# Conserved patterns across ion channels correlate with variant pathogenicity and clinical phenotypes

**DOI:** 10.1101/2022.03.23.485339

**Authors:** Tobias Brünger, Eduardo Pérez-Palma, Ludovica Montanucci, Michael Nothnagel, Rikke S. Møller, Stephanie Schorge, Sameer Zuberi, Joseph Symonds, Johannes R. Lemke, Andreas Brunklaus, Stephen F. Traynelis, Patrick May, Dennis Lal

**Author notes:** Correspondence to: Dennis Lal, 9500 Euclid Ave, NE-50, Cleveland, OH 44195.

## Abstract

Clinically identified genetic variants in ion channels can be benign or cause disease by increasing or decreasing the protein function. Consequently, therapeutic decision-making is challenging without molecular testing of each variant. Our biophysical knowledge of ion channel structures and function is just emerging, and it is currently not well understood which amino acid residues cause disease when mutated.

We sought to systematically identify biological properties associated with variant pathogenicity across all major voltage and ligand-gated ion channel families. We collected and curated 3,049 pathogenic variants from hundreds of neurodevelopmental and other disorders and 12,546 population variants for 30 ion channel or channel subunits for which a high-quality protein structure was available. Using a wide range of bioinformatics approaches, we computed 163 structural features and tested them for pathogenic variant enrichment. We developed a novel 3D spatial distance scoring approach that enables comparisons of pathogenic and population variant distribution across protein structures.

We discovered and independently replicated that several pore residue properties and proximity to the pore axis were most significantly enriched for pathogenic variants compared to population variants. Using our novel 3D scoring approach, we showed that the strongest pathogenic variant enrichment was observed for pore-lining residues and alpha-helix residues within 5Å distance from the pore axis center and not involved in gating. Within the subset of residues located at the pore, the hydrophobicity of the pore was the feature most strongly associated with variant pathogenicity. We also found an association between the identified properties and both clinical phenotypes and fucntional *in vitro* assays for voltage-gated sodium channels (*SCN1A*, *SCN2A*, *SCN8A*) and N-methyl-D-aspartate (NMDA) receptor (*GRIN1*, *GRIN2A*, *GRIN2B*) encoding genes. In an independent expert-curated dataset of 1,422 neurodevelopmental disorder pathogenic patient variants, and 679 electrophysiological experiments that pore axis distance is associated with seizure age of onset and cognitive performance as well as differential gain vs. loss-of-channel function.

In summary, we identified biological properties associated with ion-channel malfunction and show that these are correlated with *in vitro* functional read-outs and clinical phenotypes in patients with neurodevelopmental disorders. Our results suggest that clinical decision support algorithms that predict variant pathogenicity and function are feasible in the future.

## Introduction

Ion channels are membrane proteins that act as gated pathways for the movement of ions across cell membranes. They play essential roles in the physiology of all cells. At least 130 ‘channelopathies’ have been associated with genetic variants causing ion channel dysfunctions across the human body, including diseases in the nervous system (e.g., epilepsy, familial hemiplegic migraine), cardiovascular system (e.g., long QT syndrome, Brugada syndrome), respiratory system (e.g., cystic fibrosis) and endocrine system (e.g., neonatal diabetes mellitus)^1,2^.

Ion channels malfunction due to missense variants that alter the protein sequence can cause a wide spectrum of disorders^1,3^. Missense variants even within the same gene can produce different molecular effects in ion channels: increased or decreased ion permeation; dysregulation of gating elements; changes in opening and closing probabilities; altered kinetics and protein trafficking^4–9^. Not all missense variants found in human genetic studies are pathogenic^10^. The location of missense variants on protein structure has been associated with variant pathogenicity^11–13^. In addition, several recent studies showed that among pathogenic variants, the variant position in specific functional units correlates with the patient sub-phenotype^14–17,18(p1),19–21^. However, the knowledge about microdomain to phenotype associations is sparse and typically only available to experts of specific channels and cannot easily be adapted by a wider audience. Variant interpretation of less studied and recently identified channelopathies represents a major challenge.

In this study, we sought to identify biological properties associated with variant pathogenicity that are shared among all ion channels. These properties could be incorporated into future algorithms for predicting variant pathogenicity and, thus, provide insights to genotype-phenotype correlations in established and newly identified channelopathies regardless of the ion channel class. To accomplish this goal, we performed a series of association analyses for biological properties with 12,546 population and 3,049 pathogenic variants across 30 ion-channel structures. This work led to the development of a novel bioinformatic structure-based framework which showed that the distance of a residue from the central pore axis and pore hydrophobicity are the most pathogenicity-associated features shared across all ion channels. We applied our framework to two independent data sets comprising >1400 patient variants and ~700 functionally tested variants, for the NMDA receptor and voltage-gated sodium channels to demonstrate how correlations between the localization of variants and their associated molecular effect and clinical outcome can be captured.

## Material and Methods

### Collection of ion channels and protein structure selection

We collected all voltage and ligand-gated ion channels from IUPHAR^22^ (*n* = 172 genes). Out of these channels, we included those in our study that harbor at least one known pathogenic variant in the ClinVar^23^ database and whose encoded protein structures have been determined at atomic resolution and to form a pore in the protein data bank^24^. For each channel, the protein structure with the largest protein sequence coverage was chosen. All selected ion channels were divided into evolutionary derived channel families according to HGNC^25^ criteria.

### Missense variant annotation

Canonical transcripts for proteins encoded by the ion channel genes in our cohort were accessed from the UniProt^26^ database. For these transcripts, variants were collected from multiple databases. In particular, protein-coding missense variants from the general population were retrieved from the genome aggregation Database^10^ (gnomAD, https://gnomad.broadinstitute.org/, public release 2.0.2) including all ethnic backgrounds. Variant Call Format (VCFs) files were downloaded for all available exomes of ion channel genes to access the variants. These “population variants” served as control dataset in subsequent enrichment and association analyses. The extraction of missense variants (Filter = “PASS”) was performed with vcftools (version v0.1.12b) using the pre-annotated “CSQ” field. ClinVar^23^ variants were accessed from the ftp site (ftp://ftp.ncbi.nlm.nih.gov/pub/clinvar/, access July 2020). Only variants with a clinical consequence (annotated as “Pathogenic”/“Likely Pathogenic”) were extracted. Next, we obtained missense variants from the Human Gene Mutation Database (HGMD^27^) and filtered them for high-confidence calls (hgmd_confidence = “HIGH” flag) as well as disease-causing states (hgmd_variant_Type = “DM” flag) (HGMD, July 2020). Variants from ClinVar and HGMD databases were combined into a pathogenic variant data set containing unique amino acid substitutions. We applied randomized sampling to create separate subsets of the collected variants to obtain discovery and validation cohorts containing each 70% and 30% of the pathogenic and population variants, respectively.

### Collection of missense variants with detailed clinical phenotype or known molecular effect

We collected missense variants with known clinical phenotypes for three sodium channels (*SCN1A, SCN2A, SCN3A*) from the SCN Portal^28^ (https://scn-portal.broadinstitute.org, accessed 01 December 2021). Missense variants with known clinical phenotype for three genes that encode ionotropic glutamate receptors (*GRIN1A, GRIN2A, GRIN2B*) were obtained from the GRIN Portal^29^ (https://grin-portal.broadinstitute.org, access 01 December 2021). In addition, we obtained missense variants across all voltag-gated sodium channel encoding genes (*SCN1A*-*SCN11A*) with a reported molecular effect from Brunklaus et al.^21^. Variants were categorized as either gain-of-function (GoF), loss-of-function (LoF), or mixed-function (mixed) function depending on the electrophysiological readouts. In addition, missense variants with a functional effect in any of the GRIN genes had been obtained (CFERV^30^, http://functionalvariants.emory.edu/, GRIN Portal^29^).

### Collection and annotation of features

We annotated features that describe the localization of residues in functional regions of the protein as well as their protein structure (Supplementary Fig. 1). Therefore, we first mined the UniProt^26^ database (https://www.uniprot.org/, access 02.05.2021) to collect the location of the amino acids inside the protein (intracellular, membrane-spanning, or extracellular) and whether a residue is part of a specific unit (voltage-sensor, allosteric-, agonist or other binding sites). The secondary structure of amino acid residues was calculated using the DSSP^31^ (dictionary of protein secondary structure) program. Protein-protein interactions were obtained from the PDBsum^32^ database (access 17.05.2021). We further considered two differently derived pore annotations to capture residues that are predicted to be located at the pore. First, a residue was considered to be part of the protein according to its annotation in the Uniprot database. Second, we predicted the pore using the Mole2.0 webserver^33^ (https://mole.upol.cz/, access 03.04.2021) and obtained the annotation of those residues that were predicted to be pore lining. The resulting 13 features (Supplementary Fig. 1) were arranged into five main categories: whether a residue is located at the pore, whether a residue is located in the membrane, intracellular or extracellular space, the secondary structure type of the residue, the association of a residue to a functional site (e.g. a binding site), and whether a residue is reported to be involved in any protein-protein interaction. Using the features, we created different sets of residues, where all residues within one set were described by the same feature, e.g. all residues being located in the membrane. Furthermore, we created additional residue sets as combination of multiple features, e.g. a set of residues being located intracellular and simultaneously forming an alpha-helix secondary structure. To this end, we considered all possible combinations of two or more features, resulting in 163 different residue sets harboring at least one residue.

### Quantification of the residue location

To describe the localization of each channel residue with respect to its distance to the membrane center and pore axis, we predicted the membrane for each ion channel protein with the PPM^34^ server (https://opm.phar.umich.edu/ppm_server, version PPM 2.0) and obtained the location of the non-membrane spanning part (intracellular or extracellular) from Uniprot^26^. We then calculated the distance of the C-alpha atom of each protein residue to the membrane center and assigned a positive distance to all residues oriented towards the intracellular part of the protein as well as negative distance values for residues located in the extracellular space (Fig. 3A). Residues with an assigned distance of 0 were located at the membrane center. Similarly, we used the previously predicted pore dimensions (see annotation of features above) to calculate the distance of each C-alpha atom to the pore axis and normalize the distance for each gene between 0 and 1. We then divided each protein structure into 200 2D bins that group residues together based on their distance from the pore axis and from the membrane center and analyzed the enrichment of pathogenic compared to population variants in the residues of each bin.

**Figure 1.**
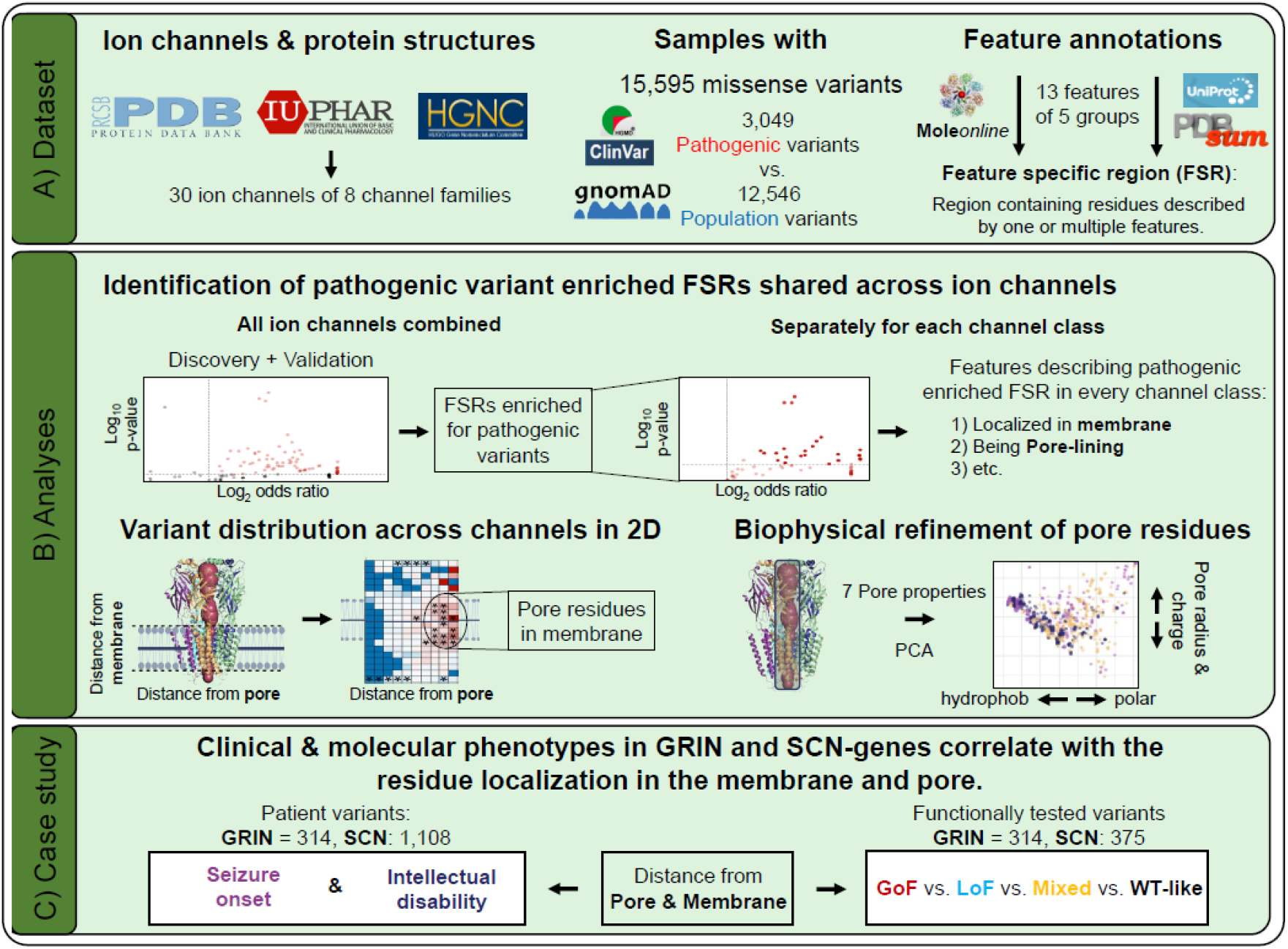
Graphical summary of the study. **(A) Dataset**. First, a subset of ion channels was selected from IUPHAR and screened for available protein structures. Second, missense variants were assembled from three different databases: gnomAD, ClinVar, and HGMD. Third, structural features were annotated to describe the location and secondary structure of all residues comprising the ion channels. **(B) Analyses**. Sets of residues described by a single feature or a combination of features that were enriched for pathogenic variants were identified across all ion channels. A subset of these features was identified to show enrichment for pathogenic variants across all considered channel families. Based on these shared features a two-dimensional reference framework was developed for describing the location of a residue with respect to the distance from the membrane and the pore. Using this framework, the highest variant burden was observed in pore residues that were located inside the membrane. Finally, within these pore residues, the impact of the biophysical pore environment on the variant burden was investigated. **(C) Case studies)** Our framework identified correlations between functional and clinical phenotypes and the localization of missense variants in the protein structure. PCA: Principal component analysis. GRIN: *GRIN1*, *GRIN2A*, *GRIN2B* genes; SCN: *SCN1A*, *SCN2A*, *SCN8A* genes. GoF: gain-of-function variant; LoF: loss-of-function variant; Mixed: Electrophysiological readouts showed conflicting functional changes; WT: wildtype.

**Figure 2.**
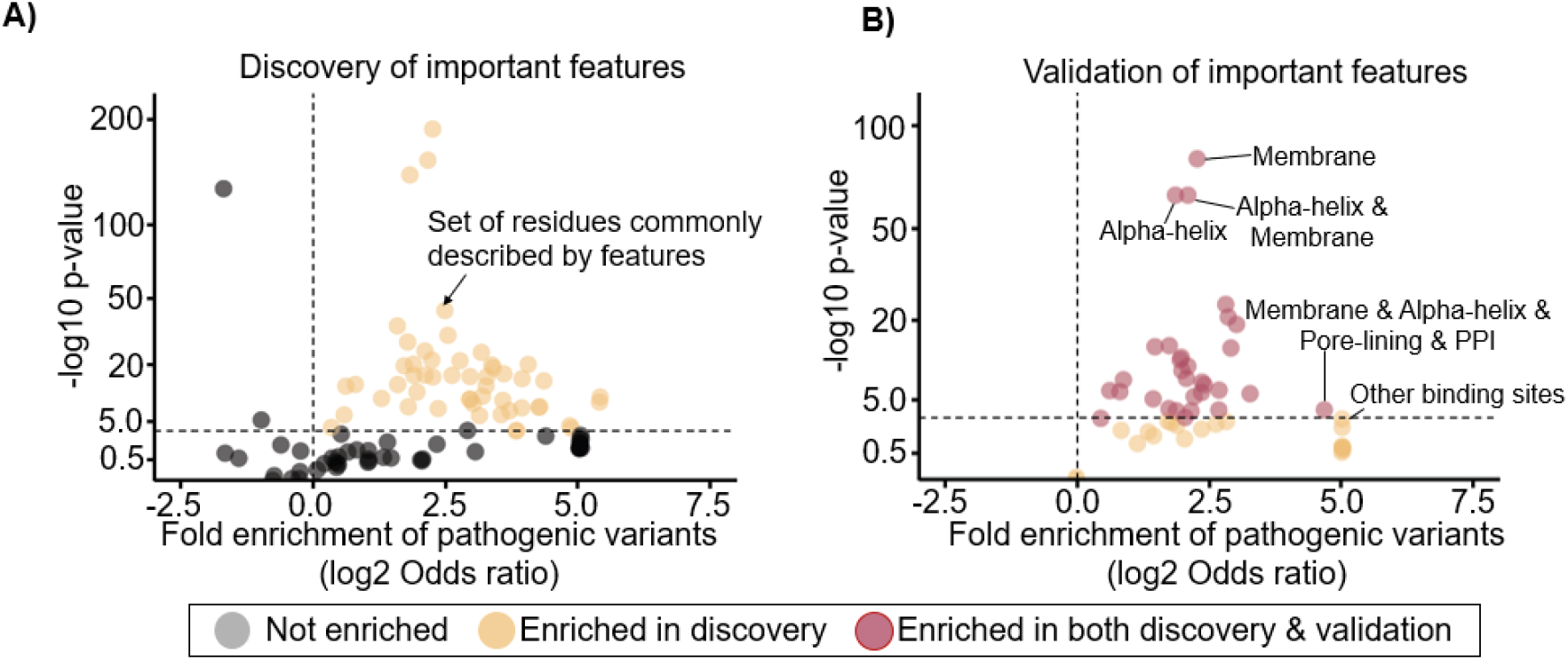
Identification of single features and feature combinations that are enriched for pathogenic variants. **(A)** Scatterplot of the odds ratio (x-axis) vs. the p-value (y-axis) of the variant burden analysis of pathogenic vs. population variants in each of the 163 different sets of residues described by one feature or a feature combination in the discovery cohort (Supplementary Fig. 1). Residue sets with significant enrichment of pathogenic variants after multiple-testing corrections (Fisher's exact test, odds ratio (*OR*) > 1, *P* < 0.05) are displayed in orange, the remaining ones in grey. (**B**) Scatterplot of variants in the validation cohort. Only the subset of the 55 features and feature combinations that were found to be enriched for pathogenic variants in the discovery cohort is shown. Residue sets enriched for pathogenic variants in both the discovery and validation cohort are displayed in red. Residue sets enriched only in the discovery cohort are displayed in orange.

**Figure 3.**
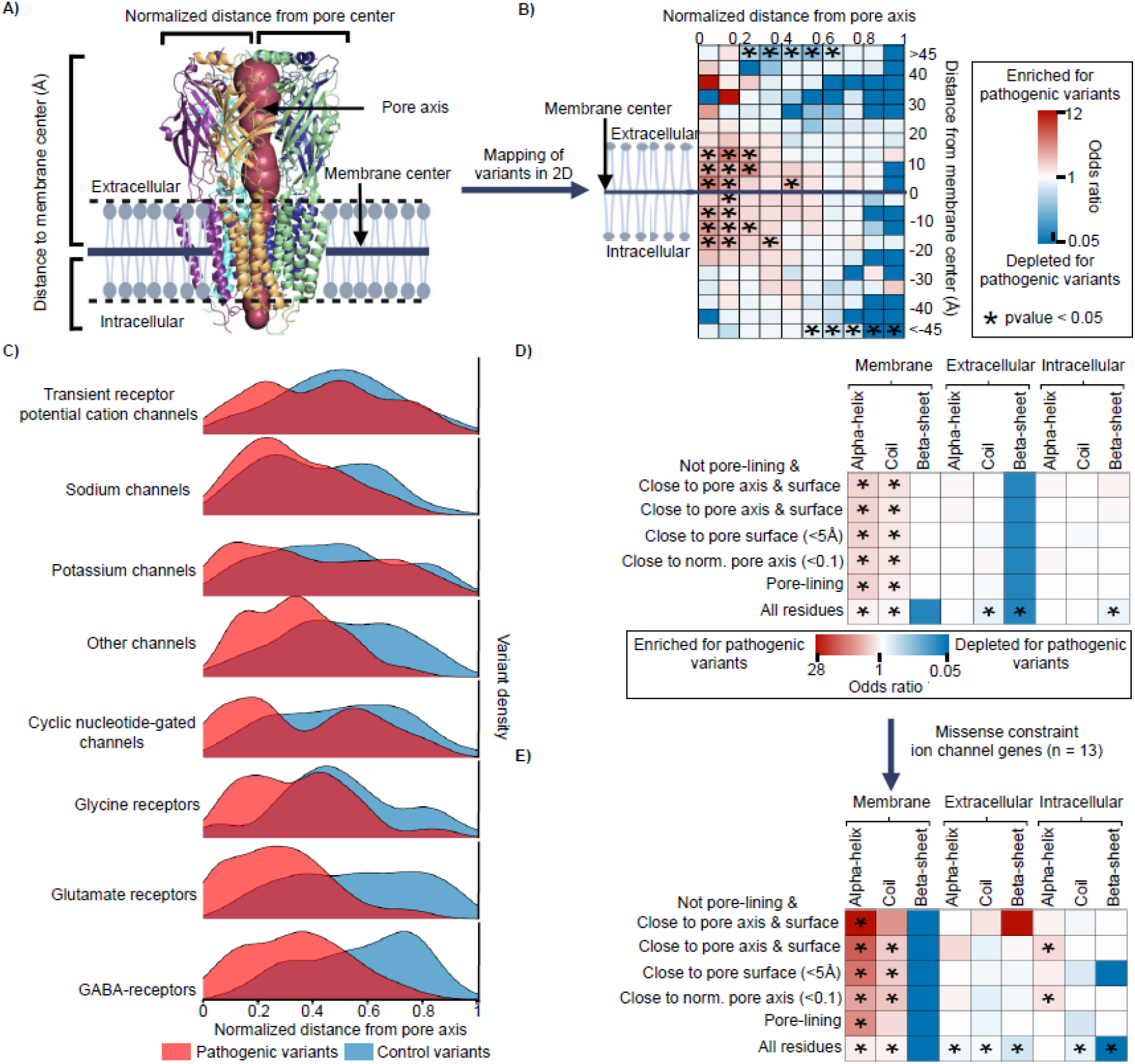
Residue distance from pore and membrane correlates with the pathogenic burden, harboring the highest burden at membrane-spanning pore residues. **(A)** Graphical representation of the framework that describes the localization of each residue in ion channel proteins on two dimensions (2D). The localization of each residue was depicted by the gene-wise normalized distance from the pore axis and the distance from the membrane center, thereby allowing a comparison of the variant distribution across ion channels. **(B)** Table of the combined enrichment or depletion of pathogenic variants across all ion channel proteins summarized over 200 different 2D regions. Each bin represents a distinct localization in the protein structure that is described by the normalized distance from the pore axis (x-axis) and the distance from the membrane center (y-axis). Bins enriched or depleted for pathogenic variants are colored in red and blue, respectively. Bins with neither depletion nor enrichment for pathogenic variants are colored white. Significant enrichments or depletions of pathogenic variants are indicated with a star. **(C)** The variant densities of pathogenic variants (red) and population variants (blue) are displayed along with the normalized distance from the pore axis, separately for each channel class (*n* = 8). **(D)** Table of the enrichment or depletion of pathogenic variants in different sets of pore residues. Each set of pore residues was defined by their location (membrane, extracellular, intracellular) and secondary structure (x-axis), together with a description of which residues were considered as pore residues (y-axis). Red and blue cells indicate an enrichment and depletion of pathogenic variants, respectively. Bins labeled with white color are neither enriched nor depleted for pathogenic variants. Significant values are indicated by a star. (**E**) Table as in D but based on a subset of channel genes that are constrained for variants in the general population (n = 13 genes).

### Annotations of different pore parts

To evaluate the variant distribution across different pore parts, we defined pore residues as: 1) residues that are pore-lining as defined by Mole2.0 webserver^33^ (https://mole.upol.cz/, access 03.04.2021), 2) residues that are close to the pore axis (normalized pore distance <0.1), and 3) residues that are close to the pore surface (pore surface distance <5A). To obtain the distance to the pore surface, we considered, as before, C-alpha atoms and calculated the closest distance of each residue to the pore surface. We then combined these pore annotations with the descriptions of the secondary structure and the general localization (membrane, intracellular, extracellular).

To study the pore properties and their correlation with the pathogenic variant burden, we assigned to each pore residue, defined as a residue that is either pore-lining, close to the pore center (normalized pore distance <0.1) or close to the pore surface (pore surface distance < 5), seven biophysical pore properties (pore radius, lipophilicity, hydropathy, hydrophobicity, polarity, water-solubility, charge) as annotated with the Mole webserver^33^. We always assigned the pore property of the closest pore segment to each pore residue.

### Statistical analysis

If not otherwise stated, we used a two-by-two contingency table to quantify the burden of pathogenic or population variants across different sets of residues, using the odds ratio (OR), augmented by the 95% confidence interval (CI), as an estimate for the enrichment. We used a two-sided Fisher’s exact test to test for the association while adjusting for multiple testing using Bonferroni correction. For the identification of important features (single features or feature combinations that describe a set of residues being enriched for pathogenic variants), a randomly subsampled set maintaining 70% of the population and pathogenic variants was considered as a discovery dataset. The remaining set consisting of each 30% population and pathogenic variants formed the validation dataset that was used to confirm the important single features and feature combinations identified in the discovery dataset.

A principal component analysis (PCA) was performed with the stats package (version 3.6.2) to aggregate the variance of the seven different physiochemical pore features across the pore residues. All statistical analyses were performed in R version 4.1.2.

### Data availability

Residue wise annotations used in this study can be obtained from the supplementary material.

## Results

### Identification of pathogenic variant associated residue features across ion channels

We sought to identify features associated with variant pathogenicity in all ion channels (Fig.1). Therefore, we collected 3,049 pathogenic and likely pathogenic classified variants from ClinVar^23^ and HGMD^27^ (term combined as ‘pathogenic’) and 12,546 population missense variants from the gnomAD database^10^. Of these, 2,543 (83.1%) pathogenic variants and 5,967 (47.6%) population variants could be mapped onto the protein structure of 30 ion channel proteins (see Methods for details, Supplementary Table 1). We calculated the relative burden of pathogenic over population variants, which served as controls in different residue sets.

For each amino acid residue, we computed 13 features that captured its structural and functional context (Supplementary Fig. 1). These features are mainly independent of each other and include the secondary protein structure, the localization within a functional site (e.g., residues located at a ligand-binding site), and within structurally defined protein regions (e.g., residues located at the pore or inside the membrane) (Supplementary Fig. 1A, Supplementary Fig. 2). We then used these features to identify residue characteristics that are associated with pathogenic genetic variants compared to population variants (Fig. 1) across all channel proteins. Groups of residues that share the same features (either one or multiple features) are here referred to as ‘residue sets’ (Supplementary Fig 1B, Methods). For example, one residue set contains all residues that form helices (10,500 residues). Furthermore, a set of residues can be also described by a combination of features. For example, 104 residues are all 1) located at the pore, 2) form helices and 3) form protein-protein interactions. The combinatorial assignment of the 13 features resulted in the identification of 163 different residue sets. We then investigated if there is an enrichment of pathogenic variants in each of the 163 residue sets (Supplementary Table 2). In the discovery screen, where we used a randomly assigned subset containing each 70% of the curated pathogenic and population variants, we found that 52 residue sets were enriched for pathogenic compared to population variants (Fisher’s exact test, Bonferroni-adjusted *P* < 0.05 for 163 tests, Fig. 2A). Of those 52 residue sets, 31 (60%) were enriched for pathogenic variants also in the validation screen, comprising the remaining 30% of variants (Fig. 2B). The highest variant enrichment of pathogenic variants was observed for the set of residues that combines ‘pore lining’, ‘located in the membrane’, ‘in an alpha-helix secondary structure’, and ‘being involved in protein-protein interactions’ (*OR* = 34.5, *P* = 3.6e-19). This residue set contains residues from 11 out of the 30 channel genes (37%), including genes encoding for gamma-aminobutyric acid (GABA) receptors, glycine receptors, N-methyl-D-aspartate (NMDA)-receptors, potassium channels, and transient receptor potential cation channels.

After identification of the residue sets most strongly associated with pathogenic variants across the full dataset of 30 ion channels, we sought to identify the subset of residue sets that are consistently associated with pathogenicity across all ion channels. Since pathogenicity of homologous residues is similar across gene families^35^, we categorized our set of ion channels into eight evolutionary derived families and assessed the residue variant burden for each residue set in every channel family. As to residue sets, the lowest number of residue sets that were enriched for pathogenic variants was found in the glycine receptor family (*n* = 9), while the highest number was found in the glutamate receptor family (*n* = 28). As to residue features, we identified four features that contributed, individually or in combination, to all residues sets enriched for pathogenic variants in all eight ion channel families, namely ‘located in the membrane’, ‘localization at the pore’, ‘alpha-helix secondary structure’ and ‘coiled secondary structure’.

### Spatial distance to pore is associated with variant pathogenicity across all ion channels

The previous analysis identified four features consistently and prominently contributing to variant pathogenicity across all ion channel families. Out of these, we observed the highest enrichment of pathogenic variants in residues that are located in the membrane (*OR* =4.8, *P* = 1.8e-273) and in residues that are pore-lining (*OR* = 3.8, *P* = 1.8e-28). To study the spatial variant localization in association with variant pathogenicity in more detail, we developed a two-dimensional framework that normalizes the localization of each residue for each ion channel relative to its distance from the pore axis and the membrane center (Fig. 3A, Supplementary Table 3 and Methods). This approach enabled us to compare the localization of variants in relation to pore and membrane distance across all ion channels, independently of their structural and sequence similarities. Using this novel approach, in the combined ion channel data set, we identified the strongest significant pathogenic variant enrichment in localizations close to the pore axis and to the membrane center, respectively (*OR* = 6.2, *P* = 8.8e-08, Fig. 3B). For each ion channel class, we observed a strong correlation between a greater relative number of pathogenic vs. population variants with closer distance to the pore axis (Pearson correlation ρ = −0.98, *P* = 8.0e-07, Fig. 3C). In contrast, no significant correlation was found between the greater relative number of pathogenic vs. population variants and the localization of the variant with respect to the membrane center (Pearson correlation ρ = −0.20, *P* = 1).

Since the highest-burden of pathogenic variants was observed closest to the pore axis, we investigated the variant burden in different subsets of residues that were located at the pore, referred to as pore residues, to identify whether one subset of residues contributed most to the pathogenic burden. Residues were considered as pore residues if they were closely located to the normalized pore axis, pore-lining, or pore surface (<5Å), to capture also pore residues of pore section with a wide pore radius. We observed enrichment for pathogenic variants in all subsets of pore residues that were located inside the membrane and that form a helix or a coiled secondary structure (Fig. 3D). Among those, the highest pathogenic burden was observed in helix forming residues that were close to the pore surface and close to the pore axis (*OR* = 5.6, *P* = 1.44e-24). However, a similar pathogenic burden was observed in membrane-located pore-lining residues forming alpha helices (*OR* = 5.5, *P* = 2.6e-13, Fig. 3D), indicating that channel residues close to the pore may be similarly prone to disease-causing variants than pore-lining residues. Interestingly, the enrichment for pathogenic variants in these pore residues differed highly between channel families, having the lowest enrichment in sodium channels (*OR* = 2.25, *P* = 2.1e-05) and the highest one in GABA receptors (*OR* = 54.5, *P* = 3.2e-11).

Notably, we observed that the fold enrichment for pathogenic compared to population variants is up to five times higher in a subset of ion channel genes that are depleted for missense variants (*n* = 13 genes, missense-z score^36^ > 3.09, Fig. 3E). This subset comprises pore residues that are close to both the pore axis and the membrane surface, but which are not pore-lining (*OR* = 27.3, *P* = 1.9e-07). Within the set of constraint ion channel genes, we identified a total of 116 pathogenic variants (7.1% out of 1,641) in pore residues that are located inside the membrane whereas only 13 population variants (0.3% out of 4,179) were found at these residue positions.

### Hydrophobic pore sections contain a higher density of pathogenic vs. population variants

Next, after observing that the enrichment of pathogenic vs. population variants correlates with the distance from the pore, we asked whether the pathogenicity of a variant also correlates with more detailed biophysical properties of the pore. Thus, we computed seven biophysical pore properties which, for each pore section, summarise its biophysical environment, namely polarity, water-solubility, hydropathy, hydrophobicity, ionizability, charge, and pore radius (Fig. 4A, Supplementary Table 3, see methods for details). Then we assigned to each pore residue these properties of the pore section in which the residue is located. Given the high correlation between these biophysical pore properties (Supplementary Fig. 3), we performed a principal component analysis to transform them into a set of equivalent but non-correlated variables (the principal components or PCs). The first two PCs discriminated majorly hydrophobic and polar pore sections (PC1, 60.4 % variance explained) and the pore radius and charge (PC2 14.7% variance explained). Among the seven biophysical properties, hydrophobicity, hydropathy, and water solubility made the highest contribution to the first two PCs (as indicated by the length of the projection of the arrow on the x and y-axis in Fig. 4B). Hence, small PC1 values describe hydrophobic pore sections whereas larger PC1 values characterize a polar and water-soluble pore environment. We mapped the pathogenic and population variants along PC1 and PC2 and observed enrichment of pathogenic variants compared to population variants at hydrophobic pore sections (median PC1 pathogenic variants: −0.12, median PC1 population variants: 0.14, P = 1.9e-06, Fig. 4C), indicating a correlation between the hydrophobicity of the pore and the pathogenicity of the pore sections.

**Figure 4.**
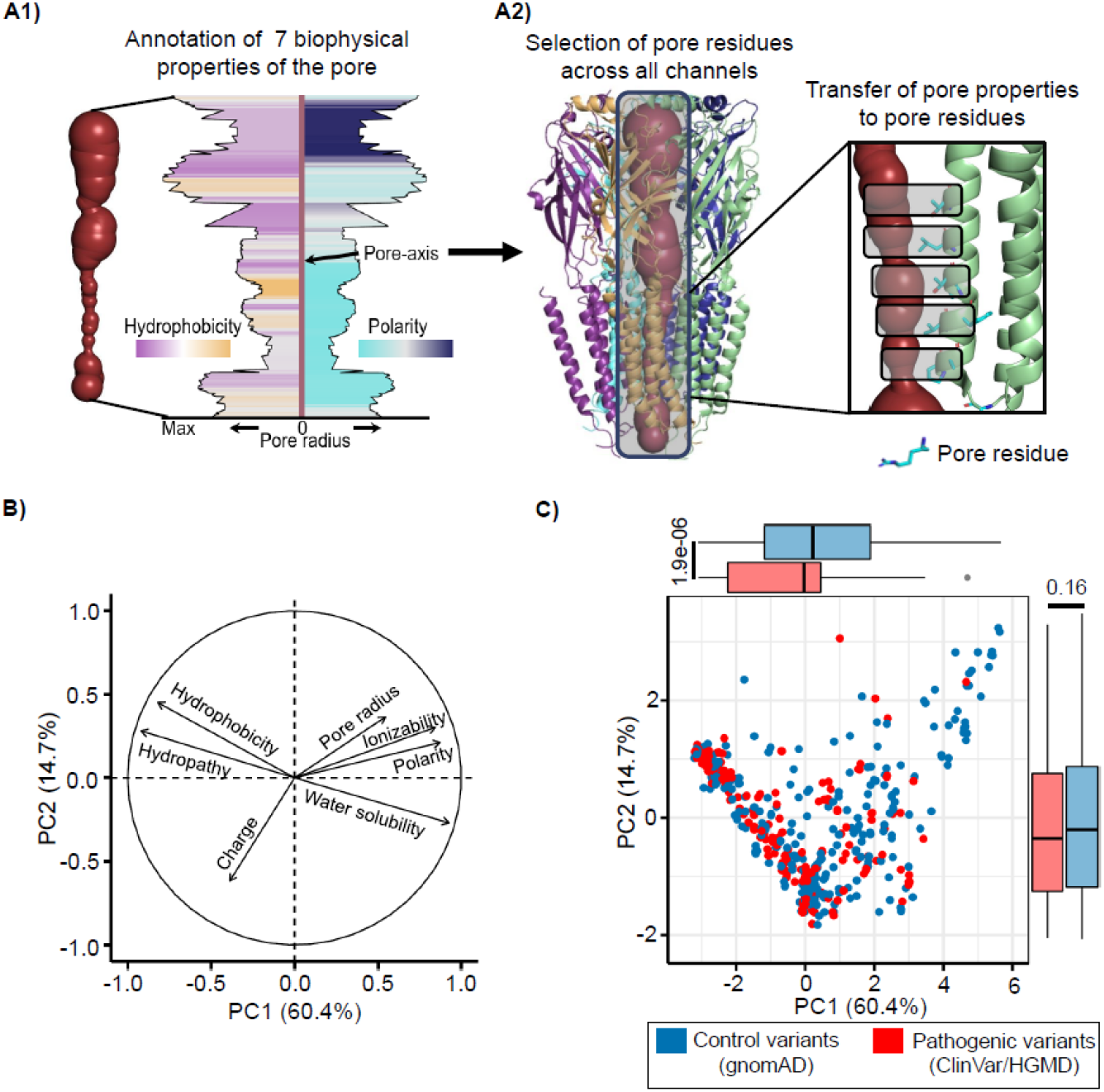
Biophysical pore properties discriminate pathogenic and population variants and identify residues that are most likely to harbor pathogenic variants in the hydrophobic pore sections. (**A1**) Graphical representation of the different biophysical pore properties for each pore section along the pore axis. (**A2**) Each pore residue is assigned to the biophysical properties of its closest pore section. (**B**) Contribution of the seven considered biophysical pore properties to the first (x-axis, PC1) and second (y-axis, PC2) dimension of the principal component analysis (PCA) that was performed on the pore properties together for all ion channel proteins. (**C**) Scatterplot along the first two dimensions of the PCA (PC1 and PC2). Each dot represents a pore residue where a pathogenic variant (ClinVar/HGMD, red) or population variant (gnomAD, blue) was observed. Pore residues where population and pathogenic variant were observed are not shown.

### Localization of missense variants correlates with functional effect and clinical phenotype

Next, we explored whether the localization of missense variants can inform about the functional effect or clinical phenotype in two independent and structurally different examples. We collected 1,104 and 314 pathogenic missense variants with curated clinical data across three voltage-gated sodium channel encoding genes (SCN genes: *SCN1A, SCN2A, SCN8A*) and three genes that encode N-methyl-D-aspartate receptor (NMDA) receptors (GRIN genes: *GRIN1, GRIN2A, GRIN2B*), respectively. Most of the residues in sodium channels are located inside the membrane, whereas NMDA receptors have two large extracellular domains on top of their membrane-spanning part. Both ion channel families have been associated with clinical and molecular heterogeneous neurodevelopmental disorders with epilepsy, intellectual disability, and other neurological disorders^20,37^. For the six selected channels, the patient variants clustered within and close to the pore whereas population variants tended to be located at the surface of the protein (Fig. 5A, D). Pathogenic variants in the selected GRIN genes that are associated with an early seizure onset were localized closer to the pore axis than variants associated with a late-onset (*P* = 0.001, Fig. 5F). Pathogenic variants in SCN genes that were observed in patients reporting intellectual disability (ID) tend to be closer to the pore (*P* = 3.8e.04, Fig. 5B) than pathogenic variants observed in patients without ID. A similar pattern was observed for the GRIN genes (*P* = 5.8e-04, Fig. 5E).

**Figure 5.**
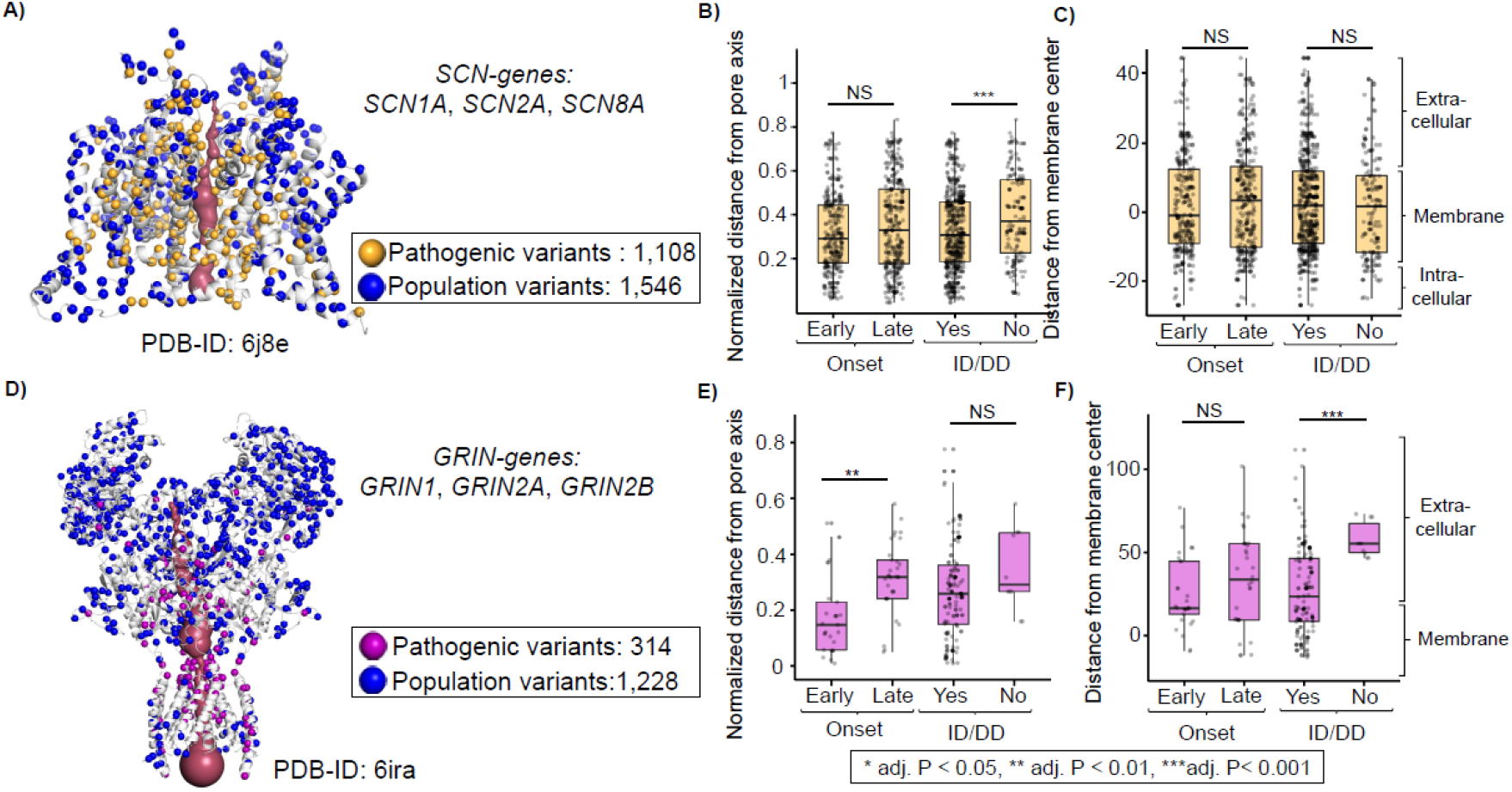
Distances to membrane and pore correlate with the clinical representation in sodium channels and NMDA-receptors. (**A**) Nav1.2 protein structure (PDB-ID: 6j8e) encoded by *SCN2A* showing patient variants and population variants observed in *SCN1A*, *SCN2A,* and *SCN8A* (SCN-genes) that were aligned to *SCN2A.* Pathogenic SCN variants were curated from the SCN Portal, population variants from gnomAD. (**B**) Boxplot of the distribution of patients’ variants grouped by their clinical phenotypes along with the normalized distance from the pore center. Boxes represent patient variants that were associated with an early seizure onset (seizure onset < median onset) or a late seizure onset (seizure onset > median onset) or patient variants associated with intellectual disability (ID) or developmental delay (DD). (**C**) Boxplots showing the same groups as in C, but along the distance to the membrane center. (**D**) Heterotetrameric protein complex consisting of two Glu1N and two Glu2NA subregions (PDB-ID: 6ira) encoded by *GRIN1* and *GRIN2A* respectively. Patient and population variants observed in *GRIN1, GRIN2A,* and *GRIN2B* (GRIN-genes) are visualized on the structure. Patient variants were recruited from our internal variant database, population variants from gnomAD. Variants in *GRIN2B* were aligned to *GRIN2A* and were visualized on the GluN2A subregions. (**E-F**) Boxplots as in B and C of the clinical phenotypes observed in the GRIN patient cohort.

Furthermore, we investigated the relationship between the functional effects of variants that were tested in 689 electrophysiological experiments^6,21^ in association with their localization in GRIN and SCN genes (Fig. 6A, D). Based on electrophysiological readouts variants were classified according to https://grin-portal.broadinstitute.org/ to have a gain of function (combined: likely/potential-LoF), loss of function (combined: likely/potential LoF), mixed (complex), or no effect. Variants in GRIN genes that were classified as GoF variants were located closer to the pore compared to variants classified as LoF (Wilcoxon rank-sum test: GRIN genes: p-value = 8.0e-06, Fig. 6B, E). In contrast, no significant difference was observed in variants classified as GoF and LoF in SCN genes (Wilcoxon rank-sum test: SCN genes: *P* = 0.075). Similarly, in both gene families, GoF-associated variants were closer located to the membrane center than variants classified as LoF (Wilcoxon Rank sum test: SCN-genes: *P* = 9.8e-11; GRIN-genes: *P* = 1.2e-05, Fig. 6C, F). In addition, functionally tested variants that showed no difference in their activity compared to the wildtype were only available for the set of GRIN genes and characterized by a larger distance from the membrane center (*P* = 6.9e-14) and pore axis (*P* = 6.3e-13) than variants that were annotated with any functional difference (Fig. 6E, F). For functionally tested variants located at the pore, we further explored whether the variants of different functional effects cluster in distinct biophysical pore environments. Whereas the variants in GRIN genes were scattered along with the PCs of the biophysical pore properties (Supplementary Fig. 4A), GoF classified variants in SCN genes cluster at low PC1 values (median = −2.7) and high PC2 values (median = 1.0) indicating a hydrophobic pore environment, whereas LoF variants were predominantly present at a less hydrophobic pore and charged pore environment (PC1 values median = −0.2; PC2 values median = −0.1, Supplementary Fig. 4B).

**Figure 6.**
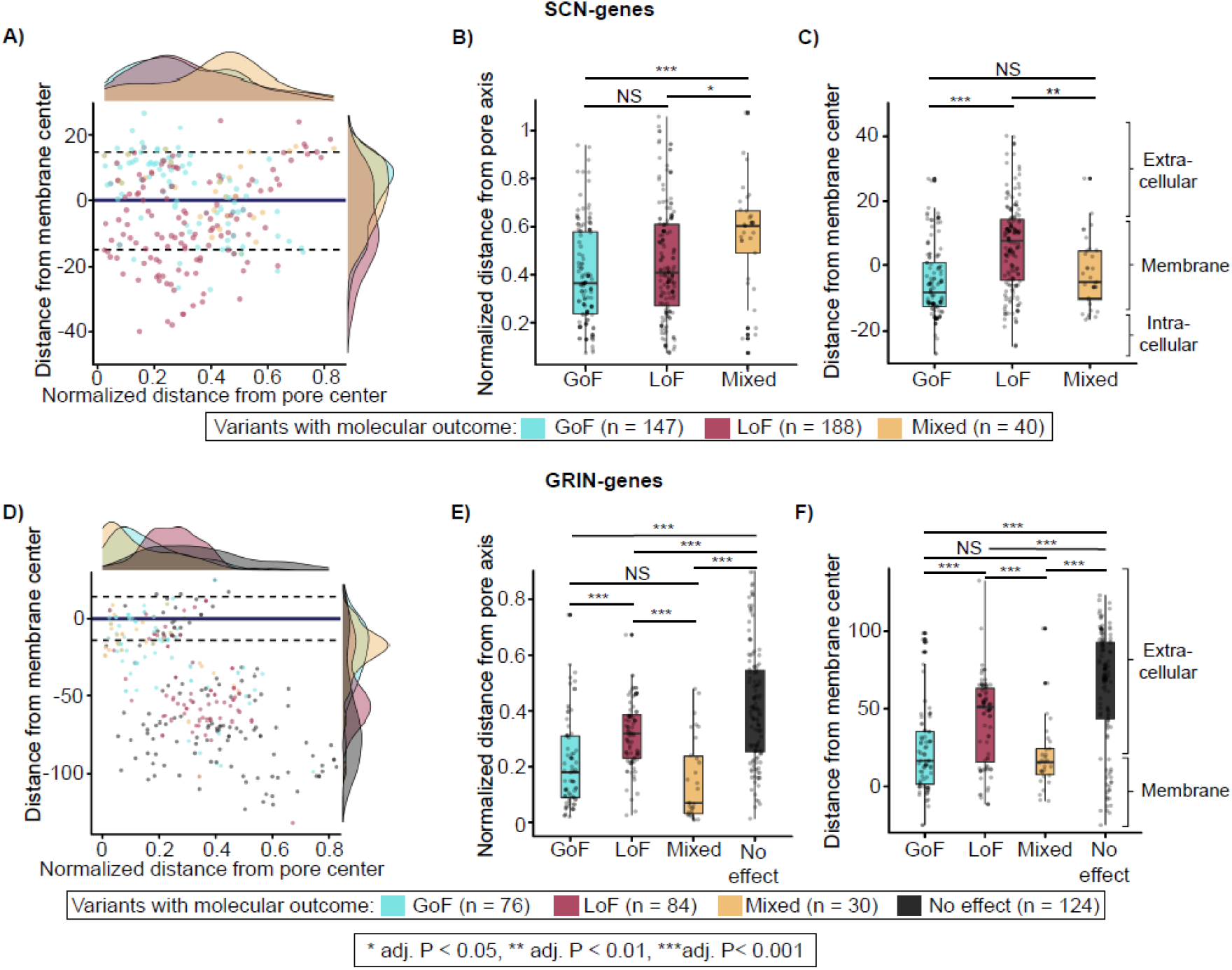
Distances to membrane and pore correlate with the molecular effect in voltage-gated sodium channels and NMDA-receptors. (**A**) Scatterplot of variants associated with a molecular effect plotted with respect to the normalized distance to the pore axis (x-axis) and the distance from the membrane center (y-axis). All readouts were assigned to the complementary *SCN2A* protein position based on a multiple sequence alignment. Variant with a mixed effect had contrary effects in different electrophysiological measurements^59^. (**B**) Boxplot of the molecular effects scattered across the normalized distance from the pore axis in SCN genes. (**C**) Boxplot of the molecular effects scattered across the distance from the membrane center in SCN genes. (**D**) Scatterplot as in A visualizing variants associated with a molecular effect in GRIN genes. (**E-F**) Boxplots as in B and C show the variants classified with a molecular effect along with the normalized distance from the pore axis and membrane center in GRIN genes.

## Discussion

We performed the first approach to systematically determine the most functionally essential structural features of residues across disease-associated ion channels. Some of the identified features are conserved across evolutionary diverse ion channels. In a proof-of-concept analysis of six neurodevelopmental disorder-associated channelopathies, we show that the newly identified conserved features are correlated with variant pathogenicity, molecular function, and clinical phenotype.

We observe that residues that are close to the pore, are located in the membrane, and form an alpha-helix, are most enriched for pathogenic variants across ion channels. Our results are in line with previous molecular biological studies, which reported ion-channels residues at the pore to be prone to disease-causing mutations^15,38,39^. For example, clusters of pathogenic variants had been found at the pore region in some specific ion channels, including voltage-gated sodium channels^40^, NMDA receptors^38^, potassium channels^15(p2)^, and GABA receptors^41^. Here, we systematically show that across 163 feature combinations characterizing amino acid residues, the pore features are most important. We were able to map these most important pore features and identify for the first time a spatially conserved pattern across eight different channel families. Although there are well characterized functional domains, such as the voltage sensor in sodium channels, that are distant from the pore and known to be enriched for pathogenic variants^39,42,43^, the overall observation that variants which are spatially more distant from the pore axis are less often pathogenic, remained stable across all eight channel families.

Structure-based bioinformatic methods that score physicochemical residue properties at the pore, such as hydrophobicity, have been shown to improve the prediction of the channel conformational states and to provide insights into the channel gating processes^44^. In our study, we observed that hydrophobicity showed the strongest association among biophysical properties with variant pathogenicity for pore residues across all investigated ion channels. Previous studies of the voltage-gated sodium channel Na_V_1.1^40^ and Na_V_1.7^45^ identified a correlation between pore hydrophobicity and variant pathogenicity. It has been suggested that a hydrophobic pore section can present an energetic barrier to ion permeation interrupting the channel gating, without having the pore physically closed due to dewetting of the hydrophobic environment, a mechanism known as hydrophobic gating^46,47^. Consequently, the variants we identified to be located at the hydrophobic pore sections may not only affect the channel activity through a possible change in the pore size but also the alterations of the hydrophobic gating^48^.

We investigated the identified pathogenicity features associated with pore axis and membrane center distance in molecular and clinical datasets. Prediction of phenotypes will guide patients and families and enable a more individualized prognosis. We showed the variant position in voltage-gated sodium channel genes (SCN genes) and NMDA receptors (GRIN genes), which are both associated with early-onset epilepsies and neurodevelopmental disorders ^43,49–52^, correlates with the functional effect and clinical phenotype. Overall, our novel spatial distance scoring approach agrees with previously reported observations that have been made on association with specific domains. For example, we find that loss-of-function variants in SCN genes are close to the pore axis in the extracellular side of the membrane, which is the region where the selectivity filter is located, whereas gain-of-function variants cluster in the intracellular site of the membrane^39^. Such knowledge of the molecular functional consequences of a variant may inform treatment decisions^53–56^. In contrast to the previous analysis, which compared the functional effect across predefined protein regions such as the voltage sensor, the S5, and S6 Linker, or the selectivity filter^39^, our novel approach has a higher spatial resolution for association analysis. Our approach can be applied to proteins in different conformational states and to ion-channel proteins where the function of domains is not well studied.

Still, our study has several limitations. First, while hundreds of genes that encode voltage- and ligand-gated ion channels have so far been identified in humans, we limited our study to only those 30 disease-associated ion channels for which 3D structural data describing proteins and protein complexes were available. Nevertheless, our study represents the most comprehensive assessment of important features across ion channels originating from eight different protein families^25^. This diversity of ion channels that are captured in our dataset suggests that our findings may also be generalizable to other ion channels, such as aquaporins, chloride channels, or piezo channels that were not part of the present study. Second, ion channels can be observed in different conformational states, namely open, closed, or inactivated conformation^57^, and this affects the location of the residues in 3D. We selected a variety of protein structures with different conformational changes, indicating that our results are likely valid for a spectrum of conformational changes. However, once human protein structures of many ion channels will be available in several conformations, our approach can be used to systematically study the structural changes and effects of the variant localization in different conformations.

We introduced a new potentially powerful approach to identify pathogenic enriched regions and identify molecular and clinical correlates across diverse ion channels. In the foreseeable future, high-quality data on protein structures and protein complexes will be available for most ion channels in the light of recent improvements in *in-silico* structure prediction tools^58^, and this will allow a wider application of our approach. Finally, our method could be expanded to other protein classes, where two features represent a horizontal and vertical axis along with the protein structure, such as membrane located transporter, to study pathogenicity and molecular and clinical phenotypes in context variant localization.

## Abbreviations

DSSP: Dictionary of Protein Secondary Structure
gnomAD: Genome aggregation Database
GoF: Gain of function
GRIN genes: *GRIN1*, *GRIN2A*. *GRIN2B*
HGMD: Human Gene Mutation Database
NMDA receptor: N-methyl-D-aspartate receptor
GABA receptor: Gamma-aminobutyric acid receptor
LoF: Loss of function
SCN genes: *SCN1A*, *SCN2A*, *SCN8A*
VCF: Variant Call Format

## Funding

Funding for this work was provided from the German Federal Ministry for Education and Research (BMBF, Treat-ION, 01GM1907D) to D.L., T.B. and P.M., the Fonds Nationale de la Recherche in Luxembourg (Research Unit FOR-2715, FNR grant INTER/DFG/17/11583046) to P.M., the Agencia Nacional de Investigación y Desarrollo (ANID, PAI77200124) of Chile to E.P., the Familie SCN2A foundation 2020 Action Potential Grant to E.P., the Dravet Syndrome Foundation (grant number, 272016) to D.L, and the NIH NINDS (Channelopathy-Associated Epilepsy Research Center, 5-U54-NS108874) to D.L.

## Competing interests

The authors report no competing interests.

## Supplementary material

Supplementary material is available.

## Notes

### Competing Interest Statement

The authors have declared no competing interest.

