## Supplementary Figures for "Conserved patterns across ion channels correlate with variant pathogenicity and clinical phenotypes"

A)

| Feature category | Feature | Source for collection or calculation |
|---|---|---|
| Protein location | Extracellular | Uniprot |
| | Intracellular | Uniprot |
| | Membrane | PPM webserver |
| Pore | Pore-lining (structure-based) | Mole 2.0 |
| | Pore (Uniprot) | Uniprot |
| Protein-Protein Interactions | Protein-Protein Interactions | PDBsum |
| Secondary protein structure | ß-strand/sheets | DSSP |
| | helices | DSSP |
| | coils | DSSP |
| Functional sites | Voltage sensor | Uniprot |
| | Allosteric binding site | Uniprot |
| | Agonist binding site | Uniprot |
| | Other binding site | Uniprot |

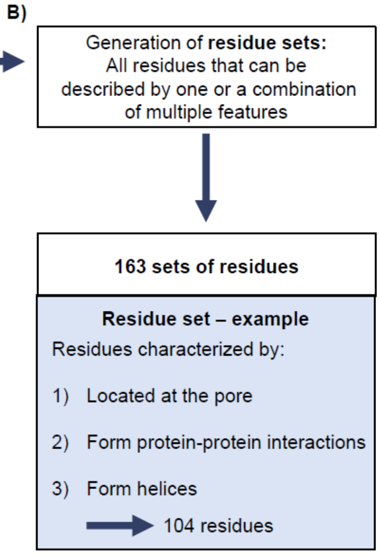


**Supplementary Figure 1: Feature-defined residue sets. A)** List of the 13 considered features along with their feature category (column one) and their source of annotation (column three). **B)** The features were used to generate sets of residues, where all residues are commonly characterized by a single feature or a combination of two or multiple features of different categories. An example for one set of residues is displayed.

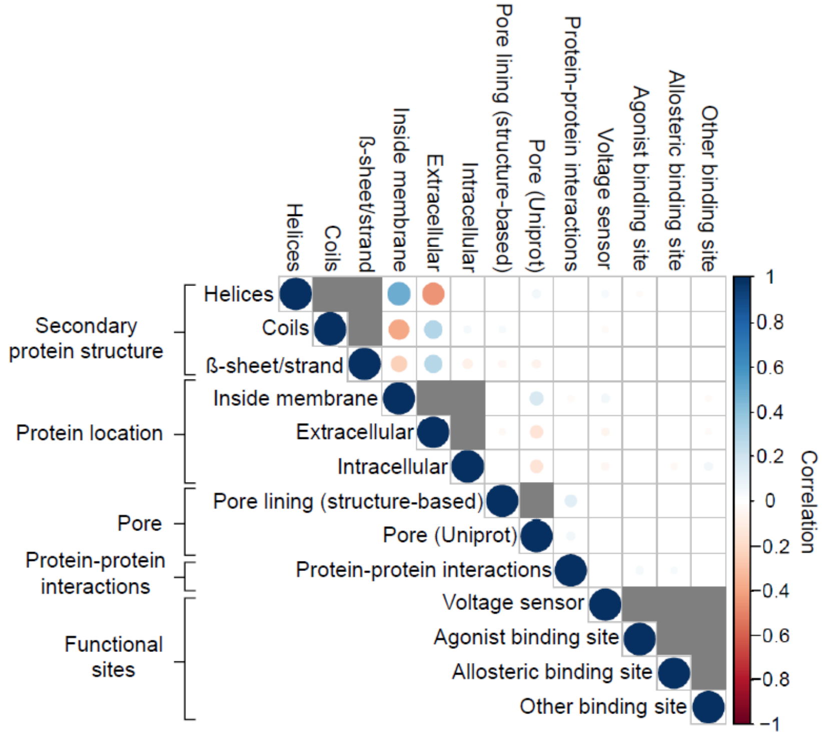


**Supplementary Figure 2: Correlation of the residue features.** The matrix depicts the correlations between the individual residue features. Correlations between features within the same feature category are not computed because they have no biological sense (for example “being in the extracellular region” and at the same time “being in the intracelluar region”) and their corresponding boxes are colored in gray (see Supplementary Figure 1, Methods). Larger spheres indicate a higher correlations. Positive correlations are shown in blue, negative correlations are shown in red. Only significant correlations (p-value < 0.05) are shown.

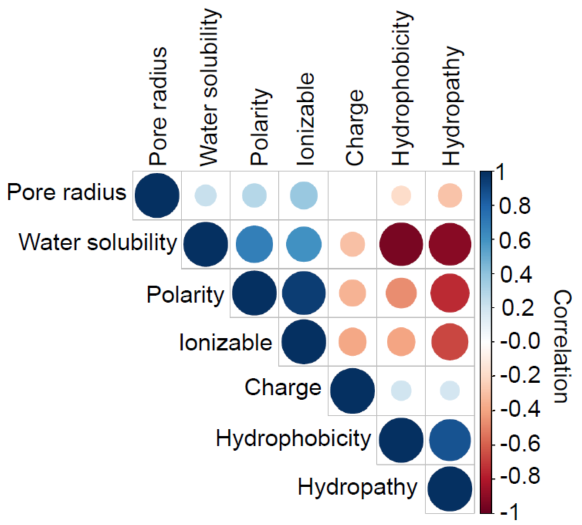


**Supplementary Figure 3: Correlation of the biophysical pore properties.** The matrix shows the correlation between the individual pore properties. Larger spheres indicate a higher correlation. Positive correlations are shown in blue, negative correlations are shown in red. Only significant correlations (p-value < 0.05) are shown.

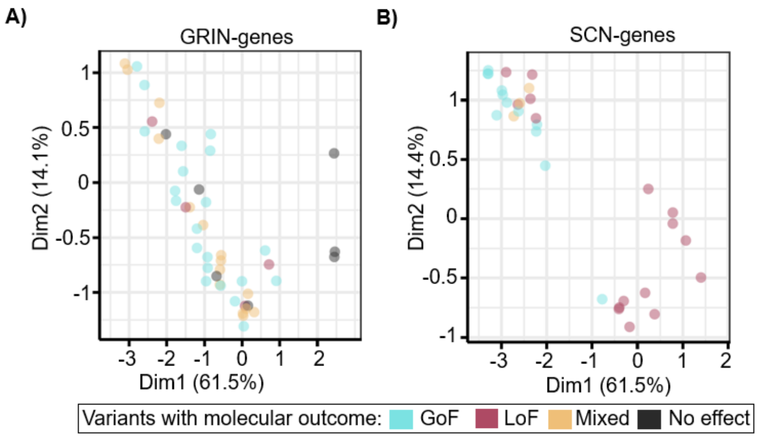


**Supplementary Figure 4: Molecular effects at pore residues differ across pore properties. A)** Scatterplot of the molecular effects in SCN genes along the first two dimensions of the PCA of the pore properties that have been performed across all ion channels (Figure 4). Variants with available electrophysiological readouts have been collected across all sodium channels and assigned to the Nav1.2 subunit, which is encoded by *SCN2A*, using a multiple sequence alignment. **B)** Scatterplot as in A, of the molecular effects in pore residues that have been collected for *GRIN2A*, *GRIN2B* and *GRIN1*.
