## Supplementary Table1 for "Conserved patterns across ion channels correlate with variant pathogenicity and clinical phenotypes"

**Supplementary Table 1:** List of all voltage-gated and ligand-gated ion channel encoding genes used in this study. Only those ion channels for which a structure of the human protein is available in the Protein Data Bank have been selected. The representative structure (PDB-ID, see methods for details) and the missense-z-score is provided for each gene.

| Genes | PDB-ID | Missense-z-score (gnomAD database) |
|---|---|---|
| CACNA1G | 6kzp | 4.64 |
| CHRNA4 | 6cnj | 0.34 |
| CHRNB2 | 6cnj | 2.1 |
| GABRA1 | 6huj | 3.15 |
| GABRA5 | 6a96 | 3.23 |
| GABRB2 | 6d6u | 3.4 |
| GABRB3 | 6i53 | 3.39 |
| GABRG2 | 6hug | 2.99 |
| GLRA1 | 2m6i | 1.42 |
| GRIN1 | 6ira | 6.22 |
| GRIN2A | 6ira | 2.83 |
| HCN1 | 5u6p | 3.66 |
| HCN4 | 6gyo | 2.75 |
| KCNK3 | 6rv3 | 3.55 |
| KCNK4 | 4wfg | 0.75 |
| KCNMA1 | 6v3g | 5.06 |
| KCNQ1 | 6uzz | 1.83 |
| KCNQ2 | 7cr7 | 4.04 |
| MCOLN1 | 5wj9 | 1.61 |
| PKD2 | 5mkf | 0.27 |
| SCN2A | 6j8e | 6.46 |
| SCN4A | 6agf | 1.56 |
| SCN9A | 6j8g | 1.06 |
| TRPA1 | 6pqo | 0.04 |
| TRPC6 | 6uza | 2.12 |
| TRPM4 | 5wp6 | 0.97 |
| TRPV3 | 6mhw | 0.17 |
| SCN1A | 7dtd | 6.04 |
| SCN5A | 7dtc | 2.75 |
| GRIN2B | 7eu8 | 5.42 |
