## Supplementary Table2 for "Conserved patterns across ion channels correlate with variant pathogenicity and clinical phenotypes"

| Supplementary Table 2: List of the 13 biological features considered in this study for each residue of each considered gene. For every residue, it is indicated whether the residue can be described (yes) or not (no) by the corresponding feature. | | | | | | | | | | | | | | | | | |
|---|---|---|---|---|---|---|---|---|---|---|---|---|---|---|---|---|---|
| | | | Collected variants | | Structural residue features | | | | | | | | | | | | |
| Gene | Protein Position | Amino acid | Population variant count (gnomAD) | Pathogenic variant count (likely-/pathogenic ClinVar/ HGMD) | Alpha helices | Coil | Beta sheet | Membrane | Extracellular | Intracellular | Pore lining | Pore (Uniprot) | Protein-Protein Interactions | Voltage sensor | Agonist binding | Allosteric binding | Unspecific binding |
| CACNA1G | 80 | N | 0 | 0 | no | yes | no | no | no | yes | no | no | no | no | no | no | no |
| CACNA1G | 81 | P | 0 | 0 | yes | no | no | no | no | no | no | no | no | no | no | no | no |
| CACNA1G | 82 | W | 0 | 0 | yes | no | no | yes | no | no | no | no | no | no | no | no | no |
| CACNA1G | 83 | F | 0 | 0 | yes | no | no | yes | no | no | no | no | no | no | no | no | no |
| CACNA1G | 84 | E | 1 | 0 | yes | no | no | yes | no | no | no | no | no | no | no | no | no |
| CACNA1G | 85 | R | 1 | 0 | yes | no | no | yes | no | no | no | no | no | no | no | no | no |
| CACNA1G | 86 | I | 0 | 0 | yes | no | no | yes | no | no | no | no | no | no | no | no | no |
| CACNA1G | 87 | S | 0 | 0 | yes | no | no | yes | no | no | no | no | no | no | no | no | no |
| CACNA1G | 88 | M | 2 | 0 | yes | no | no | yes | no | no | no | no | no | no | no | no | no |
| CACNA1G | 89 | L | 0 | 0 | yes | no | no | yes | no | no | no | no | no | no | no | no | no |
| CACNA1G | 90 | V | 0 | 0 | yes | no | no | yes | no | no | no | no | no | no | no | no | no |
| CACNA1G | 91 | I | 0 | 0 | yes | no | no | yes | no | no | no | no | no | no | no | no | no |
| CACNA1G | 92 | L | 0 | 0 | yes | no | no | yes | no | no | no | no | no | no | no | no | no |
| CACNA1G | 93 | L | 0 | 0 | yes | no | no | yes | no | no | no | no | no | no | no | no | no |
| CACNA1G | 94 | N | 0 | 0 | yes | no | no | yes | no | no | no | no | no | no | no | no | no |
| CACNA1G | 95 | C | 0 | 0 | yes | no | no | yes | no | no | no | no | no | no | no | no | no |
| CACNA1G | 96 | V | 2 | 0 | yes | no | no | yes | no | no | no | no | no | no | no | no | no |
| CACNA1G | 97 | T | 0 | 0 | yes | no | no | yes | no | no | no | no | no | no | no | no | no |
| CACNA1G | 98 | L | 0 | 0 | yes | no | no | yes | no | no | no | no | no | no | no | no | no |
| CACNA1G | 99 | G | 0 | 0 | yes | no | no | yes | no | no | no | no | no | no | no | no | no |
| CACNA1G | 100 | M | 0 | 0 | no | yes | no | yes | no | no | no | no | no | no | no | no | no |
| CACNA1G | 101 | F | 0 | 0 | no | yes | no | no | no | no | no | no | no | no | no | no | no |
| CACNA1G | 102 | R | 2 | 0 | no | yes | no | no | yes | no | no | no | no | no | no | no | no |
| CACNA1G | 103 | P | 0 | 0 | no | yes | no | no | yes | no | no | no | no | no | no | no | no |
| CACNA1G | 104 | C | 0 | 0 | no | yes | no | no | yes | no | no | no | no | no | no | no | no |
| CACNA1G | 105 | E | 1 | 0 | no | yes | no | no | yes | no | no | no | no | no | no | no | no |

| | | | | | | | | | | | | | | | | | |
|---|---|---|---|---|---|---|---|---|---|---|---|---|---|---|---|---|---|
| CACNA1G | 106 | D | 0 | 0 | no | yes | no | no | yes | no | no | no | no | no | no | no | no |
| CACNA1G | 107 | I | 0 | 0 | no | yes | no | no | yes | no | no | no | no | no | no | no | no |
| CACNA1G | 108 | A | 1 | 0 | no | yes | no | no | yes | no | no | no | no | no | no | no | no |
| CACNA1G | 109 | C | 0 | 0 | no | yes | no | no | yes | no | no | no | no | no | no | no | no |
| CACNA1G | 110 | D | 0 | 0 | no | yes | no | no | yes | no | no | no | no | no | no | no | no |
| CACNA1G | 111 | S | 0 | 0 | no | yes | no | no | yes | no | no | no | no | no | no | no | no |
| CACNA1G | 112 | Q | 1 | 0 | yes | no | no | no | yes | no | no | no | no | no | no | no | no |
| CACNA1G | 113 | R | 0 | 0 | yes | no | no | no | yes | no | no | no | no | no | no | no | no |
| CACNA1G | 114 | C | 0 | 0 | yes | no | no | no | yes | no | no | no | no | no | no | no | no |
| CACNA1G | 115 | R | 1 | 0 | yes | no | no | no | yes | no | no | no | no | no | no | no | no |
| CACNA1G | 116 | I | 0 | 0 | yes | no | no | yes | yes | no | no | no | no | no | no | no | no |
| CACNA1G | 117 | L | 0 | 0 | yes | no | no | yes | yes | no | no | no | no | no | no | no | no |
| CACNA1G | 118 | Q | 0 | 0 | yes | no | no | yes | yes | no | no | no | no | no | no | no | no |
| CACNA1G | 119 | A | 2 | 0 | yes | no | no | yes | yes | no | no | no | no | no | no | no | no |
| CACNA1G | 120 | F | 0 | 0 | yes | no | no | yes | no | no | no | no | no | no | no | no | no |
| CACNA1G | 121 | D | 0 | 0 | yes | no | no | yes | no | no | no | no | no | no | no | no | no |
| CACNA1G | 122 | D | 0 | 0 | yes | no | no | yes | no | no | no | no | no | no | no | no | no |
| CACNA1G | 123 | F | 0 | 0 | yes | no | no | yes | no | no | no | no | no | no | no | no | no |
| CACNA1G | 124 | I | 0 | 0 | yes | no | no | yes | no | no | no | no | no | no | no | no | no |
| CACNA1G | 125 | F | 0 | 0 | yes | no | no | yes | no | no | no | no | no | no | no | no | no |
| CACNA1G | 126 | A | 1 | 0 | yes | no | no | yes | no | no | no | no | no | no | no | no | no |
| CACNA1G | 127 | F | 0 | 0 | yes | no | no | yes | no | no | no | no | no | no | no | no | no |
| CACNA1G | 128 | F | 0 | 0 | yes | no | no | yes | no | no | no | no | no | no | no | no | no |
| CACNA1G | 129 | A | 0 | 0 | yes | no | no | yes | no | no | no | no | no | no | no | no | no |
| CACNA1G | 130 | V | 1 | 0 | yes | no | no | yes | no | no | no | no | no | no | no | no | no |
| CACNA1G | 131 | E | 0 | 0 | yes | no | no | yes | no | no | no | no | no | no | no | no | no |
| CACNA1G | 132 | M | 0 | 0 | yes | no | no | yes | no | no | no | no | no | no | no | no | no |
| CACNA1G | 133 | V | 0 | 0 | yes | no | no | yes | no | no | no | no | no | no | no | no | no |
| CACNA1G | 134 | V | 0 | 0 | yes | no | no | yes | no | no | no | no | no | no | no | no | no |
| CACNA1G | 135 | K | 0 | 0 | yes | no | no | yes | no | no | no | no | no | no | no | no | no |
| CACNA1G | 136 | M | 0 | 0 | no | yes | no | yes | no | no | no | no | no | no | no | no | no |
| CACNA1G | 137 | V | 0 | 0 | no | yes | no | yes | no | no | no | no | no | no | no | no | no |
| CACNA1G | 138 | A | 0 | 0 | no | yes | no | no | no | no | no | no | no | no | no | no | no |

| | | | | | | | | | | | | | | | | | |
|---|---|---|---|---|---|---|---|---|---|---|---|---|---|---|---|---|---|
| CACNA1G | 149 | G | 1 | 0 | no | yes | no | yes | no | yes | no | no | no | no | no | no | no |
| CACNA1G | 150 | D | 0 | 0 | no | yes | no | yes | no | yes | no | no | no | no | no | no | no |
| CACNA1G | 151 | T | 2 | 0 | no | yes | no | yes | no | no | no | no | no | no | no | no | no |
| CACNA1G | 152 | W | 0 | 0 | yes | no | no | yes | no | no | no | no | no | no | no | no | no |
| CACNA1G | 153 | N | 0 | 0 | yes | no | no | yes | no | no | no | no | no | no | no | no | no |
| CACNA1G | 154 | R | 2 | 0 | yes | no | no | yes | no | no | no | no | no | no | no | no | no |
| CACNA1G | 155 | L | 0 | 0 | yes | no | no | yes | no | no | no | no | no | no | no | no | no |
| CACNA1G | 156 | D | 0 | 0 | yes | no | no | yes | no | no | no | no | no | no | no | no | no |
| CACNA1G | 157 | F | 0 | 0 | yes | no | no | yes | no | no | no | no | no | no | no | no | no |
| CACNA1G | 158 | F | 1 | 0 | yes | no | no | yes | no | no | no | no | no | no | no | no | no |
| CACNA1G | 159 | I | 0 | 0 | yes | no | no | yes | no | no | no | no | no | no | no | no | no |
| CACNA1G | 160 | V | 1 | 0 | yes | no | no | yes | no | no | no | no | no | no | no | no | no |
| CACNA1G | 161 | I | 2 | 0 | yes | no | no | yes | no | no | no | no | no | no | no | no | no |
| CACNA1G | 162 | A | 0 | 0 | yes | no | no | yes | no | no | no | no | no | no | no | no | no |
| CACNA1G | 163 | G | 0 | 0 | yes | no | no | yes | no | no | no | no | no | no | no | no | no |
| CACNA1G | 164 | M | 0 | 0 | yes | no | no | yes | no | no | no | no | no | no | no | no | no |
| CACNA1G | 165 | L | 0 | 0 | yes | no | no | yes | no | no | no | no | no | no | no | no | no |
| CACNA1G | 166 | E | 0 | 0 | yes | no | no | yes | no | no | no | no | no | no | no | no | no |
| CACNA1G | 167 | Y | 0 | 0 | yes | no | no | yes | no | no | no | no | no | no | no | no | no |
| CACNA1G | 168 | S | 0 | 0 | yes | no | no | yes | no | no | no | no | no | no | no | no | no |
| CACNA1G | 169 | L | 0 | 0 | yes | no | no | yes | no | no | no | no | no | no | no | no | no |
| CACNA1G | 170 | D | 0 | 0 | yes | no | no | no | no | no | no | no | no | no | no | no | no |
| CACNA1G | 171 | L | 0 | 0 | yes | no | no | no | yes | no | no | no | no | no | no | no | no |
| CACNA1G | 172 | Q | 0 | 0 | no | yes | no | no | yes | no | no | no | no | no | no | no | no |
| CACNA1G | 173 | N | 0 | 0 | no | yes | no | no | yes | no | no | no | no | no | no | no | no |
| CACNA1G | 174 | V | 0 | 0 | no | yes | no | no | yes | no | no | no | no | no | no | no | no |
| CACNA1G | 175 | S | 0 | 0 | no | yes | no | no | yes | no | no | no | no | no | no | no | no |
| CACNA1G | 176 | F | 0 | 0 | no | yes | no | no | no | no | no | no | no | no | no | no | no |
| CACNA1G | 177 | S | 0 | 0 | no | yes | no | no | no | no | no | no | no | no | no | no | no |
| CACNA1G | 178 | A | 0 | 0 | no | yes | no | yes | no | no | no | no | no | no | no | no | no |
| CACNA1G | 179 | V | 0 | 0 | no | yes | no | yes | no | no | no | no | no | no | no | no | no |
| CACNA1G | 180 | R | 0 | 0 | yes | no | no | yes | no | no | no | no | no | no | no | no | no |
| CACNA1G | 181 | T | 0 | 0 | yes | no | no | yes | no | no | no | no | no | no | no | no | no |

| | | | | | | | | | | | | | | | | | |
|---|---|---|---|---|---|---|---|---|---|---|---|---|---|---|---|---|---|
| CACNA1G | 182 | V | 0 | 0 | yes | no | no | yes | no | no | no | no | no | no | no | no | no |
| CACNA1G | 183 | R | 2 | 0 | yes | no | no | yes | no | no | no | no | no | no | no | no | no |
| CACNA1G | 184 | V | 0 | 0 | yes | no | no | yes | no | no | no | no | no | no | no | no | no |
| CACNA1G | 185 | L | 0 | 0 | yes | no | no | yes | no | no | no | no | no | no | no | no | no |
| CACNA1G | 186 | R | 0 | 0 | yes | no | no | yes | no | no | no | no | no | no | no | no | no |
| CACNA1G | 187 | P | 0 | 0 | yes | no | no | yes | no | no | no | no | no | no | no | no | no |
| CACNA1G | 188 | L | 0 | 0 | yes | no | no | yes | no | no | no | no | no | no | no | no | no |
| CACNA1G | 189 | R | 0 | 0 | yes | no | no | yes | no | no | no | no | no | no | no | no | no |
| CACNA1G | 190 | A | 0 | 0 | yes | no | no | yes | no | no | no | no | no | no | no | no | no |
| CACNA1G | 191 | I | 0 | 0 | yes | no | no | yes | no | no | no | no | no | no | no | no | no |
| CACNA1G | 192 | N | 0 | 0 | yes | no | no | yes | no | no | no | no | no | no | no | no | no |
| CACNA1G | 193 | R | 2 | 0 | yes | no | no | yes | no | no | no | no | no | no | no | no | no |
| CACNA1G | 194 | V | 1 | 0 | no | yes | no | yes | no | yes | no | no | no | no | no | no | no |
| CACNA1G | 195 | P | 0 | 0 | yes | no | no | yes | no | yes | no | no | no | no | no | no | no |
| CACNA1G | 196 | S | 0 | 0 | yes | no | no | yes | no | yes | no | no | no | no | no | no | no |
| CACNA1G | 197 | M | 0 | 0 | yes | no | no | yes | no | yes | no | no | no | no | no | no | no |
| CACNA1G | 198 | R | 1 | 0 | yes | no | no | yes | no | yes | no | no | no | no | no | no | no |
| CACNA1G | 199 | I | 0 | 0 | yes | no | no | no | no | yes | no | no | no | no | no | no | no |
| CACNA1G | 200 | L | 0 | 0 | yes | no | no | yes | no | yes | no | no | no | no | no | no | no |
| CACNA1G | 201 | V | 0 | 0 | yes | no | no | yes | no | yes | no | no | no | no | no | no | no |
| CACNA1G | 202 | T | 1 | 0 | yes | no | no | yes | no | yes | no | no | no | no | no | no | no |
| CACNA1G | 203 | L | 0 | 0 | yes | no | no | no | no | yes | no | no | no | no | no | no | no |
| CACNA1G | 204 | L | 0 | 0 | yes | no | no | yes | no | yes | no | no | no | no | no | no | no |
| CACNA1G | 205 | L | 0 | 0 | yes | no | no | yes | no | yes | no | no | no | no | no | no | no |
| CACNA1G | 206 | D | 0 | 0 | yes | no | no | no | no | yes | no | no | no | no | no | no | no |
| CACNA1G | 207 | T | 1 | 0 | yes | no | no | yes | no | yes | no | no | no | no | no | no | no |
| CACNA1G | 208 | L | 0 | 1 | yes | no | no | yes | no | yes | no | no | no | no | no | no | no |
| CACNA1G | 209 | P | 0 | 0 | yes | no | no | yes | no | yes | no | no | no | no | no | no | no |
| CACNA1G | 210 | M | 1 | 0 | yes | no | no | yes | no | yes | no | no | no | no | no | no | no |
| CACNA1G | 211 | L | 0 | 0 | yes | no | no | yes | no | yes | no | no | no | no | no | no | no |
| CACNA1G | 212 | G | 0 | 0 | yes | no | no | yes | no | yes | no | no | no | no | no | no | no |
| CACNA1G | 213 | N | 0 | 0 | yes | no | no | yes | no | yes | no | no | no | no | no | no | no |
| CACNA1G | 214 | V | 1 | 0 | yes | no | no | yes | no | no | no | no | no | no | no | no | no |

| | | | | | | | | | | | | | | | | | |
|---|---|---|---|---|---|---|---|---|---|---|---|---|---|---|---|---|---|
| CACNA1G | 215 | L | 0 | 0 | yes | no | no | yes | no | no | no | no | no | no | no | no | no |
| CACNA1G | 216 | L | 0 | 0 | yes | no | no | yes | no | no | no | no | no | no | no | no | no |
| CACNA1G | 217 | L | 0 | 0 | yes | no | no | yes | no | no | no | no | no | no | no | no | no |
| CACNA1G | 218 | C | 0 | 0 | yes | no | no | yes | no | no | no | no | no | no | no | no | no |
| CACNA1G | 219 | F | 0 | 0 | yes | no | no | yes | no | no | no | no | no | no | no | no | no |
| CACNA1G | 220 | F | 0 | 0 | yes | no | no | yes | no | no | no | no | no | no | no | no | no |
| CACNA1G | 221 | V | 0 | 0 | yes | no | no | yes | no | no | no | no | no | no | no | no | no |
| CACNA1G | 222 | F | 0 | 0 | yes | no | no | yes | no | no | no | no | no | no | no | no | no |
| CACNA1G | 223 | F | 0 | 0 | yes | no | no | yes | no | no | no | no | no | no | no | no | no |
| CACNA1G | 224 | I | 0 | 0 | yes | no | no | yes | no | no | no | no | no | no | no | no | no |
| CACNA1G | 225 | F | 0 | 0 | yes | no | no | yes | no | no | no | no | no | no | no | no | no |
| CACNA1G | 226 | G | 0 | 0 | yes | no | no | yes | no | no | no | no | no | no | no | no | no |
| CACNA1G | 227 | I | 0 | 0 | yes | no | no | yes | no | no | no | no | no | no | no | no | no |
| CACNA1G | 228 | V | 1 | 0 | yes | no | no | yes | no | no | no | no | no | no | no | no | no |
| CACNA1G | 229 | G | 1 | 0 | yes | no | no | yes | no | no | no | no | no | no | no | no | no |
| CACNA1G | 230 | V | 1 | 0 | yes | no | no | yes | no | no | no | no | no | no | no | no | no |
| CACNA1G | 231 | Q | 0 | 0 | yes | no | no | yes | no | no | no | no | no | no | no | no | no |
| CACNA1G | 232 | L | 1 | 0 | yes | no | no | no | no | no | no | no | no | no | no | no | no |
| CACNA1G | 233 | W | 0 | 0 | yes | no | no | no | no | no | no | no | no | no | no | no | no |
| CACNA1G | 234 | A | 0 | 0 | no | yes | no | no | no | no | no | no | no | no | no | no | no |
| CACNA1G | 235 | G | 0 | 0 | no | yes | no | no | yes | no | no | no | no | no | no | no | no |
| CACNA1G | 236 | L | 0 | 0 | no | yes | no | no | yes | no | no | no | no | no | no | no | no |
| CACNA1G | 237 | L | 0 | 0 | no | yes | no | no | yes | no | no | no | no | no | no | no | no |
| CACNA1G | 238 | R | 1 | 0 | no | yes | no | no | yes | no | no | no | no | no | no | no | no |
| CACNA1G | 239 | N | 0 | 0 | no | no | yes | no | yes | no | no | no | no | no | no | no | no |
| CACNA1G | 240 | R | 0 | 0 | no | no | yes | no | yes | no | no | no | no | no | no | no | no |
| CACNA1G | 241 | C | 0 | 0 | no | no | yes | no | yes | no | no | no | no | no | no | no | no |
| CACNA1G | 242 | F | 0 | 0 | no | no | yes | no | yes | no | no | no | no | no | no | no | no |
| CACNA1G | 243 | L | 0 | 0 | no | yes | no | no | yes | no | no | no | no | no | no | no | no |
| CACNA1G | 244 | P | 1 | 0 | no | yes | no | no | yes | no | no | no | no | no | no | no | no |
| CACNA1G | 245 | E | 0 | 0 | no | yes | no | no | yes | no | no | no | no | no | no | no | no |
| CACNA1G | 246 | N | 0 | 0 | no | yes | no | no | yes | no | no | no | no | no | no | no | no |
| CACNA1G | 247 | F | 0 | 0 | no | yes | no | no | yes | no | no | no | no | no | no | no | no |

| | | | | | | | | | | | | | | | | | |
|---|---|---|---|---|---|---|---|---|---|---|---|---|---|---|---|---|---|
| CACNA1G | 248 | S | 1 | 0 | no | yes | no | no | yes | no | no | no | no | no | no | no | no |
| CACNA1G | 249 | L | 0 | 0 | no | yes | no | no | yes | no | no | no | no | no | no | no | no |
| CACNA1G | 250 | P | 0 | 0 | no | yes | no | no | yes | no | no | no | no | no | no | no | no |
| CACNA1G | 251 | L | 0 | 0 | no | yes | no | no | yes | no | no | no | no | no | no | no | no |
| CACNA1G | 252 | S | 0 | 0 | no | yes | no | no | yes | no | no | no | no | no | no | no | no |
| CACNA1G | 253 | V | 1 | 0 | no | yes | no | no | yes | no | no | no | no | no | no | no | no |
| CACNA1G | 254 | D | 1 | 0 | no | yes | no | no | yes | no | no | no | no | no | no | no | no |
| CACNA1G | 255 | L | 0 | 0 | no | yes | no | no | yes | no | no | no | no | no | no | no | no |
| CACNA1G | 256 | E | 0 | 0 | no | yes | no | no | yes | no | no | no | no | no | no | no | no |
| CACNA1G | 257 | R | 0 | 0 | no | yes | no | no | yes | no | no | no | no | no | no | no | no |
| CACNA1G | 258 | Y | 0 | 0 | no | no | yes | no | yes | no | no | no | no | no | no | no | no |
| CACNA1G | 259 | Y | 1 | 0 | no | yes | no | no | yes | no | yes | no | no | no | no | no | no |
| CACNA1G | 260 | Q | 0 | 0 | no | yes | no | no | yes | no | no | no | no | no | no | no | no |
| CACNA1G | 261 | T | 0 | 0 | no | yes | no | no | yes | no | no | no | no | no | no | no | no |
| CACNA1G | 262 | E | 0 | 0 | no | yes | no | no | yes | no | yes | no | no | no | no | no | no |
| CACNA1G | 263 | N | 0 | 0 | no | yes | no | no | yes | no | yes | no | no | no | no | no | no |
| CACNA1G | 264 | E | 0 | 0 | no | yes | no | no | yes | no | no | no | no | no | no | no | no |
| CACNA1G | 265 | D | 0 | 0 | no | yes | no | no | yes | no | no | no | no | no | no | no | no |
| CACNA1G | 266 | E | 0 | 0 | no | yes | no | no | yes | no | no | no | no | no | no | no | no |
| CACNA1G | 267 | S | 0 | 0 | no | yes | no | no | yes | no | yes | no | no | no | no | no | no |
| CACNA1G | 268 | P | 0 | 0 | no | yes | no | no | yes | no | no | no | no | no | no | no | no |
| CACNA1G | 269 | F | 0 | 0 | no | yes | no | no | yes | no | no | no | no | no | no | no | no |
| CACNA1G | 270 | I | 0 | 0 | no | yes | no | no | yes | no | no | no | no | no | no | no | no |
| CACNA1G | 271 | C | 0 | 0 | no | yes | no | no | yes | no | no | no | no | no | no | no | no |
| CACNA1G | 272 | S | 0 | 0 | no | yes | no | no | yes | no | no | no | no | no | no | no | no |
| CACNA1G | 273 | Q | 0 | 0 | no | yes | no | no | yes | no | no | no | no | no | no | no | no |
| CACNA1G | 274 | P | 0 | 0 | no | yes | no | no | yes | no | no | no | no | no | no | no | no |
| CACNA1G | 275 | R | 2 | 0 | no | yes | no | no | yes | no | no | no | no | no | no | no | no |
| CACNA1G | 276 | E | 0 | 0 | no | yes | no | no | yes | no | no | no | no | no | no | no | no |
| CACNA1G | 277 | N | 0 | 0 | no | yes | no | no | yes | no | no | no | no | no | no | no | no |
| CACNA1G | 278 | G | 0 | 0 | no | yes | no | no | yes | no | no | no | no | no | no | no | no |
| CACNA1G | 279 | M | 1 | 0 | no | yes | no | no | yes | no | no | no | no | no | no | no | no |
| CACNA1G | 280 | R | 1 | 0 | no | yes | no | no | yes | no | yes | no | no | no | no | no | no |

| | | | | | | | | | | | | | | | | | |
|---|---|---|---|---|---|---|---|---|---|---|---|---|---|---|---|---|---|
| CACNA1G | 281 | S | 0 | 0 | no | yes | no | no | yes | no | no | no | no | no | no | no | no |
| CACNA1G | 282 | C | 0 | 0 | yes | no | no | no | yes | no | no | no | no | no | no | no | no |
| CACNA1G | 283 | R | 0 | 0 | yes | no | no | no | yes | no | no | no | no | no | no | no | no |
| CACNA1G | 284 | S | 1 | 0 | yes | no | no | no | yes | no | no | no | no | no | no | no | no |
| CACNA1G | 285 | V | 1 | 0 | no | yes | no | no | yes | no | no | no | no | no | no | no | no |
| CACNA1G | 286 | P | 0 | 0 | no | yes | no | no | yes | no | yes | no | no | no | no | no | no |
| CACNA1G | 287 | T | 1 | 0 | no | yes | no | no | yes | no | yes | no | no | no | no | no | no |
| CACNA1G | 288 | L | 0 | 0 | no | yes | no | no | yes | no | yes | no | no | no | no | no | no |
| CACNA1G | 289 | R | 2 | 0 | no | yes | no | no | yes | no | yes | no | no | no | no | no | no |
| CACNA1G | 313 | C | 0 | 0 | no | yes | no | no | yes | no | no | no | no | no | no | no | no |
| CACNA1G | 314 | V | 0 | 0 | no | yes | no | no | yes | no | no | no | no | no | no | no | no |
| CACNA1G | 315 | N | 0 | 0 | no | yes | no | no | yes | no | no | no | no | no | no | no | no |
| CACNA1G | 316 | W | 0 | 0 | yes | no | no | no | yes | no | no | no | no | no | no | no | no |
| CACNA1G | 317 | N | 0 | 0 | yes | no | no | no | yes | no | yes | no | no | no | no | no | no |
| CACNA1G | 318 | Q | 1 | 0 | yes | no | no | no | yes | no | no | no | no | no | no | no | no |
| CACNA1G | 319 | Y | 1 | 0 | yes | no | no | no | yes | no | no | no | no | no | no | no | no |
| CACNA1G | 320 | Y | 0 | 0 | no | yes | no | no | yes | no | no | no | no | no | no | no | no |
| CACNA1G | 321 | T | 1 | 0 | no | yes | no | no | yes | no | no | no | no | no | no | no | no |
| CACNA1G | 322 | N | 1 | 0 | no | no | yes | no | yes | no | no | no | no | no | no | no | no |
| CACNA1G | 323 | C | 0 | 0 | no | no | yes | no | yes | no | no | no | no | no | no | no | no |
| CACNA1G | 324 | S | 0 | 0 | no | no | yes | no | yes | no | no | no | no | no | no | no | no |
| CACNA1G | 325 | A | 1 | 0 | no | no | yes | no | yes | no | no | no | no | no | no | no | no |
| CACNA1G | 326 | G | 0 | 0 | no | yes | no | no | yes | no | no | no | no | no | no | no | no |
| CACNA1G | 327 | E | 0 | 0 | no | yes | no | no | yes | no | no | no | no | no | no | no | no |
| CACNA1G | 328 | H | 0 | 0 | no | yes | no | no | yes | no | no | no | no | no | no | no | no |
| CACNA1G | 329 | N | 0 | 0 | no | yes | no | no | yes | no | no | no | no | no | no | no | no |
| CACNA1G | 330 | P | 0 | 0 | yes | no | no | no | yes | no | no | no | no | no | no | no | no |
| CACNA1G | 331 | F | 0 | 0 | yes | no | no | no | yes | no | no | no | no | no | no | no | no |
| CACNA1G | 332 | K | 0 | 0 | yes | no | no | no | yes | no | no | no | no | no | no | no | no |
| CACNA1G | 333 | G | 0 | 0 | yes | no | no | no | yes | no | no | no | no | no | no | no | no |
| CACNA1G | 334 | A | 0 | 0 | no | yes | no | no | yes | no | no | no | no | no | no | no | no |
| CACNA1G | 335 | I | 0 | 0 | no | yes | no | no | yes | no | no | no | no | no | no | no | no |
| CACNA1G | 336 | N | 0 | 0 | no | yes | no | no | yes | no | no | no | no | no | no | no | no |

| | | | | | | | | | | | | | | | | | |
|---|---|---|---|---|---|---|---|---|---|---|---|---|---|---|---|---|---|
| CACNA1G | 337 | F | 0 | 0 | no | yes | no | no | yes | no | no | no | no | no | no | no | no |
| CACNA1G | 338 | D | 0 | 0 | no | yes | no | no | yes | no | no | no | no | no | no | no | no |
| CACNA1G | 339 | N | 0 | 0 | no | yes | no | no | yes | no | no | no | no | no | no | no | no |
| CACNA1G | 340 | I | 0 | 0 | yes | no | no | yes | yes | no | no | no | no | no | no | no | no |
| CACNA1G | 341 | G | 0 | 0 | yes | no | no | yes | yes | no | no | no | no | no | no | no | no |
| CACNA1G | 342 | Y | 0 | 0 | yes | no | no | no | yes | no | no | no | no | no | no | no | no |
| CACNA1G | 343 | A | 0 | 0 | yes | no | no | yes | yes | no | no | no | no | no | no | no | no |
| CACNA1G | 344 | W | 0 | 0 | yes | no | no | yes | yes | no | no | no | no | no | no | no | no |
| CACNA1G | 345 | I | 0 | 0 | yes | no | no | yes | yes | no | no | no | no | no | no | no | no |
| CACNA1G | 346 | A | 1 | 0 | yes | no | no | yes | yes | no | no | no | no | no | no | no | no |
| CACNA1G | 347 | I | 0 | 0 | yes | no | no | yes | yes | no | no | no | no | no | no | no | no |
| CACNA1G | 348 | F | 0 | 0 | yes | no | no | yes | yes | no | no | no | no | no | no | no | no |
| CACNA1G | 349 | Q | 0 | 0 | yes | no | no | yes | yes | no | no | no | no | no | no | no | no |
| CACNA1G | 350 | V | 0 | 0 | yes | no | no | yes | yes | no | no | no | no | no | no | no | no |
| CACNA1G | 351 | I | 0 | 0 | yes | no | no | yes | yes | no | no | no | no | no | no | no | no |
| CACNA1G | 352 | T | 1 | 0 | no | yes | no | yes | yes | no | no | no | no | no | no | no | no |
| CACNA1G | 353 | L | 0 | 0 | no | yes | no | yes | yes | no | yes | no | no | no | no | no | no |
| CACNA1G | 354 | E | 0 | 0 | no | yes | no | yes | yes | no | no | yes | no | no | no | no | no |
| CACNA1G | 355 | G | 0 | 0 | no | yes | no | yes | yes | no | no | no | no | no | no | no | no |
| CACNA1G | 356 | W | 0 | 0 | yes | no | no | yes | yes | no | yes | no | no | no | no | no | no |
| CACNA1G | 357 | V | 0 | 0 | yes | no | no | no | yes | no | yes | no | no | no | no | no | no |
| CACNA1G | 358 | D | 0 | 0 | yes | no | no | no | yes | no | yes | no | no | no | no | no | no |
| CACNA1G | 359 | I | 0 | 0 | yes | no | no | no | yes | no | no | no | no | no | no | no | no |
| CACNA1G | 360 | M | 0 | 0 | yes | no | no | no | yes | no | no | no | no | no | no | no | no |
| CACNA1G | 361 | Y | 0 | 0 | yes | no | no | no | yes | no | yes | no | no | no | no | no | no |
| CACNA1G | 362 | F | 0 | 0 | yes | no | no | no | yes | no | no | no | no | no | no | no | no |
| CACNA1G | 363 | V | 1 | 0 | yes | no | no | no | yes | no | no | no | no | no | no | no | no |
| CACNA1G | 364 | M | 1 | 0 | yes | no | no | no | yes | no | no | no | no | no | no | no | no |
| CACNA1G | 365 | D | 0 | 0 | yes | no | no | no | yes | no | no | no | no | no | no | no | no |
| CACNA1G | 366 | A | 0 | 0 | yes | no | no | no | yes | no | no | no | no | no | no | no | no |
| CACNA1G | 367 | H | 0 | 0 | no | yes | no | no | yes | no | no | no | no | no | no | no | no |
| CACNA1G | 368 | S | 0 | 1 | no | yes | no | no | yes | no | no | no | no | no | no | no | no |
| CACNA1G | 369 | F | 0 | 0 | yes | no | no | no | yes | no | no | no | no | no | no | no | no |

| | | | | | | | | | | | | | | | | | |
|---|---|---|---|---|---|---|---|---|---|---|---|---|---|---|---|---|---|
| CACNA1G | 370 | Y | 0 | 0 | yes | no | no | yes | yes | no | no | no | no | no | no | no | no |
| CACNA1G | 371 | N | 1 | 0 | yes | no | no | yes | no | no | no | no | no | no | no | no | no |
| CACNA1G | 372 | F | 0 | 0 | yes | no | no | yes | no | no | no | no | no | no | no | no | no |
| CACNA1G | 373 | I | 0 | 0 | yes | no | no | yes | no | no | no | no | no | no | no | no | no |
| CACNA1G | 374 | Y | 0 | 0 | yes | no | no | yes | no | no | no | no | no | no | no | no | no |
| CACNA1G | 375 | F | 0 | 0 | yes | no | no | yes | no | no | no | no | no | no | no | no | no |
| CACNA1G | 376 | I | 0 | 0 | yes | no | no | yes | no | no | no | no | no | no | no | no | no |
| CACNA1G | 377 | L | 0 | 0 | yes | no | no | yes | no | no | no | no | no | no | no | no | no |
| CACNA1G | 378 | L | 0 | 0 | yes | no | no | yes | no | no | no | no | no | no | no | no | no |
| CACNA1G | 379 | I | 1 | 0 | yes | no | no | yes | no | no | no | no | no | no | no | no | no |
| CACNA1G | 380 | I | 0 | 0 | yes | no | no | yes | no | no | no | no | no | no | no | no | no |
| CACNA1G | 381 | V | 0 | 0 | yes | no | no | yes | no | no | no | no | no | no | no | no | no |
| CACNA1G | 382 | G | 0 | 0 | yes | no | no | yes | no | no | no | no | no | no | no | no | no |
| CACNA1G | 383 | S | 0 | 0 | yes | no | no | yes | no | no | no | no | no | no | no | no | no |
| CACNA1G | 384 | F | 0 | 0 | yes | no | no | yes | no | no | yes | no | no | no | no | no | no |
| CACNA1G | 385 | F | 0 | 0 | no | yes | no | yes | no | no | no | no | no | no | no | no | no |
| CACNA1G | 386 | M | 0 | 0 | no | yes | no | yes | no | no | no | no | no | no | no | no | no |
| CACNA1G | 387 | I | 0 | 0 | yes | no | no | yes | no | no | no | no | no | no | no | no | no |
| CACNA1G | 388 | N | 0 | 0 | yes | no | no | yes | no | no | yes | no | no | no | no | no | no |
| CACNA1G | 389 | L | 0 | 0 | yes | no | no | yes | no | no | no | no | no | no | no | no | no |
| CACNA1G | 390 | C | 0 | 0 | yes | no | no | yes | no | no | no | no | no | no | no | no | no |
| CACNA1G | 391 | L | 0 | 0 | yes | no | no | yes | no | no | no | no | no | no | no | no | no |
| CACNA1G | 392 | V | 0 | 0 | yes | no | no | yes | no | no | no | no | no | no | no | no | no |
| CACNA1G | 393 | V | 0 | 0 | yes | no | no | yes | no | no | no | no | no | no | no | no | no |
| CACNA1G | 394 | I | 0 | 0 | yes | no | no | yes | no | no | no | no | no | no | no | no | no |
| CACNA1G | 395 | A | 0 | 0 | yes | no | no | yes | no | no | no | no | no | no | no | no | no |
| CACNA1G | 396 | T | 0 | 0 | yes | no | no | no | no | yes | no | no | no | no | no | no | no |
| CACNA1G | 397 | Q | 0 | 0 | yes | no | no | no | no | yes | no | no | no | no | no | no | no |
| CACNA1G | 398 | F | 0 | 0 | yes | no | no | no | no | yes | no | no | no | no | no | no | no |
| CACNA1G | 399 | S | 0 | 0 | yes | no | no | no | no | yes | no | no | no | no | no | no | no |
| CACNA1G | 400 | E | 0 | 0 | yes | no | no | no | no | yes | no | no | no | no | no | no | no |
| CACNA1G | 401 | T | 0 | 0 | yes | no | no | no | no | yes | no | no | no | no | no | no | no |
| CACNA1G | 402 | K | 1 | 0 | yes | no | no | no | no | yes | no | no | no | no | no | no | no |

| | | | | | | | | | | | | | | | | | |
|---|---|---|---|---|---|---|---|---|---|---|---|---|---|---|---|---|---|
| CACNA1G | 403 | Q | 0 | 0 | yes | no | no | no | no | yes | no | no | no | no | no | no | no |
| CACNA1G | 404 | R | 2 | 0 | yes | no | no | no | no | yes | no | no | no | no | no | no | no |
| CACNA1G | 405 | E | 0 | 0 | no | yes | no | no | no | yes | no | no | no | no | no | no | no |
| CACNA1G | 739 | I | 2 | 0 | no | yes | no | no | no | yes | no | no | no | no | no | no | no |
| CACNA1G | 740 | V | 1 | 0 | no | yes | no | no | no | yes | no | no | no | no | no | no | no |
| CACNA1G | 741 | D | 1 | 0 | no | yes | no | no | no | yes | no | no | no | no | no | no | no |
| CACNA1G | 742 | S | 0 | 0 | no | yes | no | no | no | yes | no | no | no | no | no | no | no |
| CACNA1G | 743 | K | 0 | 0 | no | yes | no | no | no | yes | no | no | no | no | no | no | no |
| CACNA1G | 744 | Y | 0 | 0 | yes | no | no | yes | no | no | no | no | no | no | no | no | no |
| CACNA1G | 745 | F | 0 | 0 | yes | no | no | yes | no | no | no | no | no | no | no | no | no |
| CACNA1G | 746 | G | 0 | 0 | yes | no | no | yes | no | no | no | no | no | no | no | no | no |
| CACNA1G | 747 | R | 2 | 0 | yes | no | no | yes | no | no | no | no | no | no | no | no | no |
| CACNA1G | 748 | G | 0 | 0 | yes | no | no | yes | no | no | no | no | no | no | no | no | no |
| CACNA1G | 749 | I | 0 | 0 | yes | no | no | yes | no | no | no | no | no | no | no | no | no |
| CACNA1G | 750 | M | 1 | 0 | yes | no | no | yes | no | no | no | no | no | no | no | no | no |
| CACNA1G | 751 | I | 0 | 0 | yes | no | no | yes | no | no | no | no | no | no | no | no | no |
| CACNA1G | 752 | A | 0 | 0 | yes | no | no | yes | no | no | no | no | no | no | no | no | no |
| CACNA1G | 753 | I | 0 | 0 | yes | no | no | yes | no | no | no | no | no | no | no | no | no |
| CACNA1G | 754 | L | 0 | 0 | yes | no | no | yes | no | no | no | no | no | no | no | no | no |
| CACNA1G | 755 | V | 0 | 0 | yes | no | no | yes | no | no | no | no | no | no | no | no | no |
| CACNA1G | 756 | N | 0 | 0 | yes | no | no | yes | no | no | no | no | no | no | no | no | no |
| CACNA1G | 757 | T | 0 | 0 | yes | no | no | yes | no | no | no | no | no | no | no | no | no |
| CACNA1G | 758 | L | 0 | 0 | yes | no | no | yes | no | no | no | no | no | no | no | no | no |
| CACNA1G | 759 | S | 0 | 0 | yes | no | no | yes | no | no | no | no | no | no | no | no | no |
| CACNA1G | 760 | M | 1 | 0 | yes | no | no | yes | no | no | no | no | no | no | no | no | no |
| CACNA1G | 761 | G | 0 | 0 | no | yes | no | yes | no | no | no | no | no | no | no | no | no |
| CACNA1G | 762 | I | 0 | 0 | no | yes | no | yes | no | no | no | no | no | no | no | no | no |
| CACNA1G | 763 | E | 0 | 0 | no | yes | no | yes | no | no | no | no | no | no | no | no | no |
| CACNA1G | 764 | Y | 1 | 0 | no | yes | no | no | no | no | no | no | no | no | no | no | no |
| CACNA1G | 765 | H | 0 | 0 | no | yes | no | no | yes | no | no | no | no | no | no | no | no |
| CACNA1G | 766 | E | 0 | 0 | no | yes | no | no | yes | no | no | no | no | no | no | no | no |
| CACNA1G | 767 | Q | 0 | 0 | no | yes | no | no | yes | no | no | no | no | no | no | no | no |
| CACNA1G | 768 | P | 1 | 0 | no | yes | no | no | yes | no | no | no | no | no | no | no | no |

| | | | | | | | | | | | | | | | | | |
|---|---|---|---|---|---|---|---|---|---|---|---|---|---|---|---|---|---|
| CACNA1G | 769 | E | 0 | 0 | no | yes | no | no | yes | no | no | no | no | no | no | no | no |
| CACNA1G | 770 | E | 0 | 0 | yes | no | no | no | yes | no | no | no | no | no | no | no | no |
| CACNA1G | 771 | L | 0 | 0 | yes | no | no | no | yes | no | no | no | no | no | no | no | no |
| CACNA1G | 772 | T | 0 | 0 | yes | no | no | no | yes | no | no | no | no | no | no | no | no |
| CACNA1G | 773 | N | 1 | 0 | yes | no | no | no | yes | no | no | no | no | no | no | no | no |
| CACNA1G | 774 | A | 2 | 0 | yes | no | no | yes | yes | no | no | no | no | no | no | no | no |
| CACNA1G | 775 | L | 0 | 0 | yes | no | no | yes | yes | no | no | no | no | no | no | no | no |
| CACNA1G | 776 | E | 0 | 0 | yes | no | no | yes | yes | no | no | no | no | no | no | no | no |
| CACNA1G | 777 | I | 0 | 0 | yes | no | no | yes | yes | no | no | no | no | no | no | no | no |
| CACNA1G | 778 | S | 0 | 0 | yes | no | no | yes | no | no | no | no | no | no | no | no | no |
| CACNA1G | 779 | N | 0 | 0 | yes | no | no | yes | no | no | no | no | no | no | no | no | no |
| CACNA1G | 780 | I | 0 | 0 | yes | no | no | yes | no | no | no | no | no | no | no | no | no |
| CACNA1G | 781 | V | 2 | 0 | yes | no | no | yes | no | no | no | no | no | no | no | no | no |
| CACNA1G | 782 | F | 0 | 0 | yes | no | no | yes | no | no | no | no | no | no | no | no | no |
| CACNA1G | 783 | T | 0 | 0 | yes | no | no | yes | no | no | no | no | no | no | no | no | no |
| CACNA1G | 784 | S | 0 | 0 | yes | no | no | yes | no | no | no | no | no | no | no | no | no |
| CACNA1G | 785 | L | 0 | 0 | yes | no | no | yes | no | no | no | no | no | no | no | no | no |
| CACNA1G | 786 | F | 0 | 0 | yes | no | no | yes | no | no | no | no | no | no | no | no | no |
| CACNA1G | 787 | A | 2 | 0 | yes | no | no | yes | no | no | no | no | no | no | no | no | no |
| CACNA1G | 788 | L | 0 | 0 | yes | no | no | yes | no | no | no | no | no | no | no | no | no |
| CACNA1G | 789 | E | 0 | 0 | yes | no | no | yes | no | no | no | no | no | no | no | no | no |
| CACNA1G | 790 | M | 0 | 0 | yes | no | no | yes | no | no | no | no | no | no | no | no | no |
| CACNA1G | 791 | L | 0 | 0 | yes | no | no | yes | no | no | no | no | no | no | no | no | no |
| CACNA1G | 792 | L | 0 | 0 | yes | no | no | yes | no | no | no | no | no | no | no | no | no |
| CACNA1G | 793 | K | 0 | 0 | yes | no | no | yes | no | no | no | no | no | no | no | no | no |
| CACNA1G | 794 | L | 0 | 0 | yes | no | no | yes | no | no | no | no | no | no | no | no | no |
| CACNA1G | 795 | L | 0 | 0 | yes | no | no | yes | no | no | no | no | no | no | no | no | no |
| CACNA1G | 796 | V | 0 | 0 | no | yes | no | no | no | no | no | no | no | no | no | no | no |
| CACNA1G | 797 | Y | 0 | 0 | no | yes | no | no | no | no | no | no | no | no | no | no | no |
| CACNA1G | 798 | G | 0 | 0 | no | yes | no | no | no | no | no | no | no | no | no | no | no |
| CACNA1G | 799 | P | 2 | 0 | yes | no | no | no | no | no | no | no | no | no | no | no | no |
| CACNA1G | 800 | F | 0 | 0 | yes | no | no | no | no | yes | no | no | no | no | no | no | no |
| CACNA1G | 801 | G | 0 | 0 | yes | no | no | no | no | yes | no | no | no | no | no | no | no |

| | | | | | | | | | | | | | | | | | |
|---|---|---|---|---|---|---|---|---|---|---|---|---|---|---|---|---|---|
| CACNA1G | 802 | Y | 0 | 0 | yes | no | no | yes | no | yes | no | no | no | no | no | no | no |
| CACNA1G | 803 | I | 1 | 0 | no | yes | no | no | no | yes | no | no | no | no | no | no | no |
| CACNA1G | 804 | K | 0 | 0 | no | yes | no | no | no | yes | no | no | no | no | no | no | no |
| CACNA1G | 805 | N | 0 | 0 | no | yes | no | no | no | yes | no | no | no | no | no | no | no |
| CACNA1G | 806 | P | 1 | 0 | yes | no | no | no | no | no | no | no | no | no | no | no | no |
| CACNA1G | 807 | Y | 0 | 0 | yes | no | no | yes | no | no | no | no | no | no | no | no | no |
| CACNA1G | 808 | N | 0 | 0 | yes | no | no | yes | no | no | no | no | no | no | no | no | no |
| CACNA1G | 809 | I | 0 | 0 | yes | no | no | yes | no | no | no | no | no | no | no | no | no |
| CACNA1G | 810 | F | 0 | 0 | yes | no | no | yes | no | no | no | no | no | no | no | no | no |
| CACNA1G | 811 | D | 0 | 0 | yes | no | no | yes | no | no | no | no | no | no | no | no | no |
| CACNA1G | 812 | G | 0 | 0 | yes | no | no | yes | no | no | no | no | no | no | no | no | no |
| CACNA1G | 813 | V | 0 | 0 | yes | no | no | yes | no | no | no | no | no | no | no | no | no |
| CACNA1G | 814 | I | 0 | 0 | yes | no | no | yes | no | no | no | no | no | no | no | no | no |
| CACNA1G | 815 | V | 0 | 0 | yes | no | no | yes | no | no | no | no | no | no | no | no | no |
| CACNA1G | 816 | V | 0 | 0 | yes | no | no | yes | no | no | no | no | no | no | no | no | no |
| CACNA1G | 817 | I | 0 | 0 | yes | no | no | yes | no | no | no | no | no | no | no | no | no |
| CACNA1G | 818 | S | 0 | 0 | yes | no | no | yes | no | no | no | no | no | no | no | no | no |
| CACNA1G | 819 | V | 0 | 0 | yes | no | no | yes | no | no | no | no | no | no | no | no | no |
| CACNA1G | 820 | W | 0 | 0 | yes | no | no | yes | no | no | no | no | no | no | no | no | no |
| CACNA1G | 821 | E | 0 | 0 | yes | no | no | yes | no | no | no | no | no | no | no | no | no |
| CACNA1G | 822 | I | 0 | 0 | yes | no | no | yes | no | no | no | no | no | no | no | no | no |
| CACNA1G | 823 | V | 0 | 0 | no | yes | no | yes | no | no | no | no | no | no | no | no | no |
| CACNA1G | 831 | S | 0 | 0 | no | yes | no | yes | yes | no | no | no | no | no | no | no | no |
| CACNA1G | 832 | V | 0 | 0 | yes | no | no | yes | yes | no | no | no | no | no | no | no | no |
| CACNA1G | 833 | L | 0 | 0 | yes | no | no | yes | no | no | no | no | no | no | no | no | no |
| CACNA1G | 834 | R | 0 | 0 | yes | no | no | yes | no | no | no | no | no | no | no | no | no |
| CACNA1G | 835 | T | 0 | 0 | yes | no | no | yes | no | no | no | no | no | no | no | no | no |
| CACNA1G | 836 | F | 0 | 0 | yes | no | no | yes | no | no | no | no | no | no | no | no | no |
| CACNA1G | 837 | R | 1 | 0 | yes | no | no | yes | no | no | no | no | no | no | no | no | no |
| CACNA1G | 838 | L | 0 | 0 | yes | no | no | yes | no | no | no | no | no | no | no | no | no |
| CACNA1G | 839 | M | 1 | 0 | yes | no | no | yes | no | no | no | no | no | no | no | no | no |
| CACNA1G | 840 | R | 1 | 0 | yes | no | no | yes | no | no | no | no | no | no | no | no | no |
| CACNA1G | 841 | V | 0 | 0 | yes | no | no | yes | no | no | no | no | no | no | no | no | no |

| | | | | | | | | | | | | | | | | | |
|---|---|---|---|---|---|---|---|---|---|---|---|---|---|---|---|---|---|
| CACNA1G | 842 | L | 0 | 0 | yes | no | no | yes | no | no | no | no | no | no | no | no | no |
| CACNA1G | 843 | K | 0 | 0 | yes | no | no | yes | no | no | no | no | no | no | no | no | no |
| CACNA1G | 844 | L | 0 | 0 | yes | no | no | yes | no | no | no | no | no | no | no | no | no |
| CACNA1G | 845 | V | 0 | 0 | no | yes | no | yes | no | no | no | no | no | no | no | no | no |
| CACNA1G | 846 | R | 0 | 0 | no | yes | no | yes | no | no | no | no | no | no | no | no | no |
| CACNA1G | 847 | F | 0 | 0 | no | yes | no | yes | no | no | no | no | no | no | no | no | no |
| CACNA1G | 848 | L | 0 | 0 | no | yes | no | yes | no | no | no | no | no | no | no | no | no |
| CACNA1G | 849 | P | 1 | 0 | no | yes | no | no | no | no | no | no | no | no | no | no | no |
| CACNA1G | 850 | A | 1 | 0 | yes | no | no | no | no | no | no | no | no | no | no | no | no |
| CACNA1G | 851 | L | 0 | 0 | yes | no | no | yes | no | no | no | no | no | no | no | no | no |
| CACNA1G | 852 | Q | 0 | 0 | yes | no | no | no | no | no | no | no | no | no | no | no | no |
| CACNA1G | 853 | R | 2 | 1 | yes | no | no | no | no | no | no | no | no | no | no | no | no |
| CACNA1G | 854 | Q | 0 | 0 | yes | no | no | yes | no | no | no | no | no | no | no | no | no |
| CACNA1G | 855 | L | 0 | 0 | yes | no | no | yes | no | no | no | no | no | no | no | no | no |
| CACNA1G | 856 | V | 0 | 0 | yes | no | no | no | no | no | no | no | no | no | no | no | no |
| CACNA1G | 857 | V | 0 | 0 | yes | no | no | no | no | yes | no | no | no | no | no | no | no |
| CACNA1G | 858 | L | 0 | 0 | yes | no | no | yes | no | yes | no | no | no | no | no | no | no |
| CACNA1G | 859 | M | 1 | 0 | yes | no | no | yes | no | yes | no | no | no | no | no | no | no |
| CACNA1G | 860 | K | 0 | 0 | yes | no | no | no | no | yes | no | no | no | no | no | no | no |
| CACNA1G | 861 | T | 0 | 0 | yes | no | no | yes | no | yes | no | no | no | no | no | no | no |
| CACNA1G | 862 | M | 0 | 0 | yes | no | no | yes | no | yes | no | no | no | no | no | no | no |
| CACNA1G | 863 | D | 0 | 0 | yes | no | no | yes | no | yes | no | no | no | no | no | no | no |
| CACNA1G | 864 | N | 0 | 0 | yes | no | no | yes | no | yes | no | no | no | no | no | no | no |
| CACNA1G | 865 | V | 0 | 0 | no | yes | no | yes | no | yes | no | no | no | no | no | no | no |
| CACNA1G | 866 | A | 0 | 0 | yes | no | no | yes | no | yes | no | no | no | no | no | no | no |
| CACNA1G | 867 | T | 0 | 1 | yes | no | no | yes | no | yes | no | no | no | no | no | no | no |
| CACNA1G | 868 | F | 0 | 0 | yes | no | no | yes | no | no | no | no | no | no | no | no | no |
| CACNA1G | 869 | C | 0 | 0 | yes | no | no | yes | no | no | no | no | no | no | no | no | no |
| CACNA1G | 870 | M | 0 | 0 | yes | no | no | yes | no | no | no | no | no | no | no | no | no |
| CACNA1G | 871 | L | 0 | 0 | yes | no | no | yes | no | no | no | no | no | no | no | no | no |
| CACNA1G | 872 | L | 0 | 0 | yes | no | no | yes | no | no | no | no | no | no | no | no | no |
| CACNA1G | 873 | M | 0 | 0 | yes | no | no | yes | no | no | no | no | no | no | no | no | no |
| CACNA1G | 874 | L | 0 | 0 | yes | no | no | yes | no | no | no | no | no | no | no | no | no |

| | | | | | | | | | | | | | | | | | |
|---|---|---|---|---|---|---|---|---|---|---|---|---|---|---|---|---|---|
| CACNA1G | 875 | F | 0 | 0 | yes | no | no | yes | no | no | no | no | no | no | no | no | no |
| CACNA1G | 876 | I | 0 | 0 | yes | no | no | yes | no | no | no | no | no | no | no | no | no |
| CACNA1G | 877 | F | 0 | 0 | yes | no | no | yes | no | no | no | no | no | no | no | no | no |
| CACNA1G | 878 | I | 0 | 0 | yes | no | no | yes | no | no | no | no | no | no | no | no | no |
| CACNA1G | 879 | F | 0 | 0 | yes | no | no | yes | no | no | no | no | no | no | no | no | no |
| CACNA1G | 880 | S | 0 | 0 | yes | no | no | yes | no | no | no | no | no | no | no | no | no |
| CACNA1G | 881 | I | 0 | 0 | yes | no | no | yes | no | no | no | no | no | no | no | no | no |
| CACNA1G | 882 | L | 0 | 0 | yes | no | no | yes | no | no | no | no | no | no | no | no | no |
| CACNA1G | 883 | G | 0 | 0 | yes | no | no | yes | no | no | no | no | no | no | no | no | no |
| CACNA1G | 884 | M | 0 | 0 | yes | no | no | yes | no | no | no | no | no | no | no | no | no |
| CACNA1G | 885 | H | 0 | 0 | yes | no | no | yes | no | no | no | no | no | no | no | no | no |
| CACNA1G | 886 | L | 0 | 0 | yes | no | no | no | no | no | no | no | no | no | no | no | no |
| CACNA1G | 887 | F | 0 | 0 | no | yes | no | no | no | no | no | no | no | no | no | no | no |
| CACNA1G | 888 | G | 0 | 0 | no | yes | no | no | no | no | no | no | no | no | no | no | no |
| CACNA1G | 889 | C | 0 | 0 | no | yes | no | no | yes | no | no | no | no | no | no | no | no |
| CACNA1G | 890 | K | 0 | 0 | no | yes | no | no | yes | no | no | no | no | no | no | no | no |
| CACNA1G | 891 | F | 0 | 0 | no | yes | no | no | yes | no | no | no | no | no | no | no | no |
| CACNA1G | 892 | A | 0 | 0 | no | yes | no | no | yes | no | no | no | no | no | no | no | no |
| CACNA1G | 893 | S | 3 | 0 | no | yes | no | no | yes | no | no | no | no | no | no | no | no |
| CACNA1G | 900 | L | 0 | 0 | no | yes | no | no | yes | no | no | no | no | no | no | no | no |
| CACNA1G | 901 | P | 0 | 0 | no | yes | no | no | yes | no | no | no | no | no | no | no | no |
| CACNA1G | 902 | D | 0 | 0 | no | yes | no | no | yes | no | no | no | no | no | no | no | no |
| CACNA1G | 903 | R | 0 | 0 | no | yes | no | no | yes | no | no | no | no | no | no | no | no |
| CACNA1G | 904 | K | 0 | 0 | no | yes | no | no | yes | no | no | no | no | no | no | no | no |
| CACNA1G | 905 | N | 0 | 0 | no | yes | no | no | yes | no | no | no | no | no | no | no | no |
| CACNA1G | 906 | F | 0 | 0 | no | yes | no | no | yes | no | no | no | no | no | no | no | no |
| CACNA1G | 907 | D | 0 | 0 | no | yes | no | no | yes | no | no | no | no | no | no | no | no |
| CACNA1G | 908 | S | 0 | 0 | no | yes | no | no | yes | no | no | no | no | no | no | no | no |
| CACNA1G | 909 | L | 0 | 1 | no | yes | no | yes | yes | no | no | no | no | no | no | no | no |
| CACNA1G | 910 | L | 0 | 0 | yes | no | no | yes | yes | no | no | no | no | no | no | no | no |
| CACNA1G | 911 | W | 0 | 0 | yes | no | no | no | yes | no | no | no | no | no | no | no | no |
| CACNA1G | 912 | A | 1 | 0 | yes | no | no | yes | yes | no | no | no | no | no | no | no | no |
| CACNA1G | 913 | I | 1 | 0 | yes | no | no | yes | yes | no | no | no | no | no | no | no | no |

| | | | | | | | | | | | | | | | | | |
|---|---|---|---|---|---|---|---|---|---|---|---|---|---|---|---|---|---|
| CACNA1G | 914 | V | 0 | 0 | yes | no | no | yes | yes | no | no | no | no | no | no | no | no |
| CACNA1G | 915 | T | 0 | 0 | yes | no | no | yes | yes | no | no | no | no | no | no | no | no |
| CACNA1G | 916 | V | 0 | 0 | yes | no | no | yes | yes | no | no | no | no | no | no | no | no |
| CACNA1G | 917 | F | 0 | 0 | yes | no | no | yes | yes | no | no | no | no | no | no | no | no |
| CACNA1G | 918 | Q | 0 | 0 | yes | no | no | yes | yes | no | no | no | no | no | no | no | no |
| CACNA1G | 919 | I | 0 | 0 | yes | no | no | yes | yes | no | no | no | no | no | no | no | no |
| CACNA1G | 920 | L | 0 | 0 | yes | no | no | yes | yes | no | no | no | no | no | no | no | no |
| CACNA1G | 921 | T | 0 | 0 | no | yes | no | yes | yes | no | no | no | no | no | no | no | no |
| CACNA1G | 922 | Q | 0 | 0 | no | yes | no | yes | yes | no | no | no | no | no | no | no | no |
| CACNA1G | 923 | E | 0 | 0 | no | yes | no | yes | yes | no | no | no | no | no | no | no | no |
| CACNA1G | 924 | D | 0 | 0 | no | yes | no | no | yes | no | no | yes | no | no | no | no | no |
| CACNA1G | 925 | W | 0 | 0 | yes | no | no | yes | yes | no | no | no | no | no | no | no | no |
| CACNA1G | 926 | N | 0 | 0 | yes | no | no | no | yes | no | no | no | no | no | no | no | no |
| CACNA1G | 927 | K | 0 | 0 | yes | no | no | no | yes | no | no | no | no | no | no | no | no |
| CACNA1G | 928 | V | 0 | 0 | yes | no | no | no | yes | no | no | no | no | no | no | no | no |
| CACNA1G | 929 | L | 0 | 0 | yes | no | no | no | yes | no | no | no | no | no | no | no | no |
| CACNA1G | 930 | Y | 0 | 0 | yes | no | no | no | yes | no | no | no | no | no | no | no | no |
| CACNA1G | 931 | N | 0 | 0 | yes | no | no | no | yes | no | no | no | no | no | no | no | no |
| CACNA1G | 932 | G | 0 | 0 | yes | no | no | no | yes | no | no | no | no | no | no | no | no |
| CACNA1G | 933 | M | 0 | 0 | yes | no | no | no | yes | no | no | no | no | no | no | no | no |
| CACNA1G | 934 | A | 0 | 0 | yes | no | no | no | yes | no | no | no | no | no | no | no | no |
| CACNA1G | 935 | S | 0 | 0 | yes | no | no | no | yes | no | no | no | no | no | no | no | no |
| CACNA1G | 936 | T | 0 | 0 | no | yes | no | no | yes | no | no | no | no | no | no | no | no |
| CACNA1G | 937 | S | 1 | 0 | no | yes | no | no | yes | no | no | no | no | no | no | no | no |
| CACNA1G | 938 | S | 1 | 0 | yes | no | no | no | yes | no | no | no | no | no | no | no | no |
| CACNA1G | 939 | W | 0 | 0 | yes | no | no | yes | yes | no | no | no | no | no | no | no | no |
| CACNA1G | 940 | A | 0 | 0 | yes | no | no | yes | no | no | no | no | no | no | no | no | no |
| CACNA1G | 941 | A | 0 | 0 | yes | no | no | yes | no | no | no | no | no | no | no | no | no |
| CACNA1G | 942 | L | 0 | 0 | yes | no | no | yes | no | no | no | no | no | no | no | no | no |
| CACNA1G | 943 | Y | 0 | 0 | yes | no | no | yes | no | no | no | no | no | no | no | no | no |
| CACNA1G | 944 | F | 0 | 0 | yes | no | no | yes | no | no | no | no | no | no | no | no | no |
| CACNA1G | 945 | I | 0 | 0 | yes | no | no | yes | no | no | no | no | no | no | no | no | no |
| CACNA1G | 946 | A | 0 | 0 | yes | no | no | yes | no | no | no | no | no | no | no | no | no |

| | | | | | | | | | | | | | | | | | |
|---|---|---|---|---|---|---|---|---|---|---|---|---|---|---|---|---|---|
| CACNA1G | 947 | L | 0 | 0 | yes | no | no | yes | no | no | no | no | no | no | no | no | no |
| CACNA1G | 948 | M | 0 | 0 | yes | no | no | yes | no | no | no | no | no | no | no | no | no |
| CACNA1G | 949 | T | 0 | 0 | yes | no | no | yes | no | no | no | no | no | no | no | no | no |
| CACNA1G | 950 | F | 0 | 0 | yes | no | no | yes | no | no | no | no | no | no | no | no | no |
| CACNA1G | 951 | G | 1 | 0 | yes | no | no | yes | no | no | no | no | no | no | no | no | no |
| CACNA1G | 952 | N | 0 | 0 | yes | no | no | yes | no | no | no | no | no | no | no | no | no |
| CACNA1G | 953 | Y | 0 | 0 | no | yes | no | yes | no | no | no | no | no | no | no | no | no |
| CACNA1G | 954 | V | 1 | 0 | no | yes | no | yes | no | no | no | no | no | no | no | no | no |
| CACNA1G | 955 | L | 0 | 0 | yes | no | no | yes | no | no | no | no | no | no | no | no | no |
| CACNA1G | 956 | F | 0 | 0 | yes | no | no | yes | no | no | no | no | no | no | no | no | no |
| CACNA1G | 957 | N | 0 | 0 | yes | no | no | yes | no | no | no | no | no | no | no | no | no |
| CACNA1G | 958 | L | 0 | 0 | yes | no | no | yes | no | no | no | no | no | no | no | no | no |
| CACNA1G | 959 | L | 0 | 0 | yes | no | no | yes | no | no | no | no | no | no | no | no | no |
| CACNA1G | 960 | V | 0 | 0 | yes | no | no | yes | no | no | no | no | no | no | no | no | no |
| CACNA1G | 961 | A | 0 | 1 | yes | no | no | yes | no | no | no | no | no | no | no | no | no |
| CACNA1G | 962 | I | 0 | 0 | yes | no | no | yes | no | no | no | no | no | no | no | no | no |
| CACNA1G | 963 | L | 0 | 0 | yes | no | no | yes | no | no | no | no | no | no | no | no | no |
| CACNA1G | 964 | V | 0 | 0 | yes | no | no | no | no | no | no | no | no | no | no | no | no |
| CACNA1G | 965 | E | 0 | 0 | yes | no | no | no | no | yes | no | no | no | no | no | no | no |
| CACNA1G | 966 | G | 0 | 0 | yes | no | no | no | no | yes | no | no | no | no | no | no | no |
| CACNA1G | 967 | F | 0 | 0 | yes | no | no | no | no | yes | no | no | no | no | no | no | no |
| CACNA1G | 968 | Q | 0 | 0 | no | yes | no | no | no | yes | no | no | no | no | no | no | no |
| CACNA1G | 1265 | F | 0 | 0 | no | yes | no | no | no | yes | no | no | no | no | no | no | no |
| CACNA1G | 1266 | R | 2 | 0 | yes | no | no | no | no | yes | no | no | no | no | no | no | no |
| CACNA1G | 1267 | L | 0 | 0 | yes | no | no | no | no | yes | no | no | no | no | no | no | no |
| CACNA1G | 1268 | L | 0 | 0 | yes | no | no | no | no | yes | no | no | no | no | no | no | no |
| CACNA1G | 1269 | C | 0 | 0 | yes | no | no | no | no | yes | no | no | no | no | no | no | no |
| CACNA1G | 1270 | H | 1 | 0 | yes | no | no | no | no | yes | no | no | no | no | no | no | no |
| CACNA1G | 1271 | R | 1 | 0 | yes | no | no | no | no | yes | no | no | no | no | no | no | no |
| CACNA1G | 1272 | I | 0 | 0 | yes | no | no | no | no | yes | no | no | no | no | no | no | no |
| CACNA1G | 1273 | I | 1 | 0 | yes | no | no | no | no | no | no | no | no | no | no | no | no |
| CACNA1G | 1274 | T | 0 | 0 | yes | no | no | no | no | no | no | no | no | no | no | no | no |
| CACNA1G | 1275 | H | 0 | 0 | no | yes | no | no | no | no | no | no | no | no | no | no | no |

| | | | | | | | | | | | | | | | | | |
|---|---|---|---|---|---|---|---|---|---|---|---|---|---|---|---|---|---|
| CACNA1G | 1276 | K | 0 | 0 | yes | no | no | no | no | no | no | no | no | no | no | no | no |
| CACNA1G | 1277 | M | 1 | 0 | yes | no | no | yes | no | no | no | no | no | no | no | no | no |
| CACNA1G | 1278 | F | 0 | 0 | yes | no | no | yes | no | no | no | no | no | no | no | no | no |
| CACNA1G | 1279 | D | 0 | 0 | yes | no | no | yes | no | no | no | no | no | no | no | no | no |
| CACNA1G | 1280 | H | 0 | 0 | yes | no | no | yes | no | no | no | no | no | no | no | no | no |
| CACNA1G | 1281 | V | 1 | 0 | yes | no | no | yes | no | no | no | no | no | no | no | no | no |
| CACNA1G | 1282 | V | 1 | 0 | yes | no | no | yes | no | no | no | no | no | no | no | no | no |
| CACNA1G | 1283 | L | 0 | 0 | yes | no | no | yes | no | no | no | no | no | no | no | no | no |
| CACNA1G | 1284 | V | 0 | 0 | yes | no | no | yes | no | no | no | no | no | no | no | no | no |
| CACNA1G | 1285 | I | 0 | 0 | yes | no | no | yes | no | no | no | no | no | no | no | no | no |
| CACNA1G | 1286 | I | 0 | 0 | yes | no | no | yes | no | no | no | no | no | no | no | no | no |
| CACNA1G | 1287 | F | 0 | 0 | yes | no | no | yes | no | no | no | no | no | no | no | no | no |
| CACNA1G | 1288 | L | 0 | 0 | yes | no | no | yes | no | no | no | no | no | no | no | no | no |
| CACNA1G | 1289 | N | 0 | 0 | yes | no | no | yes | no | no | no | no | no | no | no | no | no |
| CACNA1G | 1290 | C | 0 | 0 | yes | no | no | yes | no | no | no | no | no | no | no | no | no |
| CACNA1G | 1291 | I | 0 | 0 | yes | no | no | yes | no | no | no | no | no | no | no | no | no |
| CACNA1G | 1292 | T | 0 | 0 | yes | no | no | yes | no | no | no | no | no | no | no | no | no |
| CACNA1G | 1293 | I | 0 | 0 | yes | no | no | yes | no | no | no | no | no | no | no | no | no |
| CACNA1G | 1294 | A | 2 | 0 | yes | no | no | yes | no | no | no | no | no | no | no | no | no |
| CACNA1G | 1295 | M | 0 | 0 | no | yes | no | yes | no | no | no | no | no | no | no | no | no |
| CACNA1G | 1296 | E | 0 | 0 | no | yes | no | yes | yes | no | no | no | no | no | no | no | no |
| CACNA1G | 1297 | R | 1 | 0 | no | yes | no | no | yes | no | no | no | no | no | no | no | no |
| CACNA1G | 1298 | P | 0 | 0 | no | yes | no | no | yes | no | no | no | no | no | no | no | no |
| CACNA1G | 1299 | K | 1 | 0 | no | yes | no | no | yes | no | no | no | no | no | no | no | no |
| CACNA1G | 1300 | I | 0 | 0 | no | yes | no | no | yes | no | no | no | no | no | no | no | no |
| CACNA1G | 1301 | D | 0 | 0 | no | yes | no | no | yes | no | no | no | no | no | no | no | no |
| CACNA1G | 1302 | P | 1 | 0 | no | yes | no | no | yes | no | no | no | no | no | no | no | no |
| CACNA1G | 1303 | H | 1 | 0 | no | yes | no | no | yes | no | no | no | no | no | no | no | no |
| CACNA1G | 1304 | S | 0 | 0 | no | yes | no | no | yes | no | no | no | no | no | no | no | no |
| CACNA1G | 1305 | A | 1 | 0 | no | yes | no | no | yes | no | no | no | no | no | no | no | no |
| CACNA1G | 1306 | E | 1 | 0 | no | yes | no | no | yes | no | no | no | no | no | no | no | no |
| CACNA1G | 1307 | R | 0 | 0 | yes | no | no | no | yes | no | no | no | no | no | no | no | no |
| CACNA1G | 1308 | I | 0 | 0 | yes | no | no | no | yes | no | no | no | no | no | no | no | no |

| | | | | | | | | | | | | | | | | | |
|---|---|---|---|---|---|---|---|---|---|---|---|---|---|---|---|---|---|
| CACNA1G | 1309 | F | 0 | 0 | yes | no | no | yes | yes | no | no | no | no | no | no | no | no |
| CACNA1G | 1310 | L | 0 | 0 | yes | no | no | yes | yes | no | no | no | no | no | no | no | no |
| CACNA1G | 1311 | T | 1 | 0 | yes | no | no | yes | yes | no | no | no | no | no | no | no | no |
| CACNA1G | 1312 | L | 0 | 0 | yes | no | no | yes | yes | no | no | no | no | no | no | no | no |
| CACNA1G | 1313 | S | 0 | 0 | yes | no | no | yes | yes | no | no | no | no | no | no | no | no |
| CACNA1G | 1314 | N | 0 | 0 | yes | no | no | yes | no | no | no | no | no | no | no | no | no |
| CACNA1G | 1315 | Y | 0 | 0 | yes | no | no | yes | no | no | no | no | no | no | no | no | no |
| CACNA1G | 1316 | I | 0 | 0 | yes | no | no | yes | no | no | no | no | no | no | no | no | no |
| CACNA1G | 1317 | F | 0 | 0 | yes | no | no | yes | no | no | no | no | no | no | no | no | no |
| CACNA1G | 1318 | T | 0 | 0 | yes | no | no | yes | no | no | no | no | no | no | no | no | no |
| CACNA1G | 1319 | A | 1 | 0 | yes | no | no | yes | no | no | no | no | no | no | no | no | no |
| CACNA1G | 1320 | V | 0 | 0 | yes | no | no | yes | no | no | no | no | no | no | no | no | no |
| CACNA1G | 1321 | F | 0 | 0 | yes | no | no | yes | no | no | no | no | no | no | no | no | no |
| CACNA1G | 1322 | L | 1 | 0 | yes | no | no | yes | no | no | no | no | no | no | no | no | no |
| CACNA1G | 1323 | A | 0 | 0 | yes | no | no | yes | no | no | no | no | no | no | no | no | no |
| CACNA1G | 1324 | E | 0 | 0 | yes | no | no | yes | no | no | no | no | no | no | no | no | no |
| CACNA1G | 1325 | M | 1 | 0 | yes | no | no | yes | no | no | no | no | no | no | no | no | no |
| CACNA1G | 1326 | T | 0 | 0 | yes | no | no | yes | no | no | no | no | no | no | no | no | no |
| CACNA1G | 1327 | V | 1 | 0 | yes | no | no | yes | no | no | no | no | no | no | no | no | no |
| CACNA1G | 1328 | K | 0 | 0 | yes | no | no | yes | no | no | no | no | no | no | no | no | no |
| CACNA1G | 1329 | V | 0 | 0 | yes | no | no | yes | no | no | no | no | no | no | no | no | no |
| CACNA1G | 1330 | V | 0 | 0 | yes | no | no | no | no | no | no | no | no | no | no | no | no |
| CACNA1G | 1331 | A | 0 | 0 | yes | no | no | no | no | no | no | no | no | no | no | no | no |
| CACNA1G | 1332 | L | 0 | 0 | no | yes | no | no | no | no | no | no | no | no | no | no | no |
| CACNA1G | 1333 | G | 0 | 0 | no | yes | no | no | no | no | no | no | no | no | no | no | no |
| CACNA1G | 1344 | S | 0 | 0 | no | yes | no | no | no | yes | no | no | no | no | no | no | no |
| CACNA1G | 1345 | S | 0 | 0 | yes | no | no | yes | no | no | no | no | no | no | no | no | no |
| CACNA1G | 1346 | W | 0 | 0 | yes | no | no | yes | no | no | no | no | no | no | no | no | no |
| CACNA1G | 1347 | N | 0 | 0 | yes | no | no | yes | no | no | no | no | no | no | no | no | no |
| CACNA1G | 1348 | V | 1 | 0 | yes | no | no | yes | no | no | no | no | no | no | no | no | no |
| CACNA1G | 1349 | L | 0 | 0 | yes | no | no | yes | no | no | no | no | no | no | no | no | no |
| CACNA1G | 1350 | D | 0 | 0 | yes | no | no | yes | no | no | no | no | no | no | no | no | no |
| CACNA1G | 1351 | G | 0 | 0 | yes | no | no | yes | no | no | no | no | no | no | no | no | no |

| | | | | | | | | | | | | | | | | | |
|---|---|---|---|---|---|---|---|---|---|---|---|---|---|---|---|---|---|
| CACNA1G | 1352 | L | 0 | 0 | yes | no | no | yes | no | no | no | no | no | no | no | no | no |
| CACNA1G | 1353 | L | 0 | 0 | yes | no | no | yes | no | no | no | no | no | no | no | no | no |
| CACNA1G | 1354 | V | 0 | 0 | yes | no | no | yes | no | no | no | no | no | no | no | no | no |
| CACNA1G | 1355 | L | 1 | 0 | yes | no | no | yes | no | no | no | no | no | no | no | no | no |
| CACNA1G | 1356 | I | 1 | 0 | yes | no | no | yes | no | no | no | no | no | no | no | no | no |
| CACNA1G | 1357 | S | 0 | 0 | yes | no | no | yes | no | no | no | no | no | no | no | no | no |
| CACNA1G | 1358 | V | 2 | 0 | yes | no | no | yes | no | no | no | no | no | no | no | no | no |
| CACNA1G | 1359 | I | 0 | 0 | yes | no | no | yes | no | no | no | no | no | no | no | no | no |
| CACNA1G | 1360 | D | 1 | 0 | yes | no | no | yes | no | no | no | no | no | no | no | no | no |
| CACNA1G | 1361 | I | 0 | 0 | yes | no | no | yes | no | no | no | no | no | no | no | no | no |
| CACNA1G | 1362 | L | 0 | 0 | yes | no | no | yes | no | no | no | no | no | no | no | no | no |
| CACNA1G | 1363 | V | 1 | 0 | yes | no | no | yes | no | no | no | no | no | no | no | no | no |
| CACNA1G | 1364 | S | 0 | 0 | yes | no | no | yes | no | no | no | no | no | no | no | no | no |
| CACNA1G | 1365 | M | 2 | 0 | yes | no | no | yes | yes | no | no | no | no | no | no | no | no |
| CACNA1G | 1366 | V | 0 | 0 | yes | no | no | no | yes | no | no | no | no | no | no | no | no |
| CACNA1G | 1367 | S | 0 | 0 | no | yes | no | no | yes | no | no | no | no | no | no | no | no |
| CACNA1G | 1372 | K | 0 | 0 | no | yes | no | no | yes | no | no | no | no | no | no | no | no |
| CACNA1G | 1373 | I | 0 | 0 | yes | no | no | no | yes | no | no | no | no | no | no | no | no |
| CACNA1G | 1374 | L | 0 | 0 | yes | no | no | no | yes | no | no | no | no | no | no | no | no |
| CACNA1G | 1375 | G | 0 | 0 | yes | no | no | no | yes | no | no | no | no | no | no | no | no |
| CACNA1G | 1376 | M | 0 | 0 | yes | no | no | yes | yes | no | no | no | no | no | no | no | no |
| CACNA1G | 1377 | L | 1 | 0 | yes | no | no | yes | yes | no | no | no | no | no | no | no | no |
| CACNA1G | 1378 | R | 0 | 0 | yes | no | no | no | yes | no | no | no | no | no | no | no | no |
| CACNA1G | 1379 | V | 0 | 0 | yes | no | no | yes | no | no | no | no | no | no | no | no | no |
| CACNA1G | 1380 | L | 0 | 0 | yes | no | no | yes | no | no | no | no | no | no | no | no | no |
| CACNA1G | 1381 | R | 1 | 0 | yes | no | no | yes | no | no | no | no | no | no | no | no | no |
| CACNA1G | 1382 | L | 0 | 0 | yes | no | no | yes | no | no | no | no | no | no | no | no | no |
| CACNA1G | 1383 | L | 0 | 0 | yes | no | no | yes | no | no | no | no | no | no | no | no | no |
| CACNA1G | 1384 | R | 1 | 0 | yes | no | no | yes | no | no | no | no | no | no | no | no | no |
| CACNA1G | 1385 | T | 0 | 0 | no | yes | no | yes | no | no | no | no | no | no | no | no | no |
| CACNA1G | 1386 | L | 0 | 0 | no | yes | no | yes | no | no | no | no | no | no | no | no | no |
| CACNA1G | 1387 | R | 0 | 0 | yes | no | no | yes | no | no | no | no | no | no | no | no | no |
| CACNA1G | 1388 | P | 2 | 0 | yes | no | no | yes | no | no | no | no | no | no | no | no | no |

| | | | | | | | | | | | | | | | | | |
|---|---|---|---|---|---|---|---|---|---|---|---|---|---|---|---|---|---|
| CACNA1G | 1389 | L | 0 | 0 | yes | no | no | yes | no | no | no | no | no | no | no | no | no |
| CACNA1G | 1390 | R | 0 | 0 | yes | no | no | yes | no | no | no | no | no | no | no | no | no |
| CACNA1G | 1391 | V | 0 | 0 | yes | no | no | yes | no | no | no | no | no | no | no | no | no |
| CACNA1G | 1392 | I | 0 | 0 | yes | no | no | yes | no | no | no | no | no | no | no | no | no |
| CACNA1G | 1393 | S | 0 | 0 | yes | no | no | yes | no | no | no | no | no | no | no | no | no |
| CACNA1G | 1394 | R | 0 | 0 | yes | no | no | yes | no | no | no | no | no | no | no | no | no |
| CACNA1G | 1395 | A | 0 | 0 | no | yes | no | yes | no | no | no | no | no | no | no | no | no |
| CACNA1G | 1396 | Q | 0 | 0 | yes | no | no | no | no | no | no | no | no | no | no | no | no |
| CACNA1G | 1397 | G | 0 | 0 | yes | no | no | yes | no | no | no | no | no | no | no | no | no |
| CACNA1G | 1398 | L | 0 | 0 | yes | no | no | yes | no | no | no | no | no | no | no | no | no |
| CACNA1G | 1399 | K | 0 | 0 | yes | no | no | yes | no | no | no | no | no | no | no | no | no |
| CACNA1G | 1400 | L | 0 | 0 | yes | no | no | no | no | no | no | no | no | no | no | no | no |
| CACNA1G | 1401 | V | 0 | 0 | yes | no | no | yes | no | yes | no | no | no | no | no | no | no |
| CACNA1G | 1402 | V | 0 | 0 | yes | no | no | yes | no | yes | no | no | no | no | no | no | no |
| CACNA1G | 1403 | E | 0 | 0 | yes | no | no | yes | no | yes | no | no | no | no | no | no | no |
| CACNA1G | 1404 | T | 0 | 0 | yes | no | no | no | no | yes | no | no | no | no | no | no | no |
| CACNA1G | 1405 | L | 0 | 0 | yes | no | no | yes | no | yes | no | no | no | no | no | no | no |
| CACNA1G | 1406 | M | 0 | 0 | yes | no | no | yes | no | yes | no | no | no | no | no | no | no |
| CACNA1G | 1407 | S | 0 | 0 | yes | no | no | no | no | yes | no | no | no | no | no | no | no |
| CACNA1G | 1408 | S | 0 | 0 | yes | no | no | yes | no | yes | no | no | no | no | no | no | no |
| CACNA1G | 1409 | L | 0 | 0 | no | yes | no | yes | no | yes | no | no | no | no | no | no | no |
| CACNA1G | 1410 | K | 0 | 0 | no | yes | no | no | no | yes | no | no | no | no | no | no | no |
| CACNA1G | 1411 | P | 0 | 0 | no | yes | no | no | no | no | no | no | no | no | no | no | no |
| CACNA1G | 1412 | I | 0 | 0 | no | yes | no | yes | no | no | no | no | no | no | no | no | no |
| CACNA1G | 1413 | G | 0 | 0 | yes | no | no | yes | no | no | no | no | no | no | no | no | no |
| CACNA1G | 1414 | N | 0 | 0 | yes | no | no | yes | no | no | no | no | no | no | no | no | no |
| CACNA1G | 1415 | I | 0 | 0 | yes | no | no | yes | no | no | no | no | no | no | no | no | no |
| CACNA1G | 1416 | V | 0 | 0 | yes | no | no | yes | no | no | no | no | no | no | no | no | no |
| CACNA1G | 1417 | V | 0 | 0 | yes | no | no | yes | no | no | no | no | no | no | no | no | no |
| CACNA1G | 1418 | I | 1 | 0 | yes | no | no | yes | no | no | no | no | no | no | no | no | no |
| CACNA1G | 1419 | C | 0 | 0 | yes | no | no | yes | no | no | no | no | no | no | no | no | no |
| CACNA1G | 1420 | C | 0 | 0 | yes | no | no | yes | no | no | no | no | no | no | no | no | no |
| CACNA1G | 1421 | A | 0 | 0 | yes | no | no | yes | no | no | no | no | no | no | no | no | no |

| | | | | | | | | | | | | | | | | | |
|---|---|---|---|---|---|---|---|---|---|---|---|---|---|---|---|---|---|
| CACNA1G | 1422 | F | 1 | 0 | yes | no | no | yes | no | no | no | no | no | no | no | no | no |
| CACNA1G | 1423 | F | 0 | 0 | yes | no | no | yes | no | no | no | no | no | no | no | no | no |
| CACNA1G | 1424 | I | 1 | 0 | yes | no | no | yes | no | no | no | no | no | no | no | no | no |
| CACNA1G | 1425 | I | 0 | 0 | yes | no | no | yes | no | no | no | no | no | no | no | no | no |
| CACNA1G | 1426 | F | 0 | 0 | yes | no | no | yes | no | no | no | no | no | no | no | no | no |
| CACNA1G | 1427 | G | 0 | 0 | yes | no | no | yes | no | no | no | no | no | no | no | no | no |
| CACNA1G | 1428 | I | 0 | 0 | yes | no | no | yes | no | no | no | no | no | no | no | no | no |
| CACNA1G | 1429 | L | 0 | 0 | yes | no | no | yes | no | no | no | no | no | no | no | no | no |
| CACNA1G | 1430 | G | 0 | 0 | yes | no | no | yes | no | no | no | no | no | no | no | no | no |
| CACNA1G | 1431 | V | 1 | 0 | yes | no | no | yes | no | no | no | no | no | no | no | no | no |
| CACNA1G | 1432 | Q | 0 | 0 | yes | no | no | yes | no | no | no | no | no | no | no | no | no |
| CACNA1G | 1433 | L | 0 | 0 | yes | no | no | yes | no | no | no | no | no | no | no | no | no |
| CACNA1G | 1434 | F | 0 | 0 | yes | no | no | no | no | no | no | no | no | no | no | no | no |
| CACNA1G | 1435 | K | 0 | 0 | no | yes | no | no | yes | no | no | no | no | no | no | no | no |
| CACNA1G | 1436 | G | 0 | 0 | no | yes | no | no | yes | no | no | no | no | no | no | no | no |
| CACNA1G | 1437 | K | 0 | 0 | no | yes | no | no | yes | no | no | no | no | no | no | no | no |
| CACNA1G | 1438 | F | 0 | 0 | no | yes | no | no | yes | no | no | no | no | no | no | no | no |
| CACNA1G | 1439 | F | 0 | 0 | no | no | yes | no | yes | no | no | no | no | no | no | no | no |
| CACNA1G | 1440 | V | 1 | 0 | no | no | yes | no | yes | no | no | no | no | no | no | no | no |
| CACNA1G | 1441 | C | 0 | 0 | no | no | yes | no | yes | no | no | no | no | no | no | no | no |
| CACNA1G | 1442 | Q | 0 | 0 | no | no | yes | no | yes | no | no | no | no | no | no | no | no |
| CACNA1G | 1443 | G | 0 | 0 | no | yes | no | no | yes | no | no | no | no | no | no | no | no |
| CACNA1G | 1444 | E | 1 | 0 | no | yes | no | no | yes | no | no | no | no | no | no | no | no |
| CACNA1G | 1445 | D | 0 | 0 | no | yes | no | no | yes | no | no | no | no | no | no | no | no |
| CACNA1G | 1446 | T | 0 | 0 | no | yes | no | no | yes | no | no | no | no | no | no | no | no |
| CACNA1G | 1447 | R | 0 | 0 | no | yes | no | no | yes | no | no | no | no | no | no | no | no |
| CACNA1G | 1448 | N | 0 | 0 | no | yes | no | no | yes | no | no | no | no | no | no | no | no |
| CACNA1G | 1449 | I | 0 | 0 | no | yes | no | no | yes | no | no | no | no | no | no | no | no |
| CACNA1G | 1450 | T | 0 | 0 | no | yes | no | no | yes | no | no | no | no | no | no | no | no |
| CACNA1G | 1451 | N | 0 | 0 | no | no | yes | no | yes | no | no | no | no | no | no | no | no |
| CACNA1G | 1452 | K | 0 | 0 | yes | no | no | no | yes | no | no | no | no | no | no | no | no |
| CACNA1G | 1453 | S | 1 | 0 | yes | no | no | no | yes | no | no | no | no | no | no | no | no |
| CACNA1G | 1454 | D | 1 | 0 | yes | no | no | no | yes | no | no | no | no | no | no | no | no |

| | | | | | | | | | | | | | | | | | |
|---|---|---|---|---|---|---|---|---|---|---|---|---|---|---|---|---|---|
| CACNA1G | 1455 | C | 0 | 0 | yes | no | no | no | yes | no | no | no | no | no | no | no | no |
| CACNA1G | 1456 | A | 1 | 0 | yes | no | no | no | yes | no | no | no | no | no | no | no | no |
| CACNA1G | 1457 | E | 3 | 0 | yes | no | no | no | yes | no | no | no | no | no | no | no | no |
| CACNA1G | 1458 | A | 0 | 0 | yes | no | no | no | yes | no | no | no | no | no | no | no | no |
| CACNA1G | 1459 | S | 1 | 0 | no | yes | no | no | yes | no | no | no | no | no | no | no | no |
| CACNA1G | 1460 | Y | 0 | 0 | no | yes | no | no | yes | no | no | no | no | no | no | no | no |
| CACNA1G | 1461 | R | 1 | 0 | no | no | yes | no | yes | no | no | no | no | no | no | no | no |
| CACNA1G | 1462 | W | 0 | 0 | no | no | yes | no | yes | no | no | no | no | no | no | no | no |
| CACNA1G | 1463 | V | 0 | 0 | no | no | yes | no | yes | no | no | no | no | no | no | no | no |
| CACNA1G | 1464 | R | 1 | 0 | no | no | yes | no | yes | no | no | no | no | no | no | no | no |
| CACNA1G | 1465 | H | 0 | 0 | no | yes | no | no | yes | no | no | yes | no | no | no | no | no |
| CACNA1G | 1466 | K | 0 | 0 | no | yes | no | no | yes | no | no | no | no | no | no | no | no |
| CACNA1G | 1467 | Y | 0 | 0 | no | yes | no | no | yes | no | no | no | no | no | no | no | no |
| CACNA1G | 1468 | N | 0 | 0 | no | yes | no | no | yes | no | no | no | no | no | no | no | no |
| CACNA1G | 1469 | F | 0 | 0 | no | yes | no | yes | yes | no | no | no | no | no | no | no | no |
| CACNA1G | 1470 | D | 0 | 0 | no | yes | no | no | yes | no | no | no | no | no | no | no | no |
| CACNA1G | 1471 | N | 0 | 0 | no | yes | no | yes | yes | no | no | no | no | no | no | no | no |
| CACNA1G | 1472 | L | 0 | 0 | yes | no | no | yes | yes | no | no | no | no | no | no | no | no |
| CACNA1G | 1473 | G | 0 | 0 | yes | no | no | yes | yes | no | no | no | no | no | no | no | no |
| CACNA1G | 1474 | Q | 0 | 0 | yes | no | no | no | yes | no | no | no | no | no | no | no | no |
| CACNA1G | 1475 | A | 0 | 0 | yes | no | no | yes | yes | no | no | no | no | no | no | no | no |
| CACNA1G | 1476 | L | 0 | 0 | yes | no | no | yes | yes | no | no | no | no | no | no | no | no |
| CACNA1G | 1477 | M | 0 | 0 | yes | no | no | yes | yes | no | no | no | no | no | no | no | no |
| CACNA1G | 1478 | S | 1 | 0 | yes | no | no | yes | yes | no | no | no | no | no | no | no | no |
| CACNA1G | 1479 | L | 0 | 0 | yes | no | no | yes | yes | no | no | no | no | no | no | no | no |
| CACNA1G | 1480 | F | 0 | 0 | yes | no | no | yes | yes | no | no | no | no | no | no | no | no |
| CACNA1G | 1481 | V | 1 | 0 | yes | no | no | yes | yes | no | no | no | no | no | no | no | no |
| CACNA1G | 1482 | L | 0 | 0 | yes | no | no | yes | yes | no | no | no | no | no | no | no | no |
| CACNA1G | 1483 | A | 0 | 0 | yes | no | no | yes | yes | no | no | no | no | no | no | no | no |
| CACNA1G | 1484 | S | 0 | 0 | no | yes | no | yes | yes | no | no | no | no | no | no | no | no |
| CACNA1G | 1485 | K | 1 | 0 | no | yes | no | yes | yes | no | no | no | no | no | no | no | no |
| CACNA1G | 1486 | D | 0 | 0 | no | yes | no | yes | yes | no | no | no | no | no | no | no | no |
| CACNA1G | 1487 | G | 0 | 0 | no | yes | no | yes | yes | no | no | no | no | no | no | no | no |

| | | | | | | | | | | | | | | | | | |
|---|---|---|---|---|---|---|---|---|---|---|---|---|---|---|---|---|---|
| CACNA1G | 1488 | W | 0 | 0 | yes | no | no | yes | yes | no | no | no | no | no | no | no | no |
| CACNA1G | 1489 | V | 0 | 0 | yes | no | no | no | yes | no | no | no | no | no | no | no | no |
| CACNA1G | 1490 | D | 0 | 0 | yes | no | no | no | yes | no | no | no | no | no | no | no | no |
| CACNA1G | 1491 | I | 0 | 0 | yes | no | no | no | yes | no | no | no | no | no | no | no | no |
| CACNA1G | 1492 | M | 1 | 0 | yes | no | no | no | yes | no | no | no | no | no | no | no | no |
| CACNA1G | 1493 | Y | 0 | 0 | yes | no | no | no | yes | no | no | no | no | no | no | no | no |
| CACNA1G | 1494 | D | 2 | 0 | yes | no | no | no | yes | no | no | no | no | no | no | no | no |
| CACNA1G | 1495 | G | 0 | 0 | yes | no | no | no | yes | no | no | no | no | no | no | no | no |
| CACNA1G | 1496 | L | 0 | 0 | yes | no | no | no | yes | no | no | no | no | no | no | no | no |
| CACNA1G | 1497 | D | 0 | 0 | yes | no | no | no | yes | no | no | no | no | no | no | no | no |
| CACNA1G | 1498 | A | 0 | 0 | no | yes | no | no | yes | no | no | no | no | no | no | no | no |
| CACNA1G | 1499 | V | 0 | 0 | no | yes | no | no | yes | no | no | no | no | no | no | no | no |
| CACNA1G | 1500 | G | 0 | 0 | no | yes | no | no | yes | no | no | no | no | no | no | no | no |
| CACNA1G | 1501 | V | 1 | 0 | no | yes | no | no | yes | no | no | no | no | no | no | no | no |
| CACNA1G | 1502 | D | 0 | 0 | no | yes | no | no | yes | no | no | no | no | no | no | no | no |
| CACNA1G | 1503 | Q | 0 | 0 | no | no | yes | no | yes | no | no | no | no | no | no | no | no |
| CACNA1G | 1504 | Q | 0 | 0 | no | yes | no | no | yes | no | no | no | no | no | no | no | no |
| CACNA1G | 1505 | P | 0 | 0 | no | yes | no | no | yes | no | no | no | no | no | no | no | no |
| CACNA1G | 1506 | I | 2 | 0 | no | yes | no | no | yes | no | no | no | no | no | no | no | no |
| CACNA1G | 1507 | M | 1 | 0 | no | yes | no | no | yes | no | no | no | no | no | no | no | no |
| CACNA1G | 1508 | N | 0 | 0 | no | yes | no | no | yes | no | no | no | no | no | no | no | no |
| CACNA1G | 1509 | H | 0 | 0 | no | yes | no | no | yes | no | no | no | no | no | no | no | no |
| CACNA1G | 1510 | N | 0 | 0 | no | yes | no | no | yes | no | no | no | no | no | no | no | no |
| CACNA1G | 1511 | P | 1 | 0 | yes | no | no | no | yes | no | no | no | no | no | no | no | no |
| CACNA1G | 1512 | W | 0 | 0 | yes | no | no | yes | no | no | no | no | no | no | no | no | no |
| CACNA1G | 1513 | M | 1 | 0 | yes | no | no | yes | no | no | no | no | no | no | no | no | no |
| CACNA1G | 1514 | L | 0 | 0 | yes | no | no | yes | no | no | no | no | no | no | no | no | no |
| CACNA1G | 1515 | L | 0 | 0 | yes | no | no | yes | no | no | no | no | no | no | no | no | no |
| CACNA1G | 1516 | Y | 0 | 0 | yes | no | no | yes | no | no | no | no | no | no | no | no | no |
| CACNA1G | 1517 | F | 0 | 0 | yes | no | no | yes | no | no | no | no | no | no | no | no | no |
| CACNA1G | 1518 | I | 0 | 0 | yes | no | no | yes | no | no | no | no | no | no | no | no | no |
| CACNA1G | 1519 | S | 1 | 0 | yes | no | no | yes | no | no | no | no | no | no | no | no | no |
| CACNA1G | 1520 | F | 0 | 0 | yes | no | no | yes | no | no | no | no | no | no | no | no | no |

| | | | | | | | | | | | | | | | | | |
|---|---|---|---|---|---|---|---|---|---|---|---|---|---|---|---|---|---|
| CACNA1G | 1521 | L | 0 | 0 | yes | no | no | yes | no | no | no | no | no | no | no | no | no |
| CACNA1G | 1522 | L | 0 | 0 | yes | no | no | yes | no | no | no | no | no | no | no | no | no |
| CACNA1G | 1523 | I | 0 | 0 | yes | no | no | yes | no | no | no | no | no | no | no | no | no |
| CACNA1G | 1524 | V | 0 | 0 | yes | no | no | yes | no | no | no | no | no | no | no | no | no |
| CACNA1G | 1525 | A | 0 | 0 | yes | no | no | yes | no | no | no | no | no | no | no | no | no |
| CACNA1G | 1526 | F | 0 | 0 | yes | no | no | yes | no | no | no | no | no | no | no | no | no |
| CACNA1G | 1527 | F | 0 | 0 | yes | no | no | yes | no | no | no | no | no | no | no | no | no |
| CACNA1G | 1528 | V | 0 | 0 | yes | no | no | yes | no | no | no | no | no | no | no | no | no |
| CACNA1G | 1529 | L | 0 | 0 | yes | no | no | yes | no | no | no | no | no | no | no | no | no |
| CACNA1G | 1530 | N | 0 | 0 | yes | no | no | yes | no | no | no | no | no | no | no | no | no |
| CACNA1G | 1531 | M | 0 | 1 | yes | no | no | yes | no | no | no | no | no | no | no | no | no |
| CACNA1G | 1532 | F | 0 | 0 | yes | no | no | yes | no | no | yes | no | no | no | no | no | no |
| CACNA1G | 1533 | V | 0 | 0 | yes | no | no | yes | no | no | no | no | no | no | no | no | no |
| CACNA1G | 1534 | G | 0 | 1 | yes | no | no | yes | no | no | no | no | no | no | no | no | no |
| CACNA1G | 1535 | V | 0 | 0 | yes | no | no | yes | no | no | no | no | no | no | no | no | no |
| CACNA1G | 1536 | V | 0 | 0 | yes | no | no | no | no | no | yes | no | no | no | no | no | no |
| CACNA1G | 1537 | V | 0 | 0 | yes | no | no | no | no | no | no | no | no | no | no | no | no |
| CACNA1G | 1538 | E | 0 | 0 | yes | no | no | no | no | yes | no | no | no | no | no | no | no |
| CACNA1G | 1539 | N | 0 | 0 | yes | no | no | no | no | yes | no | no | no | no | no | no | no |
| CACNA1G | 1540 | F | 0 | 0 | yes | no | no | no | no | yes | no | no | no | no | no | no | no |
| CACNA1G | 1541 | H | 0 | 0 | no | yes | no | no | no | yes | no | no | no | no | no | no | no |
| CACNA1G | 1612 | Y | 0 | 0 | no | yes | no | no | no | no | no | no | no | no | no | no | no |
| CACNA1G | 1613 | L | 0 | 0 | yes | no | no | yes | no | no | no | no | no | no | no | no | no |
| CACNA1G | 1614 | D | 0 | 0 | yes | no | no | yes | no | no | no | no | no | no | no | no | no |
| CACNA1G | 1615 | L | 0 | 0 | yes | no | no | yes | no | no | no | no | no | no | no | no | no |
| CACNA1G | 1616 | F | 0 | 0 | yes | no | no | yes | no | no | no | no | no | no | no | no | no |
| CACNA1G | 1617 | I | 0 | 0 | yes | no | no | yes | no | no | no | no | no | no | no | no | no |
| CACNA1G | 1618 | T | 0 | 0 | yes | no | no | yes | no | no | no | no | no | no | no | no | no |
| CACNA1G | 1619 | G | 1 | 0 | yes | no | no | yes | no | no | no | no | no | no | no | no | no |
| CACNA1G | 1620 | V | 0 | 0 | yes | no | no | yes | no | no | no | no | no | no | no | no | no |
| CACNA1G | 1621 | I | 0 | 0 | yes | no | no | yes | no | no | no | no | no | no | no | no | no |
| CACNA1G | 1622 | G | 2 | 0 | yes | no | no | yes | no | no | no | no | no | no | no | no | no |
| CACNA1G | 1623 | L | 0 | 0 | yes | no | no | yes | no | no | no | no | no | no | no | no | no |

| | | | | | | | | | | | | | | | | | |
|---|---|---|---|---|---|---|---|---|---|---|---|---|---|---|---|---|---|
| CACNA1G | 1624 | N | 0 | 0 | yes | no | no | yes | no | no | no | no | no | no | no | no | no |
| CACNA1G | 1625 | V | 1 | 0 | yes | no | no | yes | no | no | no | no | no | no | no | no | no |
| CACNA1G | 1626 | V | 1 | 0 | yes | no | no | yes | no | no | no | no | no | no | no | no | no |
| CACNA1G | 1627 | T | 0 | 0 | yes | no | no | yes | no | no | no | no | no | no | no | no | no |
| CACNA1G | 1628 | M | 0 | 0 | yes | no | no | yes | no | no | no | no | no | no | no | no | no |
| CACNA1G | 1629 | A | 0 | 0 | yes | no | no | yes | no | no | no | no | no | no | no | no | no |
| CACNA1G | 1630 | M | 0 | 0 | no | yes | no | yes | no | no | no | no | no | no | no | no | no |
| CACNA1G | 1631 | E | 0 | 0 | no | yes | no | yes | no | no | no | no | no | no | no | no | no |
| CACNA1G | 1632 | H | 0 | 0 | no | yes | no | no | yes | no | no | no | no | no | no | no | no |
| CACNA1G | 1633 | Y | 0 | 1 | no | yes | no | no | yes | no | no | no | no | no | no | no | no |
| CACNA1G | 1634 | Q | 0 | 0 | no | yes | no | no | yes | no | no | no | no | no | no | no | no |
| CACNA1G | 1635 | Q | 0 | 0 | no | yes | no | no | yes | no | no | no | no | no | no | no | no |
| CACNA1G | 1636 | P | 0 | 0 | no | yes | no | no | yes | no | no | no | no | no | no | no | no |
| CACNA1G | 1637 | Q | 0 | 0 | yes | no | no | no | yes | no | no | no | no | no | no | no | no |
| CACNA1G | 1638 | I | 0 | 0 | yes | no | no | no | yes | no | no | no | no | no | no | no | no |
| CACNA1G | 1639 | L | 0 | 0 | yes | no | no | no | yes | no | no | no | no | no | no | no | no |
| CACNA1G | 1640 | D | 0 | 0 | yes | no | no | no | yes | no | no | no | no | no | no | no | no |
| CACNA1G | 1641 | E | 0 | 0 | yes | no | no | no | yes | no | no | no | no | no | no | no | no |
| CACNA1G | 1642 | A | 0 | 0 | yes | no | no | yes | yes | no | no | no | no | no | no | no | no |
| CACNA1G | 1643 | L | 0 | 0 | yes | no | no | yes | yes | no | no | no | no | no | no | no | no |
| CACNA1G | 1644 | K | 1 | 0 | yes | no | no | yes | yes | no | no | no | no | no | no | no | no |
| CACNA1G | 1645 | I | 0 | 0 | yes | no | no | yes | yes | no | no | no | no | no | no | no | no |
| CACNA1G | 1646 | C | 0 | 0 | yes | no | no | yes | no | no | no | no | no | no | no | no | no |
| CACNA1G | 1647 | N | 1 | 0 | yes | no | no | yes | no | no | no | no | no | no | no | no | no |
| CACNA1G | 1648 | Y | 0 | 0 | yes | no | no | yes | no | no | no | no | no | no | no | no | no |
| CACNA1G | 1649 | I | 0 | 0 | yes | no | no | yes | no | no | no | no | no | no | no | no | no |
| CACNA1G | 1650 | F | 0 | 0 | yes | no | no | yes | no | no | no | no | no | no | no | no | no |
| CACNA1G | 1651 | T | 0 | 0 | yes | no | no | yes | no | no | no | no | no | no | no | no | no |
| CACNA1G | 1652 | V | 1 | 0 | yes | no | no | yes | no | no | no | no | no | no | no | no | no |
| CACNA1G | 1653 | I | 0 | 0 | yes | no | no | yes | no | no | no | no | no | no | no | no | no |
| CACNA1G | 1654 | F | 0 | 0 | yes | no | no | yes | no | no | no | no | no | no | no | no | no |
| CACNA1G | 1655 | V | 0 | 0 | yes | no | no | yes | no | no | no | no | no | no | no | no | no |
| CACNA1G | 1656 | L | 0 | 0 | yes | no | no | yes | no | no | no | no | no | no | no | no | no |

| | | | | | | | | | | | | | | | | | |
|---|---|---|---|---|---|---|---|---|---|---|---|---|---|---|---|---|---|
| CACNA1G | 1657 | E | 0 | 0 | yes | no | no | yes | no | no | no | no | no | no | no | no | no |
| CACNA1G | 1658 | S | 0 | 0 | yes | no | no | yes | no | no | no | no | no | no | no | no | no |
| CACNA1G | 1659 | V | 1 | 0 | yes | no | no | yes | no | no | no | no | no | no | no | no | no |
| CACNA1G | 1660 | F | 0 | 0 | yes | no | no | yes | no | no | no | no | no | no | no | no | no |
| CACNA1G | 1661 | K | 0 | 0 | yes | no | no | yes | no | no | no | no | no | no | no | no | no |
| CACNA1G | 1662 | L | 0 | 0 | yes | no | no | yes | no | no | no | no | no | no | no | no | no |
| CACNA1G | 1663 | V | 2 | 0 | yes | no | no | no | no | no | no | no | no | no | no | no | no |
| CACNA1G | 1664 | A | 0 | 0 | yes | no | no | no | no | no | no | no | no | no | no | no | no |
| CACNA1G | 1665 | F | 1 | 0 | yes | no | no | no | no | no | no | no | no | no | no | no | no |
| CACNA1G | 1666 | G | 0 | 0 | no | yes | no | no | no | no | no | no | no | no | no | no | no |
| CACNA1G | 1667 | F | 0 | 0 | no | yes | no | no | no | no | no | no | no | no | no | no | no |
| CACNA1G | 1668 | R | 2 | 0 | no | yes | no | no | no | yes | no | no | no | no | no | no | no |
| CACNA1G | 1669 | R | 2 | 0 | yes | no | no | no | no | yes | no | no | no | no | no | no | no |
| CACNA1G | 1670 | F | 0 | 0 | yes | no | no | yes | no | yes | no | no | no | no | no | no | no |
| CACNA1G | 1671 | F | 0 | 0 | yes | no | no | yes | no | yes | no | no | no | no | no | no | no |
| CACNA1G | 1672 | Q | 0 | 0 | yes | no | no | no | no | yes | no | no | no | no | no | no | no |
| CACNA1G | 1673 | D | 0 | 0 | no | yes | no | no | no | yes | no | no | no | no | no | no | no |
| CACNA1G | 1674 | R | 1 | 0 | no | yes | no | yes | no | yes | no | no | no | no | no | no | no |
| CACNA1G | 1675 | W | 0 | 0 | yes | no | no | yes | no | no | no | no | no | no | no | no | no |
| CACNA1G | 1676 | N | 0 | 0 | yes | no | no | yes | no | no | no | no | no | no | no | no | no |
| CACNA1G | 1677 | Q | 0 | 0 | yes | no | no | yes | no | no | no | no | no | no | no | no | no |
| CACNA1G | 1678 | L | 0 | 0 | yes | no | no | yes | no | no | no | no | no | no | no | no | no |
| CACNA1G | 1679 | D | 0 | 0 | yes | no | no | yes | no | no | no | no | no | no | no | no | no |
| CACNA1G | 1680 | L | 0 | 0 | yes | no | no | yes | no | no | no | no | no | no | no | no | no |
| CACNA1G | 1681 | A | 0 | 0 | yes | no | no | yes | no | no | no | no | no | no | no | no | no |
| CACNA1G | 1682 | I | 0 | 0 | yes | no | no | yes | no | no | no | no | no | no | no | no | no |
| CACNA1G | 1683 | V | 1 | 0 | yes | no | no | yes | no | no | no | no | no | no | no | no | no |
| CACNA1G | 1684 | L | 0 | 0 | yes | no | no | yes | no | no | no | no | no | no | no | no | no |
| CACNA1G | 1685 | L | 1 | 0 | yes | no | no | yes | no | no | no | no | no | no | no | no | no |
| CACNA1G | 1686 | S | 0 | 0 | yes | no | no | yes | no | no | no | no | no | no | no | no | no |
| CACNA1G | 1687 | I | 0 | 0 | yes | no | no | yes | no | no | no | no | no | no | no | no | no |
| CACNA1G | 1688 | M | 1 | 0 | yes | no | no | yes | no | no | no | no | no | no | no | no | no |
| CACNA1G | 1689 | G | 0 | 0 | yes | no | no | yes | no | no | no | no | no | no | no | no | no |

| | | | | | | | | | | | | | | | | | |
|---|---|---|---|---|---|---|---|---|---|---|---|---|---|---|---|---|---|
| CACNA1G | 1690 | I | 0 | 0 | yes | no | no | yes | no | no | no | no | no | no | no | no | no |
| CACNA1G | 1691 | T | 1 | 0 | no | yes | no | yes | no | no | no | no | no | no | no | no | no |
| CACNA1G | 1692 | L | 0 | 0 | no | yes | no | yes | no | no | no | no | no | no | no | no | no |
| CACNA1G | 1693 | E | 0 | 0 | yes | no | no | yes | no | no | no | no | no | no | no | no | no |
| CACNA1G | 1694 | E | 0 | 0 | yes | no | no | yes | yes | no | no | no | no | no | no | no | no |
| CACNA1G | 1695 | I | 0 | 0 | yes | no | no | no | yes | no | no | no | no | no | no | no | no |
| CACNA1G | 1696 | E | 1 | 0 | yes | no | no | no | yes | no | no | no | no | no | no | no | no |
| CACNA1G | 1697 | V | 1 | 0 | yes | no | no | no | yes | no | no | no | no | no | no | no | no |
| CACNA1G | 1698 | N | 1 | 0 | yes | no | no | no | yes | no | no | no | no | no | no | no | no |
| CACNA1G | 1699 | A | 0 | 0 | no | yes | no | no | yes | no | no | no | no | no | no | no | no |
| CACNA1G | 1700 | S | 0 | 0 | no | yes | no | no | yes | no | no | no | no | no | no | no | no |
| CACNA1G | 1701 | L | 0 | 0 | no | yes | no | no | yes | no | no | no | no | no | no | no | no |
| CACNA1G | 1702 | P | 0 | 0 | no | yes | no | no | yes | no | no | no | no | no | no | no | no |
| CACNA1G | 1703 | I | 0 | 0 | no | yes | no | no | yes | no | no | no | no | no | no | no | no |
| CACNA1G | 1704 | N | 0 | 0 | no | yes | no | no | yes | no | no | no | no | no | no | no | no |
| CACNA1G | 1705 | P | 0 | 0 | yes | no | no | no | yes | no | no | no | no | no | no | no | no |
| CACNA1G | 1706 | T | 0 | 0 | yes | no | no | no | yes | no | no | no | no | no | no | no | no |
| CACNA1G | 1707 | I | 0 | 0 | yes | no | no | yes | yes | no | no | no | no | no | no | no | no |
| CACNA1G | 1708 | I | 0 | 0 | yes | no | no | yes | no | no | no | no | no | no | no | no | no |
| CACNA1G | 1709 | R | 2 | 0 | yes | no | no | no | no | no | no | no | no | no | no | no | no |
| CACNA1G | 1710 | I | 1 | 0 | yes | no | no | yes | no | no | no | no | no | no | no | no | no |
| CACNA1G | 1711 | M | 0 | 0 | yes | no | no | yes | no | no | no | no | no | no | no | no | no |
| CACNA1G | 1712 | R | 0 | 0 | yes | no | no | yes | no | no | no | no | no | no | no | no | no |
| CACNA1G | 1713 | V | 1 | 0 | yes | no | no | yes | no | no | no | no | no | no | no | no | no |
| CACNA1G | 1714 | L | 0 | 0 | yes | no | no | yes | no | no | no | no | no | no | no | no | no |
| CACNA1G | 1715 | R | 1 | 1 | yes | no | no | yes | no | no | no | no | no | no | no | no | no |
| CACNA1G | 1716 | I | 0 | 0 | yes | no | no | yes | no | no | no | no | no | no | no | no | no |
| CACNA1G | 1717 | A | 0 | 0 | yes | no | no | yes | no | no | no | no | no | no | no | no | no |
| CACNA1G | 1718 | R | 1 | 0 | yes | no | no | yes | no | no | no | no | no | no | no | no | no |
| CACNA1G | 1719 | V | 0 | 0 | yes | no | no | yes | no | no | no | no | no | no | no | no | no |
| CACNA1G | 1720 | L | 0 | 0 | yes | no | no | yes | no | no | no | no | no | no | no | no | no |
| CACNA1G | 1721 | K | 0 | 0 | yes | no | no | yes | no | no | no | no | no | no | no | no | no |
| CACNA1G | 1722 | L | 0 | 0 | yes | no | no | yes | no | no | no | no | no | no | no | no | no |

| | | | | | | | | | | | | | | | | | |
|---|---|---|---|---|---|---|---|---|---|---|---|---|---|---|---|---|---|
| CACNA1G | 1723 | L | 0 | 0 | no | yes | no | yes | no | no | no | no | no | no | no | no | no |
| CACNA1G | 1724 | K | 0 | 0 | no | yes | no | yes | no | no | no | no | no | no | no | no | no |
| CACNA1G | 1725 | M | 0 | 0 | no | yes | no | yes | no | no | no | no | no | no | no | no | no |
| CACNA1G | 1726 | A | 0 | 0 | no | yes | no | yes | no | no | no | no | no | no | no | no | no |
| CACNA1G | 1727 | V | 0 | 0 | yes | no | no | no | no | no | no | no | no | no | no | no | no |
| CACNA1G | 1728 | G | 0 | 0 | yes | no | no | yes | no | no | no | no | no | no | no | no | no |
| CACNA1G | 1729 | M | 0 | 0 | yes | no | no | yes | no | no | no | no | no | no | no | no | no |
| CACNA1G | 1730 | R | 0 | 0 | yes | no | no | yes | no | no | no | no | no | no | no | no | no |
| CACNA1G | 1731 | A | 1 | 0 | yes | no | no | no | no | no | no | no | no | no | no | no | no |
| CACNA1G | 1732 | L | 0 | 0 | yes | no | no | yes | no | yes | no | no | no | no | no | no | no |
| CACNA1G | 1733 | L | 0 | 0 | yes | no | no | yes | no | yes | no | no | no | no | no | no | no |
| CACNA1G | 1734 | D | 0 | 0 | yes | no | no | no | no | yes | no | no | no | no | no | no | no |
| CACNA1G | 1735 | T | 1 | 0 | yes | no | no | no | no | yes | no | no | no | no | no | no | no |
| CACNA1G | 1736 | V | 0 | 0 | yes | no | no | yes | no | yes | no | no | no | no | no | no | no |
| CACNA1G | 1737 | M | 1 | 0 | yes | no | no | yes | no | yes | no | no | no | no | no | no | no |
| CACNA1G | 1738 | Q | 0 | 0 | yes | no | no | no | no | yes | no | no | no | no | no | no | no |
| CACNA1G | 1739 | A | 0 | 0 | no | yes | no | yes | no | yes | no | no | no | no | no | no | no |
| CACNA1G | 1740 | L | 0 | 0 | yes | no | no | yes | no | yes | no | no | no | no | no | no | no |
| CACNA1G | 1741 | P | 0 | 0 | yes | no | no | yes | no | yes | no | no | no | no | no | no | no |
| CACNA1G | 1742 | Q | 0 | 0 | yes | no | no | yes | no | yes | no | no | no | no | no | no | no |
| CACNA1G | 1743 | V | 1 | 0 | yes | no | no | yes | no | yes | no | no | no | no | no | no | no |
| CACNA1G | 1744 | G | 0 | 0 | yes | no | no | yes | no | yes | no | no | no | no | no | no | no |
| CACNA1G | 1745 | N | 0 | 0 | yes | no | no | yes | no | yes | no | no | no | no | no | no | no |
| CACNA1G | 1746 | L | 0 | 0 | yes | no | no | yes | no | no | no | no | no | no | no | no | no |
| CACNA1G | 1747 | G | 0 | 0 | yes | no | no | yes | no | no | no | no | no | no | no | no | no |
| CACNA1G | 1748 | L | 0 | 0 | yes | no | no | yes | no | no | no | no | no | no | no | no | no |
| CACNA1G | 1749 | L | 0 | 0 | yes | no | no | yes | no | no | no | no | no | no | no | no | no |
| CACNA1G | 1750 | F | 0 | 0 | yes | no | no | yes | no | no | no | no | no | no | no | no | no |
| CACNA1G | 1751 | M | 2 | 0 | yes | no | no | yes | no | no | no | no | no | no | no | no | no |
| CACNA1G | 1752 | L | 0 | 0 | yes | no | no | yes | no | no | no | no | no | no | no | no | no |
| CACNA1G | 1753 | L | 0 | 0 | yes | no | no | yes | no | no | no | no | no | no | no | no | no |
| CACNA1G | 1754 | F | 0 | 0 | yes | no | no | yes | no | no | no | no | no | no | no | no | no |
| CACNA1G | 1755 | F | 0 | 0 | yes | no | no | yes | no | no | no | no | no | no | no | no | no |

| | | | | | | | | | | | | | | | | | |
|---|---|---|---|---|---|---|---|---|---|---|---|---|---|---|---|---|---|
| CACNA1G | 1756 | I | 0 | 0 | yes | no | no | yes | no | no | no | no | no | no | no | no | no |
| CACNA1G | 1757 | F | 0 | 0 | yes | no | no | yes | no | no | no | no | no | no | no | no | no |
| CACNA1G | 1758 | A | 0 | 0 | yes | no | no | yes | no | no | no | no | no | no | no | no | no |
| CACNA1G | 1759 | A | 0 | 0 | yes | no | no | yes | no | no | no | no | no | no | no | no | no |
| CACNA1G | 1760 | L | 0 | 0 | yes | no | no | yes | no | no | no | no | no | no | no | no | no |
| CACNA1G | 1761 | G | 0 | 0 | yes | no | no | yes | no | no | no | no | no | no | no | no | no |
| CACNA1G | 1762 | V | 0 | 0 | yes | no | no | yes | no | no | no | no | no | no | no | no | no |
| CACNA1G | 1763 | E | 0 | 0 | yes | no | no | yes | no | no | no | no | no | no | no | no | no |
| CACNA1G | 1764 | L | 0 | 0 | yes | no | no | yes | no | no | no | no | no | no | no | no | no |
| CACNA1G | 1765 | F | 0 | 0 | yes | no | no | no | no | no | no | no | no | no | no | no | no |
| CACNA1G | 1766 | G | 0 | 0 | no | yes | no | no | no | no | no | no | no | no | no | no | no |
| CACNA1G | 1767 | D | 0 | 0 | no | yes | no | no | yes | no | no | no | no | no | no | no | no |
| CACNA1G | 1768 | L | 0 | 0 | no | yes | no | no | yes | no | no | no | no | no | no | no | no |
| CACNA1G | 1769 | E | 1 | 0 | no | yes | no | no | yes | no | no | no | no | no | no | no | no |
| CACNA1G | 1770 | C | 0 | 0 | no | yes | no | no | yes | no | no | yes | no | no | no | no | no |
| CACNA1G | 1771 | D | 0 | 0 | no | yes | no | no | yes | no | yes | no | no | no | no | no | no |
| CACNA1G | 1772 | E | 2 | 0 | no | yes | no | no | yes | no | yes | no | no | no | no | no | no |
| CACNA1G | 1773 | T | 0 | 0 | no | yes | no | no | yes | no | no | no | no | no | no | no | no |
| CACNA1G | 1774 | H | 0 | 0 | no | yes | no | no | yes | no | no | no | no | no | no | no | no |
| CACNA1G | 1775 | P | 1 | 0 | no | yes | no | no | yes | no | no | no | no | no | no | no | no |
| CACNA1G | 1776 | C | 0 | 0 | no | yes | no | no | yes | no | yes | no | no | no | no | no | no |
| CACNA1G | 1777 | E | 0 | 0 | no | yes | no | no | yes | no | no | no | no | no | no | no | no |
| CACNA1G | 1778 | G | 0 | 0 | no | yes | no | no | yes | no | yes | no | no | no | no | no | no |
| CACNA1G | 1779 | L | 0 | 0 | no | yes | no | no | yes | no | yes | no | no | no | no | no | no |
| CACNA1G | 1780 | G | 0 | 0 | no | yes | no | no | yes | no | yes | no | no | no | no | no | no |
| CACNA1G | 1781 | R | 1 | 0 | no | yes | no | no | yes | no | yes | no | no | no | no | no | no |
| CACNA1G | 1782 | H | 0 | 0 | no | yes | no | no | yes | no | no | no | no | no | no | no | no |
| CACNA1G | 1783 | A | 0 | 0 | no | yes | no | no | yes | no | no | no | no | no | no | no | no |
| CACNA1G | 1784 | T | 0 | 0 | no | yes | no | no | yes | no | no | no | no | no | no | no | no |
| CACNA1G | 1785 | F | 0 | 0 | no | yes | no | no | yes | no | no | no | no | no | no | no | no |
| CACNA1G | 1786 | R | 1 | 0 | no | yes | no | no | yes | no | no | no | no | no | no | no | no |
| CACNA1G | 1787 | N | 0 | 0 | no | yes | no | no | yes | no | no | no | no | no | no | no | no |
| CACNA1G | 1788 | F | 0 | 0 | yes | no | no | yes | yes | no | no | no | no | no | no | no | no |

| | | | | | | | | | | | | | | | | | |
|---|---|---|---|---|---|---|---|---|---|---|---|---|---|---|---|---|---|
| CACNA1G | 1789 | G | 0 | 0 | yes | no | no | yes | yes | no | no | no | no | no | no | no | no |
| CACNA1G | 1790 | M | 0 | 0 | yes | no | no | no | yes | no | no | no | no | no | no | no | no |
| CACNA1G | 1791 | A | 0 | 0 | yes | no | no | yes | yes | no | no | no | no | no | no | no | no |
| CACNA1G | 1792 | F | 0 | 0 | yes | no | no | yes | yes | no | no | no | no | no | no | no | no |
| CACNA1G | 1793 | L | 0 | 0 | yes | no | no | yes | yes | no | no | no | no | no | no | no | no |
| CACNA1G | 1794 | T | 0 | 0 | yes | no | no | yes | yes | no | no | no | no | no | no | no | no |
| CACNA1G | 1795 | L | 0 | 0 | yes | no | no | yes | yes | no | no | no | no | no | no | no | no |
| CACNA1G | 1796 | F | 0 | 0 | yes | no | no | yes | yes | no | no | no | no | no | no | no | no |
| CACNA1G | 1797 | R | 0 | 0 | yes | no | no | yes | yes | no | yes | no | no | no | no | no | no |
| CACNA1G | 1798 | V | 0 | 0 | yes | no | no | yes | yes | no | no | no | no | no | no | no | no |
| CACNA1G | 1799 | S | 0 | 0 | yes | no | no | yes | yes | no | yes | no | no | no | no | no | no |
| CACNA1G | 1800 | T | 0 | 0 | no | yes | no | yes | yes | no | yes | no | no | no | no | no | no |
| CACNA1G | 1801 | G | 0 | 0 | no | yes | no | yes | yes | no | no | no | no | no | no | no | no |
| CACNA1G | 1802 | D | 0 | 0 | no | yes | no | yes | yes | no | yes | no | no | no | no | no | no |
| CACNA1G | 1803 | N | 1 | 0 | no | yes | no | no | yes | no | no | no | no | no | no | no | no |
| CACNA1G | 1804 | W | 0 | 0 | yes | no | no | yes | yes | no | no | no | no | no | no | no | no |
| CACNA1G | 1805 | N | 0 | 0 | yes | no | no | no | yes | no | no | no | no | no | no | no | no |
| CACNA1G | 1806 | G | 0 | 0 | yes | no | no | no | yes | no | no | no | no | no | no | no | no |
| CACNA1G | 1807 | I | 0 | 0 | yes | no | no | no | yes | no | no | no | no | no | no | no | no |
| CACNA1G | 1808 | M | 0 | 0 | yes | no | no | no | yes | no | no | no | no | no | no | no | no |
| CACNA1G | 1809 | K | 0 | 0 | yes | no | no | no | yes | no | no | no | no | no | no | no | no |
| CACNA1G | 1810 | D | 0 | 0 | yes | no | no | no | yes | no | no | no | no | no | no | no | no |
| CACNA1G | 1811 | T | 1 | 0 | yes | no | no | no | yes | no | no | no | no | no | no | no | no |
| CACNA1G | 1812 | L | 0 | 0 | no | yes | no | no | yes | no | no | no | no | no | no | no | no |
| CACNA1G | 1813 | R | 0 | 0 | no | yes | no | no | yes | no | no | no | no | no | no | no | no |
| CACNA1G | 1814 | D | 0 | 0 | no | yes | no | no | yes | no | no | no | no | no | no | no | no |
| CACNA1G | 1822 | Y | 0 | 0 | no | yes | no | no | yes | no | no | no | no | no | no | no | no |
| CACNA1G | 1823 | N | 0 | 0 | yes | no | no | no | yes | no | no | no | no | no | no | no | no |
| CACNA1G | 1824 | T | 1 | 0 | yes | no | no | no | yes | no | no | no | no | no | no | no | no |
| CACNA1G | 1825 | V | 0 | 0 | yes | no | no | yes | yes | no | no | no | no | no | no | no | no |
| CACNA1G | 1826 | I | 0 | 0 | yes | no | no | yes | yes | no | no | no | no | no | no | no | no |
| CACNA1G | 1827 | S | 0 | 0 | yes | no | no | yes | no | no | no | no | no | no | no | no | no |
| CACNA1G | 1828 | P | 0 | 0 | yes | no | no | yes | no | no | no | no | no | no | no | no | no |

| | | | | | | | | | | | | | | | | | |
|---|---|---|---|---|---|---|---|---|---|---|---|---|---|---|---|---|---|
| CACNA1G | 1829 | I | 1 | 0 | yes | no | no | yes | no | no | no | no | no | no | no | no | no |
| CACNA1G | 1830 | Y | 0 | 0 | yes | no | no | yes | no | no | no | no | no | no | no | no | no |
| CACNA1G | 1831 | F | 0 | 0 | yes | no | no | yes | no | no | no | no | no | no | no | no | no |
| CACNA1G | 1832 | V | 0 | 0 | yes | no | no | yes | no | no | no | no | no | no | no | no | no |
| CACNA1G | 1833 | S | 0 | 0 | yes | no | no | yes | no | no | no | no | no | no | no | no | no |
| CACNA1G | 1834 | F | 0 | 0 | yes | no | no | yes | no | no | no | no | no | no | no | no | no |
| CACNA1G | 1835 | V | 0 | 0 | yes | no | no | yes | no | no | no | no | no | no | no | no | no |
| CACNA1G | 1836 | L | 0 | 0 | yes | no | no | yes | no | no | no | no | no | no | no | no | no |
| CACNA1G | 1837 | T | 1 | 0 | yes | no | no | yes | no | no | no | no | no | no | no | no | no |
| CACNA1G | 1838 | A | 0 | 0 | yes | no | no | yes | no | no | no | no | no | no | no | no | no |
| CACNA1G | 1839 | Q | 0 | 0 | yes | no | no | yes | no | no | yes | no | no | no | no | no | no |
| CACNA1G | 1840 | F | 0 | 0 | yes | no | no | yes | no | no | no | no | no | no | no | no | no |
| CACNA1G | 1841 | V | 0 | 0 | yes | no | no | yes | no | no | no | no | no | no | no | no | no |
| CACNA1G | 1842 | L | 0 | 0 | yes | no | no | yes | no | no | yes | no | no | no | no | no | no |
| CACNA1G | 1843 | V | 0 | 0 | yes | no | no | yes | no | no | yes | no | no | no | no | no | no |
| CACNA1G | 1844 | N | 0 | 0 | yes | no | no | yes | no | no | no | no | no | no | no | no | no |
| CACNA1G | 1845 | V | 0 | 0 | yes | no | no | yes | no | no | no | no | no | no | no | no | no |
| CACNA1G | 1846 | V | 0 | 0 | yes | no | no | yes | no | no | yes | no | no | no | no | no | no |
| CACNA1G | 1847 | I | 0 | 0 | yes | no | no | yes | no | no | yes | no | no | no | no | no | no |
| CACNA1G | 1848 | A | 0 | 0 | yes | no | no | yes | no | no | no | no | no | no | no | no | no |
| CACNA1G | 1849 | V | 0 | 0 | yes | no | no | yes | no | no | no | no | no | no | no | no | no |
| CACNA1G | 1850 | L | 0 | 0 | yes | no | no | yes | no | no | yes | no | no | no | no | no | no |
| CACNA1G | 1851 | M | 1 | 0 | yes | no | no | no | no | no | yes | no | no | no | no | no | no |
| CACNA1G | 1852 | K | 1 | 0 | yes | no | no | no | no | no | no | no | no | no | no | no | no |
| CACNA1G | 1853 | H | 0 | 0 | yes | no | no | no | no | no | no | no | no | no | no | no | no |
| CACNA1G | 1854 | L | 0 | 0 | yes | no | no | no | no | no | no | no | no | no | no | no | no |
| CACNA1G | 1855 | E | 0 | 0 | no | yes | no | no | no | yes | no | no | no | no | no | no | no |
| CACNA1G | 1856 | E | 1 | 0 | no | yes | no | no | no | yes | no | no | no | no | no | no | no |
| CACNA1G | 1857 | S | 1 | 0 | no | yes | no | no | no | yes | no | no | no | no | no | no | no |
| CACNA1G | 1858 | N | 0 | 0 | no | yes | no | no | no | yes | no | no | no | no | no | no | no |
| CACNA1G | 1859 | K | 1 | 0 | no | yes | no | no | no | yes | no | no | no | no | no | no | no |
| CHRNA4 | 31 | E | 0 | 0 | no | yes | no | no | yes | no | no | no | no | no | no | no | no |
| CHRNA4 | 32 | T | 1 | 0 | no | yes | no | no | yes | no | no | no | yes | no | no | no | no |

| | | | | | | | | | | | | | | | | | |
|---|---|---|---|---|---|---|---|---|---|---|---|---|---|---|---|---|---|
| CHRNA4 | 33 | R | 2 | 0 | yes | no | no | no | yes | no | no | no | yes | no | no | no | no |
| CHRNA4 | 34 | A | 0 | 0 | yes | no | no | no | yes | no | no | no | yes | no | no | no | no |
| CHRNA4 | 35 | H | 0 | 0 | yes | no | no | no | yes | no | no | no | no | no | no | no | no |
| CHRNA4 | 36 | A | 0 | 0 | yes | no | no | no | yes | no | no | no | no | no | no | no | no |
| CHRNA4 | 37 | E | 1 | 0 | yes | no | no | no | yes | no | no | no | no | no | no | no | no |
| CHRNA4 | 38 | E | 3 | 0 | yes | no | no | no | yes | no | no | no | yes | no | no | no | no |
| CHRNA4 | 39 | R | 2 | 0 | yes | no | no | no | yes | no | no | no | no | no | no | no | no |
| CHRNA4 | 40 | L | 1 | 0 | yes | no | no | no | yes | no | no | no | no | no | no | no | no |
| CHRNA4 | 41 | L | 0 | 0 | yes | no | no | no | yes | no | no | no | no | no | no | no | no |
| CHRNA4 | 42 | K | 0 | 0 | yes | no | no | no | yes | no | no | no | no | no | no | no | no |
| CHRNA4 | 43 | K | 0 | 0 | yes | no | no | no | yes | no | no | no | no | no | no | no | no |
| CHRNA4 | 44 | L | 0 | 0 | yes | no | no | no | yes | no | no | no | no | no | no | no | no |
| CHRNA4 | 45 | F | 0 | 0 | no | yes | no | no | yes | no | no | no | no | no | no | no | no |
| CHRNA4 | 46 | S | 0 | 0 | no | yes | no | no | yes | no | no | no | no | no | no | no | no |
| CHRNA4 | 47 | G | 2 | 0 | no | yes | no | no | yes | no | no | no | yes | no | no | no | no |
| CHRNA4 | 48 | Y | 0 | 0 | no | yes | no | no | yes | no | no | no | yes | no | no | no | no |
| CHRNA4 | 49 | N | 0 | 0 | no | yes | no | no | yes | no | no | no | yes | no | no | no | no |
| CHRNA4 | 50 | K | 0 | 0 | no | yes | no | no | yes | no | yes | no | no | no | no | no | no |
| CHRNA4 | 51 | W | 0 | 0 | no | yes | no | no | yes | no | no | no | yes | no | no | no | no |
| CHRNA4 | 52 | S | 0 | 0 | no | yes | no | no | yes | no | no | no | yes | no | no | no | no |
| CHRNA4 | 53 | R | 1 | 0 | no | yes | no | no | yes | no | no | no | yes | no | no | no | no |
| CHRNA4 | 54 | P | 1 | 0 | no | yes | no | no | yes | no | no | no | no | no | no | no | no |
| CHRNA4 | 55 | V | 1 | 0 | no | yes | no | no | yes | no | no | no | no | no | no | no | no |
| CHRNA4 | 56 | A | 1 | 0 | no | yes | no | no | yes | no | no | no | no | no | no | no | no |
| CHRNA4 | 57 | N | 0 | 0 | no | yes | no | no | yes | no | no | no | no | no | no | no | no |
| CHRNA4 | 58 | I | 1 | 0 | no | yes | no | no | yes | no | no | no | no | no | no | no | no |
| CHRNA4 | 59 | S | 1 | 0 | no | yes | no | no | yes | no | no | no | no | no | no | no | no |
| CHRNA4 | 60 | D | 1 | 0 | no | yes | no | no | yes | no | no | no | no | no | no | no | no |
| CHRNA4 | 61 | V | 1 | 0 | no | yes | no | no | yes | no | no | no | no | no | no | no | no |
| CHRNA4 | 62 | V | 0 | 0 | no | no | yes | no | yes | no | no | no | no | no | no | no | no |
| CHRNA4 | 63 | L | 0 | 0 | no | no | yes | no | yes | no | no | no | no | no | no | no | no |
| CHRNA4 | 64 | V | 1 | 0 | no | no | yes | no | yes | no | no | no | no | no | no | no | no |
| CHRNA4 | 65 | R | 2 | 0 | no | no | yes | no | yes | no | no | no | no | no | no | no | no |

| | | | | | | | | | | | | | | | | | |
|---|---|---|---|---|---|---|---|---|---|---|---|---|---|---|---|---|---|
| CHRNA4 | 66 | F | 1 | 1 | no | no | yes | no | yes | no | no | no | no | no | no | no | no |
| CHRNA4 | 67 | G | 1 | 0 | no | no | yes | no | yes | no | no | no | no | no | no | no | no |
| CHRNA4 | 68 | L | 0 | 0 | no | no | yes | no | yes | no | no | no | no | no | no | no | no |
| CHRNA4 | 69 | S | 0 | 0 | no | no | yes | no | yes | no | no | no | no | no | no | no | no |
| CHRNA4 | 70 | I | 0 | 0 | no | no | yes | no | yes | no | no | no | no | no | no | no | no |
| CHRNA4 | 71 | A | 2 | 0 | no | no | yes | no | yes | no | no | no | no | no | no | no | no |
| CHRNA4 | 72 | Q | 0 | 0 | no | no | yes | no | yes | no | no | no | no | no | no | no | no |
| CHRNA4 | 73 | L | 0 | 0 | no | no | yes | no | yes | no | no | no | no | no | no | no | no |
| CHRNA4 | 74 | I | 0 | 0 | no | no | yes | no | yes | no | no | no | no | no | no | no | no |
| CHRNA4 | 75 | D | 1 | 0 | no | no | yes | no | yes | no | no | no | no | no | no | no | no |
| CHRNA4 | 76 | V | 1 | 0 | no | no | yes | no | yes | no | no | no | no | no | no | no | no |
| CHRNA4 | 77 | D | 0 | 0 | no | no | yes | no | yes | no | no | no | no | no | no | no | no |
| CHRNA4 | 78 | E | 0 | 0 | no | yes | no | no | yes | no | no | no | no | no | no | no | no |
| CHRNA4 | 79 | K | 2 | 0 | no | yes | no | no | yes | no | no | no | no | no | no | no | no |
| CHRNA4 | 80 | N | 1 | 0 | no | yes | no | no | yes | no | no | no | no | no | no | no | no |
| CHRNA4 | 81 | Q | 0 | 0 | no | yes | no | no | yes | no | no | no | no | no | no | no | no |
| CHRNA4 | 82 | M | 0 | 0 | no | no | yes | no | yes | no | no | no | no | no | no | no | no |
| CHRNA4 | 83 | M | 1 | 0 | no | no | yes | no | yes | no | no | no | no | no | no | no | no |
| CHRNA4 | 84 | T | 1 | 0 | no | no | yes | no | yes | no | no | no | no | no | no | no | no |
| CHRNA4 | 85 | T | 1 | 0 | no | no | yes | no | yes | no | no | no | no | no | no | no | no |
| CHRNA4 | 86 | N | 1 | 0 | no | no | yes | no | yes | no | no | no | yes | no | no | no | no |
| CHRNA4 | 87 | V | 1 | 0 | no | no | yes | no | yes | no | no | no | no | no | no | no | no |
| CHRNA4 | 88 | W | 0 | 0 | no | no | yes | no | yes | no | no | no | yes | no | no | no | no |
| CHRNA4 | 89 | V | 0 | 0 | no | no | yes | no | yes | no | no | no | no | no | no | no | no |
| CHRNA4 | 90 | K | 0 | 0 | no | no | yes | no | yes | no | no | no | no | no | no | no | no |
| CHRNA4 | 91 | Q | 0 | 0 | no | no | yes | no | yes | no | no | no | no | no | no | no | no |
| CHRNA4 | 92 | E | 1 | 0 | no | no | yes | no | yes | no | no | no | no | no | no | no | no |
| CHRNA4 | 93 | W | 0 | 0 | no | no | yes | no | yes | no | no | no | no | no | no | no | no |
| CHRNA4 | 94 | H | 0 | 0 | no | no | yes | no | yes | no | no | no | no | no | no | no | no |
| CHRNA4 | 95 | D | 0 | 0 | no | yes | no | no | yes | no | no | no | no | no | no | no | no |
| CHRNA4 | 96 | Y | 0 | 0 | no | yes | no | no | yes | no | no | no | yes | no | no | no | no |
| CHRNA4 | 97 | K | 1 | 0 | no | yes | no | no | yes | no | no | no | no | no | no | no | no |
| CHRNA4 | 98 | L | 0 | 0 | no | yes | no | no | yes | no | no | no | no | no | no | no | no |

| | | | | | | | | | | | | | | | | | |
|---|---|---|---|---|---|---|---|---|---|---|---|---|---|---|---|---|---|
| CHRNA4 | 99 | R | 2 | 0 | no | yes | no | no | yes | no | no | no | no | no | no | no | no |
| CHRNA4 | 100 | W | 0 | 0 | no | yes | no | no | yes | no | no | no | no | no | no | no | no |
| CHRNA4 | 101 | D | 0 | 0 | no | yes | no | no | yes | no | no | no | no | no | no | no | no |
| CHRNA4 | 102 | P | 1 | 0 | yes | no | no | no | yes | no | no | no | no | no | no | no | no |
| CHRNA4 | 103 | A | 0 | 0 | yes | no | no | no | yes | no | no | no | no | no | no | no | no |
| CHRNA4 | 104 | D | 1 | 0 | yes | no | no | no | yes | no | no | no | no | no | no | no | no |
| CHRNA4 | 105 | Y | 1 | 0 | no | yes | no | no | yes | no | no | no | no | no | no | no | no |
| CHRNA4 | 106 | E | 0 | 0 | no | yes | no | no | yes | no | no | no | no | no | no | no | no |
| CHRNA4 | 107 | N | 0 | 0 | no | yes | no | no | yes | no | no | no | no | no | no | no | no |
| CHRNA4 | 108 | V | 0 | 0 | no | yes | no | no | yes | no | no | no | no | no | no | no | no |
| CHRNA4 | 109 | T | 2 | 0 | no | yes | no | no | yes | no | no | no | no | no | no | no | no |
| CHRNA4 | 110 | S | 0 | 0 | no | yes | no | no | yes | no | no | no | no | no | no | no | no |
| CHRNA4 | 111 | I | 2 | 0 | no | no | yes | no | yes | no | no | no | no | no | no | no | no |
| CHRNA4 | 112 | R | 1 | 0 | no | no | yes | no | yes | no | no | no | yes | no | no | no | no |
| CHRNA4 | 113 | I | 1 | 0 | no | no | yes | no | yes | no | no | no | no | no | no | no | no |
| CHRNA4 | 114 | P | 0 | 0 | no | no | yes | no | yes | no | no | no | yes | no | no | no | no |
| CHRNA4 | 115 | S | 0 | 0 | no | yes | no | no | yes | no | no | no | no | no | no | no | no |
| CHRNA4 | 116 | E | 2 | 0 | no | yes | no | no | yes | no | yes | no | no | no | no | no | no |
| CHRNA4 | 117 | L | 0 | 0 | no | yes | no | no | yes | no | no | no | no | no | no | no | no |
| CHRNA4 | 118 | I | 0 | 0 | no | yes | no | no | yes | no | no | no | no | no | no | no | no |
| CHRNA4 | 119 | W | 0 | 0 | no | yes | no | no | yes | no | no | no | no | no | no | no | no |
| CHRNA4 | 120 | R | 2 | 0 | no | yes | no | no | yes | no | no | no | no | no | no | no | no |
| CHRNA4 | 121 | P | 2 | 0 | no | yes | no | no | yes | no | no | no | no | no | no | no | no |
| CHRNA4 | 122 | D | 1 | 0 | no | yes | no | no | yes | no | no | no | yes | no | no | no | no |
| CHRNA4 | 123 | I | 1 | 0 | no | no | yes | no | yes | no | no | no | no | no | no | no | no |
| CHRNA4 | 124 | V | 1 | 0 | no | no | yes | no | yes | no | no | no | yes | no | no | no | no |
| CHRNA4 | 125 | L | 0 | 0 | no | no | yes | no | yes | no | no | no | no | no | no | no | no |
| CHRNA4 | 126 | Y | 0 | 0 | no | yes | no | no | yes | no | no | no | yes | no | no | no | no |
| CHRNA4 | 127 | N | 0 | 0 | no | yes | no | no | yes | no | no | no | yes | no | no | no | no |
| CHRNA4 | 128 | N | 0 | 0 | no | no | yes | no | yes | no | no | no | yes | no | no | no | no |
| CHRNA4 | 129 | A | 0 | 0 | no | yes | no | no | yes | no | no | no | yes | no | no | no | no |
| CHRNA4 | 130 | D | 0 | 0 | no | yes | no | no | yes | no | yes | no | yes | no | no | no | no |
| CHRNA4 | 131 | G | 1 | 0 | no | yes | no | no | yes | no | no | no | no | no | no | no | no |

| | | | | | | | | | | | | | | | | | |
|---|---|---|---|---|---|---|---|---|---|---|---|---|---|---|---|---|---|
| CHRNA4 | 132 | D | 0 | 0 | no | yes | no | no | yes | no | yes | no | no | no | no | no | no |
| CHRNA4 | 133 | F | 0 | 0 | no | yes | no | no | yes | no | no | no | yes | no | no | no | no |
| CHRNA4 | 134 | A | 1 | 0 | no | yes | no | no | yes | no | no | no | no | no | no | no | no |
| CHRNA4 | 135 | V | 0 | 0 | no | yes | no | no | yes | no | no | no | no | no | no | no | no |
| CHRNA4 | 136 | T | 0 | 0 | no | yes | no | no | yes | no | yes | no | no | no | no | no | no |
| CHRNA4 | 137 | H | 2 | 0 | no | yes | no | no | yes | no | no | no | yes | no | no | no | no |
| CHRNA4 | 138 | L | 2 | 0 | no | yes | no | no | yes | no | no | no | no | no | no | no | no |
| CHRNA4 | 139 | T | 0 | 0 | no | yes | no | no | yes | no | no | no | no | no | no | no | no |
| CHRNA4 | 140 | K | 0 | 0 | no | no | yes | no | yes | no | no | no | yes | no | no | no | no |
| CHRNA4 | 141 | A | 0 | 0 | no | no | yes | no | yes | no | no | no | no | no | no | no | no |
| CHRNA4 | 142 | H | 1 | 0 | no | no | yes | no | yes | no | no | no | no | no | no | no | no |
| CHRNA4 | 143 | L | 0 | 0 | no | no | yes | no | yes | no | no | no | no | no | no | no | no |
| CHRNA4 | 144 | F | 0 | 0 | no | yes | no | no | yes | no | no | no | no | no | no | no | no |
| CHRNA4 | 145 | H | 0 | 0 | no | yes | no | no | yes | no | no | no | no | no | no | no | no |
| CHRNA4 | 146 | D | 0 | 0 | no | yes | no | no | yes | no | no | no | no | no | no | no | no |
| CHRNA4 | 147 | G | 1 | 0 | no | yes | no | no | yes | no | no | no | no | no | no | no | no |
| CHRNA4 | 148 | R | 1 | 0 | no | no | yes | no | yes | no | no | no | no | no | no | no | no |
| CHRNA4 | 149 | V | 0 | 0 | no | no | yes | no | yes | no | no | no | no | no | no | no | no |
| CHRNA4 | 150 | Q | 1 | 0 | no | no | yes | no | yes | no | no | no | no | no | no | no | no |
| CHRNA4 | 151 | W | 1 | 0 | no | no | yes | no | yes | no | no | no | no | no | no | no | no |
| CHRNA4 | 152 | T | 1 | 0 | no | yes | no | no | yes | no | no | no | no | no | no | no | no |
| CHRNA4 | 153 | P | 0 | 0 | no | yes | no | no | yes | no | no | no | no | no | no | no | no |
| CHRNA4 | 154 | P | 1 | 0 | no | no | yes | no | yes | no | no | no | yes | no | no | no | no |
| CHRNA4 | 155 | A | 0 | 0 | no | no | yes | no | yes | no | no | no | no | no | no | no | no |
| CHRNA4 | 156 | I | 0 | 0 | no | no | yes | no | yes | no | no | no | no | no | no | no | no |
| CHRNA4 | 157 | Y | 0 | 0 | no | no | yes | no | yes | no | no | no | no | no | no | no | no |
| CHRNA4 | 158 | K | 0 | 0 | no | no | yes | no | yes | no | yes | no | no | no | no | no | no |
| CHRNA4 | 159 | S | 0 | 0 | no | no | yes | no | yes | no | no | no | no | no | no | no | no |
| CHRNA4 | 160 | S | 0 | 0 | no | yes | no | no | yes | no | no | no | yes | no | no | no | no |
| CHRNA4 | 161 | C | 0 | 0 | no | yes | no | no | yes | no | no | no | no | no | no | no | no |
| CHRNA4 | 162 | S | 1 | 0 | no | yes | no | no | yes | no | no | no | no | no | no | no | no |
| CHRNA4 | 163 | I | 0 | 0 | no | yes | no | no | yes | no | no | no | no | no | no | no | no |
| CHRNA4 | 164 | D | 1 | 0 | no | yes | no | no | yes | no | no | no | no | no | no | no | no |

| | | | | | | | | | | | | | | | | | |
|---|---|---|---|---|---|---|---|---|---|---|---|---|---|---|---|---|---|
| CHRNA4 | 165 | V | 1 | 0 | no | yes | no | no | yes | no | no | no | no | no | no | no | no |
| CHRNA4 | 166 | T | 0 | 0 | no | yes | no | no | yes | no | no | no | no | no | no | no | no |
| CHRNA4 | 167 | F | 0 | 0 | no | yes | no | no | yes | no | no | no | no | no | no | no | no |
| CHRNA4 | 168 | F | 1 | 0 | no | yes | no | no | yes | no | no | no | no | no | no | no | no |
| CHRNA4 | 169 | P | 0 | 0 | no | yes | no | no | yes | no | no | no | no | no | no | no | no |
| CHRNA4 | 170 | F | 0 | 0 | no | yes | no | no | yes | no | no | no | no | no | no | no | no |
| CHRNA4 | 171 | D | 1 | 0 | no | yes | no | no | yes | no | no | no | no | no | no | no | no |
| CHRNA4 | 172 | Q | 0 | 0 | no | no | yes | no | yes | no | no | no | no | no | no | no | no |
| CHRNA4 | 173 | Q | 0 | 0 | no | no | yes | no | yes | no | no | no | no | no | no | no | no |
| CHRNA4 | 174 | N | 0 | 0 | no | no | yes | no | yes | no | no | no | no | no | no | no | no |
| CHRNA4 | 175 | C | 1 | 0 | no | no | yes | no | yes | no | no | no | no | no | no | no | no |
| CHRNA4 | 176 | T | 1 | 0 | no | no | yes | no | yes | no | no | no | no | no | no | no | no |
| CHRNA4 | 177 | M | 0 | 0 | no | no | yes | no | yes | no | no | no | no | no | no | no | no |
| CHRNA4 | 178 | K | 0 | 0 | no | no | yes | no | yes | no | no | no | no | no | no | no | no |
| CHRNA4 | 179 | F | 0 | 0 | no | no | yes | no | yes | no | no | no | no | no | no | no | no |
| CHRNA4 | 180 | G | 1 | 0 | no | no | yes | no | yes | no | no | no | no | no | no | no | no |
| CHRNA4 | 181 | S | 0 | 0 | no | no | yes | no | yes | no | no | no | no | no | no | no | no |
| CHRNA4 | 182 | W | 0 | 1 | no | yes | no | no | yes | no | no | no | yes | no | no | no | no |
| CHRNA4 | 183 | T | 1 | 0 | no | yes | no | no | yes | no | no | no | yes | no | no | no | no |
| CHRNA4 | 184 | Y | 1 | 0 | no | yes | no | no | yes | no | no | no | no | no | no | no | no |
| CHRNA4 | 185 | D | 2 | 0 | no | yes | no | no | yes | no | no | no | no | no | no | no | no |
| CHRNA4 | 186 | K | 0 | 0 | no | yes | no | no | yes | no | no | no | no | no | no | no | no |
| CHRNA4 | 187 | A | 1 | 0 | no | yes | no | no | yes | no | no | no | no | no | no | no | no |
| CHRNA4 | 188 | K | 0 | 0 | no | yes | no | no | yes | no | no | no | yes | no | no | no | no |
| CHRNA4 | 189 | I | 1 | 0 | no | no | yes | no | yes | no | no | no | no | no | no | no | no |
| CHRNA4 | 190 | D | 2 | 0 | no | no | yes | no | yes | no | no | no | no | no | no | no | no |
| CHRNA4 | 191 | L | 0 | 0 | no | no | yes | no | yes | no | no | no | no | no | no | no | no |
| CHRNA4 | 192 | V | 0 | 0 | no | no | yes | no | yes | no | no | no | no | no | no | no | no |
| CHRNA4 | 193 | N | 0 | 0 | no | no | yes | no | yes | no | no | no | no | no | no | no | no |
| CHRNA4 | 194 | M | 0 | 0 | no | yes | no | no | yes | no | no | no | no | no | no | no | no |
| CHRNA4 | 195 | H | 1 | 0 | no | yes | no | no | yes | no | no | no | no | no | no | no | no |
| CHRNA4 | 196 | S | 1 | 0 | no | yes | no | no | yes | no | no | no | no | no | no | no | no |
| CHRNA4 | 197 | R | 2 | 0 | no | no | yes | no | yes | no | no | no | no | no | no | no | no |

| | | | | | | | | | | | | | | | | | |
|---|---|---|---|---|---|---|---|---|---|---|---|---|---|---|---|---|---|
| CHRNA4 | 198 | V | 1 | 0 | no | yes | no | no | yes | no | no | no | no | no | no | no | no |
| CHRNA4 | 199 | D | 0 | 0 | no | no | yes | no | yes | no | no | no | no | no | no | no | no |
| CHRNA4 | 200 | Q | 0 | 0 | no | yes | no | no | yes | no | no | no | no | no | no | no | no |
| CHRNA4 | 201 | L | 0 | 0 | no | yes | no | no | yes | no | no | no | no | no | no | no | no |
| CHRNA4 | 202 | D | 1 | 0 | no | yes | no | no | yes | no | no | no | no | no | no | no | no |
| CHRNA4 | 203 | F | 0 | 0 | no | yes | no | no | yes | no | no | no | no | no | no | no | no |
| CHRNA4 | 204 | W | 0 | 0 | no | no | yes | no | yes | no | no | no | yes | no | no | no | no |
| CHRNA4 | 205 | E | 0 | 0 | no | yes | no | no | yes | no | no | no | no | no | no | no | no |
| CHRNA4 | 206 | S | 0 | 0 | no | yes | no | no | yes | no | no | no | no | no | no | no | no |
| CHRNA4 | 207 | G | 0 | 0 | no | yes | no | no | yes | no | no | no | no | no | no | no | no |
| CHRNA4 | 208 | E | 1 | 0 | no | yes | no | no | yes | no | no | no | yes | no | no | no | no |
| CHRNA4 | 209 | W | 1 | 0 | no | no | yes | no | yes | no | no | no | no | no | no | no | no |
| CHRNA4 | 210 | V | 0 | 0 | no | no | yes | no | yes | no | no | no | no | no | no | no | no |
| CHRNA4 | 211 | I | 2 | 0 | no | no | yes | no | yes | no | no | no | no | no | no | no | no |
| CHRNA4 | 212 | V | 1 | 0 | no | no | yes | no | yes | no | no | no | no | no | no | no | no |
| CHRNA4 | 213 | D | 0 | 0 | no | no | yes | no | yes | no | no | no | no | no | no | no | no |
| CHRNA4 | 214 | A | 1 | 0 | no | no | yes | no | yes | no | no | no | no | no | no | no | no |
| CHRNA4 | 215 | V | 0 | 0 | no | no | yes | no | yes | no | no | no | no | no | no | no | no |
| CHRNA4 | 216 | G | 1 | 0 | no | no | yes | no | yes | no | no | no | no | no | no | no | no |
| CHRNA4 | 217 | T | 2 | 0 | no | no | yes | no | yes | no | no | no | no | no | no | no | no |
| CHRNA4 | 218 | Y | 0 | 0 | no | no | yes | no | yes | no | no | no | no | no | no | no | no |
| CHRNA4 | 219 | N | 1 | 0 | no | no | yes | no | yes | no | no | no | no | no | no | no | no |
| CHRNA4 | 220 | T | 0 | 0 | no | no | yes | no | yes | no | no | no | no | no | no | no | no |
| CHRNA4 | 221 | R | 0 | 0 | no | no | yes | no | yes | no | no | no | yes | no | no | no | no |
| CHRNA4 | 222 | K | 0 | 0 | no | yes | no | no | yes | no | no | no | no | no | no | no | no |
| CHRNA4 | 223 | Y | 0 | 0 | no | yes | no | no | yes | no | no | no | yes | no | no | no | no |
| CHRNA4 | 224 | E | 0 | 0 | no | yes | no | no | yes | no | no | no | yes | no | no | no | no |
| CHRNA4 | 225 | C | 0 | 0 | no | yes | no | no | yes | no | no | no | yes | no | no | no | no |
| CHRNA4 | 226 | C | 0 | 0 | no | yes | no | no | yes | no | no | no | no | no | no | no | no |
| CHRNA4 | 227 | A | 1 | 0 | no | yes | no | no | yes | no | no | no | no | no | no | no | no |
| CHRNA4 | 228 | E | 1 | 0 | no | yes | no | no | yes | no | no | no | no | no | no | no | no |
| CHRNA4 | 229 | I | 1 | 0 | no | yes | no | no | yes | no | no | no | no | no | no | no | no |
| CHRNA4 | 230 | Y | 0 | 0 | no | no | yes | no | yes | no | no | no | no | no | no | no | no |

| | | | | | | | | | | | | | | | | | |
|---|---|---|---|---|---|---|---|---|---|---|---|---|---|---|---|---|---|
| CHRNA4 | 231 | P | 1 | 0 | no | no | yes | no | yes | no | no | no | no | no | no | no | no |
| CHRNA4 | 232 | D | 0 | 0 | no | no | yes | no | yes | no | no | no | no | no | no | no | no |
| CHRNA4 | 233 | I | 1 | 0 | no | no | yes | no | yes | no | no | no | no | no | no | no | no |
| CHRNA4 | 234 | T | 0 | 0 | no | no | yes | no | yes | no | no | no | no | no | no | no | no |
| CHRNA4 | 235 | Y | 0 | 0 | no | no | yes | no | yes | no | no | no | no | no | no | no | no |
| CHRNA4 | 236 | A | 0 | 0 | no | no | yes | no | yes | no | no | no | no | no | no | no | no |
| CHRNA4 | 237 | F | 0 | 0 | no | no | yes | no | yes | no | no | no | no | no | no | no | no |
| CHRNA4 | 238 | V | 1 | 0 | no | no | yes | no | yes | no | no | no | no | no | no | no | no |
| CHRNA4 | 239 | I | 1 | 0 | no | no | yes | no | yes | no | no | no | no | no | no | no | no |
| CHRNA4 | 240 | R | 2 | 0 | no | no | yes | no | yes | no | no | no | no | no | no | no | no |
| CHRNA4 | 241 | R | 1 | 0 | no | no | yes | no | yes | no | no | no | no | no | no | no | no |
| CHRNA4 | 242 | L | 0 | 0 | no | yes | no | no | yes | no | no | no | yes | no | no | no | no |
| CHRNA4 | 243 | P | 1 | 0 | no | yes | no | no | no | no | no | no | no | no | no | no | no |
| CHRNA4 | 244 | L | 1 | 0 | yes | no | no | no | no | no | no | no | yes | no | no | no | no |
| CHRNA4 | 245 | F | 0 | 0 | yes | no | no | no | no | no | no | no | yes | no | no | no | no |
| CHRNA4 | 246 | Y | 1 | 0 | yes | no | no | no | no | no | no | no | no | no | no | no | no |
| CHRNA4 | 247 | T | 0 | 0 | yes | no | no | no | no | no | no | no | no | no | no | no | no |
| CHRNA4 | 248 | I | 2 | 0 | no | yes | no | yes | no | no | no | no | yes | no | no | no | no |
| CHRNA4 | 249 | N | 1 | 0 | no | yes | no | no | no | no | no | no | yes | no | no | no | no |
| CHRNA4 | 250 | L | 0 | 0 | no | yes | no | yes | no | no | no | no | no | no | no | no | no |
| CHRNA4 | 251 | I | 1 | 0 | yes | no | no | yes | no | no | no | no | no | no | no | no | no |
| CHRNA4 | 252 | I | 0 | 0 | yes | no | no | yes | no | no | no | no | yes | no | no | no | no |
| CHRNA4 | 253 | P | 1 | 0 | yes | no | no | yes | no | no | no | no | no | no | no | no | no |
| CHRNA4 | 254 | C | 0 | 0 | yes | no | no | yes | no | no | no | no | no | no | no | no | no |
| CHRNA4 | 255 | L | 0 | 0 | yes | no | no | yes | no | no | no | no | no | no | no | no | no |
| CHRNA4 | 256 | L | 0 | 0 | yes | no | no | yes | no | no | no | no | yes | no | no | no | no |
| CHRNA4 | 257 | I | 0 | 0 | yes | no | no | yes | no | no | no | no | yes | no | no | no | no |
| CHRNA4 | 258 | S | 0 | 0 | yes | no | no | yes | no | no | no | no | no | no | no | no | no |
| CHRNA4 | 259 | C | 0 | 0 | yes | no | no | yes | no | no | no | no | no | no | no | no | no |
| CHRNA4 | 260 | L | 1 | 0 | yes | no | no | yes | no | no | no | no | no | no | no | no | no |
| CHRNA4 | 261 | T | 0 | 0 | no | yes | no | yes | no | no | no | no | no | no | no | no | no |
| CHRNA4 | 262 | V | 0 | 0 | no | yes | no | yes | no | no | no | no | no | no | no | no | no |
| CHRNA4 | 263 | L | 1 | 0 | no | yes | no | yes | no | no | no | no | yes | no | no | no | no |

| | | | | | | | | | | | | | | | | | |
|---|---|---|---|---|---|---|---|---|---|---|---|---|---|---|---|---|---|
| CHRNA4 | 264 | V | 0 | 0 | yes | no | no | yes | no | no | no | no | no | no | no | no | no |
| CHRNA4 | 265 | F | 0 | 0 | yes | no | no | yes | no | no | no | no | no | no | no | no | no |
| CHRNA4 | 266 | Y | 0 | 0 | yes | no | no | yes | no | no | no | no | yes | no | no | no | no |
| CHRNA4 | 267 | L | 0 | 0 | no | yes | no | yes | no | no | no | no | yes | no | no | no | no |
| CHRNA4 | 268 | P | 1 | 0 | no | yes | no | no | no | no | no | no | yes | no | no | no | no |
| CHRNA4 | 269 | S | 0 | 0 | no | yes | no | no | no | no | no | no | no | no | no | no | no |
| CHRNA4 | 270 | E | 1 | 0 | no | yes | no | no | no | no | yes | no | yes | no | no | no | no |
| CHRNA4 | 271 | C | 0 | 0 | no | yes | no | no | no | no | yes | no | yes | no | no | no | no |
| CHRNA4 | 272 | G | 0 | 0 | no | yes | no | yes | no | no | no | no | yes | no | no | no | no |
| CHRNA4 | 273 | E | 1 | 0 | no | yes | no | yes | no | no | yes | no | yes | no | no | no | no |
| CHRNA4 | 274 | K | 0 | 0 | yes | no | no | yes | no | no | no | no | no | no | no | no | no |
| CHRNA4 | 275 | I | 1 | 1 | yes | no | no | yes | no | no | no | no | yes | no | no | no | no |
| CHRNA4 | 276 | T | 1 | 0 | yes | no | no | yes | no | no | yes | no | no | no | no | no | no |
| CHRNA4 | 277 | L | 0 | 0 | yes | no | no | yes | no | no | no | no | yes | no | no | no | no |
| CHRNA4 | 278 | C | 0 | 0 | yes | no | no | yes | no | no | no | no | no | no | no | no | no |
| CHRNA4 | 279 | I | 0 | 0 | yes | no | no | yes | no | no | no | no | yes | no | no | no | no |
| CHRNA4 | 280 | S | 0 | 1 | yes | no | no | yes | no | no | yes | no | yes | no | no | no | no |
| CHRNA4 | 281 | V | 1 | 0 | yes | no | no | yes | no | no | no | no | no | no | no | no | no |
| CHRNA4 | 282 | L | 0 | 0 | yes | no | no | yes | no | no | no | no | no | no | no | no | no |
| CHRNA4 | 283 | L | 0 | 0 | yes | no | no | yes | no | no | no | no | no | no | no | no | no |
| CHRNA4 | 284 | S | 0 | 1 | yes | no | no | yes | no | no | no | no | no | no | no | no | no |
| CHRNA4 | 285 | L | 0 | 0 | yes | no | no | yes | no | no | no | no | no | no | no | no | no |
| CHRNA4 | 286 | T | 0 | 0 | yes | no | no | yes | no | no | no | no | yes | no | no | no | no |
| CHRNA4 | 287 | V | 1 | 0 | yes | no | no | yes | no | no | yes | no | no | no | no | no | no |
| CHRNA4 | 288 | F | 0 | 0 | yes | no | no | yes | no | no | no | no | no | no | no | no | no |
| CHRNA4 | 289 | L | 0 | 0 | yes | no | no | yes | no | no | no | no | no | no | no | no | no |
| CHRNA4 | 290 | L | 0 | 1 | yes | no | no | yes | no | no | no | no | yes | no | no | no | no |
| CHRNA4 | 291 | L | 0 | 0 | yes | no | no | no | no | no | yes | no | no | no | no | no | no |
| CHRNA4 | 292 | I | 1 | 0 | yes | no | no | no | no | no | no | no | no | no | no | no | no |
| CHRNA4 | 293 | T | 0 | 1 | no | yes | no | no | no | no | no | no | yes | no | no | no | no |
| CHRNA4 | 294 | E | 1 | 0 | no | yes | no | no | no | no | yes | no | no | no | no | no | no |
| CHRNA4 | 295 | I | 0 | 0 | no | yes | no | no | no | no | no | no | no | no | no | no | no |
| CHRNA4 | 296 | I | 0 | 0 | no | yes | no | no | no | no | no | no | yes | no | no | no | no |

| | | | | | | | | | | | | | | | | | |
|---|---|---|---|---|---|---|---|---|---|---|---|---|---|---|---|---|---|
| CHRNA4 | 297 | P | 1 | 0 | no | yes | no | no | no | no | no | no | yes | no | no | no | no |
| CHRNA4 | 298 | S | 0 | 0 | no | yes | no | no | no | no | no | no | yes | no | no | no | no |
| CHRNA4 | 299 | T | 1 | 0 | no | yes | no | no | no | no | no | no | yes | no | no | no | no |
| CHRNA4 | 300 | S | 0 | 0 | no | yes | no | no | no | no | no | no | yes | no | no | no | no |
| CHRNA4 | 301 | L | 0 | 0 | no | yes | no | no | no | no | no | no | no | no | no | no | no |
| CHRNA4 | 302 | V | 1 | 0 | no | yes | no | no | no | no | no | no | no | no | no | no | no |
| CHRNA4 | 303 | I | 2 | 0 | no | yes | no | no | no | no | no | no | yes | no | no | no | no |
| CHRNA4 | 304 | P | 0 | 0 | no | yes | no | no | no | no | no | no | no | no | no | no | no |
| CHRNA4 | 305 | L | 0 | 0 | yes | no | no | yes | no | no | no | no | no | no | no | no | no |
| CHRNA4 | 306 | I | 1 | 0 | yes | no | no | yes | no | no | no | no | no | no | no | no | no |
| CHRNA4 | 307 | G | 1 | 0 | yes | no | no | yes | no | no | no | no | no | no | no | no | no |
| CHRNA4 | 308 | E | 1 | 0 | yes | no | no | yes | no | no | no | no | no | no | no | no | no |
| CHRNA4 | 309 | Y | 0 | 0 | yes | no | no | yes | no | no | no | no | no | no | no | no | no |
| CHRNA4 | 310 | L | 0 | 0 | yes | no | no | yes | no | no | no | no | no | no | no | no | no |
| CHRNA4 | 311 | L | 0 | 0 | yes | no | no | yes | no | no | no | no | yes | no | no | no | no |
| CHRNA4 | 312 | F | 0 | 0 | yes | no | no | yes | no | no | no | no | no | no | no | no | no |
| CHRNA4 | 313 | T | 1 | 0 | yes | no | no | yes | no | no | no | no | no | no | no | no | no |
| CHRNA4 | 314 | M | 2 | 0 | yes | no | no | yes | no | no | no | no | yes | no | no | no | no |
| CHRNA4 | 315 | I | 0 | 0 | yes | no | no | yes | no | no | no | no | yes | no | no | no | no |
| CHRNA4 | 316 | F | 0 | 0 | yes | no | no | yes | no | no | no | no | no | no | no | no | no |
| CHRNA4 | 317 | V | 0 | 0 | yes | no | no | yes | no | no | no | no | no | no | no | no | no |
| CHRNA4 | 318 | T | 0 | 0 | yes | no | no | yes | no | no | no | no | yes | no | no | no | no |
| CHRNA4 | 319 | L | 2 | 0 | yes | no | no | yes | no | no | no | no | no | no | no | no | no |
| CHRNA4 | 320 | S | 1 | 0 | yes | no | no | yes | no | no | no | no | no | no | no | no | no |
| CHRNA4 | 321 | I | 1 | 0 | yes | no | no | yes | no | no | no | no | yes | no | no | no | no |
| CHRNA4 | 322 | V | 1 | 0 | yes | no | no | yes | no | no | no | no | no | no | no | no | no |
| CHRNA4 | 323 | I | 1 | 0 | yes | no | no | yes | no | no | no | no | no | no | no | no | no |
| CHRNA4 | 324 | T | 1 | 0 | yes | no | no | yes | no | no | no | no | no | no | no | no | no |
| CHRNA4 | 325 | V | 0 | 0 | yes | no | no | yes | no | no | no | no | yes | no | no | no | no |
| CHRNA4 | 326 | F | 0 | 0 | yes | no | no | yes | no | no | no | no | yes | no | no | no | no |
| CHRNA4 | 327 | V | 1 | 1 | yes | no | no | yes | no | no | no | no | no | no | no | no | no |
| CHRNA4 | 328 | L | 0 | 0 | yes | no | no | yes | no | no | no | no | no | no | no | no | no |
| CHRNA4 | 329 | N | 2 | 0 | yes | no | no | no | no | no | no | no | yes | no | no | no | no |

| | | | | | | | | | | | | | | | | | |
|---|---|---|---|---|---|---|---|---|---|---|---|---|---|---|---|---|---|
| CHRNA4 | 330 | V | 1 | 0 | yes | no | no | no | no | no | no | no | no | no | no | no | no |
| CHRNA4 | 331 | H | 0 | 0 | yes | no | no | no | no | yes | no | no | no | no | no | no | no |
| CHRNA4 | 332 | H | 1 | 0 | no | yes | no | no | no | yes | yes | no | yes | no | no | no | no |
| CHRNA4 | 333 | R | 2 | 0 | no | yes | no | no | no | yes | no | no | no | no | no | no | no |
| CHRNA4 | 334 | S | 1 | 0 | no | yes | no | no | no | yes | no | no | no | no | no | no | no |
| CHRNA4 | 335 | P | 0 | 0 | no | yes | no | no | no | yes | no | no | no | no | no | no | no |
| CHRNA4 | 336 | R | 2 | 1 | no | yes | no | no | no | yes | no | no | yes | no | no | no | no |
| CHRNA4 | 337 | T | 1 | 0 | no | yes | no | no | no | yes | no | no | yes | no | no | no | no |
| CHRNA4 | 338 | H | 1 | 0 | no | yes | no | no | no | yes | no | no | yes | no | no | no | no |
| CHRNA4 | 339 | T | 0 | 0 | no | yes | no | no | no | yes | no | no | no | no | no | no | no |
| CHRNA4 | 340 | M | 0 | 0 | no | yes | no | no | no | yes | no | no | yes | no | no | no | no |
| CHRNA4 | 341 | P | 1 | 0 | no | yes | no | no | no | yes | no | no | no | no | no | no | no |
| CHRNA4 | 342 | T | 0 | 0 | yes | no | no | no | no | yes | no | no | no | no | no | no | no |
| CHRNA4 | 343 | W | 1 | 0 | yes | no | no | no | no | yes | no | no | no | no | no | no | no |
| CHRNA4 | 344 | V | 0 | 0 | yes | no | no | no | no | yes | no | no | no | no | no | no | no |
| CHRNA4 | 345 | R | 2 | 0 | yes | no | no | no | no | yes | no | no | no | no | no | no | no |
| CHRNA4 | 346 | R | 0 | 0 | yes | no | no | no | no | yes | no | no | no | no | no | no | no |
| CHRNA4 | 347 | V | 1 | 0 | yes | no | no | no | no | yes | no | no | no | no | no | no | no |
| CHRNA4 | 348 | F | 0 | 0 | yes | no | no | no | no | yes | no | no | no | no | no | no | no |
| CHRNA4 | 349 | L | 0 | 0 | no | yes | no | no | no | yes | no | no | no | no | no | no | no |
| CHRNA4 | 350 | D | 0 | 0 | no | yes | no | no | no | yes | no | no | no | no | no | no | no |
| CHRNA4 | 351 | I | 0 | 0 | yes | no | no | no | no | yes | no | no | no | no | no | no | no |
| CHRNA4 | 352 | V | 1 | 0 | yes | no | no | no | no | yes | no | no | no | no | no | no | no |
| CHRNA4 | 353 | P | 0 | 0 | yes | no | no | no | no | yes | no | no | no | no | no | no | no |
| CHRNA4 | 354 | R | 2 | 0 | yes | no | no | no | no | yes | no | no | no | no | no | no | no |
| CHRNA4 | 355 | L | 0 | 0 | yes | no | no | no | no | yes | no | no | no | no | no | no | no |
| CHRNA4 | 356 | L | 1 | 0 | no | yes | no | no | no | yes | no | no | no | no | no | no | no |
| CHRNA4 | 357 | L | 1 | 0 | no | yes | no | no | no | yes | no | no | no | no | no | no | no |
| CHRNA4 | 358 | M | 0 | 0 | no | yes | no | no | no | yes | no | no | yes | no | no | no | no |
| CHRNA4 | 359 | K | 1 | 0 | no | yes | no | no | no | yes | no | no | yes | no | no | no | no |
| CHRNA4 | 360 | R | 1 | 0 | no | yes | no | no | no | yes | no | no | yes | no | no | no | no |
| CHRNA4 | 586 | S | 1 | 0 | yes | no | no | no | no | yes | no | no | no | no | no | no | no |
| CHRNA4 | 587 | V | 1 | 0 | yes | no | no | no | no | yes | no | no | no | no | no | no | no |

| | | | | | | | | | | | | | | | | | |
|---|---|---|---|---|---|---|---|---|---|---|---|---|---|---|---|---|---|
| CHRNA4 | 588 | K | 0 | 0 | yes | no | no | no | no | yes | yes | no | no | no | no | no | no |
| CHRNA4 | 589 | E | 1 | 0 | yes | no | no | no | no | yes | no | no | yes | no | no | no | no |
| CHRNA4 | 590 | D | 0 | 0 | yes | no | no | no | no | yes | no | no | no | no | no | no | no |
| CHRNA4 | 591 | W | 0 | 0 | yes | no | no | no | no | yes | no | no | no | no | no | no | no |
| CHRNA4 | 592 | K | 0 | 0 | yes | no | no | no | no | yes | no | no | yes | no | no | no | no |
| CHRNA4 | 593 | Y | 0 | 0 | yes | no | no | no | no | yes | no | no | no | no | no | no | no |
| CHRNA4 | 594 | V | 1 | 0 | yes | no | no | no | no | yes | no | no | no | no | no | no | no |
| CHRNA4 | 595 | A | 2 | 0 | yes | no | no | no | no | yes | no | no | no | no | no | no | no |
| CHRNA4 | 596 | M | 1 | 0 | yes | no | no | no | no | yes | no | no | yes | no | no | no | no |
| CHRNA4 | 597 | V | 0 | 0 | yes | no | no | no | no | yes | no | no | no | no | no | no | no |
| CHRNA4 | 598 | I | 1 | 0 | yes | no | no | no | no | yes | no | no | no | no | no | no | no |
| CHRNA4 | 599 | D | 1 | 0 | yes | no | no | yes | no | yes | no | no | no | no | no | no | no |
| CHRNA4 | 600 | R | 2 | 0 | yes | no | no | no | no | yes | no | no | no | no | no | no | no |
| CHRNA4 | 601 | I | 2 | 0 | yes | no | no | yes | no | no | no | no | no | no | no | no | no |
| CHRNA4 | 602 | F | 0 | 0 | yes | no | no | yes | no | no | no | no | no | no | no | no | no |
| CHRNA4 | 603 | L | 0 | 0 | yes | no | no | yes | no | no | no | no | no | no | no | no | no |
| CHRNA4 | 604 | W | 0 | 0 | yes | no | no | yes | no | no | no | no | no | no | no | no | no |
| CHRNA4 | 605 | M | 1 | 0 | yes | no | no | yes | no | no | no | no | no | no | no | no | no |
| CHRNA4 | 606 | F | 0 | 0 | yes | no | no | yes | no | no | no | no | no | no | no | no | no |
| CHRNA4 | 607 | I | 0 | 0 | yes | no | no | yes | no | no | no | no | no | no | no | no | no |
| CHRNA4 | 608 | I | 2 | 0 | yes | no | no | yes | no | no | no | no | no | no | no | no | no |
| CHRNA4 | 609 | V | 2 | 0 | yes | no | no | yes | no | no | no | no | no | no | no | no | no |
| CHRNA4 | 610 | C | 0 | 0 | yes | no | no | yes | no | no | no | no | no | no | no | no | no |
| CHRNA4 | 611 | L | 0 | 0 | yes | no | no | yes | no | no | no | no | no | no | no | no | no |
| CHRNA4 | 612 | L | 0 | 0 | yes | no | no | yes | no | no | no | no | no | no | no | no | no |
| CHRNA4 | 613 | G | 0 | 0 | yes | no | no | yes | no | no | no | no | no | no | no | no | no |
| CHRNA4 | 614 | T | 1 | 0 | yes | no | no | yes | no | no | no | no | no | no | no | no | no |
| CHRNA4 | 615 | V | 0 | 0 | yes | no | no | yes | no | no | no | no | no | no | no | no | no |
| CHRNA4 | 616 | G | 0 | 0 | yes | no | no | yes | no | no | no | no | no | no | no | no | no |
| CHRNA4 | 617 | L | 0 | 0 | yes | no | no | yes | no | no | no | no | no | no | no | no | no |
| CHRNA4 | 618 | F | 0 | 0 | no | yes | no | yes | no | no | no | no | no | no | no | no | no |
| CHRNA4 | 619 | L | 0 | 0 | no | yes | no | yes | no | no | no | no | no | no | no | no | no |
| CHRNA4 | 620 | P | 2 | 0 | no | yes | no | yes | no | no | no | no | no | no | no | no | no |

| | | | | | | | | | | | | | | | | | |
|---|---|---|---|---|---|---|---|---|---|---|---|---|---|---|---|---|---|
| CHRNA4 | 621 | P | 1 | 0 | no | yes | no | no | no | no | no | no | no | no | no | no | no |
| CHRNA4 | 622 | W | 1 | 0 | no | yes | no | yes | no | no | no | no | no | no | no | no | no |
| CHRNB2 | 27 | D | 0 | 0 | no | yes | no | no | yes | no | no | no | yes | no | no | no | no |
| CHRNB2 | 28 | T | 0 | 0 | yes | no | no | no | yes | no | no | no | no | no | no | no | no |
| CHRNB2 | 29 | E | 0 | 0 | yes | no | no | no | yes | no | no | no | yes | no | no | no | no |
| CHRNB2 | 30 | E | 0 | 0 | yes | no | no | no | yes | no | no | no | yes | no | no | no | no |
| CHRNB2 | 31 | R | 2 | 0 | yes | no | no | no | yes | no | no | no | no | no | no | no | no |
| CHRNB2 | 32 | L | 0 | 0 | yes | no | no | no | yes | no | no | no | no | no | no | no | no |
| CHRNB2 | 33 | V | 0 | 0 | yes | no | no | no | yes | no | no | no | yes | no | no | no | no |
| CHRNB2 | 34 | E | 1 | 0 | yes | no | no | no | yes | no | no | no | yes | no | no | no | no |
| CHRNB2 | 35 | H | 1 | 1 | yes | no | no | no | yes | no | no | no | no | no | no | no | no |
| CHRNB2 | 36 | L | 0 | 0 | yes | no | no | no | yes | no | no | no | no | no | no | no | no |
| CHRNB2 | 37 | L | 0 | 0 | no | yes | no | no | yes | no | no | no | no | no | no | no | no |
| CHRNB2 | 38 | D | 0 | 0 | no | yes | no | no | yes | no | no | no | no | no | no | no | no |
| CHRNB2 | 39 | P | 0 | 0 | no | yes | no | no | yes | no | yes | no | no | no | no | no | no |
| CHRNB2 | 40 | S | 0 | 0 | no | yes | no | no | yes | no | no | no | no | no | no | no | no |
| CHRNB2 | 41 | R | 2 | 0 | no | yes | no | no | yes | no | no | no | yes | no | no | no | no |
| CHRNB2 | 42 | Y | 1 | 0 | no | yes | no | no | yes | no | no | no | no | no | no | no | no |
| CHRNB2 | 43 | N | 1 | 0 | no | yes | no | no | yes | no | no | no | yes | no | no | no | no |
| CHRNB2 | 44 | K | 0 | 0 | no | yes | no | no | yes | no | no | no | no | no | no | no | no |
| CHRNB2 | 45 | L | 0 | 0 | no | yes | no | no | yes | no | no | no | yes | no | no | no | no |
| CHRNB2 | 46 | I | 0 | 0 | no | yes | no | no | yes | no | no | no | no | no | no | no | no |
| CHRNB2 | 47 | R | 2 | 1 | no | yes | no | no | yes | no | no | no | no | no | no | no | no |
| CHRNB2 | 48 | P | 0 | 0 | no | yes | no | no | yes | no | no | no | no | no | no | no | no |
| CHRNB2 | 49 | A | 0 | 0 | no | yes | no | no | yes | no | no | no | no | no | no | no | no |
| CHRNB2 | 50 | T | 1 | 0 | no | yes | no | no | yes | no | no | no | no | no | no | no | no |
| CHRNB2 | 51 | N | 0 | 0 | no | yes | no | no | yes | no | no | no | no | no | no | no | no |
| CHRNB2 | 52 | G | 0 | 0 | no | yes | no | no | yes | no | no | no | no | no | no | no | no |
| CHRNB2 | 53 | S | 0 | 0 | no | yes | no | no | yes | no | no | no | no | no | no | no | no |
| CHRNB2 | 54 | E | 0 | 0 | no | yes | no | no | yes | no | no | no | no | no | no | no | no |
| CHRNB2 | 55 | L | 0 | 0 | no | yes | no | no | yes | no | no | no | no | no | no | no | no |
| CHRNB2 | 56 | V | 0 | 0 | no | no | yes | no | yes | no | no | no | no | no | no | no | no |
| CHRNB2 | 57 | T | 1 | 0 | no | no | yes | no | yes | no | no | no | no | no | no | no | no |

| | | | | | | | | | | | | | | | | | |
|---|---|---|---|---|---|---|---|---|---|---|---|---|---|---|---|---|---|
| CHRNB2 | 58 | V | 0 | 0 | no | no | yes | no | yes | no | no | no | no | no | no | no | no |
| CHRNB2 | 59 | Q | 0 | 0 | no | no | yes | no | yes | no | no | no | no | no | no | no | no |
| CHRNB2 | 60 | L | 0 | 0 | no | no | yes | no | yes | no | no | no | no | no | no | no | no |
| CHRNB2 | 61 | M | 0 | 0 | no | no | yes | no | yes | no | no | no | no | no | no | no | no |
| CHRNB2 | 62 | V | 0 | 0 | no | no | yes | no | yes | no | no | no | no | no | no | no | no |
| CHRNB2 | 63 | S | 0 | 0 | no | no | yes | no | yes | no | no | no | no | no | no | no | no |
| CHRNB2 | 64 | L | 0 | 0 | no | no | yes | no | yes | no | no | no | no | no | no | no | no |
| CHRNB2 | 65 | A | 0 | 0 | no | no | yes | no | yes | no | no | no | no | no | no | no | no |
| CHRNB2 | 66 | Q | 0 | 0 | no | no | yes | no | yes | no | no | no | yes | no | no | no | no |
| CHRNB2 | 67 | L | 0 | 0 | no | no | yes | no | yes | no | no | no | no | no | no | no | no |
| CHRNB2 | 68 | I | 1 | 0 | no | no | yes | no | yes | no | no | no | yes | no | no | no | no |
| CHRNB2 | 69 | S | 1 | 0 | no | no | yes | no | yes | no | no | no | no | no | no | no | no |
| CHRNB2 | 70 | V | 0 | 0 | no | no | yes | no | yes | no | no | no | no | no | no | no | no |
| CHRNB2 | 71 | H | 1 | 0 | no | no | yes | no | yes | no | no | no | no | no | no | no | no |
| CHRNB2 | 72 | E | 1 | 0 | no | yes | no | no | yes | no | no | no | no | no | no | no | no |
| CHRNB2 | 73 | R | 2 | 0 | no | yes | no | no | yes | no | yes | no | no | no | no | no | no |
| CHRNB2 | 74 | E | 2 | 0 | no | yes | no | no | yes | no | yes | no | no | no | no | no | no |
| CHRNB2 | 75 | Q | 0 | 0 | no | yes | no | no | yes | no | no | no | no | no | no | no | no |
| CHRNB2 | 76 | I | 0 | 0 | no | no | yes | no | yes | no | no | no | no | no | no | no | no |
| CHRNB2 | 77 | M | 0 | 0 | no | no | yes | no | yes | no | no | no | no | no | no | no | no |
| CHRNB2 | 78 | T | 1 | 0 | no | no | yes | no | yes | no | no | no | no | no | no | no | no |
| CHRNB2 | 79 | T | 1 | 0 | no | no | yes | no | yes | no | no | no | no | no | no | no | no |
| CHRNB2 | 80 | N | 0 | 0 | no | no | yes | no | yes | no | no | no | yes | no | no | no | no |
| CHRNB2 | 81 | V | 0 | 0 | no | no | yes | no | yes | no | no | no | no | no | no | no | no |
| CHRNB2 | 82 | W | 0 | 0 | no | no | yes | no | yes | no | no | no | no | no | no | no | no |
| CHRNB2 | 83 | L | 0 | 0 | no | no | yes | no | yes | no | no | no | no | no | no | no | no |
| CHRNB2 | 84 | T | 0 | 0 | no | no | yes | no | yes | no | no | no | no | no | no | no | no |
| CHRNB2 | 85 | Q | 0 | 0 | no | no | yes | no | yes | no | no | no | no | no | no | no | no |
| CHRNB2 | 86 | E | 0 | 0 | no | no | yes | no | yes | no | no | no | no | no | no | no | no |
| CHRNB2 | 87 | W | 0 | 0 | no | no | yes | no | yes | no | no | no | no | no | no | no | no |
| CHRNB2 | 88 | E | 1 | 0 | no | no | yes | no | yes | no | no | no | no | no | no | no | no |
| CHRNB2 | 89 | D | 0 | 0 | no | yes | no | no | yes | no | no | no | no | no | no | no | no |
| CHRNB2 | 90 | Y | 0 | 0 | no | yes | no | no | yes | no | no | no | no | no | no | no | no |

| | | | | | | | | | | | | | | | | | |
|---|---|---|---|---|---|---|---|---|---|---|---|---|---|---|---|---|---|
| CHRNB2 | 91 | R | 2 | 0 | no | yes | no | no | yes | no | no | no | no | no | no | no | no |
| CHRNB2 | 92 | L | 0 | 0 | no | yes | no | no | yes | no | no | no | no | no | no | no | no |
| CHRNB2 | 93 | T | 1 | 0 | no | yes | no | no | yes | no | no | no | no | no | no | no | no |
| CHRNB2 | 94 | W | 0 | 0 | no | yes | no | no | yes | no | no | no | no | no | no | no | no |
| CHRNB2 | 95 | K | 0 | 0 | no | yes | no | no | yes | no | no | no | no | no | no | no | no |
| CHRNB2 | 96 | P | 0 | 0 | no | yes | no | no | yes | no | no | no | no | no | no | no | no |
| CHRNB2 | 97 | E | 0 | 0 | no | yes | no | no | yes | no | no | no | no | no | no | no | no |
| CHRNB2 | 98 | E | 0 | 0 | no | yes | no | no | yes | no | no | no | no | no | no | no | no |
| CHRNB2 | 99 | F | 0 | 0 | no | yes | no | no | yes | no | no | no | no | no | no | no | no |
| CHRNB2 | 100 | D | 0 | 0 | no | yes | no | no | yes | no | no | no | no | no | no | no | no |
| CHRNB2 | 101 | N | 1 | 0 | no | yes | no | no | yes | no | no | no | no | no | no | no | no |
| CHRNB2 | 102 | M | 1 | 0 | no | yes | no | no | yes | no | no | no | no | no | no | no | no |
| CHRNB2 | 103 | K | 0 | 0 | no | yes | no | no | yes | no | no | no | no | no | no | no | no |
| CHRNB2 | 104 | K | 0 | 0 | no | no | yes | no | yes | no | no | no | no | no | no | no | no |
| CHRNB2 | 105 | V | 0 | 0 | no | no | yes | no | yes | no | no | no | no | no | no | no | no |
| CHRNB2 | 106 | R | 2 | 0 | no | no | yes | no | yes | no | no | no | yes | no | no | no | no |
| CHRNB2 | 107 | L | 0 | 0 | no | no | yes | no | yes | no | no | no | no | no | no | no | no |
| CHRNB2 | 108 | P | 0 | 0 | no | yes | no | no | yes | no | no | no | yes | no | no | no | no |
| CHRNB2 | 109 | S | 1 | 0 | no | yes | no | no | yes | no | no | no | no | no | no | no | no |
| CHRNB2 | 110 | K | 1 | 0 | no | yes | no | no | yes | no | yes | no | no | no | no | no | no |
| CHRNB2 | 111 | H | 1 | 0 | no | yes | no | no | yes | no | no | no | yes | no | no | no | no |
| CHRNB2 | 112 | I | 0 | 0 | no | yes | no | no | yes | no | no | no | no | no | no | no | no |
| CHRNB2 | 113 | W | 0 | 0 | no | yes | no | no | yes | no | no | no | no | no | no | no | no |
| CHRNB2 | 114 | L | 0 | 0 | no | yes | no | no | yes | no | no | no | no | no | no | no | no |
| CHRNB2 | 115 | P | 0 | 0 | no | yes | no | no | yes | no | no | no | no | no | no | no | no |
| CHRNB2 | 116 | D | 0 | 0 | no | yes | no | no | yes | no | no | no | no | no | no | no | no |
| CHRNB2 | 117 | V | 0 | 0 | no | yes | no | no | yes | no | no | no | no | no | no | no | no |
| CHRNB2 | 118 | V | 0 | 0 | no | no | yes | no | yes | no | no | no | no | no | no | no | no |
| CHRNB2 | 119 | L | 0 | 0 | no | no | yes | no | yes | no | no | no | no | no | no | no | no |
| CHRNB2 | 120 | Y | 0 | 0 | no | yes | no | no | yes | no | no | no | yes | no | no | no | no |
| CHRNB2 | 121 | N | 1 | 0 | no | yes | no | no | yes | no | no | no | no | no | no | no | no |
| CHRNB2 | 122 | N | 0 | 0 | no | no | yes | no | yes | no | no | no | no | no | no | no | no |
| CHRNB2 | 123 | A | 0 | 0 | no | yes | no | no | yes | no | no | no | no | no | no | no | no |

| | | | | | | | | | | | | | | | | | |
|---|---|---|---|---|---|---|---|---|---|---|---|---|---|---|---|---|---|
| CHRNB2 | 124 | D | 0 | 0 | no | yes | no | no | yes | no | yes | no | no | no | no | no | no |
| CHRNB2 | 125 | G | 0 | 0 | no | yes | no | no | yes | no | no | no | no | no | no | no | no |
| CHRNB2 | 126 | M | 1 | 0 | no | yes | no | no | yes | no | no | no | no | no | no | no | no |
| CHRNB2 | 127 | Y | 0 | 0 | no | yes | no | no | yes | no | no | no | yes | no | no | no | no |
| CHRNB2 | 128 | E | 0 | 0 | no | yes | no | no | yes | no | yes | no | no | no | no | no | no |
| CHRNB2 | 129 | V | 0 | 0 | no | yes | no | no | yes | no | no | no | no | no | no | no | no |
| CHRNB2 | 130 | S | 0 | 0 | no | yes | no | no | yes | no | no | no | no | no | no | no | no |
| CHRNB2 | 131 | F | 0 | 0 | no | yes | no | no | yes | no | no | no | yes | no | no | no | no |
| CHRNB2 | 132 | Y | 1 | 0 | no | yes | no | no | yes | no | no | no | no | no | no | no | no |
| CHRNB2 | 133 | S | 1 | 0 | no | yes | no | no | yes | no | no | no | yes | no | no | no | no |
| CHRNB2 | 134 | N | 0 | 0 | no | yes | no | no | yes | no | no | no | yes | no | no | no | no |
| CHRNB2 | 135 | A | 0 | 0 | no | no | yes | no | yes | no | no | no | no | no | no | no | no |
| CHRNB2 | 136 | V | 1 | 0 | no | no | yes | no | yes | no | no | no | no | no | no | no | no |
| CHRNB2 | 137 | V | 2 | 0 | no | no | yes | no | yes | no | no | no | no | no | no | no | no |
| CHRNB2 | 138 | S | 0 | 0 | no | no | yes | no | yes | no | no | no | no | no | no | no | no |
| CHRNB2 | 139 | Y | 1 | 0 | no | yes | no | no | yes | no | no | no | no | no | no | no | no |
| CHRNB2 | 140 | D | 1 | 0 | no | yes | no | no | yes | no | no | no | no | no | no | no | no |
| CHRNB2 | 141 | G | 0 | 0 | no | yes | no | no | yes | no | no | no | no | no | no | no | no |
| CHRNB2 | 142 | S | 0 | 0 | no | no | yes | no | yes | no | no | no | no | no | no | no | no |
| CHRNB2 | 143 | I | 0 | 0 | no | no | yes | no | yes | no | no | no | no | no | no | no | no |
| CHRNB2 | 144 | F | 0 | 0 | no | no | yes | no | yes | no | no | no | yes | no | no | no | no |
| CHRNB2 | 145 | W | 0 | 0 | no | no | yes | no | yes | no | no | no | no | no | no | no | no |
| CHRNB2 | 146 | L | 0 | 0 | no | yes | no | no | yes | no | no | no | yes | no | no | no | no |
| CHRNB2 | 147 | P | 1 | 0 | no | yes | no | no | yes | no | no | no | no | no | no | no | no |
| CHRNB2 | 148 | P | 1 | 0 | no | no | yes | no | yes | no | no | no | yes | no | no | no | no |
| CHRNB2 | 149 | A | 0 | 0 | no | no | yes | no | yes | no | no | no | no | no | no | no | no |
| CHRNB2 | 150 | I | 1 | 0 | no | no | yes | no | yes | no | no | no | yes | no | no | no | no |
| CHRNB2 | 151 | Y | 0 | 0 | no | no | yes | no | yes | no | no | no | no | no | no | no | no |
| CHRNB2 | 152 | K | 0 | 0 | no | no | yes | no | yes | no | no | no | yes | no | no | no | no |
| CHRNB2 | 153 | S | 1 | 0 | no | no | yes | no | yes | no | no | no | no | no | no | no | no |
| CHRNB2 | 154 | A | 2 | 0 | no | yes | no | no | yes | no | no | no | no | no | no | no | no |
| CHRNB2 | 155 | C | 0 | 0 | no | yes | no | no | yes | no | no | no | no | no | no | no | no |
| CHRNB2 | 156 | K | 0 | 0 | no | yes | no | no | yes | no | no | no | no | no | no | no | no |

| | | | | | | | | | | | | | | | | | |
|---|---|---|---|---|---|---|---|---|---|---|---|---|---|---|---|---|---|
| CHRNB2 | 157 | I | 0 | 0 | no | yes | no | no | yes | no | no | no | no | no | no | no | no |
| CHRNB2 | 158 | E | 0 | 0 | no | yes | no | no | yes | no | no | no | no | no | no | no | no |
| CHRNB2 | 159 | V | 0 | 0 | no | yes | no | no | yes | no | no | no | no | no | no | no | no |
| CHRNB2 | 160 | K | 0 | 0 | no | yes | no | no | yes | no | no | no | no | no | no | no | no |
| CHRNB2 | 161 | H | 0 | 0 | no | yes | no | no | yes | no | no | no | no | no | no | no | no |
| CHRNB2 | 162 | F | 0 | 0 | no | yes | no | no | yes | no | no | no | no | no | no | no | no |
| CHRNB2 | 163 | P | 1 | 0 | no | yes | no | no | yes | no | no | no | no | no | no | no | no |
| CHRNB2 | 164 | F | 0 | 0 | no | yes | no | no | yes | no | no | no | no | no | no | no | no |
| CHRNB2 | 165 | D | 0 | 0 | no | yes | no | no | yes | no | no | no | no | no | no | no | no |
| CHRNB2 | 166 | Q | 1 | 0 | no | no | yes | no | yes | no | no | no | no | no | no | no | no |
| CHRNB2 | 167 | Q | 0 | 0 | no | no | yes | no | yes | no | no | no | no | no | no | no | no |
| CHRNB2 | 168 | N | 0 | 0 | no | no | yes | no | yes | no | no | no | no | no | no | no | no |
| CHRNB2 | 169 | C | 0 | 0 | no | no | yes | no | yes | no | no | no | no | no | no | no | no |
| CHRNB2 | 170 | T | 0 | 0 | no | no | yes | no | yes | no | no | no | no | no | no | no | no |
| CHRNB2 | 171 | M | 0 | 0 | no | no | yes | no | yes | no | no | no | no | no | no | no | no |
| CHRNB2 | 172 | K | 1 | 0 | no | no | yes | no | yes | no | no | no | no | no | no | no | no |
| CHRNB2 | 173 | F | 0 | 0 | no | no | yes | no | yes | no | no | no | no | no | no | no | no |
| CHRNB2 | 174 | R | 0 | 0 | no | no | yes | no | yes | no | no | no | no | no | no | no | no |
| CHRNB2 | 175 | S | 1 | 0 | no | yes | no | no | yes | no | no | no | no | no | no | no | no |
| CHRNB2 | 176 | W | 0 | 0 | no | yes | no | no | yes | no | no | no | yes | no | no | no | no |
| CHRNB2 | 177 | T | 0 | 0 | no | yes | no | no | yes | no | no | no | yes | no | no | no | no |
| CHRNB2 | 178 | Y | 0 | 0 | no | yes | no | no | yes | no | no | no | no | no | no | no | no |
| CHRNB2 | 179 | D | 1 | 0 | no | yes | no | no | yes | no | no | no | no | no | no | no | no |
| CHRNB2 | 180 | R | 2 | 0 | no | yes | no | no | yes | no | no | no | no | no | no | no | no |
| CHRNB2 | 181 | T | 0 | 0 | no | yes | no | no | yes | no | no | no | no | no | no | no | no |
| CHRNB2 | 182 | E | 1 | 0 | no | yes | no | no | yes | no | no | no | no | no | no | no | no |
| CHRNB2 | 183 | I | 1 | 0 | no | no | yes | no | yes | no | no | no | no | no | no | no | no |
| CHRNB2 | 184 | D | 1 | 0 | no | no | yes | no | yes | no | no | no | no | no | no | no | no |
| CHRNB2 | 185 | L | 0 | 0 | no | no | yes | no | yes | no | no | no | no | no | no | no | no |
| CHRNB2 | 186 | V | 0 | 0 | no | no | yes | no | yes | no | no | no | no | no | no | no | no |
| CHRNB2 | 187 | L | 0 | 0 | no | yes | no | no | yes | no | no | no | no | no | no | no | no |
| CHRNB2 | 188 | K | 0 | 0 | no | yes | no | no | yes | no | no | no | no | no | no | no | no |
| CHRNB2 | 189 | S | 0 | 0 | no | yes | no | no | yes | no | no | no | no | no | no | no | no |

| | | | | | | | | | | | | | | | | | |
|---|---|---|---|---|---|---|---|---|---|---|---|---|---|---|---|---|---|
| CHRNB2 | 190 | E | 0 | 0 | no | yes | no | no | yes | no | no | no | no | no | no | no | no |
| CHRNB2 | 191 | V | 0 | 0 | no | no | yes | no | yes | no | no | no | no | no | no | no | no |
| CHRNB2 | 192 | A | 0 | 0 | no | yes | no | no | yes | no | no | no | no | no | no | no | no |
| CHRNB2 | 193 | S | 0 | 0 | no | no | yes | no | yes | no | no | no | yes | no | no | no | no |
| CHRNB2 | 194 | L | 0 | 0 | no | yes | no | no | yes | no | no | no | no | no | no | no | no |
| CHRNB2 | 195 | D | 0 | 0 | no | yes | no | no | yes | no | no | no | yes | no | no | no | no |
| CHRNB2 | 196 | D | 1 | 0 | no | yes | no | no | yes | no | no | no | yes | no | no | no | no |
| CHRNB2 | 197 | F | 0 | 0 | no | yes | no | no | yes | no | no | no | no | no | no | no | no |
| CHRNB2 | 198 | T | 0 | 0 | no | yes | no | no | yes | no | no | no | no | no | no | no | no |
| CHRNB2 | 199 | P | 0 | 0 | no | yes | no | no | yes | no | no | no | no | no | no | no | no |
| CHRNB2 | 200 | S | 1 | 0 | no | yes | no | no | yes | no | no | no | no | no | no | no | no |
| CHRNB2 | 201 | G | 1 | 0 | no | yes | no | no | yes | no | no | no | no | no | no | no | no |
| CHRNB2 | 202 | E | 1 | 0 | no | yes | no | no | yes | no | no | no | yes | no | no | no | no |
| CHRNB2 | 203 | W | 0 | 0 | no | no | yes | no | yes | no | no | no | no | no | no | no | no |
| CHRNB2 | 204 | D | 0 | 0 | no | no | yes | no | yes | no | no | no | no | no | no | no | no |
| CHRNB2 | 205 | I | 2 | 0 | no | no | yes | no | yes | no | no | no | no | no | no | no | no |
| CHRNB2 | 206 | V | 1 | 0 | no | no | yes | no | yes | no | no | no | no | no | no | no | no |
| CHRNB2 | 207 | A | 1 | 0 | no | no | yes | no | yes | no | no | no | no | no | no | no | no |
| CHRNB2 | 208 | L | 0 | 0 | no | yes | no | no | yes | no | no | no | no | no | no | no | no |
| CHRNB2 | 209 | P | 0 | 0 | no | no | yes | no | yes | no | no | no | no | no | no | no | no |
| CHRNB2 | 210 | G | 0 | 0 | no | no | yes | no | yes | no | no | no | no | no | no | no | no |
| CHRNB2 | 211 | R | 1 | 0 | no | no | yes | no | yes | no | no | no | no | no | no | no | no |
| CHRNB2 | 212 | R | 2 | 0 | no | no | yes | no | yes | no | no | no | no | no | no | no | no |
| CHRNB2 | 213 | N | 0 | 0 | no | no | yes | no | yes | no | no | no | no | no | no | no | no |
| CHRNB2 | 214 | E | 1 | 0 | no | no | yes | no | yes | no | no | no | no | no | no | no | no |
| CHRNB2 | 215 | N | 0 | 0 | no | yes | no | no | yes | no | no | no | no | no | no | no | no |
| CHRNB2 | 216 | P | 1 | 0 | no | yes | no | no | yes | no | no | no | no | no | no | no | no |
| CHRNB2 | 217 | D | 2 | 0 | no | yes | no | no | yes | no | no | no | no | no | no | no | no |
| CHRNB2 | 218 | D | 1 | 0 | no | yes | no | no | yes | no | no | no | no | no | no | no | no |
| CHRNB2 | 219 | S | 0 | 0 | no | yes | no | no | yes | no | no | no | no | no | no | no | no |
| CHRNB2 | 220 | T | 2 | 0 | no | yes | no | no | yes | no | no | no | no | no | no | no | no |
| CHRNB2 | 221 | Y | 0 | 0 | no | yes | no | no | yes | no | no | no | no | no | no | no | no |
| CHRNB2 | 222 | V | 1 | 0 | no | no | yes | no | yes | no | no | no | no | no | no | no | no |

| | | | | | | | | | | | | | | | | | |
|---|---|---|---|---|---|---|---|---|---|---|---|---|---|---|---|---|---|
| CHRNB2 | 223 | D | 1 | 0 | no | no | yes | no | yes | no | no | no | no | no | no | no | no |
| CHRNB2 | 224 | I | 0 | 0 | no | no | yes | no | yes | no | no | no | no | no | no | no | no |
| CHRNB2 | 225 | T | 2 | 0 | no | no | yes | no | yes | no | no | no | no | no | no | no | no |
| CHRNB2 | 226 | Y | 0 | 0 | no | no | yes | no | yes | no | no | no | no | no | no | no | no |
| CHRNB2 | 227 | D | 0 | 0 | no | no | yes | no | yes | no | no | no | no | no | no | no | no |
| CHRNB2 | 228 | F | 0 | 0 | no | no | yes | no | yes | no | no | no | no | no | no | no | no |
| CHRNB2 | 229 | I | 0 | 0 | no | no | yes | no | yes | no | no | no | no | no | no | no | no |
| CHRNB2 | 230 | I | 0 | 0 | no | no | yes | no | yes | no | no | no | no | no | no | no | no |
| CHRNB2 | 231 | R | 1 | 0 | no | no | yes | no | yes | no | no | no | no | no | no | no | no |
| CHRNB2 | 232 | R | 3 | 0 | no | no | yes | no | yes | no | no | no | no | no | no | no | no |
| CHRNB2 | 233 | K | 0 | 0 | no | yes | no | no | yes | no | no | no | yes | no | no | no | no |
| CHRNB2 | 234 | P | 1 | 0 | no | yes | no | no | no | no | no | no | no | no | no | no | no |
| CHRNB2 | 235 | L | 0 | 0 | yes | no | no | no | no | no | no | no | yes | no | no | no | no |
| CHRNB2 | 236 | F | 0 | 0 | yes | no | no | no | no | no | no | no | yes | no | no | no | no |
| CHRNB2 | 237 | Y | 0 | 0 | yes | no | no | no | no | no | no | no | no | no | no | no | no |
| CHRNB2 | 238 | T | 1 | 0 | yes | no | no | no | no | no | no | no | no | no | no | no | no |
| CHRNB2 | 239 | I | 0 | 0 | yes | no | no | yes | no | no | no | no | yes | no | no | no | no |
| CHRNB2 | 240 | N | 1 | 0 | yes | no | no | no | no | no | no | no | no | no | no | no | no |
| CHRNB2 | 241 | L | 1 | 0 | yes | no | no | yes | no | no | no | no | no | no | no | no | no |
| CHRNB2 | 242 | I | 0 | 0 | yes | no | no | yes | no | no | no | no | no | no | no | no | no |
| CHRNB2 | 243 | I | 0 | 0 | yes | no | no | yes | no | no | no | no | yes | no | no | no | no |
| CHRNB2 | 244 | P | 1 | 0 | yes | no | no | yes | no | no | no | no | yes | no | no | no | no |
| CHRNB2 | 245 | C | 1 | 0 | yes | no | no | yes | no | no | no | no | no | no | no | no | no |
| CHRNB2 | 246 | V | 1 | 0 | yes | no | no | yes | no | no | no | no | no | no | no | no | no |
| CHRNB2 | 247 | L | 0 | 0 | yes | no | no | yes | no | no | no | no | yes | no | no | no | no |
| CHRNB2 | 248 | I | 0 | 0 | yes | no | no | yes | no | no | no | no | no | no | no | no | no |
| CHRNB2 | 249 | T | 0 | 0 | yes | no | no | yes | no | no | no | no | no | no | no | no | no |
| CHRNB2 | 250 | S | 0 | 0 | no | yes | no | yes | no | no | no | no | no | no | no | no | no |
| CHRNB2 | 251 | L | 1 | 0 | no | yes | no | yes | no | no | no | no | yes | no | no | no | no |
| CHRNB2 | 252 | A | 0 | 0 | no | yes | no | yes | no | no | no | no | no | no | no | no | no |
| CHRNB2 | 253 | I | 0 | 0 | no | yes | no | yes | no | no | no | no | no | no | no | no | no |
| CHRNB2 | 254 | L | 0 | 0 | no | yes | no | yes | no | no | no | no | yes | no | no | no | no |
| CHRNB2 | 255 | V | 0 | 0 | no | yes | no | yes | no | no | no | no | no | no | no | no | no |

| | | | | | | | | | | | | | | | | | |
|---|---|---|---|---|---|---|---|---|---|---|---|---|---|---|---|---|---|
| CHRNB2 | 256 | F | 0 | 0 | no | yes | no | yes | no | no | no | no | no | no | no | no | no |
| CHRNB2 | 257 | Y | 0 | 0 | no | yes | no | yes | no | no | no | no | yes | no | no | no | no |
| CHRNB2 | 258 | L | 0 | 0 | no | yes | no | yes | no | no | no | no | yes | no | no | no | no |
| CHRNB2 | 259 | P | 0 | 0 | no | yes | no | no | no | yes | no | no | yes | no | no | no | no |
| CHRNB2 | 260 | S | 0 | 0 | yes | no | no | no | no | yes | no | no | no | no | no | no | no |
| CHRNB2 | 261 | D | 1 | 0 | yes | no | no | no | no | yes | yes | no | yes | no | no | no | no |
| CHRNB2 | 262 | C | 0 | 0 | yes | no | no | no | no | yes | yes | no | yes | no | no | no | no |
| CHRNB2 | 263 | G | 0 | 0 | no | yes | no | yes | no | yes | no | no | yes | no | no | no | no |
| CHRNB2 | 264 | E | 1 | 0 | no | yes | no | yes | no | yes | yes | no | yes | no | no | no | no |
| CHRNB2 | 265 | K | 0 | 0 | yes | no | no | yes | no | yes | no | no | no | no | no | no | no |
| CHRNB2 | 266 | M | 0 | 0 | yes | no | no | yes | no | no | no | no | yes | no | no | no | no |
| CHRNB2 | 267 | T | 1 | 0 | yes | no | no | yes | no | no | yes | no | no | no | no | no | no |
| CHRNB2 | 268 | L | 0 | 0 | yes | no | no | yes | no | no | no | no | no | no | no | no | no |
| CHRNB2 | 269 | C | 0 | 0 | yes | no | no | yes | no | no | no | no | no | no | no | no | no |
| CHRNB2 | 270 | I | 0 | 0 | yes | no | no | yes | no | no | no | no | yes | no | no | no | no |
| CHRNB2 | 271 | S | 0 | 0 | yes | no | no | yes | no | no | yes | no | yes | no | no | no | no |
| CHRNB2 | 272 | V | 0 | 0 | yes | no | no | yes | no | no | no | no | no | no | no | no | no |
| CHRNB2 | 273 | L | 0 | 0 | yes | no | no | yes | no | no | no | no | no | no | no | no | no |
| CHRNB2 | 274 | L | 0 | 0 | yes | no | no | yes | no | no | yes | no | no | no | no | no | no |
| CHRNB2 | 275 | A | 3 | 0 | yes | no | no | yes | no | no | no | no | no | no | no | no | no |
| CHRNB2 | 276 | L | 0 | 0 | yes | no | no | yes | no | no | no | no | no | no | no | no | no |
| CHRNB2 | 277 | T | 0 | 0 | yes | no | no | yes | no | no | no | no | no | no | no | no | no |
| CHRNB2 | 278 | V | 0 | 0 | yes | no | no | yes | no | no | yes | no | no | no | no | no | no |
| CHRNB2 | 279 | F | 0 | 0 | yes | no | no | yes | no | no | no | no | yes | no | no | no | no |
| CHRNB2 | 280 | L | 0 | 0 | yes | no | no | yes | no | no | no | no | no | no | no | no | no |
| CHRNB2 | 281 | L | 0 | 0 | yes | no | no | yes | no | no | no | no | yes | no | no | no | no |
| CHRNB2 | 282 | L | 0 | 1 | yes | no | no | no | no | no | no | no | yes | no | no | no | no |
| CHRNB2 | 283 | I | 0 | 0 | yes | no | no | no | no | no | no | no | no | no | no | no | no |
| CHRNB2 | 284 | S | 0 | 0 | yes | no | no | no | no | no | no | no | yes | no | no | no | no |
| CHRNB2 | 285 | K | 0 | 0 | yes | no | no | no | yes | no | yes | no | no | no | no | no | no |
| CHRNB2 | 286 | I | 0 | 0 | yes | no | no | no | yes | no | no | no | no | no | no | no | no |
| CHRNB2 | 287 | V | 1 | 2 | no | yes | no | no | yes | no | no | no | no | no | no | no | no |
| CHRNB2 | 288 | P | 0 | 0 | no | yes | no | no | yes | no | no | no | no | no | no | no | no |

| | | | | | | | | | | | | | | | | | |
|---|---|---|---|---|---|---|---|---|---|---|---|---|---|---|---|---|---|
| CHRNB2 | 289 | P | 0 | 0 | no | yes | no | no | yes | no | no | no | yes | no | no | no | no |
| CHRNB2 | 290 | T | 1 | 0 | no | yes | no | no | yes | no | no | no | no | no | no | no | no |
| CHRNB2 | 291 | S | 0 | 0 | no | yes | no | no | yes | no | no | no | yes | no | no | no | no |
| CHRNB2 | 292 | L | 0 | 0 | no | yes | no | no | yes | no | no | no | no | no | no | no | no |
| CHRNB2 | 293 | D | 0 | 0 | no | yes | no | no | yes | no | no | no | no | no | no | no | no |
| CHRNB2 | 294 | V | 0 | 0 | no | yes | no | no | yes | no | no | no | yes | no | no | no | no |
| CHRNB2 | 295 | P | 1 | 0 | no | yes | no | no | yes | no | no | no | no | no | no | no | no |
| CHRNB2 | 296 | L | 0 | 0 | yes | no | no | yes | yes | no | no | no | no | no | no | no | no |
| CHRNB2 | 297 | V | 1 | 0 | yes | no | no | yes | yes | no | no | no | no | no | no | no | no |
| CHRNB2 | 298 | G | 1 | 0 | yes | no | no | yes | yes | no | no | no | no | no | no | no | no |
| CHRNB2 | 299 | K | 0 | 0 | yes | no | no | yes | yes | no | no | no | no | no | no | no | no |
| CHRNB2 | 300 | Y | 0 | 0 | yes | no | no | yes | no | no | no | no | no | no | no | no | no |
| CHRNB2 | 301 | L | 0 | 1 | yes | no | no | yes | no | no | no | no | no | no | no | no | no |
| CHRNB2 | 302 | M | 0 | 0 | yes | no | no | yes | no | no | no | no | yes | no | no | no | no |
| CHRNB2 | 303 | F | 0 | 0 | yes | no | no | yes | no | no | no | no | no | no | no | no | no |
| CHRNB2 | 304 | T | 1 | 0 | yes | no | no | yes | no | no | no | no | no | no | no | no | no |
| CHRNB2 | 305 | M | 1 | 0 | yes | no | no | yes | no | no | no | no | yes | no | no | no | no |
| CHRNB2 | 306 | V | 0 | 0 | yes | no | no | yes | no | no | no | no | no | no | no | no | no |
| CHRNB2 | 307 | L | 1 | 0 | yes | no | no | yes | no | no | no | no | no | no | no | no | no |
| CHRNB2 | 308 | V | 1 | 1 | yes | no | no | yes | no | no | no | no | no | no | no | no | no |
| CHRNB2 | 309 | T | 1 | 0 | yes | no | no | yes | no | no | no | no | yes | no | no | no | no |
| CHRNB2 | 310 | F | 0 | 0 | yes | no | no | yes | no | no | no | no | no | no | no | no | no |
| CHRNB2 | 311 | S | 0 | 0 | yes | no | no | yes | no | no | no | no | no | no | no | no | no |
| CHRNB2 | 312 | I | 0 | 1 | yes | no | no | yes | no | no | no | no | no | no | no | no | no |
| CHRNB2 | 313 | V | 0 | 0 | yes | no | no | yes | no | no | no | no | no | no | no | no | no |
| CHRNB2 | 314 | T | 0 | 0 | yes | no | no | yes | no | no | no | no | no | no | no | no | no |
| CHRNB2 | 315 | S | 1 | 0 | yes | no | no | yes | no | no | no | no | no | no | no | no | no |
| CHRNB2 | 316 | V | 1 | 0 | yes | no | no | yes | no | no | no | no | yes | no | no | no | no |
| CHRNB2 | 317 | C | 0 | 0 | yes | no | no | yes | no | no | no | no | yes | no | no | no | no |
| CHRNB2 | 318 | V | 0 | 0 | yes | no | no | yes | no | no | no | no | no | no | no | no | no |
| CHRNB2 | 319 | L | 1 | 0 | yes | no | no | yes | no | no | no | no | yes | no | no | no | no |
| CHRNB2 | 320 | N | 0 | 0 | yes | no | no | no | no | no | no | no | yes | no | no | no | no |
| CHRNB2 | 321 | V | 1 | 0 | yes | no | no | no | no | no | no | no | no | no | no | no | no |

| | | | | | | | | | | | | | | | | | |
|---|---|---|---|---|---|---|---|---|---|---|---|---|---|---|---|---|---|
| CHRNB2 | 322 | H | 0 | 0 | yes | no | no | no | no | yes | no | no | no | no | no | no | no |
| CHRNB2 | 323 | H | 2 | 0 | no | yes | no | no | no | yes | no | no | yes | no | no | no | no |
| CHRNB2 | 324 | R | 3 | 0 | no | yes | no | no | no | yes | no | no | yes | no | no | no | no |
| CHRNB2 | 325 | S | 0 | 0 | no | yes | no | no | no | yes | no | no | no | no | no | no | no |
| CHRNB2 | 326 | P | 0 | 0 | no | yes | no | no | no | yes | no | no | no | no | no | no | no |
| CHRNB2 | 327 | T | 1 | 0 | no | yes | no | no | no | yes | no | no | no | no | no | no | no |
| CHRNB2 | 328 | T | 1 | 0 | no | yes | no | no | no | yes | no | no | yes | no | no | no | no |
| CHRNB2 | 329 | H | 0 | 0 | no | yes | no | no | no | yes | no | no | yes | no | no | no | no |
| CHRNB2 | 330 | T | 0 | 0 | no | yes | no | no | no | yes | no | no | no | no | no | no | no |
| CHRNB2 | 331 | M | 1 | 0 | no | yes | no | no | no | yes | no | no | no | no | no | no | no |
| CHRNB2 | 332 | A | 0 | 0 | no | yes | no | no | no | yes | no | no | no | no | no | no | no |
| CHRNB2 | 333 | P | 0 | 0 | yes | no | no | no | no | yes | no | no | no | no | no | no | no |
| CHRNB2 | 334 | W | 0 | 0 | yes | no | no | no | no | yes | no | no | no | no | no | no | no |
| CHRNB2 | 335 | V | 1 | 0 | yes | no | no | no | no | yes | no | no | no | no | no | no | no |
| CHRNB2 | 336 | K | 1 | 0 | yes | no | no | no | no | yes | no | no | no | no | no | no | no |
| CHRNB2 | 337 | V | 2 | 1 | yes | no | no | no | no | yes | no | no | no | no | no | no | no |
| CHRNB2 | 338 | V | 1 | 0 | yes | no | no | no | no | yes | no | no | no | no | no | no | no |
| CHRNB2 | 339 | F | 0 | 0 | yes | no | no | no | no | yes | no | no | no | no | no | no | no |
| CHRNB2 | 340 | L | 1 | 0 | yes | no | no | no | no | yes | no | no | no | no | no | no | no |
| CHRNB2 | 341 | E | 0 | 0 | yes | no | no | no | no | yes | no | no | no | no | no | no | no |
| CHRNB2 | 342 | K | 0 | 0 | yes | no | no | no | no | yes | no | no | no | no | no | no | no |
| CHRNB2 | 343 | L | 0 | 0 | no | yes | no | no | no | yes | no | no | no | no | no | no | no |
| CHRNB2 | 344 | P | 0 | 0 | yes | no | no | no | no | yes | no | no | no | no | no | no | no |
| CHRNB2 | 345 | A | 1 | 0 | yes | no | no | no | no | yes | no | no | no | no | no | no | no |
| CHRNB2 | 346 | L | 0 | 0 | yes | no | no | no | no | yes | no | no | no | no | no | no | no |
| CHRNB2 | 347 | L | 1 | 0 | no | yes | no | no | no | yes | no | no | no | no | no | no | no |
| CHRNB2 | 348 | F | 0 | 0 | no | yes | no | no | no | yes | no | no | yes | no | no | no | no |
| CHRNB2 | 349 | M | 3 | 0 | no | yes | no | no | no | yes | no | no | yes | no | no | no | no |
| CHRNB2 | 350 | Q | 1 | 0 | no | yes | no | no | no | yes | no | no | yes | no | no | no | no |
| CHRNB2 | 351 | Q | 0 | 0 | no | yes | no | no | no | yes | no | no | no | no | no | no | no |
| CHRNB2 | 446 | S | 0 | 0 | no | yes | no | no | no | yes | no | no | no | no | no | no | no |
| CHRNB2 | 447 | V | 1 | 0 | yes | no | no | no | no | yes | no | no | no | no | no | no | no |
| CHRNB2 | 448 | S | 2 | 0 | yes | no | no | no | no | yes | no | no | no | no | no | no | no |

| | | | | | | | | | | | | | | | | | |
|---|---|---|---|---|---|---|---|---|---|---|---|---|---|---|---|---|---|
| CHRNB2 | 449 | E | 1 | 0 | yes | no | no | no | no | yes | no | no | yes | no | no | no | no |
| CHRNB2 | 450 | D | 0 | 0 | yes | no | no | no | no | yes | no | no | no | no | no | no | no |
| CHRNB2 | 451 | W | 1 | 0 | yes | no | no | no | no | yes | no | no | no | no | no | no | no |
| CHRNB2 | 452 | K | 0 | 0 | yes | no | no | no | no | yes | no | no | no | no | no | no | no |
| CHRNB2 | 453 | Y | 1 | 0 | yes | no | no | no | no | yes | no | no | no | no | no | no | no |
| CHRNB2 | 454 | V | 1 | 0 | yes | no | no | no | no | yes | no | no | no | no | no | no | no |
| CHRNB2 | 455 | A | 2 | 0 | yes | no | no | no | no | yes | no | no | no | no | no | no | no |
| CHRNB2 | 456 | M | 1 | 0 | yes | no | no | no | no | yes | no | no | yes | no | no | no | no |
| CHRNB2 | 457 | V | 0 | 0 | yes | no | no | no | no | yes | no | no | no | no | no | no | no |
| CHRNB2 | 458 | I | 1 | 0 | yes | no | no | no | no | yes | no | no | no | no | no | no | no |
| CHRNB2 | 459 | D | 0 | 0 | yes | no | no | yes | no | no | no | no | no | no | no | no | no |
| CHRNB2 | 460 | R | 3 | 0 | yes | no | no | yes | no | no | no | no | no | no | no | no | no |
| CHRNB2 | 461 | L | 1 | 0 | yes | no | no | yes | no | no | no | no | no | no | no | no | no |
| CHRNB2 | 462 | F | 0 | 0 | yes | no | no | yes | no | no | no | no | no | no | no | no | no |
| CHRNB2 | 463 | L | 0 | 0 | yes | no | no | yes | no | no | no | no | no | no | no | no | no |
| CHRNB2 | 464 | W | 0 | 0 | yes | no | no | yes | no | no | no | no | no | no | no | no | no |
| CHRNB2 | 465 | I | 1 | 0 | yes | no | no | yes | no | no | no | no | no | no | no | no | no |
| CHRNB2 | 466 | F | 0 | 0 | yes | no | no | yes | no | no | no | no | no | no | no | no | no |
| CHRNB2 | 467 | V | 1 | 0 | yes | no | no | yes | no | no | no | no | no | no | no | no | no |
| CHRNB2 | 468 | F | 0 | 0 | yes | no | no | yes | no | no | no | no | no | no | no | no | no |
| CHRNB2 | 469 | V | 0 | 0 | yes | no | no | yes | no | no | no | no | no | no | no | no | no |
| CHRNB2 | 470 | C | 0 | 0 | yes | no | no | yes | no | no | no | no | no | no | no | no | no |
| CHRNB2 | 471 | V | 1 | 0 | yes | no | no | yes | no | no | no | no | no | no | no | no | no |
| CHRNB2 | 472 | F | 1 | 0 | yes | no | no | yes | no | no | no | no | no | no | no | no | no |
| CHRNB2 | 473 | G | 0 | 0 | yes | no | no | yes | no | no | no | no | no | no | no | no | no |
| CHRNB2 | 474 | T | 0 | 0 | yes | no | no | yes | no | no | no | no | no | no | no | no | no |
| CHRNB2 | 475 | I | 2 | 0 | yes | no | no | yes | no | no | no | no | no | no | no | no | no |
| CHRNB2 | 476 | G | 1 | 0 | no | yes | no | yes | no | no | no | no | no | no | no | no | no |
| CHRNB2 | 477 | M | 1 | 0 | no | yes | no | yes | no | no | no | no | no | no | no | no | no |
| CHRNB2 | 478 | F | 1 | 0 | no | yes | no | yes | no | no | no | no | no | no | no | no | no |
| GABRA1 | 39 | T | 0 | 0 | no | yes | no | no | yes | no | no | no | yes | no | no | no | no |
| GABRA1 | 40 | T | 1 | 0 | no | yes | no | no | yes | no | no | no | no | no | no | no | no |
| GABRA1 | 41 | V | 0 | 0 | yes | no | no | no | yes | no | no | no | no | no | no | no | no |

| | | | | | | | | | | | | | | | | | |
|---|---|---|---|---|---|---|---|---|---|---|---|---|---|---|---|---|---|
| GABRA1 | 42 | F | 0 | 0 | yes | no | no | no | yes | no | no | no | yes | no | no | no | no |
| GABRA1 | 43 | T | 0 | 0 | yes | no | no | no | yes | no | no | no | yes | no | no | no | no |
| GABRA1 | 44 | R | 0 | 0 | yes | no | no | no | yes | no | no | no | no | no | no | no | no |
| GABRA1 | 45 | I | 0 | 0 | yes | no | no | no | yes | no | no | no | no | no | no | no | no |
| GABRA1 | 46 | L | 0 | 0 | yes | no | no | no | yes | no | no | no | yes | no | no | no | no |
| GABRA1 | 47 | D | 0 | 0 | yes | no | no | no | yes | no | no | no | yes | no | no | no | no |
| GABRA1 | 48 | R | 1 | 0 | no | yes | no | no | yes | no | no | no | no | no | no | no | no |
| GABRA1 | 49 | L | 0 | 0 | no | yes | no | no | yes | no | no | no | no | no | no | no | no |
| GABRA1 | 50 | L | 0 | 0 | no | yes | no | no | yes | no | no | no | no | no | no | no | no |
| GABRA1 | 51 | D | 1 | 0 | no | yes | no | no | yes | no | no | no | no | no | no | no | no |
| GABRA1 | 52 | G | 0 | 0 | no | yes | no | no | yes | no | no | no | no | no | no | no | no |
| GABRA1 | 53 | Y | 0 | 0 | no | yes | no | no | yes | no | no | no | no | no | no | no | no |
| GABRA1 | 54 | D | 0 | 0 | no | yes | no | no | yes | no | no | no | yes | no | no | no | no |
| GABRA1 | 55 | N | 0 | 0 | no | yes | no | no | yes | no | no | no | yes | no | no | no | no |
| GABRA1 | 56 | R | 0 | 0 | no | yes | no | no | yes | no | no | no | yes | no | no | no | no |
| GABRA1 | 57 | L | 0 | 0 | no | yes | no | no | yes | no | no | no | yes | no | no | no | no |
| GABRA1 | 58 | R | 0 | 0 | no | yes | no | no | yes | no | no | no | no | no | no | no | no |
| GABRA1 | 59 | P | 1 | 0 | no | yes | no | no | yes | no | no | no | no | no | no | no | no |
| GABRA1 | 60 | G | 0 | 0 | no | yes | no | no | yes | no | no | no | no | no | no | no | no |
| GABRA1 | 61 | L | 0 | 0 | no | yes | no | no | yes | no | no | no | yes | no | no | no | no |
| GABRA1 | 62 | G | 0 | 0 | no | yes | no | no | yes | no | no | no | no | no | no | no | no |
| GABRA1 | 63 | E | 1 | 0 | no | yes | no | no | yes | no | no | no | no | no | no | no | no |
| GABRA1 | 64 | R | 0 | 0 | no | yes | no | no | yes | no | no | no | no | no | no | no | no |
| GABRA1 | 65 | V | 0 | 0 | no | yes | no | no | yes | no | no | no | no | no | no | no | no |
| GABRA1 | 66 | T | 0 | 0 | no | no | yes | no | yes | no | no | no | no | no | no | no | no |
| GABRA1 | 67 | E | 1 | 0 | no | no | yes | no | yes | no | no | no | no | no | no | no | no |
| GABRA1 | 68 | V | 0 | 0 | no | no | yes | no | yes | no | no | no | no | no | no | no | no |
| GABRA1 | 69 | K | 1 | 0 | no | no | yes | no | yes | no | no | no | no | no | no | no | no |
| GABRA1 | 70 | T | 0 | 0 | no | no | yes | no | yes | no | no | no | no | no | no | no | no |
| GABRA1 | 71 | D | 0 | 0 | no | no | yes | no | yes | no | no | no | no | no | no | no | no |
| GABRA1 | 72 | I | 0 | 0 | no | no | yes | no | yes | no | no | no | no | no | no | no | no |
| GABRA1 | 73 | F | 0 | 0 | no | no | yes | no | yes | no | no | no | yes | no | no | no | no |
| GABRA1 | 74 | V | 0 | 1 | no | no | yes | no | yes | no | no | no | no | no | no | no | no |

| | | | | | | | | | | | | | | | | | |
|---|---|---|---|---|---|---|---|---|---|---|---|---|---|---|---|---|---|
| GABRA1 | 75 | T | 0 | 0 | no | no | yes | no | yes | no | no | no | no | no | no | no | no |
| GABRA1 | 76 | S | 1 | 1 | no | no | yes | no | yes | no | no | no | no | no | no | no | no |
| GABRA1 | 77 | F | 0 | 0 | no | yes | no | no | yes | no | no | no | no | no | no | no | no |
| GABRA1 | 78 | G | 0 | 0 | no | yes | no | no | yes | no | no | no | no | no | no | no | no |
| GABRA1 | 79 | P | 0 | 0 | no | yes | no | no | yes | no | no | no | no | no | no | no | no |
| GABRA1 | 80 | V | 0 | 0 | no | no | yes | no | yes | no | no | no | no | no | no | no | no |
| GABRA1 | 81 | S | 0 | 0 | no | no | yes | no | yes | no | no | no | no | no | no | no | no |
| GABRA1 | 82 | D | 0 | 0 | no | yes | no | no | yes | no | no | no | yes | no | no | no | no |
| GABRA1 | 83 | H | 0 | 0 | no | yes | no | no | yes | no | no | no | yes | no | no | no | no |
| GABRA1 | 84 | D | 1 | 0 | no | yes | no | no | yes | no | yes | no | yes | no | no | no | no |
| GABRA1 | 85 | M | 0 | 0 | no | yes | no | no | yes | no | no | no | yes | no | no | no | no |
| GABRA1 | 86 | E | 0 | 0 | no | no | yes | no | yes | no | no | no | no | no | no | no | no |
| GABRA1 | 87 | Y | 0 | 0 | no | no | yes | no | yes | no | no | no | no | no | no | no | no |
| GABRA1 | 88 | T | 0 | 0 | no | no | yes | no | yes | no | no | no | no | no | no | no | no |
| GABRA1 | 89 | I | 1 | 0 | no | no | yes | no | yes | no | no | no | no | no | no | no | no |
| GABRA1 | 90 | D | 0 | 1 | no | no | yes | no | yes | no | no | no | no | no | no | no | no |
| GABRA1 | 91 | V | 0 | 0 | no | no | yes | no | yes | no | no | no | no | no | no | no | no |
| GABRA1 | 92 | F | 0 | 0 | no | no | yes | no | yes | no | no | no | yes | no | no | no | no |
| GABRA1 | 93 | F | 0 | 0 | no | no | yes | no | yes | no | no | no | no | no | no | no | no |
| GABRA1 | 94 | R | 1 | 0 | no | no | yes | no | yes | no | no | no | yes | no | no | no | no |
| GABRA1 | 95 | Q | 0 | 0 | no | no | yes | no | yes | no | no | no | no | no | no | no | no |
| GABRA1 | 96 | S | 0 | 0 | no | no | yes | no | yes | no | no | no | no | no | no | no | no |
| GABRA1 | 97 | W | 0 | 0 | no | no | yes | no | yes | no | no | no | no | no | no | no | no |
| GABRA1 | 98 | K | 0 | 0 | no | no | yes | no | yes | no | no | no | no | no | no | no | no |
| GABRA1 | 99 | D | 0 | 0 | no | yes | no | no | yes | no | no | no | no | no | no | no | no |
| GABRA1 | 100 | E | 0 | 0 | yes | no | no | no | yes | no | no | no | no | no | no | no | no |
| GABRA1 | 101 | R | 0 | 0 | yes | no | no | no | yes | no | no | no | no | no | no | no | no |
| GABRA1 | 102 | L | 0 | 0 | yes | no | no | no | yes | no | no | no | no | no | no | no | no |
| GABRA1 | 103 | K | 0 | 0 | no | yes | no | no | yes | no | no | no | no | no | no | no | no |
| GABRA1 | 104 | F | 0 | 1 | no | yes | no | no | yes | no | no | no | no | no | no | no | no |
| GABRA1 | 105 | K | 0 | 0 | no | yes | no | no | yes | no | no | no | no | no | no | no | no |
| GABRA1 | 106 | G | 0 | 0 | no | yes | no | no | yes | no | no | no | no | no | no | no | no |
| GABRA1 | 107 | P | 0 | 0 | no | yes | no | no | yes | no | no | no | no | no | no | no | no |

| | | | | | | | | | | | | | | | | | |
|---|---|---|---|---|---|---|---|---|---|---|---|---|---|---|---|---|---|
| GABRA1 | 108 | M | 0 | 0 | no | yes | no | no | yes | no | no | no | no | no | no | no | no |
| GABRA1 | 109 | T | 0 | 0 | no | yes | no | no | yes | no | no | no | no | no | no | no | no |
| GABRA1 | 110 | V | 1 | 0 | no | no | yes | no | yes | no | no | no | no | no | no | no | no |
| GABRA1 | 111 | L | 0 | 0 | no | no | yes | no | yes | no | no | no | no | no | no | no | no |
| GABRA1 | 112 | R | 0 | 1 | no | no | yes | no | yes | no | no | no | yes | no | no | no | no |
| GABRA1 | 113 | L | 0 | 0 | no | no | yes | no | yes | no | no | no | no | no | no | no | no |
| GABRA1 | 114 | N | 0 | 0 | no | yes | no | no | yes | no | no | no | yes | no | no | no | no |
| GABRA1 | 115 | N | 0 | 1 | no | yes | no | no | yes | no | no | no | no | no | no | no | no |
| GABRA1 | 116 | L | 0 | 0 | no | yes | no | no | yes | no | yes | no | no | no | no | no | no |
| GABRA1 | 117 | M | 1 | 0 | yes | no | no | no | yes | no | no | no | yes | no | no | no | no |
| GABRA1 | 118 | A | 0 | 0 | yes | no | no | no | yes | no | no | no | no | no | no | no | no |
| GABRA1 | 119 | S | 0 | 0 | yes | no | no | no | yes | no | yes | no | no | no | no | no | no |
| GABRA1 | 120 | K | 0 | 0 | no | yes | no | no | yes | no | no | no | no | no | no | no | no |
| GABRA1 | 121 | I | 0 | 0 | no | yes | no | no | yes | no | no | no | no | no | no | no | no |
| GABRA1 | 122 | W | 0 | 0 | no | yes | no | no | yes | no | no | no | yes | no | no | no | no |
| GABRA1 | 123 | T | 0 | 0 | no | yes | no | no | yes | no | no | no | no | no | no | no | no |
| GABRA1 | 124 | P | 0 | 0 | no | yes | no | no | yes | no | no | no | yes | no | no | no | no |
| GABRA1 | 125 | D | 0 | 0 | no | yes | no | no | yes | no | no | no | yes | no | no | no | no |
| GABRA1 | 126 | T | 0 | 0 | no | no | yes | no | yes | no | no | no | yes | no | no | no | no |
| GABRA1 | 127 | F | 0 | 0 | no | no | yes | no | yes | no | no | no | yes | no | no | no | no |
| GABRA1 | 128 | F | 0 | 0 | no | no | yes | no | yes | no | no | no | yes | no | no | no | no |
| GABRA1 | 129 | H | 2 | 0 | no | yes | no | no | yes | no | no | no | yes | no | no | no | no |
| GABRA1 | 130 | N | 0 | 0 | no | yes | no | no | yes | no | no | no | no | no | no | no | no |
| GABRA1 | 131 | G | 0 | 0 | no | no | yes | no | yes | no | no | no | yes | no | no | no | no |
| GABRA1 | 132 | K | 0 | 0 | no | yes | no | no | yes | no | no | no | yes | no | no | no | no |
| GABRA1 | 133 | K | 0 | 0 | no | yes | no | no | yes | no | yes | no | no | no | no | no | no |
| GABRA1 | 134 | S | 0 | 0 | no | yes | no | no | yes | no | no | no | yes | no | no | no | no |
| GABRA1 | 135 | V | 0 | 0 | no | no | yes | no | yes | no | no | no | no | no | no | no | no |
| GABRA1 | 136 | A | 0 | 0 | no | no | yes | no | yes | no | no | no | yes | no | no | no | no |
| GABRA1 | 137 | H | 0 | 0 | no | yes | no | no | yes | no | no | no | yes | no | no | no | no |
| GABRA1 | 138 | N | 0 | 0 | no | yes | no | no | yes | no | yes | no | no | no | no | no | no |
| GABRA1 | 139 | M | 2 | 0 | no | yes | no | no | yes | no | no | no | yes | no | no | no | no |
| GABRA1 | 140 | T | 1 | 0 | no | yes | no | no | yes | no | no | no | yes | no | no | no | no |

| | | | | | | | | | | | | | | | | | |
|---|---|---|---|---|---|---|---|---|---|---|---|---|---|---|---|---|---|
| GABRA1 | 141 | M | 0 | 0 | no | yes | no | no | yes | no | no | no | yes | no | no | no | no |
| GABRA1 | 142 | P | 1 | 0 | no | yes | no | no | yes | no | no | no | no | no | no | no | no |
| GABRA1 | 143 | N | 0 | 0 | no | yes | no | no | yes | no | no | no | yes | no | no | no | no |
| GABRA1 | 144 | K | 0 | 0 | no | no | yes | no | yes | no | no | no | yes | no | no | no | no |
| GABRA1 | 145 | L | 0 | 0 | no | no | yes | no | yes | no | no | no | yes | no | no | no | no |
| GABRA1 | 146 | L | 0 | 1 | no | no | yes | no | yes | no | no | no | no | no | no | no | no |
| GABRA1 | 147 | R | 1 | 1 | no | no | yes | no | yes | no | no | no | yes | no | no | no | no |
| GABRA1 | 148 | I | 0 | 0 | no | no | yes | no | yes | no | no | no | no | no | no | no | no |
| GABRA1 | 149 | T | 0 | 0 | no | no | yes | no | yes | no | no | no | no | no | no | no | no |
| GABRA1 | 150 | E | 0 | 0 | no | yes | no | no | yes | no | no | no | no | no | no | no | no |
| GABRA1 | 151 | D | 0 | 0 | no | yes | no | no | yes | no | no | no | no | no | no | no | no |
| GABRA1 | 152 | G | 0 | 0 | no | yes | no | no | yes | no | no | no | no | no | no | no | no |
| GABRA1 | 153 | T | 1 | 0 | no | no | yes | no | yes | no | no | no | no | no | no | no | no |
| GABRA1 | 154 | L | 0 | 0 | no | no | yes | no | yes | no | no | no | no | no | no | no | no |
| GABRA1 | 155 | L | 0 | 0 | no | no | yes | no | yes | no | no | no | no | no | no | no | no |
| GABRA1 | 156 | Y | 0 | 0 | no | no | yes | no | yes | no | no | no | no | no | no | no | no |
| GABRA1 | 157 | T | 0 | 0 | no | no | yes | no | yes | no | no | no | yes | no | no | no | no |
| GABRA1 | 158 | M | 0 | 0 | no | no | yes | no | yes | no | no | no | yes | no | no | no | no |
| GABRA1 | 159 | R | 0 | 0 | no | no | yes | no | yes | no | no | no | yes | no | no | no | no |
| GABRA1 | 160 | L | 0 | 0 | no | no | yes | no | yes | no | no | no | yes | no | no | no | no |
| GABRA1 | 161 | T | 0 | 0 | no | no | yes | no | yes | no | no | no | no | no | no | no | no |
| GABRA1 | 162 | V | 0 | 0 | no | no | yes | no | yes | no | no | no | no | no | no | no | no |
| GABRA1 | 163 | R | 0 | 0 | no | no | yes | no | yes | no | no | no | no | no | no | no | no |
| GABRA1 | 164 | A | 0 | 0 | no | no | yes | no | yes | no | no | no | no | no | no | no | no |
| GABRA1 | 165 | E | 0 | 0 | no | no | yes | no | yes | no | no | no | yes | no | no | no | no |
| GABRA1 | 166 | C | 0 | 0 | no | yes | no | no | yes | no | no | no | no | no | no | no | no |
| GABRA1 | 167 | P | 0 | 0 | no | yes | no | no | yes | no | no | no | yes | no | no | no | no |
| GABRA1 | 168 | M | 0 | 0 | no | yes | no | no | yes | no | no | no | no | no | no | no | no |
| GABRA1 | 169 | H | 0 | 0 | no | yes | no | no | yes | no | no | no | yes | no | no | no | no |
| GABRA1 | 170 | L | 0 | 0 | no | yes | no | no | yes | no | no | no | no | no | no | no | no |
| GABRA1 | 171 | E | 0 | 0 | no | yes | no | no | yes | no | no | no | no | no | no | no | no |
| GABRA1 | 172 | D | 0 | 0 | no | yes | no | no | yes | no | no | no | no | no | no | no | no |
| GABRA1 | 173 | F | 0 | 0 | no | yes | no | no | yes | no | no | no | no | no | no | no | no |

| | | | | | | | | | | | | | | | | | |
|---|---|---|---|---|---|---|---|---|---|---|---|---|---|---|---|---|---|
| GABRA1 | 174 | P | 0 | 0 | no | yes | no | no | yes | no | no | no | no | no | no | no | no |
| GABRA1 | 175 | M | 0 | 0 | no | yes | no | no | yes | no | no | no | no | no | no | no | no |
| GABRA1 | 176 | D | 0 | 0 | no | yes | no | no | yes | no | no | no | no | no | no | no | no |
| GABRA1 | 177 | A | 1 | 0 | no | no | yes | no | yes | no | no | no | no | no | no | no | no |
| GABRA1 | 178 | H | 0 | 0 | no | no | yes | no | yes | no | no | no | no | no | no | no | no |
| GABRA1 | 179 | A | 0 | 0 | no | no | yes | no | yes | no | no | no | no | no | no | no | no |
| GABRA1 | 180 | C | 0 | 0 | no | no | yes | no | yes | no | no | no | no | no | no | no | no |
| GABRA1 | 181 | P | 0 | 1 | no | no | yes | no | yes | no | no | no | no | no | no | no | no |
| GABRA1 | 182 | L | 0 | 0 | no | no | yes | no | yes | no | no | no | no | no | no | no | no |
| GABRA1 | 183 | K | 0 | 0 | no | no | yes | no | yes | no | no | no | no | no | no | no | no |
| GABRA1 | 184 | F | 0 | 0 | no | no | yes | no | yes | no | no | no | no | no | no | no | no |
| GABRA1 | 185 | G | 0 | 0 | no | no | yes | no | yes | no | no | no | no | no | no | no | no |
| GABRA1 | 186 | S | 0 | 0 | no | no | yes | no | yes | no | no | no | no | no | no | no | no |
| GABRA1 | 187 | Y | 1 | 0 | no | yes | no | no | yes | no | no | no | yes | no | no | no | no |
| GABRA1 | 188 | A | 0 | 0 | no | yes | no | no | yes | no | no | no | yes | no | no | no | no |
| GABRA1 | 189 | Y | 0 | 0 | no | yes | no | no | yes | no | no | no | yes | no | no | no | no |
| GABRA1 | 190 | T | 0 | 0 | no | yes | no | no | yes | no | no | no | yes | no | no | no | no |
| GABRA1 | 191 | R | 0 | 0 | no | yes | no | no | yes | no | no | no | no | no | no | no | no |
| GABRA1 | 192 | A | 0 | 0 | no | yes | no | no | yes | no | no | no | no | no | no | no | no |
| GABRA1 | 193 | E | 0 | 0 | no | yes | no | no | yes | no | no | no | yes | no | no | no | no |
| GABRA1 | 194 | V | 0 | 0 | no | no | yes | no | yes | no | no | no | no | no | no | no | no |
| GABRA1 | 195 | V | 0 | 0 | no | no | yes | no | yes | no | no | no | no | no | no | no | no |
| GABRA1 | 196 | Y | 0 | 0 | no | no | yes | no | yes | no | no | no | no | no | no | no | no |
| GABRA1 | 197 | E | 1 | 0 | no | no | yes | no | yes | no | no | no | no | no | no | no | no |
| GABRA1 | 198 | W | 0 | 0 | no | no | yes | no | yes | no | no | no | no | no | no | no | no |
| GABRA1 | 199 | T | 0 | 0 | no | yes | no | no | yes | no | no | no | no | no | no | no | no |
| GABRA1 | 200 | R | 0 | 0 | no | yes | no | no | yes | no | no | no | no | no | no | no | no |
| GABRA1 | 201 | E | 2 | 0 | no | yes | no | no | yes | no | no | no | no | no | no | no | no |
| GABRA1 | 202 | P | 0 | 0 | yes | no | no | no | yes | no | no | no | no | no | no | no | no |
| GABRA1 | 203 | A | 0 | 0 | yes | no | no | no | yes | no | no | no | no | no | no | no | no |
| GABRA1 | 204 | R | 1 | 0 | yes | no | no | no | yes | no | no | no | no | no | no | no | no |
| GABRA1 | 205 | S | 0 | 0 | yes | no | no | no | yes | no | no | no | no | no | no | no | no |
| GABRA1 | 206 | V | 0 | 0 | no | yes | no | no | yes | no | no | no | no | no | no | no | no |

| | | | | | | | | | | | | | | | | | |
|---|---|---|---|---|---|---|---|---|---|---|---|---|---|---|---|---|---|
| GABRA1 | 207 | V | 0 | 0 | no | no | yes | no | yes | no | no | no | no | no | no | no | no |
| GABRA1 | 208 | V | 0 | 0 | no | no | yes | no | yes | no | no | no | no | no | no | no | no |
| GABRA1 | 209 | A | 0 | 0 | no | yes | no | no | yes | no | no | no | no | no | no | no | no |
| GABRA1 | 210 | E | 0 | 0 | no | yes | no | no | yes | no | no | no | no | no | no | no | no |
| GABRA1 | 211 | D | 1 | 0 | no | yes | no | no | yes | no | no | no | no | no | no | no | no |
| GABRA1 | 212 | G | 0 | 0 | no | yes | no | no | yes | no | no | no | no | no | no | no | no |
| GABRA1 | 213 | S | 0 | 0 | no | yes | no | no | yes | no | no | no | no | no | no | no | no |
| GABRA1 | 214 | R | 0 | 3 | no | yes | no | no | yes | no | no | no | yes | no | no | no | no |
| GABRA1 | 215 | L | 0 | 2 | no | yes | no | no | yes | no | no | no | no | no | no | no | no |
| GABRA1 | 216 | N | 0 | 0 | no | yes | no | no | yes | no | no | no | yes | no | no | no | no |
| GABRA1 | 217 | Q | 0 | 0 | no | yes | no | no | yes | no | no | no | yes | no | no | no | no |
| GABRA1 | 218 | Y | 0 | 0 | no | no | yes | no | yes | no | no | no | no | no | no | no | no |
| GABRA1 | 219 | D | 0 | 1 | no | no | yes | no | yes | no | no | no | no | no | no | no | no |
| GABRA1 | 220 | L | 0 | 0 | no | no | yes | no | yes | no | no | no | no | no | no | no | no |
| GABRA1 | 221 | L | 0 | 0 | no | no | yes | no | yes | no | no | no | no | no | no | no | no |
| GABRA1 | 222 | G | 1 | 0 | no | no | yes | no | yes | no | no | no | no | no | no | no | no |
| GABRA1 | 223 | Q | 0 | 0 | no | no | yes | no | yes | no | no | no | no | no | no | no | no |
| GABRA1 | 224 | T | 0 | 0 | no | no | yes | no | yes | no | no | no | no | no | no | no | no |
| GABRA1 | 225 | V | 0 | 0 | no | no | yes | no | yes | no | no | no | no | no | no | no | no |
| GABRA1 | 226 | D | 1 | 0 | no | no | yes | no | yes | no | no | no | no | no | no | no | no |
| GABRA1 | 227 | S | 0 | 0 | no | no | yes | no | yes | no | no | no | no | no | no | no | no |
| GABRA1 | 228 | G | 0 | 0 | no | no | yes | no | yes | no | no | no | no | no | no | no | no |
| GABRA1 | 229 | I | 1 | 0 | no | no | yes | no | yes | no | no | no | no | no | no | no | no |
| GABRA1 | 230 | V | 0 | 0 | no | no | yes | no | yes | no | no | no | no | no | no | no | no |
| GABRA1 | 231 | Q | 1 | 0 | no | no | yes | no | yes | no | no | no | no | no | no | no | no |
| GABRA1 | 232 | S | 0 | 0 | no | no | yes | no | yes | no | no | no | no | no | no | no | no |
| GABRA1 | 233 | S | 0 | 0 | no | yes | no | no | yes | no | no | no | no | no | no | no | no |
| GABRA1 | 234 | T | 1 | 0 | no | yes | no | no | yes | no | no | no | yes | no | no | no | no |
| GABRA1 | 235 | G | 0 | 0 | no | no | yes | no | yes | no | no | no | no | no | no | no | no |
| GABRA1 | 236 | E | 0 | 0 | no | no | yes | no | yes | no | no | no | no | no | no | no | no |
| GABRA1 | 237 | Y | 0 | 0 | no | no | yes | no | yes | no | no | no | yes | no | no | no | no |
| GABRA1 | 238 | V | 0 | 0 | no | no | yes | no | yes | no | no | no | no | no | no | no | no |
| GABRA1 | 239 | V | 0 | 0 | no | no | yes | no | yes | no | no | no | no | no | no | no | no |

| | | | | | | | | | | | | | | | | | |
|---|---|---|---|---|---|---|---|---|---|---|---|---|---|---|---|---|---|
| GABRA1 | 240 | M | 0 | 0 | no | no | yes | no | yes | no | no | no | no | no | no | no | no |
| GABRA1 | 241 | T | 1 | 0 | no | no | yes | no | yes | no | no | no | no | no | no | no | no |
| GABRA1 | 242 | T | 1 | 0 | no | no | yes | no | yes | no | no | no | no | no | no | no | no |
| GABRA1 | 243 | H | 0 | 0 | no | no | yes | no | yes | no | no | no | no | no | no | no | no |
| GABRA1 | 244 | F | 0 | 0 | no | no | yes | no | yes | no | no | no | no | no | no | no | no |
| GABRA1 | 245 | H | 0 | 0 | no | no | yes | no | yes | no | no | no | no | no | no | no | no |
| GABRA1 | 246 | L | 0 | 0 | no | no | yes | no | yes | no | no | no | no | no | no | no | no |
| GABRA1 | 247 | K | 0 | 0 | no | no | yes | no | yes | no | no | no | no | no | no | no | no |
| GABRA1 | 248 | R | 0 | 0 | no | no | yes | no | yes | no | no | no | no | no | no | no | no |
| GABRA1 | 249 | K | 1 | 0 | no | yes | no | no | yes | no | no | no | no | no | no | no | no |
| GABRA1 | 250 | I | 0 | 0 | no | yes | no | no | yes | no | no | no | no | no | no | no | no |
| GABRA1 | 251 | G | 0 | 2 | yes | no | no | no | yes | no | no | no | yes | no | no | no | no |
| GABRA1 | 252 | Y | 0 | 0 | yes | no | no | no | no | no | no | no | yes | no | no | no | no |
| GABRA1 | 253 | F | 0 | 0 | yes | no | no | no | no | no | no | no | yes | no | no | no | no |
| GABRA1 | 254 | V | 0 | 0 | yes | no | no | yes | no | no | no | no | no | no | no | no | no |
| GABRA1 | 255 | I | 0 | 0 | yes | no | no | yes | no | no | no | no | yes | no | no | no | no |
| GABRA1 | 256 | Q | 0 | 0 | yes | no | no | yes | no | no | no | no | yes | no | no | no | no |
| GABRA1 | 257 | T | 0 | 0 | yes | no | no | yes | no | no | no | no | yes | no | no | no | no |
| GABRA1 | 258 | Y | 1 | 0 | yes | no | no | yes | no | no | no | no | no | no | no | no | no |
| GABRA1 | 259 | L | 0 | 0 | yes | no | no | yes | no | no | no | no | yes | no | no | no | no |
| GABRA1 | 260 | P | 0 | 2 | yes | no | no | yes | no | no | no | no | no | no | no | no | no |
| GABRA1 | 261 | C | 0 | 0 | yes | no | no | yes | no | no | no | no | no | no | no | no | no |
| GABRA1 | 262 | I | 0 | 0 | yes | no | no | yes | no | no | no | no | no | no | no | no | no |
| GABRA1 | 263 | M | 0 | 3 | yes | no | no | yes | no | no | no | no | yes | no | no | no | no |
| GABRA1 | 264 | T | 0 | 0 | yes | no | no | yes | no | no | no | no | no | no | no | no | no |
| GABRA1 | 265 | V | 0 | 0 | yes | no | no | yes | no | no | no | no | no | no | no | no | no |
| GABRA1 | 266 | I | 0 | 0 | yes | no | no | yes | no | no | no | no | yes | no | no | no | no |
| GABRA1 | 267 | L | 0 | 1 | yes | no | no | yes | no | no | no | no | yes | no | no | no | no |
| GABRA1 | 268 | S | 0 | 0 | yes | no | no | yes | no | no | no | no | no | no | no | no | no |
| GABRA1 | 269 | Q | 0 | 0 | yes | no | no | yes | no | no | no | no | no | no | no | no | no |
| GABRA1 | 270 | V | 0 | 0 | yes | no | no | yes | no | no | no | no | yes | no | no | no | no |
| GABRA1 | 271 | S | 0 | 0 | yes | no | no | yes | no | no | no | no | no | no | no | no | no |
| GABRA1 | 272 | F | 0 | 0 | yes | no | no | yes | no | no | no | no | no | no | no | no | no |

| | | | | | | | | | | | | | | | | | |
|---|---|---|---|---|---|---|---|---|---|---|---|---|---|---|---|---|---|
| GABRA1 | 273 | W | 0 | 0 | yes | no | no | yes | no | no | no | no | yes | no | yes | no | no |
| GABRA1 | 274 | L | 0 | 0 | no | yes | no | yes | no | no | no | no | yes | no | no | no | no |
| GABRA1 | 275 | N | 0 | 1 | no | yes | no | no | no | no | no | no | yes | no | no | no | no |
| GABRA1 | 276 | R | 0 | 0 | no | yes | no | no | no | no | no | no | no | no | no | no | no |
| GABRA1 | 277 | E | 0 | 0 | no | yes | no | no | no | no | no | no | no | no | no | no | no |
| GABRA1 | 278 | S | 0 | 0 | no | yes | no | no | no | no | no | no | yes | no | no | no | no |
| GABRA1 | 279 | V | 1 | 0 | yes | no | no | no | no | no | no | no | yes | no | no | no | no |
| GABRA1 | 280 | P | 0 | 0 | yes | no | no | yes | no | no | yes | no | no | no | no | no | no |
| GABRA1 | 281 | A | 0 | 0 | yes | no | no | yes | no | no | no | no | yes | no | no | no | no |
| GABRA1 | 282 | R | 0 | 0 | yes | no | no | yes | no | no | no | no | no | no | no | no | no |
| GABRA1 | 283 | T | 0 | 0 | yes | no | no | yes | no | no | no | no | yes | no | no | no | no |
| GABRA1 | 284 | V | 0 | 0 | yes | no | no | yes | no | no | yes | no | yes | no | no | no | no |
| GABRA1 | 285 | F | 0 | 0 | yes | no | no | yes | no | no | no | no | yes | no | no | no | no |
| GABRA1 | 286 | G | 0 | 0 | yes | no | no | yes | no | no | no | no | no | no | no | no | no |
| GABRA1 | 287 | V | 0 | 2 | yes | no | no | yes | no | no | no | no | yes | no | no | no | no |
| GABRA1 | 288 | T | 0 | 0 | yes | no | no | yes | no | no | no | no | yes | no | no | no | no |
| GABRA1 | 289 | T | 0 | 1 | yes | no | no | yes | no | no | no | no | no | no | no | no | no |
| GABRA1 | 290 | V | 0 | 1 | yes | no | no | yes | no | no | no | no | yes | no | no | no | no |
| GABRA1 | 291 | L | 0 | 0 | yes | no | no | yes | no | no | yes | no | yes | no | no | no | no |
| GABRA1 | 292 | T | 0 | 1 | yes | no | no | yes | no | no | no | no | yes | no | no | no | no |
| GABRA1 | 293 | M | 1 | 0 | yes | no | no | yes | no | no | no | no | no | no | no | no | no |
| GABRA1 | 294 | T | 1 | 1 | yes | no | no | yes | no | no | no | no | yes | no | no | no | no |
| GABRA1 | 295 | T | 0 | 1 | yes | no | no | yes | no | no | yes | no | yes | no | no | no | no |
| GABRA1 | 296 | L | 0 | 1 | yes | no | no | yes | no | no | no | no | yes | no | no | no | no |
| GABRA1 | 297 | S | 0 | 1 | yes | no | no | yes | no | no | no | no | yes | no | no | no | no |
| GABRA1 | 298 | I | 0 | 0 | yes | no | no | yes | no | no | no | no | yes | no | no | no | no |
| GABRA1 | 299 | S | 0 | 1 | yes | no | no | yes | no | no | no | no | yes | no | no | no | no |
| GABRA1 | 300 | A | 0 | 0 | no | yes | no | yes | no | no | no | no | no | no | no | no | no |
| GABRA1 | 301 | R | 0 | 1 | no | yes | no | no | no | no | no | no | yes | no | no | no | no |
| GABRA1 | 302 | N | 0 | 0 | no | yes | no | no | no | no | yes | no | yes | no | no | no | no |
| GABRA1 | 303 | S | 0 | 0 | no | yes | no | no | no | no | no | no | yes | no | no | no | no |
| GABRA1 | 304 | L | 0 | 0 | no | yes | no | no | no | no | no | no | no | no | no | no | no |
| GABRA1 | 305 | P | 0 | 0 | no | yes | no | no | no | no | no | no | no | no | no | no | no |

| | | | | | | | | | | | | | | | | | |
|---|---|---|---|---|---|---|---|---|---|---|---|---|---|---|---|---|---|
| GABRA1 | 306 | K | 0 | 1 | no | yes | no | no | no | no | yes | no | yes | no | no | no | no |
| GABRA1 | 307 | V | 0 | 0 | no | yes | no | no | no | no | no | no | yes | no | no | no | no |
| GABRA1 | 308 | A | 0 | 1 | no | yes | no | no | no | no | no | no | yes | no | no | no | no |
| GABRA1 | 309 | Y | 0 | 0 | no | yes | no | no | no | no | no | no | yes | no | no | no | no |
| GABRA1 | 310 | A | 0 | 0 | no | yes | no | no | no | no | no | no | yes | no | no | no | no |
| GABRA1 | 311 | T | 0 | 0 | no | yes | no | no | no | no | no | no | no | no | no | no | no |
| GABRA1 | 312 | A | 0 | 1 | yes | no | no | no | no | no | no | no | no | no | no | no | no |
| GABRA1 | 313 | M | 0 | 0 | yes | no | no | yes | no | no | no | no | no | no | no | no | no |
| GABRA1 | 314 | D | 0 | 1 | yes | no | no | yes | no | no | no | no | yes | no | no | no | no |
| GABRA1 | 315 | W | 0 | 1 | yes | no | no | yes | no | no | no | no | no | no | no | no | no |
| GABRA1 | 316 | F | 0 | 0 | yes | no | no | yes | no | no | no | no | no | no | no | no | no |
| GABRA1 | 317 | I | 0 | 0 | yes | no | no | yes | no | no | no | no | no | no | no | no | no |
| GABRA1 | 318 | A | 0 | 0 | yes | no | no | yes | no | no | no | no | no | no | no | no | no |
| GABRA1 | 319 | V | 0 | 0 | yes | no | no | yes | no | no | no | no | no | no | no | no | no |
| GABRA1 | 320 | C | 0 | 0 | yes | no | no | yes | no | no | no | no | no | no | no | no | no |
| GABRA1 | 321 | Y | 0 | 0 | yes | no | no | yes | no | no | no | no | yes | no | no | no | no |
| GABRA1 | 322 | A | 0 | 1 | yes | no | no | yes | no | no | no | no | no | no | no | no | no |
| GABRA1 | 323 | F | 0 | 0 | yes | no | no | yes | no | no | no | no | no | no | no | no | no |
| GABRA1 | 324 | V | 0 | 0 | yes | no | no | yes | no | no | no | no | no | no | no | no | no |
| GABRA1 | 325 | F | 0 | 1 | yes | no | no | yes | no | no | no | no | yes | no | no | no | no |
| GABRA1 | 326 | S | 0 | 0 | yes | no | no | yes | no | no | no | no | no | no | no | no | no |
| GABRA1 | 327 | A | 0 | 0 | yes | no | no | yes | no | no | no | no | no | no | no | no | no |
| GABRA1 | 328 | L | 0 | 0 | yes | no | no | yes | no | no | no | no | yes | no | no | no | no |
| GABRA1 | 329 | I | 1 | 0 | yes | no | no | yes | no | no | no | no | yes | no | no | no | no |
| GABRA1 | 330 | E | 0 | 0 | yes | no | no | yes | no | no | no | no | no | no | no | no | no |
| GABRA1 | 331 | F | 0 | 0 | yes | no | no | yes | no | no | no | no | no | no | no | no | no |
| GABRA1 | 332 | A | 0 | 0 | yes | no | no | yes | no | no | no | no | yes | no | no | no | no |
| GABRA1 | 333 | T | 0 | 0 | yes | no | no | yes | no | no | no | no | no | no | no | no | no |
| GABRA1 | 334 | V | 0 | 0 | yes | no | no | no | no | no | no | no | no | no | no | no | no |
| GABRA1 | 335 | N | 0 | 0 | yes | no | no | no | no | yes | no | no | yes | no | no | no | no |
| GABRA1 | 336 | Y | 0 | 0 | yes | no | no | no | no | yes | no | no | no | no | no | no | no |
| GABRA1 | 337 | F | 0 | 0 | yes | no | no | no | no | yes | no | no | no | no | no | no | no |
| GABRA1 | 338 | T | 1 | 0 | no | yes | no | no | no | yes | no | no | no | no | no | no | no |

| | | | | | | | | | | | | | | | | | |
|---|---|---|---|---|---|---|---|---|---|---|---|---|---|---|---|---|---|
| GABRA1 | 339 | K | 0 | 0 | no | yes | no | no | no | yes | no | no | no | no | no | no | no |
| GABRA1 | 340 | R | 1 | 0 | no | yes | no | no | no | yes | no | no | no | no | no | no | no |
| GABRA1 | 341 | G | 0 | 0 | no | yes | no | no | no | yes | yes | no | no | no | no | no | no |
| GABRA1 | 342 | Y | 0 | 0 | no | yes | no | no | no | yes | yes | no | no | no | no | no | no |
| GABRA1 | 343 | A | 0 | 0 | no | yes | no | no | no | yes | yes | no | yes | no | no | no | no |
| GABRA1 | 344 | W | 0 | 0 | no | yes | no | no | no | yes | no | no | yes | no | no | no | no |
| GABRA1 | 345 | D | 0 | 0 | no | yes | no | no | no | yes | yes | no | yes | no | no | no | no |
| GABRA1 | 346 | G | 0 | 0 | no | yes | no | no | no | yes | no | no | no | no | no | no | no |
| GABRA1 | 347 | K | 0 | 0 | no | yes | no | no | no | yes | no | no | no | no | no | no | no |
| GABRA1 | 411 | K | 0 | 0 | no | yes | no | no | no | yes | yes | no | no | no | no | no | no |
| GABRA1 | 412 | T | 1 | 0 | no | yes | no | no | no | yes | no | no | no | no | no | no | no |
| GABRA1 | 413 | F | 0 | 0 | no | yes | no | no | no | yes | no | no | no | no | no | no | no |
| GABRA1 | 414 | N | 0 | 0 | no | yes | no | no | no | yes | no | no | no | no | no | no | no |
| GABRA1 | 415 | S | 0 | 0 | no | yes | no | no | no | yes | no | no | no | no | no | no | no |
| GABRA1 | 416 | V | 0 | 0 | no | yes | no | no | no | yes | no | no | no | no | no | no | no |
| GABRA1 | 417 | S | 0 | 0 | no | yes | no | no | no | yes | no | no | no | no | no | no | no |
| GABRA1 | 418 | K | 1 | 0 | yes | no | no | no | no | yes | no | no | no | no | no | no | no |
| GABRA1 | 419 | I | 0 | 0 | yes | no | no | yes | no | yes | no | no | no | no | no | no | no |
| GABRA1 | 420 | D | 0 | 0 | yes | no | no | yes | no | yes | no | no | no | no | no | no | no |
| GABRA1 | 421 | R | 0 | 0 | yes | no | no | no | no | yes | no | no | no | no | no | no | no |
| GABRA1 | 422 | L | 0 | 0 | yes | no | no | yes | no | no | no | no | no | no | no | no | no |
| GABRA1 | 423 | S | 0 | 0 | yes | no | no | yes | no | no | no | no | no | no | no | no | no |
| GABRA1 | 424 | R | 0 | 0 | yes | no | no | yes | no | no | no | no | yes | no | no | no | no |
| GABRA1 | 425 | I | 0 | 0 | yes | no | no | yes | no | no | no | no | no | no | no | no | no |
| GABRA1 | 426 | A | 0 | 0 | yes | no | no | yes | no | no | no | no | no | no | no | no | no |
| GABRA1 | 427 | F | 0 | 0 | yes | no | no | yes | no | no | no | no | no | no | no | no | no |
| GABRA1 | 428 | P | 1 | 0 | yes | no | no | yes | no | no | no | no | no | no | no | no | no |
| GABRA1 | 429 | L | 0 | 0 | yes | no | no | yes | no | no | no | no | no | no | no | no | no |
| GABRA1 | 430 | L | 0 | 0 | yes | no | no | yes | no | no | no | no | no | no | no | no | no |
| GABRA1 | 431 | F | 0 | 0 | yes | no | no | yes | no | no | no | no | no | no | no | no | no |
| GABRA1 | 432 | G | 0 | 0 | yes | no | no | yes | no | no | no | no | no | no | no | no | no |
| GABRA1 | 433 | I | 1 | 0 | yes | no | no | yes | no | no | no | no | no | no | no | no | no |
| GABRA1 | 434 | F | 0 | 0 | yes | no | no | yes | no | no | no | no | no | no | no | no | no |

| | | | | | | | | | | | | | | | | | |
|---|---|---|---|---|---|---|---|---|---|---|---|---|---|---|---|---|---|
| GABRA1 | 435 | N | 0 | 0 | yes | no | no | yes | no | no | no | no | no | no | no | no | no |
| GABRA1 | 436 | L | 0 | 0 | yes | no | no | yes | no | no | no | no | no | no | no | no | no |
| GABRA1 | 437 | V | 0 | 0 | yes | no | no | yes | no | no | no | no | no | no | no | no | no |
| GABRA1 | 438 | Y | 0 | 0 | yes | no | no | yes | no | no | no | no | no | no | no | no | no |
| GABRA1 | 439 | W | 0 | 0 | yes | no | no | yes | no | no | no | no | no | no | no | no | no |
| GABRA1 | 440 | A | 0 | 0 | yes | no | no | yes | no | no | no | no | no | no | no | no | no |
| GABRA1 | 441 | T | 0 | 0 | yes | no | no | yes | no | no | no | no | no | no | no | no | no |
| GABRA1 | 442 | Y | 1 | 0 | no | yes | no | no | no | no | no | no | no | no | no | no | no |
| GABRA1 | 443 | L | 0 | 0 | no | yes | no | no | no | no | no | no | no | no | no | no | no |
| GABRA1 | 444 | N | 0 | 0 | no | yes | no | no | no | no | no | no | no | no | no | no | no |
| GABRA1 | 445 | R | 0 | 0 | no | yes | no | no | no | no | no | no | no | no | no | no | no |
| GABRA5 | 45 | N | 0 | 0 | no | yes | no | no | yes | no | no | no | no | no | no | no | no |
| GABRA5 | 46 | I | 0 | 0 | yes | no | no | no | yes | no | no | no | yes | no | no | no | no |
| GABRA5 | 47 | T | 0 | 0 | yes | no | no | no | yes | no | no | no | no | no | no | no | no |
| GABRA5 | 48 | I | 1 | 0 | yes | no | no | no | yes | no | no | no | no | no | no | no | no |
| GABRA5 | 49 | F | 0 | 0 | yes | no | no | no | yes | no | no | no | yes | no | no | no | no |
| GABRA5 | 50 | T | 0 | 0 | yes | no | no | no | yes | no | no | no | no | no | no | no | no |
| GABRA5 | 51 | R | 0 | 0 | yes | no | no | no | yes | no | no | no | no | no | no | no | no |
| GABRA5 | 52 | I | 0 | 0 | yes | no | no | no | yes | no | no | no | no | no | no | no | no |
| GABRA5 | 53 | L | 0 | 0 | yes | no | no | no | yes | no | no | no | yes | no | no | no | no |
| GABRA5 | 54 | D | 1 | 0 | yes | no | no | no | yes | no | no | no | yes | no | no | no | no |
| GABRA5 | 55 | G | 0 | 0 | yes | no | no | no | yes | no | no | no | no | no | no | no | no |
| GABRA5 | 56 | L | 0 | 0 | yes | no | no | no | yes | no | no | no | no | no | no | no | no |
| GABRA5 | 57 | L | 0 | 0 | yes | no | no | no | yes | no | no | no | yes | no | no | no | no |
| GABRA5 | 58 | D | 1 | 0 | no | yes | no | no | yes | no | no | no | no | no | no | no | no |
| GABRA5 | 59 | G | 1 | 0 | no | yes | no | no | yes | no | no | no | yes | no | no | no | no |
| GABRA5 | 60 | Y | 0 | 0 | no | yes | no | no | yes | no | no | no | no | no | no | no | no |
| GABRA5 | 61 | D | 1 | 0 | no | yes | no | no | yes | no | no | no | no | no | no | no | no |
| GABRA5 | 62 | N | 0 | 0 | no | yes | no | no | yes | no | no | no | yes | no | no | no | no |
| GABRA5 | 63 | R | 0 | 0 | no | yes | no | no | yes | no | no | no | yes | no | no | no | no |
| GABRA5 | 64 | L | 0 | 0 | no | yes | no | no | yes | no | no | no | yes | no | no | no | no |
| GABRA5 | 65 | R | 1 | 0 | no | yes | no | no | yes | no | no | no | no | no | no | no | no |
| GABRA5 | 66 | P | 0 | 0 | no | yes | no | no | yes | no | no | no | no | no | no | no | no |

| | | | | | | | | | | | | | | | | | |
|---|---|---|---|---|---|---|---|---|---|---|---|---|---|---|---|---|---|
| GABRA5 | 67 | G | 1 | 0 | no | yes | no | no | yes | no | no | no | no | no | no | no | no |
| GABRA5 | 68 | L | 0 | 0 | no | yes | no | no | yes | no | no | no | no | no | no | no | no |
| GABRA5 | 69 | G | 0 | 0 | no | yes | no | no | yes | no | no | no | no | no | no | no | no |
| GABRA5 | 70 | E | 1 | 0 | no | yes | no | no | yes | no | no | no | no | no | no | no | no |
| GABRA5 | 71 | R | 2 | 0 | no | yes | no | no | yes | no | no | no | no | no | no | no | no |
| GABRA5 | 72 | I | 1 | 0 | no | yes | no | no | yes | no | no | no | no | no | no | no | no |
| GABRA5 | 73 | T | 0 | 0 | no | no | yes | no | yes | no | no | no | no | no | no | no | no |
| GABRA5 | 74 | Q | 1 | 0 | no | no | yes | no | yes | no | no | no | no | no | no | no | no |
| GABRA5 | 75 | V | 0 | 0 | no | no | yes | no | yes | no | no | no | no | no | no | no | no |
| GABRA5 | 76 | R | 0 | 0 | no | no | yes | no | yes | no | no | no | no | no | no | no | no |
| GABRA5 | 77 | T | 0 | 0 | no | no | yes | no | yes | no | no | no | no | no | no | no | no |
| GABRA5 | 78 | D | 0 | 0 | no | no | yes | no | yes | no | no | no | no | no | no | no | no |
| GABRA5 | 79 | I | 0 | 0 | no | no | yes | no | yes | no | no | no | no | no | no | no | no |
| GABRA5 | 80 | Y | 1 | 0 | no | no | yes | no | yes | no | no | no | no | no | no | no | no |
| GABRA5 | 81 | V | 0 | 0 | no | no | yes | no | yes | no | no | no | no | no | no | no | no |
| GABRA5 | 82 | T | 0 | 0 | no | yes | no | no | yes | no | no | no | no | no | no | no | no |
| GABRA5 | 83 | S | 0 | 0 | no | yes | no | no | yes | no | no | no | no | no | no | no | no |
| GABRA5 | 84 | F | 1 | 0 | no | yes | no | no | yes | no | no | no | no | no | no | no | no |
| GABRA5 | 85 | G | 1 | 0 | no | yes | no | no | yes | no | no | no | no | no | no | no | no |
| GABRA5 | 86 | P | 1 | 0 | no | yes | no | no | yes | no | no | no | no | no | no | no | no |
| GABRA5 | 87 | V | 0 | 0 | no | no | yes | no | yes | no | no | no | no | no | no | no | no |
| GABRA5 | 88 | S | 1 | 0 | no | no | yes | no | yes | no | no | no | no | no | no | no | no |
| GABRA5 | 89 | D | 0 | 0 | no | yes | no | no | yes | no | no | no | no | no | no | no | no |
| GABRA5 | 90 | T | 0 | 0 | no | yes | no | no | yes | no | no | no | no | no | no | no | no |
| GABRA5 | 91 | E | 0 | 0 | no | yes | no | no | yes | no | yes | no | no | no | no | no | no |
| GABRA5 | 92 | M | 1 | 0 | no | yes | no | no | yes | no | no | no | no | no | no | no | no |
| GABRA5 | 93 | E | 0 | 0 | no | no | yes | no | yes | no | no | no | no | no | no | no | no |
| GABRA5 | 94 | Y | 0 | 0 | no | no | yes | no | yes | no | no | no | no | no | no | no | no |
| GABRA5 | 95 | T | 0 | 0 | no | no | yes | no | yes | no | no | no | no | no | no | no | no |
| GABRA5 | 96 | I | 1 | 0 | no | no | yes | no | yes | no | no | no | no | no | no | no | no |
| GABRA5 | 97 | D | 0 | 0 | no | no | yes | no | yes | no | no | no | no | no | no | no | no |
| GABRA5 | 98 | V | 1 | 0 | no | no | yes | no | yes | no | no | no | no | no | no | no | no |
| GABRA5 | 99 | F | 0 | 0 | no | no | yes | no | yes | no | no | no | yes | no | no | no | no |

| | | | | | | | | | | | | | | | | | |
|---|---|---|---|---|---|---|---|---|---|---|---|---|---|---|---|---|---|
| GABRA5 | 100 | F | 0 | 0 | no | no | yes | no | yes | no | no | no | no | no | no | no | no |
| GABRA5 | 101 | R | 0 | 0 | no | no | yes | no | yes | no | no | no | yes | no | no | no | no |
| GABRA5 | 102 | Q | 0 | 0 | no | no | yes | no | yes | no | no | no | no | no | no | no | no |
| GABRA5 | 103 | S | 0 | 0 | no | no | yes | no | yes | no | no | no | no | no | no | no | no |
| GABRA5 | 104 | W | 0 | 0 | no | no | yes | no | yes | no | no | no | no | no | no | no | no |
| GABRA5 | 105 | K | 1 | 0 | no | no | yes | no | yes | no | no | no | no | no | no | no | no |
| GABRA5 | 106 | D | 0 | 0 | no | yes | no | no | yes | no | no | no | no | no | no | no | no |
| GABRA5 | 107 | E | 0 | 0 | no | yes | no | no | yes | no | no | no | no | no | no | no | no |
| GABRA5 | 108 | R | 0 | 0 | no | yes | no | no | yes | no | no | no | yes | no | no | no | no |
| GABRA5 | 109 | L | 0 | 0 | no | yes | no | no | yes | no | no | no | no | no | no | no | no |
| GABRA5 | 110 | R | 0 | 0 | no | yes | no | no | yes | no | no | no | no | no | no | no | no |
| GABRA5 | 111 | F | 1 | 0 | no | yes | no | no | yes | no | no | no | no | no | no | no | no |
| GABRA5 | 112 | K | 0 | 0 | no | yes | no | no | yes | no | no | no | no | no | no | no | no |
| GABRA5 | 113 | G | 1 | 0 | no | yes | no | no | yes | no | no | no | no | no | no | no | no |
| GABRA5 | 114 | P | 1 | 0 | no | yes | no | no | yes | no | no | no | no | no | no | no | no |
| GABRA5 | 115 | M | 0 | 0 | no | yes | no | no | yes | no | no | no | yes | no | no | no | no |
| GABRA5 | 116 | Q | 1 | 0 | no | yes | no | no | yes | no | no | no | no | no | no | no | no |
| GABRA5 | 117 | R | 1 | 0 | no | yes | no | no | yes | no | no | no | no | no | no | no | no |
| GABRA5 | 118 | L | 0 | 0 | no | no | yes | no | yes | no | no | no | yes | no | no | no | no |
| GABRA5 | 119 | P | 0 | 0 | no | no | yes | no | yes | no | no | no | yes | no | no | no | no |
| GABRA5 | 120 | L | 0 | 0 | no | no | yes | no | yes | no | no | no | no | no | no | no | no |
| GABRA5 | 121 | N | 1 | 0 | no | yes | no | no | yes | no | no | no | yes | no | no | no | no |
| GABRA5 | 122 | N | 0 | 0 | no | yes | no | no | yes | no | no | no | no | no | no | no | no |
| GABRA5 | 123 | L | 0 | 0 | no | yes | no | no | yes | no | no | no | no | no | no | no | no |
| GABRA5 | 124 | L | 0 | 0 | yes | no | no | no | yes | no | no | no | yes | no | no | no | no |
| GABRA5 | 125 | A | 0 | 0 | yes | no | no | no | yes | no | no | no | no | no | no | no | no |
| GABRA5 | 126 | S | 1 | 0 | yes | no | no | no | yes | no | yes | no | yes | no | no | no | no |
| GABRA5 | 127 | K | 0 | 0 | yes | no | no | no | yes | no | yes | no | no | no | no | no | no |
| GABRA5 | 128 | I | 1 | 0 | no | yes | no | no | yes | no | no | no | no | no | no | no | no |
| GABRA5 | 129 | W | 0 | 0 | no | yes | no | no | yes | no | no | no | yes | no | no | no | no |
| GABRA5 | 130 | T | 0 | 0 | no | yes | no | no | yes | no | no | no | no | no | no | no | no |
| GABRA5 | 131 | P | 0 | 0 | no | yes | no | no | yes | no | no | no | no | no | no | no | no |
| GABRA5 | 132 | D | 0 | 0 | no | yes | no | no | yes | no | no | no | yes | no | no | no | no |

| | | | | | | | | | | | | | | | | | |
|---|---|---|---|---|---|---|---|---|---|---|---|---|---|---|---|---|---|
| GABRA5 | 133 | T | 1 | 0 | no | yes | no | no | yes | no | no | no | yes | no | no | no | no |
| GABRA5 | 134 | F | 0 | 0 | no | yes | no | no | yes | no | no | no | yes | no | no | no | no |
| GABRA5 | 135 | F | 0 | 0 | no | yes | no | no | yes | no | no | no | yes | no | no | no | no |
| GABRA5 | 136 | H | 0 | 0 | no | yes | no | no | yes | no | no | no | no | no | no | no | no |
| GABRA5 | 137 | N | 0 | 0 | no | yes | no | no | yes | no | no | no | no | no | no | no | no |
| GABRA5 | 138 | G | 0 | 0 | no | no | yes | no | yes | no | no | no | yes | no | no | no | no |
| GABRA5 | 139 | K | 0 | 0 | no | no | yes | no | yes | no | no | no | yes | no | no | no | no |
| GABRA5 | 140 | K | 0 | 0 | no | no | yes | no | yes | no | yes | no | no | no | no | no | no |
| GABRA5 | 141 | S | 0 | 0 | no | no | yes | no | yes | no | no | no | yes | no | no | no | no |
| GABRA5 | 142 | I | 1 | 0 | no | yes | no | no | yes | no | no | no | no | no | no | no | no |
| GABRA5 | 143 | A | 0 | 0 | no | yes | no | no | yes | no | no | no | no | no | no | no | no |
| GABRA5 | 144 | H | 0 | 0 | no | yes | no | no | yes | no | no | no | yes | no | no | no | no |
| GABRA5 | 145 | N | 0 | 0 | no | yes | no | no | yes | no | yes | no | no | no | no | no | no |
| GABRA5 | 146 | M | 0 | 0 | no | yes | no | no | yes | no | no | no | yes | no | no | no | no |
| GABRA5 | 147 | T | 0 | 0 | no | yes | no | no | yes | no | no | no | yes | no | no | no | no |
| GABRA5 | 148 | T | 0 | 0 | no | yes | no | no | yes | no | no | no | yes | no | no | no | no |
| GABRA5 | 149 | P | 0 | 0 | no | yes | no | no | yes | no | no | no | no | no | no | no | no |
| GABRA5 | 150 | N | 0 | 0 | no | yes | no | no | yes | no | no | no | yes | no | no | no | no |
| GABRA5 | 151 | K | 0 | 0 | no | no | yes | no | yes | no | no | no | yes | no | no | no | no |
| GABRA5 | 152 | L | 0 | 0 | no | yes | no | no | yes | no | no | no | yes | no | no | no | no |
| GABRA5 | 153 | L | 0 | 0 | no | no | yes | no | yes | no | no | no | no | no | no | no | no |
| GABRA5 | 154 | R | 0 | 0 | no | no | yes | no | yes | no | no | no | yes | no | no | no | no |
| GABRA5 | 155 | L | 0 | 0 | no | no | yes | no | yes | no | no | no | no | no | no | no | no |
| GABRA5 | 156 | E | 1 | 0 | no | yes | no | no | yes | no | no | no | no | no | no | no | no |
| GABRA5 | 157 | D | 1 | 0 | no | yes | no | no | yes | no | no | no | no | no | no | no | no |
| GABRA5 | 158 | D | 0 | 0 | no | yes | no | no | yes | no | no | no | no | no | no | no | no |
| GABRA5 | 159 | G | 0 | 0 | no | yes | no | no | yes | no | no | no | no | no | no | no | no |
| GABRA5 | 160 | T | 1 | 0 | no | no | yes | no | yes | no | no | no | no | no | no | no | no |
| GABRA5 | 161 | L | 0 | 0 | no | no | yes | no | yes | no | no | no | no | no | no | no | no |
| GABRA5 | 162 | L | 0 | 0 | no | no | yes | no | yes | no | no | no | no | no | no | no | no |
| GABRA5 | 163 | Y | 0 | 0 | no | no | yes | no | yes | no | no | no | no | no | no | no | no |
| GABRA5 | 164 | T | 0 | 0 | no | no | yes | no | yes | no | no | no | yes | no | no | no | no |
| GABRA5 | 165 | M | 1 | 0 | no | no | yes | no | yes | no | no | no | yes | no | no | no | no |

| | | | | | | | | | | | | | | | | | |
|---|---|---|---|---|---|---|---|---|---|---|---|---|---|---|---|---|---|
| GABRA5 | 166 | R | 0 | 0 | no | no | yes | no | yes | no | no | no | yes | no | no | no | no |
| GABRA5 | 167 | L | 0 | 0 | no | no | yes | no | yes | no | no | no | no | no | no | no | no |
| GABRA5 | 168 | T | 0 | 0 | no | no | yes | no | yes | no | no | no | no | no | no | no | no |
| GABRA5 | 169 | I | 1 | 0 | no | no | yes | no | yes | no | no | no | no | no | no | no | no |
| GABRA5 | 170 | S | 0 | 0 | no | no | yes | no | yes | no | no | no | no | no | no | no | no |
| GABRA5 | 171 | A | 0 | 0 | no | no | yes | no | yes | no | no | no | no | no | no | no | no |
| GABRA5 | 172 | E | 0 | 0 | no | no | yes | no | yes | no | no | no | yes | no | no | no | no |
| GABRA5 | 173 | C | 0 | 0 | no | yes | no | no | yes | no | no | no | no | no | no | no | no |
| GABRA5 | 174 | P | 1 | 0 | no | yes | no | no | yes | no | no | no | no | no | no | no | no |
| GABRA5 | 175 | M | 0 | 0 | no | yes | no | no | yes | no | no | no | no | no | no | no | no |
| GABRA5 | 176 | Q | 0 | 0 | no | yes | no | no | yes | no | no | no | no | no | no | no | no |
| GABRA5 | 177 | L | 0 | 0 | no | yes | no | no | yes | no | no | no | no | no | no | no | no |
| GABRA5 | 178 | E | 1 | 0 | no | yes | no | no | yes | no | no | no | no | no | no | no | no |
| GABRA5 | 179 | D | 0 | 0 | no | yes | no | no | yes | no | no | no | no | no | no | no | no |
| GABRA5 | 180 | F | 0 | 0 | no | yes | no | no | yes | no | no | no | no | no | no | no | no |
| GABRA5 | 181 | P | 1 | 0 | no | yes | no | no | yes | no | no | no | no | no | no | no | no |
| GABRA5 | 182 | M | 0 | 0 | no | yes | no | no | yes | no | no | no | no | no | no | no | no |
| GABRA5 | 183 | D | 0 | 0 | no | yes | no | no | yes | no | no | no | no | no | no | no | no |
| GABRA5 | 184 | A | 1 | 0 | no | yes | no | no | yes | no | no | no | no | no | no | no | no |
| GABRA5 | 185 | H | 0 | 0 | no | yes | no | no | yes | no | no | no | no | no | no | no | no |
| GABRA5 | 186 | A | 0 | 0 | no | no | yes | no | yes | no | no | no | no | no | no | no | no |
| GABRA5 | 187 | C | 0 | 0 | no | yes | no | no | yes | no | no | no | no | no | no | no | no |
| GABRA5 | 188 | P | 0 | 0 | no | yes | no | no | yes | no | no | no | no | no | no | no | no |
| GABRA5 | 189 | L | 0 | 0 | no | yes | no | no | yes | no | no | no | no | no | no | no | no |
| GABRA5 | 190 | K | 0 | 0 | no | no | yes | no | yes | no | no | no | no | no | no | no | no |
| GABRA5 | 191 | F | 0 | 0 | no | no | yes | no | yes | no | no | no | no | no | no | no | no |
| GABRA5 | 192 | G | 1 | 0 | no | no | yes | no | yes | no | no | no | no | no | no | no | no |
| GABRA5 | 193 | S | 0 | 0 | no | yes | no | no | yes | no | no | no | no | no | no | no | no |
| GABRA5 | 194 | Y | 0 | 0 | no | yes | no | no | yes | no | no | no | yes | no | no | no | no |
| GABRA5 | 195 | A | 1 | 0 | no | yes | no | no | yes | no | no | no | yes | no | no | no | no |
| GABRA5 | 196 | Y | 0 | 0 | no | yes | no | no | yes | no | no | no | no | no | no | no | no |
| GABRA5 | 197 | P | 0 | 0 | no | yes | no | no | yes | no | no | no | no | no | no | no | no |
| GABRA5 | 198 | N | 0 | 0 | no | yes | no | no | yes | no | no | no | no | no | no | no | no |

| | | | | | | | | | | | | | | | | | |
|---|---|---|---|---|---|---|---|---|---|---|---|---|---|---|---|---|---|
| GABRA5 | 199 | S | 1 | 0 | no | yes | no | no | yes | no | no | no | no | no | no | no | no |
| GABRA5 | 200 | E | 0 | 0 | no | yes | no | no | yes | no | no | no | yes | no | no | no | no |
| GABRA5 | 201 | V | 0 | 0 | no | no | yes | no | yes | no | no | no | no | no | no | no | no |
| GABRA5 | 202 | V | 2 | 0 | no | no | yes | no | yes | no | no | no | no | no | no | no | no |
| GABRA5 | 203 | Y | 0 | 0 | no | no | yes | no | yes | no | no | no | no | no | no | no | no |
| GABRA5 | 204 | V | 2 | 0 | no | no | yes | no | yes | no | no | no | no | no | no | no | no |
| GABRA5 | 205 | W | 0 | 0 | no | no | yes | no | yes | no | no | no | no | no | no | no | no |
| GABRA5 | 206 | T | 1 | 0 | no | yes | no | no | yes | no | no | no | no | no | no | no | no |
| GABRA5 | 207 | N | 0 | 0 | no | yes | no | no | yes | no | no | no | no | no | no | no | no |
| GABRA5 | 208 | G | 1 | 0 | no | yes | no | no | yes | no | no | no | no | no | no | no | no |
| GABRA5 | 209 | S | 0 | 0 | no | yes | no | no | yes | no | no | no | no | no | no | no | no |
| GABRA5 | 210 | T | 1 | 0 | no | yes | no | no | yes | no | no | no | no | no | no | no | no |
| GABRA5 | 211 | K | 0 | 0 | no | yes | no | no | yes | no | no | no | no | no | no | no | no |
| GABRA5 | 212 | S | 0 | 0 | no | yes | no | no | yes | no | no | no | no | no | no | no | no |
| GABRA5 | 213 | V | 0 | 0 | no | no | yes | no | yes | no | no | no | no | no | no | no | no |
| GABRA5 | 214 | V | 1 | 0 | no | no | yes | no | yes | no | no | no | no | no | no | no | no |
| GABRA5 | 215 | V | 1 | 0 | no | no | yes | no | yes | no | no | no | no | no | no | no | no |
| GABRA5 | 216 | A | 0 | 0 | no | yes | no | no | yes | no | no | no | no | no | no | no | no |
| GABRA5 | 217 | E | 0 | 0 | no | yes | no | no | yes | no | no | no | no | no | no | no | no |
| GABRA5 | 218 | D | 1 | 0 | no | yes | no | no | yes | no | no | no | no | no | no | no | no |
| GABRA5 | 219 | G | 1 | 0 | no | yes | no | no | yes | no | no | no | no | no | no | no | no |
| GABRA5 | 220 | S | 0 | 0 | no | yes | no | no | yes | no | no | no | no | no | no | no | no |
| GABRA5 | 221 | R | 0 | 0 | no | yes | no | no | yes | no | no | no | yes | no | no | no | no |
| GABRA5 | 222 | L | 0 | 0 | no | yes | no | no | yes | no | no | no | no | no | no | no | no |
| GABRA5 | 223 | N | 0 | 0 | no | yes | no | no | yes | no | no | no | yes | no | no | no | no |
| GABRA5 | 224 | Q | 1 | 0 | no | yes | no | no | yes | no | no | no | no | no | no | no | no |
| GABRA5 | 225 | Y | 1 | 0 | no | no | yes | no | yes | no | no | no | no | no | no | no | no |
| GABRA5 | 226 | H | 0 | 0 | no | no | yes | no | yes | no | no | no | no | no | no | no | no |
| GABRA5 | 227 | L | 0 | 0 | no | no | yes | no | yes | no | no | no | no | no | no | no | no |
| GABRA5 | 228 | M | 0 | 0 | no | no | yes | no | yes | no | no | no | no | no | no | no | no |
| GABRA5 | 229 | G | 1 | 0 | no | no | yes | no | yes | no | no | no | no | no | no | no | no |
| GABRA5 | 230 | Q | 0 | 0 | no | no | yes | no | yes | no | no | no | no | no | no | no | no |
| GABRA5 | 231 | T | 1 | 0 | no | no | yes | no | yes | no | no | no | no | no | no | no | no |

| | | | | | | | | | | | | | | | | | |
|---|---|---|---|---|---|---|---|---|---|---|---|---|---|---|---|---|---|
| GABRA5 | 232 | V | 0 | 0 | no | no | yes | no | yes | no | no | no | no | no | no | no | no |
| GABRA5 | 233 | G | 2 | 0 | no | no | yes | no | yes | no | no | no | no | no | no | no | no |
| GABRA5 | 234 | T | 0 | 0 | no | no | yes | no | yes | no | no | no | no | no | no | no | no |
| GABRA5 | 235 | E | 0 | 0 | no | no | yes | no | yes | no | no | no | no | no | no | no | no |
| GABRA5 | 236 | N | 0 | 0 | no | yes | no | no | yes | no | no | no | no | no | no | no | no |
| GABRA5 | 237 | I | 1 | 0 | no | yes | no | no | yes | no | no | no | no | no | no | no | no |
| GABRA5 | 238 | S | 1 | 1 | no | no | yes | no | yes | no | no | no | no | no | no | no | no |
| GABRA5 | 239 | T | 0 | 0 | no | yes | no | no | yes | no | no | no | no | no | no | no | no |
| GABRA5 | 240 | S | 1 | 0 | no | yes | no | no | yes | no | no | no | yes | no | no | no | no |
| GABRA5 | 241 | T | 0 | 0 | no | yes | no | no | yes | no | no | no | yes | no | no | no | no |
| GABRA5 | 242 | G | 0 | 0 | no | yes | no | no | yes | no | no | no | no | no | no | no | no |
| GABRA5 | 243 | E | 0 | 0 | no | no | yes | no | yes | no | no | no | no | no | no | no | no |
| GABRA5 | 244 | Y | 0 | 0 | no | yes | no | no | yes | no | no | no | no | no | no | no | no |
| GABRA5 | 245 | T | 0 | 0 | no | yes | no | no | yes | no | no | no | no | no | no | no | no |
| GABRA5 | 246 | I | 1 | 0 | no | no | yes | no | yes | no | no | no | no | no | no | no | no |
| GABRA5 | 247 | M | 0 | 0 | no | no | yes | no | yes | no | no | no | no | no | no | no | no |
| GABRA5 | 248 | T | 1 | 0 | no | no | yes | no | yes | no | no | no | no | no | no | no | no |
| GABRA5 | 249 | A | 0 | 0 | no | no | yes | no | yes | no | no | no | no | no | no | no | no |
| GABRA5 | 250 | H | 0 | 0 | no | no | yes | no | yes | no | no | no | no | no | no | no | no |
| GABRA5 | 251 | F | 0 | 0 | no | no | yes | no | yes | no | no | no | no | no | no | no | no |
| GABRA5 | 252 | H | 1 | 0 | no | no | yes | no | yes | no | no | no | no | no | no | no | no |
| GABRA5 | 253 | L | 0 | 0 | no | no | yes | no | yes | no | no | no | no | no | no | no | no |
| GABRA5 | 254 | K | 0 | 0 | no | no | yes | no | yes | no | no | no | no | no | no | no | no |
| GABRA5 | 255 | R | 0 | 0 | no | no | yes | no | yes | no | no | no | no | no | no | no | no |
| GABRA5 | 256 | K | 0 | 0 | no | yes | no | no | yes | no | no | no | no | no | no | no | no |
| GABRA5 | 257 | I | 0 | 0 | no | yes | no | no | yes | no | no | no | no | no | no | no | no |
| GABRA5 | 258 | G | 0 | 0 | no | yes | no | no | yes | no | no | no | yes | no | no | no | no |
| GABRA5 | 259 | Y | 1 | 0 | no | yes | no | no | yes | no | no | no | yes | no | no | no | no |
| GABRA5 | 260 | F | 0 | 0 | yes | no | no | no | no | no | no | no | no | no | no | no | no |
| GABRA5 | 261 | V | 1 | 0 | yes | no | no | yes | no | no | no | no | no | no | no | no | no |
| GABRA5 | 262 | I | 0 | 0 | yes | no | no | yes | no | no | no | no | yes | no | no | no | no |
| GABRA5 | 263 | Q | 1 | 0 | yes | no | no | yes | no | no | no | no | yes | no | no | no | no |
| GABRA5 | 264 | T | 0 | 0 | yes | no | no | yes | no | no | no | no | no | no | no | no | no |

| | | | | | | | | | | | | | | | | | |
|---|---|---|---|---|---|---|---|---|---|---|---|---|---|---|---|---|---|
| GABRA5 | 265 | Y | 0 | 0 | yes | no | no | yes | no | no | no | no | no | no | no | no | no |
| GABRA5 | 266 | L | 0 | 0 | yes | no | no | yes | no | no | no | no | no | no | no | no | no |
| GABRA5 | 267 | P | 0 | 0 | yes | no | no | yes | no | no | no | no | no | no | no | no | no |
| GABRA5 | 268 | C | 0 | 0 | yes | no | no | yes | no | no | no | no | no | no | no | no | no |
| GABRA5 | 269 | I | 1 | 0 | yes | no | no | yes | no | no | no | no | no | no | no | no | no |
| GABRA5 | 270 | M | 0 | 0 | yes | no | no | yes | no | no | no | no | no | no | no | no | no |
| GABRA5 | 271 | T | 0 | 0 | yes | no | no | yes | no | no | no | no | no | no | no | no | no |
| GABRA5 | 272 | V | 0 | 0 | yes | no | no | yes | no | no | no | no | no | no | no | no | no |
| GABRA5 | 273 | I | 0 | 0 | yes | no | no | yes | no | no | no | no | no | no | no | no | no |
| GABRA5 | 274 | L | 0 | 0 | yes | no | no | yes | no | no | no | no | no | no | no | no | no |
| GABRA5 | 275 | S | 0 | 0 | yes | no | no | yes | no | no | no | no | no | no | no | no | no |
| GABRA5 | 276 | Q | 0 | 0 | yes | no | no | yes | no | no | no | no | no | no | no | no | no |
| GABRA5 | 277 | V | 1 | 0 | yes | no | no | yes | no | no | no | no | no | no | no | no | no |
| GABRA5 | 278 | S | 0 | 0 | yes | no | no | yes | no | no | no | no | no | no | no | no | no |
| GABRA5 | 279 | F | 0 | 0 | yes | no | no | yes | no | no | no | no | no | no | no | no | no |
| GABRA5 | 280 | W | 0 | 0 | yes | no | no | yes | no | no | no | no | no | no | no | no | no |
| GABRA5 | 281 | L | 0 | 0 | no | yes | no | yes | no | no | no | no | no | no | no | no | no |
| GABRA5 | 282 | N | 0 | 0 | no | yes | no | no | no | no | no | no | yes | no | no | no | no |
| GABRA5 | 283 | R | 0 | 0 | no | yes | no | no | no | no | no | no | no | no | no | no | no |
| GABRA5 | 284 | E | 0 | 0 | no | yes | no | no | no | no | no | no | no | no | no | no | no |
| GABRA5 | 285 | S | 0 | 0 | no | yes | no | no | no | no | no | no | yes | no | no | no | no |
| GABRA5 | 286 | V | 0 | 0 | no | yes | no | no | no | no | no | no | no | no | no | no | no |
| GABRA5 | 287 | P | 0 | 0 | no | yes | no | no | no | no | yes | no | no | no | no | no | no |
| GABRA5 | 288 | A | 0 | 0 | yes | no | no | yes | no | no | no | no | no | no | no | no | no |
| GABRA5 | 289 | R | 0 | 0 | yes | no | no | yes | no | no | no | no | no | no | no | no | no |
| GABRA5 | 290 | T | 0 | 0 | yes | no | no | yes | no | no | no | no | no | no | no | no | no |
| GABRA5 | 291 | V | 0 | 0 | yes | no | no | yes | no | no | yes | no | no | no | no | no | no |
| GABRA5 | 292 | F | 0 | 0 | yes | no | no | yes | no | no | no | no | no | no | no | no | no |
| GABRA5 | 293 | G | 0 | 0 | yes | no | no | yes | no | no | no | no | no | no | no | no | no |
| GABRA5 | 294 | V | 0 | 2 | yes | no | no | yes | no | no | no | no | no | no | no | no | no |
| GABRA5 | 295 | T | 0 | 0 | yes | no | no | yes | no | no | yes | no | yes | no | no | no | no |
| GABRA5 | 296 | T | 1 | 0 | yes | no | no | yes | no | no | no | no | no | no | no | no | no |
| GABRA5 | 297 | V | 0 | 0 | yes | no | no | yes | no | no | no | no | no | no | no | no | no |

| | | | | | | | | | | | | | | | | | |
|---|---|---|---|---|---|---|---|---|---|---|---|---|---|---|---|---|---|
| GABRA5 | 298 | L | 0 | 0 | yes | no | no | yes | no | no | yes | no | no | no | no | no | no |
| GABRA5 | 299 | T | 0 | 0 | yes | no | no | yes | no | no | no | no | yes | no | no | no | no |
| GABRA5 | 300 | M | 0 | 0 | yes | no | no | yes | no | no | no | no | no | no | no | no | no |
| GABRA5 | 301 | T | 0 | 2 | yes | no | no | yes | no | no | no | no | no | no | no | no | no |
| GABRA5 | 302 | T | 0 | 0 | yes | no | no | yes | no | no | yes | no | no | no | no | no | no |
| GABRA5 | 303 | L | 0 | 0 | yes | no | no | yes | no | no | no | no | no | no | no | no | no |
| GABRA5 | 304 | S | 0 | 0 | yes | no | no | yes | no | no | no | no | no | no | no | no | no |
| GABRA5 | 305 | I | 0 | 0 | yes | no | no | yes | no | no | no | no | yes | no | no | no | no |
| GABRA5 | 306 | S | 0 | 0 | yes | no | no | yes | no | no | no | no | no | no | no | no | no |
| GABRA5 | 307 | A | 1 | 0 | yes | no | no | yes | no | no | no | no | no | no | no | no | no |
| GABRA5 | 308 | R | 0 | 0 | no | yes | no | yes | no | no | no | no | yes | no | no | no | no |
| GABRA5 | 309 | N | 0 | 0 | no | yes | no | no | no | no | yes | no | yes | no | no | no | no |
| GABRA5 | 310 | S | 0 | 0 | no | yes | no | no | no | no | no | no | no | no | no | no | no |
| GABRA5 | 311 | L | 0 | 0 | no | yes | no | no | no | no | no | no | no | no | no | no | no |
| GABRA5 | 312 | P | 0 | 0 | no | yes | no | no | no | no | no | no | no | no | no | no | no |
| GABRA5 | 313 | K | 0 | 0 | no | yes | no | no | no | no | no | no | no | no | no | no | no |
| GABRA5 | 314 | V | 0 | 0 | no | yes | no | no | no | no | no | no | yes | no | no | no | no |
| GABRA5 | 315 | A | 0 | 0 | no | yes | no | no | no | no | no | no | no | no | no | no | no |
| GABRA5 | 316 | Y | 0 | 0 | no | yes | no | no | no | no | no | no | no | no | no | no | no |
| GABRA5 | 317 | A | 0 | 0 | no | yes | no | no | no | no | no | no | no | no | no | no | no |
| GABRA5 | 318 | T | 0 | 0 | no | yes | no | no | no | no | no | no | no | no | no | no | no |
| GABRA5 | 319 | A | 0 | 0 | no | yes | no | yes | no | no | no | no | no | no | no | no | no |
| GABRA5 | 320 | M | 0 | 0 | no | yes | no | yes | no | no | no | no | no | no | no | no | no |
| GABRA5 | 321 | D | 0 | 0 | yes | no | no | yes | no | no | no | no | no | no | no | no | no |
| GABRA5 | 322 | W | 0 | 0 | yes | no | no | yes | no | no | no | no | no | no | no | no | no |
| GABRA5 | 323 | F | 0 | 0 | yes | no | no | yes | no | no | no | no | no | no | no | no | no |
| GABRA5 | 324 | I | 0 | 0 | yes | no | no | yes | no | no | no | no | no | no | no | no | no |
| GABRA5 | 325 | A | 0 | 0 | yes | no | no | yes | no | no | no | no | no | no | no | no | no |
| GABRA5 | 326 | V | 1 | 0 | yes | no | no | yes | no | no | no | no | no | no | no | no | no |
| GABRA5 | 327 | C | 0 | 0 | yes | no | no | yes | no | no | no | no | no | no | no | no | no |
| GABRA5 | 328 | Y | 0 | 0 | yes | no | no | yes | no | no | no | no | no | no | no | no | no |
| GABRA5 | 329 | A | 0 | 0 | yes | no | no | yes | no | no | no | no | no | no | no | no | no |
| GABRA5 | 330 | F | 0 | 0 | yes | no | no | yes | no | no | no | no | no | no | no | no | no |

| | | | | | | | | | | | | | | | | | |
|---|---|---|---|---|---|---|---|---|---|---|---|---|---|---|---|---|---|
| GABRA5 | 331 | V | 0 | 0 | yes | no | no | yes | no | no | no | no | no | no | no | no | no |
| GABRA5 | 332 | F | 0 | 0 | yes | no | no | yes | no | no | no | no | yes | no | no | no | no |
| GABRA5 | 333 | S | 1 | 0 | yes | no | no | yes | no | no | no | no | no | no | no | no | no |
| GABRA5 | 334 | A | 0 | 0 | yes | no | no | yes | no | no | no | no | no | no | no | no | no |
| GABRA5 | 335 | L | 0 | 0 | yes | no | no | yes | no | no | no | no | no | no | no | no | no |
| GABRA5 | 336 | I | 1 | 0 | yes | no | no | yes | no | no | no | no | no | no | no | no | no |
| GABRA5 | 337 | E | 0 | 0 | yes | no | no | yes | no | no | no | no | no | no | no | no | no |
| GABRA5 | 338 | F | 0 | 0 | yes | no | no | yes | no | no | no | no | no | no | no | no | no |
| GABRA5 | 339 | A | 0 | 0 | yes | no | no | yes | no | no | no | no | no | no | no | no | no |
| GABRA5 | 340 | T | 0 | 0 | yes | no | no | yes | no | no | no | no | no | no | no | no | no |
| GABRA5 | 341 | V | 0 | 0 | yes | no | no | no | no | no | no | no | no | no | no | no | no |
| GABRA5 | 342 | N | 0 | 0 | yes | no | no | no | no | yes | no | no | yes | no | no | no | no |
| GABRA5 | 343 | Y | 0 | 0 | yes | no | no | no | no | yes | no | no | yes | no | no | no | no |
| GABRA5 | 344 | F | 0 | 0 | yes | no | no | no | no | yes | no | no | no | no | no | no | no |
| GABRA5 | 345 | T | 0 | 0 | no | yes | no | no | no | yes | no | no | no | no | no | no | no |
| GABRA5 | 346 | K | 0 | 0 | no | yes | no | no | no | yes | yes | no | yes | no | no | no | no |
| GABRA5 | 424 | K | 0 | 0 | yes | no | no | no | no | yes | no | no | no | no | no | no | no |
| GABRA5 | 425 | I | 0 | 0 | yes | no | no | no | no | yes | no | no | no | no | no | no | no |
| GABRA5 | 426 | D | 1 | 0 | yes | no | no | no | no | yes | no | no | no | no | no | no | no |
| GABRA5 | 427 | K | 0 | 0 | yes | no | no | no | no | yes | no | no | no | no | no | no | no |
| GABRA5 | 428 | M | 0 | 0 | yes | no | no | yes | no | no | no | no | no | no | no | no | no |
| GABRA5 | 429 | S | 1 | 0 | yes | no | no | yes | no | no | no | no | no | no | no | no | no |
| GABRA5 | 430 | R | 1 | 0 | no | yes | no | yes | no | no | no | no | no | no | no | no | no |
| GABRA5 | 431 | I | 1 | 0 | no | yes | no | yes | no | no | no | no | no | no | no | no | no |
| GABRA5 | 432 | V | 1 | 0 | no | yes | no | yes | no | no | no | no | no | no | no | no | no |
| GABRA5 | 433 | F | 0 | 0 | yes | no | no | yes | no | no | no | no | no | no | no | no | no |
| GABRA5 | 434 | P | 0 | 0 | yes | no | no | yes | no | no | no | no | no | no | no | no | no |
| GABRA5 | 435 | V | 0 | 0 | yes | no | no | yes | no | no | no | no | no | no | no | no | no |
| GABRA5 | 436 | L | 0 | 0 | yes | no | no | yes | no | no | no | no | no | no | no | no | no |
| GABRA5 | 437 | F | 0 | 0 | yes | no | no | yes | no | no | no | no | no | no | no | no | no |
| GABRA5 | 438 | G | 1 | 0 | yes | no | no | yes | no | no | no | no | no | no | no | no | no |
| GABRA5 | 439 | T | 1 | 0 | yes | no | no | yes | no | no | no | no | no | no | no | no | no |
| GABRA5 | 440 | F | 0 | 0 | yes | no | no | yes | no | no | no | no | no | no | no | no | no |

| | | | | | | | | | | | | | | | | | |
|---|---|---|---|---|---|---|---|---|---|---|---|---|---|---|---|---|---|
| GABRA5 | 441 | N | 0 | 0 | yes | no | no | yes | no | no | no | no | no | no | no | no | no |
| GABRA5 | 442 | L | 0 | 0 | yes | no | no | yes | no | no | no | no | no | no | no | no | no |
| GABRA5 | 443 | V | 0 | 0 | yes | no | no | yes | no | no | no | no | no | no | no | no | no |
| GABRA5 | 444 | Y | 0 | 0 | yes | no | no | yes | no | no | no | no | no | no | no | no | no |
| GABRA5 | 445 | W | 1 | 0 | yes | no | no | yes | no | no | no | no | no | no | no | no | no |
| GABRA5 | 446 | A | 0 | 0 | yes | no | no | yes | no | no | no | no | no | no | no | no | no |
| GABRA5 | 447 | T | 0 | 0 | yes | no | no | yes | no | no | no | no | no | no | no | no | no |
| GABRA5 | 448 | Y | 0 | 0 | yes | no | no | no | no | no | no | no | no | no | no | no | no |
| GABRA5 | 449 | L | 2 | 0 | no | yes | no | no | no | no | no | no | no | no | no | no | no |
| GABRA5 | 450 | N | 0 | 0 | no | yes | no | no | no | no | no | no | no | no | no | no | no |
| GABRA5 | 451 | R | 1 | 0 | no | yes | no | no | no | no | no | no | no | no | no | no | no |
| GABRB2 | 32 | N | 1 | 0 | no | yes | no | no | yes | no | no | no | no | no | no | no | no |
| GABRB2 | 33 | M | 0 | 0 | yes | no | no | no | yes | no | no | no | yes | no | no | no | no |
| GABRB2 | 34 | S | 0 | 0 | yes | no | no | no | yes | no | no | no | no | no | no | no | no |
| GABRB2 | 35 | L | 0 | 0 | yes | no | no | no | yes | no | no | no | no | no | no | no | no |
| GABRB2 | 36 | V | 0 | 0 | yes | no | no | no | yes | no | no | no | yes | no | no | no | no |
| GABRB2 | 37 | K | 0 | 0 | yes | no | no | no | yes | no | no | no | no | no | no | no | no |
| GABRB2 | 38 | E | 0 | 0 | yes | no | no | no | yes | no | no | no | no | no | no | no | no |
| GABRB2 | 39 | T | 0 | 0 | yes | no | no | no | yes | no | no | no | no | no | no | no | no |
| GABRB2 | 40 | V | 0 | 0 | yes | no | no | no | yes | no | no | no | no | no | no | no | no |
| GABRB2 | 41 | D | 0 | 0 | yes | no | no | no | yes | no | no | no | no | no | no | no | no |
| GABRB2 | 42 | R | 0 | 0 | yes | no | no | no | yes | no | no | no | no | no | no | no | no |
| GABRB2 | 43 | L | 0 | 0 | yes | no | no | no | yes | no | no | no | no | no | no | no | no |
| GABRB2 | 44 | L | 0 | 0 | no | yes | no | no | yes | no | no | no | no | no | no | no | no |
| GABRB2 | 45 | K | 0 | 0 | no | yes | no | no | yes | no | no | no | no | no | no | no | no |
| GABRB2 | 46 | G | 0 | 0 | no | yes | no | no | yes | no | no | no | no | no | no | no | no |
| GABRB2 | 47 | Y | 0 | 0 | no | yes | no | no | yes | no | no | no | no | no | no | no | no |
| GABRB2 | 48 | D | 0 | 0 | no | yes | no | no | yes | no | no | no | yes | no | no | no | no |
| GABRB2 | 49 | I | 0 | 0 | no | yes | no | no | yes | no | no | no | yes | no | no | no | no |
| GABRB2 | 50 | R | 1 | 0 | no | yes | no | no | yes | no | no | no | yes | no | no | no | no |
| GABRB2 | 51 | L | 0 | 0 | no | yes | no | no | yes | no | no | no | yes | no | no | no | no |
| GABRB2 | 52 | R | 0 | 0 | no | yes | no | no | yes | no | no | no | no | no | no | no | no |
| GABRB2 | 53 | P | 0 | 0 | no | yes | no | no | yes | no | no | no | no | no | no | no | no |

| | | | | | | | | | | | | | | | | | |
|---|---|---|---|---|---|---|---|---|---|---|---|---|---|---|---|---|---|
| GABRB2 | 54 | D | 0 | 0 | no | yes | no | no | yes | no | no | no | no | no | no | no | no |
| GABRB2 | 55 | F | 0 | 0 | no | yes | no | no | yes | no | no | no | yes | no | no | no | no |
| GABRB2 | 56 | G | 0 | 0 | no | yes | no | no | yes | no | no | no | no | no | no | no | no |
| GABRB2 | 57 | G | 0 | 0 | no | yes | no | no | yes | no | no | no | no | no | no | no | no |
| GABRB2 | 58 | P | 1 | 0 | no | yes | no | no | yes | no | no | no | no | no | no | no | no |
| GABRB2 | 59 | P | 0 | 0 | no | yes | no | no | yes | no | no | no | no | no | no | no | no |
| GABRB2 | 60 | V | 0 | 0 | no | no | yes | no | yes | no | no | no | no | no | no | no | no |
| GABRB2 | 61 | A | 0 | 0 | no | no | yes | no | yes | no | no | no | no | no | no | no | no |
| GABRB2 | 62 | V | 0 | 0 | no | no | yes | no | yes | no | no | no | no | no | no | no | no |
| GABRB2 | 63 | G | 0 | 0 | no | no | yes | no | yes | no | no | no | no | no | no | no | no |
| GABRB2 | 64 | M | 0 | 0 | no | no | yes | no | yes | no | no | no | no | no | no | no | no |
| GABRB2 | 65 | N | 1 | 0 | no | no | yes | no | yes | no | no | no | yes | no | no | no | no |
| GABRB2 | 66 | I | 0 | 0 | no | no | yes | no | yes | no | no | no | no | no | no | no | no |
| GABRB2 | 67 | D | 0 | 0 | no | no | yes | no | yes | no | no | no | no | no | no | no | no |
| GABRB2 | 68 | I | 0 | 0 | no | no | yes | no | yes | no | no | no | no | no | no | no | no |
| GABRB2 | 69 | A | 0 | 0 | no | no | yes | no | yes | no | no | no | no | no | no | no | no |
| GABRB2 | 70 | S | 0 | 0 | no | no | yes | no | yes | no | no | no | no | no | no | no | no |
| GABRB2 | 71 | I | 0 | 0 | no | no | yes | no | yes | no | no | no | no | no | no | no | no |
| GABRB2 | 72 | D | 1 | 0 | no | no | yes | no | yes | no | no | no | yes | no | no | no | no |
| GABRB2 | 73 | M | 0 | 0 | no | no | yes | no | yes | no | no | no | no | no | no | no | no |
| GABRB2 | 74 | V | 0 | 0 | no | no | yes | no | yes | no | no | no | no | no | no | no | no |
| GABRB2 | 75 | S | 0 | 0 | no | yes | no | no | yes | no | no | no | no | no | no | no | no |
| GABRB2 | 76 | E | 0 | 0 | no | yes | no | no | yes | no | no | no | no | no | no | no | no |
| GABRB2 | 77 | V | 0 | 0 | no | yes | no | no | yes | no | no | no | no | no | no | no | no |
| GABRB2 | 78 | N | 0 | 0 | no | yes | no | no | yes | no | yes | no | no | no | no | no | no |
| GABRB2 | 79 | M | 0 | 1 | no | yes | no | no | yes | no | no | no | yes | no | no | no | no |
| GABRB2 | 80 | D | 0 | 0 | no | no | yes | no | yes | no | no | no | no | no | no | no | no |
| GABRB2 | 81 | Y | 0 | 0 | no | no | yes | no | yes | no | no | no | no | no | no | no | no |
| GABRB2 | 82 | T | 0 | 0 | no | no | yes | no | yes | no | no | no | no | no | no | no | no |
| GABRB2 | 83 | L | 0 | 0 | no | no | yes | no | yes | no | no | no | no | no | no | no | no |
| GABRB2 | 84 | T | 0 | 0 | no | no | yes | no | yes | no | no | no | no | no | no | no | no |
| GABRB2 | 85 | M | 1 | 0 | no | no | yes | no | yes | no | no | no | no | no | no | no | no |
| GABRB2 | 86 | Y | 0 | 0 | no | no | yes | no | yes | no | no | no | yes | no | no | no | no |

| | | | | | | | | | | | | | | | | | |
|---|---|---|---|---|---|---|---|---|---|---|---|---|---|---|---|---|---|
| GABRB2 | 87 | F | 0 | 0 | no | no | yes | no | yes | no | no | no | no | no | no | no | no |
| GABRB2 | 88 | Q | 1 | 0 | no | no | yes | no | yes | no | no | no | yes | no | no | no | no |
| GABRB2 | 89 | Q | 0 | 0 | no | no | yes | no | yes | no | no | no | no | no | no | no | no |
| GABRB2 | 90 | A | 0 | 0 | no | no | yes | no | yes | no | no | no | no | no | no | no | no |
| GABRB2 | 91 | W | 0 | 0 | no | no | yes | no | yes | no | no | no | no | no | no | no | no |
| GABRB2 | 92 | R | 0 | 0 | no | no | yes | no | yes | no | no | no | no | no | no | no | no |
| GABRB2 | 93 | D | 0 | 0 | no | yes | no | no | yes | no | no | no | no | no | no | no | no |
| GABRB2 | 94 | K | 0 | 0 | no | yes | no | no | yes | no | no | no | no | no | no | no | no |
| GABRB2 | 95 | R | 0 | 0 | no | yes | no | no | yes | no | no | no | no | no | no | no | no |
| GABRB2 | 96 | L | 0 | 0 | no | yes | no | no | yes | no | no | no | no | no | no | no | no |
| GABRB2 | 97 | S | 1 | 0 | no | yes | no | no | yes | no | no | no | no | no | no | no | no |
| GABRB2 | 98 | Y | 0 | 0 | no | yes | no | no | yes | no | no | no | no | no | no | no | no |
| GABRB2 | 99 | N | 0 | 0 | no | yes | no | no | yes | no | no | no | no | no | no | no | no |
| GABRB2 | 100 | V | 0 | 0 | no | yes | no | no | yes | no | no | no | no | no | no | no | no |
| GABRB2 | 101 | I | 0 | 0 | no | yes | no | no | yes | no | no | no | no | no | no | no | no |
| GABRB2 | 102 | P | 0 | 0 | no | yes | no | no | yes | no | no | no | no | no | no | no | no |
| GABRB2 | 103 | L | 1 | 0 | no | yes | no | no | yes | no | no | no | yes | no | no | no | no |
| GABRB2 | 104 | N | 0 | 0 | no | yes | no | no | yes | no | no | no | yes | no | no | no | no |
| GABRB2 | 105 | L | 0 | 0 | no | yes | no | no | yes | no | no | no | no | no | no | no | no |
| GABRB2 | 106 | T | 0 | 0 | no | yes | no | no | yes | no | no | no | yes | no | no | no | no |
| GABRB2 | 107 | L | 0 | 0 | no | yes | no | no | yes | no | no | no | no | no | no | no | no |
| GABRB2 | 108 | D | 0 | 0 | no | yes | no | no | yes | no | no | no | yes | no | no | no | no |
| GABRB2 | 109 | N | 0 | 0 | no | yes | no | no | yes | no | no | no | no | no | no | no | no |
| GABRB2 | 110 | R | 0 | 0 | no | yes | no | no | yes | no | no | no | yes | no | no | no | no |
| GABRB2 | 111 | V | 0 | 0 | yes | no | no | no | yes | no | no | no | no | no | no | no | no |
| GABRB2 | 112 | A | 1 | 0 | yes | no | no | no | yes | no | no | no | no | no | no | no | no |
| GABRB2 | 113 | D | 0 | 0 | yes | no | no | no | yes | no | yes | no | no | no | no | no | no |
| GABRB2 | 114 | Q | 0 | 0 | no | yes | no | no | yes | no | no | no | no | no | no | no | no |
| GABRB2 | 115 | L | 0 | 0 | no | yes | no | no | yes | no | no | no | no | no | no | no | no |
| GABRB2 | 116 | W | 0 | 0 | no | yes | no | no | yes | no | no | no | no | no | no | no | no |
| GABRB2 | 117 | V | 0 | 0 | no | yes | no | no | yes | no | no | no | yes | no | no | no | no |
| GABRB2 | 118 | P | 0 | 0 | no | yes | no | no | yes | no | no | no | no | no | no | no | no |
| GABRB2 | 119 | D | 0 | 0 | no | yes | no | no | yes | no | no | no | yes | no | yes | no | no |

| | | | | | | | | | | | | | | | | | |
|---|---|---|---|---|---|---|---|---|---|---|---|---|---|---|---|---|---|
| GABRB2 | 120 | T | 0 | 0 | no | yes | no | no | yes | no | no | no | yes | no | yes | no | no |
| GABRB2 | 121 | Y | 0 | 0 | no | no | yes | no | yes | no | no | no | yes | no | yes | no | no |
| GABRB2 | 122 | F | 0 | 0 | no | no | yes | no | yes | no | no | no | yes | no | no | no | no |
| GABRB2 | 123 | L | 0 | 0 | no | yes | no | no | yes | no | no | no | no | no | no | no | no |
| GABRB2 | 124 | N | 0 | 0 | no | yes | no | no | yes | no | no | no | no | no | no | no | no |
| GABRB2 | 125 | D | 0 | 1 | no | no | yes | no | yes | no | no | no | yes | no | no | no | no |
| GABRB2 | 126 | K | 0 | 0 | no | no | yes | no | yes | no | no | no | no | no | no | no | no |
| GABRB2 | 127 | K | 0 | 0 | no | no | yes | no | yes | no | yes | no | no | no | no | no | no |
| GABRB2 | 128 | S | 0 | 0 | no | no | yes | no | yes | no | no | no | yes | no | no | no | no |
| GABRB2 | 129 | F | 0 | 0 | no | no | yes | no | yes | no | yes | no | no | no | no | no | no |
| GABRB2 | 130 | V | 0 | 0 | no | no | yes | no | yes | no | no | no | no | no | no | no | no |
| GABRB2 | 131 | H | 0 | 0 | no | yes | no | no | yes | no | no | no | yes | no | no | no | no |
| GABRB2 | 132 | G | 0 | 0 | no | yes | no | no | yes | no | no | no | no | no | no | no | no |
| GABRB2 | 133 | V | 0 | 0 | no | yes | no | no | yes | no | no | no | yes | no | no | no | no |
| GABRB2 | 134 | T | 0 | 0 | no | yes | no | no | yes | no | no | no | yes | no | no | no | no |
| GABRB2 | 135 | V | 0 | 0 | no | yes | no | no | yes | no | no | no | yes | no | no | no | no |
| GABRB2 | 136 | K | 0 | 0 | no | yes | no | no | yes | no | yes | no | no | no | no | no | no |
| GABRB2 | 137 | N | 0 | 0 | no | yes | no | no | yes | no | no | no | yes | no | no | no | no |
| GABRB2 | 138 | R | 2 | 0 | no | yes | no | no | yes | no | no | no | yes | no | no | no | no |
| GABRB2 | 139 | M | 0 | 0 | no | no | yes | no | yes | no | no | no | yes | no | no | no | no |
| GABRB2 | 140 | I | 0 | 0 | no | no | yes | no | yes | no | no | no | no | no | no | no | no |
| GABRB2 | 141 | R | 0 | 0 | no | no | yes | no | yes | no | no | no | yes | no | no | no | no |
| GABRB2 | 142 | L | 0 | 0 | no | yes | no | no | yes | no | no | no | no | no | no | no | no |
| GABRB2 | 143 | H | 0 | 0 | no | yes | no | no | yes | no | no | no | no | no | no | no | no |
| GABRB2 | 144 | P | 0 | 0 | no | yes | no | no | yes | no | no | no | no | no | no | no | no |
| GABRB2 | 145 | D | 0 | 0 | no | yes | no | no | yes | no | no | no | no | no | no | no | no |
| GABRB2 | 146 | G | 0 | 0 | no | yes | no | no | yes | no | no | no | no | no | no | no | no |
| GABRB2 | 147 | T | 1 | 0 | no | no | yes | no | yes | no | no | no | no | no | no | no | no |
| GABRB2 | 148 | V | 0 | 0 | no | no | yes | no | yes | no | no | no | no | no | no | no | no |
| GABRB2 | 149 | L | 0 | 0 | no | no | yes | no | yes | no | no | no | no | no | no | no | no |
| GABRB2 | 150 | Y | 1 | 0 | no | no | yes | no | yes | no | no | no | no | no | no | no | no |
| GABRB2 | 151 | G | 0 | 0 | no | no | yes | no | yes | no | no | no | yes | no | no | no | no |
| GABRB2 | 152 | L | 0 | 0 | no | no | yes | no | yes | no | no | no | yes | no | no | no | no |

| | | | | | | | | | | | | | | | | | |
|---|---|---|---|---|---|---|---|---|---|---|---|---|---|---|---|---|---|
| GABRB2 | 153 | R | 0 | 0 | no | no | yes | no | yes | no | no | no | yes | no | no | no | no |
| GABRB2 | 154 | I | 0 | 0 | no | no | yes | no | yes | no | no | no | yes | no | no | no | no |
| GABRB2 | 155 | T | 0 | 0 | no | no | yes | no | yes | no | no | no | no | no | no | no | no |
| GABRB2 | 156 | T | 0 | 0 | no | no | yes | no | yes | no | no | no | no | no | no | no | no |
| GABRB2 | 157 | T | 0 | 0 | no | no | yes | no | yes | no | no | no | no | no | no | no | no |
| GABRB2 | 158 | A | 0 | 0 | no | no | yes | no | yes | no | no | no | no | no | no | no | no |
| GABRB2 | 159 | A | 0 | 0 | no | no | yes | no | yes | no | no | no | no | no | no | no | no |
| GABRB2 | 160 | C | 0 | 0 | no | yes | no | no | yes | no | no | no | no | no | no | no | no |
| GABRB2 | 161 | M | 0 | 1 | no | yes | no | no | yes | no | no | no | yes | no | no | no | no |
| GABRB2 | 162 | M | 0 | 0 | no | yes | no | no | yes | no | no | no | no | no | no | no | no |
| GABRB2 | 163 | D | 0 | 0 | no | yes | no | no | yes | no | no | no | no | no | no | no | no |
| GABRB2 | 164 | L | 0 | 0 | no | yes | no | no | yes | no | no | no | no | no | no | no | no |
| GABRB2 | 165 | R | 0 | 0 | no | yes | no | no | yes | no | no | no | no | no | no | no | no |
| GABRB2 | 166 | R | 0 | 0 | no | yes | no | no | yes | no | no | no | no | no | no | no | no |
| GABRB2 | 167 | Y | 0 | 0 | no | yes | no | no | yes | no | no | no | no | no | no | no | no |
| GABRB2 | 168 | P | 0 | 0 | no | yes | no | no | yes | no | no | no | no | no | no | no | no |
| GABRB2 | 169 | L | 0 | 0 | no | yes | no | no | yes | no | no | no | no | no | no | no | no |
| GABRB2 | 170 | D | 0 | 0 | no | yes | no | no | yes | no | no | no | no | no | no | no | no |
| GABRB2 | 171 | E | 0 | 0 | no | no | yes | no | yes | no | no | no | no | no | no | no | no |
| GABRB2 | 172 | Q | 0 | 0 | no | no | yes | no | yes | no | no | no | no | no | no | no | no |
| GABRB2 | 173 | N | 0 | 0 | no | no | yes | no | yes | no | no | no | no | no | no | no | no |
| GABRB2 | 174 | C | 0 | 0 | no | no | yes | no | yes | no | no | no | no | no | no | no | no |
| GABRB2 | 175 | T | 0 | 0 | no | no | yes | no | yes | no | no | no | no | no | no | no | no |
| GABRB2 | 176 | L | 0 | 0 | no | no | yes | no | yes | no | no | no | no | no | no | no | no |
| GABRB2 | 177 | E | 0 | 0 | no | no | yes | no | yes | no | no | no | no | no | no | no | no |
| GABRB2 | 178 | I | 0 | 0 | no | no | yes | no | yes | no | no | no | no | no | no | no | no |
| GABRB2 | 179 | E | 0 | 0 | no | no | yes | no | yes | no | no | no | no | no | yes | no | no |
| GABRB2 | 180 | S | 0 | 0 | no | yes | no | no | yes | no | no | no | no | no | yes | no | no |
| GABRB2 | 181 | Y | 0 | 0 | no | yes | no | no | yes | no | no | no | yes | no | yes | no | no |
| GABRB2 | 182 | G | 0 | 0 | no | yes | no | no | yes | no | no | no | yes | no | no | no | no |
| GABRB2 | 183 | Y | 0 | 0 | no | yes | no | no | yes | no | no | no | yes | no | no | no | no |
| GABRB2 | 184 | T | 0 | 0 | no | yes | no | no | yes | no | no | no | yes | no | no | no | no |
| GABRB2 | 185 | T | 0 | 0 | no | yes | no | no | yes | no | no | no | no | no | no | no | no |

| | | | | | | | | | | | | | | | | | |
|---|---|---|---|---|---|---|---|---|---|---|---|---|---|---|---|---|---|
| GABRB2 | 186 | D | 0 | 0 | no | yes | no | no | yes | no | no | no | no | no | no | no | no |
| GABRB2 | 187 | D | 0 | 0 | no | yes | no | no | yes | no | no | no | no | no | no | no | no |
| GABRB2 | 188 | I | 0 | 0 | no | no | yes | no | yes | no | no | no | no | no | no | no | no |
| GABRB2 | 189 | E | 0 | 0 | no | no | yes | no | yes | no | no | no | no | no | no | no | no |
| GABRB2 | 190 | F | 0 | 0 | no | no | yes | no | yes | no | no | no | no | no | no | no | no |
| GABRB2 | 191 | Y | 1 | 0 | no | no | yes | no | yes | no | no | no | no | no | no | no | no |
| GABRB2 | 192 | W | 0 | 0 | no | no | yes | no | yes | no | no | no | no | no | no | no | no |
| GABRB2 | 193 | R | 1 | 0 | no | yes | no | no | yes | no | no | no | no | no | no | no | no |
| GABRB2 | 194 | G | 0 | 0 | no | yes | no | no | yes | no | no | no | no | no | no | no | no |
| GABRB2 | 195 | D | 1 | 0 | yes | no | no | no | yes | no | no | no | no | no | no | no | no |
| GABRB2 | 196 | D | 0 | 0 | yes | no | no | no | yes | no | no | no | no | no | no | no | no |
| GABRB2 | 197 | N | 0 | 0 | yes | no | no | no | yes | no | no | no | no | no | no | no | no |
| GABRB2 | 198 | A | 2 | 0 | no | yes | no | no | yes | no | no | no | no | no | no | no | no |
| GABRB2 | 199 | V | 0 | 0 | no | no | yes | no | yes | no | no | no | no | no | no | no | no |
| GABRB2 | 200 | T | 0 | 0 | no | no | yes | no | yes | no | no | no | no | no | no | no | no |
| GABRB2 | 201 | G | 0 | 0 | no | yes | no | no | yes | no | no | no | no | no | no | no | no |
| GABRB2 | 202 | V | 0 | 0 | no | yes | no | no | yes | no | no | no | no | no | no | no | no |
| GABRB2 | 203 | T | 1 | 0 | no | yes | no | no | yes | no | no | no | no | no | no | no | no |
| GABRB2 | 204 | K | 0 | 0 | no | yes | no | no | yes | no | no | no | no | no | no | no | no |
| GABRB2 | 205 | I | 1 | 0 | no | yes | no | no | yes | no | no | no | no | no | no | no | no |
| GABRB2 | 206 | E | 0 | 0 | no | yes | no | no | yes | no | no | no | no | no | no | no | no |
| GABRB2 | 207 | L | 0 | 0 | no | yes | no | no | yes | no | no | no | no | no | no | no | no |
| GABRB2 | 208 | P | 0 | 0 | no | yes | no | no | yes | no | no | no | no | no | no | no | no |
| GABRB2 | 209 | Q | 0 | 1 | no | yes | no | no | yes | no | no | no | no | no | no | no | no |
| GABRB2 | 210 | F | 1 | 0 | no | no | yes | no | yes | no | no | no | no | no | no | no | no |
| GABRB2 | 211 | S | 0 | 0 | no | no | yes | no | yes | no | no | no | no | no | no | no | no |
| GABRB2 | 212 | I | 0 | 0 | no | no | yes | no | yes | no | no | no | no | no | no | no | no |
| GABRB2 | 213 | V | 0 | 0 | no | no | yes | no | yes | no | no | no | no | no | no | no | no |
| GABRB2 | 214 | D | 0 | 0 | no | no | yes | no | yes | no | no | no | no | no | no | no | no |
| GABRB2 | 215 | Y | 0 | 0 | no | no | yes | no | yes | no | no | no | no | no | no | no | no |
| GABRB2 | 216 | K | 0 | 0 | no | no | yes | no | yes | no | no | no | no | no | no | no | no |
| GABRB2 | 217 | L | 0 | 0 | no | no | yes | no | yes | no | no | no | no | no | no | no | no |
| GABRB2 | 218 | I | 0 | 0 | no | no | yes | no | yes | no | no | no | no | no | no | no | no |

| | | | | | | | | | | | | | | | | | |
|---|---|---|---|---|---|---|---|---|---|---|---|---|---|---|---|---|---|
| GABRB2 | 219 | T | 0 | 0 | no | no | yes | no | yes | no | no | no | no | no | no | no | no |
| GABRB2 | 220 | K | 0 | 0 | no | no | yes | no | yes | no | no | no | no | no | no | no | no |
| GABRB2 | 221 | K | 1 | 1 | no | no | yes | no | yes | no | no | no | no | no | no | no | no |
| GABRB2 | 222 | V | 0 | 0 | no | no | yes | no | yes | no | no | no | no | no | no | no | no |
| GABRB2 | 223 | V | 0 | 0 | no | no | yes | no | yes | no | no | no | no | no | no | no | no |
| GABRB2 | 224 | F | 0 | 0 | no | yes | no | no | yes | no | no | no | yes | no | yes | no | no |
| GABRB2 | 225 | S | 0 | 0 | no | yes | no | no | yes | no | no | no | yes | no | no | no | no |
| GABRB2 | 226 | T | 0 | 0 | no | yes | no | no | yes | no | no | no | no | no | no | no | no |
| GABRB2 | 227 | G | 0 | 0 | no | yes | no | no | yes | no | no | no | no | no | no | no | no |
| GABRB2 | 228 | S | 1 | 0 | no | no | yes | no | yes | no | no | no | no | no | no | no | no |
| GABRB2 | 229 | Y | 0 | 0 | no | no | yes | no | yes | no | no | no | yes | no | no | no | no |
| GABRB2 | 230 | P | 0 | 0 | no | no | yes | no | yes | no | no | no | no | no | no | no | no |
| GABRB2 | 231 | R | 0 | 0 | no | no | yes | no | yes | no | no | no | no | no | no | no | no |
| GABRB2 | 232 | L | 0 | 0 | no | no | yes | no | yes | no | no | no | no | no | no | no | no |
| GABRB2 | 233 | S | 0 | 0 | no | no | yes | no | yes | no | no | no | no | no | no | no | no |
| GABRB2 | 234 | L | 0 | 0 | no | no | yes | no | yes | no | no | no | no | no | no | no | no |
| GABRB2 | 235 | S | 0 | 0 | no | no | yes | no | yes | no | no | no | no | no | no | no | no |
| GABRB2 | 236 | F | 0 | 0 | no | no | yes | no | yes | no | no | no | no | no | no | no | no |
| GABRB2 | 237 | K | 0 | 0 | no | no | yes | no | yes | no | no | no | no | no | no | no | no |
| GABRB2 | 238 | L | 0 | 0 | no | no | yes | no | yes | no | no | no | no | no | no | no | no |
| GABRB2 | 239 | K | 0 | 0 | no | no | yes | no | yes | no | no | no | no | no | no | no | no |
| GABRB2 | 240 | R | 0 | 1 | no | no | yes | no | yes | no | no | no | no | no | no | no | no |
| GABRB2 | 241 | N | 0 | 0 | no | yes | no | no | yes | no | no | no | no | no | no | no | no |
| GABRB2 | 242 | I | 0 | 0 | no | yes | no | no | yes | no | no | no | no | no | no | no | no |
| GABRB2 | 243 | G | 0 | 0 | yes | no | no | no | yes | no | no | no | no | no | no | no | no |
| GABRB2 | 244 | Y | 0 | 1 | yes | no | no | no | yes | no | no | no | yes | no | no | no | no |
| GABRB2 | 245 | F | 0 | 1 | yes | no | no | no | no | no | no | no | yes | no | no | no | no |
| GABRB2 | 246 | I | 0 | 1 | yes | no | no | yes | no | no | no | no | no | no | no | no | no |
| GABRB2 | 247 | L | 0 | 0 | yes | no | no | yes | no | no | no | no | no | no | no | no | no |
| GABRB2 | 248 | Q | 0 | 0 | yes | no | no | yes | no | no | no | no | yes | no | no | no | no |
| GABRB2 | 249 | T | 0 | 0 | yes | no | no | yes | no | no | no | no | no | no | no | no | no |
| GABRB2 | 250 | Y | 0 | 0 | yes | no | no | yes | no | no | no | no | no | no | no | no | no |
| GABRB2 | 251 | M | 0 | 0 | yes | no | no | yes | no | no | no | no | no | no | no | no | no |

| | | | | | | | | | | | | | | | | | |
|---|---|---|---|---|---|---|---|---|---|---|---|---|---|---|---|---|---|
| GABRB2 | 252 | P | 0 | 1 | yes | no | no | yes | no | no | no | no | no | no | no | no | no |
| GABRB2 | 253 | S | 0 | 0 | yes | no | no | yes | no | no | no | no | no | no | no | no | no |
| GABRB2 | 254 | I | 0 | 0 | yes | no | no | yes | no | no | no | no | no | no | no | no | no |
| GABRB2 | 255 | L | 0 | 0 | yes | no | no | yes | no | no | no | no | no | no | no | no | no |
| GABRB2 | 256 | I | 0 | 0 | yes | no | no | yes | no | no | no | no | no | no | no | no | no |
| GABRB2 | 257 | T | 0 | 0 | yes | no | no | yes | no | no | no | no | no | no | no | no | no |
| GABRB2 | 258 | I | 0 | 0 | yes | no | no | yes | no | no | no | no | no | no | no | no | no |
| GABRB2 | 259 | L | 0 | 0 | yes | no | no | yes | no | no | no | no | no | no | no | no | no |
| GABRB2 | 260 | S | 0 | 0 | no | yes | no | yes | no | no | no | no | no | no | no | no | no |
| GABRB2 | 261 | W | 0 | 0 | no | yes | no | yes | no | no | no | no | no | no | no | no | no |
| GABRB2 | 262 | V | 0 | 0 | yes | no | no | yes | no | no | no | no | no | no | no | no | no |
| GABRB2 | 263 | S | 0 | 0 | yes | no | no | yes | no | no | no | no | no | no | no | no | no |
| GABRB2 | 264 | F | 0 | 0 | yes | no | no | yes | no | no | no | no | no | no | no | no | no |
| GABRB2 | 265 | W | 0 | 0 | no | yes | no | yes | no | no | no | no | no | no | no | no | no |
| GABRB2 | 266 | I | 0 | 0 | no | yes | no | no | no | no | no | no | no | no | no | no | no |
| GABRB2 | 267 | N | 0 | 0 | no | yes | no | no | no | no | no | no | no | no | no | no | no |
| GABRB2 | 268 | Y | 1 | 0 | no | yes | no | no | no | no | no | no | no | no | no | no | no |
| GABRB2 | 269 | D | 1 | 0 | no | yes | no | no | no | no | no | no | yes | no | no | no | no |
| GABRB2 | 270 | A | 0 | 0 | no | yes | no | no | no | no | no | no | yes | no | no | no | no |
| GABRB2 | 271 | S | 0 | 0 | no | yes | no | no | no | no | no | no | yes | no | no | no | no |
| GABRB2 | 272 | A | 0 | 0 | yes | no | no | no | no | no | no | no | yes | no | no | no | no |
| GABRB2 | 273 | A | 0 | 0 | yes | no | no | yes | no | no | no | no | no | no | no | no | no |
| GABRB2 | 274 | R | 0 | 0 | yes | no | no | yes | no | no | no | no | no | no | no | no | no |
| GABRB2 | 275 | V | 0 | 0 | yes | no | no | yes | no | no | no | no | yes | no | no | no | no |
| GABRB2 | 276 | A | 0 | 0 | yes | no | no | yes | no | no | no | no | yes | no | no | no | no |
| GABRB2 | 277 | L | 0 | 1 | yes | no | no | yes | no | no | no | no | no | no | no | no | no |
| GABRB2 | 278 | G | 0 | 0 | yes | no | no | yes | no | no | no | no | no | no | no | no | no |
| GABRB2 | 279 | I | 0 | 0 | yes | no | no | yes | no | no | no | no | yes | no | no | no | no |
| GABRB2 | 280 | T | 0 | 0 | yes | no | no | yes | no | no | no | no | yes | no | no | no | no |
| GABRB2 | 281 | T | 0 | 0 | yes | no | no | yes | no | no | no | no | no | no | no | no | no |
| GABRB2 | 282 | V | 0 | 1 | yes | no | no | yes | no | no | no | no | no | no | no | no | no |
| GABRB2 | 283 | L | 0 | 0 | yes | no | no | yes | no | no | no | no | yes | no | no | no | no |
| GABRB2 | 284 | T | 0 | 1 | yes | no | no | yes | no | no | no | no | yes | no | no | no | no |

| | | | | | | | | | | | | | | | | | |
|---|---|---|---|---|---|---|---|---|---|---|---|---|---|---|---|---|---|
| GABRB2 | 285 | M | 0 | 0 | yes | no | no | yes | no | no | no | no | no | no | no | no | no |
| GABRB2 | 286 | T | 0 | 0 | yes | no | no | yes | no | no | no | no | no | no | no | no | no |
| GABRB2 | 287 | T | 0 | 1 | yes | no | no | yes | no | no | yes | no | no | no | no | no | no |
| GABRB2 | 288 | I | 0 | 2 | yes | no | no | yes | no | no | no | no | no | no | no | no | no |
| GABRB2 | 289 | N | 1 | 1 | yes | no | no | yes | no | no | no | no | no | no | no | yes | no |
| GABRB2 | 290 | T | 0 | 0 | yes | no | no | yes | no | no | yes | no | no | no | no | yes | no |
| GABRB2 | 291 | H | 0 | 0 | yes | no | no | yes | no | no | yes | no | yes | no | no | yes | no |
| GABRB2 | 292 | L | 0 | 0 | yes | no | no | yes | no | no | no | no | yes | no | no | yes | no |
| GABRB2 | 293 | R | 1 | 1 | yes | no | no | yes | no | no | no | no | yes | no | no | yes | no |
| GABRB2 | 294 | E | 0 | 0 | yes | no | no | yes | no | no | yes | no | no | no | no | yes | no |
| GABRB2 | 295 | T | 0 | 0 | yes | no | no | no | no | no | no | no | yes | no | no | yes | no |
| GABRB2 | 296 | L | 0 | 0 | no | yes | no | no | no | no | no | no | no | no | no | yes | no |
| GABRB2 | 297 | P | 0 | 0 | no | yes | no | no | no | no | no | no | no | no | no | yes | no |
| GABRB2 | 298 | K | 0 | 0 | no | yes | no | no | no | no | no | no | yes | no | no | yes | no |
| GABRB2 | 299 | I | 0 | 2 | no | yes | no | no | no | no | no | no | yes | no | no | yes | no |
| GABRB2 | 300 | P | 0 | 1 | no | yes | no | no | no | no | no | no | yes | no | no | yes | no |
| GABRB2 | 301 | Y | 0 | 1 | no | yes | no | no | no | no | no | no | no | no | no | yes | no |
| GABRB2 | 302 | V | 0 | 1 | no | yes | no | no | no | no | no | no | no | no | no | yes | no |
| GABRB2 | 303 | K | 0 | 2 | no | yes | no | no | no | no | no | no | no | no | no | yes | no |
| GABRB2 | 304 | A | 0 | 1 | no | yes | no | no | no | no | no | no | no | no | no | yes | no |
| GABRB2 | 305 | I | 1 | 0 | yes | no | no | yes | no | no | no | no | no | no | no | yes | no |
| GABRB2 | 306 | D | 0 | 0 | yes | no | no | yes | no | no | no | no | no | no | no | yes | no |
| GABRB2 | 307 | M | 0 | 0 | yes | no | no | yes | no | no | no | no | no | no | no | yes | no |
| GABRB2 | 308 | Y | 0 | 0 | yes | no | no | yes | no | no | no | no | no | no | no | yes | no |
| GABRB2 | 309 | L | 0 | 0 | yes | no | no | yes | no | no | no | no | no | no | no | yes | no |
| GABRB2 | 310 | M | 0 | 0 | yes | no | no | yes | no | no | no | no | no | no | no | yes | no |
| GABRB2 | 311 | G | 0 | 0 | yes | no | no | yes | no | no | no | no | no | no | no | no | no |
| GABRB2 | 312 | C | 0 | 0 | yes | no | no | yes | no | no | no | no | no | no | no | no | no |
| GABRB2 | 313 | F | 0 | 0 | yes | no | no | yes | no | no | no | no | no | no | no | no | no |
| GABRB2 | 314 | V | 0 | 0 | yes | no | no | yes | no | no | no | no | no | no | no | no | no |
| GABRB2 | 315 | F | 0 | 0 | yes | no | no | yes | no | no | no | no | no | no | no | no | no |
| GABRB2 | 316 | V | 0 | 1 | yes | no | no | yes | no | no | no | no | no | no | no | no | no |
| GABRB2 | 317 | F | 0 | 0 | yes | no | no | yes | no | no | no | no | no | no | no | no | no |

| | | | | | | | | | | | | | | | | | |
|---|---|---|---|---|---|---|---|---|---|---|---|---|---|---|---|---|---|
| GABRB2 | 318 | M | 0 | 0 | yes | no | no | yes | no | no | no | no | no | no | no | no | no |
| GABRB2 | 319 | A | 0 | 0 | yes | no | no | yes | no | no | no | no | no | no | no | no | no |
| GABRB2 | 320 | L | 0 | 0 | yes | no | no | yes | no | no | no | no | no | no | no | no | no |
| GABRB2 | 321 | L | 0 | 0 | yes | no | no | yes | no | no | no | no | no | no | no | no | no |
| GABRB2 | 322 | E | 0 | 0 | yes | no | no | yes | no | no | no | no | no | no | no | no | no |
| GABRB2 | 323 | Y | 0 | 0 | yes | no | no | yes | no | no | no | no | no | no | no | no | no |
| GABRB2 | 324 | A | 0 | 0 | yes | no | no | yes | no | no | no | no | no | no | no | no | no |
| GABRB2 | 325 | L | 0 | 0 | yes | no | no | yes | no | no | no | no | no | no | no | no | no |
| GABRB2 | 326 | V | 0 | 0 | yes | no | no | no | no | no | no | no | no | no | no | no | no |
| GABRB2 | 327 | N | 0 | 0 | yes | no | no | no | no | yes | no | no | no | no | no | no | no |
| GABRB2 | 328 | Y | 0 | 0 | yes | no | no | no | no | yes | no | no | no | no | no | no | no |
| GABRB2 | 329 | I | 0 | 0 | yes | no | no | no | no | yes | no | no | no | no | no | no | no |
| GABRB2 | 330 | F | 0 | 0 | yes | no | no | no | no | yes | no | no | no | no | no | no | no |
| GABRB2 | 331 | F | 0 | 0 | yes | no | no | no | no | yes | no | no | no | no | no | no | no |
| GABRB2 | 486 | A | 1 | 0 | yes | no | no | no | no | yes | no | no | no | no | no | no | no |
| GABRB2 | 487 | I | 2 | 0 | yes | no | no | yes | no | yes | no | no | no | no | no | no | no |
| GABRB2 | 488 | D | 0 | 0 | yes | no | no | no | no | yes | no | no | no | no | no | no | no |
| GABRB2 | 489 | R | 0 | 0 | yes | no | no | no | no | yes | no | no | no | no | no | no | no |
| GABRB2 | 490 | W | 0 | 0 | yes | no | no | yes | no | no | no | no | no | no | no | no | no |
| GABRB2 | 491 | S | 0 | 0 | yes | no | no | yes | no | no | no | no | no | no | no | no | no |
| GABRB2 | 492 | R | 0 | 0 | yes | no | no | yes | no | no | no | no | no | no | no | no | no |
| GABRB2 | 493 | I | 0 | 0 | yes | no | no | yes | no | no | no | no | no | no | no | no | no |
| GABRB2 | 494 | F | 0 | 0 | yes | no | no | yes | no | no | no | no | no | no | no | no | no |
| GABRB2 | 495 | F | 0 | 0 | yes | no | no | yes | no | no | no | no | no | no | no | no | no |
| GABRB2 | 496 | P | 0 | 0 | yes | no | no | yes | no | no | no | no | no | no | no | no | no |
| GABRB2 | 497 | V | 0 | 0 | yes | no | no | yes | no | no | no | no | no | no | no | no | no |
| GABRB2 | 498 | V | 1 | 0 | yes | no | no | yes | no | no | no | no | no | no | no | no | no |
| GABRB2 | 499 | F | 0 | 0 | yes | no | no | yes | no | no | no | no | no | no | no | no | no |
| GABRB2 | 500 | S | 0 | 0 | yes | no | no | yes | no | no | no | no | no | no | no | no | no |
| GABRB2 | 501 | F | 0 | 0 | yes | no | no | yes | no | no | no | no | no | no | no | no | no |
| GABRB2 | 502 | F | 0 | 0 | yes | no | no | yes | no | no | no | no | no | no | no | no | no |
| GABRB2 | 503 | N | 0 | 0 | yes | no | no | yes | no | no | no | no | no | no | no | no | no |
| GABRB2 | 504 | I | 1 | 0 | yes | no | no | yes | no | no | no | no | no | no | no | no | no |

| | | | | | | | | | | | | | | | | | |
|---|---|---|---|---|---|---|---|---|---|---|---|---|---|---|---|---|---|
| GABRB2 | 505 | V | 1 | 0 | yes | no | no | yes | no | no | no | no | no | no | no | no | no |
| GABRB2 | 506 | Y | 0 | 0 | yes | no | no | yes | no | no | no | no | no | no | no | no | no |
| GABRB2 | 507 | W | 0 | 0 | yes | no | no | yes | no | no | no | no | no | no | no | no | no |
| GABRB2 | 508 | L | 0 | 0 | yes | no | no | yes | no | no | no | no | no | no | no | no | no |
| GABRB2 | 509 | Y | 0 | 0 | yes | no | no | yes | no | no | no | no | no | no | no | no | no |
| GABRB2 | 510 | Y | 0 | 0 | yes | no | no | no | no | no | no | no | no | no | no | no | no |
| GABRB2 | 511 | V | 0 | 0 | no | yes | no | no | no | no | no | no | no | no | no | no | no |
| GABRB3 | 35 | S | 0 | 0 | no | yes | no | no | yes | no | no | no | no | no | no | no | no |
| GABRB3 | 36 | F | 0 | 0 | yes | no | no | no | yes | no | no | no | no | no | no | no | no |
| GABRB3 | 37 | V | 0 | 1 | yes | no | no | no | yes | no | no | no | yes | no | no | no | no |
| GABRB3 | 38 | K | 0 | 0 | yes | no | no | no | yes | no | no | no | yes | no | no | no | no |
| GABRB3 | 39 | E | 1 | 0 | yes | no | no | no | yes | no | no | no | no | no | no | no | no |
| GABRB3 | 40 | T | 2 | 0 | yes | no | no | no | yes | no | no | no | no | no | no | no | no |
| GABRB3 | 41 | V | 0 | 0 | yes | no | no | no | yes | no | no | no | yes | no | no | no | no |
| GABRB3 | 42 | D | 1 | 0 | yes | no | no | no | yes | no | no | no | yes | no | no | no | no |
| GABRB3 | 43 | K | 1 | 0 | yes | no | no | no | yes | no | no | no | no | no | no | no | no |
| GABRB3 | 44 | L | 0 | 0 | yes | no | no | no | yes | no | no | no | no | no | no | no | no |
| GABRB3 | 45 | L | 0 | 0 | no | yes | no | no | yes | no | no | no | yes | no | no | no | no |
| GABRB3 | 46 | K | 1 | 0 | no | yes | no | no | yes | no | no | no | no | no | no | no | no |
| GABRB3 | 47 | G | 1 | 0 | no | yes | no | no | yes | no | no | no | no | no | no | no | no |
| GABRB3 | 48 | Y | 0 | 0 | no | yes | no | no | yes | no | no | no | no | no | no | no | no |
| GABRB3 | 49 | D | 0 | 0 | no | yes | no | no | yes | no | no | no | yes | no | no | no | no |
| GABRB3 | 50 | I | 0 | 0 | no | yes | no | no | yes | no | no | no | yes | no | no | no | no |
| GABRB3 | 51 | R | 0 | 0 | no | yes | no | no | yes | no | no | no | yes | no | no | no | no |
| GABRB3 | 52 | L | 0 | 0 | no | yes | no | no | yes | no | no | no | yes | no | no | no | no |
| GABRB3 | 53 | R | 0 | 0 | no | yes | no | no | yes | no | no | no | no | no | no | no | no |
| GABRB3 | 54 | P | 0 | 0 | no | yes | no | no | yes | no | no | no | no | no | no | no | no |
| GABRB3 | 55 | D | 0 | 0 | no | yes | no | no | yes | no | no | no | no | no | no | no | no |
| GABRB3 | 56 | F | 0 | 0 | no | yes | no | no | yes | no | no | no | yes | no | no | no | no |
| GABRB3 | 57 | G | 0 | 0 | no | yes | no | no | yes | no | no | no | no | no | no | no | no |
| GABRB3 | 58 | G | 0 | 0 | no | yes | no | no | yes | no | no | no | no | no | no | no | no |
| GABRB3 | 59 | P | 0 | 0 | no | yes | no | no | yes | no | no | no | no | no | no | no | no |
| GABRB3 | 60 | P | 0 | 0 | no | yes | no | no | yes | no | no | no | no | no | no | no | no |

| | | | | | | | | | | | | | | | | | |
|---|---|---|---|---|---|---|---|---|---|---|---|---|---|---|---|---|---|
| GABRB3 | 61 | V | 0 | 0 | no | no | yes | no | yes | no | no | no | no | no | no | no | no |
| GABRB3 | 62 | C | 0 | 0 | no | no | yes | no | yes | no | no | no | no | no | no | no | no |
| GABRB3 | 63 | V | 0 | 0 | no | no | yes | no | yes | no | no | no | no | no | no | no | no |
| GABRB3 | 64 | G | 0 | 0 | no | no | yes | no | yes | no | no | no | no | no | no | no | no |
| GABRB3 | 65 | M | 0 | 0 | no | no | yes | no | yes | no | no | no | no | no | no | no | no |
| GABRB3 | 66 | N | 1 | 0 | no | no | yes | no | yes | no | no | no | no | no | no | no | no |
| GABRB3 | 67 | I | 0 | 0 | no | no | yes | no | yes | no | no | no | no | no | no | no | no |
| GABRB3 | 68 | D | 0 | 0 | no | no | yes | no | yes | no | no | no | yes | no | no | no | no |
| GABRB3 | 69 | I | 0 | 0 | no | no | yes | no | yes | no | no | no | no | no | no | no | no |
| GABRB3 | 70 | A | 1 | 0 | no | no | yes | no | yes | no | no | no | no | no | no | no | no |
| GABRB3 | 71 | S | 1 | 0 | no | no | yes | no | yes | no | no | no | yes | no | no | no | no |
| GABRB3 | 72 | I | 0 | 0 | no | no | yes | no | yes | no | no | no | no | no | no | no | no |
| GABRB3 | 73 | D | 0 | 0 | no | no | yes | no | yes | no | no | no | yes | no | no | no | no |
| GABRB3 | 74 | M | 1 | 0 | no | no | yes | no | yes | no | no | no | yes | no | no | no | no |
| GABRB3 | 75 | V | 0 | 0 | no | no | yes | no | yes | no | no | no | no | no | no | no | no |
| GABRB3 | 76 | S | 0 | 1 | no | no | yes | no | yes | no | no | no | no | no | no | no | no |
| GABRB3 | 77 | E | 0 | 1 | no | yes | no | no | yes | no | no | no | no | no | no | no | no |
| GABRB3 | 78 | V | 0 | 0 | no | yes | no | no | yes | no | yes | no | no | no | no | no | no |
| GABRB3 | 79 | N | 0 | 0 | no | yes | no | no | yes | no | yes | no | no | no | no | no | no |
| GABRB3 | 80 | M | 0 | 1 | no | yes | no | no | yes | no | no | no | no | no | no | no | no |
| GABRB3 | 81 | D | 0 | 0 | no | no | yes | no | yes | no | no | no | no | no | no | no | no |
| GABRB3 | 82 | Y | 0 | 0 | no | no | yes | no | yes | no | no | no | no | no | no | no | no |
| GABRB3 | 83 | T | 0 | 0 | no | no | yes | no | yes | no | no | no | no | no | no | no | no |
| GABRB3 | 84 | L | 0 | 0 | no | no | yes | no | yes | no | no | no | no | no | no | no | no |
| GABRB3 | 85 | T | 0 | 0 | no | no | yes | no | yes | no | no | no | no | no | no | no | no |
| GABRB3 | 86 | M | 0 | 0 | no | no | yes | no | yes | no | no | no | no | no | no | no | no |
| GABRB3 | 87 | Y | 0 | 0 | no | no | yes | no | yes | no | no | no | yes | no | no | no | no |
| GABRB3 | 88 | F | 0 | 0 | no | no | yes | no | yes | no | no | no | no | no | no | no | no |
| GABRB3 | 89 | Q | 0 | 0 | no | no | yes | no | yes | no | no | no | yes | no | no | no | no |
| GABRB3 | 90 | Q | 0 | 0 | no | no | yes | no | yes | no | no | no | no | no | no | no | no |
| GABRB3 | 91 | Y | 0 | 0 | no | no | yes | no | yes | no | no | no | no | no | no | no | no |
| GABRB3 | 92 | W | 0 | 0 | no | no | yes | no | yes | no | no | no | no | no | no | no | no |
| GABRB3 | 93 | R | 0 | 0 | no | no | yes | no | yes | no | no | no | no | no | no | no | no |

| | | | | | | | | | | | | | | | | | |
|---|---|---|---|---|---|---|---|---|---|---|---|---|---|---|---|---|---|
| GABRB3 | 94 | D | 0 | 0 | no | yes | no | no | yes | no | no | no | no | no | no | no | no |
| GABRB3 | 95 | K | 0 | 0 | yes | no | no | no | yes | no | no | no | no | no | no | no | no |
| GABRB3 | 96 | R | 0 | 0 | yes | no | no | no | yes | no | no | no | no | no | no | no | no |
| GABRB3 | 97 | L | 0 | 0 | yes | no | no | no | yes | no | no | no | no | no | no | no | no |
| GABRB3 | 98 | A | 1 | 0 | no | yes | no | no | yes | no | no | no | no | no | no | no | no |
| GABRB3 | 99 | Y | 0 | 0 | no | yes | no | no | yes | no | no | no | no | no | no | no | no |
| GABRB3 | 100 | S | 1 | 0 | no | yes | no | no | yes | no | no | no | no | no | no | no | no |
| GABRB3 | 101 | G | 0 | 0 | no | yes | no | no | yes | no | no | no | no | no | no | no | no |
| GABRB3 | 102 | I | 0 | 0 | no | yes | no | no | yes | no | no | no | no | no | no | no | no |
| GABRB3 | 103 | P | 0 | 0 | no | yes | no | no | yes | no | no | no | no | no | no | no | no |
| GABRB3 | 104 | L | 0 | 0 | no | yes | no | no | yes | no | no | no | yes | no | no | no | no |
| GABRB3 | 105 | N | 0 | 0 | no | yes | no | no | yes | no | no | no | no | no | no | no | no |
| GABRB3 | 106 | L | 0 | 0 | no | no | yes | no | yes | no | no | no | no | no | no | no | no |
| GABRB3 | 107 | T | 0 | 0 | no | no | yes | no | yes | no | no | no | yes | no | no | no | no |
| GABRB3 | 108 | L | 0 | 0 | no | no | yes | no | yes | no | no | no | yes | no | no | no | no |
| GABRB3 | 109 | D | 0 | 0 | no | yes | no | no | yes | no | no | no | yes | no | no | no | no |
| GABRB3 | 110 | N | 1 | 1 | yes | no | no | no | yes | no | no | no | no | no | no | no | no |
| GABRB3 | 111 | R | 1 | 0 | yes | no | no | no | yes | no | no | no | yes | no | no | no | no |
| GABRB3 | 112 | V | 0 | 0 | yes | no | no | no | yes | no | no | no | yes | no | no | no | no |
| GABRB3 | 113 | A | 0 | 0 | yes | no | no | no | yes | no | no | no | no | no | no | no | no |
| GABRB3 | 114 | D | 0 | 0 | yes | no | no | no | yes | no | yes | no | no | no | no | no | no |
| GABRB3 | 115 | Q | 0 | 0 | yes | no | no | no | yes | no | yes | no | yes | no | no | no | no |
| GABRB3 | 116 | L | 0 | 0 | no | yes | no | no | yes | no | no | no | no | no | no | no | no |
| GABRB3 | 117 | W | 0 | 0 | no | yes | no | no | yes | no | no | no | no | no | no | no | no |
| GABRB3 | 118 | V | 0 | 0 | no | yes | no | no | yes | no | no | no | yes | no | no | no | no |
| GABRB3 | 119 | P | 0 | 0 | no | yes | no | no | yes | no | no | no | yes | no | no | no | no |
| GABRB3 | 120 | D | 0 | 1 | no | yes | no | no | yes | no | no | no | yes | no | yes | no | no |
| GABRB3 | 121 | T | 0 | 0 | no | no | yes | no | yes | no | no | no | yes | no | yes | no | no |
| GABRB3 | 122 | Y | 0 | 0 | no | no | yes | no | yes | no | no | no | yes | no | yes | no | no |
| GABRB3 | 123 | F | 0 | 0 | no | no | yes | no | yes | no | no | no | yes | no | no | no | no |
| GABRB3 | 124 | L | 0 | 1 | no | yes | no | no | yes | no | no | no | no | no | no | no | no |
| GABRB3 | 125 | N | 0 | 0 | no | yes | no | no | yes | no | no | no | no | no | no | no | no |
| GABRB3 | 126 | D | 0 | 0 | no | no | yes | no | yes | no | no | no | yes | no | no | no | no |

| | | | | | | | | | | | | | | | | | |
|---|---|---|---|---|---|---|---|---|---|---|---|---|---|---|---|---|---|
| GABRB3 | 127 | K | 0 | 1 | no | yes | no | no | yes | no | no | no | yes | no | no | no | no |
| GABRB3 | 128 | K | 0 | 0 | no | yes | no | no | yes | no | yes | no | no | no | no | no | no |
| GABRB3 | 129 | S | 0 | 0 | no | yes | no | no | yes | no | no | no | yes | no | no | no | no |
| GABRB3 | 130 | F | 0 | 0 | no | no | yes | no | yes | no | no | no | yes | no | no | no | no |
| GABRB3 | 131 | V | 1 | 0 | no | no | yes | no | yes | no | no | no | yes | no | no | no | no |
| GABRB3 | 132 | H | 0 | 0 | no | yes | no | no | yes | no | no | no | yes | no | no | no | no |
| GABRB3 | 133 | G | 0 | 0 | no | yes | no | no | yes | no | no | no | no | no | no | no | no |
| GABRB3 | 134 | V | 0 | 0 | no | yes | no | no | yes | no | no | no | yes | no | no | no | no |
| GABRB3 | 135 | T | 0 | 0 | no | yes | no | no | yes | no | no | no | yes | no | no | no | no |
| GABRB3 | 136 | V | 1 | 0 | no | yes | no | no | yes | no | no | no | yes | no | no | no | no |
| GABRB3 | 137 | K | 0 | 0 | no | yes | no | no | yes | no | no | no | no | no | no | no | no |
| GABRB3 | 138 | N | 0 | 0 | no | yes | no | no | yes | no | no | no | yes | no | no | no | no |
| GABRB3 | 139 | R | 2 | 0 | no | no | yes | no | yes | no | no | no | yes | no | no | no | no |
| GABRB3 | 140 | M | 0 | 0 | no | no | yes | no | yes | no | no | no | yes | no | no | no | no |
| GABRB3 | 141 | I | 0 | 0 | no | no | yes | no | yes | no | no | no | no | no | no | no | no |
| GABRB3 | 142 | R | 1 | 1 | no | no | yes | no | yes | no | no | no | yes | no | no | no | no |
| GABRB3 | 143 | L | 0 | 0 | no | no | yes | no | yes | no | no | no | no | no | no | no | no |
| GABRB3 | 144 | H | 0 | 0 | no | yes | no | no | yes | no | no | no | no | no | no | no | no |
| GABRB3 | 145 | P | 0 | 0 | no | yes | no | no | yes | no | no | no | no | no | no | no | no |
| GABRB3 | 146 | D | 0 | 0 | no | yes | no | no | yes | no | no | no | no | no | no | no | no |
| GABRB3 | 147 | G | 0 | 0 | no | yes | no | no | yes | no | no | no | no | no | no | no | no |
| GABRB3 | 148 | T | 0 | 0 | no | no | yes | no | yes | no | no | no | no | no | no | no | no |
| GABRB3 | 149 | V | 0 | 0 | no | no | yes | no | yes | no | no | no | no | no | no | no | no |
| GABRB3 | 150 | L | 0 | 0 | no | no | yes | no | yes | no | no | no | no | no | no | no | no |
| GABRB3 | 151 | Y | 0 | 0 | no | no | yes | no | yes | no | no | no | no | no | no | no | no |
| GABRB3 | 152 | G | 0 | 0 | no | no | yes | no | yes | no | no | no | yes | no | no | no | no |
| GABRB3 | 153 | L | 0 | 0 | no | no | yes | no | yes | no | no | no | yes | no | no | no | no |
| GABRB3 | 154 | R | 0 | 0 | no | no | yes | no | yes | no | no | no | yes | no | no | no | no |
| GABRB3 | 155 | I | 0 | 0 | no | no | yes | no | yes | no | no | no | yes | no | no | no | no |
| GABRB3 | 156 | T | 0 | 1 | no | no | yes | no | yes | no | no | no | no | no | no | no | no |
| GABRB3 | 157 | T | 0 | 1 | no | no | yes | no | yes | no | no | no | no | no | no | no | no |
| GABRB3 | 158 | T | 0 | 0 | no | no | yes | no | yes | no | no | no | no | no | no | no | no |
| GABRB3 | 159 | A | 0 | 0 | no | no | yes | no | yes | no | no | no | no | no | no | no | no |

| | | | | | | | | | | | | | | | | | |
|---|---|---|---|---|---|---|---|---|---|---|---|---|---|---|---|---|---|
| GABRB3 | 160 | A | 0 | 0 | no | no | yes | no | yes | no | no | no | yes | no | no | no | no |
| GABRB3 | 161 | C | 0 | 0 | no | yes | no | no | yes | no | no | no | no | no | no | no | no |
| GABRB3 | 162 | M | 0 | 1 | no | yes | no | no | yes | no | no | no | yes | no | no | no | no |
| GABRB3 | 163 | M | 0 | 0 | no | yes | no | no | yes | no | no | no | no | no | no | no | no |
| GABRB3 | 164 | D | 0 | 0 | no | yes | no | no | yes | no | no | no | no | no | no | no | no |
| GABRB3 | 165 | L | 0 | 0 | yes | no | no | no | yes | no | no | no | no | no | no | no | no |
| GABRB3 | 166 | R | 0 | 0 | yes | no | no | no | yes | no | no | no | no | no | no | no | no |
| GABRB3 | 167 | R | 0 | 0 | yes | no | no | no | yes | no | no | no | no | no | no | no | no |
| GABRB3 | 168 | Y | 0 | 0 | no | yes | no | no | yes | no | no | no | no | no | no | no | no |
| GABRB3 | 169 | P | 0 | 0 | no | yes | no | no | yes | no | no | no | no | no | no | no | no |
| GABRB3 | 170 | L | 0 | 1 | no | yes | no | no | yes | no | no | no | no | no | no | no | no |
| GABRB3 | 171 | D | 0 | 0 | no | yes | no | no | yes | no | no | no | no | no | no | no | no |
| GABRB3 | 172 | E | 0 | 0 | no | no | yes | no | yes | no | no | no | no | no | no | no | no |
| GABRB3 | 173 | Q | 0 | 0 | no | yes | no | no | yes | no | no | no | no | no | no | no | no |
| GABRB3 | 174 | N | 0 | 0 | no | yes | no | no | yes | no | no | no | no | no | no | no | no |
| GABRB3 | 175 | C | 0 | 0 | no | yes | no | no | yes | no | no | no | no | no | no | no | no |
| GABRB3 | 176 | T | 0 | 0 | no | yes | no | no | yes | no | no | no | no | no | no | no | no |
| GABRB3 | 177 | L | 0 | 0 | no | yes | no | no | yes | no | no | no | no | no | no | no | no |
| GABRB3 | 178 | E | 0 | 0 | no | no | yes | no | yes | no | no | no | no | no | no | no | no |
| GABRB3 | 179 | I | 0 | 0 | no | no | yes | no | yes | no | no | no | no | no | no | no | no |
| GABRB3 | 180 | E | 0 | 1 | no | no | yes | no | yes | no | no | no | no | no | yes | no | no |
| GABRB3 | 181 | S | 0 | 0 | no | no | yes | no | yes | no | no | no | no | no | yes | no | no |
| GABRB3 | 182 | Y | 0 | 1 | no | yes | no | no | yes | no | no | no | yes | no | yes | no | no |
| GABRB3 | 183 | G | 0 | 0 | no | yes | no | no | yes | no | no | no | yes | no | no | no | no |
| GABRB3 | 184 | Y | 0 | 1 | no | yes | no | no | yes | no | no | no | yes | no | no | no | no |
| GABRB3 | 185 | T | 0 | 1 | no | yes | no | no | yes | no | no | no | yes | no | no | no | no |
| GABRB3 | 186 | T | 1 | 0 | no | yes | no | no | yes | no | no | no | no | no | no | no | no |
| GABRB3 | 187 | D | 0 | 0 | no | yes | no | no | yes | no | no | no | no | no | no | no | no |
| GABRB3 | 188 | D | 0 | 0 | no | yes | no | no | yes | no | no | no | yes | no | no | no | no |
| GABRB3 | 189 | I | 0 | 0 | no | no | yes | no | yes | no | no | no | no | no | no | no | no |
| GABRB3 | 190 | E | 0 | 0 | no | no | yes | no | yes | no | no | no | no | no | no | no | no |
| GABRB3 | 191 | F | 0 | 0 | no | no | yes | no | yes | no | no | no | no | no | no | no | no |
| GABRB3 | 192 | Y | 0 | 0 | no | no | yes | no | yes | no | no | no | no | no | no | no | no |

| | | | | | | | | | | | | | | | | | |
|---|---|---|---|---|---|---|---|---|---|---|---|---|---|---|---|---|---|
| GABRB3 | 193 | W | 0 | 0 | no | no | yes | no | yes | no | no | no | no | no | no | no | no |
| GABRB3 | 194 | R | 0 | 0 | no | yes | no | no | yes | no | no | no | no | no | no | no | no |
| GABRB3 | 195 | G | 0 | 0 | no | yes | no | no | yes | no | no | no | no | no | no | no | no |
| GABRB3 | 196 | G | 1 | 0 | yes | no | no | no | yes | no | no | no | no | no | no | no | no |
| GABRB3 | 197 | D | 1 | 0 | yes | no | no | no | yes | no | no | no | no | no | no | no | no |
| GABRB3 | 198 | K | 0 | 0 | yes | no | no | no | yes | no | no | no | no | no | no | no | no |
| GABRB3 | 199 | A | 0 | 0 | no | yes | no | no | yes | no | no | no | no | no | no | no | no |
| GABRB3 | 200 | V | 1 | 0 | no | yes | no | no | yes | no | no | no | no | no | no | no | no |
| GABRB3 | 201 | T | 1 | 0 | no | no | yes | no | yes | no | no | no | no | no | no | no | no |
| GABRB3 | 202 | G | 0 | 0 | no | yes | no | no | yes | no | no | no | no | no | no | no | no |
| GABRB3 | 203 | V | 0 | 0 | no | yes | no | no | yes | no | no | no | no | no | no | no | no |
| GABRB3 | 204 | E | 0 | 0 | no | yes | no | no | yes | no | no | no | no | no | no | no | no |
| GABRB3 | 205 | R | 0 | 0 | no | yes | no | no | yes | no | no | no | no | no | no | no | no |
| GABRB3 | 206 | I | 0 | 0 | no | yes | no | no | yes | no | no | no | no | no | no | no | no |
| GABRB3 | 207 | E | 1 | 0 | no | yes | no | no | yes | no | no | no | no | no | no | no | no |
| GABRB3 | 208 | L | 0 | 0 | no | yes | no | no | yes | no | no | no | no | no | no | no | no |
| GABRB3 | 209 | P | 1 | 0 | no | yes | no | no | yes | no | no | no | yes | no | no | no | no |
| GABRB3 | 210 | Q | 0 | 0 | no | yes | no | no | yes | no | no | no | yes | no | no | no | no |
| GABRB3 | 211 | F | 0 | 0 | no | no | yes | no | yes | no | no | no | no | no | no | no | no |
| GABRB3 | 212 | S | 0 | 0 | no | no | yes | no | yes | no | no | no | no | no | no | no | no |
| GABRB3 | 213 | I | 1 | 0 | no | no | yes | no | yes | no | no | no | no | no | no | no | no |
| GABRB3 | 214 | V | 1 | 0 | no | no | yes | no | yes | no | no | no | no | no | no | no | no |
| GABRB3 | 215 | E | 0 | 0 | no | no | yes | no | yes | no | no | no | no | no | no | no | no |
| GABRB3 | 216 | H | 0 | 0 | no | no | yes | no | yes | no | no | no | no | no | no | no | no |
| GABRB3 | 217 | R | 1 | 0 | no | no | yes | no | yes | no | no | no | no | no | no | no | no |
| GABRB3 | 218 | L | 0 | 0 | no | no | yes | no | yes | no | no | no | no | no | no | no | no |
| GABRB3 | 219 | V | 0 | 0 | no | no | yes | no | yes | no | no | no | no | no | no | no | no |
| GABRB3 | 220 | S | 1 | 0 | no | no | yes | no | yes | no | no | no | no | no | no | no | no |
| GABRB3 | 221 | R | 1 | 0 | no | no | yes | no | yes | no | no | no | no | no | no | no | no |
| GABRB3 | 222 | N | 0 | 0 | no | no | yes | no | yes | no | no | no | no | no | no | no | no |
| GABRB3 | 223 | V | 0 | 0 | no | no | yes | no | yes | no | no | no | no | no | no | no | no |
| GABRB3 | 224 | V | 0 | 0 | no | no | yes | no | yes | no | no | no | no | no | no | no | no |
| GABRB3 | 225 | F | 0 | 0 | no | yes | no | no | yes | no | no | no | yes | no | yes | no | no |

| | | | | | | | | | | | | | | | | | |
|---|---|---|---|---|---|---|---|---|---|---|---|---|---|---|---|---|---|
| GABRB3 | 226 | A | 0 | 0 | no | yes | no | no | yes | no | no | no | no | no | no | no | no |
| GABRB3 | 227 | T | 0 | 0 | no | yes | no | no | yes | no | no | no | yes | no | no | no | no |
| GABRB3 | 228 | G | 0 | 0 | no | yes | no | no | yes | no | no | no | no | no | no | no | no |
| GABRB3 | 229 | A | 0 | 0 | no | no | yes | no | yes | no | no | no | no | no | no | no | no |
| GABRB3 | 230 | Y | 0 | 0 | no | no | yes | no | yes | no | no | no | yes | no | no | no | no |
| GABRB3 | 231 | P | 0 | 0 | no | no | yes | no | yes | no | no | no | no | no | no | no | no |
| GABRB3 | 232 | R | 0 | 1 | no | no | yes | no | yes | no | no | no | no | no | no | no | no |
| GABRB3 | 233 | L | 0 | 0 | no | no | yes | no | yes | no | no | no | no | no | no | no | no |
| GABRB3 | 234 | S | 0 | 0 | no | no | yes | no | yes | no | no | no | no | no | no | no | no |
| GABRB3 | 235 | L | 0 | 0 | no | no | yes | no | yes | no | no | no | no | no | no | no | no |
| GABRB3 | 236 | S | 0 | 0 | no | no | yes | no | yes | no | no | no | no | no | no | no | no |
| GABRB3 | 237 | F | 0 | 0 | no | no | yes | no | yes | no | no | no | no | no | no | no | no |
| GABRB3 | 238 | R | 2 | 0 | no | no | yes | no | yes | no | no | no | no | no | no | no | no |
| GABRB3 | 239 | L | 0 | 0 | no | no | yes | no | yes | no | no | no | no | no | no | no | no |
| GABRB3 | 240 | K | 0 | 0 | no | no | yes | no | yes | no | no | no | no | no | no | no | no |
| GABRB3 | 241 | R | 0 | 0 | no | no | yes | no | yes | no | no | no | no | no | no | no | no |
| GABRB3 | 242 | N | 0 | 0 | no | yes | no | no | yes | no | no | no | yes | no | no | no | no |
| GABRB3 | 243 | I | 0 | 0 | no | yes | no | no | yes | no | no | no | no | no | no | no | no |
| GABRB3 | 244 | G | 0 | 0 | yes | no | no | no | yes | no | no | no | no | no | no | no | no |
| GABRB3 | 245 | Y | 0 | 1 | yes | no | no | no | yes | no | no | no | yes | no | no | no | no |
| GABRB3 | 246 | F | 0 | 0 | yes | no | no | no | no | no | no | no | no | no | no | no | no |
| GABRB3 | 247 | I | 0 | 0 | yes | no | no | yes | no | no | no | no | no | no | no | no | no |
| GABRB3 | 248 | L | 0 | 0 | yes | no | no | yes | no | no | no | no | yes | no | no | no | no |
| GABRB3 | 249 | Q | 0 | 1 | yes | no | no | yes | no | no | no | no | yes | no | no | no | no |
| GABRB3 | 250 | T | 0 | 0 | yes | no | no | yes | no | no | no | no | no | no | no | no | no |
| GABRB3 | 251 | Y | 0 | 0 | yes | no | no | yes | no | no | no | no | no | no | no | no | no |
| GABRB3 | 252 | M | 0 | 0 | yes | no | no | yes | no | no | no | no | no | no | no | no | no |
| GABRB3 | 253 | P | 0 | 1 | yes | no | no | yes | no | no | no | no | no | no | no | no | no |
| GABRB3 | 254 | S | 0 | 1 | yes | no | no | yes | no | no | no | no | no | no | no | no | no |
| GABRB3 | 255 | I | 1 | 0 | yes | no | no | yes | no | no | no | no | no | no | no | no | no |
| GABRB3 | 256 | L | 0 | 2 | yes | no | no | yes | no | no | no | no | yes | no | no | no | no |
| GABRB3 | 257 | I | 0 | 0 | yes | no | no | yes | no | no | no | no | yes | no | no | no | no |
| GABRB3 | 258 | T | 0 | 0 | yes | no | no | yes | no | no | no | no | no | no | no | no | no |

| | | | | | | | | | | | | | | | | | |
|---|---|---|---|---|---|---|---|---|---|---|---|---|---|---|---|---|---|
| GABRB3 | 259 | I | 0 | 0 | yes | no | no | yes | no | no | no | no | no | no | no | no | no |
| GABRB3 | 260 | L | 0 | 0 | yes | no | no | yes | no | no | no | no | yes | no | no | no | no |
| GABRB3 | 261 | S | 0 | 0 | yes | no | no | yes | no | no | no | no | no | no | no | no | no |
| GABRB3 | 262 | W | 0 | 0 | yes | no | no | yes | no | no | no | no | no | no | no | no | no |
| GABRB3 | 263 | V | 0 | 0 | yes | no | no | yes | no | no | no | no | yes | no | no | no | no |
| GABRB3 | 264 | S | 0 | 0 | yes | no | no | yes | no | no | no | no | no | no | no | no | no |
| GABRB3 | 265 | F | 0 | 0 | yes | no | no | yes | no | no | no | no | no | no | no | no | no |
| GABRB3 | 266 | W | 0 | 0 | no | yes | no | yes | no | no | no | no | yes | no | no | no | no |
| GABRB3 | 267 | I | 0 | 0 | no | yes | no | yes | no | no | no | no | yes | no | no | no | no |
| GABRB3 | 268 | N | 1 | 0 | no | yes | no | no | no | yes | no | no | no | no | no | no | no |
| GABRB3 | 269 | Y | 0 | 0 | no | yes | no | no | no | yes | no | no | no | no | no | no | no |
| GABRB3 | 270 | D | 0 | 0 | no | yes | no | no | no | yes | no | no | no | no | no | no | no |
| GABRB3 | 271 | A | 0 | 0 | no | yes | no | no | no | no | no | no | no | no | no | no | no |
| GABRB3 | 272 | S | 0 | 0 | yes | no | no | no | no | no | no | no | yes | no | no | no | no |
| GABRB3 | 273 | A | 0 | 0 | yes | no | no | yes | no | no | yes | no | yes | no | no | no | no |
| GABRB3 | 274 | A | 0 | 0 | yes | no | no | yes | no | no | no | no | yes | no | no | no | no |
| GABRB3 | 275 | R | 0 | 0 | yes | no | no | yes | no | no | no | no | no | no | no | no | no |
| GABRB3 | 276 | V | 0 | 0 | yes | no | no | yes | no | no | no | no | yes | no | no | no | no |
| GABRB3 | 277 | A | 0 | 0 | yes | no | no | yes | no | no | no | no | no | no | no | no | no |
| GABRB3 | 278 | L | 0 | 0 | yes | no | no | yes | no | no | no | no | yes | no | no | no | no |
| GABRB3 | 279 | G | 0 | 0 | yes | no | no | yes | no | no | no | no | no | no | no | no | no |
| GABRB3 | 280 | I | 0 | 1 | yes | no | no | yes | no | no | no | no | yes | no | no | no | no |
| GABRB3 | 281 | T | 0 | 1 | yes | no | no | yes | no | no | yes | no | yes | no | no | no | no |
| GABRB3 | 282 | T | 0 | 0 | yes | no | no | yes | no | no | no | no | no | no | no | no | no |
| GABRB3 | 283 | V | 0 | 0 | yes | no | no | yes | no | no | no | no | yes | no | no | no | no |
| GABRB3 | 284 | L | 0 | 3 | yes | no | no | yes | no | no | yes | no | yes | no | no | no | no |
| GABRB3 | 285 | T | 0 | 0 | yes | no | no | yes | no | no | no | no | yes | no | no | no | no |
| GABRB3 | 286 | M | 0 | 0 | yes | no | no | yes | no | no | no | no | no | no | no | no | no |
| GABRB3 | 287 | T | 0 | 2 | yes | no | no | yes | no | no | no | no | no | no | no | no | no |
| GABRB3 | 288 | T | 0 | 2 | yes | no | no | yes | no | no | yes | no | no | no | no | no | no |
| GABRB3 | 289 | I | 0 | 0 | yes | no | no | yes | no | no | no | no | no | no | no | no | no |
| GABRB3 | 290 | N | 0 | 0 | yes | no | no | yes | no | no | no | no | yes | no | no | yes | no |
| GABRB3 | 291 | T | 0 | 0 | yes | no | no | yes | no | no | no | no | no | no | no | yes | no |

| | | | | | | | | | | | | | | | | | |
|---|---|---|---|---|---|---|---|---|---|---|---|---|---|---|---|---|---|
| GABRB3 | 292 | H | 0 | 0 | no | yes | no | yes | no | no | yes | no | no | no | no | yes | no |
| GABRB3 | 293 | L | 0 | 1 | no | yes | no | no | no | no | no | no | no | no | no | yes | no |
| GABRB3 | 294 | R | 0 | 1 | yes | no | no | no | yes | no | no | no | yes | no | no | yes | no |
| GABRB3 | 295 | E | 0 | 0 | yes | no | no | no | yes | no | yes | no | no | no | no | yes | no |
| GABRB3 | 296 | T | 0 | 0 | yes | no | no | no | yes | no | no | no | no | no | no | yes | no |
| GABRB3 | 297 | L | 0 | 0 | no | yes | no | no | yes | no | no | no | no | no | no | yes | no |
| GABRB3 | 298 | P | 0 | 0 | no | yes | no | no | yes | no | no | no | yes | no | no | yes | no |
| GABRB3 | 299 | K | 0 | 1 | no | yes | no | no | yes | no | no | no | yes | no | no | yes | no |
| GABRB3 | 300 | I | 0 | 1 | no | yes | no | no | yes | no | no | no | yes | no | no | yes | no |
| GABRB3 | 301 | P | 0 | 1 | no | yes | no | no | yes | no | no | no | yes | no | no | yes | no |
| GABRB3 | 302 | Y | 0 | 1 | no | yes | no | no | yes | no | no | no | no | no | no | yes | no |
| GABRB3 | 303 | V | 0 | 0 | no | yes | no | no | yes | no | no | no | yes | no | no | yes | no |
| GABRB3 | 304 | K | 0 | 0 | no | yes | no | no | yes | no | no | no | no | no | no | yes | no |
| GABRB3 | 305 | A | 0 | 3 | no | yes | no | no | no | no | no | no | no | no | no | yes | no |
| GABRB3 | 306 | I | 0 | 0 | no | yes | no | yes | no | no | no | no | no | no | no | yes | no |
| GABRB3 | 307 | D | 0 | 0 | yes | no | no | yes | no | no | no | no | yes | no | no | yes | no |
| GABRB3 | 308 | M | 1 | 0 | yes | no | no | yes | no | no | no | no | no | no | no | yes | no |
| GABRB3 | 309 | Y | 0 | 0 | yes | no | no | yes | no | no | no | no | no | no | no | yes | no |
| GABRB3 | 310 | L | 0 | 0 | yes | no | no | yes | no | no | no | no | no | no | no | yes | no |
| GABRB3 | 311 | M | 0 | 0 | yes | no | no | yes | no | no | no | no | no | no | no | yes | no |
| GABRB3 | 312 | G | 0 | 0 | yes | no | no | yes | no | no | no | no | no | no | no | no | no |
| GABRB3 | 313 | C | 0 | 0 | yes | no | no | yes | no | no | no | no | no | no | no | no | no |
| GABRB3 | 314 | F | 0 | 0 | yes | no | no | yes | no | no | no | no | yes | no | no | no | no |
| GABRB3 | 315 | V | 0 | 0 | yes | no | no | yes | no | no | no | no | no | no | no | no | no |
| GABRB3 | 316 | F | 0 | 0 | yes | no | no | yes | no | no | no | no | no | no | no | no | no |
| GABRB3 | 317 | V | 0 | 0 | yes | no | no | yes | no | no | no | no | no | no | no | no | no |
| GABRB3 | 318 | F | 0 | 0 | yes | no | no | yes | no | no | no | no | yes | no | no | no | no |
| GABRB3 | 319 | L | 0 | 0 | yes | no | no | yes | no | no | no | no | no | no | no | no | no |
| GABRB3 | 320 | A | 0 | 0 | yes | no | no | yes | no | no | no | no | no | no | no | no | no |
| GABRB3 | 321 | L | 0 | 0 | yes | no | no | yes | no | no | no | no | yes | no | no | no | no |
| GABRB3 | 322 | L | 0 | 0 | yes | no | no | yes | no | no | no | no | yes | no | no | no | no |
| GABRB3 | 323 | E | 0 | 0 | yes | no | no | yes | no | no | no | no | no | no | no | no | no |
| GABRB3 | 324 | Y | 1 | 0 | yes | no | no | yes | no | no | no | no | no | no | no | no | no |

| | | | | | | | | | | | | | | | | | |
|---|---|---|---|---|---|---|---|---|---|---|---|---|---|---|---|---|---|
| GABRB3 | 325 | A | 0 | 0 | yes | no | no | yes | no | no | no | no | yes | no | no | no | no |
| GABRB3 | 326 | F | 0 | 0 | yes | no | no | yes | no | no | no | no | no | no | no | no | no |
| GABRB3 | 327 | V | 0 | 0 | yes | no | no | no | no | no | no | no | no | no | no | no | no |
| GABRB3 | 328 | N | 0 | 1 | yes | no | no | no | no | yes | no | no | yes | no | no | no | no |
| GABRB3 | 329 | Y | 0 | 0 | yes | no | no | no | no | yes | no | no | yes | no | no | no | no |
| GABRB3 | 330 | I | 0 | 0 | yes | no | no | no | no | yes | no | no | no | no | no | no | no |
| GABRB3 | 331 | F | 1 | 0 | yes | no | no | no | no | yes | no | no | yes | no | no | no | no |
| GABRB3 | 332 | F | 0 | 0 | yes | no | no | no | no | yes | no | no | yes | no | no | no | no |
| GABRB3 | 333 | G | 0 | 0 | no | yes | no | no | no | yes | no | no | no | no | no | no | no |
| GABRB3 | 334 | R | 0 | 0 | no | yes | no | no | no | yes | no | no | no | no | no | no | no |
| GABRB3 | 335 | G | 0 | 0 | no | yes | no | no | no | yes | no | no | yes | no | no | no | no |
| GABRB3 | 336 | P | 1 | 0 | no | yes | no | no | no | yes | no | no | yes | no | no | no | no |
| GABRB3 | 337 | Q | 0 | 0 | no | yes | no | no | no | yes | no | no | no | no | no | no | no |
| GABRB3 | 440 | P | 0 | 0 | no | yes | no | yes | no | yes | no | no | no | no | no | no | no |
| GABRB3 | 441 | D | 0 | 0 | no | yes | no | no | no | yes | no | no | no | no | no | no | no |
| GABRB3 | 442 | L | 1 | 0 | no | yes | no | no | no | yes | no | no | no | no | no | no | no |
| GABRB3 | 443 | T | 1 | 0 | no | yes | no | no | no | yes | no | no | no | no | no | no | no |
| GABRB3 | 444 | D | 1 | 0 | no | yes | no | no | no | yes | no | no | no | no | no | no | no |
| GABRB3 | 445 | V | 0 | 0 | yes | no | no | no | no | yes | no | no | no | no | no | no | no |
| GABRB3 | 446 | N | 0 | 0 | yes | no | no | no | no | yes | no | no | no | no | no | no | no |
| GABRB3 | 447 | A | 1 | 0 | yes | no | no | no | no | yes | no | no | no | no | no | no | no |
| GABRB3 | 448 | I | 1 | 0 | yes | no | no | no | no | yes | no | no | no | no | no | no | no |
| GABRB3 | 449 | D | 0 | 0 | yes | no | no | no | no | yes | no | no | no | no | no | no | no |
| GABRB3 | 450 | R | 0 | 0 | yes | no | no | no | no | yes | no | no | no | no | no | no | no |
| GABRB3 | 451 | W | 0 | 0 | yes | no | no | yes | no | no | no | no | no | no | no | no | no |
| GABRB3 | 452 | S | 0 | 0 | yes | no | no | yes | no | no | no | no | no | no | no | no | no |
| GABRB3 | 453 | R | 0 | 0 | yes | no | no | yes | no | no | no | no | no | no | no | no | no |
| GABRB3 | 454 | I | 1 | 0 | yes | no | no | yes | no | no | no | no | no | no | no | no | no |
| GABRB3 | 455 | V | 0 | 0 | yes | no | no | yes | no | no | no | no | no | no | no | no | no |
| GABRB3 | 456 | F | 0 | 0 | yes | no | no | yes | no | no | no | no | no | no | no | no | no |
| GABRB3 | 457 | P | 1 | 0 | yes | no | no | yes | no | no | no | no | no | no | no | no | no |
| GABRB3 | 458 | F | 0 | 0 | yes | no | no | yes | no | no | no | no | no | no | no | no | no |
| GABRB3 | 459 | T | 0 | 0 | yes | no | no | yes | no | no | no | no | no | no | no | no | no |

| | | | | | | | | | | | | | | | | | |
|---|---|---|---|---|---|---|---|---|---|---|---|---|---|---|---|---|---|
| GABRB3 | 460 | F | 0 | 0 | yes | no | no | yes | no | no | no | no | no | no | no | no | no |
| GABRB3 | 461 | S | 0 | 0 | yes | no | no | yes | no | no | no | no | no | no | no | no | no |
| GABRB3 | 462 | L | 0 | 0 | yes | no | no | yes | no | no | no | no | no | no | no | no | no |
| GABRB3 | 463 | F | 0 | 0 | yes | no | no | yes | no | no | no | no | no | no | no | no | no |
| GABRB3 | 464 | N | 0 | 0 | yes | no | no | yes | no | no | no | no | no | no | no | no | no |
| GABRB3 | 465 | L | 0 | 0 | yes | no | no | yes | no | no | no | no | no | no | no | no | no |
| GABRB3 | 466 | V | 0 | 0 | yes | no | no | yes | no | no | no | no | no | no | no | no | no |
| GABRB3 | 467 | Y | 0 | 0 | yes | no | no | yes | no | no | no | no | no | no | no | no | no |
| GABRB3 | 468 | W | 0 | 0 | yes | no | no | yes | no | no | no | no | no | no | no | no | no |
| GABRB3 | 469 | L | 0 | 0 | yes | no | no | yes | no | no | no | no | no | no | no | no | no |
| GABRB3 | 470 | Y | 0 | 0 | yes | no | no | yes | no | no | no | no | no | no | no | no | no |
| GABRB3 | 471 | Y | 0 | 0 | no | yes | no | no | no | no | no | no | no | no | no | no | no |
| GABRB3 | 472 | V | 0 | 0 | no | yes | no | no | no | no | no | no | no | no | no | no | no |
| GABRG2 | 65 | D | 0 | 0 | no | yes | no | no | yes | no | no | no | no | no | no | no | no |
| GABRG2 | 66 | V | 0 | 0 | yes | no | no | no | yes | no | no | no | no | no | no | no | no |
| GABRG2 | 67 | T | 0 | 0 | yes | no | no | no | yes | no | no | no | no | no | no | no | no |
| GABRG2 | 68 | V | 0 | 0 | yes | no | no | no | yes | no | no | no | no | no | no | no | no |
| GABRG2 | 69 | I | 1 | 0 | yes | no | no | no | yes | no | no | no | no | no | no | no | no |
| GABRG2 | 70 | L | 0 | 0 | yes | no | no | no | yes | no | no | no | no | no | no | no | no |
| GABRG2 | 71 | N | 0 | 0 | yes | no | no | no | yes | no | no | no | no | no | no | no | no |
| GABRG2 | 72 | N | 1 | 0 | yes | no | no | no | yes | no | no | no | no | no | no | no | no |
| GABRG2 | 73 | L | 0 | 0 | yes | no | no | no | yes | no | no | no | no | no | no | no | no |
| GABRG2 | 74 | L | 0 | 1 | no | yes | no | no | yes | no | no | no | no | no | no | no | no |
| GABRG2 | 75 | E | 0 | 0 | no | yes | no | no | yes | no | no | no | no | no | no | no | no |
| GABRG2 | 76 | G | 0 | 0 | no | yes | no | no | yes | no | no | no | yes | no | no | no | no |
| GABRG2 | 77 | Y | 0 | 0 | no | yes | no | no | yes | no | no | no | no | no | no | no | no |
| GABRG2 | 78 | D | 0 | 0 | no | yes | no | no | yes | no | no | no | no | no | no | no | no |
| GABRG2 | 79 | N | 1 | 1 | no | yes | no | no | yes | no | no | no | yes | no | no | no | no |
| GABRG2 | 80 | K | 0 | 0 | no | yes | no | no | yes | no | no | no | yes | no | no | no | no |
| GABRG2 | 81 | L | 0 | 0 | no | yes | no | no | yes | no | no | no | yes | no | no | no | no |
| GABRG2 | 82 | R | 0 | 1 | no | yes | no | no | yes | no | no | no | yes | no | no | no | no |
| GABRG2 | 83 | P | 0 | 1 | no | yes | no | no | yes | no | no | no | no | no | no | no | no |
| GABRG2 | 84 | D | 0 | 0 | no | yes | no | no | yes | no | no | no | yes | no | no | no | no |

| | | | | | | | | | | | | | | | | | |
|---|---|---|---|---|---|---|---|---|---|---|---|---|---|---|---|---|---|
| GABRG2 | 85 | I | 0 | 0 | no | yes | no | no | yes | no | no | no | yes | no | no | no | no |
| GABRG2 | 86 | G | 1 | 0 | no | yes | no | no | yes | no | no | no | yes | no | no | no | no |
| GABRG2 | 87 | V | 0 | 0 | no | yes | no | no | yes | no | no | no | no | no | no | no | no |
| GABRG2 | 88 | K | 0 | 0 | no | yes | no | no | yes | no | no | no | no | no | no | no | no |
| GABRG2 | 89 | P | 0 | 0 | no | yes | no | no | yes | no | no | no | no | no | no | no | no |
| GABRG2 | 90 | T | 1 | 0 | no | no | yes | no | yes | no | no | no | no | no | no | no | no |
| GABRG2 | 91 | L | 1 | 0 | no | no | yes | no | yes | no | no | no | no | no | no | no | no |
| GABRG2 | 92 | I | 0 | 0 | no | no | yes | no | yes | no | no | no | no | no | no | no | no |
| GABRG2 | 93 | H | 0 | 0 | no | no | yes | no | yes | no | no | no | no | no | no | no | no |
| GABRG2 | 94 | T | 1 | 1 | no | no | yes | no | yes | no | no | no | no | no | no | no | no |
| GABRG2 | 95 | D | 0 | 0 | no | no | yes | no | yes | no | no | no | no | no | no | no | no |
| GABRG2 | 96 | M | 0 | 0 | no | no | yes | no | yes | no | no | no | no | no | no | no | no |
| GABRG2 | 97 | Y | 0 | 0 | no | no | yes | no | yes | no | no | no | no | no | no | no | no |
| GABRG2 | 98 | V | 1 | 0 | no | no | yes | no | yes | no | no | no | no | no | no | no | no |
| GABRG2 | 99 | N | 0 | 0 | no | no | yes | no | yes | no | no | no | no | no | no | no | no |
| GABRG2 | 100 | S | 0 | 0 | no | no | yes | no | yes | no | no | no | no | no | no | no | no |
| GABRG2 | 101 | I | 0 | 0 | no | no | yes | no | yes | no | no | no | no | no | no | no | no |
| GABRG2 | 102 | G | 0 | 0 | no | yes | no | no | yes | no | no | no | no | no | no | no | no |
| GABRG2 | 103 | P | 0 | 0 | no | yes | no | no | yes | no | no | no | no | no | no | no | no |
| GABRG2 | 104 | V | 0 | 0 | no | yes | no | no | yes | no | no | no | no | no | no | no | no |
| GABRG2 | 105 | N | 0 | 0 | no | yes | no | no | yes | no | yes | no | no | no | no | no | no |
| GABRG2 | 106 | A | 1 | 1 | no | yes | no | no | yes | no | no | no | no | no | no | no | no |
| GABRG2 | 107 | I | 0 | 1 | no | yes | no | no | yes | no | yes | no | no | no | no | no | no |
| GABRG2 | 108 | N | 0 | 0 | no | yes | no | no | yes | no | yes | no | no | no | no | no | no |
| GABRG2 | 109 | M | 0 | 0 | no | yes | no | no | yes | no | no | no | no | no | no | no | no |
| GABRG2 | 110 | E | 0 | 0 | no | no | yes | no | yes | no | no | no | no | no | no | no | no |
| GABRG2 | 111 | Y | 0 | 0 | no | no | yes | no | yes | no | no | no | no | no | no | no | no |
| GABRG2 | 112 | T | 0 | 0 | no | no | yes | no | yes | no | no | no | no | no | no | no | no |
| GABRG2 | 113 | I | 0 | 0 | no | no | yes | no | yes | no | no | no | no | no | no | no | no |
| GABRG2 | 114 | D | 0 | 0 | no | no | yes | no | yes | no | no | no | no | no | no | no | no |
| GABRG2 | 115 | I | 0 | 0 | no | no | yes | no | yes | no | no | no | no | no | no | no | no |
| GABRG2 | 116 | F | 0 | 0 | no | no | yes | no | yes | no | no | no | no | no | no | no | no |
| GABRG2 | 117 | F | 0 | 0 | no | no | yes | no | yes | no | no | no | no | no | no | no | no |

| | | | | | | | | | | | | | | | | | |
|---|---|---|---|---|---|---|---|---|---|---|---|---|---|---|---|---|---|
| GABRG2 | 118 | A | 1 | 0 | no | no | yes | no | yes | no | no | no | no | no | no | no | no |
| GABRG2 | 119 | Q | 0 | 0 | no | no | yes | no | yes | no | no | no | no | no | no | no | no |
| GABRG2 | 120 | T | 2 | 0 | no | no | yes | no | yes | no | no | no | no | no | no | no | no |
| GABRG2 | 121 | W | 0 | 0 | no | no | yes | no | yes | no | no | no | no | no | no | no | no |
| GABRG2 | 122 | Y | 0 | 0 | no | no | yes | no | yes | no | no | no | no | no | no | no | no |
| GABRG2 | 123 | D | 0 | 0 | no | yes | no | no | yes | no | no | no | no | no | no | no | no |
| GABRG2 | 124 | R | 1 | 0 | yes | no | no | no | yes | no | no | no | no | no | no | no | no |
| GABRG2 | 125 | R | 0 | 0 | yes | no | no | no | yes | no | no | no | no | no | no | no | no |
| GABRG2 | 126 | L | 0 | 0 | yes | no | no | no | yes | no | no | no | no | no | no | no | no |
| GABRG2 | 127 | K | 0 | 0 | no | yes | no | no | yes | no | no | no | no | no | no | no | no |
| GABRG2 | 128 | F | 0 | 0 | no | yes | no | no | yes | no | no | no | no | no | no | no | no |
| GABRG2 | 129 | N | 0 | 0 | no | yes | no | no | yes | no | no | no | no | no | no | no | no |
| GABRG2 | 130 | S | 1 | 0 | no | yes | no | no | yes | no | no | no | no | no | no | no | no |
| GABRG2 | 131 | T | 0 | 0 | no | yes | no | no | yes | no | no | no | no | no | no | no | no |
| GABRG2 | 132 | I | 1 | 0 | no | yes | no | no | yes | no | no | no | no | no | no | no | no |
| GABRG2 | 133 | K | 0 | 0 | no | yes | no | no | yes | no | no | no | no | no | no | no | no |
| GABRG2 | 134 | V | 0 | 0 | no | no | yes | no | yes | no | no | no | no | no | no | no | no |
| GABRG2 | 135 | L | 1 | 0 | no | no | yes | no | yes | no | no | no | no | no | no | no | no |
| GABRG2 | 136 | R | 2 | 0 | no | no | yes | no | yes | no | no | no | no | no | no | no | no |
| GABRG2 | 137 | L | 0 | 0 | no | no | yes | no | yes | no | no | no | no | no | no | no | no |
| GABRG2 | 138 | N | 1 | 0 | no | yes | no | no | yes | no | no | no | no | no | no | no | no |
| GABRG2 | 139 | S | 0 | 0 | no | yes | no | no | yes | no | no | no | no | no | no | no | no |
| GABRG2 | 140 | N | 0 | 1 | no | yes | no | no | yes | no | no | no | no | no | no | no | no |
| GABRG2 | 141 | M | 0 | 0 | no | yes | no | no | yes | no | no | no | no | no | no | no | no |
| GABRG2 | 142 | V | 1 | 0 | no | yes | no | no | yes | no | no | no | no | no | no | no | no |
| GABRG2 | 143 | G | 1 | 0 | no | yes | no | no | yes | no | no | no | yes | no | no | no | no |
| GABRG2 | 144 | K | 0 | 0 | no | yes | no | no | yes | no | no | no | no | no | no | no | no |
| GABRG2 | 145 | I | 0 | 0 | no | yes | no | no | yes | no | no | no | no | no | no | no | no |
| GABRG2 | 146 | W | 0 | 0 | no | yes | no | no | yes | no | no | no | no | no | no | no | no |
| GABRG2 | 147 | I | 0 | 0 | no | yes | no | no | yes | no | no | no | no | no | no | no | no |
| GABRG2 | 148 | P | 0 | 0 | no | yes | no | no | yes | no | no | no | yes | no | no | no | no |
| GABRG2 | 149 | D | 0 | 0 | no | yes | no | no | yes | no | no | no | yes | no | no | no | no |
| GABRG2 | 150 | T | 0 | 0 | no | no | yes | no | yes | no | no | no | yes | no | no | no | no |

| | | | | | | | | | | | | | | | | | |
|---|---|---|---|---|---|---|---|---|---|---|---|---|---|---|---|---|---|
| GABRG2 | 151 | F | 0 | 0 | no | no | yes | no | yes | no | no | no | yes | no | no | no | no |
| GABRG2 | 152 | F | 0 | 0 | no | no | yes | no | yes | no | no | no | yes | no | no | no | no |
| GABRG2 | 153 | R | 0 | 0 | no | yes | no | no | yes | no | no | no | yes | no | no | no | no |
| GABRG2 | 154 | N | 0 | 0 | no | yes | no | no | yes | no | no | no | no | no | no | no | no |
| GABRG2 | 155 | S | 0 | 0 | no | no | yes | no | yes | no | no | no | yes | no | no | no | no |
| GABRG2 | 156 | K | 1 | 0 | no | no | yes | no | yes | no | no | no | yes | no | no | no | no |
| GABRG2 | 157 | K | 1 | 0 | no | no | yes | no | yes | no | yes | no | no | no | no | no | no |
| GABRG2 | 158 | A | 0 | 0 | no | no | yes | no | yes | no | no | no | yes | no | no | no | no |
| GABRG2 | 159 | D | 0 | 0 | no | no | yes | no | yes | no | yes | no | no | no | no | no | no |
| GABRG2 | 160 | A | 0 | 0 | no | no | yes | no | yes | no | no | no | no | no | no | no | no |
| GABRG2 | 161 | H | 1 | 0 | no | yes | no | no | yes | no | no | no | no | no | no | no | no |
| GABRG2 | 162 | W | 0 | 0 | no | yes | no | no | yes | no | yes | no | no | no | no | no | no |
| GABRG2 | 163 | I | 0 | 0 | no | yes | no | no | yes | no | no | no | no | no | no | no | no |
| GABRG2 | 164 | T | 0 | 0 | no | yes | no | no | yes | no | no | no | no | no | no | no | no |
| GABRG2 | 165 | T | 0 | 0 | no | yes | no | no | yes | no | no | no | no | no | no | no | no |
| GABRG2 | 166 | P | 0 | 0 | no | yes | no | no | yes | no | no | no | no | no | no | no | no |
| GABRG2 | 167 | N | 0 | 1 | no | yes | no | no | yes | no | no | no | no | no | no | no | no |
| GABRG2 | 168 | R | 0 | 0 | no | no | yes | no | yes | no | no | no | yes | no | no | no | no |
| GABRG2 | 169 | M | 0 | 0 | no | no | yes | no | yes | no | no | no | no | no | no | no | no |
| GABRG2 | 170 | L | 0 | 0 | no | no | yes | no | yes | no | no | no | no | no | no | no | no |
| GABRG2 | 171 | R | 0 | 0 | no | no | yes | no | yes | no | no | no | no | no | no | no | no |
| GABRG2 | 172 | I | 0 | 0 | no | no | yes | no | yes | no | no | no | no | no | no | no | no |
| GABRG2 | 173 | W | 0 | 0 | no | no | yes | no | yes | no | no | no | no | no | no | no | no |
| GABRG2 | 174 | N | 0 | 0 | no | yes | no | no | yes | no | no | no | no | no | no | no | no |
| GABRG2 | 175 | D | 0 | 0 | no | yes | no | no | yes | no | no | no | no | no | no | no | no |
| GABRG2 | 176 | G | 0 | 0 | no | yes | no | no | yes | no | no | no | no | no | no | no | no |
| GABRG2 | 177 | R | 1 | 2 | no | no | yes | no | yes | no | no | no | no | no | no | no | no |
| GABRG2 | 178 | V | 0 | 0 | no | no | yes | no | yes | no | no | no | no | no | no | no | no |
| GABRG2 | 179 | L | 0 | 0 | no | no | yes | no | yes | no | no | no | no | no | no | no | no |
| GABRG2 | 180 | Y | 0 | 0 | no | no | yes | no | yes | no | no | no | no | no | no | no | no |
| GABRG2 | 181 | T | 0 | 0 | no | no | yes | no | yes | no | no | no | no | no | no | no | no |
| GABRG2 | 182 | L | 0 | 0 | no | no | yes | no | yes | no | no | no | no | no | no | no | no |
| GABRG2 | 183 | R | 0 | 0 | no | no | yes | no | yes | no | no | no | no | no | no | no | no |

| | | | | | | | | | | | | | | | | | |
|---|---|---|---|---|---|---|---|---|---|---|---|---|---|---|---|---|---|
| GABRG2 | 184 | L | 0 | 0 | no | no | yes | no | yes | no | no | no | yes | no | no | no | no |
| GABRG2 | 185 | T | 0 | 0 | no | no | yes | no | yes | no | no | no | no | no | no | no | no |
| GABRG2 | 186 | I | 1 | 0 | no | no | yes | no | yes | no | no | no | no | no | no | no | no |
| GABRG2 | 187 | D | 0 | 0 | no | no | yes | no | yes | no | no | no | no | no | no | no | no |
| GABRG2 | 188 | A | 1 | 0 | no | no | yes | no | yes | no | no | no | no | no | no | no | no |
| GABRG2 | 189 | E | 0 | 0 | no | no | yes | no | yes | no | no | no | no | no | no | no | no |
| GABRG2 | 190 | C | 0 | 0 | no | yes | no | no | yes | no | no | no | no | no | no | no | no |
| GABRG2 | 191 | Q | 0 | 0 | no | yes | no | no | yes | no | no | no | yes | no | no | no | no |
| GABRG2 | 192 | L | 0 | 0 | no | yes | no | no | yes | no | no | no | no | no | no | no | no |
| GABRG2 | 193 | Q | 0 | 0 | no | yes | no | no | yes | no | no | no | yes | no | no | no | no |
| GABRG2 | 194 | L | 0 | 0 | no | yes | no | no | yes | no | no | no | no | no | no | no | no |
| GABRG2 | 195 | H | 0 | 0 | no | yes | no | no | yes | no | no | no | no | no | no | no | no |
| GABRG2 | 196 | N | 0 | 0 | no | yes | no | no | yes | no | no | no | no | no | no | no | no |
| GABRG2 | 197 | F | 0 | 0 | no | yes | no | no | yes | no | no | no | no | no | no | no | no |
| GABRG2 | 198 | P | 0 | 0 | no | yes | no | no | yes | no | no | no | no | no | no | no | no |
| GABRG2 | 199 | M | 1 | 1 | no | yes | no | no | yes | no | no | no | no | no | no | no | no |
| GABRG2 | 200 | D | 1 | 0 | no | yes | no | no | yes | no | no | no | no | no | no | no | no |
| GABRG2 | 201 | E | 0 | 0 | no | no | yes | no | yes | no | no | no | no | no | no | no | no |
| GABRG2 | 202 | H | 0 | 0 | no | no | yes | no | yes | no | no | no | no | no | no | no | no |
| GABRG2 | 203 | S | 1 | 0 | no | no | yes | no | yes | no | no | no | no | no | no | no | no |
| GABRG2 | 204 | C | 0 | 0 | no | no | yes | no | yes | no | no | no | no | no | no | no | no |
| GABRG2 | 205 | P | 0 | 0 | no | no | yes | no | yes | no | no | no | no | no | no | no | no |
| GABRG2 | 206 | L | 0 | 0 | no | no | yes | no | yes | no | no | no | no | no | no | no | no |
| GABRG2 | 207 | E | 0 | 0 | no | no | yes | no | yes | no | no | no | no | no | no | no | no |
| GABRG2 | 208 | F | 0 | 0 | no | no | yes | no | yes | no | no | no | no | no | no | no | no |
| GABRG2 | 209 | S | 0 | 0 | no | no | yes | no | yes | no | no | no | no | no | no | no | no |
| GABRG2 | 210 | S | 0 | 0 | no | no | yes | no | yes | no | no | no | no | no | no | no | no |
| GABRG2 | 211 | Y | 0 | 0 | no | yes | no | no | yes | no | no | no | yes | no | no | no | no |
| GABRG2 | 212 | G | 0 | 0 | no | yes | no | no | yes | no | no | no | yes | no | no | no | no |
| GABRG2 | 213 | Y | 0 | 0 | no | yes | no | no | yes | no | no | no | yes | no | no | no | no |
| GABRG2 | 214 | P | 0 | 0 | no | yes | no | no | yes | no | no | no | yes | no | no | no | no |
| GABRG2 | 215 | R | 1 | 0 | no | yes | no | no | yes | no | no | no | no | no | no | no | no |
| GABRG2 | 216 | E | 0 | 0 | no | yes | no | no | yes | no | no | no | no | no | no | no | no |

| | | | | | | | | | | | | | | | | | |
|---|---|---|---|---|---|---|---|---|---|---|---|---|---|---|---|---|---|
| GABRG2 | 217 | E | 0 | 0 | no | yes | no | no | yes | no | no | no | yes | no | no | no | no |
| GABRG2 | 218 | I | 0 | 1 | no | no | yes | no | yes | no | no | no | no | no | no | no | no |
| GABRG2 | 219 | V | 1 | 0 | no | no | yes | no | yes | no | no | no | no | no | no | no | no |
| GABRG2 | 220 | Y | 0 | 0 | no | no | yes | no | yes | no | no | no | no | no | no | no | no |
| GABRG2 | 221 | Q | 0 | 0 | no | no | yes | no | yes | no | no | no | no | no | no | no | no |
| GABRG2 | 222 | W | 0 | 0 | no | no | yes | no | yes | no | no | no | no | no | no | no | no |
| GABRG2 | 223 | K | 0 | 0 | no | no | yes | no | yes | no | no | no | no | no | no | no | no |
| GABRG2 | 224 | R | 1 | 0 | no | no | yes | no | yes | no | no | no | no | no | no | no | no |
| GABRG2 | 225 | S | 0 | 0 | no | no | yes | no | yes | no | no | no | no | no | no | no | no |
| GABRG2 | 226 | S | 0 | 0 | no | no | yes | no | yes | no | no | no | no | no | no | no | no |
| GABRG2 | 227 | V | 0 | 0 | no | no | yes | no | yes | no | no | no | no | no | no | no | no |
| GABRG2 | 228 | E | 0 | 0 | no | no | yes | no | yes | no | no | no | no | no | no | no | no |
| GABRG2 | 229 | V | 0 | 0 | no | no | yes | no | yes | no | no | no | no | no | no | no | no |
| GABRG2 | 230 | G | 0 | 0 | no | yes | no | no | yes | no | no | no | no | no | no | no | no |
| GABRG2 | 231 | D | 1 | 0 | no | yes | no | no | yes | no | no | no | no | no | no | no | no |
| GABRG2 | 232 | T | 0 | 0 | no | yes | no | no | yes | no | no | no | no | no | no | no | no |
| GABRG2 | 233 | R | 0 | 0 | no | yes | no | no | yes | no | no | no | no | no | no | no | no |
| GABRG2 | 234 | S | 0 | 0 | no | yes | no | no | yes | no | no | no | no | no | no | no | no |
| GABRG2 | 235 | W | 0 | 0 | no | yes | no | no | yes | no | no | no | no | no | no | no | no |
| GABRG2 | 236 | R | 0 | 0 | no | yes | no | no | yes | no | no | no | no | no | no | no | no |
| GABRG2 | 237 | L | 0 | 0 | no | yes | no | no | yes | no | no | no | no | no | no | no | no |
| GABRG2 | 238 | Y | 0 | 0 | no | yes | no | no | yes | no | no | no | no | no | no | no | no |
| GABRG2 | 239 | Q | 0 | 0 | no | yes | no | no | yes | no | no | no | no | no | no | no | no |
| GABRG2 | 240 | F | 0 | 0 | no | no | yes | no | yes | no | no | no | no | no | no | no | no |
| GABRG2 | 241 | S | 0 | 0 | no | no | yes | no | yes | no | no | no | no | no | no | no | no |
| GABRG2 | 242 | F | 0 | 0 | no | no | yes | no | yes | no | no | no | no | no | no | no | no |
| GABRG2 | 243 | V | 2 | 0 | no | no | yes | no | yes | no | no | no | no | no | no | no | no |
| GABRG2 | 244 | G | 0 | 0 | no | no | yes | no | yes | no | no | no | no | no | no | no | no |
| GABRG2 | 245 | L | 0 | 0 | no | no | yes | no | yes | no | no | no | no | no | no | no | no |
| GABRG2 | 246 | R | 0 | 0 | no | no | yes | no | yes | no | no | no | no | no | no | no | no |
| GABRG2 | 247 | N | 0 | 0 | no | no | yes | no | yes | no | no | no | no | no | no | no | no |
| GABRG2 | 248 | T | 0 | 0 | no | no | yes | no | yes | no | no | no | no | no | no | no | no |
| GABRG2 | 249 | T | 0 | 0 | no | no | yes | no | yes | no | no | no | no | no | no | no | no |

| | | | | | | | | | | | | | | | | | |
|---|---|---|---|---|---|---|---|---|---|---|---|---|---|---|---|---|---|
| GABRG2 | 250 | E | 1 | 0 | no | no | yes | no | yes | no | no | no | no | no | no | no | no |
| GABRG2 | 251 | V | 1 | 0 | no | no | yes | no | yes | no | no | no | no | no | no | no | no |
| GABRG2 | 252 | V | 0 | 0 | no | no | yes | no | yes | no | no | no | no | no | no | no | no |
| GABRG2 | 253 | K | 0 | 0 | no | no | yes | no | yes | no | no | no | no | no | no | no | no |
| GABRG2 | 254 | T | 0 | 0 | no | yes | no | no | yes | no | no | no | yes | no | no | no | no |
| GABRG2 | 255 | T | 1 | 0 | no | yes | no | no | yes | no | no | no | yes | no | no | no | no |
| GABRG2 | 256 | S | 0 | 0 | no | yes | no | no | yes | no | no | no | yes | no | no | no | no |
| GABRG2 | 257 | G | 0 | 0 | no | yes | no | no | yes | no | no | no | no | no | no | no | no |
| GABRG2 | 258 | D | 0 | 0 | no | no | yes | no | yes | no | no | no | no | no | no | no | no |
| GABRG2 | 259 | Y | 0 | 0 | no | no | yes | no | yes | no | no | no | yes | no | no | no | no |
| GABRG2 | 260 | V | 0 | 0 | no | no | yes | no | yes | no | no | no | no | no | no | no | no |
| GABRG2 | 261 | V | 0 | 0 | no | no | yes | no | yes | no | no | no | no | no | no | no | no |
| GABRG2 | 262 | M | 0 | 0 | no | no | yes | no | yes | no | no | no | no | no | no | no | no |
| GABRG2 | 263 | S | 0 | 0 | no | no | yes | no | yes | no | no | no | no | no | no | no | no |
| GABRG2 | 264 | V | 0 | 0 | no | no | yes | no | yes | no | no | no | no | no | no | no | no |
| GABRG2 | 265 | Y | 1 | 0 | no | no | yes | no | yes | no | no | no | no | no | no | no | no |
| GABRG2 | 266 | F | 0 | 0 | no | no | yes | no | yes | no | no | no | no | no | no | no | no |
| GABRG2 | 267 | D | 0 | 0 | no | no | yes | no | yes | no | no | no | no | no | no | no | no |
| GABRG2 | 268 | L | 0 | 0 | no | no | yes | no | yes | no | no | no | no | no | no | no | no |
| GABRG2 | 269 | S | 0 | 0 | no | no | yes | no | yes | no | no | no | no | no | no | no | no |
| GABRG2 | 270 | R | 0 | 0 | no | no | yes | no | yes | no | no | no | no | no | no | no | no |
| GABRG2 | 271 | R | 0 | 0 | no | yes | no | no | yes | no | no | no | no | no | no | no | no |
| GABRG2 | 272 | M | 0 | 0 | no | yes | no | no | yes | no | no | no | no | no | no | no | no |
| GABRG2 | 273 | G | 0 | 0 | no | yes | no | no | yes | no | no | no | no | no | no | no | no |
| GABRG2 | 274 | Y | 0 | 1 | no | yes | no | no | no | no | no | no | no | no | no | no | no |
| GABRG2 | 275 | F | 0 | 0 | no | yes | no | no | no | no | no | no | no | no | no | no | no |
| GABRG2 | 276 | T | 0 | 0 | yes | no | no | yes | no | no | no | no | no | no | no | no | no |
| GABRG2 | 277 | I | 0 | 0 | yes | no | no | yes | no | no | no | no | no | no | no | no | no |
| GABRG2 | 278 | Q | 0 | 0 | yes | no | no | yes | no | no | no | no | no | no | no | no | no |
| GABRG2 | 279 | T | 1 | 0 | yes | no | no | yes | no | no | no | no | no | no | no | no | no |
| GABRG2 | 280 | Y | 0 | 0 | yes | no | no | yes | no | no | no | no | no | no | no | no | no |
| GABRG2 | 281 | I | 0 | 0 | yes | no | no | yes | no | no | no | no | no | no | no | no | no |
| GABRG2 | 282 | P | 0 | 2 | yes | no | no | yes | no | no | no | no | no | no | no | no | no |

| | | | | | | | | | | | | | | | | | |
|---|---|---|---|---|---|---|---|---|---|---|---|---|---|---|---|---|---|
| GABRG2 | 283 | C | 0 | 0 | yes | no | no | yes | no | no | no | no | no | no | no | no | no |
| GABRG2 | 284 | T | 0 | 0 | yes | no | no | yes | no | no | no | no | no | no | no | no | no |
| GABRG2 | 285 | L | 0 | 0 | yes | no | no | yes | no | no | no | no | no | no | no | no | no |
| GABRG2 | 286 | I | 0 | 0 | yes | no | no | yes | no | no | no | no | no | no | no | no | no |
| GABRG2 | 287 | V | 0 | 0 | yes | no | no | yes | no | no | no | no | no | no | no | no | no |
| GABRG2 | 288 | V | 1 | 0 | yes | no | no | yes | no | no | no | no | no | no | no | no | no |
| GABRG2 | 289 | L | 0 | 0 | yes | no | no | yes | no | no | no | no | no | no | no | no | no |
| GABRG2 | 290 | S | 0 | 0 | yes | no | no | yes | no | no | no | no | no | no | no | no | no |
| GABRG2 | 291 | W | 0 | 0 | yes | no | no | yes | no | no | no | no | no | no | no | no | no |
| GABRG2 | 292 | V | 0 | 0 | yes | no | no | yes | no | no | no | no | no | no | no | no | no |
| GABRG2 | 293 | S | 0 | 0 | yes | no | no | yes | no | no | no | no | no | no | no | no | no |
| GABRG2 | 294 | F | 0 | 0 | yes | no | no | yes | no | no | no | no | no | no | no | no | no |
| GABRG2 | 295 | W | 0 | 0 | yes | no | no | yes | no | no | no | no | no | no | no | no | no |
| GABRG2 | 296 | I | 1 | 0 | no | yes | no | yes | no | no | no | no | no | no | no | no | no |
| GABRG2 | 297 | N | 0 | 0 | no | yes | no | no | no | no | no | no | no | no | no | no | no |
| GABRG2 | 298 | K | 0 | 0 | no | yes | no | no | no | no | no | no | no | no | no | no | no |
| GABRG2 | 299 | D | 0 | 0 | no | yes | no | no | no | no | yes | no | no | no | no | no | no |
| GABRG2 | 300 | A | 1 | 0 | no | yes | no | no | no | no | no | no | no | no | no | no | no |
| GABRG2 | 301 | V | 0 | 0 | yes | no | no | no | no | no | no | no | yes | no | no | no | no |
| GABRG2 | 302 | P | 0 | 1 | yes | no | no | yes | no | no | yes | no | yes | no | no | no | no |
| GABRG2 | 303 | A | 0 | 0 | yes | no | no | yes | no | no | no | no | no | no | no | no | no |
| GABRG2 | 304 | R | 0 | 1 | yes | no | no | yes | no | no | no | no | no | no | no | no | no |
| GABRG2 | 305 | T | 0 | 0 | yes | no | no | yes | no | no | no | no | yes | no | no | no | no |
| GABRG2 | 306 | S | 0 | 1 | yes | no | no | yes | no | no | yes | no | no | no | no | no | no |
| GABRG2 | 307 | L | 0 | 1 | yes | no | no | yes | no | no | no | no | no | no | no | no | no |
| GABRG2 | 308 | G | 0 | 0 | yes | no | no | yes | no | no | no | no | no | no | no | no | no |
| GABRG2 | 309 | I | 0 | 0 | yes | no | no | yes | no | no | no | no | yes | no | no | no | no |
| GABRG2 | 310 | T | 0 | 0 | yes | no | no | yes | no | no | yes | no | no | no | no | no | no |
| GABRG2 | 311 | T | 0 | 0 | yes | no | no | yes | no | no | no | no | no | no | no | no | no |
| GABRG2 | 312 | V | 0 | 0 | yes | no | no | yes | no | no | no | no | yes | no | no | no | no |
| GABRG2 | 313 | L | 0 | 0 | yes | no | no | yes | no | no | yes | no | yes | no | no | no | no |
| GABRG2 | 314 | T | 0 | 0 | yes | no | no | yes | no | no | no | no | no | no | no | no | no |
| GABRG2 | 315 | M | 1 | 0 | yes | no | no | yes | no | no | no | no | no | no | no | no | no |

| | | | | | | | | | | | | | | | | | |
|---|---|---|---|---|---|---|---|---|---|---|---|---|---|---|---|---|---|
| GABRG2 | 316 | T | 0 | 0 | yes | no | no | yes | no | no | no | no | yes | no | no | no | no |
| GABRG2 | 317 | T | 0 | 0 | yes | no | no | yes | no | no | yes | no | no | no | no | no | no |
| GABRG2 | 318 | L | 0 | 0 | yes | no | no | yes | no | no | no | no | no | no | no | no | no |
| GABRG2 | 319 | S | 0 | 0 | yes | no | no | yes | no | no | no | no | no | no | no | no | no |
| GABRG2 | 320 | T | 0 | 0 | no | yes | no | yes | no | no | yes | no | yes | no | no | no | no |
| GABRG2 | 321 | I | 0 | 0 | yes | no | no | yes | no | no | no | no | no | no | no | no | no |
| GABRG2 | 322 | A | 0 | 0 | yes | no | no | no | no | no | no | no | no | no | no | no | no |
| GABRG2 | 323 | R | 0 | 2 | yes | no | no | no | no | no | no | no | yes | no | no | no | no |
| GABRG2 | 324 | K | 0 | 0 | no | yes | no | no | no | no | no | no | no | no | no | no | no |
| GABRG2 | 325 | S | 0 | 0 | no | yes | no | no | no | no | no | no | no | no | no | no | no |
| GABRG2 | 326 | L | 0 | 0 | no | yes | no | no | no | no | no | no | no | no | no | no | no |
| GABRG2 | 327 | P | 0 | 0 | no | yes | no | no | no | no | no | no | no | no | no | no | no |
| GABRG2 | 328 | K | 0 | 1 | no | yes | no | no | no | no | no | no | yes | no | no | no | no |
| GABRG2 | 329 | V | 1 | 1 | no | yes | no | no | no | no | no | no | yes | no | no | no | no |
| GABRG2 | 330 | S | 0 | 0 | no | yes | no | no | no | no | no | no | yes | no | no | no | no |
| GABRG2 | 331 | Y | 1 | 0 | no | yes | no | no | no | no | no | no | no | no | no | no | no |
| GABRG2 | 332 | V | 0 | 0 | no | yes | no | no | no | no | no | no | no | no | no | no | no |
| GABRG2 | 333 | T | 0 | 0 | no | yes | no | no | no | no | no | no | no | no | no | no | no |
| GABRG2 | 334 | A | 2 | 0 | yes | no | no | yes | no | no | no | no | no | no | no | no | no |
| GABRG2 | 335 | M | 0 | 0 | yes | no | no | yes | no | no | no | no | no | no | no | no | no |
| GABRG2 | 336 | D | 0 | 0 | yes | no | no | yes | no | no | no | no | yes | no | no | no | no |
| GABRG2 | 337 | L | 0 | 0 | yes | no | no | yes | no | no | no | no | no | no | no | no | no |
| GABRG2 | 338 | F | 0 | 0 | yes | no | no | yes | no | no | no | no | no | no | no | no | no |
| GABRG2 | 339 | V | 0 | 0 | yes | no | no | yes | no | no | no | no | no | no | no | no | no |
| GABRG2 | 340 | S | 0 | 0 | yes | no | no | yes | no | no | no | no | yes | no | no | no | no |
| GABRG2 | 341 | V | 0 | 0 | yes | no | no | yes | no | no | no | no | no | no | no | no | no |
| GABRG2 | 342 | C | 0 | 0 | yes | no | no | yes | no | no | no | no | no | no | no | no | no |
| GABRG2 | 343 | F | 0 | 1 | yes | no | no | yes | no | no | no | no | yes | no | no | no | no |
| GABRG2 | 344 | I | 0 | 0 | yes | no | no | yes | no | no | no | no | no | no | no | no | no |
| GABRG2 | 345 | F | 0 | 0 | yes | no | no | yes | no | no | no | no | no | no | no | no | no |
| GABRG2 | 346 | V | 0 | 0 | yes | no | no | yes | no | no | no | no | no | no | no | no | no |
| GABRG2 | 347 | F | 0 | 0 | yes | no | no | yes | no | no | no | no | yes | no | no | no | no |
| GABRG2 | 348 | S | 0 | 0 | yes | no | no | yes | no | no | no | no | no | no | no | no | no |

| | | | | | | | | | | | | | | | | | |
|---|---|---|---|---|---|---|---|---|---|---|---|---|---|---|---|---|---|
| GABRG2 | 349 | A | 1 | 0 | yes | no | no | yes | no | no | no | no | no | no | no | no | no |
| GABRG2 | 350 | L | 0 | 0 | yes | no | no | yes | no | no | no | no | yes | no | no | no | no |
| GABRG2 | 351 | V | 0 | 0 | yes | no | no | yes | no | no | no | no | no | no | no | no | no |
| GABRG2 | 352 | E | 0 | 0 | yes | no | no | yes | no | no | no | no | no | no | no | no | no |
| GABRG2 | 353 | Y | 0 | 0 | yes | no | no | yes | no | no | no | no | no | no | no | no | no |
| GABRG2 | 354 | G | 0 | 1 | yes | no | no | yes | no | no | no | no | no | no | no | no | no |
| GABRG2 | 355 | T | 0 | 0 | yes | no | no | yes | no | no | no | no | no | no | no | no | no |
| GABRG2 | 356 | L | 0 | 0 | yes | no | no | no | no | no | no | no | no | no | no | no | no |
| GABRG2 | 357 | H | 0 | 0 | yes | no | no | no | no | yes | no | no | yes | no | no | no | no |
| GABRG2 | 358 | Y | 0 | 0 | yes | no | no | no | no | yes | no | no | yes | no | no | no | no |
| GABRG2 | 359 | F | 0 | 0 | yes | no | no | no | no | yes | no | no | no | no | no | no | no |
| GABRG2 | 360 | V | 0 | 0 | yes | no | no | no | no | yes | no | no | no | no | no | no | no |
| GABRG2 | 361 | S | 1 | 0 | yes | no | no | no | no | yes | no | no | no | no | no | no | no |
| GABRG2 | 362 | N | 0 | 0 | yes | no | no | no | no | yes | no | no | yes | no | no | no | no |
| GABRG2 | 363 | R | 2 | 1 | no | yes | no | no | no | yes | no | no | no | no | no | no | no |
| GABRG2 | 437 | R | 1 | 0 | no | yes | no | no | no | yes | no | no | no | no | no | no | no |
| GABRG2 | 438 | I | 1 | 0 | no | yes | no | no | no | yes | no | no | no | no | no | no | no |
| GABRG2 | 439 | A | 0 | 0 | no | yes | no | no | no | yes | no | no | no | no | no | no | no |
| GABRG2 | 440 | K | 0 | 0 | no | yes | no | no | no | yes | no | no | no | no | no | no | no |
| GABRG2 | 441 | M | 0 | 0 | yes | no | no | yes | no | yes | no | no | no | no | no | no | no |
| GABRG2 | 442 | D | 0 | 0 | yes | no | no | yes | no | yes | no | no | no | no | no | no | no |
| GABRG2 | 443 | S | 1 | 0 | yes | no | no | no | no | yes | no | no | no | no | no | no | no |
| GABRG2 | 444 | Y | 0 | 0 | yes | no | no | yes | no | no | no | no | no | no | no | no | no |
| GABRG2 | 445 | A | 0 | 0 | yes | no | no | yes | no | no | no | no | no | no | no | no | no |
| GABRG2 | 446 | R | 1 | 1 | yes | no | no | yes | no | no | no | no | no | no | no | no | no |
| GABRG2 | 447 | I | 0 | 0 | yes | no | no | yes | no | no | no | no | no | no | no | no | no |
| GABRG2 | 448 | F | 0 | 0 | yes | no | no | yes | no | no | no | no | no | no | no | no | no |
| GABRG2 | 449 | F | 1 | 0 | yes | no | no | yes | no | no | no | no | no | no | no | no | no |
| GABRG2 | 450 | P | 0 | 0 | yes | no | no | yes | no | no | no | no | no | no | no | no | no |
| GABRG2 | 451 | T | 0 | 0 | yes | no | no | yes | no | no | no | no | no | no | no | no | no |
| GABRG2 | 452 | A | 0 | 0 | yes | no | no | yes | no | no | no | no | no | no | no | no | no |
| GABRG2 | 453 | F | 0 | 0 | yes | no | no | yes | no | no | no | no | no | no | no | no | no |
| GABRG2 | 454 | C | 0 | 0 | yes | no | no | yes | no | no | no | no | no | no | no | no | no |

| | | | | | | | | | | | | | | | | | |
|---|---|---|---|---|---|---|---|---|---|---|---|---|---|---|---|---|---|
| GABRG2 | 455 | L | 0 | 0 | yes | no | no | yes | no | no | no | no | no | no | no | no | no |
| GABRG2 | 456 | F | 0 | 0 | yes | no | no | yes | no | no | no | no | no | no | no | no | no |
| GABRG2 | 457 | N | 0 | 0 | yes | no | no | yes | no | no | no | no | no | no | no | no | no |
| GABRG2 | 458 | L | 0 | 0 | yes | no | no | yes | no | no | no | no | no | no | no | no | no |
| GABRG2 | 459 | V | 1 | 0 | yes | no | no | yes | no | no | no | no | no | no | no | no | no |
| GABRG2 | 460 | Y | 1 | 0 | yes | no | no | yes | no | no | no | no | no | no | no | no | no |
| GABRG2 | 461 | W | 0 | 0 | yes | no | no | yes | no | no | no | no | no | no | no | no | no |
| GABRG2 | 462 | V | 0 | 0 | yes | no | no | yes | no | no | no | no | no | no | no | no | no |
| GABRG2 | 463 | S | 1 | 0 | yes | no | no | yes | no | no | no | no | no | no | no | no | no |
| GABRG2 | 464 | Y | 0 | 0 | yes | no | no | yes | no | no | no | no | no | no | no | no | no |
| GABRG2 | 465 | L | 0 | 0 | yes | no | no | no | no | no | no | no | no | no | no | no | no |
| GABRG2 | 466 | Y | 0 | 0 | no | yes | no | yes | no | no | no | no | no | no | no | no | no |
| GABRG2 | 467 | L | 0 | 0 | no | yes | no | no | no | no | no | no | no | no | no | no | no |
| GLRA1 | 244 | L | 0 | 0 | no | yes | no | no | yes | no | no | no | no | no | no | no | no |
| GLRA1 | 245 | E | 0 | 0 | no | yes | no | no | yes | no | no | no | no | no | no | no | no |
| GLRA1 | 246 | R | 2 | 2 | no | yes | no | no | yes | no | no | no | no | no | no | no | no |
| GLRA1 | 247 | Q | 0 | 0 | no | yes | no | no | yes | no | no | no | no | no | no | no | no |
| GLRA1 | 249 | G | 0 | 0 | yes | no | no | no | yes | no | no | no | no | no | no | no | no |
| GLRA1 | 250 | Y | 0 | 0 | yes | no | no | no | yes | no | no | no | no | no | no | no | no |
| GLRA1 | 251 | Y | 0 | 0 | yes | no | no | no | no | no | no | no | no | no | no | no | no |
| GLRA1 | 252 | L | 0 | 0 | yes | no | no | no | no | no | no | no | no | no | no | no | no |
| GLRA1 | 253 | I | 0 | 0 | yes | no | no | no | no | no | no | no | no | no | no | no | no |
| GLRA1 | 254 | Q | 0 | 1 | yes | no | no | no | no | no | no | no | no | no | no | no | no |
| GLRA1 | 256 | Y | 0 | 1 | yes | no | no | yes | no | no | no | no | no | no | no | no | no |
| GLRA1 | 257 | I | 0 | 0 | yes | no | no | yes | no | no | no | no | no | no | no | no | no |
| GLRA1 | 258 | P | 1 | 1 | yes | no | no | yes | no | no | no | no | no | no | no | no | no |
| GLRA1 | 259 | S | 0 | 2 | yes | no | no | yes | no | no | no | no | no | no | no | no | no |
| GLRA1 | 260 | L | 0 | 0 | yes | no | no | yes | no | no | no | no | no | no | no | no | no |
| GLRA1 | 261 | L | 0 | 0 | yes | no | no | yes | no | no | no | no | no | no | no | no | no |
| GLRA1 | 262 | I | 1 | 0 | yes | no | no | yes | no | no | no | no | no | no | no | no | no |
| GLRA1 | 263 | V | 0 | 0 | yes | no | no | yes | no | no | no | no | no | no | no | no | no |
| GLRA1 | 264 | I | 0 | 0 | yes | no | no | yes | no | no | no | no | no | no | no | no | no |
| GLRA1 | 265 | L | 0 | 0 | yes | no | no | yes | no | no | no | no | yes | no | no | no | no |

| | | | | | | | | | | | | | | | | | |
|---|---|---|---|---|---|---|---|---|---|---|---|---|---|---|---|---|---|
| GLRA1 | 266 | S | 0 | 0 | yes | no | no | yes | no | no | no | no | no | no | no | no | no |
| GLRA1 | 267 | W | 0 | 1 | yes | no | no | yes | no | no | no | no | no | no | no | no | no |
| GLRA1 | 268 | I | 0 | 0 | yes | no | no | yes | no | no | no | no | yes | no | no | no | no |
| GLRA1 | 269 | S | 0 | 0 | yes | no | no | yes | no | no | no | no | yes | no | no | no | no |
| GLRA1 | 270 | F | 0 | 0 | yes | no | no | yes | no | no | no | no | no | no | no | no | no |
| GLRA1 | 271 | W | 0 | 0 | yes | no | no | yes | no | no | no | no | no | no | no | no | no |
| GLRA1 | 272 | I | 0 | 1 | yes | no | no | yes | no | no | no | no | yes | no | no | no | no |
| GLRA1 | 273 | N | 0 | 0 | yes | no | no | yes | no | yes | no | no | yes | no | no | no | no |
| GLRA1 | 275 | D | 0 | 0 | no | yes | no | no | no | yes | yes | no | no | no | no | no | no |
| GLRA1 | 276 | A | 1 | 0 | no | yes | no | yes | no | yes | no | no | no | no | no | no | no |
| GLRA1 | 277 | A | 0 | 0 | no | yes | no | yes | no | yes | no | no | yes | no | no | no | no |
| GLRA1 | 278 | P | 0 | 1 | yes | no | no | yes | no | no | yes | no | yes | no | no | no | no |
| GLRA1 | 279 | A | 1 | 0 | yes | no | no | yes | no | no | yes | no | no | no | no | no | no |
| GLRA1 | 280 | R | 1 | 2 | yes | no | no | yes | no | no | no | no | yes | no | no | no | no |
| GLRA1 | 281 | V | 0 | 0 | yes | no | no | yes | no | no | no | no | no | no | no | no | no |
| GLRA1 | 282 | G | 0 | 1 | yes | no | no | yes | no | no | no | no | yes | no | no | no | no |
| GLRA1 | 283 | L | 0 | 0 | yes | no | no | yes | no | no | yes | no | yes | no | no | no | no |
| GLRA1 | 284 | G | 0 | 0 | yes | no | no | yes | no | no | no | no | no | no | no | no | no |
| GLRA1 | 285 | I | 0 | 0 | yes | no | no | yes | no | no | no | no | no | no | no | no | no |
| GLRA1 | 286 | T | 0 | 0 | yes | no | no | yes | no | no | yes | no | no | no | no | no | no |
| GLRA1 | 287 | T | 0 | 1 | yes | no | no | yes | no | no | no | no | no | no | no | no | no |
| GLRA1 | 288 | V | 1 | 1 | yes | no | no | yes | no | no | no | no | no | no | no | no | no |
| GLRA1 | 289 | L | 0 | 0 | yes | no | no | yes | no | no | no | no | no | no | no | no | no |
| GLRA1 | 290 | T | 0 | 1 | yes | no | no | yes | no | no | yes | no | no | no | no | no | no |
| GLRA1 | 292 | T | 0 | 0 | yes | no | no | no | no | no | no | no | no | no | no | no | no |
| GLRA1 | 293 | T | 0 | 1 | yes | no | no | no | no | no | no | no | no | no | no | no | no |
| GLRA1 | 294 | Q | 0 | 1 | no | yes | no | no | no | no | no | no | no | no | no | no | no |
| GLRA1 | 295 | S | 0 | 1 | no | yes | no | no | no | no | no | no | no | no | no | no | no |
| GLRA1 | 296 | S | 0 | 0 | no | yes | no | no | no | no | no | no | no | no | no | no | no |
| GLRA1 | 297 | G | 0 | 0 | no | yes | no | no | no | no | no | no | no | no | no | no | no |
| GLRA1 | 298 | S | 0 | 1 | no | yes | no | no | no | no | no | no | no | no | no | no | no |
| GLRA1 | 299 | R | 0 | 1 | no | yes | no | no | yes | no | yes | no | no | no | no | no | no |
| GLRA1 | 300 | A | 0 | 1 | no | yes | no | no | yes | no | no | no | no | no | no | no | no |

| | | | | | | | | | | | | | | | | | |
|---|---|---|---|---|---|---|---|---|---|---|---|---|---|---|---|---|---|
| GLRA1 | 301 | S | 1 | 0 | no | yes | no | no | yes | no | no | no | no | no | no | no | no |
| GLRA1 | 302 | L | 0 | 0 | no | yes | no | no | yes | no | no | no | no | no | no | no | no |
| GLRA1 | 303 | P | 0 | 0 | yes | no | no | no | yes | no | no | no | no | no | no | no | no |
| GLRA1 | 304 | K | 0 | 2 | yes | no | no | no | yes | no | no | no | no | no | no | no | no |
| GLRA1 | 305 | V | 0 | 0 | yes | no | no | no | yes | no | no | no | no | no | no | no | no |
| GLRA1 | 306 | S | 0 | 0 | yes | no | no | no | yes | no | no | no | no | no | no | no | no |
| GLRA1 | 307 | Y | 0 | 2 | yes | no | no | no | yes | no | no | no | no | no | no | no | no |
| GLRA1 | 308 | V | 0 | 1 | yes | no | no | no | yes | no | no | no | no | no | no | no | no |
| GLRA1 | 309 | K | 0 | 0 | yes | no | no | no | yes | no | no | no | no | no | no | no | no |
| GLRA1 | 310 | A | 0 | 0 | yes | no | no | no | no | no | no | no | no | no | no | no | no |
| GLRA1 | 311 | I | 1 | 0 | yes | no | no | no | no | no | no | no | no | no | no | no | no |
| GLRA1 | 312 | D | 0 | 0 | yes | no | no | no | no | no | no | no | no | no | no | no | no |
| GLRA1 | 313 | I | 1 | 0 | yes | no | no | yes | no | no | no | no | no | no | no | no | no |
| GLRA1 | 314 | W | 0 | 0 | yes | no | no | no | no | no | no | no | no | no | no | no | no |
| GLRA1 | 316 | A | 0 | 1 | yes | no | no | yes | no | no | no | no | no | no | no | no | no |
| GLRA1 | 317 | V | 0 | 0 | yes | no | no | yes | no | no | no | no | no | no | no | no | no |
| GLRA1 | 318 | C | 0 | 0 | yes | no | no | yes | no | no | no | no | no | no | no | no | no |
| GLRA1 | 319 | L | 0 | 1 | yes | no | no | yes | no | no | no | no | no | no | no | no | no |
| GLRA1 | 320 | L | 1 | 0 | yes | no | no | yes | no | no | no | no | no | no | no | no | no |
| GLRA1 | 321 | F | 0 | 0 | yes | no | no | yes | no | no | no | no | no | no | no | no | no |
| GLRA1 | 322 | V | 0 | 0 | yes | no | no | yes | no | no | no | no | no | no | no | no | no |
| GLRA1 | 323 | F | 0 | 0 | yes | no | no | yes | no | no | no | no | no | no | no | no | no |
| GLRA1 | 324 | S | 0 | 0 | yes | no | no | yes | no | no | no | no | no | no | no | no | no |
| GLRA1 | 325 | A | 2 | 0 | yes | no | no | yes | no | no | no | no | no | no | no | no | no |
| GLRA1 | 326 | L | 0 | 0 | yes | no | no | yes | no | no | no | no | yes | no | no | no | no |
| GLRA1 | 327 | L | 0 | 0 | yes | no | no | yes | no | no | no | no | no | no | no | no | no |
| GLRA1 | 328 | E | 0 | 0 | yes | no | no | yes | no | no | no | no | no | no | no | no | no |
| GLRA1 | 329 | Y | 0 | 0 | yes | no | no | yes | no | no | no | no | yes | no | no | no | no |
| GLRA1 | 330 | A | 1 | 0 | no | yes | no | yes | no | no | no | no | yes | no | no | no | no |
| GLRA1 | 331 | A | 1 | 0 | yes | no | no | yes | no | yes | no | no | no | no | no | no | no |
| GLRA1 | 332 | V | 1 | 1 | yes | no | no | no | no | yes | no | no | yes | no | no | no | no |
| GLRA1 | 333 | N | 0 | 0 | yes | no | no | no | no | yes | no | no | yes | no | no | no | no |
| GLRA1 | 334 | F | 0 | 0 | yes | no | no | no | no | yes | no | no | yes | no | no | no | no |

| | | | | | | | | | | | | | | | | | |
|---|---|---|---|---|---|---|---|---|---|---|---|---|---|---|---|---|---|
| GLRA1 | 335 | V | 0 | 0 | yes | no | no | no | no | yes | no | no | no | no | no | no | no |
| GLRA1 | 336 | S | 0 | 0 | yes | no | no | no | no | yes | no | no | yes | no | no | no | no |
| GLRA1 | 337 | R | 2 | 0 | yes | no | no | no | no | yes | no | no | no | no | no | no | no |
| GLRA1 | 338 | Q | 0 | 0 | no | yes | no | no | no | yes | no | no | no | no | no | no | no |
| GLRA1 | 376 | G | 0 | 0 | no | yes | no | no | no | yes | no | no | no | no | no | no | no |
| GLRA1 | 378 | G | 1 | 0 | yes | no | no | no | no | yes | no | no | no | no | no | no | no |
| GLRA1 | 387 | G | 1 | 0 | yes | no | no | no | no | yes | no | no | no | no | no | no | no |
| GLRA1 | 416 | F | 0 | 0 | no | yes | no | no | no | yes | no | no | yes | no | no | no | no |
| GLRA1 | 417 | I | 0 | 0 | yes | no | no | no | no | yes | no | no | no | no | no | no | no |
| GLRA1 | 418 | Q | 1 | 0 | yes | no | no | no | no | yes | no | no | no | no | no | no | no |
| GLRA1 | 419 | R | 0 | 0 | yes | no | no | no | no | yes | no | no | no | no | no | no | no |
| GLRA1 | 420 | A | 0 | 1 | no | yes | no | no | no | yes | no | no | no | no | no | no | no |
| GLRA1 | 421 | K | 0 | 0 | no | yes | no | no | no | yes | no | no | no | no | no | no | no |
| GLRA1 | 422 | K | 1 | 0 | no | yes | no | no | no | yes | no | no | yes | no | no | no | no |
| GLRA1 | 423 | I | 1 | 0 | no | yes | no | no | no | yes | no | no | no | no | no | no | no |
| GLRA1 | 424 | D | 1 | 2 | no | yes | no | no | no | yes | no | no | no | no | no | no | no |
| GLRA1 | 425 | K | 0 | 0 | no | yes | no | no | no | yes | no | no | no | no | no | no | no |
| GLRA1 | 426 | I | 1 | 0 | no | yes | no | yes | no | no | no | no | no | no | no | no | no |
| GLRA1 | 427 | S | 0 | 0 | yes | no | no | yes | no | no | no | no | no | no | no | no | no |
| GLRA1 | 428 | R | 2 | 1 | yes | no | no | yes | no | no | no | no | no | no | no | no | no |
| GLRA1 | 429 | I | 2 | 1 | yes | no | no | yes | no | no | no | no | no | no | no | no | no |
| GLRA1 | 430 | G | 0 | 0 | yes | no | no | yes | no | no | no | no | no | no | no | no | no |
| GLRA1 | 431 | F | 0 | 0 | yes | no | no | yes | no | no | no | no | no | no | no | no | no |
| GLRA1 | 432 | P | 0 | 0 | yes | no | no | yes | no | no | no | no | no | no | no | no | no |
| GLRA1 | 434 | A | 0 | 0 | yes | no | no | yes | no | no | no | no | no | no | no | no | no |
| GLRA1 | 435 | F | 0 | 0 | yes | no | no | yes | no | no | no | no | no | no | no | no | no |
| GLRA1 | 436 | L | 0 | 0 | yes | no | no | yes | no | no | no | no | no | no | no | no | no |
| GLRA1 | 437 | I | 0 | 0 | yes | no | no | yes | no | no | no | no | no | no | no | no | no |
| GLRA1 | 438 | F | 0 | 0 | yes | no | no | yes | no | no | no | no | no | no | no | no | no |
| GLRA1 | 439 | N | 1 | 0 | yes | no | no | yes | no | no | no | no | no | no | no | no | no |
| GLRA1 | 441 | F | 0 | 0 | yes | no | no | yes | no | no | no | no | no | no | no | no | no |
| GLRA1 | 442 | Y | 0 | 0 | yes | no | no | yes | no | no | no | no | no | no | no | no | no |
| GLRA1 | 443 | W | 1 | 1 | yes | no | no | yes | no | no | no | no | no | no | no | no | no |

| | | | | | | | | | | | | | | | | | |
|---|---|---|---|---|---|---|---|---|---|---|---|---|---|---|---|---|---|
| GLRA1 | 444 | I | 0 | 0 | yes | no | no | yes | no | no | no | no | no | no | no | no | no |
| GLRA1 | 445 | I | 2 | 0 | yes | no | no | no | no | no | no | no | no | no | no | no | no |
| GLRA1 | 446 | Y | 0 | 0 | yes | no | no | no | no | no | no | no | no | no | no | no | no |
| GLRA1 | 447 | K | 2 | 0 | yes | no | no | no | yes | no | no | no | no | no | no | no | no |
| GLRA1 | 448 | I | 1 | 0 | yes | no | no | no | yes | no | no | no | no | no | no | no | no |
| GLRA1 | 449 | V | 1 | 0 | no | yes | no | no | yes | no | no | no | no | no | no | no | no |
| GLRA1 | 450 | R | 2 | 1 | no | yes | no | no | yes | no | no | no | no | no | no | no | no |
| GLRA1 | 451 | R | 0 | 0 | no | yes | no | no | yes | no | no | no | no | no | no | no | no |
| GLRA1 | 452 | E | 1 | 0 | no | yes | no | no | yes | no | no | no | no | no | no | no | no |
| GLRA1 | 453 | D | 0 | 0 | no | yes | no | no | yes | no | no | no | no | no | no | no | no |
| GRIN1 | 25 | K | 0 | 0 | no | yes | no | no | yes | no | no | no | no | no | no | no | no |
| GRIN1 | 26 | I | 0 | 0 | no | yes | no | no | yes | no | no | no | no | no | no | no | no |
| GRIN1 | 27 | V | 1 | 0 | no | no | yes | no | yes | no | no | no | no | no | no | no | no |
| GRIN1 | 28 | N | 1 | 0 | no | no | yes | no | yes | no | no | no | no | no | no | no | no |
| GRIN1 | 29 | I | 0 | 0 | no | no | yes | no | yes | no | no | no | no | no | no | no | no |
| GRIN1 | 30 | G | 0 | 0 | no | no | yes | no | yes | no | no | no | no | no | no | no | no |
| GRIN1 | 31 | A | 0 | 0 | no | no | yes | no | yes | no | no | no | no | no | no | no | no |
| GRIN1 | 32 | V | 0 | 0 | no | no | yes | no | yes | no | no | no | no | no | no | no | no |
| GRIN1 | 33 | L | 1 | 0 | no | yes | no | no | yes | no | no | no | no | no | no | no | no |
| GRIN1 | 34 | S | 0 | 0 | no | yes | no | no | yes | no | no | no | no | no | no | no | no |
| GRIN1 | 35 | T | 0 | 0 | no | yes | no | no | yes | no | no | no | no | no | no | no | no |
| GRIN1 | 36 | R | 0 | 0 | yes | no | no | no | yes | no | no | no | no | no | no | no | no |
| GRIN1 | 37 | K | 0 | 0 | yes | no | no | no | yes | no | no | no | no | no | no | no | no |
| GRIN1 | 38 | H | 1 | 0 | yes | no | no | no | yes | no | no | no | no | no | no | no | no |
| GRIN1 | 39 | E | 1 | 0 | yes | no | no | no | yes | no | no | no | no | no | no | no | no |
| GRIN1 | 40 | Q | 1 | 0 | yes | no | no | no | yes | no | no | no | no | no | no | no | no |
| GRIN1 | 41 | M | 0 | 0 | yes | no | no | no | yes | no | no | no | no | no | no | no | no |
| GRIN1 | 42 | F | 0 | 0 | yes | no | no | no | yes | no | no | no | no | no | no | no | no |
| GRIN1 | 43 | R | 0 | 0 | yes | no | no | no | yes | no | no | no | no | no | no | no | no |
| GRIN1 | 44 | E | 0 | 0 | yes | no | no | no | yes | no | no | no | no | no | no | no | no |
| GRIN1 | 45 | A | 0 | 0 | yes | no | no | no | yes | no | no | no | no | no | no | no | no |
| GRIN1 | 46 | V | 1 | 0 | yes | no | no | no | yes | no | no | no | no | no | no | no | no |
| GRIN1 | 47 | N | 0 | 0 | yes | no | no | no | yes | no | no | no | no | no | no | no | no |

| | | | | | | | | | | | | | | | | | |
|---|---|---|---|---|---|---|---|---|---|---|---|---|---|---|---|---|---|
| GRIN1 | 48 | Q | 0 | 0 | yes | no | no | no | yes | no | no | no | no | no | no | no | no |
| GRIN1 | 49 | A | 1 | 0 | yes | no | no | no | yes | no | no | no | no | no | no | no | no |
| GRIN1 | 50 | N | 0 | 0 | yes | no | no | no | yes | no | no | no | no | no | no | no | no |
| GRIN1 | 51 | K | 1 | 0 | yes | no | no | no | yes | no | no | no | no | no | no | no | no |
| GRIN1 | 52 | R | 2 | 0 | no | yes | no | no | yes | no | no | no | no | no | no | no | no |
| GRIN1 | 53 | H | 0 | 0 | no | yes | no | no | yes | no | no | no | no | no | no | no | no |
| GRIN1 | 56 | W | 0 | 0 | no | yes | no | no | yes | no | no | no | no | no | no | no | no |
| GRIN1 | 57 | K | 0 | 0 | no | yes | no | no | yes | no | no | no | no | no | no | no | no |
| GRIN1 | 58 | I | 0 | 0 | no | yes | no | no | yes | no | no | no | no | no | no | no | no |
| GRIN1 | 59 | Q | 0 | 0 | no | yes | no | no | yes | no | no | no | no | no | no | no | no |
| GRIN1 | 60 | L | 0 | 0 | no | no | yes | no | yes | no | no | no | no | no | no | no | no |
| GRIN1 | 61 | N | 0 | 0 | no | no | yes | no | yes | no | no | no | no | no | no | no | no |
| GRIN1 | 62 | A | 0 | 0 | no | no | yes | no | yes | no | no | no | no | no | no | no | no |
| GRIN1 | 63 | T | 0 | 0 | no | yes | no | no | yes | no | no | no | no | no | no | no | no |
| GRIN1 | 64 | S | 0 | 0 | no | yes | no | no | yes | no | no | no | no | no | no | no | no |
| GRIN1 | 65 | V | 0 | 0 | no | yes | no | no | yes | no | no | no | no | no | no | no | no |
| GRIN1 | 66 | T | 0 | 0 | no | yes | no | no | yes | no | no | no | no | no | no | no | no |
| GRIN1 | 67 | H | 0 | 0 | no | yes | no | no | yes | no | no | no | no | no | no | no | no |
| GRIN1 | 68 | K | 1 | 0 | no | yes | no | no | yes | no | no | no | no | no | no | no | no |
| GRIN1 | 69 | P | 0 | 0 | no | yes | no | no | yes | no | no | no | yes | no | no | no | no |
| GRIN1 | 70 | N | 2 | 0 | no | yes | no | no | yes | no | no | no | yes | no | no | no | no |
| GRIN1 | 71 | A | 1 | 0 | no | yes | no | no | yes | no | no | no | yes | no | no | no | no |
| GRIN1 | 72 | I | 0 | 0 | no | yes | no | no | yes | no | no | no | yes | no | no | no | no |
| GRIN1 | 73 | Q | 0 | 0 | no | yes | no | no | yes | no | no | no | yes | no | no | no | no |
| GRIN1 | 74 | M | 0 | 0 | no | yes | no | no | yes | no | no | no | no | no | no | no | no |
| GRIN1 | 75 | A | 1 | 0 | yes | no | no | no | yes | no | no | no | no | no | no | no | no |
| GRIN1 | 76 | L | 0 | 0 | yes | no | no | no | yes | no | no | no | no | no | no | no | no |
| GRIN1 | 77 | S | 0 | 0 | yes | no | no | no | yes | no | no | no | no | no | no | no | no |
| GRIN1 | 78 | V | 0 | 0 | yes | no | no | no | yes | no | no | no | no | no | no | no | no |
| GRIN1 | 79 | C | 0 | 0 | yes | no | no | no | yes | no | no | no | yes | no | no | no | no |
| GRIN1 | 80 | E | 1 | 0 | yes | no | no | no | yes | no | no | no | no | no | no | no | no |
| GRIN1 | 81 | D | 1 | 0 | yes | no | no | no | yes | no | no | no | no | no | no | no | no |
| GRIN1 | 82 | L | 0 | 0 | yes | no | no | no | yes | no | no | no | no | no | no | no | no |

| | | | | | | | | | | | | | | | | | |
|---|---|---|---|---|---|---|---|---|---|---|---|---|---|---|---|---|---|
| GRIN1 | 83 | I | 0 | 0 | yes | no | no | no | yes | no | no | no | no | no | no | no | no |
| GRIN1 | 84 | S | 0 | 0 | no | yes | no | no | yes | no | no | no | no | no | no | no | no |
| GRIN1 | 85 | S | 0 | 0 | no | yes | no | no | yes | no | no | no | no | no | no | no | no |
| GRIN1 | 86 | Q | 0 | 0 | no | yes | no | no | yes | no | no | no | no | no | no | no | no |
| GRIN1 | 87 | V | 0 | 0 | no | no | yes | no | yes | no | no | no | no | no | no | no | no |
| GRIN1 | 88 | Y | 0 | 0 | no | no | yes | no | yes | no | no | no | no | no | no | no | no |
| GRIN1 | 89 | A | 0 | 0 | no | no | yes | no | yes | no | no | no | no | no | no | no | no |
| GRIN1 | 90 | I | 0 | 0 | no | no | yes | no | yes | no | no | no | no | no | no | no | no |
| GRIN1 | 91 | L | 0 | 0 | no | no | yes | no | yes | no | no | no | no | no | no | no | no |
| GRIN1 | 92 | V | 0 | 0 | no | no | yes | no | yes | no | no | no | no | no | no | no | no |
| GRIN1 | 93 | S | 0 | 0 | no | yes | no | no | yes | no | no | no | no | no | no | no | no |
| GRIN1 | 94 | H | 0 | 0 | no | yes | no | no | yes | no | no | no | no | no | no | no | no |
| GRIN1 | 95 | P | 0 | 0 | no | yes | no | no | yes | no | no | no | no | no | no | no | no |
| GRIN1 | 96 | P | 0 | 0 | no | yes | no | no | yes | no | no | no | no | no | no | no | no |
| GRIN1 | 97 | T | 0 | 0 | no | yes | no | no | yes | no | no | no | no | no | no | no | no |
| GRIN1 | 98 | P | 2 | 0 | no | yes | no | no | yes | no | no | no | no | no | no | no | no |
| GRIN1 | 99 | N | 1 | 0 | no | yes | no | no | yes | no | no | no | no | no | no | no | no |
| GRIN1 | 100 | D | 0 | 0 | no | yes | no | no | yes | no | no | no | no | no | no | no | no |
| GRIN1 | 101 | H | 0 | 0 | no | yes | no | no | yes | no | no | no | no | no | no | no | no |
| GRIN1 | 102 | F | 1 | 0 | no | yes | no | no | yes | no | no | no | no | no | no | no | no |
| GRIN1 | 103 | T | 0 | 0 | no | yes | no | no | yes | no | no | no | no | no | no | no | no |
| GRIN1 | 104 | P | 0 | 0 | no | yes | no | no | yes | no | no | no | no | no | no | no | no |
| GRIN1 | 105 | T | 0 | 0 | yes | no | no | no | yes | no | no | no | yes | no | no | no | no |
| GRIN1 | 106 | P | 0 | 0 | yes | no | no | no | yes | no | no | no | no | no | no | no | no |
| GRIN1 | 107 | V | 0 | 0 | yes | no | no | no | yes | no | no | no | no | no | no | no | no |
| GRIN1 | 108 | S | 0 | 0 | yes | no | no | no | yes | no | no | no | no | no | no | no | no |
| GRIN1 | 109 | Y | 0 | 0 | yes | no | no | no | yes | no | no | no | yes | no | no | no | no |
| GRIN1 | 110 | T | 0 | 1 | yes | no | no | no | yes | no | no | no | no | no | no | no | no |
| GRIN1 | 111 | A | 0 | 0 | yes | no | no | no | yes | no | no | no | no | no | no | no | no |
| GRIN1 | 112 | G | 0 | 0 | yes | no | no | no | yes | no | no | no | no | no | no | no | no |
| GRIN1 | 113 | F | 0 | 0 | no | yes | no | no | yes | no | no | no | yes | no | no | no | no |
| GRIN1 | 114 | Y | 0 | 0 | no | yes | no | no | yes | no | no | no | no | no | no | no | no |
| GRIN1 | 115 | R | 2 | 0 | no | yes | no | no | yes | no | no | no | no | no | no | no | no |

| | | | | | | | | | | | | | | | | | |
|---|---|---|---|---|---|---|---|---|---|---|---|---|---|---|---|---|---|
| GRIN1 | 116 | I | 0 | 0 | no | yes | no | no | yes | no | no | no | no | no | no | no | no |
| GRIN1 | 117 | P | 1 | 0 | no | yes | no | no | yes | no | no | no | no | no | no | no | no |
| GRIN1 | 118 | V | 0 | 1 | no | no | yes | no | yes | no | no | no | no | no | no | no | no |
| GRIN1 | 119 | L | 0 | 0 | no | no | yes | no | yes | no | no | no | no | no | no | no | no |
| GRIN1 | 120 | G | 0 | 0 | no | no | yes | no | yes | no | no | no | no | no | no | no | no |
| GRIN1 | 121 | L | 0 | 0 | no | yes | no | no | yes | no | no | no | no | no | no | no | no |
| GRIN1 | 122 | T | 0 | 0 | no | yes | no | no | yes | no | no | no | no | no | no | no | no |
| GRIN1 | 123 | T | 0 | 0 | no | yes | no | no | yes | no | no | no | no | no | no | no | no |
| GRIN1 | 124 | R | 2 | 0 | no | yes | no | no | yes | no | no | no | no | no | no | no | no |
| GRIN1 | 125 | M | 0 | 0 | no | yes | no | no | yes | no | no | no | no | no | no | no | no |
| GRIN1 | 126 | S | 0 | 0 | yes | no | no | no | yes | no | no | no | no | no | no | no | no |
| GRIN1 | 127 | I | 0 | 0 | yes | no | no | no | yes | no | no | no | no | no | no | no | no |
| GRIN1 | 128 | Y | 0 | 0 | yes | no | no | no | yes | no | no | no | no | no | no | no | no |
| GRIN1 | 129 | S | 0 | 0 | yes | no | no | no | yes | no | no | no | no | no | no | no | no |
| GRIN1 | 130 | D | 0 | 0 | no | yes | no | no | yes | no | no | no | no | no | no | no | no |
| GRIN1 | 131 | K | 0 | 0 | no | yes | no | no | yes | no | no | no | no | no | no | no | no |
| GRIN1 | 132 | S | 0 | 0 | no | yes | no | no | yes | no | no | no | yes | no | no | no | no |
| GRIN1 | 133 | I | 0 | 0 | no | yes | no | no | yes | no | no | no | yes | no | no | no | no |
| GRIN1 | 134 | H | 0 | 0 | no | yes | no | no | yes | no | no | no | no | no | no | no | no |
| GRIN1 | 135 | L | 0 | 0 | no | yes | no | no | yes | no | no | no | no | no | no | no | no |
| GRIN1 | 136 | S | 0 | 0 | no | yes | no | no | yes | no | no | no | no | no | no | no | no |
| GRIN1 | 137 | F | 0 | 0 | no | no | yes | no | yes | no | no | no | no | no | no | no | no |
| GRIN1 | 138 | L | 0 | 0 | no | no | yes | no | yes | no | no | no | no | no | no | no | no |
| GRIN1 | 139 | R | 2 | 0 | no | no | yes | no | yes | no | no | no | no | no | no | no | no |
| GRIN1 | 140 | T | 0 | 0 | no | yes | no | no | yes | no | no | no | no | no | no | no | no |
| GRIN1 | 141 | V | 0 | 0 | no | yes | no | no | yes | no | no | no | no | no | no | no | no |
| GRIN1 | 142 | P | 1 | 0 | no | yes | no | no | yes | no | no | no | no | no | no | no | no |
| GRIN1 | 143 | P | 0 | 0 | no | yes | no | no | yes | no | no | no | no | no | no | no | no |
| GRIN1 | 144 | Y | 0 | 0 | no | yes | no | no | yes | no | no | no | no | no | no | no | no |
| GRIN1 | 145 | S | 0 | 0 | yes | no | no | no | yes | no | no | no | no | no | no | no | no |
| GRIN1 | 146 | H | 0 | 0 | yes | no | no | no | yes | no | no | no | no | no | no | no | no |
| GRIN1 | 147 | Q | 0 | 0 | yes | no | no | no | yes | no | no | no | no | no | no | no | no |
| GRIN1 | 148 | S | 0 | 0 | yes | no | no | no | yes | no | no | no | no | no | no | no | no |

| | | | | | | | | | | | | | | | | | |
|---|---|---|---|---|---|---|---|---|---|---|---|---|---|---|---|---|---|
| GRIN1 | 149 | S | 1 | 0 | yes | no | no | no | yes | no | no | no | no | no | no | no | no |
| GRIN1 | 150 | V | 0 | 0 | yes | no | no | no | yes | no | no | no | no | no | no | no | no |
| GRIN1 | 151 | W | 0 | 0 | yes | no | no | no | yes | no | no | no | no | no | no | no | no |
| GRIN1 | 152 | F | 0 | 0 | yes | no | no | no | yes | no | no | no | no | no | no | no | no |
| GRIN1 | 153 | E | 1 | 0 | yes | no | no | no | yes | no | no | no | no | no | no | no | no |
| GRIN1 | 154 | M | 0 | 0 | yes | no | no | no | yes | no | no | no | no | no | no | no | no |
| GRIN1 | 155 | M | 1 | 0 | yes | no | no | no | yes | no | no | no | no | no | no | no | no |
| GRIN1 | 156 | R | 2 | 0 | yes | no | no | no | yes | no | no | no | no | no | no | no | no |
| GRIN1 | 157 | V | 0 | 0 | yes | no | no | no | yes | no | no | no | no | no | no | no | no |
| GRIN1 | 158 | Y | 1 | 0 | yes | no | no | no | yes | no | no | no | no | no | no | no | no |
| GRIN1 | 159 | S | 0 | 0 | no | yes | no | no | yes | no | no | no | no | no | no | no | no |
| GRIN1 | 160 | W | 0 | 0 | no | yes | no | no | yes | no | no | no | no | no | no | no | no |
| GRIN1 | 161 | N | 0 | 0 | no | yes | no | no | yes | no | no | no | no | no | no | no | no |
| GRIN1 | 162 | H | 0 | 0 | no | yes | no | no | yes | no | no | no | no | no | no | no | no |
| GRIN1 | 163 | I | 0 | 0 | no | yes | no | no | yes | no | no | no | no | no | no | no | no |
| GRIN1 | 164 | I | 0 | 0 | no | no | yes | no | yes | no | no | no | no | no | no | no | no |
| GRIN1 | 165 | L | 0 | 0 | no | no | yes | no | yes | no | no | no | no | no | no | no | no |
| GRIN1 | 166 | L | 0 | 0 | no | no | yes | no | yes | no | no | no | no | no | no | no | no |
| GRIN1 | 167 | V | 0 | 0 | no | no | yes | no | yes | no | no | no | no | no | no | no | no |
| GRIN1 | 168 | S | 0 | 0 | no | no | yes | no | yes | no | no | no | no | no | no | no | no |
| GRIN1 | 169 | D | 2 | 0 | no | yes | no | no | yes | no | no | no | no | no | no | no | no |
| GRIN1 | 170 | D | 2 | 0 | no | yes | no | no | yes | no | no | no | no | no | no | no | no |
| GRIN1 | 171 | H | 0 | 0 | yes | no | no | no | yes | no | no | no | yes | no | no | no | no |
| GRIN1 | 172 | E | 0 | 0 | yes | no | no | no | yes | no | no | no | no | no | no | no | no |
| GRIN1 | 173 | G | 0 | 0 | yes | no | no | no | yes | no | no | no | no | no | no | no | no |
| GRIN1 | 174 | R | 2 | 0 | yes | no | no | no | yes | no | no | no | no | no | no | no | no |
| GRIN1 | 175 | A | 1 | 0 | yes | no | no | no | yes | no | no | no | no | no | no | no | no |
| GRIN1 | 176 | A | 0 | 0 | yes | no | no | no | yes | no | no | no | no | no | no | no | no |
| GRIN1 | 177 | Q | 0 | 0 | yes | no | no | no | yes | no | no | no | no | no | no | no | no |
| GRIN1 | 178 | K | 0 | 0 | yes | no | no | no | yes | no | no | no | yes | no | no | no | no |
| GRIN1 | 179 | R | 0 | 0 | yes | no | no | no | yes | no | no | no | no | no | no | no | no |
| GRIN1 | 180 | L | 0 | 0 | yes | no | no | no | yes | no | no | no | no | no | no | no | no |
| GRIN1 | 181 | E | 0 | 0 | yes | no | no | no | yes | no | no | no | no | no | no | no | no |

| | | | | | | | | | | | | | | | | | |
|---|---|---|---|---|---|---|---|---|---|---|---|---|---|---|---|---|---|
| GRIN1 | 182 | T | 0 | 0 | yes | no | no | no | yes | no | no | no | no | no | no | no | no |
| GRIN1 | 183 | L | 0 | 0 | yes | no | no | no | yes | no | no | no | no | no | no | no | no |
| GRIN1 | 184 | L | 0 | 0 | yes | no | no | no | yes | no | no | no | no | no | no | no | no |
| GRIN1 | 185 | E | 0 | 0 | no | yes | no | no | yes | no | no | no | no | no | no | no | no |
| GRIN1 | 186 | E | 0 | 0 | no | yes | no | no | yes | no | no | no | no | no | no | no | no |
| GRIN1 | 187 | R | 0 | 0 | no | yes | no | no | yes | no | no | no | no | no | no | no | no |
| GRIN1 | 188 | E | 0 | 0 | no | yes | no | no | yes | no | no | no | no | no | no | no | no |
| GRIN1 | 189 | S | 0 | 0 | no | yes | no | no | yes | no | no | no | no | no | no | no | no |
| GRIN1 | 190 | K | 0 | 0 | no | yes | no | no | yes | no | no | no | yes | no | no | no | no |
| GRIN1 | 191 | A | 0 | 0 | no | yes | no | no | yes | no | no | no | no | no | no | no | no |
| GRIN1 | 192 | E | 0 | 0 | no | yes | no | no | yes | no | no | no | no | no | no | no | no |
| GRIN1 | 193 | K | 0 | 0 | no | no | yes | no | yes | no | no | no | no | no | no | no | no |
| GRIN1 | 194 | V | 0 | 0 | no | no | yes | no | yes | no | no | no | no | no | no | no | no |
| GRIN1 | 195 | L | 0 | 0 | no | no | yes | no | yes | no | no | no | no | no | no | no | no |
| GRIN1 | 196 | Q | 1 | 0 | no | no | yes | no | yes | no | no | no | no | no | no | no | no |
| GRIN1 | 197 | F | 0 | 0 | no | no | yes | no | yes | no | no | no | no | no | no | no | no |
| GRIN1 | 198 | D | 2 | 0 | no | yes | no | no | yes | no | no | no | no | no | no | no | no |
| GRIN1 | 199 | P | 0 | 0 | no | yes | no | no | yes | no | no | no | no | no | no | no | no |
| GRIN1 | 200 | G | 0 | 0 | no | yes | no | no | yes | no | no | no | no | no | no | no | no |
| GRIN1 | 201 | T | 1 | 0 | no | yes | no | no | yes | no | no | no | no | no | no | no | no |
| GRIN1 | 202 | K | 0 | 0 | no | yes | no | no | yes | no | no | no | no | no | no | no | no |
| GRIN1 | 203 | N | 0 | 0 | no | yes | no | no | yes | no | no | no | no | no | no | no | no |
| GRIN1 | 204 | V | 1 | 0 | no | yes | no | no | yes | no | no | no | no | no | no | no | no |
| GRIN1 | 205 | T | 1 | 0 | yes | no | no | no | yes | no | no | no | no | no | no | no | no |
| GRIN1 | 206 | A | 1 | 0 | yes | no | no | no | yes | no | no | no | no | no | no | no | no |
| GRIN1 | 207 | L | 0 | 0 | yes | no | no | no | yes | no | no | no | no | no | no | no | no |
| GRIN1 | 208 | L | 0 | 0 | yes | no | no | no | yes | no | no | no | no | no | no | no | no |
| GRIN1 | 209 | M | 1 | 0 | yes | no | no | no | yes | no | no | no | no | no | no | no | no |
| GRIN1 | 210 | E | 1 | 0 | yes | no | no | no | yes | no | no | no | no | no | no | no | no |
| GRIN1 | 211 | A | 1 | 0 | yes | no | no | no | yes | no | no | no | no | no | no | no | no |
| GRIN1 | 212 | K | 0 | 0 | no | yes | no | no | yes | no | no | no | no | no | no | no | no |
| GRIN1 | 213 | E | 0 | 0 | no | yes | no | no | yes | no | no | no | no | no | no | no | no |
| GRIN1 | 214 | L | 0 | 0 | no | yes | no | no | yes | no | no | no | no | no | no | no | no |

| | | | | | | | | | | | | | | | | | |
|---|---|---|---|---|---|---|---|---|---|---|---|---|---|---|---|---|---|
| GRIN1 | 215 | E | 0 | 0 | no | yes | no | no | yes | no | no | no | no | no | no | no | no |
| GRIN1 | 216 | A | 0 | 0 | no | yes | no | no | yes | no | no | no | no | no | no | no | no |
| GRIN1 | 217 | R | 1 | 1 | no | yes | no | no | yes | no | no | no | no | no | no | no | no |
| GRIN1 | 218 | V | 0 | 0 | no | yes | no | no | yes | no | no | no | no | no | no | no | no |
| GRIN1 | 219 | I | 1 | 0 | no | no | yes | no | yes | no | no | no | no | no | no | no | no |
| GRIN1 | 220 | I | 0 | 0 | no | no | yes | no | yes | no | no | no | no | no | no | no | no |
| GRIN1 | 221 | L | 0 | 0 | no | no | yes | no | yes | no | no | no | no | no | no | no | no |
| GRIN1 | 222 | S | 0 | 0 | no | no | yes | no | yes | no | no | no | no | no | no | no | no |
| GRIN1 | 223 | A | 0 | 0 | no | yes | no | no | yes | no | no | no | no | no | no | no | no |
| GRIN1 | 224 | S | 0 | 0 | no | yes | no | no | yes | no | no | no | no | no | no | no | no |
| GRIN1 | 225 | E | 1 | 0 | yes | no | no | no | yes | no | no | no | no | no | no | no | no |
| GRIN1 | 226 | D | 0 | 0 | yes | no | no | no | yes | no | no | no | no | no | no | no | no |
| GRIN1 | 227 | D | 0 | 1 | yes | no | no | no | yes | no | no | no | no | no | no | no | no |
| GRIN1 | 228 | A | 0 | 0 | yes | no | no | no | yes | no | no | no | no | no | no | no | no |
| GRIN1 | 229 | A | 0 | 0 | yes | no | no | no | yes | no | no | no | no | no | no | no | no |
| GRIN1 | 230 | T | 0 | 0 | yes | no | no | no | yes | no | no | no | no | no | no | no | no |
| GRIN1 | 231 | V | 1 | 0 | yes | no | no | no | yes | no | no | no | no | no | no | no | no |
| GRIN1 | 232 | Y | 0 | 0 | yes | no | no | no | yes | no | no | no | no | no | no | no | no |
| GRIN1 | 233 | R | 0 | 0 | yes | no | no | no | yes | no | no | no | no | no | no | no | no |
| GRIN1 | 234 | A | 1 | 0 | yes | no | no | no | yes | no | no | no | no | no | no | no | no |
| GRIN1 | 235 | A | 1 | 0 | yes | no | no | no | yes | no | no | no | no | no | no | no | no |
| GRIN1 | 236 | A | 0 | 0 | yes | no | no | no | yes | no | no | no | no | no | no | no | no |
| GRIN1 | 237 | M | 0 | 0 | yes | no | no | no | yes | no | no | no | no | no | no | no | no |
| GRIN1 | 238 | L | 0 | 0 | yes | no | no | no | yes | no | no | no | no | no | no | no | no |
| GRIN1 | 239 | N | 0 | 0 | no | yes | no | no | yes | no | no | no | no | no | no | no | no |
| GRIN1 | 240 | M | 0 | 0 | no | yes | no | no | yes | no | no | no | no | no | no | no | no |
| GRIN1 | 241 | T | 0 | 0 | no | yes | no | no | yes | no | no | no | no | no | no | no | no |
| GRIN1 | 242 | G | 0 | 0 | no | yes | no | no | yes | no | no | no | no | no | no | no | no |
| GRIN1 | 243 | S | 1 | 0 | no | yes | no | no | yes | no | no | no | no | no | no | no | no |
| GRIN1 | 244 | G | 0 | 0 | no | yes | no | no | yes | no | no | no | no | no | no | no | no |
| GRIN1 | 245 | Y | 0 | 0 | no | yes | no | no | yes | no | no | no | no | no | no | no | no |
| GRIN1 | 246 | V | 0 | 0 | no | yes | no | no | yes | no | no | no | no | no | no | no | no |
| GRIN1 | 247 | W | 0 | 0 | no | yes | no | no | yes | no | no | no | no | no | no | no | no |

| | | | | | | | | | | | | | | | | | |
|---|---|---|---|---|---|---|---|---|---|---|---|---|---|---|---|---|---|
| GRIN1 | 248 | L | 0 | 0 | no | no | yes | no | yes | no | no | no | no | no | no | no | no |
| GRIN1 | 249 | V | 1 | 0 | no | no | yes | no | yes | no | no | no | no | no | no | no | no |
| GRIN1 | 250 | G | 0 | 0 | no | yes | no | no | yes | no | no | no | no | no | no | no | no |
| GRIN1 | 251 | E | 0 | 0 | yes | no | no | no | yes | no | no | no | no | no | no | no | no |
| GRIN1 | 252 | R | 1 | 0 | yes | no | no | no | yes | no | no | no | no | no | no | no | no |
| GRIN1 | 253 | E | 0 | 0 | yes | no | no | no | yes | no | no | no | no | no | no | no | no |
| GRIN1 | 254 | I | 0 | 0 | yes | no | no | no | yes | no | no | no | no | no | no | no | no |
| GRIN1 | 255 | S | 0 | 0 | no | yes | no | no | yes | no | no | no | no | no | no | no | no |
| GRIN1 | 256 | G | 0 | 0 | no | yes | no | no | yes | no | no | no | no | no | no | no | no |
| GRIN1 | 257 | N | 1 | 0 | yes | no | no | no | yes | no | no | no | no | no | no | no | no |
| GRIN1 | 258 | A | 1 | 0 | yes | no | no | no | yes | no | no | no | no | no | no | no | no |
| GRIN1 | 259 | L | 0 | 0 | yes | no | no | no | yes | no | no | no | no | no | no | no | no |
| GRIN1 | 260 | R | 0 | 0 | yes | no | no | no | yes | no | no | no | no | no | no | no | no |
| GRIN1 | 261 | Y | 0 | 0 | yes | no | no | no | yes | no | no | no | no | no | no | no | no |
| GRIN1 | 262 | A | 0 | 0 | no | yes | no | no | yes | no | no | no | no | no | no | no | no |
| GRIN1 | 263 | P | 0 | 0 | no | yes | no | no | yes | no | no | no | no | no | no | no | no |
| GRIN1 | 264 | D | 0 | 0 | no | yes | no | no | yes | no | no | no | no | no | no | no | no |
| GRIN1 | 265 | G | 0 | 0 | no | yes | no | no | yes | no | no | no | no | no | no | no | no |
| GRIN1 | 266 | I | 0 | 0 | no | yes | no | no | yes | no | no | no | no | no | no | no | no |
| GRIN1 | 267 | L | 1 | 0 | no | no | yes | no | yes | no | no | no | no | no | no | no | no |
| GRIN1 | 268 | G | 1 | 0 | no | no | yes | no | yes | no | no | no | no | no | no | no | no |
| GRIN1 | 269 | L | 0 | 0 | no | no | yes | no | yes | no | no | no | no | no | no | no | no |
| GRIN1 | 270 | Q | 0 | 0 | no | no | yes | no | yes | no | no | no | no | no | no | no | no |
| GRIN1 | 271 | L | 0 | 0 | no | no | yes | no | yes | no | no | no | no | no | no | no | no |
| GRIN1 | 272 | I | 0 | 0 | no | yes | no | no | yes | no | no | no | no | no | no | no | no |
| GRIN1 | 273 | N | 0 | 0 | no | yes | no | no | yes | no | no | no | no | no | no | no | no |
| GRIN1 | 274 | G | 0 | 0 | no | yes | no | no | yes | no | no | no | no | no | no | no | no |
| GRIN1 | 275 | K | 0 | 0 | no | yes | no | no | yes | no | no | no | no | no | no | no | no |
| GRIN1 | 276 | N | 0 | 0 | no | yes | no | no | yes | no | no | no | no | no | no | no | no |
| GRIN1 | 277 | E | 0 | 0 | yes | no | no | no | yes | no | no | no | no | no | no | no | no |
| GRIN1 | 278 | S | 0 | 0 | yes | no | no | no | yes | no | no | no | no | no | no | no | no |
| GRIN1 | 279 | A | 0 | 0 | yes | no | no | no | yes | no | no | no | no | no | no | no | no |
| GRIN1 | 280 | H | 1 | 0 | yes | no | no | no | yes | no | no | no | no | no | no | no | no |

| | | | | | | | | | | | | | | | | | |
|---|---|---|---|---|---|---|---|---|---|---|---|---|---|---|---|---|---|
| GRIN1 | 281 | I | 0 | 0 | yes | no | no | no | yes | no | no | no | no | no | no | no | no |
| GRIN1 | 282 | S | 1 | 0 | yes | no | no | no | yes | no | no | no | no | no | no | no | no |
| GRIN1 | 283 | D | 0 | 0 | yes | no | no | no | yes | no | no | no | no | no | no | no | no |
| GRIN1 | 284 | A | 1 | 0 | yes | no | no | no | yes | no | no | no | no | no | no | no | no |
| GRIN1 | 285 | V | 0 | 0 | yes | no | no | no | yes | no | no | no | no | no | no | no | no |
| GRIN1 | 286 | G | 0 | 0 | yes | no | no | no | yes | no | no | no | no | no | no | no | no |
| GRIN1 | 287 | V | 1 | 0 | yes | no | no | no | yes | no | no | no | no | no | no | no | no |
| GRIN1 | 288 | V | 1 | 0 | yes | no | no | no | yes | no | no | no | no | no | no | no | no |
| GRIN1 | 289 | A | 0 | 0 | yes | no | no | no | yes | no | no | no | no | no | no | no | no |
| GRIN1 | 290 | Q | 1 | 0 | yes | no | no | no | yes | no | no | no | no | no | no | no | no |
| GRIN1 | 291 | A | 0 | 0 | yes | no | no | no | yes | no | no | no | no | no | no | no | no |
| GRIN1 | 292 | V | 0 | 0 | yes | no | no | no | yes | no | no | no | no | no | no | no | no |
| GRIN1 | 293 | H | 0 | 0 | yes | no | no | no | yes | no | no | no | no | no | no | no | no |
| GRIN1 | 294 | E | 0 | 0 | yes | no | no | no | yes | no | no | no | no | no | no | no | no |
| GRIN1 | 295 | L | 0 | 0 | yes | no | no | no | yes | no | no | no | no | no | no | no | no |
| GRIN1 | 296 | L | 0 | 0 | yes | no | no | no | yes | no | no | no | no | no | no | no | no |
| GRIN1 | 297 | E | 0 | 0 | no | yes | no | no | yes | no | no | no | no | no | no | no | no |
| GRIN1 | 298 | K | 0 | 0 | no | yes | no | no | yes | no | no | no | no | no | no | no | no |
| GRIN1 | 299 | E | 0 | 0 | no | yes | no | no | yes | no | no | no | no | no | no | no | no |
| GRIN1 | 300 | N | 0 | 0 | no | yes | no | no | yes | no | no | no | no | no | no | no | no |
| GRIN1 | 301 | I | 1 | 0 | no | yes | no | no | yes | no | no | no | no | no | no | no | no |
| GRIN1 | 302 | T | 0 | 0 | no | yes | no | no | yes | no | no | no | no | no | no | no | no |
| GRIN1 | 303 | D | 0 | 0 | no | yes | no | no | yes | no | no | no | no | no | no | no | no |
| GRIN1 | 304 | P | 1 | 0 | no | yes | no | no | yes | no | no | no | no | no | no | no | no |
| GRIN1 | 305 | P | 0 | 0 | no | yes | no | no | yes | no | no | no | no | no | no | no | no |
| GRIN1 | 306 | R | 0 | 0 | no | yes | no | no | yes | no | no | no | no | no | no | no | no |
| GRIN1 | 307 | G | 1 | 0 | no | yes | no | no | yes | no | no | no | no | no | no | no | no |
| GRIN1 | 308 | C | 0 | 0 | no | yes | no | no | yes | no | no | no | yes | no | no | no | no |
| GRIN1 | 309 | V | 0 | 0 | no | yes | no | no | yes | no | no | no | yes | no | no | no | no |
| GRIN1 | 310 | G | 0 | 0 | no | yes | no | no | yes | no | no | no | yes | no | no | no | no |
| GRIN1 | 311 | N | 0 | 0 | no | yes | no | no | yes | no | no | no | yes | no | no | no | no |
| GRIN1 | 312 | T | 0 | 0 | no | yes | no | no | yes | no | no | no | yes | no | no | no | no |
| GRIN1 | 313 | N | 0 | 0 | no | yes | no | no | yes | no | no | no | no | no | no | no | no |

| | | | | | | | | | | | | | | | | | |
|---|---|---|---|---|---|---|---|---|---|---|---|---|---|---|---|---|---|
| GRIN1 | 314 | I | 0 | 0 | no | yes | no | no | yes | no | no | no | yes | no | no | no | no |
| GRIN1 | 315 | W | 0 | 0 | no | yes | no | no | yes | no | no | no | no | no | no | no | no |
| GRIN1 | 316 | K | 0 | 0 | no | yes | no | no | yes | no | no | no | no | no | no | no | no |
| GRIN1 | 317 | T | 0 | 0 | no | yes | no | no | yes | no | no | no | no | no | no | no | no |
| GRIN1 | 318 | G | 0 | 0 | yes | no | no | no | yes | no | no | no | no | no | no | no | no |
| GRIN1 | 319 | P | 0 | 0 | yes | no | no | no | yes | no | no | no | no | no | no | no | no |
| GRIN1 | 320 | L | 0 | 0 | yes | no | no | no | yes | no | no | no | no | no | no | no | no |
| GRIN1 | 321 | F | 0 | 0 | yes | no | no | no | yes | no | no | no | no | no | no | no | no |
| GRIN1 | 322 | K | 0 | 0 | yes | no | no | no | yes | no | no | no | no | no | no | no | no |
| GRIN1 | 323 | R | 0 | 0 | yes | no | no | no | yes | no | no | no | yes | no | no | no | no |
| GRIN1 | 324 | V | 0 | 0 | yes | no | no | no | yes | no | no | no | no | no | no | no | no |
| GRIN1 | 325 | L | 0 | 0 | yes | no | no | no | yes | no | no | no | no | no | no | no | no |
| GRIN1 | 326 | M | 1 | 0 | yes | no | no | no | yes | no | no | no | no | no | no | no | no |
| GRIN1 | 327 | S | 0 | 0 | yes | no | no | no | yes | no | no | no | no | no | no | no | no |
| GRIN1 | 328 | S | 0 | 0 | no | yes | no | no | yes | no | no | no | no | no | no | no | no |
| GRIN1 | 329 | K | 0 | 0 | no | yes | no | no | yes | no | no | no | no | no | no | no | no |
| GRIN1 | 330 | Y | 0 | 0 | no | yes | no | no | yes | no | no | no | no | no | no | no | no |
| GRIN1 | 331 | A | 1 | 0 | no | yes | no | no | yes | no | no | no | no | no | no | no | no |
| GRIN1 | 332 | D | 0 | 0 | no | yes | no | no | yes | no | no | no | no | no | no | no | no |
| GRIN1 | 333 | G | 0 | 0 | no | yes | no | no | yes | no | no | no | no | no | no | no | no |
| GRIN1 | 334 | V | 0 | 0 | no | yes | no | no | yes | no | no | no | no | no | no | no | no |
| GRIN1 | 335 | T | 0 | 0 | no | yes | no | no | yes | no | no | no | no | no | no | no | no |
| GRIN1 | 336 | G | 0 | 0 | no | yes | no | no | yes | no | no | no | no | no | no | no | no |
| GRIN1 | 337 | R | 1 | 0 | no | yes | no | no | yes | no | no | no | no | no | no | no | no |
| GRIN1 | 338 | V | 1 | 0 | no | yes | no | no | yes | no | no | no | no | no | no | no | no |
| GRIN1 | 339 | E | 0 | 0 | no | no | yes | no | yes | no | no | no | no | no | no | no | no |
| GRIN1 | 340 | F | 0 | 0 | no | no | yes | no | yes | no | no | no | no | no | no | no | no |
| GRIN1 | 341 | N | 1 | 0 | no | yes | no | no | yes | no | no | no | no | no | no | no | no |
| GRIN1 | 342 | E | 0 | 0 | no | yes | no | no | yes | no | no | no | no | no | no | no | no |
| GRIN1 | 343 | D | 0 | 0 | no | yes | no | no | yes | no | no | no | yes | no | no | no | no |
| GRIN1 | 344 | G | 1 | 0 | no | yes | no | no | yes | no | no | no | no | no | no | no | no |
| GRIN1 | 345 | D | 0 | 0 | no | yes | no | no | yes | no | no | no | no | no | no | no | no |
| GRIN1 | 346 | R | 1 | 0 | no | no | yes | no | yes | no | no | no | no | no | no | no | no |

| | | | | | | | | | | | | | | | | | |
|---|---|---|---|---|---|---|---|---|---|---|---|---|---|---|---|---|---|
| GRIN1 | 347 | K | 0 | 0 | no | no | yes | no | yes | no | no | no | no | no | no | no | no |
| GRIN1 | 348 | F | 0 | 0 | no | yes | no | no | yes | no | no | no | no | no | no | no | no |
| GRIN1 | 349 | A | 1 | 1 | no | yes | no | no | yes | no | no | no | no | no | no | no | no |
| GRIN1 | 350 | N | 1 | 0 | no | yes | no | no | yes | no | no | no | no | no | no | no | no |
| GRIN1 | 351 | Y | 2 | 0 | no | no | yes | no | yes | no | no | no | no | no | no | no | no |
| GRIN1 | 352 | S | 0 | 0 | no | no | yes | no | yes | no | no | no | no | no | no | no | no |
| GRIN1 | 353 | I | 1 | 0 | no | no | yes | no | yes | no | no | no | no | no | no | no | no |
| GRIN1 | 354 | M | 0 | 0 | no | no | yes | no | yes | no | no | no | no | no | no | no | no |
| GRIN1 | 355 | N | 0 | 0 | no | no | yes | no | yes | no | no | no | no | no | no | no | no |
| GRIN1 | 356 | L | 0 | 0 | no | no | yes | no | yes | no | no | no | no | no | no | no | no |
| GRIN1 | 357 | Q | 0 | 0 | no | yes | no | no | yes | no | no | no | no | no | no | no | no |
| GRIN1 | 358 | N | 0 | 0 | no | yes | no | no | yes | no | no | no | no | no | no | no | no |
| GRIN1 | 359 | R | 1 | 0 | no | yes | no | no | yes | no | no | no | no | no | no | no | no |
| GRIN1 | 360 | K | 0 | 0 | no | yes | no | no | yes | no | no | no | no | no | no | no | no |
| GRIN1 | 361 | L | 0 | 0 | no | no | yes | no | yes | no | no | no | no | no | no | no | no |
| GRIN1 | 362 | V | 1 | 0 | no | no | yes | no | yes | no | no | no | no | no | no | no | no |
| GRIN1 | 363 | Q | 0 | 0 | no | no | yes | no | yes | no | no | no | no | no | no | no | no |
| GRIN1 | 364 | V | 1 | 0 | no | no | yes | no | yes | no | no | no | no | no | no | no | no |
| GRIN1 | 365 | G | 1 | 0 | no | no | yes | no | yes | no | no | no | no | no | no | no | no |
| GRIN1 | 366 | I | 1 | 0 | no | no | yes | no | yes | no | no | no | no | no | no | no | no |
| GRIN1 | 367 | Y | 0 | 0 | no | no | yes | no | yes | no | no | no | no | no | no | no | no |
| GRIN1 | 368 | N | 0 | 0 | no | yes | no | no | yes | no | no | no | no | no | no | no | no |
| GRIN1 | 369 | G | 0 | 0 | no | yes | no | no | yes | no | no | no | no | no | no | no | no |
| GRIN1 | 370 | T | 1 | 0 | no | yes | no | no | yes | no | no | no | no | no | no | no | no |
| GRIN1 | 371 | H | 0 | 0 | no | yes | no | no | yes | no | no | no | no | no | no | no | no |
| GRIN1 | 372 | V | 0 | 0 | no | no | yes | no | yes | no | no | no | no | no | no | no | no |
| GRIN1 | 373 | I | 0 | 0 | no | no | yes | no | yes | no | no | no | no | no | no | no | no |
| GRIN1 | 374 | P | 1 | 0 | no | no | yes | no | yes | no | no | no | no | no | no | no | no |
| GRIN1 | 375 | N | 0 | 0 | no | yes | no | no | yes | no | no | no | no | no | no | no | no |
| GRIN1 | 376 | D | 0 | 0 | no | yes | no | no | yes | no | no | no | no | no | no | no | no |
| GRIN1 | 377 | R | 0 | 0 | no | yes | no | no | yes | no | no | no | no | no | no | no | no |
| GRIN1 | 378 | K | 0 | 0 | no | yes | no | no | yes | no | no | no | no | no | no | no | no |
| GRIN1 | 379 | I | 0 | 0 | no | yes | no | no | yes | no | no | no | no | no | no | no | no |

| | | | | | | | | | | | | | | | | | |
|---|---|---|---|---|---|---|---|---|---|---|---|---|---|---|---|---|---|
| GRIN1 | 380 | I | 0 | 0 | no | yes | no | no | yes | no | no | no | no | no | no | no | no |
| GRIN1 | 381 | W | 0 | 0 | no | yes | no | no | yes | no | no | no | no | no | no | no | no |
| GRIN1 | 382 | P | 0 | 0 | no | yes | no | no | yes | no | no | no | no | no | no | no | no |
| GRIN1 | 383 | G | 0 | 0 | no | yes | no | no | yes | no | no | no | no | no | no | no | no |
| GRIN1 | 384 | G | 1 | 0 | no | yes | no | no | yes | no | no | no | no | no | no | no | no |
| GRIN1 | 385 | E | 0 | 0 | no | yes | no | no | yes | no | no | no | no | no | no | no | no |
| GRIN1 | 386 | T | 0 | 0 | no | yes | no | no | yes | no | no | no | no | no | no | no | no |
| GRIN1 | 387 | E | 0 | 0 | no | yes | no | no | yes | no | no | no | no | no | no | no | no |
| GRIN1 | 388 | K | 0 | 0 | no | yes | no | no | yes | no | no | no | no | no | no | no | no |
| GRIN1 | 389 | P | 0 | 0 | no | yes | no | no | yes | no | no | no | no | no | no | no | no |
| GRIN1 | 390 | R | 2 | 0 | no | yes | no | no | yes | no | no | no | no | no | no | no | no |
| GRIN1 | 391 | G | 0 | 0 | no | yes | no | no | yes | no | no | no | no | no | no | no | no |
| GRIN1 | 392 | Y | 0 | 0 | no | yes | no | no | yes | no | no | no | no | no | no | no | no |
| GRIN1 | 393 | Q | 0 | 0 | no | yes | no | no | yes | no | no | no | no | no | no | no | no |
| GRIN1 | 394 | M | 1 | 0 | no | yes | no | no | yes | no | no | no | no | no | no | no | no |
| GRIN1 | 395 | S | 0 | 0 | no | yes | no | no | yes | no | no | no | no | no | no | no | no |
| GRIN1 | 396 | T | 0 | 0 | no | yes | no | no | yes | no | no | no | no | no | no | no | no |
| GRIN1 | 397 | R | 0 | 1 | no | no | yes | no | yes | no | no | no | no | no | no | no | no |
| GRIN1 | 398 | L | 0 | 0 | no | no | yes | no | yes | no | no | no | no | no | no | no | no |
| GRIN1 | 399 | K | 0 | 0 | no | no | yes | no | yes | no | no | no | no | no | no | no | no |
| GRIN1 | 400 | I | 0 | 0 | no | no | yes | no | yes | no | no | no | no | no | no | no | no |
| GRIN1 | 401 | V | 0 | 0 | no | no | yes | no | yes | no | no | no | no | no | no | no | no |
| GRIN1 | 402 | T | 0 | 0 | no | no | yes | no | yes | no | no | no | no | no | no | no | no |
| GRIN1 | 403 | I | 0 | 0 | no | yes | no | no | yes | no | no | no | no | no | no | no | no |
| GRIN1 | 404 | H | 0 | 0 | no | yes | no | no | yes | no | no | no | no | no | no | no | no |
| GRIN1 | 405 | Q | 0 | 0 | no | yes | no | no | yes | no | no | no | no | no | no | no | no |
| GRIN1 | 406 | E | 0 | 0 | no | yes | no | no | yes | no | no | no | no | no | no | no | no |
| GRIN1 | 407 | P | 0 | 0 | no | yes | no | no | yes | no | no | no | no | no | no | no | no |
| GRIN1 | 408 | F | 0 | 0 | no | yes | no | no | yes | no | no | no | no | no | no | no | no |
| GRIN1 | 409 | V | 0 | 0 | no | no | yes | no | yes | no | no | no | no | no | no | no | no |
| GRIN1 | 410 | Y | 0 | 0 | no | no | yes | no | yes | no | no | no | no | no | no | no | no |
| GRIN1 | 411 | V | 1 | 0 | no | no | yes | no | yes | no | no | no | no | no | no | no | no |
| GRIN1 | 412 | K | 0 | 0 | no | no | yes | no | yes | no | no | no | no | no | no | no | no |

| | | | | | | | | | | | | | | | | | |
|---|---|---|---|---|---|---|---|---|---|---|---|---|---|---|---|---|---|
| GRIN1 | 413 | P | 0 | 0 | no | yes | no | no | yes | no | no | no | no | no | no | no | no |
| GRIN1 | 414 | T | 0 | 0 | no | yes | no | no | yes | no | no | no | no | no | no | no | no |
| GRIN1 | 415 | L | 0 | 0 | no | yes | no | no | yes | no | no | no | no | no | no | no | no |
| GRIN1 | 416 | S | 1 | 0 | no | yes | no | no | yes | no | no | no | no | no | no | no | no |
| GRIN1 | 417 | D | 0 | 0 | no | yes | no | no | yes | no | no | no | no | no | no | no | no |
| GRIN1 | 418 | G | 0 | 0 | no | yes | no | no | yes | no | no | no | no | no | no | no | no |
| GRIN1 | 419 | T | 1 | 0 | no | yes | no | no | yes | no | no | no | no | no | no | no | no |
| GRIN1 | 420 | C | 0 | 0 | no | yes | no | no | yes | no | no | no | no | no | no | no | no |
| GRIN1 | 421 | K | 0 | 0 | no | yes | no | no | yes | no | no | no | no | no | no | no | no |
| GRIN1 | 422 | E | 0 | 0 | no | yes | no | no | yes | no | no | no | no | no | no | no | no |
| GRIN1 | 423 | E | 0 | 0 | no | yes | no | no | yes | no | no | no | no | no | no | no | no |
| GRIN1 | 424 | F | 2 | 0 | no | no | yes | no | yes | no | no | no | no | no | no | no | no |
| GRIN1 | 425 | T | 0 | 0 | no | yes | no | no | yes | no | no | no | no | no | no | no | no |
| GRIN1 | 426 | V | 0 | 0 | no | yes | no | no | yes | no | no | no | no | no | no | no | no |
| GRIN1 | 427 | N | 0 | 0 | no | yes | no | no | yes | no | no | no | no | no | no | no | no |
| GRIN1 | 428 | G | 1 | 0 | no | yes | no | no | yes | no | no | no | no | no | no | no | no |
| GRIN1 | 429 | D | 1 | 0 | no | yes | no | no | yes | no | no | no | no | no | no | no | no |
| GRIN1 | 430 | P | 1 | 0 | no | no | yes | no | yes | no | no | no | no | no | no | no | no |
| GRIN1 | 431 | V | 0 | 0 | no | yes | no | no | yes | no | no | no | no | no | no | no | no |
| GRIN1 | 432 | K | 0 | 0 | no | yes | no | no | yes | no | no | no | no | no | no | no | no |
| GRIN1 | 433 | K | 1 | 0 | no | yes | no | no | yes | no | no | no | no | no | no | no | no |
| GRIN1 | 434 | V | 0 | 0 | no | no | yes | no | yes | no | no | no | no | no | no | no | no |
| GRIN1 | 435 | I | 1 | 0 | no | no | yes | no | yes | no | no | no | no | no | no | no | no |
| GRIN1 | 436 | C | 0 | 0 | no | no | yes | no | yes | no | no | no | no | no | no | no | no |
| GRIN1 | 437 | T | 1 | 0 | no | no | yes | no | yes | no | no | no | no | no | no | no | no |
| GRIN1 | 438 | G | 0 | 0 | no | no | yes | no | yes | no | no | no | no | no | no | no | no |
| GRIN1 | 439 | P | 0 | 0 | no | no | yes | no | yes | no | no | no | no | no | no | no | no |
| GRIN1 | 440 | N | 1 | 0 | no | yes | no | no | yes | no | no | no | no | no | no | no | no |
| GRIN1 | 441 | D | 0 | 0 | no | yes | no | no | yes | no | no | no | no | no | no | no | no |
| GRIN1 | 442 | T | 0 | 0 | no | yes | no | no | yes | no | no | no | no | no | no | no | no |
| GRIN1 | 443 | S | 0 | 0 | no | yes | no | no | yes | no | no | no | no | no | no | no | no |
| GRIN1 | 444 | P | 1 | 0 | no | yes | no | no | yes | no | no | no | no | no | no | no | no |
| GRIN1 | 445 | G | 1 | 0 | no | yes | no | no | yes | no | no | no | no | no | no | no | no |

| | | | | | | | | | | | | | | | | | |
|---|---|---|---|---|---|---|---|---|---|---|---|---|---|---|---|---|---|
| GRIN1 | 446 | S | 1 | 0 | no | yes | no | no | yes | no | no | no | no | no | no | no | no |
| GRIN1 | 447 | P | 1 | 0 | no | yes | no | no | yes | no | no | no | no | no | no | no | no |
| GRIN1 | 448 | R | 1 | 0 | no | yes | no | no | yes | no | no | no | no | no | no | no | no |
| GRIN1 | 449 | H | 0 | 0 | no | yes | no | no | yes | no | no | no | no | no | no | no | no |
| GRIN1 | 450 | T | 1 | 0 | no | no | yes | no | yes | no | no | no | no | no | no | no | no |
| GRIN1 | 451 | V | 0 | 0 | no | no | yes | no | yes | no | no | no | no | no | no | no | no |
| GRIN1 | 452 | P | 0 | 0 | no | no | yes | no | yes | no | no | no | no | no | no | no | no |
| GRIN1 | 453 | Q | 1 | 0 | no | no | yes | no | yes | no | no | no | no | no | no | no | no |
| GRIN1 | 454 | C | 0 | 0 | no | no | yes | no | yes | no | no | no | no | no | no | no | no |
| GRIN1 | 455 | C | 0 | 0 | no | no | yes | no | yes | no | no | no | no | no | no | no | no |
| GRIN1 | 456 | Y | 0 | 0 | no | no | yes | no | yes | no | no | no | no | no | no | no | no |
| GRIN1 | 457 | G | 1 | 0 | no | no | yes | no | yes | no | no | no | no | no | no | no | no |
| GRIN1 | 458 | F | 0 | 0 | yes | no | no | no | yes | no | no | no | no | no | no | no | no |
| GRIN1 | 459 | C | 0 | 0 | yes | no | no | no | yes | no | no | no | no | no | no | no | no |
| GRIN1 | 460 | I | 0 | 0 | yes | no | no | no | yes | no | no | no | no | no | no | no | no |
| GRIN1 | 461 | D | 2 | 0 | yes | no | no | no | yes | no | no | no | no | no | no | no | no |
| GRIN1 | 462 | L | 0 | 0 | yes | no | no | no | yes | no | no | no | no | no | no | no | no |
| GRIN1 | 463 | L | 0 | 0 | yes | no | no | no | yes | no | no | no | no | no | no | no | no |
| GRIN1 | 464 | I | 0 | 0 | yes | no | no | no | yes | no | no | no | no | no | no | no | no |
| GRIN1 | 465 | K | 1 | 0 | yes | no | no | no | yes | no | no | no | no | no | no | no | no |
| GRIN1 | 466 | L | 0 | 0 | yes | no | no | no | yes | no | no | no | no | no | no | no | no |
| GRIN1 | 467 | A | 0 | 0 | yes | no | no | no | yes | no | no | no | no | no | no | no | no |
| GRIN1 | 468 | R | 0 | 0 | yes | no | no | no | yes | no | no | no | no | no | no | no | no |
| GRIN1 | 469 | T | 0 | 0 | yes | no | no | no | yes | no | no | no | no | no | no | no | no |
| GRIN1 | 470 | M | 0 | 0 | yes | no | no | no | yes | no | no | no | no | no | no | no | no |
| GRIN1 | 471 | N | 1 | 0 | no | yes | no | no | yes | no | no | no | no | no | no | no | no |
| GRIN1 | 472 | F | 0 | 0 | no | yes | no | no | yes | no | no | no | no | no | no | no | no |
| GRIN1 | 473 | T | 0 | 0 | no | no | yes | no | yes | no | no | no | no | no | no | no | no |
| GRIN1 | 474 | Y | 0 | 0 | no | no | yes | no | yes | no | no | no | no | no | no | no | no |
| GRIN1 | 475 | E | 0 | 0 | no | no | yes | no | yes | no | no | no | no | no | no | no | no |
| GRIN1 | 476 | V | 0 | 0 | no | no | yes | no | yes | no | no | no | no | no | no | no | no |
| GRIN1 | 477 | H | 0 | 0 | no | no | yes | no | yes | no | no | no | no | no | no | no | no |
| GRIN1 | 478 | L | 0 | 0 | no | no | yes | no | yes | no | no | no | no | no | no | no | no |

| | | | | | | | | | | | | | | | | | |
|---|---|---|---|---|---|---|---|---|---|---|---|---|---|---|---|---|---|
| GRIN1 | 479 | V | 0 | 0 | no | yes | no | no | yes | no | no | no | no | no | no | no | no |
| GRIN1 | 480 | A | 0 | 0 | no | yes | no | no | yes | no | no | no | no | no | no | no | no |
| GRIN1 | 481 | D | 0 | 0 | no | yes | no | no | yes | no | no | no | no | no | no | no | no |
| GRIN1 | 482 | G | 0 | 0 | no | yes | no | no | yes | no | no | no | no | no | no | no | no |
| GRIN1 | 483 | K | 0 | 0 | no | yes | no | no | yes | no | no | no | no | no | no | no | no |
| GRIN1 | 484 | F | 0 | 0 | no | yes | no | no | yes | no | no | no | no | no | no | no | no |
| GRIN1 | 485 | G | 0 | 0 | no | yes | no | no | yes | no | no | no | no | no | no | no | no |
| GRIN1 | 486 | T | 0 | 0 | no | yes | no | no | yes | no | no | no | no | no | no | no | no |
| GRIN1 | 487 | Q | 0 | 0 | no | no | yes | no | yes | no | no | no | yes | no | no | no | no |
| GRIN1 | 488 | E | 0 | 0 | no | no | yes | no | yes | no | no | no | no | no | no | no | no |
| GRIN1 | 489 | R | 0 | 0 | no | no | yes | no | yes | no | no | no | yes | no | no | no | no |
| GRIN1 | 490 | V | 0 | 0 | no | no | yes | no | yes | no | no | no | no | no | no | no | no |
| GRIN1 | 491 | N | 0 | 0 | no | yes | no | no | yes | no | no | no | no | no | no | no | no |
| GRIN1 | 492 | N | 0 | 0 | no | yes | no | no | yes | no | no | no | no | no | no | no | no |
| GRIN1 | 493 | S | 0 | 0 | no | yes | no | no | yes | no | no | no | no | no | no | no | no |
| GRIN1 | 494 | N | 0 | 0 | no | yes | no | no | yes | no | no | no | yes | no | no | no | no |
| GRIN1 | 495 | K | 1 | 0 | no | no | yes | no | yes | no | no | no | no | no | no | no | no |
| GRIN1 | 496 | K | 0 | 0 | no | no | yes | no | yes | no | no | no | yes | no | no | no | no |
| GRIN1 | 497 | E | 0 | 0 | no | no | yes | no | yes | no | no | no | no | no | no | no | no |
| GRIN1 | 498 | W | 0 | 0 | no | no | yes | no | yes | no | no | no | no | no | no | no | no |
| GRIN1 | 499 | N | 0 | 0 | no | yes | no | no | yes | no | no | no | no | no | no | no | no |
| GRIN1 | 500 | G | 0 | 0 | yes | no | no | no | yes | no | no | no | no | no | no | no | no |
| GRIN1 | 501 | M | 0 | 0 | yes | no | no | no | yes | no | no | no | no | no | no | no | no |
| GRIN1 | 502 | M | 0 | 0 | yes | no | no | no | yes | no | no | no | no | no | no | no | no |
| GRIN1 | 503 | G | 0 | 0 | yes | no | no | no | yes | no | no | no | no | no | no | no | no |
| GRIN1 | 504 | E | 0 | 0 | yes | no | no | no | yes | no | no | no | no | no | no | no | no |
| GRIN1 | 505 | L | 0 | 0 | yes | no | no | no | yes | no | no | no | no | no | no | no | no |
| GRIN1 | 506 | L | 0 | 0 | yes | no | no | no | yes | no | no | no | no | no | no | no | no |
| GRIN1 | 507 | S | 2 | 0 | no | yes | no | no | yes | no | no | no | no | no | no | no | no |
| GRIN1 | 508 | G | 0 | 0 | no | yes | no | no | yes | no | no | no | no | no | no | no | no |
| GRIN1 | 509 | Q | 0 | 0 | no | yes | no | no | yes | no | no | no | no | no | no | no | no |
| GRIN1 | 510 | A | 0 | 0 | no | yes | no | no | yes | no | no | no | no | no | no | no | no |
| GRIN1 | 511 | D | 0 | 0 | no | yes | no | no | yes | no | no | no | no | no | no | no | no |

| | | | | | | | | | | | | | | | | | |
|---|---|---|---|---|---|---|---|---|---|---|---|---|---|---|---|---|---|
| GRIN1 | 512 | M | 1 | 0 | no | no | yes | no | yes | no | no | no | no | no | no | no | no |
| GRIN1 | 513 | I | 0 | 0 | no | no | yes | no | yes | no | no | no | no | no | no | no | no |
| GRIN1 | 514 | V | 0 | 0 | no | no | yes | no | yes | no | no | no | no | no | no | no | no |
| GRIN1 | 515 | A | 1 | 0 | no | no | yes | no | yes | no | no | no | no | no | no | no | no |
| GRIN1 | 516 | P | 0 | 0 | no | no | yes | no | yes | no | no | no | no | no | no | no | no |
| GRIN1 | 517 | L | 0 | 0 | no | no | yes | no | yes | no | no | no | no | no | no | no | no |
| GRIN1 | 518 | T | 0 | 0 | no | yes | no | no | yes | no | no | no | no | no | no | no | no |
| GRIN1 | 519 | I | 0 | 0 | no | yes | no | no | yes | no | no | no | yes | no | no | no | no |
| GRIN1 | 520 | N | 0 | 0 | no | yes | no | no | yes | no | no | no | yes | no | no | no | no |
| GRIN1 | 521 | N | 0 | 0 | yes | no | no | no | yes | no | no | no | yes | no | no | no | no |
| GRIN1 | 522 | E | 1 | 0 | yes | no | no | no | yes | no | no | no | no | no | no | no | no |
| GRIN1 | 523 | R | 0 | 0 | yes | no | no | no | yes | no | no | no | no | no | no | no | no |
| GRIN1 | 524 | A | 1 | 0 | yes | no | no | no | yes | no | no | no | no | no | no | no | no |
| GRIN1 | 525 | Q | 0 | 0 | no | yes | no | no | yes | no | no | no | yes | no | no | no | no |
| GRIN1 | 526 | Y | 0 | 0 | no | yes | no | no | yes | no | no | no | no | no | no | no | no |
| GRIN1 | 527 | I | 0 | 0 | no | yes | no | no | yes | no | no | no | no | no | no | no | no |
| GRIN1 | 528 | E | 0 | 0 | no | no | yes | no | yes | no | no | no | yes | no | no | no | no |
| GRIN1 | 529 | F | 0 | 0 | no | no | yes | no | yes | no | no | no | no | no | no | no | no |
| GRIN1 | 530 | S | 0 | 0 | no | yes | no | no | yes | no | no | no | no | no | no | no | no |
| GRIN1 | 531 | K | 0 | 0 | no | yes | no | no | yes | no | no | no | yes | no | no | no | no |
| GRIN1 | 532 | P | 0 | 1 | no | yes | no | no | yes | no | no | no | no | no | no | no | no |
| GRIN1 | 533 | F | 0 | 0 | no | yes | no | no | yes | no | no | no | no | no | no | no | no |
| GRIN1 | 534 | K | 0 | 0 | no | no | yes | no | yes | no | no | no | no | no | no | no | no |
| GRIN1 | 535 | Y | 0 | 0 | no | no | yes | no | yes | no | no | no | yes | no | no | no | no |
| GRIN1 | 536 | Q | 0 | 1 | no | no | yes | no | yes | no | no | no | no | no | no | no | no |
| GRIN1 | 537 | G | 0 | 0 | no | yes | no | no | yes | no | no | no | no | no | no | no | no |
| GRIN1 | 538 | L | 0 | 0 | no | no | yes | no | yes | no | no | no | no | no | no | no | no |
| GRIN1 | 539 | T | 0 | 0 | no | no | yes | no | yes | no | no | no | no | no | no | no | no |
| GRIN1 | 540 | I | 0 | 0 | no | no | yes | no | yes | no | no | no | no | no | no | no | no |
| GRIN1 | 541 | L | 0 | 0 | no | no | yes | no | yes | no | no | no | no | no | no | no | no |
| GRIN1 | 542 | V | 0 | 0 | no | no | yes | no | yes | no | no | no | no | no | no | no | no |
| GRIN1 | 543 | K | 0 | 0 | no | no | yes | no | yes | no | no | no | no | no | no | no | no |
| GRIN1 | 544 | K | 1 | 0 | no | yes | no | no | yes | no | no | no | no | no | no | no | no |

| | | | | | | | | | | | | | | | | | |
|---|---|---|---|---|---|---|---|---|---|---|---|---|---|---|---|---|---|
| GRIN1 | 545 | E | 0 | 0 | no | yes | no | no | yes | no | no | no | no | no | no | no | no |
| GRIN1 | 546 | I | 0 | 0 | no | yes | no | no | yes | no | no | no | no | no | no | no | no |
| GRIN1 | 547 | P | 1 | 0 | no | yes | no | no | yes | no | no | no | no | no | no | no | no |
| GRIN1 | 548 | R | 1 | 1 | no | yes | no | no | yes | no | no | no | no | no | no | no | no |
| GRIN1 | 553 | S | 0 | 1 | no | yes | no | no | yes | no | no | no | no | no | no | no | no |
| GRIN1 | 554 | F | 0 | 0 | no | yes | no | no | yes | no | no | no | no | no | no | no | no |
| GRIN1 | 555 | M | 0 | 0 | no | yes | no | yes | yes | no | no | no | no | no | no | no | no |
| GRIN1 | 556 | Q | 0 | 0 | no | yes | no | yes | yes | no | no | no | no | no | no | no | no |
| GRIN1 | 557 | P | 0 | 1 | no | yes | no | no | yes | no | no | no | yes | no | no | no | no |
| GRIN1 | 558 | F | 0 | 0 | no | yes | no | yes | yes | no | no | no | yes | no | no | no | no |
| GRIN1 | 559 | Q | 0 | 1 | no | yes | no | no | yes | no | no | no | yes | no | no | no | no |
| GRIN1 | 560 | S | 0 | 0 | yes | no | no | yes | no | no | no | no | no | no | no | no | no |
| GRIN1 | 561 | T | 0 | 0 | yes | no | no | yes | no | no | no | no | no | no | no | no | no |
| GRIN1 | 562 | L | 0 | 0 | yes | no | no | yes | no | no | no | no | yes | no | no | no | no |
| GRIN1 | 563 | W | 0 | 0 | yes | no | no | yes | no | no | no | no | no | no | no | no | no |
| GRIN1 | 564 | L | 0 | 0 | yes | no | no | yes | no | no | no | no | no | no | no | no | no |
| GRIN1 | 565 | L | 0 | 0 | yes | no | no | yes | no | no | no | no | no | no | no | no | no |
| GRIN1 | 566 | V | 0 | 0 | yes | no | no | yes | no | no | no | no | no | no | no | no | no |
| GRIN1 | 567 | G | 0 | 0 | yes | no | no | yes | no | no | no | no | no | no | no | no | no |
| GRIN1 | 568 | L | 0 | 0 | yes | no | no | yes | no | no | no | no | no | no | no | no | no |
| GRIN1 | 569 | S | 0 | 0 | yes | no | no | yes | no | no | no | no | no | no | no | no | no |
| GRIN1 | 570 | V | 0 | 0 | yes | no | no | yes | no | no | no | no | no | no | no | no | no |
| GRIN1 | 571 | H | 0 | 0 | yes | no | no | yes | no | no | no | no | no | no | no | no | no |
| GRIN1 | 572 | V | 0 | 0 | yes | no | no | yes | no | no | no | no | no | no | no | no | no |
| GRIN1 | 573 | V | 0 | 0 | yes | no | no | yes | no | no | no | no | no | no | no | no | no |
| GRIN1 | 574 | A | 0 | 0 | yes | no | no | yes | no | no | no | no | no | no | no | no | no |
| GRIN1 | 575 | V | 0 | 0 | yes | no | no | yes | no | no | no | no | no | no | no | no | no |
| GRIN1 | 576 | M | 0 | 0 | yes | no | no | yes | no | no | no | no | no | no | no | no | no |
| GRIN1 | 577 | L | 0 | 0 | yes | no | no | yes | no | no | no | no | no | no | no | no | no |
| GRIN1 | 578 | Y | 0 | 0 | yes | no | no | yes | no | no | no | no | no | no | no | no | no |
| GRIN1 | 579 | L | 0 | 0 | yes | no | no | yes | no | no | no | no | no | no | no | no | no |
| GRIN1 | 580 | L | 0 | 0 | no | yes | no | yes | no | no | no | no | yes | no | no | no | no |
| GRIN1 | 581 | D | 0 | 0 | no | yes | no | yes | no | yes | no | no | no | no | no | no | no |

| | | | | | | | | | | | | | | | | | |
|---|---|---|---|---|---|---|---|---|---|---|---|---|---|---|---|---|---|
| GRIN1 | 582 | R | 1 | 0 | no | yes | no | no | no | yes | no | no | no | no | no | no | no |
| GRIN1 | 583 | F | 1 | 0 | no | yes | no | yes | no | yes | no | no | yes | no | no | no | no |
| GRIN1 | 584 | S | 0 | 0 | no | yes | no | no | no | yes | no | no | no | no | no | no | no |
| GRIN1 | 601 | L | 0 | 0 | no | yes | no | no | no | yes | no | no | no | no | no | no | no |
| GRIN1 | 602 | T | 0 | 0 | no | yes | no | no | no | yes | no | no | no | no | no | no | no |
| GRIN1 | 603 | L | 0 | 0 | no | yes | no | no | no | no | no | yes | no | no | no | no | no |
| GRIN1 | 604 | S | 0 | 0 | no | yes | no | yes | no | no | no | yes | no | no | no | no | no |
| GRIN1 | 605 | S | 0 | 0 | yes | no | no | yes | no | no | no | yes | no | no | no | no | no |
| GRIN1 | 606 | A | 0 | 0 | yes | no | no | yes | no | no | no | yes | no | no | no | no | no |
| GRIN1 | 607 | M | 0 | 0 | yes | no | no | yes | no | no | no | yes | no | no | no | no | no |
| GRIN1 | 608 | W | 0 | 0 | yes | no | no | yes | no | no | no | yes | yes | no | no | no | no |
| GRIN1 | 609 | F | 0 | 0 | yes | no | no | yes | no | no | no | yes | no | no | no | no | no |
| GRIN1 | 610 | S | 0 | 0 | yes | no | no | yes | no | no | no | yes | no | no | no | no | no |
| GRIN1 | 611 | W | 0 | 0 | yes | no | no | yes | no | no | no | yes | no | no | no | no | no |
| GRIN1 | 613 | V | 0 | 0 | yes | no | no | yes | no | no | no | yes | no | no | no | no | no |
| GRIN1 | 614 | L | 0 | 0 | yes | no | no | yes | no | no | no | yes | no | no | no | no | no |
| GRIN1 | 615 | L | 0 | 0 | yes | no | no | yes | no | no | no | yes | yes | no | no | no | no |
| GRIN1 | 616 | N | 0 | 1 | no | yes | no | yes | no | no | yes | yes | yes | no | no | no | no |
| GRIN1 | 617 | S | 0 | 0 | no | yes | no | yes | no | no | yes | yes | yes | no | no | no | no |
| GRIN1 | 618 | G | 0 | 1 | no | yes | no | yes | no | no | yes | yes | no | no | no | no | no |
| GRIN1 | 619 | I | 0 | 0 | no | yes | no | yes | no | no | yes | yes | yes | no | no | no | no |
| GRIN1 | 620 | G | 0 | 1 | no | yes | no | yes | no | no | yes | yes | no | no | no | no | no |
| GRIN1 | 621 | E | 0 | 0 | no | yes | no | yes | no | no | yes | yes | yes | no | no | no | no |
| GRIN1 | 622 | G | 0 | 0 | no | yes | no | yes | no | no | no | yes | no | no | no | no | no |
| GRIN1 | 626 | S | 0 | 0 | no | yes | no | yes | no | yes | no | no | no | no | no | no | no |
| GRIN1 | 627 | F | 0 | 0 | yes | no | no | yes | no | yes | no | no | yes | no | no | no | no |
| GRIN1 | 628 | S | 0 | 0 | yes | no | no | yes | no | yes | no | no | yes | no | no | no | no |
| GRIN1 | 629 | A | 1 | 0 | yes | no | no | yes | no | yes | no | no | no | no | no | no | no |
| GRIN1 | 630 | R | 0 | 0 | yes | no | no | yes | no | yes | no | no | yes | no | no | no | no |
| GRIN1 | 631 | I | 0 | 0 | yes | no | no | yes | no | no | no | no | yes | no | no | no | no |
| GRIN1 | 632 | L | 0 | 0 | yes | no | no | yes | no | no | no | no | yes | no | no | no | no |
| GRIN1 | 633 | G | 0 | 0 | yes | no | no | yes | no | no | no | no | no | no | no | no | no |
| GRIN1 | 634 | M | 0 | 0 | yes | no | no | yes | no | no | no | no | yes | no | no | no | no |

| | | | | | | | | | | | | | | | | | |
|---|---|---|---|---|---|---|---|---|---|---|---|---|---|---|---|---|---|
| GRIN1 | 635 | V | 0 | 0 | yes | no | no | yes | no | no | no | no | no | no | no | no | no |
| GRIN1 | 636 | W | 0 | 0 | yes | no | no | yes | no | no | no | no | no | no | no | no | no |
| GRIN1 | 637 | A | 0 | 1 | yes | no | no | yes | no | no | no | no | no | no | no | no | no |
| GRIN1 | 638 | G | 0 | 0 | yes | no | no | yes | no | no | no | no | no | no | no | no | no |
| GRIN1 | 639 | F | 0 | 0 | yes | no | no | yes | no | no | no | no | yes | no | no | no | no |
| GRIN1 | 640 | A | 0 | 0 | yes | no | no | yes | no | no | no | no | no | no | no | no | no |
| GRIN1 | 641 | M | 0 | 3 | yes | no | no | yes | no | no | yes | no | yes | no | no | no | no |
| GRIN1 | 642 | I | 0 | 1 | yes | no | no | yes | no | no | no | no | no | no | no | no | no |
| GRIN1 | 643 | I | 0 | 0 | yes | no | no | yes | no | no | no | no | no | no | no | no | no |
| GRIN1 | 644 | V | 0 | 1 | yes | no | no | yes | no | no | yes | no | yes | no | no | no | no |
| GRIN1 | 645 | A | 0 | 1 | yes | no | no | yes | no | no | no | no | no | no | no | no | no |
| GRIN1 | 646 | S | 0 | 0 | yes | no | no | yes | no | no | no | no | no | no | no | no | no |
| GRIN1 | 647 | Y | 0 | 2 | yes | no | no | yes | no | no | no | no | no | no | no | no | no |
| GRIN1 | 648 | T | 0 | 0 | yes | no | no | yes | yes | no | yes | no | yes | no | no | no | no |
| GRIN1 | 649 | A | 0 | 0 | yes | no | no | no | yes | no | yes | no | no | no | no | no | no |
| GRIN1 | 650 | N | 0 | 2 | yes | no | no | no | yes | no | no | no | yes | no | no | no | no |
| GRIN1 | 651 | L | 0 | 0 | yes | no | no | no | yes | no | no | no | yes | no | no | no | no |
| GRIN1 | 652 | A | 0 | 1 | yes | no | no | no | yes | no | yes | no | yes | no | no | no | no |
| GRIN1 | 653 | A | 0 | 1 | yes | no | no | no | yes | no | no | no | no | no | no | no | no |
| GRIN1 | 654 | F | 0 | 1 | yes | no | no | no | yes | no | no | no | yes | no | no | no | no |
| GRIN1 | 655 | L | 0 | 0 | yes | no | no | no | yes | no | no | no | yes | no | no | no | no |
| GRIN1 | 656 | V | 0 | 0 | yes | no | no | no | yes | no | yes | no | yes | no | no | no | no |
| GRIN1 | 657 | L | 0 | 0 | no | yes | no | no | yes | no | yes | no | yes | no | no | no | no |
| GRIN1 | 658 | D | 0 | 0 | no | yes | no | no | yes | no | no | no | no | no | no | no | no |
| GRIN1 | 659 | R | 0 | 1 | no | yes | no | no | yes | no | no | no | no | no | no | no | no |
| GRIN1 | 660 | P | 0 | 0 | no | yes | no | no | yes | no | no | no | no | no | no | no | no |
| GRIN1 | 661 | E | 0 | 0 | no | yes | no | no | yes | no | no | no | no | no | no | no | no |
| GRIN1 | 662 | E | 0 | 1 | no | yes | no | no | yes | no | no | no | no | no | no | no | no |
| GRIN1 | 663 | R | 0 | 0 | no | yes | no | no | yes | no | no | no | no | no | no | no | no |
| GRIN1 | 664 | I | 0 | 0 | no | yes | no | no | yes | no | no | no | no | no | no | no | no |
| GRIN1 | 665 | T | 0 | 0 | no | yes | no | no | yes | no | no | no | no | no | no | no | no |
| GRIN1 | 666 | G | 0 | 0 | no | yes | no | no | yes | no | no | no | no | no | no | no | no |
| GRIN1 | 667 | I | 0 | 0 | no | yes | no | no | yes | no | no | no | no | no | no | no | no |

| | | | | | | | | | | | | | | | | | |
|---|---|---|---|---|---|---|---|---|---|---|---|---|---|---|---|---|---|
| GRIN1 | 668 | N | 0 | 0 | no | yes | no | no | yes | no | yes | no | no | no | no | no | no |
| GRIN1 | 669 | D | 0 | 0 | no | yes | no | no | yes | no | no | no | no | no | no | no | no |
| GRIN1 | 670 | P | 0 | 0 | yes | no | no | no | yes | no | no | no | no | no | no | no | no |
| GRIN1 | 671 | R | 0 | 0 | yes | no | no | no | yes | no | no | no | no | no | no | no | no |
| GRIN1 | 672 | L | 0 | 0 | yes | no | no | no | yes | no | no | no | no | no | no | no | no |
| GRIN1 | 673 | R | 0 | 0 | yes | no | no | no | yes | no | no | no | no | no | no | no | no |
| GRIN1 | 674 | N | 0 | 1 | no | yes | no | no | yes | no | no | no | no | no | no | no | no |
| GRIN1 | 675 | P | 0 | 0 | no | yes | no | no | yes | no | no | no | no | no | no | no | no |
| GRIN1 | 676 | S | 0 | 0 | no | yes | no | no | yes | no | no | no | no | no | no | no | no |
| GRIN1 | 677 | D | 0 | 0 | no | yes | no | no | yes | no | no | no | no | no | no | no | no |
| GRIN1 | 678 | K | 0 | 0 | no | yes | no | no | yes | no | no | no | no | no | no | no | no |
| GRIN1 | 679 | F | 0 | 0 | no | yes | no | no | yes | no | no | no | no | no | no | no | no |
| GRIN1 | 680 | I | 0 | 0 | no | yes | no | no | yes | no | no | no | no | no | no | no | no |
| GRIN1 | 681 | Y | 0 | 0 | no | no | yes | no | yes | no | no | no | no | no | no | no | no |
| GRIN1 | 682 | A | 0 | 0 | no | yes | no | no | yes | no | no | no | no | no | no | no | no |
| GRIN1 | 683 | T | 0 | 0 | no | yes | no | no | yes | no | no | no | no | no | no | no | no |
| GRIN1 | 684 | V | 0 | 0 | no | no | yes | no | yes | no | no | no | no | no | no | no | no |
| GRIN1 | 685 | K | 1 | 0 | no | yes | no | no | yes | no | no | no | no | no | no | no | no |
| GRIN1 | 686 | Q | 0 | 0 | no | yes | no | no | yes | no | no | no | no | no | no | no | no |
| GRIN1 | 687 | S | 0 | 0 | no | yes | no | no | yes | no | no | no | no | no | no | no | no |
| GRIN1 | 688 | S | 0 | 2 | yes | no | no | no | yes | no | no | no | no | no | no | no | no |
| GRIN1 | 689 | V | 0 | 0 | yes | no | no | no | yes | no | no | no | no | no | no | no | no |
| GRIN1 | 690 | D | 0 | 0 | yes | no | no | no | yes | no | no | no | no | no | no | no | no |
| GRIN1 | 691 | I | 0 | 0 | yes | no | no | no | yes | no | no | no | no | no | no | no | no |
| GRIN1 | 692 | Y | 0 | 0 | yes | no | no | no | yes | no | no | no | no | no | no | no | no |
| GRIN1 | 693 | F | 0 | 0 | yes | no | no | no | yes | no | no | no | no | no | no | no | no |
| GRIN1 | 694 | R | 1 | 0 | yes | no | no | no | yes | no | no | no | yes | no | no | no | no |
| GRIN1 | 695 | R | 0 | 0 | yes | no | no | no | yes | no | no | no | yes | no | no | no | no |
| GRIN1 | 696 | Q | 0 | 0 | no | yes | no | no | yes | no | no | no | yes | no | no | no | no |
| GRIN1 | 697 | V | 0 | 0 | no | yes | no | no | yes | no | no | no | yes | no | no | no | no |
| GRIN1 | 698 | E | 0 | 0 | no | yes | no | no | yes | no | no | no | yes | no | no | no | no |
| GRIN1 | 699 | L | 0 | 0 | no | yes | no | no | yes | no | no | no | no | no | no | no | no |
| GRIN1 | 700 | S | 0 | 0 | yes | no | no | no | yes | no | no | no | yes | no | no | no | no |

| | | | | | | | | | | | | | | | | | |
|---|---|---|---|---|---|---|---|---|---|---|---|---|---|---|---|---|---|
| GRIN1 | 701 | T | 0 | 0 | yes | no | no | no | yes | no | no | no | no | no | no | no | no |
| GRIN1 | 702 | M | 0 | 0 | yes | no | no | no | yes | no | no | no | no | no | no | no | no |
| GRIN1 | 703 | Y | 0 | 0 | yes | no | no | no | yes | no | no | no | no | no | no | no | no |
| GRIN1 | 704 | R | 0 | 0 | yes | no | no | no | yes | no | no | no | yes | no | no | no | no |
| GRIN1 | 705 | H | 0 | 0 | yes | no | no | no | yes | no | no | no | no | no | no | no | no |
| GRIN1 | 706 | M | 0 | 1 | yes | no | no | no | yes | no | no | no | no | no | no | no | no |
| GRIN1 | 707 | E | 0 | 0 | no | yes | no | no | yes | no | no | no | no | no | no | no | no |
| GRIN1 | 708 | K | 0 | 0 | no | yes | no | no | yes | no | no | no | no | no | no | no | no |
| GRIN1 | 709 | H | 1 | 0 | no | yes | no | no | yes | no | no | no | no | no | no | no | no |
| GRIN1 | 710 | N | 0 | 0 | no | yes | no | no | yes | no | no | no | no | no | no | no | no |
| GRIN1 | 711 | Y | 0 | 0 | no | no | yes | no | yes | no | no | no | no | no | no | no | no |
| GRIN1 | 712 | E | 0 | 0 | no | yes | no | no | yes | no | no | no | no | no | no | no | no |
| GRIN1 | 713 | S | 0 | 0 | no | yes | no | no | yes | no | no | no | no | no | no | no | no |
| GRIN1 | 714 | A | 0 | 0 | yes | no | no | no | yes | no | no | no | no | no | no | no | no |
| GRIN1 | 715 | A | 0 | 0 | yes | no | no | no | yes | no | no | no | no | no | no | no | no |
| GRIN1 | 716 | E | 0 | 0 | yes | no | no | no | yes | no | no | no | no | no | no | no | no |
| GRIN1 | 717 | A | 0 | 0 | yes | no | no | no | yes | no | no | no | no | no | no | no | no |
| GRIN1 | 718 | I | 0 | 0 | yes | no | no | no | yes | no | no | no | no | no | no | no | no |
| GRIN1 | 719 | Q | 0 | 0 | yes | no | no | no | yes | no | no | no | no | no | no | no | no |
| GRIN1 | 720 | A | 0 | 0 | yes | no | no | no | yes | no | no | no | no | no | no | no | no |
| GRIN1 | 721 | V | 0 | 0 | yes | no | no | no | yes | no | no | no | no | no | no | no | no |
| GRIN1 | 722 | R | 0 | 0 | yes | no | no | no | yes | no | no | no | no | no | no | no | no |
| GRIN1 | 723 | D | 0 | 0 | no | yes | no | no | yes | no | no | no | no | no | no | no | no |
| GRIN1 | 724 | N | 0 | 0 | no | yes | no | no | yes | no | no | no | no | no | no | no | no |
| GRIN1 | 725 | K | 0 | 0 | no | yes | no | no | yes | no | no | no | no | no | no | no | no |
| GRIN1 | 726 | L | 0 | 0 | no | yes | no | no | yes | no | no | no | no | no | no | no | no |
| GRIN1 | 727 | H | 0 | 0 | no | yes | no | no | yes | no | no | no | no | no | no | no | no |
| GRIN1 | 728 | A | 0 | 0 | no | no | yes | no | yes | no | no | no | no | no | no | no | no |
| GRIN1 | 729 | F | 0 | 0 | no | no | yes | no | yes | no | no | no | no | no | no | no | no |
| GRIN1 | 730 | I | 0 | 0 | no | no | yes | no | yes | no | no | no | no | no | no | no | no |
| GRIN1 | 731 | W | 0 | 0 | no | no | yes | no | yes | no | no | no | no | no | no | no | no |
| GRIN1 | 732 | D | 0 | 0 | no | no | yes | no | yes | no | no | no | no | no | no | no | no |
| GRIN1 | 733 | S | 0 | 0 | yes | no | no | no | yes | no | no | no | no | no | no | no | no |

| | | | | | | | | | | | | | | | | | |
|---|---|---|---|---|---|---|---|---|---|---|---|---|---|---|---|---|---|
| GRIN1 | 734 | A | 0 | 0 | yes | no | no | no | yes | no | no | no | no | no | no | no | no |
| GRIN1 | 735 | V | 0 | 0 | yes | no | no | no | yes | no | no | no | no | no | no | no | no |
| GRIN1 | 736 | L | 0 | 0 | yes | no | no | no | yes | no | no | no | no | no | no | no | no |
| GRIN1 | 737 | E | 0 | 0 | yes | no | no | no | yes | no | no | no | no | no | no | no | no |
| GRIN1 | 738 | F | 0 | 0 | yes | no | no | no | yes | no | no | no | no | no | no | no | no |
| GRIN1 | 739 | E | 0 | 0 | yes | no | no | no | yes | no | no | no | no | no | no | no | no |
| GRIN1 | 740 | A | 0 | 0 | yes | no | no | no | yes | no | no | no | no | no | no | no | no |
| GRIN1 | 741 | S | 1 | 0 | yes | no | no | no | yes | no | no | no | no | no | no | no | no |
| GRIN1 | 742 | Q | 0 | 0 | yes | no | no | no | yes | no | no | no | no | no | no | no | no |
| GRIN1 | 743 | K | 0 | 0 | no | yes | no | no | yes | no | no | no | no | no | no | no | no |
| GRIN1 | 744 | C | 0 | 1 | no | yes | no | no | yes | no | no | no | no | no | no | no | no |
| GRIN1 | 745 | D | 0 | 0 | no | yes | no | no | yes | no | no | no | no | no | no | no | no |
| GRIN1 | 746 | L | 1 | 0 | no | no | yes | no | yes | no | no | no | no | no | no | no | no |
| GRIN1 | 747 | V | 0 | 0 | no | no | yes | no | yes | no | no | no | no | no | no | no | no |
| GRIN1 | 748 | T | 0 | 0 | no | no | yes | no | yes | no | no | no | no | no | no | no | no |
| GRIN1 | 749 | T | 0 | 0 | no | no | yes | no | yes | no | no | no | no | no | no | no | no |
| GRIN1 | 750 | G | 0 | 0 | no | yes | no | no | yes | no | no | no | no | no | no | no | no |
| GRIN1 | 751 | E | 0 | 0 | no | yes | no | no | yes | no | no | no | no | no | no | no | no |
| GRIN1 | 752 | L | 0 | 0 | no | yes | no | no | yes | no | no | no | no | no | no | no | no |
| GRIN1 | 753 | F | 0 | 0 | no | yes | no | no | yes | no | no | no | no | no | no | no | no |
| GRIN1 | 754 | F | 0 | 0 | no | yes | no | no | yes | no | no | no | yes | no | no | no | no |
| GRIN1 | 755 | R | 0 | 0 | no | yes | no | no | yes | no | no | no | yes | no | no | no | no |
| GRIN1 | 756 | S | 0 | 0 | no | no | yes | no | yes | no | no | no | no | no | no | no | no |
| GRIN1 | 757 | G | 0 | 0 | no | no | yes | no | yes | no | no | no | no | no | no | no | no |
| GRIN1 | 758 | F | 0 | 0 | no | no | yes | no | yes | no | no | no | no | no | no | no | no |
| GRIN1 | 759 | G | 0 | 0 | no | no | yes | no | yes | no | no | no | no | no | no | no | no |
| GRIN1 | 760 | I | 0 | 0 | no | no | yes | no | yes | no | no | no | no | no | no | no | no |
| GRIN1 | 761 | G | 0 | 0 | no | no | yes | no | yes | no | no | no | no | no | no | no | no |
| GRIN1 | 762 | M | 0 | 0 | no | no | yes | no | yes | no | no | no | no | no | no | no | no |
| GRIN1 | 763 | R | 0 | 0 | no | yes | no | no | yes | no | no | no | no | no | no | no | no |
| GRIN1 | 764 | K | 0 | 0 | no | yes | no | no | yes | no | no | no | yes | no | no | no | no |
| GRIN1 | 765 | D | 1 | 0 | no | yes | no | no | yes | no | no | no | no | no | no | no | no |
| GRIN1 | 766 | S | 0 | 0 | no | yes | no | no | yes | no | no | no | no | no | no | no | no |

| | | | | | | | | | | | | | | | | | |
|---|---|---|---|---|---|---|---|---|---|---|---|---|---|---|---|---|---|
| GRIN1 | 767 | P | 1 | 0 | no | yes | no | no | yes | no | no | no | no | no | no | no | no |
| GRIN1 | 768 | W | 0 | 0 | no | yes | no | no | yes | no | no | no | no | no | no | no | no |
| GRIN1 | 769 | K | 0 | 0 | no | yes | no | no | yes | no | no | no | no | no | no | no | no |
| GRIN1 | 770 | Q | 0 | 0 | yes | no | no | no | yes | no | no | no | no | no | no | no | no |
| GRIN1 | 771 | N | 0 | 0 | yes | no | no | no | yes | no | no | no | no | no | no | no | no |
| GRIN1 | 772 | V | 2 | 0 | yes | no | no | no | yes | no | no | no | no | no | no | no | no |
| GRIN1 | 773 | S | 0 | 0 | yes | no | no | no | yes | no | no | no | no | no | no | no | no |
| GRIN1 | 774 | L | 0 | 0 | yes | no | no | no | yes | no | no | no | no | no | no | no | no |
| GRIN1 | 775 | S | 0 | 0 | yes | no | no | no | yes | no | no | no | no | no | no | no | no |
| GRIN1 | 777 | L | 0 | 0 | yes | no | no | no | yes | no | no | no | yes | no | no | no | no |
| GRIN1 | 778 | K | 0 | 0 | yes | no | no | no | yes | no | no | no | yes | no | no | no | no |
| GRIN1 | 779 | S | 0 | 0 | yes | no | no | no | yes | no | no | no | no | no | no | no | no |
| GRIN1 | 780 | H | 0 | 0 | yes | no | no | no | yes | no | no | no | yes | no | no | no | no |
| GRIN1 | 781 | E | 0 | 0 | yes | no | no | no | yes | no | no | no | yes | no | no | no | no |
| GRIN1 | 782 | N | 0 | 0 | no | yes | no | no | yes | no | no | no | no | no | no | no | no |
| GRIN1 | 783 | G | 0 | 0 | no | yes | no | no | yes | no | no | no | no | no | no | no | no |
| GRIN1 | 784 | F | 0 | 0 | yes | no | no | no | yes | no | no | no | no | no | no | no | no |
| GRIN1 | 785 | M | 0 | 0 | yes | no | no | no | yes | no | no | no | no | no | no | no | no |
| GRIN1 | 786 | E | 0 | 0 | yes | no | no | no | yes | no | no | no | yes | no | no | no | no |
| GRIN1 | 787 | D | 0 | 0 | yes | no | no | no | yes | no | no | no | no | no | no | no | no |
| GRIN1 | 788 | L | 0 | 0 | yes | no | no | no | yes | no | no | no | no | no | no | no | no |
| GRIN1 | 789 | D | 0 | 1 | yes | no | no | no | yes | no | no | no | no | no | no | no | no |
| GRIN1 | 790 | K | 0 | 0 | yes | no | no | no | yes | no | no | no | no | no | no | no | no |
| GRIN1 | 791 | T | 0 | 0 | yes | no | no | no | yes | no | no | no | no | no | no | no | no |
| GRIN1 | 792 | W | 0 | 0 | yes | no | no | no | yes | no | no | no | no | no | no | no | no |
| GRIN1 | 793 | V | 0 | 0 | yes | no | no | no | yes | no | no | no | no | no | no | no | no |
| GRIN1 | 794 | R | 0 | 1 | no | yes | no | no | yes | no | no | no | yes | no | no | no | no |
| GRIN1 | 795 | Y | 0 | 0 | no | yes | no | no | yes | no | no | no | no | no | no | no | no |
| GRIN1 | 796 | Q | 1 | 0 | no | yes | no | no | yes | no | no | no | no | no | no | no | no |
| GRIN1 | 797 | E | 0 | 0 | no | yes | no | no | yes | no | no | no | no | no | no | no | no |
| GRIN1 | 798 | C | 0 | 0 | no | yes | no | no | yes | no | no | no | no | no | no | no | no |
| GRIN1 | 799 | D | 0 | 0 | no | yes | no | no | yes | no | no | no | no | no | no | no | no |
| GRIN1 | 800 | S | 0 | 0 | no | yes | no | no | yes | no | no | no | no | no | no | no | no |

| | | | | | | | | | | | | | | | | | |
|---|---|---|---|---|---|---|---|---|---|---|---|---|---|---|---|---|---|
| GRIN1 | 801 | R | 0 | 0 | no | yes | no | no | yes | no | no | no | no | no | no | no | no |
| GRIN1 | 802 | S | 0 | 0 | no | yes | no | no | yes | no | no | no | no | no | no | no | no |
| GRIN1 | 809 | T | 0 | 0 | no | yes | no | yes | yes | no | no | no | yes | no | no | no | no |
| GRIN1 | 810 | F | 0 | 0 | no | yes | no | no | yes | no | no | no | no | no | no | no | no |
| GRIN1 | 811 | E | 0 | 0 | yes | no | no | no | yes | no | no | no | no | no | no | no | no |
| GRIN1 | 812 | N | 0 | 0 | yes | no | no | yes | yes | no | no | no | no | no | no | no | no |
| GRIN1 | 813 | M | 0 | 0 | yes | no | no | yes | no | no | no | no | yes | no | no | no | no |
| GRIN1 | 814 | A | 0 | 1 | yes | no | no | yes | no | no | no | no | no | no | no | no | no |
| GRIN1 | 815 | G | 0 | 1 | yes | no | no | yes | no | no | no | no | no | no | no | no | no |
| GRIN1 | 816 | V | 0 | 1 | yes | no | no | yes | no | no | no | no | yes | no | no | no | no |
| GRIN1 | 817 | F | 0 | 1 | yes | no | no | yes | no | no | no | no | yes | no | no | no | no |
| GRIN1 | 818 | M | 0 | 1 | yes | no | no | yes | no | no | no | no | no | no | no | no | no |
| GRIN1 | 819 | L | 0 | 0 | yes | no | no | yes | no | no | no | no | no | no | no | no | no |
| GRIN1 | 820 | V | 0 | 0 | yes | no | no | yes | no | no | no | no | yes | no | no | no | no |
| GRIN1 | 821 | A | 0 | 0 | yes | no | no | yes | no | no | no | no | no | no | no | no | no |
| GRIN1 | 822 | G | 0 | 0 | yes | no | no | yes | no | no | no | no | no | no | no | no | no |
| GRIN1 | 823 | G | 0 | 0 | yes | no | no | yes | no | no | no | no | no | no | no | no | no |
| GRIN1 | 824 | I | 0 | 0 | yes | no | no | yes | no | no | no | no | yes | no | no | no | no |
| GRIN1 | 825 | V | 1 | 0 | yes | no | no | yes | no | no | no | no | no | no | no | no | no |
| GRIN1 | 826 | A | 0 | 0 | yes | no | no | yes | no | no | no | no | no | no | no | no | no |
| GRIN1 | 827 | G | 0 | 1 | yes | no | no | yes | no | no | no | no | no | no | no | no | no |
| GRIN1 | 828 | I | 0 | 0 | yes | no | no | yes | no | no | no | no | yes | no | no | no | no |
| GRIN1 | 829 | F | 0 | 0 | yes | no | no | yes | no | no | no | no | no | no | no | no | no |
| GRIN1 | 830 | L | 0 | 0 | yes | no | no | yes | no | no | no | no | no | no | no | no | no |
| GRIN1 | 831 | I | 0 | 0 | yes | no | no | yes | no | no | no | no | no | no | no | no | no |
| GRIN1 | 832 | F | 0 | 0 | yes | no | no | yes | no | no | no | no | no | no | no | no | no |
| GRIN1 | 833 | I | 1 | 0 | yes | no | no | yes | no | no | no | no | no | no | no | no | no |
| GRIN1 | 834 | E | 0 | 1 | yes | no | no | yes | no | yes | no | no | no | no | no | no | no |
| GRIN1 | 835 | I | 0 | 1 | yes | no | no | no | no | yes | no | no | no | no | no | no | no |
| GRIN1 | 836 | A | 0 | 0 | yes | no | no | no | no | yes | no | no | no | no | no | no | no |
| GRIN1 | 837 | Y | 0 | 0 | yes | no | no | no | no | yes | no | no | no | no | no | no | no |
| GRIN1 | 838 | K | 0 | 0 | yes | no | no | no | no | yes | no | no | no | no | no | no | no |
| GRIN1 | 839 | R | 2 | 0 | yes | no | no | no | no | yes | no | no | no | no | no | no | no |

| | | | | | | | | | | | | | | | | | |
|---|---|---|---|---|---|---|---|---|---|---|---|---|---|---|---|---|---|
| GRIN1 | 840 | H | 0 | 0 | yes | no | no | no | no | yes | no | no | no | no | no | no | no |
| GRIN1 | 841 | K | 0 | 0 | yes | no | no | no | no | yes | no | no | no | no | no | no | no |
| GRIN1 | 842 | D | 0 | 0 | yes | no | no | no | no | yes | no | no | no | no | no | no | no |
| GRIN1 | 843 | A | 0 | 0 | yes | no | no | no | no | yes | no | no | no | no | no | no | no |
| GRIN1 | 844 | R | 0 | 2 | no | yes | no | no | no | yes | no | no | no | no | no | no | no |
| GRIN2A | 34 | L | 0 | 0 | no | yes | no | no | yes | no | no | no | no | no | no | no | no |
| GRIN2A | 35 | N | 0 | 0 | no | no | yes | no | yes | no | no | no | no | no | no | no | no |
| GRIN2A | 36 | I | 0 | 0 | no | no | yes | no | yes | no | no | no | no | no | no | no | no |
| GRIN2A | 37 | A | 0 | 0 | no | no | yes | no | yes | no | no | no | no | no | no | no | no |
| GRIN2A | 38 | V | 0 | 0 | no | no | yes | no | yes | no | no | no | no | no | no | no | no |
| GRIN2A | 39 | M | 0 | 0 | no | no | yes | no | yes | no | no | no | no | no | no | no | no |
| GRIN2A | 40 | L | 1 | 0 | no | no | yes | no | yes | no | no | no | no | no | no | no | no |
| GRIN2A | 41 | G | 0 | 0 | no | yes | no | no | yes | no | no | no | no | no | no | no | no |
| GRIN2A | 42 | H | 0 | 0 | no | yes | no | no | yes | no | no | no | no | no | no | no | no |
| GRIN2A | 43 | S | 0 | 0 | no | yes | no | no | yes | no | no | no | no | no | no | no | no |
| GRIN2A | 44 | H | 0 | 0 | no | yes | no | no | yes | no | no | no | no | no | no | no | no |
| GRIN2A | 45 | D | 2 | 0 | no | yes | no | no | yes | no | no | no | no | no | no | no | no |
| GRIN2A | 46 | V | 1 | 0 | yes | no | no | no | yes | no | no | no | no | no | no | no | no |
| GRIN2A | 47 | T | 1 | 0 | yes | no | no | no | yes | no | no | no | no | no | no | no | no |
| GRIN2A | 48 | E | 0 | 0 | yes | no | no | no | yes | no | no | no | no | no | no | no | no |
| GRIN2A | 49 | R | 2 | 0 | yes | no | no | no | yes | no | no | no | no | no | no | no | no |
| GRIN2A | 50 | E | 0 | 0 | yes | no | no | no | yes | no | no | no | no | no | no | no | no |
| GRIN2A | 51 | L | 1 | 0 | yes | no | no | no | yes | no | no | no | no | no | no | no | no |
| GRIN2A | 52 | R | 0 | 0 | yes | no | no | no | yes | no | no | no | no | no | no | no | no |
| GRIN2A | 53 | T | 0 | 0 | yes | no | no | no | yes | no | no | no | no | no | no | no | no |
| GRIN2A | 54 | L | 1 | 0 | yes | no | no | no | yes | no | no | no | no | no | no | no | no |
| GRIN2A | 55 | W | 0 | 0 | yes | no | no | no | yes | no | no | no | no | no | no | no | no |
| GRIN2A | 56 | G | 1 | 0 | yes | no | no | no | yes | no | no | no | no | no | no | no | no |
| GRIN2A | 57 | P | 0 | 0 | yes | no | no | no | yes | no | no | no | no | no | no | no | no |
| GRIN2A | 58 | E | 1 | 0 | yes | no | no | no | yes | no | no | no | no | no | no | no | no |
| GRIN2A | 59 | Q | 1 | 0 | yes | no | no | no | yes | no | no | no | no | no | no | no | no |
| GRIN2A | 60 | A | 2 | 0 | no | yes | no | no | yes | no | no | no | no | no | no | no | no |
| GRIN2A | 61 | A | 2 | 0 | no | yes | no | no | yes | no | no | no | no | no | no | no | no |

| | | | | | | | | | | | | | | | | | |
|---|---|---|---|---|---|---|---|---|---|---|---|---|---|---|---|---|---|
| GRIN2A | 62 | G | 0 | 0 | no | yes | no | no | yes | no | no | no | no | no | no | no | no |
| GRIN2A | 63 | L | 0 | 0 | no | yes | no | no | yes | no | no | no | no | no | no | no | no |
| GRIN2A | 64 | P | 1 | 0 | no | yes | no | no | yes | no | no | no | no | no | no | no | no |
| GRIN2A | 65 | L | 1 | 0 | no | yes | no | no | yes | no | no | no | no | no | no | no | no |
| GRIN2A | 66 | D | 0 | 0 | no | yes | no | no | yes | no | no | no | no | no | no | no | no |
| GRIN2A | 67 | V | 0 | 0 | no | yes | no | no | yes | no | no | no | no | no | no | no | no |
| GRIN2A | 68 | N | 1 | 0 | no | no | yes | no | yes | no | no | no | no | no | no | no | no |
| GRIN2A | 69 | V | 1 | 0 | no | no | yes | no | yes | no | no | no | no | no | no | no | no |
| GRIN2A | 70 | V | 1 | 0 | no | no | yes | no | yes | no | no | no | no | no | no | no | no |
| GRIN2A | 71 | A | 0 | 0 | no | no | yes | no | yes | no | no | no | no | no | no | no | no |
| GRIN2A | 72 | L | 0 | 0 | no | no | yes | no | yes | no | no | no | no | no | no | no | no |
| GRIN2A | 73 | L | 1 | 0 | no | no | yes | no | yes | no | no | no | no | no | no | no | no |
| GRIN2A | 74 | M | 2 | 0 | no | yes | no | no | yes | no | no | no | no | no | no | no | no |
| GRIN2A | 75 | N | 0 | 0 | no | yes | no | no | yes | no | no | no | no | no | no | no | no |
| GRIN2A | 76 | R | 2 | 0 | no | yes | no | no | yes | no | no | no | yes | no | no | no | no |
| GRIN2A | 77 | T | 0 | 0 | no | yes | no | no | yes | no | no | no | yes | no | no | no | no |
| GRIN2A | 78 | D | 0 | 0 | no | yes | no | no | yes | no | no | no | yes | no | no | no | no |
| GRIN2A | 79 | P | 1 | 1 | yes | no | no | no | yes | no | no | no | yes | no | no | no | no |
| GRIN2A | 80 | K | 3 | 0 | yes | no | no | no | yes | no | no | no | yes | no | no | no | no |
| GRIN2A | 81 | S | 0 | 0 | yes | no | no | no | yes | no | no | no | no | no | no | no | no |
| GRIN2A | 82 | L | 0 | 0 | yes | no | no | no | yes | no | no | no | no | no | no | no | no |
| GRIN2A | 83 | I | 1 | 0 | yes | no | no | no | yes | no | no | no | no | no | no | no | no |
| GRIN2A | 84 | T | 1 | 0 | yes | no | no | no | yes | no | no | no | no | no | no | no | no |
| GRIN2A | 85 | H | 0 | 0 | yes | no | no | no | yes | no | no | no | no | no | no | no | no |
| GRIN2A | 86 | V | 0 | 0 | yes | no | no | no | yes | no | no | no | no | no | no | no | no |
| GRIN2A | 87 | C | 0 | 0 | yes | no | no | no | yes | no | no | no | no | no | no | no | no |
| GRIN2A | 88 | D | 1 | 0 | yes | no | no | no | yes | no | no | no | no | no | no | no | no |
| GRIN2A | 89 | L | 0 | 0 | yes | no | no | no | yes | no | no | no | no | no | no | no | no |
| GRIN2A | 90 | M | 2 | 0 | yes | no | no | no | yes | no | no | no | no | no | no | no | no |
| GRIN2A | 91 | S | 0 | 0 | yes | no | no | no | yes | no | no | no | no | no | no | no | no |
| GRIN2A | 92 | G | 1 | 0 | no | yes | no | no | yes | no | no | no | no | no | no | no | no |
| GRIN2A | 93 | A | 2 | 0 | no | yes | no | no | yes | no | no | no | no | no | no | no | no |
| GRIN2A | 94 | R | 1 | 0 | no | yes | no | no | yes | no | no | no | no | no | no | no | no |

| | | | | | | | | | | | | | | | | | |
|---|---|---|---|---|---|---|---|---|---|---|---|---|---|---|---|---|---|
| GRIN2A | 95 | I | 0 | 0 | no | yes | no | no | yes | no | no | no | no | no | no | no | no |
| GRIN2A | 96 | H | 1 | 0 | no | yes | no | no | yes | no | no | no | no | no | no | no | no |
| GRIN2A | 97 | G | 0 | 0 | no | no | yes | no | yes | no | no | no | no | no | no | no | no |
| GRIN2A | 98 | L | 0 | 1 | no | no | yes | no | yes | no | no | no | no | no | no | no | no |
| GRIN2A | 99 | V | 2 | 0 | no | no | yes | no | yes | no | no | no | no | no | no | no | no |
| GRIN2A | 100 | F | 0 | 0 | no | no | yes | no | yes | no | no | no | no | no | no | no | no |
| GRIN2A | 101 | G | 0 | 0 | no | no | yes | no | yes | no | no | no | no | no | no | no | no |
| GRIN2A | 102 | D | 1 | 0 | no | yes | no | no | yes | no | no | no | no | no | no | no | no |
| GRIN2A | 103 | D | 0 | 0 | no | yes | no | no | yes | no | no | no | no | no | no | no | no |
| GRIN2A | 104 | T | 0 | 0 | no | yes | no | no | yes | no | no | no | no | no | no | no | no |
| GRIN2A | 105 | D | 0 | 0 | no | yes | no | no | yes | no | no | no | no | no | no | no | no |
| GRIN2A | 106 | Q | 0 | 0 | no | yes | no | no | yes | no | no | no | yes | no | no | no | no |
| GRIN2A | 107 | E | 0 | 0 | no | yes | no | no | yes | no | no | no | no | no | no | no | no |
| GRIN2A | 108 | A | 0 | 0 | no | yes | no | no | yes | no | no | no | no | no | no | no | no |
| GRIN2A | 109 | V | 0 | 0 | yes | no | no | no | yes | no | no | no | no | no | no | no | no |
| GRIN2A | 110 | A | 0 | 0 | yes | no | no | no | yes | no | no | no | no | no | no | no | no |
| GRIN2A | 111 | Q | 0 | 0 | yes | no | no | no | yes | no | no | no | yes | no | no | no | no |
| GRIN2A | 112 | M | 1 | 0 | yes | no | no | no | yes | no | no | no | yes | no | no | no | no |
| GRIN2A | 113 | L | 0 | 0 | yes | no | no | no | yes | no | no | no | no | no | no | no | no |
| GRIN2A | 114 | D | 0 | 0 | yes | no | no | no | yes | no | no | no | no | no | no | no | no |
| GRIN2A | 115 | F | 0 | 0 | yes | no | no | no | yes | no | no | no | yes | no | no | no | no |
| GRIN2A | 116 | I | 1 | 0 | yes | no | no | no | yes | no | no | no | no | no | no | no | no |
| GRIN2A | 117 | S | 0 | 0 | yes | no | no | no | yes | no | no | no | no | no | no | no | no |
| GRIN2A | 118 | S | 0 | 0 | yes | no | no | no | yes | no | no | no | no | no | no | no | no |
| GRIN2A | 119 | H | 0 | 0 | yes | no | no | no | yes | no | no | no | yes | no | no | no | no |
| GRIN2A | 120 | T | 0 | 0 | no | yes | no | no | yes | no | no | no | no | no | no | no | no |
| GRIN2A | 121 | F | 0 | 0 | no | yes | no | no | yes | no | no | no | no | no | no | no | no |
| GRIN2A | 122 | V | 1 | 0 | no | yes | no | no | yes | no | no | no | no | no | no | no | no |
| GRIN2A | 123 | P | 0 | 0 | no | yes | no | no | yes | no | no | no | no | no | no | no | no |
| GRIN2A | 124 | I | 1 | 0 | no | no | yes | no | yes | no | no | no | no | no | no | no | no |
| GRIN2A | 125 | L | 0 | 0 | no | no | yes | no | yes | no | no | no | no | no | no | no | no |
| GRIN2A | 126 | G | 0 | 0 | no | no | yes | no | yes | no | no | no | no | no | no | no | no |
| GRIN2A | 127 | I | 0 | 0 | no | no | yes | no | yes | no | no | no | no | no | no | no | no |

| | | | | | | | | | | | | | | | | | |
|---|---|---|---|---|---|---|---|---|---|---|---|---|---|---|---|---|---|
| GRIN2A | 128 | H | 0 | 0 | no | no | yes | no | yes | no | no | no | no | no | no | no | no |
| GRIN2A | 129 | G | 0 | 0 | yes | no | no | no | yes | no | no | no | no | no | no | no | no |
| GRIN2A | 130 | G | 0 | 0 | yes | no | no | no | yes | no | no | no | no | no | no | no | no |
| GRIN2A | 131 | A | 0 | 0 | yes | no | no | no | yes | no | no | no | no | no | no | no | no |
| GRIN2A | 132 | S | 0 | 0 | no | yes | no | no | yes | no | no | no | no | no | no | no | no |
| GRIN2A | 133 | M | 1 | 0 | no | yes | no | no | yes | no | no | no | no | no | no | no | no |
| GRIN2A | 134 | I | 1 | 0 | no | yes | no | no | yes | no | no | no | no | no | no | no | no |
| GRIN2A | 135 | M | 0 | 0 | no | yes | no | no | yes | no | no | no | yes | no | no | no | no |
| GRIN2A | 136 | A | 0 | 0 | no | yes | no | no | yes | no | no | no | no | no | no | no | no |
| GRIN2A | 137 | D | 0 | 0 | no | yes | no | no | yes | no | no | no | yes | no | no | no | no |
| GRIN2A | 138 | K | 0 | 0 | no | yes | no | no | yes | no | no | no | yes | no | no | no | no |
| GRIN2A | 139 | D | 0 | 0 | no | yes | no | no | yes | no | no | no | no | no | no | no | no |
| GRIN2A | 140 | P | 2 | 1 | no | yes | no | no | yes | no | no | no | no | no | no | no | no |
| GRIN2A | 141 | T | 1 | 0 | no | yes | no | no | yes | no | no | no | no | no | no | no | no |
| GRIN2A | 142 | S | 0 | 0 | no | yes | no | no | yes | no | no | no | no | no | no | no | no |
| GRIN2A | 143 | T | 0 | 0 | no | yes | no | no | yes | no | no | no | no | no | no | no | no |
| GRIN2A | 144 | F | 0 | 0 | no | no | yes | no | yes | no | no | no | no | no | no | no | no |
| GRIN2A | 145 | F | 0 | 0 | no | no | yes | no | yes | no | no | no | no | no | no | no | no |
| GRIN2A | 146 | Q | 0 | 0 | no | no | yes | no | yes | no | no | no | no | no | no | no | no |
| GRIN2A | 147 | F | 1 | 0 | no | yes | no | no | yes | no | no | no | no | no | no | no | no |
| GRIN2A | 148 | G | 0 | 0 | no | yes | no | no | yes | no | no | no | no | no | no | no | no |
| GRIN2A | 149 | A | 0 | 1 | no | yes | no | no | yes | no | no | no | no | no | no | no | no |
| GRIN2A | 150 | S | 0 | 0 | no | yes | no | no | yes | no | no | no | no | no | no | no | no |
| GRIN2A | 151 | I | 0 | 0 | yes | no | no | no | yes | no | no | no | no | no | no | no | no |
| GRIN2A | 152 | Q | 0 | 0 | yes | no | no | no | yes | no | no | no | no | no | no | no | no |
| GRIN2A | 153 | Q | 1 | 0 | yes | no | no | no | yes | no | no | no | no | no | no | no | no |
| GRIN2A | 154 | Q | 0 | 0 | yes | no | no | no | yes | no | no | no | no | no | no | no | no |
| GRIN2A | 155 | A | 1 | 0 | yes | no | no | no | yes | no | no | no | no | no | no | no | no |
| GRIN2A | 156 | T | 1 | 0 | yes | no | no | no | yes | no | no | no | no | no | no | no | no |
| GRIN2A | 157 | V | 0 | 0 | yes | no | no | no | yes | no | no | no | no | no | no | no | no |
| GRIN2A | 158 | M | 0 | 0 | yes | no | no | no | yes | no | no | no | no | no | no | no | no |
| GRIN2A | 159 | L | 0 | 0 | yes | no | no | no | yes | no | no | no | no | no | no | no | no |
| GRIN2A | 160 | K | 0 | 0 | yes | no | no | no | yes | no | no | no | no | no | no | no | no |

| | | | | | | | | | | | | | | | | | |
|---|---|---|---|---|---|---|---|---|---|---|---|---|---|---|---|---|---|
| GRIN2A | 161 | I | 1 | 0 | yes | no | no | no | yes | no | no | no | no | no | no | no | no |
| GRIN2A | 162 | M | 0 | 0 | yes | no | no | no | yes | no | no | no | no | no | no | no | no |
| GRIN2A | 163 | Q | 0 | 0 | yes | no | no | no | yes | no | no | no | no | no | no | no | no |
| GRIN2A | 164 | D | 0 | 0 | yes | no | no | no | yes | no | no | no | no | no | no | no | no |
| GRIN2A | 165 | Y | 0 | 0 | no | yes | no | no | yes | no | no | no | no | no | no | no | no |
| GRIN2A | 166 | D | 2 | 0 | no | yes | no | no | yes | no | no | no | no | no | no | no | no |
| GRIN2A | 167 | W | 0 | 0 | no | yes | no | no | yes | no | no | no | no | no | no | no | no |
| GRIN2A | 168 | H | 0 | 0 | no | yes | no | no | yes | no | no | no | no | no | no | no | no |
| GRIN2A | 169 | V | 0 | 0 | no | no | yes | no | yes | no | no | no | no | no | no | no | no |
| GRIN2A | 170 | F | 0 | 0 | no | no | yes | no | yes | no | no | no | no | no | no | no | no |
| GRIN2A | 171 | S | 0 | 0 | no | no | yes | no | yes | no | no | no | no | no | no | no | no |
| GRIN2A | 172 | L | 0 | 0 | no | no | yes | no | yes | no | no | no | no | no | no | no | no |
| GRIN2A | 173 | V | 0 | 0 | no | no | yes | no | yes | no | no | no | no | no | no | no | no |
| GRIN2A | 174 | T | 0 | 0 | no | no | yes | no | yes | no | no | no | no | no | no | no | no |
| GRIN2A | 175 | T | 0 | 0 | no | yes | no | no | yes | no | no | no | no | no | no | no | no |
| GRIN2A | 176 | I | 0 | 0 | no | yes | no | no | yes | no | no | no | no | no | no | no | no |
| GRIN2A | 177 | F | 0 | 0 | no | yes | no | no | yes | no | no | no | no | no | no | no | no |
| GRIN2A | 178 | P | 0 | 0 | no | yes | no | no | yes | no | no | no | yes | no | no | no | no |
| GRIN2A | 179 | G | 0 | 0 | no | yes | no | no | yes | no | no | no | no | no | no | no | no |
| GRIN2A | 180 | Y | 0 | 0 | yes | no | no | no | yes | no | no | no | no | no | no | no | no |
| GRIN2A | 181 | R | 0 | 0 | yes | no | no | no | yes | no | no | no | yes | no | no | no | no |
| GRIN2A | 182 | E | 0 | 0 | yes | no | no | no | yes | no | no | no | yes | no | no | no | no |
| GRIN2A | 183 | F | 1 | 1 | yes | no | no | no | yes | no | no | no | no | no | no | no | no |
| GRIN2A | 184 | I | 0 | 1 | yes | no | no | no | yes | no | no | no | no | no | no | no | no |
| GRIN2A | 185 | S | 0 | 0 | yes | no | no | no | yes | no | no | no | no | no | no | no | no |
| GRIN2A | 186 | F | 0 | 0 | yes | no | no | no | yes | no | no | no | no | no | no | no | no |
| GRIN2A | 187 | V | 1 | 0 | yes | no | no | no | yes | no | no | no | no | no | no | no | no |
| GRIN2A | 188 | K | 0 | 0 | yes | no | no | no | yes | no | no | no | no | no | no | no | no |
| GRIN2A | 189 | T | 2 | 0 | yes | no | no | no | yes | no | no | no | no | no | no | no | no |
| GRIN2A | 190 | T | 1 | 0 | yes | no | no | no | yes | no | no | no | no | no | no | no | no |
| GRIN2A | 191 | V | 0 | 0 | yes | no | no | no | yes | no | no | no | no | no | no | no | no |
| GRIN2A | 192 | D | 1 | 0 | yes | no | no | no | yes | no | no | no | yes | no | no | no | no |
| GRIN2A | 193 | N | 1 | 0 | no | yes | no | no | yes | no | no | no | yes | no | no | no | no |

| | | | | | | | | | | | | | | | | | |
|---|---|---|---|---|---|---|---|---|---|---|---|---|---|---|---|---|---|
| GRIN2A | 194 | S | 0 | 0 | no | yes | no | no | yes | no | no | no | yes | no | no | no | no |
| GRIN2A | 195 | F | 1 | 0 | no | yes | no | no | yes | no | no | no | yes | no | no | no | no |
| GRIN2A | 196 | V | 0 | 0 | no | yes | no | no | yes | no | no | no | no | no | no | no | no |
| GRIN2A | 197 | G | 0 | 0 | no | yes | no | no | yes | no | no | no | no | no | no | no | no |
| GRIN2A | 198 | W | 0 | 0 | no | yes | no | no | yes | no | no | no | no | no | no | no | no |
| GRIN2A | 199 | D | 1 | 0 | no | no | yes | no | yes | no | no | no | no | no | no | no | no |
| GRIN2A | 200 | M | 2 | 0 | no | no | yes | no | yes | no | no | no | no | no | no | no | no |
| GRIN2A | 201 | Q | 2 | 0 | no | no | yes | no | yes | no | no | no | no | no | no | no | no |
| GRIN2A | 202 | N | 0 | 0 | no | no | yes | no | yes | no | no | no | no | no | no | no | no |
| GRIN2A | 203 | V | 2 | 0 | no | no | yes | no | yes | no | no | no | no | no | no | no | no |
| GRIN2A | 204 | I | 1 | 0 | no | no | yes | no | yes | no | no | no | no | no | no | no | no |
| GRIN2A | 205 | T | 0 | 0 | no | no | yes | no | yes | no | no | no | no | no | no | no | no |
| GRIN2A | 206 | L | 0 | 0 | no | yes | no | no | yes | no | no | no | no | no | no | no | no |
| GRIN2A | 207 | D | 0 | 0 | no | yes | no | no | yes | no | no | no | no | no | no | no | no |
| GRIN2A | 208 | T | 1 | 0 | no | yes | no | no | yes | no | no | no | no | no | no | no | no |
| GRIN2A | 209 | S | 1 | 0 | no | yes | no | no | yes | no | no | no | yes | no | no | no | no |
| GRIN2A | 210 | F | 0 | 0 | no | yes | no | no | yes | no | no | no | yes | no | no | no | no |
| GRIN2A | 211 | E | 1 | 0 | no | yes | no | no | yes | no | no | no | no | no | no | no | no |
| GRIN2A | 212 | D | 1 | 0 | no | yes | no | no | yes | no | no | no | no | no | no | no | no |
| GRIN2A | 213 | A | 1 | 0 | no | yes | no | no | yes | no | no | no | no | no | no | no | no |
| GRIN2A | 214 | K | 0 | 0 | yes | no | no | no | yes | no | no | no | no | no | no | no | no |
| GRIN2A | 215 | T | 0 | 0 | yes | no | no | no | yes | no | no | no | no | no | no | no | no |
| GRIN2A | 216 | Q | 0 | 0 | yes | no | no | no | yes | no | no | no | yes | no | no | no | no |
| GRIN2A | 217 | V | 0 | 0 | yes | no | no | no | yes | no | no | no | no | no | no | no | no |
| GRIN2A | 218 | Q | 0 | 0 | yes | no | no | no | yes | no | no | no | no | no | no | no | no |
| GRIN2A | 219 | L | 0 | 0 | yes | no | no | no | yes | no | no | no | no | no | no | no | no |
| GRIN2A | 220 | K | 0 | 0 | no | yes | no | no | yes | no | no | no | yes | no | no | no | no |
| GRIN2A | 221 | K | 1 | 0 | no | yes | no | no | yes | no | no | no | no | no | no | no | no |
| GRIN2A | 222 | I | 0 | 0 | no | yes | no | no | yes | no | no | no | no | no | no | no | no |
| GRIN2A | 223 | H | 1 | 0 | no | yes | no | no | yes | no | no | no | no | no | no | no | no |
| GRIN2A | 224 | S | 0 | 0 | no | yes | no | no | yes | no | no | no | no | no | no | no | no |
| GRIN2A | 225 | S | 0 | 0 | no | yes | no | no | yes | no | no | no | no | no | no | no | no |
| GRIN2A | 226 | V | 0 | 0 | no | no | yes | no | yes | no | no | no | no | no | no | no | no |

| | | | | | | | | | | | | | | | | | |
|---|---|---|---|---|---|---|---|---|---|---|---|---|---|---|---|---|---|
| GRIN2A | 227 | I | 0 | 0 | no | no | yes | no | yes | no | no | no | no | no | no | no | no |
| GRIN2A | 228 | L | 1 | 0 | no | no | yes | no | yes | no | no | no | no | no | no | no | no |
| GRIN2A | 229 | L | 0 | 0 | no | no | yes | no | yes | no | no | no | no | no | no | no | no |
| GRIN2A | 230 | Y | 1 | 0 | no | no | yes | no | yes | no | no | no | no | no | no | no | no |
| GRIN2A | 231 | C | 0 | 2 | no | yes | no | no | yes | no | no | no | no | no | no | no | no |
| GRIN2A | 232 | S | 1 | 0 | no | yes | no | no | yes | no | no | no | no | no | no | no | no |
| GRIN2A | 233 | K | 0 | 0 | yes | no | no | no | yes | no | no | no | no | no | no | no | no |
| GRIN2A | 234 | D | 1 | 0 | yes | no | no | no | yes | no | no | no | no | no | no | no | no |
| GRIN2A | 235 | E | 0 | 0 | yes | no | no | no | yes | no | no | no | no | no | no | no | no |
| GRIN2A | 236 | A | 0 | 0 | yes | no | no | no | yes | no | no | no | no | no | no | no | no |
| GRIN2A | 237 | V | 0 | 0 | yes | no | no | no | yes | no | no | no | no | no | no | no | no |
| GRIN2A | 238 | L | 1 | 0 | yes | no | no | no | yes | no | no | no | no | no | no | no | no |
| GRIN2A | 239 | I | 0 | 0 | yes | no | no | no | yes | no | no | no | no | no | no | no | no |
| GRIN2A | 240 | L | 0 | 0 | yes | no | no | no | yes | no | no | no | no | no | no | no | no |
| GRIN2A | 241 | S | 0 | 0 | yes | no | no | no | yes | no | no | no | no | no | no | no | no |
| GRIN2A | 242 | E | 0 | 0 | yes | no | no | no | yes | no | no | no | no | no | no | no | no |
| GRIN2A | 243 | A | 0 | 1 | yes | no | no | no | yes | no | no | no | no | no | no | no | no |
| GRIN2A | 244 | R | 2 | 0 | yes | no | no | no | yes | no | no | no | no | no | no | no | no |
| GRIN2A | 245 | S | 1 | 0 | yes | no | no | no | yes | no | no | no | yes | no | no | no | no |
| GRIN2A | 246 | L | 1 | 0 | yes | no | no | no | yes | no | no | no | yes | no | no | no | no |
| GRIN2A | 247 | G | 0 | 0 | no | yes | no | no | yes | no | no | no | no | no | no | no | no |
| GRIN2A | 248 | L | 0 | 0 | no | yes | no | no | yes | no | no | no | no | no | no | no | no |
| GRIN2A | 249 | T | 1 | 0 | no | yes | no | no | yes | no | no | no | no | no | no | no | no |
| GRIN2A | 250 | G | 1 | 0 | no | yes | no | no | yes | no | no | no | no | no | no | no | no |
| GRIN2A | 251 | Y | 0 | 0 | no | yes | no | no | yes | no | no | no | no | no | no | no | no |
| GRIN2A | 252 | D | 0 | 0 | no | yes | no | no | yes | no | no | no | no | no | no | no | no |
| GRIN2A | 253 | F | 1 | 0 | no | yes | no | no | yes | no | no | no | no | no | no | no | no |
| GRIN2A | 254 | F | 0 | 0 | no | no | yes | no | yes | no | no | no | no | no | no | no | no |
| GRIN2A | 255 | W | 0 | 0 | no | no | yes | no | yes | no | no | no | no | no | no | no | no |
| GRIN2A | 256 | I | 0 | 0 | no | no | yes | no | yes | no | no | no | no | no | no | no | no |
| GRIN2A | 257 | V | 0 | 0 | no | yes | no | no | yes | no | no | no | no | no | no | no | no |
| GRIN2A | 258 | P | 0 | 0 | no | yes | no | no | yes | no | no | no | no | no | no | no | no |
| GRIN2A | 259 | S | 0 | 0 | yes | no | no | no | yes | no | no | no | no | no | no | no | no |

| | | | | | | | | | | | | | | | | | |
|---|---|---|---|---|---|---|---|---|---|---|---|---|---|---|---|---|---|
| GRIN2A | 260 | L | 0 | 0 | yes | no | no | no | yes | no | no | no | no | no | no | no | no |
| GRIN2A | 261 | V | 0 | 0 | yes | no | no | no | yes | no | no | no | no | no | no | no | no |
| GRIN2A | 262 | S | 0 | 0 | yes | no | no | no | yes | no | no | no | no | no | no | no | no |
| GRIN2A | 263 | G | 0 | 0 | no | yes | no | no | yes | no | no | no | no | no | no | no | no |
| GRIN2A | 264 | N | 0 | 0 | no | yes | no | no | yes | no | no | no | no | no | no | no | no |
| GRIN2A | 265 | T | 1 | 0 | no | yes | no | no | yes | no | no | no | no | no | no | no | no |
| GRIN2A | 266 | E | 0 | 0 | no | yes | no | no | yes | no | no | no | no | no | no | no | no |
| GRIN2A | 267 | L | 1 | 0 | no | yes | no | no | yes | no | no | no | no | no | no | no | no |
| GRIN2A | 268 | I | 0 | 0 | no | yes | no | no | yes | no | no | no | no | no | no | no | no |
| GRIN2A | 269 | P | 0 | 0 | no | yes | no | no | yes | no | no | no | no | no | no | no | no |
| GRIN2A | 270 | K | 0 | 0 | no | yes | no | no | yes | no | no | no | no | no | no | no | no |
| GRIN2A | 271 | E | 0 | 0 | no | yes | no | no | yes | no | no | no | no | no | no | no | no |
| GRIN2A | 272 | F | 1 | 0 | no | yes | no | no | yes | no | no | no | no | no | no | no | no |
| GRIN2A | 273 | P | 0 | 0 | no | yes | no | no | yes | no | no | no | no | no | no | no | no |
| GRIN2A | 274 | S | 1 | 0 | no | yes | no | no | yes | no | no | no | no | no | no | no | no |
| GRIN2A | 275 | G | 0 | 0 | no | yes | no | no | yes | no | no | no | no | no | no | no | no |
| GRIN2A | 276 | L | 1 | 0 | no | yes | no | no | yes | no | no | no | no | no | no | no | no |
| GRIN2A | 277 | I | 1 | 0 | no | no | yes | no | yes | no | no | no | no | no | no | no | no |
| GRIN2A | 278 | S | 0 | 0 | no | no | yes | no | yes | no | no | no | no | no | no | no | no |
| GRIN2A | 279 | V | 1 | 0 | no | no | yes | no | yes | no | no | no | no | no | no | no | no |
| GRIN2A | 280 | S | 0 | 0 | no | no | yes | no | yes | no | no | no | no | no | no | no | no |
| GRIN2A | 281 | Y | 0 | 0 | no | yes | no | no | yes | no | no | no | no | no | no | no | no |
| GRIN2A | 282 | D | 0 | 0 | no | yes | no | no | yes | no | no | no | no | no | no | no | no |
| GRIN2A | 283 | D | 0 | 0 | no | yes | no | no | yes | no | no | no | no | no | no | no | no |
| GRIN2A | 284 | W | 0 | 0 | no | yes | no | no | yes | no | no | no | no | no | no | no | no |
| GRIN2A | 285 | D | 0 | 0 | no | yes | no | no | yes | no | no | no | no | no | no | no | no |
| GRIN2A | 286 | Y | 0 | 0 | no | yes | no | no | yes | no | no | no | no | no | no | no | no |
| GRIN2A | 287 | S | 0 | 0 | no | yes | no | no | yes | no | no | no | no | no | no | no | no |
| GRIN2A | 288 | L | 0 | 0 | yes | no | no | no | yes | no | no | no | no | no | no | no | no |
| GRIN2A | 289 | E | 1 | 0 | yes | no | no | no | yes | no | no | no | no | no | no | no | no |
| GRIN2A | 290 | A | 1 | 1 | yes | no | no | no | yes | no | no | no | no | no | no | no | no |
| GRIN2A | 291 | R | 0 | 0 | yes | no | no | no | yes | no | no | no | no | no | no | no | no |
| GRIN2A | 292 | V | 0 | 0 | yes | no | no | no | yes | no | no | no | no | no | no | no | no |

| | | | | | | | | | | | | | | | | | |
|---|---|---|---|---|---|---|---|---|---|---|---|---|---|---|---|---|---|
| GRIN2A | 293 | R | 0 | 0 | yes | no | no | no | yes | no | no | no | no | no | no | no | no |
| GRIN2A | 294 | D | 0 | 0 | yes | no | no | no | yes | no | no | no | no | no | no | no | no |
| GRIN2A | 295 | G | 1 | 1 | yes | no | no | no | yes | no | no | no | no | no | no | no | no |
| GRIN2A | 296 | I | 0 | 0 | yes | no | no | no | yes | no | no | no | no | no | no | no | no |
| GRIN2A | 297 | G | 1 | 0 | yes | no | no | no | yes | no | no | no | no | no | no | no | no |
| GRIN2A | 298 | I | 1 | 0 | yes | no | no | no | yes | no | no | no | no | no | no | no | no |
| GRIN2A | 299 | L | 0 | 0 | yes | no | no | no | yes | no | no | no | no | no | no | no | no |
| GRIN2A | 300 | T | 1 | 0 | yes | no | no | no | yes | no | no | no | no | no | no | no | no |
| GRIN2A | 301 | T | 0 | 0 | yes | no | no | no | yes | no | no | no | no | no | no | no | no |
| GRIN2A | 302 | A | 1 | 1 | yes | no | no | no | yes | no | no | no | no | no | no | no | no |
| GRIN2A | 303 | A | 0 | 0 | yes | no | no | no | yes | no | no | no | no | no | no | no | no |
| GRIN2A | 304 | S | 1 | 0 | yes | no | no | no | yes | no | no | no | no | no | no | no | no |
| GRIN2A | 305 | S | 1 | 0 | yes | no | no | no | yes | no | no | no | no | no | no | no | no |
| GRIN2A | 306 | M | 1 | 0 | yes | no | no | no | yes | no | no | no | no | no | no | no | no |
| GRIN2A | 307 | L | 0 | 0 | yes | no | no | no | yes | no | no | no | no | no | no | no | no |
| GRIN2A | 308 | E | 1 | 0 | yes | no | no | no | yes | no | no | no | no | no | no | no | no |
| GRIN2A | 309 | K | 1 | 0 | yes | no | no | no | yes | no | no | no | no | no | no | no | no |
| GRIN2A | 310 | F | 0 | 0 | no | yes | no | no | yes | no | no | no | no | no | no | no | no |
| GRIN2A | 311 | S | 1 | 0 | no | yes | no | no | yes | no | no | no | no | no | no | no | no |
| GRIN2A | 312 | Y | 0 | 0 | no | yes | no | no | yes | no | no | no | no | no | no | no | no |
| GRIN2A | 313 | I | 0 | 0 | no | yes | no | no | yes | no | no | no | no | no | no | no | no |
| GRIN2A | 314 | P | 0 | 0 | no | yes | no | no | yes | no | no | no | no | no | no | no | no |
| GRIN2A | 315 | E | 1 | 0 | no | yes | no | no | yes | no | no | no | no | no | no | no | no |
| GRIN2A | 316 | A | 1 | 0 | no | yes | no | no | yes | no | no | no | no | no | no | no | no |
| GRIN2A | 317 | K | 1 | 0 | no | yes | no | no | yes | no | no | no | no | no | no | no | no |
| GRIN2A | 318 | A | 0 | 0 | no | yes | no | no | yes | no | no | no | no | no | no | no | no |
| GRIN2A | 319 | S | 2 | 0 | no | yes | no | no | yes | no | no | no | no | no | no | no | no |
| GRIN2A | 320 | C | 0 | 0 | no | yes | no | no | yes | no | no | no | yes | no | no | no | no |
| GRIN2A | 321 | Y | 1 | 0 | no | yes | no | no | yes | no | no | no | yes | no | no | no | no |
| GRIN2A | 322 | G | 1 | 0 | no | yes | no | no | yes | no | no | no | no | no | no | no | no |
| GRIN2A | 323 | Q | 0 | 0 | no | yes | no | no | yes | no | no | no | yes | no | no | no | no |
| GRIN2A | 324 | M | 1 | 0 | no | yes | no | no | yes | no | no | no | no | no | no | no | no |
| GRIN2A | 325 | E | 0 | 0 | no | yes | no | no | yes | no | no | no | no | no | no | no | no |

| | | | | | | | | | | | | | | | | | |
|---|---|---|---|---|---|---|---|---|---|---|---|---|---|---|---|---|---|
| GRIN2A | 326 | R | 0 | 0 | no | yes | no | no | yes | no | no | no | no | no | no | no | no |
| GRIN2A | 327 | P | 2 | 0 | no | yes | no | no | yes | no | no | no | no | no | no | no | no |
| GRIN2A | 328 | E | 1 | 0 | no | yes | no | no | yes | no | no | no | no | no | no | no | no |
| GRIN2A | 329 | V | 1 | 0 | no | yes | no | no | yes | no | no | no | no | no | no | no | no |
| GRIN2A | 330 | P | 2 | 0 | no | yes | no | no | yes | no | no | no | no | no | no | no | no |
| GRIN2A | 331 | M | 0 | 0 | no | yes | no | no | yes | no | no | no | no | no | no | no | no |
| GRIN2A | 332 | H | 0 | 0 | no | yes | no | no | yes | no | no | no | no | no | no | no | no |
| GRIN2A | 333 | T | 1 | 0 | no | yes | no | no | yes | no | no | no | no | no | no | no | no |
| GRIN2A | 334 | L | 0 | 0 | no | yes | no | no | yes | no | no | no | no | no | no | no | no |
| GRIN2A | 335 | H | 1 | 0 | no | yes | no | no | yes | no | no | no | no | no | no | no | no |
| GRIN2A | 336 | P | 0 | 0 | no | yes | no | no | yes | no | no | no | no | no | no | no | no |
| GRIN2A | 337 | F | 0 | 0 | no | yes | no | no | yes | no | no | no | no | no | no | no | no |
| GRIN2A | 338 | M | 0 | 0 | no | yes | no | no | yes | no | no | no | no | no | no | no | no |
| GRIN2A | 339 | V | 0 | 0 | no | yes | no | no | yes | no | no | no | no | no | no | no | no |
| GRIN2A | 340 | N | 2 | 0 | no | yes | no | no | yes | no | no | no | no | no | no | no | no |
| GRIN2A | 341 | V | 0 | 0 | no | yes | no | no | yes | no | no | no | no | no | no | no | no |
| GRIN2A | 342 | T | 0 | 0 | no | no | yes | no | yes | no | no | no | no | no | no | no | no |
| GRIN2A | 343 | W | 2 | 0 | no | no | yes | no | yes | no | no | no | no | no | no | no | no |
| GRIN2A | 344 | D | 1 | 0 | no | yes | no | no | yes | no | no | no | no | no | no | no | no |
| GRIN2A | 345 | G | 0 | 0 | no | yes | no | no | yes | no | no | no | no | no | no | no | no |
| GRIN2A | 346 | K | 0 | 0 | no | no | yes | no | yes | no | no | no | no | no | no | no | no |
| GRIN2A | 347 | D | 1 | 0 | no | no | yes | no | yes | no | no | no | no | no | no | no | no |
| GRIN2A | 348 | L | 0 | 0 | no | yes | no | no | yes | no | no | no | no | no | no | no | no |
| GRIN2A | 349 | S | 0 | 0 | no | yes | no | no | yes | no | no | no | no | no | no | no | no |
| GRIN2A | 350 | F | 0 | 0 | no | no | yes | no | yes | no | no | no | no | no | no | no | no |
| GRIN2A | 351 | T | 0 | 0 | no | yes | no | no | yes | no | no | no | no | no | no | no | no |
| GRIN2A | 352 | E | 0 | 0 | no | yes | no | no | yes | no | no | no | no | no | no | no | no |
| GRIN2A | 353 | E | 1 | 0 | no | yes | no | no | yes | no | no | no | no | no | no | no | no |
| GRIN2A | 354 | G | 1 | 0 | no | yes | no | no | yes | no | no | no | no | no | no | no | no |
| GRIN2A | 355 | Y | 0 | 0 | no | yes | no | no | yes | no | no | no | no | no | no | no | no |
| GRIN2A | 356 | Q | 1 | 0 | no | no | yes | no | yes | no | no | no | no | no | no | no | no |
| GRIN2A | 357 | V | 1 | 0 | no | yes | no | no | yes | no | no | no | no | no | no | no | no |
| GRIN2A | 358 | H | 2 | 0 | no | yes | no | no | yes | no | no | no | no | no | no | no | no |

| | | | | | | | | | | | | | | | | | |
|---|---|---|---|---|---|---|---|---|---|---|---|---|---|---|---|---|---|
| GRIN2A | 359 | P | 0 | 0 | no | yes | no | no | yes | no | no | no | no | no | no | no | no |
| GRIN2A | 360 | R | 0 | 0 | no | yes | no | no | yes | no | no | no | no | no | no | no | no |
| GRIN2A | 361 | L | 0 | 0 | no | yes | no | no | yes | no | no | no | no | no | no | no | no |
| GRIN2A | 362 | V | 1 | 0 | no | no | yes | no | yes | no | no | no | no | no | no | no | no |
| GRIN2A | 363 | V | 0 | 0 | no | no | yes | no | yes | no | no | no | no | no | no | no | no |
| GRIN2A | 364 | I | 0 | 0 | no | no | yes | no | yes | no | no | no | no | no | no | no | no |
| GRIN2A | 365 | V | 1 | 0 | no | no | yes | no | yes | no | no | no | no | no | no | no | no |
| GRIN2A | 366 | L | 0 | 0 | no | no | yes | no | yes | no | no | no | no | no | no | no | no |
| GRIN2A | 367 | N | 1 | 0 | no | yes | no | no | yes | no | no | no | no | no | no | no | no |
| GRIN2A | 368 | K | 0 | 0 | no | yes | no | no | yes | no | no | no | no | no | no | no | no |
| GRIN2A | 369 | D | 1 | 0 | no | yes | no | no | yes | no | no | no | no | no | no | no | no |
| GRIN2A | 370 | R | 1 | 1 | no | yes | no | no | yes | no | no | no | no | no | no | no | no |
| GRIN2A | 371 | E | 0 | 0 | no | yes | no | no | yes | no | no | no | no | no | no | no | no |
| GRIN2A | 372 | W | 0 | 0 | no | no | yes | no | yes | no | no | no | no | no | no | no | no |
| GRIN2A | 373 | E | 0 | 0 | no | no | yes | no | yes | no | no | no | no | no | no | no | no |
| GRIN2A | 374 | K | 0 | 0 | no | no | yes | no | yes | no | no | no | no | no | no | no | no |
| GRIN2A | 375 | V | 1 | 0 | no | no | yes | no | yes | no | no | no | no | no | no | no | no |
| GRIN2A | 376 | G | 0 | 0 | no | no | yes | no | yes | no | no | no | no | no | no | no | no |
| GRIN2A | 377 | K | 0 | 0 | no | no | yes | no | yes | no | no | no | no | no | no | no | no |
| GRIN2A | 378 | W | 0 | 0 | no | yes | no | no | yes | no | no | no | no | no | no | no | no |
| GRIN2A | 379 | E | 0 | 0 | no | yes | no | no | yes | no | no | no | no | no | no | no | no |
| GRIN2A | 380 | N | 0 | 0 | no | yes | no | no | yes | no | no | no | no | no | no | no | no |
| GRIN2A | 381 | H | 1 | 0 | no | yes | no | no | yes | no | no | no | no | no | no | no | no |
| GRIN2A | 382 | T | 1 | 0 | no | yes | no | no | yes | no | no | no | no | no | no | no | no |
| GRIN2A | 383 | L | 0 | 0 | no | yes | no | no | yes | no | no | no | no | no | no | no | no |
| GRIN2A | 384 | S | 0 | 0 | no | yes | no | no | yes | no | no | no | no | no | no | no | no |
| GRIN2A | 385 | L | 0 | 0 | no | no | yes | no | yes | no | no | no | no | no | no | no | no |
| GRIN2A | 386 | R | 0 | 0 | no | yes | no | no | yes | no | no | no | no | no | no | no | no |
| GRIN2A | 387 | H | 1 | 0 | no | yes | no | no | yes | no | no | no | no | no | no | no | no |
| GRIN2A | 388 | A | 1 | 0 | no | yes | no | no | yes | no | no | no | no | no | no | no | no |
| GRIN2A | 389 | V | 1 | 0 | no | yes | no | no | yes | no | no | no | no | no | no | no | no |
| GRIN2A | 390 | W | 0 | 0 | no | yes | no | no | yes | no | no | no | no | no | no | no | no |
| GRIN2A | 391 | P | 0 | 0 | no | yes | no | no | yes | no | no | no | no | no | no | no | no |

| | | | | | | | | | | | | | | | | | |
|---|---|---|---|---|---|---|---|---|---|---|---|---|---|---|---|---|---|
| GRIN2A | 392 | R | 0 | 0 | no | yes | no | no | yes | no | no | no | no | no | no | no | no |
| GRIN2A | 393 | Y | 0 | 0 | no | yes | no | no | yes | no | no | no | no | no | no | no | no |
| GRIN2A | 394 | K | 0 | 0 | no | yes | no | no | yes | no | no | no | no | no | no | no | no |
| GRIN2A | 395 | S | 0 | 0 | no | yes | no | no | yes | no | no | no | no | no | no | no | no |
| GRIN2A | 396 | F | 0 | 0 | no | yes | no | no | yes | no | no | no | no | no | no | no | no |
| GRIN2A | 397 | S | 0 | 0 | no | yes | no | no | yes | no | no | no | no | no | no | no | no |
| GRIN2A | 398 | D | 0 | 0 | no | yes | no | no | yes | no | no | no | no | no | no | no | no |
| GRIN2A | 401 | P | 1 | 0 | no | yes | no | no | yes | no | no | no | no | no | no | no | no |
| GRIN2A | 402 | D | 0 | 0 | no | yes | no | no | yes | no | no | no | no | no | no | no | no |
| GRIN2A | 403 | D | 0 | 0 | no | yes | no | no | yes | no | no | no | no | no | no | no | no |
| GRIN2A | 404 | N | 0 | 0 | no | yes | no | no | yes | no | no | no | no | no | no | no | no |
| GRIN2A | 405 | H | 0 | 0 | no | no | yes | no | yes | no | no | no | no | no | no | no | no |
| GRIN2A | 406 | L | 0 | 0 | no | no | yes | no | yes | no | no | no | no | no | no | no | no |
| GRIN2A | 407 | S | 0 | 0 | no | no | yes | no | yes | no | no | no | no | no | no | no | no |
| GRIN2A | 408 | I | 0 | 0 | no | no | yes | no | yes | no | no | no | no | no | no | no | no |
| GRIN2A | 409 | V | 0 | 0 | no | no | yes | no | yes | no | no | no | no | no | no | no | no |
| GRIN2A | 410 | T | 0 | 0 | no | no | yes | no | yes | no | no | no | no | no | no | no | no |
| GRIN2A | 411 | L | 0 | 1 | no | yes | no | no | yes | no | no | no | no | no | no | no | no |
| GRIN2A | 412 | E | 0 | 0 | no | yes | no | no | yes | no | no | no | no | no | no | no | no |
| GRIN2A | 413 | E | 0 | 0 | no | yes | no | no | yes | no | no | no | no | no | no | no | no |
| GRIN2A | 414 | A | 1 | 0 | no | yes | no | no | yes | no | no | no | no | no | no | no | no |
| GRIN2A | 415 | P | 0 | 0 | no | yes | no | no | yes | no | no | no | no | no | no | no | no |
| GRIN2A | 416 | F | 0 | 0 | no | yes | no | no | yes | no | no | no | no | no | no | no | no |
| GRIN2A | 417 | V | 1 | 0 | no | yes | no | no | yes | no | no | no | no | no | no | no | no |
| GRIN2A | 418 | I | 0 | 0 | no | no | yes | no | yes | no | no | no | no | no | no | no | no |
| GRIN2A | 419 | V | 0 | 0 | no | no | yes | no | yes | no | no | no | no | no | no | no | no |
| GRIN2A | 420 | E | 0 | 0 | no | no | yes | no | yes | no | no | no | yes | no | no | no | no |
| GRIN2A | 421 | D | 0 | 0 | no | yes | no | no | yes | no | no | no | no | no | no | no | no |
| GRIN2A | 422 | I | 1 | 0 | no | yes | no | no | yes | no | no | no | no | no | no | no | no |
| GRIN2A | 423 | D | 1 | 0 | no | yes | no | no | yes | no | no | no | no | no | no | no | no |
| GRIN2A | 424 | P | 1 | 0 | no | yes | no | no | yes | no | no | no | no | no | no | no | no |
| GRIN2A | 425 | L | 1 | 0 | no | yes | no | no | yes | no | no | no | no | no | no | no | no |
| GRIN2A | 426 | T | 0 | 0 | no | yes | no | no | yes | no | no | no | yes | no | no | no | no |

| | | | | | | | | | | | | | | | | | |
|---|---|---|---|---|---|---|---|---|---|---|---|---|---|---|---|---|---|
| GRIN2A | 427 | E | 1 | 0 | no | yes | no | no | yes | no | no | no | no | no | no | no | no |
| GRIN2A | 428 | T | 0 | 0 | no | yes | no | no | yes | no | no | no | no | no | no | no | no |
| GRIN2A | 429 | C | 0 | 0 | no | yes | no | no | yes | no | no | no | no | no | no | no | no |
| GRIN2A | 430 | V | 0 | 0 | no | yes | no | no | yes | no | no | no | no | no | no | no | no |
| GRIN2A | 431 | R | 1 | 0 | no | yes | no | no | yes | no | no | no | yes | no | no | no | no |
| GRIN2A | 432 | N | 1 | 0 | no | yes | no | no | yes | no | no | no | yes | no | no | no | no |
| GRIN2A | 433 | T | 0 | 0 | no | yes | no | no | yes | no | no | no | no | no | no | no | no |
| GRIN2A | 434 | V | 0 | 0 | no | no | yes | no | yes | no | no | no | no | no | no | no | no |
| GRIN2A | 435 | P | 0 | 0 | no | no | yes | no | yes | no | no | no | no | no | no | no | no |
| GRIN2A | 436 | C | 0 | 2 | no | no | yes | no | yes | no | no | no | no | no | no | no | no |
| GRIN2A | 437 | R | 1 | 0 | no | no | yes | no | yes | no | no | no | no | no | no | no | no |
| GRIN2A | 438 | K | 1 | 0 | no | no | yes | no | yes | no | no | no | no | no | no | no | no |
| GRIN2A | 439 | F | 1 | 0 | no | no | yes | no | yes | no | no | no | no | no | no | no | no |
| GRIN2A | 440 | V | 2 | 0 | no | no | yes | no | yes | no | no | no | no | no | no | no | no |
| GRIN2A | 441 | K | 1 | 1 | no | no | yes | no | yes | no | no | no | no | no | no | no | no |
| GRIN2A | 442 | I | 0 | 0 | no | yes | no | no | yes | no | no | no | no | no | no | no | no |
| GRIN2A | 443 | N | 0 | 0 | no | yes | no | no | yes | no | no | no | no | no | no | no | no |
| GRIN2A | 444 | N | 1 | 0 | no | yes | no | no | yes | no | no | no | no | no | no | no | no |
| GRIN2A | 445 | S | 0 | 0 | no | yes | no | no | yes | no | no | no | no | no | no | no | no |
| GRIN2A | 446 | T | 0 | 0 | no | yes | no | no | yes | no | no | no | no | no | no | no | no |
| GRIN2A | 447 | N | 2 | 1 | no | yes | no | no | yes | no | no | no | no | no | no | no | no |
| GRIN2A | 448 | E | 1 | 0 | no | yes | no | no | yes | no | no | no | no | no | no | no | no |
| GRIN2A | 449 | G | 2 | 0 | no | no | yes | no | yes | no | no | no | no | no | no | no | no |
| GRIN2A | 450 | M | 0 | 0 | no | no | yes | no | yes | no | no | no | no | no | no | no | no |
| GRIN2A | 451 | N | 1 | 0 | no | no | yes | no | yes | no | no | no | no | no | no | no | no |
| GRIN2A | 452 | V | 1 | 1 | no | no | yes | no | yes | no | no | no | no | no | no | no | no |
| GRIN2A | 453 | K | 0 | 0 | no | no | yes | no | yes | no | no | no | no | no | no | no | no |
| GRIN2A | 454 | K | 0 | 0 | no | no | yes | no | yes | no | no | no | no | no | no | no | no |
| GRIN2A | 455 | C | 0 | 1 | no | no | yes | no | yes | no | no | no | no | no | no | no | no |
| GRIN2A | 456 | C | 0 | 0 | no | no | yes | no | yes | no | no | no | no | no | no | no | no |
| GRIN2A | 457 | K | 1 | 0 | no | no | yes | no | yes | no | no | no | yes | no | no | no | no |
| GRIN2A | 458 | G | 0 | 1 | no | yes | no | no | yes | no | no | no | no | no | no | no | no |
| GRIN2A | 459 | F | 1 | 0 | yes | no | no | no | yes | no | no | no | no | no | no | no | no |

| | | | | | | | | | | | | | | | | | |
|---|---|---|---|---|---|---|---|---|---|---|---|---|---|---|---|---|---|
| GRIN2A | 460 | C | 0 | 0 | yes | no | no | no | yes | no | no | no | no | no | no | no | no |
| GRIN2A | 461 | I | 0 | 0 | yes | no | no | no | yes | no | no | no | no | no | no | no | no |
| GRIN2A | 462 | D | 1 | 0 | yes | no | no | no | yes | no | no | no | no | no | no | no | no |
| GRIN2A | 463 | I | 1 | 0 | yes | no | no | no | yes | no | no | no | no | no | no | no | no |
| GRIN2A | 464 | L | 0 | 0 | yes | no | no | no | yes | no | no | no | no | no | no | no | no |
| GRIN2A | 465 | K | 0 | 0 | yes | no | no | no | yes | no | no | no | no | no | no | no | no |
| GRIN2A | 466 | K | 0 | 0 | yes | no | no | no | yes | no | no | no | no | no | no | no | no |
| GRIN2A | 467 | L | 0 | 0 | yes | no | no | no | yes | no | no | no | no | no | no | no | no |
| GRIN2A | 468 | S | 0 | 0 | yes | no | no | no | yes | no | no | no | no | no | no | no | no |
| GRIN2A | 469 | R | 0 | 0 | yes | no | no | no | yes | no | yes | no | no | no | no | no | no |
| GRIN2A | 470 | T | 1 | 0 | yes | no | no | no | yes | no | yes | no | no | no | no | no | no |
| GRIN2A | 471 | V | 1 | 0 | no | yes | no | no | yes | no | no | no | no | no | no | no | no |
| GRIN2A | 472 | K | 0 | 0 | no | yes | no | no | yes | no | yes | no | no | no | no | no | no |
| GRIN2A | 473 | F | 0 | 0 | no | yes | no | no | yes | no | no | no | no | no | no | no | no |
| GRIN2A | 474 | T | 0 | 0 | no | no | yes | no | yes | no | no | no | no | no | no | no | no |
| GRIN2A | 475 | Y | 0 | 0 | no | no | yes | no | yes | no | no | no | no | no | no | no | no |
| GRIN2A | 476 | D | 1 | 0 | no | no | yes | no | yes | no | no | no | no | no | no | no | no |
| GRIN2A | 477 | L | 0 | 0 | no | no | yes | no | yes | no | no | no | no | no | no | no | no |
| GRIN2A | 478 | Y | 1 | 0 | no | no | yes | no | yes | no | no | no | no | no | no | no | no |
| GRIN2A | 479 | L | 0 | 0 | no | no | yes | no | yes | no | no | no | no | no | no | no | no |
| GRIN2A | 480 | V | 0 | 0 | no | yes | no | no | yes | no | no | no | no | no | no | no | no |
| GRIN2A | 481 | T | 1 | 0 | no | yes | no | no | yes | no | no | no | no | no | no | no | no |
| GRIN2A | 482 | N | 1 | 0 | no | yes | no | no | yes | no | no | no | no | no | no | no | no |
| GRIN2A | 483 | G | 0 | 1 | no | yes | no | no | yes | no | no | no | no | no | no | no | no |
| GRIN2A | 484 | K | 0 | 0 | no | yes | no | no | yes | no | no | no | no | no | no | no | no |
| GRIN2A | 485 | H | 0 | 0 | no | yes | no | no | yes | no | no | no | no | no | no | no | no |
| GRIN2A | 486 | G | 0 | 0 | no | yes | no | no | yes | no | no | no | no | no | no | no | no |
| GRIN2A | 487 | K | 0 | 0 | no | yes | no | no | yes | no | no | no | no | no | no | no | no |
| GRIN2A | 488 | K | 0 | 0 | no | yes | no | no | yes | no | no | no | no | no | no | no | no |
| GRIN2A | 489 | V | 0 | 0 | no | yes | no | no | yes | no | no | no | no | no | no | no | no |
| GRIN2A | 490 | N | 0 | 0 | no | yes | no | no | yes | no | no | no | no | no | no | no | no |
| GRIN2A | 491 | N | 1 | 0 | no | yes | no | no | yes | no | no | no | no | no | no | no | no |
| GRIN2A | 492 | V | 0 | 0 | no | yes | no | no | yes | no | no | no | no | no | no | no | no |

| | | | | | | | | | | | | | | | | | |
|---|---|---|---|---|---|---|---|---|---|---|---|---|---|---|---|---|---|
| GRIN2A | 493 | W | 0 | 0 | no | yes | no | no | yes | no | no | no | no | no | no | no | no |
| GRIN2A | 494 | N | 0 | 0 | no | yes | no | no | yes | no | no | no | no | no | no | no | no |
| GRIN2A | 495 | G | 0 | 0 | yes | no | no | no | yes | no | no | no | no | no | no | no | no |
| GRIN2A | 496 | M | 0 | 0 | yes | no | no | no | yes | no | no | no | no | no | no | no | no |
| GRIN2A | 497 | I | 0 | 0 | yes | no | no | no | yes | no | no | no | no | no | no | no | no |
| GRIN2A | 498 | G | 1 | 1 | yes | no | no | no | yes | no | no | no | no | no | no | no | no |
| GRIN2A | 499 | E | 0 | 0 | yes | no | no | no | yes | no | no | no | no | no | no | no | no |
| GRIN2A | 500 | V | 0 | 0 | yes | no | no | no | yes | no | no | no | no | no | no | no | no |
| GRIN2A | 501 | V | 0 | 0 | yes | no | no | no | yes | no | no | no | no | no | no | no | no |
| GRIN2A | 502 | Y | 0 | 0 | no | yes | no | no | yes | no | no | no | no | no | no | no | no |
| GRIN2A | 503 | Q | 0 | 0 | no | yes | no | no | yes | no | no | no | no | no | no | no | no |
| GRIN2A | 504 | R | 1 | 1 | no | yes | no | no | yes | no | no | no | no | no | no | no | no |
| GRIN2A | 505 | A | 0 | 0 | no | yes | no | no | yes | no | no | no | no | no | no | no | no |
| GRIN2A | 506 | V | 1 | 0 | no | yes | no | no | yes | no | no | no | no | no | no | no | no |
| GRIN2A | 507 | M | 0 | 0 | no | no | yes | no | yes | no | no | no | no | no | no | no | no |
| GRIN2A | 508 | A | 0 | 0 | no | no | yes | no | yes | no | no | no | no | no | no | no | no |
| GRIN2A | 509 | V | 0 | 0 | no | yes | no | no | yes | no | no | no | no | no | no | no | no |
| GRIN2A | 510 | G | 0 | 0 | no | yes | no | no | yes | no | no | no | no | no | no | no | no |
| GRIN2A | 511 | S | 0 | 0 | no | yes | no | no | yes | no | no | no | no | no | no | no | no |
| GRIN2A | 512 | L | 0 | 0 | no | yes | no | no | yes | no | no | no | no | no | no | no | no |
| GRIN2A | 513 | T | 0 | 1 | no | yes | no | no | yes | no | no | no | no | no | no | no | no |
| GRIN2A | 514 | I | 0 | 0 | no | yes | no | no | yes | no | no | no | no | no | no | no | no |
| GRIN2A | 515 | N | 0 | 0 | no | yes | no | no | yes | no | no | no | yes | no | no | no | no |
| GRIN2A | 516 | E | 0 | 0 | yes | no | no | no | yes | no | no | no | yes | no | no | no | no |
| GRIN2A | 517 | E | 0 | 0 | yes | no | no | no | yes | no | no | no | no | no | no | no | no |
| GRIN2A | 518 | R | 0 | 3 | yes | no | no | no | yes | no | no | no | no | no | no | no | no |
| GRIN2A | 519 | S | 0 | 0 | yes | no | no | no | yes | no | no | no | yes | no | no | no | no |
| GRIN2A | 520 | E | 1 | 0 | yes | no | no | no | yes | no | no | no | no | no | no | no | no |
| GRIN2A | 521 | V | 1 | 0 | yes | no | no | no | yes | no | no | no | no | no | no | no | no |
| GRIN2A | 522 | V | 0 | 0 | no | yes | no | no | yes | no | no | no | no | no | no | no | no |
| GRIN2A | 523 | D | 0 | 0 | no | no | yes | no | yes | no | no | no | no | no | no | no | no |
| GRIN2A | 524 | F | 0 | 0 | no | no | yes | no | yes | no | no | no | yes | no | no | no | no |
| GRIN2A | 525 | S | 0 | 0 | no | yes | no | no | yes | no | no | no | no | no | no | no | no |

| | | | | | | | | | | | | | | | | | |
|---|---|---|---|---|---|---|---|---|---|---|---|---|---|---|---|---|---|
| GRIN2A | 526 | V | 0 | 0 | no | yes | no | no | yes | no | no | no | no | no | no | no | no |
| GRIN2A | 527 | P | 0 | 1 | no | yes | no | no | yes | no | no | no | yes | no | no | no | no |
| GRIN2A | 528 | F | 0 | 0 | no | yes | no | no | yes | no | no | no | no | no | no | no | no |
| GRIN2A | 529 | V | 0 | 0 | no | yes | no | no | yes | no | no | no | no | no | no | no | no |
| GRIN2A | 530 | E | 0 | 0 | no | yes | no | no | yes | no | no | no | yes | no | no | no | no |
| GRIN2A | 531 | T | 0 | 1 | no | yes | no | no | yes | no | no | no | no | no | no | no | no |
| GRIN2A | 532 | G | 0 | 1 | no | yes | no | no | yes | no | no | no | no | no | no | no | no |
| GRIN2A | 533 | I | 0 | 0 | no | no | yes | no | yes | no | no | no | no | no | no | no | no |
| GRIN2A | 534 | S | 0 | 0 | no | no | yes | no | yes | no | no | no | no | no | no | no | no |
| GRIN2A | 535 | V | 0 | 0 | no | no | yes | no | yes | no | no | no | no | no | no | no | no |
| GRIN2A | 536 | M | 0 | 0 | no | no | yes | no | yes | no | no | no | no | no | no | no | no |
| GRIN2A | 537 | V | 0 | 0 | no | no | yes | no | yes | no | no | no | no | no | no | no | no |
| GRIN2A | 538 | S | 0 | 0 | no | yes | no | no | yes | no | no | no | no | no | no | no | no |
| GRIN2A | 556 | S | 0 | 0 | no | yes | no | yes | no | no | no | no | yes | no | no | no | no |
| GRIN2A | 557 | V | 0 | 0 | yes | no | no | yes | no | no | no | no | yes | no | no | no | no |
| GRIN2A | 558 | W | 0 | 1 | yes | no | no | yes | no | no | no | no | no | no | no | no | no |
| GRIN2A | 559 | V | 1 | 0 | yes | no | no | yes | no | no | no | no | no | no | no | no | no |
| GRIN2A | 560 | M | 0 | 0 | yes | no | no | yes | no | no | no | no | yes | no | no | no | no |
| GRIN2A | 561 | M | 0 | 0 | yes | no | no | yes | no | no | no | no | no | no | no | no | no |
| GRIN2A | 562 | F | 0 | 0 | yes | no | no | yes | no | no | no | no | no | no | no | no | no |
| GRIN2A | 563 | V | 0 | 0 | yes | no | no | yes | no | no | no | no | no | no | no | no | no |
| GRIN2A | 564 | M | 1 | 0 | yes | no | no | yes | no | no | no | no | yes | no | no | no | no |
| GRIN2A | 565 | L | 0 | 0 | yes | no | no | yes | no | no | no | no | no | no | no | no | no |
| GRIN2A | 566 | L | 0 | 0 | yes | no | no | yes | no | no | no | no | no | no | no | no | no |
| GRIN2A | 567 | I | 0 | 0 | yes | no | no | yes | no | no | no | no | no | no | no | no | no |
| GRIN2A | 568 | V | 1 | 0 | yes | no | no | yes | no | no | no | no | no | no | no | no | no |
| GRIN2A | 569 | S | 0 | 0 | yes | no | no | yes | no | no | no | no | no | no | no | no | no |
| GRIN2A | 570 | A | 0 | 0 | yes | no | no | yes | no | no | no | no | no | no | no | no | no |
| GRIN2A | 571 | I | 1 | 0 | yes | no | no | yes | no | no | no | no | yes | no | no | no | no |
| GRIN2A | 572 | A | 0 | 0 | yes | no | no | yes | no | no | no | no | no | no | no | no | no |
| GRIN2A | 573 | V | 0 | 0 | yes | no | no | yes | no | no | no | no | no | no | no | no | no |
| GRIN2A | 574 | F | 0 | 0 | yes | no | no | yes | no | no | no | no | no | no | no | no | no |
| GRIN2A | 575 | V | 0 | 0 | yes | no | no | yes | no | no | no | no | no | no | no | no | no |

| | | | | | | | | | | | | | | | | | |
|---|---|---|---|---|---|---|---|---|---|---|---|---|---|---|---|---|---|
| GRIN2A | 576 | F | 1 | 0 | yes | no | no | no | no | no | no | no | no | no | no | no | no |
| GRIN2A | 577 | E | 0 | 0 | yes | no | no | no | no | yes | no | no | no | no | no | no | no |
| GRIN2A | 578 | Y | 0 | 0 | no | yes | no | no | no | yes | no | no | yes | no | no | no | no |
| GRIN2A | 579 | F | 0 | 0 | no | yes | no | no | no | yes | no | no | no | no | no | no | no |
| GRIN2A | 580 | S | 1 | 0 | no | yes | no | no | no | yes | no | no | no | no | no | no | no |
| GRIN2A | 581 | P | 0 | 0 | no | yes | no | no | no | yes | no | no | no | no | no | no | no |
| GRIN2A | 598 | S | 0 | 0 | no | yes | no | no | no | yes | no | no | no | no | no | no | no |
| GRIN2A | 599 | F | 0 | 0 | no | yes | no | no | no | yes | no | yes | no | no | no | no | no |
| GRIN2A | 600 | T | 0 | 0 | yes | no | no | yes | no | yes | no | yes | no | no | no | no | no |
| GRIN2A | 601 | I | 1 | 0 | yes | no | no | yes | no | no | no | yes | no | no | no | no | no |
| GRIN2A | 602 | G | 0 | 0 | yes | no | no | yes | no | no | no | yes | no | no | no | no | no |
| GRIN2A | 603 | K | 0 | 0 | yes | no | no | yes | no | no | yes | yes | no | no | no | no | no |
| GRIN2A | 604 | A | 0 | 0 | yes | no | no | yes | no | no | no | yes | no | no | no | no | no |
| GRIN2A | 605 | I | 0 | 0 | yes | no | no | yes | no | no | no | yes | no | no | no | no | no |
| GRIN2A | 606 | W | 0 | 0 | yes | no | no | yes | no | no | no | yes | yes | no | no | no | no |
| GRIN2A | 607 | L | 1 | 0 | yes | no | no | yes | no | no | no | yes | yes | no | no | no | no |
| GRIN2A | 608 | L | 0 | 0 | yes | no | no | yes | no | no | no | yes | no | no | no | no | no |
| GRIN2A | 609 | W | 0 | 0 | yes | no | no | yes | no | no | no | yes | yes | no | no | no | no |
| GRIN2A | 610 | G | 0 | 0 | yes | no | no | yes | no | no | no | yes | yes | no | no | no | no |
| GRIN2A | 611 | L | 0 | 1 | yes | no | no | yes | no | no | no | yes | no | no | no | no | no |
| GRIN2A | 612 | V | 0 | 0 | yes | no | no | yes | no | no | no | yes | no | no | no | no | no |
| GRIN2A | 613 | F | 0 | 0 | no | yes | no | yes | no | no | yes | yes | yes | no | no | no | no |
| GRIN2A | 625 | T | 0 | 0 | no | yes | no | no | no | yes | yes | no | no | no | no | no | no |
| GRIN2A | 626 | T | 0 | 0 | no | yes | no | yes | no | no | no | no | no | no | no | no | no |
| GRIN2A | 627 | S | 0 | 0 | yes | no | no | yes | no | no | no | no | no | no | no | no | no |
| GRIN2A | 628 | K | 0 | 0 | yes | no | no | yes | no | no | no | no | no | no | no | no | no |
| GRIN2A | 629 | I | 0 | 0 | yes | no | no | yes | no | no | no | no | yes | no | no | no | no |
| GRIN2A | 630 | M | 1 | 0 | yes | no | no | yes | no | no | no | no | yes | no | no | no | no |
| GRIN2A | 631 | V | 0 | 0 | yes | no | no | yes | no | no | no | no | no | no | no | no | no |
| GRIN2A | 632 | S | 0 | 1 | yes | no | no | yes | no | no | no | no | no | no | no | no | no |
| GRIN2A | 633 | V | 0 | 0 | yes | no | no | yes | no | no | no | no | no | no | no | no | no |
| GRIN2A | 634 | W | 0 | 0 | yes | no | no | yes | no | no | no | no | yes | no | no | no | no |
| GRIN2A | 635 | A | 0 | 1 | yes | no | no | yes | no | no | no | no | no | no | no | no | no |

| | | | | | | | | | | | | | | | | | |
|---|---|---|---|---|---|---|---|---|---|---|---|---|---|---|---|---|---|
| GRIN2A | 636 | F | 0 | 0 | yes | no | no | yes | no | no | no | no | yes | no | no | no | no |
| GRIN2A | 637 | F | 0 | 0 | yes | no | no | yes | no | no | no | no | yes | no | no | no | no |
| GRIN2A | 638 | A | 0 | 0 | yes | no | no | yes | no | no | no | no | no | no | no | no | no |
| GRIN2A | 639 | V | 0 | 0 | yes | no | no | yes | no | no | no | no | yes | no | no | no | no |
| GRIN2A | 640 | I | 0 | 0 | yes | no | no | yes | no | no | no | no | no | no | no | no | no |
| GRIN2A | 641 | F | 0 | 0 | yes | no | no | yes | no | no | no | no | yes | no | no | no | no |
| GRIN2A | 642 | L | 0 | 0 | yes | no | no | yes | no | no | yes | no | yes | no | no | no | no |
| GRIN2A | 643 | A | 0 | 1 | yes | no | no | yes | no | no | no | no | yes | no | no | no | no |
| GRIN2A | 644 | S | 0 | 1 | yes | no | no | yes | no | no | no | no | no | no | no | no | no |
| GRIN2A | 645 | Y | 0 | 0 | yes | no | no | yes | no | no | no | no | no | no | no | no | no |
| GRIN2A | 646 | T | 0 | 2 | yes | no | no | yes | yes | no | yes | no | no | no | no | no | no |
| GRIN2A | 647 | A | 0 | 1 | yes | no | no | no | yes | no | no | no | yes | no | no | no | no |
| GRIN2A | 648 | N | 0 | 1 | yes | no | no | no | yes | no | no | no | no | no | no | no | no |
| GRIN2A | 649 | L | 0 | 1 | yes | no | no | no | yes | no | no | no | no | no | no | no | no |
| GRIN2A | 650 | A | 0 | 0 | yes | no | no | no | yes | no | yes | no | yes | no | no | no | no |
| GRIN2A | 651 | A | 0 | 0 | yes | no | no | no | yes | no | no | no | no | no | no | no | no |
| GRIN2A | 652 | F | 0 | 1 | yes | no | no | no | yes | no | no | no | no | no | no | no | no |
| GRIN2A | 653 | M | 0 | 2 | no | yes | no | no | yes | no | no | no | no | no | no | no | no |
| GRIN2A | 654 | I | 0 | 1 | no | yes | no | no | yes | no | yes | no | yes | no | no | no | no |
| GRIN2A | 655 | Q | 0 | 0 | no | yes | no | no | yes | no | yes | no | yes | no | no | no | no |
| GRIN2A | 660 | D | 0 | 0 | no | yes | no | no | yes | no | no | no | no | no | no | no | no |
| GRIN2A | 661 | Q | 0 | 0 | no | yes | no | no | yes | no | no | no | no | no | no | no | no |
| GRIN2A | 662 | V | 0 | 0 | no | yes | no | no | yes | no | no | no | no | no | no | no | no |
| GRIN2A | 663 | T | 0 | 0 | no | yes | no | no | yes | no | no | no | no | no | no | no | no |
| GRIN2A | 664 | G | 0 | 0 | no | yes | no | no | yes | no | no | no | no | no | no | no | no |
| GRIN2A | 665 | L | 0 | 0 | no | yes | no | no | yes | no | no | no | no | no | no | no | no |
| GRIN2A | 666 | S | 0 | 0 | no | yes | no | no | yes | no | no | no | no | no | no | no | no |
| GRIN2A | 667 | D | 0 | 0 | no | yes | no | no | yes | no | no | no | no | no | no | no | no |
| GRIN2A | 668 | K | 0 | 0 | yes | no | no | no | yes | no | no | no | no | no | no | no | no |
| GRIN2A | 669 | K | 0 | 1 | yes | no | no | no | yes | no | no | no | no | no | no | no | no |
| GRIN2A | 670 | F | 0 | 0 | yes | no | no | no | yes | no | no | no | no | no | no | no | no |
| GRIN2A | 671 | Q | 1 | 0 | yes | no | no | no | yes | no | no | no | no | no | no | no | no |
| GRIN2A | 672 | R | 0 | 0 | no | yes | no | no | yes | no | no | no | no | no | no | no | no |

| | | | | | | | | | | | | | | | | | |
|---|---|---|---|---|---|---|---|---|---|---|---|---|---|---|---|---|---|
| GRIN2A | 673 | P | 1 | 0 | yes | no | no | no | yes | no | no | no | no | no | no | no | no |
| GRIN2A | 674 | H | 1 | 0 | yes | no | no | no | yes | no | no | no | no | no | no | no | no |
| GRIN2A | 675 | D | 2 | 0 | yes | no | no | no | yes | no | no | no | no | no | no | no | no |
| GRIN2A | 676 | Y | 1 | 0 | no | yes | no | no | yes | no | no | no | no | no | no | no | no |
| GRIN2A | 677 | S | 0 | 0 | no | yes | no | no | yes | no | no | no | no | no | no | no | no |
| GRIN2A | 678 | P | 1 | 0 | no | yes | no | no | yes | no | no | no | no | no | no | no | no |
| GRIN2A | 679 | P | 0 | 0 | no | yes | no | no | yes | no | no | no | no | no | no | no | no |
| GRIN2A | 680 | F | 0 | 0 | no | yes | no | no | yes | no | no | no | no | no | no | no | no |
| GRIN2A | 681 | R | 1 | 0 | no | yes | no | no | yes | no | no | no | no | no | no | no | no |
| GRIN2A | 682 | F | 1 | 0 | no | no | yes | no | yes | no | no | no | no | no | no | no | no |
| GRIN2A | 683 | G | 0 | 0 | no | no | yes | no | yes | no | no | no | no | no | no | no | no |
| GRIN2A | 684 | T | 0 | 1 | no | yes | no | no | yes | no | no | no | no | no | no | no | no |
| GRIN2A | 685 | V | 0 | 1 | no | yes | no | no | yes | no | no | no | no | no | no | no | no |
| GRIN2A | 686 | P | 0 | 0 | no | yes | no | no | yes | no | no | no | no | no | no | no | no |
| GRIN2A | 687 | N | 0 | 0 | no | yes | no | no | yes | no | no | no | no | no | no | no | no |
| GRIN2A | 688 | G | 0 | 0 | no | yes | no | no | yes | no | no | no | no | no | no | no | no |
| GRIN2A | 689 | S | 0 | 0 | yes | no | no | no | yes | no | no | no | no | no | no | no | no |
| GRIN2A | 690 | T | 1 | 1 | yes | no | no | no | yes | no | no | no | no | no | no | no | no |
| GRIN2A | 691 | E | 0 | 0 | yes | no | no | no | yes | no | no | no | no | no | no | no | no |
| GRIN2A | 692 | R | 0 | 0 | yes | no | no | no | yes | no | no | no | no | no | no | no | no |
| GRIN2A | 693 | N | 0 | 0 | yes | no | no | no | yes | no | no | no | yes | no | no | no | no |
| GRIN2A | 694 | I | 0 | 1 | yes | no | no | no | yes | no | no | no | no | no | no | no | no |
| GRIN2A | 695 | R | 0 | 1 | yes | no | no | no | yes | no | no | no | no | no | no | no | no |
| GRIN2A | 696 | N | 0 | 0 | yes | no | no | no | yes | no | no | no | no | no | no | no | no |
| GRIN2A | 697 | N | 0 | 0 | yes | no | no | no | yes | no | no | no | yes | no | no | no | no |
| GRIN2A | 698 | Y | 0 | 0 | no | yes | no | no | yes | no | no | no | no | no | no | no | no |
| GRIN2A | 699 | P | 0 | 1 | yes | no | no | no | yes | no | no | no | no | no | no | no | no |
| GRIN2A | 700 | Y | 0 | 0 | yes | no | no | no | yes | no | no | no | no | no | no | no | no |
| GRIN2A | 702 | H | 0 | 0 | yes | no | no | no | yes | no | no | no | no | no | no | no | no |
| GRIN2A | 703 | Q | 0 | 0 | yes | no | no | no | yes | no | no | no | no | no | no | no | no |
| GRIN2A | 704 | Y | 0 | 0 | yes | no | no | no | yes | no | no | no | no | no | no | no | no |
| GRIN2A | 705 | M | 0 | 1 | yes | no | no | no | yes | no | no | no | no | no | no | no | no |
| GRIN2A | 706 | T | 0 | 0 | no | yes | no | no | yes | no | no | no | no | no | no | no | no |

| | | | | | | | | | | | | | | | | | |
|---|---|---|---|---|---|---|---|---|---|---|---|---|---|---|---|---|---|
| GRIN2A | 707 | K | 0 | 0 | no | yes | no | no | yes | no | no | no | no | no | no | no | no |
| GRIN2A | 708 | F | 0 | 0 | no | yes | no | no | yes | no | no | no | no | no | no | no | no |
| GRIN2A | 709 | N | 0 | 0 | no | yes | no | no | yes | no | no | no | no | no | no | no | no |
| GRIN2A | 710 | Q | 0 | 0 | no | yes | no | no | yes | no | no | no | no | no | no | no | no |
| GRIN2A | 711 | K | 0 | 0 | no | yes | no | no | yes | no | no | no | no | no | no | no | no |
| GRIN2A | 712 | G | 0 | 0 | no | yes | no | no | yes | no | no | no | no | no | no | no | no |
| GRIN2A | 713 | V | 0 | 1 | yes | no | no | no | yes | no | no | no | no | no | no | no | no |
| GRIN2A | 714 | E | 0 | 1 | yes | no | no | no | yes | no | no | no | no | no | no | no | no |
| GRIN2A | 715 | D | 0 | 0 | yes | no | no | no | yes | no | no | no | no | no | no | no | no |
| GRIN2A | 716 | A | 0 | 1 | yes | no | no | no | yes | no | no | no | no | no | no | no | no |
| GRIN2A | 717 | L | 0 | 0 | yes | no | no | no | yes | no | no | no | no | no | no | no | no |
| GRIN2A | 718 | V | 0 | 0 | yes | no | no | no | yes | no | no | no | no | no | no | no | no |
| GRIN2A | 719 | S | 1 | 0 | yes | no | no | no | yes | no | no | no | no | no | no | no | no |
| GRIN2A | 720 | L | 0 | 0 | yes | no | no | no | yes | no | no | no | no | no | no | no | no |
| GRIN2A | 721 | K | 0 | 0 | yes | no | no | no | yes | no | no | no | no | no | no | no | no |
| GRIN2A | 722 | T | 2 | 0 | no | yes | no | no | yes | no | no | no | no | no | no | no | no |
| GRIN2A | 723 | G | 0 | 0 | no | yes | no | no | yes | no | no | no | no | no | no | no | no |
| GRIN2A | 724 | K | 0 | 0 | no | yes | no | no | yes | no | no | no | no | no | no | no | no |
| GRIN2A | 725 | L | 0 | 0 | no | yes | no | no | yes | no | no | no | no | no | no | no | no |
| GRIN2A | 726 | D | 0 | 0 | no | yes | no | no | yes | no | no | no | no | no | no | no | no |
| GRIN2A | 727 | A | 0 | 1 | no | no | yes | no | yes | no | no | no | no | no | no | no | no |
| GRIN2A | 728 | F | 0 | 0 | no | no | yes | no | yes | no | no | no | no | no | no | no | no |
| GRIN2A | 729 | I | 0 | 0 | no | no | yes | no | yes | no | no | no | no | no | no | no | no |
| GRIN2A | 730 | Y | 0 | 1 | no | no | yes | no | yes | no | no | no | no | no | no | no | no |
| GRIN2A | 731 | D | 0 | 1 | no | no | yes | no | yes | no | no | no | no | no | no | no | no |
| GRIN2A | 732 | A | 0 | 0 | yes | no | no | no | yes | no | no | no | no | no | no | no | no |
| GRIN2A | 733 | A | 0 | 1 | yes | no | no | no | yes | no | no | no | no | no | no | no | no |
| GRIN2A | 734 | V | 1 | 1 | yes | no | no | no | yes | no | no | no | no | no | no | no | no |
| GRIN2A | 735 | L | 0 | 0 | yes | no | no | no | yes | no | no | no | no | no | no | no | no |
| GRIN2A | 736 | N | 0 | 0 | yes | no | no | no | yes | no | no | no | no | no | no | no | no |
| GRIN2A | 737 | Y | 0 | 0 | yes | no | no | no | yes | no | no | no | no | no | no | no | no |
| GRIN2A | 738 | K | 0 | 0 | yes | no | no | no | yes | no | no | no | no | no | no | no | no |
| GRIN2A | 739 | A | 0 | 0 | yes | no | no | no | yes | no | no | no | no | no | no | no | no |

| | | | | | | | | | | | | | | | | | |
|---|---|---|---|---|---|---|---|---|---|---|---|---|---|---|---|---|---|
| GRIN2A | 740 | G | 0 | 0 | no | yes | no | no | yes | no | no | no | no | no | no | no | no |
| GRIN2A | 741 | R | 0 | 0 | no | yes | no | no | yes | no | no | no | no | no | no | no | no |
| GRIN2A | 742 | D | 1 | 0 | no | yes | no | no | yes | no | no | no | no | no | no | no | no |
| GRIN2A | 743 | E | 1 | 0 | no | yes | no | no | yes | no | no | no | no | no | no | no | no |
| GRIN2A | 744 | G | 0 | 0 | no | yes | no | no | yes | no | no | no | no | no | no | no | no |
| GRIN2A | 745 | C | 0 | 0 | no | yes | no | no | yes | no | no | no | no | no | no | no | no |
| GRIN2A | 746 | K | 0 | 0 | no | yes | no | no | yes | no | no | no | no | no | no | no | no |
| GRIN2A | 747 | L | 0 | 0 | no | yes | no | no | yes | no | no | no | no | no | no | no | no |
| GRIN2A | 748 | V | 0 | 0 | no | no | yes | no | yes | no | no | no | no | no | no | no | no |
| GRIN2A | 749 | T | 0 | 0 | no | no | yes | no | yes | no | no | no | no | no | no | no | no |
| GRIN2A | 750 | I | 1 | 0 | no | yes | no | no | yes | no | no | no | no | no | no | no | no |
| GRIN2A | 751 | G | 0 | 0 | no | yes | no | no | yes | no | no | no | no | no | no | no | no |
| GRIN2A | 752 | S | 0 | 0 | no | yes | no | no | yes | no | no | no | no | no | no | no | no |
| GRIN2A | 753 | G | 0 | 1 | no | yes | no | no | yes | no | no | no | no | no | no | no | no |
| GRIN2A | 754 | Y | 0 | 0 | no | yes | no | no | yes | no | no | no | yes | no | no | no | no |
| GRIN2A | 755 | I | 1 | 0 | no | yes | no | no | yes | no | no | no | no | no | no | no | no |
| GRIN2A | 756 | F | 1 | 0 | no | no | yes | no | yes | no | no | no | yes | no | no | no | no |
| GRIN2A | 757 | A | 0 | 0 | no | yes | no | no | yes | no | no | no | yes | no | no | no | no |
| GRIN2A | 758 | T | 0 | 0 | no | yes | no | no | yes | no | no | no | no | no | no | no | no |
| GRIN2A | 759 | T | 0 | 0 | no | yes | no | no | yes | no | no | no | yes | no | no | no | no |
| GRIN2A | 763 | I | 0 | 0 | no | yes | no | no | yes | no | no | no | no | no | no | no | no |
| GRIN2A | 764 | A | 0 | 0 | no | no | yes | no | yes | no | no | no | no | no | no | no | no |
| GRIN2A | 765 | L | 0 | 0 | no | no | yes | no | yes | no | no | no | no | no | no | no | no |
| GRIN2A | 766 | Q | 0 | 0 | no | yes | no | no | yes | no | no | no | no | no | no | no | no |
| GRIN2A | 767 | K | 0 | 0 | no | yes | no | no | yes | no | no | no | no | no | no | no | no |
| GRIN2A | 768 | G | 1 | 0 | no | yes | no | no | yes | no | no | no | no | no | no | no | no |
| GRIN2A | 769 | S | 0 | 0 | no | yes | no | no | yes | no | no | no | no | no | no | no | no |
| GRIN2A | 770 | P | 0 | 0 | no | yes | no | no | yes | no | no | no | no | no | no | no | no |
| GRIN2A | 771 | W | 0 | 0 | no | yes | no | no | yes | no | no | no | no | no | no | no | no |
| GRIN2A | 772 | K | 0 | 1 | yes | no | no | no | yes | no | no | no | no | no | no | no | no |
| GRIN2A | 773 | R | 1 | 0 | yes | no | no | no | yes | no | no | no | yes | no | no | no | no |
| GRIN2A | 774 | Q | 0 | 0 | yes | no | no | no | yes | no | no | no | no | no | no | no | no |
| GRIN2A | 775 | I | 1 | 0 | yes | no | no | no | yes | no | no | no | no | no | no | no | no |

| | | | | | | | | | | | | | | | | | |
|---|---|---|---|---|---|---|---|---|---|---|---|---|---|---|---|---|---|
| GRIN2A | 776 | D | 1 | 1 | yes | no | no | no | yes | no | no | no | no | no | no | no | no |
| GRIN2A | 777 | L | 0 | 0 | yes | no | no | no | yes | no | no | no | yes | no | no | no | no |
| GRIN2A | 778 | A | 0 | 0 | yes | no | no | no | yes | no | no | no | no | no | no | no | no |
| GRIN2A | 779 | L | 0 | 0 | yes | no | no | no | yes | no | no | no | no | no | no | no | no |
| GRIN2A | 780 | L | 0 | 0 | yes | no | no | no | yes | no | no | no | yes | no | no | no | no |
| GRIN2A | 781 | Q | 0 | 0 | yes | no | no | no | yes | no | yes | no | yes | no | no | no | no |
| GRIN2A | 782 | F | 0 | 0 | yes | no | no | no | yes | no | no | no | no | no | no | no | no |
| GRIN2A | 783 | V | 1 | 0 | yes | no | no | no | yes | no | no | no | yes | no | no | no | no |
| GRIN2A | 784 | G | 1 | 0 | yes | no | no | no | yes | no | no | no | yes | no | no | no | no |
| GRIN2A | 785 | D | 0 | 0 | no | yes | no | no | yes | no | yes | no | yes | no | no | no | no |
| GRIN2A | 786 | G | 0 | 0 | no | yes | no | no | yes | no | yes | no | yes | no | no | no | no |
| GRIN2A | 787 | E | 0 | 0 | yes | no | no | no | yes | no | yes | no | no | no | no | no | no |
| GRIN2A | 788 | M | 1 | 0 | yes | no | no | no | yes | no | no | no | no | no | no | no | no |
| GRIN2A | 789 | E | 0 | 0 | yes | no | no | no | yes | no | yes | no | yes | no | no | no | no |
| GRIN2A | 790 | E | 0 | 0 | yes | no | no | no | yes | no | yes | no | no | no | no | no | no |
| GRIN2A | 791 | L | 0 | 0 | yes | no | no | no | yes | no | no | no | no | no | no | no | no |
| GRIN2A | 792 | E | 0 | 0 | yes | no | no | no | yes | no | no | no | yes | no | no | no | no |
| GRIN2A | 793 | T | 0 | 0 | yes | no | no | no | yes | no | yes | no | no | no | no | no | no |
| GRIN2A | 794 | L | 0 | 0 | yes | no | no | no | yes | no | no | no | yes | no | no | no | no |
| GRIN2A | 795 | W | 0 | 0 | yes | no | no | no | yes | no | no | no | no | no | no | no | no |
| GRIN2A | 796 | L | 0 | 0 | no | yes | no | no | yes | no | no | no | no | no | no | no | no |
| GRIN2A | 797 | T | 0 | 0 | no | yes | no | no | yes | no | no | no | no | no | no | no | no |
| GRIN2A | 798 | G | 0 | 0 | no | yes | no | no | yes | no | no | no | no | no | no | no | no |
| GRIN2A | 799 | I | 0 | 0 | no | yes | no | no | yes | no | no | no | no | no | no | no | no |
| GRIN2A | 800 | C | 0 | 0 | yes | no | no | no | yes | no | no | no | no | no | no | no | no |
| GRIN2A | 801 | H | 0 | 0 | yes | no | no | no | yes | no | no | no | no | no | no | no | no |
| GRIN2A | 802 | N | 0 | 0 | yes | no | no | no | yes | no | yes | no | no | no | no | no | no |
| GRIN2A | 803 | E | 0 | 0 | no | yes | no | no | yes | no | yes | no | no | no | no | no | no |
| GRIN2A | 804 | K | 0 | 0 | no | yes | no | no | yes | no | no | no | no | no | no | no | no |
| GRIN2A | 805 | N | 0 | 0 | no | yes | no | no | yes | no | yes | no | no | no | no | no | no |
| GRIN2A | 806 | E | 0 | 0 | no | yes | no | no | yes | no | no | no | no | no | no | no | no |
| GRIN2A | 807 | V | 0 | 0 | no | yes | no | no | yes | no | no | no | no | no | no | no | no |
| GRIN2A | 808 | M | 0 | 0 | no | yes | no | no | yes | no | no | no | no | no | no | no | no |

| | | | | | | | | | | | | | | | | | |
|---|---|---|---|---|---|---|---|---|---|---|---|---|---|---|---|---|---|
| GRIN2A | 809 | S | 0 | 1 | no | yes | no | no | yes | no | no | no | yes | no | no | no | no |
| GRIN2A | 814 | I | 1 | 1 | no | yes | no | no | yes | no | no | no | yes | no | no | no | no |
| GRIN2A | 815 | D | 0 | 0 | no | yes | no | no | yes | no | no | no | yes | no | no | no | no |
| GRIN2A | 816 | N | 0 | 0 | no | yes | no | no | yes | no | no | no | yes | no | no | no | no |
| GRIN2A | 817 | M | 0 | 3 | no | yes | no | no | no | no | no | no | no | no | no | no | no |
| GRIN2A | 818 | A | 1 | 1 | no | yes | no | yes | no | no | no | no | no | no | no | no | no |
| GRIN2A | 819 | G | 0 | 0 | no | yes | no | yes | no | no | no | no | no | no | no | no | no |
| GRIN2A | 820 | V | 1 | 0 | yes | no | no | yes | no | no | no | no | no | no | no | no | no |
| GRIN2A | 821 | F | 0 | 0 | yes | no | no | yes | no | no | no | no | no | no | no | no | no |
| GRIN2A | 822 | Y | 0 | 0 | yes | no | no | yes | no | no | no | no | no | no | no | no | no |
| GRIN2A | 823 | M | 0 | 0 | yes | no | no | yes | no | no | no | no | yes | no | no | no | no |
| GRIN2A | 824 | L | 0 | 0 | yes | no | no | yes | no | no | no | no | yes | no | no | no | no |
| GRIN2A | 825 | A | 0 | 0 | yes | no | no | yes | no | no | no | no | no | no | no | no | no |
| GRIN2A | 826 | A | 0 | 0 | yes | no | no | yes | no | no | no | no | no | no | no | no | no |
| GRIN2A | 827 | A | 1 | 0 | yes | no | no | yes | no | no | no | no | yes | no | no | no | no |
| GRIN2A | 828 | M | 1 | 0 | yes | no | no | yes | no | no | no | no | no | no | no | no | no |
| GRIN2A | 829 | A | 0 | 0 | yes | no | no | yes | no | no | no | no | no | no | no | no | no |
| GRIN2A | 830 | L | 0 | 0 | yes | no | no | yes | no | no | no | no | no | no | no | no | no |
| GRIN2A | 831 | S | 0 | 0 | yes | no | no | yes | no | no | no | no | yes | no | no | no | no |
| GRIN2A | 832 | L | 2 | 0 | yes | no | no | yes | no | no | no | no | no | no | no | no | no |
| GRIN2A | 833 | I | 1 | 0 | yes | no | no | yes | no | no | no | no | no | no | no | no | no |
| GRIN2A | 834 | T | 0 | 0 | yes | no | no | yes | no | no | no | no | yes | no | no | no | no |
| GRIN2A | 835 | F | 0 | 0 | yes | no | no | yes | no | no | no | no | yes | no | no | no | no |
| GRIN2A | 836 | I | 2 | 0 | yes | no | no | yes | no | no | no | no | no | no | no | no | no |
| GRIN2A | 837 | W | 0 | 0 | no | yes | no | yes | no | no | no | no | no | no | no | no | no |
| GRIN2A | 838 | E | 0 | 0 | no | yes | no | yes | no | yes | no | no | no | no | no | no | no |
| GRIN2A | 839 | H | 2 | 0 | no | yes | no | no | no | yes | no | no | no | no | no | no | no |
| GRIN2A | 840 | L | 1 | 1 | no | yes | no | no | no | yes | no | no | no | no | no | no | no |
| GRIN2A | 841 | F | 0 | 0 | no | yes | no | no | no | yes | no | no | no | no | no | no | no |
| HCN1 | 94 | M | 3 | 0 | no | yes | no | no | no | yes | no | no | no | no | no | no | no |
| HCN1 | 95 | Q | 0 | 0 | yes | no | no | no | no | yes | no | no | no | no | no | no | no |
| HCN1 | 96 | R | 0 | 0 | yes | no | no | no | no | yes | no | no | no | no | no | no | no |
| HCN1 | 97 | Q | 0 | 0 | yes | no | no | no | no | yes | no | no | no | no | no | no | no |

| | | | | | | | | | | | | | | | | | |
|---|---|---|---|---|---|---|---|---|---|---|---|---|---|---|---|---|---|
| HCN1 | 98 | F | 0 | 0 | yes | no | no | no | no | yes | no | no | no | no | no | no | no |
| HCN1 | 99 | T | 2 | 0 | yes | no | no | no | no | yes | no | no | no | no | no | no | no |
| HCN1 | 100 | S | 1 | 1 | yes | no | no | no | no | yes | no | no | no | no | no | no | no |
| HCN1 | 101 | M | 1 | 0 | yes | no | no | no | no | yes | no | no | no | no | no | no | no |
| HCN1 | 102 | L | 0 | 0 | yes | no | no | no | no | yes | no | no | no | no | no | no | no |
| HCN1 | 103 | Q | 0 | 0 | no | yes | no | no | no | yes | no | no | no | no | no | no | no |
| HCN1 | 104 | P | 0 | 0 | no | yes | no | no | no | yes | no | no | no | no | no | no | no |
| HCN1 | 105 | G | 0 | 0 | no | yes | no | no | no | yes | no | no | no | no | no | no | no |
| HCN1 | 106 | V | 0 | 0 | no | yes | no | no | no | yes | no | no | no | no | no | no | no |
| HCN1 | 107 | N | 0 | 0 | no | yes | no | no | no | yes | no | no | no | no | no | no | no |
| HCN1 | 108 | K | 0 | 0 | yes | no | no | no | no | yes | no | no | no | no | no | no | no |
| HCN1 | 109 | F | 0 | 0 | yes | no | no | no | no | yes | no | no | no | no | no | no | no |
| HCN1 | 110 | S | 1 | 0 | yes | no | no | no | no | yes | no | no | no | no | no | no | no |
| HCN1 | 111 | L | 0 | 0 | yes | no | no | no | no | yes | no | no | no | no | no | no | no |
| HCN1 | 112 | R | 0 | 0 | yes | no | no | no | no | yes | no | no | yes | no | no | no | no |
| HCN1 | 113 | M | 0 | 0 | yes | no | no | no | no | yes | no | no | no | no | no | no | no |
| HCN1 | 114 | F | 0 | 0 | yes | no | no | no | no | yes | no | no | no | no | no | no | no |
| HCN1 | 115 | G | 0 | 0 | no | yes | no | no | no | yes | no | no | no | no | no | no | no |
| HCN1 | 116 | S | 0 | 0 | no | yes | no | no | no | yes | no | no | yes | no | no | no | no |
| HCN1 | 117 | Q | 0 | 0 | yes | no | no | no | no | yes | no | no | no | no | no | no | no |
| HCN1 | 118 | K | 0 | 0 | yes | no | no | no | no | yes | no | no | no | no | no | no | no |
| HCN1 | 119 | A | 0 | 0 | yes | no | no | no | no | yes | no | no | no | no | no | no | no |
| HCN1 | 120 | V | 0 | 0 | yes | no | no | no | no | yes | no | no | no | no | no | no | no |
| HCN1 | 121 | E | 0 | 0 | yes | no | no | no | no | yes | no | no | no | no | no | no | no |
| HCN1 | 122 | K | 0 | 0 | yes | no | no | no | no | yes | no | no | no | no | no | no | no |
| HCN1 | 123 | E | 0 | 0 | yes | no | no | no | no | yes | no | no | no | no | no | no | no |
| HCN1 | 124 | Q | 0 | 0 | yes | no | no | no | no | yes | no | no | no | no | no | no | no |
| HCN1 | 125 | E | 1 | 0 | yes | no | no | no | no | yes | no | no | no | no | no | no | no |
| HCN1 | 126 | R | 0 | 0 | yes | no | no | no | no | yes | no | no | no | no | no | no | no |
| HCN1 | 127 | V | 1 | 0 | yes | no | no | no | no | yes | no | no | no | no | no | no | no |
| HCN1 | 128 | K | 0 | 0 | no | yes | no | no | no | yes | no | no | no | no | no | no | no |
| HCN1 | 129 | T | 0 | 0 | no | yes | no | no | no | yes | no | no | no | no | no | no | no |
| HCN1 | 130 | A | 0 | 0 | no | yes | no | no | no | yes | no | no | no | no | no | no | no |

| | | | | | | | | | | | | | | | | | |
|---|---|---|---|---|---|---|---|---|---|---|---|---|---|---|---|---|---|
| HCN1 | 131 | G | 0 | 0 | no | yes | no | no | no | yes | no | no | no | no | no | no | no |
| HCN1 | 132 | F | 0 | 0 | no | yes | no | no | no | yes | no | no | no | no | no | no | no |
| HCN1 | 133 | W | 0 | 0 | no | yes | no | no | no | yes | no | no | no | no | no | no | no |
| HCN1 | 134 | I | 0 | 0 | no | yes | no | no | no | yes | no | no | no | no | no | no | no |
| HCN1 | 135 | I | 0 | 0 | no | no | yes | no | no | yes | no | no | no | no | no | no | no |
| HCN1 | 136 | H | 0 | 0 | no | yes | no | no | no | yes | no | no | no | no | no | no | no |
| HCN1 | 137 | P | 1 | 0 | no | yes | no | no | no | yes | no | no | no | no | no | no | no |
| HCN1 | 138 | Y | 0 | 0 | no | yes | no | no | no | yes | no | no | no | no | no | no | no |
| HCN1 | 139 | S | 0 | 0 | no | yes | no | no | no | yes | no | no | no | no | no | no | no |
| HCN1 | 140 | D | 0 | 0 | yes | no | no | no | no | yes | no | no | no | no | no | no | no |
| HCN1 | 141 | F | 0 | 0 | yes | no | no | no | no | yes | no | no | no | no | no | no | no |
| HCN1 | 142 | R | 0 | 0 | yes | no | no | no | no | yes | no | no | no | no | no | no | no |
| HCN1 | 143 | F | 0 | 1 | yes | no | no | no | no | no | no | no | no | no | no | no | no |
| HCN1 | 144 | Y | 0 | 0 | yes | no | no | yes | no | no | no | no | no | no | no | no | no |
| HCN1 | 145 | W | 0 | 0 | yes | no | no | yes | no | no | no | no | no | no | no | no | no |
| HCN1 | 146 | D | 0 | 0 | yes | no | no | yes | no | no | no | no | no | no | no | no | no |
| HCN1 | 147 | L | 0 | 0 | yes | no | no | yes | no | no | no | no | no | no | no | no | no |
| HCN1 | 148 | I | 0 | 0 | yes | no | no | yes | no | no | no | no | no | no | no | no | no |
| HCN1 | 149 | M | 0 | 0 | yes | no | no | yes | no | no | no | no | no | no | no | no | no |
| HCN1 | 150 | L | 0 | 0 | yes | no | no | yes | no | no | no | no | no | no | no | no | no |
| HCN1 | 151 | I | 1 | 0 | yes | no | no | yes | no | no | no | no | no | no | no | no | no |
| HCN1 | 152 | M | 0 | 0 | yes | no | no | yes | no | no | no | no | no | no | no | no | no |
| HCN1 | 153 | M | 0 | 1 | yes | no | no | yes | no | no | no | no | no | no | no | no | no |
| HCN1 | 154 | V | 0 | 0 | yes | no | no | yes | no | no | no | no | no | no | no | no | no |
| HCN1 | 155 | G | 0 | 0 | yes | no | no | yes | no | no | no | no | no | no | no | no | no |
| HCN1 | 156 | N | 0 | 0 | yes | no | no | yes | no | no | no | no | no | no | no | no | no |
| HCN1 | 157 | L | 0 | 1 | yes | no | no | yes | no | no | no | no | no | no | no | no | no |
| HCN1 | 158 | V | 0 | 0 | yes | no | no | yes | no | no | no | no | no | no | no | no | no |
| HCN1 | 159 | I | 0 | 0 | yes | no | no | yes | no | no | no | no | no | no | no | no | no |
| HCN1 | 160 | I | 0 | 0 | yes | no | no | yes | no | no | no | no | no | no | no | no | no |
| HCN1 | 161 | P | 0 | 0 | yes | no | no | yes | no | no | no | no | no | no | no | no | no |
| HCN1 | 162 | V | 1 | 0 | yes | no | no | yes | no | no | no | no | no | no | no | no | no |
| HCN1 | 163 | G | 0 | 0 | yes | no | no | yes | no | no | no | no | no | no | no | no | no |

| | | | | | | | | | | | | | | | | | |
|---|---|---|---|---|---|---|---|---|---|---|---|---|---|---|---|---|---|
| HCN1 | 164 | I | 0 | 0 | yes | no | no | yes | no | no | no | no | no | no | no | no | no |
| HCN1 | 165 | T | 0 | 0 | yes | no | no | yes | yes | no | no | no | no | no | no | no | no |
| HCN1 | 166 | F | 0 | 0 | yes | no | no | no | yes | no | no | no | no | no | no | no | no |
| HCN1 | 167 | F | 0 | 0 | no | yes | no | no | yes | no | no | no | no | no | no | no | no |
| HCN1 | 168 | T | 0 | 0 | no | yes | no | no | yes | no | no | no | no | no | no | no | no |
| HCN1 | 169 | E | 0 | 0 | no | yes | no | no | yes | no | no | no | no | no | no | no | no |
| HCN1 | 170 | Q | 0 | 0 | no | yes | no | no | yes | no | no | no | no | no | no | no | no |
| HCN1 | 171 | T | 1 | 1 | no | yes | no | no | yes | no | no | no | no | no | no | no | no |
| HCN1 | 172 | T | 0 | 1 | no | yes | no | no | yes | no | no | no | no | no | no | no | no |
| HCN1 | 173 | T | 1 | 0 | yes | no | no | yes | yes | no | no | no | no | no | no | no | no |
| HCN1 | 174 | P | 0 | 0 | yes | no | no | yes | no | no | no | no | no | no | no | no | no |
| HCN1 | 175 | W | 0 | 0 | yes | no | no | yes | no | no | no | no | no | no | no | no | no |
| HCN1 | 176 | I | 0 | 0 | yes | no | no | yes | no | no | no | no | no | no | no | no | no |
| HCN1 | 177 | I | 0 | 0 | yes | no | no | yes | no | no | no | no | no | no | no | no | no |
| HCN1 | 178 | F | 0 | 0 | yes | no | no | yes | no | no | no | no | no | no | no | no | no |
| HCN1 | 179 | N | 0 | 0 | yes | no | no | yes | no | no | no | no | no | no | no | no | no |
| HCN1 | 180 | V | 0 | 0 | yes | no | no | yes | no | no | no | no | no | no | no | no | no |
| HCN1 | 181 | A | 0 | 0 | yes | no | no | yes | no | no | no | no | no | no | no | no | no |
| HCN1 | 182 | S | 0 | 0 | yes | no | no | yes | no | no | no | no | no | no | no | no | no |
| HCN1 | 183 | D | 0 | 0 | yes | no | no | yes | no | no | no | no | no | no | no | no | no |
| HCN1 | 184 | T | 0 | 0 | yes | no | no | yes | no | no | no | no | no | no | no | no | no |
| HCN1 | 185 | V | 0 | 0 | yes | no | no | yes | no | no | no | no | no | no | no | no | no |
| HCN1 | 186 | F | 0 | 0 | yes | no | no | yes | no | no | no | no | no | no | no | no | no |
| HCN1 | 187 | L | 0 | 0 | yes | no | no | yes | no | no | no | no | no | no | no | no | no |
| HCN1 | 188 | L | 0 | 0 | yes | no | no | yes | no | no | no | no | no | no | no | no | no |
| HCN1 | 189 | D | 0 | 0 | yes | no | no | yes | no | no | no | no | no | no | no | no | no |
| HCN1 | 190 | L | 0 | 0 | yes | no | no | yes | no | no | no | no | no | no | no | no | no |
| HCN1 | 191 | I | 0 | 0 | yes | no | no | yes | no | no | no | no | no | no | no | no | no |
| HCN1 | 192 | M | 0 | 0 | yes | no | no | yes | no | no | no | no | no | no | no | no | no |
| HCN1 | 193 | N | 0 | 0 | yes | no | no | yes | no | no | no | no | no | no | no | no | no |
| HCN1 | 194 | F | 0 | 0 | yes | no | no | yes | no | no | no | no | no | no | no | no | no |
| HCN1 | 195 | R | 0 | 0 | yes | no | no | no | no | yes | no | no | no | no | no | no | no |
| HCN1 | 196 | T | 0 | 0 | no | no | yes | no | no | yes | no | no | no | no | no | no | no |

| | | | | | | | | | | | | | | | | | |
|---|---|---|---|---|---|---|---|---|---|---|---|---|---|---|---|---|---|
| HCN1 | 197 | G | 0 | 0 | no | yes | no | no | no | yes | no | no | no | no | no | no | no |
| HCN1 | 198 | T | 0 | 0 | no | no | yes | no | no | yes | no | no | no | no | no | no | no |
| HCN1 | 199 | V | 0 | 0 | no | yes | no | no | no | yes | no | no | no | no | no | no | no |
| HCN1 | 200 | N | 1 | 0 | no | yes | no | no | no | yes | no | no | no | no | no | no | no |
| HCN1 | 203 | S | 0 | 0 | no | yes | no | no | no | yes | no | no | no | no | no | no | no |
| HCN1 | 204 | S | 0 | 0 | no | yes | no | no | no | yes | no | no | no | no | no | no | no |
| HCN1 | 205 | E | 0 | 0 | no | yes | no | no | no | yes | no | no | no | no | no | no | no |
| HCN1 | 206 | I | 1 | 0 | no | no | yes | no | no | yes | no | no | no | no | no | no | no |
| HCN1 | 207 | I | 0 | 0 | no | yes | no | no | no | yes | no | no | no | no | no | no | no |
| HCN1 | 208 | L | 0 | 0 | no | yes | no | no | no | yes | no | no | no | no | no | no | no |
| HCN1 | 209 | D | 0 | 0 | yes | no | no | no | no | yes | no | no | no | no | no | no | no |
| HCN1 | 210 | P | 0 | 0 | yes | no | no | no | no | yes | no | no | no | no | no | no | no |
| HCN1 | 211 | K | 1 | 0 | yes | no | no | no | no | yes | no | no | no | no | no | no | no |
| HCN1 | 212 | V | 0 | 0 | yes | no | no | no | no | yes | no | no | no | no | no | no | no |
| HCN1 | 213 | I | 0 | 0 | yes | no | no | no | no | yes | no | no | no | no | no | no | no |
| HCN1 | 214 | K | 0 | 0 | yes | no | no | no | no | yes | no | no | no | no | no | no | no |
| HCN1 | 215 | M | 1 | 0 | yes | no | no | no | no | yes | no | no | no | no | no | no | no |
| HCN1 | 216 | N | 0 | 0 | yes | no | no | no | no | no | no | no | no | no | no | no | no |
| HCN1 | 217 | Y | 0 | 0 | yes | no | no | yes | no | no | no | no | no | no | no | no | no |
| HCN1 | 218 | L | 0 | 0 | yes | no | no | yes | no | no | no | no | no | no | no | no | no |
| HCN1 | 219 | K | 0 | 0 | no | yes | no | yes | no | no | no | no | no | no | no | no | no |
| HCN1 | 220 | S | 0 | 0 | no | yes | no | yes | no | no | no | no | no | no | no | no | no |
| HCN1 | 221 | W | 0 | 0 | no | yes | no | yes | no | no | no | no | no | no | no | no | no |
| HCN1 | 222 | F | 0 | 0 | yes | no | no | yes | no | no | no | no | no | no | no | no | no |
| HCN1 | 223 | V | 0 | 0 | yes | no | no | yes | no | no | no | no | no | no | no | no | no |
| HCN1 | 224 | V | 0 | 0 | yes | no | no | yes | no | no | no | no | no | no | no | no | no |
| HCN1 | 225 | D | 0 | 0 | yes | no | no | yes | no | no | no | no | no | no | no | no | no |
| HCN1 | 226 | F | 1 | 0 | yes | no | no | yes | no | no | no | no | no | no | no | no | no |
| HCN1 | 227 | I | 1 | 0 | yes | no | no | yes | no | no | no | no | no | no | no | no | no |
| HCN1 | 228 | S | 0 | 0 | yes | no | no | yes | no | no | no | no | no | no | no | no | no |
| HCN1 | 229 | S | 0 | 0 | yes | no | no | yes | no | no | no | no | no | no | no | no | no |
| HCN1 | 230 | I | 0 | 0 | no | yes | no | yes | no | no | no | no | no | no | no | no | no |
| HCN1 | 231 | P | 0 | 0 | no | yes | no | yes | no | no | no | no | no | no | no | no | no |

| | | | | | | | | | | | | | | | | | |
|---|---|---|---|---|---|---|---|---|---|---|---|---|---|---|---|---|---|
| HCN1 | 232 | V | 0 | 0 | yes | no | no | yes | no | no | no | no | no | no | no | no | no |
| HCN1 | 233 | D | 0 | 0 | yes | no | no | yes | no | no | no | no | no | no | no | no | no |
| HCN1 | 234 | Y | 0 | 1 | yes | no | no | yes | no | no | no | no | no | no | no | no | no |
| HCN1 | 235 | I | 1 | 0 | yes | no | no | yes | no | no | no | no | no | no | no | no | no |
| HCN1 | 236 | F | 0 | 0 | yes | no | no | yes | no | no | no | no | no | no | no | no | no |
| HCN1 | 237 | L | 0 | 0 | yes | no | no | yes | yes | no | no | no | no | no | no | no | no |
| HCN1 | 238 | I | 0 | 0 | yes | no | no | yes | yes | no | no | no | no | no | no | no | no |
| HCN1 | 239 | V | 0 | 0 | yes | no | no | yes | yes | no | no | no | no | no | no | no | no |
| HCN1 | 240 | E | 0 | 1 | yes | no | no | no | yes | no | no | no | no | no | no | no | no |
| HCN1 | 241 | K | 0 | 0 | yes | no | no | no | yes | no | no | no | no | no | no | no | no |
| HCN1 | 242 | G | 0 | 0 | no | yes | no | yes | yes | no | no | no | no | no | no | no | no |
| HCN1 | 252 | R | 0 | 0 | no | yes | no | no | yes | no | no | no | no | no | no | no | no |
| HCN1 | 253 | A | 0 | 0 | yes | no | no | no | yes | no | no | no | no | no | no | no | no |
| HCN1 | 254 | L | 0 | 0 | yes | no | no | no | yes | no | no | no | no | no | no | no | no |
| HCN1 | 255 | R | 0 | 0 | yes | no | no | no | yes | no | no | no | no | no | no | no | no |
| HCN1 | 256 | I | 0 | 0 | yes | no | no | no | yes | no | no | no | no | no | no | no | no |
| HCN1 | 257 | V | 0 | 0 | yes | no | no | no | yes | no | no | no | no | no | no | no | no |
| HCN1 | 258 | R | 0 | 0 | yes | no | no | no | yes | no | no | no | no | no | no | no | no |
| HCN1 | 259 | F | 0 | 0 | yes | no | no | yes | yes | no | no | no | no | no | no | no | no |
| HCN1 | 260 | T | 0 | 1 | yes | no | no | yes | yes | no | no | no | no | no | no | no | no |
| HCN1 | 261 | K | 0 | 0 | yes | no | no | yes | no | no | no | no | no | yes | no | no | no |
| HCN1 | 262 | I | 0 | 0 | yes | no | no | yes | no | no | no | no | no | yes | no | no | no |
| HCN1 | 263 | L | 0 | 0 | yes | no | no | yes | no | no | no | no | no | yes | no | no | no |
| HCN1 | 264 | S | 1 | 1 | yes | no | no | yes | no | no | no | no | no | yes | no | no | no |
| HCN1 | 265 | L | 0 | 0 | yes | no | no | yes | no | no | no | no | no | yes | no | no | no |
| HCN1 | 266 | L | 0 | 0 | yes | no | no | yes | no | no | no | no | no | yes | no | no | no |
| HCN1 | 267 | R | 1 | 0 | yes | no | no | yes | no | no | no | no | no | yes | no | no | no |
| HCN1 | 268 | L | 0 | 0 | yes | no | no | yes | no | no | no | no | no | yes | no | no | no |
| HCN1 | 269 | L | 0 | 0 | yes | no | no | yes | no | no | no | no | no | yes | no | no | no |
| HCN1 | 270 | R | 0 | 0 | yes | no | no | yes | no | no | no | no | no | yes | no | no | no |
| HCN1 | 271 | L | 0 | 0 | yes | no | no | yes | no | no | no | no | no | yes | no | no | no |
| HCN1 | 272 | S | 0 | 1 | yes | no | no | yes | no | no | no | no | no | yes | no | no | no |
| HCN1 | 273 | R | 0 | 0 | yes | no | no | yes | no | no | no | no | no | yes | no | no | no |

| | | | | | | | | | | | | | | | | | |
|---|---|---|---|---|---|---|---|---|---|---|---|---|---|---|---|---|---|
| HCN1 | 274 | L | 0 | 0 | yes | no | no | yes | no | no | no | no | no | yes | no | no | no |
| HCN1 | 275 | I | 0 | 1 | yes | no | no | yes | no | no | no | no | no | yes | no | no | no |
| HCN1 | 276 | R | 0 | 0 | yes | no | no | yes | no | no | no | no | no | yes | no | no | no |
| HCN1 | 277 | Y | 0 | 0 | yes | no | no | yes | no | no | no | no | no | yes | no | no | no |
| HCN1 | 278 | I | 0 | 0 | yes | no | no | yes | no | no | no | no | no | yes | no | no | no |
| HCN1 | 279 | H | 0 | 1 | yes | no | no | yes | no | no | no | no | no | yes | no | no | no |
| HCN1 | 280 | Q | 0 | 0 | yes | no | no | no | no | no | no | no | no | yes | no | no | no |
| HCN1 | 281 | W | 1 | 0 | yes | no | no | no | no | no | no | no | no | yes | no | no | no |
| HCN1 | 282 | E | 0 | 0 | yes | no | no | no | no | yes | no | no | no | no | no | no | no |
| HCN1 | 283 | E | 0 | 0 | yes | no | no | no | no | yes | no | no | no | no | no | no | no |
| HCN1 | 284 | I | 0 | 0 | yes | no | no | no | no | yes | no | no | no | no | no | no | no |
| HCN1 | 285 | F | 1 | 0 | yes | no | no | no | no | yes | no | no | no | no | no | no | no |
| HCN1 | 286 | H | 0 | 0 | yes | no | no | no | no | yes | no | no | yes | no | no | no | no |
| HCN1 | 287 | M | 0 | 0 | yes | no | no | no | no | yes | no | no | yes | no | no | no | no |
| HCN1 | 288 | T | 0 | 0 | yes | no | no | no | no | yes | no | no | no | no | no | no | no |
| HCN1 | 289 | Y | 0 | 0 | yes | no | no | no | no | yes | no | no | no | no | no | no | no |
| HCN1 | 290 | D | 0 | 0 | no | yes | no | no | no | yes | no | no | yes | no | no | no | no |
| HCN1 | 291 | L | 1 | 0 | no | yes | no | no | no | yes | no | no | yes | no | no | no | no |
| HCN1 | 292 | A | 0 | 0 | no | yes | no | no | no | yes | no | no | yes | no | no | no | no |
| HCN1 | 293 | S | 0 | 0 | yes | no | no | no | no | yes | no | no | yes | no | no | no | no |
| HCN1 | 294 | A | 0 | 0 | yes | no | no | yes | no | yes | no | no | no | no | no | no | no |
| HCN1 | 295 | V | 0 | 0 | yes | no | no | yes | no | yes | no | no | no | no | no | no | no |
| HCN1 | 296 | V | 1 | 0 | yes | no | no | yes | no | no | no | no | no | no | no | no | no |
| HCN1 | 297 | R | 1 | 1 | yes | no | no | yes | no | no | no | no | yes | no | no | no | no |
| HCN1 | 298 | I | 0 | 0 | yes | no | no | yes | no | no | no | no | no | no | no | no | no |
| HCN1 | 299 | F | 0 | 0 | yes | no | no | yes | no | no | no | no | no | no | no | no | no |
| HCN1 | 300 | N | 0 | 0 | yes | no | no | yes | no | no | no | no | no | no | no | no | no |
| HCN1 | 301 | L | 0 | 0 | yes | no | no | yes | no | no | no | no | no | no | no | no | no |
| HCN1 | 302 | I | 0 | 0 | yes | no | no | yes | no | no | no | no | no | no | no | no | no |
| HCN1 | 303 | G | 1 | 0 | yes | no | no | yes | no | no | no | no | no | no | no | no | no |
| HCN1 | 304 | M | 0 | 0 | yes | no | no | yes | no | no | no | no | no | no | no | no | no |
| HCN1 | 305 | M | 0 | 1 | yes | no | no | yes | no | no | no | no | no | no | no | no | no |
| HCN1 | 306 | L | 0 | 0 | yes | no | no | yes | no | no | no | no | no | no | no | no | no |

| | | | | | | | | | | | | | | | | | |
|---|---|---|---|---|---|---|---|---|---|---|---|---|---|---|---|---|---|
| HCN1 | 307 | L | 0 | 0 | yes | no | no | yes | no | no | no | no | no | no | no | no | no |
| HCN1 | 308 | L | 0 | 0 | yes | no | no | yes | no | no | no | no | no | no | no | no | no |
| HCN1 | 309 | C | 0 | 0 | yes | no | no | yes | no | no | no | no | no | no | no | no | no |
| HCN1 | 310 | H | 0 | 0 | yes | no | no | yes | no | no | no | no | no | no | no | no | no |
| HCN1 | 311 | W | 0 | 0 | yes | no | no | yes | no | no | no | no | no | no | no | no | no |
| HCN1 | 312 | D | 0 | 0 | yes | no | no | yes | no | no | no | no | no | no | no | no | no |
| HCN1 | 313 | G | 0 | 0 | yes | no | no | yes | no | no | no | no | no | no | no | no | no |
| HCN1 | 314 | C | 0 | 0 | yes | no | no | yes | no | no | no | no | no | no | no | no | no |
| HCN1 | 315 | L | 0 | 0 | yes | no | no | yes | no | no | no | no | no | no | no | no | no |
| HCN1 | 316 | Q | 0 | 0 | yes | no | no | yes | no | no | no | no | no | no | no | no | no |
| HCN1 | 317 | F | 0 | 0 | yes | no | no | no | no | no | no | no | no | no | no | no | no |
| HCN1 | 318 | L | 1 | 0 | yes | no | no | yes | no | no | no | no | no | no | no | no | no |
| HCN1 | 319 | V | 0 | 0 | yes | no | no | no | yes | no | no | no | no | no | no | no | no |
| HCN1 | 320 | P | 0 | 0 | yes | no | no | no | yes | no | no | no | no | no | no | no | no |
| HCN1 | 321 | L | 0 | 0 | yes | no | no | no | yes | no | no | no | no | no | no | no | no |
| HCN1 | 322 | L | 0 | 0 | yes | no | no | no | yes | no | no | no | no | no | no | no | no |
| HCN1 | 323 | Q | 0 | 0 | no | yes | no | no | yes | no | no | no | no | no | no | no | no |
| HCN1 | 324 | D | 0 | 0 | no | yes | no | no | yes | no | no | no | no | no | no | no | no |
| HCN1 | 325 | F | 0 | 0 | no | yes | no | no | yes | no | no | no | no | no | no | no | no |
| HCN1 | 326 | P | 0 | 0 | no | yes | no | no | yes | no | no | no | no | no | no | no | no |
| HCN1 | 327 | P | 0 | 0 | no | yes | no | no | yes | no | no | no | no | no | no | no | no |
| HCN1 | 328 | D | 0 | 0 | no | yes | no | no | yes | no | no | no | no | no | no | no | no |
| HCN1 | 329 | C | 0 | 1 | no | yes | no | no | yes | no | no | no | no | no | no | no | no |
| HCN1 | 330 | W | 0 | 0 | yes | no | no | no | yes | no | no | no | no | no | no | no | no |
| HCN1 | 331 | V | 0 | 0 | yes | no | no | no | yes | no | no | no | no | no | no | no | no |
| HCN1 | 332 | S | 0 | 0 | yes | no | no | no | yes | no | no | no | no | no | no | no | no |
| HCN1 | 333 | L | 0 | 0 | yes | no | no | no | yes | no | no | no | no | no | no | no | no |
| HCN1 | 334 | N | 0 | 0 | yes | no | no | no | yes | no | no | no | no | no | no | no | no |
| HCN1 | 335 | E | 0 | 0 | no | yes | no | no | yes | no | no | no | no | no | no | no | no |
| HCN1 | 336 | M | 0 | 0 | no | yes | no | no | yes | no | no | no | no | no | no | no | no |
| HCN1 | 337 | V | 0 | 0 | no | yes | no | no | yes | no | no | no | no | no | no | no | no |
| HCN1 | 338 | N | 0 | 0 | no | yes | no | no | yes | no | no | no | no | no | no | no | no |
| HCN1 | 339 | D | 0 | 0 | no | yes | no | no | yes | no | no | no | no | no | no | no | no |

| | | | | | | | | | | | | | | | | | |
|---|---|---|---|---|---|---|---|---|---|---|---|---|---|---|---|---|---|
| HCN1 | 340 | S | 0 | 0 | no | yes | no | no | yes | no | no | no | no | no | no | no | no |
| HCN1 | 341 | W | 0 | 0 | yes | no | no | no | yes | no | no | no | no | no | no | no | no |
| HCN1 | 342 | G | 1 | 0 | yes | no | no | no | yes | no | no | no | no | no | no | no | no |
| HCN1 | 343 | K | 0 | 0 | yes | no | no | no | yes | no | no | no | yes | no | no | no | no |
| HCN1 | 344 | Q | 0 | 0 | yes | no | no | no | yes | no | no | no | no | no | no | no | no |
| HCN1 | 345 | Y | 0 | 0 | yes | no | no | yes | no | no | no | yes | no | no | no | no | no |
| HCN1 | 346 | S | 0 | 0 | yes | no | no | yes | no | no | no | yes | yes | no | no | no | no |
| HCN1 | 347 | Y | 0 | 0 | yes | no | no | no | no | no | no | yes | yes | no | no | no | no |
| HCN1 | 348 | A | 0 | 0 | yes | no | no | yes | no | no | no | yes | no | no | no | no | no |
| HCN1 | 349 | L | 0 | 0 | yes | no | no | yes | no | no | no | yes | yes | no | no | no | no |
| HCN1 | 350 | F | 0 | 0 | yes | no | no | yes | no | no | no | yes | yes | no | no | no | no |
| HCN1 | 351 | K | 0 | 0 | yes | no | no | yes | no | no | no | yes | no | no | no | no | no |
| HCN1 | 352 | A | 0 | 0 | yes | no | no | yes | no | no | no | yes | no | no | no | no | no |
| HCN1 | 353 | M | 0 | 0 | yes | no | no | yes | no | no | no | yes | yes | no | no | no | no |
| HCN1 | 354 | S | 0 | 0 | yes | no | no | yes | no | no | no | yes | yes | no | no | no | no |
| HCN1 | 355 | H | 0 | 0 | yes | no | no | yes | no | no | no | yes | no | no | no | no | no |
| HCN1 | 356 | M | 0 | 0 | yes | no | no | yes | no | no | no | yes | no | no | no | no | no |
| HCN1 | 357 | L | 0 | 0 | yes | no | no | yes | no | no | no | yes | yes | no | no | no | no |
| HCN1 | 358 | C | 0 | 0 | no | yes | no | yes | no | no | yes | yes | yes | no | no | no | no |
| HCN1 | 359 | I | 0 | 0 | no | yes | no | yes | no | no | yes | yes | yes | no | no | no | no |
| HCN1 | 360 | G | 1 | 0 | no | yes | no | yes | no | no | yes | yes | yes | no | no | no | no |
| HCN1 | 361 | Y | 1 | 0 | no | yes | no | yes | no | no | yes | yes | yes | no | no | no | no |
| HCN1 | 362 | G | 0 | 0 | no | yes | no | no | no | no | no | yes | yes | no | no | no | no |
| HCN1 | 363 | A | 0 | 0 | no | yes | no | no | no | no | yes | yes | no | no | no | no | no |
| HCN1 | 364 | Q | 0 | 0 | no | yes | no | no | no | no | no | yes | no | no | no | no | no |
| HCN1 | 365 | A | 0 | 0 | no | yes | no | no | no | no | no | yes | yes | no | no | no | no |
| HCN1 | 366 | P | 0 | 0 | no | yes | no | no | no | no | no | yes | yes | no | no | no | no |
| HCN1 | 367 | V | 0 | 0 | no | yes | no | no | yes | no | no | no | no | no | no | no | no |
| HCN1 | 368 | S | 0 | 0 | no | yes | no | no | yes | no | no | no | no | no | no | no | no |
| HCN1 | 369 | M | 0 | 0 | yes | no | no | no | yes | no | no | no | yes | no | no | no | no |
| HCN1 | 370 | S | 0 | 0 | yes | no | no | no | yes | no | no | no | no | no | no | no | no |
| HCN1 | 371 | D | 0 | 0 | yes | no | no | no | yes | no | no | no | no | no | no | no | no |
| HCN1 | 372 | L | 0 | 0 | yes | no | no | yes | no | no | no | no | yes | no | no | no | no |

| | | | | | | | | | | | | | | | | | |
|---|---|---|---|---|---|---|---|---|---|---|---|---|---|---|---|---|---|
| HCN1 | 373 | W | 0 | 0 | yes | no | no | yes | no | no | no | no | yes | no | no | no | no |
| HCN1 | 374 | I | 0 | 0 | yes | no | no | yes | no | no | no | no | no | no | no | no | no |
| HCN1 | 375 | T | 0 | 0 | yes | no | no | yes | no | no | no | no | yes | no | no | no | no |
| HCN1 | 376 | M | 0 | 0 | yes | no | no | yes | no | no | no | no | yes | no | no | no | no |
| HCN1 | 377 | L | 0 | 0 | yes | no | no | yes | no | no | no | no | no | no | no | no | no |
| HCN1 | 378 | S | 0 | 0 | yes | no | no | yes | no | no | no | no | no | no | no | no | no |
| HCN1 | 379 | M | 0 | 0 | yes | no | no | yes | no | no | no | no | yes | no | no | no | no |
| HCN1 | 380 | I | 0 | 1 | yes | no | no | yes | no | no | no | no | yes | no | no | no | no |
| HCN1 | 381 | V | 0 | 0 | yes | no | no | yes | no | no | no | no | no | no | no | no | no |
| HCN1 | 382 | G | 0 | 0 | yes | no | no | yes | no | no | no | no | no | no | no | no | no |
| HCN1 | 383 | A | 0 | 0 | yes | no | no | yes | no | no | no | no | yes | no | no | no | no |
| HCN1 | 384 | T | 0 | 0 | yes | no | no | yes | no | no | no | no | no | no | no | no | no |
| HCN1 | 385 | C | 0 | 0 | yes | no | no | yes | no | no | no | no | no | no | no | no | no |
| HCN1 | 386 | Y | 0 | 0 | yes | no | no | yes | no | no | yes | no | yes | no | no | no | no |
| HCN1 | 387 | A | 0 | 0 | yes | no | no | yes | no | no | no | no | yes | no | no | no | no |
| HCN1 | 388 | M | 0 | 0 | yes | no | no | yes | no | no | no | no | yes | no | no | no | no |
| HCN1 | 389 | F | 0 | 0 | yes | no | no | yes | no | no | no | no | yes | no | no | no | no |
| HCN1 | 390 | V | 0 | 0 | yes | no | no | yes | no | no | yes | no | yes | no | no | no | no |
| HCN1 | 391 | G | 0 | 2 | yes | no | no | yes | no | no | no | no | yes | no | no | no | no |
| HCN1 | 392 | H | 1 | 0 | yes | no | no | yes | no | no | no | no | yes | no | no | no | no |
| HCN1 | 393 | A | 1 | 0 | yes | no | no | yes | no | yes | no | no | no | no | no | no | no |
| HCN1 | 394 | T | 0 | 0 | yes | no | no | yes | no | yes | yes | no | yes | no | no | no | no |
| HCN1 | 395 | A | 0 | 0 | yes | no | no | yes | no | yes | no | no | no | no | no | no | no |
| HCN1 | 396 | L | 0 | 0 | yes | no | no | yes | no | yes | no | no | no | no | no | no | no |
| HCN1 | 397 | I | 0 | 1 | yes | no | no | yes | no | yes | no | no | yes | no | no | no | no |
| HCN1 | 398 | Q | 0 | 0 | yes | no | no | no | no | yes | yes | no | yes | no | no | no | no |
| HCN1 | 399 | S | 0 | 1 | yes | no | no | no | no | yes | no | no | no | no | no | no | no |
| HCN1 | 400 | L | 0 | 0 | yes | no | no | no | no | yes | no | no | no | no | no | no | no |
| HCN1 | 401 | D | 0 | 1 | no | yes | no | no | no | yes | no | no | yes | no | no | no | no |
| HCN1 | 402 | S | 0 | 0 | yes | no | no | no | no | yes | no | no | yes | no | no | no | no |
| HCN1 | 403 | S | 0 | 0 | yes | no | no | no | no | yes | no | no | yes | no | no | no | no |
| HCN1 | 404 | R | 1 | 0 | yes | no | no | no | no | yes | no | no | yes | no | no | no | no |
| HCN1 | 405 | R | 2 | 0 | yes | no | no | no | no | yes | yes | no | yes | no | no | no | no |

| | | | | | | | | | | | | | | | | | |
|---|---|---|---|---|---|---|---|---|---|---|---|---|---|---|---|---|---|
| HCN1 | 406 | Q | 0 | 0 | yes | no | no | no | no | yes | yes | no | yes | no | no | no | no |
| HCN1 | 407 | Y | 0 | 0 | yes | no | no | no | no | yes | no | no | no | no | no | no | no |
| HCN1 | 408 | Q | 0 | 0 | yes | no | no | no | no | yes | no | no | yes | no | no | no | no |
| HCN1 | 409 | E | 0 | 0 | yes | no | no | no | no | yes | yes | no | yes | no | no | no | no |
| HCN1 | 410 | K | 0 | 0 | yes | no | no | no | no | yes | no | no | yes | no | no | no | no |
| HCN1 | 411 | Y | 0 | 0 | yes | no | no | no | no | yes | no | no | no | no | no | no | no |
| HCN1 | 412 | K | 0 | 0 | yes | no | no | no | no | yes | no | no | yes | no | no | no | no |
| HCN1 | 413 | Q | 0 | 0 | yes | no | no | no | no | yes | yes | no | yes | no | no | no | no |
| HCN1 | 414 | V | 0 | 1 | yes | no | no | no | no | yes | no | no | yes | no | no | no | no |
| HCN1 | 415 | E | 0 | 0 | yes | no | no | no | no | yes | no | no | no | no | no | no | no |
| HCN1 | 416 | Q | 0 | 0 | yes | no | no | no | no | yes | no | no | yes | no | no | no | no |
| HCN1 | 417 | Y | 0 | 0 | yes | no | no | no | no | yes | no | no | yes | no | no | no | no |
| HCN1 | 418 | M | 1 | 0 | yes | no | no | no | no | yes | no | no | no | no | no | no | no |
| HCN1 | 419 | S | 0 | 0 | yes | no | no | no | no | yes | no | no | yes | no | no | no | no |
| HCN1 | 420 | F | 0 | 0 | yes | no | no | no | no | yes | no | no | yes | no | no | no | no |
| HCN1 | 421 | H | 0 | 0 | yes | no | no | no | no | yes | no | no | yes | no | no | no | no |
| HCN1 | 422 | K | 1 | 0 | no | yes | no | no | no | yes | no | no | yes | no | no | no | no |
| HCN1 | 423 | L | 0 | 0 | no | yes | no | no | no | yes | no | no | yes | no | no | no | no |
| HCN1 | 424 | P | 0 | 0 | no | yes | no | no | no | yes | no | no | yes | no | no | no | no |
| HCN1 | 425 | A | 0 | 0 | yes | no | no | no | no | yes | no | no | no | no | no | no | no |
| HCN1 | 426 | D | 0 | 0 | yes | no | no | no | no | yes | no | no | no | no | no | no | no |
| HCN1 | 427 | M | 0 | 0 | yes | no | no | no | no | yes | no | no | yes | no | no | no | no |
| HCN1 | 428 | R | 0 | 0 | yes | no | no | no | no | yes | no | no | no | no | no | no | no |
| HCN1 | 429 | Q | 0 | 0 | yes | no | no | no | no | yes | no | no | no | no | no | no | no |
| HCN1 | 430 | K | 0 | 0 | yes | no | no | no | no | yes | no | no | yes | no | no | no | no |
| HCN1 | 431 | I | 0 | 0 | yes | no | no | no | no | yes | no | no | yes | no | no | no | no |
| HCN1 | 432 | H | 1 | 0 | yes | no | no | no | no | yes | no | no | yes | no | no | no | no |
| HCN1 | 433 | D | 1 | 0 | yes | no | no | no | no | yes | no | no | no | no | no | no | no |
| HCN1 | 434 | Y | 1 | 0 | yes | no | no | no | no | yes | no | no | yes | no | no | no | no |
| HCN1 | 435 | Y | 0 | 0 | yes | no | no | no | no | yes | no | no | yes | no | no | no | no |
| HCN1 | 436 | E | 0 | 0 | yes | no | no | no | no | yes | no | no | yes | no | no | no | no |
| HCN1 | 437 | H | 0 | 0 | yes | no | no | no | no | yes | no | no | no | no | no | no | no |
| HCN1 | 438 | R | 0 | 0 | yes | no | no | no | no | yes | no | no | yes | no | no | no | no |

| | | | | | | | | | | | | | | | | | |
|---|---|---|---|---|---|---|---|---|---|---|---|---|---|---|---|---|---|
| HCN1 | 439 | Y | 1 | 0 | yes | no | no | no | no | yes | no | no | yes | no | no | no | no |
| HCN1 | 440 | Q | 1 | 0 | no | yes | no | no | no | yes | no | no | yes | no | no | no | no |
| HCN1 | 441 | G | 0 | 0 | no | yes | no | no | no | yes | no | no | no | no | no | no | no |
| HCN1 | 442 | K | 0 | 0 | no | yes | no | no | no | yes | no | no | yes | no | no | no | no |
| HCN1 | 443 | I | 0 | 0 | no | yes | no | no | no | yes | no | no | yes | no | no | no | no |
| HCN1 | 444 | F | 0 | 0 | no | yes | no | no | no | yes | no | no | yes | no | no | no | no |
| HCN1 | 445 | D | 0 | 0 | no | yes | no | no | no | yes | yes | no | no | no | no | no | no |
| HCN1 | 446 | E | 1 | 0 | yes | no | no | no | no | yes | no | no | yes | no | no | no | no |
| HCN1 | 447 | E | 0 | 0 | yes | no | no | no | no | yes | yes | no | no | no | no | no | no |
| HCN1 | 448 | N | 1 | 0 | yes | no | no | no | no | yes | yes | no | no | no | no | no | no |
| HCN1 | 449 | I | 0 | 0 | yes | no | no | no | no | yes | no | no | yes | no | no | no | no |
| HCN1 | 450 | L | 1 | 0 | yes | no | no | no | no | yes | no | no | yes | no | no | no | no |
| HCN1 | 451 | N | 1 | 0 | yes | no | no | no | no | yes | yes | no | yes | no | no | no | no |
| HCN1 | 452 | E | 0 | 0 | yes | no | no | no | no | yes | no | no | yes | no | no | no | no |
| HCN1 | 453 | L | 0 | 0 | no | yes | no | no | no | yes | no | no | yes | no | no | no | no |
| HCN1 | 454 | N | 1 | 0 | no | yes | no | no | no | yes | no | no | yes | no | no | no | no |
| HCN1 | 455 | D | 0 | 0 | yes | no | no | no | no | yes | no | no | no | no | no | no | no |
| HCN1 | 456 | P | 0 | 0 | yes | no | no | no | no | yes | no | no | yes | no | no | no | no |
| HCN1 | 457 | L | 0 | 0 | yes | no | no | no | no | yes | no | no | yes | no | no | no | no |
| HCN1 | 458 | R | 0 | 0 | yes | no | no | no | no | yes | no | no | no | no | no | no | no |
| HCN1 | 459 | E | 1 | 0 | yes | no | no | no | no | yes | no | no | no | no | no | no | no |
| HCN1 | 460 | E | 0 | 0 | yes | no | no | no | no | yes | no | no | yes | no | no | no | no |
| HCN1 | 461 | I | 0 | 0 | yes | no | no | no | no | yes | no | no | yes | no | no | no | no |
| HCN1 | 462 | V | 0 | 0 | yes | no | no | no | no | yes | no | no | no | no | no | no | no |
| HCN1 | 463 | N | 0 | 0 | yes | no | no | no | no | yes | no | no | no | no | no | no | no |
| HCN1 | 464 | F | 0 | 0 | yes | no | no | no | no | yes | no | no | yes | no | no | no | no |
| HCN1 | 465 | N | 1 | 0 | yes | no | no | no | no | yes | no | no | yes | no | no | no | no |
| HCN1 | 466 | C | 0 | 0 | no | yes | no | no | no | yes | no | no | no | no | no | no | no |
| HCN1 | 467 | R | 1 | 0 | yes | no | no | no | no | yes | no | no | no | no | no | no | no |
| HCN1 | 468 | K | 1 | 0 | yes | no | no | no | no | yes | no | no | no | no | no | no | no |
| HCN1 | 469 | L | 0 | 0 | yes | no | no | no | no | yes | no | no | no | no | no | no | no |
| HCN1 | 470 | V | 0 | 0 | yes | no | no | no | no | yes | no | no | no | no | no | no | no |
| HCN1 | 471 | A | 0 | 0 | yes | no | no | no | no | yes | no | no | no | no | no | no | no |

| | | | | | | | | | | | | | | | | | |
|---|---|---|---|---|---|---|---|---|---|---|---|---|---|---|---|---|---|
| HCN1 | 472 | T | 0 | 0 | yes | no | no | no | no | yes | no | no | no | no | no | no | no |
| HCN1 | 473 | M | 0 | 0 | no | yes | no | no | no | yes | no | no | no | no | no | no | no |
| HCN1 | 474 | P | 0 | 0 | yes | no | no | no | no | yes | no | no | no | no | no | no | no |
| HCN1 | 475 | L | 0 | 0 | yes | no | no | no | no | yes | no | no | no | no | no | no | no |
| HCN1 | 476 | F | 0 | 0 | yes | no | no | no | no | yes | no | no | no | no | no | no | no |
| HCN1 | 477 | A | 0 | 0 | yes | no | no | no | no | yes | no | no | no | no | no | no | no |
| HCN1 | 478 | N | 0 | 0 | yes | no | no | no | no | yes | no | no | no | no | no | no | no |
| HCN1 | 479 | A | 1 | 0 | no | yes | no | no | no | yes | no | no | no | no | no | no | no |
| HCN1 | 480 | D | 1 | 0 | no | yes | no | no | no | yes | no | no | no | no | no | no | no |
| HCN1 | 481 | P | 0 | 0 | yes | no | no | no | no | yes | no | no | no | no | no | no | no |
| HCN1 | 482 | N | 0 | 0 | yes | no | no | no | no | yes | no | no | no | no | no | no | no |
| HCN1 | 483 | F | 0 | 0 | yes | no | no | no | no | yes | no | no | no | no | no | no | no |
| HCN1 | 484 | V | 0 | 0 | yes | no | no | no | no | yes | no | no | no | no | no | no | no |
| HCN1 | 485 | T | 0 | 0 | yes | no | no | no | no | yes | no | no | no | no | no | no | no |
| HCN1 | 486 | A | 0 | 0 | yes | no | no | no | no | yes | no | no | no | no | no | no | no |
| HCN1 | 487 | M | 1 | 0 | yes | no | no | no | no | yes | no | no | no | no | no | no | no |
| HCN1 | 488 | L | 0 | 0 | yes | no | no | no | no | yes | no | no | no | no | no | no | no |
| HCN1 | 489 | S | 0 | 0 | yes | no | no | no | no | yes | no | no | no | no | no | no | no |
| HCN1 | 490 | K | 0 | 0 | yes | no | no | no | no | yes | no | no | no | no | no | no | no |
| HCN1 | 491 | L | 0 | 0 | no | yes | no | no | no | yes | no | no | no | no | no | no | no |
| HCN1 | 492 | R | 1 | 0 | no | no | yes | no | no | yes | no | no | yes | no | no | no | no |
| HCN1 | 493 | F | 0 | 0 | no | no | yes | no | no | yes | no | no | no | no | no | no | no |
| HCN1 | 494 | E | 0 | 0 | no | no | yes | no | no | yes | no | no | no | no | no | no | no |
| HCN1 | 495 | V | 0 | 0 | no | no | yes | no | no | yes | no | no | yes | no | no | no | no |
| HCN1 | 496 | F | 0 | 0 | no | no | yes | no | no | yes | no | no | yes | no | no | no | no |
| HCN1 | 497 | Q | 0 | 0 | no | yes | no | no | no | yes | no | no | yes | no | no | no | no |
| HCN1 | 498 | P | 0 | 0 | no | yes | no | no | no | yes | no | no | no | no | no | no | no |
| HCN1 | 499 | G | 0 | 0 | no | yes | no | no | no | yes | no | no | no | no | no | no | no |
| HCN1 | 500 | D | 0 | 0 | no | yes | no | no | no | yes | no | no | no | no | no | no | no |
| HCN1 | 501 | Y | 1 | 0 | no | no | yes | no | no | yes | no | no | yes | no | no | no | no |
| HCN1 | 502 | I | 0 | 0 | no | no | yes | no | no | yes | no | no | no | no | no | no | no |
| HCN1 | 503 | I | 0 | 0 | no | no | yes | no | no | yes | no | no | no | no | no | no | no |
| HCN1 | 504 | R | 0 | 0 | no | yes | no | no | no | yes | no | no | no | no | no | no | no |

| | | | | | | | | | | | | | | | | | |
|---|---|---|---|---|---|---|---|---|---|---|---|---|---|---|---|---|---|
| HCN1 | 505 | E | 0 | 0 | no | yes | no | no | no | yes | no | no | no | no | no | no | no |
| HCN1 | 506 | G | 0 | 0 | no | yes | no | no | no | yes | no | no | no | no | no | no | no |
| HCN1 | 507 | A | 0 | 0 | no | yes | no | no | no | yes | no | no | no | no | no | no | no |
| HCN1 | 508 | V | 2 | 0 | no | no | yes | no | no | yes | no | no | no | no | no | no | no |
| HCN1 | 509 | G | 0 | 0 | no | yes | no | no | no | yes | no | no | no | no | no | no | no |
| HCN1 | 510 | K | 1 | 0 | no | yes | no | no | no | yes | no | no | no | no | no | no | no |
| HCN1 | 511 | K | 0 | 0 | no | no | yes | no | no | yes | no | no | no | no | no | no | no |
| HCN1 | 512 | M | 1 | 0 | no | no | yes | no | no | yes | no | no | no | no | no | no | no |
| HCN1 | 513 | Y | 0 | 0 | no | yes | no | no | no | yes | no | no | no | no | no | no | no |
| HCN1 | 514 | F | 0 | 0 | no | yes | no | no | no | yes | no | no | no | no | no | no | no |
| HCN1 | 515 | I | 0 | 0 | no | no | yes | no | no | yes | no | no | no | no | no | no | no |
| HCN1 | 516 | Q | 0 | 0 | no | no | yes | no | no | yes | no | no | no | no | no | no | no |
| HCN1 | 517 | H | 0 | 0 | no | no | yes | no | no | yes | no | no | yes | no | no | no | no |
| HCN1 | 518 | G | 0 | 0 | no | yes | no | no | no | yes | no | no | no | no | no | no | no |
| HCN1 | 519 | V | 0 | 0 | no | yes | no | no | no | yes | no | no | no | no | no | no | no |
| HCN1 | 520 | A | 1 | 0 | no | no | yes | no | no | yes | no | no | no | no | no | no | no |
| HCN1 | 521 | G | 0 | 0 | no | no | yes | no | no | yes | no | no | no | no | no | no | no |
| HCN1 | 522 | V | 0 | 0 | no | no | yes | no | no | yes | no | no | no | no | no | no | no |
| HCN1 | 523 | I | 0 | 0 | no | no | yes | no | no | yes | no | no | no | no | no | no | no |
| HCN1 | 524 | T | 1 | 0 | no | yes | no | no | no | yes | no | no | no | no | no | no | no |
| HCN1 | 525 | K | 0 | 0 | no | yes | no | no | no | yes | no | no | no | no | no | no | no |
| HCN1 | 526 | S | 1 | 0 | no | yes | no | no | no | yes | no | no | no | no | no | no | no |
| HCN1 | 527 | S | 0 | 0 | no | yes | no | no | no | yes | no | no | no | no | no | no | no |
| HCN1 | 528 | K | 1 | 0 | no | yes | no | no | no | yes | no | no | no | no | no | no | no |
| HCN1 | 529 | E | 0 | 0 | no | yes | no | no | no | yes | no | no | no | no | no | no | no |
| HCN1 | 530 | M | 0 | 0 | no | no | yes | no | no | yes | no | no | no | no | no | no | no |
| HCN1 | 531 | K | 1 | 0 | no | no | yes | no | no | yes | no | no | no | no | no | no | no |
| HCN1 | 532 | L | 1 | 0 | no | no | yes | no | no | yes | no | no | no | no | no | no | no |
| HCN1 | 533 | T | 0 | 0 | no | yes | no | no | no | yes | no | no | no | no | no | no | no |
| HCN1 | 534 | D | 0 | 0 | no | yes | no | no | no | yes | no | no | no | no | no | no | no |
| HCN1 | 535 | G | 0 | 0 | no | yes | no | no | no | yes | no | no | no | no | no | no | no |
| HCN1 | 536 | S | 0 | 0 | no | yes | no | no | no | yes | no | no | no | no | no | no | no |
| HCN1 | 537 | Y | 0 | 0 | no | yes | no | no | no | yes | no | no | no | no | no | no | no |

| | | | | | | | | | | | | | | | | | |
|---|---|---|---|---|---|---|---|---|---|---|---|---|---|---|---|---|---|
| HCN1 | 538 | F | 0 | 0 | no | yes | no | no | no | yes | no | no | no | no | no | no | no |
| HCN1 | 539 | G | 0 | 0 | no | yes | no | no | no | yes | no | no | no | no | no | no | yes |
| HCN1 | 540 | E | 0 | 0 | no | yes | no | no | no | yes | no | no | no | no | no | no | yes |
| HCN1 | 541 | I | 0 | 0 | yes | no | no | no | no | yes | no | no | no | no | no | no | yes |
| HCN1 | 542 | C | 1 | 0 | yes | no | no | no | no | yes | no | no | no | no | no | no | yes |
| HCN1 | 543 | L | 0 | 0 | yes | no | no | no | no | yes | no | no | no | no | no | no | no |
| HCN1 | 544 | L | 0 | 0 | yes | no | no | no | no | yes | no | no | no | no | no | no | no |
| HCN1 | 545 | T | 0 | 0 | yes | no | no | no | no | yes | no | no | no | no | no | no | no |
| HCN1 | 546 | K | 0 | 0 | no | yes | no | no | no | yes | no | no | no | no | no | no | no |
| HCN1 | 547 | G | 0 | 0 | no | yes | no | no | no | yes | no | no | no | no | no | no | no |
| HCN1 | 548 | R | 0 | 0 | no | no | yes | no | no | yes | no | no | yes | no | no | no | no |
| HCN1 | 549 | R | 0 | 0 | no | yes | no | no | no | yes | no | no | no | no | no | no | yes |
| HCN1 | 550 | T | 0 | 0 | no | yes | no | no | no | yes | no | no | no | no | no | no | yes |
| HCN1 | 551 | A | 0 | 0 | no | yes | no | no | no | yes | no | no | no | no | no | no | no |
| HCN1 | 552 | S | 0 | 0 | no | no | yes | no | no | yes | no | no | no | no | no | no | no |
| HCN1 | 553 | V | 1 | 0 | no | no | yes | no | no | yes | no | no | no | no | no | no | no |
| HCN1 | 554 | R | 1 | 0 | no | no | yes | no | no | yes | no | no | no | no | no | no | no |
| HCN1 | 555 | A | 0 | 0 | no | no | yes | no | no | yes | no | no | no | no | no | no | no |
| HCN1 | 556 | D | 0 | 0 | no | yes | no | no | no | yes | no | no | no | no | no | no | no |
| HCN1 | 557 | T | 0 | 0 | no | yes | no | no | no | yes | no | no | no | no | no | no | no |
| HCN1 | 558 | Y | 0 | 0 | no | yes | no | no | no | yes | no | no | no | no | no | no | no |
| HCN1 | 559 | C | 0 | 0 | no | no | yes | no | no | yes | no | no | no | no | no | no | no |
| HCN1 | 560 | R | 0 | 0 | no | no | yes | no | no | yes | no | no | no | no | no | no | no |
| HCN1 | 561 | L | 0 | 0 | no | no | yes | no | no | yes | no | no | no | no | no | no | no |
| HCN1 | 562 | Y | 0 | 0 | no | no | yes | no | no | yes | no | no | no | no | no | no | no |
| HCN1 | 563 | S | 0 | 0 | no | no | yes | no | no | yes | no | no | no | no | no | no | no |
| HCN1 | 564 | L | 0 | 0 | no | no | yes | no | no | yes | no | no | no | no | no | no | no |
| HCN1 | 565 | S | 0 | 0 | no | no | yes | no | no | yes | no | no | no | no | no | no | no |
| HCN1 | 566 | V | 1 | 0 | yes | no | no | no | no | yes | no | no | no | no | no | no | no |
| HCN1 | 567 | D | 1 | 0 | yes | no | no | no | no | yes | yes | no | no | no | no | no | no |
| HCN1 | 568 | N | 1 | 0 | yes | no | no | no | no | yes | yes | no | no | no | no | no | no |
| HCN1 | 569 | F | 0 | 0 | yes | no | no | no | no | yes | no | no | no | no | no | no | no |
| HCN1 | 570 | N | 1 | 0 | yes | no | no | no | no | yes | yes | no | no | no | no | no | no |

| | | | | | | | | | | | | | | | | | |
|---|---|---|---|---|---|---|---|---|---|---|---|---|---|---|---|---|---|
| HCN1 | 571 | E | 0 | 0 | yes | no | no | no | no | yes | no | no | no | no | no | no | no |
| HCN1 | 572 | V | 0 | 0 | yes | no | no | no | no | yes | no | no | no | no | no | no | no |
| HCN1 | 573 | L | 0 | 0 | yes | no | no | no | no | yes | no | no | no | no | no | no | no |
| HCN1 | 574 | E | 0 | 0 | yes | no | no | no | no | yes | no | no | no | no | no | no | no |
| HCN1 | 575 | E | 0 | 0 | yes | no | no | no | no | yes | no | no | yes | no | no | no | no |
| HCN1 | 576 | Y | 0 | 0 | no | yes | no | no | no | yes | no | no | no | no | no | no | no |
| HCN1 | 577 | P | 0 | 0 | yes | no | no | no | no | yes | no | no | no | no | no | no | no |
| HCN1 | 578 | M | 0 | 0 | yes | no | no | no | no | yes | no | no | no | no | no | no | no |
| HCN1 | 579 | M | 0 | 0 | yes | no | no | no | no | yes | no | no | no | no | no | no | no |
| HCN1 | 580 | R | 0 | 0 | yes | no | no | no | no | yes | no | no | no | no | no | no | no |
| HCN1 | 581 | R | 2 | 0 | yes | no | no | no | no | yes | no | no | no | no | no | no | no |
| HCN1 | 582 | A | 0 | 0 | yes | no | no | no | no | yes | no | no | no | no | no | no | no |
| HCN1 | 583 | F | 0 | 0 | yes | no | no | no | no | yes | no | no | no | no | no | no | no |
| HCN1 | 584 | E | 0 | 0 | yes | no | no | no | no | yes | no | no | no | no | no | no | no |
| HCN1 | 585 | T | 1 | 0 | yes | no | no | no | no | yes | no | no | no | no | no | no | no |
| HCN1 | 586 | V | 0 | 0 | yes | no | no | no | no | yes | no | no | no | no | no | no | no |
| HCN1 | 587 | A | 0 | 0 | yes | no | no | no | no | yes | no | no | no | no | no | no | no |
| HCN1 | 588 | I | 0 | 0 | yes | no | no | no | no | yes | no | no | no | no | no | no | no |
| HCN1 | 589 | D | 0 | 0 | yes | no | no | no | no | yes | no | no | no | no | no | no | no |
| HCN1 | 590 | R | 0 | 1 | yes | no | no | no | no | yes | no | no | no | no | no | no | yes |
| HCN1 | 591 | L | 0 | 0 | yes | no | no | no | no | yes | no | no | no | no | no | no | yes |
| HCN1 | 592 | D | 1 | 0 | yes | no | no | no | no | yes | no | no | no | no | no | no | yes |
| HCN1 | 593 | R | 2 | 0 | yes | no | no | no | no | yes | no | no | no | no | no | no | yes |
| HCN1 | 594 | I | 0 | 0 | yes | no | no | no | no | yes | no | no | no | no | no | no | no |
| HCN1 | 595 | G | 1 | 0 | no | yes | no | no | no | yes | no | no | no | no | no | no | no |
| HCN1 | 596 | K | 0 | 0 | no | yes | no | no | no | yes | no | no | no | no | no | no | no |
| HCN1 | 597 | K | 0 | 0 | no | yes | no | no | no | yes | no | no | no | no | no | no | no |
| HCN1 | 598 | N | 1 | 0 | no | yes | no | no | no | yes | no | no | no | no | no | no | no |
| HCN1 | 599 | S | 1 | 0 | yes | no | no | no | no | yes | no | no | no | no | no | no | no |
| HCN1 | 600 | I | 0 | 0 | yes | no | no | no | no | yes | no | no | no | no | no | no | no |
| HCN1 | 601 | L | 0 | 0 | yes | no | no | no | no | yes | no | no | no | no | no | no | no |
| HCN1 | 602 | L | 0 | 0 | yes | no | no | no | no | yes | no | no | no | no | no | no | no |
| HCN1 | 603 | Q | 0 | 0 | yes | no | no | no | no | yes | no | no | no | no | no | no | no |

| | | | | | | | | | | | | | | | | | |
|---|---|---|---|---|---|---|---|---|---|---|---|---|---|---|---|---|---|
| HCN1 | 604 | K | 0 | 0 | yes | no | no | no | no | yes | no | no | no | no | no | no | no |
| HCN1 | 605 | F | 1 | 0 | yes | no | no | no | no | yes | no | no | no | no | no | no | no |
| HCN1 | 606 | Q | 0 | 0 | yes | no | no | no | no | yes | no | no | no | no | no | no | no |
| HCN1 | 607 | K | 1 | 0 | yes | no | no | no | no | yes | no | no | no | no | no | no | no |
| HCN1 | 608 | D | 0 | 0 | no | yes | no | no | no | yes | no | no | no | no | no | no | no |
| HCN4 | 214 | M | 2 | 0 | no | yes | no | no | no | yes | no | no | no | no | no | no | no |
| HCN4 | 215 | Q | 0 | 0 | no | yes | no | no | no | yes | no | no | no | no | no | no | no |
| HCN4 | 216 | R | 0 | 0 | no | yes | no | no | no | yes | no | no | no | no | no | no | no |
| HCN4 | 217 | Q | 1 | 0 | no | yes | no | no | no | yes | no | no | no | no | no | no | no |
| HCN4 | 218 | F | 0 | 0 | no | yes | no | no | no | yes | no | no | no | no | no | no | no |
| HCN4 | 219 | G | 1 | 0 | no | yes | no | no | no | yes | no | no | no | no | no | no | no |
| HCN4 | 220 | A | 1 | 0 | no | yes | no | no | no | yes | no | no | no | no | no | no | no |
| HCN4 | 221 | M | 1 | 0 | no | yes | no | no | no | yes | no | no | no | no | no | no | no |
| HCN4 | 222 | L | 0 | 0 | no | yes | no | no | no | yes | no | no | no | no | no | no | no |
| HCN4 | 223 | Q | 0 | 0 | no | yes | no | no | no | yes | no | no | no | no | no | no | no |
| HCN4 | 224 | P | 1 | 0 | no | yes | no | no | no | yes | no | no | no | no | no | no | no |
| HCN4 | 225 | G | 2 | 0 | no | yes | no | no | no | yes | no | no | no | no | no | no | no |
| HCN4 | 226 | V | 1 | 0 | no | yes | no | no | no | yes | no | no | no | no | no | no | no |
| HCN4 | 227 | N | 0 | 0 | no | yes | no | no | no | yes | no | no | no | no | no | no | no |
| HCN4 | 228 | K | 0 | 0 | no | yes | no | no | no | yes | no | no | no | no | no | no | no |
| HCN4 | 229 | F | 0 | 0 | no | yes | no | no | no | yes | no | no | no | no | no | no | no |
| HCN4 | 230 | S | 0 | 0 | yes | no | no | no | no | yes | no | no | no | no | no | no | no |
| HCN4 | 231 | L | 0 | 0 | yes | no | no | no | no | yes | no | no | yes | no | no | no | no |
| HCN4 | 232 | R | 1 | 0 | yes | no | no | no | no | yes | no | no | yes | no | no | no | no |
| HCN4 | 233 | M | 1 | 0 | yes | no | no | no | no | yes | no | no | no | no | no | no | no |
| HCN4 | 234 | F | 0 | 0 | no | yes | no | no | no | yes | no | no | no | no | no | no | no |
| HCN4 | 235 | G | 0 | 0 | no | yes | no | no | no | yes | no | no | no | no | no | no | no |
| HCN4 | 236 | S | 0 | 0 | no | yes | no | no | no | yes | no | no | yes | no | no | no | no |
| HCN4 | 237 | Q | 0 | 0 | yes | no | no | no | no | yes | no | no | no | no | no | no | no |
| HCN4 | 238 | K | 0 | 0 | yes | no | no | no | no | yes | no | no | no | no | no | no | no |
| HCN4 | 239 | A | 0 | 0 | yes | no | no | no | no | yes | no | no | no | no | no | no | no |
| HCN4 | 240 | V | 1 | 0 | yes | no | no | no | no | yes | no | no | no | no | no | no | no |
| HCN4 | 241 | E | 0 | 0 | yes | no | no | no | no | yes | no | no | no | no | no | no | no |

| | | | | | | | | | | | | | | | | | |
|---|---|---|---|---|---|---|---|---|---|---|---|---|---|---|---|---|---|
| HCN4 | 242 | R | 1 | 0 | yes | no | no | no | no | yes | no | no | no | no | no | no | no |
| HCN4 | 243 | E | 1 | 0 | yes | no | no | no | no | yes | no | no | no | no | no | no | no |
| HCN4 | 244 | Q | 0 | 0 | yes | no | no | no | no | yes | no | no | no | no | no | no | no |
| HCN4 | 245 | E | 0 | 0 | yes | no | no | no | no | yes | no | no | no | no | no | no | no |
| HCN4 | 246 | R | 0 | 0 | yes | no | no | no | no | yes | no | no | no | no | no | no | no |
| HCN4 | 247 | V | 0 | 0 | yes | no | no | no | no | yes | no | no | no | no | no | no | no |
| HCN4 | 248 | K | 1 | 0 | yes | no | no | no | no | yes | no | no | no | no | no | no | no |
| HCN4 | 249 | S | 0 | 0 | no | yes | no | no | no | yes | no | no | no | no | no | no | no |
| HCN4 | 250 | A | 0 | 0 | no | yes | no | no | no | yes | no | no | no | no | no | no | no |
| HCN4 | 251 | G | 0 | 0 | no | yes | no | no | no | yes | no | no | no | no | no | no | no |
| HCN4 | 252 | F | 0 | 0 | no | yes | no | no | no | yes | no | no | no | no | no | no | no |
| HCN4 | 253 | W | 0 | 0 | no | yes | no | no | no | yes | no | no | no | no | no | no | no |
| HCN4 | 254 | I | 0 | 0 | no | yes | no | no | no | yes | no | no | no | no | no | no | no |
| HCN4 | 255 | I | 0 | 0 | no | no | yes | no | no | yes | no | no | no | no | no | no | no |
| HCN4 | 256 | H | 0 | 0 | no | yes | no | no | no | yes | no | no | no | no | no | no | no |
| HCN4 | 257 | P | 0 | 1 | no | yes | no | no | no | yes | no | no | no | no | no | no | no |
| HCN4 | 258 | Y | 0 | 0 | no | yes | no | no | no | yes | no | no | no | no | no | no | no |
| HCN4 | 259 | S | 0 | 0 | no | yes | no | no | no | yes | no | no | no | no | no | no | no |
| HCN4 | 260 | D | 0 | 0 | no | yes | no | no | no | yes | no | no | no | no | no | no | no |
| HCN4 | 261 | F | 1 | 0 | yes | no | no | no | no | yes | no | no | no | no | no | no | no |
| HCN4 | 262 | R | 0 | 0 | yes | no | no | no | no | yes | no | no | no | no | no | no | no |
| HCN4 | 263 | F | 0 | 0 | yes | no | no | no | no | yes | no | no | no | no | no | no | no |
| HCN4 | 264 | Y | 0 | 0 | yes | no | no | yes | no | yes | no | no | no | no | no | no | no |
| HCN4 | 265 | W | 0 | 0 | yes | no | no | yes | no | yes | no | no | no | no | no | no | no |
| HCN4 | 266 | D | 0 | 0 | yes | no | no | yes | no | yes | no | no | no | no | no | no | no |
| HCN4 | 267 | L | 0 | 0 | yes | no | no | yes | no | no | no | no | no | no | no | no | no |
| HCN4 | 268 | T | 0 | 0 | yes | no | no | yes | no | no | no | no | no | no | no | no | no |
| HCN4 | 269 | M | 0 | 0 | yes | no | no | yes | no | no | no | no | no | no | no | no | no |
| HCN4 | 270 | L | 0 | 0 | yes | no | no | yes | no | no | no | no | no | no | no | no | no |
| HCN4 | 271 | L | 0 | 0 | yes | no | no | yes | no | no | no | no | no | no | no | no | no |
| HCN4 | 272 | L | 0 | 0 | yes | no | no | yes | no | no | no | no | no | no | no | no | no |
| HCN4 | 273 | M | 0 | 0 | yes | no | no | yes | no | no | no | no | no | no | no | no | no |
| HCN4 | 274 | V | 1 | 0 | yes | no | no | yes | no | no | no | no | no | no | no | no | no |

| | | | | | | | | | | | | | | | | | |
|---|---|---|---|---|---|---|---|---|---|---|---|---|---|---|---|---|---|
| HCN4 | 275 | G | 0 | 0 | yes | no | no | yes | no | no | no | no | no | no | no | no | no |
| HCN4 | 276 | N | 0 | 0 | yes | no | no | yes | no | no | no | no | no | no | no | no | no |
| HCN4 | 277 | L | 0 | 0 | yes | no | no | yes | no | no | no | no | no | no | no | no | no |
| HCN4 | 278 | I | 0 | 0 | yes | no | no | yes | no | no | no | no | no | no | no | no | no |
| HCN4 | 279 | I | 0 | 0 | yes | no | no | yes | no | no | no | no | no | no | no | no | no |
| HCN4 | 280 | I | 0 | 0 | yes | no | no | yes | no | no | no | no | no | no | no | no | no |
| HCN4 | 281 | P | 0 | 0 | yes | no | no | yes | no | no | no | no | no | no | no | no | no |
| HCN4 | 282 | V | 0 | 0 | yes | no | no | yes | no | no | no | no | no | no | no | no | no |
| HCN4 | 283 | G | 1 | 0 | yes | no | no | yes | no | no | no | no | no | no | no | no | no |
| HCN4 | 284 | I | 0 | 0 | yes | no | no | yes | no | no | no | no | no | no | no | no | no |
| HCN4 | 285 | T | 0 | 0 | yes | no | no | yes | no | no | no | no | no | no | no | no | no |
| HCN4 | 286 | F | 0 | 0 | no | yes | no | no | no | no | no | no | no | no | no | no | no |
| HCN4 | 287 | F | 0 | 0 | no | yes | no | no | no | no | no | no | no | no | no | no | no |
| HCN4 | 288 | K | 0 | 0 | no | yes | no | no | yes | no | no | no | no | no | no | no | no |
| HCN4 | 289 | D | 0 | 0 | no | yes | no | no | yes | no | no | no | no | no | no | no | no |
| HCN4 | 290 | E | 0 | 0 | no | yes | no | no | yes | no | no | no | no | no | no | no | no |
| HCN4 | 291 | N | 0 | 0 | no | yes | no | no | yes | no | no | no | no | no | no | no | no |
| HCN4 | 292 | T | 0 | 0 | no | yes | no | no | yes | no | no | no | no | no | no | no | no |
| HCN4 | 293 | T | 0 | 0 | yes | no | no | yes | yes | no | no | no | no | no | no | no | no |
| HCN4 | 294 | P | 1 | 0 | yes | no | no | yes | no | no | no | no | no | no | no | no | no |
| HCN4 | 295 | W | 0 | 0 | yes | no | no | yes | no | no | no | no | no | no | no | no | no |
| HCN4 | 296 | I | 0 | 0 | yes | no | no | yes | no | no | no | no | no | no | no | no | no |
| HCN4 | 297 | V | 0 | 0 | yes | no | no | yes | no | no | no | no | no | no | no | no | no |
| HCN4 | 298 | F | 0 | 0 | yes | no | no | yes | no | no | no | no | no | no | no | no | no |
| HCN4 | 299 | N | 1 | 0 | yes | no | no | yes | no | no | no | no | no | no | no | no | no |
| HCN4 | 300 | V | 0 | 0 | yes | no | no | yes | no | no | no | no | no | no | no | no | no |
| HCN4 | 301 | V | 0 | 0 | yes | no | no | yes | no | no | no | no | no | no | no | no | no |
| HCN4 | 302 | S | 0 | 0 | yes | no | no | yes | no | no | no | no | no | no | no | no | no |
| HCN4 | 303 | D | 1 | 0 | yes | no | no | yes | no | no | no | no | no | no | no | no | no |
| HCN4 | 304 | T | 0 | 0 | yes | no | no | yes | no | no | no | no | no | no | no | no | no |
| HCN4 | 305 | F | 1 | 0 | yes | no | no | yes | no | no | no | no | no | no | no | no | no |
| HCN4 | 306 | F | 0 | 0 | yes | no | no | yes | no | no | no | no | no | no | no | no | no |
| HCN4 | 307 | L | 0 | 0 | yes | no | no | yes | no | no | no | no | no | no | no | no | no |

| | | | | | | | | | | | | | | | | | |
|---|---|---|---|---|---|---|---|---|---|---|---|---|---|---|---|---|---|
| HCN4 | 308 | I | 1 | 0 | yes | no | no | yes | no | no | no | no | no | no | no | no | no |
| HCN4 | 309 | D | 0 | 0 | yes | no | no | yes | no | no | no | no | no | no | no | no | no |
| HCN4 | 310 | L | 0 | 0 | yes | no | no | yes | no | no | no | no | no | no | no | no | no |
| HCN4 | 311 | V | 0 | 0 | yes | no | no | yes | no | no | no | no | no | no | no | no | no |
| HCN4 | 312 | L | 0 | 0 | yes | no | no | yes | no | no | no | no | no | no | no | no | no |
| HCN4 | 313 | N | 0 | 0 | no | yes | no | yes | no | no | no | no | no | no | no | no | no |
| HCN4 | 314 | F | 0 | 0 | no | yes | no | yes | no | no | no | no | no | no | no | no | no |
| HCN4 | 315 | R | 2 | 0 | no | yes | no | no | no | yes | no | no | no | no | no | no | no |
| HCN4 | 316 | T | 0 | 0 | no | no | yes | no | no | yes | no | no | no | no | no | no | no |
| HCN4 | 317 | G | 0 | 0 | no | yes | no | no | no | yes | no | no | no | no | no | no | no |
| HCN4 | 318 | I | 2 | 0 | no | no | yes | no | no | yes | no | no | no | no | no | no | no |
| HCN4 | 319 | V | 1 | 0 | no | no | yes | no | no | yes | no | no | no | no | no | no | no |
| HCN4 | 320 | V | 0 | 0 | no | yes | no | no | no | yes | no | no | yes | no | no | no | no |
| HCN4 | 321 | E | 0 | 0 | no | yes | no | no | no | yes | no | no | yes | no | no | no | no |
| HCN4 | 322 | D | 0 | 0 | no | yes | no | no | no | yes | no | no | yes | no | no | no | no |
| HCN4 | 323 | N | 0 | 0 | no | yes | no | no | no | yes | no | no | yes | no | no | no | no |
| HCN4 | 324 | T | 0 | 0 | no | yes | no | no | no | yes | no | no | yes | no | no | no | no |
| HCN4 | 325 | E | 1 | 0 | no | yes | no | no | no | yes | no | no | no | no | no | no | no |
| HCN4 | 326 | I | 0 | 0 | no | no | yes | no | no | yes | no | no | no | no | no | no | no |
| HCN4 | 327 | I | 0 | 0 | no | no | yes | no | no | yes | no | no | no | no | no | no | no |
| HCN4 | 328 | L | 0 | 0 | no | yes | no | no | no | yes | no | no | no | no | no | no | no |
| HCN4 | 329 | D | 1 | 0 | no | yes | no | no | no | yes | no | no | no | no | no | no | no |
| HCN4 | 330 | P | 1 | 0 | yes | no | no | no | no | yes | no | no | no | no | no | no | no |
| HCN4 | 331 | Q | 1 | 0 | yes | no | no | no | no | yes | no | no | no | no | no | no | no |
| HCN4 | 332 | R | 2 | 0 | yes | no | no | no | no | yes | no | no | no | no | no | no | no |
| HCN4 | 333 | I | 0 | 0 | yes | no | no | no | no | yes | no | no | no | no | no | no | no |
| HCN4 | 334 | K | 2 | 0 | yes | no | no | no | no | yes | no | no | no | no | no | no | no |
| HCN4 | 335 | M | 2 | 0 | yes | no | no | no | no | yes | no | no | no | no | no | no | no |
| HCN4 | 336 | K | 0 | 0 | yes | no | no | no | no | yes | no | no | no | no | no | no | no |
| HCN4 | 337 | Y | 0 | 0 | yes | no | no | yes | no | yes | no | no | no | no | no | no | no |
| HCN4 | 338 | L | 0 | 0 | yes | no | no | yes | no | yes | no | no | no | no | no | no | no |
| HCN4 | 339 | K | 1 | 0 | yes | no | no | yes | no | yes | no | no | no | no | no | no | no |
| HCN4 | 340 | S | 0 | 0 | no | yes | no | yes | no | yes | no | no | no | no | no | no | no |

| | | | | | | | | | | | | | | | | | |
|---|---|---|---|---|---|---|---|---|---|---|---|---|---|---|---|---|---|
| HCN4 | 341 | W | 0 | 0 | no | yes | no | yes | no | no | no | no | no | no | no | no | no |
| HCN4 | 342 | F | 0 | 0 | no | yes | no | yes | no | no | no | no | no | no | no | no | no |
| HCN4 | 343 | M | 2 | 0 | yes | no | no | yes | no | no | no | no | no | no | no | no | no |
| HCN4 | 344 | V | 0 | 0 | yes | no | no | yes | no | no | no | no | no | no | no | no | no |
| HCN4 | 345 | D | 0 | 0 | yes | no | no | yes | no | no | no | no | no | no | no | no | no |
| HCN4 | 346 | F | 0 | 0 | yes | no | no | yes | no | no | no | no | no | no | no | no | no |
| HCN4 | 347 | I | 0 | 0 | yes | no | no | yes | no | no | no | no | no | no | no | no | no |
| HCN4 | 348 | S | 0 | 0 | yes | no | no | yes | no | no | no | no | no | no | no | no | no |
| HCN4 | 349 | S | 0 | 0 | yes | no | no | yes | no | no | no | no | no | no | no | no | no |
| HCN4 | 350 | I | 0 | 0 | no | yes | no | yes | no | no | no | no | no | no | no | no | no |
| HCN4 | 351 | P | 0 | 0 | no | yes | no | yes | no | no | no | no | no | no | no | no | no |
| HCN4 | 352 | V | 1 | 0 | yes | no | no | yes | no | no | no | no | no | no | no | no | no |
| HCN4 | 353 | D | 0 | 0 | yes | no | no | yes | no | no | no | no | no | no | no | no | no |
| HCN4 | 354 | Y | 0 | 0 | yes | no | no | yes | no | no | no | no | no | no | no | no | no |
| HCN4 | 355 | I | 1 | 0 | yes | no | no | yes | no | no | no | no | no | no | no | no | no |
| HCN4 | 356 | F | 0 | 0 | yes | no | no | yes | no | no | no | no | no | no | no | no | no |
| HCN4 | 357 | L | 1 | 0 | yes | no | no | yes | no | no | no | no | no | no | no | no | no |
| HCN4 | 358 | I | 1 | 0 | yes | no | no | yes | no | no | no | no | no | no | no | no | no |
| HCN4 | 359 | V | 0 | 0 | yes | no | no | yes | no | no | no | no | no | no | no | no | no |
| HCN4 | 360 | E | 0 | 0 | yes | no | no | no | no | no | no | no | no | no | no | no | no |
| HCN4 | 361 | T | 1 | 0 | yes | no | no | no | no | no | no | no | no | no | no | no | no |
| HCN4 | 362 | R | 0 | 0 | yes | no | no | yes | yes | no | no | no | no | no | no | no | no |
| HCN4 | 363 | I | 1 | 0 | yes | no | no | no | yes | no | no | no | no | no | no | no | no |
| HCN4 | 364 | D | 1 | 0 | no | yes | no | no | yes | no | no | no | no | no | no | no | no |
| HCN4 | 365 | S | 1 | 0 | yes | no | no | no | yes | no | no | no | no | no | no | no | no |
| HCN4 | 366 | E | 0 | 0 | yes | no | no | no | yes | no | no | no | no | no | no | no | no |
| HCN4 | 367 | V | 0 | 0 | yes | no | no | no | yes | no | no | no | no | no | no | no | no |
| HCN4 | 368 | Y | 0 | 0 | no | yes | no | no | yes | no | no | no | no | no | no | no | no |
| HCN4 | 369 | K | 0 | 0 | no | yes | no | no | no | no | no | no | no | yes | no | no | no |
| HCN4 | 370 | T | 0 | 0 | no | yes | no | no | no | no | no | no | no | yes | no | no | no |
| HCN4 | 371 | A | 0 | 0 | no | yes | no | no | no | no | no | no | no | yes | no | no | no |
| HCN4 | 372 | R | 1 | 0 | yes | no | no | no | no | no | no | no | no | yes | no | no | no |
| HCN4 | 373 | A | 0 | 0 | yes | no | no | no | no | no | no | no | no | yes | no | no | no |

| | | | | | | | | | | | | | | | | | |
|---|---|---|---|---|---|---|---|---|---|---|---|---|---|---|---|---|---|
| HCN4 | 374 | L | 0 | 0 | yes | no | no | no | no | no | no | no | no | yes | no | no | no |
| HCN4 | 375 | R | 2 | 1 | yes | no | no | no | no | no | no | no | no | yes | no | no | no |
| HCN4 | 376 | I | 1 | 0 | yes | no | no | no | no | no | no | no | no | yes | no | no | no |
| HCN4 | 377 | V | 1 | 0 | yes | no | no | no | no | no | no | no | no | yes | no | no | no |
| HCN4 | 378 | R | 2 | 0 | yes | no | no | no | no | no | no | no | no | yes | no | no | no |
| HCN4 | 379 | F | 0 | 0 | yes | no | no | yes | no | no | no | no | no | yes | no | no | no |
| HCN4 | 380 | T | 0 | 0 | yes | no | no | yes | no | no | no | no | no | yes | no | no | no |
| HCN4 | 381 | K | 0 | 0 | yes | no | no | yes | no | no | no | no | no | yes | no | no | no |
| HCN4 | 382 | I | 0 | 0 | yes | no | no | yes | no | no | no | no | no | yes | no | no | no |
| HCN4 | 383 | L | 0 | 0 | yes | no | no | yes | no | no | no | no | no | yes | no | no | no |
| HCN4 | 384 | S | 0 | 0 | yes | no | no | yes | no | no | no | no | no | yes | no | no | no |
| HCN4 | 385 | L | 0 | 1 | yes | no | no | yes | no | no | no | no | no | yes | no | no | no |
| HCN4 | 386 | L | 0 | 0 | yes | no | no | yes | no | no | no | no | no | yes | no | no | no |
| HCN4 | 387 | R | 2 | 0 | yes | no | no | yes | no | no | no | no | no | yes | no | no | no |
| HCN4 | 388 | L | 0 | 0 | yes | no | no | yes | no | no | no | no | no | yes | no | no | no |
| HCN4 | 389 | L | 0 | 0 | yes | no | no | yes | no | no | no | no | no | yes | no | no | no |
| HCN4 | 390 | R | 0 | 0 | yes | no | no | yes | no | yes | no | no | no | no | no | no | no |
| HCN4 | 391 | L | 0 | 0 | yes | no | no | yes | no | yes | no | no | no | no | no | no | no |
| HCN4 | 392 | S | 0 | 0 | yes | no | no | yes | no | yes | no | no | no | no | no | no | no |
| HCN4 | 393 | R | 0 | 1 | yes | no | no | yes | no | yes | no | no | no | no | no | no | no |
| HCN4 | 394 | L | 0 | 0 | yes | no | no | yes | no | yes | no | no | no | no | no | no | no |
| HCN4 | 395 | I | 0 | 0 | yes | no | no | yes | no | yes | no | no | no | no | no | no | no |
| HCN4 | 396 | R | 0 | 0 | yes | no | no | yes | no | yes | no | no | no | no | no | no | no |
| HCN4 | 397 | Y | 0 | 0 | yes | no | no | yes | no | yes | no | no | no | no | no | no | no |
| HCN4 | 398 | I | 0 | 0 | yes | no | no | yes | no | yes | no | no | no | no | no | no | no |
| HCN4 | 399 | H | 0 | 0 | yes | no | no | yes | no | yes | no | no | no | no | no | no | no |
| HCN4 | 400 | Q | 0 | 0 | yes | no | no | no | no | yes | no | no | no | no | no | no | no |
| HCN4 | 401 | W | 0 | 0 | yes | no | no | yes | no | yes | no | no | no | no | no | no | no |
| HCN4 | 402 | E | 0 | 0 | yes | no | no | no | no | yes | no | no | no | no | no | no | no |
| HCN4 | 403 | E | 0 | 0 | yes | no | no | no | no | yes | no | no | no | no | no | no | no |
| HCN4 | 404 | I | 0 | 0 | yes | no | no | no | no | yes | no | no | no | no | no | no | no |
| HCN4 | 405 | F | 0 | 0 | no | yes | no | no | no | yes | no | no | no | no | no | no | no |
| HCN4 | 406 | H | 0 | 0 | yes | no | no | no | no | yes | no | no | yes | no | no | no | no |

| | | | | | | | | | | | | | | | | | |
|---|---|---|---|---|---|---|---|---|---|---|---|---|---|---|---|---|---|
| HCN4 | 407 | M | 0 | 0 | yes | no | no | no | no | yes | no | no | no | no | no | no | no |
| HCN4 | 408 | T | 0 | 0 | yes | no | no | no | no | yes | no | no | no | no | no | no | no |
| HCN4 | 409 | Y | 0 | 0 | yes | no | no | no | no | yes | no | no | yes | no | no | no | no |
| HCN4 | 410 | D | 0 | 0 | no | yes | no | no | no | yes | no | no | yes | no | no | no | no |
| HCN4 | 411 | L | 0 | 1 | no | yes | no | no | no | yes | no | no | yes | no | no | no | no |
| HCN4 | 412 | A | 0 | 0 | no | yes | no | no | no | yes | no | no | yes | no | no | no | no |
| HCN4 | 413 | S | 0 | 0 | yes | no | no | no | no | yes | no | no | yes | no | no | no | no |
| HCN4 | 414 | A | 1 | 1 | yes | no | no | no | no | yes | no | no | no | no | no | no | no |
| HCN4 | 415 | V | 1 | 0 | yes | no | no | yes | no | yes | no | no | no | no | no | no | no |
| HCN4 | 416 | V | 0 | 0 | yes | no | no | yes | no | yes | no | no | no | no | no | no | no |
| HCN4 | 417 | R | 1 | 0 | yes | no | no | yes | no | yes | no | no | yes | no | no | no | no |
| HCN4 | 418 | I | 1 | 0 | yes | no | no | yes | no | yes | no | no | no | no | no | no | no |
| HCN4 | 419 | V | 0 | 0 | yes | no | no | yes | no | yes | no | no | no | no | no | no | no |
| HCN4 | 420 | N | 0 | 0 | yes | no | no | yes | no | yes | no | no | no | no | no | no | no |
| HCN4 | 421 | L | 0 | 0 | yes | no | no | yes | no | no | no | no | no | no | no | no | no |
| HCN4 | 422 | I | 0 | 0 | yes | no | no | yes | no | no | no | no | no | no | no | no | no |
| HCN4 | 423 | G | 0 | 0 | yes | no | no | yes | no | no | no | no | no | no | no | no | no |
| HCN4 | 424 | M | 0 | 0 | yes | no | no | yes | no | no | no | no | no | no | no | no | no |
| HCN4 | 425 | M | 0 | 0 | yes | no | no | yes | no | no | no | no | no | no | no | no | no |
| HCN4 | 426 | L | 0 | 0 | yes | no | no | yes | no | no | no | no | no | no | no | no | no |
| HCN4 | 427 | L | 0 | 0 | yes | no | no | yes | no | no | no | no | no | no | no | no | no |
| HCN4 | 428 | L | 0 | 0 | yes | no | no | yes | no | no | no | no | no | no | no | no | no |
| HCN4 | 429 | C | 0 | 0 | yes | no | no | yes | no | no | no | no | no | no | no | no | no |
| HCN4 | 430 | H | 0 | 0 | yes | no | no | yes | no | no | no | no | no | no | no | no | no |
| HCN4 | 431 | W | 0 | 0 | yes | no | no | yes | no | no | no | no | no | no | no | no | no |
| HCN4 | 432 | D | 0 | 0 | yes | no | no | yes | no | no | no | no | no | no | no | no | no |
| HCN4 | 433 | G | 1 | 0 | yes | no | no | yes | no | no | no | no | no | no | no | no | no |
| HCN4 | 434 | C | 0 | 0 | yes | no | no | yes | no | no | no | no | no | no | no | no | no |
| HCN4 | 435 | L | 1 | 0 | yes | no | no | yes | no | no | no | no | no | no | no | no | no |
| HCN4 | 436 | Q | 0 | 0 | yes | no | no | yes | no | no | no | no | no | no | no | no | no |
| HCN4 | 437 | F | 0 | 0 | yes | no | no | no | no | no | no | no | no | no | no | no | no |
| HCN4 | 438 | L | 0 | 0 | yes | no | no | yes | no | no | no | no | no | no | no | no | no |
| HCN4 | 439 | V | 1 | 0 | yes | no | no | no | no | no | no | no | no | no | no | no | no |

| | | | | | | | | | | | | | | | | | |
|---|---|---|---|---|---|---|---|---|---|---|---|---|---|---|---|---|---|
| HCN4 | 440 | P | 1 | 0 | yes | no | no | no | no | no | no | no | no | no | no | no | no |
| HCN4 | 441 | M | 1 | 0 | yes | no | no | no | no | no | no | no | no | no | no | no | no |
| HCN4 | 442 | L | 0 | 0 | no | yes | no | no | yes | no | no | no | no | no | no | no | no |
| HCN4 | 443 | Q | 0 | 0 | no | yes | no | no | yes | no | no | no | no | no | no | no | no |
| HCN4 | 444 | D | 0 | 0 | no | yes | no | no | yes | no | no | no | no | no | no | no | no |
| HCN4 | 445 | F | 0 | 0 | no | yes | no | no | yes | no | no | no | no | no | no | no | no |
| HCN4 | 446 | P | 0 | 0 | no | yes | no | no | yes | no | no | no | no | no | no | no | no |
| HCN4 | 447 | D | 0 | 0 | no | yes | no | no | yes | no | no | no | no | no | no | no | no |
| HCN4 | 448 | D | 1 | 0 | no | yes | no | no | yes | no | no | no | no | no | no | no | no |
| HCN4 | 449 | C | 0 | 0 | no | yes | no | no | yes | no | no | no | no | no | no | no | no |
| HCN4 | 450 | W | 0 | 0 | yes | no | no | no | yes | no | no | no | no | no | no | no | no |
| HCN4 | 451 | V | 0 | 0 | yes | no | no | no | yes | no | no | no | no | no | no | no | no |
| HCN4 | 452 | S | 0 | 0 | yes | no | no | no | yes | no | no | no | no | no | no | no | no |
| HCN4 | 453 | I | 1 | 0 | yes | no | no | no | yes | no | no | no | no | no | no | no | no |
| HCN4 | 454 | N | 0 | 0 | no | yes | no | no | yes | no | no | no | no | no | no | no | no |
| HCN4 | 455 | N | 1 | 0 | no | yes | no | no | yes | no | no | no | no | no | no | no | no |
| HCN4 | 456 | M | 0 | 0 | no | yes | no | no | yes | no | no | no | no | no | no | no | no |
| HCN4 | 457 | V | 0 | 0 | no | yes | no | no | yes | no | no | no | no | no | no | no | no |
| HCN4 | 458 | N | 0 | 0 | no | yes | no | no | yes | no | no | no | no | no | no | no | no |
| HCN4 | 459 | N | 0 | 0 | no | yes | no | no | yes | no | no | no | no | no | no | no | no |
| HCN4 | 460 | S | 0 | 0 | no | yes | no | no | yes | no | no | no | no | no | no | no | no |
| HCN4 | 461 | W | 0 | 0 | yes | no | no | no | yes | no | no | no | no | no | no | no | no |
| HCN4 | 462 | G | 1 | 0 | yes | no | no | no | yes | no | no | no | no | no | no | no | no |
| HCN4 | 463 | K | 0 | 0 | yes | no | no | no | yes | no | no | no | yes | no | no | no | no |
| HCN4 | 464 | Q | 0 | 0 | yes | no | no | no | yes | no | no | no | no | no | no | no | no |
| HCN4 | 465 | Y | 0 | 0 | yes | no | no | yes | no | no | no | yes | no | no | no | no | no |
| HCN4 | 466 | S | 0 | 0 | yes | no | no | yes | no | no | no | yes | yes | no | no | no | no |
| HCN4 | 467 | Y | 0 | 0 | yes | no | no | no | no | no | no | yes | yes | no | no | no | no |
| HCN4 | 468 | A | 2 | 1 | yes | no | no | yes | no | no | no | yes | no | no | no | no | no |
| HCN4 | 469 | L | 0 | 0 | yes | no | no | yes | no | no | no | yes | yes | no | no | no | no |
| HCN4 | 470 | F | 0 | 0 | yes | no | no | yes | no | no | no | yes | yes | no | no | no | no |
| HCN4 | 471 | K | 0 | 0 | yes | no | no | yes | no | no | no | yes | no | no | no | no | no |
| HCN4 | 472 | A | 0 | 0 | yes | no | no | yes | no | no | no | yes | no | no | no | no | no |

| | | | | | | | | | | | | | | | | | |
|---|---|---|---|---|---|---|---|---|---|---|---|---|---|---|---|---|---|
| HCN4 | 473 | M | 0 | 0 | yes | no | no | yes | no | no | no | yes | yes | no | no | no | no |
| HCN4 | 474 | S | 0 | 0 | yes | no | no | yes | no | no | no | yes | yes | no | no | no | no |
| HCN4 | 475 | H | 0 | 0 | yes | no | no | yes | no | no | no | yes | no | no | no | no | no |
| HCN4 | 476 | M | 0 | 0 | yes | no | no | yes | no | no | no | yes | no | no | no | no | no |
| HCN4 | 477 | L | 0 | 0 | no | yes | no | yes | no | no | no | yes | yes | no | no | no | no |
| HCN4 | 478 | C | 0 | 0 | no | yes | no | yes | no | no | yes | yes | yes | no | no | no | no |
| HCN4 | 479 | I | 0 | 1 | no | yes | no | yes | no | no | yes | yes | yes | no | no | no | no |
| HCN4 | 480 | G | 0 | 2 | no | yes | no | yes | no | no | yes | yes | yes | no | no | no | no |
| HCN4 | 481 | Y | 0 | 1 | no | yes | no | yes | no | no | yes | yes | yes | no | no | no | no |
| HCN4 | 482 | G | 0 | 1 | no | yes | no | no | no | no | no | yes | yes | no | no | no | no |
| HCN4 | 483 | R | 2 | 0 | no | yes | no | no | no | no | yes | yes | yes | no | no | no | no |
| HCN4 | 484 | Q | 0 | 0 | no | yes | no | no | no | no | no | yes | no | no | no | no | no |
| HCN4 | 485 | A | 1 | 1 | no | yes | no | no | no | no | no | yes | yes | no | no | no | no |
| HCN4 | 486 | P | 0 | 0 | no | yes | no | no | no | no | no | yes | yes | no | no | no | no |
| HCN4 | 487 | V | 1 | 0 | no | yes | no | no | yes | no | no | no | no | no | no | no | no |
| HCN4 | 488 | G | 0 | 0 | no | yes | no | no | yes | no | no | no | no | no | no | no | no |
| HCN4 | 489 | M | 0 | 0 | yes | no | no | no | yes | no | no | no | yes | no | no | no | no |
| HCN4 | 490 | S | 0 | 0 | yes | no | no | no | yes | no | no | no | no | no | no | no | no |
| HCN4 | 491 | D | 0 | 0 | yes | no | no | no | yes | no | no | no | no | no | no | no | no |
| HCN4 | 492 | V | 1 | 1 | yes | no | no | yes | yes | no | no | no | yes | no | no | no | no |
| HCN4 | 493 | W | 0 | 0 | yes | no | no | yes | yes | no | no | no | yes | no | no | no | no |
| HCN4 | 494 | L | 0 | 0 | yes | no | no | yes | yes | no | no | no | no | no | no | no | no |
| HCN4 | 495 | T | 0 | 0 | yes | no | no | yes | yes | no | no | no | yes | no | no | no | no |
| HCN4 | 496 | M | 1 | 0 | yes | no | no | yes | yes | no | no | no | yes | no | no | no | no |
| HCN4 | 497 | L | 0 | 0 | yes | no | no | yes | no | no | no | no | no | no | no | no | no |
| HCN4 | 498 | S | 0 | 0 | yes | no | no | yes | no | no | no | no | no | no | no | no | no |
| HCN4 | 499 | M | 0 | 0 | yes | no | no | yes | no | no | no | no | yes | no | no | no | no |
| HCN4 | 500 | I | 0 | 0 | yes | no | no | yes | no | no | no | no | yes | no | no | no | no |
| HCN4 | 501 | V | 0 | 0 | yes | no | no | yes | no | no | no | no | no | no | no | no | no |
| HCN4 | 502 | G | 0 | 0 | yes | no | no | yes | no | no | no | no | no | no | no | no | no |
| HCN4 | 503 | A | 0 | 0 | yes | no | no | yes | no | no | no | no | yes | no | no | no | no |
| HCN4 | 504 | T | 0 | 0 | yes | no | no | yes | no | no | no | no | no | no | no | no | no |
| HCN4 | 505 | C | 0 | 0 | yes | no | no | yes | no | no | no | no | no | no | no | no | no |

| | | | | | | | | | | | | | | | | | |
|---|---|---|---|---|---|---|---|---|---|---|---|---|---|---|---|---|---|
| HCN4 | 506 | Y | 0 | 0 | yes | no | no | yes | no | no | yes | no | yes | no | no | no | no |
| HCN4 | 507 | A | 0 | 0 | yes | no | no | yes | no | no | no | no | yes | no | no | no | no |
| HCN4 | 508 | M | 0 | 0 | yes | no | no | yes | no | no | no | no | no | no | no | no | no |
| HCN4 | 509 | F | 0 | 0 | yes | no | no | yes | no | no | no | no | yes | no | no | no | no |
| HCN4 | 510 | I | 0 | 0 | yes | no | no | yes | no | no | yes | no | yes | no | no | no | no |
| HCN4 | 511 | G | 0 | 0 | yes | no | no | yes | no | no | no | no | yes | no | no | no | no |
| HCN4 | 512 | H | 0 | 0 | yes | no | no | yes | no | no | no | no | yes | no | no | no | no |
| HCN4 | 513 | A | 0 | 0 | yes | no | no | yes | no | no | no | no | no | no | no | no | no |
| HCN4 | 514 | T | 0 | 0 | yes | no | no | yes | no | no | yes | no | yes | no | no | no | no |
| HCN4 | 515 | A | 0 | 0 | yes | no | no | no | no | no | no | no | no | no | no | no | no |
| HCN4 | 516 | L | 1 | 0 | yes | no | no | yes | no | no | no | no | no | no | no | no | no |
| HCN4 | 517 | I | 0 | 0 | yes | no | no | yes | no | no | no | no | yes | no | no | no | no |
| HCN4 | 518 | Q | 0 | 0 | no | yes | no | no | no | yes | yes | no | yes | no | no | no | no |
| HCN4 | 519 | S | 0 | 0 | no | yes | no | no | no | yes | no | no | yes | no | no | no | no |
| HCN4 | 520 | L | 0 | 0 | no | yes | no | no | no | yes | no | no | yes | no | no | no | no |
| HCN4 | 521 | D | 0 | 0 | no | yes | no | no | no | yes | no | no | yes | no | no | no | no |
| HCN4 | 522 | S | 0 | 0 | yes | no | no | no | no | yes | no | no | yes | no | no | no | no |
| HCN4 | 523 | S | 0 | 0 | yes | no | no | no | no | yes | no | no | yes | no | no | no | no |
| HCN4 | 524 | R | 1 | 1 | yes | no | no | no | no | yes | no | no | yes | no | no | no | no |
| HCN4 | 525 | R | 1 | 0 | yes | no | no | no | no | yes | yes | no | no | no | no | no | no |
| HCN4 | 526 | Q | 1 | 0 | yes | no | no | no | no | yes | yes | no | yes | no | no | no | no |
| HCN4 | 527 | Y | 0 | 0 | yes | no | no | no | no | yes | no | no | no | no | no | no | no |
| HCN4 | 528 | Q | 1 | 0 | yes | no | no | no | no | yes | no | no | yes | no | no | no | no |
| HCN4 | 529 | E | 0 | 0 | yes | no | no | no | no | yes | yes | no | no | no | no | no | no |
| HCN4 | 530 | K | 0 | 1 | yes | no | no | no | no | yes | no | no | yes | no | no | no | no |
| HCN4 | 531 | Y | 0 | 0 | yes | no | no | no | no | yes | no | no | no | no | no | no | no |
| HCN4 | 532 | K | 0 | 0 | yes | no | no | no | no | yes | no | no | yes | no | no | no | no |
| HCN4 | 533 | Q | 0 | 0 | yes | no | no | no | no | yes | no | no | yes | no | no | no | no |
| HCN4 | 534 | V | 1 | 0 | yes | no | no | no | no | yes | no | no | no | no | no | no | no |
| HCN4 | 535 | E | 0 | 0 | yes | no | no | no | no | yes | no | no | no | no | no | no | no |
| HCN4 | 536 | Q | 0 | 0 | yes | no | no | no | no | yes | no | no | yes | no | no | no | no |
| HCN4 | 537 | Y | 0 | 0 | yes | no | no | no | no | yes | no | no | yes | no | no | no | no |
| HCN4 | 538 | M | 1 | 0 | yes | no | no | no | no | yes | no | no | no | no | no | no | no |

| | | | | | | | | | | | | | | | | | |
|---|---|---|---|---|---|---|---|---|---|---|---|---|---|---|---|---|---|
| HCN4 | 539 | S | 1 | 0 | yes | no | no | no | no | yes | no | no | no | no | no | no | no |
| HCN4 | 540 | F | 0 | 0 | yes | no | no | no | no | yes | no | no | yes | no | no | no | no |
| HCN4 | 541 | H | 1 | 0 | yes | no | no | no | no | yes | no | no | yes | no | no | no | no |
| HCN4 | 542 | K | 0 | 0 | no | yes | no | no | no | yes | no | no | yes | no | no | no | no |
| HCN4 | 543 | L | 0 | 0 | no | yes | no | no | no | yes | no | no | yes | no | no | no | no |
| HCN4 | 544 | P | 1 | 0 | no | yes | no | no | no | yes | no | no | yes | no | no | no | no |
| HCN4 | 545 | P | 1 | 0 | yes | no | no | no | no | yes | no | no | no | no | no | no | no |
| HCN4 | 546 | D | 1 | 0 | yes | no | no | no | no | yes | no | no | no | no | no | no | no |
| HCN4 | 547 | T | 1 | 0 | yes | no | no | no | no | yes | no | no | yes | no | no | no | no |
| HCN4 | 548 | R | 0 | 0 | yes | no | no | no | no | yes | no | no | no | no | no | no | no |
| HCN4 | 549 | Q | 0 | 0 | yes | no | no | no | no | yes | no | no | no | no | no | no | no |
| HCN4 | 550 | R | 2 | 1 | yes | no | no | no | no | yes | no | no | yes | no | no | no | no |
| HCN4 | 551 | I | 0 | 0 | yes | no | no | no | no | yes | no | no | yes | no | no | no | no |
| HCN4 | 552 | H | 0 | 0 | yes | no | no | no | no | yes | no | no | yes | no | no | no | no |
| HCN4 | 553 | D | 1 | 1 | yes | no | no | no | no | yes | no | no | no | no | no | no | no |
| HCN4 | 554 | Y | 1 | 0 | yes | no | no | no | no | yes | no | no | yes | no | no | no | no |
| HCN4 | 555 | Y | 0 | 0 | yes | no | no | no | no | yes | no | no | yes | no | no | no | no |
| HCN4 | 556 | E | 1 | 0 | yes | no | no | no | no | yes | no | no | yes | no | no | no | no |
| HCN4 | 557 | H | 0 | 0 | yes | no | no | no | no | yes | no | no | yes | no | no | no | no |
| HCN4 | 558 | R | 1 | 0 | yes | no | no | no | no | yes | no | no | no | no | no | no | no |
| HCN4 | 559 | Y | 0 | 0 | no | yes | no | no | no | yes | no | no | yes | no | no | no | no |
| HCN4 | 560 | Q | 2 | 0 | no | yes | no | no | no | yes | no | no | yes | no | no | no | no |
| HCN4 | 561 | G | 0 | 0 | no | yes | no | no | no | yes | no | no | yes | no | no | no | no |
| HCN4 | 562 | K | 0 | 0 | no | yes | no | no | no | yes | no | no | yes | no | no | no | no |
| HCN4 | 563 | M | 0 | 0 | no | yes | no | no | no | yes | no | no | yes | no | no | no | no |
| HCN4 | 564 | F | 0 | 0 | no | yes | no | no | no | yes | no | no | yes | no | no | no | no |
| HCN4 | 565 | D | 1 | 0 | no | yes | no | no | no | yes | yes | no | no | no | no | no | no |
| HCN4 | 566 | E | 2 | 0 | yes | no | no | no | no | yes | no | no | yes | no | no | no | no |
| HCN4 | 567 | E | 1 | 0 | yes | no | no | no | no | yes | yes | no | no | no | no | no | no |
| HCN4 | 568 | S | 3 | 0 | yes | no | no | no | no | yes | yes | no | no | no | no | no | no |
| HCN4 | 569 | I | 0 | 0 | yes | no | no | no | no | yes | no | no | yes | no | no | no | no |
| HCN4 | 570 | L | 0 | 0 | yes | no | no | no | no | yes | no | no | no | no | no | no | no |
| HCN4 | 571 | G | 1 | 0 | yes | no | no | no | no | yes | no | no | no | no | no | no | no |

| | | | | | | | | | | | | | | | | | |
|---|---|---|---|---|---|---|---|---|---|---|---|---|---|---|---|---|---|
| HCN4 | 572 | E | 0 | 0 | yes | no | no | no | no | yes | no | no | yes | no | no | no | no |
| HCN4 | 573 | L | 0 | 0 | no | yes | no | no | no | yes | no | no | yes | no | no | no | no |
| HCN4 | 574 | S | 0 | 0 | no | yes | no | no | no | yes | no | no | yes | no | no | no | no |
| HCN4 | 575 | E | 0 | 0 | yes | no | no | no | no | yes | no | no | no | no | no | no | no |
| HCN4 | 576 | P | 0 | 0 | yes | no | no | no | no | yes | no | no | yes | no | no | no | no |
| HCN4 | 577 | L | 0 | 0 | yes | no | no | no | no | yes | no | no | yes | no | no | no | no |
| HCN4 | 578 | R | 2 | 0 | yes | no | no | no | no | yes | no | no | no | no | no | no | no |
| HCN4 | 579 | E | 0 | 0 | yes | no | no | no | no | yes | no | no | no | no | no | no | no |
| HCN4 | 580 | E | 1 | 0 | yes | no | no | no | no | yes | no | no | yes | no | no | no | no |
| HCN4 | 581 | I | 0 | 0 | yes | no | no | no | no | yes | no | no | yes | no | no | no | no |
| HCN4 | 582 | I | 0 | 0 | yes | no | no | no | no | yes | no | no | no | no | no | no | no |
| HCN4 | 583 | N | 0 | 0 | yes | no | no | no | no | yes | no | no | no | no | no | no | no |
| HCN4 | 584 | F | 0 | 0 | yes | no | no | no | no | yes | no | no | yes | no | no | no | no |
| HCN4 | 585 | N | 0 | 0 | yes | no | no | no | no | yes | no | no | yes | no | no | no | no |
| HCN4 | 586 | C | 0 | 0 | no | yes | no | no | no | yes | no | no | no | no | no | no | no |
| HCN4 | 587 | R | 2 | 0 | yes | no | no | no | no | yes | no | no | no | no | no | no | no |
| HCN4 | 588 | K | 1 | 0 | yes | no | no | no | no | yes | no | no | no | no | no | no | no |
| HCN4 | 589 | L | 0 | 0 | yes | no | no | no | no | yes | no | no | no | no | no | no | no |
| HCN4 | 590 | V | 0 | 0 | yes | no | no | no | no | yes | no | no | no | no | no | no | no |
| HCN4 | 591 | A | 0 | 0 | yes | no | no | no | no | yes | no | no | no | no | no | no | no |
| HCN4 | 592 | S | 0 | 0 | yes | no | no | no | no | yes | no | no | no | no | no | no | no |
| HCN4 | 593 | M | 0 | 0 | no | yes | no | no | no | yes | no | no | no | no | no | no | no |
| HCN4 | 594 | P | 0 | 0 | no | yes | no | no | no | yes | no | no | no | no | no | no | no |
| HCN4 | 595 | L | 0 | 0 | no | yes | no | no | no | yes | no | no | no | no | no | no | no |
| HCN4 | 596 | F | 0 | 0 | yes | no | no | no | no | yes | no | no | no | no | no | no | no |
| HCN4 | 597 | A | 0 | 0 | yes | no | no | no | no | yes | no | no | no | no | no | no | no |
| HCN4 | 598 | N | 2 | 0 | yes | no | no | no | no | yes | no | no | no | no | no | no | no |
| HCN4 | 599 | A | 1 | 0 | no | yes | no | no | no | yes | no | no | no | no | no | no | no |
| HCN4 | 600 | D | 0 | 0 | no | yes | no | no | no | yes | no | no | no | no | no | no | no |
| HCN4 | 601 | P | 1 | 0 | yes | no | no | no | no | yes | no | no | no | no | no | no | no |
| HCN4 | 602 | N | 0 | 0 | yes | no | no | no | no | yes | no | no | yes | no | no | no | no |
| HCN4 | 603 | F | 0 | 0 | yes | no | no | no | no | yes | no | no | no | no | no | no | no |
| HCN4 | 604 | V | 1 | 0 | yes | no | no | no | no | yes | no | no | no | no | no | no | no |

| | | | | | | | | | | | | | | | | | |
|---|---|---|---|---|---|---|---|---|---|---|---|---|---|---|---|---|---|
| HCN4 | 605 | T | 0 | 0 | yes | no | no | no | no | yes | no | no | no | no | no | no | no |
| HCN4 | 606 | S | 0 | 0 | yes | no | no | no | no | yes | no | no | no | no | no | no | no |
| HCN4 | 607 | M | 2 | 0 | yes | no | no | no | no | yes | no | no | no | no | no | no | no |
| HCN4 | 608 | L | 0 | 0 | yes | no | no | no | no | yes | no | no | no | no | no | no | no |
| HCN4 | 609 | T | 0 | 0 | no | yes | no | no | no | yes | no | no | no | no | no | no | no |
| HCN4 | 610 | K | 0 | 0 | no | yes | no | no | no | yes | no | no | no | no | no | no | no |
| HCN4 | 611 | L | 0 | 0 | no | yes | no | no | no | yes | no | no | no | no | no | no | no |
| HCN4 | 612 | R | 2 | 0 | no | no | yes | no | no | yes | no | no | no | no | no | no | no |
| HCN4 | 613 | F | 1 | 0 | no | no | yes | no | no | yes | no | no | no | no | no | no | no |
| HCN4 | 614 | E | 1 | 0 | no | no | yes | no | no | yes | no | no | no | no | no | no | no |
| HCN4 | 615 | V | 0 | 0 | no | no | yes | no | no | yes | no | no | yes | no | no | no | no |
| HCN4 | 616 | F | 0 | 0 | no | no | yes | no | no | yes | no | no | yes | no | no | no | no |
| HCN4 | 617 | Q | 0 | 0 | no | yes | no | no | no | yes | no | no | no | no | no | no | no |
| HCN4 | 618 | P | 1 | 0 | no | yes | no | no | no | yes | no | no | no | no | no | no | no |
| HCN4 | 619 | G | 0 | 0 | no | yes | no | no | no | yes | no | no | no | no | no | no | no |
| HCN4 | 620 | D | 0 | 0 | no | yes | no | no | no | yes | no | no | yes | no | no | no | no |
| HCN4 | 621 | Y | 1 | 0 | no | no | yes | no | no | yes | no | no | no | no | no | no | no |
| HCN4 | 622 | I | 1 | 0 | no | no | yes | no | no | yes | no | no | no | no | no | no | no |
| HCN4 | 623 | I | 0 | 0 | no | no | yes | no | no | yes | no | no | no | no | no | no | no |
| HCN4 | 624 | R | 1 | 0 | no | yes | no | no | no | yes | no | no | yes | no | no | no | no |
| HCN4 | 625 | E | 0 | 0 | no | yes | no | no | no | yes | no | no | no | no | no | no | no |
| HCN4 | 626 | G | 0 | 0 | no | yes | no | no | no | yes | no | no | no | no | no | no | no |
| HCN4 | 627 | T | 0 | 0 | no | yes | no | no | no | yes | no | no | no | no | no | no | no |
| HCN4 | 628 | I | 0 | 0 | no | yes | no | no | no | yes | no | no | yes | no | no | no | no |
| HCN4 | 629 | G | 0 | 0 | no | yes | no | no | no | yes | no | no | no | no | no | no | no |
| HCN4 | 630 | K | 1 | 0 | no | yes | no | no | no | yes | no | no | yes | no | no | no | no |
| HCN4 | 631 | K | 0 | 0 | no | no | yes | no | no | yes | no | no | no | no | no | no | no |
| HCN4 | 632 | M | 1 | 0 | no | no | yes | no | no | yes | no | no | no | no | no | no | no |
| HCN4 | 633 | Y | 0 | 0 | no | no | yes | no | no | yes | no | no | no | no | no | no | no |
| HCN4 | 634 | F | 0 | 0 | no | no | yes | no | no | yes | no | no | no | no | no | no | no |
| HCN4 | 635 | I | 0 | 0 | no | no | yes | no | no | yes | no | no | no | no | no | no | no |
| HCN4 | 636 | Q | 0 | 0 | no | no | yes | no | no | yes | no | no | no | no | no | no | no |
| HCN4 | 637 | H | 0 | 0 | no | no | yes | no | no | yes | no | no | yes | no | no | no | no |

| | | | | | | | | | | | | | | | | | |
|---|---|---|---|---|---|---|---|---|---|---|---|---|---|---|---|---|---|
| HCN4 | 638 | G | 0 | 0 | no | yes | no | no | no | yes | no | no | no | no | no | no | no |
| HCN4 | 639 | V | 1 | 0 | no | yes | no | no | no | yes | no | no | no | no | no | no | no |
| HCN4 | 640 | V | 0 | 0 | no | no | yes | no | no | yes | no | no | no | no | no | no | no |
| HCN4 | 641 | S | 0 | 0 | no | no | yes | no | no | yes | no | no | no | no | no | no | no |
| HCN4 | 642 | V | 1 | 0 | no | no | yes | no | no | yes | no | no | no | no | no | no | no |
| HCN4 | 643 | L | 0 | 0 | no | no | yes | no | no | yes | no | no | no | no | no | no | no |
| HCN4 | 644 | T | 0 | 0 | no | yes | no | no | no | yes | no | no | no | no | no | no | no |
| HCN4 | 645 | K | 1 | 0 | no | yes | no | no | no | yes | no | no | no | no | no | no | no |
| HCN4 | 646 | G | 1 | 0 | no | yes | no | no | no | yes | no | no | no | no | no | no | no |
| HCN4 | 647 | N | 0 | 0 | no | yes | no | no | no | yes | no | no | no | no | no | no | no |
| HCN4 | 648 | K | 1 | 0 | no | yes | no | no | no | yes | no | no | no | no | no | no | no |
| HCN4 | 649 | E | 0 | 0 | no | yes | no | no | no | yes | no | no | no | no | no | no | no |
| HCN4 | 650 | T | 0 | 0 | no | no | yes | no | no | yes | no | no | no | no | no | no | no |
| HCN4 | 651 | K | 0 | 0 | no | no | yes | no | no | yes | no | no | no | no | no | no | no |
| HCN4 | 652 | L | 0 | 0 | no | no | yes | no | no | yes | no | no | no | no | no | no | no |
| HCN4 | 653 | A | 0 | 0 | no | yes | no | no | no | yes | no | no | no | no | no | no | no |
| HCN4 | 654 | D | 1 | 0 | no | yes | no | no | no | yes | no | no | no | no | no | no | no |
| HCN4 | 655 | G | 1 | 0 | no | yes | no | no | no | yes | no | no | no | no | no | no | no |
| HCN4 | 656 | S | 0 | 0 | no | yes | no | no | no | yes | no | no | no | no | no | no | no |
| HCN4 | 657 | Y | 0 | 0 | no | no | yes | no | no | yes | no | no | no | no | no | no | no |
| HCN4 | 658 | F | 0 | 0 | no | no | yes | no | no | yes | no | no | no | no | no | no | no |
| HCN4 | 659 | G | 0 | 0 | no | yes | no | no | no | yes | no | no | no | no | no | no | yes |
| HCN4 | 660 | E | 0 | 0 | no | yes | no | no | no | yes | no | no | no | no | no | no | yes |
| HCN4 | 661 | I | 0 | 0 | yes | no | no | no | no | yes | no | no | no | no | no | no | yes |
| HCN4 | 662 | C | 0 | 0 | yes | no | no | no | no | yes | no | no | no | no | no | no | yes |
| HCN4 | 663 | L | 0 | 0 | yes | no | no | no | no | yes | no | no | no | no | no | no | no |
| HCN4 | 664 | L | 0 | 0 | yes | no | no | no | no | yes | no | no | no | no | no | no | no |
| HCN4 | 665 | T | 0 | 0 | yes | no | no | no | no | yes | no | no | no | no | no | no | no |
| HCN4 | 666 | R | 1 | 0 | no | yes | no | no | no | yes | no | no | no | no | no | no | no |
| HCN4 | 667 | G | 0 | 0 | no | yes | no | no | no | yes | no | no | no | no | no | no | no |
| HCN4 | 668 | R | 1 | 0 | no | yes | no | no | no | yes | no | no | yes | no | no | no | no |
| HCN4 | 669 | R | 0 | 0 | no | yes | no | no | no | yes | no | no | no | no | no | no | yes |
| HCN4 | 670 | T | 0 | 0 | no | yes | no | no | no | yes | no | no | no | no | no | no | yes |

| | | | | | | | | | | | | | | | | | |
|---|---|---|---|---|---|---|---|---|---|---|---|---|---|---|---|---|---|
| HCN4 | 671 | A | 0 | 0 | no | yes | no | no | no | yes | no | no | no | no | no | no | no |
| HCN4 | 672 | S | 0 | 1 | no | no | yes | no | no | yes | no | no | no | no | no | no | no |
| HCN4 | 673 | V | 1 | 0 | no | no | yes | no | no | yes | no | no | no | no | no | no | no |
| HCN4 | 674 | R | 0 | 0 | no | no | yes | no | no | yes | no | no | no | no | no | no | no |
| HCN4 | 675 | A | 1 | 0 | no | no | yes | no | no | yes | no | no | no | no | no | no | no |
| HCN4 | 676 | D | 2 | 0 | no | yes | no | no | no | yes | no | no | no | no | no | no | no |
| HCN4 | 677 | T | 0 | 0 | no | yes | no | no | no | yes | no | no | no | no | no | no | no |
| HCN4 | 678 | Y | 1 | 0 | no | yes | no | no | no | yes | no | no | no | no | no | no | no |
| HCN4 | 679 | C | 0 | 0 | no | no | yes | no | no | yes | no | no | no | no | no | no | no |
| HCN4 | 680 | R | 1 | 0 | no | no | yes | no | no | yes | no | no | no | no | no | no | no |
| HCN4 | 681 | L | 0 | 0 | no | no | yes | no | no | yes | no | no | no | no | no | no | no |
| HCN4 | 682 | Y | 0 | 0 | no | no | yes | no | no | yes | no | no | no | no | no | no | no |
| HCN4 | 683 | S | 0 | 0 | no | no | yes | no | no | yes | no | no | no | no | no | no | no |
| HCN4 | 684 | L | 0 | 0 | no | no | yes | no | no | yes | no | no | no | no | no | no | no |
| HCN4 | 685 | S | 0 | 0 | no | no | yes | no | no | yes | no | no | no | no | no | no | no |
| HCN4 | 686 | V | 0 | 0 | yes | no | no | no | no | yes | no | no | no | no | no | no | no |
| HCN4 | 687 | D | 0 | 0 | yes | no | no | no | no | yes | yes | no | no | no | no | no | no |
| HCN4 | 688 | N | 1 | 0 | yes | no | no | no | no | yes | no | no | no | no | no | no | no |
| HCN4 | 689 | F | 0 | 0 | yes | no | no | no | no | yes | no | no | no | no | no | no | no |
| HCN4 | 690 | N | 1 | 0 | yes | no | no | no | no | yes | no | no | no | no | no | no | no |
| HCN4 | 691 | E | 0 | 0 | yes | no | no | no | no | yes | no | no | yes | no | no | no | no |
| HCN4 | 692 | V | 0 | 0 | yes | no | no | no | no | yes | no | no | no | no | no | no | no |
| HCN4 | 693 | L | 0 | 0 | yes | no | no | no | no | yes | no | no | no | no | no | no | no |
| HCN4 | 694 | E | 0 | 0 | yes | no | no | no | no | yes | no | no | yes | no | no | no | no |
| HCN4 | 695 | E | 0 | 0 | yes | no | no | no | no | yes | no | no | yes | no | no | no | no |
| HCN4 | 696 | Y | 0 | 0 | no | yes | no | no | no | yes | no | no | no | no | no | no | no |
| HCN4 | 697 | P | 0 | 0 | no | yes | no | no | no | yes | no | no | no | no | no | no | no |
| HCN4 | 698 | M | 1 | 0 | no | yes | no | no | no | yes | no | no | no | no | no | no | no |
| HCN4 | 699 | M | 0 | 0 | no | yes | no | no | no | yes | no | no | no | no | no | no | no |
| HCN4 | 700 | R | 0 | 0 | yes | no | no | no | no | yes | no | no | no | no | no | no | no |
| HCN4 | 701 | R | 0 | 0 | yes | no | no | no | no | yes | no | no | no | no | no | no | no |
| HCN4 | 702 | A | 0 | 0 | yes | no | no | no | no | yes | no | no | no | no | no | no | no |
| HCN4 | 703 | F | 0 | 0 | yes | no | no | no | no | yes | no | no | no | no | no | no | no |

| | | | | | | | | | | | | | | | | | |
|---|---|---|---|---|---|---|---|---|---|---|---|---|---|---|---|---|---|
| HCN4 | 704 | E | 0 | 0 | yes | no | no | no | no | yes | no | no | no | no | no | no | no |
| HCN4 | 705 | T | 0 | 0 | yes | no | no | no | no | yes | no | no | no | no | no | no | no |
| HCN4 | 706 | V | 1 | 0 | yes | no | no | no | no | yes | no | no | no | no | no | no | no |
| HCN4 | 707 | A | 0 | 0 | yes | no | no | no | no | yes | no | no | no | no | no | no | no |
| HCN4 | 708 | L | 0 | 0 | yes | no | no | no | no | yes | no | no | no | no | no | no | no |
| HCN4 | 709 | D | 0 | 0 | yes | no | no | no | no | yes | no | no | no | no | no | no | no |
| HCN4 | 710 | R | 0 | 0 | yes | no | no | no | no | yes | no | no | no | no | no | no | yes |
| HCN4 | 711 | L | 0 | 0 | yes | no | no | no | no | yes | no | no | no | no | no | no | yes |
| HCN4 | 712 | D | 0 | 0 | yes | no | no | no | no | yes | no | no | no | no | no | no | yes |
| HCN4 | 713 | R | 2 | 0 | no | yes | no | no | no | yes | no | no | no | no | no | no | yes |
| HCN4 | 714 | I | 1 | 0 | no | yes | no | no | no | yes | no | no | no | no | no | no | no |
| KCNK3 | 1 | M | 0 | 0 | no | yes | no | no | no | yes | no | no | no | no | no | no | no |
| KCNK3 | 2 | K | 0 | 0 | no | yes | no | no | no | yes | no | no | no | no | no | no | no |
| KCNK3 | 3 | R | 0 | 0 | yes | no | no | no | no | yes | no | no | no | no | no | no | no |
| KCNK3 | 4 | Q | 0 | 0 | yes | no | no | no | no | yes | no | no | yes | no | no | no | no |
| KCNK3 | 5 | N | 0 | 0 | yes | no | no | yes | no | yes | no | no | yes | no | no | no | no |
| KCNK3 | 6 | V | 1 | 0 | yes | no | no | yes | no | yes | no | no | no | no | no | no | no |
| KCNK3 | 7 | R | 0 | 0 | yes | no | no | yes | no | yes | no | no | no | no | no | no | no |
| KCNK3 | 8 | T | 0 | 1 | yes | no | no | yes | no | yes | no | no | yes | no | no | no | no |
| KCNK3 | 9 | L | 0 | 0 | yes | no | no | yes | no | no | no | no | no | no | no | no | no |
| KCNK3 | 10 | A | 1 | 0 | yes | no | no | yes | no | no | no | no | no | no | no | no | no |
| KCNK3 | 11 | L | 0 | 0 | yes | no | no | yes | no | no | no | no | yes | no | no | no | no |
| KCNK3 | 12 | I | 0 | 0 | yes | no | no | yes | no | no | no | no | yes | no | no | no | no |
| KCNK3 | 13 | V | 1 | 0 | yes | no | no | yes | no | no | no | no | no | no | no | no | no |
| KCNK3 | 14 | C | 1 | 0 | yes | no | no | yes | no | no | no | no | yes | no | no | no | no |
| KCNK3 | 15 | T | 0 | 0 | yes | no | no | yes | no | no | no | no | yes | no | no | no | no |
| KCNK3 | 16 | F | 0 | 0 | yes | no | no | yes | no | no | no | no | yes | no | no | no | no |
| KCNK3 | 17 | T | 0 | 0 | yes | no | no | yes | no | no | no | no | no | no | no | no | no |
| KCNK3 | 18 | Y | 0 | 0 | yes | no | no | yes | no | no | no | no | yes | no | no | no | no |
| KCNK3 | 19 | L | 0 | 0 | yes | no | no | yes | no | no | no | no | yes | no | no | no | no |
| KCNK3 | 20 | L | 0 | 0 | yes | no | no | yes | no | no | no | no | yes | no | no | no | no |
| KCNK3 | 21 | V | 0 | 0 | yes | no | no | yes | no | no | no | no | no | no | no | no | no |
| KCNK3 | 22 | G | 0 | 0 | yes | no | no | yes | no | no | no | no | yes | no | no | no | no |

| | | | | | | | | | | | | | | | | | |
|---|---|---|---|---|---|---|---|---|---|---|---|---|---|---|---|---|---|
| KCNK3 | 23 | A | 0 | 0 | yes | no | no | yes | no | no | no | no | yes | no | no | no | no |
| KCNK3 | 24 | A | 0 | 0 | yes | no | no | yes | no | no | no | no | no | no | no | no | no |
| KCNK3 | 25 | V | 0 | 0 | yes | no | no | yes | no | no | no | no | yes | no | no | no | no |
| KCNK3 | 26 | F | 0 | 0 | yes | no | no | yes | no | no | no | no | yes | no | no | no | no |
| KCNK3 | 27 | D | 0 | 0 | yes | no | no | no | no | no | no | no | yes | no | no | no | no |
| KCNK3 | 28 | A | 0 | 0 | yes | no | no | no | no | no | no | no | no | no | no | no | no |
| KCNK3 | 29 | L | 0 | 0 | yes | no | no | no | no | no | no | no | yes | no | no | no | no |
| KCNK3 | 30 | E | 0 | 0 | yes | no | no | no | no | no | no | no | yes | no | no | no | no |
| KCNK3 | 31 | S | 0 | 0 | yes | no | no | no | no | no | no | no | yes | no | no | no | no |
| KCNK3 | 32 | E | 1 | 0 | yes | no | no | no | no | no | no | no | no | no | no | no | no |
| KCNK3 | 33 | P | 0 | 0 | yes | no | no | no | no | no | no | no | yes | no | no | no | no |
| KCNK3 | 34 | E | 0 | 0 | yes | no | no | no | no | no | no | no | yes | no | no | no | no |
| KCNK3 | 35 | L | 0 | 0 | yes | no | no | no | no | no | no | no | no | no | no | no | no |
| KCNK3 | 36 | I | 0 | 0 | yes | no | no | no | no | no | no | no | no | no | no | no | no |
| KCNK3 | 37 | E | 0 | 0 | yes | no | no | no | yes | no | no | no | yes | no | no | no | no |
| KCNK3 | 38 | R | 0 | 0 | yes | no | no | no | yes | no | no | no | yes | no | no | no | no |
| KCNK3 | 39 | Q | 0 | 0 | yes | no | no | no | yes | no | no | no | no | no | no | no | no |
| KCNK3 | 40 | R | 0 | 0 | yes | no | no | no | yes | no | no | no | no | no | no | no | no |
| KCNK3 | 41 | L | 0 | 0 | yes | no | no | no | yes | no | no | no | yes | no | no | no | no |
| KCNK3 | 42 | E | 0 | 0 | yes | no | no | no | yes | no | no | no | no | no | no | no | no |
| KCNK3 | 43 | L | 0 | 0 | yes | no | no | no | yes | no | no | no | no | no | no | no | no |
| KCNK3 | 44 | R | 0 | 0 | yes | no | no | no | yes | no | no | no | yes | no | no | no | no |
| KCNK3 | 45 | Q | 1 | 0 | yes | no | no | no | yes | no | no | no | yes | no | no | no | no |
| KCNK3 | 46 | Q | 0 | 0 | yes | no | no | no | yes | no | no | no | no | no | no | no | no |
| KCNK3 | 47 | E | 0 | 0 | yes | no | no | no | no | no | no | no | no | no | no | no | no |
| KCNK3 | 48 | L | 0 | 0 | yes | no | no | no | no | no | no | no | yes | no | no | no | no |
| KCNK3 | 49 | R | 1 | 0 | yes | no | no | no | no | no | no | no | no | no | no | no | no |
| KCNK3 | 50 | A | 1 | 0 | yes | no | no | no | no | no | no | no | no | no | no | no | no |
| KCNK3 | 51 | R | 1 | 1 | yes | no | no | no | no | no | no | no | yes | no | no | no | no |
| KCNK3 | 52 | Y | 0 | 0 | no | yes | no | no | no | no | no | no | yes | no | no | no | no |
| KCNK3 | 53 | N | 0 | 0 | no | yes | no | no | no | no | no | no | no | no | no | no | no |
| KCNK3 | 54 | L | 0 | 0 | no | yes | no | no | no | no | no | no | yes | no | no | no | no |
| KCNK3 | 55 | S | 0 | 0 | no | yes | no | no | no | no | no | no | no | no | no | no | no |

| | | | | | | | | | | | | | | | | | |
|---|---|---|---|---|---|---|---|---|---|---|---|---|---|---|---|---|---|
| KCNK3 | 56 | Q | 1 | 0 | yes | no | no | no | no | no | no | no | no | no | no | no | no |
| KCNK3 | 57 | G | 0 | 0 | yes | no | no | no | no | no | no | no | no | no | no | no | no |
| KCNK3 | 58 | G | 2 | 0 | yes | no | no | no | no | no | no | no | yes | no | no | no | no |
| KCNK3 | 59 | Y | 0 | 0 | yes | no | no | no | no | no | no | no | yes | no | no | no | no |
| KCNK3 | 60 | E | 0 | 0 | yes | no | no | no | no | no | no | no | no | no | no | no | no |
| KCNK3 | 61 | E | 0 | 0 | yes | no | no | no | no | no | no | no | yes | no | no | no | no |
| KCNK3 | 62 | L | 0 | 0 | yes | no | no | no | no | no | yes | no | yes | no | no | no | no |
| KCNK3 | 63 | E | 0 | 0 | yes | no | no | no | no | no | no | no | yes | no | no | no | no |
| KCNK3 | 64 | R | 1 | 0 | yes | no | no | no | no | no | no | no | no | no | no | no | no |
| KCNK3 | 65 | V | 0 | 0 | yes | no | no | no | no | no | no | no | yes | no | no | no | no |
| KCNK3 | 66 | V | 0 | 0 | yes | no | no | no | no | no | yes | no | yes | no | no | no | no |
| KCNK3 | 67 | L | 0 | 0 | yes | no | no | no | no | no | yes | no | yes | no | no | no | no |
| KCNK3 | 68 | R | 2 | 0 | yes | no | no | no | no | no | no | no | yes | no | no | no | no |
| KCNK3 | 69 | L | 0 | 0 | yes | no | no | no | no | no | no | no | yes | no | no | no | no |
| KCNK3 | 70 | K | 0 | 0 | yes | no | no | no | no | no | yes | no | yes | no | no | no | no |
| KCNK3 | 71 | P | 0 | 0 | yes | no | no | no | no | no | no | no | no | no | no | no | no |
| KCNK3 | 72 | H | 2 | 0 | yes | no | no | no | no | no | no | no | yes | no | no | no | no |
| KCNK3 | 73 | K | 0 | 0 | yes | no | no | no | no | no | no | no | no | no | no | no | no |
| KCNK3 | 74 | A | 2 | 0 | no | yes | no | no | no | no | no | no | no | no | no | no | no |
| KCNK3 | 75 | G | 0 | 0 | no | yes | no | no | no | no | no | no | no | no | no | no | no |
| KCNK3 | 76 | V | 0 | 0 | no | yes | no | no | no | no | no | no | yes | no | no | no | no |
| KCNK3 | 77 | Q | 0 | 0 | no | yes | no | no | no | no | no | no | yes | no | no | no | no |
| KCNK3 | 78 | W | 0 | 0 | no | yes | no | no | no | no | no | yes | yes | no | no | no | no |
| KCNK3 | 79 | R | 1 | 0 | no | yes | no | no | no | no | no | yes | yes | no | no | no | no |
| KCNK3 | 80 | F | 0 | 0 | yes | no | no | no | no | no | no | yes | yes | no | no | no | no |
| KCNK3 | 81 | A | 0 | 0 | yes | no | no | no | no | no | no | yes | no | no | no | no | no |
| KCNK3 | 82 | G | 0 | 0 | yes | no | no | no | no | no | no | yes | no | no | no | no | no |
| KCNK3 | 83 | S | 0 | 0 | yes | no | no | no | no | no | no | yes | yes | no | no | no | no |
| KCNK3 | 84 | F | 0 | 0 | yes | no | no | yes | no | no | no | yes | yes | no | no | no | no |
| KCNK3 | 85 | Y | 0 | 0 | yes | no | no | yes | no | no | no | yes | no | no | no | no | no |
| KCNK3 | 86 | F | 0 | 0 | yes | no | no | yes | no | no | no | yes | yes | no | no | no | no |
| KCNK3 | 87 | A | 0 | 0 | yes | no | no | yes | no | no | no | yes | yes | no | no | no | no |
| KCNK3 | 88 | I | 0 | 0 | yes | no | no | yes | no | no | no | yes | yes | no | no | no | no |

| | | | | | | | | | | | | | | | | | |
|---|---|---|---|---|---|---|---|---|---|---|---|---|---|---|---|---|---|
| KCNK3 | 89 | T | 0 | 0 | yes | no | no | yes | no | no | no | yes | no | no | no | no | no |
| KCNK3 | 90 | V | 0 | 0 | yes | no | no | yes | no | no | no | yes | yes | no | no | no | no |
| KCNK3 | 91 | I | 0 | 0 | yes | no | no | yes | no | no | no | yes | yes | no | no | no | no |
| KCNK3 | 92 | T | 0 | 0 | no | yes | no | yes | no | no | no | yes | no | no | no | no | no |
| KCNK3 | 93 | T | 0 | 0 | no | yes | no | yes | no | no | yes | yes | yes | no | no | no | no |
| KCNK3 | 94 | I | 0 | 0 | no | yes | no | yes | no | no | yes | yes | yes | no | no | no | no |
| KCNK3 | 95 | G | 0 | 0 | no | yes | no | yes | no | no | yes | yes | yes | no | no | no | no |
| KCNK3 | 96 | Y | 0 | 0 | no | yes | no | yes | no | no | yes | yes | yes | no | no | no | no |
| KCNK3 | 97 | G | 0 | 1 | no | yes | no | no | no | no | yes | yes | yes | no | no | no | no |
| KCNK3 | 98 | H | 1 | 0 | no | yes | no | no | no | no | no | yes | no | no | no | no | no |
| KCNK3 | 99 | A | 0 | 0 | no | yes | no | no | no | no | no | yes | no | no | no | no | no |
| KCNK3 | 100 | A | 0 | 0 | no | yes | no | no | no | no | no | yes | yes | no | no | no | no |
| KCNK3 | 101 | P | 0 | 0 | no | yes | no | no | no | no | no | yes | yes | no | no | no | no |
| KCNK3 | 102 | S | 0 | 0 | no | yes | no | no | no | no | no | no | yes | no | no | no | no |
| KCNK3 | 103 | T | 0 | 0 | no | yes | no | no | no | no | no | no | yes | no | no | no | no |
| KCNK3 | 104 | D | 0 | 0 | yes | no | no | no | no | no | no | no | yes | no | no | no | no |
| KCNK3 | 105 | G | 0 | 0 | yes | no | no | no | no | no | no | no | yes | no | no | no | no |
| KCNK3 | 106 | G | 0 | 1 | yes | no | no | no | no | no | no | no | yes | no | no | no | no |
| KCNK3 | 107 | K | 0 | 0 | yes | no | no | no | no | no | no | no | yes | no | no | no | no |
| KCNK3 | 108 | V | 0 | 0 | yes | no | no | yes | no | no | no | no | yes | no | no | no | no |
| KCNK3 | 109 | F | 0 | 0 | yes | no | no | yes | no | no | no | no | yes | no | no | no | no |
| KCNK3 | 110 | C | 0 | 0 | yes | no | no | yes | no | no | no | no | yes | no | no | no | no |
| KCNK3 | 111 | M | 0 | 0 | yes | no | no | yes | no | no | no | no | yes | no | no | no | no |
| KCNK3 | 112 | F | 1 | 0 | yes | no | no | yes | no | no | no | no | no | no | no | no | no |
| KCNK3 | 113 | Y | 0 | 0 | yes | no | no | yes | no | no | no | no | yes | no | no | no | no |
| KCNK3 | 114 | A | 0 | 1 | yes | no | no | yes | no | no | no | no | yes | no | no | no | no |
| KCNK3 | 115 | L | 0 | 0 | yes | no | no | yes | no | no | no | no | no | no | no | no | no |
| KCNK3 | 116 | L | 0 | 0 | yes | no | no | yes | no | no | no | no | yes | no | no | no | no |
| KCNK3 | 117 | G | 0 | 0 | yes | no | no | yes | no | no | no | no | yes | no | no | no | no |
| KCNK3 | 118 | I | 0 | 0 | yes | no | no | yes | no | no | no | no | yes | no | no | no | no |
| KCNK3 | 119 | P | 0 | 0 | yes | no | no | yes | no | no | no | no | yes | no | no | no | no |
| KCNK3 | 120 | L | 0 | 0 | yes | no | no | yes | no | no | no | no | yes | no | no | no | no |
| KCNK3 | 121 | T | 0 | 0 | yes | no | no | yes | no | no | no | no | no | no | no | no | no |

| | | | | | | | | | | | | | | | | | |
|---|---|---|---|---|---|---|---|---|---|---|---|---|---|---|---|---|---|
| KCNK3 | 122 | L | 1 | 1 | yes | no | no | yes | no | no | yes | no | yes | no | no | no | no |
| KCNK3 | 123 | V | 0 | 1 | yes | no | no | yes | no | no | no | no | yes | no | no | no | no |
| KCNK3 | 124 | M | 0 | 0 | yes | no | no | yes | no | no | no | no | yes | no | no | no | no |
| KCNK3 | 125 | F | 0 | 0 | yes | no | no | yes | no | no | no | no | no | no | no | no | no |
| KCNK3 | 126 | Q | 0 | 0 | yes | no | no | yes | no | no | no | no | yes | no | no | no | no |
| KCNK3 | 127 | S | 0 | 0 | yes | no | no | yes | no | no | no | no | yes | no | no | no | no |
| KCNK3 | 128 | L | 0 | 0 | yes | no | no | yes | no | no | no | no | yes | no | no | no | no |
| KCNK3 | 129 | G | 0 | 0 | yes | no | no | yes | no | yes | no | no | no | no | no | no | no |
| KCNK3 | 130 | E | 1 | 0 | yes | no | no | yes | no | yes | no | no | yes | no | no | no | no |
| KCNK3 | 131 | R | 0 | 0 | yes | no | no | yes | no | yes | no | no | yes | no | no | no | no |
| KCNK3 | 132 | I | 0 | 0 | yes | no | no | yes | no | yes | no | no | no | no | no | no | no |
| KCNK3 | 133 | N | 0 | 0 | yes | no | no | yes | no | yes | no | no | no | no | no | no | no |
| KCNK3 | 134 | T | 0 | 0 | yes | no | no | yes | no | yes | no | no | no | no | no | no | no |
| KCNK3 | 135 | L | 0 | 0 | yes | no | no | yes | no | yes | no | no | no | no | no | no | no |
| KCNK3 | 136 | V | 0 | 0 | yes | no | no | yes | no | yes | no | no | no | no | no | no | no |
| KCNK3 | 137 | R | 0 | 0 | yes | no | no | no | no | yes | no | no | no | no | no | no | no |
| KCNK3 | 138 | Y | 0 | 0 | yes | no | no | no | no | yes | no | no | no | no | no | no | no |
| KCNK3 | 139 | L | 0 | 0 | yes | no | no | yes | no | yes | no | no | no | no | no | no | no |
| KCNK3 | 140 | L | 1 | 0 | yes | no | no | no | no | yes | no | no | no | no | no | no | no |
| KCNK3 | 141 | H | 1 | 0 | yes | no | no | no | no | yes | no | no | no | no | no | no | no |
| KCNK3 | 142 | R | 2 | 0 | yes | no | no | no | no | yes | no | no | no | no | no | no | no |
| KCNK3 | 143 | A | 0 | 0 | yes | no | no | no | no | yes | no | no | no | no | no | no | no |
| KCNK3 | 144 | K | 0 | 0 | yes | no | no | no | no | yes | no | no | no | no | no | no | no |
| KCNK3 | 145 | K | 0 | 0 | yes | no | no | no | no | yes | no | no | no | no | no | no | no |
| KCNK3 | 146 | G | 0 | 0 | yes | no | no | no | no | yes | no | no | no | no | no | no | no |
| KCNK3 | 147 | L | 0 | 0 | yes | no | no | no | no | yes | no | no | no | no | no | no | no |
| KCNK3 | 148 | G | 0 | 0 | no | yes | no | no | no | yes | no | no | no | no | no | no | no |
| KCNK3 | 152 | A | 0 | 0 | no | yes | no | no | no | yes | no | no | no | no | no | no | no |
| KCNK3 | 153 | D | 0 | 0 | no | yes | no | no | no | yes | no | no | no | no | no | no | no |
| KCNK3 | 154 | V | 0 | 0 | no | yes | no | no | no | yes | no | no | no | no | no | no | no |
| KCNK3 | 155 | S | 0 | 0 | no | yes | no | no | no | yes | no | no | no | no | no | no | no |
| KCNK3 | 156 | M | 0 | 0 | yes | no | no | no | no | yes | no | no | no | no | no | no | no |
| KCNK3 | 157 | A | 0 | 0 | yes | no | no | yes | no | yes | no | no | no | no | no | no | no |

| | | | | | | | | | | | | | | | | | |
|---|---|---|---|---|---|---|---|---|---|---|---|---|---|---|---|---|---|
| KCNK3 | 158 | N | 0 | 0 | yes | no | no | yes | no | yes | no | no | no | no | no | no | no |
| KCNK3 | 159 | M | 0 | 0 | yes | no | no | yes | no | no | no | no | no | no | no | no | no |
| KCNK3 | 160 | V | 0 | 0 | yes | no | no | yes | no | no | no | no | no | no | no | no | no |
| KCNK3 | 161 | L | 0 | 0 | yes | no | no | yes | no | no | no | no | no | no | no | no | no |
| KCNK3 | 162 | I | 0 | 0 | yes | no | no | yes | no | no | no | no | no | no | no | no | no |
| KCNK3 | 163 | G | 0 | 0 | yes | no | no | yes | no | no | no | no | no | no | no | no | no |
| KCNK3 | 164 | F | 0 | 0 | yes | no | no | yes | no | no | no | no | no | no | no | no | no |
| KCNK3 | 165 | F | 0 | 0 | yes | no | no | yes | no | no | no | no | no | no | no | no | no |
| KCNK3 | 166 | S | 0 | 0 | yes | no | no | yes | no | no | no | no | no | no | no | no | no |
| KCNK3 | 167 | C | 0 | 0 | yes | no | no | yes | no | no | no | no | no | no | no | no | no |
| KCNK3 | 168 | I | 1 | 0 | yes | no | no | yes | no | no | no | no | no | no | no | no | no |
| KCNK3 | 169 | S | 0 | 0 | yes | no | no | yes | no | no | no | no | no | no | no | no | no |
| KCNK3 | 170 | T | 0 | 0 | yes | no | no | yes | no | no | no | no | no | no | no | no | no |
| KCNK3 | 171 | L | 0 | 0 | yes | no | no | yes | no | no | no | no | no | no | no | no | no |
| KCNK3 | 172 | C | 0 | 0 | yes | no | no | yes | no | no | no | no | no | no | no | no | no |
| KCNK3 | 173 | I | 0 | 0 | yes | no | no | yes | no | no | no | no | no | no | no | no | no |
| KCNK3 | 174 | G | 0 | 0 | yes | no | no | yes | no | no | no | no | no | no | no | no | no |
| KCNK3 | 175 | A | 0 | 0 | yes | no | no | yes | no | no | no | no | no | no | no | no | no |
| KCNK3 | 176 | A | 1 | 0 | yes | no | no | yes | no | no | no | no | no | no | no | no | no |
| KCNK3 | 177 | A | 1 | 0 | yes | no | no | yes | no | no | no | no | no | no | no | no | no |
| KCNK3 | 178 | F | 0 | 0 | yes | no | no | yes | no | no | no | no | no | no | no | no | no |
| KCNK3 | 179 | S | 1 | 0 | yes | no | no | no | no | no | no | no | no | no | no | no | no |
| KCNK3 | 180 | H | 0 | 0 | yes | no | no | no | no | no | no | no | no | no | no | no | no |
| KCNK3 | 181 | Y | 0 | 0 | yes | no | no | no | no | no | no | no | no | no | no | no | no |
| KCNK3 | 182 | E | 1 | 2 | no | yes | no | no | no | no | no | no | no | no | no | no | no |
| KCNK3 | 183 | H | 1 | 0 | no | yes | no | no | no | no | no | no | no | no | no | no | no |
| KCNK3 | 184 | W | 0 | 0 | no | yes | no | no | no | no | no | yes | no | no | no | no | no |
| KCNK3 | 185 | T | 0 | 0 | no | yes | no | no | no | no | no | yes | no | no | no | no | no |
| KCNK3 | 186 | F | 0 | 0 | yes | no | no | yes | no | no | no | yes | no | no | no | no | no |
| KCNK3 | 187 | F | 0 | 0 | yes | no | no | yes | no | no | no | yes | yes | no | no | no | no |
| KCNK3 | 188 | Q | 0 | 0 | yes | no | no | no | no | no | no | yes | no | no | no | no | no |
| KCNK3 | 189 | A | 1 | 0 | yes | no | no | yes | no | no | no | yes | no | no | no | no | no |
| KCNK3 | 190 | Y | 0 | 0 | yes | no | no | yes | no | no | no | yes | no | no | no | no | no |

| | | | | | | | | | | | | | | | | | |
|---|---|---|---|---|---|---|---|---|---|---|---|---|---|---|---|---|---|
| KCNK3 | 191 | Y | 0 | 0 | yes | no | no | yes | no | no | no | yes | yes | no | no | no | no |
| KCNK3 | 192 | Y | 0 | 1 | yes | no | no | yes | no | no | no | yes | no | no | no | no | no |
| KCNK3 | 193 | C | 0 | 0 | yes | no | no | yes | no | no | no | yes | no | no | no | no | no |
| KCNK3 | 194 | F | 0 | 0 | yes | no | no | yes | no | no | no | yes | yes | no | no | no | no |
| KCNK3 | 195 | I | 0 | 0 | yes | no | no | yes | no | no | no | yes | no | no | no | no | no |
| KCNK3 | 196 | T | 0 | 0 | yes | no | no | yes | no | no | no | yes | no | no | no | no | no |
| KCNK3 | 197 | L | 0 | 0 | yes | no | no | yes | no | no | no | yes | no | no | no | no | no |
| KCNK3 | 198 | T | 0 | 0 | no | yes | no | yes | no | no | no | yes | yes | no | no | no | no |
| KCNK3 | 199 | T | 0 | 0 | no | yes | no | yes | no | no | yes | yes | yes | no | no | no | no |
| KCNK3 | 200 | I | 0 | 0 | no | yes | no | yes | no | no | yes | yes | yes | no | no | no | no |
| KCNK3 | 201 | G | 0 | 0 | no | yes | no | yes | no | no | yes | yes | yes | no | no | no | no |
| KCNK3 | 202 | F | 0 | 0 | no | yes | no | yes | no | no | yes | yes | yes | no | no | no | no |
| KCNK3 | 203 | G | 0 | 1 | no | yes | no | no | no | no | yes | yes | no | no | no | no | no |
| KCNK3 | 204 | D | 0 | 0 | no | yes | no | no | no | no | yes | yes | yes | no | no | no | no |
| KCNK3 | 205 | Y | 1 | 0 | no | yes | no | no | no | no | no | yes | yes | no | no | no | no |
| KCNK3 | 206 | V | 0 | 1 | no | yes | no | no | no | no | no | yes | no | no | no | no | no |
| KCNK3 | 207 | A | 0 | 0 | no | yes | no | no | no | no | no | yes | no | no | no | no | no |
| KCNK3 | 208 | L | 0 | 0 | no | yes | no | no | no | no | no | no | no | no | no | no | no |
| KCNK3 | 209 | Q | 0 | 0 | no | yes | no | no | no | no | no | no | no | no | no | no | no |
| KCNK3 | 210 | K | 0 | 0 | no | yes | no | no | no | no | no | no | no | no | no | no | no |
| KCNK3 | 211 | D | 0 | 0 | no | yes | no | no | no | no | no | no | no | no | no | no | no |
| KCNK3 | 212 | Q | 1 | 0 | no | yes | no | no | no | no | no | no | no | no | no | no | no |
| KCNK3 | 213 | A | 0 | 0 | yes | no | no | no | no | no | no | no | no | no | no | no | no |
| KCNK3 | 214 | L | 0 | 1 | yes | no | no | no | no | no | no | no | no | no | no | no | no |
| KCNK3 | 215 | Q | 0 | 0 | yes | no | no | no | no | no | no | no | no | no | no | no | no |
| KCNK3 | 216 | T | 0 | 0 | yes | no | no | no | no | no | no | no | no | no | no | no | no |
| KCNK3 | 217 | Q | 0 | 0 | no | yes | no | no | no | no | no | no | no | no | no | no | no |
| KCNK3 | 218 | P | 0 | 0 | yes | no | no | no | no | no | no | no | no | no | no | no | no |
| KCNK3 | 219 | Q | 1 | 0 | yes | no | no | no | no | no | no | no | no | no | no | no | no |
| KCNK3 | 220 | Y | 0 | 0 | yes | no | no | no | no | no | no | no | no | no | no | no | no |
| KCNK3 | 221 | V | 0 | 1 | yes | no | no | no | no | no | no | no | no | no | no | no | no |
| KCNK3 | 222 | A | 1 | 0 | yes | no | no | yes | no | no | no | no | no | no | no | no | no |
| KCNK3 | 223 | F | 0 | 0 | yes | no | no | yes | no | no | no | no | no | no | no | no | no |

| | | | | | | | | | | | | | | | | | |
|---|---|---|---|---|---|---|---|---|---|---|---|---|---|---|---|---|---|
| KCNK3 | 224 | S | 1 | 0 | yes | no | no | yes | no | no | no | no | no | no | no | no | no |
| KCNK3 | 225 | F | 0 | 0 | yes | no | no | yes | no | no | no | no | no | no | no | no | no |
| KCNK3 | 226 | V | 0 | 0 | yes | no | no | yes | no | no | no | no | no | no | no | no | no |
| KCNK3 | 227 | Y | 0 | 0 | yes | no | no | yes | no | no | no | no | no | no | no | no | no |
| KCNK3 | 228 | I | 0 | 0 | yes | no | no | yes | no | no | no | no | no | no | no | no | no |
| KCNK3 | 229 | L | 0 | 0 | yes | no | no | yes | no | no | no | no | yes | no | no | no | no |
| KCNK3 | 230 | T | 0 | 0 | yes | no | no | yes | no | no | no | no | no | no | no | no | no |
| KCNK3 | 231 | G | 0 | 0 | yes | no | no | yes | no | no | no | no | no | no | no | no | no |
| KCNK3 | 232 | L | 0 | 0 | yes | no | no | yes | no | no | no | no | no | no | no | no | no |
| KCNK3 | 233 | T | 0 | 0 | yes | no | no | yes | no | no | no | no | no | no | no | no | no |
| KCNK3 | 234 | V | 0 | 0 | yes | no | no | yes | no | no | no | no | no | no | no | no | no |
| KCNK3 | 235 | I | 0 | 0 | yes | no | no | yes | no | no | no | no | no | no | no | no | no |
| KCNK3 | 236 | G | 0 | 1 | yes | no | no | yes | no | no | no | no | no | no | no | no | no |
| KCNK3 | 237 | A | 0 | 0 | yes | no | no | yes | no | no | no | no | no | no | no | no | no |
| KCNK3 | 238 | F | 0 | 0 | yes | no | no | yes | no | no | no | no | no | no | no | no | no |
| KCNK3 | 239 | L | 0 | 0 | yes | no | no | yes | no | no | no | no | no | no | no | no | no |
| KCNK3 | 240 | N | 0 | 0 | yes | no | no | yes | no | no | no | no | no | no | no | no | no |
| KCNK3 | 241 | L | 0 | 0 | yes | no | no | yes | no | no | no | no | no | no | no | no | no |
| KCNK3 | 242 | V | 0 | 0 | no | yes | no | yes | no | no | no | no | no | no | no | no | no |
| KCNK3 | 243 | V | 1 | 0 | no | yes | no | yes | no | no | no | no | yes | no | no | no | no |
| KCNK3 | 244 | L | 0 | 0 | yes | no | no | yes | no | yes | no | no | yes | no | no | no | no |
| KCNK3 | 245 | R | 0 | 0 | yes | no | no | yes | no | yes | no | no | no | no | no | no | no |
| KCNK3 | 246 | F | 1 | 0 | yes | no | no | yes | no | yes | no | no | no | no | no | no | no |
| KCNK3 | 247 | M | 1 | 0 | yes | no | no | yes | no | yes | yes | no | yes | no | no | no | no |
| KCNK3 | 248 | T | 0 | 0 | no | yes | no | yes | no | yes | no | no | yes | no | no | no | no |
| KCNK3 | 249 | M | 0 | 0 | yes | no | no | no | no | yes | no | no | no | no | no | no | no |
| KCNK3 | 250 | N | 0 | 0 | yes | no | no | yes | no | yes | no | no | yes | no | no | no | no |
| KCNK3 | 251 | A | 1 | 0 | yes | no | no | no | no | yes | no | no | no | no | no | no | no |
| KCNK3 | 252 | E | 1 | 0 | yes | no | no | no | no | yes | no | no | no | no | no | no | no |
| KCNK3 | 253 | D | 0 | 0 | yes | no | no | no | no | yes | no | no | no | no | no | no | no |
| KCNK3 | 254 | E | 0 | 0 | yes | no | no | no | no | yes | no | no | yes | no | no | no | no |
| KCNK3 | 255 | K | 0 | 0 | yes | no | no | no | no | yes | no | no | no | no | no | no | no |
| KCNK3 | 256 | R | 0 | 0 | yes | no | no | no | no | yes | no | no | no | no | no | no | no |

| | | | | | | | | | | | | | | | | | |
|---|---|---|---|---|---|---|---|---|---|---|---|---|---|---|---|---|---|
| KCNK3 | 257 | D | 1 | 0 | yes | no | no | no | no | yes | no | no | no | no | no | no | no |
| KCNK3 | 258 | A | 0 | 0 | yes | no | no | no | no | yes | no | no | no | no | no | no | no |
| KCNK3 | 259 | E | 1 | 0 | yes | no | no | no | no | yes | no | no | no | no | no | no | no |
| KCNK4 | 2 | R | 1 | 0 | no | yes | no | no | no | yes | no | no | no | no | no | no | no |
| KCNK4 | 3 | S | 0 | 0 | yes | no | no | yes | no | yes | no | no | yes | no | no | no | no |
| KCNK4 | 4 | T | 1 | 0 | yes | no | no | yes | no | no | no | no | no | no | no | no | no |
| KCNK4 | 5 | T | 1 | 0 | yes | no | no | yes | no | no | no | no | no | no | no | no | no |
| KCNK4 | 6 | L | 1 | 0 | yes | no | no | yes | no | no | no | no | yes | no | no | no | no |
| KCNK4 | 7 | L | 0 | 0 | yes | no | no | yes | no | no | no | no | no | no | no | no | no |
| KCNK4 | 8 | A | 1 | 0 | yes | no | no | yes | no | no | no | no | no | no | no | no | no |
| KCNK4 | 9 | L | 0 | 0 | yes | no | no | yes | no | no | no | no | yes | no | no | no | no |
| KCNK4 | 10 | L | 0 | 0 | yes | no | no | yes | no | no | no | no | yes | no | no | no | no |
| KCNK4 | 11 | A | 0 | 0 | yes | no | no | yes | no | no | no | no | no | no | no | no | no |
| KCNK4 | 12 | L | 1 | 0 | yes | no | no | yes | no | no | no | no | no | no | no | no | no |
| KCNK4 | 13 | V | 0 | 0 | yes | no | no | yes | no | no | no | no | yes | no | no | no | no |
| KCNK4 | 14 | L | 0 | 0 | yes | no | no | yes | no | no | no | no | no | no | no | no | no |
| KCNK4 | 15 | L | 0 | 0 | yes | no | no | yes | no | no | no | no | no | no | no | no | no |
| KCNK4 | 16 | Y | 0 | 0 | yes | no | no | yes | no | no | no | no | yes | no | no | no | no |
| KCNK4 | 17 | L | 0 | 0 | yes | no | no | yes | no | no | no | no | yes | no | no | no | no |
| KCNK4 | 18 | V | 1 | 0 | yes | no | no | yes | no | no | no | no | no | no | no | no | no |
| KCNK4 | 19 | S | 1 | 0 | yes | no | no | yes | no | no | no | no | no | no | no | no | no |
| KCNK4 | 20 | G | 0 | 0 | yes | no | no | yes | no | no | no | no | yes | no | no | no | no |
| KCNK4 | 21 | A | 1 | 0 | yes | no | no | yes | no | no | no | no | yes | no | no | no | no |
| KCNK4 | 22 | L | 0 | 0 | yes | no | no | yes | no | no | no | no | no | no | no | no | no |
| KCNK4 | 23 | V | 0 | 0 | yes | no | no | yes | no | no | no | no | yes | no | no | no | no |
| KCNK4 | 24 | F | 0 | 0 | yes | no | no | yes | no | no | no | no | yes | no | no | no | no |
| KCNK4 | 25 | R | 0 | 0 | yes | no | no | no | yes | no | no | no | yes | no | no | no | no |
| KCNK4 | 26 | A | 2 | 0 | yes | no | no | no | yes | no | no | no | no | no | no | no | no |
| KCNK4 | 27 | L | 1 | 0 | yes | no | no | yes | yes | no | no | no | yes | no | no | no | no |
| KCNK4 | 28 | E | 0 | 0 | yes | no | no | no | yes | no | no | no | yes | no | no | no | no |
| KCNK4 | 29 | Q | 0 | 0 | yes | no | no | no | yes | no | no | no | yes | no | no | no | no |
| KCNK4 | 30 | P | 1 | 0 | yes | no | no | no | yes | no | no | no | no | no | no | no | no |
| KCNK4 | 31 | H | 1 | 0 | yes | no | no | no | yes | no | no | no | yes | no | no | no | no |

| | | | | | | | | | | | | | | | | | |
|---|---|---|---|---|---|---|---|---|---|---|---|---|---|---|---|---|---|
| KCNK4 | 32 | E | 0 | 0 | no | yes | no | no | yes | no | no | no | yes | no | no | no | no |
| KCNK4 | 33 | Q | 0 | 0 | no | yes | no | no | yes | no | no | no | yes | no | no | no | no |
| KCNK4 | 34 | Q | 0 | 0 | yes | no | no | no | yes | no | no | no | yes | no | no | no | no |
| KCNK4 | 35 | A | 0 | 0 | yes | no | no | no | yes | no | no | no | yes | no | no | no | no |
| KCNK4 | 36 | Q | 1 | 0 | yes | no | no | no | yes | no | no | no | yes | no | no | no | no |
| KCNK4 | 37 | R | 1 | 0 | yes | no | no | no | yes | no | no | no | no | no | no | no | no |
| KCNK4 | 38 | E | 2 | 0 | yes | no | no | no | yes | no | no | no | no | no | no | no | no |
| KCNK4 | 39 | L | 0 | 0 | yes | no | no | no | yes | no | no | no | yes | no | no | no | no |
| KCNK4 | 40 | G | 0 | 0 | yes | no | no | no | yes | no | no | no | no | no | no | no | no |
| KCNK4 | 41 | E | 0 | 0 | yes | no | no | no | yes | no | no | no | no | no | no | no | no |
| KCNK4 | 42 | V | 0 | 0 | yes | no | no | no | yes | no | no | no | yes | no | no | no | no |
| KCNK4 | 43 | R | 2 | 0 | yes | no | no | no | yes | no | no | no | yes | no | no | no | no |
| KCNK4 | 44 | E | 0 | 0 | yes | no | no | no | yes | no | no | no | no | no | no | no | no |
| KCNK4 | 45 | K | 0 | 0 | yes | no | no | no | yes | no | no | no | no | no | no | no | no |
| KCNK4 | 46 | F | 0 | 0 | yes | no | no | no | yes | no | no | no | no | no | no | no | no |
| KCNK4 | 47 | L | 1 | 0 | yes | no | no | no | yes | no | no | no | no | no | no | no | no |
| KCNK4 | 48 | R | 1 | 0 | yes | no | no | no | yes | no | no | no | no | no | no | no | no |
| KCNK4 | 49 | A | 0 | 0 | yes | no | no | no | yes | no | no | no | no | no | no | no | no |
| KCNK4 | 50 | H | 1 | 0 | no | yes | no | no | yes | no | no | no | yes | no | no | no | no |
| KCNK4 | 51 | P | 0 | 0 | no | yes | no | no | yes | no | no | no | no | no | no | no | no |
| KCNK4 | 52 | C | 0 | 0 | no | yes | no | no | yes | no | yes | no | yes | no | no | no | no |
| KCNK4 | 53 | V | 0 | 0 | no | yes | no | no | yes | no | yes | no | no | no | no | no | no |
| KCNK4 | 54 | S | 0 | 0 | no | yes | no | no | yes | no | yes | no | no | no | no | no | no |
| KCNK4 | 55 | D | 1 | 0 | yes | no | no | no | yes | no | no | no | yes | no | no | no | no |
| KCNK4 | 56 | Q | 1 | 0 | yes | no | no | no | yes | no | no | no | no | no | no | no | no |
| KCNK4 | 57 | E | 0 | 0 | yes | no | no | no | yes | no | yes | no | yes | no | no | no | no |
| KCNK4 | 58 | L | 0 | 0 | yes | no | no | no | yes | no | yes | no | yes | no | no | no | no |
| KCNK4 | 59 | G | 1 | 0 | yes | no | no | no | yes | no | no | no | no | no | no | no | no |
| KCNK4 | 60 | L | 0 | 0 | yes | no | no | no | yes | no | no | no | no | no | no | no | no |
| KCNK4 | 61 | L | 0 | 0 | yes | no | no | no | yes | no | yes | no | yes | no | no | no | no |
| KCNK4 | 62 | I | 0 | 0 | yes | no | no | no | yes | no | yes | no | no | no | no | no | no |
| KCNK4 | 63 | K | 0 | 0 | yes | no | no | no | yes | no | no | no | yes | no | no | no | no |
| KCNK4 | 64 | E | 0 | 0 | yes | no | no | no | yes | no | no | no | yes | no | no | no | no |

| | | | | | | | | | | | | | | | | | |
|---|---|---|---|---|---|---|---|---|---|---|---|---|---|---|---|---|---|
| KCNK4 | 65 | V | 1 | 0 | yes | no | no | no | yes | no | yes | no | yes | no | no | no | no |
| KCNK4 | 66 | A | 1 | 0 | yes | no | no | no | yes | no | yes | no | yes | no | no | no | no |
| KCNK4 | 67 | D | 0 | 0 | yes | no | no | no | yes | no | no | no | no | no | no | no | no |
| KCNK4 | 68 | A | 0 | 0 | yes | no | no | no | yes | no | no | no | yes | no | no | no | no |
| KCNK4 | 69 | L | 2 | 0 | yes | no | no | no | yes | no | yes | no | yes | no | no | no | no |
| KCNK4 | 70 | G | 1 | 0 | yes | no | no | no | yes | no | yes | no | no | no | no | no | no |
| KCNK4 | 71 | G | 1 | 0 | no | yes | no | no | yes | no | no | no | yes | no | no | no | no |
| KCNK4 | 72 | G | 1 | 0 | no | yes | no | no | yes | no | no | no | yes | no | no | no | no |
| KCNK4 | 73 | A | 0 | 0 | no | yes | no | no | yes | no | no | no | yes | no | no | no | no |
| KCNK4 | 74 | D | 1 | 0 | no | yes | no | no | yes | no | no | no | yes | no | no | no | no |
| KCNK4 | 75 | P | 0 | 0 | no | yes | no | no | yes | no | no | no | yes | no | no | no | no |
| KCNK4 | 76 | E | 0 | 0 | no | yes | no | no | yes | no | no | no | yes | no | no | no | no |
| KCNK4 | 77 | T | 0 | 0 | no | yes | no | no | yes | no | no | no | no | no | no | no | no |
| KCNK4 | 84 | S | 0 | 0 | no | yes | no | no | yes | no | no | no | no | no | no | no | no |
| KCNK4 | 85 | H | 0 | 0 | no | yes | no | no | yes | no | no | no | no | no | no | no | no |
| KCNK4 | 86 | S | 0 | 0 | no | yes | no | no | yes | no | no | no | yes | no | no | no | no |
| KCNK4 | 87 | A | 0 | 0 | no | yes | no | no | yes | no | no | no | yes | no | no | no | no |
| KCNK4 | 88 | W | 0 | 0 | no | yes | no | no | no | no | no | no | yes | no | no | no | no |
| KCNK4 | 89 | D | 0 | 0 | no | yes | no | no | no | no | no | no | yes | no | no | no | no |
| KCNK4 | 90 | L | 0 | 0 | yes | no | no | yes | no | no | no | no | yes | no | no | no | no |
| KCNK4 | 91 | G | 0 | 0 | yes | no | no | yes | no | no | no | no | no | no | no | no | no |
| KCNK4 | 92 | S | 2 | 0 | yes | no | no | yes | no | no | no | no | no | no | no | no | no |
| KCNK4 | 93 | A | 1 | 0 | yes | no | no | yes | no | no | no | no | yes | no | no | no | no |
| KCNK4 | 94 | F | 0 | 0 | yes | no | no | yes | no | no | no | no | yes | no | no | no | no |
| KCNK4 | 95 | F | 0 | 0 | yes | no | no | yes | no | no | no | no | no | no | no | no | no |
| KCNK4 | 96 | F | 1 | 0 | yes | no | no | yes | no | no | no | no | yes | no | no | no | no |
| KCNK4 | 97 | S | 0 | 0 | yes | no | no | yes | no | no | no | no | yes | no | no | no | no |
| KCNK4 | 98 | G | 0 | 0 | yes | no | no | yes | no | no | no | no | yes | no | no | no | no |
| KCNK4 | 99 | T | 0 | 0 | yes | no | no | yes | no | no | no | no | no | no | no | no | no |
| KCNK4 | 100 | I | 2 | 0 | yes | no | no | yes | no | no | no | no | yes | no | no | no | no |
| KCNK4 | 101 | I | 2 | 0 | yes | no | no | yes | no | no | no | no | yes | no | no | no | no |
| KCNK4 | 102 | T | 0 | 0 | no | yes | no | yes | no | no | no | no | no | no | no | no | no |
| KCNK4 | 103 | T | 1 | 0 | no | yes | no | yes | no | no | yes | yes | yes | no | no | no | no |

| | | | | | | | | | | | | | | | | | |
|---|---|---|---|---|---|---|---|---|---|---|---|---|---|---|---|---|---|
| KCNK4 | 104 | I | 0 | 0 | no | yes | no | yes | no | no | yes | yes | yes | no | no | no | no |
| KCNK4 | 105 | G | 0 | 0 | no | yes | no | yes | no | no | yes | yes | yes | no | no | no | no |
| KCNK4 | 106 | Y | 1 | 0 | no | yes | no | yes | no | no | yes | yes | no | no | no | no | no |
| KCNK4 | 107 | G | 0 | 0 | no | yes | no | yes | no | no | yes | yes | no | no | no | no | no |
| KCNK4 | 108 | N | 1 | 0 | no | yes | no | no | no | no | yes | yes | no | no | no | no | no |
| KCNK4 | 109 | V | 0 | 0 | no | yes | no | no | no | no | no | no | no | no | no | no | no |
| KCNK4 | 110 | A | 1 | 0 | no | yes | no | no | yes | no | no | no | no | no | no | no | no |
| KCNK4 | 111 | L | 0 | 0 | no | yes | no | no | yes | no | no | no | yes | no | no | no | no |
| KCNK4 | 112 | R | 2 | 0 | no | yes | no | no | yes | no | no | no | yes | no | no | no | no |
| KCNK4 | 113 | T | 1 | 0 | no | yes | no | no | yes | no | no | no | yes | no | no | no | no |
| KCNK4 | 114 | D | 0 | 0 | yes | no | no | no | yes | no | no | no | yes | no | no | no | no |
| KCNK4 | 115 | A | 0 | 0 | yes | no | no | no | yes | no | no | no | no | no | no | no | no |
| KCNK4 | 116 | G | 1 | 0 | yes | no | no | yes | yes | no | no | no | yes | no | no | no | no |
| KCNK4 | 117 | R | 3 | 0 | yes | no | no | no | yes | no | no | no | yes | no | no | no | no |
| KCNK4 | 118 | L | 0 | 0 | yes | no | no | yes | no | no | no | no | no | no | no | no | no |
| KCNK4 | 119 | F | 0 | 0 | yes | no | no | yes | no | no | no | no | yes | no | no | no | no |
| KCNK4 | 120 | C | 1 | 0 | yes | no | no | yes | no | no | no | no | no | no | no | no | no |
| KCNK4 | 121 | I | 2 | 0 | yes | no | no | yes | no | no | no | no | yes | no | no | no | no |
| KCNK4 | 122 | F | 0 | 0 | yes | no | no | yes | no | no | no | no | no | no | no | no | no |
| KCNK4 | 123 | Y | 0 | 0 | yes | no | no | yes | no | no | no | no | yes | no | no | no | no |
| KCNK4 | 124 | A | 1 | 0 | yes | no | no | yes | no | no | no | no | no | no | no | no | no |
| KCNK4 | 125 | L | 0 | 0 | yes | no | no | yes | no | no | no | no | yes | no | no | no | no |
| KCNK4 | 126 | V | 1 | 0 | yes | no | no | yes | no | no | no | no | yes | no | no | no | no |
| KCNK4 | 127 | G | 1 | 0 | yes | no | no | yes | no | no | no | no | yes | no | no | no | no |
| KCNK4 | 128 | I | 0 | 0 | yes | no | no | yes | no | no | no | no | yes | no | no | no | no |
| KCNK4 | 129 | P | 1 | 0 | yes | no | no | yes | no | no | no | no | yes | no | no | no | no |
| KCNK4 | 130 | L | 1 | 0 | yes | no | no | yes | no | no | no | no | yes | no | no | no | no |
| KCNK4 | 131 | F | 0 | 0 | yes | no | no | yes | no | no | no | no | no | no | no | no | no |
| KCNK4 | 132 | G | 0 | 0 | yes | no | no | yes | no | no | no | no | no | no | no | no | no |
| KCNK4 | 133 | I | 0 | 0 | yes | no | no | yes | no | no | no | no | yes | no | no | no | no |
| KCNK4 | 134 | L | 1 | 0 | yes | no | no | yes | no | no | no | no | yes | no | no | no | no |
| KCNK4 | 135 | L | 1 | 0 | yes | no | no | yes | no | no | no | no | no | no | no | no | no |
| KCNK4 | 136 | A | 0 | 0 | yes | no | no | yes | no | no | no | no | no | no | no | no | no |

| | | | | | | | | | | | | | | | | | |
|---|---|---|---|---|---|---|---|---|---|---|---|---|---|---|---|---|---|
| KCNK4 | 137 | G | 1 | 0 | yes | no | no | yes | no | no | no | no | yes | no | no | no | no |
| KCNK4 | 138 | V | 0 | 0 | yes | no | no | yes | no | no | no | no | yes | no | no | no | no |
| KCNK4 | 139 | G | 0 | 0 | yes | no | no | yes | no | no | no | no | no | no | no | no | no |
| KCNK4 | 140 | D | 1 | 0 | yes | no | no | yes | no | no | no | no | no | no | no | no | no |
| KCNK4 | 141 | R | 3 | 0 | yes | no | no | yes | no | no | no | no | yes | no | no | no | no |
| KCNK4 | 142 | L | 0 | 0 | yes | no | no | yes | no | no | no | no | no | no | no | no | no |
| KCNK4 | 143 | G | 0 | 0 | yes | no | no | yes | no | no | no | no | no | no | no | no | no |
| KCNK4 | 144 | S | 2 | 0 | yes | no | no | yes | no | no | no | no | no | no | no | no | no |
| KCNK4 | 145 | S | 0 | 0 | yes | no | no | yes | no | no | no | no | no | no | no | no | no |
| KCNK4 | 146 | L | 1 | 0 | yes | no | no | yes | no | no | no | no | no | no | no | no | no |
| KCNK4 | 147 | R | 2 | 0 | yes | no | no | yes | no | no | no | no | no | no | no | no | no |
| KCNK4 | 148 | H | 1 | 0 | yes | no | no | no | no | no | no | no | no | no | no | no | no |
| KCNK4 | 149 | G | 0 | 0 | yes | no | no | yes | no | no | no | no | no | no | no | no | no |
| KCNK4 | 150 | I | 0 | 0 | yes | no | no | yes | no | no | no | no | no | no | no | no | no |
| KCNK4 | 151 | G | 2 | 0 | yes | no | no | no | no | yes | no | no | no | no | no | no | no |
| KCNK4 | 152 | H | 0 | 0 | yes | no | no | no | no | yes | no | no | no | no | no | no | no |
| KCNK4 | 153 | I | 0 | 0 | yes | no | no | no | no | yes | no | no | no | no | no | no | no |
| KCNK4 | 154 | E | 0 | 0 | yes | no | no | no | no | yes | no | no | no | no | no | no | no |
| KCNK4 | 155 | A | 0 | 0 | yes | no | no | no | no | yes | no | no | no | no | no | no | no |
| KCNK4 | 156 | I | 1 | 0 | yes | no | no | no | no | yes | no | no | no | no | no | no | no |
| KCNK4 | 157 | F | 0 | 0 | yes | no | no | no | no | yes | no | no | no | no | no | no | no |
| KCNK4 | 158 | L | 0 | 0 | yes | no | no | no | no | yes | no | no | no | no | no | no | no |
| KCNK4 | 159 | K | 2 | 0 | yes | no | no | no | no | yes | no | no | no | no | no | no | no |
| KCNK4 | 160 | W | 0 | 0 | no | yes | no | no | no | yes | no | no | no | no | no | no | no |
| KCNK4 | 161 | H | 1 | 0 | no | yes | no | no | no | yes | no | no | no | no | no | no | no |
| KCNK4 | 162 | V | 0 | 0 | no | yes | no | no | no | yes | no | no | no | no | no | no | no |
| KCNK4 | 163 | P | 0 | 0 | no | yes | no | no | no | yes | no | no | no | no | no | no | no |
| KCNK4 | 164 | P | 3 | 0 | yes | no | no | no | no | yes | no | no | no | no | no | no | no |
| KCNK4 | 165 | E | 0 | 0 | yes | no | no | no | no | yes | no | no | no | no | no | no | no |
| KCNK4 | 166 | L | 1 | 0 | yes | no | no | no | no | yes | no | no | no | no | no | no | no |
| KCNK4 | 167 | V | 1 | 0 | yes | no | no | no | no | yes | no | no | no | no | no | no | no |
| KCNK4 | 168 | R | 0 | 0 | yes | no | no | no | no | yes | no | no | no | no | no | no | no |
| KCNK4 | 169 | V | 1 | 0 | yes | no | no | no | no | yes | no | no | no | no | no | no | no |

| | | | | | | | | | | | | | | | | | |
|---|---|---|---|---|---|---|---|---|---|---|---|---|---|---|---|---|---|
| KCNK4 | 170 | L | 0 | 0 | yes | no | no | no | no | yes | no | no | no | no | no | no | no |
| KCNK4 | 171 | S | 1 | 0 | yes | no | no | no | no | yes | no | no | no | no | no | no | no |
| KCNK4 | 172 | A | 0 | 1 | yes | no | no | yes | no | yes | no | no | no | no | no | no | no |
| KCNK4 | 173 | M | 0 | 0 | yes | no | no | yes | no | no | no | no | no | no | no | no | no |
| KCNK4 | 174 | L | 0 | 0 | yes | no | no | yes | no | no | no | no | no | no | no | no | no |
| KCNK4 | 175 | F | 0 | 0 | yes | no | no | yes | no | no | no | no | no | no | no | no | no |
| KCNK4 | 176 | L | 0 | 0 | yes | no | no | yes | no | no | no | no | no | no | no | no | no |
| KCNK4 | 177 | L | 1 | 0 | yes | no | no | yes | no | no | no | no | no | no | no | no | no |
| KCNK4 | 178 | I | 1 | 0 | yes | no | no | yes | no | no | no | no | no | no | no | no | no |
| KCNK4 | 179 | G | 1 | 0 | yes | no | no | yes | no | no | no | no | no | no | no | no | no |
| KCNK4 | 180 | C | 0 | 0 | yes | no | no | yes | no | no | no | no | no | no | no | no | no |
| KCNK4 | 181 | L | 0 | 0 | yes | no | no | yes | no | no | no | no | no | no | no | no | no |
| KCNK4 | 182 | L | 1 | 0 | yes | no | no | yes | no | no | no | no | no | no | no | no | no |
| KCNK4 | 183 | F | 0 | 0 | yes | no | no | yes | no | no | no | no | no | no | no | no | no |
| KCNK4 | 184 | V | 1 | 0 | yes | no | no | yes | no | no | no | no | no | no | no | no | no |
| KCNK4 | 185 | L | 0 | 0 | yes | no | no | yes | no | no | no | no | no | no | no | no | no |
| KCNK4 | 186 | T | 2 | 0 | yes | no | no | yes | no | no | no | no | no | no | no | no | no |
| KCNK4 | 187 | P | 0 | 0 | yes | no | no | yes | no | no | no | no | no | no | no | no | no |
| KCNK4 | 188 | T | 1 | 0 | yes | no | no | yes | no | no | no | no | no | no | no | no | no |
| KCNK4 | 189 | F | 1 | 0 | yes | no | no | yes | no | no | no | no | no | no | no | no | no |
| KCNK4 | 190 | V | 1 | 0 | yes | no | no | yes | no | no | no | no | no | no | no | no | no |
| KCNK4 | 191 | F | 0 | 0 | yes | no | no | yes | no | no | no | no | no | no | no | no | no |
| KCNK4 | 192 | C | 0 | 0 | yes | no | no | yes | no | no | no | no | no | no | no | no | no |
| KCNK4 | 193 | Y | 1 | 0 | yes | no | no | yes | no | no | no | no | no | no | no | no | no |
| KCNK4 | 194 | M | 0 | 0 | yes | no | no | yes | no | no | no | no | no | no | no | no | no |
| KCNK4 | 195 | E | 0 | 0 | yes | no | no | no | yes | no | no | no | no | no | no | no | no |
| KCNK4 | 196 | D | 2 | 0 | no | yes | no | no | yes | no | no | no | no | no | no | no | no |
| KCNK4 | 197 | W | 0 | 0 | no | yes | no | no | yes | no | no | no | no | no | no | no | no |
| KCNK4 | 198 | S | 0 | 0 | no | yes | no | yes | yes | no | no | no | no | no | no | no | no |
| KCNK4 | 199 | K | 0 | 0 | yes | no | no | yes | yes | no | no | no | no | no | no | no | no |
| KCNK4 | 200 | L | 0 | 0 | yes | no | no | yes | no | no | no | no | yes | no | no | no | no |
| KCNK4 | 201 | E | 0 | 0 | yes | no | no | yes | no | no | no | no | no | no | no | no | no |
| KCNK4 | 202 | A | 1 | 0 | yes | no | no | yes | no | no | no | no | no | no | no | no | no |

| | | | | | | | | | | | | | | | | | |
|---|---|---|---|---|---|---|---|---|---|---|---|---|---|---|---|---|---|
| KCNK4 | 203 | I | 2 | 0 | yes | no | no | yes | no | no | no | no | no | no | no | no | no |
| KCNK4 | 204 | Y | 1 | 0 | yes | no | no | yes | no | no | no | no | yes | no | no | no | no |
| KCNK4 | 205 | F | 0 | 0 | yes | no | no | yes | no | no | no | no | no | no | no | no | no |
| KCNK4 | 206 | V | 1 | 0 | yes | no | no | yes | no | no | no | no | no | no | no | no | no |
| KCNK4 | 207 | I | 2 | 0 | yes | no | no | yes | no | no | no | no | yes | no | no | no | no |
| KCNK4 | 208 | V | 1 | 0 | yes | no | no | yes | no | no | no | no | no | no | no | no | no |
| KCNK4 | 209 | T | 2 | 0 | yes | no | no | yes | no | no | no | no | no | no | no | no | no |
| KCNK4 | 210 | L | 2 | 0 | yes | no | no | yes | no | no | no | no | no | no | no | no | no |
| KCNK4 | 211 | T | 0 | 0 | no | yes | no | yes | no | no | no | no | yes | no | no | no | no |
| KCNK4 | 212 | T | 1 | 0 | no | yes | no | yes | no | no | yes | yes | yes | no | no | no | no |
| KCNK4 | 213 | V | 1 | 0 | no | yes | no | yes | no | no | yes | yes | yes | no | no | no | no |
| KCNK4 | 214 | G | 0 | 0 | no | yes | no | yes | no | no | yes | yes | yes | no | no | no | no |
| KCNK4 | 215 | F | 0 | 0 | no | yes | no | yes | no | no | yes | yes | yes | no | no | no | no |
| KCNK4 | 216 | G | 0 | 0 | no | yes | no | yes | no | no | no | yes | no | no | no | no | no |
| KCNK4 | 217 | D | 1 | 0 | no | yes | no | no | no | no | no | yes | no | no | no | no | no |
| KCNK4 | 218 | Y | 0 | 0 | no | yes | no | no | no | no | no | no | yes | no | no | no | no |
| KCNK4 | 219 | V | 0 | 0 | no | yes | no | yes | no | no | no | no | no | no | no | no | no |
| KCNK4 | 220 | A | 0 | 0 | no | yes | no | yes | yes | no | no | no | no | no | no | no | no |
| KCNK4 | 221 | G | 0 | 0 | no | yes | no | yes | yes | no | no | no | no | no | no | no | no |
| KCNK4 | 222 | A | 2 | 0 | no | yes | no | yes | yes | no | no | no | no | no | no | no | no |
| KCNK4 | 223 | D | 0 | 0 | no | yes | no | no | yes | no | no | no | no | no | no | no | no |
| KCNK4 | 224 | P | 0 | 0 | no | yes | no | no | yes | no | no | no | no | no | no | no | no |
| KCNK4 | 225 | R | 0 | 0 | no | yes | no | no | yes | no | no | no | no | no | no | no | no |
| KCNK4 | 226 | Q | 0 | 0 | no | yes | no | no | yes | no | no | no | no | no | no | no | no |
| KCNK4 | 227 | D | 1 | 0 | no | yes | no | no | yes | no | no | no | no | no | no | no | no |
| KCNK4 | 228 | S | 1 | 0 | no | yes | no | yes | yes | no | no | no | no | no | no | no | no |
| KCNK4 | 229 | P | 3 | 0 | no | yes | no | yes | yes | no | no | no | no | no | no | no | no |
| KCNK4 | 230 | A | 0 | 0 | no | yes | no | yes | yes | no | no | no | no | no | no | no | no |
| KCNK4 | 231 | Y | 0 | 0 | yes | no | no | yes | yes | no | no | no | no | no | no | no | no |
| KCNK4 | 232 | Q | 0 | 0 | yes | no | no | yes | yes | no | no | no | no | no | no | no | no |
| KCNK4 | 233 | P | 0 | 0 | yes | no | no | yes | yes | no | no | no | no | no | no | no | no |
| KCNK4 | 234 | L | 1 | 0 | yes | no | no | yes | no | no | no | no | no | no | no | no | no |
| KCNK4 | 235 | V | 0 | 0 | yes | no | no | yes | no | no | no | no | no | no | no | no | no |

| | | | | | | | | | | | | | | | | | |
|---|---|---|---|---|---|---|---|---|---|---|---|---|---|---|---|---|---|
| KCNK4 | 236 | W | 0 | 0 | yes | no | no | yes | no | no | no | no | no | no | no | no | no |
| KCNK4 | 237 | F | 1 | 0 | yes | no | no | yes | no | no | no | no | no | no | no | no | no |
| KCNK4 | 238 | W | 0 | 0 | yes | no | no | yes | no | no | no | no | no | no | no | no | no |
| KCNK4 | 239 | I | 0 | 0 | yes | no | no | yes | no | no | no | no | no | no | no | no | no |
| KCNK4 | 240 | L | 0 | 0 | yes | no | no | yes | no | no | no | no | no | no | no | no | no |
| KCNK4 | 241 | L | 0 | 0 | yes | no | no | yes | no | no | no | no | no | no | no | no | no |
| KCNK4 | 242 | G | 1 | 0 | yes | no | no | yes | no | no | no | no | no | no | no | no | no |
| KCNK4 | 243 | L | 0 | 0 | yes | no | no | yes | no | no | no | no | no | no | no | no | no |
| KCNK4 | 244 | A | 0 | 1 | yes | no | no | yes | no | no | no | no | no | no | no | no | no |
| KCNK4 | 245 | Y | 1 | 0 | yes | no | no | yes | no | no | no | no | no | no | no | no | no |
| KCNK4 | 246 | F | 0 | 0 | yes | no | no | yes | no | no | no | no | no | no | no | no | no |
| KCNK4 | 247 | A | 0 | 0 | yes | no | no | yes | no | no | yes | no | no | no | no | no | no |
| KCNK4 | 248 | S | 0 | 0 | yes | no | no | yes | no | no | no | no | no | no | no | no | no |
| KCNK4 | 249 | V | 0 | 0 | yes | no | no | yes | no | no | no | no | no | no | no | no | no |
| KCNK4 | 250 | L | 0 | 0 | yes | no | no | yes | no | no | yes | no | no | no | no | no | no |
| KCNK4 | 251 | T | 1 | 0 | yes | no | no | yes | no | no | yes | no | no | no | no | no | no |
| KCNK4 | 252 | T | 0 | 0 | yes | no | no | yes | no | no | no | no | no | no | no | no | no |
| KCNK4 | 253 | I | 0 | 0 | yes | no | no | yes | no | no | no | no | no | no | no | no | no |
| KCNK4 | 254 | G | 1 | 0 | yes | no | no | yes | no | no | no | no | yes | no | no | no | no |
| KCNK4 | 255 | N | 0 | 0 | yes | no | no | yes | no | no | yes | no | no | no | no | no | no |
| KCNK4 | 256 | W | 1 | 0 | yes | no | no | yes | no | no | no | no | no | no | no | no | no |
| KCNK4 | 257 | L | 0 | 0 | yes | no | no | yes | no | no | no | no | yes | no | no | no | no |
| KCNK4 | 258 | R | 1 | 0 | yes | no | no | yes | no | no | yes | no | no | no | no | no | no |
| KCNK4 | 259 | V | 0 | 0 | yes | no | no | no | no | no | no | no | no | no | no | no | no |
| KCNK4 | 260 | V | 2 | 0 | no | yes | no | no | no | no | no | no | no | no | no | no | no |
| KCNMA1 | 85 | R | 0 | 0 | no | yes | no | no | yes | no | no | no | no | no | no | no | no |
| KCNMA1 | 86 | M | 1 | 0 | no | yes | no | no | yes | no | no | no | no | no | no | no | no |
| KCNMA1 | 87 | W | 0 | 0 | no | yes | no | yes | no | no | no | no | no | no | no | no | no |
| KCNMA1 | 88 | W | 0 | 0 | yes | no | no | yes | no | no | no | no | no | no | no | no | no |
| KCNMA1 | 89 | A | 2 | 0 | yes | no | no | yes | no | no | no | no | no | no | no | no | no |
| KCNMA1 | 90 | F | 0 | 0 | yes | no | no | yes | no | no | no | no | no | no | no | no | no |
| KCNMA1 | 91 | L | 0 | 0 | yes | no | no | yes | no | no | no | no | no | no | no | no | no |
| KCNMA1 | 92 | A | 1 | 0 | yes | no | no | yes | no | no | no | no | no | no | no | no | no |

| | | | | | | | | | | | | | | | | | |
|---|---|---|---|---|---|---|---|---|---|---|---|---|---|---|---|---|---|
| KCNMA1 | 93 | S | 0 | 0 | yes | no | no | yes | no | no | no | no | no | no | no | no | no |
| KCNMA1 | 94 | S | 0 | 0 | yes | no | no | yes | no | no | no | no | no | no | no | no | no |
| KCNMA1 | 95 | M | 1 | 0 | yes | no | no | yes | no | no | no | no | no | no | no | no | no |
| KCNMA1 | 96 | V | 0 | 0 | yes | no | no | yes | no | no | no | no | no | no | no | no | no |
| KCNMA1 | 97 | T | 0 | 0 | yes | no | no | yes | no | no | no | no | no | no | no | no | no |
| KCNMA1 | 98 | F | 0 | 0 | yes | no | no | yes | no | no | no | no | no | no | no | no | no |
| KCNMA1 | 99 | F | 0 | 0 | no | yes | no | yes | no | no | no | no | no | no | no | no | no |
| KCNMA1 | 158 | W | 0 | 0 | no | yes | no | no | no | yes | no | no | no | no | no | no | no |
| KCNMA1 | 159 | M | 0 | 0 | yes | no | no | no | no | yes | no | no | no | no | no | no | no |
| KCNMA1 | 160 | T | 0 | 0 | yes | no | no | no | no | yes | no | no | no | no | no | no | no |
| KCNMA1 | 161 | S | 0 | 0 | yes | no | no | no | no | yes | no | no | no | no | no | no | no |
| KCNMA1 | 162 | V | 1 | 0 | yes | no | no | no | no | yes | no | no | no | no | no | no | no |
| KCNMA1 | 163 | K | 0 | 0 | no | yes | no | no | no | yes | no | no | no | no | no | no | no |
| KCNMA1 | 164 | D | 0 | 0 | no | yes | no | no | no | yes | no | no | no | no | no | no | no |
| KCNMA1 | 165 | W | 0 | 0 | no | yes | no | no | no | yes | no | no | no | no | no | no | no |
| KCNMA1 | 166 | A | 0 | 0 | no | yes | no | no | no | yes | no | no | no | no | no | no | no |
| KCNMA1 | 167 | G | 0 | 0 | yes | no | no | no | no | yes | no | no | no | no | no | no | no |
| KCNMA1 | 168 | V | 1 | 0 | yes | no | no | no | no | yes | no | no | no | no | no | no | no |
| KCNMA1 | 169 | M | 0 | 0 | yes | no | no | no | no | yes | no | no | no | no | no | no | no |
| KCNMA1 | 170 | I | 0 | 0 | yes | no | no | no | no | yes | no | no | no | no | no | no | no |
| KCNMA1 | 171 | S | 0 | 0 | no | yes | no | no | no | yes | no | no | no | no | no | no | no |
| KCNMA1 | 172 | A | 0 | 0 | no | yes | no | no | no | yes | no | no | no | no | no | no | no |
| KCNMA1 | 173 | Q | 0 | 0 | no | yes | no | no | no | yes | no | no | no | no | no | no | no |
| KCNMA1 | 174 | T | 1 | 0 | no | yes | no | no | no | yes | no | no | no | no | no | no | no |
| KCNMA1 | 175 | L | 0 | 0 | yes | no | no | no | no | yes | no | no | no | no | no | no | no |
| KCNMA1 | 176 | T | 0 | 0 | yes | no | no | no | no | yes | no | no | no | no | no | no | no |
| KCNMA1 | 177 | G | 0 | 0 | yes | no | no | no | no | yes | no | no | no | no | no | no | no |
| KCNMA1 | 178 | R | 0 | 0 | yes | no | no | no | no | yes | no | no | no | no | no | no | no |
| KCNMA1 | 179 | V | 1 | 0 | yes | no | no | yes | no | no | no | no | no | no | no | no | no |
| KCNMA1 | 180 | L | 0 | 0 | yes | no | no | yes | no | no | no | no | no | no | no | no | no |
| KCNMA1 | 181 | V | 0 | 0 | yes | no | no | yes | no | no | no | no | no | no | no | no | no |
| KCNMA1 | 182 | V | 0 | 0 | yes | no | no | yes | no | no | no | no | no | no | no | no | no |
| KCNMA1 | 183 | L | 0 | 0 | yes | no | no | yes | no | no | no | no | no | no | no | no | no |

| | | | | | | | | | | | | | | | | | |
|---|---|---|---|---|---|---|---|---|---|---|---|---|---|---|---|---|---|
| KCNMA1 | 184 | V | 0 | 0 | yes | no | no | yes | no | no | no | no | no | no | no | no | no |
| KCNMA1 | 185 | F | 0 | 0 | yes | no | no | yes | no | no | no | no | no | no | no | no | no |
| KCNMA1 | 186 | A | 0 | 0 | yes | no | no | yes | no | no | no | no | no | no | no | no | no |
| KCNMA1 | 187 | L | 0 | 0 | yes | no | no | yes | no | no | no | no | no | no | no | no | no |
| KCNMA1 | 188 | S | 0 | 0 | yes | no | no | yes | no | no | no | no | no | no | no | no | no |
| KCNMA1 | 189 | I | 0 | 0 | yes | no | no | yes | no | no | no | no | no | no | no | no | no |
| KCNMA1 | 190 | G | 1 | 0 | yes | no | no | yes | no | no | no | no | no | no | no | no | no |
| KCNMA1 | 191 | A | 1 | 0 | yes | no | no | yes | no | no | no | no | no | no | no | no | no |
| KCNMA1 | 192 | L | 0 | 0 | yes | no | no | yes | no | no | no | no | no | no | no | no | no |
| KCNMA1 | 193 | V | 0 | 0 | yes | no | no | yes | no | no | no | no | no | no | no | no | no |
| KCNMA1 | 194 | I | 1 | 0 | yes | no | no | yes | no | no | no | no | no | no | no | no | no |
| KCNMA1 | 195 | Y | 0 | 0 | yes | no | no | yes | no | no | no | no | no | no | no | no | no |
| KCNMA1 | 196 | F | 0 | 0 | yes | no | no | yes | no | no | no | no | no | no | no | no | no |
| KCNMA1 | 197 | I | 0 | 0 | yes | no | no | yes | no | no | no | no | no | no | no | no | no |
| KCNMA1 | 198 | D | 0 | 0 | yes | no | no | yes | no | no | no | no | no | no | no | no | no |
| KCNMA1 | 199 | S | 0 | 0 | no | yes | no | yes | no | no | no | no | no | no | no | no | no |
| KCNMA1 | 200 | S | 0 | 0 | no | yes | no | no | yes | no | no | no | no | no | no | no | no |
| KCNMA1 | 201 | N | 0 | 0 | no | yes | no | no | yes | no | no | no | no | no | no | no | no |
| KCNMA1 | 202 | P | 0 | 0 | no | yes | no | no | yes | no | no | no | no | no | no | no | no |
| KCNMA1 | 203 | I | 1 | 0 | no | yes | no | no | yes | no | no | no | no | no | no | no | no |
| KCNMA1 | 204 | E | 0 | 0 | no | no | yes | no | yes | no | no | no | no | no | no | no | no |
| KCNMA1 | 205 | S | 0 | 0 | no | no | yes | no | yes | no | no | no | no | no | no | no | no |
| KCNMA1 | 206 | C | 0 | 0 | no | no | yes | no | yes | no | no | no | no | no | no | no | no |
| KCNMA1 | 207 | Q | 1 | 0 | no | yes | no | no | yes | no | no | no | no | no | no | no | no |
| KCNMA1 | 208 | N | 1 | 0 | no | yes | no | no | yes | no | no | no | no | no | no | no | no |
| KCNMA1 | 209 | F | 0 | 0 | no | yes | no | yes | yes | no | no | no | no | no | no | no | no |
| KCNMA1 | 210 | Y | 0 | 0 | no | yes | no | no | yes | no | no | no | no | no | no | no | no |
| KCNMA1 | 211 | K | 1 | 0 | no | yes | no | no | yes | no | no | no | no | no | no | no | no |
| KCNMA1 | 212 | D | 1 | 0 | no | yes | no | yes | yes | no | no | no | no | no | no | no | no |
| KCNMA1 | 213 | F | 0 | 0 | yes | no | no | yes | yes | no | no | no | no | no | no | no | no |
| KCNMA1 | 214 | T | 0 | 0 | yes | no | no | yes | yes | no | no | no | no | no | no | no | no |
| KCNMA1 | 215 | L | 0 | 0 | yes | no | no | yes | no | no | no | no | no | no | no | no | no |
| KCNMA1 | 216 | Q | 0 | 0 | yes | no | no | yes | no | no | no | no | no | no | no | no | no |

| | | | | | | | | | | | | | | | | | |
|---|---|---|---|---|---|---|---|---|---|---|---|---|---|---|---|---|---|
| KCNMA1 | 217 | I | 0 | 0 | yes | no | no | yes | no | no | no | no | no | no | no | no | no |
| KCNMA1 | 218 | D | 1 | 0 | yes | no | no | yes | no | no | no | no | no | no | no | no | no |
| KCNMA1 | 219 | M | 0 | 0 | yes | no | no | yes | no | no | no | no | no | no | no | no | no |
| KCNMA1 | 220 | A | 1 | 0 | yes | no | no | yes | no | no | no | no | no | no | no | no | no |
| KCNMA1 | 221 | F | 0 | 0 | yes | no | no | yes | no | no | no | no | no | no | no | no | no |
| KCNMA1 | 222 | N | 0 | 0 | yes | no | no | yes | no | no | no | no | no | no | no | no | no |
| KCNMA1 | 223 | V | 1 | 0 | yes | no | no | yes | no | no | no | no | no | no | no | no | no |
| KCNMA1 | 224 | F | 0 | 0 | yes | no | no | yes | no | no | no | no | no | no | no | no | no |
| KCNMA1 | 225 | F | 0 | 0 | yes | no | no | yes | no | no | no | no | no | no | no | no | no |
| KCNMA1 | 226 | L | 0 | 0 | yes | no | no | yes | no | no | no | no | no | no | no | no | no |
| KCNMA1 | 227 | L | 0 | 0 | yes | no | no | yes | no | no | no | no | no | no | no | no | no |
| KCNMA1 | 228 | Y | 0 | 0 | yes | no | no | yes | no | no | no | no | no | no | no | no | no |
| KCNMA1 | 229 | F | 0 | 0 | yes | no | no | yes | no | no | no | no | no | no | no | no | no |
| KCNMA1 | 230 | G | 1 | 0 | no | yes | no | yes | no | no | no | no | no | no | no | no | no |
| KCNMA1 | 231 | L | 0 | 0 | no | yes | no | yes | no | no | no | no | no | no | no | no | no |
| KCNMA1 | 232 | R | 0 | 0 | yes | no | no | yes | no | no | no | no | no | no | no | no | no |
| KCNMA1 | 233 | F | 0 | 0 | yes | no | no | yes | no | no | no | no | no | no | no | no | no |
| KCNMA1 | 234 | I | 0 | 0 | yes | no | no | yes | no | no | no | no | no | no | no | no | no |
| KCNMA1 | 235 | A | 0 | 0 | yes | no | no | no | no | no | no | no | no | no | no | no | no |
| KCNMA1 | 236 | A | 0 | 0 | no | yes | no | no | no | yes | no | no | no | no | no | no | no |
| KCNMA1 | 237 | N | 0 | 0 | no | yes | no | no | no | yes | no | no | no | no | no | no | no |
| KCNMA1 | 238 | D | 0 | 0 | no | yes | no | no | no | yes | no | no | no | no | no | no | no |
| KCNMA1 | 239 | K | 0 | 0 | no | yes | no | no | no | yes | no | no | no | no | no | no | no |
| KCNMA1 | 240 | L | 0 | 0 | no | yes | no | no | no | no | no | no | no | no | no | no | no |
| KCNMA1 | 241 | W | 1 | 0 | yes | no | no | no | no | no | no | no | no | no | no | no | no |
| KCNMA1 | 242 | F | 0 | 0 | yes | no | no | yes | no | no | no | no | no | no | no | no | no |
| KCNMA1 | 243 | W | 1 | 0 | yes | no | no | yes | no | no | no | no | no | no | no | no | no |
| KCNMA1 | 244 | L | 0 | 0 | no | yes | no | yes | no | no | no | no | no | no | no | no | no |
| KCNMA1 | 245 | E | 0 | 0 | no | yes | no | yes | no | no | no | no | no | no | no | no | no |
| KCNMA1 | 246 | V | 0 | 0 | yes | no | no | yes | no | no | no | no | no | no | no | no | no |
| KCNMA1 | 247 | N | 0 | 0 | yes | no | no | yes | no | no | no | no | no | no | no | no | no |
| KCNMA1 | 248 | S | 0 | 0 | yes | no | no | yes | no | no | no | no | no | no | no | no | no |
| KCNMA1 | 249 | V | 0 | 0 | yes | no | no | yes | no | no | no | no | no | no | no | no | no |

| | | | | | | | | | | | | | | | | | |
|---|---|---|---|---|---|---|---|---|---|---|---|---|---|---|---|---|---|
| KCNMA1 | 250 | V | 0 | 0 | yes | no | no | yes | no | no | no | no | no | no | no | no | no |
| KCNMA1 | 251 | D | 0 | 0 | yes | no | no | yes | no | no | no | no | no | no | no | no | no |
| KCNMA1 | 252 | F | 0 | 0 | yes | no | no | yes | no | no | no | no | no | no | no | no | no |
| KCNMA1 | 253 | F | 0 | 0 | no | yes | no | yes | no | no | no | no | no | no | no | no | no |
| KCNMA1 | 254 | T | 0 | 0 | no | yes | no | yes | no | no | no | no | no | no | no | no | no |
| KCNMA1 | 255 | V | 0 | 0 | yes | no | no | yes | no | no | no | no | no | no | no | no | no |
| KCNMA1 | 256 | P | 0 | 0 | yes | no | no | yes | no | no | no | no | no | no | no | no | no |
| KCNMA1 | 257 | P | 0 | 0 | yes | no | no | yes | no | no | no | no | no | no | no | no | no |
| KCNMA1 | 258 | V | 0 | 0 | yes | no | no | yes | no | no | no | no | no | no | no | no | no |
| KCNMA1 | 259 | F | 0 | 0 | yes | no | no | yes | no | no | no | no | no | no | no | no | no |
| KCNMA1 | 260 | V | 0 | 0 | yes | no | no | yes | no | no | no | no | no | no | no | no | no |
| KCNMA1 | 261 | S | 0 | 0 | yes | no | no | yes | yes | no | no | no | no | no | no | no | no |
| KCNMA1 | 262 | V | 0 | 0 | yes | no | no | yes | yes | no | no | no | no | no | no | no | no |
| KCNMA1 | 263 | Y | 0 | 0 | yes | no | no | yes | yes | no | no | no | no | no | no | no | no |
| KCNMA1 | 264 | L | 0 | 0 | yes | no | no | no | yes | no | no | no | no | no | no | no | no |
| KCNMA1 | 265 | N | 0 | 0 | no | yes | no | no | no | no | no | no | no | no | no | no | no |
| KCNMA1 | 266 | R | 0 | 0 | no | no | yes | no | no | no | no | no | no | no | no | no | no |
| KCNMA1 | 267 | S | 0 | 0 | no | no | yes | yes | no | no | no | no | no | no | no | no | no |
| KCNMA1 | 268 | W | 1 | 0 | no | no | yes | yes | no | no | no | no | no | no | no | no | no |
| KCNMA1 | 269 | L | 1 | 0 | no | yes | no | yes | no | no | no | no | no | no | no | no | no |
| KCNMA1 | 270 | G | 0 | 1 | no | yes | no | yes | no | no | no | no | no | no | no | no | no |
| KCNMA1 | 271 | L | 0 | 0 | no | yes | no | yes | no | no | no | no | no | no | no | no | no |
| KCNMA1 | 272 | R | 0 | 0 | yes | no | no | yes | no | no | no | no | no | no | no | no | no |
| KCNMA1 | 273 | F | 0 | 0 | yes | no | no | yes | no | no | no | no | no | no | no | no | no |
| KCNMA1 | 274 | L | 0 | 0 | yes | no | no | yes | no | no | no | no | no | no | no | no | no |
| KCNMA1 | 275 | R | 0 | 0 | yes | no | no | yes | no | no | no | no | no | no | no | no | no |
| KCNMA1 | 276 | A | 0 | 0 | yes | no | no | yes | no | no | no | no | no | no | no | no | no |
| KCNMA1 | 277 | L | 0 | 0 | yes | no | no | yes | no | no | no | no | no | no | no | no | no |
| KCNMA1 | 278 | R | 0 | 0 | yes | no | no | yes | no | no | no | no | no | no | no | no | no |
| KCNMA1 | 279 | L | 0 | 0 | yes | no | no | yes | no | no | no | no | no | no | no | no | no |
| KCNMA1 | 280 | I | 0 | 0 | yes | no | no | yes | no | no | no | no | no | no | no | no | no |
| KCNMA1 | 281 | Q | 0 | 0 | yes | no | no | yes | no | no | no | no | no | no | no | no | no |
| KCNMA1 | 282 | F | 0 | 0 | yes | no | no | yes | no | no | no | no | no | no | no | no | no |

| | | | | | | | | | | | | | | | | | |
|---|---|---|---|---|---|---|---|---|---|---|---|---|---|---|---|---|---|
| KCNMA1 | 283 | S | 0 | 0 | yes | no | no | no | no | no | no | no | no | no | no | no | no |
| KCNMA1 | 284 | E | 0 | 0 | yes | no | no | no | no | no | no | no | no | no | no | no | no |
| KCNMA1 | 285 | I | 0 | 0 | yes | no | no | yes | no | no | no | no | no | no | no | no | no |
| KCNMA1 | 286 | L | 0 | 0 | yes | no | no | no | no | yes | no | no | no | no | no | no | no |
| KCNMA1 | 287 | Q | 0 | 0 | yes | no | no | no | no | yes | no | no | no | no | no | no | no |
| KCNMA1 | 288 | F | 0 | 0 | yes | no | no | no | no | yes | no | no | no | no | no | no | no |
| KCNMA1 | 289 | L | 0 | 0 | no | yes | no | no | no | yes | no | no | no | no | no | no | no |
| KCNMA1 | 290 | N | 0 | 0 | no | yes | no | no | no | yes | no | no | no | no | no | no | no |
| KCNMA1 | 291 | I | 0 | 0 | no | yes | no | no | no | yes | no | no | no | no | no | no | no |
| KCNMA1 | 292 | L | 0 | 0 | no | yes | no | no | no | yes | no | no | no | no | no | no | no |
| KCNMA1 | 293 | K | 1 | 0 | no | yes | no | no | no | yes | no | no | no | no | no | no | no |
| KCNMA1 | 294 | T | 0 | 0 | no | yes | no | no | no | yes | no | no | no | no | no | no | no |
| KCNMA1 | 295 | S | 0 | 0 | yes | no | no | no | no | yes | no | no | yes | no | no | no | no |
| KCNMA1 | 296 | N | 0 | 0 | yes | no | no | no | no | yes | no | no | yes | no | no | no | no |
| KCNMA1 | 297 | S | 0 | 0 | yes | no | no | no | no | yes | no | no | no | no | no | no | no |
| KCNMA1 | 298 | I | 0 | 0 | yes | no | no | no | no | yes | no | no | no | no | no | no | no |
| KCNMA1 | 299 | K | 0 | 0 | yes | no | no | no | no | yes | no | no | no | no | no | no | no |
| KCNMA1 | 300 | L | 0 | 0 | yes | no | no | no | no | yes | no | no | no | no | no | no | no |
| KCNMA1 | 301 | V | 0 | 0 | yes | no | no | yes | no | no | no | no | no | no | no | no | no |
| KCNMA1 | 302 | N | 0 | 0 | yes | no | no | no | no | no | no | no | no | no | no | no | no |
| KCNMA1 | 303 | L | 0 | 0 | yes | no | no | yes | no | no | no | no | no | no | no | no | no |
| KCNMA1 | 304 | L | 0 | 0 | yes | no | no | yes | no | no | no | no | no | no | no | no | no |
| KCNMA1 | 305 | S | 0 | 0 | yes | no | no | yes | no | no | no | no | no | no | no | no | no |
| KCNMA1 | 306 | I | 0 | 0 | yes | no | no | yes | no | no | no | no | no | no | no | no | no |
| KCNMA1 | 307 | F | 0 | 0 | yes | no | no | yes | no | no | no | no | no | no | no | no | no |
| KCNMA1 | 308 | I | 1 | 0 | yes | no | no | yes | no | no | no | no | no | no | no | no | no |
| KCNMA1 | 309 | S | 0 | 0 | yes | no | no | yes | no | no | no | no | no | no | no | no | no |
| KCNMA1 | 310 | T | 0 | 0 | yes | no | no | yes | no | no | no | no | no | no | no | no | no |
| KCNMA1 | 311 | W | 0 | 0 | yes | no | no | yes | no | no | no | no | no | no | no | no | no |
| KCNMA1 | 312 | L | 0 | 0 | yes | no | no | yes | no | no | no | no | no | no | no | no | no |
| KCNMA1 | 313 | T | 0 | 0 | yes | no | no | yes | no | no | no | no | no | no | no | no | no |
| KCNMA1 | 314 | A | 0 | 0 | yes | no | no | yes | no | no | no | no | no | no | no | no | no |
| KCNMA1 | 315 | A | 0 | 0 | yes | no | no | yes | no | no | no | no | no | no | no | no | no |

| | | | | | | | | | | | | | | | | | |
|---|---|---|---|---|---|---|---|---|---|---|---|---|---|---|---|---|---|
| KCNMA1 | 316 | G | 1 | 0 | yes | no | no | yes | no | no | no | no | no | no | no | no | no |
| KCNMA1 | 317 | F | 0 | 0 | yes | no | no | yes | no | no | no | no | no | no | no | no | no |
| KCNMA1 | 318 | I | 0 | 0 | yes | no | no | yes | no | no | no | no | no | no | no | no | no |
| KCNMA1 | 319 | H | 0 | 0 | yes | no | no | yes | no | no | no | no | no | no | no | no | no |
| KCNMA1 | 320 | L | 0 | 0 | yes | no | no | yes | no | no | no | no | no | no | no | no | no |
| KCNMA1 | 321 | V | 0 | 0 | yes | no | no | yes | no | no | no | no | no | no | no | no | no |
| KCNMA1 | 322 | E | 0 | 0 | yes | no | no | yes | yes | no | no | no | no | no | no | no | no |
| KCNMA1 | 323 | N | 0 | 0 | yes | no | no | yes | yes | no | no | no | no | no | no | no | no |
| KCNMA1 | 324 | S | 0 | 0 | no | yes | no | yes | yes | no | no | no | no | no | no | no | no |
| KCNMA1 | 325 | G | 0 | 0 | no | yes | no | no | yes | no | no | no | no | no | no | no | no |
| KCNMA1 | 326 | D | 0 | 0 | no | yes | no | no | yes | no | no | no | no | no | no | no | no |
| KCNMA1 | 327 | P | 1 | 0 | no | yes | no | no | yes | no | no | no | no | no | no | no | no |
| KCNMA1 | 328 | W | 0 | 0 | no | yes | no | yes | yes | no | no | no | no | no | no | no | no |
| KCNMA1 | 329 | E | 0 | 0 | no | yes | no | no | yes | no | no | no | no | no | no | no | no |
| KCNMA1 | 330 | N | 0 | 0 | no | yes | no | no | yes | no | no | no | no | no | no | no | no |
| KCNMA1 | 331 | F | 0 | 0 | no | yes | no | no | yes | no | no | no | no | no | no | no | no |
| KCNMA1 | 332 | Q | 0 | 0 | no | yes | no | no | yes | no | no | no | no | no | no | no | no |
| KCNMA1 | 333 | N | 0 | 0 | no | yes | no | no | yes | no | no | no | no | no | no | no | no |
| KCNMA1 | 334 | N | 0 | 0 | no | yes | no | no | yes | no | no | no | no | no | no | no | no |
| KCNMA1 | 335 | Q | 0 | 0 | no | yes | no | no | yes | no | no | no | no | no | no | no | no |
| KCNMA1 | 336 | A | 0 | 0 | no | yes | no | no | no | no | no | yes | no | no | no | no | no |
| KCNMA1 | 337 | L | 0 | 0 | no | yes | no | yes | no | no | no | yes | no | no | no | no | no |
| KCNMA1 | 338 | T | 0 | 0 | no | yes | no | yes | no | no | no | yes | no | no | no | no | no |
| KCNMA1 | 339 | Y | 0 | 0 | yes | no | no | yes | no | no | no | yes | no | no | no | no | no |
| KCNMA1 | 340 | W | 0 | 0 | yes | no | no | yes | no | no | no | yes | no | no | no | no | no |
| KCNMA1 | 341 | E | 1 | 0 | yes | no | no | yes | no | no | no | yes | no | no | no | no | no |
| KCNMA1 | 342 | C | 0 | 0 | yes | no | no | yes | no | no | no | yes | no | no | no | no | no |
| KCNMA1 | 343 | V | 0 | 0 | yes | no | no | yes | no | no | no | yes | no | no | no | no | no |
| KCNMA1 | 344 | Y | 0 | 0 | yes | no | no | yes | no | no | no | yes | yes | no | no | no | no |
| KCNMA1 | 345 | L | 0 | 0 | yes | no | no | yes | no | no | no | yes | yes | no | no | no | no |
| KCNMA1 | 346 | L | 0 | 0 | yes | no | no | yes | no | no | no | yes | no | no | no | no | no |
| KCNMA1 | 347 | M | 0 | 0 | yes | no | no | yes | no | no | no | yes | yes | no | no | no | no |
| KCNMA1 | 348 | V | 0 | 0 | yes | no | no | yes | no | no | no | yes | no | no | no | no | no |

| | | | | | | | | | | | | | | | | | |
|---|---|---|---|---|---|---|---|---|---|---|---|---|---|---|---|---|---|
| KCNMA1 | 349 | T | 0 | 0 | yes | no | no | yes | no | no | no | yes | yes | no | no | no | no |
| KCNMA1 | 350 | M | 0 | 0 | no | yes | no | yes | no | no | no | yes | no | no | no | no | no |
| KCNMA1 | 351 | S | 0 | 1 | no | yes | no | yes | no | no | no | yes | yes | no | no | no | no |
| KCNMA1 | 352 | T | 0 | 1 | no | yes | no | yes | no | no | yes | yes | yes | no | no | no | no |
| KCNMA1 | 353 | V | 1 | 0 | no | yes | no | yes | no | no | yes | yes | yes | no | no | no | no |
| KCNMA1 | 354 | G | 0 | 0 | no | yes | no | yes | no | no | yes | yes | yes | no | no | no | no |
| KCNMA1 | 355 | Y | 0 | 0 | no | yes | no | yes | no | no | yes | yes | yes | no | no | no | no |
| KCNMA1 | 356 | G | 0 | 1 | no | yes | no | yes | no | no | yes | yes | no | no | no | no | no |
| KCNMA1 | 357 | D | 0 | 0 | no | yes | no | yes | no | no | no | yes | yes | no | no | no | no |
| KCNMA1 | 358 | V | 0 | 0 | no | yes | no | yes | no | no | no | yes | no | no | no | no | no |
| KCNMA1 | 359 | Y | 0 | 0 | no | yes | no | yes | yes | no | yes | no | yes | no | no | no | no |
| KCNMA1 | 360 | A | 0 | 0 | no | yes | no | yes | yes | no | no | no | no | no | no | no | no |
| KCNMA1 | 361 | K | 1 | 0 | no | yes | no | yes | yes | no | no | no | no | no | no | no | no |
| KCNMA1 | 362 | T | 0 | 0 | no | yes | no | yes | yes | no | no | no | no | no | no | no | no |
| KCNMA1 | 363 | T | 0 | 0 | yes | no | no | yes | yes | no | no | no | no | no | no | no | no |
| KCNMA1 | 364 | L | 0 | 0 | yes | no | no | yes | yes | no | no | no | no | no | no | no | no |
| KCNMA1 | 365 | G | 0 | 0 | yes | no | no | yes | yes | no | no | no | no | no | no | no | no |
| KCNMA1 | 366 | R | 0 | 0 | yes | no | no | yes | yes | no | no | no | yes | no | no | no | no |
| KCNMA1 | 367 | L | 0 | 0 | yes | no | no | yes | yes | no | no | no | no | no | no | no | no |
| KCNMA1 | 368 | F | 0 | 0 | yes | no | no | yes | no | no | no | no | no | no | no | no | no |
| KCNMA1 | 369 | M | 0 | 0 | yes | no | no | yes | no | no | no | no | yes | no | no | no | no |
| KCNMA1 | 370 | V | 0 | 0 | yes | no | no | yes | no | no | no | no | yes | no | no | no | no |
| KCNMA1 | 371 | F | 0 | 0 | yes | no | no | yes | no | no | no | no | no | no | no | no | no |
| KCNMA1 | 372 | F | 0 | 0 | yes | no | no | yes | no | no | no | no | no | no | no | no | no |
| KCNMA1 | 373 | I | 0 | 0 | yes | no | no | yes | no | no | no | no | yes | no | no | no | no |
| KCNMA1 | 374 | L | 0 | 0 | yes | no | no | yes | no | no | no | no | yes | no | no | no | no |
| KCNMA1 | 375 | G | 0 | 1 | yes | no | no | yes | no | no | no | no | no | no | no | no | no |
| KCNMA1 | 376 | G | 0 | 0 | yes | no | no | yes | no | no | no | no | no | no | no | no | no |
| KCNMA1 | 377 | L | 0 | 0 | yes | no | no | yes | no | no | no | no | no | no | no | no | no |
| KCNMA1 | 378 | A | 0 | 0 | yes | no | no | yes | no | no | no | no | no | no | no | no | no |
| KCNMA1 | 379 | M | 0 | 0 | yes | no | no | yes | no | no | no | no | no | no | no | no | no |
| KCNMA1 | 380 | F | 0 | 0 | yes | no | no | yes | no | no | yes | no | no | no | no | no | no |
| KCNMA1 | 381 | A | 0 | 0 | yes | no | no | no | no | no | no | no | no | no | no | no | no |

| | | | | | | | | | | | | | | | | | |
|---|---|---|---|---|---|---|---|---|---|---|---|---|---|---|---|---|---|
| KCNMA1 | 382 | S | 0 | 0 | yes | no | no | no | no | no | no | no | no | no | no | no | no |
| KCNMA1 | 383 | Y | 0 | 0 | no | yes | no | no | no | no | no | no | no | no | no | no | no |
| KCNMA1 | 384 | V | 1 | 0 | no | yes | no | no | no | no | no | no | no | no | no | no | no |
| KCNMA1 | 385 | P | 0 | 0 | yes | no | no | no | no | no | no | no | no | no | no | no | no |
| KCNMA1 | 386 | E | 0 | 0 | yes | no | no | no | no | no | no | no | yes | no | no | no | no |
| KCNMA1 | 387 | I | 0 | 0 | yes | no | no | no | no | no | no | no | no | no | no | no | no |
| KCNMA1 | 388 | I | 0 | 0 | yes | no | no | no | no | no | no | no | no | no | no | no | no |
| KCNMA1 | 389 | E | 0 | 0 | yes | no | no | no | no | yes | yes | no | no | no | no | no | no |
| KCNMA1 | 390 | L | 0 | 0 | yes | no | no | no | no | yes | no | no | no | no | no | no | no |
| KCNMA1 | 391 | I | 0 | 0 | no | yes | no | no | no | yes | no | no | no | no | no | no | no |
| KCNMA1 | 392 | G | 0 | 0 | no | yes | no | no | no | yes | no | no | no | no | no | no | no |
| KCNMA1 | 393 | N | 0 | 0 | no | yes | no | no | no | yes | yes | no | no | no | no | no | no |
| KCNMA1 | 394 | R | 0 | 0 | no | yes | no | no | no | yes | no | no | yes | no | no | no | no |
| KCNMA1 | 395 | K | 0 | 0 | no | yes | no | no | no | yes | no | no | yes | no | no | no | no |
| KCNMA1 | 396 | K | 0 | 0 | no | yes | no | no | no | yes | no | no | no | no | no | no | no |
| KCNMA1 | 397 | Y | 0 | 0 | no | yes | no | no | no | yes | no | no | no | no | no | no | no |
| KCNMA1 | 398 | G | 0 | 0 | no | yes | no | no | no | yes | no | no | no | no | no | no | no |
| KCNMA1 | 399 | G | 0 | 0 | no | yes | no | no | no | yes | no | no | no | no | no | no | no |
| KCNMA1 | 400 | S | 0 | 0 | no | yes | no | no | no | yes | no | no | no | no | no | no | no |
| KCNMA1 | 401 | Y | 0 | 0 | no | yes | no | no | no | yes | no | no | no | no | no | no | no |
| KCNMA1 | 402 | S | 0 | 0 | no | yes | no | no | no | yes | no | no | no | no | no | no | no |
| KCNMA1 | 403 | A | 0 | 0 | no | yes | no | no | no | yes | no | no | no | no | no | no | no |
| KCNMA1 | 404 | V | 0 | 0 | no | yes | no | no | no | yes | no | no | no | no | no | no | no |
| KCNMA1 | 405 | S | 0 | 0 | no | yes | no | no | no | yes | no | no | no | no | no | no | no |
| KCNMA1 | 406 | G | 0 | 0 | no | yes | no | no | no | yes | no | no | no | no | no | no | no |
| KCNMA1 | 407 | R | 0 | 0 | no | yes | no | no | no | yes | no | no | no | no | no | no | no |
| KCNMA1 | 408 | K | 0 | 0 | no | yes | no | no | no | yes | no | no | no | no | no | no | no |
| KCNMA1 | 409 | H | 0 | 0 | no | yes | no | no | no | yes | no | no | no | no | no | no | no |
| KCNMA1 | 410 | I | 0 | 0 | no | yes | no | no | no | yes | no | no | no | no | no | no | no |
| KCNMA1 | 411 | V | 0 | 0 | no | no | yes | no | no | yes | no | no | no | no | no | no | no |
| KCNMA1 | 412 | V | 0 | 0 | no | no | yes | no | no | yes | no | no | no | no | no | no | no |
| KCNMA1 | 413 | C | 0 | 1 | no | no | yes | no | no | yes | no | no | no | no | no | no | no |
| KCNMA1 | 414 | G | 0 | 0 | no | no | yes | no | no | yes | no | no | no | no | no | no | no |

| | | | | | | | | | | | | | | | | | |
|---|---|---|---|---|---|---|---|---|---|---|---|---|---|---|---|---|---|
| KCNMA1 | 415 | H | 0 | 0 | no | yes | no | no | no | yes | no | no | no | no | no | no | no |
| KCNMA1 | 416 | I | 0 | 0 | no | yes | no | no | no | yes | no | no | no | no | no | no | no |
| KCNMA1 | 417 | T | 0 | 0 | no | yes | no | no | no | yes | no | no | no | no | no | no | no |
| KCNMA1 | 418 | L | 0 | 0 | yes | no | no | no | no | yes | no | no | no | no | no | no | no |
| KCNMA1 | 419 | E | 0 | 0 | yes | no | no | no | no | yes | no | no | no | no | no | no | no |
| KCNMA1 | 420 | S | 0 | 0 | yes | no | no | no | no | yes | no | no | no | no | no | no | no |
| KCNMA1 | 421 | V | 0 | 0 | yes | no | no | no | no | yes | no | no | no | no | no | no | no |
| KCNMA1 | 422 | S | 0 | 0 | yes | no | no | no | no | yes | no | no | no | no | no | no | no |
| KCNMA1 | 423 | N | 0 | 0 | yes | no | no | no | no | yes | no | no | no | no | no | no | no |
| KCNMA1 | 424 | F | 0 | 0 | yes | no | no | no | no | yes | no | no | no | no | no | no | no |
| KCNMA1 | 425 | L | 0 | 0 | yes | no | no | no | no | yes | no | no | no | no | no | no | no |
| KCNMA1 | 426 | K | 0 | 0 | yes | no | no | no | no | yes | no | no | no | no | no | no | no |
| KCNMA1 | 427 | D | 0 | 0 | yes | no | no | no | no | yes | no | no | no | no | no | no | no |
| KCNMA1 | 428 | F | 0 | 0 | yes | no | no | no | no | yes | no | no | no | no | no | no | no |
| KCNMA1 | 429 | L | 0 | 0 | no | yes | no | no | no | yes | no | no | no | no | no | no | no |
| KCNMA1 | 430 | H | 0 | 0 | no | yes | no | no | no | yes | no | no | no | no | no | no | no |
| KCNMA1 | 431 | K | 0 | 0 | no | yes | no | no | no | yes | no | no | no | no | no | no | no |
| KCNMA1 | 432 | D | 0 | 0 | no | yes | no | no | no | yes | no | no | no | no | no | no | no |
| KCNMA1 | 433 | R | 1 | 0 | no | yes | no | no | no | yes | no | no | no | no | no | no | no |
| KCNMA1 | 434 | D | 1 | 1 | no | yes | no | no | no | yes | no | no | no | no | no | no | no |
| KCNMA1 | 435 | D | 0 | 0 | no | yes | no | no | no | yes | no | no | no | no | no | no | no |
| KCNMA1 | 436 | V | 1 | 0 | no | yes | no | no | no | yes | no | no | no | no | no | no | no |
| KCNMA1 | 437 | N | 1 | 0 | no | yes | no | no | no | yes | no | no | no | no | no | no | no |
| KCNMA1 | 438 | V | 0 | 0 | no | yes | no | no | no | yes | no | no | no | no | no | no | no |
| KCNMA1 | 439 | E | 0 | 0 | no | yes | no | no | no | yes | no | no | no | no | no | no | no |
| KCNMA1 | 440 | I | 0 | 0 | no | no | yes | no | no | yes | no | no | no | no | no | no | no |
| KCNMA1 | 441 | V | 1 | 0 | no | no | yes | no | no | yes | no | no | no | no | no | no | no |
| KCNMA1 | 442 | F | 0 | 0 | no | no | yes | no | no | yes | no | no | no | no | no | no | no |
| KCNMA1 | 443 | L | 0 | 0 | no | no | yes | no | no | yes | no | no | no | no | no | no | no |
| KCNMA1 | 444 | H | 0 | 0 | no | yes | no | no | no | yes | no | no | no | no | no | no | no |
| KCNMA1 | 445 | N | 0 | 0 | no | yes | no | no | no | yes | no | no | no | no | no | no | no |
| KCNMA1 | 446 | I | 0 | 0 | no | yes | no | no | no | yes | yes | no | no | no | no | no | no |
| KCNMA1 | 447 | S | 0 | 0 | no | yes | no | no | no | yes | yes | no | no | no | no | no | no |

| | | | | | | | | | | | | | | | | | |
|---|---|---|---|---|---|---|---|---|---|---|---|---|---|---|---|---|---|
| KCNMA1 | 448 | P | 0 | 0 | no | yes | no | no | no | yes | no | no | no | no | no | no | no |
| KCNMA1 | 449 | N | 0 | 0 | no | yes | no | no | no | yes | yes | no | no | no | no | no | no |
| KCNMA1 | 450 | L | 0 | 0 | yes | no | no | no | no | yes | yes | no | yes | no | no | no | no |
| KCNMA1 | 451 | E | 0 | 0 | yes | no | no | no | no | yes | no | no | no | no | no | no | no |
| KCNMA1 | 452 | L | 0 | 0 | yes | no | no | no | no | yes | no | no | no | no | no | no | no |
| KCNMA1 | 453 | E | 0 | 0 | yes | no | no | no | no | yes | no | no | no | no | no | no | no |
| KCNMA1 | 454 | A | 0 | 0 | yes | no | no | no | no | yes | no | no | no | no | no | no | no |
| KCNMA1 | 455 | L | 0 | 0 | yes | no | no | no | no | yes | no | no | no | no | no | no | no |
| KCNMA1 | 456 | F | 0 | 0 | yes | no | no | no | no | yes | no | no | no | no | no | no | no |
| KCNMA1 | 457 | K | 0 | 0 | yes | no | no | no | no | yes | no | no | yes | no | no | no | no |
| KCNMA1 | 458 | R | 0 | 0 | yes | no | no | no | no | yes | no | no | no | no | no | no | no |
| KCNMA1 | 459 | H | 1 | 0 | no | yes | no | no | no | yes | no | no | no | no | no | no | no |
| KCNMA1 | 460 | F | 0 | 0 | no | yes | no | no | no | yes | no | no | yes | no | no | no | no |
| KCNMA1 | 461 | T | 0 | 0 | no | yes | no | no | no | yes | no | no | no | no | no | no | no |
| KCNMA1 | 462 | Q | 0 | 0 | no | yes | no | no | no | yes | no | no | no | no | no | no | no |
| KCNMA1 | 463 | V | 0 | 0 | no | no | yes | no | no | yes | no | no | no | no | no | no | no |
| KCNMA1 | 464 | E | 1 | 0 | no | no | yes | no | no | yes | no | no | no | no | no | no | no |
| KCNMA1 | 465 | F | 0 | 0 | no | no | yes | no | no | yes | no | no | no | no | no | no | no |
| KCNMA1 | 466 | Y | 0 | 0 | no | no | yes | no | no | yes | no | no | no | no | no | no | no |
| KCNMA1 | 467 | Q | 1 | 0 | no | yes | no | no | no | yes | no | no | no | no | no | no | no |
| KCNMA1 | 468 | G | 0 | 0 | no | yes | no | no | no | yes | no | no | no | no | no | no | no |
| KCNMA1 | 469 | S | 0 | 0 | no | yes | no | no | no | yes | no | no | no | no | no | no | no |
| KCNMA1 | 470 | V | 1 | 0 | no | yes | no | no | no | yes | no | no | no | no | no | no | no |
| KCNMA1 | 471 | L | 0 | 0 | no | yes | no | no | no | yes | no | no | yes | no | no | no | no |
| KCNMA1 | 472 | N | 0 | 0 | no | yes | no | no | no | yes | no | no | no | no | no | no | no |
| KCNMA1 | 473 | P | 0 | 0 | yes | no | no | no | no | yes | no | no | yes | no | no | no | no |
| KCNMA1 | 474 | H | 0 | 0 | yes | no | no | no | no | yes | no | no | no | no | no | no | no |
| KCNMA1 | 475 | D | 0 | 0 | yes | no | no | no | no | yes | no | no | no | no | no | no | no |
| KCNMA1 | 476 | L | 0 | 0 | yes | no | no | no | no | yes | no | no | no | no | no | no | no |
| KCNMA1 | 477 | A | 0 | 0 | yes | no | no | no | no | yes | no | no | no | no | no | no | no |
| KCNMA1 | 478 | R | 0 | 0 | yes | no | no | no | no | yes | no | no | no | no | no | no | no |
| KCNMA1 | 479 | V | 0 | 0 | no | yes | no | no | no | yes | no | no | no | no | no | no | no |
| KCNMA1 | 480 | K | 0 | 0 | no | yes | no | no | no | yes | no | no | no | no | no | no | no |

| | | | | | | | | | | | | | | | | | |
|---|---|---|---|---|---|---|---|---|---|---|---|---|---|---|---|---|---|
| KCNMA1 | 481 | I | 1 | 0 | no | yes | no | no | no | yes | no | no | no | no | no | no | no |
| KCNMA1 | 482 | E | 0 | 0 | no | yes | no | no | no | yes | no | no | no | no | no | no | no |
| KCNMA1 | 483 | S | 0 | 0 | no | yes | no | no | no | yes | no | no | no | no | no | no | no |
| KCNMA1 | 484 | A | 0 | 0 | no | yes | no | no | no | yes | no | no | no | no | no | no | no |
| KCNMA1 | 485 | D | 0 | 0 | no | yes | no | no | no | yes | no | no | no | no | no | no | no |
| KCNMA1 | 486 | A | 0 | 0 | no | no | yes | no | no | yes | no | no | no | no | no | no | no |
| KCNMA1 | 487 | C | 0 | 0 | no | no | yes | no | no | yes | no | no | no | no | no | no | no |
| KCNMA1 | 488 | L | 0 | 0 | no | no | yes | no | no | yes | no | no | no | no | no | no | no |
| KCNMA1 | 489 | I | 0 | 0 | no | no | yes | no | no | yes | no | no | no | no | no | no | no |
| KCNMA1 | 490 | L | 0 | 0 | no | no | yes | no | no | yes | no | no | no | no | no | no | no |
| KCNMA1 | 491 | A | 0 | 0 | no | yes | no | no | no | yes | no | no | no | no | no | no | no |
| KCNMA1 | 492 | N | 0 | 0 | no | yes | no | no | no | yes | no | no | no | no | no | no | no |
| KCNMA1 | 493 | K | 0 | 0 | no | yes | no | no | no | yes | no | no | no | no | no | no | no |
| KCNMA1 | 494 | Y | 0 | 0 | no | yes | no | no | no | yes | no | no | no | no | no | no | no |
| KCNMA1 | 495 | C | 0 | 0 | no | yes | no | no | no | yes | no | no | no | no | no | no | no |
| KCNMA1 | 496 | A | 1 | 0 | no | yes | no | no | no | yes | yes | no | no | no | no | no | no |
| KCNMA1 | 497 | D | 0 | 0 | no | yes | no | no | no | yes | no | no | no | no | no | no | no |
| KCNMA1 | 498 | P | 0 | 0 | yes | no | no | no | no | yes | no | no | no | no | no | no | no |
| KCNMA1 | 499 | D | 0 | 0 | yes | no | no | no | no | yes | no | no | no | no | no | no | no |
| KCNMA1 | 500 | A | 0 | 0 | yes | no | no | no | no | yes | no | no | yes | no | no | no | no |
| KCNMA1 | 501 | E | 0 | 0 | yes | no | no | no | no | yes | no | no | no | no | no | no | no |
| KCNMA1 | 502 | D | 0 | 0 | yes | no | no | no | no | yes | no | no | no | no | no | no | no |
| KCNMA1 | 503 | A | 1 | 0 | yes | no | no | no | no | yes | no | no | yes | no | no | no | no |
| KCNMA1 | 504 | S | 0 | 0 | yes | no | no | no | no | yes | no | no | yes | no | no | no | no |
| KCNMA1 | 505 | N | 0 | 0 | yes | no | no | no | no | yes | no | no | no | no | no | no | no |
| KCNMA1 | 506 | I | 0 | 0 | yes | no | no | no | no | yes | no | no | no | no | no | no | no |
| KCNMA1 | 507 | M | 0 | 0 | yes | no | no | no | no | yes | no | no | yes | no | no | no | no |
| KCNMA1 | 508 | R | 0 | 0 | yes | no | no | no | no | yes | no | no | no | no | no | no | no |
| KCNMA1 | 509 | V | 0 | 0 | yes | no | no | no | no | yes | no | no | no | no | no | no | no |
| KCNMA1 | 510 | I | 0 | 0 | yes | no | no | no | no | yes | no | no | yes | no | no | no | no |
| KCNMA1 | 511 | S | 1 | 0 | yes | no | no | no | no | yes | no | no | yes | no | no | no | no |
| KCNMA1 | 512 | I | 1 | 1 | yes | no | no | no | no | yes | no | no | no | no | no | no | no |
| KCNMA1 | 513 | K | 1 | 0 | yes | no | no | no | no | yes | no | no | no | no | no | no | no |

| | | | | | | | | | | | | | | | | | |
|---|---|---|---|---|---|---|---|---|---|---|---|---|---|---|---|---|---|
| KCNMA1 | 514 | N | 0 | 0 | yes | no | no | no | no | yes | no | no | yes | no | no | no | no |
| KCNMA1 | 515 | Y | 0 | 0 | yes | no | no | no | no | yes | no | no | yes | no | no | no | no |
| KCNMA1 | 516 | H | 2 | 0 | no | yes | no | no | no | yes | no | no | no | no | no | no | no |
| KCNMA1 | 517 | P | 1 | 0 | no | yes | no | no | no | yes | no | no | no | no | no | no | no |
| KCNMA1 | 518 | K | 1 | 0 | no | yes | no | no | no | yes | no | no | no | no | no | no | no |
| KCNMA1 | 519 | I | 0 | 0 | no | yes | no | no | no | yes | no | no | no | no | no | no | no |
| KCNMA1 | 520 | R | 1 | 0 | no | no | yes | no | no | yes | no | no | no | no | no | no | no |
| KCNMA1 | 521 | I | 0 | 0 | no | no | yes | no | no | yes | no | no | no | no | no | no | no |
| KCNMA1 | 522 | I | 1 | 0 | no | no | yes | no | no | yes | no | no | no | no | no | no | no |
| KCNMA1 | 523 | T | 0 | 0 | no | no | yes | no | no | yes | no | no | no | no | no | no | no |
| KCNMA1 | 524 | Q | 0 | 0 | no | no | yes | no | no | yes | no | no | no | no | no | no | no |
| KCNMA1 | 525 | M | 0 | 0 | no | no | yes | no | no | yes | no | no | no | no | no | no | no |
| KCNMA1 | 526 | L | 0 | 0 | no | yes | no | no | no | yes | no | no | no | no | no | no | no |
| KCNMA1 | 527 | Q | 0 | 0 | no | yes | no | no | no | yes | no | no | no | no | no | no | no |
| KCNMA1 | 528 | Y | 0 | 0 | no | yes | no | no | no | yes | no | no | no | no | no | no | no |
| KCNMA1 | 529 | H | 0 | 0 | no | yes | no | no | no | yes | no | no | no | no | no | no | no |
| KCNMA1 | 530 | N | 0 | 0 | no | yes | no | no | no | yes | no | no | no | no | no | no | no |
| KCNMA1 | 531 | K | 0 | 0 | yes | no | no | no | no | yes | no | no | no | no | no | no | no |
| KCNMA1 | 532 | A | 0 | 0 | yes | no | no | no | no | yes | no | no | no | no | no | no | no |
| KCNMA1 | 533 | H | 0 | 0 | yes | no | no | no | no | yes | no | no | yes | no | no | no | no |
| KCNMA1 | 534 | L | 0 | 0 | no | yes | no | no | no | yes | no | no | no | no | no | no | no |
| KCNMA1 | 535 | L | 0 | 0 | no | yes | no | no | no | yes | no | no | no | no | no | no | no |
| KCNMA1 | 536 | N | 0 | 1 | no | yes | no | no | no | yes | no | no | yes | no | no | no | no |
| KCNMA1 | 537 | I | 0 | 0 | no | yes | no | no | no | yes | no | no | no | no | no | no | no |
| KCNMA1 | 538 | P | 1 | 0 | no | yes | no | no | no | yes | no | no | no | no | no | no | no |
| KCNMA1 | 539 | S | 0 | 0 | no | yes | no | no | no | yes | no | no | no | no | no | no | no |
| KCNMA1 | 540 | W | 0 | 0 | no | yes | no | no | no | yes | no | no | no | no | no | no | no |
| KCNMA1 | 541 | N | 1 | 0 | no | yes | no | no | no | yes | no | no | no | no | no | no | no |
| KCNMA1 | 542 | W | 0 | 0 | no | yes | no | no | no | yes | no | no | no | no | no | no | no |
| KCNMA1 | 543 | K | 0 | 0 | no | yes | no | no | no | yes | no | no | no | no | no | no | no |
| KCNMA1 | 544 | E | 1 | 0 | no | yes | no | no | no | yes | no | no | no | no | no | no | no |
| KCNMA1 | 545 | G | 0 | 0 | no | yes | no | no | no | yes | no | no | no | no | no | no | no |
| KCNMA1 | 546 | D | 0 | 0 | no | yes | no | no | no | yes | no | no | no | no | no | no | no |

| | | | | | | | | | | | | | | | | | |
|---|---|---|---|---|---|---|---|---|---|---|---|---|---|---|---|---|---|
| KCNMA1 | 547 | D | 0 | 0 | no | no | yes | no | no | yes | no | no | no | no | no | no | no |
| KCNMA1 | 548 | A | 0 | 0 | no | no | yes | no | no | yes | no | no | no | no | no | no | no |
| KCNMA1 | 549 | I | 0 | 0 | no | no | yes | no | no | yes | no | no | no | no | no | no | no |
| KCNMA1 | 550 | C | 0 | 0 | no | no | yes | no | no | yes | no | no | no | no | no | no | no |
| KCNMA1 | 551 | L | 0 | 0 | yes | no | no | no | no | yes | no | no | no | no | no | no | no |
| KCNMA1 | 552 | A | 0 | 0 | yes | no | no | no | no | yes | no | no | no | no | no | no | no |
| KCNMA1 | 553 | E | 1 | 0 | yes | no | no | no | no | yes | no | no | no | no | no | no | no |
| KCNMA1 | 554 | L | 0 | 0 | yes | no | no | no | no | yes | no | no | no | no | no | no | no |
| KCNMA1 | 555 | K | 0 | 0 | yes | no | no | no | no | yes | no | no | no | no | no | no | no |
| KCNMA1 | 556 | L | 0 | 0 | yes | no | no | no | no | yes | no | no | no | no | no | no | no |
| KCNMA1 | 557 | G | 0 | 0 | yes | no | no | no | no | yes | no | no | no | no | no | no | no |
| KCNMA1 | 558 | F | 0 | 0 | yes | no | no | no | no | yes | no | no | no | no | no | no | no |
| KCNMA1 | 559 | I | 0 | 0 | yes | no | no | no | no | yes | no | no | no | no | no | no | no |
| KCNMA1 | 560 | A | 0 | 0 | yes | no | no | no | no | yes | no | no | no | no | no | no | no |
| KCNMA1 | 561 | Q | 0 | 0 | yes | no | no | no | no | yes | no | no | no | no | no | no | no |
| KCNMA1 | 562 | S | 0 | 0 | yes | no | no | no | no | yes | no | no | no | no | no | no | no |
| KCNMA1 | 563 | C | 0 | 0 | yes | no | no | no | no | yes | no | no | no | no | no | no | no |
| KCNMA1 | 564 | L | 0 | 0 | yes | no | no | no | no | yes | no | no | no | no | no | no | no |
| KCNMA1 | 565 | A | 0 | 0 | no | yes | no | no | no | yes | no | no | no | no | no | no | no |
| KCNMA1 | 566 | Q | 0 | 0 | no | yes | no | no | no | yes | no | no | no | no | no | no | no |
| KCNMA1 | 567 | G | 0 | 0 | no | yes | no | no | no | yes | no | no | no | no | no | no | no |
| KCNMA1 | 568 | L | 0 | 0 | yes | no | no | no | no | yes | no | no | no | no | no | no | no |
| KCNMA1 | 569 | S | 0 | 0 | yes | no | no | no | no | yes | no | no | no | no | no | no | no |
| KCNMA1 | 570 | T | 0 | 0 | yes | no | no | no | no | yes | no | no | no | no | no | no | no |
| KCNMA1 | 571 | M | 0 | 0 | yes | no | no | no | no | yes | no | no | no | no | no | no | no |
| KCNMA1 | 572 | L | 0 | 0 | yes | no | no | no | no | yes | no | no | no | no | no | no | no |
| KCNMA1 | 573 | A | 0 | 0 | yes | no | no | no | no | yes | no | no | no | no | no | no | no |
| KCNMA1 | 574 | N | 0 | 0 | yes | no | no | no | no | yes | no | no | no | no | no | no | no |
| KCNMA1 | 575 | L | 0 | 0 | no | yes | no | no | no | yes | no | no | no | no | no | no | no |
| KCNMA1 | 576 | F | 0 | 0 | no | yes | no | no | no | yes | no | no | no | no | no | no | no |
| KCNMA1 | 577 | S | 0 | 0 | no | yes | no | no | no | yes | no | no | no | no | no | no | no |
| KCNMA1 | 578 | M | 0 | 0 | no | yes | no | no | no | yes | no | no | no | no | no | no | no |
| KCNMA1 | 579 | R | 0 | 0 | no | yes | no | no | no | yes | no | no | no | no | no | no | no |

| | | | | | | | | | | | | | | | | | |
|---|---|---|---|---|---|---|---|---|---|---|---|---|---|---|---|---|---|
| KCNMA1 | 580 | S | 0 | 0 | no | yes | no | no | no | yes | no | no | no | no | no | no | no |
| KCNMA1 | 581 | F | 0 | 0 | no | yes | no | no | no | yes | no | no | no | no | no | no | no |
| KCNMA1 | 582 | I | 0 | 0 | no | yes | no | no | no | yes | no | no | no | no | no | no | no |
| KCNMA1 | 583 | K | 0 | 0 | no | yes | no | no | no | yes | no | no | no | no | no | no | no |
| KCNMA1 | 584 | I | 0 | 0 | no | yes | no | no | no | yes | no | no | no | no | no | no | no |
| KCNMA1 | 585 | E | 0 | 0 | no | yes | no | no | no | yes | no | no | no | no | no | no | no |
| KCNMA1 | 586 | E | 0 | 0 | no | yes | no | no | no | yes | no | no | no | no | no | no | no |
| KCNMA1 | 587 | D | 0 | 0 | no | yes | no | no | no | yes | no | no | no | no | no | no | no |
| KCNMA1 | 588 | T | 0 | 0 | no | yes | no | no | no | yes | no | no | no | no | no | no | no |
| KCNMA1 | 589 | W | 0 | 0 | yes | no | no | no | no | yes | no | no | no | no | no | no | no |
| KCNMA1 | 590 | Q | 0 | 0 | yes | no | no | no | no | yes | no | no | no | no | no | no | no |
| KCNMA1 | 591 | K | 0 | 0 | yes | no | no | no | no | yes | no | no | no | no | no | no | no |
| KCNMA1 | 592 | Y | 0 | 0 | yes | no | no | no | no | yes | no | no | no | no | no | no | no |
| KCNMA1 | 593 | Y | 0 | 0 | yes | no | no | no | no | yes | no | no | no | no | no | no | no |
| KCNMA1 | 594 | L | 0 | 0 | yes | no | no | no | no | yes | no | no | no | no | no | no | no |
| KCNMA1 | 595 | E | 0 | 0 | yes | no | no | no | no | yes | no | no | no | no | no | no | no |
| KCNMA1 | 596 | G | 0 | 0 | yes | no | no | no | no | yes | no | no | no | no | no | no | no |
| KCNMA1 | 597 | V | 0 | 0 | yes | no | no | no | no | yes | no | no | no | no | no | no | no |
| KCNMA1 | 598 | S | 0 | 0 | no | yes | no | no | no | yes | no | no | no | no | no | no | no |
| KCNMA1 | 599 | N | 1 | 0 | no | yes | no | no | no | yes | no | no | no | no | no | no | no |
| KCNMA1 | 600 | E | 0 | 0 | no | no | yes | no | no | yes | no | no | no | no | no | no | no |
| KCNMA1 | 601 | M | 0 | 0 | no | no | yes | no | no | yes | no | no | no | no | no | no | no |
| KCNMA1 | 602 | Y | 0 | 0 | no | no | yes | no | no | yes | no | no | no | no | no | no | no |
| KCNMA1 | 603 | T | 0 | 0 | no | no | yes | no | no | yes | no | no | no | no | no | no | no |
| KCNMA1 | 604 | E | 0 | 0 | no | no | yes | no | no | yes | no | no | no | no | no | no | no |
| KCNMA1 | 605 | Y | 0 | 0 | no | no | yes | no | no | yes | no | no | no | no | no | no | no |
| KCNMA1 | 606 | L | 0 | 0 | no | yes | no | no | no | yes | no | no | no | no | no | no | no |
| KCNMA1 | 607 | S | 0 | 0 | no | yes | no | no | no | yes | no | no | no | no | no | no | no |
| KCNMA1 | 608 | S | 1 | 0 | yes | no | no | no | no | yes | no | no | no | no | no | no | no |
| KCNMA1 | 609 | A | 0 | 0 | yes | no | no | no | no | yes | no | no | no | no | no | no | no |
| KCNMA1 | 610 | F | 0 | 0 | yes | no | no | no | no | yes | no | no | no | no | no | no | no |
| KCNMA1 | 611 | V | 1 | 0 | no | yes | no | no | no | yes | no | no | no | no | no | no | no |
| KCNMA1 | 612 | G | 0 | 0 | no | yes | no | no | no | yes | no | no | no | no | no | no | no |

| | | | | | | | | | | | | | | | | | |
|---|---|---|---|---|---|---|---|---|---|---|---|---|---|---|---|---|---|
| KCNMA1 | 613 | L | 1 | 0 | no | yes | no | no | no | yes | no | no | no | no | no | no | no |
| KCNMA1 | 614 | S | 0 | 0 | no | no | yes | no | no | yes | no | no | no | no | no | no | no |
| KCNMA1 | 615 | F | 0 | 0 | yes | no | no | no | no | yes | no | no | no | no | no | no | no |
| KCNMA1 | 616 | P | 0 | 0 | yes | no | no | no | no | yes | no | no | no | no | no | no | no |
| KCNMA1 | 617 | T | 2 | 0 | yes | no | no | no | no | yes | no | no | no | no | no | no | no |
| KCNMA1 | 618 | V | 0 | 0 | yes | no | no | no | no | yes | no | no | no | no | no | no | no |
| KCNMA1 | 619 | C | 0 | 0 | yes | no | no | no | no | yes | no | no | no | no | no | no | no |
| KCNMA1 | 620 | E | 1 | 0 | yes | no | no | no | no | yes | no | no | no | no | no | no | no |
| KCNMA1 | 621 | L | 0 | 0 | yes | no | no | no | no | yes | no | no | no | no | no | no | no |
| KCNMA1 | 622 | C | 0 | 0 | yes | no | no | no | no | yes | no | no | no | no | no | no | no |
| KCNMA1 | 623 | F | 0 | 0 | yes | no | no | no | no | yes | no | no | no | no | no | no | no |
| KCNMA1 | 624 | V | 0 | 0 | yes | no | no | no | no | yes | no | no | no | no | no | no | no |
| KCNMA1 | 625 | K | 0 | 0 | no | yes | no | no | no | yes | no | no | no | no | no | no | no |
| KCNMA1 | 626 | L | 0 | 0 | no | yes | no | no | no | yes | no | no | no | no | no | no | no |
| KCNMA1 | 627 | K | 0 | 0 | no | yes | no | no | no | yes | no | no | no | no | no | no | no |
| KCNMA1 | 628 | L | 0 | 0 | no | yes | no | no | no | yes | no | no | no | no | no | no | no |
| KCNMA1 | 629 | L | 0 | 0 | no | no | yes | no | no | yes | no | no | no | no | no | no | no |
| KCNMA1 | 630 | M | 0 | 0 | no | no | yes | no | no | yes | no | no | no | no | no | no | no |
| KCNMA1 | 631 | I | 0 | 0 | no | no | yes | no | no | yes | no | no | no | no | no | no | no |
| KCNMA1 | 632 | A | 0 | 0 | no | no | yes | no | no | yes | no | no | no | no | no | no | no |
| KCNMA1 | 633 | I | 1 | 0 | no | no | yes | no | no | yes | no | no | no | no | no | no | no |
| KCNMA1 | 634 | E | 0 | 0 | no | no | yes | no | no | yes | no | no | no | no | no | no | no |
| KCNMA1 | 635 | Y | 0 | 0 | no | no | yes | no | no | yes | no | no | no | no | no | no | no |
| KCNMA1 | 636 | K | 0 | 0 | no | yes | no | no | no | yes | no | no | no | no | no | no | no |
| KCNMA1 | 637 | S | 0 | 0 | no | yes | no | no | no | yes | no | no | no | no | no | no | no |
| KCNMA1 | 638 | A | 0 | 0 | no | yes | no | no | no | yes | no | no | no | no | no | no | no |
| KCNMA1 | 639 | N | 0 | 0 | no | yes | no | no | no | yes | no | no | no | no | no | no | no |
| KCNMA1 | 640 | R | 1 | 0 | no | yes | no | no | no | yes | no | no | no | no | no | no | no |
| KCNMA1 | 641 | E | 0 | 0 | no | yes | no | no | no | yes | no | no | no | no | no | no | no |
| KCNMA1 | 642 | S | 0 | 0 | no | yes | no | no | no | yes | no | no | no | no | no | no | no |
| KCNMA1 | 643 | R | 0 | 0 | no | no | yes | no | no | yes | no | no | no | no | no | no | no |
| KCNMA1 | 644 | I | 0 | 0 | no | no | yes | no | no | yes | no | no | no | no | no | no | no |
| KCNMA1 | 645 | L | 0 | 0 | no | no | yes | no | no | yes | no | no | no | no | no | no | no |

| | | | | | | | | | | | | | | | | | |
|---|---|---|---|---|---|---|---|---|---|---|---|---|---|---|---|---|---|
| KCNMA1 | 646 | I | 0 | 0 | no | no | yes | no | no | yes | no | no | no | no | no | no | no |
| KCNMA1 | 647 | N | 0 | 0 | no | yes | no | no | no | yes | no | no | no | no | no | no | no |
| KCNMA1 | 648 | P | 0 | 0 | no | yes | no | no | no | yes | no | no | no | no | no | no | no |
| KCNMA1 | 649 | G | 0 | 0 | no | yes | no | no | no | yes | no | no | no | no | no | no | no |
| KCNMA1 | 650 | N | 0 | 0 | no | yes | no | no | no | yes | no | no | no | no | no | no | no |
| KCNMA1 | 651 | H | 1 | 0 | no | yes | no | no | no | yes | no | no | no | no | no | no | no |
| KCNMA1 | 652 | L | 0 | 0 | no | yes | no | no | no | yes | no | no | no | no | no | no | no |
| KCNMA1 | 653 | K | 1 | 0 | no | no | yes | no | no | yes | no | no | no | no | no | no | no |
| KCNMA1 | 654 | I | 0 | 0 | no | yes | no | no | no | yes | no | no | no | no | no | no | no |
| KCNMA1 | 655 | Q | 0 | 0 | no | yes | no | no | no | yes | no | no | no | no | no | no | no |
| KCNMA1 | 656 | E | 1 | 0 | no | yes | no | no | no | yes | no | no | no | no | no | no | no |
| KCNMA1 | 657 | G | 0 | 0 | no | yes | no | no | no | yes | no | no | no | no | no | no | no |
| KCNMA1 | 658 | T | 0 | 0 | no | yes | no | no | no | yes | no | no | no | no | no | no | no |
| KCNMA1 | 659 | L | 0 | 0 | no | no | yes | no | no | yes | no | no | no | no | no | no | no |
| KCNMA1 | 660 | G | 0 | 0 | no | no | yes | no | no | yes | no | no | no | no | no | no | no |
| KCNMA1 | 661 | F | 0 | 0 | no | no | yes | no | no | yes | no | no | no | no | no | no | no |
| KCNMA1 | 662 | F | 0 | 0 | no | no | yes | no | no | yes | no | no | no | no | no | no | no |
| KCNMA1 | 663 | I | 1 | 1 | no | no | yes | no | no | yes | no | no | no | no | no | no | no |
| KCNMA1 | 664 | A | 0 | 0 | no | no | yes | no | no | yes | no | no | no | no | no | no | no |
| KCNMA1 | 665 | S | 0 | 0 | no | yes | no | no | no | yes | no | no | no | no | no | no | no |
| KCNMA1 | 666 | D | 0 | 0 | no | yes | no | no | no | yes | no | no | no | no | no | no | no |
| KCNMA1 | 667 | A | 0 | 0 | yes | no | no | no | no | yes | no | no | no | no | no | no | no |
| KCNMA1 | 668 | K | 0 | 0 | yes | no | no | no | no | yes | no | no | no | no | no | no | no |
| KCNMA1 | 669 | E | 0 | 0 | yes | no | no | no | no | yes | no | no | no | no | no | no | no |
| KCNMA1 | 670 | V | 0 | 0 | yes | no | no | no | no | yes | no | no | no | no | no | no | no |
| KCNMA1 | 671 | K | 0 | 0 | yes | no | no | no | no | yes | no | no | no | no | no | no | no |
| KCNMA1 | 672 | R | 0 | 0 | yes | no | no | no | no | yes | no | no | no | no | no | no | no |
| KCNMA1 | 673 | A | 0 | 0 | yes | no | no | no | no | yes | no | no | no | no | no | no | no |
| KCNMA1 | 674 | F | 1 | 0 | yes | no | no | no | no | yes | no | no | no | no | no | no | no |
| KCNMA1 | 675 | F | 0 | 0 | no | yes | no | no | no | yes | no | no | no | no | no | no | no |
| KCNMA1 | 676 | Y | 0 | 0 | no | yes | no | no | no | yes | no | no | no | no | no | no | no |
| KCNMA1 | 677 | C | 0 | 0 | no | yes | no | no | no | yes | no | no | no | no | no | no | no |
| KCNMA1 | 678 | K | 0 | 0 | no | yes | no | no | no | yes | no | no | no | no | no | no | no |

| | | | | | | | | | | | | | | | | | |
|---|---|---|---|---|---|---|---|---|---|---|---|---|---|---|---|---|---|
| KCNMA1 | 679 | A | 1 | 0 | no | yes | no | no | no | yes | no | no | no | no | no | no | no |
| KCNMA1 | 680 | C | 0 | 0 | no | yes | no | no | no | yes | no | no | no | no | no | no | no |
| KCNMA1 | 681 | H | 0 | 0 | no | yes | no | no | no | yes | no | no | no | no | no | no | no |
| KCNMA1 | 682 | D | 0 | 0 | no | yes | no | no | no | yes | no | no | no | no | no | no | no |
| KCNMA1 | 683 | D | 0 | 0 | no | yes | no | no | no | yes | no | no | no | no | no | no | no |
| KCNMA1 | 684 | I | 0 | 0 | no | yes | no | no | no | yes | no | no | no | no | no | no | no |
| KCNMA1 | 685 | T | 1 | 0 | no | yes | no | no | no | yes | no | no | no | no | no | no | no |
| KCNMA1 | 686 | D | 1 | 0 | no | yes | no | no | no | yes | no | no | no | no | no | no | no |
| KCNMA1 | 687 | P | 0 | 0 | yes | no | no | no | no | yes | no | no | no | no | no | no | no |
| KCNMA1 | 688 | K | 0 | 0 | yes | no | no | no | no | yes | no | no | no | no | no | no | no |
| KCNMA1 | 689 | R | 1 | 0 | yes | no | no | no | no | yes | no | no | no | no | no | no | no |
| KCNMA1 | 690 | I | 0 | 0 | yes | no | no | no | no | yes | no | no | no | no | no | no | no |
| KCNMA1 | 691 | K | 0 | 0 | no | yes | no | no | no | yes | no | no | no | no | no | no | no |
| KCNMA1 | 692 | K | 0 | 0 | no | yes | no | no | no | yes | no | no | no | no | no | no | no |
| KCNMA1 | 693 | C | 0 | 0 | no | yes | no | no | no | yes | no | no | no | no | no | no | no |
| KCNMA1 | 694 | G | 2 | 0 | no | yes | no | no | no | yes | no | no | no | no | no | no | no |
| KCNMA1 | 695 | C | 0 | 0 | no | yes | no | no | no | yes | no | no | no | no | no | no | no |
| KCNMA1 | 696 | K | 0 | 0 | no | yes | no | no | no | yes | no | no | no | no | no | no | no |
| KCNMA1 | 697 | R | 2 | 0 | no | yes | no | no | no | yes | no | no | no | no | no | no | no |
| KCNMA1 | 804 | S | 0 | 0 | no | yes | no | no | no | yes | no | no | no | no | no | no | no |
| KCNMA1 | 805 | N | 1 | 0 | no | yes | no | no | no | yes | no | no | no | no | no | no | no |
| KCNMA1 | 806 | V | 0 | 0 | no | yes | no | no | no | yes | no | no | no | no | no | no | no |
| KCNMA1 | 807 | K | 0 | 0 | no | yes | no | no | no | yes | no | no | no | no | no | no | no |
| KCNMA1 | 808 | K | 2 | 0 | no | yes | no | no | no | yes | no | no | no | no | no | no | no |
| KCNMA1 | 809 | Y | 0 | 0 | no | no | yes | no | no | yes | no | no | no | no | no | no | no |
| KCNMA1 | 810 | D | 1 | 0 | no | yes | no | no | no | yes | no | no | no | no | no | no | no |
| KCNMA1 | 811 | S | 0 | 0 | no | yes | no | no | no | yes | no | no | no | no | no | no | no |
| KCNMA1 | 812 | T | 0 | 0 | no | yes | no | no | no | yes | no | no | no | no | no | no | no |
| KCNMA1 | 813 | G | 0 | 0 | no | yes | no | no | no | yes | no | no | no | no | no | no | no |
| KCNMA1 | 814 | M | 0 | 0 | no | yes | no | no | no | yes | no | no | no | no | no | no | no |
| KCNMA1 | 815 | F | 0 | 0 | no | yes | no | no | no | yes | no | no | no | no | no | no | no |
| KCNMA1 | 816 | H | 0 | 0 | no | no | yes | no | no | yes | no | no | no | no | no | no | no |
| KCNMA1 | 817 | W | 0 | 0 | no | yes | no | no | no | yes | no | no | no | no | no | no | no |

| | | | | | | | | | | | | | | | | | |
|---|---|---|---|---|---|---|---|---|---|---|---|---|---|---|---|---|---|
| KCNMA1 | 818 | C | 0 | 0 | no | yes | no | no | no | yes | no | no | no | no | no | no | no |
| KCNMA1 | 819 | A | 0 | 0 | no | yes | no | no | no | yes | no | no | no | no | no | no | no |
| KCNMA1 | 820 | P | 1 | 0 | no | yes | no | no | no | yes | no | no | no | no | no | no | no |
| KCNMA1 | 821 | K | 1 | 0 | no | yes | no | no | no | yes | no | no | no | no | no | no | no |
| KCNMA1 | 822 | E | 0 | 0 | no | yes | no | no | no | yes | no | no | no | no | no | no | no |
| KCNMA1 | 823 | I | 0 | 0 | no | yes | no | no | no | yes | no | no | no | no | no | no | no |
| KCNMA1 | 824 | E | 0 | 0 | no | yes | no | no | no | yes | no | no | no | no | no | no | no |
| KCNMA1 | 825 | K | 0 | 0 | no | yes | no | no | no | yes | no | no | no | no | no | no | no |
| KCNMA1 | 826 | V | 0 | 0 | no | yes | no | no | no | yes | no | no | no | no | no | no | no |
| KCNMA1 | 827 | I | 3 | 0 | no | no | yes | no | no | yes | no | no | no | no | no | no | no |
| KCNMA1 | 828 | L | 0 | 0 | no | yes | no | no | no | yes | no | no | no | no | no | no | no |
| KCNMA1 | 829 | T | 0 | 0 | no | yes | no | no | no | yes | no | no | no | no | no | no | no |
| KCNMA1 | 830 | R | 1 | 0 | yes | no | no | no | no | yes | no | no | no | no | no | no | no |
| KCNMA1 | 831 | S | 1 | 0 | yes | no | no | no | no | yes | no | no | no | no | no | no | no |
| KCNMA1 | 832 | E | 0 | 0 | yes | no | no | no | no | yes | no | no | no | no | no | no | no |
| KCNMA1 | 833 | A | 1 | 0 | yes | no | no | no | no | yes | no | no | no | no | no | no | no |
| KCNMA1 | 834 | A | 2 | 0 | yes | no | no | no | no | yes | no | no | no | no | no | no | no |
| KCNMA1 | 835 | M | 0 | 0 | yes | no | no | no | no | yes | no | no | no | no | no | no | no |
| KCNMA1 | 836 | T | 1 | 0 | no | yes | no | no | no | yes | no | no | no | no | no | no | no |
| KCNMA1 | 837 | V | 1 | 0 | no | yes | no | no | no | yes | no | no | no | no | no | no | no |
| KCNMA1 | 838 | L | 0 | 0 | no | yes | no | no | no | yes | no | no | no | no | no | no | no |
| KCNMA1 | 839 | S | 0 | 0 | no | yes | no | no | no | yes | no | no | no | no | no | no | no |
| KCNMA1 | 840 | G | 0 | 0 | no | yes | no | no | no | yes | no | no | no | no | no | no | no |
| KCNMA1 | 841 | H | 0 | 0 | no | yes | no | no | no | yes | no | no | no | no | no | no | no |
| KCNMA1 | 842 | V | 0 | 0 | no | no | yes | no | no | yes | no | no | no | no | no | no | no |
| KCNMA1 | 843 | V | 1 | 0 | no | no | yes | no | no | yes | no | no | no | no | no | no | no |
| KCNMA1 | 844 | V | 0 | 0 | no | no | yes | no | no | yes | no | no | no | no | no | no | no |
| KCNMA1 | 845 | C | 0 | 0 | no | no | yes | no | no | yes | no | no | no | no | no | no | no |
| KCNMA1 | 846 | I | 0 | 0 | no | no | yes | no | no | yes | no | no | no | no | no | no | no |
| KCNMA1 | 847 | F | 0 | 0 | no | yes | no | no | no | yes | no | no | no | no | no | no | no |
| KCNMA1 | 848 | G | 0 | 0 | no | yes | no | no | no | yes | no | no | no | no | no | no | no |
| KCNMA1 | 849 | D | 2 | 0 | no | yes | no | no | no | yes | no | no | no | no | no | no | no |
| KCNMA1 | 850 | V | 1 | 0 | no | yes | no | no | no | yes | no | no | no | no | no | no | no |

| | | | | | | | | | | | | | | | | | |
|---|---|---|---|---|---|---|---|---|---|---|---|---|---|---|---|---|---|
| KCNMA1 | 851 | S | 0 | 0 | no | yes | no | no | no | yes | yes | no | no | no | no | no | no |
| KCNMA1 | 852 | S | 0 | 0 | no | yes | no | no | no | yes | no | no | no | no | no | no | no |
| KCNMA1 | 853 | A | 0 | 0 | no | yes | no | no | no | yes | no | no | no | no | no | no | no |
| KCNMA1 | 854 | L | 0 | 0 | no | yes | no | no | no | yes | no | no | no | no | no | no | no |
| KCNMA1 | 855 | I | 1 | 0 | no | yes | no | no | no | yes | no | no | no | no | no | no | no |
| KCNMA1 | 856 | G | 1 | 0 | no | yes | no | no | no | yes | no | no | no | no | no | no | no |
| KCNMA1 | 857 | L | 0 | 0 | no | yes | no | no | no | yes | no | no | no | no | no | no | no |
| KCNMA1 | 858 | R | 1 | 0 | yes | no | no | no | no | yes | no | no | no | no | no | no | no |
| KCNMA1 | 859 | N | 0 | 0 | yes | no | no | no | no | yes | no | no | no | no | no | no | no |
| KCNMA1 | 860 | L | 0 | 0 | yes | no | no | no | no | yes | no | no | no | no | no | no | no |
| KCNMA1 | 861 | V | 0 | 0 | yes | no | no | no | no | yes | no | no | no | no | no | no | no |
| KCNMA1 | 862 | M | 0 | 0 | yes | no | no | no | no | yes | no | no | no | no | no | no | no |
| KCNMA1 | 863 | P | 1 | 1 | yes | no | no | no | no | yes | no | no | no | no | no | no | no |
| KCNMA1 | 864 | L | 0 | 0 | yes | no | no | no | no | yes | no | no | no | no | no | no | no |
| KCNMA1 | 865 | R | 0 | 0 | no | yes | no | no | no | yes | no | no | no | no | no | no | no |
| KCNMA1 | 866 | A | 0 | 0 | no | yes | no | no | no | yes | no | no | no | no | no | no | no |
| KCNMA1 | 867 | S | 0 | 0 | no | yes | no | no | no | yes | no | no | no | no | no | no | no |
| KCNMA1 | 868 | N | 0 | 0 | no | yes | no | no | no | yes | no | no | no | no | no | no | no |
| KCNMA1 | 869 | F | 0 | 0 | no | yes | no | no | no | yes | no | no | no | no | no | no | no |
| KCNMA1 | 870 | H | 0 | 0 | no | yes | no | no | no | yes | no | no | no | no | no | no | no |
| KCNMA1 | 871 | Y | 0 | 0 | no | yes | no | no | no | yes | no | no | no | no | no | no | no |
| KCNMA1 | 872 | H | 1 | 0 | no | yes | no | no | no | yes | no | no | no | no | no | no | no |
| KCNMA1 | 873 | E | 0 | 0 | no | yes | no | no | no | yes | no | no | no | no | no | no | no |
| KCNMA1 | 874 | L | 0 | 0 | no | yes | no | no | no | yes | no | no | no | no | no | no | no |
| KCNMA1 | 875 | K | 0 | 0 | no | yes | no | no | no | yes | no | no | no | no | no | no | no |
| KCNMA1 | 876 | H | 0 | 0 | no | yes | no | no | no | yes | no | no | no | no | no | no | no |
| KCNMA1 | 877 | I | 1 | 0 | no | no | yes | no | no | yes | no | no | no | no | no | no | no |
| KCNMA1 | 878 | V | 0 | 0 | no | no | yes | no | no | yes | no | no | no | no | no | no | no |
| KCNMA1 | 879 | F | 0 | 0 | no | no | yes | no | no | yes | no | no | no | no | no | no | no |
| KCNMA1 | 880 | V | 0 | 0 | no | no | yes | no | no | yes | no | no | no | no | no | no | no |
| KCNMA1 | 881 | G | 0 | 0 | no | no | yes | no | no | yes | no | no | no | no | no | no | no |
| KCNMA1 | 882 | S | 1 | 0 | no | yes | no | no | no | yes | no | no | no | no | no | no | no |
| KCNMA1 | 883 | I | 1 | 0 | yes | no | no | no | no | yes | no | no | no | no | no | no | no |

| | | | | | | | | | | | | | | | | | |
|---|---|---|---|---|---|---|---|---|---|---|---|---|---|---|---|---|---|
| KCNMA1 | 884 | E | 0 | 0 | yes | no | no | no | no | yes | no | no | no | no | no | no | no |
| KCNMA1 | 885 | Y | 1 | 0 | yes | no | no | no | no | yes | no | no | no | no | no | no | no |
| KCNMA1 | 886 | L | 0 | 0 | yes | no | no | no | no | yes | no | no | no | no | no | no | no |
| KCNMA1 | 887 | K | 2 | 0 | no | yes | no | no | no | yes | no | no | no | no | no | no | no |
| KCNMA1 | 888 | R | 0 | 0 | no | yes | no | no | no | yes | no | no | no | no | no | no | no |
| KCNMA1 | 889 | E | 0 | 0 | no | yes | no | no | no | yes | no | no | no | no | no | no | no |
| KCNMA1 | 890 | W | 0 | 0 | yes | no | no | no | no | yes | no | no | no | no | no | no | no |
| KCNMA1 | 891 | E | 0 | 0 | yes | no | no | no | no | yes | no | no | no | no | no | no | no |
| KCNMA1 | 892 | T | 1 | 0 | yes | no | no | no | no | yes | no | no | no | no | no | no | no |
| KCNMA1 | 893 | L | 0 | 0 | no | yes | no | no | no | yes | no | no | no | no | no | no | no |
| KCNMA1 | 894 | H | 0 | 0 | no | yes | no | no | no | yes | no | no | no | no | no | no | no |
| KCNMA1 | 895 | N | 0 | 0 | no | yes | no | no | no | yes | no | no | no | no | no | no | no |
| KCNMA1 | 896 | F | 0 | 0 | no | yes | no | no | no | yes | no | no | no | no | no | no | no |
| KCNMA1 | 897 | P | 0 | 0 | no | yes | no | no | no | yes | no | no | no | no | no | no | no |
| KCNMA1 | 898 | K | 1 | 0 | no | yes | no | no | no | yes | no | no | no | no | no | no | no |
| KCNMA1 | 899 | V | 0 | 0 | no | no | yes | no | no | yes | no | no | no | no | no | no | no |
| KCNMA1 | 900 | S | 0 | 0 | no | no | yes | no | no | yes | no | no | no | no | no | no | no |
| KCNMA1 | 901 | I | 1 | 0 | no | no | yes | no | no | yes | no | no | no | no | no | no | no |
| KCNMA1 | 902 | L | 0 | 0 | no | no | yes | no | no | yes | no | no | no | no | no | no | no |
| KCNMA1 | 903 | P | 0 | 0 | no | no | yes | no | no | yes | no | no | no | no | no | no | no |
| KCNMA1 | 904 | G | 0 | 0 | no | yes | no | no | no | yes | no | no | no | no | no | no | no |
| KCNMA1 | 905 | T | 1 | 0 | no | yes | no | no | no | yes | no | no | no | no | no | no | no |
| KCNMA1 | 906 | P | 0 | 0 | no | yes | no | no | no | yes | no | no | no | no | no | no | no |
| KCNMA1 | 907 | L | 0 | 0 | no | yes | no | no | no | yes | no | no | no | no | no | no | no |
| KCNMA1 | 908 | S | 0 | 0 | no | yes | no | no | no | yes | no | no | no | no | no | no | no |
| KCNMA1 | 909 | R | 1 | 0 | yes | no | no | no | no | yes | no | no | yes | no | no | no | no |
| KCNMA1 | 910 | A | 0 | 0 | yes | no | no | no | no | yes | no | no | yes | no | no | no | no |
| KCNMA1 | 911 | D | 1 | 0 | yes | no | no | no | no | yes | no | no | no | no | no | no | no |
| KCNMA1 | 912 | L | 0 | 0 | yes | no | no | no | no | yes | no | no | no | no | no | no | no |
| KCNMA1 | 913 | R | 0 | 0 | yes | no | no | no | no | yes | no | no | yes | no | no | no | no |
| KCNMA1 | 914 | A | 0 | 0 | yes | no | no | no | no | yes | no | no | no | no | no | no | no |
| KCNMA1 | 915 | V | 0 | 0 | no | yes | no | no | no | yes | no | no | no | no | no | no | no |
| KCNMA1 | 916 | N | 1 | 0 | no | yes | no | no | no | yes | no | no | no | no | no | no | no |

| | | | | | | | | | | | | | | | | | |
|---|---|---|---|---|---|---|---|---|---|---|---|---|---|---|---|---|---|
| KCNMA1 | 917 | I | 0 | 0 | no | yes | no | no | no | yes | no | no | no | no | no | no | no |
| KCNMA1 | 918 | N | 1 | 0 | no | yes | no | no | no | yes | no | no | no | no | no | no | no |
| KCNMA1 | 919 | L | 0 | 0 | no | yes | no | no | no | yes | no | no | no | no | no | no | no |
| KCNMA1 | 920 | C | 0 | 0 | no | yes | no | no | no | yes | no | no | no | no | no | no | no |
| KCNMA1 | 921 | D | 0 | 0 | no | yes | no | no | no | yes | no | no | no | no | no | no | no |
| KCNMA1 | 922 | M | 0 | 0 | no | yes | no | no | no | yes | no | no | no | no | no | no | no |
| KCNMA1 | 923 | C | 0 | 0 | no | no | yes | no | no | yes | no | no | no | no | no | no | no |
| KCNMA1 | 924 | V | 1 | 0 | no | no | yes | no | no | yes | no | no | no | no | no | no | no |
| KCNMA1 | 925 | I | 0 | 0 | no | no | yes | no | no | yes | no | no | no | no | no | no | no |
| KCNMA1 | 926 | L | 0 | 0 | no | no | yes | no | no | yes | no | no | no | no | no | no | no |
| KCNMA1 | 927 | S | 1 | 0 | no | yes | no | no | no | yes | no | no | no | no | no | no | no |
| KCNMA1 | 928 | A | 0 | 0 | no | yes | no | no | no | yes | no | no | no | no | no | no | no |
| KCNMA1 | 929 | N | 1 | 0 | no | yes | no | no | no | yes | no | no | no | no | no | no | no |
| KCNMA1 | 930 | Q | 0 | 0 | no | yes | no | no | no | yes | no | no | no | no | no | no | no |
| KCNMA1 | 931 | N | 0 | 0 | no | yes | no | no | no | yes | no | no | no | no | no | no | no |
| KCNMA1 | 932 | N | 0 | 0 | no | yes | no | no | no | yes | yes | no | no | no | no | no | no |
| KCNMA1 | 933 | I | 1 | 0 | no | yes | no | no | no | yes | yes | no | no | no | no | no | no |
| KCNMA1 | 934 | D | 0 | 0 | no | yes | no | no | no | yes | no | no | no | no | no | no | no |
| KCNMA1 | 935 | D | 0 | 0 | no | yes | no | no | no | yes | no | no | no | no | no | no | no |
| KCNMA1 | 936 | T | 1 | 0 | no | yes | no | no | no | yes | no | no | no | no | no | no | no |
| KCNMA1 | 937 | S | 0 | 0 | yes | no | no | no | no | yes | no | no | yes | no | no | no | no |
| KCNMA1 | 938 | L | 0 | 0 | yes | no | no | no | no | yes | no | no | yes | no | no | no | no |
| KCNMA1 | 939 | Q | 0 | 0 | yes | no | no | no | no | yes | no | no | no | no | no | no | no |
| KCNMA1 | 940 | D | 0 | 0 | yes | no | no | no | no | yes | no | no | no | no | no | no | no |
| KCNMA1 | 941 | K | 0 | 0 | yes | no | no | no | no | yes | no | no | yes | no | no | no | no |
| KCNMA1 | 942 | E | 0 | 1 | yes | no | no | no | no | yes | no | no | no | no | no | no | no |
| KCNMA1 | 943 | C | 0 | 0 | yes | no | no | no | no | yes | no | no | no | no | no | no | no |
| KCNMA1 | 944 | I | 0 | 0 | yes | no | no | no | no | yes | no | no | yes | no | no | no | no |
| KCNMA1 | 945 | L | 0 | 0 | yes | no | no | no | no | yes | no | no | yes | no | no | no | no |
| KCNMA1 | 946 | A | 1 | 0 | yes | no | no | no | no | yes | no | no | no | no | no | no | no |
| KCNMA1 | 947 | S | 0 | 0 | yes | no | no | no | no | yes | no | no | no | no | no | no | no |
| KCNMA1 | 948 | L | 0 | 0 | yes | no | no | no | no | yes | no | no | yes | no | no | no | no |
| KCNMA1 | 949 | N | 0 | 1 | yes | no | no | no | no | yes | no | no | yes | no | no | no | no |

| | | | | | | | | | | | | | | | | | |
|---|---|---|---|---|---|---|---|---|---|---|---|---|---|---|---|---|---|
| KCNMA1 | 950 | I | 1 | 0 | yes | no | no | no | no | yes | no | no | no | no | no | no | no |
| KCNMA1 | 951 | K | 0 | 0 | yes | no | no | no | no | yes | no | no | no | no | no | no | no |
| KCNMA1 | 952 | S | 0 | 0 | yes | no | no | no | no | yes | no | no | yes | no | no | no | no |
| KCNMA1 | 953 | M | 0 | 0 | no | yes | no | no | no | yes | no | no | no | no | no | no | no |
| KCNMA1 | 954 | Q | 1 | 0 | no | no | yes | no | no | yes | no | no | no | no | no | no | no |
| KCNMA1 | 955 | F | 0 | 0 | no | yes | no | no | no | yes | no | no | no | no | no | no | no |
| KCNMA1 | 956 | D | 0 | 0 | no | yes | no | no | no | yes | no | no | no | no | no | no | no |
| KCNMA1 | 957 | D | 0 | 0 | no | yes | no | no | no | yes | no | no | no | no | no | no | no |
| KCNMA1 | 994 | T | 1 | 0 | no | yes | no | no | no | yes | no | no | no | no | no | no | no |
| KCNMA1 | 995 | T | 0 | 0 | no | no | yes | no | no | yes | no | no | no | no | no | no | no |
| KCNMA1 | 996 | G | 0 | 0 | yes | no | no | no | no | yes | no | no | no | no | no | no | no |
| KCNMA1 | 997 | V | 0 | 0 | yes | no | no | no | no | yes | no | no | no | no | no | no | no |
| KCNMA1 | 998 | N | 0 | 0 | yes | no | no | no | no | yes | no | no | no | no | no | no | no |
| KCNMA1 | 999 | I | 0 | 0 | no | yes | no | no | no | yes | no | no | no | no | no | no | no |
| KCNMA1 | 1000 | P | 1 | 0 | no | yes | no | no | no | yes | no | no | no | no | no | no | no |
| KCNMA1 | 1001 | I | 0 | 0 | no | no | yes | no | no | yes | no | no | no | no | no | no | no |
| KCNMA1 | 1002 | I | 0 | 0 | no | no | yes | no | no | yes | no | no | no | no | no | no | no |
| KCNMA1 | 1003 | T | 0 | 0 | no | no | yes | no | no | yes | no | no | no | no | no | no | no |
| KCNMA1 | 1004 | E | 0 | 0 | no | no | yes | no | no | yes | no | no | no | no | no | no | no |
| KCNMA1 | 1005 | L | 0 | 0 | no | no | yes | no | no | yes | no | no | no | no | no | no | no |
| KCNMA1 | 1006 | V | 0 | 0 | no | yes | no | no | no | yes | no | no | no | no | no | no | no |
| KCNMA1 | 1007 | N | 0 | 0 | no | yes | no | no | no | yes | no | no | no | no | no | no | no |
| KCNMA1 | 1008 | D | 0 | 1 | yes | no | no | no | no | yes | no | no | no | no | no | no | no |
| KCNMA1 | 1009 | T | 0 | 0 | yes | no | no | no | no | yes | no | no | no | no | no | no | no |
| KCNMA1 | 1010 | N | 0 | 0 | yes | no | no | no | no | yes | no | no | no | no | no | no | no |
| KCNMA1 | 1011 | V | 0 | 0 | no | yes | no | no | no | yes | no | no | no | no | no | no | no |
| KCNMA1 | 1012 | Q | 0 | 0 | no | yes | no | no | no | yes | no | no | yes | no | no | no | no |
| KCNMA1 | 1013 | F | 0 | 0 | no | yes | no | no | no | yes | no | no | yes | no | no | no | no |
| KCNMA1 | 1014 | L | 0 | 0 | no | yes | no | no | no | yes | no | no | no | no | no | no | no |
| KCNMA1 | 1015 | D | 0 | 0 | no | yes | no | no | no | yes | no | no | no | no | no | no | no |
| KCNMA1 | 1016 | Q | 0 | 0 | no | yes | no | no | no | yes | no | no | no | no | no | no | no |
| KCNMA1 | 1017 | D | 0 | 0 | no | yes | no | no | no | yes | no | no | no | no | no | no | no |
| KCNMA1 | 1018 | D | 0 | 0 | no | yes | no | no | no | yes | no | no | no | no | no | no | no |

| | | | | | | | | | | | | | | | | | |
|---|---|---|---|---|---|---|---|---|---|---|---|---|---|---|---|---|---|
| KCNMA1 | 1019 | D | 0 | 0 | no | yes | no | no | no | yes | no | no | no | no | no | no | no |
| KCNMA1 | 1020 | D | 0 | 0 | no | yes | no | no | no | yes | no | no | no | no | no | no | no |
| KCNMA1 | 1021 | D | 0 | 0 | no | yes | no | no | no | yes | no | no | no | no | no | no | no |
| KCNMA1 | 1022 | P | 0 | 0 | no | yes | no | no | no | yes | no | no | yes | no | no | no | no |
| KCNMA1 | 1023 | D | 0 | 0 | no | yes | no | no | no | yes | no | no | no | no | no | no | no |
| KCNMA1 | 1024 | T | 0 | 0 | no | yes | no | no | no | yes | no | no | no | no | no | no | no |
| KCNMA1 | 1025 | E | 0 | 0 | no | yes | no | no | no | yes | no | no | no | no | no | no | no |
| KCNMA1 | 1026 | L | 0 | 0 | no | yes | no | no | no | yes | no | no | no | no | no | no | no |
| KCNMA1 | 1027 | Y | 0 | 0 | no | yes | no | no | no | yes | no | no | no | no | no | no | no |
| KCNMA1 | 1028 | L | 0 | 0 | no | yes | no | no | no | yes | no | no | no | no | no | no | no |
| KCNMA1 | 1029 | T | 0 | 0 | no | yes | no | no | no | yes | no | no | no | no | no | no | no |
| KCNMA1 | 1030 | Q | 0 | 0 | yes | no | no | no | no | yes | no | no | no | no | no | no | no |
| KCNMA1 | 1031 | P | 0 | 0 | yes | no | no | no | no | yes | no | no | no | no | no | no | no |
| KCNMA1 | 1032 | F | 0 | 0 | yes | no | no | no | no | yes | no | no | no | no | no | no | no |
| KCNMA1 | 1033 | A | 0 | 0 | yes | no | no | no | no | yes | no | no | no | no | no | no | no |
| KCNMA1 | 1034 | C | 0 | 0 | yes | no | no | no | no | yes | no | no | no | no | no | no | no |
| KCNMA1 | 1035 | G | 0 | 0 | no | yes | no | no | no | yes | no | no | no | no | no | no | no |
| KCNMA1 | 1036 | T | 0 | 0 | no | yes | no | no | no | yes | no | no | no | no | no | no | no |
| KCNMA1 | 1037 | A | 0 | 0 | no | no | yes | no | no | yes | no | no | no | no | no | no | no |
| KCNMA1 | 1038 | F | 0 | 0 | no | no | yes | no | no | yes | no | no | no | no | no | no | no |
| KCNMA1 | 1039 | A | 0 | 0 | no | no | yes | no | no | yes | no | no | no | no | no | no | no |
| KCNMA1 | 1040 | V | 1 | 0 | no | yes | no | no | no | yes | no | no | no | no | no | no | no |
| KCNMA1 | 1041 | S | 0 | 0 | no | yes | no | no | no | yes | no | no | no | no | no | no | no |
| KCNMA1 | 1042 | V | 0 | 0 | no | yes | no | no | no | yes | no | no | no | no | no | no | no |
| KCNMA1 | 1043 | L | 0 | 0 | yes | no | no | no | no | yes | no | no | no | no | no | no | no |
| KCNMA1 | 1044 | D | 0 | 0 | yes | no | no | no | no | yes | no | no | no | no | no | no | no |
| KCNMA1 | 1045 | S | 0 | 0 | yes | no | no | no | no | yes | no | no | no | no | no | no | no |
| KCNMA1 | 1046 | L | 0 | 0 | yes | no | no | no | no | yes | no | no | no | no | no | no | no |
| KCNMA1 | 1047 | M | 0 | 0 | yes | no | no | no | no | yes | no | no | no | no | no | no | no |
| KCNMA1 | 1048 | S | 0 | 0 | yes | no | no | no | no | yes | no | no | no | no | no | no | no |
| KCNMA1 | 1049 | A | 1 | 0 | yes | no | no | no | no | yes | no | no | no | no | no | no | no |
| KCNMA1 | 1050 | T | 0 | 0 | yes | no | no | no | no | yes | no | no | no | no | no | no | no |
| KCNMA1 | 1051 | Y | 0 | 0 | yes | no | no | no | no | yes | no | no | no | no | no | no | no |

| | | | | | | | | | | | | | | | | | |
|---|---|---|---|---|---|---|---|---|---|---|---|---|---|---|---|---|---|
| KCNMA1 | 1052 | F | 0 | 0 | yes | no | no | no | no | yes | no | no | no | no | no | no | no |
| KCNMA1 | 1053 | N | 0 | 1 | no | yes | no | no | no | yes | no | no | no | no | no | no | no |
| KCNMA1 | 1054 | D | 0 | 0 | no | yes | no | no | no | yes | no | no | no | no | no | no | no |
| KCNMA1 | 1055 | N | 0 | 0 | no | yes | no | no | no | yes | no | no | no | no | no | no | no |
| KCNMA1 | 1056 | I | 0 | 0 | yes | no | no | no | no | yes | no | no | no | no | no | no | no |
| KCNMA1 | 1057 | L | 0 | 0 | yes | no | no | no | no | yes | no | no | no | no | no | no | no |
| KCNMA1 | 1058 | T | 1 | 0 | yes | no | no | no | no | yes | no | no | no | no | no | no | no |
| KCNMA1 | 1059 | L | 0 | 0 | yes | no | no | no | no | yes | no | no | no | no | no | no | no |
| KCNMA1 | 1060 | I | 0 | 0 | yes | no | no | no | no | yes | no | no | no | no | no | no | no |
| KCNMA1 | 1061 | R | 2 | 0 | yes | no | no | no | no | yes | no | no | no | no | no | no | no |
| KCNMA1 | 1062 | T | 1 | 0 | yes | no | no | no | no | yes | no | no | no | no | no | no | no |
| KCNMA1 | 1063 | L | 0 | 0 | yes | no | no | no | no | yes | no | no | no | no | no | no | no |
| KCNMA1 | 1064 | V | 0 | 0 | yes | no | no | no | no | yes | no | no | no | no | no | no | no |
| KCNMA1 | 1065 | T | 0 | 0 | no | yes | no | no | no | yes | no | no | no | no | no | no | no |
| KCNMA1 | 1066 | G | 1 | 0 | no | yes | no | no | no | yes | no | no | no | no | no | no | no |
| KCNMA1 | 1067 | G | 0 | 0 | no | yes | no | no | no | yes | no | no | no | no | no | no | no |
| KCNMA1 | 1068 | A | 0 | 0 | no | yes | no | no | no | yes | no | no | no | no | no | no | no |
| KCNMA1 | 1069 | T | 0 | 0 | no | yes | no | no | no | yes | no | no | no | no | no | no | no |
| KCNMA1 | 1070 | P | 0 | 0 | yes | no | no | no | no | yes | no | no | no | no | no | no | no |
| KCNMA1 | 1071 | E | 0 | 0 | yes | no | no | no | no | yes | no | no | no | no | no | no | no |
| KCNMA1 | 1072 | L | 0 | 0 | yes | no | no | no | no | yes | no | no | no | no | no | no | no |
| KCNMA1 | 1073 | E | 0 | 0 | yes | no | no | no | no | yes | no | no | no | no | no | no | no |
| KCNMA1 | 1074 | A | 1 | 0 | yes | no | no | no | no | yes | no | no | no | no | no | no | no |
| KCNMA1 | 1075 | L | 0 | 0 | yes | no | no | no | no | yes | no | no | no | no | no | no | no |
| KCNMA1 | 1076 | I | 0 | 0 | yes | no | no | no | no | yes | no | no | no | no | no | no | no |
| KCNMA1 | 1077 | A | 0 | 0 | no | yes | no | no | no | yes | no | no | no | no | no | no | no |
| KCNMA1 | 1078 | E | 0 | 0 | no | yes | no | no | no | yes | no | no | yes | no | no | no | no |
| KCNMA1 | 1079 | E | 0 | 0 | no | yes | no | no | no | yes | no | no | yes | no | no | no | no |
| KCNMA1 | 1080 | N | 0 | 0 | no | yes | no | no | no | yes | no | no | no | no | no | no | no |
| KCNMA1 | 1081 | A | 2 | 0 | no | yes | no | no | no | yes | no | no | no | no | no | no | no |
| KCNMA1 | 1082 | L | 0 | 0 | no | yes | no | no | no | yes | no | no | no | no | no | no | no |
| KCNMA1 | 1083 | R | 0 | 0 | no | yes | no | no | no | yes | no | no | no | no | no | no | no |
| KCNMA1 | 1084 | G | 0 | 0 | no | yes | no | no | no | yes | no | no | no | no | no | no | no |

| | | | | | | | | | | | | | | | | | |
|---|---|---|---|---|---|---|---|---|---|---|---|---|---|---|---|---|---|
| KCNMA1 | 1085 | G | 0 | 0 | no | yes | no | no | no | yes | no | no | no | no | no | no | no |
| KCNMA1 | 1086 | Y | 0 | 0 | no | yes | no | no | no | yes | no | no | no | no | no | no | no |
| KCNMA1 | 1087 | S | 0 | 0 | no | yes | no | no | no | yes | no | no | no | no | no | no | no |
| KCNMA1 | 1088 | T | 1 | 0 | no | yes | no | no | no | yes | no | no | no | no | no | no | no |
| KCNMA1 | 1089 | P | 1 | 0 | no | yes | no | no | no | yes | no | no | no | no | no | no | no |
| KCNMA1 | 1090 | Q | 0 | 0 | no | yes | no | no | no | yes | no | no | no | no | no | no | no |
| KCNMA1 | 1091 | T | 0 | 0 | no | yes | no | no | no | yes | no | no | no | no | no | no | no |
| KCNMA1 | 1092 | L | 0 | 0 | no | yes | no | no | no | yes | no | no | no | no | no | no | no |
| KCNMA1 | 1093 | A | 0 | 0 | no | yes | no | no | no | yes | no | no | no | no | no | no | no |
| KCNMA1 | 1094 | N | 1 | 0 | no | yes | no | no | no | yes | no | no | no | no | no | no | no |
| KCNMA1 | 1095 | R | 0 | 0 | no | yes | no | no | no | yes | no | no | no | no | no | no | no |
| KCNMA1 | 1096 | D | 0 | 0 | no | yes | no | no | no | yes | no | no | no | no | no | no | no |
| KCNMA1 | 1097 | R | 0 | 0 | no | yes | no | no | no | yes | no | no | no | no | no | no | no |
| KCNMA1 | 1098 | C | 0 | 0 | no | yes | no | no | no | yes | no | no | no | no | no | no | no |
| KCNMA1 | 1099 | R | 1 | 0 | no | no | yes | no | no | yes | no | no | no | no | no | no | no |
| KCNMA1 | 1100 | V | 1 | 0 | no | no | yes | no | no | yes | no | no | no | no | no | no | no |
| KCNMA1 | 1101 | A | 0 | 0 | no | no | yes | no | no | yes | no | no | no | no | no | no | no |
| KCNMA1 | 1102 | Q | 2 | 0 | no | no | yes | no | no | yes | no | no | no | no | no | no | no |
| KCNMA1 | 1103 | L | 0 | 0 | no | no | yes | no | no | yes | no | no | no | no | no | no | no |
| KCNMA1 | 1104 | A | 0 | 0 | no | yes | no | no | no | yes | no | no | no | no | no | no | no |
| KCNMA1 | 1105 | L | 0 | 0 | no | yes | no | no | no | yes | no | no | no | no | no | no | no |
| KCNMA1 | 1106 | L | 0 | 0 | no | yes | no | no | no | yes | no | no | no | no | no | no | no |
| KCNMA1 | 1107 | D | 1 | 0 | no | yes | no | no | no | yes | no | no | no | no | no | no | no |
| KCNMA1 | 1108 | G | 0 | 0 | no | yes | no | no | no | yes | no | no | no | no | no | no | no |
| KCNMA1 | 1109 | P | 0 | 0 | no | yes | no | no | no | yes | no | no | no | no | no | no | no |
| KCNMA1 | 1110 | F | 0 | 0 | no | yes | no | no | no | yes | no | no | no | no | no | no | no |
| KCNMA1 | 1111 | A | 0 | 0 | no | yes | no | no | no | yes | no | no | no | no | no | no | no |
| KCNMA1 | 1112 | D | 0 | 0 | no | yes | no | no | no | yes | no | no | no | no | no | no | no |
| KCNMA1 | 1113 | L | 0 | 0 | no | yes | no | no | no | yes | no | no | no | no | no | no | no |
| KCNMA1 | 1114 | G | 0 | 0 | no | yes | no | no | no | yes | no | no | no | no | no | no | no |
| KCNMA1 | 1115 | D | 1 | 0 | no | yes | no | no | no | yes | no | no | no | no | no | no | no |
| KCNMA1 | 1116 | G | 0 | 0 | no | yes | no | no | no | yes | no | no | no | no | no | no | no |
| KCNMA1 | 1117 | G | 0 | 0 | no | yes | no | no | no | yes | no | no | no | no | no | no | no |

| | | | | | | | | | | | | | | | | | |
|---|---|---|---|---|---|---|---|---|---|---|---|---|---|---|---|---|---|
| KCNMA1 | 1118 | C | 1 | 0 | no | no | yes | no | no | yes | no | no | no | no | no | no | no |
| KCNMA1 | 1119 | Y | 0 | 0 | yes | no | no | no | no | yes | no | no | no | no | no | no | no |
| KCNMA1 | 1120 | G | 0 | 0 | yes | no | no | no | no | yes | no | no | no | no | no | no | no |
| KCNMA1 | 1121 | D | 0 | 0 | yes | no | no | no | no | yes | no | no | no | no | no | no | no |
| KCNMA1 | 1122 | L | 0 | 0 | yes | no | no | no | no | yes | no | no | no | no | no | no | no |
| KCNMA1 | 1123 | F | 0 | 0 | yes | no | no | no | no | yes | no | no | no | no | no | no | no |
| KCNMA1 | 1124 | C | 0 | 0 | yes | no | no | no | no | yes | no | no | no | no | no | no | no |
| KCNMA1 | 1125 | K | 0 | 0 | yes | no | no | no | no | yes | no | no | no | no | no | no | no |
| KCNMA1 | 1126 | A | 0 | 0 | yes | no | no | no | no | yes | no | no | no | no | no | no | no |
| KCNMA1 | 1127 | L | 0 | 0 | yes | no | no | no | no | yes | no | no | no | no | no | no | no |
| KCNMA1 | 1128 | K | 0 | 0 | yes | no | no | no | no | yes | no | no | no | no | no | no | no |
| KCNMA1 | 1129 | T | 0 | 0 | yes | no | no | no | no | yes | no | no | no | no | no | no | no |
| KCNMA1 | 1130 | Y | 0 | 0 | yes | no | no | no | no | yes | no | no | no | no | no | no | no |
| KCNMA1 | 1131 | N | 0 | 0 | no | yes | no | no | no | yes | no | no | no | no | no | no | no |
| KCNMA1 | 1132 | M | 0 | 0 | no | yes | no | no | no | yes | no | no | no | no | no | no | no |
| KCNMA1 | 1133 | L | 0 | 0 | no | no | yes | no | no | yes | no | no | no | no | no | no | no |
| KCNMA1 | 1134 | C | 0 | 0 | no | no | yes | no | no | yes | no | no | no | no | no | no | no |
| KCNMA1 | 1135 | F | 0 | 0 | no | no | yes | no | no | yes | no | no | no | no | no | no | no |
| KCNMA1 | 1136 | G | 0 | 0 | no | no | yes | no | no | yes | no | no | no | no | no | no | no |
| KCNMA1 | 1137 | I | 0 | 0 | no | no | yes | no | no | yes | no | no | no | no | no | no | no |
| KCNMA1 | 1138 | Y | 0 | 0 | no | yes | no | no | no | yes | no | no | no | no | no | no | no |
| KCNMA1 | 1139 | R | 0 | 0 | no | no | yes | no | no | yes | no | no | no | no | no | no | no |
| KCNMA1 | 1140 | L | 0 | 0 | no | no | yes | no | no | yes | no | no | no | no | no | no | no |
| KCNMA1 | 1141 | R | 0 | 0 | yes | no | no | no | no | yes | no | no | no | no | no | no | no |
| KCNMA1 | 1142 | D | 0 | 0 | yes | no | no | no | no | yes | no | no | no | no | no | no | no |
| KCNMA1 | 1143 | A | 0 | 0 | yes | no | no | no | no | yes | no | no | no | no | no | no | no |
| KCNMA1 | 1144 | H | 1 | 0 | no | yes | no | no | no | yes | no | no | no | no | no | no | no |
| KCNMA1 | 1145 | L | 0 | 0 | no | yes | no | no | no | yes | no | no | no | no | no | no | no |
| KCNMA1 | 1146 | S | 0 | 0 | no | yes | no | no | no | yes | no | no | no | no | no | no | no |
| KCNMA1 | 1147 | T | 1 | 0 | no | yes | no | no | no | yes | no | no | no | no | no | no | no |
| KCNMA1 | 1148 | P | 1 | 0 | no | yes | no | no | no | yes | no | no | no | no | no | no | no |
| KCNMA1 | 1149 | S | 0 | 0 | no | yes | no | no | no | yes | no | no | no | no | no | no | no |
| KCNMA1 | 1150 | Q | 0 | 0 | no | yes | no | no | no | yes | no | no | no | no | no | no | no |

| | | | | | | | | | | | | | | | | | |
|---|---|---|---|---|---|---|---|---|---|---|---|---|---|---|---|---|---|
| KCNMA1 | 1151 | C | 0 | 0 | no | yes | no | no | no | yes | no | no | no | no | no | no | no |
| KCNMA1 | 1152 | T | 0 | 0 | no | yes | no | no | no | yes | no | no | no | no | no | no | no |
| KCNMA1 | 1153 | K | 0 | 0 | no | yes | no | no | no | yes | no | no | no | no | no | no | no |
| KCNMA1 | 1154 | R | 0 | 0 | no | no | yes | no | no | yes | no | no | no | no | no | no | no |
| KCNMA1 | 1155 | Y | 0 | 0 | no | yes | no | no | no | yes | no | no | no | no | no | no | no |
| KCNMA1 | 1156 | V | 0 | 0 | no | yes | no | no | no | yes | no | no | no | no | no | no | no |
| KCNMA1 | 1157 | I | 1 | 0 | no | no | yes | no | no | yes | no | no | no | no | no | no | no |
| KCNMA1 | 1158 | T | 1 | 0 | no | no | yes | no | no | yes | no | no | no | no | no | no | no |
| KCNMA1 | 1159 | N | 0 | 0 | no | yes | no | no | no | yes | no | no | no | no | no | no | no |
| KCNMA1 | 1160 | P | 0 | 0 | no | yes | no | no | no | yes | no | no | no | no | no | no | no |
| KCNMA1 | 1161 | P | 0 | 0 | no | yes | no | no | no | yes | no | no | no | no | no | no | no |
| KCNMA1 | 1162 | Y | 0 | 0 | no | yes | no | no | no | yes | no | no | no | no | no | no | no |
| KCNMA1 | 1163 | E | 0 | 1 | no | yes | no | no | no | yes | no | no | no | no | no | no | no |
| KCNMA1 | 1164 | F | 1 | 0 | no | yes | no | no | no | yes | no | no | no | no | no | no | no |
| KCNMA1 | 1165 | E | 1 | 0 | no | no | yes | no | no | yes | no | no | no | no | no | no | no |
| KCNMA1 | 1166 | L | 0 | 0 | no | yes | no | no | no | yes | no | no | no | no | no | no | no |
| KCNMA1 | 1167 | V | 1 | 0 | no | yes | no | no | no | yes | no | no | no | no | no | no | no |
| KCNMA1 | 1168 | P | 0 | 0 | no | yes | no | no | no | yes | no | no | no | no | no | no | no |
| KCNMA1 | 1169 | T | 1 | 0 | no | yes | no | no | no | yes | no | no | no | no | no | no | no |
| KCNMA1 | 1170 | D | 0 | 0 | no | no | yes | no | no | yes | no | no | no | no | no | no | no |
| KCNMA1 | 1171 | L | 0 | 0 | no | yes | no | no | no | yes | no | no | no | no | no | no | no |
| KCNMA1 | 1172 | I | 0 | 0 | no | no | yes | no | no | yes | no | no | no | no | no | no | no |
| KCNMA1 | 1173 | F | 0 | 0 | no | no | yes | no | no | yes | no | no | no | no | no | no | no |
| KCNMA1 | 1174 | C | 1 | 0 | no | no | yes | no | no | yes | no | no | no | no | no | no | no |
| KCNMA1 | 1175 | L | 0 | 0 | no | no | yes | no | no | yes | no | no | no | no | no | no | no |
| KCNMA1 | 1176 | M | 0 | 0 | no | no | yes | no | no | yes | no | no | no | no | no | no | no |
| KCNMA1 | 1177 | Q | 0 | 0 | no | yes | no | no | no | yes | no | no | no | no | no | no | no |
| KCNMA1 | 1178 | F | 0 | 0 | no | yes | no | no | no | yes | no | no | no | no | no | no | no |
| KCNMA1 | 1179 | D | 0 | 0 | no | yes | no | no | no | yes | no | no | no | no | no | no | no |
| KCNQ1 | 104 | T | 2 | 1 | no | yes | no | no | no | yes | no | no | no | no | no | no | no |
| KCNQ1 | 105 | H | 0 | 0 | yes | no | no | no | no | yes | no | no | no | no | no | no | no |
| KCNQ1 | 106 | V | 0 | 0 | yes | no | no | no | no | yes | no | no | no | no | no | no | no |
| KCNQ1 | 107 | Q | 0 | 1 | yes | no | no | no | no | yes | no | no | no | no | no | no | no |

| | | | | | | | | | | | | | | | | | |
|---|---|---|---|---|---|---|---|---|---|---|---|---|---|---|---|---|---|
| KCNQ1 | 108 | G | 1 | 0 | yes | no | no | no | no | yes | no | no | no | no | no | no | no |
| KCNQ1 | 109 | R | 2 | 1 | yes | no | no | no | no | yes | no | no | no | no | no | no | no |
| KCNQ1 | 110 | V | 1 | 0 | yes | no | no | no | no | yes | no | no | no | no | no | no | no |
| KCNQ1 | 111 | Y | 1 | 1 | yes | no | no | no | no | yes | no | no | no | no | no | no | no |
| KCNQ1 | 112 | N | 0 | 0 | yes | no | no | no | no | yes | no | no | yes | no | no | no | no |
| KCNQ1 | 113 | F | 0 | 0 | yes | no | no | yes | no | yes | no | no | no | no | no | no | no |
| KCNQ1 | 114 | L | 0 | 1 | yes | no | no | yes | no | yes | no | no | no | no | no | no | no |
| KCNQ1 | 115 | E | 0 | 2 | no | yes | no | yes | no | yes | no | no | no | no | no | no | no |
| KCNQ1 | 116 | R | 0 | 0 | no | yes | no | no | no | yes | no | no | yes | no | no | no | no |
| KCNQ1 | 117 | P | 0 | 2 | no | yes | no | yes | no | yes | no | no | no | no | no | no | no |
| KCNQ1 | 118 | T | 0 | 0 | no | yes | no | no | no | yes | no | no | no | no | no | no | no |
| KCNQ1 | 119 | G | 0 | 0 | no | yes | no | no | no | yes | no | no | no | no | no | no | no |
| KCNQ1 | 120 | W | 0 | 0 | no | yes | no | yes | no | yes | no | no | no | no | no | no | no |
| KCNQ1 | 121 | K | 0 | 0 | yes | no | no | yes | no | yes | no | no | no | no | no | no | no |
| KCNQ1 | 122 | C | 0 | 1 | yes | no | no | yes | no | no | no | no | no | no | no | no | no |
| KCNQ1 | 123 | F | 0 | 0 | yes | no | no | yes | no | no | no | no | no | no | no | no | no |
| KCNQ1 | 124 | V | 0 | 0 | yes | no | no | yes | no | no | no | no | no | no | no | no | no |
| KCNQ1 | 125 | Y | 0 | 3 | yes | no | no | yes | no | no | no | no | no | no | no | no | no |
| KCNQ1 | 126 | H | 0 | 1 | yes | no | no | yes | no | no | no | no | no | no | no | no | no |
| KCNQ1 | 127 | F | 0 | 1 | yes | no | no | yes | no | no | no | no | no | no | no | no | no |
| KCNQ1 | 128 | A | 0 | 0 | yes | no | no | yes | no | no | no | no | no | no | no | no | no |
| KCNQ1 | 129 | V | 0 | 0 | yes | no | no | yes | no | no | no | no | no | no | no | no | no |
| KCNQ1 | 130 | F | 0 | 0 | yes | no | no | yes | no | no | no | no | no | no | no | no | no |
| KCNQ1 | 131 | L | 0 | 1 | yes | no | no | yes | no | no | no | no | no | no | no | no | no |
| KCNQ1 | 132 | I | 1 | 1 | yes | no | no | yes | no | no | no | no | no | no | no | no | no |
| KCNQ1 | 133 | V | 1 | 1 | yes | no | no | yes | no | no | no | no | no | no | no | no | no |
| KCNQ1 | 134 | L | 0 | 1 | yes | no | no | yes | no | no | no | no | no | no | no | no | no |
| KCNQ1 | 135 | V | 0 | 0 | yes | no | no | yes | no | no | no | no | no | no | no | no | no |
| KCNQ1 | 136 | C | 0 | 1 | yes | no | no | yes | no | no | no | no | no | no | no | no | no |
| KCNQ1 | 137 | L | 2 | 1 | yes | no | no | yes | no | no | no | no | no | no | no | no | no |
| KCNQ1 | 138 | I | 0 | 0 | yes | no | no | yes | no | no | no | no | no | no | no | no | no |
| KCNQ1 | 139 | F | 0 | 0 | yes | no | no | yes | no | no | no | no | no | no | no | no | no |
| KCNQ1 | 140 | S | 0 | 2 | yes | no | no | yes | no | no | no | no | no | no | no | no | no |

| | | | | | | | | | | | | | | | | | |
|---|---|---|---|---|---|---|---|---|---|---|---|---|---|---|---|---|---|
| KCNQ1 | 141 | V | 0 | 1 | yes | no | no | yes | no | no | no | no | yes | no | no | no | no |
| KCNQ1 | 142 | L | 0 | 0 | yes | no | no | no | no | no | no | no | no | no | no | no | no |
| KCNQ1 | 143 | S | 1 | 0 | no | yes | no | no | yes | no | no | no | no | no | no | no | no |
| KCNQ1 | 144 | T | 0 | 1 | no | yes | no | no | yes | no | no | no | yes | no | no | no | no |
| KCNQ1 | 145 | I | 0 | 0 | no | yes | no | no | yes | no | no | no | yes | no | no | no | no |
| KCNQ1 | 146 | E | 2 | 2 | no | yes | no | no | yes | no | no | no | no | no | no | no | no |
| KCNQ1 | 147 | Q | 1 | 1 | no | yes | no | no | yes | no | no | no | no | no | no | no | no |
| KCNQ1 | 148 | Y | 0 | 0 | no | yes | no | no | no | no | no | no | no | no | no | no | no |
| KCNQ1 | 149 | A | 0 | 0 | no | yes | no | no | no | no | no | no | no | no | no | no | no |
| KCNQ1 | 150 | A | 0 | 2 | no | yes | no | no | no | no | no | no | no | no | no | no | no |
| KCNQ1 | 151 | L | 0 | 0 | yes | no | no | no | no | no | no | no | no | no | no | no | no |
| KCNQ1 | 152 | A | 0 | 0 | yes | no | no | no | no | no | no | no | no | no | no | no | no |
| KCNQ1 | 153 | T | 1 | 1 | yes | no | no | no | no | no | no | no | no | no | no | no | no |
| KCNQ1 | 154 | G | 0 | 0 | yes | no | no | no | no | no | no | no | no | no | no | no | no |
| KCNQ1 | 155 | T | 0 | 0 | no | yes | no | yes | no | no | no | no | no | no | no | no | no |
| KCNQ1 | 156 | L | 0 | 1 | no | yes | no | yes | no | no | no | no | no | no | no | no | no |
| KCNQ1 | 157 | F | 0 | 1 | yes | no | no | yes | no | no | no | no | no | no | no | no | no |
| KCNQ1 | 158 | W | 0 | 0 | yes | no | no | yes | no | no | no | no | no | no | no | no | no |
| KCNQ1 | 159 | M | 0 | 0 | yes | no | no | yes | no | no | no | no | no | no | no | no | no |
| KCNQ1 | 160 | E | 0 | 2 | yes | no | no | yes | no | no | no | no | no | no | no | no | no |
| KCNQ1 | 161 | I | 0 | 0 | yes | no | no | yes | no | no | no | no | no | no | no | no | no |
| KCNQ1 | 162 | V | 1 | 0 | yes | no | no | yes | no | no | no | no | no | no | no | no | no |
| KCNQ1 | 163 | L | 0 | 0 | yes | no | no | yes | no | no | no | no | no | no | no | no | no |
| KCNQ1 | 164 | V | 0 | 0 | yes | no | no | yes | no | no | no | no | no | no | no | no | no |
| KCNQ1 | 165 | V | 1 | 1 | yes | no | no | yes | no | no | no | no | no | no | no | no | no |
| KCNQ1 | 166 | F | 1 | 0 | yes | no | no | yes | no | no | no | no | no | no | no | no | no |
| KCNQ1 | 167 | F | 0 | 0 | yes | no | no | yes | no | no | no | no | no | no | no | no | no |
| KCNQ1 | 168 | G | 1 | 1 | yes | no | no | yes | no | no | no | no | no | no | no | no | no |
| KCNQ1 | 169 | T | 1 | 1 | yes | no | no | yes | no | yes | no | no | no | no | no | no | no |
| KCNQ1 | 170 | E | 0 | 1 | yes | no | no | yes | no | yes | no | no | no | no | no | no | no |
| KCNQ1 | 171 | Y | 0 | 0 | yes | no | no | yes | no | yes | no | no | no | no | no | no | no |
| KCNQ1 | 172 | V | 2 | 1 | yes | no | no | yes | no | yes | no | no | no | no | no | no | no |
| KCNQ1 | 173 | V | 0 | 1 | yes | no | no | yes | no | yes | no | no | no | no | no | no | no |

| | | | | | | | | | | | | | | | | | |
|---|---|---|---|---|---|---|---|---|---|---|---|---|---|---|---|---|---|
| KCNQ1 | 174 | R | 2 | 2 | yes | no | no | yes | no | yes | no | no | no | no | no | no | no |
| KCNQ1 | 175 | L | 1 | 0 | yes | no | no | yes | no | yes | no | no | no | no | no | no | no |
| KCNQ1 | 176 | W | 0 | 1 | yes | no | no | yes | no | yes | no | no | no | no | no | no | no |
| KCNQ1 | 177 | S | 0 | 0 | yes | no | no | no | no | yes | no | no | no | no | no | no | no |
| KCNQ1 | 178 | A | 1 | 2 | yes | no | no | no | no | yes | no | no | no | no | no | no | no |
| KCNQ1 | 179 | G | 0 | 2 | yes | no | no | no | no | yes | no | no | no | no | no | no | no |
| KCNQ1 | 180 | C | 0 | 0 | yes | no | no | no | no | yes | no | no | yes | no | no | no | no |
| KCNQ1 | 181 | R | 1 | 0 | no | yes | no | no | no | yes | no | no | yes | no | no | no | no |
| KCNQ1 | 182 | S | 0 | 1 | no | yes | no | no | no | yes | no | no | yes | no | no | no | no |
| KCNQ1 | 183 | K | 0 | 1 | no | yes | no | no | no | yes | no | no | no | no | no | no | no |
| KCNQ1 | 184 | Y | 0 | 4 | no | yes | no | no | no | yes | no | no | no | no | no | no | no |
| KCNQ1 | 185 | V | 1 | 0 | no | yes | no | no | no | yes | no | no | no | no | no | no | no |
| KCNQ1 | 186 | G | 0 | 3 | no | yes | no | no | no | yes | no | no | no | no | no | no | no |
| KCNQ1 | 187 | L | 2 | 1 | yes | no | no | no | no | yes | no | no | no | no | no | no | no |
| KCNQ1 | 188 | W | 3 | 0 | yes | no | no | no | no | yes | no | no | no | no | no | no | no |
| KCNQ1 | 189 | G | 0 | 2 | yes | no | no | no | no | yes | no | no | no | no | no | no | no |
| KCNQ1 | 190 | R | 2 | 2 | yes | no | no | yes | no | yes | no | no | no | no | no | no | no |
| KCNQ1 | 191 | L | 0 | 1 | yes | no | no | yes | no | yes | no | no | no | no | no | no | no |
| KCNQ1 | 192 | R | 2 | 2 | yes | no | no | no | no | yes | no | no | no | no | no | no | no |
| KCNQ1 | 193 | F | 0 | 1 | yes | no | no | yes | no | yes | no | no | no | no | no | no | no |
| KCNQ1 | 194 | A | 1 | 1 | no | yes | no | yes | no | yes | no | no | no | no | no | no | no |
| KCNQ1 | 195 | R | 2 | 2 | no | yes | no | yes | no | yes | no | no | no | no | no | no | no |
| KCNQ1 | 196 | K | 0 | 1 | no | yes | no | yes | no | yes | no | no | no | no | no | no | no |
| KCNQ1 | 197 | P | 1 | 2 | yes | no | no | yes | no | no | no | no | no | no | no | no | no |
| KCNQ1 | 198 | I | 2 | 1 | yes | no | no | yes | no | no | no | no | no | no | no | no | no |
| KCNQ1 | 199 | S | 0 | 1 | yes | no | no | yes | no | no | no | no | no | no | no | no | no |
| KCNQ1 | 200 | I | 0 | 1 | yes | no | no | yes | no | no | no | no | no | no | no | no | no |
| KCNQ1 | 201 | I | 0 | 0 | yes | no | no | yes | no | no | no | no | no | no | no | no | no |
| KCNQ1 | 202 | D | 0 | 1 | yes | no | no | yes | no | no | no | no | no | no | no | no | no |
| KCNQ1 | 203 | L | 0 | 1 | yes | no | no | yes | no | no | no | no | no | no | no | no | no |
| KCNQ1 | 204 | I | 0 | 2 | yes | no | no | yes | no | no | no | no | no | no | no | no | no |
| KCNQ1 | 205 | V | 1 | 1 | yes | no | no | yes | no | no | no | no | no | no | no | no | no |
| KCNQ1 | 206 | V | 1 | 0 | yes | no | no | yes | no | no | no | no | no | no | no | no | no |

| | | | | | | | | | | | | | | | | | |
|---|---|---|---|---|---|---|---|---|---|---|---|---|---|---|---|---|---|
| KCNQ1 | 207 | V | 2 | 0 | yes | no | no | yes | no | no | no | no | no | no | no | no | no |
| KCNQ1 | 208 | A | 1 | 0 | yes | no | no | yes | no | no | no | no | no | no | no | no | no |
| KCNQ1 | 209 | S | 0 | 2 | yes | no | no | yes | no | no | no | no | no | no | no | no | no |
| KCNQ1 | 210 | M | 1 | 0 | yes | no | no | yes | no | no | no | no | no | no | no | no | no |
| KCNQ1 | 211 | V | 0 | 0 | yes | no | no | yes | no | no | no | no | no | no | no | no | no |
| KCNQ1 | 212 | V | 0 | 0 | yes | no | no | yes | no | no | no | no | no | no | no | no | no |
| KCNQ1 | 213 | L | 0 | 0 | yes | no | no | yes | no | no | no | no | no | no | no | no | no |
| KCNQ1 | 214 | C | 1 | 0 | yes | no | no | yes | no | no | no | no | no | no | no | no | no |
| KCNQ1 | 215 | V | 2 | 1 | yes | no | no | yes | no | no | no | no | no | no | no | no | no |
| KCNQ1 | 216 | G | 0 | 0 | no | yes | no | no | no | no | no | no | no | no | no | no | no |
| KCNQ1 | 217 | S | 0 | 0 | no | yes | no | no | no | no | no | no | no | no | no | no | no |
| KCNQ1 | 218 | K | 0 | 1 | no | yes | no | no | yes | no | no | no | no | no | no | no | no |
| KCNQ1 | 223 | A | 0 | 0 | no | yes | no | no | yes | no | no | no | no | no | no | no | no |
| KCNQ1 | 224 | T | 1 | 1 | yes | no | no | no | yes | no | no | no | no | no | no | no | no |
| KCNQ1 | 225 | S | 1 | 1 | yes | no | no | no | yes | no | no | no | no | no | no | no | no |
| KCNQ1 | 226 | A | 0 | 1 | yes | no | no | no | no | no | no | no | no | yes | no | no | no |
| KCNQ1 | 227 | I | 0 | 1 | yes | no | no | no | no | no | no | no | no | yes | no | no | no |
| KCNQ1 | 228 | R | 0 | 0 | yes | no | no | no | no | no | no | no | yes | yes | no | no | no |
| KCNQ1 | 229 | G | 0 | 1 | yes | no | no | yes | no | no | no | no | no | yes | no | no | no |
| KCNQ1 | 230 | I | 0 | 0 | yes | no | no | yes | no | no | no | no | no | yes | no | no | no |
| KCNQ1 | 231 | R | 2 | 3 | yes | no | no | yes | no | no | no | no | no | yes | no | no | no |
| KCNQ1 | 232 | F | 0 | 0 | yes | no | no | yes | no | no | no | no | yes | yes | no | no | no |
| KCNQ1 | 233 | L | 1 | 0 | yes | no | no | yes | no | no | no | no | no | yes | no | no | no |
| KCNQ1 | 234 | Q | 1 | 0 | yes | no | no | yes | no | no | no | no | no | yes | no | no | no |
| KCNQ1 | 235 | I | 0 | 1 | yes | no | no | yes | no | no | no | no | yes | yes | no | no | no |
| KCNQ1 | 236 | L | 0 | 2 | yes | no | no | yes | no | no | no | no | yes | yes | no | no | no |
| KCNQ1 | 237 | R | 0 | 0 | yes | no | no | yes | no | no | no | no | no | yes | no | no | no |
| KCNQ1 | 238 | M | 1 | 0 | yes | no | no | yes | no | no | no | no | yes | yes | no | no | no |
| KCNQ1 | 239 | L | 0 | 1 | yes | no | no | yes | no | no | no | no | yes | yes | no | no | no |
| KCNQ1 | 240 | H | 1 | 1 | yes | no | no | yes | no | no | no | no | no | yes | no | no | no |
| KCNQ1 | 241 | V | 1 | 2 | no | yes | no | yes | no | no | no | no | yes | yes | no | no | no |
| KCNQ1 | 242 | D | 1 | 2 | no | yes | no | yes | no | no | no | no | yes | yes | no | no | no |
| KCNQ1 | 243 | R | 2 | 2 | no | yes | no | yes | no | no | no | no | no | yes | no | no | no |

| | | | | | | | | | | | | | | | | | |
|---|---|---|---|---|---|---|---|---|---|---|---|---|---|---|---|---|---|
| KCNQ1 | 244 | Q | 0 | 0 | no | yes | no | yes | no | no | no | no | no | yes | no | no | no |
| KCNQ1 | 245 | G | 0 | 1 | no | yes | no | yes | no | no | no | no | no | yes | no | no | no |
| KCNQ1 | 246 | G | 0 | 0 | yes | no | no | yes | no | no | no | no | no | yes | no | no | no |
| KCNQ1 | 247 | T | 0 | 0 | yes | no | no | yes | no | no | no | no | yes | yes | no | no | no |
| KCNQ1 | 248 | W | 0 | 2 | yes | no | no | yes | no | no | no | no | yes | yes | no | no | no |
| KCNQ1 | 249 | R | 1 | 0 | yes | no | no | yes | no | yes | no | no | no | no | no | no | no |
| KCNQ1 | 250 | L | 0 | 1 | yes | no | no | yes | no | yes | no | no | yes | no | no | no | no |
| KCNQ1 | 251 | L | 0 | 2 | yes | no | no | yes | no | yes | no | no | yes | no | no | no | no |
| KCNQ1 | 252 | G | 1 | 1 | yes | no | no | yes | no | yes | no | no | no | no | no | no | no |
| KCNQ1 | 253 | S | 1 | 1 | yes | no | no | no | no | yes | no | no | no | no | no | no | no |
| KCNQ1 | 254 | V | 0 | 1 | yes | no | no | yes | no | yes | no | no | no | no | no | no | no |
| KCNQ1 | 255 | V | 0 | 0 | yes | no | no | yes | no | yes | no | no | no | no | no | no | no |
| KCNQ1 | 256 | F | 0 | 0 | yes | no | no | no | no | yes | no | no | no | no | no | no | no |
| KCNQ1 | 257 | I | 1 | 0 | yes | no | no | no | no | yes | no | no | yes | no | no | no | no |
| KCNQ1 | 258 | H | 1 | 2 | no | yes | no | yes | no | yes | no | no | yes | no | no | no | no |
| KCNQ1 | 259 | R | 2 | 3 | yes | no | no | yes | no | yes | no | no | no | no | no | no | no |
| KCNQ1 | 260 | Q | 0 | 0 | yes | no | no | yes | no | yes | no | no | yes | no | no | no | no |
| KCNQ1 | 261 | E | 0 | 3 | yes | no | no | yes | no | yes | no | no | yes | no | no | no | no |
| KCNQ1 | 262 | L | 0 | 2 | yes | no | no | yes | no | no | no | no | no | no | no | no | no |
| KCNQ1 | 263 | I | 0 | 0 | yes | no | no | yes | no | no | no | no | no | no | no | no | no |
| KCNQ1 | 264 | T | 0 | 0 | yes | no | no | yes | no | no | no | no | yes | no | no | no | no |
| KCNQ1 | 265 | T | 0 | 1 | yes | no | no | yes | no | no | no | no | no | no | no | no | no |
| KCNQ1 | 266 | L | 0 | 1 | yes | no | no | yes | no | no | no | no | no | no | no | no | no |
| KCNQ1 | 267 | Y | 1 | 0 | yes | no | no | yes | no | no | no | no | yes | no | no | no | no |
| KCNQ1 | 268 | I | 1 | 1 | yes | no | no | yes | no | no | no | no | yes | no | no | no | no |
| KCNQ1 | 269 | G | 1 | 2 | yes | no | no | yes | no | no | no | no | no | no | no | no | no |
| KCNQ1 | 270 | F | 1 | 0 | yes | no | no | yes | no | no | no | no | no | no | no | no | no |
| KCNQ1 | 271 | L | 0 | 0 | yes | no | no | yes | no | no | no | no | yes | no | no | no | no |
| KCNQ1 | 272 | G | 2 | 1 | yes | no | no | yes | no | no | no | no | no | no | no | no | no |
| KCNQ1 | 273 | L | 2 | 2 | yes | no | no | yes | no | no | no | no | no | no | no | no | no |
| KCNQ1 | 274 | I | 1 | 1 | yes | no | no | yes | no | no | no | no | yes | no | no | no | no |
| KCNQ1 | 275 | F | 0 | 1 | yes | no | no | yes | no | no | no | no | yes | no | no | no | no |
| KCNQ1 | 276 | S | 0 | 1 | yes | no | no | yes | no | no | no | no | no | no | no | no | no |

| | | | | | | | | | | | | | | | | | |
|---|---|---|---|---|---|---|---|---|---|---|---|---|---|---|---|---|---|
| KCNQ1 | 277 | S | 0 | 3 | yes | no | no | yes | no | no | no | no | no | no | no | no | no |
| KCNQ1 | 278 | Y | 0 | 1 | yes | no | no | yes | no | no | no | no | yes | no | no | no | no |
| KCNQ1 | 279 | F | 0 | 1 | yes | no | no | yes | no | no | no | no | yes | no | no | no | no |
| KCNQ1 | 280 | V | 0 | 1 | yes | no | no | yes | no | no | no | no | no | no | no | no | no |
| KCNQ1 | 281 | Y | 0 | 1 | yes | no | no | no | no | no | no | no | yes | no | no | no | no |
| KCNQ1 | 282 | L | 0 | 1 | yes | no | no | no | no | no | no | no | no | no | no | no | no |
| KCNQ1 | 283 | A | 0 | 2 | yes | no | no | no | yes | no | no | no | no | no | no | no | no |
| KCNQ1 | 284 | E | 0 | 1 | yes | no | no | no | yes | no | no | no | no | no | no | no | no |
| KCNQ1 | 285 | K | 0 | 0 | no | yes | no | no | yes | no | no | no | no | no | no | no | no |
| KCNQ1 | 286 | D | 0 | 0 | no | yes | no | no | yes | no | no | no | no | no | no | no | no |
| KCNQ1 | 287 | A | 2 | 1 | no | yes | no | no | yes | no | no | no | no | no | no | no | no |
| KCNQ1 | 288 | V | 0 | 0 | no | yes | no | no | yes | no | no | no | no | no | no | no | no |
| KCNQ1 | 289 | N | 0 | 0 | no | yes | no | no | yes | no | no | no | no | no | no | no | no |
| KCNQ1 | 290 | E | 1 | 1 | no | yes | no | no | yes | no | no | no | no | no | no | no | no |
| KCNQ1 | 291 | S | 0 | 0 | no | yes | no | no | yes | no | yes | no | no | no | no | no | no |
| KCNQ1 | 292 | G | 1 | 1 | no | yes | no | no | yes | no | no | no | no | no | no | no | no |
| KCNQ1 | 293 | R | 2 | 1 | no | yes | no | no | yes | no | no | no | no | no | no | no | no |
| KCNQ1 | 294 | V | 1 | 0 | no | yes | no | no | yes | no | no | no | no | no | no | no | no |
| KCNQ1 | 295 | E | 0 | 0 | no | yes | no | no | yes | no | no | no | no | no | no | no | no |
| KCNQ1 | 296 | F | 0 | 0 | no | yes | no | no | yes | no | no | no | no | no | no | no | no |
| KCNQ1 | 297 | G | 2 | 0 | no | yes | no | no | yes | no | no | no | no | no | no | no | no |
| KCNQ1 | 298 | S | 1 | 0 | no | yes | no | no | yes | no | no | no | yes | no | no | no | no |
| KCNQ1 | 299 | Y | 0 | 0 | yes | no | no | yes | yes | no | no | no | yes | no | no | no | no |
| KCNQ1 | 300 | A | 1 | 0 | yes | no | no | no | no | no | no | yes | no | no | no | no | no |
| KCNQ1 | 301 | D | 0 | 0 | yes | no | no | no | no | no | no | yes | no | no | no | no | no |
| KCNQ1 | 302 | A | 1 | 2 | yes | no | no | yes | no | no | no | yes | no | no | no | no | no |
| KCNQ1 | 303 | L | 0 | 1 | yes | no | no | yes | no | no | no | yes | no | no | no | no | no |
| KCNQ1 | 304 | W | 0 | 1 | yes | no | no | yes | no | no | no | yes | yes | no | no | no | no |
| KCNQ1 | 305 | W | 1 | 2 | yes | no | no | yes | no | no | no | yes | yes | no | no | no | no |
| KCNQ1 | 306 | G | 0 | 2 | yes | no | no | yes | no | no | no | yes | no | no | no | no | no |
| KCNQ1 | 307 | V | 0 | 1 | yes | no | no | yes | no | no | no | yes | yes | no | no | no | no |
| KCNQ1 | 308 | V | 0 | 1 | yes | no | no | yes | no | no | no | yes | no | no | no | no | no |
| KCNQ1 | 309 | T | 0 | 1 | yes | no | no | yes | no | no | no | yes | yes | no | no | no | no |

| | | | | | | | | | | | | | | | | | |
|---|---|---|---|---|---|---|---|---|---|---|---|---|---|---|---|---|---|
| KCNQ1 | 310 | V | 0 | 1 | yes | no | no | yes | no | no | no | yes | no | no | no | no | no |
| KCNQ1 | 311 | T | 1 | 2 | no | yes | no | yes | no | no | no | yes | yes | no | no | no | no |
| KCNQ1 | 312 | T | 0 | 2 | no | yes | no | yes | no | no | yes | yes | yes | no | no | no | no |
| KCNQ1 | 313 | I | 0 | 1 | no | yes | no | yes | no | no | yes | yes | yes | no | no | no | no |
| KCNQ1 | 314 | G | 0 | 2 | no | yes | no | yes | no | no | yes | yes | yes | no | no | no | no |
| KCNQ1 | 315 | Y | 0 | 2 | no | yes | no | yes | no | no | yes | yes | yes | no | no | no | no |
| KCNQ1 | 316 | G | 0 | 2 | no | yes | no | no | no | no | yes | yes | yes | no | no | no | no |
| KCNQ1 | 317 | D | 0 | 2 | no | yes | no | no | no | no | no | yes | yes | no | no | no | no |
| KCNQ1 | 318 | K | 0 | 1 | no | yes | no | no | no | no | no | yes | no | no | no | no | no |
| KCNQ1 | 319 | V | 1 | 0 | no | yes | no | no | no | no | no | yes | yes | no | no | no | no |
| KCNQ1 | 320 | P | 0 | 1 | no | yes | no | no | no | no | no | yes | no | no | no | no | no |
| KCNQ1 | 321 | Q | 0 | 0 | no | yes | no | no | yes | no | no | no | no | no | no | no | no |
| KCNQ1 | 322 | T | 1 | 3 | no | yes | no | no | yes | no | no | no | no | no | no | no | no |
| KCNQ1 | 323 | W | 0 | 0 | yes | no | no | no | yes | no | no | no | no | no | no | no | no |
| KCNQ1 | 324 | V | 0 | 0 | yes | no | no | yes | yes | no | no | no | no | no | no | no | no |
| KCNQ1 | 325 | G | 0 | 3 | yes | no | no | yes | yes | no | no | no | no | no | no | no | no |
| KCNQ1 | 326 | K | 0 | 0 | yes | no | no | yes | yes | no | no | no | yes | no | no | no | no |
| KCNQ1 | 327 | T | 2 | 0 | yes | no | no | yes | yes | no | no | no | no | no | no | no | no |
| KCNQ1 | 328 | I | 0 | 0 | yes | no | no | yes | no | no | no | no | no | no | no | no | no |
| KCNQ1 | 329 | A | 1 | 0 | yes | no | no | yes | no | no | no | no | no | no | no | no | no |
| KCNQ1 | 330 | S | 0 | 0 | yes | no | no | yes | no | no | no | no | yes | no | no | no | no |
| KCNQ1 | 331 | C | 0 | 0 | yes | no | no | yes | no | no | no | no | no | no | no | no | no |
| KCNQ1 | 332 | F | 0 | 0 | yes | no | no | yes | no | no | no | no | no | no | no | no | no |
| KCNQ1 | 333 | S | 0 | 0 | yes | no | no | yes | no | no | no | no | yes | no | no | no | no |
| KCNQ1 | 334 | V | 0 | 0 | yes | no | no | yes | no | no | no | no | no | no | no | no | no |
| KCNQ1 | 335 | F | 0 | 0 | yes | no | no | yes | no | no | no | no | no | no | no | no | no |
| KCNQ1 | 336 | A | 1 | 0 | no | yes | no | yes | no | no | no | no | no | no | no | no | no |
| KCNQ1 | 337 | I | 0 | 0 | no | yes | no | yes | no | no | no | no | yes | no | no | no | no |
| KCNQ1 | 338 | S | 0 | 1 | yes | no | no | yes | no | no | no | no | no | no | no | no | no |
| KCNQ1 | 339 | F | 0 | 2 | yes | no | no | yes | no | no | no | no | yes | no | no | no | no |
| KCNQ1 | 340 | F | 1 | 1 | yes | no | no | yes | no | no | no | no | no | no | no | no | no |
| KCNQ1 | 341 | A | 0 | 3 | yes | no | no | yes | no | no | yes | no | yes | no | no | no | no |
| KCNQ1 | 342 | L | 0 | 1 | yes | no | no | yes | no | no | no | no | yes | no | no | no | no |

| | | | | | | | | | | | | | | | | | |
|---|---|---|---|---|---|---|---|---|---|---|---|---|---|---|---|---|---|
| KCNQ1 | 343 | P | 0 | 2 | yes | no | no | yes | no | no | no | no | no | no | no | no | no |
| KCNQ1 | 344 | A | 0 | 2 | yes | no | no | yes | no | no | no | no | yes | no | no | no | no |
| KCNQ1 | 345 | G | 0 | 3 | yes | no | no | yes | no | no | yes | no | yes | no | no | no | no |
| KCNQ1 | 346 | I | 0 | 0 | yes | no | no | yes | no | no | no | no | yes | no | no | no | no |
| KCNQ1 | 347 | L | 0 | 1 | yes | no | no | yes | no | no | no | no | yes | no | no | no | no |
| KCNQ1 | 348 | G | 0 | 0 | yes | no | no | yes | no | no | no | no | yes | no | no | no | no |
| KCNQ1 | 349 | S | 0 | 2 | yes | no | no | yes | no | yes | yes | no | yes | no | no | no | no |
| KCNQ1 | 350 | G | 0 | 2 | yes | no | no | no | no | yes | no | no | no | no | no | no | no |
| KCNQ1 | 351 | F | 1 | 2 | yes | no | no | yes | no | yes | no | no | yes | no | no | no | no |
| KCNQ1 | 352 | A | 0 | 0 | yes | no | no | no | no | yes | no | no | yes | no | no | no | no |
| KCNQ1 | 353 | L | 0 | 1 | yes | no | no | no | no | yes | yes | no | yes | no | no | no | no |
| KCNQ1 | 354 | K | 0 | 1 | yes | no | no | no | no | yes | no | no | no | no | no | no | no |
| KCNQ1 | 355 | V | 0 | 0 | yes | no | no | no | no | yes | no | no | yes | no | no | no | no |
| KCNQ1 | 356 | Q | 0 | 0 | yes | no | no | no | no | yes | yes | no | yes | no | no | no | no |
| KCNQ1 | 357 | Q | 0 | 1 | yes | no | no | no | no | yes | no | no | yes | no | no | no | no |
| KCNQ1 | 358 | K | 1 | 0 | yes | no | no | no | no | yes | no | no | no | no | no | no | no |
| KCNQ1 | 359 | Q | 0 | 1 | yes | no | no | no | no | yes | no | no | yes | no | no | no | no |
| KCNQ1 | 360 | R | 0 | 1 | yes | no | no | no | no | yes | yes | no | yes | no | no | no | no |
| KCNQ1 | 361 | Q | 0 | 0 | no | yes | no | no | no | yes | no | no | no | no | no | no | no |
| KCNQ1 | 362 | K | 1 | 1 | no | yes | no | no | no | yes | no | no | no | no | no | no | no |
| KCNQ1 | 363 | H | 0 | 1 | no | yes | no | no | no | yes | no | no | no | no | no | no | no |
| KCNQ1 | 364 | F | 1 | 0 | yes | no | no | no | no | yes | no | no | yes | no | no | no | no |
| KCNQ1 | 365 | N | 0 | 1 | yes | no | no | no | no | yes | no | no | yes | no | no | no | no |
| KCNQ1 | 366 | R | 1 | 2 | yes | no | no | no | no | yes | no | no | no | no | no | no | no |
| KCNQ1 | 367 | Q | 0 | 1 | yes | no | no | no | no | yes | no | no | no | no | no | no | no |
| KCNQ1 | 368 | I | 0 | 0 | yes | no | no | no | no | yes | no | no | yes | no | no | no | no |
| KCNQ1 | 369 | P | 0 | 0 | yes | no | no | no | no | yes | no | no | yes | no | no | no | no |
| KCNQ1 | 370 | A | 1 | 1 | yes | no | no | no | no | yes | no | no | no | no | no | no | no |
| KCNQ1 | 371 | A | 0 | 1 | yes | no | no | no | no | yes | no | no | yes | no | no | no | no |
| KCNQ1 | 372 | A | 0 | 1 | yes | no | no | no | no | yes | no | no | yes | no | no | no | no |
| KCNQ1 | 373 | S | 0 | 1 | yes | no | no | no | no | yes | no | no | no | no | no | no | no |
| KCNQ1 | 374 | L | 1 | 1 | yes | no | no | no | no | yes | no | no | yes | no | no | no | no |
| KCNQ1 | 375 | I | 0 | 1 | yes | no | no | no | no | yes | no | no | yes | no | no | no | no |

| | | | | | | | | | | | | | | | | | |
|---|---|---|---|---|---|---|---|---|---|---|---|---|---|---|---|---|---|
| KCNQ1 | 376 | Q | 0 | 0 | yes | no | no | no | no | yes | no | no | yes | no | no | no | no |
| KCNQ1 | 377 | T | 0 | 0 | yes | no | no | no | no | yes | no | no | yes | no | no | no | no |
| KCNQ1 | 378 | A | 1 | 0 | yes | no | no | no | no | yes | no | no | yes | no | no | no | no |
| KCNQ1 | 379 | W | 0 | 2 | yes | no | no | no | no | yes | no | no | yes | no | no | no | no |
| KCNQ1 | 380 | R | 0 | 2 | yes | no | no | no | no | yes | no | no | yes | no | no | no | no |
| KCNQ1 | 381 | C | 1 | 1 | yes | no | no | no | no | yes | no | no | no | no | no | no | no |
| KCNQ1 | 382 | Y | 0 | 0 | yes | no | no | no | no | yes | no | no | yes | no | no | no | no |
| KCNQ1 | 383 | A | 0 | 0 | yes | no | no | no | no | yes | no | no | no | no | no | no | no |
| KCNQ1 | 384 | A | 0 | 0 | no | yes | no | no | no | yes | no | no | no | no | no | no | no |
| KCNQ1 | 385 | E | 1 | 1 | no | yes | no | no | no | yes | no | no | no | no | no | no | no |
| KCNQ1 | 386 | N | 1 | 0 | no | yes | no | no | no | yes | no | no | no | no | no | no | no |
| KCNQ1 | 387 | P | 1 | 0 | no | yes | no | no | no | yes | no | no | no | no | no | no | no |
| KCNQ1 | 388 | D | 1 | 0 | no | yes | no | no | no | yes | no | no | no | no | no | no | no |
| KCNQ1 | 389 | S | 0 | 2 | no | yes | no | no | no | yes | no | no | yes | no | no | no | no |
| KCNQ1 | 390 | S | 0 | 0 | yes | no | no | no | no | yes | no | no | yes | no | no | no | no |
| KCNQ1 | 391 | T | 1 | 1 | yes | no | no | no | no | yes | no | no | yes | no | no | no | no |
| KCNQ1 | 392 | W | 0 | 1 | yes | no | no | no | no | yes | no | no | no | no | no | no | no |
| KCNQ1 | 393 | K | 1 | 1 | yes | no | no | no | no | yes | no | no | no | no | no | no | no |
| KCNQ1 | 394 | I | 1 | 0 | yes | no | no | no | no | yes | no | no | yes | no | no | no | no |
| KCNQ1 | 395 | Y | 1 | 0 | no | yes | no | no | no | yes | no | no | yes | no | no | no | no |
| KCNQ1 | 396 | I | 0 | 0 | no | yes | no | no | no | yes | no | no | no | no | no | no | no |
| KCNQ1 | 506 | L | 0 | 0 | no | yes | no | no | no | yes | no | no | no | no | no | no | no |
| KCNQ1 | 507 | R | 2 | 0 | no | yes | no | no | no | yes | no | no | no | no | no | no | no |
| KCNQ1 | 508 | E | 1 | 0 | no | yes | no | no | no | yes | no | no | no | no | no | no | no |
| KCNQ1 | 509 | H | 2 | 0 | no | yes | no | no | no | yes | no | no | yes | no | no | no | no |
| KCNQ1 | 510 | H | 2 | 0 | yes | no | no | no | no | yes | no | no | yes | no | no | no | no |
| KCNQ1 | 511 | R | 1 | 1 | yes | no | no | no | no | yes | no | no | no | no | no | no | no |
| KCNQ1 | 512 | A | 0 | 0 | yes | no | no | no | no | yes | no | no | no | no | no | no | no |
| KCNQ1 | 513 | T | 1 | 1 | yes | no | no | no | no | yes | no | no | yes | no | no | no | no |
| KCNQ1 | 514 | I | 0 | 1 | yes | no | no | no | no | yes | no | no | yes | no | no | no | no |
| KCNQ1 | 515 | K | 0 | 1 | yes | no | no | no | no | yes | no | no | no | no | no | no | no |
| KCNQ1 | 516 | V | 0 | 0 | yes | no | no | no | no | yes | no | no | yes | no | no | no | no |
| KCNQ1 | 517 | I | 0 | 1 | yes | no | no | no | no | yes | no | no | yes | no | no | no | no |

| | | | | | | | | | | | | | | | | | |
|---|---|---|---|---|---|---|---|---|---|---|---|---|---|---|---|---|---|
| KCNQ1 | 518 | R | 1 | 2 | yes | no | no | no | no | yes | no | no | no | no | no | no | no |
| KCNQ1 | 519 | R | 2 | 2 | yes | no | no | no | no | yes | no | no | yes | no | no | no | no |
| KCNQ1 | 520 | M | 0 | 1 | yes | no | no | no | no | yes | no | no | yes | no | no | no | no |
| KCNQ1 | 521 | Q | 0 | 0 | yes | no | no | no | no | yes | no | no | yes | no | no | no | no |
| KCNQ1 | 522 | Y | 0 | 1 | yes | no | no | no | no | yes | no | no | yes | no | no | no | no |
| KCNQ1 | 523 | F | 0 | 0 | yes | no | no | no | no | yes | no | no | yes | no | no | no | no |
| KCNQ1 | 524 | V | 0 | 1 | yes | no | no | no | no | yes | no | no | yes | no | no | no | no |
| KCNQ1 | 525 | A | 1 | 2 | yes | no | no | no | no | yes | no | no | no | no | no | no | no |
| KCNQ1 | 526 | K | 1 | 1 | yes | no | no | no | no | yes | no | no | yes | no | no | no | no |
| KCNQ1 | 527 | K | 0 | 0 | yes | no | no | no | no | yes | no | no | yes | no | no | no | no |
| KCNQ1 | 528 | K | 0 | 0 | yes | no | no | no | no | yes | no | no | yes | no | no | no | no |
| KCNQ1 | 529 | F | 0 | 0 | yes | no | no | no | no | yes | no | no | yes | no | no | no | no |
| KCNQ1 | 530 | Q | 0 | 0 | yes | no | no | no | no | yes | no | no | yes | no | no | no | no |
| KCNQ1 | 531 | Q | 0 | 0 | yes | no | no | no | no | yes | no | no | no | no | no | no | no |
| KCNQ1 | 532 | A | 0 | 0 | no | yes | no | no | no | yes | no | no | no | no | no | no | no |
| KCNQ1 | 533 | R | 1 | 1 | no | yes | no | no | no | yes | no | no | yes | no | no | no | no |
| KCNQ1 | 534 | K | 0 | 0 | no | yes | no | no | no | yes | no | no | no | no | no | no | no |
| KCNQ1 | 535 | P | 0 | 1 | no | yes | no | no | no | yes | no | no | no | no | no | no | no |
| KCNQ1 | 536 | Y | 0 | 0 | no | yes | no | no | no | yes | no | no | yes | no | no | no | no |
| KCNQ1 | 537 | D | 0 | 0 | no | yes | no | no | no | yes | no | no | yes | no | no | no | no |
| KCNQ1 | 538 | V | 0 | 0 | yes | no | no | no | no | yes | yes | no | yes | no | no | no | no |
| KCNQ1 | 539 | R | 1 | 2 | yes | no | no | no | no | yes | no | no | yes | no | no | no | no |
| KCNQ1 | 540 | D | 0 | 0 | yes | no | no | no | no | yes | no | no | yes | no | no | no | no |
| KCNQ1 | 541 | V | 1 | 1 | yes | no | no | no | no | yes | yes | no | yes | no | no | no | no |
| KCNQ1 | 542 | I | 0 | 0 | yes | no | no | no | no | yes | no | no | yes | no | no | no | no |
| KCNQ1 | 543 | E | 0 | 1 | yes | no | no | no | no | yes | no | no | no | no | no | no | no |
| KCNQ1 | 544 | Q | 1 | 0 | yes | no | no | no | no | yes | no | no | yes | no | no | no | no |
| KCNQ1 | 545 | Y | 0 | 0 | yes | no | no | no | no | yes | yes | no | yes | no | no | no | no |
| KCNQ1 | 546 | S | 0 | 1 | yes | no | no | no | no | yes | no | no | no | no | no | no | no |
| KCNQ1 | 547 | Q | 0 | 1 | yes | no | no | no | no | yes | no | no | yes | no | no | no | no |
| KCNQ1 | 548 | G | 1 | 1 | yes | no | no | no | no | yes | no | no | yes | no | no | no | no |
| KCNQ1 | 549 | H | 0 | 0 | yes | no | no | no | no | yes | no | no | yes | no | no | no | no |
| KCNQ1 | 550 | L | 0 | 0 | yes | no | no | no | no | yes | no | no | no | no | no | no | no |

| | | | | | | | | | | | | | | | | | |
|---|---|---|---|---|---|---|---|---|---|---|---|---|---|---|---|---|---|
| KCNQ1 | 551 | N | 0 | 0 | yes | no | no | no | no | yes | no | no | yes | no | no | no | no |
| KCNQ1 | 552 | L | 1 | 0 | yes | no | no | no | no | yes | yes | no | yes | no | no | no | no |
| KCNQ1 | 553 | M | 0 | 0 | yes | no | no | no | no | yes | no | no | yes | no | no | no | no |
| KCNQ1 | 554 | V | 0 | 1 | yes | no | no | no | no | yes | no | no | no | no | no | no | no |
| KCNQ1 | 555 | R | 2 | 3 | yes | no | no | no | no | yes | no | no | yes | no | no | no | no |
| KCNQ1 | 556 | I | 0 | 0 | yes | no | no | no | no | yes | yes | no | yes | no | no | no | no |
| KCNQ1 | 557 | K | 0 | 1 | yes | no | no | no | no | yes | no | no | yes | no | no | no | no |
| KCNQ1 | 558 | E | 0 | 0 | yes | no | no | no | no | yes | no | no | no | no | no | no | no |
| KCNQ1 | 559 | L | 0 | 0 | yes | no | no | no | no | yes | yes | no | yes | no | no | no | no |
| KCNQ1 | 560 | Q | 0 | 0 | yes | no | no | no | no | yes | no | no | yes | no | no | no | no |
| KCNQ1 | 561 | R | 0 | 2 | yes | no | no | no | no | yes | no | no | no | no | no | no | no |
| KCNQ1 | 562 | R | 0 | 1 | yes | no | no | no | no | yes | no | no | yes | no | no | no | no |
| KCNQ1 | 563 | L | 0 | 1 | yes | no | no | no | no | yes | yes | no | yes | no | no | no | no |
| KCNQ1 | 564 | D | 1 | 1 | yes | no | no | no | no | yes | no | no | yes | no | no | no | no |
| KCNQ1 | 565 | Q | 0 | 0 | yes | no | no | no | no | yes | no | no | no | no | no | no | no |
| KCNQ1 | 566 | S | 0 | 2 | yes | no | no | no | no | yes | no | no | yes | no | no | no | no |
| KCNQ1 | 567 | I | 0 | 2 | no | yes | no | no | no | yes | yes | no | yes | no | no | no | no |
| KCNQ1 | 568 | G | 0 | 2 | no | yes | no | no | no | yes | no | no | yes | no | no | no | no |
| KCNQ2 | 70 | R | 0 | 0 | no | yes | no | no | no | yes | no | no | no | no | no | no | no |
| KCNQ2 | 71 | N | 0 | 0 | yes | no | no | no | no | yes | no | no | no | no | no | no | no |
| KCNQ2 | 72 | A | 1 | 0 | yes | no | no | no | no | yes | no | no | no | no | no | no | no |
| KCNQ2 | 73 | F | 0 | 0 | yes | no | no | no | no | yes | no | no | no | no | no | no | no |
| KCNQ2 | 74 | Y | 0 | 0 | yes | no | no | no | no | yes | no | no | no | no | no | no | no |
| KCNQ2 | 75 | R | 0 | 0 | yes | no | no | no | no | yes | no | no | no | no | no | no | no |
| KCNQ2 | 76 | K | 0 | 0 | yes | no | no | no | no | yes | no | no | no | no | no | no | no |
| KCNQ2 | 77 | L | 0 | 0 | yes | no | no | no | no | yes | no | no | no | no | no | no | no |
| KCNQ2 | 78 | Q | 0 | 0 | yes | no | no | no | no | yes | no | no | no | no | no | no | no |
| KCNQ2 | 79 | N | 0 | 1 | yes | no | no | no | no | yes | no | no | no | no | no | no | no |
| KCNQ2 | 80 | F | 0 | 0 | yes | no | no | no | no | yes | no | no | no | no | no | no | no |
| KCNQ2 | 81 | L | 0 | 1 | yes | no | no | no | no | yes | no | no | no | no | no | no | no |
| KCNQ2 | 82 | Y | 0 | 0 | yes | no | no | no | no | yes | no | no | no | no | no | no | no |
| KCNQ2 | 83 | N | 0 | 0 | yes | no | no | no | no | yes | no | no | no | no | no | no | no |
| KCNQ2 | 84 | V | 0 | 0 | yes | no | no | no | no | yes | no | no | no | no | no | no | no |

| | | | | | | | | | | | | | | | | | |
|---|---|---|---|---|---|---|---|---|---|---|---|---|---|---|---|---|---|
| KCNQ2 | 85 | L | 0 | 0 | yes | no | no | no | no | yes | no | no | no | no | no | no | no |
| KCNQ2 | 86 | E | 0 | 0 | no | yes | no | no | no | yes | no | no | no | no | no | no | no |
| KCNQ2 | 87 | R | 0 | 0 | no | yes | no | no | no | yes | no | no | no | no | no | no | no |
| KCNQ2 | 88 | P | 0 | 0 | no | yes | no | no | no | yes | no | no | no | no | no | no | no |
| KCNQ2 | 89 | R | 0 | 0 | no | yes | no | no | no | yes | no | no | no | no | no | no | no |
| KCNQ2 | 90 | G | 0 | 0 | no | yes | no | no | no | yes | no | no | no | no | no | no | no |
| KCNQ2 | 91 | W | 0 | 0 | no | yes | no | no | no | yes | no | no | no | no | no | no | no |
| KCNQ2 | 92 | A | 0 | 0 | yes | no | no | no | no | no | no | no | no | no | no | no | no |
| KCNQ2 | 93 | F | 0 | 0 | yes | no | no | yes | no | no | no | no | no | no | no | no | no |
| KCNQ2 | 94 | I | 0 | 0 | yes | no | no | yes | no | no | no | no | no | no | no | no | no |
| KCNQ2 | 95 | Y | 0 | 0 | yes | no | no | yes | no | no | no | no | no | no | no | no | no |
| KCNQ2 | 96 | H | 0 | 1 | yes | no | no | yes | no | no | no | no | no | no | no | no | no |
| KCNQ2 | 97 | A | 1 | 0 | yes | no | no | yes | no | no | no | no | no | no | no | no | no |
| KCNQ2 | 98 | Y | 0 | 0 | yes | no | no | yes | no | no | no | no | no | no | no | no | no |
| KCNQ2 | 99 | V | 0 | 0 | yes | no | no | yes | no | no | no | no | no | no | no | no | no |
| KCNQ2 | 100 | F | 0 | 0 | yes | no | no | yes | no | no | no | no | no | no | no | no | no |
| KCNQ2 | 101 | L | 0 | 0 | yes | no | no | yes | no | no | no | no | no | no | no | no | no |
| KCNQ2 | 102 | L | 0 | 0 | yes | no | no | yes | no | no | no | no | no | no | no | no | no |
| KCNQ2 | 103 | V | 0 | 0 | yes | no | no | yes | no | no | no | no | no | no | no | no | no |
| KCNQ2 | 104 | F | 1 | 1 | yes | no | no | yes | no | no | no | no | yes | no | no | no | no |
| KCNQ2 | 105 | S | 0 | 0 | yes | no | no | yes | no | no | no | no | no | no | no | no | no |
| KCNQ2 | 106 | C | 0 | 0 | yes | no | no | yes | no | no | no | no | no | no | no | no | no |
| KCNQ2 | 107 | L | 0 | 1 | yes | no | no | yes | no | no | no | no | no | no | no | no | no |
| KCNQ2 | 108 | V | 1 | 0 | yes | no | no | yes | no | no | no | no | no | no | no | no | no |
| KCNQ2 | 109 | L | 0 | 0 | yes | no | no | yes | no | no | no | no | no | no | no | no | no |
| KCNQ2 | 110 | S | 0 | 0 | yes | no | no | yes | no | no | no | no | no | no | no | no | no |
| KCNQ2 | 111 | V | 0 | 0 | yes | no | no | yes | no | no | no | no | yes | no | no | no | no |
| KCNQ2 | 112 | F | 0 | 0 | yes | no | no | yes | no | no | no | no | no | no | no | no | no |
| KCNQ2 | 113 | S | 0 | 1 | no | yes | no | no | yes | no | no | no | no | no | no | no | no |
| KCNQ2 | 114 | T | 0 | 2 | no | yes | no | no | yes | no | no | no | yes | no | no | no | no |
| KCNQ2 | 115 | I | 1 | 0 | no | yes | no | no | yes | no | no | no | no | no | no | no | no |
| KCNQ2 | 116 | K | 1 | 0 | no | yes | no | no | yes | no | no | no | no | no | no | no | no |
| KCNQ2 | 117 | E | 1 | 0 | no | yes | no | no | yes | no | no | no | no | no | no | no | no |

| | | | | | | | | | | | | | | | | | |
|---|---|---|---|---|---|---|---|---|---|---|---|---|---|---|---|---|---|
| KCNQ2 | 118 | Y | 0 | 0 | no | yes | no | no | yes | no | no | no | no | no | no | no | no |
| KCNQ2 | 119 | E | 0 | 1 | yes | no | no | no | yes | no | no | no | no | no | no | no | no |
| KCNQ2 | 120 | K | 0 | 0 | yes | no | no | no | yes | no | no | no | no | no | no | no | no |
| KCNQ2 | 121 | S | 0 | 0 | yes | no | no | no | yes | no | no | no | no | no | no | no | no |
| KCNQ2 | 122 | S | 0 | 1 | yes | no | no | no | yes | no | no | no | no | no | no | no | no |
| KCNQ2 | 123 | E | 1 | 0 | yes | no | no | no | no | no | no | no | no | no | no | no | no |
| KCNQ2 | 124 | G | 0 | 0 | yes | no | no | no | no | no | no | no | no | no | no | no | no |
| KCNQ2 | 125 | A | 1 | 0 | yes | no | no | yes | no | no | no | no | no | no | no | no | no |
| KCNQ2 | 126 | L | 0 | 0 | yes | no | no | yes | no | no | no | no | no | no | no | no | no |
| KCNQ2 | 127 | Y | 0 | 1 | yes | no | no | yes | no | no | no | no | no | no | no | no | no |
| KCNQ2 | 128 | I | 1 | 0 | yes | no | no | yes | no | no | no | no | no | no | no | no | no |
| KCNQ2 | 129 | L | 0 | 0 | yes | no | no | yes | no | no | no | no | no | no | no | no | no |
| KCNQ2 | 130 | E | 0 | 1 | yes | no | no | yes | no | no | no | no | no | no | no | no | no |
| KCNQ2 | 131 | I | 0 | 0 | yes | no | no | yes | no | no | no | no | no | no | no | no | no |
| KCNQ2 | 132 | V | 0 | 1 | yes | no | no | yes | no | no | no | no | no | no | no | no | no |
| KCNQ2 | 133 | T | 0 | 0 | yes | no | no | yes | no | no | no | no | no | no | no | no | no |
| KCNQ2 | 134 | I | 0 | 0 | yes | no | no | yes | no | no | no | no | no | no | no | no | no |
| KCNQ2 | 135 | V | 1 | 0 | yes | no | no | yes | no | no | no | no | no | no | no | no | no |
| KCNQ2 | 136 | V | 0 | 0 | yes | no | no | yes | no | no | no | no | no | no | no | no | no |
| KCNQ2 | 137 | F | 0 | 0 | yes | no | no | yes | no | no | no | no | no | no | no | no | no |
| KCNQ2 | 138 | G | 0 | 0 | yes | no | no | yes | no | no | no | no | no | no | no | no | no |
| KCNQ2 | 139 | V | 1 | 0 | yes | no | no | yes | no | no | no | no | no | no | no | no | no |
| KCNQ2 | 140 | E | 0 | 1 | yes | no | no | yes | no | no | no | no | no | no | no | no | no |
| KCNQ2 | 141 | Y | 0 | 0 | yes | no | no | yes | no | no | no | no | no | no | no | no | no |
| KCNQ2 | 142 | F | 1 | 0 | yes | no | no | yes | no | no | no | no | no | no | no | no | no |
| KCNQ2 | 143 | V | 1 | 0 | yes | no | no | yes | no | no | no | no | no | no | no | no | no |
| KCNQ2 | 144 | R | 0 | 3 | yes | no | no | yes | no | yes | no | no | no | no | no | no | no |
| KCNQ2 | 145 | I | 0 | 0 | yes | no | no | yes | no | yes | no | no | no | no | no | no | no |
| KCNQ2 | 146 | W | 0 | 0 | yes | no | no | no | no | yes | no | no | no | no | no | no | no |
| KCNQ2 | 147 | A | 0 | 0 | yes | no | no | no | no | yes | no | no | no | no | no | no | no |
| KCNQ2 | 148 | A | 0 | 0 | yes | no | no | no | no | yes | no | no | no | no | no | no | no |
| KCNQ2 | 149 | G | 0 | 0 | yes | no | no | no | no | yes | no | no | no | no | no | no | no |
| KCNQ2 | 150 | C | 1 | 0 | yes | no | no | no | no | yes | no | no | no | no | no | no | no |

| | | | | | | | | | | | | | | | | | |
|---|---|---|---|---|---|---|---|---|---|---|---|---|---|---|---|---|---|
| KCNQ2 | 151 | C | 0 | 0 | no | yes | no | no | no | yes | no | no | no | no | no | no | no |
| KCNQ2 | 152 | C | 0 | 0 | no | yes | no | no | no | yes | no | no | no | no | no | no | no |
| KCNQ2 | 153 | R | 0 | 0 | no | yes | no | no | no | yes | no | no | yes | no | no | no | no |
| KCNQ2 | 154 | Y | 0 | 1 | no | yes | no | no | no | yes | no | no | no | no | no | no | no |
| KCNQ2 | 155 | R | 2 | 0 | no | yes | no | no | no | yes | no | no | no | no | no | no | no |
| KCNQ2 | 156 | G | 0 | 0 | no | yes | no | no | no | yes | no | no | no | no | no | no | no |
| KCNQ2 | 157 | W | 0 | 0 | yes | no | no | no | no | yes | no | no | no | no | no | no | no |
| KCNQ2 | 158 | R | 0 | 0 | yes | no | no | no | no | yes | no | no | no | no | no | no | no |
| KCNQ2 | 159 | G | 0 | 2 | yes | no | no | no | no | yes | no | no | no | no | no | no | no |
| KCNQ2 | 160 | R | 0 | 0 | yes | no | no | no | no | yes | no | no | no | no | no | no | no |
| KCNQ2 | 161 | L | 0 | 1 | no | yes | no | no | no | yes | no | no | no | no | no | no | no |
| KCNQ2 | 162 | K | 1 | 0 | no | yes | no | no | no | yes | no | no | no | no | no | no | no |
| KCNQ2 | 163 | F | 0 | 0 | yes | no | no | no | no | yes | no | no | no | no | no | no | no |
| KCNQ2 | 164 | A | 0 | 0 | yes | no | no | yes | no | yes | no | no | no | no | no | no | no |
| KCNQ2 | 165 | R | 2 | 0 | yes | no | no | yes | no | yes | no | no | no | no | no | no | no |
| KCNQ2 | 166 | K | 0 | 0 | no | yes | no | yes | no | yes | no | no | no | no | no | no | no |
| KCNQ2 | 167 | P | 1 | 0 | yes | no | no | yes | no | no | no | no | no | no | no | no | no |
| KCNQ2 | 168 | F | 0 | 0 | yes | no | no | yes | no | no | no | no | no | no | no | no | no |
| KCNQ2 | 169 | C | 0 | 1 | yes | no | no | yes | no | no | no | no | no | no | no | no | no |
| KCNQ2 | 170 | V | 0 | 0 | yes | no | no | yes | no | no | no | no | no | no | no | no | no |
| KCNQ2 | 171 | I | 0 | 0 | yes | no | no | yes | no | no | no | no | no | no | no | no | no |
| KCNQ2 | 172 | D | 0 | 1 | yes | no | no | yes | no | no | no | no | no | no | no | no | no |
| KCNQ2 | 173 | I | 1 | 0 | yes | no | no | yes | no | no | no | no | no | no | no | no | no |
| KCNQ2 | 174 | M | 0 | 0 | yes | no | no | yes | no | no | no | no | no | no | no | no | no |
| KCNQ2 | 175 | V | 0 | 1 | yes | no | no | yes | no | no | no | no | no | no | no | no | no |
| KCNQ2 | 176 | L | 0 | 0 | yes | no | no | yes | no | no | no | no | no | no | no | no | no |
| KCNQ2 | 177 | I | 1 | 0 | yes | no | no | yes | no | no | no | no | no | no | no | no | no |
| KCNQ2 | 178 | A | 0 | 0 | yes | no | no | yes | no | no | no | no | no | no | no | no | no |
| KCNQ2 | 179 | S | 0 | 0 | yes | no | no | yes | no | no | no | no | no | no | no | no | no |
| KCNQ2 | 180 | I | 1 | 0 | yes | no | no | yes | no | no | no | no | no | no | no | no | no |
| KCNQ2 | 181 | A | 0 | 0 | yes | no | no | yes | no | no | no | no | no | no | no | no | no |
| KCNQ2 | 182 | V | 0 | 1 | yes | no | no | yes | no | no | no | no | no | no | no | no | no |
| KCNQ2 | 183 | L | 0 | 0 | no | yes | no | yes | no | no | no | no | no | no | no | no | no |

| | | | | | | | | | | | | | | | | | |
|---|---|---|---|---|---|---|---|---|---|---|---|---|---|---|---|---|---|
| KCNQ2 | 184 | A | 0 | 0 | no | yes | no | yes | no | no | no | no | no | no | no | no | no |
| KCNQ2 | 195 | S | 0 | 2 | no | yes | no | no | yes | no | no | no | no | no | no | no | no |
| KCNQ2 | 196 | A | 1 | 1 | no | yes | no | yes | no | no | no | no | no | yes | no | no | no |
| KCNQ2 | 197 | L | 0 | 1 | yes | no | no | no | no | no | no | no | no | yes | no | no | no |
| KCNQ2 | 198 | R | 0 | 1 | yes | no | no | no | no | no | no | no | no | yes | no | no | no |
| KCNQ2 | 199 | S | 0 | 0 | yes | no | no | yes | no | no | no | no | no | yes | no | no | no |
| KCNQ2 | 200 | L | 0 | 1 | yes | no | no | yes | no | no | no | no | no | yes | no | no | no |
| KCNQ2 | 201 | R | 0 | 2 | yes | no | no | yes | no | no | no | no | no | yes | no | no | no |
| KCNQ2 | 202 | F | 0 | 0 | yes | no | no | yes | no | no | no | no | yes | yes | no | no | no |
| KCNQ2 | 203 | L | 0 | 1 | yes | no | no | yes | no | no | no | no | no | yes | no | no | no |
| KCNQ2 | 204 | Q | 0 | 2 | yes | no | no | yes | no | no | no | no | no | yes | no | no | no |
| KCNQ2 | 205 | I | 0 | 1 | yes | no | no | yes | no | no | no | no | no | yes | no | no | no |
| KCNQ2 | 206 | L | 0 | 1 | yes | no | no | yes | no | no | no | no | no | yes | no | no | no |
| KCNQ2 | 207 | R | 0 | 2 | yes | no | no | yes | no | no | no | no | no | yes | no | no | no |
| KCNQ2 | 208 | M | 0 | 1 | yes | no | no | yes | no | no | no | no | yes | yes | no | no | no |
| KCNQ2 | 209 | I | 0 | 1 | yes | no | no | yes | no | no | no | no | yes | yes | no | no | no |
| KCNQ2 | 210 | R | 0 | 3 | yes | no | no | yes | no | no | no | no | no | yes | no | no | no |
| KCNQ2 | 211 | M | 0 | 1 | no | yes | no | yes | no | no | no | no | no | yes | no | no | no |
| KCNQ2 | 212 | D | 0 | 3 | no | yes | no | yes | no | no | no | no | yes | yes | no | no | no |
| KCNQ2 | 213 | R | 0 | 2 | no | yes | no | yes | no | no | no | no | no | yes | no | no | no |
| KCNQ2 | 214 | R | 0 | 2 | no | yes | no | no | no | no | no | no | no | yes | no | no | no |
| KCNQ2 | 215 | G | 0 | 1 | no | yes | no | yes | no | no | no | no | no | yes | no | no | no |
| KCNQ2 | 216 | G | 0 | 0 | yes | no | no | no | no | no | no | no | no | yes | no | no | no |
| KCNQ2 | 217 | T | 0 | 2 | yes | no | no | yes | no | no | no | no | no | yes | no | no | no |
| KCNQ2 | 218 | W | 0 | 0 | yes | no | no | yes | no | no | no | no | no | yes | no | no | no |
| KCNQ2 | 219 | K | 0 | 1 | yes | no | no | no | no | yes | no | no | no | no | no | no | no |
| KCNQ2 | 220 | L | 0 | 0 | yes | no | no | no | no | yes | no | no | yes | no | no | no | no |
| KCNQ2 | 221 | L | 0 | 0 | yes | no | no | yes | no | yes | no | no | no | no | no | no | no |
| KCNQ2 | 222 | G | 0 | 0 | yes | no | no | yes | no | yes | no | no | no | no | no | no | no |
| KCNQ2 | 223 | S | 0 | 1 | yes | no | no | no | no | yes | no | no | no | no | no | no | no |
| KCNQ2 | 224 | V | 0 | 1 | yes | no | no | no | no | yes | no | no | no | no | no | no | no |
| KCNQ2 | 225 | V | 0 | 0 | yes | no | no | yes | no | yes | no | no | no | no | no | no | no |
| KCNQ2 | 226 | Y | 0 | 0 | yes | no | no | no | no | yes | no | no | no | no | no | no | no |

| | | | | | | | | | | | | | | | | | |
|---|---|---|---|---|---|---|---|---|---|---|---|---|---|---|---|---|---|
| KCNQ2 | 227 | A | 0 | 0 | yes | no | no | no | no | yes | no | no | yes | no | no | no | no |
| KCNQ2 | 228 | H | 0 | 2 | no | yes | no | no | no | yes | no | no | yes | no | no | no | no |
| KCNQ2 | 229 | S | 0 | 0 | yes | no | no | no | no | yes | no | no | no | no | no | no | no |
| KCNQ2 | 230 | K | 0 | 0 | yes | no | no | no | no | yes | no | no | no | no | no | no | no |
| KCNQ2 | 231 | E | 0 | 3 | yes | no | no | yes | no | yes | no | no | yes | no | no | no | no |
| KCNQ2 | 232 | L | 0 | 0 | yes | no | no | yes | no | no | no | no | no | no | no | no | no |
| KCNQ2 | 233 | V | 0 | 0 | yes | no | no | yes | no | no | no | no | no | no | no | no | no |
| KCNQ2 | 234 | T | 0 | 2 | yes | no | no | yes | no | no | no | no | yes | no | no | no | no |
| KCNQ2 | 235 | A | 0 | 1 | yes | no | no | yes | no | no | no | no | no | no | no | no | no |
| KCNQ2 | 236 | W | 0 | 0 | yes | no | no | yes | no | no | no | no | no | no | no | no | no |
| KCNQ2 | 237 | Y | 0 | 2 | yes | no | no | yes | no | no | no | no | yes | no | no | no | no |
| KCNQ2 | 238 | I | 1 | 0 | yes | no | no | yes | no | no | no | no | no | no | no | no | no |
| KCNQ2 | 239 | G | 0 | 1 | yes | no | no | yes | no | no | no | no | no | no | no | no | no |
| KCNQ2 | 240 | F | 0 | 0 | yes | no | no | yes | no | no | no | no | yes | no | no | no | no |
| KCNQ2 | 241 | L | 0 | 0 | yes | no | no | yes | no | no | no | no | yes | no | no | no | no |
| KCNQ2 | 242 | C | 0 | 1 | yes | no | no | yes | no | no | no | no | no | no | no | no | no |
| KCNQ2 | 243 | L | 0 | 2 | yes | no | no | yes | no | no | no | no | no | no | no | no | no |
| KCNQ2 | 244 | I | 0 | 0 | yes | no | no | yes | no | no | no | no | no | no | no | no | no |
| KCNQ2 | 245 | L | 0 | 0 | yes | no | no | yes | no | no | no | no | no | no | no | no | no |
| KCNQ2 | 246 | A | 0 | 1 | yes | no | no | yes | no | no | no | no | no | no | no | no | no |
| KCNQ2 | 247 | S | 0 | 2 | yes | no | no | yes | no | no | no | no | no | no | no | no | no |
| KCNQ2 | 248 | F | 0 | 1 | yes | no | no | yes | no | no | no | no | yes | no | no | no | no |
| KCNQ2 | 249 | L | 0 | 1 | yes | no | no | yes | no | no | no | no | no | no | no | no | no |
| KCNQ2 | 250 | V | 0 | 2 | yes | no | no | yes | no | no | no | no | no | no | no | no | no |
| KCNQ2 | 251 | Y | 0 | 0 | yes | no | no | no | no | no | no | no | no | no | no | no | no |
| KCNQ2 | 252 | L | 0 | 0 | no | yes | no | no | no | no | no | no | no | no | no | no | no |
| KCNQ2 | 253 | A | 0 | 1 | no | yes | no | no | yes | no | no | no | no | no | no | no | no |
| KCNQ2 | 254 | E | 0 | 0 | no | yes | no | no | yes | no | no | no | no | no | no | no | no |
| KCNQ2 | 255 | K | 0 | 1 | no | yes | no | no | yes | no | no | no | no | no | no | no | no |
| KCNQ2 | 256 | G | 0 | 1 | no | yes | no | no | yes | no | no | no | no | no | no | no | no |
| KCNQ2 | 257 | E | 0 | 0 | no | yes | no | no | yes | no | no | no | no | no | no | no | no |
| KCNQ2 | 258 | N | 0 | 1 | no | yes | no | no | yes | no | no | no | no | no | no | no | no |
| KCNQ2 | 259 | D | 1 | 3 | no | yes | no | no | yes | no | yes | no | no | no | no | no | no |

| | | | | | | | | | | | | | | | | | |
|---|---|---|---|---|---|---|---|---|---|---|---|---|---|---|---|---|---|
| KCNQ2 | 260 | H | 0 | 1 | no | yes | no | no | yes | no | yes | no | no | no | no | no | no |
| KCNQ2 | 261 | F | 0 | 2 | no | yes | no | no | yes | no | no | no | no | no | no | no | no |
| KCNQ2 | 262 | D | 0 | 0 | no | yes | no | no | yes | no | no | no | no | no | no | no | no |
| KCNQ2 | 263 | T | 0 | 1 | no | yes | no | no | yes | no | no | no | yes | no | no | no | no |
| KCNQ2 | 264 | Y | 0 | 1 | yes | no | no | yes | yes | no | no | no | yes | no | no | no | no |
| KCNQ2 | 265 | A | 0 | 3 | yes | no | no | no | no | no | no | yes | yes | no | no | no | no |
| KCNQ2 | 266 | D | 0 | 2 | yes | no | no | no | no | no | no | yes | yes | no | no | no | no |
| KCNQ2 | 267 | A | 0 | 0 | yes | no | no | yes | no | no | no | yes | no | no | no | no | no |
| KCNQ2 | 268 | L | 0 | 2 | yes | no | no | yes | no | no | no | yes | no | no | no | no | no |
| KCNQ2 | 269 | W | 0 | 1 | yes | no | no | yes | no | no | no | yes | yes | no | no | no | no |
| KCNQ2 | 270 | W | 0 | 1 | yes | no | no | yes | no | no | no | yes | yes | no | no | no | no |
| KCNQ2 | 271 | G | 0 | 2 | yes | no | no | yes | no | no | no | yes | no | no | no | no | no |
| KCNQ2 | 272 | L | 0 | 0 | yes | no | no | yes | no | no | no | yes | yes | no | no | no | no |
| KCNQ2 | 273 | I | 0 | 2 | yes | no | no | yes | no | no | no | yes | no | no | no | no | no |
| KCNQ2 | 274 | T | 0 | 1 | yes | no | no | yes | no | no | no | yes | yes | no | no | no | no |
| KCNQ2 | 275 | L | 0 | 0 | yes | no | no | yes | no | no | no | yes | no | no | no | no | no |
| KCNQ2 | 276 | T | 0 | 2 | no | yes | no | yes | no | no | no | yes | yes | no | no | no | no |
| KCNQ2 | 277 | T | 0 | 2 | no | yes | no | yes | no | no | yes | yes | yes | no | no | no | no |
| KCNQ2 | 278 | I | 1 | 1 | no | yes | no | yes | no | no | yes | yes | yes | no | no | no | no |
| KCNQ2 | 279 | G | 0 | 2 | no | yes | no | yes | no | no | yes | yes | yes | no | no | no | no |
| KCNQ2 | 280 | Y | 0 | 2 | no | yes | no | yes | no | no | yes | yes | yes | no | no | no | no |
| KCNQ2 | 281 | G | 0 | 2 | no | yes | no | no | no | no | yes | yes | yes | no | no | no | no |
| KCNQ2 | 282 | D | 0 | 4 | no | yes | no | no | no | no | yes | yes | yes | no | no | no | no |
| KCNQ2 | 283 | K | 0 | 2 | no | yes | no | no | no | no | no | yes | yes | no | no | no | no |
| KCNQ2 | 284 | Y | 0 | 3 | no | yes | no | no | no | no | yes | yes | yes | no | no | no | no |
| KCNQ2 | 285 | P | 0 | 2 | no | yes | no | no | no | no | no | yes | no | no | no | no | no |
| KCNQ2 | 286 | Q | 0 | 0 | no | yes | no | no | yes | no | no | no | no | no | no | no | no |
| KCNQ2 | 287 | T | 0 | 2 | no | yes | no | no | yes | no | no | no | no | no | no | no | no |
| KCNQ2 | 288 | W | 0 | 0 | yes | no | no | no | yes | no | no | no | yes | no | no | no | no |
| KCNQ2 | 289 | N | 0 | 0 | yes | no | no | no | yes | no | no | no | no | no | no | no | no |
| KCNQ2 | 290 | G | 0 | 3 | yes | no | no | no | yes | no | no | no | no | no | no | no | no |
| KCNQ2 | 291 | R | 0 | 2 | yes | no | no | no | yes | no | no | no | yes | no | no | no | no |
| KCNQ2 | 292 | L | 0 | 1 | yes | no | no | yes | no | no | no | no | no | no | no | no | no |

| | | | | | | | | | | | | | | | | | |
|---|---|---|---|---|---|---|---|---|---|---|---|---|---|---|---|---|---|
| KCNQ2 | 293 | L | 0 | 0 | yes | no | no | yes | no | no | no | no | no | no | no | no | no |
| KCNQ2 | 294 | A | 0 | 1 | yes | no | no | yes | no | no | no | no | no | no | no | no | no |
| KCNQ2 | 295 | A | 0 | 0 | yes | no | no | yes | no | no | no | no | yes | no | no | no | no |
| KCNQ2 | 296 | T | 1 | 1 | yes | no | no | yes | no | no | no | no | no | no | no | no | no |
| KCNQ2 | 297 | F | 0 | 0 | yes | no | no | yes | no | no | no | no | no | no | no | no | no |
| KCNQ2 | 298 | T | 0 | 0 | yes | no | no | yes | no | no | no | no | yes | no | no | no | no |
| KCNQ2 | 299 | L | 0 | 0 | yes | no | no | yes | no | no | no | no | yes | no | no | no | no |
| KCNQ2 | 300 | I | 0 | 0 | yes | no | no | yes | no | no | no | no | no | no | no | no | no |
| KCNQ2 | 301 | G | 0 | 3 | yes | no | no | yes | no | no | no | no | no | no | no | no | no |
| KCNQ2 | 302 | V | 0 | 0 | yes | no | no | yes | no | no | no | no | no | no | no | no | no |
| KCNQ2 | 303 | S | 0 | 0 | yes | no | no | yes | no | no | no | no | yes | no | no | no | no |
| KCNQ2 | 304 | F | 0 | 1 | yes | no | no | yes | no | no | no | no | no | no | no | no | no |
| KCNQ2 | 305 | F | 0 | 1 | yes | no | no | yes | no | no | no | no | yes | no | no | no | no |
| KCNQ2 | 306 | A | 0 | 2 | yes | no | no | yes | no | no | yes | no | yes | no | no | no | no |
| KCNQ2 | 307 | L | 0 | 0 | yes | no | no | yes | no | no | no | no | yes | no | no | no | no |
| KCNQ2 | 308 | P | 0 | 2 | yes | no | no | yes | no | no | no | no | no | no | no | no | no |
| KCNQ2 | 309 | A | 1 | 1 | yes | no | no | yes | no | no | no | no | yes | no | no | no | no |
| KCNQ2 | 310 | G | 0 | 0 | yes | no | no | yes | no | no | yes | no | no | no | no | no | no |
| KCNQ2 | 311 | I | 0 | 2 | yes | no | no | yes | no | no | no | no | yes | no | no | no | no |
| KCNQ2 | 312 | L | 0 | 1 | yes | no | no | yes | no | no | no | no | no | no | no | no | no |
| KCNQ2 | 313 | G | 0 | 1 | yes | no | no | yes | no | yes | no | no | yes | no | no | no | no |
| KCNQ2 | 314 | S | 0 | 2 | yes | no | no | no | no | yes | yes | no | yes | no | no | no | no |
| KCNQ2 | 315 | G | 0 | 2 | yes | no | no | no | no | yes | no | no | no | no | no | no | no |
| KCNQ2 | 316 | F | 0 | 0 | yes | no | no | no | no | yes | no | no | yes | no | no | no | no |
| KCNQ2 | 317 | A | 0 | 1 | yes | no | no | no | no | yes | no | no | yes | no | no | no | no |
| KCNQ2 | 318 | L | 0 | 0 | yes | no | no | no | no | yes | yes | no | yes | no | no | no | no |
| KCNQ2 | 319 | K | 0 | 2 | yes | no | no | no | no | yes | no | no | no | no | no | no | no |
| KCNQ2 | 320 | V | 0 | 1 | yes | no | no | no | no | yes | no | no | yes | no | no | no | no |
| KCNQ2 | 321 | Q | 0 | 0 | yes | no | no | no | no | yes | yes | no | yes | no | no | no | no |
| KCNQ2 | 322 | E | 0 | 0 | yes | no | no | no | no | yes | no | no | no | no | no | no | no |
| KCNQ2 | 323 | Q | 0 | 0 | yes | no | no | no | no | yes | no | no | no | no | no | no | no |
| KCNQ2 | 324 | H | 0 | 1 | yes | no | no | no | no | yes | no | no | no | no | no | no | no |
| KCNQ2 | 325 | R | 0 | 1 | yes | no | no | no | no | yes | yes | no | no | no | no | no | no |

| | | | | | | | | | | | | | | | | | |
|---|---|---|---|---|---|---|---|---|---|---|---|---|---|---|---|---|---|
| KCNQ2 | 326 | Q | 0 | 0 | yes | no | no | no | no | yes | no | no | no | no | no | no | no |
| KCNQ2 | 327 | K | 0 | 0 | yes | no | no | no | no | yes | no | no | no | no | no | no | no |
| KCNQ2 | 328 | H | 0 | 1 | yes | no | no | no | no | yes | no | no | no | no | no | no | no |
| KCNQ2 | 329 | F | 0 | 1 | yes | no | no | no | no | yes | no | no | no | no | no | no | no |
| KCNQ2 | 330 | E | 0 | 0 | yes | no | no | no | no | yes | no | no | no | no | no | no | no |
| KCNQ2 | 331 | K | 0 | 1 | yes | no | no | no | no | yes | no | no | no | no | no | no | no |
| KCNQ2 | 332 | R | 0 | 2 | yes | no | no | no | no | yes | no | no | no | no | no | no | no |
| KCNQ2 | 333 | R | 1 | 2 | yes | no | no | no | no | yes | no | no | yes | no | no | no | no |
| KCNQ2 | 334 | N | 0 | 0 | yes | no | no | no | no | yes | no | no | yes | no | no | no | no |
| KCNQ2 | 335 | P | 0 | 3 | yes | no | no | no | no | yes | no | no | no | no | no | no | no |
| KCNQ2 | 336 | A | 0 | 0 | yes | no | no | no | no | yes | no | no | yes | no | no | no | no |
| KCNQ2 | 337 | A | 0 | 2 | yes | no | no | no | no | yes | no | no | yes | no | no | no | no |
| KCNQ2 | 338 | G | 0 | 0 | yes | no | no | no | no | yes | no | no | yes | no | no | no | no |
| KCNQ2 | 339 | L | 0 | 1 | yes | no | no | no | no | yes | no | no | no | no | no | no | no |
| KCNQ2 | 340 | I | 0 | 2 | yes | no | no | no | no | yes | no | no | yes | no | no | no | no |
| KCNQ2 | 341 | Q | 0 | 1 | yes | no | no | no | no | yes | no | no | yes | no | no | no | no |
| KCNQ2 | 342 | S | 1 | 0 | yes | no | no | no | no | yes | no | no | no | no | no | no | no |
| KCNQ2 | 343 | A | 0 | 1 | yes | no | no | no | no | yes | no | no | yes | no | no | no | no |
| KCNQ2 | 344 | W | 0 | 1 | yes | no | no | no | no | yes | no | no | yes | no | no | no | no |
| KCNQ2 | 345 | R | 0 | 0 | yes | no | no | no | no | yes | no | no | yes | no | no | no | no |
| KCNQ2 | 346 | F | 0 | 0 | yes | no | no | no | no | yes | no | no | yes | no | no | no | no |
| KCNQ2 | 347 | Y | 0 | 0 | yes | no | no | no | no | yes | no | no | yes | no | no | no | no |
| KCNQ2 | 348 | A | 0 | 0 | no | yes | no | no | no | yes | no | no | no | no | no | no | no |
| KCNQ2 | 349 | T | 0 | 0 | no | yes | no | no | no | yes | no | no | no | no | no | no | no |
| KCNQ2 | 350 | N | 1 | 3 | no | yes | no | no | no | yes | no | no | no | no | no | no | no |
| KCNQ2 | 351 | L | 0 | 2 | no | yes | no | no | no | yes | no | no | no | no | no | no | no |
| KCNQ2 | 352 | S | 1 | 1 | no | yes | no | no | no | yes | no | no | no | no | no | no | no |
| KCNQ2 | 353 | R | 0 | 2 | no | yes | no | no | no | yes | no | no | yes | no | no | no | no |
| KCNQ2 | 354 | T | 0 | 0 | no | yes | no | no | no | yes | no | no | no | no | no | no | no |
| KCNQ2 | 355 | D | 0 | 1 | no | yes | no | no | no | yes | no | no | yes | no | no | no | no |
| KCNQ2 | 356 | L | 0 | 2 | no | yes | no | no | no | yes | no | no | yes | no | no | no | no |
| KCNQ2 | 357 | H | 0 | 0 | yes | no | no | no | no | yes | no | no | no | no | no | no | no |
| KCNQ2 | 358 | S | 0 | 1 | yes | no | no | no | no | yes | no | no | yes | no | no | no | no |

| | | | | | | | | | | | | | | | | | |
|---|---|---|---|---|---|---|---|---|---|---|---|---|---|---|---|---|---|
| KCNQ2 | 359 | T | 0 | 2 | yes | no | no | no | no | yes | no | no | yes | no | no | no | no |
| KCNQ2 | 360 | W | 0 | 1 | yes | no | no | no | no | yes | no | no | no | no | no | no | no |
| KCNQ2 | 361 | Q | 0 | 0 | yes | no | no | no | no | yes | no | no | no | no | no | no | no |
| KCNQ2 | 362 | Y | 0 | 1 | yes | no | no | no | no | yes | no | no | yes | no | no | no | no |
| KCNQ2 | 363 | Y | 0 | 2 | yes | no | no | no | no | yes | no | no | yes | no | no | no | no |
| KCNQ2 | 364 | E | 0 | 0 | yes | no | no | no | no | yes | no | no | no | no | no | no | no |
| KCNQ2 | 365 | R | 0 | 0 | yes | no | no | no | no | yes | no | no | no | no | no | no | no |
| KCNQ2 | 366 | T | 0 | 0 | yes | no | no | no | no | yes | no | no | yes | no | no | no | no |
| KCNQ2 | 367 | V | 1 | 0 | no | yes | no | no | no | yes | no | no | no | no | no | no | no |
| KCNQ2 | 535 | G | 0 | 0 | no | yes | no | no | no | yes | no | no | yes | no | no | no | no |
| KCNQ2 | 536 | L | 0 | 0 | yes | no | no | no | no | yes | no | no | yes | no | no | no | no |
| KCNQ2 | 537 | K | 1 | 0 | yes | no | no | no | no | yes | no | no | no | no | no | no | no |
| KCNQ2 | 538 | V | 0 | 0 | yes | no | no | no | no | yes | no | no | yes | no | no | no | no |
| KCNQ2 | 539 | S | 0 | 0 | yes | no | no | no | no | yes | no | no | yes | no | no | no | no |
| KCNQ2 | 540 | I | 0 | 0 | yes | no | no | no | no | yes | no | no | yes | no | no | no | no |
| KCNQ2 | 541 | R | 0 | 2 | yes | no | no | no | no | yes | no | no | no | no | no | no | no |
| KCNQ2 | 542 | A | 0 | 0 | yes | no | no | no | no | yes | no | no | yes | no | no | no | no |
| KCNQ2 | 543 | V | 1 | 1 | yes | no | no | no | no | yes | no | no | yes | no | no | no | no |
| KCNQ2 | 544 | C | 0 | 0 | yes | no | no | no | no | yes | no | no | no | no | no | no | no |
| KCNQ2 | 545 | V | 1 | 0 | yes | no | no | no | no | yes | no | no | no | no | no | no | no |
| KCNQ2 | 546 | M | 0 | 2 | yes | no | no | no | no | yes | no | no | yes | no | no | no | no |
| KCNQ2 | 547 | R | 1 | 1 | yes | no | no | no | no | yes | no | no | yes | no | no | no | no |
| KCNQ2 | 548 | F | 0 | 0 | yes | no | no | no | no | yes | no | no | no | no | no | no | no |
| KCNQ2 | 549 | L | 0 | 0 | yes | no | no | no | no | yes | no | no | no | no | no | no | no |
| KCNQ2 | 550 | V | 1 | 0 | yes | no | no | no | no | yes | no | no | no | no | no | no | no |
| KCNQ2 | 551 | S | 0 | 0 | yes | no | no | no | no | yes | no | no | no | no | no | no | no |
| KCNQ2 | 552 | K | 0 | 1 | yes | no | no | no | no | yes | no | no | yes | no | no | no | no |
| KCNQ2 | 553 | R | 1 | 3 | yes | no | no | no | no | yes | no | no | yes | no | no | no | no |
| KCNQ2 | 554 | K | 0 | 1 | yes | no | no | no | no | yes | no | no | no | no | no | no | no |
| KCNQ2 | 555 | F | 0 | 1 | yes | no | no | no | no | yes | no | no | yes | no | no | no | no |
| KCNQ2 | 556 | K | 0 | 1 | yes | no | no | no | no | yes | no | no | yes | no | no | no | no |
| KCNQ2 | 557 | E | 0 | 0 | no | yes | no | no | no | yes | no | no | no | no | no | no | no |
| KCNQ2 | 558 | S | 0 | 0 | no | yes | no | no | no | yes | no | no | no | no | no | no | no |

| | | | | | | | | | | | | | | | | | |
|---|---|---|---|---|---|---|---|---|---|---|---|---|---|---|---|---|---|
| KCNQ2 | 559 | L | 0 | 0 | no | yes | no | no | no | yes | no | no | no | no | no | no | no |
| KCNQ2 | 560 | R | 0 | 2 | no | yes | no | no | no | yes | no | no | no | no | no | no | no |
| KCNQ2 | 561 | P | 0 | 2 | no | yes | no | no | no | yes | no | no | no | no | no | no | no |
| KCNQ2 | 562 | Y | 0 | 0 | no | yes | no | no | no | yes | no | no | no | no | no | no | no |
| KCNQ2 | 563 | D | 0 | 4 | yes | no | no | no | no | yes | no | no | yes | no | no | no | no |
| KCNQ2 | 564 | V | 1 | 1 | yes | no | no | no | no | yes | yes | no | yes | no | no | no | no |
| KCNQ2 | 565 | M | 0 | 1 | yes | no | no | no | no | yes | no | no | no | no | no | no | no |
| KCNQ2 | 566 | D | 0 | 0 | yes | no | no | no | no | yes | no | no | no | no | no | no | no |
| KCNQ2 | 567 | V | 0 | 1 | yes | no | no | no | no | yes | yes | no | yes | no | no | no | no |
| KCNQ2 | 568 | I | 0 | 0 | yes | no | no | no | no | yes | no | no | yes | no | no | no | no |
| KCNQ2 | 569 | E | 0 | 0 | yes | no | no | no | no | yes | no | no | no | no | no | no | no |
| KCNQ2 | 570 | Q | 0 | 0 | yes | no | no | no | no | yes | no | no | no | no | no | no | no |
| KCNQ2 | 571 | Y | 0 | 0 | yes | no | no | no | no | yes | yes | no | yes | no | no | no | no |
| KCNQ2 | 572 | S | 0 | 0 | yes | no | no | no | no | yes | no | no | no | no | no | no | no |
| KCNQ2 | 573 | A | 0 | 0 | yes | no | no | no | no | yes | no | no | no | no | no | no | no |
| KCNQ2 | 574 | G | 0 | 1 | yes | no | no | no | no | yes | no | no | yes | no | no | no | no |
| KCNQ2 | 575 | H | 0 | 0 | yes | no | no | no | no | yes | yes | no | yes | no | no | no | no |
| KCNQ2 | 576 | L | 0 | 0 | yes | no | no | no | no | yes | no | no | no | no | no | no | no |
| KCNQ2 | 577 | D | 0 | 0 | yes | no | no | no | no | yes | no | no | no | no | no | no | no |
| KCNQ2 | 578 | M | 0 | 3 | yes | no | no | no | no | yes | yes | no | yes | no | no | no | no |
| KCNQ2 | 579 | L | 0 | 0 | yes | no | no | no | no | yes | no | no | yes | no | no | no | no |
| KCNQ2 | 580 | S | 0 | 0 | yes | no | no | no | no | yes | no | no | no | no | no | no | no |
| KCNQ2 | 581 | R | 0 | 2 | yes | no | no | no | no | yes | no | no | yes | no | no | no | no |
| KCNQ2 | 582 | I | 0 | 1 | yes | no | no | no | no | yes | yes | no | yes | no | no | no | no |
| KCNQ2 | 583 | K | 0 | 0 | yes | no | no | no | no | yes | no | no | no | no | no | no | no |
| KCNQ2 | 584 | S | 0 | 0 | yes | no | no | no | no | yes | no | no | no | no | no | no | no |
| KCNQ2 | 585 | L | 0 | 0 | yes | no | no | no | no | yes | yes | no | yes | no | no | no | no |
| KCNQ2 | 586 | Q | 0 | 0 | yes | no | no | no | no | yes | no | no | yes | no | no | no | no |
| KCNQ2 | 587 | S | 0 | 0 | yes | no | no | no | no | yes | no | no | no | no | no | no | no |
| KCNQ2 | 588 | R | 0 | 1 | yes | no | no | no | no | yes | no | no | yes | no | no | no | no |
| KCNQ2 | 589 | V | 0 | 1 | yes | no | no | no | no | yes | yes | no | yes | no | no | no | no |
| KCNQ2 | 590 | D | 0 | 0 | yes | no | no | no | no | yes | no | no | yes | no | no | no | no |
| KCNQ2 | 591 | Q | 0 | 0 | yes | no | no | no | no | yes | no | no | no | no | no | no | no |

| | | | | | | | | | | | | | | | | | |
|---|---|---|---|---|---|---|---|---|---|---|---|---|---|---|---|---|---|
| KCNQ2 | 592 | I | 0 | 1 | yes | no | no | no | no | yes | yes | no | no | no | no | no | no |
| KCNQ2 | 593 | V | 1 | 0 | yes | no | no | no | no | yes | no | no | no | no | no | no | no |
| KCNQ2 | 594 | G | 0 | 0 | yes | no | no | no | no | yes | no | no | no | no | no | no | no |
| KCNQ2 | 595 | R | 0 | 1 | yes | no | no | no | no | yes | no | no | no | no | no | no | no |
| KCNQ2 | 596 | G | 0 | 0 | yes | no | no | no | no | yes | yes | no | no | no | no | no | no |
| KCNQ2 | 597 | P | 1 | 0 | yes | no | no | no | no | yes | no | no | no | no | no | no | no |
| KCNQ2 | 598 | A | 2 | 0 | yes | no | no | no | no | yes | no | no | no | no | no | no | no |
| KCNQ2 | 599 | I | 0 | 0 | no | yes | no | no | no | yes | yes | no | yes | no | no | no | no |
| KCNQ2 | 600 | T | 2 | 0 | no | yes | no | no | no | yes | yes | no | yes | no | no | no | no |
| MCOLN1 | 38 | E | 1 | 0 | no | yes | no | no | no | yes | no | no | no | no | no | no | no |
| MCOLN1 | 39 | E | 1 | 0 | no | yes | no | no | no | yes | no | no | no | no | no | no | no |
| MCOLN1 | 40 | D | 4 | 0 | no | yes | no | no | no | yes | no | no | no | no | no | no | no |
| MCOLN1 | 41 | L | 0 | 0 | no | yes | no | no | no | yes | no | no | no | no | no | no | no |
| MCOLN1 | 42 | R | 3 | 0 | yes | no | no | no | no | yes | no | no | no | no | no | no | no |
| MCOLN1 | 43 | R | 2 | 0 | yes | no | no | no | no | yes | no | no | no | no | no | no | no |
| MCOLN1 | 44 | R | 1 | 0 | yes | no | no | no | no | yes | no | no | no | no | no | no | no |
| MCOLN1 | 45 | L | 0 | 0 | yes | no | no | no | no | yes | no | no | no | no | no | no | no |
| MCOLN1 | 46 | K | 1 | 0 | yes | no | no | no | no | yes | no | no | no | no | no | no | no |
| MCOLN1 | 47 | Y | 0 | 0 | yes | no | no | no | no | yes | no | no | no | no | no | no | no |
| MCOLN1 | 48 | F | 1 | 0 | yes | no | no | no | no | yes | no | no | no | no | no | no | no |
| MCOLN1 | 49 | F | 0 | 0 | yes | no | no | no | no | yes | no | no | no | no | no | no | no |
| MCOLN1 | 50 | M | 0 | 0 | no | yes | no | no | no | yes | no | no | no | no | no | no | no |
| MCOLN1 | 51 | S | 0 | 0 | no | yes | no | no | no | yes | no | no | no | no | no | no | no |
| MCOLN1 | 52 | P | 1 | 0 | yes | no | no | no | no | yes | no | no | no | no | no | no | no |
| MCOLN1 | 53 | C | 0 | 0 | yes | no | no | no | no | yes | no | no | no | no | no | no | no |
| MCOLN1 | 54 | D | 2 | 0 | yes | no | no | no | no | yes | no | no | no | no | no | no | no |
| MCOLN1 | 55 | K | 1 | 0 | yes | no | no | no | no | yes | no | no | no | no | no | no | no |
| MCOLN1 | 56 | F | 0 | 0 | yes | no | no | no | no | yes | no | no | no | no | no | no | no |
| MCOLN1 | 57 | R | 1 | 0 | yes | no | no | no | no | yes | no | no | no | no | no | no | no |
| MCOLN1 | 58 | A | 0 | 0 | yes | no | no | no | no | yes | no | no | no | no | no | no | no |
| MCOLN1 | 59 | K | 1 | 0 | no | yes | no | no | no | yes | no | no | no | no | no | no | no |
| MCOLN1 | 60 | G | 1 | 0 | no | yes | no | no | no | yes | no | no | no | no | no | no | no |
| MCOLN1 | 61 | R | 2 | 0 | no | yes | no | no | no | yes | no | no | no | no | no | no | no |

| | | | | | | | | | | | | | | | | | |
|---|---|---|---|---|---|---|---|---|---|---|---|---|---|---|---|---|---|
| MCOLN1 | 62 | K | 0 | 0 | no | yes | no | no | no | yes | no | no | no | no | no | no | no |
| MCOLN1 | 63 | P | 0 | 0 | no | yes | no | no | no | yes | no | no | no | no | no | no | no |
| MCOLN1 | 64 | C | 0 | 0 | yes | no | no | yes | no | yes | no | no | no | no | no | no | no |
| MCOLN1 | 65 | K | 0 | 0 | yes | no | no | yes | no | yes | no | no | no | no | no | no | no |
| MCOLN1 | 66 | L | 0 | 0 | yes | no | no | yes | no | no | no | no | no | no | no | no | no |
| MCOLN1 | 67 | M | 0 | 0 | yes | no | no | yes | no | no | no | no | no | no | no | no | no |
| MCOLN1 | 68 | L | 0 | 0 | yes | no | no | yes | no | no | no | no | no | no | no | no | no |
| MCOLN1 | 69 | Q | 1 | 0 | yes | no | no | yes | no | no | no | no | no | no | no | no | no |
| MCOLN1 | 70 | V | 0 | 0 | yes | no | no | yes | no | no | no | no | no | no | no | no | no |
| MCOLN1 | 71 | V | 0 | 0 | yes | no | no | yes | no | no | no | no | no | no | no | no | no |
| MCOLN1 | 72 | K | 0 | 0 | yes | no | no | yes | no | no | no | no | no | no | no | no | no |
| MCOLN1 | 73 | I | 0 | 0 | yes | no | no | yes | no | no | no | no | no | no | no | no | no |
| MCOLN1 | 74 | L | 0 | 0 | yes | no | no | yes | no | no | no | no | no | no | no | no | no |
| MCOLN1 | 75 | V | 1 | 0 | yes | no | no | yes | no | no | no | no | no | no | no | no | no |
| MCOLN1 | 76 | V | 0 | 0 | yes | no | no | yes | no | no | no | no | no | no | no | no | no |
| MCOLN1 | 77 | T | 1 | 0 | yes | no | no | yes | no | no | no | no | yes | no | no | no | no |
| MCOLN1 | 78 | V | 1 | 0 | yes | no | no | yes | no | no | no | no | no | no | no | no | no |
| MCOLN1 | 79 | Q | 0 | 0 | yes | no | no | yes | no | no | no | no | no | no | no | no | no |
| MCOLN1 | 80 | L | 2 | 0 | yes | no | no | yes | no | no | no | no | yes | no | no | no | no |
| MCOLN1 | 81 | I | 0 | 0 | yes | no | no | yes | no | no | no | no | yes | no | no | no | no |
| MCOLN1 | 82 | L | 0 | 0 | yes | no | no | yes | no | no | no | no | no | no | no | no | no |
| MCOLN1 | 83 | F | 0 | 0 | yes | no | no | yes | no | no | no | no | no | no | no | no | no |
| MCOLN1 | 84 | G | 2 | 0 | yes | no | no | yes | no | no | no | no | no | no | no | no | no |
| MCOLN1 | 85 | L | 0 | 0 | yes | no | no | no | no | no | no | no | no | no | no | no | no |
| MCOLN1 | 86 | S | 1 | 0 | yes | no | no | no | no | no | no | no | no | no | no | no | no |
| MCOLN1 | 87 | N | 0 | 0 | yes | no | no | no | yes | no | no | no | yes | no | no | no | no |
| MCOLN1 | 88 | Q | 0 | 0 | yes | no | no | no | yes | no | no | no | yes | no | no | no | no |
| MCOLN1 | 89 | L | 0 | 0 | yes | no | no | no | yes | no | no | no | no | no | no | no | no |
| MCOLN1 | 90 | A | 1 | 0 | yes | no | no | no | yes | no | no | no | no | no | no | no | no |
| MCOLN1 | 91 | V | 0 | 0 | yes | no | no | no | yes | no | no | no | yes | no | no | no | no |
| MCOLN1 | 92 | T | 0 | 0 | yes | no | no | no | yes | no | no | no | no | no | no | no | no |
| MCOLN1 | 93 | F | 0 | 0 | yes | no | no | no | yes | no | no | no | yes | no | no | no | no |
| MCOLN1 | 94 | R | 4 | 0 | yes | no | no | no | yes | no | no | no | yes | no | no | no | no |

| | | | | | | | | | | | | | | | | | |
|---|---|---|---|---|---|---|---|---|---|---|---|---|---|---|---|---|---|
| MCOLN1 | 95 | E | 0 | 0 | yes | no | no | no | yes | no | no | no | no | no | no | no | no |
| MCOLN1 | 96 | E | 1 | 0 | yes | no | no | no | yes | no | no | no | yes | no | no | no | no |
| MCOLN1 | 97 | N | 1 | 0 | yes | no | no | no | yes | no | no | no | yes | no | no | no | no |
| MCOLN1 | 98 | T | 1 | 0 | yes | no | no | no | yes | no | no | no | no | no | no | no | no |
| MCOLN1 | 99 | I | 0 | 0 | yes | no | no | no | yes | no | no | no | yes | no | no | no | no |
| MCOLN1 | 100 | A | 2 | 0 | yes | no | no | no | yes | no | no | no | yes | no | no | no | no |
| MCOLN1 | 101 | F | 0 | 0 | yes | no | no | no | yes | no | no | no | no | no | no | no | no |
| MCOLN1 | 102 | R | 1 | 0 | yes | no | no | no | yes | no | no | no | no | no | no | no | no |
| MCOLN1 | 103 | H | 0 | 0 | yes | no | no | no | yes | no | no | no | yes | no | no | no | no |
| MCOLN1 | 104 | L | 0 | 0 | yes | no | no | no | yes | no | no | no | yes | no | no | no | no |
| MCOLN1 | 105 | F | 0 | 0 | yes | no | no | no | yes | no | no | no | no | no | no | no | no |
| MCOLN1 | 106 | L | 0 | 1 | no | yes | no | no | yes | no | no | no | no | no | no | no | no |
| MCOLN1 | 107 | L | 1 | 0 | no | yes | no | no | yes | no | no | yes | no | no | no | no | no |
| MCOLN1 | 108 | G | 0 | 0 | no | yes | no | no | yes | no | no | yes | no | no | no | no | no |
| MCOLN1 | 109 | Y | 0 | 0 | no | yes | no | no | yes | no | no | yes | no | no | no | no | no |
| MCOLN1 | 110 | S | 1 | 0 | no | yes | no | no | yes | no | yes | yes | no | no | no | no | no |
| MCOLN1 | 111 | D | 0 | 0 | no | yes | no | no | yes | no | yes | yes | yes | no | no | no | no |
| MCOLN1 | 112 | G | 1 | 0 | no | yes | no | no | yes | no | yes | yes | no | no | no | no | no |
| MCOLN1 | 113 | A | 1 | 0 | no | yes | no | no | yes | no | no | yes | no | no | no | no | no |
| MCOLN1 | 114 | D | 0 | 0 | no | yes | no | no | yes | no | no | yes | no | no | no | no | no |
| MCOLN1 | 115 | D | 0 | 0 | no | yes | no | no | yes | no | no | yes | no | no | no | no | no |
| MCOLN1 | 116 | T | 1 | 0 | no | yes | no | no | yes | no | no | yes | yes | no | no | no | no |
| MCOLN1 | 117 | F | 0 | 0 | no | yes | no | no | yes | no | no | yes | no | no | no | no | no |
| MCOLN1 | 118 | A | 1 | 0 | no | no | yes | no | yes | no | no | yes | no | no | no | no | no |
| MCOLN1 | 119 | A | 0 | 0 | no | no | yes | no | yes | no | no | yes | yes | no | no | no | no |
| MCOLN1 | 120 | Y | 0 | 0 | no | yes | no | no | yes | no | no | yes | yes | no | no | no | no |
| MCOLN1 | 121 | T | 1 | 0 | no | no | yes | no | yes | no | no | yes | yes | no | no | no | no |
| MCOLN1 | 122 | R | 2 | 0 | yes | no | no | no | yes | no | no | no | yes | no | no | no | no |
| MCOLN1 | 123 | E | 2 | 0 | yes | no | no | no | yes | no | no | no | no | no | no | no | no |
| MCOLN1 | 124 | Q | 0 | 0 | yes | no | no | no | yes | no | no | no | no | no | no | no | no |
| MCOLN1 | 125 | L | 0 | 0 | yes | no | no | no | yes | no | no | no | yes | no | no | no | no |
| MCOLN1 | 126 | Y | 0 | 0 | yes | no | no | no | yes | no | no | no | no | no | no | no | no |
| MCOLN1 | 127 | Q | 1 | 0 | yes | no | no | no | yes | no | no | no | no | no | no | no | no |

| | | | | | | | | | | | | | | | | | |
|---|---|---|---|---|---|---|---|---|---|---|---|---|---|---|---|---|---|
| MCOLN1 | 128 | A | 0 | 0 | yes | no | no | no | yes | no | no | no | no | no | no | no | no |
| MCOLN1 | 129 | I | 0 | 0 | yes | no | no | no | yes | no | no | no | no | no | no | no | no |
| MCOLN1 | 130 | F | 0 | 0 | yes | no | no | no | yes | no | no | no | no | no | no | no | no |
| MCOLN1 | 131 | H | 2 | 0 | yes | no | no | no | yes | no | no | no | no | no | no | no | no |
| MCOLN1 | 132 | A | 1 | 0 | yes | no | no | no | yes | no | no | no | no | no | no | no | no |
| MCOLN1 | 133 | V | 0 | 0 | yes | no | no | no | yes | no | no | no | no | no | no | no | no |
| MCOLN1 | 134 | D | 0 | 0 | yes | no | no | no | yes | no | no | no | no | no | no | no | no |
| MCOLN1 | 135 | Q | 0 | 0 | yes | no | no | no | yes | no | no | no | no | no | no | no | no |
| MCOLN1 | 136 | Y | 0 | 0 | yes | no | no | no | yes | no | no | no | no | no | no | no | no |
| MCOLN1 | 137 | L | 1 | 1 | yes | no | no | no | yes | no | no | no | no | no | no | no | no |
| MCOLN1 | 138 | A | 2 | 0 | no | yes | no | no | yes | no | no | no | no | no | no | no | no |
| MCOLN1 | 139 | L | 0 | 0 | yes | no | no | no | yes | no | no | no | no | no | no | no | no |
| MCOLN1 | 140 | P | 0 | 0 | yes | no | no | no | yes | no | no | no | yes | no | no | no | no |
| MCOLN1 | 141 | D | 0 | 0 | yes | no | no | no | yes | no | no | no | yes | no | no | no | no |
| MCOLN1 | 142 | V | 1 | 0 | yes | no | no | no | yes | no | no | no | yes | no | no | no | no |
| MCOLN1 | 143 | S | 0 | 0 | no | no | yes | no | yes | no | no | no | yes | no | no | no | no |
| MCOLN1 | 144 | L | 0 | 0 | no | yes | no | no | yes | no | no | no | yes | no | no | no | no |
| MCOLN1 | 145 | G | 0 | 0 | no | yes | no | no | yes | no | no | no | yes | no | no | no | no |
| MCOLN1 | 146 | R | 2 | 0 | no | yes | no | no | yes | no | no | no | yes | no | no | no | no |
| MCOLN1 | 147 | Y | 0 | 0 | no | yes | no | no | yes | no | no | no | yes | no | no | no | no |
| MCOLN1 | 148 | A | 1 | 0 | no | no | yes | no | yes | no | no | no | yes | no | no | no | no |
| MCOLN1 | 149 | Y | 1 | 0 | no | yes | no | no | yes | no | no | no | no | no | no | no | no |
| MCOLN1 | 150 | V | 0 | 0 | no | yes | no | no | yes | no | no | no | no | no | no | no | no |
| MCOLN1 | 151 | R | 3 | 0 | no | yes | no | no | yes | no | no | no | no | no | no | no | no |
| MCOLN1 | 152 | G | 0 | 0 | no | yes | no | no | yes | no | no | no | no | no | no | no | no |
| MCOLN1 | 153 | G | 1 | 0 | no | yes | no | no | yes | no | no | no | no | no | no | no | no |
| MCOLN1 | 154 | G | 1 | 0 | no | yes | no | no | yes | no | no | no | no | no | no | no | no |
| MCOLN1 | 155 | D | 0 | 0 | no | yes | no | no | yes | no | no | no | no | no | no | no | no |
| MCOLN1 | 156 | P | 1 | 0 | no | yes | no | no | yes | no | no | no | no | no | no | no | no |
| MCOLN1 | 157 | W | 0 | 0 | no | yes | no | no | yes | no | no | no | no | no | no | no | no |
| MCOLN1 | 158 | T | 1 | 0 | no | yes | no | no | yes | no | no | no | no | no | no | no | no |
| MCOLN1 | 159 | N | 1 | 0 | no | yes | no | no | yes | no | no | no | no | no | no | no | no |
| MCOLN1 | 160 | G | 0 | 0 | no | yes | no | no | yes | no | no | no | no | no | no | no | no |

| | | | | | | | | | | | | | | | | | |
|---|---|---|---|---|---|---|---|---|---|---|---|---|---|---|---|---|---|
| MCOLN1 | 161 | S | 1 | 0 | no | yes | no | no | yes | no | no | no | no | no | no | no | no |
| MCOLN1 | 162 | G | 0 | 0 | no | yes | no | no | yes | no | no | no | no | no | no | no | no |
| MCOLN1 | 163 | L | 0 | 0 | no | no | yes | no | yes | no | no | no | no | no | no | no | no |
| MCOLN1 | 164 | A | 1 | 0 | no | no | yes | no | yes | no | no | no | no | no | no | no | no |
| MCOLN1 | 165 | L | 0 | 0 | no | no | yes | no | yes | no | no | no | no | no | no | no | no |
| MCOLN1 | 166 | C | 2 | 1 | no | no | yes | no | yes | no | no | no | no | no | no | no | no |
| MCOLN1 | 167 | Q | 0 | 0 | no | no | yes | no | yes | no | no | no | no | no | no | no | no |
| MCOLN1 | 168 | R | 2 | 0 | no | no | yes | no | yes | no | no | no | no | no | no | no | no |
| MCOLN1 | 169 | Y | 0 | 0 | no | no | yes | no | yes | no | no | no | no | no | no | no | no |
| MCOLN1 | 170 | Y | 0 | 0 | no | no | yes | no | yes | no | no | no | yes | no | no | no | no |
| MCOLN1 | 171 | H | 0 | 0 | no | no | yes | no | yes | no | no | no | no | no | no | no | no |
| MCOLN1 | 172 | R | 1 | 0 | no | no | yes | no | yes | no | no | no | no | no | no | no | no |
| MCOLN1 | 173 | G | 0 | 0 | no | yes | no | no | yes | no | no | no | no | no | no | no | no |
| MCOLN1 | 174 | H | 0 | 0 | no | yes | no | no | yes | no | no | no | no | no | no | no | no |
| MCOLN1 | 175 | V | 1 | 0 | no | yes | no | no | yes | no | no | no | yes | no | no | no | no |
| MCOLN1 | 176 | D | 0 | 0 | no | yes | no | no | yes | no | no | no | yes | no | no | no | no |
| MCOLN1 | 177 | P | 0 | 0 | no | yes | no | no | yes | no | no | no | yes | no | no | no | no |
| MCOLN1 | 178 | A | 1 | 0 | no | yes | no | no | yes | no | no | no | no | no | no | no | no |
| MCOLN1 | 179 | N | 0 | 0 | no | yes | no | no | yes | no | no | no | no | no | no | no | no |
| MCOLN1 | 180 | D | 2 | 0 | no | yes | no | no | yes | no | no | no | no | no | no | no | no |
| MCOLN1 | 181 | T | 0 | 0 | no | yes | no | no | yes | no | no | no | no | no | no | no | no |
| MCOLN1 | 182 | F | 0 | 0 | no | yes | no | no | yes | no | no | no | yes | no | no | no | no |
| MCOLN1 | 183 | D | 1 | 0 | no | yes | no | no | yes | no | no | no | yes | no | no | no | no |
| MCOLN1 | 184 | I | 2 | 0 | no | yes | no | no | yes | no | no | no | yes | no | no | no | no |
| MCOLN1 | 185 | D | 0 | 0 | no | no | yes | no | yes | no | no | no | yes | no | no | no | no |
| MCOLN1 | 186 | P | 1 | 0 | no | no | yes | no | yes | no | no | no | no | no | no | no | no |
| MCOLN1 | 187 | M | 2 | 0 | no | no | yes | no | yes | no | no | no | no | no | no | no | no |
| MCOLN1 | 188 | V | 1 | 0 | no | no | yes | no | yes | no | no | no | no | no | no | no | no |
| MCOLN1 | 189 | V | 1 | 0 | no | no | yes | no | yes | no | no | no | no | no | no | no | no |
| MCOLN1 | 190 | T | 0 | 0 | no | no | yes | no | yes | no | no | no | no | no | no | no | no |
| MCOLN1 | 191 | D | 0 | 0 | no | no | yes | no | yes | no | no | no | no | no | no | no | no |
| MCOLN1 | 192 | C | 0 | 0 | no | no | yes | no | yes | no | no | no | no | no | no | no | no |
| MCOLN1 | 193 | I | 0 | 0 | no | no | yes | no | yes | no | no | no | no | no | no | no | no |

| | | | | | | | | | | | | | | | | | |
|---|---|---|---|---|---|---|---|---|---|---|---|---|---|---|---|---|---|
| MCOLN1 | 194 | Q | 0 | 0 | no | no | yes | no | yes | no | no | no | no | no | no | no | no |
| MCOLN1 | 195 | V | 2 | 0 | no | no | yes | no | yes | no | no | no | no | no | no | no | no |
| MCOLN1 | 196 | D | 1 | 0 | no | yes | no | no | yes | no | no | no | no | no | no | no | no |
| MCOLN1 | 197 | P | 1 | 0 | no | yes | no | no | yes | no | no | no | no | no | no | no | no |
| MCOLN1 | 198 | P | 0 | 0 | no | yes | no | no | yes | no | no | no | no | no | no | no | no |
| MCOLN1 | 199 | E | 1 | 0 | no | yes | no | no | yes | no | no | no | no | no | no | no | no |
| MCOLN1 | 200 | R | 2 | 0 | no | yes | no | no | yes | no | no | no | no | no | no | no | no |
| MCOLN1 | 201 | P | 0 | 0 | no | yes | no | no | yes | no | no | no | no | no | no | no | no |
| MCOLN1 | 202 | P | 1 | 0 | no | yes | no | no | yes | no | no | no | no | no | no | no | no |
| MCOLN1 | 203 | P | 1 | 0 | no | yes | no | no | yes | no | no | no | no | no | no | no | no |
| MCOLN1 | 204 | P | 3 | 0 | no | yes | no | no | yes | no | no | no | no | no | no | no | no |
| MCOLN1 | 205 | P | 2 | 0 | no | yes | no | no | yes | no | no | no | no | no | no | no | no |
| MCOLN1 | 216 | S | 0 | 0 | no | yes | no | no | yes | no | no | no | no | no | no | no | no |
| MCOLN1 | 217 | S | 0 | 0 | no | yes | no | no | yes | no | no | no | no | no | no | no | no |
| MCOLN1 | 218 | Y | 1 | 0 | yes | no | no | no | yes | no | no | no | no | no | no | no | no |
| MCOLN1 | 219 | K | 2 | 0 | yes | no | no | no | yes | no | no | no | no | no | no | no | no |
| MCOLN1 | 220 | N | 0 | 0 | yes | no | no | no | yes | no | no | no | no | no | no | no | no |
| MCOLN1 | 221 | L | 0 | 0 | no | yes | no | no | yes | no | no | no | no | no | no | no | no |
| MCOLN1 | 222 | T | 1 | 0 | no | yes | no | no | yes | no | no | no | no | no | no | no | no |
| MCOLN1 | 223 | L | 0 | 0 | no | yes | no | no | yes | no | no | no | no | no | no | no | no |
| MCOLN1 | 224 | K | 0 | 0 | no | yes | no | no | yes | no | no | no | no | no | no | no | no |
| MCOLN1 | 225 | F | 0 | 0 | yes | no | no | no | yes | no | no | no | yes | no | no | no | no |
| MCOLN1 | 226 | H | 1 | 0 | yes | no | no | no | yes | no | no | no | yes | no | no | no | no |
| MCOLN1 | 227 | K | 0 | 0 | yes | no | no | no | yes | no | no | no | no | no | no | no | no |
| MCOLN1 | 228 | L | 0 | 0 | no | no | yes | no | yes | no | no | no | no | no | no | no | no |
| MCOLN1 | 229 | V | 0 | 0 | no | no | yes | no | yes | no | no | no | no | no | no | no | no |
| MCOLN1 | 230 | N | 1 | 0 | no | no | yes | no | yes | no | no | no | no | no | no | no | no |
| MCOLN1 | 231 | V | 0 | 0 | no | no | yes | no | yes | no | no | no | no | no | no | no | no |
| MCOLN1 | 232 | T | 2 | 1 | no | no | yes | no | yes | no | no | no | no | no | no | no | no |
| MCOLN1 | 233 | I | 1 | 0 | no | no | yes | no | yes | no | no | no | no | no | no | no | no |
| MCOLN1 | 234 | H | 0 | 0 | no | no | yes | no | yes | no | no | no | no | no | no | no | no |
| MCOLN1 | 235 | F | 0 | 0 | no | no | yes | no | yes | no | no | no | no | no | no | no | no |
| MCOLN1 | 236 | R | 2 | 0 | no | no | yes | no | yes | no | no | no | no | no | no | no | no |

| | | | | | | | | | | | | | | | | | |
|---|---|---|---|---|---|---|---|---|---|---|---|---|---|---|---|---|---|
| MCOLN1 | 237 | L | 0 | 0 | no | no | yes | no | yes | no | no | no | no | no | no | no | no |
| MCOLN1 | 238 | K | 0 | 0 | no | no | yes | no | yes | no | no | no | yes | no | no | no | no |
| MCOLN1 | 239 | T | 0 | 0 | no | no | yes | no | yes | no | no | no | yes | no | no | no | no |
| MCOLN1 | 240 | I | 1 | 0 | no | no | yes | no | yes | no | no | no | yes | no | no | no | no |
| MCOLN1 | 241 | N | 0 | 0 | no | yes | no | no | yes | no | no | no | no | no | no | no | no |
| MCOLN1 | 242 | L | 0 | 0 | no | yes | no | no | yes | no | no | no | yes | no | no | no | no |
| MCOLN1 | 243 | Q | 0 | 0 | no | yes | no | no | yes | no | no | no | yes | no | no | no | no |
| MCOLN1 | 244 | S | 0 | 0 | no | yes | no | no | yes | no | no | no | no | no | no | no | no |
| MCOLN1 | 245 | L | 1 | 0 | no | yes | no | no | yes | no | no | no | no | no | no | no | no |
| MCOLN1 | 246 | I | 1 | 0 | no | yes | no | no | yes | no | no | no | no | no | no | no | no |
| MCOLN1 | 247 | N | 1 | 0 | no | yes | no | no | yes | no | no | no | no | no | no | no | no |
| MCOLN1 | 248 | N | 0 | 0 | no | yes | no | no | yes | no | no | no | no | no | no | no | no |
| MCOLN1 | 249 | E | 1 | 0 | no | yes | no | no | yes | no | no | no | no | no | no | no | no |
| MCOLN1 | 250 | I | 0 | 0 | no | yes | no | no | yes | no | no | no | yes | no | no | no | no |
| MCOLN1 | 251 | P | 0 | 0 | no | yes | no | no | yes | no | no | no | no | no | no | no | no |
| MCOLN1 | 252 | D | 0 | 0 | no | yes | no | no | yes | no | no | no | no | no | no | no | no |
| MCOLN1 | 253 | C | 0 | 0 | no | no | yes | no | yes | no | no | no | no | no | no | no | no |
| MCOLN1 | 254 | Y | 0 | 0 | no | no | yes | no | yes | no | no | no | no | no | no | no | no |
| MCOLN1 | 255 | T | 0 | 0 | no | no | yes | no | yes | no | no | no | no | no | no | no | no |
| MCOLN1 | 256 | F | 0 | 0 | no | no | yes | no | yes | no | no | no | no | no | no | no | no |
| MCOLN1 | 257 | S | 1 | 0 | no | no | yes | no | yes | no | no | no | no | no | no | no | no |
| MCOLN1 | 258 | V | 1 | 0 | no | no | yes | no | yes | no | no | no | no | no | no | no | no |
| MCOLN1 | 259 | L | 0 | 0 | no | no | yes | no | yes | no | no | no | no | no | no | no | no |
| MCOLN1 | 260 | I | 0 | 0 | no | no | yes | no | yes | no | no | no | no | no | no | no | no |
| MCOLN1 | 261 | T | 1 | 0 | no | no | yes | no | yes | no | no | no | no | no | no | no | no |
| MCOLN1 | 262 | F | 0 | 0 | no | no | yes | no | yes | no | no | no | no | no | no | no | no |
| MCOLN1 | 263 | D | 0 | 0 | no | no | yes | no | yes | no | no | no | no | no | no | no | no |
| MCOLN1 | 264 | N | 2 | 0 | no | yes | no | no | yes | no | no | no | no | no | no | no | no |
| MCOLN1 | 265 | K | 0 | 0 | no | yes | no | no | yes | no | no | no | yes | no | no | no | no |
| MCOLN1 | 266 | A | 1 | 0 | no | yes | no | no | yes | no | no | no | yes | no | no | no | no |
| MCOLN1 | 267 | H | 1 | 0 | no | yes | no | no | yes | no | no | no | yes | no | no | no | no |
| MCOLN1 | 268 | S | 0 | 0 | no | yes | no | no | yes | no | no | no | yes | no | no | no | no |
| MCOLN1 | 269 | G | 0 | 0 | no | yes | no | no | yes | no | no | no | yes | no | no | no | no |

| | | | | | | | | | | | | | | | | | |
|---|---|---|---|---|---|---|---|---|---|---|---|---|---|---|---|---|---|
| MCOLN1 | 270 | R | 0 | 0 | no | yes | no | no | yes | no | no | no | yes | no | no | no | no |
| MCOLN1 | 271 | I | 0 | 0 | no | no | yes | no | yes | no | no | no | yes | no | no | no | no |
| MCOLN1 | 272 | P | 1 | 0 | no | no | yes | no | yes | no | no | no | no | no | no | no | no |
| MCOLN1 | 273 | I | 0 | 0 | no | no | yes | no | yes | no | no | no | no | no | no | no | no |
| MCOLN1 | 274 | S | 0 | 0 | no | no | yes | no | yes | no | no | no | no | no | no | no | no |
| MCOLN1 | 275 | L | 0 | 0 | no | no | yes | no | yes | no | no | no | no | no | no | no | no |
| MCOLN1 | 276 | E | 0 | 0 | no | no | yes | no | yes | no | no | no | yes | no | no | no | no |
| MCOLN1 | 277 | T | 1 | 0 | no | no | yes | no | yes | no | no | no | no | no | no | no | no |
| MCOLN1 | 278 | Q | 1 | 0 | no | no | yes | no | yes | no | no | no | no | no | no | no | no |
| MCOLN1 | 279 | A | 0 | 0 | no | yes | no | no | yes | no | no | no | no | no | no | no | no |
| MCOLN1 | 280 | H | 1 | 0 | no | yes | no | no | yes | no | no | no | no | no | no | no | no |
| MCOLN1 | 281 | I | 0 | 0 | no | no | yes | no | yes | no | no | no | no | no | no | no | no |
| MCOLN1 | 282 | Q | 0 | 0 | no | no | yes | no | yes | no | no | no | no | no | no | no | no |
| MCOLN1 | 283 | E | 0 | 0 | no | yes | no | no | yes | no | no | no | no | no | no | no | no |
| MCOLN1 | 284 | C | 1 | 0 | no | yes | no | no | yes | no | no | no | yes | no | no | no | no |
| MCOLN1 | 285 | K | 0 | 0 | no | yes | no | no | yes | no | no | no | yes | no | no | no | no |
| MCOLN1 | 286 | H | 1 | 0 | no | yes | no | no | yes | no | no | no | yes | no | no | no | no |
| MCOLN1 | 287 | P | 1 | 0 | no | yes | no | no | yes | no | no | no | no | no | no | no | no |
| MCOLN1 | 288 | S | 2 | 0 | no | yes | no | no | yes | no | no | no | no | no | no | no | no |
| MCOLN1 | 289 | V | 0 | 0 | no | yes | no | no | yes | no | no | no | yes | no | no | no | no |
| MCOLN1 | 290 | F | 0 | 0 | no | yes | no | no | yes | no | no | no | no | no | no | no | no |
| MCOLN1 | 291 | Q | 0 | 0 | no | yes | no | no | yes | no | no | no | no | no | no | no | no |
| MCOLN1 | 292 | H | 0 | 0 | yes | no | no | no | yes | no | no | no | no | no | no | no | no |
| MCOLN1 | 293 | G | 0 | 0 | yes | no | no | no | yes | no | no | no | no | no | no | no | no |
| MCOLN1 | 294 | D | 1 | 0 | yes | no | no | no | yes | no | no | no | no | no | no | no | no |
| MCOLN1 | 295 | N | 2 | 0 | yes | no | no | no | yes | no | no | no | no | no | no | no | no |
| MCOLN1 | 296 | S | 0 | 0 | yes | no | no | yes | yes | no | no | no | no | no | no | no | no |
| MCOLN1 | 297 | F | 0 | 0 | yes | no | no | yes | yes | no | no | no | no | no | no | no | no |
| MCOLN1 | 298 | R | 2 | 0 | yes | no | no | yes | yes | no | no | no | no | no | no | no | no |
| MCOLN1 | 299 | L | 0 | 0 | yes | no | no | yes | no | no | no | no | no | no | no | no | no |
| MCOLN1 | 300 | L | 0 | 0 | yes | no | no | yes | no | no | no | no | no | no | no | no | no |
| MCOLN1 | 301 | F | 1 | 0 | yes | no | no | yes | no | no | no | no | no | no | no | no | no |
| MCOLN1 | 302 | D | 0 | 0 | yes | no | no | yes | no | no | no | no | no | no | no | no | no |

| | | | | | | | | | | | | | | | | | |
|---|---|---|---|---|---|---|---|---|---|---|---|---|---|---|---|---|---|
| MCOLN1 | 303 | V | 1 | 0 | yes | no | no | yes | no | no | no | no | no | no | no | no | no |
| MCOLN1 | 304 | V | 0 | 0 | yes | no | no | yes | no | no | no | no | no | no | no | no | no |
| MCOLN1 | 305 | V | 0 | 0 | yes | no | no | yes | no | no | no | no | no | no | no | no | no |
| MCOLN1 | 306 | I | 1 | 0 | yes | no | no | yes | no | no | no | no | no | no | no | no | no |
| MCOLN1 | 307 | L | 0 | 0 | yes | no | no | yes | no | no | no | no | no | no | no | no | no |
| MCOLN1 | 308 | T | 0 | 0 | yes | no | no | yes | no | no | no | no | no | no | no | no | no |
| MCOLN1 | 309 | C | 0 | 0 | yes | no | no | yes | no | no | no | no | no | no | no | no | no |
| MCOLN1 | 310 | S | 0 | 0 | yes | no | no | yes | no | no | no | no | no | no | no | no | no |
| MCOLN1 | 311 | L | 0 | 0 | yes | no | no | yes | no | no | no | no | no | no | no | no | no |
| MCOLN1 | 312 | S | 0 | 0 | yes | no | no | yes | no | no | no | no | no | no | no | no | no |
| MCOLN1 | 313 | F | 0 | 0 | yes | no | no | yes | no | no | no | no | no | no | no | no | no |
| MCOLN1 | 314 | L | 0 | 0 | yes | no | no | yes | no | no | no | no | no | no | no | no | no |
| MCOLN1 | 315 | L | 0 | 0 | yes | no | no | yes | no | no | no | no | no | no | no | no | no |
| MCOLN1 | 316 | C | 0 | 0 | yes | no | no | yes | no | no | no | no | no | no | no | no | no |
| MCOLN1 | 317 | A | 3 | 0 | yes | no | no | yes | no | no | no | no | no | no | no | no | no |
| MCOLN1 | 318 | R | 2 | 0 | yes | no | no | no | no | no | no | no | no | no | no | no | no |
| MCOLN1 | 319 | S | 0 | 0 | yes | no | no | no | no | no | no | no | no | no | no | no | no |
| MCOLN1 | 320 | L | 0 | 0 | yes | no | no | no | no | no | no | no | no | no | no | no | no |
| MCOLN1 | 321 | L | 1 | 0 | yes | no | no | no | no | no | no | no | no | no | no | no | no |
| MCOLN1 | 322 | R | 1 | 0 | yes | no | no | no | no | yes | no | no | no | no | no | no | no |
| MCOLN1 | 323 | G | 0 | 0 | yes | no | no | no | no | yes | no | no | no | no | no | no | no |
| MCOLN1 | 324 | F | 0 | 0 | yes | no | no | no | no | yes | no | no | no | no | no | no | no |
| MCOLN1 | 325 | L | 1 | 0 | yes | no | no | no | no | yes | no | no | no | no | no | no | no |
| MCOLN1 | 326 | L | 0 | 0 | yes | no | no | no | no | yes | no | no | no | no | no | no | no |
| MCOLN1 | 327 | Q | 2 | 0 | yes | no | no | no | no | yes | no | no | no | no | no | no | no |
| MCOLN1 | 328 | N | 0 | 0 | yes | no | no | no | no | yes | no | no | no | no | no | no | no |
| MCOLN1 | 329 | E | 0 | 0 | yes | no | no | no | no | yes | no | no | no | no | no | no | no |
| MCOLN1 | 330 | F | 0 | 0 | yes | no | no | no | no | yes | no | no | no | no | no | no | no |
| MCOLN1 | 331 | V | 1 | 0 | yes | no | no | no | no | yes | no | no | no | no | no | no | no |
| MCOLN1 | 332 | G | 2 | 0 | yes | no | no | no | no | yes | no | no | no | no | no | no | no |
| MCOLN1 | 333 | F | 1 | 0 | yes | no | no | no | no | yes | no | no | no | no | no | no | no |
| MCOLN1 | 334 | M | 3 | 0 | yes | no | no | no | no | yes | no | no | no | no | no | no | no |
| MCOLN1 | 335 | W | 0 | 0 | yes | no | no | no | no | yes | no | no | no | no | no | no | no |

| | | | | | | | | | | | | | | | | | |
|---|---|---|---|---|---|---|---|---|---|---|---|---|---|---|---|---|---|
| MCOLN1 | 336 | R | 1 | 0 | yes | no | no | no | no | yes | no | no | no | no | no | no | no |
| MCOLN1 | 337 | Q | 0 | 0 | yes | no | no | no | no | yes | no | no | no | no | no | no | no |
| MCOLN1 | 338 | R | 2 | 0 | yes | no | no | no | no | yes | no | no | no | no | no | no | no |
| MCOLN1 | 339 | G | 0 | 0 | no | yes | no | no | no | yes | no | no | no | no | no | no | no |
| MCOLN1 | 340 | R | 2 | 0 | no | yes | no | no | no | yes | no | no | no | no | no | no | no |
| MCOLN1 | 341 | V | 1 | 0 | no | yes | no | no | no | yes | no | no | no | no | no | no | no |
| MCOLN1 | 342 | I | 0 | 0 | no | yes | no | no | no | yes | no | no | no | no | no | no | no |
| MCOLN1 | 343 | S | 0 | 0 | no | yes | no | no | no | yes | no | no | no | no | no | no | no |
| MCOLN1 | 344 | L | 0 | 0 | no | yes | no | no | no | yes | no | no | no | no | no | no | no |
| MCOLN1 | 345 | W | 0 | 0 | no | yes | no | no | no | yes | no | no | no | no | no | no | no |
| MCOLN1 | 346 | E | 0 | 0 | yes | no | no | no | no | yes | no | no | no | no | no | no | no |
| MCOLN1 | 347 | R | 3 | 0 | yes | no | no | no | no | yes | no | no | no | no | no | no | no |
| MCOLN1 | 348 | L | 0 | 0 | yes | no | no | no | no | yes | no | no | no | no | no | no | no |
| MCOLN1 | 349 | E | 0 | 0 | yes | no | no | no | no | yes | no | no | no | no | no | no | no |
| MCOLN1 | 350 | F | 1 | 0 | no | yes | no | no | no | yes | no | no | no | no | no | no | no |
| MCOLN1 | 351 | V | 0 | 0 | no | yes | no | no | no | no | no | no | no | no | no | no | no |
| MCOLN1 | 352 | N | 1 | 0 | no | yes | no | no | no | no | no | no | no | no | no | no | no |
| MCOLN1 | 353 | G | 0 | 0 | yes | no | no | no | no | no | no | no | no | no | no | no | no |
| MCOLN1 | 354 | W | 0 | 0 | yes | no | no | no | no | no | no | no | no | no | no | no | no |
| MCOLN1 | 355 | Y | 2 | 0 | yes | no | no | no | no | no | no | no | no | no | no | no | no |
| MCOLN1 | 356 | I | 0 | 0 | yes | no | no | yes | no | no | no | no | no | no | no | no | no |
| MCOLN1 | 357 | L | 0 | 0 | yes | no | no | yes | no | no | no | no | no | no | no | no | no |
| MCOLN1 | 358 | L | 0 | 0 | yes | no | no | yes | no | no | no | no | no | no | no | no | no |
| MCOLN1 | 359 | V | 1 | 0 | yes | no | no | yes | no | no | no | no | no | no | no | no | no |
| MCOLN1 | 360 | T | 0 | 0 | yes | no | no | yes | no | no | no | no | no | no | no | no | no |
| MCOLN1 | 361 | S | 0 | 0 | yes | no | no | yes | no | no | no | no | no | no | no | no | no |
| MCOLN1 | 362 | D | 1 | 1 | yes | no | no | yes | no | no | no | no | no | no | no | no | no |
| MCOLN1 | 363 | V | 0 | 0 | yes | no | no | yes | no | no | no | no | no | no | no | no | no |
| MCOLN1 | 364 | L | 0 | 0 | yes | no | no | yes | no | no | no | no | no | no | no | no | no |
| MCOLN1 | 365 | T | 0 | 0 | yes | no | no | yes | no | no | no | no | no | no | no | no | no |
| MCOLN1 | 366 | I | 0 | 0 | yes | no | no | yes | no | no | no | no | no | no | no | no | no |
| MCOLN1 | 367 | S | 1 | 0 | yes | no | no | yes | no | no | no | no | no | no | no | no | no |
| MCOLN1 | 368 | G | 0 | 0 | yes | no | no | yes | no | no | no | no | no | no | no | no | no |

| | | | | | | | | | | | | | | | | | |
|---|---|---|---|---|---|---|---|---|---|---|---|---|---|---|---|---|---|
| MCOLN1 | 369 | T | 0 | 0 | yes | no | no | yes | no | no | no | no | no | no | no | no | no |
| MCOLN1 | 370 | I | 1 | 0 | yes | no | no | yes | no | no | no | no | no | no | no | no | no |
| MCOLN1 | 371 | M | 1 | 0 | yes | no | no | yes | no | no | no | no | no | no | no | no | no |
| MCOLN1 | 372 | K | 0 | 0 | yes | no | no | yes | yes | no | no | no | no | no | no | no | no |
| MCOLN1 | 373 | I | 0 | 0 | yes | no | no | yes | yes | no | no | no | no | no | no | no | no |
| MCOLN1 | 374 | G | 1 | 0 | yes | no | no | yes | yes | no | no | no | no | no | no | no | no |
| MCOLN1 | 375 | I | 0 | 0 | yes | no | no | no | yes | no | no | no | no | no | no | no | no |
| MCOLN1 | 376 | E | 0 | 0 | yes | no | no | no | yes | no | no | no | no | no | no | no | no |
| MCOLN1 | 377 | A | 0 | 0 | yes | no | no | no | yes | no | no | no | no | no | no | no | no |
| MCOLN1 | 378 | K | 0 | 0 | no | yes | no | no | yes | no | no | no | no | no | no | no | no |
| MCOLN1 | 379 | N | 0 | 0 | no | yes | no | no | yes | no | no | no | no | no | no | no | no |
| MCOLN1 | 380 | L | 0 | 0 | no | yes | no | no | yes | no | no | no | no | no | no | no | no |
| MCOLN1 | 381 | A | 1 | 0 | no | yes | no | no | yes | no | no | no | no | no | no | no | no |
| MCOLN1 | 382 | S | 0 | 0 | no | yes | no | no | yes | no | no | no | no | no | no | no | no |
| MCOLN1 | 383 | Y | 0 | 0 | yes | no | no | no | no | no | no | no | no | no | no | no | no |
| MCOLN1 | 384 | D | 2 | 0 | yes | no | no | yes | no | no | no | no | yes | no | no | no | no |
| MCOLN1 | 385 | V | 1 | 0 | yes | no | no | yes | no | no | no | no | no | no | no | no | no |
| MCOLN1 | 386 | C | 1 | 0 | yes | no | no | yes | no | no | no | no | no | no | no | no | no |
| MCOLN1 | 387 | S | 1 | 0 | yes | no | no | yes | no | no | no | no | yes | no | no | no | no |
| MCOLN1 | 388 | I | 0 | 0 | yes | no | no | yes | no | no | no | no | yes | no | no | no | no |
| MCOLN1 | 389 | L | 1 | 0 | yes | no | no | yes | no | no | no | no | no | no | no | no | no |
| MCOLN1 | 390 | L | 0 | 0 | yes | no | no | yes | no | no | no | no | no | no | no | no | no |
| MCOLN1 | 391 | G | 0 | 0 | yes | no | no | yes | no | no | no | no | yes | no | no | no | no |
| MCOLN1 | 392 | T | 1 | 0 | yes | no | no | yes | no | no | no | no | yes | no | no | no | no |
| MCOLN1 | 393 | S | 1 | 0 | yes | no | no | yes | no | no | no | no | no | no | no | no | no |
| MCOLN1 | 394 | T | 1 | 0 | yes | no | no | yes | no | no | no | no | no | no | no | no | no |
| MCOLN1 | 395 | L | 1 | 0 | yes | no | no | yes | no | no | no | no | yes | no | no | no | no |
| MCOLN1 | 396 | L | 0 | 0 | yes | no | no | yes | no | no | no | no | no | no | no | no | no |
| MCOLN1 | 397 | V | 0 | 0 | yes | no | no | yes | no | no | no | no | no | no | no | no | no |
| MCOLN1 | 398 | W | 0 | 0 | yes | no | no | yes | no | no | no | no | yes | no | no | no | no |
| MCOLN1 | 399 | V | 0 | 0 | no | yes | no | yes | no | no | no | no | yes | no | no | no | no |
| MCOLN1 | 400 | G | 0 | 0 | yes | no | no | yes | no | no | no | no | no | no | no | no | no |
| MCOLN1 | 401 | V | 1 | 0 | yes | no | no | yes | no | no | no | no | yes | no | no | no | no |

| | | | | | | | | | | | | | | | | | |
|---|---|---|---|---|---|---|---|---|---|---|---|---|---|---|---|---|---|
| MCOLN1 | 402 | I | 0 | 0 | yes | no | no | yes | no | no | no | no | yes | no | no | no | no |
| MCOLN1 | 403 | R | 2 | 1 | yes | no | no | no | no | no | no | no | no | no | no | no | no |
| MCOLN1 | 404 | Y | 0 | 0 | yes | no | no | yes | no | no | no | no | no | no | no | no | no |
| MCOLN1 | 405 | L | 1 | 0 | no | yes | no | no | no | no | no | no | yes | no | no | no | no |
| MCOLN1 | 406 | T | 1 | 0 | no | yes | no | no | no | yes | no | no | no | no | no | no | no |
| MCOLN1 | 407 | F | 0 | 0 | no | yes | no | no | no | yes | no | no | no | no | no | no | no |
| MCOLN1 | 408 | F | 0 | 0 | no | yes | no | no | no | yes | no | no | no | no | no | no | no |
| MCOLN1 | 409 | H | 0 | 0 | no | yes | no | no | no | yes | no | no | no | no | no | no | no |
| MCOLN1 | 410 | N | 0 | 0 | no | yes | no | no | no | yes | no | no | yes | no | no | no | no |
| MCOLN1 | 411 | Y | 0 | 0 | yes | no | no | no | no | yes | no | no | yes | no | no | no | no |
| MCOLN1 | 412 | N | 1 | 0 | yes | no | no | no | no | yes | no | no | no | no | no | no | no |
| MCOLN1 | 413 | I | 0 | 0 | yes | no | no | no | no | yes | no | no | no | no | no | no | no |
| MCOLN1 | 414 | L | 0 | 0 | yes | no | no | no | no | yes | no | no | yes | no | no | no | no |
| MCOLN1 | 415 | I | 0 | 0 | yes | no | no | no | no | yes | no | no | yes | no | no | no | no |
| MCOLN1 | 416 | A | 2 | 0 | no | yes | no | no | no | yes | no | no | no | no | no | no | no |
| MCOLN1 | 417 | T | 0 | 0 | yes | no | no | no | no | yes | no | no | yes | no | no | no | no |
| MCOLN1 | 418 | L | 1 | 0 | yes | no | no | no | no | yes | no | no | no | no | no | no | no |
| MCOLN1 | 419 | R | 2 | 0 | yes | no | no | no | no | yes | no | no | no | no | no | no | no |
| MCOLN1 | 420 | V | 0 | 0 | yes | no | no | no | no | yes | no | no | no | no | no | no | no |
| MCOLN1 | 421 | A | 0 | 0 | yes | no | no | no | no | yes | no | no | no | no | no | no | no |
| MCOLN1 | 422 | L | 0 | 0 | yes | no | no | no | no | yes | no | no | no | no | no | no | no |
| MCOLN1 | 423 | P | 1 | 0 | yes | no | no | no | no | yes | no | no | no | no | no | no | no |
| MCOLN1 | 424 | S | 0 | 0 | yes | no | no | no | no | yes | no | no | no | no | no | no | no |
| MCOLN1 | 425 | V | 1 | 0 | yes | no | no | yes | no | yes | no | no | no | no | no | no | no |
| MCOLN1 | 426 | M | 1 | 0 | yes | no | no | yes | no | yes | no | no | no | no | no | no | no |
| MCOLN1 | 427 | R | 0 | 0 | yes | no | no | no | no | yes | no | no | yes | no | no | no | no |
| MCOLN1 | 428 | F | 0 | 0 | yes | no | no | yes | no | no | no | no | yes | no | no | no | no |
| MCOLN1 | 429 | C | 0 | 0 | yes | no | no | yes | no | no | no | no | no | no | no | no | no |
| MCOLN1 | 430 | C | 1 | 0 | yes | no | no | yes | no | no | no | no | no | no | no | no | no |
| MCOLN1 | 431 | C | 0 | 1 | yes | no | no | yes | no | no | no | no | yes | no | no | no | no |
| MCOLN1 | 432 | V | 0 | 0 | yes | no | no | yes | no | no | no | no | no | no | no | no | no |
| MCOLN1 | 433 | A | 0 | 0 | yes | no | no | yes | no | no | no | no | no | no | no | no | no |
| MCOLN1 | 434 | V | 0 | 0 | yes | no | no | yes | no | no | no | no | yes | no | no | no | no |

| | | | | | | | | | | | | | | | | | |
|---|---|---|---|---|---|---|---|---|---|---|---|---|---|---|---|---|---|
| MCOLN1 | 435 | I | 0 | 0 | yes | no | no | yes | no | no | no | no | yes | no | no | no | no |
| MCOLN1 | 436 | Y | 0 | 1 | yes | no | no | yes | no | no | no | no | no | no | no | no | no |
| MCOLN1 | 437 | L | 0 | 0 | yes | no | no | yes | no | no | no | no | no | no | no | no | no |
| MCOLN1 | 438 | G | 0 | 0 | yes | no | no | yes | no | no | no | no | yes | no | no | no | no |
| MCOLN1 | 439 | Y | 0 | 0 | yes | no | no | yes | no | no | no | no | yes | no | no | no | no |
| MCOLN1 | 440 | C | 0 | 0 | yes | no | no | yes | no | no | no | no | no | no | no | no | no |
| MCOLN1 | 441 | F | 0 | 0 | yes | no | no | yes | no | no | no | no | yes | no | no | no | no |
| MCOLN1 | 442 | C | 0 | 0 | yes | no | no | yes | no | no | no | no | yes | no | no | no | no |
| MCOLN1 | 443 | G | 0 | 0 | yes | no | no | yes | no | no | no | no | no | no | no | no | no |
| MCOLN1 | 444 | W | 0 | 0 | yes | no | no | yes | no | no | no | no | yes | no | no | no | no |
| MCOLN1 | 445 | I | 0 | 0 | yes | no | no | yes | no | no | no | no | yes | no | no | no | no |
| MCOLN1 | 446 | V | 1 | 2 | yes | no | no | yes | no | no | no | no | yes | no | no | no | no |
| MCOLN1 | 447 | L | 0 | 1 | yes | no | no | yes | no | no | no | no | no | no | no | no | no |
| MCOLN1 | 448 | G | 0 | 0 | no | yes | no | yes | no | no | no | no | no | no | no | no | no |
| MCOLN1 | 449 | P | 0 | 0 | no | yes | no | no | yes | no | no | no | yes | no | no | no | no |
| MCOLN1 | 450 | Y | 0 | 0 | no | yes | no | no | yes | no | no | no | yes | no | no | no | no |
| MCOLN1 | 451 | H | 0 | 0 | no | yes | no | yes | yes | no | no | no | no | no | no | no | no |
| MCOLN1 | 452 | V | 1 | 0 | no | yes | no | no | yes | no | no | no | no | no | no | no | no |
| MCOLN1 | 453 | K | 0 | 0 | no | yes | no | yes | yes | no | no | no | yes | no | no | no | no |
| MCOLN1 | 454 | F | 0 | 0 | no | yes | no | yes | yes | no | no | no | no | no | no | no | no |
| MCOLN1 | 455 | R | 2 | 0 | no | yes | no | no | yes | no | no | no | no | no | no | no | no |
| MCOLN1 | 456 | S | 0 | 1 | no | yes | no | yes | yes | no | no | no | no | no | no | no | no |
| MCOLN1 | 457 | L | 0 | 0 | yes | no | no | yes | no | no | no | yes | no | no | no | no | no |
| MCOLN1 | 458 | S | 0 | 0 | yes | no | no | yes | no | no | no | yes | yes | no | no | no | no |
| MCOLN1 | 459 | M | 1 | 0 | yes | no | no | yes | no | no | no | yes | yes | no | no | no | no |
| MCOLN1 | 460 | V | 1 | 0 | yes | no | no | yes | no | no | no | yes | no | no | no | no | no |
| MCOLN1 | 461 | S | 0 | 0 | yes | no | no | yes | no | no | no | yes | yes | no | no | no | no |
| MCOLN1 | 462 | E | 0 | 0 | yes | no | no | yes | no | no | no | yes | yes | no | no | no | no |
| MCOLN1 | 463 | C | 0 | 1 | yes | no | no | yes | no | no | no | yes | yes | no | no | no | no |
| MCOLN1 | 464 | L | 0 | 0 | yes | no | no | yes | no | no | no | yes | no | no | no | no | no |
| MCOLN1 | 465 | F | 0 | 1 | yes | no | no | yes | no | no | no | yes | yes | no | no | no | no |
| MCOLN1 | 466 | S | 0 | 0 | yes | no | no | yes | no | no | no | yes | yes | no | no | no | no |
| MCOLN1 | 467 | L | 0 | 0 | yes | no | no | yes | no | no | no | yes | no | no | no | no | no |

| | | | | | | | | | | | | | | | | | |
|---|---|---|---|---|---|---|---|---|---|---|---|---|---|---|---|---|---|
| MCOLN1 | 468 | I | 0 | 0 | no | yes | no | yes | no | no | no | yes | yes | no | no | no | no |
| MCOLN1 | 469 | N | 0 | 1 | no | yes | no | yes | no | no | yes | yes | yes | no | no | no | no |
| MCOLN1 | 470 | G | 1 | 0 | no | yes | no | yes | no | no | yes | yes | yes | no | no | no | no |
| MCOLN1 | 471 | D | 0 | 0 | no | yes | no | yes | no | no | yes | yes | yes | no | no | no | no |
| MCOLN1 | 472 | D | 1 | 0 | no | yes | no | yes | no | no | yes | yes | yes | no | no | no | no |
| MCOLN1 | 473 | M | 0 | 0 | yes | no | no | yes | no | no | no | yes | yes | no | no | no | no |
| MCOLN1 | 474 | F | 0 | 0 | yes | no | no | yes | no | no | no | yes | yes | no | no | no | no |
| MCOLN1 | 475 | V | 0 | 0 | yes | no | no | yes | no | no | yes | yes | no | no | no | no | no |
| MCOLN1 | 476 | T | 0 | 0 | yes | no | no | yes | no | no | no | yes | no | no | no | no | no |
| MCOLN1 | 477 | F | 0 | 0 | yes | no | no | yes | no | no | no | yes | yes | no | no | no | no |
| MCOLN1 | 478 | A | 2 | 0 | yes | no | no | yes | yes | no | no | no | no | no | no | no | no |
| MCOLN1 | 479 | A | 1 | 0 | yes | no | no | yes | yes | no | no | no | no | no | no | no | no |
| MCOLN1 | 480 | M | 0 | 0 | no | yes | no | yes | yes | no | no | no | no | no | no | no | no |
| MCOLN1 | 481 | Q | 0 | 0 | no | yes | no | yes | yes | no | no | no | yes | no | no | no | no |
| MCOLN1 | 482 | A | 2 | 0 | no | yes | no | no | yes | no | no | no | no | no | no | no | no |
| MCOLN1 | 483 | Q | 1 | 0 | yes | no | no | no | yes | no | no | no | no | no | no | no | no |
| MCOLN1 | 484 | Q | 1 | 0 | yes | no | no | yes | yes | no | no | no | no | no | no | no | no |
| MCOLN1 | 485 | G | 0 | 0 | yes | no | no | no | yes | no | no | no | no | no | no | no | no |
| MCOLN1 | 486 | R | 1 | 0 | no | yes | no | no | yes | no | no | no | yes | no | no | no | no |
| MCOLN1 | 487 | S | 0 | 0 | no | yes | no | no | yes | no | no | no | no | no | no | no | no |
| MCOLN1 | 488 | S | 0 | 0 | yes | no | no | yes | yes | no | no | no | no | no | no | no | no |
| MCOLN1 | 489 | L | 0 | 0 | yes | no | no | yes | yes | no | no | no | yes | no | no | no | no |
| MCOLN1 | 490 | V | 0 | 0 | yes | no | no | yes | yes | no | no | no | yes | no | no | no | no |
| MCOLN1 | 491 | W | 0 | 0 | yes | no | no | yes | yes | no | no | no | yes | no | no | no | no |
| MCOLN1 | 492 | L | 0 | 0 | yes | no | no | yes | no | no | no | no | no | no | no | no | no |
| MCOLN1 | 493 | F | 0 | 0 | yes | no | no | yes | no | no | no | no | yes | no | no | no | no |
| MCOLN1 | 494 | S | 0 | 0 | yes | no | no | yes | no | no | no | no | no | no | no | no | no |
| MCOLN1 | 495 | Q | 0 | 0 | yes | no | no | yes | no | no | no | no | yes | no | no | no | no |
| MCOLN1 | 496 | L | 0 | 0 | yes | no | no | yes | no | no | no | no | no | no | no | no | no |
| MCOLN1 | 497 | Y | 0 | 0 | yes | no | no | yes | no | no | no | no | no | no | no | no | no |
| MCOLN1 | 498 | L | 0 | 0 | yes | no | no | yes | no | no | no | no | no | no | no | no | no |
| MCOLN1 | 499 | Y | 0 | 0 | yes | no | no | yes | no | no | no | no | yes | no | no | no | no |
| MCOLN1 | 500 | S | 0 | 0 | yes | no | no | yes | no | no | no | no | no | no | no | no | no |

| | | | | | | | | | | | | | | | | | |
|---|---|---|---|---|---|---|---|---|---|---|---|---|---|---|---|---|---|
| MCOLN1 | 501 | F | 0 | 0 | yes | no | no | yes | no | no | no | no | no | no | no | no | no |
| MCOLN1 | 502 | I | 0 | 0 | yes | no | no | yes | no | no | no | no | yes | no | no | no | no |
| MCOLN1 | 503 | S | 0 | 0 | yes | no | no | yes | no | no | no | no | no | no | no | no | no |
| MCOLN1 | 504 | L | 1 | 0 | yes | no | no | yes | no | no | no | no | no | no | no | no | no |
| MCOLN1 | 505 | F | 0 | 0 | yes | no | no | yes | no | no | no | no | no | no | no | no | no |
| MCOLN1 | 506 | I | 1 | 0 | yes | no | no | yes | no | no | no | no | yes | no | no | no | no |
| MCOLN1 | 507 | Y | 0 | 0 | yes | no | no | yes | no | no | no | no | yes | no | no | no | no |
| MCOLN1 | 508 | M | 0 | 0 | yes | no | no | yes | no | no | no | no | yes | no | no | no | no |
| MCOLN1 | 509 | V | 0 | 0 | yes | no | no | yes | no | no | no | no | no | no | no | no | no |
| MCOLN1 | 510 | L | 0 | 0 | yes | no | no | yes | no | no | yes | no | no | no | no | no | no |
| MCOLN1 | 511 | S | 0 | 0 | yes | no | no | yes | no | no | no | no | yes | no | no | no | no |
| MCOLN1 | 512 | L | 0 | 0 | yes | no | no | no | no | no | no | no | yes | no | no | no | no |
| MCOLN1 | 513 | F | 0 | 0 | yes | no | no | yes | no | no | no | no | yes | no | no | no | no |
| MCOLN1 | 514 | I | 0 | 0 | yes | no | no | yes | no | yes | yes | no | yes | no | no | no | no |
| MCOLN1 | 515 | A | 1 | 0 | yes | no | no | no | no | yes | no | no | yes | no | no | no | no |
| MCOLN1 | 516 | L | 0 | 0 | yes | no | no | no | no | yes | no | no | yes | no | no | no | no |
| MCOLN1 | 517 | I | 0 | 0 | yes | no | no | no | no | yes | no | no | yes | no | no | no | no |
| MCOLN1 | 518 | T | 0 | 0 | yes | no | no | no | no | yes | yes | no | yes | no | no | no | no |
| MCOLN1 | 519 | G | 0 | 0 | yes | no | no | no | no | yes | no | no | no | no | no | no | no |
| MCOLN1 | 520 | A | 1 | 0 | yes | no | no | no | no | yes | no | no | no | no | no | no | no |
| MCOLN1 | 521 | Y | 1 | 0 | yes | no | no | no | no | yes | no | no | yes | no | no | no | no |
| MCOLN1 | 522 | D | 0 | 0 | yes | no | no | no | no | yes | yes | no | no | no | no | no | no |
| MCOLN1 | 523 | T | 1 | 0 | yes | no | no | no | no | yes | no | no | no | no | no | no | no |
| MCOLN1 | 524 | I | 0 | 0 | yes | no | no | no | no | yes | no | no | no | no | no | no | no |
| MCOLN1 | 525 | K | 0 | 0 | no | yes | no | no | no | yes | no | no | no | no | no | no | no |
| MCOLN1 | 526 | H | 0 | 0 | no | yes | no | no | no | yes | no | no | no | no | no | no | no |
| PKD2 | 215 | K | 0 | 0 | no | yes | no | no | no | yes | no | no | no | no | no | no | no |
| PKD2 | 216 | Y | 1 | 0 | yes | no | no | no | no | yes | no | no | no | no | no | no | no |
| PKD2 | 217 | L | 0 | 0 | yes | no | no | no | no | yes | no | no | no | no | no | no | no |
| PKD2 | 218 | K | 0 | 0 | yes | no | no | no | no | yes | no | no | no | no | no | no | no |
| PKD2 | 219 | S | 1 | 0 | yes | no | no | no | no | yes | no | no | no | no | no | no | no |
| PKD2 | 220 | V | 1 | 0 | yes | no | no | no | no | no | no | no | no | no | no | no | no |
| PKD2 | 221 | L | 0 | 0 | yes | no | no | yes | no | no | no | no | no | no | no | no | no |

| | | | | | | | | | | | | | | | | | |
|---|---|---|---|---|---|---|---|---|---|---|---|---|---|---|---|---|---|
| PKD2 | 222 | R | 2 | 0 | yes | no | no | yes | no | no | no | no | no | no | no | no | no |
| PKD2 | 223 | E | 2 | 0 | yes | no | no | yes | no | no | no | no | no | no | no | no | no |
| PKD2 | 224 | L | 0 | 0 | yes | no | no | yes | no | no | no | no | no | no | no | no | no |
| PKD2 | 225 | V | 0 | 0 | yes | no | no | yes | no | no | no | no | no | no | no | no | no |
| PKD2 | 226 | T | 0 | 0 | yes | no | no | yes | no | no | no | no | no | no | no | no | no |
| PKD2 | 227 | Y | 0 | 0 | yes | no | no | yes | no | no | no | no | no | no | no | no | no |
| PKD2 | 228 | L | 0 | 0 | yes | no | no | yes | no | no | no | no | no | no | no | no | no |
| PKD2 | 229 | L | 1 | 0 | yes | no | no | yes | no | no | no | no | no | no | no | no | no |
| PKD2 | 230 | F | 0 | 0 | yes | no | no | yes | no | no | no | no | no | no | no | no | no |
| PKD2 | 231 | L | 0 | 0 | yes | no | no | yes | no | no | no | no | no | no | no | no | no |
| PKD2 | 232 | I | 0 | 0 | yes | no | no | yes | no | no | no | no | no | no | no | no | no |
| PKD2 | 233 | V | 0 | 0 | yes | no | no | yes | no | no | no | no | no | no | no | no | no |
| PKD2 | 234 | L | 0 | 0 | yes | no | no | yes | no | no | no | no | no | no | no | no | no |
| PKD2 | 235 | C | 1 | 0 | yes | no | no | yes | no | no | no | no | no | no | no | no | no |
| PKD2 | 236 | I | 0 | 0 | yes | no | no | yes | no | no | no | no | no | no | no | no | no |
| PKD2 | 237 | L | 1 | 0 | yes | no | no | yes | no | no | no | no | no | no | no | no | no |
| PKD2 | 238 | T | 0 | 0 | yes | no | no | yes | no | no | no | no | yes | no | no | no | no |
| PKD2 | 239 | Y | 1 | 0 | yes | no | no | yes | no | no | no | no | no | no | no | no | no |
| PKD2 | 240 | G | 1 | 0 | yes | no | no | yes | no | no | no | no | no | no | no | no | no |
| PKD2 | 241 | M | 2 | 0 | yes | no | no | yes | no | no | no | no | no | no | no | no | no |
| PKD2 | 242 | M | 1 | 0 | no | yes | no | no | yes | no | no | no | no | no | no | no | no |
| PKD2 | 243 | S | 1 | 0 | no | yes | no | no | yes | no | no | no | yes | no | no | no | no |
| PKD2 | 244 | S | 0 | 0 | no | yes | no | no | yes | no | no | no | yes | no | no | no | no |
| PKD2 | 245 | N | 2 | 0 | no | yes | no | no | yes | no | no | no | no | no | no | no | no |
| PKD2 | 246 | V | 0 | 0 | yes | no | no | no | yes | no | no | no | no | no | no | no | no |
| PKD2 | 247 | Y | 0 | 0 | yes | no | no | no | yes | no | no | no | yes | no | no | no | no |
| PKD2 | 248 | Y | 1 | 0 | yes | no | no | no | yes | no | no | no | yes | no | no | no | no |
| PKD2 | 249 | Y | 0 | 0 | yes | no | no | no | yes | no | no | no | yes | no | no | no | no |
| PKD2 | 250 | T | 0 | 0 | yes | no | no | no | yes | no | no | no | yes | no | no | no | no |
| PKD2 | 251 | R | 2 | 0 | yes | no | no | no | yes | no | no | no | yes | no | no | no | no |
| PKD2 | 252 | M | 0 | 0 | yes | no | no | no | yes | no | no | no | yes | no | no | no | no |
| PKD2 | 253 | M | 0 | 0 | yes | no | no | no | yes | no | no | no | no | no | no | no | no |
| PKD2 | 254 | S | 2 | 0 | yes | no | no | no | yes | no | no | no | no | no | no | no | no |

| | | | | | | | | | | | | | | | | | |
|---|---|---|---|---|---|---|---|---|---|---|---|---|---|---|---|---|---|
| PKD2 | 255 | Q | 1 | 0 | yes | no | no | no | yes | no | no | no | no | no | no | no | no |
| PKD2 | 256 | L | 0 | 0 | yes | no | no | no | yes | no | no | no | no | no | no | no | no |
| PKD2 | 257 | F | 0 | 0 | yes | no | no | no | yes | no | no | no | no | no | no | no | no |
| PKD2 | 258 | L | 0 | 0 | no | yes | no | no | yes | no | no | no | no | no | no | no | no |
| PKD2 | 259 | D | 2 | 0 | no | yes | no | no | yes | no | no | no | no | no | no | no | no |
| PKD2 | 260 | T | 1 | 1 | no | yes | no | no | yes | no | no | no | no | no | no | no | no |
| PKD2 | 261 | P | 0 | 0 | no | yes | no | no | yes | no | no | no | no | no | no | no | no |
| PKD2 | 262 | V | 2 | 1 | no | yes | no | no | yes | no | no | no | no | no | no | no | no |
| PKD2 | 263 | S | 1 | 0 | no | yes | no | no | yes | no | no | no | no | no | no | no | no |
| PKD2 | 264 | K | 0 | 0 | no | yes | no | no | yes | no | no | no | no | no | no | no | no |
| PKD2 | 265 | T | 1 | 0 | no | yes | no | no | yes | no | yes | no | no | no | no | no | no |
| PKD2 | 266 | E | 1 | 0 | no | yes | no | no | yes | no | no | no | no | no | no | no | no |
| PKD2 | 267 | K | 1 | 0 | no | yes | no | no | yes | no | yes | no | no | no | no | no | no |
| PKD2 | 268 | T | 1 | 0 | no | yes | no | no | yes | no | no | no | no | no | no | no | no |
| PKD2 | 269 | N | 1 | 0 | no | yes | no | no | yes | no | no | no | no | no | no | no | no |
| PKD2 | 270 | F | 0 | 0 | no | yes | no | no | yes | no | no | no | no | no | no | no | no |
| PKD2 | 271 | K | 0 | 0 | no | yes | no | no | yes | no | no | no | no | no | no | no | no |
| PKD2 | 272 | T | 0 | 0 | no | yes | no | no | yes | no | no | no | no | no | no | no | no |
| PKD2 | 273 | L | 0 | 0 | no | yes | no | no | yes | no | no | no | no | no | no | no | no |
| PKD2 | 274 | S | 0 | 0 | no | yes | no | no | yes | no | no | no | no | no | no | no | no |
| PKD2 | 275 | S | 0 | 0 | no | yes | no | no | yes | no | no | no | no | no | no | no | no |
| PKD2 | 276 | M | 1 | 0 | yes | no | no | no | yes | no | no | no | yes | no | no | no | no |
| PKD2 | 277 | E | 1 | 0 | yes | no | no | no | yes | no | no | no | no | no | no | no | no |
| PKD2 | 278 | D | 0 | 0 | yes | no | no | no | yes | no | no | no | no | no | no | no | no |
| PKD2 | 279 | F | 0 | 0 | yes | no | no | no | yes | no | no | no | no | no | no | no | no |
| PKD2 | 280 | W | 0 | 0 | yes | no | no | no | yes | no | no | no | no | no | no | no | no |
| PKD2 | 281 | K | 0 | 0 | yes | no | no | no | yes | no | no | no | no | no | no | no | no |
| PKD2 | 282 | F | 0 | 0 | yes | no | no | no | yes | no | no | no | no | no | no | no | no |
| PKD2 | 283 | T | 0 | 0 | yes | no | no | no | yes | no | no | no | no | no | no | no | no |
| PKD2 | 284 | E | 0 | 0 | yes | no | no | no | yes | no | no | no | no | no | no | no | no |
| PKD2 | 285 | G | 2 | 0 | yes | no | no | no | yes | no | no | no | no | no | no | no | no |
| PKD2 | 286 | S | 1 | 0 | yes | no | no | no | yes | no | no | no | no | no | no | no | no |
| PKD2 | 287 | L | 0 | 0 | yes | no | no | no | yes | no | no | no | no | no | no | no | no |

| | | | | | | | | | | | | | | | | | |
|---|---|---|---|---|---|---|---|---|---|---|---|---|---|---|---|---|---|
| PKD2 | 288 | L | 0 | 0 | yes | no | no | no | yes | no | no | no | no | no | no | no | no |
| PKD2 | 289 | D | 1 | 0 | yes | no | no | no | yes | no | no | no | no | no | no | no | no |
| PKD2 | 290 | G | 0 | 1 | yes | no | no | no | yes | no | no | no | no | no | no | no | no |
| PKD2 | 291 | L | 0 | 0 | yes | no | no | no | yes | no | no | no | no | no | no | no | no |
| PKD2 | 292 | Y | 1 | 0 | no | yes | no | no | yes | no | no | no | no | no | no | no | no |
| PKD2 | 293 | W | 0 | 0 | no | yes | no | no | yes | no | no | no | no | no | no | no | no |
| PKD2 | 294 | K | 1 | 0 | no | yes | no | no | yes | no | no | no | no | no | no | no | no |
| PKD2 | 295 | M | 0 | 0 | no | yes | no | no | yes | no | no | no | no | no | no | no | no |
| PKD2 | 296 | Q | 0 | 0 | no | yes | no | no | yes | no | no | no | yes | no | no | no | no |
| PKD2 | 297 | P | 0 | 0 | no | yes | no | no | yes | no | no | no | no | no | no | no | no |
| PKD2 | 298 | S | 1 | 0 | no | yes | no | no | yes | no | no | no | no | no | no | no | no |
| PKD2 | 299 | N | 0 | 0 | no | yes | no | no | yes | no | no | no | no | no | no | no | no |
| PKD2 | 300 | Q | 3 | 0 | no | yes | no | no | yes | no | no | no | no | no | no | no | no |
| PKD2 | 301 | T | 1 | 0 | no | yes | no | no | yes | no | no | no | no | no | no | no | no |
| PKD2 | 302 | E | 0 | 0 | no | yes | no | no | yes | no | no | no | no | no | no | no | no |
| PKD2 | 303 | A | 1 | 0 | no | yes | no | no | yes | no | no | no | no | no | no | no | no |
| PKD2 | 304 | D | 1 | 0 | no | yes | no | no | yes | no | no | no | no | no | no | no | no |
| PKD2 | 305 | N | 1 | 0 | no | yes | no | no | yes | no | no | no | no | no | no | no | no |
| PKD2 | 306 | R | 2 | 1 | no | yes | no | no | yes | no | no | no | no | no | no | no | no |
| PKD2 | 307 | S | 1 | 0 | no | yes | no | no | yes | no | no | no | no | no | no | no | no |
| PKD2 | 308 | F | 1 | 0 | no | yes | no | no | yes | no | no | no | no | no | no | no | no |
| PKD2 | 309 | I | 0 | 0 | no | yes | no | no | yes | no | no | no | no | no | no | no | no |
| PKD2 | 310 | F | 1 | 0 | no | yes | no | no | yes | no | no | no | yes | no | no | no | no |
| PKD2 | 311 | Y | 2 | 0 | no | yes | no | no | yes | no | no | no | yes | no | no | no | no |
| PKD2 | 312 | E | 0 | 0 | no | yes | no | no | yes | no | no | no | yes | no | no | no | no |
| PKD2 | 313 | N | 0 | 0 | no | no | yes | no | yes | no | no | no | yes | no | no | no | no |
| PKD2 | 314 | L | 0 | 0 | no | no | yes | no | yes | no | no | no | no | no | no | no | no |
| PKD2 | 315 | L | 0 | 0 | no | no | yes | no | yes | no | no | no | no | no | no | no | no |
| PKD2 | 316 | L | 0 | 0 | no | no | yes | no | yes | no | no | no | no | no | no | no | no |
| PKD2 | 317 | G | 0 | 0 | no | no | yes | no | yes | no | no | no | no | no | no | no | no |
| PKD2 | 318 | V | 0 | 0 | no | no | yes | no | yes | no | no | no | no | no | no | no | no |
| PKD2 | 319 | P | 1 | 0 | no | no | yes | no | yes | no | no | no | no | no | no | no | no |
| PKD2 | 320 | R | 0 | 0 | no | no | yes | no | yes | no | no | no | no | no | no | no | no |

| | | | | | | | | | | | | | | | | | |
|---|---|---|---|---|---|---|---|---|---|---|---|---|---|---|---|---|---|
| PKD2 | 321 | I | 1 | 0 | no | no | yes | no | yes | no | no | no | no | no | no | no | no |
| PKD2 | 322 | R | 1 | 1 | no | no | yes | no | yes | no | no | no | no | no | no | no | no |
| PKD2 | 323 | Q | 0 | 0 | no | no | yes | no | yes | no | no | no | no | no | no | no | no |
| PKD2 | 324 | L | 0 | 0 | no | yes | no | no | yes | no | no | no | no | no | no | no | no |
| PKD2 | 325 | R | 0 | 2 | no | yes | no | no | yes | no | no | no | no | no | no | no | no |
| PKD2 | 326 | V | 0 | 0 | no | no | yes | no | yes | no | no | no | no | no | no | no | no |
| PKD2 | 327 | R | 0 | 0 | no | yes | no | no | yes | no | no | no | no | no | no | no | no |
| PKD2 | 328 | N | 0 | 0 | no | yes | no | no | yes | no | no | no | no | no | no | no | no |
| PKD2 | 329 | G | 2 | 0 | no | yes | no | no | yes | no | no | no | no | no | no | no | no |
| PKD2 | 330 | S | 0 | 0 | no | yes | no | no | yes | no | no | no | no | no | no | no | no |
| PKD2 | 331 | C | 0 | 1 | no | yes | no | no | yes | no | no | no | no | no | no | no | no |
| PKD2 | 332 | S | 2 | 0 | no | yes | no | no | yes | no | no | no | no | no | no | no | no |
| PKD2 | 333 | I | 1 | 0 | no | yes | no | no | yes | no | no | no | no | no | no | no | no |
| PKD2 | 334 | P | 0 | 0 | no | yes | no | no | yes | no | no | no | no | no | no | no | no |
| PKD2 | 335 | Q | 1 | 0 | yes | no | no | no | yes | no | no | no | no | no | no | no | no |
| PKD2 | 336 | D | 0 | 0 | yes | no | no | no | yes | no | no | no | yes | no | no | no | no |
| PKD2 | 337 | L | 0 | 0 | yes | no | no | no | yes | no | no | no | yes | no | no | no | no |
| PKD2 | 338 | R | 0 | 0 | yes | no | no | no | yes | no | no | no | no | no | no | no | no |
| PKD2 | 339 | D | 0 | 0 | no | yes | no | no | yes | no | no | no | no | no | no | no | no |
| PKD2 | 340 | E | 0 | 0 | no | yes | no | no | yes | no | no | no | yes | no | no | no | no |
| PKD2 | 341 | I | 0 | 0 | no | yes | no | no | yes | no | no | no | no | no | no | no | no |
| PKD2 | 342 | K | 0 | 0 | no | yes | no | no | yes | no | no | no | no | no | no | no | no |
| PKD2 | 343 | E | 0 | 0 | no | yes | no | no | yes | no | no | no | no | no | no | no | no |
| PKD2 | 344 | C | 0 | 0 | no | yes | no | no | yes | no | no | no | yes | no | no | no | no |
| PKD2 | 345 | Y | 0 | 0 | no | yes | no | no | yes | no | no | no | yes | no | no | no | no |
| PKD2 | 346 | D | 0 | 0 | no | yes | no | no | yes | no | no | no | no | no | no | no | no |
| PKD2 | 347 | V | 0 | 0 | no | yes | no | no | yes | no | no | no | no | no | no | no | no |
| PKD2 | 348 | Y | 0 | 0 | no | yes | no | no | yes | no | no | no | no | no | no | no | no |
| PKD2 | 349 | S | 0 | 1 | no | yes | no | no | yes | no | no | no | no | no | no | no | no |
| PKD2 | 350 | V | 0 | 0 | yes | no | no | no | yes | no | no | no | no | no | no | no | no |
| PKD2 | 351 | S | 0 | 0 | yes | no | no | no | yes | no | no | no | no | no | no | no | no |
| PKD2 | 352 | S | 0 | 0 | yes | no | no | no | yes | no | no | no | no | no | no | no | no |
| PKD2 | 353 | E | 0 | 0 | no | no | yes | no | yes | no | no | no | no | no | no | no | no |

| | | | | | | | | | | | | | | | | | |
|---|---|---|---|---|---|---|---|---|---|---|---|---|---|---|---|---|---|
| PKD2 | 354 | D | 0 | 0 | no | yes | no | no | yes | no | no | no | no | no | no | no | no |
| PKD2 | 355 | R | 1 | 0 | no | yes | no | no | yes | no | no | no | no | no | no | no | no |
| PKD2 | 356 | A | 1 | 1 | no | yes | no | no | yes | no | no | no | no | no | no | no | no |
| PKD2 | 357 | P | 0 | 0 | no | yes | no | no | yes | no | no | no | no | no | no | no | no |
| PKD2 | 358 | F | 0 | 0 | no | yes | no | no | yes | no | no | no | no | no | no | no | no |
| PKD2 | 359 | G | 0 | 0 | no | yes | no | no | yes | no | no | no | no | no | no | no | no |
| PKD2 | 360 | P | 0 | 0 | no | yes | no | no | yes | no | no | no | no | no | no | no | no |
| PKD2 | 361 | R | 1 | 0 | no | yes | no | no | yes | no | no | no | no | no | no | no | no |
| PKD2 | 362 | N | 1 | 0 | no | yes | no | no | yes | no | no | no | no | no | no | no | no |
| PKD2 | 363 | G | 0 | 0 | no | yes | no | no | yes | no | no | no | no | no | no | no | no |
| PKD2 | 364 | T | 1 | 0 | no | yes | no | no | yes | no | no | no | no | no | no | no | no |
| PKD2 | 365 | A | 0 | 0 | no | yes | no | no | yes | no | no | no | no | no | no | no | no |
| PKD2 | 366 | W | 0 | 0 | no | yes | no | no | yes | no | no | no | no | no | no | no | no |
| PKD2 | 367 | I | 0 | 0 | no | yes | no | no | yes | no | no | no | no | no | no | no | no |
| PKD2 | 368 | Y | 0 | 0 | no | yes | no | no | yes | no | no | no | no | no | no | no | no |
| PKD2 | 369 | T | 0 | 0 | no | yes | no | no | yes | no | no | no | no | no | no | no | no |
| PKD2 | 370 | S | 1 | 0 | no | yes | no | no | yes | no | no | no | no | no | no | no | no |
| PKD2 | 371 | E | 0 | 0 | no | yes | no | no | yes | no | no | no | no | no | no | no | no |
| PKD2 | 372 | K | 1 | 0 | no | yes | no | no | yes | no | no | no | no | no | no | no | no |
| PKD2 | 373 | D | 0 | 0 | no | yes | no | no | yes | no | no | no | no | no | no | no | no |
| PKD2 | 374 | L | 1 | 1 | no | yes | no | no | yes | no | no | no | no | no | no | no | no |
| PKD2 | 375 | N | 0 | 0 | no | yes | no | no | yes | no | no | no | no | no | no | no | no |
| PKD2 | 376 | G | 0 | 0 | no | yes | no | no | yes | no | no | no | no | no | no | no | no |
| PKD2 | 377 | S | 1 | 0 | no | yes | no | no | yes | no | no | no | no | no | no | no | no |
| PKD2 | 378 | S | 1 | 0 | no | yes | no | no | yes | no | no | no | no | no | no | no | no |
| PKD2 | 379 | H | 1 | 0 | no | no | yes | no | yes | no | no | no | no | no | no | no | no |
| PKD2 | 380 | W | 0 | 0 | no | yes | no | no | yes | no | no | no | yes | no | no | no | no |
| PKD2 | 381 | G | 0 | 1 | no | yes | no | no | yes | no | no | no | no | no | no | no | no |
| PKD2 | 382 | I | 1 | 0 | no | yes | no | no | yes | no | no | no | yes | no | no | no | no |
| PKD2 | 383 | I | 1 | 0 | no | yes | no | no | yes | no | no | no | no | no | no | no | no |
| PKD2 | 384 | A | 1 | 0 | no | yes | no | no | yes | no | no | no | no | no | no | no | no |
| PKD2 | 385 | T | 2 | 0 | no | yes | no | no | yes | no | no | no | no | no | no | no | no |
| PKD2 | 386 | Y | 0 | 0 | no | no | yes | no | yes | no | no | no | no | no | no | no | no |

| | | | | | | | | | | | | | | | | | |
|---|---|---|---|---|---|---|---|---|---|---|---|---|---|---|---|---|---|
| PKD2 | 387 | S | 0 | 0 | no | yes | no | no | yes | no | no | no | no | no | no | no | no |
| PKD2 | 388 | G | 0 | 0 | no | yes | no | no | yes | no | no | no | no | no | no | no | no |
| PKD2 | 389 | A | 0 | 1 | no | yes | no | no | yes | no | no | no | no | no | no | no | no |
| PKD2 | 390 | G | 0 | 2 | no | yes | no | no | yes | no | no | no | no | no | no | no | no |
| PKD2 | 391 | Y | 2 | 1 | no | no | yes | no | yes | no | no | no | no | no | no | no | no |
| PKD2 | 392 | Y | 0 | 1 | no | no | yes | no | yes | no | no | no | no | no | no | no | no |
| PKD2 | 393 | L | 0 | 0 | no | no | yes | no | yes | no | no | no | no | no | no | no | no |
| PKD2 | 394 | D | 0 | 0 | no | no | yes | no | yes | no | no | no | no | no | no | no | no |
| PKD2 | 395 | L | 0 | 0 | no | yes | no | no | yes | no | no | no | no | no | no | no | no |
| PKD2 | 396 | S | 2 | 0 | no | yes | no | no | yes | no | no | no | no | no | no | no | no |
| PKD2 | 397 | R | 0 | 0 | no | yes | no | no | yes | no | no | no | no | no | no | no | no |
| PKD2 | 398 | T | 0 | 0 | no | yes | no | no | yes | no | no | no | no | no | no | no | no |
| PKD2 | 399 | R | 0 | 0 | yes | no | no | no | yes | no | no | no | no | no | no | no | no |
| PKD2 | 400 | E | 0 | 0 | yes | no | no | no | yes | no | no | no | no | no | no | no | no |
| PKD2 | 401 | E | 0 | 0 | yes | no | no | no | yes | no | no | no | no | no | no | no | no |
| PKD2 | 402 | T | 0 | 0 | yes | no | no | no | yes | no | no | no | no | no | no | no | no |
| PKD2 | 403 | A | 0 | 0 | yes | no | no | no | yes | no | no | no | no | no | no | no | no |
| PKD2 | 404 | A | 1 | 0 | yes | no | no | no | yes | no | no | no | no | no | no | no | no |
| PKD2 | 405 | Q | 0 | 0 | yes | no | no | no | yes | no | no | no | no | no | no | no | no |
| PKD2 | 406 | V | 0 | 0 | yes | no | no | no | yes | no | no | no | no | no | no | no | no |
| PKD2 | 407 | A | 1 | 0 | yes | no | no | no | yes | no | no | no | no | no | no | no | no |
| PKD2 | 408 | S | 1 | 0 | yes | no | no | no | yes | no | no | no | no | no | no | no | no |
| PKD2 | 409 | L | 0 | 0 | yes | no | no | no | yes | no | no | no | no | no | no | no | no |
| PKD2 | 410 | K | 2 | 0 | yes | no | no | no | yes | no | no | no | no | no | no | no | no |
| PKD2 | 411 | K | 0 | 0 | yes | no | no | no | yes | no | no | no | no | no | no | no | no |
| PKD2 | 412 | N | 0 | 0 | yes | no | no | no | yes | no | no | no | no | no | no | no | no |
| PKD2 | 413 | V | 1 | 0 | no | yes | no | no | yes | no | no | no | no | no | no | no | no |
| PKD2 | 414 | W | 0 | 1 | no | yes | no | no | yes | no | no | no | no | no | no | no | no |
| PKD2 | 415 | L | 0 | 1 | no | yes | no | no | yes | no | no | no | no | no | no | no | no |
| PKD2 | 416 | D | 0 | 0 | no | yes | no | no | yes | no | no | no | yes | no | no | no | no |
| PKD2 | 417 | R | 1 | 0 | no | yes | no | no | yes | no | no | no | yes | no | no | no | no |
| PKD2 | 418 | G | 0 | 0 | no | yes | no | no | yes | no | no | no | yes | no | no | no | no |
| PKD2 | 419 | T | 1 | 1 | no | yes | no | no | yes | no | no | no | no | no | no | no | no |

| | | | | | | | | | | | | | | | | | |
|---|---|---|---|---|---|---|---|---|---|---|---|---|---|---|---|---|---|
| PKD2 | 420 | R | 0 | 1 | no | yes | no | no | yes | no | no | no | yes | no | no | no | no |
| PKD2 | 421 | A | 2 | 0 | no | yes | no | no | yes | no | no | no | no | no | no | no | no |
| PKD2 | 422 | T | 0 | 0 | no | no | yes | no | yes | no | no | no | no | no | no | no | no |
| PKD2 | 423 | F | 1 | 0 | no | no | yes | no | yes | no | no | no | no | no | no | no | no |
| PKD2 | 424 | I | 0 | 0 | no | no | yes | no | yes | no | no | no | no | no | no | no | no |
| PKD2 | 425 | D | 0 | 0 | no | no | yes | no | yes | no | no | no | no | no | no | no | no |
| PKD2 | 426 | F | 1 | 0 | no | no | yes | no | yes | no | no | no | no | no | no | no | no |
| PKD2 | 427 | S | 0 | 0 | no | no | yes | no | yes | no | no | no | no | no | no | no | no |
| PKD2 | 428 | V | 1 | 0 | no | no | yes | no | yes | no | no | no | no | no | no | no | no |
| PKD2 | 429 | Y | 0 | 0 | no | no | yes | no | yes | no | no | no | yes | no | no | no | no |
| PKD2 | 430 | N | 0 | 0 | no | no | yes | no | yes | no | no | no | no | no | no | no | no |
| PKD2 | 431 | A | 1 | 0 | no | yes | no | no | yes | no | no | no | no | no | no | no | no |
| PKD2 | 432 | N | 0 | 0 | no | yes | no | no | yes | no | no | no | yes | no | no | no | no |
| PKD2 | 433 | I | 1 | 0 | no | yes | no | no | yes | no | no | no | no | no | no | no | no |
| PKD2 | 434 | N | 0 | 0 | no | yes | no | no | yes | no | no | no | no | no | no | no | no |
| PKD2 | 435 | L | 0 | 0 | no | yes | no | no | yes | no | no | no | no | no | no | no | no |
| PKD2 | 436 | F | 0 | 0 | no | no | yes | no | yes | no | no | no | no | no | no | no | no |
| PKD2 | 437 | C | 0 | 0 | no | no | yes | no | yes | no | no | no | no | no | no | no | no |
| PKD2 | 438 | V | 0 | 0 | no | no | yes | no | yes | no | no | no | no | no | no | no | no |
| PKD2 | 439 | V | 0 | 0 | no | no | yes | no | yes | no | no | no | no | no | no | no | no |
| PKD2 | 440 | R | 0 | 1 | no | no | yes | no | yes | no | no | no | no | no | no | no | no |
| PKD2 | 441 | L | 0 | 1 | no | no | yes | no | yes | no | no | no | no | no | no | no | no |
| PKD2 | 442 | L | 2 | 0 | no | no | yes | no | yes | no | no | no | no | no | no | no | no |
| PKD2 | 443 | V | 0 | 1 | no | no | yes | no | yes | no | no | no | no | no | no | no | no |
| PKD2 | 444 | E | 0 | 1 | no | no | yes | no | yes | no | no | no | no | no | no | no | no |
| PKD2 | 445 | F | 0 | 0 | no | yes | no | no | yes | no | no | no | no | no | no | no | no |
| PKD2 | 446 | P | 1 | 0 | no | yes | no | no | yes | no | no | no | yes | no | no | no | no |
| PKD2 | 447 | A | 0 | 0 | no | yes | no | no | yes | no | no | no | yes | no | no | no | no |
| PKD2 | 448 | T | 1 | 0 | no | yes | no | no | yes | no | no | no | yes | no | no | no | no |
| PKD2 | 449 | G | 0 | 0 | no | yes | no | no | yes | no | no | no | yes | no | no | no | no |
| PKD2 | 450 | G | 0 | 1 | no | yes | no | no | yes | no | no | no | yes | no | no | no | no |
| PKD2 | 451 | V | 0 | 0 | no | yes | no | no | yes | no | no | no | yes | no | no | no | no |
| PKD2 | 452 | I | 1 | 0 | no | no | yes | no | yes | no | no | no | yes | no | no | no | no |

| | | | | | | | | | | | | | | | | | |
|---|---|---|---|---|---|---|---|---|---|---|---|---|---|---|---|---|---|
| PKD2 | 453 | P | 0 | 0 | no | no | yes | no | yes | no | no | no | no | no | no | no | no |
| PKD2 | 454 | S | 0 | 0 | no | no | yes | no | yes | no | no | no | yes | no | no | no | no |
| PKD2 | 455 | W | 0 | 0 | no | no | yes | no | yes | no | no | no | yes | no | no | no | no |
| PKD2 | 456 | Q | 0 | 0 | no | no | yes | no | yes | no | no | no | yes | no | no | no | no |
| PKD2 | 457 | F | 0 | 0 | no | no | yes | no | yes | no | no | no | yes | no | no | no | no |
| PKD2 | 458 | Q | 0 | 0 | no | no | yes | no | yes | no | no | no | no | no | no | no | no |
| PKD2 | 459 | P | 0 | 0 | no | no | yes | no | yes | no | no | no | no | no | no | no | no |
| PKD2 | 460 | L | 0 | 0 | no | no | yes | no | yes | no | no | no | no | no | no | no | no |
| PKD2 | 461 | K | 0 | 0 | no | yes | no | no | yes | no | no | no | no | no | no | no | no |
| PKD2 | 462 | L | 0 | 0 | no | yes | no | no | yes | no | no | no | no | no | no | no | no |
| PKD2 | 463 | I | 0 | 0 | no | yes | no | no | yes | no | no | no | no | no | no | no | no |
| PKD2 | 464 | R | 1 | 0 | no | yes | no | no | yes | no | no | no | no | no | no | no | no |
| PKD2 | 465 | Y | 1 | 0 | no | yes | no | no | yes | no | no | no | no | no | no | no | no |
| PKD2 | 466 | V | 1 | 0 | no | yes | no | no | yes | no | no | no | no | no | no | no | no |
| PKD2 | 467 | T | 1 | 0 | no | yes | no | no | yes | no | no | no | no | no | no | no | no |
| PKD2 | 468 | T | 0 | 0 | no | yes | no | no | yes | no | no | no | no | no | no | no | no |
| PKD2 | 469 | F | 0 | 0 | no | yes | no | no | no | no | no | no | no | no | no | no | no |
| PKD2 | 470 | D | 1 | 0 | yes | no | no | no | no | no | no | no | no | no | no | no | no |
| PKD2 | 471 | F | 1 | 0 | yes | no | no | no | no | no | no | no | no | no | no | no | no |
| PKD2 | 472 | F | 1 | 0 | yes | no | no | yes | no | no | no | no | no | no | no | no | no |
| PKD2 | 473 | L | 0 | 0 | yes | no | no | yes | no | no | no | no | no | no | no | no | no |
| PKD2 | 474 | A | 0 | 0 | yes | no | no | yes | no | no | no | no | no | no | no | no | no |
| PKD2 | 475 | A | 1 | 0 | yes | no | no | yes | no | no | no | no | no | no | no | no | no |
| PKD2 | 476 | C | 1 | 0 | yes | no | no | yes | no | no | no | no | no | no | no | no | no |
| PKD2 | 477 | E | 0 | 0 | yes | no | no | yes | no | no | no | no | no | no | no | no | no |
| PKD2 | 478 | I | 1 | 0 | yes | no | no | yes | no | no | no | no | no | no | no | no | no |
| PKD2 | 479 | I | 0 | 0 | yes | no | no | yes | no | no | no | no | no | no | no | no | no |
| PKD2 | 480 | F | 0 | 0 | yes | no | no | yes | no | no | no | no | no | no | no | no | no |
| PKD2 | 481 | C | 0 | 0 | yes | no | no | yes | no | no | no | no | no | no | no | no | no |
| PKD2 | 482 | F | 2 | 1 | yes | no | no | yes | no | no | no | no | no | no | no | no | no |
| PKD2 | 483 | F | 0 | 0 | yes | no | no | yes | no | no | no | no | no | no | no | no | no |
| PKD2 | 484 | I | 0 | 0 | yes | no | no | yes | no | no | no | no | no | no | no | no | no |
| PKD2 | 485 | F | 0 | 0 | yes | no | no | yes | no | no | no | no | no | no | no | no | no |

| | | | | | | | | | | | | | | | | | |
|---|---|---|---|---|---|---|---|---|---|---|---|---|---|---|---|---|---|
| PKD2 | 486 | Y | 0 | 0 | yes | no | no | yes | no | no | no | no | no | no | no | no | no |
| PKD2 | 487 | Y | 2 | 1 | yes | no | no | yes | no | no | no | no | no | no | no | no | no |
| PKD2 | 488 | V | 1 | 0 | yes | no | no | yes | no | no | no | no | no | no | no | no | no |
| PKD2 | 489 | V | 1 | 0 | yes | no | no | yes | no | no | no | no | no | no | no | no | no |
| PKD2 | 490 | E | 0 | 0 | yes | no | no | yes | no | yes | no | no | no | no | no | no | no |
| PKD2 | 491 | E | 0 | 0 | yes | no | no | yes | no | yes | no | no | no | no | no | no | no |
| PKD2 | 492 | I | 1 | 0 | yes | no | no | yes | no | yes | no | no | no | no | no | no | no |
| PKD2 | 493 | L | 0 | 0 | yes | no | no | yes | no | yes | no | no | no | no | no | no | no |
| PKD2 | 494 | E | 0 | 0 | yes | no | no | no | no | yes | no | no | no | no | no | no | no |
| PKD2 | 495 | I | 0 | 0 | yes | no | no | yes | no | yes | no | no | no | no | no | no | no |
| PKD2 | 496 | R | 1 | 0 | yes | no | no | no | no | yes | no | no | no | no | no | no | no |
| PKD2 | 497 | I | 2 | 0 | yes | no | no | no | no | yes | no | no | no | no | no | no | no |
| PKD2 | 498 | H | 0 | 0 | no | yes | no | no | no | yes | no | no | no | no | no | no | no |
| PKD2 | 499 | K | 0 | 0 | no | yes | no | no | no | yes | no | no | no | no | no | no | no |
| PKD2 | 500 | L | 1 | 0 | no | yes | no | no | no | yes | no | no | no | no | no | no | no |
| PKD2 | 501 | H | 2 | 0 | no | yes | no | no | no | yes | no | no | no | no | no | no | no |
| PKD2 | 502 | Y | 0 | 0 | no | yes | no | no | no | yes | no | no | no | no | no | no | no |
| PKD2 | 503 | F | 0 | 0 | no | yes | no | no | no | yes | no | no | no | no | no | no | no |
| PKD2 | 504 | R | 0 | 0 | no | yes | no | no | no | yes | no | no | no | no | no | no | no |
| PKD2 | 505 | S | 1 | 0 | no | yes | no | no | no | yes | no | no | no | no | no | no | no |
| PKD2 | 506 | F | 2 | 0 | yes | no | no | yes | no | no | no | no | no | no | no | no | no |
| PKD2 | 507 | W | 0 | 0 | yes | no | no | no | no | no | no | no | no | no | no | no | no |
| PKD2 | 508 | N | 1 | 0 | yes | no | no | yes | no | no | no | no | no | no | no | no | no |
| PKD2 | 509 | C | 1 | 0 | yes | no | no | yes | no | no | no | no | no | no | no | no | no |
| PKD2 | 510 | L | 0 | 0 | yes | no | no | yes | no | no | no | no | no | no | no | no | no |
| PKD2 | 511 | D | 0 | 1 | yes | no | no | yes | no | no | no | no | no | no | no | no | no |
| PKD2 | 512 | V | 2 | 0 | no | yes | no | yes | no | no | no | no | no | no | no | no | no |
| PKD2 | 513 | V | 0 | 0 | yes | no | no | yes | no | no | no | no | no | no | no | no | no |
| PKD2 | 514 | I | 0 | 0 | yes | no | no | yes | no | no | no | no | no | no | no | no | no |
| PKD2 | 515 | V | 1 | 0 | yes | no | no | yes | no | no | no | no | no | no | no | no | no |
| PKD2 | 516 | V | 1 | 0 | yes | no | no | yes | no | no | no | no | no | no | no | no | no |
| PKD2 | 517 | L | 0 | 0 | yes | no | no | yes | no | no | no | no | no | no | no | no | no |
| PKD2 | 518 | S | 0 | 0 | yes | no | no | yes | no | no | no | no | no | no | no | no | no |

| | | | | | | | | | | | | | | | | | |
|---|---|---|---|---|---|---|---|---|---|---|---|---|---|---|---|---|---|
| PKD2 | 519 | V | 1 | 0 | yes | no | no | yes | no | no | no | no | no | no | no | no | no |
| PKD2 | 520 | V | 0 | 0 | yes | no | no | yes | no | no | no | no | no | no | no | no | no |
| PKD2 | 521 | A | 1 | 0 | yes | no | no | yes | no | no | no | no | no | no | no | no | no |
| PKD2 | 522 | I | 3 | 0 | yes | no | no | yes | no | no | no | no | no | no | no | no | no |
| PKD2 | 523 | G | 0 | 0 | yes | no | no | yes | no | no | no | no | no | no | no | no | no |
| PKD2 | 524 | I | 1 | 0 | yes | no | no | yes | no | no | no | no | no | no | no | no | no |
| PKD2 | 525 | N | 0 | 0 | yes | no | no | yes | no | no | no | no | no | no | no | no | no |
| PKD2 | 526 | I | 0 | 0 | yes | no | no | yes | no | no | no | no | no | no | no | no | no |
| PKD2 | 527 | Y | 1 | 0 | no | yes | no | yes | yes | no | no | no | no | no | no | no | no |
| PKD2 | 528 | R | 1 | 0 | no | yes | no | yes | yes | no | no | no | no | no | no | no | no |
| PKD2 | 529 | T | 1 | 0 | yes | no | no | no | yes | no | no | no | no | no | no | no | no |
| PKD2 | 530 | S | 0 | 0 | yes | no | no | no | yes | no | no | no | no | no | no | no | no |
| PKD2 | 531 | N | 0 | 0 | yes | no | no | no | yes | no | no | no | no | no | no | no | no |
| PKD2 | 532 | V | 1 | 0 | yes | no | no | no | yes | no | no | no | no | no | no | no | no |
| PKD2 | 533 | E | 2 | 0 | yes | no | no | no | yes | no | no | no | no | no | no | no | no |
| PKD2 | 534 | V | 2 | 0 | yes | no | no | no | yes | no | no | no | no | no | no | no | no |
| PKD2 | 535 | L | 1 | 0 | yes | no | no | no | yes | no | no | no | no | no | no | no | no |
| PKD2 | 536 | L | 0 | 0 | no | yes | no | no | yes | no | no | no | no | no | no | no | no |
| PKD2 | 537 | Q | 0 | 0 | no | yes | no | no | yes | no | no | no | yes | no | no | no | no |
| PKD2 | 538 | F | 0 | 0 | yes | no | no | no | yes | no | no | no | no | no | no | no | no |
| PKD2 | 539 | L | 0 | 0 | yes | no | no | no | yes | no | no | no | no | no | no | no | no |
| PKD2 | 540 | E | 0 | 0 | yes | no | no | no | yes | no | no | no | no | no | no | no | no |
| PKD2 | 541 | D | 1 | 0 | yes | no | no | no | yes | no | no | no | yes | no | no | no | no |
| PKD2 | 542 | Q | 0 | 0 | no | yes | no | no | yes | no | no | no | no | no | no | no | no |
| PKD2 | 543 | N | 0 | 0 | no | yes | no | no | yes | no | no | no | no | no | no | no | no |
| PKD2 | 544 | T | 0 | 0 | no | yes | no | no | yes | no | no | no | no | no | no | no | no |
| PKD2 | 545 | F | 2 | 0 | no | yes | no | no | yes | no | no | no | no | no | no | no | no |
| PKD2 | 546 | P | 0 | 0 | no | yes | no | no | yes | no | no | no | no | no | no | no | no |
| PKD2 | 547 | N | 2 | 0 | no | yes | no | no | yes | no | no | no | no | no | no | no | no |
| PKD2 | 548 | F | 1 | 0 | no | yes | no | no | yes | no | no | no | no | no | no | no | no |
| PKD2 | 549 | E | 2 | 0 | yes | no | no | no | yes | no | no | no | no | no | no | no | no |
| PKD2 | 550 | H | 1 | 0 | yes | no | no | no | yes | no | no | no | no | no | no | no | no |
| PKD2 | 551 | L | 0 | 0 | yes | no | no | no | yes | no | no | no | no | no | no | no | no |

| | | | | | | | | | | | | | | | | | |
|---|---|---|---|---|---|---|---|---|---|---|---|---|---|---|---|---|---|
| PKD2 | 552 | A | 1 | 0 | yes | no | no | no | yes | no | no | no | no | no | no | no | no |
| PKD2 | 553 | Y | 1 | 0 | yes | no | no | no | no | no | no | no | no | no | no | no | no |
| PKD2 | 554 | W | 0 | 0 | yes | no | no | no | no | no | no | no | no | no | no | no | no |
| PKD2 | 555 | Q | 0 | 0 | yes | no | no | no | no | no | no | no | no | no | no | no | no |
| PKD2 | 556 | I | 0 | 0 | yes | no | no | no | no | no | no | no | no | no | no | no | no |
| PKD2 | 557 | Q | 0 | 0 | yes | no | no | yes | no | no | no | no | no | no | no | no | no |
| PKD2 | 558 | F | 1 | 0 | yes | no | no | yes | no | no | no | no | no | no | no | no | no |
| PKD2 | 559 | N | 0 | 0 | yes | no | no | yes | no | no | no | no | yes | no | no | no | no |
| PKD2 | 560 | N | 0 | 0 | yes | no | no | yes | no | no | no | no | yes | no | no | no | no |
| PKD2 | 561 | I | 0 | 0 | yes | no | no | yes | no | no | no | no | no | no | no | no | no |
| PKD2 | 562 | A | 0 | 0 | yes | no | no | yes | no | no | no | no | no | no | no | no | no |
| PKD2 | 563 | A | 0 | 0 | yes | no | no | yes | no | no | no | no | yes | no | no | no | no |
| PKD2 | 564 | V | 0 | 0 | yes | no | no | yes | no | no | no | no | no | no | no | no | no |
| PKD2 | 565 | T | 2 | 0 | yes | no | no | yes | no | no | no | no | no | no | no | no | no |
| PKD2 | 566 | V | 2 | 0 | yes | no | no | yes | no | no | no | no | no | no | no | no | no |
| PKD2 | 567 | F | 0 | 0 | yes | no | no | yes | no | no | no | no | yes | no | no | no | no |
| PKD2 | 568 | F | 0 | 0 | yes | no | no | yes | no | no | no | no | no | no | no | no | no |
| PKD2 | 569 | V | 0 | 0 | yes | no | no | yes | no | no | no | no | no | no | no | no | no |
| PKD2 | 570 | W | 1 | 0 | yes | no | no | yes | no | no | no | no | yes | no | no | no | no |
| PKD2 | 571 | I | 0 | 0 | yes | no | no | yes | no | no | no | no | yes | no | no | no | no |
| PKD2 | 572 | K | 0 | 0 | yes | no | no | yes | no | no | no | no | no | no | no | no | no |
| PKD2 | 573 | L | 0 | 0 | yes | no | no | yes | no | no | no | no | no | no | no | no | no |
| PKD2 | 574 | F | 0 | 0 | yes | no | no | yes | no | yes | no | no | yes | no | no | no | no |
| PKD2 | 575 | K | 1 | 0 | yes | no | no | yes | no | yes | no | no | no | no | no | no | no |
| PKD2 | 576 | F | 0 | 0 | yes | no | no | yes | no | yes | no | no | no | no | no | no | no |
| PKD2 | 577 | I | 1 | 0 | no | yes | no | yes | no | yes | no | no | no | no | no | no | no |
| PKD2 | 578 | N | 1 | 0 | no | yes | no | no | no | yes | no | no | no | no | no | no | no |
| PKD2 | 579 | F | 0 | 0 | no | yes | no | no | no | yes | no | no | no | no | no | no | no |
| PKD2 | 580 | N | 0 | 0 | no | yes | no | no | no | yes | no | no | no | no | no | no | no |
| PKD2 | 581 | R | 1 | 0 | no | yes | no | no | no | yes | no | no | no | no | no | no | no |
| PKD2 | 582 | T | 1 | 0 | yes | no | no | no | no | yes | no | no | no | no | no | no | no |
| PKD2 | 583 | M | 1 | 0 | yes | no | no | no | no | yes | no | no | no | no | no | no | no |
| PKD2 | 584 | S | 0 | 0 | yes | no | no | no | no | yes | no | no | no | no | no | no | no |

| | | | | | | | | | | | | | | | | | |
|---|---|---|---|---|---|---|---|---|---|---|---|---|---|---|---|---|---|
| PKD2 | 585 | Q | 1 | 0 | yes | no | no | no | no | yes | no | no | no | no | no | no | no |
| PKD2 | 586 | L | 0 | 0 | yes | no | no | no | no | yes | no | no | yes | no | no | no | no |
| PKD2 | 587 | S | 2 | 0 | yes | no | no | no | no | yes | no | no | no | no | no | no | no |
| PKD2 | 588 | T | 0 | 0 | yes | no | no | no | no | yes | no | no | no | no | no | no | no |
| PKD2 | 589 | T | 1 | 0 | yes | no | no | no | no | yes | no | no | no | no | no | no | no |
| PKD2 | 590 | M | 1 | 0 | yes | no | no | no | no | yes | no | no | yes | no | no | no | no |
| PKD2 | 591 | S | 0 | 0 | yes | no | no | no | no | yes | no | no | no | no | no | no | no |
| PKD2 | 592 | R | 1 | 0 | yes | no | no | no | no | yes | no | no | no | no | no | no | no |
| PKD2 | 593 | C | 2 | 0 | yes | no | no | no | no | yes | no | no | no | no | no | no | no |
| PKD2 | 594 | A | 1 | 0 | yes | no | no | no | no | yes | no | no | no | no | no | no | no |
| PKD2 | 595 | K | 0 | 0 | yes | no | no | no | no | yes | no | no | no | no | no | no | no |
| PKD2 | 596 | D | 0 | 0 | yes | no | no | no | no | yes | no | no | yes | no | no | no | no |
| PKD2 | 597 | L | 0 | 2 | yes | no | no | yes | no | yes | no | no | no | no | no | no | no |
| PKD2 | 598 | F | 1 | 0 | yes | no | no | yes | no | no | no | no | no | no | no | no | no |
| PKD2 | 599 | G | 0 | 0 | yes | no | no | yes | no | no | no | no | no | no | no | no | no |
| PKD2 | 600 | F | 1 | 0 | yes | no | no | yes | no | no | no | no | no | no | no | no | no |
| PKD2 | 601 | A | 0 | 0 | no | yes | no | yes | no | no | no | no | no | no | no | no | no |
| PKD2 | 602 | I | 1 | 0 | yes | no | no | yes | no | no | no | no | no | no | no | no | no |
| PKD2 | 603 | M | 3 | 0 | yes | no | no | yes | no | no | no | no | yes | no | no | no | no |
| PKD2 | 604 | F | 0 | 0 | yes | no | no | yes | no | no | no | no | no | no | no | no | no |
| PKD2 | 605 | F | 0 | 0 | yes | no | no | yes | no | no | no | no | no | no | no | no | no |
| PKD2 | 606 | I | 1 | 0 | yes | no | no | yes | no | no | no | no | no | no | no | no | no |
| PKD2 | 607 | I | 1 | 0 | yes | no | no | yes | no | no | no | no | yes | no | no | no | no |
| PKD2 | 608 | F | 0 | 0 | yes | no | no | yes | no | no | no | no | no | no | no | no | no |
| PKD2 | 609 | L | 0 | 0 | yes | no | no | yes | no | no | no | no | no | no | no | no | no |
| PKD2 | 610 | A | 1 | 0 | yes | no | no | yes | no | no | no | no | yes | no | no | no | no |
| PKD2 | 611 | Y | 0 | 0 | yes | no | no | yes | no | no | no | no | no | no | no | no | no |
| PKD2 | 612 | A | 0 | 0 | yes | no | no | yes | no | no | no | no | no | no | no | no | no |
| PKD2 | 613 | Q | 0 | 0 | yes | no | no | yes | no | no | no | no | yes | no | no | no | no |
| PKD2 | 614 | L | 0 | 0 | yes | no | no | yes | no | no | no | no | yes | no | no | no | no |
| PKD2 | 615 | A | 0 | 0 | yes | no | no | yes | no | no | no | no | no | no | no | no | no |
| PKD2 | 616 | Y | 0 | 0 | yes | no | no | yes | no | no | no | no | yes | no | no | no | no |
| PKD2 | 617 | L | 0 | 1 | yes | no | no | yes | no | no | no | no | yes | no | no | no | no |

| | | | | | | | | | | | | | | | | | |
|---|---|---|---|---|---|---|---|---|---|---|---|---|---|---|---|---|---|
| PKD2 | 618 | V | 0 | 0 | yes | no | no | yes | no | no | no | no | no | no | no | no | no |
| PKD2 | 619 | F | 0 | 0 | yes | no | no | yes | no | no | no | no | no | no | no | no | no |
| PKD2 | 620 | G | 1 | 0 | no | yes | no | no | yes | no | no | no | yes | no | no | no | no |
| PKD2 | 621 | T | 0 | 0 | no | yes | no | no | yes | no | no | no | yes | no | no | no | no |
| PKD2 | 622 | Q | 0 | 0 | no | yes | no | no | yes | no | no | no | yes | no | no | no | no |
| PKD2 | 624 | D | 1 | 0 | no | yes | no | no | yes | no | no | no | yes | no | no | no | no |
| PKD2 | 625 | D | 1 | 0 | no | yes | no | no | yes | no | no | no | no | no | no | no | no |
| PKD2 | 626 | F | 0 | 0 | no | yes | no | yes | yes | no | no | no | no | no | no | no | no |
| PKD2 | 627 | S | 1 | 0 | no | yes | no | no | yes | no | no | no | no | no | no | no | no |
| PKD2 | 628 | T | 1 | 0 | no | yes | no | yes | yes | no | no | no | no | no | no | no | no |
| PKD2 | 629 | F | 0 | 0 | yes | no | no | yes | yes | no | no | no | no | no | no | no | no |
| PKD2 | 630 | Q | 0 | 0 | yes | no | no | yes | yes | no | no | no | no | no | no | no | no |
| PKD2 | 631 | E | 0 | 0 | yes | no | no | yes | yes | no | no | no | no | no | no | no | no |
| PKD2 | 632 | C | 0 | 1 | yes | no | no | yes | no | no | no | yes | no | no | no | no | no |
| PKD2 | 633 | I | 0 | 0 | yes | no | no | yes | no | no | no | yes | no | no | no | no | no |
| PKD2 | 634 | F | 1 | 0 | yes | no | no | yes | no | no | no | yes | yes | no | no | no | no |
| PKD2 | 635 | T | 1 | 0 | yes | no | no | yes | no | no | no | yes | no | no | no | no | no |
| PKD2 | 636 | Q | 0 | 1 | yes | no | no | yes | no | no | no | yes | no | no | no | no | no |
| PKD2 | 637 | F | 0 | 0 | yes | no | no | yes | no | no | no | yes | no | no | no | no | no |
| PKD2 | 638 | R | 1 | 1 | yes | no | no | yes | no | no | no | yes | no | no | no | no | no |
| PKD2 | 639 | I | 0 | 0 | yes | no | no | yes | no | no | no | yes | no | no | no | no | no |
| PKD2 | 640 | I | 0 | 0 | yes | no | no | yes | no | no | no | yes | no | no | no | no | no |
| PKD2 | 641 | L | 0 | 0 | yes | no | no | yes | no | no | yes | yes | yes | no | no | no | no |
| PKD2 | 642 | G | 0 | 0 | no | yes | no | yes | no | no | yes | yes | no | no | no | no | no |
| PKD2 | 643 | D | 2 | 0 | no | yes | no | yes | no | no | yes | yes | yes | no | no | no | no |
| PKD2 | 644 | I | 1 | 0 | no | yes | no | yes | no | no | yes | yes | yes | no | no | no | no |
| PKD2 | 645 | N | 0 | 0 | no | yes | no | no | no | no | no | yes | no | no | no | no | no |
| PKD2 | 646 | F | 0 | 0 | yes | no | no | yes | no | no | no | yes | yes | no | no | no | no |
| PKD2 | 647 | A | 0 | 0 | yes | no | no | no | yes | no | no | no | no | no | no | no | no |
| PKD2 | 648 | E | 0 | 0 | yes | no | no | no | yes | no | no | no | no | no | no | no | no |
| PKD2 | 649 | I | 1 | 0 | yes | no | no | yes | yes | no | no | no | no | no | no | no | no |
| PKD2 | 650 | E | 0 | 0 | yes | no | no | no | yes | no | no | no | no | no | no | no | no |
| PKD2 | 651 | E | 1 | 0 | yes | no | no | no | yes | no | no | no | yes | no | no | no | no |

| | | | | | | | | | | | | | | | | | |
|---|---|---|---|---|---|---|---|---|---|---|---|---|---|---|---|---|---|
| PKD2 | 652 | A | 0 | 0 | yes | no | no | no | yes | no | no | no | no | no | no | no | no |
| PKD2 | 653 | N | 2 | 0 | no | yes | no | yes | yes | no | no | no | no | no | no | no | no |
| PKD2 | 654 | R | 0 | 0 | no | yes | no | no | yes | no | no | no | yes | no | no | no | no |
| PKD2 | 655 | V | 0 | 0 | no | yes | no | yes | no | no | no | no | no | no | no | no | no |
| PKD2 | 656 | L | 1 | 1 | yes | no | no | yes | no | no | no | no | yes | no | no | no | no |
| PKD2 | 657 | G | 0 | 0 | yes | no | no | yes | no | no | no | no | no | no | no | no | no |
| PKD2 | 658 | P | 0 | 0 | yes | no | no | yes | no | no | no | no | yes | no | no | no | no |
| PKD2 | 659 | I | 0 | 0 | yes | no | no | yes | no | no | no | no | no | no | no | no | no |
| PKD2 | 660 | Y | 0 | 0 | yes | no | no | yes | no | no | no | no | no | no | no | no | no |
| PKD2 | 661 | F | 0 | 0 | yes | no | no | yes | no | no | no | no | no | no | no | no | no |
| PKD2 | 662 | T | 0 | 1 | yes | no | no | yes | no | no | no | no | yes | no | no | no | no |
| PKD2 | 663 | T | 1 | 0 | yes | no | no | yes | no | no | no | no | no | no | no | no | no |
| PKD2 | 664 | F | 0 | 0 | yes | no | no | yes | no | no | no | no | no | no | no | no | no |
| PKD2 | 665 | V | 2 | 0 | yes | no | no | yes | no | no | no | no | yes | no | no | no | no |
| PKD2 | 666 | F | 0 | 0 | yes | no | no | yes | no | no | no | no | no | no | no | no | no |
| PKD2 | 667 | F | 0 | 0 | yes | no | no | yes | no | no | no | no | no | no | no | no | no |
| PKD2 | 668 | M | 1 | 0 | yes | no | no | yes | no | no | no | no | no | no | no | no | no |
| PKD2 | 669 | F | 1 | 0 | yes | no | no | yes | no | no | yes | no | no | no | no | no | no |
| PKD2 | 670 | F | 0 | 0 | yes | no | no | yes | no | no | no | no | yes | no | no | no | no |
| PKD2 | 671 | I | 0 | 0 | yes | no | no | yes | no | no | no | no | yes | no | no | no | no |
| PKD2 | 672 | L | 0 | 0 | yes | no | no | yes | no | no | no | no | no | no | no | no | no |
| PKD2 | 673 | L | 0 | 0 | yes | no | no | yes | no | no | yes | no | yes | no | no | no | no |
| PKD2 | 674 | N | 0 | 0 | yes | no | no | yes | no | no | no | no | yes | no | no | no | no |
| PKD2 | 675 | M | 2 | 0 | yes | no | no | yes | no | no | no | no | yes | no | no | no | no |
| PKD2 | 676 | F | 0 | 0 | yes | no | no | yes | no | yes | no | no | yes | no | no | no | no |
| PKD2 | 677 | L | 1 | 0 | yes | no | no | yes | no | yes | yes | no | yes | no | no | no | no |
| PKD2 | 678 | A | 1 | 0 | yes | no | no | no | no | yes | no | no | yes | no | no | no | no |
| PKD2 | 679 | I | 1 | 0 | yes | no | no | no | no | yes | no | no | yes | no | no | no | no |
| PKD2 | 680 | I | 1 | 0 | yes | no | no | no | no | yes | no | no | yes | no | no | no | no |
| PKD2 | 681 | N | 1 | 0 | yes | no | no | no | no | yes | yes | no | yes | no | no | no | no |
| PKD2 | 682 | D | 0 | 0 | yes | no | no | no | no | yes | no | no | yes | no | no | no | no |
| PKD2 | 683 | T | 1 | 0 | yes | no | no | no | no | yes | no | no | no | no | no | no | no |
| PKD2 | 684 | Y | 0 | 0 | yes | no | no | no | no | yes | no | no | yes | no | no | no | no |

| | | | | | | | | | | | | | | | | | |
|---|---|---|---|---|---|---|---|---|---|---|---|---|---|---|---|---|---|
| PKD2 | 685 | S | 0 | 0 | yes | no | no | no | no | yes | yes | no | no | no | no | no | no |
| PKD2 | 686 | E | 0 | 0 | yes | no | no | no | no | yes | no | no | no | no | no | no | no |
| PKD2 | 687 | V | 0 | 0 | yes | no | no | no | no | yes | no | no | no | no | no | no | no |
| PKD2 | 688 | K | 1 | 0 | yes | no | no | no | no | yes | no | no | no | no | no | no | no |
| PKD2 | 689 | S | 0 | 0 | yes | no | no | no | no | yes | no | no | no | no | no | no | no |
| PKD2 | 690 | D | 0 | 0 | yes | no | no | no | no | yes | no | no | no | no | no | no | no |
| PKD2 | 691 | L | 0 | 0 | yes | no | no | no | no | yes | no | no | no | no | no | no | no |
| PKD2 | 692 | A | 0 | 0 | yes | no | no | no | no | yes | no | no | no | no | no | no | no |
| PKD2 | 693 | Q | 0 | 0 | no | yes | no | no | no | yes | no | no | no | no | no | no | no |
| PKD2 | 694 | Q | 0 | 0 | no | yes | no | no | no | yes | no | no | no | no | no | no | no |
| PKD2 | 695 | K | 0 | 0 | no | yes | no | no | no | yes | no | no | no | no | no | no | no |
| SCN2A | 117 | P | 2 | 0 | no | yes | no | no | no | yes | no | no | no | no | no | no | no |
| SCN2A | 118 | I | 0 | 0 | yes | no | no | no | no | yes | no | no | no | no | no | no | no |
| SCN2A | 119 | R | 0 | 0 | yes | no | no | no | no | yes | no | no | no | no | no | no | no |
| SCN2A | 120 | K | 0 | 0 | yes | no | no | no | no | yes | no | no | no | no | no | no | no |
| SCN2A | 121 | L | 1 | 0 | yes | no | no | yes | no | yes | no | no | no | no | no | no | no |
| SCN2A | 122 | A | 1 | 0 | no | yes | no | yes | no | yes | no | no | no | no | no | no | no |
| SCN2A | 123 | I | 1 | 0 | no | yes | no | no | no | yes | no | no | no | no | no | no | no |
| SCN2A | 124 | K | 0 | 0 | yes | no | no | no | no | yes | no | no | no | no | no | no | no |
| SCN2A | 125 | I | 0 | 0 | yes | no | no | yes | no | yes | no | no | no | no | no | no | no |
| SCN2A | 126 | L | 0 | 0 | yes | no | no | yes | no | yes | no | no | no | no | no | no | no |
| SCN2A | 127 | V | 0 | 0 | yes | no | no | no | no | yes | no | no | no | no | no | no | no |
| SCN2A | 128 | H | 0 | 0 | no | yes | no | yes | no | yes | no | no | no | no | no | no | no |
| SCN2A | 129 | S | 0 | 0 | yes | no | no | yes | no | yes | no | no | no | no | no | no | no |
| SCN2A | 130 | L | 0 | 0 | yes | no | no | yes | no | no | no | no | no | no | no | no | no |
| SCN2A | 131 | F | 0 | 0 | yes | no | no | yes | no | no | no | no | no | no | no | no | no |
| SCN2A | 132 | N | 1 | 1 | yes | no | no | yes | no | no | no | no | no | no | no | no | no |
| SCN2A | 133 | M | 0 | 0 | yes | no | no | yes | no | no | no | no | no | no | no | no | no |
| SCN2A | 134 | L | 0 | 0 | yes | no | no | yes | no | no | no | no | no | no | no | no | no |
| SCN2A | 135 | I | 0 | 0 | yes | no | no | yes | no | no | no | no | no | no | no | no | no |
| SCN2A | 136 | M | 0 | 1 | yes | no | no | yes | no | no | no | no | no | no | no | no | no |
| SCN2A | 137 | C | 0 | 0 | yes | no | no | yes | no | no | no | no | no | no | no | no | no |
| SCN2A | 138 | T | 1 | 0 | yes | no | no | yes | no | no | no | no | no | no | no | no | no |

| | | | | | | | | | | | | | | | | | |
|---|---|---|---|---|---|---|---|---|---|---|---|---|---|---|---|---|---|
| SCN2A | 139 | I | 0 | 0 | yes | no | no | yes | no | no | no | no | no | no | no | no | no |
| SCN2A | 140 | L | 0 | 0 | yes | no | no | yes | no | no | no | no | no | no | no | no | no |
| SCN2A | 141 | T | 0 | 0 | yes | no | no | yes | no | no | no | no | no | no | no | no | no |
| SCN2A | 142 | N | 0 | 0 | yes | no | no | yes | no | no | no | no | no | no | no | no | no |
| SCN2A | 143 | C | 0 | 0 | yes | no | no | yes | no | no | no | no | no | no | no | no | no |
| SCN2A | 144 | V | 0 | 0 | yes | no | no | yes | no | no | no | no | no | no | no | no | no |
| SCN2A | 145 | F | 1 | 0 | yes | no | no | yes | no | no | no | no | no | no | no | no | no |
| SCN2A | 146 | M | 0 | 0 | no | yes | no | yes | no | no | no | no | no | no | no | no | no |
| SCN2A | 147 | T | 0 | 0 | no | yes | no | yes | no | no | no | no | no | no | no | no | no |
| SCN2A | 148 | M | 2 | 0 | no | yes | no | no | no | no | no | no | no | no | no | no | no |
| SCN2A | 149 | S | 0 | 0 | no | yes | no | no | yes | no | no | no | no | no | no | no | no |
| SCN2A | 150 | N | 0 | 0 | no | yes | no | no | yes | no | no | no | no | no | no | no | no |
| SCN2A | 151 | P | 1 | 0 | no | yes | no | no | yes | no | no | no | no | no | no | no | no |
| SCN2A | 152 | P | 0 | 0 | no | yes | no | no | yes | no | no | no | no | no | no | no | no |
| SCN2A | 153 | D | 0 | 0 | no | yes | no | no | yes | no | no | no | no | no | no | no | no |
| SCN2A | 154 | W | 0 | 0 | no | yes | no | no | yes | no | no | no | no | no | no | no | no |
| SCN2A | 155 | T | 0 | 0 | no | yes | no | no | yes | no | no | no | no | no | no | no | no |
| SCN2A | 156 | K | 0 | 0 | yes | no | no | no | no | no | no | no | no | no | no | no | no |
| SCN2A | 157 | N | 0 | 0 | yes | no | no | yes | no | no | no | no | no | no | no | no | no |
| SCN2A | 158 | V | 0 | 0 | yes | no | no | yes | no | no | no | no | no | no | no | no | no |
| SCN2A | 159 | E | 0 | 0 | yes | no | no | yes | no | no | no | no | no | no | no | no | no |
| SCN2A | 160 | Y | 0 | 0 | yes | no | no | yes | no | no | no | no | no | no | no | no | no |
| SCN2A | 161 | T | 0 | 0 | yes | no | no | yes | no | no | no | no | no | no | no | no | no |
| SCN2A | 162 | F | 0 | 0 | yes | no | no | yes | no | no | no | no | no | no | no | no | no |
| SCN2A | 163 | T | 0 | 0 | yes | no | no | yes | no | no | no | no | no | no | no | no | no |
| SCN2A | 164 | G | 0 | 0 | yes | no | no | yes | no | no | no | no | no | no | no | no | no |
| SCN2A | 165 | I | 0 | 0 | yes | no | no | yes | no | no | no | no | no | no | no | no | no |
| SCN2A | 166 | Y | 0 | 0 | yes | no | no | yes | no | no | no | no | no | no | no | no | no |
| SCN2A | 167 | T | 0 | 0 | yes | no | no | yes | no | no | no | no | no | no | no | no | no |
| SCN2A | 168 | F | 0 | 0 | yes | no | no | yes | no | no | no | no | no | no | no | no | no |
| SCN2A | 169 | E | 0 | 1 | yes | no | no | yes | no | no | no | no | no | no | no | no | no |
| SCN2A | 170 | S | 0 | 0 | yes | no | no | yes | no | no | no | no | no | no | no | no | no |
| SCN2A | 171 | L | 2 | 0 | yes | no | no | yes | no | no | no | no | no | no | no | no | no |

| | | | | | | | | | | | | | | | | | |
|---|---|---|---|---|---|---|---|---|---|---|---|---|---|---|---|---|---|
| SCN2A | 172 | I | 1 | 0 | yes | no | no | yes | no | no | no | no | no | no | no | no | no |
| SCN2A | 173 | K | 0 | 0 | yes | no | no | yes | no | no | no | no | no | no | no | no | no |
| SCN2A | 174 | I | 0 | 0 | yes | no | no | yes | no | no | no | no | no | no | no | no | no |
| SCN2A | 175 | L | 0 | 0 | no | yes | no | yes | no | no | no | no | no | no | no | no | no |
| SCN2A | 176 | A | 1 | 0 | no | yes | no | yes | no | no | no | no | no | no | no | no | no |
| SCN2A | 177 | R | 1 | 0 | no | yes | no | no | no | yes | no | no | no | no | no | no | no |
| SCN2A | 178 | G | 0 | 0 | no | yes | no | no | no | yes | no | no | no | no | no | no | no |
| SCN2A | 179 | F | 0 | 0 | no | yes | no | no | no | yes | no | no | no | no | no | no | no |
| SCN2A | 180 | C | 0 | 0 | no | yes | no | no | no | yes | no | no | no | no | no | no | no |
| SCN2A | 181 | L | 2 | 0 | no | yes | no | yes | no | yes | no | no | no | no | no | no | no |
| SCN2A | 182 | E | 0 | 0 | no | yes | no | no | no | yes | no | no | no | no | no | no | no |
| SCN2A | 183 | D | 0 | 0 | no | yes | no | yes | no | yes | no | no | no | no | no | no | no |
| SCN2A | 184 | F | 0 | 0 | no | yes | no | yes | no | yes | no | no | no | no | no | no | no |
| SCN2A | 185 | T | 1 | 1 | no | yes | no | yes | no | yes | no | no | no | no | no | no | no |
| SCN2A | 186 | F | 1 | 0 | no | yes | no | yes | no | yes | no | no | no | no | no | no | no |
| SCN2A | 187 | L | 0 | 0 | no | yes | no | yes | no | yes | no | no | no | no | no | no | no |
| SCN2A | 188 | R | 2 | 1 | no | yes | no | no | no | yes | no | no | no | no | no | no | no |
| SCN2A | 189 | D | 0 | 0 | no | yes | no | yes | no | yes | no | no | no | no | no | no | no |
| SCN2A | 190 | P | 0 | 0 | yes | no | no | yes | no | yes | no | no | no | no | no | no | no |
| SCN2A | 191 | W | 0 | 2 | yes | no | no | yes | no | no | no | no | no | no | no | no | no |
| SCN2A | 192 | N | 0 | 0 | yes | no | no | yes | no | no | no | no | no | no | no | no | no |
| SCN2A | 193 | W | 1 | 0 | yes | no | no | yes | no | no | no | no | no | no | no | no | no |
| SCN2A | 194 | L | 0 | 0 | yes | no | no | yes | no | no | no | no | no | no | no | no | no |
| SCN2A | 195 | D | 0 | 0 | yes | no | no | yes | no | no | no | no | no | no | no | no | no |
| SCN2A | 196 | F | 0 | 0 | yes | no | no | yes | no | no | no | no | no | no | no | no | no |
| SCN2A | 197 | T | 0 | 0 | yes | no | no | yes | no | no | no | no | no | no | no | no | no |
| SCN2A | 198 | V | 0 | 0 | yes | no | no | yes | no | no | no | no | no | no | no | no | no |
| SCN2A | 199 | I | 0 | 0 | yes | no | no | yes | no | no | no | no | no | no | no | no | no |
| SCN2A | 200 | T | 1 | 0 | yes | no | no | yes | no | no | no | no | no | no | no | no | no |
| SCN2A | 201 | F | 0 | 0 | yes | no | no | yes | no | no | no | no | no | no | no | no | no |
| SCN2A | 202 | A | 0 | 0 | yes | no | no | yes | no | no | no | no | no | no | no | no | no |
| SCN2A | 203 | Y | 0 | 0 | no | yes | no | yes | no | no | no | no | no | no | no | no | no |
| SCN2A | 204 | V | 0 | 0 | yes | no | no | yes | no | no | no | no | no | no | no | no | no |

| | | | | | | | | | | | | | | | | | |
|---|---|---|---|---|---|---|---|---|---|---|---|---|---|---|---|---|---|
| SCN2A | 205 | T | 0 | 0 | yes | no | no | yes | no | no | no | no | no | no | no | no | no |
| SCN2A | 206 | E | 0 | 0 | yes | no | no | yes | no | no | no | no | no | no | no | no | no |
| SCN2A | 207 | F | 0 | 2 | no | yes | no | yes | no | no | no | no | no | no | no | no | no |
| SCN2A | 208 | V | 0 | 1 | no | yes | no | no | no | no | no | no | no | no | no | no | no |
| SCN2A | 209 | D | 0 | 0 | no | yes | no | no | yes | no | no | no | no | no | no | no | no |
| SCN2A | 210 | L | 1 | 0 | no | yes | no | yes | yes | no | no | no | no | no | no | no | no |
| SCN2A | 211 | G | 0 | 1 | no | yes | no | no | yes | no | no | no | no | no | no | no | no |
| SCN2A | 212 | N | 0 | 1 | no | yes | no | no | yes | no | no | no | no | no | no | no | no |
| SCN2A | 213 | V | 0 | 2 | no | yes | no | no | yes | no | no | no | no | no | no | no | no |
| SCN2A | 214 | S | 1 | 1 | no | yes | no | no | yes | no | no | no | no | no | no | no | no |
| SCN2A | 215 | A | 1 | 1 | no | yes | no | no | no | no | no | no | no | no | no | no | no |
| SCN2A | 216 | L | 0 | 1 | no | yes | no | yes | no | no | no | no | no | no | no | no | no |
| SCN2A | 217 | R | 0 | 0 | no | yes | no | yes | no | no | no | no | no | no | no | no | no |
| SCN2A | 218 | T | 0 | 1 | yes | no | no | yes | no | no | no | no | no | no | no | no | no |
| SCN2A | 219 | F | 0 | 0 | yes | no | no | yes | no | no | no | no | no | no | no | no | no |
| SCN2A | 220 | R | 0 | 1 | yes | no | no | yes | no | no | no | no | no | no | no | no | no |
| SCN2A | 221 | V | 0 | 0 | yes | no | no | yes | no | no | no | no | no | no | no | no | no |
| SCN2A | 222 | L | 0 | 0 | yes | no | no | yes | no | no | no | no | no | no | no | no | no |
| SCN2A | 223 | R | 0 | 1 | yes | no | no | yes | no | no | no | no | no | no | no | no | no |
| SCN2A | 224 | A | 0 | 0 | yes | no | no | yes | no | no | no | no | no | no | no | no | no |
| SCN2A | 225 | L | 0 | 0 | no | yes | no | yes | no | no | no | no | no | no | no | no | no |
| SCN2A | 226 | K | 0 | 0 | no | yes | no | yes | no | no | no | no | no | no | no | no | no |
| SCN2A | 227 | T | 0 | 1 | yes | no | no | yes | no | no | no | no | no | no | no | no | no |
| SCN2A | 228 | I | 0 | 0 | yes | no | no | yes | no | no | no | no | no | no | no | no | no |
| SCN2A | 229 | S | 0 | 1 | yes | no | no | yes | no | no | no | no | no | no | no | no | no |
| SCN2A | 230 | V | 0 | 0 | yes | no | no | yes | no | no | no | no | no | no | no | no | no |
| SCN2A | 231 | I | 1 | 0 | no | yes | no | yes | no | no | no | no | no | no | no | no | no |
| SCN2A | 232 | P | 0 | 0 | no | yes | no | yes | no | yes | no | no | no | no | no | no | no |
| SCN2A | 233 | G | 0 | 0 | no | yes | no | yes | no | yes | no | no | no | no | no | no | no |
| SCN2A | 234 | L | 0 | 0 | yes | no | no | yes | no | yes | no | no | no | no | no | no | no |
| SCN2A | 235 | K | 0 | 0 | yes | no | no | yes | no | yes | no | no | no | no | no | no | no |
| SCN2A | 236 | T | 1 | 1 | yes | no | no | yes | no | yes | no | no | no | no | no | no | no |
| SCN2A | 237 | I | 0 | 1 | yes | no | no | yes | no | yes | no | no | no | no | no | no | no |

| | | | | | | | | | | | | | | | | | |
|---|---|---|---|---|---|---|---|---|---|---|---|---|---|---|---|---|---|
| SCN2A | 238 | V | 0 | 0 | yes | no | no | yes | no | yes | no | no | no | no | no | no | no |
| SCN2A | 239 | G | 0 | 0 | yes | no | no | yes | no | yes | no | no | no | no | no | no | no |
| SCN2A | 240 | A | 0 | 2 | yes | no | no | yes | no | yes | no | no | no | no | no | no | no |
| SCN2A | 241 | L | 0 | 0 | yes | no | no | yes | no | yes | no | no | no | no | no | no | no |
| SCN2A | 242 | I | 0 | 0 | yes | no | no | yes | no | yes | no | no | no | no | no | no | no |
| SCN2A | 243 | Q | 0 | 0 | yes | no | no | yes | no | yes | no | no | no | no | no | no | no |
| SCN2A | 244 | S | 0 | 0 | yes | no | no | yes | no | yes | no | no | no | no | no | no | no |
| SCN2A | 245 | V | 0 | 0 | yes | no | no | yes | no | yes | no | no | no | no | no | no | no |
| SCN2A | 246 | K | 0 | 0 | yes | no | no | yes | no | yes | no | no | no | no | no | no | no |
| SCN2A | 247 | K | 0 | 0 | yes | no | no | yes | no | yes | no | no | no | no | no | no | no |
| SCN2A | 248 | L | 0 | 1 | yes | no | no | yes | no | yes | no | no | no | no | no | no | no |
| SCN2A | 249 | S | 1 | 0 | yes | no | no | yes | no | yes | no | no | no | no | no | no | no |
| SCN2A | 250 | D | 0 | 0 | no | yes | no | yes | no | yes | no | no | no | no | no | no | no |
| SCN2A | 251 | V | 0 | 2 | yes | no | no | yes | no | no | no | no | no | no | no | no | no |
| SCN2A | 252 | M | 0 | 1 | yes | no | no | yes | no | no | no | no | no | no | no | no | no |
| SCN2A | 253 | I | 0 | 0 | yes | no | no | yes | no | no | no | no | no | no | no | no | no |
| SCN2A | 254 | L | 0 | 0 | yes | no | no | yes | no | no | no | no | no | no | no | no | no |
| SCN2A | 255 | T | 0 | 0 | yes | no | no | yes | no | no | no | no | no | no | no | no | no |
| SCN2A | 256 | V | 0 | 0 | yes | no | no | yes | no | no | no | no | no | no | no | no | no |
| SCN2A | 257 | F | 0 | 0 | yes | no | no | yes | no | no | no | no | no | no | no | no | no |
| SCN2A | 258 | C | 0 | 0 | yes | no | no | yes | no | no | no | no | no | no | no | no | no |
| SCN2A | 259 | L | 0 | 1 | yes | no | no | yes | no | no | no | no | no | no | no | no | no |
| SCN2A | 260 | S | 0 | 0 | yes | no | no | yes | no | no | no | no | no | no | no | no | no |
| SCN2A | 261 | V | 0 | 1 | yes | no | no | yes | no | no | no | no | no | no | no | no | no |
| SCN2A | 262 | F | 0 | 1 | yes | no | no | yes | no | no | no | no | no | no | no | no | no |
| SCN2A | 263 | A | 0 | 2 | yes | no | no | yes | no | no | no | no | no | no | no | no | no |
| SCN2A | 264 | L | 0 | 1 | yes | no | no | yes | no | no | no | no | no | no | no | no | no |
| SCN2A | 265 | I | 0 | 0 | yes | no | no | yes | no | no | no | no | no | no | no | no | no |
| SCN2A | 266 | G | 0 | 0 | yes | no | no | yes | no | no | no | no | no | no | no | no | no |
| SCN2A | 267 | L | 0 | 0 | yes | no | no | yes | no | no | no | no | no | no | no | no | no |
| SCN2A | 268 | Q | 0 | 0 | yes | no | no | yes | no | no | no | no | no | no | no | no | no |
| SCN2A | 269 | L | 0 | 0 | yes | no | no | yes | no | no | no | no | no | no | no | no | no |
| SCN2A | 270 | F | 0 | 0 | no | yes | no | yes | no | no | no | no | no | no | no | no | no |

| | | | | | | | | | | | | | | | | | |
|---|---|---|---|---|---|---|---|---|---|---|---|---|---|---|---|---|---|
| SCN2A | 271 | M | 0 | 0 | no | yes | no | no | yes | no | no | no | no | no | no | no | no |
| SCN2A | 272 | G | 0 | 0 | no | yes | no | no | yes | no | no | no | no | no | no | no | no |
| SCN2A | 273 | N | 0 | 0 | yes | no | no | no | yes | no | no | no | no | no | no | no | no |
| SCN2A | 274 | L | 0 | 0 | yes | no | no | no | yes | no | no | no | no | no | no | no | no |
| SCN2A | 275 | R | 0 | 0 | yes | no | no | no | yes | no | no | no | no | no | no | no | no |
| SCN2A | 276 | N | 0 | 0 | no | no | yes | no | yes | no | no | no | no | no | no | no | no |
| SCN2A | 277 | K | 0 | 0 | no | no | yes | no | yes | no | no | no | no | no | no | no | no |
| SCN2A | 278 | C | 0 | 0 | no | no | yes | no | yes | no | no | no | no | no | no | no | no |
| SCN2A | 279 | L | 0 | 0 | no | no | yes | no | yes | no | no | no | no | no | no | no | no |
| SCN2A | 280 | Q | 1 | 0 | no | no | yes | no | yes | no | no | no | no | no | no | no | no |
| SCN2A | 281 | W | 0 | 0 | no | yes | no | no | yes | no | no | no | no | no | no | no | no |
| SCN2A | 282 | P | 0 | 0 | no | yes | no | no | yes | no | no | no | no | no | no | no | no |
| SCN2A | 283 | P | 1 | 0 | no | yes | no | no | yes | no | no | no | no | no | no | no | no |
| SCN2A | 284 | D | 1 | 1 | no | yes | no | no | yes | no | no | no | no | no | no | no | no |
| SCN2A | 314 | F | 0 | 0 | no | yes | no | no | yes | no | no | no | no | no | no | no | no |
| SCN2A | 315 | N | 0 | 0 | no | yes | no | no | yes | no | no | no | no | no | no | no | no |
| SCN2A | 316 | W | 0 | 0 | no | yes | no | no | yes | no | no | no | no | no | no | no | no |
| SCN2A | 317 | D | 0 | 0 | no | yes | no | no | yes | no | no | no | no | no | no | no | no |
| SCN2A | 318 | E | 1 | 0 | no | yes | no | no | yes | no | no | no | no | no | no | no | no |
| SCN2A | 319 | Y | 0 | 0 | no | yes | no | no | yes | no | no | no | no | no | no | no | no |
| SCN2A | 320 | I | 0 | 0 | no | yes | no | no | yes | no | no | no | no | no | no | no | no |
| SCN2A | 321 | E | 1 | 0 | no | yes | no | no | yes | no | no | no | no | no | no | no | no |
| SCN2A | 322 | D | 0 | 1 | no | yes | no | no | yes | no | no | no | no | no | no | no | no |
| SCN2A | 323 | K | 0 | 0 | no | yes | no | no | yes | no | no | no | no | no | no | no | no |
| SCN2A | 324 | S | 0 | 0 | no | yes | no | no | yes | no | no | no | no | no | no | no | no |
| SCN2A | 325 | H | 0 | 0 | no | yes | no | no | yes | no | no | no | no | no | no | no | no |
| SCN2A | 326 | F | 0 | 0 | no | yes | no | no | yes | no | no | no | no | no | no | no | no |
| SCN2A | 327 | Y | 0 | 0 | no | yes | no | no | yes | no | no | no | no | no | no | no | no |
| SCN2A | 328 | F | 1 | 1 | no | yes | no | no | yes | no | no | no | no | no | no | no | no |
| SCN2A | 329 | L | 1 | 0 | no | yes | no | no | yes | no | no | no | no | no | no | no | no |
| SCN2A | 330 | E | 0 | 0 | no | yes | no | no | yes | no | no | no | no | no | no | no | no |
| SCN2A | 331 | G | 0 | 0 | no | yes | no | no | yes | no | no | no | no | no | no | no | no |
| SCN2A | 332 | Q | 0 | 0 | no | yes | no | no | yes | no | no | no | no | no | no | no | no |

| | | | | | | | | | | | | | | | | | |
|---|---|---|---|---|---|---|---|---|---|---|---|---|---|---|---|---|---|
| SCN2A | 333 | N | 1 | 0 | no | yes | no | no | yes | no | no | no | no | no | no | no | no |
| SCN2A | 334 | D | 1 | 0 | no | yes | no | no | yes | no | no | no | no | no | no | no | no |
| SCN2A | 335 | A | 0 | 0 | no | yes | no | no | yes | no | no | no | no | no | no | no | no |
| SCN2A | 336 | L | 0 | 0 | no | yes | no | no | yes | no | no | no | no | no | no | no | no |
| SCN2A | 337 | L | 0 | 0 | no | yes | no | no | yes | no | no | no | no | no | no | no | no |
| SCN2A | 338 | C | 0 | 0 | no | yes | no | no | yes | no | no | no | no | no | no | no | no |
| SCN2A | 339 | G | 0 | 0 | no | yes | no | no | yes | no | no | no | no | no | no | no | no |
| SCN2A | 340 | N | 0 | 0 | no | yes | no | no | yes | no | no | no | no | no | no | no | no |
| SCN2A | 341 | S | 1 | 0 | no | yes | no | no | yes | no | no | no | no | no | no | no | no |
| SCN2A | 342 | S | 0 | 0 | no | yes | no | no | yes | no | no | no | no | no | no | no | no |
| SCN2A | 343 | D | 0 | 2 | no | yes | no | no | yes | no | no | no | no | no | no | no | no |
| SCN2A | 344 | A | 0 | 1 | no | yes | no | no | yes | no | no | no | no | no | no | no | no |
| SCN2A | 345 | G | 0 | 0 | no | yes | no | no | yes | no | no | no | no | no | no | no | no |
| SCN2A | 346 | Q | 0 | 0 | no | yes | no | no | yes | no | no | no | no | no | no | no | no |
| SCN2A | 347 | C | 0 | 0 | no | yes | no | no | yes | no | no | no | no | no | no | no | no |
| SCN2A | 348 | P | 0 | 0 | no | yes | no | no | yes | no | no | no | no | no | no | no | no |
| SCN2A | 349 | E | 0 | 0 | no | yes | no | no | yes | no | no | no | no | no | no | no | no |
| SCN2A | 350 | G | 0 | 0 | no | yes | no | no | yes | no | no | no | no | no | no | no | no |
| SCN2A | 351 | Y | 0 | 0 | no | no | yes | no | yes | no | no | no | no | no | no | no | no |
| SCN2A | 352 | I | 0 | 0 | no | no | yes | no | yes | no | no | no | no | no | no | no | no |
| SCN2A | 353 | C | 0 | 1 | no | no | yes | no | yes | no | no | no | no | no | no | no | no |
| SCN2A | 354 | V | 1 | 0 | no | no | yes | no | yes | no | no | no | no | no | no | no | no |
| SCN2A | 355 | K | 0 | 0 | no | no | yes | no | yes | no | no | no | no | no | no | no | no |
| SCN2A | 356 | A | 0 | 0 | no | yes | no | no | yes | no | no | no | no | no | no | no | no |
| SCN2A | 357 | G | 0 | 0 | no | yes | no | no | yes | no | no | no | no | no | no | no | no |
| SCN2A | 358 | R | 0 | 0 | no | yes | no | no | yes | no | no | no | no | no | no | no | no |
| SCN2A | 359 | N | 1 | 0 | no | yes | no | no | yes | no | no | no | no | no | no | no | no |
| SCN2A | 360 | P | 0 | 0 | no | yes | no | no | yes | no | no | no | no | no | no | no | no |
| SCN2A | 361 | N | 0 | 0 | no | yes | no | no | yes | no | no | no | no | no | no | no | no |
| SCN2A | 362 | Y | 1 | 0 | no | yes | no | no | yes | no | no | no | no | no | no | no | no |
| SCN2A | 363 | G | 0 | 0 | no | yes | no | no | yes | no | no | no | no | no | no | no | no |
| SCN2A | 364 | Y | 0 | 0 | no | yes | no | no | yes | no | no | no | no | no | no | no | no |
| SCN2A | 365 | T | 0 | 0 | no | yes | no | no | yes | no | no | no | no | no | no | no | no |

| | | | | | | | | | | | | | | | | | |
|---|---|---|---|---|---|---|---|---|---|---|---|---|---|---|---|---|---|
| SCN2A | 366 | S | 0 | 0 | no | yes | no | no | yes | no | no | no | no | no | no | no | no |
| SCN2A | 367 | F | 0 | 0 | no | yes | no | yes | yes | no | no | no | no | no | no | no | no |
| SCN2A | 368 | D | 0 | 0 | no | yes | no | no | yes | no | no | no | no | no | no | no | no |
| SCN2A | 369 | T | 0 | 0 | no | yes | no | no | yes | no | no | no | no | no | no | no | no |
| SCN2A | 370 | F | 0 | 0 | yes | no | no | yes | no | no | no | yes | no | no | no | no | no |
| SCN2A | 371 | S | 0 | 0 | yes | no | no | yes | no | no | no | yes | no | no | no | no | no |
| SCN2A | 372 | W | 0 | 0 | yes | no | no | no | no | no | no | yes | no | no | no | no | no |
| SCN2A | 373 | A | 0 | 0 | yes | no | no | yes | no | no | no | yes | no | no | no | no | no |
| SCN2A | 374 | F | 0 | 0 | yes | no | no | yes | no | no | no | yes | no | no | no | no | no |
| SCN2A | 375 | L | 0 | 0 | yes | no | no | yes | no | no | no | yes | no | no | no | no | no |
| SCN2A | 376 | S | 0 | 0 | yes | no | no | yes | no | no | no | yes | no | no | no | no | no |
| SCN2A | 377 | L | 0 | 0 | yes | no | no | yes | no | no | no | yes | no | no | no | no | no |
| SCN2A | 378 | F | 1 | 0 | yes | no | no | yes | no | no | no | yes | no | no | no | no | no |
| SCN2A | 379 | R | 1 | 0 | yes | no | no | yes | no | no | no | yes | no | no | no | no | no |
| SCN2A | 380 | L | 0 | 0 | yes | no | no | yes | no | no | no | yes | no | no | no | no | no |
| SCN2A | 381 | M | 0 | 0 | yes | no | no | yes | no | no | no | yes | no | no | no | no | no |
| SCN2A | 382 | T | 0 | 0 | no | yes | no | yes | no | no | yes | yes | no | no | no | no | no |
| SCN2A | 383 | Q | 0 | 1 | no | yes | no | yes | no | no | yes | yes | no | no | no | no | no |
| SCN2A | 384 | D | 0 | 0 | no | yes | no | yes | no | no | no | yes | no | no | no | no | no |
| SCN2A | 385 | F | 0 | 0 | no | yes | no | no | no | no | no | yes | no | no | no | no | no |
| SCN2A | 386 | W | 0 | 0 | yes | no | no | yes | no | no | no | yes | no | no | no | no | no |
| SCN2A | 387 | E | 0 | 0 | yes | no | no | yes | no | no | no | yes | no | no | no | no | no |
| SCN2A | 388 | N | 0 | 0 | yes | no | no | no | no | no | no | yes | no | no | no | no | no |
| SCN2A | 389 | L | 1 | 0 | yes | no | no | yes | no | no | no | yes | no | no | no | no | no |
| SCN2A | 390 | Y | 0 | 0 | yes | no | no | yes | no | no | no | yes | no | no | no | no | no |
| SCN2A | 391 | Q | 0 | 0 | yes | no | no | no | no | no | no | yes | no | no | no | no | no |
| SCN2A | 392 | L | 0 | 0 | yes | no | no | no | no | no | no | yes | no | no | no | no | no |
| SCN2A | 393 | T | 0 | 1 | yes | no | no | no | no | no | no | yes | no | no | no | no | no |
| SCN2A | 394 | L | 0 | 0 | yes | no | no | no | no | no | no | yes | no | no | no | no | no |
| SCN2A | 395 | R | 0 | 1 | yes | no | no | no | yes | no | no | no | no | no | no | no | no |
| SCN2A | 396 | A | 1 | 0 | yes | no | no | no | yes | no | no | no | no | no | no | no | no |
| SCN2A | 397 | A | 0 | 0 | no | yes | no | no | yes | no | no | no | no | no | no | no | no |
| SCN2A | 398 | G | 0 | 0 | no | yes | no | no | yes | no | no | no | no | no | no | no | no |

| | | | | | | | | | | | | | | | | | |
|---|---|---|---|---|---|---|---|---|---|---|---|---|---|---|---|---|---|
| SCN2A | 399 | K | 0 | 0 | yes | no | no | no | yes | no | no | no | no | no | no | no | no |
| SCN2A | 400 | T | 1 | 0 | yes | no | no | yes | yes | no | no | no | no | no | no | no | no |
| SCN2A | 401 | Y | 0 | 0 | yes | no | no | yes | yes | no | no | no | no | no | no | no | no |
| SCN2A | 402 | M | 0 | 0 | yes | no | no | yes | no | no | no | no | no | no | no | no | no |
| SCN2A | 403 | I | 1 | 0 | yes | no | no | yes | no | no | no | no | no | no | no | no | no |
| SCN2A | 404 | F | 0 | 0 | yes | no | no | yes | no | no | no | no | no | no | no | no | no |
| SCN2A | 405 | F | 0 | 0 | yes | no | no | yes | no | no | no | no | no | no | no | no | no |
| SCN2A | 406 | V | 0 | 0 | yes | no | no | yes | no | no | no | no | no | no | no | no | no |
| SCN2A | 407 | L | 0 | 0 | yes | no | no | yes | no | no | no | no | no | no | no | no | no |
| SCN2A | 408 | V | 0 | 0 | yes | no | no | yes | no | no | no | no | no | no | no | no | no |
| SCN2A | 409 | I | 0 | 0 | yes | no | no | yes | no | no | no | no | no | no | no | no | no |
| SCN2A | 410 | F | 0 | 0 | yes | no | no | yes | no | no | no | no | no | no | no | no | no |
| SCN2A | 411 | L | 0 | 0 | yes | no | no | yes | no | no | no | no | no | no | no | no | no |
| SCN2A | 412 | G | 0 | 0 | yes | no | no | yes | no | no | no | no | no | no | no | no | no |
| SCN2A | 413 | S | 0 | 0 | yes | no | no | yes | no | no | no | no | no | no | no | no | no |
| SCN2A | 414 | F | 0 | 1 | yes | no | no | yes | no | no | no | no | no | no | no | no | no |
| SCN2A | 415 | Y | 0 | 0 | yes | no | no | yes | no | no | no | no | no | no | no | no | no |
| SCN2A | 416 | L | 0 | 0 | yes | no | no | yes | no | no | no | no | no | no | no | no | no |
| SCN2A | 417 | I | 0 | 0 | yes | no | no | yes | no | no | yes | no | no | no | no | no | no |
| SCN2A | 418 | N | 0 | 0 | yes | no | no | yes | no | no | no | no | no | no | no | no | no |
| SCN2A | 419 | L | 0 | 1 | yes | no | no | yes | no | no | no | no | no | no | no | no | no |
| SCN2A | 420 | I | 0 | 0 | yes | no | no | yes | no | no | no | no | no | no | no | no | no |
| SCN2A | 421 | L | 0 | 1 | yes | no | no | yes | no | no | yes | no | no | no | no | no | no |
| SCN2A | 422 | A | 0 | 0 | yes | no | no | yes | no | no | no | no | no | no | no | no | no |
| SCN2A | 423 | V | 0 | 1 | yes | no | no | yes | no | yes | no | no | no | no | no | no | no |
| SCN2A | 424 | V | 0 | 2 | yes | no | no | yes | no | yes | no | no | no | no | no | no | no |
| SCN2A | 425 | A | 0 | 0 | yes | no | no | yes | no | yes | no | no | no | no | no | no | no |
| SCN2A | 426 | M | 0 | 0 | yes | no | no | no | no | yes | no | no | no | no | no | no | no |
| SCN2A | 427 | A | 0 | 0 | yes | no | no | no | no | yes | no | no | no | no | no | no | no |
| SCN2A | 428 | Y | 0 | 1 | yes | no | no | yes | no | yes | no | no | no | no | no | no | no |
| SCN2A | 429 | E | 0 | 0 | yes | no | no | no | no | yes | yes | no | no | no | no | no | no |
| SCN2A | 430 | E | 0 | 1 | yes | no | no | no | no | yes | no | no | no | no | no | no | no |
| SCN2A | 431 | Q | 0 | 0 | yes | no | no | no | no | yes | no | no | no | no | no | no | no |

| | | | | | | | | | | | | | | | | | |
|---|---|---|---|---|---|---|---|---|---|---|---|---|---|---|---|---|---|
| SCN2A | 432 | N | 0 | 0 | yes | no | no | no | no | yes | no | no | no | no | no | no | no |
| SCN2A | 433 | Q | 0 | 0 | yes | no | no | no | no | yes | no | no | no | no | no | no | no |
| SCN2A | 434 | A | 0 | 2 | yes | no | no | no | no | yes | no | no | no | no | no | no | no |
| SCN2A | 435 | T | 0 | 1 | yes | no | no | no | no | yes | no | no | no | no | no | no | no |
| SCN2A | 436 | L | 0 | 1 | yes | no | no | no | no | yes | no | no | no | no | no | no | no |
| SCN2A | 437 | E | 0 | 0 | yes | no | no | no | no | yes | no | no | no | no | no | no | no |
| SCN2A | 438 | E | 0 | 2 | yes | no | no | no | no | yes | no | no | no | no | no | no | no |
| SCN2A | 439 | A | 0 | 0 | yes | no | no | no | no | yes | no | no | no | no | no | no | no |
| SCN2A | 440 | E | 0 | 0 | yes | no | no | no | no | yes | no | no | no | no | no | no | no |
| SCN2A | 441 | Q | 0 | 0 | no | yes | no | no | no | yes | no | no | no | no | no | no | no |
| SCN2A | 740 | D | 2 | 0 | no | yes | no | no | no | yes | no | no | no | no | no | no | no |
| SCN2A | 741 | C | 0 | 0 | no | yes | no | yes | no | yes | no | no | no | no | no | no | no |
| SCN2A | 742 | C | 1 | 0 | yes | no | no | no | no | yes | no | no | no | no | no | no | no |
| SCN2A | 743 | K | 0 | 0 | yes | no | no | no | no | yes | no | no | no | no | no | no | no |
| SCN2A | 744 | P | 1 | 0 | yes | no | no | no | no | yes | no | no | no | no | no | no | no |
| SCN2A | 745 | W | 0 | 0 | yes | no | no | no | no | yes | no | no | no | no | no | no | no |
| SCN2A | 746 | L | 1 | 0 | yes | no | no | no | no | yes | no | no | no | no | no | no | no |
| SCN2A | 747 | K | 0 | 0 | yes | no | no | no | no | yes | no | no | no | no | no | no | no |
| SCN2A | 748 | V | 0 | 0 | yes | no | no | yes | no | yes | no | no | no | no | no | no | no |
| SCN2A | 749 | K | 0 | 0 | yes | no | no | no | no | yes | no | no | no | no | no | no | no |
| SCN2A | 750 | H | 0 | 0 | yes | no | no | no | no | yes | no | no | no | no | no | no | no |
| SCN2A | 751 | L | 0 | 0 | yes | no | no | yes | no | yes | no | no | no | no | no | no | no |
| SCN2A | 752 | V | 0 | 0 | yes | no | no | yes | no | yes | no | no | no | no | no | no | no |
| SCN2A | 753 | N | 1 | 0 | yes | no | no | no | no | yes | no | no | no | no | no | no | no |
| SCN2A | 754 | L | 0 | 0 | no | yes | no | no | no | yes | no | no | no | no | no | no | no |
| SCN2A | 755 | V | 0 | 0 | no | yes | no | yes | no | yes | no | no | no | no | no | no | no |
| SCN2A | 756 | V | 0 | 0 | no | yes | no | yes | no | yes | no | no | no | no | no | no | no |
| SCN2A | 757 | M | 0 | 0 | no | yes | no | no | no | yes | no | no | no | no | no | no | no |
| SCN2A | 758 | D | 0 | 0 | no | yes | no | no | no | yes | no | no | no | no | no | no | no |
| SCN2A | 759 | P | 0 | 0 | no | yes | no | no | no | yes | no | no | no | no | no | no | no |
| SCN2A | 760 | F | 1 | 0 | no | yes | no | yes | no | no | no | no | no | no | no | no | no |
| SCN2A | 761 | V | 0 | 0 | no | yes | no | yes | no | no | no | no | no | no | no | no | no |
| SCN2A | 762 | D | 0 | 0 | no | yes | no | yes | no | no | no | no | no | no | no | no | no |

| | | | | | | | | | | | | | | | | | |
|---|---|---|---|---|---|---|---|---|---|---|---|---|---|---|---|---|---|
| SCN2A | 763 | L | 0 | 0 | yes | no | no | yes | no | no | no | no | no | no | no | no | no |
| SCN2A | 764 | A | 0 | 0 | yes | no | no | yes | no | no | no | no | no | no | no | no | no |
| SCN2A | 765 | I | 0 | 0 | yes | no | no | yes | no | no | no | no | no | no | no | no | no |
| SCN2A | 766 | T | 0 | 0 | yes | no | no | yes | no | no | no | no | no | no | no | no | no |
| SCN2A | 767 | I | 0 | 0 | yes | no | no | yes | no | no | no | no | no | no | no | no | no |
| SCN2A | 768 | C | 0 | 0 | yes | no | no | yes | no | no | no | no | no | no | no | no | no |
| SCN2A | 769 | I | 0 | 1 | yes | no | no | yes | no | no | no | no | no | no | no | no | no |
| SCN2A | 770 | V | 0 | 0 | yes | no | no | yes | no | no | no | no | no | no | no | no | no |
| SCN2A | 771 | L | 0 | 0 | yes | no | no | yes | no | no | no | no | no | no | no | no | no |
| SCN2A | 772 | N | 0 | 0 | yes | no | no | yes | no | no | no | no | no | no | no | no | no |
| SCN2A | 773 | T | 0 | 2 | yes | no | no | yes | no | no | no | no | no | no | no | no | no |
| SCN2A | 774 | L | 1 | 0 | yes | no | no | yes | no | no | no | no | no | no | no | no | no |
| SCN2A | 775 | F | 0 | 0 | yes | no | no | yes | no | no | no | no | no | no | no | no | no |
| SCN2A | 776 | M | 0 | 0 | yes | no | no | yes | no | no | no | no | no | no | no | no | no |
| SCN2A | 777 | A | 1 | 0 | yes | no | no | yes | no | no | no | no | no | no | no | no | no |
| SCN2A | 778 | M | 0 | 0 | no | yes | no | yes | no | no | no | no | no | no | no | no | no |
| SCN2A | 779 | E | 0 | 0 | no | yes | no | yes | yes | no | no | no | no | no | no | no | no |
| SCN2A | 780 | H | 1 | 0 | no | yes | no | no | yes | no | no | no | no | no | no | no | no |
| SCN2A | 781 | Y | 0 | 0 | no | yes | no | no | yes | no | no | no | no | no | no | no | no |
| SCN2A | 782 | P | 0 | 0 | no | yes | no | no | yes | no | no | no | no | no | no | no | no |
| SCN2A | 783 | M | 0 | 0 | no | yes | no | no | yes | no | no | no | no | no | no | no | no |
| SCN2A | 784 | T | 1 | 0 | no | yes | no | no | yes | no | no | no | no | no | no | no | no |
| SCN2A | 785 | E | 0 | 0 | no | yes | no | no | yes | no | no | no | no | no | no | no | no |
| SCN2A | 786 | Q | 0 | 0 | no | yes | no | no | yes | no | no | no | no | no | no | no | no |
| SCN2A | 787 | F | 0 | 0 | no | yes | no | no | yes | no | no | no | no | no | no | no | no |
| SCN2A | 788 | S | 1 | 0 | yes | no | no | no | yes | no | no | no | no | no | no | no | no |
| SCN2A | 789 | S | 0 | 0 | yes | no | no | no | yes | no | no | no | no | no | no | no | no |
| SCN2A | 790 | V | 0 | 0 | yes | no | no | yes | no | no | no | no | no | no | no | no | no |
| SCN2A | 791 | L | 0 | 0 | yes | no | no | yes | no | no | no | no | no | no | no | no | no |
| SCN2A | 792 | S | 0 | 0 | yes | no | no | yes | no | no | no | no | no | no | no | no | no |
| SCN2A | 793 | V | 0 | 0 | yes | no | no | yes | no | no | no | no | no | no | no | no | no |
| SCN2A | 794 | G | 0 | 0 | yes | no | no | yes | no | no | no | no | no | no | no | no | no |
| SCN2A | 795 | N | 0 | 0 | yes | no | no | yes | no | no | no | no | no | no | no | no | no |

| | | | | | | | | | | | | | | | | | |
|---|---|---|---|---|---|---|---|---|---|---|---|---|---|---|---|---|---|
| SCN2A | 796 | L | 0 | 0 | yes | no | no | yes | no | no | no | no | no | no | no | no | no |
| SCN2A | 797 | V | 1 | 0 | yes | no | no | yes | no | no | no | no | no | no | no | no | no |
| SCN2A | 798 | F | 0 | 0 | yes | no | no | yes | no | no | no | no | no | no | no | no | no |
| SCN2A | 799 | T | 0 | 0 | yes | no | no | yes | no | no | no | no | no | no | no | no | no |
| SCN2A | 800 | G | 0 | 0 | yes | no | no | yes | no | no | no | no | no | no | no | no | no |
| SCN2A | 801 | I | 0 | 0 | yes | no | no | yes | no | no | no | no | no | no | no | no | no |
| SCN2A | 802 | F | 0 | 0 | yes | no | no | yes | no | no | no | no | no | no | no | no | no |
| SCN2A | 803 | T | 0 | 0 | yes | no | no | yes | no | no | no | no | no | no | no | no | no |
| SCN2A | 804 | A | 0 | 0 | yes | no | no | yes | no | no | no | no | no | no | no | no | no |
| SCN2A | 805 | E | 0 | 0 | yes | no | no | yes | no | no | no | no | no | no | no | no | no |
| SCN2A | 806 | M | 0 | 0 | yes | no | no | yes | no | no | no | no | no | no | no | no | no |
| SCN2A | 807 | F | 0 | 0 | yes | no | no | yes | no | no | no | no | no | no | no | no | no |
| SCN2A | 808 | L | 0 | 0 | no | yes | no | yes | no | no | no | no | no | no | no | no | no |
| SCN2A | 809 | K | 0 | 0 | no | yes | no | yes | no | no | no | no | no | no | no | no | no |
| SCN2A | 810 | I | 1 | 0 | no | yes | no | yes | no | yes | no | no | no | no | no | no | no |
| SCN2A | 811 | I | 0 | 0 | no | yes | no | yes | no | yes | no | no | no | no | no | no | no |
| SCN2A | 812 | A | 0 | 0 | no | yes | no | no | no | yes | no | no | no | no | no | no | no |
| SCN2A | 813 | M | 0 | 0 | no | yes | no | no | no | yes | no | no | no | no | no | no | no |
| SCN2A | 814 | D | 0 | 0 | no | yes | no | no | no | yes | no | no | no | no | no | no | no |
| SCN2A | 815 | P | 0 | 0 | no | yes | no | yes | no | yes | no | no | no | no | no | no | no |
| SCN2A | 816 | Y | 0 | 0 | no | yes | no | yes | no | yes | no | no | no | no | no | no | no |
| SCN2A | 817 | Y | 0 | 0 | no | yes | no | no | no | yes | no | no | no | no | no | no | no |
| SCN2A | 818 | Y | 0 | 0 | no | yes | no | yes | no | yes | no | no | no | no | no | no | no |
| SCN2A | 819 | F | 0 | 0 | no | yes | no | yes | no | yes | no | no | no | no | no | no | no |
| SCN2A | 820 | Q | 0 | 0 | no | yes | no | no | no | yes | no | no | no | no | no | no | no |
| SCN2A | 821 | E | 0 | 0 | no | yes | no | no | no | yes | no | no | no | no | no | no | no |
| SCN2A | 822 | G | 1 | 0 | no | yes | no | yes | no | yes | no | no | no | no | no | no | no |
| SCN2A | 823 | W | 0 | 0 | yes | no | no | yes | no | yes | no | no | no | no | no | no | no |
| SCN2A | 824 | N | 0 | 0 | yes | no | no | yes | no | no | no | no | no | no | no | no | no |
| SCN2A | 825 | I | 0 | 0 | yes | no | no | yes | no | no | no | no | no | no | no | no | no |
| SCN2A | 826 | F | 0 | 0 | yes | no | no | yes | no | no | no | no | no | no | no | no | no |
| SCN2A | 827 | D | 0 | 1 | yes | no | no | yes | no | no | no | no | no | no | no | no | no |
| SCN2A | 828 | G | 1 | 1 | yes | no | no | yes | no | no | no | no | no | no | no | no | no |

| | | | | | | | | | | | | | | | | | |
|---|---|---|---|---|---|---|---|---|---|---|---|---|---|---|---|---|---|
| SCN2A | 829 | F | 0 | 0 | yes | no | no | yes | no | no | no | no | no | no | no | no | no |
| SCN2A | 830 | I | 0 | 0 | yes | no | no | yes | no | no | no | no | no | no | no | no | no |
| SCN2A | 831 | V | 0 | 0 | yes | no | no | yes | no | no | no | no | no | no | no | no | no |
| SCN2A | 832 | S | 1 | 0 | yes | no | no | yes | no | no | no | no | no | no | no | no | no |
| SCN2A | 833 | L | 0 | 0 | yes | no | no | yes | no | no | no | no | no | no | no | no | no |
| SCN2A | 834 | S | 0 | 1 | yes | no | no | yes | no | no | no | no | no | no | no | no | no |
| SCN2A | 835 | L | 0 | 1 | yes | no | no | yes | no | no | no | no | no | no | no | no | no |
| SCN2A | 836 | M | 0 | 0 | yes | no | no | yes | no | no | no | no | no | no | no | no | no |
| SCN2A | 837 | E | 0 | 0 | yes | no | no | yes | no | no | no | no | no | no | no | no | no |
| SCN2A | 838 | L | 0 | 0 | yes | no | no | yes | no | no | no | no | no | no | no | no | no |
| SCN2A | 839 | G | 0 | 0 | yes | no | no | yes | no | no | no | no | no | no | no | no | no |
| SCN2A | 840 | L | 0 | 0 | yes | no | no | yes | no | no | no | no | no | no | no | no | no |
| SCN2A | 841 | A | 1 | 0 | no | yes | no | no | no | no | no | no | no | no | no | no | no |
| SCN2A | 842 | N | 0 | 0 | no | yes | no | no | no | no | no | no | no | no | no | no | no |
| SCN2A | 843 | V | 0 | 1 | no | yes | no | no | no | no | no | no | no | no | no | no | no |
| SCN2A | 844 | E | 0 | 0 | no | yes | no | no | yes | no | no | no | no | no | no | no | no |
| SCN2A | 845 | G | 0 | 0 | no | yes | no | no | yes | no | no | no | no | no | no | no | no |
| SCN2A | 846 | L | 0 | 0 | no | yes | no | yes | no | no | no | no | no | no | no | no | no |
| SCN2A | 847 | S | 0 | 0 | yes | no | no | yes | no | no | no | no | no | no | no | no | no |
| SCN2A | 848 | V | 0 | 0 | yes | no | no | yes | no | no | no | no | no | no | no | no | no |
| SCN2A | 849 | L | 0 | 0 | yes | no | no | yes | no | no | no | no | no | no | no | no | no |
| SCN2A | 850 | R | 0 | 0 | yes | no | no | yes | no | no | no | no | no | no | no | no | no |
| SCN2A | 851 | S | 0 | 0 | yes | no | no | yes | no | no | no | no | no | no | no | no | no |
| SCN2A | 852 | F | 0 | 0 | yes | no | no | yes | no | no | no | no | no | no | no | no | no |
| SCN2A | 853 | R | 0 | 1 | yes | no | no | yes | no | no | no | no | no | no | no | no | no |
| SCN2A | 854 | L | 0 | 0 | yes | no | no | yes | no | no | no | no | no | no | no | no | no |
| SCN2A | 855 | L | 0 | 0 | yes | no | no | yes | no | no | no | no | no | no | no | no | no |
| SCN2A | 856 | R | 0 | 1 | yes | no | no | yes | no | no | no | no | no | no | no | no | no |
| SCN2A | 857 | V | 0 | 0 | yes | no | no | yes | no | no | no | no | no | no | no | no | no |
| SCN2A | 858 | F | 0 | 0 | yes | no | no | yes | no | no | no | no | no | no | no | no | no |
| SCN2A | 859 | K | 0 | 0 | yes | no | no | yes | no | no | no | no | no | no | no | no | no |
| SCN2A | 860 | L | 0 | 0 | yes | no | no | yes | no | no | no | no | no | no | no | no | no |
| SCN2A | 861 | A | 0 | 0 | yes | no | no | yes | no | no | no | no | no | no | no | no | no |

| | | | | | | | | | | | | | | | | | |
|---|---|---|---|---|---|---|---|---|---|---|---|---|---|---|---|---|---|
| SCN2A | 862 | K | 0 | 0 | yes | no | no | yes | no | no | no | no | no | no | no | no | no |
| SCN2A | 863 | S | 0 | 1 | no | yes | no | yes | no | no | no | no | no | no | no | no | no |
| SCN2A | 864 | W | 0 | 0 | no | yes | no | yes | no | yes | no | no | no | no | no | no | no |
| SCN2A | 865 | P | 0 | 0 | yes | no | no | yes | no | yes | no | no | no | no | no | no | no |
| SCN2A | 866 | T | 0 | 0 | yes | no | no | yes | no | yes | no | no | no | no | no | no | no |
| SCN2A | 867 | L | 0 | 0 | yes | no | no | yes | no | yes | no | no | no | no | no | no | no |
| SCN2A | 868 | N | 0 | 1 | yes | no | no | yes | no | yes | no | no | no | no | no | no | no |
| SCN2A | 869 | M | 0 | 0 | yes | no | no | yes | no | yes | no | no | no | no | no | no | no |
| SCN2A | 870 | L | 0 | 0 | yes | no | no | yes | no | yes | no | no | no | no | no | no | no |
| SCN2A | 871 | I | 0 | 0 | yes | no | no | yes | no | yes | no | no | no | no | no | no | no |
| SCN2A | 872 | K | 0 | 0 | yes | no | no | yes | no | yes | no | no | no | no | no | no | no |
| SCN2A | 873 | I | 0 | 1 | yes | no | no | yes | no | yes | no | no | no | no | no | no | no |
| SCN2A | 874 | I | 0 | 1 | yes | no | no | yes | no | yes | no | no | no | no | no | no | no |
| SCN2A | 875 | G | 0 | 0 | yes | no | no | yes | no | yes | no | no | no | no | no | no | no |
| SCN2A | 876 | N | 0 | 1 | yes | no | no | yes | no | yes | no | no | no | no | no | no | no |
| SCN2A | 877 | S | 0 | 0 | yes | no | no | yes | no | yes | no | no | no | no | no | no | no |
| SCN2A | 878 | V | 0 | 0 | yes | no | no | yes | no | yes | no | no | no | no | no | no | no |
| SCN2A | 879 | G | 0 | 2 | no | yes | no | yes | no | yes | no | no | no | no | no | no | no |
| SCN2A | 880 | A | 0 | 1 | no | yes | no | yes | no | no | no | no | no | no | no | no | no |
| SCN2A | 881 | L | 0 | 1 | no | yes | no | yes | no | no | no | no | no | no | no | no | no |
| SCN2A | 882 | G | 0 | 2 | yes | no | no | yes | no | no | no | no | no | no | no | no | no |
| SCN2A | 883 | N | 0 | 0 | yes | no | no | yes | no | no | no | no | no | no | no | no | no |
| SCN2A | 884 | L | 0 | 1 | yes | no | no | yes | no | no | no | no | no | no | no | no | no |
| SCN2A | 885 | T | 0 | 0 | yes | no | no | yes | no | no | no | no | no | no | no | no | no |
| SCN2A | 886 | L | 0 | 1 | yes | no | no | yes | no | no | no | no | no | no | no | no | no |
| SCN2A | 887 | V | 0 | 2 | yes | no | no | yes | no | no | no | no | no | no | no | no | no |
| SCN2A | 888 | L | 0 | 0 | yes | no | no | yes | no | no | no | no | no | no | no | no | no |
| SCN2A | 889 | A | 0 | 0 | yes | no | no | yes | no | no | no | no | no | no | no | no | no |
| SCN2A | 890 | I | 0 | 1 | yes | no | no | yes | no | no | no | no | no | no | no | no | no |
| SCN2A | 891 | I | 0 | 1 | yes | no | no | yes | no | no | no | no | no | no | no | no | no |
| SCN2A | 892 | V | 0 | 1 | yes | no | no | yes | no | no | no | no | no | no | no | no | no |
| SCN2A | 893 | F | 0 | 0 | yes | no | no | yes | no | no | no | no | no | no | no | no | no |
| SCN2A | 894 | I | 0 | 0 | yes | no | no | yes | no | no | no | no | no | no | no | no | no |

| | | | | | | | | | | | | | | | | | |
|---|---|---|---|---|---|---|---|---|---|---|---|---|---|---|---|---|---|
| SCN2A | 895 | F | 1 | 1 | yes | no | no | yes | no | no | no | no | no | no | no | no | no |
| SCN2A | 896 | A | 0 | 1 | yes | no | no | yes | no | no | no | no | no | no | no | no | no |
| SCN2A | 897 | V | 0 | 0 | yes | no | no | yes | no | no | no | no | no | no | no | no | no |
| SCN2A | 898 | V | 0 | 0 | yes | no | no | yes | no | no | no | no | no | no | no | no | no |
| SCN2A | 899 | G | 0 | 2 | yes | no | no | yes | yes | no | no | no | no | no | no | no | no |
| SCN2A | 900 | M | 0 | 0 | yes | no | no | yes | yes | no | no | no | no | no | no | no | no |
| SCN2A | 901 | Q | 0 | 1 | yes | no | no | yes | yes | no | no | no | no | no | no | no | no |
| SCN2A | 902 | L | 0 | 0 | yes | no | no | yes | yes | no | no | no | no | no | no | no | no |
| SCN2A | 903 | F | 0 | 0 | no | yes | no | no | yes | no | no | no | no | no | no | no | no |
| SCN2A | 904 | G | 0 | 0 | yes | no | no | no | yes | no | no | no | no | no | no | no | no |
| SCN2A | 905 | K | 0 | 2 | yes | no | no | no | yes | no | no | no | yes | no | no | no | no |
| SCN2A | 906 | S | 0 | 0 | yes | no | no | no | yes | no | no | no | no | no | no | no | no |
| SCN2A | 907 | Y | 0 | 0 | yes | no | no | no | yes | no | no | no | no | no | no | no | no |
| SCN2A | 908 | K | 1 | 1 | yes | no | no | no | yes | no | no | no | yes | no | no | no | no |
| SCN2A | 909 | E | 0 | 0 | yes | no | no | no | yes | no | no | no | yes | no | no | no | no |
| SCN2A | 910 | C | 0 | 0 | no | yes | no | no | yes | no | no | no | yes | no | no | no | no |
| SCN2A | 911 | V | 0 | 0 | no | yes | no | no | yes | no | no | no | yes | no | no | no | no |
| SCN2A | 912 | C | 0 | 0 | no | yes | no | no | yes | no | no | no | yes | no | no | no | no |
| SCN2A | 913 | K | 1 | 0 | no | yes | no | no | yes | no | no | no | yes | no | no | no | no |
| SCN2A | 914 | I | 1 | 0 | no | yes | no | no | yes | no | no | no | no | no | no | no | no |
| SCN2A | 915 | S | 0 | 0 | no | yes | no | no | yes | no | no | no | no | no | no | no | no |
| SCN2A | 916 | N | 1 | 0 | no | yes | no | no | yes | no | no | no | no | no | no | no | no |
| SCN2A | 917 | D | 0 | 0 | no | yes | no | no | yes | no | no | no | yes | no | no | no | no |
| SCN2A | 918 | C | 0 | 0 | no | yes | no | no | yes | no | no | no | yes | no | no | no | no |
| SCN2A | 919 | E | 0 | 0 | no | yes | no | no | yes | no | no | no | no | no | no | no | no |
| SCN2A | 920 | L | 1 | 0 | no | yes | no | no | yes | no | no | no | no | no | no | no | no |
| SCN2A | 921 | P | 0 | 0 | no | yes | no | no | yes | no | no | no | no | no | no | no | no |
| SCN2A | 922 | R | 0 | 1 | no | yes | no | no | yes | no | no | no | no | no | no | no | no |
| SCN2A | 923 | W | 0 | 1 | no | yes | no | no | yes | no | no | no | no | no | no | no | no |
| SCN2A | 924 | H | 0 | 0 | no | yes | no | no | yes | no | no | no | no | no | no | no | no |
| SCN2A | 925 | M | 0 | 1 | no | yes | no | no | yes | no | no | no | no | no | no | no | no |
| SCN2A | 926 | H | 1 | 1 | no | yes | no | no | yes | no | no | no | yes | no | no | no | no |
| SCN2A | 927 | D | 0 | 0 | no | yes | no | no | yes | no | no | no | no | no | no | no | no |

| | | | | | | | | | | | | | | | | | |
|---|---|---|---|---|---|---|---|---|---|---|---|---|---|---|---|---|---|
| SCN2A | 928 | F | 0 | 1 | yes | no | no | yes | no | no | no | yes | no | no | no | no | no |
| SCN2A | 929 | F | 0 | 0 | yes | no | no | yes | no | no | no | yes | no | no | no | no | no |
| SCN2A | 930 | H | 0 | 1 | yes | no | no | no | no | no | no | yes | no | no | no | no | no |
| SCN2A | 931 | S | 0 | 1 | yes | no | no | yes | no | no | no | yes | no | no | no | no | no |
| SCN2A | 932 | F | 0 | 0 | yes | no | no | yes | no | no | no | yes | no | no | no | no | no |
| SCN2A | 933 | L | 0 | 0 | yes | no | no | yes | no | no | no | yes | no | no | no | no | no |
| SCN2A | 934 | I | 0 | 0 | yes | no | no | yes | no | no | no | yes | no | no | no | no | no |
| SCN2A | 935 | V | 0 | 0 | yes | no | no | yes | no | no | no | yes | no | no | no | no | no |
| SCN2A | 936 | F | 0 | 0 | yes | no | no | yes | no | no | no | yes | no | no | no | no | no |
| SCN2A | 937 | R | 0 | 2 | yes | no | no | yes | no | no | no | yes | no | no | no | no | no |
| SCN2A | 938 | V | 0 | 0 | yes | no | no | yes | no | no | no | yes | no | no | no | no | no |
| SCN2A | 939 | L | 0 | 0 | no | yes | no | yes | no | no | no | yes | no | no | no | no | no |
| SCN2A | 940 | C | 0 | 0 | no | yes | no | yes | no | no | yes | yes | no | no | no | no | no |
| SCN2A | 941 | G | 0 | 0 | no | yes | no | yes | no | no | yes | yes | no | no | no | no | no |
| SCN2A | 942 | E | 0 | 0 | no | yes | no | yes | no | no | yes | yes | no | no | no | no | no |
| SCN2A | 943 | W | 0 | 0 | no | yes | no | yes | no | no | no | yes | no | no | no | no | no |
| SCN2A | 944 | I | 0 | 0 | yes | no | no | yes | no | no | no | yes | no | no | no | no | no |
| SCN2A | 945 | E | 0 | 0 | yes | no | no | no | no | no | no | yes | no | no | no | no | no |
| SCN2A | 946 | T | 0 | 0 | yes | no | no | no | no | no | no | yes | no | no | no | no | no |
| SCN2A | 947 | M | 0 | 0 | yes | no | no | yes | no | no | no | yes | no | no | no | no | no |
| SCN2A | 948 | W | 0 | 0 | yes | no | no | no | no | no | no | yes | no | no | no | no | no |
| SCN2A | 949 | D | 0 | 0 | yes | no | no | no | yes | no | no | no | no | no | no | no | no |
| SCN2A | 950 | C | 0 | 0 | yes | no | no | no | yes | no | no | no | no | no | no | no | no |
| SCN2A | 951 | M | 0 | 0 | yes | no | no | no | yes | no | no | no | no | no | no | no | no |
| SCN2A | 952 | E | 0 | 0 | yes | no | no | no | yes | no | no | no | no | no | no | no | no |
| SCN2A | 953 | V | 0 | 0 | yes | no | no | no | yes | no | no | no | no | no | no | no | no |
| SCN2A | 954 | A | 2 | 0 | yes | no | no | no | yes | no | no | no | no | no | no | no | no |
| SCN2A | 955 | G | 0 | 0 | no | yes | no | no | yes | no | no | no | no | no | no | no | no |
| SCN2A | 956 | Q | 0 | 0 | yes | no | no | no | yes | no | no | no | no | no | no | no | no |
| SCN2A | 957 | T | 1 | 0 | yes | no | no | yes | yes | no | no | no | no | no | no | no | no |
| SCN2A | 958 | M | 1 | 1 | yes | no | no | yes | yes | no | no | no | no | no | no | no | no |
| SCN2A | 959 | C | 0 | 1 | yes | no | no | yes | yes | no | no | no | no | no | no | no | no |
| SCN2A | 960 | L | 0 | 0 | yes | no | no | yes | yes | no | no | no | no | no | no | no | no |

| | | | | | | | | | | | | | | | | | |
|---|---|---|---|---|---|---|---|---|---|---|---|---|---|---|---|---|---|
| SCN2A | 961 | T | 0 | 0 | yes | no | no | yes | yes | no | no | no | no | no | no | no | no |
| SCN2A | 962 | V | 0 | 0 | yes | no | no | yes | no | no | no | no | no | no | no | no | no |
| SCN2A | 963 | F | 0 | 0 | yes | no | no | yes | no | no | no | no | no | no | no | no | no |
| SCN2A | 964 | M | 1 | 0 | yes | no | no | yes | no | no | no | no | no | no | no | no | no |
| SCN2A | 965 | M | 0 | 0 | yes | no | no | yes | no | no | no | no | no | no | no | no | no |
| SCN2A | 966 | V | 0 | 0 | yes | no | no | yes | no | no | no | no | no | no | no | no | no |
| SCN2A | 967 | M | 0 | 0 | yes | no | no | yes | no | no | no | no | no | no | no | no | no |
| SCN2A | 968 | V | 0 | 0 | yes | no | no | yes | no | no | no | no | no | no | no | no | no |
| SCN2A | 969 | I | 0 | 0 | yes | no | no | yes | no | no | no | no | no | no | no | no | no |
| SCN2A | 970 | G | 0 | 0 | yes | no | no | yes | no | no | no | no | no | no | no | no | no |
| SCN2A | 971 | N | 0 | 0 | yes | no | no | yes | no | no | no | no | no | no | no | no | no |
| SCN2A | 972 | L | 0 | 1 | yes | no | no | yes | no | no | no | no | no | no | no | no | no |
| SCN2A | 973 | V | 0 | 0 | yes | no | no | yes | no | no | no | no | no | no | no | no | no |
| SCN2A | 974 | V | 0 | 0 | yes | no | no | yes | no | no | no | no | no | no | no | no | no |
| SCN2A | 975 | L | 0 | 0 | yes | no | no | yes | no | no | no | no | no | no | no | no | no |
| SCN2A | 976 | N | 0 | 1 | yes | no | no | yes | no | no | no | no | no | no | no | no | no |
| SCN2A | 977 | L | 0 | 0 | yes | no | no | yes | no | no | no | no | no | no | no | no | no |
| SCN2A | 978 | F | 0 | 0 | yes | no | no | yes | no | no | no | no | no | no | no | no | no |
| SCN2A | 979 | L | 0 | 0 | yes | no | no | yes | no | no | no | no | no | no | no | no | no |
| SCN2A | 980 | A | 0 | 0 | yes | no | no | yes | no | no | no | no | no | no | no | no | no |
| SCN2A | 981 | L | 0 | 0 | yes | no | no | yes | no | no | no | no | no | no | no | no | no |
| SCN2A | 982 | L | 0 | 0 | yes | no | no | yes | no | no | no | no | no | no | no | no | no |
| SCN2A | 983 | L | 0 | 1 | yes | no | no | no | no | yes | no | no | no | no | no | no | no |
| SCN2A | 984 | S | 0 | 1 | yes | no | no | no | no | yes | no | no | no | no | no | no | no |
| SCN2A | 985 | S | 0 | 0 | yes | no | no | yes | no | yes | no | no | no | no | no | no | no |
| SCN2A | 986 | F | 0 | 0 | no | yes | no | no | no | yes | no | no | no | no | no | no | no |
| SCN2A | 987 | S | 0 | 1 | no | yes | no | no | no | yes | no | no | no | no | no | no | no |
| SCN2A | 1191 | G | 0 | 0 | no | yes | no | no | no | yes | no | no | no | no | no | no | no |
| SCN2A | 1192 | K | 0 | 0 | no | yes | no | no | no | yes | no | no | no | no | no | no | no |
| SCN2A | 1193 | L | 0 | 0 | no | yes | no | no | no | yes | no | no | no | no | no | no | no |
| SCN2A | 1194 | W | 0 | 0 | no | yes | no | no | no | yes | no | no | no | no | no | no | no |
| SCN2A | 1195 | W | 0 | 0 | yes | no | no | no | no | yes | no | no | no | no | no | no | no |
| SCN2A | 1196 | N | 1 | 0 | yes | no | no | no | no | yes | no | no | no | no | no | no | no |

| | | | | | | | | | | | | | | | | | |
|---|---|---|---|---|---|---|---|---|---|---|---|---|---|---|---|---|---|
| SCN2A | 1197 | L | 1 | 0 | yes | no | no | no | no | yes | no | no | no | no | no | no | no |
| SCN2A | 1198 | R | 0 | 0 | yes | no | no | no | no | yes | no | no | no | no | no | no | no |
| SCN2A | 1199 | K | 0 | 0 | yes | no | no | no | no | yes | no | no | no | no | no | no | no |
| SCN2A | 1200 | T | 1 | 0 | yes | no | no | no | no | yes | no | no | no | no | no | no | no |
| SCN2A | 1201 | C | 0 | 0 | yes | no | no | yes | no | yes | no | no | no | no | no | no | no |
| SCN2A | 1202 | Y | 0 | 0 | yes | no | no | no | no | yes | no | no | no | no | no | no | no |
| SCN2A | 1203 | K | 1 | 0 | yes | no | no | no | no | yes | no | no | no | no | no | no | no |
| SCN2A | 1204 | I | 0 | 0 | yes | no | no | yes | no | yes | no | no | no | no | no | no | no |
| SCN2A | 1205 | V | 0 | 0 | yes | no | no | yes | no | yes | no | no | no | no | no | no | no |
| SCN2A | 1206 | E | 0 | 0 | no | yes | no | no | no | yes | no | no | no | no | no | no | no |
| SCN2A | 1207 | H | 1 | 0 | no | yes | no | yes | no | yes | no | no | no | no | no | no | no |
| SCN2A | 1208 | N | 2 | 0 | yes | no | no | yes | no | yes | no | no | no | no | no | no | no |
| SCN2A | 1209 | W | 1 | 0 | yes | no | no | yes | no | yes | no | no | no | no | no | no | no |
| SCN2A | 1210 | F | 0 | 0 | yes | no | no | yes | no | no | no | no | no | no | no | no | no |
| SCN2A | 1211 | E | 0 | 1 | yes | no | no | yes | no | no | no | no | no | no | no | no | no |
| SCN2A | 1212 | T | 0 | 1 | yes | no | no | yes | no | no | no | no | no | no | no | no | no |
| SCN2A | 1213 | F | 0 | 0 | yes | no | no | yes | no | no | no | no | no | no | no | no | no |
| SCN2A | 1214 | I | 1 | 0 | yes | no | no | yes | no | no | no | no | no | no | no | no | no |
| SCN2A | 1215 | V | 0 | 0 | yes | no | no | yes | no | no | no | no | no | no | no | no | no |
| SCN2A | 1216 | F | 0 | 0 | yes | no | no | yes | no | no | no | no | no | no | no | no | no |
| SCN2A | 1217 | M | 0 | 0 | yes | no | no | yes | no | no | no | no | no | no | no | no | no |
| SCN2A | 1218 | I | 0 | 0 | yes | no | no | yes | no | no | no | no | no | no | no | no | no |
| SCN2A | 1219 | L | 0 | 0 | yes | no | no | yes | no | no | no | no | no | no | no | no | no |
| SCN2A | 1220 | L | 0 | 0 | yes | no | no | yes | no | no | no | no | no | no | no | no | no |
| SCN2A | 1221 | S | 0 | 0 | yes | no | no | yes | no | no | no | no | no | no | no | no | no |
| SCN2A | 1222 | S | 0 | 1 | yes | no | no | yes | no | no | no | no | no | no | no | no | no |
| SCN2A | 1223 | G | 0 | 1 | yes | no | no | yes | no | no | no | no | no | no | no | no | no |
| SCN2A | 1224 | A | 1 | 0 | yes | no | no | yes | no | no | no | no | no | no | no | no | no |
| SCN2A | 1225 | L | 0 | 0 | yes | no | no | yes | no | no | no | no | no | no | no | no | no |
| SCN2A | 1226 | A | 0 | 0 | yes | no | no | yes | no | no | no | no | no | no | no | no | no |
| SCN2A | 1227 | F | 0 | 0 | yes | no | no | yes | no | no | no | no | no | no | no | no | no |
| SCN2A | 1228 | E | 0 | 0 | no | yes | no | no | yes | no | no | no | no | no | no | no | no |
| SCN2A | 1229 | D | 0 | 0 | no | yes | no | no | yes | no | no | no | no | no | no | no | no |

| | | | | | | | | | | | | | | | | | |
|---|---|---|---|---|---|---|---|---|---|---|---|---|---|---|---|---|---|
| SCN2A | 1230 | I | 2 | 0 | no | yes | no | no | yes | no | no | no | no | no | no | no | no |
| SCN2A | 1231 | Y | 0 | 0 | no | yes | no | no | yes | no | no | no | no | no | no | no | no |
| SCN2A | 1232 | I | 0 | 0 | yes | no | no | no | yes | no | no | no | no | no | no | no | no |
| SCN2A | 1233 | E | 1 | 0 | yes | no | no | no | yes | no | no | no | no | no | no | no | no |
| SCN2A | 1234 | Q | 1 | 0 | yes | no | no | no | yes | no | no | no | no | no | no | no | no |
| SCN2A | 1235 | R | 1 | 0 | yes | no | no | no | yes | no | no | no | no | no | no | no | no |
| SCN2A | 1236 | K | 0 | 0 | no | yes | no | no | yes | no | no | no | no | no | no | no | no |
| SCN2A | 1237 | T | 1 | 0 | no | yes | no | no | yes | no | no | no | no | no | no | no | no |
| SCN2A | 1238 | I | 0 | 0 | no | yes | no | no | yes | no | no | no | no | no | no | no | no |
| SCN2A | 1239 | K | 0 | 0 | yes | no | no | no | yes | no | no | no | no | no | no | no | no |
| SCN2A | 1240 | T | 0 | 0 | yes | no | no | no | yes | no | no | no | no | no | no | no | no |
| SCN2A | 1241 | M | 0 | 0 | yes | no | no | yes | no | no | no | no | no | no | no | no | no |
| SCN2A | 1242 | L | 0 | 0 | yes | no | no | yes | no | no | no | no | no | no | no | no | no |
| SCN2A | 1243 | E | 0 | 0 | yes | no | no | yes | no | no | no | no | no | no | no | no | no |
| SCN2A | 1244 | Y | 1 | 0 | yes | no | no | yes | no | no | no | no | no | no | no | no | no |
| SCN2A | 1245 | A | 2 | 0 | yes | no | no | yes | no | no | no | no | no | no | no | no | no |
| SCN2A | 1246 | D | 0 | 0 | yes | no | no | yes | no | no | no | no | no | no | no | no | no |
| SCN2A | 1247 | K | 0 | 0 | yes | no | no | yes | no | no | no | no | no | no | no | no | no |
| SCN2A | 1248 | V | 1 | 0 | yes | no | no | yes | no | no | no | no | no | no | no | no | no |
| SCN2A | 1249 | F | 0 | 0 | yes | no | no | yes | no | no | no | no | no | no | no | no | no |
| SCN2A | 1250 | T | 0 | 0 | yes | no | no | yes | no | no | no | no | no | no | no | no | no |
| SCN2A | 1251 | Y | 0 | 0 | yes | no | no | yes | no | no | no | no | no | no | no | no | no |
| SCN2A | 1252 | I | 1 | 1 | yes | no | no | yes | no | no | no | no | no | no | no | no | no |
| SCN2A | 1253 | F | 0 | 0 | yes | no | no | yes | no | no | no | no | no | no | no | no | no |
| SCN2A | 1254 | I | 0 | 0 | yes | no | no | yes | no | no | no | no | no | no | no | no | no |
| SCN2A | 1255 | L | 0 | 0 | yes | no | no | yes | no | no | no | no | no | no | no | no | no |
| SCN2A | 1256 | E | 0 | 0 | yes | no | no | yes | no | no | no | no | no | no | no | no | no |
| SCN2A | 1257 | M | 0 | 0 | yes | no | no | yes | no | no | no | no | no | no | no | no | no |
| SCN2A | 1258 | L | 0 | 0 | yes | no | no | yes | no | no | no | no | no | no | no | no | no |
| SCN2A | 1259 | L | 0 | 0 | yes | no | no | yes | no | no | no | no | no | no | no | no | no |
| SCN2A | 1260 | K | 0 | 1 | yes | no | no | yes | no | yes | no | no | no | no | no | no | no |
| SCN2A | 1261 | W | 0 | 0 | yes | no | no | yes | no | yes | no | no | no | no | no | no | no |
| SCN2A | 1262 | V | 2 | 0 | yes | no | no | yes | no | yes | no | no | no | no | no | no | no |

| | | | | | | | | | | | | | | | | | |
|---|---|---|---|---|---|---|---|---|---|---|---|---|---|---|---|---|---|
| SCN2A | 1263 | A | 0 | 0 | yes | no | no | no | no | yes | no | no | no | no | no | no | no |
| SCN2A | 1264 | Y | 0 | 0 | yes | no | no | no | no | yes | no | no | no | no | no | no | no |
| SCN2A | 1265 | G | 0 | 0 | no | yes | no | yes | no | yes | no | no | no | no | no | no | no |
| SCN2A | 1266 | F | 0 | 0 | yes | no | no | yes | no | yes | no | no | no | no | no | no | no |
| SCN2A | 1267 | Q | 0 | 0 | yes | no | no | no | no | yes | no | no | no | no | no | no | no |
| SCN2A | 1268 | V | 1 | 0 | yes | no | no | no | no | yes | no | no | no | no | no | no | no |
| SCN2A | 1269 | Y | 0 | 0 | yes | no | no | yes | no | yes | no | no | no | no | no | no | no |
| SCN2A | 1270 | F | 0 | 0 | no | yes | no | yes | no | yes | no | no | no | no | no | no | no |
| SCN2A | 1271 | T | 0 | 0 | no | yes | no | no | no | yes | no | no | no | no | no | no | no |
| SCN2A | 1272 | N | 0 | 0 | no | yes | no | yes | no | yes | no | no | no | no | no | no | no |
| SCN2A | 1273 | A | 0 | 0 | yes | no | no | yes | no | yes | no | no | no | no | no | no | no |
| SCN2A | 1274 | W | 0 | 0 | yes | no | no | yes | no | no | no | no | no | no | no | no | no |
| SCN2A | 1275 | C | 0 | 0 | yes | no | no | yes | no | no | no | no | no | no | no | no | no |
| SCN2A | 1276 | W | 0 | 0 | yes | no | no | yes | no | no | no | no | no | no | no | no | no |
| SCN2A | 1277 | L | 2 | 0 | yes | no | no | yes | no | no | no | no | no | no | no | no | no |
| SCN2A | 1278 | D | 0 | 0 | yes | no | no | yes | no | no | no | no | no | no | no | no | no |
| SCN2A | 1279 | F | 0 | 0 | yes | no | no | yes | no | no | no | no | no | no | no | no | no |
| SCN2A | 1280 | L | 0 | 0 | yes | no | no | yes | no | no | no | no | no | no | no | no | no |
| SCN2A | 1281 | I | 0 | 1 | yes | no | no | yes | no | no | no | no | no | no | no | no | no |
| SCN2A | 1282 | V | 0 | 0 | yes | no | no | yes | no | no | no | no | no | no | no | no | no |
| SCN2A | 1283 | D | 1 | 0 | yes | no | no | yes | no | no | no | no | no | no | no | no | no |
| SCN2A | 1284 | V | 0 | 0 | yes | no | no | yes | no | no | no | no | no | no | no | no | no |
| SCN2A | 1285 | S | 0 | 0 | yes | no | no | yes | no | no | no | no | no | no | no | no | no |
| SCN2A | 1286 | L | 0 | 0 | yes | no | no | yes | no | no | no | no | no | no | no | no | no |
| SCN2A | 1287 | V | 0 | 0 | yes | no | no | yes | no | no | no | no | no | no | no | no | no |
| SCN2A | 1288 | S | 1 | 0 | yes | no | no | yes | no | no | no | no | no | no | no | no | no |
| SCN2A | 1289 | L | 0 | 0 | yes | no | no | yes | no | no | no | no | no | no | no | no | no |
| SCN2A | 1290 | T | 0 | 0 | yes | no | no | yes | no | no | no | no | no | no | no | no | no |
| SCN2A | 1291 | A | 0 | 0 | yes | no | no | yes | no | no | no | no | no | no | no | no | no |
| SCN2A | 1292 | N | 0 | 0 | yes | no | no | yes | no | no | no | no | no | no | no | no | no |
| SCN2A | 1293 | A | 0 | 0 | yes | no | no | yes | yes | no | no | no | no | no | no | no | no |
| SCN2A | 1294 | L | 0 | 0 | yes | no | no | yes | yes | no | no | no | no | no | no | no | no |
| SCN2A | 1295 | G | 1 | 0 | no | yes | no | no | yes | no | no | no | no | no | no | no | no |

| | | | | | | | | | | | | | | | | | |
|---|---|---|---|---|---|---|---|---|---|---|---|---|---|---|---|---|---|
| SCN2A | 1296 | Y | 0 | 0 | no | yes | no | no | yes | no | no | no | no | no | no | no | no |
| SCN2A | 1297 | S | 1 | 0 | no | yes | no | no | yes | no | no | no | no | no | no | no | no |
| SCN2A | 1298 | E | 0 | 0 | no | yes | no | no | yes | no | no | no | no | no | no | no | no |
| SCN2A | 1299 | L | 0 | 0 | no | yes | no | no | yes | no | no | no | no | no | no | no | no |
| SCN2A | 1300 | G | 0 | 0 | no | yes | no | no | yes | no | no | no | no | no | no | no | no |
| SCN2A | 1301 | A | 0 | 0 | no | yes | no | yes | no | no | no | no | no | no | no | no | no |
| SCN2A | 1302 | I | 0 | 0 | yes | no | no | yes | no | no | no | no | no | no | no | no | no |
| SCN2A | 1303 | K | 0 | 0 | yes | no | no | no | no | no | no | no | no | no | no | no | no |
| SCN2A | 1304 | S | 0 | 0 | yes | no | no | yes | no | no | no | no | no | no | no | no | no |
| SCN2A | 1305 | L | 1 | 0 | yes | no | no | yes | no | no | no | no | no | no | no | no | no |
| SCN2A | 1306 | R | 0 | 0 | yes | no | no | yes | no | no | no | no | no | no | no | no | no |
| SCN2A | 1307 | T | 0 | 0 | yes | no | no | yes | no | no | no | no | no | no | no | no | no |
| SCN2A | 1308 | L | 0 | 0 | yes | no | no | yes | no | no | no | no | no | no | no | no | no |
| SCN2A | 1309 | R | 0 | 0 | yes | no | no | yes | no | no | no | no | no | no | no | no | no |
| SCN2A | 1310 | A | 0 | 1 | yes | no | no | yes | no | no | no | no | no | no | no | no | no |
| SCN2A | 1311 | L | 0 | 0 | yes | no | no | yes | no | no | no | no | no | no | no | no | no |
| SCN2A | 1312 | R | 0 | 1 | yes | no | no | yes | no | no | no | no | no | no | no | no | no |
| SCN2A | 1313 | P | 0 | 0 | yes | no | no | yes | no | no | no | no | no | no | no | no | no |
| SCN2A | 1314 | L | 0 | 0 | yes | no | no | yes | no | no | no | no | no | no | no | no | no |
| SCN2A | 1315 | R | 0 | 1 | yes | no | no | yes | no | no | no | no | no | no | no | no | no |
| SCN2A | 1316 | A | 0 | 1 | yes | no | no | yes | no | no | no | no | no | no | no | no | no |
| SCN2A | 1317 | L | 0 | 0 | no | yes | no | yes | no | no | no | no | no | no | no | no | no |
| SCN2A | 1318 | S | 0 | 1 | no | yes | no | yes | no | no | no | no | no | no | no | no | no |
| SCN2A | 1319 | R | 1 | 3 | no | yes | no | yes | no | no | no | no | no | no | no | no | no |
| SCN2A | 1320 | F | 0 | 0 | no | yes | no | yes | no | yes | no | no | no | no | no | no | no |
| SCN2A | 1321 | E | 0 | 1 | yes | no | no | yes | no | yes | no | no | no | no | no | no | no |
| SCN2A | 1322 | G | 0 | 1 | yes | no | no | yes | no | yes | no | no | no | no | no | no | no |
| SCN2A | 1323 | M | 0 | 1 | yes | no | no | yes | no | yes | no | no | no | no | no | no | no |
| SCN2A | 1324 | R | 1 | 1 | yes | no | no | yes | no | yes | no | no | no | no | no | no | no |
| SCN2A | 1325 | V | 0 | 1 | yes | no | no | yes | no | yes | no | no | no | no | no | no | no |
| SCN2A | 1326 | V | 0 | 2 | yes | no | no | yes | no | yes | no | no | no | no | no | no | no |
| SCN2A | 1327 | V | 0 | 0 | yes | no | no | yes | no | yes | no | no | no | no | no | no | no |
| SCN2A | 1328 | N | 0 | 0 | yes | no | no | yes | no | yes | no | no | no | no | no | no | no |

| | | | | | | | | | | | | | | | | | |
|---|---|---|---|---|---|---|---|---|---|---|---|---|---|---|---|---|---|
| SCN2A | 1329 | A | 0 | 2 | yes | no | no | yes | no | yes | no | no | no | no | no | no | no |
| SCN2A | 1330 | L | 0 | 1 | yes | no | no | yes | no | yes | no | no | no | no | no | no | no |
| SCN2A | 1331 | L | 0 | 0 | yes | no | no | yes | no | yes | no | no | no | no | no | no | no |
| SCN2A | 1332 | G | 0 | 1 | yes | no | no | yes | no | yes | no | no | no | no | no | no | no |
| SCN2A | 1333 | A | 1 | 1 | yes | no | no | yes | no | yes | no | no | no | no | no | no | no |
| SCN2A | 1334 | I | 0 | 2 | yes | no | no | yes | no | yes | no | no | no | no | no | no | no |
| SCN2A | 1335 | P | 0 | 0 | yes | no | no | yes | no | yes | no | no | no | no | no | no | no |
| SCN2A | 1336 | S | 0 | 1 | yes | no | no | yes | no | yes | no | no | no | no | no | no | no |
| SCN2A | 1337 | I | 0 | 0 | yes | no | no | yes | no | no | no | no | no | no | no | no | no |
| SCN2A | 1338 | M | 0 | 1 | yes | no | no | yes | no | no | no | no | no | no | no | no | no |
| SCN2A | 1339 | N | 1 | 1 | yes | no | no | yes | no | no | no | no | no | no | no | no | no |
| SCN2A | 1340 | V | 0 | 0 | yes | no | no | yes | no | no | no | no | no | no | no | no | no |
| SCN2A | 1341 | L | 0 | 0 | yes | no | no | yes | no | no | no | no | no | no | no | no | no |
| SCN2A | 1342 | L | 0 | 1 | yes | no | no | yes | no | no | no | no | no | no | no | no | no |
| SCN2A | 1343 | V | 1 | 0 | yes | no | no | yes | no | no | no | no | no | no | no | no | no |
| SCN2A | 1344 | C | 0 | 1 | yes | no | no | yes | no | no | no | no | no | no | no | no | no |
| SCN2A | 1345 | L | 0 | 0 | yes | no | no | yes | no | no | no | no | no | no | no | no | no |
| SCN2A | 1346 | I | 1 | 1 | yes | no | no | yes | no | no | no | no | no | no | no | no | no |
| SCN2A | 1347 | F | 0 | 0 | yes | no | no | yes | no | no | no | no | no | no | no | no | no |
| SCN2A | 1348 | W | 0 | 0 | yes | no | no | yes | no | no | no | no | no | no | no | no | no |
| SCN2A | 1349 | L | 0 | 0 | yes | no | no | yes | no | no | no | no | no | no | no | no | no |
| SCN2A | 1350 | I | 0 | 0 | yes | no | no | yes | no | no | no | no | no | no | no | no | no |
| SCN2A | 1351 | F | 0 | 0 | yes | no | no | yes | no | no | no | no | no | no | no | no | no |
| SCN2A | 1352 | S | 0 | 0 | yes | no | no | yes | no | no | no | no | no | no | no | no | no |
| SCN2A | 1353 | I | 0 | 1 | yes | no | no | yes | no | no | no | no | no | no | no | no | no |
| SCN2A | 1354 | M | 0 | 1 | yes | no | no | yes | no | no | no | no | no | no | no | no | no |
| SCN2A | 1355 | G | 0 | 0 | yes | no | no | yes | no | no | no | no | no | no | no | no | no |
| SCN2A | 1356 | V | 0 | 0 | yes | no | no | yes | no | no | no | no | no | no | no | no | no |
| SCN2A | 1357 | N | 0 | 0 | yes | no | no | yes | yes | no | no | no | no | no | no | no | no |
| SCN2A | 1358 | L | 0 | 0 | yes | no | no | no | yes | no | no | no | no | no | no | no | no |
| SCN2A | 1359 | F | 0 | 0 | no | yes | no | no | yes | no | no | no | no | no | no | no | no |
| SCN2A | 1360 | A | 0 | 0 | no | yes | no | no | yes | no | no | no | no | no | no | no | no |
| SCN2A | 1361 | G | 0 | 0 | no | yes | no | no | yes | no | no | no | no | no | no | no | no |

| | | | | | | | | | | | | | | | | | |
|---|---|---|---|---|---|---|---|---|---|---|---|---|---|---|---|---|---|
| SCN2A | 1362 | K | 0 | 1 | no | yes | no | no | yes | no | no | no | no | no | no | no | no |
| SCN2A | 1363 | F | 0 | 0 | no | yes | no | no | yes | no | no | no | no | no | no | no | no |
| SCN2A | 1364 | Y | 1 | 0 | no | yes | no | no | yes | no | no | no | no | no | no | no | no |
| SCN2A | 1365 | H | 1 | 0 | no | no | yes | no | yes | no | no | no | no | no | no | no | no |
| SCN2A | 1366 | C | 0 | 0 | no | no | yes | no | yes | no | no | no | no | no | no | no | no |
| SCN2A | 1367 | I | 0 | 0 | no | no | yes | no | yes | no | no | no | no | no | no | no | no |
| SCN2A | 1368 | N | 1 | 0 | no | no | yes | no | yes | no | no | no | no | no | no | no | no |
| SCN2A | 1369 | Y | 0 | 0 | no | yes | no | no | yes | no | no | no | no | no | no | no | no |
| SCN2A | 1370 | T | 1 | 0 | no | yes | no | no | yes | no | no | no | no | no | no | no | no |
| SCN2A | 1371 | T | 0 | 0 | no | yes | no | no | yes | no | no | no | no | no | no | no | no |
| SCN2A | 1372 | G | 0 | 0 | no | yes | no | no | yes | no | no | no | no | no | no | no | no |
| SCN2A | 1373 | E | 1 | 0 | no | no | yes | no | yes | no | no | no | no | no | no | no | no |
| SCN2A | 1374 | M | 2 | 0 | no | no | yes | no | yes | no | no | no | no | no | no | no | no |
| SCN2A | 1375 | F | 1 | 0 | no | yes | no | no | yes | no | no | no | no | no | no | no | no |
| SCN2A | 1376 | D | 0 | 0 | no | yes | no | no | yes | no | no | no | no | no | no | no | no |
| SCN2A | 1377 | V | 1 | 0 | no | yes | no | no | yes | no | no | no | no | no | no | no | no |
| SCN2A | 1378 | S | 0 | 0 | no | yes | no | no | yes | no | no | no | no | no | no | no | no |
| SCN2A | 1379 | V | 1 | 0 | no | yes | no | no | yes | no | no | no | no | no | no | no | no |
| SCN2A | 1380 | V | 1 | 0 | no | yes | no | no | yes | no | no | no | no | no | no | no | no |
| SCN2A | 1381 | N | 0 | 0 | no | yes | no | no | yes | no | no | no | no | no | no | no | no |
| SCN2A | 1382 | N | 0 | 0 | no | no | yes | no | yes | no | no | no | no | no | no | no | no |
| SCN2A | 1383 | Y | 0 | 0 | yes | no | no | no | yes | no | no | no | no | no | no | no | no |
| SCN2A | 1384 | S | 0 | 0 | yes | no | no | no | yes | no | no | no | no | no | no | no | no |
| SCN2A | 1385 | E | 0 | 0 | yes | no | no | no | yes | no | no | no | no | no | no | no | no |
| SCN2A | 1386 | C | 0 | 1 | yes | no | no | no | yes | no | no | no | no | no | no | no | no |
| SCN2A | 1387 | K | 0 | 0 | yes | no | no | no | yes | no | no | no | no | no | no | no | no |
| SCN2A | 1388 | A | 0 | 0 | yes | no | no | no | yes | no | no | no | no | no | no | no | no |
| SCN2A | 1389 | L | 0 | 0 | yes | no | no | no | yes | no | no | no | no | no | no | no | no |
| SCN2A | 1390 | I | 1 | 0 | yes | no | no | no | yes | no | no | no | no | no | no | no | no |
| SCN2A | 1391 | E | 0 | 0 | no | yes | no | no | yes | no | no | no | no | no | no | no | no |
| SCN2A | 1392 | S | 1 | 0 | no | yes | no | no | yes | no | no | no | no | no | no | no | no |
| SCN2A | 1393 | N | 0 | 0 | no | yes | no | no | yes | no | no | no | no | no | no | no | no |
| SCN2A | 1394 | Q | 1 | 0 | no | yes | no | no | yes | no | no | no | no | no | no | no | no |

| | | | | | | | | | | | | | | | | | |
|---|---|---|---|---|---|---|---|---|---|---|---|---|---|---|---|---|---|
| SCN2A | 1395 | T | 0 | 0 | no | yes | no | no | yes | no | no | no | no | no | no | no | no |
| SCN2A | 1396 | A | 1 | 0 | no | yes | no | no | yes | no | no | no | no | no | no | no | no |
| SCN2A | 1397 | R | 0 | 0 | no | no | yes | no | yes | no | no | no | no | no | no | no | no |
| SCN2A | 1398 | W | 0 | 0 | no | no | yes | no | yes | no | no | no | no | no | no | no | no |
| SCN2A | 1399 | K | 0 | 0 | no | no | yes | no | yes | no | no | no | no | no | no | no | no |
| SCN2A | 1400 | N | 0 | 0 | no | yes | no | no | yes | no | no | no | no | no | no | no | no |
| SCN2A | 1401 | V | 1 | 0 | no | yes | no | no | yes | no | no | no | no | no | no | no | no |
| SCN2A | 1402 | K | 0 | 0 | no | yes | no | no | yes | no | no | no | no | no | no | no | no |
| SCN2A | 1403 | V | 0 | 0 | no | yes | no | no | yes | no | no | no | no | no | no | no | no |
| SCN2A | 1404 | N | 0 | 0 | no | yes | no | no | yes | no | no | no | no | no | no | no | no |
| SCN2A | 1405 | F | 0 | 0 | no | yes | no | yes | yes | no | no | no | no | no | no | no | no |
| SCN2A | 1406 | D | 0 | 0 | no | yes | no | no | yes | no | no | no | no | no | no | no | no |
| SCN2A | 1407 | N | 0 | 0 | no | yes | no | no | yes | no | no | no | no | no | no | no | no |
| SCN2A | 1408 | V | 0 | 1 | no | yes | no | yes | yes | no | no | no | no | no | no | no | no |
| SCN2A | 1409 | G | 0 | 0 | yes | no | no | yes | no | no | no | yes | no | no | no | no | no |
| SCN2A | 1410 | L | 0 | 0 | yes | no | no | yes | no | no | no | yes | no | no | no | no | no |
| SCN2A | 1411 | G | 0 | 0 | yes | no | no | yes | no | no | no | yes | no | no | no | no | no |
| SCN2A | 1412 | Y | 0 | 0 | yes | no | no | yes | no | no | no | yes | no | no | no | no | no |
| SCN2A | 1413 | L | 0 | 0 | yes | no | no | yes | no | no | no | yes | no | no | no | no | no |
| SCN2A | 1414 | S | 0 | 0 | yes | no | no | yes | no | no | no | yes | no | no | no | no | no |
| SCN2A | 1415 | L | 0 | 1 | yes | no | no | yes | no | no | no | yes | no | no | no | no | no |
| SCN2A | 1416 | L | 0 | 0 | yes | no | no | yes | no | no | no | yes | no | no | no | no | no |
| SCN2A | 1417 | Q | 0 | 0 | yes | no | no | yes | no | no | no | yes | no | no | no | no | no |
| SCN2A | 1418 | V | 0 | 0 | yes | no | no | yes | no | no | no | yes | no | no | no | no | no |
| SCN2A | 1419 | A | 0 | 0 | yes | no | no | yes | no | no | no | yes | no | no | no | no | no |
| SCN2A | 1420 | T | 0 | 0 | yes | no | no | yes | no | no | yes | yes | no | no | no | no | no |
| SCN2A | 1421 | F | 0 | 0 | no | yes | no | yes | no | no | yes | yes | no | no | no | no | no |
| SCN2A | 1422 | K | 0 | 1 | no | yes | no | yes | no | no | yes | yes | no | no | no | no | no |
| SCN2A | 1423 | G | 0 | 0 | no | yes | no | no | no | no | yes | yes | no | no | no | no | no |
| SCN2A | 1424 | W | 0 | 0 | yes | no | no | yes | no | no | yes | yes | no | no | no | no | no |
| SCN2A | 1425 | M | 0 | 0 | yes | no | no | no | no | no | yes | yes | no | no | no | no | no |
| SCN2A | 1426 | D | 0 | 0 | yes | no | no | no | no | no | yes | yes | no | no | no | no | no |
| SCN2A | 1427 | I | 0 | 0 | yes | no | no | no | no | no | no | yes | no | no | no | no | no |

| | | | | | | | | | | | | | | | | | |
|---|---|---|---|---|---|---|---|---|---|---|---|---|---|---|---|---|---|
| SCN2A | 1428 | M | 1 | 0 | yes | no | no | no | no | no | no | yes | no | no | no | no | no |
| SCN2A | 1429 | Y | 0 | 0 | yes | no | no | no | no | no | no | yes | no | no | no | no | no |
| SCN2A | 1430 | A | 1 | 0 | yes | no | no | no | no | no | no | yes | no | no | no | no | no |
| SCN2A | 1431 | A | 0 | 0 | yes | no | no | no | yes | no | no | no | no | no | no | no | no |
| SCN2A | 1432 | V | 0 | 0 | yes | no | no | no | yes | no | no | no | no | no | no | no | no |
| SCN2A | 1433 | D | 0 | 0 | no | yes | no | no | yes | no | no | no | no | no | no | no | no |
| SCN2A | 1434 | S | 0 | 0 | no | yes | no | no | yes | no | no | no | no | no | no | no | no |
| SCN2A | 1435 | R | 0 | 0 | no | yes | no | no | yes | no | no | no | no | no | no | no | no |
| SCN2A | 1436 | N | 0 | 0 | no | yes | no | no | yes | no | no | no | no | no | no | no | no |
| SCN2A | 1437 | V | 0 | 0 | no | yes | no | no | yes | no | no | no | no | no | no | no | no |
| SCN2A | 1438 | E | 0 | 0 | no | yes | no | no | yes | no | no | no | no | no | no | no | no |
| SCN2A | 1439 | L | 0 | 0 | no | no | yes | no | yes | no | no | no | no | no | no | no | no |
| SCN2A | 1440 | Q | 0 | 0 | no | yes | no | no | yes | no | no | no | no | no | no | no | no |
| SCN2A | 1441 | P | 0 | 0 | no | yes | no | no | yes | no | no | no | no | no | no | no | no |
| SCN2A | 1442 | K | 0 | 0 | no | yes | no | no | yes | no | no | no | no | no | no | no | no |
| SCN2A | 1443 | Y | 0 | 0 | no | yes | no | no | yes | no | no | no | no | no | no | no | no |
| SCN2A | 1444 | E | 0 | 0 | no | yes | no | no | yes | no | no | no | no | no | no | no | no |
| SCN2A | 1445 | D | 1 | 0 | no | yes | no | no | yes | no | no | no | no | no | no | no | no |
| SCN2A | 1446 | N | 1 | 0 | no | yes | no | no | yes | no | no | no | no | no | no | no | no |
| SCN2A | 1447 | L | 0 | 0 | yes | no | no | no | yes | no | no | no | no | no | no | no | no |
| SCN2A | 1448 | Y | 0 | 0 | yes | no | no | no | no | no | no | no | no | no | no | no | no |
| SCN2A | 1449 | M | 1 | 0 | yes | no | no | no | no | no | no | no | no | no | no | no | no |
| SCN2A | 1450 | Y | 0 | 0 | yes | no | no | no | no | no | no | no | no | no | no | no | no |
| SCN2A | 1451 | L | 0 | 0 | yes | no | no | yes | no | no | no | no | no | no | no | no | no |
| SCN2A | 1452 | Y | 0 | 0 | yes | no | no | yes | no | no | no | no | no | no | no | no | no |
| SCN2A | 1453 | F | 0 | 0 | yes | no | no | yes | no | no | no | no | no | no | no | no | no |
| SCN2A | 1454 | V | 0 | 0 | yes | no | no | yes | no | no | no | no | no | no | no | no | no |
| SCN2A | 1455 | I | 0 | 1 | yes | no | no | yes | no | no | no | no | no | no | no | no | no |
| SCN2A | 1456 | F | 0 | 0 | yes | no | no | yes | no | no | no | no | no | no | no | no | no |
| SCN2A | 1457 | I | 0 | 0 | yes | no | no | yes | no | no | no | no | no | no | no | no | no |
| SCN2A | 1458 | I | 0 | 0 | yes | no | no | yes | no | no | no | no | no | no | no | no | no |
| SCN2A | 1459 | F | 0 | 0 | yes | no | no | yes | no | no | no | no | no | no | no | no | no |
| SCN2A | 1460 | G | 0 | 1 | yes | no | no | yes | no | no | no | no | no | no | no | no | no |

| | | | | | | | | | | | | | | | | | |
|---|---|---|---|---|---|---|---|---|---|---|---|---|---|---|---|---|---|
| SCN2A | 1461 | S | 0 | 0 | yes | no | no | yes | no | no | no | no | no | no | no | no | no |
| SCN2A | 1462 | F | 0 | 0 | yes | no | no | yes | no | no | no | no | no | no | no | no | no |
| SCN2A | 1463 | F | 0 | 0 | yes | no | no | yes | no | no | no | no | no | no | no | no | no |
| SCN2A | 1464 | T | 0 | 1 | yes | no | no | yes | no | no | no | no | no | no | no | no | no |
| SCN2A | 1465 | L | 0 | 1 | yes | no | no | yes | no | no | yes | no | no | no | no | no | no |
| SCN2A | 1466 | N | 0 | 0 | yes | no | no | yes | no | no | no | no | no | no | no | no | no |
| SCN2A | 1467 | L | 0 | 0 | yes | no | no | yes | no | no | no | no | no | no | no | no | no |
| SCN2A | 1468 | F | 0 | 0 | yes | no | no | yes | no | no | no | no | no | no | no | no | no |
| SCN2A | 1469 | I | 0 | 0 | yes | no | no | yes | no | no | yes | no | no | no | no | no | no |
| SCN2A | 1470 | G | 0 | 1 | yes | no | no | yes | no | yes | no | no | no | no | no | no | no |
| SCN2A | 1471 | V | 0 | 0 | yes | no | no | yes | no | yes | no | no | no | no | no | no | no |
| SCN2A | 1472 | I | 1 | 0 | yes | no | no | yes | no | yes | no | no | no | no | no | no | no |
| SCN2A | 1473 | I | 0 | 2 | yes | no | no | yes | no | yes | yes | no | no | no | no | no | no |
| SCN2A | 1474 | D | 0 | 0 | yes | no | no | no | no | yes | no | no | no | no | no | no | no |
| SCN2A | 1475 | N | 0 | 0 | yes | no | no | no | no | yes | no | no | no | no | no | no | no |
| SCN2A | 1476 | F | 0 | 0 | yes | no | no | no | no | yes | no | no | no | no | no | no | no |
| SCN2A | 1477 | N | 1 | 0 | yes | no | no | no | no | yes | yes | no | no | no | no | no | no |
| SCN2A | 1478 | Q | 0 | 0 | yes | no | no | no | no | yes | no | no | no | no | no | no | no |
| SCN2A | 1479 | Q | 0 | 2 | yes | no | no | no | no | yes | no | no | no | no | no | no | no |
| SCN2A | 1480 | K | 0 | 1 | yes | no | no | no | no | yes | yes | no | no | no | no | no | no |
| SCN2A | 1481 | K | 0 | 0 | yes | no | no | no | no | yes | no | no | no | no | no | no | no |
| SCN2A | 1482 | K | 0 | 0 | yes | no | no | no | no | yes | no | no | no | no | no | no | no |
| SCN2A | 1483 | F | 0 | 0 | no | yes | no | no | no | yes | no | no | no | no | no | no | no |
| SCN2A | 1484 | G | 0 | 0 | no | yes | no | no | no | yes | no | no | no | no | no | no | no |
| SCN2A | 1485 | G | 0 | 0 | no | yes | no | no | no | yes | no | no | no | no | no | no | no |
| SCN2A | 1486 | Q | 0 | 0 | no | yes | no | no | no | yes | no | no | no | no | no | no | no |
| SCN2A | 1487 | D | 0 | 0 | no | yes | no | no | no | yes | no | no | no | no | no | no | no |
| SCN2A | 1488 | I | 0 | 1 | no | yes | no | no | no | yes | no | no | no | no | no | no | no |
| SCN2A | 1489 | F | 0 | 0 | no | yes | no | yes | no | yes | no | no | no | no | no | no | no |
| SCN2A | 1490 | M | 0 | 1 | no | yes | no | no | no | yes | no | no | no | no | no | no | no |
| SCN2A | 1491 | T | 0 | 0 | no | yes | no | no | no | yes | no | no | no | no | no | no | no |
| SCN2A | 1492 | E | 0 | 0 | yes | no | no | no | no | yes | no | no | no | no | no | no | no |
| SCN2A | 1493 | E | 0 | 0 | yes | no | no | no | no | yes | no | no | no | no | no | no | no |

| | | | | | | | | | | | | | | | | | |
|---|---|---|---|---|---|---|---|---|---|---|---|---|---|---|---|---|---|
| SCN2A | 1494 | Q | 0 | 0 | yes | no | no | no | no | yes | no | no | no | no | no | no | no |
| SCN2A | 1495 | K | 0 | 0 | yes | no | no | no | no | yes | no | no | no | no | no | no | no |
| SCN2A | 1496 | K | 0 | 0 | yes | no | no | no | no | yes | no | no | no | no | no | no | no |
| SCN2A | 1497 | Y | 0 | 0 | no | yes | no | no | no | yes | no | no | no | no | no | no | no |
| SCN2A | 1498 | Y | 0 | 1 | no | yes | no | no | no | yes | no | no | no | no | no | no | no |
| SCN2A | 1499 | N | 0 | 0 | no | yes | no | no | no | yes | no | no | no | no | no | no | no |
| SCN2A | 1500 | A | 0 | 1 | no | yes | no | no | no | yes | no | no | no | no | no | no | no |
| SCN2A | 1501 | M | 0 | 2 | yes | no | no | no | no | yes | no | no | no | no | no | no | no |
| SCN2A | 1502 | K | 0 | 0 | yes | no | no | no | no | yes | no | no | no | no | no | no | no |
| SCN2A | 1503 | K | 1 | 0 | yes | no | no | no | no | yes | no | no | no | no | no | no | no |
| SCN2A | 1504 | L | 0 | 0 | yes | no | no | no | no | yes | no | no | no | no | no | no | no |
| SCN2A | 1505 | G | 2 | 0 | yes | no | no | no | no | yes | no | no | no | no | no | no | no |
| SCN2A | 1506 | S | 2 | 0 | no | yes | no | no | no | yes | no | no | no | no | no | no | no |
| SCN2A | 1507 | K | 0 | 0 | no | yes | no | no | no | yes | no | no | no | no | no | no | no |
| SCN2A | 1508 | K | 0 | 0 | no | yes | no | no | no | yes | no | no | no | no | no | no | no |
| SCN2A | 1509 | P | 0 | 0 | no | yes | no | no | no | yes | no | no | no | no | no | no | no |
| SCN2A | 1510 | Q | 0 | 0 | no | yes | no | no | no | yes | no | no | no | no | no | no | no |
| SCN2A | 1511 | K | 0 | 0 | no | yes | no | no | no | yes | no | no | no | no | no | no | no |
| SCN2A | 1512 | P | 0 | 0 | no | yes | no | no | no | yes | no | no | no | no | no | no | no |
| SCN2A | 1513 | I | 0 | 0 | no | yes | no | no | no | yes | no | no | no | no | no | no | no |
| SCN2A | 1514 | P | 0 | 0 | no | yes | no | no | no | yes | no | no | no | no | no | no | no |
| SCN2A | 1515 | R | 1 | 0 | no | yes | no | no | no | yes | no | no | no | no | no | no | no |
| SCN2A | 1516 | P | 0 | 0 | no | yes | no | no | no | yes | no | no | no | no | no | no | no |
| SCN2A | 1517 | A | 0 | 0 | no | yes | no | no | no | yes | no | no | no | no | no | no | no |
| SCN2A | 1518 | N | 0 | 0 | no | yes | no | no | no | yes | no | no | no | no | no | no | no |
| SCN2A | 1519 | K | 0 | 0 | no | yes | no | no | no | yes | no | no | no | no | no | no | no |
| SCN2A | 1520 | F | 0 | 0 | yes | no | no | no | no | yes | no | no | no | no | no | no | no |
| SCN2A | 1521 | Q | 0 | 0 | yes | no | no | no | no | yes | no | no | no | no | no | no | no |
| SCN2A | 1522 | G | 1 | 1 | yes | no | no | no | no | yes | no | no | no | no | no | no | no |
| SCN2A | 1523 | M | 1 | 0 | yes | no | no | no | no | yes | no | no | no | no | no | no | no |
| SCN2A | 1524 | V | 0 | 0 | yes | no | no | no | no | yes | no | no | no | no | no | no | no |
| SCN2A | 1525 | F | 0 | 0 | yes | no | no | no | no | yes | no | no | no | no | no | no | no |
| SCN2A | 1526 | D | 0 | 0 | yes | no | no | no | no | yes | no | no | no | no | no | no | no |

| | | | | | | | | | | | | | | | | | |
|---|---|---|---|---|---|---|---|---|---|---|---|---|---|---|---|---|---|
| SCN2A | 1527 | F | 0 | 0 | yes | no | no | no | no | yes | no | no | no | no | no | no | no |
| SCN2A | 1528 | V | 0 | 0 | yes | no | no | yes | no | yes | no | no | no | no | no | no | no |
| SCN2A | 1529 | T | 0 | 0 | no | yes | no | no | no | yes | no | no | no | no | no | no | no |
| SCN2A | 1530 | K | 0 | 0 | no | yes | no | no | no | yes | no | no | no | no | no | no | no |
| SCN2A | 1531 | Q | 0 | 1 | yes | no | no | no | no | yes | no | no | no | no | no | no | no |
| SCN2A | 1532 | V | 0 | 0 | yes | no | no | yes | no | yes | no | no | no | no | no | no | no |
| SCN2A | 1533 | F | 0 | 0 | yes | no | no | yes | no | no | no | no | no | no | no | no | no |
| SCN2A | 1534 | D | 0 | 0 | yes | no | no | yes | no | no | no | no | no | no | no | no | no |
| SCN2A | 1535 | I | 0 | 0 | yes | no | no | yes | no | no | no | no | no | no | no | no | no |
| SCN2A | 1536 | S | 0 | 1 | yes | no | no | yes | no | no | no | no | no | no | no | no | no |
| SCN2A | 1537 | I | 0 | 1 | yes | no | no | yes | no | no | no | no | no | no | no | no | no |
| SCN2A | 1538 | M | 0 | 0 | yes | no | no | yes | no | no | no | no | no | no | no | no | no |
| SCN2A | 1539 | I | 2 | 0 | yes | no | no | yes | no | no | no | no | no | no | no | no | no |
| SCN2A | 1540 | L | 1 | 0 | yes | no | no | yes | no | no | no | no | no | no | no | no | no |
| SCN2A | 1541 | I | 0 | 0 | yes | no | no | yes | no | no | no | no | no | no | no | no | no |
| SCN2A | 1542 | C | 0 | 0 | yes | no | no | yes | no | no | no | no | no | no | no | no | no |
| SCN2A | 1543 | L | 0 | 0 | yes | no | no | yes | no | no | no | no | no | no | no | no | no |
| SCN2A | 1544 | N | 0 | 1 | yes | no | no | yes | no | no | no | no | no | no | no | no | no |
| SCN2A | 1545 | M | 0 | 2 | yes | no | no | yes | no | no | no | no | no | no | no | no | no |
| SCN2A | 1546 | V | 0 | 0 | yes | no | no | yes | no | no | no | no | no | no | no | no | no |
| SCN2A | 1547 | T | 0 | 0 | yes | no | no | yes | no | no | no | no | no | no | no | no | no |
| SCN2A | 1548 | M | 0 | 3 | yes | no | no | yes | no | no | no | no | no | no | no | no | no |
| SCN2A | 1549 | M | 0 | 0 | yes | no | no | yes | no | no | no | no | no | no | no | no | no |
| SCN2A | 1550 | V | 0 | 0 | no | yes | no | yes | no | no | no | no | no | no | no | no | no |
| SCN2A | 1551 | E | 0 | 0 | no | yes | no | yes | yes | no | no | no | no | no | no | no | no |
| SCN2A | 1552 | T | 0 | 0 | no | yes | no | no | yes | no | no | no | no | no | no | no | no |
| SCN2A | 1553 | D | 0 | 0 | no | yes | no | no | yes | no | no | no | no | no | no | no | no |
| SCN2A | 1554 | D | 0 | 0 | no | yes | no | no | yes | no | no | no | no | no | no | no | no |
| SCN2A | 1555 | Q | 0 | 0 | no | yes | no | no | yes | no | no | no | no | no | no | no | no |
| SCN2A | 1556 | S | 0 | 0 | no | yes | no | no | yes | no | no | no | no | no | no | no | no |
| SCN2A | 1557 | Q | 0 | 0 | yes | no | no | no | yes | no | no | no | no | no | no | no | no |
| SCN2A | 1558 | E | 1 | 0 | yes | no | no | no | yes | no | no | no | no | no | no | no | no |
| SCN2A | 1559 | M | 1 | 0 | yes | no | no | no | yes | no | no | no | no | no | no | no | no |

| | | | | | | | | | | | | | | | | | |
|---|---|---|---|---|---|---|---|---|---|---|---|---|---|---|---|---|---|
| SCN2A | 1560 | T | 0 | 0 | yes | no | no | no | yes | no | no | no | no | no | no | no | no |
| SCN2A | 1561 | N | 0 | 0 | yes | no | no | no | yes | no | no | no | no | no | no | no | no |
| SCN2A | 1562 | I | 1 | 0 | yes | no | no | yes | no | no | no | no | no | no | no | no | no |
| SCN2A | 1563 | L | 0 | 2 | yes | no | no | yes | no | no | no | no | no | no | no | no | no |
| SCN2A | 1564 | Y | 0 | 0 | yes | no | no | yes | no | no | no | no | no | no | no | no | no |
| SCN2A | 1565 | W | 0 | 0 | yes | no | no | yes | no | no | no | no | no | no | no | no | no |
| SCN2A | 1566 | I | 0 | 0 | yes | no | no | yes | no | no | no | no | no | no | no | no | no |
| SCN2A | 1567 | N | 0 | 1 | yes | no | no | yes | no | no | no | no | no | no | no | no | no |
| SCN2A | 1568 | L | 1 | 0 | yes | no | no | yes | no | no | no | no | no | no | no | no | no |
| SCN2A | 1569 | V | 0 | 0 | yes | no | no | yes | no | no | no | no | no | no | no | no | no |
| SCN2A | 1570 | F | 0 | 0 | yes | no | no | yes | no | no | no | no | no | no | no | no | no |
| SCN2A | 1571 | I | 0 | 0 | yes | no | no | yes | no | no | no | no | no | no | no | no | no |
| SCN2A | 1572 | V | 0 | 0 | yes | no | no | yes | no | no | no | no | no | no | no | no | no |
| SCN2A | 1573 | L | 0 | 1 | yes | no | no | yes | no | no | no | no | no | no | no | no | no |
| SCN2A | 1574 | F | 0 | 0 | yes | no | no | yes | no | no | no | no | no | no | no | no | no |
| SCN2A | 1575 | T | 0 | 0 | yes | no | no | yes | no | no | no | no | no | no | no | no | no |
| SCN2A | 1576 | G | 0 | 1 | yes | no | no | yes | no | no | no | no | no | no | no | no | no |
| SCN2A | 1577 | E | 0 | 0 | yes | no | no | yes | no | no | no | no | no | no | no | no | no |
| SCN2A | 1578 | C | 0 | 0 | yes | no | no | yes | no | no | no | no | no | no | no | no | no |
| SCN2A | 1579 | V | 1 | 0 | yes | no | no | yes | no | no | no | no | no | no | no | no | no |
| SCN2A | 1580 | L | 0 | 0 | yes | no | no | yes | no | no | no | no | no | no | no | no | no |
| SCN2A | 1581 | K | 0 | 0 | yes | no | no | yes | no | yes | no | no | no | no | no | no | no |
| SCN2A | 1582 | L | 0 | 1 | yes | no | no | yes | no | yes | no | no | no | no | no | no | no |
| SCN2A | 1583 | I | 1 | 0 | yes | no | no | yes | no | yes | no | no | no | no | no | no | no |
| SCN2A | 1584 | S | 0 | 1 | yes | no | no | no | no | yes | no | no | no | no | no | no | no |
| SCN2A | 1585 | L | 0 | 0 | no | yes | no | no | no | yes | no | no | no | no | no | no | no |
| SCN2A | 1586 | R | 0 | 0 | no | yes | no | no | no | yes | no | no | no | no | no | no | no |
| SCN2A | 1587 | Y | 0 | 0 | no | yes | no | no | no | yes | no | no | no | no | no | no | no |
| SCN2A | 1588 | Y | 0 | 0 | no | yes | no | no | no | yes | no | no | no | no | no | no | no |
| SCN2A | 1589 | Y | 0 | 1 | no | yes | no | yes | no | yes | no | no | no | no | no | no | no |
| SCN2A | 1590 | F | 0 | 0 | no | yes | no | yes | no | yes | no | no | no | no | no | no | no |
| SCN2A | 1591 | T | 0 | 0 | no | yes | no | no | no | yes | no | no | no | no | no | no | no |
| SCN2A | 1592 | I | 1 | 0 | no | yes | no | no | no | yes | no | no | no | no | no | no | no |

| | | | | | | | | | | | | | | | | | |
|---|---|---|---|---|---|---|---|---|---|---|---|---|---|---|---|---|---|
| SCN2A | 1593 | G | 0 | 1 | yes | no | no | yes | no | no | no | no | no | no | no | no | no |
| SCN2A | 1594 | W | 0 | 1 | yes | no | no | yes | no | no | no | no | no | no | no | no | no |
| SCN2A | 1595 | N | 0 | 0 | yes | no | no | yes | no | no | no | no | no | no | no | no | no |
| SCN2A | 1596 | I | 0 | 0 | yes | no | no | yes | no | no | no | no | no | no | no | no | no |
| SCN2A | 1597 | F | 0 | 1 | yes | no | no | yes | no | no | no | no | no | no | no | no | no |
| SCN2A | 1598 | D | 0 | 1 | yes | no | no | yes | no | no | no | no | no | no | no | no | no |
| SCN2A | 1599 | F | 0 | 0 | yes | no | no | yes | no | no | no | no | no | no | no | no | no |
| SCN2A | 1600 | V | 0 | 0 | yes | no | no | yes | no | no | no | no | no | no | no | no | no |
| SCN2A | 1601 | V | 0 | 0 | yes | no | no | yes | no | no | no | no | no | no | no | no | no |
| SCN2A | 1602 | V | 1 | 0 | yes | no | no | yes | no | no | no | no | no | no | no | no | no |
| SCN2A | 1603 | I | 0 | 0 | yes | no | no | yes | no | no | no | no | no | no | no | no | no |
| SCN2A | 1604 | L | 0 | 0 | yes | no | no | yes | no | no | no | no | no | no | no | no | no |
| SCN2A | 1605 | S | 0 | 0 | yes | no | no | yes | no | no | no | no | no | no | no | no | no |
| SCN2A | 1606 | I | 0 | 0 | yes | no | no | yes | no | no | no | no | no | no | no | no | no |
| SCN2A | 1607 | V | 0 | 1 | yes | no | no | yes | no | no | no | no | no | no | no | no | no |
| SCN2A | 1608 | G | 0 | 0 | yes | no | no | yes | no | no | no | no | no | no | no | no | no |
| SCN2A | 1609 | M | 0 | 0 | yes | no | no | yes | no | no | no | no | no | no | no | no | no |
| SCN2A | 1610 | F | 0 | 0 | yes | no | no | yes | no | no | no | no | no | no | no | no | no |
| SCN2A | 1611 | L | 0 | 0 | yes | no | no | yes | yes | no | no | no | no | no | no | no | no |
| SCN2A | 1612 | A | 0 | 1 | yes | no | no | yes | yes | no | no | no | no | no | no | no | no |
| SCN2A | 1613 | E | 0 | 0 | yes | no | no | yes | yes | no | no | no | no | no | no | no | no |
| SCN2A | 1614 | L | 0 | 1 | yes | no | no | yes | yes | no | no | no | no | no | no | no | no |
| SCN2A | 1615 | I | 1 | 0 | yes | no | no | no | yes | no | no | no | no | no | no | no | no |
| SCN2A | 1616 | E | 0 | 0 | no | yes | no | no | yes | no | no | no | no | no | no | no | no |
| SCN2A | 1617 | K | 0 | 0 | no | yes | no | no | yes | no | no | no | no | no | no | no | no |
| SCN2A | 1618 | Y | 0 | 0 | no | yes | no | no | yes | no | no | no | no | no | no | no | no |
| SCN2A | 1619 | F | 0 | 1 | no | yes | no | no | yes | no | no | no | no | no | no | no | no |
| SCN2A | 1620 | V | 0 | 0 | no | yes | no | no | yes | no | no | no | no | no | no | no | no |
| SCN2A | 1621 | S | 0 | 1 | no | yes | no | no | yes | no | no | no | no | no | no | no | no |
| SCN2A | 1622 | P | 0 | 2 | yes | no | no | no | yes | no | no | no | no | no | no | no | no |
| SCN2A | 1623 | T | 0 | 1 | yes | no | no | no | yes | no | no | no | no | no | no | no | no |
| SCN2A | 1624 | L | 0 | 0 | yes | no | no | yes | no | no | no | no | no | no | no | no | no |
| SCN2A | 1625 | F | 0 | 0 | yes | no | no | yes | no | no | no | no | no | no | no | no | no |

| | | | | | | | | | | | | | | | | | |
|---|---|---|---|---|---|---|---|---|---|---|---|---|---|---|---|---|---|
| SCN2A | 1626 | R | 0 | 1 | yes | no | no | no | no | no | no | no | no | no | no | no | no |
| SCN2A | 1627 | V | 0 | 1 | yes | no | no | yes | no | no | no | no | no | no | no | no | no |
| SCN2A | 1628 | I | 0 | 0 | yes | no | no | yes | no | no | no | no | no | no | no | no | no |
| SCN2A | 1629 | R | 0 | 1 | yes | no | no | yes | no | no | no | no | no | no | no | no | no |
| SCN2A | 1630 | L | 0 | 0 | yes | no | no | yes | no | no | no | no | no | no | no | no | no |
| SCN2A | 1631 | A | 0 | 0 | yes | no | no | yes | no | no | no | no | no | no | no | no | no |
| SCN2A | 1632 | R | 0 | 1 | yes | no | no | yes | no | no | no | no | no | no | no | no | no |
| SCN2A | 1633 | I | 0 | 0 | yes | no | no | yes | no | no | no | no | no | no | no | no | no |
| SCN2A | 1634 | G | 0 | 1 | no | yes | no | yes | no | no | no | no | no | no | no | no | no |
| SCN2A | 1635 | R | 0 | 1 | no | yes | no | yes | no | no | no | no | no | no | no | no | no |
| SCN2A | 1636 | I | 0 | 1 | yes | no | no | yes | no | no | no | no | no | no | no | no | no |
| SCN2A | 1637 | L | 0 | 0 | yes | no | no | yes | no | no | no | no | no | no | no | no | no |
| SCN2A | 1638 | R | 0 | 0 | yes | no | no | yes | no | no | no | no | no | no | no | no | no |
| SCN2A | 1639 | L | 0 | 0 | yes | no | no | yes | no | no | no | no | no | no | no | no | no |
| SCN2A | 1640 | I | 0 | 2 | yes | no | no | yes | no | no | no | no | no | no | no | no | no |
| SCN2A | 1641 | K | 1 | 1 | yes | no | no | yes | no | yes | no | no | no | no | no | no | no |
| SCN2A | 1642 | G | 1 | 0 | no | yes | no | yes | no | yes | no | no | no | no | no | no | no |
| SCN2A | 1643 | A | 0 | 0 | no | yes | no | yes | no | yes | no | no | no | no | no | no | no |
| SCN2A | 1644 | K | 0 | 0 | no | yes | no | no | no | yes | no | no | no | no | no | no | no |
| SCN2A | 1645 | G | 0 | 0 | no | yes | no | yes | no | yes | no | no | no | no | no | no | no |
| SCN2A | 1646 | I | 0 | 0 | yes | no | no | yes | no | yes | no | no | no | no | no | no | no |
| SCN2A | 1647 | R | 0 | 0 | yes | no | no | yes | no | yes | no | no | no | no | no | no | no |
| SCN2A | 1648 | T | 0 | 0 | yes | no | no | no | no | yes | no | no | no | no | no | no | no |
| SCN2A | 1649 | L | 0 | 0 | yes | no | no | yes | no | yes | no | no | no | no | no | no | no |
| SCN2A | 1650 | L | 0 | 1 | yes | no | no | yes | no | yes | no | no | no | no | no | no | no |
| SCN2A | 1651 | F | 0 | 2 | yes | no | no | no | no | yes | no | no | no | no | no | no | no |
| SCN2A | 1652 | A | 0 | 1 | yes | no | no | yes | no | yes | no | no | no | no | no | no | no |
| SCN2A | 1653 | L | 0 | 0 | yes | no | no | yes | no | yes | no | no | no | no | no | no | no |
| SCN2A | 1654 | M | 0 | 0 | yes | no | no | yes | no | yes | no | no | no | no | no | no | no |
| SCN2A | 1655 | M | 0 | 1 | no | yes | no | no | no | yes | no | no | no | no | no | no | no |
| SCN2A | 1656 | S | 0 | 1 | no | yes | no | yes | no | yes | no | no | no | no | no | no | no |
| SCN2A | 1657 | L | 0 | 0 | yes | no | no | yes | no | yes | no | no | no | no | no | no | no |
| SCN2A | 1658 | P | 0 | 0 | yes | no | no | yes | no | yes | no | no | no | no | no | no | no |

| | | | | | | | | | | | | | | | | | |
|---|---|---|---|---|---|---|---|---|---|---|---|---|---|---|---|---|---|
| SCN2A | 1659 | A | 0 | 1 | yes | no | no | yes | no | yes | no | no | no | no | no | no | no |
| SCN2A | 1660 | L | 0 | 1 | yes | no | no | yes | no | no | no | no | no | no | no | no | no |
| SCN2A | 1661 | F | 0 | 0 | yes | no | no | yes | no | no | no | no | no | no | no | no | no |
| SCN2A | 1662 | N | 0 | 0 | yes | no | no | yes | no | no | no | no | no | no | no | no | no |
| SCN2A | 1663 | I | 0 | 0 | yes | no | no | yes | no | no | no | no | no | no | no | no | no |
| SCN2A | 1664 | G | 0 | 0 | yes | no | no | yes | no | no | no | no | no | no | no | no | no |
| SCN2A | 1665 | L | 0 | 1 | yes | no | no | yes | no | no | no | no | no | no | no | no | no |
| SCN2A | 1666 | L | 0 | 0 | yes | no | no | yes | no | no | no | no | no | no | no | no | no |
| SCN2A | 1667 | L | 0 | 0 | yes | no | no | yes | no | no | no | no | no | no | no | no | no |
| SCN2A | 1668 | F | 0 | 0 | yes | no | no | yes | no | no | no | no | no | no | no | no | no |
| SCN2A | 1669 | L | 0 | 0 | yes | no | no | yes | no | no | no | no | no | no | no | no | no |
| SCN2A | 1670 | V | 0 | 0 | yes | no | no | yes | no | no | no | no | no | no | no | no | no |
| SCN2A | 1671 | M | 0 | 0 | yes | no | no | yes | no | no | no | no | no | no | no | no | no |
| SCN2A | 1672 | F | 0 | 0 | yes | no | no | yes | no | no | no | no | no | no | no | no | no |
| SCN2A | 1673 | I | 0 | 0 | yes | no | no | yes | no | no | no | no | no | no | no | no | no |
| SCN2A | 1674 | Y | 0 | 0 | yes | no | no | yes | no | no | no | no | no | no | no | no | no |
| SCN2A | 1675 | A | 0 | 0 | yes | no | no | yes | no | no | no | no | no | no | no | no | no |
| SCN2A | 1676 | I | 1 | 0 | yes | no | no | yes | no | no | no | no | no | no | no | no | no |
| SCN2A | 1677 | F | 1 | 0 | yes | no | no | yes | no | no | no | no | no | no | no | no | no |
| SCN2A | 1678 | G | 0 | 0 | yes | no | no | yes | yes | no | no | no | no | no | no | no | no |
| SCN2A | 1679 | M | 0 | 0 | yes | no | no | no | yes | no | no | no | no | no | no | no | no |
| SCN2A | 1680 | S | 0 | 0 | yes | no | no | no | yes | no | no | no | no | no | no | no | no |
| SCN2A | 1681 | N | 0 | 0 | yes | no | no | no | yes | no | no | no | no | no | no | no | no |
| SCN2A | 1682 | F | 0 | 1 | no | yes | no | no | yes | no | no | no | no | no | no | no | no |
| SCN2A | 1683 | A | 0 | 0 | no | yes | no | no | yes | no | no | no | no | no | no | no | no |
| SCN2A | 1684 | Y | 0 | 0 | no | yes | no | no | yes | no | no | no | no | no | no | no | no |
| SCN2A | 1685 | V | 0 | 0 | no | yes | no | no | yes | no | no | no | no | no | no | no | no |
| SCN2A | 1686 | K | 0 | 0 | no | yes | no | no | yes | no | no | no | no | no | no | no | no |
| SCN2A | 1687 | R | 0 | 0 | no | yes | no | no | yes | no | no | no | no | no | no | no | no |
| SCN2A | 1688 | E | 0 | 0 | no | yes | no | no | yes | no | no | no | no | no | no | no | no |
| SCN2A | 1689 | V | 1 | 0 | no | yes | no | no | yes | no | no | no | no | no | no | no | no |
| SCN2A | 1690 | G | 0 | 0 | no | yes | no | no | yes | no | no | no | no | no | no | no | no |
| SCN2A | 1691 | I | 0 | 0 | no | yes | no | no | yes | no | no | no | no | no | no | no | no |

| | | | | | | | | | | | | | | | | | |
|---|---|---|---|---|---|---|---|---|---|---|---|---|---|---|---|---|---|
| SCN2A | 1692 | D | 0 | 0 | no | yes | no | no | yes | no | no | no | no | no | no | no | no |
| SCN2A | 1693 | D | 0 | 0 | no | yes | no | no | yes | no | no | no | no | no | no | no | no |
| SCN2A | 1694 | M | 0 | 0 | no | yes | no | no | yes | no | no | no | no | no | no | no | no |
| SCN2A | 1695 | F | 0 | 0 | no | yes | no | no | yes | no | no | no | no | no | no | no | no |
| SCN2A | 1696 | N | 0 | 0 | no | yes | no | no | yes | no | no | no | no | no | no | no | no |
| SCN2A | 1697 | F | 0 | 0 | no | yes | no | no | yes | no | no | no | no | no | no | no | no |
| SCN2A | 1698 | E | 0 | 0 | no | yes | no | no | yes | no | no | no | no | no | no | no | no |
| SCN2A | 1699 | T | 0 | 0 | no | yes | no | no | yes | no | no | no | no | no | no | no | no |
| SCN2A | 1700 | F | 0 | 0 | yes | no | no | yes | no | no | no | yes | no | no | no | no | no |
| SCN2A | 1701 | G | 0 | 0 | yes | no | no | yes | no | no | no | yes | no | no | no | no | no |
| SCN2A | 1702 | N | 0 | 0 | yes | no | no | no | no | no | no | yes | no | no | no | no | no |
| SCN2A | 1703 | S | 0 | 0 | yes | no | no | yes | no | no | no | yes | no | no | no | no | no |
| SCN2A | 1704 | M | 0 | 0 | yes | no | no | yes | no | no | no | yes | no | no | no | no | no |
| SCN2A | 1705 | I | 0 | 0 | yes | no | no | yes | no | no | no | yes | no | no | no | no | no |
| SCN2A | 1706 | C | 0 | 1 | yes | no | no | yes | no | no | no | yes | no | no | no | no | no |
| SCN2A | 1707 | L | 0 | 0 | yes | no | no | yes | no | no | no | yes | no | no | no | no | no |
| SCN2A | 1708 | F | 0 | 0 | yes | no | no | yes | no | no | no | yes | no | no | no | no | no |
| SCN2A | 1709 | Q | 0 | 0 | yes | no | no | yes | no | no | no | yes | no | no | no | no | no |
| SCN2A | 1710 | I | 0 | 0 | yes | no | no | yes | no | no | no | yes | no | no | no | no | no |
| SCN2A | 1711 | T | 0 | 1 | no | yes | no | yes | no | no | no | yes | no | no | no | no | no |
| SCN2A | 1712 | T | 0 | 0 | no | yes | no | yes | no | no | no | yes | no | no | no | no | no |
| SCN2A | 1713 | S | 0 | 0 | no | yes | no | yes | no | no | no | yes | no | no | no | no | no |
| SCN2A | 1714 | A | 0 | 0 | no | yes | no | yes | no | no | no | yes | no | no | no | no | no |
| SCN2A | 1715 | G | 0 | 0 | no | yes | no | yes | no | no | no | yes | no | no | no | no | no |
| SCN2A | 1716 | W | 0 | 0 | yes | no | no | yes | no | no | no | yes | no | no | no | no | no |
| SCN2A | 1717 | D | 0 | 0 | yes | no | no | no | no | no | no | yes | no | no | no | no | no |
| SCN2A | 1718 | G | 1 | 0 | yes | no | no | no | no | no | no | yes | no | no | no | no | no |
| SCN2A | 1719 | L | 0 | 0 | yes | no | no | no | no | no | no | yes | no | no | no | no | no |
| SCN2A | 1720 | L | 0 | 0 | yes | no | no | no | no | no | no | yes | no | no | no | no | no |
| SCN2A | 1721 | A | 0 | 0 | no | yes | no | no | no | no | no | yes | no | no | no | no | no |
| SCN2A | 1722 | P | 0 | 0 | no | yes | no | no | no | no | no | yes | no | no | no | no | no |
| SCN2A | 1723 | I | 0 | 0 | no | yes | no | no | yes | no | no | no | no | no | no | no | no |
| SCN2A | 1724 | L | 0 | 0 | no | yes | no | no | yes | no | no | no | no | no | no | no | no |

| | | | | | | | | | | | | | | | | | |
|---|---|---|---|---|---|---|---|---|---|---|---|---|---|---|---|---|---|
| SCN2A | 1725 | N | 1 | 0 | no | yes | no | no | yes | no | no | no | no | no | no | no | no |
| SCN2A | 1726 | S | 0 | 0 | no | yes | no | no | yes | no | no | no | no | no | no | no | no |
| SCN2A | 1727 | G | 0 | 0 | no | yes | no | no | yes | no | no | no | no | no | no | no | no |
| SCN2A | 1728 | P | 0 | 0 | no | yes | no | no | yes | no | no | no | no | no | no | no | no |
| SCN2A | 1729 | P | 0 | 0 | no | yes | no | no | yes | no | no | no | no | no | no | no | no |
| SCN2A | 1730 | D | 1 | 0 | no | yes | no | no | yes | no | no | no | no | no | no | no | no |
| SCN2A | 1731 | C | 0 | 0 | no | yes | no | no | yes | no | no | no | no | no | no | no | no |
| SCN2A | 1732 | D | 0 | 0 | no | yes | no | no | yes | no | no | no | no | no | no | no | no |
| SCN2A | 1733 | P | 0 | 0 | no | yes | no | no | yes | no | no | no | no | no | no | no | no |
| SCN2A | 1734 | D | 0 | 0 | no | yes | no | no | yes | no | no | no | no | no | no | no | no |
| SCN2A | 1735 | K | 0 | 0 | no | yes | no | no | yes | no | no | no | no | no | no | no | no |
| SCN2A | 1736 | D | 1 | 0 | no | yes | no | no | yes | no | no | no | no | no | no | no | no |
| SCN2A | 1737 | H | 0 | 0 | no | yes | no | no | yes | no | no | no | no | no | no | no | no |
| SCN2A | 1738 | P | 0 | 0 | no | yes | no | no | yes | no | no | no | no | no | no | no | no |
| SCN2A | 1739 | G | 0 | 0 | no | yes | no | no | yes | no | no | no | no | no | no | no | no |
| SCN2A | 1740 | S | 1 | 0 | no | yes | no | no | yes | no | no | no | no | no | no | no | no |
| SCN2A | 1741 | S | 1 | 0 | no | yes | no | no | yes | no | no | no | no | no | no | no | no |
| SCN2A | 1742 | V | 0 | 0 | no | yes | no | no | yes | no | no | no | no | no | no | no | no |
| SCN2A | 1743 | K | 0 | 0 | no | yes | no | no | yes | no | no | no | no | no | no | no | no |
| SCN2A | 1744 | G | 0 | 1 | no | yes | no | no | yes | no | no | no | no | no | no | no | no |
| SCN2A | 1745 | D | 0 | 0 | no | yes | no | no | yes | no | no | no | no | no | no | no | no |
| SCN2A | 1746 | C | 0 | 0 | no | yes | no | no | yes | no | no | no | no | no | no | no | no |
| SCN2A | 1747 | G | 0 | 0 | no | yes | no | no | yes | no | no | no | no | no | no | no | no |
| SCN2A | 1748 | N | 0 | 1 | no | yes | no | no | yes | no | no | no | no | no | no | no | no |
| SCN2A | 1749 | P | 0 | 0 | yes | no | no | no | yes | no | no | no | no | no | no | no | no |
| SCN2A | 1750 | S | 0 | 0 | yes | no | no | no | yes | no | no | no | no | no | no | no | no |
| SCN2A | 1751 | V | 0 | 0 | yes | no | no | yes | yes | no | no | no | no | no | no | no | no |
| SCN2A | 1752 | G | 0 | 0 | yes | no | no | yes | yes | no | no | no | no | no | no | no | no |
| SCN2A | 1753 | I | 0 | 0 | yes | no | no | yes | no | no | no | no | no | no | no | no | no |
| SCN2A | 1754 | F | 0 | 0 | yes | no | no | yes | no | no | no | no | no | no | no | no | no |
| SCN2A | 1755 | F | 0 | 0 | yes | no | no | yes | no | no | no | no | no | no | no | no | no |
| SCN2A | 1756 | F | 0 | 0 | yes | no | no | yes | no | no | no | no | no | no | no | no | no |
| SCN2A | 1757 | V | 1 | 0 | yes | no | no | yes | no | no | no | no | no | no | no | no | no |

| | | | | | | | | | | | | | | | | | |
|---|---|---|---|---|---|---|---|---|---|---|---|---|---|---|---|---|---|
| SCN2A | 1758 | S | 1 | 1 | yes | no | no | yes | no | no | no | no | no | no | no | no | no |
| SCN2A | 1759 | Y | 0 | 0 | yes | no | no | yes | no | no | no | no | no | no | no | no | no |
| SCN2A | 1760 | I | 0 | 0 | yes | no | no | yes | no | no | no | no | no | no | no | no | no |
| SCN2A | 1761 | I | 0 | 0 | yes | no | no | yes | no | no | no | no | no | no | no | no | no |
| SCN2A | 1762 | I | 0 | 0 | yes | no | no | yes | no | no | no | no | no | no | no | no | no |
| SCN2A | 1763 | S | 0 | 0 | yes | no | no | yes | no | no | no | no | no | no | no | no | no |
| SCN2A | 1764 | F | 0 | 0 | yes | no | no | yes | no | no | no | no | no | no | no | no | no |
| SCN2A | 1765 | L | 0 | 1 | yes | no | no | yes | no | no | no | no | no | no | no | no | no |
| SCN2A | 1766 | V | 0 | 0 | yes | no | no | yes | no | no | no | no | no | no | no | no | no |
| SCN2A | 1767 | V | 0 | 0 | yes | no | no | yes | no | no | no | no | no | no | no | no | no |
| SCN2A | 1768 | V | 0 | 0 | yes | no | no | yes | no | no | yes | no | no | no | no | no | no |
| SCN2A | 1769 | N | 0 | 1 | yes | no | no | yes | no | no | no | no | no | no | no | no | no |
| SCN2A | 1770 | M | 0 | 1 | yes | no | no | yes | no | no | no | no | no | no | no | no | no |
| SCN2A | 1771 | Y | 0 | 1 | yes | no | no | yes | no | no | yes | no | no | no | no | no | no |
| SCN2A | 1772 | I | 0 | 0 | yes | no | no | yes | no | no | yes | no | no | no | no | no | no |
| SCN2A | 1773 | A | 0 | 2 | yes | no | no | yes | no | no | no | no | no | no | no | no | no |
| SCN2A | 1774 | V | 0 | 0 | yes | no | no | yes | no | no | no | no | no | no | no | no | no |
| SCN2A | 1775 | I | 0 | 0 | yes | no | no | yes | no | no | yes | no | no | no | no | no | no |
| SCN2A | 1776 | L | 0 | 0 | yes | no | no | no | no | yes | yes | no | no | no | no | no | no |
| SCN2A | 1777 | E | 0 | 0 | yes | no | no | no | no | yes | no | no | no | no | no | no | no |
| SCN2A | 1778 | N | 0 | 1 | yes | no | no | yes | no | yes | no | no | no | no | no | no | no |
| SCN2A | 1779 | F | 0 | 0 | yes | no | no | no | no | yes | yes | no | no | no | no | no | no |
| SCN2A | 1780 | S | 0 | 1 | yes | no | no | no | no | yes | no | no | no | no | no | no | no |
| SCN2A | 1781 | V | 0 | 0 | yes | no | no | no | no | yes | no | no | no | no | no | no | no |
| SCN2A | 1782 | A | 0 | 0 | yes | no | no | no | no | yes | no | no | no | no | no | no | no |
| SCN2A | 1783 | T | 0 | 0 | yes | no | no | no | no | yes | yes | no | no | no | no | no | no |
| SCN2A | 1784 | E | 0 | 0 | yes | no | no | no | no | yes | no | no | no | no | no | no | no |
| SCN2A | 1785 | E | 0 | 0 | no | yes | no | no | no | yes | no | no | no | no | no | no | no |
| SCN4A | 119 | V | 1 | 0 | no | yes | no | no | no | yes | no | no | no | no | no | no | no |
| SCN4A | 120 | V | 0 | 0 | no | yes | no | no | no | yes | no | no | no | no | no | no | no |
| SCN4A | 121 | R | 0 | 0 | yes | no | no | no | no | yes | no | no | no | no | no | no | no |
| SCN4A | 122 | R | 3 | 0 | yes | no | no | no | no | yes | no | no | no | no | no | no | no |
| SCN4A | 123 | G | 1 | 0 | yes | no | no | no | no | yes | no | no | no | no | no | no | no |

| | | | | | | | | | | | | | | | | | |
|---|---|---|---|---|---|---|---|---|---|---|---|---|---|---|---|---|---|
| SCN4A | 124 | A | 0 | 0 | yes | no | no | no | no | yes | no | no | no | no | no | no | no |
| SCN4A | 125 | I | 1 | 0 | yes | no | no | no | no | yes | no | no | no | no | no | no | no |
| SCN4A | 126 | K | 0 | 0 | yes | no | no | no | no | yes | no | no | no | no | no | no | no |
| SCN4A | 127 | V | 1 | 0 | yes | no | no | yes | no | yes | no | no | no | no | no | no | no |
| SCN4A | 128 | L | 2 | 0 | yes | no | no | no | no | yes | no | no | no | no | no | no | no |
| SCN4A | 129 | I | 1 | 0 | no | yes | no | no | no | yes | no | no | no | no | no | no | no |
| SCN4A | 130 | H | 1 | 0 | no | yes | no | no | no | yes | no | no | no | no | no | no | no |
| SCN4A | 131 | A | 0 | 0 | no | yes | no | no | no | yes | no | no | no | no | no | no | no |
| SCN4A | 132 | L | 0 | 0 | no | yes | no | yes | no | no | no | no | no | no | no | no | no |
| SCN4A | 133 | F | 0 | 0 | yes | no | no | yes | no | no | no | no | no | no | no | no | no |
| SCN4A | 134 | S | 0 | 0 | yes | no | no | yes | no | no | no | no | no | no | no | no | no |
| SCN4A | 135 | M | 2 | 0 | yes | no | no | yes | no | no | no | no | no | no | no | no | no |
| SCN4A | 136 | F | 0 | 0 | yes | no | no | yes | no | no | no | no | no | no | no | no | no |
| SCN4A | 137 | I | 1 | 0 | yes | no | no | yes | no | no | no | no | no | no | no | no | no |
| SCN4A | 138 | M | 1 | 0 | yes | no | no | yes | no | no | no | no | no | no | no | no | no |
| SCN4A | 139 | I | 1 | 0 | yes | no | no | yes | no | no | no | no | no | no | no | no | no |
| SCN4A | 140 | T | 2 | 0 | yes | no | no | yes | no | no | no | no | no | no | no | no | no |
| SCN4A | 141 | I | 0 | 1 | yes | no | no | yes | no | no | no | no | no | no | no | no | no |
| SCN4A | 142 | L | 0 | 0 | yes | no | no | yes | no | no | no | no | no | no | no | no | no |
| SCN4A | 143 | T | 0 | 0 | yes | no | no | yes | no | no | no | no | no | no | no | no | no |
| SCN4A | 144 | N | 0 | 0 | yes | no | no | yes | no | no | no | no | no | no | no | no | no |
| SCN4A | 145 | C | 0 | 0 | yes | no | no | yes | no | no | no | no | no | no | no | no | no |
| SCN4A | 146 | V | 1 | 0 | yes | no | no | yes | no | no | no | no | no | no | no | no | no |
| SCN4A | 147 | F | 0 | 0 | yes | no | no | yes | no | no | no | no | no | no | no | no | no |
| SCN4A | 148 | M | 0 | 0 | no | yes | no | yes | no | no | no | no | no | no | no | no | no |
| SCN4A | 149 | T | 0 | 0 | no | yes | no | yes | no | no | no | no | no | no | no | no | no |
| SCN4A | 150 | M | 2 | 0 | no | yes | no | yes | no | no | no | no | no | no | no | no | no |
| SCN4A | 151 | S | 0 | 0 | no | yes | no | no | yes | no | no | no | no | no | no | no | no |
| SCN4A | 152 | D | 0 | 0 | no | yes | no | no | yes | no | no | no | no | no | no | no | no |
| SCN4A | 153 | P | 0 | 0 | no | yes | no | no | yes | no | no | no | no | no | no | no | no |
| SCN4A | 154 | P | 0 | 0 | no | yes | no | no | yes | no | no | no | no | no | no | no | no |
| SCN4A | 155 | P | 0 | 0 | no | yes | no | no | yes | no | no | no | no | no | no | no | no |
| SCN4A | 156 | W | 0 | 0 | no | yes | no | no | yes | no | no | no | no | no | no | no | no |

| | | | | | | | | | | | | | | | | | |
|---|---|---|---|---|---|---|---|---|---|---|---|---|---|---|---|---|---|
| SCN4A | 157 | S | 0 | 0 | no | yes | no | yes | yes | no | no | no | no | no | no | no | no |
| SCN4A | 158 | K | 0 | 0 | yes | no | no | no | no | no | no | no | no | no | no | no | no |
| SCN4A | 159 | N | 0 | 0 | yes | no | no | yes | no | no | no | no | no | no | no | no | no |
| SCN4A | 160 | V | 0 | 0 | yes | no | no | yes | no | no | no | no | no | no | no | no | no |
| SCN4A | 161 | E | 0 | 0 | yes | no | no | yes | no | no | no | no | no | no | no | no | no |
| SCN4A | 162 | Y | 0 | 0 | yes | no | no | yes | no | no | no | no | no | no | no | no | no |
| SCN4A | 163 | T | 0 | 0 | yes | no | no | yes | no | no | no | no | no | no | no | no | no |
| SCN4A | 164 | F | 0 | 0 | yes | no | no | yes | no | no | no | no | no | no | no | no | no |
| SCN4A | 165 | T | 0 | 0 | yes | no | no | yes | no | no | no | no | no | no | no | no | no |
| SCN4A | 166 | G | 0 | 0 | yes | no | no | yes | no | no | no | no | no | no | no | no | no |
| SCN4A | 167 | I | 1 | 0 | yes | no | no | yes | no | no | no | no | no | no | no | no | no |
| SCN4A | 168 | Y | 0 | 0 | yes | no | no | yes | no | no | no | no | no | no | no | no | no |
| SCN4A | 169 | T | 1 | 0 | yes | no | no | yes | no | no | no | no | no | no | no | no | no |
| SCN4A | 170 | F | 1 | 0 | yes | no | no | yes | no | no | no | no | no | no | no | no | no |
| SCN4A | 171 | E | 0 | 0 | yes | no | no | yes | no | no | no | no | no | no | no | no | no |
| SCN4A | 172 | S | 0 | 0 | yes | no | no | yes | no | no | no | no | no | no | no | no | no |
| SCN4A | 173 | L | 1 | 0 | yes | no | no | yes | no | no | no | no | no | no | no | no | no |
| SCN4A | 174 | I | 0 | 0 | yes | no | no | yes | no | no | no | no | no | no | no | no | no |
| SCN4A | 175 | K | 1 | 0 | yes | no | no | yes | no | no | no | no | no | no | no | no | no |
| SCN4A | 176 | I | 0 | 0 | yes | no | no | yes | no | no | no | no | no | no | no | no | no |
| SCN4A | 177 | L | 0 | 0 | yes | no | no | yes | no | no | no | no | no | no | no | no | no |
| SCN4A | 178 | A | 1 | 0 | yes | no | no | no | no | no | no | no | no | no | no | no | no |
| SCN4A | 179 | R | 1 | 0 | yes | no | no | no | no | yes | no | no | no | no | no | no | no |
| SCN4A | 180 | G | 0 | 0 | no | yes | no | no | no | yes | no | no | no | no | no | no | no |
| SCN4A | 181 | F | 1 | 0 | no | yes | no | no | no | yes | no | no | no | no | no | no | no |
| SCN4A | 182 | C | 0 | 0 | no | yes | no | yes | no | yes | no | no | no | no | no | no | no |
| SCN4A | 183 | V | 0 | 0 | no | yes | no | no | no | yes | no | no | no | no | no | no | no |
| SCN4A | 184 | D | 2 | 0 | yes | no | no | no | no | yes | no | no | no | no | no | no | no |
| SCN4A | 185 | D | 1 | 0 | yes | no | no | yes | no | yes | no | no | no | no | no | no | no |
| SCN4A | 186 | F | 0 | 0 | yes | no | no | yes | no | yes | no | no | no | no | no | no | no |
| SCN4A | 187 | T | 0 | 0 | yes | no | no | no | no | yes | no | no | no | no | no | no | no |
| SCN4A | 188 | F | 0 | 0 | yes | no | no | yes | no | yes | no | no | no | no | no | no | no |
| SCN4A | 189 | L | 0 | 0 | no | yes | no | yes | no | yes | no | no | no | no | no | no | no |

| | | | | | | | | | | | | | | | | | |
|---|---|---|---|---|---|---|---|---|---|---|---|---|---|---|---|---|---|
| SCN4A | 190 | R | 2 | 0 | no | yes | no | no | no | yes | no | no | no | no | no | no | no |
| SCN4A | 191 | D | 0 | 0 | no | yes | no | yes | no | yes | no | no | no | no | no | no | no |
| SCN4A | 192 | P | 1 | 0 | no | yes | no | yes | no | yes | no | no | no | no | no | no | no |
| SCN4A | 193 | W | 0 | 0 | no | yes | no | yes | no | no | no | no | no | no | no | no | no |
| SCN4A | 194 | N | 0 | 0 | yes | no | no | yes | no | no | no | no | no | no | no | no | no |
| SCN4A | 195 | W | 0 | 0 | yes | no | no | yes | no | no | no | no | no | no | no | no | no |
| SCN4A | 196 | L | 0 | 0 | yes | no | no | yes | no | no | no | no | no | no | no | no | no |
| SCN4A | 197 | D | 0 | 0 | yes | no | no | yes | no | no | no | no | no | no | no | no | no |
| SCN4A | 198 | F | 0 | 0 | yes | no | no | yes | no | no | no | no | no | no | no | no | no |
| SCN4A | 199 | S | 1 | 0 | yes | no | no | yes | no | no | no | no | no | no | no | no | no |
| SCN4A | 200 | V | 0 | 0 | yes | no | no | yes | no | no | no | no | no | no | no | no | no |
| SCN4A | 201 | I | 0 | 0 | yes | no | no | yes | no | no | no | no | no | no | no | no | no |
| SCN4A | 202 | M | 2 | 0 | yes | no | no | yes | no | no | no | no | no | no | no | no | no |
| SCN4A | 203 | M | 1 | 1 | yes | no | no | yes | no | no | no | no | no | no | no | no | no |
| SCN4A | 204 | A | 0 | 0 | yes | no | no | yes | no | no | no | no | no | no | no | no | no |
| SCN4A | 205 | Y | 1 | 0 | yes | no | no | yes | no | no | no | no | no | no | no | no | no |
| SCN4A | 206 | L | 0 | 0 | yes | no | no | yes | no | no | no | no | no | no | no | no | no |
| SCN4A | 207 | T | 0 | 0 | yes | no | no | yes | no | no | no | no | no | no | no | no | no |
| SCN4A | 208 | E | 1 | 0 | yes | no | no | yes | no | no | no | no | no | no | no | no | no |
| SCN4A | 209 | F | 0 | 0 | yes | no | no | yes | no | no | no | no | no | no | no | no | no |
| SCN4A | 210 | V | 0 | 0 | no | yes | no | yes | no | no | no | no | no | no | no | no | no |
| SCN4A | 211 | D | 1 | 0 | no | yes | no | yes | yes | no | no | no | no | no | no | no | no |
| SCN4A | 212 | L | 1 | 0 | no | yes | no | no | yes | no | no | no | no | no | no | no | no |
| SCN4A | 213 | G | 0 | 0 | no | yes | no | no | yes | no | no | no | no | no | no | no | no |
| SCN4A | 214 | N | 0 | 0 | no | yes | no | no | yes | no | no | no | no | no | no | no | no |
| SCN4A | 215 | I | 0 | 0 | no | yes | no | no | yes | no | no | no | no | no | no | no | no |
| SCN4A | 216 | S | 0 | 0 | no | yes | no | no | yes | no | no | no | no | no | no | no | no |
| SCN4A | 217 | A | 1 | 0 | no | yes | no | yes | no | no | no | no | no | no | no | no | no |
| SCN4A | 218 | L | 0 | 0 | no | yes | no | yes | no | no | no | no | no | no | no | no | no |
| SCN4A | 219 | R | 0 | 0 | yes | no | no | yes | no | no | no | no | no | no | no | no | no |
| SCN4A | 220 | T | 1 | 0 | yes | no | no | yes | no | no | no | no | no | no | no | no | no |
| SCN4A | 221 | F | 0 | 0 | yes | no | no | yes | no | no | no | no | no | no | no | no | no |
| SCN4A | 222 | R | 0 | 2 | yes | no | no | yes | no | no | no | no | no | no | no | no | no |

| | | | | | | | | | | | | | | | | | |
|---|---|---|---|---|---|---|---|---|---|---|---|---|---|---|---|---|---|
| SCN4A | 223 | V | 0 | 0 | yes | no | no | yes | no | no | no | no | no | no | no | no | no |
| SCN4A | 224 | L | 0 | 0 | yes | no | no | yes | no | no | no | no | no | no | no | no | no |
| SCN4A | 225 | R | 2 | 1 | yes | no | no | yes | no | no | no | no | no | no | no | no | no |
| SCN4A | 226 | A | 0 | 0 | yes | no | no | yes | no | no | no | no | no | no | no | no | no |
| SCN4A | 227 | L | 1 | 0 | yes | no | no | yes | no | no | no | no | no | no | no | no | no |
| SCN4A | 228 | K | 0 | 0 | yes | no | no | yes | no | no | no | no | no | no | no | no | no |
| SCN4A | 229 | T | 0 | 0 | yes | no | no | yes | no | no | no | no | no | no | no | no | no |
| SCN4A | 230 | I | 0 | 0 | yes | no | no | yes | no | no | no | no | no | no | no | no | no |
| SCN4A | 231 | T | 1 | 1 | yes | no | no | yes | no | no | no | no | no | no | no | no | no |
| SCN4A | 232 | V | 0 | 0 | yes | no | no | yes | no | no | no | no | no | no | no | no | no |
| SCN4A | 233 | I | 0 | 0 | no | yes | no | yes | no | no | no | no | no | no | no | no | no |
| SCN4A | 234 | P | 0 | 0 | no | yes | no | yes | no | yes | no | no | no | no | no | no | no |
| SCN4A | 235 | G | 1 | 0 | no | yes | no | yes | no | yes | no | no | no | no | no | no | no |
| SCN4A | 236 | L | 0 | 0 | yes | no | no | yes | no | yes | no | no | no | no | no | no | no |
| SCN4A | 237 | K | 0 | 0 | yes | no | no | yes | no | yes | no | no | no | no | no | no | no |
| SCN4A | 238 | T | 1 | 0 | yes | no | no | no | no | yes | no | no | no | no | no | no | no |
| SCN4A | 239 | I | 0 | 0 | yes | no | no | yes | no | yes | no | no | no | no | no | no | no |
| SCN4A | 240 | V | 1 | 1 | yes | no | no | yes | no | yes | no | no | no | no | no | no | no |
| SCN4A | 241 | G | 2 | 0 | yes | no | no | yes | no | yes | no | no | no | no | no | no | no |
| SCN4A | 242 | A | 1 | 0 | yes | no | no | yes | no | yes | no | no | no | no | no | no | no |
| SCN4A | 243 | L | 0 | 0 | yes | no | no | yes | no | yes | no | no | no | no | no | no | no |
| SCN4A | 244 | I | 0 | 0 | no | yes | no | yes | no | yes | no | no | no | no | no | no | no |
| SCN4A | 245 | Q | 0 | 0 | yes | no | no | no | no | yes | no | no | no | no | no | no | no |
| SCN4A | 246 | S | 0 | 0 | yes | no | no | yes | no | yes | no | no | no | no | no | no | no |
| SCN4A | 247 | V | 1 | 0 | yes | no | no | yes | no | yes | no | no | no | no | no | no | no |
| SCN4A | 248 | K | 0 | 0 | yes | no | no | yes | no | yes | no | no | no | no | no | no | no |
| SCN4A | 249 | K | 0 | 0 | no | yes | no | no | no | yes | no | no | no | no | no | no | no |
| SCN4A | 250 | L | 0 | 1 | no | yes | no | yes | no | yes | no | no | no | no | no | no | no |
| SCN4A | 251 | S | 1 | 0 | yes | no | no | yes | no | yes | no | no | no | no | no | no | no |
| SCN4A | 252 | D | 1 | 0 | yes | no | no | yes | no | yes | no | no | no | no | no | no | no |
| SCN4A | 253 | V | 1 | 0 | yes | no | no | yes | no | no | no | no | no | no | no | no | no |
| SCN4A | 254 | M | 0 | 0 | yes | no | no | yes | no | no | no | no | no | no | no | no | no |
| SCN4A | 255 | I | 0 | 0 | yes | no | no | yes | no | no | no | no | no | no | no | no | no |

| | | | | | | | | | | | | | | | | | |
|---|---|---|---|---|---|---|---|---|---|---|---|---|---|---|---|---|---|
| SCN4A | 256 | L | 1 | 0 | yes | no | no | yes | no | no | no | no | no | no | no | no | no |
| SCN4A | 257 | T | 0 | 0 | yes | no | no | yes | no | no | no | no | no | no | no | no | no |
| SCN4A | 258 | V | 1 | 0 | yes | no | no | yes | no | no | no | no | no | no | no | no | no |
| SCN4A | 259 | F | 0 | 0 | yes | no | no | yes | no | no | no | no | no | no | no | no | no |
| SCN4A | 260 | C | 1 | 0 | yes | no | no | yes | no | no | no | no | no | no | no | no | no |
| SCN4A | 261 | L | 0 | 0 | yes | no | no | yes | no | no | no | no | no | no | no | no | no |
| SCN4A | 262 | S | 0 | 0 | yes | no | no | yes | no | no | no | no | no | no | no | no | no |
| SCN4A | 263 | V | 1 | 0 | yes | no | no | yes | no | no | no | no | no | no | no | no | no |
| SCN4A | 264 | F | 0 | 0 | yes | no | no | yes | no | no | no | no | no | no | no | no | no |
| SCN4A | 265 | A | 1 | 0 | yes | no | no | yes | no | no | no | no | no | no | no | no | no |
| SCN4A | 266 | L | 0 | 1 | yes | no | no | yes | no | no | no | no | no | no | no | no | no |
| SCN4A | 267 | V | 0 | 0 | yes | no | no | yes | no | no | no | no | no | no | no | no | no |
| SCN4A | 268 | G | 1 | 0 | yes | no | no | yes | no | no | no | no | no | no | no | no | no |
| SCN4A | 269 | L | 0 | 0 | yes | no | no | yes | no | no | no | no | no | no | no | no | no |
| SCN4A | 270 | Q | 0 | 1 | yes | no | no | yes | no | no | no | no | no | no | no | no | no |
| SCN4A | 271 | L | 0 | 0 | yes | no | no | yes | no | no | no | no | no | no | no | no | no |
| SCN4A | 272 | F | 0 | 0 | yes | no | no | yes | no | no | no | no | no | no | no | no | no |
| SCN4A | 273 | M | 3 | 0 | no | yes | no | no | yes | no | no | no | no | no | no | no | no |
| SCN4A | 274 | G | 0 | 0 | no | yes | no | no | yes | no | no | no | no | no | no | no | no |
| SCN4A | 275 | N | 0 | 0 | yes | no | no | no | yes | no | no | no | no | no | no | no | no |
| SCN4A | 276 | L | 0 | 0 | yes | no | no | no | yes | no | no | no | no | no | no | no | no |
| SCN4A | 277 | R | 1 | 0 | yes | no | no | no | yes | no | no | no | no | no | no | no | no |
| SCN4A | 278 | Q | 0 | 0 | no | yes | no | no | yes | no | no | no | no | no | no | no | no |
| SCN4A | 279 | K | 1 | 0 | no | no | yes | no | yes | no | no | no | no | no | no | no | no |
| SCN4A | 280 | C | 0 | 0 | no | no | yes | no | yes | no | no | no | no | no | no | no | no |
| SCN4A | 281 | V | 2 | 0 | no | no | yes | no | yes | no | no | no | no | no | no | no | no |
| SCN4A | 282 | R | 1 | 0 | no | no | yes | no | yes | no | no | no | yes | no | no | no | no |
| SCN4A | 283 | W | 0 | 0 | no | yes | no | no | yes | no | no | no | no | no | no | no | no |
| SCN4A | 284 | P | 0 | 0 | no | yes | no | no | yes | no | no | no | no | no | no | no | no |
| SCN4A | 285 | P | 1 | 0 | no | yes | no | no | yes | no | no | no | no | no | no | no | no |
| SCN4A | 286 | P | 1 | 0 | no | yes | no | no | yes | no | no | no | no | no | no | no | no |
| SCN4A | 336 | F | 1 | 0 | no | yes | no | no | yes | no | no | no | no | no | no | no | no |
| SCN4A | 337 | D | 2 | 0 | no | yes | no | no | yes | no | no | no | no | no | no | no | no |

| | | | | | | | | | | | | | | | | | |
|---|---|---|---|---|---|---|---|---|---|---|---|---|---|---|---|---|---|
| SCN4A | 338 | W | 0 | 0 | yes | no | no | no | yes | no | no | no | no | no | no | no | no |
| SCN4A | 339 | D | 1 | 0 | yes | no | no | no | yes | no | no | no | no | no | no | no | no |
| SCN4A | 340 | A | 1 | 0 | yes | no | no | no | yes | no | no | no | no | no | no | no | no |
| SCN4A | 341 | Y | 0 | 0 | yes | no | no | no | yes | no | no | no | no | no | no | no | no |
| SCN4A | 342 | I | 1 | 0 | yes | no | no | no | yes | no | no | no | no | no | no | no | no |
| SCN4A | 343 | S | 0 | 0 | no | yes | no | no | yes | no | no | no | no | no | no | no | no |
| SCN4A | 344 | D | 1 | 0 | no | yes | no | no | yes | no | no | no | no | no | no | no | no |
| SCN4A | 345 | E | 1 | 0 | no | yes | no | no | yes | no | no | no | no | no | no | no | no |
| SCN4A | 346 | G | 0 | 0 | no | yes | no | no | yes | no | no | no | no | no | no | no | no |
| SCN4A | 347 | N | 1 | 0 | no | yes | no | no | yes | no | no | no | no | no | no | no | no |
| SCN4A | 348 | F | 1 | 0 | no | yes | no | no | yes | no | no | no | no | no | no | no | no |
| SCN4A | 349 | Y | 0 | 0 | no | yes | no | no | yes | no | yes | no | yes | no | no | no | no |
| SCN4A | 350 | F | 0 | 0 | no | yes | no | no | yes | no | no | no | no | no | no | no | no |
| SCN4A | 351 | L | 1 | 0 | no | yes | no | no | yes | no | yes | no | yes | no | no | no | no |
| SCN4A | 352 | E | 2 | 0 | no | yes | no | no | yes | no | yes | no | no | no | no | no | no |
| SCN4A | 353 | G | 0 | 0 | no | yes | no | no | yes | no | yes | no | no | no | no | no | no |
| SCN4A | 354 | S | 0 | 0 | no | yes | no | no | yes | no | no | no | no | no | no | no | no |
| SCN4A | 355 | N | 2 | 0 | no | yes | no | no | yes | no | no | no | no | no | no | no | no |
| SCN4A | 356 | D | 2 | 0 | no | yes | no | no | yes | no | no | no | no | no | no | no | no |
| SCN4A | 357 | A | 1 | 0 | no | yes | no | no | yes | no | no | no | no | no | no | no | no |
| SCN4A | 358 | L | 0 | 0 | no | yes | no | no | yes | no | no | no | yes | no | no | no | no |
| SCN4A | 359 | L | 1 | 0 | no | yes | no | no | yes | no | no | no | no | no | no | no | no |
| SCN4A | 360 | C | 0 | 0 | no | yes | no | no | yes | no | no | no | no | no | no | no | no |
| SCN4A | 361 | G | 0 | 0 | no | yes | no | no | yes | no | no | no | no | no | no | no | no |
| SCN4A | 362 | N | 0 | 0 | no | yes | no | no | yes | no | no | no | no | no | no | no | no |
| SCN4A | 363 | S | 0 | 0 | no | yes | no | no | yes | no | no | no | no | no | no | no | no |
| SCN4A | 364 | S | 0 | 0 | no | yes | no | no | yes | no | no | no | no | no | no | no | no |
| SCN4A | 365 | D | 1 | 0 | no | yes | no | no | yes | no | no | no | no | no | no | no | no |
| SCN4A | 366 | A | 0 | 0 | no | yes | no | no | yes | no | no | no | no | no | no | no | no |
| SCN4A | 367 | G | 0 | 0 | no | yes | no | no | yes | no | no | no | no | no | no | no | no |
| SCN4A | 368 | H | 0 | 0 | no | yes | no | no | yes | no | no | no | yes | no | no | no | no |
| SCN4A | 369 | C | 0 | 0 | no | yes | no | no | yes | no | no | no | yes | no | no | no | no |
| SCN4A | 370 | P | 0 | 0 | no | yes | no | no | yes | no | no | no | yes | no | no | no | no |

| | | | | | | | | | | | | | | | | | |
|---|---|---|---|---|---|---|---|---|---|---|---|---|---|---|---|---|---|
| SCN4A | 371 | E | 0 | 0 | no | yes | no | no | yes | no | no | no | yes | no | no | no | no |
| SCN4A | 372 | G | 0 | 0 | no | yes | no | no | yes | no | no | no | yes | no | no | no | no |
| SCN4A | 373 | Y | 0 | 0 | no | no | yes | no | yes | no | no | no | yes | no | no | no | no |
| SCN4A | 374 | E | 2 | 0 | no | no | yes | no | yes | no | no | no | no | no | no | no | no |
| SCN4A | 375 | C | 0 | 1 | no | no | yes | no | yes | no | no | no | no | no | no | no | no |
| SCN4A | 376 | I | 0 | 0 | no | no | yes | no | yes | no | no | no | no | no | no | no | no |
| SCN4A | 377 | K | 0 | 0 | no | yes | no | no | yes | no | no | no | no | no | no | no | no |
| SCN4A | 378 | T | 0 | 0 | no | yes | no | no | yes | no | no | no | no | no | no | no | no |
| SCN4A | 379 | G | 1 | 0 | no | yes | no | no | yes | no | no | no | no | no | no | no | no |
| SCN4A | 380 | R | 2 | 0 | no | yes | no | no | yes | no | no | no | no | no | no | no | no |
| SCN4A | 381 | N | 0 | 0 | no | yes | no | no | yes | no | no | no | no | no | no | no | no |
| SCN4A | 382 | P | 0 | 1 | no | yes | no | no | yes | no | no | no | no | no | no | no | no |
| SCN4A | 383 | N | 0 | 0 | no | yes | no | no | yes | no | no | no | no | no | no | no | no |
| SCN4A | 384 | Y | 0 | 0 | no | yes | no | no | yes | no | no | no | no | no | no | no | no |
| SCN4A | 385 | G | 1 | 0 | no | yes | no | no | yes | no | no | no | no | no | no | no | no |
| SCN4A | 386 | Y | 1 | 0 | no | yes | no | no | yes | no | no | no | no | no | no | no | no |
| SCN4A | 387 | T | 0 | 0 | no | yes | no | no | yes | no | no | no | no | no | no | no | no |
| SCN4A | 388 | S | 0 | 0 | no | yes | no | no | yes | no | no | no | no | no | no | no | no |
| SCN4A | 389 | Y | 0 | 0 | no | yes | no | yes | yes | no | no | no | no | no | no | no | no |
| SCN4A | 390 | D | 0 | 0 | no | yes | no | no | yes | no | no | no | no | no | no | no | no |
| SCN4A | 391 | T | 1 | 0 | no | yes | no | no | yes | no | no | no | no | no | no | no | no |
| SCN4A | 392 | F | 0 | 0 | yes | no | no | yes | no | no | no | yes | no | no | no | no | no |
| SCN4A | 393 | S | 0 | 0 | yes | no | no | yes | no | no | no | yes | no | no | no | no | no |
| SCN4A | 394 | W | 0 | 0 | yes | no | no | no | no | no | no | yes | no | no | no | no | no |
| SCN4A | 395 | A | 1 | 0 | yes | no | no | yes | no | no | no | yes | no | no | no | no | no |
| SCN4A | 396 | F | 0 | 0 | yes | no | no | yes | no | no | no | yes | no | no | no | no | no |
| SCN4A | 397 | L | 0 | 0 | yes | no | no | yes | no | no | no | yes | no | no | no | no | no |
| SCN4A | 398 | A | 0 | 0 | no | yes | no | yes | no | no | no | yes | no | no | no | no | no |
| SCN4A | 399 | L | 0 | 0 | no | yes | no | yes | no | no | no | yes | no | no | no | no | no |
| SCN4A | 400 | F | 0 | 0 | yes | no | no | yes | no | no | no | yes | no | no | no | no | no |
| SCN4A | 401 | R | 0 | 1 | yes | no | no | yes | no | no | no | yes | no | no | no | no | no |
| SCN4A | 402 | L | 0 | 0 | yes | no | no | yes | no | no | no | yes | no | no | no | no | no |
| SCN4A | 403 | M | 1 | 0 | yes | no | no | yes | no | no | no | yes | no | no | no | no | no |

| | | | | | | | | | | | | | | | | | |
|---|---|---|---|---|---|---|---|---|---|---|---|---|---|---|---|---|---|
| SCN4A | 404 | T | 0 | 0 | no | yes | no | yes | no | no | yes | yes | no | no | no | no | no |
| SCN4A | 405 | Q | 2 | 0 | no | yes | no | yes | no | no | yes | yes | no | no | no | no | no |
| SCN4A | 406 | D | 0 | 0 | no | yes | no | yes | no | no | no | yes | no | no | no | no | no |
| SCN4A | 407 | Y | 1 | 0 | no | yes | no | yes | no | no | yes | yes | no | no | no | no | no |
| SCN4A | 408 | W | 0 | 0 | yes | no | no | yes | no | no | no | yes | no | no | no | no | no |
| SCN4A | 409 | E | 0 | 0 | yes | no | no | yes | no | no | yes | yes | no | no | no | no | no |
| SCN4A | 410 | N | 0 | 0 | yes | no | no | no | no | no | no | yes | no | no | no | no | no |
| SCN4A | 411 | L | 0 | 0 | yes | no | no | yes | no | no | no | yes | no | no | no | no | no |
| SCN4A | 412 | F | 0 | 0 | yes | no | no | yes | no | no | no | yes | no | no | no | no | no |
| SCN4A | 413 | Q | 0 | 0 | yes | no | no | no | no | no | no | yes | no | no | no | no | no |
| SCN4A | 414 | L | 0 | 0 | yes | no | no | no | no | no | no | yes | no | no | no | no | no |
| SCN4A | 415 | T | 0 | 0 | yes | no | no | no | no | no | no | yes | no | no | no | no | no |
| SCN4A | 416 | L | 0 | 0 | yes | no | no | no | no | no | no | yes | no | no | no | no | no |
| SCN4A | 417 | R | 1 | 0 | yes | no | no | no | yes | no | no | no | yes | no | no | no | no |
| SCN4A | 418 | A | 2 | 0 | yes | no | no | no | yes | no | no | no | no | no | no | no | no |
| SCN4A | 419 | A | 0 | 0 | yes | no | no | no | yes | no | no | no | no | no | no | no | no |
| SCN4A | 420 | G | 0 | 0 | no | yes | no | no | yes | no | no | no | no | no | no | no | no |
| SCN4A | 421 | K | 0 | 0 | yes | no | no | no | yes | no | no | no | no | no | no | no | no |
| SCN4A | 422 | T | 0 | 0 | yes | no | no | yes | yes | no | no | no | no | no | no | no | no |
| SCN4A | 423 | Y | 0 | 0 | yes | no | no | yes | yes | no | no | no | no | no | no | no | no |
| SCN4A | 424 | M | 1 | 0 | yes | no | no | yes | no | no | no | no | no | no | no | no | no |
| SCN4A | 425 | I | 0 | 0 | yes | no | no | yes | no | no | no | no | no | no | no | no | no |
| SCN4A | 426 | F | 0 | 0 | yes | no | no | yes | no | no | no | no | no | no | no | no | no |
| SCN4A | 427 | F | 1 | 0 | yes | no | no | yes | no | no | no | no | no | no | no | no | no |
| SCN4A | 428 | V | 1 | 0 | yes | no | no | yes | no | no | no | no | no | no | no | no | no |
| SCN4A | 429 | V | 0 | 0 | yes | no | no | yes | no | no | no | no | no | no | no | no | no |
| SCN4A | 430 | I | 1 | 0 | yes | no | no | yes | no | no | no | no | no | no | no | no | no |
| SCN4A | 431 | I | 1 | 0 | yes | no | no | yes | no | no | no | no | no | no | no | no | no |
| SCN4A | 432 | F | 0 | 0 | yes | no | no | yes | no | no | no | no | no | no | no | no | no |
| SCN4A | 433 | L | 1 | 0 | yes | no | no | yes | no | no | no | no | no | no | no | no | no |
| SCN4A | 434 | G | 0 | 0 | yes | no | no | yes | no | no | no | no | no | no | no | no | no |
| SCN4A | 435 | S | 0 | 0 | no | yes | no | yes | no | no | no | no | no | no | no | no | no |
| SCN4A | 436 | F | 0 | 0 | no | yes | no | yes | no | no | no | no | no | no | no | no | no |

| | | | | | | | | | | | | | | | | | |
|---|---|---|---|---|---|---|---|---|---|---|---|---|---|---|---|---|---|
| SCN4A | 437 | Y | 0 | 0 | no | yes | no | yes | no | no | no | no | no | no | no | no | no |
| SCN4A | 438 | L | 2 | 0 | no | yes | no | yes | no | no | no | no | no | no | no | no | no |
| SCN4A | 439 | I | 2 | 0 | yes | no | no | yes | no | no | yes | no | no | no | no | no | no |
| SCN4A | 440 | N | 1 | 1 | yes | no | no | yes | no | no | no | no | no | no | no | no | no |
| SCN4A | 441 | L | 0 | 0 | yes | no | no | yes | no | no | no | no | no | no | no | no | no |
| SCN4A | 442 | I | 1 | 0 | yes | no | no | yes | no | no | no | no | no | no | no | no | no |
| SCN4A | 443 | L | 0 | 0 | yes | no | no | yes | no | no | yes | no | no | no | no | no | no |
| SCN4A | 444 | A | 0 | 0 | yes | no | no | yes | no | no | no | no | no | no | no | no | no |
| SCN4A | 445 | V | 1 | 1 | yes | no | no | yes | no | yes | no | no | no | no | no | no | no |
| SCN4A | 446 | V | 0 | 0 | yes | no | no | yes | no | yes | no | no | no | no | no | no | no |
| SCN4A | 447 | A | 1 | 0 | yes | no | no | yes | no | yes | no | no | no | no | no | no | no |
| SCN4A | 448 | M | 2 | 0 | yes | no | no | no | no | yes | no | no | no | no | no | no | no |
| SCN4A | 449 | A | 1 | 0 | yes | no | no | no | no | yes | no | no | no | no | no | no | no |
| SCN4A | 450 | Y | 0 | 0 | yes | no | no | no | no | yes | no | no | no | no | no | no | no |
| SCN4A | 451 | A | 0 | 0 | yes | no | no | no | no | yes | yes | no | no | no | no | no | no |
| SCN4A | 452 | E | 2 | 1 | yes | no | no | no | no | yes | no | no | no | no | no | no | no |
| SCN4A | 453 | Q | 1 | 0 | yes | no | no | no | no | yes | no | no | no | no | no | no | no |
| SCN4A | 454 | N | 2 | 0 | no | yes | no | no | no | yes | no | no | no | no | no | no | no |
| SCN4A | 455 | E | 1 | 0 | yes | no | no | no | no | yes | yes | no | no | no | no | no | no |
| SCN4A | 456 | A | 1 | 0 | yes | no | no | no | no | yes | no | no | no | no | no | no | no |
| SCN4A | 457 | T | 0 | 0 | yes | no | no | no | no | yes | no | no | no | no | no | no | no |
| SCN4A | 458 | L | 0 | 0 | yes | no | no | no | no | yes | no | no | no | no | no | no | no |
| SCN4A | 459 | A | 2 | 0 | yes | no | no | no | no | yes | no | no | no | no | no | no | no |
| SCN4A | 460 | E | 1 | 0 | yes | no | no | no | no | yes | no | no | no | no | no | no | no |
| SCN4A | 461 | D | 0 | 0 | yes | no | no | no | no | yes | no | no | no | no | no | no | no |
| SCN4A | 462 | K | 0 | 0 | yes | no | no | no | no | yes | yes | no | no | no | no | no | no |
| SCN4A | 463 | E | 1 | 0 | no | yes | no | no | no | yes | no | no | no | no | no | no | no |
| SCN4A | 559 | N | 0 | 0 | no | yes | no | yes | no | yes | no | no | no | no | no | no | no |
| SCN4A | 560 | C | 0 | 0 | no | yes | no | no | no | yes | no | no | no | no | no | no | no |
| SCN4A | 561 | C | 0 | 0 | yes | no | no | no | no | yes | no | no | no | no | no | no | no |
| SCN4A | 562 | A | 1 | 0 | yes | no | no | no | no | yes | no | no | no | no | no | no | no |
| SCN4A | 563 | P | 1 | 0 | yes | no | no | no | no | yes | no | no | no | no | no | no | no |
| SCN4A | 564 | W | 1 | 0 | yes | no | no | no | no | yes | no | no | no | no | no | no | no |

| | | | | | | | | | | | | | | | | | |
|---|---|---|---|---|---|---|---|---|---|---|---|---|---|---|---|---|---|
| SCN4A | 565 | L | 0 | 0 | yes | no | no | no | no | yes | no | no | no | no | no | no | no |
| SCN4A | 566 | K | 1 | 0 | yes | no | no | no | no | yes | no | no | no | no | no | no | no |
| SCN4A | 567 | F | 0 | 0 | yes | no | no | no | no | yes | no | no | no | no | no | no | no |
| SCN4A | 568 | K | 0 | 0 | yes | no | no | no | no | yes | no | no | no | no | no | no | no |
| SCN4A | 569 | N | 1 | 0 | yes | no | no | no | no | yes | no | no | no | no | no | no | no |
| SCN4A | 570 | I | 2 | 0 | yes | no | no | no | no | yes | no | no | no | no | no | no | no |
| SCN4A | 571 | I | 0 | 0 | yes | no | no | no | no | yes | no | no | no | no | no | no | no |
| SCN4A | 572 | H | 0 | 0 | yes | no | no | no | no | yes | no | no | no | no | no | no | no |
| SCN4A | 573 | L | 0 | 0 | yes | no | no | no | no | yes | no | no | no | no | no | no | no |
| SCN4A | 574 | I | 0 | 0 | yes | no | no | no | no | yes | no | no | no | no | no | no | no |
| SCN4A | 575 | V | 1 | 0 | yes | no | no | no | no | yes | no | no | no | no | no | no | no |
| SCN4A | 576 | M | 0 | 0 | yes | no | no | no | no | yes | no | no | no | no | no | no | no |
| SCN4A | 577 | D | 0 | 0 | no | yes | no | no | no | yes | no | no | no | no | no | no | no |
| SCN4A | 578 | P | 2 | 0 | no | yes | no | no | no | yes | no | no | no | no | no | no | no |
| SCN4A | 579 | F | 0 | 0 | no | yes | no | no | no | no | no | no | no | no | no | no | no |
| SCN4A | 580 | V | 1 | 0 | no | yes | no | yes | no | no | no | no | no | no | no | no | no |
| SCN4A | 581 | D | 1 | 0 | yes | no | no | yes | no | no | no | no | no | no | no | no | no |
| SCN4A | 582 | L | 1 | 0 | yes | no | no | yes | no | no | no | no | no | no | no | no | no |
| SCN4A | 583 | G | 0 | 0 | yes | no | no | yes | no | no | no | no | no | no | no | no | no |
| SCN4A | 584 | I | 0 | 0 | yes | no | no | yes | no | no | no | no | no | no | no | no | no |
| SCN4A | 585 | T | 1 | 0 | yes | no | no | yes | no | no | no | no | no | no | no | no | no |
| SCN4A | 586 | I | 3 | 0 | yes | no | no | yes | no | no | no | no | no | no | no | no | no |
| SCN4A | 587 | C | 0 | 0 | yes | no | no | yes | no | no | no | no | no | no | no | no | no |
| SCN4A | 588 | I | 0 | 1 | yes | no | no | yes | no | no | no | no | no | no | no | no | no |
| SCN4A | 589 | V | 1 | 0 | yes | no | no | yes | no | no | no | no | no | no | no | no | no |
| SCN4A | 590 | L | 0 | 0 | yes | no | no | yes | no | no | no | no | no | no | no | no | no |
| SCN4A | 591 | N | 1 | 0 | yes | no | no | yes | no | no | no | no | no | no | no | no | no |
| SCN4A | 592 | T | 0 | 0 | yes | no | no | yes | no | no | no | no | no | no | no | no | no |
| SCN4A | 593 | L | 1 | 0 | yes | no | no | yes | no | no | no | no | no | no | no | no | no |
| SCN4A | 594 | F | 0 | 0 | yes | no | no | yes | no | no | no | no | no | no | no | no | no |
| SCN4A | 595 | M | 0 | 0 | yes | no | no | yes | no | no | no | no | no | no | no | no | no |
| SCN4A | 596 | A | 1 | 0 | no | yes | no | yes | no | no | no | no | no | no | no | no | no |
| SCN4A | 597 | M | 1 | 0 | no | yes | no | yes | no | no | no | no | no | no | no | no | no |

| | | | | | | | | | | | | | | | | | |
|---|---|---|---|---|---|---|---|---|---|---|---|---|---|---|---|---|---|
| SCN4A | 598 | E | 0 | 0 | no | yes | no | yes | yes | no | no | no | no | no | no | no | no |
| SCN4A | 599 | H | 1 | 0 | no | yes | no | no | yes | no | no | no | no | no | no | no | no |
| SCN4A | 600 | Y | 1 | 0 | no | yes | no | no | yes | no | no | no | no | no | no | no | no |
| SCN4A | 601 | P | 0 | 0 | no | yes | no | no | yes | no | no | no | no | no | no | no | no |
| SCN4A | 602 | M | 0 | 0 | no | yes | no | no | yes | no | no | no | no | no | no | no | no |
| SCN4A | 603 | T | 2 | 0 | no | yes | no | no | yes | no | no | no | no | no | no | no | no |
| SCN4A | 604 | E | 0 | 0 | yes | no | no | no | yes | no | no | no | no | no | no | no | no |
| SCN4A | 605 | H | 1 | 0 | yes | no | no | no | yes | no | no | no | no | no | no | no | no |
| SCN4A | 606 | F | 0 | 0 | yes | no | no | yes | yes | no | no | no | no | no | no | no | no |
| SCN4A | 607 | D | 0 | 0 | yes | no | no | no | yes | no | no | no | no | no | no | no | no |
| SCN4A | 608 | N | 1 | 0 | yes | no | no | no | yes | no | no | no | no | no | no | no | no |
| SCN4A | 609 | V | 1 | 0 | yes | no | no | yes | no | no | no | no | no | no | no | no | no |
| SCN4A | 610 | L | 1 | 0 | yes | no | no | yes | no | no | no | no | no | no | no | no | no |
| SCN4A | 611 | T | 0 | 0 | yes | no | no | yes | no | no | no | no | no | no | no | no | no |
| SCN4A | 612 | V | 1 | 0 | yes | no | no | yes | no | no | no | no | no | no | no | no | no |
| SCN4A | 613 | G | 0 | 0 | yes | no | no | yes | no | no | no | no | no | no | no | no | no |
| SCN4A | 614 | N | 1 | 0 | yes | no | no | yes | no | no | no | no | no | no | no | no | no |
| SCN4A | 615 | L | 2 | 0 | yes | no | no | yes | no | no | no | no | no | no | no | no | no |
| SCN4A | 616 | V | 0 | 0 | yes | no | no | yes | no | no | no | no | no | no | no | no | no |
| SCN4A | 617 | F | 0 | 0 | yes | no | no | yes | no | no | no | no | no | no | no | no | no |
| SCN4A | 618 | T | 1 | 0 | yes | no | no | yes | no | no | no | no | no | no | no | no | no |
| SCN4A | 619 | G | 0 | 0 | yes | no | no | yes | no | no | no | no | no | no | no | no | no |
| SCN4A | 620 | I | 1 | 0 | yes | no | no | yes | no | no | no | no | no | no | no | no | no |
| SCN4A | 621 | F | 0 | 1 | yes | no | no | yes | no | no | no | no | no | no | no | no | no |
| SCN4A | 622 | T | 1 | 0 | yes | no | no | yes | no | no | no | no | no | no | no | no | no |
| SCN4A | 623 | A | 0 | 0 | yes | no | no | yes | no | no | no | no | no | no | no | no | no |
| SCN4A | 624 | E | 0 | 0 | yes | no | no | yes | no | no | no | no | no | no | no | no | no |
| SCN4A | 625 | M | 1 | 0 | yes | no | no | yes | no | no | no | no | no | no | no | no | no |
| SCN4A | 626 | V | 1 | 0 | yes | no | no | yes | no | no | no | no | no | no | no | no | no |
| SCN4A | 627 | L | 1 | 0 | yes | no | no | yes | no | no | no | no | no | no | no | no | no |
| SCN4A | 628 | K | 0 | 0 | yes | no | no | yes | no | no | no | no | no | no | no | no | no |
| SCN4A | 629 | L | 0 | 0 | yes | no | no | yes | no | yes | no | no | no | no | no | no | no |
| SCN4A | 630 | I | 0 | 0 | yes | no | no | yes | no | yes | no | no | no | no | no | no | no |

| | | | | | | | | | | | | | | | | | |
|---|---|---|---|---|---|---|---|---|---|---|---|---|---|---|---|---|---|
| SCN4A | 631 | A | 1 | 0 | yes | no | no | no | no | yes | no | no | no | no | no | no | no |
| SCN4A | 632 | M | 1 | 0 | no | yes | no | no | no | yes | no | no | no | no | no | no | no |
| SCN4A | 633 | D | 0 | 0 | no | yes | no | no | no | yes | no | no | no | no | no | no | no |
| SCN4A | 634 | P | 0 | 0 | no | yes | no | no | no | yes | no | no | no | no | no | no | no |
| SCN4A | 635 | Y | 0 | 0 | no | yes | no | no | no | yes | no | no | no | no | no | no | no |
| SCN4A | 636 | E | 1 | 0 | no | yes | no | no | no | yes | no | no | no | no | no | no | no |
| SCN4A | 637 | Y | 0 | 0 | no | yes | no | yes | no | yes | no | no | no | no | no | no | no |
| SCN4A | 638 | F | 0 | 0 | no | yes | no | yes | no | yes | no | no | no | no | no | no | no |
| SCN4A | 639 | Q | 1 | 0 | no | yes | no | no | no | yes | no | no | no | no | no | no | no |
| SCN4A | 640 | Q | 0 | 0 | no | yes | no | no | no | yes | no | no | no | no | no | no | no |
| SCN4A | 641 | G | 0 | 0 | yes | no | no | yes | no | yes | no | no | no | no | no | no | no |
| SCN4A | 642 | W | 1 | 0 | yes | no | no | yes | no | yes | no | no | no | no | no | no | no |
| SCN4A | 643 | N | 0 | 0 | yes | no | no | yes | no | no | no | no | no | no | no | no | no |
| SCN4A | 644 | I | 0 | 0 | yes | no | no | yes | no | no | no | no | no | no | no | no | no |
| SCN4A | 645 | F | 1 | 0 | yes | no | no | yes | no | no | no | no | no | no | no | no | no |
| SCN4A | 646 | D | 0 | 0 | yes | no | no | yes | no | no | no | no | no | no | no | no | no |
| SCN4A | 647 | S | 0 | 0 | yes | no | no | yes | no | no | no | no | no | no | no | no | no |
| SCN4A | 648 | I | 1 | 0 | yes | no | no | yes | no | no | no | no | no | no | no | no | no |
| SCN4A | 649 | I | 0 | 0 | yes | no | no | yes | no | no | no | no | no | no | no | no | no |
| SCN4A | 650 | V | 1 | 0 | yes | no | no | yes | no | no | no | no | no | no | no | no | no |
| SCN4A | 651 | T | 1 | 0 | yes | no | no | yes | no | no | no | no | no | no | no | no | no |
| SCN4A | 652 | L | 1 | 0 | yes | no | no | yes | no | no | no | no | no | no | no | no | no |
| SCN4A | 653 | S | 0 | 0 | yes | no | no | yes | no | no | no | no | no | no | no | no | no |
| SCN4A | 654 | L | 1 | 0 | yes | no | no | yes | no | no | no | no | no | no | no | no | no |
| SCN4A | 655 | V | 1 | 0 | yes | no | no | yes | no | no | no | no | no | no | no | no | no |
| SCN4A | 656 | E | 0 | 0 | yes | no | no | yes | no | no | no | no | no | no | no | no | no |
| SCN4A | 657 | L | 0 | 0 | yes | no | no | yes | no | no | no | no | no | no | no | no | no |
| SCN4A | 658 | G | 1 | 0 | yes | no | no | yes | no | no | no | no | no | no | no | no | no |
| SCN4A | 659 | L | 0 | 0 | yes | no | no | yes | no | no | no | no | no | no | no | no | no |
| SCN4A | 660 | A | 0 | 0 | no | yes | no | yes | no | no | no | no | no | no | no | no | no |
| SCN4A | 661 | N | 1 | 0 | no | yes | no | no | no | no | no | no | no | no | no | no | no |
| SCN4A | 662 | V | 2 | 0 | no | yes | no | no | no | no | no | no | no | no | no | no | no |
| SCN4A | 663 | Q | 0 | 0 | no | yes | no | no | yes | no | no | no | no | no | no | no | no |

| | | | | | | | | | | | | | | | | | |
|---|---|---|---|---|---|---|---|---|---|---|---|---|---|---|---|---|---|
| SCN4A | 664 | G | 0 | 0 | no | yes | no | no | yes | no | no | no | no | no | no | no | no |
| SCN4A | 665 | L | 0 | 0 | no | yes | no | yes | no | no | no | no | no | no | no | no | no |
| SCN4A | 666 | S | 0 | 0 | no | yes | no | yes | no | no | no | no | no | no | no | no | no |
| SCN4A | 667 | V | 1 | 0 | no | yes | no | yes | no | no | no | no | no | no | no | no | no |
| SCN4A | 668 | L | 0 | 0 | yes | no | no | yes | no | no | no | no | no | no | no | no | no |
| SCN4A | 669 | R | 2 | 2 | yes | no | no | yes | no | no | no | no | no | no | no | no | no |
| SCN4A | 670 | S | 1 | 1 | yes | no | no | yes | no | no | no | no | no | no | no | no | no |
| SCN4A | 671 | F | 1 | 3 | no | yes | no | yes | no | no | no | no | no | no | no | no | no |
| SCN4A | 672 | R | 1 | 3 | yes | no | no | yes | no | no | no | no | no | no | no | no | no |
| SCN4A | 673 | L | 0 | 1 | yes | no | no | yes | no | no | no | no | no | no | no | no | no |
| SCN4A | 674 | L | 0 | 0 | yes | no | no | yes | no | no | no | no | no | no | no | no | no |
| SCN4A | 675 | R | 2 | 2 | yes | no | no | yes | no | no | no | no | no | no | no | no | no |
| SCN4A | 676 | V | 1 | 0 | yes | no | no | yes | no | no | no | no | no | no | no | no | no |
| SCN4A | 677 | F | 0 | 0 | yes | no | no | yes | no | no | no | no | no | no | no | no | no |
| SCN4A | 678 | K | 0 | 0 | yes | no | no | yes | no | no | no | no | no | no | no | no | no |
| SCN4A | 679 | L | 0 | 0 | yes | no | no | yes | no | no | no | no | no | no | no | no | no |
| SCN4A | 680 | A | 1 | 0 | yes | no | no | yes | no | no | no | no | no | no | no | no | no |
| SCN4A | 681 | K | 1 | 0 | no | yes | no | no | no | no | no | no | no | no | no | no | no |
| SCN4A | 682 | S | 1 | 1 | no | yes | no | no | no | no | no | no | no | no | no | no | no |
| SCN4A | 683 | W | 0 | 0 | no | yes | no | yes | no | yes | no | no | no | no | no | no | no |
| SCN4A | 684 | P | 1 | 0 | no | yes | no | no | no | yes | no | no | no | no | no | no | no |
| SCN4A | 685 | T | 1 | 0 | no | yes | no | yes | no | yes | no | no | no | no | no | no | no |
| SCN4A | 686 | L | 0 | 0 | yes | no | no | yes | no | yes | no | no | no | no | no | no | no |
| SCN4A | 687 | N | 0 | 0 | yes | no | no | no | no | yes | no | no | no | no | no | no | no |
| SCN4A | 688 | M | 1 | 0 | yes | no | no | yes | no | yes | no | no | no | no | no | no | no |
| SCN4A | 689 | L | 0 | 2 | yes | no | no | yes | no | yes | no | no | no | no | no | no | no |
| SCN4A | 690 | I | 0 | 0 | yes | no | no | yes | no | yes | no | no | no | no | no | no | no |
| SCN4A | 691 | K | 1 | 0 | yes | no | no | no | no | yes | no | no | no | no | no | no | no |
| SCN4A | 692 | I | 1 | 1 | yes | no | no | yes | no | yes | no | no | no | no | no | no | no |
| SCN4A | 693 | I | 0 | 3 | yes | no | no | yes | no | yes | no | no | no | no | no | no | no |
| SCN4A | 694 | G | 0 | 0 | yes | no | no | yes | no | yes | no | no | no | no | no | no | no |
| SCN4A | 695 | N | 1 | 0 | yes | no | no | yes | no | yes | no | no | no | no | no | no | no |
| SCN4A | 696 | S | 0 | 0 | yes | no | no | yes | no | yes | no | no | no | no | no | no | no |

| | | | | | | | | | | | | | | | | | |
|---|---|---|---|---|---|---|---|---|---|---|---|---|---|---|---|---|---|
| SCN4A | 697 | V | 0 | 0 | yes | no | no | yes | no | yes | no | no | no | no | no | no | no |
| SCN4A | 698 | G | 1 | 0 | no | yes | no | yes | no | yes | no | no | no | no | no | no | no |
| SCN4A | 699 | A | 1 | 0 | no | yes | no | no | no | no | no | no | no | no | no | no | no |
| SCN4A | 700 | L | 0 | 0 | no | yes | no | yes | no | no | no | no | no | no | no | no | no |
| SCN4A | 701 | G | 0 | 0 | no | yes | no | yes | no | no | no | no | no | no | no | no | no |
| SCN4A | 702 | N | 0 | 0 | no | yes | no | yes | no | no | no | no | no | no | no | no | no |
| SCN4A | 703 | L | 0 | 1 | yes | no | no | yes | no | no | no | no | no | no | no | no | no |
| SCN4A | 704 | T | 0 | 1 | yes | no | no | yes | no | no | no | no | no | no | no | no | no |
| SCN4A | 705 | L | 0 | 0 | yes | no | no | yes | no | no | no | no | no | no | no | no | no |
| SCN4A | 706 | V | 0 | 0 | yes | no | no | yes | no | no | no | no | no | no | no | no | no |
| SCN4A | 707 | L | 0 | 0 | yes | no | no | yes | no | no | no | no | no | no | no | no | no |
| SCN4A | 708 | A | 0 | 0 | yes | no | no | yes | no | no | no | no | no | no | no | no | no |
| SCN4A | 709 | I | 1 | 0 | yes | no | no | yes | no | no | no | no | no | no | no | no | no |
| SCN4A | 710 | I | 1 | 0 | yes | no | no | yes | no | no | no | no | no | no | no | no | no |
| SCN4A | 711 | V | 1 | 0 | yes | no | no | yes | no | no | no | no | no | no | no | no | no |
| SCN4A | 712 | F | 0 | 0 | yes | no | no | yes | no | no | no | no | no | no | no | no | no |
| SCN4A | 713 | I | 1 | 0 | yes | no | no | yes | no | no | no | no | no | no | no | no | no |
| SCN4A | 714 | F | 0 | 0 | yes | no | no | yes | no | no | no | no | no | no | no | no | no |
| SCN4A | 715 | A | 1 | 1 | yes | no | no | yes | no | no | no | no | no | no | no | no | no |
| SCN4A | 716 | V | 1 | 0 | yes | no | no | yes | no | no | no | no | no | no | no | no | no |
| SCN4A | 717 | V | 1 | 1 | yes | no | no | yes | no | no | no | no | no | no | no | no | no |
| SCN4A | 718 | G | 0 | 0 | yes | no | no | yes | yes | no | no | no | no | no | no | no | no |
| SCN4A | 719 | M | 0 | 0 | yes | no | no | yes | yes | no | no | no | no | no | no | no | no |
| SCN4A | 720 | Q | 0 | 0 | yes | no | no | yes | yes | no | no | no | no | no | no | no | no |
| SCN4A | 721 | L | 0 | 0 | yes | no | no | yes | yes | no | no | no | no | no | no | no | no |
| SCN4A | 722 | F | 0 | 0 | yes | no | no | no | yes | no | no | no | no | no | no | no | no |
| SCN4A | 723 | G | 0 | 0 | yes | no | no | no | yes | no | no | no | no | no | no | no | no |
| SCN4A | 724 | K | 0 | 0 | yes | no | no | no | yes | no | no | no | no | no | no | no | no |
| SCN4A | 725 | S | 1 | 0 | yes | no | no | no | yes | no | no | no | no | no | no | no | no |
| SCN4A | 726 | Y | 1 | 0 | yes | no | no | no | yes | no | no | no | no | no | no | no | no |
| SCN4A | 727 | K | 0 | 0 | no | yes | no | no | yes | no | no | no | no | no | no | no | no |
| SCN4A | 728 | E | 1 | 0 | no | yes | no | no | yes | no | no | no | no | no | no | no | no |
| SCN4A | 729 | C | 1 | 0 | no | yes | no | no | yes | no | no | no | no | no | no | no | no |

| | | | | | | | | | | | | | | | | | |
|---|---|---|---|---|---|---|---|---|---|---|---|---|---|---|---|---|---|
| SCN4A | 730 | V | 2 | 0 | yes | no | no | no | yes | no | no | no | no | no | no | no | no |
| SCN4A | 731 | C | 0 | 0 | yes | no | no | no | yes | no | no | no | no | no | no | no | no |
| SCN4A | 732 | K | 0 | 0 | yes | no | no | no | yes | no | no | no | no | no | no | no | no |
| SCN4A | 733 | I | 2 | 0 | no | yes | no | no | yes | no | no | no | no | no | no | no | no |
| SCN4A | 734 | A | 1 | 0 | no | yes | no | no | yes | no | no | no | no | no | no | no | no |
| SCN4A | 735 | L | 0 | 0 | no | yes | no | no | yes | no | no | no | no | no | no | no | no |
| SCN4A | 736 | D | 1 | 0 | no | yes | no | no | yes | no | no | no | no | no | no | no | no |
| SCN4A | 737 | C | 0 | 0 | no | yes | no | no | yes | no | no | no | no | no | no | no | no |
| SCN4A | 738 | N | 1 | 0 | no | yes | no | no | yes | no | no | no | no | no | no | no | no |
| SCN4A | 739 | L | 0 | 0 | no | yes | no | no | yes | no | no | no | no | no | no | no | no |
| SCN4A | 740 | P | 1 | 0 | no | yes | no | no | yes | no | no | no | no | no | no | no | no |
| SCN4A | 741 | R | 2 | 0 | no | yes | no | no | yes | no | no | no | no | no | no | no | no |
| SCN4A | 742 | W | 0 | 0 | no | yes | no | no | yes | no | no | no | no | no | no | no | no |
| SCN4A | 743 | H | 0 | 0 | no | yes | no | no | yes | no | no | no | no | no | no | no | no |
| SCN4A | 744 | M | 1 | 0 | no | yes | no | no | yes | no | no | no | no | no | no | no | no |
| SCN4A | 745 | H | 2 | 0 | no | yes | no | no | yes | no | no | no | no | no | no | no | no |
| SCN4A | 746 | D | 0 | 0 | no | yes | no | no | yes | no | no | no | no | no | no | no | no |
| SCN4A | 747 | F | 0 | 0 | yes | no | no | yes | no | no | no | yes | no | no | no | no | no |
| SCN4A | 748 | F | 0 | 0 | yes | no | no | yes | no | no | no | yes | no | no | no | no | no |
| SCN4A | 749 | H | 0 | 0 | yes | no | no | no | no | no | no | yes | no | no | no | no | no |
| SCN4A | 750 | S | 0 | 0 | yes | no | no | yes | no | no | no | yes | no | no | no | no | no |
| SCN4A | 751 | F | 0 | 0 | yes | no | no | yes | no | no | no | yes | no | no | no | no | no |
| SCN4A | 752 | L | 1 | 0 | yes | no | no | yes | no | no | no | yes | no | no | no | no | no |
| SCN4A | 753 | I | 0 | 0 | yes | no | no | yes | no | no | no | yes | no | no | no | no | no |
| SCN4A | 754 | V | 1 | 0 | yes | no | no | yes | no | no | no | yes | no | no | no | no | no |
| SCN4A | 755 | F | 0 | 0 | yes | no | no | yes | no | no | no | yes | no | no | no | no | no |
| SCN4A | 756 | R | 1 | 0 | yes | no | no | yes | no | no | no | yes | no | no | no | no | no |
| SCN4A | 757 | I | 0 | 0 | yes | no | no | yes | no | no | no | yes | no | no | no | no | no |
| SCN4A | 758 | L | 1 | 0 | yes | no | no | yes | no | no | yes | yes | no | no | no | no | no |
| SCN4A | 759 | C | 0 | 1 | yes | no | no | yes | no | no | yes | yes | no | no | no | no | no |
| SCN4A | 760 | G | 0 | 0 | no | yes | no | yes | no | no | yes | yes | no | no | no | no | no |
| SCN4A | 761 | E | 1 | 0 | no | yes | no | yes | no | no | yes | yes | no | no | no | no | no |
| SCN4A | 762 | W | 0 | 0 | no | yes | no | yes | no | no | no | yes | no | no | no | no | no |

| | | | | | | | | | | | | | | | | | |
|---|---|---|---|---|---|---|---|---|---|---|---|---|---|---|---|---|---|
| SCN4A | 763 | I | 0 | 0 | no | yes | no | yes | no | no | no | yes | no | no | no | no | no |
| SCN4A | 764 | E | 1 | 0 | yes | no | no | no | no | no | no | yes | no | no | no | no | no |
| SCN4A | 765 | T | 0 | 0 | yes | no | no | yes | no | no | no | yes | no | no | no | no | no |
| SCN4A | 766 | M | 1 | 0 | yes | no | no | yes | no | no | no | yes | no | no | no | no | no |
| SCN4A | 767 | W | 0 | 0 | yes | no | no | no | no | no | no | yes | no | no | no | no | no |
| SCN4A | 768 | D | 2 | 0 | yes | no | no | no | yes | no | no | no | no | no | no | no | no |
| SCN4A | 769 | C | 0 | 0 | yes | no | no | no | yes | no | no | no | no | no | no | no | no |
| SCN4A | 770 | M | 0 | 0 | yes | no | no | no | yes | no | no | no | no | no | no | no | no |
| SCN4A | 771 | E | 0 | 0 | yes | no | no | no | yes | no | no | no | no | no | no | no | no |
| SCN4A | 772 | V | 0 | 0 | yes | no | no | no | yes | no | no | no | no | no | no | no | no |
| SCN4A | 773 | A | 0 | 0 | no | yes | no | no | yes | no | no | no | no | no | no | no | no |
| SCN4A | 774 | G | 1 | 0 | no | yes | no | no | yes | no | no | no | no | no | no | no | no |
| SCN4A | 775 | Q | 0 | 0 | yes | no | no | no | yes | no | no | no | no | no | no | no | no |
| SCN4A | 776 | A | 1 | 0 | yes | no | no | yes | yes | no | no | no | no | no | no | no | no |
| SCN4A | 777 | M | 1 | 0 | yes | no | no | yes | yes | no | no | no | no | no | no | no | no |
| SCN4A | 778 | C | 0 | 0 | yes | no | no | yes | yes | no | no | no | no | no | no | no | no |
| SCN4A | 779 | L | 0 | 0 | yes | no | no | yes | no | no | no | no | no | no | no | no | no |
| SCN4A | 780 | T | 0 | 0 | yes | no | no | yes | no | no | no | no | no | no | no | no | no |
| SCN4A | 781 | V | 2 | 0 | yes | no | no | yes | no | no | no | no | no | no | no | no | no |
| SCN4A | 782 | F | 1 | 0 | yes | no | no | yes | no | no | no | no | no | no | no | no | no |
| SCN4A | 783 | L | 1 | 0 | yes | no | no | yes | no | no | no | no | no | no | no | no | no |
| SCN4A | 784 | M | 0 | 0 | yes | no | no | yes | no | no | no | no | no | no | no | no | no |
| SCN4A | 785 | V | 0 | 0 | yes | no | no | yes | no | no | no | no | no | no | no | no | no |
| SCN4A | 786 | M | 1 | 0 | yes | no | no | yes | no | no | no | no | no | no | no | no | no |
| SCN4A | 787 | V | 1 | 0 | yes | no | no | yes | no | no | no | no | no | no | no | no | no |
| SCN4A | 788 | I | 1 | 0 | yes | no | no | yes | no | no | no | no | no | no | no | no | no |
| SCN4A | 789 | G | 1 | 0 | yes | no | no | yes | no | no | no | no | no | no | no | no | no |
| SCN4A | 790 | N | 0 | 0 | yes | no | no | yes | no | no | yes | no | no | no | no | no | no |
| SCN4A | 791 | L | 0 | 0 | yes | no | no | yes | no | no | no | no | no | no | no | no | no |
| SCN4A | 792 | V | 0 | 0 | yes | no | no | yes | no | no | no | no | no | no | no | no | no |
| SCN4A | 793 | V | 0 | 0 | yes | no | no | yes | no | no | no | no | no | no | no | no | no |
| SCN4A | 794 | L | 0 | 0 | yes | no | no | yes | no | no | no | no | no | no | no | no | no |
| SCN4A | 795 | N | 0 | 0 | yes | no | no | yes | no | no | no | no | no | no | no | no | no |

| | | | | | | | | | | | | | | | | | |
|---|---|---|---|---|---|---|---|---|---|---|---|---|---|---|---|---|---|
| SCN4A | 796 | L | 0 | 1 | yes | no | no | yes | no | no | no | no | no | no | no | no | no |
| SCN4A | 797 | F | 0 | 0 | yes | no | no | yes | no | no | yes | no | no | no | no | no | no |
| SCN4A | 798 | L | 0 | 0 | yes | no | no | yes | no | yes | yes | no | no | no | no | no | no |
| SCN4A | 799 | A | 0 | 1 | yes | no | no | yes | no | yes | no | no | no | no | no | no | no |
| SCN4A | 800 | L | 0 | 0 | yes | no | no | yes | no | yes | no | no | no | no | no | no | no |
| SCN4A | 801 | L | 0 | 0 | yes | no | no | yes | no | yes | yes | no | no | no | no | no | no |
| SCN4A | 802 | L | 0 | 0 | yes | no | no | no | no | yes | yes | no | no | no | no | no | no |
| SCN4A | 803 | S | 0 | 0 | no | yes | no | no | no | yes | no | no | no | no | no | no | no |
| SCN4A | 804 | S | 1 | 1 | no | yes | no | no | no | yes | no | no | no | no | no | no | no |
| SCN4A | 805 | F | 0 | 0 | no | yes | no | no | no | yes | yes | no | no | no | no | no | no |
| SCN4A | 806 | S | 0 | 0 | no | yes | no | no | no | yes | no | no | no | no | no | no | no |
| SCN4A | 1014 | G | 0 | 0 | no | yes | no | no | no | yes | no | no | no | no | no | no | no |
| SCN4A | 1015 | K | 0 | 0 | yes | no | no | no | no | yes | no | no | no | no | no | no | no |
| SCN4A | 1016 | K | 1 | 0 | yes | no | no | no | no | yes | no | no | yes | no | no | no | no |
| SCN4A | 1017 | W | 1 | 0 | yes | no | no | no | no | yes | no | no | no | no | no | no | no |
| SCN4A | 1018 | W | 0 | 0 | yes | no | no | no | no | yes | no | no | no | no | no | no | no |
| SCN4A | 1019 | T | 0 | 0 | yes | no | no | no | no | yes | no | no | yes | no | no | no | no |
| SCN4A | 1020 | L | 1 | 0 | yes | no | no | no | no | yes | no | no | yes | no | no | no | no |
| SCN4A | 1021 | R | 2 | 0 | yes | no | no | no | no | yes | no | no | no | no | no | no | no |
| SCN4A | 1022 | R | 1 | 0 | yes | no | no | no | no | yes | no | no | yes | no | no | no | no |
| SCN4A | 1023 | A | 0 | 0 | yes | no | no | no | no | yes | no | no | no | no | no | no | no |
| SCN4A | 1024 | C | 1 | 0 | yes | no | no | no | no | yes | no | no | no | no | no | no | no |
| SCN4A | 1025 | F | 0 | 0 | yes | no | no | no | no | yes | no | no | no | no | no | no | no |
| SCN4A | 1026 | K | 0 | 0 | yes | no | no | no | no | yes | no | no | yes | no | no | no | no |
| SCN4A | 1027 | I | 1 | 0 | yes | no | no | yes | no | yes | no | no | no | no | no | no | no |
| SCN4A | 1028 | V | 0 | 0 | yes | no | no | yes | no | yes | no | no | no | no | no | no | no |
| SCN4A | 1029 | E | 2 | 0 | no | yes | no | no | no | yes | no | no | no | no | no | no | no |
| SCN4A | 1030 | H | 0 | 0 | no | yes | no | no | no | yes | no | no | no | no | no | no | no |
| SCN4A | 1031 | N | 1 | 0 | yes | no | no | yes | no | yes | no | no | no | no | no | no | no |
| SCN4A | 1032 | W | 1 | 0 | yes | no | no | yes | no | yes | no | no | no | no | no | no | no |
| SCN4A | 1033 | F | 1 | 0 | yes | no | no | yes | no | no | no | no | no | no | no | no | no |
| SCN4A | 1034 | E | 1 | 0 | yes | no | no | yes | no | no | no | no | no | no | no | no | no |
| SCN4A | 1035 | T | 0 | 0 | yes | no | no | yes | no | no | no | no | no | no | no | no | no |

| | | | | | | | | | | | | | | | | | |
|---|---|---|---|---|---|---|---|---|---|---|---|---|---|---|---|---|---|
| SCN4A | 1036 | F | 0 | 0 | yes | no | no | yes | no | no | no | no | yes | no | no | no | no |
| SCN4A | 1037 | I | 1 | 0 | yes | no | no | yes | no | no | no | no | no | no | no | no | no |
| SCN4A | 1038 | V | 0 | 0 | yes | no | no | yes | no | no | no | no | no | no | no | no | no |
| SCN4A | 1039 | F | 2 | 0 | yes | no | no | yes | no | no | no | no | no | no | no | no | no |
| SCN4A | 1040 | M | 1 | 0 | yes | no | no | yes | no | no | no | no | no | no | no | no | no |
| SCN4A | 1041 | I | 0 | 0 | yes | no | no | yes | no | no | no | no | no | no | no | no | no |
| SCN4A | 1042 | L | 0 | 0 | yes | no | no | yes | no | no | no | no | no | no | no | no | no |
| SCN4A | 1043 | L | 0 | 0 | yes | no | no | yes | no | no | no | no | no | no | no | no | no |
| SCN4A | 1044 | S | 0 | 0 | yes | no | no | yes | no | no | no | no | no | no | no | no | no |
| SCN4A | 1045 | S | 0 | 0 | yes | no | no | yes | no | no | no | no | no | no | no | no | no |
| SCN4A | 1046 | G | 1 | 0 | yes | no | no | yes | no | no | no | no | no | no | no | no | no |
| SCN4A | 1047 | A | 0 | 0 | no | yes | no | yes | no | no | no | no | no | no | no | no | no |
| SCN4A | 1048 | L | 0 | 0 | no | yes | no | yes | no | no | no | no | no | no | no | no | no |
| SCN4A | 1049 | A | 1 | 0 | no | yes | no | yes | no | no | no | no | no | no | no | no | no |
| SCN4A | 1050 | F | 0 | 0 | no | yes | no | yes | no | no | no | no | no | no | no | no | no |
| SCN4A | 1051 | E | 0 | 0 | no | yes | no | no | yes | no | no | no | no | no | no | no | no |
| SCN4A | 1052 | D | 0 | 0 | no | yes | no | no | yes | no | no | no | no | no | no | no | no |
| SCN4A | 1053 | I | 1 | 0 | yes | no | no | no | yes | no | no | no | yes | no | no | no | no |
| SCN4A | 1054 | Y | 1 | 0 | yes | no | no | no | yes | no | no | no | no | no | no | no | no |
| SCN4A | 1055 | I | 3 | 0 | yes | no | no | no | yes | no | no | no | no | no | no | no | no |
| SCN4A | 1056 | E | 0 | 0 | no | yes | no | no | yes | no | no | no | yes | no | no | no | no |
| SCN4A | 1057 | Q | 0 | 0 | no | yes | no | no | yes | no | no | no | yes | no | no | no | no |
| SCN4A | 1058 | R | 2 | 0 | no | yes | no | no | yes | no | no | no | no | no | no | no | no |
| SCN4A | 1059 | R | 1 | 0 | no | yes | no | no | yes | no | no | no | yes | no | no | no | no |
| SCN4A | 1060 | V | 1 | 0 | yes | no | no | no | yes | no | no | no | no | no | no | no | no |
| SCN4A | 1061 | I | 1 | 0 | yes | no | no | no | yes | no | no | no | no | no | no | no | no |
| SCN4A | 1062 | R | 1 | 0 | yes | no | no | no | yes | no | no | no | no | no | no | no | no |
| SCN4A | 1063 | T | 2 | 0 | yes | no | no | no | yes | no | no | no | yes | no | no | no | no |
| SCN4A | 1064 | I | 1 | 0 | yes | no | no | yes | no | no | no | no | no | no | no | no | no |
| SCN4A | 1065 | L | 0 | 0 | yes | no | no | yes | no | no | no | no | no | no | no | no | no |
| SCN4A | 1066 | E | 0 | 0 | yes | no | no | no | no | no | no | no | no | no | no | no | no |
| SCN4A | 1067 | Y | 0 | 0 | yes | no | no | yes | no | no | no | no | yes | no | no | no | no |
| SCN4A | 1068 | A | 1 | 0 | yes | no | no | yes | no | no | no | no | no | no | no | no | no |

| | | | | | | | | | | | | | | | | | |
|---|---|---|---|---|---|---|---|---|---|---|---|---|---|---|---|---|---|
| SCN4A | 1069 | D | 1 | 1 | yes | no | no | yes | no | no | no | no | no | no | no | no | no |
| SCN4A | 1070 | K | 0 | 0 | yes | no | no | yes | no | no | no | no | yes | no | no | no | no |
| SCN4A | 1071 | V | 1 | 0 | yes | no | no | yes | no | no | no | no | no | no | no | no | no |
| SCN4A | 1072 | F | 0 | 0 | yes | no | no | yes | no | no | no | no | no | no | no | no | no |
| SCN4A | 1073 | T | 1 | 0 | yes | no | no | yes | no | no | no | no | no | no | no | no | no |
| SCN4A | 1074 | Y | 1 | 0 | yes | no | no | yes | no | no | no | no | yes | no | no | no | no |
| SCN4A | 1075 | I | 0 | 0 | yes | no | no | yes | no | no | no | no | no | no | no | no | no |
| SCN4A | 1076 | F | 0 | 0 | yes | no | no | yes | no | no | no | no | no | no | no | no | no |
| SCN4A | 1077 | I | 0 | 0 | yes | no | no | yes | no | no | no | no | no | no | no | no | no |
| SCN4A | 1078 | M | 1 | 0 | yes | no | no | yes | no | no | no | no | yes | no | no | no | no |
| SCN4A | 1079 | E | 2 | 0 | no | yes | no | yes | no | no | no | no | no | no | no | no | no |
| SCN4A | 1080 | M | 1 | 1 | no | yes | no | yes | no | no | no | no | no | no | no | no | no |
| SCN4A | 1081 | L | 0 | 0 | yes | no | no | yes | no | no | no | no | no | no | no | no | no |
| SCN4A | 1082 | L | 0 | 0 | yes | no | no | yes | no | no | no | no | no | no | no | no | no |
| SCN4A | 1083 | K | 1 | 0 | yes | no | no | yes | no | yes | no | no | no | no | no | no | no |
| SCN4A | 1084 | W | 1 | 0 | yes | no | no | yes | no | yes | no | no | no | no | no | no | no |
| SCN4A | 1085 | V | 0 | 0 | yes | no | no | yes | no | yes | no | no | no | no | no | no | no |
| SCN4A | 1086 | A | 0 | 0 | yes | no | no | no | no | yes | no | no | no | no | no | no | no |
| SCN4A | 1087 | Y | 0 | 0 | yes | no | no | no | no | yes | no | no | no | no | no | no | no |
| SCN4A | 1088 | G | 1 | 0 | no | yes | no | no | no | yes | no | no | no | no | no | no | no |
| SCN4A | 1089 | F | 0 | 0 | no | yes | no | yes | no | yes | no | no | no | no | no | no | no |
| SCN4A | 1090 | K | 0 | 0 | yes | no | no | no | no | yes | no | no | no | no | no | no | no |
| SCN4A | 1091 | V | 2 | 0 | yes | no | no | no | no | yes | no | no | no | no | no | no | no |
| SCN4A | 1092 | Y | 0 | 0 | yes | no | no | yes | no | yes | no | no | no | no | no | no | no |
| SCN4A | 1093 | F | 0 | 0 | yes | no | no | yes | no | yes | no | no | no | no | no | no | no |
| SCN4A | 1094 | T | 1 | 0 | no | yes | no | no | no | yes | no | no | no | no | no | no | no |
| SCN4A | 1095 | N | 1 | 0 | no | yes | no | yes | no | yes | no | no | no | no | no | no | no |
| SCN4A | 1096 | A | 1 | 0 | no | yes | no | yes | no | yes | no | no | no | no | no | no | no |
| SCN4A | 1097 | W | 0 | 0 | no | yes | no | yes | no | no | no | no | no | no | no | no | no |
| SCN4A | 1098 | C | 0 | 0 | yes | no | no | yes | no | no | no | no | no | no | no | no | no |
| SCN4A | 1099 | W | 0 | 0 | yes | no | no | yes | no | no | no | no | no | no | no | no | no |
| SCN4A | 1100 | L | 0 | 0 | yes | no | no | yes | no | no | no | no | no | no | no | no | no |
| SCN4A | 1101 | D | 0 | 0 | yes | no | no | yes | no | no | no | no | no | no | no | no | no |

| | | | | | | | | | | | | | | | | | |
|---|---|---|---|---|---|---|---|---|---|---|---|---|---|---|---|---|---|
| SCN4A | 1102 | F | 0 | 0 | yes | no | no | yes | no | no | no | no | no | no | no | no | no |
| SCN4A | 1103 | L | 0 | 0 | yes | no | no | yes | no | no | no | no | no | no | no | no | no |
| SCN4A | 1104 | I | 1 | 0 | yes | no | no | yes | no | no | no | no | no | no | no | no | no |
| SCN4A | 1105 | V | 3 | 0 | yes | no | no | yes | no | no | no | no | no | no | no | no | no |
| SCN4A | 1106 | D | 0 | 0 | yes | no | no | yes | no | no | no | no | no | no | no | no | no |
| SCN4A | 1107 | V | 0 | 0 | yes | no | no | yes | no | no | no | no | no | no | no | no | no |
| SCN4A | 1108 | S | 1 | 0 | yes | no | no | yes | no | no | no | no | no | no | no | no | no |
| SCN4A | 1109 | I | 0 | 0 | yes | no | no | yes | no | no | no | no | no | no | no | no | no |
| SCN4A | 1110 | I | 0 | 0 | yes | no | no | yes | no | no | no | no | no | no | no | no | no |
| SCN4A | 1111 | S | 0 | 0 | yes | no | no | yes | no | no | no | no | no | no | no | no | no |
| SCN4A | 1112 | L | 1 | 0 | yes | no | no | yes | no | no | no | no | no | no | no | no | no |
| SCN4A | 1113 | V | 0 | 0 | yes | no | no | yes | no | no | no | no | no | no | no | no | no |
| SCN4A | 1114 | A | 0 | 0 | yes | no | no | yes | no | no | no | no | no | no | no | no | no |
| SCN4A | 1115 | N | 0 | 0 | yes | no | no | yes | no | no | no | no | no | no | no | no | no |
| SCN4A | 1116 | W | 0 | 0 | yes | no | no | yes | yes | no | no | no | no | no | no | no | no |
| SCN4A | 1117 | L | 0 | 0 | no | yes | no | yes | yes | no | no | no | no | no | no | no | no |
| SCN4A | 1118 | G | 0 | 0 | no | yes | no | no | yes | no | no | no | no | no | no | no | no |
| SCN4A | 1119 | Y | 0 | 0 | no | yes | no | yes | yes | no | no | no | no | no | no | no | no |
| SCN4A | 1120 | S | 1 | 1 | no | yes | no | no | yes | no | no | no | no | no | no | no | no |
| SCN4A | 1121 | E | 1 | 0 | no | yes | no | no | yes | no | no | no | no | no | no | no | no |
| SCN4A | 1122 | L | 0 | 0 | no | yes | no | no | yes | no | no | no | no | no | no | no | no |
| SCN4A | 1123 | G | 0 | 0 | no | yes | no | no | yes | no | no | no | no | no | no | no | no |
| SCN4A | 1124 | P | 1 | 0 | no | yes | no | yes | no | no | no | no | no | no | no | no | no |
| SCN4A | 1125 | I | 0 | 0 | no | yes | no | yes | no | no | no | no | no | no | no | no | no |
| SCN4A | 1126 | K | 0 | 1 | no | yes | no | no | no | no | no | no | no | no | no | no | no |
| SCN4A | 1127 | S | 0 | 0 | no | yes | no | yes | no | no | no | no | no | no | no | no | no |
| SCN4A | 1128 | L | 0 | 0 | no | yes | no | yes | no | no | no | no | no | no | no | no | no |
| SCN4A | 1129 | R | 2 | 1 | no | yes | no | yes | no | no | no | no | no | no | no | no | no |
| SCN4A | 1130 | T | 0 | 0 | yes | no | no | yes | no | no | no | no | no | no | no | no | no |
| SCN4A | 1131 | L | 2 | 0 | yes | no | no | yes | no | no | no | no | no | no | no | no | no |
| SCN4A | 1132 | R | 2 | 1 | yes | no | no | yes | no | no | no | no | no | no | no | no | no |
| SCN4A | 1133 | A | 0 | 0 | yes | no | no | yes | no | no | no | no | no | no | no | no | no |
| SCN4A | 1134 | L | 1 | 0 | yes | no | no | yes | no | no | no | no | no | no | no | no | no |

| | | | | | | | | | | | | | | | | | |
|---|---|---|---|---|---|---|---|---|---|---|---|---|---|---|---|---|---|
| SCN4A | 1135 | R | 1 | 3 | yes | no | no | yes | no | no | no | no | no | no | no | no | no |
| SCN4A | 1136 | P | 0 | 0 | yes | no | no | yes | no | no | no | no | no | no | no | no | no |
| SCN4A | 1137 | L | 0 | 0 | yes | no | no | yes | no | no | no | no | no | no | no | no | no |
| SCN4A | 1138 | R | 0 | 0 | yes | no | no | yes | no | no | no | no | no | no | no | no | no |
| SCN4A | 1139 | A | 0 | 0 | yes | no | no | yes | no | no | no | no | no | no | no | no | no |
| SCN4A | 1140 | L | 0 | 0 | yes | no | no | yes | no | no | no | no | no | no | no | no | no |
| SCN4A | 1141 | S | 0 | 0 | yes | no | no | yes | no | no | no | no | no | no | no | no | no |
| SCN4A | 1142 | R | 1 | 1 | no | yes | no | yes | no | no | no | no | no | no | no | no | no |
| SCN4A | 1143 | F | 0 | 0 | no | yes | no | yes | no | yes | no | no | no | no | no | no | no |
| SCN4A | 1144 | E | 1 | 0 | yes | no | no | yes | no | yes | no | no | no | no | no | no | no |
| SCN4A | 1145 | G | 0 | 0 | yes | no | no | yes | no | yes | no | no | no | no | no | no | no |
| SCN4A | 1146 | M | 2 | 0 | yes | no | no | yes | no | yes | no | no | no | no | no | no | no |
| SCN4A | 1147 | R | 1 | 0 | yes | no | no | yes | no | yes | no | no | no | no | no | no | no |
| SCN4A | 1148 | V | 0 | 0 | yes | no | no | no | no | yes | no | no | no | no | no | no | no |
| SCN4A | 1149 | V | 0 | 1 | yes | no | no | yes | no | yes | no | no | no | no | no | no | no |
| SCN4A | 1150 | V | 0 | 0 | yes | no | no | yes | no | yes | no | no | no | no | no | no | no |
| SCN4A | 1151 | N | 0 | 0 | yes | no | no | no | no | yes | no | no | no | no | no | no | no |
| SCN4A | 1152 | A | 0 | 1 | yes | no | no | yes | no | yes | no | no | no | no | no | no | no |
| SCN4A | 1153 | L | 0 | 0 | yes | no | no | yes | no | yes | no | no | no | no | no | no | no |
| SCN4A | 1154 | L | 0 | 0 | no | yes | no | yes | no | yes | no | no | no | no | no | no | no |
| SCN4A | 1155 | G | 0 | 0 | no | yes | no | no | no | yes | no | no | no | no | no | no | no |
| SCN4A | 1156 | A | 2 | 1 | no | yes | no | yes | no | yes | no | no | no | no | no | no | no |
| SCN4A | 1157 | I | 0 | 0 | yes | no | no | yes | no | yes | no | no | no | no | no | no | no |
| SCN4A | 1158 | P | 0 | 2 | yes | no | no | yes | no | yes | no | no | no | no | no | no | no |
| SCN4A | 1159 | S | 0 | 0 | yes | no | no | yes | no | yes | no | no | no | no | no | no | no |
| SCN4A | 1160 | I | 0 | 1 | yes | no | no | yes | no | no | no | no | no | no | no | no | no |
| SCN4A | 1161 | M | 1 | 0 | yes | no | no | yes | no | no | no | no | no | no | no | no | no |
| SCN4A | 1162 | N | 0 | 0 | yes | no | no | yes | no | no | no | no | no | no | no | no | no |
| SCN4A | 1163 | V | 0 | 0 | yes | no | no | yes | no | no | no | no | no | no | no | no | no |
| SCN4A | 1164 | L | 2 | 0 | yes | no | no | yes | no | no | no | no | no | no | no | no | no |
| SCN4A | 1165 | L | 0 | 0 | yes | no | no | yes | no | no | no | no | no | no | no | no | no |
| SCN4A | 1166 | V | 0 | 1 | yes | no | no | yes | no | no | no | no | no | no | no | no | no |
| SCN4A | 1167 | C | 0 | 0 | yes | no | no | yes | no | no | no | no | no | no | no | no | no |

| | | | | | | | | | | | | | | | | | |
|---|---|---|---|---|---|---|---|---|---|---|---|---|---|---|---|---|---|
| SCN4A | 1168 | L | 1 | 0 | yes | no | no | yes | no | no | no | no | no | no | no | no | no |
| SCN4A | 1169 | I | 0 | 0 | yes | no | no | yes | no | no | no | no | no | no | no | no | no |
| SCN4A | 1170 | F | 0 | 0 | yes | no | no | yes | no | no | no | no | no | no | no | no | no |
| SCN4A | 1171 | W | 0 | 0 | yes | no | no | yes | no | no | no | no | no | no | no | no | no |
| SCN4A | 1172 | L | 0 | 0 | yes | no | no | yes | no | no | no | no | no | no | no | no | no |
| SCN4A | 1173 | I | 0 | 0 | yes | no | no | yes | no | no | no | no | no | no | no | no | no |
| SCN4A | 1174 | F | 1 | 0 | yes | no | no | yes | no | no | no | no | no | no | no | no | no |
| SCN4A | 1175 | S | 1 | 0 | yes | no | no | yes | no | no | no | no | no | no | no | no | no |
| SCN4A | 1176 | I | 0 | 0 | yes | no | no | yes | no | no | no | no | no | no | no | no | no |
| SCN4A | 1177 | M | 1 | 0 | yes | no | no | yes | no | no | no | no | no | no | no | no | no |
| SCN4A | 1178 | G | 0 | 0 | yes | no | no | yes | no | no | no | no | no | no | no | no | no |
| SCN4A | 1179 | V | 0 | 0 | yes | no | no | yes | no | no | no | no | no | no | no | no | no |
| SCN4A | 1180 | N | 1 | 1 | yes | no | no | yes | yes | no | no | no | no | no | no | no | no |
| SCN4A | 1181 | L | 1 | 0 | yes | no | no | yes | yes | no | no | no | no | no | no | no | no |
| SCN4A | 1182 | F | 0 | 0 | yes | no | no | yes | yes | no | no | no | no | no | no | no | no |
| SCN4A | 1183 | A | 0 | 0 | no | yes | no | no | yes | no | no | no | no | no | no | no | no |
| SCN4A | 1184 | G | 1 | 0 | no | yes | no | no | yes | no | no | no | no | no | no | no | no |
| SCN4A | 1185 | K | 0 | 0 | no | yes | no | no | yes | no | no | no | no | no | no | no | no |
| SCN4A | 1186 | F | 1 | 0 | no | yes | no | no | yes | no | no | no | no | no | no | no | no |
| SCN4A | 1187 | Y | 0 | 0 | no | yes | no | no | yes | no | no | no | no | no | no | no | no |
| SCN4A | 1188 | Y | 1 | 0 | no | no | yes | no | yes | no | no | no | no | no | no | no | no |
| SCN4A | 1189 | C | 0 | 0 | no | no | yes | no | yes | no | no | no | no | no | no | no | no |
| SCN4A | 1190 | I | 0 | 0 | no | no | yes | no | yes | no | no | no | no | no | no | no | no |
| SCN4A | 1191 | N | 0 | 0 | no | no | yes | no | yes | no | no | no | no | no | no | no | no |
| SCN4A | 1192 | T | 0 | 0 | no | yes | no | no | yes | no | no | no | no | no | no | no | no |
| SCN4A | 1193 | T | 0 | 0 | no | yes | no | no | yes | no | no | no | no | no | no | no | no |
| SCN4A | 1194 | T | 0 | 0 | no | yes | no | no | yes | no | no | no | no | no | no | no | no |
| SCN4A | 1195 | S | 1 | 0 | no | yes | no | no | yes | no | no | no | no | no | no | no | no |
| SCN4A | 1196 | E | 1 | 0 | no | yes | no | no | yes | no | no | no | no | no | no | no | no |
| SCN4A | 1197 | R | 0 | 0 | no | no | yes | no | yes | no | no | no | no | no | no | no | no |
| SCN4A | 1198 | F | 2 | 0 | no | yes | no | no | yes | no | no | no | no | no | no | no | no |
| SCN4A | 1199 | D | 1 | 0 | no | yes | no | no | yes | no | no | no | no | no | no | no | no |
| SCN4A | 1200 | I | 0 | 0 | no | yes | no | no | yes | no | no | no | no | no | no | no | no |

| | | | | | | | | | | | | | | | | | |
|---|---|---|---|---|---|---|---|---|---|---|---|---|---|---|---|---|---|
| SCN4A | 1201 | S | 0 | 0 | no | yes | no | no | yes | no | no | no | no | no | no | no | no |
| SCN4A | 1202 | E | 1 | 0 | no | yes | no | no | yes | no | no | no | no | no | no | no | no |
| SCN4A | 1203 | V | 1 | 0 | no | yes | no | no | yes | no | no | no | no | no | no | no | no |
| SCN4A | 1204 | N | 0 | 0 | no | yes | no | no | yes | no | no | no | no | no | no | no | no |
| SCN4A | 1205 | N | 0 | 0 | no | no | yes | no | yes | no | no | no | no | no | no | no | no |
| SCN4A | 1206 | K | 0 | 0 | yes | no | no | no | yes | no | no | no | no | no | no | no | no |
| SCN4A | 1207 | S | 0 | 0 | yes | no | no | no | yes | no | no | no | no | no | no | no | no |
| SCN4A | 1208 | E | 2 | 0 | yes | no | no | no | yes | no | no | no | no | no | no | no | no |
| SCN4A | 1209 | C | 0 | 2 | yes | no | no | no | yes | no | no | no | no | no | no | no | no |
| SCN4A | 1210 | E | 2 | 0 | yes | no | no | no | yes | no | no | no | no | no | no | no | no |
| SCN4A | 1211 | S | 1 | 0 | yes | no | no | no | yes | no | no | no | no | no | no | no | no |
| SCN4A | 1212 | L | 0 | 0 | yes | no | no | no | yes | no | no | no | no | no | no | no | no |
| SCN4A | 1213 | M | 2 | 0 | no | yes | no | no | yes | no | no | no | no | no | no | no | no |
| SCN4A | 1214 | H | 0 | 0 | no | yes | no | no | yes | no | no | no | no | no | no | no | no |
| SCN4A | 1215 | T | 0 | 0 | no | yes | no | no | yes | no | no | no | no | no | no | no | no |
| SCN4A | 1216 | G | 0 | 0 | no | yes | no | no | yes | no | no | no | no | no | no | no | no |
| SCN4A | 1217 | Q | 2 | 0 | no | yes | no | no | yes | no | no | no | no | no | no | no | no |
| SCN4A | 1218 | V | 1 | 0 | no | no | yes | no | yes | no | no | no | no | no | no | no | no |
| SCN4A | 1219 | R | 2 | 0 | no | no | yes | no | yes | no | no | no | no | no | no | no | no |
| SCN4A | 1220 | W | 0 | 0 | no | no | yes | no | yes | no | no | no | no | no | no | no | no |
| SCN4A | 1221 | L | 0 | 0 | no | no | yes | no | yes | no | no | no | no | no | no | no | no |
| SCN4A | 1222 | N | 1 | 0 | no | yes | no | no | yes | no | no | no | no | no | no | no | no |
| SCN4A | 1223 | V | 1 | 0 | no | yes | no | no | yes | no | no | no | no | no | no | no | no |
| SCN4A | 1224 | K | 0 | 0 | no | yes | no | no | yes | no | no | no | no | no | no | no | no |
| SCN4A | 1225 | V | 0 | 0 | no | yes | no | no | yes | no | no | no | no | no | no | no | no |
| SCN4A | 1226 | N | 0 | 0 | no | yes | no | no | yes | no | no | no | no | no | no | no | no |
| SCN4A | 1227 | Y | 0 | 0 | no | yes | no | yes | yes | no | no | no | no | no | no | no | no |
| SCN4A | 1228 | D | 1 | 0 | no | yes | no | no | yes | no | no | no | no | no | no | no | no |
| SCN4A | 1229 | N | 1 | 0 | no | yes | no | yes | yes | no | no | no | no | no | no | no | no |
| SCN4A | 1230 | V | 1 | 0 | yes | no | no | yes | yes | no | no | no | no | no | no | no | no |
| SCN4A | 1231 | G | 0 | 0 | yes | no | no | yes | no | no | no | yes | no | no | no | no | no |
| SCN4A | 1232 | L | 0 | 0 | yes | no | no | yes | no | no | no | yes | no | no | no | no | no |
| SCN4A | 1233 | G | 0 | 0 | yes | no | no | yes | no | no | no | yes | no | no | no | no | no |

| | | | | | | | | | | | | | | | | | |
|---|---|---|---|---|---|---|---|---|---|---|---|---|---|---|---|---|---|
| SCN4A | 1234 | Y | 0 | 0 | yes | no | no | yes | no | no | no | yes | no | no | no | no | no |
| SCN4A | 1235 | L | 1 | 0 | yes | no | no | yes | no | no | no | yes | no | no | no | no | no |
| SCN4A | 1236 | S | 1 | 0 | yes | no | no | yes | no | no | no | yes | no | no | no | no | no |
| SCN4A | 1237 | L | 0 | 0 | yes | no | no | yes | no | no | no | yes | no | no | no | no | no |
| SCN4A | 1238 | L | 0 | 0 | yes | no | no | yes | no | no | no | yes | no | no | no | no | no |
| SCN4A | 1239 | Q | 0 | 0 | yes | no | no | yes | no | no | no | yes | no | no | no | no | no |
| SCN4A | 1240 | V | 0 | 0 | yes | no | no | yes | no | no | no | yes | no | no | no | no | no |
| SCN4A | 1241 | A | 0 | 0 | yes | no | no | yes | no | no | no | yes | no | no | no | no | no |
| SCN4A | 1242 | T | 0 | 0 | yes | no | no | yes | no | no | no | yes | no | no | no | no | no |
| SCN4A | 1243 | F | 0 | 0 | no | yes | no | yes | no | no | yes | yes | no | no | no | no | no |
| SCN4A | 1244 | K | 0 | 0 | no | yes | no | yes | no | no | yes | yes | no | no | no | no | no |
| SCN4A | 1245 | G | 0 | 0 | no | yes | no | yes | no | no | yes | yes | no | no | no | no | no |
| SCN4A | 1246 | W | 0 | 0 | yes | no | no | yes | no | no | no | yes | no | no | no | no | no |
| SCN4A | 1247 | M | 2 | 0 | yes | no | no | no | no | no | no | yes | no | no | no | no | no |
| SCN4A | 1248 | D | 1 | 0 | yes | no | no | no | no | no | no | yes | no | no | no | no | no |
| SCN4A | 1249 | I | 1 | 0 | yes | no | no | no | no | no | no | yes | no | no | no | no | no |
| SCN4A | 1250 | M | 1 | 0 | yes | no | no | yes | no | no | no | yes | no | no | no | no | no |
| SCN4A | 1251 | Y | 0 | 0 | yes | no | no | no | no | no | no | yes | no | no | no | no | no |
| SCN4A | 1252 | A | 0 | 0 | yes | no | no | no | no | no | no | yes | no | no | no | no | no |
| SCN4A | 1253 | A | 0 | 0 | yes | no | no | no | yes | no | no | no | no | no | no | no | no |
| SCN4A | 1254 | V | 1 | 0 | yes | no | no | no | yes | no | no | no | no | no | no | no | no |
| SCN4A | 1255 | D | 0 | 0 | yes | no | no | no | yes | no | no | no | no | no | no | no | no |
| SCN4A | 1256 | S | 0 | 0 | no | yes | no | no | yes | no | no | no | no | no | no | no | no |
| SCN4A | 1257 | R | 1 | 0 | no | yes | no | no | yes | no | no | no | no | no | no | no | no |
| SCN4A | 1258 | E | 0 | 0 | no | yes | no | no | yes | no | no | no | no | no | no | no | no |
| SCN4A | 1259 | K | 0 | 0 | no | yes | no | no | yes | no | no | no | no | no | no | no | no |
| SCN4A | 1260 | E | 0 | 0 | no | yes | no | no | yes | no | no | no | no | no | no | no | no |
| SCN4A | 1261 | E | 1 | 0 | no | no | yes | no | yes | no | no | no | no | no | no | no | no |
| SCN4A | 1262 | Q | 0 | 0 | no | yes | no | no | yes | no | no | no | no | no | no | no | no |
| SCN4A | 1263 | P | 1 | 0 | no | yes | no | no | yes | no | no | no | no | no | no | no | no |
| SCN4A | 1264 | Q | 0 | 0 | no | yes | no | no | yes | no | no | no | no | no | no | no | no |
| SCN4A | 1265 | Y | 1 | 0 | no | yes | no | no | yes | no | no | no | no | no | no | no | no |
| SCN4A | 1266 | E | 0 | 0 | no | yes | no | no | yes | no | no | no | no | no | no | no | no |

| | | | | | | | | | | | | | | | | | |
|---|---|---|---|---|---|---|---|---|---|---|---|---|---|---|---|---|---|
| SCN4A | 1267 | V | 0 | 0 | no | yes | no | no | yes | no | no | no | no | no | no | no | no |
| SCN4A | 1268 | N | 0 | 0 | no | yes | no | no | yes | no | no | no | no | no | no | no | no |
| SCN4A | 1269 | L | 0 | 0 | yes | no | no | no | yes | no | no | no | no | no | no | no | no |
| SCN4A | 1270 | Y | 0 | 0 | yes | no | no | yes | no | no | no | no | no | no | no | no | no |
| SCN4A | 1271 | M | 0 | 0 | yes | no | no | yes | no | no | no | no | no | no | no | no | no |
| SCN4A | 1272 | Y | 0 | 0 | yes | no | no | yes | no | no | no | no | no | no | no | no | no |
| SCN4A | 1273 | L | 0 | 0 | yes | no | no | yes | no | no | no | no | no | no | no | no | no |
| SCN4A | 1274 | Y | 1 | 0 | yes | no | no | yes | no | no | no | no | no | no | no | no | no |
| SCN4A | 1275 | F | 0 | 0 | yes | no | no | yes | no | no | no | no | no | no | no | no | no |
| SCN4A | 1276 | V | 1 | 0 | yes | no | no | yes | no | no | no | no | no | no | no | no | no |
| SCN4A | 1277 | I | 2 | 0 | yes | no | no | yes | no | no | no | no | no | no | no | no | no |
| SCN4A | 1278 | F | 0 | 0 | yes | no | no | yes | no | no | no | no | no | no | no | no | no |
| SCN4A | 1279 | I | 1 | 0 | yes | no | no | yes | no | no | no | no | no | no | no | no | no |
| SCN4A | 1280 | I | 1 | 0 | yes | no | no | yes | no | no | no | no | no | no | no | no | no |
| SCN4A | 1281 | F | 0 | 0 | yes | no | no | yes | no | no | no | no | no | no | no | no | no |
| SCN4A | 1282 | G | 0 | 0 | yes | no | no | yes | no | no | no | no | no | no | no | no | no |
| SCN4A | 1283 | S | 0 | 0 | yes | no | no | yes | no | no | no | no | no | no | no | no | no |
| SCN4A | 1284 | F | 1 | 0 | yes | no | no | yes | no | no | no | no | no | no | no | no | no |
| SCN4A | 1285 | F | 1 | 0 | yes | no | no | yes | no | no | no | no | no | no | no | no | no |
| SCN4A | 1286 | T | 0 | 0 | yes | no | no | yes | no | no | no | no | no | no | no | no | no |
| SCN4A | 1287 | L | 1 | 0 | yes | no | no | yes | no | no | yes | no | no | no | no | no | no |
| SCN4A | 1288 | N | 2 | 0 | yes | no | no | yes | no | no | no | no | no | no | no | no | no |
| SCN4A | 1289 | L | 0 | 0 | yes | no | no | yes | no | no | no | no | no | no | no | no | no |
| SCN4A | 1290 | F | 0 | 1 | yes | no | no | yes | no | no | no | no | no | no | no | no | no |
| SCN4A | 1291 | I | 2 | 0 | yes | no | no | yes | no | no | yes | no | no | no | no | no | no |
| SCN4A | 1292 | G | 1 | 1 | yes | no | no | yes | no | yes | no | no | no | no | no | no | no |
| SCN4A | 1293 | V | 0 | 1 | yes | no | no | yes | no | yes | no | no | no | no | no | no | no |
| SCN4A | 1294 | I | 0 | 0 | yes | no | no | yes | no | yes | no | no | no | no | no | no | no |
| SCN4A | 1295 | I | 1 | 0 | yes | no | no | no | no | yes | yes | no | no | no | no | no | no |
| SCN4A | 1296 | D | 0 | 0 | yes | no | no | no | no | yes | no | no | no | no | no | no | no |
| SCN4A | 1297 | N | 0 | 2 | yes | no | no | no | no | yes | no | no | no | no | no | no | no |
| SCN4A | 1298 | F | 0 | 0 | yes | no | no | no | no | yes | no | no | no | no | no | no | no |
| SCN4A | 1299 | N | 0 | 0 | yes | no | no | no | no | yes | yes | no | no | no | no | no | no |

| | | | | | | | | | | | | | | | | | |
|---|---|---|---|---|---|---|---|---|---|---|---|---|---|---|---|---|---|
| SCN4A | 1300 | Q | 1 | 0 | yes | no | no | no | no | yes | no | no | no | no | no | no | no |
| SCN4A | 1301 | Q | 0 | 0 | yes | no | no | no | no | yes | no | no | no | no | no | no | no |
| SCN4A | 1302 | K | 0 | 0 | yes | no | no | no | no | yes | no | no | no | no | no | no | no |
| SCN4A | 1303 | K | 0 | 0 | yes | no | no | no | no | yes | yes | no | no | no | no | no | no |
| SCN4A | 1304 | K | 0 | 0 | yes | no | no | no | no | yes | no | no | no | no | no | no | no |
| SCN4A | 1305 | L | 1 | 0 | no | yes | no | no | no | yes | no | no | no | no | no | no | no |
| SCN4A | 1306 | G | 1 | 1 | no | yes | no | no | no | yes | no | no | no | no | no | no | no |
| SCN4A | 1307 | G | 1 | 0 | no | yes | no | no | no | yes | no | no | no | no | no | no | no |
| SCN4A | 1308 | K | 0 | 0 | no | yes | no | no | no | yes | no | no | no | no | no | no | no |
| SCN4A | 1309 | D | 0 | 0 | no | yes | no | no | no | yes | no | no | no | no | no | no | no |
| SCN4A | 1310 | I | 0 | 0 | no | yes | no | no | no | yes | no | no | no | no | no | no | no |
| SCN4A | 1311 | F | 0 | 0 | no | yes | no | yes | no | yes | no | no | no | no | no | no | no |
| SCN4A | 1312 | M | 0 | 0 | no | yes | no | no | no | yes | no | no | no | no | no | no | no |
| SCN4A | 1313 | T | 1 | 2 | no | yes | no | no | no | yes | no | no | no | no | no | no | no |
| SCN4A | 1314 | E | 0 | 0 | no | yes | no | no | no | yes | no | no | no | no | no | no | no |
| SCN4A | 1315 | E | 0 | 0 | no | yes | no | no | no | yes | no | no | no | no | no | no | no |
| SCN4A | 1316 | Q | 0 | 0 | yes | no | no | no | no | yes | no | no | no | no | no | no | no |
| SCN4A | 1317 | K | 0 | 0 | yes | no | no | no | no | yes | no | no | no | no | no | no | no |
| SCN4A | 1318 | K | 0 | 0 | yes | no | no | no | no | yes | no | no | no | no | no | no | no |
| SCN4A | 1319 | Y | 0 | 0 | yes | no | no | no | no | yes | no | no | no | no | no | no | no |
| SCN4A | 1320 | Y | 1 | 0 | yes | no | no | no | no | yes | no | no | no | no | no | no | no |
| SCN4A | 1321 | N | 1 | 0 | yes | no | no | no | no | yes | no | no | no | no | no | no | no |
| SCN4A | 1322 | A | 0 | 0 | yes | no | no | no | no | yes | no | no | no | no | no | no | no |
| SCN4A | 1323 | M | 0 | 0 | yes | no | no | no | no | yes | no | no | no | no | no | no | no |
| SCN4A | 1324 | K | 0 | 0 | yes | no | no | no | no | yes | no | no | no | no | no | no | no |
| SCN4A | 1325 | K | 0 | 0 | no | yes | no | no | no | yes | no | no | no | no | no | no | no |
| SCN4A | 1326 | L | 1 | 0 | no | yes | no | no | no | yes | no | no | no | no | no | no | no |
| SCN4A | 1327 | G | 0 | 0 | no | yes | no | no | no | yes | no | no | no | no | no | no | no |
| SCN4A | 1328 | S | 0 | 0 | no | yes | no | no | no | yes | no | no | no | no | no | no | no |
| SCN4A | 1329 | K | 1 | 0 | no | yes | no | no | no | yes | no | no | no | no | no | no | no |
| SCN4A | 1330 | K | 1 | 0 | no | yes | no | no | no | yes | no | no | no | no | no | no | no |
| SCN4A | 1331 | P | 1 | 0 | no | yes | no | no | no | yes | no | no | no | no | no | no | no |
| SCN4A | 1332 | Q | 0 | 0 | no | yes | no | no | no | yes | no | no | no | no | no | no | no |

| | | | | | | | | | | | | | | | | | |
|---|---|---|---|---|---|---|---|---|---|---|---|---|---|---|---|---|---|
| SCN4A | 1333 | K | 1 | 0 | no | yes | no | no | no | yes | no | no | no | no | no | no | no |
| SCN4A | 1334 | P | 0 | 0 | no | yes | no | no | no | yes | no | no | no | no | no | no | no |
| SCN4A | 1335 | I | 0 | 0 | no | yes | no | no | no | yes | no | no | no | no | no | no | no |
| SCN4A | 1336 | P | 1 | 0 | no | yes | no | no | no | yes | no | no | no | no | no | no | no |
| SCN4A | 1337 | R | 2 | 1 | no | yes | no | no | no | yes | no | no | no | no | no | no | no |
| SCN4A | 1338 | P | 0 | 0 | no | yes | no | no | no | yes | no | no | no | no | no | no | no |
| SCN4A | 1339 | Q | 1 | 0 | no | yes | no | no | no | yes | no | no | no | no | no | no | no |
| SCN4A | 1340 | N | 0 | 0 | no | yes | no | no | no | yes | no | no | no | no | no | no | no |
| SCN4A | 1341 | K | 0 | 0 | no | yes | no | no | no | yes | no | no | no | no | no | no | no |
| SCN4A | 1342 | I | 0 | 0 | no | yes | no | no | no | yes | no | no | no | no | no | no | no |
| SCN4A | 1343 | Q | 0 | 0 | no | yes | no | no | no | yes | no | no | no | no | no | no | no |
| SCN4A | 1344 | G | 2 | 0 | no | yes | no | no | no | yes | no | no | no | no | no | no | no |
| SCN4A | 1345 | M | 2 | 0 | no | yes | no | no | no | yes | no | no | no | no | no | no | no |
| SCN4A | 1346 | V | 0 | 0 | no | yes | no | yes | no | yes | no | no | no | no | no | no | no |
| SCN4A | 1347 | Y | 0 | 0 | yes | no | no | no | no | yes | no | no | no | no | no | no | no |
| SCN4A | 1348 | D | 0 | 0 | yes | no | no | no | no | yes | no | no | no | no | no | no | no |
| SCN4A | 1349 | L | 0 | 0 | yes | no | no | no | no | yes | no | no | no | no | no | no | no |
| SCN4A | 1350 | V | 1 | 0 | yes | no | no | yes | no | yes | no | no | no | no | no | no | no |
| SCN4A | 1351 | T | 1 | 0 | no | yes | no | no | no | yes | no | no | no | no | no | no | no |
| SCN4A | 1352 | K | 0 | 0 | no | yes | no | no | no | yes | no | no | no | no | no | no | no |
| SCN4A | 1353 | Q | 0 | 0 | no | yes | no | no | no | yes | no | no | no | no | no | no | no |
| SCN4A | 1354 | A | 0 | 0 | yes | no | no | yes | no | yes | no | no | no | no | no | no | no |
| SCN4A | 1355 | F | 0 | 0 | yes | no | no | yes | no | no | no | no | no | no | no | no | no |
| SCN4A | 1356 | D | 1 | 0 | yes | no | no | yes | no | no | no | no | no | no | no | no | no |
| SCN4A | 1357 | I | 0 | 0 | yes | no | no | yes | no | no | no | no | no | no | no | no | no |
| SCN4A | 1358 | T | 1 | 0 | yes | no | no | yes | no | no | no | no | no | no | no | no | no |
| SCN4A | 1359 | I | 0 | 0 | yes | no | no | yes | no | no | no | no | no | no | no | no | no |
| SCN4A | 1360 | M | 1 | 1 | yes | no | no | yes | no | no | no | no | no | no | no | no | no |
| SCN4A | 1361 | I | 0 | 0 | yes | no | no | yes | no | no | no | no | no | no | no | no | no |
| SCN4A | 1362 | L | 0 | 0 | yes | no | no | yes | no | no | no | no | no | no | no | no | no |
| SCN4A | 1363 | I | 0 | 1 | no | yes | no | yes | no | no | no | no | no | no | no | no | no |
| SCN4A | 1364 | C | 0 | 0 | no | yes | no | yes | no | no | no | no | no | no | no | no | no |
| SCN4A | 1365 | L | 0 | 0 | yes | no | no | yes | no | no | no | no | no | no | no | no | no |

| | | | | | | | | | | | | | | | | | |
|---|---|---|---|---|---|---|---|---|---|---|---|---|---|---|---|---|---|
| SCN4A | 1366 | N | 0 | 1 | yes | no | no | yes | no | no | no | no | no | no | no | no | no |
| SCN4A | 1367 | M | 1 | 0 | yes | no | no | yes | no | no | no | no | no | no | no | no | no |
| SCN4A | 1368 | V | 1 | 0 | yes | no | no | yes | no | no | no | no | no | no | no | no | no |
| SCN4A | 1369 | T | 1 | 0 | yes | no | no | yes | no | no | no | no | no | no | no | no | no |
| SCN4A | 1370 | M | 1 | 1 | yes | no | no | yes | no | no | no | no | no | no | no | no | no |
| SCN4A | 1371 | M | 0 | 0 | yes | no | no | yes | no | no | no | no | no | no | no | no | no |
| SCN4A | 1372 | V | 1 | 0 | no | yes | no | yes | no | no | no | no | no | no | no | no | no |
| SCN4A | 1373 | E | 0 | 0 | no | yes | no | yes | yes | no | no | no | no | no | no | no | no |
| SCN4A | 1374 | T | 0 | 0 | no | yes | no | no | yes | no | no | no | no | no | no | no | no |
| SCN4A | 1375 | D | 0 | 0 | no | yes | no | no | yes | no | no | no | no | no | no | no | no |
| SCN4A | 1376 | N | 2 | 0 | no | yes | no | no | yes | no | no | no | no | no | no | no | no |
| SCN4A | 1377 | Q | 1 | 0 | no | yes | no | no | yes | no | no | no | no | no | no | no | no |
| SCN4A | 1378 | S | 1 | 0 | no | yes | no | no | yes | no | no | no | no | no | no | no | no |
| SCN4A | 1379 | Q | 1 | 0 | yes | no | no | no | yes | no | no | no | no | no | no | no | no |
| SCN4A | 1380 | L | 0 | 0 | yes | no | no | no | yes | no | no | no | no | no | no | no | no |
| SCN4A | 1381 | K | 0 | 0 | yes | no | no | no | yes | no | no | no | no | no | no | no | no |
| SCN4A | 1382 | V | 0 | 0 | yes | no | no | no | yes | no | no | no | no | no | no | no | no |
| SCN4A | 1383 | D | 0 | 0 | yes | no | no | no | yes | no | no | no | no | no | no | no | no |
| SCN4A | 1384 | I | 2 | 0 | yes | no | no | yes | no | no | no | no | no | no | no | no | no |
| SCN4A | 1385 | L | 0 | 0 | yes | no | no | yes | no | no | no | no | no | no | no | no | no |
| SCN4A | 1386 | Y | 1 | 0 | yes | no | no | yes | no | no | no | no | no | no | no | no | no |
| SCN4A | 1387 | N | 0 | 0 | yes | no | no | yes | no | no | no | no | no | no | no | no | no |
| SCN4A | 1388 | I | 0 | 0 | yes | no | no | yes | no | no | no | no | no | no | no | no | no |
| SCN4A | 1389 | N | 0 | 0 | yes | no | no | yes | no | no | no | no | no | no | no | no | no |
| SCN4A | 1390 | M | 1 | 0 | yes | no | no | yes | no | no | no | no | no | no | no | no | no |
| SCN4A | 1391 | I | 2 | 0 | yes | no | no | yes | no | no | no | no | no | no | no | no | no |
| SCN4A | 1392 | F | 0 | 0 | yes | no | no | yes | no | no | no | no | no | no | no | no | no |
| SCN4A | 1393 | I | 1 | 1 | yes | no | no | yes | no | no | no | no | no | no | no | no | no |
| SCN4A | 1394 | I | 2 | 0 | yes | no | no | yes | no | no | no | no | no | no | no | no | no |
| SCN4A | 1395 | I | 0 | 0 | yes | no | no | yes | no | no | no | no | no | no | no | no | no |
| SCN4A | 1396 | F | 0 | 0 | yes | no | no | yes | no | no | no | no | no | no | no | no | no |
| SCN4A | 1397 | T | 0 | 0 | yes | no | no | yes | no | no | no | no | no | no | no | no | no |
| SCN4A | 1398 | G | 0 | 0 | yes | no | no | yes | no | no | no | no | no | no | no | no | no |

| | | | | | | | | | | | | | | | | | |
|---|---|---|---|---|---|---|---|---|---|---|---|---|---|---|---|---|---|
| SCN4A | 1399 | E | 0 | 0 | yes | no | no | yes | no | no | no | no | no | no | no | no | no |
| SCN4A | 1400 | C | 0 | 0 | yes | no | no | yes | no | no | no | no | no | no | no | no | no |
| SCN4A | 1401 | V | 1 | 0 | yes | no | no | yes | no | no | no | no | no | no | no | no | no |
| SCN4A | 1402 | L | 0 | 0 | yes | no | no | yes | no | no | no | no | no | no | no | no | no |
| SCN4A | 1403 | K | 1 | 0 | yes | no | no | yes | no | yes | no | no | no | no | no | no | no |
| SCN4A | 1404 | M | 1 | 0 | yes | no | no | yes | no | yes | no | no | no | no | no | no | no |
| SCN4A | 1405 | L | 0 | 0 | no | yes | no | yes | no | yes | no | no | no | no | no | no | no |
| SCN4A | 1406 | A | 1 | 0 | no | yes | no | no | no | yes | no | no | no | no | no | no | no |
| SCN4A | 1407 | L | 0 | 0 | no | yes | no | no | no | yes | no | no | no | no | no | no | no |
| SCN4A | 1408 | R | 2 | 0 | no | yes | no | no | no | yes | no | no | no | no | no | no | no |
| SCN4A | 1409 | Q | 0 | 0 | no | yes | no | no | no | yes | no | no | no | no | no | no | no |
| SCN4A | 1410 | Y | 0 | 0 | no | yes | no | no | no | yes | no | no | no | no | no | no | no |
| SCN4A | 1411 | Y | 0 | 0 | no | yes | no | yes | no | yes | no | no | no | no | no | no | no |
| SCN4A | 1412 | F | 1 | 0 | no | yes | no | yes | no | yes | no | no | no | no | no | no | no |
| SCN4A | 1413 | T | 0 | 0 | no | yes | no | no | no | yes | no | no | no | no | no | no | no |
| SCN4A | 1414 | V | 2 | 0 | no | yes | no | no | no | yes | no | no | no | no | no | no | no |
| SCN4A | 1415 | G | 0 | 0 | yes | no | no | yes | no | no | no | no | no | no | no | no | no |
| SCN4A | 1416 | W | 0 | 0 | yes | no | no | yes | no | no | no | no | no | no | no | no | no |
| SCN4A | 1417 | N | 0 | 0 | yes | no | no | yes | no | no | no | no | no | no | no | no | no |
| SCN4A | 1418 | I | 0 | 0 | yes | no | no | yes | no | no | no | no | no | no | no | no | no |
| SCN4A | 1419 | F | 0 | 1 | yes | no | no | yes | no | no | no | no | no | no | no | no | no |
| SCN4A | 1420 | D | 0 | 0 | yes | no | no | yes | no | no | no | no | no | no | no | no | no |
| SCN4A | 1421 | F | 0 | 0 | yes | no | no | yes | no | no | no | no | no | no | no | no | no |
| SCN4A | 1422 | V | 1 | 0 | yes | no | no | yes | no | no | no | no | no | no | no | no | no |
| SCN4A | 1423 | V | 0 | 0 | yes | no | no | yes | no | no | no | no | no | no | no | no | no |
| SCN4A | 1424 | V | 2 | 0 | yes | no | no | yes | no | no | no | no | no | no | no | no | no |
| SCN4A | 1425 | I | 0 | 0 | yes | no | no | yes | no | no | no | no | no | no | no | no | no |
| SCN4A | 1426 | L | 1 | 0 | yes | no | no | yes | no | no | no | no | no | no | no | no | no |
| SCN4A | 1427 | S | 1 | 0 | yes | no | no | yes | no | no | no | no | no | no | no | no | no |
| SCN4A | 1428 | I | 1 | 0 | yes | no | no | yes | no | no | no | no | no | no | no | no | no |
| SCN4A | 1429 | V | 1 | 0 | yes | no | no | yes | no | no | no | no | no | no | no | no | no |
| SCN4A | 1430 | G | 0 | 0 | yes | no | no | yes | no | no | no | no | no | no | no | no | no |
| SCN4A | 1431 | L | 2 | 0 | yes | no | no | yes | no | no | no | no | no | no | no | no | no |

| | | | | | | | | | | | | | | | | | |
|---|---|---|---|---|---|---|---|---|---|---|---|---|---|---|---|---|---|
| SCN4A | 1432 | A | 0 | 0 | yes | no | no | yes | no | no | no | no | no | no | no | no | no |
| SCN4A | 1433 | L | 0 | 2 | yes | no | no | yes | yes | no | no | no | no | no | no | no | no |
| SCN4A | 1434 | S | 0 | 1 | yes | no | no | yes | yes | no | no | no | no | no | no | no | no |
| SCN4A | 1435 | D | 1 | 0 | yes | no | no | no | yes | no | no | no | no | no | no | no | no |
| SCN4A | 1436 | L | 0 | 1 | yes | no | no | no | yes | no | no | no | no | no | no | no | no |
| SCN4A | 1437 | I | 0 | 0 | yes | no | no | no | yes | no | no | no | no | no | no | no | no |
| SCN4A | 1438 | Q | 1 | 0 | yes | no | no | no | yes | no | no | no | no | no | no | no | no |
| SCN4A | 1439 | K | 0 | 0 | yes | no | no | no | yes | no | no | no | no | no | no | no | no |
| SCN4A | 1440 | Y | 0 | 0 | no | yes | no | no | yes | no | no | no | no | no | no | no | no |
| SCN4A | 1441 | F | 0 | 0 | no | yes | no | no | yes | no | no | no | no | no | no | no | no |
| SCN4A | 1442 | V | 1 | 2 | no | yes | no | no | yes | no | no | no | no | no | no | no | no |
| SCN4A | 1443 | S | 0 | 0 | no | yes | no | no | yes | no | no | no | no | no | no | no | no |
| SCN4A | 1444 | P | 0 | 0 | no | yes | no | no | yes | no | no | no | no | no | no | no | no |
| SCN4A | 1445 | T | 1 | 0 | no | yes | no | no | yes | no | no | no | no | no | no | no | no |
| SCN4A | 1446 | L | 0 | 0 | no | yes | no | yes | no | no | no | no | no | no | no | no | no |
| SCN4A | 1447 | F | 0 | 0 | yes | no | no | no | no | no | no | no | no | no | no | no | no |
| SCN4A | 1448 | R | 0 | 2 | yes | no | no | no | no | no | no | no | no | no | no | no | no |
| SCN4A | 1449 | V | 0 | 0 | yes | no | no | yes | no | no | no | no | no | no | no | no | no |
| SCN4A | 1450 | I | 0 | 0 | yes | no | no | yes | no | no | no | no | no | no | no | no | no |
| SCN4A | 1451 | R | 2 | 2 | yes | no | no | yes | no | no | no | no | no | no | no | no | no |
| SCN4A | 1452 | L | 1 | 0 | yes | no | no | yes | no | no | no | no | no | no | no | no | no |
| SCN4A | 1453 | A | 1 | 0 | yes | no | no | yes | no | no | no | no | no | no | no | no | no |
| SCN4A | 1454 | R | 1 | 1 | yes | no | no | yes | no | no | no | no | no | no | no | no | no |
| SCN4A | 1455 | I | 1 | 1 | yes | no | no | yes | no | no | no | no | no | no | no | no | no |
| SCN4A | 1456 | G | 0 | 1 | yes | no | no | yes | no | no | no | no | no | no | no | no | no |
| SCN4A | 1457 | R | 1 | 1 | yes | no | no | yes | no | no | no | no | no | no | no | no | no |
| SCN4A | 1458 | V | 1 | 1 | yes | no | no | yes | no | no | no | no | no | no | no | no | no |
| SCN4A | 1459 | L | 0 | 0 | yes | no | no | yes | no | no | no | no | no | no | no | no | no |
| SCN4A | 1460 | R | 2 | 2 | yes | no | no | yes | no | no | no | no | no | no | no | no | no |
| SCN4A | 1461 | L | 0 | 1 | yes | no | no | yes | no | no | no | no | no | no | no | no | no |
| SCN4A | 1462 | I | 1 | 0 | yes | no | no | yes | no | no | no | no | no | no | no | no | no |
| SCN4A | 1463 | R | 3 | 2 | yes | no | no | yes | no | yes | no | no | no | no | no | no | no |
| SCN4A | 1464 | G | 1 | 0 | no | yes | no | yes | no | yes | no | no | no | no | no | no | no |

| | | | | | | | | | | | | | | | | | |
|---|---|---|---|---|---|---|---|---|---|---|---|---|---|---|---|---|---|
| SCN4A | 1465 | A | 1 | 0 | no | yes | no | yes | no | yes | no | no | no | no | no | no | no |
| SCN4A | 1466 | K | 1 | 0 | no | yes | no | no | no | yes | no | no | no | no | no | no | no |
| SCN4A | 1467 | G | 0 | 0 | no | yes | no | yes | no | yes | no | no | no | no | no | no | no |
| SCN4A | 1468 | I | 1 | 0 | yes | no | no | yes | no | yes | no | no | no | no | no | no | no |
| SCN4A | 1469 | R | 1 | 0 | yes | no | no | no | no | yes | no | no | no | no | no | no | no |
| SCN4A | 1470 | T | 1 | 0 | yes | no | no | no | no | yes | no | no | no | no | no | no | no |
| SCN4A | 1471 | L | 0 | 0 | yes | no | no | yes | no | yes | no | no | no | no | no | no | no |
| SCN4A | 1472 | L | 0 | 0 | yes | no | no | yes | no | yes | no | no | no | no | no | no | no |
| SCN4A | 1473 | F | 0 | 1 | yes | no | no | no | no | yes | no | no | no | no | no | no | no |
| SCN4A | 1474 | A | 1 | 0 | yes | no | no | yes | no | yes | no | no | no | no | no | no | no |
| SCN4A | 1475 | L | 1 | 0 | yes | no | no | yes | no | yes | no | no | no | no | no | no | no |
| SCN4A | 1476 | M | 1 | 2 | yes | no | no | yes | no | yes | no | no | no | no | no | no | no |
| SCN4A | 1477 | M | 1 | 0 | yes | no | no | no | no | yes | no | no | no | no | no | no | no |
| SCN4A | 1478 | S | 1 | 0 | yes | no | no | yes | no | yes | no | no | no | no | no | no | no |
| SCN4A | 1479 | L | 0 | 0 | yes | no | no | yes | no | yes | no | no | no | no | no | no | no |
| SCN4A | 1480 | P | 0 | 0 | yes | no | no | yes | no | yes | no | no | no | no | no | no | no |
| SCN4A | 1481 | A | 0 | 1 | yes | no | no | yes | no | yes | no | no | no | no | no | no | no |
| SCN4A | 1482 | L | 0 | 0 | yes | no | no | yes | no | no | no | no | no | no | no | no | no |
| SCN4A | 1483 | F | 0 | 0 | yes | no | no | yes | no | no | no | no | no | no | no | no | no |
| SCN4A | 1484 | N | 0 | 0 | yes | no | no | yes | no | no | no | no | no | no | no | no | no |
| SCN4A | 1485 | I | 0 | 0 | yes | no | no | yes | no | no | no | no | no | no | no | no | no |
| SCN4A | 1486 | G | 0 | 0 | yes | no | no | yes | no | no | no | no | no | no | no | no | no |
| SCN4A | 1487 | L | 0 | 0 | yes | no | no | yes | no | no | no | no | no | no | no | no | no |
| SCN4A | 1488 | L | 0 | 1 | yes | no | no | yes | no | no | no | no | no | no | no | no | no |
| SCN4A | 1489 | L | 0 | 0 | yes | no | no | yes | no | no | no | no | no | no | no | no | no |
| SCN4A | 1490 | F | 0 | 0 | yes | no | no | yes | no | no | no | no | no | no | no | no | no |
| SCN4A | 1491 | L | 0 | 0 | yes | no | no | yes | no | no | no | no | no | no | no | no | no |
| SCN4A | 1492 | V | 0 | 0 | yes | no | no | yes | no | no | no | no | no | no | no | no | no |
| SCN4A | 1493 | M | 0 | 0 | yes | no | no | yes | no | no | no | no | no | no | no | no | no |
| SCN4A | 1494 | F | 0 | 0 | yes | no | no | yes | no | no | no | no | no | no | no | no | no |
| SCN4A | 1495 | I | 1 | 1 | yes | no | no | yes | no | no | no | no | no | no | no | no | no |
| SCN4A | 1496 | Y | 0 | 0 | yes | no | no | yes | no | no | no | no | no | no | no | no | no |
| SCN4A | 1497 | S | 0 | 0 | yes | no | no | yes | no | no | no | no | no | no | no | no | no |

| | | | | | | | | | | | | | | | | | |
|---|---|---|---|---|---|---|---|---|---|---|---|---|---|---|---|---|---|
| SCN4A | 1498 | I | 0 | 0 | yes | no | no | yes | no | no | no | no | no | no | no | no | no |
| SCN4A | 1499 | F | 0 | 0 | yes | no | no | yes | no | no | no | no | no | no | no | no | no |
| SCN4A | 1500 | G | 1 | 0 | yes | no | no | yes | yes | no | no | no | no | no | no | no | no |
| SCN4A | 1501 | M | 1 | 0 | yes | no | no | yes | yes | no | no | no | no | no | no | no | no |
| SCN4A | 1502 | S | 0 | 0 | yes | no | no | no | yes | no | no | no | no | no | no | no | no |
| SCN4A | 1503 | N | 1 | 0 | yes | no | no | no | yes | no | no | no | no | no | no | no | no |
| SCN4A | 1504 | F | 0 | 0 | no | yes | no | no | yes | no | no | no | no | no | no | no | no |
| SCN4A | 1505 | A | 0 | 0 | no | yes | no | no | yes | no | no | no | no | no | no | no | no |
| SCN4A | 1506 | Y | 1 | 0 | no | yes | no | no | yes | no | no | no | no | no | no | no | no |
| SCN4A | 1507 | V | 1 | 0 | no | yes | no | no | yes | no | no | no | no | no | no | no | no |
| SCN4A | 1508 | K | 0 | 0 | no | yes | no | no | yes | no | no | no | no | no | no | no | no |
| SCN4A | 1509 | K | 1 | 0 | no | yes | no | no | yes | no | yes | no | no | no | no | no | no |
| SCN4A | 1510 | E | 0 | 0 | no | yes | no | no | yes | no | yes | no | no | no | no | no | no |
| SCN4A | 1511 | S | 1 | 0 | no | yes | no | no | yes | no | yes | no | no | no | no | no | no |
| SCN4A | 1512 | G | 0 | 0 | no | yes | no | no | yes | no | no | no | no | no | no | no | no |
| SCN4A | 1513 | I | 1 | 0 | no | yes | no | no | yes | no | no | no | no | no | no | no | no |
| SCN4A | 1514 | D | 1 | 0 | no | yes | no | no | yes | no | no | no | no | no | no | no | no |
| SCN4A | 1515 | D | 1 | 0 | no | yes | no | no | yes | no | no | no | yes | no | no | no | no |
| SCN4A | 1516 | M | 1 | 0 | no | yes | no | no | yes | no | no | no | no | no | no | no | no |
| SCN4A | 1517 | F | 0 | 0 | no | yes | no | no | yes | no | no | no | no | no | no | no | no |
| SCN4A | 1518 | N | 0 | 0 | no | yes | no | no | yes | no | no | no | no | no | no | no | no |
| SCN4A | 1519 | F | 1 | 0 | no | yes | no | no | yes | no | no | no | no | no | no | no | no |
| SCN4A | 1520 | E | 1 | 1 | no | yes | no | no | yes | no | no | no | no | no | no | no | no |
| SCN4A | 1521 | T | 1 | 0 | no | yes | no | no | yes | no | no | no | no | no | no | no | no |
| SCN4A | 1522 | F | 0 | 0 | yes | no | no | yes | no | no | no | yes | no | no | no | no | no |
| SCN4A | 1523 | G | 1 | 0 | yes | no | no | yes | no | no | no | yes | no | no | no | no | no |
| SCN4A | 1524 | N | 1 | 0 | yes | no | no | no | no | no | no | yes | no | no | no | no | no |
| SCN4A | 1525 | S | 0 | 0 | yes | no | no | yes | no | no | no | yes | no | no | no | no | no |
| SCN4A | 1526 | I | 0 | 0 | yes | no | no | yes | no | no | no | yes | no | no | no | no | no |
| SCN4A | 1527 | I | 0 | 0 | yes | no | no | yes | no | no | no | yes | no | no | no | no | no |
| SCN4A | 1528 | C | 0 | 0 | yes | no | no | yes | no | no | no | yes | no | no | no | no | no |
| SCN4A | 1529 | L | 0 | 0 | yes | no | no | yes | no | no | no | yes | no | no | no | no | no |
| SCN4A | 1530 | F | 1 | 0 | yes | no | no | yes | no | no | no | yes | no | no | no | no | no |

| | | | | | | | | | | | | | | | | | |
|---|---|---|---|---|---|---|---|---|---|---|---|---|---|---|---|---|---|
| SCN4A | 1531 | E | 0 | 0 | yes | no | no | yes | no | no | no | yes | no | no | no | no | no |
| SCN4A | 1532 | I | 1 | 0 | yes | no | no | yes | no | no | no | yes | no | no | no | no | no |
| SCN4A | 1533 | T | 1 | 0 | yes | no | no | yes | no | no | no | yes | no | no | no | no | no |
| SCN4A | 1534 | T | 1 | 1 | yes | no | no | yes | no | no | no | yes | no | no | no | no | no |
| SCN4A | 1535 | S | 0 | 0 | no | yes | no | yes | no | no | no | yes | no | no | no | no | no |
| SCN4A | 1536 | A | 0 | 0 | no | yes | no | yes | no | no | no | yes | no | no | no | no | no |
| SCN4A | 1537 | G | 1 | 1 | no | yes | no | yes | no | no | no | yes | no | no | no | no | no |
| SCN4A | 1538 | W | 0 | 0 | yes | no | no | yes | no | no | no | yes | no | no | no | no | no |
| SCN4A | 1539 | D | 1 | 0 | yes | no | no | yes | no | no | yes | yes | no | no | no | no | no |
| SCN4A | 1540 | G | 1 | 0 | yes | no | no | no | no | no | yes | yes | no | no | no | no | no |
| SCN4A | 1541 | L | 0 | 0 | yes | no | no | no | no | no | no | yes | no | no | no | no | no |
| SCN4A | 1542 | L | 0 | 0 | yes | no | no | no | no | no | no | yes | no | no | no | no | no |
| SCN4A | 1543 | N | 1 | 0 | no | yes | no | no | no | no | no | yes | no | no | no | no | no |
| SCN4A | 1544 | P | 0 | 0 | no | yes | no | no | no | no | no | yes | no | no | no | no | no |
| SCN4A | 1545 | I | 1 | 0 | no | yes | no | no | yes | no | no | no | no | no | no | no | no |
| SCN4A | 1546 | L | 0 | 0 | no | yes | no | no | yes | no | no | no | no | no | no | no | no |
| SCN4A | 1547 | N | 1 | 0 | no | yes | no | no | yes | no | no | no | no | no | no | no | no |
| SCN4A | 1548 | S | 0 | 0 | no | yes | no | no | yes | no | no | no | no | no | no | no | no |
| SCN4A | 1549 | G | 1 | 0 | no | yes | no | no | yes | no | no | no | no | no | no | no | no |
| SCN4A | 1550 | P | 2 | 0 | no | yes | no | no | yes | no | no | no | no | no | no | no | no |
| SCN4A | 1551 | P | 0 | 0 | no | yes | no | no | yes | no | no | no | no | no | no | no | no |
| SCN4A | 1552 | D | 1 | 0 | no | yes | no | no | yes | no | no | no | no | no | no | no | no |
| SCN4A | 1553 | C | 0 | 0 | no | yes | no | no | yes | no | no | no | no | no | no | no | no |
| SCN4A | 1554 | D | 2 | 0 | no | yes | no | no | yes | no | no | no | no | no | no | no | no |
| SCN4A | 1555 | P | 0 | 0 | no | yes | no | no | yes | no | no | no | no | no | no | no | no |
| SCN4A | 1556 | N | 2 | 0 | no | yes | no | no | yes | no | no | no | no | no | no | no | no |
| SCN4A | 1557 | L | 0 | 0 | no | yes | no | no | yes | no | no | no | no | no | no | no | no |
| SCN4A | 1558 | E | 0 | 0 | no | yes | no | no | yes | no | no | no | yes | no | no | no | no |
| SCN4A | 1559 | N | 0 | 0 | no | yes | no | no | yes | no | no | no | yes | no | no | no | no |
| SCN4A | 1560 | P | 2 | 0 | no | yes | no | no | yes | no | no | no | yes | no | no | no | no |
| SCN4A | 1561 | G | 0 | 0 | no | yes | no | no | yes | no | no | no | yes | no | no | no | no |
| SCN4A | 1562 | T | 0 | 0 | no | yes | no | no | yes | no | no | no | no | no | no | no | no |
| SCN4A | 1563 | S | 1 | 0 | no | yes | no | no | yes | no | no | no | no | no | no | no | no |

| | | | | | | | | | | | | | | | | | |
|---|---|---|---|---|---|---|---|---|---|---|---|---|---|---|---|---|---|
| SCN4A | 1564 | V | 1 | 0 | no | yes | no | no | yes | no | no | no | no | no | no | no | no |
| SCN4A | 1565 | K | 0 | 0 | no | yes | no | no | yes | no | no | no | no | no | no | no | no |
| SCN4A | 1566 | G | 0 | 0 | no | yes | no | no | yes | no | no | no | no | no | no | no | no |
| SCN4A | 1567 | D | 1 | 0 | no | yes | no | no | yes | no | no | no | no | no | no | no | no |
| SCN4A | 1568 | C | 0 | 0 | no | yes | no | no | yes | no | no | no | no | no | no | no | no |
| SCN4A | 1569 | G | 2 | 0 | no | yes | no | no | yes | no | no | no | no | no | no | no | no |
| SCN4A | 1570 | N | 3 | 0 | no | yes | no | no | yes | no | no | no | no | no | no | no | no |
| SCN4A | 1571 | P | 1 | 0 | yes | no | no | no | yes | no | no | no | no | no | no | no | no |
| SCN4A | 1572 | S | 0 | 0 | yes | no | no | yes | yes | no | no | no | no | no | no | no | no |
| SCN4A | 1573 | I | 1 | 0 | yes | no | no | yes | yes | no | no | no | no | no | no | no | no |
| SCN4A | 1574 | G | 0 | 0 | yes | no | no | yes | yes | no | no | no | no | no | no | no | no |
| SCN4A | 1575 | I | 0 | 0 | yes | no | no | yes | no | no | no | no | no | no | no | no | no |
| SCN4A | 1576 | C | 1 | 0 | yes | no | no | yes | no | no | no | no | no | no | no | no | no |
| SCN4A | 1577 | F | 0 | 0 | yes | no | no | yes | no | no | no | no | no | no | no | no | no |
| SCN4A | 1578 | F | 0 | 0 | yes | no | no | yes | no | no | no | no | no | no | no | no | no |
| SCN4A | 1579 | C | 0 | 0 | yes | no | no | yes | no | no | no | no | no | no | no | no | no |
| SCN4A | 1580 | S | 0 | 0 | yes | no | no | yes | no | no | no | no | no | no | no | no | no |
| SCN4A | 1581 | Y | 1 | 0 | yes | no | no | yes | no | no | no | no | no | no | no | no | no |
| SCN4A | 1582 | I | 0 | 0 | yes | no | no | yes | no | no | no | no | no | no | no | no | no |
| SCN4A | 1583 | I | 0 | 0 | yes | no | no | yes | no | no | no | no | no | no | no | no | no |
| SCN4A | 1584 | I | 1 | 0 | yes | no | no | yes | no | no | no | no | no | no | no | no | no |
| SCN4A | 1585 | S | 1 | 0 | yes | no | no | yes | no | no | no | no | no | no | no | no | no |
| SCN4A | 1586 | F | 0 | 0 | yes | no | no | yes | no | no | no | no | no | no | no | no | no |
| SCN4A | 1587 | L | 0 | 0 | yes | no | no | yes | no | no | no | no | no | no | no | no | no |
| SCN4A | 1588 | I | 1 | 0 | yes | no | no | yes | no | no | no | no | no | no | no | no | no |
| SCN4A | 1589 | V | 1 | 1 | yes | no | no | yes | no | no | no | no | no | no | no | no | no |
| SCN4A | 1590 | V | 1 | 0 | yes | no | no | yes | no | no | no | no | no | no | no | no | no |
| SCN4A | 1591 | N | 0 | 0 | yes | no | no | yes | no | no | no | no | no | no | no | no | no |
| SCN4A | 1592 | M | 0 | 2 | yes | no | no | yes | no | no | no | no | no | no | no | no | no |
| SCN4A | 1593 | Y | 0 | 0 | yes | no | no | yes | no | no | no | no | no | no | no | no | no |
| SCN4A | 1594 | I | 1 | 0 | yes | no | no | yes | no | no | no | no | no | no | no | no | no |
| SCN4A | 1595 | A | 1 | 0 | yes | no | no | yes | no | no | no | no | no | no | no | no | no |
| SCN4A | 1596 | I | 0 | 0 | yes | no | no | yes | no | no | no | no | no | no | no | no | no |

| | | | | | | | | | | | | | | | | | |
|---|---|---|---|---|---|---|---|---|---|---|---|---|---|---|---|---|---|
| SCN4A | 1597 | I | 1 | 0 | yes | no | no | yes | no | no | yes | no | no | no | no | no | no |
| SCN4A | 1598 | L | 0 | 0 | yes | no | no | no | no | yes | no | no | no | no | no | no | no |
| SCN4A | 1599 | E | 0 | 0 | yes | no | no | no | no | yes | no | no | no | no | no | no | no |
| SCN4A | 1600 | N | 1 | 0 | yes | no | no | no | no | yes | no | no | no | no | no | no | no |
| SCN4A | 1601 | F | 0 | 0 | yes | no | no | no | no | yes | yes | no | no | no | no | no | no |
| SCN4A | 1602 | N | 1 | 0 | yes | no | no | no | no | yes | no | no | no | no | no | no | no |
| SCN4A | 1603 | V | 0 | 0 | yes | no | no | no | no | yes | no | no | no | no | no | no | no |
| SCN4A | 1604 | A | 0 | 0 | yes | no | no | no | no | yes | no | no | no | no | no | no | no |
| SCN4A | 1605 | T | 0 | 0 | yes | no | no | no | no | yes | yes | no | no | no | no | no | no |
| SCN4A | 1606 | E | 0 | 0 | no | yes | no | no | no | yes | yes | no | no | no | no | no | no |
| SCN4A | 1607 | E | 2 | 0 | no | yes | no | no | no | yes | yes | no | no | no | no | no | no |
| SCN9A | 114 | P | 0 | 0 | no | yes | no | no | no | yes | no | no | no | no | no | no | no |
| SCN9A | 115 | L | 0 | 0 | yes | no | no | no | no | yes | no | no | no | no | no | no | no |
| SCN9A | 116 | R | 0 | 0 | yes | no | no | no | no | yes | no | no | no | no | no | no | no |
| SCN9A | 117 | R | 0 | 0 | yes | no | no | no | no | yes | no | no | no | no | no | no | no |
| SCN9A | 118 | I | 3 | 0 | yes | no | no | no | no | yes | no | no | no | no | no | no | no |
| SCN9A | 119 | S | 0 | 0 | yes | no | no | no | no | yes | no | no | no | no | no | no | no |
| SCN9A | 120 | I | 2 | 1 | yes | no | no | no | no | yes | no | no | no | no | no | no | no |
| SCN9A | 121 | K | 1 | 0 | yes | no | no | no | no | yes | no | no | no | no | no | no | no |
| SCN9A | 122 | I | 0 | 0 | yes | no | no | yes | no | yes | no | no | no | no | no | no | no |
| SCN9A | 123 | L | 0 | 0 | yes | no | no | no | no | yes | no | no | no | no | no | no | no |
| SCN9A | 124 | V | 1 | 0 | no | yes | no | no | no | yes | no | no | no | no | no | no | no |
| SCN9A | 125 | H | 1 | 0 | no | yes | no | no | no | yes | no | no | no | no | no | no | no |
| SCN9A | 126 | S | 0 | 0 | yes | no | no | no | no | yes | no | no | no | no | no | no | no |
| SCN9A | 127 | L | 0 | 0 | yes | no | no | yes | no | no | no | no | no | no | no | no | no |
| SCN9A | 128 | F | 0 | 0 | yes | no | no | yes | no | no | no | no | no | no | no | no | no |
| SCN9A | 129 | S | 1 | 0 | yes | no | no | yes | no | no | no | no | no | no | no | no | no |
| SCN9A | 130 | M | 2 | 0 | yes | no | no | yes | no | no | no | no | no | no | no | no | no |
| SCN9A | 131 | L | 0 | 0 | yes | no | no | yes | no | no | no | no | no | no | no | no | no |
| SCN9A | 132 | I | 1 | 0 | yes | no | no | yes | no | no | no | no | no | no | no | no | no |
| SCN9A | 133 | M | 0 | 0 | yes | no | no | yes | no | no | no | no | no | no | no | no | no |
| SCN9A | 134 | C | 0 | 0 | yes | no | no | yes | no | no | no | no | no | no | no | no | no |
| SCN9A | 135 | T | 1 | 0 | yes | no | no | yes | no | no | no | no | no | no | no | no | no |

| | | | | | | | | | | | | | | | | | |
|---|---|---|---|---|---|---|---|---|---|---|---|---|---|---|---|---|---|
| SCN9A | 136 | I | 0 | 1 | yes | no | no | yes | no | no | no | no | no | no | no | no | no |
| SCN9A | 137 | L | 0 | 0 | yes | no | no | yes | no | no | no | no | no | no | no | no | no |
| SCN9A | 138 | T | 1 | 0 | yes | no | no | yes | no | no | no | no | no | no | no | no | no |
| SCN9A | 139 | N | 0 | 0 | yes | no | no | yes | no | no | no | no | no | no | no | no | no |
| SCN9A | 140 | C | 0 | 0 | yes | no | no | yes | no | no | no | no | no | no | no | no | no |
| SCN9A | 141 | I | 1 | 0 | yes | no | no | yes | no | no | no | no | no | no | no | no | no |
| SCN9A | 142 | F | 0 | 0 | yes | no | no | yes | no | no | no | no | no | no | no | no | no |
| SCN9A | 143 | M | 0 | 0 | yes | no | no | yes | no | no | no | no | no | no | no | no | no |
| SCN9A | 144 | T | 0 | 0 | no | yes | no | yes | no | no | no | no | no | no | no | no | no |
| SCN9A | 145 | M | 2 | 0 | no | yes | no | yes | no | no | no | no | no | no | no | no | no |
| SCN9A | 146 | N | 1 | 0 | no | yes | no | no | yes | no | no | no | no | no | no | no | no |
| SCN9A | 147 | N | 0 | 0 | no | yes | no | no | yes | no | no | no | no | no | no | no | no |
| SCN9A | 148 | P | 0 | 0 | no | yes | no | no | yes | no | no | no | no | no | no | no | no |
| SCN9A | 149 | P | 1 | 1 | no | yes | no | no | yes | no | no | no | no | no | no | no | no |
| SCN9A | 150 | D | 0 | 0 | no | yes | no | no | yes | no | no | no | no | no | no | no | no |
| SCN9A | 151 | W | 0 | 0 | no | yes | no | no | yes | no | no | no | no | no | no | no | no |
| SCN9A | 152 | T | 0 | 0 | yes | no | no | yes | yes | no | no | no | no | no | no | no | no |
| SCN9A | 153 | K | 0 | 0 | yes | no | no | yes | no | no | no | no | no | no | no | no | no |
| SCN9A | 154 | N | 1 | 0 | yes | no | no | yes | no | no | no | no | no | no | no | no | no |
| SCN9A | 155 | V | 0 | 0 | yes | no | no | yes | no | no | no | no | no | no | no | no | no |
| SCN9A | 156 | E | 0 | 0 | yes | no | no | yes | no | no | no | no | no | no | no | no | no |
| SCN9A | 157 | Y | 1 | 0 | yes | no | no | yes | no | no | no | no | no | no | no | no | no |
| SCN9A | 158 | T | 1 | 0 | yes | no | no | yes | no | no | no | no | no | no | no | no | no |
| SCN9A | 159 | F | 1 | 0 | yes | no | no | yes | no | no | no | no | no | no | no | no | no |
| SCN9A | 160 | T | 1 | 0 | yes | no | no | yes | no | no | no | no | no | no | no | no | no |
| SCN9A | 161 | G | 1 | 0 | yes | no | no | yes | no | no | no | no | no | no | no | no | no |
| SCN9A | 162 | I | 0 | 0 | yes | no | no | yes | no | no | no | no | no | no | no | no | no |
| SCN9A | 163 | Y | 0 | 0 | yes | no | no | yes | no | no | no | no | no | no | no | no | no |
| SCN9A | 164 | T | 0 | 0 | yes | no | no | yes | no | no | no | no | no | no | no | no | no |
| SCN9A | 165 | F | 2 | 0 | yes | no | no | yes | no | no | no | no | no | no | no | no | no |
| SCN9A | 166 | E | 0 | 0 | yes | no | no | yes | no | no | no | no | no | no | no | no | no |
| SCN9A | 167 | S | 0 | 0 | yes | no | no | yes | no | no | no | no | no | no | no | no | no |
| SCN9A | 168 | L | 0 | 0 | yes | no | no | yes | no | no | no | no | no | no | no | no | no |

| | | | | | | | | | | | | | | | | | |
|---|---|---|---|---|---|---|---|---|---|---|---|---|---|---|---|---|---|
| SCN9A | 169 | V | 0 | 0 | yes | no | no | yes | no | no | no | no | no | no | no | no | no |
| SCN9A | 170 | K | 0 | 0 | yes | no | no | yes | no | no | no | no | no | no | no | no | no |
| SCN9A | 171 | I | 0 | 0 | yes | no | no | yes | no | no | no | no | no | no | no | no | no |
| SCN9A | 172 | L | 0 | 0 | yes | no | no | yes | no | no | no | no | no | no | no | no | no |
| SCN9A | 173 | A | 0 | 0 | yes | no | no | no | no | no | no | no | no | no | no | no | no |
| SCN9A | 174 | R | 0 | 0 | yes | no | no | no | no | yes | no | no | no | no | no | no | no |
| SCN9A | 175 | G | 0 | 0 | yes | no | no | no | no | yes | no | no | no | no | no | no | no |
| SCN9A | 176 | F | 0 | 0 | no | yes | no | no | no | yes | no | no | no | no | no | no | no |
| SCN9A | 177 | C | 1 | 0 | no | yes | no | no | no | yes | no | no | no | no | no | no | no |
| SCN9A | 178 | V | 1 | 0 | no | yes | no | no | no | yes | no | no | no | no | no | no | no |
| SCN9A | 179 | G | 1 | 0 | no | yes | no | yes | no | yes | no | no | no | no | no | no | no |
| SCN9A | 180 | E | 1 | 0 | no | yes | no | yes | no | yes | no | no | no | no | no | no | no |
| SCN9A | 181 | F | 1 | 0 | yes | no | no | no | no | yes | no | no | no | no | no | no | no |
| SCN9A | 182 | T | 0 | 0 | yes | no | no | no | no | yes | no | no | no | no | no | no | no |
| SCN9A | 183 | F | 0 | 0 | yes | no | no | yes | no | yes | no | no | no | no | no | no | no |
| SCN9A | 184 | L | 0 | 0 | yes | no | no | yes | no | yes | no | no | no | no | no | no | no |
| SCN9A | 185 | R | 2 | 1 | yes | no | no | no | no | yes | no | no | no | no | no | no | no |
| SCN9A | 186 | D | 0 | 0 | no | yes | no | no | no | yes | no | no | no | no | no | no | no |
| SCN9A | 187 | P | 2 | 0 | yes | no | no | yes | no | yes | no | no | no | no | no | no | no |
| SCN9A | 188 | W | 0 | 0 | yes | no | no | yes | no | no | no | no | no | no | no | no | no |
| SCN9A | 189 | N | 0 | 0 | yes | no | no | yes | no | no | no | no | no | no | no | no | no |
| SCN9A | 190 | W | 0 | 0 | yes | no | no | yes | no | no | no | no | no | no | no | no | no |
| SCN9A | 191 | L | 0 | 0 | yes | no | no | yes | no | no | no | no | no | no | no | no | no |
| SCN9A | 192 | D | 0 | 0 | yes | no | no | yes | no | no | no | no | no | no | no | no | no |
| SCN9A | 193 | F | 0 | 0 | yes | no | no | yes | no | no | no | no | no | no | no | no | no |
| SCN9A | 194 | V | 0 | 0 | yes | no | no | yes | no | no | no | no | no | no | no | no | no |
| SCN9A | 195 | V | 1 | 0 | yes | no | no | yes | no | no | no | no | no | no | no | no | no |
| SCN9A | 196 | I | 1 | 0 | yes | no | no | yes | no | no | no | no | no | no | no | no | no |
| SCN9A | 197 | V | 2 | 0 | yes | no | no | yes | no | no | no | no | no | no | no | no | no |
| SCN9A | 198 | F | 0 | 0 | yes | no | no | yes | no | no | no | no | no | no | no | no | no |
| SCN9A | 199 | A | 0 | 0 | yes | no | no | yes | no | no | no | no | no | no | no | no | no |
| SCN9A | 200 | Y | 1 | 0 | yes | no | no | yes | no | no | no | no | no | no | no | no | no |
| SCN9A | 201 | L | 1 | 0 | yes | no | no | yes | no | no | no | no | no | no | no | no | no |

| | | | | | | | | | | | | | | | | | |
|---|---|---|---|---|---|---|---|---|---|---|---|---|---|---|---|---|---|
| SCN9A | 202 | T | 0 | 0 | yes | no | no | yes | no | no | no | no | no | no | no | no | no |
| SCN9A | 203 | E | 0 | 0 | yes | no | no | yes | no | no | no | no | no | no | no | no | no |
| SCN9A | 204 | F | 0 | 0 | no | yes | no | yes | no | no | no | no | no | no | no | no | no |
| SCN9A | 205 | V | 0 | 0 | no | yes | no | yes | no | no | no | no | no | no | no | no | no |
| SCN9A | 206 | N | 0 | 0 | no | yes | no | yes | yes | no | no | no | no | no | no | no | no |
| SCN9A | 207 | L | 0 | 0 | no | yes | no | yes | yes | no | no | no | no | no | no | no | no |
| SCN9A | 208 | G | 0 | 0 | no | yes | no | no | yes | no | no | no | no | no | no | no | no |
| SCN9A | 209 | N | 0 | 0 | no | yes | no | yes | yes | no | no | no | no | no | no | no | no |
| SCN9A | 210 | V | 1 | 0 | yes | no | no | no | yes | no | no | no | no | no | no | no | no |
| SCN9A | 211 | S | 0 | 1 | yes | no | no | yes | yes | no | no | no | no | no | no | no | no |
| SCN9A | 212 | A | 0 | 0 | yes | no | no | yes | no | no | no | no | no | no | no | no | no |
| SCN9A | 213 | L | 1 | 0 | yes | no | no | yes | no | no | no | no | no | no | no | no | no |
| SCN9A | 214 | R | 1 | 0 | yes | no | no | yes | no | no | no | no | no | no | no | no | no |
| SCN9A | 215 | T | 1 | 0 | yes | no | no | yes | no | no | no | no | no | no | no | no | no |
| SCN9A | 216 | F | 1 | 1 | yes | no | no | yes | no | no | no | no | no | no | no | no | no |
| SCN9A | 217 | R | 0 | 0 | yes | no | no | yes | no | no | no | no | no | no | no | no | no |
| SCN9A | 218 | V | 0 | 0 | yes | no | no | yes | no | no | no | no | no | no | no | no | no |
| SCN9A | 219 | L | 2 | 0 | yes | no | no | yes | no | no | no | no | no | no | no | no | no |
| SCN9A | 220 | R | 0 | 0 | yes | no | no | yes | no | no | no | no | no | no | no | no | no |
| SCN9A | 221 | A | 1 | 0 | yes | no | no | yes | no | no | no | no | no | no | no | no | no |
| SCN9A | 222 | L | 1 | 0 | yes | no | no | yes | no | no | no | no | no | no | no | no | no |
| SCN9A | 223 | K | 0 | 0 | yes | no | no | yes | no | no | no | no | no | no | no | no | no |
| SCN9A | 224 | T | 1 | 0 | yes | no | no | yes | no | no | no | no | no | no | no | no | no |
| SCN9A | 225 | I | 0 | 0 | yes | no | no | yes | no | no | no | no | no | no | no | no | no |
| SCN9A | 226 | S | 0 | 0 | yes | no | no | yes | no | no | no | no | no | no | no | no | no |
| SCN9A | 227 | V | 0 | 0 | yes | no | no | yes | no | no | no | no | no | no | no | no | no |
| SCN9A | 228 | I | 1 | 1 | no | yes | no | yes | no | no | no | no | no | no | no | no | no |
| SCN9A | 229 | P | 1 | 0 | no | yes | no | yes | no | yes | no | no | no | no | no | no | no |
| SCN9A | 230 | G | 0 | 0 | no | yes | no | yes | no | yes | no | no | no | no | no | no | no |
| SCN9A | 231 | L | 0 | 0 | yes | no | no | yes | no | yes | no | no | no | no | no | no | no |
| SCN9A | 232 | K | 0 | 0 | yes | no | no | yes | no | yes | no | no | no | no | no | no | no |
| SCN9A | 233 | T | 1 | 0 | yes | no | no | no | no | yes | no | no | no | no | no | no | no |
| SCN9A | 234 | I | 0 | 1 | yes | no | no | yes | no | yes | no | no | no | no | no | no | no |

| | | | | | | | | | | | | | | | | | |
|---|---|---|---|---|---|---|---|---|---|---|---|---|---|---|---|---|---|
| SCN9A | 235 | V | 0 | 0 | yes | no | no | yes | no | yes | no | no | no | no | no | no | no |
| SCN9A | 236 | G | 0 | 0 | yes | no | no | yes | no | yes | no | no | no | no | no | no | no |
| SCN9A | 237 | A | 0 | 0 | yes | no | no | yes | no | yes | no | no | no | no | no | no | no |
| SCN9A | 238 | L | 1 | 0 | yes | no | no | yes | no | yes | no | no | no | no | no | no | no |
| SCN9A | 239 | I | 0 | 0 | yes | no | no | yes | no | yes | no | no | no | no | no | no | no |
| SCN9A | 240 | Q | 1 | 0 | yes | no | no | no | no | yes | no | no | no | no | no | no | no |
| SCN9A | 241 | S | 0 | 1 | yes | no | no | yes | no | yes | no | no | no | no | no | no | no |
| SCN9A | 242 | V | 1 | 0 | yes | no | no | yes | no | yes | no | no | no | no | no | no | no |
| SCN9A | 243 | K | 0 | 0 | yes | no | no | yes | no | yes | no | no | no | no | no | no | no |
| SCN9A | 244 | K | 0 | 0 | yes | no | no | no | no | yes | no | no | no | no | no | no | no |
| SCN9A | 245 | L | 1 | 1 | no | yes | no | yes | no | yes | no | no | no | no | no | no | no |
| SCN9A | 246 | S | 1 | 0 | yes | no | no | yes | no | yes | no | no | no | no | no | no | no |
| SCN9A | 247 | D | 1 | 0 | yes | no | no | yes | no | yes | no | no | no | no | no | no | no |
| SCN9A | 248 | V | 0 | 0 | yes | no | no | yes | no | no | no | no | no | no | no | no | no |
| SCN9A | 249 | M | 0 | 0 | yes | no | no | yes | no | no | no | no | no | no | no | no | no |
| SCN9A | 250 | I | 0 | 0 | yes | no | no | yes | no | no | no | no | no | no | no | no | no |
| SCN9A | 251 | L | 0 | 0 | yes | no | no | yes | no | no | no | no | no | no | no | no | no |
| SCN9A | 252 | T | 0 | 0 | yes | no | no | yes | no | no | no | no | no | no | no | no | no |
| SCN9A | 253 | V | 0 | 0 | yes | no | no | yes | no | no | no | no | no | no | no | no | no |
| SCN9A | 254 | F | 1 | 0 | yes | no | no | yes | no | no | no | no | no | no | no | no | no |
| SCN9A | 255 | C | 1 | 0 | yes | no | no | yes | no | no | no | no | no | no | no | no | no |
| SCN9A | 256 | L | 0 | 0 | yes | no | no | yes | no | no | no | no | no | no | no | no | no |
| SCN9A | 257 | S | 2 | 0 | yes | no | no | yes | no | no | no | no | no | no | no | no | no |
| SCN9A | 258 | V | 2 | 0 | yes | no | no | yes | no | no | no | no | no | no | no | no | no |
| SCN9A | 259 | F | 0 | 0 | yes | no | no | yes | no | no | no | no | no | no | no | no | no |
| SCN9A | 260 | A | 0 | 0 | yes | no | no | yes | no | no | no | no | no | no | no | no | no |
| SCN9A | 261 | L | 1 | 0 | yes | no | no | yes | no | no | no | no | no | no | no | no | no |
| SCN9A | 262 | I | 1 | 0 | yes | no | no | yes | no | no | no | no | no | no | no | no | no |
| SCN9A | 263 | G | 0 | 0 | yes | no | no | yes | no | no | no | no | no | no | no | no | no |
| SCN9A | 264 | L | 0 | 0 | yes | no | no | yes | no | no | no | no | no | no | no | no | no |
| SCN9A | 265 | Q | 0 | 0 | yes | no | no | yes | no | no | no | no | no | no | no | no | no |
| SCN9A | 266 | L | 1 | 1 | yes | no | no | yes | no | no | no | no | no | no | no | no | no |
| SCN9A | 267 | F | 0 | 0 | yes | no | no | yes | no | no | no | no | no | no | no | no | no |

| | | | | | | | | | | | | | | | | | |
|---|---|---|---|---|---|---|---|---|---|---|---|---|---|---|---|---|---|
| SCN9A | 268 | M | 3 | 0 | no | yes | no | no | yes | no | no | no | no | no | no | no | no |
| SCN9A | 269 | G | 0 | 0 | no | yes | no | no | yes | no | no | no | no | no | no | no | no |
| SCN9A | 270 | N | 0 | 0 | yes | no | no | no | yes | no | no | no | no | no | no | no | no |
| SCN9A | 271 | L | 0 | 0 | yes | no | no | no | yes | no | no | no | no | no | no | no | no |
| SCN9A | 272 | K | 0 | 0 | yes | no | no | no | yes | no | no | no | no | no | no | no | no |
| SCN9A | 273 | H | 0 | 0 | no | no | yes | no | yes | no | no | no | no | no | no | no | no |
| SCN9A | 274 | K | 0 | 0 | no | no | yes | no | yes | no | no | no | no | no | no | no | no |
| SCN9A | 275 | C | 0 | 0 | no | no | yes | no | yes | no | no | no | no | no | no | no | no |
| SCN9A | 276 | F | 0 | 0 | no | no | yes | no | yes | no | no | no | no | no | no | no | no |
| SCN9A | 277 | R | 1 | 0 | no | no | yes | no | yes | no | no | no | yes | no | no | no | no |
| SCN9A | 278 | N | 0 | 0 | no | yes | no | no | yes | no | no | no | yes | no | no | no | no |
| SCN9A | 279 | S | 1 | 0 | no | yes | no | no | yes | no | no | no | yes | no | no | no | no |
| SCN9A | 280 | L | 1 | 0 | no | yes | no | no | yes | no | no | no | no | no | no | no | no |
| SCN9A | 281 | E | 1 | 0 | yes | no | no | no | yes | no | no | no | no | no | no | no | no |
| SCN9A | 282 | N | 0 | 0 | yes | no | no | no | yes | no | no | no | no | no | no | no | no |
| SCN9A | 283 | N | 0 | 0 | yes | no | no | no | yes | no | no | no | no | no | no | no | no |
| SCN9A | 284 | E | 0 | 0 | no | yes | no | no | yes | no | no | no | no | no | no | no | no |
| SCN9A | 285 | T | 0 | 0 | no | yes | no | no | yes | no | no | no | no | no | no | no | no |
| SCN9A | 286 | L | 1 | 0 | yes | no | no | no | yes | no | no | no | no | no | no | no | no |
| SCN9A | 287 | E | 1 | 0 | yes | no | no | no | yes | no | no | no | no | no | no | no | no |
| SCN9A | 288 | S | 0 | 0 | yes | no | no | no | yes | no | no | no | no | no | no | no | no |
| SCN9A | 289 | I | 0 | 0 | yes | no | no | no | yes | no | no | no | no | no | no | no | no |
| SCN9A | 290 | M | 0 | 0 | yes | no | no | no | yes | no | no | no | no | no | no | no | no |
| SCN9A | 291 | N | 0 | 0 | yes | no | no | no | yes | no | no | no | no | no | no | no | no |
| SCN9A | 292 | T | 1 | 0 | yes | no | no | no | yes | no | no | no | no | no | no | no | no |
| SCN9A | 293 | L | 0 | 0 | no | yes | no | no | yes | no | no | no | no | no | no | no | no |
| SCN9A | 294 | E | 0 | 0 | no | yes | no | no | yes | no | no | no | no | no | no | no | no |
| SCN9A | 295 | S | 1 | 0 | no | yes | no | no | yes | no | no | no | no | no | no | no | no |
| SCN9A | 296 | E | 0 | 0 | yes | no | no | no | yes | no | no | no | no | no | no | no | no |
| SCN9A | 297 | E | 1 | 0 | yes | no | no | no | yes | no | no | no | no | no | no | no | no |
| SCN9A | 298 | D | 2 | 0 | yes | no | no | no | yes | no | no | no | no | no | no | no | no |
| SCN9A | 299 | F | 0 | 0 | yes | no | no | no | yes | no | no | no | no | no | no | no | no |
| SCN9A | 300 | R | 0 | 0 | yes | no | no | no | yes | no | no | no | yes | no | no | no | no |

| | | | | | | | | | | | | | | | | | |
|---|---|---|---|---|---|---|---|---|---|---|---|---|---|---|---|---|---|
| SCN9A | 301 | K | 0 | 0 | yes | no | no | no | yes | no | no | no | yes | no | no | no | no |
| SCN9A | 302 | Y | 0 | 0 | yes | no | no | no | yes | no | no | no | no | no | no | no | no |
| SCN9A | 303 | F | 0 | 0 | no | no | yes | no | yes | no | no | no | no | no | no | no | no |
| SCN9A | 304 | Y | 0 | 0 | no | yes | no | no | yes | no | no | no | yes | no | no | no | no |
| SCN9A | 305 | Y | 1 | 0 | no | yes | no | no | yes | no | no | no | no | no | no | no | no |
| SCN9A | 306 | L | 0 | 0 | no | yes | no | no | yes | no | no | no | yes | no | no | no | no |
| SCN9A | 307 | E | 1 | 0 | no | yes | no | no | yes | no | yes | no | no | no | no | no | no |
| SCN9A | 308 | G | 0 | 0 | no | yes | no | no | yes | no | yes | no | no | no | no | no | no |
| SCN9A | 309 | S | 0 | 0 | no | yes | no | no | yes | no | yes | no | no | no | no | no | no |
| SCN9A | 310 | K | 1 | 0 | no | yes | no | no | yes | no | no | no | no | no | no | no | no |
| SCN9A | 311 | D | 0 | 0 | no | yes | no | no | yes | no | no | no | no | no | no | no | no |
| SCN9A | 312 | A | 0 | 0 | no | yes | no | no | yes | no | no | no | no | no | no | no | no |
| SCN9A | 313 | L | 0 | 0 | no | yes | no | no | yes | no | no | no | no | no | no | no | no |
| SCN9A | 314 | L | 1 | 0 | no | yes | no | no | yes | no | no | no | no | no | no | no | no |
| SCN9A | 315 | C | 0 | 0 | no | yes | no | no | yes | no | no | no | no | no | no | no | no |
| SCN9A | 316 | G | 0 | 0 | no | yes | no | no | yes | no | no | no | no | no | no | no | no |
| SCN9A | 317 | F | 1 | 0 | no | yes | no | no | yes | no | no | no | no | no | no | no | no |
| SCN9A | 318 | S | 0 | 0 | no | yes | no | no | yes | no | no | no | no | no | no | no | no |
| SCN9A | 319 | T | 1 | 0 | no | yes | no | no | yes | no | no | no | no | no | no | no | no |
| SCN9A | 320 | D | 1 | 0 | no | yes | no | no | yes | no | no | no | no | no | no | no | no |
| SCN9A | 321 | S | 1 | 0 | no | yes | no | no | yes | no | no | no | no | no | no | no | no |
| SCN9A | 322 | G | 0 | 0 | no | yes | no | no | yes | no | no | no | no | no | no | no | no |
| SCN9A | 323 | Q | 0 | 0 | no | yes | no | no | yes | no | no | no | yes | no | no | no | no |
| SCN9A | 324 | C | 0 | 0 | no | yes | no | no | yes | no | no | no | yes | no | no | no | no |
| SCN9A | 325 | P | 0 | 0 | no | yes | no | no | yes | no | no | no | yes | no | no | no | no |
| SCN9A | 326 | E | 1 | 0 | no | yes | no | no | yes | no | no | no | yes | no | no | no | no |
| SCN9A | 327 | G | 1 | 1 | no | yes | no | no | yes | no | no | no | yes | no | no | no | no |
| SCN9A | 328 | Y | 0 | 0 | no | no | yes | no | yes | no | no | no | yes | no | no | no | no |
| SCN9A | 329 | T | 0 | 0 | no | no | yes | no | yes | no | no | no | no | no | no | no | no |
| SCN9A | 330 | C | 0 | 0 | no | no | yes | no | yes | no | no | no | no | no | no | no | no |
| SCN9A | 331 | V | 0 | 0 | no | no | yes | no | yes | no | no | no | no | no | no | no | no |
| SCN9A | 332 | K | 0 | 0 | no | no | yes | no | yes | no | no | no | no | no | no | no | no |
| SCN9A | 333 | I | 1 | 0 | no | yes | no | no | yes | no | no | no | no | no | no | no | no |

| | | | | | | | | | | | | | | | | | |
|---|---|---|---|---|---|---|---|---|---|---|---|---|---|---|---|---|---|
| SCN9A | 334 | G | 0 | 0 | no | yes | no | no | yes | no | no | no | no | no | no | no | no |
| SCN9A | 335 | R | 0 | 0 | no | yes | no | no | yes | no | no | no | no | no | no | no | no |
| SCN9A | 336 | N | 0 | 0 | no | yes | no | no | yes | no | no | no | no | no | no | no | no |
| SCN9A | 337 | P | 1 | 0 | no | yes | no | no | yes | no | no | no | no | no | no | no | no |
| SCN9A | 338 | D | 0 | 0 | no | yes | no | no | yes | no | no | no | no | no | no | no | no |
| SCN9A | 339 | Y | 0 | 0 | no | yes | no | no | yes | no | no | no | no | no | no | no | no |
| SCN9A | 340 | G | 1 | 0 | no | yes | no | no | yes | no | no | no | no | no | no | no | no |
| SCN9A | 341 | Y | 0 | 0 | no | yes | no | no | yes | no | no | no | no | no | no | no | no |
| SCN9A | 342 | T | 1 | 0 | no | yes | no | no | yes | no | no | no | no | no | no | no | no |
| SCN9A | 343 | S | 1 | 0 | no | yes | no | no | yes | no | no | no | no | no | no | no | no |
| SCN9A | 344 | F | 0 | 0 | no | yes | no | yes | yes | no | no | no | no | no | no | no | no |
| SCN9A | 345 | D | 0 | 0 | no | yes | no | no | yes | no | no | no | no | no | no | no | no |
| SCN9A | 346 | T | 0 | 0 | no | yes | no | yes | yes | no | no | no | no | no | no | no | no |
| SCN9A | 347 | F | 0 | 0 | yes | no | no | yes | no | no | no | yes | no | no | no | no | no |
| SCN9A | 348 | S | 0 | 0 | yes | no | no | yes | no | no | no | yes | no | no | no | no | no |
| SCN9A | 349 | W | 0 | 0 | yes | no | no | yes | no | no | no | yes | no | no | no | no | no |
| SCN9A | 350 | A | 0 | 0 | yes | no | no | yes | no | no | no | yes | no | no | no | no | no |
| SCN9A | 351 | F | 1 | 0 | yes | no | no | yes | no | no | no | yes | no | no | no | no | no |
| SCN9A | 352 | L | 0 | 0 | yes | no | no | yes | no | no | no | yes | no | no | no | no | no |
| SCN9A | 353 | A | 0 | 0 | yes | no | no | yes | no | no | no | yes | no | no | no | no | no |
| SCN9A | 354 | L | 0 | 0 | yes | no | no | yes | no | no | no | yes | no | no | no | no | no |
| SCN9A | 355 | F | 0 | 0 | yes | no | no | yes | no | no | no | yes | no | no | no | no | no |
| SCN9A | 356 | R | 0 | 0 | yes | no | no | yes | no | no | no | yes | no | no | no | no | no |
| SCN9A | 357 | L | 0 | 0 | yes | no | no | yes | no | no | no | yes | no | no | no | no | no |
| SCN9A | 358 | M | 1 | 0 | yes | no | no | yes | no | no | no | yes | no | no | no | no | no |
| SCN9A | 359 | T | 1 | 0 | no | yes | no | yes | no | no | yes | yes | no | no | no | no | no |
| SCN9A | 360 | Q | 0 | 0 | no | yes | no | yes | no | no | yes | yes | no | no | no | no | no |
| SCN9A | 361 | D | 0 | 0 | no | yes | no | yes | no | no | no | yes | no | no | no | no | no |
| SCN9A | 362 | Y | 1 | 0 | no | yes | no | yes | no | no | yes | yes | no | no | no | no | no |
| SCN9A | 363 | W | 0 | 0 | yes | no | no | yes | no | no | no | yes | no | no | no | no | no |
| SCN9A | 364 | E | 0 | 0 | yes | no | no | yes | no | no | yes | yes | no | no | no | no | no |
| SCN9A | 365 | N | 0 | 0 | yes | no | no | no | no | no | no | yes | no | no | no | no | no |
| SCN9A | 366 | L | 0 | 0 | yes | no | no | yes | no | no | no | yes | no | no | no | no | no |

| | | | | | | | | | | | | | | | | | |
|---|---|---|---|---|---|---|---|---|---|---|---|---|---|---|---|---|---|
| SCN9A | 367 | Y | 1 | 0 | yes | no | no | yes | no | no | no | yes | no | no | no | no | no |
| SCN9A | 368 | Q | 0 | 0 | yes | no | no | no | no | no | no | yes | no | no | no | no | no |
| SCN9A | 369 | Q | 1 | 0 | yes | no | no | no | no | no | no | yes | no | no | no | no | no |
| SCN9A | 370 | T | 1 | 0 | yes | no | no | no | no | no | no | yes | no | no | no | no | no |
| SCN9A | 371 | L | 0 | 0 | yes | no | no | no | no | no | no | yes | no | no | no | no | no |
| SCN9A | 372 | R | 1 | 0 | yes | no | no | no | yes | no | no | no | yes | no | no | no | no |
| SCN9A | 373 | A | 0 | 0 | yes | no | no | no | yes | no | no | no | no | no | no | no | no |
| SCN9A | 374 | A | 1 | 0 | yes | no | no | no | yes | no | no | no | no | no | no | no | no |
| SCN9A | 375 | G | 0 | 0 | no | yes | no | no | yes | no | no | no | no | no | no | no | no |
| SCN9A | 376 | K | 0 | 0 | yes | no | no | no | yes | no | no | no | no | no | no | no | no |
| SCN9A | 377 | T | 1 | 0 | yes | no | no | yes | yes | no | no | no | no | no | no | no | no |
| SCN9A | 378 | Y | 0 | 0 | yes | no | no | yes | yes | no | no | no | no | no | no | no | no |
| SCN9A | 379 | M | 1 | 0 | yes | no | no | yes | no | no | no | no | no | no | no | no | no |
| SCN9A | 380 | I | 1 | 0 | yes | no | no | yes | no | no | no | no | no | no | no | no | no |
| SCN9A | 381 | F | 0 | 0 | yes | no | no | yes | no | no | no | no | no | no | no | no | no |
| SCN9A | 382 | F | 0 | 0 | yes | no | no | yes | no | no | no | no | no | no | no | no | no |
| SCN9A | 383 | V | 0 | 0 | yes | no | no | yes | no | no | no | no | no | no | no | no | no |
| SCN9A | 384 | V | 1 | 0 | yes | no | no | yes | no | no | no | no | no | no | no | no | no |
| SCN9A | 385 | V | 1 | 0 | yes | no | no | yes | no | no | no | no | no | no | no | no | no |
| SCN9A | 386 | I | 0 | 0 | yes | no | no | yes | no | no | no | no | no | no | no | no | no |
| SCN9A | 387 | F | 0 | 0 | yes | no | no | yes | no | no | no | no | no | no | no | no | no |
| SCN9A | 388 | L | 0 | 0 | yes | no | no | yes | no | no | no | no | no | no | no | no | no |
| SCN9A | 389 | G | 0 | 0 | yes | no | no | yes | no | no | no | no | no | no | no | no | no |
| SCN9A | 390 | S | 0 | 0 | yes | no | no | yes | no | no | no | no | no | no | no | no | no |
| SCN9A | 391 | F | 0 | 0 | yes | no | no | yes | no | no | no | no | no | no | no | no | no |
| SCN9A | 392 | Y | 1 | 0 | yes | no | no | yes | no | no | no | no | no | no | no | no | no |
| SCN9A | 393 | L | 0 | 0 | yes | no | no | yes | no | no | no | no | no | no | no | no | no |
| SCN9A | 394 | I | 1 | 0 | yes | no | no | yes | no | no | yes | no | no | no | no | no | no |
| SCN9A | 395 | N | 0 | 1 | yes | no | no | yes | no | no | no | no | no | no | no | no | no |
| SCN9A | 396 | L | 0 | 0 | yes | no | no | yes | no | no | no | no | no | no | no | no | no |
| SCN9A | 397 | I | 0 | 0 | yes | no | no | yes | no | no | no | no | no | no | no | no | no |
| SCN9A | 398 | L | 0 | 0 | yes | no | no | yes | no | no | yes | no | no | no | no | no | no |
| SCN9A | 399 | A | 1 | 0 | yes | no | no | yes | no | no | no | no | no | no | no | no | no |

| | | | | | | | | | | | | | | | | | |
|---|---|---|---|---|---|---|---|---|---|---|---|---|---|---|---|---|---|
| SCN9A | 400 | V | 0 | 1 | yes | no | no | yes | no | yes | no | no | no | no | no | no | no |
| SCN9A | 401 | V | 0 | 0 | yes | no | no | yes | no | yes | no | no | no | no | no | no | no |
| SCN9A | 402 | A | 1 | 0 | yes | no | no | yes | no | yes | no | no | no | no | no | no | no |
| SCN9A | 403 | M | 3 | 0 | yes | no | no | no | no | yes | no | no | no | no | no | no | no |
| SCN9A | 404 | A | 1 | 0 | yes | no | no | no | no | yes | no | no | no | no | no | no | no |
| SCN9A | 405 | Y | 0 | 0 | yes | no | no | no | no | yes | no | no | no | no | no | no | no |
| SCN9A | 407 | E | 0 | 0 | yes | no | no | no | no | yes | no | no | no | no | no | no | no |
| SCN9A | 408 | Q | 0 | 0 | yes | no | no | no | no | yes | no | no | no | no | no | no | no |
| SCN9A | 409 | N | 0 | 0 | yes | no | no | no | no | yes | no | no | no | no | no | no | no |
| SCN9A | 410 | Q | 0 | 0 | yes | no | no | no | no | yes | no | no | no | no | no | no | no |
| SCN9A | 411 | A | 0 | 0 | yes | no | no | no | no | yes | no | no | no | no | no | no | no |
| SCN9A | 412 | N | 0 | 0 | yes | no | no | no | no | yes | no | no | no | no | no | no | no |
| SCN9A | 413 | I | 2 | 0 | yes | no | no | no | no | yes | no | no | no | no | no | no | no |
| SCN9A | 414 | E | 0 | 0 | yes | no | no | no | no | yes | no | no | no | no | no | no | no |
| SCN9A | 415 | E | 0 | 0 | yes | no | no | no | no | yes | no | no | no | no | no | no | no |
| SCN9A | 416 | A | 1 | 0 | yes | no | no | no | no | yes | no | no | no | no | no | no | no |
| SCN9A | 417 | K | 0 | 0 | no | yes | no | no | no | yes | no | no | no | no | no | no | no |
| SCN9A | 726 | C | 0 | 0 | no | yes | no | no | no | yes | no | no | no | no | no | no | no |
| SCN9A | 727 | S | 1 | 0 | yes | no | no | no | no | yes | no | no | no | no | no | no | no |
| SCN9A | 728 | P | 0 | 0 | yes | no | no | no | no | yes | no | no | no | no | no | no | no |
| SCN9A | 729 | Y | 1 | 0 | yes | no | no | no | no | yes | no | no | no | no | no | no | no |
| SCN9A | 730 | W | 1 | 0 | yes | no | no | no | no | yes | no | no | no | no | no | no | no |
| SCN9A | 731 | I | 1 | 1 | yes | no | no | no | no | yes | no | no | no | no | no | no | no |
| SCN9A | 732 | K | 0 | 0 | yes | no | no | no | no | yes | no | no | no | no | no | no | no |
| SCN9A | 733 | F | 0 | 0 | yes | no | no | yes | no | yes | no | no | no | no | no | no | no |
| SCN9A | 734 | K | 0 | 0 | yes | no | no | no | no | yes | no | no | no | no | no | no | no |
| SCN9A | 735 | K | 2 | 0 | yes | no | no | no | no | yes | no | no | no | no | no | no | no |
| SCN9A | 736 | C | 0 | 0 | yes | no | no | no | no | yes | no | no | no | no | no | no | no |
| SCN9A | 737 | I | 1 | 0 | yes | no | no | no | no | yes | no | no | no | no | no | no | no |
| SCN9A | 738 | Y | 0 | 0 | yes | no | no | no | no | yes | no | no | no | no | no | no | no |
| SCN9A | 739 | F | 0 | 0 | yes | no | no | no | no | yes | no | no | no | no | no | no | no |
| SCN9A | 740 | I | 1 | 0 | no | yes | no | no | no | yes | no | no | no | no | no | no | no |
| SCN9A | 741 | V | 0 | 0 | no | yes | no | no | no | yes | no | no | no | no | no | no | no |

| | | | | | | | | | | | | | | | | | |
|---|---|---|---|---|---|---|---|---|---|---|---|---|---|---|---|---|---|
| SCN9A | 742 | M | 0 | 0 | no | yes | no | no | no | yes | no | no | no | no | no | no | no |
| SCN9A | 743 | D | 0 | 0 | no | yes | no | no | no | yes | no | no | no | no | no | no | no |
| SCN9A | 744 | P | 0 | 0 | no | yes | no | no | no | yes | no | no | no | no | no | no | no |
| SCN9A | 745 | F | 0 | 0 | no | yes | no | no | no | no | no | no | no | no | no | no | no |
| SCN9A | 746 | V | 0 | 0 | yes | no | no | yes | no | no | no | no | no | no | no | no | no |
| SCN9A | 747 | D | 2 | 0 | yes | no | no | yes | no | no | no | no | no | no | no | no | no |
| SCN9A | 748 | L | 1 | 0 | yes | no | no | yes | no | no | no | no | no | no | no | no | no |
| SCN9A | 749 | A | 0 | 0 | yes | no | no | yes | no | no | no | no | no | no | no | no | no |
| SCN9A | 750 | I | 1 | 0 | yes | no | no | yes | no | no | no | no | no | no | no | no | no |
| SCN9A | 751 | T | 0 | 0 | yes | no | no | yes | no | no | no | no | no | no | no | no | no |
| SCN9A | 752 | I | 1 | 0 | yes | no | no | yes | no | no | no | no | no | no | no | no | no |
| SCN9A | 753 | C | 1 | 0 | yes | no | no | yes | no | no | no | no | no | no | no | no | no |
| SCN9A | 754 | I | 1 | 0 | yes | no | no | yes | no | no | no | no | no | no | no | no | no |
| SCN9A | 755 | V | 1 | 0 | yes | no | no | yes | no | no | no | no | no | no | no | no | no |
| SCN9A | 756 | L | 0 | 0 | yes | no | no | yes | no | no | no | no | no | no | no | no | no |
| SCN9A | 757 | N | 0 | 0 | yes | no | no | yes | no | no | no | no | no | no | no | no | no |
| SCN9A | 758 | T | 0 | 0 | yes | no | no | yes | no | no | no | no | no | no | no | no | no |
| SCN9A | 759 | L | 0 | 0 | yes | no | no | yes | no | no | no | no | no | no | no | no | no |
| SCN9A | 760 | F | 1 | 0 | yes | no | no | yes | no | no | no | no | no | no | no | no | no |
| SCN9A | 761 | M | 0 | 0 | yes | no | no | yes | no | no | no | no | no | no | no | no | no |
| SCN9A | 762 | A | 0 | 0 | no | yes | no | yes | no | no | no | no | no | no | no | no | no |
| SCN9A | 763 | M | 1 | 0 | no | yes | no | yes | no | no | no | no | no | no | no | no | no |
| SCN9A | 764 | E | 0 | 0 | no | yes | no | yes | yes | no | no | no | no | no | no | no | no |
| SCN9A | 765 | H | 0 | 0 | no | yes | no | no | yes | no | no | no | no | no | no | no | no |
| SCN9A | 766 | H | 0 | 0 | no | yes | no | no | yes | no | no | no | no | no | no | no | no |
| SCN9A | 767 | P | 2 | 0 | no | yes | no | no | yes | no | no | no | no | no | no | no | no |
| SCN9A | 768 | M | 1 | 0 | no | yes | no | no | yes | no | no | no | no | no | no | no | no |
| SCN9A | 769 | T | 0 | 0 | no | yes | no | no | yes | no | no | no | no | no | no | no | no |
| SCN9A | 770 | E | 0 | 0 | yes | no | no | no | yes | no | no | no | no | no | no | no | no |
| SCN9A | 771 | E | 1 | 0 | yes | no | no | no | yes | no | no | no | no | no | no | no | no |
| SCN9A | 772 | F | 1 | 0 | yes | no | no | no | yes | no | no | no | no | no | no | no | no |
| SCN9A | 773 | K | 0 | 0 | yes | no | no | no | yes | no | no | no | no | no | no | no | no |
| SCN9A | 774 | N | 0 | 0 | yes | no | no | no | yes | no | no | no | no | no | no | no | no |

| | | | | | | | | | | | | | | | | | |
|---|---|---|---|---|---|---|---|---|---|---|---|---|---|---|---|---|---|
| SCN9A | 775 | V | 0 | 0 | yes | no | no | yes | no | no | no | no | no | no | no | no | no |
| SCN9A | 776 | L | 0 | 0 | yes | no | no | yes | no | no | no | no | no | no | no | no | no |
| SCN9A | 777 | A | 1 | 0 | yes | no | no | yes | no | no | no | no | no | no | no | no | no |
| SCN9A | 778 | I | 2 | 0 | yes | no | no | yes | no | no | no | no | no | no | no | no | no |
| SCN9A | 779 | G | 0 | 0 | yes | no | no | yes | no | no | no | no | no | no | no | no | no |
| SCN9A | 780 | N | 0 | 0 | yes | no | no | yes | no | no | no | no | no | no | no | no | no |
| SCN9A | 781 | L | 1 | 0 | yes | no | no | yes | no | no | no | no | no | no | no | no | no |
| SCN9A | 782 | V | 0 | 0 | yes | no | no | yes | no | no | no | no | no | no | no | no | no |
| SCN9A | 783 | F | 1 | 0 | yes | no | no | yes | no | no | no | no | no | no | no | no | no |
| SCN9A | 784 | T | 1 | 0 | yes | no | no | yes | no | no | no | no | no | no | no | no | no |
| SCN9A | 785 | G | 1 | 0 | yes | no | no | yes | no | no | no | no | no | no | no | no | no |
| SCN9A | 786 | I | 1 | 1 | yes | no | no | yes | no | no | no | no | no | no | no | no | no |
| SCN9A | 787 | F | 0 | 0 | yes | no | no | yes | no | no | no | no | no | no | no | no | no |
| SCN9A | 788 | A | 0 | 0 | yes | no | no | yes | no | no | no | no | no | no | no | no | no |
| SCN9A | 789 | A | 1 | 0 | yes | no | no | yes | no | no | no | no | no | no | no | no | no |
| SCN9A | 790 | E | 0 | 0 | yes | no | no | yes | no | no | no | no | no | no | no | no | no |
| SCN9A | 791 | M | 1 | 0 | yes | no | no | yes | no | no | no | no | no | no | no | no | no |
| SCN9A | 792 | V | 0 | 0 | yes | no | no | yes | no | no | no | no | no | no | no | no | no |
| SCN9A | 793 | L | 0 | 0 | yes | no | no | yes | no | no | no | no | no | no | no | no | no |
| SCN9A | 794 | K | 0 | 0 | yes | no | no | yes | no | no | no | no | no | no | no | no | no |
| SCN9A | 795 | L | 0 | 0 | yes | no | no | yes | no | yes | no | no | no | no | no | no | no |
| SCN9A | 796 | I | 0 | 0 | yes | no | no | yes | no | yes | no | no | no | no | no | no | no |
| SCN9A | 797 | A | 0 | 0 | yes | no | no | no | no | yes | no | no | no | no | no | no | no |
| SCN9A | 798 | M | 1 | 0 | yes | no | no | no | no | yes | no | no | no | no | no | no | no |
| SCN9A | 799 | D | 0 | 0 | no | yes | no | no | no | yes | no | no | no | no | no | no | no |
| SCN9A | 800 | P | 1 | 0 | yes | no | no | yes | no | yes | no | no | no | no | no | no | no |
| SCN9A | 801 | Y | 1 | 0 | yes | no | no | no | no | yes | no | no | no | no | no | no | no |
| SCN9A | 802 | E | 0 | 0 | yes | no | no | no | no | yes | no | no | no | no | no | no | no |
| SCN9A | 803 | Y | 0 | 0 | yes | no | no | yes | no | yes | no | no | no | no | no | no | no |
| SCN9A | 804 | F | 0 | 0 | yes | no | no | yes | no | yes | no | no | no | no | no | no | no |
| SCN9A | 805 | Q | 0 | 0 | yes | no | no | no | no | yes | no | no | no | no | no | no | no |
| SCN9A | 806 | V | 0 | 0 | no | yes | no | no | no | yes | no | no | no | no | no | no | no |
| SCN9A | 807 | G | 0 | 0 | yes | no | no | yes | no | yes | no | no | no | no | no | no | no |

| | | | | | | | | | | | | | | | | | |
|---|---|---|---|---|---|---|---|---|---|---|---|---|---|---|---|---|---|
| SCN9A | 808 | W | 1 | 0 | yes | no | no | yes | no | yes | no | no | no | no | no | no | no |
| SCN9A | 809 | N | 0 | 0 | yes | no | no | yes | no | no | no | no | no | no | no | no | no |
| SCN9A | 810 | I | 1 | 0 | yes | no | no | yes | no | no | no | no | no | no | no | no | no |
| SCN9A | 811 | F | 1 | 0 | yes | no | no | yes | no | no | no | no | no | no | no | no | no |
| SCN9A | 812 | D | 0 | 0 | yes | no | no | yes | no | no | no | no | no | no | no | no | no |
| SCN9A | 813 | S | 3 | 1 | yes | no | no | yes | no | no | no | no | no | no | no | no | no |
| SCN9A | 814 | L | 1 | 0 | yes | no | no | yes | no | no | no | no | no | no | no | no | no |
| SCN9A | 815 | I | 0 | 0 | yes | no | no | yes | no | no | no | no | no | no | no | no | no |
| SCN9A | 816 | V | 0 | 0 | yes | no | no | yes | no | no | no | no | no | no | no | no | no |
| SCN9A | 817 | T | 1 | 0 | yes | no | no | yes | no | no | no | no | no | no | no | no | no |
| SCN9A | 818 | L | 0 | 0 | yes | no | no | yes | no | no | no | no | no | no | no | no | no |
| SCN9A | 819 | S | 0 | 0 | yes | no | no | yes | no | no | no | no | no | no | no | no | no |
| SCN9A | 820 | L | 0 | 0 | yes | no | no | yes | no | no | no | no | no | no | no | no | no |
| SCN9A | 821 | V | 1 | 1 | yes | no | no | yes | no | no | no | no | no | no | no | no | no |
| SCN9A | 822 | E | 1 | 0 | yes | no | no | yes | no | no | no | no | no | no | no | no | no |
| SCN9A | 823 | L | 0 | 0 | yes | no | no | yes | no | no | no | no | no | no | no | no | no |
| SCN9A | 824 | F | 0 | 0 | yes | no | no | yes | no | no | no | no | no | no | no | no | no |
| SCN9A | 825 | L | 0 | 0 | no | yes | no | yes | no | no | no | no | no | no | no | no | no |
| SCN9A | 831 | L | 0 | 0 | no | yes | no | yes | no | no | no | no | no | no | no | no | no |
| SCN9A | 832 | S | 0 | 0 | no | yes | no | yes | no | no | no | no | no | no | no | no | no |
| SCN9A | 833 | V | 1 | 0 | yes | no | no | yes | no | no | no | no | no | no | no | no | no |
| SCN9A | 834 | L | 1 | 1 | yes | no | no | yes | no | no | no | no | no | no | no | no | no |
| SCN9A | 835 | R | 1 | 0 | yes | no | no | yes | no | no | no | no | no | no | no | no | no |
| SCN9A | 836 | S | 1 | 0 | yes | no | no | yes | no | no | no | no | no | no | no | no | no |
| SCN9A | 837 | F | 0 | 1 | yes | no | no | yes | no | no | no | no | no | no | no | no | no |
| SCN9A | 838 | R | 0 | 0 | yes | no | no | yes | no | no | no | no | no | no | no | no | no |
| SCN9A | 839 | L | 0 | 0 | yes | no | no | yes | no | no | no | no | no | no | no | no | no |
| SCN9A | 840 | L | 0 | 0 | yes | no | no | yes | no | no | no | no | no | no | no | no | no |
| SCN9A | 841 | R | 1 | 0 | yes | no | no | yes | no | no | no | no | no | no | no | no | no |
| SCN9A | 842 | V | 0 | 0 | yes | no | no | yes | no | no | no | no | no | no | no | no | no |
| SCN9A | 843 | F | 0 | 0 | yes | no | no | yes | no | no | no | no | no | no | no | no | no |
| SCN9A | 844 | K | 0 | 0 | yes | no | no | yes | no | no | no | no | no | no | no | no | no |
| SCN9A | 845 | L | 1 | 0 | yes | no | no | yes | no | no | no | no | no | no | no | no | no |

| | | | | | | | | | | | | | | | | | |
|---|---|---|---|---|---|---|---|---|---|---|---|---|---|---|---|---|---|
| SCN9A | 846 | A | 0 | 0 | yes | no | no | yes | no | no | no | no | no | no | no | no | no |
| SCN9A | 847 | K | 2 | 0 | no | yes | no | yes | no | no | no | no | no | no | no | no | no |
| SCN9A | 848 | S | 1 | 0 | no | yes | no | yes | no | no | no | no | no | no | no | no | no |
| SCN9A | 849 | W | 0 | 0 | no | yes | no | yes | no | yes | no | no | no | no | no | no | no |
| SCN9A | 850 | P | 1 | 0 | yes | no | no | no | no | yes | no | no | no | no | no | no | no |
| SCN9A | 851 | T | 1 | 0 | yes | no | no | yes | no | yes | no | no | no | no | no | no | no |
| SCN9A | 852 | L | 0 | 0 | yes | no | no | yes | no | yes | no | no | no | no | no | no | no |
| SCN9A | 853 | N | 0 | 0 | yes | no | no | yes | no | yes | no | no | no | no | no | no | no |
| SCN9A | 854 | M | 1 | 0 | yes | no | no | yes | no | yes | no | no | no | no | no | no | no |
| SCN9A | 855 | L | 0 | 0 | yes | no | no | yes | no | yes | no | no | no | no | no | no | no |
| SCN9A | 856 | I | 1 | 0 | yes | no | no | yes | no | yes | no | no | no | no | no | no | no |
| SCN9A | 857 | K | 0 | 0 | yes | no | no | yes | no | yes | no | no | no | no | no | no | no |
| SCN9A | 858 | I | 0 | 0 | yes | no | no | yes | no | yes | no | no | no | no | no | no | no |
| SCN9A | 859 | I | 0 | 1 | yes | no | no | yes | no | yes | no | no | no | no | no | no | no |
| SCN9A | 860 | G | 0 | 0 | yes | no | no | yes | no | yes | no | no | no | no | no | no | no |
| SCN9A | 861 | N | 0 | 0 | yes | no | no | yes | no | yes | no | no | no | no | no | no | no |
| SCN9A | 862 | S | 0 | 0 | yes | no | no | yes | no | yes | no | no | no | no | no | no | no |
| SCN9A | 863 | V | 1 | 0 | yes | no | no | yes | no | yes | no | no | no | no | no | no | no |
| SCN9A | 864 | G | 0 | 0 | no | yes | no | yes | no | yes | no | no | no | no | no | no | no |
| SCN9A | 865 | A | 0 | 0 | no | yes | no | no | no | no | no | no | no | no | no | no | no |
| SCN9A | 866 | L | 0 | 0 | no | yes | no | yes | no | no | no | no | no | no | no | no | no |
| SCN9A | 867 | G | 0 | 2 | yes | no | no | yes | no | no | no | no | no | no | no | no | no |
| SCN9A | 868 | N | 0 | 0 | yes | no | no | yes | no | no | no | no | no | no | no | no | no |
| SCN9A | 869 | L | 0 | 2 | yes | no | no | yes | no | no | no | no | no | no | no | no | no |
| SCN9A | 870 | T | 0 | 0 | yes | no | no | yes | no | no | no | no | no | no | no | no | no |
| SCN9A | 871 | L | 0 | 0 | yes | no | no | yes | no | no | no | no | no | no | no | no | no |
| SCN9A | 872 | V | 0 | 0 | yes | no | no | yes | no | no | no | no | no | no | no | no | no |
| SCN9A | 873 | L | 0 | 0 | yes | no | no | yes | no | no | no | no | no | no | no | no | no |
| SCN9A | 874 | A | 2 | 1 | yes | no | no | yes | no | no | no | no | no | no | no | no | no |
| SCN9A | 875 | I | 0 | 0 | yes | no | no | yes | no | no | no | no | no | no | no | no | no |
| SCN9A | 876 | I | 1 | 0 | yes | no | no | yes | no | no | no | no | no | no | no | no | no |
| SCN9A | 877 | V | 1 | 0 | yes | no | no | yes | no | no | no | no | no | no | no | no | no |
| SCN9A | 878 | F | 0 | 0 | yes | no | no | yes | no | no | no | no | no | no | no | no | no |

| | | | | | | | | | | | | | | | | | |
|---|---|---|---|---|---|---|---|---|---|---|---|---|---|---|---|---|---|
| SCN9A | 879 | I | 1 | 0 | yes | no | no | yes | no | no | no | no | no | no | no | no | no |
| SCN9A | 880 | F | 0 | 0 | yes | no | no | yes | no | no | no | no | no | no | no | no | no |
| SCN9A | 881 | A | 0 | 0 | yes | no | no | yes | no | no | no | no | no | no | no | no | no |
| SCN9A | 882 | V | 0 | 0 | yes | no | no | yes | no | no | no | no | no | no | no | no | no |
| SCN9A | 883 | V | 1 | 1 | yes | no | no | yes | no | no | no | no | no | no | no | no | no |
| SCN9A | 884 | G | 1 | 0 | yes | no | no | yes | yes | no | no | no | no | no | no | no | no |
| SCN9A | 885 | M | 2 | 0 | yes | no | no | yes | yes | no | no | no | no | no | no | no | no |
| SCN9A | 886 | Q | 0 | 1 | yes | no | no | yes | yes | no | no | no | no | no | no | no | no |
| SCN9A | 887 | L | 1 | 0 | yes | no | no | yes | yes | no | no | no | no | no | no | no | no |
| SCN9A | 888 | F | 1 | 0 | yes | no | no | no | yes | no | no | no | no | no | no | no | no |
| SCN9A | 889 | G | 1 | 0 | yes | no | no | no | yes | no | no | no | no | no | no | no | no |
| SCN9A | 890 | K | 0 | 0 | yes | no | no | no | yes | no | no | no | no | no | no | no | no |
| SCN9A | 891 | S | 1 | 0 | yes | no | no | no | yes | no | no | no | no | no | no | no | no |
| SCN9A | 892 | Y | 1 | 0 | yes | no | no | no | yes | no | no | no | no | no | no | no | no |
| SCN9A | 893 | K | 1 | 0 | yes | no | no | no | yes | no | no | no | no | no | no | no | no |
| SCN9A | 894 | E | 0 | 0 | yes | no | no | no | yes | no | no | no | yes | no | no | no | no |
| SCN9A | 895 | C | 2 | 0 | yes | no | no | no | yes | no | no | no | yes | no | no | no | no |
| SCN9A | 896 | V | 0 | 0 | no | yes | no | no | yes | no | no | no | no | no | no | no | no |
| SCN9A | 897 | C | 0 | 0 | no | yes | no | no | yes | no | no | no | yes | no | no | no | no |
| SCN9A | 898 | K | 0 | 0 | no | yes | no | no | yes | no | no | no | yes | no | no | no | no |
| SCN9A | 899 | I | 1 | 0 | no | yes | no | no | yes | no | no | no | no | no | no | no | no |
| SCN9A | 900 | N | 1 | 0 | no | yes | no | no | yes | no | no | no | no | no | no | no | no |
| SCN9A | 901 | D | 0 | 0 | no | yes | no | no | yes | no | no | no | no | no | no | no | no |
| SCN9A | 902 | D | 1 | 0 | no | yes | no | no | yes | no | no | no | no | no | no | no | no |
| SCN9A | 903 | C | 0 | 0 | no | yes | no | no | yes | no | no | no | no | no | no | no | no |
| SCN9A | 904 | T | 1 | 0 | no | yes | no | no | yes | no | no | no | no | no | no | no | no |
| SCN9A | 905 | L | 0 | 0 | no | yes | no | no | yes | no | no | no | no | no | no | no | no |
| SCN9A | 906 | P | 0 | 0 | no | yes | no | no | yes | no | no | no | no | no | no | no | no |
| SCN9A | 907 | R | 2 | 2 | no | yes | no | no | yes | no | no | no | no | no | no | no | no |
| SCN9A | 908 | W | 0 | 0 | no | yes | no | no | yes | no | no | no | no | no | no | no | no |
| SCN9A | 909 | H | 0 | 0 | no | yes | no | no | yes | no | no | no | no | no | no | no | no |
| SCN9A | 910 | M | 1 | 1 | no | yes | no | no | yes | no | no | no | no | no | no | no | no |
| SCN9A | 911 | N | 0 | 0 | no | yes | no | no | yes | no | no | no | no | no | no | no | no |

| | | | | | | | | | | | | | | | | | |
|---|---|---|---|---|---|---|---|---|---|---|---|---|---|---|---|---|---|
| SCN9A | 912 | D | 0 | 0 | no | yes | no | no | yes | no | no | no | no | no | no | no | no |
| SCN9A | 913 | F | 0 | 0 | yes | no | no | yes | no | no | no | yes | no | no | no | no | no |
| SCN9A | 914 | F | 0 | 0 | yes | no | no | yes | no | no | no | yes | no | no | no | no | no |
| SCN9A | 915 | H | 0 | 0 | yes | no | no | no | no | no | no | yes | no | no | no | no | no |
| SCN9A | 916 | S | 0 | 0 | yes | no | no | yes | no | no | no | yes | no | no | no | no | no |
| SCN9A | 917 | F | 0 | 0 | yes | no | no | yes | no | no | no | yes | no | no | no | no | no |
| SCN9A | 918 | L | 0 | 0 | yes | no | no | yes | no | no | no | yes | no | no | no | no | no |
| SCN9A | 919 | I | 0 | 0 | yes | no | no | yes | no | no | no | yes | no | no | no | no | no |
| SCN9A | 920 | V | 0 | 0 | yes | no | no | yes | no | no | no | yes | no | no | no | no | no |
| SCN9A | 921 | F | 0 | 0 | yes | no | no | yes | no | no | no | yes | no | no | no | no | no |
| SCN9A | 922 | R | 1 | 0 | yes | no | no | yes | no | no | no | yes | no | no | no | no | no |
| SCN9A | 923 | V | 1 | 0 | yes | no | no | yes | no | no | no | yes | no | no | no | no | no |
| SCN9A | 924 | L | 0 | 0 | yes | no | no | yes | no | no | no | yes | no | no | no | no | no |
| SCN9A | 925 | C | 0 | 0 | no | yes | no | yes | no | no | no | yes | no | no | no | no | no |
| SCN9A | 926 | G | 0 | 0 | no | yes | no | yes | no | no | yes | yes | no | no | no | no | no |
| SCN9A | 927 | E | 1 | 0 | no | yes | no | yes | no | no | yes | yes | no | no | no | no | no |
| SCN9A | 928 | W | 0 | 0 | yes | no | no | yes | no | no | no | yes | no | no | no | no | no |
| SCN9A | 929 | I | 2 | 0 | yes | no | no | yes | no | no | no | yes | no | no | no | no | no |
| SCN9A | 930 | E | 0 | 0 | yes | no | no | no | no | no | yes | yes | no | no | no | no | no |
| SCN9A | 931 | T | 1 | 0 | yes | no | no | yes | no | no | no | yes | no | no | no | no | no |
| SCN9A | 932 | M | 1 | 0 | yes | no | no | yes | no | no | no | yes | no | no | no | no | no |
| SCN9A | 933 | W | 0 | 0 | yes | no | no | no | no | no | no | yes | no | no | no | no | no |
| SCN9A | 934 | D | 1 | 0 | yes | no | no | no | yes | no | no | no | no | no | no | no | no |
| SCN9A | 935 | C | 0 | 0 | yes | no | no | no | yes | no | no | no | no | no | no | no | no |
| SCN9A | 936 | M | 1 | 0 | yes | no | no | no | yes | no | no | no | no | no | no | no | no |
| SCN9A | 937 | E | 1 | 0 | yes | no | no | no | yes | no | no | no | no | no | no | no | no |
| SCN9A | 938 | V | 0 | 0 | yes | no | no | no | yes | no | no | no | no | no | no | no | no |
| SCN9A | 939 | A | 1 | 0 | yes | no | no | no | yes | no | no | no | no | no | no | no | no |
| SCN9A | 940 | G | 0 | 0 | no | yes | no | no | yes | no | no | no | no | no | no | no | no |
| SCN9A | 941 | Q | 0 | 0 | yes | no | no | no | yes | no | no | no | no | no | no | no | no |
| SCN9A | 942 | A | 1 | 0 | yes | no | no | yes | yes | no | no | no | no | no | no | no | no |
| SCN9A | 943 | M | 1 | 0 | yes | no | no | yes | yes | no | no | no | no | no | no | no | no |
| SCN9A | 944 | C | 0 | 0 | yes | no | no | yes | yes | no | no | no | no | no | no | no | no |

| | | | | | | | | | | | | | | | | | |
|---|---|---|---|---|---|---|---|---|---|---|---|---|---|---|---|---|---|
| SCN9A | 945 | L | 1 | 0 | yes | no | no | yes | yes | no | no | no | no | no | no | no | no |
| SCN9A | 946 | I | 2 | 0 | yes | no | no | yes | yes | no | no | no | no | no | no | no | no |
| SCN9A | 947 | V | 0 | 0 | yes | no | no | yes | no | no | no | no | no | no | no | no | no |
| SCN9A | 948 | Y | 0 | 0 | yes | no | no | yes | no | no | no | no | no | no | no | no | no |
| SCN9A | 949 | M | 1 | 0 | yes | no | no | yes | no | no | no | no | no | no | no | no | no |
| SCN9A | 950 | M | 0 | 0 | yes | no | no | yes | no | no | no | no | no | no | no | no | no |
| SCN9A | 951 | V | 1 | 0 | yes | no | no | yes | no | no | no | no | no | no | no | no | no |
| SCN9A | 952 | M | 1 | 0 | yes | no | no | yes | no | no | no | no | no | no | no | no | no |
| SCN9A | 953 | V | 1 | 0 | yes | no | no | yes | no | no | no | no | no | no | no | no | no |
| SCN9A | 954 | I | 0 | 0 | yes | no | no | yes | no | no | no | no | no | no | no | no | no |
| SCN9A | 955 | G | 0 | 0 | yes | no | no | yes | no | no | no | no | no | no | no | no | no |
| SCN9A | 956 | N | 0 | 0 | yes | no | no | yes | no | no | no | no | no | no | no | no | no |
| SCN9A | 957 | L | 0 | 0 | yes | no | no | yes | no | no | no | no | no | no | no | no | no |
| SCN9A | 958 | V | 0 | 0 | yes | no | no | yes | no | no | no | no | no | no | no | no | no |
| SCN9A | 959 | V | 0 | 0 | yes | no | no | yes | no | no | no | no | no | no | no | no | no |
| SCN9A | 960 | L | 0 | 0 | yes | no | no | yes | no | no | no | no | no | no | no | no | no |
| SCN9A | 961 | N | 0 | 0 | yes | no | no | yes | no | no | no | no | no | no | no | no | no |
| SCN9A | 962 | L | 0 | 0 | yes | no | no | yes | no | no | no | no | no | no | no | no | no |
| SCN9A | 963 | F | 0 | 0 | yes | no | no | yes | no | no | no | no | no | no | no | no | no |
| SCN9A | 964 | L | 0 | 0 | yes | no | no | yes | no | no | no | no | no | no | no | no | no |
| SCN9A | 965 | A | 0 | 0 | yes | no | no | yes | no | no | no | no | no | no | no | no | no |
| SCN9A | 966 | L | 0 | 0 | yes | no | no | yes | no | no | no | no | no | no | no | no | no |
| SCN9A | 967 | L | 0 | 0 | yes | no | no | yes | no | no | no | no | no | no | no | no | no |
| SCN9A | 968 | L | 0 | 0 | yes | no | no | no | no | yes | no | no | no | no | no | no | no |
| SCN9A | 969 | S | 0 | 0 | yes | no | no | no | no | yes | no | no | no | no | no | no | no |
| SCN9A | 970 | S | 0 | 0 | yes | no | no | no | no | yes | no | no | no | no | no | no | no |
| SCN9A | 971 | F | 0 | 0 | yes | no | no | no | no | yes | no | no | no | no | no | no | no |
| SCN9A | 972 | S | 0 | 0 | no | yes | no | no | no | yes | no | no | no | no | no | no | no |
| SCN9A | 1175 | G | 0 | 0 | no | yes | no | no | no | yes | no | no | no | no | no | no | no |
| SCN9A | 1176 | K | 0 | 0 | no | yes | no | no | no | yes | no | no | no | no | no | no | no |
| SCN9A | 1177 | I | 0 | 0 | yes | no | no | no | no | yes | no | no | yes | no | no | no | no |
| SCN9A | 1178 | W | 1 | 0 | yes | no | no | no | no | yes | no | no | no | no | no | no | no |
| SCN9A | 1179 | W | 0 | 1 | yes | no | no | no | no | yes | no | no | no | no | no | no | no |

| | | | | | | | | | | | | | | | | | |
|---|---|---|---|---|---|---|---|---|---|---|---|---|---|---|---|---|---|
| SCN9A | 1180 | N | 1 | 0 | yes | no | no | no | no | yes | no | no | no | no | no | no | no |
| SCN9A | 1181 | I | 0 | 0 | yes | no | no | no | no | yes | no | no | no | no | no | no | no |
| SCN9A | 1182 | R | 1 | 0 | yes | no | no | no | no | yes | no | no | no | no | no | no | no |
| SCN9A | 1183 | K | 1 | 0 | yes | no | no | no | no | yes | no | no | no | no | no | no | no |
| SCN9A | 1184 | T | 0 | 0 | yes | no | no | no | no | yes | no | no | yes | no | no | no | no |
| SCN9A | 1185 | C | 0 | 0 | yes | no | no | no | no | yes | no | no | no | no | no | no | no |
| SCN9A | 1186 | Y | 1 | 0 | yes | no | no | no | no | yes | no | no | no | no | no | no | no |
| SCN9A | 1187 | K | 1 | 0 | yes | no | no | no | no | yes | no | no | no | no | no | no | no |
| SCN9A | 1188 | I | 2 | 0 | yes | no | no | yes | no | yes | no | no | no | no | no | no | no |
| SCN9A | 1189 | V | 1 | 0 | yes | no | no | yes | no | yes | no | no | no | no | no | no | no |
| SCN9A | 1190 | E | 0 | 0 | yes | no | no | no | no | yes | no | no | no | no | no | no | no |
| SCN9A | 1191 | H | 0 | 0 | no | yes | no | no | no | yes | no | no | no | no | no | no | no |
| SCN9A | 1192 | S | 0 | 0 | yes | no | no | yes | no | yes | no | no | no | no | no | no | no |
| SCN9A | 1193 | W | 1 | 0 | yes | no | no | yes | no | yes | no | no | no | no | no | no | no |
| SCN9A | 1194 | F | 0 | 0 | yes | no | no | yes | no | no | no | no | no | no | no | no | no |
| SCN9A | 1195 | E | 0 | 0 | yes | no | no | yes | no | no | no | no | no | no | no | no | no |
| SCN9A | 1196 | S | 1 | 0 | yes | no | no | yes | no | no | no | no | no | no | no | no | no |
| SCN9A | 1197 | F | 0 | 0 | yes | no | no | yes | no | no | no | no | no | no | no | no | no |
| SCN9A | 1198 | I | 1 | 0 | yes | no | no | yes | no | no | no | no | no | no | no | no | no |
| SCN9A | 1199 | V | 0 | 0 | yes | no | no | yes | no | no | no | no | no | no | no | no | no |
| SCN9A | 1200 | L | 2 | 0 | yes | no | no | yes | no | no | no | no | no | no | no | no | no |
| SCN9A | 1201 | M | 1 | 0 | yes | no | no | yes | no | no | no | no | no | no | no | no | no |
| SCN9A | 1202 | I | 0 | 0 | yes | no | no | yes | no | no | no | no | no | no | no | no | no |
| SCN9A | 1203 | L | 0 | 0 | yes | no | no | yes | no | no | no | no | no | no | no | no | no |
| SCN9A | 1204 | L | 1 | 0 | yes | no | no | yes | no | no | no | no | no | no | no | no | no |
| SCN9A | 1205 | S | 0 | 0 | yes | no | no | yes | no | no | no | no | no | no | no | no | no |
| SCN9A | 1206 | S | 0 | 0 | yes | no | no | yes | no | no | no | no | no | no | no | no | no |
| SCN9A | 1207 | G | 1 | 0 | yes | no | no | yes | no | no | no | no | no | no | no | no | no |
| SCN9A | 1208 | A | 0 | 0 | yes | no | no | yes | no | no | no | no | no | no | no | no | no |
| SCN9A | 1209 | L | 0 | 0 | yes | no | no | yes | no | no | no | no | no | no | no | no | no |
| SCN9A | 1210 | A | 0 | 0 | yes | no | no | yes | no | no | no | no | no | no | no | no | no |
| SCN9A | 1211 | F | 0 | 0 | yes | no | no | yes | no | no | no | no | no | no | no | no | no |
| SCN9A | 1212 | E | 0 | 0 | no | yes | no | no | yes | no | no | no | no | no | no | no | no |

| | | | | | | | | | | | | | | | | | |
|---|---|---|---|---|---|---|---|---|---|---|---|---|---|---|---|---|---|
| SCN9A | 1213 | D | 0 | 0 | no | yes | no | no | yes | no | no | no | no | no | no | no | no |
| SCN9A | 1214 | I | 1 | 0 | yes | no | no | no | yes | no | no | no | yes | no | no | no | no |
| SCN9A | 1215 | Y | 0 | 0 | yes | no | no | no | yes | no | no | no | yes | no | no | no | no |
| SCN9A | 1216 | I | 0 | 0 | yes | no | no | no | yes | no | no | no | no | no | no | no | no |
| SCN9A | 1217 | E | 0 | 0 | yes | no | no | no | yes | no | no | no | yes | no | no | no | no |
| SCN9A | 1218 | R | 0 | 0 | no | yes | no | no | yes | no | no | no | yes | no | no | no | no |
| SCN9A | 1219 | K | 0 | 0 | no | yes | no | no | yes | no | no | no | no | no | no | no | no |
| SCN9A | 1220 | K | 0 | 0 | yes | no | no | no | yes | no | no | no | yes | no | no | no | no |
| SCN9A | 1221 | T | 0 | 0 | yes | no | no | no | yes | no | no | no | no | no | no | no | no |
| SCN9A | 1222 | I | 0 | 0 | yes | no | no | no | yes | no | no | no | no | no | no | no | no |
| SCN9A | 1223 | K | 0 | 0 | yes | no | no | no | yes | no | no | no | no | no | no | no | no |
| SCN9A | 1224 | I | 1 | 0 | yes | no | no | no | yes | no | no | no | no | no | no | no | no |
| SCN9A | 1225 | I | 1 | 0 | yes | no | no | yes | no | no | no | no | yes | no | no | no | no |
| SCN9A | 1226 | L | 0 | 0 | yes | no | no | yes | no | no | no | no | no | no | no | no | no |
| SCN9A | 1227 | E | 0 | 0 | yes | no | no | no | no | no | no | no | no | no | no | no | no |
| SCN9A | 1228 | Y | 0 | 0 | yes | no | no | yes | no | no | no | no | yes | no | no | no | no |
| SCN9A | 1229 | A | 0 | 0 | yes | no | no | yes | no | no | no | no | no | no | no | no | no |
| SCN9A | 1230 | D | 2 | 0 | yes | no | no | yes | no | no | no | no | no | no | no | no | no |
| SCN9A | 1231 | K | 0 | 0 | yes | no | no | yes | no | no | no | no | no | no | no | no | no |
| SCN9A | 1232 | I | 0 | 0 | yes | no | no | yes | no | no | no | no | yes | no | no | no | no |
| SCN9A | 1233 | F | 0 | 0 | yes | no | no | yes | no | no | no | no | no | no | no | no | no |
| SCN9A | 1234 | T | 0 | 0 | yes | no | no | yes | no | no | no | no | no | no | no | no | no |
| SCN9A | 1235 | Y | 0 | 0 | yes | no | no | yes | no | no | no | no | yes | no | no | no | no |
| SCN9A | 1236 | I | 1 | 0 | yes | no | no | yes | no | no | no | no | yes | no | no | no | no |
| SCN9A | 1237 | F | 1 | 0 | yes | no | no | yes | no | no | no | no | no | no | no | no | no |
| SCN9A | 1238 | I | 1 | 0 | yes | no | no | yes | no | no | no | no | no | no | no | no | no |
| SCN9A | 1239 | L | 1 | 0 | yes | no | no | yes | no | no | no | no | no | no | no | no | no |
| SCN9A | 1240 | E | 0 | 0 | yes | no | no | yes | no | no | no | no | no | no | no | no | no |
| SCN9A | 1241 | M | 0 | 0 | yes | no | no | yes | no | no | no | no | no | no | no | no | no |
| SCN9A | 1242 | L | 0 | 0 | yes | no | no | yes | no | no | no | no | no | no | no | no | no |
| SCN9A | 1243 | L | 0 | 0 | yes | no | no | yes | no | no | no | no | no | no | no | no | no |
| SCN9A | 1244 | K | 0 | 0 | yes | no | no | yes | no | yes | no | no | no | no | no | no | no |
| SCN9A | 1245 | W | 0 | 0 | yes | no | no | yes | no | yes | no | no | no | no | no | no | no |

| | | | | | | | | | | | | | | | | | |
|---|---|---|---|---|---|---|---|---|---|---|---|---|---|---|---|---|---|
| SCN9A | 1246 | I | 0 | 0 | yes | no | no | yes | no | yes | no | no | no | no | no | no | no |
| SCN9A | 1247 | A | 1 | 1 | yes | no | no | no | no | yes | no | no | no | no | no | no | no |
| SCN9A | 1248 | Y | 0 | 0 | yes | no | no | no | no | yes | no | no | no | no | no | no | no |
| SCN9A | 1249 | G | 0 | 0 | yes | no | no | no | no | yes | no | no | no | no | no | no | no |
| SCN9A | 1250 | Y | 0 | 0 | yes | no | no | yes | no | yes | no | no | no | no | no | no | no |
| SCN9A | 1251 | K | 0 | 1 | yes | no | no | no | no | yes | no | no | no | no | no | no | no |
| SCN9A | 1252 | T | 1 | 0 | yes | no | no | no | no | yes | no | no | no | no | no | no | no |
| SCN9A | 1253 | Y | 0 | 0 | yes | no | no | yes | no | yes | no | no | no | no | no | no | no |
| SCN9A | 1254 | F | 0 | 0 | yes | no | no | yes | no | yes | no | no | no | no | no | no | no |
| SCN9A | 1255 | T | 0 | 0 | no | yes | no | no | no | yes | no | no | no | no | no | no | no |
| SCN9A | 1256 | N | 1 | 0 | no | yes | no | yes | no | yes | no | no | no | no | no | no | no |
| SCN9A | 1257 | A | 1 | 0 | yes | no | no | yes | no | yes | no | no | no | no | no | no | no |
| SCN9A | 1258 | W | 0 | 0 | yes | no | no | yes | no | no | no | no | no | no | no | no | no |
| SCN9A | 1259 | C | 0 | 0 | yes | no | no | yes | no | no | no | no | no | no | no | no | no |
| SCN9A | 1260 | W | 0 | 0 | yes | no | no | yes | no | no | no | no | no | no | no | no | no |
| SCN9A | 1261 | L | 0 | 0 | yes | no | no | yes | no | no | no | no | no | no | no | no | no |
| SCN9A | 1262 | D | 0 | 0 | yes | no | no | yes | no | no | no | no | no | no | no | no | no |
| SCN9A | 1263 | F | 0 | 0 | yes | no | no | yes | no | no | no | no | no | no | no | no | no |
| SCN9A | 1264 | L | 0 | 0 | yes | no | no | yes | no | no | no | no | no | no | no | no | no |
| SCN9A | 1265 | I | 1 | 0 | yes | no | no | yes | no | no | no | no | no | no | no | no | no |
| SCN9A | 1266 | V | 0 | 0 | yes | no | no | yes | no | no | no | no | no | no | no | no | no |
| SCN9A | 1267 | D | 1 | 0 | yes | no | no | yes | no | no | no | no | no | no | no | no | no |
| SCN9A | 1268 | V | 0 | 0 | yes | no | no | yes | no | no | no | no | no | no | no | no | no |
| SCN9A | 1269 | S | 0 | 0 | yes | no | no | yes | no | no | no | no | no | no | no | no | no |
| SCN9A | 1270 | L | 1 | 0 | yes | no | no | yes | no | no | no | no | no | no | no | no | no |
| SCN9A | 1271 | V | 0 | 0 | yes | no | no | yes | no | no | no | no | no | no | no | no | no |
| SCN9A | 1272 | T | 0 | 0 | yes | no | no | yes | no | no | no | no | no | no | no | no | no |
| SCN9A | 1273 | L | 0 | 0 | yes | no | no | yes | no | no | no | no | no | no | no | no | no |
| SCN9A | 1274 | V | 1 | 0 | yes | no | no | yes | no | no | no | no | no | no | no | no | no |
| SCN9A | 1275 | A | 0 | 0 | yes | no | no | yes | no | no | no | no | no | no | no | no | no |
| SCN9A | 1276 | N | 0 | 0 | yes | no | no | yes | no | no | no | no | no | no | no | no | no |
| SCN9A | 1277 | T | 1 | 0 | yes | no | no | yes | yes | no | no | no | no | no | no | no | no |
| SCN9A | 1278 | L | 1 | 0 | yes | no | no | yes | yes | no | no | no | no | no | no | no | no |

| | | | | | | | | | | | | | | | | | |
|---|---|---|---|---|---|---|---|---|---|---|---|---|---|---|---|---|---|
| SCN9A | 1279 | G | 0 | 0 | no | yes | no | no | yes | no | no | no | no | no | no | no | no |
| SCN9A | 1280 | Y | 0 | 0 | no | yes | no | no | yes | no | no | no | no | no | no | no | no |
| SCN9A | 1281 | S | 0 | 0 | no | yes | no | no | yes | no | no | no | no | no | no | no | no |
| SCN9A | 1282 | D | 0 | 0 | no | yes | no | no | yes | no | no | no | no | no | no | no | no |
| SCN9A | 1283 | L | 0 | 0 | no | yes | no | no | yes | no | no | no | no | no | no | no | no |
| SCN9A | 1284 | G | 0 | 0 | no | yes | no | no | yes | no | no | no | no | no | no | no | no |
| SCN9A | 1285 | P | 0 | 0 | no | yes | no | yes | no | no | no | no | no | no | no | no | no |
| SCN9A | 1286 | I | 0 | 0 | no | yes | no | yes | no | no | no | no | no | no | no | no | no |
| SCN9A | 1287 | K | 0 | 0 | yes | no | no | no | no | no | no | no | no | no | no | no | no |
| SCN9A | 1288 | S | 0 | 0 | yes | no | no | yes | no | no | no | no | no | no | no | no | no |
| SCN9A | 1289 | L | 0 | 0 | yes | no | no | yes | no | no | no | no | no | no | no | no | no |
| SCN9A | 1290 | R | 2 | 0 | yes | no | no | yes | no | no | no | no | no | no | no | no | no |
| SCN9A | 1291 | T | 1 | 0 | yes | no | no | yes | no | no | no | no | no | no | no | no | no |
| SCN9A | 1292 | L | 0 | 0 | yes | no | no | yes | no | no | no | no | no | no | no | no | no |
| SCN9A | 1293 | R | 0 | 0 | yes | no | no | yes | no | no | no | no | no | no | no | no | no |
| SCN9A | 1294 | A | 0 | 0 | yes | no | no | yes | no | no | no | no | no | no | no | no | no |
| SCN9A | 1295 | L | 1 | 0 | yes | no | no | yes | no | no | no | no | no | no | no | no | no |
| SCN9A | 1296 | R | 0 | 0 | yes | no | no | yes | no | no | no | no | no | no | no | no | no |
| SCN9A | 1297 | P | 0 | 0 | yes | no | no | yes | no | no | no | no | no | no | no | no | no |
| SCN9A | 1298 | L | 0 | 0 | yes | no | no | yes | no | no | no | no | no | no | no | no | no |
| SCN9A | 1299 | R | 1 | 0 | yes | no | no | yes | no | no | no | no | no | no | no | no | no |
| SCN9A | 1300 | A | 0 | 0 | yes | no | no | yes | no | no | no | no | no | no | no | no | no |
| SCN9A | 1301 | L | 0 | 0 | yes | no | no | yes | no | no | no | no | no | no | no | no | no |
| SCN9A | 1302 | S | 0 | 0 | yes | no | no | yes | no | no | no | no | no | no | no | no | no |
| SCN9A | 1303 | R | 0 | 0 | yes | no | no | yes | no | no | no | no | no | no | no | no | no |
| SCN9A | 1304 | F | 0 | 0 | no | yes | no | yes | no | yes | no | no | no | no | no | no | no |
| SCN9A | 1305 | E | 0 | 0 | yes | no | no | yes | no | yes | no | no | no | no | no | no | no |
| SCN9A | 1306 | G | 0 | 0 | yes | no | no | yes | no | yes | no | no | no | no | no | no | no |
| SCN9A | 1307 | M | 0 | 0 | yes | no | no | yes | no | yes | no | no | no | no | no | no | no |
| SCN9A | 1308 | R | 0 | 0 | yes | no | no | yes | no | yes | no | no | no | no | no | no | no |
| SCN9A | 1309 | V | 0 | 2 | yes | no | no | yes | no | yes | no | no | no | no | no | no | no |
| SCN9A | 1310 | V | 0 | 1 | yes | no | no | yes | no | yes | no | no | no | no | no | no | no |
| SCN9A | 1311 | V | 1 | 0 | yes | no | no | yes | no | yes | no | no | no | no | no | no | no |

| | | | | | | | | | | | | | | | | | |
|---|---|---|---|---|---|---|---|---|---|---|---|---|---|---|---|---|---|
| SCN9A | 1312 | N | 0 | 0 | yes | no | no | yes | no | yes | no | no | no | no | no | no | no |
| SCN9A | 1313 | A | 0 | 0 | yes | no | no | yes | no | yes | no | no | no | no | no | no | no |
| SCN9A | 1314 | L | 0 | 0 | yes | no | no | yes | no | yes | no | no | no | no | no | no | no |
| SCN9A | 1315 | I | 2 | 0 | yes | no | no | yes | no | yes | no | no | no | no | no | no | no |
| SCN9A | 1316 | G | 0 | 0 | yes | no | no | yes | no | yes | no | no | no | no | no | no | no |
| SCN9A | 1317 | A | 0 | 0 | yes | no | no | yes | no | yes | no | no | no | no | no | no | no |
| SCN9A | 1318 | I | 0 | 0 | yes | no | no | yes | no | yes | no | no | no | no | no | no | no |
| SCN9A | 1319 | P | 1 | 1 | yes | no | no | yes | no | yes | no | no | no | no | no | no | no |
| SCN9A | 1320 | S | 0 | 0 | yes | no | no | yes | no | yes | no | no | no | no | no | no | no |
| SCN9A | 1321 | I | 0 | 0 | yes | no | no | yes | no | no | no | no | no | no | no | no | no |
| SCN9A | 1322 | M | 0 | 0 | yes | no | no | yes | no | no | no | no | no | no | no | no | no |
| SCN9A | 1323 | N | 0 | 0 | yes | no | no | yes | no | no | no | no | no | no | no | no | no |
| SCN9A | 1324 | V | 2 | 0 | yes | no | no | yes | no | no | no | no | no | no | no | no | no |
| SCN9A | 1325 | L | 0 | 0 | yes | no | no | yes | no | no | no | no | no | no | no | no | no |
| SCN9A | 1326 | L | 1 | 0 | yes | no | no | yes | no | no | no | no | no | no | no | no | no |
| SCN9A | 1327 | V | 0 | 1 | yes | no | no | yes | no | no | no | no | no | no | no | no | no |
| SCN9A | 1328 | C | 0 | 0 | yes | no | no | yes | no | no | no | no | no | no | no | no | no |
| SCN9A | 1329 | L | 1 | 0 | yes | no | no | yes | no | no | no | no | no | no | no | no | no |
| SCN9A | 1330 | I | 0 | 0 | yes | no | no | yes | no | no | no | no | no | no | no | no | no |
| SCN9A | 1331 | F | 0 | 0 | yes | no | no | yes | no | no | no | no | no | no | no | no | no |
| SCN9A | 1332 | W | 0 | 0 | yes | no | no | yes | no | no | no | no | no | no | no | no | no |
| SCN9A | 1333 | L | 0 | 0 | yes | no | no | yes | no | no | no | no | no | no | no | no | no |
| SCN9A | 1334 | I | 0 | 0 | yes | no | no | yes | no | no | no | no | no | no | no | no | no |
| SCN9A | 1335 | F | 0 | 0 | yes | no | no | yes | no | no | no | no | no | no | no | no | no |
| SCN9A | 1336 | S | 0 | 0 | yes | no | no | yes | no | no | no | no | no | no | no | no | no |
| SCN9A | 1337 | I | 1 | 0 | yes | no | no | yes | no | no | no | no | no | no | no | no | no |
| SCN9A | 1338 | M | 0 | 0 | yes | no | no | yes | no | no | no | no | no | no | no | no | no |
| SCN9A | 1339 | G | 0 | 0 | yes | no | no | yes | no | no | no | no | no | no | no | no | no |
| SCN9A | 1340 | V | 0 | 0 | yes | no | no | yes | no | no | no | no | no | no | no | no | no |
| SCN9A | 1341 | N | 1 | 0 | yes | no | no | yes | yes | no | no | no | no | no | no | no | no |
| SCN9A | 1342 | L | 0 | 0 | yes | no | no | yes | yes | no | no | no | no | no | no | no | no |
| SCN9A | 1343 | F | 0 | 0 | yes | no | no | no | yes | no | no | no | no | no | no | no | no |
| SCN9A | 1344 | A | 1 | 0 | no | yes | no | no | yes | no | no | no | no | no | no | no | no |

| | | | | | | | | | | | | | | | | | |
|---|---|---|---|---|---|---|---|---|---|---|---|---|---|---|---|---|---|
| SCN9A | 1345 | G | 0 | 0 | no | yes | no | no | yes | no | no | no | no | no | no | no | no |
| SCN9A | 1346 | K | 0 | 0 | no | yes | no | no | yes | no | no | no | no | no | no | no | no |
| SCN9A | 1347 | F | 0 | 0 | no | yes | no | no | yes | no | no | no | no | no | no | no | no |
| SCN9A | 1348 | Y | 1 | 0 | no | yes | no | no | yes | no | no | no | no | no | no | no | no |
| SCN9A | 1349 | E | 0 | 0 | no | no | yes | no | yes | no | no | no | no | no | no | no | no |
| SCN9A | 1350 | C | 1 | 0 | no | no | yes | no | yes | no | no | no | no | no | no | no | no |
| SCN9A | 1351 | I | 0 | 0 | no | no | yes | no | yes | no | no | no | no | no | no | no | no |
| SCN9A | 1352 | N | 0 | 0 | no | no | yes | no | yes | no | no | no | no | no | no | no | no |
| SCN9A | 1353 | T | 0 | 0 | no | yes | no | no | yes | no | no | no | no | no | no | no | no |
| SCN9A | 1354 | T | 0 | 0 | no | yes | no | no | yes | no | no | no | no | no | no | no | no |
| SCN9A | 1355 | D | 0 | 0 | no | yes | no | no | yes | no | yes | no | no | no | no | no | no |
| SCN9A | 1356 | G | 0 | 0 | no | yes | no | no | yes | no | no | no | no | no | no | no | no |
| SCN9A | 1357 | S | 1 | 0 | no | no | yes | no | yes | no | no | no | no | no | no | no | no |
| SCN9A | 1358 | R | 2 | 0 | no | no | yes | no | yes | no | no | no | no | no | no | no | no |
| SCN9A | 1359 | F | 1 | 0 | no | yes | no | no | yes | no | no | no | no | no | no | no | no |
| SCN9A | 1360 | P | 1 | 0 | no | yes | no | no | yes | no | no | no | no | no | no | no | no |
| SCN9A | 1361 | A | 0 | 0 | no | yes | no | no | yes | no | no | no | no | no | no | no | no |
| SCN9A | 1362 | S | 1 | 0 | no | yes | no | no | yes | no | no | no | no | no | no | no | no |
| SCN9A | 1363 | Q | 0 | 0 | no | yes | no | no | yes | no | no | no | no | no | no | no | no |
| SCN9A | 1364 | V | 0 | 0 | no | yes | no | no | yes | no | no | no | no | no | no | no | no |
| SCN9A | 1365 | P | 0 | 0 | no | yes | no | no | yes | no | no | no | no | no | no | no | no |
| SCN9A | 1366 | N | 0 | 0 | no | no | yes | no | yes | no | no | no | no | no | no | no | no |
| SCN9A | 1367 | R | 1 | 0 | yes | no | no | no | yes | no | no | no | no | no | no | no | no |
| SCN9A | 1368 | S | 1 | 0 | yes | no | no | no | yes | no | no | no | no | no | no | no | no |
| SCN9A | 1369 | E | 1 | 0 | yes | no | no | no | yes | no | no | no | no | no | no | no | no |
| SCN9A | 1370 | C | 1 | 0 | yes | no | no | no | yes | no | no | no | no | no | no | no | no |
| SCN9A | 1371 | F | 0 | 0 | yes | no | no | no | yes | no | no | no | no | no | no | no | no |
| SCN9A | 1372 | A | 0 | 0 | yes | no | no | no | yes | no | no | no | no | no | no | no | no |
| SCN9A | 1373 | L | 1 | 0 | yes | no | no | no | yes | no | no | no | no | no | no | no | no |
| SCN9A | 1374 | M | 1 | 0 | yes | no | no | no | yes | no | no | no | no | no | no | no | no |
| SCN9A | 1375 | N | 1 | 0 | yes | no | no | no | yes | no | no | no | no | no | no | no | no |
| SCN9A | 1376 | V | 1 | 0 | yes | no | no | no | yes | no | no | no | no | no | no | no | no |
| SCN9A | 1377 | S | 1 | 0 | no | yes | no | no | yes | no | no | no | no | no | no | no | no |

| | | | | | | | | | | | | | | | | | |
|---|---|---|---|---|---|---|---|---|---|---|---|---|---|---|---|---|---|
| SCN9A | 1378 | Q | 0 | 0 | no | yes | no | no | yes | no | no | no | no | no | no | no | no |
| SCN9A | 1379 | N | 0 | 0 | no | yes | no | no | yes | no | no | no | no | no | no | no | no |
| SCN9A | 1380 | V | 1 | 0 | no | no | yes | no | yes | no | no | no | no | no | no | no | no |
| SCN9A | 1381 | R | 1 | 0 | no | no | yes | no | yes | no | no | no | no | no | no | no | no |
| SCN9A | 1382 | W | 1 | 0 | no | no | yes | no | yes | no | no | no | no | no | no | no | no |
| SCN9A | 1383 | K | 2 | 0 | no | no | yes | no | yes | no | no | no | no | no | no | no | no |
| SCN9A | 1384 | N | 0 | 0 | no | yes | no | no | yes | no | no | no | no | no | no | no | no |
| SCN9A | 1385 | L | 0 | 0 | no | yes | no | no | yes | no | no | no | no | no | no | no | no |
| SCN9A | 1386 | K | 2 | 0 | no | yes | no | no | yes | no | no | no | no | no | no | no | no |
| SCN9A | 1387 | V | 0 | 0 | no | yes | no | no | yes | no | no | no | no | no | no | no | no |
| SCN9A | 1388 | N | 0 | 0 | no | yes | no | no | yes | no | no | no | no | no | no | no | no |
| SCN9A | 1389 | F | 0 | 0 | no | yes | no | yes | yes | no | no | no | no | no | no | no | no |
| SCN9A | 1390 | D | 0 | 0 | no | yes | no | no | yes | no | no | no | no | no | no | no | no |
| SCN9A | 1391 | N | 0 | 0 | no | yes | no | yes | yes | no | no | no | no | no | no | no | no |
| SCN9A | 1392 | V | 0 | 0 | yes | no | no | yes | yes | no | no | no | no | no | no | no | no |
| SCN9A | 1393 | G | 1 | 0 | yes | no | no | yes | no | no | no | yes | no | no | no | no | no |
| SCN9A | 1394 | L | 0 | 0 | yes | no | no | yes | no | no | no | yes | no | no | no | no | no |
| SCN9A | 1395 | G | 1 | 0 | yes | no | no | yes | no | no | no | yes | no | no | no | no | no |
| SCN9A | 1396 | Y | 0 | 0 | yes | no | no | yes | no | no | no | yes | no | no | no | no | no |
| SCN9A | 1397 | L | 0 | 0 | yes | no | no | yes | no | no | no | yes | no | no | no | no | no |
| SCN9A | 1398 | S | 0 | 0 | yes | no | no | yes | no | no | no | yes | no | no | no | no | no |
| SCN9A | 1399 | L | 0 | 0 | yes | no | no | yes | no | no | no | yes | no | no | no | no | no |
| SCN9A | 1400 | L | 0 | 0 | yes | no | no | yes | no | no | no | yes | no | no | no | no | no |
| SCN9A | 1401 | Q | 0 | 0 | yes | no | no | yes | no | no | no | yes | no | no | no | no | no |
| SCN9A | 1402 | V | 1 | 0 | yes | no | no | yes | no | no | no | yes | no | no | no | no | no |
| SCN9A | 1403 | A | 0 | 0 | yes | no | no | yes | no | no | no | yes | no | no | no | no | no |
| SCN9A | 1404 | T | 0 | 0 | yes | no | no | yes | no | no | no | yes | no | no | no | no | no |
| SCN9A | 1405 | F | 0 | 0 | no | yes | no | yes | no | no | yes | yes | no | no | no | no | no |
| SCN9A | 1406 | K | 0 | 0 | no | yes | no | yes | no | no | yes | yes | no | no | no | no | no |
| SCN9A | 1407 | G | 0 | 0 | no | yes | no | yes | no | no | yes | yes | no | no | no | no | no |
| SCN9A | 1408 | W | 0 | 0 | yes | no | no | yes | no | no | no | yes | no | no | no | no | no |
| SCN9A | 1409 | T | 1 | 0 | yes | no | no | no | no | no | yes | yes | no | no | no | no | no |
| SCN9A | 1410 | I | 0 | 0 | yes | no | no | no | no | no | yes | yes | no | no | no | no | no |

| | | | | | | | | | | | | | | | | | |
|---|---|---|---|---|---|---|---|---|---|---|---|---|---|---|---|---|---|
| SCN9A | 1411 | I | 0 | 0 | yes | no | no | no | no | no | no | yes | no | no | no | no | no |
| SCN9A | 1412 | M | 1 | 0 | yes | no | no | no | no | no | no | yes | no | no | no | no | no |
| SCN9A | 1413 | Y | 0 | 0 | yes | no | no | no | no | no | no | yes | no | no | no | no | no |
| SCN9A | 1414 | A | 0 | 0 | yes | no | no | no | no | no | no | yes | no | no | no | no | no |
| SCN9A | 1415 | A | 0 | 0 | yes | no | no | no | yes | no | no | no | no | no | no | no | no |
| SCN9A | 1416 | V | 0 | 0 | yes | no | no | no | yes | no | no | no | no | no | no | no | no |
| SCN9A | 1417 | D | 0 | 0 | yes | no | no | no | yes | no | no | no | no | no | no | no | no |
| SCN9A | 1418 | S | 1 | 0 | no | yes | no | no | yes | no | no | no | no | no | no | no | no |
| SCN9A | 1419 | V | 0 | 0 | no | yes | no | no | yes | no | no | no | no | no | no | no | no |
| SCN9A | 1420 | N | 1 | 0 | no | yes | no | no | yes | no | no | no | no | no | no | no | no |
| SCN9A | 1421 | V | 1 | 0 | no | yes | no | no | yes | no | no | no | no | no | no | no | no |
| SCN9A | 1422 | D | 0 | 0 | no | yes | no | no | yes | no | no | no | no | no | no | no | no |
| SCN9A | 1423 | K | 0 | 0 | no | no | yes | no | yes | no | no | no | no | no | no | no | no |
| SCN9A | 1424 | Q | 0 | 0 | no | yes | no | no | yes | no | no | no | no | no | no | no | no |
| SCN9A | 1425 | P | 0 | 0 | no | yes | no | no | yes | no | no | no | no | no | no | no | no |
| SCN9A | 1426 | K | 1 | 0 | no | yes | no | no | yes | no | no | no | no | no | no | no | no |
| SCN9A | 1427 | Y | 0 | 0 | no | yes | no | no | yes | no | no | no | no | no | no | no | no |
| SCN9A | 1428 | E | 0 | 0 | no | yes | no | no | yes | no | no | no | no | no | no | no | no |
| SCN9A | 1429 | Y | 0 | 0 | no | yes | no | no | yes | no | no | no | no | no | no | no | no |
| SCN9A | 1430 | S | 1 | 0 | no | yes | no | no | yes | no | no | no | no | no | no | no | no |
| SCN9A | 1431 | L | 1 | 0 | no | yes | no | no | yes | no | no | no | no | no | no | no | no |
| SCN9A | 1432 | Y | 0 | 0 | no | yes | no | yes | no | no | no | no | no | no | no | no | no |
| SCN9A | 1433 | M | 1 | 0 | no | yes | no | yes | no | no | no | no | no | no | no | no | no |
| SCN9A | 1434 | Y | 0 | 0 | yes | no | no | yes | no | no | no | no | no | no | no | no | no |
| SCN9A | 1435 | I | 1 | 0 | yes | no | no | yes | no | no | no | no | no | no | no | no | no |
| SCN9A | 1436 | Y | 0 | 0 | yes | no | no | yes | no | no | no | no | no | no | no | no | no |
| SCN9A | 1437 | F | 0 | 0 | yes | no | no | yes | no | no | no | no | no | no | no | no | no |
| SCN9A | 1438 | V | 0 | 0 | yes | no | no | yes | no | no | no | no | no | no | no | no | no |
| SCN9A | 1439 | V | 1 | 0 | yes | no | no | yes | no | no | no | no | no | no | no | no | no |
| SCN9A | 1440 | F | 0 | 0 | yes | no | no | yes | no | no | no | no | no | no | no | no | no |
| SCN9A | 1441 | I | 1 | 0 | yes | no | no | yes | no | no | no | no | no | no | no | no | no |
| SCN9A | 1442 | I | 0 | 0 | yes | no | no | yes | no | no | no | no | no | no | no | no | no |
| SCN9A | 1443 | F | 0 | 0 | yes | no | no | yes | no | no | no | no | no | no | no | no | no |

| | | | | | | | | | | | | | | | | | |
|---|---|---|---|---|---|---|---|---|---|---|---|---|---|---|---|---|---|
| SCN9A | 1444 | G | 0 | 0 | yes | no | no | yes | no | no | no | no | no | no | no | no | no |
| SCN9A | 1445 | S | 2 | 0 | yes | no | no | yes | no | no | no | no | no | no | no | no | no |
| SCN9A | 1446 | F | 0 | 0 | yes | no | no | yes | no | no | no | no | no | no | no | no | no |
| SCN9A | 1447 | F | 0 | 0 | yes | no | no | yes | no | no | no | no | no | no | no | no | no |
| SCN9A | 1448 | T | 0 | 0 | yes | no | no | yes | no | no | no | no | no | no | no | no | no |
| SCN9A | 1449 | L | 0 | 0 | yes | no | no | yes | no | no | yes | no | no | no | no | no | no |
| SCN9A | 1450 | N | 0 | 0 | yes | no | no | yes | no | no | no | no | no | no | no | no | no |
| SCN9A | 1451 | L | 0 | 0 | yes | no | no | yes | no | no | no | no | no | no | no | no | no |
| SCN9A | 1452 | F | 0 | 0 | yes | no | no | yes | no | no | no | no | no | no | no | no | no |
| SCN9A | 1453 | I | 0 | 0 | yes | no | no | yes | no | no | yes | no | no | no | no | no | no |
| SCN9A | 1454 | G | 0 | 0 | yes | no | no | yes | no | yes | no | no | no | no | no | no | no |
| SCN9A | 1455 | V | 1 | 0 | yes | no | no | yes | no | yes | no | no | no | no | no | no | no |
| SCN9A | 1456 | I | 0 | 0 | yes | no | no | yes | no | yes | no | no | no | no | no | no | no |
| SCN9A | 1457 | I | 0 | 0 | yes | no | no | yes | no | yes | yes | no | no | no | no | no | no |
| SCN9A | 1458 | D | 0 | 0 | yes | no | no | no | no | yes | no | no | no | no | no | no | no |
| SCN9A | 1459 | N | 0 | 0 | yes | no | no | no | no | yes | no | no | no | no | no | no | no |
| SCN9A | 1460 | F | 0 | 1 | yes | no | no | yes | no | yes | no | no | no | no | no | no | no |
| SCN9A | 1461 | N | 0 | 0 | yes | no | no | no | no | yes | yes | no | no | no | no | no | no |
| SCN9A | 1462 | Q | 0 | 0 | yes | no | no | no | no | yes | no | no | no | no | no | no | no |
| SCN9A | 1463 | Q | 0 | 0 | yes | no | no | no | no | yes | no | no | no | no | no | no | no |
| SCN9A | 1464 | K | 0 | 0 | yes | no | no | no | no | yes | yes | no | no | no | no | no | no |
| SCN9A | 1465 | K | 0 | 0 | yes | no | no | no | no | yes | yes | no | no | no | no | no | no |
| SCN9A | 1466 | K | 0 | 0 | yes | no | no | no | no | yes | no | no | no | no | no | no | no |
| SCN9A | 1467 | L | 0 | 0 | no | yes | no | no | no | yes | no | no | no | no | no | no | no |
| SCN9A | 1468 | G | 0 | 0 | no | yes | no | no | no | yes | no | no | no | no | no | no | no |
| SCN9A | 1469 | G | 0 | 0 | no | yes | no | no | no | yes | no | no | no | no | no | no | no |
| SCN9A | 1470 | Q | 0 | 0 | no | yes | no | no | no | yes | no | no | no | no | no | no | no |
| SCN9A | 1471 | D | 0 | 0 | no | yes | no | no | no | yes | no | no | no | no | no | no | no |
| SCN9A | 1472 | I | 0 | 2 | no | yes | no | no | no | yes | no | no | no | no | no | no | no |
| SCN9A | 1473 | F | 0 | 1 | no | yes | no | yes | no | yes | no | no | no | no | no | no | no |
| SCN9A | 1474 | M | 0 | 0 | no | yes | no | no | no | yes | no | no | no | no | no | no | no |
| SCN9A | 1475 | T | 0 | 1 | no | yes | no | no | no | yes | no | no | no | no | no | no | no |
| SCN9A | 1476 | E | 1 | 0 | yes | no | no | no | no | yes | no | no | no | no | no | no | no |

| | | | | | | | | | | | | | | | | | |
|---|---|---|---|---|---|---|---|---|---|---|---|---|---|---|---|---|---|
| SCN9A | 1477 | E | 0 | 0 | yes | no | no | no | no | yes | no | no | no | no | no | no | no |
| SCN9A | 1478 | Q | 0 | 0 | yes | no | no | no | no | yes | no | no | no | no | no | no | no |
| SCN9A | 1479 | K | 0 | 0 | yes | no | no | no | no | yes | no | no | no | no | no | no | no |
| SCN9A | 1480 | K | 0 | 0 | yes | no | no | no | no | yes | no | no | no | no | no | no | no |
| SCN9A | 1481 | Y | 0 | 0 | yes | no | no | no | no | yes | no | no | no | no | no | no | no |
| SCN9A | 1482 | Y | 1 | 0 | yes | no | no | no | no | yes | no | no | no | no | no | no | no |
| SCN9A | 1483 | N | 0 | 0 | yes | no | no | no | no | yes | no | no | no | no | no | no | no |
| SCN9A | 1484 | A | 0 | 0 | yes | no | no | no | no | yes | no | no | no | no | no | no | no |
| SCN9A | 1485 | M | 0 | 0 | yes | no | no | no | no | yes | no | no | no | no | no | no | no |
| SCN9A | 1486 | K | 1 | 0 | yes | no | no | no | no | yes | no | no | no | no | no | no | no |
| SCN9A | 1487 | K | 1 | 0 | yes | no | no | no | no | yes | no | no | no | no | no | no | no |
| SCN9A | 1488 | L | 0 | 0 | no | yes | no | no | no | yes | no | no | no | no | no | no | no |
| SCN9A | 1489 | G | 0 | 0 | no | yes | no | no | no | yes | no | no | no | no | no | no | no |
| SCN9A | 1490 | S | 0 | 0 | no | yes | no | no | no | yes | no | no | no | no | no | no | no |
| SCN9A | 1491 | K | 0 | 0 | no | yes | no | no | no | yes | no | no | no | no | no | no | no |
| SCN9A | 1492 | K | 0 | 0 | no | yes | no | no | no | yes | no | no | no | no | no | no | no |
| SCN9A | 1493 | P | 1 | 0 | no | yes | no | no | no | yes | no | no | no | no | no | no | no |
| SCN9A | 1494 | Q | 1 | 0 | no | yes | no | no | no | yes | no | no | no | no | no | no | no |
| SCN9A | 1495 | K | 0 | 0 | no | yes | no | no | no | yes | no | no | no | no | no | no | no |
| SCN9A | 1496 | P | 1 | 0 | no | yes | no | no | no | yes | no | no | no | no | no | no | no |
| SCN9A | 1497 | I | 0 | 0 | no | yes | no | no | no | yes | no | no | no | no | no | no | no |
| SCN9A | 1498 | P | 1 | 0 | no | yes | no | no | no | yes | no | no | no | no | no | no | no |
| SCN9A | 1499 | R | 1 | 0 | no | yes | no | no | no | yes | no | no | no | no | no | no | no |
| SCN9A | 1500 | P | 0 | 0 | no | yes | no | no | no | yes | no | no | no | no | no | no | no |
| SCN9A | 1501 | G | 0 | 0 | no | yes | no | no | no | yes | no | no | no | no | no | no | no |
| SCN9A | 1502 | N | 0 | 0 | no | yes | no | no | no | yes | no | no | no | no | no | no | no |
| SCN9A | 1503 | K | 1 | 0 | yes | no | no | no | no | yes | no | no | no | no | no | no | no |
| SCN9A | 1504 | I | 1 | 0 | yes | no | no | no | no | yes | no | no | no | no | no | no | no |
| SCN9A | 1505 | Q | 0 | 0 | yes | no | no | no | no | yes | no | no | no | no | no | no | no |
| SCN9A | 1506 | G | 0 | 0 | yes | no | no | no | no | yes | no | no | no | no | no | no | no |
| SCN9A | 1507 | C | 0 | 0 | yes | no | no | no | no | yes | no | no | no | no | no | no | no |
| SCN9A | 1508 | I | 1 | 0 | yes | no | no | no | no | yes | no | no | no | no | no | no | no |
| SCN9A | 1509 | F | 0 | 0 | yes | no | no | no | no | yes | no | no | no | no | no | no | no |

| | | | | | | | | | | | | | | | | | |
|---|---|---|---|---|---|---|---|---|---|---|---|---|---|---|---|---|---|
| SCN9A | 1510 | D | 0 | 0 | yes | no | no | no | no | yes | no | no | no | no | no | no | no |
| SCN9A | 1511 | L | 0 | 0 | yes | no | no | no | no | yes | no | no | no | no | no | no | no |
| SCN9A | 1512 | V | 1 | 0 | yes | no | no | no | no | yes | no | no | no | no | no | no | no |
| SCN9A | 1513 | T | 0 | 0 | yes | no | no | no | no | yes | no | no | no | no | no | no | no |
| SCN9A | 1514 | N | 0 | 0 | no | yes | no | no | no | yes | no | no | no | no | no | no | no |
| SCN9A | 1515 | Q | 2 | 0 | yes | no | no | no | no | yes | no | no | no | no | no | no | no |
| SCN9A | 1516 | A | 2 | 0 | yes | no | no | no | no | yes | no | no | no | no | no | no | no |
| SCN9A | 1517 | F | 0 | 0 | yes | no | no | yes | no | no | no | no | no | no | no | no | no |
| SCN9A | 1518 | D | 0 | 0 | yes | no | no | yes | no | no | no | no | no | no | no | no | no |
| SCN9A | 1519 | I | 0 | 0 | yes | no | no | yes | no | no | no | no | no | no | no | no | no |
| SCN9A | 1520 | S | 0 | 0 | yes | no | no | yes | no | no | no | no | no | no | no | no | no |
| SCN9A | 1521 | I | 1 | 0 | yes | no | no | yes | no | no | no | no | no | no | no | no | no |
| SCN9A | 1522 | M | 2 | 0 | yes | no | no | yes | no | no | no | no | no | no | no | no | no |
| SCN9A | 1523 | V | 1 | 0 | yes | no | no | yes | no | no | no | no | no | no | no | no | no |
| SCN9A | 1524 | L | 1 | 0 | yes | no | no | yes | no | no | no | no | no | no | no | no | no |
| SCN9A | 1525 | I | 0 | 0 | yes | no | no | yes | no | no | no | no | no | no | no | no | no |
| SCN9A | 1526 | C | 1 | 0 | yes | no | no | yes | no | no | no | no | no | no | no | no | no |
| SCN9A | 1527 | L | 0 | 0 | yes | no | no | yes | no | no | no | no | no | no | no | no | no |
| SCN9A | 1528 | N | 0 | 0 | yes | no | no | yes | no | no | no | no | no | no | no | no | no |
| SCN9A | 1529 | M | 1 | 0 | yes | no | no | yes | no | no | no | no | no | no | no | no | no |
| SCN9A | 1530 | V | 1 | 0 | yes | no | no | yes | no | no | no | no | no | no | no | no | no |
| SCN9A | 1531 | T | 0 | 0 | yes | no | no | yes | no | no | no | no | no | no | no | no | no |
| SCN9A | 1532 | M | 0 | 0 | yes | no | no | yes | no | no | no | no | no | no | no | no | no |
| SCN9A | 1533 | M | 1 | 0 | yes | no | no | yes | no | no | no | no | no | no | no | no | no |
| SCN9A | 1534 | V | 0 | 0 | no | yes | no | yes | no | no | no | no | no | no | no | no | no |
| SCN9A | 1535 | E | 0 | 0 | no | yes | no | yes | yes | no | no | no | no | no | no | no | no |
| SCN9A | 1536 | K | 0 | 0 | no | yes | no | no | yes | no | no | no | no | no | no | no | no |
| SCN9A | 1537 | E | 0 | 0 | no | yes | no | no | yes | no | no | no | no | no | no | no | no |
| SCN9A | 1538 | G | 0 | 0 | no | yes | no | no | yes | no | no | no | no | no | no | no | no |
| SCN9A | 1539 | Q | 0 | 0 | no | yes | no | no | yes | no | no | no | no | no | no | no | no |
| SCN9A | 1540 | S | 1 | 0 | no | yes | no | no | yes | no | no | no | no | no | no | no | no |
| SCN9A | 1541 | Q | 0 | 0 | yes | no | no | no | yes | no | no | no | no | no | no | no | no |
| SCN9A | 1542 | H | 1 | 0 | yes | no | no | no | yes | no | no | no | no | no | no | no | no |

| | | | | | | | | | | | | | | | | | |
|---|---|---|---|---|---|---|---|---|---|---|---|---|---|---|---|---|---|
| SCN9A | 1543 | M | 2 | 1 | yes | no | no | no | yes | no | no | no | no | no | no | no | no |
| SCN9A | 1544 | T | 1 | 0 | yes | no | no | no | yes | no | no | no | no | no | no | no | no |
| SCN9A | 1545 | E | 1 | 0 | yes | no | no | no | yes | no | no | no | no | no | no | no | no |
| SCN9A | 1546 | V | 0 | 0 | yes | no | no | yes | no | no | no | no | no | no | no | no | no |
| SCN9A | 1547 | L | 0 | 0 | yes | no | no | yes | no | no | no | no | no | no | no | no | no |
| SCN9A | 1548 | Y | 1 | 0 | yes | no | no | yes | no | no | no | no | no | no | no | no | no |
| SCN9A | 1549 | W | 2 | 1 | yes | no | no | yes | no | no | no | no | no | no | no | no | no |
| SCN9A | 1550 | I | 0 | 0 | yes | no | no | yes | no | no | no | no | no | no | no | no | no |
| SCN9A | 1551 | N | 0 | 0 | yes | no | no | yes | no | no | no | no | no | no | no | no | no |
| SCN9A | 1552 | V | 0 | 0 | yes | no | no | yes | no | no | no | no | no | no | no | no | no |
| SCN9A | 1553 | V | 0 | 0 | yes | no | no | yes | no | no | no | no | no | no | no | no | no |
| SCN9A | 1554 | F | 0 | 0 | yes | no | no | yes | no | no | no | no | no | no | no | no | no |
| SCN9A | 1555 | I | 1 | 0 | yes | no | no | yes | no | no | no | no | no | no | no | no | no |
| SCN9A | 1556 | I | 0 | 0 | yes | no | no | yes | no | no | no | no | no | no | no | no | no |
| SCN9A | 1557 | L | 0 | 0 | yes | no | no | yes | no | no | no | no | no | no | no | no | no |
| SCN9A | 1558 | F | 0 | 0 | yes | no | no | yes | no | no | no | no | no | no | no | no | no |
| SCN9A | 1559 | T | 0 | 0 | yes | no | no | yes | no | no | no | no | no | no | no | no | no |
| SCN9A | 1560 | G | 0 | 0 | yes | no | no | yes | no | no | no | no | no | no | no | no | no |
| SCN9A | 1561 | E | 0 | 0 | yes | no | no | yes | no | no | no | no | no | no | no | no | no |
| SCN9A | 1562 | C | 1 | 0 | yes | no | no | yes | no | no | no | no | no | no | no | no | no |
| SCN9A | 1563 | V | 0 | 0 | yes | no | no | yes | no | no | no | no | no | no | no | no | no |
| SCN9A | 1564 | L | 0 | 0 | yes | no | no | yes | no | no | no | no | no | no | no | no | no |
| SCN9A | 1565 | K | 0 | 0 | yes | no | no | yes | no | yes | no | no | no | no | no | no | no |
| SCN9A | 1566 | L | 0 | 0 | yes | no | no | yes | no | yes | no | no | no | no | no | no | no |
| SCN9A | 1567 | I | 0 | 0 | yes | no | no | no | no | yes | no | no | no | no | no | no | no |
| SCN9A | 1568 | S | 1 | 0 | yes | no | no | no | no | yes | no | no | no | no | no | no | no |
| SCN9A | 1569 | L | 0 | 0 | yes | no | no | no | no | yes | no | no | no | no | no | no | no |
| SCN9A | 1570 | R | 0 | 0 | no | yes | no | no | no | yes | no | no | no | no | no | no | no |
| SCN9A | 1571 | H | 1 | 0 | yes | no | no | no | no | yes | no | no | no | no | no | no | no |
| SCN9A | 1572 | Y | 1 | 0 | yes | no | no | no | no | yes | no | no | no | no | no | no | no |
| SCN9A | 1573 | Y | 1 | 0 | yes | no | no | no | no | yes | no | no | no | no | no | no | no |
| SCN9A | 1574 | F | 0 | 0 | yes | no | no | no | no | yes | no | no | no | no | no | no | no |
| SCN9A | 1575 | T | 0 | 0 | yes | no | no | no | no | yes | no | no | no | no | no | no | no |

| | | | | | | | | | | | | | | | | | |
|---|---|---|---|---|---|---|---|---|---|---|---|---|---|---|---|---|---|
| SCN9A | 1576 | V | 1 | 0 | no | yes | no | no | no | yes | no | no | no | no | no | no | no |
| SCN9A | 1577 | G | 0 | 0 | yes | no | no | no | no | no | no | no | no | no | no | no | no |
| SCN9A | 1578 | W | 0 | 0 | yes | no | no | yes | no | no | no | no | no | no | no | no | no |
| SCN9A | 1579 | N | 2 | 0 | yes | no | no | yes | no | no | no | no | no | no | no | no | no |
| SCN9A | 1580 | I | 1 | 0 | yes | no | no | yes | no | no | no | no | no | no | no | no | no |
| SCN9A | 1581 | F | 0 | 0 | yes | no | no | yes | no | no | no | no | no | no | no | no | no |
| SCN9A | 1582 | D | 0 | 0 | yes | no | no | yes | no | no | no | no | no | no | no | no | no |
| SCN9A | 1583 | F | 1 | 0 | yes | no | no | yes | no | no | no | no | no | no | no | no | no |
| SCN9A | 1584 | V | 1 | 0 | yes | no | no | yes | no | no | no | no | no | no | no | no | no |
| SCN9A | 1585 | V | 0 | 0 | yes | no | no | yes | no | no | no | no | no | no | no | no | no |
| SCN9A | 1586 | V | 0 | 0 | yes | no | no | yes | no | no | no | no | no | no | no | no | no |
| SCN9A | 1587 | I | 2 | 0 | yes | no | no | yes | no | no | no | no | no | no | no | no | no |
| SCN9A | 1588 | I | 0 | 0 | yes | no | no | yes | no | no | no | no | no | no | no | no | no |
| SCN9A | 1589 | S | 0 | 0 | yes | no | no | yes | no | no | no | no | no | no | no | no | no |
| SCN9A | 1590 | I | 0 | 0 | yes | no | no | yes | no | no | no | no | no | no | no | no | no |
| SCN9A | 1591 | V | 0 | 0 | yes | no | no | yes | no | no | no | no | no | no | no | no | no |
| SCN9A | 1592 | G | 0 | 0 | yes | no | no | yes | no | no | no | no | no | no | no | no | no |
| SCN9A | 1593 | M | 1 | 0 | yes | no | no | yes | no | no | no | no | no | no | no | no | no |
| SCN9A | 1594 | F | 0 | 0 | yes | no | no | yes | no | no | no | no | no | no | no | no | no |
| SCN9A | 1595 | L | 0 | 0 | yes | no | no | yes | yes | no | no | no | no | no | no | no | no |
| SCN9A | 1596 | A | 2 | 0 | yes | no | no | yes | yes | no | no | no | no | no | no | no | no |
| SCN9A | 1597 | D | 2 | 0 | yes | no | no | yes | yes | no | no | no | no | no | no | no | no |
| SCN9A | 1598 | L | 0 | 0 | yes | no | no | yes | yes | no | no | no | no | no | no | no | no |
| SCN9A | 1599 | I | 0 | 0 | yes | no | no | yes | yes | no | no | no | no | no | no | no | no |
| SCN9A | 1600 | E | 0 | 0 | yes | no | no | no | yes | no | no | no | no | no | no | no | no |
| SCN9A | 1601 | T | 1 | 0 | yes | no | no | no | yes | no | no | no | no | no | no | no | no |
| SCN9A | 1602 | Y | 0 | 0 | yes | no | no | no | yes | no | no | no | no | no | no | no | no |
| SCN9A | 1603 | F | 0 | 0 | no | yes | no | no | yes | no | no | no | no | no | no | no | no |
| SCN9A | 1604 | V | 0 | 0 | no | yes | no | no | yes | no | no | no | no | no | no | no | no |
| SCN9A | 1605 | S | 0 | 0 | no | yes | no | no | yes | no | no | no | no | no | no | no | no |
| SCN9A | 1606 | P | 1 | 0 | yes | no | no | no | yes | no | no | no | no | no | no | no | no |
| SCN9A | 1607 | T | 1 | 1 | yes | no | no | no | yes | no | no | no | no | no | no | no | no |
| SCN9A | 1608 | L | 0 | 0 | yes | no | no | yes | no | no | no | no | no | no | no | no | no |

| | | | | | | | | | | | | | | | | | |
|---|---|---|---|---|---|---|---|---|---|---|---|---|---|---|---|---|---|
| SCN9A | 1609 | F | 0 | 0 | yes | no | no | yes | no | no | no | no | no | no | no | no | no |
| SCN9A | 1610 | R | 0 | 0 | yes | no | no | yes | no | no | no | no | no | no | no | no | no |
| SCN9A | 1611 | V | 0 | 0 | yes | no | no | yes | no | no | no | no | no | no | no | no | no |
| SCN9A | 1612 | I | 0 | 0 | yes | no | no | yes | no | no | no | no | no | no | no | no | no |
| SCN9A | 1613 | R | 1 | 0 | yes | no | no | yes | no | no | no | no | no | no | no | no | no |
| SCN9A | 1614 | L | 0 | 0 | yes | no | no | yes | no | no | no | no | no | no | no | no | no |
| SCN9A | 1615 | A | 0 | 0 | yes | no | no | yes | no | no | no | no | no | no | no | no | no |
| SCN9A | 1616 | R | 1 | 0 | yes | no | no | yes | no | no | no | no | no | no | no | no | no |
| SCN9A | 1617 | I | 1 | 0 | yes | no | no | yes | no | no | no | no | no | no | no | no | no |
| SCN9A | 1618 | G | 0 | 1 | yes | no | no | yes | no | no | no | no | no | no | no | no | no |
| SCN9A | 1619 | R | 0 | 0 | yes | no | no | yes | no | no | no | no | no | no | no | no | no |
| SCN9A | 1620 | I | 0 | 0 | yes | no | no | yes | no | no | no | no | no | no | no | no | no |
| SCN9A | 1621 | L | 0 | 0 | no | yes | no | yes | no | no | no | no | no | no | no | no | no |
| SCN9A | 1622 | R | 1 | 0 | no | yes | no | yes | no | no | no | no | no | no | no | no | no |
| SCN9A | 1623 | L | 0 | 1 | yes | no | no | yes | no | no | no | no | no | no | no | no | no |
| SCN9A | 1624 | V | 1 | 0 | yes | no | no | yes | no | no | no | no | no | no | no | no | no |
| SCN9A | 1625 | K | 0 | 0 | yes | no | no | yes | no | yes | no | no | no | no | no | no | no |
| SCN9A | 1626 | G | 1 | 0 | yes | no | no | yes | no | yes | no | no | no | no | no | no | no |
| SCN9A | 1627 | A | 0 | 0 | no | yes | no | yes | no | yes | no | no | no | no | no | no | no |
| SCN9A | 1628 | K | 1 | 0 | yes | no | no | no | no | yes | no | no | no | no | no | no | no |
| SCN9A | 1629 | G | 0 | 0 | yes | no | no | yes | no | yes | no | no | no | no | no | no | no |
| SCN9A | 1630 | I | 1 | 0 | yes | no | no | yes | no | yes | no | no | no | no | no | no | no |
| SCN9A | 1631 | R | 2 | 0 | yes | no | no | yes | no | yes | no | no | no | no | no | no | no |
| SCN9A | 1632 | T | 1 | 0 | yes | no | no | no | no | yes | no | no | no | no | no | no | no |
| SCN9A | 1633 | L | 0 | 0 | yes | no | no | yes | no | yes | no | no | no | no | no | no | no |
| SCN9A | 1634 | L | 1 | 0 | yes | no | no | yes | no | yes | no | no | no | no | no | no | no |
| SCN9A | 1635 | F | 0 | 0 | yes | no | no | no | no | yes | no | no | no | no | no | no | no |
| SCN9A | 1636 | A | 0 | 0 | yes | no | no | yes | no | yes | no | no | no | no | no | no | no |
| SCN9A | 1637 | L | 0 | 0 | yes | no | no | yes | no | yes | no | no | no | no | no | no | no |
| SCN9A | 1638 | M | 0 | 1 | yes | no | no | yes | no | yes | no | no | no | no | no | no | no |
| SCN9A | 1639 | M | 0 | 0 | yes | no | no | no | no | yes | no | no | no | no | no | no | no |
| SCN9A | 1640 | S | 0 | 0 | no | yes | no | yes | no | yes | no | no | no | no | no | no | no |
| SCN9A | 1641 | L | 0 | 0 | yes | no | no | yes | no | yes | no | no | no | no | no | no | no |

| | | | | | | | | | | | | | | | | | |
|---|---|---|---|---|---|---|---|---|---|---|---|---|---|---|---|---|---|
| SCN9A | 1642 | P | 0 | 0 | yes | no | no | yes | no | yes | no | no | no | no | no | no | no |
| SCN9A | 1643 | A | 1 | 3 | yes | no | no | yes | no | yes | no | no | no | no | no | no | no |
| SCN9A | 1644 | L | 1 | 0 | yes | no | no | yes | no | no | no | no | no | no | no | no | no |
| SCN9A | 1645 | F | 0 | 0 | yes | no | no | yes | no | no | no | no | no | no | no | no | no |
| SCN9A | 1646 | N | 1 | 0 | yes | no | no | yes | no | no | no | no | no | no | no | no | no |
| SCN9A | 1647 | I | 1 | 0 | yes | no | no | yes | no | no | no | no | no | no | no | no | no |
| SCN9A | 1648 | G | 1 | 0 | yes | no | no | yes | no | no | no | no | no | no | no | no | no |
| SCN9A | 1649 | L | 0 | 0 | yes | no | no | yes | no | no | no | no | no | no | no | no | no |
| SCN9A | 1650 | L | 0 | 0 | yes | no | no | yes | no | no | no | no | no | no | no | no | no |
| SCN9A | 1651 | L | 0 | 0 | yes | no | no | yes | no | no | no | no | no | no | no | no | no |
| SCN9A | 1652 | F | 0 | 0 | yes | no | no | yes | no | no | no | no | no | no | no | no | no |
| SCN9A | 1653 | L | 0 | 0 | yes | no | no | yes | no | no | no | no | no | no | no | no | no |
| SCN9A | 1654 | V | 0 | 0 | yes | no | no | yes | no | no | no | no | no | no | no | no | no |
| SCN9A | 1655 | M | 0 | 0 | yes | no | no | yes | no | no | no | no | no | no | no | no | no |
| SCN9A | 1656 | F | 1 | 0 | yes | no | no | yes | no | no | no | no | no | no | no | no | no |
| SCN9A | 1657 | I | 0 | 0 | yes | no | no | yes | no | no | no | no | no | no | no | no | no |
| SCN9A | 1658 | Y | 0 | 0 | yes | no | no | yes | no | no | no | no | no | no | no | no | no |
| SCN9A | 1659 | A | 1 | 0 | yes | no | no | yes | no | no | no | no | no | no | no | no | no |
| SCN9A | 1660 | I | 1 | 0 | yes | no | no | yes | no | no | no | no | no | no | no | no | no |
| SCN9A | 1661 | F | 0 | 0 | yes | no | no | yes | no | no | no | no | no | no | no | no | no |
| SCN9A | 1662 | G | 0 | 0 | yes | no | no | yes | yes | no | no | no | no | no | no | no | no |
| SCN9A | 1663 | M | 1 | 0 | yes | no | no | yes | yes | no | no | no | no | no | no | no | no |
| SCN9A | 1664 | S | 0 | 0 | yes | no | no | no | yes | no | no | no | no | no | no | no | no |
| SCN9A | 1665 | N | 1 | 0 | yes | no | no | no | yes | no | no | no | no | no | no | no | no |
| SCN9A | 1666 | F | 0 | 0 | yes | no | no | no | yes | no | no | no | no | no | no | no | no |
| SCN9A | 1667 | A | 0 | 0 | no | yes | no | no | yes | no | no | no | no | no | no | no | no |
| SCN9A | 1668 | Y | 1 | 0 | no | yes | no | no | yes | no | no | no | no | no | no | no | no |
| SCN9A | 1669 | V | 0 | 0 | no | yes | no | no | yes | no | no | no | no | no | no | no | no |
| SCN9A | 1670 | K | 0 | 0 | no | yes | no | no | yes | no | no | no | no | no | no | no | no |
| SCN9A | 1671 | K | 0 | 0 | no | yes | no | no | yes | no | no | no | no | no | no | no | no |
| SCN9A | 1672 | E | 1 | 0 | no | yes | no | no | yes | no | yes | no | no | no | no | no | no |
| SCN9A | 1673 | D | 0 | 0 | no | yes | no | no | yes | no | yes | no | no | no | no | no | no |
| SCN9A | 1674 | G | 0 | 0 | no | yes | no | no | yes | no | no | no | no | no | no | no | no |

| | | | | | | | | | | | | | | | | | |
|---|---|---|---|---|---|---|---|---|---|---|---|---|---|---|---|---|---|
| SCN9A | 1675 | I | 0 | 0 | no | yes | no | no | yes | no | no | no | no | no | no | no | no |
| SCN9A | 1676 | N | 0 | 0 | no | yes | no | no | yes | no | no | no | no | no | no | no | no |
| SCN9A | 1677 | D | 0 | 0 | no | yes | no | no | yes | no | no | no | yes | no | no | no | no |
| SCN9A | 1678 | M | 0 | 0 | no | yes | no | no | yes | no | no | no | no | no | no | no | no |
| SCN9A | 1679 | F | 0 | 0 | no | yes | no | no | yes | no | no | no | no | no | no | no | no |
| SCN9A | 1680 | N | 0 | 0 | no | yes | no | no | yes | no | no | no | no | no | no | no | no |
| SCN9A | 1681 | F | 0 | 0 | no | yes | no | no | yes | no | no | no | no | no | no | no | no |
| SCN9A | 1682 | E | 1 | 0 | no | yes | no | no | yes | no | no | no | yes | no | no | no | no |
| SCN9A | 1683 | T | 0 | 0 | no | yes | no | no | yes | no | no | no | no | no | no | no | no |
| SCN9A | 1684 | F | 0 | 0 | yes | no | no | yes | no | no | no | yes | no | no | no | no | no |
| SCN9A | 1685 | G | 1 | 0 | yes | no | no | yes | no | no | no | yes | no | no | no | no | no |
| SCN9A | 1686 | N | 0 | 0 | yes | no | no | no | no | no | no | yes | no | no | no | no | no |
| SCN9A | 1687 | S | 0 | 0 | yes | no | no | yes | no | no | no | yes | no | no | no | no | no |
| SCN9A | 1688 | M | 0 | 0 | yes | no | no | yes | no | no | no | yes | no | no | no | no | no |
| SCN9A | 1689 | I | 1 | 0 | yes | no | no | yes | no | no | no | yes | no | no | no | no | no |
| SCN9A | 1690 | C | 0 | 0 | yes | no | no | yes | no | no | no | yes | no | no | no | no | no |
| SCN9A | 1691 | L | 0 | 0 | yes | no | no | yes | no | no | no | yes | no | no | no | no | no |
| SCN9A | 1692 | F | 0 | 0 | yes | no | no | yes | no | no | no | yes | no | no | no | no | no |
| SCN9A | 1693 | Q | 0 | 0 | yes | no | no | yes | no | no | no | yes | no | no | no | no | no |
| SCN9A | 1694 | I | 0 | 0 | yes | no | no | yes | no | no | no | yes | no | no | no | no | no |
| SCN9A | 1695 | T | 0 | 0 | yes | no | no | yes | no | no | no | yes | no | no | no | no | no |
| SCN9A | 1696 | T | 0 | 0 | no | yes | no | yes | no | no | no | yes | no | no | no | no | no |
| SCN9A | 1697 | S | 0 | 0 | no | yes | no | yes | no | no | no | yes | no | no | no | no | no |
| SCN9A | 1698 | A | 0 | 0 | no | yes | no | yes | no | no | no | yes | no | no | no | no | no |
| SCN9A | 1699 | G | 1 | 0 | no | yes | no | yes | no | no | no | yes | no | no | no | no | no |
| SCN9A | 1700 | W | 0 | 0 | yes | no | no | yes | no | no | no | yes | no | no | no | no | no |
| SCN9A | 1701 | D | 0 | 0 | yes | no | no | yes | no | no | yes | yes | no | no | no | no | no |
| SCN9A | 1702 | G | 0 | 0 | yes | no | no | no | no | no | no | yes | no | no | no | no | no |
| SCN9A | 1703 | L | 1 | 0 | yes | no | no | no | no | no | no | yes | no | no | no | no | no |
| SCN9A | 1704 | L | 0 | 0 | yes | no | no | no | no | no | no | yes | no | no | no | no | no |
| SCN9A | 1705 | A | 0 | 0 | yes | no | no | no | no | no | no | yes | no | no | no | no | no |
| SCN9A | 1706 | P | 0 | 0 | yes | no | no | no | no | no | no | yes | no | no | no | no | no |
| SCN9A | 1707 | I | 0 | 0 | yes | no | no | no | yes | no | no | no | no | no | no | no | no |

| | | | | | | | | | | | | | | | | | |
|---|---|---|---|---|---|---|---|---|---|---|---|---|---|---|---|---|---|
| SCN9A | 1708 | L | 1 | 0 | yes | no | no | no | yes | no | no | no | no | no | no | no | no |
| SCN9A | 1709 | N | 1 | 0 | no | yes | no | no | yes | no | no | no | no | no | no | no | no |
| SCN9A | 1710 | S | 0 | 0 | no | yes | no | no | yes | no | no | no | no | no | no | no | no |
| SCN9A | 1711 | K | 0 | 0 | no | yes | no | no | yes | no | no | no | no | no | no | no | no |
| SCN9A | 1712 | P | 0 | 0 | no | yes | no | no | yes | no | no | no | no | no | no | no | no |
| SCN9A | 1713 | P | 0 | 0 | no | yes | no | no | yes | no | no | no | no | no | no | no | no |
| SCN9A | 1714 | D | 2 | 0 | no | yes | no | no | yes | no | no | no | no | no | no | no | no |
| SCN9A | 1715 | C | 1 | 0 | no | yes | no | no | yes | no | no | no | no | no | no | no | no |
| SCN9A | 1716 | D | 0 | 0 | no | yes | no | no | yes | no | no | no | no | no | no | no | no |
| SCN9A | 1717 | P | 0 | 0 | no | yes | no | no | yes | no | no | no | no | no | no | no | no |
| SCN9A | 1718 | K | 1 | 0 | no | yes | no | no | yes | no | no | no | no | no | no | no | no |
| SCN9A | 1719 | K | 1 | 0 | no | yes | no | no | yes | no | no | no | no | no | no | no | no |
| SCN9A | 1720 | V | 2 | 0 | no | yes | no | no | yes | no | no | no | no | no | no | no | no |
| SCN9A | 1721 | H | 1 | 0 | no | yes | no | no | yes | no | no | no | no | no | no | no | no |
| SCN9A | 1722 | P | 1 | 0 | no | yes | no | no | yes | no | no | no | yes | no | no | no | no |
| SCN9A | 1723 | G | 0 | 0 | no | yes | no | no | yes | no | no | no | yes | no | no | no | no |
| SCN9A | 1724 | S | 1 | 0 | no | yes | no | no | yes | no | no | no | no | no | no | no | no |
| SCN9A | 1725 | S | 0 | 0 | no | yes | no | no | yes | no | no | no | no | no | no | no | no |
| SCN9A | 1726 | V | 1 | 0 | no | yes | no | no | yes | no | no | no | no | no | no | no | no |
| SCN9A | 1727 | E | 0 | 0 | no | yes | no | no | yes | no | no | no | no | no | no | no | no |
| SCN9A | 1728 | G | 0 | 0 | no | yes | no | no | yes | no | no | no | no | no | no | no | no |
| SCN9A | 1729 | D | 1 | 0 | no | yes | no | no | yes | no | no | no | no | no | no | no | no |
| SCN9A | 1730 | C | 0 | 1 | no | yes | no | no | yes | no | no | no | no | no | no | no | no |
| SCN9A | 1731 | G | 1 | 0 | no | yes | no | no | yes | no | no | no | no | no | no | no | no |
| SCN9A | 1732 | N | 1 | 0 | no | yes | no | no | yes | no | no | no | no | no | no | no | no |
| SCN9A | 1733 | P | 2 | 0 | yes | no | no | no | yes | no | no | no | no | no | no | no | no |
| SCN9A | 1734 | S | 0 | 0 | yes | no | no | yes | yes | no | no | no | no | no | no | no | no |
| SCN9A | 1735 | V | 0 | 0 | yes | no | no | yes | yes | no | no | no | no | no | no | no | no |
| SCN9A | 1736 | G | 1 | 1 | yes | no | no | yes | yes | no | no | no | no | no | no | no | no |
| SCN9A | 1737 | I | 0 | 0 | yes | no | no | yes | no | no | no | no | no | no | no | no | no |
| SCN9A | 1738 | F | 0 | 0 | yes | no | no | yes | no | no | no | no | no | no | no | no | no |
| SCN9A | 1739 | Y | 1 | 0 | yes | no | no | yes | no | no | no | no | no | no | no | no | no |
| SCN9A | 1740 | F | 0 | 0 | yes | no | no | yes | no | no | no | no | no | no | no | no | no |

| | | | | | | | | | | | | | | | | | |
|---|---|---|---|---|---|---|---|---|---|---|---|---|---|---|---|---|---|
| SCN9A | 1741 | V | 0 | 0 | yes | no | no | yes | no | no | no | no | no | no | no | no | no |
| SCN9A | 1742 | S | 0 | 0 | yes | no | no | yes | no | no | no | no | no | no | no | no | no |
| SCN9A | 1743 | Y | 0 | 0 | yes | no | no | yes | no | no | no | no | no | no | no | no | no |
| SCN9A | 1744 | I | 1 | 0 | yes | no | no | yes | no | no | no | no | no | no | no | no | no |
| SCN9A | 1745 | I | 1 | 0 | yes | no | no | yes | no | no | no | no | no | no | no | no | no |
| SCN9A | 1746 | I | 2 | 0 | yes | no | no | yes | no | no | no | no | no | no | no | no | no |
| SCN9A | 1747 | S | 1 | 0 | yes | no | no | yes | no | no | no | no | no | no | no | no | no |
| SCN9A | 1748 | F | 0 | 0 | yes | no | no | yes | no | no | no | no | no | no | no | no | no |
| SCN9A | 1749 | L | 1 | 0 | yes | no | no | yes | no | no | no | no | no | no | no | no | no |
| SCN9A | 1750 | V | 0 | 0 | yes | no | no | yes | no | no | no | no | no | no | no | no | no |
| SCN9A | 1751 | V | 0 | 1 | yes | no | no | yes | no | no | no | no | no | no | no | no | no |
| SCN9A | 1752 | V | 0 | 0 | yes | no | no | yes | no | no | no | no | no | no | no | no | no |
| SCN9A | 1753 | N | 1 | 0 | yes | no | no | yes | no | no | no | no | no | no | no | no | no |
| SCN9A | 1754 | M | 0 | 0 | yes | no | no | yes | no | no | no | no | no | no | no | no | no |
| SCN9A | 1755 | Y | 0 | 0 | yes | no | no | yes | no | no | yes | no | no | no | no | no | no |
| SCN9A | 1756 | I | 0 | 0 | yes | no | no | yes | no | no | yes | no | no | no | no | no | no |
| SCN9A | 1757 | A | 0 | 1 | yes | no | no | yes | no | no | no | no | no | no | no | no | no |
| SCN9A | 1758 | V | 0 | 0 | yes | no | no | yes | no | no | no | no | no | no | no | no | no |
| SCN9A | 1759 | I | 0 | 0 | yes | no | no | yes | no | no | yes | no | no | no | no | no | no |
| SCN9A | 1760 | L | 1 | 0 | yes | no | no | no | no | yes | yes | no | no | no | no | no | no |
| SCN9A | 1761 | E | 0 | 0 | yes | no | no | no | no | yes | no | no | no | no | no | no | no |
| SCN9A | 1762 | N | 1 | 0 | yes | no | no | no | no | yes | no | no | no | no | no | no | no |
| SCN9A | 1763 | F | 0 | 0 | yes | no | no | no | no | yes | yes | no | no | no | no | no | no |
| SCN9A | 1764 | S | 1 | 0 | yes | no | no | no | no | yes | no | no | no | no | no | no | no |
| SCN9A | 1765 | V | 0 | 0 | yes | no | no | no | no | yes | no | no | no | no | no | no | no |
| SCN9A | 1766 | A | 0 | 0 | yes | no | no | no | no | yes | no | no | no | no | no | no | no |
| SCN9A | 1767 | T | 0 | 0 | yes | no | no | no | no | yes | yes | no | no | no | no | no | no |
| SCN9A | 1768 | E | 1 | 0 | no | yes | no | no | no | yes | no | no | no | no | no | no | no |
| TRPA1 | 446 | K | 0 | 0 | no | yes | no | no | no | yes | no | no | no | no | no | no | no |
| TRPA1 | 447 | K | 0 | 0 | yes | no | no | no | no | yes | no | no | no | no | no | no | no |
| TRPA1 | 448 | S | 0 | 0 | yes | no | no | no | no | yes | no | no | no | no | no | no | no |
| TRPA1 | 449 | P | 0 | 0 | yes | no | no | no | no | yes | no | no | no | no | no | no | no |
| TRPA1 | 450 | L | 0 | 0 | yes | no | no | no | no | yes | no | no | no | no | no | no | no |

| | | | | | | | | | | | | | | | | | |
|---|---|---|---|---|---|---|---|---|---|---|---|---|---|---|---|---|---|
| TRPA1 | 451 | H | 1 | 0 | yes | no | no | no | no | yes | no | no | no | no | no | no | no |
| TRPA1 | 452 | F | 0 | 0 | yes | no | no | no | no | yes | no | no | yes | no | no | no | no |
| TRPA1 | 453 | A | 1 | 0 | yes | no | no | no | no | yes | no | no | no | no | no | no | no |
| TRPA1 | 454 | A | 0 | 0 | yes | no | no | no | no | yes | no | no | no | no | no | no | no |
| TRPA1 | 455 | S | 0 | 0 | no | yes | no | no | no | yes | no | no | no | no | no | no | no |
| TRPA1 | 456 | Y | 1 | 0 | no | yes | no | no | no | yes | no | no | no | no | no | no | no |
| TRPA1 | 457 | G | 0 | 0 | no | yes | no | no | no | yes | no | no | yes | no | no | no | no |
| TRPA1 | 458 | R | 2 | 0 | no | yes | no | no | no | yes | no | no | yes | no | no | no | no |
| TRPA1 | 459 | I | 2 | 0 | yes | no | no | no | no | yes | no | no | no | no | no | no | no |
| TRPA1 | 460 | N | 0 | 0 | yes | no | no | no | no | yes | no | no | yes | no | no | no | no |
| TRPA1 | 461 | T | 1 | 0 | yes | no | no | no | no | yes | no | no | yes | no | no | no | no |
| TRPA1 | 462 | C | 1 | 0 | yes | no | no | no | no | yes | no | no | no | no | no | no | no |
| TRPA1 | 463 | Q | 1 | 0 | yes | no | no | no | no | yes | no | no | no | no | no | no | no |
| TRPA1 | 464 | R | 0 | 0 | yes | no | no | no | no | yes | no | no | yes | no | no | no | no |
| TRPA1 | 465 | L | 0 | 0 | yes | no | no | no | no | yes | no | no | no | no | no | no | no |
| TRPA1 | 466 | L | 0 | 0 | no | yes | no | no | no | yes | no | no | no | no | no | no | no |
| TRPA1 | 467 | Q | 0 | 0 | no | yes | no | no | no | yes | no | no | no | no | no | no | no |
| TRPA1 | 468 | D | 3 | 0 | no | yes | no | no | no | yes | no | no | no | no | no | no | no |
| TRPA1 | 469 | I | 1 | 0 | no | yes | no | no | no | yes | no | no | no | no | no | no | no |
| TRPA1 | 470 | S | 1 | 0 | no | yes | no | no | no | yes | no | no | no | no | no | no | no |
| TRPA1 | 471 | D | 1 | 0 | no | yes | no | no | no | yes | no | no | no | no | no | no | no |
| TRPA1 | 472 | T | 1 | 0 | no | yes | no | no | no | yes | no | no | no | no | no | no | no |
| TRPA1 | 473 | R | 1 | 0 | no | yes | no | no | no | yes | no | no | no | no | no | no | no |
| TRPA1 | 474 | L | 0 | 0 | no | yes | no | no | no | yes | no | no | no | no | no | no | no |
| TRPA1 | 475 | L | 0 | 0 | no | yes | no | no | no | yes | no | no | no | no | no | no | no |
| TRPA1 | 476 | N | 0 | 0 | no | yes | no | no | no | yes | no | no | no | no | no | no | no |
| TRPA1 | 477 | E | 0 | 0 | no | yes | no | no | no | yes | no | no | no | no | no | no | no |
| TRPA1 | 478 | G | 1 | 0 | no | yes | no | no | no | yes | no | no | no | no | no | no | no |
| TRPA1 | 479 | D | 1 | 0 | no | yes | no | no | no | yes | no | no | no | no | no | no | no |
| TRPA1 | 480 | L | 3 | 0 | no | yes | no | no | no | yes | no | no | no | no | no | no | no |
| TRPA1 | 481 | H | 0 | 0 | no | yes | no | no | no | yes | no | no | no | no | no | no | no |
| TRPA1 | 482 | G | 0 | 0 | no | yes | no | no | no | yes | no | no | no | no | no | no | no |
| TRPA1 | 483 | M | 0 | 0 | no | yes | no | no | no | yes | no | no | no | no | no | no | no |

| | | | | | | | | | | | | | | | | | |
|---|---|---|---|---|---|---|---|---|---|---|---|---|---|---|---|---|---|
| TRPA1 | 484 | T | 1 | 0 | no | yes | no | no | no | yes | no | no | no | no | no | no | no |
| TRPA1 | 485 | P | 2 | 0 | yes | no | no | no | no | yes | no | no | no | no | no | no | no |
| TRPA1 | 486 | L | 1 | 0 | yes | no | no | no | no | yes | no | no | no | no | no | no | no |
| TRPA1 | 487 | H | 1 | 0 | yes | no | no | no | no | yes | no | no | no | no | no | no | no |
| TRPA1 | 488 | L | 1 | 0 | yes | no | no | no | no | yes | no | no | no | no | no | no | no |
| TRPA1 | 489 | A | 1 | 0 | yes | no | no | no | no | yes | no | no | no | no | no | no | no |
| TRPA1 | 490 | A | 1 | 0 | yes | no | no | no | no | yes | no | no | no | no | no | no | no |
| TRPA1 | 491 | K | 0 | 0 | yes | no | no | no | no | yes | no | no | no | no | no | no | no |
| TRPA1 | 492 | N | 0 | 0 | no | yes | no | no | no | yes | no | no | yes | no | no | no | no |
| TRPA1 | 493 | G | 0 | 0 | no | yes | no | no | no | yes | no | no | no | no | no | no | no |
| TRPA1 | 494 | H | 0 | 0 | no | yes | no | no | no | yes | no | no | no | no | no | no | no |
| TRPA1 | 495 | D | 1 | 0 | yes | no | no | no | no | yes | no | no | no | no | no | no | no |
| TRPA1 | 496 | K | 0 | 0 | yes | no | no | no | no | yes | no | no | no | no | no | no | no |
| TRPA1 | 497 | V | 1 | 0 | yes | no | no | no | no | yes | no | no | no | no | no | no | no |
| TRPA1 | 498 | V | 0 | 0 | yes | no | no | no | no | yes | no | no | no | no | no | no | no |
| TRPA1 | 499 | Q | 0 | 0 | yes | no | no | no | no | yes | no | no | no | no | no | no | no |
| TRPA1 | 500 | L | 0 | 0 | yes | no | no | no | no | yes | no | no | no | no | no | no | no |
| TRPA1 | 501 | L | 1 | 0 | yes | no | no | no | no | yes | no | no | no | no | no | no | no |
| TRPA1 | 502 | L | 0 | 0 | yes | no | no | no | no | yes | no | no | no | no | no | no | no |
| TRPA1 | 503 | K | 0 | 0 | yes | no | no | no | no | yes | no | no | no | no | no | no | no |
| TRPA1 | 504 | K | 0 | 0 | yes | no | no | no | no | yes | no | no | no | no | no | no | no |
| TRPA1 | 505 | G | 0 | 0 | no | yes | no | no | no | yes | no | no | no | no | no | no | no |
| TRPA1 | 506 | A | 0 | 0 | no | yes | no | no | no | yes | no | no | no | no | no | no | no |
| TRPA1 | 507 | L | 0 | 0 | no | yes | no | no | no | yes | no | no | no | no | no | no | no |
| TRPA1 | 508 | F | 0 | 0 | no | yes | no | no | no | yes | no | no | no | no | no | no | no |
| TRPA1 | 509 | L | 0 | 0 | no | yes | no | no | no | yes | no | no | no | no | no | no | no |
| TRPA1 | 510 | S | 0 | 0 | no | yes | no | no | no | yes | no | no | no | no | no | no | no |
| TRPA1 | 511 | D | 0 | 0 | no | yes | no | no | no | yes | no | no | no | no | no | no | no |
| TRPA1 | 512 | H | 1 | 0 | no | yes | no | no | no | yes | no | no | no | no | no | no | no |
| TRPA1 | 513 | N | 0 | 0 | no | yes | no | no | no | yes | no | no | no | no | no | no | no |
| TRPA1 | 514 | G | 0 | 0 | no | yes | no | no | no | yes | no | no | no | no | no | no | no |
| TRPA1 | 515 | W | 0 | 0 | no | yes | no | no | no | yes | no | no | no | no | no | no | no |
| TRPA1 | 516 | T | 1 | 0 | no | yes | no | no | no | yes | no | no | no | no | no | no | no |

| | | | | | | | | | | | | | | | | | |
|---|---|---|---|---|---|---|---|---|---|---|---|---|---|---|---|---|---|
| TRPA1 | 517 | A | 0 | 0 | yes | no | no | no | no | yes | no | no | no | no | no | no | no |
| TRPA1 | 518 | L | 0 | 0 | yes | no | no | no | no | yes | no | no | no | no | no | no | no |
| TRPA1 | 519 | H | 1 | 0 | yes | no | no | no | no | yes | no | no | no | no | no | no | no |
| TRPA1 | 520 | H | 1 | 0 | yes | no | no | no | no | yes | no | no | no | no | no | no | no |
| TRPA1 | 521 | A | 3 | 0 | yes | no | no | no | no | yes | no | no | no | no | no | no | no |
| TRPA1 | 522 | S | 0 | 0 | yes | no | no | no | no | yes | no | no | no | no | no | no | no |
| TRPA1 | 523 | M | 0 | 0 | yes | no | no | no | no | yes | no | no | no | no | no | no | no |
| TRPA1 | 524 | G | 0 | 0 | yes | no | no | no | no | yes | no | no | no | no | no | no | no |
| TRPA1 | 525 | G | 2 | 0 | no | yes | no | no | no | yes | no | no | no | no | no | no | no |
| TRPA1 | 526 | Y | 0 | 0 | no | yes | no | no | no | yes | no | no | yes | no | no | no | no |
| TRPA1 | 527 | T | 1 | 0 | yes | no | no | no | no | yes | no | no | no | no | no | no | no |
| TRPA1 | 528 | Q | 0 | 0 | yes | no | no | no | no | yes | no | no | no | no | no | no | no |
| TRPA1 | 529 | T | 0 | 0 | yes | no | no | no | no | yes | no | no | no | no | no | no | no |
| TRPA1 | 530 | M | 1 | 0 | yes | no | no | no | no | yes | no | no | no | no | no | no | no |
| TRPA1 | 531 | K | 1 | 0 | yes | no | no | no | no | yes | no | no | no | no | no | no | no |
| TRPA1 | 532 | V | 1 | 0 | yes | no | no | no | no | yes | no | no | no | no | no | no | no |
| TRPA1 | 533 | I | 0 | 0 | yes | no | no | no | no | yes | no | no | no | no | no | no | no |
| TRPA1 | 534 | L | 0 | 0 | yes | no | no | no | no | yes | no | no | no | no | no | no | no |
| TRPA1 | 535 | D | 0 | 0 | yes | no | no | no | no | yes | no | no | no | no | no | no | no |
| TRPA1 | 536 | T | 0 | 0 | yes | no | no | no | no | yes | no | no | no | no | no | no | no |
| TRPA1 | 537 | N | 0 | 0 | no | yes | no | no | no | yes | no | no | no | no | no | no | no |
| TRPA1 | 538 | L | 0 | 0 | no | yes | no | no | no | yes | no | no | no | no | no | no | no |
| TRPA1 | 539 | K | 0 | 0 | no | yes | no | no | no | yes | no | no | no | no | no | no | no |
| TRPA1 | 540 | C | 0 | 0 | no | yes | no | no | no | yes | no | no | no | no | no | no | no |
| TRPA1 | 541 | T | 0 | 0 | no | yes | no | no | no | yes | no | no | no | no | no | no | no |
| TRPA1 | 542 | D | 0 | 0 | no | yes | no | no | no | yes | no | no | no | no | no | no | no |
| TRPA1 | 543 | R | 2 | 0 | no | yes | no | no | no | yes | no | no | no | no | no | no | no |
| TRPA1 | 544 | L | 0 | 0 | no | yes | no | no | no | yes | no | no | no | no | no | no | no |
| TRPA1 | 545 | D | 0 | 0 | no | yes | no | no | no | yes | no | no | no | no | no | no | no |
| TRPA1 | 546 | E | 0 | 0 | no | yes | no | no | no | yes | no | no | no | no | no | no | no |
| TRPA1 | 547 | D | 1 | 0 | no | yes | no | no | no | yes | no | no | no | no | no | no | no |
| TRPA1 | 548 | G | 2 | 0 | no | yes | no | no | no | yes | no | no | no | no | no | no | no |
| TRPA1 | 549 | N | 0 | 0 | no | yes | no | no | no | yes | no | no | no | no | no | no | no |

| | | | | | | | | | | | | | | | | | |
|---|---|---|---|---|---|---|---|---|---|---|---|---|---|---|---|---|---|
| TRPA1 | 550 | T | 1 | 0 | no | yes | no | no | no | yes | no | no | no | no | no | no | no |
| TRPA1 | 551 | A | 0 | 0 | yes | no | no | no | no | yes | no | no | no | no | no | no | no |
| TRPA1 | 552 | L | 1 | 0 | yes | no | no | no | no | yes | no | no | no | no | no | no | no |
| TRPA1 | 553 | H | 1 | 0 | yes | no | no | no | no | yes | no | no | no | no | no | no | no |
| TRPA1 | 554 | F | 0 | 0 | yes | no | no | no | no | yes | no | no | no | no | no | no | no |
| TRPA1 | 555 | A | 0 | 0 | yes | no | no | no | no | yes | no | no | no | no | no | no | no |
| TRPA1 | 556 | A | 2 | 0 | yes | no | no | no | no | yes | no | no | no | no | no | no | no |
| TRPA1 | 557 | R | 1 | 0 | yes | no | no | no | no | yes | no | no | no | no | no | no | no |
| TRPA1 | 558 | E | 0 | 0 | no | yes | no | no | no | yes | no | no | no | no | no | no | no |
| TRPA1 | 559 | G | 1 | 0 | no | yes | no | no | no | yes | no | no | no | no | no | no | no |
| TRPA1 | 560 | H | 2 | 0 | no | yes | no | no | no | yes | no | no | no | no | no | no | no |
| TRPA1 | 561 | A | 1 | 0 | yes | no | no | no | no | yes | no | no | no | no | no | no | no |
| TRPA1 | 562 | K | 1 | 0 | yes | no | no | no | no | yes | no | no | no | no | no | no | no |
| TRPA1 | 563 | A | 1 | 0 | yes | no | no | no | no | yes | no | no | no | no | no | no | no |
| TRPA1 | 564 | V | 2 | 0 | yes | no | no | no | no | yes | no | no | no | no | no | no | no |
| TRPA1 | 565 | A | 1 | 0 | yes | no | no | no | no | yes | no | no | no | no | no | no | no |
| TRPA1 | 566 | L | 0 | 0 | yes | no | no | no | no | yes | no | no | no | no | no | no | no |
| TRPA1 | 567 | L | 1 | 0 | yes | no | no | no | no | yes | no | no | no | no | no | no | no |
| TRPA1 | 568 | L | 0 | 0 | yes | no | no | no | no | yes | no | no | no | no | no | no | no |
| TRPA1 | 569 | S | 1 | 0 | yes | no | no | no | no | yes | no | no | no | no | no | no | no |
| TRPA1 | 570 | H | 1 | 0 | no | yes | no | no | no | yes | no | no | no | no | no | no | no |
| TRPA1 | 571 | N | 1 | 0 | no | yes | no | no | no | yes | no | no | no | no | no | no | no |
| TRPA1 | 572 | A | 1 | 0 | no | yes | no | no | no | yes | no | no | no | no | no | no | no |
| TRPA1 | 573 | D | 1 | 0 | no | yes | no | no | no | yes | no | no | no | no | no | no | no |
| TRPA1 | 574 | I | 1 | 0 | no | yes | no | no | no | yes | no | no | no | no | no | no | no |
| TRPA1 | 575 | V | 0 | 0 | no | yes | no | no | no | yes | no | no | no | no | no | no | no |
| TRPA1 | 576 | L | 0 | 0 | no | yes | no | no | no | yes | no | no | no | no | no | no | no |
| TRPA1 | 577 | N | 1 | 0 | no | yes | no | no | no | yes | no | no | no | no | no | no | no |
| TRPA1 | 578 | K | 0 | 0 | no | yes | no | no | no | yes | no | no | no | no | no | no | no |
| TRPA1 | 579 | Q | 1 | 0 | no | yes | no | no | no | yes | no | no | no | no | no | no | no |
| TRPA1 | 580 | Q | 1 | 0 | no | yes | no | no | no | yes | no | no | no | no | no | no | no |
| TRPA1 | 581 | A | 1 | 0 | no | yes | no | no | no | yes | no | no | no | no | no | no | no |
| TRPA1 | 582 | S | 0 | 0 | no | yes | no | no | no | yes | no | no | no | no | no | no | no |

| | | | | | | | | | | | | | | | | | |
|---|---|---|---|---|---|---|---|---|---|---|---|---|---|---|---|---|---|
| TRPA1 | 583 | F | 0 | 0 | yes | no | no | no | no | yes | no | no | no | no | no | no | no |
| TRPA1 | 584 | L | 0 | 0 | yes | no | no | no | no | yes | no | no | no | no | no | no | no |
| TRPA1 | 585 | H | 0 | 0 | yes | no | no | no | no | yes | no | no | no | no | no | no | no |
| TRPA1 | 586 | L | 0 | 0 | yes | no | no | no | no | yes | no | no | no | no | no | no | no |
| TRPA1 | 587 | A | 1 | 0 | yes | no | no | no | no | yes | no | no | no | no | no | no | no |
| TRPA1 | 588 | L | 0 | 0 | yes | no | no | no | no | yes | no | no | no | no | no | no | no |
| TRPA1 | 589 | H | 1 | 0 | no | yes | no | no | no | yes | no | no | no | no | no | no | no |
| TRPA1 | 590 | N | 0 | 0 | no | yes | no | no | no | yes | no | no | no | no | no | no | no |
| TRPA1 | 591 | K | 0 | 0 | no | yes | no | no | no | yes | no | no | no | no | no | no | no |
| TRPA1 | 592 | R | 0 | 0 | no | yes | no | no | no | yes | no | no | no | no | no | no | no |
| TRPA1 | 593 | K | 1 | 0 | yes | no | no | no | no | yes | no | no | no | no | no | no | no |
| TRPA1 | 594 | E | 1 | 0 | yes | no | no | no | no | yes | no | no | no | no | no | no | no |
| TRPA1 | 595 | V | 1 | 0 | yes | no | no | no | no | yes | no | no | no | no | no | no | no |
| TRPA1 | 596 | V | 2 | 0 | yes | no | no | no | no | yes | no | no | no | no | no | no | no |
| TRPA1 | 597 | L | 0 | 0 | yes | no | no | no | no | yes | no | no | no | no | no | no | no |
| TRPA1 | 598 | T | 2 | 0 | yes | no | no | no | no | yes | no | no | no | no | no | no | no |
| TRPA1 | 599 | I | 0 | 0 | yes | no | no | no | no | yes | no | no | no | no | no | no | no |
| TRPA1 | 600 | I | 1 | 0 | yes | no | no | no | no | yes | no | no | no | no | no | no | no |
| TRPA1 | 601 | R | 1 | 0 | no | yes | no | no | no | yes | no | no | no | no | no | no | no |
| TRPA1 | 602 | S | 0 | 0 | no | yes | no | no | no | yes | no | no | no | no | no | no | no |
| TRPA1 | 603 | K | 0 | 0 | no | yes | no | no | no | yes | no | no | no | no | no | no | no |
| TRPA1 | 604 | R | 0 | 0 | no | yes | no | no | no | yes | no | no | no | no | no | no | no |
| TRPA1 | 605 | W | 1 | 0 | yes | no | no | no | no | yes | no | no | no | no | no | no | no |
| TRPA1 | 606 | D | 1 | 0 | yes | no | no | no | no | yes | no | no | no | no | no | no | no |
| TRPA1 | 607 | E | 1 | 0 | yes | no | no | no | no | yes | no | no | no | no | no | no | no |
| TRPA1 | 608 | C | 1 | 0 | yes | no | no | no | no | yes | no | no | no | no | no | no | no |
| TRPA1 | 609 | L | 0 | 0 | yes | no | no | no | no | yes | no | no | no | no | no | no | no |
| TRPA1 | 610 | K | 0 | 0 | yes | no | no | no | no | yes | no | no | no | no | no | no | no |
| TRPA1 | 611 | I | 1 | 0 | no | yes | no | no | no | yes | no | no | no | no | no | no | no |
| TRPA1 | 612 | F | 1 | 0 | no | yes | no | no | no | yes | no | no | no | no | no | no | no |
| TRPA1 | 613 | S | 0 | 0 | no | yes | no | no | no | yes | no | no | no | no | no | no | no |
| TRPA1 | 614 | H | 0 | 0 | no | yes | no | no | no | yes | no | no | no | no | no | no | no |
| TRPA1 | 615 | N | 0 | 0 | no | yes | no | no | no | yes | no | no | no | no | no | no | no |

| | | | | | | | | | | | | | | | | | |
|---|---|---|---|---|---|---|---|---|---|---|---|---|---|---|---|---|---|
| TRPA1 | 616 | S | 1 | 0 | no | yes | no | no | no | yes | no | no | no | no | no | no | no |
| TRPA1 | 617 | P | 1 | 0 | no | yes | no | no | no | yes | no | no | no | no | no | no | no |
| TRPA1 | 618 | G | 0 | 0 | no | yes | no | no | no | yes | no | no | no | no | no | no | no |
| TRPA1 | 619 | N | 1 | 0 | no | yes | no | no | no | yes | no | no | no | no | no | no | no |
| TRPA1 | 620 | K | 1 | 0 | no | yes | no | no | no | yes | no | no | no | no | no | no | no |
| TRPA1 | 621 | C | 0 | 0 | no | yes | no | no | no | yes | no | no | no | no | yes | no | no |
| TRPA1 | 622 | P | 0 | 0 | yes | no | no | no | no | yes | no | no | no | no | no | no | no |
| TRPA1 | 623 | I | 2 | 0 | yes | no | no | no | no | yes | no | no | no | no | no | no | no |
| TRPA1 | 624 | T | 0 | 0 | yes | no | no | no | no | yes | no | no | no | no | no | no | no |
| TRPA1 | 625 | E | 0 | 0 | yes | no | no | no | no | yes | no | no | no | no | no | no | no |
| TRPA1 | 626 | M | 1 | 0 | yes | no | no | no | no | yes | no | no | no | no | no | no | no |
| TRPA1 | 627 | I | 2 | 0 | yes | no | no | no | no | yes | no | no | no | no | no | no | no |
| TRPA1 | 628 | E | 0 | 0 | yes | no | no | no | no | yes | no | no | no | no | no | no | no |
| TRPA1 | 629 | Y | 0 | 0 | no | yes | no | no | no | yes | no | no | no | no | no | no | no |
| TRPA1 | 630 | L | 0 | 0 | no | yes | no | no | no | yes | no | no | no | no | no | no | no |
| TRPA1 | 631 | P | 1 | 0 | yes | no | no | no | no | yes | no | no | no | no | no | no | no |
| TRPA1 | 632 | E | 0 | 0 | yes | no | no | no | no | yes | no | no | no | no | no | no | no |
| TRPA1 | 633 | C | 1 | 0 | yes | no | no | no | no | yes | no | no | no | no | no | no | no |
| TRPA1 | 634 | M | 3 | 0 | yes | no | no | no | no | yes | no | no | no | no | no | no | no |
| TRPA1 | 635 | K | 1 | 0 | yes | no | no | no | no | yes | no | no | no | no | no | no | no |
| TRPA1 | 636 | V | 1 | 0 | yes | no | no | no | no | yes | no | no | no | no | no | no | no |
| TRPA1 | 637 | L | 1 | 0 | yes | no | no | no | no | yes | no | no | no | no | no | no | no |
| TRPA1 | 638 | L | 0 | 0 | yes | no | no | no | no | yes | no | no | no | no | no | no | no |
| TRPA1 | 639 | D | 0 | 0 | yes | no | no | no | no | yes | no | no | no | no | no | no | no |
| TRPA1 | 640 | F | 0 | 0 | no | yes | no | no | no | yes | no | no | no | no | no | no | no |
| TRPA1 | 641 | C | 1 | 0 | no | yes | no | no | no | yes | no | no | no | no | yes | no | no |
| TRPA1 | 642 | M | 1 | 0 | no | no | yes | no | no | yes | no | no | no | no | no | no | no |
| TRPA1 | 643 | L | 0 | 0 | no | no | yes | no | no | yes | no | no | no | no | no | no | no |
| TRPA1 | 644 | H | 1 | 0 | no | no | yes | no | no | yes | no | no | no | no | no | no | no |
| TRPA1 | 645 | S | 1 | 0 | no | yes | no | no | no | yes | no | no | no | no | no | no | no |
| TRPA1 | 646 | T | 1 | 0 | no | yes | no | no | no | yes | no | no | no | no | no | no | no |
| TRPA1 | 647 | E | 1 | 0 | no | yes | no | no | no | yes | no | no | no | no | no | no | no |
| TRPA1 | 648 | D | 1 | 0 | no | yes | no | no | no | yes | no | no | no | no | no | no | no |

| | | | | | | | | | | | | | | | | | |
|---|---|---|---|---|---|---|---|---|---|---|---|---|---|---|---|---|---|
| TRPA1 | 649 | K | 0 | 0 | no | yes | no | no | no | yes | no | no | no | no | no | no | no |
| TRPA1 | 650 | S | 2 | 0 | no | yes | no | no | no | yes | no | no | no | no | no | no | no |
| TRPA1 | 651 | C | 1 | 0 | no | yes | no | no | no | yes | no | no | no | no | no | no | no |
| TRPA1 | 652 | R | 1 | 0 | no | yes | no | no | no | yes | no | no | no | no | no | no | no |
| TRPA1 | 653 | D | 1 | 0 | no | yes | no | no | no | yes | no | no | no | no | no | no | no |
| TRPA1 | 654 | Y | 0 | 0 | no | yes | no | no | no | yes | no | no | no | no | no | no | no |
| TRPA1 | 655 | Y | 0 | 0 | no | yes | no | no | no | yes | no | no | no | no | no | no | no |
| TRPA1 | 656 | I | 2 | 0 | no | no | yes | no | no | yes | no | no | no | no | no | no | no |
| TRPA1 | 657 | E | 2 | 0 | no | no | yes | no | no | yes | no | no | no | no | no | no | no |
| TRPA1 | 658 | Y | 1 | 0 | no | no | yes | no | no | yes | no | no | no | no | no | no | no |
| TRPA1 | 659 | N | 0 | 0 | no | no | yes | no | no | yes | no | no | no | no | no | no | no |
| TRPA1 | 660 | F | 1 | 0 | no | yes | no | no | no | yes | no | no | no | no | no | no | no |
| TRPA1 | 661 | K | 0 | 0 | no | yes | no | no | no | yes | no | no | no | no | no | no | no |
| TRPA1 | 662 | Y | 0 | 0 | no | yes | no | no | no | yes | no | no | no | no | no | no | no |
| TRPA1 | 663 | L | 1 | 0 | no | yes | no | no | no | yes | no | no | no | no | no | no | no |
| TRPA1 | 664 | Q | 1 | 0 | no | yes | no | no | no | yes | no | no | no | no | no | no | no |
| TRPA1 | 665 | C | 0 | 0 | no | yes | no | no | no | yes | no | no | no | no | yes | no | no |
| TRPA1 | 666 | P | 1 | 0 | no | yes | no | no | no | yes | no | no | no | no | no | no | no |
| TRPA1 | 667 | L | 1 | 0 | yes | no | no | no | no | yes | no | no | no | no | no | no | no |
| TRPA1 | 668 | E | 0 | 0 | yes | no | no | no | no | yes | no | no | no | no | no | no | no |
| TRPA1 | 669 | F | 0 | 0 | yes | no | no | no | no | yes | no | no | no | no | no | no | no |
| TRPA1 | 670 | T | 1 | 0 | no | yes | no | no | no | yes | no | no | no | no | no | no | no |
| TRPA1 | 671 | K | 0 | 0 | no | yes | no | no | no | yes | no | no | no | no | no | no | no |
| TRPA1 | 672 | K | 1 | 0 | no | yes | no | no | no | yes | no | no | no | no | no | no | no |
| TRPA1 | 673 | T | 1 | 0 | no | yes | no | no | no | yes | no | no | no | no | no | no | no |
| TRPA1 | 674 | P | 1 | 0 | no | yes | no | no | no | yes | no | no | no | no | no | no | no |
| TRPA1 | 675 | T | 0 | 0 | no | yes | no | no | no | yes | no | no | no | no | no | no | no |
| TRPA1 | 676 | Q | 0 | 0 | no | yes | no | no | no | yes | no | no | no | no | no | no | no |
| TRPA1 | 677 | D | 0 | 0 | no | yes | no | no | no | yes | no | no | no | no | no | no | no |
| TRPA1 | 678 | V | 1 | 0 | no | yes | no | no | no | yes | no | no | no | no | no | no | no |
| TRPA1 | 679 | I | 1 | 0 | no | yes | no | no | no | yes | no | no | no | no | no | no | no |
| TRPA1 | 680 | Y | 0 | 0 | no | yes | no | no | no | yes | no | no | no | no | no | no | no |
| TRPA1 | 681 | E | 0 | 0 | no | yes | no | no | no | yes | no | no | no | no | no | no | no |

| | | | | | | | | | | | | | | | | | |
|---|---|---|---|---|---|---|---|---|---|---|---|---|---|---|---|---|---|
| TRPA1 | 682 | P | 1 | 0 | no | yes | no | no | no | yes | no | no | no | no | no | no | no |
| TRPA1 | 683 | L | 0 | 0 | no | yes | no | no | no | yes | no | no | no | no | no | no | no |
| TRPA1 | 684 | T | 0 | 0 | yes | no | no | no | no | yes | no | no | no | no | no | no | no |
| TRPA1 | 685 | A | 1 | 0 | yes | no | no | no | no | yes | no | no | no | no | no | no | no |
| TRPA1 | 686 | L | 0 | 0 | yes | no | no | no | no | yes | no | no | no | no | no | no | no |
| TRPA1 | 687 | N | 1 | 0 | yes | no | no | no | no | yes | no | no | no | no | no | no | no |
| TRPA1 | 688 | A | 0 | 0 | yes | no | no | no | no | yes | no | no | no | no | no | no | no |
| TRPA1 | 689 | M | 1 | 0 | yes | no | no | no | no | yes | no | no | no | no | no | no | no |
| TRPA1 | 690 | V | 0 | 0 | yes | no | no | no | no | yes | no | no | no | no | no | no | no |
| TRPA1 | 691 | Q | 0 | 0 | no | yes | no | no | no | yes | no | no | no | no | no | no | no |
| TRPA1 | 692 | N | 0 | 0 | no | yes | no | no | no | yes | no | no | no | no | no | no | no |
| TRPA1 | 693 | N | 0 | 0 | no | yes | no | no | no | yes | no | no | no | no | no | no | no |
| TRPA1 | 694 | R | 2 | 0 | no | yes | no | no | no | yes | no | no | no | no | no | no | no |
| TRPA1 | 695 | I | 1 | 0 | yes | no | no | no | no | yes | no | no | no | no | no | no | no |
| TRPA1 | 696 | E | 0 | 0 | yes | no | no | no | no | yes | no | no | no | no | no | no | no |
| TRPA1 | 697 | L | 1 | 0 | yes | no | no | no | no | yes | no | no | no | no | no | no | no |
| TRPA1 | 698 | L | 0 | 0 | yes | no | no | no | no | yes | no | no | no | no | no | no | no |
| TRPA1 | 699 | N | 1 | 0 | no | yes | no | no | no | yes | no | no | no | no | no | no | no |
| TRPA1 | 700 | H | 3 | 0 | no | yes | no | no | no | yes | no | no | no | no | no | no | no |
| TRPA1 | 701 | P | 1 | 0 | yes | no | no | no | no | yes | no | no | no | no | no | no | no |
| TRPA1 | 702 | V | 1 | 0 | yes | no | no | no | no | yes | no | no | no | no | no | no | no |
| TRPA1 | 703 | C | 1 | 0 | yes | no | no | no | no | yes | no | no | no | no | no | no | no |
| TRPA1 | 704 | K | 0 | 0 | yes | no | no | no | no | yes | no | no | no | no | no | no | no |
| TRPA1 | 705 | E | 0 | 0 | yes | no | no | no | no | yes | no | no | no | no | no | no | no |
| TRPA1 | 706 | Y | 1 | 0 | yes | no | no | no | no | yes | no | no | no | no | no | no | no |
| TRPA1 | 707 | L | 1 | 0 | yes | no | no | no | no | yes | no | no | no | no | no | no | no |
| TRPA1 | 708 | L | 1 | 0 | yes | no | no | no | no | yes | no | no | no | no | no | no | no |
| TRPA1 | 709 | M | 0 | 0 | yes | no | no | no | no | yes | no | no | no | no | no | no | no |
| TRPA1 | 710 | K | 0 | 0 | yes | no | no | no | no | yes | no | no | no | no | yes | no | no |
| TRPA1 | 711 | W | 0 | 0 | yes | no | no | no | no | yes | no | no | no | no | no | no | no |
| TRPA1 | 712 | L | 0 | 0 | yes | no | no | no | no | yes | no | no | no | no | no | no | no |
| TRPA1 | 713 | A | 1 | 0 | yes | no | no | no | no | yes | no | no | no | no | no | no | no |
| TRPA1 | 714 | Y | 1 | 0 | yes | no | no | no | no | yes | no | no | no | no | no | no | no |

| | | | | | | | | | | | | | | | | | |
|---|---|---|---|---|---|---|---|---|---|---|---|---|---|---|---|---|---|
| TRPA1 | 715 | G | 1 | 0 | yes | no | no | no | no | yes | no | no | no | no | no | no | no |
| TRPA1 | 716 | F | 1 | 0 | yes | no | no | no | no | yes | no | no | no | no | no | no | no |
| TRPA1 | 717 | R | 0 | 0 | yes | no | no | no | no | yes | no | no | no | no | no | no | no |
| TRPA1 | 718 | A | 0 | 0 | yes | no | no | no | no | yes | no | no | no | no | no | no | no |
| TRPA1 | 719 | H | 0 | 0 | yes | no | no | no | no | yes | no | no | no | no | no | no | no |
| TRPA1 | 720 | M | 1 | 0 | yes | no | no | yes | no | no | no | no | no | no | no | no | no |
| TRPA1 | 721 | M | 0 | 0 | yes | no | no | yes | no | no | no | no | no | no | no | no | no |
| TRPA1 | 722 | N | 1 | 0 | yes | no | no | yes | no | no | no | no | no | no | no | no | no |
| TRPA1 | 723 | L | 1 | 0 | yes | no | no | yes | no | no | no | no | no | no | no | no | no |
| TRPA1 | 724 | G | 1 | 0 | yes | no | no | yes | no | no | no | no | no | no | no | no | no |
| TRPA1 | 725 | S | 0 | 0 | yes | no | no | yes | no | no | no | no | no | no | no | no | no |
| TRPA1 | 726 | Y | 0 | 0 | yes | no | no | yes | no | no | no | no | no | no | no | no | no |
| TRPA1 | 727 | C | 0 | 0 | yes | no | no | yes | no | no | no | no | no | no | no | no | no |
| TRPA1 | 728 | L | 0 | 0 | yes | no | no | yes | no | no | no | no | no | no | no | no | no |
| TRPA1 | 729 | G | 1 | 0 | yes | no | no | yes | no | no | no | no | no | no | no | no | no |
| TRPA1 | 730 | L | 1 | 0 | yes | no | no | yes | no | no | no | no | no | no | no | no | no |
| TRPA1 | 731 | I | 1 | 0 | yes | no | no | yes | no | no | no | no | no | no | no | no | no |
| TRPA1 | 732 | P | 0 | 0 | yes | no | no | yes | no | no | no | no | no | no | no | no | no |
| TRPA1 | 733 | M | 1 | 0 | yes | no | no | yes | no | no | no | no | no | no | no | no | no |
| TRPA1 | 734 | T | 1 | 0 | yes | no | no | yes | no | no | no | no | no | no | no | no | no |
| TRPA1 | 735 | I | 0 | 0 | yes | no | no | yes | no | no | no | no | no | no | no | no | no |
| TRPA1 | 736 | L | 0 | 0 | yes | no | no | yes | no | no | no | no | no | no | no | no | no |
| TRPA1 | 737 | V | 1 | 0 | yes | no | no | yes | no | no | no | no | no | no | no | no | no |
| TRPA1 | 738 | V | 0 | 0 | yes | no | no | yes | no | no | no | no | no | no | no | no | no |
| TRPA1 | 739 | N | 1 | 0 | no | yes | no | yes | no | no | no | no | no | no | no | no | no |
| TRPA1 | 740 | I | 1 | 0 | no | yes | no | no | no | no | no | no | no | no | no | no | no |
| TRPA1 | 741 | K | 0 | 0 | no | yes | no | no | yes | no | no | no | no | no | no | no | no |
| TRPA1 | 742 | P | 1 | 0 | no | yes | no | no | yes | no | no | no | yes | no | no | no | no |
| TRPA1 | 743 | G | 0 | 0 | no | yes | no | no | yes | no | no | no | no | no | no | no | no |
| TRPA1 | 744 | M | 2 | 0 | no | yes | no | no | yes | no | no | no | no | no | no | no | no |
| TRPA1 | 745 | A | 1 | 0 | no | no | yes | no | yes | no | no | no | no | no | no | no | no |
| TRPA1 | 746 | F | 0 | 0 | no | no | yes | no | yes | no | no | no | no | no | no | no | no |
| TRPA1 | 747 | N | 0 | 0 | no | no | yes | no | yes | no | no | no | no | no | no | no | no |

| | | | | | | | | | | | | | | | | | |
|---|---|---|---|---|---|---|---|---|---|---|---|---|---|---|---|---|---|
| TRPA1 | 748 | S | 0 | 0 | no | yes | no | no | yes | no | no | no | no | no | no | no | no |
| TRPA1 | 749 | T | 0 | 0 | no | yes | no | no | yes | no | no | no | no | no | no | no | no |
| TRPA1 | 750 | G | 0 | 0 | no | no | yes | no | yes | no | no | no | no | no | no | no | no |
| TRPA1 | 751 | I | 0 | 0 | no | no | yes | no | yes | no | no | no | no | no | no | no | no |
| TRPA1 | 752 | I | 1 | 0 | no | no | yes | no | yes | no | no | no | no | no | no | no | no |
| TRPA1 | 753 | N | 1 | 0 | no | yes | no | no | yes | no | no | no | no | no | no | no | no |
| TRPA1 | 761 | I | 1 | 0 | no | yes | no | no | yes | no | no | no | no | no | no | no | no |
| TRPA1 | 762 | L | 0 | 0 | no | no | yes | no | yes | no | no | no | no | no | no | no | no |
| TRPA1 | 763 | D | 1 | 0 | no | no | yes | no | yes | no | no | no | no | no | no | no | no |
| TRPA1 | 764 | T | 1 | 0 | no | yes | no | no | yes | no | no | no | no | no | no | no | no |
| TRPA1 | 765 | T | 1 | 0 | no | yes | no | no | no | no | no | no | no | no | no | no | no |
| TRPA1 | 766 | N | 0 | 0 | no | yes | no | no | no | no | no | no | no | no | no | no | no |
| TRPA1 | 767 | S | 0 | 0 | yes | no | no | no | no | no | no | no | no | no | no | no | no |
| TRPA1 | 768 | Y | 0 | 0 | yes | no | no | yes | no | no | no | no | no | no | no | no | no |
| TRPA1 | 769 | L | 1 | 0 | yes | no | no | yes | no | no | no | no | no | no | no | no | no |
| TRPA1 | 770 | I | 0 | 0 | yes | no | no | yes | no | no | no | no | no | no | no | no | no |
| TRPA1 | 771 | K | 1 | 0 | yes | no | no | yes | no | no | no | no | no | no | no | no | no |
| TRPA1 | 772 | T | 3 | 0 | yes | no | no | yes | no | no | no | no | no | no | no | no | no |
| TRPA1 | 773 | C | 1 | 0 | yes | no | no | yes | no | no | no | no | no | no | no | no | no |
| TRPA1 | 774 | M | 0 | 0 | yes | no | no | yes | no | no | no | no | no | no | no | no | no |
| TRPA1 | 775 | I | 0 | 0 | yes | no | no | yes | no | no | no | no | no | no | no | no | no |
| TRPA1 | 776 | L | 1 | 0 | yes | no | no | yes | no | no | no | no | no | no | no | no | no |
| TRPA1 | 777 | V | 1 | 0 | yes | no | no | yes | no | no | no | no | no | no | no | no | no |
| TRPA1 | 778 | F | 0 | 0 | yes | no | no | yes | no | no | no | no | no | no | no | no | no |
| TRPA1 | 779 | L | 0 | 0 | yes | no | no | yes | no | no | no | no | no | no | no | no | no |
| TRPA1 | 780 | S | 0 | 0 | yes | no | no | yes | no | no | no | no | no | no | no | no | no |
| TRPA1 | 781 | S | 0 | 0 | yes | no | no | yes | no | no | no | no | no | no | no | no | no |
| TRPA1 | 782 | I | 1 | 0 | yes | no | no | yes | no | no | no | no | no | no | no | no | no |
| TRPA1 | 783 | F | 0 | 0 | yes | no | no | yes | no | no | no | no | no | no | no | no | no |
| TRPA1 | 784 | G | 0 | 0 | yes | no | no | yes | no | no | no | no | no | no | no | no | no |
| TRPA1 | 785 | Y | 0 | 0 | yes | no | no | yes | no | no | no | no | no | no | no | no | no |
| TRPA1 | 786 | C | 0 | 0 | yes | no | no | yes | no | yes | no | no | no | no | no | no | no |
| TRPA1 | 787 | K | 0 | 0 | yes | no | no | yes | no | yes | no | no | no | no | no | no | no |

| | | | | | | | | | | | | | | | | | |
|---|---|---|---|---|---|---|---|---|---|---|---|---|---|---|---|---|---|
| TRPA1 | 788 | E | 0 | 0 | yes | no | no | yes | no | yes | no | no | no | no | no | no | no |
| TRPA1 | 789 | A | 1 | 0 | yes | no | no | yes | no | yes | no | no | no | no | no | no | no |
| TRPA1 | 790 | G | 0 | 0 | yes | no | no | yes | no | yes | no | no | no | no | no | no | no |
| TRPA1 | 791 | Q | 1 | 0 | yes | no | no | yes | no | yes | no | no | no | no | no | no | no |
| TRPA1 | 792 | I | 0 | 0 | yes | no | no | yes | no | yes | no | no | no | no | no | no | no |
| TRPA1 | 793 | F | 1 | 0 | yes | no | no | yes | no | yes | no | no | no | no | no | no | no |
| TRPA1 | 794 | Q | 0 | 0 | yes | no | no | no | no | yes | no | no | no | no | no | no | no |
| TRPA1 | 795 | Q | 0 | 0 | yes | no | no | no | no | yes | no | no | no | no | no | no | no |
| TRPA1 | 796 | K | 0 | 0 | no | yes | no | no | no | yes | no | no | no | no | no | no | no |
| TRPA1 | 797 | R | 2 | 0 | no | yes | no | no | no | yes | no | no | no | no | no | no | no |
| TRPA1 | 798 | N | 1 | 0 | no | yes | no | no | no | yes | no | no | no | no | no | no | no |
| TRPA1 | 799 | Y | 0 | 0 | no | yes | no | yes | no | yes | no | no | no | no | no | no | no |
| TRPA1 | 800 | F | 0 | 0 | no | yes | no | yes | no | yes | no | no | no | no | no | no | no |
| TRPA1 | 801 | M | 1 | 0 | no | yes | no | yes | no | yes | no | no | no | no | no | no | no |
| TRPA1 | 802 | D | 1 | 0 | no | yes | no | yes | no | yes | no | no | no | no | no | no | no |
| TRPA1 | 803 | I | 0 | 0 | yes | no | no | yes | no | yes | no | no | no | no | no | no | no |
| TRPA1 | 804 | S | 0 | 0 | yes | no | no | yes | no | no | no | no | no | no | no | no | no |
| TRPA1 | 805 | N | 1 | 0 | yes | no | no | yes | no | no | no | no | no | no | no | no | no |
| TRPA1 | 806 | V | 1 | 0 | yes | no | no | yes | no | no | no | no | no | no | no | no | no |
| TRPA1 | 807 | L | 0 | 0 | yes | no | no | yes | no | no | no | no | no | no | no | no | no |
| TRPA1 | 808 | E | 1 | 0 | yes | no | no | yes | no | no | no | no | no | no | no | no | no |
| TRPA1 | 809 | W | 0 | 0 | yes | no | no | yes | no | no | no | no | no | no | no | no | no |
| TRPA1 | 810 | I | 2 | 0 | yes | no | no | yes | no | no | no | no | no | no | no | no | no |
| TRPA1 | 811 | I | 2 | 0 | yes | no | no | yes | no | no | no | no | no | no | no | no | no |
| TRPA1 | 812 | Y | 0 | 0 | yes | no | no | yes | no | no | no | no | no | no | no | no | no |
| TRPA1 | 813 | T | 1 | 0 | yes | no | no | yes | no | no | no | no | no | no | no | no | no |
| TRPA1 | 814 | T | 1 | 0 | yes | no | no | yes | no | no | no | no | no | no | no | no | no |
| TRPA1 | 815 | G | 0 | 0 | yes | no | no | yes | no | no | no | no | no | no | no | no | no |
| TRPA1 | 816 | I | 0 | 0 | yes | no | no | yes | no | no | no | no | no | no | no | no | no |
| TRPA1 | 817 | I | 0 | 0 | yes | no | no | yes | no | no | no | no | no | no | no | no | no |
| TRPA1 | 818 | F | 0 | 0 | yes | no | no | yes | no | no | no | no | no | no | no | no | no |
| TRPA1 | 819 | V | 1 | 0 | yes | no | no | yes | no | no | no | no | no | no | no | no | no |
| TRPA1 | 820 | L | 0 | 0 | yes | no | no | yes | no | no | no | no | no | no | no | no | no |

| | | | | | | | | | | | | | | | | | |
|---|---|---|---|---|---|---|---|---|---|---|---|---|---|---|---|---|---|
| TRPA1 | 821 | P | 1 | 0 | yes | no | no | yes | no | no | no | no | no | no | no | no | no |
| TRPA1 | 822 | L | 0 | 0 | yes | no | no | yes | no | no | no | no | no | no | no | no | no |
| TRPA1 | 823 | F | 1 | 0 | no | yes | no | yes | no | no | no | no | no | no | no | no | no |
| TRPA1 | 824 | V | 1 | 0 | no | yes | no | yes | no | no | no | no | no | no | no | no | no |
| TRPA1 | 825 | E | 0 | 0 | no | yes | no | no | yes | no | no | no | no | no | no | no | no |
| TRPA1 | 826 | I | 0 | 0 | no | yes | no | no | yes | no | no | no | no | no | no | no | no |
| TRPA1 | 827 | P | 0 | 0 | no | yes | no | no | yes | no | no | no | no | no | no | no | no |
| TRPA1 | 828 | A | 0 | 0 | yes | no | no | no | yes | no | no | no | no | no | no | no | no |
| TRPA1 | 829 | H | 0 | 0 | yes | no | no | no | yes | no | no | no | yes | no | no | no | no |
| TRPA1 | 830 | L | 0 | 0 | yes | no | no | yes | no | no | no | no | no | no | no | no | no |
| TRPA1 | 831 | Q | 1 | 0 | yes | no | no | yes | no | no | no | no | no | no | no | no | no |
| TRPA1 | 832 | W | 2 | 0 | yes | no | no | yes | no | no | no | no | no | no | no | no | no |
| TRPA1 | 833 | Q | 0 | 0 | yes | no | no | yes | no | no | no | no | yes | no | no | no | no |
| TRPA1 | 834 | C | 1 | 0 | yes | no | no | yes | no | no | no | no | no | no | no | no | no |
| TRPA1 | 835 | G | 0 | 0 | yes | no | no | yes | no | no | no | no | no | no | no | no | no |
| TRPA1 | 836 | A | 2 | 0 | yes | no | no | yes | no | no | no | no | yes | no | no | no | no |
| TRPA1 | 837 | I | 0 | 0 | yes | no | no | yes | no | no | no | no | no | no | no | no | no |
| TRPA1 | 838 | A | 0 | 0 | yes | no | no | yes | no | no | no | no | no | no | no | no | no |
| TRPA1 | 839 | V | 1 | 0 | yes | no | no | yes | no | no | no | no | no | no | no | no | no |
| TRPA1 | 840 | Y | 1 | 0 | yes | no | no | yes | no | no | no | no | yes | no | no | no | no |
| TRPA1 | 841 | F | 1 | 0 | yes | no | no | yes | no | no | no | no | no | no | no | no | no |
| TRPA1 | 842 | Y | 2 | 0 | yes | no | no | yes | no | no | no | no | no | no | no | no | no |
| TRPA1 | 843 | W | 0 | 0 | yes | no | no | yes | no | no | no | no | yes | no | no | no | no |
| TRPA1 | 844 | M | 0 | 0 | yes | no | no | yes | no | no | no | no | yes | no | no | no | no |
| TRPA1 | 845 | N | 1 | 0 | yes | no | no | yes | no | no | no | no | no | no | no | no | no |
| TRPA1 | 846 | F | 0 | 0 | yes | no | no | yes | no | no | no | no | yes | no | no | no | no |
| TRPA1 | 847 | L | 0 | 0 | yes | no | no | yes | no | no | no | no | yes | no | no | no | no |
| TRPA1 | 848 | L | 0 | 0 | yes | no | no | yes | no | no | no | no | no | no | no | no | no |
| TRPA1 | 849 | Y | 0 | 0 | yes | no | no | yes | no | no | no | no | no | no | no | no | no |
| TRPA1 | 850 | L | 0 | 0 | yes | no | no | yes | no | no | no | no | yes | no | no | no | no |
| TRPA1 | 851 | Q | 0 | 0 | yes | no | no | no | no | yes | no | no | no | no | no | no | no |
| TRPA1 | 852 | R | 0 | 0 | yes | no | no | no | no | yes | no | no | no | no | no | no | no |
| TRPA1 | 853 | F | 1 | 0 | no | yes | no | no | no | yes | no | no | no | no | no | no | no |

| | | | | | | | | | | | | | | | | | |
|---|---|---|---|---|---|---|---|---|---|---|---|---|---|---|---|---|---|
| TRPA1 | 854 | E | 0 | 0 | no | yes | no | no | no | yes | no | no | no | no | no | no | no |
| TRPA1 | 855 | N | 0 | 1 | no | yes | no | no | no | yes | no | no | no | no | no | no | no |
| TRPA1 | 856 | C | 0 | 0 | no | yes | no | yes | no | yes | no | no | no | no | no | no | no |
| TRPA1 | 857 | G | 0 | 0 | yes | no | no | no | no | yes | no | no | no | no | no | no | no |
| TRPA1 | 858 | I | 2 | 0 | yes | no | no | no | no | yes | no | no | no | no | no | no | no |
| TRPA1 | 859 | F | 0 | 0 | yes | no | no | yes | no | yes | no | no | yes | no | no | no | no |
| TRPA1 | 860 | I | 0 | 0 | yes | no | no | yes | no | yes | no | no | no | no | no | no | no |
| TRPA1 | 861 | V | 0 | 0 | yes | no | no | no | no | yes | no | no | no | no | no | no | no |
| TRPA1 | 862 | M | 1 | 0 | yes | no | no | no | no | yes | no | no | yes | no | no | no | no |
| TRPA1 | 863 | L | 0 | 0 | yes | no | no | yes | no | yes | no | no | no | no | no | no | no |
| TRPA1 | 864 | E | 1 | 0 | yes | no | no | yes | no | yes | no | no | no | no | no | no | no |
| TRPA1 | 865 | V | 0 | 0 | yes | no | no | no | no | yes | no | no | no | no | no | no | no |
| TRPA1 | 866 | I | 0 | 0 | yes | no | no | yes | no | yes | no | no | yes | no | no | no | no |
| TRPA1 | 867 | L | 0 | 0 | yes | no | no | yes | no | yes | no | no | yes | no | no | no | no |
| TRPA1 | 868 | K | 0 | 0 | yes | no | no | yes | no | yes | no | no | no | no | no | no | no |
| TRPA1 | 869 | T | 0 | 0 | yes | no | no | yes | no | yes | no | no | no | no | no | no | no |
| TRPA1 | 870 | L | 0 | 0 | yes | no | no | yes | no | yes | no | no | yes | no | no | no | no |
| TRPA1 | 871 | L | 0 | 0 | yes | no | no | yes | no | yes | no | no | no | no | no | no | no |
| TRPA1 | 872 | R | 0 | 0 | yes | no | no | yes | no | yes | no | no | yes | no | no | no | no |
| TRPA1 | 873 | S | 1 | 0 | yes | no | no | yes | no | yes | no | no | no | no | no | no | no |
| TRPA1 | 874 | T | 0 | 0 | yes | no | no | yes | no | no | no | no | no | no | no | no | no |
| TRPA1 | 875 | V | 0 | 0 | yes | no | no | yes | no | no | no | no | no | no | no | no | no |
| TRPA1 | 876 | V | 0 | 0 | yes | no | no | yes | no | no | no | no | no | no | no | no | no |
| TRPA1 | 877 | F | 0 | 0 | yes | no | no | yes | no | no | no | no | no | no | no | no | no |
| TRPA1 | 878 | I | 0 | 0 | yes | no | no | yes | no | no | no | no | no | no | no | no | no |
| TRPA1 | 879 | F | 0 | 0 | yes | no | no | yes | no | no | no | no | yes | no | no | no | no |
| TRPA1 | 880 | L | 1 | 0 | yes | no | no | yes | no | no | no | no | no | no | no | no | no |
| TRPA1 | 881 | L | 0 | 0 | yes | no | no | yes | no | no | no | no | no | no | no | no | no |
| TRPA1 | 882 | L | 0 | 0 | yes | no | no | yes | no | no | no | no | no | no | no | no | no |
| TRPA1 | 883 | A | 0 | 0 | yes | no | no | yes | no | no | no | no | no | no | no | no | no |
| TRPA1 | 884 | F | 0 | 0 | yes | no | no | yes | no | no | no | no | no | no | no | no | no |
| TRPA1 | 885 | G | 2 | 0 | yes | no | no | yes | no | no | no | no | no | no | no | no | no |
| TRPA1 | 886 | L | 0 | 0 | yes | no | no | yes | no | no | no | no | no | no | no | no | no |

| | | | | | | | | | | | | | | | | | |
|---|---|---|---|---|---|---|---|---|---|---|---|---|---|---|---|---|---|
| TRPA1 | 887 | S | 0 | 0 | yes | no | no | yes | no | no | no | no | no | no | no | no | no |
| TRPA1 | 888 | F | 0 | 0 | yes | no | no | yes | no | no | no | no | no | no | no | no | no |
| TRPA1 | 889 | Y | 0 | 0 | yes | no | no | yes | no | no | no | no | yes | no | no | no | no |
| TRPA1 | 890 | I | 1 | 0 | yes | no | no | yes | no | no | no | no | no | no | no | no | no |
| TRPA1 | 891 | L | 1 | 0 | yes | no | no | yes | no | no | no | no | no | no | no | no | no |
| TRPA1 | 892 | L | 0 | 0 | no | yes | no | no | no | no | no | no | no | no | no | no | no |
| TRPA1 | 893 | N | 0 | 0 | no | yes | no | no | no | no | no | no | yes | no | no | no | no |
| TRPA1 | 894 | L | 0 | 0 | no | yes | no | no | no | no | no | no | no | no | no | no | no |
| TRPA1 | 895 | Q | 1 | 0 | no | yes | no | no | yes | no | no | no | no | no | no | no | no |
| TRPA1 | 896 | D | 1 | 0 | yes | no | no | no | yes | no | no | no | no | no | no | no | no |
| TRPA1 | 897 | P | 1 | 0 | yes | no | no | no | yes | no | no | no | yes | no | no | no | no |
| TRPA1 | 898 | F | 1 | 0 | yes | no | no | yes | yes | no | no | no | no | no | no | no | no |
| TRPA1 | 899 | S | 0 | 0 | no | yes | no | no | yes | no | no | no | no | no | no | no | no |
| TRPA1 | 900 | S | 1 | 0 | no | yes | no | no | yes | no | no | no | no | no | no | no | no |
| TRPA1 | 901 | P | 1 | 0 | yes | no | no | yes | yes | no | no | no | no | no | no | no | no |
| TRPA1 | 902 | L | 1 | 0 | yes | no | no | yes | no | no | no | yes | no | no | no | no | no |
| TRPA1 | 903 | L | 0 | 0 | yes | no | no | yes | no | no | no | yes | no | no | no | no | no |
| TRPA1 | 904 | S | 0 | 0 | yes | no | no | yes | no | no | no | yes | no | no | no | no | no |
| TRPA1 | 905 | I | 0 | 0 | yes | no | no | yes | no | no | no | yes | no | no | no | no | no |
| TRPA1 | 906 | I | 1 | 0 | yes | no | no | yes | no | no | no | yes | yes | no | no | no | no |
| TRPA1 | 907 | Q | 0 | 0 | yes | no | no | yes | no | no | no | yes | yes | no | no | no | no |
| TRPA1 | 908 | T | 1 | 0 | yes | no | no | yes | no | no | no | yes | no | no | no | no | no |
| TRPA1 | 909 | F | 0 | 0 | yes | no | no | yes | no | no | no | yes | yes | no | yes | no | no |
| TRPA1 | 910 | S | 1 | 0 | yes | no | no | yes | no | no | no | yes | yes | no | no | no | no |
| TRPA1 | 911 | M | 2 | 0 | yes | no | no | yes | no | no | no | yes | no | no | no | no | no |
| TRPA1 | 912 | M | 1 | 0 | no | yes | no | yes | no | no | no | yes | no | no | no | no | no |
| TRPA1 | 913 | L | 2 | 0 | no | yes | no | yes | no | no | yes | yes | yes | no | no | no | no |
| TRPA1 | 914 | G | 0 | 0 | no | yes | no | yes | no | no | yes | yes | no | no | no | no | no |
| TRPA1 | 915 | D | 0 | 0 | no | yes | no | yes | no | no | yes | yes | no | no | no | no | no |
| TRPA1 | 916 | I | 0 | 0 | no | yes | no | yes | no | no | no | yes | no | no | no | no | no |
| TRPA1 | 917 | N | 1 | 0 | no | yes | no | yes | no | no | yes | yes | yes | no | no | no | no |
| TRPA1 | 918 | Y | 0 | 0 | yes | no | no | yes | no | no | no | yes | yes | no | no | no | no |
| TRPA1 | 919 | R | 1 | 0 | yes | no | no | no | no | no | no | yes | yes | no | no | no | no |

| | | | | | | | | | | | | | | | | | |
|---|---|---|---|---|---|---|---|---|---|---|---|---|---|---|---|---|---|
| TRPA1 | 920 | E | 0 | 0 | yes | no | no | no | no | no | yes | yes | no | no | no | no | no |
| TRPA1 | 921 | S | 0 | 0 | yes | no | no | no | no | no | no | yes | yes | no | no | no | no |
| TRPA1 | 922 | F | 0 | 0 | yes | no | no | yes | no | no | no | yes | no | no | no | no | no |
| TRPA1 | 923 | L | 0 | 0 | yes | no | no | no | yes | no | no | no | yes | no | no | no | no |
| TRPA1 | 924 | E | 0 | 0 | yes | no | no | no | yes | no | no | no | no | no | no | no | no |
| TRPA1 | 925 | P | 1 | 0 | yes | no | no | no | yes | no | no | no | no | no | no | no | no |
| TRPA1 | 926 | Y | 0 | 0 | yes | no | no | no | yes | no | no | no | no | no | no | no | no |
| TRPA1 | 927 | L | 1 | 0 | no | yes | no | no | yes | no | no | no | yes | no | no | no | no |
| TRPA1 | 928 | R | 1 | 0 | no | yes | no | no | yes | no | no | no | no | no | no | no | no |
| TRPA1 | 929 | N | 0 | 0 | no | yes | no | no | yes | no | no | no | no | no | no | no | no |
| TRPA1 | 930 | E | 0 | 0 | no | yes | no | no | yes | no | no | no | no | no | no | no | no |
| TRPA1 | 931 | L | 0 | 0 | no | yes | no | no | yes | no | no | no | no | no | no | no | no |
| TRPA1 | 932 | A | 0 | 0 | no | yes | no | no | yes | no | no | no | no | no | no | no | no |
| TRPA1 | 933 | H | 1 | 0 | no | yes | no | no | yes | no | no | no | yes | no | no | no | no |
| TRPA1 | 934 | P | 0 | 0 | yes | no | no | no | yes | no | no | no | no | no | no | no | no |
| TRPA1 | 935 | V | 0 | 0 | yes | no | no | yes | no | no | no | no | no | no | no | no | no |
| TRPA1 | 936 | L | 0 | 0 | yes | no | no | yes | no | no | no | no | no | no | no | no | no |
| TRPA1 | 937 | S | 1 | 0 | yes | no | no | yes | no | no | no | no | no | no | no | no | no |
| TRPA1 | 938 | F | 0 | 0 | yes | no | no | yes | no | no | no | no | no | no | no | no | no |
| TRPA1 | 939 | A | 0 | 0 | yes | no | no | yes | no | no | no | no | no | no | no | no | no |
| TRPA1 | 940 | Q | 1 | 0 | yes | no | no | yes | no | no | no | no | yes | no | no | no | no |
| TRPA1 | 941 | L | 1 | 0 | yes | no | no | yes | no | no | no | no | no | no | no | no | no |
| TRPA1 | 942 | V | 0 | 0 | yes | no | no | yes | no | no | no | no | yes | no | no | no | no |
| TRPA1 | 943 | S | 1 | 0 | yes | no | no | yes | no | no | no | no | yes | no | no | no | no |
| TRPA1 | 944 | F | 0 | 0 | yes | no | no | yes | no | no | no | no | no | no | yes | no | no |
| TRPA1 | 945 | T | 0 | 0 | yes | no | no | yes | no | no | no | no | yes | no | no | no | no |
| TRPA1 | 946 | I | 0 | 0 | yes | no | no | yes | no | no | no | no | yes | no | no | no | no |
| TRPA1 | 947 | F | 0 | 0 | yes | no | no | yes | no | no | no | no | no | no | no | no | no |
| TRPA1 | 948 | V | 0 | 0 | yes | no | no | yes | no | no | no | no | no | no | no | no | no |
| TRPA1 | 949 | P | 1 | 0 | yes | no | no | yes | no | no | no | no | yes | no | no | no | no |
| TRPA1 | 950 | I | 0 | 0 | yes | no | no | yes | no | no | no | no | yes | no | no | no | no |
| TRPA1 | 951 | V | 0 | 0 | yes | no | no | yes | no | no | no | no | yes | no | no | no | no |
| TRPA1 | 952 | L | 0 | 0 | yes | no | no | yes | no | no | no | no | no | no | no | no | no |

| | | | | | | | | | | | | | | | | | |
|---|---|---|---|---|---|---|---|---|---|---|---|---|---|---|---|---|---|
| TRPA1 | 953 | M | 2 | 0 | yes | no | no | yes | no | no | no | no | yes | no | no | no | no |
| TRPA1 | 954 | N | 1 | 0 | yes | no | no | yes | no | no | no | no | yes | no | no | no | no |
| TRPA1 | 955 | L | 0 | 0 | yes | no | no | yes | no | no | no | no | yes | no | no | no | no |
| TRPA1 | 956 | L | 0 | 0 | yes | no | no | yes | no | no | no | no | no | no | no | no | no |
| TRPA1 | 957 | I | 0 | 0 | yes | no | no | yes | no | yes | yes | no | yes | no | no | no | no |
| TRPA1 | 958 | G | 0 | 0 | yes | no | no | yes | no | yes | no | no | no | no | no | no | no |
| TRPA1 | 959 | L | 0 | 0 | yes | no | no | yes | no | yes | no | no | yes | no | no | no | no |
| TRPA1 | 960 | A | 0 | 0 | yes | no | no | yes | no | yes | no | no | yes | no | no | no | no |
| TRPA1 | 961 | V | 0 | 0 | yes | no | no | no | no | yes | yes | no | no | no | no | no | no |
| TRPA1 | 962 | G | 0 | 0 | yes | no | no | no | no | yes | no | no | no | no | no | no | no |
| TRPA1 | 963 | D | 1 | 0 | yes | no | no | no | no | yes | no | no | no | no | no | no | no |
| TRPA1 | 964 | I | 0 | 0 | yes | no | no | no | no | yes | no | no | yes | no | no | no | no |
| TRPA1 | 965 | A | 0 | 0 | yes | no | no | no | no | yes | yes | no | no | no | no | no | no |
| TRPA1 | 966 | E | 1 | 0 | yes | no | no | no | no | yes | no | no | no | no | no | no | no |
| TRPA1 | 967 | V | 1 | 0 | yes | no | no | no | no | yes | no | no | no | no | no | no | no |
| TRPA1 | 968 | Q | 0 | 0 | yes | no | no | no | no | yes | no | no | no | no | no | no | no |
| TRPA1 | 969 | K | 0 | 0 | no | yes | no | no | no | yes | yes | no | no | no | no | no | no |
| TRPA1 | 970 | H | 0 | 0 | no | yes | no | no | no | yes | no | no | no | no | no | no | no |
| TRPA1 | 971 | A | 0 | 0 | yes | no | no | no | no | yes | no | no | no | no | no | no | no |
| TRPA1 | 972 | S | 0 | 0 | yes | no | no | no | no | yes | no | no | no | no | no | no | no |
| TRPA1 | 973 | L | 2 | 0 | yes | no | no | no | no | yes | no | no | no | no | no | no | no |
| TRPA1 | 974 | K | 0 | 0 | yes | no | no | no | no | yes | no | no | no | no | no | no | no |
| TRPA1 | 975 | R | 0 | 0 | yes | no | no | no | no | yes | no | no | no | no | no | no | no |
| TRPA1 | 976 | I | 0 | 0 | yes | no | no | no | no | yes | no | no | no | no | no | no | no |
| TRPA1 | 977 | A | 0 | 0 | yes | no | no | no | no | yes | no | no | no | no | no | no | no |
| TRPA1 | 978 | M | 1 | 0 | yes | no | no | no | no | yes | no | no | no | no | no | no | no |
| TRPA1 | 979 | Q | 0 | 0 | yes | no | no | no | no | yes | no | no | no | no | no | no | no |
| TRPA1 | 980 | V | 0 | 0 | yes | no | no | no | no | yes | no | no | no | no | no | no | no |
| TRPA1 | 981 | E | 0 | 0 | yes | no | no | no | no | yes | no | no | no | no | no | no | no |
| TRPA1 | 982 | L | 0 | 0 | yes | no | no | no | no | yes | no | no | no | no | no | no | no |
| TRPA1 | 983 | H | 0 | 0 | yes | no | no | no | no | yes | no | no | no | no | no | no | no |
| TRPA1 | 984 | T | 0 | 0 | yes | no | no | no | no | yes | no | no | no | no | no | no | no |
| TRPA1 | 985 | S | 1 | 0 | yes | no | no | no | no | yes | no | no | no | no | no | no | no |

| | | | | | | | | | | | | | | | | | |
|---|---|---|---|---|---|---|---|---|---|---|---|---|---|---|---|---|---|
| TRPA1 | 986 | L | 0 | 0 | yes | no | no | no | no | yes | no | no | no | no | no | no | no |
| TRPA1 | 987 | E | 0 | 0 | no | yes | no | no | no | yes | no | no | no | no | no | no | no |
| TRPA1 | 988 | K | 0 | 0 | no | yes | no | no | no | yes | no | no | no | no | no | no | no |
| TRPA1 | 989 | K | 0 | 0 | no | yes | no | no | no | yes | no | no | no | no | no | no | no |
| TRPA1 | 990 | L | 0 | 0 | no | yes | no | no | no | yes | no | no | no | no | no | no | no |
| TRPA1 | 991 | P | 0 | 0 | no | yes | no | no | no | yes | no | no | no | no | no | no | no |
| TRPA1 | 992 | L | 1 | 0 | yes | no | no | no | no | yes | no | no | no | no | no | no | no |
| TRPA1 | 993 | W | 0 | 0 | yes | no | no | no | no | yes | no | no | no | no | no | no | no |
| TRPA1 | 994 | F | 1 | 0 | yes | no | no | no | no | yes | no | no | no | no | no | no | no |
| TRPA1 | 995 | L | 0 | 0 | yes | no | no | no | no | yes | no | no | no | no | no | no | no |
| TRPA1 | 996 | R | 2 | 0 | yes | no | no | no | no | yes | no | no | no | no | no | no | no |
| TRPA1 | 997 | K | 0 | 0 | yes | no | no | no | no | yes | no | no | no | no | no | no | no |
| TRPA1 | 998 | V | 1 | 0 | yes | no | no | no | no | yes | no | no | no | no | no | no | no |
| TRPA1 | 999 | D | 1 | 0 | no | yes | no | no | no | yes | no | no | no | no | no | no | no |
| TRPA1 | 1000 | Q | 0 | 0 | no | yes | no | no | no | yes | no | no | no | no | no | no | no |
| TRPA1 | 1001 | K | 0 | 0 | no | yes | no | no | no | yes | no | no | no | no | no | no | no |
| TRPA1 | 1002 | S | 1 | 0 | no | no | yes | no | no | yes | no | no | no | no | no | no | no |
| TRPA1 | 1003 | T | 1 | 0 | no | no | yes | no | no | yes | no | no | no | no | no | no | no |
| TRPA1 | 1004 | I | 0 | 0 | no | no | yes | no | no | yes | no | no | no | no | no | no | no |
| TRPA1 | 1005 | V | 1 | 0 | no | no | yes | no | no | yes | no | no | no | no | no | no | no |
| TRPA1 | 1006 | Y | 1 | 0 | no | yes | no | no | no | yes | no | no | no | no | no | no | no |
| TRPA1 | 1007 | P | 1 | 0 | no | yes | no | no | no | yes | no | no | no | no | no | no | no |
| TRPA1 | 1008 | N | 0 | 0 | no | yes | no | no | no | yes | no | no | no | no | no | no | no |
| TRPA1 | 1009 | K | 0 | 0 | no | yes | no | no | no | yes | no | no | no | no | no | no | no |
| TRPA1 | 1010 | P | 0 | 0 | no | yes | no | no | no | yes | no | no | no | no | no | no | no |
| TRPA1 | 1011 | R | 0 | 0 | no | yes | no | no | no | yes | no | no | no | no | no | no | no |
| TRPA1 | 1012 | S | 0 | 0 | no | yes | no | no | no | yes | no | no | no | no | no | no | no |
| TRPA1 | 1015 | M | 2 | 0 | no | yes | no | yes | no | yes | no | no | no | no | no | no | no |
| TRPA1 | 1016 | L | 1 | 0 | yes | no | no | yes | no | yes | no | no | no | no | no | no | no |
| TRPA1 | 1017 | F | 0 | 0 | yes | no | no | no | no | yes | no | no | no | no | no | no | no |
| TRPA1 | 1018 | H | 2 | 0 | yes | no | no | no | no | yes | no | no | no | no | no | no | no |
| TRPA1 | 1019 | I | 2 | 0 | yes | no | no | yes | no | yes | no | no | no | no | no | no | no |
| TRPA1 | 1020 | F | 0 | 0 | yes | no | no | yes | no | yes | no | no | no | no | no | no | no |

| | | | | | | | | | | | | | | | | | |
|---|---|---|---|---|---|---|---|---|---|---|---|---|---|---|---|---|---|
| TRPA1 | 1021 | C | 1 | 0 | yes | no | no | no | no | yes | no | no | no | no | no | no | no |
| TRPA1 | 1022 | F | 0 | 0 | yes | no | no | yes | no | yes | no | no | no | no | no | no | no |
| TRPA1 | 1023 | L | 0 | 0 | yes | no | no | yes | no | yes | no | no | no | no | no | no | no |
| TRPA1 | 1024 | F | 0 | 0 | no | yes | no | no | no | yes | no | no | no | no | no | no | no |
| TRPA1 | 1025 | C | 0 | 0 | no | yes | no | no | no | yes | no | no | no | no | no | no | no |
| TRPA1 | 1039 | S | 0 | 0 | no | yes | no | no | no | yes | no | no | no | no | no | no | no |
| TRPA1 | 1040 | L | 0 | 0 | yes | no | no | no | no | yes | yes | no | no | no | no | no | no |
| TRPA1 | 1041 | E | 0 | 0 | yes | no | no | no | no | yes | no | no | no | no | no | no | no |
| TRPA1 | 1042 | M | 2 | 0 | yes | no | no | no | no | yes | no | no | no | no | no | no | no |
| TRPA1 | 1043 | E | 0 | 0 | yes | no | no | no | no | yes | no | no | yes | no | no | no | no |
| TRPA1 | 1044 | I | 0 | 0 | yes | no | no | no | no | yes | yes | no | no | no | no | no | no |
| TRPA1 | 1045 | L | 0 | 0 | yes | no | no | no | no | yes | no | no | no | no | no | no | no |
| TRPA1 | 1046 | K | 1 | 0 | yes | no | no | no | no | yes | no | no | no | no | no | no | no |
| TRPA1 | 1047 | Q | 0 | 0 | yes | no | no | no | no | yes | yes | no | yes | no | no | no | no |
| TRPA1 | 1048 | K | 0 | 0 | yes | no | no | no | no | yes | no | no | yes | no | no | no | no |
| TRPA1 | 1049 | Y | 0 | 0 | yes | no | no | no | no | yes | no | no | no | no | no | no | no |
| TRPA1 | 1050 | R | 0 | 0 | yes | no | no | no | no | yes | no | no | no | no | no | no | no |
| TRPA1 | 1051 | L | 2 | 0 | yes | no | no | no | no | yes | yes | no | yes | no | no | no | no |
| TRPA1 | 1052 | K | 0 | 0 | yes | no | no | no | no | yes | no | no | no | no | no | no | no |
| TRPA1 | 1053 | D | 0 | 0 | yes | no | no | no | no | yes | no | no | no | no | no | no | no |
| TRPA1 | 1054 | L | 0 | 0 | yes | no | no | no | no | yes | yes | no | yes | no | no | no | no |
| TRPA1 | 1055 | T | 0 | 0 | yes | no | no | no | no | yes | no | no | yes | no | no | no | no |
| TRPA1 | 1056 | F | 1 | 0 | yes | no | no | no | no | yes | no | no | no | no | no | no | no |
| TRPA1 | 1057 | L | 0 | 0 | yes | no | no | no | no | yes | no | no | yes | no | no | no | no |
| TRPA1 | 1058 | L | 1 | 0 | yes | no | no | no | no | yes | yes | no | yes | no | no | no | no |
| TRPA1 | 1059 | E | 0 | 0 | yes | no | no | no | no | yes | no | no | yes | no | no | no | no |
| TRPA1 | 1060 | K | 1 | 0 | yes | no | no | no | no | yes | no | no | no | no | no | no | no |
| TRPA1 | 1061 | Q | 1 | 0 | yes | no | no | no | no | yes | yes | no | yes | no | no | no | no |
| TRPA1 | 1062 | H | 2 | 0 | yes | no | no | no | no | yes | no | no | yes | no | no | no | no |
| TRPA1 | 1063 | E | 0 | 0 | yes | no | no | no | no | yes | no | no | no | no | no | no | no |
| TRPA1 | 1064 | L | 0 | 0 | yes | no | no | no | no | yes | no | no | yes | no | no | no | no |
| TRPA1 | 1065 | I | 0 | 0 | yes | no | no | no | no | yes | yes | no | yes | no | no | no | no |
| TRPA1 | 1066 | K | 0 | 0 | yes | no | no | no | no | yes | no | no | yes | no | no | no | no |

| | | | | | | | | | | | | | | | | | |
|---|---|---|---|---|---|---|---|---|---|---|---|---|---|---|---|---|---|
| TRPA1 | 1067 | L | 0 | 0 | yes | no | no | no | no | yes | no | no | no | no | no | no | no |
| TRPA1 | 1068 | I | 1 | 0 | yes | no | no | no | no | yes | yes | no | yes | no | no | no | no |
| TRPA1 | 1069 | I | 0 | 0 | yes | no | no | no | no | yes | no | no | yes | no | no | no | no |
| TRPA1 | 1070 | Q | 1 | 0 | yes | no | no | no | no | yes | no | no | yes | no | no | no | no |
| TRPA1 | 1071 | K | 0 | 0 | yes | no | no | no | no | yes | no | no | yes | no | no | no | no |
| TRPA1 | 1072 | M | 2 | 0 | no | yes | no | no | no | yes | yes | no | yes | no | no | no | no |
| TRPA1 | 1073 | E | 0 | 0 | no | yes | no | no | no | yes | yes | no | yes | no | no | no | no |
| TRPA1 | 1074 | I | 0 | 0 | no | yes | no | no | no | yes | no | no | yes | no | no | no | no |
| TRPA1 | 1075 | I | 1 | 0 | no | yes | no | no | no | yes | no | no | yes | no | no | no | no |
| TRPA1 | 1076 | S | 0 | 0 | no | yes | no | no | no | yes | no | no | yes | no | no | no | no |
| TRPA1 | 1077 | E | 0 | 0 | no | yes | no | no | no | yes | no | no | yes | no | no | no | no |
| TRPA1 | 1078 | T | 1 | 0 | no | yes | no | no | no | yes | no | no | yes | no | no | no | no |
| TRPA1 | 1079 | E | 0 | 0 | no | yes | no | no | no | yes | no | no | no | no | no | no | no |
| TRPC6 | 85 | A | 0 | 0 | no | yes | no | no | no | yes | no | no | no | no | no | no | no |
| TRPC6 | 86 | Y | 0 | 0 | no | yes | no | no | no | yes | no | no | yes | no | no | no | no |
| TRPC6 | 87 | M | 1 | 0 | no | yes | no | no | no | yes | no | no | yes | no | no | no | no |
| TRPC6 | 88 | F | 0 | 0 | no | yes | no | no | no | yes | no | no | yes | no | no | no | no |
| TRPC6 | 89 | S | 0 | 1 | no | yes | no | no | no | yes | no | no | yes | no | no | no | no |
| TRPC6 | 90 | D | 1 | 0 | no | yes | no | no | no | yes | no | no | no | no | no | no | no |
| TRPC6 | 91 | R | 2 | 0 | no | yes | no | no | no | yes | no | no | no | no | no | no | no |
| TRPC6 | 92 | S | 1 | 0 | no | yes | no | no | no | yes | no | no | no | no | no | no | no |
| TRPC6 | 93 | T | 1 | 0 | no | yes | no | no | no | yes | no | no | no | no | no | no | no |
| TRPC6 | 94 | S | 0 | 0 | no | yes | no | no | no | yes | no | no | no | no | no | no | no |
| TRPC6 | 95 | L | 1 | 0 | no | yes | no | no | no | yes | no | no | no | no | no | no | no |
| TRPC6 | 96 | S | 0 | 0 | no | yes | no | no | no | yes | no | no | no | no | no | no | no |
| TRPC6 | 97 | I | 2 | 0 | no | yes | no | no | no | yes | no | no | no | no | no | no | no |
| TRPC6 | 98 | E | 1 | 0 | no | yes | no | no | no | yes | no | no | no | no | no | no | no |
| TRPC6 | 99 | E | 0 | 0 | no | yes | no | no | no | yes | no | no | no | no | no | no | no |
| TRPC6 | 100 | E | 0 | 0 | yes | no | no | no | no | yes | no | no | no | no | no | no | no |
| TRPC6 | 101 | R | 2 | 0 | yes | no | no | no | no | yes | no | no | no | no | no | no | no |
| TRPC6 | 102 | F | 2 | 0 | yes | no | no | no | no | yes | no | no | no | no | no | no | no |
| TRPC6 | 103 | L | 0 | 0 | yes | no | no | no | no | yes | no | no | no | no | no | no | no |
| TRPC6 | 104 | D | 0 | 0 | yes | no | no | no | no | yes | no | no | no | no | no | no | no |

| | | | | | | | | | | | | | | | | | |
|---|---|---|---|---|---|---|---|---|---|---|---|---|---|---|---|---|---|
| TRPC6 | 105 | A | 1 | 0 | yes | no | no | no | no | yes | no | no | no | no | no | no | no |
| TRPC6 | 106 | A | 0 | 0 | yes | no | no | no | no | yes | no | no | no | no | no | no | no |
| TRPC6 | 107 | E | 0 | 0 | yes | no | no | no | no | yes | no | no | no | no | no | no | no |
| TRPC6 | 108 | Y | 0 | 0 | yes | no | no | no | no | yes | no | no | yes | no | no | no | no |
| TRPC6 | 109 | G | 0 | 0 | no | yes | no | no | no | yes | no | no | no | no | no | no | no |
| TRPC6 | 110 | N | 0 | 1 | no | yes | no | no | no | yes | no | no | no | no | no | no | no |
| TRPC6 | 111 | I | 1 | 0 | yes | no | no | no | no | yes | no | no | no | no | no | no | no |
| TRPC6 | 112 | P | 0 | 1 | yes | no | no | no | no | yes | no | no | no | no | no | no | no |
| TRPC6 | 113 | V | 0 | 0 | yes | no | no | no | no | yes | no | no | no | no | no | no | no |
| TRPC6 | 114 | V | 0 | 0 | yes | no | no | no | no | yes | no | no | no | no | no | no | no |
| TRPC6 | 115 | R | 2 | 0 | yes | no | no | no | no | yes | no | no | no | no | no | no | no |
| TRPC6 | 116 | K | 1 | 0 | yes | no | no | no | no | yes | no | no | no | no | no | no | no |
| TRPC6 | 117 | M | 1 | 0 | yes | no | no | no | no | yes | no | no | no | no | no | no | no |
| TRPC6 | 118 | L | 0 | 0 | no | yes | no | no | no | yes | no | no | no | no | no | no | no |
| TRPC6 | 119 | E | 0 | 0 | no | yes | no | no | no | yes | no | no | no | no | no | no | no |
| TRPC6 | 120 | E | 0 | 0 | no | yes | no | no | no | yes | no | no | no | no | no | no | no |
| TRPC6 | 121 | C | 0 | 0 | no | yes | no | no | no | yes | no | no | no | no | no | no | no |
| TRPC6 | 122 | H | 1 | 0 | no | yes | no | no | no | yes | no | no | no | no | no | no | no |
| TRPC6 | 123 | S | 0 | 0 | no | yes | no | no | no | yes | no | no | no | no | no | no | no |
| TRPC6 | 124 | L | 0 | 0 | no | yes | no | no | no | yes | no | no | no | no | no | no | no |
| TRPC6 | 125 | N | 3 | 1 | no | yes | no | no | no | yes | no | no | no | no | no | no | no |
| TRPC6 | 126 | V | 2 | 0 | no | yes | no | no | no | yes | no | no | no | no | no | no | no |
| TRPC6 | 127 | N | 0 | 0 | no | yes | no | no | no | yes | no | no | no | no | no | no | no |
| TRPC6 | 128 | C | 1 | 0 | no | yes | no | no | no | yes | no | no | no | no | no | no | no |
| TRPC6 | 129 | V | 0 | 0 | no | yes | no | no | no | yes | no | no | no | no | no | no | no |
| TRPC6 | 130 | D | 0 | 0 | no | yes | no | no | no | yes | no | no | no | no | no | no | no |
| TRPC6 | 131 | Y | 0 | 0 | no | yes | no | no | no | yes | no | no | yes | no | no | no | no |
| TRPC6 | 132 | M | 1 | 1 | no | yes | no | no | no | yes | no | no | yes | no | no | no | no |
| TRPC6 | 133 | G | 0 | 0 | no | yes | no | no | no | yes | no | no | no | no | no | no | no |
| TRPC6 | 134 | Q | 0 | 0 | no | yes | no | no | no | yes | no | no | no | no | no | no | no |
| TRPC6 | 135 | N | 1 | 0 | no | yes | no | no | no | yes | no | no | no | no | no | no | no |
| TRPC6 | 136 | A | 1 | 0 | yes | no | no | no | no | yes | no | no | no | no | no | no | no |
| TRPC6 | 137 | L | 0 | 0 | yes | no | no | no | no | yes | no | no | no | no | no | no | no |

| | | | | | | | | | | | | | | | | | |
|---|---|---|---|---|---|---|---|---|---|---|---|---|---|---|---|---|---|
| TRPC6 | 138 | Q | 0 | 0 | yes | no | no | no | no | yes | no | no | no | no | no | no | no |
| TRPC6 | 139 | L | 0 | 0 | yes | no | no | no | no | yes | no | no | no | no | no | no | no |
| TRPC6 | 140 | A | 0 | 0 | yes | no | no | no | no | yes | no | no | no | no | no | no | no |
| TRPC6 | 141 | V | 1 | 0 | no | yes | no | no | no | yes | no | no | no | no | no | no | no |
| TRPC6 | 142 | A | 0 | 0 | no | yes | no | no | no | yes | no | no | no | no | no | no | no |
| TRPC6 | 143 | N | 1 | 0 | no | yes | no | no | no | yes | no | no | no | no | no | no | no |
| TRPC6 | 144 | E | 0 | 0 | no | yes | no | no | no | yes | no | no | yes | no | no | no | no |
| TRPC6 | 145 | H | 0 | 1 | no | yes | no | no | no | yes | no | no | no | no | no | no | no |
| TRPC6 | 146 | L | 1 | 0 | yes | no | no | no | no | yes | no | no | no | no | no | no | no |
| TRPC6 | 147 | E | 0 | 0 | yes | no | no | no | no | yes | no | no | yes | no | no | no | no |
| TRPC6 | 148 | I | 0 | 0 | yes | no | no | no | no | yes | no | no | no | no | no | no | no |
| TRPC6 | 149 | T | 0 | 0 | yes | no | no | no | no | yes | no | no | no | no | no | no | no |
| TRPC6 | 150 | E | 0 | 0 | yes | no | no | no | no | yes | no | no | no | no | no | no | no |
| TRPC6 | 151 | L | 1 | 0 | yes | no | no | no | no | yes | no | no | no | no | no | no | no |
| TRPC6 | 152 | L | 0 | 0 | yes | no | no | no | no | yes | no | no | no | no | no | no | no |
| TRPC6 | 153 | L | 1 | 0 | no | yes | no | no | no | yes | no | no | no | no | no | no | no |
| TRPC6 | 154 | K | 0 | 0 | no | yes | no | no | no | yes | no | no | no | no | no | no | no |
| TRPC6 | 155 | K | 0 | 0 | no | yes | no | no | no | yes | no | no | no | no | no | no | no |
| TRPC6 | 156 | E | 0 | 0 | no | yes | no | no | no | yes | no | no | no | no | no | no | no |
| TRPC6 | 157 | N | 0 | 0 | no | yes | no | no | no | yes | no | no | no | no | no | no | no |
| TRPC6 | 158 | L | 1 | 0 | no | yes | no | no | no | yes | no | no | no | no | no | no | no |
| TRPC6 | 159 | S | 0 | 0 | no | yes | no | no | no | yes | no | no | no | no | no | no | no |
| TRPC6 | 160 | R | 1 | 0 | no | yes | no | no | no | yes | no | no | no | no | no | no | no |
| TRPC6 | 161 | V | 0 | 0 | no | yes | no | no | no | yes | no | no | no | no | no | no | no |
| TRPC6 | 162 | G | 0 | 0 | yes | no | no | no | no | yes | no | no | no | no | no | no | no |
| TRPC6 | 163 | D | 1 | 0 | yes | no | no | no | no | yes | no | no | no | no | no | no | no |
| TRPC6 | 164 | A | 1 | 0 | yes | no | no | no | no | yes | no | no | no | no | no | no | no |
| TRPC6 | 165 | L | 0 | 0 | yes | no | no | no | no | yes | no | no | no | no | no | no | no |
| TRPC6 | 166 | L | 1 | 0 | yes | no | no | no | no | yes | no | no | no | no | no | no | no |
| TRPC6 | 167 | L | 0 | 0 | yes | no | no | no | no | yes | no | no | no | no | no | no | no |
| TRPC6 | 168 | A | 0 | 0 | yes | no | no | no | no | yes | no | no | no | no | no | no | no |
| TRPC6 | 169 | I | 0 | 0 | yes | no | no | no | no | yes | no | no | no | no | no | no | no |
| TRPC6 | 170 | S | 1 | 0 | yes | no | no | no | no | yes | no | no | no | no | no | no | no |

| | | | | | | | | | | | | | | | | | |
|---|---|---|---|---|---|---|---|---|---|---|---|---|---|---|---|---|---|
| TRPC6 | 171 | K | 0 | 0 | yes | no | no | no | no | yes | no | no | no | no | no | no | no |
| TRPC6 | 172 | G | 0 | 0 | no | yes | no | no | no | yes | no | no | no | no | no | no | no |
| TRPC6 | 173 | Y | 0 | 1 | no | yes | no | no | no | yes | no | no | no | no | no | no | no |
| TRPC6 | 174 | V | 1 | 0 | yes | no | no | no | no | yes | no | no | no | no | no | no | no |
| TRPC6 | 175 | R | 0 | 2 | yes | no | no | no | no | yes | no | no | yes | no | no | no | no |
| TRPC6 | 176 | I | 0 | 0 | yes | no | no | no | no | yes | no | no | no | no | no | no | no |
| TRPC6 | 177 | V | 1 | 0 | yes | no | no | no | no | yes | no | no | no | no | no | no | no |
| TRPC6 | 178 | E | 0 | 0 | yes | no | no | no | no | yes | no | no | no | no | no | no | no |
| TRPC6 | 179 | A | 0 | 0 | yes | no | no | no | no | yes | no | no | no | no | no | no | no |
| TRPC6 | 180 | I | 1 | 0 | yes | no | no | no | no | yes | no | no | no | no | no | no | no |
| TRPC6 | 181 | L | 1 | 0 | no | yes | no | no | no | yes | no | no | no | no | no | no | no |
| TRPC6 | 182 | S | 2 | 0 | no | yes | no | no | no | yes | no | no | no | no | no | no | no |
| TRPC6 | 183 | H | 1 | 0 | no | yes | no | no | no | yes | no | no | no | no | no | no | no |
| TRPC6 | 184 | P | 2 | 0 | no | yes | no | no | no | yes | no | no | no | no | no | no | no |
| TRPC6 | 185 | A | 0 | 0 | no | yes | no | no | no | yes | no | no | no | no | no | no | no |
| TRPC6 | 186 | F | 0 | 0 | no | yes | no | no | no | yes | no | no | no | no | no | no | no |
| TRPC6 | 187 | A | 0 | 0 | no | yes | no | no | no | yes | no | no | no | no | no | no | no |
| TRPC6 | 188 | E | 0 | 0 | no | yes | no | no | no | yes | no | no | no | no | no | no | no |
| TRPC6 | 189 | G | 1 | 0 | no | yes | no | no | no | yes | no | no | no | no | no | no | no |
| TRPC6 | 190 | K | 2 | 0 | no | yes | no | no | no | yes | no | no | no | no | no | no | no |
| TRPC6 | 191 | R | 1 | 0 | no | yes | no | no | no | yes | no | no | no | no | no | no | no |
| TRPC6 | 192 | L | 0 | 0 | no | yes | no | no | no | yes | no | no | no | no | no | no | no |
| TRPC6 | 193 | A | 1 | 0 | no | yes | no | no | no | yes | no | no | no | no | no | no | no |
| TRPC6 | 194 | T | 0 | 0 | no | yes | no | no | no | yes | no | no | no | no | no | no | no |
| TRPC6 | 203 | Q | 1 | 0 | no | yes | no | no | no | yes | no | no | no | no | no | no | no |
| TRPC6 | 204 | D | 0 | 0 | no | yes | no | no | no | yes | no | no | no | no | no | no | no |
| TRPC6 | 205 | D | 2 | 0 | no | yes | no | no | no | yes | no | no | no | no | no | no | no |
| TRPC6 | 206 | F | 0 | 0 | no | yes | no | no | no | yes | no | no | no | no | no | no | no |
| TRPC6 | 207 | Y | 0 | 0 | no | yes | no | no | no | yes | no | no | no | no | no | no | no |
| TRPC6 | 208 | A | 0 | 0 | no | no | yes | no | no | yes | no | no | no | no | no | no | no |
| TRPC6 | 209 | Y | 2 | 0 | no | yes | no | no | no | yes | no | no | yes | no | no | no | no |
| TRPC6 | 210 | D | 1 | 0 | no | yes | no | no | no | yes | no | no | yes | no | no | no | no |
| TRPC6 | 211 | E | 0 | 0 | no | yes | no | no | no | yes | no | no | no | no | no | no | no |

| | | | | | | | | | | | | | | | | | |
|---|---|---|---|---|---|---|---|---|---|---|---|---|---|---|---|---|---|
| TRPC6 | 212 | D | 1 | 0 | no | yes | no | no | no | yes | no | no | yes | no | no | no | no |
| TRPC6 | 213 | G | 0 | 0 | no | yes | no | no | no | yes | no | no | no | no | no | no | no |
| TRPC6 | 214 | T | 0 | 0 | no | no | yes | no | no | yes | no | no | yes | no | no | no | no |
| TRPC6 | 215 | R | 1 | 0 | no | yes | no | no | no | yes | no | no | yes | no | no | no | no |
| TRPC6 | 216 | F | 0 | 0 | no | yes | no | no | no | yes | no | no | yes | no | no | no | no |
| TRPC6 | 217 | S | 0 | 0 | no | yes | no | no | no | yes | no | no | yes | no | no | no | no |
| TRPC6 | 218 | H | 1 | 1 | no | yes | no | no | no | yes | no | no | no | no | no | no | no |
| TRPC6 | 219 | D | 0 | 0 | no | yes | no | no | no | yes | no | no | no | no | no | no | no |
| TRPC6 | 220 | V | 0 | 0 | no | yes | no | no | no | yes | no | no | no | no | no | no | no |
| TRPC6 | 221 | T | 0 | 0 | no | yes | no | no | no | yes | no | no | no | no | no | no | no |
| TRPC6 | 222 | P | 0 | 0 | yes | no | no | no | no | yes | no | no | no | no | no | no | no |
| TRPC6 | 223 | I | 1 | 0 | yes | no | no | no | no | yes | no | no | no | no | no | no | no |
| TRPC6 | 224 | I | 0 | 0 | yes | no | no | no | no | yes | no | no | no | no | no | no | no |
| TRPC6 | 225 | L | 1 | 0 | yes | no | no | no | no | yes | no | no | no | no | no | no | no |
| TRPC6 | 226 | A | 0 | 0 | yes | no | no | no | no | yes | no | no | no | no | no | no | no |
| TRPC6 | 227 | A | 1 | 0 | yes | no | no | no | no | yes | no | no | no | no | no | no | no |
| TRPC6 | 228 | H | 0 | 0 | yes | no | no | no | no | yes | no | no | no | no | no | no | no |
| TRPC6 | 229 | C | 0 | 0 | no | yes | no | no | no | yes | no | no | no | no | no | no | no |
| TRPC6 | 230 | Q | 0 | 0 | no | yes | no | no | no | yes | no | no | no | no | no | no | no |
| TRPC6 | 231 | E | 1 | 0 | no | yes | no | no | no | yes | no | no | no | no | no | no | no |
| TRPC6 | 232 | Y | 0 | 0 | yes | no | no | no | no | yes | no | no | no | no | no | no | no |
| TRPC6 | 233 | E | 0 | 0 | yes | no | no | no | no | yes | no | no | yes | no | no | no | no |
| TRPC6 | 234 | I | 0 | 0 | yes | no | no | no | no | yes | no | no | no | no | no | no | no |
| TRPC6 | 235 | V | 1 | 0 | yes | no | no | no | no | yes | no | no | no | no | no | no | no |
| TRPC6 | 236 | H | 0 | 0 | yes | no | no | no | no | yes | no | no | yes | no | no | no | no |
| TRPC6 | 237 | T | 0 | 0 | yes | no | no | no | no | yes | no | no | no | no | no | no | no |
| TRPC6 | 238 | L | 0 | 0 | yes | no | no | no | no | yes | no | no | no | no | no | no | no |
| TRPC6 | 239 | L | 0 | 0 | yes | no | no | no | no | yes | no | no | yes | no | no | no | no |
| TRPC6 | 240 | R | 2 | 0 | yes | no | no | no | no | yes | no | no | yes | no | no | no | no |
| TRPC6 | 241 | K | 1 | 0 | no | yes | no | no | no | yes | no | no | no | no | no | no | no |
| TRPC6 | 242 | G | 0 | 0 | no | yes | no | no | no | yes | no | no | no | no | no | no | no |
| TRPC6 | 243 | A | 2 | 0 | no | yes | no | no | no | yes | no | no | no | no | no | no | no |
| TRPC6 | 244 | R | 1 | 0 | no | yes | no | no | no | yes | no | no | yes | no | no | no | no |

| | | | | | | | | | | | | | | | | | |
|---|---|---|---|---|---|---|---|---|---|---|---|---|---|---|---|---|---|
| TRPC6 | 245 | I | 0 | 0 | no | yes | no | no | no | yes | no | no | no | no | no | no | no |
| TRPC6 | 246 | E | 0 | 0 | no | yes | no | no | no | yes | no | no | no | no | no | no | no |
| TRPC6 | 247 | R | 2 | 0 | no | yes | no | no | no | yes | no | no | no | no | no | no | no |
| TRPC6 | 248 | P | 0 | 0 | no | yes | no | no | no | yes | no | no | no | no | no | no | no |
| TRPC6 | 249 | H | 0 | 0 | no | yes | no | no | no | yes | no | no | no | no | no | no | no |
| TRPC6 | 250 | D | 2 | 0 | no | yes | no | no | no | yes | no | no | no | no | no | no | no |
| TRPC6 | 251 | Y | 1 | 0 | no | yes | no | no | no | yes | no | no | no | no | no | no | no |
| TRPC6 | 252 | F | 0 | 0 | no | yes | no | no | no | yes | no | no | no | no | no | no | no |
| TRPC6 | 253 | C | 1 | 0 | no | yes | no | no | no | yes | no | no | no | no | no | no | no |
| TRPC6 | 254 | K | 0 | 0 | no | yes | no | no | no | yes | no | no | no | no | no | no | no |
| TRPC6 | 255 | C | 0 | 0 | no | yes | no | no | no | yes | no | no | no | no | no | no | no |
| TRPC6 | 256 | N | 1 | 0 | no | yes | no | no | no | yes | no | no | no | no | no | no | no |
| TRPC6 | 257 | D | 0 | 0 | no | yes | no | no | no | yes | no | no | no | no | no | no | no |
| TRPC6 | 258 | C | 0 | 0 | yes | no | no | no | no | yes | no | no | no | no | no | no | no |
| TRPC6 | 259 | N | 0 | 0 | yes | no | no | no | no | yes | no | no | no | no | no | no | no |
| TRPC6 | 260 | Q | 0 | 0 | yes | no | no | no | no | yes | no | no | no | no | no | no | no |
| TRPC6 | 261 | K | 0 | 0 | yes | no | no | no | no | yes | no | no | no | no | no | no | no |
| TRPC6 | 262 | Q | 0 | 0 | yes | no | no | no | no | yes | no | no | no | no | no | no | no |
| TRPC6 | 263 | K | 0 | 0 | yes | no | no | no | no | yes | no | no | no | no | no | no | no |
| TRPC6 | 264 | H | 1 | 0 | yes | no | no | no | no | yes | no | no | no | no | no | no | no |
| TRPC6 | 265 | D | 0 | 0 | no | yes | no | no | no | yes | no | no | yes | no | no | no | no |
| TRPC6 | 266 | S | 0 | 0 | no | yes | no | no | no | yes | no | no | no | no | no | no | no |
| TRPC6 | 267 | F | 0 | 0 | no | yes | no | no | no | yes | no | no | yes | no | no | no | no |
| TRPC6 | 268 | S | 0 | 0 | yes | no | no | no | no | yes | no | no | yes | no | no | no | no |
| TRPC6 | 269 | H | 0 | 0 | yes | no | no | no | no | yes | no | no | no | no | no | no | no |
| TRPC6 | 270 | S | 1 | 1 | yes | no | no | no | no | yes | no | no | no | no | no | no | no |
| TRPC6 | 271 | R | 0 | 0 | yes | no | no | no | no | yes | no | no | yes | no | no | no | no |
| TRPC6 | 272 | S | 0 | 0 | yes | no | no | no | no | yes | no | no | no | no | no | no | no |
| TRPC6 | 273 | R | 2 | 0 | yes | no | no | no | no | yes | no | no | no | no | no | no | no |
| TRPC6 | 274 | I | 0 | 0 | yes | no | no | no | no | yes | no | no | no | no | no | no | no |
| TRPC6 | 275 | N | 0 | 0 | yes | no | no | no | no | yes | no | no | no | no | no | no | no |
| TRPC6 | 276 | A | 0 | 0 | yes | no | no | no | no | yes | no | no | no | no | no | no | no |
| TRPC6 | 277 | Y | 1 | 0 | yes | no | no | no | no | yes | no | no | no | no | no | no | no |

| | | | | | | | | | | | | | | | | | |
|---|---|---|---|---|---|---|---|---|---|---|---|---|---|---|---|---|---|
| TRPC6 | 278 | K | 0 | 0 | yes | no | no | no | no | yes | no | no | no | no | no | no | no |
| TRPC6 | 279 | G | 1 | 0 | yes | no | no | no | no | yes | no | no | no | no | no | no | no |
| TRPC6 | 280 | L | 1 | 0 | yes | no | no | no | no | yes | no | no | no | no | no | no | no |
| TRPC6 | 281 | A | 0 | 0 | no | yes | no | no | no | yes | no | no | no | no | no | no | no |
| TRPC6 | 282 | S | 0 | 0 | no | yes | no | no | no | yes | no | no | no | no | no | no | no |
| TRPC6 | 283 | P | 1 | 0 | yes | no | no | no | no | yes | no | no | no | no | no | no | no |
| TRPC6 | 284 | A | 1 | 0 | yes | no | no | no | no | yes | no | no | no | no | no | no | no |
| TRPC6 | 285 | Y | 0 | 0 | yes | no | no | no | no | yes | no | no | no | no | no | no | no |
| TRPC6 | 286 | L | 0 | 0 | yes | no | no | no | no | yes | no | no | no | no | no | no | no |
| TRPC6 | 287 | S | 0 | 0 | yes | no | no | no | no | yes | no | no | no | no | no | no | no |
| TRPC6 | 288 | L | 0 | 0 | yes | no | no | no | no | yes | no | no | yes | no | no | no | no |
| TRPC6 | 289 | S | 1 | 0 | yes | no | no | no | no | yes | no | no | yes | no | no | no | no |
| TRPC6 | 290 | S | 1 | 0 | no | yes | no | no | no | yes | no | no | yes | no | no | no | no |
| TRPC6 | 291 | E | 1 | 0 | no | yes | no | no | no | yes | no | no | yes | no | no | no | no |
| TRPC6 | 292 | D | 0 | 0 | no | yes | no | no | no | yes | no | no | no | no | no | no | no |
| TRPC6 | 293 | P | 0 | 0 | yes | no | no | no | no | yes | no | no | no | no | no | no | no |
| TRPC6 | 294 | V | 0 | 0 | yes | no | no | no | no | yes | no | no | no | no | no | no | no |
| TRPC6 | 295 | M | 0 | 0 | yes | no | no | no | no | yes | no | no | no | no | no | no | no |
| TRPC6 | 296 | T | 2 | 0 | yes | no | no | no | no | yes | no | no | no | no | no | no | no |
| TRPC6 | 297 | A | 0 | 0 | yes | no | no | no | no | yes | no | no | no | no | no | no | no |
| TRPC6 | 298 | L | 0 | 0 | yes | no | no | no | no | yes | no | no | no | no | no | no | no |
| TRPC6 | 299 | E | 0 | 0 | yes | no | no | no | no | yes | no | no | no | no | no | no | no |
| TRPC6 | 300 | L | 0 | 0 | yes | no | no | no | no | yes | no | no | no | no | no | no | no |
| TRPC6 | 301 | S | 0 | 0 | yes | no | no | no | no | yes | no | no | no | no | no | no | no |
| TRPC6 | 302 | N | 0 | 0 | yes | no | no | no | no | yes | no | no | no | no | no | no | no |
| TRPC6 | 303 | E | 0 | 0 | yes | no | no | no | no | yes | no | no | no | no | no | no | no |
| TRPC6 | 304 | L | 0 | 0 | yes | no | no | no | no | yes | no | no | no | no | no | no | no |
| TRPC6 | 305 | A | 0 | 0 | yes | no | no | no | no | yes | no | no | no | no | no | no | no |
| TRPC6 | 306 | V | 0 | 0 | yes | no | no | no | no | yes | no | no | no | no | no | no | no |
| TRPC6 | 307 | L | 0 | 0 | yes | no | no | no | no | yes | no | no | no | no | no | no | no |
| TRPC6 | 308 | A | 0 | 0 | yes | no | no | no | no | yes | no | no | no | no | no | no | no |
| TRPC6 | 309 | N | 0 | 0 | yes | no | no | no | no | yes | no | no | no | no | no | no | no |
| TRPC6 | 310 | I | 0 | 0 | yes | no | no | no | no | yes | no | no | no | no | no | no | no |

| | | | | | | | | | | | | | | | | | |
|---|---|---|---|---|---|---|---|---|---|---|---|---|---|---|---|---|---|
| TRPC6 | 311 | E | 0 | 0 | no | yes | no | no | no | yes | no | no | no | no | no | no | no |
| TRPC6 | 312 | K | 0 | 0 | no | yes | no | no | no | yes | no | no | no | no | no | no | no |
| TRPC6 | 313 | E | 0 | 0 | no | yes | no | no | no | yes | no | no | no | no | no | no | no |
| TRPC6 | 314 | F | 0 | 0 | no | yes | no | no | no | yes | no | no | no | no | no | no | no |
| TRPC6 | 315 | K | 0 | 0 | yes | no | no | no | no | yes | no | no | no | no | no | no | no |
| TRPC6 | 316 | N | 1 | 0 | yes | no | no | no | no | yes | no | no | no | no | no | no | no |
| TRPC6 | 317 | D | 1 | 0 | yes | no | no | no | no | yes | no | no | no | no | no | no | no |
| TRPC6 | 318 | Y | 1 | 0 | yes | no | no | no | no | yes | no | no | no | no | no | no | no |
| TRPC6 | 319 | K | 1 | 0 | yes | no | no | no | no | yes | no | no | no | no | no | no | no |
| TRPC6 | 320 | K | 0 | 0 | yes | no | no | no | no | yes | no | no | no | no | no | no | no |
| TRPC6 | 321 | L | 0 | 0 | yes | no | no | no | no | yes | no | no | no | no | no | no | no |
| TRPC6 | 322 | S | 1 | 0 | yes | no | no | no | no | yes | no | no | no | no | no | no | no |
| TRPC6 | 323 | M | 1 | 0 | yes | no | no | no | no | yes | no | no | no | no | no | no | no |
| TRPC6 | 324 | Q | 1 | 0 | yes | no | no | no | no | yes | no | no | no | no | no | no | no |
| TRPC6 | 325 | C | 1 | 0 | yes | no | no | no | no | yes | no | no | no | no | no | no | no |
| TRPC6 | 326 | K | 0 | 0 | yes | no | no | no | no | yes | no | no | no | no | no | no | no |
| TRPC6 | 327 | D | 1 | 0 | yes | no | no | no | no | yes | no | no | no | no | no | no | no |
| TRPC6 | 328 | F | 0 | 0 | yes | no | no | no | no | yes | no | no | no | no | no | no | no |
| TRPC6 | 329 | V | 0 | 0 | yes | no | no | no | no | yes | no | no | no | no | no | no | no |
| TRPC6 | 330 | V | 0 | 0 | yes | no | no | no | no | yes | no | no | no | no | no | no | no |
| TRPC6 | 331 | G | 1 | 0 | yes | no | no | no | no | yes | no | no | no | no | no | no | no |
| TRPC6 | 332 | L | 1 | 0 | yes | no | no | no | no | yes | no | no | no | no | no | no | no |
| TRPC6 | 333 | L | 0 | 0 | yes | no | no | no | no | yes | no | no | no | no | no | no | no |
| TRPC6 | 334 | D | 0 | 0 | no | yes | no | no | no | yes | no | no | no | no | no | no | no |
| TRPC6 | 335 | L | 0 | 0 | no | yes | no | no | no | yes | no | no | no | no | no | no | no |
| TRPC6 | 336 | C | 0 | 0 | no | yes | no | no | no | yes | no | no | yes | no | no | no | no |
| TRPC6 | 337 | R | 1 | 0 | no | yes | no | no | no | yes | no | no | yes | no | no | no | no |
| TRPC6 | 338 | N | 1 | 0 | no | yes | no | no | no | yes | no | no | yes | no | no | no | no |
| TRPC6 | 339 | T | 0 | 0 | no | yes | no | no | no | yes | no | no | yes | no | no | no | no |
| TRPC6 | 340 | E | 0 | 0 | yes | no | no | no | no | yes | no | no | no | no | no | no | no |
| TRPC6 | 341 | E | 0 | 0 | yes | no | no | no | no | yes | no | no | no | no | no | no | no |
| TRPC6 | 342 | V | 0 | 0 | yes | no | no | no | no | yes | no | no | no | no | no | no | no |
| TRPC6 | 343 | E | 0 | 0 | yes | no | no | no | no | yes | no | no | no | no | no | no | no |

| | | | | | | | | | | | | | | | | | |
|---|---|---|---|---|---|---|---|---|---|---|---|---|---|---|---|---|---|
| TRPC6 | 344 | A | 1 | 0 | yes | no | no | no | no | yes | no | no | no | no | no | no | no |
| TRPC6 | 345 | I | 0 | 0 | yes | no | no | no | no | yes | no | no | no | no | no | no | no |
| TRPC6 | 346 | L | 0 | 0 | yes | no | no | no | no | yes | no | no | no | no | no | no | no |
| TRPC6 | 347 | N | 0 | 0 | no | yes | no | no | no | yes | no | no | no | no | no | no | no |
| TRPC6 | 348 | G | 0 | 0 | no | yes | no | no | no | yes | no | no | no | no | no | no | no |
| TRPC6 | 349 | D | 0 | 0 | no | yes | no | no | no | yes | no | no | no | no | no | no | no |
| TRPC6 | 350 | V | 0 | 0 | yes | no | no | no | no | yes | no | no | no | no | no | no | no |
| TRPC6 | 351 | E | 0 | 0 | yes | no | no | no | no | yes | no | no | no | no | no | no | no |
| TRPC6 | 352 | T | 1 | 0 | yes | no | no | no | no | yes | no | no | no | no | no | no | no |
| TRPC6 | 353 | L | 1 | 0 | no | yes | no | no | no | yes | no | no | no | no | no | no | no |
| TRPC6 | 354 | Q | 0 | 0 | no | yes | no | no | no | yes | no | no | no | no | no | no | no |
| TRPC6 | 355 | S | 1 | 0 | no | yes | no | no | no | yes | no | no | no | no | no | no | no |
| TRPC6 | 356 | G | 0 | 0 | no | yes | no | no | no | yes | no | no | no | no | no | no | no |
| TRPC6 | 357 | D | 0 | 0 | no | yes | no | no | no | yes | no | no | no | no | no | no | no |
| TRPC6 | 358 | H | 0 | 0 | no | yes | no | no | no | yes | no | no | no | no | no | no | no |
| TRPC6 | 359 | G | 1 | 0 | no | yes | no | no | no | yes | no | no | no | no | no | no | no |
| TRPC6 | 360 | R | 2 | 1 | no | yes | no | no | no | yes | no | no | no | no | no | no | no |
| TRPC6 | 361 | P | 0 | 0 | no | yes | no | no | no | yes | no | no | no | no | no | no | no |
| TRPC6 | 362 | N | 0 | 0 | no | yes | no | no | no | yes | no | no | no | no | no | no | no |
| TRPC6 | 363 | L | 0 | 0 | no | yes | no | no | no | yes | no | no | no | no | no | no | no |
| TRPC6 | 364 | S | 1 | 0 | no | yes | no | no | no | yes | no | no | no | no | no | no | no |
| TRPC6 | 365 | R | 2 | 0 | yes | no | no | no | no | yes | no | no | no | no | no | no | no |
| TRPC6 | 366 | L | 0 | 0 | yes | no | no | no | no | yes | no | no | no | no | no | no | no |
| TRPC6 | 367 | K | 0 | 0 | yes | no | no | no | no | yes | no | no | no | no | no | no | no |
| TRPC6 | 368 | L | 0 | 0 | yes | no | no | no | no | yes | no | no | no | no | no | no | no |
| TRPC6 | 369 | A | 0 | 0 | yes | no | no | no | no | yes | no | no | no | no | no | no | no |
| TRPC6 | 370 | I | 1 | 0 | yes | no | no | no | no | yes | no | no | no | no | no | no | no |
| TRPC6 | 371 | K | 0 | 0 | yes | no | no | no | no | yes | no | no | no | no | no | no | no |
| TRPC6 | 372 | Y | 0 | 0 | no | yes | no | no | no | yes | no | no | no | no | no | no | no |
| TRPC6 | 373 | E | 1 | 0 | no | yes | no | no | no | yes | no | no | no | no | no | no | no |
| TRPC6 | 374 | V | 1 | 0 | no | yes | no | no | no | yes | no | no | no | no | no | no | no |
| TRPC6 | 375 | K | 0 | 0 | yes | no | no | no | no | yes | no | no | no | no | no | no | no |
| TRPC6 | 376 | K | 1 | 0 | yes | no | no | no | no | yes | no | no | no | no | no | no | no |

| | | | | | | | | | | | | | | | | | |
|---|---|---|---|---|---|---|---|---|---|---|---|---|---|---|---|---|---|
| TRPC6 | 377 | F | 0 | 0 | yes | no | no | no | no | yes | no | no | no | no | no | no | no |
| TRPC6 | 378 | V | 1 | 0 | yes | no | no | no | no | yes | no | no | no | no | no | no | no |
| TRPC6 | 379 | A | 0 | 0 | no | yes | no | no | no | yes | no | no | no | no | no | no | no |
| TRPC6 | 380 | H | 0 | 0 | no | yes | no | no | no | yes | no | no | no | no | no | no | no |
| TRPC6 | 381 | P | 0 | 0 | yes | no | no | no | no | yes | no | no | no | no | no | no | no |
| TRPC6 | 382 | N | 0 | 0 | yes | no | no | no | no | yes | no | no | yes | no | no | no | no |
| TRPC6 | 383 | C | 0 | 0 | yes | no | no | no | no | yes | no | no | no | no | no | no | no |
| TRPC6 | 384 | Q | 0 | 0 | yes | no | no | no | no | yes | no | no | no | no | no | no | no |
| TRPC6 | 385 | Q | 1 | 0 | yes | no | no | no | no | yes | no | no | no | no | no | no | no |
| TRPC6 | 386 | Q | 0 | 0 | yes | no | no | no | no | yes | no | no | no | no | no | no | no |
| TRPC6 | 387 | L | 0 | 0 | yes | no | no | no | no | yes | no | no | no | no | no | no | no |
| TRPC6 | 388 | L | 0 | 0 | yes | no | no | no | no | yes | no | no | no | no | no | no | no |
| TRPC6 | 389 | S | 0 | 0 | yes | no | no | no | no | yes | no | no | no | no | no | no | no |
| TRPC6 | 390 | I | 0 | 0 | yes | no | no | no | no | yes | no | no | no | no | no | no | no |
| TRPC6 | 391 | W | 0 | 0 | yes | no | no | no | no | yes | no | no | no | no | no | no | no |
| TRPC6 | 392 | Y | 1 | 0 | no | yes | no | no | no | yes | no | no | no | no | no | no | no |
| TRPC6 | 393 | E | 1 | 0 | no | yes | no | no | no | yes | no | no | no | no | no | no | no |
| TRPC6 | 394 | N | 1 | 0 | no | yes | no | no | no | yes | no | no | no | no | no | no | no |
| TRPC6 | 395 | L | 0 | 0 | no | yes | no | no | no | yes | no | no | no | no | no | no | no |
| TRPC6 | 396 | S | 1 | 0 | no | yes | no | no | no | yes | no | no | no | no | no | no | no |
| TRPC6 | 397 | G | 0 | 0 | no | yes | no | no | no | yes | no | no | no | no | no | no | no |
| TRPC6 | 398 | L | 0 | 0 | no | yes | no | no | no | yes | no | no | no | no | no | no | no |
| TRPC6 | 399 | R | 1 | 0 | no | yes | no | no | no | yes | no | no | no | no | no | no | no |
| TRPC6 | 400 | Q | 0 | 0 | no | yes | no | no | no | yes | no | no | no | no | no | no | no |
| TRPC6 | 401 | Q | 0 | 0 | no | yes | no | no | no | yes | no | no | no | no | no | no | no |
| TRPC6 | 402 | T | 1 | 0 | no | yes | no | no | no | yes | no | no | no | no | no | no | no |
| TRPC6 | 403 | M | 2 | 0 | no | yes | no | no | no | yes | no | no | no | no | no | no | no |
| TRPC6 | 404 | A | 1 | 0 | no | yes | no | no | no | yes | no | no | no | no | no | no | no |
| TRPC6 | 405 | V | 0 | 0 | no | yes | no | no | no | yes | no | no | no | no | no | no | no |
| TRPC6 | 406 | K | 0 | 0 | yes | no | no | no | no | yes | no | no | no | no | no | no | no |
| TRPC6 | 407 | F | 0 | 0 | yes | no | no | yes | no | yes | no | no | no | no | no | no | no |
| TRPC6 | 408 | L | 1 | 0 | yes | no | no | yes | no | yes | no | no | no | no | no | no | no |
| TRPC6 | 409 | V | 1 | 0 | yes | no | no | yes | no | yes | no | no | no | no | no | no | no |

| | | | | | | | | | | | | | | | | | |
|---|---|---|---|---|---|---|---|---|---|---|---|---|---|---|---|---|---|
| TRPC6 | 410 | V | 0 | 0 | yes | no | no | yes | no | yes | no | no | no | no | no | no | no |
| TRPC6 | 411 | L | 1 | 0 | yes | no | no | yes | no | yes | no | no | no | no | no | no | no |
| TRPC6 | 412 | A | 0 | 0 | yes | no | no | yes | no | yes | no | no | no | no | no | no | no |
| TRPC6 | 413 | V | 1 | 0 | yes | no | no | yes | no | yes | no | no | no | no | no | no | no |
| TRPC6 | 414 | A | 0 | 0 | yes | no | no | yes | no | yes | no | no | no | no | no | no | no |
| TRPC6 | 415 | I | 2 | 0 | yes | no | no | yes | no | yes | no | no | no | no | no | no | no |
| TRPC6 | 416 | G | 0 | 0 | no | yes | no | yes | no | yes | no | no | no | no | no | no | no |
| TRPC6 | 417 | L | 0 | 0 | yes | no | no | yes | no | yes | no | no | no | no | no | no | no |
| TRPC6 | 418 | P | 0 | 0 | yes | no | no | yes | no | yes | no | no | no | no | no | no | no |
| TRPC6 | 419 | F | 0 | 0 | yes | no | no | yes | no | yes | no | no | no | no | no | no | no |
| TRPC6 | 420 | L | 0 | 0 | yes | no | no | yes | no | yes | no | no | no | no | no | no | no |
| TRPC6 | 421 | A | 0 | 0 | yes | no | no | yes | no | yes | no | no | no | no | no | no | no |
| TRPC6 | 422 | L | 0 | 0 | yes | no | no | yes | no | yes | no | no | no | no | no | no | no |
| TRPC6 | 423 | I | 0 | 0 | yes | no | no | yes | no | yes | no | no | no | no | no | no | no |
| TRPC6 | 424 | Y | 0 | 0 | yes | no | no | no | no | yes | no | no | no | no | no | no | no |
| TRPC6 | 425 | W | 0 | 0 | yes | no | no | yes | no | yes | no | no | no | no | no | no | no |
| TRPC6 | 426 | F | 1 | 0 | yes | no | no | yes | no | yes | no | no | no | no | no | no | no |
| TRPC6 | 427 | A | 0 | 0 | no | yes | no | no | no | yes | no | no | no | no | no | no | no |
| TRPC6 | 428 | P | 0 | 0 | no | yes | no | no | no | yes | no | no | no | no | no | no | no |
| TRPC6 | 429 | C | 0 | 0 | no | yes | no | no | no | yes | no | no | no | no | no | no | no |
| TRPC6 | 430 | S | 1 | 0 | no | yes | no | no | no | yes | no | no | no | no | no | no | no |
| TRPC6 | 431 | K | 0 | 0 | no | yes | no | no | no | yes | no | no | no | no | no | no | no |
| TRPC6 | 432 | M | 1 | 0 | no | yes | no | no | no | yes | no | no | no | no | no | no | no |
| TRPC6 | 433 | G | 0 | 0 | no | yes | no | no | no | yes | no | no | no | no | no | no | no |
| TRPC6 | 434 | K | 0 | 0 | no | yes | no | no | no | yes | no | no | no | no | no | no | no |
| TRPC6 | 435 | I | 1 | 0 | no | yes | no | no | no | yes | no | no | no | no | no | no | no |
| TRPC6 | 436 | M | 0 | 0 | no | yes | no | no | no | yes | no | no | no | no | no | no | no |
| TRPC6 | 437 | R | 2 | 0 | no | yes | no | no | no | yes | no | no | no | no | no | no | no |
| TRPC6 | 438 | G | 1 | 0 | no | yes | no | no | no | yes | no | no | no | no | no | no | no |
| TRPC6 | 439 | P | 2 | 0 | yes | no | no | no | no | no | no | no | no | no | no | no | no |
| TRPC6 | 440 | F | 0 | 0 | yes | no | no | no | no | no | no | no | no | no | no | no | no |
| TRPC6 | 441 | M | 0 | 0 | yes | no | no | no | no | no | no | no | no | no | no | no | no |
| TRPC6 | 442 | K | 0 | 0 | yes | no | no | no | no | no | no | no | no | no | no | no | no |

| | | | | | | | | | | | | | | | | | |
|---|---|---|---|---|---|---|---|---|---|---|---|---|---|---|---|---|---|
| TRPC6 | 443 | F | 0 | 0 | yes | no | no | no | no | no | no | no | no | no | no | no | no |
| TRPC6 | 444 | V | 2 | 0 | yes | no | no | no | no | no | no | no | no | no | no | no | no |
| TRPC6 | 445 | A | 0 | 0 | yes | no | no | yes | no | no | no | no | no | no | no | no | no |
| TRPC6 | 446 | H | 0 | 0 | yes | no | no | yes | no | no | no | no | no | no | no | no | no |
| TRPC6 | 447 | A | 1 | 0 | yes | no | no | yes | no | no | no | no | no | no | no | no | no |
| TRPC6 | 448 | A | 0 | 0 | yes | no | no | yes | no | no | no | no | no | no | no | no | no |
| TRPC6 | 449 | S | 1 | 0 | yes | no | no | yes | no | no | no | no | no | no | no | no | no |
| TRPC6 | 450 | F | 0 | 0 | yes | no | no | yes | no | no | no | no | no | no | no | no | no |
| TRPC6 | 451 | T | 0 | 0 | yes | no | no | yes | no | no | no | no | no | no | no | no | no |
| TRPC6 | 452 | I | 1 | 0 | yes | no | no | yes | no | no | no | no | no | no | no | no | no |
| TRPC6 | 453 | F | 1 | 0 | yes | no | no | yes | no | no | no | no | no | no | no | no | no |
| TRPC6 | 454 | L | 0 | 0 | yes | no | no | yes | no | no | no | no | no | no | no | no | no |
| TRPC6 | 455 | G | 0 | 0 | yes | no | no | yes | no | no | no | no | no | no | no | no | no |
| TRPC6 | 456 | L | 0 | 0 | yes | no | no | yes | no | no | no | no | no | no | no | no | no |
| TRPC6 | 457 | L | 0 | 0 | yes | no | no | yes | no | no | no | no | no | no | no | no | no |
| TRPC6 | 458 | V | 0 | 0 | yes | no | no | yes | no | no | no | no | yes | no | no | no | no |
| TRPC6 | 459 | M | 0 | 0 | yes | no | no | yes | no | no | no | no | no | no | no | no | no |
| TRPC6 | 460 | N | 0 | 0 | yes | no | no | yes | yes | no | no | no | no | no | no | no | no |
| TRPC6 | 461 | A | 0 | 0 | no | yes | no | yes | yes | no | no | no | yes | no | no | no | no |
| TRPC6 | 462 | A | 0 | 0 | no | yes | no | yes | yes | no | no | no | no | no | no | no | no |
| TRPC6 | 463 | D | 0 | 0 | no | yes | no | yes | yes | no | no | no | no | no | no | no | no |
| TRPC6 | 464 | R | 0 | 0 | no | yes | no | yes | yes | no | no | no | yes | no | no | no | no |
| TRPC6 | 465 | F | 0 | 0 | no | yes | no | yes | yes | no | no | no | yes | no | no | no | no |
| TRPC6 | 466 | E | 0 | 0 | no | yes | no | yes | yes | no | no | no | yes | no | no | no | no |
| TRPC6 | 467 | G | 0 | 0 | no | yes | no | no | yes | no | no | no | yes | no | no | no | no |
| TRPC6 | 468 | T | 0 | 0 | no | yes | no | no | yes | no | no | no | yes | no | no | no | no |
| TRPC6 | 469 | K | 1 | 0 | no | yes | no | no | yes | no | no | no | no | no | no | no | no |
| TRPC6 | 470 | L | 0 | 0 | no | yes | no | no | yes | no | no | no | no | no | no | no | no |
| TRPC6 | 471 | L | 0 | 0 | no | yes | no | no | yes | no | no | no | no | no | no | no | no |
| TRPC6 | 472 | P | 0 | 0 | no | yes | no | no | yes | no | no | no | yes | no | no | no | no |
| TRPC6 | 473 | N | 0 | 0 | no | yes | no | no | yes | no | no | no | no | no | no | no | no |
| TRPC6 | 474 | E | 1 | 0 | no | yes | no | no | yes | no | no | no | no | no | no | no | no |
| TRPC6 | 475 | T | 0 | 0 | no | yes | no | no | yes | no | no | no | no | no | no | no | no |

| | | | | | | | | | | | | | | | | | |
|---|---|---|---|---|---|---|---|---|---|---|---|---|---|---|---|---|---|
| TRPC6 | 476 | S | 2 | 0 | no | yes | no | no | yes | no | no | no | no | no | no | no | no |
| TRPC6 | 477 | T | 0 | 0 | no | yes | no | no | yes | no | no | no | no | no | no | no | no |
| TRPC6 | 478 | D | 1 | 0 | no | yes | no | no | yes | no | no | no | no | no | no | no | no |
| TRPC6 | 479 | N | 0 | 0 | no | yes | no | no | yes | no | no | no | no | no | no | no | no |
| TRPC6 | 480 | A | 0 | 0 | no | yes | no | no | yes | no | no | no | no | no | no | no | no |
| TRPC6 | 481 | K | 0 | 0 | no | yes | no | no | yes | no | no | no | no | no | no | no | no |
| TRPC6 | 482 | Q | 0 | 0 | no | yes | no | no | yes | no | no | no | no | no | no | no | no |
| TRPC6 | 483 | L | 0 | 0 | no | yes | no | no | yes | no | no | no | no | no | no | no | no |
| TRPC6 | 484 | F | 1 | 0 | no | yes | no | no | yes | no | no | no | no | no | no | no | no |
| TRPC6 | 485 | R | 0 | 0 | no | yes | no | no | yes | no | no | no | no | no | no | no | no |
| TRPC6 | 486 | M | 0 | 0 | no | yes | no | no | yes | no | no | no | no | no | no | no | no |
| TRPC6 | 487 | K | 1 | 0 | no | yes | no | no | yes | no | no | no | no | no | no | no | no |
| TRPC6 | 488 | T | 0 | 0 | no | yes | no | no | no | no | no | no | no | no | no | no | no |
| TRPC6 | 489 | S | 0 | 0 | no | yes | no | no | no | no | no | no | no | no | no | no | no |
| TRPC6 | 490 | C | 0 | 0 | no | yes | no | no | no | no | no | no | no | no | no | no | no |
| TRPC6 | 491 | F | 0 | 0 | no | yes | no | yes | no | no | no | no | no | no | no | no | no |
| TRPC6 | 492 | S | 1 | 0 | no | yes | no | yes | no | no | no | no | no | no | no | no | no |
| TRPC6 | 493 | W | 0 | 0 | no | yes | no | yes | no | no | no | no | no | no | no | no | no |
| TRPC6 | 494 | M | 0 | 0 | no | yes | no | yes | no | no | no | no | no | no | no | no | no |
| TRPC6 | 495 | E | 0 | 0 | no | yes | no | yes | no | no | no | no | no | no | no | no | no |
| TRPC6 | 496 | M | 0 | 0 | no | yes | no | yes | no | no | no | no | no | no | no | no | no |
| TRPC6 | 497 | L | 0 | 0 | yes | no | no | yes | no | no | no | no | no | no | no | no | no |
| TRPC6 | 498 | I | 0 | 0 | yes | no | no | yes | no | no | no | no | no | no | no | no | no |
| TRPC6 | 499 | I | 1 | 0 | yes | no | no | yes | no | no | no | no | no | no | no | no | no |
| TRPC6 | 500 | S | 0 | 0 | yes | no | no | yes | no | no | no | no | no | no | no | no | no |
| TRPC6 | 501 | W | 0 | 0 | yes | no | no | yes | no | no | no | no | no | no | no | no | no |
| TRPC6 | 502 | V | 0 | 0 | yes | no | no | yes | no | no | no | no | no | no | no | no | no |
| TRPC6 | 503 | I | 0 | 0 | yes | no | no | yes | no | no | no | no | no | no | no | no | no |
| TRPC6 | 504 | G | 0 | 0 | yes | no | no | yes | no | no | no | no | no | no | no | no | no |
| TRPC6 | 505 | M | 1 | 0 | yes | no | no | yes | no | no | no | no | no | no | no | no | no |
| TRPC6 | 506 | I | 1 | 0 | yes | no | no | yes | no | no | no | no | no | no | no | no | no |
| TRPC6 | 507 | W | 0 | 0 | yes | no | no | yes | no | no | no | no | no | no | no | no | no |
| TRPC6 | 508 | A | 0 | 0 | yes | no | no | yes | no | no | no | no | no | no | no | no | no |

| | | | | | | | | | | | | | | | | | |
|---|---|---|---|---|---|---|---|---|---|---|---|---|---|---|---|---|---|
| TRPC6 | 509 | E | 0 | 0 | yes | no | no | yes | no | yes | no | no | no | no | no | no | no |
| TRPC6 | 510 | C | 1 | 0 | yes | no | no | yes | no | yes | no | no | no | no | no | no | no |
| TRPC6 | 511 | K | 0 | 0 | yes | no | no | yes | no | yes | no | no | no | no | no | no | no |
| TRPC6 | 512 | E | 0 | 0 | yes | no | no | no | no | yes | no | no | no | no | no | no | no |
| TRPC6 | 513 | I | 1 | 0 | yes | no | no | yes | no | yes | no | no | no | no | no | no | no |
| TRPC6 | 514 | W | 0 | 0 | yes | no | no | yes | no | yes | no | no | no | no | no | no | no |
| TRPC6 | 515 | T | 0 | 0 | yes | no | no | no | no | yes | no | no | no | no | no | no | no |
| TRPC6 | 516 | Q | 0 | 0 | yes | no | no | no | no | yes | no | no | no | no | no | no | no |
| TRPC6 | 517 | G | 0 | 0 | no | yes | no | no | no | yes | no | no | no | no | no | no | no |
| TRPC6 | 518 | P | 1 | 0 | no | yes | no | no | no | yes | no | no | no | no | no | no | no |
| TRPC6 | 519 | K | 2 | 0 | no | yes | no | no | no | yes | no | no | no | no | no | no | no |
| TRPC6 | 520 | E | 0 | 0 | no | yes | no | no | no | yes | no | no | no | no | no | no | no |
| TRPC6 | 521 | Y | 0 | 0 | no | yes | no | yes | no | yes | no | no | no | no | no | no | no |
| TRPC6 | 522 | L | 1 | 0 | no | yes | no | yes | no | no | no | no | no | no | no | no | no |
| TRPC6 | 523 | F | 0 | 0 | no | yes | no | no | no | no | no | no | no | no | no | no | no |
| TRPC6 | 524 | E | 1 | 0 | no | yes | no | no | no | no | no | no | no | no | no | no | no |
| TRPC6 | 525 | L | 0 | 0 | yes | no | no | yes | no | no | no | no | no | no | no | no | no |
| TRPC6 | 526 | W | 1 | 0 | yes | no | no | yes | no | no | no | no | no | no | no | no | no |
| TRPC6 | 527 | N | 0 | 0 | yes | no | no | yes | no | no | no | no | no | no | no | no | no |
| TRPC6 | 528 | M | 1 | 0 | yes | no | no | yes | no | no | no | no | no | no | no | no | no |
| TRPC6 | 529 | L | 0 | 0 | yes | no | no | yes | no | no | no | no | no | no | no | no | no |
| TRPC6 | 530 | D | 0 | 0 | yes | no | no | yes | no | no | no | no | no | no | no | no | no |
| TRPC6 | 531 | F | 1 | 0 | yes | no | no | yes | no | no | no | no | no | no | no | no | no |
| TRPC6 | 532 | G | 0 | 0 | yes | no | no | yes | no | no | no | no | no | no | no | no | no |
| TRPC6 | 533 | M | 1 | 0 | yes | no | no | yes | no | no | no | no | no | no | no | no | no |
| TRPC6 | 534 | L | 1 | 0 | yes | no | no | yes | no | no | no | no | no | no | no | no | no |
| TRPC6 | 535 | A | 0 | 0 | yes | no | no | yes | no | no | no | no | no | no | no | no | no |
| TRPC6 | 536 | I | 1 | 0 | yes | no | no | yes | no | no | no | no | no | no | no | no | no |
| TRPC6 | 537 | F | 0 | 0 | yes | no | no | yes | no | no | no | no | no | no | no | no | no |
| TRPC6 | 538 | A | 1 | 0 | yes | no | no | yes | no | no | no | no | no | no | no | no | no |
| TRPC6 | 539 | A | 0 | 0 | yes | no | no | yes | no | no | no | no | no | no | no | no | no |
| TRPC6 | 540 | S | 0 | 0 | yes | no | no | yes | no | no | no | no | no | no | no | no | no |
| TRPC6 | 541 | F | 0 | 0 | yes | no | no | yes | no | no | no | no | no | no | no | no | no |

| | | | | | | | | | | | | | | | | | |
|---|---|---|---|---|---|---|---|---|---|---|---|---|---|---|---|---|---|
| TRPC6 | 542 | I | 0 | 0 | yes | no | no | yes | no | no | no | no | no | no | no | no | no |
| TRPC6 | 543 | A | 1 | 0 | yes | no | no | yes | yes | no | no | no | no | no | no | no | no |
| TRPC6 | 544 | R | 0 | 0 | yes | no | no | yes | yes | no | no | no | no | no | no | no | no |
| TRPC6 | 545 | F | 0 | 0 | yes | no | no | yes | yes | no | no | no | no | no | no | no | no |
| TRPC6 | 546 | M | 0 | 0 | yes | no | no | yes | yes | no | no | no | no | no | no | no | no |
| TRPC6 | 547 | A | 0 | 0 | yes | no | no | no | yes | no | no | no | no | no | no | no | no |
| TRPC6 | 548 | F | 0 | 0 | yes | no | no | no | yes | no | no | no | no | no | no | no | no |
| TRPC6 | 549 | W | 0 | 0 | yes | no | no | no | yes | no | no | no | no | no | no | no | no |
| TRPC6 | 550 | H | 0 | 0 | yes | no | no | no | yes | no | no | no | no | no | no | no | no |
| TRPC6 | 551 | A | 0 | 0 | yes | no | no | no | yes | no | no | no | no | no | no | no | no |
| TRPC6 | 552 | S | 0 | 0 | yes | no | no | no | yes | no | no | no | no | no | no | no | no |
| TRPC6 | 553 | K | 0 | 0 | yes | no | no | no | yes | no | no | no | no | no | no | no | no |
| TRPC6 | 554 | A | 0 | 0 | yes | no | no | no | yes | no | no | no | no | no | no | no | no |
| TRPC6 | 555 | Q | 0 | 0 | yes | no | no | no | yes | no | no | no | no | no | no | no | no |
| TRPC6 | 556 | S | 1 | 0 | no | yes | no | no | yes | no | no | no | no | no | no | no | no |
| TRPC6 | 557 | I | 1 | 0 | no | yes | no | no | yes | no | no | no | no | no | no | no | no |
| TRPC6 | 558 | I | 0 | 0 | no | yes | no | no | yes | no | no | no | no | no | no | no | no |
| TRPC6 | 559 | D | 0 | 0 | no | yes | no | no | yes | no | no | no | no | no | no | no | no |
| TRPC6 | 569 | K | 0 | 0 | no | yes | no | no | yes | no | no | no | no | no | no | no | no |
| TRPC6 | 570 | V | 1 | 0 | no | yes | no | no | yes | no | no | no | no | no | no | no | no |
| TRPC6 | 571 | T | 1 | 0 | no | yes | no | no | yes | no | no | no | no | no | no | no | no |
| TRPC6 | 572 | L | 0 | 0 | no | yes | no | no | yes | no | no | no | no | no | no | no | no |
| TRPC6 | 573 | G | 0 | 0 | no | yes | no | no | yes | no | no | no | no | no | no | no | no |
| TRPC6 | 574 | D | 0 | 0 | no | yes | no | no | yes | no | no | no | no | no | no | no | no |
| TRPC6 | 575 | N | 1 | 0 | no | yes | no | no | yes | no | no | no | no | no | no | no | no |
| TRPC6 | 576 | V | 0 | 0 | no | yes | no | no | yes | no | no | no | no | no | no | no | no |
| TRPC6 | 577 | K | 0 | 0 | no | yes | no | no | yes | no | no | no | no | no | no | no | no |
| TRPC6 | 578 | Y | 0 | 0 | no | yes | no | no | yes | no | no | no | no | no | no | no | no |
| TRPC6 | 579 | Y | 1 | 0 | no | yes | no | no | yes | no | no | no | no | no | no | no | no |
| TRPC6 | 580 | N | 1 | 0 | no | yes | no | no | yes | no | no | no | no | no | no | no | no |
| TRPC6 | 581 | L | 0 | 0 | no | yes | no | no | yes | no | no | no | no | no | no | no | no |
| TRPC6 | 582 | A | 0 | 0 | no | yes | no | no | yes | no | no | no | no | no | no | no | no |
| TRPC6 | 583 | R | 0 | 0 | yes | no | no | no | yes | no | no | no | yes | no | no | no | no |

| | | | | | | | | | | | | | | | | | |
|---|---|---|---|---|---|---|---|---|---|---|---|---|---|---|---|---|---|
| TRPC6 | 584 | I | 1 | 0 | yes | no | no | no | yes | no | no | no | yes | no | no | no | no |
| TRPC6 | 585 | K | 0 | 0 | yes | no | no | no | yes | no | no | no | no | no | no | no | no |
| TRPC6 | 586 | W | 0 | 0 | no | yes | no | no | yes | no | no | no | no | no | no | no | no |
| TRPC6 | 587 | D | 1 | 0 | no | yes | no | no | yes | no | no | no | no | no | no | no | no |
| TRPC6 | 588 | P | 1 | 0 | no | yes | no | no | yes | no | no | no | yes | no | no | no | no |
| TRPC6 | 589 | S | 0 | 0 | no | yes | no | no | yes | no | no | no | no | no | no | no | no |
| TRPC6 | 590 | D | 0 | 0 | no | yes | no | no | yes | no | no | no | no | no | no | no | no |
| TRPC6 | 591 | P | 0 | 0 | no | yes | no | no | yes | no | no | no | no | no | no | no | no |
| TRPC6 | 592 | Q | 0 | 0 | yes | no | no | no | yes | no | no | no | no | no | no | no | no |
| TRPC6 | 593 | I | 0 | 0 | yes | no | no | no | no | no | no | no | yes | no | no | no | no |
| TRPC6 | 594 | I | 1 | 0 | yes | no | no | yes | no | no | no | no | yes | no | no | no | no |
| TRPC6 | 595 | S | 0 | 0 | yes | no | no | yes | no | no | no | no | no | no | no | no | no |
| TRPC6 | 596 | E | 0 | 0 | yes | no | no | yes | no | no | no | no | yes | no | no | no | no |
| TRPC6 | 597 | G | 0 | 0 | yes | no | no | yes | no | no | no | no | yes | no | no | no | no |
| TRPC6 | 598 | L | 0 | 0 | yes | no | no | yes | no | no | no | no | no | no | no | no | no |
| TRPC6 | 599 | Y | 0 | 0 | yes | no | no | yes | no | no | no | no | yes | no | no | no | no |
| TRPC6 | 600 | A | 0 | 0 | yes | no | no | yes | no | no | no | no | yes | no | no | no | no |
| TRPC6 | 601 | I | 0 | 0 | yes | no | no | yes | no | no | no | no | yes | no | no | no | no |
| TRPC6 | 602 | A | 0 | 0 | yes | no | no | yes | no | no | no | no | no | no | no | no | no |
| TRPC6 | 603 | V | 1 | 0 | yes | no | no | yes | no | no | no | no | yes | no | no | no | no |
| TRPC6 | 604 | V | 0 | 0 | yes | no | no | yes | no | no | no | no | yes | no | no | no | no |
| TRPC6 | 605 | L | 0 | 0 | yes | no | no | yes | no | no | no | no | no | no | no | no | no |
| TRPC6 | 606 | S | 0 | 0 | yes | no | no | yes | no | no | no | no | no | no | no | no | no |
| TRPC6 | 607 | F | 0 | 0 | yes | no | no | yes | no | no | no | no | yes | no | no | no | no |
| TRPC6 | 608 | S | 0 | 0 | yes | no | no | yes | no | no | no | no | no | no | no | no | no |
| TRPC6 | 609 | R | 0 | 0 | yes | no | no | yes | no | no | no | no | no | no | no | no | no |
| TRPC6 | 610 | I | 0 | 0 | yes | no | no | yes | no | no | no | no | yes | no | no | no | no |
| TRPC6 | 611 | A | 1 | 0 | yes | no | no | yes | no | no | no | no | no | no | no | no | no |
| TRPC6 | 612 | Y | 0 | 0 | yes | no | no | yes | no | no | no | no | no | no | no | no | no |
| TRPC6 | 613 | I | 1 | 0 | no | yes | no | yes | no | no | no | no | no | no | no | no | no |
| TRPC6 | 614 | L | 0 | 0 | yes | no | no | no | no | yes | no | no | no | no | no | no | no |
| TRPC6 | 615 | P | 1 | 0 | yes | no | no | no | no | yes | no | no | no | no | no | no | no |
| TRPC6 | 616 | A | 0 | 0 | yes | no | no | no | no | yes | no | no | no | no | no | no | no |

| | | | | | | | | | | | | | | | | | |
|---|---|---|---|---|---|---|---|---|---|---|---|---|---|---|---|---|---|
| TRPC6 | 617 | N | 0 | 0 | no | yes | no | no | no | yes | no | no | no | no | no | no | no |
| TRPC6 | 618 | E | 0 | 0 | no | yes | no | no | no | yes | no | no | no | no | no | no | no |
| TRPC6 | 619 | S | 0 | 0 | no | yes | no | no | no | yes | no | no | no | no | no | no | no |
| TRPC6 | 620 | F | 0 | 0 | no | yes | no | no | no | yes | no | no | yes | no | no | no | no |
| TRPC6 | 621 | G | 0 | 0 | yes | no | no | no | no | yes | no | no | no | no | no | no | no |
| TRPC6 | 622 | P | 1 | 0 | yes | no | no | no | no | yes | no | no | no | no | no | no | no |
| TRPC6 | 623 | L | 1 | 0 | yes | no | no | no | no | yes | no | no | yes | no | no | no | no |
| TRPC6 | 624 | Q | 0 | 0 | yes | no | no | no | no | yes | no | no | no | no | no | no | no |
| TRPC6 | 625 | I | 0 | 0 | yes | no | no | no | no | yes | no | no | no | no | no | no | no |
| TRPC6 | 626 | S | 1 | 0 | yes | no | no | no | no | yes | no | no | yes | no | no | no | no |
| TRPC6 | 627 | L | 0 | 0 | yes | no | no | no | no | yes | no | no | yes | no | no | no | no |
| TRPC6 | 628 | G | 1 | 0 | yes | no | no | no | no | yes | no | no | no | no | no | no | no |
| TRPC6 | 629 | R | 1 | 0 | no | yes | no | no | no | yes | no | no | no | no | no | no | no |
| TRPC6 | 630 | T | 0 | 0 | no | yes | no | no | no | yes | no | no | yes | no | no | no | no |
| TRPC6 | 631 | V | 1 | 0 | no | yes | no | yes | no | yes | no | no | no | no | no | no | no |
| TRPC6 | 632 | K | 0 | 0 | no | yes | no | no | no | yes | no | no | no | no | no | no | no |
| TRPC6 | 633 | D | 0 | 0 | no | yes | no | no | no | yes | no | no | no | no | no | no | no |
| TRPC6 | 634 | I | 0 | 0 | no | yes | no | yes | no | yes | no | no | yes | no | no | no | no |
| TRPC6 | 635 | F | 0 | 0 | no | yes | no | yes | no | yes | no | no | no | no | no | no | no |
| TRPC6 | 636 | K | 0 | 0 | no | yes | no | no | no | yes | no | no | yes | no | no | no | no |
| TRPC6 | 637 | F | 0 | 0 | no | yes | no | yes | no | no | no | no | no | no | no | no | no |
| TRPC6 | 638 | M | 0 | 0 | yes | no | no | yes | no | no | no | no | no | no | no | no | no |
| TRPC6 | 639 | V | 0 | 0 | yes | no | no | yes | no | no | no | no | yes | no | no | no | no |
| TRPC6 | 640 | I | 0 | 0 | yes | no | no | yes | no | no | no | no | no | no | no | no | no |
| TRPC6 | 641 | F | 0 | 0 | yes | no | no | yes | no | no | no | no | no | no | no | no | no |
| TRPC6 | 642 | I | 0 | 0 | yes | no | no | yes | no | no | no | no | no | no | no | no | no |
| TRPC6 | 643 | M | 1 | 0 | yes | no | no | yes | no | no | no | no | yes | no | no | no | no |
| TRPC6 | 644 | V | 0 | 0 | yes | no | no | yes | no | no | no | no | no | no | no | no | no |
| TRPC6 | 645 | F | 0 | 0 | yes | no | no | yes | no | no | no | no | no | no | no | no | no |
| TRPC6 | 646 | V | 0 | 0 | yes | no | no | yes | no | no | no | no | no | no | no | no | no |
| TRPC6 | 647 | A | 1 | 0 | yes | no | no | yes | no | no | no | no | yes | no | no | no | no |
| TRPC6 | 648 | F | 0 | 0 | yes | no | no | yes | no | no | no | no | yes | no | no | no | no |
| TRPC6 | 649 | M | 0 | 0 | yes | no | no | yes | no | no | no | no | no | no | no | no | no |

| | | | | | | | | | | | | | | | | | |
|---|---|---|---|---|---|---|---|---|---|---|---|---|---|---|---|---|---|
| TRPC6 | 650 | I | 0 | 0 | yes | no | no | yes | no | no | no | no | yes | no | no | no | no |
| TRPC6 | 651 | G | 0 | 0 | yes | no | no | yes | no | no | no | no | yes | no | no | no | no |
| TRPC6 | 652 | M | 1 | 0 | yes | no | no | yes | no | no | no | no | no | no | no | no | no |
| TRPC6 | 653 | F | 0 | 0 | yes | no | no | yes | no | no | no | no | yes | no | no | no | no |
| TRPC6 | 654 | N | 0 | 0 | no | yes | no | yes | no | no | no | no | yes | no | no | no | no |
| TRPC6 | 655 | L | 0 | 0 | no | yes | no | yes | no | no | no | no | yes | no | no | no | no |
| TRPC6 | 656 | Y | 0 | 0 | no | yes | no | yes | no | no | no | no | no | no | no | no | no |
| TRPC6 | 657 | S | 0 | 0 | no | yes | no | yes | no | no | no | no | yes | no | no | no | no |
| TRPC6 | 658 | Y | 1 | 0 | no | yes | no | no | yes | no | no | no | yes | no | no | no | no |
| TRPC6 | 659 | Y | 1 | 0 | no | yes | no | no | yes | no | no | no | yes | no | no | no | no |
| TRPC6 | 660 | I | 1 | 0 | no | yes | no | no | yes | no | no | no | yes | no | no | no | no |
| TRPC6 | 661 | G | 0 | 0 | no | yes | no | no | yes | no | no | no | no | no | no | no | no |
| TRPC6 | 662 | A | 0 | 0 | no | yes | no | no | yes | no | no | no | no | no | no | no | no |
| TRPC6 | 663 | K | 0 | 0 | no | no | yes | no | yes | no | no | no | no | no | no | no | no |
| TRPC6 | 664 | Q | 0 | 0 | no | yes | no | no | yes | no | no | no | no | no | no | no | no |
| TRPC6 | 665 | N | 0 | 0 | no | yes | no | no | yes | no | no | no | no | no | no | no | no |
| TRPC6 | 666 | E | 0 | 0 | no | yes | no | no | yes | no | no | no | no | no | no | no | no |
| TRPC6 | 667 | A | 0 | 0 | no | yes | no | yes | yes | no | no | no | no | no | no | no | no |
| TRPC6 | 668 | F | 0 | 0 | no | yes | no | yes | yes | no | no | no | no | no | no | no | no |
| TRPC6 | 669 | T | 0 | 0 | no | yes | no | yes | yes | no | no | no | no | no | no | no | no |
| TRPC6 | 670 | T | 0 | 0 | no | yes | no | yes | yes | no | no | no | yes | no | no | no | no |
| TRPC6 | 671 | V | 0 | 0 | yes | no | no | yes | yes | no | no | no | yes | no | no | no | no |
| TRPC6 | 672 | E | 0 | 0 | yes | no | no | yes | yes | no | no | no | no | no | no | no | no |
| TRPC6 | 673 | E | 0 | 0 | yes | no | no | yes | yes | no | no | no | no | no | no | no | no |
| TRPC6 | 674 | S | 0 | 0 | yes | no | no | yes | yes | no | no | no | no | no | no | no | no |
| TRPC6 | 675 | F | 0 | 0 | yes | no | no | yes | yes | no | no | no | no | no | no | no | no |
| TRPC6 | 676 | K | 0 | 0 | yes | no | no | yes | yes | no | no | no | no | no | no | no | no |
| TRPC6 | 677 | T | 0 | 0 | yes | no | no | yes | yes | no | no | no | no | no | no | no | no |
| TRPC6 | 678 | L | 0 | 0 | yes | no | no | yes | yes | no | no | no | no | no | no | no | no |
| TRPC6 | 679 | F | 0 | 0 | yes | no | no | yes | yes | no | no | no | yes | no | no | no | no |
| TRPC6 | 680 | W | 0 | 0 | no | yes | no | yes | yes | no | no | no | yes | no | no | no | no |
| TRPC6 | 681 | A | 0 | 0 | no | yes | no | yes | yes | no | no | no | no | no | no | no | no |
| TRPC6 | 682 | I | 0 | 0 | no | yes | no | yes | yes | no | yes | no | no | no | no | no | no |

| | | | | | | | | | | | | | | | | | |
|---|---|---|---|---|---|---|---|---|---|---|---|---|---|---|---|---|---|
| TRPC6 | 683 | F | 0 | 0 | no | yes | no | yes | yes | no | no | no | yes | no | no | no | no |
| TRPC6 | 684 | G | 0 | 0 | no | yes | no | yes | yes | no | no | no | yes | no | no | no | no |
| TRPC6 | 685 | L | 0 | 0 | no | yes | no | yes | yes | no | no | no | yes | no | no | no | no |
| TRPC6 | 686 | S | 1 | 0 | no | yes | no | yes | yes | no | no | no | yes | no | no | no | no |
| TRPC6 | 687 | E | 0 | 0 | no | yes | no | yes | yes | no | no | no | no | no | no | no | no |
| TRPC6 | 688 | V | 0 | 0 | yes | no | no | yes | yes | no | no | no | yes | no | no | no | no |
| TRPC6 | 689 | K | 0 | 0 | yes | no | no | yes | yes | no | no | no | no | no | no | no | no |
| TRPC6 | 690 | S | 0 | 0 | yes | no | no | yes | yes | no | no | no | no | no | no | no | no |
| TRPC6 | 691 | V | 0 | 0 | yes | no | no | yes | yes | no | no | no | no | no | no | no | no |
| TRPC6 | 692 | V | 1 | 0 | no | yes | no | no | yes | no | no | no | no | no | no | no | no |
| TRPC6 | 693 | I | 2 | 0 | no | no | yes | no | yes | no | no | no | no | no | no | no | no |
| TRPC6 | 694 | N | 1 | 0 | no | yes | no | no | yes | no | no | no | no | no | no | no | no |
| TRPC6 | 695 | Y | 1 | 0 | no | yes | no | no | yes | no | no | no | yes | no | no | no | no |
| TRPC6 | 696 | N | 0 | 0 | no | yes | no | no | yes | no | no | no | yes | no | no | no | no |
| TRPC6 | 697 | H | 0 | 0 | no | yes | no | no | yes | no | no | no | yes | no | no | no | no |
| TRPC6 | 698 | K | 0 | 0 | yes | no | no | no | yes | no | no | no | no | no | no | no | no |
| TRPC6 | 699 | F | 1 | 0 | yes | no | no | yes | yes | no | no | no | yes | no | no | no | no |
| TRPC6 | 700 | I | 1 | 0 | yes | no | no | yes | yes | no | no | no | no | no | no | no | no |
| TRPC6 | 701 | E | 0 | 0 | yes | no | no | yes | yes | no | no | no | no | no | no | no | no |
| TRPC6 | 702 | N | 1 | 0 | yes | no | no | yes | yes | no | no | no | no | no | no | no | no |
| TRPC6 | 703 | I | 1 | 0 | yes | no | no | yes | yes | no | no | no | yes | no | no | no | no |
| TRPC6 | 704 | G | 0 | 0 | yes | no | no | yes | yes | no | no | no | no | no | no | no | no |
| TRPC6 | 705 | Y | 0 | 0 | yes | no | no | yes | yes | no | no | no | no | no | no | no | no |
| TRPC6 | 706 | V | 1 | 0 | yes | no | no | yes | yes | no | no | no | no | no | no | no | no |
| TRPC6 | 707 | L | 2 | 0 | yes | no | no | yes | no | no | no | no | no | no | no | no | no |
| TRPC6 | 708 | Y | 0 | 0 | yes | no | no | yes | no | no | no | no | yes | no | no | no | no |
| TRPC6 | 709 | G | 0 | 0 | yes | no | no | yes | no | no | no | no | yes | no | no | no | no |
| TRPC6 | 710 | V | 0 | 0 | yes | no | no | yes | no | no | no | no | no | no | no | no | no |
| TRPC6 | 711 | Y | 0 | 0 | yes | no | no | yes | no | no | no | no | no | no | no | no | no |
| TRPC6 | 712 | N | 0 | 0 | yes | no | no | yes | no | no | no | no | yes | no | no | no | no |
| TRPC6 | 713 | V | 1 | 0 | yes | no | no | yes | no | no | no | no | no | no | no | no | no |
| TRPC6 | 714 | T | 1 | 0 | yes | no | no | yes | no | no | no | no | no | no | no | no | no |
| TRPC6 | 715 | M | 0 | 0 | yes | no | no | yes | no | no | no | no | no | no | no | no | no |

| | | | | | | | | | | | | | | | | | |
|---|---|---|---|---|---|---|---|---|---|---|---|---|---|---|---|---|---|
| TRPC6 | 716 | V | 0 | 0 | yes | no | no | yes | no | no | yes | no | no | no | no | no | no |
| TRPC6 | 717 | I | 0 | 0 | yes | no | no | yes | no | no | no | no | yes | no | no | no | no |
| TRPC6 | 718 | V | 0 | 0 | yes | no | no | yes | no | no | no | no | no | no | no | no | no |
| TRPC6 | 719 | L | 0 | 0 | yes | no | no | yes | no | no | no | no | no | no | no | no | no |
| TRPC6 | 720 | L | 0 | 0 | yes | no | no | yes | no | no | no | no | yes | no | no | no | no |
| TRPC6 | 721 | N | 0 | 0 | yes | no | no | no | no | no | no | no | yes | no | no | no | no |
| TRPC6 | 722 | M | 2 | 0 | yes | no | no | yes | no | no | no | no | no | no | no | no | no |
| TRPC6 | 723 | L | 0 | 0 | yes | no | no | yes | no | no | yes | no | no | no | no | no | no |
| TRPC6 | 724 | I | 0 | 0 | yes | no | no | no | no | no | no | no | yes | no | no | no | no |
| TRPC6 | 725 | A | 0 | 0 | yes | no | no | no | no | no | no | no | no | no | no | no | no |
| TRPC6 | 726 | M | 1 | 0 | yes | no | no | no | no | no | no | no | yes | no | no | no | no |
| TRPC6 | 727 | I | 0 | 0 | yes | no | no | no | no | no | yes | no | yes | no | no | no | no |
| TRPC6 | 728 | N | 0 | 0 | yes | no | no | no | no | yes | no | no | yes | no | no | no | no |
| TRPC6 | 729 | S | 0 | 0 | no | yes | no | no | no | yes | no | no | no | no | no | no | no |
| TRPC6 | 730 | S | 0 | 0 | no | yes | no | no | no | yes | no | no | yes | no | no | no | no |
| TRPC6 | 731 | F | 0 | 0 | no | yes | no | no | no | yes | yes | no | yes | no | no | no | no |
| TRPC6 | 732 | Q | 0 | 0 | no | yes | no | no | no | yes | no | no | no | no | no | no | no |
| TRPC6 | 733 | E | 0 | 0 | no | yes | no | no | no | yes | no | no | no | no | no | no | no |
| TRPC6 | 734 | I | 0 | 0 | no | yes | no | no | no | yes | no | no | no | no | no | no | no |
| TRPC6 | 735 | E | 0 | 0 | no | yes | no | no | no | yes | no | no | no | no | no | no | no |
| TRPC6 | 736 | D | 0 | 0 | no | yes | no | no | no | yes | no | no | no | no | no | no | no |
| TRPC6 | 737 | D | 0 | 0 | yes | no | no | no | no | yes | no | no | no | no | no | no | no |
| TRPC6 | 738 | A | 0 | 0 | yes | no | no | no | no | yes | no | no | no | no | no | no | no |
| TRPC6 | 739 | D | 0 | 0 | yes | no | no | no | no | yes | no | no | no | no | no | no | no |
| TRPC6 | 740 | V | 0 | 0 | yes | no | no | no | no | yes | no | no | no | no | no | no | no |
| TRPC6 | 741 | E | 0 | 0 | yes | no | no | no | no | yes | no | no | no | no | no | no | no |
| TRPC6 | 742 | W | 0 | 0 | yes | no | no | no | no | yes | no | no | no | no | no | no | no |
| TRPC6 | 743 | K | 0 | 0 | yes | no | no | no | no | yes | no | no | no | no | no | no | no |
| TRPC6 | 744 | F | 0 | 0 | yes | no | no | no | no | yes | no | no | no | no | no | no | no |
| TRPC6 | 745 | A | 1 | 0 | yes | no | no | no | no | yes | no | no | no | no | no | no | no |
| TRPC6 | 746 | R | 0 | 0 | yes | no | no | no | no | yes | no | no | no | no | no | no | no |
| TRPC6 | 747 | A | 1 | 0 | yes | no | no | no | no | yes | no | no | no | no | no | no | no |
| TRPC6 | 748 | K | 1 | 0 | yes | no | no | no | no | yes | no | no | no | no | no | no | no |

| | | | | | | | | | | | | | | | | | |
|---|---|---|---|---|---|---|---|---|---|---|---|---|---|---|---|---|---|
| TRPC6 | 749 | L | 0 | 0 | yes | no | no | no | no | yes | no | no | no | no | no | no | no |
| TRPC6 | 750 | W | 0 | 0 | yes | no | no | no | no | yes | no | no | no | no | no | no | no |
| TRPC6 | 751 | F | 0 | 0 | no | yes | no | no | no | yes | no | no | no | no | no | no | no |
| TRPC6 | 752 | S | 0 | 0 | no | yes | no | no | no | yes | no | no | no | no | no | no | no |
| TRPC6 | 753 | Y | 1 | 0 | no | yes | no | no | no | yes | no | no | no | no | no | no | no |
| TRPC6 | 754 | F | 0 | 0 | no | yes | no | no | no | yes | no | no | no | no | no | no | no |
| TRPC6 | 755 | E | 1 | 0 | no | yes | no | no | no | yes | no | no | no | no | no | no | no |
| TRPC6 | 756 | E | 1 | 0 | no | yes | no | no | no | yes | no | no | no | no | no | no | no |
| TRPC6 | 757 | G | 0 | 1 | no | yes | no | no | no | yes | no | no | no | no | no | no | no |
| TRPC6 | 758 | R | 0 | 0 | no | yes | no | no | no | yes | no | no | no | no | no | no | no |
| TRPC6 | 759 | T | 0 | 0 | no | yes | no | no | no | yes | no | no | no | no | no | no | no |
| TRPC6 | 760 | L | 1 | 0 | no | yes | no | yes | no | yes | no | no | no | no | no | no | no |
| TRPC6 | 761 | P | 2 | 0 | no | yes | no | yes | no | yes | no | no | no | no | no | no | no |
| TRPC6 | 762 | V | 0 | 1 | no | yes | no | yes | no | yes | no | no | no | no | no | no | no |
| TRPC6 | 763 | P | 0 | 0 | no | yes | no | yes | no | yes | no | no | no | no | no | no | no |
| TRPC6 | 764 | F | 0 | 0 | no | yes | no | yes | no | yes | no | no | no | no | no | no | no |
| TRPC6 | 765 | N | 1 | 0 | no | yes | no | yes | no | yes | no | no | no | no | no | no | no |
| TRPC6 | 766 | L | 1 | 0 | no | yes | no | yes | no | yes | no | no | no | no | no | no | no |
| TRPC6 | 767 | V | 1 | 0 | no | yes | no | yes | no | yes | no | no | no | no | no | no | no |
| TRPC6 | 853 | R | 0 | 0 | no | yes | no | no | no | yes | no | no | no | no | no | no | no |
| TRPC6 | 854 | Q | 0 | 0 | yes | no | no | no | no | yes | no | no | no | no | no | no | no |
| TRPC6 | 855 | Y | 0 | 0 | yes | no | no | no | no | yes | no | no | no | no | no | no | no |
| TRPC6 | 856 | Q | 0 | 0 | yes | no | no | no | no | yes | no | no | no | no | no | no | no |
| TRPC6 | 857 | K | 0 | 0 | yes | no | no | no | no | yes | no | no | no | no | no | no | no |
| TRPC6 | 858 | I | 0 | 0 | yes | no | no | no | no | yes | no | no | no | no | no | no | no |
| TRPC6 | 859 | M | 1 | 0 | yes | no | no | no | no | yes | no | no | no | no | no | no | no |
| TRPC6 | 860 | K | 0 | 0 | yes | no | no | no | no | yes | no | no | yes | no | no | no | no |
| TRPC6 | 861 | R | 0 | 0 | yes | no | no | no | no | yes | no | no | yes | no | no | no | no |
| TRPC6 | 862 | L | 0 | 0 | yes | no | no | no | no | yes | no | no | no | no | no | no | no |
| TRPC6 | 863 | I | 1 | 0 | yes | no | no | no | no | yes | no | no | yes | no | no | no | no |
| TRPC6 | 864 | K | 1 | 0 | yes | no | no | no | no | yes | no | no | yes | no | no | no | no |
| TRPC6 | 865 | R | 0 | 0 | yes | no | no | no | no | yes | no | no | no | no | no | no | no |
| TRPC6 | 866 | Y | 0 | 0 | yes | no | no | no | no | yes | no | no | no | no | no | no | no |

| | | | | | | | | | | | | | | | | | |
|---|---|---|---|---|---|---|---|---|---|---|---|---|---|---|---|---|---|
| TRPC6 | 867 | V | 0 | 0 | yes | no | no | no | no | yes | no | no | yes | no | no | no | no |
| TRPC6 | 868 | L | 1 | 0 | yes | no | no | no | no | yes | no | no | yes | no | no | no | no |
| TRPC6 | 869 | Q | 0 | 0 | yes | no | no | no | no | yes | no | no | no | no | no | no | no |
| TRPC6 | 870 | A | 0 | 0 | yes | no | no | no | no | yes | no | no | no | no | no | no | no |
| TRPC6 | 871 | Q | 0 | 0 | yes | no | no | no | no | yes | no | no | no | no | no | no | no |
| TRPC6 | 872 | I | 2 | 0 | yes | no | no | no | no | yes | no | no | no | no | no | no | no |
| TRPC6 | 873 | D | 0 | 0 | yes | no | no | no | no | yes | no | no | no | no | no | no | no |
| TRPC6 | 874 | K | 0 | 0 | yes | no | no | no | no | yes | no | no | yes | no | no | no | no |
| TRPC6 | 875 | E | 0 | 0 | no | yes | no | no | no | yes | no | no | yes | no | no | no | no |
| TRPC6 | 876 | S | 0 | 0 | no | yes | no | no | no | yes | no | no | yes | no | no | no | no |
| TRPC6 | 877 | D | 0 | 0 | no | yes | no | no | no | yes | no | no | yes | no | no | no | no |
| TRPC6 | 878 | E | 2 | 0 | no | no | yes | no | no | yes | yes | no | no | no | no | no | no |
| TRPC6 | 879 | V | 0 | 0 | no | yes | no | no | no | yes | yes | no | yes | no | no | no | no |
| TRPC6 | 880 | N | 1 | 0 | no | no | yes | no | no | yes | no | no | yes | no | no | no | no |
| TRPC6 | 881 | E | 0 | 0 | yes | no | no | no | no | yes | no | no | yes | no | no | no | no |
| TRPC6 | 882 | G | 0 | 0 | yes | no | no | no | no | yes | no | no | yes | no | no | no | no |
| TRPC6 | 883 | E | 0 | 0 | yes | no | no | no | no | yes | no | no | yes | no | no | no | no |
| TRPC6 | 884 | L | 0 | 0 | yes | no | no | no | no | yes | yes | no | yes | no | no | no | no |
| TRPC6 | 885 | K | 0 | 0 | yes | no | no | no | no | yes | no | no | yes | no | no | no | no |
| TRPC6 | 886 | E | 0 | 1 | yes | no | no | no | no | yes | no | no | yes | no | no | no | no |
| TRPC6 | 887 | I | 1 | 0 | yes | no | no | no | no | yes | yes | no | yes | no | no | no | no |
| TRPC6 | 888 | K | 0 | 0 | yes | no | no | no | no | yes | no | no | yes | no | no | no | no |
| TRPC6 | 889 | Q | 0 | 1 | yes | no | no | no | no | yes | no | no | no | no | no | no | no |
| TRPC6 | 890 | D | 0 | 1 | yes | no | no | no | no | yes | no | no | yes | no | no | no | no |
| TRPC6 | 891 | I | 1 | 0 | yes | no | no | no | no | yes | yes | no | yes | no | no | no | no |
| TRPC6 | 892 | S | 0 | 0 | yes | no | no | no | no | yes | no | no | no | no | no | no | no |
| TRPC6 | 893 | S | 0 | 0 | yes | no | no | no | no | yes | no | no | no | no | no | no | no |
| TRPC6 | 894 | L | 0 | 0 | yes | no | no | no | no | yes | yes | no | yes | no | no | no | no |
| TRPC6 | 895 | R | 0 | 2 | yes | no | no | no | no | yes | no | no | yes | no | no | no | no |
| TRPC6 | 896 | Y | 0 | 0 | yes | no | no | no | no | yes | no | no | yes | no | no | no | no |
| TRPC6 | 897 | E | 0 | 2 | yes | no | no | no | no | yes | no | no | yes | no | no | no | no |
| TRPC6 | 898 | L | 0 | 0 | yes | no | no | no | no | yes | yes | no | yes | no | no | no | no |
| TRPC6 | 899 | L | 0 | 0 | yes | no | no | no | no | yes | no | no | yes | no | no | no | no |

| | | | | | | | | | | | | | | | | | |
|---|---|---|---|---|---|---|---|---|---|---|---|---|---|---|---|---|---|
| TRPC6 | 900 | E | 0 | 0 | yes | no | no | no | no | yes | no | no | no | no | no | no | no |
| TRPC6 | 901 | E | 1 | 0 | yes | no | no | no | no | yes | no | no | no | no | no | no | no |
| TRPC6 | 902 | K | 1 | 0 | yes | no | no | no | no | yes | yes | no | no | no | no | no | no |
| TRPC6 | 903 | S | 1 | 0 | yes | no | no | no | no | yes | no | no | no | no | no | no | no |
| TRPC6 | 904 | Q | 1 | 0 | yes | no | no | no | no | yes | no | no | no | no | no | no | no |
| TRPC6 | 905 | N | 0 | 0 | yes | no | no | no | no | yes | yes | no | no | no | no | no | no |
| TRPC6 | 906 | T | 0 | 0 | yes | no | no | no | no | yes | no | no | no | no | no | no | no |
| TRPC6 | 907 | E | 0 | 0 | yes | no | no | no | no | yes | no | no | no | no | no | no | no |
| TRPC6 | 908 | D | 1 | 0 | yes | no | no | no | no | yes | no | no | no | no | no | no | no |
| TRPC6 | 909 | L | 0 | 0 | yes | no | no | no | no | yes | yes | no | no | no | no | no | no |
| TRPC6 | 910 | A | 0 | 0 | yes | no | no | no | no | yes | no | no | no | no | no | no | no |
| TRPC6 | 911 | E | 1 | 0 | yes | no | no | no | no | yes | no | no | no | no | no | no | no |
| TRPC6 | 912 | L | 0 | 0 | yes | no | no | no | no | yes | no | no | yes | no | no | no | no |
| TRPC6 | 913 | I | 2 | 0 | yes | no | no | no | no | yes | yes | no | yes | no | no | no | no |
| TRPC6 | 914 | R | 0 | 0 | yes | no | no | no | no | yes | no | no | no | no | no | no | no |
| TRPC6 | 915 | E | 0 | 0 | yes | no | no | no | no | yes | no | no | no | no | no | no | no |
| TRPC6 | 916 | L | 0 | 0 | yes | no | no | no | no | yes | yes | no | yes | no | no | no | no |
| TRPC6 | 917 | G | 0 | 0 | yes | no | no | no | no | yes | no | no | no | no | no | no | no |
| TRPC6 | 918 | E | 0 | 0 | no | yes | no | no | no | yes | no | no | no | no | no | no | no |
| TRPC6 | 919 | K | 0 | 0 | no | yes | no | no | no | yes | no | no | yes | no | no | no | no |
| TRPC6 | 920 | L | 0 | 0 | no | yes | no | no | no | yes | yes | no | yes | no | no | no | no |
| TRPM4 | 7 | E | 0 | 1 | no | yes | no | no | no | yes | no | no | no | no | no | no | no |
| TRPM4 | 8 | Q | 0 | 0 | no | yes | no | no | no | yes | no | no | no | no | no | no | no |
| TRPM4 | 9 | S | 1 | 0 | no | yes | no | no | no | yes | no | no | no | no | no | no | no |
| TRPM4 | 10 | W | 1 | 0 | no | yes | no | no | no | yes | no | no | no | no | no | no | no |
| TRPM4 | 11 | I | 1 | 0 | yes | no | no | no | no | yes | no | no | no | no | no | no | no |
| TRPM4 | 12 | P | 1 | 0 | yes | no | no | no | no | yes | no | no | no | no | no | no | no |
| TRPM4 | 13 | K | 2 | 0 | yes | no | no | no | no | yes | no | no | no | no | no | no | no |
| TRPM4 | 14 | I | 1 | 0 | yes | no | no | no | no | yes | no | no | no | no | no | no | no |
| TRPM4 | 15 | F | 0 | 0 | no | yes | no | no | no | yes | no | no | no | no | no | no | no |
| TRPM4 | 16 | K | 1 | 0 | no | no | yes | no | no | yes | no | no | no | no | no | no | no |
| TRPM4 | 17 | K | 0 | 0 | no | no | yes | no | no | yes | no | no | no | no | no | no | no |
| TRPM4 | 18 | K | 0 | 0 | no | yes | no | no | no | yes | no | no | no | no | no | no | no |

| | | | | | | | | | | | | | | | | | |
|---|---|---|---|---|---|---|---|---|---|---|---|---|---|---|---|---|---|
| TRPM4 | 19 | T | 0 | 0 | no | yes | no | no | no | yes | no | no | no | no | no | no | no |
| TRPM4 | 66 | H | 0 | 0 | no | yes | no | no | no | yes | no | no | no | no | no | no | no |
| TRPM4 | 67 | T | 1 | 0 | no | yes | no | no | no | yes | no | no | no | no | no | no | no |
| TRPM4 | 68 | T | 1 | 0 | no | yes | no | no | no | yes | no | no | no | no | no | no | no |
| TRPM4 | 69 | E | 1 | 0 | no | yes | no | no | no | yes | no | no | no | no | no | no | no |
| TRPM4 | 70 | K | 1 | 0 | no | no | yes | no | no | yes | no | no | no | no | no | no | no |
| TRPM4 | 71 | P | 2 | 0 | no | no | yes | no | no | yes | no | no | no | no | no | no | no |
| TRPM4 | 72 | T | 1 | 0 | no | yes | no | no | no | yes | no | no | no | no | no | no | no |
| TRPM4 | 73 | D | 3 | 0 | no | yes | no | no | no | yes | no | no | no | no | no | no | no |
| TRPM4 | 74 | A | 0 | 0 | no | yes | no | no | no | yes | no | no | no | no | no | no | no |
| TRPM4 | 75 | Y | 1 | 0 | no | no | yes | no | no | yes | no | no | no | no | no | no | no |
| TRPM4 | 76 | G | 1 | 0 | no | no | yes | no | no | yes | no | no | no | no | no | no | no |
| TRPM4 | 77 | E | 0 | 0 | no | no | yes | no | no | yes | no | no | no | no | no | no | no |
| TRPM4 | 78 | L | 0 | 0 | no | no | yes | no | no | yes | no | no | no | no | no | no | no |
| TRPM4 | 79 | D | 1 | 0 | no | yes | no | no | no | yes | no | no | no | no | no | no | no |
| TRPM4 | 80 | F | 1 | 0 | no | yes | no | no | no | yes | no | no | no | no | no | no | no |
| TRPM4 | 81 | T | 0 | 0 | no | yes | no | no | no | yes | no | no | yes | no | no | no | no |
| TRPM4 | 82 | G | 0 | 0 | no | yes | no | no | no | yes | no | no | no | no | no | no | no |
| TRPM4 | 83 | A | 2 | 0 | no | yes | no | no | no | yes | no | no | no | no | no | no | no |
| TRPM4 | 84 | G | 1 | 0 | no | yes | no | no | no | yes | no | no | no | no | no | no | no |
| TRPM4 | 85 | R | 2 | 0 | no | yes | no | no | no | yes | no | no | no | no | no | no | no |
| TRPM4 | 86 | K | 0 | 0 | no | yes | no | no | no | yes | no | no | no | no | no | no | no |
| TRPM4 | 87 | H | 0 | 0 | no | yes | no | no | no | yes | no | no | no | no | no | no | no |
| TRPM4 | 88 | S | 0 | 0 | no | no | yes | no | no | yes | no | no | no | no | no | no | no |
| TRPM4 | 89 | N | 0 | 0 | no | no | yes | no | no | yes | no | no | no | no | no | no | no |
| TRPM4 | 90 | F | 0 | 0 | no | no | yes | no | no | yes | no | no | no | no | no | no | no |
| TRPM4 | 91 | L | 0 | 0 | no | no | yes | no | no | yes | no | no | no | no | no | no | no |
| TRPM4 | 92 | R | 1 | 0 | no | no | yes | no | no | yes | no | no | no | no | no | no | no |
| TRPM4 | 93 | L | 0 | 0 | no | no | yes | no | no | yes | no | no | no | no | no | no | no |
| TRPM4 | 94 | S | 0 | 0 | no | no | yes | no | no | yes | no | no | no | no | no | no | no |
| TRPM4 | 95 | D | 0 | 0 | no | yes | no | no | no | yes | no | no | no | no | no | no | no |
| TRPM4 | 96 | R | 1 | 0 | no | yes | no | no | no | yes | no | no | no | no | no | no | no |
| TRPM4 | 97 | T | 2 | 0 | no | yes | no | no | no | yes | no | no | no | no | no | no | no |

| | | | | | | | | | | | | | | | | | |
|---|---|---|---|---|---|---|---|---|---|---|---|---|---|---|---|---|---|
| TRPM4 | 98 | D | 0 | 0 | no | yes | no | no | no | yes | no | no | no | no | no | no | no |
| TRPM4 | 99 | P | 0 | 0 | yes | no | no | no | no | yes | no | no | no | no | no | no | no |
| TRPM4 | 100 | A | 1 | 0 | yes | no | no | no | no | yes | no | no | no | no | no | no | no |
| TRPM4 | 101 | A | 1 | 0 | yes | no | no | no | no | yes | no | no | no | no | no | no | no |
| TRPM4 | 102 | V | 0 | 0 | yes | no | no | no | no | yes | no | no | no | no | no | no | no |
| TRPM4 | 103 | Y | 1 | 0 | yes | no | no | no | no | yes | no | no | no | no | no | no | no |
| TRPM4 | 104 | S | 0 | 0 | yes | no | no | no | no | yes | no | no | no | no | no | no | no |
| TRPM4 | 105 | L | 0 | 0 | yes | no | no | no | no | yes | no | no | no | no | no | no | no |
| TRPM4 | 106 | V | 1 | 0 | yes | no | no | no | no | yes | no | no | no | no | no | no | no |
| TRPM4 | 107 | T | 0 | 0 | yes | no | no | no | no | yes | no | no | no | no | no | no | no |
| TRPM4 | 108 | R | 2 | 0 | yes | no | no | no | no | yes | no | no | no | no | no | no | no |
| TRPM4 | 109 | T | 1 | 0 | yes | no | no | no | no | yes | no | no | no | no | no | no | no |
| TRPM4 | 110 | W | 1 | 0 | no | yes | no | no | no | yes | no | no | no | no | no | no | no |
| TRPM4 | 111 | G | 1 | 0 | no | yes | no | no | no | yes | no | no | no | no | no | no | no |
| TRPM4 | 112 | F | 0 | 0 | no | yes | no | no | no | yes | no | no | no | no | no | no | no |
| TRPM4 | 113 | R | 3 | 0 | no | yes | no | no | no | yes | no | no | no | no | no | no | no |
| TRPM4 | 114 | A | 2 | 0 | no | yes | no | no | no | yes | no | no | no | no | no | no | no |
| TRPM4 | 115 | P | 2 | 0 | no | yes | no | no | no | yes | no | no | no | no | no | no | no |
| TRPM4 | 116 | N | 0 | 0 | no | yes | no | no | no | yes | no | no | no | no | no | no | no |
| TRPM4 | 117 | L | 0 | 0 | no | no | yes | no | no | yes | no | no | no | no | no | no | no |
| TRPM4 | 118 | V | 1 | 0 | no | no | yes | no | no | yes | no | no | no | no | no | no | no |
| TRPM4 | 119 | V | 0 | 0 | no | no | yes | no | no | yes | no | no | no | no | no | no | no |
| TRPM4 | 120 | S | 0 | 0 | no | no | yes | no | no | yes | no | no | no | no | no | no | no |
| TRPM4 | 121 | V | 1 | 0 | no | no | yes | no | no | yes | no | no | no | no | no | no | no |
| TRPM4 | 122 | L | 1 | 0 | no | yes | no | no | no | yes | no | no | no | no | no | no | no |
| TRPM4 | 123 | G | 0 | 0 | no | yes | no | no | no | yes | no | no | no | no | no | no | no |
| TRPM4 | 124 | G | 1 | 0 | no | yes | no | no | no | yes | no | no | no | no | no | no | no |
| TRPM4 | 125 | S | 1 | 0 | no | yes | no | no | no | yes | no | no | no | no | no | no | no |
| TRPM4 | 126 | G | 0 | 0 | no | yes | no | no | no | yes | no | no | no | no | no | no | no |
| TRPM4 | 127 | G | 2 | 0 | no | yes | no | no | no | yes | no | no | no | no | no | no | no |
| TRPM4 | 128 | P | 1 | 0 | no | yes | no | no | no | yes | no | no | no | no | no | no | no |
| TRPM4 | 129 | V | 1 | 0 | no | yes | no | no | no | yes | no | no | yes | no | no | no | no |
| TRPM4 | 130 | L | 0 | 0 | no | yes | no | no | no | yes | no | no | no | no | no | no | no |

| | | | | | | | | | | | | | | | | | |
|---|---|---|---|---|---|---|---|---|---|---|---|---|---|---|---|---|---|
| TRPM4 | 131 | Q | 2 | 1 | no | yes | no | no | no | yes | no | no | no | no | no | no | no |
| TRPM4 | 132 | T | 1 | 0 | yes | no | no | no | no | yes | no | no | no | no | no | no | no |
| TRPM4 | 133 | W | 0 | 0 | yes | no | no | no | no | yes | no | no | no | no | no | no | no |
| TRPM4 | 134 | L | 0 | 0 | yes | no | no | no | no | yes | no | no | no | no | no | no | no |
| TRPM4 | 135 | Q | 0 | 0 | yes | no | no | no | no | yes | no | no | no | no | no | no | no |
| TRPM4 | 136 | D | 2 | 0 | yes | no | no | no | no | yes | no | no | no | no | no | no | no |
| TRPM4 | 137 | L | 0 | 0 | yes | no | no | no | no | yes | no | no | no | no | no | no | no |
| TRPM4 | 138 | L | 0 | 0 | yes | no | no | no | no | yes | no | no | no | no | no | no | no |
| TRPM4 | 139 | R | 2 | 0 | yes | no | no | no | no | yes | no | no | no | no | no | no | no |
| TRPM4 | 140 | R | 1 | 0 | yes | no | no | no | no | yes | no | no | no | no | no | no | no |
| TRPM4 | 141 | G | 1 | 0 | yes | no | no | no | no | yes | no | no | no | no | no | no | no |
| TRPM4 | 142 | L | 0 | 0 | yes | no | no | no | no | yes | no | no | no | no | no | no | no |
| TRPM4 | 143 | V | 0 | 0 | yes | no | no | no | no | yes | no | no | no | no | no | no | no |
| TRPM4 | 144 | R | 2 | 0 | yes | no | no | no | no | yes | no | no | no | no | no | no | no |
| TRPM4 | 145 | A | 2 | 0 | yes | no | no | no | no | yes | no | no | no | no | no | no | no |
| TRPM4 | 146 | A | 2 | 0 | yes | no | no | no | no | yes | no | no | no | no | no | no | no |
| TRPM4 | 147 | Q | 1 | 0 | yes | no | no | no | no | yes | no | no | no | no | no | no | no |
| TRPM4 | 148 | S | 1 | 0 | yes | no | no | no | no | yes | no | no | no | no | no | no | no |
| TRPM4 | 149 | T | 0 | 0 | no | yes | no | no | no | yes | no | no | no | no | no | no | no |
| TRPM4 | 150 | G | 0 | 0 | no | yes | no | no | no | yes | no | no | no | no | no | no | no |
| TRPM4 | 151 | A | 0 | 0 | no | no | yes | no | no | yes | no | no | no | no | no | no | no |
| TRPM4 | 152 | W | 0 | 0 | no | no | yes | no | no | yes | no | no | no | no | no | no | no |
| TRPM4 | 153 | I | 0 | 0 | no | no | yes | no | no | yes | no | no | no | no | no | no | no |
| TRPM4 | 154 | V | 0 | 0 | no | no | yes | no | no | yes | no | no | no | no | no | no | no |
| TRPM4 | 155 | T | 1 | 0 | no | no | yes | no | no | yes | no | no | no | no | no | no | no |
| TRPM4 | 156 | G | 0 | 0 | no | no | yes | no | no | yes | no | no | no | no | no | no | no |
| TRPM4 | 157 | G | 0 | 0 | no | yes | no | no | no | yes | no | no | no | no | no | no | no |
| TRPM4 | 158 | L | 1 | 0 | no | no | yes | no | no | yes | no | no | no | no | no | no | no |
| TRPM4 | 159 | H | 1 | 0 | no | yes | no | no | no | yes | no | no | no | no | no | no | no |
| TRPM4 | 160 | T | 2 | 0 | no | yes | no | no | no | yes | no | no | no | no | no | no | no |
| TRPM4 | 161 | G | 0 | 0 | no | yes | no | no | no | yes | no | no | no | no | no | no | no |
| TRPM4 | 162 | I | 0 | 0 | no | yes | no | no | no | yes | no | no | no | no | no | no | no |
| TRPM4 | 163 | G | 2 | 0 | yes | no | no | no | no | yes | no | no | no | no | no | no | no |

| | | | | | | | | | | | | | | | | | |
|---|---|---|---|---|---|---|---|---|---|---|---|---|---|---|---|---|---|
| TRPM4 | 164 | R | 2 | 1 | yes | no | no | no | no | yes | no | no | no | no | no | no | no |
| TRPM4 | 165 | H | 0 | 0 | yes | no | no | no | no | yes | no | no | no | no | no | no | no |
| TRPM4 | 166 | V | 0 | 0 | yes | no | no | no | no | yes | no | no | no | no | no | no | no |
| TRPM4 | 167 | G | 1 | 0 | yes | no | no | no | no | yes | no | no | no | no | no | no | no |
| TRPM4 | 168 | V | 1 | 1 | yes | no | no | no | no | yes | no | no | no | no | no | no | no |
| TRPM4 | 169 | A | 0 | 0 | yes | no | no | no | no | yes | no | no | no | no | no | no | no |
| TRPM4 | 170 | V | 1 | 0 | yes | no | no | no | no | yes | no | no | no | no | no | no | no |
| TRPM4 | 171 | R | 0 | 0 | yes | no | no | no | no | yes | no | no | yes | no | no | no | yes |
| TRPM4 | 172 | D | 0 | 0 | yes | no | no | no | no | yes | no | no | no | no | no | no | no |
| TRPM4 | 173 | H | 0 | 0 | yes | no | no | no | no | yes | no | no | no | no | no | no | no |
| TRPM4 | 174 | Q | 0 | 0 | yes | no | no | no | no | yes | no | no | no | no | no | no | no |
| TRPM4 | 175 | M | 0 | 0 | yes | no | no | no | no | yes | no | no | no | no | no | no | no |
| TRPM4 | 176 | A | 1 | 0 | yes | no | no | no | no | yes | no | no | no | no | no | no | no |
| TRPM4 | 177 | S | 1 | 0 | no | yes | no | no | no | yes | no | no | yes | no | no | no | no |
| TRPM4 | 178 | T | 0 | 0 | no | yes | no | no | no | yes | no | no | no | no | no | no | no |
| TRPM4 | 179 | G | 1 | 0 | no | yes | no | no | no | yes | no | no | no | no | no | no | no |
| TRPM4 | 180 | G | 1 | 0 | no | yes | no | no | no | yes | no | no | no | no | no | no | no |
| TRPM4 | 181 | T | 0 | 0 | no | yes | no | no | no | yes | no | no | no | no | no | no | no |
| TRPM4 | 182 | K | 1 | 0 | no | yes | no | no | no | yes | no | no | no | no | no | no | no |
| TRPM4 | 183 | V | 0 | 0 | no | yes | no | no | no | yes | no | no | no | no | no | no | no |
| TRPM4 | 184 | V | 0 | 0 | no | yes | no | no | no | yes | no | no | no | no | no | no | no |
| TRPM4 | 185 | A | 0 | 0 | no | yes | no | no | no | yes | no | no | no | no | no | no | no |
| TRPM4 | 186 | M | 1 | 0 | no | no | yes | no | no | yes | no | no | no | no | no | no | no |
| TRPM4 | 187 | G | 0 | 0 | no | no | yes | no | no | yes | no | no | no | no | no | no | no |
| TRPM4 | 188 | V | 0 | 0 | no | no | yes | no | no | yes | no | no | no | no | no | no | no |
| TRPM4 | 189 | A | 1 | 0 | no | no | yes | no | no | yes | no | no | no | no | no | no | no |
| TRPM4 | 190 | P | 0 | 0 | no | no | yes | no | no | yes | no | no | no | no | no | no | no |
| TRPM4 | 191 | W | 0 | 0 | no | yes | no | no | no | yes | no | no | no | no | no | no | no |
| TRPM4 | 192 | G | 2 | 0 | no | yes | no | no | no | yes | no | no | no | no | no | no | no |
| TRPM4 | 193 | V | 0 | 0 | no | yes | no | no | no | yes | no | no | no | no | no | no | no |
| TRPM4 | 194 | V | 1 | 0 | no | no | yes | no | no | yes | no | no | no | no | no | no | no |
| TRPM4 | 195 | R | 2 | 0 | no | yes | no | no | no | yes | no | no | no | no | no | no | no |
| TRPM4 | 196 | N | 0 | 0 | no | yes | no | no | no | yes | no | no | no | no | no | no | no |

| | | | | | | | | | | | | | | | | | |
|---|---|---|---|---|---|---|---|---|---|---|---|---|---|---|---|---|---|
| TRPM4 | 197 | R | 1 | 0 | no | yes | no | no | no | yes | no | no | no | no | no | no | no |
| TRPM4 | 198 | D | 1 | 0 | no | yes | no | no | no | yes | no | no | no | no | no | no | no |
| TRPM4 | 199 | T | 1 | 0 | no | yes | no | no | no | yes | no | no | no | no | no | no | no |
| TRPM4 | 200 | L | 1 | 0 | no | yes | no | no | no | yes | no | no | no | no | no | no | no |
| TRPM4 | 201 | I | 0 | 0 | no | yes | no | no | no | yes | no | no | no | no | no | no | no |
| TRPM4 | 202 | N | 0 | 0 | no | yes | no | no | no | yes | no | no | no | no | no | no | no |
| TRPM4 | 203 | P | 0 | 0 | no | yes | no | no | no | yes | no | no | no | no | no | no | no |
| TRPM4 | 204 | K | 0 | 0 | no | yes | no | no | no | yes | no | no | no | no | no | no | no |
| TRPM4 | 205 | G | 0 | 0 | no | yes | no | no | no | yes | no | no | no | no | no | no | no |
| TRPM4 | 206 | S | 1 | 0 | no | yes | no | no | no | yes | no | no | no | no | no | no | no |
| TRPM4 | 207 | F | 1 | 0 | no | yes | no | no | no | yes | no | no | yes | no | no | no | no |
| TRPM4 | 208 | P | 2 | 0 | no | yes | no | no | no | yes | no | no | no | no | no | no | no |
| TRPM4 | 209 | A | 1 | 0 | no | yes | no | no | no | yes | no | no | no | no | no | no | no |
| TRPM4 | 210 | R | 0 | 0 | no | yes | no | no | no | yes | no | no | no | no | no | no | no |
| TRPM4 | 211 | Y | 0 | 0 | no | yes | no | no | no | yes | no | no | no | no | no | no | no |
| TRPM4 | 212 | R | 2 | 0 | no | yes | no | no | no | yes | no | no | yes | no | no | no | no |
| TRPM4 | 213 | W | 0 | 0 | no | yes | no | no | no | yes | no | no | yes | no | no | no | no |
| TRPM4 | 214 | R | 2 | 0 | no | yes | no | no | no | yes | no | no | yes | no | no | no | no |
| TRPM4 | 215 | G | 1 | 0 | no | yes | no | no | no | yes | no | no | no | no | no | no | no |
| TRPM4 | 221 | V | 0 | 0 | no | yes | no | no | no | yes | no | no | no | no | no | no | no |
| TRPM4 | 222 | Q | 0 | 0 | no | yes | no | no | no | yes | no | no | no | no | no | no | no |
| TRPM4 | 223 | F | 1 | 0 | no | no | yes | no | no | yes | no | no | no | no | no | no | no |
| TRPM4 | 224 | P | 2 | 0 | no | yes | no | no | no | yes | no | no | no | no | no | no | no |
| TRPM4 | 225 | L | 0 | 0 | no | no | yes | no | no | yes | no | no | no | no | no | no | no |
| TRPM4 | 226 | D | 2 | 0 | no | yes | no | no | no | yes | no | no | no | no | no | no | no |
| TRPM4 | 227 | Y | 1 | 0 | no | yes | no | no | no | yes | no | no | yes | no | no | no | no |
| TRPM4 | 228 | N | 0 | 0 | no | yes | no | no | no | yes | no | no | no | no | no | no | no |
| TRPM4 | 229 | Y | 0 | 0 | no | yes | no | no | no | yes | no | no | no | no | no | no | no |
| TRPM4 | 230 | S | 1 | 0 | no | yes | no | no | no | yes | no | no | no | no | no | no | no |
| TRPM4 | 231 | A | 1 | 0 | no | no | yes | no | no | yes | no | no | no | no | no | no | no |
| TRPM4 | 232 | F | 0 | 0 | no | no | yes | no | no | yes | no | no | no | no | no | no | no |
| TRPM4 | 233 | F | 1 | 0 | no | no | yes | no | no | yes | no | no | no | no | no | no | no |
| TRPM4 | 234 | L | 0 | 0 | no | no | yes | no | no | yes | no | no | no | no | no | no | no |

| | | | | | | | | | | | | | | | | | |
|---|---|---|---|---|---|---|---|---|---|---|---|---|---|---|---|---|---|
| TRPM4 | 235 | V | 0 | 0 | no | no | yes | no | no | yes | no | no | no | no | no | no | no |
| TRPM4 | 236 | D | 2 | 0 | no | no | yes | no | no | yes | no | no | no | no | no | no | no |
| TRPM4 | 237 | D | 0 | 0 | no | yes | no | no | no | yes | no | no | no | no | no | no | no |
| TRPM4 | 238 | G | 0 | 0 | no | yes | no | no | no | yes | no | no | no | no | no | no | no |
| TRPM4 | 239 | T | 0 | 0 | no | yes | no | no | no | yes | no | no | no | no | no | no | no |
| TRPM4 | 240 | H | 0 | 0 | no | yes | no | no | no | yes | no | no | no | no | no | no | no |
| TRPM4 | 241 | G | 1 | 0 | no | yes | no | no | no | yes | no | no | no | no | no | no | no |
| TRPM4 | 242 | C | 0 | 0 | no | yes | no | no | no | yes | no | no | no | no | no | no | no |
| TRPM4 | 243 | L | 0 | 0 | no | yes | no | no | no | yes | no | no | no | no | no | no | no |
| TRPM4 | 244 | G | 0 | 0 | no | yes | no | no | no | yes | no | no | no | no | no | no | no |
| TRPM4 | 245 | G | 1 | 0 | no | yes | no | no | no | yes | no | no | no | no | no | no | no |
| TRPM4 | 246 | E | 1 | 0 | no | yes | no | no | no | yes | no | no | no | no | no | no | no |
| TRPM4 | 247 | N | 1 | 0 | no | yes | no | no | no | yes | no | no | no | no | no | no | no |
| TRPM4 | 248 | R | 2 | 0 | yes | no | no | no | no | yes | no | no | no | no | no | no | no |
| TRPM4 | 249 | F | 0 | 0 | yes | no | no | no | no | yes | no | no | no | no | no | no | no |
| TRPM4 | 250 | R | 2 | 0 | yes | no | no | no | no | yes | no | no | no | no | no | no | no |
| TRPM4 | 251 | L | 0 | 0 | yes | no | no | no | no | yes | no | no | no | no | no | no | no |
| TRPM4 | 252 | R | 3 | 0 | yes | no | no | no | no | yes | no | no | no | no | no | no | no |
| TRPM4 | 253 | L | 0 | 0 | yes | no | no | no | no | yes | no | no | no | no | no | no | no |
| TRPM4 | 254 | E | 1 | 0 | yes | no | no | no | no | yes | no | no | no | no | no | no | no |
| TRPM4 | 255 | S | 1 | 0 | yes | no | no | no | no | yes | no | no | no | no | no | no | no |
| TRPM4 | 256 | Y | 0 | 0 | yes | no | no | no | no | yes | no | no | no | no | no | no | no |
| TRPM4 | 257 | I | 0 | 0 | yes | no | no | no | no | yes | no | no | no | no | no | no | no |
| TRPM4 | 258 | S | 0 | 0 | no | yes | no | no | no | yes | no | no | no | no | no | no | no |
| TRPM4 | 259 | Q | 0 | 0 | no | yes | no | no | no | yes | no | no | no | no | no | no | no |
| TRPM4 | 260 | Q | 0 | 0 | no | yes | no | no | no | yes | no | no | no | no | no | no | no |
| TRPM4 | 261 | K | 0 | 0 | no | no | yes | no | no | yes | no | no | no | no | no | no | no |
| TRPM4 | 262 | T | 0 | 0 | no | yes | no | no | no | yes | no | no | no | no | no | no | no |
| TRPM4 | 263 | G | 0 | 0 | no | yes | no | no | no | yes | no | no | no | no | no | no | no |
| TRPM4 | 264 | V | 2 | 0 | no | yes | no | no | no | yes | no | no | no | no | no | no | no |
| TRPM4 | 265 | G | 0 | 0 | no | yes | no | no | no | yes | no | no | no | no | no | no | no |
| TRPM4 | 266 | G | 0 | 0 | no | yes | no | no | no | yes | no | no | no | no | no | no | no |
| TRPM4 | 267 | T | 0 | 0 | no | yes | no | no | no | yes | no | no | no | no | no | no | no |

| | | | | | | | | | | | | | | | | | |
|---|---|---|---|---|---|---|---|---|---|---|---|---|---|---|---|---|---|
| TRPM4 | 268 | G | 0 | 0 | no | yes | no | no | no | yes | no | no | no | no | no | no | no |
| TRPM4 | 269 | I | 0 | 0 | no | yes | no | no | no | yes | no | no | no | no | no | no | no |
| TRPM4 | 270 | D | 0 | 0 | no | no | yes | no | no | yes | no | no | no | no | no | no | no |
| TRPM4 | 271 | I | 1 | 0 | no | yes | no | no | no | yes | no | no | no | no | no | no | no |
| TRPM4 | 272 | P | 1 | 0 | no | yes | no | no | no | yes | no | no | no | no | no | no | no |
| TRPM4 | 273 | V | 1 | 0 | no | no | yes | no | no | yes | no | no | no | no | no | no | no |
| TRPM4 | 274 | L | 0 | 0 | no | no | yes | no | no | yes | no | no | no | no | no | no | no |
| TRPM4 | 275 | L | 1 | 0 | no | no | yes | no | no | yes | no | no | no | no | no | no | no |
| TRPM4 | 276 | L | 1 | 0 | no | no | yes | no | no | yes | no | no | no | no | no | no | no |
| TRPM4 | 277 | L | 0 | 0 | no | no | yes | no | no | yes | no | no | no | no | no | no | no |
| TRPM4 | 278 | I | 0 | 0 | no | yes | no | no | no | yes | no | no | no | no | no | no | no |
| TRPM4 | 279 | D | 0 | 0 | no | yes | no | no | no | yes | no | no | no | no | no | no | no |
| TRPM4 | 280 | G | 0 | 0 | no | yes | no | no | no | yes | no | no | no | no | no | no | no |
| TRPM4 | 281 | D | 1 | 0 | no | yes | no | no | no | yes | no | no | no | no | no | no | no |
| TRPM4 | 282 | E | 0 | 0 | no | yes | no | no | no | yes | no | no | no | no | no | no | no |
| TRPM4 | 283 | K | 0 | 0 | no | yes | no | no | no | yes | no | no | no | no | no | no | no |
| TRPM4 | 284 | M | 3 | 0 | yes | no | no | no | no | yes | no | no | no | no | no | no | no |
| TRPM4 | 285 | L | 0 | 0 | yes | no | no | no | no | yes | no | no | no | no | no | no | no |
| TRPM4 | 286 | T | 2 | 0 | yes | no | no | no | no | yes | no | no | no | no | no | no | no |
| TRPM4 | 287 | R | 2 | 0 | yes | no | no | no | no | yes | no | no | no | no | no | no | no |
| TRPM4 | 288 | I | 0 | 0 | yes | no | no | no | no | yes | no | no | no | no | no | no | no |
| TRPM4 | 289 | E | 2 | 0 | yes | no | no | no | no | yes | no | no | no | no | no | no | no |
| TRPM4 | 290 | N | 0 | 0 | yes | no | no | no | no | yes | no | no | no | no | no | no | no |
| TRPM4 | 291 | A | 1 | 0 | yes | no | no | no | no | yes | no | no | no | no | no | no | no |
| TRPM4 | 292 | T | 1 | 0 | yes | no | no | no | no | yes | no | no | no | no | no | no | no |
| TRPM4 | 293 | Q | 1 | 1 | yes | no | no | no | no | yes | no | no | no | no | no | no | no |
| TRPM4 | 294 | A | 0 | 0 | yes | no | no | no | no | yes | no | no | no | no | no | no | no |
| TRPM4 | 295 | Q | 1 | 0 | no | yes | no | no | no | yes | no | no | no | no | no | no | no |
| TRPM4 | 296 | L | 1 | 0 | no | yes | no | no | no | yes | no | no | no | no | no | no | no |
| TRPM4 | 297 | P | 0 | 0 | no | yes | no | no | no | yes | no | no | no | no | no | no | no |
| TRPM4 | 298 | C | 1 | 0 | no | no | yes | no | no | yes | no | no | no | no | no | no | no |
| TRPM4 | 299 | L | 0 | 0 | no | no | yes | no | no | yes | no | no | no | no | no | no | no |
| TRPM4 | 300 | L | 0 | 0 | no | no | yes | no | no | yes | no | no | no | no | no | no | no |

| | | | | | | | | | | | | | | | | | |
|---|---|---|---|---|---|---|---|---|---|---|---|---|---|---|---|---|---|
| TRPM4 | 301 | V | 1 | 0 | no | no | yes | no | no | yes | no | no | no | no | no | no | no |
| TRPM4 | 302 | A | 0 | 0 | no | yes | no | no | no | yes | no | no | no | no | no | no | no |
| TRPM4 | 303 | G | 1 | 0 | no | yes | no | no | no | yes | no | no | no | no | no | no | no |
| TRPM4 | 304 | S | 1 | 0 | no | yes | no | no | no | yes | no | no | no | no | no | no | no |
| TRPM4 | 305 | G | 0 | 0 | no | yes | no | no | no | yes | no | no | no | no | no | no | no |
| TRPM4 | 306 | G | 0 | 0 | no | yes | no | no | no | yes | no | no | no | no | no | no | no |
| TRPM4 | 307 | A | 1 | 0 | no | yes | no | no | no | yes | no | no | no | no | no | no | no |
| TRPM4 | 308 | A | 1 | 0 | no | yes | no | no | no | yes | no | no | no | no | no | no | no |
| TRPM4 | 309 | D | 0 | 0 | yes | no | no | no | no | yes | no | no | no | no | no | no | no |
| TRPM4 | 310 | C | 1 | 0 | yes | no | no | no | no | yes | no | no | no | no | no | no | no |
| TRPM4 | 311 | L | 0 | 0 | yes | no | no | no | no | yes | no | no | no | no | no | no | no |
| TRPM4 | 312 | A | 1 | 0 | yes | no | no | no | no | yes | no | no | no | no | no | no | no |
| TRPM4 | 313 | E | 0 | 0 | yes | no | no | no | no | yes | no | no | no | no | no | no | no |
| TRPM4 | 314 | T | 0 | 0 | yes | no | no | no | no | yes | no | no | no | no | no | no | no |
| TRPM4 | 315 | L | 1 | 0 | yes | no | no | no | no | yes | no | no | no | no | no | no | no |
| TRPM4 | 316 | E | 0 | 0 | no | yes | no | no | no | yes | no | no | no | no | no | no | no |
| TRPM4 | 317 | D | 0 | 0 | no | yes | no | no | no | yes | no | no | no | no | no | no | no |
| TRPM4 | 327 | R | 2 | 0 | no | yes | no | no | no | yes | no | no | no | no | no | no | no |
| TRPM4 | 328 | Q | 1 | 0 | yes | no | no | no | no | yes | no | no | no | no | no | no | no |
| TRPM4 | 329 | G | 0 | 0 | yes | no | no | no | no | yes | no | no | no | no | no | no | no |
| TRPM4 | 330 | E | 1 | 0 | yes | no | no | no | no | yes | no | no | no | no | no | no | no |
| TRPM4 | 331 | A | 0 | 0 | yes | no | no | no | no | yes | no | no | no | no | no | no | no |
| TRPM4 | 332 | R | 1 | 0 | yes | no | no | no | no | yes | no | no | no | no | no | no | no |
| TRPM4 | 333 | D | 0 | 0 | no | yes | no | no | no | yes | no | no | no | no | no | no | no |
| TRPM4 | 334 | R | 1 | 0 | no | yes | no | no | no | yes | no | no | no | no | no | no | no |
| TRPM4 | 335 | I | 1 | 0 | no | yes | no | no | no | yes | no | no | no | no | no | no | no |
| TRPM4 | 336 | R | 2 | 0 | no | yes | no | no | no | yes | no | no | no | no | no | no | no |
| TRPM4 | 337 | R | 2 | 0 | no | yes | no | no | no | yes | no | no | no | no | no | no | no |
| TRPM4 | 338 | F | 0 | 0 | no | yes | no | no | no | yes | no | no | no | no | no | no | no |
| TRPM4 | 339 | F | 0 | 0 | no | yes | no | no | no | yes | no | no | no | no | no | no | no |
| TRPM4 | 340 | P | 0 | 0 | no | yes | no | no | no | yes | no | no | no | no | no | no | no |
| TRPM4 | 341 | K | 0 | 0 | no | yes | no | no | no | yes | no | no | no | no | no | no | no |
| TRPM4 | 342 | G | 0 | 0 | no | yes | no | no | no | yes | no | no | no | no | no | no | no |

| | | | | | | | | | | | | | | | | | |
|---|---|---|---|---|---|---|---|---|---|---|---|---|---|---|---|---|---|
| TRPM4 | 343 | D | 1 | 0 | no | yes | no | no | no | yes | no | no | no | no | no | no | no |
| TRPM4 | 344 | L | 0 | 0 | no | yes | no | no | no | yes | no | no | no | no | no | no | no |
| TRPM4 | 345 | E | 2 | 0 | no | yes | no | no | no | yes | no | no | no | no | no | no | no |
| TRPM4 | 346 | V | 0 | 0 | no | yes | no | no | no | yes | no | no | no | no | no | no | no |
| TRPM4 | 347 | L | 0 | 0 | no | yes | no | no | no | yes | no | no | no | no | no | no | no |
| TRPM4 | 348 | Q | 0 | 0 | no | yes | no | no | no | yes | no | no | no | no | no | no | no |
| TRPM4 | 349 | A | 1 | 0 | yes | no | no | no | no | yes | no | no | no | no | no | no | no |
| TRPM4 | 350 | Q | 0 | 0 | yes | no | no | no | no | yes | no | no | no | no | no | no | no |
| TRPM4 | 351 | V | 2 | 0 | yes | no | no | no | no | yes | no | no | no | no | no | no | no |
| TRPM4 | 352 | E | 0 | 0 | yes | no | no | no | no | yes | no | no | no | no | no | no | no |
| TRPM4 | 353 | R | 0 | 0 | no | yes | no | no | no | yes | no | no | no | no | no | no | no |
| TRPM4 | 354 | I | 0 | 0 | no | yes | no | no | no | yes | no | no | no | no | no | no | no |
| TRPM4 | 355 | M | 0 | 0 | no | yes | no | no | no | yes | no | no | no | no | no | no | no |
| TRPM4 | 356 | T | 0 | 0 | no | yes | no | no | no | yes | no | no | no | no | no | no | no |
| TRPM4 | 357 | R | 2 | 0 | no | yes | no | no | no | yes | no | no | no | no | no | no | no |
| TRPM4 | 358 | K | 0 | 0 | no | yes | no | no | no | yes | no | no | no | no | no | no | no |
| TRPM4 | 359 | E | 0 | 0 | no | yes | no | no | no | yes | no | no | no | no | no | no | no |
| TRPM4 | 360 | L | 0 | 0 | no | yes | no | no | no | yes | no | no | no | no | no | no | no |
| TRPM4 | 361 | L | 1 | 0 | no | yes | no | no | no | yes | no | no | no | no | no | no | no |
| TRPM4 | 362 | T | 0 | 0 | no | yes | no | no | no | yes | no | no | no | no | no | no | no |
| TRPM4 | 363 | V | 1 | 0 | no | no | yes | no | no | yes | no | no | no | no | no | no | no |
| TRPM4 | 364 | Y | 0 | 0 | no | yes | no | no | no | yes | no | no | no | no | no | no | no |
| TRPM4 | 365 | S | 0 | 0 | no | yes | no | no | no | yes | no | no | no | no | no | no | no |
| TRPM4 | 366 | S | 0 | 0 | no | yes | no | no | no | yes | no | no | no | no | no | no | no |
| TRPM4 | 367 | E | 1 | 0 | no | yes | no | no | no | yes | no | no | no | no | no | no | no |
| TRPM4 | 368 | D | 1 | 0 | no | yes | no | no | no | yes | no | no | no | no | no | no | no |
| TRPM4 | 369 | G | 1 | 0 | no | yes | no | no | no | yes | no | no | no | no | no | no | no |
| TRPM4 | 370 | S | 1 | 0 | no | yes | no | no | no | yes | no | no | no | no | no | no | no |
| TRPM4 | 371 | E | 0 | 0 | no | yes | no | no | no | yes | no | no | no | no | no | no | no |
| TRPM4 | 372 | E | 1 | 0 | no | yes | no | no | no | yes | no | no | no | no | no | no | no |
| TRPM4 | 373 | F | 1 | 0 | yes | no | no | no | no | yes | no | no | no | no | no | no | no |
| TRPM4 | 374 | E | 1 | 0 | yes | no | no | no | no | yes | no | no | no | no | no | no | no |
| TRPM4 | 375 | T | 0 | 0 | yes | no | no | no | no | yes | no | no | no | no | no | no | no |

| | | | | | | | | | | | | | | | | | |
|---|---|---|---|---|---|---|---|---|---|---|---|---|---|---|---|---|---|
| TRPM4 | 376 | I | 0 | 1 | yes | no | no | no | no | yes | no | no | no | no | no | no | no |
| TRPM4 | 377 | V | 0 | 0 | yes | no | no | no | no | yes | no | no | no | no | no | no | no |
| TRPM4 | 378 | L | 0 | 0 | yes | no | no | no | no | yes | no | no | no | no | no | no | no |
| TRPM4 | 379 | K | 0 | 0 | yes | no | no | no | no | yes | no | no | no | no | no | no | no |
| TRPM4 | 380 | A | 2 | 0 | yes | no | no | no | no | yes | no | no | no | no | no | no | no |
| TRPM4 | 381 | L | 2 | 0 | yes | no | no | no | no | yes | no | no | no | no | no | no | no |
| TRPM4 | 382 | V | 0 | 0 | yes | no | no | no | no | yes | no | no | no | no | no | no | no |
| TRPM4 | 383 | K | 0 | 0 | yes | no | no | no | no | yes | no | no | no | no | no | no | no |
| TRPM4 | 384 | A | 2 | 0 | no | yes | no | no | no | yes | no | no | no | no | no | no | no |
| TRPM4 | 385 | C | 0 | 0 | no | yes | no | no | no | yes | no | no | no | no | no | no | no |
| TRPM4 | 386 | G | 0 | 0 | no | yes | no | no | no | yes | no | no | no | no | no | no | no |
| TRPM4 | 393 | Y | 0 | 0 | no | yes | no | no | no | yes | no | no | no | no | no | no | no |
| TRPM4 | 394 | L | 1 | 0 | no | yes | no | no | no | yes | no | no | no | no | no | no | no |
| TRPM4 | 395 | D | 2 | 0 | no | yes | no | no | no | yes | no | no | no | no | no | no | no |
| TRPM4 | 396 | E | 0 | 0 | yes | no | no | no | no | yes | no | no | no | no | no | no | no |
| TRPM4 | 397 | L | 0 | 0 | yes | no | no | no | no | yes | no | no | no | no | no | no | no |
| TRPM4 | 398 | R | 1 | 0 | yes | no | no | no | no | yes | no | no | no | no | no | no | no |
| TRPM4 | 399 | L | 0 | 0 | yes | no | no | no | no | yes | no | no | no | no | no | no | no |
| TRPM4 | 400 | A | 0 | 0 | yes | no | no | no | no | yes | no | no | no | no | no | no | no |
| TRPM4 | 401 | V | 1 | 0 | yes | no | no | no | no | yes | no | no | no | no | no | no | no |
| TRPM4 | 402 | A | 0 | 0 | yes | no | no | no | no | yes | no | no | no | no | no | no | no |
| TRPM4 | 403 | W | 0 | 0 | no | yes | no | no | no | yes | no | no | no | no | no | no | no |
| TRPM4 | 404 | N | 1 | 0 | no | yes | no | no | no | yes | no | no | no | no | no | no | no |
| TRPM4 | 405 | R | 3 | 0 | no | yes | no | no | no | yes | no | no | no | no | no | no | no |
| TRPM4 | 406 | V | 1 | 0 | no | yes | no | no | no | yes | no | no | no | no | no | no | no |
| TRPM4 | 407 | D | 0 | 0 | yes | no | no | no | no | yes | no | no | no | no | no | no | no |
| TRPM4 | 408 | I | 0 | 0 | yes | no | no | no | no | yes | no | no | no | no | no | no | no |
| TRPM4 | 409 | A | 0 | 0 | yes | no | no | no | no | yes | no | no | no | no | no | no | no |
| TRPM4 | 410 | Q | 1 | 0 | yes | no | no | no | no | yes | no | no | no | no | no | no | no |
| TRPM4 | 411 | S | 0 | 0 | yes | no | no | no | no | yes | no | no | no | no | no | no | no |
| TRPM4 | 412 | E | 1 | 0 | yes | no | no | no | no | yes | no | no | no | no | no | no | no |
| TRPM4 | 413 | L | 1 | 0 | yes | no | no | no | no | yes | no | no | no | no | no | no | no |
| TRPM4 | 414 | F | 0 | 0 | yes | no | no | no | no | yes | no | no | no | no | no | no | no |

| | | | | | | | | | | | | | | | | | |
|---|---|---|---|---|---|---|---|---|---|---|---|---|---|---|---|---|---|
| TRPM4 | 415 | R | 1 | 0 | no | yes | no | no | no | yes | no | no | no | no | no | no | no |
| TRPM4 | 416 | G | 1 | 0 | no | yes | no | no | no | yes | no | no | yes | no | no | no | no |
| TRPM4 | 417 | D | 1 | 0 | no | yes | no | no | no | yes | no | no | no | no | no | no | no |
| TRPM4 | 418 | I | 2 | 0 | no | yes | no | no | no | yes | no | no | no | no | no | no | no |
| TRPM4 | 419 | Q | 0 | 0 | no | yes | no | no | no | yes | no | no | no | no | no | no | no |
| TRPM4 | 420 | W | 0 | 0 | no | yes | no | no | no | yes | no | no | no | no | no | no | no |
| TRPM4 | 421 | R | 1 | 0 | no | yes | no | no | no | yes | no | no | no | no | no | no | yes |
| TRPM4 | 422 | S | 1 | 0 | no | yes | no | no | no | yes | no | no | no | no | no | no | no |
| TRPM4 | 423 | F | 0 | 0 | yes | no | no | no | no | yes | no | no | no | no | no | no | no |
| TRPM4 | 424 | H | 0 | 0 | yes | no | no | no | no | yes | no | no | no | no | no | no | no |
| TRPM4 | 425 | L | 0 | 0 | yes | no | no | no | no | yes | no | no | no | no | no | no | no |
| TRPM4 | 426 | E | 0 | 0 | yes | no | no | no | no | yes | no | no | no | no | no | no | no |
| TRPM4 | 427 | A | 1 | 0 | yes | no | no | no | no | yes | no | no | no | no | no | no | no |
| TRPM4 | 428 | S | 1 | 0 | yes | no | no | no | no | yes | no | no | no | no | no | no | no |
| TRPM4 | 429 | L | 0 | 0 | yes | no | no | no | no | yes | no | no | no | no | no | no | no |
| TRPM4 | 430 | M | 1 | 0 | yes | no | no | no | no | yes | no | no | no | no | no | no | no |
| TRPM4 | 431 | D | 0 | 0 | yes | no | no | no | no | yes | no | no | no | no | no | no | no |
| TRPM4 | 432 | A | 2 | 1 | yes | no | no | no | no | yes | no | no | no | no | no | no | no |
| TRPM4 | 433 | L | 1 | 0 | yes | no | no | no | no | yes | no | no | no | no | no | no | no |
| TRPM4 | 434 | L | 0 | 0 | yes | no | no | no | no | yes | no | no | no | no | no | no | no |
| TRPM4 | 435 | N | 0 | 0 | no | yes | no | no | no | yes | no | no | no | no | no | no | no |
| TRPM4 | 436 | D | 1 | 0 | no | yes | no | no | no | yes | no | no | no | no | no | no | no |
| TRPM4 | 437 | R | 2 | 0 | no | yes | no | no | no | yes | no | no | no | no | no | no | no |
| TRPM4 | 438 | P | 0 | 0 | yes | no | no | no | no | yes | no | no | no | no | no | no | no |
| TRPM4 | 439 | E | 0 | 0 | yes | no | no | no | no | yes | no | no | no | no | no | no | no |
| TRPM4 | 440 | F | 0 | 0 | yes | no | no | no | no | yes | no | no | no | no | no | no | no |
| TRPM4 | 441 | V | 2 | 1 | yes | no | no | no | no | yes | no | no | no | no | no | no | no |
| TRPM4 | 442 | R | 2 | 0 | yes | no | no | no | no | yes | no | no | no | no | no | no | no |
| TRPM4 | 443 | L | 0 | 0 | yes | no | no | no | no | yes | no | no | no | no | no | no | no |
| TRPM4 | 444 | L | 0 | 0 | yes | no | no | no | no | yes | no | no | no | no | no | no | no |
| TRPM4 | 445 | I | 0 | 0 | yes | no | no | no | no | yes | no | no | no | no | no | no | no |
| TRPM4 | 446 | S | 1 | 0 | yes | no | no | no | no | yes | no | no | no | no | no | no | no |
| TRPM4 | 447 | H | 0 | 0 | no | yes | no | no | no | yes | no | no | yes | no | no | no | no |

| | | | | | | | | | | | | | | | | | |
|---|---|---|---|---|---|---|---|---|---|---|---|---|---|---|---|---|---|
| TRPM4 | 448 | G | 1 | 0 | no | yes | no | no | no | yes | no | no | yes | no | no | no | yes |
| TRPM4 | 449 | L | 0 | 0 | no | yes | no | no | no | yes | no | no | no | no | no | no | no |
| TRPM4 | 450 | S | 0 | 0 | no | yes | no | no | no | yes | no | no | yes | no | no | no | no |
| TRPM4 | 451 | L | 0 | 0 | yes | no | no | no | no | yes | no | no | yes | no | no | no | no |
| TRPM4 | 452 | G | 0 | 0 | yes | no | no | no | no | yes | no | no | yes | no | no | no | no |
| TRPM4 | 453 | H | 0 | 0 | yes | no | no | no | no | yes | no | no | yes | no | no | no | no |
| TRPM4 | 454 | F | 0 | 0 | yes | no | no | no | no | yes | no | no | no | no | no | no | no |
| TRPM4 | 455 | L | 0 | 0 | no | yes | no | no | no | yes | no | no | no | no | no | no | no |
| TRPM4 | 456 | T | 0 | 0 | no | yes | no | no | no | yes | no | no | no | no | no | no | no |
| TRPM4 | 457 | P | 1 | 0 | yes | no | no | no | no | yes | no | no | no | no | no | no | no |
| TRPM4 | 458 | M | 2 | 0 | yes | no | no | no | no | yes | no | no | no | no | no | no | no |
| TRPM4 | 459 | R | 2 | 0 | yes | no | no | no | no | yes | no | no | no | no | no | no | no |
| TRPM4 | 460 | L | 0 | 0 | yes | no | no | no | no | yes | no | no | no | no | no | no | no |
| TRPM4 | 461 | A | 0 | 0 | yes | no | no | no | no | yes | no | no | no | no | no | no | no |
| TRPM4 | 462 | Q | 1 | 0 | yes | no | no | no | no | yes | no | no | no | no | no | no | no |
| TRPM4 | 463 | L | 0 | 0 | yes | no | no | no | no | yes | no | no | no | no | no | no | no |
| TRPM4 | 464 | Y | 2 | 0 | yes | no | no | no | no | yes | no | no | no | no | no | no | no |
| TRPM4 | 465 | S | 0 | 0 | yes | no | no | no | no | yes | no | no | no | no | no | no | no |
| TRPM4 | 466 | A | 0 | 0 | no | yes | no | no | no | yes | no | no | no | no | no | no | no |
| TRPM4 | 467 | A | 0 | 0 | no | yes | no | no | no | yes | no | no | no | no | no | no | no |
| TRPM4 | 468 | P | 2 | 0 | no | yes | no | no | no | yes | no | no | no | no | no | no | no |
| TRPM4 | 469 | S | 0 | 0 | no | yes | no | no | no | yes | no | no | no | no | no | no | no |
| TRPM4 | 470 | N | 2 | 0 | no | yes | no | no | no | yes | no | no | no | no | no | no | no |
| TRPM4 | 471 | S | 1 | 0 | no | yes | no | no | no | yes | no | no | no | no | no | no | no |
| TRPM4 | 472 | L | 0 | 0 | no | yes | no | no | no | yes | no | no | no | no | no | no | no |
| TRPM4 | 473 | I | 1 | 0 | yes | no | no | no | no | yes | no | no | no | no | no | no | no |
| TRPM4 | 474 | R | 2 | 0 | yes | no | no | no | no | yes | no | no | no | no | no | no | no |
| TRPM4 | 475 | N | 1 | 0 | yes | no | no | no | no | yes | no | no | no | no | no | no | no |
| TRPM4 | 476 | L | 0 | 0 | yes | no | no | no | no | yes | no | no | no | no | no | no | no |
| TRPM4 | 477 | L | 1 | 0 | yes | no | no | no | no | yes | no | no | no | no | no | no | no |
| TRPM4 | 478 | D | 1 | 0 | yes | no | no | no | no | yes | no | no | no | no | no | no | no |
| TRPM4 | 479 | Q | 0 | 0 | no | yes | no | no | no | yes | no | no | no | no | no | no | no |
| TRPM4 | 480 | A | 0 | 0 | no | yes | no | no | no | yes | no | no | no | no | no | no | no |

| | | | | | | | | | | | | | | | | | |
|---|---|---|---|---|---|---|---|---|---|---|---|---|---|---|---|---|---|
| TRPM4 | 558 | P | 1 | 0 | no | yes | no | no | no | yes | no | no | no | no | no | no | no |
| TRPM4 | 559 | W | 0 | 0 | yes | no | no | no | no | yes | no | no | no | no | no | no | no |
| TRPM4 | 560 | S | 0 | 0 | yes | no | no | no | no | yes | no | no | no | no | no | no | no |
| TRPM4 | 561 | D | 3 | 0 | yes | no | no | no | no | yes | no | no | no | no | no | no | no |
| TRPM4 | 562 | L | 0 | 0 | yes | no | no | no | no | yes | no | no | no | no | no | no | no |
| TRPM4 | 563 | L | 0 | 0 | yes | no | no | no | no | yes | no | no | no | no | no | no | no |
| TRPM4 | 564 | L | 1 | 0 | yes | no | no | no | no | yes | no | no | no | no | no | no | no |
| TRPM4 | 565 | W | 0 | 0 | yes | no | no | no | no | yes | no | no | no | no | no | no | no |
| TRPM4 | 566 | A | 0 | 0 | yes | no | no | no | no | yes | no | no | no | no | no | no | no |
| TRPM4 | 567 | L | 0 | 0 | yes | no | no | no | no | yes | no | no | no | no | no | no | no |
| TRPM4 | 568 | L | 1 | 0 | yes | no | no | no | no | yes | no | no | no | no | no | no | no |
| TRPM4 | 569 | L | 2 | 0 | no | yes | no | no | no | yes | no | no | no | no | no | no | no |
| TRPM4 | 570 | N | 1 | 0 | no | yes | no | no | no | yes | no | no | no | no | no | no | no |
| TRPM4 | 571 | R | 1 | 0 | no | yes | no | no | no | yes | no | no | no | no | no | no | no |
| TRPM4 | 572 | A | 2 | 0 | yes | no | no | no | no | yes | no | no | no | no | no | no | no |
| TRPM4 | 573 | Q | 1 | 0 | yes | no | no | no | no | yes | no | no | no | no | no | no | no |
| TRPM4 | 574 | M | 0 | 0 | yes | no | no | no | no | yes | no | no | no | no | no | no | no |
| TRPM4 | 575 | A | 1 | 0 | yes | no | no | no | no | yes | no | no | no | no | no | no | no |
| TRPM4 | 576 | M | 1 | 0 | yes | no | no | no | no | yes | no | no | no | no | no | no | no |
| TRPM4 | 577 | Y | 0 | 0 | yes | no | no | no | no | yes | no | no | no | no | no | no | no |
| TRPM4 | 578 | F | 0 | 0 | yes | no | no | no | no | yes | no | no | no | no | no | no | no |
| TRPM4 | 579 | W | 0 | 0 | yes | no | no | no | no | yes | no | no | no | no | no | no | no |
| TRPM4 | 580 | E | 0 | 0 | yes | no | no | no | no | yes | no | no | no | no | no | no | no |
| TRPM4 | 581 | M | 1 | 0 | no | yes | no | no | no | yes | no | no | no | no | no | no | no |
| TRPM4 | 582 | G | 1 | 1 | no | yes | no | no | no | yes | no | no | no | no | no | no | no |
| TRPM4 | 583 | S | 0 | 0 | no | yes | no | no | no | yes | no | no | no | no | no | no | no |
| TRPM4 | 584 | N | 2 | 0 | no | yes | no | no | no | yes | no | no | no | no | no | no | no |
| TRPM4 | 585 | A | 0 | 0 | yes | no | no | no | no | yes | no | no | no | no | no | no | no |
| TRPM4 | 586 | V | 2 | 0 | yes | no | no | no | no | yes | no | no | no | no | no | no | no |
| TRPM4 | 587 | S | 1 | 0 | yes | no | no | no | no | yes | no | no | no | no | no | no | no |
| TRPM4 | 588 | S | 0 | 0 | yes | no | no | no | no | yes | no | no | no | no | no | no | no |
| TRPM4 | 589 | A | 1 | 0 | yes | no | no | no | no | yes | no | no | no | no | no | no | no |
| TRPM4 | 590 | L | 1 | 0 | yes | no | no | no | no | yes | no | no | no | no | no | no | no |

| | | | | | | | | | | | | | | | | | |
|---|---|---|---|---|---|---|---|---|---|---|---|---|---|---|---|---|---|
| TRPM4 | 591 | G | 1 | 0 | yes | no | no | no | no | yes | no | no | no | no | no | no | no |
| TRPM4 | 592 | A | 2 | 0 | yes | no | no | no | no | yes | no | no | no | no | no | no | no |
| TRPM4 | 593 | C | 1 | 0 | yes | no | no | no | no | yes | no | no | no | no | no | no | no |
| TRPM4 | 594 | L | 0 | 0 | yes | no | no | no | no | yes | no | no | no | no | no | no | no |
| TRPM4 | 595 | L | 2 | 0 | yes | no | no | no | no | yes | no | no | no | no | no | no | no |
| TRPM4 | 596 | L | 0 | 0 | yes | no | no | no | no | yes | no | no | no | no | no | no | no |
| TRPM4 | 597 | R | 2 | 0 | yes | no | no | no | no | yes | no | no | no | no | no | no | no |
| TRPM4 | 598 | V | 0 | 0 | yes | no | no | no | no | yes | no | no | no | no | no | no | no |
| TRPM4 | 599 | M | 3 | 0 | yes | no | no | no | no | yes | no | no | no | no | no | no | no |
| TRPM4 | 600 | A | 0 | 0 | no | yes | no | no | no | yes | no | no | no | no | no | no | no |
| TRPM4 | 601 | R | 2 | 0 | no | yes | no | no | no | yes | no | no | no | no | no | no | no |
| TRPM4 | 602 | L | 0 | 0 | no | yes | no | no | no | yes | no | no | no | no | no | no | no |
| TRPM4 | 603 | E | 0 | 0 | no | yes | no | no | no | yes | no | no | no | no | no | no | no |
| TRPM4 | 604 | P | 1 | 0 | no | yes | no | no | no | yes | no | no | no | no | no | no | no |
| TRPM4 | 605 | D | 1 | 0 | no | yes | no | no | no | yes | no | no | no | no | no | no | no |
| TRPM4 | 606 | A | 1 | 0 | no | yes | no | no | no | yes | no | no | no | no | no | no | no |
| TRPM4 | 607 | E | 0 | 0 | no | yes | no | no | no | yes | no | no | no | no | no | no | no |
| TRPM4 | 608 | E | 0 | 0 | no | yes | no | no | no | yes | no | no | no | no | no | no | no |
| TRPM4 | 609 | A | 2 | 0 | yes | no | no | no | no | yes | no | no | no | no | no | no | no |
| TRPM4 | 610 | A | 1 | 0 | yes | no | no | no | no | yes | no | no | no | no | no | no | no |
| TRPM4 | 611 | R | 2 | 0 | yes | no | no | no | no | yes | no | no | no | no | no | no | no |
| TRPM4 | 612 | R | 0 | 0 | yes | no | no | no | no | yes | no | no | no | no | no | no | no |
| TRPM4 | 613 | K | 0 | 0 | yes | no | no | no | no | yes | no | no | no | no | no | no | no |
| TRPM4 | 614 | D | 2 | 0 | yes | no | no | no | no | yes | no | no | no | no | no | no | no |
| TRPM4 | 615 | L | 0 | 0 | yes | no | no | no | no | yes | no | no | no | no | no | no | no |
| TRPM4 | 616 | A | 1 | 0 | yes | no | no | no | no | yes | no | no | no | no | no | no | no |
| TRPM4 | 617 | F | 0 | 0 | yes | no | no | no | no | yes | no | no | no | no | no | no | no |
| TRPM4 | 618 | K | 0 | 0 | yes | no | no | no | no | yes | no | no | no | no | no | no | no |
| TRPM4 | 619 | F | 0 | 0 | yes | no | no | no | no | yes | no | no | no | no | no | no | no |
| TRPM4 | 620 | E | 0 | 0 | yes | no | no | no | no | yes | no | no | no | no | no | no | no |
| TRPM4 | 621 | G | 1 | 0 | yes | no | no | no | no | yes | no | no | no | no | no | no | no |
| TRPM4 | 622 | M | 0 | 0 | yes | no | no | no | no | yes | no | no | no | no | no | no | no |
| TRPM4 | 623 | G | 0 | 0 | yes | no | no | no | no | yes | no | no | no | no | no | no | no |

| | | | | | | | | | | | | | | | | | |
|---|---|---|---|---|---|---|---|---|---|---|---|---|---|---|---|---|---|
| TRPM4 | 624 | V | 0 | 0 | yes | no | no | no | no | yes | no | no | no | no | no | no | no |
| TRPM4 | 625 | D | 0 | 0 | yes | no | no | no | no | yes | no | no | no | no | no | no | no |
| TRPM4 | 626 | L | 1 | 0 | yes | no | no | no | no | yes | no | no | no | no | no | no | no |
| TRPM4 | 627 | F | 1 | 0 | yes | no | no | no | no | yes | no | no | no | no | no | no | no |
| TRPM4 | 628 | G | 0 | 0 | yes | no | no | no | no | yes | no | no | no | no | no | no | no |
| TRPM4 | 629 | E | 1 | 0 | yes | no | no | no | no | yes | no | no | no | no | no | no | no |
| TRPM4 | 630 | C | 0 | 0 | yes | no | no | no | no | yes | no | no | no | no | no | no | no |
| TRPM4 | 631 | Y | 1 | 0 | yes | no | no | no | no | yes | no | no | no | no | no | no | no |
| TRPM4 | 632 | R | 1 | 0 | no | yes | no | no | no | yes | no | no | no | no | no | no | no |
| TRPM4 | 633 | S | 1 | 0 | no | yes | no | no | no | yes | no | no | no | no | no | no | no |
| TRPM4 | 634 | S | 1 | 0 | no | yes | no | no | no | yes | no | no | no | no | no | no | no |
| TRPM4 | 635 | E | 0 | 0 | yes | no | no | no | no | yes | no | no | no | no | no | no | no |
| TRPM4 | 636 | V | 0 | 0 | yes | no | no | no | no | yes | no | no | no | no | no | no | no |
| TRPM4 | 637 | R | 1 | 0 | yes | no | no | no | no | yes | no | no | no | no | no | no | no |
| TRPM4 | 638 | A | 0 | 0 | yes | no | no | no | no | yes | no | no | no | no | no | no | no |
| TRPM4 | 639 | A | 0 | 0 | yes | no | no | no | no | yes | no | no | no | no | no | no | no |
| TRPM4 | 640 | R | 1 | 0 | yes | no | no | no | no | yes | no | no | no | no | no | no | no |
| TRPM4 | 641 | L | 0 | 0 | yes | no | no | no | no | yes | no | no | no | no | no | no | no |
| TRPM4 | 642 | L | 0 | 0 | yes | no | no | no | no | yes | no | no | no | no | no | no | no |
| TRPM4 | 643 | L | 1 | 0 | yes | no | no | no | no | yes | no | no | no | no | no | no | no |
| TRPM4 | 644 | R | 2 | 0 | no | yes | no | no | no | yes | no | no | no | no | no | no | no |
| TRPM4 | 645 | R | 1 | 0 | no | no | yes | no | no | yes | no | no | no | no | no | no | no |
| TRPM4 | 646 | C | 0 | 0 | no | no | yes | no | no | yes | no | no | no | no | no | no | no |
| TRPM4 | 647 | P | 1 | 0 | yes | no | no | no | no | yes | no | no | no | no | no | no | no |
| TRPM4 | 648 | L | 0 | 0 | yes | no | no | no | no | yes | no | no | no | no | no | no | no |
| TRPM4 | 649 | W | 0 | 0 | yes | no | no | no | no | yes | no | no | no | no | no | no | no |
| TRPM4 | 650 | G | 0 | 0 | no | yes | no | no | no | yes | no | no | no | no | no | no | no |
| TRPM4 | 651 | D | 1 | 0 | no | yes | no | no | no | yes | no | no | no | no | no | no | no |
| TRPM4 | 652 | A | 0 | 0 | no | no | yes | no | no | yes | no | no | no | no | no | no | no |
| TRPM4 | 653 | T | 1 | 0 | no | no | yes | no | no | yes | no | no | no | no | no | no | no |
| TRPM4 | 654 | C | 0 | 0 | yes | no | no | no | no | yes | no | no | no | no | no | no | no |
| TRPM4 | 655 | L | 0 | 0 | yes | no | no | no | no | yes | no | no | no | no | no | no | no |
| TRPM4 | 656 | Q | 0 | 0 | yes | no | no | no | no | yes | no | no | no | no | no | no | no |

| | | | | | | | | | | | | | | | | | |
|---|---|---|---|---|---|---|---|---|---|---|---|---|---|---|---|---|---|
| TRPM4 | 657 | L | 2 | 0 | yes | no | no | no | no | yes | no | no | no | no | no | no | no |
| TRPM4 | 658 | A | 0 | 0 | yes | no | no | no | no | yes | no | no | no | no | no | no | no |
| TRPM4 | 659 | M | 0 | 0 | yes | no | no | no | no | yes | no | no | no | no | no | no | no |
| TRPM4 | 660 | Q | 0 | 0 | no | yes | no | no | no | yes | no | no | no | no | no | no | no |
| TRPM4 | 661 | A | 0 | 0 | no | yes | no | no | no | yes | no | no | no | no | no | no | no |
| TRPM4 | 662 | D | 0 | 0 | no | yes | no | no | no | yes | no | no | no | no | no | no | no |
| TRPM4 | 663 | A | 1 | 0 | no | yes | no | no | no | yes | no | no | no | no | no | no | no |
| TRPM4 | 664 | R | 2 | 0 | yes | no | no | no | no | yes | no | no | no | no | no | no | no |
| TRPM4 | 665 | A | 0 | 0 | yes | no | no | no | no | yes | no | no | no | no | no | no | no |
| TRPM4 | 666 | F | 0 | 0 | yes | no | no | no | no | yes | no | no | no | no | no | no | no |
| TRPM4 | 667 | F | 0 | 0 | yes | no | no | no | no | yes | no | no | no | no | no | no | no |
| TRPM4 | 668 | A | 0 | 0 | yes | no | no | no | no | yes | no | no | no | no | no | no | no |
| TRPM4 | 669 | Q | 0 | 0 | no | yes | no | no | no | yes | no | no | no | no | no | no | no |
| TRPM4 | 670 | D | 0 | 0 | yes | no | no | no | no | yes | no | no | no | no | no | no | no |
| TRPM4 | 671 | G | 1 | 0 | yes | no | no | no | no | yes | no | no | no | no | no | no | no |
| TRPM4 | 672 | V | 0 | 0 | yes | no | no | no | no | yes | no | no | no | no | no | no | no |
| TRPM4 | 673 | Q | 1 | 0 | yes | no | no | no | no | yes | no | no | no | no | no | no | no |
| TRPM4 | 674 | S | 0 | 0 | yes | no | no | no | no | yes | no | no | no | no | no | no | no |
| TRPM4 | 675 | L | 0 | 0 | yes | no | no | no | no | yes | no | no | no | no | no | no | no |
| TRPM4 | 676 | L | 0 | 0 | yes | no | no | no | no | yes | no | no | no | no | no | no | no |
| TRPM4 | 677 | T | 0 | 0 | yes | no | no | no | no | yes | no | no | no | no | no | no | no |
| TRPM4 | 678 | Q | 0 | 0 | yes | no | no | no | no | yes | no | no | no | no | no | no | no |
| TRPM4 | 679 | K | 0 | 0 | yes | no | no | no | no | yes | no | no | no | no | no | no | no |
| TRPM4 | 680 | W | 1 | 0 | yes | no | no | no | no | yes | no | no | no | no | no | no | no |
| TRPM4 | 681 | W | 0 | 0 | no | yes | no | no | no | yes | no | no | no | no | no | no | no |
| TRPM4 | 682 | G | 0 | 0 | no | yes | no | no | no | yes | no | no | no | no | no | no | no |
| TRPM4 | 683 | D | 1 | 0 | no | yes | no | no | no | yes | no | no | no | no | no | no | no |
| TRPM4 | 684 | M | 1 | 0 | no | yes | no | no | no | yes | no | no | no | no | no | no | no |
| TRPM4 | 685 | A | 1 | 0 | no | yes | no | no | no | yes | no | no | no | no | no | no | no |
| TRPM4 | 686 | S | 0 | 0 | no | yes | no | no | no | yes | no | no | no | no | no | no | no |
| TRPM4 | 687 | T | 0 | 0 | no | yes | no | no | no | yes | no | no | no | no | no | no | no |
| TRPM4 | 688 | T | 1 | 0 | no | yes | no | no | no | yes | no | no | no | no | no | no | no |
| TRPM4 | 689 | P | 0 | 0 | no | yes | no | no | no | yes | no | no | no | no | no | no | no |

| | | | | | | | | | | | | | | | | | |
|---|---|---|---|---|---|---|---|---|---|---|---|---|---|---|---|---|---|
| TRPM4 | 690 | I | 0 | 0 | yes | no | no | yes | no | yes | no | no | no | no | no | no | no |
| TRPM4 | 691 | W | 1 | 0 | yes | no | no | yes | no | yes | no | no | no | no | no | no | no |
| TRPM4 | 692 | A | 1 | 0 | yes | no | no | yes | no | yes | no | no | no | no | no | no | no |
| TRPM4 | 693 | L | 0 | 0 | yes | no | no | yes | no | yes | no | no | no | no | no | no | no |
| TRPM4 | 694 | V | 0 | 0 | yes | no | no | yes | no | yes | no | no | no | no | no | no | no |
| TRPM4 | 695 | L | 0 | 0 | yes | no | no | yes | no | yes | no | no | no | no | no | no | no |
| TRPM4 | 696 | A | 1 | 0 | yes | no | no | yes | no | yes | no | no | no | no | no | no | no |
| TRPM4 | 697 | F | 0 | 0 | yes | no | no | yes | no | yes | no | no | no | no | no | no | no |
| TRPM4 | 698 | F | 0 | 0 | yes | no | no | yes | no | yes | no | no | no | no | no | no | no |
| TRPM4 | 699 | C | 0 | 0 | no | yes | no | yes | no | yes | no | no | no | no | no | no | no |
| TRPM4 | 700 | P | 1 | 0 | yes | no | no | yes | no | yes | no | no | no | no | no | no | no |
| TRPM4 | 701 | P | 0 | 0 | yes | no | no | yes | no | yes | no | no | no | no | no | no | no |
| TRPM4 | 702 | L | 1 | 0 | yes | no | no | yes | no | yes | no | no | no | no | no | no | no |
| TRPM4 | 703 | I | 0 | 0 | no | yes | no | yes | no | yes | no | no | no | no | no | no | no |
| TRPM4 | 704 | Y | 0 | 0 | no | yes | no | yes | no | yes | no | no | no | no | no | no | no |
| TRPM4 | 705 | T | 1 | 0 | no | yes | no | yes | no | yes | no | no | no | no | no | no | no |
| TRPM4 | 706 | R | 2 | 0 | no | yes | no | no | no | yes | no | no | no | no | no | no | no |
| TRPM4 | 707 | L | 0 | 0 | no | yes | no | no | no | yes | no | no | no | no | no | no | no |
| TRPM4 | 708 | I | 2 | 0 | no | yes | no | no | no | yes | no | no | no | no | no | no | no |
| TRPM4 | 709 | T | 1 | 0 | no | yes | no | no | no | yes | no | no | no | no | no | no | no |
| TRPM4 | 710 | F | 0 | 0 | no | yes | no | no | no | yes | no | no | no | no | no | no | no |
| TRPM4 | 766 | R | 0 | 0 | no | yes | no | no | no | yes | no | no | no | no | no | no | no |
| TRPM4 | 767 | R | 1 | 0 | no | yes | no | no | no | yes | no | no | no | no | no | no | no |
| TRPM4 | 768 | C | 0 | 0 | yes | no | no | no | no | yes | no | no | no | no | no | no | no |
| TRPM4 | 769 | L | 0 | 0 | yes | no | no | no | no | yes | no | no | no | no | no | no | no |
| TRPM4 | 770 | R | 2 | 0 | yes | no | no | no | no | yes | no | no | no | no | no | no | no |
| TRPM4 | 771 | R | 1 | 0 | yes | no | no | no | no | yes | no | no | no | no | no | no | no |
| TRPM4 | 772 | W | 0 | 0 | yes | no | no | yes | no | yes | no | no | no | no | no | no | no |
| TRPM4 | 773 | F | 0 | 1 | yes | no | no | no | no | yes | no | no | no | no | no | no | no |
| TRPM4 | 774 | H | 1 | 0 | yes | no | no | no | no | yes | no | no | no | no | no | no | no |
| TRPM4 | 775 | F | 0 | 0 | yes | no | no | no | no | yes | no | no | no | no | no | no | no |
| TRPM4 | 776 | W | 0 | 0 | no | yes | no | yes | no | yes | no | no | no | no | no | no | no |
| TRPM4 | 777 | G | 0 | 0 | no | yes | no | no | no | yes | no | no | no | no | no | no | no |

| | | | | | | | | | | | | | | | | | |
|---|---|---|---|---|---|---|---|---|---|---|---|---|---|---|---|---|---|
| TRPM4 | 778 | A | 2 | 0 | no | yes | no | no | no | yes | no | no | no | no | no | no | no |
| TRPM4 | 779 | P | 1 | 1 | yes | no | no | no | no | yes | no | no | no | no | no | no | no |
| TRPM4 | 780 | V | 0 | 0 | yes | no | no | no | no | yes | no | no | no | no | no | no | no |
| TRPM4 | 781 | T | 0 | 0 | yes | no | no | yes | no | yes | no | no | no | no | no | no | no |
| TRPM4 | 782 | I | 0 | 0 | yes | no | no | yes | no | yes | no | no | no | no | no | no | no |
| TRPM4 | 783 | F | 0 | 0 | yes | no | no | yes | no | no | no | no | no | no | no | no | no |
| TRPM4 | 784 | M | 1 | 0 | yes | no | no | yes | no | no | no | no | no | no | no | no | no |
| TRPM4 | 785 | G | 0 | 0 | yes | no | no | yes | no | no | no | no | no | no | no | no | no |
| TRPM4 | 786 | N | 0 | 0 | yes | no | no | yes | no | no | no | no | no | no | no | no | no |
| TRPM4 | 787 | V | 0 | 0 | yes | no | no | yes | no | no | no | no | no | no | no | no | no |
| TRPM4 | 788 | V | 1 | 0 | yes | no | no | yes | no | no | no | no | no | no | no | no | no |
| TRPM4 | 789 | S | 0 | 0 | yes | no | no | yes | no | no | no | no | no | no | no | no | no |
| TRPM4 | 790 | Y | 0 | 1 | yes | no | no | yes | no | no | no | no | no | no | no | no | no |
| TRPM4 | 791 | L | 0 | 0 | yes | no | no | yes | no | no | no | no | no | no | no | no | no |
| TRPM4 | 792 | L | 1 | 0 | yes | no | no | yes | no | no | no | no | no | no | no | no | no |
| TRPM4 | 793 | F | 0 | 0 | yes | no | no | yes | no | no | no | no | no | no | no | no | no |
| TRPM4 | 794 | L | 0 | 0 | yes | no | no | yes | no | no | no | no | no | no | no | no | no |
| TRPM4 | 795 | L | 1 | 0 | yes | no | no | yes | no | no | no | no | no | no | no | no | no |
| TRPM4 | 796 | L | 1 | 0 | yes | no | no | yes | no | no | no | no | no | no | no | no | no |
| TRPM4 | 797 | F | 0 | 0 | yes | no | no | yes | no | no | no | no | no | no | no | no | no |
| TRPM4 | 798 | S | 1 | 0 | yes | no | no | yes | no | no | no | no | no | no | no | no | no |
| TRPM4 | 799 | R | 0 | 0 | yes | no | no | yes | no | no | no | no | no | no | no | no | no |
| TRPM4 | 800 | V | 1 | 0 | yes | no | no | yes | no | no | no | no | no | no | no | no | no |
| TRPM4 | 801 | L | 0 | 0 | yes | no | no | yes | no | no | no | no | no | no | no | no | no |
| TRPM4 | 802 | L | 1 | 0 | yes | no | no | yes | no | no | no | no | yes | no | no | no | no |
| TRPM4 | 803 | V | 0 | 0 | no | yes | no | no | no | no | no | no | no | no | no | no | no |
| TRPM4 | 804 | D | 1 | 0 | no | yes | no | no | yes | no | no | no | no | no | no | no | no |
| TRPM4 | 805 | F | 0 | 0 | no | yes | no | no | yes | no | no | no | no | no | no | no | no |
| TRPM4 | 806 | Q | 0 | 0 | no | yes | no | no | yes | no | no | no | no | no | no | no | no |
| TRPM4 | 807 | P | 1 | 0 | no | yes | no | no | yes | no | no | no | no | no | no | no | no |
| TRPM4 | 808 | A | 0 | 0 | no | yes | no | no | yes | no | no | no | no | no | no | no | no |
| TRPM4 | 809 | P | 2 | 0 | no | yes | no | no | yes | no | no | no | no | no | no | no | no |
| TRPM4 | 810 | P | 1 | 0 | no | yes | no | no | yes | no | no | no | no | no | no | no | no |

| | | | | | | | | | | | | | | | | | |
|---|---|---|---|---|---|---|---|---|---|---|---|---|---|---|---|---|---|
| TRPM4 | 811 | G | 0 | 0 | no | yes | no | yes | yes | no | no | no | no | no | no | no | no |
| TRPM4 | 812 | S | 0 | 0 | yes | no | no | yes | yes | no | no | no | no | no | no | no | no |
| TRPM4 | 813 | L | 1 | 0 | yes | no | no | yes | yes | no | no | no | no | no | no | no | no |
| TRPM4 | 814 | E | 0 | 0 | yes | no | no | yes | yes | no | no | no | no | no | no | no | no |
| TRPM4 | 815 | L | 0 | 0 | yes | no | no | yes | no | no | no | no | no | no | no | no | no |
| TRPM4 | 816 | L | 1 | 0 | yes | no | no | yes | no | no | no | no | no | no | no | no | no |
| TRPM4 | 817 | L | 0 | 0 | yes | no | no | yes | no | no | no | no | no | no | no | no | no |
| TRPM4 | 818 | Y | 1 | 0 | yes | no | no | yes | no | no | no | no | no | no | no | no | no |
| TRPM4 | 819 | F | 0 | 0 | yes | no | no | yes | no | no | no | no | no | no | no | no | no |
| TRPM4 | 820 | W | 0 | 0 | yes | no | no | yes | no | no | no | no | no | no | no | no | no |
| TRPM4 | 821 | A | 1 | 0 | yes | no | no | yes | no | no | no | no | no | no | no | no | no |
| TRPM4 | 822 | F | 1 | 0 | yes | no | no | yes | no | no | no | no | no | no | no | no | no |
| TRPM4 | 823 | T | 1 | 0 | yes | no | no | yes | no | no | no | no | no | no | no | no | no |
| TRPM4 | 824 | L | 0 | 0 | yes | no | no | yes | no | no | no | no | no | no | no | no | no |
| TRPM4 | 825 | L | 0 | 0 | yes | no | no | yes | no | no | no | no | no | no | no | no | no |
| TRPM4 | 826 | C | 0 | 0 | yes | no | no | yes | no | no | no | no | no | no | no | no | no |
| TRPM4 | 827 | E | 1 | 0 | yes | no | no | yes | no | no | no | no | no | no | no | no | no |
| TRPM4 | 828 | E | 0 | 0 | yes | no | no | yes | no | no | no | no | no | no | no | no | no |
| TRPM4 | 829 | L | 1 | 0 | yes | no | no | yes | no | no | no | no | no | no | no | no | no |
| TRPM4 | 830 | R | 1 | 0 | yes | no | no | yes | no | no | no | no | no | no | no | no | no |
| TRPM4 | 831 | Q | 0 | 0 | yes | no | no | yes | no | no | no | no | no | no | no | no | no |
| TRPM4 | 832 | G | 0 | 0 | yes | no | no | yes | no | no | no | no | no | no | no | no | no |
| TRPM4 | 833 | L | 1 | 0 | yes | no | no | no | no | no | no | no | no | no | no | no | no |
| TRPM4 | 834 | S | 1 | 0 | yes | no | no | no | no | no | no | no | no | no | no | no | no |
| TRPM4 | 835 | G | 0 | 0 | no | yes | no | no | no | no | no | no | no | no | no | no | no |
| TRPM4 | 838 | G | 1 | 0 | no | yes | no | no | no | yes | no | no | no | no | no | no | no |
| TRPM4 | 839 | S | 1 | 0 | no | yes | no | no | no | yes | no | no | no | no | no | no | no |
| TRPM4 | 840 | L | 0 | 0 | no | yes | no | no | no | yes | no | no | no | no | no | no | no |
| TRPM4 | 841 | A | 0 | 0 | no | yes | no | no | no | yes | no | no | no | no | no | no | no |
| TRPM4 | 842 | S | 1 | 0 | no | yes | no | no | no | yes | no | no | no | no | no | no | no |
| TRPM4 | 843 | G | 1 | 0 | no | yes | no | no | no | yes | no | no | no | no | no | no | no |
| TRPM4 | 844 | G | 2 | 1 | no | yes | no | no | no | yes | no | no | no | no | no | no | no |
| TRPM4 | 845 | P | 2 | 0 | no | yes | no | no | no | yes | no | no | no | no | no | no | no |

| | | | | | | | | | | | | | | | | | |
|---|---|---|---|---|---|---|---|---|---|---|---|---|---|---|---|---|---|
| TRPM4 | 846 | G | 2 | 0 | no | yes | no | no | no | yes | no | no | no | no | no | no | no |
| TRPM4 | 847 | P | 0 | 0 | no | yes | no | no | no | yes | no | no | no | no | no | no | no |
| TRPM4 | 848 | G | 0 | 0 | no | yes | no | no | no | yes | no | no | no | no | no | no | no |
| TRPM4 | 849 | H | 2 | 0 | no | yes | no | no | no | yes | no | no | no | no | no | no | no |
| TRPM4 | 850 | A | 0 | 0 | no | yes | no | no | no | yes | no | no | no | no | no | no | no |
| TRPM4 | 851 | S | 0 | 0 | no | yes | no | no | no | yes | no | no | no | no | no | no | no |
| TRPM4 | 852 | L | 0 | 0 | yes | no | no | no | no | yes | no | no | no | no | no | no | no |
| TRPM4 | 853 | S | 1 | 0 | yes | no | no | no | no | yes | no | no | no | no | no | no | no |
| TRPM4 | 854 | Q | 1 | 0 | yes | no | no | no | no | yes | no | no | no | no | no | no | no |
| TRPM4 | 855 | R | 2 | 0 | yes | no | no | no | no | yes | no | no | no | no | no | no | no |
| TRPM4 | 856 | L | 0 | 0 | yes | no | no | no | no | yes | no | no | no | no | no | no | no |
| TRPM4 | 857 | R | 1 | 0 | yes | no | no | no | no | yes | no | no | no | no | no | no | no |
| TRPM4 | 858 | L | 1 | 0 | yes | no | no | no | no | yes | no | no | no | no | no | no | no |
| TRPM4 | 859 | Y | 1 | 0 | yes | no | no | yes | no | yes | no | no | no | no | no | no | no |
| TRPM4 | 860 | L | 0 | 0 | no | yes | no | yes | no | yes | no | no | no | no | no | no | no |
| TRPM4 | 861 | A | 0 | 0 | no | yes | no | no | no | yes | no | no | no | no | no | no | no |
| TRPM4 | 862 | D | 0 | 0 | no | yes | no | no | no | yes | no | no | no | no | no | no | no |
| TRPM4 | 863 | S | 1 | 0 | yes | no | no | yes | no | yes | no | no | no | no | no | no | no |
| TRPM4 | 864 | W | 0 | 0 | yes | no | no | yes | no | no | no | no | no | no | no | no | no |
| TRPM4 | 865 | N | 2 | 0 | yes | no | no | yes | no | no | no | no | no | no | no | no | no |
| TRPM4 | 866 | Q | 0 | 0 | yes | no | no | yes | no | no | no | no | no | no | no | no | no |
| TRPM4 | 867 | C | 0 | 0 | yes | no | no | yes | no | no | no | no | no | no | no | no | no |
| TRPM4 | 868 | D | 0 | 0 | yes | no | no | yes | no | no | no | no | no | no | no | no | no |
| TRPM4 | 869 | L | 0 | 0 | yes | no | no | yes | no | no | no | no | no | no | no | no | no |
| TRPM4 | 870 | V | 0 | 0 | yes | no | no | yes | no | no | no | no | no | no | no | no | no |
| TRPM4 | 871 | A | 0 | 0 | yes | no | no | yes | no | no | no | no | no | no | no | no | no |
| TRPM4 | 872 | L | 1 | 0 | yes | no | no | yes | no | no | no | no | no | no | no | no | no |
| TRPM4 | 873 | T | 0 | 1 | yes | no | no | yes | no | no | no | no | no | no | no | no | no |
| TRPM4 | 874 | C | 0 | 0 | yes | no | no | yes | no | no | no | no | no | no | no | no | no |
| TRPM4 | 875 | F | 0 | 0 | yes | no | no | yes | no | no | no | no | no | no | no | no | no |
| TRPM4 | 876 | L | 1 | 0 | yes | no | no | yes | no | no | no | no | no | no | no | no | no |
| TRPM4 | 877 | L | 0 | 0 | yes | no | no | yes | no | no | no | no | no | no | no | no | no |
| TRPM4 | 878 | G | 0 | 0 | yes | no | no | yes | no | no | no | no | no | no | no | no | no |

| | | | | | | | | | | | | | | | | | |
|---|---|---|---|---|---|---|---|---|---|---|---|---|---|---|---|---|---|
| TRPM4 | 879 | V | 0 | 0 | yes | no | no | yes | no | no | no | no | no | no | no | no | no |
| TRPM4 | 880 | G | 1 | 0 | yes | no | no | yes | no | no | no | no | no | no | no | no | no |
| TRPM4 | 881 | C | 0 | 0 | yes | no | no | yes | no | no | no | no | no | no | no | no | no |
| TRPM4 | 882 | R | 0 | 0 | yes | no | no | yes | no | no | no | no | no | no | no | no | no |
| TRPM4 | 883 | L | 0 | 0 | yes | no | no | yes | no | no | no | no | no | no | no | no | no |
| TRPM4 | 884 | T | 0 | 0 | no | yes | no | no | no | no | no | no | no | no | no | no | no |
| TRPM4 | 885 | P | 2 | 0 | no | yes | no | no | yes | no | no | no | no | no | no | no | no |
| TRPM4 | 886 | G | 0 | 0 | no | yes | no | no | yes | no | no | no | no | no | no | no | no |
| TRPM4 | 887 | L | 0 | 0 | no | yes | no | no | no | no | no | no | no | no | no | no | no |
| TRPM4 | 888 | Y | 0 | 0 | yes | no | no | no | no | no | no | no | no | no | no | no | no |
| TRPM4 | 889 | H | 0 | 0 | yes | no | no | no | no | no | no | no | yes | no | no | no | no |
| TRPM4 | 890 | L | 0 | 0 | yes | no | no | yes | no | no | no | no | no | no | no | no | no |
| TRPM4 | 891 | G | 1 | 0 | yes | no | no | yes | no | no | no | no | no | no | no | no | no |
| TRPM4 | 892 | R | 2 | 0 | yes | no | no | yes | no | no | no | no | yes | no | no | no | no |
| TRPM4 | 893 | T | 0 | 0 | yes | no | no | yes | no | no | no | no | yes | no | no | no | no |
| TRPM4 | 894 | V | 1 | 0 | yes | no | no | yes | no | no | no | no | no | no | no | no | no |
| TRPM4 | 895 | L | 1 | 0 | yes | no | no | yes | no | no | no | no | no | no | no | no | no |
| TRPM4 | 896 | C | 0 | 0 | yes | no | no | yes | no | no | no | no | no | no | no | no | no |
| TRPM4 | 897 | I | 1 | 0 | yes | no | no | yes | no | no | no | no | yes | no | no | no | no |
| TRPM4 | 898 | D | 2 | 0 | yes | no | no | yes | no | no | no | no | no | no | no | no | no |
| TRPM4 | 899 | F | 1 | 0 | yes | no | no | yes | no | no | no | no | yes | no | no | no | no |
| TRPM4 | 900 | M | 2 | 0 | yes | no | no | yes | no | no | no | no | yes | no | no | no | no |
| TRPM4 | 901 | V | 0 | 0 | yes | no | no | yes | no | no | no | no | no | no | no | no | no |
| TRPM4 | 902 | F | 1 | 0 | yes | no | no | yes | no | no | no | no | no | no | no | no | no |
| TRPM4 | 903 | T | 1 | 0 | yes | no | no | yes | no | no | no | no | yes | no | no | no | no |
| TRPM4 | 904 | V | 1 | 0 | yes | no | no | yes | no | no | no | no | no | no | no | no | no |
| TRPM4 | 905 | R | 1 | 0 | yes | no | no | yes | no | no | no | no | no | no | no | no | no |
| TRPM4 | 906 | L | 0 | 0 | yes | no | no | yes | no | no | no | no | no | no | no | no | no |
| TRPM4 | 907 | L | 0 | 0 | yes | no | no | yes | no | no | no | no | yes | no | no | no | no |
| TRPM4 | 908 | H | 0 | 0 | no | yes | no | yes | no | no | no | no | no | no | no | no | no |
| TRPM4 | 909 | I | 0 | 0 | yes | no | no | yes | no | no | no | no | no | no | no | no | no |
| TRPM4 | 910 | F | 0 | 0 | yes | no | no | yes | no | no | no | no | yes | no | no | no | no |
| TRPM4 | 911 | T | 1 | 0 | yes | no | no | no | no | yes | no | no | no | no | no | no | no |

| | | | | | | | | | | | | | | | | | |
|---|---|---|---|---|---|---|---|---|---|---|---|---|---|---|---|---|---|
| TRPM4 | 912 | V | 0 | 0 | no | yes | no | no | no | yes | no | no | no | no | no | no | no |
| TRPM4 | 913 | N | 2 | 0 | no | yes | no | no | no | yes | no | no | no | no | no | no | no |
| TRPM4 | 914 | K | 1 | 1 | no | yes | no | no | no | yes | no | no | no | no | no | no | no |
| TRPM4 | 915 | Q | 0 | 0 | no | yes | no | no | no | yes | no | no | no | no | no | no | no |
| TRPM4 | 916 | L | 0 | 0 | no | yes | no | yes | no | yes | no | no | no | no | no | no | no |
| TRPM4 | 917 | G | 1 | 0 | yes | no | no | no | no | yes | no | no | no | no | no | no | no |
| TRPM4 | 918 | P | 0 | 0 | yes | no | no | no | no | yes | no | no | no | no | no | no | no |
| TRPM4 | 919 | K | 0 | 0 | yes | no | no | no | no | yes | no | no | yes | no | no | no | no |
| TRPM4 | 920 | I | 1 | 0 | yes | no | no | yes | no | yes | no | no | no | no | no | no | no |
| TRPM4 | 921 | V | 2 | 0 | yes | no | no | no | no | yes | no | no | no | no | no | no | no |
| TRPM4 | 922 | I | 0 | 0 | yes | no | no | no | no | yes | no | no | no | no | no | no | no |
| TRPM4 | 923 | V | 2 | 0 | yes | no | no | yes | no | yes | no | no | no | no | no | no | no |
| TRPM4 | 924 | S | 0 | 0 | yes | no | no | yes | no | yes | no | no | no | no | no | no | no |
| TRPM4 | 925 | K | 1 | 0 | no | yes | no | no | no | yes | no | no | no | no | no | no | no |
| TRPM4 | 926 | M | 1 | 0 | no | yes | no | yes | no | yes | no | no | no | no | no | no | no |
| TRPM4 | 927 | M | 0 | 0 | no | yes | no | yes | no | yes | no | no | no | no | no | no | no |
| TRPM4 | 928 | K | 0 | 0 | no | yes | no | no | no | yes | no | no | no | no | no | no | no |
| TRPM4 | 929 | D | 0 | 0 | no | yes | no | yes | no | yes | no | no | no | no | no | no | no |
| TRPM4 | 930 | V | 1 | 0 | yes | no | no | yes | no | yes | no | no | no | no | no | no | no |
| TRPM4 | 931 | F | 0 | 0 | yes | no | no | yes | no | no | no | no | no | no | no | no | no |
| TRPM4 | 932 | F | 0 | 0 | yes | no | no | yes | no | no | no | no | no | no | no | no | no |
| TRPM4 | 933 | F | 0 | 0 | yes | no | no | yes | no | no | no | no | no | no | no | no | no |
| TRPM4 | 934 | L | 0 | 0 | yes | no | no | yes | no | no | no | no | no | no | no | no | no |
| TRPM4 | 935 | F | 0 | 0 | yes | no | no | yes | no | no | no | no | yes | no | no | no | no |
| TRPM4 | 936 | F | 3 | 0 | yes | no | no | yes | no | no | no | no | yes | no | no | no | no |
| TRPM4 | 937 | L | 0 | 0 | yes | no | no | yes | no | no | no | no | no | no | no | no | no |
| TRPM4 | 938 | G | 1 | 0 | yes | no | no | yes | no | no | no | no | no | no | no | no | no |
| TRPM4 | 939 | V | 1 | 0 | yes | no | no | yes | no | no | no | no | no | no | no | no | no |
| TRPM4 | 940 | W | 1 | 0 | yes | no | no | yes | no | no | no | no | no | no | no | no | no |
| TRPM4 | 941 | L | 1 | 0 | yes | no | no | yes | no | no | no | no | no | no | no | no | no |
| TRPM4 | 942 | V | 0 | 0 | yes | no | no | yes | no | no | no | no | no | no | no | no | no |
| TRPM4 | 943 | A | 0 | 0 | yes | no | no | yes | no | no | no | no | yes | no | no | no | no |
| TRPM4 | 944 | Y | 1 | 0 | yes | no | no | yes | no | no | no | no | no | no | no | no | no |

| | | | | | | | | | | | | | | | | | |
|---|---|---|---|---|---|---|---|---|---|---|---|---|---|---|---|---|---|
| TRPM4 | 945 | G | 1 | 0 | yes | no | no | yes | no | no | no | no | no | no | no | no | no |
| TRPM4 | 946 | V | 0 | 0 | yes | no | no | yes | no | no | no | no | yes | no | no | no | no |
| TRPM4 | 947 | A | 2 | 0 | yes | no | no | yes | no | no | no | no | no | no | no | no | no |
| TRPM4 | 948 | T | 1 | 0 | yes | no | no | yes | no | no | no | no | no | no | no | no | no |
| TRPM4 | 949 | E | 0 | 0 | yes | no | no | yes | no | no | no | no | yes | no | no | no | no |
| TRPM4 | 950 | G | 1 | 0 | yes | no | no | yes | no | no | no | no | no | no | no | no | no |
| TRPM4 | 951 | L | 1 | 0 | yes | no | no | yes | no | no | no | no | no | no | no | no | no |
| TRPM4 | 952 | L | 1 | 0 | yes | no | no | no | yes | no | no | no | no | no | no | no | no |
| TRPM4 | 953 | R | 0 | 0 | no | yes | no | no | yes | no | no | no | yes | no | no | no | no |
| TRPM4 | 954 | P | 1 | 0 | no | yes | no | no | yes | no | no | no | no | no | no | no | no |
| TRPM4 | 955 | R | 1 | 0 | no | yes | no | no | yes | no | no | no | no | no | no | no | no |
| TRPM4 | 956 | D | 0 | 0 | no | yes | no | no | yes | no | no | no | no | no | no | no | no |
| TRPM4 | 957 | S | 0 | 0 | no | yes | no | no | yes | no | no | no | no | no | no | no | no |
| TRPM4 | 958 | D | 0 | 0 | no | yes | no | no | yes | no | no | no | no | no | no | no | no |
| TRPM4 | 959 | F | 0 | 0 | yes | no | no | no | yes | no | no | no | no | no | no | no | no |
| TRPM4 | 960 | P | 0 | 0 | yes | no | no | no | yes | no | no | no | no | no | no | no | no |
| TRPM4 | 961 | S | 0 | 0 | yes | no | no | no | yes | no | no | no | no | no | no | no | no |
| TRPM4 | 962 | I | 1 | 0 | yes | no | no | no | yes | no | no | no | no | no | no | no | no |
| TRPM4 | 963 | L | 1 | 0 | yes | no | no | yes | yes | no | no | no | no | no | no | no | no |
| TRPM4 | 964 | R | 2 | 0 | yes | no | no | yes | no | no | no | yes | yes | no | no | no | no |
| TRPM4 | 965 | R | 2 | 0 | yes | no | no | no | no | no | no | yes | no | no | no | no | no |
| TRPM4 | 966 | V | 1 | 0 | yes | no | no | yes | no | no | no | yes | no | no | no | no | no |
| TRPM4 | 967 | F | 0 | 0 | yes | no | no | yes | no | no | no | yes | no | no | no | no | no |
| TRPM4 | 968 | Y | 0 | 0 | yes | no | no | yes | no | no | no | yes | yes | no | no | no | no |
| TRPM4 | 969 | R | 2 | 0 | yes | no | no | yes | no | no | no | yes | no | no | no | no | no |
| TRPM4 | 970 | P | 1 | 1 | yes | no | no | yes | no | no | no | yes | no | no | no | no | no |
| TRPM4 | 971 | Y | 1 | 0 | yes | no | no | yes | no | no | no | yes | yes | no | no | no | no |
| TRPM4 | 972 | L | 0 | 0 | no | yes | no | yes | no | no | no | yes | yes | no | no | no | no |
| TRPM4 | 973 | Q | 1 | 0 | no | yes | no | yes | no | no | no | yes | no | no | no | no | no |
| TRPM4 | 974 | I | 1 | 0 | no | yes | no | yes | no | no | no | yes | no | no | no | no | no |
| TRPM4 | 975 | F | 0 | 0 | no | yes | no | yes | no | no | yes | yes | yes | no | no | no | no |
| TRPM4 | 976 | G | 1 | 0 | no | yes | no | yes | no | no | yes | yes | no | no | no | no | no |
| TRPM4 | 977 | Q | 0 | 0 | no | yes | no | yes | no | no | yes | yes | no | no | no | no | no |

| | | | | | | | | | | | | | | | | | |
|---|---|---|---|---|---|---|---|---|---|---|---|---|---|---|---|---|---|
| TRPM4 | 978 | I | 2 | 0 | no | yes | no | yes | no | no | no | yes | no | no | no | no | no |
| TRPM4 | 979 | P | 1 | 0 | no | yes | no | yes | no | no | no | yes | no | no | no | no | no |
| TRPM4 | 980 | Q | 1 | 0 | no | yes | no | no | no | no | no | yes | yes | no | no | no | no |
| TRPM4 | 981 | E | 0 | 0 | no | yes | no | no | no | no | yes | yes | no | no | no | no | no |
| TRPM4 | 982 | D | 1 | 0 | no | yes | no | no | no | no | no | yes | no | no | no | no | no |
| TRPM4 | 983 | M | 1 | 0 | no | yes | no | no | no | no | no | yes | no | no | no | no | no |
| TRPM4 | 984 | D | 1 | 0 | no | yes | no | no | no | no | no | yes | yes | no | no | no | no |
| TRPM4 | 985 | V | 0 | 0 | no | yes | no | no | yes | no | no | no | no | no | no | no | no |
| TRPM4 | 986 | A | 0 | 0 | no | yes | no | no | yes | no | no | no | yes | no | no | no | no |
| TRPM4 | 987 | L | 0 | 0 | no | yes | no | no | yes | no | no | no | yes | no | no | no | no |
| TRPM4 | 988 | M | 0 | 0 | no | yes | no | no | yes | no | no | no | no | no | no | no | no |
| TRPM4 | 989 | E | 0 | 0 | no | no | yes | no | yes | no | no | no | no | no | no | no | no |
| TRPM4 | 990 | H | 1 | 0 | no | yes | no | no | yes | no | no | no | no | no | no | no | no |
| TRPM4 | 991 | S | 2 | 0 | no | yes | no | no | yes | no | no | no | no | no | no | no | no |
| TRPM4 | 992 | N | 0 | 0 | no | yes | no | no | yes | no | no | no | no | no | no | no | no |
| TRPM4 | 993 | C | 1 | 0 | no | yes | no | no | yes | no | no | no | no | no | no | no | no |
| TRPM4 | 994 | S | 1 | 0 | no | yes | no | no | yes | no | no | no | no | no | no | no | no |
| TRPM4 | 995 | S | 1 | 0 | no | yes | no | no | yes | no | no | no | no | no | no | no | no |
| TRPM4 | 996 | E | 1 | 0 | no | yes | no | no | yes | no | no | no | no | no | no | no | no |
| TRPM4 | 997 | P | 1 | 0 | no | yes | no | no | yes | no | no | no | no | no | no | no | no |
| TRPM4 | 998 | G | 1 | 0 | no | yes | no | no | yes | no | no | no | no | no | no | no | no |
| TRPM4 | 999 | F | 0 | 0 | no | yes | no | no | yes | no | no | no | no | no | no | no | no |
| TRPM4 | 1000 | W | 0 | 0 | no | no | yes | no | yes | no | no | no | no | no | no | no | no |
| TRPM4 | 1001 | A | 1 | 0 | no | no | yes | no | yes | no | no | no | no | no | no | no | no |
| TRPM4 | 1002 | H | 1 | 0 | no | no | yes | no | yes | no | no | no | no | no | no | no | no |
| TRPM4 | 1003 | P | 2 | 0 | no | yes | no | no | yes | no | no | no | no | no | no | no | no |
| TRPM4 | 1004 | P | 0 | 0 | no | yes | no | no | yes | no | yes | no | no | no | no | no | no |
| TRPM4 | 1005 | G | 0 | 0 | no | yes | no | no | yes | no | no | no | no | no | no | no | no |
| TRPM4 | 1006 | A | 0 | 0 | no | yes | no | no | yes | no | no | no | no | no | no | no | no |
| TRPM4 | 1007 | Q | 0 | 0 | no | yes | no | no | yes | no | no | no | yes | no | no | no | no |
| TRPM4 | 1008 | A | 1 | 0 | no | yes | no | no | yes | no | no | no | no | no | no | no | no |
| TRPM4 | 1009 | G | 1 | 0 | no | yes | no | no | yes | no | no | no | no | no | no | no | no |
| TRPM4 | 1010 | T | 0 | 0 | no | yes | no | no | yes | no | no | no | no | no | no | no | no |

| | | | | | | | | | | | | | | | | | |
|---|---|---|---|---|---|---|---|---|---|---|---|---|---|---|---|---|---|
| TRPM4 | 1011 | C | 0 | 0 | no | yes | no | no | yes | no | no | no | no | no | no | no | no |
| TRPM4 | 1012 | V | 1 | 0 | no | no | yes | no | yes | no | no | no | no | no | no | no | no |
| TRPM4 | 1013 | S | 0 | 0 | no | no | yes | no | yes | no | no | no | no | no | no | no | no |
| TRPM4 | 1014 | Q | 0 | 0 | no | yes | no | no | yes | no | no | no | no | no | no | no | no |
| TRPM4 | 1015 | Y | 1 | 0 | no | yes | no | no | yes | no | no | no | yes | no | no | no | no |
| TRPM4 | 1016 | A | 0 | 0 | no | yes | no | no | yes | no | no | no | no | no | no | no | no |
| TRPM4 | 1017 | N | 1 | 0 | no | yes | no | no | yes | no | no | no | no | no | no | no | no |
| TRPM4 | 1018 | W | 0 | 0 | yes | no | no | yes | yes | no | no | no | no | no | no | no | no |
| TRPM4 | 1019 | L | 1 | 0 | yes | no | no | yes | yes | no | no | no | yes | no | no | no | no |
| TRPM4 | 1020 | V | 1 | 0 | yes | no | no | yes | no | no | no | no | no | no | no | no | no |
| TRPM4 | 1021 | V | 1 | 0 | yes | no | no | yes | no | no | no | no | yes | no | no | no | no |
| TRPM4 | 1022 | L | 0 | 0 | yes | no | no | yes | no | no | no | no | no | no | no | no | no |
| TRPM4 | 1023 | L | 0 | 0 | yes | no | no | yes | no | no | no | no | no | no | no | no | no |
| TRPM4 | 1024 | L | 0 | 0 | yes | no | no | yes | no | no | no | no | no | no | no | no | no |
| TRPM4 | 1025 | V | 2 | 0 | yes | no | no | yes | no | no | no | no | yes | no | no | no | no |
| TRPM4 | 1026 | I | 1 | 0 | yes | no | no | yes | no | no | no | no | no | no | no | no | no |
| TRPM4 | 1027 | F | 0 | 0 | yes | no | no | yes | no | no | no | no | no | no | no | no | no |
| TRPM4 | 1028 | L | 0 | 0 | yes | no | no | yes | no | no | no | no | yes | no | no | no | no |
| TRPM4 | 1029 | L | 1 | 0 | yes | no | no | yes | no | no | no | no | yes | no | no | no | no |
| TRPM4 | 1030 | V | 0 | 0 | yes | no | no | yes | no | no | no | no | no | no | no | no | no |
| TRPM4 | 1031 | A | 0 | 0 | yes | no | no | yes | no | no | no | no | no | no | no | no | no |
| TRPM4 | 1032 | N | 0 | 0 | yes | no | no | yes | no | no | yes | no | yes | no | no | no | no |
| TRPM4 | 1033 | I | 0 | 1 | yes | no | no | yes | no | no | no | no | yes | no | no | no | no |
| TRPM4 | 1034 | L | 0 | 0 | yes | no | no | yes | no | no | no | no | yes | no | no | no | no |
| TRPM4 | 1035 | L | 0 | 0 | yes | no | no | yes | no | no | no | no | no | no | no | no | no |
| TRPM4 | 1036 | V | 0 | 0 | yes | no | no | yes | no | no | yes | no | no | no | no | no | no |
| TRPM4 | 1037 | N | 0 | 0 | yes | no | no | yes | no | no | no | no | yes | no | no | no | no |
| TRPM4 | 1038 | L | 0 | 0 | yes | no | no | yes | no | no | no | no | no | no | no | no | no |
| TRPM4 | 1039 | L | 0 | 0 | yes | no | no | yes | no | no | no | no | yes | no | no | no | no |
| TRPM4 | 1040 | I | 0 | 1 | yes | no | no | yes | no | no | yes | no | yes | no | no | no | no |
| TRPM4 | 1041 | A | 0 | 0 | yes | no | no | no | no | yes | no | no | no | no | no | no | no |
| TRPM4 | 1042 | M | 2 | 0 | yes | no | no | no | no | yes | no | no | yes | no | no | no | no |
| TRPM4 | 1043 | F | 0 | 0 | yes | no | no | yes | no | yes | no | no | yes | no | no | no | no |

| | | | | | | | | | | | | | | | | | |
|---|---|---|---|---|---|---|---|---|---|---|---|---|---|---|---|---|---|
| TRPM4 | 1044 | S | 0 | 0 | yes | no | no | no | no | yes | yes | no | no | no | no | no | no |
| TRPM4 | 1045 | Y | 0 | 0 | yes | no | no | no | no | yes | no | no | yes | no | no | no | no |
| TRPM4 | 1046 | T | 0 | 0 | yes | no | no | no | no | yes | no | no | no | no | no | no | no |
| TRPM4 | 1047 | F | 0 | 0 | yes | no | no | no | no | yes | no | no | no | no | no | no | no |
| TRPM4 | 1048 | G | 1 | 0 | yes | no | no | no | no | yes | yes | no | no | no | no | no | no |
| TRPM4 | 1049 | K | 1 | 0 | yes | no | no | no | no | yes | no | no | no | no | no | no | no |
| TRPM4 | 1050 | V | 1 | 0 | yes | no | no | no | no | yes | no | no | no | no | no | no | no |
| TRPM4 | 1051 | Q | 1 | 0 | yes | no | no | no | no | yes | no | no | yes | no | no | no | no |
| TRPM4 | 1052 | G | 0 | 0 | yes | no | no | no | no | yes | no | no | no | no | no | no | no |
| TRPM4 | 1053 | N | 0 | 0 | yes | no | no | no | no | yes | no | no | no | no | no | no | no |
| TRPM4 | 1054 | S | 2 | 0 | yes | no | no | no | no | yes | no | no | no | no | no | no | no |
| TRPM4 | 1055 | D | 0 | 0 | yes | no | no | no | no | yes | no | no | no | no | no | no | no |
| TRPM4 | 1056 | L | 0 | 0 | yes | no | no | no | no | yes | no | no | no | no | no | no | no |
| TRPM4 | 1057 | Y | 1 | 0 | yes | no | no | no | no | yes | no | no | no | no | no | no | no |
| TRPM4 | 1058 | W | 1 | 0 | yes | no | no | no | no | yes | no | no | no | no | no | no | no |
| TRPM4 | 1059 | K | 1 | 0 | yes | no | no | no | no | yes | no | no | no | no | no | no | no |
| TRPM4 | 1060 | A | 1 | 0 | yes | no | no | no | no | yes | no | no | no | no | no | no | no |
| TRPM4 | 1061 | Q | 0 | 0 | no | yes | no | no | no | yes | no | no | no | no | no | no | no |
| TRPM4 | 1062 | R | 1 | 0 | yes | no | no | no | no | yes | no | no | no | no | no | no | no |
| TRPM4 | 1063 | Y | 0 | 0 | yes | no | no | no | no | yes | no | no | no | no | no | no | no |
| TRPM4 | 1064 | R | 1 | 0 | yes | no | no | no | no | yes | no | no | no | no | no | no | no |
| TRPM4 | 1065 | L | 0 | 0 | yes | no | no | no | no | yes | no | no | no | no | no | no | no |
| TRPM4 | 1066 | I | 0 | 0 | yes | no | no | no | no | yes | no | no | no | no | no | no | no |
| TRPM4 | 1067 | R | 1 | 0 | yes | no | no | no | no | yes | no | no | no | no | no | no | no |
| TRPM4 | 1068 | E | 0 | 0 | yes | no | no | no | no | yes | no | no | no | no | no | no | no |
| TRPM4 | 1069 | F | 0 | 0 | yes | no | no | no | no | yes | no | no | no | no | no | no | no |
| TRPM4 | 1070 | H | 0 | 0 | yes | no | no | no | no | yes | no | no | no | no | no | no | no |
| TRPM4 | 1071 | S | 0 | 0 | yes | no | no | no | no | yes | no | no | no | no | no | no | no |
| TRPM4 | 1072 | R | 1 | 0 | no | yes | no | no | no | yes | no | no | no | no | no | no | no |
| TRPM4 | 1073 | P | 2 | 0 | no | yes | no | yes | no | yes | no | no | no | no | no | no | no |
| TRPM4 | 1074 | A | 1 | 0 | no | yes | no | yes | no | yes | no | no | no | no | no | no | no |
| TRPM4 | 1075 | L | 2 | 1 | no | yes | no | yes | no | yes | no | no | no | no | no | no | no |
| TRPM4 | 1076 | A | 0 | 0 | no | yes | no | yes | no | yes | no | no | no | no | no | no | no |

| | | | | | | | | | | | | | | | | | |
|---|---|---|---|---|---|---|---|---|---|---|---|---|---|---|---|---|---|
| TRPM4 | 1077 | P | 0 | 0 | no | yes | no | yes | no | yes | no | no | no | no | no | no | no |
| TRPM4 | 1078 | P | 0 | 0 | no | yes | no | yes | no | yes | no | no | no | no | no | no | no |
| TRPM4 | 1079 | F | 0 | 0 | no | yes | no | yes | no | yes | no | no | no | no | no | no | no |
| TRPM4 | 1080 | I | 1 | 0 | yes | no | no | yes | no | yes | no | no | no | no | no | no | no |
| TRPM4 | 1081 | V | 1 | 0 | yes | no | no | yes | no | yes | no | no | no | no | no | no | no |
| TRPM4 | 1082 | I | 0 | 0 | yes | no | no | yes | no | yes | no | no | no | no | no | no | no |
| TRPM4 | 1083 | S | 1 | 0 | yes | no | no | yes | no | yes | no | no | no | no | no | no | no |
| TRPM4 | 1084 | H | 2 | 0 | yes | no | no | yes | no | yes | no | no | no | no | no | no | no |
| TRPM4 | 1085 | L | 1 | 0 | yes | no | no | yes | no | yes | no | no | no | no | no | no | no |
| TRPM4 | 1086 | R | 2 | 0 | yes | no | no | yes | no | yes | no | no | no | no | no | no | no |
| TRPM4 | 1087 | L | 2 | 0 | yes | no | no | yes | no | yes | no | no | no | no | no | no | no |
| TRPM4 | 1088 | L | 2 | 0 | no | yes | no | yes | no | yes | no | no | no | no | no | no | no |
| TRPM4 | 1089 | L | 0 | 0 | no | yes | no | yes | no | yes | no | no | no | no | no | no | no |
| TRPM4 | 1090 | R | 1 | 0 | no | yes | no | no | no | yes | no | no | no | no | no | no | no |
| TRPM4 | 1105 | A | 1 | 0 | no | yes | no | no | no | yes | no | no | no | no | no | no | no |
| TRPM4 | 1106 | L | 1 | 0 | no | yes | no | no | no | yes | no | no | no | no | no | no | no |
| TRPM4 | 1107 | E | 1 | 0 | no | yes | no | no | no | yes | no | no | no | no | no | no | no |
| TRPM4 | 1108 | H | 0 | 0 | no | yes | no | no | no | yes | no | no | no | no | no | no | no |
| TRPM4 | 1109 | F | 0 | 0 | no | yes | no | no | no | yes | no | no | no | no | no | no | no |
| TRPM4 | 1110 | R | 0 | 0 | no | yes | no | no | no | yes | no | no | no | no | no | no | no |
| TRPM4 | 1111 | V | 1 | 0 | no | yes | no | no | no | yes | no | no | no | no | no | no | no |
| TRPM4 | 1112 | Y | 1 | 0 | no | yes | no | no | no | yes | no | no | no | no | no | no | no |
| TRPM4 | 1113 | L | 2 | 0 | no | yes | no | no | no | yes | no | no | no | no | no | no | no |
| TRPM4 | 1114 | S | 0 | 0 | no | yes | no | no | no | yes | no | no | no | no | no | no | no |
| TRPM4 | 1115 | K | 0 | 0 | yes | no | no | no | no | yes | no | no | no | no | no | no | no |
| TRPM4 | 1116 | E | 0 | 0 | yes | no | no | no | no | yes | no | no | no | no | no | no | no |
| TRPM4 | 1117 | A | 0 | 0 | yes | no | no | no | no | yes | no | no | no | no | no | no | no |
| TRPM4 | 1118 | E | 1 | 0 | yes | no | no | no | no | yes | no | no | no | no | no | no | no |
| TRPM4 | 1119 | R | 0 | 0 | yes | no | no | no | no | yes | no | no | yes | no | no | no | no |
| TRPM4 | 1120 | K | 1 | 0 | yes | no | no | no | no | yes | no | no | no | no | no | no | no |
| TRPM4 | 1121 | L | 1 | 0 | yes | no | no | no | no | yes | no | no | no | no | no | no | no |
| TRPM4 | 1122 | L | 0 | 0 | yes | no | no | no | no | yes | no | no | no | no | no | no | no |
| TRPM4 | 1123 | T | 1 | 0 | yes | no | no | no | no | yes | no | no | no | no | no | no | no |

| | | | | | | | | | | | | | | | | | |
|---|---|---|---|---|---|---|---|---|---|---|---|---|---|---|---|---|---|
| TRPM4 | 1124 | W | 0 | 0 | yes | no | no | no | no | yes | no | no | no | no | no | no | no |
| TRPM4 | 1125 | E | 0 | 0 | yes | no | no | no | no | yes | no | no | no | no | no | no | no |
| TRPM4 | 1126 | S | 1 | 0 | yes | no | no | no | no | yes | no | no | no | no | no | no | no |
| TRPM4 | 1127 | V | 1 | 0 | yes | no | no | no | no | yes | no | no | no | no | no | no | no |
| TRPM4 | 1128 | H | 2 | 0 | yes | no | no | no | no | yes | no | no | no | no | no | no | no |
| TRPM4 | 1129 | K | 2 | 0 | yes | no | no | no | no | yes | no | no | no | no | no | no | no |
| TRPM4 | 1130 | E | 0 | 0 | yes | no | no | no | no | yes | no | no | no | no | no | no | no |
| TRPM4 | 1131 | N | 1 | 0 | yes | no | no | no | no | yes | no | no | no | no | no | no | no |
| TRPM4 | 1132 | F | 1 | 0 | yes | no | no | no | no | yes | no | no | no | no | no | no | no |
| TRPM4 | 1133 | L | 1 | 0 | yes | no | no | no | no | yes | no | no | no | no | no | no | no |
| TRPM4 | 1134 | L | 0 | 0 | yes | no | no | no | no | yes | no | no | no | no | no | no | no |
| TRPM4 | 1135 | A | 1 | 0 | yes | no | no | no | no | yes | no | no | no | no | no | no | no |
| TRPM4 | 1136 | R | 2 | 0 | yes | no | no | no | no | yes | no | no | no | no | no | no | no |
| TRPM4 | 1137 | A | 1 | 0 | yes | no | no | no | no | yes | no | no | no | no | no | no | no |
| TRPM4 | 1138 | R | 0 | 0 | yes | no | no | no | no | yes | no | no | no | no | no | no | no |
| TRPM4 | 1139 | D | 0 | 0 | yes | no | no | no | no | yes | no | no | no | no | no | no | no |
| TRPM4 | 1140 | K | 0 | 0 | yes | no | no | no | no | yes | no | no | no | no | no | no | no |
| TRPM4 | 1141 | R | 1 | 0 | yes | no | no | no | no | yes | no | no | yes | no | no | no | no |
| TRPM4 | 1142 | E | 1 | 0 | no | yes | no | no | no | yes | no | no | no | no | no | no | no |
| TRPM4 | 1143 | S | 1 | 0 | no | yes | no | no | no | yes | no | no | no | no | no | no | no |
| TRPM4 | 1144 | D | 0 | 0 | no | yes | no | no | no | yes | yes | no | no | no | no | no | no |
| TRPM4 | 1145 | S | 2 | 0 | no | yes | no | no | no | yes | no | no | yes | no | no | no | no |
| TRPM4 | 1146 | E | 0 | 0 | no | yes | no | no | no | yes | no | no | no | no | no | no | no |
| TRPM4 | 1147 | R | 1 | 0 | yes | no | no | no | no | yes | no | no | yes | no | no | no | no |
| TRPM4 | 1148 | L | 1 | 0 | yes | no | no | no | no | yes | yes | no | yes | no | no | no | no |
| TRPM4 | 1149 | K | 1 | 0 | yes | no | no | no | no | yes | no | no | yes | no | no | no | no |
| TRPM4 | 1150 | R | 2 | 0 | yes | no | no | no | no | yes | no | no | no | no | no | no | no |
| TRPM4 | 1151 | T | 1 | 0 | yes | no | no | no | no | yes | yes | no | yes | no | no | no | no |
| TRPM4 | 1152 | S | 1 | 0 | yes | no | no | no | no | yes | no | no | yes | no | no | no | no |
| TRPM4 | 1153 | Q | 1 | 0 | yes | no | no | no | no | yes | no | no | no | no | no | no | no |
| TRPM4 | 1154 | K | 0 | 0 | yes | no | no | no | no | yes | no | no | yes | no | no | no | no |
| TRPM4 | 1155 | V | 0 | 0 | yes | no | no | no | no | yes | yes | no | yes | no | no | no | no |
| TRPM4 | 1156 | D | 1 | 0 | yes | no | no | no | no | yes | no | no | yes | no | no | no | no |

| | | | | | | | | | | | | | | | | | |
|---|---|---|---|---|---|---|---|---|---|---|---|---|---|---|---|---|---|
| TRPM4 | 1157 | L | 1 | 0 | yes | no | no | no | no | yes | no | no | no | no | no | no | no |
| TRPM4 | 1158 | A | 1 | 0 | yes | no | no | no | no | yes | yes | no | yes | no | no | no | no |
| TRPM4 | 1159 | L | 0 | 0 | yes | no | no | no | no | yes | no | no | yes | no | no | no | no |
| TRPM4 | 1160 | K | 0 | 0 | yes | no | no | no | no | yes | no | no | no | no | no | no | no |
| TRPM4 | 1161 | Q | 0 | 0 | yes | no | no | no | no | yes | no | no | yes | no | no | no | no |
| TRPM4 | 1162 | L | 0 | 0 | yes | no | no | no | no | yes | yes | no | yes | no | no | no | no |
| TRPM4 | 1163 | G | 1 | 0 | no | yes | no | no | no | yes | no | no | yes | no | no | no | no |
| TRPM4 | 1164 | H | 0 | 0 | no | yes | no | no | no | yes | no | no | yes | no | no | no | no |
| TRPM4 | 1165 | I | 1 | 0 | no | yes | no | no | no | yes | yes | no | no | no | no | no | no |
| TRPM4 | 1166 | R | 1 | 0 | yes | no | no | no | no | yes | no | no | yes | no | no | no | no |
| TRPM4 | 1167 | E | 1 | 0 | yes | no | no | no | no | yes | no | no | no | no | no | no | no |
| TRPM4 | 1168 | Y | 0 | 0 | yes | no | no | no | no | yes | no | no | yes | no | no | no | no |
| TRPM4 | 1169 | E | 1 | 0 | yes | no | no | no | no | yes | yes | no | yes | no | no | no | no |
| TRPM4 | 1170 | Q | 1 | 0 | yes | no | no | no | no | yes | no | no | no | no | no | no | no |
| TRPM4 | 1171 | R | 3 | 0 | yes | no | no | no | no | yes | no | no | no | no | no | no | no |
| TRPM4 | 1172 | L | 0 | 0 | yes | no | no | no | no | yes | yes | no | no | no | no | no | no |
| TRPM4 | 1173 | K | 0 | 0 | yes | no | no | no | no | yes | no | no | no | no | no | no | no |
| TRPM4 | 1174 | V | 0 | 0 | yes | no | no | no | no | yes | no | no | no | no | no | no | no |
| TRPM4 | 1175 | L | 0 | 0 | yes | no | no | no | no | yes | no | no | no | no | no | no | no |
| TRPM4 | 1176 | E | 0 | 0 | yes | no | no | no | no | yes | yes | no | no | no | no | no | no |
| TRPM4 | 1177 | R | 1 | 0 | yes | no | no | no | no | yes | no | no | no | no | no | no | no |
| TRPM4 | 1178 | E | 0 | 0 | yes | no | no | no | no | yes | no | no | no | no | no | no | no |
| TRPM4 | 1179 | V | 0 | 0 | yes | no | no | no | no | yes | yes | no | no | no | no | no | no |
| TRPM4 | 1180 | Q | 0 | 0 | yes | no | no | no | no | yes | no | no | no | no | no | no | no |
| TRPM4 | 1181 | Q | 1 | 0 | yes | no | no | no | no | yes | no | no | no | no | no | no | no |
| TRPM4 | 1182 | C | 2 | 0 | yes | no | no | no | no | yes | no | no | no | no | no | no | no |
| TRPM4 | 1183 | S | 0 | 0 | yes | no | no | no | no | yes | yes | no | no | no | no | no | no |
| TRPM4 | 1184 | R | 1 | 0 | yes | no | no | no | no | yes | no | no | no | no | no | no | no |
| TRPM4 | 1185 | V | 1 | 0 | yes | no | no | no | no | yes | no | no | no | no | no | no | no |
| TRPM4 | 1186 | L | 1 | 0 | yes | no | no | no | no | yes | yes | no | no | no | no | no | no |
| TRPM4 | 1187 | G | 1 | 0 | yes | no | no | no | no | yes | no | no | no | no | no | no | no |
| TRPM4 | 1188 | W | 1 | 0 | yes | no | no | no | no | yes | no | no | no | no | no | no | no |
| TRPM4 | 1189 | V | 0 | 0 | yes | no | no | no | no | yes | no | no | no | no | no | no | no |

| | | | | | | | | | | | | | | | | | |
|---|---|---|---|---|---|---|---|---|---|---|---|---|---|---|---|---|---|
| TRPM4 | 1190 | A | 1 | 0 | yes | no | no | no | no | yes | no | no | no | no | no | no | no |
| TRPM4 | 1191 | E | 0 | 0 | yes | no | no | no | no | yes | no | no | no | no | no | no | no |
| TRPM4 | 1192 | A | 1 | 0 | no | yes | no | no | no | yes | no | no | no | no | no | no | no |
| TRPV3 | 114 | R | 1 | 0 | no | yes | no | no | no | yes | no | no | no | no | no | no | no |
| TRPV3 | 115 | K | 0 | 0 | yes | no | no | no | no | yes | no | no | no | no | no | no | no |
| TRPV3 | 116 | K | 1 | 0 | yes | no | no | no | no | yes | no | no | no | no | no | no | no |
| TRPV3 | 117 | R | 1 | 0 | yes | no | no | no | no | yes | no | no | no | no | no | no | no |
| TRPV3 | 118 | R | 2 | 0 | yes | no | no | no | no | yes | no | no | no | no | no | no | no |
| TRPV3 | 119 | L | 0 | 0 | yes | no | no | no | no | yes | no | no | no | no | no | no | no |
| TRPV3 | 120 | K | 1 | 0 | yes | no | no | no | no | yes | no | no | no | no | no | no | no |
| TRPV3 | 121 | K | 0 | 0 | yes | no | no | no | no | yes | no | no | no | no | no | no | no |
| TRPV3 | 122 | R | 3 | 0 | yes | no | no | no | no | yes | no | no | no | no | no | no | no |
| TRPV3 | 123 | I | 2 | 0 | yes | no | no | no | no | yes | no | no | no | no | no | no | no |
| TRPV3 | 124 | F | 0 | 0 | yes | no | no | no | no | yes | no | no | no | no | no | no | no |
| TRPV3 | 125 | A | 0 | 0 | yes | no | no | no | no | yes | no | no | no | no | no | no | no |
| TRPV3 | 126 | A | 0 | 0 | yes | no | no | no | no | yes | no | no | no | no | no | no | no |
| TRPV3 | 127 | V | 2 | 0 | yes | no | no | no | no | yes | no | no | no | no | no | no | no |
| TRPV3 | 128 | S | 0 | 0 | yes | no | no | no | no | yes | no | no | no | no | no | no | no |
| TRPV3 | 129 | E | 0 | 0 | yes | no | no | no | no | yes | no | no | no | no | no | no | no |
| TRPV3 | 130 | G | 1 | 0 | no | yes | no | no | no | yes | no | no | no | no | no | no | no |
| TRPV3 | 131 | C | 0 | 0 | no | yes | no | no | no | yes | no | no | no | no | no | no | no |
| TRPV3 | 132 | V | 2 | 0 | no | yes | no | no | no | yes | no | no | no | no | no | no | no |
| TRPV3 | 133 | E | 0 | 0 | yes | no | no | no | no | yes | no | no | no | no | no | no | no |
| TRPV3 | 134 | E | 2 | 0 | yes | no | no | no | no | yes | no | no | no | no | no | no | no |
| TRPV3 | 135 | L | 0 | 0 | yes | no | no | no | no | yes | no | no | no | no | no | no | no |
| TRPV3 | 136 | V | 0 | 0 | yes | no | no | no | no | yes | no | no | no | no | no | no | no |
| TRPV3 | 137 | E | 0 | 0 | yes | no | no | no | no | yes | no | no | no | no | no | no | no |
| TRPV3 | 138 | L | 0 | 0 | yes | no | no | no | no | yes | no | no | no | no | no | no | no |
| TRPV3 | 139 | L | 0 | 0 | yes | no | no | no | no | yes | no | no | no | no | no | no | no |
| TRPV3 | 140 | V | 0 | 0 | yes | no | no | no | no | yes | no | no | no | no | no | no | no |
| TRPV3 | 141 | E | 0 | 0 | yes | no | no | no | no | yes | no | no | no | no | no | no | no |
| TRPV3 | 142 | L | 0 | 0 | yes | no | no | no | no | yes | no | no | no | no | no | no | no |
| TRPV3 | 143 | Q | 1 | 0 | yes | no | no | no | no | yes | no | no | no | no | no | no | no |

| | | | | | | | | | | | | | | | | | |
|---|---|---|---|---|---|---|---|---|---|---|---|---|---|---|---|---|---|
| TRPV3 | 144 | E | 0 | 0 | yes | no | no | no | no | yes | no | no | no | no | no | no | no |
| TRPV3 | 145 | L | 0 | 0 | yes | no | no | no | no | yes | no | no | no | no | no | no | no |
| TRPV3 | 146 | C | 0 | 0 | yes | no | no | no | no | yes | no | no | no | no | no | no | no |
| TRPV3 | 147 | R | 0 | 0 | yes | no | no | no | no | yes | no | no | no | no | no | no | no |
| TRPV3 | 148 | R | 3 | 0 | no | yes | no | no | no | yes | no | no | no | no | no | no | no |
| TRPV3 | 149 | R | 2 | 0 | no | yes | no | no | no | yes | no | no | no | no | no | no | no |
| TRPV3 | 150 | H | 0 | 0 | no | yes | no | no | no | yes | no | no | no | no | no | no | no |
| TRPV3 | 151 | D | 0 | 0 | no | yes | no | no | no | yes | no | no | no | no | no | no | no |
| TRPV3 | 152 | E | 0 | 0 | no | yes | no | no | no | yes | no | no | no | no | no | no | no |
| TRPV3 | 153 | D | 0 | 0 | no | yes | no | no | no | yes | no | no | no | no | no | no | no |
| TRPV3 | 154 | V | 0 | 0 | yes | no | no | no | no | yes | no | no | no | no | no | no | no |
| TRPV3 | 155 | P | 1 | 0 | yes | no | no | no | no | yes | no | no | no | no | no | no | no |
| TRPV3 | 156 | D | 0 | 0 | yes | no | no | no | no | yes | no | no | no | no | no | no | no |
| TRPV3 | 157 | F | 1 | 0 | yes | no | no | no | no | yes | no | no | no | no | no | no | no |
| TRPV3 | 158 | L | 0 | 0 | yes | no | no | no | no | yes | no | no | no | no | no | no | no |
| TRPV3 | 159 | M | 3 | 0 | yes | no | no | no | no | yes | no | no | no | no | no | no | no |
| TRPV3 | 160 | H | 0 | 0 | yes | no | no | no | no | yes | no | no | no | no | no | no | no |
| TRPV3 | 161 | K | 0 | 0 | yes | no | no | no | no | yes | no | no | no | no | no | no | no |
| TRPV3 | 162 | L | 0 | 0 | yes | no | no | no | no | yes | no | no | no | no | no | no | no |
| TRPV3 | 163 | T | 1 | 0 | no | no | yes | no | no | yes | no | no | no | no | no | no | no |
| TRPV3 | 164 | A | 1 | 0 | no | no | yes | no | no | yes | no | no | no | no | no | no | no |
| TRPV3 | 165 | S | 1 | 0 | no | yes | no | no | no | yes | no | no | no | no | no | no | no |
| TRPV3 | 166 | D | 1 | 0 | no | yes | no | no | no | yes | no | no | no | no | no | no | no |
| TRPV3 | 167 | T | 1 | 0 | no | yes | no | no | no | yes | no | no | no | no | no | no | no |
| TRPV3 | 168 | G | 0 | 0 | no | yes | no | no | no | yes | no | no | no | no | no | no | no |
| TRPV3 | 169 | K | 0 | 0 | no | no | yes | no | no | yes | no | no | yes | no | no | no | no |
| TRPV3 | 170 | T | 3 | 0 | no | no | yes | no | no | yes | no | no | no | no | no | no | no |
| TRPV3 | 171 | C | 0 | 0 | yes | no | no | no | no | yes | no | no | no | no | no | no | no |
| TRPV3 | 172 | L | 0 | 0 | yes | no | no | no | no | yes | no | no | no | no | no | no | no |
| TRPV3 | 173 | M | 1 | 0 | yes | no | no | no | no | yes | no | no | no | no | no | no | no |
| TRPV3 | 174 | K | 0 | 0 | yes | no | no | no | no | yes | no | no | no | no | no | no | no |
| TRPV3 | 175 | A | 0 | 0 | yes | no | no | no | no | yes | no | no | no | no | no | no | no |
| TRPV3 | 176 | L | 0 | 0 | yes | no | no | no | no | yes | no | no | no | no | no | no | no |

| | | | | | | | | | | | | | | | | | |
|---|---|---|---|---|---|---|---|---|---|---|---|---|---|---|---|---|---|
| TRPV3 | 177 | L | 0 | 0 | yes | no | no | no | no | yes | no | no | yes | no | no | no | no |
| TRPV3 | 178 | N | 0 | 0 | no | yes | no | no | no | yes | no | no | yes | no | no | no | no |
| TRPV3 | 179 | I | 1 | 0 | no | yes | no | no | no | yes | no | no | no | no | no | no | no |
| TRPV3 | 180 | N | 2 | 0 | no | yes | no | no | no | yes | no | no | no | no | no | no | no |
| TRPV3 | 181 | P | 2 | 0 | no | yes | no | no | no | yes | no | no | no | no | no | no | no |
| TRPV3 | 182 | N | 1 | 0 | no | yes | no | no | no | yes | no | no | no | no | no | no | no |
| TRPV3 | 183 | T | 0 | 0 | yes | no | no | no | no | yes | no | no | no | no | no | no | no |
| TRPV3 | 184 | K | 0 | 0 | yes | no | no | no | no | yes | no | no | no | no | no | no | no |
| TRPV3 | 185 | E | 1 | 0 | yes | no | no | no | no | yes | no | no | no | no | no | no | no |
| TRPV3 | 186 | I | 2 | 0 | yes | no | no | no | no | yes | no | no | no | no | no | no | no |
| TRPV3 | 187 | V | 2 | 0 | yes | no | no | no | no | yes | no | no | no | no | no | no | no |
| TRPV3 | 188 | R | 2 | 0 | yes | no | no | no | no | yes | no | no | no | no | no | no | no |
| TRPV3 | 189 | I | 0 | 0 | yes | no | no | no | no | yes | no | no | no | no | no | no | no |
| TRPV3 | 190 | L | 0 | 0 | yes | no | no | no | no | yes | no | no | no | no | no | no | no |
| TRPV3 | 191 | L | 0 | 0 | yes | no | no | no | no | yes | no | no | no | no | no | no | no |
| TRPV3 | 192 | A | 0 | 0 | yes | no | no | no | no | yes | no | no | no | no | no | no | no |
| TRPV3 | 193 | F | 1 | 0 | yes | no | no | no | no | yes | no | no | no | no | no | no | no |
| TRPV3 | 194 | A | 0 | 0 | yes | no | no | no | no | yes | no | no | no | no | no | no | no |
| TRPV3 | 195 | E | 0 | 0 | yes | no | no | no | no | yes | no | no | no | no | no | no | no |
| TRPV3 | 196 | E | 1 | 0 | yes | no | no | no | no | yes | no | no | no | no | no | no | no |
| TRPV3 | 197 | N | 1 | 0 | no | yes | no | no | no | yes | no | no | no | no | no | no | no |
| TRPV3 | 198 | D | 1 | 0 | no | yes | no | no | no | yes | no | no | no | no | no | no | no |
| TRPV3 | 199 | I | 1 | 0 | no | yes | no | no | no | yes | no | no | no | no | no | no | no |
| TRPV3 | 200 | L | 0 | 0 | yes | no | no | no | no | yes | no | no | no | no | no | no | no |
| TRPV3 | 201 | G | 1 | 0 | yes | no | no | no | no | yes | no | no | no | no | no | no | no |
| TRPV3 | 202 | R | 2 | 0 | yes | no | no | no | no | yes | no | no | no | no | no | no | no |
| TRPV3 | 203 | F | 0 | 0 | yes | no | no | no | no | yes | no | no | no | no | no | no | no |
| TRPV3 | 204 | I | 0 | 0 | yes | no | no | no | no | yes | no | no | no | no | no | no | no |
| TRPV3 | 205 | N | 1 | 0 | no | yes | no | no | no | yes | no | no | no | no | no | no | no |
| TRPV3 | 206 | A | 2 | 0 | no | yes | no | no | no | yes | no | no | no | no | no | no | no |
| TRPV3 | 207 | E | 1 | 0 | no | yes | no | no | no | yes | no | no | no | no | no | no | no |
| TRPV3 | 208 | Y | 0 | 0 | no | no | yes | no | no | yes | no | no | yes | no | no | no | no |
| TRPV3 | 209 | T | 0 | 0 | no | yes | no | no | no | yes | no | no | no | no | no | no | no |

| | | | | | | | | | | | | | | | | | |
|---|---|---|---|---|---|---|---|---|---|---|---|---|---|---|---|---|---|
| TRPV3 | 210 | E | 0 | 0 | no | yes | no | no | no | yes | no | no | no | no | no | no | no |
| TRPV3 | 211 | E | 1 | 0 | no | yes | no | no | no | yes | no | no | no | no | no | no | no |
| TRPV3 | 212 | A | 0 | 0 | no | yes | no | no | no | yes | no | no | no | no | no | no | no |
| TRPV3 | 213 | Y | 1 | 0 | no | yes | no | no | no | yes | no | no | yes | no | no | no | no |
| TRPV3 | 214 | E | 1 | 0 | no | no | yes | no | no | yes | no | no | no | no | no | no | no |
| TRPV3 | 215 | G | 0 | 0 | no | yes | no | no | no | yes | no | no | no | no | no | no | no |
| TRPV3 | 216 | Q | 0 | 0 | no | yes | no | no | no | yes | no | no | no | no | no | no | no |
| TRPV3 | 217 | T | 1 | 0 | no | yes | no | no | no | yes | no | no | no | no | no | no | no |
| TRPV3 | 218 | A | 1 | 0 | yes | no | no | no | no | yes | no | no | no | no | no | no | no |
| TRPV3 | 219 | L | 0 | 0 | yes | no | no | no | no | yes | no | no | no | no | no | no | no |
| TRPV3 | 220 | N | 0 | 0 | yes | no | no | no | no | yes | no | no | no | no | no | no | no |
| TRPV3 | 221 | I | 2 | 0 | yes | no | no | no | no | yes | no | no | no | no | no | no | no |
| TRPV3 | 222 | A | 0 | 0 | yes | no | no | no | no | yes | no | no | no | no | no | no | no |
| TRPV3 | 223 | I | 1 | 0 | yes | no | no | no | no | yes | no | no | no | no | no | no | no |
| TRPV3 | 224 | E | 0 | 0 | yes | no | no | no | no | yes | no | no | yes | no | no | no | no |
| TRPV3 | 225 | R | 0 | 0 | no | yes | no | no | no | yes | no | no | yes | no | no | no | no |
| TRPV3 | 226 | R | 0 | 0 | no | yes | no | no | no | yes | no | no | yes | no | no | no | no |
| TRPV3 | 227 | Q | 0 | 0 | no | yes | no | no | no | yes | no | no | yes | no | no | no | no |
| TRPV3 | 228 | G | 1 | 0 | no | yes | no | no | no | yes | no | no | no | no | no | no | no |
| TRPV3 | 229 | D | 0 | 0 | yes | no | no | no | no | yes | no | no | no | no | no | no | no |
| TRPV3 | 230 | I | 1 | 0 | yes | no | no | no | no | yes | no | no | no | no | no | no | no |
| TRPV3 | 231 | A | 0 | 0 | yes | no | no | no | no | yes | no | no | no | no | no | no | no |
| TRPV3 | 232 | A | 1 | 0 | yes | no | no | no | no | yes | no | no | no | no | no | no | no |
| TRPV3 | 233 | L | 0 | 0 | yes | no | no | no | no | yes | no | no | no | no | no | no | no |
| TRPV3 | 234 | L | 0 | 0 | yes | no | no | no | no | yes | no | no | no | no | no | no | no |
| TRPV3 | 235 | I | 0 | 0 | yes | no | no | no | no | yes | no | no | no | no | no | no | no |
| TRPV3 | 236 | A | 1 | 0 | yes | no | no | no | no | yes | no | no | no | no | no | no | no |
| TRPV3 | 237 | A | 0 | 0 | no | yes | no | no | no | yes | no | no | no | no | no | no | no |
| TRPV3 | 238 | G | 2 | 0 | no | yes | no | no | no | yes | no | no | no | no | no | no | no |
| TRPV3 | 239 | A | 0 | 0 | no | yes | no | no | no | yes | no | no | no | no | no | no | no |
| TRPV3 | 240 | D | 2 | 0 | no | yes | no | no | no | yes | no | no | no | no | no | no | no |
| TRPV3 | 241 | V | 0 | 0 | no | yes | no | no | no | yes | no | no | no | no | no | no | no |
| TRPV3 | 242 | N | 1 | 0 | no | yes | no | no | no | yes | no | no | no | no | no | no | no |

| | | | | | | | | | | | | | | | | | |
|---|---|---|---|---|---|---|---|---|---|---|---|---|---|---|---|---|---|
| TRPV3 | 243 | A | 1 | 0 | no | yes | no | no | no | yes | no | no | no | no | no | no | no |
| TRPV3 | 244 | H | 1 | 0 | no | yes | no | no | no | yes | no | no | no | no | no | no | no |
| TRPV3 | 245 | A | 0 | 0 | no | yes | no | no | no | yes | no | no | no | no | no | no | no |
| TRPV3 | 246 | K | 1 | 0 | no | yes | no | no | no | yes | no | no | no | no | no | no | no |
| TRPV3 | 247 | G | 0 | 0 | no | yes | no | no | no | yes | no | no | no | no | no | no | no |
| TRPV3 | 248 | A | 0 | 0 | no | yes | no | no | no | yes | no | no | no | no | no | no | no |
| TRPV3 | 249 | F | 0 | 0 | no | yes | no | no | no | yes | no | no | yes | no | no | no | no |
| TRPV3 | 250 | F | 0 | 0 | no | yes | no | no | no | yes | no | no | yes | no | no | no | no |
| TRPV3 | 251 | N | 1 | 0 | no | yes | no | no | no | yes | no | no | no | no | no | no | no |
| TRPV3 | 252 | P | 1 | 0 | no | yes | no | no | no | yes | no | no | no | no | no | no | no |
| TRPV3 | 253 | K | 0 | 0 | no | yes | no | no | no | yes | no | no | no | no | no | no | no |
| TRPV3 | 254 | Y | 1 | 0 | no | yes | no | no | no | yes | no | no | no | no | no | no | no |
| TRPV3 | 255 | Q | 1 | 0 | no | yes | no | no | no | yes | no | no | no | no | no | no | no |
| TRPV3 | 256 | H | 0 | 0 | no | yes | no | no | no | yes | no | no | no | no | no | no | no |
| TRPV3 | 257 | E | 0 | 0 | no | yes | no | no | no | yes | no | no | no | no | no | no | no |
| TRPV3 | 258 | G | 1 | 0 | no | yes | no | no | no | yes | no | no | no | no | no | no | no |
| TRPV3 | 259 | F | 0 | 0 | no | yes | no | no | no | yes | no | no | yes | no | no | no | no |
| TRPV3 | 260 | Y | 0 | 0 | no | yes | no | no | no | yes | no | no | no | no | no | no | no |
| TRPV3 | 261 | F | 0 | 0 | no | yes | no | no | no | yes | no | no | yes | no | no | no | no |
| TRPV3 | 262 | G | 0 | 0 | no | yes | no | no | no | yes | no | no | no | no | no | no | no |
| TRPV3 | 263 | E | 0 | 0 | no | yes | no | no | no | yes | no | no | no | no | no | no | no |
| TRPV3 | 264 | T | 1 | 0 | no | yes | no | no | no | yes | no | no | no | no | no | no | no |
| TRPV3 | 265 | P | 1 | 0 | yes | no | no | no | no | yes | no | no | no | no | no | no | no |
| TRPV3 | 266 | L | 0 | 0 | yes | no | no | no | no | yes | no | no | no | no | no | no | no |
| TRPV3 | 267 | A | 1 | 0 | yes | no | no | no | no | yes | no | no | no | no | no | no | no |
| TRPV3 | 268 | L | 0 | 0 | yes | no | no | no | no | yes | no | no | yes | no | no | no | no |
| TRPV3 | 269 | A | 0 | 0 | yes | no | no | no | no | yes | no | no | no | no | no | no | no |
| TRPV3 | 270 | A | 1 | 0 | yes | no | no | no | no | yes | no | no | no | no | no | no | no |
| TRPV3 | 271 | C | 1 | 0 | yes | no | no | no | no | yes | no | no | no | no | no | no | no |
| TRPV3 | 272 | T | 0 | 0 | no | yes | no | no | no | yes | no | no | no | no | no | no | no |
| TRPV3 | 273 | N | 1 | 0 | no | yes | no | no | no | yes | no | no | no | no | no | no | no |
| TRPV3 | 274 | Q | 0 | 0 | no | yes | no | no | no | yes | no | no | no | no | no | no | no |
| TRPV3 | 275 | P | 0 | 0 | yes | no | no | no | no | yes | no | no | no | no | no | no | no |

| | | | | | | | | | | | | | | | | | |
|---|---|---|---|---|---|---|---|---|---|---|---|---|---|---|---|---|---|
| TRPV3 | 276 | E | 1 | 0 | yes | no | no | no | no | yes | no | no | no | no | no | no | no |
| TRPV3 | 277 | I | 1 | 0 | yes | no | no | no | no | yes | no | no | no | no | no | no | no |
| TRPV3 | 278 | V | 0 | 0 | yes | no | no | no | no | yes | no | no | no | no | no | no | no |
| TRPV3 | 279 | Q | 2 | 0 | yes | no | no | no | no | yes | no | no | no | no | no | no | no |
| TRPV3 | 280 | L | 0 | 0 | yes | no | no | no | no | yes | no | no | no | no | no | no | no |
| TRPV3 | 281 | L | 0 | 0 | yes | no | no | no | no | yes | no | no | no | no | no | no | no |
| TRPV3 | 282 | M | 1 | 0 | yes | no | no | no | no | yes | no | no | no | no | no | no | no |
| TRPV3 | 283 | E | 0 | 0 | yes | no | no | no | no | yes | no | no | no | no | no | no | no |
| TRPV3 | 284 | H | 2 | 0 | no | yes | no | no | no | yes | no | no | no | no | no | no | no |
| TRPV3 | 285 | E | 1 | 0 | no | yes | no | no | no | yes | no | no | no | no | no | no | no |
| TRPV3 | 286 | Q | 1 | 0 | no | yes | no | no | no | yes | no | no | no | no | no | no | no |
| TRPV3 | 287 | T | 3 | 0 | no | yes | no | no | no | yes | no | no | no | no | no | no | no |
| TRPV3 | 288 | D | 0 | 0 | no | yes | no | no | no | yes | no | no | no | no | no | no | no |
| TRPV3 | 289 | I | 1 | 0 | no | yes | no | no | no | yes | no | no | no | no | no | no | no |
| TRPV3 | 290 | T | 1 | 0 | no | yes | no | no | no | yes | no | no | no | no | no | no | no |
| TRPV3 | 291 | S | 1 | 0 | no | yes | no | no | no | yes | no | no | no | no | no | no | no |
| TRPV3 | 292 | R | 2 | 0 | no | yes | no | no | no | yes | no | no | no | no | no | no | no |
| TRPV3 | 293 | D | 1 | 0 | no | yes | no | no | no | yes | no | no | no | no | no | no | no |
| TRPV3 | 294 | S | 0 | 0 | no | yes | no | no | no | yes | no | no | no | no | no | no | no |
| TRPV3 | 295 | R | 1 | 0 | no | yes | no | no | no | yes | no | no | no | no | no | no | no |
| TRPV3 | 296 | G | 1 | 0 | no | yes | no | no | no | yes | no | no | no | no | no | no | no |
| TRPV3 | 297 | N | 2 | 0 | no | yes | no | no | no | yes | no | no | no | no | no | no | no |
| TRPV3 | 298 | N | 1 | 0 | no | yes | no | no | no | yes | no | no | no | no | no | no | no |
| TRPV3 | 299 | I | 0 | 0 | yes | no | no | no | no | yes | no | no | no | no | no | no | no |
| TRPV3 | 300 | L | 0 | 0 | yes | no | no | no | no | yes | no | no | no | no | no | no | no |
| TRPV3 | 301 | H | 1 | 0 | yes | no | no | no | no | yes | no | no | no | no | no | no | no |
| TRPV3 | 302 | A | 1 | 0 | yes | no | no | no | no | yes | no | no | no | no | no | no | no |
| TRPV3 | 303 | L | 0 | 0 | yes | no | no | no | no | yes | no | no | no | no | no | no | no |
| TRPV3 | 304 | V | 0 | 0 | yes | no | no | no | no | yes | no | no | no | no | no | no | no |
| TRPV3 | 305 | T | 1 | 0 | no | yes | no | no | no | yes | no | no | no | no | no | no | no |
| TRPV3 | 306 | V | 1 | 0 | no | yes | no | no | no | yes | no | no | yes | no | no | no | no |
| TRPV3 | 307 | A | 0 | 0 | no | yes | no | no | no | yes | no | no | no | no | no | no | no |
| TRPV3 | 308 | E | 1 | 0 | no | yes | no | no | no | yes | no | no | no | no | no | no | no |

| | | | | | | | | | | | | | | | | | |
|---|---|---|---|---|---|---|---|---|---|---|---|---|---|---|---|---|---|
| TRPV3 | 309 | D | 0 | 0 | no | yes | no | no | no | yes | no | no | no | no | no | no | no |
| TRPV3 | 310 | F | 1 | 0 | yes | no | no | no | no | yes | yes | no | no | no | no | no | no |
| TRPV3 | 311 | K | 0 | 0 | yes | no | no | no | no | yes | yes | no | no | no | no | no | no |
| TRPV3 | 312 | T | 2 | 0 | yes | no | no | no | no | yes | no | no | yes | no | no | no | no |
| TRPV3 | 313 | Q | 0 | 0 | yes | no | no | no | no | yes | no | no | no | no | no | no | no |
| TRPV3 | 314 | N | 0 | 0 | no | yes | no | no | no | yes | yes | no | no | no | no | no | no |
| TRPV3 | 315 | D | 0 | 0 | no | yes | no | no | no | yes | no | no | no | no | no | no | no |
| TRPV3 | 316 | F | 1 | 0 | no | yes | no | no | no | yes | no | no | no | no | no | no | no |
| TRPV3 | 317 | V | 1 | 0 | yes | no | no | no | no | yes | no | no | no | no | no | no | no |
| TRPV3 | 318 | K | 0 | 0 | yes | no | no | no | no | yes | no | no | no | no | no | no | no |
| TRPV3 | 319 | R | 3 | 0 | yes | no | no | no | no | yes | no | no | no | no | no | no | no |
| TRPV3 | 320 | M | 2 | 0 | yes | no | no | no | no | yes | no | no | no | no | no | no | no |
| TRPV3 | 321 | Y | 0 | 0 | yes | no | no | no | no | yes | no | no | no | no | no | no | no |
| TRPV3 | 322 | D | 1 | 0 | yes | no | no | no | no | yes | no | no | no | no | no | no | no |
| TRPV3 | 323 | M | 1 | 0 | yes | no | no | no | no | yes | no | no | no | no | no | no | no |
| TRPV3 | 324 | I | 3 | 0 | yes | no | no | no | no | yes | no | no | no | no | no | no | no |
| TRPV3 | 325 | L | 0 | 0 | yes | no | no | no | no | yes | no | no | no | no | no | no | no |
| TRPV3 | 326 | L | 1 | 0 | yes | no | no | no | no | yes | no | no | no | no | no | no | no |
| TRPV3 | 327 | R | 2 | 0 | yes | no | no | no | no | yes | no | no | no | no | no | no | no |
| TRPV3 | 328 | S | 1 | 0 | no | yes | no | no | no | yes | no | no | no | no | no | no | no |
| TRPV3 | 329 | G | 0 | 0 | no | yes | no | no | no | yes | no | no | no | no | no | no | no |
| TRPV3 | 330 | N | 2 | 0 | no | yes | no | no | no | yes | no | no | no | no | no | no | no |
| TRPV3 | 331 | W | 0 | 0 | no | yes | no | no | no | yes | no | no | no | no | no | no | no |
| TRPV3 | 332 | E | 1 | 0 | yes | no | no | no | no | yes | no | no | no | no | no | no | no |
| TRPV3 | 333 | L | 0 | 0 | yes | no | no | no | no | yes | no | no | no | no | no | no | no |
| TRPV3 | 334 | E | 0 | 0 | yes | no | no | no | no | yes | no | no | no | no | no | no | no |
| TRPV3 | 335 | T | 0 | 0 | yes | no | no | no | no | yes | no | no | no | no | no | no | no |
| TRPV3 | 336 | T | 1 | 0 | no | yes | no | no | no | yes | no | no | no | no | no | no | no |
| TRPV3 | 337 | R | 2 | 0 | no | yes | no | no | no | yes | no | no | no | no | no | no | no |
| TRPV3 | 338 | N | 0 | 0 | no | yes | no | no | no | yes | no | no | no | no | no | no | no |
| TRPV3 | 339 | N | 2 | 0 | no | yes | no | no | no | yes | no | no | no | no | no | no | no |
| TRPV3 | 340 | D | 3 | 0 | no | yes | no | no | no | yes | no | no | no | no | no | no | no |
| TRPV3 | 341 | G | 1 | 0 | no | yes | no | no | no | yes | no | no | no | no | no | no | no |

| | | | | | | | | | | | | | | | | | |
|---|---|---|---|---|---|---|---|---|---|---|---|---|---|---|---|---|---|
| TRPV3 | 342 | L | 0 | 0 | no | yes | no | no | no | yes | no | no | no | no | no | no | no |
| TRPV3 | 343 | T | 1 | 0 | no | yes | no | no | no | yes | no | no | no | no | no | no | no |
| TRPV3 | 344 | P | 1 | 0 | yes | no | no | no | no | yes | no | no | no | no | no | no | no |
| TRPV3 | 345 | L | 1 | 0 | yes | no | no | no | no | yes | no | no | no | no | no | no | no |
| TRPV3 | 346 | Q | 0 | 0 | yes | no | no | no | no | yes | no | no | no | no | no | no | no |
| TRPV3 | 347 | L | 0 | 0 | yes | no | no | no | no | yes | no | no | no | no | no | no | no |
| TRPV3 | 348 | A | 0 | 0 | yes | no | no | no | no | yes | no | no | no | no | no | no | no |
| TRPV3 | 349 | A | 1 | 0 | yes | no | no | no | no | yes | no | no | no | no | no | no | no |
| TRPV3 | 350 | K | 1 | 0 | yes | no | no | no | no | yes | no | no | no | no | no | no | no |
| TRPV3 | 351 | M | 0 | 0 | no | yes | no | no | no | yes | no | no | no | no | no | no | no |
| TRPV3 | 352 | G | 0 | 0 | no | yes | no | no | no | yes | no | no | no | no | no | no | no |
| TRPV3 | 353 | K | 0 | 0 | no | yes | no | no | no | yes | no | no | no | no | no | no | no |
| TRPV3 | 354 | A | 1 | 0 | no | yes | no | no | no | yes | no | no | no | no | no | no | no |
| TRPV3 | 355 | E | 1 | 0 | yes | no | no | no | no | yes | no | no | no | no | no | no | no |
| TRPV3 | 356 | I | 1 | 0 | yes | no | no | no | no | yes | no | no | no | no | no | no | no |
| TRPV3 | 357 | L | 0 | 0 | yes | no | no | no | no | yes | no | no | no | no | no | no | no |
| TRPV3 | 358 | K | 1 | 0 | yes | no | no | no | no | yes | no | no | no | no | no | no | no |
| TRPV3 | 359 | Y | 0 | 0 | yes | no | no | no | no | yes | no | no | no | no | no | no | no |
| TRPV3 | 360 | I | 2 | 0 | yes | no | no | no | no | yes | no | no | no | no | no | no | no |
| TRPV3 | 361 | L | 0 | 0 | yes | no | no | no | no | yes | no | no | no | no | no | no | no |
| TRPV3 | 362 | S | 1 | 0 | no | yes | no | no | no | yes | no | no | no | no | no | no | no |
| TRPV3 | 363 | R | 3 | 0 | no | yes | no | no | no | yes | no | no | no | no | no | no | no |
| TRPV3 | 364 | E | 1 | 0 | no | yes | no | no | no | yes | no | no | no | no | no | no | no |
| TRPV3 | 365 | I | 0 | 0 | no | yes | no | no | no | yes | no | no | no | no | no | no | no |
| TRPV3 | 366 | K | 1 | 0 | no | yes | no | no | no | yes | no | no | no | no | no | no | no |
| TRPV3 | 367 | E | 0 | 0 | no | yes | no | no | no | yes | no | no | no | no | no | no | no |
| TRPV3 | 368 | K | 0 | 0 | no | yes | no | no | no | yes | no | no | no | no | no | no | no |
| TRPV3 | 369 | R | 2 | 0 | no | yes | no | no | no | yes | no | no | no | no | no | no | no |
| TRPV3 | 370 | L | 0 | 0 | yes | no | no | no | no | yes | no | no | no | no | no | no | no |
| TRPV3 | 371 | R | 2 | 0 | yes | no | no | no | no | yes | no | no | no | no | no | no | no |
| TRPV3 | 372 | S | 0 | 0 | yes | no | no | no | no | yes | no | no | no | no | no | no | no |
| TRPV3 | 373 | L | 1 | 0 | yes | no | no | no | no | yes | no | no | no | no | no | no | no |
| TRPV3 | 374 | S | 0 | 0 | no | yes | no | no | no | yes | no | no | no | no | no | no | no |

| | | | | | | | | | | | | | | | | | |
|---|---|---|---|---|---|---|---|---|---|---|---|---|---|---|---|---|---|
| TRPV3 | 375 | R | 0 | 0 | no | yes | no | no | no | yes | no | no | no | no | no | no | no |
| TRPV3 | 376 | K | 0 | 0 | no | yes | no | no | no | yes | no | no | no | no | no | no | no |
| TRPV3 | 377 | F | 1 | 0 | no | no | yes | no | no | yes | no | no | no | no | no | no | no |
| TRPV3 | 378 | T | 0 | 0 | no | no | yes | no | no | yes | no | no | no | no | no | no | no |
| TRPV3 | 379 | D | 1 | 0 | no | yes | no | no | no | yes | no | no | no | no | no | no | no |
| TRPV3 | 380 | W | 0 | 0 | no | yes | no | no | no | yes | no | no | no | no | no | no | no |
| TRPV3 | 381 | A | 1 | 0 | no | yes | no | no | no | yes | no | no | yes | no | no | no | no |
| TRPV3 | 382 | Y | 0 | 0 | no | yes | no | no | no | yes | no | no | yes | no | no | no | no |
| TRPV3 | 383 | G | 1 | 0 | no | yes | no | no | no | yes | no | no | no | no | no | no | no |
| TRPV3 | 384 | P | 0 | 0 | no | yes | no | no | no | yes | no | no | yes | no | no | no | no |
| TRPV3 | 385 | V | 1 | 0 | no | no | yes | no | no | yes | no | no | yes | no | no | no | no |
| TRPV3 | 386 | S | 0 | 0 | no | no | yes | no | no | yes | no | no | no | no | no | no | no |
| TRPV3 | 387 | S | 1 | 0 | no | no | yes | no | no | yes | no | no | no | no | no | no | no |
| TRPV3 | 388 | S | 0 | 0 | no | no | yes | no | no | yes | no | no | no | no | no | no | no |
| TRPV3 | 389 | L | 1 | 0 | no | no | yes | no | no | yes | no | no | no | no | no | no | no |
| TRPV3 | 390 | Y | 0 | 0 | no | no | yes | no | no | yes | no | no | no | no | no | no | no |
| TRPV3 | 391 | D | 3 | 0 | no | no | yes | no | no | yes | no | no | no | no | no | no | no |
| TRPV3 | 392 | L | 0 | 0 | no | yes | no | no | no | yes | no | no | no | no | no | no | no |
| TRPV3 | 393 | T | 0 | 0 | no | yes | no | no | no | yes | no | no | no | no | no | no | no |
| TRPV3 | 394 | N | 2 | 0 | no | yes | no | no | no | yes | no | no | no | no | no | no | no |
| TRPV3 | 395 | V | 1 | 0 | no | yes | no | no | no | yes | no | no | no | no | no | no | no |
| TRPV3 | 396 | D | 0 | 0 | no | yes | no | no | no | yes | no | no | no | no | no | no | no |
| TRPV3 | 397 | T | 1 | 0 | no | yes | no | no | no | yes | no | no | no | no | no | no | no |
| TRPV3 | 398 | T | 1 | 0 | no | yes | no | no | no | yes | no | no | no | no | no | no | no |
| TRPV3 | 399 | T | 1 | 0 | no | yes | no | no | no | yes | no | no | no | no | no | no | no |
| TRPV3 | 400 | D | 0 | 0 | no | yes | no | no | no | yes | no | no | no | no | no | no | no |
| TRPV3 | 401 | N | 1 | 0 | no | yes | no | no | no | yes | no | no | no | no | no | no | no |
| TRPV3 | 402 | S | 0 | 0 | no | yes | no | no | no | yes | no | no | no | no | no | no | no |
| TRPV3 | 403 | V | 0 | 0 | yes | no | no | no | no | yes | no | no | no | no | no | no | no |
| TRPV3 | 404 | L | 0 | 0 | yes | no | no | no | no | yes | no | no | no | no | no | no | no |
| TRPV3 | 405 | E | 0 | 0 | yes | no | no | no | no | yes | no | no | no | no | no | no | no |
| TRPV3 | 406 | I | 2 | 0 | yes | no | no | no | no | yes | no | no | no | no | no | no | no |
| TRPV3 | 407 | T | 1 | 0 | yes | no | no | no | no | yes | no | no | no | no | no | no | no |

| | | | | | | | | | | | | | | | | | |
|---|---|---|---|---|---|---|---|---|---|---|---|---|---|---|---|---|---|
| TRPV3 | 408 | V | 0 | 0 | yes | no | no | no | no | yes | no | no | no | no | no | no | no |
| TRPV3 | 409 | Y | 0 | 0 | yes | no | no | no | no | yes | no | no | no | no | no | no | no |
| TRPV3 | 410 | N | 0 | 0 | no | yes | no | no | no | yes | no | no | no | no | no | no | no |
| TRPV3 | 411 | T | 0 | 0 | no | yes | no | no | no | yes | no | no | no | no | no | no | no |
| TRPV3 | 412 | N | 0 | 0 | no | yes | no | no | no | yes | no | no | no | no | no | no | no |
| TRPV3 | 413 | I | 0 | 0 | no | yes | no | no | no | yes | no | no | no | no | no | no | no |
| TRPV3 | 414 | D | 1 | 0 | no | yes | no | no | no | yes | no | no | no | no | no | no | no |
| TRPV3 | 415 | N | 0 | 0 | no | yes | no | no | no | yes | no | no | no | no | no | no | no |
| TRPV3 | 416 | R | 2 | 0 | yes | no | no | no | no | yes | no | no | no | no | no | no | no |
| TRPV3 | 417 | H | 1 | 0 | yes | no | no | no | no | yes | no | no | no | no | no | no | no |
| TRPV3 | 418 | E | 0 | 0 | yes | no | no | no | no | yes | no | no | no | no | no | no | no |
| TRPV3 | 419 | M | 0 | 0 | yes | no | no | no | no | yes | no | no | no | no | no | no | no |
| TRPV3 | 420 | L | 0 | 0 | yes | no | no | no | no | yes | no | no | no | no | no | no | no |
| TRPV3 | 421 | T | 2 | 0 | no | yes | no | no | no | yes | no | no | no | no | no | no | no |
| TRPV3 | 422 | L | 1 | 0 | no | yes | no | no | no | yes | no | no | no | no | no | no | no |
| TRPV3 | 423 | E | 1 | 0 | yes | no | no | no | no | yes | no | no | no | no | no | no | no |
| TRPV3 | 424 | P | 1 | 0 | yes | no | no | no | no | yes | no | no | no | no | no | no | no |
| TRPV3 | 425 | L | 0 | 0 | yes | no | no | no | no | yes | no | no | no | no | no | no | no |
| TRPV3 | 426 | H | 1 | 0 | yes | no | no | no | no | yes | no | no | no | no | no | no | no |
| TRPV3 | 427 | T | 2 | 0 | yes | no | no | no | no | yes | no | no | no | no | no | no | no |
| TRPV3 | 428 | L | 1 | 0 | yes | no | no | no | no | yes | no | no | no | no | no | no | no |
| TRPV3 | 429 | L | 1 | 0 | yes | no | no | no | no | yes | no | no | no | no | no | no | no |
| TRPV3 | 430 | H | 1 | 0 | yes | no | no | no | no | yes | no | no | no | no | no | no | no |
| TRPV3 | 431 | M | 2 | 0 | yes | no | no | no | no | yes | no | no | no | no | no | no | no |
| TRPV3 | 432 | K | 0 | 0 | yes | no | no | no | no | yes | no | no | no | no | no | no | no |
| TRPV3 | 433 | W | 0 | 0 | yes | no | no | no | no | yes | no | no | no | no | no | no | no |
| TRPV3 | 434 | K | 1 | 0 | yes | no | no | no | no | yes | no | no | no | no | no | no | no |
| TRPV3 | 435 | K | 0 | 0 | yes | no | no | no | no | yes | no | no | no | no | no | no | no |
| TRPV3 | 436 | F | 0 | 0 | yes | no | no | no | no | yes | no | no | no | no | no | no | no |
| TRPV3 | 437 | A | 0 | 0 | yes | no | no | no | no | yes | no | no | no | no | no | no | no |
| TRPV3 | 438 | K | 0 | 0 | yes | no | no | no | no | yes | no | no | no | no | no | no | no |
| TRPV3 | 439 | H | 0 | 0 | yes | no | no | no | no | yes | no | no | no | no | no | no | no |
| TRPV3 | 440 | M | 1 | 0 | yes | no | no | no | no | no | no | no | no | no | no | no | no |

| | | | | | | | | | | | | | | | | | |
|---|---|---|---|---|---|---|---|---|---|---|---|---|---|---|---|---|---|
| TRPV3 | 441 | F | 0 | 0 | yes | no | no | no | no | no | no | no | no | no | no | no | no |
| TRPV3 | 442 | F | 0 | 0 | yes | no | no | yes | no | no | no | no | no | no | no | no | no |
| TRPV3 | 443 | L | 0 | 0 | yes | no | no | yes | no | no | no | no | no | no | no | no | no |
| TRPV3 | 444 | S | 1 | 0 | yes | no | no | yes | no | no | no | no | no | no | no | no | no |
| TRPV3 | 445 | F | 0 | 0 | yes | no | no | yes | no | no | no | no | no | no | no | no | no |
| TRPV3 | 446 | C | 1 | 0 | yes | no | no | yes | no | no | no | no | no | no | no | no | no |
| TRPV3 | 447 | F | 1 | 0 | yes | no | no | yes | no | no | no | no | no | no | no | no | no |
| TRPV3 | 448 | Y | 0 | 0 | yes | no | no | yes | no | no | no | no | no | no | no | no | no |
| TRPV3 | 449 | F | 0 | 0 | yes | no | no | yes | no | no | no | no | no | no | no | no | no |
| TRPV3 | 450 | F | 0 | 0 | yes | no | no | yes | no | no | no | no | no | no | no | no | no |
| TRPV3 | 451 | Y | 0 | 0 | yes | no | no | yes | no | no | no | no | no | no | no | no | no |
| TRPV3 | 452 | N | 1 | 0 | yes | no | no | yes | no | no | no | no | no | no | no | no | no |
| TRPV3 | 453 | I | 0 | 0 | yes | no | no | yes | no | no | no | no | no | no | no | no | no |
| TRPV3 | 454 | T | 2 | 0 | yes | no | no | yes | no | no | no | no | no | no | no | no | no |
| TRPV3 | 455 | L | 1 | 0 | yes | no | no | yes | no | no | no | no | no | no | no | no | no |
| TRPV3 | 456 | T | 0 | 0 | yes | no | no | yes | no | no | no | no | no | no | no | no | no |
| TRPV3 | 457 | L | 0 | 0 | yes | no | no | yes | no | no | no | no | no | no | no | no | no |
| TRPV3 | 458 | V | 1 | 0 | yes | no | no | yes | no | no | no | no | no | no | no | no | no |
| TRPV3 | 459 | S | 1 | 0 | yes | no | no | yes | no | no | no | no | yes | no | no | no | no |
| TRPV3 | 460 | Y | 0 | 0 | yes | no | no | yes | no | no | no | no | no | no | no | no | no |
| TRPV3 | 461 | Y | 1 | 0 | no | yes | no | yes | yes | no | no | no | no | no | no | no | no |
| TRPV3 | 462 | R | 2 | 0 | no | yes | no | yes | yes | no | no | no | no | no | no | no | no |
| TRPV3 | 484 | L | 2 | 0 | no | yes | no | yes | yes | no | no | no | no | no | no | no | no |
| TRPV3 | 485 | L | 0 | 0 | yes | no | no | yes | yes | no | no | no | no | no | no | no | no |
| TRPV3 | 486 | G | 1 | 0 | yes | no | no | yes | yes | no | no | no | no | no | no | no | no |
| TRPV3 | 487 | R | 2 | 0 | yes | no | no | yes | yes | no | no | no | no | no | no | no | no |
| TRPV3 | 488 | M | 2 | 0 | yes | no | no | yes | no | no | no | no | no | no | no | no | no |
| TRPV3 | 489 | F | 1 | 0 | yes | no | no | yes | no | no | no | no | no | no | no | no | no |
| TRPV3 | 490 | V | 2 | 0 | yes | no | no | yes | no | no | no | no | no | no | no | no | no |
| TRPV3 | 491 | L | 1 | 0 | yes | no | no | yes | no | no | no | no | no | no | no | no | no |
| TRPV3 | 492 | I | 0 | 0 | yes | no | no | yes | no | no | no | no | no | no | no | no | no |
| TRPV3 | 493 | W | 0 | 0 | yes | no | no | yes | no | no | no | no | no | no | no | no | no |
| TRPV3 | 494 | A | 0 | 0 | yes | no | no | yes | no | no | no | no | no | no | no | no | no |

| | | | | | | | | | | | | | | | | | |
|---|---|---|---|---|---|---|---|---|---|---|---|---|---|---|---|---|---|
| TRPV3 | 495 | M | 1 | 0 | yes | no | no | yes | no | no | no | no | no | no | no | no | no |
| TRPV3 | 496 | C | 1 | 0 | yes | no | no | yes | no | no | no | no | no | no | no | no | no |
| TRPV3 | 497 | I | 1 | 0 | yes | no | no | yes | no | no | no | no | no | no | no | no | no |
| TRPV3 | 498 | S | 1 | 0 | yes | no | no | yes | no | no | no | no | no | no | no | no | no |
| TRPV3 | 499 | V | 2 | 0 | yes | no | no | yes | no | no | no | no | no | no | no | no | no |
| TRPV3 | 500 | K | 0 | 0 | yes | no | no | yes | no | no | no | no | no | no | no | no | no |
| TRPV3 | 501 | E | 0 | 0 | yes | no | no | yes | no | no | no | no | no | no | no | no | no |
| TRPV3 | 502 | G | 1 | 0 | yes | no | no | yes | no | no | no | no | no | no | no | no | no |
| TRPV3 | 503 | I | 2 | 0 | yes | no | no | yes | no | no | no | no | no | no | no | no | no |
| TRPV3 | 504 | A | 0 | 0 | yes | no | no | yes | no | no | no | no | no | no | no | no | no |
| TRPV3 | 505 | I | 1 | 0 | yes | no | no | yes | no | no | no | no | no | no | no | no | no |
| TRPV3 | 506 | F | 0 | 0 | yes | no | no | no | no | no | no | no | no | no | no | no | no |
| TRPV3 | 507 | L | 0 | 0 | yes | no | no | no | no | no | no | no | no | no | no | no | no |
| TRPV3 | 508 | L | 1 | 0 | no | yes | no | no | no | no | no | no | no | no | no | no | no |
| TRPV3 | 518 | S | 1 | 0 | no | yes | no | no | no | yes | no | no | no | no | no | no | no |
| TRPV3 | 519 | D | 0 | 0 | no | yes | no | no | no | yes | no | no | no | no | no | no | no |
| TRPV3 | 520 | A | 0 | 0 | no | yes | no | no | no | yes | no | no | no | no | no | no | no |
| TRPV3 | 521 | W | 0 | 1 | yes | no | no | yes | no | yes | no | no | no | no | no | no | no |
| TRPV3 | 522 | F | 0 | 0 | yes | no | no | yes | no | yes | no | no | no | no | no | no | no |
| TRPV3 | 523 | H | 0 | 0 | yes | no | no | yes | no | yes | no | no | no | no | no | no | no |
| TRPV3 | 524 | F | 1 | 0 | yes | no | no | yes | no | no | no | no | no | no | no | no | no |
| TRPV3 | 525 | V | 0 | 0 | yes | no | no | yes | no | no | no | no | no | no | no | no | no |
| TRPV3 | 526 | F | 0 | 0 | yes | no | no | yes | no | no | no | no | no | no | no | no | no |
| TRPV3 | 527 | F | 0 | 0 | yes | no | no | yes | no | no | no | no | no | no | no | no | no |
| TRPV3 | 528 | I | 0 | 0 | yes | no | no | yes | no | no | no | no | no | no | no | no | no |
| TRPV3 | 529 | Q | 0 | 0 | yes | no | no | yes | no | no | no | no | no | no | no | no | no |
| TRPV3 | 530 | A | 0 | 0 | yes | no | no | yes | no | no | no | no | no | no | no | no | no |
| TRPV3 | 531 | V | 0 | 0 | yes | no | no | yes | no | no | no | no | no | no | no | no | no |
| TRPV3 | 532 | L | 0 | 0 | yes | no | no | yes | no | no | no | no | no | no | no | no | no |
| TRPV3 | 533 | V | 1 | 0 | yes | no | no | yes | no | no | no | no | no | no | no | no | no |
| TRPV3 | 534 | I | 1 | 0 | yes | no | no | yes | no | no | no | no | no | no | no | no | no |
| TRPV3 | 535 | L | 1 | 0 | yes | no | no | yes | no | no | no | no | no | no | no | no | no |
| TRPV3 | 536 | S | 1 | 0 | yes | no | no | yes | no | no | no | no | no | no | no | no | no |

| | | | | | | | | | | | | | | | | | |
|---|---|---|---|---|---|---|---|---|---|---|---|---|---|---|---|---|---|
| TRPV3 | 537 | V | 1 | 0 | yes | no | no | yes | no | no | no | no | no | no | no | no | no |
| TRPV3 | 538 | F | 0 | 0 | yes | no | no | yes | no | no | no | no | no | no | no | no | no |
| TRPV3 | 539 | L | 1 | 0 | yes | no | no | yes | no | no | no | no | no | no | no | no | no |
| TRPV3 | 540 | Y | 0 | 0 | yes | no | no | yes | no | no | no | no | no | no | no | no | no |
| TRPV3 | 541 | L | 1 | 0 | yes | no | no | yes | no | no | no | no | no | no | no | no | no |
| TRPV3 | 542 | F | 0 | 0 | no | yes | no | yes | no | no | no | no | no | no | no | no | no |
| TRPV3 | 543 | A | 1 | 0 | no | yes | no | no | no | no | no | no | no | no | no | no | no |
| TRPV3 | 544 | Y | 1 | 0 | no | yes | no | no | no | no | no | no | no | no | no | no | no |
| TRPV3 | 545 | K | 1 | 0 | no | yes | no | no | yes | no | no | no | no | no | no | no | no |
| TRPV3 | 546 | E | 0 | 0 | no | yes | no | yes | no | no | no | no | yes | no | no | no | no |
| TRPV3 | 547 | Y | 1 | 0 | yes | no | no | yes | no | no | no | no | no | no | no | no | no |
| TRPV3 | 548 | L | 0 | 0 | yes | no | no | yes | no | no | no | no | no | no | no | no | no |
| TRPV3 | 549 | A | 2 | 0 | yes | no | no | yes | no | no | no | no | yes | no | no | no | no |
| TRPV3 | 550 | C | 2 | 0 | yes | no | no | yes | no | no | no | no | no | no | no | no | no |
| TRPV3 | 551 | L | 0 | 0 | yes | no | no | yes | no | no | no | no | no | no | no | no | no |
| TRPV3 | 552 | V | 1 | 0 | yes | no | no | yes | no | no | no | no | yes | no | no | no | no |
| TRPV3 | 553 | L | 1 | 0 | yes | no | no | yes | no | no | no | no | yes | no | no | no | no |
| TRPV3 | 554 | A | 1 | 0 | yes | no | no | yes | no | no | no | no | no | no | no | no | no |
| TRPV3 | 555 | M | 2 | 0 | yes | no | no | yes | no | no | no | no | yes | no | no | no | no |
| TRPV3 | 556 | A | 2 | 0 | yes | no | no | yes | no | no | no | no | no | no | no | no | no |
| TRPV3 | 557 | L | 1 | 0 | yes | no | no | yes | no | no | no | no | no | no | no | no | no |
| TRPV3 | 558 | G | 1 | 0 | yes | no | no | yes | no | no | no | no | no | no | no | no | no |
| TRPV3 | 559 | W | 0 | 0 | yes | no | no | yes | no | no | no | no | yes | no | no | no | no |
| TRPV3 | 560 | A | 1 | 0 | yes | no | no | yes | no | no | no | no | no | no | no | no | no |
| TRPV3 | 561 | N | 0 | 0 | yes | no | no | yes | no | no | no | no | no | no | no | no | no |
| TRPV3 | 562 | M | 3 | 0 | yes | no | no | yes | no | no | no | no | no | no | no | no | no |
| TRPV3 | 563 | L | 1 | 0 | yes | no | no | yes | no | no | no | no | no | no | no | no | no |
| TRPV3 | 564 | Y | 0 | 0 | yes | no | no | yes | no | no | no | no | no | no | no | no | no |
| TRPV3 | 565 | Y | 2 | 0 | yes | no | no | yes | no | no | no | no | no | no | no | no | no |
| TRPV3 | 566 | T | 0 | 0 | yes | no | no | yes | no | no | no | no | no | no | no | no | no |
| TRPV3 | 567 | R | 2 | 0 | no | yes | no | no | no | yes | no | no | no | no | no | no | no |
| TRPV3 | 568 | G | 1 | 2 | no | yes | no | no | no | yes | no | no | no | no | no | no | no |
| TRPV3 | 569 | F | 2 | 0 | no | yes | no | no | no | yes | no | no | no | no | no | no | no |

| | | | | | | | | | | | | | | | | | |
|---|---|---|---|---|---|---|---|---|---|---|---|---|---|---|---|---|---|
| TRPV3 | 570 | Q | 1 | 0 | yes | no | no | no | no | yes | no | no | no | no | no | no | no |
| TRPV3 | 571 | S | 2 | 0 | yes | no | no | no | no | yes | no | no | no | no | no | no | no |
| TRPV3 | 572 | M | 2 | 0 | yes | no | no | no | no | yes | no | no | no | no | no | no | no |
| TRPV3 | 573 | G | 0 | 4 | yes | no | no | no | no | yes | no | no | no | no | no | no | no |
| TRPV3 | 574 | M | 0 | 0 | yes | no | no | no | no | yes | no | no | no | no | no | no | no |
| TRPV3 | 575 | Y | 0 | 0 | yes | no | no | yes | no | yes | no | no | yes | no | no | no | no |
| TRPV3 | 576 | S | 1 | 0 | yes | no | no | yes | no | yes | no | no | no | no | no | no | no |
| TRPV3 | 577 | V | 1 | 0 | yes | no | no | no | no | yes | no | no | no | no | no | no | no |
| TRPV3 | 578 | M | 1 | 0 | yes | no | no | no | no | yes | no | no | yes | no | no | no | no |
| TRPV3 | 579 | I | 0 | 0 | yes | no | no | yes | no | yes | no | no | yes | no | no | no | no |
| TRPV3 | 580 | Q | 0 | 1 | yes | no | no | yes | no | yes | no | no | no | no | no | no | no |
| TRPV3 | 581 | K | 0 | 0 | yes | no | no | no | no | yes | no | no | no | no | no | no | no |
| TRPV3 | 582 | V | 0 | 0 | yes | no | no | yes | no | yes | no | no | yes | no | no | no | no |
| TRPV3 | 583 | I | 0 | 0 | yes | no | no | yes | no | yes | no | no | yes | no | no | no | no |
| TRPV3 | 584 | L | 0 | 0 | yes | no | no | yes | no | yes | no | no | no | no | no | no | no |
| TRPV3 | 585 | H | 2 | 0 | yes | no | no | no | no | yes | no | no | no | no | no | no | no |
| TRPV3 | 586 | D | 0 | 0 | no | yes | no | yes | no | yes | no | no | no | no | no | no | no |
| TRPV3 | 587 | V | 1 | 0 | yes | no | no | yes | no | yes | no | no | no | no | no | no | no |
| TRPV3 | 588 | L | 1 | 0 | yes | no | no | yes | no | yes | no | no | no | no | no | no | no |
| TRPV3 | 589 | K | 0 | 0 | yes | no | no | yes | no | yes | no | no | yes | no | no | no | no |
| TRPV3 | 590 | F | 1 | 0 | yes | no | no | yes | no | no | no | no | no | no | no | no | no |
| TRPV3 | 591 | L | 0 | 0 | yes | no | no | yes | no | no | no | no | no | no | no | no | no |
| TRPV3 | 592 | F | 0 | 0 | yes | no | no | yes | no | no | no | no | no | no | no | no | no |
| TRPV3 | 593 | V | 0 | 0 | yes | no | no | yes | no | no | no | no | no | no | no | no | no |
| TRPV3 | 594 | Y | 0 | 0 | yes | no | no | yes | no | no | no | no | no | no | no | no | no |
| TRPV3 | 595 | I | 1 | 0 | yes | no | no | yes | no | no | no | no | no | no | no | no | no |
| TRPV3 | 596 | V | 1 | 0 | yes | no | no | yes | no | no | no | no | yes | no | no | no | no |
| TRPV3 | 597 | F | 0 | 0 | yes | no | no | yes | no | no | no | no | no | no | no | no | no |
| TRPV3 | 598 | L | 0 | 0 | yes | no | no | yes | no | no | no | no | no | no | no | no | no |
| TRPV3 | 599 | L | 0 | 0 | yes | no | no | yes | no | no | no | no | no | no | no | no | no |
| TRPV3 | 600 | G | 0 | 0 | yes | no | no | yes | no | no | no | no | yes | no | no | no | no |
| TRPV3 | 601 | F | 0 | 0 | yes | no | no | yes | no | no | no | no | no | no | no | no | no |
| TRPV3 | 602 | G | 0 | 0 | yes | no | no | yes | no | no | no | no | no | no | no | no | no |

| | | | | | | | | | | | | | | | | | |
|---|---|---|---|---|---|---|---|---|---|---|---|---|---|---|---|---|---|
| TRPV3 | 603 | V | 2 | 0 | yes | no | no | yes | no | no | no | no | yes | no | no | no | no |
| TRPV3 | 604 | A | 1 | 0 | yes | no | no | yes | no | no | no | no | yes | no | no | no | no |
| TRPV3 | 605 | L | 0 | 0 | yes | no | no | yes | no | no | no | no | no | no | no | no | no |
| TRPV3 | 606 | A | 0 | 0 | yes | no | no | yes | no | no | no | no | yes | no | no | no | no |
| TRPV3 | 607 | S | 1 | 0 | yes | no | no | yes | no | no | no | no | yes | no | no | no | no |
| TRPV3 | 608 | L | 0 | 0 | yes | no | no | yes | no | no | no | no | no | no | no | no | no |
| TRPV3 | 609 | I | 0 | 0 | no | yes | no | no | no | no | no | no | no | no | no | no | no |
| TRPV3 | 610 | E | 1 | 0 | no | yes | no | no | no | no | no | no | no | no | no | no | no |
| TRPV3 | 611 | K | 0 | 0 | no | yes | no | no | no | no | no | no | no | no | no | no | no |
| TRPV3 | 612 | C | 1 | 0 | no | yes | no | no | no | no | no | no | no | no | no | no | no |
| TRPV3 | 613 | P | 0 | 0 | no | yes | no | no | no | no | no | no | no | no | no | no | no |
| TRPV3 | 614 | K | 0 | 0 | no | yes | no | no | no | no | no | no | no | no | no | no | no |
| TRPV3 | 615 | D | 0 | 0 | no | yes | no | no | no | no | no | no | no | no | no | no | no |
| TRPV3 | 616 | N | 0 | 0 | no | yes | no | no | no | no | no | no | no | no | no | no | no |
| TRPV3 | 617 | K | 0 | 0 | no | yes | no | no | no | no | no | no | no | no | no | no | no |
| TRPV3 | 618 | D | 0 | 0 | no | yes | no | no | no | no | no | no | no | no | no | no | no |
| TRPV3 | 619 | C | 0 | 0 | no | yes | no | no | no | no | no | no | no | no | no | no | no |
| TRPV3 | 620 | S | 0 | 0 | no | yes | no | no | no | no | no | no | no | no | no | no | no |
| TRPV3 | 621 | S | 1 | 0 | no | yes | no | no | no | no | no | yes | no | no | no | no | no |
| TRPV3 | 622 | Y | 1 | 0 | no | yes | no | yes | no | no | no | yes | no | no | no | no | no |
| TRPV3 | 623 | G | 2 | 0 | no | yes | no | no | no | no | no | yes | no | no | no | no | no |
| TRPV3 | 624 | S | 0 | 0 | no | yes | no | no | no | no | no | yes | no | no | no | no | no |
| TRPV3 | 625 | F | 0 | 0 | yes | no | no | yes | no | no | no | yes | no | no | no | no | no |
| TRPV3 | 626 | S | 1 | 0 | yes | no | no | yes | no | no | no | yes | no | no | no | no | no |
| TRPV3 | 627 | D | 1 | 0 | yes | no | no | yes | no | no | no | yes | no | no | no | no | no |
| TRPV3 | 628 | A | 1 | 0 | yes | no | no | yes | no | no | no | yes | no | no | no | no | no |
| TRPV3 | 629 | V | 0 | 0 | yes | no | no | yes | no | no | no | yes | no | no | no | no | no |
| TRPV3 | 630 | L | 0 | 0 | yes | no | no | yes | no | no | no | yes | no | no | no | no | no |
| TRPV3 | 631 | E | 0 | 0 | yes | no | no | yes | no | no | no | yes | no | no | no | no | no |
| TRPV3 | 632 | L | 0 | 0 | yes | no | no | yes | no | no | no | yes | no | no | no | no | no |
| TRPV3 | 633 | F | 0 | 0 | yes | no | no | yes | no | no | no | yes | yes | no | no | no | no |
| TRPV3 | 634 | K | 0 | 0 | yes | no | no | yes | no | no | no | yes | yes | no | no | no | no |
| TRPV3 | 635 | L | 0 | 0 | yes | no | no | yes | no | no | no | yes | yes | no | no | no | no |

| | | | | | | | | | | | | | | | | | |
|---|---|---|---|---|---|---|---|---|---|---|---|---|---|---|---|---|---|
| TRPV3 | 636 | T | 2 | 0 | yes | no | no | yes | no | no | no | yes | no | no | no | no | no |
| TRPV3 | 637 | I | 1 | 0 | yes | no | no | yes | no | no | yes | yes | no | no | no | no | no |
| TRPV3 | 638 | G | 0 | 0 | no | yes | no | yes | no | no | yes | no | yes | no | no | no | no |
| TRPV3 | 639 | L | 1 | 0 | no | yes | no | yes | no | no | yes | no | yes | no | no | no | no |
| TRPV3 | 640 | G | 0 | 0 | no | yes | no | yes | no | no | yes | no | yes | no | no | no | no |
| TRPV3 | 641 | D | 1 | 0 | no | yes | no | no | no | no | yes | no | no | no | no | no | no |
| TRPV3 | 642 | L | 1 | 0 | no | yes | no | yes | no | no | no | no | yes | no | no | no | no |
| TRPV3 | 643 | N | 1 | 0 | no | yes | no | no | no | no | no | no | no | no | no | no | no |
| TRPV3 | 644 | I | 0 | 0 | no | yes | no | no | no | no | no | no | no | no | no | no | no |
| TRPV3 | 645 | Q | 2 | 0 | no | yes | no | no | no | no | no | no | no | no | no | no | no |
| TRPV3 | 646 | Q | 1 | 0 | no | yes | no | no | no | no | no | no | no | no | no | no | no |
| TRPV3 | 647 | N | 1 | 0 | no | yes | no | no | no | no | no | no | no | no | no | no | no |
| TRPV3 | 648 | S | 0 | 0 | no | yes | no | no | no | no | no | no | no | no | no | no | no |
| TRPV3 | 649 | K | 1 | 0 | no | yes | no | no | no | no | no | no | no | no | no | no | no |
| TRPV3 | 650 | Y | 0 | 0 | no | yes | no | no | no | no | no | no | yes | no | no | no | no |
| TRPV3 | 651 | P | 0 | 0 | yes | no | no | no | no | no | no | no | no | no | no | no | no |
| TRPV3 | 652 | I | 1 | 0 | yes | no | no | yes | no | no | no | no | no | no | no | no | no |
| TRPV3 | 653 | L | 0 | 0 | yes | no | no | yes | no | no | no | no | yes | no | no | no | no |
| TRPV3 | 654 | F | 0 | 0 | yes | no | no | yes | no | no | no | no | no | no | no | no | no |
| TRPV3 | 655 | L | 1 | 0 | yes | no | no | yes | no | no | no | no | yes | no | no | no | no |
| TRPV3 | 656 | F | 0 | 0 | yes | no | no | yes | no | no | no | no | no | no | no | no | no |
| TRPV3 | 657 | L | 0 | 0 | yes | no | no | yes | no | no | no | no | no | no | no | no | no |
| TRPV3 | 658 | L | 0 | 0 | yes | no | no | yes | no | no | no | no | no | no | no | no | no |
| TRPV3 | 659 | I | 1 | 0 | yes | no | no | yes | no | no | no | no | no | no | no | no | no |
| TRPV3 | 660 | T | 0 | 0 | yes | no | no | yes | no | no | no | no | no | no | no | no | no |
| TRPV3 | 661 | Y | 1 | 0 | yes | no | no | yes | no | no | no | no | no | no | no | no | no |
| TRPV3 | 662 | V | 0 | 0 | yes | no | no | yes | no | no | no | no | yes | no | no | no | no |
| TRPV3 | 663 | I | 0 | 0 | yes | no | no | yes | no | no | no | no | no | no | no | no | no |
| TRPV3 | 664 | L | 1 | 0 | yes | no | no | yes | no | no | no | no | no | no | no | no | no |
| TRPV3 | 665 | T | 0 | 0 | yes | no | no | yes | no | no | no | no | no | no | no | no | no |
| TRPV3 | 666 | F | 0 | 0 | yes | no | no | yes | no | no | no | no | yes | no | no | no | no |
| TRPV3 | 667 | V | 1 | 0 | yes | no | no | yes | no | no | no | no | yes | no | no | no | no |
| TRPV3 | 668 | L | 0 | 0 | yes | no | no | yes | no | no | no | no | yes | no | no | no | no |

| | | | | | | | | | | | | | | | | | |
|---|---|---|---|---|---|---|---|---|---|---|---|---|---|---|---|---|---|
| TRPV3 | 669 | L | 1 | 0 | yes | no | no | yes | no | no | no | no | yes | no | no | no | no |
| TRPV3 | 670 | L | 0 | 0 | yes | no | no | yes | no | no | no | no | yes | no | no | no | no |
| TRPV3 | 671 | N | 1 | 0 | yes | no | no | yes | no | yes | no | no | yes | no | no | no | no |
| TRPV3 | 672 | M | 2 | 1 | yes | no | no | yes | no | yes | no | no | no | no | no | no | no |
| TRPV3 | 673 | L | 0 | 1 | yes | no | no | yes | no | yes | yes | no | yes | no | no | no | no |
| TRPV3 | 674 | I | 0 | 0 | yes | no | no | yes | no | yes | no | no | yes | no | no | no | no |
| TRPV3 | 675 | A | 1 | 0 | yes | no | no | no | no | yes | no | no | no | no | no | no | no |
| TRPV3 | 676 | L | 0 | 0 | yes | no | no | no | no | yes | no | no | yes | no | no | no | no |
| TRPV3 | 677 | M | 0 | 0 | yes | no | no | no | no | yes | yes | no | yes | no | no | no | no |
| TRPV3 | 678 | G | 0 | 0 | yes | no | no | no | no | yes | no | no | yes | no | no | no | no |
| TRPV3 | 679 | E | 2 | 0 | yes | no | no | no | no | yes | no | no | no | no | no | no | no |
| TRPV3 | 680 | T | 0 | 0 | yes | no | no | no | no | yes | no | no | yes | no | no | no | no |
| TRPV3 | 681 | V | 0 | 0 | no | yes | no | no | no | yes | yes | no | no | no | no | no | no |
| TRPV3 | 682 | E | 0 | 0 | yes | no | no | no | no | yes | yes | no | no | no | no | no | no |
| TRPV3 | 683 | N | 0 | 0 | yes | no | no | no | no | yes | no | no | no | no | no | no | no |
| TRPV3 | 684 | V | 1 | 0 | yes | no | no | no | no | yes | no | no | no | no | no | no | no |
| TRPV3 | 685 | S | 1 | 0 | yes | no | no | no | no | yes | no | no | no | no | no | no | no |
| TRPV3 | 686 | K | 0 | 0 | yes | no | no | no | no | yes | no | no | no | no | no | no | no |
| TRPV3 | 687 | E | 1 | 0 | yes | no | no | no | no | yes | no | no | no | no | no | no | no |
| TRPV3 | 688 | S | 1 | 0 | yes | no | no | no | no | yes | no | no | no | no | no | no | no |
| TRPV3 | 689 | E | 1 | 0 | yes | no | no | no | no | yes | no | no | no | no | no | no | no |
| TRPV3 | 690 | R | 2 | 0 | yes | no | no | no | no | yes | no | no | no | no | no | no | no |
| TRPV3 | 691 | I | 1 | 0 | yes | no | no | no | no | yes | no | no | no | no | no | no | no |
| TRPV3 | 692 | W | 0 | 2 | yes | no | no | no | no | yes | no | no | no | no | no | no | no |
| TRPV3 | 693 | R | 2 | 0 | yes | no | no | no | no | yes | no | no | no | no | no | no | no |
| TRPV3 | 694 | L | 0 | 0 | yes | no | no | no | no | yes | no | no | no | no | no | no | no |
| TRPV3 | 695 | Q | 0 | 0 | yes | no | no | no | no | yes | no | no | no | no | no | no | no |
| TRPV3 | 696 | R | 0 | 0 | yes | no | no | no | no | yes | no | no | no | no | no | no | no |
| TRPV3 | 697 | A | 0 | 0 | yes | no | no | no | no | yes | no | no | no | no | no | no | no |
| TRPV3 | 698 | R | 0 | 0 | yes | no | no | no | no | yes | no | no | no | no | no | no | no |
| TRPV3 | 699 | T | 0 | 0 | yes | no | no | no | no | yes | no | no | no | no | no | no | no |
| TRPV3 | 700 | I | 1 | 0 | yes | no | no | no | no | yes | no | no | no | no | no | no | no |
| TRPV3 | 701 | L | 0 | 0 | yes | no | no | no | no | yes | no | no | no | no | no | no | no |

| | | | | | | | | | | | | | | | | | |
|---|---|---|---|---|---|---|---|---|---|---|---|---|---|---|---|---|---|
| TRPV3 | 702 | E | 0 | 0 | yes | no | no | no | no | yes | no | no | no | no | no | no | no |
| TRPV3 | 703 | F | 0 | 0 | yes | no | no | no | no | yes | no | no | no | no | no | no | no |
| TRPV3 | 704 | E | 0 | 0 | yes | no | no | no | no | yes | no | no | no | no | no | no | no |
| TRPV3 | 705 | K | 1 | 0 | yes | no | no | no | no | yes | no | no | no | no | no | no | no |
| TRPV3 | 706 | M | 0 | 0 | yes | no | no | no | no | yes | no | no | no | no | no | no | no |
| TRPV3 | 707 | L | 0 | 0 | no | yes | no | no | no | yes | no | no | no | no | no | no | no |
| TRPV3 | 708 | P | 1 | 0 | no | yes | no | no | no | yes | no | no | no | no | no | no | no |
| TRPV3 | 709 | E | 1 | 0 | yes | no | no | no | no | yes | no | no | no | no | no | no | no |
| TRPV3 | 710 | W | 0 | 0 | yes | no | no | no | no | yes | no | no | no | no | no | no | no |
| TRPV3 | 711 | L | 0 | 0 | yes | no | no | no | no | yes | no | no | no | no | no | no | no |
| TRPV3 | 712 | R | 0 | 0 | yes | no | no | no | no | yes | no | no | no | no | no | no | no |
| TRPV3 | 713 | S | 0 | 0 | yes | no | no | no | no | yes | no | no | no | no | no | no | no |
| TRPV3 | 714 | R | 0 | 0 | yes | no | no | no | no | yes | no | no | no | no | no | no | no |
| TRPV3 | 715 | F | 0 | 0 | no | yes | no | no | no | yes | no | no | no | no | no | no | no |
| TRPV3 | 716 | R | 3 | 0 | no | yes | no | no | no | yes | no | no | no | no | no | no | no |
| TRPV3 | 717 | M | 3 | 0 | no | yes | no | no | no | yes | no | no | no | no | no | no | no |
| TRPV3 | 718 | G | 1 | 0 | no | yes | no | no | no | yes | no | no | no | no | no | no | no |
| TRPV3 | 719 | E | 0 | 0 | no | no | yes | no | no | yes | no | no | no | no | no | no | no |
| TRPV3 | 720 | L | 0 | 0 | no | no | yes | no | no | yes | no | no | no | no | no | no | no |
| TRPV3 | 721 | C | 1 | 0 | no | no | yes | no | no | yes | no | no | no | no | no | no | no |
| TRPV3 | 722 | K | 1 | 0 | no | no | yes | no | no | yes | no | no | no | no | no | no | no |
| TRPV3 | 723 | V | 0 | 0 | no | yes | no | no | no | yes | no | no | no | no | no | no | no |
| TRPV3 | 724 | A | 0 | 0 | no | yes | no | no | no | yes | no | no | no | no | no | no | no |
| TRPV3 | 725 | E | 1 | 0 | no | yes | no | no | no | yes | no | no | no | no | no | no | no |
| TRPV3 | 726 | D | 1 | 0 | no | yes | no | no | no | yes | no | no | no | no | no | no | no |
| TRPV3 | 727 | D | 0 | 0 | no | yes | no | no | no | yes | no | no | no | no | no | no | no |
| TRPV3 | 728 | F | 0 | 0 | no | no | yes | no | no | yes | no | no | no | no | no | no | no |
| TRPV3 | 729 | R | 1 | 0 | no | no | yes | no | no | yes | no | no | no | no | no | no | no |
| TRPV3 | 730 | L | 0 | 0 | no | no | yes | no | no | yes | no | no | no | no | no | no | no |
| TRPV3 | 731 | C | 0 | 0 | no | no | yes | no | no | yes | no | no | no | no | no | no | no |
| TRPV3 | 732 | L | 1 | 0 | no | no | yes | no | no | yes | no | no | no | no | no | no | no |
| TRPV3 | 733 | R | 0 | 0 | no | no | yes | no | no | yes | no | no | no | no | no | no | no |
| TRPV3 | 734 | I | 0 | 0 | no | no | yes | no | no | yes | no | no | no | no | no | no | no |

| | | | | | | | | | | | | | | | | | |
|---|---|---|---|---|---|---|---|---|---|---|---|---|---|---|---|---|---|
| TRPV3 | 735 | N | 2 | 0 | no | no | yes | no | no | yes | no | no | no | no | no | no | no |
| TRPV3 | 736 | E | 0 | 0 | no | no | yes | no | no | yes | no | no | no | no | no | no | no |
| TRPV3 | 737 | V | 1 | 0 | no | no | yes | no | no | yes | no | no | no | no | no | no | no |
| TRPV3 | 738 | K | 0 | 0 | no | yes | no | no | no | yes | no | no | no | no | no | no | no |
| TRPV3 | 739 | W | 1 | 0 | no | yes | no | no | no | yes | no | no | yes | no | no | no | no |
| TRPV3 | 740 | T | 0 | 0 | no | yes | no | no | no | yes | no | no | yes | no | no | no | no |
| TRPV3 | 741 | E | 0 | 0 | no | yes | no | no | no | yes | no | no | no | no | no | no | no |
| TRPV3 | 742 | W | 0 | 0 | no | yes | no | no | no | yes | no | no | yes | no | no | no | no |
| TRPV3 | 743 | K | 0 | 0 | no | yes | no | no | no | yes | no | no | no | no | no | no | no |
| TRPV3 | 744 | T | 1 | 0 | no | yes | no | no | no | yes | no | no | yes | no | no | no | no |
| TRPV3 | 745 | H | 0 | 0 | no | yes | no | no | no | yes | no | no | no | no | no | no | no |
| TRPV3 | 746 | V | 1 | 0 | no | yes | no | no | no | yes | no | no | no | no | no | no | no |
| TRPV3 | 747 | S | 1 | 0 | no | yes | no | no | no | yes | no | no | no | no | no | no | no |
| TRPV3 | 748 | F | 0 | 0 | no | yes | no | no | no | yes | no | no | yes | no | no | no | no |
| TRPV3 | 749 | L | 1 | 0 | no | yes | no | no | no | yes | no | no | no | no | no | no | no |
| TRPV3 | 750 | N | 2 | 0 | no | yes | no | no | no | yes | no | no | no | no | no | no | no |
| TRPV3 | 751 | E | 1 | 0 | no | yes | no | no | no | yes | no | no | yes | no | no | no | no |
| TRPV3 | 752 | D | 2 | 0 | no | yes | no | no | no | yes | no | no | yes | no | no | no | no |
| TRPV3 | 753 | P | 1 | 0 | no | yes | no | no | no | yes | no | no | yes | no | no | no | no |
| TRPV3 | 754 | G | 2 | 0 | no | yes | no | no | no | yes | no | no | yes | no | no | no | no |
| TRPV3 | 755 | P | 1 | 0 | no | yes | no | no | no | yes | no | no | yes | no | no | no | no |
| TRPV3 | 756 | V | 0 | 0 | no | yes | no | no | no | yes | no | no | no | no | no | no | no |
| SCN1A | 116 | P | 2 | 0 | no | yes | no | no | no | yes | no | no | no | no | no | no | no |
| SCN1A | 117 | L | 0 | 1 | yes | no | no | yes | no | yes | no | no | no | no | no | no | no |
| SCN1A | 118 | R | 0 | 1 | yes | no | no | no | no | yes | no | no | no | no | no | no | no |
| SCN1A | 119 | K | 0 | 0 | yes | no | no | no | no | yes | no | no | no | no | no | no | no |
| SCN1A | 120 | I | 0 | 0 | yes | no | no | yes | no | yes | no | no | no | no | no | no | no |
| SCN1A | 121 | A | 0 | 1 | yes | no | no | yes | no | yes | no | no | no | no | no | no | no |
| SCN1A | 122 | I | 1 | 1 | yes | no | no | no | no | yes | no | no | no | no | no | no | no |
| SCN1A | 123 | K | 0 | 0 | yes | no | no | no | no | yes | no | no | no | no | no | no | no |
| SCN1A | 124 | I | 0 | 1 | yes | no | no | yes | no | yes | no | no | no | no | no | no | no |
| SCN1A | 125 | L | 0 | 0 | yes | no | no | yes | no | yes | no | no | no | no | no | no | no |
| SCN1A | 126 | V | 0 | 0 | no | yes | no | no | no | yes | no | no | no | no | no | no | no |

| | | | | | | | | | | | | | | | | | |
|---|---|---|---|---|---|---|---|---|---|---|---|---|---|---|---|---|---|
| SCN1A | 127 | H | 1 | 2 | no | yes | no | yes | no | yes | no | no | no | no | no | no | no |
| SCN1A | 128 | S | 0 | 0 | yes | no | no | yes | no | yes | no | no | no | no | no | no | no |
| SCN1A | 129 | L | 1 | 0 | yes | no | no | yes | no | no | no | no | no | no | no | no | no |
| SCN1A | 130 | F | 0 | 0 | yes | no | no | yes | no | no | no | no | no | no | no | no | no |
| SCN1A | 131 | S | 0 | 1 | yes | no | no | yes | no | no | no | no | no | no | no | no | no |
| SCN1A | 132 | M | 1 | 0 | yes | no | no | yes | no | no | no | no | no | no | no | no | no |
| SCN1A | 133 | L | 0 | 0 | yes | no | no | yes | no | no | no | no | no | no | no | no | no |
| SCN1A | 134 | I | 0 | 0 | yes | no | no | yes | no | no | no | no | no | no | no | no | no |
| SCN1A | 135 | M | 0 | 1 | yes | no | no | yes | no | no | no | no | no | no | no | no | no |
| SCN1A | 136 | C | 0 | 0 | yes | no | no | yes | no | no | no | no | no | no | no | no | no |
| SCN1A | 137 | T | 0 | 0 | yes | no | no | yes | no | no | no | no | no | no | no | no | no |
| SCN1A | 138 | I | 0 | 2 | yes | no | no | yes | no | no | no | no | no | no | no | no | no |
| SCN1A | 139 | L | 0 | 0 | yes | no | no | yes | no | no | no | no | no | no | no | no | no |
| SCN1A | 140 | T | 0 | 1 | yes | no | no | yes | no | no | no | no | no | no | no | no | no |
| SCN1A | 141 | N | 0 | 1 | yes | no | no | yes | no | no | no | no | no | no | no | no | no |
| SCN1A | 142 | C | 0 | 0 | yes | no | no | yes | no | no | no | no | no | no | no | no | no |
| SCN1A | 143 | V | 0 | 1 | yes | no | no | yes | no | no | no | no | no | no | no | no | no |
| SCN1A | 144 | F | 0 | 0 | yes | no | no | yes | no | no | no | no | no | no | no | no | no |
| SCN1A | 145 | M | 0 | 2 | yes | no | no | yes | no | no | no | no | no | no | no | no | no |
| SCN1A | 146 | T | 0 | 1 | no | yes | no | yes | no | no | no | no | no | no | no | no | no |
| SCN1A | 147 | M | 0 | 0 | no | yes | no | yes | no | no | no | no | no | no | no | no | no |
| SCN1A | 148 | S | 0 | 0 | no | yes | no | no | yes | no | no | no | no | no | no | no | no |
| SCN1A | 149 | N | 0 | 0 | no | yes | no | no | yes | no | no | no | no | no | no | no | no |
| SCN1A | 150 | P | 0 | 0 | no | yes | no | no | yes | no | no | no | no | no | no | no | no |
| SCN1A | 151 | P | 0 | 0 | no | yes | no | no | yes | no | no | no | no | no | no | no | no |
| SCN1A | 152 | D | 0 | 0 | yes | no | no | no | yes | no | no | no | no | no | no | no | no |
| SCN1A | 153 | W | 0 | 0 | yes | no | no | no | yes | no | no | no | no | no | no | no | no |
| SCN1A | 154 | T | 0 | 0 | yes | no | no | yes | yes | no | no | no | no | no | no | no | no |
| SCN1A | 155 | K | 0 | 0 | yes | no | no | yes | no | no | no | no | no | no | no | no | no |
| SCN1A | 156 | N | 0 | 0 | yes | no | no | yes | no | no | no | no | no | no | no | no | no |
| SCN1A | 157 | V | 1 | 0 | yes | no | no | yes | no | no | no | no | no | no | no | no | no |
| SCN1A | 158 | E | 0 | 0 | yes | no | no | yes | no | no | no | no | no | no | no | no | no |
| SCN1A | 159 | Y | 0 | 0 | yes | no | no | yes | no | no | no | no | no | no | no | no | no |

| | | | | | | | | | | | | | | | | | |
|---|---|---|---|---|---|---|---|---|---|---|---|---|---|---|---|---|---|
| SCN1A | 160 | T | 0 | 0 | yes | no | no | yes | no | no | no | no | no | no | no | no | no |
| SCN1A | 161 | F | 0 | 0 | yes | no | no | yes | no | no | no | no | no | no | no | no | no |
| SCN1A | 162 | T | 0 | 2 | yes | no | no | yes | no | no | no | no | no | no | no | no | no |
| SCN1A | 163 | G | 0 | 1 | yes | no | no | yes | no | no | no | no | no | no | no | no | no |
| SCN1A | 164 | I | 0 | 1 | yes | no | no | yes | no | no | no | no | no | no | no | no | no |
| SCN1A | 165 | Y | 0 | 1 | yes | no | no | yes | no | no | no | no | no | no | no | no | no |
| SCN1A | 166 | T | 0 | 0 | yes | no | no | yes | no | no | no | no | no | no | no | no | no |
| SCN1A | 167 | F | 1 | 0 | yes | no | no | yes | no | no | no | no | no | no | no | no | no |
| SCN1A | 168 | E | 0 | 0 | yes | no | no | yes | no | no | no | no | no | no | no | no | no |
| SCN1A | 169 | S | 0 | 1 | yes | no | no | yes | no | no | no | no | no | no | no | no | no |
| SCN1A | 170 | L | 0 | 1 | yes | no | no | yes | no | no | no | no | no | no | no | no | no |
| SCN1A | 171 | I | 1 | 1 | yes | no | no | yes | no | no | no | no | no | no | no | no | no |
| SCN1A | 172 | K | 0 | 0 | yes | no | no | yes | no | no | no | no | no | no | no | no | no |
| SCN1A | 173 | I | 1 | 0 | yes | no | no | yes | no | no | no | no | no | no | no | no | no |
| SCN1A | 174 | I | 0 | 0 | yes | no | no | yes | no | no | no | no | no | no | no | no | no |
| SCN1A | 175 | A | 0 | 2 | yes | no | no | yes | no | no | no | no | no | no | no | no | no |
| SCN1A | 176 | R | 0 | 0 | yes | no | no | no | no | yes | no | no | no | no | no | no | no |
| SCN1A | 177 | G | 0 | 2 | no | yes | no | yes | no | yes | no | no | no | no | no | no | no |
| SCN1A | 178 | F | 0 | 1 | no | yes | no | no | no | yes | no | no | no | no | no | no | no |
| SCN1A | 179 | C | 0 | 1 | no | yes | no | yes | no | yes | no | no | no | no | no | no | no |
| SCN1A | 180 | L | 0 | 0 | no | yes | no | yes | no | yes | no | no | no | no | no | no | no |
| SCN1A | 181 | E | 0 | 0 | no | yes | no | no | no | yes | no | no | no | no | no | no | no |
| SCN1A | 182 | D | 0 | 0 | no | yes | no | yes | no | yes | no | no | no | no | no | no | no |
| SCN1A | 183 | F | 0 | 0 | no | yes | no | yes | no | yes | no | no | no | no | no | no | no |
| SCN1A | 184 | T | 1 | 0 | no | yes | no | yes | no | yes | no | no | no | no | no | no | no |
| SCN1A | 185 | F | 1 | 0 | no | yes | no | yes | no | yes | no | no | no | no | no | no | no |
| SCN1A | 186 | L | 0 | 1 | no | yes | no | yes | no | yes | no | no | no | no | no | no | no |
| SCN1A | 187 | R | 2 | 0 | no | yes | no | no | no | yes | no | no | no | no | no | no | no |
| SCN1A | 188 | D | 0 | 1 | no | yes | no | yes | no | yes | no | no | no | no | no | no | no |
| SCN1A | 189 | P | 0 | 0 | yes | no | no | yes | no | yes | no | no | no | no | no | no | no |
| SCN1A | 190 | W | 0 | 1 | yes | no | no | yes | no | no | no | no | no | no | no | no | no |
| SCN1A | 191 | N | 0 | 3 | yes | no | no | yes | no | no | no | no | no | no | no | no | no |
| SCN1A | 192 | W | 0 | 0 | yes | no | no | yes | no | no | no | no | no | no | no | no | no |

| | | | | | | | | | | | | | | | | | |
|---|---|---|---|---|---|---|---|---|---|---|---|---|---|---|---|---|---|
| SCN1A | 193 | L | 0 | 1 | yes | no | no | yes | no | no | no | no | no | no | no | no | no |
| SCN1A | 194 | D | 0 | 3 | yes | no | no | yes | no | no | no | no | no | no | no | no | no |
| SCN1A | 195 | F | 0 | 0 | yes | no | no | yes | no | no | no | no | no | no | no | no | no |
| SCN1A | 196 | T | 0 | 0 | yes | no | no | yes | no | no | no | no | no | no | no | no | no |
| SCN1A | 197 | V | 0 | 0 | yes | no | no | yes | no | no | no | no | no | no | no | no | no |
| SCN1A | 198 | I | 0 | 0 | yes | no | no | yes | no | no | no | no | no | no | no | no | no |
| SCN1A | 199 | T | 0 | 1 | yes | no | no | yes | no | no | no | no | no | no | no | no | no |
| SCN1A | 200 | F | 0 | 0 | yes | no | no | yes | no | no | no | no | no | no | no | no | no |
| SCN1A | 201 | A | 0 | 0 | yes | no | no | yes | no | no | no | no | no | no | no | no | no |
| SCN1A | 202 | Y | 0 | 0 | yes | no | no | yes | no | no | no | no | no | no | no | no | no |
| SCN1A | 203 | V | 1 | 0 | yes | no | no | yes | no | no | no | no | no | no | no | no | no |
| SCN1A | 204 | T | 0 | 0 | yes | no | no | yes | no | no | no | no | no | no | no | no | no |
| SCN1A | 205 | E | 0 | 0 | no | yes | no | yes | no | no | no | no | no | no | no | no | no |
| SCN1A | 206 | F | 0 | 0 | no | yes | no | yes | no | no | no | no | no | no | no | no | no |
| SCN1A | 207 | V | 0 | 1 | no | yes | no | no | no | no | no | no | no | no | no | no | no |
| SCN1A | 208 | D | 0 | 0 | no | yes | no | no | yes | no | no | no | no | no | no | no | no |
| SCN1A | 209 | L | 0 | 1 | no | yes | no | yes | yes | no | no | no | no | no | no | no | no |
| SCN1A | 210 | G | 0 | 1 | no | yes | no | no | yes | no | no | no | no | no | no | no | no |
| SCN1A | 211 | N | 0 | 1 | no | yes | no | no | yes | no | no | no | no | no | no | no | no |
| SCN1A | 212 | V | 1 | 1 | no | yes | no | no | yes | no | no | no | no | no | no | no | no |
| SCN1A | 213 | S | 1 | 0 | no | yes | no | no | yes | no | no | no | no | no | no | no | no |
| SCN1A | 214 | A | 0 | 0 | no | yes | no | no | no | no | no | no | no | no | no | no | no |
| SCN1A | 215 | L | 0 | 0 | yes | no | no | yes | no | no | no | no | no | no | no | no | no |
| SCN1A | 216 | R | 0 | 0 | yes | no | no | yes | no | no | no | no | no | no | no | no | no |
| SCN1A | 217 | T | 1 | 1 | yes | no | no | yes | no | no | no | no | no | no | no | no | no |
| SCN1A | 218 | F | 0 | 1 | yes | no | no | yes | no | no | no | no | no | no | no | no | no |
| SCN1A | 219 | R | 0 | 2 | yes | no | no | yes | no | no | no | no | no | no | no | no | no |
| SCN1A | 220 | V | 1 | 1 | yes | no | no | yes | no | no | no | no | no | no | no | no | no |
| SCN1A | 221 | L | 0 | 2 | yes | no | no | yes | no | no | no | no | no | no | no | no | no |
| SCN1A | 222 | R | 0 | 0 | yes | no | no | yes | no | no | no | no | no | no | no | no | no |
| SCN1A | 223 | A | 0 | 2 | yes | no | no | yes | no | no | no | no | no | no | no | no | no |
| SCN1A | 224 | L | 1 | 1 | no | yes | no | yes | no | no | no | no | no | no | no | no | no |
| SCN1A | 225 | K | 0 | 2 | no | yes | no | yes | no | no | no | no | no | no | no | no | no |

| | | | | | | | | | | | | | | | | | |
|---|---|---|---|---|---|---|---|---|---|---|---|---|---|---|---|---|---|
| SCN1A | 226 | T | 0 | 2 | yes | no | no | yes | no | no | no | no | no | no | no | no | no |
| SCN1A | 227 | I | 0 | 1 | yes | no | no | yes | no | no | no | no | no | no | no | no | no |
| SCN1A | 228 | S | 0 | 2 | yes | no | no | yes | no | no | no | no | no | no | no | no | no |
| SCN1A | 229 | V | 1 | 0 | yes | no | no | yes | no | no | no | no | no | no | no | no | no |
| SCN1A | 230 | I | 0 | 0 | no | yes | no | yes | no | no | no | no | no | no | no | no | no |
| SCN1A | 231 | P | 0 | 0 | no | yes | no | yes | no | yes | no | no | no | no | no | no | no |
| SCN1A | 232 | G | 1 | 0 | no | yes | no | yes | no | yes | no | no | no | no | no | no | no |
| SCN1A | 233 | L | 0 | 0 | no | yes | no | yes | no | yes | no | no | no | no | no | no | no |
| SCN1A | 234 | K | 0 | 0 | yes | no | no | yes | no | yes | no | no | no | no | no | no | no |
| SCN1A | 235 | T | 0 | 1 | yes | no | no | yes | no | yes | no | no | no | no | no | no | no |
| SCN1A | 236 | I | 0 | 1 | yes | no | no | yes | no | yes | no | no | no | no | no | no | no |
| SCN1A | 237 | V | 0 | 0 | yes | no | no | yes | no | yes | no | no | no | no | no | no | no |
| SCN1A | 238 | G | 1 | 0 | yes | no | no | yes | no | yes | no | no | no | no | no | no | no |
| SCN1A | 239 | A | 0 | 2 | yes | no | no | yes | no | yes | no | no | no | no | no | no | no |
| SCN1A | 240 | L | 0 | 1 | yes | no | no | yes | no | yes | no | no | no | no | no | no | no |
| SCN1A | 241 | I | 0 | 0 | yes | no | no | yes | no | yes | no | no | no | no | no | no | no |
| SCN1A | 242 | Q | 0 | 0 | yes | no | no | yes | no | yes | no | no | no | no | no | no | no |
| SCN1A | 243 | S | 0 | 1 | yes | no | no | yes | no | yes | no | no | no | no | no | no | no |
| SCN1A | 244 | V | 0 | 1 | yes | no | no | yes | no | yes | no | no | no | no | no | no | no |
| SCN1A | 245 | K | 1 | 0 | yes | no | no | yes | no | yes | no | no | no | no | no | no | no |
| SCN1A | 246 | K | 0 | 0 | no | yes | no | yes | no | yes | no | no | no | no | no | no | no |
| SCN1A | 247 | L | 0 | 1 | no | yes | no | yes | no | yes | no | no | no | no | no | no | no |
| SCN1A | 248 | S | 0 | 0 | yes | no | no | yes | no | yes | no | no | no | no | no | no | no |
| SCN1A | 249 | D | 1 | 1 | yes | no | no | yes | no | yes | no | no | no | no | no | no | no |
| SCN1A | 250 | V | 0 | 2 | yes | no | no | yes | no | no | no | no | no | no | no | no | no |
| SCN1A | 251 | M | 0 | 1 | yes | no | no | yes | no | no | no | no | no | no | no | no | no |
| SCN1A | 252 | I | 0 | 2 | yes | no | no | yes | no | no | no | no | no | no | no | no | no |
| SCN1A | 253 | L | 0 | 0 | yes | no | no | yes | no | no | no | no | no | no | no | no | no |
| SCN1A | 254 | T | 0 | 1 | yes | no | no | yes | no | no | no | no | no | no | no | no | no |
| SCN1A | 255 | V | 1 | 0 | yes | no | no | yes | no | no | no | no | no | no | no | no | no |
| SCN1A | 256 | F | 0 | 0 | yes | no | no | yes | no | no | no | no | no | no | no | no | no |
| SCN1A | 257 | C | 0 | 1 | yes | no | no | yes | no | no | no | no | no | no | no | no | no |
| SCN1A | 258 | L | 0 | 0 | yes | no | no | yes | no | no | no | no | no | no | no | no | no |

| | | | | | | | | | | | | | | | | | |
|---|---|---|---|---|---|---|---|---|---|---|---|---|---|---|---|---|---|
| SCN1A | 259 | S | 0 | 1 | yes | no | no | yes | no | no | no | no | no | no | no | no | no |
| SCN1A | 260 | V | 1 | 0 | yes | no | no | yes | no | no | no | no | no | no | no | no | no |
| SCN1A | 261 | F | 0 | 0 | yes | no | no | yes | no | no | no | no | no | no | no | no | no |
| SCN1A | 262 | A | 0 | 0 | yes | no | no | yes | no | no | no | no | no | no | no | no | no |
| SCN1A | 263 | L | 0 | 1 | yes | no | no | yes | no | no | no | no | no | no | no | no | no |
| SCN1A | 264 | I | 1 | 1 | yes | no | no | yes | no | no | no | no | no | no | no | no | no |
| SCN1A | 265 | G | 0 | 1 | yes | no | no | yes | no | no | no | no | no | no | no | no | no |
| SCN1A | 266 | L | 0 | 1 | yes | no | no | yes | no | no | no | no | no | no | no | no | no |
| SCN1A | 267 | Q | 0 | 1 | yes | no | no | yes | no | no | no | no | no | no | no | no | no |
| SCN1A | 268 | L | 0 | 0 | yes | no | no | yes | no | no | no | no | no | no | no | no | no |
| SCN1A | 269 | F | 0 | 0 | no | yes | no | no | no | no | no | no | no | no | no | no | no |
| SCN1A | 270 | M | 0 | 0 | no | yes | no | no | yes | no | no | no | no | no | no | no | no |
| SCN1A | 271 | G | 0 | 1 | no | yes | no | no | yes | no | no | no | no | no | no | no | no |
| SCN1A | 272 | N | 0 | 0 | yes | no | no | no | yes | no | no | no | no | no | no | no | no |
| SCN1A | 273 | L | 0 | 1 | yes | no | no | no | yes | no | no | no | no | no | no | no | no |
| SCN1A | 274 | R | 1 | 0 | yes | no | no | no | yes | no | no | no | no | no | no | no | no |
| SCN1A | 275 | N | 0 | 0 | no | no | yes | no | yes | no | no | no | no | no | no | no | no |
| SCN1A | 276 | K | 0 | 2 | no | yes | no | no | yes | no | no | no | no | no | no | no | no |
| SCN1A | 277 | C | 0 | 2 | no | yes | no | no | yes | no | no | no | no | no | no | no | no |
| SCN1A | 278 | I | 0 | 0 | no | no | yes | no | yes | no | no | no | no | no | no | no | no |
| SCN1A | 279 | Q | 0 | 0 | no | yes | no | no | yes | no | no | no | no | no | no | no | no |
| SCN1A | 280 | W | 0 | 2 | no | yes | no | no | yes | no | no | no | no | no | no | no | no |
| SCN1A | 281 | P | 0 | 4 | no | yes | no | no | yes | no | no | no | no | no | no | no | no |
| SCN1A | 282 | P | 0 | 0 | no | yes | no | no | yes | no | no | no | no | no | no | no | no |
| SCN1A | 312 | F | 0 | 1 | no | yes | no | no | yes | no | no | no | no | no | no | no | no |
| SCN1A | 313 | D | 1 | 0 | no | yes | no | no | yes | no | no | no | no | no | no | no | no |
| SCN1A | 314 | W | 0 | 1 | no | yes | no | no | yes | no | no | no | no | no | no | no | no |
| SCN1A | 315 | K | 0 | 0 | no | yes | no | no | yes | no | no | no | no | no | no | no | no |
| SCN1A | 316 | S | 0 | 0 | no | yes | no | no | yes | no | no | no | no | no | no | no | no |
| SCN1A | 317 | Y | 0 | 2 | no | yes | no | no | yes | no | no | no | no | no | no | no | no |
| SCN1A | 318 | I | 0 | 0 | no | yes | no | no | yes | no | no | no | no | no | no | no | no |
| SCN1A | 319 | Q | 0 | 0 | no | yes | no | no | yes | no | no | no | no | no | no | no | no |
| SCN1A | 320 | D | 1 | 0 | no | yes | no | no | yes | no | no | no | no | no | no | no | no |

| | | | | | | | | | | | | | | | | | |
|---|---|---|---|---|---|---|---|---|---|---|---|---|---|---|---|---|---|
| SCN1A | 321 | S | 0 | 0 | no | yes | no | no | yes | no | no | no | no | no | no | no | no |
| SCN1A | 322 | R | 0 | 1 | no | yes | no | no | yes | no | no | no | no | no | no | no | no |
| SCN1A | 323 | Y | 0 | 0 | no | yes | no | no | yes | no | no | no | no | no | no | no | no |
| SCN1A | 324 | H | 0 | 2 | no | no | yes | no | yes | no | no | no | no | no | no | no | no |
| SCN1A | 325 | Y | 0 | 1 | no | yes | no | no | yes | no | no | no | no | no | no | no | no |
| SCN1A | 326 | F | 0 | 0 | no | yes | no | no | yes | no | no | no | no | no | no | no | no |
| SCN1A | 327 | L | 0 | 1 | no | yes | no | no | yes | no | no | no | no | no | no | no | no |
| SCN1A | 328 | E | 1 | 0 | no | yes | no | no | yes | no | no | no | no | no | no | no | no |
| SCN1A | 329 | G | 1 | 2 | no | yes | no | no | yes | no | no | no | no | no | no | no | no |
| SCN1A | 330 | F | 0 | 0 | no | yes | no | no | yes | no | no | no | no | no | no | no | no |
| SCN1A | 331 | L | 1 | 0 | no | yes | no | no | yes | no | no | no | no | no | no | no | no |
| SCN1A | 332 | D | 0 | 1 | no | yes | no | no | yes | no | no | no | no | no | no | no | no |
| SCN1A | 333 | A | 0 | 0 | no | yes | no | no | yes | no | no | no | no | no | no | no | no |
| SCN1A | 334 | L | 1 | 0 | no | yes | no | no | yes | no | no | no | no | no | no | no | no |
| SCN1A | 335 | L | 0 | 0 | no | yes | no | no | yes | no | no | no | no | no | no | no | no |
| SCN1A | 336 | C | 0 | 3 | no | yes | no | no | yes | no | no | no | no | no | no | no | no |
| SCN1A | 337 | G | 0 | 1 | no | yes | no | no | yes | no | no | no | no | no | no | no | no |
| SCN1A | 338 | N | 0 | 0 | no | yes | no | no | yes | no | no | no | no | no | no | no | no |
| SCN1A | 339 | S | 0 | 0 | no | yes | no | no | yes | no | no | no | no | no | no | no | no |
| SCN1A | 340 | S | 0 | 1 | no | yes | no | no | yes | no | no | no | no | no | no | no | no |
| SCN1A | 341 | D | 0 | 0 | no | yes | no | no | yes | no | no | no | no | no | no | no | no |
| SCN1A | 342 | A | 0 | 2 | no | yes | no | no | yes | no | no | no | no | no | no | no | no |
| SCN1A | 343 | G | 0 | 0 | no | yes | no | no | yes | no | no | no | no | no | no | no | no |
| SCN1A | 344 | Q | 0 | 0 | no | yes | no | no | yes | no | no | no | no | no | no | no | no |
| SCN1A | 345 | C | 0 | 2 | no | yes | no | no | yes | no | no | no | no | no | no | no | no |
| SCN1A | 346 | P | 0 | 0 | no | yes | no | no | yes | no | no | no | no | no | no | no | no |
| SCN1A | 347 | E | 0 | 0 | no | yes | no | no | yes | no | no | no | no | no | no | no | no |
| SCN1A | 348 | G | 0 | 1 | no | yes | no | no | yes | no | no | no | no | no | no | no | no |
| SCN1A | 349 | Y | 0 | 1 | no | yes | no | no | yes | no | no | no | no | no | no | no | no |
| SCN1A | 350 | M | 1 | 1 | no | no | yes | no | yes | no | no | no | no | no | no | no | no |
| SCN1A | 351 | C | 0 | 1 | no | yes | no | no | yes | no | no | no | no | no | no | no | no |
| SCN1A | 352 | V | 1 | 0 | no | yes | no | no | yes | no | no | no | no | no | no | no | no |
| SCN1A | 353 | K | 0 | 0 | no | no | yes | no | yes | no | no | no | no | no | no | no | no |

| | | | | | | | | | | | | | | | | | |
|---|---|---|---|---|---|---|---|---|---|---|---|---|---|---|---|---|---|
| SCN1A | 354 | A | 0 | 1 | no | yes | no | no | yes | no | no | no | no | no | no | no | no |
| SCN1A | 355 | G | 0 | 1 | no | yes | no | no | yes | no | no | no | no | no | no | no | no |
| SCN1A | 356 | R | 0 | 1 | no | yes | no | no | yes | no | no | no | no | no | no | no | no |
| SCN1A | 357 | N | 0 | 2 | no | yes | no | no | yes | no | no | no | no | no | no | no | no |
| SCN1A | 358 | P | 0 | 3 | no | yes | no | no | yes | no | no | no | no | no | no | no | no |
| SCN1A | 359 | N | 0 | 2 | no | yes | no | no | yes | no | no | no | no | no | no | no | no |
| SCN1A | 360 | Y | 0 | 0 | no | yes | no | no | yes | no | no | no | no | no | no | no | no |
| SCN1A | 361 | G | 0 | 0 | no | yes | no | no | yes | no | no | no | no | no | no | no | no |
| SCN1A | 362 | Y | 0 | 0 | no | yes | no | no | yes | no | no | no | no | no | no | no | no |
| SCN1A | 363 | T | 0 | 1 | no | yes | no | no | yes | no | no | no | no | no | no | no | no |
| SCN1A | 364 | S | 0 | 1 | no | yes | no | no | yes | no | no | no | no | no | no | no | no |
| SCN1A | 365 | F | 0 | 1 | no | yes | no | no | yes | no | no | no | no | no | no | no | no |
| SCN1A | 366 | D | 0 | 2 | no | yes | no | no | yes | no | no | no | no | no | no | no | no |
| SCN1A | 367 | T | 0 | 0 | no | yes | no | no | yes | no | no | no | no | no | no | no | no |
| SCN1A | 368 | F | 0 | 0 | no | yes | no | yes | no | no | no | yes | no | no | no | no | no |
| SCN1A | 369 | S | 1 | 0 | yes | no | no | yes | no | no | no | yes | no | no | no | no | no |
| SCN1A | 370 | W | 0 | 0 | yes | no | no | no | no | no | no | yes | no | no | no | no | no |
| SCN1A | 371 | A | 0 | 2 | yes | no | no | yes | no | no | no | yes | no | no | no | no | no |
| SCN1A | 372 | F | 0 | 0 | yes | no | no | yes | no | no | no | yes | no | no | no | no | no |
| SCN1A | 373 | L | 0 | 1 | yes | no | no | yes | no | no | no | yes | no | no | no | no | no |
| SCN1A | 374 | S | 0 | 1 | yes | no | no | yes | no | no | no | yes | no | no | no | no | no |
| SCN1A | 375 | L | 0 | 1 | yes | no | no | yes | no | no | no | yes | no | no | no | no | no |
| SCN1A | 376 | F | 0 | 0 | yes | no | no | yes | no | no | no | yes | no | no | no | no | no |
| SCN1A | 377 | R | 1 | 1 | yes | no | no | yes | no | no | no | yes | no | no | no | no | no |
| SCN1A | 378 | L | 0 | 1 | yes | no | no | yes | no | no | no | yes | no | no | no | no | no |
| SCN1A | 379 | M | 0 | 2 | yes | no | no | yes | no | no | no | yes | no | no | no | no | no |
| SCN1A | 380 | T | 0 | 0 | no | yes | no | yes | no | no | no | yes | no | no | no | no | no |
| SCN1A | 381 | Q | 0 | 0 | no | yes | no | yes | no | no | no | yes | no | no | no | no | no |
| SCN1A | 382 | D | 0 | 2 | no | yes | no | yes | no | no | no | yes | no | no | no | no | no |
| SCN1A | 383 | F | 0 | 1 | no | yes | no | no | no | no | no | yes | no | no | no | no | no |
| SCN1A | 384 | W | 0 | 1 | yes | no | no | yes | no | no | no | yes | no | no | no | no | no |
| SCN1A | 385 | E | 0 | 3 | yes | no | no | yes | no | no | no | yes | no | no | no | no | no |
| SCN1A | 386 | N | 0 | 0 | yes | no | no | no | no | no | no | yes | no | no | no | no | no |

| | | | | | | | | | | | | | | | | | |
|---|---|---|---|---|---|---|---|---|---|---|---|---|---|---|---|---|---|
| SCN1A | 387 | L | 0 | 0 | yes | no | no | no | no | no | no | yes | no | no | no | no | no |
| SCN1A | 388 | Y | 0 | 1 | yes | no | no | yes | no | no | no | yes | no | no | no | no | no |
| SCN1A | 389 | Q | 0 | 0 | yes | no | no | no | no | no | no | yes | no | no | no | no | no |
| SCN1A | 390 | L | 1 | 0 | yes | no | no | no | no | no | no | yes | no | no | no | no | no |
| SCN1A | 391 | T | 0 | 1 | yes | no | no | no | no | no | no | yes | no | no | no | no | no |
| SCN1A | 392 | L | 0 | 1 | yes | no | no | no | no | no | no | yes | no | no | no | no | no |
| SCN1A | 393 | R | 0 | 4 | yes | no | no | no | yes | no | no | no | no | no | no | no | no |
| SCN1A | 394 | A | 0 | 1 | yes | no | no | no | yes | no | no | no | no | no | no | no | no |
| SCN1A | 395 | A | 2 | 1 | yes | no | no | no | yes | no | no | no | no | no | no | no | no |
| SCN1A | 396 | G | 0 | 1 | no | yes | no | no | yes | no | no | no | no | no | no | no | no |
| SCN1A | 397 | K | 0 | 0 | yes | no | no | no | yes | no | no | no | no | no | no | no | no |
| SCN1A | 398 | T | 1 | 0 | yes | no | no | yes | yes | no | no | no | no | no | no | no | no |
| SCN1A | 399 | Y | 0 | 2 | yes | no | no | yes | yes | no | no | no | no | no | no | no | no |
| SCN1A | 400 | M | 1 | 1 | yes | no | no | yes | no | no | no | no | no | no | no | no | no |
| SCN1A | 401 | I | 0 | 0 | yes | no | no | yes | no | no | no | no | no | no | no | no | no |
| SCN1A | 402 | F | 0 | 2 | yes | no | no | yes | no | no | no | no | no | no | no | no | no |
| SCN1A | 403 | F | 0 | 1 | yes | no | no | yes | no | no | no | no | no | no | no | no | no |
| SCN1A | 404 | V | 0 | 0 | yes | no | no | yes | no | no | no | no | no | no | no | no | no |
| SCN1A | 405 | L | 0 | 0 | yes | no | no | yes | no | no | no | no | no | no | no | no | no |
| SCN1A | 406 | V | 0 | 1 | yes | no | no | yes | no | no | no | no | no | no | no | no | no |
| SCN1A | 407 | I | 1 | 0 | yes | no | no | yes | no | no | no | no | no | no | no | no | no |
| SCN1A | 408 | F | 0 | 1 | yes | no | no | yes | no | no | no | no | no | no | no | no | no |
| SCN1A | 409 | L | 0 | 1 | yes | no | no | yes | no | no | no | no | no | no | no | no | no |
| SCN1A | 410 | G | 0 | 0 | yes | no | no | yes | no | no | no | no | no | no | no | no | no |
| SCN1A | 411 | S | 0 | 0 | yes | no | no | yes | no | no | no | no | no | no | no | no | no |
| SCN1A | 412 | F | 0 | 1 | yes | no | no | yes | no | no | no | no | no | no | no | no | no |
| SCN1A | 413 | Y | 0 | 1 | yes | no | no | yes | no | no | no | no | no | no | no | no | no |
| SCN1A | 414 | L | 0 | 0 | yes | no | no | yes | no | no | no | no | no | no | no | no | no |
| SCN1A | 415 | I | 0 | 1 | yes | no | no | yes | no | no | no | no | no | no | no | no | no |
| SCN1A | 416 | N | 0 | 1 | yes | no | no | yes | no | no | no | no | no | no | no | no | no |
| SCN1A | 417 | L | 0 | 0 | yes | no | no | yes | no | no | no | no | no | no | no | no | no |
| SCN1A | 418 | I | 0 | 1 | yes | no | no | yes | no | no | no | no | no | no | no | no | no |
| SCN1A | 419 | L | 0 | 0 | yes | no | no | yes | no | no | yes | no | no | no | no | no | no |

| | | | | | | | | | | | | | | | | | |
|---|---|---|---|---|---|---|---|---|---|---|---|---|---|---|---|---|---|
| SCN1A | 420 | A | 0 | 1 | yes | no | no | yes | no | no | no | no | no | no | no | no | no |
| SCN1A | 421 | V | 1 | 1 | yes | no | no | yes | no | yes | no | no | no | no | no | no | no |
| SCN1A | 422 | V | 0 | 2 | yes | no | no | yes | no | yes | no | no | no | no | no | no | no |
| SCN1A | 423 | A | 0 | 0 | yes | no | no | yes | no | yes | yes | no | no | no | no | no | no |
| SCN1A | 424 | M | 0 | 0 | yes | no | no | no | no | yes | no | no | no | no | no | no | no |
| SCN1A | 425 | A | 0 | 0 | yes | no | no | no | no | yes | no | no | no | no | no | no | no |
| SCN1A | 426 | Y | 0 | 2 | yes | no | no | yes | no | yes | no | no | no | no | no | no | no |
| SCN1A | 427 | E | 1 | 0 | yes | no | no | no | no | yes | yes | no | no | no | no | no | no |
| SCN1A | 428 | E | 0 | 0 | yes | no | no | no | no | yes | no | no | no | no | no | no | no |
| SCN1A | 429 | Q | 1 | 0 | yes | no | no | no | no | yes | no | no | no | no | no | no | no |
| SCN1A | 430 | N | 0 | 0 | yes | no | no | no | no | yes | no | no | no | no | no | no | no |
| SCN1A | 431 | Q | 0 | 0 | yes | no | no | no | no | yes | yes | no | no | no | no | no | no |
| SCN1A | 432 | A | 0 | 1 | yes | no | no | no | no | yes | no | no | no | no | no | no | no |
| SCN1A | 433 | T | 0 | 0 | yes | no | no | no | no | yes | no | no | no | no | no | no | no |
| SCN1A | 434 | L | 2 | 0 | yes | no | no | no | no | yes | no | no | no | no | no | no | no |
| SCN1A | 435 | E | 2 | 0 | yes | no | no | no | no | yes | no | no | no | no | no | no | no |
| SCN1A | 436 | E | 0 | 0 | yes | no | no | no | no | yes | no | no | no | no | no | no | no |
| SCN1A | 437 | A | 0 | 0 | yes | no | no | no | no | yes | no | no | no | no | no | no | no |
| SCN1A | 438 | E | 0 | 0 | yes | no | no | no | no | yes | yes | no | no | no | no | no | no |
| SCN1A | 439 | Q | 0 | 0 | no | yes | no | no | no | yes | no | no | no | no | no | no | no |
| SCN1A | 749 | D | 1 | 0 | no | yes | no | no | no | yes | no | no | no | no | no | no | no |
| SCN1A | 750 | C | 1 | 0 | yes | no | no | no | no | yes | no | no | no | no | no | no | no |
| SCN1A | 751 | S | 0 | 0 | yes | no | no | no | no | yes | no | no | no | no | no | no | no |
| SCN1A | 752 | P | 0 | 0 | yes | no | no | no | no | yes | no | no | no | no | no | no | no |
| SCN1A | 753 | Y | 0 | 0 | yes | no | no | no | no | yes | no | no | no | no | no | no | no |
| SCN1A | 754 | W | 0 | 1 | yes | no | no | no | no | yes | no | no | no | no | no | no | no |
| SCN1A | 755 | L | 0 | 0 | yes | no | no | no | no | yes | no | no | no | no | no | no | no |
| SCN1A | 756 | K | 1 | 0 | yes | no | no | no | no | yes | no | no | no | no | no | no | no |
| SCN1A | 757 | V | 0 | 0 | yes | no | no | no | no | yes | no | no | no | no | no | no | no |
| SCN1A | 758 | K | 0 | 0 | yes | no | no | no | no | yes | no | no | no | no | no | no | no |
| SCN1A | 759 | H | 0 | 0 | yes | no | no | no | no | yes | no | no | no | no | no | no | no |
| SCN1A | 760 | V | 0 | 0 | yes | no | no | no | no | yes | no | no | no | no | no | no | no |
| SCN1A | 761 | V | 0 | 0 | yes | no | no | no | no | yes | no | no | no | no | no | no | no |

| | | | | | | | | | | | | | | | | | |
|---|---|---|---|---|---|---|---|---|---|---|---|---|---|---|---|---|---|
| SCN1A | 762 | N | 0 | 0 | yes | no | no | no | no | yes | no | no | no | no | no | no | no |
| SCN1A | 763 | L | 1 | 0 | no | yes | no | no | no | yes | no | no | no | no | no | no | no |
| SCN1A | 764 | V | 0 | 0 | no | yes | no | no | no | yes | no | no | no | no | no | no | no |
| SCN1A | 765 | V | 2 | 0 | no | yes | no | no | no | yes | no | no | no | no | no | no | no |
| SCN1A | 766 | M | 0 | 0 | no | yes | no | no | no | yes | no | no | no | no | no | no | no |
| SCN1A | 767 | D | 1 | 0 | no | yes | no | no | no | yes | no | no | no | no | no | no | no |
| SCN1A | 768 | P | 1 | 1 | no | yes | no | no | no | yes | no | no | no | no | no | no | no |
| SCN1A | 769 | F | 0 | 1 | no | yes | no | yes | no | no | no | no | no | no | no | no | no |
| SCN1A | 770 | V | 0 | 0 | yes | no | no | yes | no | no | no | no | no | no | no | no | no |
| SCN1A | 771 | D | 0 | 0 | yes | no | no | yes | no | no | no | no | no | no | no | no | no |
| SCN1A | 772 | L | 0 | 0 | yes | no | no | yes | no | no | no | no | no | no | no | no | no |
| SCN1A | 773 | A | 0 | 0 | yes | no | no | yes | no | no | no | no | no | no | no | no | no |
| SCN1A | 774 | I | 0 | 0 | yes | no | no | yes | no | no | no | no | no | no | no | no | no |
| SCN1A | 775 | T | 0 | 0 | yes | no | no | yes | no | no | no | no | no | no | no | no | no |
| SCN1A | 776 | I | 0 | 0 | yes | no | no | yes | no | no | no | no | no | no | no | no | no |
| SCN1A | 777 | C | 0 | 0 | yes | no | no | yes | no | no | no | no | no | no | no | no | no |
| SCN1A | 778 | I | 0 | 0 | yes | no | no | yes | no | no | no | no | no | no | no | no | no |
| SCN1A | 779 | V | 0 | 0 | yes | no | no | yes | no | no | no | no | no | no | no | no | no |
| SCN1A | 780 | L | 0 | 0 | yes | no | no | yes | no | no | no | no | no | no | no | no | no |
| SCN1A | 781 | N | 0 | 0 | yes | no | no | yes | no | no | no | no | no | no | no | no | no |
| SCN1A | 782 | T | 0 | 0 | yes | no | no | yes | no | no | no | no | no | no | no | no | no |
| SCN1A | 783 | L | 0 | 1 | yes | no | no | yes | no | no | no | no | no | no | no | no | no |
| SCN1A | 784 | F | 0 | 0 | yes | no | no | yes | no | no | no | no | no | no | no | no | no |
| SCN1A | 785 | M | 1 | 1 | yes | no | no | yes | no | no | no | no | no | no | no | no | no |
| SCN1A | 786 | A | 1 | 1 | yes | no | no | yes | no | no | no | no | no | no | no | no | no |
| SCN1A | 787 | M | 1 | 1 | no | yes | no | yes | no | no | no | no | no | no | no | no | no |
| SCN1A | 788 | E | 0 | 1 | no | yes | no | yes | yes | no | no | no | no | no | no | no | no |
| SCN1A | 789 | H | 0 | 0 | no | yes | no | no | yes | no | no | no | no | no | no | no | no |
| SCN1A | 790 | Y | 1 | 1 | no | yes | no | no | yes | no | no | no | no | no | no | no | no |
| SCN1A | 791 | P | 0 | 0 | no | yes | no | no | yes | no | no | no | no | no | no | no | no |
| SCN1A | 792 | M | 2 | 0 | no | yes | no | no | yes | no | no | no | no | no | no | no | no |
| SCN1A | 793 | T | 1 | 0 | no | yes | no | no | yes | no | no | no | no | no | no | no | no |
| SCN1A | 794 | D | 1 | 0 | yes | no | no | no | yes | no | no | no | no | no | no | no | no |

| | | | | | | | | | | | | | | | | | |
|---|---|---|---|---|---|---|---|---|---|---|---|---|---|---|---|---|---|
| SCN1A | 795 | H | 0 | 0 | yes | no | no | no | yes | no | no | no | no | no | no | no | no |
| SCN1A | 796 | F | 0 | 0 | yes | no | no | yes | yes | no | no | no | no | no | no | no | no |
| SCN1A | 797 | N | 1 | 0 | yes | no | no | no | yes | no | no | no | no | no | no | no | no |
| SCN1A | 798 | N | 0 | 0 | yes | no | no | no | yes | no | no | no | no | no | no | no | no |
| SCN1A | 799 | V | 0 | 0 | yes | no | no | yes | no | no | no | no | no | no | no | no | no |
| SCN1A | 800 | L | 0 | 0 | yes | no | no | yes | no | no | no | no | no | no | no | no | no |
| SCN1A | 801 | T | 0 | 0 | yes | no | no | yes | no | no | no | no | no | no | no | no | no |
| SCN1A | 802 | V | 1 | 0 | yes | no | no | yes | no | no | no | no | no | no | no | no | no |
| SCN1A | 803 | G | 0 | 0 | yes | no | no | yes | no | no | no | no | no | no | no | no | no |
| SCN1A | 804 | N | 0 | 1 | yes | no | no | yes | no | no | no | no | no | no | no | no | no |
| SCN1A | 805 | L | 0 | 0 | yes | no | no | yes | no | no | no | no | no | no | no | no | no |
| SCN1A | 806 | V | 0 | 0 | yes | no | no | yes | no | no | no | no | no | no | no | no | no |
| SCN1A | 807 | F | 0 | 0 | yes | no | no | yes | no | no | no | no | no | no | no | no | no |
| SCN1A | 808 | T | 0 | 1 | yes | no | no | yes | no | no | no | no | no | no | no | no | no |
| SCN1A | 809 | G | 0 | 0 | yes | no | no | yes | no | no | no | no | no | no | no | no | no |
| SCN1A | 810 | I | 0 | 0 | yes | no | no | yes | no | no | no | no | no | no | no | no | no |
| SCN1A | 811 | F | 0 | 0 | yes | no | no | yes | no | no | no | no | no | no | no | no | no |
| SCN1A | 812 | T | 0 | 1 | yes | no | no | yes | no | no | no | no | no | no | no | no | no |
| SCN1A | 813 | A | 1 | 0 | yes | no | no | yes | no | no | no | no | no | no | no | no | no |
| SCN1A | 814 | E | 0 | 0 | yes | no | no | yes | no | no | no | no | no | no | no | no | no |
| SCN1A | 815 | M | 0 | 0 | yes | no | no | yes | no | no | no | no | no | no | no | no | no |
| SCN1A | 816 | F | 0 | 0 | yes | no | no | yes | no | no | no | no | no | no | no | no | no |
| SCN1A | 817 | L | 0 | 0 | yes | no | no | yes | no | no | no | no | no | no | no | no | no |
| SCN1A | 818 | K | 0 | 0 | yes | no | no | yes | no | no | no | no | no | no | no | no | no |
| SCN1A | 819 | I | 0 | 0 | no | yes | no | yes | no | yes | no | no | no | no | no | no | no |
| SCN1A | 820 | I | 0 | 0 | no | yes | no | yes | no | yes | no | no | no | no | no | no | no |
| SCN1A | 821 | A | 0 | 0 | no | yes | no | no | no | yes | no | no | no | no | no | no | no |
| SCN1A | 822 | M | 0 | 0 | no | yes | no | no | no | yes | no | no | no | no | no | no | no |
| SCN1A | 823 | D | 0 | 0 | no | yes | no | no | no | yes | no | no | no | no | no | no | no |
| SCN1A | 824 | P | 1 | 0 | no | yes | no | yes | no | yes | no | no | no | no | no | no | no |
| SCN1A | 825 | Y | 1 | 0 | no | yes | no | no | no | yes | no | no | no | no | no | no | no |
| SCN1A | 826 | Y | 1 | 0 | no | yes | no | no | no | yes | no | no | no | no | no | no | no |
| SCN1A | 827 | Y | 0 | 1 | no | yes | no | yes | no | yes | no | no | no | no | no | no | no |

| | | | | | | | | | | | | | | | | | |
|---|---|---|---|---|---|---|---|---|---|---|---|---|---|---|---|---|---|
| SCN1A | 828 | F | 0 | 0 | no | yes | no | yes | no | yes | no | no | no | no | no | no | no |
| SCN1A | 829 | Q | 1 | 0 | no | yes | no | no | no | yes | no | no | no | no | no | no | no |
| SCN1A | 830 | E | 0 | 0 | no | yes | no | no | no | yes | no | no | no | no | no | no | no |
| SCN1A | 831 | G | 0 | 0 | no | yes | no | yes | no | yes | no | no | no | no | no | no | no |
| SCN1A | 832 | W | 0 | 0 | yes | no | no | yes | no | yes | no | no | no | no | no | no | no |
| SCN1A | 833 | N | 0 | 0 | yes | no | no | yes | no | no | no | no | no | no | no | no | no |
| SCN1A | 834 | I | 0 | 0 | yes | no | no | yes | no | no | no | no | no | no | no | no | no |
| SCN1A | 835 | F | 0 | 1 | yes | no | no | yes | no | no | no | no | no | no | no | no | no |
| SCN1A | 836 | D | 0 | 0 | yes | no | no | yes | no | no | no | no | no | no | no | no | no |
| SCN1A | 837 | G | 1 | 0 | yes | no | no | yes | no | no | no | no | no | no | no | no | no |
| SCN1A | 838 | F | 0 | 0 | yes | no | no | yes | no | no | no | no | no | no | no | no | no |
| SCN1A | 839 | I | 0 | 0 | yes | no | no | yes | no | no | no | no | no | no | no | no | no |
| SCN1A | 840 | V | 0 | 0 | yes | no | no | yes | no | no | no | no | no | no | no | no | no |
| SCN1A | 841 | T | 1 | 0 | yes | no | no | yes | no | no | no | no | no | no | no | no | no |
| SCN1A | 842 | L | 0 | 0 | yes | no | no | yes | no | no | no | no | no | no | no | no | no |
| SCN1A | 843 | S | 0 | 1 | yes | no | no | yes | no | no | no | no | no | no | no | no | no |
| SCN1A | 844 | L | 0 | 0 | yes | no | no | yes | no | no | no | no | no | no | no | no | no |
| SCN1A | 845 | V | 0 | 0 | yes | no | no | yes | no | no | no | no | no | no | no | no | no |
| SCN1A | 846 | E | 0 | 1 | yes | no | no | yes | no | no | no | no | no | no | no | no | no |
| SCN1A | 847 | L | 0 | 0 | yes | no | no | yes | no | no | no | no | no | no | no | no | no |
| SCN1A | 848 | G | 1 | 0 | no | yes | no | yes | no | no | no | no | no | no | no | no | no |
| SCN1A | 849 | L | 0 | 1 | no | yes | no | yes | no | no | no | no | no | no | no | no | no |
| SCN1A | 850 | A | 1 | 0 | no | yes | no | no | no | no | no | no | no | no | no | no | no |
| SCN1A | 851 | N | 2 | 0 | no | yes | no | no | no | no | no | no | no | no | no | no | no |
| SCN1A | 852 | V | 0 | 0 | no | yes | no | no | no | no | no | no | no | no | no | no | no |
| SCN1A | 853 | E | 1 | 1 | no | yes | no | no | yes | no | no | no | no | no | no | no | no |
| SCN1A | 854 | G | 0 | 0 | no | yes | no | no | yes | no | no | no | no | no | no | no | no |
| SCN1A | 855 | L | 0 | 0 | no | yes | no | yes | no | no | no | no | no | no | no | no | no |
| SCN1A | 856 | S | 0 | 0 | yes | no | no | yes | no | no | no | no | no | no | no | no | no |
| SCN1A | 857 | V | 0 | 1 | yes | no | no | yes | no | no | no | no | no | no | no | no | no |
| SCN1A | 858 | L | 0 | 1 | yes | no | no | yes | no | no | no | no | no | no | no | no | no |
| SCN1A | 859 | R | 1 | 2 | yes | no | no | yes | no | no | no | no | no | no | no | no | no |
| SCN1A | 860 | S | 0 | 0 | yes | no | no | yes | no | no | no | no | no | no | no | no | no |

| | | | | | | | | | | | | | | | | | |
|---|---|---|---|---|---|---|---|---|---|---|---|---|---|---|---|---|---|
| SCN1A | 861 | F | 0 | 0 | yes | no | no | yes | no | no | no | no | no | no | no | no | no |
| SCN1A | 862 | R | 0 | 2 | yes | no | no | yes | no | no | no | no | no | no | no | no | no |
| SCN1A | 863 | L | 0 | 1 | yes | no | no | yes | no | no | no | no | no | no | no | no | no |
| SCN1A | 864 | L | 1 | 0 | yes | no | no | yes | no | no | no | no | no | no | no | no | no |
| SCN1A | 865 | R | 0 | 1 | yes | no | no | yes | no | no | no | no | no | no | no | no | no |
| SCN1A | 866 | V | 0 | 0 | yes | no | no | yes | no | no | no | no | no | no | no | no | no |
| SCN1A | 867 | F | 1 | 0 | yes | no | no | yes | no | no | no | no | no | no | no | no | no |
| SCN1A | 868 | K | 0 | 1 | yes | no | no | yes | no | no | no | no | no | no | no | no | no |
| SCN1A | 869 | L | 0 | 1 | yes | no | no | yes | no | no | no | no | no | no | no | no | no |
| SCN1A | 870 | A | 0 | 0 | yes | no | no | yes | no | no | no | no | no | no | no | no | no |
| SCN1A | 871 | K | 0 | 0 | yes | no | no | yes | no | no | no | no | no | no | no | no | no |
| SCN1A | 872 | S | 0 | 1 | no | yes | no | yes | no | no | no | no | no | no | no | no | no |
| SCN1A | 873 | W | 0 | 1 | no | yes | no | yes | no | yes | no | no | no | no | no | no | no |
| SCN1A | 874 | P | 0 | 0 | yes | no | no | yes | no | yes | no | no | no | no | no | no | no |
| SCN1A | 875 | T | 0 | 1 | yes | no | no | yes | no | yes | no | no | no | no | no | no | no |
| SCN1A | 876 | L | 0 | 1 | yes | no | no | yes | no | yes | no | no | no | no | no | no | no |
| SCN1A | 877 | N | 1 | 0 | yes | no | no | yes | no | yes | no | no | no | no | no | no | no |
| SCN1A | 878 | M | 0 | 0 | yes | no | no | yes | no | yes | no | no | no | no | no | no | no |
| SCN1A | 879 | L | 0 | 2 | yes | no | no | yes | no | yes | no | no | no | no | no | no | no |
| SCN1A | 880 | I | 0 | 0 | yes | no | no | yes | no | yes | no | no | no | no | no | no | no |
| SCN1A | 881 | K | 0 | 0 | yes | no | no | yes | no | yes | no | no | no | no | no | no | no |
| SCN1A | 882 | I | 0 | 1 | yes | no | no | yes | no | yes | no | no | no | no | no | no | no |
| SCN1A | 883 | I | 0 | 0 | yes | no | no | yes | no | yes | no | no | no | no | no | no | no |
| SCN1A | 884 | G | 1 | 1 | yes | no | no | yes | no | yes | no | no | no | no | no | no | no |
| SCN1A | 885 | N | 0 | 0 | yes | no | no | yes | no | yes | no | no | no | no | no | no | no |
| SCN1A | 886 | S | 0 | 1 | no | yes | no | yes | no | yes | no | no | no | no | no | no | no |
| SCN1A | 887 | V | 1 | 0 | no | yes | no | yes | no | yes | no | no | no | no | no | no | no |
| SCN1A | 888 | G | 0 | 0 | no | yes | no | yes | no | yes | no | no | no | no | no | no | no |
| SCN1A | 889 | A | 1 | 1 | no | yes | no | yes | no | no | no | no | no | no | no | no | no |
| SCN1A | 890 | L | 0 | 1 | no | yes | no | yes | no | no | no | no | no | no | no | no | no |
| SCN1A | 891 | G | 0 | 0 | yes | no | no | yes | no | no | no | no | no | no | no | no | no |
| SCN1A | 892 | N | 0 | 0 | yes | no | no | yes | no | no | no | no | no | no | no | no | no |
| SCN1A | 893 | L | 0 | 0 | yes | no | no | yes | no | no | no | no | no | no | no | no | no |

| | | | | | | | | | | | | | | | | | |
|---|---|---|---|---|---|---|---|---|---|---|---|---|---|---|---|---|---|
| SCN1A | 894 | T | 0 | 1 | yes | no | no | yes | no | no | no | no | no | no | no | no | no |
| SCN1A | 895 | L | 1 | 0 | yes | no | no | yes | no | no | no | no | no | no | no | no | no |
| SCN1A | 896 | V | 1 | 1 | yes | no | no | yes | no | no | no | no | no | no | no | no | no |
| SCN1A | 897 | L | 0 | 2 | yes | no | no | yes | no | no | no | no | no | no | no | no | no |
| SCN1A | 898 | A | 0 | 0 | yes | no | no | yes | no | no | no | no | no | no | no | no | no |
| SCN1A | 899 | I | 1 | 1 | yes | no | no | yes | no | no | no | no | no | no | no | no | no |
| SCN1A | 900 | I | 0 | 0 | yes | no | no | yes | no | no | no | no | no | no | no | no | no |
| SCN1A | 901 | V | 1 | 0 | yes | no | no | yes | no | no | no | no | no | no | no | no | no |
| SCN1A | 902 | F | 0 | 1 | yes | no | no | yes | no | no | no | no | no | no | no | no | no |
| SCN1A | 903 | I | 0 | 0 | yes | no | no | yes | no | no | no | no | no | no | no | no | no |
| SCN1A | 904 | F | 0 | 0 | yes | no | no | yes | no | no | no | no | no | no | no | no | no |
| SCN1A | 905 | A | 0 | 2 | yes | no | no | yes | no | no | no | no | no | no | no | no | no |
| SCN1A | 906 | V | 1 | 0 | yes | no | no | yes | no | no | no | no | no | no | no | no | no |
| SCN1A | 907 | V | 0 | 1 | yes | no | no | yes | no | no | no | no | no | no | no | no | no |
| SCN1A | 908 | G | 0 | 1 | yes | no | no | yes | yes | no | no | no | no | no | no | no | no |
| SCN1A | 909 | M | 0 | 0 | yes | no | no | yes | yes | no | no | no | no | no | no | no | no |
| SCN1A | 910 | Q | 0 | 2 | yes | no | no | yes | yes | no | no | no | no | no | no | no | no |
| SCN1A | 911 | L | 0 | 0 | yes | no | no | yes | yes | no | no | no | no | no | no | no | no |
| SCN1A | 912 | F | 0 | 0 | no | yes | no | no | yes | no | no | no | no | no | no | no | no |
| SCN1A | 913 | G | 0 | 0 | no | yes | no | no | yes | no | no | no | no | no | no | no | no |
| SCN1A | 914 | K | 0 | 0 | no | yes | no | no | yes | no | no | no | no | no | no | no | no |
| SCN1A | 915 | S | 0 | 0 | yes | no | no | no | yes | no | no | no | no | no | no | no | no |
| SCN1A | 916 | Y | 0 | 0 | yes | no | no | no | yes | no | no | no | no | no | no | no | no |
| SCN1A | 917 | K | 0 | 0 | yes | no | no | no | yes | no | no | no | no | no | no | no | no |
| SCN1A | 918 | D | 1 | 0 | yes | no | no | no | yes | no | no | no | yes | no | no | no | no |
| SCN1A | 919 | C | 0 | 0 | no | yes | no | no | yes | no | no | no | yes | no | no | no | no |
| SCN1A | 920 | V | 0 | 0 | yes | no | no | no | yes | no | no | no | no | no | no | no | no |
| SCN1A | 921 | C | 0 | 0 | yes | no | no | no | yes | no | no | no | yes | no | no | no | no |
| SCN1A | 922 | K | 0 | 0 | yes | no | no | no | yes | no | no | no | yes | no | no | no | no |
| SCN1A | 923 | I | 0 | 0 | no | yes | no | no | yes | no | no | no | no | no | no | no | no |
| SCN1A | 924 | A | 1 | 0 | no | yes | no | no | yes | no | no | no | no | no | no | no | no |
| SCN1A | 925 | S | 0 | 0 | no | yes | no | no | yes | no | no | no | no | no | no | no | no |
| SCN1A | 926 | D | 0 | 0 | no | yes | no | no | yes | no | no | no | no | no | no | no | no |

| | | | | | | | | | | | | | | | | | |
|---|---|---|---|---|---|---|---|---|---|---|---|---|---|---|---|---|---|
| SCN1A | 927 | C | 0 | 1 | no | yes | no | no | yes | no | no | no | no | no | no | no | no |
| SCN1A | 928 | Q | 1 | 0 | no | yes | no | no | yes | no | no | no | no | no | no | no | no |
| SCN1A | 929 | L | 0 | 0 | no | yes | no | no | yes | no | no | no | no | no | no | no | no |
| SCN1A | 930 | P | 0 | 0 | no | yes | no | no | yes | no | no | no | no | no | no | no | no |
| SCN1A | 931 | R | 0 | 3 | no | yes | no | no | yes | no | no | no | no | no | no | no | no |
| SCN1A | 932 | W | 0 | 1 | no | yes | no | no | yes | no | no | no | no | no | no | no | no |
| SCN1A | 933 | H | 0 | 2 | no | yes | no | no | yes | no | no | no | no | no | no | no | no |
| SCN1A | 934 | M | 0 | 3 | no | yes | no | no | yes | no | no | no | no | no | no | no | no |
| SCN1A | 935 | N | 1 | 1 | no | yes | no | no | yes | no | no | no | no | no | no | no | no |
| SCN1A | 936 | D | 0 | 2 | no | yes | no | no | yes | no | no | no | no | no | no | no | no |
| SCN1A | 937 | F | 0 | 0 | yes | no | no | yes | no | no | no | yes | no | no | no | no | no |
| SCN1A | 938 | F | 0 | 0 | yes | no | no | yes | no | no | no | yes | no | no | no | no | no |
| SCN1A | 939 | H | 0 | 2 | yes | no | no | no | no | no | no | yes | no | no | no | no | no |
| SCN1A | 940 | S | 0 | 1 | yes | no | no | yes | no | no | no | yes | no | no | no | no | no |
| SCN1A | 941 | F | 0 | 0 | yes | no | no | yes | no | no | no | yes | no | no | no | no | no |
| SCN1A | 942 | L | 0 | 2 | yes | no | no | yes | no | no | no | yes | no | no | no | no | no |
| SCN1A | 943 | I | 0 | 1 | yes | no | no | yes | no | no | no | yes | no | no | no | no | no |
| SCN1A | 944 | V | 0 | 1 | yes | no | no | yes | no | no | no | yes | no | no | no | no | no |
| SCN1A | 945 | F | 0 | 1 | yes | no | no | yes | no | no | no | yes | no | no | no | no | no |
| SCN1A | 946 | R | 0 | 2 | yes | no | no | yes | no | no | no | yes | no | no | no | no | no |
| SCN1A | 947 | V | 0 | 1 | yes | no | no | yes | no | no | no | yes | no | no | no | no | no |
| SCN1A | 948 | L | 0 | 0 | yes | no | no | yes | no | no | no | yes | no | no | no | no | no |
| SCN1A | 949 | C | 0 | 2 | yes | no | no | yes | no | no | no | yes | no | no | no | no | no |
| SCN1A | 950 | G | 0 | 1 | no | yes | no | yes | no | no | no | yes | no | no | no | no | no |
| SCN1A | 951 | E | 0 | 0 | no | yes | no | yes | no | no | no | yes | no | no | no | no | no |
| SCN1A | 952 | W | 0 | 2 | no | yes | no | yes | no | no | no | yes | no | no | no | no | no |
| SCN1A | 953 | I | 0 | 0 | no | yes | no | yes | no | no | no | yes | no | no | no | no | no |
| SCN1A | 954 | E | 0 | 1 | yes | no | no | no | no | no | no | yes | no | no | no | no | no |
| SCN1A | 955 | T | 0 | 1 | yes | no | no | no | no | no | no | yes | no | no | no | no | no |
| SCN1A | 956 | M | 0 | 1 | yes | no | no | yes | no | no | no | yes | no | no | no | no | no |
| SCN1A | 957 | W | 0 | 2 | yes | no | no | no | no | no | no | yes | no | no | no | no | no |
| SCN1A | 958 | D | 0 | 0 | yes | no | no | no | yes | no | no | no | no | no | no | no | no |
| SCN1A | 959 | C | 0 | 3 | yes | no | no | no | yes | no | no | no | no | no | no | no | no |

| | | | | | | | | | | | | | | | | | |
|---|---|---|---|---|---|---|---|---|---|---|---|---|---|---|---|---|---|
| SCN1A | 960 | M | 0 | 3 | yes | no | no | no | yes | no | no | no | no | no | no | no | no |
| SCN1A | 961 | E | 0 | 0 | yes | no | no | no | yes | no | no | no | no | no | no | no | no |
| SCN1A | 962 | V | 1 | 0 | yes | no | no | no | yes | no | no | no | no | no | no | no | no |
| SCN1A | 963 | A | 0 | 0 | yes | no | no | no | yes | no | no | no | no | no | no | no | no |
| SCN1A | 964 | G | 1 | 0 | no | yes | no | no | yes | no | no | no | no | no | no | no | no |
| SCN1A | 965 | Q | 0 | 0 | yes | no | no | no | yes | no | no | no | no | no | no | no | no |
| SCN1A | 966 | A | 0 | 0 | yes | no | no | yes | yes | no | no | no | no | no | no | no | no |
| SCN1A | 967 | M | 1 | 0 | yes | no | no | yes | yes | no | no | no | no | no | no | no | no |
| SCN1A | 968 | C | 0 | 2 | yes | no | no | yes | yes | no | no | no | no | no | no | no | no |
| SCN1A | 969 | L | 0 | 1 | yes | no | no | yes | yes | no | no | no | no | no | no | no | no |
| SCN1A | 970 | T | 1 | 0 | yes | no | no | yes | yes | no | no | no | no | no | no | no | no |
| SCN1A | 971 | V | 1 | 1 | yes | no | no | yes | no | no | no | no | no | no | no | no | no |
| SCN1A | 972 | F | 0 | 0 | yes | no | no | yes | no | no | no | no | no | no | no | no | no |
| SCN1A | 973 | M | 0 | 2 | yes | no | no | yes | no | no | no | no | no | no | no | no | no |
| SCN1A | 974 | M | 1 | 0 | yes | no | no | yes | no | no | no | no | no | no | no | no | no |
| SCN1A | 975 | V | 0 | 0 | yes | no | no | yes | no | no | no | no | no | no | no | no | no |
| SCN1A | 976 | M | 1 | 3 | yes | no | no | yes | no | no | no | no | no | no | no | no | no |
| SCN1A | 977 | V | 0 | 1 | yes | no | no | yes | no | no | no | no | no | no | no | no | no |
| SCN1A | 978 | I | 0 | 0 | yes | no | no | yes | no | no | no | no | no | no | no | no | no |
| SCN1A | 979 | G | 0 | 2 | yes | no | no | yes | no | no | no | no | no | no | no | no | no |
| SCN1A | 980 | N | 0 | 1 | yes | no | no | yes | no | no | no | no | no | no | no | no | no |
| SCN1A | 981 | L | 0 | 1 | yes | no | no | yes | no | no | no | no | no | no | no | no | no |
| SCN1A | 982 | V | 0 | 1 | yes | no | no | yes | no | no | no | no | no | no | no | no | no |
| SCN1A | 983 | V | 0 | 2 | yes | no | no | yes | no | no | no | no | no | no | no | no | no |
| SCN1A | 984 | L | 0 | 1 | yes | no | no | yes | no | no | no | no | no | no | no | no | no |
| SCN1A | 985 | N | 0 | 1 | yes | no | no | yes | no | no | no | no | no | no | no | no | no |
| SCN1A | 986 | L | 0 | 2 | yes | no | no | yes | no | no | no | no | no | no | no | no | no |
| SCN1A | 987 | F | 0 | 1 | yes | no | no | yes | no | no | yes | no | no | no | no | no | no |
| SCN1A | 988 | L | 0 | 1 | yes | no | no | yes | no | no | yes | no | no | no | no | no | no |
| SCN1A | 989 | A | 0 | 2 | yes | no | no | yes | no | no | no | no | no | no | no | no | no |
| SCN1A | 990 | L | 0 | 1 | yes | no | no | yes | no | no | no | no | no | no | no | no | no |
| SCN1A | 991 | L | 0 | 0 | yes | no | no | yes | no | no | yes | no | no | no | no | no | no |
| SCN1A | 992 | L | 0 | 1 | yes | no | no | no | no | yes | yes | no | no | no | no | no | no |

| | | | | | | | | | | | | | | | | | |
|---|---|---|---|---|---|---|---|---|---|---|---|---|---|---|---|---|---|
| SCN1A | 993 | S | 0 | 0 | yes | no | no | no | no | yes | no | no | no | no | no | no | no |
| SCN1A | 994 | S | 0 | 1 | yes | no | no | no | no | yes | no | no | no | no | no | no | no |
| SCN1A | 995 | F | 0 | 1 | no | yes | no | no | no | yes | no | no | no | no | no | no | no |
| SCN1A | 996 | S | 0 | 0 | no | yes | no | no | no | yes | no | no | no | no | no | no | no |
| SCN1A | 1201 | G | 0 | 0 | no | yes | no | no | no | yes | no | no | no | no | no | no | no |
| SCN1A | 1202 | K | 0 | 0 | no | yes | no | no | no | yes | no | no | no | no | no | no | no |
| SCN1A | 1203 | Q | 0 | 0 | no | yes | no | no | no | yes | no | no | no | no | no | no | no |
| SCN1A | 1204 | W | 0 | 1 | yes | no | no | no | no | yes | no | no | no | no | no | no | no |
| SCN1A | 1205 | W | 0 | 0 | yes | no | no | no | no | yes | no | no | no | no | no | no | no |
| SCN1A | 1206 | N | 1 | 0 | yes | no | no | no | no | yes | no | no | no | no | no | no | no |
| SCN1A | 1207 | L | 0 | 1 | yes | no | no | yes | no | yes | no | no | no | no | no | no | no |
| SCN1A | 1208 | R | 0 | 0 | yes | no | no | no | no | yes | no | no | no | no | no | no | no |
| SCN1A | 1209 | R | 0 | 0 | yes | no | no | no | no | yes | no | no | no | no | no | no | no |
| SCN1A | 1210 | T | 1 | 1 | yes | no | no | no | no | yes | no | no | no | no | no | no | no |
| SCN1A | 1211 | C | 0 | 0 | yes | no | no | yes | no | yes | no | no | no | no | no | no | no |
| SCN1A | 1212 | F | 0 | 0 | yes | no | no | no | no | yes | no | no | no | no | no | no | no |
| SCN1A | 1213 | R | 1 | 1 | yes | no | no | no | no | yes | no | no | no | no | no | no | no |
| SCN1A | 1214 | I | 0 | 0 | yes | no | no | yes | no | yes | no | no | no | no | no | no | no |
| SCN1A | 1215 | V | 1 | 0 | yes | no | no | yes | no | yes | no | no | no | no | no | no | no |
| SCN1A | 1216 | E | 0 | 1 | no | yes | no | no | no | yes | no | no | no | no | no | no | no |
| SCN1A | 1217 | H | 0 | 0 | no | yes | no | yes | no | yes | no | no | no | no | no | no | no |
| SCN1A | 1218 | N | 0 | 0 | yes | no | no | yes | no | yes | no | no | no | no | no | no | no |
| SCN1A | 1219 | W | 0 | 0 | yes | no | no | yes | no | yes | no | no | no | no | no | no | no |
| SCN1A | 1220 | F | 0 | 0 | yes | no | no | yes | no | no | no | no | no | no | no | no | no |
| SCN1A | 1221 | E | 0 | 1 | yes | no | no | yes | no | no | no | no | no | no | no | no | no |
| SCN1A | 1222 | T | 0 | 0 | yes | no | no | yes | no | no | no | no | no | no | no | no | no |
| SCN1A | 1223 | F | 0 | 0 | yes | no | no | yes | no | no | no | no | no | no | no | no | no |
| SCN1A | 1224 | I | 0 | 0 | yes | no | no | yes | no | no | no | no | no | no | no | no | no |
| SCN1A | 1225 | V | 0 | 0 | yes | no | no | yes | no | no | no | no | no | no | no | no | no |
| SCN1A | 1226 | F | 0 | 0 | yes | no | no | yes | no | no | no | no | no | no | no | no | no |
| SCN1A | 1227 | M | 0 | 0 | yes | no | no | yes | no | no | no | no | no | no | no | no | no |
| SCN1A | 1228 | I | 0 | 1 | yes | no | no | yes | no | no | no | no | no | no | no | no | no |
| SCN1A | 1229 | L | 0 | 0 | yes | no | no | yes | no | no | no | no | no | no | no | no | no |

| | | | | | | | | | | | | | | | | | |
|---|---|---|---|---|---|---|---|---|---|---|---|---|---|---|---|---|---|
| SCN1A | 1230 | L | 0 | 0 | yes | no | no | yes | no | no | no | no | no | no | no | no | no |
| SCN1A | 1231 | S | 0 | 2 | yes | no | no | yes | no | no | no | no | no | no | no | no | no |
| SCN1A | 1232 | S | 0 | 0 | yes | no | no | yes | no | no | no | no | no | no | no | no | no |
| SCN1A | 1233 | G | 0 | 1 | yes | no | no | yes | no | no | no | no | no | no | no | no | no |
| SCN1A | 1234 | A | 0 | 0 | no | yes | no | yes | no | no | no | no | no | no | no | no | no |
| SCN1A | 1235 | L | 0 | 0 | no | yes | no | yes | no | no | no | no | no | no | no | no | no |
| SCN1A | 1236 | A | 0 | 1 | no | yes | no | yes | no | no | no | no | no | no | no | no | no |
| SCN1A | 1237 | F | 0 | 0 | no | yes | no | yes | no | no | no | no | no | no | no | no | no |
| SCN1A | 1238 | E | 2 | 1 | no | yes | no | no | yes | no | no | no | no | no | no | no | no |
| SCN1A | 1239 | D | 0 | 2 | no | yes | no | no | yes | no | no | no | no | no | no | no | no |
| SCN1A | 1240 | I | 1 | 0 | yes | no | no | no | yes | no | no | no | no | no | no | no | no |
| SCN1A | 1241 | Y | 0 | 1 | yes | no | no | no | yes | no | no | no | no | no | no | no | no |
| SCN1A | 1242 | I | 1 | 0 | yes | no | no | no | yes | no | no | no | no | no | no | no | no |
| SCN1A | 1243 | D | 0 | 0 | yes | no | no | no | yes | no | no | no | no | no | no | no | no |
| SCN1A | 1244 | Q | 0 | 0 | yes | no | no | no | yes | no | no | no | no | no | no | no | no |
| SCN1A | 1245 | R | 0 | 1 | no | yes | no | no | yes | no | no | no | no | no | no | no | no |
| SCN1A | 1246 | K | 0 | 0 | no | yes | no | no | yes | no | no | no | no | no | no | no | no |
| SCN1A | 1247 | T | 1 | 0 | no | yes | no | no | yes | no | no | no | no | no | no | no | no |
| SCN1A | 1248 | I | 0 | 0 | no | yes | no | no | yes | no | no | no | no | no | no | no | no |
| SCN1A | 1249 | K | 0 | 1 | yes | no | no | no | yes | no | no | no | no | no | no | no | no |
| SCN1A | 1250 | T | 1 | 0 | yes | no | no | no | yes | no | no | no | no | no | no | no | no |
| SCN1A | 1251 | M | 1 | 0 | yes | no | no | no | no | no | no | no | no | no | no | no | no |
| SCN1A | 1252 | L | 0 | 0 | yes | no | no | yes | no | no | no | no | no | no | no | no | no |
| SCN1A | 1253 | E | 0 | 0 | yes | no | no | no | no | no | no | no | no | no | no | no | no |
| SCN1A | 1254 | Y | 1 | 1 | yes | no | no | yes | no | no | no | no | no | no | no | no | no |
| SCN1A | 1255 | A | 2 | 2 | yes | no | no | yes | no | no | no | no | no | no | no | no | no |
| SCN1A | 1256 | D | 0 | 0 | yes | no | no | yes | no | no | no | no | no | no | no | no | no |
| SCN1A | 1257 | K | 0 | 0 | yes | no | no | yes | no | no | no | no | no | no | no | no | no |
| SCN1A | 1258 | V | 0 | 0 | yes | no | no | yes | no | no | no | no | no | no | no | no | no |
| SCN1A | 1259 | F | 1 | 1 | yes | no | no | yes | no | no | no | no | no | no | no | no | no |
| SCN1A | 1260 | T | 0 | 1 | yes | no | no | yes | no | no | no | no | no | no | no | no | no |
| SCN1A | 1261 | Y | 1 | 0 | yes | no | no | yes | no | no | no | no | no | no | no | no | no |
| SCN1A | 1262 | I | 0 | 0 | yes | no | no | yes | no | no | no | no | no | no | no | no | no |

| | | | | | | | | | | | | | | | | | |
|---|---|---|---|---|---|---|---|---|---|---|---|---|---|---|---|---|---|
| SCN1A | 1263 | F | 0 | 1 | yes | no | no | yes | no | no | no | no | no | no | no | no | no |
| SCN1A | 1264 | I | 0 | 0 | yes | no | no | yes | no | no | no | no | no | no | no | no | no |
| SCN1A | 1265 | L | 0 | 1 | yes | no | no | yes | no | no | no | no | no | no | no | no | no |
| SCN1A | 1266 | E | 0 | 0 | yes | no | no | yes | no | no | no | no | no | no | no | no | no |
| SCN1A | 1267 | M | 0 | 2 | yes | no | no | yes | no | no | no | no | no | no | no | no | no |
| SCN1A | 1268 | L | 0 | 0 | yes | no | no | yes | no | no | no | no | no | no | no | no | no |
| SCN1A | 1269 | L | 0 | 0 | yes | no | no | yes | no | no | no | no | no | no | no | no | no |
| SCN1A | 1270 | K | 0 | 1 | yes | no | no | yes | no | yes | no | no | no | no | no | no | no |
| SCN1A | 1271 | W | 1 | 0 | yes | no | no | yes | no | yes | no | no | no | no | no | no | no |
| SCN1A | 1272 | V | 0 | 0 | yes | no | no | yes | no | yes | no | no | no | no | no | no | no |
| SCN1A | 1273 | A | 0 | 1 | yes | no | no | yes | no | yes | no | no | no | no | no | no | no |
| SCN1A | 1274 | Y | 0 | 2 | yes | no | no | yes | no | yes | no | no | no | no | no | no | no |
| SCN1A | 1275 | G | 0 | 1 | yes | no | no | yes | no | yes | no | no | no | no | no | no | no |
| SCN1A | 1276 | Y | 0 | 0 | yes | no | no | yes | no | yes | no | no | no | no | no | no | no |
| SCN1A | 1277 | Q | 0 | 0 | yes | no | no | no | no | yes | no | no | no | no | no | no | no |
| SCN1A | 1278 | T | 0 | 0 | yes | no | no | no | no | yes | no | no | no | no | no | no | no |
| SCN1A | 1279 | Y | 0 | 0 | yes | no | no | yes | no | yes | no | no | no | no | no | no | no |
| SCN1A | 1280 | F | 0 | 0 | yes | no | no | yes | no | yes | no | no | no | no | no | no | no |
| SCN1A | 1281 | T | 0 | 0 | no | yes | no | no | no | yes | no | no | no | no | no | no | no |
| SCN1A | 1282 | N | 0 | 0 | no | yes | no | yes | no | yes | no | no | no | no | no | no | no |
| SCN1A | 1283 | A | 0 | 0 | yes | no | no | yes | no | yes | no | no | no | no | no | no | no |
| SCN1A | 1284 | W | 0 | 2 | yes | no | no | yes | no | no | no | no | no | no | no | no | no |
| SCN1A | 1285 | C | 0 | 0 | yes | no | no | yes | no | no | no | no | no | no | no | no | no |
| SCN1A | 1286 | W | 0 | 0 | yes | no | no | yes | no | no | no | no | no | no | no | no | no |
| SCN1A | 1287 | L | 0 | 1 | yes | no | no | yes | no | no | no | no | no | no | no | no | no |
| SCN1A | 1288 | D | 0 | 2 | yes | no | no | yes | no | no | no | no | no | no | no | no | no |
| SCN1A | 1289 | F | 0 | 0 | yes | no | no | yes | no | no | no | no | no | no | no | no | no |
| SCN1A | 1290 | L | 0 | 0 | yes | no | no | yes | no | no | no | no | no | no | no | no | no |
| SCN1A | 1291 | I | 0 | 0 | yes | no | no | yes | no | no | no | no | no | no | no | no | no |
| SCN1A | 1292 | V | 0 | 0 | yes | no | no | yes | no | no | no | no | no | no | no | no | no |
| SCN1A | 1293 | D | 0 | 0 | yes | no | no | yes | no | no | no | no | no | no | no | no | no |
| SCN1A | 1294 | V | 0 | 0 | yes | no | no | yes | no | no | no | no | no | no | no | no | no |
| SCN1A | 1295 | S | 0 | 0 | yes | no | no | yes | no | no | no | no | no | no | no | no | no |

| | | | | | | | | | | | | | | | | | |
|---|---|---|---|---|---|---|---|---|---|---|---|---|---|---|---|---|---|
| SCN1A | 1296 | L | 1 | 0 | yes | no | no | yes | no | no | no | no | no | no | no | no | no |
| SCN1A | 1297 | V | 2 | 0 | yes | no | no | yes | no | no | no | no | no | no | no | no | no |
| SCN1A | 1298 | S | 0 | 0 | yes | no | no | yes | no | no | no | no | no | no | no | no | no |
| SCN1A | 1299 | L | 1 | 0 | yes | no | no | yes | no | no | no | no | no | no | no | no | no |
| SCN1A | 1300 | T | 1 | 1 | yes | no | no | yes | no | no | no | no | no | no | no | no | no |
| SCN1A | 1301 | A | 1 | 0 | yes | no | no | yes | no | no | no | no | no | no | no | no | no |
| SCN1A | 1302 | N | 1 | 0 | yes | no | no | no | no | no | no | no | no | no | no | no | no |
| SCN1A | 1303 | A | 1 | 0 | yes | no | no | yes | yes | no | no | no | no | no | no | no | no |
| SCN1A | 1304 | L | 0 | 0 | yes | no | no | yes | yes | no | no | no | no | no | no | no | no |
| SCN1A | 1305 | G | 0 | 0 | no | yes | no | no | yes | no | no | no | no | no | no | no | no |
| SCN1A | 1306 | Y | 0 | 0 | no | yes | no | no | yes | no | no | no | no | no | no | no | no |
| SCN1A | 1307 | S | 0 | 0 | no | yes | no | no | yes | no | no | no | no | no | no | no | no |
| SCN1A | 1308 | E | 1 | 1 | no | yes | no | no | yes | no | no | no | no | no | no | no | no |
| SCN1A | 1309 | L | 1 | 2 | no | yes | no | no | yes | no | no | no | no | no | no | no | no |
| SCN1A | 1310 | G | 0 | 0 | no | yes | no | no | yes | no | no | no | no | no | no | no | no |
| SCN1A | 1311 | A | 1 | 0 | no | yes | no | no | no | no | no | no | no | no | no | no | no |
| SCN1A | 1312 | I | 0 | 0 | yes | no | no | no | no | no | no | no | no | no | no | no | no |
| SCN1A | 1313 | K | 1 | 2 | yes | no | no | no | no | no | no | no | no | no | no | no | no |
| SCN1A | 1314 | S | 0 | 0 | yes | no | no | no | no | no | no | no | no | no | no | no | no |
| SCN1A | 1315 | L | 0 | 0 | yes | no | no | yes | no | no | no | no | no | no | no | no | no |
| SCN1A | 1316 | R | 0 | 2 | yes | no | no | no | no | no | no | no | no | no | no | no | no |
| SCN1A | 1317 | T | 0 | 0 | yes | no | no | yes | no | no | no | no | no | no | no | no | no |
| SCN1A | 1318 | L | 0 | 1 | yes | no | no | yes | no | no | no | no | no | no | no | no | no |
| SCN1A | 1319 | R | 0 | 0 | yes | no | no | yes | no | no | no | no | no | no | no | no | no |
| SCN1A | 1320 | A | 0 | 0 | yes | no | no | yes | no | no | no | no | no | no | no | no | no |
| SCN1A | 1321 | L | 0 | 0 | yes | no | no | yes | no | no | no | no | no | no | no | no | no |
| SCN1A | 1322 | R | 0 | 1 | yes | no | no | yes | no | no | no | no | no | no | no | no | no |
| SCN1A | 1323 | P | 0 | 1 | yes | no | no | yes | no | no | no | no | no | no | no | no | no |
| SCN1A | 1324 | L | 0 | 0 | yes | no | no | yes | no | no | no | no | no | no | no | no | no |
| SCN1A | 1325 | R | 0 | 1 | yes | no | no | yes | no | no | no | no | no | no | no | no | no |
| SCN1A | 1326 | A | 0 | 2 | yes | no | no | yes | no | no | no | no | no | no | no | no | no |
| SCN1A | 1327 | L | 0 | 0 | no | yes | no | yes | no | no | no | no | no | no | no | no | no |
| SCN1A | 1328 | S | 0 | 1 | no | yes | no | yes | no | no | no | no | no | no | no | no | no |

| | | | | | | | | | | | | | | | | | |
|---|---|---|---|---|---|---|---|---|---|---|---|---|---|---|---|---|---|
| SCN1A | 1329 | R | 0 | 1 | no | yes | no | yes | no | no | no | no | no | no | no | no | no |
| SCN1A | 1330 | F | 0 | 0 | no | yes | no | yes | no | yes | no | no | no | no | no | no | no |
| SCN1A | 1331 | E | 0 | 0 | yes | no | no | yes | no | yes | no | no | no | no | no | no | no |
| SCN1A | 1332 | G | 0 | 1 | yes | no | no | yes | no | yes | no | no | no | no | no | no | no |
| SCN1A | 1333 | M | 0 | 1 | yes | no | no | yes | no | yes | no | no | no | no | no | no | no |
| SCN1A | 1334 | R | 2 | 0 | yes | no | no | yes | no | yes | no | no | no | no | no | no | no |
| SCN1A | 1335 | V | 0 | 1 | yes | no | no | yes | no | yes | no | no | no | no | no | no | no |
| SCN1A | 1336 | V | 0 | 0 | yes | no | no | yes | no | yes | no | no | no | no | no | no | no |
| SCN1A | 1337 | V | 0 | 1 | yes | no | no | yes | no | yes | no | no | no | no | no | no | no |
| SCN1A | 1338 | N | 1 | 1 | yes | no | no | yes | no | yes | no | no | no | no | no | no | no |
| SCN1A | 1339 | A | 0 | 1 | yes | no | no | yes | no | yes | no | no | no | no | no | no | no |
| SCN1A | 1340 | L | 0 | 1 | yes | no | no | yes | no | yes | no | no | no | no | no | no | no |
| SCN1A | 1341 | L | 0 | 0 | yes | no | no | yes | no | yes | no | no | no | no | no | no | no |
| SCN1A | 1342 | G | 0 | 0 | no | yes | no | yes | no | yes | no | no | no | no | no | no | no |
| SCN1A | 1343 | A | 0 | 1 | no | yes | no | yes | no | yes | no | no | no | no | no | no | no |
| SCN1A | 1344 | I | 0 | 1 | yes | no | no | yes | no | yes | no | no | no | no | no | no | no |
| SCN1A | 1345 | P | 0 | 2 | yes | no | no | yes | no | yes | no | no | no | no | no | no | no |
| SCN1A | 1346 | S | 1 | 0 | yes | no | no | yes | no | yes | no | no | no | no | no | no | no |
| SCN1A | 1347 | I | 0 | 1 | yes | no | no | yes | no | no | no | no | no | no | no | no | no |
| SCN1A | 1348 | M | 0 | 2 | yes | no | no | yes | no | no | no | no | no | no | no | no | no |
| SCN1A | 1349 | N | 0 | 0 | yes | no | no | yes | no | no | no | no | no | no | no | no | no |
| SCN1A | 1350 | V | 0 | 0 | yes | no | no | yes | no | no | no | no | no | no | no | no | no |
| SCN1A | 1351 | L | 0 | 0 | yes | no | no | yes | no | no | no | no | no | no | no | no | no |
| SCN1A | 1352 | L | 0 | 2 | yes | no | no | yes | no | no | no | no | no | no | no | no | no |
| SCN1A | 1353 | V | 0 | 2 | yes | no | no | yes | no | no | no | no | no | no | no | no | no |
| SCN1A | 1354 | C | 0 | 3 | yes | no | no | yes | no | no | no | no | no | no | no | no | no |
| SCN1A | 1355 | L | 0 | 1 | yes | no | no | yes | no | no | no | no | no | no | no | no | no |
| SCN1A | 1356 | I | 0 | 0 | yes | no | no | yes | no | no | no | no | no | no | no | no | no |
| SCN1A | 1357 | F | 0 | 1 | yes | no | no | yes | no | no | no | no | no | no | no | no | no |
| SCN1A | 1358 | W | 0 | 1 | yes | no | no | yes | no | no | no | no | no | no | no | no | no |
| SCN1A | 1359 | L | 0 | 2 | yes | no | no | yes | no | no | no | no | no | no | no | no | no |
| SCN1A | 1360 | I | 1 | 0 | yes | no | no | yes | no | no | no | no | no | no | no | no | no |
| SCN1A | 1361 | F | 0 | 1 | yes | no | no | yes | no | no | no | no | no | no | no | no | no |

| | | | | | | | | | | | | | | | | | |
|---|---|---|---|---|---|---|---|---|---|---|---|---|---|---|---|---|---|
| SCN1A | 1362 | S | 0 | 1 | yes | no | no | yes | no | no | no | no | no | no | no | no | no |
| SCN1A | 1363 | I | 0 | 1 | yes | no | no | yes | no | no | no | no | no | no | no | no | no |
| SCN1A | 1364 | M | 0 | 1 | yes | no | no | yes | no | no | no | no | no | no | no | no | no |
| SCN1A | 1365 | G | 0 | 2 | yes | no | no | yes | no | no | no | no | no | no | no | no | no |
| SCN1A | 1366 | V | 1 | 1 | yes | no | no | yes | no | no | no | no | no | no | no | no | no |
| SCN1A | 1367 | N | 0 | 1 | yes | no | no | yes | yes | no | no | no | no | no | no | no | no |
| SCN1A | 1368 | L | 0 | 0 | yes | no | no | yes | yes | no | no | no | no | no | no | no | no |
| SCN1A | 1369 | F | 0 | 0 | no | yes | no | no | yes | no | no | no | no | no | no | no | no |
| SCN1A | 1370 | A | 0 | 1 | no | yes | no | no | yes | no | no | no | no | no | no | no | no |
| SCN1A | 1371 | G | 0 | 1 | no | yes | no | no | yes | no | no | no | no | no | no | no | no |
| SCN1A | 1372 | K | 0 | 1 | no | yes | no | no | yes | no | no | no | no | no | no | no | no |
| SCN1A | 1373 | F | 0 | 1 | no | yes | no | no | yes | no | no | no | no | no | no | no | no |
| SCN1A | 1374 | Y | 1 | 0 | no | yes | no | no | yes | no | no | no | no | no | no | no | no |
| SCN1A | 1375 | H | 0 | 0 | no | no | yes | no | yes | no | no | no | no | no | no | no | no |
| SCN1A | 1376 | C | 0 | 1 | no | no | yes | no | yes | no | no | no | no | no | no | no | no |
| SCN1A | 1377 | I | 1 | 1 | no | no | yes | no | yes | no | yes | no | no | no | no | no | no |
| SCN1A | 1378 | N | 0 | 1 | no | no | yes | no | yes | no | no | no | no | no | no | no | no |
| SCN1A | 1379 | T | 0 | 0 | no | yes | no | no | yes | no | yes | no | no | no | no | no | no |
| SCN1A | 1380 | T | 0 | 0 | no | yes | no | no | yes | no | no | no | no | no | no | no | no |
| SCN1A | 1381 | T | 0 | 0 | no | yes | no | no | yes | no | no | no | no | no | no | no | no |
| SCN1A | 1382 | G | 0 | 1 | no | yes | no | no | yes | no | yes | no | no | no | no | no | no |
| SCN1A | 1383 | D | 0 | 0 | no | yes | no | no | yes | no | no | no | no | no | no | no | no |
| SCN1A | 1384 | R | 0 | 0 | no | yes | no | no | yes | no | no | no | no | no | no | no | no |
| SCN1A | 1385 | F | 0 | 1 | no | yes | no | no | yes | no | no | no | no | no | no | no | no |
| SCN1A | 1386 | D | 0 | 0 | no | yes | no | no | yes | no | no | no | no | no | no | no | no |
| SCN1A | 1387 | I | 1 | 0 | no | yes | no | no | yes | no | no | no | no | no | no | no | no |
| SCN1A | 1388 | E | 1 | 0 | no | yes | no | no | yes | no | no | no | no | no | no | no | no |
| SCN1A | 1389 | D | 0 | 0 | no | yes | no | no | yes | no | no | no | no | no | no | no | no |
| SCN1A | 1390 | V | 0 | 1 | no | yes | no | no | yes | no | no | no | no | no | no | no | no |
| SCN1A | 1391 | N | 1 | 1 | no | yes | no | no | yes | no | no | no | no | no | no | no | no |
| SCN1A | 1392 | N | 0 | 1 | no | no | yes | no | yes | no | no | no | no | no | no | no | no |
| SCN1A | 1393 | H | 0 | 2 | yes | no | no | no | yes | no | no | no | no | no | no | no | no |
| SCN1A | 1394 | T | 0 | 0 | yes | no | no | no | yes | no | no | no | no | no | no | no | no |

| | | | | | | | | | | | | | | | | | |
|---|---|---|---|---|---|---|---|---|---|---|---|---|---|---|---|---|---|
| SCN1A | 1395 | D | 0 | 1 | yes | no | no | no | yes | no | no | no | no | no | no | no | no |
| SCN1A | 1396 | C | 0 | 1 | yes | no | no | no | yes | no | no | no | no | no | no | no | no |
| SCN1A | 1397 | L | 0 | 0 | yes | no | no | no | yes | no | no | no | no | no | no | no | no |
| SCN1A | 1398 | K | 0 | 0 | yes | no | no | no | yes | no | no | no | no | no | no | no | no |
| SCN1A | 1399 | L | 1 | 0 | yes | no | no | no | yes | no | no | no | no | no | no | no | no |
| SCN1A | 1400 | I | 0 | 0 | yes | no | no | no | yes | no | no | no | no | no | no | no | no |
| SCN1A | 1401 | E | 0 | 0 | yes | no | no | no | yes | no | no | no | no | no | no | no | no |
| SCN1A | 1402 | R | 1 | 0 | no | yes | no | no | yes | no | no | no | no | no | no | no | no |
| SCN1A | 1403 | N | 0 | 0 | no | yes | no | no | yes | no | no | no | no | no | no | no | no |
| SCN1A | 1404 | E | 0 | 1 | no | yes | no | no | yes | no | no | no | no | no | no | no | no |
| SCN1A | 1405 | T | 0 | 0 | no | yes | no | no | yes | no | no | no | no | no | no | no | no |
| SCN1A | 1406 | A | 0 | 1 | no | no | yes | no | yes | no | no | no | no | no | no | no | no |
| SCN1A | 1407 | R | 0 | 0 | no | no | yes | no | yes | no | no | no | no | no | no | no | no |
| SCN1A | 1408 | W | 0 | 0 | no | no | yes | no | yes | no | no | no | no | no | no | no | no |
| SCN1A | 1409 | K | 0 | 0 | no | no | yes | no | yes | no | yes | no | no | no | no | no | no |
| SCN1A | 1410 | N | 0 | 0 | no | yes | no | no | yes | no | no | no | no | no | no | no | no |
| SCN1A | 1411 | V | 0 | 0 | no | yes | no | no | yes | no | no | no | no | no | no | no | no |
| SCN1A | 1412 | K | 0 | 0 | no | yes | no | no | yes | no | no | no | no | no | no | no | no |
| SCN1A | 1413 | V | 0 | 0 | no | yes | no | no | yes | no | no | no | no | no | no | no | no |
| SCN1A | 1414 | N | 0 | 1 | no | yes | no | no | yes | no | no | no | no | no | no | no | no |
| SCN1A | 1415 | F | 0 | 2 | no | yes | no | yes | yes | no | no | no | no | no | no | no | no |
| SCN1A | 1416 | D | 0 | 2 | no | yes | no | no | yes | no | no | no | no | no | no | no | no |
| SCN1A | 1417 | N | 0 | 0 | no | yes | no | yes | yes | no | no | no | no | no | no | no | no |
| SCN1A | 1418 | V | 0 | 1 | yes | no | no | yes | yes | no | no | no | no | no | no | no | no |
| SCN1A | 1419 | G | 1 | 0 | yes | no | no | yes | no | no | no | yes | no | no | no | no | no |
| SCN1A | 1420 | F | 0 | 0 | yes | no | no | yes | no | no | no | yes | no | no | no | no | no |
| SCN1A | 1421 | G | 0 | 2 | yes | no | no | yes | no | no | no | yes | no | no | no | no | no |
| SCN1A | 1422 | Y | 1 | 1 | yes | no | no | yes | no | no | no | yes | no | no | no | no | no |
| SCN1A | 1423 | L | 0 | 1 | yes | no | no | yes | no | no | no | yes | no | no | no | no | no |
| SCN1A | 1424 | S | 0 | 0 | yes | no | no | yes | no | no | no | yes | no | no | no | no | no |
| SCN1A | 1425 | L | 0 | 0 | yes | no | no | yes | no | no | no | yes | no | no | no | no | no |
| SCN1A | 1426 | L | 0 | 1 | yes | no | no | yes | no | no | no | yes | no | no | no | no | no |
| SCN1A | 1427 | Q | 0 | 2 | yes | no | no | yes | no | no | no | yes | no | no | no | no | no |

| | | | | | | | | | | | | | | | | | |
|---|---|---|---|---|---|---|---|---|---|---|---|---|---|---|---|---|---|
| SCN1A | 1428 | V | 0 | 2 | yes | no | no | yes | no | no | no | yes | no | no | no | no | no |
| SCN1A | 1429 | A | 0 | 2 | yes | no | no | yes | no | no | no | yes | no | no | no | no | no |
| SCN1A | 1430 | T | 0 | 0 | yes | no | no | yes | no | no | yes | yes | no | no | no | no | no |
| SCN1A | 1431 | F | 0 | 1 | no | yes | no | yes | no | no | yes | yes | no | no | no | no | no |
| SCN1A | 1432 | K | 0 | 3 | no | yes | no | yes | no | no | yes | yes | no | no | no | no | no |
| SCN1A | 1433 | G | 0 | 2 | no | yes | no | yes | no | no | no | yes | no | no | no | no | no |
| SCN1A | 1434 | W | 0 | 1 | yes | no | no | yes | no | no | no | yes | no | no | no | no | no |
| SCN1A | 1435 | M | 0 | 1 | yes | no | no | no | no | no | no | yes | no | no | no | no | no |
| SCN1A | 1436 | D | 0 | 0 | yes | no | no | no | no | no | yes | yes | no | no | no | no | no |
| SCN1A | 1437 | I | 0 | 1 | yes | no | no | no | no | no | no | yes | no | no | no | no | no |
| SCN1A | 1438 | M | 0 | 1 | yes | no | no | no | no | no | no | yes | no | no | no | no | no |
| SCN1A | 1439 | Y | 0 | 0 | yes | no | no | no | no | no | no | yes | no | no | no | no | no |
| SCN1A | 1440 | A | 0 | 1 | yes | no | no | no | no | no | no | yes | no | no | no | no | no |
| SCN1A | 1441 | A | 0 | 1 | yes | no | no | no | yes | no | no | no | no | no | no | no | no |
| SCN1A | 1442 | V | 0 | 1 | yes | no | no | no | yes | no | no | no | no | no | no | no | no |
| SCN1A | 1443 | D | 0 | 2 | no | yes | no | no | yes | no | no | no | no | no | no | no | no |
| SCN1A | 1444 | S | 0 | 1 | no | yes | no | no | yes | no | no | no | no | no | no | no | no |
| SCN1A | 1445 | R | 0 | 0 | no | yes | no | no | yes | no | no | no | no | no | no | no | no |
| SCN1A | 1446 | N | 0 | 0 | no | yes | no | no | yes | no | no | no | no | no | no | no | no |
| SCN1A | 1447 | V | 0 | 0 | no | yes | no | no | yes | no | no | no | no | no | no | no | no |
| SCN1A | 1448 | E | 1 | 0 | no | yes | no | no | yes | no | no | no | no | no | no | no | no |
| SCN1A | 1449 | L | 1 | 0 | no | no | yes | no | yes | no | no | no | no | no | no | no | no |
| SCN1A | 1450 | Q | 0 | 2 | no | yes | no | no | yes | no | no | no | no | no | no | no | no |
| SCN1A | 1451 | P | 0 | 2 | no | yes | no | no | yes | no | no | no | no | no | no | no | no |
| SCN1A | 1452 | K | 0 | 0 | no | yes | no | no | yes | no | no | no | no | no | no | no | no |
| SCN1A | 1453 | Y | 0 | 1 | no | yes | no | no | yes | no | no | no | no | no | no | no | no |
| SCN1A | 1454 | E | 0 | 0 | no | yes | no | no | yes | no | no | no | no | no | no | no | no |
| SCN1A | 1455 | E | 1 | 1 | no | yes | no | no | yes | no | no | no | no | no | no | no | no |
| SCN1A | 1456 | S | 0 | 0 | no | yes | no | no | yes | no | no | no | no | no | no | no | no |
| SCN1A | 1457 | L | 0 | 0 | yes | no | no | no | yes | no | no | no | no | no | no | no | no |
| SCN1A | 1458 | Y | 0 | 0 | yes | no | no | yes | no | no | no | no | no | no | no | no | no |
| SCN1A | 1459 | M | 0 | 0 | yes | no | no | yes | no | no | no | no | no | no | no | no | no |
| SCN1A | 1460 | Y | 0 | 0 | yes | no | no | yes | no | no | no | no | no | no | no | no | no |

| | | | | | | | | | | | | | | | | | |
|---|---|---|---|---|---|---|---|---|---|---|---|---|---|---|---|---|---|
| SCN1A | 1461 | L | 1 | 1 | yes | no | no | yes | no | no | no | no | no | no | no | no | no |
| SCN1A | 1462 | Y | 0 | 2 | yes | no | no | yes | no | no | no | no | no | no | no | no | no |
| SCN1A | 1463 | F | 0 | 2 | yes | no | no | yes | no | no | no | no | no | no | no | no | no |
| SCN1A | 1464 | V | 0 | 0 | yes | no | no | yes | no | no | no | no | no | no | no | no | no |
| SCN1A | 1465 | I | 1 | 0 | yes | no | no | yes | no | no | no | no | no | no | no | no | no |
| SCN1A | 1466 | F | 0 | 1 | yes | no | no | yes | no | no | no | no | no | no | no | no | no |
| SCN1A | 1467 | I | 0 | 1 | yes | no | no | yes | no | no | no | no | no | no | no | no | no |
| SCN1A | 1468 | I | 0 | 0 | yes | no | no | yes | no | no | no | no | no | no | no | no | no |
| SCN1A | 1469 | F | 0 | 0 | yes | no | no | yes | no | no | no | no | no | no | no | no | no |
| SCN1A | 1470 | G | 0 | 1 | yes | no | no | yes | no | no | no | no | no | no | no | no | no |
| SCN1A | 1471 | S | 0 | 1 | yes | no | no | yes | no | no | no | no | no | no | no | no | no |
| SCN1A | 1472 | F | 0 | 1 | yes | no | no | yes | no | no | no | no | no | no | no | no | no |
| SCN1A | 1473 | F | 0 | 0 | yes | no | no | yes | no | no | no | no | no | no | no | no | no |
| SCN1A | 1474 | T | 0 | 0 | yes | no | no | yes | no | no | no | no | no | no | no | no | no |
| SCN1A | 1475 | L | 0 | 1 | yes | no | no | yes | no | no | yes | no | no | no | no | no | no |
| SCN1A | 1476 | N | 0 | 1 | yes | no | no | yes | no | no | no | no | no | no | no | no | no |
| SCN1A | 1477 | L | 0 | 0 | yes | no | no | yes | no | no | no | no | no | no | no | no | no |
| SCN1A | 1478 | F | 0 | 1 | yes | no | no | yes | no | no | no | no | no | no | no | no | no |
| SCN1A | 1479 | I | 0 | 0 | yes | no | no | yes | no | no | yes | no | no | no | no | no | no |
| SCN1A | 1480 | G | 0 | 2 | yes | no | no | yes | no | yes | no | no | no | no | no | no | no |
| SCN1A | 1481 | V | 0 | 1 | yes | no | no | yes | no | yes | no | no | no | no | no | no | no |
| SCN1A | 1482 | I | 1 | 1 | yes | no | no | yes | no | yes | no | no | no | no | no | no | no |
| SCN1A | 1483 | I | 0 | 0 | yes | no | no | no | no | yes | yes | no | no | no | no | no | no |
| SCN1A | 1484 | D | 0 | 1 | yes | no | no | no | no | yes | no | no | no | no | no | no | no |
| SCN1A | 1485 | N | 0 | 1 | yes | no | no | no | no | yes | no | no | no | no | no | no | no |
| SCN1A | 1486 | F | 0 | 1 | yes | no | no | no | no | yes | no | no | no | no | no | no | no |
| SCN1A | 1487 | N | 0 | 0 | yes | no | no | no | no | yes | yes | no | no | no | no | no | no |
| SCN1A | 1488 | Q | 0 | 0 | yes | no | no | no | no | yes | no | no | no | no | no | no | no |
| SCN1A | 1489 | Q | 0 | 2 | yes | no | no | no | no | yes | no | no | no | no | no | no | no |
| SCN1A | 1490 | K | 0 | 0 | yes | no | no | no | no | yes | yes | no | no | no | no | no | no |
| SCN1A | 1491 | K | 0 | 0 | yes | no | no | no | no | yes | yes | no | no | no | no | no | no |
| SCN1A | 1492 | K | 0 | 0 | yes | no | no | no | no | yes | no | no | no | no | no | no | no |
| SCN1A | 1493 | F | 0 | 0 | yes | no | no | no | no | yes | no | no | no | no | no | no | no |

| | | | | | | | | | | | | | | | | | |
|---|---|---|---|---|---|---|---|---|---|---|---|---|---|---|---|---|---|
| SCN1A | 1494 | G | 0 | 0 | no | yes | no | no | no | yes | no | no | no | no | no | no | no |
| SCN1A | 1495 | G | 0 | 0 | no | yes | no | no | no | yes | no | no | no | no | no | no | no |
| SCN1A | 1496 | Q | 0 | 0 | no | yes | no | no | no | yes | no | no | no | no | no | no | no |
| SCN1A | 1497 | D | 0 | 0 | no | yes | no | no | no | yes | no | no | no | no | no | no | no |
| SCN1A | 1498 | I | 0 | 2 | no | yes | no | no | no | yes | no | no | no | no | no | no | no |
| SCN1A | 1499 | F | 0 | 1 | no | yes | no | yes | no | yes | no | no | no | no | no | no | no |
| SCN1A | 1500 | M | 1 | 1 | no | yes | no | no | no | yes | no | no | no | no | no | no | no |
| SCN1A | 1501 | T | 0 | 0 | no | yes | no | no | no | yes | no | no | no | no | no | no | no |
| SCN1A | 1502 | E | 0 | 0 | no | yes | no | no | no | yes | no | no | no | no | no | no | no |
| SCN1A | 1503 | E | 0 | 2 | no | yes | no | no | no | yes | no | no | no | no | no | no | no |
| SCN1A | 1504 | Q | 0 | 2 | yes | no | no | yes | no | yes | no | no | no | no | no | no | no |
| SCN1A | 1505 | K | 0 | 0 | yes | no | no | no | no | yes | no | no | no | no | no | no | no |
| SCN1A | 1506 | K | 0 | 0 | yes | no | no | no | no | yes | no | no | no | no | no | no | no |
| SCN1A | 1507 | Y | 0 | 1 | yes | no | no | no | no | yes | no | no | no | no | no | no | no |
| SCN1A | 1508 | Y | 0 | 0 | yes | no | no | no | no | yes | no | no | no | no | no | no | no |
| SCN1A | 1509 | N | 1 | 1 | yes | no | no | no | no | yes | no | no | no | no | no | no | no |
| SCN1A | 1510 | A | 0 | 1 | yes | no | no | no | no | yes | no | no | no | no | no | no | no |
| SCN1A | 1511 | M | 0 | 2 | yes | no | no | no | no | yes | no | no | no | no | no | no | no |
| SCN1A | 1512 | K | 0 | 0 | yes | no | no | no | no | yes | no | no | no | no | no | no | no |
| SCN1A | 1513 | K | 0 | 0 | yes | no | no | no | no | yes | no | no | no | no | no | no | no |
| SCN1A | 1514 | L | 1 | 1 | yes | no | no | no | no | yes | no | no | no | no | no | no | no |
| SCN1A | 1515 | G | 0 | 1 | no | yes | no | no | no | yes | no | no | no | no | no | no | no |
| SCN1A | 1516 | S | 1 | 1 | no | yes | no | no | no | yes | no | no | no | no | no | no | no |
| SCN1A | 1517 | K | 0 | 0 | no | yes | no | no | no | yes | no | no | no | no | no | no | no |
| SCN1A | 1518 | K | 0 | 0 | no | yes | no | no | no | yes | no | no | no | no | no | no | no |
| SCN1A | 1519 | P | 1 | 2 | no | yes | no | no | no | yes | no | no | no | no | no | no | no |
| SCN1A | 1520 | Q | 0 | 0 | no | yes | no | no | no | yes | no | no | no | no | no | no | no |
| SCN1A | 1521 | K | 0 | 0 | no | yes | no | no | no | yes | no | no | no | no | no | no | no |
| SCN1A | 1522 | P | 0 | 0 | no | yes | no | no | no | yes | no | no | no | no | no | no | no |
| SCN1A | 1523 | I | 1 | 1 | no | yes | no | no | no | yes | no | no | no | no | no | no | no |
| SCN1A | 1524 | P | 1 | 0 | no | yes | no | no | no | yes | no | no | no | no | no | no | no |
| SCN1A | 1525 | R | 1 | 1 | no | yes | no | no | no | yes | no | no | no | no | no | no | no |
| SCN1A | 1526 | P | 0 | 1 | no | yes | no | no | no | yes | no | no | no | no | no | no | no |

| | | | | | | | | | | | | | | | | | |
|---|---|---|---|---|---|---|---|---|---|---|---|---|---|---|---|---|---|
| SCN1A | 1527 | G | 1 | 0 | no | yes | no | no | no | yes | no | no | no | no | no | no | no |
| SCN1A | 1528 | N | 0 | 1 | no | yes | no | no | no | yes | no | no | no | no | no | no | no |
| SCN1A | 1529 | K | 0 | 0 | no | yes | no | no | no | yes | no | no | no | no | no | no | no |
| SCN1A | 1530 | F | 0 | 0 | yes | no | no | no | no | yes | no | no | no | no | no | no | no |
| SCN1A | 1531 | Q | 0 | 0 | yes | no | no | no | no | yes | no | no | no | no | no | no | no |
| SCN1A | 1532 | G | 0 | 0 | yes | no | no | no | no | yes | no | no | no | no | no | no | no |
| SCN1A | 1533 | M | 0 | 0 | yes | no | no | no | no | yes | no | no | no | no | no | no | no |
| SCN1A | 1534 | V | 0 | 0 | yes | no | no | yes | no | yes | no | no | no | no | no | no | no |
| SCN1A | 1535 | F | 0 | 0 | yes | no | no | no | no | yes | no | no | no | no | no | no | no |
| SCN1A | 1536 | D | 0 | 0 | yes | no | no | no | no | yes | no | no | no | no | no | no | no |
| SCN1A | 1537 | F | 1 | 0 | yes | no | no | yes | no | yes | no | no | no | no | no | no | no |
| SCN1A | 1538 | V | 1 | 1 | yes | no | no | yes | no | yes | no | no | no | no | no | no | no |
| SCN1A | 1539 | T | 0 | 1 | yes | no | no | no | no | yes | no | no | no | no | no | no | no |
| SCN1A | 1540 | R | 1 | 0 | no | yes | no | no | no | yes | no | no | no | no | no | no | no |
| SCN1A | 1541 | Q | 1 | 0 | yes | no | no | no | no | yes | no | no | no | no | no | no | no |
| SCN1A | 1542 | V | 1 | 0 | yes | no | no | yes | no | yes | no | no | no | no | no | no | no |
| SCN1A | 1543 | F | 1 | 1 | yes | no | no | yes | no | no | no | no | no | no | no | no | no |
| SCN1A | 1544 | D | 0 | 3 | yes | no | no | yes | no | no | no | no | no | no | no | no | no |
| SCN1A | 1545 | I | 0 | 1 | yes | no | no | yes | no | no | no | no | no | no | no | no | no |
| SCN1A | 1546 | S | 0 | 0 | yes | no | no | yes | no | no | no | no | no | no | no | no | no |
| SCN1A | 1547 | I | 0 | 0 | yes | no | no | yes | no | no | no | no | no | no | no | no | no |
| SCN1A | 1548 | M | 0 | 1 | yes | no | no | yes | no | no | no | no | no | no | no | no | no |
| SCN1A | 1549 | I | 0 | 0 | yes | no | no | yes | no | no | no | no | no | no | no | no | no |
| SCN1A | 1550 | L | 0 | 0 | yes | no | no | yes | no | no | no | no | no | no | no | no | no |
| SCN1A | 1551 | I | 0 | 0 | yes | no | no | yes | no | no | no | no | no | no | no | no | no |
| SCN1A | 1552 | C | 0 | 1 | yes | no | no | yes | no | no | no | no | no | no | no | no | no |
| SCN1A | 1553 | L | 0 | 1 | yes | no | no | yes | no | no | no | no | no | no | no | no | no |
| SCN1A | 1554 | N | 0 | 0 | yes | no | no | yes | no | no | no | no | no | no | no | no | no |
| SCN1A | 1555 | M | 0 | 1 | yes | no | no | yes | no | no | no | no | no | no | no | no | no |
| SCN1A | 1556 | V | 1 | 0 | yes | no | no | yes | no | no | no | no | no | no | no | no | no |
| SCN1A | 1557 | T | 0 | 0 | yes | no | no | yes | no | no | no | no | no | no | no | no | no |
| SCN1A | 1558 | M | 0 | 1 | yes | no | no | yes | no | no | no | no | no | no | no | no | no |
| SCN1A | 1559 | M | 1 | 1 | yes | no | no | yes | no | no | no | no | no | no | no | no | no |

| | | | | | | | | | | | | | | | | | |
|---|---|---|---|---|---|---|---|---|---|---|---|---|---|---|---|---|---|
| SCN1A | 1560 | V | 0 | 0 | no | yes | no | yes | no | no | no | no | no | no | no | no | no |
| SCN1A | 1561 | E | 0 | 1 | no | yes | no | yes | yes | no | no | no | no | no | no | no | no |
| SCN1A | 1562 | T | 1 | 0 | no | yes | no | no | yes | no | no | no | no | no | no | no | no |
| SCN1A | 1563 | D | 0 | 0 | no | yes | no | no | yes | no | no | no | no | no | no | no | no |
| SCN1A | 1564 | D | 1 | 0 | no | yes | no | no | yes | no | no | no | no | no | no | no | no |
| SCN1A | 1565 | Q | 1 | 0 | no | yes | no | no | yes | no | no | no | no | no | no | no | no |
| SCN1A | 1566 | S | 2 | 0 | no | yes | no | no | yes | no | no | no | no | no | no | no | no |
| SCN1A | 1567 | E | 0 | 0 | yes | no | no | no | yes | no | no | no | no | no | no | no | no |
| SCN1A | 1568 | Y | 1 | 0 | yes | no | no | no | yes | no | no | no | no | no | no | no | no |
| SCN1A | 1569 | V | 0 | 0 | yes | no | no | no | yes | no | no | no | no | no | no | no | no |
| SCN1A | 1570 | T | 1 | 0 | yes | no | no | no | yes | no | no | no | no | no | no | no | no |
| SCN1A | 1571 | T | 1 | 0 | yes | no | no | no | yes | no | no | no | no | no | no | no | no |
| SCN1A | 1572 | I | 0 | 0 | yes | no | no | yes | no | no | no | no | no | no | no | no | no |
| SCN1A | 1573 | L | 0 | 0 | yes | no | no | yes | no | no | no | no | no | no | no | no | no |
| SCN1A | 1574 | S | 0 | 0 | yes | no | no | yes | no | no | no | no | no | no | no | no | no |
| SCN1A | 1575 | R | 2 | 1 | yes | no | no | yes | no | no | no | no | no | no | no | no | no |
| SCN1A | 1576 | I | 0 | 0 | yes | no | no | yes | no | no | no | no | no | no | no | no | no |
| SCN1A | 1577 | N | 0 | 0 | yes | no | no | yes | no | no | no | no | no | no | no | no | no |
| SCN1A | 1578 | L | 0 | 0 | yes | no | no | yes | no | no | no | no | no | no | no | no | no |
| SCN1A | 1579 | V | 1 | 1 | yes | no | no | yes | no | no | no | no | no | no | no | no | no |
| SCN1A | 1580 | F | 0 | 0 | yes | no | no | yes | no | no | no | no | no | no | no | no | no |
| SCN1A | 1581 | I | 1 | 0 | yes | no | no | yes | no | no | no | no | no | no | no | no | no |
| SCN1A | 1582 | V | 1 | 0 | yes | no | no | yes | no | no | no | no | no | no | no | no | no |
| SCN1A | 1583 | L | 0 | 0 | yes | no | no | yes | no | no | no | no | no | no | no | no | no |
| SCN1A | 1584 | F | 0 | 0 | yes | no | no | yes | no | no | no | no | no | no | no | no | no |
| SCN1A | 1585 | T | 0 | 0 | yes | no | no | yes | no | no | no | no | no | no | no | no | no |
| SCN1A | 1586 | G | 0 | 1 | yes | no | no | yes | no | no | no | no | no | no | no | no | no |
| SCN1A | 1587 | E | 0 | 0 | yes | no | no | yes | no | no | no | no | no | no | no | no | no |
| SCN1A | 1588 | C | 0 | 1 | yes | no | no | yes | no | no | no | no | no | no | no | no | no |
| SCN1A | 1589 | V | 1 | 1 | yes | no | no | yes | no | no | no | no | no | no | no | no | no |
| SCN1A | 1590 | L | 0 | 1 | yes | no | no | yes | no | no | no | no | no | no | no | no | no |
| SCN1A | 1591 | K | 0 | 1 | yes | no | no | yes | no | yes | no | no | no | no | no | no | no |
| SCN1A | 1592 | L | 0 | 0 | yes | no | no | yes | no | yes | no | no | no | no | no | no | no |

| | | | | | | | | | | | | | | | | | |
|---|---|---|---|---|---|---|---|---|---|---|---|---|---|---|---|---|---|
| SCN1A | 1593 | I | 0 | 0 | yes | no | no | yes | no | yes | no | no | no | no | no | no | no |
| SCN1A | 1594 | S | 0 | 1 | yes | no | no | no | no | yes | no | no | no | no | no | no | no |
| SCN1A | 1595 | L | 1 | 0 | no | yes | no | no | no | yes | no | no | no | no | no | no | no |
| SCN1A | 1596 | R | 1 | 2 | no | yes | no | no | no | yes | no | no | no | no | no | no | no |
| SCN1A | 1597 | H | 0 | 1 | no | yes | no | no | no | yes | no | no | no | no | no | no | no |
| SCN1A | 1598 | Y | 1 | 1 | no | yes | no | no | no | yes | no | no | no | no | no | no | no |
| SCN1A | 1599 | Y | 0 | 0 | no | yes | no | yes | no | yes | no | no | no | no | no | no | no |
| SCN1A | 1600 | F | 0 | 0 | no | yes | no | yes | no | yes | no | no | no | no | no | no | no |
| SCN1A | 1601 | T | 0 | 0 | no | yes | no | no | no | yes | no | no | no | no | no | no | no |
| SCN1A | 1602 | I | 1 | 0 | no | yes | no | no | no | yes | no | no | no | no | no | no | no |
| SCN1A | 1603 | G | 1 | 0 | yes | no | no | yes | no | no | no | no | no | no | no | no | no |
| SCN1A | 1604 | W | 0 | 0 | yes | no | no | yes | no | no | no | no | no | no | no | no | no |
| SCN1A | 1605 | N | 0 | 1 | yes | no | no | yes | no | no | no | no | no | no | no | no | no |
| SCN1A | 1606 | I | 0 | 0 | yes | no | no | yes | no | no | no | no | no | no | no | no | no |
| SCN1A | 1607 | F | 0 | 0 | yes | no | no | yes | no | no | no | no | no | no | no | no | no |
| SCN1A | 1608 | D | 0 | 2 | yes | no | no | yes | no | no | no | no | no | no | no | no | no |
| SCN1A | 1609 | F | 1 | 0 | yes | no | no | yes | no | no | no | no | no | no | no | no | no |
| SCN1A | 1610 | V | 0 | 0 | yes | no | no | yes | no | no | no | no | no | no | no | no | no |
| SCN1A | 1611 | V | 1 | 1 | yes | no | no | yes | no | no | no | no | no | no | no | no | no |
| SCN1A | 1612 | V | 1 | 1 | yes | no | no | yes | no | no | no | no | no | no | no | no | no |
| SCN1A | 1613 | I | 1 | 0 | yes | no | no | yes | no | no | no | no | no | no | no | no | no |
| SCN1A | 1614 | L | 0 | 0 | yes | no | no | yes | no | no | no | no | no | no | no | no | no |
| SCN1A | 1615 | S | 0 | 0 | yes | no | no | yes | no | no | no | no | no | no | no | no | no |
| SCN1A | 1616 | I | 0 | 1 | yes | no | no | yes | no | no | no | no | no | no | no | no | no |
| SCN1A | 1617 | V | 0 | 0 | yes | no | no | yes | no | no | no | no | no | no | no | no | no |
| SCN1A | 1618 | G | 0 | 0 | yes | no | no | yes | no | no | no | no | no | no | no | no | no |
| SCN1A | 1619 | M | 1 | 1 | yes | no | no | yes | no | no | no | no | no | no | no | no | no |
| SCN1A | 1620 | F | 0 | 1 | yes | no | no | yes | no | no | no | no | no | no | no | no | no |
| SCN1A | 1621 | L | 0 | 1 | no | yes | no | yes | yes | no | no | no | no | no | no | no | no |
| SCN1A | 1622 | A | 0 | 0 | no | yes | no | yes | yes | no | no | no | no | no | no | no | no |
| SCN1A | 1623 | E | 0 | 0 | no | yes | no | no | yes | no | no | no | no | no | no | no | no |
| SCN1A | 1624 | L | 0 | 1 | no | yes | no | no | yes | no | no | no | no | no | no | no | no |
| SCN1A | 1625 | I | 1 | 0 | no | yes | no | no | yes | no | no | no | no | no | no | no | no |

| | | | | | | | | | | | | | | | | | |
|---|---|---|---|---|---|---|---|---|---|---|---|---|---|---|---|---|---|
| SCN1A | 1626 | E | 0 | 0 | no | yes | no | no | yes | no | no | no | no | no | no | no | no |
| SCN1A | 1627 | K | 1 | 0 | no | yes | no | no | yes | no | no | no | no | no | no | no | no |
| SCN1A | 1628 | Y | 0 | 0 | no | yes | no | no | yes | no | no | no | no | no | no | no | no |
| SCN1A | 1629 | F | 0 | 0 | no | yes | no | no | yes | no | no | no | no | no | no | no | no |
| SCN1A | 1630 | V | 0 | 2 | no | yes | no | no | yes | no | no | no | no | no | no | no | no |
| SCN1A | 1631 | S | 0 | 0 | no | yes | no | no | yes | no | no | no | no | no | no | no | no |
| SCN1A | 1632 | P | 0 | 1 | yes | no | no | no | yes | no | no | no | no | no | no | no | no |
| SCN1A | 1633 | T | 1 | 0 | yes | no | no | no | yes | no | no | no | no | no | no | no | no |
| SCN1A | 1634 | L | 1 | 0 | yes | no | no | no | no | no | no | no | no | no | no | no | no |
| SCN1A | 1635 | F | 0 | 0 | yes | no | no | no | no | no | no | no | no | no | no | no | no |
| SCN1A | 1636 | R | 0 | 2 | yes | no | no | no | no | no | no | no | no | no | no | no | no |
| SCN1A | 1637 | V | 0 | 1 | yes | no | no | yes | no | no | no | no | no | no | no | no | no |
| SCN1A | 1638 | I | 0 | 1 | yes | no | no | yes | no | no | no | no | no | no | no | no | no |
| SCN1A | 1639 | R | 0 | 2 | yes | no | no | yes | no | no | no | no | no | no | no | no | no |
| SCN1A | 1640 | L | 0 | 0 | yes | no | no | yes | no | no | no | no | no | no | no | no | no |
| SCN1A | 1641 | A | 0 | 1 | yes | no | no | yes | no | no | no | no | no | no | no | no | no |
| SCN1A | 1642 | R | 0 | 2 | yes | no | no | yes | no | no | no | no | no | no | no | no | no |
| SCN1A | 1643 | I | 0 | 0 | yes | no | no | yes | no | no | no | no | no | no | no | no | no |
| SCN1A | 1644 | G | 1 | 1 | yes | no | no | yes | no | no | no | no | no | no | no | no | no |
| SCN1A | 1645 | R | 0 | 2 | yes | no | no | yes | no | no | no | no | no | no | no | no | no |
| SCN1A | 1646 | I | 0 | 0 | yes | no | no | yes | no | no | no | no | no | no | no | no | no |
| SCN1A | 1647 | L | 0 | 0 | yes | no | no | yes | no | no | no | no | no | no | no | no | no |
| SCN1A | 1648 | R | 1 | 3 | yes | no | no | yes | no | no | no | no | no | no | no | no | no |
| SCN1A | 1649 | L | 0 | 1 | yes | no | no | yes | no | no | no | no | no | no | no | no | no |
| SCN1A | 1650 | I | 0 | 0 | yes | no | no | yes | no | no | no | no | no | no | no | no | no |
| SCN1A | 1651 | K | 0 | 0 | yes | no | no | yes | no | yes | no | no | no | no | no | no | no |
| SCN1A | 1652 | G | 0 | 0 | no | yes | no | yes | no | yes | no | no | no | no | no | no | no |
| SCN1A | 1653 | A | 0 | 0 | no | yes | no | yes | no | yes | no | no | no | no | no | no | no |
| SCN1A | 1654 | K | 1 | 0 | no | yes | no | yes | no | yes | no | no | no | no | no | no | no |
| SCN1A | 1655 | G | 0 | 1 | no | yes | no | yes | no | yes | no | no | no | no | no | no | no |
| SCN1A | 1656 | I | 0 | 2 | yes | no | no | yes | no | yes | no | no | no | no | no | no | no |
| SCN1A | 1657 | R | 0 | 2 | yes | no | no | yes | no | yes | no | no | no | no | no | no | no |
| SCN1A | 1658 | T | 0 | 2 | yes | no | no | yes | no | yes | no | no | no | no | no | no | no |

| | | | | | | | | | | | | | | | | | |
|---|---|---|---|---|---|---|---|---|---|---|---|---|---|---|---|---|---|
| SCN1A | 1659 | L | 0 | 2 | yes | no | no | yes | no | yes | no | no | no | no | no | no | no |
| SCN1A | 1660 | L | 0 | 1 | yes | no | no | yes | no | yes | no | no | no | no | no | no | no |
| SCN1A | 1661 | F | 0 | 2 | yes | no | no | yes | no | yes | no | no | no | no | no | no | no |
| SCN1A | 1662 | A | 0 | 1 | yes | no | no | yes | no | yes | no | no | no | no | no | no | no |
| SCN1A | 1663 | L | 0 | 1 | yes | no | no | yes | no | yes | no | no | no | no | no | no | no |
| SCN1A | 1664 | M | 0 | 1 | yes | no | no | yes | no | yes | no | no | no | no | no | no | no |
| SCN1A | 1665 | M | 0 | 1 | yes | no | no | yes | no | yes | no | no | no | no | no | no | no |
| SCN1A | 1666 | S | 0 | 1 | yes | no | no | yes | no | yes | no | no | no | no | no | no | no |
| SCN1A | 1667 | L | 0 | 2 | yes | no | no | yes | no | yes | no | no | no | no | no | no | no |
| SCN1A | 1668 | P | 0 | 2 | yes | no | no | yes | no | yes | no | no | no | no | no | no | no |
| SCN1A | 1669 | A | 0 | 2 | yes | no | no | yes | no | yes | no | no | no | no | no | no | no |
| SCN1A | 1670 | L | 0 | 1 | yes | no | no | yes | no | no | no | no | no | no | no | no | no |
| SCN1A | 1671 | F | 0 | 0 | yes | no | no | yes | no | no | no | no | no | no | no | no | no |
| SCN1A | 1672 | N | 0 | 1 | yes | no | no | yes | no | no | no | no | no | no | no | no | no |
| SCN1A | 1673 | I | 0 | 1 | yes | no | no | yes | no | no | no | no | no | no | no | no | no |
| SCN1A | 1674 | G | 0 | 1 | yes | no | no | yes | no | no | no | no | no | no | no | no | no |
| SCN1A | 1675 | L | 0 | 1 | yes | no | no | yes | no | no | no | no | no | no | no | no | no |
| SCN1A | 1676 | L | 0 | 0 | yes | no | no | yes | no | no | no | no | no | no | no | no | no |
| SCN1A | 1677 | L | 0 | 1 | yes | no | no | yes | no | no | no | no | no | no | no | no | no |
| SCN1A | 1678 | F | 1 | 0 | yes | no | no | yes | no | no | no | no | no | no | no | no | no |
| SCN1A | 1679 | L | 0 | 0 | yes | no | no | yes | no | no | no | no | no | no | no | no | no |
| SCN1A | 1680 | V | 0 | 1 | yes | no | no | yes | no | no | no | no | no | no | no | no | no |
| SCN1A | 1681 | M | 0 | 0 | yes | no | no | yes | no | no | no | no | no | no | no | no | no |
| SCN1A | 1682 | F | 0 | 0 | yes | no | no | yes | no | no | no | no | no | no | no | no | no |
| SCN1A | 1683 | I | 0 | 0 | yes | no | no | yes | no | no | no | no | no | no | no | no | no |
| SCN1A | 1684 | Y | 0 | 1 | yes | no | no | yes | no | no | no | no | no | no | no | no | no |
| SCN1A | 1685 | A | 1 | 2 | yes | no | no | yes | no | no | no | no | no | no | no | no | no |
| SCN1A | 1686 | I | 0 | 0 | yes | no | no | yes | no | no | no | no | no | no | no | no | no |
| SCN1A | 1687 | F | 0 | 1 | yes | no | no | yes | no | no | no | no | no | no | no | no | no |
| SCN1A | 1688 | G | 0 | 1 | yes | no | no | no | yes | no | no | no | no | no | no | no | no |
| SCN1A | 1689 | M | 0 | 1 | yes | no | no | no | yes | no | no | no | no | no | no | no | no |
| SCN1A | 1690 | S | 0 | 0 | yes | no | no | no | yes | no | no | no | no | no | no | no | no |
| SCN1A | 1691 | N | 1 | 0 | yes | no | no | no | yes | no | no | no | no | no | no | no | no |

| | | | | | | | | | | | | | | | | | |
|---|---|---|---|---|---|---|---|---|---|---|---|---|---|---|---|---|---|
| SCN1A | 1692 | F | 0 | 1 | no | yes | no | no | yes | no | no | no | no | no | no | no | no |
| SCN1A | 1693 | A | 0 | 0 | no | yes | no | no | yes | no | no | no | no | no | no | no | no |
| SCN1A | 1694 | Y | 1 | 1 | no | yes | no | no | yes | no | no | no | no | no | no | no | no |
| SCN1A | 1695 | V | 0 | 0 | no | yes | no | no | yes | no | no | no | no | no | no | no | no |
| SCN1A | 1696 | K | 0 | 0 | no | yes | no | no | yes | no | no | no | no | no | no | no | no |
| SCN1A | 1697 | R | 0 | 0 | no | yes | no | no | yes | no | no | no | no | no | no | no | no |
| SCN1A | 1698 | E | 2 | 0 | no | yes | no | no | yes | no | yes | no | no | no | no | no | no |
| SCN1A | 1699 | V | 2 | 0 | no | yes | no | no | yes | no | yes | no | no | no | no | no | no |
| SCN1A | 1700 | G | 0 | 0 | no | yes | no | no | yes | no | no | no | no | no | no | no | no |
| SCN1A | 1701 | I | 0 | 0 | no | yes | no | no | yes | no | no | no | no | no | no | no | no |
| SCN1A | 1702 | D | 1 | 2 | no | yes | no | no | yes | no | no | no | no | no | no | no | no |
| SCN1A | 1703 | D | 0 | 2 | no | yes | no | no | yes | no | no | no | no | no | no | no | no |
| SCN1A | 1704 | M | 0 | 0 | no | yes | no | no | yes | no | no | no | no | no | no | no | no |
| SCN1A | 1705 | F | 0 | 0 | no | yes | no | no | yes | no | no | no | no | no | no | no | no |
| SCN1A | 1706 | N | 0 | 0 | no | yes | no | no | yes | no | no | no | no | no | no | no | no |
| SCN1A | 1707 | F | 0 | 2 | no | yes | no | no | yes | no | no | no | no | no | no | no | no |
| SCN1A | 1708 | E | 0 | 0 | no | yes | no | no | yes | no | no | no | no | no | no | no | no |
| SCN1A | 1709 | T | 0 | 1 | no | yes | no | no | yes | no | no | no | no | no | no | no | no |
| SCN1A | 1710 | F | 0 | 0 | yes | no | no | yes | no | no | no | yes | no | no | no | no | no |
| SCN1A | 1711 | G | 0 | 1 | yes | no | no | no | no | no | no | yes | no | no | no | no | no |
| SCN1A | 1712 | N | 0 | 1 | yes | no | no | no | no | no | no | yes | no | no | no | no | no |
| SCN1A | 1713 | S | 0 | 1 | yes | no | no | yes | no | no | no | yes | no | no | no | no | no |
| SCN1A | 1714 | M | 1 | 1 | yes | no | no | yes | no | no | no | yes | no | no | no | no | no |
| SCN1A | 1715 | I | 0 | 0 | yes | no | no | yes | no | no | no | yes | no | no | no | no | no |
| SCN1A | 1716 | C | 0 | 1 | yes | no | no | yes | no | no | no | yes | no | no | no | no | no |
| SCN1A | 1717 | L | 0 | 1 | yes | no | no | yes | no | no | no | yes | no | no | no | no | no |
| SCN1A | 1718 | F | 0 | 1 | yes | no | no | yes | no | no | no | yes | no | no | no | no | no |
| SCN1A | 1719 | Q | 0 | 0 | yes | no | no | yes | no | no | no | yes | no | no | no | no | no |
| SCN1A | 1720 | I | 0 | 0 | yes | no | no | yes | no | no | no | yes | no | no | no | no | no |
| SCN1A | 1721 | T | 0 | 1 | yes | no | no | yes | no | no | no | yes | no | no | no | no | no |
| SCN1A | 1722 | T | 0 | 1 | no | yes | no | yes | no | no | no | yes | no | no | no | no | no |
| SCN1A | 1723 | S | 0 | 2 | no | yes | no | yes | no | no | yes | yes | no | no | no | no | no |
| SCN1A | 1724 | A | 0 | 2 | no | yes | no | yes | no | no | no | yes | no | no | no | no | no |

| | | | | | | | | | | | | | | | | | |
|---|---|---|---|---|---|---|---|---|---|---|---|---|---|---|---|---|---|
| SCN1A | 1725 | G | 0 | 3 | no | yes | no | yes | no | no | no | yes | no | no | no | no | no |
| SCN1A | 1726 | W | 0 | 1 | yes | no | no | yes | no | no | yes | yes | no | no | no | no | no |
| SCN1A | 1727 | D | 0 | 2 | yes | no | no | no | no | no | yes | yes | no | no | no | no | no |
| SCN1A | 1728 | G | 1 | 1 | yes | no | no | no | no | no | yes | yes | no | no | no | no | no |
| SCN1A | 1729 | L | 1 | 1 | yes | no | no | no | no | no | no | yes | no | no | no | no | no |
| SCN1A | 1730 | L | 0 | 1 | yes | no | no | no | no | no | no | yes | no | no | no | no | no |
| SCN1A | 1731 | A | 0 | 1 | yes | no | no | no | no | no | yes | yes | no | no | no | no | no |
| SCN1A | 1732 | P | 0 | 1 | yes | no | no | no | no | no | no | yes | no | no | no | no | no |
| SCN1A | 1733 | I | 0 | 0 | yes | no | no | no | yes | no | no | no | no | no | no | no | no |
| SCN1A | 1734 | L | 0 | 1 | no | yes | no | no | yes | no | no | no | no | no | no | no | no |
| SCN1A | 1735 | N | 0 | 0 | no | yes | no | no | yes | no | no | no | no | no | no | no | no |
| SCN1A | 1736 | S | 0 | 0 | no | yes | no | no | yes | no | no | no | no | no | no | no | no |
| SCN1A | 1737 | K | 0 | 0 | no | yes | no | no | yes | no | no | no | no | no | no | no | no |
| SCN1A | 1738 | P | 0 | 0 | no | yes | no | no | yes | no | no | no | no | no | no | no | no |
| SCN1A | 1739 | P | 0 | 0 | no | yes | no | no | yes | no | no | no | no | no | no | no | no |
| SCN1A | 1740 | D | 0 | 1 | no | yes | no | no | yes | no | no | no | no | no | no | no | no |
| SCN1A | 1741 | C | 0 | 2 | no | yes | no | no | yes | no | no | no | no | no | no | no | no |
| SCN1A | 1742 | D | 2 | 1 | no | yes | no | no | yes | no | no | no | no | no | no | no | no |
| SCN1A | 1743 | P | 0 | 0 | no | yes | no | no | yes | no | no | no | no | no | no | no | no |
| SCN1A | 1744 | N | 0 | 1 | no | yes | no | no | yes | no | no | no | no | no | no | no | no |
| SCN1A | 1745 | K | 0 | 0 | no | yes | no | no | yes | no | no | no | no | no | no | no | no |
| SCN1A | 1746 | V | 1 | 0 | no | yes | no | no | yes | no | no | no | no | no | no | no | no |
| SCN1A | 1747 | N | 0 | 0 | no | yes | no | no | yes | no | no | no | no | no | no | no | no |
| SCN1A | 1748 | P | 0 | 0 | no | yes | no | no | yes | no | no | no | no | no | no | no | no |
| SCN1A | 1749 | G | 0 | 2 | no | yes | no | no | yes | no | no | no | no | no | no | no | no |
| SCN1A | 1750 | S | 1 | 0 | no | yes | no | no | yes | no | no | no | no | no | no | no | no |
| SCN1A | 1751 | S | 0 | 0 | no | yes | no | no | yes | no | no | no | no | no | no | no | no |
| SCN1A | 1752 | V | 1 | 0 | no | yes | no | no | yes | no | no | no | no | no | no | no | no |
| SCN1A | 1753 | K | 0 | 0 | no | yes | no | no | yes | no | no | no | no | no | no | no | no |
| SCN1A | 1754 | G | 0 | 3 | no | yes | no | no | yes | no | no | no | no | no | no | no | no |
| SCN1A | 1755 | D | 0 | 2 | no | yes | no | no | yes | no | no | no | no | no | no | no | no |
| SCN1A | 1756 | C | 0 | 1 | no | yes | no | no | yes | no | no | no | no | no | no | no | no |
| SCN1A | 1757 | G | 0 | 3 | no | yes | no | no | yes | no | no | no | no | no | no | no | no |

| | | | | | | | | | | | | | | | | | |
|---|---|---|---|---|---|---|---|---|---|---|---|---|---|---|---|---|---|
| SCN1A | 1758 | N | 0 | 0 | no | yes | no | no | yes | no | no | no | no | no | no | no | no |
| SCN1A | 1759 | P | 0 | 1 | yes | no | no | no | yes | no | no | no | no | no | no | no | no |
| SCN1A | 1760 | S | 0 | 0 | yes | no | no | yes | yes | no | no | no | no | no | no | no | no |
| SCN1A | 1761 | V | 0 | 1 | yes | no | no | yes | yes | no | no | no | no | no | no | no | no |
| SCN1A | 1762 | G | 0 | 1 | yes | no | no | yes | yes | no | no | no | no | no | no | no | no |
| SCN1A | 1763 | I | 0 | 0 | yes | no | no | yes | no | no | no | no | no | no | no | no | no |
| SCN1A | 1764 | F | 0 | 1 | yes | no | no | yes | no | no | no | no | no | no | no | no | no |
| SCN1A | 1765 | F | 0 | 4 | yes | no | no | yes | no | no | no | no | no | no | no | no | no |
| SCN1A | 1766 | F | 0 | 0 | yes | no | no | yes | no | no | no | no | no | no | no | no | no |
| SCN1A | 1767 | V | 0 | 0 | yes | no | no | yes | no | no | no | no | no | no | no | no | no |
| SCN1A | 1768 | S | 0 | 1 | yes | no | no | yes | no | no | no | no | no | no | no | no | no |
| SCN1A | 1769 | Y | 0 | 1 | yes | no | no | yes | no | no | no | no | no | no | no | no | no |
| SCN1A | 1770 | I | 0 | 0 | yes | no | no | yes | no | no | no | no | no | no | no | no | no |
| SCN1A | 1771 | I | 0 | 3 | yes | no | no | yes | no | no | no | no | no | no | no | no | no |
| SCN1A | 1772 | I | 1 | 0 | yes | no | no | yes | no | no | no | no | no | no | no | no | no |
| SCN1A | 1773 | S | 0 | 1 | yes | no | no | yes | no | no | no | no | no | no | no | no | no |
| SCN1A | 1774 | F | 0 | 1 | yes | no | no | yes | no | no | yes | no | no | no | no | no | no |
| SCN1A | 1775 | L | 0 | 1 | yes | no | no | yes | no | no | no | no | no | no | no | no | no |
| SCN1A | 1776 | V | 0 | 1 | yes | no | no | yes | no | no | no | no | no | no | no | no | no |
| SCN1A | 1777 | V | 0 | 0 | yes | no | no | yes | no | no | no | no | no | no | no | no | no |
| SCN1A | 1778 | V | 0 | 0 | yes | no | no | yes | no | no | no | no | no | no | no | no | no |
| SCN1A | 1779 | N | 0 | 0 | yes | no | no | yes | no | no | no | no | no | no | no | no | no |
| SCN1A | 1780 | M | 0 | 2 | yes | no | no | yes | no | no | no | no | no | no | no | no | no |
| SCN1A | 1781 | Y | 0 | 2 | yes | no | no | yes | no | no | yes | no | no | no | no | no | no |
| SCN1A | 1782 | I | 0 | 2 | yes | no | no | yes | no | no | no | no | no | no | no | no | no |
| SCN1A | 1783 | A | 0 | 2 | yes | no | no | yes | no | no | no | no | no | no | no | no | no |
| SCN1A | 1784 | V | 1 | 0 | yes | no | no | yes | no | no | no | no | no | no | no | no | no |
| SCN1A | 1785 | I | 0 | 0 | yes | no | no | yes | no | no | yes | no | no | no | no | no | no |
| SCN1A | 1786 | L | 0 | 1 | yes | no | no | no | no | yes | no | no | no | no | no | no | no |
| SCN1A | 1787 | E | 0 | 2 | yes | no | no | yes | no | yes | no | no | no | no | no | no | no |
| SCN1A | 1788 | N | 0 | 0 | yes | no | no | yes | no | yes | no | no | no | no | no | no | no |
| SCN1A | 1789 | F | 0 | 0 | yes | no | no | no | no | yes | yes | no | no | no | no | no | no |
| SCN1A | 1790 | S | 2 | 0 | yes | no | no | no | no | yes | no | no | no | no | no | no | no |

| | | | | | | | | | | | | | | | | | |
|---|---|---|---|---|---|---|---|---|---|---|---|---|---|---|---|---|---|
| SCN1A | 1791 | V | 0 | 0 | yes | no | no | no | no | yes | no | no | no | no | no | no | no |
| SCN1A | 1792 | A | 0 | 3 | yes | no | no | no | no | yes | no | no | no | no | no | no | no |
| SCN1A | 1793 | T | 0 | 1 | no | yes | no | no | no | yes | yes | no | no | no | no | no | no |
| SCN1A | 1794 | E | 0 | 0 | no | yes | no | no | no | yes | yes | no | no | no | no | no | no |
| SCN1A | 1795 | E | 0 | 2 | no | yes | no | no | no | yes | no | no | no | no | no | no | no |
| SCN5A | 119 | P | 2 | 0 | no | yes | no | no | no | yes | no | no | no | no | no | no | no |
| SCN5A | 120 | I | 0 | 0 | no | yes | no | no | no | yes | no | no | no | no | no | no | no |
| SCN5A | 121 | R | 1 | 2 | yes | no | no | no | no | yes | no | no | no | no | no | no | no |
| SCN5A | 122 | R | 0 | 0 | yes | no | no | no | no | yes | no | no | no | no | no | no | no |
| SCN5A | 123 | A | 1 | 0 | yes | no | no | no | no | yes | no | no | no | no | no | no | no |
| SCN5A | 124 | A | 0 | 1 | yes | no | no | no | no | yes | no | no | no | no | no | no | no |
| SCN5A | 125 | V | 1 | 1 | yes | no | no | no | no | yes | no | no | no | no | no | no | no |
| SCN5A | 126 | K | 0 | 1 | yes | no | no | no | no | yes | no | no | no | no | no | no | no |
| SCN5A | 127 | I | 0 | 0 | yes | no | no | yes | no | yes | no | no | no | no | no | no | no |
| SCN5A | 128 | L | 0 | 0 | yes | no | no | no | no | yes | no | no | no | no | no | no | no |
| SCN5A | 129 | V | 0 | 0 | no | yes | no | no | no | yes | no | no | no | no | no | no | no |
| SCN5A | 130 | H | 0 | 0 | no | yes | no | no | no | yes | no | no | no | no | no | no | no |
| SCN5A | 131 | S | 0 | 0 | yes | no | no | yes | no | yes | no | no | no | no | no | no | no |
| SCN5A | 132 | L | 0 | 0 | yes | no | no | yes | no | no | no | no | no | no | no | no | no |
| SCN5A | 133 | F | 0 | 0 | yes | no | no | yes | no | no | no | no | no | no | no | no | no |
| SCN5A | 134 | N | 1 | 0 | yes | no | no | yes | no | no | no | no | no | no | no | no | no |
| SCN5A | 135 | M | 1 | 0 | yes | no | no | yes | no | no | no | no | no | no | no | no | no |
| SCN5A | 136 | L | 0 | 1 | yes | no | no | yes | no | no | no | no | no | no | no | no | no |
| SCN5A | 137 | I | 1 | 1 | yes | no | no | yes | no | no | no | no | no | no | no | no | no |
| SCN5A | 138 | M | 1 | 1 | yes | no | no | yes | no | no | no | no | no | no | no | no | no |
| SCN5A | 139 | C | 0 | 0 | yes | no | no | yes | no | no | no | no | no | no | no | no | no |
| SCN5A | 140 | T | 0 | 0 | yes | no | no | yes | no | no | no | no | no | no | no | no | no |
| SCN5A | 141 | I | 1 | 1 | yes | no | no | yes | no | no | no | no | no | no | no | no | no |
| SCN5A | 142 | L | 0 | 0 | yes | no | no | yes | no | no | no | no | no | no | no | no | no |
| SCN5A | 143 | T | 0 | 0 | yes | no | no | yes | no | no | no | no | no | no | no | no | no |
| SCN5A | 144 | N | 0 | 0 | yes | no | no | yes | no | no | no | no | no | no | no | no | no |
| SCN5A | 145 | C | 0 | 0 | yes | no | no | yes | no | no | no | no | no | no | no | no | no |
| SCN5A | 146 | V | 2 | 1 | yes | no | no | yes | no | no | no | no | no | no | no | no | no |

| | | | | | | | | | | | | | | | | | |
|---|---|---|---|---|---|---|---|---|---|---|---|---|---|---|---|---|---|
| SCN5A | 147 | F | 0 | 0 | yes | no | no | yes | no | no | no | no | no | no | no | no | no |
| SCN5A | 148 | M | 0 | 0 | yes | no | no | yes | no | no | no | no | no | no | no | no | no |
| SCN5A | 149 | A | 0 | 1 | no | yes | no | yes | no | no | no | no | no | no | no | no | no |
| SCN5A | 150 | Q | 0 | 0 | no | yes | no | yes | no | no | no | no | no | no | no | no | no |
| SCN5A | 151 | H | 0 | 0 | no | yes | no | no | yes | no | no | no | no | no | no | no | no |
| SCN5A | 152 | D | 1 | 0 | no | yes | no | no | yes | no | no | no | no | no | no | no | no |
| SCN5A | 153 | P | 0 | 0 | no | yes | no | no | yes | no | no | no | no | no | no | no | no |
| SCN5A | 154 | P | 1 | 0 | no | yes | no | no | yes | no | no | no | no | no | no | no | no |
| SCN5A | 155 | P | 0 | 0 | no | yes | no | no | yes | no | no | no | no | no | no | no | no |
| SCN5A | 156 | W | 1 | 0 | no | yes | no | no | yes | no | no | no | no | no | no | no | no |
| SCN5A | 157 | T | 1 | 0 | no | yes | no | yes | yes | no | no | no | no | no | no | no | no |
| SCN5A | 158 | K | 1 | 0 | yes | no | no | yes | no | no | no | no | no | no | no | no | no |
| SCN5A | 159 | Y | 1 | 0 | yes | no | no | yes | no | no | no | no | no | no | no | no | no |
| SCN5A | 160 | V | 0 | 0 | yes | no | no | yes | no | no | no | no | no | no | no | no | no |
| SCN5A | 161 | E | 0 | 0 | yes | no | no | yes | no | no | no | no | no | no | no | no | no |
| SCN5A | 162 | Y | 2 | 0 | yes | no | no | yes | no | no | no | no | no | no | no | no | no |
| SCN5A | 163 | T | 0 | 0 | yes | no | no | yes | no | no | no | no | no | no | no | no | no |
| SCN5A | 164 | F | 1 | 0 | yes | no | no | yes | no | no | no | no | no | no | no | no | no |
| SCN5A | 165 | T | 0 | 0 | yes | no | no | yes | no | no | no | no | no | no | no | no | no |
| SCN5A | 166 | A | 1 | 0 | yes | no | no | yes | no | no | no | no | no | no | no | no | no |
| SCN5A | 167 | I | 0 | 0 | yes | no | no | yes | no | no | no | no | no | no | no | no | no |
| SCN5A | 168 | Y | 0 | 0 | yes | no | no | yes | no | no | no | no | no | no | no | no | no |
| SCN5A | 169 | T | 0 | 0 | yes | no | no | yes | no | no | no | no | no | no | no | no | no |
| SCN5A | 170 | F | 1 | 0 | yes | no | no | yes | no | no | no | no | no | no | no | no | no |
| SCN5A | 171 | E | 0 | 0 | yes | no | no | yes | no | no | no | no | no | no | no | no | no |
| SCN5A | 172 | S | 0 | 0 | yes | no | no | yes | no | no | no | no | no | no | no | no | no |
| SCN5A | 173 | L | 0 | 0 | yes | no | no | yes | no | no | no | no | no | no | no | no | no |
| SCN5A | 174 | V | 1 | 0 | yes | no | no | yes | no | no | no | no | no | no | no | no | no |
| SCN5A | 175 | K | 0 | 1 | yes | no | no | yes | no | no | no | no | no | no | no | no | no |
| SCN5A | 176 | I | 0 | 1 | yes | no | no | yes | no | no | no | no | no | no | no | no | no |
| SCN5A | 177 | L | 0 | 0 | no | yes | no | yes | no | no | no | no | no | no | no | no | no |
| SCN5A | 178 | A | 0 | 1 | no | yes | no | no | no | no | no | no | no | no | no | no | no |
| SCN5A | 179 | R | 1 | 0 | no | yes | no | no | no | yes | no | no | no | no | no | no | no |

| | | | | | | | | | | | | | | | | | |
|---|---|---|---|---|---|---|---|---|---|---|---|---|---|---|---|---|---|
| SCN5A | 180 | G | 0 | 1 | no | yes | no | no | no | yes | no | no | no | no | no | no | no |
| SCN5A | 181 | F | 0 | 0 | no | yes | no | no | no | yes | no | no | no | no | no | no | no |
| SCN5A | 182 | C | 1 | 1 | no | yes | no | no | no | yes | no | no | no | no | no | no | no |
| SCN5A | 183 | L | 0 | 0 | no | yes | no | no | no | yes | no | no | no | no | no | no | no |
| SCN5A | 184 | H | 1 | 0 | no | yes | no | yes | no | yes | no | no | no | no | no | no | no |
| SCN5A | 185 | A | 2 | 2 | no | yes | no | no | no | yes | no | no | no | no | no | no | no |
| SCN5A | 186 | F | 0 | 0 | no | yes | no | no | no | yes | no | no | no | no | no | no | no |
| SCN5A | 187 | T | 1 | 1 | no | yes | no | no | no | yes | no | no | no | no | no | no | no |
| SCN5A | 188 | F | 0 | 0 | no | yes | no | yes | no | yes | no | no | no | no | no | no | no |
| SCN5A | 189 | L | 0 | 0 | no | yes | no | yes | no | yes | no | no | no | no | no | no | no |
| SCN5A | 190 | R | 4 | 2 | no | yes | no | no | no | yes | no | no | no | no | no | no | no |
| SCN5A | 191 | D | 0 | 0 | no | yes | no | yes | no | yes | no | no | no | no | no | no | no |
| SCN5A | 192 | P | 0 | 0 | yes | no | no | yes | no | yes | no | no | no | no | no | no | no |
| SCN5A | 193 | W | 1 | 0 | yes | no | no | yes | no | no | no | no | no | no | no | no | no |
| SCN5A | 194 | N | 0 | 0 | yes | no | no | yes | no | no | no | no | no | no | no | no | no |
| SCN5A | 195 | W | 0 | 0 | yes | no | no | yes | no | no | no | no | no | no | no | no | no |
| SCN5A | 196 | L | 0 | 0 | yes | no | no | yes | no | no | no | no | no | no | no | no | no |
| SCN5A | 197 | D | 0 | 2 | yes | no | no | yes | no | no | no | no | no | no | no | no | no |
| SCN5A | 198 | F | 0 | 0 | yes | no | no | yes | no | no | no | no | no | no | no | no | no |
| SCN5A | 199 | S | 1 | 0 | yes | no | no | yes | no | no | no | no | no | no | no | no | no |
| SCN5A | 200 | V | 0 | 0 | yes | no | no | yes | no | no | no | no | no | no | no | no | no |
| SCN5A | 201 | I | 0 | 0 | yes | no | no | yes | no | no | no | no | no | no | no | no | no |
| SCN5A | 202 | I | 1 | 0 | yes | no | no | yes | no | no | no | no | no | no | no | no | no |
| SCN5A | 203 | M | 0 | 0 | yes | no | no | yes | no | no | no | no | no | no | no | no | no |
| SCN5A | 204 | A | 0 | 0 | yes | no | no | yes | no | no | no | no | no | no | no | no | no |
| SCN5A | 205 | Y | 0 | 0 | yes | no | no | yes | no | no | no | no | no | no | no | no | no |
| SCN5A | 206 | T | 0 | 0 | yes | no | no | yes | no | no | no | no | no | no | no | no | no |
| SCN5A | 207 | T | 0 | 0 | no | yes | no | yes | no | no | no | no | no | no | no | no | no |
| SCN5A | 208 | E | 0 | 0 | no | yes | no | yes | no | no | no | no | no | no | no | no | no |
| SCN5A | 209 | F | 0 | 0 | no | yes | no | yes | no | no | no | no | no | no | no | no | no |
| SCN5A | 210 | V | 0 | 0 | no | yes | no | yes | no | no | no | no | no | no | no | no | no |
| SCN5A | 211 | D | 0 | 0 | no | yes | no | no | yes | no | no | no | no | no | no | no | no |
| SCN5A | 212 | L | 0 | 1 | no | yes | no | no | yes | no | no | no | no | no | no | no | no |

| | | | | | | | | | | | | | | | | | |
|---|---|---|---|---|---|---|---|---|---|---|---|---|---|---|---|---|---|
| SCN5A | 213 | G | 0 | 0 | no | yes | no | no | yes | no | no | no | no | no | no | no | no |
| SCN5A | 214 | N | 0 | 0 | no | yes | no | no | yes | no | no | no | no | no | no | no | no |
| SCN5A | 215 | V | 0 | 0 | no | yes | no | no | yes | no | no | no | no | no | no | no | no |
| SCN5A | 216 | S | 1 | 0 | no | yes | no | no | yes | no | no | no | no | no | no | no | no |
| SCN5A | 217 | A | 0 | 0 | no | yes | no | yes | no | no | no | no | no | no | no | no | no |
| SCN5A | 218 | L | 0 | 0 | no | yes | no | yes | no | no | no | no | no | no | no | no | no |
| SCN5A | 219 | R | 1 | 1 | yes | no | no | yes | no | no | no | no | no | no | no | no | no |
| SCN5A | 220 | T | 1 | 1 | yes | no | no | yes | no | no | no | no | no | no | no | no | no |
| SCN5A | 221 | F | 0 | 0 | yes | no | no | yes | no | no | no | no | no | no | no | no | no |
| SCN5A | 222 | R | 1 | 2 | yes | no | no | yes | no | no | no | no | no | no | no | no | no |
| SCN5A | 223 | V | 0 | 2 | yes | no | no | yes | no | no | no | no | no | no | no | no | no |
| SCN5A | 224 | L | 0 | 0 | yes | no | no | yes | no | no | no | no | no | no | no | no | no |
| SCN5A | 225 | R | 2 | 2 | yes | no | no | yes | no | no | no | no | no | no | no | no | no |
| SCN5A | 226 | A | 1 | 1 | yes | no | no | yes | no | no | no | no | no | no | no | no | no |
| SCN5A | 227 | L | 1 | 0 | yes | no | no | yes | no | no | no | no | no | no | no | no | no |
| SCN5A | 228 | K | 0 | 0 | yes | no | no | yes | no | no | no | no | no | no | no | no | no |
| SCN5A | 229 | T | 0 | 0 | yes | no | no | yes | no | no | no | no | no | no | no | no | no |
| SCN5A | 230 | I | 2 | 2 | yes | no | no | yes | no | no | no | no | no | no | no | no | no |
| SCN5A | 231 | S | 0 | 0 | no | yes | no | yes | no | no | no | no | no | no | no | no | no |
| SCN5A | 232 | V | 1 | 0 | no | yes | no | yes | no | no | no | no | no | no | no | no | no |
| SCN5A | 233 | I | 0 | 0 | no | yes | no | yes | no | no | no | no | no | no | no | no | no |
| SCN5A | 234 | S | 0 | 0 | no | yes | no | yes | no | yes | no | no | no | no | no | no | no |
| SCN5A | 235 | G | 0 | 0 | no | yes | no | yes | no | yes | no | no | no | no | no | no | no |
| SCN5A | 236 | L | 0 | 0 | yes | no | no | yes | no | yes | no | no | no | no | no | no | no |
| SCN5A | 237 | K | 0 | 0 | yes | no | no | yes | no | yes | no | no | no | no | no | no | no |
| SCN5A | 238 | T | 0 | 0 | yes | no | no | yes | no | yes | no | no | no | no | no | no | no |
| SCN5A | 239 | I | 0 | 1 | yes | no | no | yes | no | yes | no | no | no | no | no | no | no |
| SCN5A | 240 | V | 1 | 1 | yes | no | no | yes | no | yes | no | no | no | no | no | no | no |
| SCN5A | 241 | G | 0 | 0 | yes | no | no | yes | no | yes | no | no | no | no | no | no | no |
| SCN5A | 242 | A | 0 | 0 | yes | no | no | yes | no | yes | no | no | no | no | no | no | no |
| SCN5A | 243 | L | 0 | 0 | yes | no | no | yes | no | yes | no | no | no | no | no | no | no |
| SCN5A | 244 | I | 0 | 0 | yes | no | no | yes | no | yes | no | no | no | no | no | no | no |
| SCN5A | 245 | Q | 1 | 1 | yes | no | no | yes | no | yes | no | no | no | no | no | no | no |

| | | | | | | | | | | | | | | | | | |
|---|---|---|---|---|---|---|---|---|---|---|---|---|---|---|---|---|---|
| SCN5A | 246 | S | 0 | 0 | yes | no | no | yes | no | yes | no | no | no | no | no | no | no |
| SCN5A | 247 | V | 1 | 1 | yes | no | no | yes | no | yes | no | no | no | no | no | no | no |
| SCN5A | 248 | K | 0 | 0 | yes | no | no | yes | no | yes | no | no | no | no | no | no | no |
| SCN5A | 249 | K | 0 | 0 | yes | no | no | yes | no | yes | no | no | no | no | no | no | no |
| SCN5A | 250 | L | 0 | 1 | yes | no | no | yes | no | yes | no | no | no | no | no | no | no |
| SCN5A | 251 | A | 0 | 0 | yes | no | no | yes | no | yes | no | no | no | no | no | no | no |
| SCN5A | 252 | D | 0 | 0 | yes | no | no | yes | no | yes | no | no | no | no | no | no | no |
| SCN5A | 253 | V | 0 | 0 | yes | no | no | yes | no | no | no | no | no | no | no | no | no |
| SCN5A | 254 | M | 0 | 0 | yes | no | no | yes | no | no | no | no | no | no | no | no | no |
| SCN5A | 255 | V | 0 | 0 | yes | no | no | yes | no | no | no | no | no | no | no | no | no |
| SCN5A | 256 | L | 0 | 0 | yes | no | no | yes | no | no | no | no | no | no | no | no | no |
| SCN5A | 257 | T | 0 | 0 | yes | no | no | yes | no | no | no | no | no | no | no | no | no |
| SCN5A | 258 | V | 0 | 1 | yes | no | no | yes | no | no | no | no | no | no | no | no | no |
| SCN5A | 259 | F | 0 | 0 | yes | no | no | yes | no | no | no | no | no | no | no | no | no |
| SCN5A | 260 | C | 0 | 0 | yes | no | no | yes | no | no | no | no | no | no | no | no | no |
| SCN5A | 261 | L | 0 | 0 | yes | no | no | yes | no | no | no | no | no | no | no | no | no |
| SCN5A | 262 | S | 1 | 0 | yes | no | no | yes | no | no | no | no | no | no | no | no | no |
| SCN5A | 263 | V | 1 | 0 | yes | no | no | yes | no | no | no | no | no | no | no | no | no |
| SCN5A | 264 | F | 0 | 0 | yes | no | no | yes | no | no | no | no | no | no | no | no | no |
| SCN5A | 265 | A | 1 | 0 | yes | no | no | yes | no | no | no | no | no | no | no | no | no |
| SCN5A | 266 | L | 1 | 0 | yes | no | no | yes | no | no | no | no | no | no | no | no | no |
| SCN5A | 267 | I | 0 | 0 | yes | no | no | yes | no | no | no | no | no | no | no | no | no |
| SCN5A | 268 | G | 1 | 0 | yes | no | no | yes | no | no | no | no | no | no | no | no | no |
| SCN5A | 269 | L | 0 | 0 | yes | no | no | yes | no | no | no | no | no | no | no | no | no |
| SCN5A | 270 | Q | 0 | 1 | yes | no | no | yes | no | no | no | no | no | no | no | no | no |
| SCN5A | 271 | L | 1 | 0 | yes | no | no | yes | no | no | no | no | no | no | no | no | no |
| SCN5A | 272 | F | 0 | 0 | yes | no | no | no | no | no | no | no | no | no | no | no | no |
| SCN5A | 273 | M | 0 | 0 | no | yes | no | no | yes | no | no | no | no | no | no | no | no |
| SCN5A | 274 | G | 1 | 0 | no | yes | no | no | yes | no | no | no | no | no | no | no | no |
| SCN5A | 275 | N | 0 | 1 | no | yes | no | no | yes | no | no | no | no | no | no | no | no |
| SCN5A | 276 | L | 0 | 1 | no | yes | no | no | yes | no | no | no | no | no | no | no | no |
| SCN5A | 277 | R | 0 | 0 | no | yes | no | no | yes | no | no | no | no | no | no | no | no |
| SCN5A | 278 | H | 0 | 2 | no | no | yes | no | yes | no | no | no | no | no | no | no | no |

| | | | | | | | | | | | | | | | | | |
|---|---|---|---|---|---|---|---|---|---|---|---|---|---|---|---|---|---|
| SCN5A | 279 | K | 0 | 0 | no | no | yes | no | yes | no | no | no | no | no | no | no | no |
| SCN5A | 280 | C | 0 | 1 | no | no | yes | no | yes | no | no | no | no | no | no | no | no |
| SCN5A | 281 | V | 1 | 1 | no | no | yes | no | yes | no | no | no | no | no | no | no | no |
| SCN5A | 282 | R | 1 | 2 | no | yes | no | no | yes | no | no | no | no | no | no | no | no |
| SCN5A | 283 | N | 0 | 0 | no | yes | no | no | yes | no | no | no | no | no | no | no | no |
| SCN5A | 284 | F | 0 | 0 | no | yes | no | no | yes | no | no | no | no | no | no | no | no |
| SCN5A | 285 | T | 1 | 0 | no | no | yes | no | yes | no | no | no | no | no | no | no | no |
| SCN5A | 286 | A | 2 | 0 | no | yes | no | no | yes | no | no | no | no | no | no | no | no |
| SCN5A | 287 | L | 0 | 0 | no | yes | no | no | yes | no | no | no | no | no | no | no | no |
| SCN5A | 288 | N | 1 | 0 | no | yes | no | no | yes | no | no | no | no | no | no | no | no |
| SCN5A | 289 | G | 1 | 1 | no | yes | no | no | yes | no | no | no | no | no | no | no | no |
| SCN5A | 290 | T | 0 | 0 | no | yes | no | no | yes | no | no | no | no | no | no | no | no |
| SCN5A | 291 | N | 3 | 0 | no | yes | no | no | yes | no | no | no | no | no | no | no | no |
| SCN5A | 292 | G | 1 | 1 | no | yes | no | no | yes | no | no | no | no | no | no | no | no |
| SCN5A | 293 | S | 0 | 0 | no | yes | no | no | yes | no | no | no | no | no | no | no | no |
| SCN5A | 294 | V | 1 | 1 | no | yes | no | no | yes | no | no | no | no | no | no | no | no |
| SCN5A | 295 | E | 1 | 0 | no | yes | no | no | yes | no | no | no | no | no | no | no | no |
| SCN5A | 296 | A | 0 | 0 | no | no | yes | no | yes | no | no | no | no | no | no | no | no |
| SCN5A | 297 | D | 0 | 0 | no | yes | no | no | yes | no | no | no | no | no | no | no | no |
| SCN5A | 298 | G | 2 | 1 | no | yes | no | no | yes | no | no | no | no | no | no | no | no |
| SCN5A | 299 | L | 2 | 0 | no | yes | no | no | yes | no | no | no | no | no | no | no | no |
| SCN5A | 300 | V | 1 | 1 | no | no | yes | no | yes | no | no | no | no | no | no | no | no |
| SCN5A | 301 | W | 0 | 0 | no | yes | no | no | yes | no | no | no | no | no | no | no | no |
| SCN5A | 302 | E | 0 | 0 | no | yes | no | no | yes | no | no | no | no | no | no | no | no |
| SCN5A | 303 | S | 0 | 0 | no | yes | no | no | yes | no | no | no | no | no | no | no | no |
| SCN5A | 304 | L | 0 | 0 | no | yes | no | no | yes | no | no | no | no | no | no | no | no |
| SCN5A | 305 | D | 1 | 0 | yes | no | no | no | yes | no | no | no | no | no | no | no | no |
| SCN5A | 306 | L | 1 | 0 | yes | no | no | no | yes | no | no | no | no | no | no | no | no |
| SCN5A | 307 | Y | 0 | 0 | yes | no | no | no | yes | no | no | no | no | no | no | no | no |
| SCN5A | 308 | L | 0 | 0 | yes | no | no | no | yes | no | no | no | no | no | no | no | no |
| SCN5A | 309 | S | 2 | 0 | yes | no | no | no | yes | no | no | no | no | no | no | no | no |
| SCN5A | 310 | D | 1 | 0 | no | yes | no | no | yes | no | no | no | no | no | no | no | no |
| SCN5A | 311 | P | 0 | 0 | no | yes | no | no | yes | no | no | no | no | no | no | no | no |

| | | | | | | | | | | | | | | | | | |
|---|---|---|---|---|---|---|---|---|---|---|---|---|---|---|---|---|---|
| SCN5A | 312 | E | 0 | 0 | no | yes | no | no | yes | no | yes | no | no | no | no | no | no |
| SCN5A | 313 | N | 0 | 0 | no | yes | no | no | yes | no | no | no | no | no | no | no | no |
| SCN5A | 314 | Y | 0 | 0 | no | yes | no | no | yes | no | no | no | no | no | no | no | no |
| SCN5A | 315 | L | 0 | 1 | no | yes | no | no | yes | no | no | no | no | no | no | no | no |
| SCN5A | 316 | L | 0 | 0 | no | no | yes | no | yes | no | yes | no | no | no | no | no | no |
| SCN5A | 317 | K | 2 | 1 | no | yes | no | no | yes | no | no | no | no | no | no | no | no |
| SCN5A | 318 | N | 0 | 0 | no | yes | no | no | yes | no | no | no | no | no | no | no | no |
| SCN5A | 319 | G | 2 | 1 | no | yes | no | no | yes | no | no | no | no | no | no | no | no |
| SCN5A | 320 | T | 1 | 1 | no | yes | no | no | yes | no | no | no | no | no | no | no | no |
| SCN5A | 321 | S | 0 | 0 | no | yes | no | no | yes | no | yes | no | no | no | no | no | no |
| SCN5A | 322 | D | 0 | 0 | no | yes | no | no | yes | no | yes | no | no | no | no | no | no |
| SCN5A | 323 | V | 0 | 0 | no | no | yes | no | yes | no | no | no | no | no | no | no | no |
| SCN5A | 324 | L | 0 | 0 | no | yes | no | no | yes | no | no | no | no | no | no | no | no |
| SCN5A | 325 | L | 0 | 1 | no | yes | no | no | yes | no | no | no | no | no | no | no | no |
| SCN5A | 326 | C | 0 | 0 | no | yes | no | no | yes | no | no | no | no | no | no | no | no |
| SCN5A | 327 | G | 0 | 0 | no | yes | no | no | yes | no | no | no | no | no | no | no | no |
| SCN5A | 328 | N | 0 | 0 | no | yes | no | no | yes | no | no | no | no | no | no | no | no |
| SCN5A | 329 | S | 0 | 0 | no | yes | no | no | yes | no | no | no | no | no | no | no | no |
| SCN5A | 330 | S | 1 | 0 | no | yes | no | no | yes | no | no | no | no | no | no | no | no |
| SCN5A | 331 | D | 0 | 0 | no | yes | no | no | yes | no | no | no | no | no | no | no | no |
| SCN5A | 332 | A | 1 | 0 | no | yes | no | no | yes | no | no | no | no | no | no | no | no |
| SCN5A | 333 | G | 0 | 0 | no | yes | no | no | yes | no | no | no | no | no | no | no | no |
| SCN5A | 334 | T | 0 | 0 | no | yes | no | no | yes | no | no | no | no | no | no | no | no |
| SCN5A | 335 | C | 0 | 1 | no | yes | no | no | yes | no | no | no | no | no | no | no | no |
| SCN5A | 336 | P | 0 | 1 | no | yes | no | no | yes | no | no | no | no | no | no | no | no |
| SCN5A | 337 | E | 0 | 0 | no | yes | no | no | yes | no | no | no | no | no | no | no | no |
| SCN5A | 338 | G | 0 | 0 | no | yes | no | no | yes | no | no | no | no | no | no | no | no |
| SCN5A | 339 | Y | 0 | 0 | no | yes | no | no | yes | no | no | no | no | no | no | no | no |
| SCN5A | 340 | R | 2 | 2 | no | no | yes | no | yes | no | no | no | no | no | no | no | no |
| SCN5A | 341 | C | 0 | 0 | no | no | yes | no | yes | no | no | no | no | no | no | no | no |
| SCN5A | 342 | L | 0 | 0 | no | no | yes | no | yes | no | no | no | no | no | no | no | no |
| SCN5A | 343 | K | 0 | 0 | no | no | yes | no | yes | no | no | no | no | no | no | no | no |
| SCN5A | 344 | A | 1 | 0 | no | yes | no | no | yes | no | no | no | no | no | no | no | no |

| | | | | | | | | | | | | | | | | | |
|---|---|---|---|---|---|---|---|---|---|---|---|---|---|---|---|---|---|
| SCN5A | 345 | G | 0 | 0 | no | yes | no | no | yes | no | no | no | no | no | no | no | no |
| SCN5A | 346 | E | 2 | 0 | no | yes | no | no | yes | no | no | no | no | no | no | no | no |
| SCN5A | 347 | N | 0 | 0 | no | yes | no | no | yes | no | no | no | no | no | no | no | no |
| SCN5A | 348 | P | 1 | 0 | no | yes | no | no | yes | no | no | no | no | no | no | no | no |
| SCN5A | 349 | D | 1 | 1 | yes | no | no | no | yes | no | no | no | no | no | no | no | no |
| SCN5A | 350 | H | 1 | 0 | yes | no | no | no | yes | no | no | no | no | no | no | no | no |
| SCN5A | 351 | G | 1 | 1 | yes | no | no | no | yes | no | no | no | no | no | no | no | no |
| SCN5A | 352 | Y | 0 | 1 | no | yes | no | no | yes | no | no | no | no | no | no | no | no |
| SCN5A | 353 | T | 0 | 2 | no | yes | no | no | yes | no | no | no | no | no | no | no | no |
| SCN5A | 354 | S | 0 | 0 | no | yes | no | no | yes | no | no | no | no | no | no | no | no |
| SCN5A | 355 | F | 0 | 1 | no | yes | no | no | yes | no | no | no | no | no | no | no | no |
| SCN5A | 356 | D | 1 | 2 | no | yes | no | no | yes | no | no | no | no | no | no | no | no |
| SCN5A | 357 | S | 0 | 0 | no | yes | no | no | yes | no | no | no | no | no | no | no | no |
| SCN5A | 358 | F | 0 | 0 | yes | no | no | yes | no | no | no | yes | no | no | no | no | no |
| SCN5A | 359 | A | 1 | 0 | yes | no | no | yes | no | no | no | yes | no | no | no | no | no |
| SCN5A | 360 | W | 0 | 0 | yes | no | no | no | no | no | no | yes | no | no | no | no | no |
| SCN5A | 361 | A | 0 | 0 | yes | no | no | yes | no | no | no | yes | no | no | no | no | no |
| SCN5A | 362 | F | 0 | 0 | yes | no | no | yes | no | no | no | yes | no | no | no | no | no |
| SCN5A | 363 | L | 0 | 0 | yes | no | no | yes | no | no | no | yes | no | no | no | no | no |
| SCN5A | 364 | A | 0 | 0 | yes | no | no | yes | no | no | no | yes | no | no | no | no | no |
| SCN5A | 365 | L | 0 | 0 | yes | no | no | yes | no | no | no | yes | no | no | no | no | no |
| SCN5A | 366 | F | 0 | 0 | yes | no | no | yes | no | no | no | yes | no | no | no | no | no |
| SCN5A | 367 | R | 1 | 3 | yes | no | no | yes | no | no | no | yes | no | no | no | no | no |
| SCN5A | 368 | L | 0 | 0 | yes | no | no | yes | no | no | no | yes | no | no | no | no | no |
| SCN5A | 369 | M | 0 | 1 | yes | no | no | yes | no | no | no | yes | no | no | no | no | no |
| SCN5A | 370 | T | 0 | 1 | no | yes | no | yes | no | no | yes | yes | no | no | no | no | no |
| SCN5A | 371 | Q | 0 | 0 | no | yes | no | yes | no | no | yes | yes | no | no | no | no | no |
| SCN5A | 372 | D | 0 | 0 | no | yes | no | yes | no | no | yes | yes | no | no | no | no | no |
| SCN5A | 373 | C | 0 | 0 | no | yes | no | yes | no | no | yes | yes | no | no | no | no | no |
| SCN5A | 374 | W | 0 | 1 | yes | no | no | yes | no | no | no | yes | no | no | no | no | no |
| SCN5A | 375 | E | 0 | 0 | yes | no | no | no | no | no | no | yes | no | no | no | no | no |
| SCN5A | 376 | R | 1 | 2 | yes | no | no | no | no | no | yes | yes | no | no | no | no | no |
| SCN5A | 377 | L | 0 | 0 | yes | no | no | no | no | no | no | yes | no | no | no | no | no |

| | | | | | | | | | | | | | | | | | |
|---|---|---|---|---|---|---|---|---|---|---|---|---|---|---|---|---|---|
| SCN5A | 378 | Y | 0 | 0 | yes | no | no | yes | no | no | no | yes | no | no | no | no | no |
| SCN5A | 379 | Q | 0 | 0 | yes | no | no | no | no | no | no | yes | no | no | no | no | no |
| SCN5A | 380 | Q | 0 | 0 | yes | no | no | no | no | no | no | yes | no | no | no | no | no |
| SCN5A | 381 | T | 0 | 0 | yes | no | no | no | no | no | no | yes | no | no | no | no | no |
| SCN5A | 382 | L | 0 | 0 | yes | no | no | no | no | no | no | yes | no | no | no | no | no |
| SCN5A | 383 | R | 0 | 0 | yes | no | no | no | yes | no | no | no | no | no | no | no | no |
| SCN5A | 384 | S | 1 | 0 | yes | no | no | no | yes | no | no | no | no | no | no | no | no |
| SCN5A | 385 | A | 1 | 1 | yes | no | no | no | yes | no | no | no | no | no | no | no | no |
| SCN5A | 386 | G | 0 | 2 | no | yes | no | no | yes | no | no | no | no | no | no | no | no |
| SCN5A | 387 | K | 0 | 0 | yes | no | no | no | yes | no | no | no | no | no | no | no | no |
| SCN5A | 388 | I | 1 | 0 | yes | no | no | yes | yes | no | no | no | no | no | no | no | no |
| SCN5A | 389 | Y | 1 | 0 | yes | no | no | yes | yes | no | no | no | no | no | no | no | no |
| SCN5A | 390 | M | 0 | 0 | yes | no | no | yes | no | no | no | no | no | no | no | no | no |
| SCN5A | 391 | I | 0 | 0 | yes | no | no | yes | no | no | no | no | no | no | no | no | no |
| SCN5A | 392 | F | 0 | 0 | yes | no | no | yes | no | no | no | no | no | no | no | no | no |
| SCN5A | 393 | F | 0 | 0 | yes | no | no | yes | no | no | no | no | no | no | no | no | no |
| SCN5A | 394 | M | 0 | 0 | yes | no | no | yes | no | no | no | no | no | no | no | no | no |
| SCN5A | 395 | L | 0 | 0 | yes | no | no | yes | no | no | no | no | no | no | no | no | no |
| SCN5A | 396 | V | 0 | 2 | yes | no | no | yes | no | no | no | no | no | no | no | no | no |
| SCN5A | 397 | I | 1 | 2 | yes | no | no | yes | no | no | no | no | no | no | no | no | no |
| SCN5A | 398 | F | 0 | 0 | yes | no | no | yes | no | no | no | no | no | no | no | no | no |
| SCN5A | 399 | L | 0 | 0 | yes | no | no | yes | no | no | no | no | no | no | no | no | no |
| SCN5A | 400 | G | 2 | 2 | yes | no | no | yes | no | no | no | no | no | no | no | no | no |
| SCN5A | 401 | S | 0 | 0 | yes | no | no | yes | no | no | no | no | no | no | no | no | no |
| SCN5A | 402 | F | 1 | 0 | yes | no | no | yes | no | no | no | no | no | no | no | no | no |
| SCN5A | 403 | Y | 0 | 0 | yes | no | no | yes | no | no | no | no | no | no | no | no | no |
| SCN5A | 404 | L | 1 | 1 | yes | no | no | yes | no | no | no | no | no | no | no | no | no |
| SCN5A | 405 | V | 0 | 0 | yes | no | no | yes | no | no | yes | no | no | no | no | no | no |
| SCN5A | 406 | N | 0 | 2 | yes | no | no | yes | no | no | no | no | no | no | no | no | no |
| SCN5A | 407 | L | 0 | 0 | yes | no | no | yes | no | no | no | no | no | no | no | no | no |
| SCN5A | 408 | I | 0 | 0 | yes | no | no | yes | no | no | no | no | no | no | no | no | no |
| SCN5A | 409 | L | 0 | 2 | yes | no | no | yes | no | no | yes | no | no | no | no | no | no |
| SCN5A | 410 | A | 1 | 0 | yes | no | no | yes | no | no | no | no | no | no | no | no | no |

| | | | | | | | | | | | | | | | | | |
|---|---|---|---|---|---|---|---|---|---|---|---|---|---|---|---|---|---|
| SCN5A | 411 | V | 0 | 1 | yes | no | no | yes | no | yes | no | no | no | no | no | no | no |
| SCN5A | 412 | V | 0 | 0 | yes | no | no | yes | no | yes | no | no | no | no | no | no | no |
| SCN5A | 413 | A | 1 | 2 | yes | no | no | yes | no | yes | yes | no | no | no | no | no | no |
| SCN5A | 414 | M | 1 | 0 | yes | no | no | no | no | yes | no | no | no | no | no | no | no |
| SCN5A | 415 | A | 1 | 0 | yes | no | no | no | no | yes | no | no | no | no | no | no | no |
| SCN5A | 416 | Y | 1 | 1 | yes | no | no | yes | no | yes | no | no | no | no | no | no | no |
| SCN5A | 417 | E | 0 | 0 | yes | no | no | no | no | yes | yes | no | no | no | no | no | no |
| SCN5A | 418 | E | 1 | 0 | yes | no | no | no | no | yes | no | no | no | no | no | no | no |
| SCN5A | 419 | Q | 0 | 0 | yes | no | no | no | no | yes | no | no | no | no | no | no | no |
| SCN5A | 420 | N | 0 | 0 | yes | no | no | no | no | yes | no | no | no | no | no | no | no |
| SCN5A | 421 | Q | 0 | 0 | yes | no | no | no | no | yes | yes | no | no | no | no | no | no |
| SCN5A | 422 | A | 0 | 0 | yes | no | no | no | no | yes | no | no | no | no | no | no | no |
| SCN5A | 423 | T | 0 | 0 | yes | no | no | no | no | yes | no | no | no | no | no | no | no |
| SCN5A | 424 | I | 1 | 0 | yes | no | no | no | no | yes | no | no | no | no | no | no | no |
| SCN5A | 425 | A | 1 | 0 | yes | no | no | no | no | yes | no | no | no | no | no | no | no |
| SCN5A | 426 | E | 0 | 0 | yes | no | no | no | no | yes | no | no | no | no | no | no | no |
| SCN5A | 427 | T | 0 | 0 | yes | no | no | no | no | yes | no | no | no | no | no | no | no |
| SCN5A | 428 | E | 1 | 0 | yes | no | no | no | no | yes | no | no | no | no | no | no | no |
| SCN5A | 429 | E | 1 | 0 | no | yes | no | no | no | yes | no | no | no | no | no | no | no |
| SCN5A | 699 | C | 0 | 0 | no | yes | no | no | no | yes | no | no | no | no | no | no | no |
| SCN5A | 700 | C | 0 | 0 | yes | no | no | no | no | yes | no | no | no | no | no | no | no |
| SCN5A | 701 | P | 1 | 1 | yes | no | no | no | no | yes | no | no | no | no | no | no | no |
| SCN5A | 702 | L | 0 | 0 | yes | no | no | no | no | yes | no | no | no | no | no | no | no |
| SCN5A | 703 | W | 0 | 0 | yes | no | no | no | no | yes | no | no | no | no | no | no | no |
| SCN5A | 704 | M | 0 | 0 | yes | no | no | no | no | yes | no | no | no | no | no | no | no |
| SCN5A | 705 | S | 2 | 0 | yes | no | no | no | no | yes | no | no | no | no | no | no | no |
| SCN5A | 706 | I | 0 | 0 | yes | no | no | no | no | yes | no | no | no | no | no | no | no |
| SCN5A | 707 | K | 0 | 0 | yes | no | no | no | no | yes | no | no | no | no | no | no | no |
| SCN5A | 708 | Q | 0 | 0 | yes | no | no | no | no | yes | no | no | no | no | no | no | no |
| SCN5A | 709 | G | 0 | 1 | yes | no | no | no | no | yes | no | no | no | no | no | no | no |
| SCN5A | 710 | V | 0 | 0 | yes | no | no | no | no | yes | no | no | no | no | no | no | no |
| SCN5A | 711 | K | 0 | 0 | yes | no | no | no | no | yes | no | no | no | no | no | no | no |
| SCN5A | 712 | L | 0 | 0 | yes | no | no | no | no | yes | no | no | no | no | no | no | no |

| | | | | | | | | | | | | | | | | | |
|---|---|---|---|---|---|---|---|---|---|---|---|---|---|---|---|---|---|
| SCN5A | 713 | V | 0 | 0 | yes | no | no | no | no | yes | no | no | no | no | no | no | no |
| SCN5A | 714 | V | 3 | 0 | yes | no | no | yes | no | yes | no | no | no | no | no | no | no |
| SCN5A | 715 | M | 2 | 0 | no | yes | no | no | no | yes | no | no | no | no | no | no | no |
| SCN5A | 716 | D | 0 | 0 | no | yes | no | no | no | yes | no | no | no | no | no | no | no |
| SCN5A | 717 | P | 1 | 0 | yes | no | no | no | no | yes | no | no | no | no | no | no | no |
| SCN5A | 718 | F | 1 | 0 | yes | no | no | yes | no | no | no | no | no | no | no | no | no |
| SCN5A | 719 | T | 0 | 0 | yes | no | no | yes | no | no | no | no | no | no | no | no | no |
| SCN5A | 720 | D | 0 | 0 | yes | no | no | yes | no | no | no | no | no | no | no | no | no |
| SCN5A | 721 | L | 0 | 0 | yes | no | no | yes | no | no | no | no | no | no | no | no | no |
| SCN5A | 722 | T | 0 | 0 | yes | no | no | yes | no | no | no | no | no | no | no | no | no |
| SCN5A | 723 | I | 1 | 0 | yes | no | no | yes | no | no | no | no | no | no | no | no | no |
| SCN5A | 724 | T | 1 | 0 | yes | no | no | yes | no | no | no | no | no | no | no | no | no |
| SCN5A | 725 | M | 0 | 0 | yes | no | no | yes | no | no | no | no | no | no | no | no | no |
| SCN5A | 726 | C | 0 | 0 | yes | no | no | yes | no | no | no | no | no | no | no | no | no |
| SCN5A | 727 | I | 0 | 0 | yes | no | no | yes | no | no | no | no | no | no | no | no | no |
| SCN5A | 728 | V | 1 | 1 | yes | no | no | yes | no | no | no | no | no | no | no | no | no |
| SCN5A | 729 | L | 0 | 0 | yes | no | no | yes | no | no | no | no | no | no | no | no | no |
| SCN5A | 730 | N | 1 | 0 | yes | no | no | yes | no | no | no | no | no | no | no | no | no |
| SCN5A | 731 | T | 0 | 1 | yes | no | no | yes | no | no | no | no | no | no | no | no | no |
| SCN5A | 732 | L | 0 | 0 | yes | no | no | yes | no | no | no | no | no | no | no | no | no |
| SCN5A | 733 | F | 1 | 0 | yes | no | no | yes | no | no | no | no | no | no | no | no | no |
| SCN5A | 734 | M | 0 | 0 | yes | no | no | yes | no | no | no | no | no | no | no | no | no |
| SCN5A | 735 | A | 1 | 3 | yes | no | no | yes | no | no | no | no | no | no | no | no | no |
| SCN5A | 736 | L | 0 | 0 | no | yes | no | yes | no | no | no | no | no | no | no | no | no |
| SCN5A | 737 | E | 0 | 0 | no | yes | no | yes | yes | no | no | no | no | no | no | no | no |
| SCN5A | 738 | H | 0 | 0 | no | no | yes | no | yes | no | no | no | no | no | no | no | no |
| SCN5A | 739 | Y | 0 | 0 | no | yes | no | no | yes | no | no | no | no | no | no | no | no |
| SCN5A | 740 | N | 0 | 0 | no | yes | no | no | yes | no | no | no | no | no | no | no | no |
| SCN5A | 741 | M | 0 | 0 | no | yes | no | no | yes | no | no | no | no | no | no | no | no |
| SCN5A | 742 | T | 1 | 0 | no | yes | no | no | yes | no | no | no | no | no | no | no | no |
| SCN5A | 743 | S | 0 | 0 | no | yes | no | no | yes | no | no | no | no | no | no | no | no |
| SCN5A | 744 | E | 0 | 0 | no | yes | no | no | yes | no | no | no | no | no | no | no | no |
| SCN5A | 745 | F | 0 | 0 | yes | no | no | no | yes | no | no | no | no | no | no | no | no |

| | | | | | | | | | | | | | | | | | |
|---|---|---|---|---|---|---|---|---|---|---|---|---|---|---|---|---|---|
| SCN5A | 746 | E | 1 | 1 | yes | no | no | no | yes | no | no | no | no | no | no | no | no |
| SCN5A | 747 | E | 1 | 0 | yes | no | no | no | yes | no | no | no | no | no | no | no | no |
| SCN5A | 748 | M | 1 | 0 | yes | no | no | yes | no | no | no | no | no | no | no | no | no |
| SCN5A | 749 | L | 0 | 0 | yes | no | no | yes | no | no | no | no | no | no | no | no | no |
| SCN5A | 750 | Q | 0 | 1 | yes | no | no | yes | no | no | no | no | no | no | no | no | no |
| SCN5A | 751 | V | 2 | 0 | yes | no | no | yes | no | no | no | no | no | no | no | no | no |
| SCN5A | 752 | G | 1 | 1 | yes | no | no | yes | no | no | no | no | no | no | no | no | no |
| SCN5A | 753 | N | 0 | 0 | yes | no | no | yes | no | no | no | no | no | no | no | no | no |
| SCN5A | 754 | L | 0 | 0 | yes | no | no | yes | no | no | no | no | no | no | no | no | no |
| SCN5A | 755 | V | 0 | 0 | yes | no | no | yes | no | no | no | no | no | no | no | no | no |
| SCN5A | 756 | F | 0 | 0 | yes | no | no | yes | no | no | no | no | no | no | no | no | no |
| SCN5A | 757 | T | 0 | 0 | yes | no | no | yes | no | no | no | no | no | no | no | no | no |
| SCN5A | 758 | G | 0 | 1 | yes | no | no | yes | no | no | no | no | no | no | no | no | no |
| SCN5A | 759 | I | 1 | 1 | yes | no | no | yes | no | no | no | no | no | no | no | no | no |
| SCN5A | 760 | F | 0 | 0 | yes | no | no | yes | no | no | no | no | no | no | no | no | no |
| SCN5A | 761 | T | 0 | 0 | yes | no | no | yes | no | no | no | no | no | no | no | no | no |
| SCN5A | 762 | A | 0 | 0 | yes | no | no | yes | no | no | no | no | no | no | no | no | no |
| SCN5A | 763 | E | 2 | 0 | yes | no | no | yes | no | no | no | no | no | no | no | no | no |
| SCN5A | 764 | M | 0 | 1 | yes | no | no | yes | no | no | no | no | no | no | no | no | no |
| SCN5A | 765 | T | 0 | 0 | yes | no | no | yes | no | no | no | no | no | no | no | no | no |
| SCN5A | 766 | F | 0 | 0 | yes | no | no | yes | no | no | no | no | no | no | no | no | no |
| SCN5A | 767 | K | 0 | 0 | yes | no | no | yes | no | no | no | no | no | no | no | no | no |
| SCN5A | 768 | I | 0 | 0 | yes | no | no | yes | no | yes | no | no | no | no | no | no | no |
| SCN5A | 769 | I | 0 | 0 | no | yes | no | yes | no | yes | no | no | no | no | no | no | no |
| SCN5A | 770 | A | 0 | 0 | no | yes | no | no | no | yes | no | no | no | no | no | no | no |
| SCN5A | 771 | L | 0 | 0 | no | yes | no | no | no | yes | no | no | no | no | no | no | no |
| SCN5A | 772 | D | 1 | 0 | no | yes | no | no | no | yes | no | no | no | no | no | no | no |
| SCN5A | 773 | P | 0 | 1 | yes | no | no | no | no | yes | no | no | no | no | no | no | no |
| SCN5A | 774 | Y | 2 | 0 | yes | no | no | no | no | yes | no | no | no | no | no | no | no |
| SCN5A | 775 | Y | 0 | 0 | yes | no | no | no | no | yes | no | no | no | no | no | no | no |
| SCN5A | 776 | Y | 0 | 0 | yes | no | no | yes | no | yes | no | no | no | no | no | no | no |
| SCN5A | 777 | F | 0 | 1 | no | yes | no | yes | no | yes | no | no | no | no | no | no | no |
| SCN5A | 778 | Q | 0 | 0 | no | yes | no | no | no | yes | no | no | no | no | no | no | no |

| | | | | | | | | | | | | | | | | | |
|---|---|---|---|---|---|---|---|---|---|---|---|---|---|---|---|---|---|
| SCN5A | 779 | Q | 0 | 1 | no | yes | no | no | no | yes | no | no | no | no | no | no | no |
| SCN5A | 780 | G | 0 | 0 | no | yes | no | yes | no | yes | no | no | no | no | no | no | no |
| SCN5A | 781 | W | 0 | 0 | no | yes | no | yes | no | yes | no | no | no | no | no | no | no |
| SCN5A | 782 | N | 0 | 1 | yes | no | no | yes | no | no | no | no | no | no | no | no | no |
| SCN5A | 783 | I | 1 | 0 | yes | no | no | yes | no | no | no | no | no | no | no | no | no |
| SCN5A | 784 | F | 1 | 0 | yes | no | no | yes | no | no | no | no | no | no | no | no | no |
| SCN5A | 785 | D | 0 | 1 | yes | no | no | yes | no | no | no | no | no | no | no | no | no |
| SCN5A | 786 | S | 0 | 0 | yes | no | no | yes | no | no | no | no | no | no | no | no | no |
| SCN5A | 787 | I | 0 | 0 | yes | no | no | yes | no | no | no | no | no | no | no | no | no |
| SCN5A | 788 | I | 1 | 0 | yes | no | no | yes | no | no | no | no | no | no | no | no | no |
| SCN5A | 789 | V | 2 | 1 | yes | no | no | yes | no | no | no | no | no | no | no | no | no |
| SCN5A | 790 | I | 0 | 0 | yes | no | no | yes | no | no | no | no | no | no | no | no | no |
| SCN5A | 791 | L | 1 | 0 | yes | no | no | yes | no | no | no | no | no | no | no | no | no |
| SCN5A | 792 | S | 0 | 0 | yes | no | no | yes | no | no | no | no | no | no | no | no | no |
| SCN5A | 793 | L | 1 | 0 | yes | no | no | yes | no | no | no | no | no | no | no | no | no |
| SCN5A | 794 | M | 0 | 0 | yes | no | no | yes | no | no | no | no | no | no | no | no | no |
| SCN5A | 795 | E | 0 | 0 | yes | no | no | yes | no | no | no | no | no | no | no | no | no |
| SCN5A | 796 | L | 0 | 0 | yes | no | no | yes | no | no | no | no | no | no | no | no | no |
| SCN5A | 797 | G | 0 | 0 | yes | no | no | yes | no | no | no | no | no | no | no | no | no |
| SCN5A | 798 | L | 0 | 0 | no | yes | no | yes | no | no | no | no | no | no | no | no | no |
| SCN5A | 799 | S | 0 | 0 | no | yes | no | yes | no | no | no | no | no | no | no | no | no |
| SCN5A | 800 | R | 3 | 1 | no | yes | no | no | no | no | no | no | no | no | no | no | no |
| SCN5A | 801 | M | 1 | 0 | no | yes | no | no | no | no | no | no | no | no | no | no | no |
| SCN5A | 802 | S | 0 | 0 | no | yes | no | no | yes | no | no | no | no | no | no | no | no |
| SCN5A | 803 | N | 0 | 0 | no | yes | no | yes | yes | no | no | no | no | no | no | no | no |
| SCN5A | 804 | L | 0 | 0 | no | yes | no | no | no | no | no | no | no | no | no | no | no |
| SCN5A | 805 | S | 1 | 0 | no | yes | no | yes | no | no | no | no | no | no | no | no | no |
| SCN5A | 806 | V | 1 | 0 | no | yes | no | yes | no | no | no | no | no | no | no | no | no |
| SCN5A | 807 | L | 0 | 0 | no | yes | no | yes | no | no | no | no | no | no | no | no | no |
| SCN5A | 808 | R | 2 | 2 | no | yes | no | yes | no | no | no | no | no | no | no | no | no |
| SCN5A | 809 | S | 0 | 0 | yes | no | no | yes | no | no | no | no | no | no | no | no | no |
| SCN5A | 810 | F | 0 | 0 | yes | no | no | yes | no | no | no | no | no | no | no | no | no |
| SCN5A | 811 | R | 2 | 1 | yes | no | no | yes | no | no | no | no | no | no | no | no | no |

| | | | | | | | | | | | | | | | | | |
|---|---|---|---|---|---|---|---|---|---|---|---|---|---|---|---|---|---|
| SCN5A | 812 | L | 0 | 1 | yes | no | no | yes | no | no | no | no | no | no | no | no | no |
| SCN5A | 813 | L | 0 | 0 | yes | no | no | yes | no | no | no | no | no | no | no | no | no |
| SCN5A | 814 | R | 1 | 2 | yes | no | no | yes | no | no | no | no | no | no | no | no | no |
| SCN5A | 815 | V | 0 | 0 | yes | no | no | yes | no | no | no | no | no | no | no | no | no |
| SCN5A | 816 | F | 1 | 1 | yes | no | no | yes | no | no | no | no | no | no | no | no | no |
| SCN5A | 817 | K | 0 | 1 | yes | no | no | yes | no | no | no | no | no | no | no | no | no |
| SCN5A | 818 | L | 0 | 0 | yes | no | no | yes | no | no | no | no | no | no | no | no | no |
| SCN5A | 819 | A | 0 | 0 | yes | no | no | yes | no | no | no | no | no | no | no | no | no |
| SCN5A | 820 | K | 0 | 0 | no | yes | no | yes | no | no | no | no | no | no | no | no | no |
| SCN5A | 821 | S | 0 | 0 | no | yes | no | yes | no | no | no | no | no | no | no | no | no |
| SCN5A | 822 | W | 0 | 1 | no | yes | no | yes | no | yes | no | no | no | no | no | no | no |
| SCN5A | 823 | P | 1 | 0 | no | yes | no | yes | no | yes | no | no | no | no | no | no | no |
| SCN5A | 824 | T | 0 | 0 | no | yes | no | yes | no | yes | no | no | no | no | no | no | no |
| SCN5A | 825 | L | 0 | 0 | yes | no | no | yes | no | yes | no | no | no | no | no | no | no |
| SCN5A | 826 | N | 1 | 0 | yes | no | no | yes | no | yes | no | no | no | no | no | no | no |
| SCN5A | 827 | T | 0 | 0 | yes | no | no | yes | no | yes | no | no | no | no | no | no | no |
| SCN5A | 828 | L | 0 | 1 | yes | no | no | yes | no | yes | no | no | no | no | no | no | no |
| SCN5A | 829 | I | 0 | 0 | yes | no | no | yes | no | yes | no | no | no | no | no | no | no |
| SCN5A | 830 | K | 0 | 0 | yes | no | no | yes | no | yes | no | no | no | no | no | no | no |
| SCN5A | 831 | I | 0 | 0 | yes | no | no | yes | no | yes | no | no | no | no | no | no | no |
| SCN5A | 832 | I | 0 | 0 | yes | no | no | yes | no | yes | no | no | no | no | no | no | no |
| SCN5A | 833 | G | 1 | 1 | yes | no | no | yes | no | yes | no | no | no | no | no | no | no |
| SCN5A | 834 | N | 0 | 1 | yes | no | no | yes | no | yes | no | no | no | no | no | no | no |
| SCN5A | 835 | S | 1 | 1 | yes | no | no | yes | no | yes | no | no | no | no | no | no | no |
| SCN5A | 836 | V | 1 | 0 | yes | no | no | yes | no | yes | no | no | no | no | no | no | no |
| SCN5A | 837 | G | 0 | 0 | yes | no | no | yes | no | yes | no | no | no | no | no | no | no |
| SCN5A | 838 | A | 0 | 0 | yes | no | no | yes | no | no | no | no | no | no | no | no | no |
| SCN5A | 839 | L | 0 | 1 | no | yes | no | yes | no | no | no | no | no | no | no | no | no |
| SCN5A | 840 | G | 0 | 1 | no | yes | no | yes | no | no | no | no | no | no | no | no | no |
| SCN5A | 841 | N | 1 | 0 | no | yes | no | yes | no | no | no | no | no | no | no | no | no |
| SCN5A | 842 | L | 0 | 0 | yes | no | no | yes | no | no | no | no | no | no | no | no | no |
| SCN5A | 843 | T | 0 | 1 | yes | no | no | yes | no | no | no | no | no | no | no | no | no |
| SCN5A | 844 | L | 0 | 0 | yes | no | no | yes | no | no | no | no | no | no | no | no | no |

| | | | | | | | | | | | | | | | | | |
|---|---|---|---|---|---|---|---|---|---|---|---|---|---|---|---|---|---|
| SCN5A | 845 | V | 0 | 0 | yes | no | no | yes | no | no | no | no | no | no | no | no | no |
| SCN5A | 846 | L | 0 | 1 | yes | no | no | yes | no | no | no | no | no | no | no | no | no |
| SCN5A | 847 | A | 0 | 0 | yes | no | no | yes | no | no | no | no | no | no | no | no | no |
| SCN5A | 848 | I | 1 | 1 | yes | no | no | yes | no | no | no | no | no | no | no | no | no |
| SCN5A | 849 | I | 0 | 1 | yes | no | no | yes | no | no | no | no | no | no | no | no | no |
| SCN5A | 850 | V | 1 | 0 | yes | no | no | yes | no | no | no | no | no | no | no | no | no |
| SCN5A | 851 | F | 1 | 1 | yes | no | no | yes | no | no | no | no | no | no | no | no | no |
| SCN5A | 852 | I | 0 | 0 | yes | no | no | yes | no | no | no | no | no | no | no | no | no |
| SCN5A | 853 | F | 0 | 0 | yes | no | no | yes | no | no | no | no | no | no | no | no | no |
| SCN5A | 854 | A | 0 | 0 | yes | no | no | yes | no | no | no | no | no | no | no | no | no |
| SCN5A | 855 | V | 0 | 0 | yes | no | no | yes | no | no | no | no | no | no | no | no | no |
| SCN5A | 856 | V | 1 | 0 | yes | no | no | yes | no | no | no | no | no | no | no | no | no |
| SCN5A | 857 | G | 1 | 0 | yes | no | no | yes | yes | no | no | no | no | no | no | no | no |
| SCN5A | 858 | M | 1 | 1 | yes | no | no | yes | yes | no | no | no | no | no | no | no | no |
| SCN5A | 859 | Q | 0 | 0 | yes | no | no | yes | yes | no | no | no | no | no | no | no | no |
| SCN5A | 860 | L | 0 | 0 | yes | no | no | yes | yes | no | no | no | no | no | no | no | no |
| SCN5A | 861 | F | 0 | 0 | yes | no | no | no | yes | no | no | no | no | no | no | no | no |
| SCN5A | 862 | G | 0 | 0 | yes | no | no | no | yes | no | no | no | no | no | no | no | no |
| SCN5A | 863 | K | 0 | 0 | yes | no | no | no | yes | no | no | no | no | no | no | no | no |
| SCN5A | 864 | N | 0 | 0 | yes | no | no | no | yes | no | no | no | no | no | no | no | no |
| SCN5A | 865 | Y | 0 | 0 | yes | no | no | no | yes | no | no | no | no | no | no | no | no |
| SCN5A | 866 | S | 2 | 0 | yes | no | no | no | yes | no | no | no | no | no | no | no | no |
| SCN5A | 867 | E | 1 | 1 | yes | no | no | no | yes | no | no | no | no | no | no | no | no |
| SCN5A | 868 | L | 0 | 0 | yes | no | no | no | yes | no | no | no | no | no | no | no | no |
| SCN5A | 869 | R | 1 | 0 | yes | no | no | no | yes | no | no | no | no | no | no | no | no |
| SCN5A | 870 | D | 0 | 0 | no | yes | no | no | yes | no | yes | no | no | no | no | no | no |
| SCN5A | 871 | S | 0 | 0 | no | yes | no | no | yes | no | yes | no | no | no | no | no | no |
| SCN5A | 872 | D | 1 | 0 | no | yes | no | no | yes | no | yes | no | no | no | no | no | no |
| SCN5A | 873 | S | 1 | 0 | no | yes | no | no | yes | no | no | no | no | no | no | no | no |
| SCN5A | 874 | G | 1 | 0 | no | yes | no | no | yes | no | no | no | no | no | no | no | no |
| SCN5A | 875 | L | 0 | 0 | no | yes | no | no | yes | no | no | no | no | no | no | no | no |
| SCN5A | 876 | L | 0 | 0 | no | yes | no | no | yes | no | no | no | no | no | no | no | no |
| SCN5A | 877 | P | 0 | 0 | no | yes | no | no | yes | no | yes | no | no | no | no | no | no |

| | | | | | | | | | | | | | | | | | |
|---|---|---|---|---|---|---|---|---|---|---|---|---|---|---|---|---|---|
| SCN5A | 878 | R | 0 | 2 | no | yes | no | no | yes | no | yes | no | no | no | no | no | no |
| SCN5A | 879 | W | 0 | 1 | no | yes | no | no | yes | no | no | no | no | no | no | no | no |
| SCN5A | 880 | H | 0 | 0 | no | yes | no | no | yes | no | no | no | no | no | no | no | no |
| SCN5A | 881 | M | 0 | 0 | no | yes | no | no | yes | no | no | no | no | no | no | no | no |
| SCN5A | 882 | M | 0 | 0 | no | yes | no | no | yes | no | no | no | no | no | no | no | no |
| SCN5A | 883 | D | 0 | 0 | no | yes | no | no | yes | no | no | no | no | no | no | no | no |
| SCN5A | 884 | F | 0 | 0 | no | yes | no | yes | no | no | no | yes | no | no | no | no | no |
| SCN5A | 885 | F | 0 | 0 | yes | no | no | yes | no | no | no | yes | no | no | no | no | no |
| SCN5A | 886 | H | 1 | 1 | yes | no | no | no | no | no | no | yes | no | no | no | no | no |
| SCN5A | 887 | A | 0 | 0 | yes | no | no | yes | no | no | no | yes | no | no | no | no | no |
| SCN5A | 888 | F | 0 | 0 | yes | no | no | yes | no | no | no | yes | no | no | no | no | no |
| SCN5A | 889 | L | 0 | 0 | yes | no | no | yes | no | no | no | yes | no | no | no | no | no |
| SCN5A | 890 | I | 0 | 1 | yes | no | no | yes | no | no | no | yes | no | no | no | no | no |
| SCN5A | 891 | I | 2 | 0 | yes | no | no | yes | no | no | no | yes | no | no | no | no | no |
| SCN5A | 892 | F | 0 | 1 | yes | no | no | yes | no | no | no | yes | no | no | no | no | no |
| SCN5A | 893 | R | 2 | 2 | yes | no | no | yes | no | no | yes | yes | no | no | no | no | no |
| SCN5A | 894 | I | 1 | 0 | yes | no | no | yes | no | no | no | yes | no | no | no | no | no |
| SCN5A | 895 | L | 1 | 0 | yes | no | no | yes | no | no | no | yes | no | no | no | no | no |
| SCN5A | 896 | C | 0 | 2 | yes | no | no | yes | no | no | yes | yes | no | no | no | no | no |
| SCN5A | 897 | G | 0 | 1 | no | yes | no | yes | no | no | yes | yes | no | no | no | no | no |
| SCN5A | 898 | E | 0 | 0 | no | yes | no | yes | no | no | yes | yes | no | no | no | no | no |
| SCN5A | 899 | W | 0 | 0 | no | yes | no | yes | no | no | no | yes | no | no | no | no | no |
| SCN5A | 900 | I | 0 | 0 | no | yes | no | yes | no | no | no | yes | no | no | no | no | no |
| SCN5A | 901 | E | 0 | 1 | yes | no | no | no | no | no | yes | yes | no | no | no | no | no |
| SCN5A | 902 | T | 0 | 0 | yes | no | no | no | no | no | no | yes | no | no | no | no | no |
| SCN5A | 903 | M | 0 | 0 | yes | no | no | no | no | no | no | yes | no | no | no | no | no |
| SCN5A | 904 | W | 0 | 0 | yes | no | no | no | no | no | no | yes | no | no | no | no | no |
| SCN5A | 905 | D | 0 | 0 | yes | no | no | no | yes | no | yes | no | no | no | no | no | no |
| SCN5A | 906 | C | 0 | 0 | yes | no | no | no | yes | no | no | no | no | no | no | no | no |
| SCN5A | 907 | M | 0 | 0 | yes | no | no | no | yes | no | no | no | no | no | no | no | no |
| SCN5A | 908 | E | 0 | 0 | yes | no | no | no | yes | no | no | no | no | no | no | no | no |
| SCN5A | 909 | V | 0 | 0 | yes | no | no | no | yes | no | no | no | no | no | no | no | no |
| SCN5A | 910 | S | 1 | 1 | yes | no | no | no | yes | no | no | no | no | no | no | no | no |

| | | | | | | | | | | | | | | | | | |
|---|---|---|---|---|---|---|---|---|---|---|---|---|---|---|---|---|---|
| SCN5A | 911 | G | 1 | 0 | no | yes | no | no | yes | no | no | no | no | no | no | no | no |
| SCN5A | 912 | Q | 1 | 1 | yes | no | no | no | yes | no | no | no | no | no | no | no | no |
| SCN5A | 913 | S | 0 | 0 | yes | no | no | yes | yes | no | no | no | no | no | no | no | no |
| SCN5A | 914 | L | 0 | 0 | yes | no | no | yes | yes | no | no | no | no | no | no | no | no |
| SCN5A | 915 | C | 0 | 1 | yes | no | no | yes | yes | no | no | no | no | no | no | no | no |
| SCN5A | 916 | L | 0 | 0 | yes | no | no | yes | yes | no | no | no | no | no | no | no | no |
| SCN5A | 917 | L | 1 | 1 | yes | no | no | yes | yes | no | no | no | no | no | no | no | no |
| SCN5A | 918 | V | 0 | 0 | yes | no | no | yes | no | no | no | no | no | no | no | no | no |
| SCN5A | 919 | F | 0 | 1 | yes | no | no | yes | no | no | no | no | no | no | no | no | no |
| SCN5A | 920 | L | 0 | 0 | yes | no | no | yes | no | no | no | no | no | no | no | no | no |
| SCN5A | 921 | L | 0 | 0 | yes | no | no | yes | no | no | no | no | no | no | no | no | no |
| SCN5A | 922 | V | 1 | 0 | yes | no | no | yes | no | no | no | no | no | no | no | no | no |
| SCN5A | 923 | M | 0 | 0 | yes | no | no | yes | no | no | no | no | no | no | no | no | no |
| SCN5A | 924 | V | 1 | 0 | yes | no | no | yes | no | no | no | no | no | no | no | no | no |
| SCN5A | 925 | I | 1 | 0 | yes | no | no | yes | no | no | no | no | no | no | no | no | no |
| SCN5A | 926 | G | 0 | 0 | yes | no | no | yes | no | no | no | no | no | no | no | no | no |
| SCN5A | 927 | N | 1 | 1 | yes | no | no | yes | no | no | no | no | no | no | no | no | no |
| SCN5A | 928 | L | 0 | 1 | yes | no | no | yes | no | no | no | no | no | no | no | no | no |
| SCN5A | 929 | V | 0 | 0 | yes | no | no | yes | no | no | no | no | no | no | no | no | no |
| SCN5A | 930 | V | 0 | 0 | yes | no | no | yes | no | no | no | no | no | no | no | no | no |
| SCN5A | 931 | L | 0 | 0 | yes | no | no | yes | no | no | no | no | no | no | no | no | no |
| SCN5A | 932 | N | 0 | 0 | yes | no | no | yes | no | no | no | no | no | no | no | no | no |
| SCN5A | 933 | L | 0 | 0 | yes | no | no | yes | no | no | no | no | no | no | no | no | no |
| SCN5A | 934 | F | 0 | 0 | yes | no | no | yes | no | no | yes | no | no | no | no | no | no |
| SCN5A | 935 | L | 0 | 1 | yes | no | no | yes | no | no | yes | no | no | no | no | no | no |
| SCN5A | 936 | A | 0 | 0 | yes | no | no | yes | no | no | no | no | no | no | no | no | no |
| SCN5A | 937 | L | 0 | 0 | yes | no | no | yes | no | no | no | no | no | no | no | no | no |
| SCN5A | 938 | L | 0 | 0 | yes | no | no | yes | no | no | yes | no | no | no | no | no | no |
| SCN5A | 939 | L | 1 | 0 | yes | no | no | no | no | yes | yes | no | no | no | no | no | no |
| SCN5A | 940 | S | 1 | 0 | yes | no | no | no | no | yes | no | no | no | no | no | no | no |
| SCN5A | 941 | S | 0 | 1 | yes | no | no | yes | no | yes | no | no | no | no | no | no | no |
| SCN5A | 942 | F | 0 | 0 | yes | no | no | no | no | yes | yes | no | no | no | no | no | no |
| SCN5A | 943 | S | 1 | 0 | yes | no | no | no | no | yes | no | no | no | no | no | no | no |

| | | | | | | | | | | | | | | | | | |
|---|---|---|---|---|---|---|---|---|---|---|---|---|---|---|---|---|---|
| SCN5A | 944 | A | 0 | 0 | no | yes | no | no | no | yes | no | no | no | no | no | no | no |
| SCN5A | 1188 | G | 0 | 0 | no | yes | no | no | no | yes | no | no | no | no | no | no | no |
| SCN5A | 1189 | K | 1 | 0 | no | yes | no | no | no | yes | no | no | no | no | no | no | no |
| SCN5A | 1190 | V | 1 | 0 | no | yes | no | no | no | yes | no | no | no | no | no | no | no |
| SCN5A | 1191 | W | 0 | 0 | yes | no | no | no | no | yes | no | no | no | no | no | no | no |
| SCN5A | 1192 | W | 0 | 0 | yes | no | no | no | no | yes | no | no | no | no | no | no | no |
| SCN5A | 1193 | R | 2 | 0 | yes | no | no | no | no | yes | no | no | no | no | no | no | no |
| SCN5A | 1194 | L | 0 | 0 | yes | no | no | yes | no | yes | no | no | no | no | no | no | no |
| SCN5A | 1195 | R | 1 | 1 | yes | no | no | no | no | yes | no | no | no | no | no | no | no |
| SCN5A | 1196 | K | 0 | 0 | yes | no | no | no | no | yes | no | no | no | no | no | no | no |
| SCN5A | 1197 | T | 0 | 0 | yes | no | no | yes | no | yes | no | no | no | no | no | no | no |
| SCN5A | 1198 | C | 0 | 0 | yes | no | no | yes | no | yes | no | no | no | no | no | no | no |
| SCN5A | 1199 | Y | 0 | 1 | yes | no | no | no | no | yes | no | no | no | no | no | no | no |
| SCN5A | 1200 | H | 1 | 0 | yes | no | no | no | no | yes | no | no | no | no | no | no | no |
| SCN5A | 1201 | I | 1 | 0 | yes | no | no | yes | no | yes | no | no | no | no | no | no | no |
| SCN5A | 1202 | V | 1 | 0 | yes | no | no | yes | no | yes | no | no | no | no | no | no | no |
| SCN5A | 1203 | E | 0 | 0 | yes | no | no | no | no | yes | no | no | no | no | no | no | no |
| SCN5A | 1204 | H | 0 | 0 | no | yes | no | yes | no | yes | no | no | no | no | no | no | no |
| SCN5A | 1205 | S | 0 | 0 | yes | no | no | yes | no | yes | no | no | no | no | no | no | no |
| SCN5A | 1206 | W | 0 | 1 | yes | no | no | yes | no | yes | no | no | no | no | no | no | no |
| SCN5A | 1207 | F | 0 | 0 | yes | no | no | yes | no | no | no | no | no | no | no | no | no |
| SCN5A | 1208 | E | 1 | 1 | yes | no | no | yes | no | no | no | no | no | no | no | no | no |
| SCN5A | 1209 | T | 0 | 0 | yes | no | no | yes | no | no | no | no | no | no | no | no | no |
| SCN5A | 1210 | F | 0 | 0 | yes | no | no | yes | no | no | no | no | no | no | no | no | no |
| SCN5A | 1211 | I | 0 | 0 | yes | no | no | yes | no | no | no | no | no | no | no | no | no |
| SCN5A | 1212 | I | 0 | 0 | yes | no | no | yes | no | no | no | no | no | no | no | no | no |
| SCN5A | 1213 | F | 0 | 0 | yes | no | no | yes | no | no | no | no | no | no | no | no | no |
| SCN5A | 1214 | M | 1 | 0 | yes | no | no | yes | no | no | no | no | no | no | no | no | no |
| SCN5A | 1215 | I | 1 | 0 | yes | no | no | yes | no | no | no | no | no | no | no | no | no |
| SCN5A | 1216 | L | 1 | 0 | yes | no | no | yes | no | no | no | no | no | no | no | no | no |
| SCN5A | 1217 | L | 0 | 1 | yes | no | no | yes | no | no | no | no | no | no | no | no | no |
| SCN5A | 1218 | S | 1 | 1 | yes | no | no | yes | no | no | no | no | no | no | no | no | no |
| SCN5A | 1219 | S | 0 | 1 | yes | no | no | yes | no | no | no | no | no | no | no | no | no |

| | | | | | | | | | | | | | | | | | |
|---|---|---|---|---|---|---|---|---|---|---|---|---|---|---|---|---|---|
| SCN5A | 1220 | G | 1 | 0 | yes | no | no | yes | no | no | no | no | no | no | no | no | no |
| SCN5A | 1221 | A | 1 | 0 | no | yes | no | yes | no | no | no | no | no | no | no | no | no |
| SCN5A | 1222 | L | 0 | 0 | no | yes | no | yes | no | no | no | no | no | no | no | no | no |
| SCN5A | 1223 | A | 0 | 0 | no | yes | no | yes | no | no | no | no | no | no | no | no | no |
| SCN5A | 1224 | F | 0 | 0 | no | yes | no | no | no | no | no | no | no | no | no | no | no |
| SCN5A | 1225 | E | 1 | 1 | no | yes | no | no | yes | no | no | no | no | no | no | no | no |
| SCN5A | 1226 | D | 0 | 0 | no | yes | no | no | yes | no | no | no | no | no | no | no | no |
| SCN5A | 1227 | I | 0 | 0 | no | yes | no | no | yes | no | no | no | no | no | no | no | no |
| SCN5A | 1228 | Y | 2 | 1 | no | yes | no | no | yes | no | no | no | no | no | no | no | no |
| SCN5A | 1229 | L | 0 | 0 | no | yes | no | no | yes | no | no | no | no | no | no | no | no |
| SCN5A | 1230 | E | 1 | 0 | no | yes | no | no | yes | no | no | no | no | no | no | no | no |
| SCN5A | 1231 | E | 1 | 1 | no | yes | no | no | yes | no | no | no | no | no | no | no | no |
| SCN5A | 1232 | R | 1 | 1 | no | yes | no | no | yes | no | no | no | no | no | no | no | no |
| SCN5A | 1233 | K | 1 | 0 | no | yes | no | no | yes | no | no | no | no | no | no | no | no |
| SCN5A | 1234 | T | 0 | 0 | no | yes | no | no | yes | no | no | no | no | no | no | no | no |
| SCN5A | 1235 | I | 0 | 0 | yes | no | no | no | yes | no | no | no | no | no | no | no | no |
| SCN5A | 1236 | K | 1 | 1 | yes | no | no | no | yes | no | no | no | no | no | no | no | no |
| SCN5A | 1237 | V | 1 | 0 | yes | no | no | no | yes | no | no | no | no | no | no | no | no |
| SCN5A | 1238 | L | 0 | 0 | yes | no | no | no | no | no | no | no | no | no | no | no | no |
| SCN5A | 1239 | L | 1 | 1 | yes | no | no | no | no | no | no | no | no | no | no | no | no |
| SCN5A | 1240 | E | 1 | 0 | yes | no | no | no | no | no | no | no | no | no | no | no | no |
| SCN5A | 1241 | Y | 0 | 1 | yes | no | no | no | no | no | no | no | no | no | no | no | no |
| SCN5A | 1242 | A | 0 | 0 | yes | no | no | yes | no | no | no | no | no | no | no | no | no |
| SCN5A | 1243 | D | 1 | 1 | yes | no | no | yes | no | no | no | no | no | no | no | no | no |
| SCN5A | 1244 | K | 1 | 0 | yes | no | no | yes | no | no | no | no | no | no | no | no | no |
| SCN5A | 1245 | M | 1 | 0 | yes | no | no | yes | no | no | no | no | no | no | no | no | no |
| SCN5A | 1246 | F | 0 | 0 | yes | no | no | yes | no | no | no | no | no | no | no | no | no |
| SCN5A | 1247 | T | 1 | 0 | yes | no | no | yes | no | no | no | no | no | no | no | no | no |
| SCN5A | 1248 | Y | 0 | 0 | yes | no | no | yes | no | no | no | no | no | no | no | no | no |
| SCN5A | 1249 | V | 0 | 1 | yes | no | no | yes | no | no | no | no | no | no | no | no | no |
| SCN5A | 1250 | F | 0 | 1 | yes | no | no | yes | no | no | no | no | no | no | no | no | no |
| SCN5A | 1251 | V | 1 | 0 | yes | no | no | yes | no | no | no | no | no | no | no | no | no |
| SCN5A | 1252 | L | 0 | 0 | yes | no | no | yes | no | no | no | no | no | no | no | no | no |

| | | | | | | | | | | | | | | | | | |
|---|---|---|---|---|---|---|---|---|---|---|---|---|---|---|---|---|---|
| SCN5A | 1253 | E | 0 | 1 | yes | no | no | yes | no | no | no | no | no | no | no | no | no |
| SCN5A | 1254 | M | 0 | 0 | yes | no | no | yes | no | no | no | no | no | no | no | no | no |
| SCN5A | 1255 | L | 1 | 0 | yes | no | no | yes | no | no | no | no | no | no | no | no | no |
| SCN5A | 1256 | L | 0 | 0 | yes | no | no | yes | no | no | no | no | no | no | no | no | no |
| SCN5A | 1257 | K | 0 | 0 | yes | no | no | yes | no | yes | no | no | no | no | no | no | no |
| SCN5A | 1258 | W | 0 | 0 | yes | no | no | yes | no | yes | no | no | no | no | no | no | no |
| SCN5A | 1259 | V | 0 | 0 | yes | no | no | yes | no | yes | no | no | no | no | no | no | no |
| SCN5A | 1260 | A | 0 | 0 | yes | no | no | yes | no | yes | no | no | no | no | no | no | no |
| SCN5A | 1261 | Y | 0 | 0 | yes | no | no | yes | no | yes | no | no | no | no | no | no | no |
| SCN5A | 1262 | G | 1 | 1 | no | yes | no | yes | no | yes | no | no | no | no | no | no | no |
| SCN5A | 1263 | F | 0 | 0 | no | yes | no | yes | no | yes | no | no | no | no | no | no | no |
| SCN5A | 1264 | K | 2 | 0 | yes | no | no | no | no | yes | no | no | no | no | no | no | no |
| SCN5A | 1265 | K | 0 | 0 | yes | no | no | no | no | yes | no | no | no | no | no | no | no |
| SCN5A | 1266 | Y | 0 | 0 | yes | no | no | yes | no | yes | no | no | no | no | no | no | no |
| SCN5A | 1267 | F | 0 | 0 | yes | no | no | yes | no | yes | no | no | no | no | no | no | no |
| SCN5A | 1268 | T | 2 | 0 | no | yes | no | yes | no | yes | no | no | no | no | no | no | no |
| SCN5A | 1269 | N | 1 | 1 | no | yes | no | yes | no | yes | no | no | no | no | no | no | no |
| SCN5A | 1270 | A | 0 | 0 | no | yes | no | yes | no | yes | no | no | no | no | no | no | no |
| SCN5A | 1271 | W | 0 | 1 | no | yes | no | yes | no | no | no | no | no | no | no | no | no |
| SCN5A | 1272 | C | 0 | 0 | yes | no | no | yes | no | no | no | no | no | no | no | no | no |
| SCN5A | 1273 | W | 1 | 0 | yes | no | no | yes | no | no | no | no | no | no | no | no | no |
| SCN5A | 1274 | L | 0 | 0 | yes | no | no | yes | no | no | no | no | no | no | no | no | no |
| SCN5A | 1275 | D | 1 | 1 | yes | no | no | yes | no | no | no | no | no | no | no | no | no |
| SCN5A | 1276 | F | 0 | 0 | yes | no | no | yes | no | no | no | no | no | no | no | no | no |
| SCN5A | 1277 | L | 0 | 0 | yes | no | no | yes | no | no | no | no | no | no | no | no | no |
| SCN5A | 1278 | I | 0 | 1 | yes | no | no | yes | no | no | no | no | no | no | no | no | no |
| SCN5A | 1279 | V | 1 | 0 | yes | no | no | yes | no | no | no | no | no | no | no | no | no |
| SCN5A | 1280 | D | 0 | 0 | yes | no | no | yes | no | no | no | no | no | no | no | no | no |
| SCN5A | 1281 | V | 0 | 0 | yes | no | no | yes | no | no | no | no | no | no | no | no | no |
| SCN5A | 1282 | S | 1 | 0 | yes | no | no | yes | no | no | no | no | no | no | no | no | no |
| SCN5A | 1283 | L | 0 | 1 | yes | no | no | yes | no | no | no | no | no | no | no | no | no |
| SCN5A | 1284 | V | 0 | 0 | yes | no | no | yes | no | no | no | no | no | no | no | no | no |
| SCN5A | 1285 | S | 0 | 1 | yes | no | no | yes | no | no | no | no | no | no | no | no | no |

| | | | | | | | | | | | | | | | | | |
|---|---|---|---|---|---|---|---|---|---|---|---|---|---|---|---|---|---|
| SCN5A | 1286 | L | 0 | 0 | yes | no | no | yes | no | no | no | no | no | no | no | no | no |
| SCN5A | 1287 | V | 0 | 0 | yes | no | no | yes | no | no | no | no | no | no | no | no | no |
| SCN5A | 1288 | A | 0 | 1 | yes | no | no | yes | no | no | no | no | no | no | no | no | no |
| SCN5A | 1289 | N | 0 | 0 | yes | no | no | no | no | no | no | no | no | no | no | no | no |
| SCN5A | 1290 | T | 0 | 0 | yes | no | no | no | yes | no | no | no | no | no | no | no | no |
| SCN5A | 1291 | L | 0 | 0 | yes | no | no | yes | yes | no | no | no | no | no | no | no | no |
| SCN5A | 1292 | G | 0 | 0 | yes | no | no | no | yes | no | no | no | no | no | no | no | no |
| SCN5A | 1293 | F | 1 | 0 | no | yes | no | no | yes | no | no | no | no | no | no | no | no |
| SCN5A | 1294 | A | 1 | 1 | no | yes | no | no | yes | no | no | no | no | no | no | no | no |
| SCN5A | 1295 | E | 1 | 1 | no | yes | no | no | yes | no | no | no | no | no | no | no | no |
| SCN5A | 1296 | M | 1 | 0 | no | yes | no | no | yes | no | no | no | no | no | no | no | no |
| SCN5A | 1297 | G | 0 | 0 | no | yes | no | no | yes | no | no | no | no | no | no | no | no |
| SCN5A | 1298 | P | 0 | 1 | no | yes | no | no | no | no | no | no | no | no | no | no | no |
| SCN5A | 1299 | I | 0 | 0 | no | yes | no | no | no | no | no | no | no | no | no | no | no |
| SCN5A | 1300 | K | 0 | 0 | yes | no | no | no | no | no | no | no | no | no | no | no | no |
| SCN5A | 1301 | S | 0 | 0 | yes | no | no | yes | no | no | no | no | no | no | no | no | no |
| SCN5A | 1302 | L | 0 | 0 | yes | no | no | yes | no | no | no | no | no | no | no | no | no |
| SCN5A | 1303 | R | 2 | 1 | yes | no | no | no | no | no | no | no | no | no | no | no | no |
| SCN5A | 1304 | T | 1 | 1 | no | yes | no | yes | no | no | no | no | no | no | no | no | no |
| SCN5A | 1305 | L | 0 | 0 | no | yes | no | yes | no | no | no | no | no | no | no | no | no |
| SCN5A | 1306 | R | 2 | 0 | yes | no | no | yes | no | no | no | no | no | no | no | no | no |
| SCN5A | 1307 | A | 0 | 0 | yes | no | no | yes | no | no | no | no | no | no | no | no | no |
| SCN5A | 1308 | L | 1 | 0 | yes | no | no | yes | no | no | no | no | no | no | no | no | no |
| SCN5A | 1309 | R | 2 | 1 | yes | no | no | yes | no | no | no | no | no | no | no | no | no |
| SCN5A | 1310 | P | 0 | 0 | yes | no | no | yes | no | no | no | no | no | no | no | no | no |
| SCN5A | 1311 | L | 0 | 1 | yes | no | no | yes | no | no | no | no | no | no | no | no | no |
| SCN5A | 1312 | R | 0 | 0 | yes | no | no | yes | no | no | no | no | no | no | no | no | no |
| SCN5A | 1313 | A | 0 | 0 | yes | no | no | yes | no | no | no | no | no | no | no | no | no |
| SCN5A | 1314 | L | 0 | 0 | no | yes | no | yes | no | no | no | no | no | no | no | no | no |
| SCN5A | 1315 | S | 0 | 0 | no | yes | no | yes | no | no | no | no | no | no | no | no | no |
| SCN5A | 1316 | R | 1 | 1 | no | yes | no | yes | no | no | no | no | no | no | no | no | no |
| SCN5A | 1317 | F | 1 | 0 | no | yes | no | yes | no | yes | no | no | no | no | no | no | no |
| SCN5A | 1318 | E | 0 | 0 | yes | no | no | yes | no | yes | no | no | no | no | no | no | no |

| | | | | | | | | | | | | | | | | | |
|---|---|---|---|---|---|---|---|---|---|---|---|---|---|---|---|---|---|
| SCN5A | 1319 | G | 1 | 1 | yes | no | no | yes | no | yes | no | no | no | no | no | no | no |
| SCN5A | 1320 | M | 1 | 0 | yes | no | no | yes | no | yes | no | no | no | no | no | no | no |
| SCN5A | 1321 | R | 1 | 0 | yes | no | no | yes | no | yes | no | no | no | no | no | no | no |
| SCN5A | 1322 | V | 0 | 0 | yes | no | no | yes | no | yes | no | no | no | no | no | no | no |
| SCN5A | 1323 | V | 0 | 1 | yes | no | no | yes | no | yes | no | no | no | no | no | no | no |
| SCN5A | 1324 | V | 0 | 0 | yes | no | no | yes | no | yes | no | no | no | no | no | no | no |
| SCN5A | 1325 | N | 0 | 1 | yes | no | no | yes | no | yes | no | no | no | no | no | no | no |
| SCN5A | 1326 | A | 0 | 1 | yes | no | no | yes | no | yes | no | no | no | no | no | no | no |
| SCN5A | 1327 | L | 0 | 0 | yes | no | no | yes | no | yes | no | no | no | no | no | no | no |
| SCN5A | 1328 | V | 0 | 0 | yes | no | no | yes | no | yes | no | no | no | no | no | no | no |
| SCN5A | 1329 | G | 1 | 1 | yes | no | no | yes | no | yes | no | no | no | no | no | no | no |
| SCN5A | 1330 | A | 0 | 2 | no | yes | no | yes | no | yes | no | no | no | no | no | no | no |
| SCN5A | 1331 | I | 1 | 0 | no | yes | no | yes | no | yes | no | no | no | no | no | no | no |
| SCN5A | 1332 | P | 0 | 3 | yes | no | no | yes | no | yes | no | no | no | no | no | no | no |
| SCN5A | 1333 | S | 0 | 1 | yes | no | no | yes | no | yes | no | no | no | no | no | no | no |
| SCN5A | 1334 | I | 1 | 1 | yes | no | no | yes | no | no | no | no | no | no | no | no | no |
| SCN5A | 1335 | M | 1 | 0 | yes | no | no | yes | no | no | no | no | no | no | no | no | no |
| SCN5A | 1336 | N | 0 | 0 | yes | no | no | yes | no | no | no | no | no | no | no | no | no |
| SCN5A | 1337 | V | 0 | 0 | yes | no | no | yes | no | no | no | no | no | no | no | no | no |
| SCN5A | 1338 | L | 1 | 1 | yes | no | no | yes | no | no | no | no | no | no | no | no | no |
| SCN5A | 1339 | L | 1 | 0 | yes | no | no | yes | no | no | no | no | no | no | no | no | no |
| SCN5A | 1340 | V | 1 | 1 | yes | no | no | yes | no | no | no | no | no | no | no | no | no |
| SCN5A | 1341 | C | 0 | 0 | yes | no | no | yes | no | no | no | no | no | no | no | no | no |
| SCN5A | 1342 | L | 0 | 0 | yes | no | no | yes | no | no | no | no | no | no | no | no | no |
| SCN5A | 1343 | I | 0 | 1 | yes | no | no | yes | no | no | no | no | no | no | no | no | no |
| SCN5A | 1344 | F | 0 | 2 | yes | no | no | yes | no | no | no | no | no | no | no | no | no |
| SCN5A | 1345 | W | 0 | 1 | yes | no | no | yes | no | no | no | no | no | no | no | no | no |
| SCN5A | 1346 | L | 0 | 2 | yes | no | no | yes | no | no | no | no | no | no | no | no | no |
| SCN5A | 1347 | I | 0 | 0 | yes | no | no | yes | no | no | no | no | no | no | no | no | no |
| SCN5A | 1348 | F | 1 | 0 | yes | no | no | yes | no | no | no | no | no | no | no | no | no |
| SCN5A | 1349 | S | 0 | 0 | yes | no | no | yes | no | no | no | no | no | no | no | no | no |
| SCN5A | 1350 | I | 1 | 1 | yes | no | no | yes | no | no | no | no | no | no | no | no | no |
| SCN5A | 1351 | M | 1 | 1 | yes | no | no | yes | no | no | no | no | no | no | no | no | no |

| | | | | | | | | | | | | | | | | | |
|---|---|---|---|---|---|---|---|---|---|---|---|---|---|---|---|---|---|
| SCN5A | 1352 | G | 0 | 0 | yes | no | no | yes | no | no | no | no | no | no | no | no | no |
| SCN5A | 1353 | V | 1 | 1 | yes | no | no | yes | no | no | no | no | no | no | no | no | no |
| SCN5A | 1354 | N | 0 | 1 | yes | no | no | yes | yes | no | no | no | no | no | no | no | no |
| SCN5A | 1355 | L | 0 | 0 | yes | no | no | no | yes | no | no | no | no | no | no | no | no |
| SCN5A | 1356 | F | 0 | 0 | yes | no | no | no | yes | no | no | no | no | no | no | no | no |
| SCN5A | 1357 | A | 1 | 1 | no | yes | no | no | yes | no | no | no | no | no | no | no | no |
| SCN5A | 1358 | G | 1 | 1 | no | yes | no | no | yes | no | no | no | no | no | no | no | no |
| SCN5A | 1359 | K | 1 | 1 | no | yes | no | no | yes | no | no | no | no | no | no | no | no |
| SCN5A | 1360 | F | 0 | 1 | no | yes | no | no | yes | no | no | no | no | no | no | no | no |
| SCN5A | 1361 | G | 0 | 0 | no | no | yes | no | yes | no | no | no | no | no | no | no | no |
| SCN5A | 1362 | R | 1 | 0 | no | no | yes | no | yes | no | no | no | no | no | no | no | no |
| SCN5A | 1363 | C | 0 | 1 | no | yes | no | no | yes | no | no | no | no | no | no | no | no |
| SCN5A | 1364 | I | 1 | 0 | no | yes | no | no | yes | no | no | no | no | no | no | no | no |
| SCN5A | 1365 | N | 0 | 1 | no | no | yes | no | yes | no | no | no | no | no | no | no | no |
| SCN5A | 1366 | Q | 2 | 0 | no | yes | no | no | yes | no | no | no | no | no | no | no | no |
| SCN5A | 1367 | T | 0 | 0 | no | yes | no | no | yes | no | no | no | no | no | no | no | no |
| SCN5A | 1368 | E | 0 | 0 | no | yes | no | no | yes | no | yes | no | no | no | no | no | no |
| SCN5A | 1369 | G | 1 | 0 | no | yes | no | no | yes | no | no | no | no | no | no | no | no |
| SCN5A | 1370 | D | 1 | 0 | no | yes | no | no | yes | no | no | no | no | no | no | no | no |
| SCN5A | 1371 | L | 0 | 0 | no | yes | no | no | yes | no | yes | no | no | no | no | no | no |
| SCN5A | 1372 | P | 0 | 1 | no | yes | no | no | yes | no | no | no | no | no | no | no | no |
| SCN5A | 1373 | L | 0 | 0 | no | yes | no | no | yes | no | no | no | no | no | no | no | no |
| SCN5A | 1374 | N | 0 | 0 | no | yes | no | no | yes | no | no | no | no | no | no | no | no |
| SCN5A | 1375 | Y | 1 | 0 | no | yes | no | no | yes | no | no | no | no | no | no | no | no |
| SCN5A | 1376 | T | 0 | 0 | no | yes | no | no | yes | no | no | no | no | no | no | no | no |
| SCN5A | 1377 | I | 1 | 0 | no | yes | no | no | yes | no | no | no | no | no | no | no | no |
| SCN5A | 1378 | V | 1 | 1 | no | yes | no | no | yes | no | no | no | no | no | no | no | no |
| SCN5A | 1379 | N | 0 | 0 | no | yes | no | no | yes | no | no | no | no | no | no | no | no |
| SCN5A | 1380 | N | 0 | 1 | no | no | yes | no | yes | no | no | no | no | no | no | no | no |
| SCN5A | 1381 | K | 0 | 0 | yes | no | no | no | yes | no | no | no | no | no | no | no | no |
| SCN5A | 1382 | S | 0 | 1 | yes | no | no | no | yes | no | no | no | no | no | no | no | no |
| SCN5A | 1383 | Q | 0 | 0 | yes | no | no | no | yes | no | no | no | no | no | no | no | no |
| SCN5A | 1384 | C | 1 | 0 | yes | no | no | no | yes | no | no | no | no | no | no | no | no |

| | | | | | | | | | | | | | | | | | |
|---|---|---|---|---|---|---|---|---|---|---|---|---|---|---|---|---|---|
| SCN5A | 1385 | E | 0 | 0 | no | yes | no | no | yes | no | no | no | no | no | no | no | no |
| SCN5A | 1386 | S | 0 | 0 | no | yes | no | no | yes | no | no | no | no | no | no | no | no |
| SCN5A | 1387 | L | 1 | 0 | no | yes | no | no | yes | no | no | no | no | no | no | no | no |
| SCN5A | 1388 | N | 0 | 0 | no | yes | no | no | yes | no | no | no | no | no | no | no | no |
| SCN5A | 1389 | L | 0 | 0 | no | yes | no | no | yes | no | no | no | no | no | no | no | no |
| SCN5A | 1390 | T | 0 | 0 | no | yes | no | no | yes | no | no | no | no | no | no | no | no |
| SCN5A | 1391 | G | 1 | 1 | no | yes | no | no | yes | no | no | no | no | no | no | no | no |
| SCN5A | 1392 | E | 0 | 0 | no | yes | no | no | yes | no | no | no | no | no | no | no | no |
| SCN5A | 1393 | L | 0 | 0 | no | no | yes | no | yes | no | no | no | no | no | no | no | no |
| SCN5A | 1394 | Y | 0 | 0 | no | yes | no | no | yes | no | no | no | no | no | no | no | no |
| SCN5A | 1395 | W | 0 | 0 | no | yes | no | no | yes | no | no | no | no | no | no | no | no |
| SCN5A | 1396 | T | 0 | 0 | no | no | yes | no | yes | no | no | no | no | no | no | no | no |
| SCN5A | 1397 | K | 0 | 0 | no | no | yes | no | yes | no | no | no | no | no | no | no | no |
| SCN5A | 1398 | V | 1 | 0 | no | yes | no | no | yes | no | no | no | no | no | no | no | no |
| SCN5A | 1399 | K | 0 | 0 | no | yes | no | no | yes | no | no | no | no | no | no | no | no |
| SCN5A | 1400 | V | 1 | 0 | no | yes | no | no | yes | no | no | no | no | no | no | no | no |
| SCN5A | 1401 | N | 0 | 0 | no | yes | no | no | yes | no | no | no | no | no | no | no | no |
| SCN5A | 1402 | F | 0 | 0 | no | yes | no | no | yes | no | no | no | no | no | no | no | no |
| SCN5A | 1403 | D | 0 | 0 | no | yes | no | no | yes | no | no | no | no | no | no | no | no |
| SCN5A | 1404 | N | 0 | 0 | no | no | yes | no | yes | no | no | no | no | no | no | no | no |
| SCN5A | 1405 | V | 0 | 1 | yes | no | no | yes | yes | no | no | no | no | no | no | no | no |
| SCN5A | 1406 | G | 0 | 2 | yes | no | no | yes | no | no | no | yes | no | no | no | no | no |
| SCN5A | 1407 | A | 0 | 0 | yes | no | no | yes | no | no | no | yes | no | no | no | no | no |
| SCN5A | 1408 | G | 0 | 1 | yes | no | no | yes | no | no | no | yes | no | no | no | no | no |
| SCN5A | 1409 | Y | 0 | 1 | yes | no | no | yes | no | no | no | yes | no | no | no | no | no |
| SCN5A | 1410 | L | 0 | 0 | yes | no | no | yes | no | no | no | yes | no | no | no | no | no |
| SCN5A | 1411 | A | 0 | 0 | yes | no | no | yes | no | no | no | yes | no | no | no | no | no |
| SCN5A | 1412 | L | 0 | 1 | yes | no | no | yes | no | no | no | yes | no | no | no | no | no |
| SCN5A | 1413 | L | 0 | 0 | yes | no | no | yes | no | no | no | yes | no | no | no | no | no |
| SCN5A | 1414 | Q | 1 | 0 | yes | no | no | yes | no | no | no | yes | no | no | no | no | no |
| SCN5A | 1415 | V | 0 | 0 | yes | no | no | yes | no | no | no | yes | no | no | no | no | no |
| SCN5A | 1416 | A | 0 | 0 | yes | no | no | yes | no | no | no | yes | no | no | no | no | no |
| SCN5A | 1417 | T | 0 | 0 | yes | no | no | yes | no | no | no | yes | no | no | no | no | no |

| | | | | | | | | | | | | | | | | | |
|---|---|---|---|---|---|---|---|---|---|---|---|---|---|---|---|---|---|
| SCN5A | 1418 | F | 0 | 0 | no | yes | no | yes | no | no | yes | yes | no | no | no | no | no |
| SCN5A | 1419 | K | 0 | 1 | no | yes | no | yes | no | no | yes | yes | no | no | no | no | no |
| SCN5A | 1420 | G | 0 | 1 | no | yes | no | no | no | no | no | yes | no | no | no | no | no |
| SCN5A | 1421 | W | 0 | 0 | yes | no | no | yes | no | no | no | yes | no | no | no | no | no |
| SCN5A | 1422 | M | 1 | 0 | yes | no | no | no | no | no | yes | yes | no | no | no | no | no |
| SCN5A | 1423 | D | 1 | 0 | yes | no | no | no | no | no | no | yes | no | no | no | no | no |
| SCN5A | 1424 | I | 1 | 0 | yes | no | no | no | no | no | no | yes | no | no | no | no | no |
| SCN5A | 1425 | M | 0 | 0 | yes | no | no | no | no | no | no | yes | no | no | no | no | no |
| SCN5A | 1426 | Y | 0 | 0 | yes | no | no | no | no | no | yes | yes | no | no | no | no | no |
| SCN5A | 1427 | A | 1 | 1 | yes | no | no | no | no | no | no | yes | no | no | no | no | no |
| SCN5A | 1428 | A | 1 | 2 | yes | no | no | no | yes | no | no | no | no | no | no | no | no |
| SCN5A | 1429 | V | 0 | 0 | no | yes | no | no | yes | no | no | no | no | no | no | no | no |
| SCN5A | 1430 | D | 0 | 1 | no | yes | no | no | yes | no | no | no | no | no | no | no | no |
| SCN5A | 1431 | S | 1 | 0 | no | yes | no | no | yes | no | no | no | no | no | no | no | no |
| SCN5A | 1432 | R | 0 | 2 | no | yes | no | no | yes | no | no | no | no | no | no | no | no |
| SCN5A | 1433 | G | 0 | 2 | no | yes | no | no | yes | no | no | no | no | no | no | no | no |
| SCN5A | 1434 | Y | 0 | 0 | no | yes | no | no | yes | no | no | no | no | no | no | no | no |
| SCN5A | 1435 | E | 0 | 0 | no | yes | no | no | yes | no | no | no | no | no | no | no | no |
| SCN5A | 1436 | E | 0 | 0 | no | no | yes | no | yes | no | no | no | no | no | no | no | no |
| SCN5A | 1437 | Q | 0 | 0 | no | yes | no | no | yes | no | no | no | no | no | no | no | no |
| SCN5A | 1438 | P | 0 | 2 | no | yes | no | no | yes | no | no | no | no | no | no | no | no |
| SCN5A | 1439 | Q | 2 | 0 | no | yes | no | no | yes | no | no | no | no | no | no | no | no |
| SCN5A | 1440 | W | 0 | 0 | no | yes | no | no | yes | no | yes | no | no | no | no | no | no |
| SCN5A | 1441 | E | 0 | 1 | no | yes | no | no | yes | no | no | no | no | no | no | no | no |
| SCN5A | 1442 | Y | 2 | 0 | no | yes | no | no | yes | no | no | no | no | no | no | no | no |
| SCN5A | 1443 | N | 1 | 1 | no | yes | no | no | yes | no | no | no | no | no | no | no | no |
| SCN5A | 1444 | L | 1 | 0 | yes | no | no | no | yes | no | no | no | no | no | no | no | no |
| SCN5A | 1445 | Y | 1 | 0 | yes | no | no | no | no | no | no | no | no | no | no | no | no |
| SCN5A | 1446 | M | 0 | 1 | yes | no | no | no | no | no | no | no | no | no | no | no | no |
| SCN5A | 1447 | Y | 0 | 1 | yes | no | no | no | no | no | no | no | no | no | no | no | no |
| SCN5A | 1448 | I | 1 | 2 | yes | no | no | yes | no | no | no | no | no | no | no | no | no |
| SCN5A | 1449 | Y | 0 | 2 | yes | no | no | yes | no | no | no | no | no | no | no | no | no |
| SCN5A | 1450 | F | 0 | 0 | yes | no | no | yes | no | no | no | no | no | no | no | no | no |

| | | | | | | | | | | | | | | | | | |
|---|---|---|---|---|---|---|---|---|---|---|---|---|---|---|---|---|---|
| SCN5A | 1451 | V | 1 | 1 | yes | no | no | yes | no | no | no | no | no | no | no | no | no |
| SCN5A | 1452 | I | 0 | 0 | yes | no | no | yes | no | no | no | no | no | no | no | no | no |
| SCN5A | 1453 | F | 0 | 0 | yes | no | no | yes | no | no | no | no | no | no | no | no | no |
| SCN5A | 1454 | I | 0 | 0 | yes | no | no | yes | no | no | no | no | no | no | no | no | no |
| SCN5A | 1455 | I | 0 | 0 | yes | no | no | yes | no | no | no | no | no | no | no | no | no |
| SCN5A | 1456 | F | 0 | 0 | yes | no | no | yes | no | no | no | no | no | no | no | no | no |
| SCN5A | 1457 | G | 0 | 0 | yes | no | no | yes | no | no | no | no | no | no | no | no | no |
| SCN5A | 1458 | S | 0 | 1 | yes | no | no | yes | no | no | no | no | no | no | no | no | no |
| SCN5A | 1459 | F | 0 | 0 | yes | no | no | yes | no | no | no | no | no | no | no | no | no |
| SCN5A | 1460 | F | 0 | 0 | yes | no | no | yes | no | no | no | no | no | no | no | no | no |
| SCN5A | 1461 | T | 0 | 1 | yes | no | no | yes | no | no | no | no | no | no | no | no | no |
| SCN5A | 1462 | L | 0 | 0 | yes | no | no | yes | no | no | yes | no | no | no | no | no | no |
| SCN5A | 1463 | N | 0 | 1 | yes | no | no | yes | no | no | no | no | no | no | no | no | no |
| SCN5A | 1464 | L | 1 | 0 | yes | no | no | yes | no | no | no | no | no | no | no | no | no |
| SCN5A | 1465 | F | 0 | 0 | yes | no | no | yes | no | no | no | no | no | no | no | no | no |
| SCN5A | 1466 | I | 0 | 0 | yes | no | no | yes | no | no | yes | no | no | no | no | no | no |
| SCN5A | 1467 | G | 0 | 0 | yes | no | no | yes | no | yes | no | no | no | no | no | no | no |
| SCN5A | 1468 | V | 1 | 1 | yes | no | no | yes | no | yes | no | no | no | no | no | no | no |
| SCN5A | 1469 | I | 1 | 0 | yes | no | no | yes | no | yes | no | no | no | no | no | no | no |
| SCN5A | 1470 | I | 0 | 0 | yes | no | no | yes | no | yes | yes | no | no | no | no | no | no |
| SCN5A | 1471 | D | 0 | 0 | yes | no | no | no | no | yes | no | no | no | no | no | no | no |
| SCN5A | 1472 | N | 0 | 1 | yes | no | no | yes | no | yes | no | no | no | no | no | no | no |
| SCN5A | 1473 | F | 0 | 2 | yes | no | no | yes | no | yes | no | no | no | no | no | no | no |
| SCN5A | 1474 | N | 0 | 0 | yes | no | no | no | no | yes | yes | no | no | no | no | no | no |
| SCN5A | 1475 | Q | 0 | 1 | yes | no | no | no | no | yes | no | no | no | no | no | no | no |
| SCN5A | 1476 | Q | 0 | 1 | yes | no | no | no | no | yes | no | no | no | no | no | no | no |
| SCN5A | 1477 | K | 0 | 0 | yes | no | no | no | no | yes | yes | no | no | no | no | no | no |
| SCN5A | 1478 | K | 1 | 0 | yes | no | no | no | no | yes | yes | no | no | no | no | no | no |
| SCN5A | 1479 | K | 0 | 0 | yes | no | no | no | no | yes | no | no | no | no | no | no | no |
| SCN5A | 1480 | L | 0 | 0 | no | yes | no | no | no | yes | no | no | no | no | no | no | no |
| SCN5A | 1481 | G | 0 | 2 | no | yes | no | no | no | yes | no | no | no | no | no | no | no |
| SCN5A | 1482 | G | 0 | 0 | no | yes | no | no | no | yes | no | no | no | no | no | no | no |
| SCN5A | 1483 | Q | 1 | 0 | no | yes | no | no | no | yes | yes | no | no | no | no | no | no |

| | | | | | | | | | | | | | | | | | |
|---|---|---|---|---|---|---|---|---|---|---|---|---|---|---|---|---|---|
| SCN5A | 1484 | D | 0 | 0 | no | yes | no | no | no | yes | no | no | no | no | no | no | no |
| SCN5A | 1485 | I | 0 | 1 | no | yes | no | no | no | yes | no | no | no | no | no | no | no |
| SCN5A | 1486 | F | 0 | 1 | no | yes | no | yes | no | yes | no | no | no | no | no | no | no |
| SCN5A | 1487 | M | 1 | 2 | no | yes | no | yes | no | yes | no | no | no | no | no | no | no |
| SCN5A | 1488 | T | 0 | 1 | no | yes | no | yes | no | yes | no | no | no | no | no | no | no |
| SCN5A | 1489 | E | 1 | 1 | yes | no | no | no | no | yes | no | no | no | no | no | no | no |
| SCN5A | 1490 | E | 0 | 0 | yes | no | no | no | no | yes | no | no | no | no | no | no | no |
| SCN5A | 1491 | Q | 0 | 0 | yes | no | no | yes | no | yes | no | no | no | no | no | no | no |
| SCN5A | 1492 | K | 0 | 0 | yes | no | no | no | no | yes | no | no | no | no | no | no | no |
| SCN5A | 1493 | K | 1 | 1 | yes | no | no | no | no | yes | no | no | no | no | no | no | no |
| SCN5A | 1494 | Y | 0 | 1 | yes | no | no | no | no | yes | no | no | no | no | no | no | no |
| SCN5A | 1495 | Y | 0 | 1 | yes | no | no | no | no | yes | no | no | no | no | no | no | no |
| SCN5A | 1496 | N | 0 | 0 | yes | no | no | no | no | yes | no | no | no | no | no | no | no |
| SCN5A | 1497 | A | 0 | 0 | yes | no | no | no | no | yes | no | no | no | no | no | no | no |
| SCN5A | 1498 | M | 0 | 2 | yes | no | no | no | no | yes | no | no | no | no | no | no | no |
| SCN5A | 1499 | K | 0 | 0 | no | yes | no | no | no | yes | no | no | no | no | no | no | no |
| SCN5A | 1500 | K | 0 | 0 | no | yes | no | no | no | yes | no | no | no | no | no | no | no |
| SCN5A | 1501 | L | 1 | 1 | no | yes | no | no | no | yes | no | no | no | no | no | no | no |
| SCN5A | 1502 | G | 2 | 1 | no | yes | no | no | no | yes | no | no | no | no | no | no | no |
| SCN5A | 1503 | S | 1 | 0 | no | yes | no | no | no | yes | no | no | no | no | no | no | no |
| SCN5A | 1504 | K | 0 | 0 | no | yes | no | no | no | yes | no | no | no | no | no | no | no |
| SCN5A | 1505 | K | 0 | 1 | no | yes | no | no | no | yes | no | no | no | no | no | no | no |
| SCN5A | 1506 | P | 0 | 1 | no | yes | no | no | no | yes | no | no | no | no | no | no | no |
| SCN5A | 1507 | Q | 0 | 0 | no | yes | no | no | no | yes | no | no | no | no | no | no | no |
| SCN5A | 1508 | K | 0 | 0 | no | yes | no | no | no | yes | no | no | no | no | no | no | no |
| SCN5A | 1509 | P | 1 | 0 | no | yes | no | no | no | yes | no | no | no | no | no | no | no |
| SCN5A | 1510 | I | 0 | 0 | no | yes | no | no | no | yes | no | no | no | no | no | no | no |
| SCN5A | 1511 | P | 0 | 0 | no | yes | no | no | no | yes | no | no | no | no | no | no | no |
| SCN5A | 1512 | R | 2 | 1 | no | yes | no | no | no | yes | no | no | no | no | no | no | no |
| SCN5A | 1513 | P | 0 | 0 | no | yes | no | no | no | yes | no | no | no | no | no | no | no |
| SCN5A | 1514 | L | 1 | 0 | no | yes | no | no | no | yes | no | no | no | no | no | no | no |
| SCN5A | 1515 | N | 1 | 0 | no | yes | no | no | no | yes | no | no | no | no | no | no | no |
| SCN5A | 1516 | K | 0 | 0 | yes | no | no | no | no | yes | no | no | no | no | no | no | no |

| | | | | | | | | | | | | | | | | | |
|---|---|---|---|---|---|---|---|---|---|---|---|---|---|---|---|---|---|
| SCN5A | 1517 | Y | 0 | 0 | yes | no | no | no | no | yes | no | no | no | no | no | no | no |
| SCN5A | 1518 | Q | 0 | 0 | yes | no | no | no | no | yes | no | no | no | no | no | no | no |
| SCN5A | 1519 | G | 0 | 0 | yes | no | no | no | no | yes | no | no | no | no | no | no | no |
| SCN5A | 1520 | F | 0 | 1 | yes | no | no | no | no | yes | no | no | no | no | no | no | no |
| SCN5A | 1521 | I | 1 | 1 | yes | no | no | no | no | yes | no | no | no | no | no | no | no |
| SCN5A | 1522 | F | 0 | 0 | yes | no | no | no | no | yes | no | no | no | no | no | no | no |
| SCN5A | 1523 | D | 1 | 0 | yes | no | no | no | no | yes | no | no | no | no | no | no | no |
| SCN5A | 1524 | I | 1 | 0 | yes | no | no | no | no | yes | no | no | no | no | no | no | no |
| SCN5A | 1525 | V | 2 | 0 | yes | no | no | yes | no | yes | no | no | no | no | no | no | no |
| SCN5A | 1526 | T | 0 | 0 | no | yes | no | no | no | yes | no | no | no | no | no | no | no |
| SCN5A | 1527 | K | 0 | 0 | no | yes | no | no | no | yes | no | no | no | no | no | no | no |
| SCN5A | 1528 | Q | 0 | 0 | no | yes | no | no | no | yes | no | no | no | no | no | no | no |
| SCN5A | 1529 | A | 0 | 0 | no | yes | no | yes | no | yes | no | no | no | no | no | no | no |
| SCN5A | 1530 | F | 0 | 0 | yes | no | no | yes | no | no | no | no | no | no | no | no | no |
| SCN5A | 1531 | D | 0 | 0 | yes | no | no | yes | no | no | no | no | no | no | no | no | no |
| SCN5A | 1532 | V | 1 | 1 | yes | no | no | yes | no | no | no | no | no | no | no | no | no |
| SCN5A | 1533 | T | 1 | 0 | yes | no | no | yes | no | no | no | no | no | no | no | no | no |
| SCN5A | 1534 | I | 0 | 0 | yes | no | no | yes | no | no | no | no | no | no | no | no | no |
| SCN5A | 1535 | M | 0 | 0 | yes | no | no | yes | no | no | no | no | no | no | no | no | no |
| SCN5A | 1536 | F | 0 | 0 | yes | no | no | yes | no | no | no | no | no | no | no | no | no |
| SCN5A | 1537 | L | 0 | 0 | yes | no | no | yes | no | no | no | no | no | no | no | no | no |
| SCN5A | 1538 | I | 0 | 0 | yes | no | no | yes | no | no | no | no | no | no | no | no | no |
| SCN5A | 1539 | C | 1 | 0 | yes | no | no | yes | no | no | no | no | no | no | no | no | no |
| SCN5A | 1540 | L | 0 | 0 | yes | no | no | yes | no | no | no | no | no | no | no | no | no |
| SCN5A | 1541 | N | 0 | 0 | yes | no | no | yes | no | no | no | no | no | no | no | no | no |
| SCN5A | 1542 | M | 0 | 0 | yes | no | no | yes | no | no | no | no | no | no | no | no | no |
| SCN5A | 1543 | V | 2 | 0 | yes | no | no | yes | no | no | no | no | no | no | no | no | no |
| SCN5A | 1544 | T | 0 | 1 | yes | no | no | yes | no | no | no | no | no | no | no | no | no |
| SCN5A | 1545 | M | 0 | 0 | yes | no | no | yes | no | no | no | no | no | no | no | no | no |
| SCN5A | 1546 | M | 1 | 0 | yes | no | no | yes | no | no | no | no | no | no | no | no | no |
| SCN5A | 1547 | V | 1 | 0 | no | yes | no | yes | no | no | no | no | no | no | no | no | no |
| SCN5A | 1548 | E | 1 | 1 | no | yes | no | yes | yes | no | no | no | no | no | no | no | no |
| SCN5A | 1549 | T | 0 | 0 | no | yes | no | no | yes | no | no | no | no | no | no | no | no |

| | | | | | | | | | | | | | | | | | |
|---|---|---|---|---|---|---|---|---|---|---|---|---|---|---|---|---|---|
| SCN5A | 1550 | D | 0 | 0 | no | yes | no | no | yes | no | no | no | no | no | no | no | no |
| SCN5A | 1551 | D | 1 | 0 | no | yes | no | no | yes | no | no | no | no | no | no | no | no |
| SCN5A | 1552 | Q | 1 | 0 | no | yes | no | no | yes | no | no | no | no | no | no | no | no |
| SCN5A | 1553 | S | 0 | 1 | no | yes | no | no | yes | no | no | no | no | no | no | no | no |
| SCN5A | 1554 | P | 0 | 0 | yes | no | no | no | yes | no | no | no | no | no | no | no | no |
| SCN5A | 1555 | E | 1 | 0 | yes | no | no | no | yes | no | no | no | no | no | no | no | no |
| SCN5A | 1556 | K | 0 | 0 | yes | no | no | no | yes | no | no | no | no | no | no | no | no |
| SCN5A | 1557 | I | 1 | 0 | yes | no | no | no | yes | no | no | no | no | no | no | no | no |
| SCN5A | 1558 | N | 0 | 0 | yes | no | no | no | yes | no | no | no | no | no | no | no | no |
| SCN5A | 1559 | I | 1 | 0 | yes | no | no | yes | no | no | no | no | no | no | no | no | no |
| SCN5A | 1560 | L | 1 | 1 | yes | no | no | yes | no | no | no | no | no | no | no | no | no |
| SCN5A | 1561 | A | 0 | 0 | yes | no | no | yes | no | no | no | no | no | no | no | no | no |
| SCN5A | 1562 | K | 0 | 0 | yes | no | no | yes | no | no | no | no | no | no | no | no | no |
| SCN5A | 1563 | I | 0 | 0 | yes | no | no | yes | no | no | no | no | no | no | no | no | no |
| SCN5A | 1564 | N | 0 | 0 | yes | no | no | yes | no | no | no | no | no | no | no | no | no |
| SCN5A | 1565 | L | 1 | 0 | yes | no | no | yes | no | no | no | no | no | no | no | no | no |
| SCN5A | 1566 | L | 0 | 0 | yes | no | no | yes | no | no | no | no | no | no | no | no | no |
| SCN5A | 1567 | F | 1 | 0 | yes | no | no | yes | no | no | no | no | no | no | no | no | no |
| SCN5A | 1568 | V | 0 | 0 | yes | no | no | yes | no | no | no | no | no | no | no | no | no |
| SCN5A | 1569 | A | 0 | 0 | yes | no | no | yes | no | no | no | no | no | no | no | no | no |
| SCN5A | 1570 | I | 1 | 0 | yes | no | no | yes | no | no | no | no | no | no | no | no | no |
| SCN5A | 1571 | F | 0 | 2 | yes | no | no | yes | no | no | no | no | no | no | no | no | no |
| SCN5A | 1572 | T | 0 | 0 | yes | no | no | yes | no | no | no | no | no | no | no | no | no |
| SCN5A | 1573 | G | 0 | 0 | yes | no | no | yes | no | no | no | no | no | no | no | no | no |
| SCN5A | 1574 | E | 0 | 1 | yes | no | no | yes | no | no | no | no | no | no | no | no | no |
| SCN5A | 1575 | C | 1 | 0 | yes | no | no | yes | no | no | no | no | no | no | no | no | no |
| SCN5A | 1576 | I | 0 | 0 | yes | no | no | yes | no | no | no | no | no | no | no | no | no |
| SCN5A | 1577 | V | 0 | 0 | yes | no | no | yes | no | no | no | no | no | no | no | no | no |
| SCN5A | 1578 | K | 0 | 1 | yes | no | no | yes | no | yes | no | no | no | no | no | no | no |
| SCN5A | 1579 | L | 0 | 0 | yes | no | no | yes | no | yes | no | no | no | no | no | no | no |
| SCN5A | 1580 | A | 0 | 0 | yes | no | no | yes | no | yes | no | no | no | no | no | no | no |
| SCN5A | 1581 | A | 1 | 0 | yes | no | no | no | no | yes | no | no | no | no | no | no | no |
| SCN5A | 1582 | L | 0 | 1 | no | yes | no | no | no | yes | no | no | no | no | no | no | no |

| | | | | | | | | | | | | | | | | | |
|---|---|---|---|---|---|---|---|---|---|---|---|---|---|---|---|---|---|
| SCN5A | 1583 | R | 2 | 2 | no | yes | no | no | no | yes | no | no | no | no | no | no | no |
| SCN5A | 1584 | H | 0 | 0 | no | yes | no | no | no | yes | no | no | no | no | no | no | no |
| SCN5A | 1585 | Y | 1 | 0 | no | yes | no | no | no | yes | no | no | no | no | no | no | no |
| SCN5A | 1586 | Y | 0 | 0 | no | yes | no | yes | no | yes | no | no | no | no | no | no | no |
| SCN5A | 1587 | F | 1 | 0 | no | yes | no | yes | no | yes | no | no | no | no | no | no | no |
| SCN5A | 1588 | T | 1 | 0 | no | yes | no | no | no | yes | no | no | no | no | no | no | no |
| SCN5A | 1589 | N | 0 | 0 | no | yes | no | no | no | yes | no | no | no | no | no | no | no |
| SCN5A | 1590 | S | 0 | 0 | no | yes | no | yes | no | no | no | no | no | no | no | no | no |
| SCN5A | 1591 | W | 0 | 0 | no | yes | no | yes | no | no | no | no | no | no | no | no | no |
| SCN5A | 1592 | N | 0 | 0 | yes | no | no | yes | no | no | no | no | no | no | no | no | no |
| SCN5A | 1593 | I | 1 | 1 | yes | no | no | yes | no | no | no | no | no | no | no | no | no |
| SCN5A | 1594 | F | 0 | 1 | yes | no | no | yes | no | no | no | no | no | no | no | no | no |
| SCN5A | 1595 | D | 0 | 2 | yes | no | no | yes | no | no | no | no | no | no | no | no | no |
| SCN5A | 1596 | F | 2 | 0 | yes | no | no | yes | no | no | no | no | no | no | no | no | no |
| SCN5A | 1597 | V | 1 | 1 | yes | no | no | yes | no | no | no | no | no | no | no | no | no |
| SCN5A | 1598 | V | 1 | 0 | yes | no | no | yes | no | no | no | no | no | no | no | no | no |
| SCN5A | 1599 | V | 0 | 0 | yes | no | no | yes | no | no | no | no | no | no | no | no | no |
| SCN5A | 1600 | I | 0 | 0 | yes | no | no | yes | no | no | no | no | no | no | no | no | no |
| SCN5A | 1601 | L | 1 | 0 | yes | no | no | yes | no | no | no | no | no | no | no | no | no |
| SCN5A | 1602 | S | 0 | 0 | yes | no | no | yes | no | no | no | no | no | no | no | no | no |
| SCN5A | 1603 | I | 1 | 0 | yes | no | no | yes | no | no | no | no | no | no | no | no | no |
| SCN5A | 1604 | V | 1 | 1 | yes | no | no | yes | no | no | no | no | no | no | no | no | no |
| SCN5A | 1605 | G | 1 | 0 | yes | no | no | yes | no | no | no | no | no | no | no | no | no |
| SCN5A | 1606 | T | 1 | 0 | yes | no | no | yes | no | no | no | no | no | no | no | no | no |
| SCN5A | 1607 | V | 0 | 0 | yes | no | no | yes | no | no | no | no | no | no | no | no | no |
| SCN5A | 1608 | L | 0 | 0 | yes | no | no | yes | yes | no | no | no | no | no | no | no | no |
| SCN5A | 1609 | S | 1 | 1 | no | yes | no | yes | yes | no | no | no | no | no | no | no | no |
| SCN5A | 1610 | D | 1 | 0 | no | yes | no | no | yes | no | no | no | no | no | no | no | no |
| SCN5A | 1611 | I | 1 | 0 | no | yes | no | no | yes | no | no | no | no | no | no | no | no |
| SCN5A | 1612 | I | 0 | 0 | no | yes | no | no | yes | no | no | no | no | no | no | no | no |
| SCN5A | 1613 | Q | 1 | 1 | no | yes | no | no | yes | no | no | no | no | no | no | no | no |
| SCN5A | 1614 | K | 0 | 0 | no | yes | no | no | yes | no | no | no | no | no | no | no | no |
| SCN5A | 1615 | Y | 0 | 0 | no | yes | no | no | yes | no | no | no | no | no | no | no | no |

| | | | | | | | | | | | | | | | | | |
|---|---|---|---|---|---|---|---|---|---|---|---|---|---|---|---|---|---|
| SCN5A | 1616 | F | 0 | 0 | no | yes | no | no | yes | no | no | no | no | no | no | no | no |
| SCN5A | 1617 | F | 0 | 0 | no | yes | no | no | yes | no | no | no | no | no | no | no | no |
| SCN5A | 1618 | S | 0 | 0 | no | yes | no | no | yes | no | no | no | no | no | no | no | no |
| SCN5A | 1619 | P | 1 | 0 | no | yes | no | no | yes | no | no | no | no | no | no | no | no |
| SCN5A | 1620 | T | 1 | 1 | no | yes | no | no | yes | no | no | no | no | no | no | no | no |
| SCN5A | 1621 | L | 0 | 0 | no | yes | no | yes | no | no | no | no | no | no | no | no | no |
| SCN5A | 1622 | F | 0 | 0 | yes | no | no | no | no | no | no | no | no | no | no | no | no |
| SCN5A | 1623 | R | 0 | 1 | yes | no | no | no | no | no | no | no | no | no | no | no | no |
| SCN5A | 1624 | V | 1 | 0 | yes | no | no | yes | no | no | no | no | no | no | no | no | no |
| SCN5A | 1625 | I | 0 | 0 | yes | no | no | yes | no | no | no | no | no | no | no | no | no |
| SCN5A | 1626 | R | 2 | 2 | yes | no | no | yes | no | no | no | no | no | no | no | no | no |
| SCN5A | 1627 | L | 0 | 0 | yes | no | no | yes | no | no | no | no | no | no | no | no | no |
| SCN5A | 1628 | A | 0 | 0 | yes | no | no | yes | no | no | no | no | no | no | no | no | no |
| SCN5A | 1629 | R | 1 | 2 | yes | no | no | yes | no | no | no | no | no | no | no | no | no |
| SCN5A | 1630 | I | 2 | 0 | yes | no | no | yes | no | no | no | no | no | no | no | no | no |
| SCN5A | 1631 | G | 0 | 1 | yes | no | no | yes | no | no | no | no | no | no | no | no | no |
| SCN5A | 1632 | R | 2 | 2 | yes | no | no | yes | no | no | no | no | no | no | no | no | no |
| SCN5A | 1633 | I | 0 | 0 | yes | no | no | yes | no | no | no | no | no | no | no | no | no |
| SCN5A | 1634 | L | 0 | 0 | yes | no | no | yes | no | no | no | no | no | no | no | no | no |
| SCN5A | 1635 | R | 0 | 0 | yes | no | no | yes | no | no | no | no | no | no | no | no | no |
| SCN5A | 1636 | L | 0 | 0 | yes | no | no | yes | no | no | no | no | no | no | no | no | no |
| SCN5A | 1637 | I | 0 | 0 | yes | no | no | yes | no | no | no | no | no | no | no | no | no |
| SCN5A | 1638 | R | 1 | 1 | no | yes | no | yes | no | yes | no | no | no | no | no | no | no |
| SCN5A | 1639 | G | 1 | 0 | no | yes | no | yes | no | yes | no | no | no | no | no | no | no |
| SCN5A | 1640 | A | 0 | 0 | no | yes | no | yes | no | yes | no | no | no | no | no | no | no |
| SCN5A | 1641 | K | 0 | 0 | yes | no | no | no | no | yes | no | no | no | no | no | no | no |
| SCN5A | 1642 | G | 0 | 1 | yes | no | no | yes | no | yes | no | no | no | no | no | no | no |
| SCN5A | 1643 | I | 1 | 0 | yes | no | no | yes | no | yes | no | no | no | no | no | no | no |
| SCN5A | 1644 | R | 2 | 2 | yes | no | no | yes | no | yes | no | no | no | no | no | no | no |
| SCN5A | 1645 | T | 1 | 1 | yes | no | no | yes | no | yes | no | no | no | no | no | no | no |
| SCN5A | 1646 | L | 0 | 1 | yes | no | no | yes | no | yes | no | no | no | no | no | no | no |
| SCN5A | 1647 | L | 0 | 0 | yes | no | no | yes | no | yes | no | no | no | no | no | no | no |
| SCN5A | 1648 | F | 0 | 0 | yes | no | no | yes | no | yes | no | no | no | no | no | no | no |

| | | | | | | | | | | | | | | | | | |
|---|---|---|---|---|---|---|---|---|---|---|---|---|---|---|---|---|---|
| SCN5A | 1649 | A | 0 | 1 | yes | no | no | yes | no | yes | no | no | no | no | no | no | no |
| SCN5A | 1650 | L | 0 | 2 | yes | no | no | yes | no | yes | no | no | no | no | no | no | no |
| SCN5A | 1651 | M | 0 | 1 | yes | no | no | yes | no | yes | no | no | no | no | no | no | no |
| SCN5A | 1652 | M | 0 | 1 | no | yes | no | yes | no | yes | no | no | no | no | no | no | no |
| SCN5A | 1653 | S | 0 | 0 | no | yes | no | yes | no | yes | no | no | no | no | no | no | no |
| SCN5A | 1654 | L | 0 | 0 | yes | no | no | yes | no | yes | no | no | no | no | no | no | no |
| SCN5A | 1655 | P | 0 | 0 | yes | no | no | yes | no | yes | no | no | no | no | no | no | no |
| SCN5A | 1656 | A | 0 | 1 | yes | no | no | yes | no | yes | no | no | no | no | no | no | no |
| SCN5A | 1657 | L | 0 | 0 | yes | no | no | yes | no | no | no | no | no | no | no | no | no |
| SCN5A | 1658 | F | 0 | 0 | yes | no | no | yes | no | no | no | no | no | no | no | no | no |
| SCN5A | 1659 | N | 0 | 0 | yes | no | no | yes | no | no | no | no | no | no | no | no | no |
| SCN5A | 1660 | I | 1 | 1 | yes | no | no | yes | no | no | no | no | no | no | no | no | no |
| SCN5A | 1661 | G | 1 | 1 | yes | no | no | yes | no | no | no | no | no | no | no | no | no |
| SCN5A | 1662 | L | 0 | 0 | yes | no | no | yes | no | no | no | no | no | no | no | no | no |
| SCN5A | 1663 | L | 0 | 0 | yes | no | no | yes | no | no | no | no | no | no | no | no | no |
| SCN5A | 1664 | L | 0 | 0 | yes | no | no | yes | no | no | no | no | no | no | no | no | no |
| SCN5A | 1665 | F | 0 | 0 | yes | no | no | yes | no | no | no | no | no | no | no | no | no |
| SCN5A | 1666 | L | 0 | 0 | yes | no | no | yes | no | no | no | no | no | no | no | no | no |
| SCN5A | 1667 | V | 1 | 1 | yes | no | no | yes | no | no | no | no | no | no | no | no | no |
| SCN5A | 1668 | M | 1 | 0 | yes | no | no | yes | no | no | no | no | no | no | no | no | no |
| SCN5A | 1669 | F | 0 | 0 | yes | no | no | yes | no | no | no | no | no | no | no | no | no |
| SCN5A | 1670 | I | 0 | 0 | yes | no | no | yes | no | no | no | no | no | no | no | no | no |
| SCN5A | 1671 | Y | 0 | 0 | yes | no | no | yes | no | no | no | no | no | no | no | no | no |
| SCN5A | 1672 | S | 0 | 1 | yes | no | no | yes | no | no | no | no | no | no | no | no | no |
| SCN5A | 1673 | I | 0 | 0 | yes | no | no | yes | no | no | no | no | no | no | no | no | no |
| SCN5A | 1674 | F | 1 | 0 | yes | no | no | yes | no | no | no | no | no | no | no | no | no |
| SCN5A | 1675 | G | 0 | 0 | yes | no | no | no | yes | no | no | no | no | no | no | no | no |
| SCN5A | 1676 | M | 2 | 2 | yes | no | no | no | yes | no | no | no | no | no | no | no | no |
| SCN5A | 1677 | A | 0 | 0 | yes | no | no | no | yes | no | no | no | no | no | no | no | no |
| SCN5A | 1678 | N | 1 | 0 | yes | no | no | no | yes | no | no | no | no | no | no | no | no |
| SCN5A | 1679 | F | 0 | 0 | no | yes | no | no | yes | no | no | no | no | no | no | no | no |
| SCN5A | 1680 | A | 1 | 1 | no | yes | no | no | yes | no | no | no | no | no | no | no | no |
| SCN5A | 1681 | Y | 1 | 0 | no | yes | no | no | yes | no | no | no | no | no | no | no | no |

| | | | | | | | | | | | | | | | | | |
|---|---|---|---|---|---|---|---|---|---|---|---|---|---|---|---|---|---|
| SCN5A | 1682 | V | 0 | 0 | no | no | yes | no | yes | no | no | no | no | no | no | no | no |
| SCN5A | 1683 | K | 0 | 0 | no | yes | no | no | yes | no | no | no | no | no | no | no | no |
| SCN5A | 1684 | W | 1 | 0 | no | yes | no | no | yes | no | no | no | no | no | no | no | no |
| SCN5A | 1685 | E | 0 | 0 | no | no | yes | no | yes | no | no | no | no | no | no | no | no |
| SCN5A | 1686 | A | 0 | 0 | no | yes | no | no | yes | no | no | no | no | no | no | no | no |
| SCN5A | 1687 | G | 0 | 0 | no | yes | no | no | yes | no | no | no | no | no | no | no | no |
| SCN5A | 1688 | I | 0 | 0 | no | no | yes | no | yes | no | no | no | no | no | no | no | no |
| SCN5A | 1689 | D | 1 | 0 | no | yes | no | no | yes | no | no | no | no | no | no | no | no |
| SCN5A | 1690 | D | 1 | 1 | no | yes | no | no | yes | no | no | no | no | no | no | no | no |
| SCN5A | 1691 | M | 0 | 1 | no | yes | no | no | yes | no | no | no | no | no | no | no | no |
| SCN5A | 1692 | F | 0 | 0 | no | yes | no | no | yes | no | no | no | no | no | no | no | no |
| SCN5A | 1693 | N | 0 | 0 | no | yes | no | no | yes | no | no | no | no | no | no | no | no |
| SCN5A | 1694 | F | 0 | 0 | no | yes | no | no | yes | no | no | no | no | no | no | no | no |
| SCN5A | 1695 | Q | 0 | 0 | no | yes | no | no | yes | no | no | no | no | no | no | no | no |
| SCN5A | 1696 | T | 0 | 0 | no | yes | no | no | yes | no | no | no | no | no | no | no | no |
| SCN5A | 1697 | F | 0 | 0 | yes | no | no | yes | no | no | no | yes | no | no | no | no | no |
| SCN5A | 1698 | A | 1 | 1 | yes | no | no | no | no | no | no | yes | no | no | no | no | no |
| SCN5A | 1699 | N | 0 | 0 | yes | no | no | no | no | no | no | yes | no | no | no | no | no |
| SCN5A | 1700 | S | 0 | 1 | yes | no | no | no | no | no | no | yes | no | no | no | no | no |
| SCN5A | 1701 | M | 1 | 0 | yes | no | no | yes | no | no | no | yes | no | no | no | no | no |
| SCN5A | 1702 | L | 0 | 0 | yes | no | no | yes | no | no | no | yes | no | no | no | no | no |
| SCN5A | 1703 | C | 0 | 1 | yes | no | no | yes | no | no | no | yes | no | no | no | no | no |
| SCN5A | 1704 | L | 0 | 0 | yes | no | no | yes | no | no | no | yes | no | no | no | no | no |
| SCN5A | 1705 | F | 0 | 1 | yes | no | no | yes | no | no | no | yes | no | no | no | no | no |
| SCN5A | 1706 | Q | 0 | 1 | yes | no | no | yes | no | no | no | yes | no | no | no | no | no |
| SCN5A | 1707 | I | 0 | 0 | yes | no | no | yes | no | no | no | yes | no | no | no | no | no |
| SCN5A | 1708 | T | 1 | 1 | yes | no | no | yes | no | no | no | yes | no | no | no | no | no |
| SCN5A | 1709 | T | 1 | 1 | no | yes | no | yes | no | no | no | yes | no | no | no | no | no |
| SCN5A | 1710 | S | 1 | 1 | no | yes | no | yes | no | no | no | yes | no | no | no | no | no |
| SCN5A | 1711 | A | 0 | 0 | no | yes | no | yes | no | no | no | yes | no | no | no | no | no |
| SCN5A | 1712 | G | 1 | 1 | no | yes | no | yes | no | no | no | yes | no | no | no | no | no |
| SCN5A | 1713 | W | 0 | 0 | yes | no | no | yes | no | no | no | yes | no | no | no | no | no |
| SCN5A | 1714 | D | 0 | 2 | yes | no | no | no | no | no | no | yes | no | no | no | no | no |

| | | | | | | | | | | | | | | | | | |
|---|---|---|---|---|---|---|---|---|---|---|---|---|---|---|---|---|---|
| SCN5A | 1715 | G | 0 | 0 | yes | no | no | no | no | no | no | yes | no | no | no | no | no |
| SCN5A | 1716 | L | 0 | 0 | yes | no | no | no | no | no | no | yes | no | no | no | no | no |
| SCN5A | 1717 | L | 0 | 1 | yes | no | no | no | no | no | no | yes | no | no | no | no | no |
| SCN5A | 1718 | S | 1 | 0 | no | yes | no | no | no | no | no | yes | no | no | no | no | no |
| SCN5A | 1719 | P | 0 | 0 | no | yes | no | no | no | no | no | yes | no | no | no | no | no |
| SCN5A | 1720 | I | 0 | 0 | no | yes | no | no | yes | no | no | no | no | no | no | no | no |
| SCN5A | 1721 | L | 0 | 0 | no | yes | no | no | yes | no | no | no | no | no | no | no | no |
| SCN5A | 1722 | N | 0 | 1 | no | yes | no | no | yes | no | no | no | no | no | no | no | no |
| SCN5A | 1723 | T | 1 | 1 | no | yes | no | no | yes | no | no | no | no | no | no | no | no |
| SCN5A | 1724 | G | 0 | 0 | no | yes | no | no | yes | no | no | no | no | no | no | no | no |
| SCN5A | 1725 | P | 1 | 1 | no | yes | no | no | yes | no | no | no | no | no | no | no | no |
| SCN5A | 1726 | P | 0 | 0 | no | yes | no | no | yes | no | no | no | no | no | no | no | no |
| SCN5A | 1727 | Y | 0 | 0 | no | yes | no | no | yes | no | no | no | no | no | no | no | no |
| SCN5A | 1728 | C | 0 | 2 | no | yes | no | no | yes | no | no | no | no | no | no | no | no |
| SCN5A | 1729 | D | 1 | 0 | no | yes | no | no | yes | no | no | no | no | no | no | no | no |
| SCN5A | 1730 | P | 2 | 1 | no | yes | no | no | yes | no | no | no | no | no | no | no | no |
| SCN5A | 1731 | T | 0 | 0 | no | yes | no | no | yes | no | no | no | no | no | no | no | no |
| SCN5A | 1732 | L | 0 | 0 | no | yes | no | no | yes | no | no | no | no | no | no | no | no |
| SCN5A | 1733 | P | 0 | 0 | no | yes | no | no | yes | no | no | no | no | no | no | no | no |
| SCN5A | 1734 | N | 0 | 0 | no | yes | no | no | yes | no | no | no | no | no | no | no | no |
| SCN5A | 1735 | S | 0 | 0 | no | yes | no | no | yes | no | no | no | no | no | no | no | no |
| SCN5A | 1736 | N | 0 | 0 | no | yes | no | no | yes | no | no | no | no | no | no | no | no |
| SCN5A | 1737 | G | 1 | 0 | no | yes | no | no | yes | no | no | no | no | no | no | no | no |
| SCN5A | 1738 | S | 2 | 0 | no | yes | no | no | yes | no | no | no | no | no | no | no | no |
| SCN5A | 1739 | R | 2 | 1 | no | yes | no | no | yes | no | no | no | no | no | no | no | no |
| SCN5A | 1740 | G | 0 | 1 | no | yes | no | no | yes | no | no | no | no | no | no | no | no |
| SCN5A | 1741 | D | 2 | 0 | no | no | yes | no | yes | no | no | no | no | no | no | no | no |
| SCN5A | 1742 | C | 0 | 0 | no | yes | no | no | yes | no | no | no | no | no | no | no | no |
| SCN5A | 1743 | G | 0 | 2 | no | yes | no | no | yes | no | no | no | no | no | no | no | no |
| SCN5A | 1744 | S | 1 | 1 | no | yes | no | no | yes | no | no | no | no | no | no | no | no |
| SCN5A | 1745 | P | 0 | 0 | yes | no | no | no | yes | no | no | no | no | no | no | no | no |
| SCN5A | 1746 | A | 2 | 1 | yes | no | no | no | yes | no | no | no | no | no | no | no | no |
| SCN5A | 1747 | V | 1 | 1 | yes | no | no | no | yes | no | no | no | no | no | no | no | no |

| | | | | | | | | | | | | | | | | | |
|---|---|---|---|---|---|---|---|---|---|---|---|---|---|---|---|---|---|
| SCN5A | 1748 | G | 0 | 1 | yes | no | no | yes | yes | no | no | no | no | no | no | no | no |
| SCN5A | 1749 | I | 0 | 0 | yes | no | no | yes | no | no | no | no | no | no | no | no | no |
| SCN5A | 1750 | L | 1 | 0 | yes | no | no | yes | no | no | no | no | no | no | no | no | no |
| SCN5A | 1751 | F | 0 | 0 | yes | no | no | yes | no | no | no | no | no | no | no | no | no |
| SCN5A | 1752 | F | 0 | 0 | yes | no | no | yes | no | no | no | no | no | no | no | no | no |
| SCN5A | 1753 | T | 1 | 0 | yes | no | no | yes | no | no | no | no | no | no | no | no | no |
| SCN5A | 1754 | T | 0 | 0 | yes | no | no | yes | no | no | no | no | no | no | no | no | no |
| SCN5A | 1755 | Y | 0 | 0 | yes | no | no | yes | no | no | no | no | no | no | no | no | no |
| SCN5A | 1756 | I | 1 | 0 | yes | no | no | yes | no | no | no | no | no | no | no | no | no |
| SCN5A | 1757 | I | 0 | 0 | yes | no | no | yes | no | no | no | no | no | no | no | no | no |
| SCN5A | 1758 | I | 1 | 1 | yes | no | no | yes | no | no | no | no | no | no | no | no | no |
| SCN5A | 1759 | S | 1 | 0 | yes | no | no | yes | no | no | no | no | no | no | no | no | no |
| SCN5A | 1760 | F | 0 | 1 | yes | no | no | yes | no | no | no | no | no | no | no | no | no |
| SCN5A | 1761 | L | 0 | 2 | yes | no | no | yes | no | no | no | no | no | no | no | no | no |
| SCN5A | 1762 | I | 1 | 0 | yes | no | no | yes | no | no | no | no | no | no | no | no | no |
| SCN5A | 1763 | V | 0 | 2 | yes | no | no | yes | no | no | no | no | no | no | no | no | no |
| SCN5A | 1764 | V | 0 | 1 | yes | no | no | yes | no | no | no | no | no | no | no | no | no |
| SCN5A | 1765 | N | 0 | 0 | yes | no | no | yes | no | no | no | no | no | no | no | no | no |
| SCN5A | 1766 | M | 1 | 2 | yes | no | no | yes | no | no | no | no | no | no | no | no | no |
| SCN5A | 1767 | Y | 0 | 1 | yes | no | no | yes | no | no | yes | no | no | no | no | no | no |
| SCN5A | 1768 | I | 0 | 1 | yes | no | no | yes | no | no | no | no | no | no | no | no | no |
| SCN5A | 1769 | A | 0 | 0 | yes | no | no | yes | no | no | no | no | no | no | no | no | no |
| SCN5A | 1770 | I | 1 | 0 | yes | no | no | yes | no | no | no | no | no | no | no | no | no |
| SCN5A | 1771 | I | 1 | 0 | yes | no | no | yes | no | no | yes | no | no | no | no | no | no |
| SCN5A | 1772 | L | 0 | 1 | yes | no | no | no | no | yes | no | no | no | no | no | no | no |
| SCN5A | 1773 | E | 0 | 0 | yes | no | no | yes | no | yes | no | no | no | no | no | no | no |
| SCN5A | 1774 | N | 0 | 2 | yes | no | no | yes | no | yes | no | no | no | no | no | no | no |
| SCN5A | 1775 | F | 1 | 0 | yes | no | no | no | no | yes | yes | no | no | no | no | no | no |
| SCN5A | 1776 | S | 0 | 0 | yes | no | no | no | no | yes | no | no | no | no | no | no | no |
| SCN5A | 1777 | V | 1 | 1 | yes | no | no | no | no | yes | no | no | no | no | no | no | no |
| SCN5A | 1778 | A | 0 | 0 | yes | no | no | no | no | yes | no | no | no | no | no | no | no |
| SCN5A | 1779 | T | 1 | 1 | no | yes | no | no | no | yes | yes | no | no | no | no | no | no |
| SCN5A | 1780 | E | 0 | 0 | no | yes | no | no | no | yes | no | no | no | no | no | no | no |

| | | | | | | | | | | | | | | | | | |
|---|---|---|---|---|---|---|---|---|---|---|---|---|---|---|---|---|---|
| SCN5A | 1781 | E | 1 | 1 | no | yes | no | no | no | yes | no | no | no | no | no | no | no |
| GRIN2B | 34 | S | 1 | 0 | no | yes | no | no | yes | no | no | no | no | no | no | no | no |
| GRIN2B | 35 | I | 2 | 0 | no | yes | no | no | yes | no | no | no | no | no | no | no | no |
| GRIN2B | 36 | G | 0 | 0 | no | no | yes | no | yes | no | no | no | no | no | no | no | no |
| GRIN2B | 37 | I | 0 | 0 | no | no | yes | no | yes | no | no | no | no | no | no | no | no |
| GRIN2B | 38 | A | 0 | 0 | no | no | yes | no | yes | no | no | no | no | no | no | no | no |
| GRIN2B | 39 | V | 0 | 0 | no | no | yes | no | yes | no | no | no | no | no | no | no | no |
| GRIN2B | 40 | I | 0 | 0 | no | no | yes | no | yes | no | no | no | no | no | no | no | no |
| GRIN2B | 41 | L | 0 | 0 | no | no | yes | no | yes | no | no | no | no | no | no | no | no |
| GRIN2B | 42 | V | 0 | 0 | no | yes | no | no | yes | no | no | no | no | no | no | no | no |
| GRIN2B | 45 | S | 0 | 0 | no | yes | no | no | yes | no | no | no | no | no | no | no | no |
| GRIN2B | 46 | D | 1 | 0 | no | yes | no | no | yes | no | no | no | no | no | no | no | no |
| GRIN2B | 47 | E | 1 | 0 | no | yes | no | no | yes | no | no | no | no | no | no | no | no |
| GRIN2B | 48 | V | 1 | 0 | yes | no | no | no | yes | no | no | no | no | no | no | no | no |
| GRIN2B | 49 | A | 0 | 0 | yes | no | no | no | yes | no | no | no | no | no | no | no | no |
| GRIN2B | 50 | I | 0 | 0 | yes | no | no | no | yes | no | no | no | no | no | no | no | no |
| GRIN2B | 51 | K | 0 | 0 | yes | no | no | no | yes | no | no | no | no | no | no | no | no |
| GRIN2B | 52 | D | 0 | 0 | no | yes | no | no | yes | no | no | no | no | no | no | no | no |
| GRIN2B | 53 | A | 1 | 0 | no | yes | no | no | yes | no | no | no | no | no | no | no | no |
| GRIN2B | 54 | H | 0 | 0 | no | yes | no | no | yes | no | no | no | no | no | no | no | no |
| GRIN2B | 55 | E | 1 | 0 | no | yes | no | no | yes | no | no | no | no | no | no | no | no |
| GRIN2B | 56 | K | 0 | 0 | no | yes | no | no | yes | no | no | no | no | no | no | no | no |
| GRIN2B | 57 | D | 1 | 0 | no | yes | no | no | yes | no | no | no | no | no | no | no | no |
| GRIN2B | 58 | D | 0 | 0 | no | yes | no | no | yes | no | no | no | no | no | no | no | no |
| GRIN2B | 59 | F | 0 | 0 | no | yes | no | no | yes | no | no | no | no | no | no | no | no |
| GRIN2B | 60 | H | 0 | 0 | no | yes | no | no | yes | no | no | no | no | no | no | no | no |
| GRIN2B | 61 | H | 1 | 0 | no | yes | no | no | yes | no | no | no | no | no | no | no | no |
| GRIN2B | 62 | L | 1 | 0 | no | yes | no | no | yes | no | no | no | no | no | no | no | no |
| GRIN2B | 63 | S | 0 | 0 | no | yes | no | no | yes | no | no | no | no | no | no | no | no |
| GRIN2B | 64 | V | 2 | 0 | no | yes | no | no | yes | no | no | no | no | no | no | no | no |
| GRIN2B | 65 | V | 1 | 0 | no | yes | no | no | yes | no | no | no | no | no | no | no | no |
| GRIN2B | 66 | P | 0 | 0 | no | yes | no | no | yes | no | no | no | no | no | no | no | no |
| GRIN2B | 67 | R | 1 | 0 | no | no | yes | no | yes | no | no | no | no | no | no | no | no |

| | | | | | | | | | | | | | | | | | |
|---|---|---|---|---|---|---|---|---|---|---|---|---|---|---|---|---|---|
| GRIN2B | 68 | V | 0 | 0 | no | no | yes | no | yes | no | no | no | no | no | no | no | no |
| GRIN2B | 69 | E | 0 | 0 | no | no | yes | no | yes | no | no | no | no | no | no | no | no |
| GRIN2B | 70 | L | 0 | 0 | no | no | yes | no | yes | no | no | no | no | no | no | no | no |
| GRIN2B | 71 | V | 0 | 0 | no | no | yes | no | yes | no | no | no | no | no | no | no | no |
| GRIN2B | 72 | A | 0 | 0 | no | no | yes | no | yes | no | no | no | no | no | no | no | no |
| GRIN2B | 73 | M | 0 | 0 | no | yes | no | no | yes | no | no | no | no | no | no | no | no |
| GRIN2B | 74 | N | 1 | 0 | no | yes | no | no | yes | no | no | no | no | no | no | no | no |
| GRIN2B | 75 | E | 0 | 0 | no | yes | no | no | yes | no | no | no | no | no | no | no | no |
| GRIN2B | 76 | T | 0 | 0 | no | yes | no | no | yes | no | no | no | yes | no | no | no | no |
| GRIN2B | 77 | D | 0 | 0 | no | yes | no | no | yes | no | no | no | yes | no | no | no | no |
| GRIN2B | 78 | P | 0 | 0 | yes | no | no | no | yes | no | no | no | no | no | no | no | no |
| GRIN2B | 79 | K | 0 | 0 | yes | no | no | no | yes | no | no | no | no | no | no | no | no |
| GRIN2B | 80 | S | 0 | 0 | yes | no | no | no | yes | no | no | no | no | no | no | no | no |
| GRIN2B | 81 | I | 1 | 0 | yes | no | no | no | yes | no | no | no | no | no | no | no | no |
| GRIN2B | 82 | I | 0 | 0 | yes | no | no | no | yes | no | no | no | no | no | no | no | no |
| GRIN2B | 83 | T | 0 | 0 | yes | no | no | no | yes | no | no | no | no | no | no | no | no |
| GRIN2B | 84 | R | 2 | 0 | yes | no | no | no | yes | no | no | no | no | no | no | no | no |
| GRIN2B | 85 | I | 0 | 0 | yes | no | no | no | yes | no | no | no | no | no | no | no | no |
| GRIN2B | 86 | C | 0 | 0 | yes | no | no | no | yes | no | no | no | no | no | no | no | no |
| GRIN2B | 87 | D | 0 | 0 | yes | no | no | no | yes | no | no | no | no | no | no | no | no |
| GRIN2B | 88 | L | 1 | 0 | yes | no | no | no | yes | no | no | no | no | no | no | no | no |
| GRIN2B | 89 | M | 1 | 0 | yes | no | no | no | yes | no | no | no | no | no | no | no | no |
| GRIN2B | 90 | S | 0 | 0 | yes | no | no | no | yes | no | no | no | no | no | no | no | no |
| GRIN2B | 91 | D | 0 | 0 | no | yes | no | no | yes | no | no | no | no | no | no | no | no |
| GRIN2B | 92 | R | 2 | 0 | no | yes | no | no | yes | no | no | no | no | no | no | no | no |
| GRIN2B | 93 | K | 0 | 0 | no | yes | no | no | yes | no | no | no | no | no | no | no | no |
| GRIN2B | 94 | I | 0 | 0 | no | yes | no | no | yes | no | no | no | no | no | no | no | no |
| GRIN2B | 95 | Q | 0 | 0 | no | yes | no | no | yes | no | no | no | no | no | no | no | no |
| GRIN2B | 96 | G | 0 | 0 | no | yes | no | no | yes | no | no | no | no | no | no | no | no |
| GRIN2B | 97 | V | 1 | 0 | no | no | yes | no | yes | no | no | no | no | no | no | no | no |
| GRIN2B | 98 | V | 0 | 0 | no | no | yes | no | yes | no | no | no | no | no | no | no | no |
| GRIN2B | 99 | F | 0 | 0 | no | no | yes | no | yes | no | no | no | no | no | no | no | no |
| GRIN2B | 100 | A | 0 | 0 | no | no | yes | no | yes | no | no | no | no | no | no | no | no |

| | | | | | | | | | | | | | | | | | |
|---|---|---|---|---|---|---|---|---|---|---|---|---|---|---|---|---|---|
| GRIN2B | 101 | D | 0 | 0 | no | yes | no | no | yes | no | no | no | no | no | no | no | no |
| GRIN2B | 102 | D | 0 | 0 | no | yes | no | no | yes | no | no | no | no | no | no | no | no |
| GRIN2B | 103 | T | 0 | 0 | no | yes | no | no | yes | no | no | no | no | no | no | no | no |
| GRIN2B | 104 | D | 0 | 0 | no | yes | no | no | yes | no | no | no | no | no | no | no | no |
| GRIN2B | 105 | Q | 2 | 0 | no | yes | no | no | yes | no | no | no | no | no | no | no | no |
| GRIN2B | 106 | E | 0 | 0 | no | yes | no | no | yes | no | no | no | no | no | no | no | no |
| GRIN2B | 107 | A | 0 | 0 | yes | no | no | no | yes | no | no | no | yes | no | no | no | no |
| GRIN2B | 108 | I | 0 | 0 | yes | no | no | no | yes | no | no | no | no | no | no | no | no |
| GRIN2B | 109 | A | 2 | 0 | yes | no | no | no | yes | no | no | no | no | no | no | no | no |
| GRIN2B | 110 | Q | 0 | 0 | yes | no | no | no | yes | no | no | no | no | no | no | no | no |
| GRIN2B | 111 | I | 0 | 0 | yes | no | no | no | yes | no | no | no | yes | no | no | no | no |
| GRIN2B | 112 | L | 0 | 0 | yes | no | no | no | yes | no | no | no | no | no | no | no | no |
| GRIN2B | 113 | D | 0 | 0 | yes | no | no | no | yes | no | no | no | no | no | no | no | no |
| GRIN2B | 114 | F | 0 | 0 | yes | no | no | no | yes | no | no | no | yes | no | no | no | no |
| GRIN2B | 115 | I | 0 | 0 | yes | no | no | no | yes | no | no | no | no | no | no | no | no |
| GRIN2B | 116 | S | 0 | 0 | yes | no | no | no | yes | no | no | no | no | no | no | no | no |
| GRIN2B | 117 | A | 0 | 0 | yes | no | no | no | yes | no | no | no | no | no | no | no | no |
| GRIN2B | 118 | Q | 0 | 0 | yes | no | no | no | yes | no | no | no | yes | no | no | no | no |
| GRIN2B | 119 | T | 0 | 0 | yes | no | no | no | yes | no | no | no | no | no | no | no | no |
| GRIN2B | 120 | L | 0 | 0 | no | yes | no | no | yes | no | no | no | no | no | no | no | no |
| GRIN2B | 121 | T | 0 | 0 | no | yes | no | no | yes | no | no | no | no | no | no | no | no |
| GRIN2B | 122 | P | 0 | 0 | no | yes | no | no | yes | no | no | no | no | no | no | no | no |
| GRIN2B | 123 | I | 0 | 0 | no | no | yes | no | yes | no | no | no | no | no | no | no | no |
| GRIN2B | 124 | L | 0 | 0 | no | no | yes | no | yes | no | no | no | no | no | no | no | no |
| GRIN2B | 125 | G | 0 | 0 | no | no | yes | no | yes | no | no | no | no | no | no | no | no |
| GRIN2B | 126 | I | 0 | 0 | no | yes | no | no | yes | no | no | no | no | no | no | no | no |
| GRIN2B | 127 | H | 0 | 0 | no | yes | no | no | yes | no | no | no | no | no | no | no | no |
| GRIN2B | 128 | G | 0 | 0 | no | yes | no | no | yes | no | no | no | no | no | no | no | no |
| GRIN2B | 129 | G | 0 | 0 | no | yes | no | no | yes | no | no | no | no | no | no | no | no |
| GRIN2B | 130 | S | 0 | 0 | no | yes | no | no | yes | no | no | no | no | no | no | no | no |
| GRIN2B | 131 | S | 0 | 0 | no | yes | no | no | yes | no | no | no | no | no | no | no | no |
| GRIN2B | 132 | M | 0 | 0 | no | yes | no | no | yes | no | no | no | no | no | no | no | no |
| GRIN2B | 133 | I | 0 | 0 | no | yes | no | no | yes | no | no | no | no | no | no | no | no |

| | | | | | | | | | | | | | | | | | |
|---|---|---|---|---|---|---|---|---|---|---|---|---|---|---|---|---|---|
| GRIN2B | 134 | M | 0 | 0 | no | yes | no | no | yes | no | no | no | no | no | no | no | no |
| GRIN2B | 135 | A | 0 | 0 | no | yes | no | no | yes | no | no | no | no | no | no | no | no |
| GRIN2B | 136 | D | 0 | 0 | no | yes | no | no | yes | no | no | no | yes | no | no | no | no |
| GRIN2B | 137 | K | 0 | 0 | no | yes | no | no | yes | no | no | no | no | no | no | no | no |
| GRIN2B | 138 | D | 0 | 0 | no | yes | no | no | yes | no | no | no | no | no | no | no | no |
| GRIN2B | 139 | E | 0 | 0 | no | yes | no | no | yes | no | no | no | no | no | no | no | no |
| GRIN2B | 140 | S | 1 | 0 | no | yes | no | no | yes | no | no | no | no | no | no | no | no |
| GRIN2B | 141 | S | 0 | 0 | no | yes | no | no | yes | no | no | no | no | no | no | no | no |
| GRIN2B | 142 | M | 2 | 0 | no | yes | no | no | yes | no | no | no | no | no | no | no | no |
| GRIN2B | 143 | F | 2 | 0 | no | yes | no | no | yes | no | no | no | no | no | no | no | no |
| GRIN2B | 144 | F | 0 | 0 | no | yes | no | no | yes | no | no | no | no | no | no | no | no |
| GRIN2B | 145 | Q | 0 | 0 | no | yes | no | no | yes | no | no | no | no | no | no | no | no |
| GRIN2B | 146 | F | 0 | 0 | no | yes | no | no | yes | no | no | no | no | no | no | no | no |
| GRIN2B | 147 | G | 0 | 0 | no | yes | no | no | yes | no | no | no | no | no | no | no | no |
| GRIN2B | 148 | P | 0 | 0 | no | yes | no | no | yes | no | no | no | no | no | no | no | no |
| GRIN2B | 149 | S | 0 | 0 | no | yes | no | no | yes | no | no | no | no | no | no | no | no |
| GRIN2B | 150 | I | 0 | 1 | yes | no | no | no | yes | no | no | no | no | no | no | no | no |
| GRIN2B | 151 | E | 0 | 0 | yes | no | no | no | yes | no | no | no | no | no | no | no | no |
| GRIN2B | 152 | Q | 0 | 0 | yes | no | no | no | yes | no | no | no | no | no | no | no | no |
| GRIN2B | 153 | Q | 0 | 0 | yes | no | no | no | yes | no | no | no | no | no | no | no | no |
| GRIN2B | 154 | A | 0 | 0 | yes | no | no | no | yes | no | no | no | no | no | no | no | no |
| GRIN2B | 155 | S | 0 | 0 | yes | no | no | no | yes | no | no | no | no | no | no | no | no |
| GRIN2B | 156 | V | 0 | 0 | yes | no | no | no | yes | no | no | no | no | no | no | no | no |
| GRIN2B | 157 | M | 0 | 0 | yes | no | no | no | yes | no | no | no | no | no | no | no | no |
| GRIN2B | 158 | L | 1 | 0 | yes | no | no | no | yes | no | no | no | no | no | no | no | no |
| GRIN2B | 159 | N | 1 | 0 | yes | no | no | no | yes | no | no | no | no | no | no | no | no |
| GRIN2B | 160 | I | 0 | 0 | yes | no | no | no | yes | no | no | no | no | no | no | no | no |
| GRIN2B | 161 | M | 0 | 0 | yes | no | no | no | yes | no | no | no | no | no | no | no | no |
| GRIN2B | 162 | E | 0 | 0 | yes | no | no | no | yes | no | no | no | no | no | no | no | no |
| GRIN2B | 163 | E | 0 | 0 | yes | no | no | no | yes | no | no | no | no | no | no | no | no |
| GRIN2B | 164 | Y | 0 | 0 | yes | no | no | no | yes | no | no | no | no | no | no | no | no |
| GRIN2B | 165 | D | 0 | 0 | no | yes | no | no | yes | no | no | no | no | no | no | no | no |
| GRIN2B | 166 | W | 0 | 0 | no | yes | no | no | yes | no | no | no | no | no | no | no | no |

| | | | | | | | | | | | | | | | | | |
|---|---|---|---|---|---|---|---|---|---|---|---|---|---|---|---|---|---|
| GRIN2B | 167 | Y | 0 | 0 | no | yes | no | no | yes | no | no | no | no | no | no | no | no |
| GRIN2B | 168 | I | 0 | 0 | no | yes | no | no | yes | no | no | no | no | no | no | no | no |
| GRIN2B | 169 | F | 0 | 0 | no | yes | no | no | yes | no | no | no | no | no | no | no | no |
| GRIN2B | 170 | S | 0 | 0 | no | yes | no | no | yes | no | no | no | no | no | no | no | no |
| GRIN2B | 171 | I | 0 | 0 | no | no | yes | no | yes | no | no | no | no | no | no | no | no |
| GRIN2B | 172 | V | 1 | 0 | no | yes | no | no | yes | no | no | no | no | no | no | no | no |
| GRIN2B | 173 | T | 0 | 0 | no | yes | no | no | yes | no | no | no | no | no | no | no | no |
| GRIN2B | 174 | T | 0 | 0 | no | yes | no | no | yes | no | no | no | no | no | no | no | no |
| GRIN2B | 175 | Y | 0 | 0 | no | yes | no | no | yes | no | no | no | yes | no | no | no | no |
| GRIN2B | 176 | F | 0 | 0 | no | yes | no | no | yes | no | no | no | no | no | no | no | no |
| GRIN2B | 177 | P | 1 | 0 | yes | no | no | no | yes | no | no | no | yes | no | no | no | no |
| GRIN2B | 178 | G | 0 | 0 | yes | no | no | no | yes | no | no | no | no | no | no | no | no |
| GRIN2B | 179 | Y | 1 | 0 | yes | no | no | no | yes | no | no | no | no | no | no | no | no |
| GRIN2B | 180 | Q | 1 | 0 | yes | no | no | no | yes | no | no | no | no | no | no | no | no |
| GRIN2B | 181 | D | 0 | 0 | yes | no | no | no | yes | no | no | no | no | no | no | no | no |
| GRIN2B | 182 | F | 0 | 0 | yes | no | no | no | yes | no | no | no | no | no | no | no | no |
| GRIN2B | 183 | V | 0 | 0 | yes | no | no | no | yes | no | no | no | no | no | no | no | no |
| GRIN2B | 184 | N | 0 | 0 | yes | no | no | no | yes | no | no | no | no | no | no | no | no |
| GRIN2B | 185 | K | 0 | 0 | yes | no | no | no | yes | no | no | no | no | no | no | no | no |
| GRIN2B | 186 | I | 0 | 0 | yes | no | no | no | yes | no | no | no | no | no | no | no | no |
| GRIN2B | 187 | R | 1 | 0 | yes | no | no | no | yes | no | no | no | no | no | no | no | no |
| GRIN2B | 188 | S | 0 | 0 | yes | no | no | no | yes | no | no | no | no | no | no | no | no |
| GRIN2B | 189 | T | 0 | 0 | yes | no | no | no | yes | no | no | no | no | no | no | no | no |
| GRIN2B | 190 | I | 0 | 0 | yes | no | no | no | yes | no | no | no | no | no | no | no | no |
| GRIN2B | 191 | E | 0 | 0 | yes | no | no | no | yes | no | no | no | yes | no | no | no | no |
| GRIN2B | 192 | N | 0 | 0 | yes | no | no | no | yes | no | no | no | yes | no | no | no | no |
| GRIN2B | 193 | S | 0 | 0 | no | yes | no | no | yes | no | no | no | no | no | no | no | no |
| GRIN2B | 194 | F | 0 | 0 | no | yes | no | no | yes | no | no | no | no | no | no | no | no |
| GRIN2B | 195 | V | 0 | 0 | no | yes | no | no | yes | no | no | no | no | no | no | no | no |
| GRIN2B | 196 | G | 0 | 0 | no | yes | no | no | yes | no | no | no | no | no | no | no | no |
| GRIN2B | 197 | W | 0 | 0 | no | yes | no | no | yes | no | no | no | no | no | no | no | no |
| GRIN2B | 198 | E | 0 | 0 | no | yes | no | no | yes | no | yes | no | no | no | no | no | no |
| GRIN2B | 199 | L | 0 | 0 | no | yes | no | no | yes | no | no | no | no | no | no | no | no |

| | | | | | | | | | | | | | | | | | |
|---|---|---|---|---|---|---|---|---|---|---|---|---|---|---|---|---|---|
| GRIN2B | 200 | E | 0 | 0 | no | yes | no | no | yes | no | no | no | no | no | no | no | no |
| GRIN2B | 215 | K | 0 | 0 | no | yes | no | no | yes | no | no | no | no | no | no | no | no |
| GRIN2B | 216 | I | 2 | 0 | yes | no | no | no | yes | no | no | no | no | no | no | no | no |
| GRIN2B | 217 | Q | 0 | 0 | yes | no | no | no | yes | no | no | no | yes | no | no | no | no |
| GRIN2B | 218 | N | 0 | 0 | yes | no | no | no | yes | no | no | no | yes | no | no | no | no |
| GRIN2B | 219 | Q | 0 | 0 | yes | no | no | no | yes | no | no | no | no | no | no | no | no |
| GRIN2B | 220 | L | 0 | 0 | yes | no | no | no | yes | no | no | no | no | no | no | no | no |
| GRIN2B | 221 | K | 0 | 0 | yes | no | no | no | yes | no | yes | no | yes | no | no | no | no |
| GRIN2B | 222 | K | 0 | 0 | yes | no | no | no | yes | no | yes | no | no | no | no | no | no |
| GRIN2B | 223 | L | 0 | 0 | no | yes | no | no | yes | no | no | no | no | no | no | no | no |
| GRIN2B | 224 | Q | 0 | 0 | no | yes | no | no | yes | no | yes | no | no | no | no | no | no |
| GRIN2B | 225 | S | 1 | 0 | no | yes | no | no | yes | no | no | no | no | no | no | no | no |
| GRIN2B | 226 | P | 1 | 0 | no | yes | no | no | yes | no | no | no | no | no | no | no | no |
| GRIN2B | 227 | I | 1 | 0 | no | no | yes | no | yes | no | no | no | no | no | no | no | no |
| GRIN2B | 228 | I | 0 | 0 | no | no | yes | no | yes | no | no | no | no | no | no | no | no |
| GRIN2B | 229 | L | 1 | 0 | no | no | yes | no | yes | no | no | no | no | no | no | no | no |
| GRIN2B | 230 | L | 1 | 0 | no | yes | no | no | yes | no | no | no | no | no | no | no | no |
| GRIN2B | 231 | Y | 0 | 0 | no | yes | no | no | yes | no | no | no | no | no | no | no | no |
| GRIN2B | 232 | C | 0 | 0 | no | yes | no | no | yes | no | no | no | no | no | no | no | no |
| GRIN2B | 233 | T | 0 | 0 | no | yes | no | no | yes | no | no | no | no | no | no | no | no |
| GRIN2B | 234 | K | 0 | 0 | yes | no | no | no | yes | no | no | no | no | no | no | no | no |
| GRIN2B | 235 | E | 0 | 0 | yes | no | no | no | yes | no | no | no | no | no | no | no | no |
| GRIN2B | 236 | E | 0 | 0 | yes | no | no | no | yes | no | no | no | no | no | no | no | no |
| GRIN2B | 237 | A | 0 | 0 | yes | no | no | no | yes | no | no | no | no | no | no | no | no |
| GRIN2B | 238 | T | 0 | 0 | yes | no | no | no | yes | no | no | no | no | no | no | no | no |
| GRIN2B | 239 | Y | 0 | 0 | yes | no | no | no | yes | no | yes | no | no | no | no | no | no |
| GRIN2B | 240 | I | 1 | 0 | yes | no | no | no | yes | no | no | no | no | no | no | no | no |
| GRIN2B | 241 | F | 0 | 0 | yes | no | no | no | yes | no | no | no | no | no | no | no | no |
| GRIN2B | 242 | E | 0 | 0 | yes | no | no | no | yes | no | yes | no | no | no | no | no | no |
| GRIN2B | 243 | V | 0 | 0 | yes | no | no | no | yes | no | yes | no | no | no | no | no | no |
| GRIN2B | 244 | A | 0 | 0 | yes | no | no | no | yes | no | no | no | no | no | no | no | no |
| GRIN2B | 245 | N | 0 | 0 | yes | no | no | no | yes | no | no | no | no | no | no | no | no |
| GRIN2B | 246 | S | 0 | 0 | yes | no | no | no | yes | no | yes | no | no | no | no | no | no |

| | | | | | | | | | | | | | | | | | |
|---|---|---|---|---|---|---|---|---|---|---|---|---|---|---|---|---|---|
| GRIN2B | 247 | V | 0 | 0 | yes | no | no | no | yes | no | yes | no | yes | no | no | no | no |
| GRIN2B | 248 | G | 0 | 0 | no | yes | no | no | yes | no | no | no | no | no | no | no | no |
| GRIN2B | 253 | G | 0 | 0 | no | yes | no | no | yes | no | no | no | no | no | no | no | no |
| GRIN2B | 254 | Y | 0 | 0 | no | yes | no | no | yes | no | no | no | no | no | no | no | no |
| GRIN2B | 255 | T | 1 | 0 | no | no | yes | no | yes | no | no | no | no | no | no | no | no |
| GRIN2B | 256 | W | 0 | 0 | no | no | yes | no | yes | no | no | no | no | no | no | no | no |
| GRIN2B | 257 | I | 0 | 0 | no | no | yes | no | yes | no | no | no | no | no | no | no | no |
| GRIN2B | 258 | V | 0 | 0 | no | yes | no | no | yes | no | no | no | no | no | no | no | no |
| GRIN2B | 259 | P | 0 | 0 | no | yes | no | no | yes | no | no | no | no | no | no | no | no |
| GRIN2B | 260 | S | 0 | 0 | no | yes | no | no | yes | no | no | no | no | no | no | no | no |
| GRIN2B | 261 | L | 0 | 0 | no | yes | no | no | yes | no | no | no | no | no | no | no | no |
| GRIN2B | 262 | V | 0 | 0 | no | yes | no | no | yes | no | no | no | no | no | no | no | no |
| GRIN2B | 263 | A | 0 | 0 | no | yes | no | no | yes | no | no | no | no | no | no | no | no |
| GRIN2B | 264 | G | 0 | 0 | no | yes | no | no | yes | no | no | no | no | no | no | no | no |
| GRIN2B | 265 | D | 0 | 0 | no | yes | no | no | yes | no | no | no | no | no | no | no | no |
| GRIN2B | 266 | T | 1 | 0 | no | yes | no | no | yes | no | no | no | no | no | no | no | no |
| GRIN2B | 267 | D | 0 | 0 | no | yes | no | no | yes | no | no | no | no | no | no | no | no |
| GRIN2B | 268 | T | 1 | 0 | no | yes | no | no | yes | no | no | no | no | no | no | no | no |
| GRIN2B | 269 | V | 1 | 0 | no | yes | no | no | yes | no | no | no | no | no | no | no | no |
| GRIN2B | 270 | P | 0 | 0 | no | yes | no | no | yes | no | no | no | no | no | no | no | no |
| GRIN2B | 271 | A | 1 | 0 | no | yes | no | no | yes | no | no | no | no | no | no | no | no |
| GRIN2B | 272 | E | 1 | 0 | no | yes | no | no | yes | no | no | no | no | no | no | no | no |
| GRIN2B | 273 | F | 1 | 0 | no | yes | no | no | yes | no | no | no | no | no | no | no | no |
| GRIN2B | 274 | P | 0 | 0 | no | yes | no | no | yes | no | no | no | no | no | no | no | no |
| GRIN2B | 275 | T | 1 | 0 | no | yes | no | no | yes | no | no | no | no | no | no | no | no |
| GRIN2B | 276 | G | 0 | 0 | no | yes | no | no | yes | no | no | no | no | no | no | no | no |
| GRIN2B | 277 | L | 0 | 0 | no | yes | no | no | yes | no | no | no | no | no | no | no | no |
| GRIN2B | 278 | I | 0 | 0 | no | no | yes | no | yes | no | no | no | no | no | no | no | no |
| GRIN2B | 279 | S | 0 | 0 | no | no | yes | no | yes | no | no | no | no | no | no | no | no |
| GRIN2B | 280 | V | 1 | 0 | no | no | yes | no | yes | no | no | no | no | no | no | no | no |
| GRIN2B | 281 | S | 0 | 0 | no | no | yes | no | yes | no | no | no | no | no | no | no | no |
| GRIN2B | 282 | Y | 0 | 0 | no | no | yes | no | yes | no | no | no | no | no | no | no | no |
| GRIN2B | 283 | D | 0 | 0 | no | yes | no | no | yes | no | no | no | no | no | no | no | no |

| | | | | | | | | | | | | | | | | | |
|---|---|---|---|---|---|---|---|---|---|---|---|---|---|---|---|---|---|
| GRIN2B | 284 | E | 0 | 0 | no | yes | no | no | yes | no | no | no | no | no | no | no | no |
| GRIN2B | 285 | W | 0 | 0 | no | yes | no | no | yes | no | no | no | no | no | no | no | no |
| GRIN2B | 286 | D | 0 | 0 | no | yes | no | no | yes | no | no | no | no | no | no | no | no |
| GRIN2B | 287 | Y | 0 | 0 | no | yes | no | no | yes | no | no | no | no | no | no | no | no |
| GRIN2B | 288 | G | 0 | 0 | no | yes | no | no | yes | no | no | no | no | no | no | no | no |
| GRIN2B | 289 | L | 0 | 0 | yes | no | no | no | yes | no | no | no | no | no | no | no | no |
| GRIN2B | 290 | P | 0 | 0 | yes | no | no | no | yes | no | no | no | no | no | no | no | no |
| GRIN2B | 291 | A | 1 | 0 | yes | no | no | no | yes | no | no | no | no | no | no | no | no |
| GRIN2B | 292 | R | 0 | 0 | yes | no | no | no | yes | no | no | no | no | no | no | no | no |
| GRIN2B | 293 | V | 0 | 0 | yes | no | no | no | yes | no | no | no | no | no | no | no | no |
| GRIN2B | 294 | R | 0 | 0 | yes | no | no | no | yes | no | no | no | no | no | no | no | no |
| GRIN2B | 295 | D | 0 | 0 | yes | no | no | no | yes | no | no | no | no | no | no | no | no |
| GRIN2B | 296 | G | 0 | 0 | yes | no | no | no | yes | no | no | no | no | no | no | no | no |
| GRIN2B | 297 | I | 0 | 0 | yes | no | no | no | yes | no | no | no | no | no | no | no | no |
| GRIN2B | 298 | A | 0 | 0 | yes | no | no | no | yes | no | no | no | no | no | no | no | no |
| GRIN2B | 299 | I | 1 | 0 | yes | no | no | no | yes | no | no | no | no | no | no | no | no |
| GRIN2B | 300 | I | 0 | 0 | yes | no | no | no | yes | no | no | no | no | no | no | no | no |
| GRIN2B | 301 | T | 0 | 0 | yes | no | no | no | yes | no | no | no | no | no | no | no | no |
| GRIN2B | 302 | T | 1 | 0 | yes | no | no | no | yes | no | no | no | no | no | no | no | no |
| GRIN2B | 303 | A | 0 | 0 | yes | no | no | no | yes | no | no | no | no | no | no | no | no |
| GRIN2B | 304 | A | 0 | 0 | yes | no | no | no | yes | no | no | no | no | no | no | no | no |
| GRIN2B | 305 | S | 0 | 0 | yes | no | no | no | yes | no | no | no | no | no | no | no | no |
| GRIN2B | 306 | D | 0 | 0 | yes | no | no | no | yes | no | no | no | no | no | no | no | no |
| GRIN2B | 307 | M | 0 | 0 | yes | no | no | no | yes | no | no | no | no | no | no | no | no |
| GRIN2B | 308 | L | 0 | 0 | yes | no | no | no | yes | no | no | no | no | no | no | no | no |
| GRIN2B | 309 | S | 0 | 0 | yes | no | no | no | yes | no | no | no | no | no | no | no | no |
| GRIN2B | 310 | E | 0 | 0 | yes | no | no | no | yes | no | no | no | no | no | no | no | no |
| GRIN2B | 311 | H | 0 | 0 | yes | no | no | no | yes | no | no | no | no | no | no | no | no |
| GRIN2B | 312 | S | 1 | 0 | no | yes | no | no | yes | no | no | no | no | no | no | no | no |
| GRIN2B | 313 | F | 0 | 0 | no | yes | no | no | yes | no | no | no | no | no | no | no | no |
| GRIN2B | 314 | I | 0 | 0 | no | yes | no | no | yes | no | no | no | no | no | no | no | no |
| GRIN2B | 315 | P | 0 | 0 | no | yes | no | no | yes | no | no | no | no | no | no | no | no |
| GRIN2B | 316 | E | 0 | 0 | no | yes | no | no | yes | no | no | no | no | no | no | no | no |

| | | | | | | | | | | | | | | | | | |
|---|---|---|---|---|---|---|---|---|---|---|---|---|---|---|---|---|---|
| GRIN2B | 317 | P | 0 | 0 | no | yes | no | no | yes | no | no | no | no | no | no | no | no |
| GRIN2B | 318 | K | 0 | 0 | no | yes | no | no | yes | no | no | no | no | no | no | no | no |
| GRIN2B | 319 | S | 0 | 0 | no | yes | no | no | yes | no | no | no | no | no | no | no | no |
| GRIN2B | 320 | S | 1 | 0 | no | yes | no | no | yes | no | no | no | no | no | no | no | no |
| GRIN2B | 321 | C | 0 | 0 | yes | no | no | no | yes | no | no | no | no | no | no | no | no |
| GRIN2B | 322 | Y | 0 | 0 | yes | no | no | no | yes | no | no | no | yes | no | no | no | no |
| GRIN2B | 323 | N | 0 | 0 | yes | no | no | no | yes | no | no | no | no | no | no | no | no |
| GRIN2B | 324 | T | 0 | 0 | yes | no | no | no | yes | no | no | no | no | no | no | no | no |
| GRIN2B | 325 | H | 1 | 0 | no | yes | no | no | yes | no | no | no | no | no | no | no | no |
| GRIN2B | 326 | E | 0 | 0 | no | yes | no | no | yes | no | no | no | no | no | no | no | no |
| GRIN2B | 327 | K | 0 | 0 | no | yes | no | no | yes | no | no | no | no | no | no | no | no |
| GRIN2B | 328 | R | 0 | 0 | no | yes | no | no | yes | no | no | no | no | no | no | no | no |
| GRIN2B | 329 | I | 0 | 0 | no | yes | no | no | yes | no | no | no | no | no | no | no | no |
| GRIN2B | 330 | Y | 0 | 0 | no | yes | no | no | yes | no | no | no | no | no | no | no | no |
| GRIN2B | 331 | Q | 2 | 0 | no | yes | no | no | yes | no | no | no | no | no | no | no | no |
| GRIN2B | 332 | S | 0 | 0 | yes | no | no | no | yes | no | no | no | no | no | no | no | no |
| GRIN2B | 333 | N | 1 | 0 | yes | no | no | no | yes | no | no | no | no | no | no | no | no |
| GRIN2B | 334 | M | 0 | 0 | yes | no | no | no | yes | no | no | no | no | no | no | no | no |
| GRIN2B | 335 | L | 0 | 0 | yes | no | no | no | yes | no | no | no | no | no | no | no | no |
| GRIN2B | 336 | N | 1 | 0 | yes | no | no | no | yes | no | no | no | no | no | no | no | no |
| GRIN2B | 337 | R | 1 | 0 | yes | no | no | no | yes | no | no | no | no | no | no | no | no |
| GRIN2B | 338 | Y | 1 | 0 | yes | no | no | no | yes | no | no | no | no | no | no | no | no |
| GRIN2B | 339 | L | 0 | 0 | yes | no | no | no | yes | no | no | no | no | no | no | no | no |
| GRIN2B | 340 | I | 0 | 0 | yes | no | no | no | yes | no | no | no | no | no | no | no | no |
| GRIN2B | 341 | N | 0 | 0 | no | yes | no | no | yes | no | no | no | no | no | no | no | no |
| GRIN2B | 342 | V | 0 | 0 | no | yes | no | no | yes | no | no | no | no | no | no | no | no |
| GRIN2B | 343 | T | 0 | 0 | no | no | yes | no | yes | no | no | no | no | no | no | no | no |
| GRIN2B | 344 | F | 0 | 0 | no | no | yes | no | yes | no | no | no | no | no | no | no | no |
| GRIN2B | 345 | E | 0 | 0 | no | yes | no | no | yes | no | no | no | no | no | no | no | no |
| GRIN2B | 346 | G | 0 | 0 | no | yes | no | no | yes | no | no | no | no | no | no | no | no |
| GRIN2B | 347 | R | 1 | 0 | no | no | yes | no | yes | no | no | no | no | no | no | no | no |
| GRIN2B | 348 | N | 0 | 0 | no | no | yes | no | yes | no | no | no | no | no | no | no | no |
| GRIN2B | 349 | L | 0 | 0 | no | yes | no | no | yes | no | no | no | no | no | no | no | no |

| | | | | | | | | | | | | | | | | | |
|---|---|---|---|---|---|---|---|---|---|---|---|---|---|---|---|---|---|
| GRIN2B | 350 | S | 0 | 0 | no | yes | no | no | yes | no | no | no | no | no | no | no | no |
| GRIN2B | 351 | F | 0 | 0 | no | no | yes | no | yes | no | no | no | no | no | no | no | no |
| GRIN2B | 352 | S | 0 | 0 | no | yes | no | no | yes | no | no | no | no | no | no | no | no |
| GRIN2B | 353 | E | 0 | 0 | no | yes | no | no | yes | no | no | no | no | no | no | no | no |
| GRIN2B | 354 | D | 1 | 0 | no | yes | no | no | yes | no | no | no | no | no | no | no | no |
| GRIN2B | 355 | G | 0 | 0 | no | yes | no | no | yes | no | no | no | no | no | no | no | no |
| GRIN2B | 356 | Y | 0 | 0 | no | yes | no | no | yes | no | no | no | no | no | no | no | no |
| GRIN2B | 357 | Q | 0 | 0 | no | no | yes | no | yes | no | no | no | no | no | no | no | no |
| GRIN2B | 358 | M | 0 | 0 | no | yes | no | no | yes | no | no | no | no | no | no | no | no |
| GRIN2B | 359 | H | 1 | 0 | no | yes | no | no | yes | no | no | no | no | no | no | no | no |
| GRIN2B | 360 | P | 0 | 0 | no | yes | no | no | yes | no | no | no | no | no | no | no | no |
| GRIN2B | 361 | K | 0 | 0 | no | yes | no | no | yes | no | no | no | no | no | no | no | no |
| GRIN2B | 362 | L | 0 | 0 | no | no | yes | no | yes | no | no | no | no | no | no | no | no |
| GRIN2B | 363 | V | 0 | 0 | no | no | yes | no | yes | no | no | no | no | no | no | no | no |
| GRIN2B | 364 | I | 0 | 0 | no | no | yes | no | yes | no | no | no | no | no | no | no | no |
| GRIN2B | 365 | I | 0 | 0 | no | no | yes | no | yes | no | no | no | no | no | no | no | no |
| GRIN2B | 366 | L | 1 | 0 | no | no | yes | no | yes | no | no | no | no | no | no | no | no |
| GRIN2B | 367 | L | 0 | 0 | no | yes | no | no | yes | no | no | no | no | no | no | no | no |
| GRIN2B | 368 | N | 1 | 0 | no | yes | no | no | yes | no | no | no | no | no | no | no | no |
| GRIN2B | 369 | K | 0 | 0 | no | yes | no | no | yes | no | no | no | no | no | no | no | no |
| GRIN2B | 370 | E | 0 | 0 | no | yes | no | no | yes | no | no | no | no | no | no | no | no |
| GRIN2B | 371 | R | 1 | 0 | no | yes | no | no | yes | no | no | no | no | no | no | no | no |
| GRIN2B | 372 | K | 0 | 0 | no | yes | no | no | yes | no | no | no | no | no | no | no | no |
| GRIN2B | 373 | W | 0 | 0 | no | yes | no | no | yes | no | no | no | no | no | no | no | no |
| GRIN2B | 374 | E | 0 | 0 | no | yes | no | no | yes | no | no | no | no | no | no | no | no |
| GRIN2B | 375 | R | 1 | 0 | no | yes | no | no | yes | no | no | no | no | no | no | no | no |
| GRIN2B | 376 | V | 0 | 0 | no | yes | no | no | yes | no | no | no | no | no | no | no | no |
| GRIN2B | 377 | G | 0 | 0 | no | yes | no | no | yes | no | no | no | no | no | no | no | no |
| GRIN2B | 378 | K | 0 | 0 | no | no | yes | no | yes | no | no | no | no | no | no | no | no |
| GRIN2B | 379 | W | 0 | 0 | no | yes | no | no | yes | no | no | no | no | no | no | no | no |
| GRIN2B | 380 | K | 0 | 0 | no | yes | no | no | yes | no | no | no | no | no | no | no | no |
| GRIN2B | 381 | D | 0 | 0 | no | yes | no | no | yes | no | no | no | no | no | no | no | no |
| GRIN2B | 382 | K | 1 | 0 | no | yes | no | no | yes | no | no | no | no | no | no | no | no |

| | | | | | | | | | | | | | | | | | |
|---|---|---|---|---|---|---|---|---|---|---|---|---|---|---|---|---|---|
| GRIN2B | 383 | S | 2 | 0 | no | yes | no | no | yes | no | no | no | no | no | no | no | no |
| GRIN2B | 384 | L | 0 | 0 | no | yes | no | no | yes | no | no | no | no | no | no | no | no |
| GRIN2B | 385 | Q | 0 | 0 | no | no | yes | no | yes | no | no | no | no | no | no | no | no |
| GRIN2B | 386 | M | 0 | 0 | no | yes | no | no | yes | no | no | no | no | no | no | no | no |
| GRIN2B | 387 | K | 0 | 0 | no | yes | no | no | yes | no | no | no | no | no | no | no | no |
| GRIN2B | 388 | Y | 0 | 0 | no | yes | no | no | yes | no | no | no | no | no | no | no | no |
| GRIN2B | 389 | Y | 0 | 0 | no | yes | no | no | yes | no | no | no | no | no | no | no | no |
| GRIN2B | 390 | V | 0 | 0 | no | yes | no | no | yes | no | no | no | no | no | no | no | no |
| GRIN2B | 391 | W | 0 | 0 | no | yes | no | no | yes | no | no | no | no | no | no | no | no |
| GRIN2B | 392 | P | 0 | 0 | no | yes | no | no | yes | no | no | no | no | no | no | no | no |
| GRIN2B | 406 | L | 0 | 0 | no | yes | no | no | yes | no | no | no | no | no | no | no | no |
| GRIN2B | 407 | S | 1 | 0 | no | yes | no | no | yes | no | no | no | no | no | no | no | no |
| GRIN2B | 408 | I | 0 | 0 | no | no | yes | no | yes | no | no | no | no | no | no | no | no |
| GRIN2B | 409 | V | 0 | 0 | no | no | yes | no | yes | no | no | no | no | no | no | no | no |
| GRIN2B | 410 | T | 0 | 0 | no | no | yes | no | yes | no | no | no | no | no | no | no | no |
| GRIN2B | 411 | L | 0 | 0 | no | yes | no | no | yes | no | no | no | no | no | no | no | no |
| GRIN2B | 412 | E | 0 | 0 | no | yes | no | no | yes | no | no | no | no | no | no | no | no |
| GRIN2B | 413 | E | 0 | 1 | no | yes | no | no | yes | no | no | no | no | no | no | no | no |
| GRIN2B | 414 | A | 0 | 0 | yes | no | no | no | yes | no | no | no | no | no | no | no | no |
| GRIN2B | 415 | P | 0 | 0 | yes | no | no | no | yes | no | no | no | no | no | no | no | no |
| GRIN2B | 416 | F | 0 | 0 | yes | no | no | no | yes | no | no | no | no | no | no | no | no |
| GRIN2B | 417 | V | 0 | 0 | no | yes | no | no | yes | no | no | no | no | no | no | no | no |
| GRIN2B | 418 | I | 0 | 0 | no | yes | no | no | yes | no | no | no | no | no | no | no | no |
| GRIN2B | 419 | V | 0 | 0 | no | no | yes | no | yes | no | no | no | no | no | no | no | no |
| GRIN2B | 420 | E | 0 | 0 | no | yes | no | no | yes | no | no | no | no | no | no | no | no |
| GRIN2B | 421 | S | 0 | 0 | no | yes | no | no | yes | no | no | no | no | no | no | no | no |
| GRIN2B | 422 | V | 0 | 0 | no | yes | no | no | yes | no | no | no | no | no | no | no | no |
| GRIN2B | 423 | D | 0 | 0 | no | no | yes | no | yes | no | no | no | no | no | no | no | no |
| GRIN2B | 424 | P | 1 | 0 | no | yes | no | no | yes | no | no | no | no | no | no | no | no |
| GRIN2B | 425 | L | 0 | 0 | no | yes | no | no | yes | no | no | no | no | no | no | no | no |
| GRIN2B | 426 | S | 0 | 0 | no | yes | no | no | yes | no | no | no | no | no | no | no | no |
| GRIN2B | 427 | G | 0 | 1 | no | yes | no | no | yes | no | no | no | no | no | no | no | no |
| GRIN2B | 428 | T | 0 | 0 | no | no | yes | no | yes | no | no | no | no | no | no | no | no |

| | | | | | | | | | | | | | | | | | |
|---|---|---|---|---|---|---|---|---|---|---|---|---|---|---|---|---|---|
| GRIN2B | 429 | C | 0 | 0 | no | yes | no | no | yes | no | no | no | no | no | no | no | no |
| GRIN2B | 430 | M | 1 | 0 | no | yes | no | no | yes | no | no | no | no | no | no | no | no |
| GRIN2B | 431 | R | 0 | 0 | no | yes | no | no | yes | no | no | no | no | no | no | no | no |
| GRIN2B | 432 | N | 0 | 0 | no | yes | no | no | yes | no | no | no | no | no | no | no | no |
| GRIN2B | 433 | T | 0 | 0 | no | yes | no | no | yes | no | no | no | no | no | no | no | no |
| GRIN2B | 434 | V | 0 | 0 | no | no | yes | no | yes | no | no | no | no | no | no | no | no |
| GRIN2B | 435 | P | 0 | 0 | no | no | yes | no | yes | no | no | no | no | no | no | no | no |
| GRIN2B | 436 | C | 0 | 1 | no | yes | no | no | yes | no | no | no | no | no | no | no | no |
| GRIN2B | 437 | Q | 0 | 0 | no | no | yes | no | yes | no | no | no | no | no | no | no | no |
| GRIN2B | 438 | K | 0 | 0 | no | yes | no | no | yes | no | no | no | no | no | no | no | no |
| GRIN2B | 451 | G | 0 | 0 | no | yes | no | no | yes | no | no | no | no | no | no | no | no |
| GRIN2B | 452 | Y | 0 | 0 | no | yes | no | no | yes | no | no | no | no | no | no | no | no |
| GRIN2B | 453 | I | 0 | 0 | no | yes | no | no | yes | no | no | no | no | no | no | no | no |
| GRIN2B | 454 | K | 0 | 0 | no | yes | no | no | yes | no | no | no | no | no | no | no | no |
| GRIN2B | 455 | K | 0 | 0 | no | yes | no | no | yes | no | no | no | no | no | no | no | no |
| GRIN2B | 456 | C | 0 | 1 | no | no | yes | no | yes | no | no | no | no | no | no | no | no |
| GRIN2B | 457 | C | 0 | 0 | no | no | yes | no | yes | no | no | no | no | no | no | no | no |
| GRIN2B | 458 | K | 0 | 0 | no | no | yes | no | yes | no | no | no | no | no | no | no | no |
| GRIN2B | 459 | G | 0 | 0 | no | yes | no | no | yes | no | no | no | no | no | no | no | no |
| GRIN2B | 460 | F | 0 | 0 | yes | no | no | no | yes | no | no | no | no | no | no | no | no |
| GRIN2B | 461 | C | 0 | 1 | yes | no | no | no | yes | no | no | no | no | no | no | no | no |
| GRIN2B | 462 | I | 1 | 0 | yes | no | no | no | yes | no | no | no | no | no | no | no | no |
| GRIN2B | 463 | D | 0 | 0 | yes | no | no | no | yes | no | no | no | no | no | no | no | no |
| GRIN2B | 464 | I | 0 | 0 | yes | no | no | no | yes | no | no | no | no | no | no | no | no |
| GRIN2B | 465 | L | 0 | 0 | yes | no | no | no | yes | no | no | no | no | no | no | no | no |
| GRIN2B | 466 | K | 0 | 0 | yes | no | no | no | yes | no | no | no | no | no | no | no | no |
| GRIN2B | 467 | K | 0 | 0 | yes | no | no | no | yes | no | no | no | no | no | no | no | no |
| GRIN2B | 468 | I | 0 | 0 | yes | no | no | no | yes | no | no | no | no | no | no | no | no |
| GRIN2B | 469 | S | 0 | 0 | yes | no | no | no | yes | no | no | no | no | no | no | no | no |
| GRIN2B | 470 | K | 0 | 0 | yes | no | no | no | yes | no | yes | no | no | no | no | no | no |
| GRIN2B | 471 | S | 0 | 0 | yes | no | no | no | yes | no | no | no | no | no | no | no | no |
| GRIN2B | 472 | V | 0 | 0 | no | yes | no | no | yes | no | no | no | no | no | no | no | no |
| GRIN2B | 473 | K | 0 | 0 | no | yes | no | no | yes | no | no | no | no | no | no | no | no |

| | | | | | | | | | | | | | | | | | |
|---|---|---|---|---|---|---|---|---|---|---|---|---|---|---|---|---|---|
| GRIN2B | 474 | F | 0 | 0 | no | yes | no | no | yes | no | no | no | no | no | no | no | no |
| GRIN2B | 475 | T | 0 | 0 | no | yes | no | no | yes | no | no | no | no | no | no | no | no |
| GRIN2B | 476 | Y | 0 | 0 | no | yes | no | no | yes | no | no | no | no | no | no | no | no |
| GRIN2B | 477 | D | 0 | 0 | no | yes | no | no | yes | no | no | no | no | no | no | no | no |
| GRIN2B | 478 | L | 0 | 0 | no | no | yes | no | yes | no | no | no | no | no | no | no | no |
| GRIN2B | 479 | Y | 0 | 0 | no | no | yes | no | yes | no | no | no | no | no | no | no | no |
| GRIN2B | 480 | L | 0 | 0 | no | no | yes | no | yes | no | no | no | no | no | no | no | no |
| GRIN2B | 481 | V | 0 | 0 | no | yes | no | no | yes | no | no | no | no | no | no | no | no |
| GRIN2B | 482 | T | 0 | 0 | no | yes | no | no | yes | no | no | no | no | no | no | no | no |
| GRIN2B | 483 | N | 0 | 0 | no | yes | no | no | yes | no | no | no | no | no | no | no | no |
| GRIN2B | 484 | G | 0 | 0 | no | yes | no | no | yes | no | no | no | no | no | no | no | no |
| GRIN2B | 485 | K | 0 | 0 | no | yes | no | no | yes | no | no | no | no | no | no | no | no |
| GRIN2B | 486 | H | 0 | 1 | no | yes | no | no | yes | no | no | no | no | no | no | no | no |
| GRIN2B | 487 | G | 0 | 0 | no | yes | no | no | yes | no | no | no | no | no | no | no | no |
| GRIN2B | 488 | K | 0 | 0 | no | yes | no | no | yes | no | no | no | no | no | no | no | no |
| GRIN2B | 489 | K | 0 | 0 | no | yes | no | no | yes | no | no | no | no | no | no | no | no |
| GRIN2B | 490 | I | 1 | 0 | no | no | yes | no | yes | no | no | no | no | no | no | no | no |
| GRIN2B | 491 | N | 0 | 0 | no | yes | no | no | yes | no | no | no | no | no | no | no | no |
| GRIN2B | 492 | G | 0 | 0 | no | yes | no | no | yes | no | no | no | no | no | no | no | no |
| GRIN2B | 493 | T | 0 | 0 | no | no | yes | no | yes | no | no | no | no | no | no | no | no |
| GRIN2B | 494 | W | 0 | 0 | no | yes | no | no | yes | no | no | no | no | no | no | no | no |
| GRIN2B | 495 | N | 0 | 0 | no | yes | no | no | yes | no | no | no | no | no | no | no | no |
| GRIN2B | 496 | G | 0 | 0 | yes | no | no | no | yes | no | no | no | no | no | no | no | no |
| GRIN2B | 497 | M | 0 | 0 | yes | no | no | no | yes | no | no | no | no | no | no | no | no |
| GRIN2B | 498 | I | 0 | 0 | yes | no | no | no | yes | no | no | no | no | no | no | no | no |
| GRIN2B | 499 | G | 0 | 1 | yes | no | no | no | yes | no | no | no | no | no | no | no | no |
| GRIN2B | 500 | E | 0 | 0 | yes | no | no | no | yes | no | no | no | no | no | no | no | no |
| GRIN2B | 501 | V | 0 | 0 | yes | no | no | no | yes | no | no | no | no | no | no | no | no |
| GRIN2B | 502 | V | 0 | 0 | yes | no | no | no | yes | no | no | no | no | no | no | no | no |
| GRIN2B | 503 | M | 1 | 0 | no | yes | no | no | yes | no | no | no | no | no | no | no | no |
| GRIN2B | 504 | K | 0 | 0 | no | yes | no | no | yes | no | no | no | no | no | no | no | no |
| GRIN2B | 505 | R | 0 | 0 | no | yes | no | no | yes | no | no | no | no | no | no | no | no |
| GRIN2B | 506 | A | 0 | 0 | no | yes | no | no | yes | no | no | no | no | no | no | no | no |

| | | | | | | | | | | | | | | | | | |
|---|---|---|---|---|---|---|---|---|---|---|---|---|---|---|---|---|---|
| GRIN2B | 507 | Y | 0 | 0 | no | yes | no | no | yes | no | no | no | no | no | no | no | no |
| GRIN2B | 508 | M | 0 | 0 | no | no | yes | no | yes | no | no | no | no | no | no | no | no |
| GRIN2B | 509 | A | 0 | 0 | no | no | yes | no | yes | no | no | no | no | no | no | no | no |
| GRIN2B | 510 | V | 0 | 0 | no | no | yes | no | yes | no | no | no | no | no | no | no | no |
| GRIN2B | 511 | G | 0 | 0 | no | yes | no | no | yes | no | no | no | no | no | no | no | no |
| GRIN2B | 512 | S | 0 | 0 | no | yes | no | no | yes | no | no | no | no | no | no | no | no |
| GRIN2B | 513 | L | 0 | 0 | no | yes | no | no | yes | no | no | no | no | no | no | no | no |
| GRIN2B | 514 | T | 0 | 1 | no | yes | no | no | yes | no | no | no | no | no | no | no | no |
| GRIN2B | 515 | I | 0 | 0 | no | yes | no | no | yes | no | no | no | yes | no | no | no | no |
| GRIN2B | 516 | N | 0 | 1 | no | yes | no | no | yes | no | no | no | yes | no | no | no | no |
| GRIN2B | 517 | E | 0 | 0 | yes | no | no | no | yes | no | no | no | yes | no | no | no | no |
| GRIN2B | 518 | E | 0 | 0 | yes | no | no | no | yes | no | no | no | no | no | no | no | no |
| GRIN2B | 519 | R | 0 | 0 | yes | no | no | no | yes | no | no | no | no | no | no | no | no |
| GRIN2B | 520 | S | 0 | 0 | yes | no | no | no | yes | no | no | no | yes | no | no | no | no |
| GRIN2B | 521 | E | 0 | 0 | yes | no | no | no | yes | no | no | no | no | no | no | no | no |
| GRIN2B | 522 | V | 0 | 0 | yes | no | no | no | yes | no | no | no | no | no | no | no | no |
| GRIN2B | 523 | V | 0 | 0 | no | yes | no | no | yes | no | no | no | no | no | no | no | no |
| GRIN2B | 524 | D | 0 | 0 | no | no | yes | no | yes | no | no | no | no | no | no | no | no |
| GRIN2B | 525 | F | 0 | 1 | no | yes | no | no | yes | no | no | no | no | no | no | no | no |
| GRIN2B | 526 | S | 0 | 1 | no | yes | no | no | yes | no | no | no | yes | no | no | no | no |
| GRIN2B | 527 | V | 0 | 0 | no | yes | no | no | yes | no | no | no | no | no | no | no | no |
| GRIN2B | 528 | P | 0 | 0 | no | yes | no | no | yes | no | no | no | yes | no | no | no | no |
| GRIN2B | 529 | F | 0 | 0 | no | yes | no | no | yes | no | no | no | no | no | no | no | no |
| GRIN2B | 530 | I | 0 | 0 | no | yes | no | no | yes | no | no | no | no | no | no | no | no |
| GRIN2B | 531 | E | 0 | 0 | no | no | yes | no | yes | no | no | no | yes | no | no | no | no |
| GRIN2B | 532 | T | 0 | 0 | no | no | yes | no | yes | no | no | no | no | no | no | no | no |
| GRIN2B | 533 | G | 0 | 1 | no | no | yes | no | yes | no | no | no | no | no | no | no | no |
| GRIN2B | 534 | I | 0 | 0 | no | no | yes | no | yes | no | no | no | no | no | no | no | no |
| GRIN2B | 535 | S | 0 | 0 | no | yes | no | no | yes | no | no | no | no | no | no | no | no |
| GRIN2B | 536 | V | 0 | 0 | no | yes | no | no | yes | no | no | no | no | no | no | no | no |
| GRIN2B | 537 | M | 0 | 0 | no | yes | no | no | yes | no | no | no | no | no | no | no | no |
| GRIN2B | 538 | V | 0 | 0 | no | yes | no | no | yes | no | no | no | no | no | no | no | no |
| GRIN2B | 539 | S | 0 | 0 | no | yes | no | no | yes | no | no | no | no | no | no | no | no |

| | | | | | | | | | | | | | | | | | |
|---|---|---|---|---|---|---|---|---|---|---|---|---|---|---|---|---|---|
| GRIN2B | 540 | R | 0 | 1 | no | yes | no | no | yes | no | no | no | no | no | no | no | no |
| GRIN2B | 541 | S | 0 | 1 | no | yes | no | no | yes | no | no | no | no | no | no | no | no |
| GRIN2B | 542 | N | 0 | 0 | no | yes | no | no | yes | no | no | no | no | no | no | no | no |
| GRIN2B | 543 | G | 0 | 0 | no | yes | no | no | yes | no | no | no | no | no | no | no | no |
| GRIN2B | 544 | T | 0 | 0 | no | yes | no | no | yes | no | no | no | no | no | no | no | no |
| GRIN2B | 545 | V | 0 | 0 | no | yes | no | no | yes | no | no | no | no | no | no | no | no |
| GRIN2B | 546 | S | 0 | 0 | no | yes | no | no | yes | no | no | no | no | no | no | no | no |
| GRIN2B | 547 | P | 0 | 0 | no | yes | no | no | yes | no | no | no | no | no | no | no | no |
| GRIN2B | 548 | S | 0 | 0 | no | yes | no | no | yes | no | no | no | no | no | no | no | no |
| GRIN2B | 549 | A | 0 | 0 | no | yes | no | yes | yes | no | no | no | no | no | no | no | no |
| GRIN2B | 550 | F | 0 | 0 | no | yes | no | yes | yes | no | no | no | no | no | no | no | no |
| GRIN2B | 551 | L | 0 | 0 | no | yes | no | yes | yes | no | no | no | no | no | no | no | no |
| GRIN2B | 552 | E | 0 | 0 | no | yes | no | yes | yes | no | no | no | no | no | no | no | no |
| GRIN2B | 553 | P | 0 | 1 | no | yes | no | no | yes | no | no | no | no | no | no | no | no |
| GRIN2B | 554 | F | 0 | 0 | no | yes | no | no | yes | no | no | no | yes | no | no | no | no |
| GRIN2B | 555 | S | 0 | 1 | no | yes | no | yes | yes | no | no | no | yes | no | no | no | no |
| GRIN2B | 556 | A | 0 | 0 | yes | no | no | yes | yes | no | no | no | no | no | no | no | no |
| GRIN2B | 557 | D | 0 | 0 | yes | no | no | yes | yes | no | no | no | yes | no | no | no | no |
| GRIN2B | 558 | V | 0 | 0 | yes | no | no | yes | no | no | no | no | yes | no | no | no | no |
| GRIN2B | 559 | W | 0 | 1 | yes | no | no | yes | no | no | no | no | no | no | no | no | no |
| GRIN2B | 560 | V | 0 | 0 | yes | no | no | yes | no | no | no | no | no | no | no | no | no |
| GRIN2B | 561 | M | 0 | 0 | yes | no | no | yes | no | no | no | no | yes | no | no | no | no |
| GRIN2B | 562 | M | 0 | 0 | yes | no | no | yes | no | no | no | no | yes | no | no | no | no |
| GRIN2B | 563 | F | 0 | 0 | yes | no | no | yes | no | no | no | no | no | no | no | no | no |
| GRIN2B | 564 | V | 0 | 0 | yes | no | no | yes | no | no | no | no | no | no | no | no | no |
| GRIN2B | 565 | M | 0 | 0 | yes | no | no | yes | no | no | no | no | yes | no | no | no | no |
| GRIN2B | 566 | L | 0 | 0 | yes | no | no | yes | no | no | no | no | no | no | no | no | no |
| GRIN2B | 567 | L | 0 | 0 | yes | no | no | yes | no | no | no | no | no | no | no | no | no |
| GRIN2B | 568 | I | 1 | 0 | yes | no | no | yes | no | no | no | no | no | no | no | no | no |
| GRIN2B | 569 | V | 0 | 0 | yes | no | no | yes | no | no | no | no | no | no | no | no | no |
| GRIN2B | 570 | S | 0 | 0 | yes | no | no | yes | no | no | no | no | no | no | no | no | no |
| GRIN2B | 571 | A | 0 | 0 | yes | no | no | yes | no | no | no | no | no | no | no | no | no |
| GRIN2B | 572 | V | 1 | 0 | yes | no | no | yes | no | no | no | no | no | no | no | no | no |

| | | | | | | | | | | | | | | | | | |
|---|---|---|---|---|---|---|---|---|---|---|---|---|---|---|---|---|---|
| GRIN2B | 573 | A | 0 | 0 | yes | no | no | yes | no | no | no | no | no | no | no | no | no |
| GRIN2B | 574 | V | 0 | 0 | yes | no | no | yes | no | no | no | no | no | no | no | no | no |
| GRIN2B | 575 | F | 1 | 0 | yes | no | no | yes | no | no | no | no | no | no | no | no | no |
| GRIN2B | 576 | V | 1 | 0 | yes | no | no | yes | no | no | no | no | no | no | no | no | no |
| GRIN2B | 577 | F | 0 | 0 | yes | no | no | no | no | yes | no | no | no | no | no | no | no |
| GRIN2B | 578 | E | 0 | 0 | yes | no | no | no | no | yes | no | no | no | no | no | no | no |
| GRIN2B | 579 | Y | 0 | 0 | no | yes | no | no | no | yes | no | no | no | no | no | no | no |
| GRIN2B | 600 | F | 0 | 0 | no | yes | no | no | no | yes | no | no | no | no | no | no | no |
| GRIN2B | 601 | T | 0 | 0 | yes | no | no | no | no | yes | no | no | no | no | no | no | no |
| GRIN2B | 602 | I | 0 | 0 | yes | no | no | yes | no | yes | no | no | no | no | no | no | no |
| GRIN2B | 603 | G | 1 | 0 | yes | no | no | yes | no | yes | no | no | no | no | no | no | no |
| GRIN2B | 604 | K | 0 | 0 | yes | no | no | yes | no | no | no | yes | no | no | no | no | no |
| GRIN2B | 605 | A | 0 | 0 | yes | no | no | yes | no | no | no | yes | no | no | no | no | no |
| GRIN2B | 606 | I | 0 | 0 | yes | no | no | yes | no | no | no | yes | no | no | no | no | no |
| GRIN2B | 607 | W | 0 | 1 | yes | no | no | yes | no | no | no | yes | yes | no | no | no | no |
| GRIN2B | 608 | L | 0 | 0 | yes | no | no | yes | no | no | no | yes | no | no | no | no | no |
| GRIN2B | 609 | L | 0 | 0 | yes | no | no | yes | no | no | no | yes | no | no | no | no | no |
| GRIN2B | 610 | W | 0 | 0 | yes | no | no | yes | no | no | no | yes | yes | no | no | no | no |
| GRIN2B | 611 | G | 0 | 1 | yes | no | no | yes | no | no | no | yes | no | no | no | no | no |
| GRIN2B | 612 | L | 0 | 0 | yes | no | no | yes | no | no | no | yes | no | no | no | no | no |
| GRIN2B | 613 | V | 0 | 0 | yes | no | no | yes | no | no | no | yes | no | no | no | no | no |
| GRIN2B | 614 | F | 0 | 0 | yes | no | no | yes | no | no | no | yes | yes | no | no | no | no |
| GRIN2B | 615 | N | 0 | 2 | no | yes | no | yes | no | no | no | yes | no | no | no | no | no |
| GRIN2B | 616 | N | 0 | 2 | no | yes | no | yes | no | no | no | yes | yes | no | no | no | no |
| GRIN2B | 617 | S | 0 | 0 | no | yes | no | yes | no | no | no | yes | yes | no | no | no | no |
| GRIN2B | 618 | V | 0 | 1 | no | yes | no | yes | no | no | no | yes | no | no | no | no | no |
| GRIN2B | 619 | P | 0 | 0 | no | yes | no | yes | no | no | no | yes | no | no | no | no | no |
| GRIN2B | 620 | V | 0 | 1 | no | yes | no | yes | no | no | no | yes | no | no | no | no | no |
| GRIN2B | 621 | Q | 0 | 0 | no | yes | no | yes | no | no | no | yes | no | no | no | no | no |
| GRIN2B | 622 | N | 0 | 0 | no | yes | no | yes | no | no | no | yes | no | no | no | no | no |
| GRIN2B | 623 | P | 0 | 0 | no | yes | no | yes | no | no | no | yes | no | no | no | no | no |
| GRIN2B | 624 | K | 0 | 0 | no | yes | no | no | no | yes | no | no | no | no | no | no | no |
| GRIN2B | 625 | G | 0 | 0 | no | yes | no | no | no | yes | no | no | no | no | no | no | no |

| | | | | | | | | | | | | | | | | | |
|---|---|---|---|---|---|---|---|---|---|---|---|---|---|---|---|---|---|
| GRIN2B | 626 | T | 0 | 0 | no | yes | no | yes | no | yes | no | no | no | no | no | no | no |
| GRIN2B | 627 | T | 0 | 0 | no | yes | no | yes | no | yes | no | no | yes | no | no | no | no |
| GRIN2B | 628 | S | 0 | 2 | no | yes | no | yes | no | yes | no | no | no | no | no | no | no |
| GRIN2B | 629 | K | 0 | 0 | yes | no | no | yes | no | yes | no | no | yes | no | no | no | no |
| GRIN2B | 630 | I | 0 | 0 | yes | no | no | yes | no | yes | no | no | no | no | no | no | no |
| GRIN2B | 631 | M | 0 | 0 | yes | no | no | yes | no | no | no | no | no | no | no | no | no |
| GRIN2B | 632 | V | 0 | 0 | yes | no | no | yes | no | no | no | no | no | no | no | no | no |
| GRIN2B | 633 | S | 0 | 0 | yes | no | no | yes | no | no | no | no | yes | no | no | no | no |
| GRIN2B | 634 | V | 0 | 0 | yes | no | no | yes | no | no | no | no | no | no | no | no | no |
| GRIN2B | 635 | W | 0 | 0 | yes | no | no | yes | no | no | no | no | yes | no | no | no | no |
| GRIN2B | 636 | A | 0 | 2 | yes | no | no | yes | no | no | no | no | yes | no | no | no | no |
| GRIN2B | 637 | F | 0 | 0 | yes | no | no | yes | no | no | no | no | yes | no | no | no | no |
| GRIN2B | 638 | F | 0 | 0 | yes | no | no | yes | no | no | no | no | no | no | no | no | no |
| GRIN2B | 639 | A | 1 | 1 | yes | no | no | yes | no | no | no | no | no | no | no | no | no |
| GRIN2B | 640 | V | 0 | 0 | yes | no | no | yes | no | no | no | no | yes | no | no | no | no |
| GRIN2B | 641 | I | 0 | 1 | yes | no | no | yes | no | no | no | no | yes | no | no | no | no |
| GRIN2B | 642 | F | 0 | 0 | yes | no | no | yes | no | no | no | no | no | no | no | no | no |
| GRIN2B | 643 | L | 0 | 1 | yes | no | no | yes | no | no | no | no | no | no | no | no | no |
| GRIN2B | 644 | A | 0 | 0 | yes | no | no | yes | no | no | no | no | yes | no | no | no | no |
| GRIN2B | 645 | S | 0 | 0 | yes | no | no | yes | no | no | no | no | yes | no | no | no | no |
| GRIN2B | 646 | Y | 0 | 0 | yes | no | no | yes | no | no | no | no | yes | no | no | no | no |
| GRIN2B | 647 | T | 0 | 0 | yes | no | no | yes | yes | no | yes | no | no | no | no | no | no |
| GRIN2B | 648 | A | 0 | 0 | yes | no | no | yes | yes | no | no | no | yes | no | no | no | no |
| GRIN2B | 649 | N | 0 | 0 | yes | no | no | no | yes | no | no | no | yes | no | no | no | no |
| GRIN2B | 650 | L | 0 | 0 | yes | no | no | yes | yes | no | no | no | yes | no | no | no | no |
| GRIN2B | 651 | A | 0 | 0 | yes | no | no | no | yes | no | yes | no | yes | no | no | no | no |
| GRIN2B | 652 | A | 0 | 2 | yes | no | no | no | yes | no | no | no | yes | no | no | no | no |
| GRIN2B | 653 | F | 0 | 0 | yes | no | no | no | yes | no | no | no | no | no | no | no | no |
| GRIN2B | 654 | M | 0 | 0 | yes | no | no | no | yes | no | no | no | yes | no | no | no | no |
| GRIN2B | 655 | I | 0 | 1 | no | yes | no | no | yes | no | yes | no | yes | no | no | no | no |
| GRIN2B | 656 | Q | 0 | 0 | no | yes | no | no | yes | no | no | no | no | no | no | no | no |
| GRIN2B | 657 | E | 0 | 1 | no | yes | no | no | yes | no | no | no | no | no | no | no | no |
| GRIN2B | 658 | E | 0 | 0 | no | yes | no | no | yes | no | no | no | no | no | no | no | no |

| | | | | | | | | | | | | | | | | | |
|---|---|---|---|---|---|---|---|---|---|---|---|---|---|---|---|---|---|
| GRIN2B | 659 | Y | 0 | 0 | no | yes | no | no | yes | no | no | no | no | no | no | no | no |
| GRIN2B | 660 | V | 0 | 0 | no | yes | no | no | yes | no | no | no | no | no | no | no | no |
| GRIN2B | 661 | D | 1 | 0 | no | yes | no | no | yes | no | no | no | no | no | no | no | no |
| GRIN2B | 662 | Q | 0 | 1 | no | yes | no | no | yes | no | no | no | no | no | no | no | no |
| GRIN2B | 663 | V | 0 | 0 | no | yes | no | no | yes | no | no | no | no | no | no | no | no |
| GRIN2B | 664 | S | 0 | 0 | no | yes | no | no | yes | no | no | no | no | no | no | no | no |
| GRIN2B | 665 | G | 0 | 0 | no | yes | no | no | yes | no | no | no | no | no | no | no | no |
| GRIN2B | 666 | L | 0 | 0 | no | yes | no | no | yes | no | no | no | no | no | no | no | no |
| GRIN2B | 667 | S | 0 | 0 | no | yes | no | no | yes | no | no | no | no | no | no | no | no |
| GRIN2B | 668 | D | 0 | 1 | no | yes | no | no | yes | no | no | no | no | no | no | no | no |
| GRIN2B | 669 | K | 0 | 0 | yes | no | no | no | yes | no | no | no | no | no | no | no | no |
| GRIN2B | 670 | K | 0 | 0 | yes | no | no | no | yes | no | no | no | no | no | no | no | no |
| GRIN2B | 671 | F | 0 | 0 | yes | no | no | no | yes | no | no | no | no | no | no | no | no |
| GRIN2B | 672 | Q | 0 | 0 | yes | no | no | no | yes | no | no | no | no | no | no | no | no |
| GRIN2B | 673 | R | 0 | 0 | no | yes | no | no | yes | no | no | no | no | no | no | no | no |
| GRIN2B | 674 | P | 0 | 0 | yes | no | no | no | yes | no | no | no | no | no | no | no | no |
| GRIN2B | 675 | N | 0 | 0 | yes | no | no | no | yes | no | no | no | no | no | no | no | no |
| GRIN2B | 676 | D | 0 | 1 | yes | no | no | no | yes | no | no | no | no | no | no | no | no |
| GRIN2B | 677 | F | 0 | 0 | yes | no | no | no | yes | no | no | no | no | no | no | no | no |
| GRIN2B | 678 | S | 0 | 0 | no | yes | no | no | yes | no | no | no | no | no | no | no | no |
| GRIN2B | 679 | P | 0 | 0 | no | yes | no | no | yes | no | no | no | no | no | no | no | no |
| GRIN2B | 680 | P | 0 | 0 | no | yes | no | no | yes | no | no | no | no | no | no | no | no |
| GRIN2B | 681 | F | 0 | 0 | no | yes | no | no | yes | no | no | no | no | no | no | no | no |
| GRIN2B | 682 | R | 0 | 2 | no | yes | no | no | yes | no | no | no | no | no | no | no | no |
| GRIN2B | 683 | F | 0 | 0 | no | yes | no | no | yes | no | no | no | no | no | no | no | no |
| GRIN2B | 684 | G | 0 | 0 | no | yes | no | no | yes | no | no | no | no | no | no | no | no |
| GRIN2B | 685 | T | 0 | 0 | no | yes | no | no | yes | no | no | no | no | no | no | no | no |
| GRIN2B | 686 | V | 0 | 0 | no | yes | no | no | yes | no | no | no | no | no | no | no | no |
| GRIN2B | 687 | P | 0 | 2 | no | yes | no | no | yes | no | no | no | no | no | no | no | no |
| GRIN2B | 688 | N | 0 | 0 | no | yes | no | no | yes | no | no | no | no | no | no | no | no |
| GRIN2B | 689 | G | 0 | 1 | no | yes | no | no | yes | no | no | no | no | no | no | no | no |
| GRIN2B | 690 | S | 0 | 0 | yes | no | no | no | yes | no | no | no | no | no | no | no | no |
| GRIN2B | 691 | T | 0 | 0 | yes | no | no | no | yes | no | no | no | no | no | no | no | no |

| | | | | | | | | | | | | | | | | | |
|---|---|---|---|---|---|---|---|---|---|---|---|---|---|---|---|---|---|
| GRIN2B | 692 | E | 0 | 0 | yes | no | no | no | yes | no | no | no | no | no | no | no | no |
| GRIN2B | 693 | R | 0 | 1 | yes | no | no | no | yes | no | no | no | no | no | no | no | no |
| GRIN2B | 694 | N | 0 | 1 | yes | no | no | no | yes | no | no | no | no | no | no | no | no |
| GRIN2B | 695 | I | 0 | 2 | yes | no | no | no | yes | no | no | no | no | no | no | no | no |
| GRIN2B | 696 | R | 0 | 1 | yes | no | no | no | yes | no | no | no | no | no | no | no | no |
| GRIN2B | 697 | N | 0 | 0 | yes | no | no | no | yes | no | no | no | no | no | no | no | no |
| GRIN2B | 698 | N | 0 | 0 | yes | no | no | no | yes | no | no | no | no | no | no | no | no |
| GRIN2B | 699 | Y | 0 | 0 | no | yes | no | no | yes | no | no | no | no | no | no | no | no |
| GRIN2B | 700 | A | 1 | 0 | yes | no | no | no | yes | no | no | no | no | no | no | no | no |
| GRIN2B | 701 | E | 0 | 0 | yes | no | no | no | yes | no | no | no | no | no | no | no | no |
| GRIN2B | 702 | M | 0 | 0 | yes | no | no | no | yes | no | no | no | no | no | no | no | no |
| GRIN2B | 703 | H | 0 | 0 | yes | no | no | no | yes | no | no | no | no | no | no | no | no |
| GRIN2B | 704 | A | 0 | 0 | yes | no | no | no | yes | no | no | no | no | no | no | no | no |
| GRIN2B | 705 | Y | 0 | 0 | yes | no | no | no | yes | no | no | no | no | no | no | no | no |
| GRIN2B | 706 | M | 0 | 1 | yes | no | no | no | yes | no | no | no | no | no | no | no | no |
| GRIN2B | 707 | G | 0 | 0 | yes | no | no | no | yes | no | no | no | no | no | no | no | no |
| GRIN2B | 708 | K | 0 | 0 | yes | no | no | no | yes | no | no | no | no | no | no | no | no |
| GRIN2B | 709 | F | 0 | 0 | yes | no | no | no | yes | no | no | no | no | no | no | no | no |
| GRIN2B | 710 | N | 0 | 0 | no | yes | no | no | yes | no | no | no | no | no | no | no | no |
| GRIN2B | 711 | Q | 0 | 0 | no | yes | no | no | yes | no | no | no | no | no | no | no | no |
| GRIN2B | 712 | R | 0 | 0 | no | yes | no | no | yes | no | no | no | no | no | no | no | no |
| GRIN2B | 713 | G | 0 | 0 | no | yes | no | no | yes | no | no | no | no | no | no | no | no |
| GRIN2B | 714 | V | 0 | 0 | no | yes | no | no | yes | no | no | no | no | no | no | no | no |
| GRIN2B | 715 | D | 1 | 0 | yes | no | no | no | yes | no | no | no | no | no | no | no | no |
| GRIN2B | 716 | D | 0 | 0 | yes | no | no | no | yes | no | no | no | no | no | no | no | no |
| GRIN2B | 717 | A | 0 | 0 | yes | no | no | no | yes | no | no | no | no | no | no | no | no |
| GRIN2B | 718 | L | 0 | 0 | yes | no | no | no | yes | no | no | no | no | no | no | no | no |
| GRIN2B | 719 | L | 0 | 0 | yes | no | no | no | yes | no | no | no | no | no | no | no | no |
| GRIN2B | 720 | S | 0 | 0 | yes | no | no | no | yes | no | no | no | no | no | no | no | no |
| GRIN2B | 721 | L | 0 | 0 | yes | no | no | no | yes | no | no | no | no | no | no | no | no |
| GRIN2B | 722 | K | 0 | 0 | yes | no | no | no | yes | no | no | no | no | no | no | no | no |
| GRIN2B | 723 | T | 0 | 0 | yes | no | no | no | yes | no | no | no | no | no | no | no | no |
| GRIN2B | 724 | G | 0 | 0 | no | yes | no | no | yes | no | no | no | no | no | no | no | no |

| | | | | | | | | | | | | | | | | | |
|---|---|---|---|---|---|---|---|---|---|---|---|---|---|---|---|---|---|
| GRIN2B | 725 | K | 0 | 0 | no | yes | no | no | yes | no | no | no | no | no | no | no | no |
| GRIN2B | 726 | L | 0 | 0 | no | yes | no | no | yes | no | no | no | no | no | no | no | no |
| GRIN2B | 727 | D | 0 | 0 | no | yes | no | no | yes | no | no | no | no | no | no | no | no |
| GRIN2B | 728 | A | 0 | 0 | no | yes | no | no | yes | no | no | no | no | no | no | no | no |
| GRIN2B | 729 | F | 0 | 0 | no | yes | no | no | yes | no | no | no | no | no | no | no | no |
| GRIN2B | 730 | I | 0 | 0 | no | yes | no | no | yes | no | no | no | no | no | no | no | no |
| GRIN2B | 731 | Y | 0 | 0 | no | yes | no | no | yes | no | no | no | no | no | no | no | no |
| GRIN2B | 732 | D | 0 | 1 | no | no | yes | no | yes | no | no | no | no | no | no | no | no |
| GRIN2B | 733 | A | 0 | 0 | yes | no | no | no | yes | no | no | no | no | no | no | no | no |
| GRIN2B | 734 | A | 0 | 1 | yes | no | no | no | yes | no | no | no | no | no | no | no | no |
| GRIN2B | 735 | V | 0 | 0 | yes | no | no | no | yes | no | no | no | no | no | no | no | no |
| GRIN2B | 736 | L | 0 | 0 | yes | no | no | no | yes | no | no | no | no | no | no | no | no |
| GRIN2B | 737 | N | 0 | 0 | yes | no | no | no | yes | no | no | no | no | no | no | no | no |
| GRIN2B | 738 | Y | 0 | 0 | yes | no | no | no | yes | no | no | no | no | no | no | no | no |
| GRIN2B | 739 | M | 0 | 1 | yes | no | no | no | yes | no | no | no | no | no | no | no | no |
| GRIN2B | 740 | A | 0 | 0 | yes | no | no | no | yes | no | no | no | no | no | no | no | no |
| GRIN2B | 741 | G | 0 | 0 | no | yes | no | no | yes | no | no | no | no | no | no | no | no |
| GRIN2B | 742 | R | 0 | 1 | no | yes | no | no | yes | no | no | no | no | no | no | no | no |
| GRIN2B | 743 | D | 1 | 0 | yes | no | no | no | yes | no | no | no | no | no | no | no | no |
| GRIN2B | 744 | E | 0 | 0 | yes | no | no | no | yes | no | no | no | no | no | no | no | no |
| GRIN2B | 745 | G | 0 | 0 | yes | no | no | no | yes | no | no | no | no | no | no | no | no |
| GRIN2B | 746 | C | 0 | 0 | no | yes | no | no | yes | no | no | no | no | no | no | no | no |
| GRIN2B | 747 | K | 0 | 0 | no | yes | no | no | yes | no | no | no | no | no | no | no | no |
| GRIN2B | 748 | L | 0 | 0 | no | yes | no | no | yes | no | no | no | no | no | no | no | no |
| GRIN2B | 749 | V | 0 | 0 | no | yes | no | no | yes | no | no | no | no | no | no | no | no |
| GRIN2B | 750 | T | 0 | 0 | no | yes | no | no | yes | no | no | no | no | no | no | no | no |
| GRIN2B | 751 | I | 0 | 2 | no | yes | no | no | yes | no | no | no | no | no | no | no | no |
| GRIN2B | 752 | G | 0 | 0 | no | yes | no | no | yes | no | no | no | no | no | no | no | no |
| GRIN2B | 753 | S | 0 | 0 | no | yes | no | no | yes | no | no | no | no | no | no | no | no |
| GRIN2B | 754 | G | 0 | 0 | no | yes | no | no | yes | no | no | no | no | no | no | no | no |
| GRIN2B | 755 | K | 0 | 0 | no | yes | no | no | yes | no | no | no | no | no | no | no | no |
| GRIN2B | 756 | V | 0 | 0 | no | yes | no | no | yes | no | no | no | no | no | no | no | no |
| GRIN2B | 757 | F | 0 | 0 | no | no | yes | no | yes | no | no | no | no | no | no | no | no |

| | | | | | | | | | | | | | | | | | |
|---|---|---|---|---|---|---|---|---|---|---|---|---|---|---|---|---|---|
| GRIN2B | 758 | A | 0 | 0 | no | no | yes | no | yes | no | no | no | yes | no | no | no | no |
| GRIN2B | 759 | S | 0 | 0 | no | no | yes | no | yes | no | no | no | yes | no | no | no | no |
| GRIN2B | 760 | T | 0 | 0 | no | no | yes | no | yes | no | no | no | yes | no | no | no | no |
| GRIN2B | 761 | G | 0 | 0 | no | no | yes | no | yes | no | no | no | no | no | no | no | no |
| GRIN2B | 762 | Y | 0 | 0 | no | yes | no | no | yes | no | no | no | no | no | no | no | no |
| GRIN2B | 763 | G | 0 | 0 | no | yes | no | no | yes | no | no | no | no | no | no | no | no |
| GRIN2B | 764 | I | 0 | 0 | no | yes | no | no | yes | no | no | no | no | no | no | no | no |
| GRIN2B | 765 | A | 0 | 0 | no | no | yes | no | yes | no | no | no | no | no | no | no | no |
| GRIN2B | 766 | I | 0 | 0 | no | no | yes | no | yes | no | no | no | no | no | no | no | no |
| GRIN2B | 767 | Q | 0 | 0 | no | yes | no | no | yes | no | no | no | no | no | no | no | no |
| GRIN2B | 768 | K | 0 | 0 | no | yes | no | no | yes | no | no | no | no | no | no | no | no |
| GRIN2B | 769 | D | 0 | 0 | no | yes | no | no | yes | no | no | no | no | no | no | no | no |
| GRIN2B | 770 | S | 0 | 0 | no | yes | no | no | yes | no | no | no | no | no | no | no | no |
| GRIN2B | 771 | G | 0 | 0 | no | yes | no | no | yes | no | no | no | no | no | no | no | no |
| GRIN2B | 772 | W | 0 | 0 | no | yes | no | no | yes | no | no | no | no | no | no | no | no |
| GRIN2B | 773 | K | 0 | 0 | yes | no | no | no | yes | no | no | no | no | no | no | no | no |
| GRIN2B | 774 | R | 0 | 0 | yes | no | no | no | yes | no | no | no | no | no | no | no | no |
| GRIN2B | 775 | Q | 2 | 0 | yes | no | no | no | yes | no | no | no | no | no | no | no | no |
| GRIN2B | 776 | V | 0 | 0 | yes | no | no | no | yes | no | no | no | no | no | no | no | no |
| GRIN2B | 777 | D | 0 | 0 | yes | no | no | no | yes | no | no | no | no | no | no | no | no |
| GRIN2B | 778 | L | 0 | 0 | yes | no | no | no | yes | no | no | no | yes | no | no | no | no |
| GRIN2B | 779 | A | 0 | 0 | yes | no | no | no | yes | no | no | no | no | no | no | no | no |
| GRIN2B | 780 | I | 0 | 0 | yes | no | no | no | yes | no | no | no | no | no | no | no | no |
| GRIN2B | 781 | L | 0 | 1 | yes | no | no | no | yes | no | no | no | yes | no | no | no | no |
| GRIN2B | 782 | Q | 0 | 0 | yes | no | no | no | yes | no | no | no | no | no | no | no | no |
| GRIN2B | 783 | L | 0 | 0 | yes | no | no | no | yes | no | no | no | no | no | no | no | no |
| GRIN2B | 784 | F | 0 | 0 | yes | no | no | no | yes | no | no | no | yes | no | no | no | no |
| GRIN2B | 785 | G | 0 | 0 | yes | no | no | no | yes | no | no | no | yes | no | no | no | no |
| GRIN2B | 786 | D | 0 | 0 | no | yes | no | no | yes | no | no | no | no | no | no | no | no |
| GRIN2B | 787 | G | 0 | 0 | no | yes | no | no | yes | no | no | no | no | no | no | no | no |
| GRIN2B | 788 | E | 0 | 0 | yes | no | no | no | yes | no | no | no | no | no | no | no | no |
| GRIN2B | 789 | M | 0 | 0 | yes | no | no | no | yes | no | no | no | no | no | no | no | no |
| GRIN2B | 790 | E | 1 | 0 | yes | no | no | no | yes | no | no | no | no | no | no | no | no |

| | | | | | | | | | | | | | | | | | |
|---|---|---|---|---|---|---|---|---|---|---|---|---|---|---|---|---|---|
| GRIN2B | 791 | E | 0 | 0 | yes | no | no | no | yes | no | yes | no | no | no | no | no | no |
| GRIN2B | 792 | L | 0 | 0 | yes | no | no | no | yes | no | no | no | no | no | no | no | no |
| GRIN2B | 793 | E | 0 | 0 | yes | no | no | no | yes | no | no | no | no | no | no | no | no |
| GRIN2B | 794 | A | 0 | 0 | yes | no | no | no | yes | no | yes | no | no | no | no | no | no |
| GRIN2B | 795 | L | 0 | 0 | yes | no | no | no | yes | no | yes | no | yes | no | no | no | no |
| GRIN2B | 796 | W | 0 | 0 | yes | no | no | no | yes | no | no | no | no | no | no | no | no |
| GRIN2B | 797 | L | 0 | 0 | no | yes | no | no | yes | no | no | no | no | no | no | no | no |
| GRIN2B | 798 | T | 0 | 1 | no | yes | no | no | yes | no | yes | no | yes | no | no | no | no |
| GRIN2B | 799 | G | 0 | 1 | no | yes | no | no | yes | no | no | no | no | no | no | no | no |
| GRIN2B | 800 | I | 0 | 0 | no | yes | no | no | yes | no | no | no | no | no | no | no | no |
| GRIN2B | 801 | C | 0 | 0 | no | yes | no | no | yes | no | no | no | no | no | no | no | no |
| GRIN2B | 802 | H | 0 | 0 | no | yes | no | no | yes | no | yes | no | no | no | no | no | no |
| GRIN2B | 803 | N | 1 | 0 | no | yes | no | no | yes | no | yes | no | no | no | no | no | no |
| GRIN2B | 809 | M | 0 | 0 | no | yes | no | no | yes | no | yes | no | no | no | no | no | no |
| GRIN2B | 810 | S | 0 | 2 | no | yes | no | no | yes | no | no | no | no | no | no | no | no |
| GRIN2B | 811 | S | 0 | 0 | no | yes | no | no | yes | no | no | no | no | no | no | no | no |
| GRIN2B | 812 | Q | 0 | 0 | no | yes | no | no | yes | no | no | no | no | no | no | no | no |
| GRIN2B | 813 | L | 0 | 1 | no | yes | no | no | yes | no | no | no | yes | no | no | no | no |
| GRIN2B | 814 | D | 0 | 0 | no | yes | no | no | yes | no | no | no | no | no | no | no | no |
| GRIN2B | 815 | I | 0 | 0 | yes | no | no | yes | yes | no | no | no | no | no | no | no | no |
| GRIN2B | 816 | D | 0 | 0 | yes | no | no | no | yes | no | no | no | no | no | no | no | no |
| GRIN2B | 817 | N | 0 | 1 | yes | no | no | no | yes | no | no | no | no | no | no | no | no |
| GRIN2B | 818 | M | 0 | 2 | yes | no | no | yes | no | no | no | no | no | no | no | no | no |
| GRIN2B | 819 | A | 0 | 1 | yes | no | no | yes | no | no | no | no | no | no | no | no | no |
| GRIN2B | 820 | G | 1 | 2 | yes | no | no | yes | no | no | no | no | no | no | no | no | no |
| GRIN2B | 821 | V | 0 | 0 | yes | no | no | yes | no | no | no | no | yes | no | no | no | no |
| GRIN2B | 822 | F | 0 | 0 | yes | no | no | yes | no | no | no | no | yes | no | no | no | no |
| GRIN2B | 823 | Y | 0 | 0 | yes | no | no | yes | no | no | no | no | no | no | no | no | no |
| GRIN2B | 824 | M | 0 | 0 | yes | no | no | yes | no | no | no | no | no | no | no | no | no |
| GRIN2B | 825 | L | 0 | 1 | yes | no | no | yes | no | no | no | no | yes | no | no | no | no |
| GRIN2B | 826 | G | 0 | 1 | yes | no | no | yes | no | no | no | no | no | no | no | no | no |
| GRIN2B | 827 | A | 0 | 0 | yes | no | no | yes | no | no | no | no | no | no | no | no | no |
| GRIN2B | 828 | A | 0 | 0 | yes | no | no | yes | no | no | no | no | yes | no | no | no | no |

| | | | | | | | | | | | | | | | | | |
|---|---|---|---|---|---|---|---|---|---|---|---|---|---|---|---|---|---|
| GRIN2B | 829 | M | 0 | 0 | yes | no | no | yes | no | no | no | no | no | no | no | no | no |
| GRIN2B | 830 | A | 0 | 0 | yes | no | no | yes | no | no | no | no | no | no | no | no | no |
| GRIN2B | 831 | L | 0 | 0 | yes | no | no | yes | no | no | no | no | no | no | no | no | no |
| GRIN2B | 832 | S | 0 | 0 | yes | no | no | yes | no | no | no | no | no | no | no | no | no |
| GRIN2B | 833 | L | 0 | 0 | yes | no | no | yes | no | no | no | no | no | no | no | no | no |
| GRIN2B | 834 | I | 0 | 0 | yes | no | no | yes | no | no | no | no | no | no | no | no | no |
| GRIN2B | 835 | T | 0 | 0 | yes | no | no | yes | no | no | no | no | no | no | no | no | no |
| GRIN2B | 836 | F | 0 | 0 | yes | no | no | yes | no | no | no | no | no | no | no | no | no |
| GRIN2B | 837 | I | 0 | 0 | no | yes | no | yes | no | no | no | no | no | no | no | no | no |
