## Supplementary Table3 for "Conserved patterns across ion channels correlate with variant pathogenicity and clinical phenotypes"

| Supplementary Table 3: List of all the pore-specific features for each residue. For each residue the distances from the pore axis and membrane center along with the seven biophysical pore properties (only for pore residues, see methods for details) are reported. | | | | | | | | | | | | | |
|---|---|---|---|---|---|---|---|---|---|---|---|---|---|
| | | | Distances from membrane center and pore axis | | | | Biophysical pore properties | | | | | | |
| Gene | Protein Position | Amino acid | Distance from membrane center (in Å) | Distance to pore axis (in Å) | Distance to pore axis (normalized) | Distance to pore surface (in Å) | Pore radius (in Å) | Hydrophobicity | Hydropathy | Polarity | Water solubility | Ionizability | Charge |
| CACNA1G | 80 | N | -16.4995 | 43.80896141 | 0.827972434 | 42.59347891 | NA | NA | NA | NA | NA | NA | NA |
| CACNA1G | 81 | P | -15.95142857 | 42.30741015 | 0.798202725 | 41.09165613 | NA | NA | NA | NA | NA | NA | NA |
| CACNA1G | 82 | W | -12.92528571 | 44.53647113 | 0.842396019 | 43.32063994 | NA | NA | NA | NA | NA | NA | NA |
| CACNA1G | 83 | F | -10.38045455 | 42.56338452 | 0.803277665 | 41.33785271 | NA | NA | NA | NA | NA | NA | NA |
| CACNA1G | 84 | E | -11.19066667 | 36.82711411 | 0.689550545 | 35.60388079 | NA | NA | NA | NA | NA | NA | NA |
| CACNA1G | 85 | R | -11.28654545 | 39.16389548 | 0.7358795 | 37.95238498 | NA | NA | NA | NA | NA | NA | NA |
| CACNA1G | 86 | I | -6.34625 | 42.27544413 | 0.797568968 | 41.04868825 | NA | NA | NA | NA | NA | NA | NA |
| CACNA1G | 87 | S | -5.4915 | 40.01392911 | 0.752732241 | 38.77606309 | NA | NA | NA | NA | NA | NA | NA |
| CACNA1G | 88 | M | -4.716 | 34.05333495 | 0.634557685 | 32.92800843 | NA | NA | NA | NA | NA | NA | NA |
| CACNA1G | 89 | L | -2.15475 | 40.1286639 | 0.755006969 | 38.98764897 | NA | NA | NA | NA | NA | NA | NA |
| CACNA1G | 90 | V | -0.502142857 | 40.4206734 | 0.76079634 | 39.23549836 | NA | NA | NA | NA | NA | NA | NA |
| CACNA1G | 91 | I | -1.07625 | 35.80060069 | 0.669198921 | 34.61449912 | NA | NA | NA | NA | NA | NA | NA |
| CACNA1G | 92 | L | 0.61975 | 35.65343031 | 0.666281126 | 34.48663964 | NA | NA | NA | NA | NA | NA | NA |
| CACNA1G | 93 | L | 3.306625 | 39.63700657 | 0.745259386 | 37.14029811 | NA | NA | NA | NA | NA | NA | NA |
| CACNA1G | 94 | N | 3.74675 | 37.61893861 | 0.705249233 | 34.98188932 | NA | NA | NA | NA | NA | NA | NA |
| CACNA1G | 95 | C | 3.900333333 | 34.04978511 | 0.634487306 | 31.36329709 | NA | NA | NA | NA | NA | NA | NA |
| CACNA1G | 96 | V | 6.581857143 | 37.44171868 | 0.701735676 | 36.67433552 | NA | NA | NA | NA | NA | NA | NA |
| CACNA1G | 97 | T | 8.363142857 | 37.97511371 | 0.712310749 | 37.11908171 | NA | NA | NA | NA | NA | NA | NA |
| CACNA1G | 98 | L | 8.13475 | 34.26892607 | 0.638831987 | 33.38305905 | NA | NA | NA | NA | NA | NA | NA |
| CACNA1G | 99 | G | 10.35825 | 33.62031911 | 0.625972726 | 32.77235104 | NA | NA | NA | NA | NA | NA | NA |
| CACNA1G | 100 | M | 12.359125 | 35.74486907 | 0.668093988 | 34.88257544 | NA | NA | NA | NA | NA | NA | NA |
| CACNA1G | 101 | F | 15.04563636 | 34.0089485 | 0.63367768 | 33.05008757 | NA | NA | NA | NA | NA | NA | NA |
| CACNA1G | 102 | R | 18.87254545 | 35.33801886 | 0.660027788 | 34.36857618 | NA | NA | NA | NA | NA | NA | NA |
| CACNA1G | 103 | P | 18.92271429 | 31.88730006 | 0.591613944 | 30.91644891 | NA | NA | NA | NA | NA | NA | NA |
| CACNA1G | 104 | C | 22.74583333 | 32.89252698 | 0.611543542 | 32.14901473 | NA | NA | NA | NA | NA | NA | NA |
| CACNA1G | 105 | E | 23.15377778 | 37.33595944 | 0.699638896 | 36.35681853 | NA | NA | NA | NA | NA | NA | NA |
| CACNA1G | 106 | D | 19.399375 | 38.52091766 | 0.723131842 | 37.55460083 | NA | NA | NA | NA | NA | NA | NA |
| CACNA1G | 107 | I | 22.95 | 39.15939527 | 0.735790279 | 38.18572632 | NA | NA | NA | NA | NA | NA | NA |

| | | | | | | | | | | | | | |
|---|---|---|---|---|---|---|---|---|---|---|---|---|---|
| CACNA1G | 108 | A | 23.3442 | 41.81428203 | 0.788425982 | 40.84010362 | NA | NA | NA | NA | NA | NA | NA |
| CACNA1G | 109 | C | 21.7088 | 44.71794076 | 0.84599383 | 43.74852953 | NA | NA | NA | NA | NA | NA | NA |
| CACNA1G | 110 | D | 24.3092 | 46.40110616 | 0.879364216 | 45.42948255 | NA | NA | NA | NA | NA | NA | NA |
| CACNA1G | 111 | S | 22.1464 | 45.02036843 | 0.851989752 | 44.04743716 | NA | NA | NA | NA | NA | NA | NA |
| CACNA1G | 112 | Q | 20.1236 | 47.46230524 | 0.900403516 | 46.49697163 | NA | NA | NA | NA | NA | NA | NA |
| CACNA1G | 113 | R | 17.30427273 | 39.35294091 | 0.739627509 | 38.44204043 | NA | NA | NA | NA | NA | NA | NA |
| CACNA1G | 114 | C | 17.7265 | 43.20918964 | 0.816081377 | 42.24704669 | NA | NA | NA | NA | NA | NA | NA |
| CACNA1G | 115 | R | 18.76045455 | 46.26141565 | 0.876594716 | 45.30100453 | NA | NA | NA | NA | NA | NA | NA |
| CACNA1G | 116 | I | 13.929375 | 46.67795795 | 0.884853071 | 45.74507828 | NA | NA | NA | NA | NA | NA | NA |
| CACNA1G | 117 | L | 12.625 | 41.24423859 | 0.777124318 | 40.3105987 | NA | NA | NA | NA | NA | NA | NA |
| CACNA1G | 118 | Q | 13.72611111 | 44.97225741 | 0.851035905 | 44.02454951 | NA | NA | NA | NA | NA | NA | NA |
| CACNA1G | 119 | A | 10.6482 | 47.3302525 | 0.897785442 | 46.41361191 | NA | NA | NA | NA | NA | NA | NA |
| CACNA1G | 120 | F | 8.073181818 | 45.40271114 | 0.859570067 | 44.54373804 | NA | NA | NA | NA | NA | NA | NA |
| CACNA1G | 121 | D | 7.8835 | 41.53222554 | 0.782833939 | 40.59905273 | NA | NA | NA | NA | NA | NA | NA |
| CACNA1G | 122 | D | 8.326125 | 46.35153708 | 0.878381461 | 45.42078524 | NA | NA | NA | NA | NA | NA | NA |
| CACNA1G | 123 | F | 3.517454545 | 47.77345357 | 0.906572333 | 47.06121419 | NA | NA | NA | NA | NA | NA | NA |
| CACNA1G | 124 | I | 3.002875 | 42.8586839 | 0.809132261 | 40.09811613 | NA | NA | NA | NA | NA | NA | NA |
| CACNA1G | 125 | F | 3.190545455 | 44.78105918 | 0.847245214 | 42.2565129 | NA | NA | NA | NA | NA | NA | NA |
| CACNA1G | 126 | A | 1.3012 | 47.87250346 | 0.908536093 | 45.11283646 | NA | NA | NA | NA | NA | NA | NA |
| CACNA1G | 127 | F | -1.804818182 | 44.55476283 | 0.84275867 | 43.31574027 | NA | NA | NA | NA | NA | NA | NA |
| CACNA1G | 128 | F | -1.037090909 | 40.56827974 | 0.763722779 | 39.56925762 | NA | NA | NA | NA | NA | NA | NA |
| CACNA1G | 129 | A | -1.3056 | 46.39076777 | 0.879159247 | 45.18463224 | NA | NA | NA | NA | NA | NA | NA |
| CACNA1G | 130 | V | -3.460714286 | 48.33859016 | 0.917776714 | 47.10113118 | NA | NA | NA | NA | NA | NA | NA |
| CACNA1G | 131 | E | -6.061 | 42.26244225 | 0.797311193 | 41.02951065 | NA | NA | NA | NA | NA | NA | NA |
| CACNA1G | 132 | M | -6.39725 | 42.72218778 | 0.806426093 | 41.49536208 | NA | NA | NA | NA | NA | NA | NA |
| CACNA1G | 133 | V | -6.942285714 | 47.90126222 | 0.909106263 | 46.66851343 | NA | NA | NA | NA | NA | NA | NA |
| CACNA1G | 134 | V | -9.923285714 | 47.81736072 | 0.907442835 | 46.59493771 | NA | NA | NA | NA | NA | NA | NA |
| CACNA1G | 135 | K | -11.41644444 | 41.5969811 | 0.784117781 | 40.38152847 | NA | NA | NA | NA | NA | NA | NA |
| CACNA1G | 136 | M | -10.416125 | 45.56066808 | 0.862701716 | 44.33976068 | NA | NA | NA | NA | NA | NA | NA |
| CACNA1G | 137 | V | -14.11714286 | 49.22901727 | 0.935430294 | 48.01510718 | NA | NA | NA | NA | NA | NA | NA |
| CACNA1G | 138 | A | -16.039 | 46.52032934 | 0.881727931 | 45.44061488 | NA | NA | NA | NA | NA | NA | NA |
| CACNA1G | 149 | G | -12.72575 | 38.69914281 | 0.726665328 | 37.50079043 | NA | NA | NA | NA | NA | NA | NA |
| CACNA1G | 150 | D | -13.9995 | 35.18330471 | 0.65696043 | 34.12007111 | NA | NA | NA | NA | NA | NA | NA |

| | | | | | | | | | | | | | |
|---|---|---|---|---|---|---|---|---|---|---|---|---|---|
| CACNA1G | 151 | T | -11.086 | 31.18432869 | 0.577676855 | 29.97382834 | NA | NA | NA | NA | NA | NA | NA |
| CACNA1G | 152 | W | -9.347714286 | 28.50539817 | 0.524564461 | 27.27942879 | NA | NA | NA | NA | NA | NA | NA |
| CACNA1G | 153 | N | -9.659875 | 36.34497063 | 0.679991583 | 35.12403858 | NA | NA | NA | NA | NA | NA | NA |
| CACNA1G | 154 | R | -8.980363636 | 36.40769914 | 0.681235237 | 35.21676823 | NA | NA | NA | NA | NA | NA | NA |
| CACNA1G | 155 | L | -4.149125 | 32.30197087 | 0.599835194 | 31.05548603 | NA | NA | NA | NA | NA | NA | NA |
| CACNA1G | 156 | D | -4.661875 | 36.95570181 | 0.69209992 | 35.7159032 | NA | NA | NA | NA | NA | NA | NA |
| CACNA1G | 157 | F | -3.804 | 39.53792196 | 0.743294938 | 38.29890858 | NA | NA | NA | NA | NA | NA | NA |
| CACNA1G | 158 | F | -0.638181818 | 36.475144 | 0.682572396 | 36.17454988 | NA | NA | NA | NA | NA | NA | NA |
| CACNA1G | 159 | I | 0.373125 | 36.25010379 | 0.678110756 | 33.57727304 | NA | NA | NA | NA | NA | NA | NA |
| CACNA1G | 160 | V | 0.391857143 | 41.64166761 | 0.785003734 | 38.83817976 | NA | NA | NA | NA | NA | NA | NA |
| CACNA1G | 161 | I | 1.01375 | 43.06984033 | 0.813318642 | 40.46797052 | NA | NA | NA | NA | NA | NA | NA |
| CACNA1G | 162 | A | 4.1322 | 39.51476213 | 0.742835771 | 38.58843757 | NA | NA | NA | NA | NA | NA | NA |
| CACNA1G | 163 | G | 5.0825 | 42.15987406 | 0.795277679 | 41.22830814 | NA | NA | NA | NA | NA | NA | NA |
| CACNA1G | 164 | M | 3.913875 | 44.91939817 | 0.849987919 | 43.9929257 | NA | NA | NA | NA | NA | NA | NA |
| CACNA1G | 165 | L | 7.485625 | 42.49353138 | 0.801892759 | 41.61180815 | NA | NA | NA | NA | NA | NA | NA |
| CACNA1G | 166 | E | 9.941888889 | 41.19402061 | 0.776128698 | 40.28548726 | NA | NA | NA | NA | NA | NA | NA |
| CACNA1G | 167 | Y | 8.892 | 45.49404104 | 0.861380771 | 44.56787187 | NA | NA | NA | NA | NA | NA | NA |
| CACNA1G | 168 | S | 9.913333333 | 47.91731801 | 0.909424585 | 47.0236616 | NA | NA | NA | NA | NA | NA | NA |
| CACNA1G | 169 | L | 13.209625 | 44.15366633 | 0.834806543 | 43.23954436 | NA | NA | NA | NA | NA | NA | NA |
| CACNA1G | 170 | D | 14.798 | 46.76905595 | 0.886659177 | 45.83520589 | NA | NA | NA | NA | NA | NA | NA |
| CACNA1G | 171 | L | 14.7645 | 50.04456431 | 0.951599305 | 49.11797843 | NA | NA | NA | NA | NA | NA | NA |
| CACNA1G | 172 | Q | 16.46977778 | 50.12070855 | 0.953108938 | 49.20185608 | NA | NA | NA | NA | NA | NA | NA |
| CACNA1G | 173 | N | 20.020125 | 48.18243716 | 0.914680829 | 47.22983011 | NA | NA | NA | NA | NA | NA | NA |
| CACNA1G | 174 | V | 19.53571429 | 46.65685892 | 0.884434762 | 45.71549942 | NA | NA | NA | NA | NA | NA | NA |
| CACNA1G | 175 | S | 17.9 | 43.03965189 | 0.812720127 | 42.0968023 | NA | NA | NA | NA | NA | NA | NA |
| CACNA1G | 176 | F | 17.65363636 | 39.83786594 | 0.749241618 | 38.89975588 | NA | NA | NA | NA | NA | NA | NA |
| CACNA1G | 177 | S | 16.93966667 | 35.62543021 | 0.665725997 | 34.68261661 | NA | NA | NA | NA | NA | NA | NA |
| CACNA1G | 178 | A | 13.4374 | 34.94741998 | 0.652283787 | 34.03009388 | NA | NA | NA | NA | NA | NA | NA |
| CACNA1G | 179 | V | 10.81471429 | 37.14659101 | 0.695884484 | 36.22659728 | NA | NA | NA | NA | NA | NA | NA |
| CACNA1G | 180 | R | 13.56790909 | 35.91029882 | 0.671373793 | 34.9899606 | NA | NA | NA | NA | NA | NA | NA |
| CACNA1G | 181 | T | 10.44785714 | 31.09155942 | 0.575837614 | 30.17265759 | NA | NA | NA | NA | NA | NA | NA |
| CACNA1G | 182 | V | 6.672142857 | 34.62889421 | 0.645968705 | 33.72433921 | NA | NA | NA | NA | NA | NA | NA |
| CACNA1G | 183 | R | 8.624272727 | 34.93674588 | 0.652072163 | 33.99417003 | NA | NA | NA | NA | NA | NA | NA |

| | | | | | | | | | | | | | |
|---|---|---|---|---|---|---|---|---|---|---|---|---|---|
| CACNA1G | 184 | V | 5.852857143 | 29.99731998 | 0.554143257 | 29.05006134 | NA | NA | NA | NA | NA | NA | NA |
| CACNA1G | 185 | L | 2.052875 | 31.03888223 | 0.574793238 | 28.42783952 | NA | NA | NA | NA | NA | NA | NA |
| CACNA1G | 186 | R | 3.235454545 | 34.799408 | 0.649349306 | 31.99447206 | NA | NA | NA | NA | NA | NA | NA |
| CACNA1G | 187 | P | 1.130285714 | 31.69795191 | 0.587859933 | 29.25874976 | NA | NA | NA | NA | NA | NA | NA |
| CACNA1G | 188 | L | -1.474625 | 29.06341152 | 0.535627617 | 27.80728239 | NA | NA | NA | NA | NA | NA | NA |
| CACNA1G | 189 | R | -2.172 | 33.61213 | 0.625810369 | 32.49474987 | NA | NA | NA | NA | NA | NA | NA |
| CACNA1G | 190 | A | -4.8606 | 31.68552635 | 0.587613584 | 30.45082414 | NA | NA | NA | NA | NA | NA | NA |
| CACNA1G | 191 | I | -6.843375 | 27.48215066 | 0.504277588 | 26.23565335 | NA | NA | NA | NA | NA | NA | NA |
| CACNA1G | 192 | N | -10.0015 | 31.65508929 | 0.58701014 | 30.42268505 | NA | NA | NA | NA | NA | NA | NA |
| CACNA1G | 193 | R | -8.045454545 | 33.38394861 | 0.621286452 | 32.15860563 | NA | NA | NA | NA | NA | NA | NA |
| CACNA1G | 194 | V | -9.374 | 29.69393785 | 0.548128412 | 28.46868556 | NA | NA | NA | NA | NA | NA | NA |
| CACNA1G | 195 | P | -13.38885714 | 28.52825873 | 0.525017694 | 27.31570457 | NA | NA | NA | NA | NA | NA | NA |
| CACNA1G | 196 | S | -12.60716667 | 25.0518061 | 0.456093651 | 23.83453902 | NA | NA | NA | NA | NA | NA | NA |
| CACNA1G | 197 | M | -8.703875 | 25.22908752 | 0.459608427 | 24.00363235 | NA | NA | NA | NA | NA | NA | NA |
| CACNA1G | 198 | R | -13.30127273 | 26.19202018 | 0.4786995 | 24.97874628 | NA | NA | NA | NA | NA | NA | NA |
| CACNA1G | 199 | I | -15.42375 | 23.16834673 | 0.418752244 | 22.08718832 | NA | NA | NA | NA | NA | NA | NA |
| CACNA1G | 200 | L | -11.558875 | 19.40018088 | 0.344044704 | 18.18088844 | NA | NA | NA | NA | NA | NA | NA |
| CACNA1G | 201 | V | -10.00557143 | 22.98455836 | 0.415108461 | 21.76317492 | NA | NA | NA | NA | NA | NA | NA |
| CACNA1G | 202 | T | -13.98442857 | 22.70780912 | 0.409621639 | 21.6145182 | NA | NA | NA | NA | NA | NA | NA |
| CACNA1G | 203 | L | -14.919375 | 18.92382662 | 0.334600519 | 17.83239948 | NA | NA | NA | NA | NA | NA | NA |
| CACNA1G | 204 | L | -9.5865 | 19.20582633 | 0.340191437 | 17.96308944 | NA | NA | NA | NA | NA | NA | NA |
| CACNA1G | 205 | L | -10.513125 | 21.47496778 | 0.385179365 | 20.26767661 | NA | NA | NA | NA | NA | NA | NA |
| CACNA1G | 206 | D | -15.490375 | 20.18014016 | 0.359508153 | 19.07489244 | NA | NA | NA | NA | NA | NA | NA |
| CACNA1G | 207 | T | -12.61614286 | 15.50041306 | 0.266728027 | 14.40826435 | NA | NA | NA | NA | NA | NA | NA |
| CACNA1G | 208 | L | -9.04825 | 18.5874172 | 0.327930876 | 17.36406864 | NA | NA | NA | NA | NA | NA | NA |
| CACNA1G | 209 | P | -12.19957143 | 20.09545148 | 0.357829117 | 19.02709098 | NA | NA | NA | NA | NA | NA | NA |
| CACNA1G | 210 | M | -13.883875 | 16.57244638 | 0.287982127 | 15.51067536 | NA | NA | NA | NA | NA | NA | NA |
| CACNA1G | 211 | L | -8.443125 | 14.455122 | 0.246004119 | 13.45923994 | NA | NA | NA | NA | NA | NA | NA |
| CACNA1G | 212 | G | -8.29575 | 18.91018162 | 0.334329994 | 17.88239858 | NA | NA | NA | NA | NA | NA | NA |
| CACNA1G | 213 | N | -9.98775 | 18.07633771 | 0.31779823 | 17.06429051 | NA | NA | NA | NA | NA | NA | NA |
| CACNA1G | 214 | V | -6.127 | 14.38646235 | 0.244642875 | 13.18969217 | NA | NA | NA | NA | NA | NA | NA |
| CACNA1G | 215 | L | -3.437125 | 18.49071067 | 0.326013576 | 16.38996769 | NA | NA | NA | NA | NA | NA | NA |
| CACNA1G | 216 | L | -5.607375 | 20.43604184 | 0.364581651 | 18.29328652 | NA | NA | NA | NA | NA | NA | NA |

| | | | | | | | | | | | | | |
|---|---|---|---|---|---|---|---|---|---|---|---|---|---|
| CACNA1G | 217 | L | -3.99175 | 15.62388534 | 0.269175985 | 13.14271667 | NA | NA | NA | NA | NA | NA | NA |
| CACNA1G | 218 | C | -0.656333333 | 15.93061876 | 0.275257272 | 13.39408597 | NA | NA | NA | NA | NA | NA | NA |
| CACNA1G | 219 | F | -0.253545455 | 19.35939475 | 0.343236079 | 16.82409052 | NA | NA | NA | NA | NA | NA | NA |
| CACNA1G | 220 | F | -1.977727273 | 19.61924711 | 0.348387904 | 17.05996195 | NA | NA | NA | NA | NA | NA | NA |
| CACNA1G | 221 | V | 1.255571429 | 14.70431263 | 0.250944565 | 13.96435226 | NA | NA | NA | NA | NA | NA | NA |
| CACNA1G | 222 | F | 3.805545455 | 14.34556583 | 0.243832062 | 13.5649819 | NA | NA | NA | NA | NA | NA | NA |
| CACNA1G | 223 | F | 3.671727273 | 19.67721233 | 0.349537121 | 18.94408714 | NA | NA | NA | NA | NA | NA | NA |
| CACNA1G | 224 | I | 3.065 | 19.35167075 | 0.343082944 | 18.6627699 | NA | NA | NA | NA | NA | NA | NA |
| CACNA1G | 225 | F | 5.247454545 | 14.98101886 | 0.256430534 | 14.30915952 | NA | NA | NA | NA | NA | NA | NA |
| CACNA1G | 226 | G | 8.53475 | 18.21622604 | 0.320571652 | 17.4915188 | NA | NA | NA | NA | NA | NA | NA |
| CACNA1G | 227 | I | 8.532625 | 20.82738075 | 0.372340325 | 20.07823934 | NA | NA | NA | NA | NA | NA | NA |
| CACNA1G | 228 | V | 9.526285714 | 20.11220341 | 0.358161241 | 19.35141396 | NA | NA | NA | NA | NA | NA | NA |
| CACNA1G | 229 | G | 12.64125 | 18.78309445 | 0.331810367 | 18.30148928 | NA | NA | NA | NA | NA | NA | NA |
| CACNA1G | 230 | V | 13.82628571 | 20.7342751 | 0.370494415 | 19.88187023 | NA | NA | NA | NA | NA | NA | NA |
| CACNA1G | 231 | Q | 13.30022222 | 23.10451592 | 0.417486736 | 22.01874101 | NA | NA | NA | NA | NA | NA | NA |
| CACNA1G | 232 | L | 14.821375 | 19.1307337 | 0.338702653 | 19.09894114 | NA | NA | NA | NA | NA | NA | NA |
| CACNA1G | 233 | W | 15.92764286 | 15.32057734 | 0.26316261 | 15.30723915 | NA | NA | NA | NA | NA | NA | NA |
| CACNA1G | 234 | A | 19.4474 | 20.85568783 | 0.37290154 | 19.82348556 | NA | NA | NA | NA | NA | NA | NA |
| CACNA1G | 235 | G | 22.934 | 19.731426 | 0.35061196 | 16.08390229 | NA | NA | NA | NA | NA | NA | NA |
| CACNA1G | 236 | L | 22.533 | 15.76168855 | 0.271908067 | 12.11843787 | NA | NA | NA | NA | NA | NA | NA |
| CACNA1G | 237 | L | 22.67425 | 13.6870057 | 0.230775469 | 10.00240489 | NA | NA | NA | NA | NA | NA | NA |
| CACNA1G | 238 | R | 26.16345455 | 16.83840934 | 0.2932551 | 13.05665382 | NA | NA | NA | NA | NA | NA | NA |
| CACNA1G | 239 | N | 27.91425 | 15.15598532 | 0.259899414 | 11.36657569 | NA | NA | NA | NA | NA | NA | NA |
| CACNA1G | 240 | R | 30.69127273 | 13.63868245 | 0.229817414 | 11.68625276 | NA | NA | NA | NA | NA | NA | NA |
| CACNA1G | 241 | C | 34.80433333 | 12.81004329 | 0.213388839 | 10.29071482 | NA | NA | NA | NA | NA | NA | NA |
| CACNA1G | 242 | F | 36.24790909 | 14.9388533 | 0.255594561 | 12.41791652 | NA | NA | NA | NA | NA | NA | NA |
| CACNA1G | 243 | L | 41.54775 | 13.65972983 | 0.230234698 | 11.21309945 | NA | NA | NA | NA | NA | NA | NA |
| CACNA1G | 244 | P | 42.30542857 | 17.24933167 | 0.301402034 | 14.81852662 | NA | NA | NA | NA | NA | NA | NA |
| CACNA1G | 245 | E | 42.20377778 | 21.17261271 | 0.379184883 | 18.71664943 | NA | NA | NA | NA | NA | NA | NA |
| CACNA1G | 246 | N | 46.456375 | 21.56481115 | 0.386960597 | 21.3351672 | NA | NA | NA | NA | NA | NA | NA |
| CACNA1G | 247 | F | 46.73627273 | 14.68515267 | 0.2505647 | 14.51161526 | NA | NA | NA | NA | NA | NA | NA |
| CACNA1G | 248 | S | 49.69733333 | 16.99009723 | 0.29626246 | 16.81672091 | NA | NA | NA | NA | NA | NA | NA |
| CACNA1G | 249 | L | 48.306875 | 12.5133764 | 0.207507131 | 12.25479065 | NA | NA | NA | NA | NA | NA | NA |

| | | | | | | | | | | | | | |
|---|---|---|---|---|---|---|---|---|---|---|---|---|---|
| CACNA1G | 250 | P | 51.43028571 | 14.28852074 | 0.242701088 | 14.11852679 | NA | NA | NA | NA | NA | NA | NA |
| CACNA1G | 251 | L | 52.646375 | 15.72009141 | 0.271083364 | 15.33614116 | NA | NA | NA | NA | NA | NA | NA |
| CACNA1G | 252 | S | 51.89566667 | 12.12761403 | 0.199859018 | 11.68818626 | NA | NA | NA | NA | NA | NA | NA |
| CACNA1G | 253 | V | 48.76628571 | 9.064009819 | 0.139120095 | 8.548844657 | NA | NA | NA | NA | NA | NA | NA |
| CACNA1G | 254 | D | 45.40625 | 9.536295612 | 0.148483619 | 8.908737205 | NA | NA | NA | NA | NA | NA | NA |
| CACNA1G | 255 | L | 43.745 | 7.275757349 | 0.103666257 | 6.669906426 | NA | NA | NA | NA | NA | NA | NA |
| CACNA1G | 256 | E | 39.49877778 | 10.90581634 | 0.175635695 | 8.472593227 | NA | NA | NA | NA | NA | NA | NA |
| CACNA1G | 257 | R | 41.56563636 | 12.58312195 | 0.208889904 | 10.14840859 | NA | NA | NA | NA | NA | NA | NA |
| CACNA1G | 258 | Y | 36.14341667 | 9.92627211 | 0.156215281 | 7.481670201 | NA | NA | NA | NA | NA | NA | NA |
| CACNA1G | 259 | Y | 36.00766667 | 4.143044533 | 0.041557189 | 1.80146859 | 2.61 | -0.3 | -1.93 | 26.35 | 0.65 | 2 | 0 |
| CACNA1G | 260 | Q | 36.08644444 | 5.888550586 | 0.076163539 | 3.361357093 | 2.61 | -0.3 | -1.93 | 26.35 | 0.65 | 2 | 0 |
| CACNA1G | 261 | T | 33.191 | 6.4563801 | 0.087421309 | 4.450191638 | 2.5 | -0.02 | -1.8 | 21.4 | 0.23 | 2 | 0 |
| CACNA1G | 262 | E | 34.58366667 | 4.194999881 | 0.042587254 | 1.869375851 | 3.37 | -0.32 | -2.48 | 25.75 | 0.43 | 2 | -2 |
| CACNA1G | 263 | N | 29.812625 | 6.476571006 | 0.087821613 | 1.819667299 | 4.66 | -0.83 | -1.92 | 22.07 | 1.32 | 2 | 0 |
| CACNA1G | 264 | E | 31.42633333 | 10.24566801 | 0.162547614 | 5.601425767 | NA | NA | NA | NA | NA | NA | NA |
| CACNA1G | 265 | D | 27.75025 | 10.76645206 | 0.172872663 | 6.956274613 | NA | NA | NA | NA | NA | NA | NA |
| CACNA1G | 266 | E | 24.92277778 | 7.192253958 | 0.102010721 | 3.385837448 | 3.94 | -0.38 | -1.48 | 12.41 | 0.58 | 1 | 1 |
| CACNA1G | 267 | S | 27.04983333 | 5.424076235 | 0.066954884 | 1.589624498 | 3.94 | -0.38 | -1.48 | 12.41 | 0.58 | 1 | 1 |
| CACNA1G | 268 | P | 27.88728571 | 8.420678951 | 0.126365437 | 4.576175632 | 3.88 | -0.75 | -1.53 | 15.11 | 1.09 | 1 | 1 |
| CACNA1G | 269 | F | 31.02872727 | 7.637523879 | 0.110838629 | 3.662144469 | 4.22 | -0.4 | -3.2 | 38.35 | 1.16 | 3 | -1 |
| CACNA1G | 270 | I | 26.703375 | 10.68570077 | 0.17127169 | 6.901101515 | NA | NA | NA | NA | NA | NA | NA |
| CACNA1G | 271 | C | 28.72 | 13.4332368 | 0.225744255 | 9.472980221 | NA | NA | NA | NA | NA | NA | NA |
| CACNA1G | 272 | S | 27.30116667 | 15.88584206 | 0.274369531 | 12.05041668 | NA | NA | NA | NA | NA | NA | NA |
| CACNA1G | 273 | Q | 27.52511111 | 19.16572675 | 0.339396424 | 15.33948428 | NA | NA | NA | NA | NA | NA | NA |
| CACNA1G | 274 | P | 28.91285714 | 22.0211131 | 0.396007225 | 18.05836366 | NA | NA | NA | NA | NA | NA | NA |
| CACNA1G | 275 | R | 26.108 | 23.57555473 | 0.426825537 | 19.75141682 | NA | NA | NA | NA | NA | NA | NA |
| CACNA1G | 276 | E | 23.40955556 | 20.35189217 | 0.362913303 | 16.67818451 | NA | NA | NA | NA | NA | NA | NA |
| CACNA1G | 277 | N | 22.1825 | 18.14845208 | 0.319227967 | 14.46766578 | NA | NA | NA | NA | NA | NA | NA |
| CACNA1G | 278 | G | 22.88075 | 15.38070262 | 0.264354652 | 11.6841351 | NA | NA | NA | NA | NA | NA | NA |
| CACNA1G | 279 | M | 22.86725 | 11.5183142 | 0.187779058 | 7.634899018 | NA | NA | NA | NA | NA | NA | NA |
| CACNA1G | 280 | R | 27.73154545 | 5.876779816 | 0.075930172 | 2.313072383 | 4.1 | -0.75 | -1.53 | 15.11 | 1.09 | 1 | 1 |
| CACNA1G | 281 | S | 30.39283333 | 12.58419171 | 0.208911113 | 8.598718884 | NA | NA | NA | NA | NA | NA | NA |
| CACNA1G | 282 | C | 34.08566667 | 12.96991207 | 0.216558393 | 9.498724782 | NA | NA | NA | NA | NA | NA | NA |

| | | | | | | | | | | | | | |
|---|---|---|---|---|---|---|---|---|---|---|---|---|---|
| CACNA1G | 283 | R | 33.75645455 | 12.45585136 | 0.206366641 | 9.204128111 | NA | NA | NA | NA | NA | NA | NA |
| CACNA1G | 284 | S | 33.02083333 | 8.350812535 | 0.124980268 | 5.236116249 | NA | NA | NA | NA | NA | NA | NA |
| CACNA1G | 285 | V | 36.40714286 | 7.64363042 | 0.110959697 | 5.358595758 | NA | NA | NA | NA | NA | NA | NA |
| CACNA1G | 286 | P | 37.41371429 | 4.464621149 | 0.047932757 | 1.718330293 | 2.74 | -0.14 | -2.73 | 26.27 | 0.28 | 2 | 0 |
| CACNA1G | 287 | T | 41.38385714 | 4.576724921 | 0.050155323 | 2.259843302 | 2.45 | -0.31 | -3.2 | 26.72 | 0.74 | 2 | 0 |
| CACNA1G | 288 | L | 43.74925 | 2.250595699 | 0.004037557 | 1.707128562 | 1.19 | 0.36 | -0.35 | 26.07 | -0.08 | 1 | 1 |
| CACNA1G | 289 | R | 42.62018182 | 2.250608362 | 0.004037808 | 1.781167981 | 0.64 | 0.17 | 1.7 | 1.76 | -0.49 | 0 | 0 |
| CACNA1G | 313 | C | 51.66383333 | 6.752958537 | 0.093301264 | 6.792884223 | 0.64 | 0.17 | 1.7 | 1.76 | -0.49 | 0 | 0 |
| CACNA1G | 314 | V | 50.46171429 | 6.782995282 | 0.093896771 | 6.852271057 | 0.64 | 0.17 | 1.7 | 1.76 | -0.49 | 0 | 0 |
| CACNA1G | 315 | N | 46.259625 | 6.295539056 | 0.084232479 | 6.322291779 | 0.64 | 0.17 | 1.7 | 1.76 | -0.49 | 0 | 0 |
| CACNA1G | 316 | W | 47.24492857 | 7.034075561 | 0.098874681 | 6.971559596 | 0.64 | 0.17 | 1.7 | 1.76 | -0.49 | 0 | 0 |
| CACNA1G | 317 | N | 41.256125 | 4.033983887 | 0.039394956 | 1.894253011 | 2.33 | -0.02 | -1.4 | 18.5 | 0.13 | 1 | 1 |
| CACNA1G | 318 | Q | 43.94077778 | 9.692584433 | 0.151582196 | 7.567908154 | NA | NA | NA | NA | NA | NA | NA |
| CACNA1G | 319 | Y | 45.15225 | 11.92336152 | 0.195809514 | 9.783085925 | NA | NA | NA | NA | NA | NA | NA |
| CACNA1G | 320 | Y | 39.803 | 8.46766544 | 0.12729699 | 5.983612708 | NA | NA | NA | NA | NA | NA | NA |
| CACNA1G | 321 | T | 38.16342857 | 14.42139581 | 0.245335465 | 12.08740862 | NA | NA | NA | NA | NA | NA | NA |
| CACNA1G | 322 | N | 36.374875 | 16.6929988 | 0.290372196 | 14.19769697 | NA | NA | NA | NA | NA | NA | NA |
| CACNA1G | 323 | C | 32.17916667 | 17.39073739 | 0.304205539 | 15.66615597 | NA | NA | NA | NA | NA | NA | NA |
| CACNA1G | 324 | S | 32.28183333 | 18.72273231 | 0.330613629 | 16.18671263 | NA | NA | NA | NA | NA | NA | NA |
| CACNA1G | 325 | A | 30.1624 | 20.67421718 | 0.369303708 | 16.9396322 | NA | NA | NA | NA | NA | NA | NA |
| CACNA1G | 326 | G | 30.37275 | 19.47122433 | 0.345453209 | 16.95249623 | NA | NA | NA | NA | NA | NA | NA |
| CACNA1G | 327 | E | 30.81777778 | 20.13454268 | 0.358604138 | 17.62047675 | NA | NA | NA | NA | NA | NA | NA |
| CACNA1G | 328 | H | 29.5011 | 16.66206677 | 0.289758938 | 13.03202977 | NA | NA | NA | NA | NA | NA | NA |
| CACNA1G | 329 | N | 24.306 | 15.06963742 | 0.258187483 | 11.42828789 | NA | NA | NA | NA | NA | NA | NA |
| CACNA1G | 330 | P | 23.60371429 | 11.55394569 | 0.188485487 | 7.90059531 | NA | NA | NA | NA | NA | NA | NA |
| CACNA1G | 331 | F | 21.81418182 | 8.878102782 | 0.135434308 | 5.354578646 | NA | NA | NA | NA | NA | NA | NA |
| CACNA1G | 332 | K | 24.68666667 | 15.12113954 | 0.259208562 | 11.44846422 | NA | NA | NA | NA | NA | NA | NA |
| CACNA1G | 333 | G | 22.361 | 16.94269866 | 0.295322737 | 13.27156448 | NA | NA | NA | NA | NA | NA | NA |
| CACNA1G | 334 | A | 19.1182 | 16.28589577 | 0.282300983 | 15.2758899 | NA | NA | NA | NA | NA | NA | NA |
| CACNA1G | 335 | I | 18.54 | 12.53700116 | 0.207975514 | 11.54196735 | NA | NA | NA | NA | NA | NA | NA |
| CACNA1G | 336 | N | 18.841625 | 16.21950647 | 0.280984751 | 15.21023696 | NA | NA | NA | NA | NA | NA | NA |
| CACNA1G | 337 | F | 15.77136364 | 15.02311572 | 0.257265145 | 14.07942018 | NA | NA | NA | NA | NA | NA | NA |
| CACNA1G | 338 | D | 19.113625 | 21.03161703 | 0.376389507 | 20.02448004 | NA | NA | NA | NA | NA | NA | NA |

| | | | | | | | | | | | | | |
|---|---|---|---|---|---|---|---|---|---|---|---|---|---|
| CACNA1G | 339 | N | 15.7195 | 21.42687007 | 0.384225781 | 20.53543897 | NA | NA | NA | NA | NA | NA | NA |
| CACNA1G | 340 | I | 10.6365 | 20.69039731 | 0.369624495 | 19.8549582 | NA | NA | NA | NA | NA | NA | NA |
| CACNA1G | 341 | G | 12.27175 | 19.36048026 | 0.343257601 | 18.47361208 | NA | NA | NA | NA | NA | NA | NA |
| CACNA1G | 342 | Y | 16.49483333 | 17.06488764 | 0.297745252 | 16.13150117 | NA | NA | NA | NA | NA | NA | NA |
| CACNA1G | 343 | A | 11.8544 | 15.50108128 | 0.266741276 | 14.64833542 | NA | NA | NA | NA | NA | NA | NA |
| CACNA1G | 344 | W | 7.883071429 | 14.77221507 | 0.252290797 | 13.94097251 | NA | NA | NA | NA | NA | NA | NA |
| CACNA1G | 345 | I | 12.228875 | 12.82818343 | 0.213748485 | 11.91007732 | NA | NA | NA | NA | NA | NA | NA |
| CACNA1G | 346 | A | 12.0214 | 10.70648873 | 0.171683832 | 9.808885265 | NA | NA | NA | NA | NA | NA | NA |
| CACNA1G | 347 | I | 8.258625 | 10.22785256 | 0.162194406 | 9.468523425 | NA | NA | NA | NA | NA | NA | NA |
| CACNA1G | 348 | F | 6.94 | 9.644467741 | 0.150628236 | 8.814983188 | NA | NA | NA | NA | NA | NA | NA |
| CACNA1G | 349 | Q | 11.573 | 6.535907971 | 0.088998026 | 5.636755366 | 1.03 | 0.26 | -0.33 | 13.4 | -0.22 | 1 | -1 |
| CACNA1G | 350 | V | 8.276285714 | 5.770854183 | 0.073830093 | 5.105770131 | 1.03 | 0.26 | -0.33 | 13.4 | -0.22 | 1 | -1 |
| CACNA1G | 351 | I | 4.3305 | 6.618664971 | 0.090638764 | 6.086379388 | 1.01 | 0.69 | 0.73 | 1.3 | -1.17 | 0 | 0 |
| CACNA1G | 352 | T | 6.275428571 | 5.107597772 | 0.060680392 | 4.283017656 | 1.01 | 0.69 | 0.73 | 1.3 | -1.17 | 0 | 0 |
| CACNA1G | 353 | L | 6.64925 | 2.210915647 | 0.003250862 | 1.489965719 | 1.01 | 0.69 | 0.73 | 1.3 | -1.17 | 0 | 0 |
| CACNA1G | 354 | E | 11.07877778 | 4.228718364 | 0.043255756 | 3.28831176 | 1.03 | 0.26 | -0.33 | 13.4 | -0.22 | 1 | -1 |
| CACNA1G | 355 | G | 13.60225 | 5.323605827 | 0.064962961 | 4.39113527 | 0.97 | -0.23 | -1.38 | 14.21 | 0.43 | 1 | -1 |
| CACNA1G | 356 | W | 10.99214286 | 2.252822674 | 0.004081709 | 1.468938926 | 0.97 | -0.23 | -1.38 | 14.21 | 0.43 | 1 | -1 |
| CACNA1G | 357 | V | 14.82 | 2.144314576 | 0.001930431 | 1.649445171 | 0.51 | 0.36 | -0.15 | 26.07 | 0.17 | 1 | 1 |
| CACNA1G | 358 | D | 18.2185 | 3.243172829 | 0.023716361 | 2.199081676 | 1.31 | 0.2 | -1.28 | 25.86 | 0.38 | 2 | 0 |
| CACNA1G | 359 | I | 15.861125 | 7.148402409 | 0.101141322 | 6.829775952 | NA | NA | NA | NA | NA | NA | NA |
| CACNA1G | 360 | M | 15.131 | 7.0600017 | 0.099388692 | 6.938927781 | 0.67 | 0.61 | -0.53 | 17.91 | -0.37 | 1 | 1 |
| CACNA1G | 361 | Y | 18.43808333 | 2.717638865 | 0.013297141 | 1.908169522 | 1.31 | 0.2 | -1.28 | 25.86 | 0.38 | 2 | 0 |
| CACNA1G | 362 | F | 21.30772727 | 7.386551225 | 0.105862853 | 3.721982679 | 3.79 | -0.42 | -2.1 | 21.67 | 0.95 | 2 | 0 |
| CACNA1G | 363 | V | 18.19914286 | 10.96611148 | 0.176831104 | 10.60527505 | NA | NA | NA | NA | NA | NA | NA |
| CACNA1G | 364 | M | 18.61 | 9.129598951 | 0.140420463 | 8.69987889 | NA | NA | NA | NA | NA | NA | NA |
| CACNA1G | 365 | D | 23.4325 | 7.804025692 | 0.114139689 | 4.14426441 | 3.79 | -0.42 | -2.1 | 21.67 | 0.95 | 2 | 0 |
| CACNA1G | 366 | A | 22.9298 | 13.15431826 | 0.220214424 | 9.481777388 | NA | NA | NA | NA | NA | NA | NA |
| CACNA1G | 367 | H | 19.0381 | 15.65504216 | 0.269793699 | 11.9664197 | NA | NA | NA | NA | NA | NA | NA |
| CACNA1G | 368 | S | 18.2055 | 14.33232776 | 0.243569604 | 13.84452708 | NA | NA | NA | NA | NA | NA | NA |
| CACNA1G | 369 | F | 16.50409091 | 12.53364847 | 0.207909044 | 12.20959905 | NA | NA | NA | NA | NA | NA | NA |
| CACNA1G | 370 | Y | 14.20916667 | 15.62082626 | 0.269115336 | 15.26081308 | NA | NA | NA | NA | NA | NA | NA |
| CACNA1G | 371 | N | 14.170125 | 12.73974635 | 0.211995134 | 11.47299407 | NA | NA | NA | NA | NA | NA | NA |

| | | | | | | | | | | | | | |
|---|---|---|---|---|---|---|---|---|---|---|---|---|---|
| CACNA1G | 372 | F | 11.05627273 | 10.81122135 | 0.173760257 | 9.549557778 | NA | NA | NA | NA | NA | NA | NA |
| CACNA1G | 373 | I | 8.281875 | 12.93196145 | 0.215805985 | 11.67487664 | NA | NA | NA | NA | NA | NA | NA |
| CACNA1G | 374 | Y | 9.036833333 | 12.39567045 | 0.205173496 | 11.10589957 | NA | NA | NA | NA | NA | NA | NA |
| CACNA1G | 375 | F | 10.41863636 | 8.989672853 | 0.137646293 | 7.702226864 | NA | NA | NA | NA | NA | NA | NA |
| CACNA1G | 376 | I | 5.99525 | 8.633848331 | 0.130591727 | 7.374500498 | NA | NA | NA | NA | NA | NA | NA |
| CACNA1G | 377 | L | 3.326125 | 11.96546969 | 0.196644349 | 11.84410534 | NA | NA | NA | NA | NA | NA | NA |
| CACNA1G | 378 | L | 4.78475 | 9.836237899 | 0.154430265 | 9.147437776 | NA | NA | NA | NA | NA | NA | NA |
| CACNA1G | 379 | I | 4.11375 | 5.47491936 | 0.067962899 | 5.405689923 | 0.53 | 0.19 | 1.55 | 0.9 | -0.51 | 0 | 0 |
| CACNA1G | 380 | I | 0.575 | 9.173006759 | 0.141281065 | 7.146677008 | NA | NA | NA | NA | NA | NA | NA |
| CACNA1G | 381 | V | -0.104428571 | 10.98631631 | 0.177231684 | 8.406658545 | NA | NA | NA | NA | NA | NA | NA |
| CACNA1G | 382 | G | 0.2855 | 9.343634678 | 0.144663929 | 6.78922906 | NA | NA | NA | NA | NA | NA | NA |
| CACNA1G | 383 | S | 1.359833333 | 5.406538726 | 0.066607186 | 2.851997461 | 2.77 | 0.63 | 1.73 | 1.04 | -1.32 | 0 | 0 |
| CACNA1G | 384 | F | -1.830909091 | 3.969803773 | 0.038122523 | 1.843565484 | 2.77 | 0.63 | 1.73 | 1.04 | -1.32 | 0 | 0 |
| CACNA1G | 385 | F | -4.214909091 | 8.501799692 | 0.127973735 | 6.305810218 | NA | NA | NA | NA | NA | NA | NA |
| CACNA1G | 386 | M | -1.99225 | 9.942808155 | 0.156543124 | 7.757739157 | NA | NA | NA | NA | NA | NA | NA |
| CACNA1G | 387 | I | -1.377625 | 7.347620159 | 0.105091007 | 5.251556168 | NA | NA | NA | NA | NA | NA | NA |
| CACNA1G | 388 | N | -4.547875 | 4.358107502 | 0.045821021 | 3.150809894 | 1.43 | 0.49 | 2.67 | 1.21 | -0.6 | 0 | 0 |
| CACNA1G | 389 | L | -7.48625 | 9.120040241 | 0.140230953 | 8.221583869 | NA | NA | NA | NA | NA | NA | NA |
| CACNA1G | 390 | C | -6.288333333 | 10.1237456 | 0.160130384 | 9.090641061 | NA | NA | NA | NA | NA | NA | NA |
| CACNA1G | 391 | L | -8.007125 | 4.772882358 | 0.054044335 | 3.494473532 | 1.31 | 0.22 | 1.45 | 1.81 | -0.5 | 0 | 0 |
| CACNA1G | 392 | V | -11.27257143 | 5.559846131 | 0.069646654 | 4.542830148 | 1.17 | -0.15 | 1 | 2.3 | -0.06 | 0 | 0 |
| CACNA1G | 393 | V | -11.73828571 | 9.342799687 | 0.144647375 | 8.290521541 | NA | NA | NA | NA | NA | NA | NA |
| CACNA1G | 394 | I | -10.5825 | 10.6253783 | 0.170075739 | 9.533686881 | NA | NA | NA | NA | NA | NA | NA |
| CACNA1G | 395 | A | -13.545 | 7.291297415 | 0.103974354 | 6.170863033 | NA | NA | NA | NA | NA | NA | NA |
| CACNA1G | 396 | T | -16.36285714 | 7.797164998 | 0.114003669 | 7.347768289 | NA | NA | NA | NA | NA | NA | NA |
| CACNA1G | 397 | Q | -16.49855556 | 11.7484267 | 0.192341262 | 11.28171253 | NA | NA | NA | NA | NA | NA | NA |
| CACNA1G | 398 | F | -15.50209091 | 13.31032956 | 0.2233075 | 12.81653762 | NA | NA | NA | NA | NA | NA | NA |
| CACNA1G | 399 | S | -19.3565 | 10.67170881 | 0.170994286 | 10.232072 | NA | NA | NA | NA | NA | NA | NA |
| CACNA1G | 400 | E | -21.55444444 | 13.25150667 | 0.222141279 | 12.83154043 | NA | NA | NA | NA | NA | NA | NA |
| CACNA1G | 401 | T | -19.43914286 | 15.87057497 | 0.274066846 | 15.44132007 | NA | NA | NA | NA | NA | NA | NA |
| CACNA1G | 402 | K | -20.47366667 | 17.18032348 | 0.30003388 | 16.73159766 | NA | NA | NA | NA | NA | NA | NA |
| CACNA1G | 403 | Q | -24.59366667 | 15.24058401 | 0.261576665 | 14.84694877 | NA | NA | NA | NA | NA | NA | NA |
| CACNA1G | 404 | R | -23.37690909 | 18.61272095 | 0.328432548 | 18.22295707 | NA | NA | NA | NA | NA | NA | NA |

| | | | | | | | | | | | | | |
|---|---|---|---|---|---|---|---|---|---|---|---|---|---|
| CACNA1G | 405 | E | -23.69011111 | 20.96251295 | 0.375019451 | 20.55356643 | NA | NA | NA | NA | NA | NA | NA |
| CACNA1G | 739 | I | -18.991875 | 48.22932284 | 0.915610383 | 47.26080115 | NA | NA | NA | NA | NA | NA | NA |
| CACNA1G | 740 | V | -16.46885714 | 44.89139056 | 0.849432641 | 43.90201533 | NA | NA | NA | NA | NA | NA | NA |
| CACNA1G | 741 | D | -18.793875 | 42.86556654 | 0.809268716 | 41.8997872 | NA | NA | NA | NA | NA | NA | NA |
| CACNA1G | 742 | S | -18.78283333 | 44.2718302 | 0.837149256 | 43.33571834 | NA | NA | NA | NA | NA | NA | NA |
| CACNA1G | 743 | K | -18.07855556 | 39.0768353 | 0.734153448 | 38.17711225 | NA | NA | NA | NA | NA | NA | NA |
| CACNA1G | 744 | Y | -13.62316667 | 43.74238117 | 0.826652416 | 43.08571562 | NA | NA | NA | NA | NA | NA | NA |
| CACNA1G | 745 | F | -12.33645455 | 42.62109658 | 0.804421862 | 41.7232251 | NA | NA | NA | NA | NA | NA | NA |
| CACNA1G | 746 | G | -13.071 | 39.0329097 | 0.73328258 | 38.13146279 | NA | NA | NA | NA | NA | NA | NA |
| CACNA1G | 747 | R | -11.73790909 | 36.63141795 | 0.685670679 | 35.76391334 | NA | NA | NA | NA | NA | NA | NA |
| CACNA1G | 748 | G | -8.87175 | 41.2101427 | 0.776448334 | 40.35696193 | NA | NA | NA | NA | NA | NA | NA |
| CACNA1G | 749 | I | -8.12275 | 38.16573706 | 0.716090042 | 37.30741476 | NA | NA | NA | NA | NA | NA | NA |
| CACNA1G | 750 | M | -8.352125 | 32.39676629 | 0.601714605 | 31.53236424 | NA | NA | NA | NA | NA | NA | NA |
| CACNA1G | 751 | I | -5.49 | 37.47228108 | 0.702341605 | 36.62551291 | NA | NA | NA | NA | NA | NA | NA |
| CACNA1G | 752 | A | -3.508 | 38.5211409 | 0.723136267 | 37.69619121 | NA | NA | NA | NA | NA | NA | NA |
| CACNA1G | 753 | I | -3.49875 | 34.73357104 | 0.648044024 | 33.9049603 | NA | NA | NA | NA | NA | NA | NA |
| CACNA1G | 754 | L | -2.41 | 31.14423338 | 0.576881927 | 30.33875659 | NA | NA | NA | NA | NA | NA | NA |
| CACNA1G | 755 | V | -0.518 | 36.1475082 | 0.676076699 | 34.9044764 | NA | NA | NA | NA | NA | NA | NA |
| CACNA1G | 756 | N | 1.2695 | 36.54130596 | 0.683884121 | 35.23809195 | NA | NA | NA | NA | NA | NA | NA |
| CACNA1G | 757 | T | 0.993571429 | 31.64487383 | 0.586807609 | 30.34997097 | NA | NA | NA | NA | NA | NA | NA |
| CACNA1G | 758 | L | 1.91025 | 31.57806911 | 0.58548314 | 30.24279102 | NA | NA | NA | NA | NA | NA | NA |
| CACNA1G | 759 | S | 5.6845 | 35.89240506 | 0.671019032 | 34.50542464 | NA | NA | NA | NA | NA | NA | NA |
| CACNA1G | 760 | M | 5.7705 | 33.93601325 | 0.632231668 | 31.68814445 | NA | NA | NA | NA | NA | NA | NA |
| CACNA1G | 761 | G | 7.83 | 31.6275736 | 0.586464615 | 30.01935035 | NA | NA | NA | NA | NA | NA | NA |
| CACNA1G | 762 | I | 10.03675 | 33.58378027 | 0.625248308 | 32.23508157 | NA | NA | NA | NA | NA | NA | NA |
| CACNA1G | 763 | E | 12.90466667 | 33.83273879 | 0.630184152 | 32.40021776 | NA | NA | NA | NA | NA | NA | NA |
| CACNA1G | 764 | Y | 16.91583333 | 33.30715491 | 0.619763942 | 33.12397394 | NA | NA | NA | NA | NA | NA | NA |
| CACNA1G | 765 | H | 20.4041 | 30.5719561 | 0.565535975 | 27.62072324 | NA | NA | NA | NA | NA | NA | NA |
| CACNA1G | 766 | E | 22.80577778 | 33.82221508 | 0.629975509 | 30.86059147 | NA | NA | NA | NA | NA | NA | NA |
| CACNA1G | 767 | Q | 18.44388889 | 36.4538999 | 0.682151211 | 33.51472382 | NA | NA | NA | NA | NA | NA | NA |
| CACNA1G | 768 | P | 20.18471429 | 39.46784191 | 0.741905533 | 36.53790122 | NA | NA | NA | NA | NA | NA | NA |
| CACNA1G | 769 | E | 20.35855556 | 41.96823314 | 0.791478212 | 39.02878296 | NA | NA | NA | NA | NA | NA | NA |
| CACNA1G | 770 | E | 19.135 | 44.37067772 | 0.839109004 | 41.42380718 | NA | NA | NA | NA | NA | NA | NA |

| | | | | | | | | | | | | | |
|---|---|---|---|---|---|---|---|---|---|---|---|---|---|
| CACNA1G | 771 | L | 14.63775 | 39.38805855 | 0.74032375 | 39.18278635 | NA | NA | NA | NA | NA | NA | NA |
| CACNA1G | 772 | T | 15.438 | 41.32816818 | 0.778788304 | 40.95325243 | NA | NA | NA | NA | NA | NA | NA |
| CACNA1G | 773 | N | 15.54275 | 45.33439961 | 0.858215725 | 44.89237957 | NA | NA | NA | NA | NA | NA | NA |
| CACNA1G | 774 | A | 11.5868 | 44.86469468 | 0.848903369 | 43.51732205 | NA | NA | NA | NA | NA | NA | NA |
| CACNA1G | 775 | L | 10.54275 | 39.70670164 | 0.746641158 | 38.32349179 | NA | NA | NA | NA | NA | NA | NA |
| CACNA1G | 776 | E | 10.54555556 | 44.47114189 | 0.841100804 | 43.12471239 | NA | NA | NA | NA | NA | NA | NA |
| CACNA1G | 777 | I | 9.215375 | 46.87593927 | 0.888778242 | 45.70818548 | NA | NA | NA | NA | NA | NA | NA |
| CACNA1G | 778 | S | 5.822 | 42.81040072 | 0.808175 | 41.45456681 | NA | NA | NA | NA | NA | NA | NA |
| CACNA1G | 779 | N | 5.590375 | 43.02108662 | 0.812352053 | 41.7236257 | NA | NA | NA | NA | NA | NA | NA |
| CACNA1G | 780 | I | 5.589625 | 46.48802797 | 0.881087525 | 45.17618516 | NA | NA | NA | NA | NA | NA | NA |
| CACNA1G | 781 | V | 2.259285714 | 46.26603278 | 0.876686255 | 45.01538828 | NA | NA | NA | NA | NA | NA | NA |
| CACNA1G | 782 | F | 0.539272727 | 41.46742511 | 0.781549207 | 40.19398901 | NA | NA | NA | NA | NA | NA | NA |
| CACNA1G | 783 | T | 0.912428571 | 43.9333279 | 0.83043812 | 42.6891972 | NA | NA | NA | NA | NA | NA | NA |
| CACNA1G | 784 | S | -0.401166667 | 47.11887738 | 0.893594726 | 46.35985067 | NA | NA | NA | NA | NA | NA | NA |
| CACNA1G | 785 | L | -3.336875 | 44.54265759 | 0.842518672 | 43.69390889 | NA | NA | NA | NA | NA | NA | NA |
| CACNA1G | 786 | F | -3.564363636 | 40.51284069 | 0.762623646 | 39.68661655 | NA | NA | NA | NA | NA | NA | NA |
| CACNA1G | 787 | A | -3.7496 | 46.44511999 | 0.880236833 | 45.62497103 | NA | NA | NA | NA | NA | NA | NA |
| CACNA1G | 788 | L | -6.013375 | 48.46668336 | 0.920316286 | 47.61979463 | NA | NA | NA | NA | NA | NA | NA |
| CACNA1G | 789 | E | -8.621333333 | 42.02752704 | 0.792653771 | 41.12202284 | NA | NA | NA | NA | NA | NA | NA |
| CACNA1G | 790 | M | -7.76375 | 46.75541916 | 0.886388814 | 45.85957339 | NA | NA | NA | NA | NA | NA | NA |
| CACNA1G | 791 | L | -8.734625 | 49.41848951 | 0.939186765 | 48.51956774 | NA | NA | NA | NA | NA | NA | NA |
| CACNA1G | 792 | L | -11.662875 | 50.03694036 | 0.951448153 | 49.1231632 | NA | NA | NA | NA | NA | NA | NA |
| CACNA1G | 793 | K | -13.41922222 | 43.67165226 | 0.825250147 | 42.7851916 | NA | NA | NA | NA | NA | NA | NA |
| CACNA1G | 794 | L | -12.657125 | 49.88217752 | 0.948379829 | 48.94112578 | NA | NA | NA | NA | NA | NA | NA |
| CACNA1G | 795 | L | -14.117375 | 52.48584196 | 1 | 51.54179724 | NA | NA | NA | NA | NA | NA | NA |
| CACNA1G | 796 | V | -17.75671429 | 51.44929268 | 0.979449406 | 50.48789058 | NA | NA | NA | NA | NA | NA | NA |
| CACNA1G | 797 | Y | -18.75325 | 45.77337026 | 0.866918743 | 44.81865961 | NA | NA | NA | NA | NA | NA | NA |
| CACNA1G | 798 | G | -18.20875 | 49.28732301 | 0.936586262 | 48.29520249 | NA | NA | NA | NA | NA | NA | NA |
| CACNA1G | 799 | P | -16.10214286 | 49.25100589 | 0.93586624 | 48.22815094 | NA | NA | NA | NA | NA | NA | NA |
| CACNA1G | 800 | F | -19.37309091 | 47.77151693 | 0.906533937 | 46.79886186 | NA | NA | NA | NA | NA | NA | NA |
| CACNA1G | 801 | G | -18.68425 | 45.02467316 | 0.852075098 | 44.0464322 | NA | NA | NA | NA | NA | NA | NA |
| CACNA1G | 802 | Y | -13.34608333 | 44.17233907 | 0.835176748 | 43.13599418 | NA | NA | NA | NA | NA | NA | NA |
| CACNA1G | 803 | I | -14.497125 | 43.03262764 | 0.812580865 | 42.0029908 | NA | NA | NA | NA | NA | NA | NA |

| | | | | | | | | | | | | | |
|---|---|---|---|---|---|---|---|---|---|---|---|---|---|
| CACNA1G | 804 | K | -19.87622222 | 41.2840092 | 0.777912809 | 40.31525281 | NA | NA | NA | NA | NA | NA | NA |
| CACNA1G | 805 | N | -16.60875 | 39.07133568 | 0.734044412 | 38.08821001 | NA | NA | NA | NA | NA | NA | NA |
| CACNA1G | 806 | P | -15.11028571 | 36.96538597 | 0.692291918 | 35.9628726 | NA | NA | NA | NA | NA | NA | NA |
| CACNA1G | 807 | Y | -13.70583333 | 31.85045915 | 0.590883537 | 31.64999312 | NA | NA | NA | NA | NA | NA | NA |
| CACNA1G | 808 | N | -12.43 | 39.92469763 | 0.75096314 | 38.95773645 | NA | NA | NA | NA | NA | NA | NA |
| CACNA1G | 809 | I | -9.896625 | 40.07848408 | 0.754012105 | 39.18876434 | NA | NA | NA | NA | NA | NA | NA |
| CACNA1G | 810 | F | -6.483272727 | 34.53534514 | 0.644114004 | 33.72044226 | NA | NA | NA | NA | NA | NA | NA |
| CACNA1G | 811 | D | -6.93575 | 39.28186402 | 0.738218341 | 38.40557841 | NA | NA | NA | NA | NA | NA | NA |
| CACNA1G | 812 | G | -5.596 | 41.94648239 | 0.791046982 | 41.08410397 | NA | NA | NA | NA | NA | NA | NA |
| CACNA1G | 813 | V | -3.478428571 | 40.73047541 | 0.766938465 | 39.96557846 | NA | NA | NA | NA | NA | NA | NA |
| CACNA1G | 814 | I | -1.9345 | 37.41216805 | 0.701149806 | 36.65706448 | NA | NA | NA | NA | NA | NA | NA |
| CACNA1G | 815 | V | -1.785 | 42.61964773 | 0.804393138 | 41.84154567 | NA | NA | NA | NA | NA | NA | NA |
| CACNA1G | 816 | V | -0.751714286 | 44.74378619 | 0.846506241 | 43.9864255 | NA | NA | NA | NA | NA | NA | NA |
| CACNA1G | 817 | I | 1.278375 | 41.44753227 | 0.781154813 | 40.24813331 | NA | NA | NA | NA | NA | NA | NA |
| CACNA1G | 818 | S | 2.419166667 | 42.6029249 | 0.804061591 | 41.39233029 | NA | NA | NA | NA | NA | NA | NA |
| CACNA1G | 819 | V | 2.188571429 | 45.57622359 | 0.863010119 | 44.37278193 | NA | NA | NA | NA | NA | NA | NA |
| CACNA1G | 820 | W | 5.337571429 | 45.58137372 | 0.863112226 | 44.36339509 | NA | NA | NA | NA | NA | NA | NA |
| CACNA1G | 821 | E | 6.955222222 | 41.67305808 | 0.78562608 | 40.38184038 | NA | NA | NA | NA | NA | NA | NA |
| CACNA1G | 822 | I | 6.892375 | 47.05306955 | 0.892290022 | 45.77293804 | NA | NA | NA | NA | NA | NA | NA |
| CACNA1G | 823 | V | 7.051428571 | 49.8334353 | 0.947413467 | 48.55825988 | NA | NA | NA | NA | NA | NA | NA |
| CACNA1G | 831 | S | 12.9425 | 38.76214243 | 0.727914357 | 37.36904268 | NA | NA | NA | NA | NA | NA | NA |
| CACNA1G | 832 | V | 9.572857143 | 38.80661754 | 0.728796119 | 37.5861943 | NA | NA | NA | NA | NA | NA | NA |
| CACNA1G | 833 | L | 7.62575 | 39.87834426 | 0.75004414 | 38.56762 | NA | NA | NA | NA | NA | NA | NA |
| CACNA1G | 834 | R | 9.881545455 | 37.8537472 | 0.709904541 | 35.39542909 | NA | NA | NA | NA | NA | NA | NA |
| CACNA1G | 835 | T | 7.937857143 | 34.28018107 | 0.639055129 | 31.88410464 | NA | NA | NA | NA | NA | NA | NA |
| CACNA1G | 836 | F | 4.403636364 | 36.37424794 | 0.680572034 | 35.10568541 | NA | NA | NA | NA | NA | NA | NA |
| CACNA1G | 837 | R | 5.702363636 | 35.22557816 | 0.657798542 | 33.95574666 | NA | NA | NA | NA | NA | NA | NA |
| CACNA1G | 838 | L | 3.04325 | 31.42955981 | 0.5825388 | 30.06607631 | NA | NA | NA | NA | NA | NA | NA |
| CACNA1G | 839 | M | 0.30575 | 32.94338528 | 0.612551857 | 31.75673523 | NA | NA | NA | NA | NA | NA | NA |
| CACNA1G | 840 | R | -1.072272727 | 33.81385404 | 0.629809743 | 33.04588571 | NA | NA | NA | NA | NA | NA | NA |
| CACNA1G | 841 | V | -2.083571429 | 29.49057354 | 0.544096518 | 28.75116592 | NA | NA | NA | NA | NA | NA | NA |
| CACNA1G | 842 | L | -4.86725 | 28.66043168 | 0.527638151 | 27.82959938 | NA | NA | NA | NA | NA | NA | NA |
| CACNA1G | 843 | K | -6.295444444 | 32.91688887 | 0.61202654 | 32.06024221 | NA | NA | NA | NA | NA | NA | NA |

| | | | | | | | | | | | | | |
|---|---|---|---|---|---|---|---|---|---|---|---|---|---|
| CACNA1G | 844 | L | -7.582125 | 30.59935021 | 0.56607909 | 29.72062263 | NA | NA | NA | NA | NA | NA | NA |
| CACNA1G | 845 | V | -10.88528571 | 29.37999268 | 0.541904145 | 28.45192331 | NA | NA | NA | NA | NA | NA | NA |
| CACNA1G | 846 | R | -12.89172727 | 32.91414116 | 0.611972064 | 31.97339944 | NA | NA | NA | NA | NA | NA | NA |
| CACNA1G | 847 | F | -12.03381818 | 32.76098463 | 0.608935587 | 31.95230961 | NA | NA | NA | NA | NA | NA | NA |
| CACNA1G | 848 | L | -13.149625 | 27.98552608 | 0.514257493 | 27.15013216 | NA | NA | NA | NA | NA | NA | NA |
| CACNA1G | 849 | P | -17.31428571 | 26.33192143 | 0.481473178 | 25.40326859 | NA | NA | NA | NA | NA | NA | NA |
| CACNA1G | 850 | A | -15.6782 | 23.17829832 | 0.418949544 | 22.20444094 | NA | NA | NA | NA | NA | NA | NA |
| CACNA1G | 851 | L | -11.993875 | 22.63581562 | 0.408194299 | 21.89411098 | NA | NA | NA | NA | NA | NA | NA |
| CACNA1G | 852 | Q | -15.47622222 | 25.34231313 | 0.461853235 | 24.35313148 | NA | NA | NA | NA | NA | NA | NA |
| CACNA1G | 853 | R | -18.49081818 | 21.72149445 | 0.390066995 | 20.77665589 | NA | NA | NA | NA | NA | NA | NA |
| CACNA1G | 854 | Q | -13.517 | 17.41820668 | 0.304750145 | 16.35944071 | NA | NA | NA | NA | NA | NA | NA |
| CACNA1G | 855 | L | -11.212625 | 22.64978382 | 0.408471232 | 21.58699404 | NA | NA | NA | NA | NA | NA | NA |
| CACNA1G | 856 | V | -15.57842857 | 22.48757268 | 0.405255239 | 21.4686319 | NA | NA | NA | NA | NA | NA | NA |
| CACNA1G | 857 | V | -15.72557143 | 17.92547037 | 0.314807139 | 16.98562094 | NA | NA | NA | NA | NA | NA | NA |
| CACNA1G | 858 | L | -10.72 | 17.36869039 | 0.303768436 | 16.67173504 | NA | NA | NA | NA | NA | NA | NA |
| CACNA1G | 859 | M | -11.02025 | 21.56186212 | 0.38690213 | 20.38176336 | NA | NA | NA | NA | NA | NA | NA |
| CACNA1G | 860 | K | -16.21877778 | 20.44650293 | 0.364789053 | 19.35618782 | NA | NA | NA | NA | NA | NA | NA |
| CACNA1G | 861 | T | -12.12257143 | 16.65431539 | 0.289605259 | 15.4475957 | NA | NA | NA | NA | NA | NA | NA |
| CACNA1G | 862 | M | -9.046375 | 20.30681359 | 0.362019576 | 19.09942263 | NA | NA | NA | NA | NA | NA | NA |
| CACNA1G | 863 | D | -13.692875 | 22.20650655 | 0.39968283 | 20.99929232 | NA | NA | NA | NA | NA | NA | NA |
| CACNA1G | 864 | N | -14.0625 | 18.90758559 | 0.334278525 | 17.77802834 | NA | NA | NA | NA | NA | NA | NA |
| CACNA1G | 865 | V | -9.229428571 | 16.91227782 | 0.294719615 | 15.69769714 | NA | NA | NA | NA | NA | NA | NA |
| CACNA1G | 866 | A | -8.5984 | 21.64349131 | 0.388520507 | 20.42706294 | NA | NA | NA | NA | NA | NA | NA |
| CACNA1G | 867 | T | -9.590142857 | 22.12069759 | 0.397981584 | 20.90313576 | NA | NA | NA | NA | NA | NA | NA |
| CACNA1G | 868 | F | -6.162818182 | 16.45220669 | 0.285598259 | 15.79877922 | NA | NA | NA | NA | NA | NA | NA |
| CACNA1G | 869 | C | -4.133833333 | 20.0800881 | 0.357524523 | 19.34859938 | NA | NA | NA | NA | NA | NA | NA |
| CACNA1G | 870 | M | -5.026125 | 23.58049868 | 0.426923556 | 22.83504376 | NA | NA | NA | NA | NA | NA | NA |
| CACNA1G | 871 | L | -4.56775 | 21.90271458 | 0.39365986 | 21.11920393 | NA | NA | NA | NA | NA | NA | NA |
| CACNA1G | 872 | L | -0.860125 | 18.79073724 | 0.331961893 | 17.52871656 | NA | NA | NA | NA | NA | NA | NA |
| CACNA1G | 873 | M | 0.22075 | 23.25159889 | 0.420402799 | 22.03768484 | NA | NA | NA | NA | NA | NA | NA |
| CACNA1G | 874 | L | -0.562 | 25.39874198 | 0.462971991 | 22.88787033 | NA | NA | NA | NA | NA | NA | NA |
| CACNA1G | 875 | F | 0.789272727 | 19.89976445 | 0.353949432 | 17.12650903 | NA | NA | NA | NA | NA | NA | NA |
| CACNA1G | 876 | I | 3.824 | 21.12915524 | 0.378323296 | 18.31163057 | NA | NA | NA | NA | NA | NA | NA |

| | | | | | | | | | | | | | |
|---|---|---|---|---|---|---|---|---|---|---|---|---|---|
| CACNA1G | 877 | F | 4.877818182 | 25.9790586 | 0.474477331 | 23.17820518 | NA | NA | NA | NA | NA | NA | NA |
| CACNA1G | 878 | I | 3.649625 | 26.88330607 | 0.492404913 | 25.99241135 | NA | NA | NA | NA | NA | NA | NA |
| CACNA1G | 879 | F | 5.627181818 | 22.54885853 | 0.40647029 | 21.6081971 | NA | NA | NA | NA | NA | NA | NA |
| CACNA1G | 880 | S | 9.046833333 | 24.68756971 | 0.448872312 | 23.86650113 | NA | NA | NA | NA | NA | NA | NA |
| CACNA1G | 881 | I | 8.944 | 27.91272128 | 0.512814067 | 27.00536804 | NA | NA | NA | NA | NA | NA | NA |
| CACNA1G | 882 | L | 9.6005 | 25.87440964 | 0.472402564 | 24.95560206 | NA | NA | NA | NA | NA | NA | NA |
| CACNA1G | 883 | G | 13.0525 | 25.48363618 | 0.464655101 | 24.53455024 | NA | NA | NA | NA | NA | NA | NA |
| CACNA1G | 884 | M | 14.264375 | 27.54538869 | 0.505531343 | 26.58888024 | NA | NA | NA | NA | NA | NA | NA |
| CACNA1G | 885 | H | 13.6234 | 29.73162925 | 0.548875681 | 28.77741801 | NA | NA | NA | NA | NA | NA | NA |
| CACNA1G | 886 | L | 14.48125 | 25.91973459 | 0.473301175 | 24.98980811 | NA | NA | NA | NA | NA | NA | NA |
| CACNA1G | 887 | F | 16.52009091 | 22.6572444 | 0.408619145 | 21.69869001 | NA | NA | NA | NA | NA | NA | NA |
| CACNA1G | 888 | G | 19.31975 | 27.98924874 | 0.514331298 | 27.01649747 | NA | NA | NA | NA | NA | NA | NA |
| CACNA1G | 889 | C | 23.133 | 29.22525849 | 0.53883639 | 25.47665759 | NA | NA | NA | NA | NA | NA | NA |
| CACNA1G | 890 | K | 22.49922222 | 25.72914062 | 0.469522465 | 21.99967204 | NA | NA | NA | NA | NA | NA | NA |
| CACNA1G | 891 | F | 21.78 | 22.30782793 | 0.401691625 | 19.53081575 | NA | NA | NA | NA | NA | NA | NA |
| CACNA1G | 892 | A | 26.9428 | 21.28550479 | 0.381423078 | 17.55580345 | NA | NA | NA | NA | NA | NA | NA |
| CACNA1G | 893 | S | 28.2285 | 24.55210586 | 0.44618661 | 20.6678842 | NA | NA | NA | NA | NA | NA | NA |
| CACNA1G | 900 | L | 29.5485 | 22.22664727 | 0.40008214 | 17.62936271 | NA | NA | NA | NA | NA | NA | NA |
| CACNA1G | 901 | P | 26.496 | 23.48491452 | 0.425028507 | 19.71804761 | NA | NA | NA | NA | NA | NA | NA |
| CACNA1G | 902 | D | 23.144375 | 20.65254878 | 0.368874111 | 16.87374489 | NA | NA | NA | NA | NA | NA | NA |
| CACNA1G | 903 | R | 21.06027273 | 20.74850968 | 0.370776629 | 18.62954441 | NA | NA | NA | NA | NA | NA | NA |
| CACNA1G | 904 | K | 18.07633333 | 15.13022221 | 0.259388635 | 14.70038819 | NA | NA | NA | NA | NA | NA | NA |
| CACNA1G | 905 | N | 19.1315 | 23.14805951 | 0.41835003 | 22.17274292 | NA | NA | NA | NA | NA | NA | NA |
| CACNA1G | 906 | F | 15.28681818 | 21.67649566 | 0.389174851 | 20.71599877 | NA | NA | NA | NA | NA | NA | NA |
| CACNA1G | 907 | D | 19.46475 | 27.68977929 | 0.508394026 | 26.71136141 | NA | NA | NA | NA | NA | NA | NA |
| CACNA1G | 908 | S | 17.20733333 | 29.11907317 | 0.536731163 | 28.1636766 | NA | NA | NA | NA | NA | NA | NA |
| CACNA1G | 909 | L | 11.945875 | 27.38534504 | 0.502358322 | 26.53836563 | NA | NA | NA | NA | NA | NA | NA |
| CACNA1G | 910 | L | 13.28025 | 25.7389133 | 0.469716217 | 24.92383874 | NA | NA | NA | NA | NA | NA | NA |
| CACNA1G | 911 | W | 16.28021429 | 23.81025882 | 0.431478773 | 22.83634046 | NA | NA | NA | NA | NA | NA | NA |
| CACNA1G | 912 | A | 12.4212 | 23.30791938 | 0.421519407 | 22.42436904 | NA | NA | NA | NA | NA | NA | NA |
| CACNA1G | 913 | I | 9.107 | 21.92495425 | 0.394100783 | 21.19118065 | NA | NA | NA | NA | NA | NA | NA |
| CACNA1G | 914 | V | 11.352 | 19.40610468 | 0.344162149 | 18.62686498 | NA | NA | NA | NA | NA | NA | NA |
| CACNA1G | 915 | T | 12.50157143 | 18.01772491 | 0.316636175 | 17.11765232 | NA | NA | NA | NA | NA | NA | NA |

| | | | | | | | | | | | | | |
|---|---|---|---|---|---|---|---|---|---|---|---|---|---|
| CACNA1G | 916 | V | 8.392285714 | 17.89495261 | 0.314202095 | 17.11761973 | NA | NA | NA | NA | NA | NA | NA |
| CACNA1G | 917 | F | 6.527909091 | 16.22007873 | 0.280996097 | 15.55786401 | NA | NA | NA | NA | NA | NA | NA |
| CACNA1G | 918 | Q | 11.28533333 | 13.6370723 | 0.229785491 | 12.88759563 | NA | NA | NA | NA | NA | NA | NA |
| CACNA1G | 919 | I | 9.25975 | 12.91973827 | 0.215563649 | 12.08456424 | NA | NA | NA | NA | NA | NA | NA |
| CACNA1G | 920 | L | 4.60725 | 12.91489156 | 0.215467558 | 12.3697702 | NA | NA | NA | NA | NA | NA | NA |
| CACNA1G | 921 | T | 6.255285714 | 10.06116946 | 0.158889752 | 9.929492262 | NA | NA | NA | NA | NA | NA | NA |
| CACNA1G | 922 | Q | 6.351666667 | 7.236010088 | 0.102878229 | 6.787247346 | NA | NA | NA | NA | NA | NA | NA |
| CACNA1G | 923 | E | 11.54788889 | 8.833587097 | 0.134551742 | 8.012523315 | NA | NA | NA | NA | NA | NA | NA |
| CACNA1G | 924 | D | 14.99875 | 10.33988486 | 0.164415555 | 9.454260261 | NA | NA | NA | NA | NA | NA | NA |
| CACNA1G | 925 | W | 11.1885 | 9.047640024 | 0.138795548 | 8.1104217 | NA | NA | NA | NA | NA | NA | NA |
| CACNA1G | 926 | N | 15.257 | 9.487042268 | 0.147507124 | 8.766288227 | NA | NA | NA | NA | NA | NA | NA |
| CACNA1G | 927 | K | 18.43666667 | 9.963128876 | 0.156946002 | 8.944787061 | NA | NA | NA | NA | NA | NA | NA |
| CACNA1G | 928 | V | 16.02628571 | 15.70694073 | 0.270822639 | 14.72257376 | NA | NA | NA | NA | NA | NA | NA |
| CACNA1G | 929 | L | 15.26875 | 14.58396664 | 0.248558589 | 13.61266666 | NA | NA | NA | NA | NA | NA | NA |
| CACNA1G | 930 | Y | 18.56366667 | 11.52517462 | 0.187915073 | 10.54861354 | NA | NA | NA | NA | NA | NA | NA |
| CACNA1G | 931 | N | 20.86725 | 18.34747628 | 0.323173815 | 17.32164623 | NA | NA | NA | NA | NA | NA | NA |
| CACNA1G | 932 | G | 18.588 | 20.9714603 | 0.375196841 | 19.98796725 | NA | NA | NA | NA | NA | NA | NA |
| CACNA1G | 933 | M | 18.131125 | 18.37392459 | 0.323698179 | 17.3935367 | NA | NA | NA | NA | NA | NA | NA |
| CACNA1G | 934 | A | 22.7298 | 20.55222708 | 0.366885136 | 16.84794982 | NA | NA | NA | NA | NA | NA | NA |
| CACNA1G | 935 | S | 22.51166667 | 23.69899789 | 0.429272917 | 19.98091026 | NA | NA | NA | NA | NA | NA | NA |
| CACNA1G | 936 | T | 19.78671429 | 25.50029753 | 0.464985428 | 24.5254924 | NA | NA | NA | NA | NA | NA | NA |
| CACNA1G | 937 | S | 18.33366667 | 23.99176913 | 0.435077391 | 23.02031028 | NA | NA | NA | NA | NA | NA | NA |
| CACNA1G | 938 | S | 17.11533333 | 22.06251391 | 0.396828036 | 21.11624065 | NA | NA | NA | NA | NA | NA | NA |
| CACNA1G | 939 | W | 13.38342857 | 23.66585728 | 0.428615873 | 22.73602853 | NA | NA | NA | NA | NA | NA | NA |
| CACNA1G | 940 | A | 13.9378 | 20.81826491 | 0.372159594 | 19.88044227 | NA | NA | NA | NA | NA | NA | NA |
| CACNA1G | 941 | A | 11.5764 | 18.50650353 | 0.326326684 | 17.60959215 | NA | NA | NA | NA | NA | NA | NA |
| CACNA1G | 942 | L | 8.62875 | 20.6495506 | 0.36881467 | 19.75336834 | NA | NA | NA | NA | NA | NA | NA |
| CACNA1G | 943 | Y | 9.56275 | 19.6849032 | 0.3496896 | 18.79694795 | NA | NA | NA | NA | NA | NA | NA |
| CACNA1G | 944 | F | 10.81154545 | 15.85629913 | 0.273783814 | 14.98032315 | NA | NA | NA | NA | NA | NA | NA |
| CACNA1G | 945 | I | 6.30325 | 16.70711767 | 0.290652116 | 15.8525501 | NA | NA | NA | NA | NA | NA | NA |
| CACNA1G | 946 | A | 4.0838 | 19.03216735 | 0.336748479 | 18.11614651 | NA | NA | NA | NA | NA | NA | NA |
| CACNA1G | 947 | L | 5.286625 | 17.48910712 | 0.306155815 | 16.67848876 | NA | NA | NA | NA | NA | NA | NA |
| CACNA1G | 948 | M | 4.69 | 11.33417814 | 0.184128382 | 10.38842379 | NA | NA | NA | NA | NA | NA | NA |

| | | | | | | | | | | | | | |
|---|---|---|---|---|---|---|---|---|---|---|---|---|---|
| CACNA1G | 949 | T | 0.917428571 | 15.74568303 | 0.271590742 | 13.1784676 | NA | NA | NA | NA | NA | NA | NA |
| CACNA1G | 950 | F | -0.014818182 | 17.45851892 | 0.305549374 | 14.81940231 | NA | NA | NA | NA | NA | NA | NA |
| CACNA1G | 951 | G | 0 | 14.89081603 | 0.254642176 | 12.06262746 | NA | NA | NA | NA | NA | NA | NA |
| CACNA1G | 952 | N | 1.054375 | 11.41453017 | 0.185721439 | 8.58194739 | NA | NA | NA | NA | NA | NA | NA |
| CACNA1G | 953 | Y | -1.2665 | 9.432751719 | 0.146430761 | 7.475031737 | NA | NA | NA | NA | NA | NA | NA |
| CACNA1G | 954 | V | -4.267714286 | 14.62719997 | 0.249415732 | 13.27491135 | NA | NA | NA | NA | NA | NA | NA |
| CACNA1G | 955 | L | -3.4095 | 14.64246294 | 0.249718335 | 13.43971891 | NA | NA | NA | NA | NA | NA | NA |
| CACNA1G | 956 | F | -3.342 | 8.550261867 | 0.128934544 | 7.337740062 | NA | NA | NA | NA | NA | NA | NA |
| CACNA1G | 957 | N | -6.299375 | 8.85351834 | 0.134946898 | 8.317938097 | NA | NA | NA | NA | NA | NA | NA |
| CACNA1G | 958 | L | -8.88475 | 12.65370416 | 0.210289265 | 11.40255336 | NA | NA | NA | NA | NA | NA | NA |
| CACNA1G | 959 | L | -7.551875 | 11.8318616 | 0.193995439 | 10.60279761 | NA | NA | NA | NA | NA | NA | NA |
| CACNA1G | 960 | V | -9.442 | 6.926416462 | 0.096740235 | 5.696224677 | 1.31 | 0.22 | 1.45 | 1.81 | -0.5 | 0 | 0 |
| CACNA1G | 961 | A | -12.6276 | 10.15922792 | 0.160833856 | 8.939213245 | NA | NA | NA | NA | NA | NA | NA |
| CACNA1G | 962 | I | -12.588875 | 12.44034477 | 0.206059208 | 11.22962171 | NA | NA | NA | NA | NA | NA | NA |
| CACNA1G | 963 | L | -12.41675 | 11.00666816 | 0.17763518 | 9.887906977 | NA | NA | NA | NA | NA | NA | NA |
| CACNA1G | 964 | V | -15.52885714 | 7.079336339 | 0.09977202 | 5.925050247 | 1.25 | 0.49 | 2.53 | 1.21 | -0.76 | 0 | 0 |
| CACNA1G | 965 | E | -18.86322222 | 11.02368695 | 0.177972594 | 10.11986131 | NA | NA | NA | NA | NA | NA | NA |
| CACNA1G | 966 | G | -18.09325 | 13.19181644 | 0.220957862 | 12.3186327 | NA | NA | NA | NA | NA | NA | NA |
| CACNA1G | 967 | F | -16.96081818 | 11.33211304 | 0.18408744 | 10.59014137 | NA | NA | NA | NA | NA | NA | NA |
| CACNA1G | 968 | Q | -21.45622222 | 9.768717572 | 0.15309161 | 9.312068671 | NA | NA | NA | NA | NA | NA | NA |
| CACNA1G | 1265 | F | -16.75290909 | 47.9622066 | 0.910314545 | 47.46188664 | NA | NA | NA | NA | NA | NA | NA |
| CACNA1G | 1266 | R | -20.35381818 | 44.81630335 | 0.847943964 | 44.31115073 | NA | NA | NA | NA | NA | NA | NA |
| CACNA1G | 1267 | L | -23.24875 | 45.27052278 | 0.856949305 | 44.79640103 | NA | NA | NA | NA | NA | NA | NA |
| CACNA1G | 1268 | L | -20.1105 | 44.83245241 | 0.848264135 | 44.34655547 | NA | NA | NA | NA | NA | NA | NA |
| CACNA1G | 1269 | C | -17.16866667 | 42.16817934 | 0.795442339 | 41.66732086 | NA | NA | NA | NA | NA | NA | NA |
| CACNA1G | 1270 | H | -21.0929 | 39.04857597 | 0.733593179 | 38.5694198 | NA | NA | NA | NA | NA | NA | NA |
| CACNA1G | 1271 | R | -22.26681818 | 40.27612285 | 0.757930486 | 39.80798083 | NA | NA | NA | NA | NA | NA | NA |
| CACNA1G | 1272 | I | -16.371125 | 38.53580762 | 0.72342705 | 38.04460291 | NA | NA | NA | NA | NA | NA | NA |
| CACNA1G | 1273 | I | -16.92275 | 35.61784657 | 0.665575644 | 35.13123561 | NA | NA | NA | NA | NA | NA | NA |
| CACNA1G | 1274 | T | -20.786 | 35.47512822 | 0.662746114 | 35.0169805 | NA | NA | NA | NA | NA | NA | NA |
| CACNA1G | 1275 | H | -17.7529 | 35.54700646 | 0.66417117 | 35.06819404 | NA | NA | NA | NA | NA | NA | NA |
| CACNA1G | 1276 | K | -17.909 | 31.2502843 | 0.578984489 | 30.82719745 | NA | NA | NA | NA | NA | NA | NA |
| CACNA1G | 1277 | M | -13.613625 | 35.53460198 | 0.663925239 | 35.05861246 | NA | NA | NA | NA | NA | NA | NA |

| | | | | | | | | | | | | | |
|---|---|---|---|---|---|---|---|---|---|---|---|---|---|
| CACNA1G | 1278 | F | -12.32063636 | 34.76887011 | 0.648743863 | 34.67427879 | NA | NA | NA | NA | NA | NA | NA |
| CACNA1G | 1279 | D | -12.575125 | 29.61103061 | 0.546484696 | 28.92244476 | NA | NA | NA | NA | NA | NA | NA |
| CACNA1G | 1280 | H | -11.8087 | 30.00501741 | 0.554295866 | 29.58298072 | NA | NA | NA | NA | NA | NA | NA |
| CACNA1G | 1281 | V | -8.192571429 | 34.42376599 | 0.641901839 | 32.3679518 | NA | NA | NA | NA | NA | NA | NA |
| CACNA1G | 1282 | V | -6.917857143 | 31.66220111 | 0.587151139 | 29.52230411 | NA | NA | NA | NA | NA | NA | NA |
| CACNA1G | 1283 | L | -6.336625 | 27.40343019 | 0.502716878 | 25.53025116 | NA | NA | NA | NA | NA | NA | NA |
| CACNA1G | 1284 | V | -3.743285714 | 31.10145588 | 0.576033821 | 28.96097496 | NA | NA | NA | NA | NA | NA | NA |
| CACNA1G | 1285 | I | -2.3495 | 31.25494351 | 0.579076862 | 29.1068884 | NA | NA | NA | NA | NA | NA | NA |
| CACNA1G | 1286 | I | -2.15625 | 27.09073131 | 0.49651732 | 24.94487106 | NA | NA | NA | NA | NA | NA | NA |
| CACNA1G | 1287 | F | -0.311545455 | 25.06121988 | 0.456280289 | 23.08196805 | NA | NA | NA | NA | NA | NA | NA |
| CACNA1G | 1288 | L | 1.1375 | 29.39188965 | 0.542140014 | 28.15401509 | NA | NA | NA | NA | NA | NA | NA |
| CACNA1G | 1289 | N | 2.45875 | 26.98285904 | 0.494378647 | 25.74213575 | NA | NA | NA | NA | NA | NA | NA |
| CACNA1G | 1290 | C | 2.880833333 | 23.74883618 | 0.43026101 | 22.51137667 | NA | NA | NA | NA | NA | NA | NA |
| CACNA1G | 1291 | I | 5.4545 | 25.22406599 | 0.459508871 | 23.99451178 | NA | NA | NA | NA | NA | NA | NA |
| CACNA1G | 1292 | T | 7.215571429 | 26.33214249 | 0.481477561 | 25.09920351 | NA | NA | NA | NA | NA | NA | NA |
| CACNA1G | 1293 | I | 6.944625 | 22.792046 | 0.411291717 | 21.55968626 | NA | NA | NA | NA | NA | NA | NA |
| CACNA1G | 1294 | A | 9.4464 | 21.38145423 | 0.383325368 | 20.15172284 | NA | NA | NA | NA | NA | NA | NA |
| CACNA1G | 1295 | M | 12.063625 | 23.5732117 | 0.426779084 | 22.34692643 | NA | NA | NA | NA | NA | NA | NA |
| CACNA1G | 1296 | E | 13.36033333 | 21.58488047 | 0.387358491 | 21.41989522 | NA | NA | NA | NA | NA | NA | NA |
| CACNA1G | 1297 | R | 17.36781818 | 21.08302751 | 0.377408769 | 20.91327322 | NA | NA | NA | NA | NA | NA | NA |
| CACNA1G | 1298 | P | 19.58771429 | 20.25395174 | 0.360971539 | 17.85231112 | NA | NA | NA | NA | NA | NA | NA |
| CACNA1G | 1299 | K | 23.08544444 | 18.60947788 | 0.328368251 | 14.39089802 | NA | NA | NA | NA | NA | NA | NA |
| CACNA1G | 1300 | I | 19.63625 | 25.26125684 | 0.460246215 | 21.13722706 | NA | NA | NA | NA | NA | NA | NA |
| CACNA1G | 1301 | D | 23.368 | 27.80539782 | 0.510686276 | 23.58633329 | NA | NA | NA | NA | NA | NA | NA |
| CACNA1G | 1302 | P | 22.28042857 | 30.04857025 | 0.555159343 | 25.82012684 | NA | NA | NA | NA | NA | NA | NA |
| CACNA1G | 1303 | H | 23.4514 | 33.2634479 | 0.618897408 | 29.0125714 | NA | NA | NA | NA | NA | NA | NA |
| CACNA1G | 1304 | S | 20.7315 | 32.12910509 | 0.596407963 | 27.83823901 | NA | NA | NA | NA | NA | NA | NA |
| CACNA1G | 1305 | A | 18.1782 | 33.58616803 | 0.625295648 | 33.40823952 | NA | NA | NA | NA | NA | NA | NA |
| CACNA1G | 1306 | E | 16.44233333 | 26.73238254 | 0.489412708 | 26.5564467 | NA | NA | NA | NA | NA | NA | NA |
| CACNA1G | 1307 | R | 16.76854545 | 27.89949616 | 0.512551867 | 27.72592929 | NA | NA | NA | NA | NA | NA | NA |
| CACNA1G | 1308 | I | 15.418 | 34.22261308 | 0.637913788 | 34.05326891 | NA | NA | NA | NA | NA | NA | NA |
| CACNA1G | 1309 | F | 10.94818182 | 31.83118597 | 0.590501427 | 30.60572187 | NA | NA | NA | NA | NA | NA | NA |
| CACNA1G | 1310 | L | 11.9995 | 28.2031314 | 0.51857173 | 27.55922443 | NA | NA | NA | NA | NA | NA | NA |

| | | | | | | | | | | | | | |
|---|---|---|---|---|---|---|---|---|---|---|---|---|---|
| CACNA1G | 1311 | T | 12.25857143 | 33.7977559 | 0.629490582 | 32.94033033 | NA | NA | NA | NA | NA | NA | NA |
| CACNA1G | 1312 | L | 9.88125 | 36.34512337 | 0.679994611 | 35.24354519 | NA | NA | NA | NA | NA | NA | NA |
| CACNA1G | 1313 | S | 6.758333333 | 33.33813476 | 0.620378148 | 32.10491973 | NA | NA | NA | NA | NA | NA | NA |
| CACNA1G | 1314 | N | 6.96975 | 33.54984693 | 0.624575547 | 32.30836182 | NA | NA | NA | NA | NA | NA | NA |
| CACNA1G | 1315 | Y | 7.28975 | 36.76299659 | 0.688279353 | 35.52563404 | NA | NA | NA | NA | NA | NA | NA |
| CACNA1G | 1316 | I | 2.96925 | 37.00577817 | 0.693092733 | 35.76925437 | NA | NA | NA | NA | NA | NA | NA |
| CACNA1G | 1317 | F | 1.627454545 | 32.47719486 | 0.60330918 | 31.23893154 | NA | NA | NA | NA | NA | NA | NA |
| CACNA1G | 1318 | T | 1.545142857 | 35.48604583 | 0.662962566 | 34.23988573 | NA | NA | NA | NA | NA | NA | NA |
| CACNA1G | 1319 | A | -0.5528 | 38.88796201 | 0.730408851 | 37.64083743 | NA | NA | NA | NA | NA | NA | NA |
| CACNA1G | 1320 | V | -2.321 | 37.13339027 | 0.695622766 | 34.98145684 | NA | NA | NA | NA | NA | NA | NA |
| CACNA1G | 1321 | F | -2.554636364 | 32.13474671 | 0.596519813 | 29.98369647 | NA | NA | NA | NA | NA | NA | NA |
| CACNA1G | 1322 | L | -3.293125 | 38.18888104 | 0.716548894 | 36.03011154 | NA | NA | NA | NA | NA | NA | NA |
| CACNA1G | 1323 | A | -5.9804 | 40.5243682 | 0.76285219 | 38.36883417 | NA | NA | NA | NA | NA | NA | NA |
| CACNA1G | 1324 | E | -8.044444444 | 34.94999648 | 0.652334868 | 33.82415706 | NA | NA | NA | NA | NA | NA | NA |
| CACNA1G | 1325 | M | -7.728375 | 37.58783923 | 0.704632657 | 36.97571776 | NA | NA | NA | NA | NA | NA | NA |
| CACNA1G | 1326 | T | -9.026 | 40.78103416 | 0.767940842 | 40.00040517 | NA | NA | NA | NA | NA | NA | NA |
| CACNA1G | 1327 | V | -12.03685714 | 41.30674359 | 0.778363541 | 40.41833856 | NA | NA | NA | NA | NA | NA | NA |
| CACNA1G | 1328 | K | -13.27177778 | 34.61629948 | 0.645719002 | 33.74755776 | NA | NA | NA | NA | NA | NA | NA |
| CACNA1G | 1329 | V | -13.10028571 | 40.25642617 | 0.75753998 | 39.35348766 | NA | NA | NA | NA | NA | NA | NA |
| CACNA1G | 1330 | V | -15.16828571 | 43.30517596 | 0.817984399 | 42.41835893 | NA | NA | NA | NA | NA | NA | NA |
| CACNA1G | 1331 | A | -17.9756 | 41.81271077 | 0.78839483 | 41.08777841 | NA | NA | NA | NA | NA | NA | NA |
| CACNA1G | 1332 | L | -18.440625 | 37.27160058 | 0.69836292 | 36.56051729 | NA | NA | NA | NA | NA | NA | NA |
| CACNA1G | 1333 | G | -16.857 | 41.13605144 | 0.774979403 | 40.26696399 | NA | NA | NA | NA | NA | NA | NA |
| CACNA1G | 1344 | S | -14.7485 | 28.95322588 | 0.53344308 | 27.89283696 | NA | NA | NA | NA | NA | NA | NA |
| CACNA1G | 1345 | S | -11.5735 | 26.7425553 | 0.489614393 | 25.83826772 | NA | NA | NA | NA | NA | NA | NA |
| CACNA1G | 1346 | W | -10.51278571 | 22.14476801 | 0.398458804 | 21.23922803 | NA | NA | NA | NA | NA | NA | NA |
| CACNA1G | 1347 | N | -11.492875 | 30.27388368 | 0.5596264 | 29.37377368 | NA | NA | NA | NA | NA | NA | NA |
| CACNA1G | 1348 | V | -8.632428571 | 31.29782275 | 0.579926985 | 30.46135838 | NA | NA | NA | NA | NA | NA | NA |
| CACNA1G | 1349 | L | -5.6425 | 25.44504217 | 0.463889937 | 24.30103013 | NA | NA | NA | NA | NA | NA | NA |
| CACNA1G | 1350 | D | -6.248 | 29.25265485 | 0.539379549 | 28.12078553 | NA | NA | NA | NA | NA | NA | NA |
| CACNA1G | 1351 | G | -4.5745 | 33.16175306 | 0.61688121 | 32.01910787 | NA | NA | NA | NA | NA | NA | NA |
| CACNA1G | 1352 | L | -2.49475 | 32.31310938 | 0.600056026 | 30.26303035 | NA | NA | NA | NA | NA | NA | NA |
| CACNA1G | 1353 | L | -0.75375 | 28.40825604 | 0.522638524 | 26.24128657 | NA | NA | NA | NA | NA | NA | NA |

| | | | | | | | | | | | | | |
|---|---|---|---|---|---|---|---|---|---|---|---|---|---|
| CACNA1G | 1354 | V | -0.978142857 | 33.70941711 | 0.62773918 | 31.53976486 | NA | NA | NA | NA | NA | NA | NA |
| CACNA1G | 1355 | L | -0.60075 | 35.68154417 | 0.66683851 | 33.50968947 | NA | NA | NA | NA | NA | NA | NA |
| CACNA1G | 1356 | I | 2.519 | 30.97464232 | 0.57351962 | 28.80156756 | NA | NA | NA | NA | NA | NA | NA |
| CACNA1G | 1357 | S | 3.084666667 | 32.92621287 | 0.612211397 | 31.67107865 | NA | NA | NA | NA | NA | NA | NA |
| CACNA1G | 1358 | V | 2.745571429 | 37.01664091 | 0.693308097 | 35.76113832 | NA | NA | NA | NA | NA | NA | NA |
| CACNA1G | 1359 | I | 5.079 | 37.74278754 | 0.707704658 | 36.48828267 | NA | NA | NA | NA | NA | NA | NA |
| CACNA1G | 1360 | D | 8.504125 | 33.23192408 | 0.618272418 | 31.98246862 | NA | NA | NA | NA | NA | NA | NA |
| CACNA1G | 1361 | I | 7.72775 | 37.04500572 | 0.693870457 | 35.80210395 | NA | NA | NA | NA | NA | NA | NA |
| CACNA1G | 1362 | L | 7.9785 | 40.02088372 | 0.752870123 | 38.77244266 | NA | NA | NA | NA | NA | NA | NA |
| CACNA1G | 1363 | V | 11.47442857 | 37.48819195 | 0.702657054 | 37.30689071 | NA | NA | NA | NA | NA | NA | NA |
| CACNA1G | 1364 | S | 13.25033333 | 36.89608697 | 0.690917999 | 36.89710981 | NA | NA | NA | NA | NA | NA | NA |
| CACNA1G | 1365 | M | 12.614 | 40.33042748 | 0.759007128 | 40.27911735 | NA | NA | NA | NA | NA | NA | NA |
| CACNA1G | 1366 | V | 15.10314286 | 41.85046653 | 0.789143375 | 41.65579569 | NA | NA | NA | NA | NA | NA | NA |
| CACNA1G | 1367 | S | 18.55933333 | 38.81202537 | 0.728903334 | 36.04030941 | NA | NA | NA | NA | NA | NA | NA |
| CACNA1G | 1372 | K | 19.67933333 | 36.09554384 | 0.675046455 | 33.33491427 | NA | NA | NA | NA | NA | NA | NA |
| CACNA1G | 1373 | I | 16.255875 | 35.21061961 | 0.657501974 | 32.73315451 | NA | NA | NA | NA | NA | NA | NA |
| CACNA1G | 1374 | L | 18.371625 | 31.78839905 | 0.589653135 | 29.26145801 | NA | NA | NA | NA | NA | NA | NA |
| CACNA1G | 1375 | G | 15.96825 | 31.58736105 | 0.585667362 | 29.17522381 | NA | NA | NA | NA | NA | NA | NA |
| CACNA1G | 1376 | M | 12.312625 | 33.10762201 | 0.615808009 | 32.78243895 | NA | NA | NA | NA | NA | NA | NA |
| CACNA1G | 1377 | L | 13.135875 | 31.15477763 | 0.577090977 | 30.8996933 | NA | NA | NA | NA | NA | NA | NA |
| CACNA1G | 1378 | R | 15.33963636 | 27.32929847 | 0.501247145 | 27.17691657 | NA | NA | NA | NA | NA | NA | NA |
| CACNA1G | 1379 | V | 10.04685714 | 28.77124041 | 0.529835041 | 27.50691482 | NA | NA | NA | NA | NA | NA | NA |
| CACNA1G | 1380 | L | 8.356125 | 29.39895367 | 0.542280065 | 28.14153209 | NA | NA | NA | NA | NA | NA | NA |
| CACNA1G | 1381 | R | 12.18018182 | 26.11216692 | 0.477116332 | 24.99059 | NA | NA | NA | NA | NA | NA | NA |
| CACNA1G | 1382 | L | 9.1915 | 23.35970792 | 0.422546165 | 22.09916474 | NA | NA | NA | NA | NA | NA | NA |
| CACNA1G | 1383 | L | 4.913875 | 26.98737025 | 0.494468086 | 25.72372899 | NA | NA | NA | NA | NA | NA | NA |
| CACNA1G | 1384 | R | 6.854727273 | 26.37476138 | 0.482322522 | 25.11953674 | NA | NA | NA | NA | NA | NA | NA |
| CACNA1G | 1385 | T | 5.464571429 | 22.74065349 | 0.410272811 | 21.47909357 | NA | NA | NA | NA | NA | NA | NA |
| CACNA1G | 1386 | L | 1.441125 | 23.0512671 | 0.416431027 | 20.88437242 | NA | NA | NA | NA | NA | NA | NA |
| CACNA1G | 1387 | R | 2.367454545 | 25.78247934 | 0.470579956 | 23.62020491 | NA | NA | NA | NA | NA | NA | NA |
| CACNA1G | 1388 | P | 0.910857143 | 22.55787466 | 0.406649043 | 20.40159037 | NA | NA | NA | NA | NA | NA | NA |
| CACNA1G | 1389 | L | -2.615625 | 22.53163574 | 0.406128831 | 20.37029824 | NA | NA | NA | NA | NA | NA | NA |
| CACNA1G | 1390 | R | -3.109909091 | 25.44778045 | 0.463944227 | 23.29337737 | NA | NA | NA | NA | NA | NA | NA |

| | | | | | | | | | | | | | |
|---|---|---|---|---|---|---|---|---|---|---|---|---|---|
| CACNA1G | 1391 | V | -5.741428571 | 23.15546506 | 0.418496852 | 22.1500101 | NA | NA | NA | NA | NA | NA | NA |
| CACNA1G | 1392 | I | -7.65225 | 19.96526947 | 0.355248133 | 19.47326944 | NA | NA | NA | NA | NA | NA | NA |
| CACNA1G | 1393 | S | -10.65883333 | 24.67462091 | 0.448615589 | 23.77082961 | NA | NA | NA | NA | NA | NA | NA |
| CACNA1G | 1394 | R | -9.307727273 | 26.32648495 | 0.481365395 | 25.45302341 | NA | NA | NA | NA | NA | NA | NA |
| CACNA1G | 1395 | A | -10.94 | 22.91127253 | 0.413655499 | 22.05706929 | NA | NA | NA | NA | NA | NA | NA |
| CACNA1G | 1396 | Q | -15.06777778 | 20.66104535 | 0.369042564 | 20.1521586 | NA | NA | NA | NA | NA | NA | NA |
| CACNA1G | 1397 | G | -12.789 | 17.9260424 | 0.31481848 | 17.60172328 | NA | NA | NA | NA | NA | NA | NA |
| CACNA1G | 1398 | L | -9.4115 | 17.80245256 | 0.312368191 | 17.03152274 | NA | NA | NA | NA | NA | NA | NA |
| CACNA1G | 1399 | K | -13.63933333 | 19.50912592 | 0.346204645 | 18.69264695 | NA | NA | NA | NA | NA | NA | NA |
| CACNA1G | 1400 | L | -15.91075 | 16.2758726 | 0.282102264 | 15.35351713 | NA | NA | NA | NA | NA | NA | NA |
| CACNA1G | 1401 | V | -12.01971429 | 13.63125801 | 0.229670217 | 13.68123907 | NA | NA | NA | NA | NA | NA | NA |
| CACNA1G | 1402 | V | -10.14757143 | 15.81705238 | 0.273005709 | 14.93078863 | NA | NA | NA | NA | NA | NA | NA |
| CACNA1G | 1403 | E | -14.381 | 16.15564341 | 0.279718604 | 15.17699956 | NA | NA | NA | NA | NA | NA | NA |
| CACNA1G | 1404 | T | -14.68942857 | 11.77338932 | 0.19283617 | 10.6721151 | NA | NA | NA | NA | NA | NA | NA |
| CACNA1G | 1405 | L | -10.425375 | 9.959744274 | 0.156878899 | 9.06640163 | NA | NA | NA | NA | NA | NA | NA |
| CACNA1G | 1406 | M | -10.5265 | 14.56114484 | 0.248106125 | 13.67022422 | NA | NA | NA | NA | NA | NA | NA |
| CACNA1G | 1407 | S | -15.27233333 | 14.15225706 | 0.239999528 | 13.20346993 | NA | NA | NA | NA | NA | NA | NA |
| CACNA1G | 1408 | S | -13.46016667 | 10.48492771 | 0.16729117 | 9.722541423 | NA | NA | NA | NA | NA | NA | NA |
| CACNA1G | 1409 | L | -10.550375 | 11.68480749 | 0.191079949 | 10.82262597 | NA | NA | NA | NA | NA | NA | NA |
| CACNA1G | 1410 | K | -14.86133333 | 15.038209 | 0.257564384 | 14.05358343 | NA | NA | NA | NA | NA | NA | NA |
| CACNA1G | 1411 | P | -15.26814286 | 12.69178746 | 0.211044303 | 11.82746104 | NA | NA | NA | NA | NA | NA | NA |
| CACNA1G | 1412 | I | -11.143625 | 12.06407962 | 0.198599387 | 11.18739641 | NA | NA | NA | NA | NA | NA | NA |
| CACNA1G | 1413 | G | -10.7815 | 16.28417407 | 0.282266849 | 15.34451417 | NA | NA | NA | NA | NA | NA | NA |
| CACNA1G | 1414 | N | -12.016 | 18.28494733 | 0.321934118 | 17.37118608 | NA | NA | NA | NA | NA | NA | NA |
| CACNA1G | 1415 | I | -9.136125 | 14.60673793 | 0.249010052 | 13.76631887 | NA | NA | NA | NA | NA | NA | NA |
| CACNA1G | 1416 | V | -6.117857143 | 14.5840217 | 0.248559681 | 13.74689328 | NA | NA | NA | NA | NA | NA | NA |
| CACNA1G | 1417 | V | -6.745428571 | 18.79705559 | 0.332087161 | 17.94598758 | NA | NA | NA | NA | NA | NA | NA |
| CACNA1G | 1418 | I | -6.288 | 19.82635145 | 0.352493949 | 19.00998617 | NA | NA | NA | NA | NA | NA | NA |
| CACNA1G | 1419 | C | -2.559333333 | 15.7465547 | 0.271608024 | 14.47007224 | NA | NA | NA | NA | NA | NA | NA |
| CACNA1G | 1420 | C | -1.5835 | 18.95696663 | 0.335257552 | 17.66710387 | NA | NA | NA | NA | NA | NA | NA |
| CACNA1G | 1421 | A | -1.2848 | 21.83605381 | 0.392338246 | 20.58316706 | NA | NA | NA | NA | NA | NA | NA |
| CACNA1G | 1422 | F | -0.321545455 | 20.30387463 | 0.361961308 | 19.00722748 | NA | NA | NA | NA | NA | NA | NA |
| CACNA1G | 1423 | F | 1.965090909 | 16.75315161 | 0.291564783 | 14.04937936 | NA | NA | NA | NA | NA | NA | NA |

| | | | | | | | | | | | | | |
|---|---|---|---|---|---|---|---|---|---|---|---|---|---|
| CACNA1G | 1424 | I | 2.47975 | 23.83677581 | 0.432004498 | 22.43075752 | NA | NA | NA | NA | NA | NA | NA |
| CACNA1G | 1425 | I | 1.906625 | 25.6887045 | 0.468720779 | 24.32490087 | NA | NA | NA | NA | NA | NA | NA |
| CACNA1G | 1426 | F | 4.317909091 | 23.25287528 | 0.420428104 | 20.55367237 | NA | NA | NA | NA | NA | NA | NA |
| CACNA1G | 1427 | G | 7.209 | 25.0938035 | 0.45692629 | 23.80327492 | NA | NA | NA | NA | NA | NA | NA |
| CACNA1G | 1428 | I | 6.927375 | 27.66888075 | 0.507979693 | 26.35613857 | NA | NA | NA | NA | NA | NA | NA |
| CACNA1G | 1429 | L | 7.84125 | 28.50737357 | 0.524603625 | 27.19564626 | NA | NA | NA | NA | NA | NA | NA |
| CACNA1G | 1430 | G | 11.04225 | 27.15210404 | 0.497734094 | 25.72191669 | NA | NA | NA | NA | NA | NA | NA |
| CACNA1G | 1431 | V | 12.48885714 | 28.51063442 | 0.524668275 | 27.06802909 | NA | NA | NA | NA | NA | NA | NA |
| CACNA1G | 1432 | Q | 11.26344444 | 31.24941791 | 0.578967312 | 29.80978309 | NA | NA | NA | NA | NA | NA | NA |
| CACNA1G | 1433 | L | 13.46275 | 29.85883208 | 0.5513976 | 29.49931984 | NA | NA | NA | NA | NA | NA | NA |
| CACNA1G | 1434 | F | 15.56254545 | 26.38561936 | 0.482537792 | 26.23144539 | NA | NA | NA | NA | NA | NA | NA |
| CACNA1G | 1435 | K | 17.439 | 30.12694795 | 0.556713257 | 27.5272793 | NA | NA | NA | NA | NA | NA | NA |
| CACNA1G | 1436 | G | 21.385 | 29.11624911 | 0.536675173 | 26.16860276 | NA | NA | NA | NA | NA | NA | NA |
| CACNA1G | 1437 | K | 21.10255556 | 26.75774376 | 0.489915519 | 23.79658696 | NA | NA | NA | NA | NA | NA | NA |
| CACNA1G | 1438 | F | 20.10918182 | 22.90674019 | 0.413565641 | 20.16996106 | NA | NA | NA | NA | NA | NA | NA |
| CACNA1G | 1439 | F | 25.49336364 | 24.37643801 | 0.442703824 | 21.52421734 | NA | NA | NA | NA | NA | NA | NA |
| CACNA1G | 1440 | V | 28.04057143 | 20.31230989 | 0.362128546 | 15.63548188 | NA | NA | NA | NA | NA | NA | NA |
| CACNA1G | 1441 | C | 32.285 | 20.33398802 | 0.362558336 | 15.6811709 | NA | NA | NA | NA | NA | NA | NA |
| CACNA1G | 1442 | Q | 31.34377778 | 15.50860032 | 0.266890348 | 10.85842294 | NA | NA | NA | NA | NA | NA | NA |
| CACNA1G | 1443 | G | 35.504 | 16.1522712 | 0.279651747 | 12.98453396 | NA | NA | NA | NA | NA | NA | NA |
| CACNA1G | 1444 | E | 38.964 | 12.57378431 | 0.208704776 | 10.95539959 | NA | NA | NA | NA | NA | NA | NA |
| CACNA1G | 1445 | D | 38.560375 | 17.58754016 | 0.308107345 | 17.55777557 | NA | NA | NA | NA | NA | NA | NA |
| CACNA1G | 1446 | T | 35.03514286 | 20.13321579 | 0.358577831 | 16.98945449 | NA | NA | NA | NA | NA | NA | NA |
| CACNA1G | 1447 | R | 36.45245455 | 20.94923259 | 0.374756155 | 19.28676788 | NA | NA | NA | NA | NA | NA | NA |
| CACNA1G | 1448 | N | 38.831375 | 24.99755358 | 0.455018042 | 24.98666774 | NA | NA | NA | NA | NA | NA | NA |
| CACNA1G | 1449 | I | 35.689125 | 24.39821215 | 0.443135518 | 21.57574073 | NA | NA | NA | NA | NA | NA | NA |
| CACNA1G | 1450 | T | 33.856 | 28.94039504 | 0.533188696 | 24.47514511 | NA | NA | NA | NA | NA | NA | NA |
| CACNA1G | 1451 | N | 33.764 | 28.49982928 | 0.524454052 | 23.94674406 | NA | NA | NA | NA | NA | NA | NA |
| CACNA1G | 1452 | K | 32.57877778 | 28.22003255 | 0.518906811 | 23.90157248 | NA | NA | NA | NA | NA | NA | NA |
| CACNA1G | 1453 | S | 37.0605 | 29.57922524 | 0.545854124 | 26.51853407 | NA | NA | NA | NA | NA | NA | NA |
| CACNA1G | 1454 | D | 38.089625 | 26.48625653 | 0.484533021 | 23.75939284 | NA | NA | NA | NA | NA | NA | NA |
| CACNA1G | 1455 | C | 36.89866667 | 24.74345376 | 0.449980267 | 21.54947414 | NA | NA | NA | NA | NA | NA | NA |
| CACNA1G | 1456 | A | 39.5864 | 26.76597912 | 0.490078793 | 23.70127433 | NA | NA | NA | NA | NA | NA | NA |

| | | | | | | | | | | | | | |
|---|---|---|---|---|---|---|---|---|---|---|---|---|---|
| CACNA1G | 1457 | E | 42.76111111 | 24.81868673 | 0.451471834 | 22.91476754 | NA | NA | NA | NA | NA | NA | NA |
| CACNA1G | 1458 | A | 41.427 | 21.09591975 | 0.37766437 | 21.39271605 | NA | NA | NA | NA | NA | NA | NA |
| CACNA1G | 1459 | S | 41.06033333 | 20.49723913 | 0.365794947 | 17.43047325 | NA | NA | NA | NA | NA | NA | NA |
| CACNA1G | 1460 | Y | 38.24508333 | 18.47998566 | 0.325800942 | 16.18187636 | NA | NA | NA | NA | NA | NA | NA |
| CACNA1G | 1461 | R | 34.158 | 14.71214345 | 0.251099819 | 11.59521673 | NA | NA | NA | NA | NA | NA | NA |
| CACNA1G | 1462 | W | 30.5245 | 22.54790815 | 0.406451448 | 17.89171914 | NA | NA | NA | NA | NA | NA | NA |
| CACNA1G | 1463 | V | 28.40714286 | 18.78420437 | 0.331832373 | 14.18793472 | NA | NA | NA | NA | NA | NA | NA |
| CACNA1G | 1464 | R | 22.93409091 | 19.67026045 | 0.349399293 | 16.69730521 | NA | NA | NA | NA | NA | NA | NA |
| CACNA1G | 1465 | H | 22.3408 | 18.90873169 | 0.334301247 | 15.94201847 | NA | NA | NA | NA | NA | NA | NA |
| CACNA1G | 1466 | K | 20.58788889 | 15.59684946 | 0.268639973 | 12.64537809 | NA | NA | NA | NA | NA | NA | NA |
| CACNA1G | 1467 | Y | 15.98975 | 13.34783001 | 0.224050982 | 13.16885035 | NA | NA | NA | NA | NA | NA | NA |
| CACNA1G | 1468 | N | 17.035375 | 21.07166144 | 0.377183426 | 20.91668898 | NA | NA | NA | NA | NA | NA | NA |
| CACNA1G | 1469 | F | 14.03263636 | 22.88260547 | 0.413087146 | 22.55112314 | NA | NA | NA | NA | NA | NA | NA |
| CACNA1G | 1470 | D | 17.64225 | 26.47295429 | 0.484269292 | 23.54270464 | NA | NA | NA | NA | NA | NA | NA |
| CACNA1G | 1471 | N | 13.902875 | 27.1170522 | 0.497039157 | 26.63559521 | NA | NA | NA | NA | NA | NA | NA |
| CACNA1G | 1472 | L | 8.909875 | 25.40804329 | 0.463156399 | 23.99304816 | NA | NA | NA | NA | NA | NA | NA |
| CACNA1G | 1473 | G | 11.11425 | 23.69579796 | 0.429209476 | 22.30024217 | NA | NA | NA | NA | NA | NA | NA |
| CACNA1G | 1474 | Q | 14.56844444 | 22.08637639 | 0.397301133 | 21.5528965 | NA | NA | NA | NA | NA | NA | NA |
| CACNA1G | 1475 | A | 10.6928 | 21.76560082 | 0.390941447 | 20.30972957 | NA | NA | NA | NA | NA | NA | NA |
| CACNA1G | 1476 | L | 7.29 | 19.15096345 | 0.339103727 | 17.74337534 | NA | NA | NA | NA | NA | NA | NA |
| CACNA1G | 1477 | M | 10.593875 | 16.89671879 | 0.294411142 | 15.85618662 | NA | NA | NA | NA | NA | NA | NA |
| CACNA1G | 1478 | S | 11.64416667 | 18.07978169 | 0.31786651 | 18.04367722 | NA | NA | NA | NA | NA | NA | NA |
| CACNA1G | 1479 | L | 7.526625 | 16.49660847 | 0.286478567 | 15.12309658 | NA | NA | NA | NA | NA | NA | NA |
| CACNA1G | 1480 | F | 6.219090909 | 12.15304373 | 0.200363186 | 10.80516694 | NA | NA | NA | NA | NA | NA | NA |
| CACNA1G | 1481 | V | 10.33271429 | 13.10759085 | 0.219288008 | 13.00916874 | NA | NA | NA | NA | NA | NA | NA |
| CACNA1G | 1482 | L | 8.724375 | 13.75693051 | 0.232161796 | 13.60034737 | NA | NA | NA | NA | NA | NA | NA |
| CACNA1G | 1483 | A | 4.7176 | 11.30766103 | 0.183602655 | 10.02652544 | NA | NA | NA | NA | NA | NA | NA |
| CACNA1G | 1484 | S | 6.204833333 | 8.698163772 | 0.131866843 | 7.158128119 | NA | NA | NA | NA | NA | NA | NA |
| CACNA1G | 1485 | K | 6.035777778 | 9.880325197 | 0.155304339 | 9.984449367 | NA | NA | NA | NA | NA | NA | NA |
| CACNA1G | 1486 | D | 11.00225 | 6.910588036 | 0.096426421 | 6.245994063 | 0.97 | -0.23 | -1.38 | 14.21 | 0.43 | 1 | -1 |
| CACNA1G | 1487 | G | 13.406 | 10.75911455 | 0.17272719 | 10.31847683 | NA | NA | NA | NA | NA | NA | NA |
| CACNA1G | 1488 | W | 10.80285714 | 12.85662047 | 0.214312277 | 12.45632667 | NA | NA | NA | NA | NA | NA | NA |
| CACNA1G | 1489 | V | 14.64714286 | 14.71983386 | 0.251252288 | 13.86337087 | NA | NA | NA | NA | NA | NA | NA |

| | | | | | | | | | | | | | |
|---|---|---|---|---|---|---|---|---|---|---|---|---|---|
| CACNA1G | 1490 | D | 17.79025 | 12.98281067 | 0.21681412 | 10.91988577 | NA | NA | NA | NA | NA | NA | NA |
| CACNA1G | 1491 | I | 14.974625 | 16.65759842 | 0.289670349 | 16.16846443 | NA | NA | NA | NA | NA | NA | NA |
| CACNA1G | 1492 | M | 14.45575 | 19.41620535 | 0.344362405 | 18.75280287 | NA | NA | NA | NA | NA | NA | NA |
| CACNA1G | 1493 | Y | 18.49016667 | 18.46111505 | 0.325426814 | 16.4165739 | NA | NA | NA | NA | NA | NA | NA |
| CACNA1G | 1494 | D | 19.595625 | 19.03289623 | 0.33676293 | 16.99611868 | NA | NA | NA | NA | NA | NA | NA |
| CACNA1G | 1495 | G | 17.455 | 23.45417423 | 0.424419051 | 21.67501914 | NA | NA | NA | NA | NA | NA | NA |
| CACNA1G | 1496 | L | 17.925125 | 24.7844031 | 0.450792128 | 23.31379369 | NA | NA | NA | NA | NA | NA | NA |
| CACNA1G | 1497 | D | 22.246375 | 22.86583112 | 0.412754579 | 19.91149408 | NA | NA | NA | NA | NA | NA | NA |
| CACNA1G | 1498 | A | 21.067 | 27.04789687 | 0.495668085 | 24.12491386 | NA | NA | NA | NA | NA | NA | NA |
| CACNA1G | 1499 | V | 24.08485714 | 29.73334132 | 0.548909624 | 26.7802354 | NA | NA | NA | NA | NA | NA | NA |
| CACNA1G | 1500 | G | 24.08325 | 30.77367009 | 0.56953515 | 27.80835731 | NA | NA | NA | NA | NA | NA | NA |
| CACNA1G | 1501 | V | 24.61614286 | 30.62473224 | 0.566582313 | 27.65192433 | NA | NA | NA | NA | NA | NA | NA |
| CACNA1G | 1502 | D | 28.57775 | 30.98568783 | 0.573738607 | 26.27550612 | NA | NA | NA | NA | NA | NA | NA |
| CACNA1G | 1503 | Q | 28.55622222 | 28.6461699 | 0.527355397 | 24.00766962 | NA | NA | NA | NA | NA | NA | NA |
| CACNA1G | 1504 | Q | 29.932 | 23.67658713 | 0.428828602 | 19.01939629 | NA | NA | NA | NA | NA | NA | NA |
| CACNA1G | 1505 | P | 26.67785714 | 25.19715835 | 0.458975401 | 22.22246343 | NA | NA | NA | NA | NA | NA | NA |
| CACNA1G | 1506 | I | 26.67925 | 27.27082217 | 0.500087795 | 24.294917 | NA | NA | NA | NA | NA | NA | NA |
| CACNA1G | 1507 | M | 25.00375 | 24.81578089 | 0.451414223 | 21.84174985 | NA | NA | NA | NA | NA | NA | NA |
| CACNA1G | 1508 | N | 21.410625 | 26.12946678 | 0.477459319 | 24.03584809 | NA | NA | NA | NA | NA | NA | NA |
| CACNA1G | 1509 | H | 22.4158 | 29.03790471 | 0.535121919 | 26.11363877 | NA | NA | NA | NA | NA | NA | NA |
| CACNA1G | 1510 | N | 17.710125 | 29.05127689 | 0.535387036 | 27.69339582 | NA | NA | NA | NA | NA | NA | NA |
| CACNA1G | 1511 | P | 16.56942857 | 28.54018698 | 0.525254183 | 27.85601786 | NA | NA | NA | NA | NA | NA | NA |
| CACNA1G | 1512 | W | 12.73371429 | 29.08911594 | 0.536137232 | 28.54384754 | NA | NA | NA | NA | NA | NA | NA |
| CACNA1G | 1513 | M | 13.425875 | 25.29454281 | 0.460906142 | 24.74487894 | NA | NA | NA | NA | NA | NA | NA |
| CACNA1G | 1514 | L | 11.48575 | 23.90414462 | 0.43334015 | 23.21256713 | NA | NA | NA | NA | NA | NA | NA |
| CACNA1G | 1515 | L | 7.71125 | 25.79107113 | 0.470750297 | 24.4133035 | NA | NA | NA | NA | NA | NA | NA |
| CACNA1G | 1516 | Y | 8.699166667 | 23.16984234 | 0.418781896 | 21.77899185 | NA | NA | NA | NA | NA | NA | NA |
| CACNA1G | 1517 | F | 10.60636364 | 19.94394926 | 0.354825439 | 19.70899011 | NA | NA | NA | NA | NA | NA | NA |
| CACNA1G | 1518 | I | 6.655125 | 20.49733907 | 0.365796928 | 19.32726576 | NA | NA | NA | NA | NA | NA | NA |
| CACNA1G | 1519 | S | 3.348166667 | 20.59725877 | 0.367777933 | 17.93928287 | NA | NA | NA | NA | NA | NA | NA |
| CACNA1G | 1520 | F | 4.313454545 | 17.72729683 | 0.310878156 | 15.17550907 | NA | NA | NA | NA | NA | NA | NA |
| CACNA1G | 1521 | L | 4.56725 | 15.65447936 | 0.269782541 | 12.87574709 | NA | NA | NA | NA | NA | NA | NA |
| CACNA1G | 1522 | L | 0.98925 | 16.3438378 | 0.28344974 | 14.18449966 | NA | NA | NA | NA | NA | NA | NA |

| | | | | | | | | | | | | | |
|---|---|---|---|---|---|---|---|---|---|---|---|---|---|
| CACNA1G | 1523 | I | -0.35 | 15.94067812 | 0.275456709 | 14.74714521 | NA | NA | NA | NA | NA | NA | NA |
| CACNA1G | 1524 | V | -0.469 | 12.88502255 | 0.214875376 | 11.60739349 | NA | NA | NA | NA | NA | NA | NA |
| CACNA1G | 1525 | A | -1.1308 | 11.62570295 | 0.189908144 | 10.472681 | NA | NA | NA | NA | NA | NA | NA |
| CACNA1G | 1526 | F | -5.455363636 | 13.04979513 | 0.218142152 | 12.23862482 | NA | NA | NA | NA | NA | NA | NA |
| CACNA1G | 1527 | F | -4.702727273 | 11.64231188 | 0.190237433 | 10.78768565 | NA | NA | NA | NA | NA | NA | NA |
| CACNA1G | 1528 | V | -3.683285714 | 8.176456262 | 0.121523486 | 6.919586651 | NA | NA | NA | NA | NA | NA | NA |
| CACNA1G | 1529 | L | -6.436125 | 8.179056914 | 0.121575046 | 7.360986944 | NA | NA | NA | NA | NA | NA | NA |
| CACNA1G | 1530 | N | -9.131875 | 10.86694856 | 0.174865103 | 9.921343136 | NA | NA | NA | NA | NA | NA | NA |
| CACNA1G | 1531 | M | -7.574 | 8.810204595 | 0.134088161 | 7.955986795 | NA | NA | NA | NA | NA | NA | NA |
| CACNA1G | 1532 | F | -9.029363636 | 2.584874078 | 0.01066495 | 1.556480775 | 1.02 | 0.22 | 1.55 | 1.81 | -0.37 | 0 | 0 |
| CACNA1G | 1533 | V | -12.51228571 | 7.480987167 | 0.107735137 | 6.469949687 | NA | NA | NA | NA | NA | NA | NA |
| CACNA1G | 1534 | G | -13.91125 | 8.735561917 | 0.132608297 | 7.762620192 | NA | NA | NA | NA | NA | NA | NA |
| CACNA1G | 1535 | V | -13.08628571 | 6.147984141 | 0.08130706 | 5.236888654 | 1.25 | 0.49 | 2.53 | 1.21 | -0.76 | 0 | 0 |
| CACNA1G | 1536 | V | -14.86742857 | 3.004155122 | 0.018977603 | 2.546003658 | 1.12 | 0.45 | 1.77 | 1.65 | -0.57 | 0 | 0 |
| CACNA1G | 1537 | V | -18.16014286 | 6.270223361 | 0.083730571 | 5.421308514 | 1.12 | 0.45 | 1.77 | 1.65 | -0.57 | 0 | 0 |
| CACNA1G | 1538 | E | -18.05888889 | 8.27482024 | 0.123473647 | 7.359469212 | NA | NA | NA | NA | NA | NA | NA |
| CACNA1G | 1539 | N | -17.78625 | 5.859966297 | 0.075596827 | 5.384486819 | 0.51 | 1 | 1.9 | 1.43 | -0.72 | 0 | 0 |
| CACNA1G | 1540 | F | -21.85545455 | 5.799332893 | 0.074394711 | 5.48138633 | 0.51 | 1 | 1.9 | 1.43 | -0.72 | 0 | 0 |
| CACNA1G | 1541 | H | -24.0534 | 8.697779487 | 0.131859224 | 8.291984708 | NA | NA | NA | NA | NA | NA | NA |
| CACNA1G | 1612 | Y | -14.6335 | 40.31657908 | 0.75873257 | 39.92961142 | NA | NA | NA | NA | NA | NA | NA |
| CACNA1G | 1613 | L | -11.15475 | 39.74492683 | 0.74739901 | 39.33192861 | NA | NA | NA | NA | NA | NA | NA |
| CACNA1G | 1614 | D | -11.894875 | 35.05769202 | 0.654470037 | 34.66511597 | NA | NA | NA | NA | NA | NA | NA |
| CACNA1G | 1615 | L | -10.415125 | 37.239123 | 0.69771902 | 36.82476923 | NA | NA | NA | NA | NA | NA | NA |
| CACNA1G | 1616 | F | -6.099090909 | 38.98746309 | 0.732381557 | 38.18806694 | NA | NA | NA | NA | NA | NA | NA |
| CACNA1G | 1617 | I | -6.167625 | 35.59329693 | 0.665088923 | 33.04939518 | NA | NA | NA | NA | NA | NA | NA |
| CACNA1G | 1618 | T | -6.218857143 | 33.26266889 | 0.618881964 | 30.86530864 | NA | NA | NA | NA | NA | NA | NA |
| CACNA1G | 1619 | G | -2.984 | 36.08127649 | 0.674763591 | 33.63795838 | NA | NA | NA | NA | NA | NA | NA |
| CACNA1G | 1620 | V | -1.316571429 | 34.92541451 | 0.651847507 | 34.19924095 | NA | NA | NA | NA | NA | NA | NA |
| CACNA1G | 1621 | I | -1.30475 | 31.19818126 | 0.577951496 | 30.49037515 | NA | NA | NA | NA | NA | NA | NA |
| CACNA1G | 1622 | G | -0.449 | 32.25722485 | 0.598948061 | 31.50811993 | NA | NA | NA | NA | NA | NA | NA |
| CACNA1G | 1623 | L | 2.07875 | 33.28897597 | 0.619403527 | 32.56502119 | NA | NA | NA | NA | NA | NA | NA |
| CACNA1G | 1624 | N | 2.97875 | 30.48966889 | 0.563904551 | 29.7792536 | NA | NA | NA | NA | NA | NA | NA |
| CACNA1G | 1625 | V | 2.878857143 | 27.39168434 | 0.502484005 | 26.65264088 | NA | NA | NA | NA | NA | NA | NA |

| | | | | | | | | | | | | | |
|---|---|---|---|---|---|---|---|---|---|---|---|---|---|
| CACNA1G | 1626 | V | 5.561285714 | 30.16796848 | 0.557526529 | 29.43834646 | NA | NA | NA | NA | NA | NA | NA |
| CACNA1G | 1627 | T | 7.565428571 | 31.07176039 | 0.575445079 | 30.31183815 | NA | NA | NA | NA | NA | NA | NA |
| CACNA1G | 1628 | M | 7.44425 | 27.46157084 | 0.503869573 | 26.67584896 | NA | NA | NA | NA | NA | NA | NA |
| CACNA1G | 1629 | A | 9.4068 | 26.38299138 | 0.48248569 | 25.61078682 | NA | NA | NA | NA | NA | NA | NA |
| CACNA1G | 1630 | M | 11.684625 | 28.83160651 | 0.531031858 | 28.02456195 | NA | NA | NA | NA | NA | NA | NA |
| CACNA1G | 1631 | E | 13.12977778 | 26.82474839 | 0.49124395 | 25.95762222 | NA | NA | NA | NA | NA | NA | NA |
| CACNA1G | 1632 | H | 17.5658 | 27.4932461 | 0.504497565 | 26.47486811 | NA | NA | NA | NA | NA | NA | NA |
| CACNA1G | 1633 | Y | 22.58491667 | 23.81664244 | 0.431605335 | 20.191669 | NA | NA | NA | NA | NA | NA | NA |
| CACNA1G | 1634 | Q | 24.35233333 | 27.01004993 | 0.494917733 | 23.38945134 | NA | NA | NA | NA | NA | NA | NA |
| CACNA1G | 1635 | Q | 19.26833333 | 30.68502355 | 0.567777647 | 27.0260574 | NA | NA | NA | NA | NA | NA | NA |
| CACNA1G | 1636 | P | 21.23128571 | 34.19089616 | 0.637284969 | 30.53305676 | NA | NA | NA | NA | NA | NA | NA |
| CACNA1G | 1637 | Q | 22.16588889 | 35.80257517 | 0.669238067 | 32.15432936 | NA | NA | NA | NA | NA | NA | NA |
| CACNA1G | 1638 | I | 19.497 | 39.31972534 | 0.738968978 | 35.66100368 | NA | NA | NA | NA | NA | NA | NA |
| CACNA1G | 1639 | L | 16.401125 | 33.85298702 | 0.630585593 | 32.98464855 | NA | NA | NA | NA | NA | NA | NA |
| CACNA1G | 1640 | D | 17.50075 | 34.77502646 | 0.648865918 | 33.73354887 | NA | NA | NA | NA | NA | NA | NA |
| CACNA1G | 1641 | E | 17.10488889 | 38.70920057 | 0.726864733 | 38.01039583 | NA | NA | NA | NA | NA | NA | NA |
| CACNA1G | 1642 | A | 13.1898 | 38.66200059 | 0.725928948 | 37.85470339 | NA | NA | NA | NA | NA | NA | NA |
| CACNA1G | 1643 | L | 11.835375 | 33.16431006 | 0.616931905 | 32.34352202 | NA | NA | NA | NA | NA | NA | NA |
| CACNA1G | 1644 | K | 13.05711111 | 37.79580202 | 0.708755721 | 36.99615516 | NA | NA | NA | NA | NA | NA | NA |
| CACNA1G | 1645 | I | 10.332625 | 40.20838725 | 0.756587562 | 39.42526141 | NA | NA | NA | NA | NA | NA | NA |
| CACNA1G | 1646 | C | 7.471833333 | 37.85366051 | 0.709902822 | 37.10347653 | NA | NA | NA | NA | NA | NA | NA |
| CACNA1G | 1647 | N | 7.13625 | 35.20650536 | 0.657420406 | 34.46338332 | NA | NA | NA | NA | NA | NA | NA |
| CACNA1G | 1648 | Y | 8.47625 | 39.88975233 | 0.750270316 | 39.15607438 | NA | NA | NA | NA | NA | NA | NA |
| CACNA1G | 1649 | I | 3.660125 | 40.85321776 | 0.769371951 | 40.15605285 | NA | NA | NA | NA | NA | NA | NA |
| CACNA1G | 1650 | F | 2.375363636 | 36.07682716 | 0.674675379 | 35.37112143 | NA | NA | NA | NA | NA | NA | NA |
| CACNA1G | 1651 | T | 1.948285714 | 38.08106917 | 0.714411419 | 37.40255097 | NA | NA | NA | NA | NA | NA | NA |
| CACNA1G | 1652 | V | 0.903571429 | 41.74311609 | 0.787015048 | 41.05697841 | NA | NA | NA | NA | NA | NA | NA |
| CACNA1G | 1653 | I | -1.39275 | 40.99247415 | 0.772132844 | 40.27979834 | NA | NA | NA | NA | NA | NA | NA |
| CACNA1G | 1654 | F | -1.591454545 | 35.70669877 | 0.667337225 | 35.01017195 | NA | NA | NA | NA | NA | NA | NA |
| CACNA1G | 1655 | V | -2.525714286 | 41.94875582 | 0.791092055 | 41.25906198 | NA | NA | NA | NA | NA | NA | NA |
| CACNA1G | 1656 | L | -4.983875 | 44.42008841 | 0.840088619 | 42.04630156 | NA | NA | NA | NA | NA | NA | NA |
| CACNA1G | 1657 | E | -7.021888889 | 39.10233882 | 0.73465908 | 36.53856072 | NA | NA | NA | NA | NA | NA | NA |
| CACNA1G | 1658 | S | -7.833 | 42.23264459 | 0.796720425 | 39.65458147 | NA | NA | NA | NA | NA | NA | NA |

| | | | | | | | | | | | | | |
|---|---|---|---|---|---|---|---|---|---|---|---|---|---|
| CACNA1G | 1659 | V | -8.305428571 | 45.29557746 | 0.857446038 | 45.1472528 | NA | NA | NA | NA | NA | NA | NA |
| CACNA1G | 1660 | F | -11.44254545 | 45.72464851 | 0.865952787 | 45.32724975 | NA | NA | NA | NA | NA | NA | NA |
| CACNA1G | 1661 | K | -13.06233333 | 38.36923055 | 0.720124498 | 37.98633155 | NA | NA | NA | NA | NA | NA | NA |
| CACNA1G | 1662 | L | -12.848875 | 44.55310078 | 0.842725718 | 44.1610816 | NA | NA | NA | NA | NA | NA | NA |
| CACNA1G | 1663 | V | -14.87914286 | 46.79598228 | 0.887193018 | 46.41256415 | NA | NA | NA | NA | NA | NA | NA |
| CACNA1G | 1664 | A | -17.9246 | 45.14541757 | 0.854468973 | 44.78160142 | NA | NA | NA | NA | NA | NA | NA |
| CACNA1G | 1665 | F | -17.69554545 | 39.5745583 | 0.744021289 | 39.20534218 | NA | NA | NA | NA | NA | NA | NA |
| CACNA1G | 1666 | G | -17.66 | 45.50976737 | 0.86169256 | 45.14558993 | NA | NA | NA | NA | NA | NA | NA |
| CACNA1G | 1667 | F | -14.95454545 | 44.1932461 | 0.83559125 | 43.82286453 | NA | NA | NA | NA | NA | NA | NA |
| CACNA1G | 1668 | R | -20.96418182 | 40.46664544 | 0.761707781 | 40.1086288 | NA | NA | NA | NA | NA | NA | NA |
| CACNA1G | 1669 | R | -20.398 | 38.53016661 | 0.723315211 | 38.20215162 | NA | NA | NA | NA | NA | NA | NA |
| CACNA1G | 1670 | F | -12.90636364 | 38.82075972 | 0.729076501 | 38.43300957 | NA | NA | NA | NA | NA | NA | NA |
| CACNA1G | 1671 | F | -13.69545455 | 37.09632204 | 0.694887853 | 36.72038079 | NA | NA | NA | NA | NA | NA | NA |
| CACNA1G | 1672 | Q | -19.23444444 | 34.54173315 | 0.644240652 | 34.1837696 | NA | NA | NA | NA | NA | NA | NA |
| CACNA1G | 1673 | D | -15.715 | 32.71997931 | 0.608122617 | 32.34795172 | NA | NA | NA | NA | NA | NA | NA |
| CACNA1G | 1674 | R | -13.54754545 | 29.85241218 | 0.551270319 | 29.45574424 | NA | NA | NA | NA | NA | NA | NA |
| CACNA1G | 1675 | W | -11.66657143 | 26.91869835 | 0.493106599 | 26.5084055 | NA | NA | NA | NA | NA | NA | NA |
| CACNA1G | 1676 | N | -11.5855 | 34.66401474 | 0.646665003 | 34.25994012 | NA | NA | NA | NA | NA | NA | NA |
| CACNA1G | 1677 | Q | -10.69977778 | 34.31461971 | 0.639737908 | 33.91140588 | NA | NA | NA | NA | NA | NA | NA |
| CACNA1G | 1678 | L | -6.722625 | 30.42732657 | 0.562668554 | 27.83622631 | NA | NA | NA | NA | NA | NA | NA |
| CACNA1G | 1679 | D | -6.517 | 34.66680385 | 0.6467203 | 32.09231999 | NA | NA | NA | NA | NA | NA | NA |
| CACNA1G | 1680 | L | -5.639875 | 37.09886012 | 0.694938173 | 34.51153988 | NA | NA | NA | NA | NA | NA | NA |
| CACNA1G | 1681 | A | -3.201 | 34.76525441 | 0.648672178 | 32.14591749 | NA | NA | NA | NA | NA | NA | NA |
| CACNA1G | 1682 | I | -1.5245 | 31.73577294 | 0.588609772 | 31.29744798 | NA | NA | NA | NA | NA | NA | NA |
| CACNA1G | 1683 | V | -1.323857143 | 36.4437191 | 0.681949367 | 35.76212884 | NA | NA | NA | NA | NA | NA | NA |
| CACNA1G | 1684 | L | -0.4895 | 37.42644863 | 0.701432932 | 36.7578628 | NA | NA | NA | NA | NA | NA | NA |
| CACNA1G | 1685 | L | 1.5855 | 32.31860494 | 0.600164981 | 31.27099918 | NA | NA | NA | NA | NA | NA | NA |
| CACNA1G | 1686 | S | 2.953833333 | 35.70925946 | 0.667387993 | 35.04069562 | NA | NA | NA | NA | NA | NA | NA |
| CACNA1G | 1687 | I | 2.877875 | 38.22493609 | 0.71726372 | 37.28882126 | NA | NA | NA | NA | NA | NA | NA |
| CACNA1G | 1688 | M | 5.0465 | 36.30473765 | 0.679193925 | 35.02859113 | NA | NA | NA | NA | NA | NA | NA |
| CACNA1G | 1689 | G | 8.0795 | 36.37240354 | 0.680535467 | 35.08377378 | NA | NA | NA | NA | NA | NA | NA |
| CACNA1G | 1690 | I | 7.87025 | 38.00147794 | 0.712833446 | 36.70837462 | NA | NA | NA | NA | NA | NA | NA |
| CACNA1G | 1691 | T | 8.741714286 | 40.4600448 | 0.761576916 | 39.16948125 | NA | NA | NA | NA | NA | NA | NA |

| | | | | | | | | | | | | | |
|---|---|---|---|---|---|---|---|---|---|---|---|---|---|
| CACNA1G | 1692 | L | 11.350875 | 36.81334834 | 0.689277625 | 35.5376208 | NA | NA | NA | NA | NA | NA | NA |
| CACNA1G | 1693 | E | 12.97677778 | 34.03005323 | 0.634096102 | 32.73757038 | NA | NA | NA | NA | NA | NA | NA |
| CACNA1G | 1694 | E | 14.117 | 40.68992289 | 0.766134472 | 39.40301822 | NA | NA | NA | NA | NA | NA | NA |
| CACNA1G | 1695 | I | 15.253875 | 41.93861985 | 0.7908911 | 41.41036824 | NA | NA | NA | NA | NA | NA | NA |
| CACNA1G | 1696 | E | 18.89677778 | 35.50058197 | 0.663250759 | 35.16035429 | NA | NA | NA | NA | NA | NA | NA |
| CACNA1G | 1697 | V | 18.85457143 | 40.27205419 | 0.757849821 | 36.79697845 | NA | NA | NA | NA | NA | NA | NA |
| CACNA1G | 1698 | N | 18.6745 | 43.65262669 | 0.824872947 | 40.06124888 | NA | NA | NA | NA | NA | NA | NA |
| CACNA1G | 1699 | A | 21.471 | 42.87993289 | 0.809553543 | 39.2167115 | NA | NA | NA | NA | NA | NA | NA |
| CACNA1G | 1700 | S | 19.4705 | 44.76084152 | 0.84684438 | 41.0852623 | NA | NA | NA | NA | NA | NA | NA |
| CACNA1G | 1701 | L | 19.21825 | 41.22867329 | 0.776815721 | 37.55965854 | NA | NA | NA | NA | NA | NA | NA |
| CACNA1G | 1702 | P | 21.98414286 | 38.40369638 | 0.720807816 | 34.72071779 | NA | NA | NA | NA | NA | NA | NA |
| CACNA1G | 1703 | I | 17.681 | 36.04425764 | 0.674029656 | 35.54368239 | NA | NA | NA | NA | NA | NA | NA |
| CACNA1G | 1704 | N | 17.62475 | 31.84900284 | 0.590854664 | 31.34446559 | NA | NA | NA | NA | NA | NA | NA |
| CACNA1G | 1705 | P | 18.79628571 | 30.59909257 | 0.566073982 | 30.06217811 | NA | NA | NA | NA | NA | NA | NA |
| CACNA1G | 1706 | T | 15.91728571 | 28.33264043 | 0.521139372 | 27.79263647 | NA | NA | NA | NA | NA | NA | NA |
| CACNA1G | 1707 | I | 12.543875 | 31.53227951 | 0.584575317 | 30.28597404 | NA | NA | NA | NA | NA | NA | NA |
| CACNA1G | 1708 | I | 14.24025 | 31.41263149 | 0.582203179 | 30.12814933 | NA | NA | NA | NA | NA | NA | NA |
| CACNA1G | 1709 | R | 15.45118182 | 25.4146505 | 0.463287393 | 24.97266635 | NA | NA | NA | NA | NA | NA | NA |
| CACNA1G | 1710 | I | 10.185625 | 27.00079645 | 0.494734274 | 25.72937443 | NA | NA | NA | NA | NA | NA | NA |
| CACNA1G | 1711 | M | 8.401375 | 30.1022307 | 0.556223214 | 28.81929406 | NA | NA | NA | NA | NA | NA | NA |
| CACNA1G | 1712 | R | 12.01654545 | 28.41997011 | 0.522870767 | 27.31410071 | NA | NA | NA | NA | NA | NA | NA |
| CACNA1G | 1713 | V | 8.574571429 | 23.9888378 | 0.435019275 | 22.69453121 | NA | NA | NA | NA | NA | NA | NA |
| CACNA1G | 1714 | L | 4.609875 | 27.47613681 | 0.504158357 | 26.24933114 | NA | NA | NA | NA | NA | NA | NA |
| CACNA1G | 1715 | R | 6.780363636 | 28.88480196 | 0.532086509 | 28.20984995 | NA | NA | NA | NA | NA | NA | NA |
| CACNA1G | 1716 | I | 3.682375 | 23.1092671 | 0.417580933 | 22.42773724 | NA | NA | NA | NA | NA | NA | NA |
| CACNA1G | 1717 | A | 0.5686 | 27.65150936 | 0.507635288 | 26.97193122 | NA | NA | NA | NA | NA | NA | NA |
| CACNA1G | 1718 | R | -1.8 | 29.59822302 | 0.546230773 | 28.9095534 | NA | NA | NA | NA | NA | NA | NA |
| CACNA1G | 1719 | V | -0.788714286 | 25.96644113 | 0.474227177 | 25.26040967 | NA | NA | NA | NA | NA | NA | NA |
| CACNA1G | 1720 | L | -3.416 | 25.3114293 | 0.461240933 | 22.72522067 | NA | NA | NA | NA | NA | NA | NA |
| CACNA1G | 1721 | K | -5.023888889 | 29.6532004 | 0.547320753 | 27.0932017 | NA | NA | NA | NA | NA | NA | NA |
| CACNA1G | 1722 | L | -5.625125 | 27.55802431 | 0.505781856 | 25.56830499 | NA | NA | NA | NA | NA | NA | NA |
| CACNA1G | 1723 | L | -7.95125 | 24.62031765 | 0.447538974 | 24.11847656 | NA | NA | NA | NA | NA | NA | NA |
| CACNA1G | 1724 | K | -11.47533333 | 27.48953454 | 0.50442398 | 27.08489249 | NA | NA | NA | NA | NA | NA | NA |

| | | | | | | | | | | | | | |
|---|---|---|---|---|---|---|---|---|---|---|---|---|---|
| CACNA1G | 1725 | M | -10.964 | 26.91782824 | 0.493089349 | 26.50168699 | NA | NA | NA | NA | NA | NA | NA |
| CACNA1G | 1726 | A | -11.0252 | 23.34847751 | 0.422323511 | 22.92468602 | NA | NA | NA | NA | NA | NA | NA |
| CACNA1G | 1727 | V | -14.98085714 | 21.38260915 | 0.383348266 | 20.99660247 | NA | NA | NA | NA | NA | NA | NA |
| CACNA1G | 1728 | G | -13.7765 | 18.5028774 | 0.326254793 | 18.08358541 | NA | NA | NA | NA | NA | NA | NA |
| CACNA1G | 1729 | M | -9.864625 | 18.14340795 | 0.319127963 | 17.13310805 | NA | NA | NA | NA | NA | NA | NA |
| CACNA1G | 1730 | R | -13.18890909 | 18.3128586 | 0.322487486 | 17.91306082 | NA | NA | NA | NA | NA | NA | NA |
| CACNA1G | 1731 | A | -15.5312 | 15.10996592 | 0.258987034 | 14.73129929 | NA | NA | NA | NA | NA | NA | NA |
| CACNA1G | 1732 | L | -12.34975 | 13.10119972 | 0.219161298 | 12.65680912 | NA | NA | NA | NA | NA | NA | NA |
| CACNA1G | 1733 | L | -10.675875 | 15.12710478 | 0.259326829 | 14.69136413 | NA | NA | NA | NA | NA | NA | NA |
| CACNA1G | 1734 | D | -15.8945 | 14.26375862 | 0.242210155 | 13.89423065 | NA | NA | NA | NA | NA | NA | NA |
| CACNA1G | 1735 | T | -15.39171429 | 9.787349488 | 0.153461005 | 9.408351689 | NA | NA | NA | NA | NA | NA | NA |
| CACNA1G | 1736 | V | -11.91671429 | 11.65937108 | 0.190575648 | 11.27162715 | NA | NA | NA | NA | NA | NA | NA |
| CACNA1G | 1737 | M | -12.9965 | 13.54390815 | 0.227938421 | 13.11511397 | NA | NA | NA | NA | NA | NA | NA |
| CACNA1G | 1738 | Q | -17.82022222 | 11.77411385 | 0.192850534 | 11.39581592 | NA | NA | NA | NA | NA | NA | NA |
| CACNA1G | 1739 | A | -14.2634 | 8.782939087 | 0.133547596 | 8.352003338 | NA | NA | NA | NA | NA | NA | NA |
| CACNA1G | 1740 | L | -11.404 | 12.26435273 | 0.202569995 | 11.81221184 | NA | NA | NA | NA | NA | NA | NA |
| CACNA1G | 1741 | P | -13.81985714 | 13.91916406 | 0.235378233 | 13.44923424 | NA | NA | NA | NA | NA | NA | NA |
| CACNA1G | 1742 | Q | -12.82933333 | 9.968139897 | 0.15704535 | 9.472713814 | NA | NA | NA | NA | NA | NA | NA |
| CACNA1G | 1743 | V | -8.894571429 | 10.01974067 | 0.158068386 | 9.114357347 | NA | NA | NA | NA | NA | NA | NA |
| CACNA1G | 1744 | G | -8.33325 | 13.95169355 | 0.236023162 | 13.10665789 | NA | NA | NA | NA | NA | NA | NA |
| CACNA1G | 1745 | N | -10.180375 | 14.30592933 | 0.24304623 | 13.41322362 | NA | NA | NA | NA | NA | NA | NA |
| CACNA1G | 1746 | L | -7.569625 | 8.677908216 | 0.131465257 | 7.814355433 | NA | NA | NA | NA | NA | NA | NA |
| CACNA1G | 1747 | G | -4.1915 | 12.52731651 | 0.207783507 | 10.36361129 | NA | NA | NA | NA | NA | NA | NA |
| CACNA1G | 1748 | L | -4.85725 | 14.81723719 | 0.253183404 | 12.55280469 | NA | NA | NA | NA | NA | NA | NA |
| CACNA1G | 1749 | L | -4.479625 | 13.92498916 | 0.235493722 | 11.70581804 | NA | NA | NA | NA | NA | NA | NA |
| CACNA1G | 1750 | F | -0.651818182 | 7.662538548 | 0.111334569 | 5.519658742 | NA | NA | NA | NA | NA | NA | NA |
| CACNA1G | 1751 | M | 0.20525 | 12.61850249 | 0.209591357 | 10.48303137 | NA | NA | NA | NA | NA | NA | NA |
| CACNA1G | 1752 | L | -1.146125 | 16.0734603 | 0.278089244 | 13.92839223 | NA | NA | NA | NA | NA | NA | NA |
| CACNA1G | 1753 | L | 0.714625 | 12.1404126 | 0.200112762 | 9.976258539 | NA | NA | NA | NA | NA | NA | NA |
| CACNA1G | 1754 | F | 3.912818182 | 9.619108742 | 0.150125469 | 7.704320355 | NA | NA | NA | NA | NA | NA | NA |
| CACNA1G | 1755 | F | 3.896272727 | 16.38540466 | 0.284273844 | 15.12567235 | NA | NA | NA | NA | NA | NA | NA |
| CACNA1G | 1756 | I | 3.171 | 18.08572202 | 0.317984283 | 16.82355597 | NA | NA | NA | NA | NA | NA | NA |
| CACNA1G | 1757 | F | 5.337090909 | 14.94177315 | 0.25565245 | 13.67054928 | NA | NA | NA | NA | NA | NA | NA |

| | | | | | | | | | | | | | |
|---|---|---|---|---|---|---|---|---|---|---|---|---|---|
| CACNA1G | 1758 | A | 8.6866 | 15.03083664 | 0.25741822 | 13.78907773 | NA | NA | NA | NA | NA | NA | NA |
| CACNA1G | 1759 | A | 8.853 | 17.67010914 | 0.309744355 | 16.42455226 | NA | NA | NA | NA | NA | NA | NA |
| CACNA1G | 1760 | L | 8.90725 | 18.38570045 | 0.323931646 | 17.13250618 | NA | NA | NA | NA | NA | NA | NA |
| CACNA1G | 1761 | G | 12.54875 | 16.41302638 | 0.284821471 | 15.16878242 | NA | NA | NA | NA | NA | NA | NA |
| CACNA1G | 1762 | V | 14.06585714 | 17.31727513 | 0.302749079 | 17.11474188 | NA | NA | NA | NA | NA | NA | NA |
| CACNA1G | 1763 | E | 13.52255556 | 19.71785716 | 0.350342944 | 19.57038625 | NA | NA | NA | NA | NA | NA | NA |
| CACNA1G | 1764 | L | 14.174625 | 18.42120596 | 0.324635577 | 18.30356165 | NA | NA | NA | NA | NA | NA | NA |
| CACNA1G | 1765 | F | 15.81909091 | 14.80896144 | 0.253019329 | 14.58715311 | NA | NA | NA | NA | NA | NA | NA |
| CACNA1G | 1766 | G | 19.39275 | 16.91629348 | 0.294799229 | 13.96357171 | NA | NA | NA | NA | NA | NA | NA |
| CACNA1G | 1767 | D | 22.238125 | 16.64963831 | 0.289512532 | 12.40374805 | NA | NA | NA | NA | NA | NA | NA |
| CACNA1G | 1768 | L | 22.222125 | 12.61940252 | 0.209609201 | 9.047156389 | NA | NA | NA | NA | NA | NA | NA |
| CACNA1G | 1769 | E | 26.68566667 | 10.58391426 | 0.169253674 | 6.365374848 | NA | NA | NA | NA | NA | NA | NA |
| CACNA1G | 1770 | C | 26.09066667 | 7.238079994 | 0.102919267 | 3.07515961 | 4.37 | -0.65 | -2.55 | 27.26 | 1.3 | 2 | 2 |
| CACNA1G | 1771 | D | 30.065625 | 6.196868725 | 0.082276244 | 1.740940842 | 4.59 | -0.68 | -2.12 | 22.49 | 1.2 | 2 | 2 |
| CACNA1G | 1772 | E | 34.69666667 | 5.096116463 | 0.060452764 | 2.000640389 | 3.37 | -0.32 | -2.48 | 25.75 | 0.43 | 2 | -2 |
| CACNA1G | 1773 | T | 33.79328571 | 7.87150208 | 0.115477474 | 4.650931364 | 3.37 | -0.32 | -2.48 | 25.75 | 0.43 | 2 | -2 |
| CACNA1G | 1774 | H | 30.3239 | 7.607507213 | 0.11024352 | 3.197986077 | 4.68 | -0.68 | -2.74 | 22.49 | 1.14 | 2 | 2 |
| CACNA1G | 1775 | P | 28.55771429 | 7.050956318 | 0.099209359 | 2.61006601 | 4.59 | -0.68 | -2.12 | 22.49 | 1.2 | 2 | 2 |
| CACNA1G | 1776 | C | 24.6765 | 5.532894089 | 0.069112304 | 1.608182837 | 4.1 | -0.75 | -1.53 | 15.11 | 1.09 | 1 | 1 |
| CACNA1G | 1777 | E | 22.70311111 | 4.808419699 | 0.054748897 | 1.855025313 | 2.98 | -0.24 | -1.73 | 18.23 | 0.67 | 1 | -1 |
| CACNA1G | 1778 | G | 19.63125 | 4.163432959 | 0.04196141 | 1.437617552 | 2.98 | -0.24 | -1.73 | 18.23 | 0.67 | 1 | -1 |
| CACNA1G | 1779 | L | 20.234375 | 4.78304516 | 0.054245822 | 1.922632665 | 2.98 | -0.24 | -1.73 | 18.23 | 0.67 | 1 | -1 |
| CACNA1G | 1780 | G | 22.44025 | 5.507930374 | 0.068617374 | 2.434905819 | 3.13 | -0.38 | -1.4 | 14.52 | 0.7 | 1 | -1 |
| CACNA1G | 1781 | R | 24.38109091 | 5.479508098 | 0.068053875 | 1.575523592 | 3.88 | -0.75 | -1.53 | 15.11 | 1.09 | 1 | 1 |
| CACNA1G | 1782 | H | 20.036 | 6.164043721 | 0.081625457 | 5.886383542 | 0.67 | 0.61 | -0.53 | 17.91 | -0.37 | 1 | 1 |
| CACNA1G | 1783 | A | 18.1684 | 5.09426452 | 0.060416047 | 4.923389983 | 0.67 | 0.61 | -0.53 | 17.91 | -0.37 | 1 | 1 |
| CACNA1G | 1784 | T | 18.42171429 | 8.622027894 | 0.130357375 | 8.447059572 | NA | NA | NA | NA | NA | NA | NA |
| CACNA1G | 1785 | F | 15.39181818 | 11.0633189 | 0.178758335 | 10.86520633 | NA | NA | NA | NA | NA | NA | NA |
| CACNA1G | 1786 | R | 20.00627273 | 13.26914771 | 0.222491029 | 13.14578997 | NA | NA | NA | NA | NA | NA | NA |
| CACNA1G | 1787 | N | 15.87075 | 15.3062007 | 0.262877579 | 15.14197928 | NA | NA | NA | NA | NA | NA | NA |
| CACNA1G | 1788 | F | 9.992727273 | 15.19277072 | 0.26062872 | 13.96822691 | NA | NA | NA | NA | NA | NA | NA |
| CACNA1G | 1789 | G | 12.40575 | 12.9987369 | 0.217129873 | 11.77984051 | NA | NA | NA | NA | NA | NA | NA |
| CACNA1G | 1790 | M | 15.378 | 10.03428214 | 0.158356684 | 9.827245972 | NA | NA | NA | NA | NA | NA | NA |

| | | | | | | | | | | | | | |
|---|---|---|---|---|---|---|---|---|---|---|---|---|---|
| CACNA1G | 1791 | A | 11.9342 | 11.19735388 | 0.181415709 | 9.963737575 | NA | NA | NA | NA | NA | NA | NA |
| CACNA1G | 1792 | F | 7.560727273 | 9.702194288 | 0.151772721 | 8.467400861 | NA | NA | NA | NA | NA | NA | NA |
| CACNA1G | 1793 | L | 11.3985 | 6.336916758 | 0.085052832 | 5.114895374 | 1.32 | 0.41 | -0.48 | 13.97 | -0.35 | 1 | 1 |
| CACNA1G | 1794 | T | 12.541 | 6.301236387 | 0.084345434 | 5.055481216 | 1.32 | 0.41 | -0.48 | 13.97 | -0.35 | 1 | 1 |
| CACNA1G | 1795 | L | 8.43725 | 7.545160436 | 0.109007434 | 6.286541457 | NA | NA | NA | NA | NA | NA | NA |
| CACNA1G | 1796 | F | 6.892090909 | 4.661617852 | 0.051838408 | 3.382643417 | 1.32 | 0.41 | -0.48 | 13.97 | -0.35 | 1 | 1 |
| CACNA1G | 1797 | R | 12.58490909 | 2.12918623 | 0.001630497 | 1.452864189 | 1.32 | 0.41 | -0.48 | 13.97 | -0.35 | 1 | 1 |
| CACNA1G | 1798 | V | 8.850857143 | 4.768609336 | 0.053959618 | 3.485204085 | 1.32 | 0.41 | -0.48 | 13.97 | -0.35 | 1 | 1 |
| CACNA1G | 1799 | S | 5.0635 | 3.543576865 | 0.029672162 | 1.848326286 | 1.7 | -0.15 | 1 | 2.3 | -0.06 | 0 | 0 |
| CACNA1G | 1800 | T | 6.378857143 | 2.139969392 | 0.001844283 | 1.480372588 | 0.53 | 0.19 | 1.55 | 0.9 | -0.51 | 0 | 0 |
| CACNA1G | 1801 | G | 8.32125 | 3.888301043 | 0.036506653 | 3.223856358 | 0.53 | 0.19 | 1.55 | 0.9 | -0.51 | 0 | 0 |
| CACNA1G | 1802 | D | 11.723875 | 2.046945774 | 0 | 1.556416882 | 0.67 | -0.03 | -1.7 | 17.82 | 0.31 | 1 | -1 |
| CACNA1G | 1803 | N | 15.05375 | 3.829240264 | 0.035335716 | 3.689827491 | 0.8 | -0.11 | -1.27 | 33.94 | 0.99 | 2 | 0 |
| CACNA1G | 1804 | W | 10.70035714 | 5.804541842 | 0.074497984 | 5.193770777 | 0.8 | -0.11 | -1.27 | 33.94 | 0.99 | 2 | 0 |
| CACNA1G | 1805 | N | 14.495625 | 8.283635253 | 0.123648413 | 7.72703782 | NA | NA | NA | NA | NA | NA | NA |
| CACNA1G | 1806 | G | 17.5445 | 8.368345476 | 0.125327876 | 6.349923653 | NA | NA | NA | NA | NA | NA | NA |
| CACNA1G | 1807 | I | 15.968375 | 6.145090154 | 0.081249684 | 5.644907662 | 0.51 | 0.36 | -0.15 | 26.07 | 0.17 | 1 | 1 |
| CACNA1G | 1808 | M | 14.986625 | 10.48939665 | 0.167379771 | 10.01564612 | NA | NA | NA | NA | NA | NA | NA |
| CACNA1G | 1809 | K | 19.374 | 11.76012113 | 0.192573115 | 8.857157345 | NA | NA | NA | NA | NA | NA | NA |
| CACNA1G | 1810 | D | 20.96025 | 7.712381539 | 0.112322755 | 4.747192013 | 2.98 | -0.24 | -1.73 | 18.23 | 0.67 | 1 | -1 |
| CACNA1G | 1811 | T | 18.36414286 | 13.07529805 | 0.218647772 | 10.14241706 | NA | NA | NA | NA | NA | NA | NA |
| CACNA1G | 1812 | L | 19.31025 | 15.65575996 | 0.26980793 | 12.71911829 | NA | NA | NA | NA | NA | NA | NA |
| CACNA1G | 1813 | R | 24.82763636 | 10.31827045 | 0.163987028 | 8.832151615 | NA | NA | NA | NA | NA | NA | NA |
| CACNA1G | 1814 | D | 25.32825 | 17.15803351 | 0.299591959 | 12.90228896 | NA | NA | NA | NA | NA | NA | NA |
| CACNA1G | 1822 | Y | 17.96958333 | 23.22369912 | 0.419849659 | 20.31955107 | NA | NA | NA | NA | NA | NA | NA |
| CACNA1G | 1823 | N | 17.26875 | 19.26930512 | 0.341449965 | 16.32304791 | NA | NA | NA | NA | NA | NA | NA |
| CACNA1G | 1824 | T | 16.57971429 | 21.97096944 | 0.395013079 | 21.56385909 | NA | NA | NA | NA | NA | NA | NA |
| CACNA1G | 1825 | V | 12.63228571 | 21.98273109 | 0.395246265 | 21.84488897 | NA | NA | NA | NA | NA | NA | NA |
| CACNA1G | 1826 | I | 12.5225 | 19.36786207 | 0.343403952 | 19.25902248 | NA | NA | NA | NA | NA | NA | NA |
| CACNA1G | 1827 | S | 13.25833333 | 16.27837157 | 0.282151809 | 16.20601386 | NA | NA | NA | NA | NA | NA | NA |
| CACNA1G | 1828 | P | 10.89442857 | 17.70076388 | 0.310352115 | 17.26674715 | NA | NA | NA | NA | NA | NA | NA |
| CACNA1G | 1829 | I | 7.4775 | 17.72560775 | 0.310844669 | 16.20882722 | NA | NA | NA | NA | NA | NA | NA |
| CACNA1G | 1830 | Y | 9.112166667 | 14.03290754 | 0.237633308 | 12.75836857 | NA | NA | NA | NA | NA | NA | NA |

| | | | | | | | | | | | | | |
|---|---|---|---|---|---|---|---|---|---|---|---|---|---|
| CACNA1G | 1831 | F | 10.11127273 | 10.84377139 | 0.174405593 | 10.14171217 | NA | NA | NA | NA | NA | NA | NA |
| CACNA1G | 1832 | V | 5.356428571 | 13.45167636 | 0.226109837 | 11.71938876 | NA | NA | NA | NA | NA | NA | NA |
| CACNA1G | 1833 | S | 3.365166667 | 13.57213123 | 0.228497971 | 11.75637093 | NA | NA | NA | NA | NA | NA | NA |
| CACNA1G | 1834 | F | 4.715636364 | 9.815838731 | 0.154025832 | 7.925015033 | NA | NA | NA | NA | NA | NA | NA |
| CACNA1G | 1835 | V | 3.894285714 | 8.407128047 | 0.126096778 | 6.713048074 | NA | NA | NA | NA | NA | NA | NA |
| CACNA1G | 1836 | L | 0.714 | 10.57976488 | 0.169171408 | 8.776530974 | NA | NA | NA | NA | NA | NA | NA |
| CACNA1G | 1837 | T | -0.190714286 | 9.799369163 | 0.153699307 | 7.351737151 | NA | NA | NA | NA | NA | NA | NA |
| CACNA1G | 1838 | A | 0.2862 | 6.466642405 | 0.087624769 | 4.642589756 | 2.22 | -0.08 | 0.66 | 2.11 | -0.37 | 0 | 0 |
| CACNA1G | 1839 | Q | -0.807111111 | 3.5311593 | 0.029425972 | 1.615610733 | 2.57 | 0.16 | -0.1 | 1.85 | -0.73 | 0 | 0 |
| CACNA1G | 1840 | F | -5.147181818 | 7.087071398 | 0.099925375 | 6.028832008 | 1.43 | 0.49 | 2.67 | 1.21 | -0.6 | 0 | 0 |
| CACNA1G | 1841 | V | -3.820142857 | 6.632200314 | 0.090907115 | 5.504271865 | 1.43 | 0.49 | 2.67 | 1.21 | -0.6 | 0 | 0 |
| CACNA1G | 1842 | L | -2.21625 | 3.324777887 | 0.02533426 | 2.205389644 | 1.43 | 0.49 | 2.67 | 1.21 | -0.6 | 0 | 0 |
| CACNA1G | 1843 | V | -5.655714286 | 2.558473959 | 0.010141542 | 1.614415276 | 1.52 | 1.13 | 4.2 | 0.13 | -1.3 | 0 | 0 |
| CACNA1G | 1844 | N | -8.391 | 4.692859363 | 0.052457801 | 3.848711278 | 1.02 | 0.22 | 1.55 | 1.81 | -0.37 | 0 | 0 |
| CACNA1G | 1845 | V | -7.218428571 | 5.046145559 | 0.059462042 | 4.160000867 | 1.02 | 0.22 | 1.55 | 1.81 | -0.37 | 0 | 0 |
| CACNA1G | 1846 | V | -7.925142857 | 2.604963339 | 0.011063239 | 1.640909966 | 1.01 | 0.56 | 2.2 | 1.29 | -0.77 | 0 | 0 |
| CACNA1G | 1847 | I | -11.31425 | 2.490170476 | 0.008787359 | 1.557756328 | 1.19 | 0.17 | 1.7 | 1.76 | -0.49 | 0 | 0 |
| CACNA1G | 1848 | A | -12.645 | 4.678918037 | 0.052181401 | 4.222691302 | 0.93 | 0.1 | 0.75 | 2.41 | 0.05 | 0 | 0 |
| CACNA1G | 1849 | V | -11.79271429 | 5.219273992 | 0.062894481 | 4.315022123 | 1.19 | 0.17 | 1.7 | 1.76 | -0.49 | 0 | 0 |
| CACNA1G | 1850 | L | -13.757375 | 2.689165856 | 0.012732636 | 2.176041779 | 0.73 | 0.1 | 0.75 | 2.41 | 0.05 | 0 | 0 |
| CACNA1G | 1851 | M | -17.137625 | 2.115492614 | 0.001359008 | 1.666308753 | 0.51 | 1 | 1.9 | 1.43 | -0.72 | 0 | 0 |
| CACNA1G | 1852 | K | -17.89122222 | 4.454122809 | 0.047724618 | 4.127455115 | 0.51 | 1 | 1.9 | 1.43 | -0.72 | 0 | 0 |
| CACNA1G | 1853 | H | -16.4492 | 5.753588619 | 0.073487787 | 5.424785873 | 0.51 | 1 | 1.9 | 1.43 | -0.72 | 0 | 0 |
| CACNA1G | 1854 | L | -18.880875 | 4.448720041 | 0.047617503 | 4.118219505 | 0.51 | 1 | 1.9 | 1.43 | -0.72 | 0 | 0 |
| CACNA1G | 1855 | E | -21.79811111 | 5.673114577 | 0.071892311 | 5.410872392 | 0.51 | 1 | 1.9 | 1.43 | -0.72 | 0 | 0 |
| CACNA1G | 1856 | E | -21.94744444 | 8.394320163 | 0.125842849 | 8.151884392 | NA | NA | NA | NA | NA | NA | NA |
| CACNA1G | 1857 | S | -22.32433333 | 9.825116793 | 0.154209779 | 9.555342065 | NA | NA | NA | NA | NA | NA | NA |
| CACNA1G | 1858 | N | -25.8795 | 10.52577194 | 0.168100946 | 10.27530723 | NA | NA | NA | NA | NA | NA | NA |
| CACNA1G | 1859 | K | -27.66822222 | 12.19060347 | 0.201107845 | 11.95550789 | NA | NA | NA | NA | NA | NA | NA |
| CHRNA4 | 31 | E | 83.20583333 | 31.08634612 | 0.76466826 | 21.05882262 | NA | NA | NA | NA | NA | NA | NA |
| CHRNA4 | 32 | T | 79.02488095 | 28.28359726 | 0.687782679 | 18.31902912 | NA | NA | NA | NA | NA | NA | NA |
| CHRNA4 | 33 | R | 74.31516667 | 26.94788324 | 0.651141099 | 18.07334679 | NA | NA | NA | NA | NA | NA | NA |
| CHRNA4 | 34 | A | 75.97296667 | 23.86505841 | 0.566572416 | 15.05582949 | NA | NA | NA | NA | NA | NA | NA |

| | | | | | | | | | | | | | |
|---|---|---|---|---|---|---|---|---|---|---|---|---|---|
| CHRNA4 | 35 | H | 81.00966667 | 24.39198944 | 0.581027296 | 14.37355299 | NA | NA | NA | NA | NA | NA | NA |
| CHRNA4 | 36 | A | 79.18636667 | 24.90476551 | 0.595093874 | 14.93823009 | NA | NA | NA | NA | NA | NA | NA |
| CHRNA4 | 37 | E | 74.8665 | 22.10620264 | 0.518323124 | 12.14863658 | NA | NA | NA | NA | NA | NA | NA |
| CHRNA4 | 38 | E | 78.6205 | 18.23115079 | 0.412021908 | 8.30585555 | NA | NA | NA | NA | NA | NA | NA |
| CHRNA4 | 39 | R | 82.57925758 | 21.74239198 | 0.508342996 | 12.02856059 | NA | NA | NA | NA | NA | NA | NA |
| CHRNA4 | 40 | L | 77.67441667 | 21.42913748 | 0.499749734 | 11.71137687 | NA | NA | NA | NA | NA | NA | NA |
| CHRNA4 | 41 | L | 75.59679167 | 16.73725369 | 0.371041017 | 6.95744291 | NA | NA | NA | NA | NA | NA | NA |
| CHRNA4 | 42 | K | 81.04516667 | 17.47443604 | 0.391263554 | 7.774501836 | NA | NA | NA | NA | NA | NA | NA |
| CHRNA4 | 43 | K | 81.56872222 | 19.8714932 | 0.457020117 | 10.16220402 | NA | NA | NA | NA | NA | NA | NA |
| CHRNA4 | 44 | L | 76.90554167 | 19.34406395 | 0.44255157 | 9.62817743 | NA | NA | NA | NA | NA | NA | NA |
| CHRNA4 | 45 | F | 75.01562121 | 15.14364646 | 0.327324857 | 5.434795372 | NA | NA | NA | NA | NA | NA | NA |
| CHRNA4 | 46 | S | 80.6115 | 15.53704805 | 0.33811673 | 5.856250072 | NA | NA | NA | NA | NA | NA | NA |
| CHRNA4 | 47 | G | 79.57991667 | 17.4813494 | 0.391453203 | 7.791160194 | NA | NA | NA | NA | NA | NA | NA |
| CHRNA4 | 48 | Y | 74.324 | 15.75353211 | 0.344055365 | 6.038953332 | NA | NA | NA | NA | NA | NA | NA |
| CHRNA4 | 49 | N | 73.92729167 | 16.43470891 | 0.362741546 | 6.717116642 | NA | NA | NA | NA | NA | NA | NA |
| CHRNA4 | 50 | K | 70.05561111 | 11.02691788 | 0.214393915 | 2.270778372 | 8.88 | -0.67 | -3.4 | 50.47 | 1.42 | 2 | -2 |
| CHRNA4 | 51 | W | 69.33959524 | 12.87717628 | 0.265150583 | 3.992880653 | 8.88 | -0.67 | -3.4 | 50.47 | 1.42 | 2 | -2 |
| CHRNA4 | 52 | S | 71.7755 | 20.33065299 | 0.469615883 | 10.66117797 | NA | NA | NA | NA | NA | NA | NA |
| CHRNA4 | 53 | R | 68.88380303 | 23.6596 | 0.560936239 | 14.7765487 | NA | NA | NA | NA | NA | NA | NA |
| CHRNA4 | 54 | P | 70.47430952 | 24.17434479 | 0.575056824 | 15.35760947 | NA | NA | NA | NA | NA | NA | NA |
| CHRNA4 | 55 | V | 71.89802381 | 27.9013047 | 0.677295551 | 19.03042793 | NA | NA | NA | NA | NA | NA | NA |
| CHRNA4 | 56 | A | 75.14736667 | 30.93922722 | 0.760632464 | 21.22190333 | NA | NA | NA | NA | NA | NA | NA |
| CHRNA4 | 57 | N | 75.23254167 | 32.48742214 | 0.803102865 | 22.76977304 | NA | NA | NA | NA | NA | NA | NA |
| CHRNA4 | 58 | I | 73.05266667 | 32.16413933 | 0.794234504 | 22.45266937 | NA | NA | NA | NA | NA | NA | NA |
| CHRNA4 | 59 | S | 70.97983333 | 35.19934864 | 0.877496988 | 25.48155633 | NA | NA | NA | NA | NA | NA | NA |
| CHRNA4 | 60 | D | 70.18116667 | 33.50293053 | 0.830960499 | 24.65868669 | NA | NA | NA | NA | NA | NA | NA |
| CHRNA4 | 61 | V | 66.5477381 | 33.05304121 | 0.818619043 | 25.1288596 | NA | NA | NA | NA | NA | NA | NA |
| CHRNA4 | 62 | V | 64.95202381 | 29.58854728 | 0.723580336 | 21.72513301 | NA | NA | NA | NA | NA | NA | NA |
| CHRNA4 | 63 | L | 62.53129167 | 30.80437834 | 0.756933262 | 22.77035427 | NA | NA | NA | NA | NA | NA | NA |
| CHRNA4 | 64 | V | 59.47288095 | 27.72510209 | 0.672461925 | 19.69054489 | NA | NA | NA | NA | NA | NA | NA |
| CHRNA4 | 65 | R | 58.21625758 | 29.35740719 | 0.717239653 | 21.3457605 | NA | NA | NA | NA | NA | NA | NA |
| CHRNA4 | 66 | F | 53.24089394 | 25.461593 | 0.61036888 | 17.16675448 | NA | NA | NA | NA | NA | NA | NA |
| CHRNA4 | 67 | G | 51.67316667 | 26.71740212 | 0.644818493 | 18.50393129 | NA | NA | NA | NA | NA | NA | NA |

| | | | | | | | | | | | | | |
|---|---|---|---|---|---|---|---|---|---|---|---|---|---|
| CHRNA4 | 68 | L | 48.18066667 | 24.85537047 | 0.593738859 | 16.66712651 | NA | NA | NA | NA | NA | NA | NA |
| CHRNA4 | 69 | S | 47.20483333 | 23.92601783 | 0.568244667 | 16.80362702 | NA | NA | NA | NA | NA | NA | NA |
| CHRNA4 | 70 | I | 42.85254167 | 23.41998851 | 0.554363168 | 16.61679217 | NA | NA | NA | NA | NA | NA | NA |
| CHRNA4 | 71 | A | 43.14116667 | 21.63668433 | 0.505443202 | 14.70125916 | NA | NA | NA | NA | NA | NA | NA |
| CHRNA4 | 72 | Q | 39.9885 | 17.66277965 | 0.396430235 | 10.35802339 | NA | NA | NA | NA | NA | NA | NA |
| CHRNA4 | 73 | L | 36.65004167 | 18.93981441 | 0.431462114 | 10.92295175 | NA | NA | NA | NA | NA | NA | NA |
| CHRNA4 | 74 | I | 37.41754167 | 14.09224787 | 0.298482675 | 6.090560385 | NA | NA | NA | NA | NA | NA | NA |
| CHRNA4 | 75 | D | 33.68391667 | 12.59432753 | 0.257391418 | 5.521240402 | NA | NA | NA | NA | NA | NA | NA |
| CHRNA4 | 76 | V | 32.14359524 | 16.04601966 | 0.352078943 | 9.02267621 | NA | NA | NA | NA | NA | NA | NA |
| CHRNA4 | 77 | D | 31.22541667 | 10.99980154 | 0.213650054 | 3.846291178 | 7.22 | -1.14 | -3.5 | 49.9 | 2.23 | 5 | -5 |
| CHRNA4 | 78 | E | 27.92638889 | 15.66491341 | 0.341624358 | 8.460194121 | NA | NA | NA | NA | NA | NA | NA |
| CHRNA4 | 79 | K | 27.16661111 | 10.7236761 | 0.206075324 | 3.869216863 | 7.28 | -1.14 | -3.5 | 49.9 | 2.23 | 5 | -5 |
| CHRNA4 | 80 | N | 31.80029167 | 10.49057458 | 0.199680836 | 3.375035162 | 7.22 | -1.14 | -3.5 | 49.9 | 2.23 | 5 | -5 |
| CHRNA4 | 81 | Q | 32.34127778 | 16.126653 | 0.354290894 | 9.040340755 | NA | NA | NA | NA | NA | NA | NA |
| CHRNA4 | 82 | M | 35.71966667 | 11.95989319 | 0.239987485 | 4.068143373 | 8.03 | -0.81 | -3.66 | 49.66 | 2.08 | 5 | -1 |
| CHRNA4 | 83 | M | 37.02554167 | 16.59232741 | 0.36706537 | 8.65781849 | NA | NA | NA | NA | NA | NA | NA |
| CHRNA4 | 84 | T | 38.3757381 | 13.03831454 | 0.26957096 | 5.724312638 | NA | NA | NA | NA | NA | NA | NA |
| CHRNA4 | 85 | T | 41.5637381 | 16.66265735 | 0.368994675 | 9.825583301 | NA | NA | NA | NA | NA | NA | NA |
| CHRNA4 | 86 | N | 44.22279167 | 17.82771284 | 0.400954716 | 10.98290917 | NA | NA | NA | NA | NA | NA | NA |
| CHRNA4 | 87 | V | 46.81888095 | 19.43579666 | 0.445068001 | 12.31140864 | NA | NA | NA | NA | NA | NA | NA |
| CHRNA4 | 88 | W | 49.93966667 | 21.80168239 | 0.509969463 | 13.62123874 | NA | NA | NA | NA | NA | NA | NA |
| CHRNA4 | 89 | V | 52.38330952 | 21.83447201 | 0.510868954 | 13.5506192 | NA | NA | NA | NA | NA | NA | NA |
| CHRNA4 | 90 | K | 53.7295 | 25.42240917 | 0.609293981 | 17.1245927 | NA | NA | NA | NA | NA | NA | NA |
| CHRNA4 | 91 | Q | 57.42705556 | 22.92419248 | 0.540762389 | 14.91373576 | NA | NA | NA | NA | NA | NA | NA |
| CHRNA4 | 92 | E | 59.75661111 | 27.41621055 | 0.663988349 | 19.40355106 | NA | NA | NA | NA | NA | NA | NA |
| CHRNA4 | 93 | W | 63.56502381 | 22.30498635 | 0.5237762 | 14.28218975 | NA | NA | NA | NA | NA | NA | NA |
| CHRNA4 | 94 | H | 67.35036667 | 28.3587286 | 0.689843698 | 20.45818823 | NA | NA | NA | NA | NA | NA | NA |
| CHRNA4 | 95 | D | 70.66829167 | 24.06428318 | 0.572037591 | 15.23894082 | NA | NA | NA | NA | NA | NA | NA |
| CHRNA4 | 96 | Y | 75.2285 | 26.62315657 | 0.64223313 | 16.91033827 | NA | NA | NA | NA | NA | NA | NA |
| CHRNA4 | 97 | K | 76.80038889 | 22.74927397 | 0.535963989 | 13.04382248 | NA | NA | NA | NA | NA | NA | NA |
| CHRNA4 | 98 | L | 73.24691667 | 21.88946438 | 0.512377516 | 12.44596569 | NA | NA | NA | NA | NA | NA | NA |
| CHRNA4 | 99 | R | 76.06234848 | 26.86780387 | 0.648944345 | 17.0788334 | NA | NA | NA | NA | NA | NA | NA |
| CHRNA4 | 100 | W | 76.01802381 | 25.68891907 | 0.616604936 | 16.12992852 | NA | NA | NA | NA | NA | NA | NA |

| | | | | | | | | | | | | | |
|---|---|---|---|---|---|---|---|---|---|---|---|---|---|
| CHRNA4 | 101 | D | 79.95829167 | 30.72171527 | 0.754665632 | 21.00482465 | NA | NA | NA | NA | NA | NA | NA |
| CHRNA4 | 102 | P | 77.2997381 | 32.54927302 | 0.804799571 | 22.75857318 | NA | NA | NA | NA | NA | NA | NA |
| CHRNA4 | 103 | A | 80.60636667 | 34.39531961 | 0.855440699 | 24.60156913 | NA | NA | NA | NA | NA | NA | NA |
| CHRNA4 | 104 | D | 83.21916667 | 30.87137234 | 0.758771055 | 21.16212777 | NA | NA | NA | NA | NA | NA | NA |
| CHRNA4 | 105 | Y | 80.80366667 | 26.55132217 | 0.640262554 | 16.80726902 | NA | NA | NA | NA | NA | NA | NA |
| CHRNA4 | 106 | E | 79.59383333 | 31.32228276 | 0.771140522 | 21.53790299 | NA | NA | NA | NA | NA | NA | NA |
| CHRNA4 | 107 | N | 75.55466667 | 32.57367876 | 0.805469074 | 23.72745578 | NA | NA | NA | NA | NA | NA | NA |
| CHRNA4 | 108 | V | 74.6607381 | 28.69523032 | 0.699074682 | 19.81950379 | NA | NA | NA | NA | NA | NA | NA |
| CHRNA4 | 109 | T | 71.1557381 | 28.9693022 | 0.706593078 | 21.1136193 | NA | NA | NA | NA | NA | NA | NA |
| CHRNA4 | 110 | S | 68.47083333 | 26.38286179 | 0.635641314 | 18.55668824 | NA | NA | NA | NA | NA | NA | NA |
| CHRNA4 | 111 | I | 69.88829167 | 22.06888747 | 0.517299487 | 14.24550066 | NA | NA | NA | NA | NA | NA | NA |
| CHRNA4 | 112 | R | 65.4505303 | 19.30616578 | 0.44151194 | 11.5011743 | NA | NA | NA | NA | NA | NA | NA |
| CHRNA4 | 113 | I | 67.51129167 | 16.74395467 | 0.37122484 | 9.020154059 | NA | NA | NA | NA | NA | NA | NA |
| CHRNA4 | 114 | P | 66.2107381 | 13.14222664 | 0.272421498 | 5.325770826 | NA | NA | NA | NA | NA | NA | NA |
| CHRNA4 | 115 | S | 63.53816667 | 14.43173548 | 0.307795569 | 6.469305737 | NA | NA | NA | NA | NA | NA | NA |
| CHRNA4 | 116 | E | 66.08938889 | 9.501790463 | 0.172556307 | 1.874305824 | 8.08 | -0.9 | -3.63 | 49.77 | 1.97 | 3 | -1 |
| CHRNA4 | 117 | L | 69.41779167 | 11.26314814 | 0.220874232 | 3.61003821 | 7.94 | -1.14 | -3.5 | 49.9 | 2.23 | 2 | -2 |
| CHRNA4 | 118 | I | 67.67079167 | 15.84991763 | 0.346699432 | 7.961161282 | NA | NA | NA | NA | NA | NA | NA |
| CHRNA4 | 119 | W | 66.11338095 | 18.53849392 | 0.420453008 | 10.63173557 | NA | NA | NA | NA | NA | NA | NA |
| CHRNA4 | 120 | R | 62.11007576 | 14.22155863 | 0.302029955 | 6.201999477 | NA | NA | NA | NA | NA | NA | NA |
| CHRNA4 | 121 | P | 61.01688095 | 21.18609039 | 0.493082417 | 13.12429024 | NA | NA | NA | NA | NA | NA | NA |
| CHRNA4 | 122 | D | 58.74566667 | 20.41742672 | 0.471996277 | 12.40342328 | NA | NA | NA | NA | NA | NA | NA |
| CHRNA4 | 123 | I | 55.07829167 | 20.77111672 | 0.481698774 | 12.8202691 | NA | NA | NA | NA | NA | NA | NA |
| CHRNA4 | 124 | V | 52.34716667 | 19.95670947 | 0.459357787 | 11.66027205 | NA | NA | NA | NA | NA | NA | NA |
| CHRNA4 | 125 | L | 47.85604167 | 18.85903017 | 0.429246024 | 10.96819741 | NA | NA | NA | NA | NA | NA | NA |
| CHRNA4 | 126 | Y | 48.63408333 | 21.98029003 | 0.514869064 | 14.87841295 | NA | NA | NA | NA | NA | NA | NA |
| CHRNA4 | 127 | N | 42.36516667 | 19.36954359 | 0.443250533 | 12.46520872 | NA | NA | NA | NA | NA | NA | NA |
| CHRNA4 | 128 | N | 44.32041667 | 14.83026385 | 0.318728081 | 7.914076432 | NA | NA | NA | NA | NA | NA | NA |
| CHRNA4 | 129 | A | 41.06436667 | 13.2599382 | 0.275650586 | 6.363701911 | NA | NA | NA | NA | NA | NA | NA |
| CHRNA4 | 130 | D | 41.64291667 | 8.678873775 | 0.149981888 | 2.063022153 | 6.94 | -0.88 | -3.6 | 49.65 | 2.34 | 4 | -2 |
| CHRNA4 | 131 | G | 45.34241667 | 11.59696413 | 0.23003154 | 4.504193647 | 7.14 | -0.88 | -3.6 | 49.65 | 2.34 | 4 | -2 |
| CHRNA4 | 132 | D | 48.20666667 | 9.759795592 | 0.179633957 | 1.749352862 | 8.02 | -0.63 | -2.42 | 40.05 | 1.96 | 4 | -4 |
| CHRNA4 | 133 | F | 49.52307576 | 14.19254649 | 0.301234088 | 5.970335931 | NA | NA | NA | NA | NA | NA | NA |

| | | | | | | | | | | | | | |
|---|---|---|---|---|---|---|---|---|---|---|---|---|---|
| CHRNA4 | 134 | A | 52.65376667 | 14.71572248 | 0.315585958 | 6.623028867 | NA | NA | NA | NA | NA | NA | NA |
| CHRNA4 | 135 | V | 54.76345238 | 14.38522944 | 0.306519806 | 6.137735546 | NA | NA | NA | NA | NA | NA | NA |
| CHRNA4 | 136 | T | 54.46859524 | 9.516720969 | 0.172965884 | 1.394263636 | 8.11 | -0.92 | -1.76 | 21.3 | 1.36 | 2 | -2 |
| CHRNA4 | 137 | H | 53.41436667 | 12.38616087 | 0.251680947 | 4.21221795 | 8.26 | -0.4 | -0.75 | 13.61 | 0.86 | 1 | -1 |
| CHRNA4 | 138 | L | 58.95116667 | 10.87571483 | 0.210246082 | 2.871280704 | 8.25 | -0.5 | -3.1 | 37.54 | 1.29 | 3 | 3 |
| CHRNA4 | 139 | T | 58.69130952 | 15.49726637 | 0.337025431 | 7.191281747 | NA | NA | NA | NA | NA | NA | NA |
| CHRNA4 | 140 | K | 62.21916667 | 16.01322822 | 0.351179402 | 8.127171993 | NA | NA | NA | NA | NA | NA | NA |
| CHRNA4 | 141 | A | 63.74156667 | 18.9385666 | 0.431427884 | 10.99390777 | NA | NA | NA | NA | NA | NA | NA |
| CHRNA4 | 142 | H | 63.48736667 | 22.31904286 | 0.524161801 | 14.39312706 | NA | NA | NA | NA | NA | NA | NA |
| CHRNA4 | 143 | L | 68.65054167 | 21.94816359 | 0.513987765 | 14.09744688 | NA | NA | NA | NA | NA | NA | NA |
| CHRNA4 | 144 | F | 67.55171212 | 27.21990266 | 0.658603191 | 19.36593157 | NA | NA | NA | NA | NA | NA | NA |
| CHRNA4 | 145 | H | 72.85876667 | 29.89571223 | 0.732006548 | 21.20150102 | NA | NA | NA | NA | NA | NA | NA |
| CHRNA4 | 146 | D | 71.21716667 | 30.89032575 | 0.759290989 | 22.07189274 | NA | NA | NA | NA | NA | NA | NA |
| CHRNA4 | 147 | G | 70.35916667 | 27.77205666 | 0.673749993 | 19.1788438 | NA | NA | NA | NA | NA | NA | NA |
| CHRNA4 | 148 | R | 65.18598485 | 27.11250079 | 0.655656921 | 19.26833111 | NA | NA | NA | NA | NA | NA | NA |
| CHRNA4 | 149 | V | 64.9947381 | 23.22266337 | 0.548950104 | 15.30696281 | NA | NA | NA | NA | NA | NA | NA |
| CHRNA4 | 150 | Q | 60.82927778 | 23.18514214 | 0.547920814 | 15.21215141 | NA | NA | NA | NA | NA | NA | NA |
| CHRNA4 | 151 | W | 59.10916667 | 17.93018118 | 0.403765648 | 9.933810976 | NA | NA | NA | NA | NA | NA | NA |
| CHRNA4 | 152 | T | 56.20216667 | 21.11716506 | 0.491191643 | 12.87616057 | NA | NA | NA | NA | NA | NA | NA |
| CHRNA4 | 153 | P | 54.6177381 | 18.74704267 | 0.42617396 | 10.53575959 | NA | NA | NA | NA | NA | NA | NA |
| CHRNA4 | 154 | P | 51.76516667 | 16.33237659 | 0.359934345 | 8.125966432 | NA | NA | NA | NA | NA | NA | NA |
| CHRNA4 | 155 | A | 48.54456667 | 16.00260607 | 0.350888013 | 7.937952568 | NA | NA | NA | NA | NA | NA | NA |
| CHRNA4 | 156 | I | 45.37591667 | 12.81464432 | 0.263435193 | 5.774552783 | NA | NA | NA | NA | NA | NA | NA |
| CHRNA4 | 157 | Y | 46.263 | 15.43296543 | 0.335261514 | 8.501177841 | NA | NA | NA | NA | NA | NA | NA |
| CHRNA4 | 158 | K | 41.39694444 | 8.747842363 | 0.151873848 | 1.847005773 | 6.91 | -0.79 | -3.66 | 49.62 | 2.16 | 5 | -1 |
| CHRNA4 | 159 | S | 39.91583333 | 17.14205664 | 0.382145654 | 10.24711763 | NA | NA | NA | NA | NA | NA | NA |
| CHRNA4 | 160 | S | 38.0885 | 18.96101316 | 0.432043642 | 11.66547861 | NA | NA | NA | NA | NA | NA | NA |
| CHRNA4 | 161 | C | 35.012 | 20.88921162 | 0.484938377 | 12.80830822 | NA | NA | NA | NA | NA | NA | NA |
| CHRNA4 | 162 | S | 32.46183333 | 23.69799538 | 0.561989509 | 16.63581864 | NA | NA | NA | NA | NA | NA | NA |
| CHRNA4 | 163 | I | 29.52854167 | 20.45127727 | 0.472924873 | 13.24032406 | NA | NA | NA | NA | NA | NA | NA |
| CHRNA4 | 164 | D | 26.51741667 | 24.55164557 | 0.585407016 | 17.29007831 | NA | NA | NA | NA | NA | NA | NA |
| CHRNA4 | 165 | V | 23.39302381 | 21.86985041 | 0.511839462 | 16.62794231 | NA | NA | NA | NA | NA | NA | NA |
| CHRNA4 | 166 | T | 20.94716667 | 25.76661578 | 0.618736328 | 20.31855049 | NA | NA | NA | NA | NA | NA | NA |

| | | | | | | | | | | | | | |
|---|---|---|---|---|---|---|---|---|---|---|---|---|---|
| CHRNA4 | 167 | F | 19.52098485 | 25.79765179 | 0.619587714 | 20.30568122 | NA | NA | NA | NA | NA | NA | NA |
| CHRNA4 | 168 | F | 19.61516667 | 18.69958195 | 0.424872008 | 13.22138526 | NA | NA | NA | NA | NA | NA | NA |
| CHRNA4 | 169 | P | 19.88802381 | 22.45236315 | 0.52781907 | 16.96535664 | NA | NA | NA | NA | NA | NA | NA |
| CHRNA4 | 170 | F | 19.27734848 | 25.65704638 | 0.615730597 | 20.23203532 | NA | NA | NA | NA | NA | NA | NA |
| CHRNA4 | 171 | D | 24.84779167 | 24.41688696 | 0.58171029 | 17.16756148 | NA | NA | NA | NA | NA | NA | NA |
| CHRNA4 | 172 | Q | 27.49205556 | 27.37395302 | 0.662829132 | 20.13370681 | NA | NA | NA | NA | NA | NA | NA |
| CHRNA4 | 173 | Q | 29.98016667 | 23.5117662 | 0.556880832 | 16.30369611 | NA | NA | NA | NA | NA | NA | NA |
| CHRNA4 | 174 | N | 33.85754167 | 26.66624364 | 0.643415103 | 18.67301258 | NA | NA | NA | NA | NA | NA | NA |
| CHRNA4 | 175 | C | 36.21666667 | 23.06144217 | 0.544527451 | 15.06519116 | NA | NA | NA | NA | NA | NA | NA |
| CHRNA4 | 176 | T | 39.57016667 | 23.7317398 | 0.562915193 | 16.84675032 | NA | NA | NA | NA | NA | NA | NA |
| CHRNA4 | 177 | M | 43.24241667 | 21.41506145 | 0.499363598 | 14.48089553 | NA | NA | NA | NA | NA | NA | NA |
| CHRNA4 | 178 | K | 46.30961111 | 24.58682267 | 0.586372001 | 17.49894631 | NA | NA | NA | NA | NA | NA | NA |
| CHRNA4 | 179 | F | 48.80571212 | 22.24575353 | 0.522151313 | 14.32645945 | NA | NA | NA | NA | NA | NA | NA |
| CHRNA4 | 180 | G | 52.92466667 | 24.81155757 | 0.592536975 | 16.56383483 | NA | NA | NA | NA | NA | NA | NA |
| CHRNA4 | 181 | S | 56.1795 | 25.81295665 | 0.62000756 | 17.80401608 | NA | NA | NA | NA | NA | NA | NA |
| CHRNA4 | 182 | W | 55.12509524 | 22.50374371 | 0.529228553 | 14.5612049 | NA | NA | NA | NA | NA | NA | NA |
| CHRNA4 | 183 | T | 60.44830952 | 23.16676415 | 0.547416665 | 15.15758947 | NA | NA | NA | NA | NA | NA | NA |
| CHRNA4 | 184 | Y | 62.36533333 | 23.92344175 | 0.568174 | 17.21847378 | NA | NA | NA | NA | NA | NA | NA |
| CHRNA4 | 185 | D | 61.24929167 | 30.88634917 | 0.759181903 | 22.86972618 | NA | NA | NA | NA | NA | NA | NA |
| CHRNA4 | 186 | K | 58.81316667 | 33.82392387 | 0.839766054 | 25.80633824 | NA | NA | NA | NA | NA | NA | NA |
| CHRNA4 | 187 | A | 63.56856667 | 34.49029026 | 0.858045953 | 26.42814126 | NA | NA | NA | NA | NA | NA | NA |
| CHRNA4 | 188 | K | 64.9615 | 30.28719066 | 0.742745664 | 22.27038595 | NA | NA | NA | NA | NA | NA | NA |
| CHRNA4 | 189 | I | 61.16441667 | 27.275416 | 0.660126044 | 19.22862015 | NA | NA | NA | NA | NA | NA | NA |
| CHRNA4 | 190 | D | 59.05304167 | 32.6222981 | 0.80680281 | 24.56996504 | NA | NA | NA | NA | NA | NA | NA |
| CHRNA4 | 191 | L | 55.78366667 | 29.43676738 | 0.719416678 | 21.36951162 | NA | NA | NA | NA | NA | NA | NA |
| CHRNA4 | 192 | V | 55.97230952 | 33.47507863 | 0.83019646 | 25.533697 | NA | NA | NA | NA | NA | NA | NA |
| CHRNA4 | 193 | N | 51.92979167 | 33.9402636 | 0.842957509 | 25.6461567 | NA | NA | NA | NA | NA | NA | NA |
| CHRNA4 | 194 | M | 53.92679167 | 32.68897748 | 0.808631972 | 24.54957828 | NA | NA | NA | NA | NA | NA | NA |
| CHRNA4 | 195 | H | 50.39006667 | 33.36193425 | 0.82709266 | 25.47497971 | NA | NA | NA | NA | NA | NA | NA |
| CHRNA4 | 196 | S | 46.80166667 | 35.36817024 | 0.882128137 | 28.26746281 | NA | NA | NA | NA | NA | NA | NA |
| CHRNA4 | 197 | R | 43.11462121 | 33.23088699 | 0.823497745 | 26.30098906 | NA | NA | NA | NA | NA | NA | NA |
| CHRNA4 | 198 | V | 44.15645238 | 28.52892671 | 0.694512607 | 21.58987345 | NA | NA | NA | NA | NA | NA | NA |
| CHRNA4 | 199 | D | 46.01241667 | 29.3111618 | 0.71597104 | 22.37264349 | NA | NA | NA | NA | NA | NA | NA |

| | | | | | | | | | | | | | |
|---|---|---|---|---|---|---|---|---|---|---|---|---|---|
| CHRNA4 | 200 | Q | 41.38683333 | 29.8063148 | 0.729554179 | 23.00280901 | NA | NA | NA | NA | NA | NA | NA |
| CHRNA4 | 201 | L | 45.22691667 | 30.96262552 | 0.761274331 | 24.02320034 | NA | NA | NA | NA | NA | NA | NA |
| CHRNA4 | 202 | D | 45.11979167 | 26.59631963 | 0.641496934 | 19.6762684 | NA | NA | NA | NA | NA | NA | NA |
| CHRNA4 | 203 | F | 39.24434848 | 25.2587025 | 0.604803146 | 18.56878329 | NA | NA | NA | NA | NA | NA | NA |
| CHRNA4 | 204 | W | 38.20388095 | 22.47292978 | 0.528383259 | 15.19853836 | NA | NA | NA | NA | NA | NA | NA |
| CHRNA4 | 205 | E | 35.11105556 | 23.9479391 | 0.568846016 | 15.95892443 | NA | NA | NA | NA | NA | NA | NA |
| CHRNA4 | 206 | S | 32.97216667 | 21.37156868 | 0.498170495 | 14.28719913 | NA | NA | NA | NA | NA | NA | NA |
| CHRNA4 | 207 | G | 29.64566667 | 23.24621599 | 0.549596204 | 16.04762407 | NA | NA | NA | NA | NA | NA | NA |
| CHRNA4 | 208 | E | 27.63461111 | 18.09644866 | 0.408326732 | 10.97576978 | NA | NA | NA | NA | NA | NA | NA |
| CHRNA4 | 209 | W | 30.12195238 | 19.71261269 | 0.452661674 | 12.50756593 | NA | NA | NA | NA | NA | NA | NA |
| CHRNA4 | 210 | V | 31.40088095 | 25.88688975 | 0.622035708 | 18.69842246 | NA | NA | NA | NA | NA | NA | NA |
| CHRNA4 | 211 | I | 35.53241667 | 26.30065395 | 0.633386172 | 18.32217713 | NA | NA | NA | NA | NA | NA | NA |
| CHRNA4 | 212 | V | 33.1787381 | 30.29927408 | 0.743077139 | 22.28690999 | NA | NA | NA | NA | NA | NA | NA |
| CHRNA4 | 213 | D | 36.19579167 | 30.59545166 | 0.751201943 | 22.59816606 | NA | NA | NA | NA | NA | NA | NA |
| CHRNA4 | 214 | A | 40.27816667 | 30.04509228 | 0.736104372 | 22.78354265 | NA | NA | NA | NA | NA | NA | NA |
| CHRNA4 | 215 | V | 41.50788095 | 31.6283437 | 0.779536449 | 24.73125532 | NA | NA | NA | NA | NA | NA | NA |
| CHRNA4 | 216 | G | 45.19841667 | 32.4836353 | 0.802998983 | 25.5562539 | NA | NA | NA | NA | NA | NA | NA |
| CHRNA4 | 217 | T | 45.71345238 | 33.10525052 | 0.820051259 | 25.99587806 | NA | NA | NA | NA | NA | NA | NA |
| CHRNA4 | 218 | Y | 51.57875 | 34.86643575 | 0.868364453 | 26.97332808 | NA | NA | NA | NA | NA | NA | NA |
| CHRNA4 | 219 | N | 48.38254167 | 34.37424638 | 0.854862614 | 27.26913372 | NA | NA | NA | NA | NA | NA | NA |
| CHRNA4 | 220 | T | 51.30802381 | 35.96592036 | 0.89852574 | 27.96035915 | NA | NA | NA | NA | NA | NA | NA |
| CHRNA4 | 221 | R | 48.66343939 | 31.130281 | 0.765873491 | 23.3940696 | NA | NA | NA | NA | NA | NA | NA |
| CHRNA4 | 222 | K | 52.54538889 | 35.34432459 | 0.881473998 | 27.33039962 | NA | NA | NA | NA | NA | NA | NA |
| CHRNA4 | 223 | Y | 50.98183333 | 29.58227863 | 0.723408373 | 21.91067335 | NA | NA | NA | NA | NA | NA | NA |
| CHRNA4 | 224 | E | 51.18461111 | 34.00149128 | 0.844637119 | 25.94629368 | NA | NA | NA | NA | NA | NA | NA |
| CHRNA4 | 225 | C | 55.41683333 | 31.02930855 | 0.763103594 | 23.02314002 | NA | NA | NA | NA | NA | NA | NA |
| CHRNA4 | 226 | C | 57.0535 | 31.44545298 | 0.774519353 | 23.50544595 | NA | NA | NA | NA | NA | NA | NA |
| CHRNA4 | 227 | A | 58.38236667 | 36.09269771 | 0.902003523 | 28.14239222 | NA | NA | NA | NA | NA | NA | NA |
| CHRNA4 | 228 | E | 59.81616667 | 35.00931445 | 0.872283931 | 27.01901367 | NA | NA | NA | NA | NA | NA | NA |
| CHRNA4 | 229 | I | 56.13904167 | 34.69065582 | 0.863542422 | 26.68719635 | NA | NA | NA | NA | NA | NA | NA |
| CHRNA4 | 230 | Y | 55.99641667 | 30.0865801 | 0.737242474 | 22.05360256 | NA | NA | NA | NA | NA | NA | NA |
| CHRNA4 | 231 | P | 54.60259524 | 30.66380598 | 0.753077052 | 22.58278465 | NA | NA | NA | NA | NA | NA | NA |
| CHRNA4 | 232 | D | 50.19604167 | 30.14013958 | 0.738711729 | 22.12692374 | NA | NA | NA | NA | NA | NA | NA |

| | | | | | | | | | | | | | |
|---|---|---|---|---|---|---|---|---|---|---|---|---|---|
| CHRNA4 | 233 | I | 49.40966667 | 28.37271178 | 0.690227287 | 20.49708073 | NA | NA | NA | NA | NA | NA | NA |
| CHRNA4 | 234 | T | 44.73745238 | 29.69922576 | 0.72661649 | 22.78047155 | NA | NA | NA | NA | NA | NA | NA |
| CHRNA4 | 235 | Y | 45.34658333 | 27.26197973 | 0.659757458 | 20.40050718 | NA | NA | NA | NA | NA | NA | NA |
| CHRNA4 | 236 | A | 39.71136667 | 29.05891426 | 0.709051334 | 22.24504272 | NA | NA | NA | NA | NA | NA | NA |
| CHRNA4 | 237 | F | 38.81207576 | 25.0033173 | 0.597797367 | 17.810954 | NA | NA | NA | NA | NA | NA | NA |
| CHRNA4 | 238 | V | 33.92002381 | 29.27270992 | 0.71491622 | 21.2591144 | NA | NA | NA | NA | NA | NA | NA |
| CHRNA4 | 239 | I | 32.20629167 | 23.99686723 | 0.570188223 | 16.89927402 | NA | NA | NA | NA | NA | NA | NA |
| CHRNA4 | 240 | R | 27.94516667 | 25.75130533 | 0.618316328 | 18.55302128 | NA | NA | NA | NA | NA | NA | NA |
| CHRNA4 | 241 | R | 25.37980303 | 20.69011527 | 0.479476726 | 13.43705741 | NA | NA | NA | NA | NA | NA | NA |
| CHRNA4 | 242 | L | 25.25766667 | 23.73315112 | 0.562953908 | 16.80736194 | NA | NA | NA | NA | NA | NA | NA |
| CHRNA4 | 243 | P | 20.94488095 | 23.60468473 | 0.559429792 | 18.21913457 | NA | NA | NA | NA | NA | NA | NA |
| CHRNA4 | 244 | L | 19.15966667 | 22.70977765 | 0.534880518 | 17.24383271 | NA | NA | NA | NA | NA | NA | NA |
| CHRNA4 | 245 | F | 20.68034848 | 16.68075193 | 0.369491049 | 11.25669075 | NA | NA | NA | NA | NA | NA | NA |
| CHRNA4 | 246 | Y | 20.03216667 | 16.22287801 | 0.356930558 | 10.88382255 | NA | NA | NA | NA | NA | NA | NA |
| CHRNA4 | 247 | T | 15.30859524 | 21.46471116 | 0.500725599 | 14.89483686 | NA | NA | NA | NA | NA | NA | NA |
| CHRNA4 | 248 | I | 14.50116667 | 19.43147565 | 0.444949466 | 12.82837829 | NA | NA | NA | NA | NA | NA | NA |
| CHRNA4 | 249 | N | 15.19166667 | 15.46095844 | 0.336029424 | 8.845761398 | NA | NA | NA | NA | NA | NA | NA |
| CHRNA4 | 250 | L | 14.57754167 | 15.32375375 | 0.332265597 | 8.78379689 | NA | NA | NA | NA | NA | NA | NA |
| CHRNA4 | 251 | I | 12.42691667 | 18.74507605 | 0.426120011 | 13.88617898 | NA | NA | NA | NA | NA | NA | NA |
| CHRNA4 | 252 | I | 9.503041667 | 18.8874512 | 0.430025676 | 13.78876668 | NA | NA | NA | NA | NA | NA | NA |
| CHRNA4 | 253 | P | 8.797738095 | 15.3650603 | 0.333398727 | 10.29703432 | NA | NA | NA | NA | NA | NA | NA |
| CHRNA4 | 254 | C | 7.987666667 | 16.41549932 | 0.362214585 | 11.95314251 | NA | NA | NA | NA | NA | NA | NA |
| CHRNA4 | 255 | L | 6.429041667 | 18.82504191 | 0.428313651 | 14.99764709 | NA | NA | NA | NA | NA | NA | NA |
| CHRNA4 | 256 | L | 3.686041667 | 17.26223395 | 0.385442383 | 13.61572512 | NA | NA | NA | NA | NA | NA | NA |
| CHRNA4 | 257 | I | 3.400791667 | 13.86861687 | 0.292347984 | 10.21948783 | NA | NA | NA | NA | NA | NA | NA |
| CHRNA4 | 258 | S | 2.471166667 | 17.28206591 | 0.385986417 | 13.64973505 | NA | NA | NA | NA | NA | NA | NA |
| CHRNA4 | 259 | C | 0.412333333 | 18.91355918 | 0.430741875 | 14.71846168 | NA | NA | NA | NA | NA | NA | NA |
| CHRNA4 | 260 | L | -0.843791667 | 14.07789604 | 0.298088973 | 10.1282526 | NA | NA | NA | NA | NA | NA | NA |
| CHRNA4 | 261 | T | -0.998595238 | 17.28197816 | 0.38598401 | 13.33412889 | NA | NA | NA | NA | NA | NA | NA |
| CHRNA4 | 262 | V | -3.940595238 | 19.35441216 | 0.442835444 | 16.63169178 | NA | NA | NA | NA | NA | NA | NA |
| CHRNA4 | 263 | L | -5.414666667 | 15.80143088 | 0.345369334 | 13.13720972 | NA | NA | NA | NA | NA | NA | NA |
| CHRNA4 | 264 | V | -6.881595238 | 13.39508358 | 0.279357922 | 10.75567486 | NA | NA | NA | NA | NA | NA | NA |
| CHRNA4 | 265 | F | -8.346166667 | 17.25036771 | 0.385116866 | 14.71741025 | NA | NA | NA | NA | NA | NA | NA |

| | | | | | | | | | | | | | |
|---|---|---|---|---|---|---|---|---|---|---|---|---|---|
| CHRNA4 | 266 | Y | -12.28616667 | 17.18313129 | 0.383272423 | 13.71279449 | NA | NA | NA | NA | NA | NA | NA |
| CHRNA4 | 267 | L | -11.59416667 | 11.65761862 | 0.231695427 | 8.555819101 | NA | NA | NA | NA | NA | NA | NA |
| CHRNA4 | 268 | P | -15.05788095 | 11.51134215 | 0.227682741 | 7.852311227 | NA | NA | NA | NA | NA | NA | NA |
| CHRNA4 | 269 | S | -16.13283333 | 9.295198008 | 0.166889021 | 5.350454372 | NA | NA | NA | NA | NA | NA | NA |
| CHRNA4 | 270 | E | -18.7755 | 6.432277124 | 0.088352788 | 1.631472954 | 4.88 | -0.8 | -0.4 | 3.38 | 0.81 | 0 | 0 |
| CHRNA4 | 271 | C | -15.103 | 5.119958301 | 0.05235299 | 1.302912913 | 3.95 | -0.87 | -1.02 | 12.68 | 1.09 | 1 | -1 |
| CHRNA4 | 272 | G | -12.60341667 | 5.178814343 | 0.053967541 | 2.375454178 | 3.48 | -0.94 | -1.64 | 21.99 | 1.38 | 2 | -2 |
| CHRNA4 | 273 | E | -9.481388889 | 3.253006148 | 0.001138376 | 1.598022635 | 1.83 | -1.14 | -3.5 | 49.9 | 2.23 | 4 | -4 |
| CHRNA4 | 274 | K | -9.668833333 | 9.095003793 | 0.161397252 | 6.638711076 | NA | NA | NA | NA | NA | NA | NA |
| CHRNA4 | 275 | I | -6.679291667 | 6.793394512 | 0.098259035 | 4.391119749 | 2.55 | -0.92 | -1.82 | 20.96 | 1.35 | 2 | -2 |
| CHRNA4 | 276 | T | -4.756309524 | 4.456221942 | 0.034145238 | 1.849123116 | 2.81 | -0.77 | -0.7 | 1.66 | 0.77 | 0 | 0 |
| CHRNA4 | 277 | L | -3.631041667 | 8.240854385 | 0.13796605 | 5.404487807 | NA | NA | NA | NA | NA | NA | NA |
| CHRNA4 | 278 | C | -2.648166667 | 10.57938188 | 0.202117016 | 7.741246272 | NA | NA | NA | NA | NA | NA | NA |
| CHRNA4 | 279 | I | -0.706041667 | 7.777954037 | 0.125267673 | 3.821044772 | 3.96 | -0.85 | -0.74 | 1.66 | 0.91 | 0 | 0 |
| CHRNA4 | 280 | S | 0.257333333 | 6.195722637 | 0.081863577 | 2.110798492 | 4.34 | -0.97 | -0.8 | 1.67 | 1.11 | 0 | 0 |
| CHRNA4 | 281 | V | 0.587452381 | 10.15503402 | 0.190476218 | 5.60272596 | NA | NA | NA | NA | NA | NA | NA |
| CHRNA4 | 282 | L | 2.048041667 | 10.98770886 | 0.213318325 | 7.140513251 | NA | NA | NA | NA | NA | NA | NA |
| CHRNA4 | 283 | L | 3.666541667 | 6.596196328 | 0.092849454 | 3.020934007 | 3.71 | 1.14 | 3.8 | 0.13 | -1.79 | 0 | 0 |
| CHRNA4 | 284 | S | 5.394666667 | 8.735543372 | 0.15153646 | 5.096829805 | NA | NA | NA | NA | NA | NA | NA |
| CHRNA4 | 285 | L | 6.461541667 | 11.26839336 | 0.22101812 | 7.47141897 | NA | NA | NA | NA | NA | NA | NA |
| CHRNA4 | 286 | T | 7.89802381 | 10.87180693 | 0.21013888 | 6.449711932 | NA | NA | NA | NA | NA | NA | NA |
| CHRNA4 | 287 | V | 9.947738095 | 6.706608905 | 0.095878314 | 1.594839683 | 5.12 | 1.13 | 4.1 | 0.13 | -1.42 | 0 | 0 |
| CHRNA4 | 288 | F | 11.40471212 | 10.35703901 | 0.19601766 | 5.238863677 | NA | NA | NA | NA | NA | NA | NA |
| CHRNA4 | 289 | L | 12.27179167 | 12.8721915 | 0.265013839 | 7.754730078 | NA | NA | NA | NA | NA | NA | NA |
| CHRNA4 | 290 | L | 14.19166667 | 8.872079181 | 0.155281938 | 2.282685097 | 6.65 | 0.21 | -0.74 | 29.75 | 0.26 | 3 | 3 |
| CHRNA4 | 291 | L | 15.96841667 | 8.117360532 | 0.134578342 | 1.583764436 | 6.65 | 0.21 | -0.74 | 29.75 | 0.26 | 3 | 3 |
| CHRNA4 | 292 | I | 16.73254167 | 12.33935707 | 0.250397016 | 6.186664805 | NA | NA | NA | NA | NA | NA | NA |
| CHRNA4 | 293 | T | 18.94188095 | 12.75317968 | 0.261749082 | 6.882962807 | NA | NA | NA | NA | NA | NA | NA |
| CHRNA4 | 294 | E | 21.48772222 | 6.91594238 | 0.101620793 | 1.51858101 | 5.43 | -0.65 | -3.77 | 49.63 | 1.72 | 3 | 1 |
| CHRNA4 | 295 | I | 22.15941667 | 12.9901642 | 0.26825009 | 7.566845124 | NA | NA | NA | NA | NA | NA | NA |
| CHRNA4 | 296 | I | 22.03416667 | 15.73004542 | 0.343411073 | 10.30512317 | NA | NA | NA | NA | NA | NA | NA |
| CHRNA4 | 297 | P | 24.99488095 | 17.74256363 | 0.398618885 | 10.43575549 | NA | NA | NA | NA | NA | NA | NA |
| CHRNA4 | 298 | S | 24.96133333 | 17.90009209 | 0.402940238 | 12.64082193 | NA | NA | NA | NA | NA | NA | NA |

| | | | | | | | | | | | | | |
|---|---|---|---|---|---|---|---|---|---|---|---|---|---|
| CHRNA4 | 299 | T | 25.01902381 | 20.9153874 | 0.485656437 | 13.59656083 | NA | NA | NA | NA | NA | NA | NA |
| CHRNA4 | 300 | S | 23.2245 | 24.18088282 | 0.575236176 | 18.83845652 | NA | NA | NA | NA | NA | NA | NA |
| CHRNA4 | 301 | L | 24.89754167 | 26.70146768 | 0.644381376 | 19.38092681 | NA | NA | NA | NA | NA | NA | NA |
| CHRNA4 | 302 | V | 22.18159524 | 24.97788346 | 0.597099661 | 19.52936111 | NA | NA | NA | NA | NA | NA | NA |
| CHRNA4 | 303 | I | 17.96241667 | 24.664034 | 0.588490078 | 18.79475168 | NA | NA | NA | NA | NA | NA | NA |
| CHRNA4 | 304 | P | 17.91659524 | 21.38879831 | 0.498643142 | 15.53557628 | NA | NA | NA | NA | NA | NA | NA |
| CHRNA4 | 305 | L | 14.39341667 | 23.94018456 | 0.568633292 | 17.32488601 | NA | NA | NA | NA | NA | NA | NA |
| CHRNA4 | 306 | I | 13.83204167 | 19.77871384 | 0.454474975 | 13.16590995 | NA | NA | NA | NA | NA | NA | NA |
| CHRNA4 | 307 | G | 13.13191667 | 20.46771958 | 0.473375922 | 13.864041 | NA | NA | NA | NA | NA | NA | NA |
| CHRNA4 | 308 | E | 11.70705556 | 22.76145977 | 0.536298272 | 17.87715201 | NA | NA | NA | NA | NA | NA | NA |
| CHRNA4 | 309 | Y | 7.793833333 | 21.63023398 | 0.505266255 | 16.60292024 | NA | NA | NA | NA | NA | NA | NA |
| CHRNA4 | 310 | L | 9.328916667 | 17.08927386 | 0.380697706 | 12.0401169 | NA | NA | NA | NA | NA | NA | NA |
| CHRNA4 | 311 | L | 8.483291667 | 20.24823958 | 0.467355101 | 15.17916793 | NA | NA | NA | NA | NA | NA | NA |
| CHRNA4 | 312 | F | 4.901530303 | 22.11713483 | 0.518623018 | 18.38881438 | NA | NA | NA | NA | NA | NA | NA |
| CHRNA4 | 313 | T | 3.910738095 | 18.86801659 | 0.429492541 | 15.23849744 | NA | NA | NA | NA | NA | NA | NA |
| CHRNA4 | 314 | M | 4.983541667 | 15.64684345 | 0.34112866 | 12.0555293 | NA | NA | NA | NA | NA | NA | NA |
| CHRNA4 | 315 | I | 2.824291667 | 20.32812601 | 0.469546562 | 16.69944598 | NA | NA | NA | NA | NA | NA | NA |
| CHRNA4 | 316 | F | 0.224348485 | 20.37116121 | 0.470727112 | 16.41705891 | NA | NA | NA | NA | NA | NA | NA |
| CHRNA4 | 317 | V | -0.040452381 | 15.84789011 | 0.346643813 | 11.8897508 | NA | NA | NA | NA | NA | NA | NA |
| CHRNA4 | 318 | T | -0.16702381 | 17.97905365 | 0.405106328 | 14.02045792 | NA | NA | NA | NA | NA | NA | NA |
| CHRNA4 | 319 | L | -2.432291667 | 20.07756823 | 0.46267321 | 16.12217922 | NA | NA | NA | NA | NA | NA | NA |
| CHRNA4 | 320 | S | -3.763166667 | 17.41031203 | 0.389504491 | 14.70310862 | NA | NA | NA | NA | NA | NA | NA |
| CHRNA4 | 321 | I | -3.679416667 | 14.57633514 | 0.311762256 | 11.94689911 | NA | NA | NA | NA | NA | NA | NA |
| CHRNA4 | 322 | V | -5.503309524 | 17.74847444 | 0.398781032 | 15.3586147 | NA | NA | NA | NA | NA | NA | NA |
| CHRNA4 | 323 | I | -8.321291667 | 18.66093315 | 0.423811786 | 16.17162305 | NA | NA | NA | NA | NA | NA | NA |
| CHRNA4 | 324 | T | -9.07602381 | 14.71366498 | 0.315529517 | 13.04486904 | NA | NA | NA | NA | NA | NA | NA |
| CHRNA4 | 325 | V | -9.510738095 | 14.46643996 | 0.308747589 | 12.66088519 | NA | NA | NA | NA | NA | NA | NA |
| CHRNA4 | 326 | F | -12.47625758 | 17.37253721 | 0.388468244 | 15.45977816 | NA | NA | NA | NA | NA | NA | NA |
| CHRNA4 | 327 | V | -13.91616667 | 15.8313411 | 0.346189837 | 12.18562133 | NA | NA | NA | NA | NA | NA | NA |
| CHRNA4 | 328 | L | -13.76779167 | 11.34935945 | 0.223239198 | 7.769520369 | NA | NA | NA | NA | NA | NA | NA |
| CHRNA4 | 329 | N | -15.54179167 | 15.00569629 | 0.323540579 | 11.07330181 | NA | NA | NA | NA | NA | NA | NA |
| CHRNA4 | 330 | V | -17.87545238 | 16.65157437 | 0.368690644 | 11.79610117 | NA | NA | NA | NA | NA | NA | NA |
| CHRNA4 | 331 | H | -19.66386667 | 11.38209506 | 0.224137208 | 6.3706395 | NA | NA | NA | NA | NA | NA | NA |

| | | | | | | | | | | | | | |
|---|---|---|---|---|---|---|---|---|---|---|---|---|---|
| CHRNA4 | 332 | H | -19.57396667 | 7.955556612 | 0.130139703 | 2.805889543 | 5.22 | -0.8 | -0.4 | 3.38 | 0.81 | 0 | 0 |
| CHRNA4 | 333 | R | -21.30016667 | 13.2328135 | 0.274906495 | 5.974461668 | NA | NA | NA | NA | NA | NA | NA |
| CHRNA4 | 334 | S | -25.304 | 13.82868765 | 0.291252637 | 6.486726904 | NA | NA | NA | NA | NA | NA | NA |
| CHRNA4 | 335 | P | -28.67345238 | 13.70999376 | 0.287996602 | 6.030108644 | NA | NA | NA | NA | NA | NA | NA |
| CHRNA4 | 336 | R | -31.35207576 | 15.15659609 | 0.327680094 | 7.195314948 | NA | NA | NA | NA | NA | NA | NA |
| CHRNA4 | 337 | T | -26.01916667 | 17.62134598 | 0.395293618 | 9.749091867 | NA | NA | NA | NA | NA | NA | NA |
| CHRNA4 | 338 | H | -23.37176667 | 19.54587141 | 0.448087593 | 12.21860071 | NA | NA | NA | NA | NA | NA | NA |
| CHRNA4 | 339 | T | -28.05859524 | 21.02418574 | 0.488641015 | 13.34553738 | NA | NA | NA | NA | NA | NA | NA |
| CHRNA4 | 340 | M | -25.39329167 | 23.44409668 | 0.555024508 | 15.76541275 | NA | NA | NA | NA | NA | NA | NA |
| CHRNA4 | 341 | P | -28.29488095 | 23.23254254 | 0.549221111 | 15.59212123 | NA | NA | NA | NA | NA | NA | NA |
| CHRNA4 | 342 | T | -28.54916667 | 26.62289652 | 0.642225996 | 18.97554925 | NA | NA | NA | NA | NA | NA | NA |
| CHRNA4 | 343 | W | -22.95266667 | 28.52316357 | 0.694354511 | 20.89954383 | NA | NA | NA | NA | NA | NA | NA |
| CHRNA4 | 344 | V | -24.00030952 | 24.3215886 | 0.579096046 | 16.70009355 | NA | NA | NA | NA | NA | NA | NA |
| CHRNA4 | 345 | R | -29.16398485 | 21.53490019 | 0.502651038 | 14.22157816 | NA | NA | NA | NA | NA | NA | NA |
| CHRNA4 | 346 | R | -27.62489394 | 27.96896485 | 0.679151619 | 20.67418662 | NA | NA | NA | NA | NA | NA | NA |
| CHRNA4 | 347 | V | -22.83616667 | 28.40882912 | 0.691218065 | 21.12458046 | NA | NA | NA | NA | NA | NA | NA |
| CHRNA4 | 348 | F | -21.07262121 | 24.55582483 | 0.585521662 | 17.27443759 | NA | NA | NA | NA | NA | NA | NA |
| CHRNA4 | 349 | L | -26.30166667 | 23.11594337 | 0.546022539 | 15.86821304 | NA | NA | NA | NA | NA | NA | NA |
| CHRNA4 | 350 | D | -25.53291667 | 28.16055752 | 0.684407428 | 21.26595995 | NA | NA | NA | NA | NA | NA | NA |
| CHRNA4 | 351 | I | -20.44254167 | 29.02981788 | 0.708253156 | 21.76721694 | NA | NA | NA | NA | NA | NA | NA |
| CHRNA4 | 352 | V | -22.50945238 | 25.09107433 | 0.600204736 | 18.18620619 | NA | NA | NA | NA | NA | NA | NA |
| CHRNA4 | 353 | P | -23.76302381 | 29.01293522 | 0.707790028 | 22.05439562 | NA | NA | NA | NA | NA | NA | NA |
| CHRNA4 | 354 | R | -18.65507576 | 31.4721287 | 0.775251127 | 24.53215929 | NA | NA | NA | NA | NA | NA | NA |
| CHRNA4 | 355 | L | -17.76004167 | 27.10587545 | 0.655475173 | 20.288164 | NA | NA | NA | NA | NA | NA | NA |
| CHRNA4 | 356 | L | -19.76416667 | 28.37056364 | 0.690168359 | 21.4089412 | NA | NA | NA | NA | NA | NA | NA |
| CHRNA4 | 357 | L | -21.90541667 | 23.21124751 | 0.548636942 | 16.27099914 | NA | NA | NA | NA | NA | NA | NA |
| CHRNA4 | 358 | M | -25.62254167 | 22.99521004 | 0.542710558 | 16.47047368 | NA | NA | NA | NA | NA | NA | NA |
| CHRNA4 | 359 | K | -28.54327778 | 21.72879095 | 0.50796989 | 15.28484992 | NA | NA | NA | NA | NA | NA | NA |
| CHRNA4 | 360 | R | -30.34425758 | 18.01248223 | 0.406023347 | 11.48568703 | NA | NA | NA | NA | NA | NA | NA |
| CHRNA4 | 586 | S | -32.84166667 | 14.14769384 | 0.30000368 | 7.647612272 | NA | NA | NA | NA | NA | NA | NA |
| CHRNA4 | 587 | V | -30.35916667 | 15.22973197 | 0.329686372 | 7.921803675 | NA | NA | NA | NA | NA | NA | NA |
| CHRNA4 | 588 | K | -27.36527778 | 9.502026836 | 0.172562791 | 2.324579566 | 7.35 | -0.4 | -3.8 | 50.46 | 1.4 | 4 | 2 |
| CHRNA4 | 589 | E | -28.28983333 | 10.24054867 | 0.192822074 | 3.290920317 | 6.98 | -0.41 | -4.1 | 50.33 | 1.52 | 3 | 3 |

| | | | | | | | | | | | | | |
|---|---|---|---|---|---|---|---|---|---|---|---|---|---|
| CHRNA4 | 590 | D | -27.51029167 | 15.87956205 | 0.347512644 | 8.601747462 | NA | NA | NA | NA | NA | NA | NA |
| CHRNA4 | 591 | W | -23.56245238 | 15.98960978 | 0.350531496 | 8.722085533 | NA | NA | NA | NA | NA | NA | NA |
| CHRNA4 | 592 | K | -23.42172222 | 10.97729789 | 0.213032729 | 3.784584666 | 7.35 | -0.4 | -3.8 | 50.46 | 1.4 | 4 | 2 |
| CHRNA4 | 593 | Y | -24.09816667 | 16.41786228 | 0.362279406 | 9.138129688 | NA | NA | NA | NA | NA | NA | NA |
| CHRNA4 | 594 | V | -21.52588095 | 19.01528914 | 0.433532552 | 11.73582785 | NA | NA | NA | NA | NA | NA | NA |
| CHRNA4 | 595 | A | -18.67456667 | 17.24032671 | 0.384841419 | 10.00375755 | NA | NA | NA | NA | NA | NA | NA |
| CHRNA4 | 596 | M | -19.03529167 | 16.36324946 | 0.360781256 | 9.259586171 | NA | NA | NA | NA | NA | NA | NA |
| CHRNA4 | 597 | V | -18.90259524 | 21.00177678 | 0.488026288 | 13.68670798 | NA | NA | NA | NA | NA | NA | NA |
| CHRNA4 | 598 | I | -15.50291667 | 21.17219748 | 0.492701303 | 17.22896036 | NA | NA | NA | NA | NA | NA | NA |
| CHRNA4 | 599 | D | -13.36416667 | 17.38656749 | 0.388853126 | 13.91047858 | NA | NA | NA | NA | NA | NA | NA |
| CHRNA4 | 600 | R | -14.8515303 | 22.1557231 | 0.51968158 | 18.47903686 | NA | NA | NA | NA | NA | NA | NA |
| CHRNA4 | 601 | I | -13.57854167 | 24.89450907 | 0.594812518 | 21.42343311 | NA | NA | NA | NA | NA | NA | NA |
| CHRNA4 | 602 | F | -8.808075758 | 22.58794293 | 0.531538323 | 20.98368905 | NA | NA | NA | NA | NA | NA | NA |
| CHRNA4 | 603 | L | -9.230291667 | 21.74926015 | 0.508531405 | 19.22037092 | NA | NA | NA | NA | NA | NA | NA |
| CHRNA4 | 604 | W | -10.90959524 | 26.24618715 | 0.631892028 | 23.71366663 | NA | NA | NA | NA | NA | NA | NA |
| CHRNA4 | 605 | M | -6.409416667 | 27.6919801 | 0.671553316 | 25.16754722 | NA | NA | NA | NA | NA | NA | NA |
| CHRNA4 | 606 | F | -3.622075758 | 21.55935651 | 0.503321929 | 19.07543407 | NA | NA | NA | NA | NA | NA | NA |
| CHRNA4 | 607 | I | -4.157666667 | 25.49007546 | 0.611150217 | 22.68154257 | NA | NA | NA | NA | NA | NA | NA |
| CHRNA4 | 608 | I | -3.827291667 | 28.59343248 | 0.696282143 | 25.79320948 | NA | NA | NA | NA | NA | NA | NA |
| CHRNA4 | 609 | V | -0.910166667 | 28.31944309 | 0.688766009 | 24.36598638 | NA | NA | NA | NA | NA | NA | NA |
| CHRNA4 | 610 | C | 0 | 24.38646916 | 0.580875863 | 20.42980341 | NA | NA | NA | NA | NA | NA | NA |
| CHRNA4 | 611 | L | 0.097916667 | 28.20930084 | 0.685744565 | 24.25674183 | NA | NA | NA | NA | NA | NA | NA |
| CHRNA4 | 612 | L | 0.661666667 | 30.84209638 | 0.757967951 | 26.30144977 | NA | NA | NA | NA | NA | NA | NA |
| CHRNA4 | 613 | G | 3.769666667 | 29.09278692 | 0.709980536 | 25.44600731 | NA | NA | NA | NA | NA | NA | NA |
| CHRNA4 | 614 | T | 4.310595238 | 26.13091414 | 0.628729835 | 22.48910297 | NA | NA | NA | NA | NA | NA | NA |
| CHRNA4 | 615 | V | 4.711595238 | 30.70529914 | 0.754215301 | 27.05511253 | NA | NA | NA | NA | NA | NA | NA |
| CHRNA4 | 616 | G | 7.268666667 | 32.36796741 | 0.799825958 | 28.55685835 | NA | NA | NA | NA | NA | NA | NA |
| CHRNA4 | 617 | L | 9.241666667 | 27.05908894 | 0.654191716 | 21.99066975 | NA | NA | NA | NA | NA | NA | NA |
| CHRNA4 | 618 | F | 10.43407576 | 25.98788221 | 0.624806154 | 20.88875784 | NA | NA | NA | NA | NA | NA | NA |
| CHRNA4 | 619 | L | 10.40266667 | 33.38059595 | 0.827604592 | 28.28068458 | NA | NA | NA | NA | NA | NA | NA |
| CHRNA4 | 620 | P | 14.21659524 | 34.08258425 | 0.846861678 | 27.46171499 | NA | NA | NA | NA | NA | NA | NA |
| CHRNA4 | 621 | P | 15.20345238 | 32.17091968 | 0.794420504 | 26.02638732 | NA | NA | NA | NA | NA | NA | NA |
| CHRNA4 | 622 | W | 13.47602381 | 33.58827416 | 0.833301663 | 26.95559306 | NA | NA | NA | NA | NA | NA | NA |

| | | | | | | | | | | | | | |
|---|---|---|---|---|---|---|---|---|---|---|---|---|---|
| CHRNB2 | 27 | D | 80.12557143 | 25.10763091 | 0.60065892 | 15.4197354 | NA | NA | NA | NA | NA | NA | NA |
| CHRNB2 | 28 | T | 79.28071429 | 25.4899257 | 0.611146109 | 15.80393108 | NA | NA | NA | NA | NA | NA | NA |
| CHRNB2 | 29 | E | 74.72901587 | 23.11538739 | 0.546007287 | 13.41533506 | NA | NA | NA | NA | NA | NA | NA |
| CHRNB2 | 30 | E | 77.38257143 | 19.4270594 | 0.444828318 | 9.728455463 | NA | NA | NA | NA | NA | NA | NA |
| CHRNB2 | 31 | R | 81.91738961 | 21.6871849 | 0.506828544 | 12.0198796 | NA | NA | NA | NA | NA | NA | NA |
| CHRNB2 | 32 | L | 77.61682143 | 21.97017665 | 0.514591632 | 12.29148911 | NA | NA | NA | NA | NA | NA | NA |
| CHRNB2 | 33 | V | 74.954 | 18.76531857 | 0.426675308 | 9.078212334 | NA | NA | NA | NA | NA | NA | NA |
| CHRNB2 | 34 | E | 77.84390476 | 15.29117265 | 0.331371825 | 5.617516962 | NA | NA | NA | NA | NA | NA | NA |
| CHRNB2 | 35 | H | 80.64467143 | 19.79932416 | 0.455040361 | 10.13597212 | NA | NA | NA | NA | NA | NA | NA |
| CHRNB2 | 36 | L | 76.00407143 | 19.09513493 | 0.435722898 | 9.423340038 | NA | NA | NA | NA | NA | NA | NA |
| CHRNB2 | 37 | L | 74.15532143 | 14.58228782 | 0.311925552 | 4.907958719 | 9.72 | -0.53 | -2.95 | 38.25 | 0.74 | 2 | -2 |
| CHRNB2 | 38 | D | 79.12719643 | 14.66531929 | 0.314203288 | 5.038251427 | NA | NA | NA | NA | NA | NA | NA |
| CHRNB2 | 39 | P | 77.24042857 | 11.22923159 | 0.219943826 | 1.577335077 | 9.72 | -0.53 | -2.95 | 38.25 | 0.74 | 2 | -2 |
| CHRNB2 | 40 | S | 80.69707143 | 13.74498541 | 0.2889565 | 4.170721304 | 9.72 | -0.53 | -2.95 | 38.25 | 0.74 | 2 | -2 |
| CHRNB2 | 41 | R | 79.82102597 | 16.72576937 | 0.370725977 | 7.09818326 | NA | NA | NA | NA | NA | NA | NA |
| CHRNB2 | 42 | Y | 73.84015476 | 15.588898 | 0.339539088 | 5.928831482 | NA | NA | NA | NA | NA | NA | NA |
| CHRNB2 | 43 | N | 73.86432143 | 15.46577877 | 0.336161656 | 5.79361781 | NA | NA | NA | NA | NA | NA | NA |
| CHRNB2 | 44 | K | 68.87712698 | 13.0212524 | 0.269102908 | 4.015994205 | 9.69 | -0.32 | -2.77 | 34.36 | 0.24 | 1 | -1 |
| CHRNB2 | 45 | L | 69.58482143 | 14.72337461 | 0.315795873 | 5.089557312 | NA | NA | NA | NA | NA | NA | NA |
| CHRNB2 | 46 | I | 71.54907143 | 19.52993049 | 0.447650299 | 9.859689896 | NA | NA | NA | NA | NA | NA | NA |
| CHRNB2 | 47 | R | 68.85548052 | 22.90760007 | 0.540307223 | 13.23170743 | NA | NA | NA | NA | NA | NA | NA |
| CHRNB2 | 48 | P | 70.13785714 | 25.0776207 | 0.599835673 | 15.37890349 | NA | NA | NA | NA | NA | NA | NA |
| CHRNB2 | 49 | A | 72.81937143 | 27.75662721 | 0.673326729 | 18.07739138 | NA | NA | NA | NA | NA | NA | NA |
| CHRNB2 | 50 | T | 74.44371429 | 30.93792904 | 0.760596852 | 21.26680861 | NA | NA | NA | NA | NA | NA | NA |
| CHRNB2 | 51 | N | 75.81619643 | 33.25157623 | 0.824065296 | 23.59166538 | NA | NA | NA | NA | NA | NA | NA |
| CHRNB2 | 52 | G | 73.35732143 | 33.56533176 | 0.832672302 | 23.90296371 | NA | NA | NA | NA | NA | NA | NA |
| CHRNB2 | 53 | S | 70.1897381 | 32.58077362 | 0.805663702 | 22.90097831 | NA | NA | NA | NA | NA | NA | NA |
| CHRNB2 | 54 | E | 70.78168254 | 32.72069009 | 0.809501919 | 23.05284884 | NA | NA | NA | NA | NA | NA | NA |
| CHRNB2 | 55 | L | 65.43657143 | 33.0136646 | 0.817538856 | 25.19592678 | NA | NA | NA | NA | NA | NA | NA |
| CHRNB2 | 56 | V | 65.60328571 | 28.98319027 | 0.706974058 | 21.1547055 | NA | NA | NA | NA | NA | NA | NA |
| CHRNB2 | 57 | T | 62.81285714 | 29.9301629 | 0.732951605 | 22.12454582 | NA | NA | NA | NA | NA | NA | NA |
| CHRNB2 | 58 | V | 59.63028571 | 26.54746574 | 0.640156764 | 18.78208893 | NA | NA | NA | NA | NA | NA | NA |
| CHRNB2 | 59 | Q | 58.41812698 | 28.7591277 | 0.700827528 | 20.99633598 | NA | NA | NA | NA | NA | NA | NA |

| | | | | | | | | | | | | | |
|---|---|---|---|---|---|---|---|---|---|---|---|---|---|
| CHRNB2 | 60 | L | 53.97257143 | 26.21715257 | 0.631095545 | 18.32630789 | NA | NA | NA | NA | NA | NA | NA |
| CHRNB2 | 61 | M | 51.69769643 | 27.67079157 | 0.670972068 | 19.51665861 | NA | NA | NA | NA | NA | NA | NA |
| CHRNB2 | 62 | V | 47.94957143 | 25.78342797 | 0.619197523 | 17.92368652 | NA | NA | NA | NA | NA | NA | NA |
| CHRNB2 | 63 | S | 47.18790476 | 25.41576347 | 0.609111675 | 18.31584493 | NA | NA | NA | NA | NA | NA | NA |
| CHRNB2 | 64 | L | 42.63019643 | 23.98673857 | 0.569910371 | 17.0821109 | NA | NA | NA | NA | NA | NA | NA |
| CHRNB2 | 65 | A | 43.20497143 | 23.67028663 | 0.561229397 | 16.75150871 | NA | NA | NA | NA | NA | NA | NA |
| CHRNB2 | 66 | Q | 39.94868254 | 20.91664108 | 0.485690828 | 14.04086914 | NA | NA | NA | NA | NA | NA | NA |
| CHRNB2 | 67 | L | 36.49344643 | 20.83355987 | 0.483411727 | 13.53385337 | NA | NA | NA | NA | NA | NA | NA |
| CHRNB2 | 68 | I | 37.78382143 | 16.40764587 | 0.361999148 | 9.114993187 | NA | NA | NA | NA | NA | NA | NA |
| CHRNB2 | 69 | S | 34.85357143 | 15.4664709 | 0.336180643 | 7.394394135 | NA | NA | NA | NA | NA | NA | NA |
| CHRNB2 | 70 | V | 32.15885714 | 16.16866816 | 0.355443462 | 9.191323574 | NA | NA | NA | NA | NA | NA | NA |
| CHRNB2 | 71 | H | 32.19607143 | 11.62905744 | 0.230911931 | 5.284533142 | NA | NA | NA | NA | NA | NA | NA |
| CHRNB2 | 72 | E | 27.98734921 | 14.9847983 | 0.322967301 | 7.998093572 | NA | NA | NA | NA | NA | NA | NA |
| CHRNB2 | 73 | R | 26.85593506 | 8.37525528 | 0.141652963 | 1.584410688 | 7.01 | -0.96 | -3.75 | 50.43 | 2.08 | 4 | -2 |
| CHRNB2 | 74 | E | 32.30590476 | 8.761103013 | 0.152237617 | 2.043444411 | 7.01 | -0.96 | -3.75 | 50.43 | 2.08 | 4 | -2 |
| CHRNB2 | 75 | Q | 32.3572381 | 15.33943591 | 0.332695793 | 8.368881247 | NA | NA | NA | NA | NA | NA | NA |
| CHRNB2 | 76 | I | 35.74369643 | 12.35231189 | 0.250752395 | 4.306732071 | 8.05 | -0.95 | -3.58 | 49.74 | 2.24 | 5 | -3 |
| CHRNB2 | 77 | M | 37.31082143 | 16.94783895 | 0.376817835 | 8.908117094 | NA | NA | NA | NA | NA | NA | NA |
| CHRNB2 | 78 | T | 38.35142857 | 13.67715076 | 0.287095646 | 6.962574194 | NA | NA | NA | NA | NA | NA | NA |
| CHRNB2 | 79 | T | 41.17414286 | 17.73168771 | 0.398320535 | 10.83405137 | NA | NA | NA | NA | NA | NA | NA |
| CHRNB2 | 80 | N | 44.12132143 | 19.70386954 | 0.452421831 | 12.8170516 | NA | NA | NA | NA | NA | NA | NA |
| CHRNB2 | 81 | V | 46.75685714 | 20.81506303 | 0.482904318 | 13.71029674 | NA | NA | NA | NA | NA | NA | NA |
| CHRNB2 | 82 | W | 49.29578571 | 22.97217436 | 0.542078638 | 14.97442593 | NA | NA | NA | NA | NA | NA | NA |
| CHRNB2 | 83 | L | 52.31294643 | 20.97640296 | 0.487330228 | 12.82256481 | NA | NA | NA | NA | NA | NA | NA |
| CHRNB2 | 84 | T | 54.59214286 | 25.72793738 | 0.617675294 | 17.76803468 | NA | NA | NA | NA | NA | NA | NA |
| CHRNB2 | 85 | Q | 57.63079365 | 21.99155356 | 0.515178047 | 14.25529976 | NA | NA | NA | NA | NA | NA | NA |
| CHRNB2 | 86 | E | 60.26579365 | 27.58598331 | 0.66864559 | 19.78901594 | NA | NA | NA | NA | NA | NA | NA |
| CHRNB2 | 87 | W | 63.71321429 | 23.09389647 | 0.545417744 | 15.25102242 | NA | NA | NA | NA | NA | NA | NA |
| CHRNB2 | 88 | E | 68.31168254 | 29.68232346 | 0.726152823 | 20.03754617 | NA | NA | NA | NA | NA | NA | NA |
| CHRNB2 | 89 | D | 70.47757143 | 24.52645857 | 0.584716081 | 14.82555486 | NA | NA | NA | NA | NA | NA | NA |
| CHRNB2 | 90 | Y | 74.91032143 | 27.04965798 | 0.653933004 | 17.39009463 | NA | NA | NA | NA | NA | NA | NA |
| CHRNB2 | 91 | R | 74.1602987 | 21.4994873 | 0.501679585 | 11.8090089 | NA | NA | NA | NA | NA | NA | NA |
| CHRNB2 | 92 | L | 73.23907143 | 21.88888179 | 0.512361534 | 12.19742657 | NA | NA | NA | NA | NA | NA | NA |

| | | | | | | | | | | | | | |
|---|---|---|---|---|---|---|---|---|---|---|---|---|---|
| CHRNB2 | 93 | T | 76.09514286 | 26.8363749 | 0.648082179 | 17.15817735 | NA | NA | NA | NA | NA | NA | NA |
| CHRNB2 | 94 | W | 75.91928571 | 26.71035161 | 0.644625082 | 17.01265726 | NA | NA | NA | NA | NA | NA | NA |
| CHRNB2 | 95 | K | 81.20190476 | 30.36285066 | 0.744821184 | 20.69768689 | NA | NA | NA | NA | NA | NA | NA |
| CHRNB2 | 96 | P | 78.50085714 | 32.87779483 | 0.813811648 | 23.20380876 | NA | NA | NA | NA | NA | NA | NA |
| CHRNB2 | 97 | E | 83.02890476 | 34.70005916 | 0.863800377 | 25.02838194 | NA | NA | NA | NA | NA | NA | NA |
| CHRNB2 | 98 | E | 83.6972381 | 30.16671455 | 0.739440739 | 20.51762473 | NA | NA | NA | NA | NA | NA | NA |
| CHRNB2 | 99 | F | 80.7282987 | 27.29865933 | 0.66076366 | 17.6394346 | NA | NA | NA | NA | NA | NA | NA |
| CHRNB2 | 100 | D | 79.61932143 | 32.3518893 | 0.7993849 | 22.66492871 | NA | NA | NA | NA | NA | NA | NA |
| CHRNB2 | 101 | N | 77.75157143 | 34.29739567 | 0.852754429 | 24.60147281 | NA | NA | NA | NA | NA | NA | NA |
| CHRNB2 | 102 | M | 74.97032143 | 28.23811219 | 0.686534923 | 18.53127743 | NA | NA | NA | NA | NA | NA | NA |
| CHRNB2 | 103 | K | 72.22734921 | 31.71432252 | 0.781895038 | 22.00468191 | NA | NA | NA | NA | NA | NA | NA |
| CHRNB2 | 104 | K | 68.03901587 | 29.31052263 | 0.715953507 | 19.59262066 | NA | NA | NA | NA | NA | NA | NA |
| CHRNB2 | 105 | V | 70.16928571 | 24.75928614 | 0.591103054 | 15.13850129 | NA | NA | NA | NA | NA | NA | NA |
| CHRNB2 | 106 | R | 66.10902597 | 22.6645497 | 0.533639815 | 13.77944288 | NA | NA | NA | NA | NA | NA | NA |
| CHRNB2 | 107 | L | 68.18857143 | 20.24145432 | 0.467168966 | 11.49532244 | NA | NA | NA | NA | NA | NA | NA |
| CHRNB2 | 108 | P | 66.52457143 | 16.70264775 | 0.3700917 | 8.730720116 | NA | NA | NA | NA | NA | NA | NA |
| CHRNB2 | 109 | S | 63.2647381 | 15.77547115 | 0.344657201 | 7.856178765 | NA | NA | NA | NA | NA | NA | NA |
| CHRNB2 | 110 | K | 63.52146032 | 9.607145101 | 0.175446417 | 1.640699981 | 8.53 | -0.9 | -3.63 | 49.77 | 1.97 | 3 | -1 |
| CHRNB2 | 111 | H | 68.64437143 | 11.53950818 | 0.228455397 | 2.653439335 | 8.88 | -0.67 | -3.4 | 50.47 | 1.42 | 2 | -2 |
| CHRNB2 | 112 | I | 67.45207143 | 17.42009225 | 0.389772784 | 8.533402917 | NA | NA | NA | NA | NA | NA | NA |
| CHRNB2 | 113 | W | 65.99107143 | 19.63218521 | 0.450455371 | 10.7449894 | NA | NA | NA | NA | NA | NA | NA |
| CHRNB2 | 114 | L | 62.37869643 | 16.82446689 | 0.373433467 | 8.957194299 | NA | NA | NA | NA | NA | NA | NA |
| CHRNB2 | 115 | P | 60.67485714 | 19.90409352 | 0.457914416 | 12.13402439 | NA | NA | NA | NA | NA | NA | NA |
| CHRNB2 | 116 | D | 58.53832143 | 16.73625627 | 0.371013655 | 8.962812372 | NA | NA | NA | NA | NA | NA | NA |
| CHRNB2 | 117 | V | 54.76571429 | 19.74902099 | 0.453660434 | 11.89377981 | NA | NA | NA | NA | NA | NA | NA |
| CHRNB2 | 118 | V | 51.991 | 16.38602398 | 0.361406012 | 8.218823086 | NA | NA | NA | NA | NA | NA | NA |
| CHRNB2 | 119 | L | 47.92607143 | 18.92631721 | 0.431091856 | 10.75063163 | NA | NA | NA | NA | NA | NA | NA |
| CHRNB2 | 120 | Y | 48.6312381 | 19.9809788 | 0.460023549 | 12.28281021 | NA | NA | NA | NA | NA | NA | NA |
| CHRNB2 | 121 | N | 43.10694643 | 19.12277318 | 0.436481076 | 12.18820703 | NA | NA | NA | NA | NA | NA | NA |
| CHRNB2 | 122 | N | 44.49507143 | 15.13598077 | 0.32711457 | 8.040959963 | NA | NA | NA | NA | NA | NA | NA |
| CHRNB2 | 123 | A | 42.18997143 | 13.30317992 | 0.276836802 | 6.392057131 | NA | NA | NA | NA | NA | NA | NA |
| CHRNB2 | 124 | D | 42.78307143 | 8.398757587 | 0.142297683 | 1.566327417 | 6.94 | -0.88 | -3.6 | 49.65 | 2.34 | 4 | -2 |
| CHRNB2 | 125 | G | 46.40282143 | 11.62902051 | 0.230910918 | 3.899247422 | 7.78 | -0.88 | -3.6 | 49.65 | 2.34 | 4 | -2 |

| | | | | | | | | | | | | | |
|---|---|---|---|---|---|---|---|---|---|---|---|---|---|
| CHRNB2 | 126 | M | 47.61594643 | 10.73081609 | 0.20627119 | 2.713390118 | 8.05 | -0.63 | -2.42 | 40.05 | 1.96 | 4 | -4 |
| CHRNB2 | 127 | Y | 49.4937381 | 12.77607557 | 0.262377167 | 4.608706861 | 8.2 | -0.56 | -1.3 | 20.91 | 1.06 | 2 | -2 |
| CHRNB2 | 128 | E | 53.17612698 | 9.419888163 | 0.170309547 | 1.353483886 | 8.11 | -0.92 | -1.76 | 21.3 | 1.36 | 2 | -2 |
| CHRNB2 | 129 | V | 53.39414286 | 12.52392646 | 0.255460161 | 4.450472943 | 8.11 | -0.92 | -1.76 | 21.3 | 1.36 | 2 | -2 |
| CHRNB2 | 130 | S | 52.31140476 | 12.00347708 | 0.241183087 | 4.053281045 | 8.2 | -0.56 | -1.3 | 20.91 | 1.06 | 2 | -2 |
| CHRNB2 | 131 | F | 55.51893506 | 14.16669817 | 0.300525011 | 6.116401108 | NA | NA | NA | NA | NA | NA | NA |
| CHRNB2 | 132 | Y | 58.1557381 | 11.11683772 | 0.216860615 | 3.222089957 | 8.02 | -0.4 | -1.96 | 21.21 | 0.77 | 2 | 0 |
| CHRNB2 | 133 | S | 59.01207143 | 17.22081499 | 0.38430617 | 9.266397764 | NA | NA | NA | NA | NA | NA | NA |
| CHRNB2 | 134 | N | 62.23357143 | 19.47023372 | 0.446012685 | 11.83700743 | NA | NA | NA | NA | NA | NA | NA |
| CHRNB2 | 135 | A | 63.91637143 | 22.21207034 | 0.521227309 | 14.28007075 | NA | NA | NA | NA | NA | NA | NA |
| CHRNB2 | 136 | V | 64.58671429 | 25.60897116 | 0.614411788 | 17.30925609 | NA | NA | NA | NA | NA | NA | NA |
| CHRNB2 | 137 | V | 68.60457143 | 25.2422268 | 0.604351182 | 16.36606939 | NA | NA | NA | NA | NA | NA | NA |
| CHRNB2 | 138 | S | 69.51590476 | 29.57566654 | 0.723226988 | 19.87133706 | NA | NA | NA | NA | NA | NA | NA |
| CHRNB2 | 139 | Y | 73.54165476 | 31.00030564 | 0.76230798 | 21.30509032 | NA | NA | NA | NA | NA | NA | NA |
| CHRNB2 | 140 | D | 71.83769643 | 32.09766736 | 0.792411032 | 22.40352114 | NA | NA | NA | NA | NA | NA | NA |
| CHRNB2 | 141 | G | 70.30857143 | 29.31373449 | 0.716041615 | 19.60719661 | NA | NA | NA | NA | NA | NA | NA |
| CHRNB2 | 142 | S | 67.16240476 | 29.73528564 | 0.727605692 | 20.08886156 | NA | NA | NA | NA | NA | NA | NA |
| CHRNB2 | 143 | I | 65.21544643 | 25.43506243 | 0.609641088 | 17.54380658 | NA | NA | NA | NA | NA | NA | NA |
| CHRNB2 | 144 | F | 60.31620779 | 25.90391247 | 0.622502678 | 18.03479779 | NA | NA | NA | NA | NA | NA | NA |
| CHRNB2 | 145 | W | 59.27764286 | 18.02403975 | 0.406340396 | 10.20282078 | NA | NA | NA | NA | NA | NA | NA |
| CHRNB2 | 146 | L | 55.79219643 | 22.02569175 | 0.516114533 | 14.04990074 | NA | NA | NA | NA | NA | NA | NA |
| CHRNB2 | 147 | P | 54.63228571 | 18.6060086 | 0.422305084 | 10.62783398 | NA | NA | NA | NA | NA | NA | NA |
| CHRNB2 | 148 | P | 51.70842857 | 17.55082827 | 0.393359161 | 9.361322336 | NA | NA | NA | NA | NA | NA | NA |
| CHRNB2 | 149 | A | 48.36937143 | 17.22188544 | 0.384335534 | 9.305229793 | NA | NA | NA | NA | NA | NA | NA |
| CHRNB2 | 150 | I | 45.38332143 | 12.27236118 | 0.248559171 | 5.172140108 | NA | NA | NA | NA | NA | NA | NA |
| CHRNB2 | 151 | Y | 46.08382143 | 15.96874287 | 0.349959071 | 9.029649252 | NA | NA | NA | NA | NA | NA | NA |
| CHRNB2 | 152 | K | 41.46990476 | 9.017157313 | 0.159261751 | 2.119427274 | 6.91 | -0.79 | -3.66 | 49.62 | 2.16 | 5 | -1 |
| CHRNB2 | 153 | S | 40.05507143 | 17.09296139 | 0.380798863 | 10.19332905 | NA | NA | NA | NA | NA | NA | NA |
| CHRNB2 | 154 | A | 38.43697143 | 18.44439015 | 0.417871534 | 11.17630972 | NA | NA | NA | NA | NA | NA | NA |
| CHRNB2 | 155 | C | 35.29640476 | 20.57593036 | 0.476344382 | 12.52778654 | NA | NA | NA | NA | NA | NA | NA |
| CHRNB2 | 156 | K | 32.6012381 | 23.0361444 | 0.543833477 | 16.05662185 | NA | NA | NA | NA | NA | NA | NA |
| CHRNB2 | 157 | I | 29.50132143 | 19.97361412 | 0.459821519 | 12.9909204 | NA | NA | NA | NA | NA | NA | NA |
| CHRNB2 | 158 | E | 27.05746032 | 23.88193334 | 0.567035333 | 17.06940868 | NA | NA | NA | NA | NA | NA | NA |

| | | | | | | | | | | | | | |
|---|---|---|---|---|---|---|---|---|---|---|---|---|---|
| CHRNB2 | 159 | V | 23.51028571 | 21.33763469 | 0.497239611 | 14.73125878 | NA | NA | NA | NA | NA | NA | NA |
| CHRNB2 | 160 | K | 20.49612698 | 25.52881419 | 0.612212906 | 20.25963428 | NA | NA | NA | NA | NA | NA | NA |
| CHRNB2 | 161 | H | 19.77427143 | 26.44089955 | 0.637233418 | 21.06282868 | NA | NA | NA | NA | NA | NA | NA |
| CHRNB2 | 162 | F | 19.71575325 | 19.48166166 | 0.446326179 | 14.0895908 | NA | NA | NA | NA | NA | NA | NA |
| CHRNB2 | 163 | P | 20.283 | 23.57989296 | 0.558749699 | 18.25777956 | NA | NA | NA | NA | NA | NA | NA |
| CHRNB2 | 164 | F | 19.46075325 | 26.6459432 | 0.642858218 | 21.24901251 | NA | NA | NA | NA | NA | NA | NA |
| CHRNB2 | 165 | D | 24.81982143 | 23.68418607 | 0.561610689 | 17.08463696 | NA | NA | NA | NA | NA | NA | NA |
| CHRNB2 | 166 | Q | 27.87990476 | 27.6177721 | 0.669517627 | 20.62698232 | NA | NA | NA | NA | NA | NA | NA |
| CHRNB2 | 167 | Q | 29.89990476 | 23.22467261 | 0.549005222 | 16.23689108 | NA | NA | NA | NA | NA | NA | NA |
| CHRNB2 | 168 | N | 33.98907143 | 26.72767713 | 0.64510036 | 19.95919356 | NA | NA | NA | NA | NA | NA | NA |
| CHRNB2 | 169 | C | 36.3312381 | 23.20885551 | 0.548571324 | 15.18376249 | NA | NA | NA | NA | NA | NA | NA |
| CHRNB2 | 170 | T | 39.54442857 | 23.86902072 | 0.566681111 | 16.96695418 | NA | NA | NA | NA | NA | NA | NA |
| CHRNB2 | 171 | M | 43.44707143 | 22.66850593 | 0.533748343 | 15.79183749 | NA | NA | NA | NA | NA | NA | NA |
| CHRNB2 | 172 | K | 45.87734921 | 23.34319151 | 0.552256457 | 16.28342529 | NA | NA | NA | NA | NA | NA | NA |
| CHRNB2 | 173 | F | 49.33938961 | 23.0662541 | 0.544659453 | 14.95941359 | NA | NA | NA | NA | NA | NA | NA |
| CHRNB2 | 174 | R | 52.62038961 | 22.37022847 | 0.525565935 | 14.36402992 | NA | NA | NA | NA | NA | NA | NA |
| CHRNB2 | 175 | S | 56.12540476 | 22.94124323 | 0.541230129 | 15.24616282 | NA | NA | NA | NA | NA | NA | NA |
| CHRNB2 | 176 | W | 55.27121429 | 15.77322684 | 0.344595635 | 8.124083931 | NA | NA | NA | NA | NA | NA | NA |
| CHRNB2 | 177 | T | 59.60057143 | 18.85408118 | 0.429110262 | 11.1088703 | NA | NA | NA | NA | NA | NA | NA |
| CHRNB2 | 178 | Y | 61.9242381 | 21.9972631 | 0.515334673 | 14.16231653 | NA | NA | NA | NA | NA | NA | NA |
| CHRNB2 | 179 | D | 60.04807143 | 27.02433637 | 0.653238376 | 19.28482922 | NA | NA | NA | NA | NA | NA | NA |
| CHRNB2 | 180 | R | 58.93057143 | 30.30379252 | 0.74320109 | 22.57541689 | NA | NA | NA | NA | NA | NA | NA |
| CHRNB2 | 181 | T | 63.41885714 | 30.29696708 | 0.743013853 | 22.52086055 | NA | NA | NA | NA | NA | NA | NA |
| CHRNB2 | 182 | E | 65.32812698 | 26.86445657 | 0.648852521 | 19.04071166 | NA | NA | NA | NA | NA | NA | NA |
| CHRNB2 | 183 | I | 61.43107143 | 25.5023564 | 0.61148711 | 17.7454586 | NA | NA | NA | NA | NA | NA | NA |
| CHRNB2 | 184 | D | 59.31419643 | 30.4833604 | 0.748127032 | 22.72315613 | NA | NA | NA | NA | NA | NA | NA |
| CHRNB2 | 185 | L | 56.18432143 | 27.31831785 | 0.661302937 | 19.61923039 | NA | NA | NA | NA | NA | NA | NA |
| CHRNB2 | 186 | V | 56.13742857 | 31.80167419 | 0.784291286 | 24.07821677 | NA | NA | NA | NA | NA | NA | NA |
| CHRNB2 | 187 | L | 52.13394643 | 31.79098493 | 0.783998057 | 23.7682203 | NA | NA | NA | NA | NA | NA | NA |
| CHRNB2 | 188 | K | 54.52534921 | 32.69214808 | 0.808718949 | 24.85072566 | NA | NA | NA | NA | NA | NA | NA |
| CHRNB2 | 189 | S | 50.0412381 | 35.14547715 | 0.876019174 | 26.95774294 | NA | NA | NA | NA | NA | NA | NA |
| CHRNB2 | 190 | E | 47.39657143 | 35.81533775 | 0.894394927 | 28.70886188 | NA | NA | NA | NA | NA | NA | NA |
| CHRNB2 | 191 | V | 44.74814286 | 34.0661142 | 0.846409869 | 27.18587976 | NA | NA | NA | NA | NA | NA | NA |

| | | | | | | | | | | | | | |
|---|---|---|---|---|---|---|---|---|---|---|---|---|---|
| CHRNB2 | 192 | A | 44.57417143 | 30.55742316 | 0.750158737 | 23.46784163 | NA | NA | NA | NA | NA | NA | NA |
| CHRNB2 | 193 | S | 46.1167381 | 30.39441996 | 0.7456872 | 23.29889973 | NA | NA | NA | NA | NA | NA | NA |
| CHRNB2 | 194 | L | 42.23469643 | 29.81159241 | 0.729698955 | 22.91776042 | NA | NA | NA | NA | NA | NA | NA |
| CHRNB2 | 195 | D | 45.46382143 | 32.35799824 | 0.799552482 | 25.43204432 | NA | NA | NA | NA | NA | NA | NA |
| CHRNB2 | 196 | D | 44.81657143 | 28.19301603 | 0.685297836 | 21.2564835 | NA | NA | NA | NA | NA | NA | NA |
| CHRNB2 | 197 | F | 39.26238961 | 26.72123975 | 0.644923768 | 19.91515334 | NA | NA | NA | NA | NA | NA | NA |
| CHRNB2 | 198 | T | 38.315 | 26.0126624 | 0.625485929 | 18.74085179 | NA | NA | NA | NA | NA | NA | NA |
| CHRNB2 | 199 | P | 34.56 | 26.4217599 | 0.636708375 | 18.34722469 | NA | NA | NA | NA | NA | NA | NA |
| CHRNB2 | 200 | S | 33.3282381 | 23.32236787 | 0.551685218 | 15.38672574 | NA | NA | NA | NA | NA | NA | NA |
| CHRNB2 | 201 | G | 29.90957143 | 24.80341325 | 0.592313559 | 17.61620379 | NA | NA | NA | NA | NA | NA | NA |
| CHRNB2 | 202 | E | 27.60290476 | 19.15866368 | 0.437465632 | 11.82870097 | NA | NA | NA | NA | NA | NA | NA |
| CHRNB2 | 203 | W | 29.90564286 | 20.54172695 | 0.475406107 | 13.19381581 | NA | NA | NA | NA | NA | NA | NA |
| CHRNB2 | 204 | D | 30.89707143 | 27.1547573 | 0.65681611 | 19.97032544 | NA | NA | NA | NA | NA | NA | NA |
| CHRNB2 | 205 | I | 34.81282143 | 27.92188548 | 0.677860128 | 19.83794335 | NA | NA | NA | NA | NA | NA | NA |
| CHRNB2 | 206 | V | 33.289 | 31.56324704 | 0.777750704 | 23.48425411 | NA | NA | NA | NA | NA | NA | NA |
| CHRNB2 | 207 | A | 36.78717143 | 31.87467631 | 0.786293896 | 23.87620095 | NA | NA | NA | NA | NA | NA | NA |
| CHRNB2 | 208 | L | 40.28144643 | 28.89949346 | 0.70467807 | 21.99997706 | NA | NA | NA | NA | NA | NA | NA |
| CHRNB2 | 209 | P | 41.12585714 | 31.97619016 | 0.789078644 | 25.04044954 | NA | NA | NA | NA | NA | NA | NA |
| CHRNB2 | 210 | G | 45.09082143 | 32.78492021 | 0.811263893 | 25.67809451 | NA | NA | NA | NA | NA | NA | NA |
| CHRNB2 | 211 | R | 44.84275325 | 32.83347708 | 0.812595915 | 25.7371476 | NA | NA | NA | NA | NA | NA | NA |
| CHRNB2 | 212 | R | 51.16738961 | 33.92640943 | 0.842577459 | 25.80181286 | NA | NA | NA | NA | NA | NA | NA |
| CHRNB2 | 213 | N | 47.72632143 | 32.82226668 | 0.812288389 | 25.00161073 | NA | NA | NA | NA | NA | NA | NA |
| CHRNB2 | 214 | E | 52.07279365 | 34.51589485 | 0.858748344 | 26.63961098 | NA | NA | NA | NA | NA | NA | NA |
| CHRNB2 | 215 | N | 50.46019643 | 35.34663504 | 0.881537379 | 27.17613917 | NA | NA | NA | NA | NA | NA | NA |
| CHRNB2 | 216 | P | 51.45857143 | 38.56482399 | 0.969819397 | 30.5425079 | NA | NA | NA | NA | NA | NA | NA |
| CHRNB2 | 217 | D | 51.06457143 | 38.60137342 | 0.970822029 | 30.67171799 | NA | NA | NA | NA | NA | NA | NA |
| CHRNB2 | 218 | D | 52.81569643 | 33.98274171 | 0.844122777 | 25.97186946 | NA | NA | NA | NA | NA | NA | NA |
| CHRNB2 | 219 | S | 56.52357143 | 34.98794312 | 0.871697669 | 27.28708342 | NA | NA | NA | NA | NA | NA | NA |
| CHRNB2 | 220 | T | 56.53514286 | 30.42701857 | 0.746581452 | 22.73282487 | NA | NA | NA | NA | NA | NA | NA |
| CHRNB2 | 221 | Y | 50.97007143 | 28.95516579 | 0.706205285 | 21.15338797 | NA | NA | NA | NA | NA | NA | NA |
| CHRNB2 | 222 | V | 53.98128571 | 28.6060473 | 0.696628195 | 20.64417011 | NA | NA | NA | NA | NA | NA | NA |
| CHRNB2 | 223 | D | 49.82694643 | 28.53555097 | 0.694694325 | 20.36020044 | NA | NA | NA | NA | NA | NA | NA |
| CHRNB2 | 224 | I | 49.41969643 | 27.96686647 | 0.679094055 | 19.82094021 | NA | NA | NA | NA | NA | NA | NA |

| | | | | | | | | | | | | | |
|---|---|---|---|---|---|---|---|---|---|---|---|---|---|
| CHRNB2 | 225 | T | 44.897 | 29.35432794 | 0.717155183 | 22.24282771 | NA | NA | NA | NA | NA | NA | NA |
| CHRNB2 | 226 | Y | 45.25682143 | 27.53170451 | 0.667156603 | 20.60929707 | NA | NA | NA | NA | NA | NA | NA |
| CHRNB2 | 227 | D | 39.73757143 | 29.29685104 | 0.715578465 | 22.41497704 | NA | NA | NA | NA | NA | NA | NA |
| CHRNB2 | 228 | F | 38.73520779 | 24.74809142 | 0.590795959 | 17.12447197 | NA | NA | NA | NA | NA | NA | NA |
| CHRNB2 | 229 | I | 33.81307143 | 29.33294862 | 0.716568701 | 21.25468769 | NA | NA | NA | NA | NA | NA | NA |
| CHRNB2 | 230 | I | 32.28669643 | 25.5621452 | 0.613127249 | 18.59616598 | NA | NA | NA | NA | NA | NA | NA |
| CHRNB2 | 231 | R | 27.89248052 | 26.81091854 | 0.647383855 | 19.81302984 | NA | NA | NA | NA | NA | NA | NA |
| CHRNB2 | 232 | R | 25.16293506 | 20.59014388 | 0.47673429 | 13.7523197 | NA | NA | NA | NA | NA | NA | NA |
| CHRNB2 | 233 | K | 25.36146032 | 24.70259707 | 0.589547948 | 18.16435515 | NA | NA | NA | NA | NA | NA | NA |
| CHRNB2 | 234 | P | 20.55442857 | 24.58413938 | 0.586298393 | 19.24751422 | NA | NA | NA | NA | NA | NA | NA |
| CHRNB2 | 235 | L | 18.95257143 | 24.24472611 | 0.576987538 | 18.39343062 | NA | NA | NA | NA | NA | NA | NA |
| CHRNB2 | 236 | F | 20.36311688 | 18.25874547 | 0.412778891 | 12.8249037 | NA | NA | NA | NA | NA | NA | NA |
| CHRNB2 | 237 | Y | 19.8064881 | 15.67546341 | 0.341913768 | 10.27902669 | NA | NA | NA | NA | NA | NA | NA |
| CHRNB2 | 238 | T | 15.38371429 | 21.32404933 | 0.496866935 | 15.29151976 | NA | NA | NA | NA | NA | NA | NA |
| CHRNB2 | 239 | I | 13.79194643 | 19.98914871 | 0.460247667 | 13.41526341 | NA | NA | NA | NA | NA | NA | NA |
| CHRNB2 | 240 | N | 15.56794643 | 16.37976758 | 0.361234385 | 9.798976423 | NA | NA | NA | NA | NA | NA | NA |
| CHRNB2 | 241 | L | 14.18932143 | 16.04112456 | 0.35194466 | 9.45352824 | NA | NA | NA | NA | NA | NA | NA |
| CHRNB2 | 242 | I | 12.22857143 | 19.81308113 | 0.455417745 | 14.69725139 | NA | NA | NA | NA | NA | NA | NA |
| CHRNB2 | 243 | I | 9.243821429 | 19.80058204 | 0.455074868 | 14.7432774 | NA | NA | NA | NA | NA | NA | NA |
| CHRNB2 | 244 | P | 8.727857143 | 16.28608553 | 0.35866448 | 11.21632098 | NA | NA | NA | NA | NA | NA | NA |
| CHRNB2 | 245 | C | 8.210738095 | 17.69396439 | 0.397285701 | 12.63515653 | NA | NA | NA | NA | NA | NA | NA |
| CHRNB2 | 246 | V | 5.707714286 | 19.91596412 | 0.458240053 | 16.0714071 | NA | NA | NA | NA | NA | NA | NA |
| CHRNB2 | 247 | L | 3.655821429 | 17.021699 | 0.378843979 | 13.4055068 | NA | NA | NA | NA | NA | NA | NA |
| CHRNB2 | 248 | I | 3.841821429 | 14.16179847 | 0.300390602 | 10.52354706 | NA | NA | NA | NA | NA | NA | NA |
| CHRNB2 | 249 | T | 3.011857143 | 17.61541353 | 0.395130878 | 13.9899132 | NA | NA | NA | NA | NA | NA | NA |
| CHRNB2 | 250 | S | 0.434404762 | 19.59202708 | 0.449353745 | 15.13018593 | NA | NA | NA | NA | NA | NA | NA |
| CHRNB2 | 251 | L | -0.499946429 | 15.01943364 | 0.323917425 | 11.06243627 | NA | NA | NA | NA | NA | NA | NA |
| CHRNB2 | 252 | A | -1.252371429 | 18.057482 | 0.40725779 | 13.95818997 | NA | NA | NA | NA | NA | NA | NA |
| CHRNB2 | 253 | I | -3.773071429 | 19.34417827 | 0.442554706 | 16.8617967 | NA | NA | NA | NA | NA | NA | NA |
| CHRNB2 | 254 | L | -5.684196429 | 15.25965809 | 0.330507312 | 12.77070087 | NA | NA | NA | NA | NA | NA | NA |
| CHRNB2 | 255 | V | -6.742428571 | 13.04094682 | 0.26964317 | 10.56940735 | NA | NA | NA | NA | NA | NA | NA |
| CHRNB2 | 256 | F | -8.302662338 | 16.49956175 | 0.364520603 | 14.74301753 | NA | NA | NA | NA | NA | NA | NA |
| CHRNB2 | 257 | Y | -11.93190476 | 16.3809171 | 0.361265918 | 14.46363665 | NA | NA | NA | NA | NA | NA | NA |

| | | | | | | | | | | | | | |
|---|---|---|---|---|---|---|---|---|---|---|---|---|---|
| CHRNB2 | 258 | L | -11.36044643 | 10.41201988 | 0.197525907 | 8.623401248 | NA | NA | NA | NA | NA | NA | NA |
| CHRNB2 | 259 | P | -14.87171429 | 10.96613195 | 0.212726423 | 7.063061319 | NA | NA | NA | NA | NA | NA | NA |
| CHRNB2 | 260 | S | -15.7887381 | 9.536170877 | 0.173499438 | 5.586336828 | NA | NA | NA | NA | NA | NA | NA |
| CHRNB2 | 261 | D | -18.84332143 | 6.520072546 | 0.09076121 | 1.682487871 | 4.88 | -0.8 | -0.4 | 3.38 | 0.81 | 0 | 0 |
| CHRNB2 | 262 | C | -15.0022381 | 5.125156485 | 0.052495587 | 1.292624036 | 3.95 | -0.87 | -1.02 | 12.68 | 1.09 | 1 | -1 |
| CHRNB2 | 263 | G | -12.64732143 | 5.045064321 | 0.050298483 | 2.968106283 | 2.51 | -1.14 | -3.5 | 49.9 | 2.23 | 5 | -5 |
| CHRNB2 | 264 | E | -9.963349206 | 3.260856483 | 0.001353728 | 1.783900269 | 1.83 | -1.14 | -3.5 | 49.9 | 2.23 | 4 | -4 |
| CHRNB2 | 265 | K | -9.600571429 | 8.355631574 | 0.141114642 | 5.96182386 | NA | NA | NA | NA | NA | NA | NA |
| CHRNB2 | 266 | M | -6.470571429 | 7.656529501 | 0.12193673 | 5.145324547 | NA | NA | NA | NA | NA | NA | NA |
| CHRNB2 | 267 | T | -4.590142857 | 3.962742106 | 0.020607998 | 1.610401903 | 2.88 | -0.77 | -0.7 | 1.66 | 0.77 | 0 | 0 |
| CHRNB2 | 268 | L | -3.711946429 | 8.000465799 | 0.131371661 | 5.199790362 | NA | NA | NA | NA | NA | NA | NA |
| CHRNB2 | 269 | C | -2.211904762 | 10.34128005 | 0.195585357 | 7.329551835 | NA | NA | NA | NA | NA | NA | NA |
| CHRNB2 | 270 | I | -0.741571429 | 7.895010196 | 0.128478782 | 3.708110119 | 4.19 | -0.89 | -0.76 | 1.67 | 0.97 | 0 | 0 |
| CHRNB2 | 271 | S | 0.255404762 | 5.741532287 | 0.069404134 | 1.523262239 | 4.34 | -0.97 | -0.8 | 1.67 | 1.11 | 0 | 0 |
| CHRNB2 | 272 | V | 0.710714286 | 10.00420697 | 0.1863387 | 5.448097097 | NA | NA | NA | NA | NA | NA | NA |
| CHRNB2 | 273 | L | 2.370321429 | 10.96660335 | 0.212739354 | 7.321328832 | NA | NA | NA | NA | NA | NA | NA |
| CHRNB2 | 274 | L | 4.394071429 | 5.355531253 | 0.058815275 | 1.711141047 | 3.71 | 1.14 | 3.8 | 0.13 | -1.79 | 0 | 0 |
| CHRNB2 | 275 | A | 5.787371429 | 8.876057514 | 0.155391073 | 5.224744219 | NA | NA | NA | NA | NA | NA | NA |
| CHRNB2 | 276 | L | 6.333946429 | 11.47970583 | 0.226814887 | 7.690691623 | NA | NA | NA | NA | NA | NA | NA |
| CHRNB2 | 277 | T | 8.196428571 | 9.973676002 | 0.185501168 | 4.976620607 | 5.08 | 1.13 | 4.1 | 0.13 | -1.42 | 0 | 0 |
| CHRNB2 | 278 | V | 10.24342857 | 6.844370387 | 0.099657415 | 1.877558362 | 5.09 | 1.13 | 4.2 | 0.13 | -1.3 | 0 | 0 |
| CHRNB2 | 279 | F | 10.30075325 | 11.06440857 | 0.215422367 | 5.929811629 | NA | NA | NA | NA | NA | NA | NA |
| CHRNB2 | 280 | L | 12.39469643 | 13.14527854 | 0.272505219 | 7.908652781 | NA | NA | NA | NA | NA | NA | NA |
| CHRNB2 | 281 | L | 14.28569643 | 9.417454911 | 0.170242797 | 3.151235752 | 6.65 | 0.21 | -0.74 | 29.75 | 0.26 | 3 | 3 |
| CHRNB2 | 282 | L | 16.28082143 | 9.098737605 | 0.161499679 | 2.93302024 | 6.17 | -0.25 | -2.28 | 39.71 | 0.96 | 4 | 2 |
| CHRNB2 | 283 | I | 16.66644643 | 13.01976455 | 0.269062093 | 6.889354438 | NA | NA | NA | NA | NA | NA | NA |
| CHRNB2 | 284 | S | 18.41457143 | 12.98747069 | 0.268176201 | 7.159357773 | NA | NA | NA | NA | NA | NA | NA |
| CHRNB2 | 285 | K | 20.70990476 | 6.919961561 | 0.101731048 | 1.724769085 | 5.26 | -0.78 | -3.7 | 49.7 | 1.85 | 4 | 0 |
| CHRNB2 | 286 | I | 21.67094643 | 12.51951513 | 0.255339148 | 7.176847254 | NA | NA | NA | NA | NA | NA | NA |
| CHRNB2 | 287 | V | 21.92457143 | 14.69671001 | 0.315064404 | 9.423689728 | NA | NA | NA | NA | NA | NA | NA |
| CHRNB2 | 288 | P | 25.09885714 | 15.93208354 | 0.348953424 | 9.097323737 | NA | NA | NA | NA | NA | NA | NA |
| CHRNB2 | 289 | P | 25.67628571 | 14.29235803 | 0.303972138 | 7.730103709 | NA | NA | NA | NA | NA | NA | NA |
| CHRNB2 | 290 | T | 25.20057143 | 18.57340424 | 0.421410675 | 11.73918533 | NA | NA | NA | NA | NA | NA | NA |

| | | | | | | | | | | | | | |
|---|---|---|---|---|---|---|---|---|---|---|---|---|---|
| CHRNB2 | 291 | S | 23.72490476 | 21.34251262 | 0.497373423 | 14.74924306 | NA | NA | NA | NA | NA | NA | NA |
| CHRNB2 | 292 | L | 25.45094643 | 23.90764325 | 0.567740612 | 17.07253927 | NA | NA | NA | NA | NA | NA | NA |
| CHRNB2 | 293 | D | 22.10419643 | 23.28832053 | 0.550751225 | 16.71457463 | NA | NA | NA | NA | NA | NA | NA |
| CHRNB2 | 294 | V | 18.45685714 | 23.84177774 | 0.565933776 | 18.91221724 | NA | NA | NA | NA | NA | NA | NA |
| CHRNB2 | 295 | P | 17.972 | 20.92394239 | 0.485891119 | 15.0631139 | NA | NA | NA | NA | NA | NA | NA |
| CHRNB2 | 296 | L | 14.67757143 | 23.84382536 | 0.565989947 | 17.25944886 | NA | NA | NA | NA | NA | NA | NA |
| CHRNB2 | 297 | V | 13.69571429 | 20.00991894 | 0.46081744 | 13.43835699 | NA | NA | NA | NA | NA | NA | NA |
| CHRNB2 | 298 | G | 13.26482143 | 20.06976534 | 0.462459159 | 13.49101688 | NA | NA | NA | NA | NA | NA | NA |
| CHRNB2 | 299 | K | 12.16868254 | 22.32908664 | 0.524437324 | 17.12440389 | NA | NA | NA | NA | NA | NA | NA |
| CHRNB2 | 300 | Y | 7.786488095 | 21.9297287 | 0.513482055 | 16.86409934 | NA | NA | NA | NA | NA | NA | NA |
| CHRNB2 | 301 | L | 9.248446429 | 17.27913878 | 0.38590612 | 12.25399659 | NA | NA | NA | NA | NA | NA | NA |
| CHRNB2 | 302 | M | 8.792946429 | 20.10253579 | 0.463358125 | 15.03225919 | NA | NA | NA | NA | NA | NA | NA |
| CHRNB2 | 303 | F | 5.175662338 | 22.28920732 | 0.523343346 | 18.51550607 | NA | NA | NA | NA | NA | NA | NA |
| CHRNB2 | 304 | T | 3.974571429 | 19.44779682 | 0.445397191 | 15.81841918 | NA | NA | NA | NA | NA | NA | NA |
| CHRNB2 | 305 | M | 5.008696429 | 15.2487769 | 0.330208817 | 11.77095208 | NA | NA | NA | NA | NA | NA | NA |
| CHRNB2 | 306 | V | 2.919142857 | 20.12265529 | 0.463910047 | 16.51535128 | NA | NA | NA | NA | NA | NA | NA |
| CHRNB2 | 307 | L | 0.185571429 | 20.44171837 | 0.472662651 | 15.92645803 | NA | NA | NA | NA | NA | NA | NA |
| CHRNB2 | 308 | V | 0 | 15.56631305 | 0.338919533 | 11.3028479 | NA | NA | NA | NA | NA | NA | NA |
| CHRNB2 | 309 | T | -0.372 | 17.97392083 | 0.404965523 | 13.98048736 | NA | NA | NA | NA | NA | NA | NA |
| CHRNB2 | 310 | F | -3.086298701 | 19.90161514 | 0.457846428 | 17.25195 | NA | NA | NA | NA | NA | NA | NA |
| CHRNB2 | 311 | S | -3.962571429 | 16.92647728 | 0.376231837 | 14.41877808 | NA | NA | NA | NA | NA | NA | NA |
| CHRNB2 | 312 | I | -4.018321429 | 12.64787417 | 0.25886032 | 10.1372045 | NA | NA | NA | NA | NA | NA | NA |
| CHRNB2 | 313 | V | -5.533142857 | 17.29412996 | 0.386317361 | 14.78293668 | NA | NA | NA | NA | NA | NA | NA |
| CHRNB2 | 314 | T | -8.235857143 | 17.77254557 | 0.399441356 | 15.27766299 | NA | NA | NA | NA | NA | NA | NA |
| CHRNB2 | 315 | S | -9.043904762 | 13.38787888 | 0.279160281 | 11.59730908 | NA | NA | NA | NA | NA | NA | NA |
| CHRNB2 | 316 | V | -9.542857143 | 14.04798882 | 0.297268552 | 12.25374096 | NA | NA | NA | NA | NA | NA | NA |
| CHRNB2 | 317 | C | -12.1042381 | 16.77905909 | 0.372187831 | 14.98259691 | NA | NA | NA | NA | NA | NA | NA |
| CHRNB2 | 318 | V | -13.92057143 | 15.6495691 | 0.34120343 | 13.89675777 | NA | NA | NA | NA | NA | NA | NA |
| CHRNB2 | 319 | L | -13.87082143 | 9.950718416 | 0.184871391 | 7.607287598 | NA | NA | NA | NA | NA | NA | NA |
| CHRNB2 | 320 | N | -15.55632143 | 14.72918185 | 0.315955179 | 10.92036479 | NA | NA | NA | NA | NA | NA | NA |
| CHRNB2 | 321 | V | -18.09985714 | 16.24547078 | 0.357550327 | 11.07787812 | NA | NA | NA | NA | NA | NA | NA |
| CHRNB2 | 322 | H | -19.39717143 | 10.93719626 | 0.211932653 | 5.853920001 | NA | NA | NA | NA | NA | NA | NA |
| CHRNB2 | 323 | H | -19.52737143 | 8.488866002 | 0.144769556 | 3.120491788 | 5.58 | -0.96 | -2.47 | 34.26 | 2.02 | 2 | -2 |

| | | | | | | | | | | | | | |
|---|---|---|---|---|---|---|---|---|---|---|---|---|---|
| CHRNB2 | 324 | R | -21.06011688 | 14.44031153 | 0.308030829 | 8.992682941 | NA | NA | NA | NA | NA | NA | NA |
| CHRNB2 | 325 | S | -26.11840476 | 14.73542687 | 0.316126493 | 7.936716464 | NA | NA | NA | NA | NA | NA | NA |
| CHRNB2 | 326 | P | -29.27257143 | 15.69856761 | 0.342547567 | 7.730058144 | NA | NA | NA | NA | NA | NA | NA |
| CHRNB2 | 327 | T | -29.6 | 17.65941013 | 0.396337801 | 9.743725151 | NA | NA | NA | NA | NA | NA | NA |
| CHRNB2 | 328 | T | -25.86885714 | 18.2520947 | 0.412596446 | 12.73473824 | NA | NA | NA | NA | NA | NA | NA |
| CHRNB2 | 329 | H | -23.61977143 | 19.58939494 | 0.44928154 | 14.22800739 | NA | NA | NA | NA | NA | NA | NA |
| CHRNB2 | 330 | T | -28.52814286 | 22.76682775 | 0.536445528 | 16.00283032 | NA | NA | NA | NA | NA | NA | NA |
| CHRNB2 | 331 | M | -25.72769643 | 24.95463474 | 0.596461897 | 19.47605903 | NA | NA | NA | NA | NA | NA | NA |
| CHRNB2 | 332 | A | -29.10717143 | 25.825934 | 0.620363557 | 17.84994157 | NA | NA | NA | NA | NA | NA | NA |
| CHRNB2 | 333 | P | -29.26042857 | 28.52959204 | 0.694530859 | 20.55548289 | NA | NA | NA | NA | NA | NA | NA |
| CHRNB2 | 334 | W | -22.95678571 | 27.86654692 | 0.676342069 | 22.58470505 | NA | NA | NA | NA | NA | NA | NA |
| CHRNB2 | 335 | V | -24.93642857 | 24.60968714 | 0.586999224 | 19.18705227 | NA | NA | NA | NA | NA | NA | NA |
| CHRNB2 | 336 | K | -28.61168254 | 23.51714619 | 0.557028417 | 15.54136192 | NA | NA | NA | NA | NA | NA | NA |
| CHRNB2 | 337 | V | -27.25985714 | 30.16009029 | 0.73925902 | 22.20876518 | NA | NA | NA | NA | NA | NA | NA |
| CHRNB2 | 338 | V | -23.251 | 29.08589643 | 0.709791514 | 23.72279187 | NA | NA | NA | NA | NA | NA | NA |
| CHRNB2 | 339 | F | -22.53166234 | 24.35030353 | 0.57988376 | 19.0258649 | NA | NA | NA | NA | NA | NA | NA |
| CHRNB2 | 340 | L | -27.59757143 | 27.35492133 | 0.662307051 | 20.17237045 | NA | NA | NA | NA | NA | NA | NA |
| CHRNB2 | 341 | E | -25.30112698 | 31.27993211 | 0.769978751 | 25.84467112 | NA | NA | NA | NA | NA | NA | NA |
| CHRNB2 | 342 | K | -19.76757143 | 27.98192522 | 0.67950715 | 22.80214231 | NA | NA | NA | NA | NA | NA | NA |
| CHRNB2 | 343 | L | -22.83419643 | 26.32490543 | 0.634051444 | 20.92842419 | NA | NA | NA | NA | NA | NA | NA |
| CHRNB2 | 344 | P | -22.95085714 | 31.08722485 | 0.764692366 | 25.68725973 | NA | NA | NA | NA | NA | NA | NA |
| CHRNB2 | 345 | A | -18.92437143 | 32.59305317 | 0.806000557 | 27.42621168 | NA | NA | NA | NA | NA | NA | NA |
| CHRNB2 | 346 | L | -17.13894643 | 26.87989991 | 0.649276166 | 21.7253149 | NA | NA | NA | NA | NA | NA | NA |
| CHRNB2 | 347 | L | -18.53119643 | 29.58513921 | 0.723486845 | 24.4586732 | NA | NA | NA | NA | NA | NA | NA |
| CHRNB2 | 348 | F | -20.86175325 | 22.50223302 | 0.529187111 | 17.40519729 | NA | NA | NA | NA | NA | NA | NA |
| CHRNB2 | 349 | M | -24.38907143 | 25.39138574 | 0.60844294 | 18.04735423 | NA | NA | NA | NA | NA | NA | NA |
| CHRNB2 | 350 | Q | -27.77834921 | 24.65436763 | 0.588224908 | 16.67699777 | NA | NA | NA | NA | NA | NA | NA |
| CHRNB2 | 351 | Q | -28.92212698 | 23.11602118 | 0.546024673 | 15.13981718 | NA | NA | NA | NA | NA | NA | NA |
| CHRNB2 | 446 | S | -32.57540476 | 15.33606521 | 0.332603327 | 7.358300939 | NA | NA | NA | NA | NA | NA | NA |
| CHRNB2 | 447 | V | -30.10357143 | 16.70972693 | 0.370285897 | 8.733145034 | NA | NA | NA | NA | NA | NA | NA |
| CHRNB2 | 448 | S | -28.0047381 | 12.65654815 | 0.259098266 | 4.678932151 | 7.98 | -0.53 | -3.6 | 50 | 1.6 | 4 | 0 |
| CHRNB2 | 449 | E | -28.14679365 | 12.4611814 | 0.253738926 | 4.488187292 | 7.98 | -0.53 | -3.6 | 50 | 1.6 | 4 | 0 |
| CHRNB2 | 450 | D | -26.96532143 | 17.10883944 | 0.381234433 | 9.13360552 | NA | NA | NA | NA | NA | NA | NA |

| | | | | | | | | | | | | | |
|---|---|---|---|---|---|---|---|---|---|---|---|---|---|
| CHRNB2 | 451 | W | -24.19442857 | 16.55456028 | 0.366029334 | 11.10952224 | NA | NA | NA | NA | NA | NA | NA |
| CHRNB2 | 452 | K | -23.08368254 | 9.643743723 | 0.176450398 | 4.352071134 | 5.56 | -0.96 | -2.47 | 34.26 | 2.02 | 2 | -2 |
| CHRNB2 | 453 | Y | -23.8002381 | 17.34056441 | 0.38759116 | 11.91328079 | NA | NA | NA | NA | NA | NA | NA |
| CHRNB2 | 454 | V | -21.13614286 | 19.46541233 | 0.445880423 | 14.09494568 | NA | NA | NA | NA | NA | NA | NA |
| CHRNB2 | 455 | A | -18.16217143 | 16.51507763 | 0.364946238 | 11.3505782 | NA | NA | NA | NA | NA | NA | NA |
| CHRNB2 | 456 | M | -18.73769643 | 16.88830927 | 0.375184804 | 11.80600557 | NA | NA | NA | NA | NA | NA | NA |
| CHRNB2 | 457 | V | -18.56071429 | 20.92336402 | 0.485875253 | 15.75599662 | NA | NA | NA | NA | NA | NA | NA |
| CHRNB2 | 458 | I | -15.57444643 | 20.59620909 | 0.476900672 | 16.65991776 | NA | NA | NA | NA | NA | NA | NA |
| CHRNB2 | 459 | D | -13.62294643 | 17.14863852 | 0.38232621 | 15.26018672 | NA | NA | NA | NA | NA | NA | NA |
| CHRNB2 | 460 | R | -14.74466234 | 21.54414737 | 0.502904709 | 19.6291634 | NA | NA | NA | NA | NA | NA | NA |
| CHRNB2 | 461 | L | -12.69694643 | 24.18513349 | 0.575352781 | 22.26305622 | NA | NA | NA | NA | NA | NA | NA |
| CHRNB2 | 462 | F | -9.859662338 | 22.03642387 | 0.516408939 | 20.28378549 | NA | NA | NA | NA | NA | NA | NA |
| CHRNB2 | 463 | L | -8.979946429 | 20.90059516 | 0.485250653 | 19.10814952 | NA | NA | NA | NA | NA | NA | NA |
| CHRNB2 | 464 | W | -11.088 | 25.23833231 | 0.604244347 | 23.44937472 | NA | NA | NA | NA | NA | NA | NA |
| CHRNB2 | 465 | I | -6.900821429 | 25.85928185 | 0.621278362 | 23.35485377 | NA | NA | NA | NA | NA | NA | NA |
| CHRNB2 | 466 | F | -3.590207792 | 21.35003656 | 0.497579822 | 18.92740756 | NA | NA | NA | NA | NA | NA | NA |
| CHRNB2 | 467 | V | -4.206714286 | 25.57976546 | 0.613610611 | 23.10491561 | NA | NA | NA | NA | NA | NA | NA |
| CHRNB2 | 468 | F | -3.398116883 | 28.32852649 | 0.689015187 | 25.85322028 | NA | NA | NA | NA | NA | NA | NA |
| CHRNB2 | 469 | V | -0.712428571 | 27.2043843 | 0.658177488 | 23.03837558 | NA | NA | NA | NA | NA | NA | NA |
| CHRNB2 | 470 | C | 0.091238095 | 24.88806061 | 0.594635622 | 20.69713457 | NA | NA | NA | NA | NA | NA | NA |
| CHRNB2 | 471 | V | 0.319857143 | 28.92560993 | 0.705394502 | 24.72656438 | NA | NA | NA | NA | NA | NA | NA |
| CHRNB2 | 472 | F | 0.912844156 | 30.76819444 | 0.755940658 | 26.56506671 | NA | NA | NA | NA | NA | NA | NA |
| CHRNB2 | 473 | G | 4.196821429 | 28.25044895 | 0.686873348 | 24.58905382 | NA | NA | NA | NA | NA | NA | NA |
| CHRNB2 | 474 | T | 4.420142857 | 27.18286392 | 0.657587137 | 23.54778993 | NA | NA | NA | NA | NA | NA | NA |
| CHRNB2 | 475 | I | 4.746071429 | 31.16971803 | 0.766955336 | 27.5195188 | NA | NA | NA | NA | NA | NA | NA |
| CHRNB2 | 476 | G | 8.303821429 | 32.09735587 | 0.792402487 | 27.09134828 | NA | NA | NA | NA | NA | NA | NA |
| CHRNB2 | 477 | M | 10.26932143 | 27.04180049 | 0.653717456 | 21.96978227 | NA | NA | NA | NA | NA | NA | NA |
| CHRNB2 | 478 | F | 8.355662338 | 32.50241419 | 0.80351413 | 27.43657742 | NA | NA | NA | NA | NA | NA | NA |
| GABRA1 | 39 | T | 76.59614286 | 24.90089263 | 0.649151808 | 15.91504369 | NA | NA | NA | NA | NA | NA | NA |
| GABRA1 | 40 | T | 80.51371429 | 24.94755529 | 0.650539884 | 15.95695629 | NA | NA | NA | NA | NA | NA | NA |
| GABRA1 | 41 | V | 79.87557143 | 26.12889592 | 0.685681259 | 17.07739242 | NA | NA | NA | NA | NA | NA | NA |
| GABRA1 | 42 | F | 75.46728571 | 23.35896612 | 0.603284073 | 14.27261456 | NA | NA | NA | NA | NA | NA | NA |
| GABRA1 | 43 | T | 77.38714286 | 20.42418001 | 0.515982901 | 11.43262441 | NA | NA | NA | NA | NA | NA | NA |

| | | | | | | | | | | | | | |
|---|---|---|---|---|---|---|---|---|---|---|---|---|---|
| GABRA1 | 44 | R | 82.1101039 | 22.95712752 | 0.591330568 | 13.90094002 | NA | NA | NA | NA | NA | NA | NA |
| GABRA1 | 45 | I | 77.54566071 | 22.55716689 | 0.579432927 | 13.42218162 | NA | NA | NA | NA | NA | NA | NA |
| GABRA1 | 46 | L | 74.61478571 | 18.94608419 | 0.472013939 | 9.80660061 | NA | NA | NA | NA | NA | NA | NA |
| GABRA1 | 47 | D | 78.28866071 | 17.25575492 | 0.421731663 | 8.225420033 | NA | NA | NA | NA | NA | NA | NA |
| GABRA1 | 48 | R | 81.56519481 | 19.31584456 | 0.483013212 | 10.31378614 | NA | NA | NA | NA | NA | NA | NA |
| GABRA1 | 49 | L | 75.96166071 | 20.2247797 | 0.510051334 | 11.01772469 | NA | NA | NA | NA | NA | NA | NA |
| GABRA1 | 50 | L | 75.40228571 | 15.54113468 | 0.370726802 | 6.296619551 | NA | NA | NA | NA | NA | NA | NA |
| GABRA1 | 51 | D | 80.25203571 | 18.13160128 | 0.447785491 | 9.037922806 | NA | NA | NA | NA | NA | NA | NA |
| GABRA1 | 52 | G | 78.87553571 | 19.23597333 | 0.48063728 | 10.02366065 | NA | NA | NA | NA | NA | NA | NA |
| GABRA1 | 53 | Y | 73.54845238 | 16.87112477 | 0.410290058 | 7.562648415 | NA | NA | NA | NA | NA | NA | NA |
| GABRA1 | 54 | D | 73.52016071 | 16.33154585 | 0.394239187 | 7.016437209 | NA | NA | NA | NA | NA | NA | NA |
| GABRA1 | 55 | N | 69.68053571 | 13.11534067 | 0.298566633 | 4.057945201 | 9.09 | 0.19 | 0.05 | 13.32 | -0.5 | 1 | 1 |
| GABRA1 | 56 | R | 71.2561039 | 15.20074663 | 0.360601268 | 6.173871663 | NA | NA | NA | NA | NA | NA | NA |
| GABRA1 | 57 | L | 71.14316071 | 19.67971303 | 0.493837219 | 10.63379307 | NA | NA | NA | NA | NA | NA | NA |
| GABRA1 | 58 | R | 67.38192208 | 22.96011903 | 0.591419556 | 14.01022289 | NA | NA | NA | NA | NA | NA | NA |
| GABRA1 | 59 | P | 68.81314286 | 24.4054423 | 0.634413632 | 15.35724416 | NA | NA | NA | NA | NA | NA | NA |
| GABRA1 | 60 | G | 70.44828571 | 27.77184895 | 0.734554233 | 18.48305522 | NA | NA | NA | NA | NA | NA | NA |
| GABRA1 | 61 | L | 70.91203571 | 25.25130329 | 0.659575485 | 15.96406444 | NA | NA | NA | NA | NA | NA | NA |
| GABRA1 | 62 | G | 73.01803571 | 29.6380107 | 0.790067004 | 20.34206876 | NA | NA | NA | NA | NA | NA | NA |
| GABRA1 | 63 | E | 73.18761905 | 31.07436767 | 0.832794359 | 21.75586274 | NA | NA | NA | NA | NA | NA | NA |
| GABRA1 | 64 | R | 69.24101299 | 30.08767962 | 0.803443319 | 21.02492547 | NA | NA | NA | NA | NA | NA | NA |
| GABRA1 | 65 | V | 64.707 | 31.56455465 | 0.847375966 | 22.59928667 | NA | NA | NA | NA | NA | NA | NA |
| GABRA1 | 66 | T | 64.09671429 | 27.2150741 | 0.717991835 | 18.2574131 | NA | NA | NA | NA | NA | NA | NA |
| GABRA1 | 67 | E | 61.46106349 | 29.50047503 | 0.785975726 | 20.55495692 | NA | NA | NA | NA | NA | NA | NA |
| GABRA1 | 68 | V | 59.01728571 | 26.20491912 | 0.687942724 | 17.26457341 | NA | NA | NA | NA | NA | NA | NA |
| GABRA1 | 69 | K | 58.04928571 | 28.62837825 | 0.760033437 | 19.69184694 | NA | NA | NA | NA | NA | NA | NA |
| GABRA1 | 70 | T | 53.69328571 | 26.9018851 | 0.708675392 | 19.58466567 | NA | NA | NA | NA | NA | NA | NA |
| GABRA1 | 71 | D | 51.75266071 | 26.33402472 | 0.691783232 | 22.26839078 | NA | NA | NA | NA | NA | NA | NA |
| GABRA1 | 72 | I | 48.06128571 | 24.36772088 | 0.633291532 | 20.29481581 | NA | NA | NA | NA | NA | NA | NA |
| GABRA1 | 73 | F | 46.78628571 | 24.35540897 | 0.632925289 | 20.25079351 | NA | NA | NA | NA | NA | NA | NA |
| GABRA1 | 74 | V | 43.28142857 | 23.59304853 | 0.610247329 | 19.47222617 | NA | NA | NA | NA | NA | NA | NA |
| GABRA1 | 75 | T | 43.45342857 | 21.92793654 | 0.560715192 | 18.12816148 | NA | NA | NA | NA | NA | NA | NA |
| GABRA1 | 76 | S | 40.38078571 | 20.20039962 | 0.509326099 | 16.34535351 | NA | NA | NA | NA | NA | NA | NA |

| | | | | | | | | | | | | | |
|---|---|---|---|---|---|---|---|---|---|---|---|---|---|
| GABRA1 | 77 | F | 37.08983117 | 20.61902117 | 0.521778847 | 16.79386081 | NA | NA | NA | NA | NA | NA | NA |
| GABRA1 | 78 | G | 36.26303571 | 17.88079061 | 0.440324618 | 10.88850967 | NA | NA | NA | NA | NA | NA | NA |
| GABRA1 | 79 | P | 34.37642857 | 14.46564181 | 0.338734082 | 7.459687827 | NA | NA | NA | NA | NA | NA | NA |
| GABRA1 | 80 | V | 32.07685714 | 15.97075283 | 0.383506666 | 8.11885869 | NA | NA | NA | NA | NA | NA | NA |
| GABRA1 | 81 | S | 31.97645238 | 12.70983544 | 0.286504057 | 4.654424682 | 8.12 | -0.88 | -3.5 | 21.91 | 1.38 | 2 | -2 |
| GABRA1 | 82 | D | 28.12953571 | 14.16497886 | 0.329790252 | 8.313208399 | NA | NA | NA | NA | NA | NA | NA |
| GABRA1 | 83 | H | 27.85718571 | 9.378879784 | 0.187418017 | 3.578292398 | 5.91 | 0.05 | -0.93 | 25.63 | 0.47 | 2 | 2 |
| GABRA1 | 84 | D | 33.22491071 | 10.24757518 | 0.213259125 | 2.31886855 | 7.99 | -0.75 | -3.58 | 21.87 | 1.14 | 2 | 0 |
| GABRA1 | 85 | M | 32.87178571 | 15.72760497 | 0.376273739 | 7.738139326 | NA | NA | NA | NA | NA | NA | NA |
| GABRA1 | 86 | E | 35.85173016 | 11.62174746 | 0.25413667 | 7.768989367 | NA | NA | NA | NA | NA | NA | NA |
| GABRA1 | 87 | Y | 37.98770238 | 16.08334284 | 0.386855884 | 12.34487006 | NA | NA | NA | NA | NA | NA | NA |
| GABRA1 | 88 | T | 38.64542857 | 13.82572739 | 0.319698529 | 9.955702634 | NA | NA | NA | NA | NA | NA | NA |
| GABRA1 | 89 | I | 41.80591071 | 16.98789993 | 0.413763772 | 12.87285328 | NA | NA | NA | NA | NA | NA | NA |
| GABRA1 | 90 | D | 43.99828571 | 18.22099114 | 0.450444574 | 14.09925618 | NA | NA | NA | NA | NA | NA | NA |
| GABRA1 | 91 | V | 46.88185714 | 19.77758744 | 0.496748692 | 15.67003371 | NA | NA | NA | NA | NA | NA | NA |
| GABRA1 | 92 | F | 49.84455844 | 22.4589325 | 0.576510745 | 18.3763928 | NA | NA | NA | NA | NA | NA | NA |
| GABRA1 | 93 | F | 52.11301299 | 21.61833106 | 0.551505349 | 14.21190663 | NA | NA | NA | NA | NA | NA | NA |
| GABRA1 | 94 | R | 53.32755844 | 24.50447102 | 0.637359442 | 17.14118932 | NA | NA | NA | NA | NA | NA | NA |
| GABRA1 | 95 | Q | 57.22373016 | 21.06790533 | 0.535131817 | 12.13455898 | NA | NA | NA | NA | NA | NA | NA |
| GABRA1 | 96 | S | 60.17928571 | 26.55983874 | 0.698500529 | 17.61900772 | NA | NA | NA | NA | NA | NA | NA |
| GABRA1 | 97 | W | 63.30707143 | 21.65716503 | 0.552660544 | 12.71026535 | NA | NA | NA | NA | NA | NA | NA |
| GABRA1 | 98 | K | 67.92273016 | 27.79728471 | 0.735310872 | 18.81771995 | NA | NA | NA | NA | NA | NA | NA |
| GABRA1 | 99 | D | 69.98553571 | 22.92150416 | 0.590270879 | 13.85123053 | NA | NA | NA | NA | NA | NA | NA |
| GABRA1 | 100 | E | 74.10973016 | 27.34563956 | 0.72187577 | 18.0268671 | NA | NA | NA | NA | NA | NA | NA |
| GABRA1 | 101 | R | 73.60546753 | 21.42430202 | 0.545733561 | 12.11050199 | NA | NA | NA | NA | NA | NA | NA |
| GABRA1 | 102 | L | 72.36953571 | 21.51075619 | 0.548305316 | 12.19338742 | NA | NA | NA | NA | NA | NA | NA |
| GABRA1 | 103 | K | 75.67584127 | 26.76959236 | 0.704740076 | 17.47111136 | NA | NA | NA | NA | NA | NA | NA |
| GABRA1 | 104 | F | 74.69964935 | 27.03193426 | 0.712543969 | 17.7850333 | NA | NA | NA | NA | NA | NA | NA |
| GABRA1 | 105 | K | 79.4321746 | 29.97327068 | 0.800039993 | 20.79085333 | NA | NA | NA | NA | NA | NA | NA |
| GABRA1 | 106 | G | 77.41178571 | 31.69306973 | 0.851198908 | 22.55489376 | NA | NA | NA | NA | NA | NA | NA |
| GABRA1 | 107 | P | 76.47528571 | 32.16153414 | 0.865134333 | 23.05223685 | NA | NA | NA | NA | NA | NA | NA |
| GABRA1 | 108 | M | 73.41153571 | 29.17846605 | 0.776396915 | 20.03510716 | NA | NA | NA | NA | NA | NA | NA |
| GABRA1 | 109 | T | 70.28942857 | 30.76603662 | 0.823622426 | 21.97911912 | NA | NA | NA | NA | NA | NA | NA |

| | | | | | | | | | | | | | |
|---|---|---|---|---|---|---|---|---|---|---|---|---|---|
| GABRA1 | 110 | V | 67.51842857 | 28.4387162 | 0.754391554 | 19.43790739 | NA | NA | NA | NA | NA | NA | NA |
| GABRA1 | 111 | L | 69.37103571 | 23.76653892 | 0.615408153 | 14.97027257 | NA | NA | NA | NA | NA | NA | NA |
| GABRA1 | 112 | R | 64.76283117 | 22.31362841 | 0.57218838 | 13.35659622 | NA | NA | NA | NA | NA | NA | NA |
| GABRA1 | 113 | L | 66.40341071 | 19.13247538 | 0.477558523 | 10.12713649 | NA | NA | NA | NA | NA | NA | NA |
| GABRA1 | 114 | N | 62.54178571 | 16.23466837 | 0.391357369 | 7.27703195 | NA | NA | NA | NA | NA | NA | NA |
| GABRA1 | 115 | N | 63.18091071 | 11.10618152 | 0.238800114 | 2.231830488 | 9.01 | -0.48 | -1 | 25.3 | 1.15 | 2 | -2 |
| GABRA1 | 116 | L | 67.84353571 | 10.46217783 | 0.219642917 | 1.722280066 | 8.83 | -0.48 | -1 | 25.3 | 1.15 | 2 | -2 |
| GABRA1 | 117 | M | 69.39066071 | 14.50015824 | 0.339760844 | 5.703723731 | NA | NA | NA | NA | NA | NA | NA |
| GABRA1 | 118 | A | 64.96008571 | 14.48928846 | 0.3394375 | 5.49772918 | NA | NA | NA | NA | NA | NA | NA |
| GABRA1 | 119 | S | 65.77195238 | 10.48451668 | 0.220307431 | 1.530823085 | 9.01 | -0.48 | -1 | 25.3 | 1.15 | 2 | -2 |
| GABRA1 | 120 | K | 70.06973016 | 11.41315473 | 0.247931656 | 2.551269056 | 9.01 | -0.05 | -0.66 | 20.6 | 0.13 | 2 | 0 |
| GABRA1 | 121 | I | 67.94891071 | 15.79718032 | 0.378343399 | 6.814122683 | NA | NA | NA | NA | NA | NA | NA |
| GABRA1 | 122 | W | 65.33807143 | 18.3383198 | 0.453934753 | 9.383501156 | NA | NA | NA | NA | NA | NA | NA |
| GABRA1 | 123 | T | 62.54157143 | 16.70411309 | 0.405321956 | 7.758894695 | NA | NA | NA | NA | NA | NA | NA |
| GABRA1 | 124 | P | 60.046 | 19.60060971 | 0.49148413 | 10.66154237 | NA | NA | NA | NA | NA | NA | NA |
| GABRA1 | 125 | D | 58.44578571 | 19.34221981 | 0.483797797 | 10.39915929 | NA | NA | NA | NA | NA | NA | NA |
| GABRA1 | 126 | T | 54.77614286 | 19.11153173 | 0.476935512 | 11.82138798 | NA | NA | NA | NA | NA | NA | NA |
| GABRA1 | 127 | F | 51.56592208 | 18.59033077 | 0.461431331 | 14.58013409 | NA | NA | NA | NA | NA | NA | NA |
| GABRA1 | 128 | F | 48.23864935 | 17.41672997 | 0.426520192 | 13.37123924 | NA | NA | NA | NA | NA | NA | NA |
| GABRA1 | 129 | H | 47.67658571 | 18.80346495 | 0.46777144 | 14.7582089 | NA | NA | NA | NA | NA | NA | NA |
| GABRA1 | 130 | N | 42.81366071 | 16.53246712 | 0.400215998 | 12.45831996 | NA | NA | NA | NA | NA | NA | NA |
| GABRA1 | 131 | G | 44.38728571 | 13.95980566 | 0.323686959 | 9.899396388 | NA | NA | NA | NA | NA | NA | NA |
| GABRA1 | 132 | K | 41.24428571 | 10.26209048 | 0.213690912 | 6.213567797 | NA | NA | NA | NA | NA | NA | NA |
| GABRA1 | 133 | K | 45.20706349 | 5.605740005 | 0.075178312 | 1.591726415 | 4.13 | -0.57 | -3.8 | 49.55 | 1.75 | 4 | 2 |
| GABRA1 | 134 | S | 46.96578571 | 9.940638863 | 0.204128681 | 5.895483557 | NA | NA | NA | NA | NA | NA | NA |
| GABRA1 | 135 | V | 47.72742857 | 8.735027247 | 0.168265316 | 4.641095731 | 4.13 | -0.57 | -3.8 | 49.55 | 1.75 | 4 | 2 |
| GABRA1 | 136 | A | 51.26868571 | 10.87962614 | 0.232060765 | 3.40669207 | 7.5 | -0.06 | -2.95 | 14.59 | 0.02 | 1 | 1 |
| GABRA1 | 137 | H | 49.32708571 | 12.61004703 | 0.283535648 | 5.1533397 | NA | NA | NA | NA | NA | NA | NA |
| GABRA1 | 138 | N | 53.99428571 | 8.983750943 | 0.175664107 | 1.736861078 | 7.5 | -0.06 | -2.95 | 14.59 | 0.02 | 1 | 1 |
| GABRA1 | 139 | M | 51.57466071 | 13.29597495 | 0.303939967 | 5.838414437 | NA | NA | NA | NA | NA | NA | NA |
| GABRA1 | 140 | T | 54.76814286 | 14.16013065 | 0.329646032 | 6.793185918 | NA | NA | NA | NA | NA | NA | NA |
| GABRA1 | 141 | M | 57.91166071 | 13.70467636 | 0.31609762 | 7.022548345 | NA | NA | NA | NA | NA | NA | NA |
| GABRA1 | 142 | P | 57.65628571 | 11.72109611 | 0.257091997 | 4.908228565 | 7.41 | 0.06 | -2.63 | 2.95 | -0.47 | 0 | 0 |

| | | | | | | | | | | | | | |
|---|---|---|---|---|---|---|---|---|---|---|---|---|---|
| GABRA1 | 143 | N | 54.27491071 | 15.47089467 | 0.36863737 | 8.104412549 | NA | NA | NA | NA | NA | NA | NA |
| GABRA1 | 144 | K | 58.02495238 | 13.30410392 | 0.304181779 | 4.376275993 | 9.01 | -0.34 | -2.18 | 25.79 | 0.97 | 2 | -2 |
| GABRA1 | 145 | L | 59.49278571 | 19.3591943 | 0.484302738 | 10.41599754 | NA | NA | NA | NA | NA | NA | NA |
| GABRA1 | 146 | L | 64.07716071 | 19.25181981 | 0.481108666 | 10.35446132 | NA | NA | NA | NA | NA | NA | NA |
| GABRA1 | 147 | R | 62.47401299 | 23.06027446 | 0.594398883 | 14.08918142 | NA | NA | NA | NA | NA | NA | NA |
| GABRA1 | 148 | I | 68.55953571 | 22.54411715 | 0.579044736 | 13.6525702 | NA | NA | NA | NA | NA | NA | NA |
| GABRA1 | 149 | T | 68.49114286 | 27.89071268 | 0.738090077 | 18.92002607 | NA | NA | NA | NA | NA | NA | NA |
| GABRA1 | 150 | E | 72.71961905 | 30.72104858 | 0.822284165 | 21.44861226 | NA | NA | NA | NA | NA | NA | NA |
| GABRA1 | 151 | D | 70.72178571 | 30.86696551 | 0.826624761 | 21.5469688 | NA | NA | NA | NA | NA | NA | NA |
| GABRA1 | 152 | G | 69.59978571 | 27.53918105 | 0.727633054 | 18.51536278 | NA | NA | NA | NA | NA | NA | NA |
| GABRA1 | 153 | T | 66.26371429 | 27.77580856 | 0.73467202 | 18.78481655 | NA | NA | NA | NA | NA | NA | NA |
| GABRA1 | 154 | L | 65.21428571 | 23.05806484 | 0.594333153 | 14.07408649 | NA | NA | NA | NA | NA | NA | NA |
| GABRA1 | 155 | L | 60.74341071 | 23.2250496 | 0.599300454 | 14.27536243 | NA | NA | NA | NA | NA | NA | NA |
| GABRA1 | 156 | Y | 59.67395238 | 15.73861195 | 0.376601164 | 6.800702349 | NA | NA | NA | NA | NA | NA | NA |
| GABRA1 | 157 | T | 56.44471429 | 19.87173533 | 0.499549312 | 10.93913663 | NA | NA | NA | NA | NA | NA | NA |
| GABRA1 | 158 | M | 54.01316071 | 17.24981052 | 0.421554834 | 9.900571284 | NA | NA | NA | NA | NA | NA | NA |
| GABRA1 | 159 | R | 49.58719481 | 16.30547426 | 0.393463635 | 8.830670154 | NA | NA | NA | NA | NA | NA | NA |
| GABRA1 | 160 | L | 49.17366071 | 15.72056201 | 0.376064232 | 11.89514909 | NA | NA | NA | NA | NA | NA | NA |
| GABRA1 | 161 | T | 44.94871429 | 12.80008816 | 0.289188807 | 8.700465379 | NA | NA | NA | NA | NA | NA | NA |
| GABRA1 | 162 | V | 44.23485714 | 14.55855041 | 0.341497837 | 10.45675705 | NA | NA | NA | NA | NA | NA | NA |
| GABRA1 | 163 | R | 41.1721039 | 7.625634728 | 0.135264182 | 3.903885795 | 4.13 | -0.57 | -3.8 | 49.55 | 1.75 | 4 | 2 |
| GABRA1 | 164 | A | 40.53388571 | 15.32351193 | 0.364253171 | 11.22468671 | NA | NA | NA | NA | NA | NA | NA |
| GABRA1 | 165 | E | 38.52295238 | 16.97977824 | 0.413522176 | 12.88626777 | NA | NA | NA | NA | NA | NA | NA |
| GABRA1 | 166 | C | 35.07695238 | 19.29251295 | 0.482319166 | 15.18797491 | NA | NA | NA | NA | NA | NA | NA |
| GABRA1 | 167 | P | 34.34671429 | 22.1398341 | 0.567018515 | 18.03666744 | NA | NA | NA | NA | NA | NA | NA |
| GABRA1 | 168 | M | 30.22278571 | 18.68750208 | 0.464321889 | 12.84442454 | NA | NA | NA | NA | NA | NA | NA |
| GABRA1 | 169 | H | 28.34348571 | 22.61061885 | 0.581022964 | 16.92585843 | NA | NA | NA | NA | NA | NA | NA |
| GABRA1 | 170 | L | 24.46841071 | 17.43518838 | 0.427069275 | 12.25226639 | NA | NA | NA | NA | NA | NA | NA |
| GABRA1 | 171 | E | 22.0121746 | 22.3363405 | 0.572863997 | 18.71723875 | NA | NA | NA | NA | NA | NA | NA |
| GABRA1 | 172 | D | 20.63353571 | 22.61021099 | 0.581010831 | 18.94625266 | NA | NA | NA | NA | NA | NA | NA |
| GABRA1 | 173 | F | 20.12783117 | 15.95202467 | 0.382949559 | 12.29095099 | NA | NA | NA | NA | NA | NA | NA |
| GABRA1 | 174 | P | 19.38957143 | 19.93941123 | 0.501562469 | 16.26023475 | NA | NA | NA | NA | NA | NA | NA |
| GABRA1 | 175 | M | 18.80703571 | 23.06829367 | 0.594637431 | 19.39426193 | NA | NA | NA | NA | NA | NA | NA |

| | | | | | | | | | | | | | |
|---|---|---|---|---|---|---|---|---|---|---|---|---|---|
| GABRA1 | 176 | D | 23.83516071 | 21.87834674 | 0.559240043 | 18.27972763 | NA | NA | NA | NA | NA | NA | NA |
| GABRA1 | 177 | A | 27.07108571 | 26.23922718 | 0.688963287 | 21.26873891 | NA | NA | NA | NA | NA | NA | NA |
| GABRA1 | 178 | H | 29.41278571 | 22.20713984 | 0.569020661 | 16.42672353 | NA | NA | NA | NA | NA | NA | NA |
| GABRA1 | 179 | A | 34.07748571 | 26.09056605 | 0.68454106 | 22.46701337 | NA | NA | NA | NA | NA | NA | NA |
| GABRA1 | 180 | C | 36.16645238 | 22.89879818 | 0.589595443 | 19.02393539 | NA | NA | NA | NA | NA | NA | NA |
| GABRA1 | 181 | P | 39.53428571 | 21.5954253 | 0.55082397 | 17.5070673 | NA | NA | NA | NA | NA | NA | NA |
| GABRA1 | 182 | L | 43.48766071 | 20.00697391 | 0.503572258 | 15.92249595 | NA | NA | NA | NA | NA | NA | NA |
| GABRA1 | 183 | K | 44.45895238 | 21.37223453 | 0.544184708 | 17.32335708 | NA | NA | NA | NA | NA | NA | NA |
| GABRA1 | 184 | F | 49.70137662 | 22.21425313 | 0.56923226 | 18.17196223 | NA | NA | NA | NA | NA | NA | NA |
| GABRA1 | 185 | G | 52.73803571 | 22.90668472 | 0.589830044 | 18.89148444 | NA | NA | NA | NA | NA | NA | NA |
| GABRA1 | 186 | S | 55.93011905 | 23.9626866 | 0.621242964 | 16.61750087 | NA | NA | NA | NA | NA | NA | NA |
| GABRA1 | 187 | Y | 55.39411905 | 20.4225232 | 0.515933615 | 13.27343302 | NA | NA | NA | NA | NA | NA | NA |
| GABRA1 | 188 | A | 59.61048571 | 22.16792904 | 0.567854256 | 13.21637591 | NA | NA | NA | NA | NA | NA | NA |
| GABRA1 | 189 | Y | 61.87003571 | 21.58563332 | 0.550532688 | 12.67096654 | NA | NA | NA | NA | NA | NA | NA |
| GABRA1 | 190 | T | 59.30228571 | 28.25004851 | 0.74877925 | 19.30073067 | NA | NA | NA | NA | NA | NA | NA |
| GABRA1 | 191 | R | 57.55428571 | 31.50953954 | 0.84573943 | 22.56193266 | NA | NA | NA | NA | NA | NA | NA |
| GABRA1 | 192 | A | 62.44668571 | 32.06492685 | 0.862260553 | 23.11941398 | NA | NA | NA | NA | NA | NA | NA |
| GABRA1 | 193 | E | 63.24584127 | 28.52481174 | 0.756952641 | 19.58110466 | NA | NA | NA | NA | NA | NA | NA |
| GABRA1 | 194 | V | 60.37014286 | 26.88680407 | 0.708226776 | 17.94143091 | NA | NA | NA | NA | NA | NA | NA |
| GABRA1 | 195 | V | 58.67 | 30.11544406 | 0.804269229 | 21.17038806 | NA | NA | NA | NA | NA | NA | NA |
| GABRA1 | 196 | Y | 55.79878571 | 24.25506021 | 0.629940211 | 15.31788695 | NA | NA | NA | NA | NA | NA | NA |
| GABRA1 | 197 | E | 55.78284127 | 31.57025591 | 0.847545561 | 27.53694835 | NA | NA | NA | NA | NA | NA | NA |
| GABRA1 | 198 | W | 50.56471429 | 28.00160219 | 0.74138871 | 23.93634774 | NA | NA | NA | NA | NA | NA | NA |
| GABRA1 | 199 | T | 54.31257143 | 31.58669175 | 0.848034479 | 22.65247863 | NA | NA | NA | NA | NA | NA | NA |
| GABRA1 | 200 | R | 51.9861039 | 33.28778494 | 0.89863695 | 29.20702751 | NA | NA | NA | NA | NA | NA | NA |
| GABRA1 | 201 | E | 49.08061905 | 34.81323089 | 0.944014438 | 30.74058579 | NA | NA | NA | NA | NA | NA | NA |
| GABRA1 | 202 | P | 47.609 | 33.19194313 | 0.895785941 | 29.11110047 | NA | NA | NA | NA | NA | NA | NA |
| GABRA1 | 203 | A | 44.57088571 | 33.9035325 | 0.916953612 | 29.80706064 | NA | NA | NA | NA | NA | NA | NA |
| GABRA1 | 204 | R | 47.03583117 | 33.59991622 | 0.907921929 | 29.50994539 | NA | NA | NA | NA | NA | NA | NA |
| GABRA1 | 205 | S | 48.34745238 | 30.29876587 | 0.809722508 | 26.2118029 | NA | NA | NA | NA | NA | NA | NA |
| GABRA1 | 206 | V | 45.02528571 | 27.36462806 | 0.722440621 | 23.26747035 | NA | NA | NA | NA | NA | NA | NA |
| GABRA1 | 207 | V | 44.63071429 | 27.97860384 | 0.740704578 | 23.86729558 | NA | NA | NA | NA | NA | NA | NA |
| GABRA1 | 208 | V | 41.00585714 | 27.44828803 | 0.724929257 | 23.32836945 | NA | NA | NA | NA | NA | NA | NA |

| | | | | | | | | | | | | | |
|---|---|---|---|---|---|---|---|---|---|---|---|---|---|
| GABRA1 | 209 | A | 40.64408571 | 27.79006133 | 0.735095998 | 23.96829633 | NA | NA | NA | NA | NA | NA | NA |
| GABRA1 | 210 | E | 39.28828571 | 29.88049297 | 0.797280132 | 22.90613027 | NA | NA | NA | NA | NA | NA | NA |
| GABRA1 | 211 | D | 37.11216071 | 25.48712438 | 0.666590462 | 18.50956321 | NA | NA | NA | NA | NA | NA | NA |
| GABRA1 | 212 | G | 37.39528571 | 24.41191916 | 0.634606299 | 17.43433051 | NA | NA | NA | NA | NA | NA | NA |
| GABRA1 | 213 | S | 34.09795238 | 22.68378042 | 0.583199303 | 15.68193979 | NA | NA | NA | NA | NA | NA | NA |
| GABRA1 | 214 | R | 36.15637662 | 18.3368316 | 0.453890483 | 11.48095702 | NA | NA | NA | NA | NA | NA | NA |
| GABRA1 | 215 | L | 31.13878571 | 21.10784274 | 0.536319837 | 14.08162378 | NA | NA | NA | NA | NA | NA | NA |
| GABRA1 | 216 | N | 27.66203571 | 16.69184307 | 0.404956959 | 8.092208748 | NA | NA | NA | NA | NA | NA | NA |
| GABRA1 | 217 | Q | 25.69939683 | 16.42225965 | 0.396937653 | 11.46255117 | NA | NA | NA | NA | NA | NA | NA |
| GABRA1 | 218 | Y | 28.21345238 | 19.29815424 | 0.482486977 | 11.02287984 | NA | NA | NA | NA | NA | NA | NA |
| GABRA1 | 219 | D | 27.88916071 | 25.44162298 | 0.66523693 | 16.83852634 | NA | NA | NA | NA | NA | NA | NA |
| GABRA1 | 220 | L | 33.44228571 | 27.13838466 | 0.715710552 | 20.18185083 | NA | NA | NA | NA | NA | NA | NA |
| GABRA1 | 221 | L | 30.76266071 | 29.97776634 | 0.800173726 | 21.39914419 | NA | NA | NA | NA | NA | NA | NA |
| GABRA1 | 222 | G | 35.21778571 | 30.92281011 | 0.828285971 | 27.07700545 | NA | NA | NA | NA | NA | NA | NA |
| GABRA1 | 223 | Q | 39.23728571 | 29.46317278 | 0.784866095 | 25.35447993 | NA | NA | NA | NA | NA | NA | NA |
| GABRA1 | 224 | T | 39.50214286 | 30.24188237 | 0.808030393 | 26.15074029 | NA | NA | NA | NA | NA | NA | NA |
| GABRA1 | 225 | V | 43.712 | 30.20921995 | 0.807058783 | 26.12359607 | NA | NA | NA | NA | NA | NA | NA |
| GABRA1 | 226 | D | 44.90616071 | 30.66356832 | 0.820574298 | 26.60150577 | NA | NA | NA | NA | NA | NA | NA |
| GABRA1 | 227 | S | 48.32611905 | 32.4492383 | 0.873692677 | 28.39685784 | NA | NA | NA | NA | NA | NA | NA |
| GABRA1 | 228 | G | 49.17678571 | 31.54340277 | 0.84674676 | 27.50764222 | NA | NA | NA | NA | NA | NA | NA |
| GABRA1 | 229 | I | 50.60216071 | 31.50172976 | 0.845507112 | 27.46896112 | NA | NA | NA | NA | NA | NA | NA |
| GABRA1 | 230 | V | 49.59442857 | 27.95766607 | 0.740081741 | 23.92950799 | NA | NA | NA | NA | NA | NA | NA |
| GABRA1 | 231 | Q | 50.58395238 | 31.05991378 | 0.832364399 | 27.03968249 | NA | NA | NA | NA | NA | NA | NA |
| GABRA1 | 232 | S | 51.75828571 | 27.23514725 | 0.718588952 | 23.22350263 | NA | NA | NA | NA | NA | NA | NA |
| GABRA1 | 233 | S | 53.20211905 | 28.59546656 | 0.759054412 | 24.58847978 | NA | NA | NA | NA | NA | NA | NA |
| GABRA1 | 234 | T | 56.26171429 | 27.94346373 | 0.739659263 | 23.9464321 | NA | NA | NA | NA | NA | NA | NA |
| GABRA1 | 235 | G | 55.53878571 | 31.12294441 | 0.834239373 | 27.11479599 | NA | NA | NA | NA | NA | NA | NA |
| GABRA1 | 236 | E | 55.11828571 | 31.53008603 | 0.846350627 | 27.52011924 | NA | NA | NA | NA | NA | NA | NA |
| GABRA1 | 237 | Y | 54.68436905 | 27.41873166 | 0.724050043 | 23.61982995 | NA | NA | NA | NA | NA | NA | NA |
| GABRA1 | 238 | V | 53.82942857 | 27.96375402 | 0.740262839 | 23.94028603 | NA | NA | NA | NA | NA | NA | NA |
| GABRA1 | 239 | V | 49.57914286 | 26.55803671 | 0.698446924 | 22.5147944 | NA | NA | NA | NA | NA | NA | NA |
| GABRA1 | 240 | M | 49.68266071 | 25.87072548 | 0.678001455 | 21.81614653 | NA | NA | NA | NA | NA | NA | NA |
| GABRA1 | 241 | T | 45.00471429 | 27.37126396 | 0.72263802 | 23.30510111 | NA | NA | NA | NA | NA | NA | NA |

| | | | | | | | | | | | | | |
|---|---|---|---|---|---|---|---|---|---|---|---|---|---|
| GABRA1 | 242 | T | 42.88471429 | 25.49171407 | 0.666726991 | 21.40150594 | NA | NA | NA | NA | NA | NA | NA |
| GABRA1 | 243 | H | 39.08078571 | 27.56797274 | 0.728489522 | 23.47060195 | NA | NA | NA | NA | NA | NA | NA |
| GABRA1 | 244 | F | 38.27501299 | 24.0317144 | 0.623296336 | 19.92647052 | NA | NA | NA | NA | NA | NA | NA |
| GABRA1 | 245 | H | 33.90078571 | 28.36745216 | 0.75227166 | 24.65213678 | NA | NA | NA | NA | NA | NA | NA |
| GABRA1 | 246 | L | 31.21816071 | 23.33162472 | 0.602470747 | 14.85778129 | NA | NA | NA | NA | NA | NA | NA |
| GABRA1 | 247 | K | 25.57139683 | 25.05577526 | 0.653759106 | 20.07806957 | NA | NA | NA | NA | NA | NA | NA |
| GABRA1 | 248 | R | 24.14874026 | 18.75985109 | 0.466474057 | 13.78908574 | NA | NA | NA | NA | NA | NA | NA |
| GABRA1 | 249 | K | 23.17039683 | 21.48603812 | 0.547570026 | 16.79369139 | NA | NA | NA | NA | NA | NA | NA |
| GABRA1 | 250 | I | 17.76841071 | 21.77367936 | 0.556126499 | 18.21772191 | NA | NA | NA | NA | NA | NA | NA |
| GABRA1 | 251 | G | 16.80578571 | 21.67561905 | 0.553209496 | 18.82762935 | NA | NA | NA | NA | NA | NA | NA |
| GABRA1 | 252 | Y | 17.89203571 | 13.81537698 | 0.319390634 | 10.82566794 | NA | NA | NA | NA | NA | NA | NA |
| GABRA1 | 253 | F | 17.39328571 | 15.48182115 | 0.3689624 | 13.36795333 | NA | NA | NA | NA | NA | NA | NA |
| GABRA1 | 254 | V | 12.49042857 | 19.46790749 | 0.487536633 | 17.08424888 | NA | NA | NA | NA | NA | NA | NA |
| GABRA1 | 255 | I | 12.12103571 | 17.17748634 | 0.419403405 | 14.89325268 | NA | NA | NA | NA | NA | NA | NA |
| GABRA1 | 256 | Q | 13.29973016 | 13.71973619 | 0.316545605 | 11.37174853 | NA | NA | NA | NA | NA | NA | NA |
| GABRA1 | 257 | T | 11.71828571 | 14.37062462 | 0.335907603 | 11.9566397 | NA | NA | NA | NA | NA | NA | NA |
| GABRA1 | 258 | Y | 11.32920238 | 17.74118003 | 0.436171618 | 15.10854014 | NA | NA | NA | NA | NA | NA | NA |
| GABRA1 | 259 | L | 7.351410714 | 17.65087165 | 0.433485212 | 14.83045797 | NA | NA | NA | NA | NA | NA | NA |
| GABRA1 | 260 | P | 6.741 | 14.10333563 | 0.327956549 | 11.34581979 | NA | NA | NA | NA | NA | NA | NA |
| GABRA1 | 261 | C | 5.860119048 | 14.54654743 | 0.341140784 | 11.96866064 | NA | NA | NA | NA | NA | NA | NA |
| GABRA1 | 262 | I | 3.940285714 | 17.8170183 | 0.438427581 | 14.88843525 | NA | NA | NA | NA | NA | NA | NA |
| GABRA1 | 263 | M | 1.886785714 | 15.83417841 | 0.379443983 | 14.06717386 | NA | NA | NA | NA | NA | NA | NA |
| GABRA1 | 264 | T | 1.132857143 | 12.29599764 | 0.274193611 | 10.48595088 | NA | NA | NA | NA | NA | NA | NA |
| GABRA1 | 265 | V | 0.520285714 | 16.29600678 | 0.393182005 | 14.48708007 | NA | NA | NA | NA | NA | NA | NA |
| GABRA1 | 266 | I | -0.751785714 | 17.66797314 | 0.433993931 | 15.59570194 | NA | NA | NA | NA | NA | NA | NA |
| GABRA1 | 267 | L | -2.349035714 | 12.89413607 | 0.291986453 | 10.11944918 | NA | NA | NA | NA | NA | NA | NA |
| GABRA1 | 268 | S | -3.060285714 | 14.51548993 | 0.340216916 | 11.53642798 | NA | NA | NA | NA | NA | NA | NA |
| GABRA1 | 269 | Q | -4.049730159 | 16.66747161 | 0.404231981 | 13.68335008 | NA | NA | NA | NA | NA | NA | NA |
| GABRA1 | 270 | V | -6.852428571 | 16.10888122 | 0.387615575 | 14.18095774 | NA | NA | NA | NA | NA | NA | NA |
| GABRA1 | 271 | S | -9.049952381 | 13.34038942 | 0.305261165 | 11.28141144 | NA | NA | NA | NA | NA | NA | NA |
| GABRA1 | 272 | F | -9.387194805 | 16.48327486 | 0.398752674 | 14.28056739 | NA | NA | NA | NA | NA | NA | NA |
| GABRA1 | 273 | W | -11.24135714 | 16.13297967 | 0.388332433 | 12.08187353 | NA | NA | NA | NA | NA | NA | NA |
| GABRA1 | 274 | L | -13.54753571 | 11.69741206 | 0.256387467 | 8.4854682 | NA | NA | NA | NA | NA | NA | NA |

| | | | | | | | | | | | | | |
|---|---|---|---|---|---|---|---|---|---|---|---|---|---|
| GABRA1 | 275 | N | -17.71578571 | 9.444129446 | 0.189359 | 5.404355637 | NA | NA | NA | NA | NA | NA | NA |
| GABRA1 | 276 | R | -19.08792208 | 11.01941673 | 0.236219119 | 6.946216258 | NA | NA | NA | NA | NA | NA | NA |
| GABRA1 | 277 | E | -20.38728571 | 6.788632116 | 0.11036584 | 2.715357696 | 4.09 | -0.28 | -0.55 | 1.21 | -0.23 | 0 | 0 |
| GABRA1 | 278 | S | -15.80811905 | 7.071307729 | 0.1187746 | 3.037028347 | 4.09 | -0.28 | -0.55 | 1.21 | -0.23 | 0 | 0 |
| GABRA1 | 279 | V | -15.07885714 | 6.676271489 | 0.107023445 | 4.545697902 | 2.78 | -0.05 | -0.47 | 1.05 | -0.67 | 0 | 0 |
| GABRA1 | 280 | P | -13.30485714 | 3.12619801 | 0.001419301 | 1.589973671 | 1.63 | -0.04 | 0.1 | 0.79 | -0.36 | 0 | 0 |
| GABRA1 | 281 | A | -11.55688571 | 5.689581004 | 0.077672333 | 3.452877463 | 2.34 | -0.04 | 0.1 | 0.79 | -0.36 | 0 | 0 |
| GABRA1 | 282 | R | -11.73937662 | 8.073515343 | 0.148587301 | 5.908027091 | NA | NA | NA | NA | NA | NA | NA |
| GABRA1 | 283 | T | -8.659714286 | 7.394825488 | 0.128398293 | 5.407385014 | NA | NA | NA | NA | NA | NA | NA |
| GABRA1 | 284 | V | -6.633142857 | 3.560084269 | 0.014326128 | 1.864502025 | 1.96 | 0.33 | 2.35 | 0.48 | -0.23 | 0 | 0 |
| GABRA1 | 285 | F | -6.314285714 | 8.113302287 | 0.149770844 | 5.090234594 | NA | NA | NA | NA | NA | NA | NA |
| GABRA1 | 286 | G | -3.922785714 | 9.650734946 | 0.1955049 | 6.621352714 | NA | NA | NA | NA | NA | NA | NA |
| GABRA1 | 287 | V | -2.533571429 | 7.73782198 | 0.13860142 | 4.705900322 | 3.07 | 0.18 | 1.75 | 0.9 | -0.27 | 0 | 0 |
| GABRA1 | 288 | T | -0.931142857 | 4.879289395 | 0.053568564 | 2.711946833 | 2.18 | 0.66 | 2.68 | 0.51 | -1.15 | 0 | 0 |
| GABRA1 | 289 | T | -0.478428571 | 8.731624248 | 0.168164086 | 6.91294458 | NA | NA | NA | NA | NA | NA | NA |
| GABRA1 | 290 | V | 0.561714286 | 9.020977663 | 0.176771491 | 7.214039871 | NA | NA | NA | NA | NA | NA | NA |
| GABRA1 | 291 | L | 1.868160714 | 3.087448947 | 2.67E-04 | 1.415803483 | 1.81 | 1.14 | 3.8 | 0.13 | -1.79 | 0 | 0 |
| GABRA1 | 292 | T | 2.789571429 | 7.211616254 | 0.122948362 | 4.906052414 | 2.41 | 1.14 | 3.8 | 0.13 | -1.79 | 0 | 0 |
| GABRA1 | 293 | M | 4.724285714 | 9.470375758 | 0.19013975 | 6.538527706 | NA | NA | NA | NA | NA | NA | NA |
| GABRA1 | 294 | T | 6.247285714 | 7.411600165 | 0.12889729 | 4.718024639 | 2.82 | -0.77 | -0.7 | 1.66 | 0.77 | 0 | 0 |
| GABRA1 | 295 | T | 7.661285714 | 4.375114284 | 0.038570851 | 1.552386534 | 2.93 | -0.51 | -1.33 | 14.15 | 0.53 | 0 | 0 |
| GABRA1 | 296 | L | 8.336035714 | 8.312094922 | 0.155684335 | 5.456587932 | NA | NA | NA | NA | NA | NA | NA |
| GABRA1 | 297 | S | 10.66545238 | 9.332128642 | 0.186027309 | 6.820339905 | NA | NA | NA | NA | NA | NA | NA |
| GABRA1 | 298 | I | 12.42653571 | 4.714582272 | 0.048669016 | 2.178662063 | 2.59 | -0.7 | -2.73 | 38.27 | 1.26 | 2 | -2 |
| GABRA1 | 299 | S | 13.93495238 | 7.033268017 | 0.117643032 | 4.749655653 | 2.43 | -0.67 | -3.4 | 50.47 | 1.42 | 2 | -2 |
| GABRA1 | 300 | A | 14.87908571 | 9.604300807 | 0.194123623 | 7.207179942 | NA | NA | NA | NA | NA | NA | NA |
| GABRA1 | 301 | R | 17.83146753 | 9.471983636 | 0.19018758 | 7.167898382 | NA | NA | NA | NA | NA | NA | NA |
| GABRA1 | 302 | N | 18.44416071 | 4.63148799 | 0.046197208 | 2.06585951 | 3 | -1.02 | -3.5 | 34.39 | 1.67 | 2 | -2 |
| GABRA1 | 303 | S | 20.05695238 | 8.507117197 | 0.161485668 | 4.825115085 | 3.69 | -1.02 | -3.5 | 34.39 | 1.67 | 2 | -2 |
| GABRA1 | 304 | L | 21.46591071 | 10.0190016 | 0.20645974 | 6.4302643 | NA | NA | NA | NA | NA | NA | NA |
| GABRA1 | 305 | P | 24.61257143 | 11.81571441 | 0.259906611 | 6.289967215 | NA | NA | NA | NA | NA | NA | NA |
| GABRA1 | 306 | K | 23.53673016 | 7.188250204 | 0.122253291 | 1.696597848 | 5.53 | -0.59 | -3.8 | 49.6 | 1.65 | 4 | 2 |
| GABRA1 | 307 | V | 23.37657143 | 14.41438427 | 0.337209323 | 8.89118069 | NA | NA | NA | NA | NA | NA | NA |

| | | | | | | | | | | | | | |
|---|---|---|---|---|---|---|---|---|---|---|---|---|---|
| GABRA1 | 308 | A | 22.11328571 | 17.21263051 | 0.420448839 | 13.6068244 | NA | NA | NA | NA | NA | NA | NA |
| GABRA1 | 309 | Y | 21.97470238 | 18.50830865 | 0.458991417 | 14.84195431 | NA | NA | NA | NA | NA | NA | NA |
| GABRA1 | 310 | A | 17.34568571 | 20.30138729 | 0.512330182 | 17.31392222 | NA | NA | NA | NA | NA | NA | NA |
| GABRA1 | 311 | T | 17.86928571 | 17.92925545 | 0.441766303 | 14.9279167 | NA | NA | NA | NA | NA | NA | NA |
| GABRA1 | 312 | A | 15.86588571 | 21.06661748 | 0.535093508 | 18.52098181 | NA | NA | NA | NA | NA | NA | NA |
| GABRA1 | 313 | M | 14.78928571 | 15.62094815 | 0.373101015 | 13.07388136 | NA | NA | NA | NA | NA | NA | NA |
| GABRA1 | 314 | D | 14.33928571 | 15.53819645 | 0.370639398 | 13.01905071 | NA | NA | NA | NA | NA | NA | NA |
| GABRA1 | 315 | W | 12.473 | 20.06023235 | 0.505156538 | 17.53179268 | NA | NA | NA | NA | NA | NA | NA |
| GABRA1 | 316 | F | 9.031922078 | 19.39531745 | 0.485377295 | 17.06342251 | NA | NA | NA | NA | NA | NA | NA |
| GABRA1 | 317 | I | 9.427035714 | 14.45714419 | 0.338481303 | 11.94820903 | NA | NA | NA | NA | NA | NA | NA |
| GABRA1 | 318 | A | 7.591485714 | 17.93799827 | 0.442026376 | 15.2127452 | NA | NA | NA | NA | NA | NA | NA |
| GABRA1 | 319 | V | 5.417285714 | 20.08928344 | 0.506020722 | 17.31941934 | NA | NA | NA | NA | NA | NA | NA |
| GABRA1 | 320 | C | 4.373619048 | 16.81895627 | 0.4087382 | 13.9059345 | NA | NA | NA | NA | NA | NA | NA |
| GABRA1 | 321 | Y | 3.789285714 | 12.59577108 | 0.28311098 | 9.741677872 | NA | NA | NA | NA | NA | NA | NA |
| GABRA1 | 322 | A | 1.165485714 | 18.27656439 | 0.452097713 | 15.9208568 | NA | NA | NA | NA | NA | NA | NA |
| GABRA1 | 323 | F | 0.16374026 | 18.35272517 | 0.45436327 | 16.50516668 | NA | NA | NA | NA | NA | NA | NA |
| GABRA1 | 324 | V | 0 | 13.46616252 | 0.309002541 | 11.63684688 | NA | NA | NA | NA | NA | NA | NA |
| GABRA1 | 325 | F | -0.507012987 | 16.26232582 | 0.392180097 | 14.41495224 | NA | NA | NA | NA | NA | NA | NA |
| GABRA1 | 326 | S | -2.784952381 | 18.36804478 | 0.454818983 | 15.34066721 | NA | NA | NA | NA | NA | NA | NA |
| GABRA1 | 327 | A | -5.123485714 | 15.90134576 | 0.381442012 | 13.02971633 | NA | NA | NA | NA | NA | NA | NA |
| GABRA1 | 328 | L | -6.112410714 | 11.61504012 | 0.253937147 | 9.802146767 | NA | NA | NA | NA | NA | NA | NA |
| GABRA1 | 329 | I | -5.994285714 | 16.82713487 | 0.408981489 | 14.90146187 | NA | NA | NA | NA | NA | NA | NA |
| GABRA1 | 330 | E | -9.701952381 | 17.64020527 | 0.433167919 | 15.63745633 | NA | NA | NA | NA | NA | NA | NA |
| GABRA1 | 331 | F | -11.5241039 | 10.9495093 | 0.234139581 | 8.89607168 | NA | NA | NA | NA | NA | NA | NA |
| GABRA1 | 332 | A | -11.23488571 | 15.87619255 | 0.380693778 | 13.55904843 | NA | NA | NA | NA | NA | NA | NA |
| GABRA1 | 333 | T | -12.862 | 18.25481076 | 0.451450607 | 16.71845807 | NA | NA | NA | NA | NA | NA | NA |
| GABRA1 | 334 | V | -15.10928571 | 15.80791533 | 0.378662734 | 11.74189281 | NA | NA | NA | NA | NA | NA | NA |
| GABRA1 | 335 | N | -15.96841071 | 11.69737941 | 0.256386496 | 9.034048824 | NA | NA | NA | NA | NA | NA | NA |
| GABRA1 | 336 | Y | -17.21570238 | 16.99944746 | 0.414107277 | 12.96729156 | NA | NA | NA | NA | NA | NA | NA |
| GABRA1 | 337 | F | -19.01555844 | 17.36190036 | 0.424889174 | 12.96231704 | NA | NA | NA | NA | NA | NA | NA |
| GABRA1 | 338 | T | -21.20771429 | 12.57947495 | 0.282626219 | 7.51598623 | NA | NA | NA | NA | NA | NA | NA |
| GABRA1 | 339 | K | -25.4311746 | 14.25741625 | 0.33253999 | 9.595836384 | NA | NA | NA | NA | NA | NA | NA |
| GABRA1 | 340 | R | -26.56646753 | 10.99568374 | 0.235513134 | 5.506106191 | NA | NA | NA | NA | NA | NA | NA |

| | | | | | | | | | | | | | |
|---|---|---|---|---|---|---|---|---|---|---|---|---|---|
| GABRA1 | 341 | G | -24.08303571 | 6.664696692 | 0.106679129 | 1.589121159 | 5.11 | -0.67 | -0.8 | 2.5 | 0.36 | 0 | 0 |
| GABRA1 | 342 | Y | -27.36970238 | 6.914284996 | 0.11410364 | 1.586682005 | 5.35 | -0.47 | -1.2 | 12.29 | 0.72 | 1 | -1 |
| GABRA1 | 343 | A | -24.65648571 | 6.979380488 | 0.116040038 | 1.655618921 | 5.35 | -0.47 | -1.2 | 12.29 | 0.72 | 1 | -1 |
| GABRA1 | 344 | W | -26.899 | 8.520997477 | 0.161898565 | 3.086918631 | 5.45 | -0.16 | -0.7 | 2.79 | 0.06 | 0 | 0 |
| GABRA1 | 345 | D | -29.23066071 | 7.024476493 | 0.11738151 | 1.607238767 | 5.52 | 0.1 | -1.63 | 14.08 | 0.14 | 1 | -1 |
| GABRA1 | 346 | G | -29.43453571 | 10.25557414 | 0.213497071 | 4.728132372 | 5.53 | 0.1 | -1.63 | 14.08 | 0.14 | 1 | -1 |
| GABRA1 | 347 | K | -33.18806349 | 12.62907831 | 0.284101772 | 7.261267518 | NA | NA | NA | NA | NA | NA | NA |
| GABRA1 | 411 | K | -33.64450794 | 7.030099146 | 0.117548767 | 1.503954122 | 5.54 | 0.6 | -2.17 | 17.57 | -0.47 | 1 | 1 |
| GABRA1 | 412 | T | -32.13957143 | 12.67088509 | 0.285345399 | 7.245049477 | NA | NA | NA | NA | NA | NA | NA |
| GABRA1 | 413 | F | -29.25064935 | 10.19320568 | 0.211641794 | 4.85409831 | 5.39 | 0.19 | -2.5 | 25.61 | 0.3 | 2 | 0 |
| GABRA1 | 414 | N | -24.86928571 | 12.72124121 | 0.286843344 | 7.381834258 | NA | NA | NA | NA | NA | NA | NA |
| GABRA1 | 415 | S | -23.24361905 | 16.99799312 | 0.414064014 | 11.70058542 | NA | NA | NA | NA | NA | NA | NA |
| GABRA1 | 416 | V | -20.21471429 | 18.81450448 | 0.468099833 | 14.78908085 | NA | NA | NA | NA | NA | NA | NA |
| GABRA1 | 417 | S | -18.12578571 | 17.9169011 | 0.441398798 | 13.84458312 | NA | NA | NA | NA | NA | NA | NA |
| GABRA1 | 418 | K | -18.70573016 | 21.96922249 | 0.561943327 | 17.90657169 | NA | NA | NA | NA | NA | NA | NA |
| GABRA1 | 419 | I | -14.22528571 | 20.82284594 | 0.527842028 | 16.75547484 | NA | NA | NA | NA | NA | NA | NA |
| GABRA1 | 420 | D | -13.54478571 | 17.28661578 | 0.422649681 | 13.2328871 | NA | NA | NA | NA | NA | NA | NA |
| GABRA1 | 421 | R | -17.26001299 | 22.19337602 | 0.568611228 | 18.12273699 | NA | NA | NA | NA | NA | NA | NA |
| GABRA1 | 422 | L | -12.12291071 | 24.80202802 | 0.646210879 | 20.73959118 | NA | NA | NA | NA | NA | NA | NA |
| GABRA1 | 423 | S | -9.320952381 | 21.25529264 | 0.540706033 | 19.05811223 | NA | NA | NA | NA | NA | NA | NA |
| GABRA1 | 424 | R | -11.91646753 | 18.5307002 | 0.459657499 | 18.02899012 | NA | NA | NA | NA | NA | NA | NA |
| GABRA1 | 425 | I | -8.701410714 | 25.6856794 | 0.672496884 | 23.4829382 | NA | NA | NA | NA | NA | NA | NA |
| GABRA1 | 426 | A | -5.608885714 | 26.34727375 | 0.692177351 | 24.61078987 | NA | NA | NA | NA | NA | NA | NA |
| GABRA1 | 427 | F | -4.272285714 | 21.29376721 | 0.541850538 | 18.26464565 | NA | NA | NA | NA | NA | NA | NA |
| GABRA1 | 428 | P | -3.432 | 23.95211577 | 0.620928514 | 20.96343517 | NA | NA | NA | NA | NA | NA | NA |
| GABRA1 | 429 | L | -2.291660714 | 27.18793253 | 0.717184454 | 24.1998527 | NA | NA | NA | NA | NA | NA | NA |
| GABRA1 | 430 | L | -0.783660714 | 26.97465985 | 0.710840225 | 25.17607418 | NA | NA | NA | NA | NA | NA | NA |
| GABRA1 | 431 | F | 1.121012987 | 21.45808962 | 0.546738641 | 19.76841665 | NA | NA | NA | NA | NA | NA | NA |
| GABRA1 | 432 | G | 0.913535714 | 26.92857861 | 0.709469445 | 25.11941779 | NA | NA | NA | NA | NA | NA | NA |
| GABRA1 | 433 | I | 1.419035714 | 29.37367954 | 0.782203937 | 26.54175222 | NA | NA | NA | NA | NA | NA | NA |
| GABRA1 | 434 | F | 5.429194805 | 23.83943317 | 0.617576541 | 21.02142568 | NA | NA | NA | NA | NA | NA | NA |
| GABRA1 | 435 | N | 5.411410714 | 24.3345806 | 0.632305707 | 21.41059571 | NA | NA | NA | NA | NA | NA | NA |
| GABRA1 | 436 | L | 5.419285714 | 29.60213295 | 0.788999748 | 26.67662824 | NA | NA | NA | NA | NA | NA | NA |

| | | | | | | | | | | | | | |
|---|---|---|---|---|---|---|---|---|---|---|---|---|---|
| GABRA1 | 437 | V | 7.907285714 | 30.14610985 | 0.805181445 | 27.47956968 | NA | NA | NA | NA | NA | NA | NA |
| GABRA1 | 438 | Y | 11.13761905 | 22.20017502 | 0.568813479 | 19.83021005 | NA | NA | NA | NA | NA | NA | NA |
| GABRA1 | 439 | W | 11.10835714 | 23.83229114 | 0.617364087 | 21.44782638 | NA | NA | NA | NA | NA | NA | NA |
| GABRA1 | 440 | A | 12.16788571 | 30.97012678 | 0.829693502 | 28.59439111 | NA | NA | NA | NA | NA | NA | NA |
| GABRA1 | 441 | T | 14.04414286 | 30.68978863 | 0.821354274 | 28.33214958 | NA | NA | NA | NA | NA | NA | NA |
| GABRA1 | 442 | Y | 15.79061905 | 25.62112179 | 0.670576487 | 23.15672238 | NA | NA | NA | NA | NA | NA | NA |
| GABRA1 | 443 | L | 15.94978571 | 28.58608744 | 0.758775411 | 25.60647343 | NA | NA | NA | NA | NA | NA | NA |
| GABRA1 | 444 | N | 17.80628571 | 32.08008033 | 0.862711324 | 28.39924135 | NA | NA | NA | NA | NA | NA | NA |
| GABRA1 | 445 | R | 20.2591039 | 28.68087643 | 0.761595102 | 25.01507234 | NA | NA | NA | NA | NA | NA | NA |
| GABRA5 | 45 | N | 80.50421591 | 28.15765583 | 0.68287684 | 16.68154108 | NA | NA | NA | NA | NA | NA | NA |
| GABRA5 | 46 | I | 77.23209091 | 24.25261145 | 0.565927271 | 14.67894143 | NA | NA | NA | NA | NA | NA | NA |
| GABRA5 | 47 | T | 81.56109091 | 23.37892421 | 0.539761795 | 11.93934789 | NA | NA | NA | NA | NA | NA | NA |
| GABRA5 | 48 | I | 81.55334091 | 25.12969329 | 0.59219441 | 13.55345624 | NA | NA | NA | NA | NA | NA | NA |
| GABRA5 | 49 | F | 77.17618182 | 22.89480216 | 0.525263147 | 13.34682895 | NA | NA | NA | NA | NA | NA | NA |
| GABRA5 | 50 | T | 79.29251948 | 19.10551078 | 0.411780187 | 7.593771688 | NA | NA | NA | NA | NA | NA | NA |
| GABRA5 | 51 | R | 84.56063636 | 21.07956589 | 0.470899846 | 7.976621459 | NA | NA | NA | NA | NA | NA | NA |
| GABRA5 | 52 | I | 80.14071591 | 22.42195125 | 0.51110205 | 10.89956952 | NA | NA | NA | NA | NA | NA | NA |
| GABRA5 | 53 | L | 76.72571591 | 19.10935054 | 0.411895181 | 9.642075976 | NA | NA | NA | NA | NA | NA | NA |
| GABRA5 | 54 | D | 79.49146591 | 16.22073278 | 0.325385895 | 4.850886627 | 11.59 | -0.93 | -3.6 | 15.02 | 0.46 | 1 | 1 |
| GABRA5 | 55 | G | 81.66709091 | 18.98225608 | 0.408088914 | 5.866711889 | NA | NA | NA | NA | NA | NA | NA |
| GABRA5 | 56 | L | 78.04971591 | 19.69348024 | 0.429388892 | 8.210664444 | NA | NA | NA | NA | NA | NA | NA |
| GABRA5 | 57 | L | 76.93134091 | 16.14395227 | 0.323086447 | 6.598859654 | NA | NA | NA | NA | NA | NA | NA |
| GABRA5 | 58 | D | 81.78521591 | 15.13637357 | 0.292911145 | 3.723489067 | 11.59 | -0.93 | -3.6 | 15.02 | 0.46 | 1 | 1 |
| GABRA5 | 59 | G | 79.83784091 | 16.62017371 | 0.337348485 | 5.028234804 | NA | NA | NA | NA | NA | NA | NA |
| GABRA5 | 60 | Y | 74.58684091 | 16.35752564 | 0.329482613 | 6.80689608 | NA | NA | NA | NA | NA | NA | NA |
| GABRA5 | 61 | D | 74.11934091 | 16.46073665 | 0.332573611 | 7.042567892 | NA | NA | NA | NA | NA | NA | NA |
| GABRA5 | 62 | N | 70.50896591 | 13.61350701 | 0.24730383 | 4.278574765 | 9.37 | -0.94 | -3.04 | 12.35 | 0.59 | 1 | 1 |
| GABRA5 | 63 | R | 71.79863636 | 14.35548205 | 0.269524746 | 5.361762647 | NA | NA | NA | NA | NA | NA | NA |
| GABRA5 | 64 | L | 71.69346591 | 20.40433077 | 0.45067768 | 11.10527643 | NA | NA | NA | NA | NA | NA | NA |
| GABRA5 | 65 | R | 68.38972727 | 23.4918535 | 0.543143839 | 14.79163787 | NA | NA | NA | NA | NA | NA | NA |
| GABRA5 | 66 | P | 70.07437662 | 25.42743135 | 0.601111168 | 16.17075481 | NA | NA | NA | NA | NA | NA | NA |
| GABRA5 | 67 | G | 71.28584091 | 28.80178503 | 0.702167436 | 19.43724987 | NA | NA | NA | NA | NA | NA | NA |
| GABRA5 | 68 | L | 71.35096591 | 26.34552353 | 0.628606499 | 16.98622645 | NA | NA | NA | NA | NA | NA | NA |

| | | | | | | | | | | | | | |
|---|---|---|---|---|---|---|---|---|---|---|---|---|---|
| GABRA5 | 69 | G | 72.94784091 | 30.44914069 | 0.751502991 | 21.0296309 | NA | NA | NA | NA | NA | NA | NA |
| GABRA5 | 70 | E | 73.77909091 | 32.55832313 | 0.814669489 | 23.13879703 | NA | NA | NA | NA | NA | NA | NA |
| GABRA5 | 71 | R | 70.08954545 | 31.8898802 | 0.794650738 | 23.1916718 | NA | NA | NA | NA | NA | NA | NA |
| GABRA5 | 72 | I | 65.52134091 | 33.05083492 | 0.829419396 | 24.60499324 | NA | NA | NA | NA | NA | NA | NA |
| GABRA5 | 73 | T | 64.99594805 | 28.96507209 | 0.707057611 | 20.52570854 | NA | NA | NA | NA | NA | NA | NA |
| GABRA5 | 74 | Q | 62.5399798 | 30.54317019 | 0.754319018 | 22.15160462 | NA | NA | NA | NA | NA | NA | NA |
| GABRA5 | 75 | V | 59.77337662 | 27.49894751 | 0.663149624 | 19.09707412 | NA | NA | NA | NA | NA | NA | NA |
| GABRA5 | 76 | R | 59.48454545 | 28.6642235 | 0.698047697 | 20.30317176 | NA | NA | NA | NA | NA | NA | NA |
| GABRA5 | 77 | T | 54.42994805 | 28.22543346 | 0.684906667 | 20.35465156 | NA | NA | NA | NA | NA | NA | NA |
| GABRA5 | 78 | D | 52.71934091 | 27.31251049 | 0.657566146 | 19.6633321 | NA | NA | NA | NA | NA | NA | NA |
| GABRA5 | 79 | I | 48.83784091 | 26.46231993 | 0.632104356 | 21.37492619 | NA | NA | NA | NA | NA | NA | NA |
| GABRA5 | 80 | Y | 47.43067424 | 25.40467408 | 0.600429626 | 20.56606679 | NA | NA | NA | NA | NA | NA | NA |
| GABRA5 | 81 | V | 43.79180519 | 25.0122048 | 0.588675826 | 20.49207751 | NA | NA | NA | NA | NA | NA | NA |
| GABRA5 | 82 | T | 43.95851948 | 23.4643767 | 0.542320954 | 18.92660386 | NA | NA | NA | NA | NA | NA | NA |
| GABRA5 | 83 | S | 40.66642424 | 20.74330678 | 0.460829447 | 16.01165063 | NA | NA | NA | NA | NA | NA | NA |
| GABRA5 | 84 | F | 37.17218182 | 20.79608357 | 0.462410024 | 16.03421246 | NA | NA | NA | NA | NA | NA | NA |
| GABRA5 | 85 | G | 36.11209091 | 18.4446123 | 0.391987379 | 10.28292208 | NA | NA | NA | NA | NA | NA | NA |
| GABRA5 | 86 | P | 33.90880519 | 15.2680764 | 0.296855425 | 6.31805896 | NA | NA | NA | NA | NA | NA | NA |
| GABRA5 | 87 | V | 31.55337662 | 16.37447874 | 0.32999033 | 7.676437934 | NA | NA | NA | NA | NA | NA | NA |
| GABRA5 | 88 | S | 30.59109091 | 12.82817236 | 0.223784367 | 3.747205742 | 9.15 | -0.08 | -0.42 | 11.38 | 0.14 | 1 | -1 |
| GABRA5 | 89 | D | 27.31434091 | 15.65208919 | 0.308355968 | 6.521559497 | NA | NA | NA | NA | NA | NA | NA |
| GABRA5 | 90 | T | 28.08723377 | 11.99177059 | 0.198735528 | 2.870873239 | 9.15 | -0.08 | -0.42 | 11.38 | 0.14 | 1 | -1 |
| GABRA5 | 91 | E | 32.23442424 | 10.09521763 | 0.141936929 | 1.374287899 | 8.73 | -0.86 | -3.5 | 15.01 | 0.96 | 1 | -1 |
| GABRA5 | 92 | M | 32.01359091 | 16.93568508 | 0.346797525 | 7.974410877 | NA | NA | NA | NA | NA | NA | NA |
| GABRA5 | 93 | E | 35.37542424 | 12.19364593 | 0.204781358 | 4.074104164 | 8.17 | -0.56 | -3.82 | 49.58 | 1.61 | 5 | 3 |
| GABRA5 | 94 | Y | 37.60984091 | 17.1934922 | 0.354518418 | 9.27847133 | NA | NA | NA | NA | NA | NA | NA |
| GABRA5 | 95 | T | 38.49651948 | 14.55829544 | 0.275598668 | 9.791717952 | NA | NA | NA | NA | NA | NA | NA |
| GABRA5 | 96 | I | 41.95609091 | 17.87284625 | 0.374863939 | 13.32139796 | NA | NA | NA | NA | NA | NA | NA |
| GABRA5 | 97 | D | 44.03446591 | 19.0213464 | 0.409259604 | 14.54898647 | NA | NA | NA | NA | NA | NA | NA |
| GABRA5 | 98 | V | 47.08494805 | 20.88773923 | 0.465154958 | 16.0415953 | NA | NA | NA | NA | NA | NA | NA |
| GABRA5 | 99 | F | 49.67590909 | 22.6472618 | 0.517849725 | 15.27751589 | NA | NA | NA | NA | NA | NA | NA |
| GABRA5 | 100 | F | 52.47690909 | 22.46509217 | 0.512394049 | 14.8133373 | NA | NA | NA | NA | NA | NA | NA |
| GABRA5 | 101 | R | 54.36554545 | 25.19649093 | 0.594194888 | 17.30416038 | NA | NA | NA | NA | NA | NA | NA |

| | | | | | | | | | | | | | |
|---|---|---|---|---|---|---|---|---|---|---|---|---|---|
| GABRA5 | 102 | Q | 57.86542424 | 22.5478717 | 0.514873157 | 14.46816124 | NA | NA | NA | NA | NA | NA | NA |
| GABRA5 | 103 | S | 60.72292424 | 27.15327962 | 0.652797447 | 18.74891423 | NA | NA | NA | NA | NA | NA | NA |
| GABRA5 | 104 | W | 63.94859091 | 22.75452614 | 0.521062114 | 14.36174359 | NA | NA | NA | NA | NA | NA | NA |
| GABRA5 | 105 | K | 68.66831313 | 29.0852464 | 0.710656631 | 20.72617736 | NA | NA | NA | NA | NA | NA | NA |
| GABRA5 | 106 | D | 70.94246591 | 24.35174583 | 0.568896181 | 15.17430443 | NA | NA | NA | NA | NA | NA | NA |
| GABRA5 | 107 | E | 74.91986869 | 28.0279072 | 0.678991085 | 18.65059383 | NA | NA | NA | NA | NA | NA | NA |
| GABRA5 | 108 | R | 74.76927273 | 23.63665122 | 0.547480289 | 14.1710775 | NA | NA | NA | NA | NA | NA | NA |
| GABRA5 | 109 | L | 73.57596591 | 22.67424702 | 0.518657888 | 13.25431358 | NA | NA | NA | NA | NA | NA | NA |
| GABRA5 | 110 | R | 76.754 | 27.42721924 | 0.661001482 | 17.86381119 | NA | NA | NA | NA | NA | NA | NA |
| GABRA5 | 111 | F | 76.718 | 27.68463939 | 0.668710786 | 18.11617178 | NA | NA | NA | NA | NA | NA | NA |
| GABRA5 | 112 | K | 81.0559798 | 30.04250732 | 0.739325 | 18.49518246 | NA | NA | NA | NA | NA | NA | NA |
| GABRA5 | 113 | G | 77.86584091 | 30.86073178 | 0.763829458 | 21.30179674 | NA | NA | NA | NA | NA | NA | NA |
| GABRA5 | 114 | P | 77.00337662 | 31.13799022 | 0.772132886 | 21.57742134 | NA | NA | NA | NA | NA | NA | NA |
| GABRA5 | 115 | M | 72.55959091 | 31.13789084 | 0.77212991 | 21.71924003 | NA | NA | NA | NA | NA | NA | NA |
| GABRA5 | 116 | Q | 72.59342424 | 31.58724531 | 0.785587327 | 22.24590097 | NA | NA | NA | NA | NA | NA | NA |
| GABRA5 | 117 | R | 67.25645455 | 29.22767753 | 0.714922206 | 20.1242192 | NA | NA | NA | NA | NA | NA | NA |
| GABRA5 | 118 | L | 69.91209091 | 25.20341995 | 0.5944024 | 16.18600808 | NA | NA | NA | NA | NA | NA | NA |
| GABRA5 | 119 | P | 66.64166234 | 23.70041875 | 0.54939002 | 15.26628294 | NA | NA | NA | NA | NA | NA | NA |
| GABRA5 | 120 | L | 67.68159091 | 20.20841575 | 0.444810352 | 11.75580336 | NA | NA | NA | NA | NA | NA | NA |
| GABRA5 | 121 | N | 65.90459091 | 15.58690312 | 0.306403754 | 7.159697783 | NA | NA | NA | NA | NA | NA | NA |
| GABRA5 | 122 | N | 62.73034091 | 14.57446747 | 0.276082994 | 6.284583636 | NA | NA | NA | NA | NA | NA | NA |
| GABRA5 | 123 | L | 67.24546591 | 12.15460279 | 0.203612081 | 3.753401649 | 8.47 | -1.04 | -3.5 | 49.7 | 2.63 | 4 | -4 |
| GABRA5 | 124 | L | 68.80834091 | 14.65021897 | 0.278351625 | 6.065210902 | NA | NA | NA | NA | NA | NA | NA |
| GABRA5 | 125 | A | 65.34989091 | 14.37977003 | 0.27025213 | 5.93115488 | NA | NA | NA | NA | NA | NA | NA |
| GABRA5 | 126 | S | 66.93942424 | 10.72436152 | 0.160778739 | 2.340387457 | 8.47 | -1.04 | -3.5 | 49.7 | 2.63 | 4 | -4 |
| GABRA5 | 127 | K | 70.91709091 | 11.05297182 | 0.17062007 | 2.186301145 | 9.12 | -1.03 | -2.96 | 40.09 | 2.33 | 4 | -4 |
| GABRA5 | 128 | I | 68.53209091 | 15.70100666 | 0.309820964 | 7.000236423 | NA | NA | NA | NA | NA | NA | NA |
| GABRA5 | 129 | W | 66.40616234 | 18.45518978 | 0.392304157 | 10.17889095 | NA | NA | NA | NA | NA | NA | NA |
| GABRA5 | 130 | T | 63.36337662 | 17.81799736 | 0.373221307 | 9.409417591 | NA | NA | NA | NA | NA | NA | NA |
| GABRA5 | 131 | P | 61.23923377 | 19.99922523 | 0.438545445 | 11.59376439 | NA | NA | NA | NA | NA | NA | NA |
| GABRA5 | 132 | D | 58.23784091 | 20.08589371 | 0.441141021 | 12.01023497 | NA | NA | NA | NA | NA | NA | NA |
| GABRA5 | 133 | T | 54.26609091 | 19.61357354 | 0.42699582 | 12.12299362 | NA | NA | NA | NA | NA | NA | NA |
| GABRA5 | 134 | F | 51.384 | 20.35918903 | 0.449325761 | 12.70555655 | NA | NA | NA | NA | NA | NA | NA |

| | | | | | | | | | | | | | |
|---|---|---|---|---|---|---|---|---|---|---|---|---|---|
| GABRA5 | 135 | F | 48.16336364 | 19.74489081 | 0.430928553 | 14.96663579 | NA | NA | NA | NA | NA | NA | NA |
| GABRA5 | 136 | H | 46.78039091 | 22.17176087 | 0.503609265 | 17.38100757 | NA | NA | NA | NA | NA | NA | NA |
| GABRA5 | 137 | N | 42.40684091 | 18.52383152 | 0.394359863 | 14.01488234 | NA | NA | NA | NA | NA | NA | NA |
| GABRA5 | 138 | G | 44.46134091 | 16.15890816 | 0.323534351 | 11.65957328 | NA | NA | NA | NA | NA | NA | NA |
| GABRA5 | 139 | K | 40.8979798 | 12.6184108 | 0.217502357 | 8.06931611 | NA | NA | NA | NA | NA | NA | NA |
| GABRA5 | 140 | K | 43.95475758 | 6.565114089 | 0.036216213 | 2.174407681 | 4.57 | -0.41 | -3.9 | 49.5 | 1.46 | 5 | 5 |
| GABRA5 | 141 | S | 47.22125758 | 10.74850264 | 0.161501726 | 6.060977105 | NA | NA | NA | NA | NA | NA | NA |
| GABRA5 | 142 | I | 47.84246591 | 9.362434406 | 0.119991293 | 4.351143709 | 5.18 | -0.41 | -3.9 | 49.5 | 1.46 | 4 | 4 |
| GABRA5 | 143 | A | 52.02009091 | 11.42668832 | 0.181812256 | 3.808616556 | 7.66 | 0.57 | 0.2 | 10.79 | -0.69 | 1 | 1 |
| GABRA5 | 144 | H | 50.10879091 | 12.29922327 | 0.207943223 | 4.96217808 | 7.66 | 0.57 | 0.2 | 10.79 | -0.69 | 1 | 1 |
| GABRA5 | 145 | N | 54.31009091 | 8.810635051 | 0.103465822 | 1.419556728 | 7.62 | 0.64 | 0.7 | 1.36 | -1.03 | 0 | 0 |
| GABRA5 | 146 | M | 51.89846591 | 13.61343032 | 0.247301533 | 5.952438181 | NA | NA | NA | NA | NA | NA | NA |
| GABRA5 | 147 | T | 55.85451948 | 14.97328731 | 0.288026983 | 6.957883735 | NA | NA | NA | NA | NA | NA | NA |
| GABRA5 | 148 | T | 57.72951948 | 14.52805827 | 0.274693116 | 6.546109824 | NA | NA | NA | NA | NA | NA | NA |
| GABRA5 | 149 | P | 57.99980519 | 11.66873554 | 0.189061167 | 3.707244785 | 8.04 | -0.73 | -3.66 | 40.36 | 1.74 | 4 | 0 |
| GABRA5 | 150 | N | 54.85284091 | 15.86444304 | 0.314715612 | 7.908874698 | NA | NA | NA | NA | NA | NA | NA |
| GABRA5 | 151 | K | 59.00353535 | 13.30705978 | 0.238126247 | 5.221593595 | NA | NA | NA | NA | NA | NA | NA |
| GABRA5 | 152 | L | 60.34121591 | 19.72373758 | 0.430295049 | 11.33330135 | NA | NA | NA | NA | NA | NA | NA |
| GABRA5 | 153 | L | 65.29284091 | 20.01240048 | 0.438940022 | 11.63963657 | NA | NA | NA | NA | NA | NA | NA |
| GABRA5 | 154 | R | 63.77336364 | 24.01527733 | 0.55881951 | 15.58265696 | NA | NA | NA | NA | NA | NA | NA |
| GABRA5 | 155 | L | 68.85234091 | 23.66664095 | 0.548378432 | 15.02207222 | NA | NA | NA | NA | NA | NA | NA |
| GABRA5 | 156 | E | 68.65653535 | 28.92140088 | 0.705749731 | 19.45587827 | NA | NA | NA | NA | NA | NA | NA |
| GABRA5 | 157 | D | 73.57209091 | 31.37731558 | 0.779300282 | 21.96943085 | NA | NA | NA | NA | NA | NA | NA |
| GABRA5 | 158 | D | 70.89646591 | 30.96211118 | 0.766865602 | 21.50847463 | NA | NA | NA | NA | NA | NA | NA |
| GABRA5 | 159 | G | 69.96834091 | 28.52800046 | 0.693968044 | 19.09788376 | NA | NA | NA | NA | NA | NA | NA |
| GABRA5 | 160 | T | 66.77409091 | 28.1651607 | 0.683101598 | 19.7332213 | NA | NA | NA | NA | NA | NA | NA |
| GABRA5 | 161 | L | 65.53396591 | 23.1826363 | 0.533883299 | 14.75460731 | NA | NA | NA | NA | NA | NA | NA |
| GABRA5 | 162 | L | 61.35859091 | 23.57199463 | 0.545543932 | 15.15616749 | NA | NA | NA | NA | NA | NA | NA |
| GABRA5 | 163 | Y | 60.27267424 | 16.32502723 | 0.32850934 | 7.926653264 | NA | NA | NA | NA | NA | NA | NA |
| GABRA5 | 164 | T | 56.93580519 | 19.96648519 | 0.437564935 | 11.89111974 | NA | NA | NA | NA | NA | NA | NA |
| GABRA5 | 165 | M | 54.48859091 | 16.43670965 | 0.331854042 | 8.908754454 | NA | NA | NA | NA | NA | NA | NA |
| GABRA5 | 166 | R | 49.92718182 | 16.2151268 | 0.325218005 | 8.843674381 | NA | NA | NA | NA | NA | NA | NA |
| GABRA5 | 167 | L | 49.22021591 | 16.79457216 | 0.342571428 | 11.70133416 | NA | NA | NA | NA | NA | NA | NA |

| | | | | | | | | | | | | | |
|---|---|---|---|---|---|---|---|---|---|---|---|---|---|
| GABRA5 | 168 | T | 45.34337662 | 13.84753715 | 0.254312643 | 9.011100165 | NA | NA | NA | NA | NA | NA | NA |
| GABRA5 | 169 | I | 44.30021591 | 15.72184681 | 0.310445092 | 11.1850198 | NA | NA | NA | NA | NA | NA | NA |
| GABRA5 | 170 | S | 40.98092424 | 12.70711525 | 0.220158908 | 8.176013083 | NA | NA | NA | NA | NA | NA | NA |
| GABRA5 | 171 | A | 40.09829091 | 16.90412757 | 0.34585243 | 12.27896423 | NA | NA | NA | NA | NA | NA | NA |
| GABRA5 | 172 | E | 38.34553535 | 18.27273195 | 0.38683985 | 13.50478374 | NA | NA | NA | NA | NA | NA | NA |
| GABRA5 | 173 | C | 35.04425758 | 20.99806203 | 0.468458942 | 12.92193203 | NA | NA | NA | NA | NA | NA | NA |
| GABRA5 | 174 | P | 33.73123377 | 23.65643794 | 0.548072868 | 14.94273017 | NA | NA | NA | NA | NA | NA | NA |
| GABRA5 | 175 | M | 29.61409091 | 20.92635011 | 0.466311289 | 11.8259329 | NA | NA | NA | NA | NA | NA | NA |
| GABRA5 | 176 | Q | 27.41520202 | 25.86439323 | 0.614197448 | 16.28641134 | NA | NA | NA | NA | NA | NA | NA |
| GABRA5 | 177 | L | 23.87271591 | 20.87670647 | 0.464824545 | 13.81197827 | NA | NA | NA | NA | NA | NA | NA |
| GABRA5 | 178 | E | 21.7659798 | 26.59929033 | 0.636206391 | 21.70086459 | NA | NA | NA | NA | NA | NA | NA |
| GABRA5 | 179 | D | 20.36259091 | 26.62890801 | 0.637093392 | 21.98108997 | NA | NA | NA | NA | NA | NA | NA |
| GABRA5 | 180 | F | 19.96072727 | 19.61875233 | 0.427150916 | 15.1295385 | NA | NA | NA | NA | NA | NA | NA |
| GABRA5 | 181 | P | 18.84837662 | 22.70474369 | 0.519571212 | 18.04699393 | NA | NA | NA | NA | NA | NA | NA |
| GABRA5 | 182 | M | 19.69896591 | 26.21784995 | 0.624782888 | 21.56770094 | NA | NA | NA | NA | NA | NA | NA |
| GABRA5 | 183 | D | 24.44309091 | 24.21219529 | 0.564716875 | 17.17630497 | NA | NA | NA | NA | NA | NA | NA |
| GABRA5 | 184 | A | 27.09029091 | 28.98545778 | 0.707668128 | 19.43801419 | NA | NA | NA | NA | NA | NA | NA |
| GABRA5 | 185 | H | 29.88959091 | 24.32154091 | 0.567991593 | 15.20682382 | NA | NA | NA | NA | NA | NA | NA |
| GABRA5 | 186 | A | 33.90349091 | 27.69491697 | 0.669018583 | 19.52819438 | NA | NA | NA | NA | NA | NA | NA |
| GABRA5 | 187 | C | 35.80575758 | 23.75755263 | 0.551101085 | 15.59176671 | NA | NA | NA | NA | NA | NA | NA |
| GABRA5 | 188 | P | 39.16080519 | 24.52269696 | 0.574015882 | 19.79966802 | NA | NA | NA | NA | NA | NA | NA |
| GABRA5 | 189 | L | 43.43971591 | 22.27360604 | 0.506659358 | 17.72057785 | NA | NA | NA | NA | NA | NA | NA |
| GABRA5 | 190 | K | 44.5419798 | 24.34664055 | 0.568743286 | 19.50638705 | NA | NA | NA | NA | NA | NA | NA |
| GABRA5 | 191 | F | 49.71736364 | 24.77802006 | 0.581662383 | 19.99751173 | NA | NA | NA | NA | NA | NA | NA |
| GABRA5 | 192 | G | 52.41609091 | 25.05208824 | 0.589870268 | 17.40519871 | NA | NA | NA | NA | NA | NA | NA |
| GABRA5 | 193 | S | 55.93325758 | 25.6710067 | 0.608405844 | 17.60398476 | NA | NA | NA | NA | NA | NA | NA |
| GABRA5 | 194 | Y | 56.42559091 | 20.93645935 | 0.466614044 | 12.86196069 | NA | NA | NA | NA | NA | NA | NA |
| GABRA5 | 195 | A | 59.74289091 | 22.90079252 | 0.525442548 | 14.48852956 | NA | NA | NA | NA | NA | NA | NA |
| GABRA5 | 196 | Y | 61.59200758 | 22.45966469 | 0.512231505 | 14.18440104 | NA | NA | NA | NA | NA | NA | NA |
| GABRA5 | 197 | P | 60.44637662 | 29.59736453 | 0.725993715 | 21.17512105 | NA | NA | NA | NA | NA | NA | NA |
| GABRA5 | 198 | N | 58.53509091 | 32.91148728 | 0.825246167 | 24.50037702 | NA | NA | NA | NA | NA | NA | NA |
| GABRA5 | 199 | S | 63.39259091 | 33.43070729 | 0.84079594 | 25.07887251 | NA | NA | NA | NA | NA | NA | NA |
| GABRA5 | 200 | E | 64.35009091 | 28.33890321 | 0.688304897 | 19.91901366 | NA | NA | NA | NA | NA | NA | NA |

| | | | | | | | | | | | | | |
|---|---|---|---|---|---|---|---|---|---|---|---|---|---|
| GABRA5 | 201 | V | 60.99709091 | 27.66985907 | 0.66826814 | 19.26063202 | NA | NA | NA | NA | NA | NA | NA |
| GABRA5 | 202 | V | 59.24994805 | 31.47217293 | 0.782141101 | 23.04966291 | NA | NA | NA | NA | NA | NA | NA |
| GABRA5 | 203 | Y | 56.06959091 | 25.39041876 | 0.600002703 | 17.31778943 | NA | NA | NA | NA | NA | NA | NA |
| GABRA5 | 204 | V | 56.26023377 | 32.61477049 | 0.816359993 | 24.53904608 | NA | NA | NA | NA | NA | NA | NA |
| GABRA5 | 205 | W | 51.19923377 | 31.06617125 | 0.769982028 | 23.66712957 | NA | NA | NA | NA | NA | NA | NA |
| GABRA5 | 206 | T | 55.32966234 | 31.99781652 | 0.79788325 | 23.920299 | NA | NA | NA | NA | NA | NA | NA |
| GABRA5 | 207 | N | 53.05309091 | 36.15030868 | 0.922243468 | 28.07805233 | NA | NA | NA | NA | NA | NA | NA |
| GABRA5 | 208 | G | 50.91784091 | 38.09091997 | 0.980361541 | 30.5319841 | NA | NA | NA | NA | NA | NA | NA |
| GABRA5 | 209 | S | 47.88309091 | 37.46833507 | 0.961716161 | 32.63111659 | NA | NA | NA | NA | NA | NA | NA |
| GABRA5 | 210 | T | 46.19337662 | 36.71479339 | 0.939148844 | 31.88362877 | NA | NA | NA | NA | NA | NA | NA |
| GABRA5 | 211 | K | 48.75420202 | 35.57968394 | 0.905154208 | 30.75310665 | NA | NA | NA | NA | NA | NA | NA |
| GABRA5 | 212 | S | 49.52625758 | 32.25862914 | 0.805694154 | 27.45072664 | NA | NA | NA | NA | NA | NA | NA |
| GABRA5 | 213 | V | 45.93966234 | 30.39320329 | 0.749827759 | 25.59220517 | NA | NA | NA | NA | NA | NA | NA |
| GABRA5 | 214 | V | 45.59623377 | 29.56332326 | 0.724974236 | 24.75271951 | NA | NA | NA | NA | NA | NA | NA |
| GABRA5 | 215 | V | 41.41366234 | 29.20032769 | 0.714103124 | 24.87933221 | NA | NA | NA | NA | NA | NA | NA |
| GABRA5 | 216 | A | 40.99649091 | 28.58633957 | 0.695715203 | 24.0681204 | NA | NA | NA | NA | NA | NA | NA |
| GABRA5 | 217 | E | 39.21053535 | 30.23984988 | 0.745235081 | 25.51520771 | NA | NA | NA | NA | NA | NA | NA |
| GABRA5 | 218 | D | 37.60659091 | 26.86552482 | 0.64417967 | 18.70580781 | NA | NA | NA | NA | NA | NA | NA |
| GABRA5 | 219 | G | 37.62409091 | 23.87604486 | 0.554649729 | 19.13600441 | NA | NA | NA | NA | NA | NA | NA |
| GABRA5 | 220 | S | 34.86392424 | 25.00988824 | 0.588606449 | 16.84779219 | NA | NA | NA | NA | NA | NA | NA |
| GABRA5 | 221 | R | 34.72609091 | 17.35125958 | 0.359243288 | 9.151066768 | NA | NA | NA | NA | NA | NA | NA |
| GABRA5 | 222 | L | 30.99909091 | 23.16365578 | 0.533314864 | 14.44067847 | NA | NA | NA | NA | NA | NA | NA |
| GABRA5 | 223 | N | 27.75934091 | 19.51637133 | 0.424084776 | 9.959287498 | NA | NA | NA | NA | NA | NA | NA |
| GABRA5 | 224 | Q | 25.99653535 | 17.8301274 | 0.373584581 | 9.619579565 | NA | NA | NA | NA | NA | NA | NA |
| GABRA5 | 225 | Y | 27.99309091 | 20.59452279 | 0.456373614 | 11.41565067 | NA | NA | NA | NA | NA | NA | NA |
| GABRA5 | 226 | H | 28.82559091 | 26.41710745 | 0.630750318 | 17.27740285 | NA | NA | NA | NA | NA | NA | NA |
| GABRA5 | 227 | L | 33.28121591 | 28.78479962 | 0.701658751 | 20.0660325 | NA | NA | NA | NA | NA | NA | NA |
| GABRA5 | 228 | M | 31.43071591 | 31.20069784 | 0.774010875 | 22.47344053 | NA | NA | NA | NA | NA | NA | NA |
| GABRA5 | 229 | G | 35.84609091 | 32.04742258 | 0.799368869 | 23.94793857 | NA | NA | NA | NA | NA | NA | NA |
| GABRA5 | 230 | Q | 39.86686869 | 31.49479979 | 0.782818738 | 26.77050088 | NA | NA | NA | NA | NA | NA | NA |
| GABRA5 | 231 | T | 40.48523377 | 32.94152196 | 0.826145655 | 28.32506476 | NA | NA | NA | NA | NA | NA | NA |
| GABRA5 | 232 | V | 44.63980519 | 33.50412794 | 0.842994766 | 28.96154368 | NA | NA | NA | NA | NA | NA | NA |
| GABRA5 | 233 | G | 46.37759091 | 33.6700131 | 0.84796275 | 28.82673838 | NA | NA | NA | NA | NA | NA | NA |

| | | | | | | | | | | | | | |
|---|---|---|---|---|---|---|---|---|---|---|---|---|---|
| GABRA5 | 234 | T | 49.15237662 | 35.48054364 | 0.902185121 | 30.6428201 | NA | NA | NA | NA | NA | NA | NA |
| GABRA5 | 235 | E | 47.58909091 | 33.67831022 | 0.848211235 | 28.8670065 | NA | NA | NA | NA | NA | NA | NA |
| GABRA5 | 236 | N | 51.54459091 | 35.50518264 | 0.902923018 | 27.85281625 | NA | NA | NA | NA | NA | NA | NA |
| GABRA5 | 237 | I | 49.27084091 | 31.22157001 | 0.774635962 | 24.07305273 | NA | NA | NA | NA | NA | NA | NA |
| GABRA5 | 238 | S | 50.43175758 | 34.23270289 | 0.864814371 | 26.57962152 | NA | NA | NA | NA | NA | NA | NA |
| GABRA5 | 239 | T | 51.76037662 | 30.73505772 | 0.76006573 | 23.08459317 | NA | NA | NA | NA | NA | NA | NA |
| GABRA5 | 240 | S | 53.43559091 | 30.80160952 | 0.762058845 | 23.14561177 | NA | NA | NA | NA | NA | NA | NA |
| GABRA5 | 241 | T | 56.63609091 | 28.96191178 | 0.706962965 | 20.88275956 | NA | NA | NA | NA | NA | NA | NA |
| GABRA5 | 242 | G | 55.95834091 | 32.03803248 | 0.799087651 | 23.95786066 | NA | NA | NA | NA | NA | NA | NA |
| GABRA5 | 243 | E | 55.1599798 | 33.24565681 | 0.835253987 | 25.16855756 | NA | NA | NA | NA | NA | NA | NA |
| GABRA5 | 244 | Y | 55.13634091 | 28.04434433 | 0.679483349 | 19.96536525 | NA | NA | NA | NA | NA | NA | NA |
| GABRA5 | 245 | T | 54.32951948 | 30.13439585 | 0.742076908 | 22.48311471 | NA | NA | NA | NA | NA | NA | NA |
| GABRA5 | 246 | I | 49.93646591 | 29.74091177 | 0.730292716 | 22.08913571 | NA | NA | NA | NA | NA | NA | NA |
| GABRA5 | 247 | M | 49.71996591 | 29.19346367 | 0.713897558 | 24.349973 | NA | NA | NA | NA | NA | NA | NA |
| GABRA5 | 248 | T | 45.03466234 | 30.33243225 | 0.748007768 | 25.52037713 | NA | NA | NA | NA | NA | NA | NA |
| GABRA5 | 249 | A | 42.74149091 | 28.15148923 | 0.682692161 | 23.67926753 | NA | NA | NA | NA | NA | NA | NA |
| GABRA5 | 250 | H | 39.41739091 | 29.78818976 | 0.731708613 | 25.01426538 | NA | NA | NA | NA | NA | NA | NA |
| GABRA5 | 251 | F | 38.24054545 | 26.59931379 | 0.636207094 | 21.91268508 | NA | NA | NA | NA | NA | NA | NA |
| GABRA5 | 252 | H | 33.69739091 | 29.60356958 | 0.726179546 | 20.85575786 | NA | NA | NA | NA | NA | NA | NA |
| GABRA5 | 253 | L | 31.08246591 | 24.57950996 | 0.575717337 | 15.85301163 | NA | NA | NA | NA | NA | NA | NA |
| GABRA5 | 254 | K | 25.84331313 | 26.91334457 | 0.645611792 | 17.32968367 | NA | NA | NA | NA | NA | NA | NA |
| GABRA5 | 255 | R | 23.661 | 20.5601197 | 0.455343299 | 13.48355589 | NA | NA | NA | NA | NA | NA | NA |
| GABRA5 | 256 | K | 23.60053535 | 24.32524263 | 0.568102454 | 19.4130538 | NA | NA | NA | NA | NA | NA | NA |
| GABRA5 | 257 | I | 18.45396591 | 24.78292126 | 0.581809166 | 20.12945236 | NA | NA | NA | NA | NA | NA | NA |
| GABRA5 | 258 | G | 17.59034091 | 24.54920058 | 0.574809621 | 19.9326834 | NA | NA | NA | NA | NA | NA | NA |
| GABRA5 | 259 | Y | 17.63025758 | 16.29829 | 0.327708605 | 11.68587164 | NA | NA | NA | NA | NA | NA | NA |
| GABRA5 | 260 | F | 16.90245455 | 17.94592202 | 0.377052437 | 13.77823632 | NA | NA | NA | NA | NA | NA | NA |
| GABRA5 | 261 | V | 12.28251948 | 21.55003761 | 0.48498969 | 17.68233914 | NA | NA | NA | NA | NA | NA | NA |
| GABRA5 | 262 | I | 12.16359091 | 19.1995688 | 0.414597068 | 15.37409431 | NA | NA | NA | NA | NA | NA | NA |
| GABRA5 | 263 | Q | 13.64442424 | 14.38115646 | 0.270293651 | 10.4709385 | NA | NA | NA | NA | NA | NA | NA |
| GABRA5 | 264 | T | 11.21209091 | 17.01535195 | 0.349183415 | 13.20702344 | NA | NA | NA | NA | NA | NA | NA |
| GABRA5 | 265 | Y | 10.57842424 | 20.02067344 | 0.439187783 | 16.08062571 | NA | NA | NA | NA | NA | NA | NA |
| GABRA5 | 266 | L | 6.722090909 | 19.81074017 | 0.432900631 | 14.56099562 | NA | NA | NA | NA | NA | NA | NA |

| | | | | | | | | | | | | | |
|---|---|---|---|---|---|---|---|---|---|---|---|---|---|
| GABRA5 | 267 | P | 5.538948052 | 16.21872159 | 0.325325663 | 10.86084942 | NA | NA | NA | NA | NA | NA | NA |
| GABRA5 | 268 | C | 4.498757576 | 18.08966606 | 0.381357331 | 12.73181933 | NA | NA | NA | NA | NA | NA | NA |
| GABRA5 | 269 | I | 2.962465909 | 20.69165885 | 0.459282677 | 15.64523019 | NA | NA | NA | NA | NA | NA | NA |
| GABRA5 | 270 | M | 1.281340909 | 18.86135557 | 0.404468146 | 14.48554641 | NA | NA | NA | NA | NA | NA | NA |
| GABRA5 | 271 | T | 0.289233766 | 15.23456442 | 0.295851797 | 10.86194045 | NA | NA | NA | NA | NA | NA | NA |
| GABRA5 | 272 | V | -0.187805195 | 19.23045777 | 0.415522141 | 14.85785598 | NA | NA | NA | NA | NA | NA | NA |
| GABRA5 | 273 | I | -1.379090909 | 20.62601937 | 0.457316884 | 15.9950157 | NA | NA | NA | NA | NA | NA | NA |
| GABRA5 | 274 | L | -2.857090909 | 14.56172236 | 0.275701299 | 9.776314081 | NA | NA | NA | NA | NA | NA | NA |
| GABRA5 | 275 | S | -4.205257576 | 18.39325844 | 0.390449417 | 13.18346119 | NA | NA | NA | NA | NA | NA | NA |
| GABRA5 | 276 | Q | -4.323313131 | 20.86683026 | 0.464528769 | 15.68107852 | NA | NA | NA | NA | NA | NA | NA |
| GABRA5 | 277 | V | -7.299376623 | 21.59564468 | 0.486355546 | 16.50883449 | NA | NA | NA | NA | NA | NA | NA |
| GABRA5 | 278 | S | -9.163924242 | 17.20215225 | 0.354777772 | 12.91705192 | NA | NA | NA | NA | NA | NA | NA |
| GABRA5 | 279 | F | -9.975727273 | 20.77160507 | 0.461676933 | 16.50788623 | NA | NA | NA | NA | NA | NA | NA |
| GABRA5 | 280 | W | -12.65394805 | 20.83792557 | 0.463663122 | 16.76731733 | NA | NA | NA | NA | NA | NA | NA |
| GABRA5 | 281 | L | -13.49709091 | 16.28550982 | 0.32732586 | 12.2172655 | NA | NA | NA | NA | NA | NA | NA |
| GABRA5 | 282 | N | -18.42546591 | 15.52597375 | 0.304579021 | 11.72007422 | NA | NA | NA | NA | NA | NA | NA |
| GABRA5 | 283 | R | -19.864 | 14.9157032 | 0.286302435 | 11.25007547 | NA | NA | NA | NA | NA | NA | NA |
| GABRA5 | 284 | E | -22.2069798 | 11.80650537 | 0.193187144 | 7.879974197 | NA | NA | NA | NA | NA | NA | NA |
| GABRA5 | 285 | S | -17.22125758 | 11.06795415 | 0.171068766 | 7.283264636 | NA | NA | NA | NA | NA | NA | NA |
| GABRA5 | 286 | V | -18.41894805 | 8.921845437 | 0.106796388 | 5.133520452 | NA | NA | NA | NA | NA | NA | NA |
| GABRA5 | 287 | P | -16.18551948 | 5.381857858 | 7.80E-04 | 1.580613976 | 3.82 | -0.01 | 0.95 | 0.4 | 0.12 | 0 | 0 |
| GABRA5 | 288 | A | -13.34389091 | 7.965930015 | 0.078168315 | 3.91777801 | 4.14 | -0.02 | 0.67 | 0.53 | -0.04 | 0 | 0 |
| GABRA5 | 289 | R | -11.80063636 | 10.15351555 | 0.143682854 | 5.949532509 | NA | NA | NA | NA | NA | NA | NA |
| GABRA5 | 290 | T | -11.34180519 | 6.95794553 | 0.04798086 | 2.716738993 | 4.3 | 0.06 | 1 | 1.27 | -0.3 | 0 | 0 |
| GABRA5 | 291 | V | -8.315519481 | 5.574098133 | 0.006536937 | 1.72563944 | 4.03 | 0.39 | 2.6 | 0.04 | -0.04 | 0 | 0 |
| GABRA5 | 292 | F | -6.963545455 | 9.705453776 | 0.130264151 | 5.709967824 | NA | NA | NA | NA | NA | NA | NA |
| GABRA5 | 293 | G | -5.648340909 | 12.01830479 | 0.199530183 | 6.679855214 | NA | NA | NA | NA | NA | NA | NA |
| GABRA5 | 294 | V | -4.150805195 | 8.112962098 | 0.08257168 | 2.777097652 | 5.34 | -0.01 | 1.18 | 1.05 | -0.16 | 0 | 0 |
| GABRA5 | 295 | T | -1.734376623 | 6.773758632 | 0.042464769 | 2.180447897 | 4.63 | 0.38 | 2 | 0.74 | -0.77 | 0 | 0 |
| GABRA5 | 296 | T | -1.655948052 | 11.08989991 | 0.171726005 | 6.475796486 | NA | NA | NA | NA | NA | NA | NA |
| GABRA5 | 297 | V | -0.239376623 | 11.46381996 | 0.182924287 | 7.101146834 | NA | NA | NA | NA | NA | NA | NA |
| GABRA5 | 298 | L | 0.340215909 | 5.714224182 | 0.010733479 | 1.413045106 | 4.39 | 1.14 | 3.8 | 0.13 | -1.79 | 0 | 0 |
| GABRA5 | 299 | T | 1.805805195 | 9.745613475 | 0.131466867 | 5.056221336 | NA | NA | NA | NA | NA | NA | NA |

| | | | | | | | | | | | | | |
|---|---|---|---|---|---|---|---|---|---|---|---|---|---|
| GABRA5 | 300 | M | 2.877590909 | 12.24320518 | 0.206265575 | 7.296120545 | NA | NA | NA | NA | NA | NA | NA |
| GABRA5 | 301 | T | 4.684233766 | 10.44083909 | 0.152287715 | 5.579035501 | NA | NA | NA | NA | NA | NA | NA |
| GABRA5 | 302 | T | 6.109233766 | 6.811696852 | 0.043600956 | 1.773577964 | 5.26 | 0.05 | -2.7 | 41.61 | -0.01 | 0 | 0 |
| GABRA5 | 303 | L | 7.680090909 | 10.69846779 | 0.160003265 | 5.443375633 | NA | NA | NA | NA | NA | NA | NA |
| GABRA5 | 304 | S | 9.296757576 | 11.91956568 | 0.196573111 | 7.991063399 | NA | NA | NA | NA | NA | NA | NA |
| GABRA5 | 305 | I | 10.38034091 | 7.0963857 | 0.052126912 | 3.247083022 | 3.85 | 0.26 | -3.2 | 51.6 | -0.2 | 0 | 0 |
| GABRA5 | 306 | S | 12.55875758 | 9.614747579 | 0.127547652 | 5.814907786 | NA | NA | NA | NA | NA | NA | NA |
| GABRA5 | 307 | A | 13.38749091 | 12.06023972 | 0.200786064 | 8.146845035 | NA | NA | NA | NA | NA | NA | NA |
| GABRA5 | 308 | R | 13.51127273 | 10.8021284 | 0.163107728 | 6.62952527 | NA | NA | NA | NA | NA | NA | NA |
| GABRA5 | 309 | N | 16.82834091 | 7.090315931 | 0.051945132 | 2.450998266 | 4.66 | -0.44 | -3.35 | 50.75 | 1.02 | 2 | -2 |
| GABRA5 | 310 | S | 19.20759091 | 10.94983119 | 0.16753118 | 7.066604649 | NA | NA | NA | NA | NA | NA | NA |
| GABRA5 | 311 | L | 18.95184091 | 13.26598108 | 0.236896008 | 9.246131729 | NA | NA | NA | NA | NA | NA | NA |
| GABRA5 | 312 | P | 22.39537662 | 14.73471157 | 0.280882037 | 9.858646421 | NA | NA | NA | NA | NA | NA | NA |
| GABRA5 | 313 | K | 22.60664646 | 12.84561439 | 0.224306726 | 7.958147042 | NA | NA | NA | NA | NA | NA | NA |
| GABRA5 | 314 | V | 21.98709091 | 18.59825083 | 0.396588597 | 13.68999053 | NA | NA | NA | NA | NA | NA | NA |
| GABRA5 | 315 | A | 21.46369091 | 21.99965213 | 0.498454896 | 17.08832262 | NA | NA | NA | NA | NA | NA | NA |
| GABRA5 | 316 | Y | 21.91134091 | 23.58741618 | 0.546005782 | 19.02394644 | NA | NA | NA | NA | NA | NA | NA |
| GABRA5 | 317 | A | 16.79209091 | 25.19055672 | 0.594017168 | 20.53227958 | NA | NA | NA | NA | NA | NA | NA |
| GABRA5 | 318 | T | 16.58923377 | 21.3339059 | 0.478516906 | 16.67633595 | NA | NA | NA | NA | NA | NA | NA |
| GABRA5 | 319 | A | 14.73089091 | 24.06812143 | 0.560402102 | 19.8106904 | NA | NA | NA | NA | NA | NA | NA |
| GABRA5 | 320 | M | 13.98496591 | 17.40268318 | 0.360783339 | 13.35325864 | NA | NA | NA | NA | NA | NA | NA |
| GABRA5 | 321 | D | 12.07334091 | 19.89370056 | 0.435385157 | 16.01509862 | NA | NA | NA | NA | NA | NA | NA |
| GABRA5 | 322 | W | 10.67666234 | 23.24762941 | 0.535829734 | 19.41017492 | NA | NA | NA | NA | NA | NA | NA |
| GABRA5 | 323 | F | 8.548090909 | 22.2556848 | 0.506122647 | 18.32989949 | NA | NA | NA | NA | NA | NA | NA |
| GABRA5 | 324 | I | 8.712465909 | 17.61925461 | 0.367269293 | 13.69428016 | NA | NA | NA | NA | NA | NA | NA |
| GABRA5 | 325 | A | 5.958290909 | 21.00017259 | 0.46852215 | 15.75249622 | NA | NA | NA | NA | NA | NA | NA |
| GABRA5 | 326 | V | 4.049805195 | 22.98239 | 0.527886256 | 17.70848989 | NA | NA | NA | NA | NA | NA | NA |
| GABRA5 | 327 | C | 3.532924242 | 19.67446614 | 0.428819452 | 14.69404361 | NA | NA | NA | NA | NA | NA | NA |
| GABRA5 | 328 | Y | 2.998757576 | 15.44531469 | 0.302163416 | 10.40581048 | NA | NA | NA | NA | NA | NA | NA |
| GABRA5 | 329 | A | 0.522690909 | 20.77508734 | 0.461781222 | 16.5606795 | NA | NA | NA | NA | NA | NA | NA |
| GABRA5 | 330 | F | 0 | 20.62747791 | 0.457360565 | 16.3602166 | NA | NA | NA | NA | NA | NA | NA |
| GABRA5 | 331 | V | -1.124090909 | 16.54426058 | 0.335075013 | 12.23970395 | NA | NA | NA | NA | NA | NA | NA |
| GABRA5 | 332 | F | -1.069363636 | 18.77051701 | 0.401747682 | 14.26156284 | NA | NA | NA | NA | NA | NA | NA |

| | | | | | | | | | | | | | |
|---|---|---|---|---|---|---|---|---|---|---|---|---|---|
| GABRA5 | 333 | S | -3.724924242 | 21.12840103 | 0.472362377 | 15.79669842 | NA | NA | NA | NA | NA | NA | NA |
| GABRA5 | 334 | A | -6.038890909 | 19.9251531 | 0.436327108 | 14.58576084 | NA | NA | NA | NA | NA | NA | NA |
| GABRA5 | 335 | L | -6.651340909 | 15.43388292 | 0.301821054 | 10.09469485 | NA | NA | NA | NA | NA | NA | NA |
| GABRA5 | 336 | I | -8.173590909 | 20.18227195 | 0.444027389 | 16.21649393 | NA | NA | NA | NA | NA | NA | NA |
| GABRA5 | 337 | E | -11.19553535 | 20.37610181 | 0.44983227 | 16.17540943 | NA | NA | NA | NA | NA | NA | NA |
| GABRA5 | 338 | F | -12.58363636 | 13.44220298 | 0.24217356 | 9.394207289 | NA | NA | NA | NA | NA | NA | NA |
| GABRA5 | 339 | A | -11.93229091 | 17.36334835 | 0.359605327 | 13.31530902 | NA | NA | NA | NA | NA | NA | NA |
| GABRA5 | 340 | T | -14.45194805 | 19.39570244 | 0.420470943 | 15.35005282 | NA | NA | NA | NA | NA | NA | NA |
| GABRA5 | 341 | V | -17.06437662 | 17.90153633 | 0.375723159 | 14.11795962 | NA | NA | NA | NA | NA | NA | NA |
| GABRA5 | 342 | N | -17.44659091 | 13.97214028 | 0.258044299 | 10.28315636 | NA | NA | NA | NA | NA | NA | NA |
| GABRA5 | 343 | Y | -18.50250758 | 18.4863963 | 0.39323874 | 14.69703691 | NA | NA | NA | NA | NA | NA | NA |
| GABRA5 | 344 | F | -20.29854545 | 20.1690478 | 0.443631348 | 16.32734838 | NA | NA | NA | NA | NA | NA | NA |
| GABRA5 | 345 | T | -22.41223377 | 17.30335728 | 0.357808694 | 10.58817983 | NA | NA | NA | NA | NA | NA | NA |
| GABRA5 | 346 | K | -24.21253535 | 13.36055167 | 0.23972824 | 3.982179729 | 9.71 | -0.64 | -1.83 | 18.22 | 0.71 | 1 | -1 |
| GABRA5 | 424 | K | -20.43131313 | 28.32573662 | 0.687910579 | 24.59179807 | NA | NA | NA | NA | NA | NA | NA |
| GABRA5 | 425 | I | -15.58271591 | 27.21339475 | 0.654597795 | 23.17012857 | NA | NA | NA | NA | NA | NA | NA |
| GABRA5 | 426 | D | -15.47996591 | 22.51652125 | 0.513934264 | 18.4820646 | NA | NA | NA | NA | NA | NA | NA |
| GABRA5 | 427 | K | -17.91231313 | 26.55196878 | 0.63478919 | 22.50892647 | NA | NA | NA | NA | NA | NA | NA |
| GABRA5 | 428 | M | -13.70109091 | 28.8353038 | 0.703171267 | 24.78640037 | NA | NA | NA | NA | NA | NA | NA |
| GABRA5 | 429 | S | -11.12192424 | 25.57880752 | 0.605644632 | 21.52523867 | NA | NA | NA | NA | NA | NA | NA |
| GABRA5 | 430 | R | -12.71809091 | 22.3604906 | 0.509261406 | 18.24966509 | NA | NA | NA | NA | NA | NA | NA |
| GABRA5 | 431 | I | -10.21584091 | 29.58945768 | 0.725756918 | 25.36189476 | NA | NA | NA | NA | NA | NA | NA |
| GABRA5 | 432 | V | -7.156090909 | 29.79004671 | 0.731764225 | 24.60851893 | NA | NA | NA | NA | NA | NA | NA |
| GABRA5 | 433 | F | -5.405363636 | 25.02970815 | 0.589200022 | 20.25729663 | NA | NA | NA | NA | NA | NA | NA |
| GABRA5 | 434 | P | -4.272376623 | 26.95103961 | 0.646740696 | 22.16890021 | NA | NA | NA | NA | NA | NA | NA |
| GABRA5 | 435 | V | -3.401805195 | 30.32854967 | 0.747891491 | 25.71011668 | NA | NA | NA | NA | NA | NA | NA |
| GABRA5 | 436 | L | -1.635965909 | 30.21193772 | 0.744399158 | 25.85012781 | NA | NA | NA | NA | NA | NA | NA |
| GABRA5 | 437 | F | 0.553909091 | 24.67526296 | 0.578584979 | 20.44590903 | NA | NA | NA | NA | NA | NA | NA |
| GABRA5 | 438 | G | 0.197340909 | 29.8628731 | 0.733945254 | 25.49250901 | NA | NA | NA | NA | NA | NA | NA |
| GABRA5 | 439 | T | 0.640662338 | 32.39416447 | 0.809753211 | 28.03981013 | NA | NA | NA | NA | NA | NA | NA |
| GABRA5 | 440 | F | 4.336090909 | 27.45459049 | 0.661821205 | 22.69326048 | NA | NA | NA | NA | NA | NA | NA |
| GABRA5 | 441 | N | 4.938465909 | 27.17495305 | 0.65344653 | 21.92126538 | NA | NA | NA | NA | NA | NA | NA |
| GABRA5 | 442 | L | 5.488215909 | 32.55191454 | 0.814477562 | 27.29561809 | NA | NA | NA | NA | NA | NA | NA |

| | | | | | | | | | | | | | |
|---|---|---|---|---|---|---|---|---|---|---|---|---|---|
| GABRA5 | 443 | V | 7.009519481 | 33.2921467 | 0.836646282 | 28.07402445 | NA | NA | NA | NA | NA | NA | NA |
| GABRA5 | 444 | Y | 10.37942424 | 25.40647378 | 0.600483524 | 21.77151127 | NA | NA | NA | NA | NA | NA | NA |
| GABRA5 | 445 | W | 11.14801948 | 26.95763797 | 0.646938306 | 23.13076451 | NA | NA | NA | NA | NA | NA | NA |
| GABRA5 | 446 | A | 11.84009091 | 34.02580397 | 0.858618094 | 30.18757026 | NA | NA | NA | NA | NA | NA | NA |
| GABRA5 | 447 | T | 13.75609091 | 33.27736026 | 0.836203452 | 29.20236592 | NA | NA | NA | NA | NA | NA | NA |
| GABRA5 | 448 | Y | 15.90125758 | 29.42039247 | 0.720693697 | 24.88723901 | NA | NA | NA | NA | NA | NA | NA |
| GABRA5 | 449 | L | 16.37609091 | 31.85768129 | 0.793686434 | 27.31366574 | NA | NA | NA | NA | NA | NA | NA |
| GABRA5 | 450 | N | 18.00271591 | 35.59357664 | 0.905570271 | 30.93682208 | NA | NA | NA | NA | NA | NA | NA |
| GABRA5 | 451 | R | 20.50945455 | 34.48202075 | 0.872281026 | 29.82472773 | NA | NA | NA | NA | NA | NA | NA |
| GABRB2 | 32 | N | 79.267875 | 32.32409788 | 0.741210924 | 23.4030364 | NA | NA | NA | NA | NA | NA | NA |
| GABRB2 | 33 | M | 77.086375 | 26.4214845 | 0.591075511 | 17.50461796 | NA | NA | NA | NA | NA | NA | NA |
| GABRB2 | 34 | S | 81.05329167 | 30.39340198 | 0.692102875 | 21.33241477 | NA | NA | NA | NA | NA | NA | NA |
| GABRB2 | 35 | L | 80.750375 | 30.71165178 | 0.70019769 | 21.64662756 | NA | NA | NA | NA | NA | NA | NA |
| GABRB2 | 36 | V | 76.44441071 | 27.49693572 | 0.618430058 | 18.57329129 | NA | NA | NA | NA | NA | NA | NA |
| GABRB2 | 37 | K | 78.94468056 | 25.20406733 | 0.560110002 | 16.14776857 | NA | NA | NA | NA | NA | NA | NA |
| GABRB2 | 38 | E | 82.12879167 | 28.2117621 | 0.636611962 | 19.14742624 | NA | NA | NA | NA | NA | NA | NA |
| GABRB2 | 39 | T | 78.53383929 | 27.53532685 | 0.619406552 | 18.46115932 | NA | NA | NA | NA | NA | NA | NA |
| GABRB2 | 40 | V | 75.84726786 | 23.13962052 | 0.507599943 | 14.06935236 | NA | NA | NA | NA | NA | NA | NA |
| GABRB2 | 41 | D | 79.161375 | 23.01078375 | 0.504322927 | 13.94365082 | NA | NA | NA | NA | NA | NA | NA |
| GABRB2 | 42 | R | 81.84930682 | 26.46851913 | 0.592271856 | 17.39454564 | NA | NA | NA | NA | NA | NA | NA |
| GABRB2 | 43 | L | 76.488875 | 24.85189234 | 0.551152285 | 15.77225749 | NA | NA | NA | NA | NA | NA | NA |
| GABRB2 | 44 | L | 75.360625 | 19.68691456 | 0.419778939 | 10.60856291 | NA | NA | NA | NA | NA | NA | NA |
| GABRB2 | 45 | K | 81.56434722 | 23.70031597 | 0.521861464 | 14.6211969 | NA | NA | NA | NA | NA | NA | NA |
| GABRB2 | 46 | G | 79.129625 | 24.03722268 | 0.530430825 | 14.95727917 | NA | NA | NA | NA | NA | NA | NA |
| GABRB2 | 47 | Y | 73.854375 | 21.31536246 | 0.461199185 | 12.25426596 | NA | NA | NA | NA | NA | NA | NA |
| GABRB2 | 48 | D | 74.834625 | 21.95823558 | 0.477550929 | 12.93698149 | NA | NA | NA | NA | NA | NA | NA |
| GABRB2 | 49 | I | 70.27625 | 16.75422735 | 0.345184827 | 8.476268887 | NA | NA | NA | NA | NA | NA | NA |
| GABRB2 | 50 | R | 72.311125 | 20.09255867 | 0.430096665 | 11.90269401 | NA | NA | NA | NA | NA | NA | NA |
| GABRB2 | 51 | L | 72.1045 | 23.2659591 | 0.510813417 | 14.99195116 | NA | NA | NA | NA | NA | NA | NA |
| GABRB2 | 52 | R | 68.49985227 | 25.31265354 | 0.562871937 | 17.04310707 | NA | NA | NA | NA | NA | NA | NA |
| GABRB2 | 53 | P | 69.48841071 | 26.5887584 | 0.595330192 | 18.31786427 | NA | NA | NA | NA | NA | NA | NA |
| GABRB2 | 54 | D | 71.696375 | 30.49286641 | 0.694632794 | 22.21910161 | NA | NA | NA | NA | NA | NA | NA |
| GABRB2 | 55 | F | 71.70185227 | 28.36705316 | 0.640561855 | 20.09723281 | NA | NA | NA | NA | NA | NA | NA |

| | | | | | | | | | | | | | |
|---|---|---|---|---|---|---|---|---|---|---|---|---|---|
| GABRB2 | 56 | G | 72.660375 | 32.95853465 | 0.757348085 | 24.68786562 | NA | NA | NA | NA | NA | NA | NA |
| GABRB2 | 57 | G | 71.005625 | 35.12096886 | 0.812350494 | 26.85159477 | NA | NA | NA | NA | NA | NA | NA |
| GABRB2 | 58 | P | 69.16698214 | 34.28911528 | 0.791191954 | 27.05041763 | NA | NA | NA | NA | NA | NA | NA |
| GABRB2 | 59 | P | 65.38855357 | 33.44609098 | 0.769749282 | 26.04124766 | NA | NA | NA | NA | NA | NA | NA |
| GABRB2 | 60 | V | 64.58641071 | 29.31612242 | 0.664701824 | 21.90474076 | NA | NA | NA | NA | NA | NA | NA |
| GABRB2 | 61 | A | 62.371925 | 31.0560359 | 0.708957243 | 23.66246777 | NA | NA | NA | NA | NA | NA | NA |
| GABRB2 | 62 | V | 59.53883929 | 26.88739999 | 0.602926264 | 19.47023405 | NA | NA | NA | NA | NA | NA | NA |
| GABRB2 | 63 | G | 57.368625 | 28.49610393 | 0.643844314 | 21.09528809 | NA | NA | NA | NA | NA | NA | NA |
| GABRB2 | 64 | M | 53.717 | 24.98675051 | 0.554582459 | 18.39523249 | NA | NA | NA | NA | NA | NA | NA |
| GABRB2 | 65 | N | 52.248375 | 25.82224524 | 0.575833612 | 19.22940575 | NA | NA | NA | NA | NA | NA | NA |
| GABRB2 | 66 | I | 48.441 | 24.4086243 | 0.539877579 | 17.84396347 | NA | NA | NA | NA | NA | NA | NA |
| GABRB2 | 67 | D | 47.316875 | 23.89632493 | 0.526847033 | 17.33334983 | NA | NA | NA | NA | NA | NA | NA |
| GABRB2 | 68 | I | 43.23925 | 24.24593642 | 0.535739546 | 17.89396355 | NA | NA | NA | NA | NA | NA | NA |
| GABRB2 | 69 | A | 42.684525 | 22.31262867 | 0.486565063 | 15.89538768 | NA | NA | NA | NA | NA | NA | NA |
| GABRB2 | 70 | S | 39.95295833 | 20.79183794 | 0.447883122 | 16.4435056 | NA | NA | NA | NA | NA | NA | NA |
| GABRB2 | 71 | I | 37.058375 | 19.67646785 | 0.419513223 | 11.61080218 | NA | NA | NA | NA | NA | NA | NA |
| GABRB2 | 72 | D | 36.680625 | 16.33288477 | 0.334467804 | 8.144609002 | NA | NA | NA | NA | NA | NA | NA |
| GABRB2 | 73 | M | 33.704375 | 11.63354675 | 0.214938198 | 3.440380981 | 8.21 | -0.59 | -3.85 | 27.07 | 1.04 | 2 | 2 |
| GABRB2 | 74 | V | 31.38169643 | 16.41768632 | 0.336624766 | 8.429450511 | NA | NA | NA | NA | NA | NA | NA |
| GABRB2 | 75 | S | 30.149625 | 12.72991677 | 0.242824823 | 4.868983516 | 7.92 | -0.4 | -2.15 | 14.47 | 0.75 | 1 | -1 |
| GABRB2 | 76 | E | 26.99134722 | 14.9087915 | 0.298245403 | 6.523373506 | NA | NA | NA | NA | NA | NA | NA |
| GABRB2 | 77 | V | 27.963125 | 10.53360152 | 0.186960636 | 2.157075933 | 8.41 | -0.27 | -1.8 | 27.49 | 0.31 | 0 | 0 |
| GABRB2 | 78 | N | 32.098375 | 9.487311632 | 0.16034782 | 1.658998388 | 7.92 | -0.4 | -2.15 | 14.47 | 0.75 | 1 | -1 |
| GABRB2 | 79 | M | 31.810125 | 15.00920418 | 0.300799441 | 7.155456326 | NA | NA | NA | NA | NA | NA | NA |
| GABRB2 | 80 | D | 34.96625 | 12.13050951 | 0.227578651 | 4.262010749 | 7.88 | -0.53 | -3.97 | 34.96 | 1.21 | 2 | 2 |
| GABRB2 | 81 | Y | 37.42479167 | 16.26974742 | 0.332861879 | 8.395976612 | NA | NA | NA | NA | NA | NA | NA |
| GABRB2 | 82 | T | 38.65155357 | 13.58424455 | 0.264555003 | 5.718847934 | NA | NA | NA | NA | NA | NA | NA |
| GABRB2 | 83 | L | 41.797125 | 17.13555281 | 0.354883998 | 12.81022605 | NA | NA | NA | NA | NA | NA | NA |
| GABRB2 | 84 | T | 43.58755357 | 16.96933095 | 0.350656076 | 10.72327188 | NA | NA | NA | NA | NA | NA | NA |
| GABRB2 | 85 | M | 46.8865 | 19.09933195 | 0.404833532 | 12.70194212 | NA | NA | NA | NA | NA | NA | NA |
| GABRB2 | 86 | Y | 49.17445833 | 21.03842226 | 0.454155096 | 14.45379689 | NA | NA | NA | NA | NA | NA | NA |
| GABRB2 | 87 | F | 52.42094318 | 20.44332823 | 0.439018634 | 13.85036525 | NA | NA | NA | NA | NA | NA | NA |
| GABRB2 | 88 | Q | 54.68334722 | 23.5153166 | 0.517155929 | 16.28790744 | NA | NA | NA | NA | NA | NA | NA |

| | | | | | | | | | | | | | |
|---|---|---|---|---|---|---|---|---|---|---|---|---|---|
| GABRB2 | 89 | Q | 57.552125 | 21.41810718 | 0.463812539 | 14.008253 | NA | NA | NA | NA | NA | NA | NA |
| GABRB2 | 90 | A | 60.446525 | 27.09532035 | 0.608214805 | 19.67795903 | NA | NA | NA | NA | NA | NA | NA |
| GABRB2 | 91 | W | 63.81783929 | 23.00475257 | 0.504169521 | 15.60564543 | NA | NA | NA | NA | NA | NA | NA |
| GABRB2 | 92 | R | 68.68985227 | 30.13946327 | 0.685643839 | 22.80380528 | NA | NA | NA | NA | NA | NA | NA |
| GABRB2 | 93 | D | 70.18525 | 25.61215657 | 0.57048992 | 17.32535593 | NA | NA | NA | NA | NA | NA | NA |
| GABRB2 | 94 | K | 73.53556944 | 30.55660672 | 0.696254055 | 22.26824922 | NA | NA | NA | NA | NA | NA | NA |
| GABRB2 | 95 | R | 74.15239773 | 25.41994426 | 0.565600921 | 17.14003834 | NA | NA | NA | NA | NA | NA | NA |
| GABRB2 | 96 | L | 72.3775 | 24.48553602 | 0.541833861 | 16.18327687 | NA | NA | NA | NA | NA | NA | NA |
| GABRB2 | 97 | S | 75.65379167 | 30.02846984 | 0.682820675 | 21.72025185 | NA | NA | NA | NA | NA | NA | NA |
| GABRB2 | 98 | Y | 74.673875 | 29.96803107 | 0.68128339 | 21.13828322 | NA | NA | NA | NA | NA | NA | NA |
| GABRB2 | 99 | N | 78.44875 | 34.87295651 | 0.806042197 | 26.02018883 | NA | NA | NA | NA | NA | NA | NA |
| GABRB2 | 100 | V | 79.666125 | 35.23186656 | 0.815171223 | 26.16113588 | NA | NA | NA | NA | NA | NA | NA |
| GABRB2 | 101 | I | 76.062625 | 32.64187305 | 0.749293667 | 23.71347243 | NA | NA | NA | NA | NA | NA | NA |
| GABRB2 | 102 | P | 73.61155357 | 35.24601246 | 0.81553103 | 26.89109234 | NA | NA | NA | NA | NA | NA | NA |
| GABRB2 | 103 | L | 71.136 | 32.28113669 | 0.740118188 | 23.92957746 | NA | NA | NA | NA | NA | NA | NA |
| GABRB2 | 104 | N | 67.0935 | 29.09260557 | 0.65901658 | 20.72673397 | NA | NA | NA | NA | NA | NA | NA |
| GABRB2 | 105 | L | 69.9645 | 26.4669141 | 0.592231032 | 18.1159721 | NA | NA | NA | NA | NA | NA | NA |
| GABRB2 | 106 | T | 65.68183929 | 23.69091476 | 0.52162234 | 15.31132662 | NA | NA | NA | NA | NA | NA | NA |
| GABRB2 | 107 | L | 67.837625 | 20.70275866 | 0.445617354 | 12.33058743 | NA | NA | NA | NA | NA | NA | NA |
| GABRB2 | 108 | D | 65.35225 | 16.7514094 | 0.345113151 | 8.333928491 | NA | NA | NA | NA | NA | NA | NA |
| GABRB2 | 109 | N | 62.43825 | 14.32875968 | 0.283492053 | 5.988755859 | NA | NA | NA | NA | NA | NA | NA |
| GABRB2 | 110 | R | 65.95385227 | 12.43018757 | 0.235201086 | 4.077086181 | 8.49 | -0.13 | -2.95 | 26.17 | 0.54 | 2 | 0 |
| GABRB2 | 111 | V | 67.67526786 | 15.61665617 | 0.316250233 | 7.273619706 | NA | NA | NA | NA | NA | NA | NA |
| GABRB2 | 112 | A | 64.744325 | 15.04700648 | 0.301760958 | 6.747509488 | NA | NA | NA | NA | NA | NA | NA |
| GABRB2 | 113 | D | 65.706875 | 10.10377974 | 0.176027941 | 1.79168524 | 8.49 | -0.13 | -2.95 | 26.17 | 0.54 | 2 | 0 |
| GABRB2 | 114 | Q | 70.51834722 | 14.67218791 | 0.292227292 | 6.557703621 | NA | NA | NA | NA | NA | NA | NA |
| GABRB2 | 115 | L | 68.46925 | 17.7435229 | 0.370347968 | 9.448762308 | NA | NA | NA | NA | NA | NA | NA |
| GABRB2 | 116 | W | 65.91641071 | 20.16975848 | 0.432060274 | 11.88706142 | NA | NA | NA | NA | NA | NA | NA |
| GABRB2 | 117 | V | 62.96383929 | 17.73272585 | 0.370073341 | 10.3305719 | NA | NA | NA | NA | NA | NA | NA |
| GABRB2 | 118 | P | 60.712125 | 20.60573634 | 0.443149551 | 13.18746877 | NA | NA | NA | NA | NA | NA | NA |
| GABRB2 | 119 | D | 58.816125 | 19.68480757 | 0.419725347 | 12.26794978 | NA | NA | NA | NA | NA | NA | NA |
| GABRB2 | 120 | T | 54.81169643 | 18.99023407 | 0.402058583 | 11.86451293 | NA | NA | NA | NA | NA | NA | NA |
| GABRB2 | 121 | Y | 51.669625 | 18.87838025 | 0.399213534 | 12.28493639 | NA | NA | NA | NA | NA | NA | NA |

| | | | | | | | | | | | | | |
|---|---|---|---|---|---|---|---|---|---|---|---|---|---|
| GABRB2 | 122 | F | 48.48921591 | 17.86659313 | 0.37347831 | 11.30860018 | NA | NA | NA | NA | NA | NA | NA |
| GABRB2 | 123 | L | 46.3225 | 20.28865893 | 0.435084556 | 15.9817634 | NA | NA | NA | NA | NA | NA | NA |
| GABRB2 | 124 | N | 42.995875 | 17.90915978 | 0.374561011 | 13.85781758 | NA | NA | NA | NA | NA | NA | NA |
| GABRB2 | 125 | D | 45.191 | 14.09011671 | 0.277422071 | 10.07689668 | NA | NA | NA | NA | NA | NA | NA |
| GABRB2 | 126 | K | 40.67656944 | 10.55646267 | 0.187542118 | 6.497132577 | NA | NA | NA | NA | NA | NA | NA |
| GABRB2 | 127 | K | 43.32590278 | 5.649191535 | 0.062723647 | 1.675629055 | 4.35 | -0.41 | -3.9 | 49.5 | 1.46 | 3 | 3 |
| GABRB2 | 128 | S | 47.09745833 | 9.193133307 | 0.152865272 | 2.78675651 | 6.44 | -0.18 | -2.48 | 39.71 | 1.04 | 4 | 2 |
| GABRB2 | 129 | F | 47.527125 | 8.147799089 | 0.126276764 | 1.583832537 | 6.6 | -0.18 | -2.48 | 39.71 | 1.04 | 4 | 2 |
| GABRB2 | 130 | V | 51.96341071 | 9.902846358 | 0.170917117 | 3.32090955 | 6.6 | -0.18 | -2.48 | 39.71 | 1.04 | 4 | 2 |
| GABRB2 | 131 | H | 49.057925 | 11.78580651 | 0.218810988 | 5.191717859 | NA | NA | NA | NA | NA | NA | NA |
| GABRB2 | 132 | G | 52.333875 | 11.24532623 | 0.205063648 | 3.884450543 | 7.4 | 0.06 | -2.63 | 2.95 | -0.47 | 0 | 0 |
| GABRB2 | 133 | V | 51.46241071 | 13.26510437 | 0.256437541 | 6.638338888 | NA | NA | NA | NA | NA | NA | NA |
| GABRB2 | 134 | T | 55.19455357 | 13.62069381 | 0.265482105 | 6.46212096 | NA | NA | NA | NA | NA | NA | NA |
| GABRB2 | 135 | V | 57.37869643 | 13.05126546 | 0.250998459 | 5.694981795 | NA | NA | NA | NA | NA | NA | NA |
| GABRB2 | 136 | K | 57.23823611 | 9.019204289 | 0.148441316 | 1.602249998 | 7.42 | -0.06 | -2.95 | 14.59 | 0.02 | 1 | 1 |
| GABRB2 | 137 | N | 54.411875 | 14.10938773 | 0.277912237 | 6.716846563 | NA | NA | NA | NA | NA | NA | NA |
| GABRB2 | 138 | R | 59.298125 | 12.37498687 | 0.233797034 | 5.506020314 | NA | NA | NA | NA | NA | NA | NA |
| GABRB2 | 139 | M | 59.415 | 19.2486133 | 0.408630565 | 11.99024979 | NA | NA | NA | NA | NA | NA | NA |
| GABRB2 | 140 | I | 63.926875 | 19.3254741 | 0.410585551 | 12.12522341 | NA | NA | NA | NA | NA | NA | NA |
| GABRB2 | 141 | R | 62.99457955 | 23.95634747 | 0.528373731 | 16.68265688 | NA | NA | NA | NA | NA | NA | NA |
| GABRB2 | 142 | L | 68.264375 | 25.15994708 | 0.558987785 | 16.83041278 | NA | NA | NA | NA | NA | NA | NA |
| GABRB2 | 143 | H | 67.223025 | 29.40747079 | 0.667025307 | 21.07984424 | NA | NA | NA | NA | NA | NA | NA |
| GABRB2 | 144 | P | 71.46369643 | 32.85580731 | 0.754735173 | 24.5271087 | NA | NA | NA | NA | NA | NA | NA |
| GABRB2 | 145 | D | 70.151375 | 33.28422278 | 0.765632098 | 24.97531643 | NA | NA | NA | NA | NA | NA | NA |
| GABRB2 | 146 | G | 69.757625 | 29.70387074 | 0.674564363 | 21.40358935 | NA | NA | NA | NA | NA | NA | NA |
| GABRB2 | 147 | T | 66.20969643 | 29.08979103 | 0.658944991 | 20.78403737 | NA | NA | NA | NA | NA | NA | NA |
| GABRB2 | 148 | V | 65.27226786 | 24.23922002 | 0.535568711 | 17.01512752 | NA | NA | NA | NA | NA | NA | NA |
| GABRB2 | 149 | L | 61.136375 | 23.7275163 | 0.522553316 | 16.34694421 | NA | NA | NA | NA | NA | NA | NA |
| GABRB2 | 150 | Y | 59.70320833 | 16.07646049 | 0.327945546 | 8.687505992 | NA | NA | NA | NA | NA | NA | NA |
| GABRB2 | 151 | G | 57.066375 | 18.915313 | 0.400152934 | 11.49879975 | NA | NA | NA | NA | NA | NA | NA |
| GABRB2 | 152 | L | 54.489125 | 15.7851936 | 0.320537053 | 8.475391529 | NA | NA | NA | NA | NA | NA | NA |
| GABRB2 | 153 | R | 49.53930682 | 14.79250516 | 0.295287612 | 8.199666945 | NA | NA | NA | NA | NA | NA | NA |
| GABRB2 | 154 | I | 48.94875 | 15.2792886 | 0.30766915 | 8.700031 | NA | NA | NA | NA | NA | NA | NA |

| | | | | | | | | | | | | | |
|---|---|---|---|---|---|---|---|---|---|---|---|---|---|
| GABRB2 | 155 | T | 45.15641071 | 12.84327201 | 0.24570806 | 6.434412114 | NA | NA | NA | NA | NA | NA | NA |
| GABRB2 | 156 | T | 43.957125 | 14.87044176 | 0.297269961 | 10.52750484 | NA | NA | NA | NA | NA | NA | NA |
| GABRB2 | 157 | T | 40.69526786 | 12.00697352 | 0.224436462 | 7.964343812 | NA | NA | NA | NA | NA | NA | NA |
| GABRB2 | 158 | A | 40.110525 | 15.97854981 | 0.325455148 | 11.93094025 | NA | NA | NA | NA | NA | NA | NA |
| GABRB2 | 159 | A | 38.411725 | 17.10046824 | 0.353991607 | 12.60995619 | NA | NA | NA | NA | NA | NA | NA |
| GABRB2 | 160 | C | 35.28145833 | 19.68077016 | 0.419622654 | 11.86571033 | NA | NA | NA | NA | NA | NA | NA |
| GABRB2 | 161 | M | 33.258875 | 21.57827697 | 0.467886524 | 13.63023392 | NA | NA | NA | NA | NA | NA | NA |
| GABRB2 | 162 | M | 29.85375 | 19.59278883 | 0.417384812 | 10.9774785 | NA | NA | NA | NA | NA | NA | NA |
| GABRB2 | 163 | D | 26.4475 | 24.0813048 | 0.531552072 | 15.71996698 | NA | NA | NA | NA | NA | NA | NA |
| GABRB2 | 164 | L | 23.762375 | 19.16899943 | 0.406605553 | 15.70237542 | NA | NA | NA | NA | NA | NA | NA |
| GABRB2 | 165 | R | 21.80530682 | 23.45324127 | 0.515577017 | 19.90656108 | NA | NA | NA | NA | NA | NA | NA |
| GABRB2 | 166 | R | 20.87030682 | 23.85640377 | 0.525831622 | 20.30888945 | NA | NA | NA | NA | NA | NA | NA |
| GABRB2 | 167 | Y | 19.84395833 | 16.77284845 | 0.345658462 | 13.2478465 | NA | NA | NA | NA | NA | NA | NA |
| GABRB2 | 168 | P | 18.90998214 | 20.52391042 | 0.441068275 | 17.39994538 | NA | NA | NA | NA | NA | NA | NA |
| GABRB2 | 169 | L | 18.8845 | 23.67978199 | 0.521339174 | 20.61130944 | NA | NA | NA | NA | NA | NA | NA |
| GABRB2 | 170 | D | 23.85475 | 23.50336986 | 0.516852058 | 19.99696341 | NA | NA | NA | NA | NA | NA | NA |
| GABRB2 | 171 | E | 26.75479167 | 27.96755917 | 0.63040056 | 19.57994497 | NA | NA | NA | NA | NA | NA | NA |
| GABRB2 | 172 | Q | 29.31023611 | 23.88587216 | 0.526581162 | 16.03810977 | NA | NA | NA | NA | NA | NA | NA |
| GABRB2 | 173 | N | 34.13175 | 26.22124549 | 0.585982349 | 18.20760105 | NA | NA | NA | NA | NA | NA | NA |
| GABRB2 | 174 | C | 35.56195833 | 22.39344875 | 0.488620756 | 14.51618928 | NA | NA | NA | NA | NA | NA | NA |
| GABRB2 | 175 | T | 39.69198214 | 22.94757338 | 0.502715145 | 18.91003332 | NA | NA | NA | NA | NA | NA | NA |
| GABRB2 | 176 | L | 43.691125 | 20.94184056 | 0.451698501 | 16.5984483 | NA | NA | NA | NA | NA | NA | NA |
| GABRB2 | 177 | E | 44.77001389 | 22.81004996 | 0.49921718 | 18.5512416 | NA | NA | NA | NA | NA | NA | NA |
| GABRB2 | 178 | I | 49.40725 | 23.41834945 | 0.514689529 | 16.84530378 | NA | NA | NA | NA | NA | NA | NA |
| GABRB2 | 179 | E | 51.33223611 | 23.30243803 | 0.511741274 | 16.70963651 | NA | NA | NA | NA | NA | NA | NA |
| GABRB2 | 180 | S | 55.85329167 | 24.59938613 | 0.544729685 | 17.22451918 | NA | NA | NA | NA | NA | NA | NA |
| GABRB2 | 181 | Y | 55.81620833 | 20.83779924 | 0.449052167 | 13.46274678 | NA | NA | NA | NA | NA | NA | NA |
| GABRB2 | 182 | G | 60.170625 | 23.73387785 | 0.522715124 | 16.32491517 | NA | NA | NA | NA | NA | NA | NA |
| GABRB2 | 183 | Y | 62.238625 | 23.41776057 | 0.51467455 | 16.38971073 | NA | NA | NA | NA | NA | NA | NA |
| GABRB2 | 184 | T | 60.52526786 | 29.52900909 | 0.670116684 | 22.11116478 | NA | NA | NA | NA | NA | NA | NA |
| GABRB2 | 185 | T | 59.09669643 | 32.39492962 | 0.743012558 | 24.98740518 | NA | NA | NA | NA | NA | NA | NA |
| GABRB2 | 186 | D | 62.936 | 33.88407973 | 0.780889707 | 26.48124943 | NA | NA | NA | NA | NA | NA | NA |
| GABRB2 | 187 | D | 64.13275 | 28.92591088 | 0.654776632 | 21.50400257 | NA | NA | NA | NA | NA | NA | NA |

| | | | | | | | | | | | | | |
|---|---|---|---|---|---|---|---|---|---|---|---|---|---|
| GABRB2 | 188 | I | 60.757 | 26.77693765 | 0.600116609 | 19.36253175 | NA | NA | NA | NA | NA | NA | NA |
| GABRB2 | 189 | E | 59.307125 | 30.98318542 | 0.707104261 | 23.57070482 | NA | NA | NA | NA | NA | NA | NA |
| GABRB2 | 190 | F | 55.77439773 | 25.99266983 | 0.580168433 | 18.6149941 | NA | NA | NA | NA | NA | NA | NA |
| GABRB2 | 191 | Y | 56.693125 | 31.9302663 | 0.731193655 | 25.33831327 | NA | NA | NA | NA | NA | NA | NA |
| GABRB2 | 192 | W | 50.30376786 | 28.03154254 | 0.632028003 | 21.45765623 | NA | NA | NA | NA | NA | NA | NA |
| GABRB2 | 193 | R | 55.56421591 | 30.28686796 | 0.689393138 | 24.15248396 | NA | NA | NA | NA | NA | NA | NA |
| GABRB2 | 194 | G | 51.584125 | 35.19989035 | 0.814357895 | 28.61347069 | NA | NA | NA | NA | NA | NA | NA |
| GABRB2 | 195 | D | 49.537 | 35.66510617 | 0.826190852 | 29.09298837 | NA | NA | NA | NA | NA | NA | NA |
| GABRB2 | 196 | D | 46.95675 | 36.08037983 | 0.836753509 | 29.51616143 | NA | NA | NA | NA | NA | NA | NA |
| GABRB2 | 197 | N | 48.42025 | 32.75398518 | 0.752145285 | 26.16951736 | NA | NA | NA | NA | NA | NA | NA |
| GABRB2 | 198 | A | 48.618925 | 29.78339982 | 0.676587218 | 23.20242705 | NA | NA | NA | NA | NA | NA | NA |
| GABRB2 | 199 | V | 45.41498214 | 28.73431384 | 0.649903282 | 22.31943373 | NA | NA | NA | NA | NA | NA | NA |
| GABRB2 | 200 | T | 45.44826786 | 28.61584676 | 0.646890023 | 22.19982943 | NA | NA | NA | NA | NA | NA | NA |
| GABRB2 | 201 | G | 41.764375 | 28.93206541 | 0.654933175 | 22.52351324 | NA | NA | NA | NA | NA | NA | NA |
| GABRB2 | 202 | V | 39.17098214 | 29.70477898 | 0.674587464 | 25.50839008 | NA | NA | NA | NA | NA | NA | NA |
| GABRB2 | 203 | T | 37.296125 | 31.12519873 | 0.710716428 | 22.9286846 | NA | NA | NA | NA | NA | NA | NA |
| GABRB2 | 204 | K | 38.562125 | 28.29785068 | 0.638801661 | 20.10079444 | NA | NA | NA | NA | NA | NA | NA |
| GABRB2 | 205 | I | 36.342625 | 25.31369606 | 0.562898454 | 17.11994055 | NA | NA | NA | NA | NA | NA | NA |
| GABRB2 | 206 | E | 31.324125 | 24.02516822 | 0.530124215 | 15.84202474 | NA | NA | NA | NA | NA | NA | NA |
| GABRB2 | 207 | L | 31.14175 | 21.32164492 | 0.461358982 | 13.12662579 | NA | NA | NA | NA | NA | NA | NA |
| GABRB2 | 208 | P | 27.96598214 | 19.98324208 | 0.427316152 | 11.58590257 | NA | NA | NA | NA | NA | NA | NA |
| GABRB2 | 209 | Q | 25.18934722 | 18.61288339 | 0.392460512 | 10.28872456 | NA | NA | NA | NA | NA | NA | NA |
| GABRB2 | 210 | F | 27.97948864 | 21.36210828 | 0.462388184 | 12.98093498 | NA | NA | NA | NA | NA | NA | NA |
| GABRB2 | 211 | S | 29.047625 | 26.67735791 | 0.597583757 | 18.81798338 | NA | NA | NA | NA | NA | NA | NA |
| GABRB2 | 212 | I | 33.305625 | 26.36031741 | 0.589519701 | 18.17264349 | NA | NA | NA | NA | NA | NA | NA |
| GABRB2 | 213 | V | 31.455125 | 30.84062918 | 0.703478284 | 22.64884272 | NA | NA | NA | NA | NA | NA | NA |
| GABRB2 | 214 | D | 35.288375 | 31.6741637 | 0.724679579 | 23.59940409 | NA | NA | NA | NA | NA | NA | NA |
| GABRB2 | 215 | Y | 39.83295833 | 31.56603322 | 0.721929236 | 23.49796609 | NA | NA | NA | NA | NA | NA | NA |
| GABRB2 | 216 | K | 39.39301389 | 31.96564634 | 0.732093561 | 27.61670028 | NA | NA | NA | NA | NA | NA | NA |
| GABRB2 | 217 | L | 44.633875 | 31.37164704 | 0.716984943 | 24.96091601 | NA | NA | NA | NA | NA | NA | NA |
| GABRB2 | 218 | I | 44.988375 | 32.22994944 | 0.738816219 | 25.82733399 | NA | NA | NA | NA | NA | NA | NA |
| GABRB2 | 219 | T | 49.323125 | 33.81577824 | 0.779152431 | 27.23980374 | NA | NA | NA | NA | NA | NA | NA |
| GABRB2 | 220 | K | 47.82590278 | 31.47740078 | 0.719674833 | 25.38579063 | NA | NA | NA | NA | NA | NA | NA |

| | | | | | | | | | | | | | |
|---|---|---|---|---|---|---|---|---|---|---|---|---|---|
| GABRB2 | 221 | K | 52.906125 | 33.97024117 | 0.783081259 | 27.38572863 | NA | NA | NA | NA | NA | NA | NA |
| GABRB2 | 222 | V | 50.29726786 | 30.93639916 | 0.705914233 | 24.40547375 | NA | NA | NA | NA | NA | NA | NA |
| GABRB2 | 223 | V | 51.35569643 | 32.98868138 | 0.75811488 | 26.40353392 | NA | NA | NA | NA | NA | NA | NA |
| GABRB2 | 224 | F | 51.03748864 | 27.66585262 | 0.622726529 | 21.1939821 | NA | NA | NA | NA | NA | NA | NA |
| GABRB2 | 225 | S | 53.52195833 | 32.19933915 | 0.738037634 | 25.60794235 | NA | NA | NA | NA | NA | NA | NA |
| GABRB2 | 226 | T | 56.85955357 | 29.60381426 | 0.672019385 | 22.2267912 | NA | NA | NA | NA | NA | NA | NA |
| GABRB2 | 227 | G | 56.160625 | 32.53303437 | 0.74652531 | 25.15633732 | NA | NA | NA | NA | NA | NA | NA |
| GABRB2 | 228 | S | 56.04995833 | 32.4327077 | 0.74397346 | 26.00336941 | NA | NA | NA | NA | NA | NA | NA |
| GABRB2 | 229 | Y | 55.436125 | 27.70943713 | 0.623835119 | 21.47291381 | NA | NA | NA | NA | NA | NA | NA |
| GABRB2 | 230 | P | 54.60398214 | 28.59961133 | 0.646477068 | 22.00776994 | NA | NA | NA | NA | NA | NA | NA |
| GABRB2 | 231 | R | 48.96794318 | 27.365791 | 0.615094337 | 20.91081337 | NA | NA | NA | NA | NA | NA | NA |
| GABRB2 | 232 | L | 49.8125 | 27.27447415 | 0.612771656 | 20.70987123 | NA | NA | NA | NA | NA | NA | NA |
| GABRB2 | 233 | S | 45.01395833 | 28.68353836 | 0.648611787 | 22.27029637 | NA | NA | NA | NA | NA | NA | NA |
| GABRB2 | 234 | L | 42.97025 | 26.52747834 | 0.593771508 | 22.20557007 | NA | NA | NA | NA | NA | NA | NA |
| GABRB2 | 235 | S | 39.51245833 | 28.94358647 | 0.655226218 | 24.89499599 | NA | NA | NA | NA | NA | NA | NA |
| GABRB2 | 236 | F | 36.97485227 | 24.55609598 | 0.543628583 | 16.68899475 | NA | NA | NA | NA | NA | NA | NA |
| GABRB2 | 237 | K | 32.05323611 | 28.28420444 | 0.638454563 | 20.21164203 | NA | NA | NA | NA | NA | NA | NA |
| GABRB2 | 238 | L | 30.794625 | 23.43990243 | 0.515237738 | 15.6099651 | NA | NA | NA | NA | NA | NA | NA |
| GABRB2 | 239 | K | 25.75823611 | 26.59907775 | 0.595592669 | 18.22870012 | NA | NA | NA | NA | NA | NA | NA |
| GABRB2 | 240 | R | 23.82721591 | 20.3910473 | 0.437688847 | 16.94833572 | NA | NA | NA | NA | NA | NA | NA |
| GABRB2 | 241 | N | 22.292 | 24.07396183 | 0.531365301 | 20.55452952 | NA | NA | NA | NA | NA | NA | NA |
| GABRB2 | 242 | I | 18.063625 | 23.09537257 | 0.506474478 | 20.01423723 | NA | NA | NA | NA | NA | NA | NA |
| GABRB2 | 243 | G | 16.496375 | 23.22618477 | 0.509801741 | 20.13350791 | NA | NA | NA | NA | NA | NA | NA |
| GABRB2 | 244 | Y | 18.25429167 | 18.59181707 | 0.391924681 | 16.71443524 | NA | NA | NA | NA | NA | NA | NA |
| GABRB2 | 245 | F | 17.28521591 | 18.44296749 | 0.388138631 | 15.35586266 | NA | NA | NA | NA | NA | NA | NA |
| GABRB2 | 246 | I | 12.488375 | 18.90413949 | 0.399868731 | 15.81564439 | NA | NA | NA | NA | NA | NA | NA |
| GABRB2 | 247 | L | 10.6925 | 17.32502035 | 0.359703183 | 14.23005546 | NA | NA | NA | NA | NA | NA | NA |
| GABRB2 | 248 | Q | 12.28556944 | 14.13096692 | 0.278461113 | 11.03334105 | NA | NA | NA | NA | NA | NA | NA |
| GABRB2 | 249 | T | 11.363125 | 14.73333292 | 0.293782541 | 11.67925807 | NA | NA | NA | NA | NA | NA | NA |
| GABRB2 | 250 | Y | 10.669375 | 16.87123439 | 0.348160949 | 13.97268337 | NA | NA | NA | NA | NA | NA | NA |
| GABRB2 | 251 | M | 6.253125 | 16.92707931 | 0.349581388 | 14.60746261 | NA | NA | NA | NA | NA | NA | NA |
| GABRB2 | 252 | P | 4.864982143 | 13.36807574 | 0.25905666 | 11.04837242 | NA | NA | NA | NA | NA | NA | NA |
| GABRB2 | 253 | S | 3.786791667 | 13.4270779 | 0.260557404 | 11.17667769 | NA | NA | NA | NA | NA | NA | NA |

| | | | | | | | | | | | | | |
|---|---|---|---|---|---|---|---|---|---|---|---|---|---|
| GABRB2 | 254 | I | 3.145 | 17.11696276 | 0.354411152 | 15.8320235 | NA | NA | NA | NA | NA | NA | NA |
| GABRB2 | 255 | L | 0.65575 | 16.3309662 | 0.334419004 | 14.94643373 | NA | NA | NA | NA | NA | NA | NA |
| GABRB2 | 256 | I | 0 | 12.03512131 | 0.225152413 | 10.67480795 | NA | NA | NA | NA | NA | NA | NA |
| GABRB2 | 257 | T | -0.308410714 | 16.08374944 | 0.328130944 | 14.62605775 | NA | NA | NA | NA | NA | NA | NA |
| GABRB2 | 258 | I | -1.54 | 18.55422254 | 0.390968449 | 17.12637293 | NA | NA | NA | NA | NA | NA | NA |
| GABRB2 | 259 | L | -4.432375 | 14.22685397 | 0.28090004 | 12.93254918 | NA | NA | NA | NA | NA | NA | NA |
| GABRB2 | 260 | S | -4.962458333 | 16.5871805 | 0.340935921 | 15.13051962 | NA | NA | NA | NA | NA | NA | NA |
| GABRB2 | 261 | W | -4.490482143 | 19.4350027 | 0.413371457 | 18.02232526 | NA | NA | NA | NA | NA | NA | NA |
| GABRB2 | 262 | V | -9.296410714 | 20.70965565 | 0.445792782 | 19.37125631 | NA | NA | NA | NA | NA | NA | NA |
| GABRB2 | 263 | S | -10.95395833 | 18.12195191 | 0.379973467 | 16.73069519 | NA | NA | NA | NA | NA | NA | NA |
| GABRB2 | 264 | F | -11.10294318 | 22.53777261 | 0.492291693 | 21.06012098 | NA | NA | NA | NA | NA | NA | NA |
| GABRB2 | 265 | W | -12.56576786 | 24.70438676 | 0.54740042 | 23.3383559 | NA | NA | NA | NA | NA | NA | NA |
| GABRB2 | 266 | I | -15.802125 | 21.28688361 | 0.460474814 | 19.98284401 | NA | NA | NA | NA | NA | NA | NA |
| GABRB2 | 267 | N | -20.55225 | 25.79538077 | 0.575150304 | 24.43539857 | NA | NA | NA | NA | NA | NA | NA |
| GABRB2 | 268 | Y | -21.31854167 | 26.43222111 | 0.591348601 | 25.01929982 | NA | NA | NA | NA | NA | NA | NA |
| GABRB2 | 269 | D | -23.179 | 26.32979292 | 0.588743298 | 24.95151301 | NA | NA | NA | NA | NA | NA | NA |
| GABRB2 | 270 | A | -19.645925 | 22.82002553 | 0.499470912 | 21.45595067 | NA | NA | NA | NA | NA | NA | NA |
| GABRB2 | 271 | S | -18.17495833 | 19.78214207 | 0.42220109 | 18.38670793 | NA | NA | NA | NA | NA | NA | NA |
| GABRB2 | 272 | A | -16.313525 | 17.4855317 | 0.363785856 | 16.14508315 | NA | NA | NA | NA | NA | NA | NA |
| GABRB2 | 273 | A | -14.781925 | 18.01826387 | 0.377336119 | 16.69884581 | NA | NA | NA | NA | NA | NA | NA |
| GABRB2 | 274 | R | -13.76230682 | 16.61058843 | 0.341531311 | 15.20218522 | NA | NA | NA | NA | NA | NA | NA |
| GABRB2 | 275 | V | -12.50283929 | 13.98870527 | 0.274842629 | 12.56060633 | NA | NA | NA | NA | NA | NA | NA |
| GABRB2 | 276 | A | -10.456125 | 12.11740191 | 0.227245254 | 10.770029 | NA | NA | NA | NA | NA | NA | NA |
| GABRB2 | 277 | L | -8.824125 | 12.7254229 | 0.242710519 | 11.33345556 | NA | NA | NA | NA | NA | NA | NA |
| GABRB2 | 278 | G | -6.994625 | 11.80798158 | 0.21937502 | 10.30878428 | NA | NA | NA | NA | NA | NA | NA |
| GABRB2 | 279 | I | -7.044375 | 8.604469071 | 0.137892354 | 7.118358953 | NA | NA | NA | NA | NA | NA | NA |
| GABRB2 | 280 | T | -4.844696429 | 8.145984041 | 0.126230598 | 6.755385086 | NA | NA | NA | NA | NA | NA | NA |
| GABRB2 | 281 | T | -3.013125 | 9.937291834 | 0.171793252 | 8.422951723 | NA | NA | NA | NA | NA | NA | NA |
| GABRB2 | 282 | V | -1.728125 | 8.282791377 | 0.129710349 | 6.800610923 | NA | NA | NA | NA | NA | NA | NA |
| GABRB2 | 283 | L | -1.386 | 5.256251231 | 0.052729048 | 3.725678368 | 1.57 | 1.35 | 2.8 | 0.35 | -1.81 | 0 | 0 |
| GABRB2 | 284 | T | 0.286553571 | 6.644881113 | 0.088049424 | 5.154393378 | 1.57 | 1.35 | 2.8 | 0.35 | -1.81 | 0 | 0 |
| GABRB2 | 285 | M | 1.728375 | 8.723346204 | 0.140916043 | 7.173966293 | NA | NA | NA | NA | NA | NA | NA |
| GABRB2 | 286 | T | 2.353267857 | 6.054349924 | 0.073029019 | 4.461382434 | 1.79 | 0.24 | 1.3 | 0.95 | -0.52 | 0 | 0 |

| | | | | | | | | | | | | | |
|---|---|---|---|---|---|---|---|---|---|---|---|---|---|
| GABRB2 | 287 | T | 3.417410714 | 3.489238456 | 0.007784348 | 1.853325419 | 1.85 | 0.57 | 1.97 | 0.71 | -0.94 | 0 | 0 |
| GABRB2 | 288 | I | 5.326375 | 6.50713616 | 0.084545824 | 4.293410684 | 2.38 | 0.24 | 1.3 | 0.95 | -0.52 | 0 | 0 |
| GABRB2 | 289 | N | 7.569625 | 8.018086181 | 0.122977463 | 5.740620037 | NA | NA | NA | NA | NA | NA | NA |
| GABRB2 | 290 | T | 7.997267857 | 4.018625885 | 0.021249536 | 1.773819971 | 2.38 | 0.24 | 1.3 | 0.95 | -0.52 | 0 | 0 |
| GABRB2 | 291 | H | 10.251925 | 4.758695725 | 0.040073518 | 1.679812598 | 3.13 | -0.57 | -2.52 | 31.36 | 1.04 | 2 | 0 |
| GABRB2 | 292 | L | 11.5665 | 7.668578877 | 0.1140876 | 4.602976094 | 3.13 | -0.57 | -2.52 | 31.36 | 1.04 | 2 | 0 |
| GABRB2 | 293 | R | 12.75894318 | 9.30391509 | 0.155683053 | 6.255550104 | NA | NA | NA | NA | NA | NA | NA |
| GABRB2 | 294 | E | 13.25134722 | 4.067535618 | 0.022493575 | 2.209886473 | 3.57 | -0.69 | -3.85 | 38.7 | 1.47 | 3 | 1 |
| GABRB2 | 295 | T | 15.33326786 | 8.280569546 | 0.129653836 | 4.740609374 | 3.57 | -0.69 | -3.85 | 38.7 | 1.47 | 3 | 1 |
| GABRB2 | 296 | L | 17.178125 | 9.994107114 | 0.173238372 | 6.460335191 | NA | NA | NA | NA | NA | NA | NA |
| GABRB2 | 297 | P | 21.03483929 | 11.66147581 | 0.215648585 | 8.646815289 | NA | NA | NA | NA | NA | NA | NA |
| GABRB2 | 298 | K | 22.66945833 | 10.87585744 | 0.195666057 | 7.766668576 | NA | NA | NA | NA | NA | NA | NA |
| GABRB2 | 299 | I | 21.977875 | 13.79514563 | 0.269919359 | 10.27560012 | NA | NA | NA | NA | NA | NA | NA |
| GABRB2 | 300 | P | 22.42483929 | 16.08205096 | 0.328087742 | 12.57417601 | NA | NA | NA | NA | NA | NA | NA |
| GABRB2 | 301 | Y | 22.47345833 | 18.11123745 | 0.37970094 | 14.55286439 | NA | NA | NA | NA | NA | NA | NA |
| GABRB2 | 302 | V | 17.83741071 | 20.206606 | 0.432997506 | 16.65774622 | NA | NA | NA | NA | NA | NA | NA |
| GABRB2 | 303 | K | 17.46923611 | 15.46033664 | 0.312274181 | 11.91065626 | NA | NA | NA | NA | NA | NA | NA |
| GABRB2 | 304 | A | 14.959925 | 20.05217218 | 0.429069418 | 17.01448863 | NA | NA | NA | NA | NA | NA | NA |
| GABRB2 | 305 | I | 13.29575 | 15.92774093 | 0.324162803 | 12.86056741 | NA | NA | NA | NA | NA | NA | NA |
| GABRB2 | 306 | D | 12.512375 | 14.68122726 | 0.292457212 | 12.51985082 | NA | NA | NA | NA | NA | NA | NA |
| GABRB2 | 307 | M | 11.3275 | 18.71748362 | 0.395121062 | 16.46002639 | NA | NA | NA | NA | NA | NA | NA |
| GABRB2 | 308 | Y | 7.947041667 | 17.68903937 | 0.368962157 | 15.47882884 | NA | NA | NA | NA | NA | NA | NA |
| GABRB2 | 309 | L | 7.793875 | 12.97321737 | 0.249013274 | 10.68529104 | NA | NA | NA | NA | NA | NA | NA |
| GABRB2 | 310 | M | 6.243375 | 16.10089081 | 0.328566942 | 14.38253084 | NA | NA | NA | NA | NA | NA | NA |
| GABRB2 | 311 | G | 4.132625 | 18.32142882 | 0.385047244 | 16.58926369 | NA | NA | NA | NA | NA | NA | NA |
| GABRB2 | 312 | C | 2.785125 | 16.3891372 | 0.335898608 | 14.75831728 | NA | NA | NA | NA | NA | NA | NA |
| GABRB2 | 313 | F | 1.321306818 | 11.59846119 | 0.214045782 | 9.80583643 | NA | NA | NA | NA | NA | NA | NA |
| GABRB2 | 314 | V | 0.782267857 | 17.01611278 | 0.351845991 | 15.24230511 | NA | NA | NA | NA | NA | NA | NA |
| GABRB2 | 315 | F | -0.398306818 | 18.15174554 | 0.38073128 | 16.65416595 | NA | NA | NA | NA | NA | NA | NA |
| GABRB2 | 316 | V | -1.478553571 | 13.95182085 | 0.273904458 | 12.44830908 | NA | NA | NA | NA | NA | NA | NA |
| GABRB2 | 317 | F | -2.324670455 | 14.63404021 | 0.29125699 | 13.23343372 | NA | NA | NA | NA | NA | NA | NA |
| GABRB2 | 318 | M | -3.667625 | 18.1739808 | 0.381296843 | 16.71690315 | NA | NA | NA | NA | NA | NA | NA |
| GABRB2 | 319 | A | -6.351325 | 18.82290063 | 0.397802387 | 17.33454356 | NA | NA | NA | NA | NA | NA | NA |

| | | | | | | | | | | | | | |
|---|---|---|---|---|---|---|---|---|---|---|---|---|---|
| GABRB2 | 320 | L | -7.37675 | 13.83414096 | 0.270911222 | 12.37619055 | NA | NA | NA | NA | NA | NA | NA |
| GABRB2 | 321 | L | -7.327625 | 18.51179997 | 0.389889413 | 17.05430267 | NA | NA | NA | NA | NA | NA | NA |
| GABRB2 | 322 | E | -10.40101389 | 21.16326943 | 0.457330636 | 19.69784018 | NA | NA | NA | NA | NA | NA | NA |
| GABRB2 | 323 | Y | -13.87829167 | 20.2520198 | 0.434152625 | 18.76639569 | NA | NA | NA | NA | NA | NA | NA |
| GABRB2 | 324 | A | -12.553325 | 20.0844283 | 0.429889865 | 18.63522712 | NA | NA | NA | NA | NA | NA | NA |
| GABRB2 | 325 | L | -13.47 | 23.1416913 | 0.507652614 | 21.68648087 | NA | NA | NA | NA | NA | NA | NA |
| GABRB2 | 326 | V | -15.999125 | 25.16268308 | 0.559057376 | 23.69561356 | NA | NA | NA | NA | NA | NA | NA |
| GABRB2 | 327 | N | -17.91725 | 22.79378409 | 0.498803451 | 21.32368608 | NA | NA | NA | NA | NA | NA | NA |
| GABRB2 | 328 | Y | -18.269375 | 26.07779972 | 0.582333746 | 24.62006061 | NA | NA | NA | NA | NA | NA | NA |
| GABRB2 | 329 | I | -18.7945 | 28.63929727 | 0.647486496 | 27.18160314 | NA | NA | NA | NA | NA | NA | NA |
| GABRB2 | 330 | F | -22.76548864 | 29.30640619 | 0.664454688 | 27.82879812 | NA | NA | NA | NA | NA | NA | NA |
| GABRB2 | 331 | F | -24.81421591 | 30.55488329 | 0.696210219 | 29.08677288 | NA | NA | NA | NA | NA | NA | NA |
| GABRB2 | 486 | A | -17.464125 | 32.15938011 | 0.737021259 | 30.65813646 | NA | NA | NA | NA | NA | NA | NA |
| GABRB2 | 487 | I | -14.50275 | 28.5896515 | 0.646223736 | 27.1077393 | NA | NA | NA | NA | NA | NA | NA |
| GABRB2 | 488 | D | -15.25925 | 26.60326362 | 0.595699138 | 25.10990301 | NA | NA | NA | NA | NA | NA | NA |
| GABRB2 | 489 | R | -16.75967045 | 30.27912867 | 0.689196286 | 28.79938407 | NA | NA | NA | NA | NA | NA | NA |
| GABRB2 | 490 | W | -13.05098214 | 30.15728531 | 0.68609715 | 28.64741583 | NA | NA | NA | NA | NA | NA | NA |
| GABRB2 | 491 | S | -9.709625 | 26.55713849 | 0.594525926 | 25.05190959 | NA | NA | NA | NA | NA | NA | NA |
| GABRB2 | 492 | R | -12.78567045 | 27.81420376 | 0.626499902 | 26.3698335 | NA | NA | NA | NA | NA | NA | NA |
| GABRB2 | 493 | I | -9.6655 | 30.03148678 | 0.682897412 | 28.54783215 | NA | NA | NA | NA | NA | NA | NA |
| GABRB2 | 494 | F | -6.913306818 | 28.57380389 | 0.645820645 | 27.04780914 | NA | NA | NA | NA | NA | NA | NA |
| GABRB2 | 495 | F | -4.978670455 | 23.12207755 | 0.50715373 | 21.61469822 | NA | NA | NA | NA | NA | NA | NA |
| GABRB2 | 496 | P | -3.570267857 | 26.81687816 | 0.601132512 | 25.31568834 | NA | NA | NA | NA | NA | NA | NA |
| GABRB2 | 497 | V | -1.707267857 | 29.04194052 | 0.657727894 | 27.5271258 | NA | NA | NA | NA | NA | NA | NA |
| GABRB2 | 498 | V | -0.516410714 | 26.99558907 | 0.605678098 | 25.48866202 | NA | NA | NA | NA | NA | NA | NA |
| GABRB2 | 499 | F | 0.598579545 | 21.84418579 | 0.474650025 | 20.30100727 | NA | NA | NA | NA | NA | NA | NA |
| GABRB2 | 500 | S | 1.264458333 | 27.33486159 | 0.614307635 | 25.79602264 | NA | NA | NA | NA | NA | NA | NA |
| GABRB2 | 501 | F | 3.237943182 | 29.21603101 | 0.662155957 | 27.90098555 | NA | NA | NA | NA | NA | NA | NA |
| GABRB2 | 502 | F | 5.092579545 | 22.80081819 | 0.498982366 | 21.09573781 | NA | NA | NA | NA | NA | NA | NA |
| GABRB2 | 503 | N | 5.562 | 23.57057636 | 0.518561483 | 21.35162502 | NA | NA | NA | NA | NA | NA | NA |
| GABRB2 | 504 | I | 6.851875 | 28.71397745 | 0.649386017 | 26.50058733 | NA | NA | NA | NA | NA | NA | NA |
| GABRB2 | 505 | V | 8.709553571 | 29.18694299 | 0.661416092 | 26.9549352 | NA | NA | NA | NA | NA | NA | NA |
| GABRB2 | 506 | Y | 11.03779167 | 21.91624744 | 0.476482943 | 19.62424461 | NA | NA | NA | NA | NA | NA | NA |

| | | | | | | | | | | | | | |
|---|---|---|---|---|---|---|---|---|---|---|---|---|---|
| GABRB2 | 507 | W | 11.375625 | 23.5700287 | 0.518547553 | 21.26991585 | NA | NA | NA | NA | NA | NA | NA |
| GABRB2 | 508 | L | 12.620875 | 30.47827995 | 0.694261781 | 28.19290781 | NA | NA | NA | NA | NA | NA | NA |
| GABRB2 | 509 | Y | 13.90729167 | 30.79133286 | 0.702224411 | 28.49953082 | NA | NA | NA | NA | NA | NA | NA |
| GABRB2 | 510 | Y | 16.30520833 | 25.0286187 | 0.555647393 | 21.96757695 | NA | NA | NA | NA | NA | NA | NA |
| GABRB2 | 511 | V | 17.33569643 | 29.00151656 | 0.656699694 | 25.9291885 | NA | NA | NA | NA | NA | NA | NA |
| GABRB3 | 35 | S | 78.40916667 | 26.51759603 | 0.661783249 | 17.2169169 | NA | NA | NA | NA | NA | NA | NA |
| GABRB3 | 36 | F | 79.39356061 | 27.16278942 | 0.680045877 | 17.84998468 | NA | NA | NA | NA | NA | NA | NA |
| GABRB3 | 37 | V | 75.1672619 | 24.11193858 | 0.593689518 | 14.78840439 | NA | NA | NA | NA | NA | NA | NA |
| GABRB3 | 38 | K | 77.30794444 | 20.17097018 | 0.48213779 | 10.8901647 | NA | NA | NA | NA | NA | NA | NA |
| GABRB3 | 39 | E | 80.92361111 | 23.7796832 | 0.584284808 | 14.47493826 | NA | NA | NA | NA | NA | NA | NA |
| GABRB3 | 40 | T | 77.09040476 | 23.69608999 | 0.581918647 | 14.36982818 | NA | NA | NA | NA | NA | NA | NA |
| GABRB3 | 41 | V | 74.26711905 | 19.86492592 | 0.473475004 | 10.52426203 | NA | NA | NA | NA | NA | NA | NA |
| GABRB3 | 42 | D | 77.47145833 | 18.15519741 | 0.425080003 | 8.850812142 | NA | NA | NA | NA | NA | NA | NA |
| GABRB3 | 43 | K | 80.07827778 | 21.99910525 | 0.533884367 | 12.68869089 | NA | NA | NA | NA | NA | NA | NA |
| GABRB3 | 44 | L | 74.84208333 | 20.63176277 | 0.49518083 | 11.28718756 | NA | NA | NA | NA | NA | NA | NA |
| GABRB3 | 45 | L | 74.01595833 | 16.76140883 | 0.385627892 | 7.442854248 | NA | NA | NA | NA | NA | NA | NA |
| GABRB3 | 46 | K | 80.33861111 | 18.88594422 | 0.445764277 | 9.577731294 | NA | NA | NA | NA | NA | NA | NA |
| GABRB3 | 47 | G | 77.35633333 | 18.23223442 | 0.427260587 | 8.918578409 | NA | NA | NA | NA | NA | NA | NA |
| GABRB3 | 48 | Y | 72.23858333 | 16.26807604 | 0.37166378 | 6.90783865 | NA | NA | NA | NA | NA | NA | NA |
| GABRB3 | 49 | D | 72.69583333 | 16.88860995 | 0.389228404 | 7.505197749 | NA | NA | NA | NA | NA | NA | NA |
| GABRB3 | 50 | I | 68.64208333 | 12.89653136 | 0.276229969 | 3.635662777 | 9.31 | -0.49 | -2.12 | 30.51 | 1.01 | 3 | -1 |
| GABRB3 | 51 | R | 70.35110606 | 15.56284123 | 0.35170164 | 6.921750356 | NA | NA | NA | NA | NA | NA | NA |
| GABRB3 | 52 | L | 70.41395833 | 19.63471798 | 0.466958815 | 10.4499965 | NA | NA | NA | NA | NA | NA | NA |
| GABRB3 | 53 | R | 66.60265152 | 22.62136055 | 0.551497717 | 13.93609573 | NA | NA | NA | NA | NA | NA | NA |
| GABRB3 | 54 | P | 67.59297619 | 24.06835859 | 0.592455957 | 14.97587935 | NA | NA | NA | NA | NA | NA | NA |
| GABRB3 | 55 | D | 69.88045833 | 27.62907083 | 0.693244282 | 18.42956611 | NA | NA | NA | NA | NA | NA | NA |
| GABRB3 | 56 | F | 69.88610606 | 24.30216608 | 0.599074033 | 15.63205349 | NA | NA | NA | NA | NA | NA | NA |
| GABRB3 | 57 | G | 71.17933333 | 29.29715592 | 0.740460538 | 20.10064108 | NA | NA | NA | NA | NA | NA | NA |
| GABRB3 | 58 | G | 69.17408333 | 31.68201717 | 0.807965619 | 22.99916652 | NA | NA | NA | NA | NA | NA | NA |
| GABRB3 | 59 | P | 67.55983333 | 31.40520993 | 0.800130406 | 22.76338092 | NA | NA | NA | NA | NA | NA | NA |
| GABRB3 | 60 | P | 63.59411905 | 31.18548941 | 0.793911071 | 22.49257835 | NA | NA | NA | NA | NA | NA | NA |
| GABRB3 | 61 | V | 62.78697619 | 27.10735194 | 0.678476682 | 18.41721682 | NA | NA | NA | NA | NA | NA | NA |
| GABRB3 | 62 | C | 60.65933333 | 30.07271256 | 0.762413184 | 21.39632291 | NA | NA | NA | NA | NA | NA | NA |

| | | | | | | | | | | | | | |
|---|---|---|---|---|---|---|---|---|---|---|---|---|---|
| GABRB3 | 63 | V | 58.01354762 | 26.1859451 | 0.65239565 | 17.68584647 | NA | NA | NA | NA | NA | NA | NA |
| GABRB3 | 64 | G | 56.13108333 | 28.83242061 | 0.727305896 | 20.30956938 | NA | NA | NA | NA | NA | NA | NA |
| GABRB3 | 65 | M | 52.81545833 | 25.41543724 | 0.630585913 | 18.03339072 | NA | NA | NA | NA | NA | NA | NA |
| GABRB3 | 66 | N | 51.12145833 | 26.19342501 | 0.652607373 | 18.80628767 | NA | NA | NA | NA | NA | NA | NA |
| GABRB3 | 67 | I | 47.26833333 | 25.27542075 | 0.626622653 | 19.1960158 | NA | NA | NA | NA | NA | NA | NA |
| GABRB3 | 68 | D | 46.47958333 | 24.70806421 | 0.610563249 | 18.63332415 | NA | NA | NA | NA | NA | NA | NA |
| GABRB3 | 69 | I | 42.39195833 | 24.77597354 | 0.612485468 | 18.98844002 | NA | NA | NA | NA | NA | NA | NA |
| GABRB3 | 70 | A | 41.81983333 | 22.73174435 | 0.554622203 | 16.63444917 | NA | NA | NA | NA | NA | NA | NA |
| GABRB3 | 71 | S | 39.10866667 | 20.6122132 | 0.494627467 | 13.72809607 | NA | NA | NA | NA | NA | NA | NA |
| GABRB3 | 72 | I | 36.07258333 | 19.03157634 | 0.449886491 | 12.16016348 | NA | NA | NA | NA | NA | NA | NA |
| GABRB3 | 73 | D | 36.10395833 | 15.92739919 | 0.362020695 | 9.053289194 | NA | NA | NA | NA | NA | NA | NA |
| GABRB3 | 74 | M | 33.15658333 | 11.10108504 | 0.225408668 | 4.249415467 | 6.95 | -0.5 | -3.95 | 38.6 | 1.27 | 3 | 3 |
| GABRB3 | 75 | V | 30.84540476 | 15.60108493 | 0.352784154 | 8.745635633 | NA | NA | NA | NA | NA | NA | NA |
| GABRB3 | 76 | S | 29.824 | 11.71604562 | 0.242815536 | 4.865129817 | 6.95 | -0.5 | -3.95 | 38.6 | 1.27 | 3 | 3 |
| GABRB3 | 77 | E | 26.49783333 | 14.20557524 | 0.313283325 | 6.324220525 | NA | NA | NA | NA | NA | NA | NA |
| GABRB3 | 78 | V | 27.25383333 | 9.620741396 | 0.183506559 | 1.719903193 | 7.91 | 0.61 | 0.5 | 13.81 | -0.7 | 0 | 0 |
| GABRB3 | 79 | N | 31.38295833 | 8.181466678 | 0.142766932 | 1.314133516 | 6.95 | -0.5 | -3.95 | 38.6 | 1.27 | 3 | 3 |
| GABRB3 | 80 | M | 30.63733333 | 13.57096588 | 0.295320286 | 6.689170707 | NA | NA | NA | NA | NA | NA | NA |
| GABRB3 | 81 | D | 34.01458333 | 11.21752852 | 0.228704678 | 4.316506219 | 6.95 | -0.5 | -3.95 | 38.6 | 1.27 | 3 | 3 |
| GABRB3 | 82 | Y | 36.68566667 | 15.38610344 | 0.34669896 | 8.494182264 | NA | NA | NA | NA | NA | NA | NA |
| GABRB3 | 83 | T | 37.63697619 | 13.11719398 | 0.282475971 | 6.221545203 | NA | NA | NA | NA | NA | NA | NA |
| GABRB3 | 84 | L | 40.91395833 | 17.50236661 | 0.406601194 | 10.69901555 | NA | NA | NA | NA | NA | NA | NA |
| GABRB3 | 85 | T | 42.52840476 | 18.09100721 | 0.423263057 | 11.96792014 | NA | NA | NA | NA | NA | NA | NA |
| GABRB3 | 86 | M | 45.89108333 | 20.19316382 | 0.482765996 | 14.12765578 | NA | NA | NA | NA | NA | NA | NA |
| GABRB3 | 87 | Y | 48.257 | 22.44907448 | 0.546621045 | 16.38416976 | NA | NA | NA | NA | NA | NA | NA |
| GABRB3 | 88 | F | 51.03674242 | 21.08379726 | 0.507975967 | 13.69698332 | NA | NA | NA | NA | NA | NA | NA |
| GABRB3 | 89 | Q | 53.29483333 | 23.9382504 | 0.588773158 | 16.54863682 | NA | NA | NA | NA | NA | NA | NA |
| GABRB3 | 90 | Q | 56.40283333 | 22.52351866 | 0.548728237 | 14.00818841 | NA | NA | NA | NA | NA | NA | NA |
| GABRB3 | 91 | Y | 58.24208333 | 27.45932037 | 0.688439382 | 18.93699026 | NA | NA | NA | NA | NA | NA | NA |
| GABRB3 | 92 | W | 62.24140476 | 22.13794437 | 0.537814301 | 13.45222953 | NA | NA | NA | NA | NA | NA | NA |
| GABRB3 | 93 | R | 66.85274242 | 28.06414506 | 0.705559347 | 19.36832532 | NA | NA | NA | NA | NA | NA | NA |
| GABRB3 | 94 | D | 69.08695833 | 23.76793819 | 0.583952358 | 14.41194232 | NA | NA | NA | NA | NA | NA | NA |
| GABRB3 | 95 | K | 72.08394444 | 27.3398002 | 0.685056284 | 17.96675898 | NA | NA | NA | NA | NA | NA | NA |

| | | | | | | | | | | | | | |
|---|---|---|---|---|---|---|---|---|---|---|---|---|---|
| GABRB3 | 96 | R | 72.60856061 | 21.27629284 | 0.513424682 | 11.89180417 | NA | NA | NA | NA | NA | NA | NA |
| GABRB3 | 97 | L | 70.94883333 | 21.61509491 | 0.5230147 | 12.24281837 | NA | NA | NA | NA | NA | NA | NA |
| GABRB3 | 98 | A | 73.97383333 | 26.93514266 | 0.673602184 | 17.57076914 | NA | NA | NA | NA | NA | NA | NA |
| GABRB3 | 99 | Y | 72.939 | 27.46149697 | 0.688500992 | 18.10097105 | NA | NA | NA | NA | NA | NA | NA |
| GABRB3 | 100 | S | 77.49983333 | 31.0350464 | 0.789652682 | 21.69276463 | NA | NA | NA | NA | NA | NA | NA |
| GABRB3 | 101 | G | 77.06333333 | 33.71836924 | 0.865605917 | 24.38063134 | NA | NA | NA | NA | NA | NA | NA |
| GABRB3 | 102 | I | 74.28595833 | 28.80852225 | 0.726629437 | 19.47171076 | NA | NA | NA | NA | NA | NA | NA |
| GABRB3 | 103 | P | 71.91583333 | 33.90007351 | 0.870749177 | 24.5361378 | NA | NA | NA | NA | NA | NA | NA |
| GABRB3 | 104 | L | 69.38208333 | 30.6243679 | 0.778028154 | 21.25585753 | NA | NA | NA | NA | NA | NA | NA |
| GABRB3 | 105 | N | 65.95083333 | 28.67777463 | 0.722928539 | 19.32446088 | NA | NA | NA | NA | NA | NA | NA |
| GABRB3 | 106 | L | 68.66208333 | 24.785886 | 0.612766047 | 15.40607072 | NA | NA | NA | NA | NA | NA | NA |
| GABRB3 | 107 | T | 64.15911905 | 23.94707333 | 0.589022897 | 14.5828743 | NA | NA | NA | NA | NA | NA | NA |
| GABRB3 | 108 | L | 66.24495833 | 20.51011953 | 0.491737638 | 11.17453079 | NA | NA | NA | NA | NA | NA | NA |
| GABRB3 | 109 | D | 64.04895833 | 16.35595213 | 0.374151171 | 6.989367704 | NA | NA | NA | NA | NA | NA | NA |
| GABRB3 | 110 | N | 60.98520833 | 15.42995953 | 0.347940335 | 6.923955081 | NA | NA | NA | NA | NA | NA | NA |
| GABRB3 | 111 | R | 64.19165152 | 12.53472178 | 0.265988708 | 3.845625042 | 8.7 | -0.48 | -1 | 25.3 | 1.15 | 2 | -2 |
| GABRB3 | 112 | V | 66.35083333 | 14.97664629 | 0.335109003 | 5.611530077 | NA | NA | NA | NA | NA | NA | NA |
| GABRB3 | 113 | A | 63.34723333 | 14.40160061 | 0.318831953 | 5.811729982 | NA | NA | NA | NA | NA | NA | NA |
| GABRB3 | 114 | D | 64.01258333 | 10.15179881 | 0.198538491 | 1.7364126 | 8.7 | -0.48 | -1 | 25.3 | 1.15 | 2 | -2 |
| GABRB3 | 115 | Q | 68.8955 | 10.94597739 | 0.221018243 | 1.664580015 | 9.37 | -0.49 | -2.12 | 30.51 | 1.01 | 3 | -1 |
| GABRB3 | 116 | L | 66.69420833 | 15.3978166 | 0.347030508 | 6.198379544 | NA | NA | NA | NA | NA | NA | NA |
| GABRB3 | 117 | W | 64.35319048 | 18.10387069 | 0.423627166 | 9.407996175 | NA | NA | NA | NA | NA | NA | NA |
| GABRB3 | 118 | V | 61.45011905 | 16.63575072 | 0.382071056 | 7.954836416 | NA | NA | NA | NA | NA | NA | NA |
| GABRB3 | 119 | P | 59.0362619 | 19.14661539 | 0.453142748 | 10.49693652 | NA | NA | NA | NA | NA | NA | NA |
| GABRB3 | 120 | D | 57.28933333 | 18.66990611 | 0.439649175 | 10.20879132 | NA | NA | NA | NA | NA | NA | NA |
| GABRB3 | 121 | T | 53.85597619 | 18.86112534 | 0.445061762 | 11.85144133 | NA | NA | NA | NA | NA | NA | NA |
| GABRB3 | 122 | Y | 50.2455 | 18.90927912 | 0.446424787 | 11.51206337 | NA | NA | NA | NA | NA | NA | NA |
| GABRB3 | 123 | F | 47.1754697 | 18.32691147 | 0.429940484 | 12.24064653 | NA | NA | NA | NA | NA | NA | NA |
| GABRB3 | 124 | L | 44.81283333 | 19.51372517 | 0.463534034 | 15.88005943 | NA | NA | NA | NA | NA | NA | NA |
| GABRB3 | 125 | N | 41.19833333 | 17.01999783 | 0.392947425 | 11.91465082 | NA | NA | NA | NA | NA | NA | NA |
| GABRB3 | 126 | D | 44.18820833 | 14.54734732 | 0.322957411 | 11.32122118 | NA | NA | NA | NA | NA | NA | NA |
| GABRB3 | 127 | K | 39.69616667 | 11.59901802 | 0.239502992 | 7.718368184 | NA | NA | NA | NA | NA | NA | NA |
| GABRB3 | 128 | K | 42.61827778 | 5.371928332 | 0.063241083 | 1.72744896 | 3.68 | -0.41 | -3.9 | 49.5 | 1.46 | 3 | 3 |

| | | | | | | | | | | | | | |
|---|---|---|---|---|---|---|---|---|---|---|---|---|---|
| GABRB3 | 129 | S | 46.22766667 | 9.893410838 | 0.191224648 | 3.801480179 | 6.15 | -0.41 | -3.9 | 49.5 | 1.46 | 4 | 4 |
| GABRB3 | 130 | F | 46.34237879 | 9.361070558 | 0.176156403 | 3.286841421 | 6.15 | -0.41 | -3.9 | 49.5 | 1.46 | 4 | 4 |
| GABRB3 | 131 | V | 51.06183333 | 10.93714515 | 0.220768241 | 3.547086119 | 7.52 | -0.06 | -2.95 | 14.59 | 0.02 | 1 | 1 |
| GABRB3 | 132 | H | 48.28683333 | 12.9491819 | 0.277720277 | 5.542697509 | NA | NA | NA | NA | NA | NA | NA |
| GABRB3 | 133 | G | 51.87408333 | 11.98673091 | 0.250477463 | 4.554542578 | 7.52 | -0.06 | -2.95 | 14.59 | 0.02 | 1 | 1 |
| GABRB3 | 134 | V | 50.50897619 | 13.41023658 | 0.290770736 | 6.05679074 | NA | NA | NA | NA | NA | NA | NA |
| GABRB3 | 135 | T | 53.72040476 | 15.31320574 | 0.344635542 | 7.888674477 | NA | NA | NA | NA | NA | NA | NA |
| GABRB3 | 136 | V | 55.89997619 | 14.61260117 | 0.324804464 | 7.395965645 | NA | NA | NA | NA | NA | NA | NA |
| GABRB3 | 137 | K | 55.9075 | 11.09715464 | 0.225297416 | 4.051755169 | 7.2 | 0.06 | -2.63 | 2.95 | -0.47 | 0 | 0 |
| GABRB3 | 138 | N | 53.28645833 | 15.3160651 | 0.344716478 | 7.911411886 | NA | NA | NA | NA | NA | NA | NA |
| GABRB3 | 139 | R | 58.29065152 | 12.64877789 | 0.269217142 | 4.892005853 | 8.05 | -0.26 | -3.06 | 21.61 | 0.54 | 2 | 0 |
| GABRB3 | 140 | M | 58.45270833 | 20.04072496 | 0.478451113 | 11.96933931 | NA | NA | NA | NA | NA | NA | NA |
| GABRB3 | 141 | I | 62.92495833 | 19.48375462 | 0.462685697 | 10.88727257 | NA | NA | NA | NA | NA | NA | NA |
| GABRB3 | 142 | R | 61.50728788 | 24.20416729 | 0.596300112 | 15.50063317 | NA | NA | NA | NA | NA | NA | NA |
| GABRB3 | 143 | L | 67.12120833 | 22.92840921 | 0.560188933 | 13.59185943 | NA | NA | NA | NA | NA | NA | NA |
| GABRB3 | 144 | H | 65.55643333 | 28.04577694 | 0.705039425 | 18.68166183 | NA | NA | NA | NA | NA | NA | NA |
| GABRB3 | 145 | P | 70.35611905 | 30.73015758 | 0.781022601 | 21.35325039 | NA | NA | NA | NA | NA | NA | NA |
| GABRB3 | 146 | D | 69.01508333 | 30.88741807 | 0.785473964 | 21.50569029 | NA | NA | NA | NA | NA | NA | NA |
| GABRB3 | 147 | G | 68.34558333 | 27.47089161 | 0.688766914 | 18.10090733 | NA | NA | NA | NA | NA | NA | NA |
| GABRB3 | 148 | T | 64.99040476 | 27.72062492 | 0.695835781 | 18.52113782 | NA | NA | NA | NA | NA | NA | NA |
| GABRB3 | 149 | V | 63.51769048 | 23.35022743 | 0.572128778 | 14.66762678 | NA | NA | NA | NA | NA | NA | NA |
| GABRB3 | 150 | L | 59.58295833 | 23.41148705 | 0.573862772 | 14.88175884 | NA | NA | NA | NA | NA | NA | NA |
| GABRB3 | 151 | Y | 58.26183333 | 16.25848053 | 0.371392172 | 7.718766872 | NA | NA | NA | NA | NA | NA | NA |
| GABRB3 | 152 | G | 55.70808333 | 19.91766113 | 0.474967709 | 12.74943597 | NA | NA | NA | NA | NA | NA | NA |
| GABRB3 | 153 | L | 53.12520833 | 16.78297751 | 0.386238408 | 9.417275081 | NA | NA | NA | NA | NA | NA | NA |
| GABRB3 | 154 | R | 48.26701515 | 15.94555019 | 0.362534472 | 8.570040188 | NA | NA | NA | NA | NA | NA | NA |
| GABRB3 | 155 | I | 47.97295833 | 16.02133737 | 0.364679678 | 9.949152842 | NA | NA | NA | NA | NA | NA | NA |
| GABRB3 | 156 | T | 44.08297619 | 13.60204485 | 0.296199996 | 7.48771185 | NA | NA | NA | NA | NA | NA | NA |
| GABRB3 | 157 | T | 42.94940476 | 15.60670782 | 0.352943313 | 11.34502834 | NA | NA | NA | NA | NA | NA | NA |
| GABRB3 | 158 | T | 39.83354762 | 12.33246297 | 0.260263639 | 7.339835174 | NA | NA | NA | NA | NA | NA | NA |
| GABRB3 | 159 | A | 38.93543333 | 16.27939468 | 0.371984161 | 9.851555887 | NA | NA | NA | NA | NA | NA | NA |
| GABRB3 | 160 | A | 37.05963333 | 16.89772165 | 0.389486317 | 9.990060631 | NA | NA | NA | NA | NA | NA | NA |
| GABRB3 | 161 | C | 33.67583333 | 18.07164873 | 0.422715103 | 11.17192073 | NA | NA | NA | NA | NA | NA | NA |

| | | | | | | | | | | | | | |
|---|---|---|---|---|---|---|---|---|---|---|---|---|---|
| GABRB3 | 162 | M | 31.62070833 | 18.75011717 | 0.441919602 | 11.85318986 | NA | NA | NA | NA | NA | NA | NA |
| GABRB3 | 163 | M | 28.49570833 | 18.34708631 | 0.430511546 | 11.47708382 | NA | NA | NA | NA | NA | NA | NA |
| GABRB3 | 164 | D | 25.19745833 | 23.64600391 | 0.580500928 | 15.87588463 | NA | NA | NA | NA | NA | NA | NA |
| GABRB3 | 165 | L | 22.91420833 | 18.58288164 | 0.437185889 | 16.37064884 | NA | NA | NA | NA | NA | NA | NA |
| GABRB3 | 166 | R | 21.23819697 | 23.84792465 | 0.586216428 | 21.32780895 | NA | NA | NA | NA | NA | NA | NA |
| GABRB3 | 167 | R | 19.26028788 | 24.35928566 | 0.600690841 | 21.78173977 | NA | NA | NA | NA | NA | NA | NA |
| GABRB3 | 168 | Y | 19.75366667 | 16.65399448 | 0.382587457 | 14.20554207 | NA | NA | NA | NA | NA | NA | NA |
| GABRB3 | 169 | P | 18.31240476 | 21.23720351 | 0.512318233 | 18.65613582 | NA | NA | NA | NA | NA | NA | NA |
| GABRB3 | 170 | L | 18.04945833 | 24.51946663 | 0.60522487 | 21.94707908 | NA | NA | NA | NA | NA | NA | NA |
| GABRB3 | 171 | D | 22.87483333 | 23.62441383 | 0.579889806 | 21.32838027 | NA | NA | NA | NA | NA | NA | NA |
| GABRB3 | 172 | E | 25.91616667 | 26.72474885 | 0.667646847 | 18.9016716 | NA | NA | NA | NA | NA | NA | NA |
| GABRB3 | 173 | Q | 28.46761111 | 23.18661918 | 0.567497738 | 16.31449132 | NA | NA | NA | NA | NA | NA | NA |
| GABRB3 | 174 | N | 33.21295833 | 24.26805163 | 0.598108401 | 17.37617956 | NA | NA | NA | NA | NA | NA | NA |
| GABRB3 | 175 | C | 35.1795 | 20.97260246 | 0.504828524 | 14.08644079 | NA | NA | NA | NA | NA | NA | NA |
| GABRB3 | 176 | T | 38.75254762 | 22.47551345 | 0.547369418 | 15.56480585 | NA | NA | NA | NA | NA | NA | NA |
| GABRB3 | 177 | L | 42.99908333 | 21.47935348 | 0.519172448 | 16.58573141 | NA | NA | NA | NA | NA | NA | NA |
| GABRB3 | 178 | E | 43.80927778 | 23.35889289 | 0.572374059 | 19.26605984 | NA | NA | NA | NA | NA | NA | NA |
| GABRB3 | 179 | I | 48.28145833 | 24.14828729 | 0.594718392 | 18.21242069 | NA | NA | NA | NA | NA | NA | NA |
| GABRB3 | 180 | E | 50.31983333 | 22.82294911 | 0.557203815 | 15.43198787 | NA | NA | NA | NA | NA | NA | NA |
| GABRB3 | 181 | S | 54.46483333 | 24.0038303 | 0.590629441 | 16.87371403 | NA | NA | NA | NA | NA | NA | NA |
| GABRB3 | 182 | Y | 54.37341667 | 20.308652 | 0.486034965 | 11.86779176 | NA | NA | NA | NA | NA | NA | NA |
| GABRB3 | 183 | G | 58.60608333 | 22.08453171 | 0.53630242 | 13.44807212 | NA | NA | NA | NA | NA | NA | NA |
| GABRB3 | 184 | Y | 60.799 | 20.98844839 | 0.505277054 | 12.2933885 | NA | NA | NA | NA | NA | NA | NA |
| GABRB3 | 185 | T | 58.58683333 | 27.41830779 | 0.687278494 | 18.76358788 | NA | NA | NA | NA | NA | NA | NA |
| GABRB3 | 186 | T | 57.74654762 | 30.74141477 | 0.781341244 | 22.09038006 | NA | NA | NA | NA | NA | NA | NA |
| GABRB3 | 187 | D | 61.50208333 | 31.25035406 | 0.795747108 | 22.57260039 | NA | NA | NA | NA | NA | NA | NA |
| GABRB3 | 188 | D | 62.66395833 | 27.27443099 | 0.683205965 | 18.59326579 | NA | NA | NA | NA | NA | NA | NA |
| GABRB3 | 189 | I | 59.16420833 | 25.32485414 | 0.628021898 | 16.66194333 | NA | NA | NA | NA | NA | NA | NA |
| GABRB3 | 190 | E | 58.34261111 | 30.07162014 | 0.762382262 | 21.40827304 | NA | NA | NA | NA | NA | NA | NA |
| GABRB3 | 191 | F | 54.32656061 | 26.15702625 | 0.651577082 | 17.66946144 | NA | NA | NA | NA | NA | NA | NA |
| GABRB3 | 192 | Y | 55.23116667 | 31.6072037 | 0.805847974 | 24.22252574 | NA | NA | NA | NA | NA | NA | NA |
| GABRB3 | 193 | W | 48.88090476 | 28.39509607 | 0.714927135 | 21.11405459 | NA | NA | NA | NA | NA | NA | NA |
| GABRB3 | 194 | R | 54.45874242 | 32.11144254 | 0.820120789 | 24.73218754 | NA | NA | NA | NA | NA | NA | NA |

| | | | | | | | | | | | | | |
|---|---|---|---|---|---|---|---|---|---|---|---|---|---|
| GABRB3 | 195 | G | 50.40558333 | 36.02599486 | 0.930924792 | 28.63881958 | NA | NA | NA | NA | NA | NA | NA |
| GABRB3 | 196 | G | 48.23883333 | 35.63447571 | 0.919842583 | 28.28365134 | NA | NA | NA | NA | NA | NA | NA |
| GABRB3 | 197 | D | 44.37408333 | 35.77281188 | 0.92375828 | 29.68785902 | NA | NA | NA | NA | NA | NA | NA |
| GABRB3 | 198 | K | 46.9655 | 34.38515361 | 0.884479691 | 28.31974005 | NA | NA | NA | NA | NA | NA | NA |
| GABRB3 | 199 | A | 47.53743333 | 31.08705832 | 0.791124914 | 25.01995962 | NA | NA | NA | NA | NA | NA | NA |
| GABRB3 | 200 | V | 44.37511905 | 29.20467505 | 0.737842805 | 23.10975846 | NA | NA | NA | NA | NA | NA | NA |
| GABRB3 | 201 | T | 44.3372619 | 29.11204186 | 0.735220761 | 23.02085532 | NA | NA | NA | NA | NA | NA | NA |
| GABRB3 | 202 | G | 40.80258333 | 28.31094207 | 0.7125451 | 23.08132717 | NA | NA | NA | NA | NA | NA | NA |
| GABRB3 | 203 | V | 38.11069048 | 28.95265572 | 0.730709231 | 22.07978358 | NA | NA | NA | NA | NA | NA | NA |
| GABRB3 | 204 | E | 37.02205556 | 30.92260225 | 0.786469876 | 24.04645053 | NA | NA | NA | NA | NA | NA | NA |
| GABRB3 | 205 | R | 39.2964697 | 28.08723174 | 0.70621283 | 21.22429493 | NA | NA | NA | NA | NA | NA | NA |
| GABRB3 | 206 | I | 35.28658333 | 24.49963222 | 0.604663444 | 17.63375479 | NA | NA | NA | NA | NA | NA | NA |
| GABRB3 | 207 | E | 30.4245 | 24.12640027 | 0.594098865 | 17.28147877 | NA | NA | NA | NA | NA | NA | NA |
| GABRB3 | 208 | L | 30.33433333 | 21.06535616 | 0.507453979 | 14.2108597 | NA | NA | NA | NA | NA | NA | NA |
| GABRB3 | 209 | P | 27.04540476 | 20.49583965 | 0.491333436 | 12.65278705 | NA | NA | NA | NA | NA | NA | NA |
| GABRB3 | 210 | Q | 24.5995 | 19.11498164 | 0.452247333 | 11.42206444 | NA | NA | NA | NA | NA | NA | NA |
| GABRB3 | 211 | F | 27.37665152 | 20.21355468 | 0.483343173 | 12.35861282 | NA | NA | NA | NA | NA | NA | NA |
| GABRB3 | 212 | S | 27.97516667 | 26.13851478 | 0.651053103 | 19.29042598 | NA | NA | NA | NA | NA | NA | NA |
| GABRB3 | 213 | I | 32.19270833 | 26.07758171 | 0.649328352 | 19.20689505 | NA | NA | NA | NA | NA | NA | NA |
| GABRB3 | 214 | V | 30.51397619 | 30.06705674 | 0.762253092 | 23.20093398 | NA | NA | NA | NA | NA | NA | NA |
| GABRB3 | 215 | E | 33.7075 | 30.19650625 | 0.765917246 | 23.3148148 | NA | NA | NA | NA | NA | NA | NA |
| GABRB3 | 216 | H | 38.60093333 | 30.47618936 | 0.773833862 | 23.58406342 | NA | NA | NA | NA | NA | NA | NA |
| GABRB3 | 217 | R | 37.77492424 | 31.38974165 | 0.799692566 | 24.48228338 | NA | NA | NA | NA | NA | NA | NA |
| GABRB3 | 218 | L | 43.83870833 | 31.58481247 | 0.805214176 | 25.49197024 | NA | NA | NA | NA | NA | NA | NA |
| GABRB3 | 219 | V | 44.12369048 | 32.25342348 | 0.824139654 | 26.15983672 | NA | NA | NA | NA | NA | NA | NA |
| GABRB3 | 220 | S | 48.2585 | 34.06165342 | 0.875322803 | 28.03485128 | NA | NA | NA | NA | NA | NA | NA |
| GABRB3 | 221 | R | 46.33019697 | 30.67872416 | 0.779566744 | 24.45040158 | NA | NA | NA | NA | NA | NA | NA |
| GABRB3 | 222 | N | 50.89983333 | 33.75399772 | 0.866614404 | 26.35468373 | NA | NA | NA | NA | NA | NA | NA |
| GABRB3 | 223 | V | 49.01283333 | 30.17556265 | 0.765324423 | 22.7733337 | NA | NA | NA | NA | NA | NA | NA |
| GABRB3 | 224 | V | 49.8822619 | 32.7287679 | 0.837594593 | 25.31432313 | NA | NA | NA | NA | NA | NA | NA |
| GABRB3 | 225 | F | 49.70983333 | 27.73860007 | 0.69634458 | 20.32790248 | NA | NA | NA | NA | NA | NA | NA |
| GABRB3 | 226 | A | 52.53183333 | 31.57542961 | 0.804948587 | 24.52586603 | NA | NA | NA | NA | NA | NA | NA |
| GABRB3 | 227 | T | 55.44283333 | 28.10433853 | 0.706697049 | 19.4806545 | NA | NA | NA | NA | NA | NA | NA |

| | | | | | | | | | | | | | |
|---|---|---|---|---|---|---|---|---|---|---|---|---|---|
| GABRB3 | 228 | G | 54.71408333 | 30.93579546 | 0.786843318 | 22.30590378 | NA | NA | NA | NA | NA | NA | NA |
| GABRB3 | 229 | A | 54.64843333 | 31.38396078 | 0.799528935 | 22.74519423 | NA | NA | NA | NA | NA | NA | NA |
| GABRB3 | 230 | Y | 53.91816667 | 26.63738962 | 0.665174087 | 18.21409167 | NA | NA | NA | NA | NA | NA | NA |
| GABRB3 | 231 | P | 53.25583333 | 28.2811032 | 0.711700491 | 20.88792718 | NA | NA | NA | NA | NA | NA | NA |
| GABRB3 | 232 | R | 47.54501515 | 27.97231876 | 0.702960142 | 20.69245302 | NA | NA | NA | NA | NA | NA | NA |
| GABRB3 | 233 | L | 48.51808333 | 27.62417796 | 0.693105786 | 21.52867772 | NA | NA | NA | NA | NA | NA | NA |
| GABRB3 | 234 | S | 43.8225 | 28.7199382 | 0.724122007 | 22.61594716 | NA | NA | NA | NA | NA | NA | NA |
| GABRB3 | 235 | L | 41.94183333 | 26.25428363 | 0.654330017 | 19.34460141 | NA | NA | NA | NA | NA | NA | NA |
| GABRB3 | 236 | S | 38.427 | 27.77743496 | 0.697443827 | 20.87474296 | NA | NA | NA | NA | NA | NA | NA |
| GABRB3 | 237 | F | 36.56774242 | 24.84713217 | 0.61449966 | 17.94948322 | NA | NA | NA | NA | NA | NA | NA |
| GABRB3 | 238 | R | 30.93156061 | 27.62118053 | 0.693020941 | 20.75066675 | NA | NA | NA | NA | NA | NA | NA |
| GABRB3 | 239 | L | 30.19270833 | 22.16459257 | 0.538568596 | 15.29410461 | NA | NA | NA | NA | NA | NA | NA |
| GABRB3 | 240 | K | 24.93638889 | 26.76957712 | 0.668915741 | 18.90424577 | NA | NA | NA | NA | NA | NA | NA |
| GABRB3 | 241 | R | 23.32892424 | 19.95806666 | 0.476111415 | 12.13175302 | NA | NA | NA | NA | NA | NA | NA |
| GABRB3 | 242 | N | 22.32958333 | 24.01441963 | 0.590929179 | 17.09142061 | NA | NA | NA | NA | NA | NA | NA |
| GABRB3 | 243 | I | 17.34158333 | 24.48247782 | 0.604177877 | 21.87880721 | NA | NA | NA | NA | NA | NA | NA |
| GABRB3 | 244 | G | 16.87158333 | 23.87139259 | 0.586880704 | 21.24428995 | NA | NA | NA | NA | NA | NA | NA |
| GABRB3 | 245 | Y | 18.07875 | 15.39992834 | 0.347090283 | 12.82506578 | NA | NA | NA | NA | NA | NA | NA |
| GABRB3 | 246 | F | 17.00601515 | 17.85319109 | 0.416531514 | 15.62894394 | NA | NA | NA | NA | NA | NA | NA |
| GABRB3 | 247 | I | 12.31408333 | 20.18837027 | 0.482630311 | 16.50321547 | NA | NA | NA | NA | NA | NA | NA |
| GABRB3 | 248 | L | 12.24508333 | 18.0069508 | 0.420883784 | 14.32567503 | NA | NA | NA | NA | NA | NA | NA |
| GABRB3 | 249 | Q | 13.67761111 | 14.00694924 | 0.307661084 | 10.67325363 | NA | NA | NA | NA | NA | NA | NA |
| GABRB3 | 250 | T | 11.59154762 | 15.95674296 | 0.36285129 | 12.82060252 | NA | NA | NA | NA | NA | NA | NA |
| GABRB3 | 251 | Y | 10.81575 | 18.9124537 | 0.446514645 | 15.22713274 | NA | NA | NA | NA | NA | NA | NA |
| GABRB3 | 252 | M | 6.645458333 | 18.55050649 | 0.436269489 | 15.15522656 | NA | NA | NA | NA | NA | NA | NA |
| GABRB3 | 253 | P | 6.558547619 | 15.08020961 | 0.338040432 | 11.70059419 | NA | NA | NA | NA | NA | NA | NA |
| GABRB3 | 254 | S | 5.042666667 | 16.14385738 | 0.368147688 | 12.7405653 | NA | NA | NA | NA | NA | NA | NA |
| GABRB3 | 255 | I | 3.393208333 | 18.2352442 | 0.427345781 | 15.03801398 | NA | NA | NA | NA | NA | NA | NA |
| GABRB3 | 256 | L | 1.377458333 | 16.27872262 | 0.371965138 | 13.94265779 | NA | NA | NA | NA | NA | NA | NA |
| GABRB3 | 257 | I | 1.061458333 | 12.36686173 | 0.261237318 | 10.09898135 | NA | NA | NA | NA | NA | NA | NA |
| GABRB3 | 258 | T | 0.198404762 | 16.14977158 | 0.368315093 | 13.88262845 | NA | NA | NA | NA | NA | NA | NA |
| GABRB3 | 259 | I | -0.983958333 | 17.43496226 | 0.404693269 | 14.26224689 | NA | NA | NA | NA | NA | NA | NA |
| GABRB3 | 260 | L | -2.292708333 | 13.53535149 | 0.294312197 | 10.13816908 | NA | NA | NA | NA | NA | NA | NA |

| | | | | | | | | | | | | | |
|---|---|---|---|---|---|---|---|---|---|---|---|---|---|
| GABRB3 | 261 | S | -3.726833333 | 14.67895095 | 0.326682539 | 11.27698103 | NA | NA | NA | NA | NA | NA | NA |
| GABRB3 | 262 | W | -4.177404762 | 16.62263198 | 0.381699721 | 13.22293607 | NA | NA | NA | NA | NA | NA | NA |
| GABRB3 | 263 | V | -8.182404762 | 14.84241722 | 0.33130956 | 12.51575556 | NA | NA | NA | NA | NA | NA | NA |
| GABRB3 | 264 | S | -10.06716667 | 13.82455862 | 0.302498397 | 11.62688984 | NA | NA | NA | NA | NA | NA | NA |
| GABRB3 | 265 | F | -10.27828788 | 16.61434937 | 0.381465276 | 14.41916908 | NA | NA | NA | NA | NA | NA | NA |
| GABRB3 | 266 | W | -12.79133333 | 16.35229858 | 0.374047755 | 14.51698544 | NA | NA | NA | NA | NA | NA | NA |
| GABRB3 | 267 | I | -13.88920833 | 11.40079497 | 0.233892157 | 9.576237787 | NA | NA | NA | NA | NA | NA | NA |
| GABRB3 | 268 | N | -18.77458333 | 11.70192471 | 0.242415834 | 9.80029264 | NA | NA | NA | NA | NA | NA | NA |
| GABRB3 | 269 | Y | -20.01491667 | 11.42149491 | 0.234478083 | 9.529411932 | NA | NA | NA | NA | NA | NA | NA |
| GABRB3 | 270 | D | -20.54258333 | 7.933249902 | 0.135740991 | 5.566566704 | NA | NA | NA | NA | NA | NA | NA |
| GABRB3 | 271 | A | -16.19483333 | 7.69734032 | 0.129063414 | 5.826455741 | NA | NA | NA | NA | NA | NA | NA |
| GABRB3 | 272 | S | -14.8265 | 6.248205262 | 0.088044684 | 4.345339997 | 1.91 | -0.06 | -0.75 | 1.19 | -0.83 | 0 | 0 |
| GABRB3 | 273 | A | -12.51783333 | 3.197674311 | 0.001697379 | 1.532414026 | 1.72 | -0.04 | 0.1 | 0.79 | -0.36 | 0 | 0 |
| GABRB3 | 274 | A | -11.41503333 | 6.12760157 | 0.084630916 | 3.889453619 | 2.31 | -0.33 | 0.24 | 1.67 | 0.3 | 0 | 0 |
| GABRB3 | 275 | R | -12.01783333 | 8.571154531 | 0.153797305 | 6.348705991 | NA | NA | NA | NA | NA | NA | NA |
| GABRB3 | 276 | V | -8.694833333 | 7.8965766 | 0.134702929 | 5.650727554 | NA | NA | NA | NA | NA | NA | NA |
| GABRB3 | 277 | A | -6.634633333 | 5.036356223 | 0.053742492 | 2.094821218 | 3.05 | 0.43 | 2.53 | 0.64 | -0.5 | 0 | 0 |
| GABRB3 | 278 | L | -6.103833333 | 8.07206473 | 0.139670237 | 4.758381688 | 3.41 | -0.01 | 1.26 | 1.05 | -0.06 | 0 | 0 |
| GABRB3 | 279 | G | -3.780583333 | 9.703520289 | 0.18584967 | 6.3413611 | NA | NA | NA | NA | NA | NA | NA |
| GABRB3 | 280 | I | -3.071208333 | 6.099265857 | 0.083828855 | 2.787217904 | 3.35 | -0.39 | 0.28 | 1.35 | 0.36 | 0 | 0 |
| GABRB3 | 281 | T | -1.011833333 | 4.911152309 | 0.050198512 | 1.705927656 | 3.28 | -0.01 | 1.1 | 1.05 | -0.25 | 0 | 0 |
| GABRB3 | 282 | T | -0.31797619 | 8.699841148 | 0.157439865 | 6.739929381 | NA | NA | NA | NA | NA | NA | NA |
| GABRB3 | 283 | V | 0.029261905 | 9.145509609 | 0.170054807 | 6.929785298 | NA | NA | NA | NA | NA | NA | NA |
| GABRB3 | 284 | L | 1.165708333 | 4.006943723 | 0.024604288 | 1.87774998 | 2.53 | 1.14 | 3.8 | 0.13 | -1.79 | 0 | 0 |
| GABRB3 | 285 | T | 2.638404762 | 7.604390705 | 0.126432413 | 4.676912607 | 2.96 | 1.14 | 3.8 | 0.13 | -1.79 | 0 | 0 |
| GABRB3 | 286 | M | 4.108458333 | 9.987447572 | 0.19388642 | 6.784669668 | NA | NA | NA | NA | NA | NA | NA |
| GABRB3 | 287 | T | 5.912261905 | 8.513354568 | 0.152161239 | 4.860383981 | 3.68 | -0.77 | -0.7 | 1.66 | 0.77 | 0 | 0 |
| GABRB3 | 288 | T | 7.295547619 | 5.123906615 | 0.056220664 | 1.737911507 | 3.7 | -0.43 | -1.53 | 18.31 | 0.45 | 0 | 0 |
| GABRB3 | 289 | I | 8.635833333 | 9.358768402 | 0.176091239 | 5.665209095 | NA | NA | NA | NA | NA | NA | NA |
| GABRB3 | 290 | N | 10.62308333 | 9.42851351 | 0.178065421 | 6.121240074 | NA | NA | NA | NA | NA | NA | NA |
| GABRB3 | 291 | T | 11.48169048 | 5.84725628 | 0.076695557 | 2.535318778 | 3.42 | -0.16 | -0.74 | 20.99 | 0.41 | 1 | -1 |
| GABRB3 | 292 | H | 12.66393333 | 4.949388144 | 0.051280803 | 1.879198267 | 3.42 | -0.16 | -0.74 | 20.99 | 0.41 | 1 | -1 |
| GABRB3 | 293 | L | 15.31995833 | 9.604624147 | 0.183050349 | 7.063520162 | NA | NA | NA | NA | NA | NA | NA |

| | | | | | | | | | | | | | |
|---|---|---|---|---|---|---|---|---|---|---|---|---|---|
| GABRB3 | 294 | R | 15.55237879 | 7.00688754 | 0.109519689 | 4.85418118 | 2.57 | -0.82 | -2.93 | 26.11 | 1.34 | 2 | 0 |
| GABRB3 | 295 | E | 18.1205 | 3.946902203 | 0.022904772 | 2.237965723 | 2.78 | -0.77 | -3.63 | 34.26 | 1.41 | 2 | 0 |
| GABRB3 | 296 | T | 20.94069048 | 8.611614715 | 0.154942558 | 6.148408659 | NA | NA | NA | NA | NA | NA | NA |
| GABRB3 | 297 | L | 20.79745833 | 10.52500437 | 0.209102322 | 8.244615526 | NA | NA | NA | NA | NA | NA | NA |
| GABRB3 | 298 | P | 23.36869048 | 12.88537582 | 0.275914204 | 7.760009049 | NA | NA | NA | NA | NA | NA | NA |
| GABRB3 | 299 | K | 22.56327778 | 10.57518596 | 0.210522746 | 8.155500855 | NA | NA | NA | NA | NA | NA | NA |
| GABRB3 | 300 | I | 21.78133333 | 16.31984822 | 0.373129226 | 13.83232006 | NA | NA | NA | NA | NA | NA | NA |
| GABRB3 | 301 | P | 20.22597619 | 19.41447988 | 0.46072483 | 16.81415285 | NA | NA | NA | NA | NA | NA | NA |
| GABRB3 | 302 | Y | 20.43766667 | 20.25683847 | 0.484568349 | 17.77094211 | NA | NA | NA | NA | NA | NA | NA |
| GABRB3 | 303 | V | 15.07311905 | 21.59856361 | 0.52254677 | 19.03430955 | NA | NA | NA | NA | NA | NA | NA |
| GABRB3 | 304 | K | 16.91361111 | 16.60535727 | 0.381210749 | 14.17672159 | NA | NA | NA | NA | NA | NA | NA |
| GABRB3 | 305 | A | 14.61323333 | 22.13308047 | 0.537676625 | 19.64873573 | NA | NA | NA | NA | NA | NA | NA |
| GABRB3 | 306 | I | 13.79570833 | 17.9947155 | 0.420537456 | 14.65727197 | NA | NA | NA | NA | NA | NA | NA |
| GABRB3 | 307 | D | 13.02458333 | 16.12553894 | 0.367629172 | 12.85189297 | NA | NA | NA | NA | NA | NA | NA |
| GABRB3 | 308 | M | 10.32233333 | 20.77204056 | 0.499151486 | 17.47099313 | NA | NA | NA | NA | NA | NA | NA |
| GABRB3 | 309 | Y | 7.943416667 | 20.15040677 | 0.481555729 | 16.92038686 | NA | NA | NA | NA | NA | NA | NA |
| GABRB3 | 310 | L | 8.499583333 | 15.17830603 | 0.340817116 | 11.9830439 | NA | NA | NA | NA | NA | NA | NA |
| GABRB3 | 311 | M | 7.464958333 | 18.30261069 | 0.429252634 | 15.01991514 | NA | NA | NA | NA | NA | NA | NA |
| GABRB3 | 312 | G | 4.346333333 | 20.72877985 | 0.497926963 | 17.52363432 | NA | NA | NA | NA | NA | NA | NA |
| GABRB3 | 313 | C | 3.3115 | 17.42844359 | 0.404508754 | 14.49567271 | NA | NA | NA | NA | NA | NA | NA |
| GABRB3 | 314 | F | 2.275015152 | 13.64550791 | 0.297430247 | 10.71123833 | NA | NA | NA | NA | NA | NA | NA |
| GABRB3 | 315 | V | 0.428261905 | 18.9085523 | 0.446404214 | 16.38747915 | NA | NA | NA | NA | NA | NA | NA |
| GABRB3 | 316 | F | -0.014560606 | 18.53033702 | 0.435698579 | 16.4849106 | NA | NA | NA | NA | NA | NA | NA |
| GABRB3 | 317 | V | -0.251119048 | 13.87671589 | 0.303974743 | 10.72299235 | NA | NA | NA | NA | NA | NA | NA |
| GABRB3 | 318 | F | -1.200469697 | 16.32242412 | 0.373202138 | 12.96632574 | NA | NA | NA | NA | NA | NA | NA |
| GABRB3 | 319 | L | -3.381583333 | 18.71009107 | 0.440786637 | 15.38148553 | NA | NA | NA | NA | NA | NA | NA |
| GABRB3 | 320 | A | -5.517433333 | 16.1252142 | 0.367619981 | 12.76936084 | NA | NA | NA | NA | NA | NA | NA |
| GABRB3 | 321 | L | -5.974958333 | 12.04423717 | 0.252105216 | 8.700039478 | NA | NA | NA | NA | NA | NA | NA |
| GABRB3 | 322 | L | -6.492083333 | 16.99624915 | 0.392275203 | 13.63339387 | NA | NA | NA | NA | NA | NA | NA |
| GABRB3 | 323 | E | -10.40383333 | 17.91858237 | 0.418382457 | 15.65159532 | NA | NA | NA | NA | NA | NA | NA |
| GABRB3 | 324 | Y | -12.69266667 | 10.64492344 | 0.212496711 | 9.473704367 | NA | NA | NA | NA | NA | NA | NA |
| GABRB3 | 325 | A | -11.67763333 | 16.01390858 | 0.364469401 | 13.72360706 | NA | NA | NA | NA | NA | NA | NA |
| GABRB3 | 326 | F | -13.25237879 | 18.62809193 | 0.438465597 | 16.93476222 | NA | NA | NA | NA | NA | NA | NA |

| | | | | | | | | | | | | | |
|---|---|---|---|---|---|---|---|---|---|---|---|---|---|
| GABRB3 | 327 | V | -15.67097619 | 16.76795423 | 0.385813164 | 14.88379208 | NA | NA | NA | NA | NA | NA | NA |
| GABRB3 | 328 | N | -16.21133333 | 12.14454602 | 0.254944524 | 10.28017125 | NA | NA | NA | NA | NA | NA | NA |
| GABRB3 | 329 | Y | -16.72708333 | 17.50290822 | 0.406616525 | 15.61772091 | NA | NA | NA | NA | NA | NA | NA |
| GABRB3 | 330 | I | -18.99120833 | 18.92760238 | 0.446943439 | 14.69552187 | NA | NA | NA | NA | NA | NA | NA |
| GABRB3 | 331 | F | -22.63965152 | 12.15792293 | 0.255323167 | 8.011631849 | NA | NA | NA | NA | NA | NA | NA |
| GABRB3 | 332 | F | -21.36992424 | 11.13084076 | 0.226250924 | 6.808351515 | NA | NA | NA | NA | NA | NA | NA |
| GABRB3 | 333 | G | -21.20083333 | 17.98878565 | 0.420369608 | 13.69681785 | NA | NA | NA | NA | NA | NA | NA |
| GABRB3 | 334 | R | -21.01083333 | 20.09544819 | 0.48000009 | 15.95037862 | NA | NA | NA | NA | NA | NA | NA |
| GABRB3 | 335 | G | -25.24158333 | 17.89944633 | 0.417840799 | 13.63951063 | NA | NA | NA | NA | NA | NA | NA |
| GABRB3 | 336 | P | -26.91654762 | 16.38439886 | 0.374956374 | 12.09445025 | NA | NA | NA | NA | NA | NA | NA |
| GABRB3 | 337 | Q | -26.25538889 | 20.30408966 | 0.485905825 | 16.02534534 | NA | NA | NA | NA | NA | NA | NA |
| GABRB3 | 440 | P | -13.91483333 | 30.89996561 | 0.78582913 | 29.04773811 | NA | NA | NA | NA | NA | NA | NA |
| GABRB3 | 441 | D | -16.11908333 | 26.19458832 | 0.652640301 | 22.70810491 | NA | NA | NA | NA | NA | NA | NA |
| GABRB3 | 442 | L | -20.73708333 | 25.70969037 | 0.638914943 | 21.50027376 | NA | NA | NA | NA | NA | NA | NA |
| GABRB3 | 443 | T | -20.58769048 | 21.97561874 | 0.533219566 | 17.756391 | NA | NA | NA | NA | NA | NA | NA |
| GABRB3 | 444 | D | -22.14420833 | 21.24185124 | 0.51244979 | 17.03062475 | NA | NA | NA | NA | NA | NA | NA |
| GABRB3 | 445 | V | -20.14469048 | 17.11810124 | 0.395724307 | 12.89666471 | NA | NA | NA | NA | NA | NA | NA |
| GABRB3 | 446 | N | -20.26183333 | 19.44571706 | 0.461609019 | 15.30376213 | NA | NA | NA | NA | NA | NA | NA |
| GABRB3 | 447 | A | -18.33943333 | 22.14517324 | 0.538018919 | 18.00251973 | NA | NA | NA | NA | NA | NA | NA |
| GABRB3 | 448 | I | -15.46708333 | 22.32131022 | 0.543004593 | 20.48185862 | NA | NA | NA | NA | NA | NA | NA |
| GABRB3 | 449 | D | -14.87083333 | 17.709528 | 0.412465035 | 15.83468957 | NA | NA | NA | NA | NA | NA | NA |
| GABRB3 | 450 | R | -17.92319697 | 21.9620665 | 0.532835961 | 21.03569225 | NA | NA | NA | NA | NA | NA | NA |
| GABRB3 | 451 | W | -11.73654762 | 25.16848923 | 0.623595885 | 23.32683786 | NA | NA | NA | NA | NA | NA | NA |
| GABRB3 | 452 | S | -10.29 | 22.29310214 | 0.542206144 | 20.06948666 | NA | NA | NA | NA | NA | NA | NA |
| GABRB3 | 453 | R | -12.59428788 | 19.67607199 | 0.468129368 | 18.24645061 | NA | NA | NA | NA | NA | NA | NA |
| GABRB3 | 454 | I | -9.777958333 | 26.01663675 | 0.647603264 | 23.81954563 | NA | NA | NA | NA | NA | NA | NA |
| GABRB3 | 455 | V | -6.938404762 | 26.62745209 | 0.664892798 | 23.22448782 | NA | NA | NA | NA | NA | NA | NA |
| GABRB3 | 456 | F | -5.00919697 | 21.38694078 | 0.516556645 | 17.98619932 | NA | NA | NA | NA | NA | NA | NA |
| GABRB3 | 457 | P | -4.301547619 | 24.51175306 | 0.605006532 | 21.11222065 | NA | NA | NA | NA | NA | NA | NA |
| GABRB3 | 458 | F | -3.235742424 | 27.59172853 | 0.692187283 | 24.18924346 | NA | NA | NA | NA | NA | NA | NA |
| GABRB3 | 459 | T | -1.261119048 | 27.34535502 | 0.685213517 | 24.16777133 | NA | NA | NA | NA | NA | NA | NA |
| GABRB3 | 460 | F | -0.377015152 | 21.81177194 | 0.528581773 | 19.60186128 | NA | NA | NA | NA | NA | NA | NA |
| GABRB3 | 461 | S | 0 | 27.17441904 | 0.680375061 | 24.71831297 | NA | NA | NA | NA | NA | NA | NA |

| | | | | | | | | | | | | | |
|---|---|---|---|---|---|---|---|---|---|---|---|---|---|
| GABRB3 | 462 | L | 0.354083333 | 29.48282057 | 0.745715899 | 26.98437814 | NA | NA | NA | NA | NA | NA | NA |
| GABRB3 | 463 | F | 3.818833333 | 24.00066143 | 0.590539744 | 21.08761331 | NA | NA | NA | NA | NA | NA | NA |
| GABRB3 | 464 | N | 4.614083333 | 24.86184649 | 0.614916159 | 21.66346278 | NA | NA | NA | NA | NA | NA | NA |
| GABRB3 | 465 | L | 4.681583333 | 30.0334942 | 0.761303082 | 26.82850921 | NA | NA | NA | NA | NA | NA | NA |
| GABRB3 | 466 | V | 6.47797619 | 31.08831411 | 0.79116046 | 27.68416326 | NA | NA | NA | NA | NA | NA | NA |
| GABRB3 | 467 | Y | 9.971833333 | 23.61274118 | 0.579559404 | 21.46739147 | NA | NA | NA | NA | NA | NA | NA |
| GABRB3 | 468 | W | 10.19104762 | 25.47183492 | 0.632182287 | 21.78348335 | NA | NA | NA | NA | NA | NA | NA |
| GABRB3 | 469 | L | 10.28745833 | 32.29903901 | 0.825430832 | 28.97552768 | NA | NA | NA | NA | NA | NA | NA |
| GABRB3 | 470 | Y | 13.33108333 | 30.78869301 | 0.782679485 | 27.47895272 | NA | NA | NA | NA | NA | NA | NA |
| GABRB3 | 471 | Y | 14.54091667 | 26.66719157 | 0.66601765 | 24.17903719 | NA | NA | NA | NA | NA | NA | NA |
| GABRB3 | 472 | V | 15.34411905 | 30.38228989 | 0.771175975 | 28.00832325 | NA | NA | NA | NA | NA | NA | NA |
| GABRG2 | 65 | D | 78.76980682 | 29.54555224 | 0.782816624 | 20.83956703 | NA | NA | NA | NA | NA | NA | NA |
| GABRG2 | 66 | V | 75.28875325 | 26.11914526 | 0.681058279 | 17.11551511 | NA | NA | NA | NA | NA | NA | NA |
| GABRG2 | 67 | T | 77.49246753 | 24.16313566 | 0.622968187 | 15.48621616 | NA | NA | NA | NA | NA | NA | NA |
| GABRG2 | 68 | V | 80.47775325 | 27.04747406 | 0.708628035 | 18.3683988 | NA | NA | NA | NA | NA | NA | NA |
| GABRG2 | 69 | I | 76.97793182 | 26.21514497 | 0.683909304 | 17.5221911 | NA | NA | NA | NA | NA | NA | NA |
| GABRG2 | 70 | L | 74.43155682 | 22.64694198 | 0.577939864 | 13.96299336 | NA | NA | NA | NA | NA | NA | NA |
| GABRG2 | 71 | N | 78.39768182 | 21.13189433 | 0.532945576 | 12.49722863 | NA | NA | NA | NA | NA | NA | NA |
| GABRG2 | 72 | N | 80.12368182 | 25.28737916 | 0.656356269 | 16.63936837 | NA | NA | NA | NA | NA | NA | NA |
| GABRG2 | 73 | L | 75.42005682 | 23.74784588 | 0.6106348 | 15.07909303 | NA | NA | NA | NA | NA | NA | NA |
| GABRG2 | 74 | L | 75.14543182 | 18.94906098 | 0.468119211 | 10.26468377 | NA | NA | NA | NA | NA | NA | NA |
| GABRG2 | 75 | E | 80.9719596 | 22.93236713 | 0.586416496 | 14.31351971 | NA | NA | NA | NA | NA | NA | NA |
| GABRG2 | 76 | G | 78.54668182 | 23.32160102 | 0.597976067 | 14.73852787 | NA | NA | NA | NA | NA | NA | NA |
| GABRG2 | 77 | Y | 73.18759848 | 20.2415198 | 0.506502997 | 11.62891953 | NA | NA | NA | NA | NA | NA | NA |
| GABRG2 | 78 | D | 73.21918182 | 19.8274393 | 0.494205524 | 11.2163392 | NA | NA | NA | NA | NA | NA | NA |
| GABRG2 | 79 | N | 69.52693182 | 15.27298366 | 0.35894609 | 6.586336088 | NA | NA | NA | NA | NA | NA | NA |
| GABRG2 | 80 | K | 69.84929293 | 17.79164593 | 0.433745992 | 9.173802432 | NA | NA | NA | NA | NA | NA | NA |
| GABRG2 | 81 | L | 70.72930682 | 21.83083713 | 0.553702965 | 13.15214614 | NA | NA | NA | NA | NA | NA | NA |
| GABRG2 | 82 | R | 66.45363636 | 24.52130362 | 0.633605154 | 15.59690524 | NA | NA | NA | NA | NA | NA | NA |
| GABRG2 | 83 | P | 68.00718182 | 25.78244538 | 0.671058877 | 16.8675264 | NA | NA | NA | NA | NA | NA | NA |
| GABRG2 | 84 | D | 69.40130682 | 29.96468925 | 0.795264267 | 21.12483144 | NA | NA | NA | NA | NA | NA | NA |
| GABRG2 | 85 | I | 69.70230682 | 26.74088447 | 0.699522856 | 18.06191277 | NA | NA | NA | NA | NA | NA | NA |
| GABRG2 | 86 | G | 70.89168182 | 32.06233794 | 0.857560795 | 23.37604545 | NA | NA | NA | NA | NA | NA | NA |

| | | | | | | | | | | | | | |
|---|---|---|---|---|---|---|---|---|---|---|---|---|---|
| GABRG2 | 87 | V | 70.33661039 | 34.34732632 | 0.925420987 | 25.64672256 | NA | NA | NA | NA | NA | NA | NA |
| GABRG2 | 88 | K | 67.05940404 | 32.96709658 | 0.884430557 | 23.91106637 | NA | NA | NA | NA | NA | NA | NA |
| GABRG2 | 89 | P | 63.37203896 | 32.50601621 | 0.87073727 | 23.5476632 | NA | NA | NA | NA | NA | NA | NA |
| GABRG2 | 90 | T | 62.94275325 | 28.46652506 | 0.750771388 | 19.58037479 | NA | NA | NA | NA | NA | NA | NA |
| GABRG2 | 91 | L | 60.31118182 | 30.85456028 | 0.821691893 | 22.34667549 | NA | NA | NA | NA | NA | NA | NA |
| GABRG2 | 92 | I | 58.37943182 | 26.50600153 | 0.69254724 | 18.90586998 | NA | NA | NA | NA | NA | NA | NA |
| GABRG2 | 93 | H | 57.26388182 | 29.07659119 | 0.768889294 | 21.56391781 | NA | NA | NA | NA | NA | NA | NA |
| GABRG2 | 94 | T | 53.00646753 | 26.91709589 | 0.704756029 | 23.07629319 | NA | NA | NA | NA | NA | NA | NA |
| GABRG2 | 95 | D | 51.35355682 | 25.01885141 | 0.64838146 | 21.17981955 | NA | NA | NA | NA | NA | NA | NA |
| GABRG2 | 96 | M | 47.50043182 | 22.50556102 | 0.573741095 | 18.70413489 | NA | NA | NA | NA | NA | NA | NA |
| GABRG2 | 97 | Y | 46.09334848 | 22.22998246 | 0.565556889 | 18.57780113 | NA | NA | NA | NA | NA | NA | NA |
| GABRG2 | 98 | V | 42.74875325 | 22.06142763 | 0.560551103 | 18.39815235 | NA | NA | NA | NA | NA | NA | NA |
| GABRG2 | 99 | N | 43.32118182 | 19.40205804 | 0.481572438 | 15.74746693 | NA | NA | NA | NA | NA | NA | NA |
| GABRG2 | 100 | S | 39.53984848 | 17.97301149 | 0.439132234 | 14.3556243 | NA | NA | NA | NA | NA | NA | NA |
| GABRG2 | 101 | I | 36.84518182 | 18.78181338 | 0.463152248 | 15.32237489 | NA | NA | NA | NA | NA | NA | NA |
| GABRG2 | 102 | G | 35.33893182 | 16.7505105 | 0.402826075 | 13.28925079 | NA | NA | NA | NA | NA | NA | NA |
| GABRG2 | 103 | P | 33.25332468 | 13.91483618 | 0.318611465 | 6.12988 | NA | NA | NA | NA | NA | NA | NA |
| GABRG2 | 104 | V | 31.27832468 | 15.09534567 | 0.35367055 | 7.265822377 | NA | NA | NA | NA | NA | NA | NA |
| GABRG2 | 105 | N | 31.22218182 | 9.956697896 | 0.201061623 | 2.034027991 | 8.04 | -0.88 | -3.5 | 21.91 | 1.38 | 2 | -2 |
| GABRG2 | 106 | A | 28.18158182 | 14.16092998 | 0.325920025 | 7.694052737 | NA | NA | NA | NA | NA | NA | NA |
| GABRG2 | 107 | I | 27.83130682 | 9.064243818 | 0.174557284 | 2.68895279 | 6.47 | -0.7 | -1.28 | 14.91 | 0.97 | 1 | 1 |
| GABRG2 | 108 | N | 32.20830682 | 9.434546571 | 0.185554634 | 1.564675312 | 8.04 | -0.88 | -3.5 | 21.91 | 1.38 | 2 | -2 |
| GABRG2 | 109 | M | 31.72918182 | 15.07120941 | 0.352953745 | 7.303960353 | NA | NA | NA | NA | NA | NA | NA |
| GABRG2 | 110 | E | 35.20618182 | 11.70772023 | 0.253063949 | 4.002234195 | 7.89 | -0.81 | -3.58 | 31.13 | 1.56 | 3 | -1 |
| GABRG2 | 111 | Y | 37.21918182 | 15.42822414 | 0.363556463 | 12.03943702 | NA | NA | NA | NA | NA | NA | NA |
| GABRG2 | 112 | T | 38.14461039 | 12.74206082 | 0.283782071 | 9.431955403 | NA | NA | NA | NA | NA | NA | NA |
| GABRG2 | 113 | I | 41.39968182 | 15.0704127 | 0.352930084 | 11.39481072 | NA | NA | NA | NA | NA | NA | NA |
| GABRG2 | 114 | D | 43.89343182 | 15.9321669 | 0.378522689 | 12.27947007 | NA | NA | NA | NA | NA | NA | NA |
| GABRG2 | 115 | I | 46.58305682 | 17.83768118 | 0.435113159 | 14.18233963 | NA | NA | NA | NA | NA | NA | NA |
| GABRG2 | 116 | F | 49.37063636 | 20.34819611 | 0.509671098 | 16.50907042 | NA | NA | NA | NA | NA | NA | NA |
| GABRG2 | 117 | F | 51.89009091 | 20.58774609 | 0.516785317 | 16.7473357 | NA | NA | NA | NA | NA | NA | NA |
| GABRG2 | 118 | A | 54.22298182 | 23.83379689 | 0.613187396 | 19.99265519 | NA | NA | NA | NA | NA | NA | NA |
| GABRG2 | 119 | Q | 57.04029293 | 22.86682354 | 0.584469965 | 15.26323299 | NA | NA | NA | NA | NA | NA | NA |

| | | | | | | | | | | | | | |
|---|---|---|---|---|---|---|---|---|---|---|---|---|---|
| GABRG2 | 120 | T | 59.58718182 | 27.68415485 | 0.72753635 | 18.77694431 | NA | NA | NA | NA | NA | NA | NA |
| GABRG2 | 121 | W | 62.89068182 | 22.84295876 | 0.583761222 | 13.8460145 | NA | NA | NA | NA | NA | NA | NA |
| GABRG2 | 122 | Y | 65.82776515 | 29.51277806 | 0.781843288 | 20.46937853 | NA | NA | NA | NA | NA | NA | NA |
| GABRG2 | 123 | D | 69.37030682 | 24.79723577 | 0.64179986 | 15.74608871 | NA | NA | NA | NA | NA | NA | NA |
| GABRG2 | 124 | R | 72.92345455 | 29.77953544 | 0.78976552 | 20.84011453 | NA | NA | NA | NA | NA | NA | NA |
| GABRG2 | 125 | R | 73.04554545 | 23.93500973 | 0.616193242 | 15.25952675 | NA | NA | NA | NA | NA | NA | NA |
| GABRG2 | 126 | L | 71.85805682 | 24.10525571 | 0.621249252 | 15.17383897 | NA | NA | NA | NA | NA | NA | NA |
| GABRG2 | 127 | K | 74.60118182 | 29.54892128 | 0.782916679 | 20.60057092 | NA | NA | NA | NA | NA | NA | NA |
| GABRG2 | 128 | F | 74.63781818 | 29.58700642 | 0.784047742 | 20.91455202 | NA | NA | NA | NA | NA | NA | NA |
| GABRG2 | 129 | N | 79.08030682 | 33.82248397 | 0.909834079 | 24.98461327 | NA | NA | NA | NA | NA | NA | NA |
| GABRG2 | 130 | S | 77.19001515 | 33.8124682 | 0.909536628 | 24.7834375 | NA | NA | NA | NA | NA | NA | NA |
| GABRG2 | 131 | T | 75.90246753 | 35.69462381 | 0.965433386 | 26.64875553 | NA | NA | NA | NA | NA | NA | NA |
| GABRG2 | 132 | I | 73.09605682 | 31.139087 | 0.830141843 | 22.10010085 | NA | NA | NA | NA | NA | NA | NA |
| GABRG2 | 133 | K | 70.07984848 | 32.66231472 | 0.875379065 | 23.636118 | NA | NA | NA | NA | NA | NA | NA |
| GABRG2 | 134 | V | 68.09075325 | 29.3246785 | 0.776257057 | 20.30386073 | NA | NA | NA | NA | NA | NA | NA |
| GABRG2 | 135 | L | 69.33555682 | 25.16746322 | 0.652794973 | 16.15728163 | NA | NA | NA | NA | NA | NA | NA |
| GABRG2 | 136 | R | 64.34081818 | 24.50937784 | 0.633250979 | 15.51642773 | NA | NA | NA | NA | NA | NA | NA |
| GABRG2 | 137 | L | 66.23318182 | 21.18826187 | 0.534619594 | 12.20261251 | NA | NA | NA | NA | NA | NA | NA |
| GABRG2 | 138 | N | 63.97768182 | 18.12674433 | 0.443697833 | 9.152396299 | NA | NA | NA | NA | NA | NA | NA |
| GABRG2 | 139 | S | 62.17984848 | 14.01043086 | 0.321450462 | 5.042596738 | NA | NA | NA | NA | NA | NA | NA |
| GABRG2 | 140 | N | 65.88893182 | 13.66444529 | 0.31117529 | 4.660650402 | 9.06 | 0 | -1.52 | 21 | 0.31 | 2 | -2 |
| GABRG2 | 141 | M | 68.00693182 | 15.99733537 | 0.38045808 | 6.96885334 | NA | NA | NA | NA | NA | NA | NA |
| GABRG2 | 142 | V | 63.63275325 | 14.89070962 | 0.347593214 | 5.863356489 | NA | NA | NA | NA | NA | NA | NA |
| GABRG2 | 143 | G | 66.21418182 | 13.08312826 | 0.293911182 | 4.033076429 | 9.06 | 0 | -1.52 | 21 | 0.31 | 2 | -2 |
| GABRG2 | 144 | K | 69.99062626 | 14.46996866 | 0.335097937 | 5.617205265 | NA | NA | NA | NA | NA | NA | NA |
| GABRG2 | 145 | I | 67.74330682 | 17.45853611 | 0.423853207 | 8.409546064 | NA | NA | NA | NA | NA | NA | NA |
| GABRG2 | 146 | W | 64.95575325 | 19.67274882 | 0.489611485 | 10.61864725 | NA | NA | NA | NA | NA | NA | NA |
| GABRG2 | 147 | I | 62.18805682 | 17.59221612 | 0.427823272 | 8.583659128 | NA | NA | NA | NA | NA | NA | NA |
| GABRG2 | 148 | P | 59.83818182 | 19.64706731 | 0.488848789 | 11.22113993 | NA | NA | NA | NA | NA | NA | NA |
| GABRG2 | 149 | D | 57.90218182 | 19.63234968 | 0.488411701 | 12.01741455 | NA | NA | NA | NA | NA | NA | NA |
| GABRG2 | 150 | T | 54.37503896 | 18.43085701 | 0.452729452 | 11.6552614 | NA | NA | NA | NA | NA | NA | NA |
| GABRG2 | 151 | F | 51.03545455 | 18.35236146 | 0.45039827 | 11.73990685 | NA | NA | NA | NA | NA | NA | NA |
| GABRG2 | 152 | F | 48.07563636 | 16.73672354 | 0.402416626 | 12.89975941 | NA | NA | NA | NA | NA | NA | NA |

| | | | | | | | | | | | | | |
|---|---|---|---|---|---|---|---|---|---|---|---|---|---|
| GABRG2 | 153 | R | 46.93427273 | 19.26499097 | 0.477501784 | 15.46274909 | NA | NA | NA | NA | NA | NA | NA |
| GABRG2 | 154 | N | 42.30093182 | 17.11541963 | 0.413663243 | 13.44769173 | NA | NA | NA | NA | NA | NA | NA |
| GABRG2 | 155 | S | 44.24001515 | 14.32508949 | 0.330795277 | 10.63222068 | NA | NA | NA | NA | NA | NA | NA |
| GABRG2 | 156 | K | 40.29618182 | 10.78109336 | 0.225544738 | 7.087827252 | NA | NA | NA | NA | NA | NA | NA |
| GABRG2 | 157 | K | 42.56273737 | 5.508943093 | 0.068971019 | 2.618800182 | 3.72 | -0.41 | -3.9 | 49.5 | 1.46 | 4 | 4 |
| GABRG2 | 158 | A | 46.86838182 | 9.417319842 | 0.18504303 | 5.571723982 | NA | NA | NA | NA | NA | NA | NA |
| GABRG2 | 159 | D | 47.42880682 | 6.301126645 | 0.092497496 | 2.507962966 | 3.88 | -0.57 | -3.8 | 49.55 | 1.75 | 4 | 2 |
| GABRG2 | 160 | A | 50.96938182 | 10.06299091 | 0.204218341 | 5.940383362 | NA | NA | NA | NA | NA | NA | NA |
| GABRG2 | 161 | H | 49.92878182 | 12.26083786 | 0.269490584 | 8.137100978 | NA | NA | NA | NA | NA | NA | NA |
| GABRG2 | 162 | W | 54.6693961 | 9.10872642 | 0.175878341 | 1.575266545 | 7.58 | 0.06 | -2.63 | 2.95 | -0.47 | 0 | 0 |
| GABRG2 | 163 | I | 51.85580682 | 12.13593552 | 0.265781201 | 4.63163077 | 7.57 | 0.06 | -2.63 | 2.95 | -0.47 | 0 | 0 |
| GABRG2 | 164 | T | 54.82103896 | 14.4138737 | 0.333432014 | 6.860475986 | NA | NA | NA | NA | NA | NA | NA |
| GABRG2 | 165 | T | 57.14503896 | 14.29984301 | 0.3300455 | 6.780570847 | NA | NA | NA | NA | NA | NA | NA |
| GABRG2 | 166 | P | 58.00518182 | 12.00779668 | 0.261975699 | 5.043065834 | NA | NA | NA | NA | NA | NA | NA |
| GABRG2 | 167 | N | 54.37280682 | 15.17956475 | 0.35617171 | 7.626757559 | NA | NA | NA | NA | NA | NA | NA |
| GABRG2 | 168 | R | 57.748 | 12.3454441 | 0.272003242 | 4.847839233 | 7.66 | 0.06 | -2.63 | 2.95 | -0.47 | 0 | 0 |
| GABRG2 | 169 | M | 59.37493182 | 19.62014692 | 0.4880493 | 10.64542111 | NA | NA | NA | NA | NA | NA | NA |
| GABRG2 | 170 | L | 63.79118182 | 20.32977848 | 0.509124126 | 11.30991626 | NA | NA | NA | NA | NA | NA | NA |
| GABRG2 | 171 | R | 62.26181818 | 24.11247275 | 0.621463586 | 15.12382913 | NA | NA | NA | NA | NA | NA | NA |
| GABRG2 | 172 | I | 68.09655682 | 23.59942688 | 0.606227013 | 14.55647809 | NA | NA | NA | NA | NA | NA | NA |
| GABRG2 | 173 | W | 66.10682468 | 29.18519178 | 0.772114544 | 20.16040668 | NA | NA | NA | NA | NA | NA | NA |
| GABRG2 | 174 | N | 72.05668182 | 32.14178285 | 0.859920171 | 23.10153272 | NA | NA | NA | NA | NA | NA | NA |
| GABRG2 | 175 | D | 69.99030682 | 32.7925342 | 0.879246357 | 23.74342165 | NA | NA | NA | NA | NA | NA | NA |
| GABRG2 | 176 | G | 69.15693182 | 28.95832112 | 0.765376878 | 19.91720849 | NA | NA | NA | NA | NA | NA | NA |
| GABRG2 | 177 | R | 64.39390909 | 28.45303817 | 0.750370851 | 19.43418684 | NA | NA | NA | NA | NA | NA | NA |
| GABRG2 | 178 | V | 64.44918182 | 23.94355228 | 0.616446941 | 14.92989052 | NA | NA | NA | NA | NA | NA | NA |
| GABRG2 | 179 | L | 60.49468182 | 23.72138316 | 0.609848903 | 14.73792533 | NA | NA | NA | NA | NA | NA | NA |
| GABRG2 | 180 | Y | 58.89301515 | 16.69273614 | 0.401110277 | 8.025441252 | NA | NA | NA | NA | NA | NA | NA |
| GABRG2 | 181 | T | 55.8858961 | 20.22659378 | 0.50605972 | 12.71329304 | NA | NA | NA | NA | NA | NA | NA |
| GABRG2 | 182 | L | 53.86118182 | 16.25233894 | 0.388031243 | 8.686861686 | NA | NA | NA | NA | NA | NA | NA |
| GABRG2 | 183 | R | 48.949 | 14.81789786 | 0.345430831 | 10.98538037 | NA | NA | NA | NA | NA | NA | NA |
| GABRG2 | 184 | L | 48.94580682 | 14.20573789 | 0.327250742 | 10.39345482 | NA | NA | NA | NA | NA | NA | NA |
| GABRG2 | 185 | T | 45.0558961 | 10.49686091 | 0.217103528 | 6.844976631 | NA | NA | NA | NA | NA | NA | NA |

| | | | | | | | | | | | | | |
|---|---|---|---|---|---|---|---|---|---|---|---|---|---|
| GABRG2 | 186 | I | 43.67230682 | 12.84196597 | 0.286749081 | 9.18093852 | NA | NA | NA | NA | NA | NA | NA |
| GABRG2 | 187 | D | 40.48055682 | 8.924206631 | 0.170398423 | 5.480442544 | NA | NA | NA | NA | NA | NA | NA |
| GABRG2 | 188 | A | 39.69078182 | 14.63389111 | 0.33996615 | 11.02616753 | NA | NA | NA | NA | NA | NA | NA |
| GABRG2 | 189 | E | 37.67673737 | 17.25355946 | 0.417765756 | 13.68864812 | NA | NA | NA | NA | NA | NA | NA |
| GABRG2 | 190 | C | 34.62218182 | 19.73146153 | 0.491355151 | 16.16449223 | NA | NA | NA | NA | NA | NA | NA |
| GABRG2 | 191 | Q | 32.78929293 | 20.90258008 | 0.52613534 | 13.15520728 | NA | NA | NA | NA | NA | NA | NA |
| GABRG2 | 192 | L | 29.29130682 | 18.54402367 | 0.456090306 | 12.68474497 | NA | NA | NA | NA | NA | NA | NA |
| GABRG2 | 193 | Q | 27.29773737 | 23.10960956 | 0.591680288 | 17.27355077 | NA | NA | NA | NA | NA | NA | NA |
| GABRG2 | 194 | L | 24.05430682 | 17.73344059 | 0.432017394 | 11.89605744 | NA | NA | NA | NA | NA | NA | NA |
| GABRG2 | 195 | H | 21.80968182 | 23.16288009 | 0.593262331 | 21.13897295 | NA | NA | NA | NA | NA | NA | NA |
| GABRG2 | 196 | N | 19.51030682 | 22.43681299 | 0.571699397 | 20.23859464 | NA | NA | NA | NA | NA | NA | NA |
| GABRG2 | 197 | F | 19.47936364 | 15.43272539 | 0.363690142 | 13.27570293 | NA | NA | NA | NA | NA | NA | NA |
| GABRG2 | 198 | P | 18.60646753 | 18.39737055 | 0.451734962 | 16.16886107 | NA | NA | NA | NA | NA | NA | NA |
| GABRG2 | 199 | M | 18.38768182 | 21.92392344 | 0.556467467 | 19.72546422 | NA | NA | NA | NA | NA | NA | NA |
| GABRG2 | 200 | D | 23.30793182 | 21.27624819 | 0.537232635 | 15.46752465 | NA | NA | NA | NA | NA | NA | NA |
| GABRG2 | 201 | E | 26.36740404 | 25.32874298 | 0.657584702 | 19.51611852 | NA | NA | NA | NA | NA | NA | NA |
| GABRG2 | 202 | H | 28.94338182 | 22.03682743 | 0.55982052 | 16.20653782 | NA | NA | NA | NA | NA | NA | NA |
| GABRG2 | 203 | S | 33.66101515 | 26.33155738 | 0.687366551 | 18.61940801 | NA | NA | NA | NA | NA | NA | NA |
| GABRG2 | 204 | C | 35.48818182 | 22.05707095 | 0.560421717 | 19.2794653 | NA | NA | NA | NA | NA | NA | NA |
| GABRG2 | 205 | P | 38.75561039 | 21.51482087 | 0.54431783 | 17.87358579 | NA | NA | NA | NA | NA | NA | NA |
| GABRG2 | 206 | L | 43.07330682 | 19.01546302 | 0.470091237 | 15.33410358 | NA | NA | NA | NA | NA | NA | NA |
| GABRG2 | 207 | E | 44.26629293 | 21.14121619 | 0.533222419 | 17.49748848 | NA | NA | NA | NA | NA | NA | NA |
| GABRG2 | 208 | F | 49.44227273 | 22.13359191 | 0.562694257 | 18.2885148 | NA | NA | NA | NA | NA | NA | NA |
| GABRG2 | 209 | S | 51.87768182 | 22.52994696 | 0.574465315 | 15.91467135 | NA | NA | NA | NA | NA | NA | NA |
| GABRG2 | 210 | S | 55.30868182 | 23.29049285 | 0.597052208 | 16.51607588 | NA | NA | NA | NA | NA | NA | NA |
| GABRG2 | 211 | Y | 54.73134848 | 19.82799135 | 0.494221919 | 13.06246529 | NA | NA | NA | NA | NA | NA | NA |
| GABRG2 | 212 | G | 58.44718182 | 23.16799924 | 0.593414361 | 15.59016718 | NA | NA | NA | NA | NA | NA | NA |
| GABRG2 | 213 | Y | 60.88084848 | 22.65186873 | 0.57808618 | 14.17415873 | NA | NA | NA | NA | NA | NA | NA |
| GABRG2 | 214 | P | 58.55118182 | 28.87778213 | 0.76298501 | 22.28377975 | NA | NA | NA | NA | NA | NA | NA |
| GABRG2 | 215 | R | 56.28654545 | 31.80715941 | 0.849982435 | 25.03655107 | NA | NA | NA | NA | NA | NA | NA |
| GABRG2 | 216 | E | 60.50240404 | 33.04725019 | 0.88681098 | 24.47112278 | NA | NA | NA | NA | NA | NA | NA |
| GABRG2 | 217 | E | 62.5889596 | 28.19980376 | 0.742850228 | 19.38386874 | NA | NA | NA | NA | NA | NA | NA |
| GABRG2 | 218 | I | 59.46080682 | 25.7621432 | 0.670455937 | 17.18911418 | NA | NA | NA | NA | NA | NA | NA |

| | | | | | | | | | | | | | |
|---|---|---|---|---|---|---|---|---|---|---|---|---|---|
| GABRG2 | 219 | V | 57.5848961 | 31.11023505 | 0.82928499 | 23.49419017 | NA | NA | NA | NA | NA | NA | NA |
| GABRG2 | 220 | Y | 54.76384848 | 24.34158261 | 0.628267751 | 17.57365104 | NA | NA | NA | NA | NA | NA | NA |
| GABRG2 | 221 | Q | 54.55951515 | 31.21070747 | 0.832268847 | 27.36813602 | NA | NA | NA | NA | NA | NA | NA |
| GABRG2 | 222 | W | 49.59553896 | 26.8584133 | 0.703013258 | 23.04802154 | NA | NA | NA | NA | NA | NA | NA |
| GABRG2 | 223 | K | 53.43018182 | 31.79393096 | 0.849589573 | 27.95336085 | NA | NA | NA | NA | NA | NA | NA |
| GABRG2 | 224 | R | 51.93009091 | 34.98236626 | 0.944280572 | 31.17306384 | NA | NA | NA | NA | NA | NA | NA |
| GABRG2 | 225 | S | 49.16601515 | 31.18547364 | 0.831519446 | 27.36976028 | NA | NA | NA | NA | NA | NA | NA |
| GABRG2 | 226 | S | 47.42234848 | 28.50161438 | 0.75181348 | 24.69737091 | NA | NA | NA | NA | NA | NA | NA |
| GABRG2 | 227 | V | 44.72161039 | 26.45422962 | 0.691009704 | 22.78187311 | NA | NA | NA | NA | NA | NA | NA |
| GABRG2 | 228 | E | 45.59307071 | 26.37702409 | 0.688716833 | 22.70697796 | NA | NA | NA | NA | NA | NA | NA |
| GABRG2 | 229 | V | 40.95361039 | 27.67222698 | 0.727182113 | 23.99929874 | NA | NA | NA | NA | NA | NA | NA |
| GABRG2 | 230 | G | 40.23843182 | 26.99849368 | 0.707173402 | 23.33022305 | NA | NA | NA | NA | NA | NA | NA |
| GABRG2 | 231 | D | 37.40243182 | 26.29614622 | 0.686314901 | 22.67876022 | NA | NA | NA | NA | NA | NA | NA |
| GABRG2 | 232 | T | 35.0208961 | 26.49712605 | 0.692283653 | 23.26996836 | NA | NA | NA | NA | NA | NA | NA |
| GABRG2 | 233 | R | 32.19572727 | 25.44164991 | 0.660937842 | 17.62781732 | NA | NA | NA | NA | NA | NA | NA |
| GABRG2 | 234 | S | 35.74518182 | 22.11651105 | 0.562186985 | 18.63845215 | NA | NA | NA | NA | NA | NA | NA |
| GABRG2 | 235 | W | 36.14818182 | 21.22308149 | 0.535653677 | 18.04526919 | NA | NA | NA | NA | NA | NA | NA |
| GABRG2 | 236 | R | 35.19645455 | 14.7146562 | 0.342364733 | 8.324392513 | NA | NA | NA | NA | NA | NA | NA |
| GABRG2 | 237 | L | 31.10080682 | 19.21326107 | 0.475965496 | 13.37378721 | NA | NA | NA | NA | NA | NA | NA |
| GABRG2 | 238 | Y | 28.04751515 | 13.68273792 | 0.31171855 | 7.828931199 | NA | NA | NA | NA | NA | NA | NA |
| GABRG2 | 239 | Q | 24.80107071 | 13.45262264 | 0.304884525 | 7.66139626 | NA | NA | NA | NA | NA | NA | NA |
| GABRG2 | 240 | F | 27.85254545 | 18.27999568 | 0.448249132 | 12.44584239 | NA | NA | NA | NA | NA | NA | NA |
| GABRG2 | 241 | S | 28.13018182 | 23.36467218 | 0.599255206 | 17.54532674 | NA | NA | NA | NA | NA | NA | NA |
| GABRG2 | 242 | F | 33.67363636 | 25.94494269 | 0.675884765 | 20.11160701 | NA | NA | NA | NA | NA | NA | NA |
| GABRG2 | 243 | V | 30.79132468 | 28.79913202 | 0.760649238 | 22.964409 | NA | NA | NA | NA | NA | NA | NA |
| GABRG2 | 244 | G | 34.62543182 | 29.06250913 | 0.768471082 | 25.48097308 | NA | NA | NA | NA | NA | NA | NA |
| GABRG2 | 245 | L | 38.84768182 | 27.74833609 | 0.729442422 | 24.08545998 | NA | NA | NA | NA | NA | NA | NA |
| GABRG2 | 246 | R | 38.13181818 | 29.25036323 | 0.774050023 | 25.57919012 | NA | NA | NA | NA | NA | NA | NA |
| GABRG2 | 247 | N | 43.44068182 | 30.72099565 | 0.817725255 | 27.04163592 | NA | NA | NA | NA | NA | NA | NA |
| GABRG2 | 248 | T | 44.10175325 | 30.59218464 | 0.813899791 | 26.92842861 | NA | NA | NA | NA | NA | NA | NA |
| GABRG2 | 249 | T | 47.63932468 | 32.4975472 | 0.870485755 | 28.68809785 | NA | NA | NA | NA | NA | NA | NA |
| GABRG2 | 250 | E | 46.06218182 | 28.98662838 | 0.766217555 | 25.22194476 | NA | NA | NA | NA | NA | NA | NA |
| GABRG2 | 251 | V | 49.72575325 | 32.60943132 | 0.873808519 | 28.76189373 | NA | NA | NA | NA | NA | NA | NA |

| | | | | | | | | | | | | | |
|---|---|---|---|---|---|---|---|---|---|---|---|---|---|
| GABRG2 | 252 | V | 48.52818182 | 28.89518617 | 0.76350188 | 25.04056959 | NA | NA | NA | NA | NA | NA | NA |
| GABRG2 | 253 | K | 49.51240404 | 31.72295582 | 0.847481734 | 25.40172679 | NA | NA | NA | NA | NA | NA | NA |
| GABRG2 | 254 | T | 50.76103896 | 27.61301688 | 0.725423676 | 21.29568259 | NA | NA | NA | NA | NA | NA | NA |
| GABRG2 | 255 | T | 52.53475325 | 29.52196724 | 0.782116191 | 22.9122083 | NA | NA | NA | NA | NA | NA | NA |
| GABRG2 | 256 | S | 55.46251515 | 27.04874189 | 0.708665687 | 20.28127244 | NA | NA | NA | NA | NA | NA | NA |
| GABRG2 | 257 | G | 54.33518182 | 30.52916499 | 0.812028217 | 23.76396275 | NA | NA | NA | NA | NA | NA | NA |
| GABRG2 | 258 | D | 53.87418182 | 31.32058962 | 0.835532156 | 24.70669073 | NA | NA | NA | NA | NA | NA | NA |
| GABRG2 | 259 | Y | 53.57918182 | 25.85873477 | 0.673324539 | 19.24546307 | NA | NA | NA | NA | NA | NA | NA |
| GABRG2 | 260 | V | 52.76575325 | 28.35501949 | 0.747459866 | 21.78228533 | NA | NA | NA | NA | NA | NA | NA |
| GABRG2 | 261 | V | 48.5848961 | 26.6133995 | 0.695736773 | 22.78880612 | NA | NA | NA | NA | NA | NA | NA |
| GABRG2 | 262 | M | 48.84455682 | 25.98625764 | 0.677111747 | 22.17801838 | NA | NA | NA | NA | NA | NA | NA |
| GABRG2 | 263 | S | 44.07034848 | 26.69779877 | 0.698243285 | 23.02539042 | NA | NA | NA | NA | NA | NA | NA |
| GABRG2 | 264 | V | 42.48946753 | 24.85991724 | 0.643661391 | 21.19300306 | NA | NA | NA | NA | NA | NA | NA |
| GABRG2 | 265 | Y | 38.93351515 | 26.97010934 | 0.706330437 | 23.3063991 | NA | NA | NA | NA | NA | NA | NA |
| GABRG2 | 266 | F | 37.74263636 | 22.87293529 | 0.584651473 | 19.21647222 | NA | NA | NA | NA | NA | NA | NA |
| GABRG2 | 267 | D | 33.03755682 | 27.52740424 | 0.722881129 | 19.83541691 | NA | NA | NA | NA | NA | NA | NA |
| GABRG2 | 268 | L | 30.79543182 | 22.54222602 | 0.574829981 | 17.08243958 | NA | NA | NA | NA | NA | NA | NA |
| GABRG2 | 269 | S | 26.61934848 | 23.78931939 | 0.611866492 | 17.97691811 | NA | NA | NA | NA | NA | NA | NA |
| GABRG2 | 270 | R | 23.32863636 | 17.61704771 | 0.428560727 | 11.83614413 | NA | NA | NA | NA | NA | NA | NA |
| GABRG2 | 271 | R | 23.19581818 | 20.43517737 | 0.512254291 | 18.29446035 | NA | NA | NA | NA | NA | NA | NA |
| GABRG2 | 272 | M | 17.43043182 | 19.99260636 | 0.499110699 | 17.75967401 | NA | NA | NA | NA | NA | NA | NA |
| GABRG2 | 273 | G | 16.49368182 | 20.03633647 | 0.500409407 | 17.80039109 | NA | NA | NA | NA | NA | NA | NA |
| GABRG2 | 274 | Y | 17.09284848 | 11.42438278 | 0.244649319 | 9.225808202 | NA | NA | NA | NA | NA | NA | NA |
| GABRG2 | 275 | F | 16.34327273 | 13.8129132 | 0.31558453 | 11.60653699 | NA | NA | NA | NA | NA | NA | NA |
| GABRG2 | 276 | T | 11.83946753 | 16.96316085 | 0.409141421 | 14.56677008 | NA | NA | NA | NA | NA | NA | NA |
| GABRG2 | 277 | I | 11.25055682 | 14.16428872 | 0.326019773 | 11.99384415 | NA | NA | NA | NA | NA | NA | NA |
| GABRG2 | 278 | Q | 12.19240404 | 10.80423755 | 0.226232081 | 8.472521642 | NA | NA | NA | NA | NA | NA | NA |
| GABRG2 | 279 | T | 10.68503896 | 12.08507195 | 0.264270641 | 9.431463952 | NA | NA | NA | NA | NA | NA | NA |
| GABRG2 | 280 | Y | 10.66518182 | 15.35345492 | 0.361335947 | 12.90945471 | NA | NA | NA | NA | NA | NA | NA |
| GABRG2 | 281 | I | 6.552056818 | 15.88407687 | 0.377094499 | 13.64974007 | NA | NA | NA | NA | NA | NA | NA |
| GABRG2 | 282 | P | 6.035324675 | 12.255574 | 0.269334256 | 10.01719334 | NA | NA | NA | NA | NA | NA | NA |
| GABRG2 | 283 | C | 5.048348485 | 13.2473934 | 0.298789572 | 10.86931988 | NA | NA | NA | NA | NA | NA | NA |
| GABRG2 | 284 | T | 3.009038961 | 16.93528326 | 0.408313505 | 14.68040696 | NA | NA | NA | NA | NA | NA | NA |

| | | | | | | | | | | | | | |
|---|---|---|---|---|---|---|---|---|---|---|---|---|---|
| GABRG2 | 285 | L | 0.929806818 | 16.45459091 | 0.394037776 | 13.90991163 | NA | NA | NA | NA | NA | NA | NA |
| GABRG2 | 286 | I | 0.683806818 | 11.61535066 | 0.250320733 | 10.19250015 | NA | NA | NA | NA | NA | NA | NA |
| GABRG2 | 287 | V | -0.17561039 | 16.32095429 | 0.390069 | 14.57418103 | NA | NA | NA | NA | NA | NA | NA |
| GABRG2 | 288 | V | -1.74661039 | 17.23662783 | 0.417262916 | 14.2614079 | NA | NA | NA | NA | NA | NA | NA |
| GABRG2 | 289 | L | -2.595306818 | 12.17308593 | 0.266884504 | 9.418854856 | NA | NA | NA | NA | NA | NA | NA |
| GABRG2 | 290 | S | -3.918348485 | 14.34576356 | 0.331409261 | 11.38286614 | NA | NA | NA | NA | NA | NA | NA |
| GABRG2 | 291 | W | -4.150824675 | 16.39086822 | 0.392145323 | 13.4287959 | NA | NA | NA | NA | NA | NA | NA |
| GABRG2 | 292 | V | -8.004467532 | 14.49270054 | 0.335773035 | 12.50710586 | NA | NA | NA | NA | NA | NA | NA |
| GABRG2 | 293 | S | -9.999515152 | 13.46942259 | 0.305383454 | 10.96660519 | NA | NA | NA | NA | NA | NA | NA |
| GABRG2 | 294 | F | -10.15381818 | 16.30781362 | 0.389678745 | 13.73529609 | NA | NA | NA | NA | NA | NA | NA |
| GABRG2 | 295 | W | -12.01396753 | 16.46852887 | 0.394451709 | 14.52618683 | NA | NA | NA | NA | NA | NA | NA |
| GABRG2 | 296 | I | -13.99268182 | 11.55773788 | 0.248609734 | 9.615611128 | NA | NA | NA | NA | NA | NA | NA |
| GABRG2 | 297 | N | -18.81805682 | 11.2093019 | 0.238261789 | 8.551350049 | NA | NA | NA | NA | NA | NA | NA |
| GABRG2 | 298 | K | -19.23640404 | 12.30990991 | 0.270947939 | 5.974424151 | NA | NA | NA | NA | NA | NA | NA |
| GABRG2 | 299 | D | -20.77143182 | 7.908502576 | 0.140233774 | 1.56710515 | 6.37 | -0.62 | -0.7 | 2.93 | 0.28 | 0 | 0 |
| GABRG2 | 300 | A | -16.67338182 | 7.98656741 | 0.142552164 | 5.499588921 | NA | NA | NA | NA | NA | NA | NA |
| GABRG2 | 301 | V | -15.33503896 | 6.97218194 | 0.112426675 | 5.10077576 | NA | NA | NA | NA | NA | NA | NA |
| GABRG2 | 302 | P | -13.69861039 | 3.368498033 | 0.005403513 | 1.652517484 | 1.99 | -0.05 | -0.24 | 0.95 | -0.54 | 0 | 0 |
| GABRG2 | 303 | A | -11.92038182 | 5.757498155 | 0.076352673 | 3.785980645 | 1.99 | -0.05 | -0.24 | 0.95 | -0.54 | 0 | 0 |
| GABRG2 | 304 | R | -12.072 | 8.333966643 | 0.152869319 | 5.764601489 | NA | NA | NA | NA | NA | NA | NA |
| GABRG2 | 305 | T | -9.069753247 | 7.249794825 | 0.120671296 | 5.509955195 | NA | NA | NA | NA | NA | NA | NA |
| GABRG2 | 306 | S | -7.471848485 | 3.553260615 | 0.010890641 | 1.835558715 | 2.04 | 0.43 | 2.53 | 0.64 | -0.5 | 0 | 0 |
| GABRG2 | 307 | L | -6.788306818 | 7.475513494 | 0.127374749 | 4.595815432 | 2.93 | 0.13 | 1.73 | 0.9 | -0.18 | 0 | 0 |
| GABRG2 | 308 | G | -4.528931818 | 9.125105205 | 0.176364762 | 6.242642369 | NA | NA | NA | NA | NA | NA | NA |
| GABRG2 | 309 | I | -2.937931818 | 6.283799249 | 0.091982903 | 3.423399639 | 2.93 | 0.13 | 1.73 | 0.9 | -0.18 | 0 | 0 |
| GABRG2 | 310 | T | -1.49561039 | 4.556203024 | 0.040676291 | 1.621379101 | 3.21 | 0.38 | 2 | 0.74 | -0.77 | 0 | 0 |
| GABRG2 | 311 | T | -0.721753247 | 8.5587044 | 0.15954364 | 5.943659578 | NA | NA | NA | NA | NA | NA | NA |
| GABRG2 | 312 | V | 0.194324675 | 9.118394157 | 0.176165456 | 7.375411279 | NA | NA | NA | NA | NA | NA | NA |
| GABRG2 | 313 | L | 1.479431818 | 3.186550957 | 0 | 1.464936112 | 1.74 | 1.14 | 3.8 | 0.13 | -1.79 | 0 | 0 |
| GABRG2 | 314 | T | 2.527038961 | 7.060516553 | 0.11505006 | 4.348916197 | 2.85 | 0.66 | 2.68 | 0.51 | -1.15 | 0 | 0 |
| GABRG2 | 315 | M | 3.897681818 | 9.329043949 | 0.182421389 | 6.454869117 | NA | NA | NA | NA | NA | NA | NA |
| GABRG2 | 316 | T | 5.840467532 | 7.032924143 | 0.114230613 | 4.757838392 | 2.93 | -0.77 | -0.7 | 1.66 | 0.77 | 0 | 0 |
| GABRG2 | 317 | T | 7.068324675 | 3.95556039 | 0.022838246 | 1.682473725 | 2.72 | 0.13 | -0.03 | 13.76 | -0.13 | 0 | 0 |

| | | | | | | | | | | | | | |
|---|---|---|---|---|---|---|---|---|---|---|---|---|---|
| GABRG2 | 318 | L | 7.988806818 | 7.154535205 | 0.117842251 | 4.487792905 | 2.72 | 0.13 | -0.03 | 13.76 | -0.13 | 0 | 0 |
| GABRG2 | 319 | S | 10.25734848 | 8.371232705 | 0.153976056 | 5.805269593 | NA | NA | NA | NA | NA | NA | NA |
| GABRG2 | 320 | T | 12.44675325 | 4.15195472 | 0.028670818 | 1.734372189 | 2.57 | 0.43 | 0.2 | 17.8 | -0.43 | 0 | 0 |
| GABRG2 | 321 | I | 13.57243182 | 7.060532983 | 0.115050548 | 4.979852322 | 2.4 | -1.02 | -2.57 | 33.82 | 1.74 | 2 | -2 |
| GABRG2 | 322 | A | 14.87558182 | 9.798641743 | 0.196367629 | 7.588833587 | NA | NA | NA | NA | NA | NA | NA |
| GABRG2 | 323 | R | 18.15590909 | 9.631041948 | 0.191390206 | 7.775257785 | NA | NA | NA | NA | NA | NA | NA |
| GABRG2 | 324 | K | 19.44351515 | 7.088000917 | 0.115866298 | 5.102876449 | NA | NA | NA | NA | NA | NA | NA |
| GABRG2 | 325 | S | 22.60451515 | 10.34170126 | 0.212495555 | 4.639456657 | 5.85 | -0.66 | -2.6 | 18.75 | 0.94 | 1 | 1 |
| GABRG2 | 326 | L | 22.21380682 | 11.61502583 | 0.250311086 | 5.777071653 | NA | NA | NA | NA | NA | NA | NA |
| GABRG2 | 327 | P | 24.5688961 | 13.57699926 | 0.308578295 | 7.727396176 | NA | NA | NA | NA | NA | NA | NA |
| GABRG2 | 328 | K | 22.53662626 | 11.80749067 | 0.256026958 | 6.360682443 | NA | NA | NA | NA | NA | NA | NA |
| GABRG2 | 329 | V | 22.23575325 | 17.3513562 | 0.42067015 | 11.5462551 | NA | NA | NA | NA | NA | NA | NA |
| GABRG2 | 330 | S | 20.18651515 | 20.50152572 | 0.514224722 | 18.32227092 | NA | NA | NA | NA | NA | NA | NA |
| GABRG2 | 331 | Y | 20.49476515 | 20.26556878 | 0.50721721 | 18.04775964 | NA | NA | NA | NA | NA | NA | NA |
| GABRG2 | 332 | V | 15.3598961 | 21.1919077 | 0.534727869 | 18.99226237 | NA | NA | NA | NA | NA | NA | NA |
| GABRG2 | 333 | T | 16.6688961 | 17.93131746 | 0.437893994 | 15.69603112 | NA | NA | NA | NA | NA | NA | NA |
| GABRG2 | 334 | A | 14.62538182 | 19.81154956 | 0.493733626 | 17.66482306 | NA | NA | NA | NA | NA | NA | NA |
| GABRG2 | 335 | M | 13.96255682 | 15.12031666 | 0.354412145 | 12.89586609 | NA | NA | NA | NA | NA | NA | NA |
| GABRG2 | 336 | D | 13.67318182 | 15.06829858 | 0.352867298 | 12.77618125 | NA | NA | NA | NA | NA | NA | NA |
| GABRG2 | 337 | L | 10.45005682 | 19.02410284 | 0.470347824 | 16.41284294 | NA | NA | NA | NA | NA | NA | NA |
| GABRG2 | 338 | F | 8.231090909 | 17.53502372 | 0.426124757 | 14.86226296 | NA | NA | NA | NA | NA | NA | NA |
| GABRG2 | 339 | V | 8.467324675 | 12.89820425 | 0.28841926 | 10.26483685 | NA | NA | NA | NA | NA | NA | NA |
| GABRG2 | 340 | S | 7.185348485 | 16.63743379 | 0.399467893 | 14.02491315 | NA | NA | NA | NA | NA | NA | NA |
| GABRG2 | 341 | V | 4.58661039 | 19.07391276 | 0.471827093 | 16.45145578 | NA | NA | NA | NA | NA | NA | NA |
| GABRG2 | 342 | C | 3.511181818 | 16.88665938 | 0.40686946 | 14.32187427 | NA | NA | NA | NA | NA | NA | NA |
| GABRG2 | 343 | F | 2.854818182 | 12.99021509 | 0.291151822 | 10.06172372 | NA | NA | NA | NA | NA | NA | NA |
| GABRG2 | 344 | I | 1.424431818 | 18.2943394 | 0.448675116 | 15.38832206 | NA | NA | NA | NA | NA | NA | NA |
| GABRG2 | 345 | F | 0 | 18.29555665 | 0.448711266 | 16.76522819 | NA | NA | NA | NA | NA | NA | NA |
| GABRG2 | 346 | V | -0.069324675 | 13.49895659 | 0.306260563 | 10.9579266 | NA | NA | NA | NA | NA | NA | NA |
| GABRG2 | 347 | F | -0.661545455 | 16.4298052 | 0.393301684 | 13.83901707 | NA | NA | NA | NA | NA | NA | NA |
| GABRG2 | 348 | S | -3.123848485 | 18.40823245 | 0.452057542 | 15.534278 | NA | NA | NA | NA | NA | NA | NA |
| GABRG2 | 349 | A | -5.463181818 | 15.93797054 | 0.378695047 | 13.05208211 | NA | NA | NA | NA | NA | NA | NA |
| GABRG2 | 350 | L | -6.027931818 | 11.72474469 | 0.253569546 | 8.823275223 | NA | NA | NA | NA | NA | NA | NA |

| | | | | | | | | | | | | | |
|---|---|---|---|---|---|---|---|---|---|---|---|---|---|
| GABRG2 | 351 | V | -6.464038961 | 16.43055026 | 0.393323811 | 13.53976242 | NA | NA | NA | NA | NA | NA | NA |
| GABRG2 | 352 | E | -9.931181818 | 18.20712264 | 0.446084929 | 15.43887089 | NA | NA | NA | NA | NA | NA | NA |
| GABRG2 | 353 | Y | -12.59626515 | 11.35822116 | 0.242684433 | 9.532806357 | NA | NA | NA | NA | NA | NA | NA |
| GABRG2 | 354 | G | -11.56068182 | 16.22683315 | 0.387273766 | 13.71235875 | NA | NA | NA | NA | NA | NA | NA |
| GABRG2 | 355 | T | -12.19761039 | 18.76786576 | 0.462738027 | 16.54128745 | NA | NA | NA | NA | NA | NA | NA |
| GABRG2 | 356 | L | -15.34818182 | 17.30343544 | 0.419246987 | 15.4717738 | NA | NA | NA | NA | NA | NA | NA |
| GABRG2 | 357 | H | -16.54128182 | 12.21018042 | 0.267986146 | 10.50630886 | NA | NA | NA | NA | NA | NA | NA |
| GABRG2 | 358 | Y | -16.71043182 | 17.77747356 | 0.433325096 | 10.4164461 | NA | NA | NA | NA | NA | NA | NA |
| GABRG2 | 359 | F | -16.849 | 18.67682987 | 0.460034419 | 11.01390198 | NA | NA | NA | NA | NA | NA | NA |
| GABRG2 | 360 | V | -20.90818182 | 15.07546795 | 0.353080216 | 7.440337434 | NA | NA | NA | NA | NA | NA | NA |
| GABRG2 | 361 | S | -21.32951515 | 12.58229892 | 0.27903742 | 4.902129675 | 7.76 | 0.04 | -2.4 | 25.66 | 0.6 | 2 | -2 |
| GABRG2 | 362 | N | -19.72043182 | 14.89302565 | 0.347661996 | 7.218205769 | NA | NA | NA | NA | NA | NA | NA |
| GABRG2 | 363 | R | -23.31181818 | 16.55281662 | 0.396954909 | 8.676436634 | NA | NA | NA | NA | NA | NA | NA |
| GABRG2 | 437 | R | -20.57318182 | 24.56199336 | 0.634813569 | 16.89632799 | NA | NA | NA | NA | NA | NA | NA |
| GABRG2 | 438 | I | -16.80343182 | 21.10910045 | 0.532268637 | 13.42540145 | NA | NA | NA | NA | NA | NA | NA |
| GABRG2 | 439 | A | -19.98158182 | 19.80050739 | 0.493405693 | 12.15248963 | NA | NA | NA | NA | NA | NA | NA |
| GABRG2 | 440 | K | -19.48451515 | 21.95840682 | 0.557491564 | 14.28336197 | NA | NA | NA | NA | NA | NA | NA |
| GABRG2 | 441 | M | -13.81568182 | 21.7367821 | 0.550909694 | 19.87899288 | NA | NA | NA | NA | NA | NA | NA |
| GABRG2 | 442 | D | -14.60430682 | 17.9261615 | 0.43774087 | 16.08836869 | NA | NA | NA | NA | NA | NA | NA |
| GABRG2 | 443 | S | -15.33318182 | 23.07852911 | 0.590757253 | 21.19318049 | NA | NA | NA | NA | NA | NA | NA |
| GABRG2 | 444 | Y | -13.94459848 | 25.22191369 | 0.654412058 | 23.24423561 | NA | NA | NA | NA | NA | NA | NA |
| GABRG2 | 445 | A | -9.793181818 | 21.92132042 | 0.556390162 | 19.41067995 | NA | NA | NA | NA | NA | NA | NA |
| GABRG2 | 446 | R | -12.56563636 | 19.68922218 | 0.490100715 | 17.65770341 | NA | NA | NA | NA | NA | NA | NA |
| GABRG2 | 447 | I | -9.593556818 | 25.87263833 | 0.673737451 | 23.3625983 | NA | NA | NA | NA | NA | NA | NA |
| GABRG2 | 448 | F | -5.703636364 | 26.01868317 | 0.67807473 | 23.14402141 | NA | NA | NA | NA | NA | NA | NA |
| GABRG2 | 449 | F | -4.681181818 | 20.67954973 | 0.519511726 | 17.73765324 | NA | NA | NA | NA | NA | NA | NA |
| GABRG2 | 450 | P | -4.127038961 | 24.00562851 | 0.618290497 | 21.04907665 | NA | NA | NA | NA | NA | NA | NA |
| GABRG2 | 451 | T | -3.171038961 | 27.20413399 | 0.713280563 | 24.24519839 | NA | NA | NA | NA | NA | NA | NA |
| GABRG2 | 452 | A | -0.675181818 | 27.21177741 | 0.713507559 | 24.26612316 | NA | NA | NA | NA | NA | NA | NA |
| GABRG2 | 453 | F | 0.690090909 | 21.30162855 | 0.537986388 | 20.26176659 | NA | NA | NA | NA | NA | NA | NA |
| GABRG2 | 454 | C | 0.156348485 | 27.36503442 | 0.718059027 | 24.67688602 | NA | NA | NA | NA | NA | NA | NA |
| GABRG2 | 455 | L | 0.659431818 | 29.16926225 | 0.771641464 | 26.49762797 | NA | NA | NA | NA | NA | NA | NA |
| GABRG2 | 456 | F | 4.638909091 | 22.73047613 | 0.580420683 | 20.07245501 | NA | NA | NA | NA | NA | NA | NA |

| | | | | | | | | | | | | | |
|---|---|---|---|---|---|---|---|---|---|---|---|---|---|
| GABRG2 | 457 | N | 4.629056818 | 23.42576244 | 0.601069481 | 20.74051833 | NA | NA | NA | NA | NA | NA | NA |
| GABRG2 | 458 | L | 4.921806818 | 28.57133077 | 0.753883936 | 25.89076801 | NA | NA | NA | NA | NA | NA | NA |
| GABRG2 | 459 | V | 7.056753247 | 28.67855268 | 0.757068241 | 26.01550124 | NA | NA | NA | NA | NA | NA | NA |
| GABRG2 | 460 | Y | 10.25326515 | 20.95085323 | 0.527568969 | 18.29519383 | NA | NA | NA | NA | NA | NA | NA |
| GABRG2 | 461 | W | 9.977467532 | 21.8455617 | 0.554140259 | 19.16179484 | NA | NA | NA | NA | NA | NA | NA |
| GABRG2 | 462 | V | 11.00761039 | 29.32160477 | 0.776165773 | 26.64871829 | NA | NA | NA | NA | NA | NA | NA |
| GABRG2 | 463 | S | 12.89968182 | 29.56372867 | 0.783356433 | 27.02287852 | NA | NA | NA | NA | NA | NA | NA |
| GABRG2 | 464 | Y | 14.56734848 | 25.19347427 | 0.653567456 | 23.04181915 | NA | NA | NA | NA | NA | NA | NA |
| GABRG2 | 465 | L | 15.49205682 | 26.46443143 | 0.69131268 | 24.3491033 | NA | NA | NA | NA | NA | NA | NA |
| GABRG2 | 466 | Y | 14.45434848 | 31.33130661 | 0.835850432 | 29.0997601 | NA | NA | NA | NA | NA | NA | NA |
| GABRG2 | 467 | L | 16.10243182 | 31.23916294 | 0.833113925 | 29.00752947 | NA | NA | NA | NA | NA | NA | NA |
| GLRA1 | 244 | L | 28.84213636 | 24.64911516 | 0.553263673 | 16.68802046 | NA | NA | NA | NA | NA | NA | NA |
| GLRA1 | 245 | E | 28.00808081 | 23.58897047 | 0.524575058 | 15.6305156 | NA | NA | NA | NA | NA | NA | NA |
| GLRA1 | 246 | R | 27.05590909 | 26.63973455 | 0.607131899 | 18.67279443 | NA | NA | NA | NA | NA | NA | NA |
| GLRA1 | 247 | Q | 26.54219192 | 26.09207276 | 0.592311603 | 18.10864658 | NA | NA | NA | NA | NA | NA | NA |
| GLRA1 | 249 | G | 19.46413636 | 25.05059945 | 0.564128255 | 16.73642682 | NA | NA | NA | NA | NA | NA | NA |
| GLRA1 | 250 | Y | 21.4652197 | 26.09791522 | 0.592469706 | 17.78201713 | NA | NA | NA | NA | NA | NA | NA |
| GLRA1 | 251 | Y | 18.4174697 | 18.3402211 | 0.382538453 | 10.02708846 | NA | NA | NA | NA | NA | NA | NA |
| GLRA1 | 252 | L | 16.30713636 | 19.85390773 | 0.423500384 | 11.51620428 | NA | NA | NA | NA | NA | NA | NA |
| GLRA1 | 253 | I | 14.85388636 | 25.25170565 | 0.569570397 | 18.08043738 | NA | NA | NA | NA | NA | NA | NA |
| GLRA1 | 254 | Q | 14.9239697 | 24.82649057 | 0.558063636 | 17.66207307 | NA | NA | NA | NA | NA | NA | NA |
| GLRA1 | 256 | Y | 11.26463636 | 20.38047463 | 0.437749831 | 13.74880409 | NA | NA | NA | NA | NA | NA | NA |
| GLRA1 | 257 | I | 8.391761364 | 24.50760188 | 0.549434177 | 18.0445782 | NA | NA | NA | NA | NA | NA | NA |
| GLRA1 | 258 | P | 7.939636364 | 21.09775858 | 0.457160279 | 15.09459396 | NA | NA | NA | NA | NA | NA | NA |
| GLRA1 | 259 | S | 7.326969697 | 18.77586009 | 0.394327296 | 12.70517099 | NA | NA | NA | NA | NA | NA | NA |
| GLRA1 | 260 | L | 5.206636364 | 21.84697018 | 0.477434722 | 16.85981454 | NA | NA | NA | NA | NA | NA | NA |
| GLRA1 | 261 | L | 2.925761364 | 22.04272463 | 0.482732041 | 17.7794562 | NA | NA | NA | NA | NA | NA | NA |
| GLRA1 | 262 | I | 2.540511364 | 16.77251114 | 0.340114595 | 12.46736508 | NA | NA | NA | NA | NA | NA | NA |
| GLRA1 | 263 | V | 1.427779221 | 18.09105044 | 0.375795636 | 13.98323743 | NA | NA | NA | NA | NA | NA | NA |
| GLRA1 | 264 | I | -0.329761364 | 21.10478574 | 0.457350441 | 16.5168871 | NA | NA | NA | NA | NA | NA | NA |
| GLRA1 | 265 | L | -1.146886364 | 18.45934349 | 0.385762028 | 14.03340768 | NA | NA | NA | NA | NA | NA | NA |
| GLRA1 | 266 | S | -1.486969697 | 15.41531647 | 0.303387499 | 10.78337902 | NA | NA | NA | NA | NA | NA | NA |
| GLRA1 | 267 | W | -4.055207792 | 17.49603161 | 0.359693809 | 14.21134587 | NA | NA | NA | NA | NA | NA | NA |

| | | | | | | | | | | | | | |
|---|---|---|---|---|---|---|---|---|---|---|---|---|---|
| GLRA1 | 268 | I | -6.061511364 | 18.48950256 | 0.386578164 | 13.92484463 | NA | NA | NA | NA | NA | NA | NA |
| GLRA1 | 269 | S | -6.29130303 | 14.29209887 | 0.272992065 | 9.864257642 | NA | NA | NA | NA | NA | NA | NA |
| GLRA1 | 270 | F | -6.444363636 | 13.62809286 | 0.255023373 | 9.227712527 | NA | NA | NA | NA | NA | NA | NA |
| GLRA1 | 271 | W | -9.536136364 | 14.36746178 | 0.275031463 | 9.616863418 | NA | NA | NA | NA | NA | NA | NA |
| GLRA1 | 272 | I | -11.37626136 | 10.92182526 | 0.181788963 | 6.267259337 | NA | NA | NA | NA | NA | NA | NA |
| GLRA1 | 273 | N | -7.733386364 | 9.527405785 | 0.144054526 | 4.937538554 | 4.59 | -0.09 | -1.6 | 1.58 | -1.3 | 0 | 0 |
| GLRA1 | 275 | D | -12.63851136 | 6.194640103 | 0.053866428 | 1.539360723 | 4.68 | -0.8 | -0.4 | 3.38 | 0.81 | 0 | 0 |
| GLRA1 | 276 | A | -11.93923636 | 6.263664981 | 0.055734313 | 1.628230192 | 4.68 | -0.8 | -0.4 | 3.38 | 0.81 | 0 | 0 |
| GLRA1 | 277 | A | -8.332836364 | 6.112160665 | 0.051634449 | 1.521679778 | 4.6 | -0.09 | -1.6 | 1.58 | -1.3 | 0 | 0 |
| GLRA1 | 278 | P | -6.373207792 | 5.286646007 | 0.029295165 | 0.750730657 | 4.59 | -0.09 | -1.6 | 1.58 | -1.3 | 0 | 0 |
| GLRA1 | 279 | A | -3.440836364 | 4.204115365 | 7.63E-07 | 0.924479992 | 3.28 | 0.02 | 1.8 | 0 | 0.59 | 0 | 0 |
| GLRA1 | 280 | R | -1.869909091 | 7.852389063 | 0.098726835 | 4.572075752 | 3.28 | 0.02 | 1.8 | 0 | 0.59 | 0 | 0 |
| GLRA1 | 281 | V | -2.384922078 | 9.360331511 | 0.139533322 | 6.073951079 | NA | NA | NA | NA | NA | NA | NA |
| GLRA1 | 282 | G | -0.339636364 | 7.128597688 | 0.079140289 | 2.489205534 | 4.64 | 0.47 | 2.6 | 0.05 | -0.36 | 0 | 0 |
| GLRA1 | 283 | L | 0.498636364 | 5.044523169 | 0.02274307 | 0.897807843 | 4.15 | 1.14 | 3.8 | 0.13 | -1.79 | 0 | 0 |
| GLRA1 | 284 | G | 1.307386364 | 10.34952699 | 0.166301979 | 6.336185431 | NA | NA | NA | NA | NA | NA | NA |
| GLRA1 | 285 | I | 1.467386364 | 11.17918714 | 0.188753443 | 7.031944892 | NA | NA | NA | NA | NA | NA | NA |
| GLRA1 | 286 | T | 4.651207792 | 6.795035246 | 0.070113743 | 1.814711983 | 4.99 | 1.14 | 3.8 | 0.13 | -1.79 | 0 | 0 |
| GLRA1 | 287 | T | 5.590207792 | 9.985302099 | 0.156445675 | 4.461619879 | 5.69 | 1.14 | 3.8 | 0.13 | -1.79 | 0 | 0 |
| GLRA1 | 288 | V | 6.467493506 | 12.9227381 | 0.235935741 | 7.041603724 | NA | NA | NA | NA | NA | NA | NA |
| GLRA1 | 289 | L | 8.313261364 | 12.03314709 | 0.211862486 | 5.58877793 | NA | NA | NA | NA | NA | NA | NA |
| GLRA1 | 290 | T | 10.08163636 | 7.692111804 | 0.094389566 | 0.901917534 | 6.81 | -0.77 | -0.7 | 1.66 | 0.77 | 0 | 0 |
| GLRA1 | 292 | T | 12.90963636 | 13.88894114 | 0.262082198 | 6.994541522 | NA | NA | NA | NA | NA | NA | NA |
| GLRA1 | 293 | T | 14.77863636 | 9.929096434 | 0.154924691 | 2.745014294 | 7.21 | -0.77 | -0.7 | 1.66 | 0.77 | 0 | 0 |
| GLRA1 | 294 | Q | 15.54219192 | 9.263048148 | 0.136900734 | 0.953691036 | 8.34 | -0.7 | -1.46 | 11.73 | 0.94 | 1 | 1 |
| GLRA1 | 295 | S | 16.88713636 | 13.42749254 | 0.24959492 | 5.090179387 | NA | NA | NA | NA | NA | NA | NA |
| GLRA1 | 296 | S | 19.7519697 | 13.51847932 | 0.252057117 | 5.50734841 | NA | NA | NA | NA | NA | NA | NA |
| GLRA1 | 297 | G | 21.60738636 | 10.89330574 | 0.181017196 | 2.923192617 | 8.05 | -0.96 | -3.7 | 13.22 | 0.43 | 1 | 1 |
| GLRA1 | 298 | S | 24.02330303 | 13.43785976 | 0.249875468 | 5.486148708 | NA | NA | NA | NA | NA | NA | NA |
| GLRA1 | 299 | R | 24.59827273 | 8.647734328 | 0.120249703 | 0.714032913 | 8.05 | -0.96 | -3.7 | 13.22 | 0.43 | 1 | 1 |
| GLRA1 | 300 | A | 27.89163636 | 17.30552354 | 0.354538463 | 9.372662371 | NA | NA | NA | NA | NA | NA | NA |
| GLRA1 | 301 | S | 26.5134697 | 19.19683776 | 0.405719389 | 11.25519161 | NA | NA | NA | NA | NA | NA | NA |
| GLRA1 | 302 | L | 28.45051136 | 19.95439415 | 0.426219651 | 12.02210998 | NA | NA | NA | NA | NA | NA | NA |

| | | | | | | | | | | | | | |
|---|---|---|---|---|---|---|---|---|---|---|---|---|---|
| GLRA1 | 303 | P | 23.92606494 | 16.51270871 | 0.333084071 | 8.552880727 | NA | NA | NA | NA | NA | NA | NA |
| GLRA1 | 304 | K | 25.42285859 | 18.02215009 | 0.373931122 | 10.06733593 | NA | NA | NA | NA | NA | NA | NA |
| GLRA1 | 305 | V | 25.11906494 | 22.79602487 | 0.503117127 | 14.83135907 | NA | NA | NA | NA | NA | NA | NA |
| GLRA1 | 306 | S | 21.1304697 | 21.02615103 | 0.455222504 | 13.03433356 | NA | NA | NA | NA | NA | NA | NA |
| GLRA1 | 307 | Y | 19.95630303 | 13.93749497 | 0.263396115 | 5.92880223 | NA | NA | NA | NA | NA | NA | NA |
| GLRA1 | 308 | V | 21.15292208 | 22.49178968 | 0.494884207 | 14.50022512 | NA | NA | NA | NA | NA | NA | NA |
| GLRA1 | 309 | K | 19.21563636 | 24.59098138 | 0.551690512 | 16.31978929 | NA | NA | NA | NA | NA | NA | NA |
| GLRA1 | 310 | A | 16.30983636 | 20.9454295 | 0.453038096 | 12.61563538 | NA | NA | NA | NA | NA | NA | NA |
| GLRA1 | 311 | I | 16.55751136 | 20.45656445 | 0.4398089 | 12.16905726 | NA | NA | NA | NA | NA | NA | NA |
| GLRA1 | 312 | D | 14.73676136 | 25.03559712 | 0.563722276 | 17.90763289 | NA | NA | NA | NA | NA | NA | NA |
| GLRA1 | 313 | I | 12.29001136 | 24.04260136 | 0.53685078 | 17.20304561 | NA | NA | NA | NA | NA | NA | NA |
| GLRA1 | 314 | W | 12.75599351 | 16.52705642 | 0.333472335 | 10.13914276 | NA | NA | NA | NA | NA | NA | NA |
| GLRA1 | 316 | A | 6.950436364 | 23.31467757 | 0.517152407 | 17.24562497 | NA | NA | NA | NA | NA | NA | NA |
| GLRA1 | 317 | V | 6.461779221 | 20.75907659 | 0.447995193 | 14.9693492 | NA | NA | NA | NA | NA | NA | NA |
| GLRA1 | 318 | C | 5.85630303 | 18.01093251 | 0.373627562 | 12.33594622 | NA | NA | NA | NA | NA | NA | NA |
| GLRA1 | 319 | L | 3.638886364 | 20.98718335 | 0.454167998 | 16.36680184 | NA | NA | NA | NA | NA | NA | NA |
| GLRA1 | 320 | L | 1.802886364 | 22.35313548 | 0.49113208 | 18.20341443 | NA | NA | NA | NA | NA | NA | NA |
| GLRA1 | 321 | F | 1.119090909 | 15.48313311 | 0.305222687 | 11.42762645 | NA | NA | NA | NA | NA | NA | NA |
| GLRA1 | 322 | V | 0.389636364 | 15.38262647 | 0.302502873 | 10.9353965 | NA | NA | NA | NA | NA | NA | NA |
| GLRA1 | 323 | F | -1.063636364 | 20.35078097 | 0.436946289 | 15.73024881 | NA | NA | NA | NA | NA | NA | NA |
| GLRA1 | 324 | S | -2.99630303 | 20.23189596 | 0.433729138 | 16.9819476 | NA | NA | NA | NA | NA | NA | NA |
| GLRA1 | 325 | A | -3.512836364 | 15.84116631 | 0.314911437 | 12.58786659 | NA | NA | NA | NA | NA | NA | NA |
| GLRA1 | 326 | L | -3.589386364 | 15.40373844 | 0.303074185 | 12.13539869 | NA | NA | NA | NA | NA | NA | NA |
| GLRA1 | 327 | L | -7.185886364 | 19.77470933 | 0.421357193 | 15.3165298 | NA | NA | NA | NA | NA | NA | NA |
| GLRA1 | 328 | E | -9.265858586 | 16.73639205 | 0.339137175 | 12.24749577 | NA | NA | NA | NA | NA | NA | NA |
| GLRA1 | 329 | Y | -7.176969697 | 8.546707963 | 0.117515825 | 4.185278484 | 4.54 | -0.05 | -0.24 | 0.95 | -0.54 | 0 | 0 |
| GLRA1 | 330 | A | -9.704036364 | 16.52045708 | 0.33329375 | 11.78090518 | NA | NA | NA | NA | NA | NA | NA |
| GLRA1 | 331 | A | -12.42463636 | 18.21686016 | 0.379200178 | 13.64354884 | NA | NA | NA | NA | NA | NA | NA |
| GLRA1 | 332 | V | -14.45092208 | 12.91834684 | 0.235816909 | 7.639684636 | NA | NA | NA | NA | NA | NA | NA |
| GLRA1 | 333 | N | -14.73551136 | 14.28117495 | 0.272696452 | 9.00343333 | NA | NA | NA | NA | NA | NA | NA |
| GLRA1 | 334 | F | -13.73436364 | 18.74281476 | 0.393433055 | 13.46420971 | NA | NA | NA | NA | NA | NA | NA |
| GLRA1 | 335 | V | -18.59620779 | 17.9047695 | 0.370754681 | 12.41078641 | NA | NA | NA | NA | NA | NA | NA |
| GLRA1 | 336 | S | -19.65130303 | 20.2170485 | 0.43332735 | 12.18766968 | NA | NA | NA | NA | NA | NA | NA |

| | | | | | | | | | | | | | |
|---|---|---|---|---|---|---|---|---|---|---|---|---|---|
| GLRA1 | 337 | R | -17.28318182 | 22.75008547 | 0.501873959 | 17.24102425 | NA | NA | NA | NA | NA | NA | NA |
| GLRA1 | 338 | Q | -22.10452525 | 24.17922083 | 0.540547845 | 13.49165414 | NA | NA | NA | NA | NA | NA | NA |
| GLRA1 | 376 | G | -24.93313636 | 24.73086804 | 0.555475991 | 13.99250652 | NA | NA | NA | NA | NA | NA | NA |
| GLRA1 | 378 | G | -27.94988636 | 20.46565154 | 0.440054806 | 9.753066307 | NA | NA | NA | NA | NA | NA | NA |
| GLRA1 | 387 | G | -25.37813636 | 19.20677167 | 0.405988211 | 8.475536194 | NA | NA | NA | NA | NA | NA | NA |
| GLRA1 | 416 | F | -24.00118182 | 14.56133692 | 0.280277926 | 3.786745487 | 10.8 | 0.67 | 1.3 | 19.96 | -0.06 | 2 | -2 |
| GLRA1 | 417 | I | -26.37938636 | 13.25088397 | 0.244815709 | 2.526119759 | 10.8 | 0.67 | 1.3 | 19.96 | -0.06 | 2 | -2 |
| GLRA1 | 418 | Q | -26.78708081 | 15.97811622 | 0.318617444 | 5.270194618 | NA | NA | NA | NA | NA | NA | NA |
| GLRA1 | 419 | R | -19.55490909 | 16.42861984 | 0.330808539 | 5.697150356 | NA | NA | NA | NA | NA | NA | NA |
| GLRA1 | 420 | A | -21.04463636 | 12.6199815 | 0.227742833 | 1.872739563 | 10.8 | 0.67 | 1.3 | 19.96 | -0.06 | 2 | -2 |
| GLRA1 | 421 | K | -23.17230303 | 13.45614339 | 0.250370242 | 2.759531338 | 10.8 | 0.67 | 1.3 | 19.96 | -0.06 | 2 | -2 |
| GLRA1 | 422 | K | -17.18730303 | 14.61575417 | 0.281750513 | 9.183137734 | NA | NA | NA | NA | NA | NA | NA |
| GLRA1 | 423 | I | -18.77488636 | 20.4212328 | 0.438852789 | 15.00405741 | NA | NA | NA | NA | NA | NA | NA |
| GLRA1 | 424 | D | -14.51213636 | 20.53302778 | 0.441878077 | 15.33179208 | NA | NA | NA | NA | NA | NA | NA |
| GLRA1 | 425 | K | -14.71819192 | 22.2620831 | 0.488668108 | 17.00916015 | NA | NA | NA | NA | NA | NA | NA |
| GLRA1 | 426 | I | -10.42176136 | 19.21131057 | 0.406111038 | 14.55710712 | NA | NA | NA | NA | NA | NA | NA |
| GLRA1 | 427 | S | -10.18813636 | 23.11427775 | 0.51172938 | 18.30186631 | NA | NA | NA | NA | NA | NA | NA |
| GLRA1 | 428 | R | -12.34918182 | 25.37002473 | 0.572772234 | 20.54956805 | NA | NA | NA | NA | NA | NA | NA |
| GLRA1 | 429 | I | -6.908386364 | 23.2907502 | 0.516504908 | 18.74705601 | NA | NA | NA | NA | NA | NA | NA |
| GLRA1 | 430 | G | -4.878636364 | 23.65936039 | 0.526479882 | 20.45005322 | NA | NA | NA | NA | NA | NA | NA |
| GLRA1 | 431 | F | -6.535909091 | 24.5464098 | 0.550484359 | 21.60011751 | NA | NA | NA | NA | NA | NA | NA |
| GLRA1 | 432 | P | -4.205779221 | 27.71947925 | 0.636350911 | 24.55081173 | NA | NA | NA | NA | NA | NA | NA |
| GLRA1 | 434 | A | -0.134636364 | 24.75402367 | 0.556102606 | 20.11479168 | NA | NA | NA | NA | NA | NA | NA |
| GLRA1 | 435 | F | 0 | 28.78116394 | 0.665081201 | 24.43840918 | NA | NA | NA | NA | NA | NA | NA |
| GLRA1 | 436 | L | 0.403511364 | 29.93831158 | 0.696394817 | 25.85016194 | NA | NA | NA | NA | NA | NA | NA |
| GLRA1 | 437 | I | 3.082636364 | 24.42603971 | 0.54722702 | 20.12800598 | NA | NA | NA | NA | NA | NA | NA |
| GLRA1 | 438 | F | 4.086454545 | 28.14024801 | 0.647737351 | 23.55114845 | NA | NA | NA | NA | NA | NA | NA |
| GLRA1 | 439 | N | 4.370386364 | 31.06570302 | 0.726903199 | 26.13416618 | NA | NA | NA | NA | NA | NA | NA |
| GLRA1 | 441 | F | 9.238363636 | 24.61948761 | 0.552461921 | 17.87467154 | NA | NA | NA | NA | NA | NA | NA |
| GLRA1 | 442 | Y | 9.06130303 | 31.20662217 | 0.730716617 | 24.43926004 | NA | NA | NA | NA | NA | NA | NA |
| GLRA1 | 443 | W | 7.744922078 | 33.51621069 | 0.793216479 | 26.70668428 | NA | NA | NA | NA | NA | NA | NA |
| GLRA1 | 444 | I | 11.99588636 | 30.72069749 | 0.717566991 | 23.92028716 | NA | NA | NA | NA | NA | NA | NA |
| GLRA1 | 445 | I | 14.00888636 | 27.81893409 | 0.639042263 | 20.63184017 | NA | NA | NA | NA | NA | NA | NA |

| | | | | | | | | | | | | | |
|---|---|---|---|---|---|---|---|---|---|---|---|---|---|
| GLRA1 | 446 | Y | 14.2359697 | 33.4562893 | 0.791594945 | 26.2832584 | NA | NA | NA | NA | NA | NA | NA |
| GLRA1 | 447 | K | 13.7019697 | 35.02163663 | 0.833954867 | 27.81904044 | NA | NA | NA | NA | NA | NA | NA |
| GLRA1 | 448 | I | 18.11551136 | 30.84267119 | 0.720867726 | 22.50800552 | NA | NA | NA | NA | NA | NA | NA |
| GLRA1 | 449 | V | 20.11077922 | 32.9154314 | 0.776958768 | 24.60216057 | NA | NA | NA | NA | NA | NA | NA |
| GLRA1 | 450 | R | 18.91672727 | 36.89491916 | 0.884647837 | 28.56562119 | NA | NA | NA | NA | NA | NA | NA |
| GLRA1 | 451 | R | 20.501 | 37.02313127 | 0.888117389 | 28.72014017 | NA | NA | NA | NA | NA | NA | NA |
| GLRA1 | 452 | E | 20.40085859 | 40.81908016 | 0.990839705 | 32.50592137 | NA | NA | NA | NA | NA | NA | NA |
| GLRA1 | 453 | D | 16.99363636 | 41.15231233 | 0.999857314 | 32.81949597 | NA | NA | NA | NA | NA | NA | NA |
| GRIN1 | 25 | K | 138.3893651 | 52.8835153 | 0.682323885 | 51.86692312 | NA | NA | NA | NA | NA | NA | NA |
| GRIN1 | 26 | I | 136.7901429 | 54.41945448 | 0.703130109 | 53.35734701 | NA | NA | NA | NA | NA | NA | NA |
| GRIN1 | 27 | V | 132.6891429 | 52.13287066 | 0.672155461 | 51.08078708 | NA | NA | NA | NA | NA | NA | NA |
| GRIN1 | 28 | N | 132.1321429 | 54.67054244 | 0.706531411 | 53.5540807 | NA | NA | NA | NA | NA | NA | NA |
| GRIN1 | 29 | I | 129.1053929 | 54.69525328 | 0.70686615 | 53.60537748 | NA | NA | NA | NA | NA | NA | NA |
| GRIN1 | 30 | G | 126.7433929 | 56.98808765 | 0.73792547 | 55.84302756 | NA | NA | NA | NA | NA | NA | NA |
| GRIN1 | 31 | A | 125.6553429 | 59.16852292 | 0.767462202 | 57.8036783 | NA | NA | NA | NA | NA | NA | NA |
| GRIN1 | 32 | V | 122.854 | 60.43481042 | 0.784615656 | 59.10325385 | NA | NA | NA | NA | NA | NA | NA |
| GRIN1 | 33 | L | 124.5865179 | 62.48339716 | 0.812366335 | 61.16952885 | NA | NA | NA | NA | NA | NA | NA |
| GRIN1 | 34 | S | 124.0928095 | 65.98682543 | 0.859824669 | 64.57171294 | NA | NA | NA | NA | NA | NA | NA |
| GRIN1 | 35 | T | 126.6885714 | 67.80805057 | 0.884495449 | 66.48548032 | NA | NA | NA | NA | NA | NA | NA |
| GRIN1 | 36 | R | 131.1462338 | 68.98015675 | 0.900373099 | 67.62821722 | NA | NA | NA | NA | NA | NA | NA |
| GRIN1 | 37 | K | 130.8098095 | 67.51392178 | 0.880511106 | 66.17086355 | NA | NA | NA | NA | NA | NA | NA |
| GRIN1 | 38 | H | 127.0482429 | 64.18781073 | 0.835454756 | 62.84083256 | NA | NA | NA | NA | NA | NA | NA |
| GRIN1 | 39 | E | 130.9498095 | 65.06296669 | 0.847309842 | 63.76421753 | NA | NA | NA | NA | NA | NA | NA |
| GRIN1 | 40 | Q | 133.9454762 | 65.60274218 | 0.854621779 | 64.51821428 | NA | NA | NA | NA | NA | NA | NA |
| GRIN1 | 41 | M | 130.0975179 | 62.41924666 | 0.811497336 | 61.12749924 | NA | NA | NA | NA | NA | NA | NA |
| GRIN1 | 42 | F | 128.9205065 | 58.68362248 | 0.760893617 | 57.58190546 | NA | NA | NA | NA | NA | NA | NA |
| GRIN1 | 43 | R | 135.6357792 | 62.79536268 | 0.8165923 | 61.74076419 | NA | NA | NA | NA | NA | NA | NA |
| GRIN1 | 44 | E | 135.3104762 | 62.06887199 | 0.806751071 | 61.02689148 | NA | NA | NA | NA | NA | NA | NA |
| GRIN1 | 45 | A | 131.9385429 | 58.498719 | 0.758388868 | 57.45090071 | NA | NA | NA | NA | NA | NA | NA |
| GRIN1 | 46 | V | 134.025 | 58.77862452 | 0.762180539 | 57.75067076 | NA | NA | NA | NA | NA | NA | NA |
| GRIN1 | 47 | N | 137.2115179 | 60.15359039 | 0.780806178 | 59.1501958 | NA | NA | NA | NA | NA | NA | NA |
| GRIN1 | 48 | Q | 133.3166984 | 57.44193009 | 0.744073335 | 56.43743976 | NA | NA | NA | NA | NA | NA | NA |
| GRIN1 | 49 | A | 133.2733429 | 55.09662354 | 0.712303214 | 54.09668372 | NA | NA | NA | NA | NA | NA | NA |

| | | | | | | | | | | | | | |
|---|---|---|---|---|---|---|---|---|---|---|---|---|---|
| GRIN1 | 50 | N | 136.9545179 | 56.93993721 | 0.737273212 | 55.96835373 | NA | NA | NA | NA | NA | NA | NA |
| GRIN1 | 51 | K | 137.4100317 | 56.24404102 | 0.727846424 | 55.28190181 | NA | NA | NA | NA | NA | NA | NA |
| GRIN1 | 52 | R | 132.5295974 | 53.58044735 | 0.691764704 | 52.59214915 | NA | NA | NA | NA | NA | NA | NA |
| GRIN1 | 53 | H | 132.2181429 | 50.36309675 | 0.648181652 | 49.39351993 | NA | NA | NA | NA | NA | NA | NA |
| GRIN1 | 56 | W | 140.0293429 | 52.68221426 | 0.679597009 | 51.74337961 | NA | NA | NA | NA | NA | NA | NA |
| GRIN1 | 57 | K | 141.6793429 | 55.35210042 | 0.715763969 | 54.4083615 | NA | NA | NA | NA | NA | NA | NA |
| GRIN1 | 58 | I | 141.3613929 | 56.19347927 | 0.727161502 | 55.20439883 | NA | NA | NA | NA | NA | NA | NA |
| GRIN1 | 59 | Q | 136.2351429 | 51.29002182 | 0.660738015 | 50.31006749 | NA | NA | NA | NA | NA | NA | NA |
| GRIN1 | 60 | L | 135.3353929 | 54.75124369 | 0.70762461 | 53.7072424 | NA | NA | NA | NA | NA | NA | NA |
| GRIN1 | 61 | N | 136.3061429 | 58.58301059 | 0.759530703 | 57.52766108 | NA | NA | NA | NA | NA | NA | NA |
| GRIN1 | 62 | A | 134.0187429 | 60.8773652 | 0.790610616 | 59.79664757 | NA | NA | NA | NA | NA | NA | NA |
| GRIN1 | 63 | T | 131.2988571 | 59.72716657 | 0.775029732 | 58.59152553 | NA | NA | NA | NA | NA | NA | NA |
| GRIN1 | 64 | S | 129.8393095 | 62.8174232 | 0.816891137 | 61.67204617 | NA | NA | NA | NA | NA | NA | NA |
| GRIN1 | 65 | V | 127.5151429 | 63.54464697 | 0.826742296 | 62.20362457 | NA | NA | NA | NA | NA | NA | NA |
| GRIN1 | 66 | T | 125.8501429 | 65.23152195 | 0.849593135 | 63.90785355 | NA | NA | NA | NA | NA | NA | NA |
| GRIN1 | 67 | H | 121.2346429 | 62.68006963 | 0.815030511 | 61.24039847 | NA | NA | NA | NA | NA | NA | NA |
| GRIN1 | 68 | K | 122.1219206 | 65.96567775 | 0.859538197 | 64.52015476 | NA | NA | NA | NA | NA | NA | NA |
| GRIN1 | 69 | P | 120.0375714 | 69.09741388 | 0.901961494 | 67.62692312 | NA | NA | NA | NA | NA | NA | NA |
| GRIN1 | 70 | N | 118.9051429 | 66.08205955 | 0.861114734 | 64.61686674 | NA | NA | NA | NA | NA | NA | NA |
| GRIN1 | 71 | A | 116.1557429 | 63.90671581 | 0.831646973 | 62.42769727 | NA | NA | NA | NA | NA | NA | NA |
| GRIN1 | 72 | I | 117.9130179 | 64.81198763 | 0.843910016 | 63.33485435 | NA | NA | NA | NA | NA | NA | NA |
| GRIN1 | 73 | Q | 121.8049206 | 63.31179256 | 0.823587991 | 61.8492639 | NA | NA | NA | NA | NA | NA | NA |
| GRIN1 | 74 | M | 120.3191429 | 62.24429321 | 0.809127372 | 60.78885375 | NA | NA | NA | NA | NA | NA | NA |
| GRIN1 | 75 | A | 122.0273429 | 60.35185977 | 0.783491985 | 59.00690202 | NA | NA | NA | NA | NA | NA | NA |
| GRIN1 | 76 | L | 124.8128929 | 63.04727531 | 0.820004772 | 61.68816521 | NA | NA | NA | NA | NA | NA | NA |
| GRIN1 | 77 | S | 125.8093095 | 62.38875823 | 0.811084332 | 61.00292455 | NA | NA | NA | NA | NA | NA | NA |
| GRIN1 | 78 | V | 124.1544286 | 57.79708495 | 0.748884354 | 56.43115218 | NA | NA | NA | NA | NA | NA | NA |
| GRIN1 | 79 | C | 125.7734762 | 58.67416045 | 0.760765442 | 57.46055198 | NA | NA | NA | NA | NA | NA | NA |
| GRIN1 | 80 | E | 128.9540317 | 62.07856527 | 0.806882379 | 60.85845288 | NA | NA | NA | NA | NA | NA | NA |
| GRIN1 | 81 | D | 129.8873929 | 61.4212243 | 0.79797787 | 60.25359441 | NA | NA | NA | NA | NA | NA | NA |
| GRIN1 | 82 | L | 127.8355179 | 57.70792835 | 0.747676616 | 56.54068829 | NA | NA | NA | NA | NA | NA | NA |
| GRIN1 | 83 | I | 126.5285179 | 55.87703398 | 0.722874853 | 54.67002684 | NA | NA | NA | NA | NA | NA | NA |
| GRIN1 | 84 | S | 131.0591429 | 58.91749047 | 0.764061653 | 57.73521886 | NA | NA | NA | NA | NA | NA | NA |

| | | | | | | | | | | | | | |
|---|---|---|---|---|---|---|---|---|---|---|---|---|---|
| GRIN1 | 85 | S | 131.7299762 | 57.52714039 | 0.745227616 | 56.40147907 | NA | NA | NA | NA | NA | NA | NA |
| GRIN1 | 86 | Q | 131.1791429 | 53.99607265 | 0.697394871 | 52.86836606 | NA | NA | NA | NA | NA | NA | NA |
| GRIN1 | 87 | V | 127.566 | 52.93947458 | 0.683081923 | 51.77770391 | NA | NA | NA | NA | NA | NA | NA |
| GRIN1 | 88 | Y | 127.2465595 | 49.25369599 | 0.633153426 | 48.14152213 | NA | NA | NA | NA | NA | NA | NA |
| GRIN1 | 89 | A | 124.4525429 | 51.7597803 | 0.667101484 | 50.60580131 | NA | NA | NA | NA | NA | NA | NA |
| GRIN1 | 90 | I | 122.8912679 | 54.05148184 | 0.698145458 | 52.6874785 | NA | NA | NA | NA | NA | NA | NA |
| GRIN1 | 91 | L | 122.2002679 | 53.30305427 | 0.688007067 | 51.97328782 | NA | NA | NA | NA | NA | NA | NA |
| GRIN1 | 92 | V | 119.1655714 | 56.8816432 | 0.736483546 | 55.46387953 | NA | NA | NA | NA | NA | NA | NA |
| GRIN1 | 93 | S | 118.4311429 | 59.19426006 | 0.767810844 | 57.76865936 | NA | NA | NA | NA | NA | NA | NA |
| GRIN1 | 94 | H | 114.5251429 | 58.73553528 | 0.761596841 | 57.2235228 | NA | NA | NA | NA | NA | NA | NA |
| GRIN1 | 95 | P | 116.9961429 | 63.27314885 | 0.823064513 | 61.82590754 | NA | NA | NA | NA | NA | NA | NA |
| GRIN1 | 96 | P | 118.0735429 | 66.87230848 | 0.871819648 | 65.51663591 | NA | NA | NA | NA | NA | NA | NA |
| GRIN1 | 97 | T | 114.2047429 | 67.09346613 | 0.874815506 | 65.58151027 | NA | NA | NA | NA | NA | NA | NA |
| GRIN1 | 98 | P | 112.6215429 | 63.63589774 | 0.827978402 | 62.12439097 | NA | NA | NA | NA | NA | NA | NA |
| GRIN1 | 99 | N | 109.3247429 | 63.93701232 | 0.832057377 | 62.42757563 | NA | NA | NA | NA | NA | NA | NA |
| GRIN1 | 100 | D | 109.9649429 | 62.63698571 | 0.814446885 | 61.12511857 | NA | NA | NA | NA | NA | NA | NA |
| GRIN1 | 101 | H | 109.1329429 | 62.49821057 | 0.812567001 | 60.98562865 | NA | NA | NA | NA | NA | NA | NA |
| GRIN1 | 102 | F | 112.7475429 | 61.93228693 | 0.804900855 | 60.50693659 | NA | NA | NA | NA | NA | NA | NA |
| GRIN1 | 103 | T | 110.9542857 | 58.66277346 | 0.760611191 | 57.15020164 | NA | NA | NA | NA | NA | NA | NA |
| GRIN1 | 104 | P | 114.053 | 56.20976787 | 0.727382151 | 54.74638351 | NA | NA | NA | NA | NA | NA | NA |
| GRIN1 | 105 | T | 112.5081429 | 55.41032074 | 0.716552636 | 53.94024405 | NA | NA | NA | NA | NA | NA | NA |
| GRIN1 | 106 | P | 115.529 | 56.24571868 | 0.72786915 | 54.78107739 | NA | NA | NA | NA | NA | NA | NA |
| GRIN1 | 107 | V | 118.4438571 | 54.60772558 | 0.705680477 | 53.16031853 | NA | NA | NA | NA | NA | NA | NA |
| GRIN1 | 108 | S | 115.2559762 | 51.57968758 | 0.664661902 | 50.12307618 | NA | NA | NA | NA | NA | NA | NA |
| GRIN1 | 109 | Y | 112.5284762 | 51.48174625 | 0.663335163 | 49.98204796 | NA | NA | NA | NA | NA | NA | NA |
| GRIN1 | 110 | T | 118.7367143 | 53.09534271 | 0.685193353 | 51.64893641 | NA | NA | NA | NA | NA | NA | NA |
| GRIN1 | 111 | A | 118.3585429 | 49.68885559 | 0.639048209 | 48.32781526 | NA | NA | NA | NA | NA | NA | NA |
| GRIN1 | 112 | G | 115.8471429 | 48.33076845 | 0.620651214 | 46.86837715 | NA | NA | NA | NA | NA | NA | NA |
| GRIN1 | 113 | F | 117.5035065 | 49.06673967 | 0.630620868 | 47.68171189 | NA | NA | NA | NA | NA | NA | NA |
| GRIN1 | 114 | Y | 120.7750595 | 47.17833784 | 0.605040094 | 45.94844726 | NA | NA | NA | NA | NA | NA | NA |
| GRIN1 | 115 | R | 115.8032338 | 41.17627223 | 0.523734581 | 39.80259892 | NA | NA | NA | NA | NA | NA | NA |
| GRIN1 | 116 | I | 120.2870179 | 47.9134032 | 0.614997477 | 46.5439083 | NA | NA | NA | NA | NA | NA | NA |
| GRIN1 | 117 | P | 120.124 | 46.96277474 | 0.602120022 | 45.5971391 | NA | NA | NA | NA | NA | NA | NA |

| | | | | | | | | | | | | | |
|---|---|---|---|---|---|---|---|---|---|---|---|---|---|
| GRIN1 | 118 | V | 118.155 | 48.99808925 | 0.629690912 | 47.69384395 | NA | NA | NA | NA | NA | NA | NA |
| GRIN1 | 119 | L | 118.1607679 | 47.81401426 | 0.61365113 | 46.4111751 | NA | NA | NA | NA | NA | NA | NA |
| GRIN1 | 120 | G | 115.4331429 | 51.20435583 | 0.659577562 | 49.77730567 | NA | NA | NA | NA | NA | NA | NA |
| GRIN1 | 121 | L | 117.8343929 | 53.99426041 | 0.697370322 | 52.56893308 | NA | NA | NA | NA | NA | NA | NA |
| GRIN1 | 122 | T | 114.7091429 | 55.65138741 | 0.719818187 | 54.30576427 | NA | NA | NA | NA | NA | NA | NA |
| GRIN1 | 123 | T | 111.5352857 | 53.33031544 | 0.688376353 | 51.81830741 | NA | NA | NA | NA | NA | NA | NA |
| GRIN1 | 124 | R | 107.7575065 | 52.44527301 | 0.676387343 | 50.93685473 | NA | NA | NA | NA | NA | NA | NA |
| GRIN1 | 125 | M | 106.2232679 | 51.63124003 | 0.665360244 | 50.11933592 | NA | NA | NA | NA | NA | NA | NA |
| GRIN1 | 126 | S | 103.8248095 | 48.10900287 | 0.61764712 | 46.59729854 | NA | NA | NA | NA | NA | NA | NA |
| GRIN1 | 127 | I | 104.0586429 | 48.79966133 | 0.627002956 | 47.28717718 | NA | NA | NA | NA | NA | NA | NA |
| GRIN1 | 128 | Y | 109.1541429 | 47.75558554 | 0.612859639 | 46.24261104 | NA | NA | NA | NA | NA | NA | NA |
| GRIN1 | 129 | S | 105.7088095 | 44.17324502 | 0.56433234 | 42.66062567 | NA | NA | NA | NA | NA | NA | NA |
| GRIN1 | 130 | D | 104.3290179 | 43.13238805 | 0.550232626 | 41.62027834 | NA | NA | NA | NA | NA | NA | NA |
| GRIN1 | 131 | K | 108.0529206 | 39.40440211 | 0.499732377 | 37.95587233 | NA | NA | NA | NA | NA | NA | NA |
| GRIN1 | 132 | S | 105.8559762 | 42.31707629 | 0.539188205 | 40.80503947 | NA | NA | NA | NA | NA | NA | NA |
| GRIN1 | 133 | I | 106.2888929 | 44.9440024 | 0.574773216 | 43.44023375 | NA | NA | NA | NA | NA | NA | NA |
| GRIN1 | 134 | H | 109.0047429 | 45.28984922 | 0.579458145 | 43.81772002 | NA | NA | NA | NA | NA | NA | NA |
| GRIN1 | 135 | L | 111.9050179 | 43.41583602 | 0.554072284 | 41.96062764 | NA | NA | NA | NA | NA | NA | NA |
| GRIN1 | 136 | S | 114.8956429 | 45.19498793 | 0.57817313 | 43.75553322 | NA | NA | NA | NA | NA | NA | NA |
| GRIN1 | 137 | F | 113.7286883 | 46.55129449 | 0.596546005 | 45.10836167 | NA | NA | NA | NA | NA | NA | NA |
| GRIN1 | 138 | L | 113.7917679 | 43.36455305 | 0.553377592 | 41.94632338 | NA | NA | NA | NA | NA | NA | NA |
| GRIN1 | 139 | R | 111.279961 | 47.72045603 | 0.612383766 | 46.2911808 | NA | NA | NA | NA | NA | NA | NA |
| GRIN1 | 140 | T | 115.0724286 | 50.09335711 | 0.64452769 | 48.67408405 | NA | NA | NA | NA | NA | NA | NA |
| GRIN1 | 141 | V | 111.6994286 | 50.46440976 | 0.649554063 | 48.95240559 | NA | NA | NA | NA | NA | NA | NA |
| GRIN1 | 142 | P | 108.3121429 | 48.65337771 | 0.625021361 | 47.14370178 | NA | NA | NA | NA | NA | NA | NA |
| GRIN1 | 143 | P | 105.4008571 | 47.73866269 | 0.612630398 | 46.23030419 | NA | NA | NA | NA | NA | NA | NA |
| GRIN1 | 144 | Y | 103.2555595 | 49.60414689 | 0.637900723 | 48.14357832 | NA | NA | NA | NA | NA | NA | NA |
| GRIN1 | 145 | S | 100.9783095 | 46.02558537 | 0.589424615 | 44.57021407 | NA | NA | NA | NA | NA | NA | NA |
| GRIN1 | 146 | H | 102.9265429 | 43.48452869 | 0.555002813 | 42.03326131 | NA | NA | NA | NA | NA | NA | NA |
| GRIN1 | 147 | Q | 101.0764762 | 48.46050056 | 0.622408598 | 46.9966805 | NA | NA | NA | NA | NA | NA | NA |
| GRIN1 | 148 | S | 97.24080952 | 45.34571292 | 0.580214889 | 43.87583915 | NA | NA | NA | NA | NA | NA | NA |
| GRIN1 | 149 | S | 98.48497619 | 44.07690707 | 0.563027321 | 42.60700078 | NA | NA | NA | NA | NA | NA | NA |
| GRIN1 | 150 | V | 99.10942857 | 47.05078502 | 0.603312231 | 45.58070569 | NA | NA | NA | NA | NA | NA | NA |

| | | | | | | | | | | | | | |
|---|---|---|---|---|---|---|---|---|---|---|---|---|---|
| GRIN1 | 151 | W | 95.90421429 | 47.77274741 | 0.613092118 | 46.30198641 | NA | NA | NA | NA | NA | NA | NA |
| GRIN1 | 152 | F | 93.6585974 | 42.50391541 | 0.541719175 | 41.03478081 | NA | NA | NA | NA | NA | NA | NA |
| GRIN1 | 153 | E | 98.33114286 | 45.74449781 | 0.585616931 | 44.274834 | NA | NA | NA | NA | NA | NA | NA |
| GRIN1 | 154 | M | 95.06964286 | 48.708279 | 0.625765068 | 47.23738582 | NA | NA | NA | NA | NA | NA | NA |
| GRIN1 | 155 | M | 91.02901786 | 45.79217393 | 0.586262764 | 44.94625938 | NA | NA | NA | NA | NA | NA | NA |
| GRIN1 | 156 | R | 94.31514286 | 40.64179688 | 0.516494442 | 39.23979583 | NA | NA | NA | NA | NA | NA | NA |
| GRIN1 | 157 | V | 93.99 | 48.94875453 | 0.629022611 | 47.53391776 | NA | NA | NA | NA | NA | NA | NA |
| GRIN1 | 158 | Y | 92.55522619 | 50.85752157 | 0.654879256 | 49.43877631 | NA | NA | NA | NA | NA | NA | NA |
| GRIN1 | 159 | S | 87.49680952 | 48.79548023 | 0.626946318 | 44.82794358 | NA | NA | NA | NA | NA | NA | NA |
| GRIN1 | 160 | W | 86.95828571 | 45.9176923 | 0.587963068 | 41.93911238 | NA | NA | NA | NA | NA | NA | NA |
| GRIN1 | 161 | N | 83.44126786 | 43.71927233 | 0.55818271 | 39.72853864 | NA | NA | NA | NA | NA | NA | NA |
| GRIN1 | 162 | H | 81.83504286 | 44.95888127 | 0.574974769 | 40.92035338 | NA | NA | NA | NA | NA | NA | NA |
| GRIN1 | 163 | I | 86.00901786 | 45.82631861 | 0.586725297 | 41.83330052 | NA | NA | NA | NA | NA | NA | NA |
| GRIN1 | 164 | I | 85.46739286 | 50.35537045 | 0.648076989 | 46.33408224 | NA | NA | NA | NA | NA | NA | NA |
| GRIN1 | 165 | L | 88.30051786 | 47.37939003 | 0.607763599 | 43.36513433 | NA | NA | NA | NA | NA | NA | NA |
| GRIN1 | 166 | L | 89.05614286 | 52.89686793 | 0.682504763 | 48.87090911 | NA | NA | NA | NA | NA | NA | NA |
| GRIN1 | 167 | V | 91.28928571 | 50.84716032 | 0.6547389 | 49.65500445 | NA | NA | NA | NA | NA | NA | NA |
| GRIN1 | 168 | S | 94.21864286 | 53.10924158 | 0.685381631 | 51.63912286 | NA | NA | NA | NA | NA | NA | NA |
| GRIN1 | 169 | D | 92.25451786 | 53.12962543 | 0.685657755 | 52.20590173 | NA | NA | NA | NA | NA | NA | NA |
| GRIN1 | 170 | D | 96.68064286 | 50.4314834 | 0.649108035 | 48.96553947 | NA | NA | NA | NA | NA | NA | NA |
| GRIN1 | 171 | H | 98.79234286 | 48.17747094 | 0.618574606 | 46.71606183 | NA | NA | NA | NA | NA | NA | NA |
| GRIN1 | 172 | E | 100.0865873 | 49.05935939 | 0.630520893 | 47.59450077 | NA | NA | NA | NA | NA | NA | NA |
| GRIN1 | 173 | G | 95.35789286 | 48.62797037 | 0.624677187 | 47.15787342 | NA | NA | NA | NA | NA | NA | NA |
| GRIN1 | 174 | R | 93.26441558 | 45.31902465 | 0.579853363 | 43.84895943 | NA | NA | NA | NA | NA | NA | NA |
| GRIN1 | 175 | A | 96.86014286 | 44.43821369 | 0.567921673 | 42.97027709 | NA | NA | NA | NA | NA | NA | NA |
| GRIN1 | 176 | A | 95.62114286 | 45.48947885 | 0.582162379 | 44.01936306 | NA | NA | NA | NA | NA | NA | NA |
| GRIN1 | 177 | Q | 91.54603175 | 43.58851595 | 0.556411451 | 42.11802234 | NA | NA | NA | NA | NA | NA | NA |
| GRIN1 | 178 | K | 93.64936508 | 40.56873911 | 0.515504783 | 39.09852113 | NA | NA | NA | NA | NA | NA | NA |
| GRIN1 | 179 | R | 98.28105195 | 41.27681694 | 0.525096586 | 39.80662263 | NA | NA | NA | NA | NA | NA | NA |
| GRIN1 | 180 | L | 92.65901786 | 41.74322168 | 0.531414623 | 40.2724791 | NA | NA | NA | NA | NA | NA | NA |
| GRIN1 | 181 | E | 89.62625397 | 38.77506668 | 0.491207238 | 37.34868758 | NA | NA | NA | NA | NA | NA | NA |
| GRIN1 | 182 | T | 93.09371429 | 36.92810644 | 0.466187843 | 35.45745366 | NA | NA | NA | NA | NA | NA | NA |
| GRIN1 | 183 | L | 94.15551786 | 38.78160132 | 0.491295758 | 37.31144361 | NA | NA | NA | NA | NA | NA | NA |

| | | | | | | | | | | | | | |
|---|---|---|---|---|---|---|---|---|---|---|---|---|---|
| GRIN1 | 184 | L | 88.97589286 | 37.96008368 | 0.48016727 | 37.01261289 | NA | NA | NA | NA | NA | NA | NA |
| GRIN1 | 185 | E | 87.73880952 | 34.11781297 | 0.42811889 | 30.1541144 | NA | NA | NA | NA | NA | NA | NA |
| GRIN1 | 186 | E | 92.84692063 | 33.9897024 | 0.426383472 | 32.51962668 | NA | NA | NA | NA | NA | NA | NA |
| GRIN1 | 187 | R | 91.66596104 | 36.16837794 | 0.455896367 | 34.75127766 | NA | NA | NA | NA | NA | NA | NA |
| GRIN1 | 188 | E | 85.43536508 | 31.23711053 | 0.38909616 | 27.26945996 | NA | NA | NA | NA | NA | NA | NA |
| GRIN1 | 189 | S | 86.37130952 | 38.21590253 | 0.483632658 | 34.23643798 | NA | NA | NA | NA | NA | NA | NA |
| GRIN1 | 190 | K | 82.58892063 | 34.27097273 | 0.430193632 | 30.23607082 | NA | NA | NA | NA | NA | NA | NA |
| GRIN1 | 191 | A | 83.57734286 | 41.99267404 | 0.534793769 | 37.967304 | NA | NA | NA | NA | NA | NA | NA |
| GRIN1 | 192 | E | 80.1255873 | 44.76920406 | 0.572405353 | 40.7225784 | NA | NA | NA | NA | NA | NA | NA |
| GRIN1 | 193 | K | 80.76592063 | 46.67540286 | 0.598227209 | 42.61416594 | NA | NA | NA | NA | NA | NA | NA |
| GRIN1 | 194 | V | 85.12028571 | 46.28158582 | 0.592892463 | 42.23496343 | NA | NA | NA | NA | NA | NA | NA |
| GRIN1 | 195 | L | 84.83814286 | 50.59075929 | 0.651265627 | 46.52837702 | NA | NA | NA | NA | NA | NA | NA |
| GRIN1 | 196 | Q | 86.80247619 | 49.04642214 | 0.630345641 | 44.95832555 | NA | NA | NA | NA | NA | NA | NA |
| GRIN1 | 197 | F | 88.97050649 | 55.16573862 | 0.713239464 | 51.09935909 | NA | NA | NA | NA | NA | NA | NA |
| GRIN1 | 198 | D | 90.43276786 | 57.24181807 | 0.741362567 | 53.15462934 | NA | NA | NA | NA | NA | NA | NA |
| GRIN1 | 199 | P | 94.28328571 | 59.31768597 | 0.769482803 | 57.84755 | NA | NA | NA | NA | NA | NA | NA |
| GRIN1 | 200 | G | 93.19164286 | 63.42394358 | 0.825107217 | 61.95377345 | NA | NA | NA | NA | NA | NA | NA |
| GRIN1 | 201 | T | 89.95071429 | 62.10084449 | 0.807184179 | 58.0249429 | NA | NA | NA | NA | NA | NA | NA |
| GRIN1 | 202 | K | 89.84236508 | 65.76099021 | 0.856765447 | 61.68249936 | NA | NA | NA | NA | NA | NA | NA |
| GRIN1 | 203 | N | 84.57301786 | 63.74959459 | 0.829518569 | 59.65261409 | NA | NA | NA | NA | NA | NA | NA |
| GRIN1 | 204 | V | 85.01385714 | 61.04743422 | 0.792914415 | 56.97703072 | NA | NA | NA | NA | NA | NA | NA |
| GRIN1 | 205 | T | 81.42514286 | 62.1160287 | 0.807389868 | 58.02207601 | NA | NA | NA | NA | NA | NA | NA |
| GRIN1 | 206 | A | 79.27294286 | 59.02281202 | 0.765488365 | 54.91649243 | NA | NA | NA | NA | NA | NA | NA |
| GRIN1 | 207 | L | 81.94826786 | 56.71263462 | 0.734194112 | 52.60978391 | NA | NA | NA | NA | NA | NA | NA |
| GRIN1 | 208 | L | 82.93726786 | 59.35585531 | 0.769999855 | 55.27780364 | NA | NA | NA | NA | NA | NA | NA |
| GRIN1 | 209 | M | 77.80676786 | 58.47326786 | 0.7580441 | 54.37530455 | NA | NA | NA | NA | NA | NA | NA |
| GRIN1 | 210 | E | 79.1085873 | 53.37747313 | 0.689015164 | 49.27868982 | NA | NA | NA | NA | NA | NA | NA |
| GRIN1 | 211 | A | 79.86874286 | 55.63433195 | 0.719587149 | 51.55749351 | NA | NA | NA | NA | NA | NA | NA |
| GRIN1 | 212 | K | 77.86254286 | 57.61134413 | 0.746368261 | 53.52629794 | NA | NA | NA | NA | NA | NA | NA |
| GRIN1 | 213 | E | 75.10080952 | 54.49988079 | 0.704219584 | 50.40069047 | NA | NA | NA | NA | NA | NA | NA |
| GRIN1 | 214 | L | 77.52764286 | 51.34114534 | 0.661430548 | 47.24719587 | NA | NA | NA | NA | NA | NA | NA |
| GRIN1 | 215 | E | 76.65725397 | 48.63861086 | 0.624821325 | 44.56883271 | NA | NA | NA | NA | NA | NA | NA |
| GRIN1 | 216 | A | 81.18654286 | 50.98475046 | 0.656602731 | 46.95175863 | NA | NA | NA | NA | NA | NA | NA |

| | | | | | | | | | | | | | |
|---|---|---|---|---|---|---|---|---|---|---|---|---|---|
| GRIN1 | 217 | R | 81.43614286 | 54.0107174 | 0.697593252 | 49.98329275 | NA | NA | NA | NA | NA | NA | NA |
| GRIN1 | 218 | V | 86.81542857 | 52.64787472 | 0.679131837 | 48.63660594 | NA | NA | NA | NA | NA | NA | NA |
| GRIN1 | 219 | I | 88.10376786 | 54.63123068 | 0.705998883 | 50.62627561 | NA | NA | NA | NA | NA | NA | NA |
| GRIN1 | 220 | I | 91.50226786 | 50.75129249 | 0.65344025 | 49.29714674 | NA | NA | NA | NA | NA | NA | NA |
| GRIN1 | 221 | L | 93.13714286 | 55.32712121 | 0.715425594 | 53.85691233 | NA | NA | NA | NA | NA | NA | NA |
| GRIN1 | 222 | S | 95.00280952 | 52.92519647 | 0.682888508 | 51.4550229 | NA | NA | NA | NA | NA | NA | NA |
| GRIN1 | 223 | A | 97.06634286 | 56.56733846 | 0.732225894 | 55.09738484 | NA | NA | NA | NA | NA | NA | NA |
| GRIN1 | 224 | S | 98.67114286 | 59.35659176 | 0.770009831 | 57.88838677 | NA | NA | NA | NA | NA | NA | NA |
| GRIN1 | 225 | E | 101.4010317 | 62.43947336 | 0.811771332 | 60.973789 | NA | NA | NA | NA | NA | NA | NA |
| GRIN1 | 226 | D | 98.55839286 | 64.90184395 | 0.845127233 | 63.43374395 | NA | NA | NA | NA | NA | NA | NA |
| GRIN1 | 227 | D | 94.62376786 | 59.82300949 | 0.776328045 | 58.35287013 | NA | NA | NA | NA | NA | NA | NA |
| GRIN1 | 228 | A | 94.97334286 | 62.37263636 | 0.810865941 | 60.90253069 | NA | NA | NA | NA | NA | NA | NA |
| GRIN1 | 229 | A | 94.96254286 | 65.3357973 | 0.851005676 | 63.86564311 | NA | NA | NA | NA | NA | NA | NA |
| GRIN1 | 230 | T | 92.66571429 | 65.79458453 | 0.857220525 | 64.58211935 | NA | NA | NA | NA | NA | NA | NA |
| GRIN1 | 231 | V | 91.09342857 | 61.53419056 | 0.79950814 | 57.50385154 | NA | NA | NA | NA | NA | NA | NA |
| GRIN1 | 232 | Y | 93.93605952 | 61.12654574 | 0.793986079 | 57.86715244 | NA | NA | NA | NA | NA | NA | NA |
| GRIN1 | 233 | R | 91.66587013 | 67.37725834 | 0.878659828 | 63.35572565 | NA | NA | NA | NA | NA | NA | NA |
| GRIN1 | 234 | A | 87.77554286 | 66.23815342 | 0.863229222 | 62.18798953 | NA | NA | NA | NA | NA | NA | NA |
| GRIN1 | 235 | A | 86.69274286 | 63.64287006 | 0.828072851 | 59.60671829 | NA | NA | NA | NA | NA | NA | NA |
| GRIN1 | 236 | A | 87.12974286 | 66.48715545 | 0.866602267 | 62.4553562 | NA | NA | NA | NA | NA | NA | NA |
| GRIN1 | 237 | M | 84.93864286 | 68.27470291 | 0.890816841 | 64.21981414 | NA | NA | NA | NA | NA | NA | NA |
| GRIN1 | 238 | L | 82.61989286 | 63.71451423 | 0.829043361 | 59.64644866 | NA | NA | NA | NA | NA | NA | NA |
| GRIN1 | 239 | N | 82.42314286 | 64.69522877 | 0.842328371 | 60.64913339 | NA | NA | NA | NA | NA | NA | NA |
| GRIN1 | 240 | M | 84.12964286 | 59.63218522 | 0.77374309 | 55.57291108 | NA | NA | NA | NA | NA | NA | NA |
| GRIN1 | 241 | T | 87.68571429 | 64.3848898 | 0.83812444 | 60.36218419 | NA | NA | NA | NA | NA | NA | NA |
| GRIN1 | 242 | G | 85.77064286 | 63.62504778 | 0.827831426 | 59.61502301 | NA | NA | NA | NA | NA | NA | NA |
| GRIN1 | 243 | S | 84.27114286 | 63.04790875 | 0.820013353 | 59.0346648 | NA | NA | NA | NA | NA | NA | NA |
| GRIN1 | 244 | G | 83.86014286 | 58.94955264 | 0.764495975 | 54.92994859 | NA | NA | NA | NA | NA | NA | NA |
| GRIN1 | 245 | Y | 84.21539286 | 58.91706266 | 0.764055858 | 54.91802184 | NA | NA | NA | NA | NA | NA | NA |
| GRIN1 | 246 | V | 88.62042857 | 56.01970152 | 0.724807464 | 52.02525069 | NA | NA | NA | NA | NA | NA | NA |
| GRIN1 | 247 | W | 90.89485714 | 57.90912154 | 0.75040203 | 56.43832708 | NA | NA | NA | NA | NA | NA | NA |
| GRIN1 | 248 | L | 93.70164286 | 52.82263098 | 0.68149913 | 51.35187418 | NA | NA | NA | NA | NA | NA | NA |
| GRIN1 | 249 | V | 97.26971429 | 56.18792791 | 0.727086302 | 54.71758727 | NA | NA | NA | NA | NA | NA | NA |

| | | | | | | | | | | | | | |
|---|---|---|---|---|---|---|---|---|---|---|---|---|---|
| GRIN1 | 250 | G | 99.95189286 | 55.99547477 | 0.724479282 | 54.52587444 | NA | NA | NA | NA | NA | NA | NA |
| GRIN1 | 251 | E | 104.5120317 | 55.71350812 | 0.72065969 | 54.25604313 | NA | NA | NA | NA | NA | NA | NA |
| GRIN1 | 252 | R | 103.7217792 | 57.1918493 | 0.740685677 | 55.74195172 | NA | NA | NA | NA | NA | NA | NA |
| GRIN1 | 253 | E | 98.72103175 | 57.80756737 | 0.749026351 | 56.33753493 | NA | NA | NA | NA | NA | NA | NA |
| GRIN1 | 254 | I | 102.4897679 | 59.562458 | 0.772798548 | 58.09349457 | NA | NA | NA | NA | NA | NA | NA |
| GRIN1 | 255 | S | 104.0364762 | 63.83419627 | 0.830664605 | 62.37306576 | NA | NA | NA | NA | NA | NA | NA |
| GRIN1 | 256 | G | 104.1176429 | 67.35468702 | 0.878354071 | 65.89236724 | NA | NA | NA | NA | NA | NA | NA |
| GRIN1 | 257 | N | 101.7801429 | 68.7392444 | 0.897109639 | 67.2694923 | NA | NA | NA | NA | NA | NA | NA |
| GRIN1 | 258 | A | 99.86914286 | 65.08065449 | 0.847549446 | 63.61082103 | NA | NA | NA | NA | NA | NA | NA |
| GRIN1 | 259 | L | 102.2046429 | 66.1418895 | 0.861925206 | 64.673204 | NA | NA | NA | NA | NA | NA | NA |
| GRIN1 | 260 | R | 101.8926883 | 69.87416636 | 0.912483581 | 68.40421747 | NA | NA | NA | NA | NA | NA | NA |
| GRIN1 | 261 | Y | 96.92580952 | 69.34401645 | 0.905302035 | 67.87948361 | NA | NA | NA | NA | NA | NA | NA |
| GRIN1 | 262 | A | 95.42454286 | 66.2828463 | 0.863834643 | 64.81245873 | NA | NA | NA | NA | NA | NA | NA |
| GRIN1 | 263 | P | 93.45728571 | 65.97535674 | 0.859669311 | 64.80674354 | NA | NA | NA | NA | NA | NA | NA |
| GRIN1 | 264 | D | 96.11376786 | 66.61180965 | 0.868290865 | 65.14089502 | NA | NA | NA | NA | NA | NA | NA |
| GRIN1 | 265 | G | 95.39814286 | 62.95180026 | 0.818711443 | 61.48087337 | NA | NA | NA | NA | NA | NA | NA |
| GRIN1 | 266 | I | 95.75414286 | 61.24940219 | 0.795650324 | 59.77886148 | NA | NA | NA | NA | NA | NA | NA |
| GRIN1 | 267 | L | 96.50414286 | 57.05040435 | 0.738769628 | 55.57943728 | NA | NA | NA | NA | NA | NA | NA |
| GRIN1 | 268 | G | 100.4678929 | 57.79660814 | 0.748877895 | 56.32649422 | NA | NA | NA | NA | NA | NA | NA |
| GRIN1 | 269 | L | 102.4207679 | 53.49122378 | 0.690556059 | 52.02132072 | NA | NA | NA | NA | NA | NA | NA |
| GRIN1 | 270 | Q | 107.5258095 | 56.03711151 | 0.725043304 | 54.57915569 | NA | NA | NA | NA | NA | NA | NA |
| GRIN1 | 271 | L | 108.6888929 | 52.36401564 | 0.67528661 | 50.85994447 | NA | NA | NA | NA | NA | NA | NA |
| GRIN1 | 272 | I | 111.6041429 | 55.86597652 | 0.722725066 | 54.35940042 | NA | NA | NA | NA | NA | NA | NA |
| GRIN1 | 273 | N | 115.5848929 | 57.73483448 | 0.748041093 | 56.22233838 | NA | NA | NA | NA | NA | NA | NA |
| GRIN1 | 274 | G | 113.8453929 | 59.01289623 | 0.765354043 | 57.50123936 | NA | NA | NA | NA | NA | NA | NA |
| GRIN1 | 275 | K | 114.6340317 | 60.93970845 | 0.791455134 | 59.42777972 | NA | NA | NA | NA | NA | NA | NA |
| GRIN1 | 276 | N | 119.7402679 | 60.77496987 | 0.789223543 | 59.46383202 | NA | NA | NA | NA | NA | NA | NA |
| GRIN1 | 277 | E | 120.6819206 | 59.64540437 | 0.77392216 | 58.25207456 | NA | NA | NA | NA | NA | NA | NA |
| GRIN1 | 278 | S | 123.9911429 | 60.06631454 | 0.779623917 | 58.75644534 | NA | NA | NA | NA | NA | NA | NA |
| GRIN1 | 279 | A | 122.5081429 | 57.13967472 | 0.739978907 | 55.83799821 | NA | NA | NA | NA | NA | NA | NA |
| GRIN1 | 280 | H | 119.1969429 | 55.36492103 | 0.71593764 | 54.061061 | NA | NA | NA | NA | NA | NA | NA |
| GRIN1 | 281 | I | 123.9757679 | 56.53866433 | 0.731837467 | 55.20385716 | NA | NA | NA | NA | NA | NA | NA |
| GRIN1 | 282 | S | 125.4688095 | 55.49079053 | 0.717642701 | 54.14222121 | NA | NA | NA | NA | NA | NA | NA |

| | | | | | | | | | | | | | |
|---|---|---|---|---|---|---|---|---|---|---|---|---|---|
| GRIN1 | 283 | D | 122.6643929 | 52.21338053 | 0.673246068 | 50.87581735 | NA | NA | NA | NA | NA | NA | NA |
| GRIN1 | 284 | A | 122.9379429 | 52.05439583 | 0.671092421 | 50.7009892 | NA | NA | NA | NA | NA | NA | NA |
| GRIN1 | 285 | V | 126.7895714 | 52.99607536 | 0.683848652 | 51.75766792 | NA | NA | NA | NA | NA | NA | NA |
| GRIN1 | 286 | G | 126.1418929 | 50.40911339 | 0.648805005 | 49.34974923 | NA | NA | NA | NA | NA | NA | NA |
| GRIN1 | 287 | V | 122.974 | 47.79495359 | 0.613392929 | 46.44769901 | NA | NA | NA | NA | NA | NA | NA |
| GRIN1 | 288 | V | 125.4738571 | 49.39869582 | 0.63511763 | 48.32358158 | NA | NA | NA | NA | NA | NA | NA |
| GRIN1 | 289 | A | 128.3667429 | 49.24296496 | 0.63300806 | 48.20683523 | NA | NA | NA | NA | NA | NA | NA |
| GRIN1 | 290 | Q | 126.8463651 | 45.6798097 | 0.58474065 | 44.64624636 | NA | NA | NA | NA | NA | NA | NA |
| GRIN1 | 291 | A | 125.2059429 | 45.13574026 | 0.577370546 | 44.07710602 | NA | NA | NA | NA | NA | NA | NA |
| GRIN1 | 292 | V | 128.6155714 | 46.58931777 | 0.597061078 | 45.55697016 | NA | NA | NA | NA | NA | NA | NA |
| GRIN1 | 293 | H | 130.8974429 | 44.81717734 | 0.573055212 | 43.82009888 | NA | NA | NA | NA | NA | NA | NA |
| GRIN1 | 294 | E | 125.5686984 | 41.45600596 | 0.527523926 | 40.43410484 | NA | NA | NA | NA | NA | NA | NA |
| GRIN1 | 295 | L | 126.0941429 | 42.642444 | 0.543595718 | 41.55684449 | NA | NA | NA | NA | NA | NA | NA |
| GRIN1 | 296 | L | 130.8892679 | 43.84350858 | 0.559865646 | 42.85230825 | NA | NA | NA | NA | NA | NA | NA |
| GRIN1 | 297 | E | 128.2321429 | 40.18150487 | 0.510259209 | 39.21481662 | NA | NA | NA | NA | NA | NA | NA |
| GRIN1 | 298 | K | 125.480254 | 37.36041914 | 0.472044061 | 36.2873612 | NA | NA | NA | NA | NA | NA | NA |
| GRIN1 | 299 | E | 127.4404762 | 35.91403139 | 0.452450924 | 34.93669388 | NA | NA | NA | NA | NA | NA | NA |
| GRIN1 | 300 | N | 127.4038929 | 37.40555555 | 0.472655491 | 36.36269758 | NA | NA | NA | NA | NA | NA | NA |
| GRIN1 | 301 | I | 129.4662679 | 42.55806391 | 0.542452684 | 41.51445317 | NA | NA | NA | NA | NA | NA | NA |
| GRIN1 | 302 | T | 127.8165714 | 43.53279874 | 0.555656691 | 42.42126312 | NA | NA | NA | NA | NA | NA | NA |
| GRIN1 | 303 | D | 131.1236429 | 47.79087522 | 0.613337682 | 46.69185001 | NA | NA | NA | NA | NA | NA | NA |
| GRIN1 | 304 | P | 130.0157143 | 50.6008876 | 0.651402827 | 49.48544603 | NA | NA | NA | NA | NA | NA | NA |
| GRIN1 | 305 | P | 127.9601429 | 51.07932014 | 0.657883797 | 49.91433407 | NA | NA | NA | NA | NA | NA | NA |
| GRIN1 | 306 | R | 132.6741429 | 55.47558989 | 0.717436789 | 54.30836027 | NA | NA | NA | NA | NA | NA | NA |
| GRIN1 | 307 | G | 127.9141429 | 56.03530971 | 0.725018896 | 54.81755236 | NA | NA | NA | NA | NA | NA | NA |
| GRIN1 | 308 | C | 124.8793095 | 56.8851866 | 0.736531546 | 55.63974005 | NA | NA | NA | NA | NA | NA | NA |
| GRIN1 | 309 | V | 126.7777143 | 58.55598663 | 0.759164629 | 57.3212757 | NA | NA | NA | NA | NA | NA | NA |
| GRIN1 | 310 | G | 125.7518929 | 54.75238987 | 0.707640137 | 53.52420919 | NA | NA | NA | NA | NA | NA | NA |
| GRIN1 | 311 | N | 124.7313929 | 53.14141123 | 0.685817409 | 51.89668345 | NA | NA | NA | NA | NA | NA | NA |
| GRIN1 | 312 | T | 120.1601429 | 48.88882515 | 0.628210792 | 47.63352006 | NA | NA | NA | NA | NA | NA | NA |
| GRIN1 | 313 | N | 122.1215179 | 47.77691494 | 0.613148573 | 46.54152209 | NA | NA | NA | NA | NA | NA | NA |
| GRIN1 | 314 | I | 119.1325179 | 44.30286883 | 0.566088257 | 43.05648092 | NA | NA | NA | NA | NA | NA | NA |
| GRIN1 | 315 | W | 123.3306429 | 42.69331688 | 0.544284856 | 41.48632204 | NA | NA | NA | NA | NA | NA | NA |

| | | | | | | | | | | | | | |
|---|---|---|---|---|---|---|---|---|---|---|---|---|---|
| GRIN1 | 316 | K | 123.0770317 | 39.67411423 | 0.503385966 | 38.5067567 | NA | NA | NA | NA | NA | NA | NA |
| GRIN1 | 317 | T | 122.1841429 | 41.31149008 | 0.525566277 | 40.15882116 | NA | NA | NA | NA | NA | NA | NA |
| GRIN1 | 318 | G | 118.8953929 | 41.40445098 | 0.526825549 | 40.20891175 | NA | NA | NA | NA | NA | NA | NA |
| GRIN1 | 319 | P | 116.7777143 | 37.93703822 | 0.479855091 | 36.72575505 | NA | NA | NA | NA | NA | NA | NA |
| GRIN1 | 320 | L | 120.1100179 | 36.80203759 | 0.464480082 | 35.68232811 | NA | NA | NA | NA | NA | NA | NA |
| GRIN1 | 321 | F | 121.6528701 | 41.45491383 | 0.527509132 | 40.30999568 | NA | NA | NA | NA | NA | NA | NA |
| GRIN1 | 322 | K | 115.6786984 | 39.96453689 | 0.507320105 | 38.60105848 | NA | NA | NA | NA | NA | NA | NA |
| GRIN1 | 323 | R | 115.9333247 | 32.68108519 | 0.408656609 | 31.31615576 | NA | NA | NA | NA | NA | NA | NA |
| GRIN1 | 324 | V | 120.4635714 | 39.57643608 | 0.502062793 | 38.4552611 | NA | NA | NA | NA | NA | NA | NA |
| GRIN1 | 325 | L | 118.7582679 | 41.75089994 | 0.531518635 | 40.40924814 | NA | NA | NA | NA | NA | NA | NA |
| GRIN1 | 326 | M | 114.1497679 | 37.96989062 | 0.480300118 | 36.69574613 | NA | NA | NA | NA | NA | NA | NA |
| GRIN1 | 327 | S | 117.0193095 | 36.83610711 | 0.464941597 | 35.49785965 | NA | NA | NA | NA | NA | NA | NA |
| GRIN1 | 328 | S | 118.7721429 | 40.21537176 | 0.510717978 | 38.86064017 | NA | NA | NA | NA | NA | NA | NA |
| GRIN1 | 329 | K | 118.8333651 | 38.07362324 | 0.481705306 | 36.74520751 | NA | NA | NA | NA | NA | NA | NA |
| GRIN1 | 330 | Y | 121.6033929 | 45.1117912 | 0.577046126 | 43.7746353 | NA | NA | NA | NA | NA | NA | NA |
| GRIN1 | 331 | A | 120.9203429 | 47.29335447 | 0.606598139 | 45.95894934 | NA | NA | NA | NA | NA | NA | NA |
| GRIN1 | 332 | D | 123.6857679 | 49.99672471 | 0.643218682 | 48.66464777 | NA | NA | NA | NA | NA | NA | NA |
| GRIN1 | 333 | G | 122.3068929 | 51.95331198 | 0.669723113 | 50.63396741 | NA | NA | NA | NA | NA | NA | NA |
| GRIN1 | 334 | V | 122.0617143 | 54.88240479 | 0.709401352 | 53.57787446 | NA | NA | NA | NA | NA | NA | NA |
| GRIN1 | 335 | T | 118.0115714 | 52.38741519 | 0.675603586 | 51.12137205 | NA | NA | NA | NA | NA | NA | NA |
| GRIN1 | 336 | G | 118.5741429 | 50.07783655 | 0.644317444 | 48.78862165 | NA | NA | NA | NA | NA | NA | NA |
| GRIN1 | 337 | R | 115.5994156 | 43.66623242 | 0.557464218 | 42.41340407 | NA | NA | NA | NA | NA | NA | NA |
| GRIN1 | 338 | V | 116.8372857 | 45.07430875 | 0.576538379 | 43.78000786 | NA | NA | NA | NA | NA | NA | NA |
| GRIN1 | 339 | E | 113.7476984 | 40.28597216 | 0.511674349 | 38.91608241 | NA | NA | NA | NA | NA | NA | NA |
| GRIN1 | 340 | F | 113.6364156 | 40.18603979 | 0.51032064 | 38.77731228 | NA | NA | NA | NA | NA | NA | NA |
| GRIN1 | 341 | N | 108.1221429 | 39.19155294 | 0.496849067 | 37.70808404 | NA | NA | NA | NA | NA | NA | NA |
| GRIN1 | 342 | E | 106.0700317 | 33.04409105 | 0.413573979 | 31.53208483 | NA | NA | NA | NA | NA | NA | NA |
| GRIN1 | 343 | D | 106.4807679 | 39.09463699 | 0.495536219 | 37.58250528 | NA | NA | NA | NA | NA | NA | NA |
| GRIN1 | 344 | G | 110.9323929 | 41.28120654 | 0.525156048 | 39.84246485 | NA | NA | NA | NA | NA | NA | NA |
| GRIN1 | 345 | D | 110.1427679 | 43.74889903 | 0.558584041 | 42.23646124 | NA | NA | NA | NA | NA | NA | NA |
| GRIN1 | 346 | R | 113.3395974 | 45.20755721 | 0.578343397 | 43.695346 | NA | NA | NA | NA | NA | NA | NA |
| GRIN1 | 347 | K | 109.144254 | 37.40463204 | 0.472642981 | 35.89210806 | NA | NA | NA | NA | NA | NA | NA |
| GRIN1 | 348 | F | 108.0457792 | 40.54170426 | 0.515138562 | 39.03497934 | NA | NA | NA | NA | NA | NA | NA |

| | | | | | | | | | | | | | |
|---|---|---|---|---|---|---|---|---|---|---|---|---|---|
| GRIN1 | 349 | A | 109.5429429 | 47.51120611 | 0.609549213 | 46.00099443 | NA | NA | NA | NA | NA | NA | NA |
| GRIN1 | 350 | N | 110.5758929 | 50.94344116 | 0.656043145 | 49.43502707 | NA | NA | NA | NA | NA | NA | NA |
| GRIN1 | 351 | Y | 105.9853095 | 47.23764044 | 0.605843423 | 45.78217358 | NA | NA | NA | NA | NA | NA | NA |
| GRIN1 | 352 | S | 106.0088095 | 55.10721083 | 0.712446632 | 53.64539761 | NA | NA | NA | NA | NA | NA | NA |
| GRIN1 | 353 | I | 102.0617679 | 54.6503885 | 0.7062584 | 53.17991361 | NA | NA | NA | NA | NA | NA | NA |
| GRIN1 | 354 | M | 103.5740179 | 59.62953122 | 0.773707139 | 58.15939376 | NA | NA | NA | NA | NA | NA | NA |
| GRIN1 | 355 | N | 99.99601786 | 61.78844222 | 0.802952298 | 60.31780274 | NA | NA | NA | NA | NA | NA | NA |
| GRIN1 | 356 | L | 100.4276429 | 65.4336173 | 0.85233077 | 63.96329741 | NA | NA | NA | NA | NA | NA | NA |
| GRIN1 | 357 | Q | 98.97403175 | 65.26445738 | 0.850039287 | 63.79350241 | NA | NA | NA | NA | NA | NA | NA |
| GRIN1 | 358 | N | 99.10751786 | 71.10751869 | 0.929190887 | 69.63687599 | NA | NA | NA | NA | NA | NA | NA |
| GRIN1 | 359 | R | 100.083961 | 71.38189427 | 0.932907649 | 69.91172413 | NA | NA | NA | NA | NA | NA | NA |
| GRIN1 | 360 | K | 104.5514762 | 68.55911319 | 0.894669536 | 67.08886198 | NA | NA | NA | NA | NA | NA | NA |
| GRIN1 | 361 | L | 104.9683929 | 64.79065798 | 0.843621079 | 63.32070369 | NA | NA | NA | NA | NA | NA | NA |
| GRIN1 | 362 | V | 104.0291429 | 63.27300275 | 0.823062534 | 61.80273459 | NA | NA | NA | NA | NA | NA | NA |
| GRIN1 | 363 | Q | 107.4575873 | 61.87015814 | 0.804059243 | 60.39986245 | NA | NA | NA | NA | NA | NA | NA |
| GRIN1 | 364 | V | 102.4814286 | 58.48384353 | 0.758187361 | 57.01327287 | NA | NA | NA | NA | NA | NA | NA |
| GRIN1 | 365 | G | 104.1931429 | 55.97261111 | 0.724169565 | 54.50319323 | NA | NA | NA | NA | NA | NA | NA |
| GRIN1 | 366 | I | 106.4288929 | 52.58198304 | 0.678239252 | 51.11359247 | NA | NA | NA | NA | NA | NA | NA |
| GRIN1 | 367 | Y | 103.9462262 | 50.10892192 | 0.644738534 | 48.63862098 | NA | NA | NA | NA | NA | NA | NA |
| GRIN1 | 368 | N | 108.8332679 | 48.1137312 | 0.617711172 | 46.65893634 | NA | NA | NA | NA | NA | NA | NA |
| GRIN1 | 369 | G | 105.5916429 | 45.90615831 | 0.587806826 | 44.44495643 | NA | NA | NA | NA | NA | NA | NA |
| GRIN1 | 370 | T | 102.8447143 | 42.99060109 | 0.548311943 | 41.52408657 | NA | NA | NA | NA | NA | NA | NA |
| GRIN1 | 371 | H | 104.5350429 | 44.70793242 | 0.571575352 | 43.24346267 | NA | NA | NA | NA | NA | NA | NA |
| GRIN1 | 372 | V | 101.292 | 48.55813424 | 0.623731169 | 47.08727317 | NA | NA | NA | NA | NA | NA | NA |
| GRIN1 | 373 | I | 104.3147679 | 52.54113814 | 0.677685956 | 51.07042586 | NA | NA | NA | NA | NA | NA | NA |
| GRIN1 | 374 | P | 101.4725714 | 54.93184993 | 0.710071149 | 53.46073236 | NA | NA | NA | NA | NA | NA | NA |
| GRIN1 | 375 | N | 104.1363929 | 58.96154758 | 0.764658462 | 57.4907293 | NA | NA | NA | NA | NA | NA | NA |
| GRIN1 | 376 | D | 102.7270179 | 60.9654715 | 0.791804127 | 59.4941031 | NA | NA | NA | NA | NA | NA | NA |
| GRIN1 | 377 | R | 104.0985974 | 62.960775 | 0.818833017 | 61.48970705 | NA | NA | NA | NA | NA | NA | NA |
| GRIN1 | 378 | K | 97.7545873 | 64.04878925 | 0.833571536 | 62.57749743 | NA | NA | NA | NA | NA | NA | NA |
| GRIN1 | 379 | I | 95.51589286 | 59.01366158 | 0.765364411 | 57.54296341 | NA | NA | NA | NA | NA | NA | NA |
| GRIN1 | 380 | I | 94.60414286 | 64.74482528 | 0.843000218 | 63.2767877 | NA | NA | NA | NA | NA | NA | NA |
| GRIN1 | 381 | W | 91.40235714 | 57.97803299 | 0.751335522 | 56.56248954 | NA | NA | NA | NA | NA | NA | NA |

| | | | | | | | | | | | | | |
|---|---|---|---|---|---|---|---|---|---|---|---|---|---|
| GRIN1 | 382 | P | 90.98157143 | 62.73799108 | 0.815815129 | 58.76436015 | NA | NA | NA | NA | NA | NA | NA |
| GRIN1 | 383 | G | 88.02564286 | 65.09683535 | 0.847768636 | 61.10796154 | NA | NA | NA | NA | NA | NA | NA |
| GRIN1 | 384 | G | 87.28814286 | 66.17000929 | 0.862306124 | 62.17397183 | NA | NA | NA | NA | NA | NA | NA |
| GRIN1 | 385 | E | 86.51992063 | 64.47489755 | 0.839343708 | 60.49899645 | NA | NA | NA | NA | NA | NA | NA |
| GRIN1 | 386 | T | 90.91514286 | 62.93793569 | 0.81852363 | 61.7588549 | NA | NA | NA | NA | NA | NA | NA |
| GRIN1 | 387 | E | 86.92325397 | 60.80256326 | 0.78959733 | 56.83194899 | NA | NA | NA | NA | NA | NA | NA |
| GRIN1 | 388 | K | 89.25692063 | 56.83566417 | 0.735860702 | 53.25817336 | NA | NA | NA | NA | NA | NA | NA |
| GRIN1 | 389 | P | 85.44728571 | 55.16263233 | 0.713197386 | 51.17570311 | NA | NA | NA | NA | NA | NA | NA |
| GRIN1 | 390 | R | 81.36005195 | 52.87986178 | 0.682274393 | 48.86674843 | NA | NA | NA | NA | NA | NA | NA |
| GRIN1 | 391 | G | 81.11964286 | 51.32373572 | 0.661194712 | 47.29120769 | NA | NA | NA | NA | NA | NA | NA |
| GRIN1 | 392 | Y | 79.80039286 | 47.47041307 | 0.60899662 | 43.47254032 | NA | NA | NA | NA | NA | NA | NA |
| GRIN1 | 393 | Q | 76.87880952 | 53.38822572 | 0.689160821 | 49.34008036 | NA | NA | NA | NA | NA | NA | NA |
| GRIN1 | 394 | M | 72.41026786 | 50.21769053 | 0.646211942 | 46.25392367 | NA | NA | NA | NA | NA | NA | NA |
| GRIN1 | 395 | S | 69.64364286 | 53.34721126 | 0.688605229 | 49.34446965 | NA | NA | NA | NA | NA | NA | NA |
| GRIN1 | 396 | T | 66.64014286 | 51.41878926 | 0.662482332 | 47.53354767 | NA | NA | NA | NA | NA | NA | NA |
| GRIN1 | 397 | R | 65.96850649 | 50.93921855 | 0.655985945 | 46.97216003 | NA | NA | NA | NA | NA | NA | NA |
| GRIN1 | 398 | L | 64.94101786 | 47.1526219 | 0.60469174 | 43.17097826 | NA | NA | NA | NA | NA | NA | NA |
| GRIN1 | 399 | K | 65.38603175 | 45.13562522 | 0.577368988 | 41.1194387 | NA | NA | NA | NA | NA | NA | NA |
| GRIN1 | 400 | I | 60.65001786 | 42.43617698 | 0.540801573 | 37.43795051 | NA | NA | NA | NA | NA | NA | NA |
| GRIN1 | 401 | V | 60.06442857 | 39.92143618 | 0.506736252 | 34.91176257 | NA | NA | NA | NA | NA | NA | NA |
| GRIN1 | 402 | T | 55.92514286 | 39.3344399 | 0.49878465 | 34.31491971 | NA | NA | NA | NA | NA | NA | NA |
| GRIN1 | 403 | I | 55.11826786 | 35.29040309 | 0.444003095 | 30.26752247 | NA | NA | NA | NA | NA | NA | NA |
| GRIN1 | 404 | H | 51.32784286 | 35.25255368 | 0.443490378 | 30.21571557 | NA | NA | NA | NA | NA | NA | NA |
| GRIN1 | 405 | Q | 50.74947619 | 30.73687904 | 0.382319896 | 25.70572064 | NA | NA | NA | NA | NA | NA | NA |
| GRIN1 | 406 | E | 45.1135873 | 31.81683954 | 0.396949317 | 26.79444797 | NA | NA | NA | NA | NA | NA | NA |
| GRIN1 | 407 | P | 45.69471429 | 31.35162699 | 0.390647429 | 26.33819712 | NA | NA | NA | NA | NA | NA | NA |
| GRIN1 | 408 | F | 49.42768831 | 29.30536746 | 0.362928275 | 24.3077722 | NA | NA | NA | NA | NA | NA | NA |
| GRIN1 | 409 | V | 49.70128571 | 35.88331095 | 0.452034777 | 30.87310863 | NA | NA | NA | NA | NA | NA | NA |
| GRIN1 | 410 | Y | 44.96439286 | 35.78720847 | 0.450732948 | 30.77222385 | NA | NA | NA | NA | NA | NA | NA |
| GRIN1 | 411 | V | 47.80328571 | 41.25243182 | 0.524766259 | 36.2294066 | NA | NA | NA | NA | NA | NA | NA |
| GRIN1 | 412 | K | 43.72980952 | 43.17296291 | 0.550782263 | 38.14983723 | NA | NA | NA | NA | NA | NA | NA |
| GRIN1 | 413 | P | 44.574 | 47.36600098 | 0.607582227 | 42.33275011 | NA | NA | NA | NA | NA | NA | NA |
| GRIN1 | 414 | T | 45.58542857 | 51.01933213 | 0.657071184 | 45.99270516 | NA | NA | NA | NA | NA | NA | NA |

| | | | | | | | | | | | | | |
|---|---|---|---|---|---|---|---|---|---|---|---|---|---|
| GRIN1 | 415 | L | 42.65201786 | 53.84461928 | 0.695343245 | 48.82200449 | NA | NA | NA | NA | NA | NA | NA |
| GRIN1 | 416 | S | 42.91730952 | 57.01160579 | 0.738244052 | 51.98509335 | NA | NA | NA | NA | NA | NA | NA |
| GRIN1 | 417 | D | 44.66876786 | 58.44296356 | 0.75763359 | 53.42197038 | NA | NA | NA | NA | NA | NA | NA |
| GRIN1 | 418 | G | 47.57714286 | 55.0026798 | 0.711030628 | 49.98556555 | NA | NA | NA | NA | NA | NA | NA |
| GRIN1 | 419 | T | 45.99671429 | 53.75713307 | 0.694158134 | 48.74069281 | NA | NA | NA | NA | NA | NA | NA |
| GRIN1 | 420 | C | 45.21247619 | 50.5374993 | 0.650544153 | 45.52757584 | NA | NA | NA | NA | NA | NA | NA |
| GRIN1 | 421 | K | 40.61492063 | 51.00385826 | 0.656861571 | 45.99810078 | NA | NA | NA | NA | NA | NA | NA |
| GRIN1 | 422 | E | 40.33414286 | 51.25689259 | 0.660289238 | 46.25313381 | NA | NA | NA | NA | NA | NA | NA |
| GRIN1 | 423 | E | 38.45480952 | 46.84168461 | 0.600479704 | 41.24319901 | NA | NA | NA | NA | NA | NA | NA |
| GRIN1 | 424 | F | 36.13577922 | 44.92561583 | 0.574524147 | 39.35327302 | NA | NA | NA | NA | NA | NA | NA |
| GRIN1 | 425 | T | 37.89071429 | 43.30787609 | 0.552609831 | 37.81106716 | NA | NA | NA | NA | NA | NA | NA |
| GRIN1 | 426 | V | 34.86471429 | 39.95068105 | 0.50713241 | 33.08223069 | NA | NA | NA | NA | NA | NA | NA |
| GRIN1 | 427 | N | 35.28326786 | 41.95993796 | 0.534350318 | 36.40622492 | NA | NA | NA | NA | NA | NA | NA |
| GRIN1 | 428 | G | 36.29289286 | 45.75811465 | 0.585801388 | 40.15873919 | NA | NA | NA | NA | NA | NA | NA |
| GRIN1 | 429 | D | 39.56376786 | 44.78170807 | 0.572574736 | 39.79928048 | NA | NA | NA | NA | NA | NA | NA |
| GRIN1 | 430 | P | 41.50914286 | 47.37733304 | 0.607735734 | 42.38868261 | NA | NA | NA | NA | NA | NA | NA |
| GRIN1 | 431 | V | 43.29185714 | 44.27625425 | 0.565727729 | 39.27898223 | NA | NA | NA | NA | NA | NA | NA |
| GRIN1 | 432 | K | 47.70692063 | 45.11171472 | 0.57704509 | 40.11714264 | NA | NA | NA | NA | NA | NA | NA |
| GRIN1 | 433 | K | 46.14914286 | 47.0838623 | 0.603760305 | 42.08508258 | NA | NA | NA | NA | NA | NA | NA |
| GRIN1 | 434 | V | 51.61085714 | 46.48201061 | 0.595607468 | 41.48643751 | NA | NA | NA | NA | NA | NA | NA |
| GRIN1 | 435 | I | 53.02201786 | 48.61387558 | 0.624486255 | 43.60672844 | NA | NA | NA | NA | NA | NA | NA |
| GRIN1 | 436 | C | 53.74964286 | 45.50668511 | 0.58239546 | 40.49560805 | NA | NA | NA | NA | NA | NA | NA |
| GRIN1 | 437 | T | 56.78828571 | 47.45434545 | 0.608778964 | 42.42938448 | NA | NA | NA | NA | NA | NA | NA |
| GRIN1 | 438 | G | 55.53439286 | 47.81243961 | 0.613629799 | 42.77908623 | NA | NA | NA | NA | NA | NA | NA |
| GRIN1 | 439 | P | 55.144 | 46.32704357 | 0.593508245 | 41.28037761 | NA | NA | NA | NA | NA | NA | NA |
| GRIN1 | 440 | N | 51.29426786 | 46.32769225 | 0.593517032 | 41.27090439 | NA | NA | NA | NA | NA | NA | NA |
| GRIN1 | 441 | D | 52.15451786 | 46.49611203 | 0.595798489 | 41.43157476 | NA | NA | NA | NA | NA | NA | NA |
| GRIN1 | 442 | T | 53.18257143 | 42.30177085 | 0.538980873 | 37.23298529 | NA | NA | NA | NA | NA | NA | NA |
| GRIN1 | 443 | S | 54.16064286 | 43.92499416 | 0.56096947 | 38.84749208 | NA | NA | NA | NA | NA | NA | NA |
| GRIN1 | 444 | P | 56.483 | 42.38142994 | 0.540059955 | 37.29223679 | NA | NA | NA | NA | NA | NA | NA |
| GRIN1 | 445 | G | 57.93064286 | 46.51602208 | 0.596068196 | 42.47859433 | NA | NA | NA | NA | NA | NA | NA |
| GRIN1 | 446 | S | 56.62197619 | 47.79463465 | 0.613388608 | 42.71933546 | NA | NA | NA | NA | NA | NA | NA |
| GRIN1 | 447 | P | 56.73371429 | 50.91901251 | 0.655712228 | 45.85267312 | NA | NA | NA | NA | NA | NA | NA |

| | | | | | | | | | | | | | |
|---|---|---|---|---|---|---|---|---|---|---|---|---|---|
| GRIN1 | 448 | R | 58.49268831 | 44.91185472 | 0.574337735 | 39.84339379 | NA | NA | NA | NA | NA | NA | NA |
| GRIN1 | 449 | H | 55.45214286 | 51.3457563 | 0.661493009 | 46.30093091 | NA | NA | NA | NA | NA | NA | NA |
| GRIN1 | 450 | T | 57.10128571 | 52.47891082 | 0.676843009 | 47.44059151 | NA | NA | NA | NA | NA | NA | NA |
| GRIN1 | 451 | V | 52.991 | 50.33346458 | 0.647780247 | 45.30441643 | NA | NA | NA | NA | NA | NA | NA |
| GRIN1 | 452 | P | 52.34228571 | 51.05986495 | 0.657620252 | 46.03573232 | NA | NA | NA | NA | NA | NA | NA |
| GRIN1 | 453 | Q | 49.87525397 | 46.62300596 | 0.597517427 | 41.58889908 | NA | NA | NA | NA | NA | NA | NA |
| GRIN1 | 454 | C | 47.65397619 | 45.51731641 | 0.582539474 | 40.49914376 | NA | NA | NA | NA | NA | NA | NA |
| GRIN1 | 455 | C | 49.30397619 | 43.86118963 | 0.560105158 | 38.85416353 | NA | NA | NA | NA | NA | NA | NA |
| GRIN1 | 456 | Y | 45.46289286 | 40.12984945 | 0.509559471 | 35.12414077 | NA | NA | NA | NA | NA | NA | NA |
| GRIN1 | 457 | G | 48.63264286 | 38.51335309 | 0.487661999 | 33.51918176 | NA | NA | NA | NA | NA | NA | NA |
| GRIN1 | 458 | F | 49.55105195 | 32.72081226 | 0.409194762 | 27.73295022 | NA | NA | NA | NA | NA | NA | NA |
| GRIN1 | 459 | C | 53.88980952 | 36.5806647 | 0.461481309 | 31.58719944 | NA | NA | NA | NA | NA | NA | NA |
| GRIN1 | 460 | I | 52.01564286 | 40.6546825 | 0.516668994 | 35.6645333 | NA | NA | NA | NA | NA | NA | NA |
| GRIN1 | 461 | D | 50.41726786 | 40.68099725 | 0.51702546 | 35.69558523 | NA | NA | NA | NA | NA | NA | NA |
| GRIN1 | 462 | L | 54.22851786 | 36.38649263 | 0.458851004 | 31.40586368 | NA | NA | NA | NA | NA | NA | NA |
| GRIN1 | 463 | L | 57.20001786 | 41.73651113 | 0.531323721 | 36.75663699 | NA | NA | NA | NA | NA | NA | NA |
| GRIN1 | 464 | I | 53.91551786 | 44.23237889 | 0.565133382 | 39.25362108 | NA | NA | NA | NA | NA | NA | NA |
| GRIN1 | 465 | K | 53.81914286 | 39.44330931 | 0.500259423 | 34.47368998 | NA | NA | NA | NA | NA | NA | NA |
| GRIN1 | 466 | L | 59.04976786 | 41.24924616 | 0.524723105 | 36.27561542 | NA | NA | NA | NA | NA | NA | NA |
| GRIN1 | 467 | A | 58.52594286 | 46.46858199 | 0.59542556 | 41.49489926 | NA | NA | NA | NA | NA | NA | NA |
| GRIN1 | 468 | R | 54.62532468 | 48.24898528 | 0.619543358 | 43.27840367 | NA | NA | NA | NA | NA | NA | NA |
| GRIN1 | 469 | T | 59.83371429 | 45.30467483 | 0.579658977 | 40.33927973 | NA | NA | NA | NA | NA | NA | NA |
| GRIN1 | 470 | M | 63.30514286 | 45.97679215 | 0.58876365 | 41.01049463 | NA | NA | NA | NA | NA | NA | NA |
| GRIN1 | 471 | N | 62.98114286 | 50.19307778 | 0.645878531 | 45.22350208 | NA | NA | NA | NA | NA | NA | NA |
| GRIN1 | 472 | F | 63.75305195 | 47.21356497 | 0.60551729 | 42.24046568 | NA | NA | NA | NA | NA | NA | NA |
| GRIN1 | 473 | T | 62.06328571 | 49.71280094 | 0.639372579 | 44.72632442 | NA | NA | NA | NA | NA | NA | NA |
| GRIN1 | 474 | Y | 59.69680952 | 46.18151826 | 0.591536922 | 41.1994685 | NA | NA | NA | NA | NA | NA | NA |
| GRIN1 | 475 | E | 61.8005873 | 47.68867205 | 0.611953212 | 42.68625907 | NA | NA | NA | NA | NA | NA | NA |
| GRIN1 | 476 | V | 57.83042857 | 44.73585208 | 0.571953559 | 39.72924184 | NA | NA | NA | NA | NA | NA | NA |
| GRIN1 | 477 | H | 60.13754286 | 43.77620364 | 0.558953916 | 38.75924266 | NA | NA | NA | NA | NA | NA | NA |
| GRIN1 | 478 | L | 56.35676786 | 43.08364535 | 0.549572345 | 38.05822595 | NA | NA | NA | NA | NA | NA | NA |
| GRIN1 | 479 | V | 58.63528571 | 39.58788351 | 0.502217862 | 34.55667107 | NA | NA | NA | NA | NA | NA | NA |
| GRIN1 | 480 | A | 60.45674286 | 43.52535946 | 0.555555917 | 39.55094187 | NA | NA | NA | NA | NA | NA | NA |

| | | | | | | | | | | | | | |
|---|---|---|---|---|---|---|---|---|---|---|---|---|---|
| GRIN1 | 481 | D | 60.20001786 | 40.42284279 | 0.513528434 | 36.49348126 | NA | NA | NA | NA | NA | NA | NA |
| GRIN1 | 482 | G | 56.40439286 | 37.1959625 | 0.46981629 | 32.14296229 | NA | NA | NA | NA | NA | NA | NA |
| GRIN1 | 483 | K | 58.06069841 | 35.97623245 | 0.453293515 | 30.93236886 | NA | NA | NA | NA | NA | NA | NA |
| GRIN1 | 484 | F | 58.33223377 | 30.48257681 | 0.378875053 | 26.64445678 | NA | NA | NA | NA | NA | NA | NA |
| GRIN1 | 485 | G | 63.05339286 | 33.50490522 | 0.419816286 | 29.49503499 | NA | NA | NA | NA | NA | NA | NA |
| GRIN1 | 486 | T | 64.92042857 | 31.27195966 | 0.389568235 | 27.26308512 | NA | NA | NA | NA | NA | NA | NA |
| GRIN1 | 487 | Q | 69.63880952 | 33.54953658 | 0.420420873 | 29.54600138 | NA | NA | NA | NA | NA | NA | NA |
| GRIN1 | 488 | E | 67.32780952 | 35.32708621 | 0.444500014 | 31.32739504 | NA | NA | NA | NA | NA | NA | NA |
| GRIN1 | 489 | R | 72.63914286 | 33.6577895 | 0.421887295 | 31.94382737 | NA | NA | NA | NA | NA | NA | NA |
| GRIN1 | 490 | V | 72.323 | 40.74399237 | 0.517878808 | 36.74805505 | NA | NA | NA | NA | NA | NA | NA |
| GRIN1 | 491 | N | 74.12094286 | 43.66615285 | 0.55746314 | 39.66485758 | NA | NA | NA | NA | NA | NA | NA |
| GRIN1 | 492 | N | 76.82054286 | 40.98550817 | 0.521150443 | 36.72277539 | NA | NA | NA | NA | NA | NA | NA |
| GRIN1 | 493 | S | 75.08554286 | 37.83959235 | 0.478535064 | 33.56131336 | NA | NA | NA | NA | NA | NA | NA |
| GRIN1 | 494 | N | 77.67626786 | 36.17154586 | 0.45593928 | 31.91657461 | NA | NA | NA | NA | NA | NA | NA |
| GRIN1 | 495 | K | 77.19847619 | 38.92152362 | 0.493191181 | 34.68563213 | NA | NA | NA | NA | NA | NA | NA |
| GRIN1 | 496 | K | 74.12380952 | 31.41379905 | 0.391489628 | 27.17708451 | NA | NA | NA | NA | NA | NA | NA |
| GRIN1 | 497 | E | 71.86392063 | 37.57360355 | 0.474931912 | 33.57578791 | NA | NA | NA | NA | NA | NA | NA |
| GRIN1 | 498 | W | 69.38307143 | 32.7395657 | 0.409448801 | 28.77495422 | NA | NA | NA | NA | NA | NA | NA |
| GRIN1 | 499 | N | 65.67001786 | 36.59905435 | 0.461730419 | 32.59151797 | NA | NA | NA | NA | NA | NA | NA |
| GRIN1 | 500 | G | 64.12464286 | 38.88697495 | 0.492723176 | 34.88707752 | NA | NA | NA | NA | NA | NA | NA |
| GRIN1 | 501 | M | 62.78876786 | 37.1001109 | 0.46851786 | 33.20637727 | NA | NA | NA | NA | NA | NA | NA |
| GRIN1 | 502 | M | 66.74351786 | 34.37458166 | 0.431597145 | 30.38401614 | NA | NA | NA | NA | NA | NA | NA |
| GRIN1 | 503 | G | 68.61264286 | 39.27525824 | 0.497982961 | 35.26861869 | NA | NA | NA | NA | NA | NA | NA |
| GRIN1 | 504 | E | 65.83169841 | 41.59872283 | 0.529457205 | 37.58851165 | NA | NA | NA | NA | NA | NA | NA |
| GRIN1 | 505 | L | 68.49276786 | 39.31380771 | 0.498505162 | 35.31540148 | NA | NA | NA | NA | NA | NA | NA |
| GRIN1 | 506 | L | 72.23014286 | 39.15016988 | 0.496288482 | 35.20440122 | NA | NA | NA | NA | NA | NA | NA |
| GRIN1 | 507 | S | 71.61047619 | 44.37560299 | 0.567073533 | 40.38626938 | NA | NA | NA | NA | NA | NA | NA |
| GRIN1 | 508 | G | 71.22814286 | 46.48311088 | 0.595622373 | 42.4912014 | NA | NA | NA | NA | NA | NA | NA |
| GRIN1 | 509 | Q | 68.3575873 | 47.27567463 | 0.606358643 | 43.26339379 | NA | NA | NA | NA | NA | NA | NA |
| GRIN1 | 510 | A | 67.02094286 | 44.08174567 | 0.563092866 | 40.07051558 | NA | NA | NA | NA | NA | NA | NA |
| GRIN1 | 511 | D | 67.88139286 | 43.68826902 | 0.557762731 | 39.68372969 | NA | NA | NA | NA | NA | NA | NA |
| GRIN1 | 512 | M | 64.05526786 | 41.27916848 | 0.52512844 | 37.24271505 | NA | NA | NA | NA | NA | NA | NA |
| GRIN1 | 513 | I | 62.50289286 | 36.54986048 | 0.461064027 | 32.52467077 | NA | NA | NA | NA | NA | NA | NA |

| | | | | | | | | | | | | | |
|---|---|---|---|---|---|---|---|---|---|---|---|---|---|
| GRIN1 | 514 | V | 58.675 | 37.21556531 | 0.470081834 | 32.21907269 | NA | NA | NA | NA | NA | NA | NA |
| GRIN1 | 515 | A | 57.23114286 | 33.38151885 | 0.418144862 | 28.38349743 | NA | NA | NA | NA | NA | NA | NA |
| GRIN1 | 516 | P | 55.03957143 | 30.4685176 | 0.378684604 | 25.45773667 | NA | NA | NA | NA | NA | NA | NA |
| GRIN1 | 517 | L | 59.30001786 | 29.15494762 | 0.360890649 | 25.18546323 | NA | NA | NA | NA | NA | NA | NA |
| GRIN1 | 518 | T | 59.29642857 | 25.99677588 | 0.318109249 | 21.99432278 | NA | NA | NA | NA | NA | NA | NA |
| GRIN1 | 519 | I | 62.03876786 | 24.85039955 | 0.302580142 | 20.85414291 | NA | NA | NA | NA | NA | NA | NA |
| GRIN1 | 520 | N | 64.32351786 | 25.0396736 | 0.305144097 | 21.03831215 | NA | NA | NA | NA | NA | NA | NA |
| GRIN1 | 521 | N | 68.27401786 | 23.37418247 | 0.282582929 | 19.22803135 | NA | NA | NA | NA | NA | NA | NA |
| GRIN1 | 522 | E | 69.39125397 | 27.75244236 | 0.341891955 | 23.89148656 | NA | NA | NA | NA | NA | NA | NA |
| GRIN1 | 523 | R | 65.38750649 | 29.67018473 | 0.367870183 | 25.6597129 | NA | NA | NA | NA | NA | NA | NA |
| GRIN1 | 524 | A | 69.88074286 | 29.94462965 | 0.371587884 | 26.1060603 | NA | NA | NA | NA | NA | NA | NA |
| GRIN1 | 525 | Q | 73.02325397 | 27.31405239 | 0.335953413 | 23.11899413 | NA | NA | NA | NA | NA | NA | NA |
| GRIN1 | 526 | Y | 73.61097619 | 31.80396499 | 0.396774915 | 27.63443921 | NA | NA | NA | NA | NA | NA | NA |
| GRIN1 | 527 | I | 69.53439286 | 33.86151724 | 0.424647043 | 29.97648312 | NA | NA | NA | NA | NA | NA | NA |
| GRIN1 | 528 | E | 70.51880952 | 34.27791039 | 0.430287611 | 30.66765933 | NA | NA | NA | NA | NA | NA | NA |
| GRIN1 | 529 | F | 65.09841558 | 30.02262044 | 0.372644367 | 26.05399742 | NA | NA | NA | NA | NA | NA | NA |
| GRIN1 | 530 | S | 63.74314286 | 31.75396479 | 0.3960976 | 27.79061175 | NA | NA | NA | NA | NA | NA | NA |
| GRIN1 | 531 | K | 62.93436508 | 31.58302169 | 0.393781961 | 27.58204662 | NA | NA | NA | NA | NA | NA | NA |
| GRIN1 | 532 | P | 58.95985714 | 28.04212093 | 0.345816015 | 23.07512992 | NA | NA | NA | NA | NA | NA | NA |
| GRIN1 | 533 | F | 55.70105195 | 28.89236247 | 0.357333604 | 23.92423164 | NA | NA | NA | NA | NA | NA | NA |
| GRIN1 | 534 | K | 51.91114286 | 25.87127049 | 0.316409121 | 20.89680606 | NA | NA | NA | NA | NA | NA | NA |
| GRIN1 | 535 | Y | 55.05047619 | 21.98724933 | 0.263795179 | 17.01413883 | NA | NA | NA | NA | NA | NA | NA |
| GRIN1 | 536 | Q | 51.39869841 | 21.0256183 | 0.250768679 | 16.03804514 | NA | NA | NA | NA | NA | NA | NA |
| GRIN1 | 537 | G | 52.00139286 | 20.17337696 | 0.239224 | 15.16280738 | NA | NA | NA | NA | NA | NA | NA |
| GRIN1 | 538 | L | 51.54739286 | 17.0280922 | 0.19661717 | 12.00604973 | NA | NA | NA | NA | NA | NA | NA |
| GRIN1 | 539 | T | 46.95228571 | 18.19364213 | 0.212406007 | 13.15885362 | NA | NA | NA | NA | NA | NA | NA |
| GRIN1 | 540 | I | 45.13189286 | 15.09803391 | 0.170472108 | 9.425816076 | NA | NA | NA | NA | NA | NA | NA |
| GRIN1 | 541 | L | 42.88114286 | 20.281473 | 0.240688297 | 14.18841968 | NA | NA | NA | NA | NA | NA | NA |
| GRIN1 | 542 | V | 39.48014286 | 17.57292975 | 0.203997679 | 10.79666945 | NA | NA | NA | NA | NA | NA | NA |
| GRIN1 | 543 | K | 36.55947619 | 19.9141506 | 0.235712454 | 13.13093425 | NA | NA | NA | NA | NA | NA | NA |
| GRIN1 | 544 | K | 32.2185873 | 21.43326083 | 0.256290709 | 15.30788689 | NA | NA | NA | NA | NA | NA | NA |
| GRIN1 | 545 | E | 30.97154286 | 20.23010136 | 0.239992404 | 14.29017446 | NA | NA | NA | NA | NA | NA | NA |
| GRIN1 | 546 | I | 27.86754286 | 19.67029489 | 0.232409122 | 18.01355792 | NA | NA | NA | NA | NA | NA | NA |

| | | | | | | | | | | | | | |
|---|---|---|---|---|---|---|---|---|---|---|---|---|---|
| GRIN1 | 547 | P | 24.73854286 | 18.32516922 | 0.214187707 | 16.67049413 | NA | NA | NA | NA | NA | NA | NA |
| GRIN1 | 548 | R | 22.40614286 | 16.19207646 | 0.185292287 | 14.57684724 | NA | NA | NA | NA | NA | NA | NA |
| GRIN1 | 553 | S | 17.47097619 | 13.63681755 | 0.150678098 | 11.72345617 | NA | NA | NA | NA | NA | NA | NA |
| GRIN1 | 554 | F | 17.67005195 | 14.87359103 | 0.167431747 | 12.97339923 | NA | NA | NA | NA | NA | NA | NA |
| GRIN1 | 555 | M | 12.81689286 | 16.57976396 | 0.190544001 | 14.6609337 | NA | NA | NA | NA | NA | NA | NA |
| GRIN1 | 556 | Q | 10.40336508 | 15.41674392 | 0.174789435 | 14.41804703 | NA | NA | NA | NA | NA | NA | NA |
| GRIN1 | 557 | P | 15.286 | 13.85907951 | 0.153688915 | 11.93972712 | NA | NA | NA | NA | NA | NA | NA |
| GRIN1 | 558 | F | 13.19077922 | 11.32063214 | 0.119302459 | 9.233012783 | NA | NA | NA | NA | NA | NA | NA |
| GRIN1 | 559 | Q | 14.78425397 | 18.14530683 | 0.211751245 | 16.05073096 | NA | NA | NA | NA | NA | NA | NA |
| GRIN1 | 560 | S | 11.54714286 | 21.1031405 | 0.251818815 | 18.66954755 | NA | NA | NA | NA | NA | NA | NA |
| GRIN1 | 561 | T | 12.26985714 | 22.0735052 | 0.264963623 | 19.98325527 | NA | NA | NA | NA | NA | NA | NA |
| GRIN1 | 562 | L | 11.23064286 | 17.47246102 | 0.202636704 | 15.68247771 | NA | NA | NA | NA | NA | NA | NA |
| GRIN1 | 563 | W | 8.343 | 14.26951912 | 0.159248835 | 11.33535679 | NA | NA | NA | NA | NA | NA | NA |
| GRIN1 | 564 | L | 6.806892857 | 20.79417 | 0.247633421 | 17.89232164 | NA | NA | NA | NA | NA | NA | NA |
| GRIN1 | 565 | L | 7.490517857 | 19.59998349 | 0.231456666 | 16.69565414 | NA | NA | NA | NA | NA | NA | NA |
| GRIN1 | 566 | V | 4.614571429 | 15.00302573 | 0.169185103 | 12.13707544 | NA | NA | NA | NA | NA | NA | NA |
| GRIN1 | 567 | G | 1.861642857 | 18.47753585 | 0.216251704 | 15.48328317 | NA | NA | NA | NA | NA | NA | NA |
| GRIN1 | 568 | L | 2.141142857 | 20.79431014 | 0.24763532 | 17.76567684 | NA | NA | NA | NA | NA | NA | NA |
| GRIN1 | 569 | S | 1.240142857 | 17.29177599 | 0.200189098 | 14.27167788 | NA | NA | NA | NA | NA | NA | NA |
| GRIN1 | 570 | V | -0.209285714 | 16.10696927 | 0.184139404 | 13.00142965 | NA | NA | NA | NA | NA | NA | NA |
| GRIN1 | 571 | H | -0.923342857 | 20.38072374 | 0.242032773 | 17.25517149 | NA | NA | NA | NA | NA | NA | NA |
| GRIN1 | 572 | V | -0.467285714 | 21.253132 | 0.253850638 | 18.10428664 | NA | NA | NA | NA | NA | NA | NA |
| GRIN1 | 573 | V | -2.73 | 17.03076258 | 0.196653344 | 13.87649788 | NA | NA | NA | NA | NA | NA | NA |
| GRIN1 | 574 | A | -5.554342857 | 20.31557951 | 0.241150313 | 17.66608751 | NA | NA | NA | NA | NA | NA | NA |
| GRIN1 | 575 | V | -5.308428571 | 23.08730043 | 0.278696752 | 20.42774556 | NA | NA | NA | NA | NA | NA | NA |
| GRIN1 | 576 | M | -4.889017857 | 22.928419 | 0.276544503 | 20.23984772 | NA | NA | NA | NA | NA | NA | NA |
| GRIN1 | 577 | L | -7.764517857 | 17.3904357 | 0.201525568 | 14.63042004 | NA | NA | NA | NA | NA | NA | NA |
| GRIN1 | 578 | Y | -11.10247619 | 19.04759914 | 0.223973927 | 15.71894399 | NA | NA | NA | NA | NA | NA | NA |
| GRIN1 | 579 | L | -9.265142857 | 25.10009181 | 0.305962538 | 21.56118808 | NA | NA | NA | NA | NA | NA | NA |
| GRIN1 | 580 | L | -9.566517857 | 23.43871522 | 0.283457106 | 20.05710806 | NA | NA | NA | NA | NA | NA | NA |
| GRIN1 | 581 | D | -13.91451786 | 21.25168993 | 0.253831103 | 14.89030399 | NA | NA | NA | NA | NA | NA | NA |
| GRIN1 | 582 | R | -16.5425974 | 25.47855196 | 0.311089255 | 19.07080989 | NA | NA | NA | NA | NA | NA | NA |
| GRIN1 | 583 | F | -13.43041558 | 26.38294589 | 0.323340406 | 19.97896449 | NA | NA | NA | NA | NA | NA | NA |

| | | | | | | | | | | | | | |
|---|---|---|---|---|---|---|---|---|---|---|---|---|---|
| GRIN1 | 584 | S | -17.71430952 | 23.1108047 | 0.279015147 | 15.68640474 | NA | NA | NA | NA | NA | NA | NA |
| GRIN1 | 601 | L | -17.67164286 | 15.40683008 | 0.17465514 | 8.999511156 | NA | NA | NA | NA | NA | NA | NA |
| GRIN1 | 602 | T | -16.65242857 | 20.50790687 | 0.243755628 | 14.10138035 | NA | NA | NA | NA | NA | NA | NA |
| GRIN1 | 603 | L | -15.72901786 | 21.18009927 | 0.252861318 | 14.83996216 | NA | NA | NA | NA | NA | NA | NA |
| GRIN1 | 604 | S | -13.36130952 | 17.03244064 | 0.196676075 | 11.66495033 | NA | NA | NA | NA | NA | NA | NA |
| GRIN1 | 605 | S | -13.52380952 | 14.97947666 | 0.168866101 | 9.65913263 | NA | NA | NA | NA | NA | NA | NA |
| GRIN1 | 606 | A | -10.34574286 | 16.15557786 | 0.184797868 | 12.46182627 | NA | NA | NA | NA | NA | NA | NA |
| GRIN1 | 607 | M | -8.420017857 | 15.64134949 | 0.177831999 | 12.09647078 | NA | NA | NA | NA | NA | NA | NA |
| GRIN1 | 608 | W | -9.433857143 | 12.09278276 | 0.129762209 | 8.288398507 | NA | NA | NA | NA | NA | NA | NA |
| GRIN1 | 609 | F | -10.48841558 | 10.7982869 | 0.112226637 | 7.730969499 | NA | NA | NA | NA | NA | NA | NA |
| GRIN1 | 610 | S | -5.294642857 | 12.21170807 | 0.131373201 | 10.01806137 | NA | NA | NA | NA | NA | NA | NA |
| GRIN1 | 611 | W | -3.605857143 | 10.63725308 | 0.110045232 | 7.988342646 | NA | NA | NA | NA | NA | NA | NA |
| GRIN1 | 613 | V | -3.189142857 | 7.749748706 | 0.070930361 | 5.199439607 | 2.69 | 0.51 | 2.05 | 1.76 | -0.52 | 0 | 0 |
| GRIN1 | 614 | L | -0.312892857 | 8.458257799 | 0.080528006 | 5.318053665 | 3.23 | 0.63 | 1.6 | 1.27 | -0.82 | 0 | 0 |
| GRIN1 | 615 | L | -0.356017857 | 6.846840877 | 0.058699341 | 3.681705737 | 3.23 | 0.63 | 1.6 | 1.27 | -0.82 | 0 | 0 |
| GRIN1 | 616 | N | -0.537267857 | 4.286003616 | 0.024009585 | 1.610840215 | 2.76 | 0.51 | 2.05 | 1.76 | -0.52 | 0 | 0 |
| GRIN1 | 617 | S | -4.594142857 | 4.054119017 | 0.020868417 | 1.5678071 | 2.69 | 0.51 | 2.05 | 1.76 | -0.52 | 0 | 0 |
| GRIN1 | 618 | G | -5.417392857 | 4.430951252 | 0.025973083 | 1.626459828 | 2.81 | -0.8 | -0.4 | 3.38 | 0.81 | 0 | 0 |
| GRIN1 | 619 | I | -4.534767857 | 4.277675654 | 0.023896773 | 1.58894558 | 2.72 | -0.15 | 0.83 | 2.57 | 0.15 | 0 | 0 |
| GRIN1 | 620 | G | -8.777892857 | 5.190104912 | 0.036256772 | 1.618133582 | 3.6 | -0.8 | -0.4 | 3.38 | 0.81 | 0 | 0 |
| GRIN1 | 621 | E | -12.26580952 | 6.903569077 | 0.059467795 | 2.469056452 | 4.44 | -0.67 | -1.77 | 19.59 | 1.08 | 1 | 1 |
| GRIN1 | 622 | G | -12.30064286 | 10.05830935 | 0.102202712 | 5.191197984 | NA | NA | NA | NA | NA | NA | NA |
| GRIN1 | 626 | S | -13.92547619 | 19.21019979 | 0.226176557 | 14.33498856 | NA | NA | NA | NA | NA | NA | NA |
| GRIN1 | 627 | F | -12.20114286 | 19.14427123 | 0.225283472 | 15.0102276 | NA | NA | NA | NA | NA | NA | NA |
| GRIN1 | 628 | S | -8.49697619 | 20.39129456 | 0.242175968 | 16.95485417 | NA | NA | NA | NA | NA | NA | NA |
| GRIN1 | 629 | A | -8.495742857 | 17.72148157 | 0.20601 | 14.24751795 | NA | NA | NA | NA | NA | NA | NA |
| GRIN1 | 630 | R | -11.01187013 | 15.47585468 | 0.175590164 | 12.12257882 | NA | NA | NA | NA | NA | NA | NA |
| GRIN1 | 631 | I | -5.300392857 | 17.0974713 | 0.197556997 | 14.43813463 | NA | NA | NA | NA | NA | NA | NA |
| GRIN1 | 632 | L | -3.641267857 | 17.04067733 | 0.196787652 | 14.39068523 | NA | NA | NA | NA | NA | NA | NA |
| GRIN1 | 633 | G | -3.819892857 | 13.5272471 | 0.149193829 | 10.87938292 | NA | NA | NA | NA | NA | NA | NA |
| GRIN1 | 634 | M | -3.120392857 | 11.16284686 | 0.117165059 | 8.502370919 | NA | NA | NA | NA | NA | NA | NA |
| GRIN1 | 635 | V | 0 | 14.36107642 | 0.160489094 | 11.20237344 | NA | NA | NA | NA | NA | NA | NA |
| GRIN1 | 636 | W | -0.313571429 | 11.85755325 | 0.12657573 | 8.69953257 | NA | NA | NA | NA | NA | NA | NA |

| | | | | | | | | | | | | | |
|---|---|---|---|---|---|---|---|---|---|---|---|---|---|
| GRIN1 | 637 | A | -0.163742857 | 9.52268292 | 0.09494698 | 6.35877693 | 3.23 | 0.63 | 1.6 | 1.27 | -0.82 | 0 | 0 |
| GRIN1 | 638 | G | 1.935642857 | 11.13968985 | 0.116851369 | 8.060618131 | NA | NA | NA | NA | NA | NA | NA |
| GRIN1 | 639 | F | 2.802051948 | 10.9656951 | 0.114494391 | 7.938859506 | NA | NA | NA | NA | NA | NA | NA |
| GRIN1 | 640 | A | 2.529142857 | 7.717493699 | 0.070493426 | 4.71933751 | 3.12 | 1.1 | 3.53 | 0.46 | -1.28 | 0 | 0 |
| GRIN1 | 641 | M | 3.491767857 | 4.713443964 | 0.029799801 | 1.623226788 | 3.12 | 1.1 | 3.53 | 0.46 | -1.28 | 0 | 0 |
| GRIN1 | 642 | I | 7.175517857 | 9.338255619 | 0.092448681 | 6.244742447 | 3.19 | 1.11 | 3.58 | 0.39 | -1.38 | 0 | 0 |
| GRIN1 | 643 | I | 6.775892857 | 9.360583796 | 0.092751144 | 6.271800988 | 3.13 | 1.11 | 3.58 | 0.39 | -1.38 | 0 | 0 |
| GRIN1 | 644 | V | 6.576 | 4.765972514 | 0.030511367 | 1.684029178 | 3.19 | 1.11 | 3.58 | 0.39 | -1.38 | 0 | 0 |
| GRIN1 | 645 | A | 9.815742857 | 6.18398108 | 0.049720073 | 3.92881968 | 2.71 | -0.77 | -0.7 | 1.66 | 0.77 | 0 | 0 |
| GRIN1 | 646 | S | 11.66897619 | 8.060853553 | 0.075144666 | 5.986909434 | 2.11 | -0.18 | 1.18 | 0.42 | 0.64 | 0 | 0 |
| GRIN1 | 647 | Y | 10.02939286 | 7.667651727 | 0.069818254 | 5.209948645 | 2.67 | -0.77 | -0.7 | 1.66 | 0.77 | 0 | 0 |
| GRIN1 | 648 | T | 11.64942857 | 4.182871502 | 0.022612531 | 1.623091593 | 2.69 | -0.77 | -0.7 | 1.66 | 0.77 | 0 | 0 |
| GRIN1 | 649 | A | 15.21714286 | 4.624831456 | 0.028599434 | 2.53931143 | 2.11 | -0.18 | 1.18 | 0.42 | 0.64 | 0 | 0 |
| GRIN1 | 650 | N | 16.65839286 | 6.704843771 | 0.056775812 | 5.048359585 | 1.82 | -0.19 | 1.25 | 0.85 | 0.65 | 0 | 0 |
| GRIN1 | 651 | L | 14.64476786 | 6.551780369 | 0.054702376 | 5.097541286 | 1.93 | -0.31 | 0.92 | 1.35 | 0.68 | 0 | 0 |
| GRIN1 | 652 | A | 17.75834286 | 3.499872141 | 0.013360448 | 1.60782906 | 2.01 | 0.99 | 3.5 | 0.09 | -0.85 | 0 | 0 |
| GRIN1 | 653 | A | 20.41114286 | 4.22904079 | 0.023237952 | 2.304175952 | 2.01 | 0.99 | 3.5 | 0.09 | -0.85 | 0 | 0 |
| GRIN1 | 654 | F | 21.37041558 | 6.393567862 | 0.052559189 | 4.488057315 | 2.01 | 0.99 | 3.5 | 0.09 | -0.85 | 0 | 0 |
| GRIN1 | 655 | L | 19.94026786 | 6.36347468 | 0.052151539 | 4.480306178 | 2.01 | 0.99 | 3.5 | 0.09 | -0.85 | 0 | 0 |
| GRIN1 | 656 | V | 23.25028571 | 3.31047338 | 0.010794804 | 1.874802304 | 1.53 | 1.58 | 4.4 | 0.13 | -1.67 | 0 | 0 |
| GRIN1 | 657 | L | 25.12064286 | 3.59107825 | 0.014595949 | 2.152790343 | 1.55 | 1.47 | 4.25 | 0.13 | -1.7 | 0 | 0 |
| GRIN1 | 658 | D | 27.59834286 | 7.685814531 | 0.070064292 | 5.125550225 | 2.74 | 1.31 | 4.08 | 0.13 | -1.68 | 0 | 0 |
| GRIN1 | 659 | R | 28.47274286 | 11.00806005 | 0.115068278 | 8.268210788 | NA | NA | NA | NA | NA | NA | NA |
| GRIN1 | 660 | P | 28.90294286 | 10.97280976 | 0.114590768 | 9.173305957 | NA | NA | NA | NA | NA | NA | NA |
| GRIN1 | 661 | E | 32.24054286 | 12.79406382 | 0.139261941 | 6.845514419 | NA | NA | NA | NA | NA | NA | NA |
| GRIN1 | 662 | E | 35.79214286 | 14.1074659 | 0.157053621 | 7.220016799 | NA | NA | NA | NA | NA | NA | NA |
| GRIN1 | 663 | R | 35.47932468 | 11.71190787 | 0.12460278 | 4.923950855 | 6.89 | -0.78 | -2.03 | 2.95 | 0.67 | 0 | 0 |
| GRIN1 | 664 | I | 41.98289286 | 12.82245456 | 0.139646529 | 6.27356102 | NA | NA | NA | NA | NA | NA | NA |
| GRIN1 | 665 | T | 43.18085714 | 8.789389114 | 0.085013595 | 2.241363445 | 6.57 | -0.77 | -3.5 | 3.38 | 0.54 | 0 | 0 |
| GRIN1 | 666 | G | 46.62964286 | 11.31347718 | 0.119205536 | 5.235079133 | NA | NA | NA | NA | NA | NA | NA |
| GRIN1 | 667 | I | 50.79301786 | 11.53851208 | 0.122253916 | 6.454214083 | NA | NA | NA | NA | NA | NA | NA |
| GRIN1 | 668 | N | 49.44176786 | 6.877899098 | 0.059120063 | 1.779923096 | 5.14 | -0.68 | -3.05 | 15.11 | 0.87 | 1 | 1 |
| GRIN1 | 669 | D | 46.04364286 | 9.525974806 | 0.094991573 | 2.980408369 | 6.57 | -0.77 | -3.5 | 3.38 | 0.54 | 0 | 0 |

| | | | | | | | | | | | | |
|---|---|---|---|---|---|---|---|---|---|---|---|---|
| GRIN1 | 670 | P | 46.62214286 | 11.40662303 | 0.120467314 | 4.879797713 | 6.57 | -0.77 | -3.5 | 3.38 | 0.54 | 0 | 0 |
| GRIN1 | 671 | R | 43.27050649 | 14.60768681 | 0.163829741 | 8.070277835 | NA | NA | NA | NA | NA | NA | NA |
| GRIN1 | 672 | L | 49.42076786 | 15.85878618 | 0.180777452 | 9.310767391 | NA | NA | NA | NA | NA | NA | NA |
| GRIN1 | 673 | R | 51.32187013 | 8.047059214 | 0.074957805 | 4.167863128 | 4.08 | -0.65 | -2.9 | 19.01 | 0.98 | 1 | 1 |
| GRIN1 | 674 | N | 49.93551786 | 16.36160915 | 0.187588821 | 9.843500519 | NA | NA | NA | NA | NA | NA | NA |
| GRIN1 | 675 | P | 50.39214286 | 19.68456913 | 0.232602485 | 13.14300182 | NA | NA | NA | NA | NA | NA | NA |
| GRIN1 | 676 | S | 46.77897619 | 21.63747721 | 0.259057076 | 15.10720479 | NA | NA | NA | NA | NA | NA | NA |
| GRIN1 | 677 | D | 46.77839286 | 24.91329553 | 0.303432147 | 18.38123135 | NA | NA | NA | NA | NA | NA | NA |
| GRIN1 | 678 | K | 41.65125397 | 24.15856225 | 0.293208338 | 17.62505178 | NA | NA | NA | NA | NA | NA | NA |
| GRIN1 | 679 | F | 43.26450649 | 19.42266115 | 0.229054612 | 13.03730227 | NA | NA | NA | NA | NA | NA | NA |
| GRIN1 | 680 | I | 46.13776786 | 23.99431656 | 0.290983424 | 17.44890606 | NA | NA | NA | NA | NA | NA | NA |
| GRIN1 | 681 | Y | 47.45355952 | 18.62948741 | 0.218310078 | 12.08358766 | NA | NA | NA | NA | NA | NA | NA |
| GRIN1 | 682 | A | 49.92954286 | 25.02514226 | 0.304947252 | 19.92714722 | NA | NA | NA | NA | NA | NA | NA |
| GRIN1 | 683 | T | 52.55214286 | 24.55714081 | 0.298607585 | 19.46968764 | NA | NA | NA | NA | NA | NA | NA |
| GRIN1 | 684 | V | 53.939 | 28.64436044 | 0.353974105 | 23.56203893 | NA | NA | NA | NA | NA | NA | NA |
| GRIN1 | 685 | K | 57.45725397 | 31.23787824 | 0.389106559 | 27.16745639 | NA | NA | NA | NA | NA | NA | NA |
| GRIN1 | 686 | Q | 60.76525397 | 31.30139633 | 0.389966992 | 27.33348761 | NA | NA | NA | NA | NA | NA | NA |
| GRIN1 | 687 | S | 58.26897619 | 27.89267664 | 0.343791604 | 23.78615126 | NA | NA | NA | NA | NA | NA | NA |
| GRIN1 | 688 | S | 58.47880952 | 24.44895795 | 0.297142112 | 20.46708593 | NA | NA | NA | NA | NA | NA | NA |
| GRIN1 | 689 | V | 56.38514286 | 22.85851152 | 0.275597519 | 18.78668588 | NA | NA | NA | NA | NA | NA | NA |
| GRIN1 | 690 | D | 59.27489286 | 24.6599453 | 0.3000002 | 20.65342805 | NA | NA | NA | NA | NA | NA | NA |
| GRIN1 | 691 | I | 62.28464286 | 23.00496281 | 0.277581385 | 18.99985024 | NA | NA | NA | NA | NA | NA | NA |
| GRIN1 | 692 | Y | 59.72639286 | 19.45161788 | 0.229446868 | 15.44120786 | NA | NA | NA | NA | NA | NA | NA |
| GRIN1 | 693 | F | 57.77696104 | 18.69792259 | 0.219237119 | 14.79912765 | NA | NA | NA | NA | NA | NA | NA |
| GRIN1 | 694 | R | 63.21223377 | 22.54605642 | 0.271364922 | 18.5362064 | NA | NA | NA | NA | NA | NA | NA |
| GRIN1 | 695 | R | 65.15314286 | 20.2232217 | 0.23989921 | 16.33059576 | NA | NA | NA | NA | NA | NA | NA |
| GRIN1 | 696 | Q | 61.53614286 | 14.74467016 | 0.165685352 | 10.76063075 | NA | NA | NA | NA | NA | NA | NA |
| GRIN1 | 697 | V | 63.244 | 16.80299595 | 0.193567959 | 12.94601588 | NA | NA | NA | NA | NA | NA | NA |
| GRIN1 | 698 | E | 60.04947619 | 11.3031285 | 0.11906535 | 7.34301518 | NA | NA | NA | NA | NA | NA | NA |
| GRIN1 | 699 | L | 57.39226786 | 14.81495545 | 0.166637455 | 10.88322186 | NA | NA | NA | NA | NA | NA | NA |
| GRIN1 | 700 | S | 60.19030952 | 18.85277147 | 0.221334741 | 14.84659188 | NA | NA | NA | NA | NA | NA | NA |
| GRIN1 | 701 | T | 56.90357143 | 18.90538032 | 0.222047394 | 14.97488278 | NA | NA | NA | NA | NA | NA | NA |
| GRIN1 | 702 | M | 54.93239286 | 18.65793158 | 0.21869539 | 14.78704588 | NA | NA | NA | NA | NA | NA | NA |

| | | | | | | | | | | | | | |
|---|---|---|---|---|---|---|---|---|---|---|---|---|---|
| GRIN1 | 703 | Y | 59.56672619 | 21.06176992 | 0.251258398 | 17.05298166 | NA | NA | NA | NA | NA | NA | NA |
| GRIN1 | 704 | R | 59.29941558 | 25.29713361 | 0.308631716 | 21.35704215 | NA | NA | NA | NA | NA | NA | NA |
| GRIN1 | 705 | H | 52.73494286 | 24.3296801 | 0.295526344 | 20.41526558 | NA | NA | NA | NA | NA | NA | NA |
| GRIN1 | 706 | M | 54.02114286 | 22.79815104 | 0.27477986 | 18.78953874 | NA | NA | NA | NA | NA | NA | NA |
| GRIN1 | 707 | E | 58.4385873 | 26.7484862 | 0.328292109 | 22.75475933 | NA | NA | NA | NA | NA | NA | NA |
| GRIN1 | 708 | K | 55.42525397 | 29.53989472 | 0.366105241 | 25.58611533 | NA | NA | NA | NA | NA | NA | NA |
| GRIN1 | 709 | H | 51.09674286 | 29.08971461 | 0.360006986 | 24.18382562 | NA | NA | NA | NA | NA | NA | NA |
| GRIN1 | 710 | N | 53.59376786 | 28.36594035 | 0.350202556 | 23.29371165 | NA | NA | NA | NA | NA | NA | NA |
| GRIN1 | 711 | Y | 49.74280952 | 30.67243489 | 0.381446919 | 25.70797937 | NA | NA | NA | NA | NA | NA | NA |
| GRIN1 | 712 | E | 53.49736508 | 34.02615396 | 0.426877254 | 28.94150658 | NA | NA | NA | NA | NA | NA | NA |
| GRIN1 | 713 | S | 49.95464286 | 32.52413352 | 0.406530502 | 27.45353758 | NA | NA | NA | NA | NA | NA | NA |
| GRIN1 | 714 | A | 47.58534286 | 31.42910711 | 0.391696994 | 26.3644311 | NA | NA | NA | NA | NA | NA | NA |
| GRIN1 | 715 | A | 44.92734286 | 33.51489658 | 0.419951631 | 27.54892758 | NA | NA | NA | NA | NA | NA | NA |
| GRIN1 | 716 | E | 47.3165873 | 32.33106792 | 0.403915186 | 26.26616562 | NA | NA | NA | NA | NA | NA | NA |
| GRIN1 | 717 | A | 45.10854286 | 29.15287125 | 0.360862522 | 23.04520725 | NA | NA | NA | NA | NA | NA | NA |
| GRIN1 | 718 | I | 42.24014286 | 30.51065106 | 0.379255354 | 24.43545701 | NA | NA | NA | NA | NA | NA | NA |
| GRIN1 | 719 | Q | 40.81980952 | 32.23851689 | 0.402661466 | 25.70550738 | NA | NA | NA | NA | NA | NA | NA |
| GRIN1 | 720 | A | 41.78314286 | 29.47871425 | 0.365276475 | 22.92994562 | NA | NA | NA | NA | NA | NA | NA |
| GRIN1 | 721 | V | 39.62628571 | 26.5420068 | 0.325495086 | 20.00090296 | NA | NA | NA | NA | NA | NA | NA |
| GRIN1 | 722 | R | 37.5735974 | 30.11914278 | 0.373951883 | 23.33907482 | NA | NA | NA | NA | NA | NA | NA |
| GRIN1 | 723 | D | 37.90376786 | 31.54371242 | 0.393249467 | 24.75630583 | NA | NA | NA | NA | NA | NA | NA |
| GRIN1 | 724 | N | 36.36614286 | 27.95719977 | 0.344665651 | 21.1668038 | NA | NA | NA | NA | NA | NA | NA |
| GRIN1 | 725 | K | 41.23525397 | 28.31078966 | 0.34945547 | 21.76411953 | NA | NA | NA | NA | NA | NA | NA |
| GRIN1 | 726 | L | 43.42801786 | 25.41410295 | 0.310216213 | 18.86423826 | NA | NA | NA | NA | NA | NA | NA |
| GRIN1 | 727 | H | 41.34524286 | 22.18435523 | 0.266465225 | 15.63523889 | NA | NA | NA | NA | NA | NA | NA |
| GRIN1 | 728 | A | 44.23274286 | 19.86071502 | 0.234988602 | 13.31314293 | NA | NA | NA | NA | NA | NA | NA |
| GRIN1 | 729 | F | 46.19723377 | 20.20166553 | 0.239607205 | 14.11046669 | NA | NA | NA | NA | NA | NA | NA |
| GRIN1 | 730 | I | 49.99614286 | 17.79629136 | 0.207023392 | 12.70183862 | NA | NA | NA | NA | NA | NA | NA |
| GRIN1 | 731 | W | 50.41007143 | 21.44769186 | 0.256486195 | 16.39164513 | NA | NA | NA | NA | NA | NA | NA |
| GRIN1 | 732 | D | 50.11501786 | 23.73857799 | 0.287519124 | 18.71467875 | NA | NA | NA | NA | NA | NA | NA |
| GRIN1 | 733 | S | 47.81047619 | 21.09547018 | 0.251714911 | 16.08702552 | NA | NA | NA | NA | NA | NA | NA |
| GRIN1 | 734 | A | 46.45194286 | 24.10951325 | 0.292543907 | 19.10150846 | NA | NA | NA | NA | NA | NA | NA |
| GRIN1 | 735 | V | 45.798 | 26.3794989 | 0.323293712 | 21.36376053 | NA | NA | NA | NA | NA | NA | NA |

| | | | | | | | | | | | | | |
|---|---|---|---|---|---|---|---|---|---|---|---|---|---|
| GRIN1 | 736 | L | 44.16664286 | 22.57132307 | 0.271707191 | 17.02883448 | NA | NA | NA | NA | NA | NA | NA |
| GRIN1 | 737 | E | 42.43869841 | 20.83407997 | 0.248174052 | 15.34828225 | NA | NA | NA | NA | NA | NA | NA |
| GRIN1 | 738 | F | 41.67105195 | 26.8531398 | 0.329709773 | 21.18838515 | NA | NA | NA | NA | NA | NA | NA |
| GRIN1 | 739 | E | 40.47480952 | 27.41153153 | 0.33727389 | 21.51309504 | NA | NA | NA | NA | NA | NA | NA |
| GRIN1 | 740 | A | 37.43054286 | 23.99529731 | 0.290996709 | 17.17460715 | NA | NA | NA | NA | NA | NA | NA |
| GRIN1 | 741 | S | 36.33930952 | 26.59320221 | 0.326188592 | 19.74981253 | NA | NA | NA | NA | NA | NA | NA |
| GRIN1 | 742 | Q | 36.81947619 | 29.15295344 | 0.360863636 | 22.29963881 | NA | NA | NA | NA | NA | NA | NA |
| GRIN1 | 743 | K | 35.02536508 | 27.43544139 | 0.337597779 | 20.56820925 | NA | NA | NA | NA | NA | NA | NA |
| GRIN1 | 744 | C | 31.49614286 | 25.47630534 | 0.311058822 | 19.34948025 | NA | NA | NA | NA | NA | NA | NA |
| GRIN1 | 745 | D | 34.16726786 | 24.41560284 | 0.296690275 | 17.57533049 | NA | NA | NA | NA | NA | NA | NA |
| GRIN1 | 746 | L | 37.68164286 | 22.65776092 | 0.2728781 | 15.88402338 | NA | NA | NA | NA | NA | NA | NA |
| GRIN1 | 747 | V | 38.05985714 | 17.99602276 | 0.209729005 | 11.22316932 | NA | NA | NA | NA | NA | NA | NA |
| GRIN1 | 748 | T | 41.25557143 | 17.17965547 | 0.198670285 | 11.13877638 | NA | NA | NA | NA | NA | NA | NA |
| GRIN1 | 749 | T | 42.23585714 | 14.46868221 | 0.161946749 | 8.390718489 | NA | NA | NA | NA | NA | NA | NA |
| GRIN1 | 750 | G | 44.39014286 | 12.41639195 | 0.134145901 | 6.75642816 | NA | NA | NA | NA | NA | NA | NA |
| GRIN1 | 751 | E | 47.90314286 | 8.115972277 | 0.075891319 | 3.138536927 | 5.14 | -0.68 | -3.05 | 15.11 | 0.87 | 1 | 1 |
| GRIN1 | 752 | L | 50.08164286 | 13.18812625 | 0.144600011 | 8.180776086 | NA | NA | NA | NA | NA | NA | NA |
| GRIN1 | 753 | F | 51.98941558 | 11.95899887 | 0.127949938 | 6.908988122 | NA | NA | NA | NA | NA | NA | NA |
| GRIN1 | 754 | F | 57.21241558 | 16.53692635 | 0.189963712 | 12.38210979 | NA | NA | NA | NA | NA | NA | NA |
| GRIN1 | 755 | R | 54.25623377 | 13.35315266 | 0.146835501 | 10.02778504 | NA | NA | NA | NA | NA | NA | NA |
| GRIN1 | 756 | S | 57.34780952 | 20.97048769 | 0.250021866 | 16.95725611 | NA | NA | NA | NA | NA | NA | NA |
| GRIN1 | 757 | G | 57.84264286 | 24.51595293 | 0.298049643 | 19.5360762 | NA | NA | NA | NA | NA | NA | NA |
| GRIN1 | 758 | F | 55.61332468 | 25.4879426 | 0.311216463 | 20.49118139 | NA | NA | NA | NA | NA | NA | NA |
| GRIN1 | 759 | G | 59.56814286 | 30.73386878 | 0.382279118 | 26.84112891 | NA | NA | NA | NA | NA | NA | NA |
| GRIN1 | 760 | I | 61.93251786 | 34.08502903 | 0.427674791 | 30.09102661 | NA | NA | NA | NA | NA | NA | NA |
| GRIN1 | 761 | G | 64.69464286 | 36.33533623 | 0.458158027 | 32.32529663 | NA | NA | NA | NA | NA | NA | NA |
| GRIN1 | 762 | M | 67.74114286 | 37.94304101 | 0.479936406 | 33.96200265 | NA | NA | NA | NA | NA | NA | NA |
| GRIN1 | 763 | R | 71.77577922 | 39.96381727 | 0.507310357 | 36.56977403 | NA | NA | NA | NA | NA | NA | NA |
| GRIN1 | 764 | K | 74.5665873 | 32.88714962 | 0.41144801 | 28.77452661 | NA | NA | NA | NA | NA | NA | NA |
| GRIN1 | 765 | D | 76.92089286 | 40.26366373 | 0.511372154 | 36.21346553 | NA | NA | NA | NA | NA | NA | NA |
| GRIN1 | 766 | S | 72.40264286 | 42.11477629 | 0.536447797 | 38.04549523 | NA | NA | NA | NA | NA | NA | NA |
| GRIN1 | 767 | P | 72.793 | 44.11846092 | 0.563590221 | 40.03659226 | NA | NA | NA | NA | NA | NA | NA |
| GRIN1 | 768 | W | 69.04835714 | 41.78577549 | 0.531991068 | 37.91136908 | NA | NA | NA | NA | NA | NA | NA |

| | | | | | | | | | | | | | |
|---|---|---|---|---|---|---|---|---|---|---|---|---|---|
| GRIN1 | 769 | K | 70.31914286 | 37.37480195 | 0.472238895 | 35.82974222 | NA | NA | NA | NA | NA | NA | NA |
| GRIN1 | 770 | Q | 69.89580952 | 39.73875338 | 0.504261584 | 36.87207803 | NA | NA | NA | NA | NA | NA | NA |
| GRIN1 | 771 | N | 66.71651786 | 42.39122264 | 0.54019261 | 38.74614272 | NA | NA | NA | NA | NA | NA | NA |
| GRIN1 | 772 | V | 63.86414286 | 39.4004315 | 0.499678589 | 35.50665735 | NA | NA | NA | NA | NA | NA | NA |
| GRIN1 | 773 | S | 64.00030952 | 37.63603479 | 0.475777621 | 33.71497078 | NA | NA | NA | NA | NA | NA | NA |
| GRIN1 | 774 | L | 63.64301786 | 39.65213192 | 0.503088188 | 35.78648979 | NA | NA | NA | NA | NA | NA | NA |
| GRIN1 | 775 | S | 59.96664286 | 38.99797843 | 0.494226858 | 34.03773402 | NA | NA | NA | NA | NA | NA | NA |
| GRIN1 | 777 | L | 59.36601786 | 33.82641338 | 0.424171517 | 28.87299039 | NA | NA | NA | NA | NA | NA | NA |
| GRIN1 | 778 | K | 57.9075873 | 38.00188908 | 0.480733577 | 33.04633127 | NA | NA | NA | NA | NA | NA | NA |
| GRIN1 | 779 | S | 54.45980952 | 35.52144915 | 0.447132904 | 30.56130133 | NA | NA | NA | NA | NA | NA | NA |
| GRIN1 | 780 | H | 54.82094286 | 31.18052334 | 0.388329615 | 26.21849797 | NA | NA | NA | NA | NA | NA | NA |
| GRIN1 | 781 | E | 54.78969841 | 35.17948644 | 0.44250059 | 30.22720651 | NA | NA | NA | NA | NA | NA | NA |
| GRIN1 | 782 | N | 51.98326786 | 37.13622213 | 0.469007031 | 32.17987981 | NA | NA | NA | NA | NA | NA | NA |
| GRIN1 | 783 | G | 48.55764286 | 35.96342511 | 0.453120024 | 31.00315898 | NA | NA | NA | NA | NA | NA | NA |
| GRIN1 | 784 | F | 50.11396104 | 35.75371223 | 0.450279199 | 30.78775768 | NA | NA | NA | NA | NA | NA | NA |
| GRIN1 | 785 | M | 50.23076786 | 31.23637457 | 0.38908619 | 26.2687749 | NA | NA | NA | NA | NA | NA | NA |
| GRIN1 | 786 | E | 45.75492063 | 31.60662369 | 0.394101679 | 26.64551789 | NA | NA | NA | NA | NA | NA | NA |
| GRIN1 | 787 | D | 45.52751786 | 35.39361878 | 0.445401282 | 30.42457866 | NA | NA | NA | NA | NA | NA | NA |
| GRIN1 | 788 | L | 46.81351786 | 34.04259818 | 0.427100012 | 29.05993964 | NA | NA | NA | NA | NA | NA | NA |
| GRIN1 | 789 | D | 44.70489286 | 29.92667267 | 0.371344634 | 24.95592116 | NA | NA | NA | NA | NA | NA | NA |
| GRIN1 | 790 | K | 40.78747619 | 33.78889952 | 0.423663345 | 28.80933059 | NA | NA | NA | NA | NA | NA | NA |
| GRIN1 | 791 | T | 42.09657143 | 36.14220886 | 0.455541874 | 31.15906864 | NA | NA | NA | NA | NA | NA | NA |
| GRIN1 | 792 | W | 43.52907143 | 33.51125853 | 0.419902349 | 28.51425908 | NA | NA | NA | NA | NA | NA | NA |
| GRIN1 | 793 | V | 42.438 | 29.12706394 | 0.36051293 | 24.13772034 | NA | NA | NA | NA | NA | NA | NA |
| GRIN1 | 794 | R | 40.34650649 | 25.60889541 | 0.312854921 | 20.25892019 | NA | NA | NA | NA | NA | NA | NA |
| GRIN1 | 795 | Y | 36.65889286 | 32.48587091 | 0.406012187 | 25.59856607 | NA | NA | NA | NA | NA | NA | NA |
| GRIN1 | 796 | Q | 35.57114286 | 29.04353687 | 0.359381451 | 22.15791027 | NA | NA | NA | NA | NA | NA | NA |
| GRIN1 | 797 | E | 31.86892063 | 27.79850893 | 0.342515985 | 21.75615775 | NA | NA | NA | NA | NA | NA | NA |
| GRIN1 | 798 | C | 30.60547619 | 25.39215332 | 0.309918877 | 23.67244258 | NA | NA | NA | NA | NA | NA | NA |
| GRIN1 | 799 | D | 26.71964286 | 22.95686882 | 0.276929892 | 21.2543707 | NA | NA | NA | NA | NA | NA | NA |
| GRIN1 | 800 | S | 27.25380952 | 18.4499932 | 0.215878604 | 16.76202305 | NA | NA | NA | NA | NA | NA | NA |
| GRIN1 | 801 | R | 25.32323377 | 15.64735137 | 0.177913302 | 14.03582858 | NA | NA | NA | NA | NA | NA | NA |
| GRIN1 | 802 | S | 22.25080952 | 18.89321521 | 0.221882603 | 17.18894854 | NA | NA | NA | NA | NA | NA | NA |

| | | | | | | | | | | | | | |
|---|---|---|---|---|---|---|---|---|---|---|---|---|---|
| GRIN1 | 809 | T | 12.65971429 | 19.37589474 | 0.228421103 | 17.02488133 | NA | NA | NA | NA | NA | NA | NA |
| GRIN1 | 810 | F | 14.5265974 | 15.79674194 | 0.179936985 | 13.85431726 | NA | NA | NA | NA | NA | NA | NA |
| GRIN1 | 811 | E | 16.29925397 | 18.48222433 | 0.216315215 | 16.55918502 | NA | NA | NA | NA | NA | NA | NA |
| GRIN1 | 812 | N | 13.23301786 | 13.65825948 | 0.150968556 | 11.73780882 | NA | NA | NA | NA | NA | NA | NA |
| GRIN1 | 813 | M | 9.955267857 | 15.99138096 | 0.182573615 | 13.37460689 | NA | NA | NA | NA | NA | NA | NA |
| GRIN1 | 814 | A | 10.37034286 | 20.48888921 | 0.24349801 | 17.8255695 | NA | NA | NA | NA | NA | NA | NA |
| GRIN1 | 815 | G | 9.462392857 | 20.27745351 | 0.240633848 | 18.06093702 | NA | NA | NA | NA | NA | NA | NA |
| GRIN1 | 816 | V | 6.928857143 | 15.91817797 | 0.181581989 | 12.99450376 | NA | NA | NA | NA | NA | NA | NA |
| GRIN1 | 817 | F | 5.215142857 | 18.87292627 | 0.221607763 | 15.73663079 | NA | NA | NA | NA | NA | NA | NA |
| GRIN1 | 818 | M | 6.720892857 | 22.01373246 | 0.264153926 | 18.96604628 | NA | NA | NA | NA | NA | NA | NA |
| GRIN1 | 819 | L | 5.452642857 | 19.24045579 | 0.226586412 | 16.29601649 | NA | NA | NA | NA | NA | NA | NA |
| GRIN1 | 820 | V | 1.619 | 16.70464013 | 0.192235606 | 13.5906471 | NA | NA | NA | NA | NA | NA | NA |
| GRIN1 | 821 | A | 0.710942857 | 21.03915597 | 0.250952064 | 17.8877397 | NA | NA | NA | NA | NA | NA | NA |
| GRIN1 | 822 | G | 0.373142857 | 23.44647679 | 0.283562246 | 20.30300189 | NA | NA | NA | NA | NA | NA | NA |
| GRIN1 | 823 | G | 0.217392857 | 22.21490846 | 0.266879107 | 19.04977103 | NA | NA | NA | NA | NA | NA | NA |
| GRIN1 | 824 | I | -1.849142857 | 17.9869868 | 0.209606601 | 15.28162357 | NA | NA | NA | NA | NA | NA | NA |
| GRIN1 | 825 | V | -2.178285714 | 23.77275924 | 0.287982152 | 21.07349417 | NA | NA | NA | NA | NA | NA | NA |
| GRIN1 | 826 | A | -2.537142857 | 25.64039105 | 0.313281569 | 22.94416848 | NA | NA | NA | NA | NA | NA | NA |
| GRIN1 | 827 | G | -5.302142857 | 23.7981777 | 0.288326477 | 21.17574381 | NA | NA | NA | NA | NA | NA | NA |
| GRIN1 | 828 | I | -6.820892857 | 24.23479682 | 0.294241031 | 21.5845408 | NA | NA | NA | NA | NA | NA | NA |
| GRIN1 | 829 | F | -6.438597403 | 27.30179346 | 0.33578735 | 24.65771537 | NA | NA | NA | NA | NA | NA | NA |
| GRIN1 | 830 | L | -7.367267857 | 26.55713857 | 0.325700065 | 23.939912 | NA | NA | NA | NA | NA | NA | NA |
| GRIN1 | 831 | I | -10.85251786 | 22.97858805 | 0.277224106 | 17.67728658 | NA | NA | NA | NA | NA | NA | NA |
| GRIN1 | 832 | F | -12.67032468 | 27.53905104 | 0.339001302 | 21.98814682 | NA | NA | NA | NA | NA | NA | NA |
| GRIN1 | 833 | I | -12.44101786 | 29.79531644 | 0.369565249 | 24.30848687 | NA | NA | NA | NA | NA | NA | NA |
| GRIN1 | 834 | E | -11.75914286 | 26.39554701 | 0.323511104 | 21.10901882 | NA | NA | NA | NA | NA | NA | NA |
| GRIN1 | 835 | I | -15.84914286 | 22.90572125 | 0.276237034 | 16.8936912 | NA | NA | NA | NA | NA | NA | NA |
| GRIN1 | 836 | A | -17.90954286 | 28.09956587 | 0.346594179 | 22.08960304 | NA | NA | NA | NA | NA | NA | NA |
| GRIN1 | 837 | Y | -16.82205952 | 29.70609611 | 0.368356648 | 23.73129026 | NA | NA | NA | NA | NA | NA | NA |
| GRIN1 | 838 | K | -17.86603175 | 24.01584746 | 0.291275087 | 18.03148545 | NA | NA | NA | NA | NA | NA | NA |
| GRIN1 | 839 | R | -20.50605195 | 24.86178632 | 0.30273439 | 18.5445458 | NA | NA | NA | NA | NA | NA | NA |
| GRIN1 | 840 | H | -22.06984286 | 30.44302919 | 0.378339331 | 24.02453944 | NA | NA | NA | NA | NA | NA | NA |
| GRIN1 | 841 | K | -21.2815873 | 29.0113665 | 0.358945663 | 22.63143064 | NA | NA | NA | NA | NA | NA | NA |

| | | | | | | | | | | | | | |
|---|---|---|---|---|---|---|---|---|---|---|---|---|---|
| GRIN1 | 842 | D | -24.72664286 | 25.24692839 | 0.307951623 | 15.50497552 | NA | NA | NA | NA | NA | NA | NA |
| GRIN1 | 843 | A | -27.15934286 | 30.03448841 | 0.372805133 | 20.22733258 | NA | NA | NA | NA | NA | NA | NA |
| GRIN1 | 844 | R | -25.01841558 | 31.78986412 | 0.396583901 | 21.98130254 | NA | NA | NA | NA | NA | NA | NA |
| GRIN2A | 34 | L | 102.93475 | 72.97532279 | 0.954492638 | 71.45893427 | NA | NA | NA | NA | NA | NA | NA |
| GRIN2A | 35 | N | 106.311 | 69.90443663 | 0.91289363 | 68.36129217 | NA | NA | NA | NA | NA | NA | NA |
| GRIN2A | 36 | I | 104.89025 | 66.14602763 | 0.861981262 | 64.60123452 | NA | NA | NA | NA | NA | NA | NA |
| GRIN2A | 37 | A | 108.3507 | 64.29281171 | 0.836877126 | 62.75556952 | NA | NA | NA | NA | NA | NA | NA |
| GRIN2A | 38 | V | 108.7185 | 61.8849131 | 0.804259117 | 60.34868645 | NA | NA | NA | NA | NA | NA | NA |
| GRIN2A | 39 | M | 112.0115 | 59.25152571 | 0.768586579 | 57.72601961 | NA | NA | NA | NA | NA | NA | NA |
| GRIN2A | 40 | L | 111.263375 | 58.15118718 | 0.753681113 | 56.61857416 | NA | NA | NA | NA | NA | NA | NA |
| GRIN2A | 41 | G | 113.61475 | 57.32938989 | 0.742548837 | 55.81495723 | NA | NA | NA | NA | NA | NA | NA |
| GRIN2A | 42 | H | 116.7829 | 55.88388057 | 0.722967599 | 54.62154781 | NA | NA | NA | NA | NA | NA | NA |
| GRIN2A | 43 | S | 112.3926667 | 55.80145659 | 0.721851062 | 54.61762344 | NA | NA | NA | NA | NA | NA | NA |
| GRIN2A | 44 | H | 109.7341 | 55.7795046 | 0.721553695 | 54.24418857 | NA | NA | NA | NA | NA | NA | NA |
| GRIN2A | 45 | D | 112.753125 | 61.41083423 | 0.797837124 | 59.87968935 | NA | NA | NA | NA | NA | NA | NA |
| GRIN2A | 46 | V | 111.6977857 | 64.32570778 | 0.837322745 | 62.79514851 | NA | NA | NA | NA | NA | NA | NA |
| GRIN2A | 47 | T | 111.0073571 | 66.59759859 | 0.868098358 | 65.06437886 | NA | NA | NA | NA | NA | NA | NA |
| GRIN2A | 48 | E | 107.5422778 | 62.29503563 | 0.809814742 | 60.75704098 | NA | NA | NA | NA | NA | NA | NA |
| GRIN2A | 49 | R | 107.0707 | 63.98145385 | 0.832659393 | 62.44260937 | NA | NA | NA | NA | NA | NA | NA |
| GRIN2A | 50 | E | 107.2567 | 67.10525399 | 0.874975188 | 65.56530232 | NA | NA | NA | NA | NA | NA | NA |
| GRIN2A | 51 | L | 104.5201 | 67.77434693 | 0.884038891 | 66.23099147 | NA | NA | NA | NA | NA | NA | NA |
| GRIN2A | 52 | R | 102.7481 | 64.47259792 | 0.839312556 | 62.92863326 | NA | NA | NA | NA | NA | NA | NA |
| GRIN2A | 53 | T | 103.1379 | 67.25448442 | 0.876996701 | 65.71007281 | NA | NA | NA | NA | NA | NA | NA |
| GRIN2A | 54 | L | 102.0599 | 69.9664416 | 0.913733565 | 68.42143748 | NA | NA | NA | NA | NA | NA | NA |
| GRIN2A | 55 | W | 99.2045 | 68.20515244 | 0.889874693 | 66.66232921 | NA | NA | NA | NA | NA | NA | NA |
| GRIN2A | 56 | G | 98.0095 | 68.11960042 | 0.888715783 | 66.60041535 | NA | NA | NA | NA | NA | NA | NA |
| GRIN2A | 57 | P | 98.4453 | 70.51117963 | 0.921112726 | 68.9649334 | NA | NA | NA | NA | NA | NA | NA |
| GRIN2A | 58 | E | 96.2783 | 70.96939643 | 0.927319848 | 67.85173142 | NA | NA | NA | NA | NA | NA | NA |
| GRIN2A | 59 | Q | 93.6403 | 67.39092531 | 0.878844964 | 64.27378888 | NA | NA | NA | NA | NA | NA | NA |
| GRIN2A | 60 | A | 91.7619 | 67.46542654 | 0.879854177 | 64.34049324 | NA | NA | NA | NA | NA | NA | NA |
| GRIN2A | 61 | A | 90.9741 | 69.37855999 | 0.905769971 | 66.25977101 | NA | NA | NA | NA | NA | NA | NA |
| GRIN2A | 62 | G | 93.29175 | 72.18643148 | 0.943806115 | 69.07457498 | NA | NA | NA | NA | NA | NA | NA |
| GRIN2A | 63 | L | 96.01575 | 70.08468847 | 0.915335368 | 66.99429975 | NA | NA | NA | NA | NA | NA | NA |

| | | | | | | | | | | | | | |
|---|---|---|---|---|---|---|---|---|---|---|---|---|---|
| GRIN2A | 64 | P | 95.0975 | 74.26187093 | 0.971920548 | 71.1665764 | NA | NA | NA | NA | NA | NA | NA |
| GRIN2A | 65 | L | 97.790125 | 73.67243884 | 0.96393595 | 72.36761808 | NA | NA | NA | NA | NA | NA | NA |
| GRIN2A | 66 | D | 100.7015 | 75.55240823 | 0.989402495 | 74.08973602 | NA | NA | NA | NA | NA | NA | NA |
| GRIN2A | 67 | V | 101.8772143 | 70.71974328 | 0.923937982 | 69.17092847 | NA | NA | NA | NA | NA | NA | NA |
| GRIN2A | 68 | N | 106.06575 | 71.509402 | 0.934634901 | 69.96711377 | NA | NA | NA | NA | NA | NA | NA |
| GRIN2A | 69 | V | 106.6196429 | 67.383343 | 0.878742252 | 65.84135845 | NA | NA | NA | NA | NA | NA | NA |
| GRIN2A | 70 | V | 110.9195 | 67.14517672 | 0.875515991 | 65.6128764 | NA | NA | NA | NA | NA | NA | NA |
| GRIN2A | 71 | A | 111.6585 | 65.1503843 | 0.848494024 | 63.61757185 | NA | NA | NA | NA | NA | NA | NA |
| GRIN2A | 72 | L | 114.913375 | 62.93377555 | 0.818467276 | 61.41452219 | NA | NA | NA | NA | NA | NA | NA |
| GRIN2A | 73 | L | 116.200875 | 61.23270675 | 0.795424164 | 59.71320872 | NA | NA | NA | NA | NA | NA | NA |
| GRIN2A | 74 | M | 119.208625 | 58.84967549 | 0.763143014 | 57.62243679 | NA | NA | NA | NA | NA | NA | NA |
| GRIN2A | 75 | N | 119.076625 | 55.72576159 | 0.720825679 | 54.46193906 | NA | NA | NA | NA | NA | NA | NA |
| GRIN2A | 76 | R | 121.5827727 | 56.59542491 | 0.73260636 | 55.38079368 | NA | NA | NA | NA | NA | NA | NA |
| GRIN2A | 77 | T | 117.2336429 | 53.30806248 | 0.688074909 | 51.82658015 | NA | NA | NA | NA | NA | NA | NA |
| GRIN2A | 78 | D | 120.627625 | 56.31901325 | 0.728862017 | 55.05882439 | NA | NA | NA | NA | NA | NA | NA |
| GRIN2A | 79 | P | 119.2857857 | 57.54060134 | 0.745409961 | 56.07441986 | NA | NA | NA | NA | NA | NA | NA |
| GRIN2A | 80 | K | 123.2217222 | 60.90513835 | 0.790986838 | 59.5393428 | NA | NA | NA | NA | NA | NA | NA |
| GRIN2A | 81 | S | 119.3298333 | 60.22753366 | 0.781807832 | 58.74103039 | NA | NA | NA | NA | NA | NA | NA |
| GRIN2A | 82 | L | 116.150625 | 57.82468913 | 0.749258287 | 56.33082779 | NA | NA | NA | NA | NA | NA | NA |
| GRIN2A | 83 | I | 118.676125 | 63.18391924 | 0.821855786 | 61.69390314 | NA | NA | NA | NA | NA | NA | NA |
| GRIN2A | 84 | T | 120.1575 | 65.76515766 | 0.856821901 | 64.27707553 | NA | NA | NA | NA | NA | NA | NA |
| GRIN2A | 85 | H | 117.9551 | 64.29926458 | 0.836964539 | 62.80432861 | NA | NA | NA | NA | NA | NA | NA |
| GRIN2A | 86 | V | 113.9476429 | 63.44362607 | 0.825373841 | 61.93144088 | NA | NA | NA | NA | NA | NA | NA |
| GRIN2A | 87 | C | 116.3911667 | 67.64814339 | 0.882329306 | 66.14348051 | NA | NA | NA | NA | NA | NA | NA |
| GRIN2A | 88 | D | 117.819125 | 69.78614629 | 0.911291239 | 68.28135583 | NA | NA | NA | NA | NA | NA | NA |
| GRIN2A | 89 | L | 112.963625 | 66.10314251 | 0.86140033 | 64.58143559 | NA | NA | NA | NA | NA | NA | NA |
| GRIN2A | 90 | M | 112.38675 | 69.65901721 | 0.909569116 | 68.14075032 | NA | NA | NA | NA | NA | NA | NA |
| GRIN2A | 91 | S | 115.6121667 | 72.97042142 | 0.954426243 | 71.45499504 | NA | NA | NA | NA | NA | NA | NA |
| GRIN2A | 92 | G | 113.28875 | 73.93597783 | 0.967505917 | 72.40838267 | NA | NA | NA | NA | NA | NA | NA |
| GRIN2A | 93 | A | 110.4991 | 75.39704013 | 0.987297839 | 73.86096188 | NA | NA | NA | NA | NA | NA | NA |
| GRIN2A | 94 | R | 109.4133182 | 72.80089093 | 0.95212974 | 71.26620501 | NA | NA | NA | NA | NA | NA | NA |
| GRIN2A | 95 | I | 108.702375 | 68.75239613 | 0.897287796 | 67.2168811 | NA | NA | NA | NA | NA | NA | NA |
| GRIN2A | 96 | H | 104.5334 | 66.73286257 | 0.869930678 | 65.21735508 | NA | NA | NA | NA | NA | NA | NA |

| | | | | | | | | | | | | | |
|---|---|---|---|---|---|---|---|---|---|---|---|---|---|
| GRIN2A | 97 | G | 105.4795 | 64.81621958 | 0.843967343 | 63.275455 | NA | NA | NA | NA | NA | NA | NA |
| GRIN2A | 98 | L | 106.949875 | 60.79176596 | 0.789451067 | 59.25046763 | NA | NA | NA | NA | NA | NA | NA |
| GRIN2A | 99 | V | 106.2276429 | 59.62526828 | 0.773649392 | 58.08938393 | NA | NA | NA | NA | NA | NA | NA |
| GRIN2A | 100 | F | 108.7207727 | 55.25872107 | 0.714499028 | 53.72239492 | NA | NA | NA | NA | NA | NA | NA |
| GRIN2A | 101 | G | 108.22025 | 54.47341229 | 0.703861035 | 52.94417301 | NA | NA | NA | NA | NA | NA | NA |
| GRIN2A | 102 | D | 110.216125 | 50.89243351 | 0.655352183 | 49.36796698 | NA | NA | NA | NA | NA | NA | NA |
| GRIN2A | 103 | D | 111.382625 | 49.99632653 | 0.643213289 | 48.48907428 | NA | NA | NA | NA | NA | NA | NA |
| GRIN2A | 104 | T | 113.3122143 | 47.5575944 | 0.610177601 | 46.04782274 | NA | NA | NA | NA | NA | NA | NA |
| GRIN2A | 105 | D | 112.270875 | 44.23679287 | 0.565193175 | 42.74018914 | NA | NA | NA | NA | NA | NA | NA |
| GRIN2A | 106 | Q | 114.5807222 | 47.15470657 | 0.604719979 | 45.66259347 | NA | NA | NA | NA | NA | NA | NA |
| GRIN2A | 107 | E | 109.1321667 | 42.36316826 | 0.539812578 | 40.85596123 | NA | NA | NA | NA | NA | NA | NA |
| GRIN2A | 108 | A | 110.5633 | 48.39584945 | 0.621532817 | 46.8952805 | NA | NA | NA | NA | NA | NA | NA |
| GRIN2A | 109 | V | 112.1200714 | 50.94499004 | 0.656064127 | 49.44689881 | NA | NA | NA | NA | NA | NA | NA |
| GRIN2A | 110 | A | 108.3647 | 51.83301425 | 0.668093529 | 50.30976995 | NA | NA | NA | NA | NA | NA | NA |
| GRIN2A | 111 | Q | 107.9547222 | 48.29124852 | 0.620115867 | 46.77204163 | NA | NA | NA | NA | NA | NA | NA |
| GRIN2A | 112 | M | 112.783375 | 53.97955067 | 0.69717106 | 52.47617812 | NA | NA | NA | NA | NA | NA | NA |
| GRIN2A | 113 | L | 109.633375 | 56.12567781 | 0.726243046 | 54.60410186 | NA | NA | NA | NA | NA | NA | NA |
| GRIN2A | 114 | D | 107.302375 | 53.73376574 | 0.693841594 | 52.21225425 | NA | NA | NA | NA | NA | NA | NA |
| GRIN2A | 115 | F | 111.2608636 | 55.73483913 | 0.720948645 | 54.24037463 | NA | NA | NA | NA | NA | NA | NA |
| GRIN2A | 116 | I | 110.456625 | 60.68516367 | 0.788007005 | 59.17536093 | NA | NA | NA | NA | NA | NA | NA |
| GRIN2A | 117 | S | 106.7311667 | 59.35405135 | 0.769975418 | 57.84000059 | NA | NA | NA | NA | NA | NA | NA |
| GRIN2A | 118 | S | 109.1466667 | 61.73616134 | 0.802244088 | 60.22213718 | NA | NA | NA | NA | NA | NA | NA |
| GRIN2A | 119 | H | 112.0339 | 63.99522237 | 0.832845905 | 62.49743126 | NA | NA | NA | NA | NA | NA | NA |
| GRIN2A | 120 | T | 108.6576429 | 64.72076307 | 0.842674265 | 63.19522041 | NA | NA | NA | NA | NA | NA | NA |
| GRIN2A | 121 | F | 105.2536818 | 64.18686477 | 0.835441942 | 62.67367462 | NA | NA | NA | NA | NA | NA | NA |
| GRIN2A | 122 | V | 104.9033571 | 62.31211716 | 0.810046132 | 60.79730936 | NA | NA | NA | NA | NA | NA | NA |
| GRIN2A | 123 | P | 101.9646429 | 61.72237788 | 0.802057374 | 60.20710036 | NA | NA | NA | NA | NA | NA | NA |
| GRIN2A | 124 | I | 104.08425 | 57.73298463 | 0.748016034 | 56.21751583 | NA | NA | NA | NA | NA | NA | NA |
| GRIN2A | 125 | L | 101.562625 | 56.46230288 | 0.730803055 | 55.07375656 | NA | NA | NA | NA | NA | NA | NA |
| GRIN2A | 126 | G | 102.5845 | 52.8791536 | 0.6822648 | 51.33194073 | NA | NA | NA | NA | NA | NA | NA |
| GRIN2A | 127 | I | 103.402125 | 50.83762686 | 0.654609758 | 49.2935803 | NA | NA | NA | NA | NA | NA | NA |
| GRIN2A | 128 | H | 104.9519 | 50.05211577 | 0.643969024 | 48.51963652 | NA | NA | NA | NA | NA | NA | NA |
| GRIN2A | 129 | G | 107.14525 | 47.34567402 | 0.607306873 | 45.81774107 | NA | NA | NA | NA | NA | NA | NA |

| | | | | | | | | | | | | | |
|---|---|---|---|---|---|---|---|---|---|---|---|---|---|
| GRIN2A | 130 | G | 107.6685 | 48.61616275 | 0.624517238 | 47.09673302 | NA | NA | NA | NA | NA | NA | NA |
| GRIN2A | 131 | A | 105.4541 | 50.11025324 | 0.644756569 | 48.57365296 | NA | NA | NA | NA | NA | NA | NA |
| GRIN2A | 132 | S | 103.5011667 | 46.67099009 | 0.598167432 | 45.12683398 | NA | NA | NA | NA | NA | NA | NA |
| GRIN2A | 133 | M | 104.244375 | 41.52424035 | 0.528448246 | 39.99147087 | NA | NA | NA | NA | NA | NA | NA |
| GRIN2A | 134 | I | 100.685125 | 43.72355985 | 0.55824079 | 42.20843504 | NA | NA | NA | NA | NA | NA | NA |
| GRIN2A | 135 | M | 101.891625 | 40.65251168 | 0.516639587 | 39.13838399 | NA | NA | NA | NA | NA | NA | NA |
| GRIN2A | 136 | A | 101.7817 | 46.77652715 | 0.599597064 | 45.262181 | NA | NA | NA | NA | NA | NA | NA |
| GRIN2A | 137 | D | 100.37075 | 45.69611767 | 0.584961562 | 44.18413823 | NA | NA | NA | NA | NA | NA | NA |
| GRIN2A | 138 | K | 99.12172222 | 50.9172629 | 0.655688528 | 49.40434838 | NA | NA | NA | NA | NA | NA | NA |
| GRIN2A | 139 | D | 102.002875 | 54.36716286 | 0.702421753 | 52.85431669 | NA | NA | NA | NA | NA | NA | NA |
| GRIN2A | 140 | P | 98.80064286 | 56.61025067 | 0.732807193 | 55.10280542 | NA | NA | NA | NA | NA | NA | NA |
| GRIN2A | 141 | T | 101.6187857 | 59.02597213 | 0.765531173 | 57.51643883 | NA | NA | NA | NA | NA | NA | NA |
| GRIN2A | 142 | S | 100.8643333 | 55.81036958 | 0.7219718 | 54.29707583 | NA | NA | NA | NA | NA | NA | NA |
| GRIN2A | 143 | T | 101.4596429 | 58.31230077 | 0.755863599 | 56.79788789 | NA | NA | NA | NA | NA | NA | NA |
| GRIN2A | 144 | F | 102.6625 | 54.75455944 | 0.707669526 | 53.24008524 | NA | NA | NA | NA | NA | NA | NA |
| GRIN2A | 145 | F | 97.71286364 | 54.31626514 | 0.70173228 | 52.80072515 | NA | NA | NA | NA | NA | NA | NA |
| GRIN2A | 146 | Q | 100.3673889 | 50.59596856 | 0.651336193 | 49.09142406 | NA | NA | NA | NA | NA | NA | NA |
| GRIN2A | 147 | F | 97.31713636 | 48.23240817 | 0.6193188 | 46.71526123 | NA | NA | NA | NA | NA | NA | NA |
| GRIN2A | 148 | G | 97.9945 | 46.00865734 | 0.589195304 | 44.49245793 | NA | NA | NA | NA | NA | NA | NA |
| GRIN2A | 149 | A | 97.4895 | 42.40251872 | 0.540345629 | 40.88635526 | NA | NA | NA | NA | NA | NA | NA |
| GRIN2A | 150 | S | 94.0085 | 41.61930872 | 0.529736067 | 40.19894509 | NA | NA | NA | NA | NA | NA | NA |
| GRIN2A | 151 | I | 94.5335 | 38.76605559 | 0.491085172 | 37.41277662 | NA | NA | NA | NA | NA | NA | NA |
| GRIN2A | 152 | Q | 89.91616667 | 39.02353745 | 0.494573087 | 35.94854937 | NA | NA | NA | NA | NA | NA | NA |
| GRIN2A | 153 | Q | 89.95705556 | 39.12692448 | 0.495973594 | 36.04496844 | NA | NA | NA | NA | NA | NA | NA |
| GRIN2A | 154 | Q | 93.5825 | 36.48150349 | 0.460138046 | 35.58135609 | NA | NA | NA | NA | NA | NA | NA |
| GRIN2A | 155 | A | 89.3865 | 34.4674871 | 0.432855665 | 31.38461926 | NA | NA | NA | NA | NA | NA | NA |
| GRIN2A | 156 | T | 86.56064286 | 35.63039612 | 0.448608728 | 32.52422838 | NA | NA | NA | NA | NA | NA | NA |
| GRIN2A | 157 | V | 88.05035714 | 34.6402372 | 0.435195782 | 31.53508898 | NA | NA | NA | NA | NA | NA | NA |
| GRIN2A | 158 | M | 89.663125 | 31.30762401 | 0.390051353 | 28.24500407 | NA | NA | NA | NA | NA | NA | NA |
| GRIN2A | 159 | L | 85.2815 | 30.31293828 | 0.376577086 | 27.22169038 | NA | NA | NA | NA | NA | NA | NA |
| GRIN2A | 160 | K | 82.8545 | 32.48785652 | 0.406039084 | 29.34373029 | NA | NA | NA | NA | NA | NA | NA |
| GRIN2A | 161 | I | 86.7385 | 29.75526325 | 0.369022678 | 26.61981656 | NA | NA | NA | NA | NA | NA | NA |
| GRIN2A | 162 | M | 85.034 | 25.67977574 | 0.313815084 | 22.58143423 | NA | NA | NA | NA | NA | NA | NA |

| | | | | | | | | | | | | | |
|---|---|---|---|---|---|---|---|---|---|---|---|---|---|
| GRIN2A | 163 | Q | 80.55772222 | 28.95724548 | 0.358212526 | 24.64403253 | NA | NA | NA | NA | NA | NA | NA |
| GRIN2A | 164 | D | 80.76325 | 29.4774849 | 0.365259822 | 25.43550471 | NA | NA | NA | NA | NA | NA | NA |
| GRIN2A | 165 | Y | 83.73916667 | 25.40458943 | 0.31008734 | 22.22835173 | NA | NA | NA | NA | NA | NA | NA |
| GRIN2A | 166 | D | 79.97625 | 23.00094748 | 0.277526993 | 18.92184142 | NA | NA | NA | NA | NA | NA | NA |
| GRIN2A | 167 | W | 84.56685714 | 21.94602634 | 0.263236761 | 18.77928716 | NA | NA | NA | NA | NA | NA | NA |
| GRIN2A | 168 | H | 80.0694 | 21.68031434 | 0.259637359 | 17.44866113 | NA | NA | NA | NA | NA | NA | NA |
| GRIN2A | 169 | V | 84.03864286 | 17.83749814 | 0.20758159 | 14.71504823 | NA | NA | NA | NA | NA | NA | NA |
| GRIN2A | 170 | F | 85.81059091 | 20.85542495 | 0.248463197 | 17.80058491 | NA | NA | NA | NA | NA | NA | NA |
| GRIN2A | 171 | S | 90.64233333 | 20.89106804 | 0.248946027 | 19.42390487 | NA | NA | NA | NA | NA | NA | NA |
| GRIN2A | 172 | L | 90.944375 | 23.57984779 | 0.285368924 | 22.11163389 | NA | NA | NA | NA | NA | NA | NA |
| GRIN2A | 173 | V | 95.81921429 | 21.94721171 | 0.263252819 | 20.43407404 | NA | NA | NA | NA | NA | NA | NA |
| GRIN2A | 174 | T | 97.52392857 | 26.06644638 | 0.319053023 | 24.55360671 | NA | NA | NA | NA | NA | NA | NA |
| GRIN2A | 175 | T | 101.2995 | 28.31174235 | 0.349468376 | 26.97229855 | NA | NA | NA | NA | NA | NA | NA |
| GRIN2A | 176 | I | 102.080875 | 30.61038028 | 0.380606312 | 29.09840556 | NA | NA | NA | NA | NA | NA | NA |
| GRIN2A | 177 | F | 104.7470455 | 33.7605896 | 0.423279851 | 32.37978574 | NA | NA | NA | NA | NA | NA | NA |
| GRIN2A | 178 | P | 102.6205 | 37.37551629 | 0.472248571 | 35.86234045 | NA | NA | NA | NA | NA | NA | NA |
| GRIN2A | 179 | G | 98.2435 | 36.52726961 | 0.460758005 | 35.01815878 | NA | NA | NA | NA | NA | NA | NA |
| GRIN2A | 180 | Y | 99.29466667 | 33.08651704 | 0.414148692 | 31.58273151 | NA | NA | NA | NA | NA | NA | NA |
| GRIN2A | 181 | R | 97.84704545 | 34.38387096 | 0.43172298 | 32.92946537 | NA | NA | NA | NA | NA | NA | NA |
| GRIN2A | 182 | E | 95.03372222 | 36.11743803 | 0.455206322 | 34.65302464 | NA | NA | NA | NA | NA | NA | NA |
| GRIN2A | 183 | F | 94.71340909 | 33.3411444 | 0.41759794 | 31.87091014 | NA | NA | NA | NA | NA | NA | NA |
| GRIN2A | 184 | I | 92.175125 | 30.7908377 | 0.383050834 | 29.32778082 | NA | NA | NA | NA | NA | NA | NA |
| GRIN2A | 185 | S | 89.78383333 | 34.3745864 | 0.431597209 | 32.90395176 | NA | NA | NA | NA | NA | NA | NA |
| GRIN2A | 186 | F | 89.25086364 | 33.87446042 | 0.424822375 | 30.82701951 | NA | NA | NA | NA | NA | NA | NA |
| GRIN2A | 187 | V | 87.74664286 | 29.16547606 | 0.36103327 | 26.10566486 | NA | NA | NA | NA | NA | NA | NA |
| GRIN2A | 188 | K | 86.61616667 | 30.63090443 | 0.380884337 | 26.60455453 | NA | NA | NA | NA | NA | NA | NA |
| GRIN2A | 189 | T | 84.23164286 | 33.09488803 | 0.414262088 | 28.98424799 | NA | NA | NA | NA | NA | NA | NA |
| GRIN2A | 190 | T | 83.53435714 | 32.13965829 | 0.401322302 | 27.95141421 | NA | NA | NA | NA | NA | NA | NA |
| GRIN2A | 191 | V | 81.71778571 | 27.53136555 | 0.338897192 | 23.37262519 | NA | NA | NA | NA | NA | NA | NA |
| GRIN2A | 192 | D | 80.01475 | 29.28127123 | 0.362601861 | 25.14631397 | NA | NA | NA | NA | NA | NA | NA |
| GRIN2A | 193 | N | 78.885125 | 32.83652529 | 0.41076224 | 28.65365757 | NA | NA | NA | NA | NA | NA | NA |
| GRIN2A | 194 | S | 77.3545 | 29.9774833 | 0.372032928 | 25.75395079 | NA | NA | NA | NA | NA | NA | NA |
| GRIN2A | 195 | F | 72.45122727 | 28.95206882 | 0.358142401 | 24.64828208 | NA | NA | NA | NA | NA | NA | NA |

| | | | | | | | | | | | | | |
|---|---|---|---|---|---|---|---|---|---|---|---|---|---|
| GRIN2A | 196 | V | 75.92835714 | 25.89126511 | 0.316679973 | 21.58918272 | NA | NA | NA | NA | NA | NA | NA |
| GRIN2A | 197 | G | 76.77575 | 23.3306693 | 0.281993488 | 19.07005511 | NA | NA | NA | NA | NA | NA | NA |
| GRIN2A | 198 | W | 80.2695 | 23.53750441 | 0.28479533 | 19.3358007 | NA | NA | NA | NA | NA | NA | NA |
| GRIN2A | 199 | D | 79.78825 | 20.03320024 | 0.237325131 | 15.79573194 | NA | NA | NA | NA | NA | NA | NA |
| GRIN2A | 200 | M | 83.68975 | 19.87828999 | 0.235226677 | 15.78897527 | NA | NA | NA | NA | NA | NA | NA |
| GRIN2A | 201 | Q | 87.34838889 | 16.42992009 | 0.188514178 | 14.58971825 | NA | NA | NA | NA | NA | NA | NA |
| GRIN2A | 202 | N | 87.936 | 22.44303623 | 0.269969384 | 20.64229561 | NA | NA | NA | NA | NA | NA | NA |
| GRIN2A | 203 | V | 92.0055 | 24.47309288 | 0.29746905 | 23.0029339 | NA | NA | NA | NA | NA | NA | NA |
| GRIN2A | 204 | I | 94.40525 | 19.67200086 | 0.232432232 | 18.20183953 | NA | NA | NA | NA | NA | NA | NA |
| GRIN2A | 205 | T | 97.83764286 | 24.21898049 | 0.294026779 | 22.71340583 | NA | NA | NA | NA | NA | NA | NA |
| GRIN2A | 206 | L | 101.144625 | 22.56821307 | 0.271665062 | 21.14369601 | NA | NA | NA | NA | NA | NA | NA |
| GRIN2A | 207 | D | 102.76875 | 26.54502055 | 0.325535911 | 25.03185767 | NA | NA | NA | NA | NA | NA | NA |
| GRIN2A | 208 | T | 107.0470714 | 26.94301592 | 0.330927258 | 25.51043158 | NA | NA | NA | NA | NA | NA | NA |
| GRIN2A | 209 | S | 110.0368333 | 27.79861473 | 0.342517418 | 26.44715992 | NA | NA | NA | NA | NA | NA | NA |
| GRIN2A | 210 | F | 112.1911364 | 25.62405747 | 0.31306031 | 24.46792434 | NA | NA | NA | NA | NA | NA | NA |
| GRIN2A | 211 | E | 107.2576111 | 22.80800932 | 0.274913403 | 21.43748133 | NA | NA | NA | NA | NA | NA | NA |
| GRIN2A | 212 | D | 109.309625 | 18.88332995 | 0.221748694 | 17.55472965 | NA | NA | NA | NA | NA | NA | NA |
| GRIN2A | 213 | A | 106.1469 | 18.27722307 | 0.213538216 | 16.99950988 | NA | NA | NA | NA | NA | NA | NA |
| GRIN2A | 214 | K | 105.2896111 | 20.29213712 | 0.240832756 | 18.84973603 | NA | NA | NA | NA | NA | NA | NA |
| GRIN2A | 215 | T | 103.8710714 | 18.47435244 | 0.216208581 | 17.00757195 | NA | NA | NA | NA | NA | NA | NA |
| GRIN2A | 216 | Q | 103.6331667 | 15.49747508 | 0.17588304 | 14.11284909 | NA | NA | NA | NA | NA | NA | NA |
| GRIN2A | 217 | V | 100.3490714 | 16.5293937 | 0.189861673 | 15.01732494 | NA | NA | NA | NA | NA | NA | NA |
| GRIN2A | 218 | Q | 98.7595 | 17.00103317 | 0.196250622 | 15.48893664 | NA | NA | NA | NA | NA | NA | NA |
| GRIN2A | 219 | L | 98.9435 | 15.12175893 | 0.170793493 | 13.60725986 | NA | NA | NA | NA | NA | NA | NA |
| GRIN2A | 220 | K | 98.27572222 | 11.71600038 | 0.124658218 | 10.60769149 | NA | NA | NA | NA | NA | NA | NA |
| GRIN2A | 221 | K | 94.39116667 | 14.26817287 | 0.159230598 | 12.8225011 | NA | NA | NA | NA | NA | NA | NA |
| GRIN2A | 222 | I | 93.74725 | 13.21496285 | 0.144963546 | 11.70766568 | NA | NA | NA | NA | NA | NA | NA |
| GRIN2A | 223 | H | 89.9914 | 11.10752421 | 0.116415645 | 9.636915117 | NA | NA | NA | NA | NA | NA | NA |
| GRIN2A | 224 | S | 89.008 | 15.57676029 | 0.176957058 | 13.66425888 | NA | NA | NA | NA | NA | NA | NA |
| GRIN2A | 225 | S | 88.24966667 | 17.15861681 | 0.19838529 | 15.41822868 | NA | NA | NA | NA | NA | NA | NA |
| GRIN2A | 226 | V | 88.82078571 | 19.97289423 | 0.23650821 | 16.89934897 | NA | NA | NA | NA | NA | NA | NA |
| GRIN2A | 227 | I | 92.939375 | 19.71134524 | 0.232965201 | 18.19360578 | NA | NA | NA | NA | NA | NA | NA |
| GRIN2A | 228 | L | 92.582125 | 24.77333094 | 0.301536151 | 23.25775636 | NA | NA | NA | NA | NA | NA | NA |

| | | | | | | | | | | | | | |
|---|---|---|---|---|---|---|---|---|---|---|---|---|---|
| GRIN2A | 229 | L | 97.685625 | 24.98097256 | 0.304348918 | 23.4654245 | NA | NA | NA | NA | NA | NA | NA |
| GRIN2A | 230 | Y | 96.06658333 | 29.19255045 | 0.361400027 | 27.67773317 | NA | NA | NA | NA | NA | NA | NA |
| GRIN2A | 231 | C | 101.7953333 | 27.7676855 | 0.342098443 | 26.23887069 | NA | NA | NA | NA | NA | NA | NA |
| GRIN2A | 232 | S | 105.5675 | 30.60528838 | 0.380537336 | 29.09886624 | NA | NA | NA | NA | NA | NA | NA |
| GRIN2A | 233 | K | 107.9251667 | 30.14883847 | 0.374354149 | 28.6590696 | NA | NA | NA | NA | NA | NA | NA |
| GRIN2A | 234 | D | 110.437375 | 28.71893053 | 0.354984251 | 27.32454244 | NA | NA | NA | NA | NA | NA | NA |
| GRIN2A | 235 | E | 106.4362778 | 26.22473058 | 0.321197181 | 24.74687014 | NA | NA | NA | NA | NA | NA | NA |
| GRIN2A | 236 | A | 104.9669 | 25.391529 | 0.30991042 | 23.89197548 | NA | NA | NA | NA | NA | NA | NA |
| GRIN2A | 237 | V | 108.2950714 | 24.44618574 | 0.297104559 | 23.18017219 | NA | NA | NA | NA | NA | NA | NA |
| GRIN2A | 238 | L | 108.461125 | 22.06670929 | 0.264871564 | 20.86982358 | NA | NA | NA | NA | NA | NA | NA |
| GRIN2A | 239 | I | 103.479375 | 20.53751502 | 0.244156707 | 19.04098918 | NA | NA | NA | NA | NA | NA | NA |
| GRIN2A | 240 | L | 102.973625 | 20.27632733 | 0.240618592 | 18.77592183 | NA | NA | NA | NA | NA | NA | NA |
| GRIN2A | 241 | S | 106.7156667 | 18.61046711 | 0.218052425 | 17.35411163 | NA | NA | NA | NA | NA | NA | NA |
| GRIN2A | 242 | E | 105.0742778 | 16.15955238 | 0.184851708 | 15.02071986 | NA | NA | NA | NA | NA | NA | NA |
| GRIN2A | 243 | A | 101.3699 | 16.05831305 | 0.183480294 | 14.53069496 | NA | NA | NA | NA | NA | NA | NA |
| GRIN2A | 244 | R | 103.8003182 | 14.4397635 | 0.161555009 | 13.02438177 | NA | NA | NA | NA | NA | NA | NA |
| GRIN2A | 245 | S | 103.3056667 | 12.03047476 | 0.128918169 | 10.76985704 | NA | NA | NA | NA | NA | NA | NA |
| GRIN2A | 246 | L | 99.843125 | 11.75773507 | 0.125223567 | 10.2177208 | NA | NA | NA | NA | NA | NA | NA |
| GRIN2A | 247 | G | 97.581 | 12.76612541 | 0.13888348 | 11.2261413 | NA | NA | NA | NA | NA | NA | NA |
| GRIN2A | 248 | L | 96.136 | 14.17776417 | 0.158005899 | 12.74974013 | NA | NA | NA | NA | NA | NA | NA |
| GRIN2A | 249 | T | 98.38821429 | 16.87188946 | 0.194501208 | 15.32971874 | NA | NA | NA | NA | NA | NA | NA |
| GRIN2A | 250 | G | 94.94425 | 17.08999216 | 0.197455683 | 15.5681305 | NA | NA | NA | NA | NA | NA | NA |
| GRIN2A | 251 | Y | 95.59775 | 19.02778623 | 0.223705536 | 17.54831578 | NA | NA | NA | NA | NA | NA | NA |
| GRIN2A | 252 | D | 89.63125 | 17.76052888 | 0.206538944 | 15.97841486 | NA | NA | NA | NA | NA | NA | NA |
| GRIN2A | 253 | F | 92.20995455 | 15.63807952 | 0.177787703 | 14.12150009 | NA | NA | NA | NA | NA | NA | NA |
| GRIN2A | 254 | F | 90.32377273 | 22.38210334 | 0.269143972 | 20.86122172 | NA | NA | NA | NA | NA | NA | NA |
| GRIN2A | 255 | W | 95.90214286 | 20.67402801 | 0.246005947 | 19.51363824 | NA | NA | NA | NA | NA | NA | NA |
| GRIN2A | 256 | I | 93.0375 | 27.46082725 | 0.337941662 | 25.94237167 | NA | NA | NA | NA | NA | NA | NA |
| GRIN2A | 257 | V | 97.45464286 | 28.48089003 | 0.351759693 | 27.146055 | NA | NA | NA | NA | NA | NA | NA |
| GRIN2A | 258 | P | 97.44864286 | 32.38084423 | 0.404589468 | 30.864577 | NA | NA | NA | NA | NA | NA | NA |
| GRIN2A | 259 | S | 100.1398333 | 36.24981194 | 0.456999493 | 34.69881614 | NA | NA | NA | NA | NA | NA | NA |
| GRIN2A | 260 | L | 102.438375 | 35.56876193 | 0.447773815 | 34.03869004 | NA | NA | NA | NA | NA | NA | NA |
| GRIN2A | 261 | V | 103.3557857 | 31.25141022 | 0.389289867 | 29.72633785 | NA | NA | NA | NA | NA | NA | NA |

| | | | | | | | | | | | | | |
|---|---|---|---|---|---|---|---|---|---|---|---|---|---|
| GRIN2A | 262 | S | 102.8791667 | 34.29761756 | 0.430554569 | 32.7618182 | NA | NA | NA | NA | NA | NA | NA |
| GRIN2A | 263 | G | 104.3295 | 37.17328783 | 0.469509133 | 35.63791603 | NA | NA | NA | NA | NA | NA | NA |
| GRIN2A | 264 | N | 107.889625 | 37.97781896 | 0.480407517 | 36.4537943 | NA | NA | NA | NA | NA | NA | NA |
| GRIN2A | 265 | T | 105.0950714 | 38.43669653 | 0.48662359 | 36.9067855 | NA | NA | NA | NA | NA | NA | NA |
| GRIN2A | 266 | E | 108.9199444 | 39.74172978 | 0.504301903 | 38.48147433 | NA | NA | NA | NA | NA | NA | NA |
| GRIN2A | 267 | L | 110.596125 | 36.15723824 | 0.455745466 | 34.89609858 | NA | NA | NA | NA | NA | NA | NA |
| GRIN2A | 268 | I | 108.0895 | 33.49432264 | 0.419672931 | 32.23765643 | NA | NA | NA | NA | NA | NA | NA |
| GRIN2A | 269 | P | 108.4343571 | 30.25360093 | 0.375773288 | 28.99378164 | NA | NA | NA | NA | NA | NA | NA |
| GRIN2A | 270 | K | 110.1939444 | 27.28989676 | 0.335626194 | 26.03227678 | NA | NA | NA | NA | NA | NA | NA |
| GRIN2A | 271 | E | 107.5258333 | 23.85751358 | 0.289130256 | 22.62275681 | NA | NA | NA | NA | NA | NA | NA |
| GRIN2A | 272 | F | 104.1198636 | 26.18895616 | 0.320712572 | 24.80646629 | NA | NA | NA | NA | NA | NA | NA |
| GRIN2A | 273 | P | 100.7247857 | 23.7300689 | 0.287403857 | 22.19096233 | NA | NA | NA | NA | NA | NA | NA |
| GRIN2A | 274 | S | 98.20366667 | 26.24225869 | 0.321434622 | 24.70694446 | NA | NA | NA | NA | NA | NA | NA |
| GRIN2A | 275 | G | 95.693 | 28.49639263 | 0.351969695 | 26.96953142 | NA | NA | NA | NA | NA | NA | NA |
| GRIN2A | 276 | L | 97.527625 | 26.12679605 | 0.319870535 | 24.58537144 | NA | NA | NA | NA | NA | NA | NA |
| GRIN2A | 277 | I | 93.854625 | 29.63679647 | 0.367417897 | 28.805696 | NA | NA | NA | NA | NA | NA | NA |
| GRIN2A | 278 | S | 96.08683333 | 33.5632342 | 0.420606425 | 32.02342568 | NA | NA | NA | NA | NA | NA | NA |
| GRIN2A | 279 | V | 94.7905 | 36.54720747 | 0.461028089 | 35.02863538 | NA | NA | NA | NA | NA | NA | NA |
| GRIN2A | 280 | S | 97.724 | 39.86780279 | 0.50600972 | 38.3197889 | NA | NA | NA | NA | NA | NA | NA |
| GRIN2A | 281 | Y | 97.24008333 | 43.1835045 | 0.550925062 | 41.63448842 | NA | NA | NA | NA | NA | NA | NA |
| GRIN2A | 282 | D | 100.21775 | 40.73364431 | 0.517738631 | 39.18960062 | NA | NA | NA | NA | NA | NA | NA |
| GRIN2A | 283 | D | 101.235875 | 46.30159802 | 0.593163553 | 44.75744415 | NA | NA | NA | NA | NA | NA | NA |
| GRIN2A | 284 | W | 100.3180714 | 42.69078423 | 0.544250548 | 41.15041113 | NA | NA | NA | NA | NA | NA | NA |
| GRIN2A | 285 | D | 96.336 | 47.7448473 | 0.612714176 | 46.32208727 | NA | NA | NA | NA | NA | NA | NA |
| GRIN2A | 286 | Y | 96.46333333 | 49.3757933 | 0.634807387 | 48.07563925 | NA | NA | NA | NA | NA | NA | NA |
| GRIN2A | 287 | S | 100.7328333 | 54.11897639 | 0.699059756 | 52.57706768 | NA | NA | NA | NA | NA | NA | NA |
| GRIN2A | 288 | L | 104.167375 | 53.46886003 | 0.690253114 | 51.92819308 | NA | NA | NA | NA | NA | NA | NA |
| GRIN2A | 289 | E | 103.1610556 | 57.92559008 | 0.750625117 | 56.38203915 | NA | NA | NA | NA | NA | NA | NA |
| GRIN2A | 290 | A | 98.7305 | 57.4468886 | 0.744140504 | 55.90433334 | NA | NA | NA | NA | NA | NA | NA |
| GRIN2A | 291 | R | 99.11440909 | 49.91207037 | 0.642071933 | 48.36632479 | NA | NA | NA | NA | NA | NA | NA |
| GRIN2A | 292 | V | 102.0965 | 58.51553706 | 0.758616689 | 56.96854594 | NA | NA | NA | NA | NA | NA | NA |
| GRIN2A | 293 | R | 99.04040909 | 61.17937111 | 0.794701665 | 59.6312642 | NA | NA | NA | NA | NA | NA | NA |
| GRIN2A | 294 | D | 96.29925 | 56.90229247 | 0.736763266 | 55.6420644 | NA | NA | NA | NA | NA | NA | NA |

| | | | | | | | | | | | | | |
|---|---|---|---|---|---|---|---|---|---|---|---|---|---|
| GRIN2A | 295 | G | 98.49675 | 59.87928604 | 0.777090381 | 58.36180823 | NA | NA | NA | NA | NA | NA | NA |
| GRIN2A | 296 | I | 99.912375 | 62.40649711 | 0.811324627 | 60.89823545 | NA | NA | NA | NA | NA | NA | NA |
| GRIN2A | 297 | G | 95.61675 | 63.50703307 | 0.826232768 | 60.41814514 | NA | NA | NA | NA | NA | NA | NA |
| GRIN2A | 298 | I | 95.117375 | 58.28855467 | 0.755541928 | 55.20591165 | NA | NA | NA | NA | NA | NA | NA |
| GRIN2A | 299 | L | 98.4575 | 63.66062884 | 0.828313416 | 62.21234189 | NA | NA | NA | NA | NA | NA | NA |
| GRIN2A | 300 | T | 95.45664286 | 66.20444252 | 0.862772565 | 63.12277866 | NA | NA | NA | NA | NA | NA | NA |
| GRIN2A | 301 | T | 92.2615 | 63.20918789 | 0.822198082 | 60.11887556 | NA | NA | NA | NA | NA | NA | NA |
| GRIN2A | 302 | A | 94.0559 | 65.23774372 | 0.849677417 | 62.16150051 | NA | NA | NA | NA | NA | NA | NA |
| GRIN2A | 303 | A | 94.6563 | 68.0289721 | 0.887488109 | 64.94816118 | NA | NA | NA | NA | NA | NA | NA |
| GRIN2A | 304 | S | 91.07166667 | 67.9079177 | 0.885848275 | 64.80897765 | NA | NA | NA | NA | NA | NA | NA |
| GRIN2A | 305 | S | 90.3105 | 66.54312925 | 0.867360502 | 62.61241865 | NA | NA | NA | NA | NA | NA | NA |
| GRIN2A | 306 | M | 93.154125 | 69.51110203 | 0.907565419 | 66.42458651 | NA | NA | NA | NA | NA | NA | NA |
| GRIN2A | 307 | L | 92.27075 | 72.4085348 | 0.946814783 | 69.32029581 | NA | NA | NA | NA | NA | NA | NA |
| GRIN2A | 308 | E | 87.03461111 | 69.07189771 | 0.901615845 | 64.8417122 | NA | NA | NA | NA | NA | NA | NA |
| GRIN2A | 309 | K | 86.76994444 | 66.36760406 | 0.864982793 | 62.154321 | NA | NA | NA | NA | NA | NA | NA |
| GRIN2A | 310 | F | 91.2485 | 71.43429621 | 0.933617499 | 67.29479549 | NA | NA | NA | NA | NA | NA | NA |
| GRIN2A | 311 | S | 92.74316667 | 76.3347281 | 1 | 72.28043133 | NA | NA | NA | NA | NA | NA | NA |
| GRIN2A | 312 | Y | 96.14466667 | 74.81604873 | 0.979427582 | 73.57396079 | NA | NA | NA | NA | NA | NA | NA |
| GRIN2A | 313 | I | 99.5525 | 74.50950875 | 0.975275113 | 72.99459208 | NA | NA | NA | NA | NA | NA | NA |
| GRIN2A | 314 | P | 100.3807857 | 71.38178816 | 0.932906211 | 69.86758999 | NA | NA | NA | NA | NA | NA | NA |
| GRIN2A | 315 | E | 104.3492778 | 73.66997759 | 0.963902609 | 72.15571022 | NA | NA | NA | NA | NA | NA | NA |
| GRIN2A | 316 | A | 107.2597 | 72.61805217 | 0.949652959 | 71.14180491 | NA | NA | NA | NA | NA | NA | NA |
| GRIN2A | 317 | K | 109.3389444 | 68.59751002 | 0.895189669 | 67.08365618 | NA | NA | NA | NA | NA | NA | NA |
| GRIN2A | 318 | A | 112.8977 | 74.97708111 | 0.981608967 | 73.45680963 | NA | NA | NA | NA | NA | NA | NA |
| GRIN2A | 319 | S | 115.2135 | 72.22135286 | 0.944279169 | 70.71396854 | NA | NA | NA | NA | NA | NA | NA |
| GRIN2A | 320 | C | 116.0981667 | 69.29057844 | 0.904578151 | 67.78864886 | NA | NA | NA | NA | NA | NA | NA |
| GRIN2A | 321 | Y | 120.19425 | 70.41841942 | 0.919856172 | 68.94152328 | NA | NA | NA | NA | NA | NA | NA |
| GRIN2A | 322 | G | 116.3965 | 72.8535398 | 0.952842935 | 71.35576396 | NA | NA | NA | NA | NA | NA | NA |
| GRIN2A | 323 | Q | 115.7475 | 69.82986048 | 0.911883403 | 68.34675159 | NA | NA | NA | NA | NA | NA | NA |
| GRIN2A | 324 | M | 113.5017 | 75.68645122 | 0.991218276 | 74.19173761 | NA | NA | NA | NA | NA | NA | NA |
| GRIN2A | 325 | E | 109.9693 | 75.8347368 | 0.99322699 | 74.32228079 | NA | NA | NA | NA | NA | NA | NA |
| GRIN2A | 326 | R | 107.5725 | 72.81340916 | 0.952299315 | 71.3006672 | NA | NA | NA | NA | NA | NA | NA |
| GRIN2A | 327 | P | 105.0845 | 69.55806469 | 0.908201588 | 68.04530263 | NA | NA | NA | NA | NA | NA | NA |

| | | | | | | | | | | | | | |
|---|---|---|---|---|---|---|---|---|---|---|---|---|---|
| GRIN2A | 328 | E | 102.2113 | 71.65013469 | 0.936541302 | 70.14079452 | NA | NA | NA | NA | NA | NA | NA |
| GRIN2A | 329 | V | 99.5115 | 68.94329099 | 0.899873706 | 67.43463397 | NA | NA | NA | NA | NA | NA | NA |
| GRIN2A | 330 | P | 98.0675 | 72.55158798 | 0.948752618 | 71.09687446 | NA | NA | NA | NA | NA | NA | NA |
| GRIN2A | 331 | M | 97.3331 | 71.51655318 | 0.934731773 | 70.11042298 | NA | NA | NA | NA | NA | NA | NA |
| GRIN2A | 332 | H | 94.5654 | 68.53202993 | 0.894302659 | 67.25901808 | NA | NA | NA | NA | NA | NA | NA |
| GRIN2A | 333 | T | 97.52178571 | 65.70895688 | 0.85606059 | 64.2268058 | NA | NA | NA | NA | NA | NA | NA |
| GRIN2A | 334 | L | 96.066125 | 63.05808688 | 0.820151229 | 61.85820667 | NA | NA | NA | NA | NA | NA | NA |
| GRIN2A | 335 | H | 95.3676 | 60.35236144 | 0.783498781 | 58.89768024 | NA | NA | NA | NA | NA | NA | NA |
| GRIN2A | 336 | P | 90.83021429 | 62.84159712 | 0.817218603 | 58.7546288 | NA | NA | NA | NA | NA | NA | NA |
| GRIN2A | 337 | F | 91.75095455 | 62.85616347 | 0.817415923 | 59.77424757 | NA | NA | NA | NA | NA | NA | NA |
| GRIN2A | 338 | M | 93.22125 | 58.48976775 | 0.758267612 | 55.40796223 | NA | NA | NA | NA | NA | NA | NA |
| GRIN2A | 339 | V | 88.36164286 | 57.35961207 | 0.742958234 | 53.30862616 | NA | NA | NA | NA | NA | NA | NA |
| GRIN2A | 340 | N | 86.04875 | 57.40456095 | 0.743567123 | 53.17887715 | NA | NA | NA | NA | NA | NA | NA |
| GRIN2A | 341 | V | 89.53678571 | 57.83408123 | 0.749385515 | 54.72804183 | NA | NA | NA | NA | NA | NA | NA |
| GRIN2A | 342 | T | 86.83364286 | 58.57532754 | 0.759426626 | 55.45200444 | NA | NA | NA | NA | NA | NA | NA |
| GRIN2A | 343 | W | 90.93078571 | 58.25986904 | 0.755153345 | 55.13642769 | NA | NA | NA | NA | NA | NA | NA |
| GRIN2A | 344 | D | 88.866125 | 58.18489282 | 0.754137698 | 55.04627335 | NA | NA | NA | NA | NA | NA | NA |
| GRIN2A | 345 | G | 85.21725 | 55.32072829 | 0.715338994 | 52.17699178 | NA | NA | NA | NA | NA | NA | NA |
| GRIN2A | 346 | K | 88.2155 | 54.04428333 | 0.698047945 | 50.91679824 | NA | NA | NA | NA | NA | NA | NA |
| GRIN2A | 347 | D | 85.621 | 53.02233064 | 0.684204313 | 49.91152725 | NA | NA | NA | NA | NA | NA | NA |
| GRIN2A | 348 | L | 90.543 | 54.0258455 | 0.697798181 | 50.92599045 | NA | NA | NA | NA | NA | NA | NA |
| GRIN2A | 349 | S | 88.5465 | 53.45333977 | 0.690042873 | 50.36477965 | NA | NA | NA | NA | NA | NA | NA |
| GRIN2A | 350 | F | 91.58913636 | 53.16500105 | 0.686136963 | 50.07973085 | NA | NA | NA | NA | NA | NA | NA |
| GRIN2A | 351 | T | 89.64735714 | 49.59975328 | 0.637841206 | 45.5730196 | NA | NA | NA | NA | NA | NA | NA |
| GRIN2A | 352 | E | 89.16283333 | 52.75673543 | 0.680606492 | 48.65761016 | NA | NA | NA | NA | NA | NA | NA |
| GRIN2A | 353 | E | 92.55583333 | 47.24126417 | 0.60589251 | 46.02285254 | NA | NA | NA | NA | NA | NA | NA |
| GRIN2A | 354 | G | 93.924 | 52.00203259 | 0.670383095 | 50.67012534 | NA | NA | NA | NA | NA | NA | NA |
| GRIN2A | 355 | Y | 93.98333333 | 46.43910862 | 0.595026307 | 44.92770026 | NA | NA | NA | NA | NA | NA | NA |
| GRIN2A | 356 | Q | 92.95761111 | 47.95019274 | 0.615495838 | 44.86545013 | NA | NA | NA | NA | NA | NA | NA |
| GRIN2A | 357 | V | 87.48292857 | 47.19355629 | 0.605246247 | 44.0903241 | NA | NA | NA | NA | NA | NA | NA |
| GRIN2A | 358 | H | 86.6684 | 47.78506814 | 0.613259018 | 44.66155982 | NA | NA | NA | NA | NA | NA | NA |
| GRIN2A | 359 | P | 91.12992857 | 46.19952438 | 0.591780837 | 43.11295196 | NA | NA | NA | NA | NA | NA | NA |
| GRIN2A | 360 | R | 93.31068182 | 44.86357838 | 0.573683772 | 41.75726711 | NA | NA | NA | NA | NA | NA | NA |

| | | | | | | | | | | | | | |
|---|---|---|---|---|---|---|---|---|---|---|---|---|---|
| GRIN2A | 361 | L | 91.945625 | 41.75619635 | 0.531590382 | 38.65268065 | NA | NA | NA | NA | NA | NA | NA |
| GRIN2A | 362 | V | 94.56764286 | 40.14680703 | 0.509789183 | 38.66478389 | NA | NA | NA | NA | NA | NA | NA |
| GRIN2A | 363 | V | 92.16835714 | 36.14854221 | 0.455627667 | 33.03919761 | NA | NA | NA | NA | NA | NA | NA |
| GRIN2A | 364 | I | 96.13275 | 34.54105315 | 0.43385221 | 33.00988622 | NA | NA | NA | NA | NA | NA | NA |
| GRIN2A | 365 | V | 95.26035714 | 32.07555222 | 0.400453905 | 30.56261684 | NA | NA | NA | NA | NA | NA | NA |
| GRIN2A | 366 | L | 99.356125 | 31.40624088 | 0.391387243 | 29.86798937 | NA | NA | NA | NA | NA | NA | NA |
| GRIN2A | 367 | N | 97.63625 | 31.96880694 | 0.399007906 | 30.43637463 | NA | NA | NA | NA | NA | NA | NA |
| GRIN2A | 368 | K | 98.83705556 | 30.44186223 | 0.378323523 | 29.16914631 | NA | NA | NA | NA | NA | NA | NA |
| GRIN2A | 369 | D | 102.9935 | 33.55327278 | 0.420471485 | 32.13219145 | NA | NA | NA | NA | NA | NA | NA |
| GRIN2A | 370 | R | 104.7361364 | 27.26531441 | 0.335293196 | 25.88095689 | NA | NA | NA | NA | NA | NA | NA |
| GRIN2A | 371 | E | 101.8589444 | 35.24690764 | 0.443413895 | 33.71140473 | NA | NA | NA | NA | NA | NA | NA |
| GRIN2A | 372 | W | 101.2441429 | 33.10638236 | 0.414417793 | 31.56539742 | NA | NA | NA | NA | NA | NA | NA |
| GRIN2A | 373 | E | 97.33561111 | 37.56942307 | 0.474875282 | 36.05353082 | NA | NA | NA | NA | NA | NA | NA |
| GRIN2A | 374 | K | 94.60438889 | 39.49399652 | 0.500946045 | 38.17723348 | NA | NA | NA | NA | NA | NA | NA |
| GRIN2A | 375 | V | 90.95235714 | 35.69269812 | 0.449452687 | 32.55174382 | NA | NA | NA | NA | NA | NA | NA |
| GRIN2A | 376 | G | 89.88325 | 38.55328471 | 0.488202923 | 35.41962046 | NA | NA | NA | NA | NA | NA | NA |
| GRIN2A | 377 | K | 88.91083333 | 40.63308813 | 0.516376471 | 37.49501528 | NA | NA | NA | NA | NA | NA | NA |
| GRIN2A | 378 | W | 86.65085714 | 38.47055152 | 0.487082198 | 35.34504732 | NA | NA | NA | NA | NA | NA | NA |
| GRIN2A | 379 | E | 87.77061111 | 45.44213875 | 0.581521098 | 42.30580022 | NA | NA | NA | NA | NA | NA | NA |
| GRIN2A | 380 | N | 82.920875 | 44.94363898 | 0.574768293 | 41.78656906 | NA | NA | NA | NA | NA | NA | NA |
| GRIN2A | 381 | H | 80.4857 | 44.79837644 | 0.57280053 | 40.50372437 | NA | NA | NA | NA | NA | NA | NA |
| GRIN2A | 382 | T | 82.96235714 | 41.77697459 | 0.531871849 | 37.54496567 | NA | NA | NA | NA | NA | NA | NA |
| GRIN2A | 383 | L | 81.39325 | 40.90362167 | 0.520041187 | 36.70146585 | NA | NA | NA | NA | NA | NA | NA |
| GRIN2A | 384 | S | 83.29866667 | 36.98843503 | 0.46700507 | 33.82807986 | NA | NA | NA | NA | NA | NA | NA |
| GRIN2A | 385 | L | 84.225875 | 39.22943591 | 0.49736224 | 36.06029265 | NA | NA | NA | NA | NA | NA | NA |
| GRIN2A | 386 | R | 84.76668182 | 32.855879 | 0.411024411 | 29.70329648 | NA | NA | NA | NA | NA | NA | NA |
| GRIN2A | 387 | H | 87.4183 | 36.06899242 | 0.454550065 | 32.87112642 | NA | NA | NA | NA | NA | NA | NA |
| GRIN2A | 388 | A | 84.1683 | 32.6477853 | 0.40820552 | 29.43240031 | NA | NA | NA | NA | NA | NA | NA |
| GRIN2A | 389 | V | 87.2875 | 31.23608423 | 0.389082257 | 28.06083709 | NA | NA | NA | NA | NA | NA | NA |
| GRIN2A | 390 | W | 86.5817 | 27.97591704 | 0.3449192 | 24.80673445 | NA | NA | NA | NA | NA | NA | NA |
| GRIN2A | 391 | P | 88.5401 | 26.10744126 | 0.31960835 | 22.91828634 | NA | NA | NA | NA | NA | NA | NA |
| GRIN2A | 392 | R | 92.3845 | 28.31512363 | 0.349514179 | 26.84198029 | NA | NA | NA | NA | NA | NA | NA |
| GRIN2A | 393 | Y | 93.5301 | 24.84036163 | 0.302444166 | 23.3791827 | NA | NA | NA | NA | NA | NA | NA |

| | | | | | | | | | | | | | |
|---|---|---|---|---|---|---|---|---|---|---|---|---|---|
| GRIN2A | 394 | K | 95.7021 | 24.79324807 | 0.301805954 | 23.29346882 | NA | NA | NA | NA | NA | NA | NA |
| GRIN2A | 395 | S | 98.2513 | 22.2961248 | 0.267979285 | 20.86524513 | NA | NA | NA | NA | NA | NA | NA |
| GRIN2A | 396 | F | 101.1729 | 23.85521872 | 0.289099169 | 22.44731982 | NA | NA | NA | NA | NA | NA | NA |
| GRIN2A | 397 | S | 101.9817 | 19.55898006 | 0.230901223 | 18.30737597 | NA | NA | NA | NA | NA | NA | NA |
| GRIN2A | 398 | D | 102.0267 | 19.91234205 | 0.235687955 | 18.66399221 | NA | NA | NA | NA | NA | NA | NA |
| GRIN2A | 401 | P | 94.6323 | 18.60459623 | 0.217972896 | 17.3716171 | NA | NA | NA | NA | NA | NA | NA |
| GRIN2A | 402 | D | 91.3313 | 19.51623598 | 0.230322201 | 18.13048646 | NA | NA | NA | NA | NA | NA | NA |
| GRIN2A | 403 | D | 87.6427 | 19.4218807 | 0.22904404 | 16.18570462 | NA | NA | NA | NA | NA | NA | NA |
| GRIN2A | 404 | N | 85.675625 | 16.08860271 | 0.183890606 | 12.88686526 | NA | NA | NA | NA | NA | NA | NA |
| GRIN2A | 405 | H | 85.4325 | 20.17531078 | 0.239250196 | 16.94035909 | NA | NA | NA | NA | NA | NA | NA |
| GRIN2A | 406 | L | 80.7515 | 19.37149496 | 0.228361502 | 16.09927652 | NA | NA | NA | NA | NA | NA | NA |
| GRIN2A | 407 | S | 79.1795 | 24.4358273 | 0.296964241 | 19.34968979 | NA | NA | NA | NA | NA | NA | NA |
| GRIN2A | 408 | I | 75.178875 | 21.11437468 | 0.251970996 | 16.07883501 | NA | NA | NA | NA | NA | NA | NA |
| GRIN2A | 409 | V | 73.34764286 | 26.7715223 | 0.328604162 | 21.79091487 | NA | NA | NA | NA | NA | NA | NA |
| GRIN2A | 410 | T | 69.48678571 | 24.91787056 | 0.303494122 | 20.12314871 | NA | NA | NA | NA | NA | NA | NA |
| GRIN2A | 411 | L | 65.74 | 27.77081504 | 0.342140836 | 24.34845232 | NA | NA | NA | NA | NA | NA | NA |
| GRIN2A | 412 | E | 63.32272222 | 28.28599 | 0.349119528 | 24.46002735 | NA | NA | NA | NA | NA | NA | NA |
| GRIN2A | 413 | E | 60.66072222 | 24.13569788 | 0.292898611 | 20.17000224 | NA | NA | NA | NA | NA | NA | NA |
| GRIN2A | 414 | A | 58.1137 | 22.43897524 | 0.269914373 | 18.4299725 | NA | NA | NA | NA | NA | NA | NA |
| GRIN2A | 415 | P | 58.27678571 | 18.87825696 | 0.221679974 | 14.87395743 | NA | NA | NA | NA | NA | NA | NA |
| GRIN2A | 416 | F | 60.82740909 | 19.56490608 | 0.230981499 | 16.00144292 | NA | NA | NA | NA | NA | NA | NA |
| GRIN2A | 417 | V | 63.54207143 | 22.92846137 | 0.276545077 | 19.08940131 | NA | NA | NA | NA | NA | NA | NA |
| GRIN2A | 418 | I | 62.10625 | 21.62873048 | 0.258938591 | 17.62027371 | NA | NA | NA | NA | NA | NA | NA |
| GRIN2A | 419 | V | 65.17392857 | 26.74141354 | 0.328196301 | 22.74227891 | NA | NA | NA | NA | NA | NA | NA |
| GRIN2A | 420 | E | 64.73427778 | 25.06840312 | 0.305533274 | 21.05992238 | NA | NA | NA | NA | NA | NA | NA |
| GRIN2A | 421 | D | 67.037 | 30.44528238 | 0.378369853 | 26.45877926 | NA | NA | NA | NA | NA | NA | NA |
| GRIN2A | 422 | I | 70.704875 | 32.34643796 | 0.404123392 | 28.57072822 | NA | NA | NA | NA | NA | NA | NA |
| GRIN2A | 423 | D | 68.6811 | 30.23154429 | 0.375474503 | 26.23191466 | NA | NA | NA | NA | NA | NA | NA |
| GRIN2A | 424 | P | 69.1465 | 33.17831703 | 0.415392239 | 29.18550034 | NA | NA | NA | NA | NA | NA | NA |
| GRIN2A | 425 | L | 70.0667 | 30.0194921 | 0.37260199 | 26.04296484 | NA | NA | NA | NA | NA | NA | NA |
| GRIN2A | 426 | T | 73.4545 | 32.14186821 | 0.401352238 | 27.74565534 | NA | NA | NA | NA | NA | NA | NA |
| GRIN2A | 427 | E | 73.8819 | 29.54232613 | 0.366138178 | 25.08177412 | NA | NA | NA | NA | NA | NA | NA |
| GRIN2A | 428 | T | 74.0989 | 27.69278877 | 0.341083872 | 22.99989659 | NA | NA | NA | NA | NA | NA | NA |

| | | | | | | | | | | | | | |
|---|---|---|---|---|---|---|---|---|---|---|---|---|---|
| GRIN2A | 429 | C | 72.8457 | 25.3585844 | 0.309464144 | 20.62446191 | NA | NA | NA | NA | NA | NA | NA |
| GRIN2A | 430 | V | 69.27021429 | 22.47650182 | 0.270422718 | 19.80632988 | NA | NA | NA | NA | NA | NA | NA |
| GRIN2A | 431 | R | 67.82731818 | 18.99727417 | 0.223292212 | 16.03264535 | NA | NA | NA | NA | NA | NA | NA |
| GRIN2A | 432 | N | 67.087625 | 17.47028852 | 0.202607274 | 14.46691408 | NA | NA | NA | NA | NA | NA | NA |
| GRIN2A | 433 | T | 69.43192857 | 21.90641758 | 0.262700211 | 17.08067395 | NA | NA | NA | NA | NA | NA | NA |
| GRIN2A | 434 | V | 72.37435714 | 20.34123595 | 0.241497861 | 15.48199061 | NA | NA | NA | NA | NA | NA | NA |
| GRIN2A | 435 | P | 74.35735714 | 24.68524551 | 0.300342924 | 19.73109099 | NA | NA | NA | NA | NA | NA | NA |
| GRIN2A | 436 | C | 73.08816667 | 25.93457904 | 0.317266715 | 21.22327904 | NA | NA | NA | NA | NA | NA | NA |
| GRIN2A | 437 | R | 78.19922727 | 29.78314735 | 0.369400403 | 24.81244194 | NA | NA | NA | NA | NA | NA | NA |
| GRIN2A | 438 | K | 71.50094444 | 32.79947599 | 0.410260361 | 27.9153355 | NA | NA | NA | NA | NA | NA | NA |
| GRIN2A | 439 | F | 75.69331818 | 35.46272707 | 0.44633744 | 30.37520474 | NA | NA | NA | NA | NA | NA | NA |
| GRIN2A | 440 | V | 72.78035714 | 38.55520839 | 0.488228982 | 33.71399996 | NA | NA | NA | NA | NA | NA | NA |
| GRIN2A | 441 | K | 71.55894444 | 42.03670771 | 0.53539026 | 37.1840287 | NA | NA | NA | NA | NA | NA | NA |
| GRIN2A | 442 | I | 74.461125 | 45.32377196 | 0.579917672 | 40.4480748 | NA | NA | NA | NA | NA | NA | NA |
| GRIN2A | 443 | N | 71.61 | 47.44890492 | 0.608705265 | 42.59588869 | NA | NA | NA | NA | NA | NA | NA |
| GRIN2A | 444 | N | 70.00475 | 47.76511461 | 0.612988722 | 42.96281023 | NA | NA | NA | NA | NA | NA | NA |
| GRIN2A | 445 | S | 74.2735 | 50.83546968 | 0.654580536 | 46.00025554 | NA | NA | NA | NA | NA | NA | NA |
| GRIN2A | 446 | T | 76.22792857 | 47.20840875 | 0.605447443 | 42.26764298 | NA | NA | NA | NA | NA | NA | NA |
| GRIN2A | 447 | N | 75.507625 | 43.69527831 | 0.557857681 | 38.6919327 | NA | NA | NA | NA | NA | NA | NA |
| GRIN2A | 448 | E | 78.97105556 | 43.02933489 | 0.548836641 | 37.99513824 | NA | NA | NA | NA | NA | NA | NA |
| GRIN2A | 449 | G | 77.22275 | 39.7951243 | 0.505025199 | 34.70505068 | NA | NA | NA | NA | NA | NA | NA |
| GRIN2A | 450 | M | 77.28675 | 37.64795288 | 0.475939067 | 32.672537 | NA | NA | NA | NA | NA | NA | NA |
| GRIN2A | 451 | N | 78.342375 | 35.14628333 | 0.442050812 | 30.0573343 | NA | NA | NA | NA | NA | NA | NA |
| GRIN2A | 452 | V | 74.0305 | 32.85526273 | 0.411016063 | 27.96979962 | NA | NA | NA | NA | NA | NA | NA |
| GRIN2A | 453 | K | 76.10638889 | 30.69929208 | 0.381810734 | 25.96981031 | NA | NA | NA | NA | NA | NA | NA |
| GRIN2A | 454 | K | 70.18661111 | 28.03388724 | 0.345704479 | 23.20498988 | NA | NA | NA | NA | NA | NA | NA |
| GRIN2A | 455 | C | 68.795 | 26.92610952 | 0.33069824 | 22.17201477 | NA | NA | NA | NA | NA | NA | NA |
| GRIN2A | 456 | C | 68.60783333 | 22.94422204 | 0.276758575 | 18.21674914 | NA | NA | NA | NA | NA | NA | NA |
| GRIN2A | 457 | K | 64.54883333 | 21.31532207 | 0.25469308 | 17.8157297 | NA | NA | NA | NA | NA | NA | NA |
| GRIN2A | 458 | G | 63.89475 | 18.55113563 | 0.217248705 | 14.60251082 | NA | NA | NA | NA | NA | NA | NA |
| GRIN2A | 459 | F | 61.80077273 | 14.62734911 | 0.164096091 | 11.15776363 | NA | NA | NA | NA | NA | NA | NA |
| GRIN2A | 460 | C | 66.6535 | 17.78668853 | 0.20689331 | 15.88013305 | NA | NA | NA | NA | NA | NA | NA |
| GRIN2A | 461 | I | 68.705375 | 16.49551903 | 0.189402798 | 11.75492407 | NA | NA | NA | NA | NA | NA | NA |

| | | | | | | | | | | | | | |
|---|---|---|---|---|---|---|---|---|---|---|---|---|---|
| GRIN2A | 462 | D | 66.81225 | 13.47216293 | 0.148447644 | 11.40242192 | NA | NA | NA | NA | NA | NA | NA |
| GRIN2A | 463 | I | 68.019375 | 13.14585357 | 0.144027375 | 8.392207998 | NA | NA | NA | NA | NA | NA | NA |
| GRIN2A | 464 | L | 72.0735 | 14.08910285 | 0.15680487 | 9.188279236 | NA | NA | NA | NA | NA | NA | NA |
| GRIN2A | 465 | K | 71.78127778 | 11.37252342 | 0.120005392 | 6.479456339 | NA | NA | NA | NA | NA | NA | NA |
| GRIN2A | 466 | K | 69.75038889 | 9.158651102 | 0.090015712 | 4.266264605 | 5.15 | -0.83 | -2.52 | 22.05 | 1.19 | 2 | 0 |
| GRIN2A | 467 | L | 73.7195 | 11.33407433 | 0.119484551 | 6.456404935 | NA | NA | NA | NA | NA | NA | NA |
| GRIN2A | 468 | S | 76.31866667 | 10.75877544 | 0.111691405 | 5.735388444 | NA | NA | NA | NA | NA | NA | NA |
| GRIN2A | 469 | R | 74.67686364 | 6.552396356 | 0.05471072 | 1.494202289 | 5.12 | -0.69 | -2.6 | 31.25 | 1.46 | 3 | 1 |
| GRIN2A | 470 | T | 75.6375 | 6.587441233 | 0.055185447 | 1.655966001 | 5.12 | -0.69 | -2.6 | 31.25 | 1.46 | 3 | 1 |
| GRIN2A | 471 | V | 78.09507143 | 9.881131514 | 0.099802616 | 4.795886561 | 5.12 | -0.69 | -2.6 | 31.25 | 1.46 | 3 | 1 |
| GRIN2A | 472 | K | 80.29683333 | 5.024607447 | 0.034014901 | 1.616382022 | 3.43 | -0.3 | -0.9 | 14.14 | 0.28 | 1 | 1 |
| GRIN2A | 473 | F | 82.27568182 | 13.17276547 | 0.14439193 | 9.884919215 | NA | NA | NA | NA | NA | NA | NA |
| GRIN2A | 474 | T | 82.3245 | 16.26018023 | 0.186214839 | 12.99467004 | NA | NA | NA | NA | NA | NA | NA |
| GRIN2A | 475 | Y | 78.35883333 | 14.47740225 | 0.162064873 | 9.393444984 | NA | NA | NA | NA | NA | NA | NA |
| GRIN2A | 476 | D | 79.192 | 21.88595282 | 0.26242299 | 16.79928748 | NA | NA | NA | NA | NA | NA | NA |
| GRIN2A | 477 | L | 74.6755 | 22.29849795 | 0.268011432 | 17.48928926 | NA | NA | NA | NA | NA | NA | NA |
| GRIN2A | 478 | Y | 76.4005 | 27.12381142 | 0.33337636 | 22.18174399 | NA | NA | NA | NA | NA | NA | NA |
| GRIN2A | 479 | L | 70.74825 | 28.95443968 | 0.358174518 | 24.17725039 | NA | NA | NA | NA | NA | NA | NA |
| GRIN2A | 480 | V | 69.60207143 | 31.5744682 | 0.393666093 | 26.76190408 | NA | NA | NA | NA | NA | NA | NA |
| GRIN2A | 481 | T | 69.79807143 | 35.71264136 | 0.449722843 | 30.896724 | NA | NA | NA | NA | NA | NA | NA |
| GRIN2A | 482 | N | 70.48775 | 37.48294667 | 0.473703851 | 32.66760464 | NA | NA | NA | NA | NA | NA | NA |
| GRIN2A | 483 | G | 66.42275 | 36.29743408 | 0.457644594 | 31.53802106 | NA | NA | NA | NA | NA | NA | NA |
| GRIN2A | 484 | K | 63.62161111 | 33.0440419 | 0.413573313 | 30.33475046 | NA | NA | NA | NA | NA | NA | NA |
| GRIN2A | 485 | H | 64.0791 | 30.17432566 | 0.374699405 | 26.75417684 | NA | NA | NA | NA | NA | NA | NA |
| GRIN2A | 486 | G | 65.72225 | 35.13531117 | 0.441902181 | 30.36838128 | NA | NA | NA | NA | NA | NA | NA |
| GRIN2A | 487 | K | 64.7475 | 38.11013364 | 0.482199885 | 33.32113052 | NA | NA | NA | NA | NA | NA | NA |
| GRIN2A | 488 | K | 66.55938889 | 40.32960945 | 0.512265471 | 35.48937532 | NA | NA | NA | NA | NA | NA | NA |
| GRIN2A | 489 | V | 68.3075 | 43.06012301 | 0.549253705 | 38.22915311 | NA | NA | NA | NA | NA | NA | NA |
| GRIN2A | 490 | N | 68.5835 | 45.66189345 | 0.584497952 | 40.79048297 | NA | NA | NA | NA | NA | NA | NA |
| GRIN2A | 491 | N | 69.955375 | 42.51364509 | 0.541850976 | 37.60330616 | NA | NA | NA | NA | NA | NA | NA |
| GRIN2A | 492 | V | 71.66321429 | 41.14397255 | 0.523297042 | 36.22463093 | NA | NA | NA | NA | NA | NA | NA |
| GRIN2A | 493 | W | 69.73571429 | 37.26209406 | 0.470712125 | 32.43385791 | NA | NA | NA | NA | NA | NA | NA |
| GRIN2A | 494 | N | 68.958 | 35.73928758 | 0.450083799 | 30.90282745 | NA | NA | NA | NA | NA | NA | NA |

| | | | | | | | | | | | | | |
|---|---|---|---|---|---|---|---|---|---|---|---|---|---|
| GRIN2A | 495 | G | 69.702 | 33.68086295 | 0.422199854 | 28.85375258 | NA | NA | NA | NA | NA | NA | NA |
| GRIN2A | 496 | M | 69.5185 | 27.49650034 | 0.338424899 | 22.70692292 | NA | NA | NA | NA | NA | NA | NA |
| GRIN2A | 497 | I | 70.031125 | 32.34913639 | 0.404159946 | 27.50810949 | NA | NA | NA | NA | NA | NA | NA |
| GRIN2A | 498 | G | 73.38725 | 34.47735538 | 0.432989344 | 29.55461013 | NA | NA | NA | NA | NA | NA | NA |
| GRIN2A | 499 | E | 74.01461111 | 31.05773717 | 0.386666322 | 26.2365191 | NA | NA | NA | NA | NA | NA | NA |
| GRIN2A | 500 | V | 75.00578571 | 29.35389838 | 0.363585687 | 24.45569458 | NA | NA | NA | NA | NA | NA | NA |
| GRIN2A | 501 | V | 76.38278571 | 33.32290885 | 0.417350916 | 28.27272744 | NA | NA | NA | NA | NA | NA | NA |
| GRIN2A | 502 | Y | 77.56933333 | 34.98927963 | 0.439924 | 29.91463411 | NA | NA | NA | NA | NA | NA | NA |
| GRIN2A | 503 | Q | 81.27883333 | 32.18813976 | 0.401979044 | 27.22128674 | NA | NA | NA | NA | NA | NA | NA |
| GRIN2A | 504 | R | 78.19995455 | 30.83477694 | 0.383646046 | 25.75022065 | NA | NA | NA | NA | NA | NA | NA |
| GRIN2A | 505 | A | 77.9161 | 28.37434396 | 0.350316393 | 23.29897836 | NA | NA | NA | NA | NA | NA | NA |
| GRIN2A | 506 | V | 78.7125 | 24.98967373 | 0.304466786 | 19.91130489 | NA | NA | NA | NA | NA | NA | NA |
| GRIN2A | 507 | M | 75.249125 | 21.24776445 | 0.253777928 | 16.26325392 | NA | NA | NA | NA | NA | NA | NA |
| GRIN2A | 508 | A | 71.4725 | 23.84566701 | 0.288969779 | 19.00958704 | NA | NA | NA | NA | NA | NA | NA |
| GRIN2A | 509 | V | 68.63864286 | 21.12015864 | 0.252049347 | 16.3840329 | NA | NA | NA | NA | NA | NA | NA |
| GRIN2A | 510 | G | 65.4675 | 23.15974473 | 0.279678101 | 19.98083672 | NA | NA | NA | NA | NA | NA | NA |
| GRIN2A | 511 | S | 61.72183333 | 23.86549788 | 0.289238413 | 20.4529669 | NA | NA | NA | NA | NA | NA | NA |
| GRIN2A | 512 | L | 64.53825 | 25.60094283 | 0.312747193 | 22.1797223 | NA | NA | NA | NA | NA | NA | NA |
| GRIN2A | 513 | T | 61.15378571 | 27.55472393 | 0.339213611 | 24.13168607 | NA | NA | NA | NA | NA | NA | NA |
| GRIN2A | 514 | I | 62.623 | 24.29476034 | 0.295053312 | 21.15320673 | NA | NA | NA | NA | NA | NA | NA |
| GRIN2A | 515 | N | 61.77025 | 30.34807877 | 0.377053109 | 27.11983419 | NA | NA | NA | NA | NA | NA | NA |
| GRIN2A | 516 | E | 62.67127778 | 33.60139862 | 0.42112341 | 30.44767541 | NA | NA | NA | NA | NA | NA | NA |
| GRIN2A | 517 | E | 64.99472222 | 35.33564407 | 0.444615941 | 30.51545808 | NA | NA | NA | NA | NA | NA | NA |
| GRIN2A | 518 | R | 65.29431818 | 30.5695357 | 0.380053021 | 26.53209714 | NA | NA | NA | NA | NA | NA | NA |
| GRIN2A | 519 | S | 67.8035 | 31.55592391 | 0.393414887 | 26.81830254 | NA | NA | NA | NA | NA | NA | NA |
| GRIN2A | 520 | E | 68.79583333 | 33.99225179 | 0.426418007 | 29.11138305 | NA | NA | NA | NA | NA | NA | NA |
| GRIN2A | 521 | V | 72.31178571 | 32.44099237 | 0.40540425 | 27.35694197 | NA | NA | NA | NA | NA | NA | NA |
| GRIN2A | 522 | V | 72.2645 | 28.19566644 | 0.347895982 | 23.16138994 | NA | NA | NA | NA | NA | NA | NA |
| GRIN2A | 523 | D | 73.09125 | 26.1808827 | 0.320603207 | 21.22565329 | NA | NA | NA | NA | NA | NA | NA |
| GRIN2A | 524 | F | 68.91977273 | 23.19411447 | 0.280143682 | 18.29219951 | NA | NA | NA | NA | NA | NA | NA |
| GRIN2A | 525 | S | 68.58633333 | 20.0348434 | 0.237347389 | 15.19573269 | NA | NA | NA | NA | NA | NA | NA |
| GRIN2A | 526 | V | 66.60878571 | 18.2591298 | 0.21329312 | 14.47151281 | NA | NA | NA | NA | NA | NA | NA |
| GRIN2A | 527 | P | 63.43307143 | 18.37615724 | 0.214878403 | 15.59856256 | NA | NA | NA | NA | NA | NA | NA |

| | | | | | | | | | | | | | |
|---|---|---|---|---|---|---|---|---|---|---|---|---|---|
| GRIN2A | 528 | F | 63.31895455 | 16.35065142 | 0.187440384 | 13.30817575 | NA | NA | NA | NA | NA | NA | NA |
| GRIN2A | 529 | V | 59.23521429 | 16.32984988 | 0.187158601 | 12.90885954 | NA | NA | NA | NA | NA | NA | NA |
| GRIN2A | 530 | E | 55.96138889 | 16.94923464 | 0.195548946 | 13.46049524 | NA | NA | NA | NA | NA | NA | NA |
| GRIN2A | 531 | T | 55.29564286 | 22.15859781 | 0.266116309 | 18.65260516 | NA | NA | NA | NA | NA | NA | NA |
| GRIN2A | 532 | G | 53.06025 | 25.4824726 | 0.311142365 | 21.98778866 | NA | NA | NA | NA | NA | NA | NA |
| GRIN2A | 533 | I | 50.424375 | 28.42108358 | 0.35094954 | 24.87316174 | NA | NA | NA | NA | NA | NA | NA |
| GRIN2A | 534 | S | 47.62166667 | 25.34472318 | 0.309276376 | 18.69741169 | NA | NA | NA | NA | NA | NA | NA |
| GRIN2A | 535 | V | 43.8895 | 25.97194611 | 0.317772898 | 19.38261305 | NA | NA | NA | NA | NA | NA | NA |
| GRIN2A | 536 | M | 43.881125 | 23.66876397 | 0.286573405 | 17.1353382 | NA | NA | NA | NA | NA | NA | NA |
| GRIN2A | 537 | V | 39.56578571 | 24.65574012 | 0.299943236 | 17.84493121 | NA | NA | NA | NA | NA | NA | NA |
| GRIN2A | 538 | S | 36.77166667 | 24.36379671 | 0.295988496 | 18.59983837 | NA | NA | NA | NA | NA | NA | NA |
| GRIN2A | 556 | S | 10.6305 | 22.79546179 | 0.274743431 | 20.75615423 | NA | NA | NA | NA | NA | NA | NA |
| GRIN2A | 557 | V | 9.575071429 | 18.01316638 | 0.209961236 | 16.00125806 | NA | NA | NA | NA | NA | NA | NA |
| GRIN2A | 558 | W | 7.096357143 | 16.62652282 | 0.191177409 | 13.46634774 | NA | NA | NA | NA | NA | NA | NA |
| GRIN2A | 559 | V | 5.859214286 | 22.87939575 | 0.275880422 | 19.72591372 | NA | NA | NA | NA | NA | NA | NA |
| GRIN2A | 560 | M | 6.212 | 21.56939512 | 0.258134819 | 18.42500211 | NA | NA | NA | NA | NA | NA | NA |
| GRIN2A | 561 | M | 3.401625 | 15.80136368 | 0.179999592 | 12.69888089 | NA | NA | NA | NA | NA | NA | NA |
| GRIN2A | 562 | F | 0.7235 | 21.47875336 | 0.256906962 | 18.40070176 | NA | NA | NA | NA | NA | NA | NA |
| GRIN2A | 563 | V | 1.226357143 | 23.86015834 | 0.289166082 | 20.78660551 | NA | NA | NA | NA | NA | NA | NA |
| GRIN2A | 564 | M | 1.13475 | 21.35625086 | 0.255247512 | 18.70792887 | NA | NA | NA | NA | NA | NA | NA |
| GRIN2A | 565 | L | -0.84325 | 17.16341429 | 0.198450278 | 14.52068146 | NA | NA | NA | NA | NA | NA | NA |
| GRIN2A | 566 | L | -2.477625 | 22.18503897 | 0.266474488 | 19.57217658 | NA | NA | NA | NA | NA | NA | NA |
| GRIN2A | 567 | I | -1.90025 | 23.0250377 | 0.277853325 | 20.38203333 | NA | NA | NA | NA | NA | NA | NA |
| GRIN2A | 568 | V | -3.949071429 | 18.35931279 | 0.214650224 | 15.70842985 | NA | NA | NA | NA | NA | NA | NA |
| GRIN2A | 569 | S | -6.718166667 | 20.54097622 | 0.244203594 | 16.33286851 | NA | NA | NA | NA | NA | NA | NA |
| GRIN2A | 570 | A | -7.6883 | 22.7523919 | 0.274159995 | 18.54203703 | NA | NA | NA | NA | NA | NA | NA |
| GRIN2A | 571 | I | -7.078 | 21.56853507 | 0.258123168 | 17.35236349 | NA | NA | NA | NA | NA | NA | NA |
| GRIN2A | 572 | A | -10.4875 | 18.34730956 | 0.214487625 | 14.13204622 | NA | NA | NA | NA | NA | NA | NA |
| GRIN2A | 573 | V | -12.34907143 | 21.22712296 | 0.253498313 | 17.01693714 | NA | NA | NA | NA | NA | NA | NA |
| GRIN2A | 574 | F | -11.33222727 | 23.41556724 | 0.283143538 | 19.21590275 | NA | NA | NA | NA | NA | NA | NA |
| GRIN2A | 575 | V | -13.39335714 | 19.7612306 | 0.233640961 | 15.5631363 | NA | NA | NA | NA | NA | NA | NA |
| GRIN2A | 576 | F | -15.81540909 | 16.59069899 | 0.19069213 | 7.282739861 | NA | NA | NA | NA | NA | NA | NA |
| GRIN2A | 577 | E | -17.17883333 | 22.56026429 | 0.271557386 | 12.68199388 | NA | NA | NA | NA | NA | NA | NA |

| | | | | | | | | | | | | | |
|---|---|---|---|---|---|---|---|---|---|---|---|---|---|
| GRIN2A | 578 | Y | -17.18158333 | 22.29935793 | 0.268023081 | 12.40494644 | NA | NA | NA | NA | NA | NA | NA |
| GRIN2A | 579 | F | -18.13168182 | 17.54697324 | 0.203646065 | 7.725453081 | NA | NA | NA | NA | NA | NA | NA |
| GRIN2A | 580 | S | -22.2525 | 20.9171473 | 0.249299303 | 11.03657201 | NA | NA | NA | NA | NA | NA | NA |
| GRIN2A | 581 | P | -23.91978571 | 21.07256543 | 0.251404637 | 11.28137709 | NA | NA | NA | NA | NA | NA | NA |
| GRIN2A | 598 | S | -17.937 | 13.16490505 | 0.144285451 | 5.005022182 | NA | NA | NA | NA | NA | NA | NA |
| GRIN2A | 599 | F | -17.29840909 | 15.47152139 | 0.175531465 | 7.31490417 | NA | NA | NA | NA | NA | NA | NA |
| GRIN2A | 600 | T | -13.0825 | 17.84330869 | 0.207660301 | 13.63383965 | NA | NA | NA | NA | NA | NA | NA |
| GRIN2A | 601 | I | -12.834625 | 19.38234483 | 0.228508477 | 15.1832971 | NA | NA | NA | NA | NA | NA | NA |
| GRIN2A | 602 | G | -12.3965 | 16.29693747 | 0.186712762 | 11.91828692 | NA | NA | NA | NA | NA | NA | NA |
| GRIN2A | 603 | K | -13.83794444 | 10.44538381 | 0.107446122 | 8.045721799 | 4.24 | -0.71 | -1.43 | 15.54 | 1.02 | 1 | 1 |
| GRIN2A | 604 | A | -9.5611 | 14.48981508 | 0.16223302 | 10.27817402 | NA | NA | NA | NA | NA | NA | NA |
| GRIN2A | 605 | I | -7.10375 | 14.41224518 | 0.161182239 | 11.02527611 | NA | NA | NA | NA | NA | NA | NA |
| GRIN2A | 606 | W | -8.637785714 | 11.22096769 | 0.117952379 | 7.723567571 | NA | NA | NA | NA | NA | NA | NA |
| GRIN2A | 607 | L | -8.271125 | 8.414282501 | 0.079932305 | 5.161274456 | 3.55 | -0.8 | -0.4 | 3.38 | 0.81 | 0 | 0 |
| GRIN2A | 608 | L | -4.281875 | 11.67005681 | 0.124035855 | 9.07821693 | NA | NA | NA | NA | NA | NA | NA |
| GRIN2A | 609 | W | -1.619571429 | 10.57317403 | 0.109177201 | 7.956836175 | NA | NA | NA | NA | NA | NA | NA |
| GRIN2A | 610 | G | -3.44225 | 6.964308796 | 0.060290591 | 4.409675048 | 2.72 | -0.15 | 0.83 | 2.57 | 0.15 | 0 | 0 |
| GRIN2A | 611 | L | -2.610125 | 5.570534086 | 0.041410163 | 2.92086001 | 2.71 | 0.07 | 1.23 | 2.3 | -0.08 | 0 | 0 |
| GRIN2A | 612 | V | 0.112071429 | 9.253924249 | 0.091306306 | 5.43601699 | 3.9 | -0.14 | 0.05 | 2.57 | 0.08 | 0 | 0 |
| GRIN2A | 613 | F | 0.231863636 | 5.515076246 | 0.040658917 | 1.688609692 | 3.85 | 0.14 | 0.45 | 2.08 | -0.29 | 0 | 0 |
| GRIN2A | 625 | T | -14.87907143 | 11.4567261 | 0.121146023 | 2.008728832 | 9.49 | -0.78 | -3.85 | 50.33 | 1.89 | 4 | 0 |
| GRIN2A | 626 | T | -12.39178571 | 15.32315571 | 0.173521665 | 11.13305374 | NA | NA | NA | NA | NA | NA | NA |
| GRIN2A | 627 | S | -9.976166667 | 15.55468978 | 0.176658085 | 11.34291147 | NA | NA | NA | NA | NA | NA | NA |
| GRIN2A | 628 | K | -10.9345 | 10.75253589 | 0.111606882 | 6.665296229 | NA | NA | NA | NA | NA | NA | NA |
| GRIN2A | 629 | I | -7.638 | 13.35454518 | 0.146854364 | 9.147889267 | NA | NA | NA | NA | NA | NA | NA |
| GRIN2A | 630 | M | -5.716 | 15.95382992 | 0.182064939 | 11.92162973 | NA | NA | NA | NA | NA | NA | NA |
| GRIN2A | 631 | V | -4.462357143 | 14.43566864 | 0.161499539 | 11.78719043 | NA | NA | NA | NA | NA | NA | NA |
| GRIN2A | 632 | S | -3.554833333 | 11.53131783 | 0.122156461 | 8.877973314 | NA | NA | NA | NA | NA | NA | NA |
| GRIN2A | 633 | V | -0.892642857 | 14.24036028 | 0.158853842 | 11.65244744 | NA | NA | NA | NA | NA | NA | NA |
| GRIN2A | 634 | W | 0.340428571 | 13.48736939 | 0.148653635 | 11.17668757 | NA | NA | NA | NA | NA | NA | NA |
| GRIN2A | 635 | A | -0.2337 | 10.2850753 | 0.105274542 | 7.653578535 | NA | NA | NA | NA | NA | NA | NA |
| GRIN2A | 636 | F | 2.2895 | 11.00529445 | 0.115030814 | 7.924453216 | NA | NA | NA | NA | NA | NA | NA |
| GRIN2A | 637 | F | 3.827681818 | 11.74662701 | 0.125073094 | 8.105135475 | NA | NA | NA | NA | NA | NA | NA |

| | | | | | | | | | | | | | |
|---|---|---|---|---|---|---|---|---|---|---|---|---|---|
| GRIN2A | 638 | A | 3.2657 | 9.077490017 | 0.088916283 | 5.926804007 | 3.19 | 1.11 | 3.58 | 0.39 | -1.38 | 0 | 0 |
| GRIN2A | 639 | V | 4.682785714 | 6.821490013 | 0.058355931 | 3.245110167 | 3.69 | -0.01 | 1.1 | 1.05 | -0.25 | 0 | 0 |
| GRIN2A | 640 | I | 7.710125 | 9.235391167 | 0.091055253 | 6.018162157 | 3.31 | -0.77 | -0.7 | 1.66 | 0.77 | 0 | 0 |
| GRIN2A | 641 | F | 6.485863636 | 9.492081805 | 0.09453245 | 6.200876478 | 3.31 | -0.77 | -0.7 | 1.66 | 0.77 | 0 | 0 |
| GRIN2A | 642 | L | 6.627875 | 4.836091811 | 0.03146122 | 2.625360253 | 3.31 | -0.77 | -0.7 | 1.66 | 0.77 | 0 | 0 |
| GRIN2A | 643 | A | 10.7371 | 6.319502591 | 0.051555882 | 3.66429064 | 2.67 | -0.77 | -0.7 | 1.66 | 0.77 | 0 | 0 |
| GRIN2A | 644 | S | 12.05316667 | 8.16677972 | 0.076579569 | 6.097978535 | 2.11 | -0.18 | 1.18 | 0.42 | 0.64 | 0 | 0 |
| GRIN2A | 645 | Y | 11.37583333 | 7.331274446 | 0.065261601 | 5.322180724 | 2.11 | -0.18 | 1.18 | 0.42 | 0.64 | 0 | 0 |
| GRIN2A | 646 | T | 12.87221429 | 4.041627148 | 0.020699199 | 2.262536254 | 2.11 | -0.18 | 1.18 | 0.42 | 0.64 | 0 | 0 |
| GRIN2A | 647 | A | 16.1977 | 5.268661785 | 0.037320924 | 3.558337644 | 1.93 | -0.31 | 0.92 | 1.35 | 0.68 | 0 | 0 |
| GRIN2A | 648 | N | 16.4315 | 7.477653308 | 0.067244487 | 5.571584612 | 1.93 | -0.31 | 0.92 | 1.35 | 0.68 | 0 | 0 |
| GRIN2A | 649 | L | 15.999625 | 6.521863691 | 0.054297117 | 4.643956658 | 1.93 | -0.31 | 0.92 | 1.35 | 0.68 | 0 | 0 |
| GRIN2A | 650 | A | 18.9025 | 3.314995777 | 0.010856065 | 1.515704432 | 1.82 | -0.19 | 1.25 | 0.85 | 0.65 | 0 | 0 |
| GRIN2A | 651 | A | 21.0563 | 6.476501293 | 0.053682626 | 4.698525735 | 1.81 | 0.47 | 2.48 | 0.03 | -0.02 | 0 | 0 |
| GRIN2A | 652 | F | 20.83386364 | 8.778725819 | 0.084869147 | 7.003596338 | 1.81 | 0.47 | 2.48 | 0.03 | -0.02 | 0 | 0 |
| GRIN2A | 653 | M | 20.441875 | 7.479535413 | 0.067269982 | 6.03813409 | 1.81 | 0.47 | 2.48 | 0.03 | -0.02 | 0 | 0 |
| GRIN2A | 654 | I | 23.827125 | 3.156117235 | 0.008703856 | 1.7183665 | 1.55 | 1.47 | 4.25 | 0.13 | -1.7 | 0 | 0 |
| GRIN2A | 655 | Q | 26.35938889 | 5.795172991 | 0.044453179 | 3.071150225 | 2.74 | 1.31 | 4.08 | 0.13 | -1.68 | 0 | 0 |
| GRIN2A | 660 | D | 33.4087 | 22.52391454 | 0.271064982 | 17.3714766 | NA | NA | NA | NA | NA | NA | NA |
| GRIN2A | 661 | Q | 32.67927778 | 25.90519118 | 0.316868619 | 20.75594251 | NA | NA | NA | NA | NA | NA | NA |
| GRIN2A | 662 | V | 36.8735 | 27.55284891 | 0.339188211 | 21.92463009 | NA | NA | NA | NA | NA | NA | NA |
| GRIN2A | 663 | T | 37.69021429 | 28.54883719 | 0.352680123 | 21.41676477 | NA | NA | NA | NA | NA | NA | NA |
| GRIN2A | 664 | G | 40.6495 | 30.55514998 | 0.379858148 | 23.54607662 | NA | NA | NA | NA | NA | NA | NA |
| GRIN2A | 665 | L | 43.96025 | 32.93742498 | 0.412129053 | 26.32526621 | NA | NA | NA | NA | NA | NA | NA |
| GRIN2A | 666 | S | 40.0195 | 33.86646405 | 0.424714054 | 26.73520359 | NA | NA | NA | NA | NA | NA | NA |
| GRIN2A | 667 | D | 37.67675 | 32.02267415 | 0.399737605 | 24.89058133 | NA | NA | NA | NA | NA | NA | NA |
| GRIN2A | 668 | K | 36.43072222 | 37.62888384 | 0.475680753 | 30.49862458 | NA | NA | NA | NA | NA | NA | NA |
| GRIN2A | 669 | K | 35.48761111 | 31.49533164 | 0.392594089 | 25.77141401 | NA | NA | NA | NA | NA | NA | NA |
| GRIN2A | 670 | F | 40.62477273 | 33.52029317 | 0.420024735 | 26.70271599 | NA | NA | NA | NA | NA | NA | NA |
| GRIN2A | 671 | Q | 40.38594444 | 38.81237684 | 0.491712651 | 31.83429055 | NA | NA | NA | NA | NA | NA | NA |
| GRIN2A | 672 | R | 36.06268182 | 41.74008899 | 0.531372187 | 35.99838875 | NA | NA | NA | NA | NA | NA | NA |
| GRIN2A | 673 | P | 37.39107143 | 42.17295975 | 0.537235965 | 36.4179512 | NA | NA | NA | NA | NA | NA | NA |
| GRIN2A | 674 | H | 35.8622 | 44.23489043 | 0.565167404 | 39.09248883 | NA | NA | NA | NA | NA | NA | NA |

| | | | | | | | | | | | | | |
|---|---|---|---|---|---|---|---|---|---|---|---|---|---|
| GRIN2A | 675 | D | 32.610125 | 42.83718803 | 0.546233771 | 37.73014052 | NA | NA | NA | NA | NA | NA | NA |
| GRIN2A | 676 | Y | 31.64358333 | 38.78350814 | 0.491321588 | 33.68461487 | NA | NA | NA | NA | NA | NA | NA |
| GRIN2A | 677 | S | 29.32916667 | 39.82214395 | 0.505391214 | 36.06832214 | NA | NA | NA | NA | NA | NA | NA |
| GRIN2A | 678 | P | 31.8005 | 39.2853377 | 0.4981195 | 34.14490302 | NA | NA | NA | NA | NA | NA | NA |
| GRIN2A | 679 | P | 35.27878571 | 40.63763395 | 0.51643805 | 34.8794543 | NA | NA | NA | NA | NA | NA | NA |
| GRIN2A | 680 | F | 36.75295455 | 35.82510089 | 0.451246248 | 30.07456701 | NA | NA | NA | NA | NA | NA | NA |
| GRIN2A | 681 | R | 39.02904545 | 35.96986751 | 0.453207294 | 29.1790814 | NA | NA | NA | NA | NA | NA | NA |
| GRIN2A | 682 | F | 42.23240909 | 33.52933538 | 0.420147223 | 26.90607728 | NA | NA | NA | NA | NA | NA | NA |
| GRIN2A | 683 | G | 47.24425 | 34.89534912 | 0.438651594 | 28.29659521 | NA | NA | NA | NA | NA | NA | NA |
| GRIN2A | 684 | T | 50.83964286 | 36.00683781 | 0.453708103 | 29.4093336 | NA | NA | NA | NA | NA | NA | NA |
| GRIN2A | 685 | V | 54.49878571 | 34.3112178 | 0.430738802 | 30.35407614 | NA | NA | NA | NA | NA | NA | NA |
| GRIN2A | 686 | P | 56.10407143 | 38.67230954 | 0.489815264 | 35.10773308 | NA | NA | NA | NA | NA | NA | NA |
| GRIN2A | 687 | N | 59.21925 | 37.39573881 | 0.472522511 | 33.88843023 | NA | NA | NA | NA | NA | NA | NA |
| GRIN2A | 688 | G | 57.675 | 35.63845946 | 0.448717956 | 32.13295549 | NA | NA | NA | NA | NA | NA | NA |
| GRIN2A | 689 | S | 57.82033333 | 31.94866202 | 0.398735018 | 28.44256635 | NA | NA | NA | NA | NA | NA | NA |
| GRIN2A | 690 | T | 54.1165 | 32.26672597 | 0.403043593 | 28.79240915 | NA | NA | NA | NA | NA | NA | NA |
| GRIN2A | 691 | E | 54.48294444 | 36.6601039 | 0.462557412 | 33.165749 | NA | NA | NA | NA | NA | NA | NA |
| GRIN2A | 692 | R | 59.57431818 | 37.79151945 | 0.477883856 | 34.46849105 | NA | NA | NA | NA | NA | NA | NA |
| GRIN2A | 693 | N | 55.15825 | 33.43533984 | 0.418873935 | 29.93114024 | NA | NA | NA | NA | NA | NA | NA |
| GRIN2A | 694 | I | 51.04025 | 37.08312584 | 0.468287776 | 33.65728234 | NA | NA | NA | NA | NA | NA | NA |
| GRIN2A | 695 | R | 55.09259091 | 40.84664233 | 0.519269331 | 37.34245613 | NA | NA | NA | NA | NA | NA | NA |
| GRIN2A | 696 | N | 55.14625 | 39.21306878 | 0.497140526 | 35.72888407 | NA | NA | NA | NA | NA | NA | NA |
| GRIN2A | 697 | N | 50.566375 | 37.5295033 | 0.474334519 | 34.02738336 | NA | NA | NA | NA | NA | NA | NA |
| GRIN2A | 698 | Y | 47.14716667 | 39.60604829 | 0.502463927 | 36.69595319 | NA | NA | NA | NA | NA | NA | NA |
| GRIN2A | 699 | P | 49.5285 | 45.37019494 | 0.580546529 | 41.87993866 | NA | NA | NA | NA | NA | NA | NA |
| GRIN2A | 700 | Y | 46.26625 | 46.14443708 | 0.591034611 | 39.52851909 | NA | NA | NA | NA | NA | NA | NA |
| GRIN2A | 702 | H | 50.1836 | 41.09179023 | 0.522590167 | 34.96648074 | NA | NA | NA | NA | NA | NA | NA |
| GRIN2A | 703 | Q | 47.55016667 | 46.24494469 | 0.592396112 | 39.60266095 | NA | NA | NA | NA | NA | NA | NA |
| GRIN2A | 704 | Y | 42.97316667 | 44.1439742 | 0.56393583 | 37.52626841 | NA | NA | NA | NA | NA | NA | NA |
| GRIN2A | 705 | M | 46.75975 | 37.91761022 | 0.479591914 | 31.27756591 | NA | NA | NA | NA | NA | NA | NA |
| GRIN2A | 706 | T | 49.25164286 | 43.88818791 | 0.560470884 | 37.29010269 | NA | NA | NA | NA | NA | NA | NA |
| GRIN2A | 707 | K | 44.97661111 | 43.4486426 | 0.554516691 | 36.84897134 | NA | NA | NA | NA | NA | NA | NA |
| GRIN2A | 708 | F | 44.97786364 | 39.16477532 | 0.496486331 | 32.56626543 | NA | NA | NA | NA | NA | NA | NA |

| | | | | | | | | | | | | | |
|---|---|---|---|---|---|---|---|---|---|---|---|---|---|
| GRIN2A | 709 | N | 50.6055 | 39.86443394 | 0.505964085 | 33.26791633 | NA | NA | NA | NA | NA | NA | NA |
| GRIN2A | 710 | Q | 50.17916667 | 34.2616342 | 0.430067129 | 27.6667428 | NA | NA | NA | NA | NA | NA | NA |
| GRIN2A | 711 | K | 53.73627778 | 36.5354619 | 0.46086898 | 32.58147405 | NA | NA | NA | NA | NA | NA | NA |
| GRIN2A | 712 | G | 52.84275 | 33.46929357 | 0.419333881 | 29.61001222 | NA | NA | NA | NA | NA | NA | NA |
| GRIN2A | 713 | V | 51.90578571 | 28.63070373 | 0.353789108 | 22.04549899 | NA | NA | NA | NA | NA | NA | NA |
| GRIN2A | 714 | E | 49.89161111 | 30.00606009 | 0.372420036 | 23.4377492 | NA | NA | NA | NA | NA | NA | NA |
| GRIN2A | 715 | D | 49.073875 | 33.3585433 | 0.41783363 | 26.77128846 | NA | NA | NA | NA | NA | NA | NA |
| GRIN2A | 716 | A | 46.7599 | 30.73681345 | 0.382319008 | 24.13996173 | NA | NA | NA | NA | NA | NA | NA |
| GRIN2A | 717 | L | 46.323875 | 26.53526397 | 0.325403746 | 19.94488505 | NA | NA | NA | NA | NA | NA | NA |
| GRIN2A | 718 | V | 43.88635714 | 31.05623668 | 0.386645996 | 24.46541921 | NA | NA | NA | NA | NA | NA | NA |
| GRIN2A | 719 | S | 42.8115 | 32.91033313 | 0.41176206 | 26.27761211 | NA | NA | NA | NA | NA | NA | NA |
| GRIN2A | 720 | L | 41.2445 | 28.54511496 | 0.352629701 | 21.78742065 | NA | NA | NA | NA | NA | NA | NA |
| GRIN2A | 721 | K | 40.8845 | 25.81439252 | 0.315638637 | 19.23284713 | NA | NA | NA | NA | NA | NA | NA |
| GRIN2A | 722 | T | 38.93007143 | 32.27597684 | 0.403168908 | 25.46414017 | NA | NA | NA | NA | NA | NA | NA |
| GRIN2A | 723 | G | 37.7235 | 33.34204773 | 0.417610176 | 26.53324504 | NA | NA | NA | NA | NA | NA | NA |
| GRIN2A | 724 | K | 40.7485 | 35.37811224 | 0.445191226 | 28.56273731 | NA | NA | NA | NA | NA | NA | NA |
| GRIN2A | 725 | L | 41.739125 | 33.3906958 | 0.418269176 | 26.57680585 | NA | NA | NA | NA | NA | NA | NA |
| GRIN2A | 726 | D | 38.629375 | 31.02520662 | 0.386225655 | 24.21226118 | NA | NA | NA | NA | NA | NA | NA |
| GRIN2A | 727 | A | 41.9529 | 29.87009422 | 0.370578208 | 23.24454405 | NA | NA | NA | NA | NA | NA | NA |
| GRIN2A | 728 | F | 45.49322727 | 29.41369637 | 0.364395726 | 22.79106855 | NA | NA | NA | NA | NA | NA | NA |
| GRIN2A | 729 | I | 47.362875 | 30.89930645 | 0.384520179 | 24.26359984 | NA | NA | NA | NA | NA | NA | NA |
| GRIN2A | 730 | Y | 50.73066667 | 28.82113346 | 0.356368718 | 22.44228951 | NA | NA | NA | NA | NA | NA | NA |
| GRIN2A | 731 | D | 54.240375 | 26.10208923 | 0.31953585 | 22.82488411 | NA | NA | NA | NA | NA | NA | NA |
| GRIN2A | 732 | A | 50.8451 | 23.03556403 | 0.277995917 | 16.43860591 | NA | NA | NA | NA | NA | NA | NA |
| GRIN2A | 733 | A | 52.7205 | 20.88016087 | 0.248798276 | 16.93819482 | NA | NA | NA | NA | NA | NA | NA |
| GRIN2A | 734 | V | 53.24535714 | 22.81753098 | 0.275042386 | 18.92710022 | NA | NA | NA | NA | NA | NA | NA |
| GRIN2A | 735 | L | 49.13825 | 21.58853552 | 0.2583941 | 14.99070776 | NA | NA | NA | NA | NA | NA | NA |
| GRIN2A | 736 | N | 48.959 | 18.16814834 | 0.212060662 | 11.57177025 | NA | NA | NA | NA | NA | NA | NA |
| GRIN2A | 737 | Y | 53.08575 | 17.39432948 | 0.201578314 | 12.38684641 | NA | NA | NA | NA | NA | NA | NA |
| GRIN2A | 738 | K | 50.05038889 | 20.94306389 | 0.249650376 | 14.35042779 | NA | NA | NA | NA | NA | NA | NA |
| GRIN2A | 739 | A | 45.7587 | 18.48519705 | 0.216355484 | 11.88846205 | NA | NA | NA | NA | NA | NA | NA |
| GRIN2A | 740 | G | 46.1895 | 16.08733176 | 0.183873389 | 9.504768382 | NA | NA | NA | NA | NA | NA | NA |
| GRIN2A | 741 | R | 49.49177273 | 18.16852385 | 0.212065748 | 11.58237823 | NA | NA | NA | NA | NA | NA | NA |

| | | | | | | | | | | | | | |
|---|---|---|---|---|---|---|---|---|---|---|---|---|---|
| GRIN2A | 742 | D | 43.559125 | 19.66948594 | 0.232398164 | 13.08439784 | NA | NA | NA | NA | NA | NA | NA |
| GRIN2A | 743 | E | 41.54838889 | 21.4277906 | 0.256216608 | 14.62996711 | NA | NA | NA | NA | NA | NA | NA |
| GRIN2A | 744 | G | 38.56925 | 18.87786633 | 0.221674683 | 12.10509465 | NA | NA | NA | NA | NA | NA | NA |
| GRIN2A | 745 | C | 39.75 | 16.76967635 | 0.193116603 | 9.952887293 | NA | NA | NA | NA | NA | NA | NA |
| GRIN2A | 746 | K | 36.0965 | 20.88046372 | 0.248802378 | 14.078807 | NA | NA | NA | NA | NA | NA | NA |
| GRIN2A | 747 | L | 40.422625 | 21.7378075 | 0.260416176 | 14.9521789 | NA | NA | NA | NA | NA | NA | NA |
| GRIN2A | 748 | V | 40.24064286 | 19.72702694 | 0.233177629 | 12.9139341 | NA | NA | NA | NA | NA | NA | NA |
| GRIN2A | 749 | T | 43.13978571 | 22.45001425 | 0.27006391 | 15.89709643 | NA | NA | NA | NA | NA | NA | NA |
| GRIN2A | 750 | I | 42.453125 | 23.30024848 | 0.2815814 | 16.68575614 | NA | NA | NA | NA | NA | NA | NA |
| GRIN2A | 751 | G | 40.50125 | 21.36252207 | 0.255332464 | 14.59271603 | NA | NA | NA | NA | NA | NA | NA |
| GRIN2A | 752 | S | 41.382 | 19.37411823 | 0.228397038 | 12.81787997 | NA | NA | NA | NA | NA | NA | NA |
| GRIN2A | 753 | G | 44.99075 | 20.71616338 | 0.246576724 | 14.12476608 | NA | NA | NA | NA | NA | NA | NA |
| GRIN2A | 754 | Y | 42.68516667 | 22.93064144 | 0.276574609 | 16.53265658 | NA | NA | NA | NA | NA | NA | NA |
| GRIN2A | 755 | I | 48.627375 | 22.66613158 | 0.272991491 | 17.01095814 | NA | NA | NA | NA | NA | NA | NA |
| GRIN2A | 756 | F | 46.54122727 | 28.38362465 | 0.350442112 | 24.99459537 | NA | NA | NA | NA | NA | NA | NA |
| GRIN2A | 757 | A | 51.9253 | 28.76850747 | 0.355655832 | 25.26613916 | NA | NA | NA | NA | NA | NA | NA |
| GRIN2A | 758 | T | 53.23292857 | 26.70324087 | 0.327679204 | 23.19970566 | NA | NA | NA | NA | NA | NA | NA |
| GRIN2A | 759 | T | 56.89692857 | 24.66758257 | 0.300103657 | 21.24693924 | NA | NA | NA | NA | NA | NA | NA |
| GRIN2A | 763 | I | 69.16475 | 18.31808396 | 0.214091728 | 13.53648254 | NA | NA | NA | NA | NA | NA | NA |
| GRIN2A | 764 | A | 71.4839 | 23.73202985 | 0.287430421 | 18.88677176 | NA | NA | NA | NA | NA | NA | NA |
| GRIN2A | 765 | L | 74.447125 | 21.88548743 | 0.262416686 | 16.77186394 | NA | NA | NA | NA | NA | NA | NA |
| GRIN2A | 766 | Q | 77.85083333 | 26.56235242 | 0.325770693 | 21.51683161 | NA | NA | NA | NA | NA | NA | NA |
| GRIN2A | 767 | K | 76.5595 | 28.8907051 | 0.357311153 | 23.83633039 | NA | NA | NA | NA | NA | NA | NA |
| GRIN2A | 768 | G | 80.56975 | 26.88802027 | 0.330182273 | 22.06355415 | NA | NA | NA | NA | NA | NA | NA |
| GRIN2A | 769 | S | 79.77816667 | 23.85980878 | 0.289161347 | 18.80223887 | NA | NA | NA | NA | NA | NA | NA |
| GRIN2A | 770 | P | 81.81864286 | 20.02815104 | 0.237256733 | 16.73337458 | NA | NA | NA | NA | NA | NA | NA |
| GRIN2A | 771 | W | 80.01621429 | 17.40000006 | 0.201655129 | 12.8221769 | NA | NA | NA | NA | NA | NA | NA |
| GRIN2A | 772 | K | 75.2095 | 18.14506966 | 0.211748032 | 13.1318967 | NA | NA | NA | NA | NA | NA | NA |
| GRIN2A | 773 | R | 77.42877273 | 15.7018184 | 0.178651127 | 10.685954 | NA | NA | NA | NA | NA | NA | NA |
| GRIN2A | 774 | Q | 78.23438889 | 13.24285124 | 0.14534133 | 8.221080508 | NA | NA | NA | NA | NA | NA | NA |
| GRIN2A | 775 | I | 73.8005 | 14.25160377 | 0.159006149 | 9.305462758 | NA | NA | NA | NA | NA | NA | NA |
| GRIN2A | 776 | D | 71.355875 | 14.2481244 | 0.158959017 | 9.341532968 | NA | NA | NA | NA | NA | NA | NA |
| GRIN2A | 777 | L | 73.209625 | 11.08869064 | 0.11616052 | 6.161257581 | NA | NA | NA | NA | NA | NA | NA |

| | | | | | | | | | | | | | |
|---|---|---|---|---|---|---|---|---|---|---|---|---|---|
| GRIN2A | 778 | A | 71.5373 | 10.10466199 | 0.102830617 | 5.312107099 | NA | NA | NA | NA | NA | NA | NA |
| GRIN2A | 779 | L | 68.466875 | 11.62354116 | 0.123405742 | 7.390881302 | NA | NA | NA | NA | NA | NA | NA |
| GRIN2A | 780 | L | 67.339375 | 10.28309355 | 0.105247697 | 7.139649762 | NA | NA | NA | NA | NA | NA | NA |
| GRIN2A | 781 | Q | 69.79705556 | 6.591244116 | 0.055236962 | 1.894310971 | 4.99 | -0.89 | -3.14 | 31.36 | 1.48 | 3 | -1 |
| GRIN2A | 782 | F | 67.69622727 | 7.706269461 | 0.07034138 | 6.37188684 | 2.3 | -0.35 | -0.93 | 33.24 | 1.19 | 2 | -2 |
| GRIN2A | 783 | V | 63.17407143 | 9.177835311 | 0.090275586 | 6.139794821 | 3.12 | -1.05 | -2.73 | 26.68 | 1.35 | 2 | -2 |
| GRIN2A | 784 | G | 63.695 | 7.199231487 | 0.063472914 | 4.93258907 | 3.12 | -1.05 | -2.73 | 26.68 | 1.35 | 2 | -2 |
| GRIN2A | 785 | D | 64.972375 | 3.912649614 | 0.018952037 | 1.633215965 | 3.04 | -0.27 | 0.1 | 17.8 | 0.58 | 1 | -1 |
| GRIN2A | 786 | G | 61.1145 | 4.869272122 | 0.031910689 | 1.722027427 | 3.21 | -1 | -2.26 | 22.02 | 1.24 | 2 | -2 |
| GRIN2A | 787 | E | 64.15472222 | 4.199340543 | 0.022835625 | 2.745838003 | 3.03 | -1.03 | -2.47 | 34.39 | 1.76 | 2 | -2 |
| GRIN2A | 788 | M | 61.120625 | 8.081130181 | 0.075419339 | 4.71464643 | 3.43 | -1.05 | -2.73 | 26.68 | 1.35 | 2 | -2 |
| GRIN2A | 789 | E | 56.81327778 | 5.175773565 | 0.036062636 | 1.641739063 | 3.58 | -1.06 | -2.73 | 38.27 | 1.88 | 3 | -3 |
| GRIN2A | 790 | E | 60.33972222 | 4.517567266 | 0.027146406 | 1.673350239 | 3.84 | -1.05 | -3.5 | 38.27 | 1.81 | 3 | -3 |
| GRIN2A | 791 | L | 60.560875 | 9.218258458 | 0.090823169 | 5.641264508 | 3.58 | -1.06 | -2.73 | 38.27 | 1.88 | 3 | -3 |
| GRIN2A | 792 | E | 55.69538889 | 9.588006362 | 0.095831868 | 5.888876696 | 3.84 | -1.05 | -3.5 | 38.27 | 1.81 | 3 | -3 |
| GRIN2A | 793 | T | 55.33464286 | 6.059527704 | 0.048034195 | 2.048716791 | 4.02 | -0.68 | -2.35 | 14.68 | 0.93 | 1 | 1 |
| GRIN2A | 794 | L | 58.165 | 9.876332973 | 0.099737614 | 5.875496481 | 4.02 | -0.68 | -2.35 | 14.68 | 0.93 | 1 | 1 |
| GRIN2A | 795 | W | 58.61078571 | 12.57748759 | 0.136328144 | 8.565621285 | NA | NA | NA | NA | NA | NA | NA |
| GRIN2A | 796 | L | 53.523 | 13.44299896 | 0.148052582 | 9.45675827 | NA | NA | NA | NA | NA | NA | NA |
| GRIN2A | 797 | T | 50.2495 | 10.67639419 | 0.110575447 | 4.421411689 | 6.63 | -0.77 | -3.5 | 3.38 | 0.54 | 0 | 0 |
| GRIN2A | 798 | G | 47.37925 | 11.54663232 | 0.122363915 | 4.96822908 | 6.63 | -0.77 | -3.5 | 3.38 | 0.54 | 0 | 0 |
| GRIN2A | 799 | I | 43.549 | 11.50807499 | 0.121841608 | 5.006285342 | NA | NA | NA | NA | NA | NA | NA |
| GRIN2A | 800 | C | 43.12083333 | 12.31123727 | 0.132721449 | 5.790250114 | NA | NA | NA | NA | NA | NA | NA |
| GRIN2A | 801 | H | 44.3367 | 8.8620558 | 0.085997956 | 2.249635184 | 6.63 | -0.77 | -2.57 | 2.81 | 0.62 | 0 | 0 |
| GRIN2A | 802 | N | 42.2321 | 8.25716392 | 0.077803937 | 1.688817106 | 6.65 | -0.77 | -2.57 | 2.81 | 0.62 | 0 | 0 |
| GRIN2A | 803 | E | 39.1939 | 8.540747157 | 0.081645428 | 1.836147202 | 6.82 | -0.78 | -2.03 | 2.95 | 0.67 | 0 | 0 |
| GRIN2A | 804 | K | 35.7219 | 9.817817578 | 0.098944949 | 3.185675758 | 6.96 | -0.79 | -1.08 | 3.04 | 0.75 | 0 | 0 |
| GRIN2A | 805 | N | 33.0805 | 7.634344635 | 0.069367068 | 1.870013202 | 5.77 | 0.44 | 1.02 | 1.46 | -0.95 | 0 | 0 |
| GRIN2A | 806 | E | 32.3825 | 11.26038059 | 0.118486276 | 5.499286191 | NA | NA | NA | NA | NA | NA | NA |
| GRIN2A | 807 | V | 29.1781 | 11.36049673 | 0.119842475 | 7.651992876 | NA | NA | NA | NA | NA | NA | NA |
| GRIN2A | 808 | M | 26.5575 | 12.5532585 | 0.13599993 | 10.07897477 | NA | NA | NA | NA | NA | NA | NA |
| GRIN2A | 809 | S | 24.9057 | 13.55957617 | 0.149631767 | 11.91583337 | NA | NA | NA | NA | NA | NA | NA |
| GRIN2A | 814 | I | 17.503375 | 12.23034836 | 0.131625707 | 10.82572607 | NA | NA | NA | NA | NA | NA | NA |

| | | | | | | | | | | | | | |
|---|---|---|---|---|---|---|---|---|---|---|---|---|---|
| GRIN2A | 815 | D | 17.163125 | 11.06718054 | 0.115869139 | 9.157471713 | NA | NA | NA | NA | NA | NA | NA |
| GRIN2A | 816 | N | 17.212125 | 16.61013131 | 0.190955365 | 14.69624784 | NA | NA | NA | NA | NA | NA | NA |
| GRIN2A | 817 | M | 16.51025 | 15.68293764 | 0.178395363 | 13.76951524 | NA | NA | NA | NA | NA | NA | NA |
| GRIN2A | 818 | A | 13.7983 | 19.56114692 | 0.230930576 | 17.49380614 | NA | NA | NA | NA | NA | NA | NA |
| GRIN2A | 819 | G | 11.16225 | 18.600539 | 0.217917936 | 16.53036037 | NA | NA | NA | NA | NA | NA | NA |
| GRIN2A | 820 | V | 8.132928571 | 16.87872237 | 0.194593768 | 13.84535947 | NA | NA | NA | NA | NA | NA | NA |
| GRIN2A | 821 | F | 9.833681818 | 21.18001629 | 0.252860194 | 18.14012807 | NA | NA | NA | NA | NA | NA | NA |
| GRIN2A | 822 | Y | 10.7565 | 22.0948184 | 0.265252337 | 19.02116767 | NA | NA | NA | NA | NA | NA | NA |
| GRIN2A | 823 | M | 6.022625 | 17.47516752 | 0.202673367 | 14.40988615 | NA | NA | NA | NA | NA | NA | NA |
| GRIN2A | 824 | L | 4.04875 | 19.96999224 | 0.236468899 | 17.02294678 | NA | NA | NA | NA | NA | NA | NA |
| GRIN2A | 825 | A | 4.3341 | 23.39482979 | 0.282862623 | 20.36922741 | NA | NA | NA | NA | NA | NA | NA |
| GRIN2A | 826 | A | 2.8125 | 22.53813579 | 0.271257627 | 19.4612559 | NA | NA | NA | NA | NA | NA | NA |
| GRIN2A | 827 | A | 0.3529 | 20.79891812 | 0.247697741 | 17.67045872 | NA | NA | NA | NA | NA | NA | NA |
| GRIN2A | 828 | M | 0 | 23.9250922 | 0.290045693 | 20.79911372 | NA | NA | NA | NA | NA | NA | NA |
| GRIN2A | 829 | A | 0.3069 | 26.18544315 | 0.320664984 | 23.05711736 | NA | NA | NA | NA | NA | NA | NA |
| GRIN2A | 830 | L | -0.843625 | 22.21972723 | 0.266944384 | 19.03709339 | NA | NA | NA | NA | NA | NA | NA |
| GRIN2A | 831 | S | -3.508 | 25.59208389 | 0.312627188 | 22.93504218 | NA | NA | NA | NA | NA | NA | NA |
| GRIN2A | 832 | L | -2.5375 | 28.23082989 | 0.348372315 | 25.56692099 | NA | NA | NA | NA | NA | NA | NA |
| GRIN2A | 833 | I | -3.063 | 28.69531294 | 0.354664321 | 26.16180779 | NA | NA | NA | NA | NA | NA | NA |
| GRIN2A | 834 | T | -6.876357143 | 25.2066441 | 0.307405922 | 22.52248403 | NA | NA | NA | NA | NA | NA | NA |
| GRIN2A | 835 | F | -9.030772727 | 24.97269136 | 0.304236739 | 21.43293279 | NA | NA | NA | NA | NA | NA | NA |
| GRIN2A | 836 | I | -7.83225 | 31.25451635 | 0.389331943 | 27.71659639 | NA | NA | NA | NA | NA | NA | NA |
| GRIN2A | 837 | W | -9.259857143 | 30.80976159 | 0.383307182 | 27.32489297 | NA | NA | NA | NA | NA | NA | NA |
| GRIN2A | 838 | E | -13.90016667 | 32.38567747 | 0.404654941 | 27.51924322 | NA | NA | NA | NA | NA | NA | NA |
| GRIN2A | 839 | H | -16.6232 | 29.24669925 | 0.36213354 | 21.79928789 | NA | NA | NA | NA | NA | NA | NA |
| GRIN2A | 840 | L | -17.272125 | 35.40106349 | 0.445502129 | 27.94687108 | NA | NA | NA | NA | NA | NA | NA |
| GRIN2A | 841 | F | -17.4375 | 35.15834719 | 0.442214233 | 27.67458658 | NA | NA | NA | NA | NA | NA | NA |
| HCN1 | 94 | M | -18.69833333 | 53.11999191 | 0.996813968 | 51.48167666 | NA | NA | NA | NA | NA | NA | NA |
| HCN1 | 95 | Q | -20.66593333 | 52.00080071 | 0.974914658 | 50.10587258 | NA | NA | NA | NA | NA | NA | NA |
| HCN1 | 96 | R | -23.39593333 | 50.25022582 | 0.940661009 | 46.21881494 | NA | NA | NA | NA | NA | NA | NA |
| HCN1 | 97 | Q | -21.30553333 | 48.10635133 | 0.898711634 | 44.73735528 | NA | NA | NA | NA | NA | NA | NA |
| HCN1 | 98 | F | -19.83433333 | 47.20338211 | 0.881043158 | 45.58207294 | NA | NA | NA | NA | NA | NA | NA |
| HCN1 | 99 | T | -22.94193333 | 46.33957111 | 0.864140895 | 42.93537953 | NA | NA | NA | NA | NA | NA | NA |

| | | | | | | | | | | | | | |
|---|---|---|---|---|---|---|---|---|---|---|---|---|---|
| HCN1 | 100 | S | -23.82293333 | 44.10637276 | 0.820443712 | 40.0750518 | NA | NA | NA | NA | NA | NA | NA |
| HCN1 | 101 | M | -20.09233333 | 42.38342089 | 0.786730566 | 39.01570434 | NA | NA | NA | NA | NA | NA | NA |
| HCN1 | 102 | L | -21.57213333 | 41.9530003 | 0.778308489 | 38.58678704 | NA | NA | NA | NA | NA | NA | NA |
| HCN1 | 103 | Q | -24.57273333 | 41.33513707 | 0.766218707 | 37.30071898 | NA | NA | NA | NA | NA | NA | NA |
| HCN1 | 104 | P | -26.17604762 | 38.10082403 | 0.702932627 | 34.07312037 | NA | NA | NA | NA | NA | NA | NA |
| HCN1 | 105 | G | -26.51308333 | 37.13372707 | 0.684009358 | 32.91375283 | NA | NA | NA | NA | NA | NA | NA |
| HCN1 | 106 | V | -29.46647619 | 36.90530806 | 0.679539864 | 32.22968838 | NA | NA | NA | NA | NA | NA | NA |
| HCN1 | 107 | N | -26.15920833 | 33.73940813 | 0.61759243 | 29.18636841 | NA | NA | NA | NA | NA | NA | NA |
| HCN1 | 108 | K | -27.78011111 | 28.34572994 | 0.512053868 | 23.71807214 | NA | NA | NA | NA | NA | NA | NA |
| HCN1 | 109 | F | -25.57342424 | 28.31525276 | 0.511457518 | 23.88789647 | NA | NA | NA | NA | NA | NA | NA |
| HCN1 | 110 | S | -28.53883333 | 31.62383718 | 0.576196872 | 26.94637281 | NA | NA | NA | NA | NA | NA | NA |
| HCN1 | 111 | L | -31.65413333 | 30.06657303 | 0.545725752 | 23.25418918 | NA | NA | NA | NA | NA | NA | NA |
| HCN1 | 112 | R | -29.86978788 | 24.71193908 | 0.440951172 | 18.51234982 | NA | NA | NA | NA | NA | NA | NA |
| HCN1 | 113 | M | -29.58295833 | 27.65844847 | 0.498605773 | 22.79157957 | NA | NA | NA | NA | NA | NA | NA |
| HCN1 | 114 | F | -31.53660606 | 30.34685391 | 0.551210032 | 23.53440249 | NA | NA | NA | NA | NA | NA | NA |
| HCN1 | 115 | G | -35.32508333 | 29.96489756 | 0.543736259 | 22.94563554 | NA | NA | NA | NA | NA | NA | NA |
| HCN1 | 116 | S | -37.39833333 | 31.98139314 | 0.5831932 | 26.18005918 | NA | NA | NA | NA | NA | NA | NA |
| HCN1 | 117 | Q | -35.66388889 | 35.33080541 | 0.648731436 | 28.31961848 | NA | NA | NA | NA | NA | NA | NA |
| HCN1 | 118 | K | -37.90253333 | 37.45540815 | 0.69030372 | 31.64844231 | NA | NA | NA | NA | NA | NA | NA |
| HCN1 | 119 | A | -35.80613333 | 34.64406264 | 0.635293882 | 27.62449837 | NA | NA | NA | NA | NA | NA | NA |
| HCN1 | 120 | V | -32.56419048 | 35.5999708 | 0.653998218 | 28.64916772 | NA | NA | NA | NA | NA | NA | NA |
| HCN1 | 121 | E | -34.21233333 | 39.38620534 | 0.728083793 | 32.42841516 | NA | NA | NA | NA | NA | NA | NA |
| HCN1 | 122 | K | -35.66955556 | 38.14404632 | 0.703778361 | 31.30241939 | NA | NA | NA | NA | NA | NA | NA |
| HCN1 | 123 | E | -30.50011111 | 36.80357739 | 0.677549291 | 29.99222023 | NA | NA | NA | NA | NA | NA | NA |
| HCN1 | 124 | Q | -30.17166667 | 41.07302824 | 0.761090001 | 34.26148953 | NA | NA | NA | NA | NA | NA | NA |
| HCN1 | 125 | E | -33.76311111 | 43.52554503 | 0.809078606 | 36.71275152 | NA | NA | NA | NA | NA | NA | NA |
| HCN1 | 126 | R | -30.53178788 | 37.46773647 | 0.690544949 | 30.70315916 | NA | NA | NA | NA | NA | NA | NA |
| HCN1 | 127 | V | -26.97533333 | 41.61614702 | 0.771717252 | 36.94668449 | NA | NA | NA | NA | NA | NA | NA |
| HCN1 | 128 | K | -28.84822222 | 45.83829342 | 0.854332352 | 41.16144306 | NA | NA | NA | NA | NA | NA | NA |
| HCN1 | 129 | T | -28.44090476 | 46.39397948 | 0.865205508 | 41.71755457 | NA | NA | NA | NA | NA | NA | NA |
| HCN1 | 130 | A | -24.60293333 | 47.72688376 | 0.89128656 | 43.69525922 | NA | NA | NA | NA | NA | NA | NA |
| HCN1 | 131 | G | -22.55883333 | 50.37314051 | 0.943066092 | 46.34149499 | NA | NA | NA | NA | NA | NA | NA |
| HCN1 | 132 | F | -19.65233333 | 49.97445446 | 0.935264968 | 48.05353458 | NA | NA | NA | NA | NA | NA | NA |

| | | | | | | | | | | | | | |
|---|---|---|---|---|---|---|---|---|---|---|---|---|---|
| HCN1 | 133 | W | -19.00233333 | 46.03687545 | 0.858218023 | 44.40280053 | NA | NA | NA | NA | NA | NA | NA |
| HCN1 | 134 | I | -21.12195833 | 43.23630287 | 0.80341898 | 39.87111358 | NA | NA | NA | NA | NA | NA | NA |
| HCN1 | 135 | I | -19.09795833 | 40.09082575 | 0.741871161 | 38.18335507 | NA | NA | NA | NA | NA | NA | NA |
| HCN1 | 136 | H | -23.73953333 | 36.1055912 | 0.663891736 | 32.87200508 | NA | NA | NA | NA | NA | NA | NA |
| HCN1 | 137 | P | -21.32404762 | 34.43381572 | 0.631179962 | 31.0655574 | NA | NA | NA | NA | NA | NA | NA |
| HCN1 | 138 | Y | -23.46475 | 29.34656808 | 0.531637353 | 25.99530383 | NA | NA | NA | NA | NA | NA | NA |
| HCN1 | 139 | S | -20.76966667 | 36.68287633 | 0.675187523 | 33.31305891 | NA | NA | NA | NA | NA | NA | NA |
| HCN1 | 140 | D | -19.36133333 | 38.23951964 | 0.705646496 | 36.31224251 | NA | NA | NA | NA | NA | NA | NA |
| HCN1 | 141 | F | -15.76878788 | 39.55971017 | 0.731478777 | 38.44469931 | NA | NA | NA | NA | NA | NA | NA |
| HCN1 | 142 | R | -15.49606061 | 36.05998744 | 0.662999403 | 35.20728766 | NA | NA | NA | NA | NA | NA | NA |
| HCN1 | 143 | F | -15.67778788 | 32.49072078 | 0.593159258 | 31.60245207 | NA | NA | NA | NA | NA | NA | NA |
| HCN1 | 144 | Y | -14.09383333 | 37.17036379 | 0.684726232 | 35.89136831 | NA | NA | NA | NA | NA | NA | NA |
| HCN1 | 145 | W | -10.20440476 | 36.83827773 | 0.678228276 | 35.81147059 | NA | NA | NA | NA | NA | NA | NA |
| HCN1 | 146 | D | -11.78533333 | 33.49893842 | 0.612887138 | 32.48090473 | NA | NA | NA | NA | NA | NA | NA |
| HCN1 | 147 | L | -9.779208333 | 32.95101718 | 0.602165917 | 31.58198032 | NA | NA | NA | NA | NA | NA | NA |
| HCN1 | 148 | I | -7.269583333 | 35.91649634 | 0.660191701 | 34.80230244 | NA | NA | NA | NA | NA | NA | NA |
| HCN1 | 149 | M | -6.356083333 | 33.55191158 | 0.613923669 | 32.47393931 | NA | NA | NA | NA | NA | NA | NA |
| HCN1 | 150 | L | -6.820958333 | 29.62083372 | 0.537003932 | 28.54202945 | NA | NA | NA | NA | NA | NA | NA |
| HCN1 | 151 | I | -3.372083333 | 32.1751819 | 0.586985081 | 29.85501821 | NA | NA | NA | NA | NA | NA | NA |
| HCN1 | 152 | M | -1.100708333 | 33.21734855 | 0.607377245 | 31.04981596 | NA | NA | NA | NA | NA | NA | NA |
| HCN1 | 153 | M | -2.799083333 | 30.26840237 | 0.549674964 | 28.02856825 | NA | NA | NA | NA | NA | NA | NA |
| HCN1 | 154 | V | -0.817333333 | 27.85807106 | 0.502511805 | 25.69593506 | NA | NA | NA | NA | NA | NA | NA |
| HCN1 | 155 | G | 1.205333333 | 30.63446298 | 0.556837703 | 27.75847914 | NA | NA | NA | NA | NA | NA | NA |
| HCN1 | 156 | N | 1.559333333 | 30.22400023 | 0.548806143 | 27.37449369 | NA | NA | NA | NA | NA | NA | NA |
| HCN1 | 157 | L | 1.030333333 | 26.83008955 | 0.482397203 | 24.00105254 | NA | NA | NA | NA | NA | NA | NA |
| HCN1 | 158 | V | 3.862761905 | 26.43242647 | 0.474616096 | 24.61008833 | NA | NA | NA | NA | NA | NA | NA |
| HCN1 | 159 | I | 6.391083333 | 29.17623106 | 0.528304354 | 27.80709094 | NA | NA | NA | NA | NA | NA | NA |
| HCN1 | 160 | I | 6.138208333 | 30.00068916 | 0.544436596 | 28.82447631 | NA | NA | NA | NA | NA | NA | NA |
| HCN1 | 161 | P | 7.781619048 | 26.46993389 | 0.475350006 | 25.25671299 | NA | NA | NA | NA | NA | NA | NA |
| HCN1 | 162 | V | 10.71590476 | 26.24244581 | 0.470898728 | 25.47401952 | NA | NA | NA | NA | NA | NA | NA |
| HCN1 | 163 | G | 12.05733333 | 29.73544365 | 0.539246515 | 28.76599913 | NA | NA | NA | NA | NA | NA | NA |
| HCN1 | 164 | I | 11.55070833 | 29.75120705 | 0.539554959 | 28.79364862 | NA | NA | NA | NA | NA | NA | NA |
| HCN1 | 165 | T | 14.06161905 | 25.81008016 | 0.462438592 | 22.46041459 | NA | NA | NA | NA | NA | NA | NA |

| | | | | | | | | | | | | | |
|---|---|---|---|---|---|---|---|---|---|---|---|---|---|
| HCN1 | 166 | F | 15.52124242 | 22.96000531 | 0.406670934 | 19.63818175 | NA | NA | NA | NA | NA | NA | NA |
| HCN1 | 167 | F | 15.83706061 | 29.95799838 | 0.543601262 | 26.61104881 | NA | NA | NA | NA | NA | NA | NA |
| HCN1 | 168 | T | 19.32233333 | 32.02312397 | 0.584009751 | 27.64343945 | NA | NA | NA | NA | NA | NA | NA |
| HCN1 | 169 | E | 18.38213333 | 34.77855814 | 0.637925567 | 30.39957251 | NA | NA | NA | NA | NA | NA | NA |
| HCN1 | 170 | Q | 15.23033333 | 35.98053892 | 0.661444827 | 32.63078647 | NA | NA | NA | NA | NA | NA | NA |
| HCN1 | 171 | T | 16.17033333 | 39.15703204 | 0.723599539 | 35.8088907 | NA | NA | NA | NA | NA | NA | NA |
| HCN1 | 172 | T | 15.83933333 | 39.49765969 | 0.73026463 | 36.14818044 | NA | NA | NA | NA | NA | NA | NA |
| HCN1 | 173 | T | 13.64193333 | 43.07880041 | 0.800337116 | 41.06900953 | NA | NA | NA | NA | NA | NA | NA |
| HCN1 | 174 | P | 12.21261905 | 41.25847652 | 0.764718683 | 40.28316337 | NA | NA | NA | NA | NA | NA | NA |
| HCN1 | 175 | W | 10.84240476 | 33.5806551 | 0.614486096 | 32.63086553 | NA | NA | NA | NA | NA | NA | NA |
| HCN1 | 176 | I | 9.835533333 | 41.13506313 | 0.762303843 | 40.36683947 | NA | NA | NA | NA | NA | NA | NA |
| HCN1 | 177 | I | 7.524333333 | 42.81561404 | 0.795187326 | 41.71169133 | NA | NA | NA | NA | NA | NA | NA |
| HCN1 | 178 | F | 5.432969697 | 35.91171788 | 0.6600982 | 34.56770822 | NA | NA | NA | NA | NA | NA | NA |
| HCN1 | 179 | N | 5.761208333 | 38.66234852 | 0.713920025 | 37.3122861 | NA | NA | NA | NA | NA | NA | NA |
| HCN1 | 180 | V | 3.862933333 | 41.92913853 | 0.777841584 | 40.23844889 | NA | NA | NA | NA | NA | NA | NA |
| HCN1 | 181 | A | 1.550133333 | 41.9483033 | 0.778216583 | 39.05244976 | NA | NA | NA | NA | NA | NA | NA |
| HCN1 | 182 | S | 0.9475 | 38.22459089 | 0.705354384 | 35.32773506 | NA | NA | NA | NA | NA | NA | NA |
| HCN1 | 183 | D | 0.398533333 | 40.31773787 | 0.74631117 | 37.42272083 | NA | NA | NA | NA | NA | NA | NA |
| HCN1 | 184 | T | -0.638619048 | 42.62243627 | 0.7914074 | 40.44795472 | NA | NA | NA | NA | NA | NA | NA |
| HCN1 | 185 | V | -2.897333333 | 39.85705496 | 0.737296948 | 37.57688386 | NA | NA | NA | NA | NA | NA | NA |
| HCN1 | 186 | F | -2.268333333 | 35.12887113 | 0.644780171 | 32.84641811 | NA | NA | NA | NA | NA | NA | NA |
| HCN1 | 187 | L | -3.601458333 | 41.31840207 | 0.765891252 | 39.03505059 | NA | NA | NA | NA | NA | NA | NA |
| HCN1 | 188 | L | -6.318458333 | 42.35005909 | 0.786077772 | 41.2208918 | NA | NA | NA | NA | NA | NA | NA |
| HCN1 | 189 | D | -8.283083333 | 36.34639149 | 0.668603496 | 34.93293839 | NA | NA | NA | NA | NA | NA | NA |
| HCN1 | 190 | L | -7.772708333 | 39.70205469 | 0.734264044 | 38.28839957 | NA | NA | NA | NA | NA | NA | NA |
| HCN1 | 191 | I | -9.629208333 | 42.58094739 | 0.790595584 | 41.21276225 | NA | NA | NA | NA | NA | NA | NA |
| HCN1 | 192 | M | -12.47033333 | 41.22599898 | 0.764083192 | 40.12136998 | NA | NA | NA | NA | NA | NA | NA |
| HCN1 | 193 | N | -12.77420833 | 36.96535863 | 0.680714879 | 35.68560857 | NA | NA | NA | NA | NA | NA | NA |
| HCN1 | 194 | F | -13.73451515 | 41.48578322 | 0.769166413 | 40.60773428 | NA | NA | NA | NA | NA | NA | NA |
| HCN1 | 195 | R | -17.10933333 | 42.10795615 | 0.781340523 | 40.98400732 | NA | NA | NA | NA | NA | NA | NA |
| HCN1 | 196 | T | -17.63253333 | 38.59483178 | 0.712598919 | 37.42030864 | NA | NA | NA | NA | NA | NA | NA |
| HCN1 | 197 | G | -20.29433333 | 36.04594708 | 0.662724674 | 32.69346733 | NA | NA | NA | NA | NA | NA | NA |
| HCN1 | 198 | T | -21.49253333 | 34.23774001 | 0.627343332 | 30.87110802 | NA | NA | NA | NA | NA | NA | NA |

| | | | | | | | | | | | | | |
|---|---|---|---|---|---|---|---|---|---|---|---|---|---|
| HCN1 | 199 | V | -22.67913333 | 30.5063329 | 0.554330571 | 27.0872367 | NA | NA | NA | NA | NA | NA | NA |
| HCN1 | 200 | N | -24.84413333 | 28.6354345 | 0.517722542 | 24.60154265 | NA | NA | NA | NA | NA | NA | NA |
| HCN1 | 203 | S | -29.29013333 | 29.0243746 | 0.525332966 | 24.35752559 | NA | NA | NA | NA | NA | NA | NA |
| HCN1 | 204 | S | -29.41253333 | 32.15398215 | 0.586570264 | 27.50136279 | NA | NA | NA | NA | NA | NA | NA |
| HCN1 | 205 | E | -26.74813333 | 34.15452405 | 0.625715038 | 29.70119369 | NA | NA | NA | NA | NA | NA | NA |
| HCN1 | 206 | I | -24.67693333 | 36.37950596 | 0.669251449 | 32.34490773 | NA | NA | NA | NA | NA | NA | NA |
| HCN1 | 207 | I | -25.65593333 | 39.64426124 | 0.733133195 | 35.6088076 | NA | NA | NA | NA | NA | NA | NA |
| HCN1 | 208 | L | -24.55433333 | 42.18665793 | 0.782880488 | 38.15209456 | NA | NA | NA | NA | NA | NA | NA |
| HCN1 | 209 | D | -21.68433333 | 44.96095486 | 0.837165393 | 41.63699255 | NA | NA | NA | NA | NA | NA | NA |
| HCN1 | 210 | P | -21.13019048 | 46.55122959 | 0.868282434 | 43.51588777 | NA | NA | NA | NA | NA | NA | NA |
| HCN1 | 211 | K | -22.05593333 | 43.46341453 | 0.807862893 | 40.09790701 | NA | NA | NA | NA | NA | NA | NA |
| HCN1 | 212 | V | -22.19613333 | 40.23580465 | 0.744707975 | 36.86339501 | NA | NA | NA | NA | NA | NA | NA |
| HCN1 | 213 | I | -19.21433333 | 40.36456621 | 0.747227464 | 38.50235692 | NA | NA | NA | NA | NA | NA | NA |
| HCN1 | 214 | K | -16.87513333 | 42.3959006 | 0.786974757 | 41.46325233 | NA | NA | NA | NA | NA | NA | NA |
| HCN1 | 215 | M | -17.58433333 | 41.10161372 | 0.761649335 | 39.98059659 | NA | NA | NA | NA | NA | NA | NA |
| HCN1 | 216 | N | -17.51313333 | 37.65970465 | 0.694301206 | 36.55262254 | NA | NA | NA | NA | NA | NA | NA |
| HCN1 | 217 | Y | -12.96916667 | 36.66530766 | 0.674843756 | 35.89230555 | NA | NA | NA | NA | NA | NA | NA |
| HCN1 | 218 | L | -12.62313333 | 40.0735445 | 0.741533017 | 38.78527183 | NA | NA | NA | NA | NA | NA | NA |
| HCN1 | 219 | K | -13.44553333 | 36.84060073 | 0.67827373 | 35.5509458 | NA | NA | NA | NA | NA | NA | NA |
| HCN1 | 220 | S | -12.58513333 | 34.1526867 | 0.625679087 | 32.92193449 | NA | NA | NA | NA | NA | NA | NA |
| HCN1 | 221 | W | -10.35833333 | 28.91558792 | 0.523204328 | 27.53940426 | NA | NA | NA | NA | NA | NA | NA |
| HCN1 | 222 | F | -8.14269697 | 36.99727982 | 0.681339483 | 35.17754732 | NA | NA | NA | NA | NA | NA | NA |
| HCN1 | 223 | V | -5.340133333 | 37.15057806 | 0.684339083 | 36.05236657 | NA | NA | NA | NA | NA | NA | NA |
| HCN1 | 224 | V | -3.979133333 | 33.49893894 | 0.612887149 | 32.50533974 | NA | NA | NA | NA | NA | NA | NA |
| HCN1 | 225 | D | -5.708458333 | 34.15661031 | 0.62575586 | 33.26684687 | NA | NA | NA | NA | NA | NA | NA |
| HCN1 | 226 | F | -2.595878788 | 38.37555475 | 0.708308306 | 36.10760596 | NA | NA | NA | NA | NA | NA | NA |
| HCN1 | 227 | I | -0.216708333 | 37.1319147 | 0.683973895 | 34.95337085 | NA | NA | NA | NA | NA | NA | NA |
| HCN1 | 228 | S | -0.459333333 | 34.94162209 | 0.641116253 | 32.77233723 | NA | NA | NA | NA | NA | NA | NA |
| HCN1 | 229 | S | -0.9955 | 38.03267671 | 0.701599183 | 35.84355092 | NA | NA | NA | NA | NA | NA | NA |
| HCN1 | 230 | I | 0.429583333 | 39.70532515 | 0.734328037 | 36.80637031 | NA | NA | NA | NA | NA | NA | NA |
| HCN1 | 231 | P | 3.841047619 | 41.16180251 | 0.762827054 | 39.29296467 | NA | NA | NA | NA | NA | NA | NA |
| HCN1 | 232 | V | 5.475904762 | 39.86109891 | 0.737376076 | 38.3528378 | NA | NA | NA | NA | NA | NA | NA |
| HCN1 | 233 | D | 9.008208333 | 38.27838009 | 0.706406882 | 37.29298848 | NA | NA | NA | NA | NA | NA | NA |

| | | | | | | | | | | | | | |
|---|---|---|---|---|---|---|---|---|---|---|---|---|---|
| HCN1 | 234 | Y | 9.6055 | 41.07534103 | 0.761135255 | 40.30636451 | NA | NA | NA | NA | NA | NA | NA |
| HCN1 | 235 | I | 6.965583333 | 44.29198945 | 0.82407569 | 43.06523056 | NA | NA | NA | NA | NA | NA | NA |
| HCN1 | 236 | F | 11.64278788 | 41.8035374 | 0.775383936 | 41.20014673 | NA | NA | NA | NA | NA | NA | NA |
| HCN1 | 237 | L | 13.81745833 | 43.99954663 | 0.818353436 | 42.00494585 | NA | NA | NA | NA | NA | NA | NA |
| HCN1 | 238 | I | 11.13608333 | 46.7039133 | 0.87127001 | 45.71608766 | NA | NA | NA | NA | NA | NA | NA |
| HCN1 | 239 | V | 11.65033333 | 47.95874424 | 0.895823394 | 46.97427839 | NA | NA | NA | NA | NA | NA | NA |
| HCN1 | 240 | E | 15.55433333 | 46.03270412 | 0.858136402 | 42.68495101 | NA | NA | NA | NA | NA | NA | NA |
| HCN1 | 241 | K | 16.32293333 | 48.70505105 | 0.910426443 | 45.35718494 | NA | NA | NA | NA | NA | NA | NA |
| HCN1 | 242 | G | 15.02583333 | 51.45899408 | 0.964313081 | 48.10996575 | NA | NA | NA | NA | NA | NA | NA |
| HCN1 | 252 | R | 23.96313333 | 40.92504164 | 0.758194334 | 36.65502504 | NA | NA | NA | NA | NA | NA | NA |
| HCN1 | 253 | A | 22.61153333 | 38.70016605 | 0.714660004 | 34.44217617 | NA | NA | NA | NA | NA | NA | NA |
| HCN1 | 254 | L | 22.28593333 | 35.61732759 | 0.65433784 | 31.38075178 | NA | NA | NA | NA | NA | NA | NA |
| HCN1 | 255 | R | 19.81833333 | 36.79483659 | 0.677378259 | 32.58918023 | NA | NA | NA | NA | NA | NA | NA |
| HCN1 | 256 | I | 18.09195833 | 37.94136272 | 0.699812434 | 33.70821644 | NA | NA | NA | NA | NA | NA | NA |
| HCN1 | 257 | V | 18.28433333 | 35.04059506 | 0.643052865 | 30.75410229 | NA | NA | NA | NA | NA | NA | NA |
| HCN1 | 258 | R | 18.61642424 | 32.04631033 | 0.584463441 | 28.97454361 | NA | NA | NA | NA | NA | NA | NA |
| HCN1 | 259 | F | 12.98006061 | 35.33559436 | 0.648825142 | 31.91946447 | NA | NA | NA | NA | NA | NA | NA |
| HCN1 | 260 | T | 13.17761905 | 35.57176186 | 0.653446252 | 33.5648831 | NA | NA | NA | NA | NA | NA | NA |
| HCN1 | 261 | K | 13.50433333 | 31.39552231 | 0.571729416 | 30.59956538 | NA | NA | NA | NA | NA | NA | NA |
| HCN1 | 262 | I | 10.23620833 | 31.40431074 | 0.57190138 | 30.67129911 | NA | NA | NA | NA | NA | NA | NA |
| HCN1 | 263 | L | 8.336708333 | 34.65058987 | 0.635421601 | 33.43680024 | NA | NA | NA | NA | NA | NA | NA |
| HCN1 | 264 | S | 7.764333333 | 33.58561601 | 0.614583166 | 32.39018595 | NA | NA | NA | NA | NA | NA | NA |
| HCN1 | 265 | L | 6.704958333 | 28.59913247 | 0.517012217 | 27.35321627 | NA | NA | NA | NA | NA | NA | NA |
| HCN1 | 266 | L | 3.099458333 | 32.87806524 | 0.60073846 | 30.90430236 | NA | NA | NA | NA | NA | NA | NA |
| HCN1 | 267 | R | 3.941060606 | 32.36774676 | 0.590753015 | 29.62445912 | NA | NA | NA | NA | NA | NA | NA |
| HCN1 | 268 | L | 2.241458333 | 27.72335119 | 0.49987573 | 24.99843796 | NA | NA | NA | NA | NA | NA | NA |
| HCN1 | 269 | L | -0.223083333 | 29.81752025 | 0.540852515 | 27.63862749 | NA | NA | NA | NA | NA | NA | NA |
| HCN1 | 270 | R | -1.248424242 | 30.00164919 | 0.544455381 | 27.71780851 | NA | NA | NA | NA | NA | NA | NA |
| HCN1 | 271 | L | -2.673458333 | 24.04681927 | 0.427936716 | 21.7674611 | NA | NA | NA | NA | NA | NA | NA |
| HCN1 | 272 | S | -5.832333333 | 27.75622406 | 0.500518956 | 26.6367092 | NA | NA | NA | NA | NA | NA | NA |
| HCN1 | 273 | R | -6.452606061 | 29.56736052 | 0.535957618 | 28.42852594 | NA | NA | NA | NA | NA | NA | NA |
| HCN1 | 274 | L | -6.204958333 | 25.73383899 | 0.460946775 | 24.6323819 | NA | NA | NA | NA | NA | NA | NA |
| HCN1 | 275 | I | -8.103208333 | 23.67796733 | 0.420719358 | 21.84888992 | NA | NA | NA | NA | NA | NA | NA |

| | | | | | | | | | | | | | |
|---|---|---|---|---|---|---|---|---|---|---|---|---|---|
| HCN1 | 276 | R | -10.96324242 | 26.75510587 | 0.480929991 | 25.71850838 | NA | NA | NA | NA | NA | NA | NA |
| HCN1 | 277 | Y | -11.11166667 | 27.75607202 | 0.500515981 | 26.63647181 | NA | NA | NA | NA | NA | NA | NA |
| HCN1 | 278 | I | -11.09620833 | 23.43452566 | 0.415955915 | 22.37111875 | NA | NA | NA | NA | NA | NA | NA |
| HCN1 | 279 | H | -13.51103333 | 20.37077811 | 0.356007305 | 19.40248603 | NA | NA | NA | NA | NA | NA | NA |
| HCN1 | 280 | Q | -16.27711111 | 26.08461069 | 0.467810355 | 25.16816161 | NA | NA | NA | NA | NA | NA | NA |
| HCN1 | 281 | W | -16.3282619 | 26.23390118 | 0.470731534 | 25.29673586 | NA | NA | NA | NA | NA | NA | NA |
| HCN1 | 282 | E | -15.14133333 | 20.7881452 | 0.364173962 | 20.08195142 | NA | NA | NA | NA | NA | NA | NA |
| HCN1 | 283 | E | -19.906 | 22.80372928 | 0.403613068 | 20.876406 | NA | NA | NA | NA | NA | NA | NA |
| HCN1 | 284 | I | -20.67958333 | 25.57661078 | 0.457870277 | 22.23221712 | NA | NA | NA | NA | NA | NA | NA |
| HCN1 | 285 | F | -17.78869697 | 24.25618424 | 0.432033378 | 22.65128123 | NA | NA | NA | NA | NA | NA | NA |
| HCN1 | 286 | H | -20.72593333 | 18.60250327 | 0.32140732 | 15.39947951 | NA | NA | NA | NA | NA | NA | NA |
| HCN1 | 287 | M | -25.45745833 | 22.67606463 | 0.401115043 | 18.66938487 | NA | NA | NA | NA | NA | NA | NA |
| HCN1 | 288 | T | -22.98290476 | 25.86960956 | 0.463603409 | 22.44661992 | NA | NA | NA | NA | NA | NA | NA |
| HCN1 | 289 | Y | -20.77991667 | 23.96709611 | 0.426376766 | 20.59946305 | NA | NA | NA | NA | NA | NA | NA |
| HCN1 | 290 | D | -23.46458333 | 19.49018735 | 0.33877671 | 16.13931888 | NA | NA | NA | NA | NA | NA | NA |
| HCN1 | 291 | L | -18.95433333 | 18.2527673 | 0.314564006 | 16.33015789 | NA | NA | NA | NA | NA | NA | NA |
| HCN1 | 292 | A | -18.01813333 | 16.36674412 | 0.277660029 | 14.75535075 | NA | NA | NA | NA | NA | NA | NA |
| HCN1 | 293 | S | -17.4565 | 14.26122723 | 0.236461201 | 13.10928252 | NA | NA | NA | NA | NA | NA | NA |
| HCN1 | 294 | A | -14.70393333 | 11.79884405 | 0.188279539 | 10.91082661 | NA | NA | NA | NA | NA | NA | NA |
| HCN1 | 295 | V | -13.07004762 | 14.60252588 | 0.24313942 | 13.32527533 | NA | NA | NA | NA | NA | NA | NA |
| HCN1 | 296 | V | -13.41219048 | 16.80660016 | 0.28626673 | 15.51679526 | NA | NA | NA | NA | NA | NA | NA |
| HCN1 | 297 | R | -13.14460606 | 12.98403423 | 0.211470256 | 11.99661989 | NA | NA | NA | NA | NA | NA | NA |
| HCN1 | 298 | I | -8.659458333 | 11.942964 | 0.191099546 | 10.16371706 | NA | NA | NA | NA | NA | NA | NA |
| HCN1 | 299 | F | -8.129878788 | 16.36038034 | 0.277535509 | 14.97316516 | NA | NA | NA | NA | NA | NA | NA |
| HCN1 | 300 | N | -8.570208333 | 17.68353658 | 0.30342582 | 16.28676041 | NA | NA | NA | NA | NA | NA | NA |
| HCN1 | 301 | L | -5.449208333 | 11.72444886 | 0.186823842 | 10.62164927 | NA | NA | NA | NA | NA | NA | NA |
| HCN1 | 302 | I | -3.448833333 | 15.08094539 | 0.252500696 | 12.78172819 | NA | NA | NA | NA | NA | NA | NA |
| HCN1 | 303 | G | -2.407333333 | 18.28523596 | 0.315199323 | 15.91105927 | NA | NA | NA | NA | NA | NA | NA |
| HCN1 | 304 | M | -2.652458333 | 17.8251455 | 0.306196694 | 15.54807839 | NA | NA | NA | NA | NA | NA | NA |
| HCN1 | 305 | M | -0.557958333 | 11.3161379 | 0.178834386 | 9.14650066 | NA | NA | NA | NA | NA | NA | NA |
| HCN1 | 306 | L | 0.561958333 | 17.02318164 | 0.290504598 | 14.73417338 | NA | NA | NA | NA | NA | NA | NA |
| HCN1 | 307 | L | 0.628458333 | 19.4458718 | 0.337909584 | 16.56411747 | NA | NA | NA | NA | NA | NA | NA |
| HCN1 | 308 | L | 1.234208333 | 16.95228695 | 0.289117396 | 14.23594793 | NA | NA | NA | NA | NA | NA | NA |

| | | | | | | | | | | | | | |
|---|---|---|---|---|---|---|---|---|---|---|---|---|---|
| HCN1 | 309 | C | 4.856833333 | 14.99453517 | 0.2508099 | 13.36782128 | NA | NA | NA | NA | NA | NA | NA |
| HCN1 | 310 | H | 6.106533333 | 18.77778552 | 0.324837082 | 17.41496132 | NA | NA | NA | NA | NA | NA | NA |
| HCN1 | 311 | W | 3.662119048 | 20.44581811 | 0.357475619 | 19.0938033 | NA | NA | NA | NA | NA | NA | NA |
| HCN1 | 312 | D | 7.097833333 | 15.62007061 | 0.263049805 | 14.54994254 | NA | NA | NA | NA | NA | NA | NA |
| HCN1 | 313 | G | 10.56858333 | 18.28845157 | 0.315262243 | 17.54282522 | NA | NA | NA | NA | NA | NA | NA |
| HCN1 | 314 | C | 10.58166667 | 20.74305354 | 0.36329165 | 19.96567077 | NA | NA | NA | NA | NA | NA | NA |
| HCN1 | 315 | L | 10.25508333 | 20.95863225 | 0.367509897 | 20.26153295 | NA | NA | NA | NA | NA | NA | NA |
| HCN1 | 316 | Q | 12.94955556 | 14.45266003 | 0.240206983 | 12.63268179 | NA | NA | NA | NA | NA | NA | NA |
| HCN1 | 317 | F | 16.99760606 | 20.15194177 | 0.351725315 | 16.81089968 | NA | NA | NA | NA | NA | NA | NA |
| HCN1 | 318 | L | 14.04320833 | 22.79358254 | 0.403414526 | 19.23356876 | NA | NA | NA | NA | NA | NA | NA |
| HCN1 | 319 | V | 15.77004762 | 22.49185413 | 0.39751058 | 19.14328543 | NA | NA | NA | NA | NA | NA | NA |
| HCN1 | 320 | P | 19.01261905 | 20.63283623 | 0.361135018 | 16.25588688 | NA | NA | NA | NA | NA | NA | NA |
| HCN1 | 321 | L | 19.06283333 | 24.64905394 | 0.439720693 | 20.26974607 | NA | NA | NA | NA | NA | NA | NA |
| HCN1 | 322 | L | 17.00295833 | 26.95100553 | 0.484763176 | 22.56086407 | NA | NA | NA | NA | NA | NA | NA |
| HCN1 | 323 | Q | 20.40833333 | 23.27850137 | 0.412902974 | 19.02029902 | NA | NA | NA | NA | NA | NA | NA |
| HCN1 | 324 | D | 23.77793333 | 25.60935813 | 0.458511047 | 21.36326184 | NA | NA | NA | NA | NA | NA | NA |
| HCN1 | 325 | F | 23.55760606 | 22.63124897 | 0.400238131 | 18.37059115 | NA | NA | NA | NA | NA | NA | NA |
| HCN1 | 326 | P | 24.43190476 | 19.89931642 | 0.346782173 | 15.67413674 | NA | NA | NA | NA | NA | NA | NA |
| HCN1 | 327 | P | 27.73090476 | 17.19787586 | 0.293922855 | 13.0180125 | NA | NA | NA | NA | NA | NA | NA |
| HCN1 | 328 | D | 24.72753333 | 16.11050946 | 0.272646264 | 11.85703219 | NA | NA | NA | NA | NA | NA | NA |
| HCN1 | 329 | C | 22.78966667 | 15.64960894 | 0.263627784 | 11.40712469 | NA | NA | NA | NA | NA | NA | NA |
| HCN1 | 330 | W | 19.20847619 | 14.1210177 | 0.233717709 | 9.84508878 | NA | NA | NA | NA | NA | NA | NA |
| HCN1 | 331 | V | 22.39847619 | 16.572745 | 0.281690866 | 12.30398393 | NA | NA | NA | NA | NA | NA | NA |
| HCN1 | 332 | S | 26.12416667 | 16.60263964 | 0.282275817 | 12.4181015 | NA | NA | NA | NA | NA | NA | NA |
| HCN1 | 333 | L | 26.09733333 | 12.70971569 | 0.206102642 | 8.499549853 | NA | NA | NA | NA | NA | NA | NA |
| HCN1 | 334 | N | 24.55345833 | 11.57981658 | 0.18399381 | 7.34604348 | NA | NA | NA | NA | NA | NA | NA |
| HCN1 | 335 | E | 27.65593333 | 16.228285 | 0.274950788 | 12.05049361 | NA | NA | NA | NA | NA | NA | NA |
| HCN1 | 336 | M | 23.50308333 | 16.4567313 | 0.279420816 | 12.18888777 | NA | NA | NA | NA | NA | NA | NA |
| HCN1 | 337 | V | 25.32404762 | 20.89021067 | 0.366171086 | 16.66889262 | NA | NA | NA | NA | NA | NA | NA |
| HCN1 | 338 | N | 27.66845833 | 23.46677792 | 0.416586997 | 19.25484604 | NA | NA | NA | NA | NA | NA | NA |
| HCN1 | 339 | D | 24.18733333 | 22.0288847 | 0.388451618 | 17.78671278 | NA | NA | NA | NA | NA | NA | NA |
| HCN1 | 340 | S | 21.55333333 | 24.1885815 | 0.430710589 | 19.96960117 | NA | NA | NA | NA | NA | NA | NA |
| HCN1 | 341 | W | 16.6537619 | 24.23439622 | 0.43160705 | 19.90425849 | NA | NA | NA | NA | NA | NA | NA |

| | | | | | | | | | | | | | |
|---|---|---|---|---|---|---|---|---|---|---|---|---|---|
| HCN1 | 342 | G | 16.75658333 | 22.74342617 | 0.402433112 | 19.39451243 | NA | NA | NA | NA | NA | NA | NA |
| HCN1 | 343 | K | 20.07088889 | 19.11478823 | 0.331431243 | 15.11113035 | NA | NA | NA | NA | NA | NA | NA |
| HCN1 | 344 | Q | 19.13944444 | 19.56067614 | 0.34015597 | 15.21079409 | NA | NA | NA | NA | NA | NA | NA |
| HCN1 | 345 | Y | 12.20625 | 19.55116503 | 0.339969865 | 16.18737971 | NA | NA | NA | NA | NA | NA | NA |
| HCN1 | 346 | S | 14.31783333 | 17.06154304 | 0.291255219 | 15.03624718 | NA | NA | NA | NA | NA | NA | NA |
| HCN1 | 347 | Y | 17.47816667 | 11.04789523 | 0.173585659 | 8.465720436 | NA | NA | NA | NA | NA | NA | NA |
| HCN1 | 348 | A | 13.72873333 | 15.79271237 | 0.266427901 | 13.83288022 | NA | NA | NA | NA | NA | NA | NA |
| HCN1 | 349 | L | 9.999833333 | 14.90143986 | 0.248988296 | 14.12895947 | NA | NA | NA | NA | NA | NA | NA |
| HCN1 | 350 | F | 12.95687879 | 10.23277406 | 0.157636114 | 9.44481493 | NA | NA | NA | NA | NA | NA | NA |
| HCN1 | 351 | K | 13.54866667 | 11.25007831 | 0.177541793 | 10.3100045 | NA | NA | NA | NA | NA | NA | NA |
| HCN1 | 352 | A | 9.000533333 | 11.9785995 | 0.191796829 | 11.03987508 | NA | NA | NA | NA | NA | NA | NA |
| HCN1 | 353 | M | 7.111833333 | 9.98362324 | 0.152760958 | 8.816515573 | NA | NA | NA | NA | NA | NA | NA |
| HCN1 | 354 | S | 9.206 | 7.060078541 | 0.095555709 | 6.184357878 | 1 | -0.8 | -0.4 | 3.38 | 0.81 | 0 | 0 |
| HCN1 | 355 | H | 9.059333333 | 8.101977104 | 0.115942628 | 7.335179983 | NA | NA | NA | NA | NA | NA | NA |
| HCN1 | 356 | M | 3.437333333 | 7.879331 | 0.111586092 | 6.274124489 | NA | NA | NA | NA | NA | NA | NA |
| HCN1 | 357 | L | 3.856083333 | 4.84658911 | 0.052244173 | 3.155434364 | 1.7 | -0.02 | 1.05 | 2.43 | 0.49 | 0 | 0 |
| HCN1 | 358 | C | 5.4615 | 2.256866855 | 0.001570856 | 1.131550014 | 1.38 | -0.41 | 0.33 | 2.91 | 0.65 | 0 | 0 |
| HCN1 | 359 | I | 8.947083333 | 2.199415832 | 4.47E-04 | 1.791196584 | 1.08 | -0.8 | -0.4 | 3.38 | 0.81 | 0 | 0 |
| HCN1 | 360 | G | 11.74883333 | 4.035583105 | 0.036375149 | 3.045477395 | 0.99 | -0.8 | -0.4 | 3.38 | 0.81 | 0 | 0 |
| HCN1 | 361 | Y | 14.4285 | 5.274846917 | 0.06062393 | 1.890286586 | 3.43 | -0.8 | -0.4 | 3.38 | 0.81 | 0 | 0 |
| HCN1 | 362 | G | 17.34183333 | 6.858249704 | 0.091606507 | 2.831113162 | 4.15 | 1.11 | -1.3 | 1.61 | -1.44 | 0 | 0 |
| HCN1 | 363 | A | 20.79073333 | 5.590022809 | 0.066791003 | 1.291368224 | 4.31 | -0.8 | -0.4 | 3.38 | 0.81 | 0 | 0 |
| HCN1 | 364 | Q | 20.57673333 | 6.843729247 | 0.091322384 | 2.533513649 | 4.31 | -0.8 | -0.4 | 3.38 | 0.81 | 0 | 0 |
| HCN1 | 365 | A | 19.35353333 | 7.369366119 | 0.101607566 | 3.02538211 | 4.38 | -0.42 | -0.58 | 3.03 | 0.36 | 0 | 0 |
| HCN1 | 366 | P | 18.18861905 | 10.65692554 | 0.165935522 | 6.345539587 | NA | NA | NA | NA | NA | NA | NA |
| HCN1 | 367 | V | 22.34233333 | 12.30368327 | 0.198157771 | 8.065376151 | NA | NA | NA | NA | NA | NA | NA |
| HCN1 | 368 | S | 20.54483333 | 16.16414047 | 0.273695667 | 11.86465327 | NA | NA | NA | NA | NA | NA | NA |
| HCN1 | 369 | M | 17.84645833 | 19.0336993 | 0.329844569 | 14.71395132 | NA | NA | NA | NA | NA | NA | NA |
| HCN1 | 370 | S | 16.85916667 | 19.24377848 | 0.333955206 | 15.89501168 | NA | NA | NA | NA | NA | NA | NA |
| HCN1 | 371 | D | 17.28295833 | 15.9283265 | 0.269081474 | 12.58112313 | NA | NA | NA | NA | NA | NA | NA |
| HCN1 | 372 | L | 15.11020833 | 14.03749728 | 0.232083458 | 10.69378784 | NA | NA | NA | NA | NA | NA | NA |
| HCN1 | 373 | W | 11.73019048 | 17.38453672 | 0.297575264 | 16.4758569 | NA | NA | NA | NA | NA | NA | NA |
| HCN1 | 374 | I | 11.93095833 | 16.1178685 | 0.272790259 | 15.2498236 | NA | NA | NA | NA | NA | NA | NA |

| | | | | | | | | | | | | | |
|---|---|---|---|---|---|---|---|---|---|---|---|---|---|
| HCN1 | 375 | T | 11.64147619 | 11.50951945 | 0.1826183 | 10.52759217 | NA | NA | NA | NA | NA | NA | NA |
| HCN1 | 376 | M | 9.283958333 | 14.02490367 | 0.231837037 | 13.14944405 | NA | NA | NA | NA | NA | NA | NA |
| HCN1 | 377 | L | 6.371583333 | 15.29491298 | 0.256687418 | 14.2693899 | NA | NA | NA | NA | NA | NA | NA |
| HCN1 | 378 | S | 6.925333333 | 12.69184975 | 0.205753057 | 11.6907032 | NA | NA | NA | NA | NA | NA | NA |
| HCN1 | 379 | M | 6.699583333 | 9.780643077 | 0.148789228 | 8.802558027 | NA | NA | NA | NA | NA | NA | NA |
| HCN1 | 380 | I | 2.950958333 | 12.55749852 | 0.203124195 | 10.64441596 | NA | NA | NA | NA | NA | NA | NA |
| HCN1 | 381 | V | 1.811333333 | 12.59443258 | 0.203846887 | 9.710620324 | NA | NA | NA | NA | NA | NA | NA |
| HCN1 | 382 | G | 1.729333333 | 9.513745109 | 0.143566813 | 7.748126085 | NA | NA | NA | NA | NA | NA | NA |
| HCN1 | 383 | A | 0.695733333 | 7.726803026 | 0.108601564 | 5.267315557 | NA | NA | NA | NA | NA | NA | NA |
| HCN1 | 384 | T | -0.713904762 | 9.642505224 | 0.146086273 | 7.529275167 | NA | NA | NA | NA | NA | NA | NA |
| HCN1 | 385 | C | -0.982666667 | 9.620708082 | 0.145659766 | 7.474652672 | NA | NA | NA | NA | NA | NA | NA |
| HCN1 | 386 | Y | 0 | 3.727437994 | 0.030345647 | 1.645858616 | 2.18 | 1.11 | -1.3 | 1.61 | -1.44 | 0 | 0 |
| HCN1 | 387 | A | -3.621133333 | 5.920925857 | 0.073265812 | 3.831583993 | 2.38 | 1.13 | 4.2 | 0.13 | -1.3 | 0 | 0 |
| HCN1 | 388 | M | -6.129458333 | 8.023979935 | 0.11441645 | 7.025308852 | NA | NA | NA | NA | NA | NA | NA |
| HCN1 | 389 | F | -4.253242424 | 7.523605851 | 0.104625588 | 6.594115324 | NA | NA | NA | NA | NA | NA | NA |
| HCN1 | 390 | V | -5.580333333 | 2.598422598 | 0.008254107 | 1.760282958 | 1 | 1.13 | 4.2 | 0.13 | -1.3 | 0 | 0 |
| HCN1 | 391 | G | -9.184333333 | 5.192908048 | 0.059020625 | 3.356038176 | 1.84 | -0.77 | -0.7 | 1.66 | 0.77 | 0 | 0 |
| HCN1 | 392 | H | -9.818233333 | 7.245007522 | 0.099174231 | 5.877886523 | 1.39 | -0.77 | -0.7 | 1.66 | 0.77 | 0 | 0 |
| HCN1 | 393 | A | -9.273733333 | 7.175442913 | 0.097813054 | 5.461349165 | 1.84 | -0.77 | -0.7 | 1.66 | 0.77 | 0 | 0 |
| HCN1 | 394 | T | -11.34647619 | 2.676487437 | 0.009781608 | 1.669553348 | 1.08 | -0.77 | -0.7 | 1.66 | 0.77 | 0 | 0 |
| HCN1 | 395 | A | -14.12633333 | 5.825454317 | 0.071397712 | 4.952822115 | 1.02 | -1.03 | -2.94 | 3.16 | 0.26 | 0 | 0 |
| HCN1 | 396 | L | -13.03145833 | 8.421946271 | 0.122203492 | 7.265153021 | NA | NA | NA | NA | NA | NA | NA |
| HCN1 | 397 | I | -12.93395833 | 8.521801511 | 0.124157368 | 7.293782343 | NA | NA | NA | NA | NA | NA | NA |
| HCN1 | 398 | Q | -16.67366667 | 2.398161171 | 0.004335574 | 1.573229535 | 1.18 | -1.1 | -3.5 | 3.53 | 0.13 | 0 | 0 |
| HCN1 | 399 | S | -18.2465 | 8.403101868 | 0.121834761 | 6.865050783 | NA | NA | NA | NA | NA | NA | NA |
| HCN1 | 400 | L | -16.65958333 | 11.03829158 | 0.173397744 | 9.855617146 | NA | NA | NA | NA | NA | NA | NA |
| HCN1 | 401 | D | -18.56883333 | 9.705142451 | 0.147311901 | 8.130383287 | NA | NA | NA | NA | NA | NA | NA |
| HCN1 | 402 | S | -22.35933333 | 9.07225424 | 0.134928123 | 5.734857505 | NA | NA | NA | NA | NA | NA | NA |
| HCN1 | 403 | S | -24.37433333 | 11.40775337 | 0.180627034 | 7.824863444 | NA | NA | NA | NA | NA | NA | NA |
| HCN1 | 404 | R | -22.36206061 | 12.53752727 | 0.202733416 | 9.116482407 | NA | NA | NA | NA | NA | NA | NA |
| HCN1 | 405 | R | -22.09987879 | 4.986153929 | 0.054975049 | 1.69003175 | 3.34 | -0.69 | -4.1 | 32.61 | 1.03 | 3 | 3 |
| HCN1 | 406 | Q | -26.54422222 | 6.409512462 | 0.082826027 | 2.592815483 | 4.1 | -0.85 | -3.9 | 50.74 | 1.99 | 5 | -1 |
| HCN1 | 407 | Y | -28.07308333 | 11.24346713 | 0.177412431 | 6.389495194 | NA | NA | NA | NA | NA | NA | NA |

| | | | | | | | | | | | | | |
|---|---|---|---|---|---|---|---|---|---|---|---|---|---|
| HCN1 | 408 | Q | -23.92655556 | 12.37335985 | 0.199521138 | 8.738198003 | NA | NA | NA | NA | NA | NA | NA |
| HCN1 | 409 | E | -25.55088889 | 5.996384327 | 0.074742314 | 1.930834615 | 4.1 | -0.85 | -3.9 | 50.74 | 1.99 | 5 | -1 |
| HCN1 | 410 | K | -31.21711111 | 10.38894595 | 0.160691942 | 5.435236881 | NA | NA | NA | NA | NA | NA | NA |
| HCN1 | 411 | Y | -29.64116667 | 12.96633973 | 0.211124026 | 7.908818108 | NA | NA | NA | NA | NA | NA | NA |
| HCN1 | 412 | K | -25.89577778 | 13.20191463 | 0.215733541 | 8.722212979 | NA | NA | NA | NA | NA | NA | NA |
| HCN1 | 413 | Q | -31.11011111 | 8.283414876 | 0.119492836 | 3.245366663 | 5.07 | -1.11 | -3.5 | 12.8 | 0.55 | 1 | -1 |
| HCN1 | 414 | V | -32.88861905 | 14.50450361 | 0.24122141 | 7.774707135 | NA | NA | NA | NA | NA | NA | NA |
| HCN1 | 415 | E | -28.79388889 | 17.11159484 | 0.292234587 | 12.0704623 | NA | NA | NA | NA | NA | NA | NA |
| HCN1 | 416 | Q | -29.54066667 | 14.57452208 | 0.242591468 | 9.558696894 | NA | NA | NA | NA | NA | NA | NA |
| HCN1 | 417 | Y | -36.53683333 | 16.80902433 | 0.286314164 | 10.14645213 | NA | NA | NA | NA | NA | NA | NA |
| HCN1 | 418 | M | -33.73470833 | 19.54008032 | 0.33975297 | 12.81287529 | NA | NA | NA | NA | NA | NA | NA |
| HCN1 | 419 | S | -31.2795 | 21.51400321 | 0.378376888 | 15.56843134 | NA | NA | NA | NA | NA | NA | NA |
| HCN1 | 420 | F | -35.22469697 | 20.36368751 | 0.355868562 | 13.64964042 | NA | NA | NA | NA | NA | NA | NA |
| HCN1 | 421 | H | -37.87763333 | 19.97147558 | 0.348194118 | 14.27284421 | NA | NA | NA | NA | NA | NA | NA |
| HCN1 | 422 | K | -34.07755556 | 24.99612978 | 0.446511956 | 18.28049092 | NA | NA | NA | NA | NA | NA | NA |
| HCN1 | 423 | L | -35.91420833 | 23.40475238 | 0.415373338 | 17.59038311 | NA | NA | NA | NA | NA | NA | NA |
| HCN1 | 424 | P | -35.59261905 | 28.19080871 | 0.509022511 | 21.19985561 | NA | NA | NA | NA | NA | NA | NA |
| HCN1 | 425 | A | -32.23993333 | 28.70992753 | 0.519180153 | 22.77549512 | NA | NA | NA | NA | NA | NA | NA |
| HCN1 | 426 | D | -34.66033333 | 29.06388751 | 0.526106118 | 22.35337259 | NA | NA | NA | NA | NA | NA | NA |
| HCN1 | 427 | M | -37.56383333 | 25.87008763 | 0.463612764 | 20.0752472 | NA | NA | NA | NA | NA | NA | NA |
| HCN1 | 428 | R | -32.1179697 | 23.80221406 | 0.423150505 | 17.2876249 | NA | NA | NA | NA | NA | NA | NA |
| HCN1 | 429 | Q | -32.01144444 | 26.27683683 | 0.47157166 | 19.53561707 | NA | NA | NA | NA | NA | NA | NA |
| HCN1 | 430 | K | -37.83244444 | 25.15131444 | 0.449548467 | 19.35504627 | NA | NA | NA | NA | NA | NA | NA |
| HCN1 | 431 | I | -36.20945833 | 20.81557864 | 0.364710754 | 15.0398315 | NA | NA | NA | NA | NA | NA | NA |
| HCN1 | 432 | H | -30.85023333 | 21.98301456 | 0.387554073 | 15.59391885 | NA | NA | NA | NA | NA | NA | NA |
| HCN1 | 433 | D | -33.71020833 | 24.44572701 | 0.435742178 | 17.72024089 | NA | NA | NA | NA | NA | NA | NA |
| HCN1 | 434 | Y | -38.462 | 18.70941688 | 0.323499307 | 13.29489301 | NA | NA | NA | NA | NA | NA | NA |
| HCN1 | 435 | Y | -34.9105 | 15.16458585 | 0.254137296 | 8.346306834 | NA | NA | NA | NA | NA | NA | NA |
| HCN1 | 436 | E | -31.25366667 | 21.64583371 | 0.380956426 | 15.74455525 | NA | NA | NA | NA | NA | NA | NA |
| HCN1 | 437 | H | -35.40843333 | 22.84107548 | 0.404343824 | 15.88172215 | NA | NA | NA | NA | NA | NA | NA |
| HCN1 | 438 | R | -38.42587879 | 16.40390871 | 0.278387232 | 11.04652437 | NA | NA | NA | NA | NA | NA | NA |
| HCN1 | 439 | Y | -35.275 | 14.14700039 | 0.234226114 | 7.753970943 | NA | NA | NA | NA | NA | NA | NA |
| HCN1 | 440 | Q | -32.06577778 | 20.38626741 | 0.356310385 | 14.44873614 | NA | NA | NA | NA | NA | NA | NA |

| | | | | | | | | | | | | | |
|---|---|---|---|---|---|---|---|---|---|---|---|---|---|
| HCN1 | 441 | G | -28.73233333 | 17.16208737 | 0.293222579 | 12.3035715 | NA | NA | NA | NA | NA | NA | NA |
| HCN1 | 442 | K | -30.31411111 | 15.31483856 | 0.257077304 | 10.2762742 | NA | NA | NA | NA | NA | NA | NA |
| HCN1 | 443 | I | -30.71733333 | 9.36456566 | 0.140647806 | 3.505154419 | 5.95 | -1.1 | -3.5 | 3.53 | 0.13 | 0 | 0 |
| HCN1 | 444 | F | -34.98987879 | 11.70374244 | 0.186418677 | 4.753035938 | 7.04 | -1.06 | -3.5 | 31.23 | 1.63 | 3 | -3 |
| HCN1 | 445 | D | -38.13483333 | 7.127470659 | 0.096874377 | 1.502235209 | 5.81 | -0.99 | -3.5 | 40.44 | 2.21 | 4 | -4 |
| HCN1 | 446 | E | -40.71277778 | 11.978555 | 0.191795958 | 5.949937338 | NA | NA | NA | NA | NA | NA | NA |
| HCN1 | 447 | E | -42.38033333 | 8.781085696 | 0.129230803 | 2.388782894 | 6.4 | -1.06 | -3.5 | 49.74 | 2.55 | 5 | -5 |
| HCN1 | 448 | N | -39.31945833 | 8.14443976 | 0.116773498 | 2.45992081 | 6.03 | -0.99 | -3.5 | 40.44 | 2.21 | 4 | -4 |
| HCN1 | 449 | I | -38.42908333 | 11.44078446 | 0.181273356 | 5.797947805 | NA | NA | NA | NA | NA | NA | NA |
| HCN1 | 450 | L | -42.81908333 | 13.53145166 | 0.22218162 | 7.190682929 | NA | NA | NA | NA | NA | NA | NA |
| HCN1 | 451 | N | -44.02508333 | 8.623047373 | 0.126138454 | 1.901232185 | 6.78 | -0.99 | -3.5 | 40.44 | 2.21 | 4 | -4 |
| HCN1 | 452 | E | -39.308 | 11.18565769 | 0.176281269 | 6.112366888 | NA | NA | NA | NA | NA | NA | NA |
| HCN1 | 453 | L | -41.49145833 | 15.75616911 | 0.265712856 | 9.726241408 | NA | NA | NA | NA | NA | NA | NA |
| HCN1 | 454 | N | -44.34658333 | 18.76018819 | 0.324492754 | 11.7094912 | NA | NA | NA | NA | NA | NA | NA |
| HCN1 | 455 | D | -47.77713333 | 19.99393716 | 0.348633625 | 12.36574189 | NA | NA | NA | NA | NA | NA | NA |
| HCN1 | 456 | P | -46.6507619 | 22.58719141 | 0.399376053 | 14.96961899 | NA | NA | NA | NA | NA | NA | NA |
| HCN1 | 457 | L | -42.61145833 | 22.07694841 | 0.389392084 | 15.30262165 | NA | NA | NA | NA | NA | NA | NA |
| HCN1 | 458 | R | -46.06751515 | 15.67827609 | 0.264188717 | 8.666407565 | NA | NA | NA | NA | NA | NA | NA |
| HCN1 | 459 | E | -48.66988889 | 22.04053686 | 0.388679617 | 14.41178684 | NA | NA | NA | NA | NA | NA | NA |
| HCN1 | 460 | E | -43.49655556 | 24.53691333 | 0.437526428 | 17.41738634 | NA | NA | NA | NA | NA | NA | NA |
| HCN1 | 461 | I | -42.25445833 | 20.7312653 | 0.363060988 | 14.4277716 | NA | NA | NA | NA | NA | NA | NA |
| HCN1 | 462 | V | -45.95113333 | 22.138196 | 0.390590521 | 15.1472284 | NA | NA | NA | NA | NA | NA | NA |
| HCN1 | 463 | N | -47.19183333 | 25.12541586 | 0.449041707 | 17.49488541 | NA | NA | NA | NA | NA | NA | NA |
| HCN1 | 464 | F | -41.15242424 | 26.8561495 | 0.48290712 | 20.09075141 | NA | NA | NA | NA | NA | NA | NA |
| HCN1 | 465 | N | -42.92620833 | 21.91509199 | 0.386225026 | 15.16009928 | NA | NA | NA | NA | NA | NA | NA |
| HCN1 | 466 | C | -47.024 | 24.5161675 | 0.437120493 | 16.90677388 | NA | NA | NA | NA | NA | NA | NA |
| HCN1 | 467 | R | -43.79951515 | 29.0451046 | 0.525738592 | 21.93011357 | NA | NA | NA | NA | NA | NA | NA |
| HCN1 | 468 | K | -46.34893333 | 31.312114 | 0.570097358 | 24.19952772 | NA | NA | NA | NA | NA | NA | NA |
| HCN1 | 469 | L | -49.69733333 | 26.96737707 | 0.48508352 | 17.8664211 | NA | NA | NA | NA | NA | NA | NA |
| HCN1 | 470 | V | -51.14933333 | 29.32501291 | 0.531215581 | 18.3909825 | NA | NA | NA | NA | NA | NA | NA |
| HCN1 | 471 | A | -50.20973333 | 33.45033554 | 0.611936122 | 24.3495953 | NA | NA | NA | NA | NA | NA | NA |
| HCN1 | 472 | T | -51.42433333 | 34.47688002 | 0.632022605 | 23.55811161 | NA | NA | NA | NA | NA | NA | NA |
| HCN1 | 473 | M | -54.58945833 | 29.37976028 | 0.532286828 | 18.50020479 | NA | NA | NA | NA | NA | NA | NA |

| | | | | | | | | | | | | | |
|---|---|---|---|---|---|---|---|---|---|---|---|---|---|
| HCN1 | 474 | P | -57.71119048 | 34.40876535 | 0.6306898 | 24.78345924 | NA | NA | NA | NA | NA | NA | NA |
| HCN1 | 475 | L | -59.90558333 | 28.85117552 | 0.521943965 | 19.28387277 | NA | NA | NA | NA | NA | NA | NA |
| HCN1 | 476 | F | -55.9029697 | 26.48460825 | 0.475637141 | 16.71967736 | NA | NA | NA | NA | NA | NA | NA |
| HCN1 | 477 | A | -56.43433333 | 33.24155184 | 0.607850833 | 23.49386382 | NA | NA | NA | NA | NA | NA | NA |
| HCN1 | 478 | N | -60.10108333 | 34.11154659 | 0.624874095 | 24.56977007 | NA | NA | NA | NA | NA | NA | NA |
| HCN1 | 479 | A | -60.09893333 | 30.450432 | 0.553236753 | 20.88524429 | NA | NA | NA | NA | NA | NA | NA |
| HCN1 | 480 | D | -59.65713333 | 29.07294889 | 0.526283423 | 19.71816267 | NA | NA | NA | NA | NA | NA | NA |
| HCN1 | 481 | P | -56.37790476 | 27.54175018 | 0.496322327 | 17.77707904 | NA | NA | NA | NA | NA | NA | NA |
| HCN1 | 482 | N | -57.82970833 | 24.60411464 | 0.438841362 | 14.91318186 | NA | NA | NA | NA | NA | NA | NA |
| HCN1 | 483 | F | -59.44787879 | 24.06708983 | 0.428333352 | 14.78605621 | NA | NA | NA | NA | NA | NA | NA |
| HCN1 | 484 | V | -55.10319048 | 24.43569964 | 0.435545971 | 14.68415833 | NA | NA | NA | NA | NA | NA | NA |
| HCN1 | 485 | T | -53.65704762 | 21.92157618 | 0.386351903 | 11.03177733 | NA | NA | NA | NA | NA | NA | NA |
| HCN1 | 486 | A | -55.99533333 | 19.65187614 | 0.341940488 | 9.885756819 | NA | NA | NA | NA | NA | NA | NA |
| HCN1 | 487 | M | -55.92183333 | 20.48367865 | 0.358216439 | 10.75151111 | NA | NA | NA | NA | NA | NA | NA |
| HCN1 | 488 | L | -51.31870833 | 20.40429418 | 0.356663116 | 9.655836524 | NA | NA | NA | NA | NA | NA | NA |
| HCN1 | 489 | S | -51.34133333 | 17.03284016 | 0.290693588 | 7.934626095 | NA | NA | NA | NA | NA | NA | NA |
| HCN1 | 490 | K | -54.05077778 | 15.40659878 | 0.258872784 | 4.622113227 | 10.97 | -0.93 | -3.5 | 31.17 | 1.79 | 3 | -3 |
| HCN1 | 491 | L | -50.75595833 | 17.27486961 | 0.295429398 | 8.202157422 | NA | NA | NA | NA | NA | NA | NA |
| HCN1 | 492 | R | -47.79333333 | 13.97734481 | 0.230906449 | 5.137364112 | NA | NA | NA | NA | NA | NA | NA |
| HCN1 | 493 | F | -43.56687879 | 18.10972893 | 0.311765162 | 11.00534482 | NA | NA | NA | NA | NA | NA | NA |
| HCN1 | 494 | E | -46.28688889 | 18.23329446 | 0.314182979 | 11.50882101 | NA | NA | NA | NA | NA | NA | NA |
| HCN1 | 495 | V | -42.04333333 | 21.95862168 | 0.387076775 | 15.59140438 | NA | NA | NA | NA | NA | NA | NA |
| HCN1 | 496 | F | -45.25106061 | 23.39597703 | 0.41520163 | 16.8647635 | NA | NA | NA | NA | NA | NA | NA |
| HCN1 | 497 | Q | -40.58455556 | 26.17021951 | 0.46948547 | 20.21498261 | NA | NA | NA | NA | NA | NA | NA |
| HCN1 | 498 | P | -41.2567619 | 30.76437545 | 0.559379711 | 24.37178812 | NA | NA | NA | NA | NA | NA | NA |
| HCN1 | 499 | G | -44.57208333 | 31.79229386 | 0.579493079 | 25.04083449 | NA | NA | NA | NA | NA | NA | NA |
| HCN1 | 500 | D | -45.24033333 | 26.97366126 | 0.485206483 | 19.89231852 | NA | NA | NA | NA | NA | NA | NA |
| HCN1 | 501 | Y | -50.24716667 | 27.11937916 | 0.488057758 | 18.01645516 | NA | NA | NA | NA | NA | NA | NA |
| HCN1 | 502 | I | -50.57158333 | 25.07295134 | 0.44801513 | 16.1884366 | NA | NA | NA | NA | NA | NA | NA |
| HCN1 | 503 | I | -54.87658333 | 25.97360774 | 0.46563835 | 15.11358828 | NA | NA | NA | NA | NA | NA | NA |
| HCN1 | 504 | R | -53.96042424 | 27.62052268 | 0.497863676 | 17.89763202 | NA | NA | NA | NA | NA | NA | NA |
| HCN1 | 505 | E | -59.24753333 | 29.47297815 | 0.53411083 | 19.95399046 | NA | NA | NA | NA | NA | NA | NA |
| HCN1 | 506 | G | -61.77958333 | 26.24953601 | 0.471037462 | 16.64896542 | NA | NA | NA | NA | NA | NA | NA |

| | | | | | | | | | | | | | |
|---|---|---|---|---|---|---|---|---|---|---|---|---|---|
| HCN1 | 507 | A | -60.26153333 | 23.777608 | 0.422669036 | 14.21891093 | NA | NA | NA | NA | NA | NA | NA |
| HCN1 | 508 | V | -60.56153333 | 20.29901539 | 0.354603117 | 10.71963226 | NA | NA | NA | NA | NA | NA | NA |
| HCN1 | 509 | G | -59.98033333 | 19.00075838 | 0.329200011 | 9.471576655 | NA | NA | NA | NA | NA | NA | NA |
| HCN1 | 510 | K | -57.88573333 | 16.47421883 | 0.279762996 | 6.851766083 | NA | NA | NA | NA | NA | NA | NA |
| HCN1 | 511 | K | -56.07953333 | 18.38753072 | 0.317200933 | 8.620026298 | NA | NA | NA | NA | NA | NA | NA |
| HCN1 | 512 | M | -56.61545833 | 20.68865027 | 0.362227136 | 10.96671294 | NA | NA | NA | NA | NA | NA | NA |
| HCN1 | 513 | Y | -52.78108333 | 21.8550159 | 0.385049512 | 10.90907326 | NA | NA | NA | NA | NA | NA | NA |
| HCN1 | 514 | F | -52.87151515 | 23.73475389 | 0.421830506 | 12.79898878 | NA | NA | NA | NA | NA | NA | NA |
| HCN1 | 515 | I | -49.32345833 | 28.31996446 | 0.511549713 | 19.22000876 | NA | NA | NA | NA | NA | NA | NA |
| HCN1 | 516 | Q | -44.87677778 | 25.82936813 | 0.462816002 | 18.74054882 | NA | NA | NA | NA | NA | NA | NA |
| HCN1 | 517 | H | -42.28813333 | 29.77304339 | 0.539982232 | 23.01979617 | NA | NA | NA | NA | NA | NA | NA |
| HCN1 | 518 | G | -44.56908333 | 32.01419165 | 0.583834972 | 25.24823813 | NA | NA | NA | NA | NA | NA | NA |
| HCN1 | 519 | V | -46.00961905 | 33.63061345 | 0.615463635 | 26.53049406 | NA | NA | NA | NA | NA | NA | NA |
| HCN1 | 520 | A | -49.37973333 | 32.03198555 | 0.584183146 | 22.93706112 | NA | NA | NA | NA | NA | NA | NA |
| HCN1 | 521 | G | -51.41233333 | 34.00522986 | 0.622793786 | 23.11091064 | NA | NA | NA | NA | NA | NA | NA |
| HCN1 | 522 | V | -55.2597619 | 34.23718993 | 0.627332569 | 24.47041455 | NA | NA | NA | NA | NA | NA | NA |
| HCN1 | 523 | I | -55.28695833 | 34.84425417 | 0.639211047 | 25.1578446 | NA | NA | NA | NA | NA | NA | NA |
| HCN1 | 524 | T | -59.56213333 | 36.51992438 | 0.671999029 | 26.95590132 | NA | NA | NA | NA | NA | NA | NA |
| HCN1 | 525 | K | -62.10693333 | 38.01454361 | 0.701244371 | 28.43385005 | NA | NA | NA | NA | NA | NA | NA |
| HCN1 | 526 | S | -62.49333333 | 41.36872322 | 0.76687589 | 31.77296425 | NA | NA | NA | NA | NA | NA | NA |
| HCN1 | 527 | S | -60.18913333 | 39.52033515 | 0.730708322 | 29.95347406 | NA | NA | NA | NA | NA | NA | NA |
| HCN1 | 528 | K | -57.44013333 | 41.43010909 | 0.768077032 | 31.67767967 | NA | NA | NA | NA | NA | NA | NA |
| HCN1 | 529 | E | -54.76813333 | 41.24754806 | 0.764504845 | 31.47863136 | NA | NA | NA | NA | NA | NA | NA |
| HCN1 | 530 | M | -56.18595833 | 37.73093968 | 0.695695068 | 28.00761963 | NA | NA | NA | NA | NA | NA | NA |
| HCN1 | 531 | K | -50.37422222 | 37.68402298 | 0.694777045 | 26.72795223 | NA | NA | NA | NA | NA | NA | NA |
| HCN1 | 532 | L | -52.18033333 | 32.91994333 | 0.601557892 | 21.95301591 | NA | NA | NA | NA | NA | NA | NA |
| HCN1 | 533 | T | -49.53713333 | 34.00361725 | 0.622762232 | 24.90189819 | NA | NA | NA | NA | NA | NA | NA |
| HCN1 | 534 | D | -48.24413333 | 33.84915656 | 0.619739887 | 26.25481927 | NA | NA | NA | NA | NA | NA | NA |
| HCN1 | 535 | G | -51.30733333 | 32.6422222 | 0.596123699 | 23.65896956 | NA | NA | NA | NA | NA | NA | NA |
| HCN1 | 536 | S | -52.85666667 | 30.09768254 | 0.546334474 | 19.13819903 | NA | NA | NA | NA | NA | NA | NA |
| HCN1 | 537 | Y | -57.44091667 | 27.99838719 | 0.505257383 | 18.39885389 | NA | NA | NA | NA | NA | NA | NA |
| HCN1 | 538 | F | -54.47642424 | 25.96418229 | 0.465453922 | 16.21015326 | NA | NA | NA | NA | NA | NA | NA |
| HCN1 | 539 | G | -58.69208333 | 23.70296492 | 0.421208488 | 14.33528486 | NA | NA | NA | NA | NA | NA | NA |

| | | | | | | | | | | | | | |
|---|---|---|---|---|---|---|---|---|---|---|---|---|---|
| HCN1 | 540 | E | -61.42173333 | 24.05411058 | 0.428079386 | 14.47073323 | NA | NA | NA | NA | NA | NA | NA |
| HCN1 | 541 | I | -64.23158333 | 24.14198877 | 0.429798906 | 14.76025668 | NA | NA | NA | NA | NA | NA | NA |
| HCN1 | 542 | C | -63.64393333 | 21.03868943 | 0.369076382 | 11.59089668 | NA | NA | NA | NA | NA | NA | NA |
| HCN1 | 543 | L | -62.70283333 | 19.25853956 | 0.334244038 | 9.710914914 | NA | NA | NA | NA | NA | NA | NA |
| HCN1 | 544 | L | -66.55608333 | 21.15294107 | 0.371311954 | 8.087461083 | NA | NA | NA | NA | NA | NA | NA |
| HCN1 | 545 | T | -68.68793333 | 21.26709077 | 0.373545531 | 8.20591355 | NA | NA | NA | NA | NA | NA | NA |
| HCN1 | 546 | K | -68.07733333 | 18.51228187 | 0.31964195 | 5.448841079 | NA | NA | NA | NA | NA | NA | NA |
| HCN1 | 547 | G | -66.42708333 | 21.4079365 | 0.376301471 | 8.418819145 | NA | NA | NA | NA | NA | NA | NA |
| HCN1 | 548 | R | -64.97678788 | 23.09113269 | 0.409236715 | 13.71808106 | NA | NA | NA | NA | NA | NA | NA |
| HCN1 | 549 | R | -60.15306061 | 22.27028828 | 0.393175182 | 13.22355439 | NA | NA | NA | NA | NA | NA | NA |
| HCN1 | 550 | T | -63.29047619 | 28.66914533 | 0.518382165 | 19.07607439 | NA | NA | NA | NA | NA | NA | NA |
| HCN1 | 551 | A | -59.73933333 | 30.24032832 | 0.549125636 | 20.86114006 | NA | NA | NA | NA | NA | NA | NA |
| HCN1 | 552 | S | -56.35666667 | 31.6232251 | 0.576184896 | 21.88203498 | NA | NA | NA | NA | NA | NA | NA |
| HCN1 | 553 | V | -53.43190476 | 30.10362495 | 0.54645075 | 19.15348839 | NA | NA | NA | NA | NA | NA | NA |
| HCN1 | 554 | R | -48.80087879 | 32.13963513 | 0.586289535 | 23.14585917 | NA | NA | NA | NA | NA | NA | NA |
| HCN1 | 555 | A | -46.50213333 | 30.81591269 | 0.560388145 | 23.71192044 | NA | NA | NA | NA | NA | NA | NA |
| HCN1 | 556 | D | -44.64853333 | 34.04290534 | 0.623530986 | 26.93995877 | NA | NA | NA | NA | NA | NA | NA |
| HCN1 | 557 | T | -42.28753333 | 31.81323077 | 0.579902753 | 25.42400823 | NA | NA | NA | NA | NA | NA | NA |
| HCN1 | 558 | Y | -38.83116667 | 28.5822822 | 0.516682506 | 22.75318727 | NA | NA | NA | NA | NA | NA | NA |
| HCN1 | 559 | C | -44.11283333 | 26.51887179 | 0.476307579 | 19.7528273 | NA | NA | NA | NA | NA | NA | NA |
| HCN1 | 560 | R | -41.27742424 | 24.18980056 | 0.430734443 | 17.808175 | NA | NA | NA | NA | NA | NA | NA |
| HCN1 | 561 | L | -47.28620833 | 22.76738114 | 0.402901841 | 15.1619961 | NA | NA | NA | NA | NA | NA | NA |
| HCN1 | 562 | Y | -47.439 | 21.79684551 | 0.383911287 | 12.93529257 | NA | NA | NA | NA | NA | NA | NA |
| HCN1 | 563 | S | -51.02683333 | 18.6439023 | 0.322217378 | 7.918310829 | NA | NA | NA | NA | NA | NA | NA |
| HCN1 | 564 | L | -55.27408333 | 17.92577153 | 0.308165652 | 7.093910368 | NA | NA | NA | NA | NA | NA | NA |
| HCN1 | 565 | S | -56.37816667 | 13.67459122 | 0.224982444 | 3.927501102 | 9.77 | -0.99 | -3.5 | 40.44 | 2.21 | 4 | -4 |
| HCN1 | 566 | V | -60.43461905 | 14.25396668 | 0.236319133 | 4.75285311 | 9.6 | -1.04 | -3.5 | 49.7 | 2.63 | 4 | -4 |
| HCN1 | 567 | D | -60.21473333 | 11.06337128 | 0.17388848 | 1.723016565 | 9.6 | -1.04 | -3.5 | 49.7 | 2.63 | 4 | -4 |
| HCN1 | 568 | N | -57.42195833 | 12.82650623 | 0.208387892 | 3.088021464 | 9.77 | -0.99 | -3.5 | 40.44 | 2.21 | 4 | -4 |
| HCN1 | 569 | F | -61.24751515 | 16.17662069 | 0.273939868 | 6.811389105 | NA | NA | NA | NA | NA | NA | NA |
| HCN1 | 570 | N | -63.89508333 | 14.29372439 | 0.237097075 | 4.8530376 | 9.6 | -1.04 | -3.5 | 49.7 | 2.63 | 4 | -4 |
| HCN1 | 571 | E | -62.13753333 | 13.97035168 | 0.230769614 | 4.423000185 | 9.6 | -1.04 | -3.5 | 49.7 | 2.63 | 4 | -4 |
| HCN1 | 572 | V | -60.98804762 | 16.9726492 | 0.289515826 | 7.387778787 | NA | NA | NA | NA | NA | NA | NA |

| | | | | | | | | | | | | | |
|---|---|---|---|---|---|---|---|---|---|---|---|---|---|
| HCN1 | 573 | L | -64.63245833 | 18.8861288 | 0.326957045 | 9.315529303 | NA | NA | NA | NA | NA | NA | NA |
| HCN1 | 574 | E | -66.66273333 | 16.11617678 | 0.272757157 | 3.257825257 | 13.07 | -0.99 | -3.5 | 40.44 | 2.21 | 4 | -4 |
| HCN1 | 575 | E | -65.22493333 | 18.94517596 | 0.328112425 | 9.617092084 | NA | NA | NA | NA | NA | NA | NA |
| HCN1 | 576 | Y | -63.652 | 21.69488202 | 0.381916159 | 12.3215083 | NA | NA | NA | NA | NA | NA | NA |
| HCN1 | 577 | P | -69.02947619 | 22.01767077 | 0.388232194 | 8.954505961 | NA | NA | NA | NA | NA | NA | NA |
| HCN1 | 578 | M | -68.13753333 | 25.50609806 | 0.456490548 | 12.44472434 | NA | NA | NA | NA | NA | NA | NA |
| HCN1 | 579 | M | -65.19913333 | 26.21181598 | 0.470299391 | 16.67063505 | NA | NA | NA | NA | NA | NA | NA |
| HCN1 | 580 | R | -67.68560606 | 18.5400353 | 0.320185004 | 5.559549534 | NA | NA | NA | NA | NA | NA | NA |
| HCN1 | 581 | R | -69.00193333 | 26.5902554 | 0.477704348 | 13.52989565 | NA | NA | NA | NA | NA | NA | NA |
| HCN1 | 582 | A | -66.13393333 | 29.11023038 | 0.527012913 | 16.09617132 | NA | NA | NA | NA | NA | NA | NA |
| HCN1 | 583 | F | -63.25087879 | 25.32345446 | 0.452916746 | 15.73432065 | NA | NA | NA | NA | NA | NA | NA |
| HCN1 | 584 | E | -68.20673333 | 27.47190905 | 0.49495574 | 14.40195724 | NA | NA | NA | NA | NA | NA | NA |
| HCN1 | 585 | T | -68.32333333 | 30.32758225 | 0.550832941 | 17.27003586 | NA | NA | NA | NA | NA | NA | NA |
| HCN1 | 586 | V | -64.29347619 | 31.89360974 | 0.581475535 | 22.32276716 | NA | NA | NA | NA | NA | NA | NA |
| HCN1 | 587 | A | -65.99033333 | 29.16834923 | 0.52815013 | 19.82182742 | NA | NA | NA | NA | NA | NA | NA |
| HCN1 | 588 | I | -69.63608333 | 31.00354039 | 0.564059472 | 18.0373401 | NA | NA | NA | NA | NA | NA | NA |
| HCN1 | 589 | D | -66.71233333 | 34.59269994 | 0.634288863 | 21.55925499 | NA | NA | NA | NA | NA | NA | NA |
| HCN1 | 590 | R | -62.18451515 | 31.03141352 | 0.564604868 | 21.58611843 | NA | NA | NA | NA | NA | NA | NA |
| HCN1 | 591 | L | -67.42520833 | 30.51670895 | 0.5545336 | 17.45230562 | NA | NA | NA | NA | NA | NA | NA |
| HCN1 | 592 | D | -68.43973333 | 36.02066937 | 0.662230063 | 22.96604436 | NA | NA | NA | NA | NA | NA | NA |
| HCN1 | 593 | R | -63.91069697 | 38.32744637 | 0.707366966 | 28.77000848 | NA | NA | NA | NA | NA | NA | NA |
| HCN1 | 594 | I | -65.26393333 | 36.16663209 | 0.665086128 | 26.6552685 | NA | NA | NA | NA | NA | NA | NA |
| HCN1 | 595 | G | -68.62183333 | 37.1412069 | 0.684155717 | 24.07088283 | NA | NA | NA | NA | NA | NA | NA |
| HCN1 | 596 | K | -69.00073333 | 33.04336455 | 0.603972886 | 19.97220135 | NA | NA | NA | NA | NA | NA | NA |
| HCN1 | 597 | K | -72.34813333 | 31.387751 | 0.571577354 | 16.84416954 | NA | NA | NA | NA | NA | NA | NA |
| HCN1 | 598 | N | -72.32013333 | 28.49373603 | 0.514949916 | 13.96792159 | NA | NA | NA | NA | NA | NA | NA |
| HCN1 | 599 | S | -75.76513333 | 27.2023319 | 0.489680901 | 12.72416521 | NA | NA | NA | NA | NA | NA | NA |
| HCN1 | 600 | I | -74.94093333 | 23.626019 | 0.419702881 | 9.123115329 | NA | NA | NA | NA | NA | NA | NA |
| HCN1 | 601 | L | -73.29073333 | 25.84573857 | 0.463136323 | 11.32276128 | NA | NA | NA | NA | NA | NA | NA |
| HCN1 | 602 | L | -76.04113333 | 28.2323391 | 0.50983514 | 13.72718444 | NA | NA | NA | NA | NA | NA | NA |
| HCN1 | 603 | Q | -78.18453333 | 26.13244302 | 0.468746294 | 11.72011181 | NA | NA | NA | NA | NA | NA | NA |
| HCN1 | 604 | K | -76.13933333 | 24.66260574 | 0.439985862 | 10.16919939 | NA | NA | NA | NA | NA | NA | NA |
| HCN1 | 605 | F | -75.09160606 | 27.62631342 | 0.497976983 | 13.14594418 | NA | NA | NA | NA | NA | NA | NA |

| | | | | | | | | | | | | | |
|---|---|---|---|---|---|---|---|---|---|---|---|---|---|
| HCN1 | 606 | Q | -79.64213333 | 29.53169433 | 0.535259735 | 15.09934318 | NA | NA | NA | NA | NA | NA | NA |
| HCN1 | 607 | K | -79.86213333 | 25.94427145 | 0.465064325 | 11.55531275 | NA | NA | NA | NA | NA | NA | NA |
| HCN1 | 608 | D | -77.74053333 | 27.8616618 | 0.502582065 | 13.43295176 | NA | NA | NA | NA | NA | NA | NA |
| HCN4 | 214 | M | -18.790575 | 53.997461 | 0.999751833 | 51.68471037 | NA | NA | NA | NA | NA | NA | NA |
| HCN4 | 215 | Q | -19.95759722 | 53.14633464 | 0.983239338 | 50.84805243 | NA | NA | NA | NA | NA | NA | NA |
| HCN4 | 216 | R | -24.87010227 | 51.23570325 | 0.946171643 | 46.61657998 | NA | NA | NA | NA | NA | NA | NA |
| HCN4 | 217 | Q | -20.78948611 | 49.57854469 | 0.914021512 | 45.84731537 | NA | NA | NA | NA | NA | NA | NA |
| HCN4 | 218 | F | -18.690175 | 49.5291477 | 0.913063173 | 48.23867803 | NA | NA | NA | NA | NA | NA | NA |
| HCN4 | 219 | G | -21.830875 | 47.69179143 | 0.877417071 | 43.94444181 | NA | NA | NA | NA | NA | NA | NA |
| HCN4 | 220 | A | -23.158375 | 45.02139969 | 0.825609449 | 40.40158097 | NA | NA | NA | NA | NA | NA | NA |
| HCN4 | 221 | M | -19.819625 | 42.77855838 | 0.78209663 | 40.4596803 | NA | NA | NA | NA | NA | NA | NA |
| HCN4 | 222 | L | -20.378375 | 42.77867269 | 0.782098848 | 40.51938026 | NA | NA | NA | NA | NA | NA | NA |
| HCN4 | 223 | Q | -25.36781944 | 41.68659487 | 0.760911712 | 36.74687986 | NA | NA | NA | NA | NA | NA | NA |
| HCN4 | 224 | P | -26.43908929 | 38.46711388 | 0.698451343 | 32.95807476 | NA | NA | NA | NA | NA | NA | NA |
| HCN4 | 225 | G | -26.884375 | 38.3832511 | 0.696824342 | 32.77991186 | NA | NA | NA | NA | NA | NA | NA |
| HCN4 | 226 | V | -29.61523214 | 38.49929881 | 0.699075755 | 31.6493206 | NA | NA | NA | NA | NA | NA | NA |
| HCN4 | 227 | N | -27.2635 | 34.82987374 | 0.627886131 | 28.99952635 | NA | NA | NA | NA | NA | NA | NA |
| HCN4 | 228 | K | -28.26270833 | 31.22926592 | 0.558031614 | 25.38088027 | NA | NA | NA | NA | NA | NA | NA |
| HCN4 | 229 | F | -25.717375 | 29.63287667 | 0.527060454 | 24.00484833 | NA | NA | NA | NA | NA | NA | NA |
| HCN4 | 230 | S | -29.19804167 | 33.43766895 | 0.600876305 | 26.59556895 | NA | NA | NA | NA | NA | NA | NA |
| HCN4 | 231 | L | -32.5675 | 30.86713032 | 0.551005909 | 23.7610646 | NA | NA | NA | NA | NA | NA | NA |
| HCN4 | 232 | R | -29.15364773 | 25.31501357 | 0.443290641 | 18.5254512 | NA | NA | NA | NA | NA | NA | NA |
| HCN4 | 233 | M | -29.31825 | 27.87056975 | 0.492870366 | 21.00760228 | NA | NA | NA | NA | NA | NA | NA |
| HCN4 | 234 | F | -31.50601136 | 31.72374133 | 0.567624811 | 24.60601464 | NA | NA | NA | NA | NA | NA | NA |
| HCN4 | 235 | G | -35.432375 | 31.45447819 | 0.562400902 | 23.43535394 | NA | NA | NA | NA | NA | NA | NA |
| HCN4 | 236 | S | -37.271875 | 33.41580599 | 0.600452147 | 25.48679277 | NA | NA | NA | NA | NA | NA | NA |
| HCN4 | 237 | Q | -35.85304167 | 36.63513771 | 0.662909619 | 28.6180015 | NA | NA | NA | NA | NA | NA | NA |
| HCN4 | 238 | K | -39.61970833 | 38.500362 | 0.699096382 | 30.60779358 | NA | NA | NA | NA | NA | NA | NA |
| HCN4 | 239 | A | -35.478975 | 35.85123236 | 0.647701262 | 27.83163981 | NA | NA | NA | NA | NA | NA | NA |
| HCN4 | 240 | V | -32.51723214 | 36.86195158 | 0.66730998 | 29.74182771 | NA | NA | NA | NA | NA | NA | NA |
| HCN4 | 241 | E | -34.17204167 | 40.7009381 | 0.741789224 | 33.26289635 | NA | NA | NA | NA | NA | NA | NA |
| HCN4 | 242 | R | -36.48882955 | 36.94964001 | 0.669011202 | 29.71348214 | NA | NA | NA | NA | NA | NA | NA |
| HCN4 | 243 | E | -30.44248611 | 37.82951114 | 0.686081367 | 30.83209356 | NA | NA | NA | NA | NA | NA | NA |

| | | | | | | | | | | | | | |
|---|---|---|---|---|---|---|---|---|---|---|---|---|---|
| HCN4 | 244 | Q | -29.94670833 | 42.15380799 | 0.769976 | 35.29455364 | NA | NA | NA | NA | NA | NA | NA |
| HCN4 | 245 | E | -32.77604167 | 44.68119408 | 0.819009203 | 37.56612516 | NA | NA | NA | NA | NA | NA | NA |
| HCN4 | 246 | R | -30.17619318 | 38.75950207 | 0.704123896 | 31.80727518 | NA | NA | NA | NA | NA | NA | NA |
| HCN4 | 247 | V | -26.80066071 | 42.39178537 | 0.774592941 | 36.47902921 | NA | NA | NA | NA | NA | NA | NA |
| HCN4 | 248 | K | -28.05859722 | 46.7652372 | 0.859441217 | 40.90570809 | NA | NA | NA | NA | NA | NA | NA |
| HCN4 | 249 | S | -29.25854167 | 47.80263066 | 0.879567436 | 40.94160436 | NA | NA | NA | NA | NA | NA | NA |
| HCN4 | 250 | A | -25.961375 | 46.44122329 | 0.853155102 | 40.84206022 | NA | NA | NA | NA | NA | NA | NA |
| HCN4 | 251 | G | -23.414875 | 49.35726113 | 0.909728443 | 44.74916186 | NA | NA | NA | NA | NA | NA | NA |
| HCN4 | 252 | F | -20.03155682 | 49.75739552 | 0.917491354 | 47.45053857 | NA | NA | NA | NA | NA | NA | NA |
| HCN4 | 253 | W | -18.81716071 | 47.16748191 | 0.867245069 | 44.87396741 | NA | NA | NA | NA | NA | NA | NA |
| HCN4 | 254 | I | -21.190375 | 44.15662313 | 0.808832129 | 41.87862564 | NA | NA | NA | NA | NA | NA | NA |
| HCN4 | 255 | I | -18.671625 | 40.86645134 | 0.745000306 | 38.54777826 | NA | NA | NA | NA | NA | NA | NA |
| HCN4 | 256 | H | -23.516475 | 37.15640763 | 0.67302265 | 33.56540889 | NA | NA | NA | NA | NA | NA | NA |
| HCN4 | 257 | P | -21.16780357 | 35.11996135 | 0.63351405 | 32.79939299 | NA | NA | NA | NA | NA | NA | NA |
| HCN4 | 258 | Y | -23.75654167 | 30.50041149 | 0.543891286 | 26.16242319 | NA | NA | NA | NA | NA | NA | NA |
| HCN4 | 259 | S | -21.03304167 | 37.34557121 | 0.676692567 | 33.61816803 | NA | NA | NA | NA | NA | NA | NA |
| HCN4 | 260 | D | -20.07175 | 38.80817607 | 0.705068208 | 36.51376245 | NA | NA | NA | NA | NA | NA | NA |
| HCN4 | 261 | F | -15.31255682 | 39.87300929 | 0.725726778 | 39.01694001 | NA | NA | NA | NA | NA | NA | NA |
| HCN4 | 262 | R | -15.36692045 | 36.56034231 | 0.661458532 | 35.76938749 | NA | NA | NA | NA | NA | NA | NA |
| HCN4 | 263 | F | -16.38055682 | 32.3171528 | 0.579137443 | 31.4322617 | NA | NA | NA | NA | NA | NA | NA |
| HCN4 | 264 | Y | -14.68345833 | 37.2305166 | 0.67446042 | 36.09270746 | NA | NA | NA | NA | NA | NA | NA |
| HCN4 | 265 | W | -10.18966071 | 37.08055018 | 0.671550958 | 35.88006458 | NA | NA | NA | NA | NA | NA | NA |
| HCN4 | 266 | D | -12.088625 | 33.5657814 | 0.603361783 | 32.37613305 | NA | NA | NA | NA | NA | NA | NA |
| HCN4 | 267 | L | -10.205 | 32.4288785 | 0.581305006 | 30.78885485 | NA | NA | NA | NA | NA | NA | NA |
| HCN4 | 268 | T | -7.503517857 | 35.83462094 | 0.647378988 | 34.49417795 | NA | NA | NA | NA | NA | NA | NA |
| HCN4 | 269 | M | -6.400375 | 33.42781424 | 0.600685116 | 32.35019691 | NA | NA | NA | NA | NA | NA | NA |
| HCN4 | 270 | L | -7.232 | 29.71082887 | 0.528572786 | 28.65569872 | NA | NA | NA | NA | NA | NA | NA |
| HCN4 | 271 | L | -4.379625 | 32.13396865 | 0.575583532 | 30.90669593 | NA | NA | NA | NA | NA | NA | NA |
| HCN4 | 272 | L | -1.51425 | 33.38150994 | 0.599786777 | 31.47485776 | NA | NA | NA | NA | NA | NA | NA |
| HCN4 | 273 | M | -2.931 | 29.99299458 | 0.534047014 | 28.08641053 | NA | NA | NA | NA | NA | NA | NA |
| HCN4 | 274 | V | -1.128946429 | 27.78880253 | 0.49128402 | 25.86990141 | NA | NA | NA | NA | NA | NA | NA |
| HCN4 | 275 | G | 0.775625 | 30.82273174 | 0.550144543 | 28.05625847 | NA | NA | NA | NA | NA | NA | NA |
| HCN4 | 276 | N | 1.14425 | 30.47890223 | 0.54347399 | 28.49403618 | NA | NA | NA | NA | NA | NA | NA |

| | | | | | | | | | | | | | |
|---|---|---|---|---|---|---|---|---|---|---|---|---|---|
| HCN4 | 277 | L | 0.81025 | 26.62274178 | 0.468661559 | 23.97947334 | NA | NA | NA | NA | NA | NA | NA |
| HCN4 | 278 | I | 3.09 | 26.30463406 | 0.462490028 | 24.28387761 | NA | NA | NA | NA | NA | NA | NA |
| HCN4 | 279 | I | 5.854125 | 29.30833535 | 0.520764107 | 27.88328222 | NA | NA | NA | NA | NA | NA | NA |
| HCN4 | 280 | I | 5.89225 | 29.92768608 | 0.53277998 | 28.57584211 | NA | NA | NA | NA | NA | NA | NA |
| HCN4 | 281 | P | 7.583375 | 26.45527613 | 0.465412598 | 25.09154664 | NA | NA | NA | NA | NA | NA | NA |
| HCN4 | 282 | V | 10.45908929 | 25.98813412 | 0.45634969 | 24.55997826 | NA | NA | NA | NA | NA | NA | NA |
| HCN4 | 283 | G | 11.723125 | 29.7929204 | 0.530165424 | 28.38340603 | NA | NA | NA | NA | NA | NA | NA |
| HCN4 | 284 | I | 11.424375 | 29.72705397 | 0.528887565 | 28.25674063 | NA | NA | NA | NA | NA | NA | NA |
| HCN4 | 285 | T | 13.625375 | 25.77255713 | 0.452167333 | 23.65898524 | NA | NA | NA | NA | NA | NA | NA |
| HCN4 | 286 | F | 15.04628409 | 22.74755895 | 0.393480078 | 19.27340513 | NA | NA | NA | NA | NA | NA | NA |
| HCN4 | 287 | F | 15.973375 | 29.42360333 | 0.523000393 | 25.00176418 | NA | NA | NA | NA | NA | NA | NA |
| HCN4 | 288 | K | 21.28970833 | 30.8285934 | 0.550258264 | 26.55348279 | NA | NA | NA | NA | NA | NA | NA |
| HCN4 | 289 | D | 19.124175 | 34.45323686 | 0.620579091 | 30.16144648 | NA | NA | NA | NA | NA | NA | NA |
| HCN4 | 290 | E | 15.266775 | 34.62963087 | 0.624001268 | 31.08332741 | NA | NA | NA | NA | NA | NA | NA |
| HCN4 | 291 | N | 15.51425 | 38.1854552 | 0.692986952 | 36.02905507 | NA | NA | NA | NA | NA | NA | NA |
| HCN4 | 292 | T | 15.51323214 | 39.35933295 | 0.715761069 | 35.79203829 | NA | NA | NA | NA | NA | NA | NA |
| HCN4 | 293 | T | 13.28094643 | 42.77045048 | 0.781939331 | 40.59072506 | NA | NA | NA | NA | NA | NA | NA |
| HCN4 | 294 | P | 11.61023214 | 40.51547228 | 0.738191047 | 39.14883825 | NA | NA | NA | NA | NA | NA | NA |
| HCN4 | 295 | W | 10.20744643 | 33.64989358 | 0.604993623 | 32.22091992 | NA | NA | NA | NA | NA | NA | NA |
| HCN4 | 296 | I | 10.250125 | 40.88499693 | 0.745360105 | 39.55591959 | NA | NA | NA | NA | NA | NA | NA |
| HCN4 | 297 | V | 7.301517857 | 42.40151871 | 0.774781775 | 40.84876291 | NA | NA | NA | NA | NA | NA | NA |
| HCN4 | 298 | F | 4.933465909 | 35.61990048 | 0.643213248 | 34.19272794 | NA | NA | NA | NA | NA | NA | NA |
| HCN4 | 299 | N | 5.749625 | 36.89057573 | 0.66786531 | 35.45527426 | NA | NA | NA | NA | NA | NA | NA |
| HCN4 | 300 | V | 4.071803571 | 41.89627742 | 0.764979712 | 40.12857137 | NA | NA | NA | NA | NA | NA | NA |
| HCN4 | 301 | V | 1.358660714 | 41.96520809 | 0.766317019 | 39.97986931 | NA | NA | NA | NA | NA | NA | NA |
| HCN4 | 302 | S | 0.525041667 | 37.98842247 | 0.689164368 | 35.19544512 | NA | NA | NA | NA | NA | NA | NA |
| HCN4 | 303 | D | 0.93375 | 38.97857973 | 0.708374168 | 36.3815828 | NA | NA | NA | NA | NA | NA | NA |
| HCN4 | 304 | T | -0.566517857 | 42.94414271 | 0.785309092 | 40.71123015 | NA | NA | NA | NA | NA | NA | NA |
| HCN4 | 305 | F | -2.825193182 | 40.83995943 | 0.744486343 | 39.88340495 | NA | NA | NA | NA | NA | NA | NA |
| HCN4 | 306 | F | -2.546920455 | 35.25989173 | 0.636228806 | 34.32517013 | NA | NA | NA | NA | NA | NA | NA |
| HCN4 | 307 | L | -3.08475 | 41.91291603 | 0.765302513 | 40.9522487 | NA | NA | NA | NA | NA | NA | NA |
| HCN4 | 308 | I | -6.388375 | 42.91457181 | 0.784735394 | 41.81270305 | NA | NA | NA | NA | NA | NA | NA |
| HCN4 | 309 | D | -8.448375 | 37.35645252 | 0.676903672 | 35.92175217 | NA | NA | NA | NA | NA | NA | NA |

| | | | | | | | | | | | | | |
|---|---|---|---|---|---|---|---|---|---|---|---|---|---|
| HCN4 | 310 | L | -7.265375 | 40.60214185 | 0.739872502 | 39.17817316 | NA | NA | NA | NA | NA | NA | NA |
| HCN4 | 311 | V | -9.542232143 | 43.39407062 | 0.794038034 | 41.58987665 | NA | NA | NA | NA | NA | NA | NA |
| HCN4 | 312 | L | -12.19025 | 41.86754306 | 0.764422243 | 40.82950699 | NA | NA | NA | NA | NA | NA | NA |
| HCN4 | 313 | N | -12.39125 | 37.87111226 | 0.686888461 | 36.72828769 | NA | NA | NA | NA | NA | NA | NA |
| HCN4 | 314 | F | -12.95610227 | 42.521756 | 0.77711447 | 41.19752877 | NA | NA | NA | NA | NA | NA | NA |
| HCN4 | 315 | R | -16.81864773 | 42.78587726 | 0.782238622 | 41.91063626 | NA | NA | NA | NA | NA | NA | NA |
| HCN4 | 316 | T | -17.64851786 | 38.12773127 | 0.691867064 | 36.82853082 | NA | NA | NA | NA | NA | NA | NA |
| HCN4 | 317 | G | -20.658625 | 36.87472602 | 0.667557813 | 33.15594183 | NA | NA | NA | NA | NA | NA | NA |
| HCN4 | 318 | I | -20.859 | 35.08126413 | 0.632763295 | 31.32245209 | NA | NA | NA | NA | NA | NA | NA |
| HCN4 | 319 | V | -22.90808929 | 30.81707931 | 0.550034882 | 26.20649977 | NA | NA | NA | NA | NA | NA | NA |
| HCN4 | 320 | V | -24.39951786 | 30.8870621 | 0.551392601 | 26.0793234 | NA | NA | NA | NA | NA | NA | NA |
| HCN4 | 321 | E | -25.484375 | 27.45110187 | 0.484732372 | 21.98401105 | NA | NA | NA | NA | NA | NA | NA |
| HCN4 | 322 | D | -28.279375 | 30.2218936 | 0.538487829 | 24.38379583 | NA | NA | NA | NA | NA | NA | NA |
| HCN4 | 323 | N | -29.74425 | 31.44140819 | 0.562147335 | 24.57665451 | NA | NA | NA | NA | NA | NA | NA |
| HCN4 | 324 | T | -32.126375 | 30.22942067 | 0.538633859 | 23.34854438 | NA | NA | NA | NA | NA | NA | NA |
| HCN4 | 325 | E | -30.260375 | 34.01862851 | 0.61214736 | 27.04082621 | NA | NA | NA | NA | NA | NA | NA |
| HCN4 | 326 | I | -26.626 | 35.04584428 | 0.632076123 | 29.43638354 | NA | NA | NA | NA | NA | NA | NA |
| HCN4 | 327 | I | -24.612625 | 37.77460082 | 0.685016066 | 32.83834756 | NA | NA | NA | NA | NA | NA | NA |
| HCN4 | 328 | L | -25.04175 | 41.15217351 | 0.750543533 | 36.21651371 | NA | NA | NA | NA | NA | NA | NA |
| HCN4 | 329 | D | -24.32875 | 43.5523572 | 0.797108913 | 38.61283381 | NA | NA | NA | NA | NA | NA | NA |
| HCN4 | 330 | P | -20.96866071 | 45.61963842 | 0.837215734 | 43.30807249 | NA | NA | NA | NA | NA | NA | NA |
| HCN4 | 331 | Q | -20.29870833 | 44.91503955 | 0.823545982 | 42.59628499 | NA | NA | NA | NA | NA | NA | NA |
| HCN4 | 332 | R | -23.90228409 | 40.79444796 | 0.743603386 | 37.41013842 | NA | NA | NA | NA | NA | NA | NA |
| HCN4 | 333 | I | -19.09575 | 41.20047364 | 0.751480592 | 38.99012876 | NA | NA | NA | NA | NA | NA | NA |
| HCN4 | 334 | K | -15.78715278 | 42.86039247 | 0.783684274 | 41.80675554 | NA | NA | NA | NA | NA | NA | NA |
| HCN4 | 335 | M | -17.10925 | 41.08390543 | 0.74921908 | 39.98628301 | NA | NA | NA | NA | NA | NA | NA |
| HCN4 | 336 | K | -18.02248611 | 34.42538822 | 0.620038806 | 33.17893977 | NA | NA | NA | NA | NA | NA | NA |
| HCN4 | 337 | Y | -12.39820833 | 37.7572262 | 0.684678985 | 36.50947595 | NA | NA | NA | NA | NA | NA | NA |
| HCN4 | 338 | L | -12.04325 | 40.18328106 | 0.731746286 | 39.11371562 | NA | NA | NA | NA | NA | NA | NA |
| HCN4 | 339 | K | -13.476375 | 37.50445602 | 0.679775052 | 36.36317103 | NA | NA | NA | NA | NA | NA | NA |
| HCN4 | 340 | S | -12.249875 | 34.43422799 | 0.620210304 | 33.37362596 | NA | NA | NA | NA | NA | NA | NA |
| HCN4 | 341 | W | -10.35866071 | 29.56330391 | 0.52571069 | 27.9233612 | NA | NA | NA | NA | NA | NA | NA |
| HCN4 | 342 | F | -7.623193182 | 37.23692912 | 0.674584828 | 35.80780348 | NA | NA | NA | NA | NA | NA | NA |

| | | | | | | | | | | | | | |
|---|---|---|---|---|---|---|---|---|---|---|---|---|---|
| HCN4 | 343 | M | -4.309875 | 37.37352916 | 0.677234972 | 36.16587706 | NA | NA | NA | NA | NA | NA | NA |
| HCN4 | 344 | V | -4.017803571 | 32.69557215 | 0.586479065 | 31.73551038 | NA | NA | NA | NA | NA | NA | NA |
| HCN4 | 345 | D | -5.493875 | 34.53653916 | 0.622195218 | 33.3345644 | NA | NA | NA | NA | NA | NA | NA |
| HCN4 | 346 | F | -2.333193182 | 38.71062388 | 0.703175622 | 37.75780361 | NA | NA | NA | NA | NA | NA | NA |
| HCN4 | 347 | I | 0 | 36.57148884 | 0.661674783 | 34.28934473 | NA | NA | NA | NA | NA | NA | NA |
| HCN4 | 348 | S | -0.417708333 | 34.89596852 | 0.62916842 | 32.60260119 | NA | NA | NA | NA | NA | NA | NA |
| HCN4 | 349 | S | -0.917708333 | 38.13892733 | 0.692084276 | 35.88933456 | NA | NA | NA | NA | NA | NA | NA |
| HCN4 | 350 | I | 0.40375 | 39.91575452 | 0.726556068 | 37.13410826 | NA | NA | NA | NA | NA | NA | NA |
| HCN4 | 351 | P | 3.823375 | 41.09363723 | 0.749407885 | 39.31347138 | NA | NA | NA | NA | NA | NA | NA |
| HCN4 | 352 | V | 5.317089286 | 39.60929359 | 0.720610494 | 38.2301638 | NA | NA | NA | NA | NA | NA | NA |
| HCN4 | 353 | D | 8.7325 | 38.03550327 | 0.690077771 | 36.68646438 | NA | NA | NA | NA | NA | NA | NA |
| HCN4 | 354 | Y | 9.235791667 | 40.05329442 | 0.729224446 | 38.71056351 | NA | NA | NA | NA | NA | NA | NA |
| HCN4 | 355 | I | 6.87525 | 44.33446471 | 0.812282391 | 42.77875585 | NA | NA | NA | NA | NA | NA | NA |
| HCN4 | 356 | F | 11.49473864 | 41.46626511 | 0.756637147 | 40.06846086 | NA | NA | NA | NA | NA | NA | NA |
| HCN4 | 357 | L | 13.4555 | 41.8689705 | 0.764449937 | 40.41388309 | NA | NA | NA | NA | NA | NA | NA |
| HCN4 | 358 | I | 11.389625 | 46.61832409 | 0.856590992 | 45.13914371 | NA | NA | NA | NA | NA | NA | NA |
| HCN4 | 359 | V | 11.454375 | 48.07415407 | 0.884835195 | 46.66046989 | NA | NA | NA | NA | NA | NA | NA |
| HCN4 | 360 | E | 16.01870833 | 45.228276 | 0.829623006 | 43.10279633 | NA | NA | NA | NA | NA | NA | NA |
| HCN4 | 361 | T | 16.48380357 | 47.91098851 | 0.88166966 | 44.33146852 | NA | NA | NA | NA | NA | NA | NA |
| HCN4 | 362 | R | 13.37355682 | 51.00193987 | 0.941636456 | 48.82887138 | NA | NA | NA | NA | NA | NA | NA |
| HCN4 | 363 | I | 15.815375 | 49.80383982 | 0.918392408 | 46.30109462 | NA | NA | NA | NA | NA | NA | NA |
| HCN4 | 364 | D | 20.557125 | 49.6318543 | 0.915055759 | 45.2377115 | NA | NA | NA | NA | NA | NA | NA |
| HCN4 | 365 | S | 21.336175 | 50.05061832 | 0.923180098 | 45.66344215 | NA | NA | NA | NA | NA | NA | NA |
| HCN4 | 366 | E | 24.410575 | 49.38741738 | 0.910313497 | 45.10388453 | NA | NA | NA | NA | NA | NA | NA |
| HCN4 | 367 | V | 22.42880357 | 45.0815635 | 0.826776673 | 40.79854761 | NA | NA | NA | NA | NA | NA | NA |
| HCN4 | 368 | Y | 24.563575 | 44.87767225 | 0.822821028 | 40.60016877 | NA | NA | NA | NA | NA | NA | NA |
| HCN4 | 369 | K | 27.950575 | 44.63598091 | 0.818132033 | 40.3635759 | NA | NA | NA | NA | NA | NA | NA |
| HCN4 | 370 | T | 26.78923214 | 41.32029115 | 0.753805142 | 37.04573217 | NA | NA | NA | NA | NA | NA | NA |
| HCN4 | 371 | A | 25.708175 | 39.07818585 | 0.710306602 | 34.79823246 | NA | NA | NA | NA | NA | NA | NA |
| HCN4 | 372 | R | 21.07892045 | 40.32898066 | 0.734572968 | 36.04069811 | NA | NA | NA | NA | NA | NA | NA |
| HCN4 | 373 | A | 22.173775 | 39.06467517 | 0.710044485 | 34.7759268 | NA | NA | NA | NA | NA | NA | NA |
| HCN4 | 374 | L | 22.59475 | 34.51305824 | 0.621739671 | 30.23731919 | NA | NA | NA | NA | NA | NA | NA |
| HCN4 | 375 | R | 19.53710227 | 36.14349265 | 0.653371333 | 31.74855423 | NA | NA | NA | NA | NA | NA | NA |

| | | | | | | | | | | | | | |
|---|---|---|---|---|---|---|---|---|---|---|---|---|---|
| HCN4 | 376 | I | 17.422125 | 38.30181103 | 0.695244343 | 33.92287093 | NA | NA | NA | NA | NA | NA | NA |
| HCN4 | 377 | V | 18.02123214 | 35.34389049 | 0.637858445 | 30.96502981 | NA | NA | NA | NA | NA | NA | NA |
| HCN4 | 378 | R | 18.71864773 | 30.16513075 | 0.537386586 | 25.77313277 | NA | NA | NA | NA | NA | NA | NA |
| HCN4 | 379 | F | 12.56964773 | 35.43261403 | 0.639579749 | 33.26125556 | NA | NA | NA | NA | NA | NA | NA |
| HCN4 | 380 | T | 12.96894643 | 35.670537 | 0.644195635 | 34.20845588 | NA | NA | NA | NA | NA | NA | NA |
| HCN4 | 381 | K | 13.86581944 | 31.88497046 | 0.570752778 | 30.43425202 | NA | NA | NA | NA | NA | NA | NA |
| HCN4 | 382 | I | 10.66225 | 31.60660146 | 0.565352209 | 30.15798138 | NA | NA | NA | NA | NA | NA | NA |
| HCN4 | 383 | L | 8.249375 | 34.67192983 | 0.6248219 | 33.3555083 | NA | NA | NA | NA | NA | NA | NA |
| HCN4 | 384 | S | 7.949208333 | 33.43779749 | 0.600878798 | 31.89128385 | NA | NA | NA | NA | NA | NA | NA |
| HCN4 | 385 | L | 6.531125 | 28.78793904 | 0.510668024 | 27.48072557 | NA | NA | NA | NA | NA | NA | NA |
| HCN4 | 386 | L | 3.003625 | 33.04776823 | 0.593311935 | 31.07919202 | NA | NA | NA | NA | NA | NA | NA |
| HCN4 | 387 | R | 3.358375 | 31.92000891 | 0.571432551 | 29.9495996 | NA | NA | NA | NA | NA | NA | NA |
| HCN4 | 388 | L | 1.837625 | 27.56725612 | 0.486985852 | 25.09133747 | NA | NA | NA | NA | NA | NA | NA |
| HCN4 | 389 | L | -0.372 | 30.55105689 | 0.544873845 | 28.28147798 | NA | NA | NA | NA | NA | NA | NA |
| HCN4 | 390 | R | -1.072375 | 30.07675932 | 0.535672113 | 28.20024886 | NA | NA | NA | NA | NA | NA | NA |
| HCN4 | 391 | L | -3.0515 | 24.2261717 | 0.422166285 | 23.30736017 | NA | NA | NA | NA | NA | NA | NA |
| HCN4 | 392 | S | -6.147875 | 28.59978871 | 0.507017766 | 27.55670875 | NA | NA | NA | NA | NA | NA | NA |
| HCN4 | 393 | R | -6.233102273 | 30.1931598 | 0.537930371 | 29.09388779 | NA | NA | NA | NA | NA | NA | NA |
| HCN4 | 394 | L | -5.970625 | 26.32294894 | 0.462845351 | 25.35746651 | NA | NA | NA | NA | NA | NA | NA |
| HCN4 | 395 | I | -8.46475 | 23.72956748 | 0.412531787 | 21.93318133 | NA | NA | NA | NA | NA | NA | NA |
| HCN4 | 396 | R | -10.95801136 | 27.5514108 | 0.486678441 | 26.35071059 | NA | NA | NA | NA | NA | NA | NA |
| HCN4 | 397 | Y | -10.806875 | 28.75481848 | 0.510025461 | 27.51473483 | NA | NA | NA | NA | NA | NA | NA |
| HCN4 | 398 | I | -10.699375 | 24.24930816 | 0.42261515 | 22.85958701 | NA | NA | NA | NA | NA | NA | NA |
| HCN4 | 399 | H | -13.744475 | 21.75925185 | 0.374306172 | 20.51524645 | NA | NA | NA | NA | NA | NA | NA |
| HCN4 | 400 | Q | -16.08626389 | 27.33345739 | 0.48244998 | 26.47539661 | NA | NA | NA | NA | NA | NA | NA |
| HCN4 | 401 | W | -13.97708929 | 27.15209023 | 0.478931319 | 26.33658441 | NA | NA | NA | NA | NA | NA | NA |
| HCN4 | 402 | E | -15.05293056 | 22.1883731 | 0.38263145 | 21.50548481 | NA | NA | NA | NA | NA | NA | NA |
| HCN4 | 403 | E | -19.70570833 | 24.55372129 | 0.428520995 | 22.2355724 | NA | NA | NA | NA | NA | NA | NA |
| HCN4 | 404 | I | -20.080125 | 27.75335146 | 0.490596242 | 25.43649667 | NA | NA | NA | NA | NA | NA | NA |
| HCN4 | 405 | F | -17.46201136 | 26.80032033 | 0.472106717 | 25.54365571 | NA | NA | NA | NA | NA | NA | NA |
| HCN4 | 406 | H | -22.758175 | 25.85222014 | 0.453712856 | 22.10087539 | NA | NA | NA | NA | NA | NA | NA |
| HCN4 | 407 | M | -18.562875 | 24.3342162 | 0.42426243 | 22.0679205 | NA | NA | NA | NA | NA | NA | NA |
| HCN4 | 408 | T | -17.27494643 | 21.99781644 | 0.378934506 | 20.71595122 | NA | NA | NA | NA | NA | NA | NA |

| | | | | | | | | | | | | | |
|---|---|---|---|---|---|---|---|---|---|---|---|---|---|
| HCN4 | 409 | Y | -23.09029167 | 21.38565515 | 0.367058114 | 19.17338415 | NA | NA | NA | NA | NA | NA | NA |
| HCN4 | 410 | D | -22.8995 | 20.51378905 | 0.350143251 | 16.13680483 | NA | NA | NA | NA | NA | NA | NA |
| HCN4 | 411 | L | -18.482625 | 18.65887022 | 0.314156422 | 16.34269299 | NA | NA | NA | NA | NA | NA | NA |
| HCN4 | 412 | A | -18.841975 | 15.39297531 | 0.250795589 | 13.09193307 | NA | NA | NA | NA | NA | NA | NA |
| HCN4 | 413 | S | -18.16154167 | 13.84513366 | 0.220766289 | 12.65999757 | NA | NA | NA | NA | NA | NA | NA |
| HCN4 | 414 | A | -15.358775 | 11.84761774 | 0.182012968 | 10.82978525 | NA | NA | NA | NA | NA | NA | NA |
| HCN4 | 415 | V | -14.06880357 | 14.9345624 | 0.241902031 | 13.626584 | NA | NA | NA | NA | NA | NA | NA |
| HCN4 | 416 | V | -14.09908929 | 17.33121118 | 0.288398832 | 16.00149515 | NA | NA | NA | NA | NA | NA | NA |
| HCN4 | 417 | R | -13.75419318 | 13.43719993 | 0.212852066 | 12.48769073 | NA | NA | NA | NA | NA | NA | NA |
| HCN4 | 418 | I | -9.535875 | 11.7062989 | 0.179271276 | 9.900769888 | NA | NA | NA | NA | NA | NA | NA |
| HCN4 | 419 | V | -8.832946429 | 16.73806817 | 0.276891408 | 14.83289478 | NA | NA | NA | NA | NA | NA | NA |
| HCN4 | 420 | N | -9.1775 | 17.89275879 | 0.29929328 | 15.98456393 | NA | NA | NA | NA | NA | NA | NA |
| HCN4 | 421 | L | -6.103625 | 11.95158287 | 0.18402997 | 10.84147577 | NA | NA | NA | NA | NA | NA | NA |
| HCN4 | 422 | I | -4.193875 | 14.96685518 | 0.242528535 | 13.738578 | NA | NA | NA | NA | NA | NA | NA |
| HCN4 | 423 | G | -3.023375 | 18.44865136 | 0.310078017 | 17.51466123 | NA | NA | NA | NA | NA | NA | NA |
| HCN4 | 424 | M | -2.78875 | 17.76484559 | 0.296811668 | 16.87540697 | NA | NA | NA | NA | NA | NA | NA |
| HCN4 | 425 | M | -0.92475 | 11.46817592 | 0.174651509 | 9.22050404 | NA | NA | NA | NA | NA | NA | NA |
| HCN4 | 426 | L | 0.063125 | 17.31559901 | 0.288095944 | 14.7293778 | NA | NA | NA | NA | NA | NA | NA |
| HCN4 | 427 | L | 0.206125 | 19.64955972 | 0.333376548 | 17.03151101 | NA | NA | NA | NA | NA | NA | NA |
| HCN4 | 428 | L | 0.83875 | 17.36702847 | 0.289093714 | 14.75636336 | NA | NA | NA | NA | NA | NA | NA |
| HCN4 | 429 | C | 4.240708333 | 15.20895585 | 0.247225472 | 13.46891527 | NA | NA | NA | NA | NA | NA | NA |
| HCN4 | 430 | H | 5.504375 | 18.92677226 | 0.319353924 | 17.52299293 | NA | NA | NA | NA | NA | NA | NA |
| HCN4 | 431 | W | 3.099089286 | 20.59036073 | 0.3516288 | 19.15990402 | NA | NA | NA | NA | NA | NA | NA |
| HCN4 | 432 | D | 7.089375 | 15.65045322 | 0.255790855 | 14.32680761 | NA | NA | NA | NA | NA | NA | NA |
| HCN4 | 433 | G | 9.888625 | 18.30907002 | 0.307370033 | 16.98179529 | NA | NA | NA | NA | NA | NA | NA |
| HCN4 | 434 | C | 9.992875 | 20.77711217 | 0.355251919 | 19.33431721 | NA | NA | NA | NA | NA | NA | NA |
| HCN4 | 435 | L | 9.858375 | 20.69864616 | 0.353729619 | 19.22924236 | NA | NA | NA | NA | NA | NA | NA |
| HCN4 | 436 | Q | 12.37481944 | 15.43665705 | 0.251643047 | 13.96304895 | NA | NA | NA | NA | NA | NA | NA |
| HCN4 | 437 | F | 16.39682955 | 20.08658532 | 0.341855176 | 17.91776629 | NA | NA | NA | NA | NA | NA | NA |
| HCN4 | 438 | L | 13.55175 | 22.69704236 | 0.392500018 | 20.52105576 | NA | NA | NA | NA | NA | NA | NA |
| HCN4 | 439 | V | 14.93566071 | 22.62571221 | 0.391116159 | 20.44814011 | NA | NA | NA | NA | NA | NA | NA |
| HCN4 | 440 | P | 18.24694643 | 19.89261737 | 0.338092051 | 15.49467612 | NA | NA | NA | NA | NA | NA | NA |
| HCN4 | 441 | M | 18.3995 | 23.91971614 | 0.416220815 | 19.52436394 | NA | NA | NA | NA | NA | NA | NA |

| | | | | | | | | | | | | | |
|---|---|---|---|---|---|---|---|---|---|---|---|---|---|
| HCN4 | 442 | L | 17.24625 | 26.42857007 | 0.464894481 | 22.03214234 | NA | NA | NA | NA | NA | NA | NA |
| HCN4 | 443 | Q | 20.31559722 | 23.15010041 | 0.401289687 | 18.86358816 | NA | NA | NA | NA | NA | NA | NA |
| HCN4 | 444 | D | 24.45525 | 25.81227206 | 0.452937833 | 21.54310519 | NA | NA | NA | NA | NA | NA | NA |
| HCN4 | 445 | F | 23.04810227 | 22.42410353 | 0.387204798 | 18.14756963 | NA | NA | NA | NA | NA | NA | NA |
| HCN4 | 446 | P | 24.12908929 | 20.57875451 | 0.35140363 | 16.31963919 | NA | NA | NA | NA | NA | NA | NA |
| HCN4 | 447 | D | 27.93625 | 18.52776066 | 0.311612797 | 14.28425332 | NA | NA | NA | NA | NA | NA | NA |
| HCN4 | 448 | D | 23.513 | 16.14383043 | 0.265362746 | 11.8780904 | NA | NA | NA | NA | NA | NA | NA |
| HCN4 | 449 | C | 21.90570833 | 16.4210859 | 0.270741712 | 12.15862035 | NA | NA | NA | NA | NA | NA | NA |
| HCN4 | 450 | W | 18.614875 | 13.94639921 | 0.222730917 | 9.662871253 | NA | NA | NA | NA | NA | NA | NA |
| HCN4 | 451 | V | 21.93294643 | 16.7905734 | 0.277910049 | 12.51730964 | NA | NA | NA | NA | NA | NA | NA |
| HCN4 | 452 | S | 25.62570833 | 17.77220687 | 0.296954482 | 13.52598966 | NA | NA | NA | NA | NA | NA | NA |
| HCN4 | 453 | I | 24.971875 | 13.2056054 | 0.208358957 | 8.967763554 | NA | NA | NA | NA | NA | NA | NA |
| HCN4 | 454 | N | 23.29925 | 12.12784062 | 0.187449504 | 7.874113847 | NA | NA | NA | NA | NA | NA | NA |
| HCN4 | 455 | N | 27.403 | 17.26553996 | 0.28712476 | 13.01967778 | NA | NA | NA | NA | NA | NA | NA |
| HCN4 | 456 | M | 22.47875 | 15.48542315 | 0.252589147 | 11.20940454 | NA | NA | NA | NA | NA | NA | NA |
| HCN4 | 457 | V | 25.06780357 | 21.34454415 | 0.366260529 | 17.0828538 | NA | NA | NA | NA | NA | NA | NA |
| HCN4 | 458 | N | 27.0135 | 23.79067094 | 0.413717241 | 19.52972698 | NA | NA | NA | NA | NA | NA | NA |
| HCN4 | 459 | N | 23.643375 | 20.20948505 | 0.344239523 | 15.94244976 | NA | NA | NA | NA | NA | NA | NA |
| HCN4 | 460 | S | 20.79954167 | 23.95727213 | 0.41694943 | 19.66926967 | NA | NA | NA | NA | NA | NA | NA |
| HCN4 | 461 | W | 16.07973214 | 24.20047421 | 0.421667734 | 19.92404163 | NA | NA | NA | NA | NA | NA | NA |
| HCN4 | 462 | G | 16.425625 | 22.28212256 | 0.38445026 | 18.70958865 | NA | NA | NA | NA | NA | NA | NA |
| HCN4 | 463 | K | 19.70315278 | 19.3571648 | 0.327703865 | 15.05843965 | NA | NA | NA | NA | NA | NA | NA |
| HCN4 | 464 | Q | 18.50259722 | 19.54752816 | 0.331397059 | 15.30160934 | NA | NA | NA | NA | NA | NA | NA |
| HCN4 | 465 | Y | 11.577625 | 19.49835267 | 0.330443017 | 17.41069262 | NA | NA | NA | NA | NA | NA | NA |
| HCN4 | 466 | S | 13.840875 | 16.87260742 | 0.279501571 | 14.75047158 | NA | NA | NA | NA | NA | NA | NA |
| HCN4 | 467 | Y | 16.63254167 | 10.63307712 | 0.158449961 | 7.196992912 | NA | NA | NA | NA | NA | NA | NA |
| HCN4 | 468 | A | 13.055375 | 15.8694516 | 0.260039589 | 13.95722249 | NA | NA | NA | NA | NA | NA | NA |
| HCN4 | 469 | L | 9.33075 | 14.70319986 | 0.237413423 | 13.31528156 | NA | NA | NA | NA | NA | NA | NA |
| HCN4 | 470 | F | 12.14410227 | 9.816031021 | 0.142598648 | 8.537496372 | NA | NA | NA | NA | NA | NA | NA |
| HCN4 | 471 | K | 12.99493056 | 11.51383885 | 0.175537405 | 10.16597474 | NA | NA | NA | NA | NA | NA | NA |
| HCN4 | 472 | A | 8.348775 | 11.93152199 | 0.183640774 | 10.79353546 | NA | NA | NA | NA | NA | NA | NA |
| HCN4 | 473 | M | 6.493375 | 9.726191958 | 0.140855702 | 7.963438729 | NA | NA | NA | NA | NA | NA | NA |
| HCN4 | 474 | S | 8.735208333 | 6.703790197 | 0.08221882 | 5.565856527 | 1.35 | -0.8 | -0.4 | 3.38 | 0.81 | 0 | 0 |

| | | | | | | | | | | | | | |
|---|---|---|---|---|---|---|---|---|---|---|---|---|---|
| HCN4 | 475 | H | 7.982975 | 8.576609178 | 0.118552926 | 7.295259312 | NA | NA | NA | NA | NA | NA | NA |
| HCN4 | 476 | M | 2.53625 | 8.101954024 | 0.109344257 | 6.568437027 | NA | NA | NA | NA | NA | NA | NA |
| HCN4 | 477 | L | 2.632125 | 5.62984165 | 0.061383405 | 3.636804792 | 2.04 | 0.77 | 2.5 | 1.48 | 0.16 | 0 | 0 |
| HCN4 | 478 | C | 3.921541667 | 2.74898163 | 0.00549254 | 1.294969873 | 1.78 | -0.41 | 0.33 | 2.91 | 0.65 | 0 | 0 |
| HCN4 | 479 | I | 7.817125 | 2.465872057 | 0 | 1.662249742 | 1.37 | -0.8 | -0.4 | 3.38 | 0.81 | 0 | 0 |
| HCN4 | 480 | G | 11.015375 | 3.184602644 | 0.013943917 | 1.903737618 | 1.48 | -0.8 | -0.4 | 3.38 | 0.81 | 0 | 0 |
| HCN4 | 481 | Y | 13.26645833 | 4.852921388 | 0.046310564 | 3.441371267 | 1.48 | -0.8 | -0.4 | 3.38 | 0.81 | 0 | 0 |
| HCN4 | 482 | G | 16.132375 | 6.725984538 | 0.082649407 | 3.176272152 | 3.58 | -0.42 | -0.58 | 3.03 | 0.36 | 0 | 0 |
| HCN4 | 483 | R | 21.77173864 | 5.754057872 | 0.063793294 | 1.502862815 | 4.3 | -0.42 | -0.58 | 3.03 | 0.36 | 0 | 0 |
| HCN4 | 484 | Q | 19.270575 | 6.631487918 | 0.080816101 | 2.372244856 | 4.3 | -0.42 | -0.58 | 3.03 | 0.36 | 0 | 0 |
| HCN4 | 485 | A | 18.386775 | 7.063269286 | 0.089192986 | 2.689882428 | 4.4 | -0.42 | -0.58 | 3.03 | 0.36 | 0 | 0 |
| HCN4 | 486 | P | 17.01408929 | 9.997556001 | 0.14612037 | 5.70101148 | NA | NA | NA | NA | NA | NA | NA |
| HCN4 | 487 | V | 21.05994643 | 12.64286854 | 0.197441436 | 8.385753507 | NA | NA | NA | NA | NA | NA | NA |
| HCN4 | 488 | G | 19.300625 | 16.11419613 | 0.264787818 | 11.83735909 | NA | NA | NA | NA | NA | NA | NA |
| HCN4 | 489 | M | 17.339625 | 18.51126646 | 0.311292797 | 14.11878587 | NA | NA | NA | NA | NA | NA | NA |
| HCN4 | 490 | S | 16.06054167 | 18.43732085 | 0.309858196 | 14.97057035 | NA | NA | NA | NA | NA | NA | NA |
| HCN4 | 491 | D | 16.524375 | 15.0393064 | 0.243934144 | 11.64613587 | NA | NA | NA | NA | NA | NA | NA |
| HCN4 | 492 | V | 14.06723214 | 15.20337883 | 0.247117273 | 13.11075443 | NA | NA | NA | NA | NA | NA | NA |
| HCN4 | 493 | W | 11.04944643 | 17.40877371 | 0.289903603 | 16.04898477 | NA | NA | NA | NA | NA | NA | NA |
| HCN4 | 494 | L | 11.0655 | 15.7049752 | 0.256848623 | 14.3067162 | NA | NA | NA | NA | NA | NA | NA |
| HCN4 | 495 | T | 10.79180357 | 11.2732714 | 0.170870214 | 9.812015836 | NA | NA | NA | NA | NA | NA | NA |
| HCN4 | 496 | M | 8.297625 | 14.19964806 | 0.227644137 | 12.85378401 | NA | NA | NA | NA | NA | NA | NA |
| HCN4 | 497 | L | 5.659625 | 15.23435765 | 0.247718286 | 14.05031688 | NA | NA | NA | NA | NA | NA | NA |
| HCN4 | 498 | S | 6.179708333 | 12.21708001 | 0.189180816 | 10.82131451 | NA | NA | NA | NA | NA | NA | NA |
| HCN4 | 499 | M | 5.80075 | 9.531698799 | 0.137082387 | 8.288140662 | NA | NA | NA | NA | NA | NA | NA |
| HCN4 | 500 | I | 1.8665 | 12.39381225 | 0.192609555 | 10.50077687 | NA | NA | NA | NA | NA | NA | NA |
| HCN4 | 501 | V | 0.962517857 | 12.58967811 | 0.196409501 | 9.900244407 | NA | NA | NA | NA | NA | NA | NA |
| HCN4 | 502 | G | 0.902125 | 9.224557117 | 0.131123606 | 7.426033846 | NA | NA | NA | NA | NA | NA | NA |
| HCN4 | 503 | A | 0.380975 | 8.068018654 | 0.108685885 | 5.760753775 | NA | NA | NA | NA | NA | NA | NA |
| HCN4 | 504 | T | -1.770089286 | 10.10613052 | 0.148226798 | 8.352631776 | NA | NA | NA | NA | NA | NA | NA |
| HCN4 | 505 | C | -1.739541667 | 9.840663545 | 0.143076537 | 8.015331245 | NA | NA | NA | NA | NA | NA | NA |
| HCN4 | 506 | Y | -0.818541667 | 3.975620077 | 0.029290254 | 2.190239825 | 1.92 | 1.67 | 3.34 | 0.43 | -1.77 | 0 | 0 |
| HCN4 | 507 | A | -4.361775 | 6.145417642 | 0.07138597 | 5.389554896 | 0.96 | 1.81 | 4.5 | 0.13 | -1.85 | 0 | 0 |

| | | | | | | | | | | | | |
|---|---|---|---|---|---|---|---|---|---|---|---|---|
| HCN4 | 508 | M | -6.567875 | 8.166044146 | 0.110587653 | 7.167862629 | NA | NA | NA | NA | NA | NA | NA |
| HCN4 | 509 | F | -4.973556818 | 7.40653583 | 0.095852617 | 6.26740394 | 1.22 | 1.81 | 4.5 | 0.13 | -1.85 | 0 | 0 |
| HCN4 | 510 | I | -6.028625 | 2.619016037 | 0.002971109 | 1.523395534 | 1.11 | 1.81 | 4.5 | 0.13 | -1.85 | 0 | 0 |
| HCN4 | 511 | G | -9.930625 | 5.39992324 | 0.056922814 | 3.672729143 | 1.82 | -0.77 | -0.7 | 1.66 | 0.77 | 0 | 0 |
| HCN4 | 512 | H | -10.569075 | 7.509461832 | 0.097849459 | 5.882181483 | 1.64 | -0.77 | -0.7 | 1.66 | 0.77 | 0 | 0 |
| HCN4 | 513 | A | -10.053375 | 7.096300445 | 0.089833816 | 5.385242719 | 1.82 | -0.77 | -0.7 | 1.66 | 0.77 | 0 | 0 |
| HCN4 | 514 | T | -12.15223214 | 2.676617455 | 0.00408862 | 1.63892713 | 1.08 | -0.77 | -0.7 | 1.66 | 0.77 | 0 | 0 |
| HCN4 | 515 | A | -14.967975 | 5.799147524 | 0.064668067 | 4.881647852 | 1.07 | -1.1 | -3.5 | 3.53 | 0.13 | 0 | 0 |
| HCN4 | 516 | L | -14.13425 | 8.341069596 | 0.11398328 | 7.185477002 | NA | NA | NA | NA | NA | NA | NA |
| HCN4 | 517 | I | -13.68575 | 8.467375686 | 0.116433713 | 7.390733494 | NA | NA | NA | NA | NA | NA | NA |
| HCN4 | 518 | Q | -17.25148611 | 2.521074374 | 0.001070967 | 1.293262722 | 1.3 | -1.1 | -3.5 | 3.53 | 0.13 | 0 | 0 |
| HCN4 | 519 | S | -19.19620833 | 8.542574202 | 0.117892622 | 6.372432703 | NA | NA | NA | NA | NA | NA | NA |
| HCN4 | 520 | L | -17.846125 | 11.20802115 | 0.16960431 | 9.987631983 | NA | NA | NA | NA | NA | NA | NA |
| HCN4 | 521 | D | -21.340775 | 12.77197127 | 0.199946126 | 9.037177969 | NA | NA | NA | NA | NA | NA | NA |
| HCN4 | 522 | S | -24.53504167 | 11.02547981 | 0.16606287 | 6.364779597 | NA | NA | NA | NA | NA | NA | NA |
| HCN4 | 523 | S | -27.002375 | 12.93030379 | 0.203017897 | 7.137239267 | NA | NA | NA | NA | NA | NA | NA |
| HCN4 | 524 | R | -23.89319318 | 13.8999082 | 0.221828956 | 8.9594431 | NA | NA | NA | NA | NA | NA | NA |
| HCN4 | 525 | R | -23.28082955 | 6.330625956 | 0.074979151 | 1.505836198 | 4.84 | -0.42 | -4.5 | 52 | 1.63 | 4 | 4 |
| HCN4 | 526 | Q | -28.82126389 | 8.55984147 | 0.11822762 | 3.626766615 | 5.92 | -1 | -3.7 | 50.32 | 2.11 | 5 | -3 |
| HCN4 | 527 | Y | -28.684875 | 12.80357231 | 0.20055921 | 6.946048151 | NA | NA | NA | NA | NA | NA | NA |
| HCN4 | 528 | Q | -24.27893056 | 13.51912338 | 0.214441443 | 8.015952554 | NA | NA | NA | NA | NA | NA | NA |
| HCN4 | 529 | E | -26.05281944 | 7.207027959 | 0.091982013 | 1.842572237 | 5.51 | -0.56 | -4.3 | 51.58 | 1.75 | 5 | 3 |
| HCN4 | 530 | K | -30.83708929 | 11.35270576 | 0.172411301 | 4.766455606 | 6.82 | -1.13 | -3.5 | 40.63 | 1.81 | 4 | -4 |
| HCN4 | 531 | Y | -28.868875 | 14.71575092 | 0.237656923 | 7.853318995 | NA | NA | NA | NA | NA | NA | NA |
| HCN4 | 532 | K | -25.14904167 | 13.56083699 | 0.215250718 | 8.890859208 | NA | NA | NA | NA | NA | NA | NA |
| HCN4 | 533 | Q | -29.99815278 | 9.75587669 | 0.141431608 | 2.889342457 | 6.87 | -1.13 | -3.5 | 40.63 | 1.81 | 4 | -4 |
| HCN4 | 534 | V | -31.87008929 | 16.09518496 | 0.264418987 | 9.039303778 | NA | NA | NA | NA | NA | NA | NA |
| HCN4 | 535 | E | -27.46604167 | 18.64419719 | 0.313871754 | 12.85485397 | NA | NA | NA | NA | NA | NA | NA |
| HCN4 | 536 | Q | -27.53426389 | 15.7196742 | 0.257133794 | 9.965553458 | NA | NA | NA | NA | NA | NA | NA |
| HCN4 | 537 | Y | -34.81204167 | 18.18118847 | 0.304889034 | 11.28926193 | NA | NA | NA | NA | NA | NA | NA |
| HCN4 | 538 | M | -31.940625 | 21.154626 | 0.362575973 | 14.0345237 | NA | NA | NA | NA | NA | NA | NA |
| HCN4 | 539 | S | -29.15304167 | 22.05071895 | 0.379960855 | 15.26913257 | NA | NA | NA | NA | NA | NA | NA |
| HCN4 | 540 | F | -31.97619318 | 21.552583 | 0.37029664 | 14.73518372 | NA | NA | NA | NA | NA | NA | NA |

| | | | | | | | | | | | | | |
|---|---|---|---|---|---|---|---|---|---|---|---|---|---|
| HCN4 | 541 | H | -35.385175 | 20.96451798 | 0.358887734 | 13.63790503 | NA | NA | NA | NA | NA | NA | NA |
| HCN4 | 542 | K | -31.43504167 | 26.14500346 | 0.459393074 | 19.03076136 | NA | NA | NA | NA | NA | NA | NA |
| HCN4 | 543 | L | -34.42 | 24.28787562 | 0.423363388 | 16.87671052 | NA | NA | NA | NA | NA | NA | NA |
| HCN4 | 544 | P | -34.40551786 | 29.50815955 | 0.524640848 | 22.08034759 | NA | NA | NA | NA | NA | NA | NA |
| HCN4 | 545 | P | -31.16851786 | 30.56979179 | 0.545237317 | 23.75656351 | NA | NA | NA | NA | NA | NA | NA |
| HCN4 | 546 | D | -34.40975 | 30.2200248 | 0.538451572 | 22.78091267 | NA | NA | NA | NA | NA | NA | NA |
| HCN4 | 547 | T | -35.81108929 | 26.65369342 | 0.469262044 | 18.63505045 | NA | NA | NA | NA | NA | NA | NA |
| HCN4 | 548 | R | -31.05173864 | 26.13302964 | 0.459160772 | 19.01774991 | NA | NA | NA | NA | NA | NA | NA |
| HCN4 | 549 | Q | -31.14459722 | 27.96096967 | 0.494624193 | 20.84234294 | NA | NA | NA | NA | NA | NA | NA |
| HCN4 | 550 | R | -37.55664773 | 26.12011886 | 0.458910293 | 18.09302167 | NA | NA | NA | NA | NA | NA | NA |
| HCN4 | 551 | I | -34.70175 | 22.052898 | 0.38000313 | 14.61216145 | NA | NA | NA | NA | NA | NA | NA |
| HCN4 | 552 | H | -29.580675 | 24.21570668 | 0.421963256 | 17.51619541 | NA | NA | NA | NA | NA | NA | NA |
| HCN4 | 553 | D | -32.8465 | 25.72629272 | 0.451269768 | 18.6581134 | NA | NA | NA | NA | NA | NA | NA |
| HCN4 | 554 | Y | -37.45279167 | 20.18122268 | 0.343691212 | 12.56105005 | NA | NA | NA | NA | NA | NA | NA |
| HCN4 | 555 | Y | -33.75695833 | 16.84937135 | 0.279050774 | 9.463561419 | NA | NA | NA | NA | NA | NA | NA |
| HCN4 | 556 | E | -29.89193056 | 22.99862328 | 0.398350917 | 16.20118519 | NA | NA | NA | NA | NA | NA | NA |
| HCN4 | 557 | H | -34.544775 | 24.04582145 | 0.418667354 | 16.61792664 | NA | NA | NA | NA | NA | NA | NA |
| HCN4 | 558 | R | -37.86355682 | 17.53423999 | 0.292337744 | 10.01670068 | NA | NA | NA | NA | NA | NA | NA |
| HCN4 | 559 | Y | -34.79379167 | 15.45155866 | 0.25193215 | 8.369185701 | NA | NA | NA | NA | NA | NA | NA |
| HCN4 | 560 | Q | -31.68959722 | 21.46644111 | 0.368625423 | 14.67498474 | NA | NA | NA | NA | NA | NA | NA |
| HCN4 | 561 | G | -28.254125 | 17.89353679 | 0.299308374 | 12.06209921 | NA | NA | NA | NA | NA | NA | NA |
| HCN4 | 562 | K | -28.68326389 | 16.73638963 | 0.276858843 | 9.897958101 | NA | NA | NA | NA | NA | NA | NA |
| HCN4 | 563 | M | -32.126 | 12.67984235 | 0.198158755 | 5.586729081 | NA | NA | NA | NA | NA | NA | NA |
| HCN4 | 564 | F | -33.11501136 | 12.75916349 | 0.199697646 | 5.333558342 | NA | NA | NA | NA | NA | NA | NA |
| HCN4 | 565 | D | -36.229975 | 9.635518979 | 0.139096578 | 1.621803151 | 8.02 | -1.05 | -3.5 | 40.47 | 2.13 | 4 | -4 |
| HCN4 | 566 | E | -39.24670833 | 12.80683997 | 0.200622605 | 4.923475813 | 7.9 | -0.97 | -0.8 | 1.67 | 1.11 | 0 | 0 |
| HCN4 | 567 | E | -41.462875 | 9.724523125 | 0.140823325 | 1.64125117 | 8.22 | -1.11 | -2.96 | 40.25 | 2.01 | 4 | -4 |
| HCN4 | 568 | S | -38.63004167 | 9.458697743 | 0.135666112 | 1.591214833 | 7.94 | -1 | -1.34 | 11.32 | 1.33 | 1 | -1 |
| HCN4 | 569 | I | -36.936 | 12.64072154 | 0.197399782 | 4.750956721 | 7.93 | -0.98 | -1.34 | 11.28 | 1.41 | 1 | -1 |
| HCN4 | 570 | L | -41.262 | 14.92435463 | 0.241703993 | 6.855608818 | NA | NA | NA | NA | NA | NA | NA |
| HCN4 | 571 | G | -41.853375 | 13.50164864 | 0.214102419 | 5.325101106 | NA | NA | NA | NA | NA | NA | NA |
| HCN4 | 572 | E | -38.02704167 | 11.8842834 | 0.18272431 | 4.495116136 | 7.94 | -1 | -1.34 | 11.32 | 1.33 | 1 | -1 |
| HCN4 | 573 | L | -40.0415 | 16.98733146 | 0.281727305 | 8.969296814 | NA | NA | NA | NA | NA | NA | NA |

| | | | | | | | | | | | | | |
|---|---|---|---|---|---|---|---|---|---|---|---|---|---|
| HCN4 | 574 | S | -43.06320833 | 20.06365774 | 0.341410363 | 12.00351949 | NA | NA | NA | NA | NA | NA | NA |
| HCN4 | 575 | E | -47.56204167 | 19.29604234 | 0.326518043 | 10.5608761 | NA | NA | NA | NA | NA | NA | NA |
| HCN4 | 576 | P | -45.268375 | 23.63342775 | 0.410666604 | 15.55035383 | NA | NA | NA | NA | NA | NA | NA |
| HCN4 | 577 | L | -41.291 | 22.80570501 | 0.394608156 | 14.63195474 | NA | NA | NA | NA | NA | NA | NA |
| HCN4 | 578 | R | -44.83819318 | 16.84683116 | 0.279001493 | 8.717755312 | NA | NA | NA | NA | NA | NA | NA |
| HCN4 | 579 | E | -46.379375 | 22.82436967 | 0.394970265 | 14.14111926 | NA | NA | NA | NA | NA | NA | NA |
| HCN4 | 580 | E | -42.26093056 | 25.2960726 | 0.442923172 | 17.14003278 | NA | NA | NA | NA | NA | NA | NA |
| HCN4 | 581 | I | -40.651375 | 22.09123471 | 0.380746891 | 14.0770139 | NA | NA | NA | NA | NA | NA | NA |
| HCN4 | 582 | I | -44.698 | 19.98313892 | 0.339848237 | 11.8812894 | NA | NA | NA | NA | NA | NA | NA |
| HCN4 | 583 | N | -45.87675 | 25.76090255 | 0.451941225 | 17.62259945 | NA | NA | NA | NA | NA | NA | NA |
| HCN4 | 584 | F | -39.70764773 | 27.55248602 | 0.486699301 | 19.38885956 | NA | NA | NA | NA | NA | NA | NA |
| HCN4 | 585 | N | -40.72675 | 23.08778718 | 0.400080764 | 14.92150721 | NA | NA | NA | NA | NA | NA | NA |
| HCN4 | 586 | C | -44.92004167 | 24.90244968 | 0.43528659 | 16.78162984 | NA | NA | NA | NA | NA | NA | NA |
| HCN4 | 587 | R | -41.80819318 | 29.73671767 | 0.529075048 | 21.57701533 | NA | NA | NA | NA | NA | NA | NA |
| HCN4 | 588 | K | -42.59981944 | 31.70195226 | 0.567202087 | 23.54213451 | NA | NA | NA | NA | NA | NA | NA |
| HCN4 | 589 | L | -47.833625 | 27.60182249 | 0.487656466 | 18.85592056 | NA | NA | NA | NA | NA | NA | NA |
| HCN4 | 590 | V | -49.09694643 | 29.81980644 | 0.530687033 | 19.12233949 | NA | NA | NA | NA | NA | NA | NA |
| HCN4 | 591 | A | -47.893175 | 33.91893679 | 0.610213265 | 25.14108824 | NA | NA | NA | NA | NA | NA | NA |
| HCN4 | 592 | S | -48.79104167 | 33.49851986 | 0.602056858 | 24.72102567 | NA | NA | NA | NA | NA | NA | NA |
| HCN4 | 593 | M | -52.2365 | 29.83313327 | 0.530945584 | 19.97338856 | NA | NA | NA | NA | NA | NA | NA |
| HCN4 | 594 | P | -55.067375 | 35.14285092 | 0.633958126 | 25.44108544 | NA | NA | NA | NA | NA | NA | NA |
| HCN4 | 595 | L | -57.11475 | 29.3975195 | 0.522494347 | 20.19772117 | NA | NA | NA | NA | NA | NA | NA |
| HCN4 | 596 | F | -53.52955682 | 27.0652164 | 0.477245901 | 17.22896037 | NA | NA | NA | NA | NA | NA | NA |
| HCN4 | 597 | A | -53.598575 | 34.14704857 | 0.614638806 | 24.30744897 | NA | NA | NA | NA | NA | NA | NA |
| HCN4 | 598 | N | -57.125125 | 34.82162964 | 0.62772619 | 25.64157784 | NA | NA | NA | NA | NA | NA | NA |
| HCN4 | 599 | A | -57.044175 | 31.64379942 | 0.566073878 | 22.44463234 | NA | NA | NA | NA | NA | NA | NA |
| HCN4 | 600 | D | -57.50825 | 29.89507133 | 0.532147229 | 20.75343929 | NA | NA | NA | NA | NA | NA | NA |
| HCN4 | 601 | P | -53.83380357 | 28.57735805 | 0.506582594 | 18.72258094 | NA | NA | NA | NA | NA | NA | NA |
| HCN4 | 602 | N | -55.321 | 25.75890209 | 0.451902415 | 16.02589079 | NA | NA | NA | NA | NA | NA | NA |
| HCN4 | 603 | F | -57.49328409 | 25.13536206 | 0.439805266 | 15.98735741 | NA | NA | NA | NA | NA | NA | NA |
| HCN4 | 604 | V | -52.902375 | 25.52070943 | 0.447281297 | 15.66317132 | NA | NA | NA | NA | NA | NA | NA |
| HCN4 | 605 | T | -51.19008929 | 22.90160453 | 0.396468679 | 11.88410713 | NA | NA | NA | NA | NA | NA | NA |
| HCN4 | 606 | S | -54.33520833 | 20.68973789 | 0.353556792 | 10.88675044 | NA | NA | NA | NA | NA | NA | NA |

| | | | | | | | | | | | | | |
|---|---|---|---|---|---|---|---|---|---|---|---|---|---|
| HCN4 | 607 | M | -54.200625 | 21.33191304 | 0.366015476 | 11.47270517 | NA | NA | NA | NA | NA | NA | NA |
| HCN4 | 608 | L | -49.45225 | 21.02753407 | 0.360110294 | 10.38036965 | NA | NA | NA | NA | NA | NA | NA |
| HCN4 | 609 | T | -49.86808929 | 17.42909992 | 0.290297947 | 6.731301228 | NA | NA | NA | NA | NA | NA | NA |
| HCN4 | 610 | K | -52.44526389 | 16.15333396 | 0.265547122 | 5.509736098 | NA | NA | NA | NA | NA | NA | NA |
| HCN4 | 611 | L | -49.58625 | 18.10448083 | 0.303400848 | 7.574847912 | NA | NA | NA | NA | NA | NA | NA |
| HCN4 | 612 | R | -47.34373864 | 14.46061676 | 0.232707127 | 5.680643349 | NA | NA | NA | NA | NA | NA | NA |
| HCN4 | 613 | F | -42.38319318 | 19.23452295 | 0.325324521 | 11.07534865 | NA | NA | NA | NA | NA | NA | NA |
| HCN4 | 614 | E | -45.45226389 | 18.93548484 | 0.319522955 | 11.1805564 | NA | NA | NA | NA | NA | NA | NA |
| HCN4 | 615 | V | -41.27294643 | 22.91462356 | 0.396721258 | 14.74583296 | NA | NA | NA | NA | NA | NA | NA |
| HCN4 | 616 | F | -44.70873864 | 23.56869409 | 0.409410722 | 15.51973819 | NA | NA | NA | NA | NA | NA | NA |
| HCN4 | 617 | Q | -39.93226389 | 26.89919006 | 0.474024865 | 19.07200764 | NA | NA | NA | NA | NA | NA | NA |
| HCN4 | 618 | P | -41.23423214 | 31.40667875 | 0.561473557 | 23.23434436 | NA | NA | NA | NA | NA | NA | NA |
| HCN4 | 619 | G | -44.780125 | 32.40193786 | 0.580782337 | 24.26684433 | NA | NA | NA | NA | NA | NA | NA |
| HCN4 | 620 | D | -45.042375 | 27.85602831 | 0.492588251 | 19.71343118 | NA | NA | NA | NA | NA | NA | NA |
| HCN4 | 621 | Y | -49.89795833 | 27.17616296 | 0.479398348 | 16.45494237 | NA | NA | NA | NA | NA | NA | NA |
| HCN4 | 622 | I | -50.339375 | 25.24473527 | 0.441927189 | 14.55777652 | NA | NA | NA | NA | NA | NA | NA |
| HCN4 | 623 | I | -54.579875 | 25.97346096 | 0.456065019 | 16.20987942 | NA | NA | NA | NA | NA | NA | NA |
| HCN4 | 624 | R | -53.82692045 | 27.40250458 | 0.483789547 | 18.27897082 | NA | NA | NA | NA | NA | NA | NA |
| HCN4 | 625 | E | -60.06215278 | 29.46128383 | 0.523731423 | 20.34289638 | NA | NA | NA | NA | NA | NA | NA |
| HCN4 | 626 | G | -61.329875 | 26.77020444 | 0.471522446 | 16.81180312 | NA | NA | NA | NA | NA | NA | NA |
| HCN4 | 627 | T | -59.71251786 | 23.51753442 | 0.408418185 | 14.53682035 | NA | NA | NA | NA | NA | NA | NA |
| HCN4 | 628 | I | -61.557375 | 18.8042183 | 0.316976285 | 8.827952197 | NA | NA | NA | NA | NA | NA | NA |
| HCN4 | 629 | G | -59.861625 | 18.85333291 | 0.317929145 | 9.840828901 | NA | NA | NA | NA | NA | NA | NA |
| HCN4 | 630 | K | -57.69648611 | 14.61756043 | 0.235751953 | 5.423467244 | NA | NA | NA | NA | NA | NA | NA |
| HCN4 | 631 | K | -53.97370833 | 18.10302433 | 0.303372591 | 9.375982269 | NA | NA | NA | NA | NA | NA | NA |
| HCN4 | 632 | M | -55.902125 | 21.154214 | 0.36256798 | 12.07345097 | NA | NA | NA | NA | NA | NA | NA |
| HCN4 | 633 | Y | -52.137875 | 21.91863137 | 0.377398256 | 12.08959027 | NA | NA | NA | NA | NA | NA | NA |
| HCN4 | 634 | F | -52.23428409 | 24.41282751 | 0.425787549 | 13.62817006 | NA | NA | NA | NA | NA | NA | NA |
| HCN4 | 635 | I | -48.8345 | 28.7918538 | 0.510743974 | 20.10148431 | NA | NA | NA | NA | NA | NA | NA |
| HCN4 | 636 | Q | -44.08870833 | 26.59709437 | 0.46816398 | 18.45601883 | NA | NA | NA | NA | NA | NA | NA |
| HCN4 | 637 | H | -41.820475 | 30.54484449 | 0.54475332 | 22.38755938 | NA | NA | NA | NA | NA | NA | NA |
| HCN4 | 638 | G | -44.606875 | 33.01335194 | 0.592644233 | 24.85410653 | NA | NA | NA | NA | NA | NA | NA |
| HCN4 | 639 | V | -46.23408929 | 34.3174891 | 0.617945481 | 25.55986499 | NA | NA | NA | NA | NA | NA | NA |

| | | | | | | | | | | | | | |
|---|---|---|---|---|---|---|---|---|---|---|---|---|---|
| HCN4 | 640 | V | -49.61966071 | 31.70251995 | 0.5672131 | 21.01254399 | NA | NA | NA | NA | NA | NA | NA |
| HCN4 | 641 | S | -51.16704167 | 34.63064282 | 0.624020901 | 23.73172999 | NA | NA | NA | NA | NA | NA | NA |
| HCN4 | 642 | V | -55.49894643 | 34.06867786 | 0.613118355 | 24.34847943 | NA | NA | NA | NA | NA | NA | NA |
| HCN4 | 643 | L | -55.211375 | 34.8433755 | 0.628148076 | 25.64206385 | NA | NA | NA | NA | NA | NA | NA |
| HCN4 | 644 | T | -60.09466071 | 36.07023945 | 0.651950165 | 26.95740164 | NA | NA | NA | NA | NA | NA | NA |
| HCN4 | 645 | K | -61.64081944 | 37.25621449 | 0.674958978 | 27.32896825 | NA | NA | NA | NA | NA | NA | NA |
| HCN4 | 646 | G | -62.503875 | 40.46541133 | 0.737219826 | 30.49306912 | NA | NA | NA | NA | NA | NA | NA |
| HCN4 | 647 | N | -61.971 | 39.19714298 | 0.71261446 | 29.22187551 | NA | NA | NA | NA | NA | NA | NA |
| HCN4 | 648 | K | -58.431575 | 40.64269351 | 0.740659235 | 31.44746134 | NA | NA | NA | NA | NA | NA | NA |
| HCN4 | 649 | E | -53.87893056 | 40.26139681 | 0.73326179 | 31.14874788 | NA | NA | NA | NA | NA | NA | NA |
| HCN4 | 650 | T | -55.46608929 | 37.58378487 | 0.681314092 | 28.48243691 | NA | NA | NA | NA | NA | NA | NA |
| HCN4 | 651 | K | -50.87070833 | 38.03484914 | 0.69006508 | 28.17505563 | NA | NA | NA | NA | NA | NA | NA |
| HCN4 | 652 | L | -52.36175 | 33.62444199 | 0.604499843 | 22.8681665 | NA | NA | NA | NA | NA | NA | NA |
| HCN4 | 653 | A | -49.321175 | 34.3248037 | 0.61808739 | 23.59837709 | NA | NA | NA | NA | NA | NA | NA |
| HCN4 | 654 | D | -46.941125 | 34.11968397 | 0.614107912 | 25.40743402 | NA | NA | NA | NA | NA | NA | NA |
| HCN4 | 655 | G | -51.073125 | 32.74466508 | 0.587431505 | 22.09804434 | NA | NA | NA | NA | NA | NA | NA |
| HCN4 | 656 | S | -52.745375 | 31.30432796 | 0.559487874 | 21.44625901 | NA | NA | NA | NA | NA | NA | NA |
| HCN4 | 657 | Y | -56.545625 | 28.70176448 | 0.508996173 | 19.17929217 | NA | NA | NA | NA | NA | NA | NA |
| HCN4 | 658 | F | -54.27992045 | 26.58955678 | 0.468017745 | 17.45102055 | NA | NA | NA | NA | NA | NA | NA |
| HCN4 | 659 | G | -58.041375 | 24.26187878 | 0.422859029 | 15.06996278 | NA | NA | NA | NA | NA | NA | NA |
| HCN4 | 660 | E | -60.71326389 | 24.51010041 | 0.427674717 | 14.52101918 | NA | NA | NA | NA | NA | NA | NA |
| HCN4 | 661 | I | -64.141625 | 23.92803567 | 0.41638222 | 14.04686696 | NA | NA | NA | NA | NA | NA | NA |
| HCN4 | 662 | C | -63.12370833 | 20.47474791 | 0.349385824 | 10.50578054 | NA | NA | NA | NA | NA | NA | NA |
| HCN4 | 663 | L | -61.7555 | 19.83732371 | 0.337019312 | 9.913430949 | NA | NA | NA | NA | NA | NA | NA |
| HCN4 | 664 | L | -65.945875 | 21.23813881 | 0.364196185 | 10.64504299 | NA | NA | NA | NA | NA | NA | NA |
| HCN4 | 665 | T | -68.09123214 | 20.68558346 | 0.353476193 | 10.14157695 | NA | NA | NA | NA | NA | NA | NA |
| HCN4 | 666 | R | -68.11382955 | 15.88144102 | 0.260272193 | 5.29554942 | NA | NA | NA | NA | NA | NA | NA |
| HCN4 | 667 | G | -66.105375 | 21.37010225 | 0.366756376 | 10.70781728 | NA | NA | NA | NA | NA | NA | NA |
| HCN4 | 668 | R | -64.48219318 | 23.42160938 | 0.406557167 | 12.8533272 | NA | NA | NA | NA | NA | NA | NA |
| HCN4 | 669 | R | -59.944375 | 22.56229467 | 0.389885811 | 13.33260939 | NA | NA | NA | NA | NA | NA | NA |
| HCN4 | 670 | T | -63.03451786 | 28.62629122 | 0.507531934 | 18.64148997 | NA | NA | NA | NA | NA | NA | NA |
| HCN4 | 671 | A | -59.300775 | 30.31898132 | 0.540371404 | 21.1211721 | NA | NA | NA | NA | NA | NA | NA |
| HCN4 | 672 | S | -55.82720833 | 31.55090343 | 0.564271625 | 22.38044786 | NA | NA | NA | NA | NA | NA | NA |

| | | | | | | | | | | | | | |
|---|---|---|---|---|---|---|---|---|---|---|---|---|---|
| HCN4 | 673 | V | -53.06508929 | 30.42027674 | 0.542336612 | 20.57097583 | NA | NA | NA | NA | NA | NA | NA |
| HCN4 | 674 | R | -49.17173864 | 32.46573649 | 0.582020079 | 21.74954917 | NA | NA | NA | NA | NA | NA | NA |
| HCN4 | 675 | A | -46.390975 | 31.25261669 | 0.558484637 | 22.53538628 | NA | NA | NA | NA | NA | NA | NA |
| HCN4 | 676 | D | -45.296625 | 34.1562524 | 0.614817367 | 26.03114837 | NA | NA | NA | NA | NA | NA | NA |
| HCN4 | 677 | T | -41.886375 | 31.99732373 | 0.572932517 | 23.81922104 | NA | NA | NA | NA | NA | NA | NA |
| HCN4 | 678 | Y | -38.08820833 | 29.84725096 | 0.531219478 | 21.95653687 | NA | NA | NA | NA | NA | NA | NA |
| HCN4 | 679 | C | -43.42504167 | 27.27847331 | 0.481383247 | 19.14607784 | NA | NA | NA | NA | NA | NA | NA |
| HCN4 | 680 | R | -40.51573864 | 25.54392425 | 0.447731682 | 18.12837604 | NA | NA | NA | NA | NA | NA | NA |
| HCN4 | 681 | L | -46.673875 | 23.63611984 | 0.410718832 | 14.93438681 | NA | NA | NA | NA | NA | NA | NA |
| HCN4 | 682 | Y | -46.88929167 | 22.72743492 | 0.393089657 | 14.01858788 | NA | NA | NA | NA | NA | NA | NA |
| HCN4 | 683 | S | -50.19820833 | 18.88609144 | 0.318564686 | 8.180343662 | NA | NA | NA | NA | NA | NA | NA |
| HCN4 | 684 | L | -54.368125 | 18.44494663 | 0.310006142 | 8.694230946 | NA | NA | NA | NA | NA | NA | NA |
| HCN4 | 685 | S | -55.64320833 | 14.4222968 | 0.231963691 | 5.423827483 | NA | NA | NA | NA | NA | NA | NA |
| HCN4 | 686 | V | -59.65251786 | 15.17364459 | 0.246540406 | 6.07758591 | NA | NA | NA | NA | NA | NA | NA |
| HCN4 | 687 | D | -59.111 | 10.29813818 | 0.151951892 | 1.125272938 | 9.2 | -1.04 | -3.5 | 49.7 | 2.63 | 4 | -4 |
| HCN4 | 688 | N | -56.260875 | 13.74076042 | 0.218741369 | 4.821743018 | 9.2 | -1.04 | -3.5 | 49.7 | 2.63 | 4 | -4 |
| HCN4 | 689 | F | -60.22564773 | 17.1349677 | 0.28459156 | 7.94909147 | NA | NA | NA | NA | NA | NA | NA |
| HCN4 | 690 | N | -62.87425 | 15.23877374 | 0.247803961 | 5.25625628 | NA | NA | NA | NA | NA | NA | NA |
| HCN4 | 691 | E | -59.48093056 | 13.74925096 | 0.218906092 | 4.818405736 | 9.99 | -1.04 | -3.5 | 49.7 | 2.63 | 4 | -4 |
| HCN4 | 692 | V | -59.34308929 | 18.2491237 | 0.306207029 | 9.185535245 | NA | NA | NA | NA | NA | NA | NA |
| HCN4 | 693 | L | -63.301625 | 20.17107863 | 0.34349441 | 10.21770158 | NA | NA | NA | NA | NA | NA | NA |
| HCN4 | 694 | E | -64.718875 | 16.49387968 | 0.272153967 | 5.951922969 | NA | NA | NA | NA | NA | NA | NA |
| HCN4 | 695 | E | -61.64415278 | 18.97285527 | 0.32024797 | 9.184562035 | NA | NA | NA | NA | NA | NA | NA |
| HCN4 | 696 | Y | -61.52229167 | 22.80963623 | 0.394684425 | 12.81876644 | NA | NA | NA | NA | NA | NA | NA |
| HCN4 | 697 | P | -66.94823214 | 23.48749499 | 0.407835398 | 12.87253399 | NA | NA | NA | NA | NA | NA | NA |
| HCN4 | 698 | M | -65.717 | 26.39918862 | 0.464324458 | 15.71059495 | NA | NA | NA | NA | NA | NA | NA |
| HCN4 | 699 | M | -62.768875 | 27.07057851 | 0.47734993 | 17.09488608 | NA | NA | NA | NA | NA | NA | NA |
| HCN4 | 700 | R | -66.68046591 | 20.67056903 | 0.353184902 | 10.00571127 | NA | NA | NA | NA | NA | NA | NA |
| HCN4 | 701 | R | -68.90101136 | 25.94008647 | 0.455417529 | 15.40154822 | NA | NA | NA | NA | NA | NA | NA |
| HCN4 | 702 | A | -63.829775 | 29.8957754 | 0.532160889 | 19.32827822 | NA | NA | NA | NA | NA | NA | NA |
| HCN4 | 703 | F | -61.84646591 | 25.87093698 | 0.454075976 | 16.02147859 | NA | NA | NA | NA | NA | NA | NA |
| HCN4 | 704 | E | -68.02915278 | 24.95833929 | 0.43637089 | 14.40121035 | NA | NA | NA | NA | NA | NA | NA |
| HCN4 | 705 | T | -66.59666071 | 30.91887149 | 0.552009727 | 20.24150607 | NA | NA | NA | NA | NA | NA | NA |

| | | | | | | | | | | | | | |
|---|---|---|---|---|---|---|---|---|---|---|---|---|---|
| HCN4 | 706 | V | -63.046375 | 32.08245664 | 0.57458416 | 22.10951226 | NA | NA | NA | NA | NA | NA | NA |
| HCN4 | 707 | A | -65.431975 | 28.88661135 | 0.512582342 | 18.17607023 | NA | NA | NA | NA | NA | NA | NA |
| HCN4 | 708 | L | -69.264625 | 31.49338872 | 0.563155796 | 20.82758813 | NA | NA | NA | NA | NA | NA | NA |
| HCN4 | 709 | D | -65.49 | 34.24611547 | 0.616560779 | 23.56777067 | NA | NA | NA | NA | NA | NA | NA |
| HCN4 | 710 | R | -61.75155682 | 31.32823728 | 0.559951733 | 21.34601225 | NA | NA | NA | NA | NA | NA | NA |
| HCN4 | 711 | L | -67.732 | 30.13726572 | 0.536845983 | 19.53179433 | NA | NA | NA | NA | NA | NA | NA |
| HCN4 | 712 | D | -68.77475 | 35.63069505 | 0.643422671 | 24.98718898 | NA | NA | NA | NA | NA | NA | NA |
| HCN4 | 713 | R | -62.99982955 | 37.72552245 | 0.684063908 | 27.03216032 | NA | NA | NA | NA | NA | NA | NA |
| HCN4 | 714 | I | -66.104125 | 34.93362405 | 0.629898966 | 24.25699438 | NA | NA | NA | NA | NA | NA | NA |
| KCNK3 | 1 | M | -16.332 | 21.78422057 | 0.540899583 | 21.75424429 | NA | NA | NA | NA | NA | NA | NA |
| KCNK3 | 2 | K | -18.04621429 | 19.82509052 | 0.486991246 | 19.90856641 | NA | NA | NA | NA | NA | NA | NA |
| KCNK3 | 3 | R | -17.92440909 | 17.07252755 | 0.411250439 | 17.16277285 | NA | NA | NA | NA | NA | NA | NA |
| KCNK3 | 4 | Q | -15.72127778 | 15.03151027 | 0.355088855 | 15.07218809 | NA | NA | NA | NA | NA | NA | NA |
| KCNK3 | 5 | N | -13.245625 | 17.09329418 | 0.411821863 | 17.14611452 | NA | NA | NA | NA | NA | NA | NA |
| KCNK3 | 6 | V | -13.14592857 | 17.68093428 | 0.427991642 | 17.65205723 | NA | NA | NA | NA | NA | NA | NA |
| KCNK3 | 7 | R | -12.50868182 | 12.48449707 | 0.285004051 | 12.5479035 | NA | NA | NA | NA | NA | NA | NA |
| KCNK3 | 8 | T | -8.637357143 | 12.95126361 | 0.297847817 | 12.9139099 | NA | NA | NA | NA | NA | NA | NA |
| KCNK3 | 9 | L | -7.93925 | 16.24144467 | 0.388381974 | 16.08130815 | NA | NA | NA | NA | NA | NA | NA |
| KCNK3 | 10 | A | -7.3087 | 16.53458348 | 0.396448119 | 16.24660301 | NA | NA | NA | NA | NA | NA | NA |
| KCNK3 | 11 | L | -5.67025 | 11.3423625 | 0.253576543 | 11.17565103 | NA | NA | NA | NA | NA | NA | NA |
| KCNK3 | 12 | I | -3.065125 | 14.92589877 | 0.352182799 | 14.49412097 | NA | NA | NA | NA | NA | NA | NA |
| KCNK3 | 13 | V | -2.9495 | 17.79682528 | 0.431180553 | 17.33851537 | NA | NA | NA | NA | NA | NA | NA |
| KCNK3 | 14 | C | -2.343 | 16.72516613 | 0.40169228 | 16.22009831 | NA | NA | NA | NA | NA | NA | NA |
| KCNK3 | 15 | T | -0.061928571 | 13.65366508 | 0.317175424 | 12.96098124 | NA | NA | NA | NA | NA | NA | NA |
| KCNK3 | 16 | F | 0.6175 | 17.90482337 | 0.434152279 | 15.51007894 | NA | NA | NA | NA | NA | NA | NA |
| KCNK3 | 17 | T | 0.162785714 | 20.37048858 | 0.501998673 | 17.82487248 | NA | NA | NA | NA | NA | NA | NA |
| KCNK3 | 18 | Y | 1.110083333 | 15.49277154 | 0.367781135 | 14.3200157 | NA | NA | NA | NA | NA | NA | NA |
| KCNK3 | 19 | L | 4.315625 | 14.76360942 | 0.34771717 | 13.8764967 | NA | NA | NA | NA | NA | NA | NA |
| KCNK3 | 20 | L | 4.817375 | 19.93835725 | 0.490107946 | 19.05173961 | NA | NA | NA | NA | NA | NA | NA |
| KCNK3 | 21 | V | 4.409357143 | 20.87341192 | 0.515837347 | 20.021289 | NA | NA | NA | NA | NA | NA | NA |
| KCNK3 | 22 | G | 7.34 | 18.55747736 | 0.452111011 | 17.63922874 | NA | NA | NA | NA | NA | NA | NA |
| KCNK3 | 23 | A | 9.7051 | 18.88539512 | 0.461134149 | 18.0792065 | NA | NA | NA | NA | NA | NA | NA |
| KCNK3 | 24 | A | 9.7171 | 21.3388218 | 0.528643782 | 20.46286201 | NA | NA | NA | NA | NA | NA | NA |

| | | | | | | | | | | | | | |
|---|---|---|---|---|---|---|---|---|---|---|---|---|---|
| KCNK3 | 25 | V | 9.707214286 | 20.72484441 | 0.511749294 | 19.8695733 | NA | NA | NA | NA | NA | NA | NA |
| KCNK3 | 26 | F | 11.62059091 | 14.98475215 | 0.353802237 | 14.23243135 | NA | NA | NA | NA | NA | NA | NA |
| KCNK3 | 27 | D | 14.945125 | 20.02632757 | 0.492528578 | 18.98789415 | NA | NA | NA | NA | NA | NA | NA |
| KCNK3 | 28 | A | 14.5763 | 22.07493928 | 0.548899134 | 18.13103789 | NA | NA | NA | NA | NA | NA | NA |
| KCNK3 | 29 | L | 14.550625 | 18.38819355 | 0.447452918 | 14.46188875 | NA | NA | NA | NA | NA | NA | NA |
| KCNK3 | 30 | E | 17.78505556 | 14.06148075 | 0.32839707 | 10.12665753 | NA | NA | NA | NA | NA | NA | NA |
| KCNK3 | 31 | S | 19.715 | 18.446231 | 0.449049904 | 14.49967564 | NA | NA | NA | NA | NA | NA | NA |
| KCNK3 | 32 | E | 22.50561111 | 18.91998422 | 0.462085919 | 15.01533454 | NA | NA | NA | NA | NA | NA | NA |
| KCNK3 | 33 | P | 22.39407143 | 15.94714376 | 0.380283853 | 12.04577399 | NA | NA | NA | NA | NA | NA | NA |
| KCNK3 | 34 | E | 23.44827778 | 13.38661399 | 0.309827121 | 9.810478418 | NA | NA | NA | NA | NA | NA | NA |
| KCNK3 | 35 | L | 26.502625 | 16.83332198 | 0.404668347 | 15.64623112 | NA | NA | NA | NA | NA | NA | NA |
| KCNK3 | 36 | I | 27.3925 | 16.75987052 | 0.402647222 | 15.55113325 | NA | NA | NA | NA | NA | NA | NA |
| KCNK3 | 37 | E | 27.94592857 | 11.21337862 | 0.250027363 | 10.23854052 | NA | NA | NA | NA | NA | NA | NA |
| KCNK3 | 38 | R | 29.56422727 | 13.50730003 | 0.313147975 | 12.35217466 | NA | NA | NA | NA | NA | NA | NA |
| KCNK3 | 39 | Q | 31.78516667 | 15.64638559 | 0.372008051 | 15.18196751 | NA | NA | NA | NA | NA | NA | NA |
| KCNK3 | 40 | R | 31.51259091 | 13.86579871 | 0.323012592 | 13.18819998 | NA | NA | NA | NA | NA | NA | NA |
| KCNK3 | 41 | L | 32.766125 | 9.532716349 | 0.203781475 | 8.842311847 | NA | NA | NA | NA | NA | NA | NA |
| KCNK3 | 42 | E | 34.87605556 | 12.58586374 | 0.287793304 | 11.92505319 | NA | NA | NA | NA | NA | NA | NA |
| KCNK3 | 43 | L | 37.88825 | 13.7584353 | 0.32005833 | 12.90735383 | NA | NA | NA | NA | NA | NA | NA |
| KCNK3 | 44 | R | 36.63422727 | 9.21367055 | 0.195002462 | 8.387559232 | NA | NA | NA | NA | NA | NA | NA |
| KCNK3 | 45 | Q | 38.69516667 | 9.245243642 | 0.195871242 | 8.420511838 | NA | NA | NA | NA | NA | NA | NA |
| KCNK3 | 46 | Q | 41.37238889 | 12.18157453 | 0.276668693 | 11.0167684 | NA | NA | NA | NA | NA | NA | NA |
| KCNK3 | 47 | E | 43.69105556 | 11.3384431 | 0.253468695 | 10.05254906 | NA | NA | NA | NA | NA | NA | NA |
| KCNK3 | 48 | L | 42.2805 | 7.769022912 | 0.155250864 | 6.557023466 | NA | NA | NA | NA | NA | NA | NA |
| KCNK3 | 49 | R | 43.53513636 | 8.993245299 | 0.188937138 | 7.76040079 | NA | NA | NA | NA | NA | NA | NA |
| KCNK3 | 50 | A | 47.5331 | 10.93193162 | 0.242282936 | 9.675364525 | NA | NA | NA | NA | NA | NA | NA |
| KCNK3 | 51 | R | 46.70913636 | 9.934400083 | 0.214834392 | 8.753011026 | NA | NA | NA | NA | NA | NA | NA |
| KCNK3 | 52 | Y | 46.51208333 | 6.478931393 | 0.119752104 | 5.256094201 | NA | NA | NA | NA | NA | NA | NA |
| KCNK3 | 53 | N | 49.5967 | 9.699846391 | 0.208380304 | 8.564665782 | NA | NA | NA | NA | NA | NA | NA |
| KCNK3 | 54 | L | 45.925125 | 5.4538546 | 0.091545612 | 4.223983398 | 1.52 | 1.13 | 2.1 | 0.62 | -1.67 | 0 | 0 |
| KCNK3 | 55 | S | 45.93783333 | 10.38988032 | 0.227367599 | 9.257293271 | NA | NA | NA | NA | NA | NA | NA |
| KCNK3 | 56 | Q | 42.8699 | 11.77040717 | 0.26535482 | 10.3013204 | NA | NA | NA | NA | NA | NA | NA |
| KCNK3 | 57 | G | 43.01575 | 10.69139542 | 0.235664229 | 9.212489969 | NA | NA | NA | NA | NA | NA | NA |

| | | | | | | | | | | | | | |
|---|---|---|---|---|---|---|---|---|---|---|---|---|---|
| KCNK3 | 58 | G | 43.49775 | 7.227741694 | 0.140356717 | 5.763507767 | NA | NA | NA | NA | NA | NA | NA |
| KCNK3 | 59 | Y | 39.26083333 | 6.286063156 | 0.114445051 | 5.658420896 | NA | NA | NA | NA | NA | NA | NA |
| KCNK3 | 60 | E | 38.81783333 | 8.915597007 | 0.186800532 | 8.127975483 | NA | NA | NA | NA | NA | NA | NA |
| KCNK3 | 61 | E | 40.0199 | 7.780959581 | 0.155579319 | 6.407762813 | NA | NA | NA | NA | NA | NA | NA |
| KCNK3 | 62 | L | 40.072625 | 3.514852628 | 0.038191129 | 3.045016321 | 1.5 | 1.13 | 2.23 | 0.62 | -1.51 | 0 | 0 |
| KCNK3 | 63 | E | 35.29816667 | 4.837103059 | 0.074574789 | 4.561578035 | 0.95 | 0.17 | 1.9 | 1.76 | -0.25 | 0 | 0 |
| KCNK3 | 64 | R | 35.93404545 | 7.107586792 | 0.137050479 | 5.961347683 | NA | NA | NA | NA | NA | NA | NA |
| KCNK3 | 65 | V | 35.98207143 | 6.556292092 | 0.121880797 | 5.823344247 | NA | NA | NA | NA | NA | NA | NA |
| KCNK3 | 66 | V | 33.92792857 | 2.260154198 | 0.003666262 | 1.706373324 | 1.07 | 1.13 | 4.07 | 0.13 | -1.46 | 0 | 0 |
| KCNK3 | 67 | L | 30.52375 | 2.565183229 | 0.012059583 | 1.699494144 | 1.22 | 1.14 | 4 | 0.13 | -1.55 | 0 | 0 |
| KCNK3 | 68 | R | 31.1595 | 6.299161055 | 0.114805459 | 5.09796057 | NA | NA | NA | NA | NA | NA | NA |
| KCNK3 | 69 | L | 32.255625 | 7.388331476 | 0.144775581 | 6.314861628 | NA | NA | NA | NA | NA | NA | NA |
| KCNK3 | 70 | K | 27.91133333 | 4.55335327 | 0.066766997 | 4.785620301 | 1.17 | 1.13 | 4.07 | 0.13 | -1.46 | 0 | 0 |
| KCNK3 | 71 | P | 26.06978571 | 7.394348856 | 0.144941158 | 3.925994667 | 3.63 | -0.61 | -2.15 | 26.44 | 1.14 | 2 | 2 |
| KCNK3 | 72 | H | 28.7188 | 10.04465455 | 0.217868206 | 6.580168514 | NA | NA | NA | NA | NA | NA | NA |
| KCNK3 | 73 | K | 29.0119 | 9.43281432 | 0.201032525 | 8.391477248 | NA | NA | NA | NA | NA | NA | NA |
| KCNK3 | 74 | A | 25.8307 | 10.16703738 | 0.221235749 | 6.577872048 | NA | NA | NA | NA | NA | NA | NA |
| KCNK3 | 75 | G | 26.13525 | 12.80659728 | 0.293867111 | 9.249736612 | NA | NA | NA | NA | NA | NA | NA |
| KCNK3 | 76 | V | 24.7185 | 14.95462393 | 0.352973215 | 11.49213993 | NA | NA | NA | NA | NA | NA | NA |
| KCNK3 | 77 | Q | 21.76561111 | 9.312071091 | 0.197710098 | 5.838568289 | NA | NA | NA | NA | NA | NA | NA |
| KCNK3 | 78 | W | 19.94078571 | 12.76010478 | 0.292587801 | 9.638896993 | NA | NA | NA | NA | NA | NA | NA |
| KCNK3 | 79 | R | 20.31433333 | 18.15502586 | 0.441036967 | 14.71452981 | NA | NA | NA | NA | NA | NA | NA |
| KCNK3 | 80 | F | 14.93240909 | 19.47819791 | 0.477445987 | 18.57863558 | NA | NA | NA | NA | NA | NA | NA |
| KCNK3 | 81 | A | 16.5525 | 17.13008082 | 0.412834101 | 16.20791611 | NA | NA | NA | NA | NA | NA | NA |
| KCNK3 | 82 | G | 17.20925 | 14.35771225 | 0.336548314 | 13.54276693 | NA | NA | NA | NA | NA | NA | NA |
| KCNK3 | 83 | S | 15.47716667 | 15.54487716 | 0.369214898 | 14.79741278 | NA | NA | NA | NA | NA | NA | NA |
| KCNK3 | 84 | F | 10.87922727 | 14.77633628 | 0.348067368 | 13.59533305 | NA | NA | NA | NA | NA | NA | NA |
| KCNK3 | 85 | Y | 14.04908333 | 11.36367947 | 0.254163111 | 10.45390688 | NA | NA | NA | NA | NA | NA | NA |
| KCNK3 | 86 | F | 13.99722727 | 10.01426148 | 0.217031896 | 8.999821623 | NA | NA | NA | NA | NA | NA | NA |
| KCNK3 | 87 | A | 10.1005 | 11.81246465 | 0.266512093 | 10.84945561 | NA | NA | NA | NA | NA | NA | NA |
| KCNK3 | 88 | I | 8.724375 | 9.507945204 | 0.203099861 | 8.655766733 | NA | NA | NA | NA | NA | NA | NA |
| KCNK3 | 89 | T | 10.65507143 | 6.474277257 | 0.119624038 | 5.443324425 | NA | NA | NA | NA | NA | NA | NA |
| KCNK3 | 90 | V | 8.8165 | 7.484600791 | 0.147424572 | 6.912368727 | NA | NA | NA | NA | NA | NA | NA |

| | | | | | | | | | | | | | |
|---|---|---|---|---|---|---|---|---|---|---|---|---|---|
| KCNK3 | 91 | I | 4.954875 | 7.860247579 | 0.157761044 | 7.262008647 | NA | NA | NA | NA | NA | NA | NA |
| KCNK3 | 92 | T | 5.076642857 | 4.920546819 | 0.076870866 | 3.71853286 | 1.25 | -0.79 | -0.55 | 2.52 | 0.79 | 0 | 0 |
| KCNK3 | 93 | T | 5.630071429 | 2.157625778 | 8.45E-04 | 1.679242116 | 1.03 | -0.78 | -0.6 | 2.23 | 0.78 | 0 | 0 |
| KCNK3 | 94 | I | 8.753 | 2.423879948 | 0.008171416 | 1.782083483 | 1.12 | -0.8 | -0.4 | 3.38 | 0.81 | 0 | 0 |
| KCNK3 | 95 | G | 11.9615 | 2.480835545 | 0.009738633 | 1.525153096 | 1.23 | -0.8 | -0.4 | 3.38 | 0.81 | 0 | 0 |
| KCNK3 | 96 | Y | 13.87558333 | 2.29729406 | 0.004688219 | 1.619353978 | 1.16 | -0.8 | -0.4 | 3.38 | 0.81 | 0 | 0 |
| KCNK3 | 97 | G | 17.668 | 4.089560123 | 0.054005049 | 3.257897003 | 0.9 | -0.8 | -0.4 | 3.38 | 0.81 | 0 | 0 |
| KCNK3 | 98 | H | 18.6961 | 5.222588822 | 0.085181995 | 2.367431172 | 3.32 | -0.8 | -0.4 | 3.38 | 0.81 | 0 | 0 |
| KCNK3 | 99 | A | 18.9097 | 7.605769784 | 0.150758715 | 5.545598466 | NA | NA | NA | NA | NA | NA | NA |
| KCNK3 | 100 | A | 19.0579 | 8.296117887 | 0.169754655 | 7.414062726 | NA | NA | NA | NA | NA | NA | NA |
| KCNK3 | 101 | P | 18.09407143 | 11.22453037 | 0.250334219 | 10.43884759 | NA | NA | NA | NA | NA | NA | NA |
| KCNK3 | 102 | S | 22.5245 | 13.57110946 | 0.314903785 | 10.36698384 | NA | NA | NA | NA | NA | NA | NA |
| KCNK3 | 103 | T | 20.91664286 | 16.17393817 | 0.386524434 | 14.13567985 | NA | NA | NA | NA | NA | NA | NA |
| KCNK3 | 104 | D | 18.622 | 18.79446131 | 0.458631972 | 17.92337158 | NA | NA | NA | NA | NA | NA | NA |
| KCNK3 | 105 | G | 16.8035 | 19.44181625 | 0.476444892 | 18.51909477 | NA | NA | NA | NA | NA | NA | NA |
| KCNK3 | 106 | G | 16.11875 | 16.03664432 | 0.382746592 | 15.17455999 | NA | NA | NA | NA | NA | NA | NA |
| KCNK3 | 107 | K | 16.6975 | 13.60192545 | 0.315751732 | 12.90937938 | NA | NA | NA | NA | NA | NA | NA |
| KCNK3 | 108 | V | 12.86007143 | 17.53743428 | 0.424043029 | 16.39841545 | NA | NA | NA | NA | NA | NA | NA |
| KCNK3 | 109 | F | 11.04468182 | 16.79194593 | 0.403529824 | 15.88248866 | NA | NA | NA | NA | NA | NA | NA |
| KCNK3 | 110 | C | 11.18783333 | 11.86020021 | 0.267825607 | 10.91369862 | NA | NA | NA | NA | NA | NA | NA |
| KCNK3 | 111 | M | 9.59225 | 14.20275132 | 0.332284337 | 13.66902074 | NA | NA | NA | NA | NA | NA | NA |
| KCNK3 | 112 | F | 7.016772727 | 15.93047978 | 0.379825319 | 15.01569506 | NA | NA | NA | NA | NA | NA | NA |
| KCNK3 | 113 | Y | 7.317166667 | 13.6975788 | 0.318383774 | 12.76527513 | NA | NA | NA | NA | NA | NA | NA |
| KCNK3 | 114 | A | 5.2915 | 10.86093886 | 0.240329466 | 9.83777927 | NA | NA | NA | NA | NA | NA | NA |
| KCNK3 | 115 | L | 2.51325 | 11.80434822 | 0.266288758 | 9.674821337 | NA | NA | NA | NA | NA | NA | NA |
| KCNK3 | 116 | L | 0.706875 | 12.40833143 | 0.282908242 | 9.914299591 | NA | NA | NA | NA | NA | NA | NA |
| KCNK3 | 117 | G | 1.86175 | 10.69496741 | 0.235762518 | 7.919457214 | NA | NA | NA | NA | NA | NA | NA |
| KCNK3 | 118 | I | 1.887625 | 6.13527644 | 0.110295934 | 3.338891981 | 2.8 | 0.5 | 2.3 | 0.64 | -0.94 | 0 | 0 |
| KCNK3 | 119 | P | -0.1855 | 8.175611231 | 0.166438738 | 5.674795518 | NA | NA | NA | NA | NA | NA | NA |
| KCNK3 | 120 | L | 0 | 10.23393429 | 0.223076515 | 7.723582143 | NA | NA | NA | NA | NA | NA | NA |
| KCNK3 | 121 | T | 0.265642857 | 9.056873909 | 0.190687973 | 6.84548408 | NA | NA | NA | NA | NA | NA | NA |
| KCNK3 | 122 | L | -0.861125 | 3.808940798 | 0.046283396 | 1.469793266 | 2.51 | 1.09 | 3.17 | 0.56 | -1.43 | 0 | 0 |
| KCNK3 | 123 | V | -4.3235 | 8.232095602 | 0.167992988 | 5.452818788 | NA | NA | NA | NA | NA | NA | NA |

| | | | | | | | | | | | | | |
|---|---|---|---|---|---|---|---|---|---|---|---|---|---|
| KCNK3 | 124 | M | -1.74325 | 10.90123016 | 0.24143814 | 8.424413565 | NA | NA | NA | NA | NA | NA | NA |
| KCNK3 | 125 | F | -1.073772727 | 9.219530411 | 0.195163705 | 6.539290153 | NA | NA | NA | NA | NA | NA | NA |
| KCNK3 | 126 | Q | -5.4925 | 7.285117089 | 0.141935485 | 4.981639983 | 2.33 | 0.55 | 1.5 | 1.31 | -1.04 | 0 | 0 |
| KCNK3 | 127 | S | -7.1375 | 11.69107856 | 0.263171977 | 9.437562455 | NA | NA | NA | NA | NA | NA | NA |
| KCNK3 | 128 | L | -4.549 | 14.23424483 | 0.333150927 | 11.6041046 | NA | NA | NA | NA | NA | NA | NA |
| KCNK3 | 129 | G | -6.3605 | 13.95328381 | 0.325419872 | 11.6583271 | NA | NA | NA | NA | NA | NA | NA |
| KCNK3 | 130 | E | -10.92072222 | 14.21854834 | 0.332719015 | 11.91975111 | NA | NA | NA | NA | NA | NA | NA |
| KCNK3 | 131 | R | -10.81259091 | 16.82129181 | 0.404337319 | 14.51656478 | NA | NA | NA | NA | NA | NA | NA |
| KCNK3 | 132 | I | -6.74175 | 17.54839186 | 0.424344543 | 16.00133153 | NA | NA | NA | NA | NA | NA | NA |
| KCNK3 | 133 | N | -9.84275 | 14.99140207 | 0.353985219 | 14.04883047 | NA | NA | NA | NA | NA | NA | NA |
| KCNK3 | 134 | T | -13.32564286 | 19.17211944 | 0.469023789 | 18.23391061 | NA | NA | NA | NA | NA | NA | NA |
| KCNK3 | 135 | L | -10.644625 | 22.03332916 | 0.547754171 | 19.76432993 | NA | NA | NA | NA | NA | NA | NA |
| KCNK3 | 136 | V | -10.51507143 | 22.10685407 | 0.549777316 | 21.40023109 | NA | NA | NA | NA | NA | NA | NA |
| KCNK3 | 137 | R | -15.02231818 | 18.44307279 | 0.448963001 | 17.71686041 | NA | NA | NA | NA | NA | NA | NA |
| KCNK3 | 138 | Y | -16.22616667 | 25.15554607 | 0.633666548 | 24.22285975 | NA | NA | NA | NA | NA | NA | NA |
| KCNK3 | 139 | L | -12.300875 | 26.78463358 | 0.67849328 | 26.08947561 | NA | NA | NA | NA | NA | NA | NA |
| KCNK3 | 140 | L | -14.31975 | 26.28242183 | 0.664674187 | 25.43062427 | NA | NA | NA | NA | NA | NA | NA |
| KCNK3 | 141 | H | -19.6283 | 28.63970061 | 0.72953817 | 27.77274652 | NA | NA | NA | NA | NA | NA | NA |
| KCNK3 | 142 | R | -17.70913636 | 30.9305153 | 0.792573296 | 30.03674184 | NA | NA | NA | NA | NA | NA | NA |
| KCNK3 | 143 | A | -15.9045 | 32.24004673 | 0.828606974 | 31.29630022 | NA | NA | NA | NA | NA | NA | NA |
| KCNK3 | 144 | K | -18.3565 | 28.97195012 | 0.738680503 | 28.28147292 | NA | NA | NA | NA | NA | NA | NA |
| KCNK3 | 145 | K | -21.45264286 | 34.46373099 | 0.889794908 | 33.64980721 | NA | NA | NA | NA | NA | NA | NA |
| KCNK3 | 146 | G | -19.11275 | 36.46739679 | 0.944928711 | 35.59371784 | NA | NA | NA | NA | NA | NA | NA |
| KCNK3 | 147 | L | -18.4731 | 37.54880942 | 0.974685365 | 36.65466712 | NA | NA | NA | NA | NA | NA | NA |
| KCNK3 | 148 | G | -21.451 | 38.46879071 | 1 | 37.72330452 | NA | NA | NA | NA | NA | NA | NA |
| KCNK3 | 152 | A | -22.1019 | 30.12039598 | 0.770281675 | 29.43078203 | NA | NA | NA | NA | NA | NA | NA |
| KCNK3 | 153 | D | -22.292125 | 27.94660854 | 0.710466725 | 27.33247027 | NA | NA | NA | NA | NA | NA | NA |
| KCNK3 | 154 | V | -17.79664286 | 24.13911798 | 0.605698039 | 23.40404451 | NA | NA | NA | NA | NA | NA | NA |
| KCNK3 | 155 | S | -16.96783333 | 22.51799571 | 0.561090482 | 21.86928011 | NA | NA | NA | NA | NA | NA | NA |
| KCNK3 | 156 | M | -14.382375 | 15.91968602 | 0.379528313 | 15.38572153 | NA | NA | NA | NA | NA | NA | NA |
| KCNK3 | 157 | A | -13.3143 | 21.85102428 | 0.542737784 | 21.26110298 | NA | NA | NA | NA | NA | NA | NA |
| KCNK3 | 158 | N | -13.523125 | 22.99275105 | 0.574154071 | 22.18006423 | NA | NA | NA | NA | NA | NA | NA |
| KCNK3 | 159 | M | -11.492375 | 19.65880655 | 0.482415699 | 18.75708269 | NA | NA | NA | NA | NA | NA | NA |

| | | | | | | | | | | | | | |
|---|---|---|---|---|---|---|---|---|---|---|---|---|---|
| KCNK3 | 160 | V | -8.890214286 | 17.96954716 | 0.435933249 | 17.25919633 | NA | NA | NA | NA | NA | NA | NA |
| KCNK3 | 161 | L | -7.837875 | 21.63290924 | 0.536736029 | 19.2664506 | NA | NA | NA | NA | NA | NA | NA |
| KCNK3 | 162 | I | -7.536 | 20.73790165 | 0.512108583 | 18.105009 | NA | NA | NA | NA | NA | NA | NA |
| KCNK3 | 163 | G | -4.8275 | 17.59739893 | 0.425693044 | 14.95732755 | NA | NA | NA | NA | NA | NA | NA |
| KCNK3 | 164 | F | -2.263318182 | 18.15193998 | 0.440952055 | 15.50519395 | NA | NA | NA | NA | NA | NA | NA |
| KCNK3 | 165 | F | -2.445136364 | 20.56662687 | 0.507395706 | 17.92061833 | NA | NA | NA | NA | NA | NA | NA |
| KCNK3 | 166 | S | -1.376333333 | 18.76157158 | 0.457726962 | 16.10078055 | NA | NA | NA | NA | NA | NA | NA |
| KCNK3 | 167 | C | -0.120666667 | 14.96954993 | 0.353383925 | 12.12585414 | NA | NA | NA | NA | NA | NA | NA |
| KCNK3 | 168 | I | 0.678625 | 18.8942034 | 0.461376521 | 16.35263098 | NA | NA | NA | NA | NA | NA | NA |
| KCNK3 | 169 | S | 0.8645 | 20.84241598 | 0.514984448 | 18.39798253 | NA | NA | NA | NA | NA | NA | NA |
| KCNK3 | 170 | T | 2.448214286 | 17.13661778 | 0.413013975 | 15.9978596 | NA | NA | NA | NA | NA | NA | NA |
| KCNK3 | 171 | L | 4.964125 | 14.76611862 | 0.347786214 | 13.58442423 | NA | NA | NA | NA | NA | NA | NA |
| KCNK3 | 172 | C | 5.880833333 | 20.10771618 | 0.494768105 | 19.4169618 | NA | NA | NA | NA | NA | NA | NA |
| KCNK3 | 173 | I | 5.663875 | 21.38883515 | 0.530019973 | 20.67569752 | NA | NA | NA | NA | NA | NA | NA |
| KCNK3 | 174 | G | 9.00825 | 19.10168079 | 0.467085566 | 17.88317674 | NA | NA | NA | NA | NA | NA | NA |
| KCNK3 | 175 | A | 10.8131 | 19.36595273 | 0.474357396 | 18.16170631 | NA | NA | NA | NA | NA | NA | NA |
| KCNK3 | 176 | A | 11.4959 | 21.92541699 | 0.544784809 | 18.36072555 | NA | NA | NA | NA | NA | NA | NA |
| KCNK3 | 177 | A | 12.5733 | 20.72152236 | 0.511657883 | 20.42349886 | NA | NA | NA | NA | NA | NA | NA |
| KCNK3 | 178 | F | 13.3775 | 16.36178704 | 0.391693371 | 13.25418433 | NA | NA | NA | NA | NA | NA | NA |
| KCNK3 | 179 | S | 16.13716667 | 18.52621896 | 0.45125089 | 14.61934429 | NA | NA | NA | NA | NA | NA | NA |
| KCNK3 | 180 | H | 17.5809 | 20.06601819 | 0.493620724 | 16.15388962 | NA | NA | NA | NA | NA | NA | NA |
| KCNK3 | 181 | Y | 17.09583333 | 17.62906864 | 0.426564483 | 13.70464202 | NA | NA | NA | NA | NA | NA | NA |
| KCNK3 | 182 | E | 19.58127778 | 13.1913577 | 0.304454358 | 9.257561229 | NA | NA | NA | NA | NA | NA | NA |
| KCNK3 | 183 | H | 22.3586 | 15.59452856 | 0.370581129 | 11.69715157 | NA | NA | NA | NA | NA | NA | NA |
| KCNK3 | 184 | W | 18.49985714 | 11.28616015 | 0.252030053 | 7.367266795 | NA | NA | NA | NA | NA | NA | NA |
| KCNK3 | 185 | T | 17.75435714 | 16.37538598 | 0.392067566 | 12.48251272 | NA | NA | NA | NA | NA | NA | NA |
| KCNK3 | 186 | F | 11.96613636 | 18.88868013 | 0.461224541 | 14.99357936 | NA | NA | NA | NA | NA | NA | NA |
| KCNK3 | 187 | F | 13.65913636 | 16.53855353 | 0.396557361 | 12.65547283 | NA | NA | NA | NA | NA | NA | NA |
| KCNK3 | 188 | Q | 17.1665 | 12.96090282 | 0.298113054 | 9.077453347 | NA | NA | NA | NA | NA | NA | NA |
| KCNK3 | 189 | A | 13.3053 | 15.612468 | 0.371074759 | 11.68220314 | NA | NA | NA | NA | NA | NA | NA |
| KCNK3 | 190 | Y | 8.313833333 | 15.08759802 | 0.356632192 | 14.01167979 | NA | NA | NA | NA | NA | NA | NA |
| KCNK3 | 191 | Y | 12.61208333 | 10.41506937 | 0.228060713 | 9.877360506 | NA | NA | NA | NA | NA | NA | NA |
| KCNK3 | 192 | Y | 12.75458333 | 10.4928479 | 0.230200903 | 9.594735474 | NA | NA | NA | NA | NA | NA | NA |

| | | | | | | | | | | | | | |
|---|---|---|---|---|---|---|---|---|---|---|---|---|---|
| KCNK3 | 193 | C | 8.745833333 | 11.88534711 | 0.268517561 | 10.91594678 | NA | NA | NA | NA | NA | NA | NA |
| KCNK3 | 194 | F | 6.050681818 | 9.955301552 | 0.215409527 | 9.232894088 | NA | NA | NA | NA | NA | NA | NA |
| KCNK3 | 195 | I | 9.539375 | 6.759909467 | 0.127483627 | 5.707412944 | NA | NA | NA | NA | NA | NA | NA |
| KCNK3 | 196 | T | 8.510928571 | 7.353896178 | 0.143828043 | 6.397788359 | NA | NA | NA | NA | NA | NA | NA |
| KCNK3 | 197 | L | 4.510375 | 8.465441867 | 0.174413853 | 7.315835934 | NA | NA | NA | NA | NA | NA | NA |
| KCNK3 | 198 | T | 4.2905 | 4.947008591 | 0.077599001 | 3.817694046 | 1.19 | -0.77 | -0.7 | 1.66 | 0.77 | 0 | 0 |
| KCNK3 | 199 | T | 5.345071429 | 2.137984331 | 3.05E-04 | 1.518075968 | 0.89 | -0.79 | -0.48 | 2.95 | 0.8 | 0 | 0 |
| KCNK3 | 200 | I | 8.224125 | 2.421704359 | 0.008111551 | 1.769351734 | 1.12 | -0.8 | -0.4 | 3.38 | 0.81 | 0 | 0 |
| KCNK3 | 201 | G | 11.58575 | 2.263447371 | 0.003756878 | 1.510127659 | 1.24 | -0.8 | -0.4 | 3.38 | 0.81 | 0 | 0 |
| KCNK3 | 202 | F | 13.4655 | 2.467749987 | 0.009378564 | 2.01116138 | 1.01 | -0.8 | -0.4 | 3.38 | 0.81 | 0 | 0 |
| KCNK3 | 203 | G | 17.15 | 4.283669222 | 0.059346245 | 3.458824143 | 0.9 | -0.8 | -0.4 | 3.38 | 0.81 | 0 | 0 |
| KCNK3 | 204 | D | 17.844875 | 5.725377979 | 0.099016976 | 1.736984593 | 3.99 | -0.64 | -1.8 | 21.83 | 1.07 | 2 | 2 |
| KCNK3 | 205 | Y | 17.76866667 | 7.508878345 | 0.148092605 | 3.521132317 | 3.99 | -0.64 | -1.8 | 21.83 | 1.07 | 2 | 2 |
| KCNK3 | 206 | V | 17.60607143 | 7.155947317 | 0.13838119 | 6.280954964 | NA | NA | NA | NA | NA | NA | NA |
| KCNK3 | 207 | A | 16.6031 | 11.85621672 | 0.267715996 | 8.476978122 | NA | NA | NA | NA | NA | NA | NA |
| KCNK3 | 208 | L | 17.83825 | 13.44012876 | 0.311299659 | 9.839071159 | NA | NA | NA | NA | NA | NA | NA |
| KCNK3 | 209 | Q | 21.5375 | 8.684397964 | 0.180438751 | 5.168588803 | NA | NA | NA | NA | NA | NA | NA |
| KCNK3 | 210 | K | 23.945 | 12.62893143 | 0.288978374 | 8.710313177 | NA | NA | NA | NA | NA | NA | NA |
| KCNK3 | 211 | D | 26.184375 | 15.41890794 | 0.36574867 | 11.50809466 | NA | NA | NA | NA | NA | NA | NA |
| KCNK3 | 212 | Q | 26.47338889 | 16.56024278 | 0.397154172 | 13.09435167 | NA | NA | NA | NA | NA | NA | NA |
| KCNK3 | 213 | A | 21.2567 | 17.21060266 | 0.415049778 | 13.61674141 | NA | NA | NA | NA | NA | NA | NA |
| KCNK3 | 214 | L | 19.7785 | 14.23628389 | 0.333207035 | 10.83668748 | NA | NA | NA | NA | NA | NA | NA |
| KCNK3 | 215 | Q | 22.50861111 | 19.16081193 | 0.468712646 | 15.7510069 | NA | NA | NA | NA | NA | NA | NA |
| KCNK3 | 216 | T | 23.55692857 | 20.9036802 | 0.516670223 | 17.30836826 | NA | NA | NA | NA | NA | NA | NA |
| KCNK3 | 217 | Q | 20.44738889 | 19.54989138 | 0.479418738 | 16.16796128 | NA | NA | NA | NA | NA | NA | NA |
| KCNK3 | 218 | P | 17.1465 | 23.65052615 | 0.592253718 | 20.15082941 | NA | NA | NA | NA | NA | NA | NA |
| KCNK3 | 219 | Q | 14.93572222 | 23.76863212 | 0.595503577 | 22.65018227 | NA | NA | NA | NA | NA | NA | NA |
| KCNK3 | 220 | Y | 15.89 | 16.9496569 | 0.407869473 | 16.51927272 | NA | NA | NA | NA | NA | NA | NA |
| KCNK3 | 221 | V | 14.71578571 | 19.21465173 | 0.470194127 | 18.23396282 | NA | NA | NA | NA | NA | NA | NA |
| KCNK3 | 222 | A | 11.0837 | 20.75410675 | 0.51255449 | 19.79311087 | NA | NA | NA | NA | NA | NA | NA |
| KCNK3 | 223 | F | 10.26222727 | 19.28529748 | 0.472138049 | 18.33378221 | NA | NA | NA | NA | NA | NA | NA |
| KCNK3 | 224 | S | 11.35516667 | 15.75983242 | 0.375129707 | 14.77969302 | NA | NA | NA | NA | NA | NA | NA |
| KCNK3 | 225 | F | 10.42186364 | 16.23804529 | 0.388288436 | 15.18111425 | NA | NA | NA | NA | NA | NA | NA |

| | | | | | | | | | | | | | |
|---|---|---|---|---|---|---|---|---|---|---|---|---|---|
| KCNK3 | 226 | V | 5.8355 | 17.5308386 | 0.423861539 | 16.30182648 | NA | NA | NA | NA | NA | NA | NA |
| KCNK3 | 227 | Y | 6.599 | 15.33719182 | 0.363500131 | 14.51818833 | NA | NA | NA | NA | NA | NA | NA |
| KCNK3 | 228 | I | 7.082 | 11.50551959 | 0.25806605 | 10.66291672 | NA | NA | NA | NA | NA | NA | NA |
| KCNK3 | 229 | L | 3.589 | 13.82979595 | 0.322021923 | 12.61537318 | NA | NA | NA | NA | NA | NA | NA |
| KCNK3 | 230 | T | 1.4875 | 15.23877062 | 0.360791927 | 12.73319317 | NA | NA | NA | NA | NA | NA | NA |
| KCNK3 | 231 | G | 1.711 | 12.96556493 | 0.298241339 | 12.01901305 | NA | NA | NA | NA | NA | NA | NA |
| KCNK3 | 232 | L | 1.67075 | 8.268649587 | 0.168998825 | 5.896539554 | NA | NA | NA | NA | NA | NA | NA |
| KCNK3 | 233 | T | -0.461214286 | 12.35174696 | 0.281351237 | 9.673678456 | NA | NA | NA | NA | NA | NA | NA |
| KCNK3 | 234 | V | -0.461785714 | 13.3030681 | 0.307528234 | 10.64999405 | NA | NA | NA | NA | NA | NA | NA |
| KCNK3 | 235 | I | -0.25125 | 9.995991697 | 0.216529176 | 7.341880849 | NA | NA | NA | NA | NA | NA | NA |
| KCNK3 | 236 | G | -2.6485 | 9.59701167 | 0.205550655 | 6.974956216 | NA | NA | NA | NA | NA | NA | NA |
| KCNK3 | 237 | A | -5.1357 | 11.88639563 | 0.268546413 | 9.275926403 | NA | NA | NA | NA | NA | NA | NA |
| KCNK3 | 238 | F | -4.579590909 | 12.16559041 | 0.276228867 | 9.628856541 | NA | NA | NA | NA | NA | NA | NA |
| KCNK3 | 239 | L | -3.99425 | 7.03709926 | 0.135110911 | 7.79840131 | NA | NA | NA | NA | NA | NA | NA |
| KCNK3 | 240 | N | -8.435 | 9.864743129 | 0.212917679 | 8.934447643 | NA | NA | NA | NA | NA | NA | NA |
| KCNK3 | 241 | L | -9.92075 | 11.87794044 | 0.268313756 | 10.99715942 | NA | NA | NA | NA | NA | NA | NA |
| KCNK3 | 242 | V | -8.379785714 | 10.98392466 | 0.243713601 | 10.37264588 | NA | NA | NA | NA | NA | NA | NA |
| KCNK3 | 243 | V | -6.594642857 | 7.520133709 | 0.148402312 | 6.799064464 | NA | NA | NA | NA | NA | NA | NA |
| KCNK3 | 244 | L | -8.8435 | 5.161635497 | 0.083504775 | 4.338361516 | 0.97 | 0.35 | 0.73 | 1.7 | -0.79 | 0 | 0 |
| KCNK3 | 245 | R | -12.97604545 | 8.307382801 | 0.170064626 | 8.020074476 | NA | NA | NA | NA | NA | NA | NA |
| KCNK3 | 246 | F | -10.7675 | 7.598053567 | 0.150546392 | 7.630719967 | NA | NA | NA | NA | NA | NA | NA |
| KCNK3 | 247 | M | -8.107625 | 2.742221909 | 0.016931062 | 2.320504464 | 0.97 | 0.35 | 0.73 | 1.7 | -0.79 | 0 | 0 |
| KCNK3 | 248 | T | -12.96335714 | 6.623421019 | 0.123727948 | 6.759767823 | NA | NA | NA | NA | NA | NA | NA |
| KCNK3 | 249 | M | -14.60975 | 9.212482076 | 0.19496976 | 9.368603313 | NA | NA | NA | NA | NA | NA | NA |
| KCNK3 | 250 | N | -13.358625 | 10.55044696 | 0.231785826 | 10.72715164 | NA | NA | NA | NA | NA | NA | NA |
| KCNK3 | 251 | A | -15.9517 | 10.59194128 | 0.232927603 | 10.67357459 | NA | NA | NA | NA | NA | NA | NA |
| KCNK3 | 252 | E | -18.47838889 | 12.3959017 | 0.28256622 | 12.41324222 | NA | NA | NA | NA | NA | NA | NA |
| KCNK3 | 253 | D | -18.786875 | 14.67803192 | 0.34536238 | 14.84710376 | NA | NA | NA | NA | NA | NA | NA |
| KCNK3 | 254 | E | -18.55738889 | 15.48978567 | 0.367698974 | 15.59345361 | NA | NA | NA | NA | NA | NA | NA |
| KCNK3 | 255 | K | -21.36907143 | 15.40631189 | 0.365402071 | 15.41761347 | NA | NA | NA | NA | NA | NA | NA |
| KCNK3 | 256 | R | -23.8799 | 18.62170223 | 0.453878252 | 18.75461171 | NA | NA | NA | NA | NA | NA | NA |
| KCNK3 | 257 | D | -22.93925 | 20.17371629 | 0.496584195 | 20.31698945 | NA | NA | NA | NA | NA | NA | NA |
| KCNK3 | 258 | A | -25.3309 | 21.64143362 | 0.53697059 | 21.75419003 | NA | NA | NA | NA | NA | NA | NA |

| | | | | | | | | | | | | | |
|---|---|---|---|---|---|---|---|---|---|---|---|---|---|
| KCNK3 | 259 | E | -27.8047 | 22.74897261 | 0.56744615 | 22.85193453 | NA | NA | NA | NA | NA | NA | NA |
| KCNK4 | 2 | R | -17.16276623 | 16.06971257 | 0.401790875 | 11.6996996 | NA | NA | NA | NA | NA | NA | NA |
| KCNK4 | 3 | S | -12.49335714 | 18.20316646 | 0.46318884 | 13.8111421 | NA | NA | NA | NA | NA | NA | NA |
| KCNK4 | 4 | T | -12.564 | 19.96787275 | 0.513974739 | 15.74327391 | NA | NA | NA | NA | NA | NA | NA |
| KCNK4 | 5 | T | -12.27328571 | 16.56799101 | 0.416130665 | 12.36077866 | NA | NA | NA | NA | NA | NA | NA |
| KCNK4 | 6 | L | -9.530482143 | 13.01159998 | 0.313782462 | 8.795767555 | NA | NA | NA | NA | NA | NA | NA |
| KCNK4 | 7 | L | -6.995982143 | 18.05039041 | 0.458792148 | 13.89467445 | NA | NA | NA | NA | NA | NA | NA |
| KCNK4 | 8 | A | -7.399457143 | 18.57413013 | 0.473864681 | 14.49219526 | NA | NA | NA | NA | NA | NA | NA |
| KCNK4 | 9 | L | -6.697857143 | 12.58265683 | 0.301438049 | 8.471117025 | NA | NA | NA | NA | NA | NA | NA |
| KCNK4 | 10 | L | -2.764982143 | 16.71494631 | 0.420359843 | 12.72332396 | NA | NA | NA | NA | NA | NA | NA |
| KCNK4 | 11 | A | -2.376657143 | 19.2128584 | 0.492246431 | 15.22355769 | NA | NA | NA | NA | NA | NA | NA |
| KCNK4 | 12 | L | -3.193857143 | 16.81576311 | 0.423261217 | 12.98958029 | NA | NA | NA | NA | NA | NA | NA |
| KCNK4 | 13 | V | -0.776857143 | 14.50689719 | 0.356815127 | 10.55240682 | NA | NA | NA | NA | NA | NA | NA |
| KCNK4 | 14 | L | 0.937982143 | 18.8314256 | 0.481269302 | 18.75979278 | NA | NA | NA | NA | NA | NA | NA |
| KCNK4 | 15 | L | 0.804607143 | 21.13178085 | 0.547470466 | 20.75057104 | NA | NA | NA | NA | NA | NA | NA |
| KCNK4 | 16 | Y | 1.05002381 | 15.84366097 | 0.39528541 | 15.14376854 | NA | NA | NA | NA | NA | NA | NA |
| KCNK4 | 17 | L | 4.205107143 | 15.05889654 | 0.372700934 | 14.08474142 | NA | NA | NA | NA | NA | NA | NA |
| KCNK4 | 18 | V | 5.914428571 | 20.11219802 | 0.518128228 | 19.11432466 | NA | NA | NA | NA | NA | NA | NA |
| KCNK4 | 19 | S | 5.24852381 | 20.81893057 | 0.538467051 | 19.79755121 | NA | NA | NA | NA | NA | NA | NA |
| KCNK4 | 20 | G | 7.608607143 | 18.52307971 | 0.472395518 | 17.49260395 | NA | NA | NA | NA | NA | NA | NA |
| KCNK4 | 21 | A | 10.05025714 | 18.86367181 | 0.482197305 | 18.15510886 | NA | NA | NA | NA | NA | NA | NA |
| KCNK4 | 22 | L | 10.15235714 | 21.24910963 | 0.550847032 | 20.36497987 | NA | NA | NA | NA | NA | NA | NA |
| KCNK4 | 23 | V | 10.44857143 | 20.19458603 | 0.520499246 | 19.24132152 | NA | NA | NA | NA | NA | NA | NA |
| KCNK4 | 24 | F | 11.54058442 | 14.90979416 | 0.368409966 | 14.08718141 | NA | NA | NA | NA | NA | NA | NA |
| KCNK4 | 25 | R | 15.30431169 | 19.07359258 | 0.488238546 | 16.53768203 | NA | NA | NA | NA | NA | NA | NA |
| KCNK4 | 26 | A | 15.58645714 | 21.37603469 | 0.554499767 | 18.83863662 | NA | NA | NA | NA | NA | NA | NA |
| KCNK4 | 27 | L | 15.02373214 | 19.32301198 | 0.495416504 | 16.81507148 | NA | NA | NA | NA | NA | NA | NA |
| KCNK4 | 28 | E | 17.77719048 | 14.89108515 | 0.367871545 | 12.3912923 | NA | NA | NA | NA | NA | NA | NA |
| KCNK4 | 29 | Q | 20.52174603 | 18.42077732 | 0.469451391 | 15.36641253 | NA | NA | NA | NA | NA | NA | NA |
| KCNK4 | 30 | P | 22.14928571 | 19.83149361 | 0.510049929 | 15.65423709 | NA | NA | NA | NA | NA | NA | NA |
| KCNK4 | 31 | H | 22.77095714 | 15.43645727 | 0.383566629 | 10.9568281 | NA | NA | NA | NA | NA | NA | NA |
| KCNK4 | 32 | E | 23.39107937 | 13.69219099 | 0.333368966 | 9.253259581 | NA | NA | NA | NA | NA | NA | NA |
| KCNK4 | 33 | Q | 27.88052381 | 15.1594681 | 0.37559525 | 10.76820387 | NA | NA | NA | NA | NA | NA | NA |

| | | | | | | | | | | | | | |
|---|---|---|---|---|---|---|---|---|---|---|---|---|---|
| KCNK4 | 34 | Q | 28.15407937 | 14.31736851 | 0.351360744 | 13.17132482 | NA | NA | NA | NA | NA | NA | NA |
| KCNK4 | 35 | A | 28.72125714 | 10.89387452 | 0.25283714 | 9.677155803 | NA | NA | NA | NA | NA | NA | NA |
| KCNK4 | 36 | Q | 31.38952381 | 10.60034363 | 0.244389712 | 9.62330477 | NA | NA | NA | NA | NA | NA | NA |
| KCNK4 | 37 | R | 32.11667532 | 13.44221678 | 0.326175041 | 12.57691461 | NA | NA | NA | NA | NA | NA | NA |
| KCNK4 | 38 | E | 31.27230159 | 10.8480277 | 0.25151773 | 10.2967771 | NA | NA | NA | NA | NA | NA | NA |
| KCNK4 | 39 | L | 34.34860714 | 6.136680699 | 0.11593143 | 4.902882769 | 1.24 | 0.49 | 2.67 | 1.21 | -0.6 | 0 | 0 |
| KCNK4 | 40 | G | 37.16285714 | 10.53863378 | 0.242613785 | 9.68990125 | NA | NA | NA | NA | NA | NA | NA |
| KCNK4 | 41 | E | 36.46730159 | 11.11667077 | 0.25924892 | 10.38426491 | NA | NA | NA | NA | NA | NA | NA |
| KCNK4 | 42 | V | 37.59057143 | 7.323860731 | 0.150096892 | 6.784620312 | NA | NA | NA | NA | NA | NA | NA |
| KCNK4 | 43 | R | 40.09576623 | 6.321145308 | 0.121240076 | 5.61644808 | NA | NA | NA | NA | NA | NA | NA |
| KCNK4 | 44 | E | 42.07852381 | 9.504719301 | 0.212859101 | 9.146372097 | NA | NA | NA | NA | NA | NA | NA |
| KCNK4 | 45 | K | 41.73596825 | 10.68398133 | 0.246796694 | 10.14919747 | NA | NA | NA | NA | NA | NA | NA |
| KCNK4 | 46 | F | 43.47303896 | 5.092955331 | 0.085894403 | 4.372210042 | 0.84 | 1.21 | 3.47 | 0.2 | -1.8 | 0 | 0 |
| KCNK4 | 47 | L | 46.23760714 | 6.058806566 | 0.113690316 | 5.503756183 | NA | NA | NA | NA | NA | NA | NA |
| KCNK4 | 48 | R | 46.86194805 | 10.34798773 | 0.237127245 | 9.646850613 | NA | NA | NA | NA | NA | NA | NA |
| KCNK4 | 49 | A | 47.71825714 | 10.37126511 | 0.237797137 | 9.635171874 | NA | NA | NA | NA | NA | NA | NA |
| KCNK4 | 50 | H | 47.66005714 | 5.89632165 | 0.109014217 | 5.187696888 | NA | NA | NA | NA | NA | NA | NA |
| KCNK4 | 51 | P | 52.34428571 | 3.791482296 | 0.04843974 | 2.179708903 | 1.65 | -0.79 | -1.18 | 3.38 | 0.74 | 0 | 0 |
| KCNK4 | 52 | C | 51.76735714 | 2.114833563 | 1.88E-04 | 1.573525223 | 1.65 | -0.79 | -1.18 | 3.38 | 0.74 | 0 | 0 |
| KCNK4 | 53 | V | 48.50671429 | 2.184777792 | 0.002200922 | 1.449867796 | 0.74 | -0.89 | -1.18 | 15.01 | 1.17 | 1 | -1 |
| KCNK4 | 54 | S | 48.22752381 | 2.108300263 | 0 | 1.598754798 | 0.66 | -0.86 | -1.64 | 12.68 | 1.04 | 1 | -1 |
| KCNK4 | 55 | D | 44.77785714 | 5.457006505 | 0.096371311 | 4.618016566 | 1.12 | 0.14 | 0.78 | 13.44 | -0.02 | 1 | -1 |
| KCNK4 | 56 | Q | 46.09196825 | 5.416625241 | 0.095209192 | 4.281296666 | 1.16 | 0.45 | 1.17 | 16.79 | -0.29 | 1 | -1 |
| KCNK4 | 57 | E | 46.85474603 | 2.208273081 | 0.002877085 | 1.634438887 | 1.16 | 0.45 | 1.17 | 16.79 | -0.29 | 1 | -1 |
| KCNK4 | 58 | L | 42.33035714 | 2.260915301 | 0.004392058 | 1.636484023 | 0.84 | -0.08 | 0.67 | 2.37 | -0.06 | 0 | 0 |
| KCNK4 | 59 | G | 40.44435714 | 4.58473707 | 0.071268557 | 4.067021662 | 0.96 | 0.72 | 2.63 | 1.21 | -0.94 | 0 | 0 |
| KCNK4 | 60 | L | 41.36548214 | 4.949842018 | 0.081775792 | 4.110022723 | 0.96 | 0.72 | 2.63 | 1.21 | -0.94 | 0 | 0 |
| KCNK4 | 61 | L | 39.71548214 | 2.274370682 | 0.004779286 | 1.607859882 | 0.96 | 0.72 | 2.63 | 1.21 | -0.94 | 0 | 0 |
| KCNK4 | 62 | I | 36.79260714 | 2.430296072 | 0.009266611 | 1.499283275 | 1.24 | 0.49 | 2.67 | 1.21 | -0.6 | 0 | 0 |
| KCNK4 | 63 | K | 36.54019048 | 4.406984797 | 0.066153083 | 3.833144352 | 1.07 | 0.36 | 2.28 | 1.43 | -0.55 | 0 | 0 |
| KCNK4 | 64 | E | 37.50063492 | 5.298498089 | 0.09180965 | 4.421693705 | 1.03 | 0.49 | 2.67 | 1.21 | -0.6 | 0 | 0 |
| KCNK4 | 65 | V | 34.106 | 2.438308635 | 0.009497202 | 1.620394146 | 0.93 | 0.17 | 1.8 | 1.76 | -0.37 | 0 | 0 |
| KCNK4 | 66 | A | 31.31145714 | 2.40847296 | 0.008638571 | 1.564514993 | 0.88 | 0.17 | 1.7 | 1.76 | -0.49 | 0 | 0 |

| | | | | | | | | | | | | | |
|---|---|---|---|---|---|---|---|---|---|---|---|---|---|
| KCNK4 | 67 | D | 31.60023214 | 4.667720429 | 0.073656708 | 3.654156873 | 1.02 | 0.77 | 3.13 | 0.09 | -1 | 0 | 0 |
| KCNK4 | 68 | A | 30.46965714 | 5.787193275 | 0.105873647 | 4.774338113 | 1.02 | 0.77 | 3.13 | 0.09 | -1 | 0 | 0 |
| KCNK4 | 69 | L | 28.66098214 | 2.443428943 | 0.009644558 | 1.589233051 | 1.36 | 0.49 | 2.4 | 1.21 | -0.92 | 0 | 0 |
| KCNK4 | 70 | G | 26.13135714 | 4.133361949 | 0.058278582 | 1.896804711 | 2.38 | 0.17 | 1.7 | 1.76 | -0.49 | 0 | 0 |
| KCNK4 | 71 | G | 25.92785714 | 6.805314173 | 0.135173812 | 4.449835125 | 2.38 | 0.17 | 1.7 | 1.76 | -0.49 | 0 | 0 |
| KCNK4 | 72 | G | 25.40235714 | 7.635272687 | 0.159058914 | 4.374114296 | 3.29 | -0.32 | 0.65 | 2.57 | 0.16 | 0 | 0 |
| KCNK4 | 73 | A | 27.90085714 | 7.349042387 | 0.150821587 | 6.005344646 | NA | NA | NA | NA | NA | NA | NA |
| KCNK4 | 74 | D | 27.25885714 | 8.781357184 | 0.192041702 | 8.067320844 | NA | NA | NA | NA | NA | NA | NA |
| KCNK4 | 75 | P | 30.21142857 | 7.397882738 | 0.152227147 | 6.675666168 | NA | NA | NA | NA | NA | NA | NA |
| KCNK4 | 76 | E | 33.24074603 | 11.08400568 | 0.258308863 | 10.40806114 | NA | NA | NA | NA | NA | NA | NA |
| KCNK4 | 77 | T | 33.10785714 | 12.5080552 | 0.299291113 | 10.96363355 | NA | NA | NA | NA | NA | NA | NA |
| KCNK4 | 84 | S | 23.76185714 | 17.66471616 | 0.447692956 | 14.40930055 | NA | NA | NA | NA | NA | NA | NA |
| KCNK4 | 85 | H | 23.81285714 | 13.31894425 | 0.322627422 | 10.04613779 | NA | NA | NA | NA | NA | NA | NA |
| KCNK4 | 86 | S | 21.76569048 | 14.79114526 | 0.364995408 | 11.52441768 | NA | NA | NA | NA | NA | NA | NA |
| KCNK4 | 87 | A | 19.36105714 | 11.74754196 | 0.277404554 | 7.716464644 | NA | NA | NA | NA | NA | NA | NA |
| KCNK4 | 88 | W | 18.10364286 | 11.98370531 | 0.284201021 | 7.972428384 | NA | NA | NA | NA | NA | NA | NA |
| KCNK4 | 89 | D | 18.10698214 | 17.46375361 | 0.441909521 | 13.4332009 | NA | NA | NA | NA | NA | NA | NA |
| KCNK4 | 90 | L | 14.12398214 | 19.48076859 | 0.49995653 | 18.25515489 | NA | NA | NA | NA | NA | NA | NA |
| KCNK4 | 91 | G | 13.67185714 | 17.36018989 | 0.438929095 | 16.19139383 | NA | NA | NA | NA | NA | NA | NA |
| KCNK4 | 92 | S | 15.20502381 | 14.3408529 | 0.352036593 | 13.02309388 | NA | NA | NA | NA | NA | NA | NA |
| KCNK4 | 93 | A | 13.05965714 | 15.18553476 | 0.376345414 | 14.18054154 | NA | NA | NA | NA | NA | NA | NA |
| KCNK4 | 94 | F | 8.563402597 | 14.60896728 | 0.359752568 | 13.5077035 | NA | NA | NA | NA | NA | NA | NA |
| KCNK4 | 95 | F | 10.88167532 | 10.92778015 | 0.253812899 | 10.03522927 | NA | NA | NA | NA | NA | NA | NA |
| KCNK4 | 96 | F | 12.01458442 | 9.499277709 | 0.212702499 | 8.915897952 | NA | NA | NA | NA | NA | NA | NA |
| KCNK4 | 97 | S | 8.187690476 | 11.52415871 | 0.270975881 | 10.66874095 | NA | NA | NA | NA | NA | NA | NA |
| KCNK4 | 98 | G | 6.276857143 | 9.453190414 | 0.211376168 | 8.44326012 | NA | NA | NA | NA | NA | NA | NA |
| KCNK4 | 99 | T | 8.299285714 | 6.462887977 | 0.125319242 | 5.672186051 | NA | NA | NA | NA | NA | NA | NA |
| KCNK4 | 100 | I | 6.614357143 | 7.171979573 | 0.145725955 | 6.462352093 | NA | NA | NA | NA | NA | NA | NA |
| KCNK4 | 101 | I | 2.563732143 | 7.732713495 | 0.161863131 | 7.4157406 | NA | NA | NA | NA | NA | NA | NA |
| KCNK4 | 102 | T | 2.917285714 | 4.697291241 | 0.074507716 | 4.213759879 | 0.65 | -0.78 | -0.6 | 2.23 | 0.78 | 0 | 0 |
| KCNK4 | 103 | T | 3.262285714 | 2.192984496 | 0.0024371 | 1.780432818 | 0.65 | -0.78 | -0.6 | 2.23 | 0.78 | 0 | 0 |
| KCNK4 | 104 | I | 6.237607143 | 2.349968936 | 0.006954903 | 1.818821304 | 1.15 | -0.8 | -0.4 | 3.38 | 0.81 | 0 | 0 |
| KCNK4 | 105 | G | 9.612357143 | 2.318500809 | 0.006049292 | 1.618784441 | 1.11 | -0.8 | -0.4 | 3.38 | 0.81 | 0 | 0 |

| | | | | | | | | | | | | | |
|---|---|---|---|---|---|---|---|---|---|---|---|---|---|
| KCNK4 | 106 | Y | 11.12869048 | 2.466719887 | 0.01031484 | 1.802113395 | 0.99 | -0.8 | -0.4 | 3.38 | 0.81 | 0 | 0 |
| KCNK4 | 107 | G | 15.21260714 | 3.805875589 | 0.04885396 | 2.728581646 | 1.37 | -0.8 | -0.4 | 3.38 | 0.81 | 0 | 0 |
| KCNK4 | 108 | N | 17.15573214 | 4.51459821 | 0.069250054 | 2.112936563 | 2.46 | -0.8 | -0.4 | 3.38 | 0.81 | 0 | 0 |
| KCNK4 | 109 | V | 17.93185714 | 6.436974755 | 0.124573494 | 3.037490487 | 4.26 | -0.8 | -0.4 | 3.38 | 0.81 | 0 | 0 |
| KCNK4 | 110 | A | 18.59885714 | 6.671047444 | 0.131309794 | 2.477773304 | 4.26 | -0.8 | -0.4 | 3.38 | 0.81 | 0 | 0 |
| KCNK4 | 111 | L | 16.69435714 | 9.769176117 | 0.220469817 | 5.564459278 | NA | NA | NA | NA | NA | NA | NA |
| KCNK4 | 112 | R | 21.81803896 | 8.014023521 | 0.16995886 | 3.779262625 | 4.26 | -0.8 | -0.4 | 3.38 | 0.81 | 0 | 0 |
| KCNK4 | 113 | T | 19.15857143 | 13.8396681 | 0.337613161 | 9.60689032 | NA | NA | NA | NA | NA | NA | NA |
| KCNK4 | 114 | D | 17.92010714 | 16.61228949 | 0.417405517 | 12.38262566 | NA | NA | NA | NA | NA | NA | NA |
| KCNK4 | 115 | A | 15.50505714 | 18.73857489 | 0.478597182 | 14.51282084 | NA | NA | NA | NA | NA | NA | NA |
| KCNK4 | 116 | G | 14.41535714 | 15.62492157 | 0.388990381 | 13.40542354 | NA | NA | NA | NA | NA | NA | NA |
| KCNK4 | 117 | R | 15.7014026 | 11.3407979 | 0.265699001 | 13.0758697 | NA | NA | NA | NA | NA | NA | NA |
| KCNK4 | 118 | L | 11.09560714 | 17.30958235 | 0.437472678 | 16.38571903 | NA | NA | NA | NA | NA | NA | NA |
| KCNK4 | 119 | F | 9.46812987 | 16.7535678 | 0.421471319 | 15.66309144 | NA | NA | NA | NA | NA | NA | NA |
| KCNK4 | 120 | C | 9.216690476 | 11.62308513 | 0.273822852 | 10.52377969 | NA | NA | NA | NA | NA | NA | NA |
| KCNK4 | 121 | I | 7.261607143 | 14.17113154 | 0.347152238 | 13.34808085 | NA | NA | NA | NA | NA | NA | NA |
| KCNK4 | 122 | F | 5.240584416 | 15.87920483 | 0.396308315 | 15.23608135 | NA | NA | NA | NA | NA | NA | NA |
| KCNK4 | 123 | Y | 5.498440476 | 13.87664408 | 0.338677281 | 13.22884677 | NA | NA | NA | NA | NA | NA | NA |
| KCNK4 | 124 | A | 3.374257143 | 10.88184212 | 0.252490864 | 10.24855344 | NA | NA | NA | NA | NA | NA | NA |
| KCNK4 | 125 | L | 1.322107143 | 12.81067812 | 0.308000199 | 12.19180486 | NA | NA | NA | NA | NA | NA | NA |
| KCNK4 | 126 | V | 0 | 13.94847214 | 0.340744397 | 13.38122966 | NA | NA | NA | NA | NA | NA | NA |
| KCNK4 | 127 | G | 0.230357143 | 11.69419937 | 0.275869425 | 10.96760918 | NA | NA | NA | NA | NA | NA | NA |
| KCNK4 | 128 | I | 0.640482143 | 7.100197251 | 0.143660155 | 6.474198414 | NA | NA | NA | NA | NA | NA | NA |
| KCNK4 | 129 | P | -2.833571429 | 9.598197435 | 0.215549278 | 4.196307117 | 5.47 | -0.77 | -0.7 | 1.66 | 0.77 | 0 | 0 |
| KCNK4 | 130 | L | -2.669232143 | 11.51951622 | 0.270842276 | 6.129407577 | NA | NA | NA | NA | NA | NA | NA |
| KCNK4 | 131 | F | -0.838038961 | 9.16432327 | 0.203062956 | 3.984071785 | 5.6 | -0.77 | -0.7 | 1.66 | 0.77 | 0 | 0 |
| KCNK4 | 132 | G | -4.715357143 | 7.783758796 | 0.163332147 | 2.390539746 | 5.47 | -0.77 | -0.7 | 1.66 | 0.77 | 0 | 0 |
| KCNK4 | 133 | I | -7.397732143 | 9.735656526 | 0.219505167 | 4.315542679 | 5.47 | -0.77 | -0.7 | 1.66 | 0.77 | 0 | 0 |
| KCNK4 | 134 | L | -4.757857143 | 11.95698198 | 0.283431959 | 6.549202999 | NA | NA | NA | NA | NA | NA | NA |
| KCNK4 | 135 | L | -4.637732143 | 7.485702439 | 0.154754482 | 2.120571528 | 5.47 | -0.77 | -0.7 | 1.66 | 0.77 | 0 | 0 |
| KCNK4 | 136 | A | -8.916857143 | 9.891059094 | 0.223977447 | 4.452942435 | 5.47 | -0.77 | -0.7 | 1.66 | 0.77 | 0 | 0 |
| KCNK4 | 137 | G | -9.518607143 | 12.56399821 | 0.300901079 | 8.851843513 | NA | NA | NA | NA | NA | NA | NA |
| KCNK4 | 138 | V | -6.867142857 | 13.98719672 | 0.341858839 | 8.558061014 | NA | NA | NA | NA | NA | NA | NA |

| | | | | | | | | | | | | | |
|---|---|---|---|---|---|---|---|---|---|---|---|---|---|
| KCNK4 | 139 | G | -9.145107143 | 12.78454794 | 0.307248207 | 7.382266572 | NA | NA | NA | NA | NA | NA | NA |
| KCNK4 | 140 | D | -12.67460714 | 9.890125429 | 0.223950577 | 5.886445136 | NA | NA | NA | NA | NA | NA | NA |
| KCNK4 | 141 | R | -11.57476623 | 14.92564498 | 0.368866131 | 10.90399511 | NA | NA | NA | NA | NA | NA | NA |
| KCNK4 | 142 | L | -8.396107143 | 16.29658449 | 0.408319947 | 12.46811926 | NA | NA | NA | NA | NA | NA | NA |
| KCNK4 | 143 | G | -12.06260714 | 14.96325185 | 0.369948407 | 10.91494885 | NA | NA | NA | NA | NA | NA | NA |
| KCNK4 | 144 | S | -14.29752381 | 15.82487475 | 0.394744768 | 11.81696203 | NA | NA | NA | NA | NA | NA | NA |
| KCNK4 | 145 | S | -11.88735714 | 18.72254491 | 0.478135861 | 14.7120301 | NA | NA | NA | NA | NA | NA | NA |
| KCNK4 | 146 | L | -10.70285714 | 18.12011231 | 0.460798652 | 14.01074204 | NA | NA | NA | NA | NA | NA | NA |
| KCNK4 | 147 | R | -14.80803896 | 11.53059465 | 0.271161099 | 7.472415179 | NA | NA | NA | NA | NA | NA | NA |
| KCNK4 | 148 | H | -16.70705714 | 19.85037597 | 0.510593338 | 15.86065662 | NA | NA | NA | NA | NA | NA | NA |
| KCNK4 | 149 | G | -14.74110714 | 22.52234069 | 0.587488929 | 18.51855496 | NA | NA | NA | NA | NA | NA | NA |
| KCNK4 | 150 | I | -14.89223214 | 19.78411459 | 0.508686424 | 15.72162763 | NA | NA | NA | NA | NA | NA | NA |
| KCNK4 | 151 | G | -18.73410714 | 22.86865186 | 0.597455304 | 18.87518947 | NA | NA | NA | NA | NA | NA | NA |
| KCNK4 | 152 | H | -18.10315714 | 25.65188868 | 0.677553157 | 21.6656237 | NA | NA | NA | NA | NA | NA | NA |
| KCNK4 | 153 | I | -16.39885714 | 26.49519741 | 0.70182246 | 22.48276962 | NA | NA | NA | NA | NA | NA | NA |
| KCNK4 | 154 | E | -19.70296825 | 20.30596425 | 0.523704563 | 16.8500627 | NA | NA | NA | NA | NA | NA | NA |
| KCNK4 | 155 | A | -22.37545714 | 26.46298915 | 0.70089555 | 22.96506304 | NA | NA | NA | NA | NA | NA | NA |
| KCNK4 | 156 | I | -20.22248214 | 29.02148004 | 0.774525514 | 25.55058846 | NA | NA | NA | NA | NA | NA | NA |
| KCNK4 | 157 | F | -19.44894805 | 25.5575527 | 0.674838293 | 22.52086169 | NA | NA | NA | NA | NA | NA | NA |
| KCNK4 | 158 | L | -25.10610714 | 24.71664957 | 0.650638219 | 21.34555396 | NA | NA | NA | NA | NA | NA | NA |
| KCNK4 | 159 | K | -25.67252381 | 29.98062709 | 0.80212849 | 26.53909688 | NA | NA | NA | NA | NA | NA | NA |
| KCNK4 | 160 | W | -21.48107143 | 31.73440611 | 0.852599917 | 28.25475479 | NA | NA | NA | NA | NA | NA | NA |
| KCNK4 | 161 | H | -27.90605714 | 29.95354627 | 0.80134914 | 26.51319542 | NA | NA | NA | NA | NA | NA | NA |
| KCNK4 | 162 | V | -23.39685714 | 26.66609388 | 0.706740634 | 23.21828277 | NA | NA | NA | NA | NA | NA | NA |
| KCNK4 | 163 | P | -24.78185714 | 24.04271251 | 0.631243207 | 20.59104157 | NA | NA | NA | NA | NA | NA | NA |
| KCNK4 | 164 | P | -25.67571429 | 20.76157607 | 0.536816465 | 17.30235111 | NA | NA | NA | NA | NA | NA | NA |
| KCNK4 | 165 | E | -24.60030159 | 19.79499758 | 0.508999622 | 16.45402344 | NA | NA | NA | NA | NA | NA | NA |
| KCNK4 | 166 | L | -20.62010714 | 21.83504509 | 0.567709476 | 18.36594922 | NA | NA | NA | NA | NA | NA | NA |
| KCNK4 | 167 | V | -20.673 | 21.08669403 | 0.546172927 | 17.91517322 | NA | NA | NA | NA | NA | NA | NA |
| KCNK4 | 168 | R | -21.52358442 | 17.58200381 | 0.445312605 | 14.10372882 | NA | NA | NA | NA | NA | NA | NA |
| KCNK4 | 169 | V | -16.77014286 | 18.89211542 | 0.483015874 | 15.41508515 | NA | NA | NA | NA | NA | NA | NA |
| KCNK4 | 170 | L | -15.72535714 | 20.97679709 | 0.54301024 | 17.41424339 | NA | NA | NA | NA | NA | NA | NA |
| KCNK4 | 171 | S | -15.35419048 | 18.61705715 | 0.475100063 | 15.07276564 | NA | NA | NA | NA | NA | NA | NA |

| | | | | | | | | | | | | | |
|---|---|---|---|---|---|---|---|---|---|---|---|---|---|
| KCNK4 | 172 | A | -12.92325714 | 16.30910013 | 0.40868013 | 12.7102852 | NA | NA | NA | NA | NA | NA | NA |
| KCNK4 | 173 | M | -10.84110714 | 18.99396162 | 0.485946872 | 15.39011551 | NA | NA | NA | NA | NA | NA | NA |
| KCNK4 | 174 | L | -11.02773214 | 20.72584158 | 0.535788074 | 17.12123135 | NA | NA | NA | NA | NA | NA | NA |
| KCNK4 | 175 | F | -10.3954026 | 13.85049523 | 0.337924752 | 10.25114952 | NA | NA | NA | NA | NA | NA | NA |
| KCNK4 | 176 | L | -7.209732143 | 17.56541295 | 0.444835142 | 13.9697334 | NA | NA | NA | NA | NA | NA | NA |
| KCNK4 | 177 | L | -5.986982143 | 20.62881094 | 0.532995661 | 17.03604624 | NA | NA | NA | NA | NA | NA | NA |
| KCNK4 | 178 | I | -6.404732143 | 21.64095019 | 0.562123682 | 17.49773057 | NA | NA | NA | NA | NA | NA | NA |
| KCNK4 | 179 | G | -3.790107143 | 19.52073833 | 0.501106806 | 15.08082197 | NA | NA | NA | NA | NA | NA | NA |
| KCNK4 | 180 | C | -1.536190476 | 19.56201137 | 0.502294589 | 15.10188991 | NA | NA | NA | NA | NA | NA | NA |
| KCNK4 | 181 | L | -0.977732143 | 21.82785876 | 0.567502663 | 21.22262457 | NA | NA | NA | NA | NA | NA | NA |
| KCNK4 | 182 | L | -1.394732143 | 20.54424012 | 0.530561825 | 19.83698 | NA | NA | NA | NA | NA | NA | NA |
| KCNK4 | 183 | F | 1.322857143 | 16.88841949 | 0.425352171 | 16.30360813 | NA | NA | NA | NA | NA | NA | NA |
| KCNK4 | 184 | V | 1.568 | 16.76136194 | 0.421695623 | 15.90484772 | NA | NA | NA | NA | NA | NA | NA |
| KCNK4 | 185 | L | 2.178607143 | 20.50994525 | 0.529574865 | 19.48805698 | NA | NA | NA | NA | NA | NA | NA |
| KCNK4 | 186 | T | 3.156857143 | 22.06054872 | 0.57419917 | 21.00725488 | NA | NA | NA | NA | NA | NA | NA |
| KCNK4 | 187 | P | 5.134 | 18.56085674 | 0.47348269 | 17.64313899 | NA | NA | NA | NA | NA | NA | NA |
| KCNK4 | 188 | T | 7.216571429 | 18.61020376 | 0.474902832 | 17.56328563 | NA | NA | NA | NA | NA | NA | NA |
| KCNK4 | 189 | F | 7.583493506 | 22.1505495 | 0.576789273 | 21.04060284 | NA | NA | NA | NA | NA | NA | NA |
| KCNK4 | 190 | V | 9.006142857 | 21.89293772 | 0.569375548 | 20.78723485 | NA | NA | NA | NA | NA | NA | NA |
| KCNK4 | 191 | F | 10.44312987 | 16.53691395 | 0.415236309 | 15.91680518 | NA | NA | NA | NA | NA | NA | NA |
| KCNK4 | 192 | C | 13.33985714 | 20.81941815 | 0.538481083 | 19.74090176 | NA | NA | NA | NA | NA | NA | NA |
| KCNK4 | 193 | Y | 14.76377381 | 23.17171968 | 0.606177192 | 21.80055877 | NA | NA | NA | NA | NA | NA | NA |
| KCNK4 | 194 | M | 14.37773214 | 20.7085662 | 0.535290912 | 18.57206032 | NA | NA | NA | NA | NA | NA | NA |
| KCNK4 | 195 | E | 16.22596825 | 16.30372798 | 0.408525527 | 13.87775334 | NA | NA | NA | NA | NA | NA | NA |
| KCNK4 | 196 | D | 19.37535714 | 19.14374702 | 0.490257497 | 16.79163487 | NA | NA | NA | NA | NA | NA | NA |
| KCNK4 | 197 | W | 15.40614286 | 13.77240524 | 0.335677426 | 11.33512692 | NA | NA | NA | NA | NA | NA | NA |
| KCNK4 | 198 | S | 14.97785714 | 19.49235802 | 0.500290058 | 17.10233872 | NA | NA | NA | NA | NA | NA | NA |
| KCNK4 | 199 | K | 11.64796825 | 19.80484552 | 0.509283032 | 18.79413564 | NA | NA | NA | NA | NA | NA | NA |
| KCNK4 | 200 | L | 11.88398214 | 17.16511471 | 0.433315091 | 16.13458812 | NA | NA | NA | NA | NA | NA | NA |
| KCNK4 | 201 | E | 14.87507937 | 14.9086817 | 0.368377951 | 13.73405158 | NA | NA | NA | NA | NA | NA | NA |
| KCNK4 | 202 | A | 10.43265714 | 15.7054934 | 0.391309132 | 14.92758184 | NA | NA | NA | NA | NA | NA | NA |
| KCNK4 | 203 | I | 7.400232143 | 14.74313003 | 0.363613594 | 13.7304242 | NA | NA | NA | NA | NA | NA | NA |
| KCNK4 | 204 | Y | 10.69827381 | 9.814288257 | 0.221768084 | 9.493195159 | NA | NA | NA | NA | NA | NA | NA |

| | | | | | | | | | | | | | |
|---|---|---|---|---|---|---|---|---|---|---|---|---|---|
| KCNK4 | 205 | F | 9.840857143 | 10.90641696 | 0.253198095 | 9.833390755 | NA | NA | NA | NA | NA | NA | NA |
| KCNK4 | 206 | V | 5.859571429 | 11.60035913 | 0.273168828 | 10.72428261 | NA | NA | NA | NA | NA | NA | NA |
| KCNK4 | 207 | I | 4.272107143 | 9.736147133 | 0.219519286 | 8.681915817 | NA | NA | NA | NA | NA | NA | NA |
| KCNK4 | 208 | V | 7.327 | 6.770335073 | 0.13416716 | 5.849344838 | NA | NA | NA | NA | NA | NA | NA |
| KCNK4 | 209 | T | 5.888285714 | 7.274401212 | 0.148673513 | 6.6183159 | NA | NA | NA | NA | NA | NA | NA |
| KCNK4 | 210 | L | 1.852482143 | 7.743283025 | 0.162167308 | 7.127803306 | NA | NA | NA | NA | NA | NA | NA |
| KCNK4 | 211 | T | 2.151571429 | 4.760739648 | 0.076333677 | 3.942284681 | 1.06 | -0.77 | -0.7 | 1.66 | 0.77 | 0 | 0 |
| KCNK4 | 212 | T | 3.019714286 | 2.187322793 | 0.002274163 | 1.504254688 | 1.06 | -0.77 | -0.7 | 1.66 | 0.77 | 0 | 0 |
| KCNK4 | 213 | V | 6.320857143 | 2.332155655 | 0.00644226 | 1.623081417 | 1.03 | -0.8 | -0.4 | 3.38 | 0.81 | 0 | 0 |
| KCNK4 | 214 | G | 9.288107143 | 2.308912081 | 0.005773341 | 1.607512975 | 1.11 | -0.8 | -0.4 | 3.38 | 0.81 | 0 | 0 |
| KCNK4 | 215 | F | 11.44422078 | 2.471846274 | 0.010462371 | 1.843706783 | 0.99 | -0.8 | -0.4 | 3.38 | 0.81 | 0 | 0 |
| KCNK4 | 216 | G | 14.77660714 | 4.373334197 | 0.065184664 | 3.087682297 | 1.37 | -0.8 | -0.4 | 3.38 | 0.81 | 0 | 0 |
| KCNK4 | 217 | D | 15.68023214 | 5.799387726 | 0.106224587 | 3.419723798 | 2.46 | -0.8 | -0.4 | 3.38 | 0.81 | 0 | 0 |
| KCNK4 | 218 | Y | 15.36044048 | 8.30475972 | 0.178325861 | 6.956303553 | NA | NA | NA | NA | NA | NA | NA |
| KCNK4 | 219 | V | 14.46528571 | 7.93940892 | 0.167811551 | 6.564472264 | NA | NA | NA | NA | NA | NA | NA |
| KCNK4 | 220 | A | 13.03545714 | 12.41753458 | 0.29668605 | 11.3387721 | NA | NA | NA | NA | NA | NA | NA |
| KCNK4 | 221 | G | 12.37635714 | 14.1205172 | 0.345695625 | 13.14555128 | NA | NA | NA | NA | NA | NA | NA |
| KCNK4 | 222 | A | 14.85745714 | 12.76053678 | 0.306557197 | 11.41719473 | NA | NA | NA | NA | NA | NA | NA |
| KCNK4 | 223 | D | 17.30648214 | 15.97293692 | 0.3990058 | 13.52860316 | NA | NA | NA | NA | NA | NA | NA |
| KCNK4 | 224 | P | 16.45628571 | 19.62966311 | 0.504241516 | 17.41962565 | NA | NA | NA | NA | NA | NA | NA |
| KCNK4 | 225 | R | 17.14612987 | 21.29979103 | 0.552305576 | 18.87330512 | NA | NA | NA | NA | NA | NA | NA |
| KCNK4 | 226 | Q | 20.62563492 | 24.61882008 | 0.647822817 | 21.3338817 | NA | NA | NA | NA | NA | NA | NA |
| KCNK4 | 227 | D | 18.61835714 | 27.57114044 | 0.732786669 | 25.55023545 | NA | NA | NA | NA | NA | NA | NA |
| KCNK4 | 228 | S | 14.34802381 | 25.3602274 | 0.669159533 | 23.9949491 | NA | NA | NA | NA | NA | NA | NA |
| KCNK4 | 229 | P | 12.16385714 | 25.03286921 | 0.6597386 | 23.93101869 | NA | NA | NA | NA | NA | NA | NA |
| KCNK4 | 230 | A | 8.965457143 | 23.62431218 | 0.619202202 | 22.51693707 | NA | NA | NA | NA | NA | NA | NA |
| KCNK4 | 231 | Y | 11.62827381 | 18.2389029 | 0.464217287 | 17.56755974 | NA | NA | NA | NA | NA | NA | NA |
| KCNK4 | 232 | Q | 11.39152381 | 18.0153562 | 0.45778391 | 17.21934773 | NA | NA | NA | NA | NA | NA | NA |
| KCNK4 | 233 | P | 7.353285714 | 19.80211931 | 0.509204576 | 18.8758891 | NA | NA | NA | NA | NA | NA | NA |
| KCNK4 | 234 | L | 5.198232143 | 19.35705783 | 0.496396299 | 18.3125746 | NA | NA | NA | NA | NA | NA | NA |
| KCNK4 | 235 | V | 7.129285714 | 15.74327491 | 0.392396433 | 14.63577888 | NA | NA | NA | NA | NA | NA | NA |
| KCNK4 | 236 | W | 7.080428571 | 15.67088632 | 0.390313186 | 15.02265288 | NA | NA | NA | NA | NA | NA | NA |
| KCNK4 | 237 | F | 1.063220779 | 16.97415612 | 0.427819557 | 16.46148854 | NA | NA | NA | NA | NA | NA | NA |

| | | | | | | | | | | | | | |
|---|---|---|---|---|---|---|---|---|---|---|---|---|---|
| KCNK4 | 238 | W | 3.446714286 | 13.93309854 | 0.340301965 | 13.4410445 | NA | NA | NA | NA | NA | NA | NA |
| KCNK4 | 239 | I | 3.287107143 | 11.09922601 | 0.258746883 | 10.45301215 | NA | NA | NA | NA | NA | NA | NA |
| KCNK4 | 240 | L | 0.566357143 | 12.90933844 | 0.310839511 | 12.36421388 | NA | NA | NA | NA | NA | NA | NA |
| KCNK4 | 241 | L | -1.679482143 | 13.82169823 | 0.337096012 | 9.352146431 | NA | NA | NA | NA | NA | NA | NA |
| KCNK4 | 242 | G | -0.561107143 | 11.95707364 | 0.283434597 | 11.44672809 | NA | NA | NA | NA | NA | NA | NA |
| KCNK4 | 243 | L | -0.908482143 | 7.471386819 | 0.154342497 | 6.763967902 | NA | NA | NA | NA | NA | NA | NA |
| KCNK4 | 244 | A | -4.422457143 | 10.91993361 | 0.253587086 | 5.348027165 | NA | NA | NA | NA | NA | NA | NA |
| KCNK4 | 245 | Y | -4.439690476 | 12.1738818 | 0.289674047 | 6.631339112 | NA | NA | NA | NA | NA | NA | NA |
| KCNK4 | 246 | F | -1.646220779 | 9.315823367 | 0.207422927 | 4.873144606 | 4.52 | -0.77 | -0.7 | 1.66 | 0.77 | 0 | 0 |
| KCNK4 | 247 | A | -5.613457143 | 7.334789568 | 0.15041141 | 1.749953429 | 5.63 | -0.61 | -0.2 | 1.33 | 0.73 | 0 | 0 |
| KCNK4 | 248 | S | -8.52102381 | 9.135636759 | 0.202237397 | 5.305562772 | NA | NA | NA | NA | NA | NA | NA |
| KCNK4 | 249 | V | -7.084142857 | 9.597199279 | 0.215520552 | 5.466944607 | NA | NA | NA | NA | NA | NA | NA |
| KCNK4 | 250 | L | -5.886857143 | 6.35115769 | 0.122103793 | 2.267935169 | 4.25 | -0.38 | 0.55 | 0.83 | 0.68 | 0 | 0 |
| KCNK4 | 251 | T | -10.07085714 | 5.404002036 | 0.094845913 | 1.684364764 | 3.94 | -0.6 | -2.6 | 26.83 | 1.2 | 1 | 1 |
| KCNK4 | 252 | T | -11.87271429 | 7.094965257 | 0.143509585 | 3.509913572 | 3.62 | -0.66 | -2.8 | 19.59 | 0.99 | 1 | 1 |
| KCNK4 | 253 | I | -9.047232143 | 8.604857349 | 0.186962271 | 4.433012028 | 4.25 | -0.38 | 0.55 | 0.83 | 0.68 | 0 | 0 |
| KCNK4 | 254 | G | -10.87185714 | 6.336822705 | 0.121691251 | 2.159651146 | 4.25 | -0.38 | 0.55 | 0.83 | 0.68 | 0 | 0 |
| KCNK4 | 255 | N | -14.77448214 | 5.331268611 | 0.092752742 | 1.86063504 | 3.62 | -0.66 | -2.8 | 19.59 | 0.99 | 1 | 1 |
| KCNK4 | 256 | W | -13.3035 | 8.376670938 | 0.18039537 | 4.809133128 | 3.62 | -0.66 | -2.8 | 19.59 | 0.99 | 1 | 1 |
| KCNK4 | 257 | L | -11.20673214 | 9.576630932 | 0.214928622 | 5.395091718 | NA | NA | NA | NA | NA | NA | NA |
| KCNK4 | 258 | R | -15.12976623 | 5.357702026 | 0.09351346 | 1.742293016 | 4.41 | -0.39 | -1.13 | 17.89 | 1 | 1 | 1 |
| KCNK4 | 259 | V | -17.51914286 | 8.830115175 | 0.193444892 | 5.749879836 | NA | NA | NA | NA | NA | NA | NA |
| KCNK4 | 260 | V | -15.34814286 | 12.2691397 | 0.292415443 | 8.780907772 | NA | NA | NA | NA | NA | NA | NA |
| KCNMA1 | 85 | R | 18.59075 | 28.80742429 | 0.363932374 | 23.1771229 | NA | NA | NA | NA | NA | NA | NA |
| KCNMA1 | 86 | M | 16.265875 | 35.23390567 | 0.451370956 | 29.60326715 | NA | NA | NA | NA | NA | NA | NA |
| KCNMA1 | 87 | W | 12.60167857 | 31.32072818 | 0.398128338 | 27.51771137 | NA | NA | NA | NA | NA | NA | NA |
| KCNMA1 | 88 | W | 12.96153571 | 34.64075626 | 0.443300576 | 31.63767166 | NA | NA | NA | NA | NA | NA | NA |
| KCNMA1 | 89 | A | 11.13535 | 37.30073625 | 0.479492213 | 34.2920361 | NA | NA | NA | NA | NA | NA | NA |
| KCNMA1 | 90 | F | 9.790204545 | 36.24637794 | 0.465146634 | 35.48162657 | NA | NA | NA | NA | NA | NA | NA |
| KCNMA1 | 91 | L | 6.362125 | 31.89585968 | 0.405953566 | 31.21694887 | NA | NA | NA | NA | NA | NA | NA |
| KCNMA1 | 92 | A | 6.38075 | 36.78479036 | 0.472472263 | 36.10825245 | NA | NA | NA | NA | NA | NA | NA |
| KCNMA1 | 93 | S | 5.43075 | 39.27893289 | 0.506407518 | 38.4395546 | NA | NA | NA | NA | NA | NA | NA |
| KCNMA1 | 94 | S | 2.820583333 | 36.49706972 | 0.468557542 | 35.47791239 | NA | NA | NA | NA | NA | NA | NA |

| | | | | | | | | | | | | | |
|---|---|---|---|---|---|---|---|---|---|---|---|---|---|
| KCNMA1 | 95 | M | 1.626375 | 32.97845024 | 0.420683273 | 32.1722228 | NA | NA | NA | NA | NA | NA | NA |
| KCNMA1 | 96 | V | 1.787892857 | 40.0267585 | 0.516582419 | 39.22282755 | NA | NA | NA | NA | NA | NA | NA |
| KCNMA1 | 97 | T | 0.439321429 | 40.93589653 | 0.528952134 | 39.98604719 | NA | NA | NA | NA | NA | NA | NA |
| KCNMA1 | 98 | F | -2.319386364 | 35.84484351 | 0.459683364 | 35.12334578 | NA | NA | NA | NA | NA | NA | NA |
| KCNMA1 | 99 | F | -0.491386364 | 38.4380666 | 0.494966708 | 37.59233047 | NA | NA | NA | NA | NA | NA | NA |
| KCNMA1 | 158 | W | -15.10782143 | 57.4834423 | 0.754097722 | 52.15891612 | NA | NA | NA | NA | NA | NA | NA |
| KCNMA1 | 159 | M | -15.29025 | 53.7118708 | 0.702781793 | 47.70258068 | NA | NA | NA | NA | NA | NA | NA |
| KCNMA1 | 160 | T | -20.37317857 | 53.07158104 | 0.694070023 | 48.26266541 | NA | NA | NA | NA | NA | NA | NA |
| KCNMA1 | 161 | S | -19.96075 | 54.19769445 | 0.7093919 | 49.4098536 | NA | NA | NA | NA | NA | NA | NA |
| KCNMA1 | 162 | V | -15.67317857 | 51.97490054 | 0.679148609 | 45.9450837 | NA | NA | NA | NA | NA | NA | NA |
| KCNMA1 | 163 | K | -17.85830556 | 48.24221837 | 0.628361808 | 42.39494904 | NA | NA | NA | NA | NA | NA | NA |
| KCNMA1 | 164 | D | -20.156625 | 48.43234053 | 0.630948606 | 43.62141354 | NA | NA | NA | NA | NA | NA | NA |
| KCNMA1 | 165 | W | -15.83682143 | 50.37506259 | 0.657381245 | 45.06799091 | NA | NA | NA | NA | NA | NA | NA |
| KCNMA1 | 166 | A | -16.34715 | 46.01646743 | 0.598078283 | 40.00313334 | NA | NA | NA | NA | NA | NA | NA |
| KCNMA1 | 167 | G | -19.4395 | 44.700135 | 0.58016829 | 39.88907864 | NA | NA | NA | NA | NA | NA | NA |
| KCNMA1 | 168 | V | -21.06746429 | 44.68389313 | 0.579947303 | 39.87378399 | NA | NA | NA | NA | NA | NA | NA |
| KCNMA1 | 169 | M | -16.57825 | 43.20506721 | 0.559826426 | 37.86665884 | NA | NA | NA | NA | NA | NA | NA |
| KCNMA1 | 170 | I | -17.292 | 40.2396744 | 0.519479349 | 34.94767055 | NA | NA | NA | NA | NA | NA | NA |
| KCNMA1 | 171 | S | -20.98125 | 40.19534452 | 0.518876197 | 35.45561998 | NA | NA | NA | NA | NA | NA | NA |
| KCNMA1 | 172 | A | -21.80055 | 38.76098262 | 0.499360297 | 33.95339631 | NA | NA | NA | NA | NA | NA | NA |
| KCNMA1 | 173 | Q | -23.77408333 | 42.23191918 | 0.546585793 | 37.11098077 | NA | NA | NA | NA | NA | NA | NA |
| KCNMA1 | 174 | T | -20.61989286 | 44.15200558 | 0.572710451 | 39.3521677 | NA | NA | NA | NA | NA | NA | NA |
| KCNMA1 | 175 | L | -18.177625 | 45.11367816 | 0.58579495 | 40.37099515 | NA | NA | NA | NA | NA | NA | NA |
| KCNMA1 | 176 | T | -15.95160714 | 45.60565207 | 0.592488737 | 39.58601066 | NA | NA | NA | NA | NA | NA | NA |
| KCNMA1 | 177 | G | -16.1245 | 42.07907728 | 0.544506229 | 36.04944849 | NA | NA | NA | NA | NA | NA | NA |
| KCNMA1 | 178 | R | -14.99684091 | 40.91558688 | 0.528675801 | 34.89690747 | NA | NA | NA | NA | NA | NA | NA |
| KCNMA1 | 179 | V | -11.78832143 | 42.73377383 | 0.553414017 | 37.48267034 | NA | NA | NA | NA | NA | NA | NA |
| KCNMA1 | 180 | L | -10.875625 | 41.15413983 | 0.531921548 | 35.81146053 | NA | NA | NA | NA | NA | NA | NA |
| KCNMA1 | 181 | V | -11.47175 | 36.89652226 | 0.473992485 | 31.59460146 | NA | NA | NA | NA | NA | NA | NA |
| KCNMA1 | 182 | V | -9.815178571 | 38.7471256 | 0.499171758 | 33.46837809 | NA | NA | NA | NA | NA | NA | NA |
| KCNMA1 | 183 | L | -6.566 | 40.49159273 | 0.522906945 | 38.07221118 | NA | NA | NA | NA | NA | NA | NA |
| KCNMA1 | 184 | V | -6.31075 | 37.06513032 | 0.476286563 | 34.66766054 | NA | NA | NA | NA | NA | NA | NA |
| KCNMA1 | 185 | F | -6.889386364 | 31.51661849 | 0.400793618 | 29.12639012 | NA | NA | NA | NA | NA | NA | NA |

| | | | | | | | | | | | | | |
|---|---|---|---|---|---|---|---|---|---|---|---|---|---|
| KCNMA1 | 186 | A | -3.73035 | 36.67393502 | 0.470963967 | 35.1298795 | NA | NA | NA | NA | NA | NA | NA |
| KCNMA1 | 187 | L | -1.3455 | 36.7362214 | 0.471811435 | 35.90423973 | NA | NA | NA | NA | NA | NA | NA |
| KCNMA1 | 188 | S | -1.966416667 | 33.20105976 | 0.423712094 | 32.35738213 | NA | NA | NA | NA | NA | NA | NA |
| KCNMA1 | 189 | I | -1.582375 | 32.71093889 | 0.417043518 | 31.92187892 | NA | NA | NA | NA | NA | NA | NA |
| KCNMA1 | 190 | G | 0.883 | 34.92214743 | 0.447129179 | 34.0659678 | NA | NA | NA | NA | NA | NA | NA |
| KCNMA1 | 191 | A | 1.75055 | 33.00116909 | 0.420992385 | 32.23379432 | NA | NA | NA | NA | NA | NA | NA |
| KCNMA1 | 192 | L | 1.071 | 28.30168599 | 0.357051309 | 27.60399317 | NA | NA | NA | NA | NA | NA | NA |
| KCNMA1 | 193 | V | 3.80975 | 31.10739102 | 0.395225677 | 30.48019494 | NA | NA | NA | NA | NA | NA | NA |
| KCNMA1 | 194 | I | 6.28725 | 31.561661 | 0.401406466 | 30.90570719 | NA | NA | NA | NA | NA | NA | NA |
| KCNMA1 | 195 | Y | 5.901166667 | 24.90888368 | 0.310888906 | 24.20031431 | NA | NA | NA | NA | NA | NA | NA |
| KCNMA1 | 196 | F | 4.717931818 | 25.93655685 | 0.324871407 | 25.54009213 | NA | NA | NA | NA | NA | NA | NA |
| KCNMA1 | 197 | I | 9.658125 | 29.30385144 | 0.370686752 | 27.02757528 | NA | NA | NA | NA | NA | NA | NA |
| KCNMA1 | 198 | D | 10.951375 | 28.48540549 | 0.359550992 | 25.59198777 | NA | NA | NA | NA | NA | NA | NA |
| KCNMA1 | 199 | S | 11.28691667 | 25.08495168 | 0.313284484 | 22.16424355 | NA | NA | NA | NA | NA | NA | NA |
| KCNMA1 | 200 | S | 14.57691667 | 26.24221197 | 0.329030145 | 21.5952437 | NA | NA | NA | NA | NA | NA | NA |
| KCNMA1 | 201 | N | 15.72525 | 27.44649096 | 0.345415541 | 21.80933975 | NA | NA | NA | NA | NA | NA | NA |
| KCNMA1 | 202 | P | 17.41775 | 28.11861129 | 0.354560398 | 22.48241413 | NA | NA | NA | NA | NA | NA | NA |
| KCNMA1 | 203 | I | 14.71475 | 25.64823013 | 0.320948439 | 20.0152282 | NA | NA | NA | NA | NA | NA | NA |
| KCNMA1 | 204 | E | 17.88030556 | 30.49031896 | 0.386829806 | 24.85625042 | NA | NA | NA | NA | NA | NA | NA |
| KCNMA1 | 205 | S | 17.79125 | 33.71746902 | 0.430738348 | 28.08282742 | NA | NA | NA | NA | NA | NA | NA |
| KCNMA1 | 206 | C | 19.40591667 | 36.7276537 | 0.471694863 | 31.09227648 | NA | NA | NA | NA | NA | NA | NA |
| KCNMA1 | 207 | Q | 16.16241667 | 36.1476074 | 0.463802764 | 30.5105753 | NA | NA | NA | NA | NA | NA | NA |
| KCNMA1 | 208 | N | 17.79 | 42.29027132 | 0.547379731 | 36.65362526 | NA | NA | NA | NA | NA | NA | NA |
| KCNMA1 | 209 | F | 13.70784091 | 45.64667705 | 0.593046923 | 40.00969744 | NA | NA | NA | NA | NA | NA | NA |
| KCNMA1 | 210 | Y | 16.93558333 | 45.51244012 | 0.591220497 | 39.87884105 | NA | NA | NA | NA | NA | NA | NA |
| KCNMA1 | 211 | K | 17.26352778 | 40.32820574 | 0.520683904 | 34.69096895 | NA | NA | NA | NA | NA | NA | NA |
| KCNMA1 | 212 | D | 14.116875 | 38.80308417 | 0.49993313 | 34.1919904 | NA | NA | NA | NA | NA | NA | NA |
| KCNMA1 | 213 | F | 10.71565909 | 42.03806009 | 0.54394815 | 39.04989187 | NA | NA | NA | NA | NA | NA | NA |
| KCNMA1 | 214 | T | 9.33375 | 37.31925494 | 0.479744178 | 35.00641312 | NA | NA | NA | NA | NA | NA | NA |
| KCNMA1 | 215 | L | 11.138625 | 38.8832494 | 0.501023856 | 35.96121159 | NA | NA | NA | NA | NA | NA | NA |
| KCNMA1 | 216 | Q | 9.703416667 | 42.27295975 | 0.54714419 | 39.94314807 | NA | NA | NA | NA | NA | NA | NA |
| KCNMA1 | 217 | I | 5.822 | 42.11800614 | 0.545035894 | 40.85064553 | NA | NA | NA | NA | NA | NA | NA |
| KCNMA1 | 218 | D | 5.573125 | 37.0184625 | 0.475651602 | 36.18428815 | NA | NA | NA | NA | NA | NA | NA |

| | | | | | | | | | | | | | |
|---|---|---|---|---|---|---|---|---|---|---|---|---|---|
| KCNMA1 | 219 | M | 6.23475 | 41.94193818 | 0.542640317 | 41.10235742 | NA | NA | NA | NA | NA | NA | NA |
| KCNMA1 | 220 | A | 2.72175 | 44.14689536 | 0.572640922 | 43.13703671 | NA | NA | NA | NA | NA | NA | NA |
| KCNMA1 | 221 | F | 1.187386364 | 39.74630717 | 0.512766604 | 38.94317959 | NA | NA | NA | NA | NA | NA | NA |
| KCNMA1 | 222 | N | 1.881875 | 38.95046179 | 0.501938347 | 38.15368251 | NA | NA | NA | NA | NA | NA | NA |
| KCNMA1 | 223 | V | 0.740321429 | 44.02734166 | 0.571014276 | 43.11906841 | NA | NA | NA | NA | NA | NA | NA |
| KCNMA1 | 224 | F | -2.678204545 | 43.3204614 | 0.561396477 | 42.50953793 | NA | NA | NA | NA | NA | NA | NA |
| KCNMA1 | 225 | F | -1.466659091 | 37.92549612 | 0.487992684 | 37.02409341 | NA | NA | NA | NA | NA | NA | NA |
| KCNMA1 | 226 | L | -1.647625 | 43.04460511 | 0.557643182 | 42.07671594 | NA | NA | NA | NA | NA | NA | NA |
| KCNMA1 | 227 | L | -3.49325 | 45.54488541 | 0.591661947 | 44.03531834 | NA | NA | NA | NA | NA | NA | NA |
| KCNMA1 | 228 | Y | -7.94175 | 40.35437565 | 0.521039972 | 38.03712342 | NA | NA | NA | NA | NA | NA | NA |
| KCNMA1 | 229 | F | -5.404568182 | 40.49337183 | 0.522931151 | 37.78194955 | NA | NA | NA | NA | NA | NA | NA |
| KCNMA1 | 230 | G | -7.253 | 44.40446889 | 0.576145462 | 41.98676348 | NA | NA | NA | NA | NA | NA | NA |
| KCNMA1 | 231 | L | -9.974375 | 46.27047834 | 0.601534351 | 40.92485342 | NA | NA | NA | NA | NA | NA | NA |
| KCNMA1 | 232 | R | -11.43602273 | 39.68280405 | 0.511902581 | 34.4833219 | NA | NA | NA | NA | NA | NA | NA |
| KCNMA1 | 233 | F | -10.08820455 | 44.97869728 | 0.583958403 | 39.6299706 | NA | NA | NA | NA | NA | NA | NA |
| KCNMA1 | 234 | I | -13.672625 | 47.13991651 | 0.61336391 | 41.77687561 | NA | NA | NA | NA | NA | NA | NA |
| KCNMA1 | 235 | A | -15.68695 | 44.67911106 | 0.579882238 | 38.67778636 | NA | NA | NA | NA | NA | NA | NA |
| KCNMA1 | 236 | A | -16.19615 | 42.50453539 | 0.550295004 | 36.47702851 | NA | NA | NA | NA | NA | NA | NA |
| KCNMA1 | 237 | N | -20.099875 | 41.97293453 | 0.543062053 | 37.17594697 | NA | NA | NA | NA | NA | NA | NA |
| KCNMA1 | 238 | D | -17.568875 | 38.90660495 | 0.501341631 | 33.62750898 | NA | NA | NA | NA | NA | NA | NA |
| KCNMA1 | 239 | K | -14.76908333 | 42.75057041 | 0.553642551 | 37.22264886 | NA | NA | NA | NA | NA | NA | NA |
| KCNMA1 | 240 | L | -14.50325 | 39.44450787 | 0.508660328 | 33.98492198 | NA | NA | NA | NA | NA | NA | NA |
| KCNMA1 | 241 | W | -16.46525 | 34.8658658 | 0.446363412 | 30.08952356 | NA | NA | NA | NA | NA | NA | NA |
| KCNMA1 | 242 | F | -10.53465909 | 38.64562044 | 0.497790681 | 33.30414235 | NA | NA | NA | NA | NA | NA | NA |
| KCNMA1 | 243 | W | -7.353821429 | 37.68465015 | 0.484715738 | 32.76388621 | NA | NA | NA | NA | NA | NA | NA |
| KCNMA1 | 244 | L | -10.068375 | 33.22478305 | 0.424034872 | 28.25498732 | NA | NA | NA | NA | NA | NA | NA |
| KCNMA1 | 245 | E | -10.76797222 | 32.98379393 | 0.420755979 | 27.71726088 | NA | NA | NA | NA | NA | NA | NA |
| KCNMA1 | 246 | V | -8.475035714 | 28.91964905 | 0.365459302 | 26.02042319 | NA | NA | NA | NA | NA | NA | NA |
| KCNMA1 | 247 | N | -8.810875 | 32.20436264 | 0.410151052 | 29.34052515 | NA | NA | NA | NA | NA | NA | NA |
| KCNMA1 | 248 | S | -6.501583333 | 35.09259452 | 0.449448279 | 32.69338973 | NA | NA | NA | NA | NA | NA | NA |
| KCNMA1 | 249 | V | -3.923035714 | 32.6376984 | 0.41604701 | 31.00567983 | NA | NA | NA | NA | NA | NA | NA |
| KCNMA1 | 250 | V | -2.896607143 | 31.26905747 | 0.397425308 | 29.80055466 | NA | NA | NA | NA | NA | NA | NA |
| KCNMA1 | 251 | D | -2.4505 | 35.12245228 | 0.449854523 | 34.25139488 | NA | NA | NA | NA | NA | NA | NA |

| | | | | | | | | | | | | | |
|---|---|---|---|---|---|---|---|---|---|---|---|---|---|
| KCNMA1 | 252 | F | -2.231477273 | 36.90582953 | 0.47411912 | 35.94162744 | NA | NA | NA | NA | NA | NA | NA |
| KCNMA1 | 253 | F | 0.274840909 | 32.48575751 | 0.413979705 | 31.59438537 | NA | NA | NA | NA | NA | NA | NA |
| KCNMA1 | 254 | T | 1.494607143 | 33.70236039 | 0.43053278 | 32.93812359 | NA | NA | NA | NA | NA | NA | NA |
| KCNMA1 | 255 | V | 1.419607143 | 38.03229681 | 0.489445812 | 37.24586505 | NA | NA | NA | NA | NA | NA | NA |
| KCNMA1 | 256 | P | 2.769892857 | 39.72805002 | 0.512518197 | 38.75226663 | NA | NA | NA | NA | NA | NA | NA |
| KCNMA1 | 257 | P | 5.957321429 | 36.91439966 | 0.474235725 | 36.21216834 | NA | NA | NA | NA | NA | NA | NA |
| KCNMA1 | 258 | V | 6.535035714 | 38.58797753 | 0.497006393 | 37.91562786 | NA | NA | NA | NA | NA | NA | NA |
| KCNMA1 | 259 | F | 4.547840909 | 42.36633694 | 0.548414678 | 41.09685953 | NA | NA | NA | NA | NA | NA | NA |
| KCNMA1 | 260 | V | 8.811607143 | 42.68969534 | 0.552814286 | 41.87017201 | NA | NA | NA | NA | NA | NA | NA |
| KCNMA1 | 261 | S | 11.27825 | 39.95256094 | 0.515572889 | 37.0375527 | NA | NA | NA | NA | NA | NA | NA |
| KCNMA1 | 262 | V | 10.75817857 | 43.61185222 | 0.565361135 | 40.68330544 | NA | NA | NA | NA | NA | NA | NA |
| KCNMA1 | 263 | Y | 13.24291667 | 45.90830845 | 0.596606674 | 42.58890186 | NA | NA | NA | NA | NA | NA | NA |
| KCNMA1 | 264 | L | 14.780625 | 41.75485238 | 0.540094831 | 36.12847358 | NA | NA | NA | NA | NA | NA | NA |
| KCNMA1 | 265 | N | 16.17675 | 41.55615075 | 0.5373913 | 35.92123994 | NA | NA | NA | NA | NA | NA | NA |
| KCNMA1 | 266 | R | 16.00484091 | 38.28829224 | 0.492928881 | 32.65554422 | NA | NA | NA | NA | NA | NA | NA |
| KCNMA1 | 267 | S | 13.69275 | 34.58563098 | 0.442550543 | 30.80915347 | NA | NA | NA | NA | NA | NA | NA |
| KCNMA1 | 268 | W | 11.55067857 | 30.06732331 | 0.381074536 | 26.31932676 | NA | NA | NA | NA | NA | NA | NA |
| KCNMA1 | 269 | L | 10.363375 | 27.23289454 | 0.342509353 | 24.30298337 | NA | NA | NA | NA | NA | NA | NA |
| KCNMA1 | 270 | G | 7.36675 | 29.62927216 | 0.37511442 | 28.84753179 | NA | NA | NA | NA | NA | NA | NA |
| KCNMA1 | 271 | L | 3.637 | 29.02271419 | 0.366861604 | 28.18246654 | NA | NA | NA | NA | NA | NA | NA |
| KCNMA1 | 272 | R | 5.661659091 | 30.01162348 | 0.380316685 | 29.26525881 | NA | NA | NA | NA | NA | NA | NA |
| KCNMA1 | 273 | F | 2.309113636 | 25.60505079 | 0.320360942 | 24.80394149 | NA | NA | NA | NA | NA | NA | NA |
| KCNMA1 | 274 | L | 0 | 27.70192055 | 0.348890911 | 26.63859716 | NA | NA | NA | NA | NA | NA | NA |
| KCNMA1 | 275 | R | -0.85475 | 30.04596249 | 0.380783901 | 29.21193453 | NA | NA | NA | NA | NA | NA | NA |
| KCNMA1 | 276 | A | -2.50795 | 26.70921028 | 0.335384115 | 25.74688212 | NA | NA | NA | NA | NA | NA | NA |
| KCNMA1 | 277 | L | -4.811625 | 25.64970596 | 0.320968519 | 23.98006067 | NA | NA | NA | NA | NA | NA | NA |
| KCNMA1 | 278 | R | -5.211659091 | 29.79444019 | 0.377361693 | 27.38040571 | NA | NA | NA | NA | NA | NA | NA |
| KCNMA1 | 279 | L | -6.446375 | 26.71543391 | 0.335468793 | 24.07119957 | NA | NA | NA | NA | NA | NA | NA |
| KCNMA1 | 280 | I | -10.9915 | 25.12234784 | 0.313793295 | 19.84141872 | NA | NA | NA | NA | NA | NA | NA |
| KCNMA1 | 281 | Q | -10.91408333 | 29.7697765 | 0.37702612 | 24.42213078 | NA | NA | NA | NA | NA | NA | NA |
| KCNMA1 | 282 | F | -10.67138636 | 28.19445488 | 0.355592324 | 22.84696888 | NA | NA | NA | NA | NA | NA | NA |
| KCNMA1 | 283 | S | -14.96825 | 27.66936246 | 0.348447927 | 21.670488 | NA | NA | NA | NA | NA | NA | NA |
| KCNMA1 | 284 | E | -16.25519444 | 30.87470662 | 0.392059778 | 24.8505413 | NA | NA | NA | NA | NA | NA | NA |

| | | | | | | | | | | | | | |
|---|---|---|---|---|---|---|---|---|---|---|---|---|---|
| KCNMA1 | 285 | I | -13.088125 | 33.36762569 | 0.425978386 | 28.02121521 | NA | NA | NA | NA | NA | NA | NA |
| KCNMA1 | 286 | L | -14.621125 | 29.95095533 | 0.379491235 | 24.00246657 | NA | NA | NA | NA | NA | NA | NA |
| KCNMA1 | 287 | Q | -19.29075 | 29.86834803 | 0.378367282 | 25.05635412 | NA | NA | NA | NA | NA | NA | NA |
| KCNMA1 | 288 | F | -17.87575 | 35.50473632 | 0.455055872 | 30.21071285 | NA | NA | NA | NA | NA | NA | NA |
| KCNMA1 | 289 | L | -15.475875 | 36.93148129 | 0.474468137 | 30.91906846 | NA | NA | NA | NA | NA | NA | NA |
| KCNMA1 | 290 | N | -20.257125 | 35.36362023 | 0.45313585 | 30.57211748 | NA | NA | NA | NA | NA | NA | NA |
| KCNMA1 | 291 | I | -17.698375 | 31.53485819 | 0.401041787 | 26.74332678 | NA | NA | NA | NA | NA | NA | NA |
| KCNMA1 | 292 | L | -18.744625 | 28.28412808 | 0.356812416 | 23.67669741 | NA | NA | NA | NA | NA | NA | NA |
| KCNMA1 | 293 | K | -23.56308333 | 29.0991144 | 0.367901104 | 23.86407544 | NA | NA | NA | NA | NA | NA | NA |
| KCNMA1 | 294 | T | -23.90446429 | 25.05140679 | 0.312828072 | 19.82479286 | NA | NA | NA | NA | NA | NA | NA |
| KCNMA1 | 295 | S | -23.99208333 | 23.37561796 | 0.290027322 | 18.35907568 | NA | NA | NA | NA | NA | NA | NA |
| KCNMA1 | 296 | N | -23.044375 | 20.00780073 | 0.244204865 | 15.56328299 | NA | NA | NA | NA | NA | NA | NA |
| KCNMA1 | 297 | S | -19.50058333 | 23.71025392 | 0.294580372 | 18.90143305 | NA | NA | NA | NA | NA | NA | NA |
| KCNMA1 | 298 | I | -19.687 | 24.18610316 | 0.301054768 | 19.4534095 | NA | NA | NA | NA | NA | NA | NA |
| KCNMA1 | 299 | K | -20.06941667 | 18.36076657 | 0.22179535 | 13.56619508 | NA | NA | NA | NA | NA | NA | NA |
| KCNMA1 | 300 | L | -15.8975 | 19.39332929 | 0.235844379 | 14.05563206 | NA | NA | NA | NA | NA | NA | NA |
| KCNMA1 | 301 | V | -14.23960714 | 23.02941743 | 0.285316924 | 17.03136419 | NA | NA | NA | NA | NA | NA | NA |
| KCNMA1 | 302 | N | -15.30625 | 21.92371045 | 0.270272696 | 15.96730508 | NA | NA | NA | NA | NA | NA | NA |
| KCNMA1 | 303 | L | -13.85875 | 17.01265135 | 0.203452921 | 11.68920689 | NA | NA | NA | NA | NA | NA | NA |
| KCNMA1 | 304 | L | -10.400875 | 19.30664202 | 0.234664914 | 14.01840432 | NA | NA | NA | NA | NA | NA | NA |
| KCNMA1 | 305 | S | -9.514083333 | 22.37390163 | 0.276397989 | 17.42215406 | NA | NA | NA | NA | NA | NA | NA |
| KCNMA1 | 306 | I | -9.836625 | 20.06956312 | 0.245045203 | 15.20281138 | NA | NA | NA | NA | NA | NA | NA |
| KCNMA1 | 307 | F | -6.827568182 | 14.52654522 | 0.169627009 | 11.69354725 | NA | NA | NA | NA | NA | NA | NA |
| KCNMA1 | 308 | I | -4.90875 | 20.12670333 | 0.245822652 | 18.51121781 | NA | NA | NA | NA | NA | NA | NA |
| KCNMA1 | 309 | S | -5.043916667 | 22.13885349 | 0.273199928 | 20.47392231 | NA | NA | NA | NA | NA | NA | NA |
| KCNMA1 | 310 | T | -4.616607143 | 18.90408384 | 0.229187715 | 17.23687299 | NA | NA | NA | NA | NA | NA | NA |
| KCNMA1 | 311 | W | -1.07525 | 16.15884677 | 0.191836072 | 15.39071587 | NA | NA | NA | NA | NA | NA | NA |
| KCNMA1 | 312 | L | -0.083 | 20.42974877 | 0.249945882 | 19.39815488 | NA | NA | NA | NA | NA | NA | NA |
| KCNMA1 | 313 | T | -0.571178571 | 21.98168087 | 0.271061441 | 20.92408535 | NA | NA | NA | NA | NA | NA | NA |
| KCNMA1 | 314 | A | 0.29795 | 18.72202655 | 0.226710647 | 17.66526291 | NA | NA | NA | NA | NA | NA | NA |
| KCNMA1 | 315 | A | 2.49195 | 18.86319684 | 0.228631407 | 18.0574583 | NA | NA | NA | NA | NA | NA | NA |
| KCNMA1 | 316 | G | 3.8415 | 21.40858748 | 0.263263943 | 20.66701085 | NA | NA | NA | NA | NA | NA | NA |
| KCNMA1 | 317 | F | 3.118659091 | 21.60895236 | 0.265990104 | 20.89572513 | NA | NA | NA | NA | NA | NA | NA |

| | | | | | | | | | | | | | |
|---|---|---|---|---|---|---|---|---|---|---|---|---|---|
| KCNMA1 | 318 | I | 5.43325 | 16.99513654 | 0.203214615 | 15.94773767 | NA | NA | NA | NA | NA | NA | NA |
| KCNMA1 | 319 | H | 8.62975 | 19.93687222 | 0.243239814 | 19.17899175 | NA | NA | NA | NA | NA | NA | NA |
| KCNMA1 | 320 | L | 8.483875 | 22.35536835 | 0.276145825 | 21.53533416 | NA | NA | NA | NA | NA | NA | NA |
| KCNMA1 | 321 | V | 8.763607143 | 20.82868618 | 0.255373817 | 20.01004647 | NA | NA | NA | NA | NA | NA | NA |
| KCNMA1 | 322 | E | 11.02197222 | 15.94171901 | 0.188881836 | 13.03259446 | NA | NA | NA | NA | NA | NA | NA |
| KCNMA1 | 323 | N | 13.113125 | 20.59741739 | 0.252227178 | 16.77591532 | NA | NA | NA | NA | NA | NA | NA |
| KCNMA1 | 324 | S | 13.75275 | 21.28686114 | 0.261607737 | 15.66088279 | NA | NA | NA | NA | NA | NA | NA |
| KCNMA1 | 325 | G | 14.83675 | 22.0033295 | 0.271355991 | 16.3745425 | NA | NA | NA | NA | NA | NA | NA |
| KCNMA1 | 326 | D | 14.612375 | 20.38965605 | 0.249400382 | 14.76352948 | NA | NA | NA | NA | NA | NA | NA |
| KCNMA1 | 327 | P | 14.32403571 | 25.88509975 | 0.324171283 | 20.25829299 | NA | NA | NA | NA | NA | NA | NA |
| KCNMA1 | 328 | W | 13.82182143 | 22.98857808 | 0.284761265 | 17.6440069 | NA | NA | NA | NA | NA | NA | NA |
| KCNMA1 | 329 | E | 18.45841667 | 22.34025575 | 0.275940204 | 16.71561713 | NA | NA | NA | NA | NA | NA | NA |
| KCNMA1 | 330 | N | 19.9045 | 25.95934102 | 0.325181408 | 20.33230146 | NA | NA | NA | NA | NA | NA | NA |
| KCNMA1 | 331 | F | 17.21702273 | 23.41481655 | 0.290560657 | 17.7861498 | NA | NA | NA | NA | NA | NA | NA |
| KCNMA1 | 332 | Q | 22.03819444 | 19.87392684 | 0.24238338 | 14.24641202 | NA | NA | NA | NA | NA | NA | NA |
| KCNMA1 | 333 | N | 18.125375 | 18.05009753 | 0.217568394 | 12.42431296 | NA | NA | NA | NA | NA | NA | NA |
| KCNMA1 | 334 | N | 18.33675 | 16.63768713 | 0.198351165 | 11.0073739 | NA | NA | NA | NA | NA | NA | NA |
| KCNMA1 | 335 | Q | 14.86408333 | 12.29560987 | 0.139272945 | 6.844182677 | NA | NA | NA | NA | NA | NA | NA |
| KCNMA1 | 336 | A | 16.19755 | 16.08473621 | 0.190827725 | 10.44785295 | NA | NA | NA | NA | NA | NA | NA |
| KCNMA1 | 337 | L | 12.345375 | 14.55390552 | 0.169999272 | 10.73066882 | NA | NA | NA | NA | NA | NA | NA |
| KCNMA1 | 338 | T | 11.50246429 | 18.5928476 | 0.224953041 | 15.27448163 | NA | NA | NA | NA | NA | NA | NA |
| KCNMA1 | 339 | Y | 5.796083333 | 19.42917729 | 0.236332126 | 18.69345546 | NA | NA | NA | NA | NA | NA | NA |
| KCNMA1 | 340 | W | 6.33525 | 17.33064837 | 0.207779582 | 16.65385201 | NA | NA | NA | NA | NA | NA | NA |
| KCNMA1 | 341 | E | 9.969416667 | 14.45404559 | 0.16864058 | 13.66367502 | NA | NA | NA | NA | NA | NA | NA |
| KCNMA1 | 342 | C | 6.10125 | 15.45306015 | 0.182233153 | 14.90470494 | NA | NA | NA | NA | NA | NA | NA |
| KCNMA1 | 343 | V | 2.995035714 | 15.05380354 | 0.176800875 | 14.34753416 | NA | NA | NA | NA | NA | NA | NA |
| KCNMA1 | 344 | Y | 5.12125 | 10.13156281 | 0.109828963 | 9.47420612 | NA | NA | NA | NA | NA | NA | NA |
| KCNMA1 | 345 | L | 4.65625 | 10.196802 | 0.110716606 | 9.437719025 | NA | NA | NA | NA | NA | NA | NA |
| KCNMA1 | 346 | L | 1.143375 | 13.00635879 | 0.148943381 | 12.24401248 | NA | NA | NA | NA | NA | NA | NA |
| KCNMA1 | 347 | M | 0.303875 | 10.35722926 | 0.112899376 | 9.320331591 | NA | NA | NA | NA | NA | NA | NA |
| KCNMA1 | 348 | V | 1.305321429 | 7.137896819 | 0.069097201 | 6.492550011 | 0.81 | -0.8 | -0.4 | 3.38 | 0.81 | 0 | 0 |
| KCNMA1 | 349 | T | 0.392464286 | 7.475047425 | 0.073684466 | 6.794024684 | 0.97 | -0.8 | -0.4 | 3.38 | 0.81 | 0 | 0 |
| KCNMA1 | 350 | M | -2.672 | 8.661214234 | 0.089823428 | 7.733674556 | 0.97 | -0.79 | -0.48 | 2.95 | 0.8 | 0 | 0 |

| | | | | | | | | | | | | | |
|---|---|---|---|---|---|---|---|---|---|---|---|---|---|
| KCNMA1 | 351 | S | -2.260583333 | 5.091264479 | 0.041250761 | 3.622627746 | 1.47 | -0.77 | -0.7 | 1.66 | 0.77 | 0 | 0 |
| KCNMA1 | 352 | T | -2.070892857 | 2.147763022 | 0.001201537 | 1.486378505 | 0.88 | -0.79 | -0.48 | 2.95 | 0.8 | 0 | 0 |
| KCNMA1 | 353 | V | 0.309035714 | 2.072561941 | 1.78E-04 | 1.523769492 | 0.97 | -0.8 | -0.4 | 3.38 | 0.81 | 0 | 0 |
| KCNMA1 | 354 | G | 3.1375 | 2.060926733 | 2.00E-05 | 1.541869608 | 1.02 | -0.8 | -0.4 | 3.38 | 0.81 | 0 | 0 |
| KCNMA1 | 355 | Y | 5.08475 | 2.34607502 | 0.003899766 | 1.940466515 | 0.84 | -0.8 | -0.4 | 3.38 | 0.81 | 0 | 0 |
| KCNMA1 | 356 | G | 9.1055 | 4.423827528 | 0.032169627 | 3.629268492 | 0.82 | -0.8 | -0.4 | 3.38 | 0.81 | 0 | 0 |
| KCNMA1 | 357 | D | 10.224625 | 5.688395732 | 0.049375317 | 2.856499536 | 2.93 | -0.8 | -0.4 | 3.38 | 0.81 | 0 | 0 |
| KCNMA1 | 358 | V | 9.877464286 | 8.163329835 | 0.083049223 | 5.253351312 | 2.93 | -0.8 | -0.4 | 3.38 | 0.81 | 0 | 0 |
| KCNMA1 | 359 | Y | 11.48525 | 6.644549797 | 0.062384731 | 4.862151115 | 2.33 | -0.8 | -0.4 | 3.38 | 0.81 | 0 | 0 |
| KCNMA1 | 360 | A | 9.57715 | 11.32740835 | 0.126099614 | 9.024979265 | NA | NA | NA | NA | NA | NA | NA |
| KCNMA1 | 361 | K | 13.84741667 | 9.803032082 | 0.10535898 | 6.101051622 | NA | NA | NA | NA | NA | NA | NA |
| KCNMA1 | 362 | T | 11.47517857 | 15.90972511 | 0.188446528 | 12.59291962 | NA | NA | NA | NA | NA | NA | NA |
| KCNMA1 | 363 | T | 10.02703571 | 18.09235844 | 0.218143395 | 15.80709554 | NA | NA | NA | NA | NA | NA | NA |
| KCNMA1 | 364 | L | 7.65825 | 19.71100348 | 0.240166648 | 18.90326437 | NA | NA | NA | NA | NA | NA | NA |
| KCNMA1 | 365 | G | 6.806 | 16.4787289 | 0.196188382 | 15.87123829 | NA | NA | NA | NA | NA | NA | NA |
| KCNMA1 | 366 | R | 8.180568182 | 11.52165075 | 0.128742472 | 11.38811765 | NA | NA | NA | NA | NA | NA | NA |
| KCNMA1 | 367 | L | 3.558125 | 17.57123798 | 0.21105304 | 16.75453492 | NA | NA | NA | NA | NA | NA | NA |
| KCNMA1 | 368 | F | 1.52275 | 16.98264376 | 0.203044638 | 16.26900223 | NA | NA | NA | NA | NA | NA | NA |
| KCNMA1 | 369 | M | 2.13775 | 12.23529166 | 0.138452257 | 11.44955175 | NA | NA | NA | NA | NA | NA | NA |
| KCNMA1 | 370 | V | 0.583464286 | 14.92081854 | 0.174991484 | 14.04909019 | NA | NA | NA | NA | NA | NA | NA |
| KCNMA1 | 371 | F | -0.51775 | 16.80695204 | 0.20065418 | 15.85608657 | NA | NA | NA | NA | NA | NA | NA |
| KCNMA1 | 372 | F | -1.30475 | 14.41173782 | 0.168064941 | 13.65583413 | NA | NA | NA | NA | NA | NA | NA |
| KCNMA1 | 373 | I | -1.78675 | 10.81360412 | 0.119108804 | 9.98019066 | NA | NA | NA | NA | NA | NA | NA |
| KCNMA1 | 374 | L | -4.655125 | 13.66596897 | 0.157918025 | 12.05699476 | NA | NA | NA | NA | NA | NA | NA |
| KCNMA1 | 375 | G | -6.2165 | 15.27934717 | 0.179869618 | 12.86208728 | NA | NA | NA | NA | NA | NA | NA |
| KCNMA1 | 376 | G | -6.76875 | 12.32577661 | 0.139683393 | 10.07013725 | NA | NA | NA | NA | NA | NA | NA |
| KCNMA1 | 377 | L | -7.9415 | 8.455168656 | 0.087019976 | 5.574025091 | 2.9 | -0.77 | -0.7 | 1.66 | 0.77 | 0 | 0 |
| KCNMA1 | 378 | A | -10.56575 | 13.39952197 | 0.154292752 | 8.075713054 | NA | NA | NA | NA | NA | NA | NA |
| KCNMA1 | 379 | M | -10.445375 | 13.594979 | 0.156952137 | 8.304056137 | NA | NA | NA | NA | NA | NA | NA |
| KCNMA1 | 380 | F | -11.44329545 | 6.810538525 | 0.06464317 | 1.695912116 | 5.24 | 0.93 | 2.1 | 0.61 | -1.29 | 0 | 0 |
| KCNMA1 | 381 | A | -14.32755 | 10.45209472 | 0.114190114 | 5.091641368 | NA | NA | NA | NA | NA | NA | NA |
| KCNMA1 | 382 | S | -16.13108333 | 12.61366644 | 0.143600417 | 7.289819413 | NA | NA | NA | NA | NA | NA | NA |
| KCNMA1 | 383 | Y | -15.52408333 | 14.2147248 | 0.165384386 | 8.18481708 | NA | NA | NA | NA | NA | NA | NA |

| | | | | | | | | | | | | | |
|---|---|---|---|---|---|---|---|---|---|---|---|---|---|
| KCNMA1 | 384 | V | -15.21517857 | 11.53711194 | 0.128952837 | 5.673414486 | NA | NA | NA | NA | NA | NA | NA |
| KCNMA1 | 385 | P | -17.50689286 | 8.660430128 | 0.08981276 | 3.386267554 | 5.35 | 0.85 | 1.54 | 10.26 | -1 | 1 | -1 |
| KCNMA1 | 386 | E | -20.61408333 | 10.54946544 | 0.115514938 | 5.800201414 | NA | NA | NA | NA | NA | NA | NA |
| KCNMA1 | 387 | I | -19.72875 | 12.5516967 | 0.142757258 | 7.757455118 | NA | NA | NA | NA | NA | NA | NA |
| KCNMA1 | 388 | I | -17.713125 | 10.31563905 | 0.112333501 | 6.196538804 | NA | NA | NA | NA | NA | NA | NA |
| KCNMA1 | 389 | E | -21.94908333 | 5.441555384 | 0.046016812 | 0.66430872 | 4.81 | -1.14 | -3.5 | 49.9 | 2.23 | 4 | -4 |
| KCNMA1 | 390 | L | -23.64975 | 12.04235812 | 0.135827207 | 7.154736118 | NA | NA | NA | NA | NA | NA | NA |
| KCNMA1 | 391 | I | -20.380125 | 13.75819861 | 0.1591729 | 8.955708451 | NA | NA | NA | NA | NA | NA | NA |
| KCNMA1 | 392 | G | -22.85275 | 11.38709976 | 0.126911774 | 6.957048333 | NA | NA | NA | NA | NA | NA | NA |
| KCNMA1 | 393 | N | -26.355625 | 10.36947583 | 0.113066003 | 5.248679199 | 5.24 | -1.14 | -3.5 | 49.9 | 2.23 | 4 | -4 |
| KCNMA1 | 394 | R | -23.63538636 | 13.97410587 | 0.16211053 | 8.733936157 | NA | NA | NA | NA | NA | NA | NA |
| KCNMA1 | 395 | K | -29.27141667 | 15.4541377 | 0.182247814 | 7.726569186 | NA | NA | NA | NA | NA | NA | NA |
| KCNMA1 | 396 | K | -29.91175 | 15.73011252 | 0.186002722 | 7.44948329 | NA | NA | NA | NA | NA | NA | NA |
| KCNMA1 | 397 | Y | -32.55566667 | 18.24275456 | 0.220189681 | 8.177197401 | NA | NA | NA | NA | NA | NA | NA |
| KCNMA1 | 398 | G | -31.26475 | 21.32282939 | 0.26209712 | 12.01354519 | NA | NA | NA | NA | NA | NA | NA |
| KCNMA1 | 399 | G | -30.92475 | 24.40840419 | 0.304079391 | 16.12040705 | NA | NA | NA | NA | NA | NA | NA |
| KCNMA1 | 400 | S | -30.69341667 | 27.72075522 | 0.349147176 | 18.42620495 | NA | NA | NA | NA | NA | NA | NA |
| KCNMA1 | 401 | Y | -31.915 | 30.55409082 | 0.387697485 | 21.24478624 | NA | NA | NA | NA | NA | NA | NA |
| KCNMA1 | 402 | S | -28.36491667 | 32.63723282 | 0.416040675 | 25.70189031 | NA | NA | NA | NA | NA | NA | NA |
| KCNMA1 | 403 | A | -29.79815 | 35.9927538 | 0.461695829 | 27.7098946 | NA | NA | NA | NA | NA | NA | NA |
| KCNMA1 | 404 | V | -28.56803571 | 36.73210088 | 0.471755371 | 29.78528101 | NA | NA | NA | NA | NA | NA | NA |
| KCNMA1 | 405 | S | -29.12825 | 41.02888047 | 0.530217271 | 33.19227637 | NA | NA | NA | NA | NA | NA | NA |
| KCNMA1 | 406 | G | -31.7015 | 42.09986961 | 0.544789129 | 32.79351159 | NA | NA | NA | NA | NA | NA | NA |
| KCNMA1 | 407 | R | -31.17147727 | 36.8947325 | 0.473968134 | 26.98447812 | NA | NA | NA | NA | NA | NA | NA |
| KCNMA1 | 408 | K | -37.27119444 | 38.11437591 | 0.490562578 | 26.01336693 | NA | NA | NA | NA | NA | NA | NA |
| KCNMA1 | 409 | H | -36.68245 | 34.75790053 | 0.444894439 | 22.64824476 | NA | NA | NA | NA | NA | NA | NA |
| KCNMA1 | 410 | I | -40.586625 | 32.491453 | 0.414057198 | 19.94548717 | NA | NA | NA | NA | NA | NA | NA |
| KCNMA1 | 411 | V | -41.59603571 | 30.56825324 | 0.387890179 | 18.11848245 | NA | NA | NA | NA | NA | NA | NA |
| KCNMA1 | 412 | V | -43.09446429 | 27.23313287 | 0.342512595 | 14.81429986 | NA | NA | NA | NA | NA | NA | NA |
| KCNMA1 | 413 | C | -45.72158333 | 25.79072378 | 0.322887205 | 13.29267392 | NA | NA | NA | NA | NA | NA | NA |
| KCNMA1 | 414 | G | -47.07425 | 22.71580835 | 0.281049965 | 9.831561846 | NA | NA | NA | NA | NA | NA | NA |
| KCNMA1 | 415 | H | -48.55265 | 19.27083968 | 0.234177788 | 6.376627505 | NA | NA | NA | NA | NA | NA | NA |
| KCNMA1 | 416 | I | -43.983125 | 19.55537185 | 0.238049127 | 7.131130711 | NA | NA | NA | NA | NA | NA | NA |

| | | | | | | | | | | | | | |
|---|---|---|---|---|---|---|---|---|---|---|---|---|---|
| KCNMA1 | 417 | T | -45.01803571 | 18.66282157 | 0.225905105 | 6.454776121 | NA | NA | NA | NA | NA | NA | NA |
| KCNMA1 | 418 | L | -41.951625 | 21.0086967 | 0.257823037 | 8.586309791 | NA | NA | NA | NA | NA | NA | NA |
| KCNMA1 | 419 | E | -46.83730556 | 22.58666155 | 0.279292796 | 10.11758537 | NA | NA | NA | NA | NA | NA | NA |
| KCNMA1 | 420 | S | -46.70158333 | 23.36011291 | 0.28981636 | 10.45573798 | NA | NA | NA | NA | NA | NA | NA |
| KCNMA1 | 421 | V | -43.12817857 | 24.54626898 | 0.305955177 | 12.11774543 | NA | NA | NA | NA | NA | NA | NA |
| KCNMA1 | 422 | S | -42.47191667 | 25.80617271 | 0.323097403 | 13.39881109 | NA | NA | NA | NA | NA | NA | NA |
| KCNMA1 | 423 | N | -45.902375 | 28.91961494 | 0.365458838 | 16.45527712 | NA | NA | NA | NA | NA | NA | NA |
| KCNMA1 | 424 | F | -44.35320455 | 30.39310961 | 0.385507178 | 18.05277022 | NA | NA | NA | NA | NA | NA | NA |
| KCNMA1 | 425 | L | -40.341125 | 28.04691887 | 0.353584952 | 15.61826138 | NA | NA | NA | NA | NA | NA | NA |
| KCNMA1 | 426 | K | -42.05863889 | 32.25499597 | 0.410839968 | 19.78081816 | NA | NA | NA | NA | NA | NA | NA |
| KCNMA1 | 427 | D | -44.356875 | 34.6291587 | 0.44314278 | 22.20350671 | NA | NA | NA | NA | NA | NA | NA |
| KCNMA1 | 428 | F | -41.35320455 | 35.01383946 | 0.448376739 | 22.5650194 | NA | NA | NA | NA | NA | NA | NA |
| KCNMA1 | 429 | L | -37.59125 | 33.15446638 | 0.423078145 | 20.72284874 | NA | NA | NA | NA | NA | NA | NA |
| KCNMA1 | 430 | H | -39.25285 | 37.70931128 | 0.485051277 | 25.22950738 | NA | NA | NA | NA | NA | NA | NA |
| KCNMA1 | 431 | K | -35.30930556 | 39.09546589 | 0.50391127 | 27.12639932 | NA | NA | NA | NA | NA | NA | NA |
| KCNMA1 | 432 | D | -38.640625 | 42.56915313 | 0.551174191 | 30.10125235 | NA | NA | NA | NA | NA | NA | NA |
| KCNMA1 | 433 | R | -39.74775 | 40.277773 | 0.519997718 | 27.98150649 | NA | NA | NA | NA | NA | NA | NA |
| KCNMA1 | 434 | D | -36.987 | 45.24367989 | 0.587563751 | 32.94595493 | NA | NA | NA | NA | NA | NA | NA |
| KCNMA1 | 435 | D | -36.072875 | 44.10408717 | 0.572058474 | 32.18477334 | NA | NA | NA | NA | NA | NA | NA |
| KCNMA1 | 436 | V | -35.33060714 | 40.64266193 | 0.524962389 | 28.72493223 | NA | NA | NA | NA | NA | NA | NA |
| KCNMA1 | 437 | N | -32.89675 | 40.19301987 | 0.518844568 | 30.07558463 | NA | NA | NA | NA | NA | NA | NA |
| KCNMA1 | 438 | V | -36.17917857 | 36.22692907 | 0.464882013 | 24.30760814 | NA | NA | NA | NA | NA | NA | NA |
| KCNMA1 | 439 | E | -33.29130556 | 32.32137585 | 0.411743131 | 21.32710852 | NA | NA | NA | NA | NA | NA | NA |
| KCNMA1 | 440 | I | -37.614625 | 30.24208414 | 0.383452328 | 17.97732479 | NA | NA | NA | NA | NA | NA | NA |
| KCNMA1 | 441 | V | -37.47303571 | 28.73482462 | 0.362944584 | 16.45606822 | NA | NA | NA | NA | NA | NA | NA |
| KCNMA1 | 442 | F | -39.46502273 | 25.5166151 | 0.319157688 | 12.95995527 | NA | NA | NA | NA | NA | NA | NA |
| KCNMA1 | 443 | L | -40.624125 | 23.84755233 | 0.296448452 | 11.32452382 | NA | NA | NA | NA | NA | NA | NA |
| KCNMA1 | 444 | H | -42.70965 | 18.21077848 | 0.219754615 | 5.96347191 | NA | NA | NA | NA | NA | NA | NA |
| KCNMA1 | 445 | N | -42.604 | 16.85650361 | 0.201328378 | 4.463819947 | 12.45 | 1.81 | 4.5 | 0.13 | -1.85 | 0 | 0 |
| KCNMA1 | 446 | I | -42.026625 | 14.01557145 | 0.162674709 | 1.580997016 | 12.45 | 1.81 | 4.5 | 0.13 | -1.85 | 0 | 0 |
| KCNMA1 | 447 | S | -37.89258333 | 14.1956163 | 0.165124396 | 1.765822646 | 12.47 | -0.39 | -2.04 | 2.73 | 0.07 | 0 | 0 |
| KCNMA1 | 448 | P | -37.24517857 | 14.31866017 | 0.166798529 | 1.906917991 | 12.5 | -0.78 | -2.88 | 3.38 | 0.59 | 0 | 0 |
| KCNMA1 | 449 | N | -38.132625 | 14.11951224 | 0.164088926 | 1.874658066 | 12.3 | 1.14 | 3.8 | 0.13 | -1.79 | 0 | 0 |

| | | | | | | | | | | | | | |
|---|---|---|---|---|---|---|---|---|---|---|---|---|---|
| KCNMA1 | 450 | L | -34.15525 | 13.27198595 | 0.152557499 | 2.068736181 | 11.33 | -0.39 | -2.04 | 2.73 | 0.07 | 0 | 0 |
| KCNMA1 | 451 | E | -38.03063889 | 17.43914017 | 0.209255719 | 5.332713798 | NA | NA | NA | NA | NA | NA | NA |
| KCNMA1 | 452 | L | -37.677125 | 19.16853633 | 0.23278585 | 7.063354656 | NA | NA | NA | NA | NA | NA | NA |
| KCNMA1 | 453 | E | -32.59775 | 14.87731273 | 0.174399545 | 4.786878536 | 10.13 | -0.84 | -3.5 | 12.68 | 0.88 | 1 | -1 |
| KCNMA1 | 454 | A | -33.00195 | 20.575723 | 0.251932005 | 10.49137913 | NA | NA | NA | NA | NA | NA | NA |
| KCNMA1 | 455 | L | -36.0735 | 22.95112097 | 0.284251624 | 11.04726032 | NA | NA | NA | NA | NA | NA | NA |
| KCNMA1 | 456 | F | -34.83184091 | 22.20140529 | 0.274051007 | 12.31139691 | NA | NA | NA | NA | NA | NA | NA |
| KCNMA1 | 457 | K | -29.25986111 | 20.18002968 | 0.246548209 | 12.10073278 | NA | NA | NA | NA | NA | NA | NA |
| KCNMA1 | 458 | R | -32.92265909 | 26.39928929 | 0.331167336 | 17.09110005 | NA | NA | NA | NA | NA | NA | NA |
| KCNMA1 | 459 | H | -33.70265 | 28.49921846 | 0.359738932 | 18.37945321 | NA | NA | NA | NA | NA | NA | NA |
| KCNMA1 | 460 | F | -27.33965909 | 26.54020377 | 0.333084615 | 18.87992657 | NA | NA | NA | NA | NA | NA | NA |
| KCNMA1 | 461 | T | -28.87246429 | 31.06533653 | 0.394653485 | 23.59168657 | NA | NA | NA | NA | NA | NA | NA |
| KCNMA1 | 462 | Q | -32.63986111 | 31.70065269 | 0.403297584 | 21.59688405 | NA | NA | NA | NA | NA | NA | NA |
| KCNMA1 | 463 | V | -33.94046429 | 27.4209424 | 0.345067928 | 17.38594384 | NA | NA | NA | NA | NA | NA | NA |
| KCNMA1 | 464 | E | -32.57486111 | 26.47378016 | 0.332180857 | 16.37174929 | NA | NA | NA | NA | NA | NA | NA |
| KCNMA1 | 465 | F | -36.19493182 | 20.08863641 | 0.245304714 | 8.221742494 | NA | NA | NA | NA | NA | NA | NA |
| KCNMA1 | 466 | Y | -34.51258333 | 22.55627642 | 0.278879377 | 10.68044932 | NA | NA | NA | NA | NA | NA | NA |
| KCNMA1 | 467 | Q | -36.94330556 | 16.25569211 | 0.193153748 | 3.956643132 | 12.3 | 1.14 | 3.8 | 0.13 | -1.79 | 0 | 0 |
| KCNMA1 | 468 | G | -40.38925 | 21.3942256 | 0.263068536 | 8.841343463 | NA | NA | NA | NA | NA | NA | NA |
| KCNMA1 | 469 | S | -43.44891667 | 22.97043885 | 0.284514463 | 10.58139945 | NA | NA | NA | NA | NA | NA | NA |
| KCNMA1 | 470 | V | -45.38960714 | 25.3565863 | 0.316980339 | 12.88906328 | NA | NA | NA | NA | NA | NA | NA |
| KCNMA1 | 471 | L | -47.189 | 24.41527131 | 0.304172825 | 12.13991682 | NA | NA | NA | NA | NA | NA | NA |
| KCNMA1 | 472 | N | -42.261 | 26.7060484 | 0.335341094 | 14.46295541 | NA | NA | NA | NA | NA | NA | NA |
| KCNMA1 | 473 | P | -42.33503571 | 30.26810523 | 0.383806371 | 17.84074164 | NA | NA | NA | NA | NA | NA | NA |
| KCNMA1 | 474 | H | -37.12515 | 28.44861643 | 0.359050441 | 16.05357774 | NA | NA | NA | NA | NA | NA | NA |
| KCNMA1 | 475 | D | -39.357125 | 25.0247575 | 0.312465483 | 12.53304316 | NA | NA | NA | NA | NA | NA | NA |
| KCNMA1 | 476 | L | -41.400125 | 29.9107629 | 0.378944378 | 17.36268393 | NA | NA | NA | NA | NA | NA | NA |
| KCNMA1 | 477 | A | -37.00035 | 32.14956737 | 0.409405508 | 19.87809665 | NA | NA | NA | NA | NA | NA | NA |
| KCNMA1 | 478 | R | -35.26520455 | 24.67992563 | 0.307773707 | 12.79109169 | NA | NA | NA | NA | NA | NA | NA |
| KCNMA1 | 479 | V | -37.19832143 | 28.86174655 | 0.364671482 | 16.7641138 | NA | NA | NA | NA | NA | NA | NA |
| KCNMA1 | 480 | K | -35.17930556 | 32.9661918 | 0.420516485 | 21.10043045 | NA | NA | NA | NA | NA | NA | NA |
| KCNMA1 | 481 | I | -41.166 | 33.99357813 | 0.434495083 | 21.53818155 | NA | NA | NA | NA | NA | NA | NA |
| KCNMA1 | 482 | E | -39.04986111 | 35.8127467 | 0.459246656 | 23.33155365 | NA | NA | NA | NA | NA | NA | NA |

| | | | | | | | | | | | | | |
|---|---|---|---|---|---|---|---|---|---|---|---|---|---|
| KCNMA1 | 483 | S | -37.86608333 | 38.39891647 | 0.494434032 | 25.92566393 | NA | NA | NA | NA | NA | NA | NA |
| KCNMA1 | 484 | A | -40.46775 | 37.12661512 | 0.477123124 | 24.56963855 | NA | NA | NA | NA | NA | NA | NA |
| KCNMA1 | 485 | D | -42.2185 | 39.4640624 | 0.508926387 | 27.03055255 | NA | NA | NA | NA | NA | NA | NA |
| KCNMA1 | 486 | A | -44.11395 | 35.8342078 | 0.459538655 | 23.47269935 | NA | NA | NA | NA | NA | NA | NA |
| KCNMA1 | 487 | C | -45.84758333 | 31.84911573 | 0.405317569 | 19.35170746 | NA | NA | NA | NA | NA | NA | NA |
| KCNMA1 | 488 | L | -46.556875 | 29.92606558 | 0.379152586 | 17.01865869 | NA | NA | NA | NA | NA | NA | NA |
| KCNMA1 | 489 | I | -49.607625 | 27.36981998 | 0.344372358 | 14.50635339 | NA | NA | NA | NA | NA | NA | NA |
| KCNMA1 | 490 | L | -48.6375 | 25.38493155 | 0.317366004 | 12.6730172 | NA | NA | NA | NA | NA | NA | NA |
| KCNMA1 | 491 | A | -52.30835 | 23.46404042 | 0.291230396 | 10.99819425 | NA | NA | NA | NA | NA | NA | NA |
| KCNMA1 | 492 | N | -53.65625 | 18.25539359 | 0.220361648 | 6.130881127 | NA | NA | NA | NA | NA | NA | NA |
| KCNMA1 | 493 | K | -54.18563889 | 21.41527039 | 0.263354871 | 10.87288718 | NA | NA | NA | NA | NA | NA | NA |
| KCNMA1 | 494 | Y | -54.20733333 | 16.60991577 | 0.197973308 | 6.233799473 | NA | NA | NA | NA | NA | NA | NA |
| KCNMA1 | 495 | C | -57.80591667 | 15.2112815 | 0.178943517 | 4.737111326 | 10.48 | -0.14 | 0.74 | 0.68 | 0.58 | 0 | 0 |
| KCNMA1 | 496 | A | -59.89535 | 12.08882302 | 0.136459407 | 1.75311482 | 10.34 | -0.18 | 1.28 | 0.33 | 0.69 | 0 | 0 |
| KCNMA1 | 497 | D | -61.646375 | 15.00297507 | 0.176109304 | 5.137859102 | NA | NA | NA | NA | NA | NA | NA |
| KCNMA1 | 498 | P | -61.53375 | 18.30510557 | 0.221038028 | 8.479215862 | NA | NA | NA | NA | NA | NA | NA |
| KCNMA1 | 499 | D | -62.78125 | 19.92567665 | 0.243087487 | 10.11142031 | NA | NA | NA | NA | NA | NA | NA |
| KCNMA1 | 500 | A | -58.36195 | 17.33772745 | 0.2078759 | 7.008472881 | NA | NA | NA | NA | NA | NA | NA |
| KCNMA1 | 501 | E | -55.09086111 | 18.44822889 | 0.222985361 | 8.060927719 | NA | NA | NA | NA | NA | NA | NA |
| KCNMA1 | 502 | D | -59.1545 | 22.52697394 | 0.278480688 | 12.04927532 | NA | NA | NA | NA | NA | NA | NA |
| KCNMA1 | 503 | A | -58.16135 | 23.50735317 | 0.291819709 | 13.63113316 | NA | NA | NA | NA | NA | NA | NA |
| KCNMA1 | 504 | S | -54.65291667 | 20.92754085 | 0.256718832 | 10.38677721 | NA | NA | NA | NA | NA | NA | NA |
| KCNMA1 | 505 | N | -53.903625 | 24.67643931 | 0.307726272 | 14.15610955 | NA | NA | NA | NA | NA | NA | NA |
| KCNMA1 | 506 | I | -57.088125 | 27.28268957 | 0.343186863 | 16.73393626 | NA | NA | NA | NA | NA | NA | NA |
| KCNMA1 | 507 | M | -53.8565 | 26.2952205 | 0.329751378 | 15.74526763 | NA | NA | NA | NA | NA | NA | NA |
| KCNMA1 | 508 | R | -50.45711364 | 21.63727053 | 0.2663754 | 9.105943534 | NA | NA | NA | NA | NA | NA | NA |
| KCNMA1 | 509 | V | -52.71975 | 30.10981586 | 0.381652689 | 17.64130003 | NA | NA | NA | NA | NA | NA | NA |
| KCNMA1 | 510 | I | -53.75775 | 30.94473542 | 0.393012588 | 19.0719816 | NA | NA | NA | NA | NA | NA | NA |
| KCNMA1 | 511 | S | -49.08225 | 30.32449304 | 0.384573582 | 17.60023008 | NA | NA | NA | NA | NA | NA | NA |
| KCNMA1 | 512 | I | -47.6365 | 30.96724889 | 0.393318906 | 18.03156139 | NA | NA | NA | NA | NA | NA | NA |
| KCNMA1 | 513 | K | -50.50486111 | 35.15385069 | 0.450281729 | 22.42423834 | NA | NA | NA | NA | NA | NA | NA |
| KCNMA1 | 514 | N | -49.393875 | 35.4326094 | 0.454074515 | 22.70449952 | NA | NA | NA | NA | NA | NA | NA |
| KCNMA1 | 515 | Y | -43.33791667 | 35.38027668 | 0.453362477 | 23.01429239 | NA | NA | NA | NA | NA | NA | NA |

| | | | | | | | | | | | | | |
|---|---|---|---|---|---|---|---|---|---|---|---|---|---|
| KCNMA1 | 516 | H | -44.87275 | 38.09901848 | 0.490353625 | 25.59904604 | NA | NA | NA | NA | NA | NA | NA |
| KCNMA1 | 517 | P | -48.63917857 | 40.73252058 | 0.526185004 | 27.83268137 | NA | NA | NA | NA | NA | NA | NA |
| KCNMA1 | 518 | K | -47.78819444 | 41.99871015 | 0.543412755 | 29.0606526 | NA | NA | NA | NA | NA | NA | NA |
| KCNMA1 | 519 | I | -46.672125 | 37.10686434 | 0.476854395 | 24.32985964 | NA | NA | NA | NA | NA | NA | NA |
| KCNMA1 | 520 | R | -48.07520455 | 37.37675835 | 0.480526568 | 24.47945485 | NA | NA | NA | NA | NA | NA | NA |
| KCNMA1 | 521 | I | -50.31 | 33.77442566 | 0.431513299 | 21.06096148 | NA | NA | NA | NA | NA | NA | NA |
| KCNMA1 | 522 | I | -50.443875 | 32.73878374 | 0.417422375 | 20.17908746 | NA | NA | NA | NA | NA | NA | NA |
| KCNMA1 | 523 | T | -53.36403571 | 29.82514109 | 0.377779409 | 17.65517719 | NA | NA | NA | NA | NA | NA | NA |
| KCNMA1 | 524 | Q | -53.86486111 | 27.31725801 | 0.3436572 | 16.91376252 | NA | NA | NA | NA | NA | NA | NA |
| KCNMA1 | 525 | M | -58.572875 | 26.90464956 | 0.338043258 | 16.42796679 | NA | NA | NA | NA | NA | NA | NA |
| KCNMA1 | 526 | L | -59.574375 | 21.93897653 | 0.270480406 | 11.64529022 | NA | NA | NA | NA | NA | NA | NA |
| KCNMA1 | 527 | Q | -63.99063889 | 23.0016304 | 0.284938854 | 13.2431494 | NA | NA | NA | NA | NA | NA | NA |
| KCNMA1 | 528 | Y | -66.59141667 | 28.47510142 | 0.359410796 | 18.71210238 | NA | NA | NA | NA | NA | NA | NA |
| KCNMA1 | 529 | H | -67.42325 | 24.6915586 | 0.307931985 | 18.20417209 | NA | NA | NA | NA | NA | NA | NA |
| KCNMA1 | 530 | N | -63.81 | 24.38842022 | 0.30380749 | 16.0902919 | NA | NA | NA | NA | NA | NA | NA |
| KCNMA1 | 531 | K | -62.66486111 | 29.96150744 | 0.379634807 | 20.15137909 | NA | NA | NA | NA | NA | NA | NA |
| KCNMA1 | 532 | A | -64.79855 | 32.19755466 | 0.410058422 | 23.81608753 | NA | NA | NA | NA | NA | NA | NA |
| KCNMA1 | 533 | H | -63.21675 | 27.48392177 | 0.345924824 | 19.22031853 | NA | NA | NA | NA | NA | NA | NA |
| KCNMA1 | 534 | L | -59.312 | 32.07564879 | 0.408399774 | 22.18225223 | NA | NA | NA | NA | NA | NA | NA |
| KCNMA1 | 535 | L | -63.073875 | 35.7684451 | 0.458643889 | 27.36276796 | NA | NA | NA | NA | NA | NA | NA |
| KCNMA1 | 536 | N | -62.320625 | 32.88202745 | 0.419371346 | 26.3921065 | NA | NA | NA | NA | NA | NA | NA |
| KCNMA1 | 537 | I | -58.039375 | 34.26017246 | 0.438122361 | 26.00304272 | NA | NA | NA | NA | NA | NA | NA |
| KCNMA1 | 538 | P | -58.08032143 | 39.01309917 | 0.50279059 | 30.58529988 | NA | NA | NA | NA | NA | NA | NA |
| KCNMA1 | 539 | S | -54.36525 | 40.37360203 | 0.521301565 | 29.860343 | NA | NA | NA | NA | NA | NA | NA |
| KCNMA1 | 540 | W | -56.27532143 | 35.79139519 | 0.458956148 | 25.26249484 | NA | NA | NA | NA | NA | NA | NA |
| KCNMA1 | 541 | N | -56.147 | 41.40622072 | 0.535351355 | 30.85763069 | NA | NA | NA | NA | NA | NA | NA |
| KCNMA1 | 542 | W | -59.68339286 | 41.49860665 | 0.536608357 | 31.02010646 | NA | NA | NA | NA | NA | NA | NA |
| KCNMA1 | 543 | K | -57.59341667 | 44.36961902 | 0.575671296 | 33.89407749 | NA | NA | NA | NA | NA | NA | NA |
| KCNMA1 | 544 | E | -52.59208333 | 44.70395357 | 0.580220245 | 32.51858959 | NA | NA | NA | NA | NA | NA | NA |
| KCNMA1 | 545 | G | -52.77575 | 41.01516897 | 0.530030713 | 28.83343204 | NA | NA | NA | NA | NA | NA | NA |
| KCNMA1 | 546 | D | -53.245125 | 39.22770943 | 0.505710573 | 27.08574724 | NA | NA | NA | NA | NA | NA | NA |
| KCNMA1 | 547 | D | -54.825625 | 35.88631525 | 0.460247628 | 25.36157523 | NA | NA | NA | NA | NA | NA | NA |
| KCNMA1 | 548 | A | -57.05795 | 33.92133716 | 0.433512174 | 23.44501213 | NA | NA | NA | NA | NA | NA | NA |

| | | | | | | | | | | | | | |
|---|---|---|---|---|---|---|---|---|---|---|---|---|---|
| KCNMA1 | 549 | I | -55.8535 | 31.23580058 | 0.396972815 | 20.76648532 | NA | NA | NA | NA | NA | NA | NA |
| KCNMA1 | 550 | C | -60.10675 | 30.15091234 | 0.382211847 | 19.7836443 | NA | NA | NA | NA | NA | NA | NA |
| KCNMA1 | 551 | L | -57.989375 | 26.43858442 | 0.331701984 | 16.00940851 | NA | NA | NA | NA | NA | NA | NA |
| KCNMA1 | 552 | A | -62.61595 | 27.75429642 | 0.349603537 | 17.93755264 | NA | NA | NA | NA | NA | NA | NA |
| KCNMA1 | 553 | E | -63.88619444 | 30.82781343 | 0.39142175 | 20.99397775 | NA | NA | NA | NA | NA | NA | NA |
| KCNMA1 | 554 | L | -60.434125 | 33.04284941 | 0.421559487 | 22.75232467 | NA | NA | NA | NA | NA | NA | NA |
| KCNMA1 | 555 | K | -61.26730556 | 27.65686591 | 0.348277899 | 17.32708771 | NA | NA | NA | NA | NA | NA | NA |
| KCNMA1 | 556 | L | -65.515 | 29.7771571 | 0.37712654 | 20.04502665 | NA | NA | NA | NA | NA | NA | NA |
| KCNMA1 | 557 | G | -64.865 | 33.76299451 | 0.431357767 | 23.95128258 | NA | NA | NA | NA | NA | NA | NA |
| KCNMA1 | 558 | F | -61.08247727 | 35.81659497 | 0.459299015 | 25.9726922 | NA | NA | NA | NA | NA | NA | NA |
| KCNMA1 | 559 | I | -64.90425 | 31.87341375 | 0.405648168 | 22.04186835 | NA | NA | NA | NA | NA | NA | NA |
| KCNMA1 | 560 | A | -68.03695 | 36.29578407 | 0.465818853 | 26.52364636 | NA | NA | NA | NA | NA | NA | NA |
| KCNMA1 | 561 | Q | -66.95097222 | 38.99662224 | 0.502566405 | 29.21909213 | NA | NA | NA | NA | NA | NA | NA |
| KCNMA1 | 562 | S | -65.49791667 | 38.98606511 | 0.502422765 | 29.1318297 | NA | NA | NA | NA | NA | NA | NA |
| KCNMA1 | 563 | C | -69.43675 | 37.60354774 | 0.48361226 | 27.88182693 | NA | NA | NA | NA | NA | NA | NA |
| KCNMA1 | 564 | L | -70.39875 | 42.1592339 | 0.545596838 | 32.41602169 | NA | NA | NA | NA | NA | NA | NA |
| KCNMA1 | 565 | A | -66.41135 | 43.2224272 | 0.560062626 | 33.39222692 | NA | NA | NA | NA | NA | NA | NA |
| KCNMA1 | 566 | Q | -64.92708333 | 41.18999086 | 0.532409336 | 31.34428059 | NA | NA | NA | NA | NA | NA | NA |
| KCNMA1 | 567 | G | -61.39375 | 42.75434736 | 0.55369394 | 32.33043513 | NA | NA | NA | NA | NA | NA | NA |
| KCNMA1 | 568 | L | -61.74 | 41.83837355 | 0.541231218 | 31.36529013 | NA | NA | NA | NA | NA | NA | NA |
| KCNMA1 | 569 | S | -59.30225 | 37.91750026 | 0.487883892 | 27.44574377 | NA | NA | NA | NA | NA | NA | NA |
| KCNMA1 | 570 | T | -56.34517857 | 40.69344876 | 0.525653394 | 30.22664932 | NA | NA | NA | NA | NA | NA | NA |
| KCNMA1 | 571 | M | -57.060125 | 43.10585097 | 0.558476492 | 32.64126769 | NA | NA | NA | NA | NA | NA | NA |
| KCNMA1 | 572 | L | -57.6585 | 40.51721166 | 0.523255515 | 30.07744763 | NA | NA | NA | NA | NA | NA | NA |
| KCNMA1 | 573 | A | -53.63315 | 38.70480938 | 0.498596005 | 28.17513123 | NA | NA | NA | NA | NA | NA | NA |
| KCNMA1 | 574 | N | -51.616625 | 41.39770228 | 0.535235454 | 28.90357191 | NA | NA | NA | NA | NA | NA | NA |
| KCNMA1 | 575 | L | -52.976125 | 43.61045969 | 0.565342188 | 31.14290792 | NA | NA | NA | NA | NA | NA | NA |
| KCNMA1 | 576 | F | -52.33011364 | 40.96508236 | 0.529349235 | 28.4640335 | NA | NA | NA | NA | NA | NA | NA |
| KCNMA1 | 577 | S | -48.15191667 | 38.41012932 | 0.494586594 | 25.49388672 | NA | NA | NA | NA | NA | NA | NA |
| KCNMA1 | 578 | M | -44.131625 | 40.69537071 | 0.525679544 | 28.33280935 | NA | NA | NA | NA | NA | NA | NA |
| KCNMA1 | 579 | R | -46.20029545 | 38.15750215 | 0.491149353 | 26.21594206 | NA | NA | NA | NA | NA | NA | NA |
| KCNMA1 | 580 | S | -41.37408333 | 42.43057052 | 0.549288639 | 30.00596505 | NA | NA | NA | NA | NA | NA | NA |
| KCNMA1 | 581 | F | -41.77456818 | 46.1077932 | 0.59932086 | 33.65758808 | NA | NA | NA | NA | NA | NA | NA |

| | | | | | | | | | | | | | |
|---|---|---|---|---|---|---|---|---|---|---|---|---|---|
| KCNMA1 | 582 | I | -40.687625 | 44.01486192 | 0.570844477 | 31.46608773 | NA | NA | NA | NA | NA | NA | NA |
| KCNMA1 | 583 | K | -39.92919444 | 49.81494842 | 0.649760342 | 37.26531112 | NA | NA | NA | NA | NA | NA | NA |
| KCNMA1 | 584 | I | -42.38325 | 49.11502504 | 0.640237198 | 36.66457541 | NA | NA | NA | NA | NA | NA | NA |
| KCNMA1 | 585 | E | -40.67919444 | 53.97702054 | 0.706389415 | 41.42942551 | NA | NA | NA | NA | NA | NA | NA |
| KCNMA1 | 586 | E | -42.40797222 | 53.14558939 | 0.695076979 | 40.77449716 | NA | NA | NA | NA | NA | NA | NA |
| KCNMA1 | 587 | D | -47.904 | 55.54567539 | 0.727732503 | 42.60624032 | NA | NA | NA | NA | NA | NA | NA |
| KCNMA1 | 588 | T | -48.16217857 | 51.50996745 | 0.672822739 | 38.61571475 | NA | NA | NA | NA | NA | NA | NA |
| KCNMA1 | 589 | W | -51.26853571 | 45.37630558 | 0.589368253 | 32.88011828 | NA | NA | NA | NA | NA | NA | NA |
| KCNMA1 | 590 | Q | -47.75641667 | 46.07044268 | 0.59881267 | 33.17272013 | NA | NA | NA | NA | NA | NA | NA |
| KCNMA1 | 591 | K | -46.71908333 | 51.53895172 | 0.673217098 | 38.59966755 | NA | NA | NA | NA | NA | NA | NA |
| KCNMA1 | 592 | Y | -51.07966667 | 52.04168412 | 0.680057265 | 39.47179601 | NA | NA | NA | NA | NA | NA | NA |
| KCNMA1 | 593 | Y | -51.11775 | 43.52144437 | 0.564131048 | 30.95677314 | NA | NA | NA | NA | NA | NA | NA |
| KCNMA1 | 594 | L | -46.834375 | 46.54490134 | 0.605268145 | 33.61706571 | NA | NA | NA | NA | NA | NA | NA |
| KCNMA1 | 595 | E | -48.50875 | 51.34030633 | 0.670514333 | 38.43086946 | NA | NA | NA | NA | NA | NA | NA |
| KCNMA1 | 596 | G | -50.48275 | 48.85111724 | 0.636646474 | 36.28113514 | NA | NA | NA | NA | NA | NA | NA |
| KCNMA1 | 597 | V | -48.60389286 | 45.12259898 | 0.585916326 | 32.21938705 | NA | NA | NA | NA | NA | NA | NA |
| KCNMA1 | 598 | S | -46.48008333 | 48.44441109 | 0.631112838 | 35.97651032 | NA | NA | NA | NA | NA | NA | NA |
| KCNMA1 | 599 | N | -49.7845 | 47.74330694 | 0.621573628 | 35.01516588 | NA | NA | NA | NA | NA | NA | NA |
| KCNMA1 | 600 | E | -48.60563889 | 46.42367719 | 0.603618771 | 33.73215972 | NA | NA | NA | NA | NA | NA | NA |
| KCNMA1 | 601 | M | -52.7545 | 47.37265243 | 0.61653051 | 34.90798925 | NA | NA | NA | NA | NA | NA | NA |
| KCNMA1 | 602 | Y | -48.8145 | 49.39303645 | 0.644019816 | 37.1549733 | NA | NA | NA | NA | NA | NA | NA |
| KCNMA1 | 603 | T | -51.67732143 | 49.23740942 | 0.641902358 | 36.77174102 | NA | NA | NA | NA | NA | NA | NA |
| KCNMA1 | 604 | E | -50.71130556 | 53.85098539 | 0.704674584 | 41.3535505 | NA | NA | NA | NA | NA | NA | NA |
| KCNMA1 | 605 | Y | -52.15116667 | 54.81394445 | 0.717776586 | 42.34439136 | NA | NA | NA | NA | NA | NA | NA |
| KCNMA1 | 606 | L | -55.314625 | 58.64282232 | 0.769872225 | 48.12807853 | NA | NA | NA | NA | NA | NA | NA |
| KCNMA1 | 607 | S | -53.60558333 | 63.07800162 | 0.830217189 | 50.62308112 | NA | NA | NA | NA | NA | NA | NA |
| KCNMA1 | 608 | S | -54.74091667 | 66.17767568 | 0.872391294 | 55.64875374 | NA | NA | NA | NA | NA | NA | NA |
| KCNMA1 | 609 | A | -56.20295 | 68.14284891 | 0.899129403 | 57.59501986 | NA | NA | NA | NA | NA | NA | NA |
| KCNMA1 | 610 | F | -57.11593182 | 63.80768556 | 0.840145254 | 53.33909175 | NA | NA | NA | NA | NA | NA | NA |
| KCNMA1 | 611 | V | -60.06317857 | 65.55413033 | 0.863907348 | 55.07966608 | NA | NA | NA | NA | NA | NA | NA |
| KCNMA1 | 612 | G | -64.064 | 66.69273017 | 0.879399116 | 56.3813853 | NA | NA | NA | NA | NA | NA | NA |
| KCNMA1 | 613 | L | -63.1655 | 65.1794018 | 0.858808799 | 54.90199217 | NA | NA | NA | NA | NA | NA | NA |
| KCNMA1 | 614 | S | -65.84058333 | 63.87768996 | 0.841097733 | 54.02022048 | NA | NA | NA | NA | NA | NA | NA |

| | | | | | | | | | | | | | |
|---|---|---|---|---|---|---|---|---|---|---|---|---|---|
| KCNMA1 | 615 | F | -64.37093182 | 56.74900018 | 0.744104917 | 46.87298976 | NA | NA | NA | NA | NA | NA | NA |
| KCNMA1 | 616 | P | -65.70089286 | 62.24799532 | 0.818924139 | 52.3960055 | NA | NA | NA | NA | NA | NA | NA |
| KCNMA1 | 617 | T | -63.32332143 | 64.61883345 | 0.851181717 | 54.32857469 | NA | NA | NA | NA | NA | NA | NA |
| KCNMA1 | 618 | V | -60.24832143 | 62.664376 | 0.824589406 | 52.19083787 | NA | NA | NA | NA | NA | NA | NA |
| KCNMA1 | 619 | C | -60.69675 | 58.97674744 | 0.774415603 | 48.50987542 | NA | NA | NA | NA | NA | NA | NA |
| KCNMA1 | 620 | E | -61.19686111 | 63.38544469 | 0.834400253 | 52.9110139 | NA | NA | NA | NA | NA | NA | NA |
| KCNMA1 | 621 | L | -57.567875 | 64.77776094 | 0.853344081 | 54.31008535 | NA | NA | NA | NA | NA | NA | NA |
| KCNMA1 | 622 | C | -55.65575 | 61.31080291 | 0.806172717 | 50.77520537 | NA | NA | NA | NA | NA | NA | NA |
| KCNMA1 | 623 | F | -56.79393182 | 59.62132629 | 0.783185731 | 49.15401767 | NA | NA | NA | NA | NA | NA | NA |
| KCNMA1 | 624 | V | -55.62560714 | 64.41893028 | 0.848461838 | 53.89601214 | NA | NA | NA | NA | NA | NA | NA |
| KCNMA1 | 625 | K | -53.49041667 | 65.66761873 | 0.865451469 | 53.19792126 | NA | NA | NA | NA | NA | NA | NA |
| KCNMA1 | 626 | L | -51.643 | 61.40155146 | 0.80740744 | 48.83785749 | NA | NA | NA | NA | NA | NA | NA |
| KCNMA1 | 627 | K | -50.87375 | 58.90250984 | 0.773405528 | 46.43354616 | NA | NA | NA | NA | NA | NA | NA |
| KCNMA1 | 628 | L | -52.573125 | 57.83483002 | 0.758878697 | 45.66922052 | NA | NA | NA | NA | NA | NA | NA |
| KCNMA1 | 629 | L | -57.1245 | 54.2783477 | 0.710489267 | 43.80818419 | NA | NA | NA | NA | NA | NA | NA |
| KCNMA1 | 630 | M | -59.02025 | 54.86139598 | 0.718422211 | 44.38813753 | NA | NA | NA | NA | NA | NA | NA |
| KCNMA1 | 631 | I | -60.06975 | 50.16942668 | 0.654583367 | 39.69453348 | NA | NA | NA | NA | NA | NA | NA |
| KCNMA1 | 632 | A | -60.76995 | 51.22682799 | 0.668970349 | 40.80068444 | NA | NA | NA | NA | NA | NA | NA |
| KCNMA1 | 633 | I | -61.852375 | 52.77301048 | 0.690007678 | 42.36541021 | NA | NA | NA | NA | NA | NA | NA |
| KCNMA1 | 634 | E | -61.44641667 | 50.20135274 | 0.655017752 | 39.89000044 | NA | NA | NA | NA | NA | NA | NA |
| KCNMA1 | 635 | Y | -65.21016667 | 54.06901048 | 0.707641029 | 44.22321627 | NA | NA | NA | NA | NA | NA | NA |
| KCNMA1 | 636 | K | -62.14152778 | 55.61732824 | 0.72870741 | 45.75396978 | NA | NA | NA | NA | NA | NA | NA |
| KCNMA1 | 637 | S | -66.67458333 | 55.59135756 | 0.728354054 | 45.76384924 | NA | NA | NA | NA | NA | NA | NA |
| KCNMA1 | 638 | A | -67.41175 | 55.13820963 | 0.722188532 | 49.88983158 | NA | NA | NA | NA | NA | NA | NA |
| KCNMA1 | 639 | N | -70.70075 | 52.71562981 | 0.689226957 | 47.46651731 | NA | NA | NA | NA | NA | NA | NA |
| KCNMA1 | 640 | R | -68.95820455 | 48.13951468 | 0.626964423 | 42.88957945 | NA | NA | NA | NA | NA | NA | NA |
| KCNMA1 | 641 | E | -64.18097222 | 48.47071206 | 0.631470688 | 38.59592142 | NA | NA | NA | NA | NA | NA | NA |
| KCNMA1 | 642 | S | -63.16691667 | 49.32835453 | 0.643139755 | 39.47015947 | NA | NA | NA | NA | NA | NA | NA |
| KCNMA1 | 643 | R | -67.69175 | 48.02967868 | 0.625469997 | 38.28942617 | NA | NA | NA | NA | NA | NA | NA |
| KCNMA1 | 644 | I | -63.0105 | 47.04139866 | 0.612023478 | 37.13366818 | NA | NA | NA | NA | NA | NA | NA |
| KCNMA1 | 645 | L | -66.40225 | 51.10560708 | 0.667321019 | 41.26023946 | NA | NA | NA | NA | NA | NA | NA |
| KCNMA1 | 646 | I | -65.111125 | 48.42044974 | 0.63078682 | 38.55476396 | NA | NA | NA | NA | NA | NA | NA |
| KCNMA1 | 647 | N | -66.82075 | 53.16444956 | 0.69533359 | 43.32115131 | NA | NA | NA | NA | NA | NA | NA |

| | | | | | | | | | | | | | |
|---|---|---|---|---|---|---|---|---|---|---|---|---|---|
| KCNMA1 | 648 | P | -66.99846429 | 55.17756042 | 0.722723938 | 45.33478353 | NA | NA | NA | NA | NA | NA | NA |
| KCNMA1 | 649 | G | -70.079 | 57.69824758 | 0.757020359 | 47.91172846 | NA | NA | NA | NA | NA | NA | NA |
| KCNMA1 | 650 | N | -71.395875 | 61.29191362 | 0.80591571 | 51.50753021 | NA | NA | NA | NA | NA | NA | NA |
| KCNMA1 | 651 | H | -73.00695 | 58.87183959 | 0.772988229 | 49.13445244 | NA | NA | NA | NA | NA | NA | NA |
| KCNMA1 | 652 | L | -69.861 | 57.28202688 | 0.751357268 | 47.49930801 | NA | NA | NA | NA | NA | NA | NA |
| KCNMA1 | 653 | K | -68.32897222 | 61.52283967 | 0.809057685 | 51.68313399 | NA | NA | NA | NA | NA | NA | NA |
| KCNMA1 | 654 | I | -63.1415 | 61.38164946 | 0.807136654 | 51.49812049 | NA | NA | NA | NA | NA | NA | NA |
| KCNMA1 | 655 | Q | -63.55086111 | 59.77968976 | 0.785340421 | 49.50382599 | NA | NA | NA | NA | NA | NA | NA |
| KCNMA1 | 656 | E | -58.82063889 | 62.11208225 | 0.817074908 | 51.63557754 | NA | NA | NA | NA | NA | NA | NA |
| KCNMA1 | 657 | G | -57.039 | 58.81590414 | 0.772227172 | 48.3433298 | NA | NA | NA | NA | NA | NA | NA |
| KCNMA1 | 658 | T | -58.32360714 | 57.2602518 | 0.751060997 | 46.78800603 | NA | NA | NA | NA | NA | NA | NA |
| KCNMA1 | 659 | L | -56.3175 | 52.28196535 | 0.683326527 | 41.90911426 | NA | NA | NA | NA | NA | NA | NA |
| KCNMA1 | 660 | G | -56.066 | 53.7849906 | 0.70377666 | 43.37514642 | NA | NA | NA | NA | NA | NA | NA |
| KCNMA1 | 661 | F | -56.94329545 | 49.15775118 | 0.640818529 | 38.73176055 | NA | NA | NA | NA | NA | NA | NA |
| KCNMA1 | 662 | F | -54.07275 | 51.66974469 | 0.674996665 | 41.18564615 | NA | NA | NA | NA | NA | NA | NA |
| KCNMA1 | 663 | I | -53.96375 | 50.75845257 | 0.662597642 | 38.7529525 | NA | NA | NA | NA | NA | NA | NA |
| KCNMA1 | 664 | A | -49.65835 | 51.89067664 | 0.678002661 | 39.16296957 | NA | NA | NA | NA | NA | NA | NA |
| KCNMA1 | 665 | S | -46.14075 | 51.95914433 | 0.678934231 | 39.12386445 | NA | NA | NA | NA | NA | NA | NA |
| KCNMA1 | 666 | D | -43.300375 | 52.52545423 | 0.686639432 | 40.12933927 | NA | NA | NA | NA | NA | NA | NA |
| KCNMA1 | 667 | A | -43.66735 | 52.23890532 | 0.682740653 | 39.81097182 | NA | NA | NA | NA | NA | NA | NA |
| KCNMA1 | 668 | K | -41.51819444 | 55.227458 | 0.723402843 | 42.79745865 | NA | NA | NA | NA | NA | NA | NA |
| KCNMA1 | 669 | E | -45.53675 | 56.98656341 | 0.747337198 | 44.50907314 | NA | NA | NA | NA | NA | NA | NA |
| KCNMA1 | 670 | V | -47.73017857 | 54.42515772 | 0.712486761 | 41.48803808 | NA | NA | NA | NA | NA | NA | NA |
| KCNMA1 | 671 | K | -43.44263889 | 55.67424466 | 0.729481814 | 43.40194832 | NA | NA | NA | NA | NA | NA | NA |
| KCNMA1 | 672 | R | -46.12711364 | 60.17159367 | 0.790672658 | 47.67163067 | NA | NA | NA | NA | NA | NA | NA |
| KCNMA1 | 673 | A | -50.23675 | 59.44412073 | 0.780774675 | 46.71911621 | NA | NA | NA | NA | NA | NA | NA |
| KCNMA1 | 674 | F | -48.74475 | 60.10922966 | 0.789824135 | 47.43266608 | NA | NA | NA | NA | NA | NA | NA |
| KCNMA1 | 675 | F | -46.19702273 | 63.24990319 | 0.832556078 | 50.44706403 | NA | NA | NA | NA | NA | NA | NA |
| KCNMA1 | 676 | Y | -50.83266667 | 64.0144622 | 0.842958653 | 51.45782684 | NA | NA | NA | NA | NA | NA | NA |
| KCNMA1 | 677 | C | -51.56725 | 70.32349398 | 0.928799218 | 57.82505372 | NA | NA | NA | NA | NA | NA | NA |
| KCNMA1 | 678 | K | -53.52215 | 72.96015955 | 0.964673639 | 60.77822104 | NA | NA | NA | NA | NA | NA | NA |
| KCNMA1 | 679 | A | -50.69235 | 75.55654432 | 1 | 62.99336108 | NA | NA | NA | NA | NA | NA | NA |
| KCNMA1 | 680 | C | -48.09108333 | 72.97285898 | 0.964846427 | 60.03367127 | NA | NA | NA | NA | NA | NA | NA |

| | | | | | | | | | | | | | |
|---|---|---|---|---|---|---|---|---|---|---|---|---|---|
| KCNMA1 | 681 | H | -49.00725 | 69.78587497 | 0.921484384 | 57.12366419 | NA | NA | NA | NA | NA | NA | NA |
| KCNMA1 | 682 | D | -53.312 | 73.56469076 | 0.972898879 | 61.10060718 | NA | NA | NA | NA | NA | NA | NA |
| KCNMA1 | 683 | D | -51.439375 | 74.37101711 | 0.98386974 | 61.81168208 | NA | NA | NA | NA | NA | NA | NA |
| KCNMA1 | 684 | I | -48.28975 | 70.42940473 | 0.930240238 | 57.52746641 | NA | NA | NA | NA | NA | NA | NA |
| KCNMA1 | 685 | T | -49.02889286 | 68.69205242 | 0.906601856 | 55.78284796 | NA | NA | NA | NA | NA | NA | NA |
| KCNMA1 | 686 | D | -45.179875 | 67.58717107 | 0.891568861 | 55.0882936 | NA | NA | NA | NA | NA | NA | NA |
| KCNMA1 | 687 | P | -45.16046429 | 63.92518313 | 0.841743924 | 51.42616104 | NA | NA | NA | NA | NA | NA | NA |
| KCNMA1 | 688 | K | -41.33030556 | 61.34152723 | 0.806590752 | 48.91093739 | NA | NA | NA | NA | NA | NA | NA |
| KCNMA1 | 689 | R | -42.06684091 | 67.95141274 | 0.896524726 | 55.521619 | NA | NA | NA | NA | NA | NA | NA |
| KCNMA1 | 690 | I | -46.17 | 66.15993398 | 0.872149901 | 53.7108371 | NA | NA | NA | NA | NA | NA | NA |
| KCNMA1 | 691 | K | -41.56708333 | 60.50700559 | 0.795236266 | 48.19637748 | NA | NA | NA | NA | NA | NA | NA |
| KCNMA1 | 692 | K | -43.55408333 | 67.66767567 | 0.892664205 | 55.23838484 | NA | NA | NA | NA | NA | NA | NA |
| KCNMA1 | 693 | C | -42.10908333 | 66.04402823 | 0.87057289 | 53.62538628 | NA | NA | NA | NA | NA | NA | NA |
| KCNMA1 | 694 | G | -43.96 | 70.80573213 | 0.935360541 | 58.37573448 | NA | NA | NA | NA | NA | NA | NA |
| KCNMA1 | 695 | C | -46.17458333 | 69.72268017 | 0.920624557 | 57.24711145 | NA | NA | NA | NA | NA | NA | NA |
| KCNMA1 | 696 | K | -46.39130556 | 73.55207291 | 0.972727201 | 61.05488596 | NA | NA | NA | NA | NA | NA | NA |
| KCNMA1 | 697 | R | -49.59575 | 69.54882063 | 0.918259027 | 56.7080443 | NA | NA | NA | NA | NA | NA | NA |
| KCNMA1 | 804 | S | -95.51291667 | 48.50925595 | 0.631995116 | 43.26641169 | NA | NA | NA | NA | NA | NA | NA |
| KCNMA1 | 805 | N | -94.47075 | 44.17588891 | 0.573035407 | 38.93423439 | NA | NA | NA | NA | NA | NA | NA |
| KCNMA1 | 806 | V | -92.28875 | 43.30860322 | 0.561235135 | 38.06014614 | NA | NA | NA | NA | NA | NA | NA |
| KCNMA1 | 807 | K | -89.19430556 | 38.5534473 | 0.496536575 | 33.31317946 | NA | NA | NA | NA | NA | NA | NA |
| KCNMA1 | 808 | K | -86.97630556 | 42.30963297 | 0.547643165 | 37.05170386 | NA | NA | NA | NA | NA | NA | NA |
| KCNMA1 | 809 | Y | -83.28425 | 41.29037121 | 0.533775109 | 36.03705955 | NA | NA | NA | NA | NA | NA | NA |
| KCNMA1 | 810 | D | -80.116375 | 39.55227116 | 0.510126553 | 34.29394022 | NA | NA | NA | NA | NA | NA | NA |
| KCNMA1 | 811 | S | -79.99891667 | 39.6980568 | 0.51211011 | 34.44693857 | NA | NA | NA | NA | NA | NA | NA |
| KCNMA1 | 812 | T | -77.24603571 | 42.11685261 | 0.545020199 | 36.85964956 | NA | NA | NA | NA | NA | NA | NA |
| KCNMA1 | 813 | G | -79.0355 | 43.87347233 | 0.568920733 | 38.61650991 | NA | NA | NA | NA | NA | NA | NA |
| KCNMA1 | 814 | M | -75.5685 | 43.64606418 | 0.565826622 | 38.37872614 | NA | NA | NA | NA | NA | NA | NA |
| KCNMA1 | 815 | F | -76.23502273 | 40.50697605 | 0.52311625 | 35.35150597 | NA | NA | NA | NA | NA | NA | NA |
| KCNMA1 | 816 | H | -80.95895 | 39.23170083 | 0.50576488 | 33.96553454 | NA | NA | NA | NA | NA | NA | NA |
| KCNMA1 | 817 | W | -83.71189286 | 35.12646907 | 0.449909175 | 29.87038798 | NA | NA | NA | NA | NA | NA | NA |
| KCNMA1 | 818 | C | -85.75325 | 36.19468454 | 0.464443295 | 30.93998749 | NA | NA | NA | NA | NA | NA | NA |
| KCNMA1 | 819 | A | -87.41075 | 33.84578912 | 0.432484269 | 28.58971365 | NA | NA | NA | NA | NA | NA | NA |

| | | | | | | | | | | | | | |
|---|---|---|---|---|---|---|---|---|---|---|---|---|---|
| KCNMA1 | 820 | P | -86.37689286 | 30.72597706 | 0.390036166 | 25.47434765 | NA | NA | NA | NA | NA | NA | NA |
| KCNMA1 | 821 | K | -86.18708333 | 28.37505352 | 0.358049546 | 23.11730198 | NA | NA | NA | NA | NA | NA | NA |
| KCNMA1 | 822 | E | -87.30730556 | 24.00729025 | 0.298621843 | 18.75118275 | NA | NA | NA | NA | NA | NA | NA |
| KCNMA1 | 823 | I | -83.7385 | 24.61144988 | 0.306842027 | 19.34147363 | NA | NA | NA | NA | NA | NA | NA |
| KCNMA1 | 824 | E | -87.22486111 | 24.31974474 | 0.302873093 | 19.04995512 | NA | NA | NA | NA | NA | NA | NA |
| KCNMA1 | 825 | K | -89.04063889 | 29.91460911 | 0.378996709 | 24.64937188 | NA | NA | NA | NA | NA | NA | NA |
| KCNMA1 | 826 | V | -84.51375 | 30.83371643 | 0.391502066 | 25.56011641 | NA | NA | NA | NA | NA | NA | NA |
| KCNMA1 | 827 | I | -84.5375 | 28.13035846 | 0.354720229 | 22.1047013 | NA | NA | NA | NA | NA | NA | NA |
| KCNMA1 | 828 | L | -84.148625 | 31.7089412 | 0.403410357 | 25.68291993 | NA | NA | NA | NA | NA | NA | NA |
| KCNMA1 | 829 | T | -85.13446429 | 32.09850173 | 0.40871071 | 26.10984554 | NA | NA | NA | NA | NA | NA | NA |
| KCNMA1 | 830 | R | -81.26875 | 31.57881564 | 0.401639871 | 25.91307385 | NA | NA | NA | NA | NA | NA | NA |
| KCNMA1 | 831 | S | -86.37758333 | 36.31743394 | 0.466113421 | 30.29882417 | NA | NA | NA | NA | NA | NA | NA |
| KCNMA1 | 832 | E | -87.93319444 | 34.96948992 | 0.44777332 | 29.01389593 | NA | NA | NA | NA | NA | NA | NA |
| KCNMA1 | 833 | A | -84.44775 | 37.19914459 | 0.478109959 | 31.17983401 | NA | NA | NA | NA | NA | NA | NA |
| KCNMA1 | 834 | A | -85.34055 | 39.37031824 | 0.507650905 | 33.35145557 | NA | NA | NA | NA | NA | NA | NA |
| KCNMA1 | 835 | M | -89.390625 | 41.51321484 | 0.536807115 | 35.4928236 | NA | NA | NA | NA | NA | NA | NA |
| KCNMA1 | 836 | T | -87.30489286 | 42.08187669 | 0.544544318 | 36.05659243 | NA | NA | NA | NA | NA | NA | NA |
| KCNMA1 | 837 | V | -85.98803571 | 44.62593253 | 0.579158692 | 38.60464854 | NA | NA | NA | NA | NA | NA | NA |
| KCNMA1 | 838 | L | -82.751125 | 40.47121072 | 0.522629627 | 34.45045491 | NA | NA | NA | NA | NA | NA | NA |
| KCNMA1 | 839 | S | -80.97358333 | 45.22220324 | 0.58727154 | 35.68506406 | NA | NA | NA | NA | NA | NA | NA |
| KCNMA1 | 840 | G | -78.24225 | 43.60742154 | 0.565300851 | 33.99676263 | NA | NA | NA | NA | NA | NA | NA |
| KCNMA1 | 841 | H | -77.89385 | 39.46475781 | 0.508935848 | 29.89782964 | NA | NA | NA | NA | NA | NA | NA |
| KCNMA1 | 842 | V | -74.54675 | 36.66818616 | 0.470885748 | 27.08052177 | NA | NA | NA | NA | NA | NA | NA |
| KCNMA1 | 843 | V | -73.60032143 | 34.16937333 | 0.436886949 | 24.54378024 | NA | NA | NA | NA | NA | NA | NA |
| KCNMA1 | 844 | V | -71.67589286 | 30.19228295 | 0.382774734 | 20.59608458 | NA | NA | NA | NA | NA | NA | NA |
| KCNMA1 | 845 | C | -69.56925 | 27.73591178 | 0.349353395 | 18.06350152 | NA | NA | NA | NA | NA | NA | NA |
| KCNMA1 | 846 | I | -69.74975 | 23.83446127 | 0.296270335 | 14.23599884 | NA | NA | NA | NA | NA | NA | NA |
| KCNMA1 | 847 | F | -66.23920455 | 20.20702863 | 0.246915556 | 10.52636552 | NA | NA | NA | NA | NA | NA | NA |
| KCNMA1 | 848 | G | -68.64375 | 19.22964321 | 0.233617269 | 9.680407873 | NA | NA | NA | NA | NA | NA | NA |
| KCNMA1 | 849 | D | -68.0955 | 14.14863078 | 0.164485112 | 4.643285314 | 9.92 | -0.97 | -0.8 | 1.67 | 1.11 | 0 | 0 |
| KCNMA1 | 850 | V | -70.39517857 | 14.79729414 | 0.173310813 | 8.768126611 | NA | NA | NA | NA | NA | NA | NA |
| KCNMA1 | 851 | S | -66.29125 | 11.58456982 | 0.129598548 | 3.62841144 | 8.43 | -0.64 | 0.07 | 1.11 | 0.94 | 0 | 0 |
| KCNMA1 | 852 | S | -65.79391667 | 14.4411244 | 0.168464775 | 4.880863587 | 9.92 | -0.97 | -0.8 | 1.67 | 1.11 | 0 | 0 |

| | | | | | | | | | | | | | |
|---|---|---|---|---|---|---|---|---|---|---|---|---|---|
| KCNMA1 | 853 | A | -64.42795 | 16.1339502 | 0.19149733 | 6.450016638 | NA | NA | NA | NA | NA | NA | NA |
| KCNMA1 | 854 | L | -66.6505 | 19.01991998 | 0.230763779 | 9.351236074 | NA | NA | NA | NA | NA | NA | NA |
| KCNMA1 | 855 | I | -66.077875 | 22.41560807 | 0.276965446 | 12.73901398 | NA | NA | NA | NA | NA | NA | NA |
| KCNMA1 | 856 | G | -68.22525 | 25.04288693 | 0.312712151 | 15.37909725 | NA | NA | NA | NA | NA | NA | NA |
| KCNMA1 | 857 | L | -70.541375 | 26.2804984 | 0.329551069 | 16.66004293 | NA | NA | NA | NA | NA | NA | NA |
| KCNMA1 | 858 | R | -72.34465909 | 29.80384844 | 0.377489702 | 23.71480454 | NA | NA | NA | NA | NA | NA | NA |
| KCNMA1 | 859 | N | -68.574 | 30.30810342 | 0.384350585 | 20.58408478 | NA | NA | NA | NA | NA | NA | NA |
| KCNMA1 | 860 | L | -70.268125 | 32.15999 | 0.409547319 | 22.4931824 | NA | NA | NA | NA | NA | NA | NA |
| KCNMA1 | 861 | V | -74.26746429 | 33.52247788 | 0.428085302 | 23.94943019 | NA | NA | NA | NA | NA | NA | NA |
| KCNMA1 | 862 | M | -73.007875 | 35.56080777 | 0.455818779 | 25.92328395 | NA | NA | NA | NA | NA | NA | NA |
| KCNMA1 | 863 | P | -71.08646429 | 36.8982245 | 0.474015646 | 27.21098496 | NA | NA | NA | NA | NA | NA | NA |
| KCNMA1 | 864 | L | -73.847375 | 37.91265016 | 0.487817902 | 28.26905195 | NA | NA | NA | NA | NA | NA | NA |
| KCNMA1 | 865 | R | -77.85738636 | 39.31174728 | 0.50685399 | 33.28634232 | NA | NA | NA | NA | NA | NA | NA |
| KCNMA1 | 866 | A | -75.36195 | 42.60159948 | 0.551615656 | 32.97998937 | NA | NA | NA | NA | NA | NA | NA |
| KCNMA1 | 867 | S | -78.41408333 | 44.88193802 | 0.582641898 | 39.6728842 | NA | NA | NA | NA | NA | NA | NA |
| KCNMA1 | 868 | N | -75.664125 | 47.24161947 | 0.614747678 | 37.60457381 | NA | NA | NA | NA | NA | NA | NA |
| KCNMA1 | 869 | F | -76.76365909 | 44.64642402 | 0.579437499 | 34.98105243 | NA | NA | NA | NA | NA | NA | NA |
| KCNMA1 | 870 | H | -82.03075 | 47.88585114 | 0.623513082 | 42.66121776 | NA | NA | NA | NA | NA | NA | NA |
| KCNMA1 | 871 | Y | -85.6715 | 42.63826555 | 0.552114534 | 37.37349455 | NA | NA | NA | NA | NA | NA | NA |
| KCNMA1 | 872 | H | -84.54155 | 48.71807605 | 0.634836318 | 43.44724047 | NA | NA | NA | NA | NA | NA | NA |
| KCNMA1 | 873 | E | -79.81530556 | 47.17005154 | 0.613773926 | 41.63569682 | NA | NA | NA | NA | NA | NA | NA |
| KCNMA1 | 874 | L | -80.250125 | 43.81016162 | 0.568059329 | 37.78592493 | NA | NA | NA | NA | NA | NA | NA |
| KCNMA1 | 875 | K | -76.39430556 | 40.76055224 | 0.526566403 | 31.19120139 | NA | NA | NA | NA | NA | NA | NA |
| KCNMA1 | 876 | H | -80.66025 | 38.25250511 | 0.492441962 | 32.22834976 | NA | NA | NA | NA | NA | NA | NA |
| KCNMA1 | 877 | I | -77.128875 | 34.60534152 | 0.442818724 | 29.10926967 | NA | NA | NA | NA | NA | NA | NA |
| KCNMA1 | 878 | V | -77.60989286 | 33.48948713 | 0.427636431 | 28.035961 | NA | NA | NA | NA | NA | NA | NA |
| KCNMA1 | 879 | F | -75.26065909 | 30.1953434 | 0.382816375 | 20.68056431 | NA | NA | NA | NA | NA | NA | NA |
| KCNMA1 | 880 | V | -74.14317857 | 27.77995009 | 0.34995258 | 18.21674253 | NA | NA | NA | NA | NA | NA | NA |
| KCNMA1 | 881 | G | -74.117 | 25.35945492 | 0.317019369 | 15.8820947 | NA | NA | NA | NA | NA | NA | NA |
| KCNMA1 | 882 | S | -73.83475 | 21.24088859 | 0.260982235 | 15.23339049 | NA | NA | NA | NA | NA | NA | NA |
| KCNMA1 | 883 | I | -77.90725 | 22.06897426 | 0.272249153 | 16.04687986 | NA | NA | NA | NA | NA | NA | NA |
| KCNMA1 | 884 | E | -75.90997222 | 17.46604234 | 0.20962175 | 11.45012685 | NA | NA | NA | NA | NA | NA | NA |
| KCNMA1 | 885 | Y | -71.21708333 | 20.16439349 | 0.246335464 | 14.43394092 | NA | NA | NA | NA | NA | NA | NA |

| | | | | | | | | | | | | | |
|---|---|---|---|---|---|---|---|---|---|---|---|---|---|
| KCNMA1 | 886 | L | -75.105375 | 22.63406358 | 0.279937747 | 16.61812713 | NA | NA | NA | NA | NA | NA | NA |
| KCNMA1 | 887 | K | -78.40608333 | 19.86033094 | 0.242198394 | 14.05682494 | NA | NA | NA | NA | NA | NA | NA |
| KCNMA1 | 888 | R | -71.86747727 | 17.21559601 | 0.206214182 | 11.20024668 | NA | NA | NA | NA | NA | NA | NA |
| KCNMA1 | 889 | E | -72.52075 | 22.52074808 | 0.278395979 | 16.49614343 | NA | NA | NA | NA | NA | NA | NA |
| KCNMA1 | 890 | W | -80.06575 | 24.66345623 | 0.307549624 | 18.63263134 | NA | NA | NA | NA | NA | NA | NA |
| KCNMA1 | 891 | E | -77.06763889 | 21.44110172 | 0.263706331 | 16.17313176 | NA | NA | NA | NA | NA | NA | NA |
| KCNMA1 | 892 | T | -75.17603571 | 26.77325608 | 0.336255521 | 21.56224184 | NA | NA | NA | NA | NA | NA | NA |
| KCNMA1 | 893 | L | -76.798625 | 28.94926598 | 0.365862269 | 22.92604217 | NA | NA | NA | NA | NA | NA | NA |
| KCNMA1 | 894 | H | -81.43355 | 27.49036817 | 0.346012534 | 22.23033881 | NA | NA | NA | NA | NA | NA | NA |
| KCNMA1 | 895 | N | -81.503375 | 31.74790936 | 0.403940557 | 26.4896301 | NA | NA | NA | NA | NA | NA | NA |
| KCNMA1 | 896 | F | -79.36456818 | 34.11222464 | 0.436109385 | 28.83773586 | NA | NA | NA | NA | NA | NA | NA |
| KCNMA1 | 897 | P | -83.44817857 | 37.73035505 | 0.485337598 | 32.46254484 | NA | NA | NA | NA | NA | NA | NA |
| KCNMA1 | 898 | K | -83.25208333 | 37.71868737 | 0.485178848 | 32.55556294 | NA | NA | NA | NA | NA | NA | NA |
| KCNMA1 | 899 | V | -81.12932143 | 33.79725894 | 0.431823968 | 27.76650142 | NA | NA | NA | NA | NA | NA | NA |
| KCNMA1 | 900 | S | -80.89058333 | 32.42700541 | 0.413180325 | 26.40600838 | NA | NA | NA | NA | NA | NA | NA |
| KCNMA1 | 901 | I | -79.740125 | 28.04492375 | 0.353557806 | 22.02352279 | NA | NA | NA | NA | NA | NA | NA |
| KCNMA1 | 902 | L | -78.707875 | 29.0417645 | 0.367120802 | 23.02901846 | NA | NA | NA | NA | NA | NA | NA |
| KCNMA1 | 903 | P | -79.10075 | 26.77586686 | 0.336291043 | 20.75731867 | NA | NA | NA | NA | NA | NA | NA |
| KCNMA1 | 904 | G | -75.205 | 25.81634447 | 0.3232358 | 16.32715432 | NA | NA | NA | NA | NA | NA | NA |
| KCNMA1 | 905 | T | -72.05060714 | 25.06444911 | 0.313005526 | 15.53243624 | NA | NA | NA | NA | NA | NA | NA |
| KCNMA1 | 906 | P | -70.08703571 | 26.41213185 | 0.331342071 | 16.80426625 | NA | NA | NA | NA | NA | NA | NA |
| KCNMA1 | 907 | L | -67.96075 | 25.33428464 | 0.316676903 | 15.59797332 | NA | NA | NA | NA | NA | NA | NA |
| KCNMA1 | 908 | S | -72.82191667 | 29.20142438 | 0.369293132 | 19.58080592 | NA | NA | NA | NA | NA | NA | NA |
| KCNMA1 | 909 | R | -71.51056818 | 32.44032833 | 0.413361596 | 22.82395848 | NA | NA | NA | NA | NA | NA | NA |
| KCNMA1 | 910 | A | -76.85195 | 34.38067028 | 0.439761852 | 24.80592485 | NA | NA | NA | NA | NA | NA | NA |
| KCNMA1 | 911 | D | -76.88375 | 30.46023626 | 0.386420502 | 20.96173522 | NA | NA | NA | NA | NA | NA | NA |
| KCNMA1 | 912 | L | -74.134625 | 33.36441203 | 0.425934661 | 23.70913414 | NA | NA | NA | NA | NA | NA | NA |
| KCNMA1 | 913 | R | -76.53293182 | 37.27407839 | 0.479129507 | 27.69725379 | NA | NA | NA | NA | NA | NA | NA |
| KCNMA1 | 914 | A | -80.26895 | 37.53788505 | 0.482718855 | 28.05171236 | NA | NA | NA | NA | NA | NA | NA |
| KCNMA1 | 915 | V | -77.96189286 | 36.00823314 | 0.46190644 | 26.47271911 | NA | NA | NA | NA | NA | NA | NA |
| KCNMA1 | 916 | N | -79.611125 | 39.77378936 | 0.513140526 | 30.22357242 | NA | NA | NA | NA | NA | NA | NA |
| KCNMA1 | 917 | I | -74.06225 | 37.75178381 | 0.485629157 | 28.088245 | NA | NA | NA | NA | NA | NA | NA |
| KCNMA1 | 918 | N | -75.72125 | 41.47795735 | 0.536327403 | 31.83916667 | NA | NA | NA | NA | NA | NA | NA |

| | | | | | | | | | | | | | |
|---|---|---|---|---|---|---|---|---|---|---|---|---|---|
| KCNMA1 | 919 | L | -78.111875 | 43.8185622 | 0.568173627 | 34.1984265 | NA | NA | NA | NA | NA | NA | NA |
| KCNMA1 | 920 | C | -74.77191667 | 39.61552349 | 0.510987163 | 29.98334304 | NA | NA | NA | NA | NA | NA | NA |
| KCNMA1 | 921 | D | -72.778125 | 39.94664738 | 0.515492429 | 30.26024282 | NA | NA | NA | NA | NA | NA | NA |
| KCNMA1 | 922 | M | -68.82425 | 37.82190231 | 0.486583188 | 28.12290384 | NA | NA | NA | NA | NA | NA | NA |
| KCNMA1 | 923 | C | -69.07675 | 34.06940694 | 0.435526809 | 24.36239637 | NA | NA | NA | NA | NA | NA | NA |
| KCNMA1 | 924 | V | -67.96146429 | 31.17642476 | 0.396164949 | 21.47204019 | NA | NA | NA | NA | NA | NA | NA |
| KCNMA1 | 925 | I | -65.229375 | 28.64946354 | 0.361783163 | 18.85446885 | NA | NA | NA | NA | NA | NA | NA |
| KCNMA1 | 926 | L | -65.429375 | 25.15009791 | 0.314170862 | 15.41112426 | NA | NA | NA | NA | NA | NA | NA |
| KCNMA1 | 927 | S | -62.40925 | 22.29425188 | 0.275314276 | 12.46845396 | NA | NA | NA | NA | NA | NA | NA |
| KCNMA1 | 928 | A | -64.31195 | 19.3594464 | 0.235383369 | 9.58011342 | NA | NA | NA | NA | NA | NA | NA |
| KCNMA1 | 929 | N | -61.76425 | 14.93956187 | 0.175246505 | 5.167909226 | NA | NA | NA | NA | NA | NA | NA |
| KCNMA1 | 930 | Q | -59.04408333 | 17.84526136 | 0.214781397 | 7.376043501 | NA | NA | NA | NA | NA | NA | NA |
| KCNMA1 | 931 | N | -59.47525 | 14.1578261 | 0.164610224 | 3.872024968 | 10.48 | -0.14 | 0.74 | 0.68 | 0.58 | 0 | 0 |
| KCNMA1 | 932 | N | -58.257875 | 12.18948338 | 0.13782899 | 1.739801498 | 10.48 | -0.14 | 0.74 | 0.68 | 0.58 | 0 | 0 |
| KCNMA1 | 933 | I | -53.348875 | 14.3068219 | 0.166637458 | 4.058241426 | 10.55 | -0.14 | 0.74 | 0.68 | 0.58 | 0 | 0 |
| KCNMA1 | 934 | D | -53.293875 | 14.28660569 | 0.166362396 | 2.281095862 | 12.19 | -0.14 | 0.74 | 0.68 | 0.58 | 0 | 0 |
| KCNMA1 | 935 | D | -48.668875 | 16.63903771 | 0.198369541 | 3.806711587 | 12.91 | 1.81 | 4.5 | 0.13 | -1.85 | 0 | 0 |
| KCNMA1 | 936 | T | -49.44603571 | 19.72693481 | 0.240383409 | 6.998744435 | NA | NA | NA | NA | NA | NA | NA |
| KCNMA1 | 937 | S | -51.56208333 | 22.49688943 | 0.278071358 | 9.935397435 | NA | NA | NA | NA | NA | NA | NA |
| KCNMA1 | 938 | L | -52.853875 | 19.93901688 | 0.243268994 | 7.791728478 | NA | NA | NA | NA | NA | NA | NA |
| KCNMA1 | 939 | Q | -55.34530556 | 20.10815469 | 0.24557028 | 9.564377216 | NA | NA | NA | NA | NA | NA | NA |
| KCNMA1 | 940 | D | -56.37125 | 22.84980186 | 0.282873078 | 12.44027208 | NA | NA | NA | NA | NA | NA | NA |
| KCNMA1 | 941 | K | -57.78119444 | 19.33383886 | 0.235034953 | 8.87258359 | NA | NA | NA | NA | NA | NA | NA |
| KCNMA1 | 942 | E | -60.84941667 | 20.06223838 | 0.244945543 | 9.629253106 | NA | NA | NA | NA | NA | NA | NA |
| KCNMA1 | 943 | C | -60.96941667 | 25.59924151 | 0.320281901 | 15.16217157 | NA | NA | NA | NA | NA | NA | NA |
| KCNMA1 | 944 | I | -58.417625 | 28.17245224 | 0.355292957 | 17.702525 | NA | NA | NA | NA | NA | NA | NA |
| KCNMA1 | 945 | L | -61.75325 | 26.69990423 | 0.335257497 | 16.2619272 | NA | NA | NA | NA | NA | NA | NA |
| KCNMA1 | 946 | A | -64.41575 | 29.03270676 | 0.366997563 | 19.24774292 | NA | NA | NA | NA | NA | NA | NA |
| KCNMA1 | 947 | S | -63.00058333 | 31.421973 | 0.399505873 | 21.58800255 | NA | NA | NA | NA | NA | NA | NA |
| KCNMA1 | 948 | L | -61.4365 | 33.30960843 | 0.425189005 | 22.9052293 | NA | NA | NA | NA | NA | NA | NA |
| KCNMA1 | 949 | N | -66.564625 | 31.86291087 | 0.405505266 | 22.13378109 | NA | NA | NA | NA | NA | NA | NA |
| KCNMA1 | 950 | I | -67.76425 | 33.75086067 | 0.431192674 | 24.01939047 | NA | NA | NA | NA | NA | NA | NA |
| KCNMA1 | 951 | K | -64.38475 | 37.06219046 | 0.476246563 | 27.26314357 | NA | NA | NA | NA | NA | NA | NA |

| | | | | | | | | | | | | | |
|---|---|---|---|---|---|---|---|---|---|---|---|---|---|
| KCNMA1 | 952 | S | -66.53058333 | 36.99918748 | 0.475389346 | 27.20638899 | NA | NA | NA | NA | NA | NA | NA |
| KCNMA1 | 953 | M | -70.47875 | 36.79701579 | 0.472638602 | 27.11383731 | NA | NA | NA | NA | NA | NA | NA |
| KCNMA1 | 954 | Q | -71.06419444 | 41.94359781 | 0.542662898 | 32.22932917 | NA | NA | NA | NA | NA | NA | NA |
| KCNMA1 | 955 | F | -74.35229545 | 41.34553929 | 0.534525725 | 31.66581425 | NA | NA | NA | NA | NA | NA | NA |
| KCNMA1 | 956 | D | -77.676125 | 47.75377707 | 0.621716085 | 38.10763375 | NA | NA | NA | NA | NA | NA | NA |
| KCNMA1 | 957 | D | -79.27275 | 51.15504062 | 0.667993611 | 41.52001216 | NA | NA | NA | NA | NA | NA | NA |
| KCNMA1 | 994 | T | -72.83903571 | 46.13444043 | 0.599683422 | 36.41577185 | NA | NA | NA | NA | NA | NA | NA |
| KCNMA1 | 995 | T | -69.00103571 | 44.66825054 | 0.57973447 | 34.90521798 | NA | NA | NA | NA | NA | NA | NA |
| KCNMA1 | 996 | G | -68.38625 | 41.13683097 | 0.531686044 | 31.35487431 | NA | NA | NA | NA | NA | NA | NA |
| KCNMA1 | 997 | V | -65.11632143 | 42.62442035 | 0.551926156 | 32.79937739 | NA | NA | NA | NA | NA | NA | NA |
| KCNMA1 | 998 | N | -67.5205 | 43.79562195 | 0.567861503 | 34.0078053 | NA | NA | NA | NA | NA | NA | NA |
| KCNMA1 | 999 | I | -68.834625 | 40.02326511 | 0.516534888 | 30.29225895 | NA | NA | NA | NA | NA | NA | NA |
| KCNMA1 | 1000 | P | -68.15703571 | 39.11551696 | 0.504184084 | 29.35741887 | NA | NA | NA | NA | NA | NA | NA |
| KCNMA1 | 1001 | I | -65.146875 | 35.97947205 | 0.461515118 | 26.14251037 | NA | NA | NA | NA | NA | NA | NA |
| KCNMA1 | 1002 | I | -63.774 | 33.13960738 | 0.422875974 | 23.32837377 | NA | NA | NA | NA | NA | NA | NA |
| KCNMA1 | 1003 | T | -61.37303571 | 29.938892 | 0.379327102 | 19.51619345 | NA | NA | NA | NA | NA | NA | NA |
| KCNMA1 | 1004 | E | -60.43352778 | 26.50121243 | 0.3325541 | 16.02769691 | NA | NA | NA | NA | NA | NA | NA |
| KCNMA1 | 1005 | L | -56.50425 | 27.66198729 | 0.34834758 | 17.20364087 | NA | NA | NA | NA | NA | NA | NA |
| KCNMA1 | 1006 | V | -54.46689286 | 23.91201802 | 0.297325571 | 13.3743423 | NA | NA | NA | NA | NA | NA | NA |
| KCNMA1 | 1007 | N | -51.195375 | 24.28070232 | 0.302341882 | 11.71839594 | NA | NA | NA | NA | NA | NA | NA |
| KCNMA1 | 1008 | D | -48.82125 | 28.93153492 | 0.365621021 | 16.20137206 | NA | NA | NA | NA | NA | NA | NA |
| KCNMA1 | 1009 | T | -47.42175 | 27.99362633 | 0.352859855 | 15.06912491 | NA | NA | NA | NA | NA | NA | NA |
| KCNMA1 | 1010 | N | -52.055625 | 26.7620545 | 0.336103112 | 14.25481351 | NA | NA | NA | NA | NA | NA | NA |
| KCNMA1 | 1011 | V | -52.74475 | 32.16331348 | 0.409592538 | 19.9858446 | NA | NA | NA | NA | NA | NA | NA |
| KCNMA1 | 1012 | Q | -50.28852778 | 33.51711172 | 0.42801229 | 21.13310195 | NA | NA | NA | NA | NA | NA | NA |
| KCNMA1 | 1013 | F | -52.80902273 | 29.8690049 | 0.378376219 | 17.69977145 | NA | NA | NA | NA | NA | NA | NA |
| KCNMA1 | 1014 | L | -57.374375 | 33.16374475 | 0.423204386 | 22.69756362 | NA | NA | NA | NA | NA | NA | NA |
| KCNMA1 | 1015 | D | -55.427625 | 37.52227405 | 0.482506452 | 27.06466937 | NA | NA | NA | NA | NA | NA | NA |
| KCNMA1 | 1016 | Q | -57.92730556 | 40.02702654 | 0.516586066 | 29.56775114 | NA | NA | NA | NA | NA | NA | NA |
| KCNMA1 | 1017 | D | -54.965 | 42.33623188 | 0.548005069 | 31.86950232 | NA | NA | NA | NA | NA | NA | NA |
| KCNMA1 | 1018 | D | -50.333875 | 43.74877607 | 0.567224118 | 31.17951114 | NA | NA | NA | NA | NA | NA | NA |
| KCNMA1 | 1019 | D | -47.952125 | 40.5028002 | 0.523059433 | 27.60099046 | NA | NA | NA | NA | NA | NA | NA |
| KCNMA1 | 1020 | D | -43.803 | 42.90618373 | 0.555759823 | 30.44517373 | NA | NA | NA | NA | NA | NA | NA |

| | | | | | | | | | | | | | |
|---|---|---|---|---|---|---|---|---|---|---|---|---|---|
| KCNMA1 | 1021 | D | -43.221625 | 39.23220312 | 0.505771714 | 26.8410501 | NA | NA | NA | NA | NA | NA | NA |
| KCNMA1 | 1022 | P | -43.76117857 | 35.86467623 | 0.459953208 | 23.51226293 | NA | NA | NA | NA | NA | NA | NA |
| KCNMA1 | 1023 | D | -41.2605 | 35.91696026 | 0.460664583 | 23.53512907 | NA | NA | NA | NA | NA | NA | NA |
| KCNMA1 | 1024 | T | -45.13575 | 36.82011886 | 0.472952942 | 24.4979927 | NA | NA | NA | NA | NA | NA | NA |
| KCNMA1 | 1025 | E | -44.34475 | 37.08483282 | 0.476554635 | 24.71503809 | NA | NA | NA | NA | NA | NA | NA |
| KCNMA1 | 1026 | L | -49.278125 | 34.1976714 | 0.437271972 | 21.33404232 | NA | NA | NA | NA | NA | NA | NA |
| KCNMA1 | 1027 | Y | -46.93758333 | 38.83584738 | 0.500378905 | 26.12490561 | NA | NA | NA | NA | NA | NA | NA |
| KCNMA1 | 1028 | L | -47.35225 | 41.12294966 | 0.531497175 | 28.18309205 | NA | NA | NA | NA | NA | NA | NA |
| KCNMA1 | 1029 | T | -50.70889286 | 37.97980008 | 0.488731542 | 25.25349485 | NA | NA | NA | NA | NA | NA | NA |
| KCNMA1 | 1030 | Q | -53.69841667 | 41.13636524 | 0.531679707 | 28.96037986 | NA | NA | NA | NA | NA | NA | NA |
| KCNMA1 | 1031 | P | -55.09746429 | 37.79553504 | 0.486224436 | 27.27468273 | NA | NA | NA | NA | NA | NA | NA |
| KCNMA1 | 1032 | F | -53.91493182 | 37.49107773 | 0.482081995 | 26.98308891 | NA | NA | NA | NA | NA | NA | NA |
| KCNMA1 | 1033 | A | -55.02555 | 41.36426235 | 0.534780471 | 30.83078076 | NA | NA | NA | NA | NA | NA | NA |
| KCNMA1 | 1034 | C | -58.16475 | 42.44581061 | 0.549495995 | 31.97817828 | NA | NA | NA | NA | NA | NA | NA |
| KCNMA1 | 1035 | G | -59.89625 | 39.53479951 | 0.509888834 | 29.06217081 | NA | NA | NA | NA | NA | NA | NA |
| KCNMA1 | 1036 | T | -60.45946429 | 36.0525855 | 0.462509898 | 25.5810981 | NA | NA | NA | NA | NA | NA | NA |
| KCNMA1 | 1037 | A | -58.14715 | 34.96385118 | 0.4476966 | 24.49762955 | NA | NA | NA | NA | NA | NA | NA |
| KCNMA1 | 1038 | F | -58.50347727 | 31.41299876 | 0.39938377 | 20.94685271 | NA | NA | NA | NA | NA | NA | NA |
| KCNMA1 | 1039 | A | -55.22655 | 31.20565247 | 0.39656262 | 20.65823345 | NA | NA | NA | NA | NA | NA | NA |
| KCNMA1 | 1040 | V | -55.32089286 | 27.06018987 | 0.340159536 | 16.51612737 | NA | NA | NA | NA | NA | NA | NA |
| KCNMA1 | 1041 | S | -51.25758333 | 29.39821175 | 0.371970617 | 16.97041586 | NA | NA | NA | NA | NA | NA | NA |
| KCNMA1 | 1042 | V | -52.32875 | 32.28171636 | 0.411203525 | 19.81343113 | NA | NA | NA | NA | NA | NA | NA |
| KCNMA1 | 1043 | L | -54.421875 | 31.86069532 | 0.405475121 | 19.81307434 | NA | NA | NA | NA | NA | NA | NA |
| KCNMA1 | 1044 | D | -50.206375 | 28.20631755 | 0.355753727 | 15.65058708 | NA | NA | NA | NA | NA | NA | NA |
| KCNMA1 | 1045 | S | -49.32241667 | 33.67485996 | 0.43015861 | 20.78454135 | NA | NA | NA | NA | NA | NA | NA |
| KCNMA1 | 1046 | L | -52.363375 | 35.62730976 | 0.456723604 | 23.13531413 | NA | NA | NA | NA | NA | NA | NA |
| KCNMA1 | 1047 | M | -49.58525 | 31.06757461 | 0.394683936 | 18.34214935 | NA | NA | NA | NA | NA | NA | NA |
| KCNMA1 | 1048 | S | -46.86825 | 36.02118252 | 0.462082629 | 23.11369779 | NA | NA | NA | NA | NA | NA | NA |
| KCNMA1 | 1049 | A | -49.17935 | 38.74875515 | 0.49919393 | 25.83961081 | NA | NA | NA | NA | NA | NA | NA |
| KCNMA1 | 1050 | T | -50.96146429 | 38.32772414 | 0.49346539 | 25.77987387 | NA | NA | NA | NA | NA | NA | NA |
| KCNMA1 | 1051 | Y | -44.99833333 | 38.73745074 | 0.499040123 | 25.83113354 | NA | NA | NA | NA | NA | NA | NA |
| KCNMA1 | 1052 | F | -44.50984091 | 40.21419263 | 0.519132644 | 27.81313787 | NA | NA | NA | NA | NA | NA | NA |
| KCNMA1 | 1053 | N | -49.324625 | 43.22258828 | 0.560064817 | 30.51224167 | NA | NA | NA | NA | NA | NA | NA |

| | | | | | | | | | | | | | |
|---|---|---|---|---|---|---|---|---|---|---|---|---|---|
| KCNMA1 | 1054 | D | -51.69225 | 43.86450834 | 0.568798769 | 31.42573842 | NA | NA | NA | NA | NA | NA | NA |
| KCNMA1 | 1055 | N | -54.738 | 45.18412088 | 0.586753392 | 34.63659686 | NA | NA | NA | NA | NA | NA | NA |
| KCNMA1 | 1056 | I | -54.067125 | 42.39694181 | 0.548831088 | 31.88983154 | NA | NA | NA | NA | NA | NA | NA |
| KCNMA1 | 1057 | L | -56.9165 | 37.91482382 | 0.487847476 | 27.44473931 | NA | NA | NA | NA | NA | NA | NA |
| KCNMA1 | 1058 | T | -59.58089286 | 42.40639514 | 0.548959709 | 31.93287601 | NA | NA | NA | NA | NA | NA | NA |
| KCNMA1 | 1059 | L | -59.194875 | 43.94436512 | 0.569885299 | 33.4686891 | NA | NA | NA | NA | NA | NA | NA |
| KCNMA1 | 1060 | I | -59.218 | 39.87837523 | 0.514563519 | 29.404168 | NA | NA | NA | NA | NA | NA | NA |
| KCNMA1 | 1061 | R | -63.10311364 | 38.10769205 | 0.490471638 | 28.26222692 | NA | NA | NA | NA | NA | NA | NA |
| KCNMA1 | 1062 | T | -64.38075 | 42.53819909 | 0.550753031 | 32.6844548 | NA | NA | NA | NA | NA | NA | NA |
| KCNMA1 | 1063 | L | -63.703 | 43.18188126 | 0.559510958 | 33.35185883 | NA | NA | NA | NA | NA | NA | NA |
| KCNMA1 | 1064 | V | -64.65632143 | 38.61150374 | 0.49732649 | 28.7762251 | NA | NA | NA | NA | NA | NA | NA |
| KCNMA1 | 1065 | T | -67.16760714 | 37.75860751 | 0.485722 | 28.00148815 | NA | NA | NA | NA | NA | NA | NA |
| KCNMA1 | 1066 | G | -69.631 | 41.59006214 | 0.537852698 | 31.83091817 | NA | NA | NA | NA | NA | NA | NA |
| KCNMA1 | 1067 | G | -69.463 | 42.74169808 | 0.553521835 | 32.99921218 | NA | NA | NA | NA | NA | NA | NA |
| KCNMA1 | 1068 | A | -67.42735 | 39.63063677 | 0.511192794 | 29.8540688 | NA | NA | NA | NA | NA | NA | NA |
| KCNMA1 | 1069 | T | -69.38432143 | 41.20863436 | 0.532662999 | 31.47272792 | NA | NA | NA | NA | NA | NA | NA |
| KCNMA1 | 1070 | P | -67.23875 | 40.96735624 | 0.529380174 | 32.72545754 | NA | NA | NA | NA | NA | NA | NA |
| KCNMA1 | 1071 | E | -71.82563889 | 40.17269403 | 0.518568015 | 34.9170891 | NA | NA | NA | NA | NA | NA | NA |
| KCNMA1 | 1072 | L | -72.471375 | 37.56910157 | 0.483143586 | 32.30645969 | NA | NA | NA | NA | NA | NA | NA |
| KCNMA1 | 1073 | E | -67.51275 | 36.15964857 | 0.463966596 | 30.03763528 | NA | NA | NA | NA | NA | NA | NA |
| KCNMA1 | 1074 | A | -71.06755 | 35.8095116 | 0.459202639 | 30.55822648 | NA | NA | NA | NA | NA | NA | NA |
| KCNMA1 | 1075 | L | -74.805 | 34.46017577 | 0.440843602 | 29.2075617 | NA | NA | NA | NA | NA | NA | NA |
| KCNMA1 | 1076 | I | -72.611125 | 31.81902949 | 0.404908216 | 26.56038849 | NA | NA | NA | NA | NA | NA | NA |
| KCNMA1 | 1077 | A | -71.14175 | 31.69680523 | 0.403245235 | 26.44742858 | NA | NA | NA | NA | NA | NA | NA |
| KCNMA1 | 1078 | E | -74.18819444 | 30.15272913 | 0.382236566 | 24.91141346 | NA | NA | NA | NA | NA | NA | NA |
| KCNMA1 | 1079 | E | -76.66819444 | 27.45444962 | 0.345523827 | 22.20174439 | NA | NA | NA | NA | NA | NA | NA |
| KCNMA1 | 1080 | N | -73.31025 | 26.43508158 | 0.331654325 | 21.1642648 | NA | NA | NA | NA | NA | NA | NA |
| KCNMA1 | 1081 | A | -76.65475 | 29.54498585 | 0.373967622 | 24.2889638 | NA | NA | NA | NA | NA | NA | NA |
| KCNMA1 | 1082 | L | -77.57825 | 32.30384429 | 0.411504597 | 27.04336352 | NA | NA | NA | NA | NA | NA | NA |
| KCNMA1 | 1083 | R | -80.60856818 | 34.01514054 | 0.434788461 | 28.76712628 | NA | NA | NA | NA | NA | NA | NA |
| KCNMA1 | 1084 | G | -83.586 | 36.60520721 | 0.470028858 | 31.35546967 | NA | NA | NA | NA | NA | NA | NA |
| KCNMA1 | 1085 | G | -85.984 | 39.77702359 | 0.513184531 | 34.53045739 | NA | NA | NA | NA | NA | NA | NA |
| KCNMA1 | 1086 | Y | -89.71741667 | 42.67496094 | 0.55261381 | 37.43132646 | NA | NA | NA | NA | NA | NA | NA |

| | | | | | | | | | | | | | |
|---|---|---|---|---|---|---|---|---|---|---|---|---|---|
| KCNMA1 | 1087 | S | -87.42525 | 46.05807015 | 0.598644329 | 40.81182912 | NA | NA | NA | NA | NA | NA | NA |
| KCNMA1 | 1088 | T | -85.40060714 | 48.69538999 | 0.634527652 | 43.45210304 | NA | NA | NA | NA | NA | NA | NA |
| KCNMA1 | 1089 | P | -85.36460714 | 52.35534832 | 0.684324975 | 47.10805121 | NA | NA | NA | NA | NA | NA | NA |
| KCNMA1 | 1090 | Q | -80.61963889 | 50.78905853 | 0.663014066 | 45.53725152 | NA | NA | NA | NA | NA | NA | NA |
| KCNMA1 | 1091 | T | -82.06903571 | 47.11413131 | 0.613013077 | 41.86010801 | NA | NA | NA | NA | NA | NA | NA |
| KCNMA1 | 1092 | L | -83.44975 | 50.0971284 | 0.653599678 | 44.84377778 | NA | NA | NA | NA | NA | NA | NA |
| KCNMA1 | 1093 | A | -79.44715 | 52.57519029 | 0.68731614 | 47.31799001 | NA | NA | NA | NA | NA | NA | NA |
| KCNMA1 | 1094 | N | -76.607375 | 47.7668971 | 0.621894596 | 42.5069858 | NA | NA | NA | NA | NA | NA | NA |
| KCNMA1 | 1095 | R | -79.90475 | 49.44270386 | 0.64469559 | 44.17749768 | NA | NA | NA | NA | NA | NA | NA |
| KCNMA1 | 1096 | D | -75.838 | 52.25881764 | 0.68301158 | 42.62751478 | NA | NA | NA | NA | NA | NA | NA |
| KCNMA1 | 1097 | R | -72.85847727 | 45.89943578 | 0.596485953 | 36.19069434 | NA | NA | NA | NA | NA | NA | NA |
| KCNMA1 | 1098 | C | -70.00641667 | 48.86415399 | 0.636823852 | 39.09574227 | NA | NA | NA | NA | NA | NA | NA |
| KCNMA1 | 1099 | R | -72.88311364 | 48.48626925 | 0.631682359 | 38.76353648 | NA | NA | NA | NA | NA | NA | NA |
| KCNMA1 | 1100 | V | -68.32475 | 45.96815108 | 0.597420892 | 36.16600111 | NA | NA | NA | NA | NA | NA | NA |
| KCNMA1 | 1101 | A | -69.17875 | 48.7147555 | 0.634791139 | 38.93624332 | NA | NA | NA | NA | NA | NA | NA |
| KCNMA1 | 1102 | Q | -68.30852778 | 46.90206673 | 0.61012773 | 37.10200765 | NA | NA | NA | NA | NA | NA | NA |
| KCNMA1 | 1103 | L | -70.245375 | 52.56642504 | 0.68719688 | 42.77905138 | NA | NA | NA | NA | NA | NA | NA |
| KCNMA1 | 1104 | A | -70.45475 | 54.83511825 | 0.718064676 | 45.05960131 | NA | NA | NA | NA | NA | NA | NA |
| KCNMA1 | 1105 | L | -68.198 | 57.59223363 | 0.755577935 | 47.7947702 | NA | NA | NA | NA | NA | NA | NA |
| KCNMA1 | 1106 | L | -72.030375 | 58.97416366 | 0.774380448 | 49.19813529 | NA | NA | NA | NA | NA | NA | NA |
| KCNMA1 | 1107 | D | -75.579 | 58.13418954 | 0.762951777 | 48.43169387 | NA | NA | NA | NA | NA | NA | NA |
| KCNMA1 | 1108 | G | -74.003 | 61.1270447 | 0.803672506 | 51.380335 | NA | NA | NA | NA | NA | NA | NA |
| KCNMA1 | 1109 | P | -73.22032143 | 62.03898312 | 0.816080323 | 52.28061866 | NA | NA | NA | NA | NA | NA | NA |
| KCNMA1 | 1110 | F | -69.26775 | 59.90745812 | 0.787078835 | 50.09465672 | NA | NA | NA | NA | NA | NA | NA |
| KCNMA1 | 1111 | A | -70.54115 | 64.82650298 | 0.854007264 | 55.03010989 | NA | NA | NA | NA | NA | NA | NA |
| KCNMA1 | 1112 | D | -68.568375 | 66.98863747 | 0.883425225 | 57.17606073 | NA | NA | NA | NA | NA | NA | NA |
| KCNMA1 | 1113 | L | -65.51425 | 64.20250651 | 0.84551718 | 53.90884499 | NA | NA | NA | NA | NA | NA | NA |
| KCNMA1 | 1114 | G | -65.57925 | 62.42640823 | 0.821351622 | 52.12933441 | NA | NA | NA | NA | NA | NA | NA |
| KCNMA1 | 1115 | D | -63.181375 | 64.6588064 | 0.851725588 | 54.22189434 | NA | NA | NA | NA | NA | NA | NA |
| KCNMA1 | 1116 | G | -60.4055 | 65.24938876 | 0.85976104 | 54.77795452 | NA | NA | NA | NA | NA | NA | NA |
| KCNMA1 | 1117 | G | -60.7045 | 63.51987346 | 0.836229288 | 53.04996933 | NA | NA | NA | NA | NA | NA | NA |
| KCNMA1 | 1118 | C | -58.54508333 | 63.38784963 | 0.834432975 | 52.91883089 | NA | NA | NA | NA | NA | NA | NA |
| KCNMA1 | 1119 | Y | -59.1525 | 55.98265435 | 0.73367803 | 45.51040227 | NA | NA | NA | NA | NA | NA | NA |

| | | | | | | | | | | | | | |
|---|---|---|---|---|---|---|---|---|---|---|---|---|---|
| KCNMA1 | 1120 | G | -59.4955 | 62.78263159 | 0.82619839 | 52.3141177 | NA | NA | NA | NA | NA | NA | NA |
| KCNMA1 | 1121 | D | -61.44 | 64.78889023 | 0.853495506 | 54.32034435 | NA | NA | NA | NA | NA | NA | NA |
| KCNMA1 | 1122 | L | -64.13875 | 59.94652145 | 0.78761033 | 49.53989247 | NA | NA | NA | NA | NA | NA | NA |
| KCNMA1 | 1123 | F | -61.80756818 | 58.48383744 | 0.767709079 | 48.01020268 | NA | NA | NA | NA | NA | NA | NA |
| KCNMA1 | 1124 | C | -63.83425 | 63.59956333 | 0.837313547 | 53.17470354 | NA | NA | NA | NA | NA | NA | NA |
| KCNMA1 | 1125 | K | -67.12063889 | 64.35073644 | 0.847533994 | 54.51979728 | NA | NA | NA | NA | NA | NA | NA |
| KCNMA1 | 1126 | A | -68.00495 | 60.72900105 | 0.798256732 | 50.89841215 | NA | NA | NA | NA | NA | NA | NA |
| KCNMA1 | 1127 | L | -66.6645 | 60.27945638 | 0.792140236 | 50.43811764 | NA | NA | NA | NA | NA | NA | NA |
| KCNMA1 | 1128 | K | -68.15730556 | 64.22906829 | 0.84587858 | 54.40024436 | NA | NA | NA | NA | NA | NA | NA |
| KCNMA1 | 1129 | T | -72.15875 | 63.99240234 | 0.842658507 | 54.22216285 | NA | NA | NA | NA | NA | NA | NA |
| KCNMA1 | 1130 | Y | -73.59125 | 59.75786764 | 0.78504351 | 49.99930173 | NA | NA | NA | NA | NA | NA | NA |
| KCNMA1 | 1131 | N | -70.489375 | 57.71600224 | 0.757261928 | 47.92128364 | NA | NA | NA | NA | NA | NA | NA |
| KCNMA1 | 1132 | M | -69.40875 | 57.00663637 | 0.74761031 | 47.24026223 | NA | NA | NA | NA | NA | NA | NA |
| KCNMA1 | 1133 | L | -64.30525 | 54.01799295 | 0.706946885 | 44.13404403 | NA | NA | NA | NA | NA | NA | NA |
| KCNMA1 | 1134 | C | -63.24925 | 55.00437365 | 0.720367562 | 44.57385304 | NA | NA | NA | NA | NA | NA | NA |
| KCNMA1 | 1135 | F | -61.15065909 | 48.09461936 | 0.626353578 | 37.62252503 | NA | NA | NA | NA | NA | NA | NA |
| KCNMA1 | 1136 | G | -60.701 | 51.95369305 | 0.678860061 | 41.48217472 | NA | NA | NA | NA | NA | NA | NA |
| KCNMA1 | 1137 | I | -59.58825 | 51.23505054 | 0.669082225 | 40.76696948 | NA | NA | NA | NA | NA | NA | NA |
| KCNMA1 | 1138 | Y | -62.01866667 | 45.19377101 | 0.586884692 | 34.74264708 | NA | NA | NA | NA | NA | NA | NA |
| KCNMA1 | 1139 | R | -58.23402273 | 50.40375657 | 0.657771655 | 39.9363103 | NA | NA | NA | NA | NA | NA | NA |
| KCNMA1 | 1140 | L | -60.576625 | 51.65562882 | 0.674804604 | 41.18599111 | NA | NA | NA | NA | NA | NA | NA |
| KCNMA1 | 1141 | R | -56.18902273 | 47.1787632 | 0.613892457 | 36.7478716 | NA | NA | NA | NA | NA | NA | NA |
| KCNMA1 | 1142 | D | -57.55525 | 51.37615183 | 0.671002046 | 40.91325559 | NA | NA | NA | NA | NA | NA | NA |
| KCNMA1 | 1143 | A | -57.67675 | 54.99680641 | 0.720264602 | 44.53335156 | NA | NA | NA | NA | NA | NA | NA |
| KCNMA1 | 1144 | H | -53.33955 | 53.93054116 | 0.705757018 | 43.46063628 | NA | NA | NA | NA | NA | NA | NA |
| KCNMA1 | 1145 | L | -54.295125 | 54.60616021 | 0.714949478 | 44.07813218 | NA | NA | NA | NA | NA | NA | NA |
| KCNMA1 | 1146 | S | -55.22875 | 60.08392872 | 0.789479891 | 49.54468956 | NA | NA | NA | NA | NA | NA | NA |
| KCNMA1 | 1147 | T | -56.53946429 | 58.11213384 | 0.762651687 | 47.64921864 | NA | NA | NA | NA | NA | NA | NA |
| KCNMA1 | 1148 | P | -60.41989286 | 57.24010198 | 0.750786839 | 46.77240724 | NA | NA | NA | NA | NA | NA | NA |
| KCNMA1 | 1149 | S | -60.51808333 | 52.55444697 | 0.687033907 | 42.08848783 | NA | NA | NA | NA | NA | NA | NA |
| KCNMA1 | 1150 | Q | -63.95786111 | 50.55835117 | 0.659875066 | 40.12484455 | NA | NA | NA | NA | NA | NA | NA |
| KCNMA1 | 1151 | C | -61.47075 | 47.72508604 | 0.621325715 | 37.25694124 | NA | NA | NA | NA | NA | NA | NA |
| KCNMA1 | 1152 | T | -64.28860714 | 47.06149435 | 0.612296899 | 36.69678125 | NA | NA | NA | NA | NA | NA | NA |

| | | | | | | | | | | | | | |
|---|---|---|---|---|---|---|---|---|---|---|---|---|---|
| KCNMA1 | 1153 | K | -60.31408333 | 41.65031679 | 0.538672522 | 31.18449888 | NA | NA | NA | NA | NA | NA | NA |
| KCNMA1 | 1154 | R | -62.65665909 | 45.94267222 | 0.597074227 | 35.47365354 | NA | NA | NA | NA | NA | NA | NA |
| KCNMA1 | 1155 | Y | -56.05433333 | 47.21223375 | 0.614347857 | 36.74828533 | NA | NA | NA | NA | NA | NA | NA |
| KCNMA1 | 1156 | V | -56.99603571 | 45.63966445 | 0.592951509 | 35.17204056 | NA | NA | NA | NA | NA | NA | NA |
| KCNMA1 | 1157 | I | -55.443 | 49.76402522 | 0.649067482 | 39.26165023 | NA | NA | NA | NA | NA | NA | NA |
| KCNMA1 | 1158 | T | -54.75689286 | 48.20606299 | 0.627869878 | 37.65974581 | NA | NA | NA | NA | NA | NA | NA |
| KCNMA1 | 1159 | N | -54.048 | 52.65912294 | 0.688458126 | 42.11221972 | NA | NA | NA | NA | NA | NA | NA |
| KCNMA1 | 1160 | P | -55.39017857 | 55.34239108 | 0.724966621 | 44.84199906 | NA | NA | NA | NA | NA | NA | NA |
| KCNMA1 | 1161 | P | -52.54117857 | 57.86047056 | 0.759227562 | 45.3907054 | NA | NA | NA | NA | NA | NA | NA |
| KCNMA1 | 1162 | Y | -54.344 | 62.13935556 | 0.817445988 | 49.95571927 | NA | NA | NA | NA | NA | NA | NA |
| KCNMA1 | 1163 | E | -50.78697222 | 62.54906589 | 0.823020499 | 50.05417487 | NA | NA | NA | NA | NA | NA | NA |
| KCNMA1 | 1164 | F | -55.00647727 | 56.47977048 | 0.740441783 | 45.94554336 | NA | NA | NA | NA | NA | NA | NA |
| KCNMA1 | 1165 | E | -56.35330556 | 60.27721195 | 0.792109698 | 49.81311167 | NA | NA | NA | NA | NA | NA | NA |
| KCNMA1 | 1166 | L | -60.808375 | 58.90638985 | 0.773458319 | 48.43419078 | NA | NA | NA | NA | NA | NA | NA |
| KCNMA1 | 1167 | V | -61.35460714 | 58.25012791 | 0.764529232 | 47.77947009 | NA | NA | NA | NA | NA | NA | NA |
| KCNMA1 | 1168 | P | -65.24732143 | 58.87949243 | 0.773092354 | 48.51908099 | NA | NA | NA | NA | NA | NA | NA |
| KCNMA1 | 1169 | T | -64.80746429 | 55.15292752 | 0.722388783 | 44.85802124 | NA | NA | NA | NA | NA | NA | NA |
| KCNMA1 | 1170 | D | -63.818 | 54.40827244 | 0.712257021 | 44.03572352 | NA | NA | NA | NA | NA | NA | NA |
| KCNMA1 | 1171 | L | -66.430625 | 50.36235349 | 0.657208325 | 40.55036437 | NA | NA | NA | NA | NA | NA | NA |
| KCNMA1 | 1172 | I | -65.0285 | 52.42676211 | 0.685296629 | 42.15818165 | NA | NA | NA | NA | NA | NA | NA |
| KCNMA1 | 1173 | F | -63.31438636 | 46.62947683 | 0.606418877 | 36.32436529 | NA | NA | NA | NA | NA | NA | NA |
| KCNMA1 | 1174 | C | -66.84858333 | 50.8720282 | 0.66414295 | 41.06394955 | NA | NA | NA | NA | NA | NA | NA |
| KCNMA1 | 1175 | L | -66.6155 | 49.24604574 | 0.642019864 | 39.41730684 | NA | NA | NA | NA | NA | NA | NA |
| KCNMA1 | 1176 | M | -71.84175 | 52.42451016 | 0.685265989 | 42.66024904 | NA | NA | NA | NA | NA | NA | NA |
| KCNMA1 | 1177 | Q | -72.58119444 | 54.12140902 | 0.708353962 | 44.41383774 | NA | NA | NA | NA | NA | NA | NA |
| KCNMA1 | 1178 | F | -78.14820455 | 52.86844087 | 0.691306102 | 43.24198156 | NA | NA | NA | NA | NA | NA | NA |
| KCNMA1 | 1179 | D | -80.253625 | 56.42606059 | 0.739711007 | 46.79051716 | NA | NA | NA | NA | NA | NA | NA |
| KCNQ1 | 104 | T | -23.62171429 | 45.30826601 | 0.9226226 | 42.93873601 | NA | NA | NA | NA | NA | NA | NA |
| KCNQ1 | 105 | H | -23.8795 | 44.57043851 | 0.906821999 | 41.94784617 | NA | NA | NA | NA | NA | NA | NA |
| KCNQ1 | 106 | V | -19.32014286 | 44.85985757 | 0.913019918 | 43.57885759 | NA | NA | NA | NA | NA | NA | NA |
| KCNQ1 | 107 | Q | -19.14888889 | 42.75192433 | 0.867878457 | 41.4718224 | NA | NA | NA | NA | NA | NA | NA |
| KCNQ1 | 108 | G | -20.34425 | 40.06106169 | 0.810253543 | 38.7238242 | NA | NA | NA | NA | NA | NA | NA |
| KCNQ1 | 109 | R | -19.18881818 | 40.44132158 | 0.818396821 | 39.18272261 | NA | NA | NA | NA | NA | NA | NA |

| | | | | | | | | | | | | | |
|---|---|---|---|---|---|---|---|---|---|---|---|---|---|
| KCNQ1 | 110 | V | -15.49728571 | 39.90222868 | 0.806852129 | 38.77500382 | NA | NA | NA | NA | NA | NA | NA |
| KCNQ1 | 111 | Y | -17.043 | 35.89554942 | 0.721048957 | 34.76523119 | NA | NA | NA | NA | NA | NA | NA |
| KCNQ1 | 112 | N | -18.747875 | 35.59940519 | 0.714707019 | 33.63796583 | NA | NA | NA | NA | NA | NA | NA |
| KCNQ1 | 113 | F | -13.192 | 36.08779751 | 0.725165957 | 35.04087837 | NA | NA | NA | NA | NA | NA | NA |
| KCNQ1 | 114 | L | -11.671875 | 34.6626311 | 0.69464597 | 32.61517588 | NA | NA | NA | NA | NA | NA | NA |
| KCNQ1 | 115 | E | -13.78322222 | 32.15352946 | 0.640913474 | 30.75675605 | NA | NA | NA | NA | NA | NA | NA |
| KCNQ1 | 116 | R | -18.68318182 | 31.96789554 | 0.636938117 | 30.86614346 | NA | NA | NA | NA | NA | NA | NA |
| KCNQ1 | 117 | P | -14.57571429 | 30.28983263 | 0.601002344 | 29.24342806 | NA | NA | NA | NA | NA | NA | NA |
| KCNQ1 | 118 | T | -17.413 | 30.49639184 | 0.605425816 | 28.98525056 | NA | NA | NA | NA | NA | NA | NA |
| KCNQ1 | 119 | G | -15.71775 | 31.8054188 | 0.633458672 | 30.34936402 | NA | NA | NA | NA | NA | NA | NA |
| KCNQ1 | 120 | W | -13.64807143 | 33.92918794 | 0.67893926 | 32.6569412 | NA | NA | NA | NA | NA | NA | NA |
| KCNQ1 | 121 | K | -13.76388889 | 36.12175552 | 0.725893169 | 34.55631115 | NA | NA | NA | NA | NA | NA | NA |
| KCNQ1 | 122 | C | -12.42733333 | 33.72807786 | 0.674632481 | 31.72744037 | NA | NA | NA | NA | NA | NA | NA |
| KCNQ1 | 123 | F | -9.298272727 | 28.74361595 | 0.567890061 | 26.12121832 | NA | NA | NA | NA | NA | NA | NA |
| KCNQ1 | 124 | V | -7.717285714 | 34.35288898 | 0.688012832 | 31.95585424 | NA | NA | NA | NA | NA | NA | NA |
| KCNQ1 | 125 | Y | -7.0005 | 35.0998059 | 0.704008083 | 32.64249977 | NA | NA | NA | NA | NA | NA | NA |
| KCNQ1 | 126 | H | -7.8417 | 31.5738736 | 0.628500124 | 29.19748431 | NA | NA | NA | NA | NA | NA | NA |
| KCNQ1 | 127 | F | -3.623636364 | 29.01957772 | 0.573799792 | 25.7426278 | NA | NA | NA | NA | NA | NA | NA |
| KCNQ1 | 128 | A | -2.1428 | 33.84094235 | 0.677049478 | 30.6954212 | NA | NA | NA | NA | NA | NA | NA |
| KCNQ1 | 129 | V | -2.064714286 | 32.79558284 | 0.654663069 | 29.6297515 | NA | NA | NA | NA | NA | NA | NA |
| KCNQ1 | 130 | F | -0.824818182 | 25.47919 | 0.49798227 | 22.31183045 | NA | NA | NA | NA | NA | NA | NA |
| KCNQ1 | 131 | L | 0.326625 | 30.23674541 | 0.599865479 | 27.69579105 | NA | NA | NA | NA | NA | NA | NA |
| KCNQ1 | 132 | I | 1.532 | 33.21656558 | 0.663678428 | 31.68504038 | NA | NA | NA | NA | NA | NA | NA |
| KCNQ1 | 133 | V | 1.875428571 | 30.2224659 | 0.599559683 | 28.70035851 | NA | NA | NA | NA | NA | NA | NA |
| KCNQ1 | 134 | L | 3.497875 | 25.89047923 | 0.506790043 | 24.33368082 | NA | NA | NA | NA | NA | NA | NA |
| KCNQ1 | 135 | V | 6.240428571 | 30.48984456 | 0.605285606 | 29.62779568 | NA | NA | NA | NA | NA | NA | NA |
| KCNQ1 | 136 | C | 7.2395 | 31.08839441 | 0.618103571 | 30.20038696 | NA | NA | NA | NA | NA | NA | NA |
| KCNQ1 | 137 | L | 7.378625 | 25.33624441 | 0.494921085 | 24.47332466 | NA | NA | NA | NA | NA | NA | NA |
| KCNQ1 | 138 | I | 9.615625 | 27.1430774 | 0.533614475 | 26.30602009 | NA | NA | NA | NA | NA | NA | NA |
| KCNQ1 | 139 | F | 12.089 | 29.97102069 | 0.594174975 | 29.10137703 | NA | NA | NA | NA | NA | NA | NA |
| KCNQ1 | 140 | S | 12.00866667 | 27.74411655 | 0.546485748 | 26.90117821 | NA | NA | NA | NA | NA | NA | NA |
| KCNQ1 | 141 | V | 13.31542857 | 24.00771661 | 0.466470618 | 23.01858414 | NA | NA | NA | NA | NA | NA | NA |
| KCNQ1 | 142 | L | 15.910625 | 27.01034715 | 0.530772052 | 25.41813506 | NA | NA | NA | NA | NA | NA | NA |

| | | | | | | | | | | | | | |
|---|---|---|---|---|---|---|---|---|---|---|---|---|---|
| KCNQ1 | 143 | S | 17.20183333 | 28.81288236 | 0.569373404 | 26.84394662 | NA | NA | NA | NA | NA | NA | NA |
| KCNQ1 | 144 | T | 18.62585714 | 24.80725728 | 0.483592808 | 22.56845154 | NA | NA | NA | NA | NA | NA | NA |
| KCNQ1 | 145 | I | 20.452625 | 24.23124563 | 0.471257499 | 20.65703123 | NA | NA | NA | NA | NA | NA | NA |
| KCNQ1 | 146 | E | 25.58844444 | 25.99944657 | 0.509123582 | 20.09358361 | NA | NA | NA | NA | NA | NA | NA |
| KCNQ1 | 147 | Q | 24.38477778 | 25.21541009 | 0.492333414 | 19.30276921 | NA | NA | NA | NA | NA | NA | NA |
| KCNQ1 | 148 | Y | 20.52283333 | 28.24420544 | 0.557195168 | 24.45865963 | NA | NA | NA | NA | NA | NA | NA |
| KCNQ1 | 149 | A | 21.562 | 30.56182573 | 0.606827085 | 26.71258407 | NA | NA | NA | NA | NA | NA | NA |
| KCNQ1 | 150 | A | 22.178 | 33.28696623 | 0.665186061 | 28.3896182 | NA | NA | NA | NA | NA | NA | NA |
| KCNQ1 | 151 | L | 19.136 | 34.76885453 | 0.696920748 | 31.36566199 | NA | NA | NA | NA | NA | NA | NA |
| KCNQ1 | 152 | A | 16.838 | 32.22841897 | 0.642517235 | 30.62778849 | NA | NA | NA | NA | NA | NA | NA |
| KCNQ1 | 153 | T | 17.75442857 | 34.14436359 | 0.683547254 | 32.18268476 | NA | NA | NA | NA | NA | NA | NA |
| KCNQ1 | 154 | G | 17.01075 | 37.64410207 | 0.758494271 | 35.67387617 | NA | NA | NA | NA | NA | NA | NA |
| KCNQ1 | 155 | T | 13.96371429 | 36.36201327 | 0.731038296 | 34.99156304 | NA | NA | NA | NA | NA | NA | NA |
| KCNQ1 | 156 | L | 12.424375 | 33.6258505 | 0.672443279 | 32.63699217 | NA | NA | NA | NA | NA | NA | NA |
| KCNQ1 | 157 | F | 13.58427273 | 38.78539743 | 0.78293515 | 37.79623881 | NA | NA | NA | NA | NA | NA | NA |
| KCNQ1 | 158 | W | 12.73221429 | 41.08336097 | 0.832146116 | 40.02254808 | NA | NA | NA | NA | NA | NA | NA |
| KCNQ1 | 159 | M | 8.184875 | 37.42363048 | 0.753772865 | 36.5475751 | NA | NA | NA | NA | NA | NA | NA |
| KCNQ1 | 160 | E | 7.505111111 | 36.32675214 | 0.730283178 | 35.37834074 | NA | NA | NA | NA | NA | NA | NA |
| KCNQ1 | 161 | I | 7.71375 | 41.67092183 | 0.844728752 | 40.72077809 | NA | NA | NA | NA | NA | NA | NA |
| KCNQ1 | 162 | V | 4.532857143 | 41.93102089 | 0.850298782 | 40.73580944 | NA | NA | NA | NA | NA | NA | NA |
| KCNQ1 | 163 | L | 3.044125 | 36.12153471 | 0.72588844 | 34.57255433 | NA | NA | NA | NA | NA | NA | NA |
| KCNQ1 | 164 | V | 2.986285714 | 41.02663443 | 0.830931316 | 39.47702862 | NA | NA | NA | NA | NA | NA | NA |
| KCNQ1 | 165 | V | 1.230857143 | 43.50021808 | 0.883903193 | 41.94034094 | NA | NA | NA | NA | NA | NA | NA |
| KCNQ1 | 166 | F | -0.874 | 39.17475082 | 0.791273166 | 35.99155375 | NA | NA | NA | NA | NA | NA | NA |
| KCNQ1 | 167 | F | -0.547636364 | 35.69660396 | 0.716788534 | 32.536064 | NA | NA | NA | NA | NA | NA | NA |
| KCNQ1 | 168 | G | -1.805 | 42.07883965 | 0.853464326 | 38.89063012 | NA | NA | NA | NA | NA | NA | NA |
| KCNQ1 | 169 | T | -3.540714286 | 43.81931443 | 0.890736652 | 40.48045227 | NA | NA | NA | NA | NA | NA | NA |
| KCNQ1 | 170 | E | -5.607666667 | 37.11065456 | 0.747070475 | 33.99366758 | NA | NA | NA | NA | NA | NA | NA |
| KCNQ1 | 171 | Y | -5.946 | 39.99704844 | 0.808882697 | 36.55965263 | NA | NA | NA | NA | NA | NA | NA |
| KCNQ1 | 172 | V | -7.046142857 | 43.6367404 | 0.886826823 | 41.18334313 | NA | NA | NA | NA | NA | NA | NA |
| KCNQ1 | 173 | V | -9.660571429 | 43.34928275 | 0.880670907 | 40.77675383 | NA | NA | NA | NA | NA | NA | NA |
| KCNQ1 | 174 | R | -9.843545455 | 36.32415556 | 0.730227572 | 33.93074206 | NA | NA | NA | NA | NA | NA | NA |
| KCNQ1 | 175 | L | -10.992875 | 42.24768002 | 0.857080048 | 39.65834978 | NA | NA | NA | NA | NA | NA | NA |

| | | | | | | | | | | | | | |
|---|---|---|---|---|---|---|---|---|---|---|---|---|---|
| KCNQ1 | 176 | W | -14.44428571 | 44.66979223 | 0.908949662 | 43.39205005 | NA | NA | NA | NA | NA | NA | NA |
| KCNQ1 | 177 | S | -15.39583333 | 41.70218747 | 0.845398307 | 40.5600994 | NA | NA | NA | NA | NA | NA | NA |
| KCNQ1 | 178 | A | -16.5336 | 40.65325073 | 0.822935291 | 39.50844575 | NA | NA | NA | NA | NA | NA | NA |
| KCNQ1 | 179 | G | -19.47825 | 43.04195676 | 0.874089511 | 41.77642641 | NA | NA | NA | NA | NA | NA | NA |
| KCNQ1 | 180 | C | -21.51833333 | 40.24848383 | 0.814267194 | 38.89356782 | NA | NA | NA | NA | NA | NA | NA |
| KCNQ1 | 181 | R | -21.545 | 34.18045133 | 0.684320074 | 31.69265828 | NA | NA | NA | NA | NA | NA | NA |
| KCNQ1 | 182 | S | -24.71316667 | 40.12691588 | 0.811663812 | 37.13233845 | NA | NA | NA | NA | NA | NA | NA |
| KCNQ1 | 183 | K | -21.49111111 | 34.2296927 | 0.685374579 | 31.82521506 | NA | NA | NA | NA | NA | NA | NA |
| KCNQ1 | 184 | Y | -18.24483333 | 35.6343334 | 0.715455007 | 34.39797876 | NA | NA | NA | NA | NA | NA | NA |
| KCNQ1 | 185 | V | -21.51385714 | 43.64272107 | 0.886954899 | 42.28463795 | NA | NA | NA | NA | NA | NA | NA |
| KCNQ1 | 186 | G | -18.71925 | 46.83212979 | 0.955256194 | 45.57571646 | NA | NA | NA | NA | NA | NA | NA |
| KCNQ1 | 187 | L | -15.189375 | 47.09566403 | 0.960899789 | 45.958931 | NA | NA | NA | NA | NA | NA | NA |
| KCNQ1 | 188 | W | -16.4435 | 44.29372694 | 0.900896212 | 42.73772252 | NA | NA | NA | NA | NA | NA | NA |
| KCNQ1 | 189 | G | -17.03375 | 42.31683748 | 0.858561057 | 40.31641833 | NA | NA | NA | NA | NA | NA | NA |
| KCNQ1 | 190 | R | -15.12563636 | 42.84887602 | 0.86995468 | 41.80499446 | NA | NA | NA | NA | NA | NA | NA |
| KCNQ1 | 191 | L | -12.351375 | 41.87788229 | 0.849160817 | 40.31066478 | NA | NA | NA | NA | NA | NA | NA |
| KCNQ1 | 192 | R | -17.01172727 | 38.74870452 | 0.78214937 | 37.67280129 | NA | NA | NA | NA | NA | NA | NA |
| KCNQ1 | 193 | F | -12.98572727 | 37.44396552 | 0.75420834 | 36.14422499 | NA | NA | NA | NA | NA | NA | NA |
| KCNQ1 | 194 | A | -9.735 | 39.14279589 | 0.79058885 | 36.53564699 | NA | NA | NA | NA | NA | NA | NA |
| KCNQ1 | 195 | R | -12.38818182 | 36.16197742 | 0.726754522 | 33.51655464 | NA | NA | NA | NA | NA | NA | NA |
| KCNQ1 | 196 | K | -11.631 | 33.52232529 | 0.670226283 | 31.01790249 | NA | NA | NA | NA | NA | NA | NA |
| KCNQ1 | 197 | P | -8.330142857 | 31.28237921 | 0.622257762 | 28.84114563 | NA | NA | NA | NA | NA | NA | NA |
| KCNQ1 | 198 | I | -7.241375 | 30.18479811 | 0.598753026 | 27.73952884 | NA | NA | NA | NA | NA | NA | NA |
| KCNQ1 | 199 | S | -7.189166667 | 34.54555718 | 0.692138828 | 32.08847278 | NA | NA | NA | NA | NA | NA | NA |
| KCNQ1 | 200 | I | -4.739 | 35.63935238 | 0.715562489 | 32.44228028 | NA | NA | NA | NA | NA | NA | NA |
| KCNQ1 | 201 | I | -2.201 | 31.45317734 | 0.62591541 | 28.2539001 | NA | NA | NA | NA | NA | NA | NA |
| KCNQ1 | 202 | D | -2.843125 | 35.01482746 | 0.702188267 | 31.65578909 | NA | NA | NA | NA | NA | NA | NA |
| KCNQ1 | 203 | L | -1.904875 | 37.75376319 | 0.760842668 | 34.56398183 | NA | NA | NA | NA | NA | NA | NA |
| KCNQ1 | 204 | I | 0.30375 | 35.95119649 | 0.722240641 | 32.66767314 | NA | NA | NA | NA | NA | NA | NA |
| KCNQ1 | 205 | V | 1.154142857 | 34.00404595 | 0.680542347 | 32.49389721 | NA | NA | NA | NA | NA | NA | NA |
| KCNQ1 | 206 | V | 0.697285714 | 38.56443979 | 0.778203334 | 35.99989691 | NA | NA | NA | NA | NA | NA | NA |
| KCNQ1 | 207 | V | 2.134428571 | 40.88183769 | 0.827830488 | 39.33537633 | NA | NA | NA | NA | NA | NA | NA |
| KCNQ1 | 208 | A | 5.2846 | 38.33457242 | 0.773280717 | 37.2874597 | NA | NA | NA | NA | NA | NA | NA |

| | | | | | | | | | | | | | |
|---|---|---|---|---|---|---|---|---|---|---|---|---|---|
| KCNQ1 | 209 | S | 5.84 | 38.08349507 | 0.767903887 | 37.2084003 | NA | NA | NA | NA | NA | NA | NA |
| KCNQ1 | 210 | M | 4.937625 | 42.22176086 | 0.856524988 | 41.32148363 | NA | NA | NA | NA | NA | NA | NA |
| KCNQ1 | 211 | V | 8.273714286 | 42.90121364 | 0.871075492 | 41.88924159 | NA | NA | NA | NA | NA | NA | NA |
| KCNQ1 | 212 | V | 10.85985714 | 38.86178427 | 0.784570977 | 37.95548443 | NA | NA | NA | NA | NA | NA | NA |
| KCNQ1 | 213 | L | 10.499625 | 41.27347315 | 0.836217375 | 40.07401543 | NA | NA | NA | NA | NA | NA | NA |
| KCNQ1 | 214 | C | 11.25983333 | 44.96585351 | 0.915289825 | 43.89856732 | NA | NA | NA | NA | NA | NA | NA |
| KCNQ1 | 215 | V | 14.12 | 43.89135319 | 0.892279364 | 42.52285791 | NA | NA | NA | NA | NA | NA | NA |
| KCNQ1 | 216 | G | 15.86 | 42.84631955 | 0.869899934 | 41.23821357 | NA | NA | NA | NA | NA | NA | NA |
| KCNQ1 | 217 | S | 16.18433333 | 44.82490436 | 0.912271394 | 43.21711164 | NA | NA | NA | NA | NA | NA | NA |
| KCNQ1 | 218 | K | 19.98622222 | 46.30759901 | 0.94402335 | 43.9778603 | NA | NA | NA | NA | NA | NA | NA |
| KCNQ1 | 223 | A | 19.631 | 37.70890359 | 0.759881998 | 35.38861899 | NA | NA | NA | NA | NA | NA | NA |
| KCNQ1 | 224 | T | 20.65057143 | 34.02133697 | 0.680912635 | 30.63738239 | NA | NA | NA | NA | NA | NA | NA |
| KCNQ1 | 225 | S | 17.73433333 | 35.80054969 | 0.719014535 | 33.9172387 | NA | NA | NA | NA | NA | NA | NA |
| KCNQ1 | 226 | A | 15.284 | 36.11995826 | 0.72585468 | 34.54747691 | NA | NA | NA | NA | NA | NA | NA |
| KCNQ1 | 227 | I | 16.3375 | 32.70675915 | 0.652760906 | 31.1197234 | NA | NA | NA | NA | NA | NA | NA |
| KCNQ1 | 228 | R | 17.41881818 | 31.32076867 | 0.623079874 | 30.30103177 | NA | NA | NA | NA | NA | NA | NA |
| KCNQ1 | 229 | G | 11.52875 | 33.93355403 | 0.67903276 | 33.07168234 | NA | NA | NA | NA | NA | NA | NA |
| KCNQ1 | 230 | I | 10.842375 | 32.6213176 | 0.650931173 | 31.74635497 | NA | NA | NA | NA | NA | NA | NA |
| KCNQ1 | 231 | R | 13.50127273 | 28.63660429 | 0.565598403 | 28.25806943 | NA | NA | NA | NA | NA | NA | NA |
| KCNQ1 | 232 | F | 7.253727273 | 27.83333607 | 0.548396387 | 26.93070845 | NA | NA | NA | NA | NA | NA | NA |
| KCNQ1 | 233 | L | 6.010625 | 31.68797849 | 0.630943684 | 30.78703432 | NA | NA | NA | NA | NA | NA | NA |
| KCNQ1 | 234 | Q | 6.898111111 | 29.31881321 | 0.58020793 | 28.46405257 | NA | NA | NA | NA | NA | NA | NA |
| KCNQ1 | 235 | I | 5.54875 | 25.57812257 | 0.500100914 | 24.529766 | NA | NA | NA | NA | NA | NA | NA |
| KCNQ1 | 236 | L | 2.006875 | 28.82141152 | 0.569556056 | 27.29166136 | NA | NA | NA | NA | NA | NA | NA |
| KCNQ1 | 237 | R | 2.081727273 | 29.89207475 | 0.592484345 | 27.32311055 | NA | NA | NA | NA | NA | NA | NA |
| KCNQ1 | 238 | M | 0.83325 | 26.2661782 | 0.514835648 | 23.70321675 | NA | NA | NA | NA | NA | NA | NA |
| KCNQ1 | 239 | L | -1.778875 | 26.00301638 | 0.509200029 | 22.81122593 | NA | NA | NA | NA | NA | NA | NA |
| KCNQ1 | 240 | H | -3.5442 | 29.45808409 | 0.58319042 | 26.10761739 | NA | NA | NA | NA | NA | NA | NA |
| KCNQ1 | 241 | V | -4.688285714 | 26.66472642 | 0.523370572 | 23.49358271 | NA | NA | NA | NA | NA | NA | NA |
| KCNQ1 | 242 | D | -7.792625 | 24.24433884 | 0.471537891 | 21.87773774 | NA | NA | NA | NA | NA | NA | NA |
| KCNQ1 | 243 | R | -9.217454545 | 28.29732858 | 0.558332802 | 25.72412545 | NA | NA | NA | NA | NA | NA | NA |
| KCNQ1 | 244 | Q | -14.06144444 | 27.38810327 | 0.538861712 | 25.82778541 | NA | NA | NA | NA | NA | NA | NA |
| KCNQ1 | 245 | G | -11.713 | 23.93335635 | 0.46487819 | 21.37613217 | NA | NA | NA | NA | NA | NA | NA |

| | | | | | | | | | | | | | |
|---|---|---|---|---|---|---|---|---|---|---|---|---|---|
| KCNQ1 | 246 | G | -12.817 | 20.7862423 | 0.397482637 | 19.49231404 | NA | NA | NA | NA | NA | NA | NA |
| KCNQ1 | 247 | T | -9.667714286 | 19.29715331 | 0.365593746 | 16.62736822 | NA | NA | NA | NA | NA | NA | NA |
| KCNQ1 | 248 | W | -7.332785714 | 20.56564176 | 0.392758469 | 18.02915846 | NA | NA | NA | NA | NA | NA | NA |
| KCNQ1 | 249 | R | -14.91490909 | 19.46225586 | 0.369129422 | 18.18773346 | NA | NA | NA | NA | NA | NA | NA |
| KCNQ1 | 250 | L | -13.94375 | 15.94508692 | 0.293809131 | 14.64560521 | NA | NA | NA | NA | NA | NA | NA |
| KCNQ1 | 251 | L | -9.918 | 14.67615897 | 0.266634997 | 12.27596522 | NA | NA | NA | NA | NA | NA | NA |
| KCNQ1 | 252 | G | -12.573 | 18.24066537 | 0.342969021 | 16.19469286 | NA | NA | NA | NA | NA | NA | NA |
| KCNQ1 | 253 | S | -15.8765 | 15.99579585 | 0.294895065 | 14.84549529 | NA | NA | NA | NA | NA | NA | NA |
| KCNQ1 | 254 | V | -14.16728571 | 11.58004788 | 0.200331673 | 10.5247456 | NA | NA | NA | NA | NA | NA | NA |
| KCNQ1 | 255 | V | -11.30471429 | 14.94521157 | 0.272396767 | 12.90681127 | NA | NA | NA | NA | NA | NA | NA |
| KCNQ1 | 256 | F | -15.62436364 | 16.91102995 | 0.314494834 | 15.85882267 | NA | NA | NA | NA | NA | NA | NA |
| KCNQ1 | 257 | I | -17.4995 | 13.57946774 | 0.243149317 | 12.12764681 | NA | NA | NA | NA | NA | NA | NA |
| KCNQ1 | 258 | H | -14.5828 | 9.615873595 | 0.158268815 | 8.68603909 | NA | NA | NA | NA | NA | NA | NA |
| KCNQ1 | 259 | R | -14.47981818 | 15.81124872 | 0.290942982 | 14.26133196 | NA | NA | NA | NA | NA | NA | NA |
| KCNQ1 | 260 | Q | -13.37344444 | 16.33263114 | 0.302108404 | 14.77180606 | NA | NA | NA | NA | NA | NA | NA |
| KCNQ1 | 261 | E | -12.72166667 | 12.6992275 | 0.224298943 | 10.47274915 | NA | NA | NA | NA | NA | NA | NA |
| KCNQ1 | 262 | L | -8.54825 | 11.89874304 | 0.207156541 | 9.559728491 | NA | NA | NA | NA | NA | NA | NA |
| KCNQ1 | 263 | I | -8.11125 | 16.50059578 | 0.305705372 | 14.10967513 | NA | NA | NA | NA | NA | NA | NA |
| KCNQ1 | 264 | T | -8.340285714 | 16.18470889 | 0.298940644 | 13.9571062 | NA | NA | NA | NA | NA | NA | NA |
| KCNQ1 | 265 | T | -5.638428571 | 12.28608404 | 0.215451461 | 8.953449091 | NA | NA | NA | NA | NA | NA | NA |
| KCNQ1 | 266 | L | -2.993375 | 13.95796138 | 0.251254771 | 10.59993508 | NA | NA | NA | NA | NA | NA | NA |
| KCNQ1 | 267 | Y | -3.468833333 | 17.64078575 | 0.330122579 | 14.27346794 | NA | NA | NA | NA | NA | NA | NA |
| KCNQ1 | 268 | I | -3.22125 | 16.94778204 | 0.315281881 | 13.67954561 | NA | NA | NA | NA | NA | NA | NA |
| KCNQ1 | 269 | G | -0.442 | 15.1326797 | 0.276411403 | 11.90455637 | NA | NA | NA | NA | NA | NA | NA |
| KCNQ1 | 270 | F | 1.117454545 | 16.81237975 | 0.312382237 | 14.25337446 | NA | NA | NA | NA | NA | NA | NA |
| KCNQ1 | 271 | L | 0.355125 | 19.16046842 | 0.362666634 | 16.59207888 | NA | NA | NA | NA | NA | NA | NA |
| KCNQ1 | 272 | G | 2.2345 | 17.69093793 | 0.33119659 | 16.13709061 | NA | NA | NA | NA | NA | NA | NA |
| KCNQ1 | 273 | L | 4.912375 | 15.3847713 | 0.281809954 | 14.24896219 | NA | NA | NA | NA | NA | NA | NA |
| KCNQ1 | 274 | I | 5.429125 | 19.29522814 | 0.365552518 | 18.25562736 | NA | NA | NA | NA | NA | NA | NA |
| KCNQ1 | 275 | F | 4.147363636 | 21.03376158 | 0.40278327 | 20.01673161 | NA | NA | NA | NA | NA | NA | NA |
| KCNQ1 | 276 | S | 7.448166667 | 16.99067094 | 0.316200348 | 16.10247302 | NA | NA | NA | NA | NA | NA | NA |
| KCNQ1 | 277 | S | 9.888833333 | 16.89253806 | 0.314098829 | 16.03193305 | NA | NA | NA | NA | NA | NA | NA |
| KCNQ1 | 278 | Y | 11.30633333 | 20.83917726 | 0.398616241 | 19.94290062 | NA | NA | NA | NA | NA | NA | NA |

| | | | | | | | | | | | | | |
|---|---|---|---|---|---|---|---|---|---|---|---|---|---|
| KCNQ1 | 279 | F | 9.375 | 21.28379762 | 0.408137801 | 20.41584911 | NA | NA | NA | NA | NA | NA | NA |
| KCNQ1 | 280 | V | 13.30628571 | 16.73361924 | 0.310695578 | 15.75268078 | NA | NA | NA | NA | NA | NA | NA |
| KCNQ1 | 281 | Y | 16.556 | 20.0429628 | 0.381565281 | 18.46922476 | NA | NA | NA | NA | NA | NA | NA |
| KCNQ1 | 282 | L | 15.336375 | 22.56621209 | 0.43560075 | 20.95643319 | NA | NA | NA | NA | NA | NA | NA |
| KCNQ1 | 283 | A | 16.993 | 21.05540952 | 0.403246862 | 19.55221615 | NA | NA | NA | NA | NA | NA | NA |
| KCNQ1 | 284 | E | 18.825 | 16.40933061 | 0.303750926 | 14.08124005 | NA | NA | NA | NA | NA | NA | NA |
| KCNQ1 | 285 | K | 21.003 | 20.77078492 | 0.397151616 | 17.16625799 | NA | NA | NA | NA | NA | NA | NA |
| KCNQ1 | 286 | D | 24.3355 | 20.13428978 | 0.383521052 | 14.73972874 | NA | NA | NA | NA | NA | NA | NA |
| KCNQ1 | 287 | A | 24.228 | 16.99751394 | 0.316346891 | 11.64950316 | NA | NA | NA | NA | NA | NA | NA |
| KCNQ1 | 288 | V | 27.45828571 | 14.21595027 | 0.256779612 | 8.17591571 | NA | NA | NA | NA | NA | NA | NA |
| KCNQ1 | 289 | N | 26.40825 | 8.785075811 | 0.140477252 | 2.666173342 | 6.12 | -0.94 | -0.72 | 2.01 | 1.05 | 0 | 0 |
| KCNQ1 | 290 | E | 27.709 | 9.214101638 | 0.149664855 | 3.044496376 | 6.17 | -0.97 | -0.8 | 1.67 | 1.11 | 0 | 0 |
| KCNQ1 | 291 | S | 29.979 | 6.384416496 | 0.089067052 | 0.41555481 | 6.17 | -0.97 | -0.8 | 1.67 | 1.11 | 0 | 0 |
| KCNQ1 | 292 | G | 30.829 | 10.59703756 | 0.179280474 | 4.426925863 | 6.17 | -0.97 | -0.8 | 1.67 | 1.11 | 0 | 0 |
| KCNQ1 | 293 | R | 28.20481818 | 10.11327959 | 0.168920781 | 4.027087711 | 6.17 | -0.97 | -0.8 | 1.67 | 1.11 | 0 | 0 |
| KCNQ1 | 294 | V | 25.61042857 | 14.57877862 | 0.264549593 | 8.571259646 | NA | NA | NA | NA | NA | NA | NA |
| KCNQ1 | 295 | E | 22.05377778 | 9.7540996 | 0.161228929 | 4.855185173 | 4.9 | -0.8 | -0.4 | 3.38 | 0.81 | 0 | 0 |
| KCNQ1 | 296 | F | 18.63754545 | 13.65195048 | 0.244701537 | 10.48283149 | NA | NA | NA | NA | NA | NA | NA |
| KCNQ1 | 297 | G | 21.473 | 18.06281166 | 0.339160278 | 14.20391454 | NA | NA | NA | NA | NA | NA | NA |
| KCNQ1 | 298 | S | 18.823 | 20.04953877 | 0.381706106 | 17.80491065 | NA | NA | NA | NA | NA | NA | NA |
| KCNQ1 | 299 | Y | 13.09383333 | 20.2402772 | 0.385790776 | 18.68768643 | NA | NA | NA | NA | NA | NA | NA |
| KCNQ1 | 300 | A | 15.2146 | 17.81612228 | 0.333877417 | 16.25951942 | NA | NA | NA | NA | NA | NA | NA |
| KCNQ1 | 301 | D | 17.636875 | 15.19197272 | 0.277681165 | 13.63646137 | NA | NA | NA | NA | NA | NA | NA |
| KCNQ1 | 302 | A | 13.946 | 15.98648057 | 0.294695578 | 14.77398591 | NA | NA | NA | NA | NA | NA | NA |
| KCNQ1 | 303 | L | 10.759625 | 15.45580169 | 0.283331072 | 14.41817319 | NA | NA | NA | NA | NA | NA | NA |
| KCNQ1 | 304 | W | 13.378 | 11.43409957 | 0.197206185 | 10.4631659 | NA | NA | NA | NA | NA | NA | NA |
| KCNQ1 | 305 | W | 12.19092857 | 11.11621149 | 0.190398602 | 10.12645924 | NA | NA | NA | NA | NA | NA | NA |
| KCNQ1 | 306 | G | 8.958 | 12.08796927 | 0.211208827 | 11.39786606 | NA | NA | NA | NA | NA | NA | NA |
| KCNQ1 | 307 | V | 7.893428571 | 10.22402372 | 0.17129237 | 9.299413934 | NA | NA | NA | NA | NA | NA | NA |
| KCNQ1 | 308 | V | 10.05228571 | 7.112716921 | 0.104663631 | 6.316773965 | NA | NA | NA | NA | NA | NA | NA |
| KCNQ1 | 309 | T | 8.297714286 | 7.725376366 | 0.117783753 | 7.020485433 | NA | NA | NA | NA | NA | NA | NA |
| KCNQ1 | 310 | V | 4.205571429 | 8.481238353 | 0.133970563 | 7.478011021 | NA | NA | NA | NA | NA | NA | NA |
| KCNQ1 | 311 | T | 4.341714286 | 5.145182796 | 0.062528821 | 3.932991149 | 1.23 | -0.79 | -0.48 | 2.95 | 0.8 | 0 | 0 |

| | | | | | | | | | | | | | |
|---|---|---|---|---|---|---|---|---|---|---|---|---|---|
| KCNQ1 | 312 | T | 5.115 | 2.355757628 | 0.002793187 | 1.649664711 | 1.05 | -0.79 | -0.48 | 2.95 | 0.8 | 0 | 0 |
| KCNQ1 | 313 | I | 8.64175 | 2.229450605 | 8.83E-05 | 1.782938539 | 0.96 | -0.8 | -0.4 | 3.38 | 0.81 | 0 | 0 |
| KCNQ1 | 314 | G | 11.7415 | 2.303786883 | 0.001680232 | 1.647694248 | 1.07 | -0.8 | -0.4 | 3.38 | 0.81 | 0 | 0 |
| KCNQ1 | 315 | Y | 13.72425 | 2.887143052 | 0.014172824 | 2.05660189 | 1.37 | -0.8 | -0.4 | 3.38 | 0.81 | 0 | 0 |
| KCNQ1 | 316 | G | 17.2535 | 4.322290712 | 0.044906559 | 2.728574941 | 1.6 | -0.8 | -0.4 | 3.38 | 0.81 | 0 | 0 |
| KCNQ1 | 317 | D | 18.452125 | 5.684118753 | 0.074070153 | 3.568824877 | 2.33 | -0.8 | -0.4 | 3.38 | 0.81 | 0 | 0 |
| KCNQ1 | 318 | K | 17.94133333 | 8.237675461 | 0.128754656 | 6.267708505 | NA | NA | NA | NA | NA | NA | NA |
| KCNQ1 | 319 | V | 17.71228571 | 7.486672158 | 0.112671895 | 5.550615304 | NA | NA | NA | NA | NA | NA | NA |
| KCNQ1 | 320 | P | 16.63828571 | 12.02917562 | 0.209949759 | 10.48791926 | NA | NA | NA | NA | NA | NA | NA |
| KCNQ1 | 321 | Q | 21.015 | 11.23726363 | 0.192990937 | 7.836630772 | NA | NA | NA | NA | NA | NA | NA |
| KCNQ1 | 322 | T | 19.00928571 | 17.23828965 | 0.321503111 | 14.90861299 | NA | NA | NA | NA | NA | NA | NA |
| KCNQ1 | 323 | W | 16.50514286 | 19.74252233 | 0.375131339 | 18.19213949 | NA | NA | NA | NA | NA | NA | NA |
| KCNQ1 | 324 | V | 14.706 | 20.10176122 | 0.382824451 | 18.513917 | NA | NA | NA | NA | NA | NA | NA |
| KCNQ1 | 325 | G | 14.0525 | 16.52679948 | 0.306266525 | 15.2398388 | NA | NA | NA | NA | NA | NA | NA |
| KCNQ1 | 326 | K | 14.89588889 | 14.40546133 | 0.260837997 | 14.06699757 | NA | NA | NA | NA | NA | NA | NA |
| KCNQ1 | 327 | T | 10.63457143 | 18.3659434 | 0.345651854 | 17.51094708 | NA | NA | NA | NA | NA | NA | NA |
| KCNQ1 | 328 | I | 9.528 | 16.8852984 | 0.313943792 | 16.1073129 | NA | NA | NA | NA | NA | NA | NA |
| KCNQ1 | 329 | A | 9.3766 | 13.46268294 | 0.240648367 | 12.68874496 | NA | NA | NA | NA | NA | NA | NA |
| KCNQ1 | 330 | S | 7.7355 | 14.55145453 | 0.263964447 | 13.78190775 | NA | NA | NA | NA | NA | NA | NA |
| KCNQ1 | 331 | C | 4.760333333 | 15.6478518 | 0.287443831 | 14.52515804 | NA | NA | NA | NA | NA | NA | NA |
| KCNQ1 | 332 | F | 5.115909091 | 13.15101886 | 0.23397407 | 11.99040905 | NA | NA | NA | NA | NA | NA | NA |
| KCNQ1 | 333 | S | 4.760833333 | 9.761896383 | 0.161395898 | 8.604400618 | NA | NA | NA | NA | NA | NA | NA |
| KCNQ1 | 334 | V | 1.840571429 | 11.66789973 | 0.202213024 | 10.3515344 | NA | NA | NA | NA | NA | NA | NA |
| KCNQ1 | 335 | F | 0 | 12.31655106 | 0.216103914 | 9.056865665 | NA | NA | NA | NA | NA | NA | NA |
| KCNQ1 | 336 | A | 0.4108 | 11.66811604 | 0.202217656 | 9.10230774 | NA | NA | NA | NA | NA | NA | NA |
| KCNQ1 | 337 | I | 1.206625 | 7.055569857 | 0.103439824 | 4.58609906 | 2.57 | -0.61 | -0.2 | 1.33 | 0.73 | 0 | 0 |
| KCNQ1 | 338 | S | -1.957666667 | 8.41860612 | 0.132629292 | 5.364123704 | NA | NA | NA | NA | NA | NA | NA |
| KCNQ1 | 339 | F | -2.851181818 | 10.26803043 | 0.172234776 | 6.988247848 | NA | NA | NA | NA | NA | NA | NA |
| KCNQ1 | 340 | F | -0.417090909 | 7.638967797 | 0.115933311 | 4.509559974 | 3.23 | -0.14 | 1.3 | 0.33 | 0.63 | 0 | 0 |
| KCNQ1 | 341 | A | -2.6252 | 4.764495881 | 0.054376398 | 1.565868084 | 3.2 | 0.02 | 1.8 | 0 | 0.59 | 0 | 0 |
| KCNQ1 | 342 | L | -6.222625 | 6.971842439 | 0.101646799 | 3.712416678 | 3.4 | -0.8 | -0.4 | 3.38 | 0.81 | 0 | 0 |
| KCNQ1 | 343 | P | -5.649428571 | 7.75135672 | 0.118340123 | 4.34523935 | 3.44 | -0.8 | -0.4 | 3.38 | 0.81 | 0 | 0 |
| KCNQ1 | 344 | A | -6.2414 | 4.952943872 | 0.058412018 | 1.789047199 | 3.44 | -0.8 | -0.4 | 3.38 | 0.81 | 0 | 0 |

| | | | | | | | | | | | | | |
|---|---|---|---|---|---|---|---|---|---|---|---|---|---|
| KCNQ1 | 345 | G | -9.0865 | 3.907567658 | 0.036025251 | 1.582392828 | 2.36 | -0.8 | -0.4 | 3.38 | 0.81 | 0 | 0 |
| KCNQ1 | 346 | I | -10.90675 | 5.996826161 | 0.080766792 | 3.517011217 | 2.59 | -0.83 | -0.48 | 3.04 | 0.87 | 0 | 0 |
| KCNQ1 | 347 | L | -10.244375 | 7.455129912 | 0.111996416 | 4.964616113 | 2.5 | -0.94 | -0.72 | 2.01 | 1.05 | 0 | 0 |
| KCNQ1 | 348 | G | -11.475 | 5.23428744 | 0.064437 | 2.899203263 | 2.5 | -0.94 | -0.72 | 2.01 | 1.05 | 0 | 0 |
| KCNQ1 | 349 | S | -14.39483333 | 2.532073854 | 0.006569005 | 1.544788553 | 1.4 | -0.97 | -0.8 | 1.67 | 1.11 | 0 | 0 |
| KCNQ1 | 350 | G | -15.54575 | 6.19134525 | 0.084932425 | 5.232693487 | 1.15 | -0.97 | -0.8 | 1.67 | 1.11 | 0 | 0 |
| KCNQ1 | 351 | F | -13.40663636 | 8.161743135 | 0.127128562 | 7.084469579 | NA | NA | NA | NA | NA | NA | NA |
| KCNQ1 | 352 | A | -16.7774 | 5.784047891 | 0.076210139 | 4.644026839 | 1.15 | -0.97 | -0.8 | 1.67 | 1.11 | 0 | 0 |
| KCNQ1 | 353 | L | -19.5865 | 2.827333196 | 0.012891994 | 1.673832558 | 1.29 | 1.14 | 3.8 | 0.13 | -1.79 | 0 | 0 |
| KCNQ1 | 354 | K | -19.69522222 | 8.184045088 | 0.127606159 | 6.897869441 | NA | NA | NA | NA | NA | NA | NA |
| KCNQ1 | 355 | V | -19.61128571 | 9.866266822 | 0.163630994 | 8.582628938 | NA | NA | NA | NA | NA | NA | NA |
| KCNQ1 | 356 | Q | -22.99733333 | 4.037569194 | 0.038809238 | 1.540779133 | 2.5 | -0.65 | -2.04 | 2.85 | -0.25 | 0 | 0 |
| KCNQ1 | 357 | Q | -24.97088889 | 8.478058445 | 0.133902465 | 5.537785632 | NA | NA | NA | NA | NA | NA | NA |
| KCNQ1 | 358 | K | -23.15122222 | 11.14693384 | 0.191056522 | 8.150056435 | NA | NA | NA | NA | NA | NA | NA |
| KCNQ1 | 359 | Q | -26.31633333 | 9.345024131 | 0.152468565 | 5.515923578 | NA | NA | NA | NA | NA | NA | NA |
| KCNQ1 | 360 | R | -28.86909091 | 5.499746267 | 0.07012181 | 2.236078769 | 3.53 | 0.82 | 2.46 | 10.5 | -0.71 | 1 | 1 |
| KCNQ1 | 361 | Q | -29.19855556 | 12.80277021 | 0.226516313 | 9.944159748 | NA | NA | NA | NA | NA | NA | NA |
| KCNQ1 | 362 | K | -34.37711111 | 12.96272772 | 0.229941809 | 11.28412414 | NA | NA | NA | NA | NA | NA | NA |
| KCNQ1 | 363 | H | -35.2089 | 16.87382956 | 0.313698186 | 15.07144808 | NA | NA | NA | NA | NA | NA | NA |
| KCNQ1 | 364 | F | -33.87190909 | 20.43489432 | 0.389958508 | 18.61635212 | NA | NA | NA | NA | NA | NA | NA |
| KCNQ1 | 365 | N | -38.77425 | 20.24412875 | 0.385873257 | 18.40799125 | NA | NA | NA | NA | NA | NA | NA |
| KCNQ1 | 366 | R | -39.33781818 | 13.32941176 | 0.23779436 | 12.15768119 | NA | NA | NA | NA | NA | NA | NA |
| KCNQ1 | 367 | Q | -35.84955556 | 20.72045456 | 0.39607379 | 18.63230519 | NA | NA | NA | NA | NA | NA | NA |
| KCNQ1 | 368 | I | -39.168375 | 23.3360601 | 0.452087071 | 22.10707975 | NA | NA | NA | NA | NA | NA | NA |
| KCNQ1 | 369 | P | -42.391 | 22.11300332 | 0.425895269 | 20.50132125 | NA | NA | NA | NA | NA | NA | NA |
| KCNQ1 | 370 | A | -40.3922 | 21.69614586 | 0.416968252 | 20.34946693 | NA | NA | NA | NA | NA | NA | NA |
| KCNQ1 | 371 | A | -38.704 | 24.74539123 | 0.482267944 | 22.9234311 | NA | NA | NA | NA | NA | NA | NA |
| KCNQ1 | 372 | A | -41.716 | 27.22567867 | 0.535383383 | 25.8625719 | NA | NA | NA | NA | NA | NA | NA |
| KCNQ1 | 373 | S | -43.32466667 | 25.39859856 | 0.496256401 | 23.26666526 | NA | NA | NA | NA | NA | NA | NA |
| KCNQ1 | 374 | L | -39.36375 | 27.05915368 | 0.531817245 | 25.73472768 | NA | NA | NA | NA | NA | NA | NA |
| KCNQ1 | 375 | I | -39.794125 | 29.71045274 | 0.588594904 | 28.48567276 | NA | NA | NA | NA | NA | NA | NA |
| KCNQ1 | 376 | Q | -44.66488889 | 29.23425677 | 0.578397151 | 26.96507226 | NA | NA | NA | NA | NA | NA | NA |
| KCNQ1 | 377 | T | -44.19914286 | 28.40981501 | 0.560741703 | 25.87621996 | NA | NA | NA | NA | NA | NA | NA |

| | | | | | | | | | | | | | |
|---|---|---|---|---|---|---|---|---|---|---|---|---|---|
| KCNQ1 | 378 | A | -41.4974 | 31.74241785 | 0.632109505 | 30.40172779 | NA | NA | NA | NA | NA | NA | NA |
| KCNQ1 | 379 | W | -44.31964286 | 34.40850622 | 0.689203877 | 32.25183556 | NA | NA | NA | NA | NA | NA | NA |
| KCNQ1 | 380 | R | -48.204 | 31.36561171 | 0.624040189 | 30.32645656 | NA | NA | NA | NA | NA | NA | NA |
| KCNQ1 | 381 | C | -44.46833333 | 33.11591524 | 0.661522998 | 30.56652893 | NA | NA | NA | NA | NA | NA | NA |
| KCNQ1 | 382 | Y | -41.95133333 | 37.30968392 | 0.751332695 | 35.14044943 | NA | NA | NA | NA | NA | NA | NA |
| KCNQ1 | 383 | A | -46.6818 | 39.2780471 | 0.793485259 | 38.03456466 | NA | NA | NA | NA | NA | NA | NA |
| KCNQ1 | 384 | A | -48.0786 | 38.63374779 | 0.779687567 | 37.64910505 | NA | NA | NA | NA | NA | NA | NA |
| KCNQ1 | 385 | E | -45.562 | 39.91566707 | 0.807139912 | 38.72023794 | NA | NA | NA | NA | NA | NA | NA |
| KCNQ1 | 386 | N | -48.075375 | 44.01924481 | 0.895018168 | 43.0319882 | NA | NA | NA | NA | NA | NA | NA |
| KCNQ1 | 387 | P | -51.48271429 | 45.62230543 | 0.929347765 | 43.01538663 | NA | NA | NA | NA | NA | NA | NA |
| KCNQ1 | 388 | D | -52.31325 | 46.24260782 | 0.942631561 | 44.32649548 | NA | NA | NA | NA | NA | NA | NA |
| KCNQ1 | 389 | S | -51.1985 | 43.21347188 | 0.877762513 | 41.03216965 | NA | NA | NA | NA | NA | NA | NA |
| KCNQ1 | 390 | S | -52.88933333 | 40.18768355 | 0.812965154 | 38.42833377 | NA | NA | NA | NA | NA | NA | NA |
| KCNQ1 | 391 | T | -51.84985714 | 37.33702175 | 0.751918136 | 34.9027951 | NA | NA | NA | NA | NA | NA | NA |
| KCNQ1 | 392 | W | -54.04471429 | 38.13759063 | 0.769062345 | 36.59909076 | NA | NA | NA | NA | NA | NA | NA |
| KCNQ1 | 393 | K | -57.88411111 | 37.21573863 | 0.749320854 | 35.55789229 | NA | NA | NA | NA | NA | NA | NA |
| KCNQ1 | 394 | I | -56.296 | 32.94195047 | 0.657797536 | 31.34869641 | NA | NA | NA | NA | NA | NA | NA |
| KCNQ1 | 395 | Y | -56.80741667 | 32.2588783 | 0.643169523 | 30.13669627 | NA | NA | NA | NA | NA | NA | NA |
| KCNQ1 | 396 | I | -62.799375 | 34.51508299 | 0.691486222 | 32.80315536 | NA | NA | NA | NA | NA | NA | NA |
| KCNQ1 | 506 | L | -63.94875 | 40.69228336 | 0.823771176 | 38.9208716 | NA | NA | NA | NA | NA | NA | NA |
| KCNQ1 | 507 | R | -60.18172727 | 38.41187387 | 0.77493613 | 36.76298591 | NA | NA | NA | NA | NA | NA | NA |
| KCNQ1 | 508 | E | -59.60444444 | 42.78181186 | 0.868518499 | 40.70677071 | NA | NA | NA | NA | NA | NA | NA |
| KCNQ1 | 509 | H | -59.2917 | 39.13753843 | 0.790476261 | 36.83697153 | NA | NA | NA | NA | NA | NA | NA |
| KCNQ1 | 510 | H | -59.3307 | 36.56322651 | 0.735347285 | 35.0228038 | NA | NA | NA | NA | NA | NA | NA |
| KCNQ1 | 511 | R | -54.96709091 | 38.76878608 | 0.782579417 | 37.2647048 | NA | NA | NA | NA | NA | NA | NA |
| KCNQ1 | 512 | A | -52.861 | 36.81264302 | 0.740688548 | 34.90442374 | NA | NA | NA | NA | NA | NA | NA |
| KCNQ1 | 513 | T | -54.66971429 | 33.57566221 | 0.671368495 | 31.81768433 | NA | NA | NA | NA | NA | NA | NA |
| KCNQ1 | 514 | I | -53.48125 | 33.99424925 | 0.68033255 | 32.33456206 | NA | NA | NA | NA | NA | NA | NA |
| KCNQ1 | 515 | K | -49.25577778 | 34.4041382 | 0.689110336 | 33.36878853 | NA | NA | NA | NA | NA | NA | NA |
| KCNQ1 | 516 | V | -49.671 | 31.36986066 | 0.62413118 | 30.32030304 | NA | NA | NA | NA | NA | NA | NA |
| KCNQ1 | 517 | I | -51.53125 | 29.1912842 | 0.577476892 | 27.0292017 | NA | NA | NA | NA | NA | NA | NA |
| KCNQ1 | 518 | R | -49.271 | 30.82839551 | 0.612535686 | 29.86914906 | NA | NA | NA | NA | NA | NA | NA |
| KCNQ1 | 519 | R | -44.02463636 | 29.54406968 | 0.585031805 | 28.01925959 | NA | NA | NA | NA | NA | NA | NA |

| | | | | | | | | | | | | | |
|---|---|---|---|---|---|---|---|---|---|---|---|---|---|
| KCNQ1 | 520 | M | -47.234 | 26.25157757 | 0.514522976 | 25.00574129 | NA | NA | NA | NA | NA | NA | NA |
| KCNQ1 | 521 | Q | -47.90555556 | 26.32503953 | 0.516096166 | 25.12614181 | NA | NA | NA | NA | NA | NA | NA |
| KCNQ1 | 522 | Y | -40.88 | 26.94886725 | 0.529455457 | 24.84170972 | NA | NA | NA | NA | NA | NA | NA |
| KCNQ1 | 523 | F | -41.26454545 | 24.1871249 | 0.470312652 | 22.60294848 | NA | NA | NA | NA | NA | NA | NA |
| KCNQ1 | 524 | V | -43.74728571 | 21.58339443 | 0.414553676 | 19.49250238 | NA | NA | NA | NA | NA | NA | NA |
| KCNQ1 | 525 | A | -41.635 | 23.51542979 | 0.455928279 | 21.99373037 | NA | NA | NA | NA | NA | NA | NA |
| KCNQ1 | 526 | K | -37.50122222 | 22.72903909 | 0.439087695 | 20.89994978 | NA | NA | NA | NA | NA | NA | NA |
| KCNQ1 | 527 | K | -39.54544444 | 17.32534156 | 0.323367331 | 15.99349198 | NA | NA | NA | NA | NA | NA | NA |
| KCNQ1 | 528 | K | -41.50988889 | 17.21800378 | 0.321068689 | 16.77114519 | NA | NA | NA | NA | NA | NA | NA |
| KCNQ1 | 529 | F | -35.86545455 | 20.63833736 | 0.394315248 | 18.48397785 | NA | NA | NA | NA | NA | NA | NA |
| KCNQ1 | 530 | Q | -34.158 | 17.90332142 | 0.335744789 | 16.0714105 | NA | NA | NA | NA | NA | NA | NA |
| KCNQ1 | 531 | Q | -36.20266667 | 13.92366633 | 0.250520341 | 11.91236053 | NA | NA | NA | NA | NA | NA | NA |
| KCNQ1 | 532 | A | -35.6996 | 17.39285293 | 0.324813089 | 15.05265372 | NA | NA | NA | NA | NA | NA | NA |
| KCNQ1 | 533 | R | -31.23809091 | 16.75304205 | 0.311111518 | 15.07163571 | NA | NA | NA | NA | NA | NA | NA |
| KCNQ1 | 534 | K | -31.56711111 | 13.52745349 | 0.24203543 | 10.64351375 | NA | NA | NA | NA | NA | NA | NA |
| KCNQ1 | 535 | P | -30.49628571 | 10.61371363 | 0.179637593 | 7.77960682 | NA | NA | NA | NA | NA | NA | NA |
| KCNQ1 | 536 | Y | -34.6915 | 7.388485298 | 0.11056922 | 5.789381382 | NA | NA | NA | NA | NA | NA | NA |
| KCNQ1 | 537 | D | -33.35925 | 5.934232722 | 0.079426352 | 4.190873056 | 1.83 | 1.13 | 4.2 | 0.13 | -1.3 | 0 | 0 |
| KCNQ1 | 538 | V | -35.56728571 | 3.485416761 | 0.026984876 | 1.974196661 | 2.02 | 1.13 | 4.2 | 0.13 | -1.3 | 0 | 0 |
| KCNQ1 | 539 | R | -36.73409091 | 7.327991539 | 0.109273744 | 4.699742513 | 2.66 | 1.13 | 4.2 | 0.13 | -1.3 | 0 | 0 |
| KCNQ1 | 540 | D | -38.174 | 7.417561931 | 0.111191897 | 6.201860124 | NA | NA | NA | NA | NA | NA | NA |
| KCNQ1 | 541 | V | -40.52742857 | 2.651896303 | 0.009135007 | 1.283791565 | 1.37 | 1.13 | 4.2 | 0.13 | -1.3 | 0 | 0 |
| KCNQ1 | 542 | I | -41.195 | 6.130475838 | 0.083628905 | 4.657330999 | 1.62 | 1.13 | 4.2 | 0.13 | -1.3 | 0 | 0 |
| KCNQ1 | 543 | E | -42.33288889 | 8.572366884 | 0.135922084 | 6.49181001 | NA | NA | NA | NA | NA | NA | NA |
| KCNQ1 | 544 | Q | -44.68277778 | 6.262717461 | 0.086460863 | 3.739680792 | 2.55 | 1.13 | 3.1 | 0.43 | -1.33 | 0 | 0 |
| KCNQ1 | 545 | Y | -47.55158333 | 2.631032877 | 0.008688216 | 1.812572105 | 1.28 | 1.11 | -1.3 | 1.61 | -1.44 | 0 | 0 |
| KCNQ1 | 546 | S | -47.65233333 | 7.807005636 | 0.119531847 | 6.864570773 | NA | NA | NA | NA | NA | NA | NA |
| KCNQ1 | 547 | Q | -48.21333333 | 9.284335787 | 0.151168922 | 8.254209883 | NA | NA | NA | NA | NA | NA | NA |
| KCNQ1 | 548 | G | -51.27875 | 6.542808724 | 0.092459027 | 4.464792461 | 2.19 | 1.11 | -1.3 | 1.61 | -1.44 | 0 | 0 |
| KCNQ1 | 549 | H | -52.5568 | 5.735359971 | 0.075167485 | 3.464788468 | 2.29 | 1.11 | -1.3 | 1.61 | -1.44 | 0 | 0 |
| KCNQ1 | 550 | L | -53.550875 | 8.568465965 | 0.135838545 | 6.874148438 | NA | NA | NA | NA | NA | NA | NA |
| KCNQ1 | 551 | N | -54.9485 | 8.516171499 | 0.134718658 | 6.994018933 | NA | NA | NA | NA | NA | NA | NA |
| KCNQ1 | 552 | L | -56.825875 | 2.615148371 | 0.008348049 | 1.195820745 | 1.6 | 1.14 | 3.8 | 0.13 | -1.79 | 0 | 0 |

| | | | | | | | | | | | | | |
|---|---|---|---|---|---|---|---|---|---|---|---|---|---|
| KCNQ1 | 553 | M | -57.918125 | 6.937428198 | 0.100909817 | 4.823781389 | 2.2 | 1.68 | 4.36 | 0.13 | -1.84 | 0 | 0 |
| KCNQ1 | 554 | V | -59.70342857 | 9.239263012 | 0.150203687 | 7.166094221 | NA | NA | NA | NA | NA | NA | NA |
| KCNQ1 | 555 | R | -61.29127273 | 6.972607403 | 0.10166318 | 5.497417616 | NA | NA | NA | NA | NA | NA | NA |
| KCNQ1 | 556 | I | -62.5415 | 2.853619631 | 0.013454919 | 1.460187105 | 1.72 | 1.68 | 4.36 | 0.13 | -1.84 | 0 | 0 |
| KCNQ1 | 557 | K | -63.37766667 | 8.053353153 | 0.124807387 | 6.297803183 | NA | NA | NA | NA | NA | NA | NA |
| KCNQ1 | 558 | E | -66.35077778 | 8.643173781 | 0.137438416 | 7.27117007 | NA | NA | NA | NA | NA | NA | NA |
| KCNQ1 | 559 | L | -67.656625 | 2.665063977 | 0.009416993 | 1.580309617 | 1.42 | 1.14 | 3.8 | 0.13 | -1.79 | 0 | 0 |
| KCNQ1 | 560 | Q | -68.46933333 | 6.813836951 | 0.098263106 | 5.404773849 | 1.42 | 1.14 | 3.8 | 0.13 | -1.79 | 0 | 0 |
| KCNQ1 | 561 | R | -70.27436364 | 9.257459425 | 0.150593363 | 6.556170523 | NA | NA | NA | NA | NA | NA | NA |
| KCNQ1 | 562 | R | -71.70681818 | 7.891405515 | 0.121339273 | 5.810066174 | NA | NA | NA | NA | NA | NA | NA |
| KCNQ1 | 563 | L | -73.278875 | 3.100707177 | 0.018746307 | 1.088123868 | 2.01 | 1.14 | 3.8 | 0.13 | -1.79 | 0 | 0 |
| KCNQ1 | 564 | D | -73.74475 | 8.12304555 | 0.126299852 | 6.285662218 | NA | NA | NA | NA | NA | NA | NA |
| KCNQ1 | 565 | Q | -76.01066667 | 9.971214069 | 0.165878443 | 8.096511177 | NA | NA | NA | NA | NA | NA | NA |
| KCNQ1 | 566 | S | -78.5335 | 7.548983309 | 0.11400629 | 5.73612847 | NA | NA | NA | NA | NA | NA | NA |
| KCNQ1 | 567 | I | -78.77775 | 3.254267506 | 0.022034806 | 1.464706061 | 1.9 | 1.54 | 4.22 | 0.13 | -1.83 | 0 | 0 |
| KCNQ1 | 568 | G | -78.469 | 8.520832354 | 0.13481847 | 6.677600504 | NA | NA | NA | NA | NA | NA | NA |
| KCNQ2 | 70 | R | -21.64471591 | 52.17973364 | 1 | 49.15455454 | NA | NA | NA | NA | NA | NA | NA |
| KCNQ2 | 71 | N | -25.581375 | 50.85998619 | 0.973787522 | 47.8238143 | NA | NA | NA | NA | NA | NA | NA |
| KCNQ2 | 72 | A | -26.829425 | 48.55116242 | 0.927930268 | 45.53847909 | NA | NA | NA | NA | NA | NA | NA |
| KCNQ2 | 73 | F | -24.44844318 | 48.09394448 | 0.918849125 | 45.08279433 | NA | NA | NA | NA | NA | NA | NA |
| KCNQ2 | 74 | Y | -21.607625 | 47.76329621 | 0.912281876 | 45.71785594 | NA | NA | NA | NA | NA | NA | NA |
| KCNQ2 | 75 | R | -25.57380682 | 44.94231986 | 0.856252382 | 41.90690269 | NA | NA | NA | NA | NA | NA | NA |
| KCNQ2 | 76 | K | -24.328625 | 43.19595225 | 0.821566486 | 40.21404884 | NA | NA | NA | NA | NA | NA | NA |
| KCNQ2 | 77 | L | -20.14725 | 42.94352926 | 0.816552926 | 41.02818365 | NA | NA | NA | NA | NA | NA | NA |
| KCNQ2 | 78 | Q | -20.64695833 | 41.93965229 | 0.796614185 | 40.02421997 | NA | NA | NA | NA | NA | NA | NA |
| KCNQ2 | 79 | N | -24.0105 | 39.48659866 | 0.747892276 | 36.73944242 | NA | NA | NA | NA | NA | NA | NA |
| KCNQ2 | 80 | F | -22.19626136 | 37.08893435 | 0.700270495 | 34.46313465 | NA | NA | NA | NA | NA | NA | NA |
| KCNQ2 | 81 | L | -18.5845 | 38.63122979 | 0.730903163 | 36.52362666 | NA | NA | NA | NA | NA | NA | NA |
| KCNQ2 | 82 | Y | -18.08920833 | 36.73470186 | 0.693234822 | 34.62862188 | NA | NA | NA | NA | NA | NA | NA |
| KCNQ2 | 83 | N | -21.49575 | 33.86040642 | 0.636146318 | 31.95042364 | NA | NA | NA | NA | NA | NA | NA |
| KCNQ2 | 84 | V | -17.82119643 | 33.72223534 | 0.633402001 | 31.6302653 | NA | NA | NA | NA | NA | NA | NA |
| KCNQ2 | 85 | L | -14.773875 | 34.34719839 | 0.645814853 | 32.33963624 | NA | NA | NA | NA | NA | NA | NA |
| KCNQ2 | 86 | E | -17.37851389 | 32.55984903 | 0.610314988 | 30.45153941 | NA | NA | NA | NA | NA | NA | NA |

| | | | | | | | | | | | | | |
|---|---|---|---|---|---|---|---|---|---|---|---|---|---|
| KCNQ2 | 87 | R | -19.02153409 | 26.0124614 | 0.480272489 | 23.90673678 | NA | NA | NA | NA | NA | NA | NA |
| KCNQ2 | 88 | P | -18.569625 | 28.09267008 | 0.521589049 | 26.1775146 | NA | NA | NA | NA | NA | NA | NA |
| KCNQ2 | 89 | R | -22.84398864 | 26.95468349 | 0.498986657 | 24.95434137 | NA | NA | NA | NA | NA | NA | NA |
| KCNQ2 | 90 | G | -19.253625 | 29.05321087 | 0.540667059 | 27.19387604 | NA | NA | NA | NA | NA | NA | NA |
| KCNQ2 | 91 | W | -16.78491071 | 30.59199153 | 0.571229918 | 28.36944031 | NA | NA | NA | NA | NA | NA | NA |
| KCNQ2 | 92 | A | -14.574825 | 31.38113883 | 0.586903755 | 29.43422652 | NA | NA | NA | NA | NA | NA | NA |
| KCNQ2 | 93 | F | -13.33744318 | 25.90683705 | 0.478174606 | 23.69951588 | NA | NA | NA | NA | NA | NA | NA |
| KCNQ2 | 94 | I | -11.323625 | 30.19970566 | 0.563438438 | 27.80499778 | NA | NA | NA | NA | NA | NA | NA |
| KCNQ2 | 95 | Y | -9.061958333 | 33.05884143 | 0.620225845 | 30.63805581 | NA | NA | NA | NA | NA | NA | NA |
| KCNQ2 | 96 | H | -10.161025 | 30.36907516 | 0.566802411 | 28.32979251 | NA | NA | NA | NA | NA | NA | NA |
| KCNQ2 | 97 | A | -6.612425 | 28.13299915 | 0.522390055 | 25.83336664 | NA | NA | NA | NA | NA | NA | NA |
| KCNQ2 | 98 | Y | -5.977625 | 30.99145051 | 0.579163867 | 27.67361378 | NA | NA | NA | NA | NA | NA | NA |
| KCNQ2 | 99 | V | -4.219053571 | 32.11321144 | 0.601443989 | 28.80033904 | NA | NA | NA | NA | NA | NA | NA |
| KCNQ2 | 100 | F | -2.819534091 | 24.83781991 | 0.456942067 | 21.41587368 | NA | NA | NA | NA | NA | NA | NA |
| KCNQ2 | 101 | L | -1.43075 | 27.67641539 | 0.513321508 | 23.97418504 | NA | NA | NA | NA | NA | NA | NA |
| KCNQ2 | 102 | L | 0.057625 | 31.73973124 | 0.594026023 | 28.012562 | NA | NA | NA | NA | NA | NA | NA |
| KCNQ2 | 103 | V | 0.481482143 | 30.44241822 | 0.568259132 | 26.88021828 | NA | NA | NA | NA | NA | NA | NA |
| KCNQ2 | 104 | F | 1.562715909 | 23.92176557 | 0.438747636 | 22.19917285 | NA | NA | NA | NA | NA | NA | NA |
| KCNQ2 | 105 | S | 4.024958333 | 28.88927417 | 0.537410991 | 28.06291139 | NA | NA | NA | NA | NA | NA | NA |
| KCNQ2 | 106 | C | 5.225125 | 30.59655577 | 0.571320571 | 29.95554918 | NA | NA | NA | NA | NA | NA | NA |
| KCNQ2 | 107 | L | 5.584625 | 25.66315133 | 0.473334584 | 25.00136215 | NA | NA | NA | NA | NA | NA | NA |
| KCNQ2 | 108 | V | 7.284339286 | 25.19614504 | 0.464059027 | 24.44376597 | NA | NA | NA | NA | NA | NA | NA |
| KCNQ2 | 109 | L | 9.8965 | 28.55831868 | 0.53083764 | 27.80711502 | NA | NA | NA | NA | NA | NA | NA |
| KCNQ2 | 110 | S | 10.21129167 | 28.64117381 | 0.532483287 | 27.80301606 | NA | NA | NA | NA | NA | NA | NA |
| KCNQ2 | 111 | V | 11.68891071 | 23.81178607 | 0.436563252 | 22.76494791 | NA | NA | NA | NA | NA | NA | NA |
| KCNQ2 | 112 | F | 12.53535227 | 26.25740379 | 0.485137471 | 25.26935998 | NA | NA | NA | NA | NA | NA | NA |
| KCNQ2 | 113 | S | 15.56329167 | 28.46538515 | 0.528991818 | 27.41868751 | NA | NA | NA | NA | NA | NA | NA |
| KCNQ2 | 114 | T | 16.10005357 | 24.24801456 | 0.445227508 | 22.07917033 | NA | NA | NA | NA | NA | NA | NA |
| KCNQ2 | 115 | I | 18.16525 | 23.99977404 | 0.440297019 | 21.839897 | NA | NA | NA | NA | NA | NA | NA |
| KCNQ2 | 116 | K | 22.66018056 | 25.85694183 | 0.4771836 | 19.95841504 | NA | NA | NA | NA | NA | NA | NA |
| KCNQ2 | 117 | E | 22.97795833 | 22.77469231 | 0.415964767 | 16.94721006 | NA | NA | NA | NA | NA | NA | NA |
| KCNQ2 | 118 | Y | 19.68629167 | 26.05942121 | 0.481205192 | 21.95972328 | NA | NA | NA | NA | NA | NA | NA |
| KCNQ2 | 119 | E | 20.02951389 | 29.57288748 | 0.55098874 | 27.43444962 | NA | NA | NA | NA | NA | NA | NA |

| | | | | | | | | | | | | | |
|---|---|---|---|---|---|---|---|---|---|---|---|---|---|
| KCNQ2 | 120 | K | 21.46784722 | 31.74273237 | 0.594085631 | 27.73174489 | NA | NA | NA | NA | NA | NA | NA |
| KCNQ2 | 121 | S | 17.15895833 | 34.05456118 | 0.640002569 | 31.89552742 | NA | NA | NA | NA | NA | NA | NA |
| KCNQ2 | 122 | S | 15.30079167 | 30.04806711 | 0.560426633 | 28.98538098 | NA | NA | NA | NA | NA | NA | NA |
| KCNQ2 | 123 | E | 14.93773611 | 33.99684743 | 0.638856274 | 32.92331725 | NA | NA | NA | NA | NA | NA | NA |
| KCNQ2 | 124 | G | 14.689875 | 36.55254381 | 0.689616847 | 35.47901277 | NA | NA | NA | NA | NA | NA | NA |
| KCNQ2 | 125 | A | 11.606625 | 36.29001649 | 0.684402598 | 35.20011126 | NA | NA | NA | NA | NA | NA | NA |
| KCNQ2 | 126 | L | 10.59175 | 32.60159496 | 0.611144135 | 31.76740024 | NA | NA | NA | NA | NA | NA | NA |
| KCNQ2 | 127 | Y | 10.58395833 | 37.50599111 | 0.708553967 | 36.6673953 | NA | NA | NA | NA | NA | NA | NA |
| KCNQ2 | 128 | I | 8.64075 | 39.18204474 | 0.741843306 | 38.42983805 | NA | NA | NA | NA | NA | NA | NA |
| KCNQ2 | 129 | L | 5.94325 | 35.38627805 | 0.666452781 | 34.71747723 | NA | NA | NA | NA | NA | NA | NA |
| KCNQ2 | 130 | E | 6.730958333 | 34.75150535 | 0.653845092 | 34.1090511 | NA | NA | NA | NA | NA | NA | NA |
| KCNQ2 | 131 | I | 5.4625 | 40.23639001 | 0.762784435 | 39.60078801 | NA | NA | NA | NA | NA | NA | NA |
| KCNQ2 | 132 | V | 2.057910714 | 39.33186257 | 0.744818948 | 37.62698113 | NA | NA | NA | NA | NA | NA | NA |
| KCNQ2 | 133 | T | 1.493196429 | 35.6424367 | 0.671540537 | 33.93619673 | NA | NA | NA | NA | NA | NA | NA |
| KCNQ2 | 134 | I | 1.588875 | 39.51499399 | 0.748456256 | 35.83293824 | NA | NA | NA | NA | NA | NA | NA |
| KCNQ2 | 135 | V | -0.511339286 | 41.83980347 | 0.794631014 | 38.12672922 | NA | NA | NA | NA | NA | NA | NA |
| KCNQ2 | 136 | V | -2.386053571 | 37.89782155 | 0.716336401 | 34.28821949 | NA | NA | NA | NA | NA | NA | NA |
| KCNQ2 | 137 | F | -3.156806818 | 34.15061617 | 0.641910389 | 30.69874481 | NA | NA | NA | NA | NA | NA | NA |
| KCNQ2 | 138 | G | -3.852375 | 41.46339578 | 0.787154903 | 38.01535984 | NA | NA | NA | NA | NA | NA | NA |
| KCNQ2 | 139 | V | -5.527339286 | 42.68751309 | 0.811468 | 39.10050343 | NA | NA | NA | NA | NA | NA | NA |
| KCNQ2 | 140 | E | -8.188736111 | 37.4007713 | 0.706464119 | 35.22212355 | NA | NA | NA | NA | NA | NA | NA |
| KCNQ2 | 141 | Y | -7.649791667 | 41.39599605 | 0.785816227 | 39.09542638 | NA | NA | NA | NA | NA | NA | NA |
| KCNQ2 | 142 | F | -9.941625 | 44.09335144 | 0.839390394 | 41.72075769 | NA | NA | NA | NA | NA | NA | NA |
| KCNQ2 | 143 | V | -11.23605357 | 42.24229793 | 0.802625253 | 39.86143623 | NA | NA | NA | NA | NA | NA | NA |
| KCNQ2 | 144 | R | -13.058625 | 35.36188686 | 0.66596833 | 33.07509328 | NA | NA | NA | NA | NA | NA | NA |
| KCNQ2 | 145 | I | -13.166875 | 43.1512698 | 0.820679015 | 40.86267674 | NA | NA | NA | NA | NA | NA | NA |
| KCNQ2 | 146 | W | -17.061125 | 45.18258886 | 0.861024542 | 42.73629769 | NA | NA | NA | NA | NA | NA | NA |
| KCNQ2 | 147 | A | -17.399025 | 42.18036562 | 0.80139517 | 40.0894117 | NA | NA | NA | NA | NA | NA | NA |
| KCNQ2 | 148 | A | -18.611625 | 41.99057463 | 0.797625591 | 39.88241261 | NA | NA | NA | NA | NA | NA | NA |
| KCNQ2 | 149 | G | -21.293625 | 44.26236244 | 0.842747246 | 42.35254962 | NA | NA | NA | NA | NA | NA | NA |
| KCNQ2 | 150 | C | -22.775625 | 39.6049138 | 0.75024222 | 36.98076002 | NA | NA | NA | NA | NA | NA | NA |
| KCNQ2 | 151 | C | -23.52729167 | 40.52216134 | 0.768460351 | 37.49802583 | NA | NA | NA | NA | NA | NA | NA |
| KCNQ2 | 152 | C | -27.128625 | 41.24718364 | 0.782860554 | 38.25831938 | NA | NA | NA | NA | NA | NA | NA |

| | | | | | | | | | | | | | |
|---|---|---|---|---|---|---|---|---|---|---|---|---|---|
| KCNQ2 | 153 | R | -26.41753409 | 37.66162644 | 0.711645155 | 34.76694661 | NA | NA | NA | NA | NA | NA | NA |
| KCNQ2 | 154 | Y | -21.97954167 | 41.9246656 | 0.796316523 | 39.30684062 | NA | NA | NA | NA | NA | NA | NA |
| KCNQ2 | 155 | R | -25.52780682 | 44.54192507 | 0.848299846 | 42.14122185 | NA | NA | NA | NA | NA | NA | NA |
| KCNQ2 | 156 | G | -20.412375 | 49.87362015 | 0.954196578 | 47.96848255 | NA | NA | NA | NA | NA | NA | NA |
| KCNQ2 | 157 | W | -18.02419643 | 51.16141082 | 0.979774339 | 49.28178699 | NA | NA | NA | NA | NA | NA | NA |
| KCNQ2 | 158 | R | -22.77580682 | 49.01835839 | 0.937209592 | 47.11438279 | NA | NA | NA | NA | NA | NA | NA |
| KCNQ2 | 159 | G | -19.932125 | 46.16553759 | 0.880547612 | 44.27439764 | NA | NA | NA | NA | NA | NA | NA |
| KCNQ2 | 160 | R | -17.90935227 | 46.68973466 | 0.890959077 | 44.30851659 | NA | NA | NA | NA | NA | NA | NA |
| KCNQ2 | 161 | L | -15.551625 | 48.76407413 | 0.932159065 | 46.31574861 | NA | NA | NA | NA | NA | NA | NA |
| KCNQ2 | 162 | K | -18.88084722 | 45.3302433 | 0.863957216 | 42.88255803 | NA | NA | NA | NA | NA | NA | NA |
| KCNQ2 | 163 | F | -16.62244318 | 38.59741813 | 0.730231605 | 36.15216158 | NA | NA | NA | NA | NA | NA | NA |
| KCNQ2 | 164 | A | -12.424625 | 43.21235562 | 0.821892285 | 40.96390191 | NA | NA | NA | NA | NA | NA | NA |
| KCNQ2 | 165 | R | -13.39207955 | 41.8128565 | 0.7940958 | 39.41486636 | NA | NA | NA | NA | NA | NA | NA |
| KCNQ2 | 166 | K | -12.06895833 | 34.71424113 | 0.65310496 | 32.33897196 | NA | NA | NA | NA | NA | NA | NA |
| KCNQ2 | 167 | P | -9.462910714 | 36.61625449 | 0.690882251 | 34.51214213 | NA | NA | NA | NA | NA | NA | NA |
| KCNQ2 | 168 | F | -8.591443182 | 31.75034905 | 0.594236911 | 29.62429611 | NA | NA | NA | NA | NA | NA | NA |
| KCNQ2 | 169 | C | -8.508791667 | 38.08606067 | 0.720075157 | 35.94277821 | NA | NA | NA | NA | NA | NA | NA |
| KCNQ2 | 170 | V | -5.896625 | 40.14392827 | 0.760947984 | 37.19938477 | NA | NA | NA | NA | NA | NA | NA |
| KCNQ2 | 171 | I | -3.50525 | 36.0182171 | 0.679004189 | 32.58166728 | NA | NA | NA | NA | NA | NA | NA |
| KCNQ2 | 172 | D | -4.50025 | 37.7507839 | 0.713415977 | 34.4293538 | NA | NA | NA | NA | NA | NA | NA |
| KCNQ2 | 173 | I | -3.169375 | 40.72312218 | 0.772451782 | 37.2686501 | NA | NA | NA | NA | NA | NA | NA |
| KCNQ2 | 174 | M | -0.22 | 38.46311657 | 0.727564142 | 34.7372535 | NA | NA | NA | NA | NA | NA | NA |
| KCNQ2 | 175 | V | 0.173625 | 36.63017109 | 0.691158659 | 32.91055213 | NA | NA | NA | NA | NA | NA | NA |
| KCNQ2 | 176 | L | 0.5505 | 40.05130264 | 0.759108278 | 36.32924587 | NA | NA | NA | NA | NA | NA | NA |
| KCNQ2 | 177 | I | 0.89725 | 42.75696521 | 0.81284744 | 39.04154298 | NA | NA | NA | NA | NA | NA | NA |
| KCNQ2 | 178 | A | 4.095425 | 40.92774443 | 0.776515936 | 40.08327481 | NA | NA | NA | NA | NA | NA | NA |
| KCNQ2 | 179 | S | 3.715791667 | 40.08872441 | 0.75985154 | 39.24537634 | NA | NA | NA | NA | NA | NA | NA |
| KCNQ2 | 180 | I | 3.338125 | 43.06162395 | 0.818898492 | 42.21548204 | NA | NA | NA | NA | NA | NA | NA |
| KCNQ2 | 181 | A | 6.916425 | 44.66337585 | 0.850712069 | 44.01311136 | NA | NA | NA | NA | NA | NA | NA |
| KCNQ2 | 182 | V | 9.229053571 | 40.70666762 | 0.772124966 | 39.98759217 | NA | NA | NA | NA | NA | NA | NA |
| KCNQ2 | 183 | L | 7.353125 | 40.9336773 | 0.776633773 | 40.18196495 | NA | NA | NA | NA | NA | NA | NA |
| KCNQ2 | 184 | A | 9.313025 | 45.97339867 | 0.876731399 | 45.22111403 | NA | NA | NA | NA | NA | NA | NA |
| KCNQ2 | 195 | S | 16.66045833 | 36.49489864 | 0.688471913 | 34.47888264 | NA | NA | NA | NA | NA | NA | NA |

| | | | | | | | | | | | | | |
|---|---|---|---|---|---|---|---|---|---|---|---|---|---|
| KCNQ2 | 196 | A | 12.520025 | 35.36063916 | 0.665943548 | 34.28949121 | NA | NA | NA | NA | NA | NA | NA |
| KCNQ2 | 197 | L | 13.802125 | 30.84396208 | 0.576234491 | 29.75489683 | NA | NA | NA | NA | NA | NA | NA |
| KCNQ2 | 198 | R | 13.76089773 | 30.22880835 | 0.564016468 | 29.64576666 | NA | NA | NA | NA | NA | NA | NA |
| KCNQ2 | 199 | S | 8.716291667 | 33.35770652 | 0.626161825 | 32.6259561 | NA | NA | NA | NA | NA | NA | NA |
| KCNQ2 | 200 | L | 8.458 | 32.95116242 | 0.618087152 | 32.19881064 | NA | NA | NA | NA | NA | NA | NA |
| KCNQ2 | 201 | R | 9.822352273 | 26.5665466 | 0.491277584 | 26.29753576 | NA | NA | NA | NA | NA | NA | NA |
| KCNQ2 | 202 | F | 6.236625 | 28.77238423 | 0.535089354 | 28.1901898 | NA | NA | NA | NA | NA | NA | NA |
| KCNQ2 | 203 | L | 3.659375 | 32.89981508 | 0.617067305 | 32.05735089 | NA | NA | NA | NA | NA | NA | NA |
| KCNQ2 | 204 | Q | 3.986513889 | 30.40964125 | 0.567608124 | 28.7060164 | NA | NA | NA | NA | NA | NA | NA |
| KCNQ2 | 205 | I | 2.084875 | 27.64604207 | 0.51271824 | 26.02179391 | NA | NA | NA | NA | NA | NA | NA |
| KCNQ2 | 206 | L | -0.256625 | 30.66148299 | 0.572610139 | 26.95700416 | NA | NA | NA | NA | NA | NA | NA |
| KCNQ2 | 207 | R | -0.364261364 | 30.76383945 | 0.574643116 | 27.05747874 | NA | NA | NA | NA | NA | NA | NA |
| KCNQ2 | 208 | M | -1.40325 | 24.57364942 | 0.451695182 | 20.99509881 | NA | NA | NA | NA | NA | NA | NA |
| KCNQ2 | 209 | I | -2.88425 | 27.85068152 | 0.516782736 | 24.50914981 | NA | NA | NA | NA | NA | NA | NA |
| KCNQ2 | 210 | R | -6.591625 | 30.12474642 | 0.561949617 | 26.54481314 | NA | NA | NA | NA | NA | NA | NA |
| KCNQ2 | 211 | M | -7.326875 | 24.92219126 | 0.458617829 | 22.63885934 | NA | NA | NA | NA | NA | NA | NA |
| KCNQ2 | 212 | D | -8.92325 | 25.32434631 | 0.466605327 | 23.00898571 | NA | NA | NA | NA | NA | NA | NA |
| KCNQ2 | 213 | R | -12.29289773 | 29.44791351 | 0.548506539 | 27.09372684 | NA | NA | NA | NA | NA | NA | NA |
| KCNQ2 | 214 | R | -15.08371591 | 27.10571182 | 0.501986342 | 24.83032503 | NA | NA | NA | NA | NA | NA | NA |
| KCNQ2 | 215 | G | -12.018375 | 23.73333533 | 0.435005084 | 21.33494475 | NA | NA | NA | NA | NA | NA | NA |
| KCNQ2 | 216 | G | -13.604875 | 20.75875926 | 0.375924832 | 18.68811925 | NA | NA | NA | NA | NA | NA | NA |
| KCNQ2 | 217 | T | -10.62805357 | 19.35935763 | 0.348130284 | 16.78552163 | NA | NA | NA | NA | NA | NA | NA |
| KCNQ2 | 218 | W | -8.287839286 | 20.59949431 | 0.372761554 | 18.25245274 | NA | NA | NA | NA | NA | NA | NA |
| KCNQ2 | 219 | K | -15.112625 | 19.64691225 | 0.353841618 | 17.61486932 | NA | NA | NA | NA | NA | NA | NA |
| KCNQ2 | 220 | L | -14.6475 | 15.51912823 | 0.271856652 | 13.51621661 | NA | NA | NA | NA | NA | NA | NA |
| KCNQ2 | 221 | L | -11.055125 | 15.64325174 | 0.274321961 | 13.42034111 | NA | NA | NA | NA | NA | NA | NA |
| KCNQ2 | 222 | G | -13.572375 | 18.37422839 | 0.328563904 | 16.1779962 | NA | NA | NA | NA | NA | NA | NA |
| KCNQ2 | 223 | S | -16.74979167 | 16.19082002 | 0.285197619 | 13.74401411 | NA | NA | NA | NA | NA | NA | NA |
| KCNQ2 | 224 | V | -14.95633929 | 11.80002953 | 0.197988888 | 9.801497714 | NA | NA | NA | NA | NA | NA | NA |
| KCNQ2 | 225 | V | -12.50419643 | 15.25639984 | 0.26663841 | 12.99471518 | NA | NA | NA | NA | NA | NA | NA |
| KCNQ2 | 226 | Y | -17.15904167 | 17.09838065 | 0.303223351 | 15.14735635 | NA | NA | NA | NA | NA | NA | NA |
| KCNQ2 | 227 | A | -17.521825 | 14.17928863 | 0.245245109 | 12.07298768 | NA | NA | NA | NA | NA | NA | NA |
| KCNQ2 | 228 | H | -15.118325 | 9.511476279 | 0.152534242 | 7.569621371 | NA | NA | NA | NA | NA | NA | NA |

| | | | | | | | | | | | | | |
|---|---|---|---|---|---|---|---|---|---|---|---|---|---|
| KCNQ2 | 229 | S | -13.74529167 | 16.01356731 | 0.281677072 | 13.96218185 | NA | NA | NA | NA | NA | NA | NA |
| KCNQ2 | 230 | K | -15.22229167 | 16.50179078 | 0.291374039 | 14.44082891 | NA | NA | NA | NA | NA | NA | NA |
| KCNQ2 | 231 | E | -13.58895833 | 10.86929404 | 0.179502863 | 9.174794079 | NA | NA | NA | NA | NA | NA | NA |
| KCNQ2 | 232 | L | -9.469375 | 12.44903213 | 0.210879207 | 10.05024836 | NA | NA | NA | NA | NA | NA | NA |
| KCNQ2 | 233 | V | -8.891767857 | 16.67394569 | 0.294793335 | 14.53671758 | NA | NA | NA | NA | NA | NA | NA |
| KCNQ2 | 234 | T | -9.143625 | 16.30173469 | 0.287400577 | 14.18559284 | NA | NA | NA | NA | NA | NA | NA |
| KCNQ2 | 235 | A | -6.046425 | 12.85067212 | 0.218856476 | 10.55096132 | NA | NA | NA | NA | NA | NA | NA |
| KCNQ2 | 236 | W | -4.705053571 | 14.17846642 | 0.245228779 | 10.76098904 | NA | NA | NA | NA | NA | NA | NA |
| KCNQ2 | 237 | Y | -4.154541667 | 17.97536442 | 0.320641773 | 14.4548822 | NA | NA | NA | NA | NA | NA | NA |
| KCNQ2 | 238 | I | -3.908125 | 17.19174468 | 0.305077722 | 13.69844162 | NA | NA | NA | NA | NA | NA | NA |
| KCNQ2 | 239 | G | -0.707125 | 15.3852 | 0.269196605 | 11.65834649 | NA | NA | NA | NA | NA | NA | NA |
| KCNQ2 | 240 | F | 0.661443182 | 17.13989714 | 0.30404794 | 13.79920301 | NA | NA | NA | NA | NA | NA | NA |
| KCNQ2 | 241 | L | 0.448625 | 19.38698226 | 0.348678957 | 16.03747394 | NA | NA | NA | NA | NA | NA | NA |
| KCNQ2 | 242 | C | 1.436458333 | 16.65035717 | 0.294324826 | 14.96182328 | NA | NA | NA | NA | NA | NA | NA |
| KCNQ2 | 243 | L | 4.517875 | 15.84732435 | 0.278375198 | 14.83255187 | NA | NA | NA | NA | NA | NA | NA |
| KCNQ2 | 244 | I | 4.9075 | 19.11881427 | 0.343352674 | 18.38837967 | NA | NA | NA | NA | NA | NA | NA |
| KCNQ2 | 245 | L | 4.85175 | 20.84196577 | 0.377577458 | 20.11348869 | NA | NA | NA | NA | NA | NA | NA |
| KCNQ2 | 246 | A | 7.202025 | 17.68340041 | 0.31484286 | 16.65719439 | NA | NA | NA | NA | NA | NA | NA |
| KCNQ2 | 247 | S | 9.464958333 | 17.90561448 | 0.319256418 | 17.05402624 | NA | NA | NA | NA | NA | NA | NA |
| KCNQ2 | 248 | F | 10.30989773 | 20.58105337 | 0.372395285 | 19.58754679 | NA | NA | NA | NA | NA | NA | NA |
| KCNQ2 | 249 | L | 9.751 | 21.60735331 | 0.392779385 | 20.59574072 | NA | NA | NA | NA | NA | NA | NA |
| KCNQ2 | 250 | V | 13.01933929 | 17.28047884 | 0.306840137 | 16.32605224 | NA | NA | NA | NA | NA | NA | NA |
| KCNQ2 | 251 | Y | 16.056375 | 19.95436717 | 0.359948207 | 19.38830716 | NA | NA | NA | NA | NA | NA | NA |
| KCNQ2 | 252 | L | 14.941375 | 22.51422 | 0.410791334 | 21.92166097 | NA | NA | NA | NA | NA | NA | NA |
| KCNQ2 | 253 | A | 16.055825 | 21.14972986 | 0.383690188 | 20.56660529 | NA | NA | NA | NA | NA | NA | NA |
| KCNQ2 | 254 | E | 18.94773611 | 16.04187673 | 0.282239346 | 12.07277537 | NA | NA | NA | NA | NA | NA | NA |
| KCNQ2 | 255 | K | 20.53129167 | 21.15544899 | 0.38380378 | 17.18687364 | NA | NA | NA | NA | NA | NA | NA |
| KCNQ2 | 256 | G | 23.936125 | 19.76836592 | 0.356253899 | 13.06639802 | NA | NA | NA | NA | NA | NA | NA |
| KCNQ2 | 257 | E | 23.97706944 | 16.5987605 | 0.293300026 | 9.795144291 | NA | NA | NA | NA | NA | NA | NA |
| KCNQ2 | 258 | N | 22.584375 | 12.9237587 | 0.220308102 | 7.136763962 | NA | NA | NA | NA | NA | NA | NA |
| KCNQ2 | 259 | D | 24.728125 | 12.29537909 | 0.207827391 | 5.566556713 | 6.8 | -0.8 | -0.4 | 3.38 | 0.81 | 0 | 0 |
| KCNQ2 | 260 | H | 21.157125 | 9.206743507 | 0.14648172 | 4.097302916 | 5.11 | -0.8 | -0.4 | 3.38 | 0.81 | 0 | 0 |
| KCNQ2 | 261 | F | 18.29553409 | 13.38420031 | 0.229453273 | 9.540047592 | NA | NA | NA | NA | NA | NA | NA |

| | | | | | | | | | | | | | |
|---|---|---|---|---|---|---|---|---|---|---|---|---|---|
| KCNQ2 | 262 | D | 22.0485 | 18.2057851 | 0.325218328 | 13.12516304 | NA | NA | NA | NA | NA | NA | NA |
| KCNQ2 | 263 | T | 18.55219643 | 19.79924486 | 0.356867209 | 15.68680866 | NA | NA | NA | NA | NA | NA | NA |
| KCNQ2 | 264 | Y | 12.46554167 | 19.9914584 | 0.360684904 | 19.42366203 | NA | NA | NA | NA | NA | NA | NA |
| KCNQ2 | 265 | A | 15.016025 | 17.84580161 | 0.31806843 | 17.25208784 | NA | NA | NA | NA | NA | NA | NA |
| KCNQ2 | 266 | D | 17.23975 | 15.43242278 | 0.270134532 | 14.70986675 | NA | NA | NA | NA | NA | NA | NA |
| KCNQ2 | 267 | A | 13.312425 | 15.36198428 | 0.2687355 | 14.97221623 | NA | NA | NA | NA | NA | NA | NA |
| KCNQ2 | 268 | L | 10.394 | 15.18016657 | 0.265124285 | 14.16629407 | NA | NA | NA | NA | NA | NA | NA |
| KCNQ2 | 269 | W | 12.93498214 | 11.66170832 | 0.195241588 | 10.82353362 | NA | NA | NA | NA | NA | NA | NA |
| KCNQ2 | 270 | W | 12.13233929 | 10.62478899 | 0.174646568 | 9.604226551 | NA | NA | NA | NA | NA | NA | NA |
| KCNQ2 | 271 | G | 8.691875 | 11.67151717 | 0.195436409 | 10.80497123 | NA | NA | NA | NA | NA | NA | NA |
| KCNQ2 | 272 | L | 6.955875 | 9.991057251 | 0.162059554 | 8.981726333 | NA | NA | NA | NA | NA | NA | NA |
| KCNQ2 | 273 | I | 10.065125 | 6.993095595 | 0.102514824 | 6.150296611 | NA | NA | NA | NA | NA | NA | NA |
| KCNQ2 | 274 | T | 8.298910714 | 7.246903684 | 0.107555894 | 6.56806496 | NA | NA | NA | NA | NA | NA | NA |
| KCNQ2 | 275 | L | 4.15325 | 8.059721273 | 0.123699864 | 7.587183513 | NA | NA | NA | NA | NA | NA | NA |
| KCNQ2 | 276 | T | 4.383196429 | 4.788395243 | 0.058725643 | 3.920532056 | 0.88 | -0.79 | -0.48 | 2.95 | 0.8 | 0 | 0 |
| KCNQ2 | 277 | T | 4.822482143 | 2.126345927 | 0.005852716 | 1.712897813 | 0.75 | -0.79 | -0.48 | 2.95 | 0.8 | 0 | 0 |
| KCNQ2 | 278 | I | 8.251 | 2.154861713 | 0.006419089 | 2.001291632 | 0.76 | -0.8 | -0.4 | 3.38 | 0.81 | 0 | 0 |
| KCNQ2 | 279 | G | 11.362375 | 1.838097386 | 1.28E-04 | 1.531498412 | 1.02 | -0.8 | -0.4 | 3.38 | 0.81 | 0 | 0 |
| KCNQ2 | 280 | Y | 13.536375 | 2.480415086 | 0.012885145 | 1.906054482 | 1.11 | -0.8 | -0.4 | 3.38 | 0.81 | 0 | 0 |
| KCNQ2 | 281 | G | 16.788875 | 3.977072416 | 0.042611361 | 3.236584401 | 0.77 | -0.8 | -0.4 | 3.38 | 0.81 | 0 | 0 |
| KCNQ2 | 282 | D | 18.21425 | 5.425165988 | 0.071373017 | 3.395130916 | 2.18 | -0.8 | -0.4 | 3.38 | 0.81 | 0 | 0 |
| KCNQ2 | 283 | K | 17.77451389 | 7.861832738 | 0.119769454 | 5.757353825 | NA | NA | NA | NA | NA | NA | NA |
| KCNQ2 | 284 | Y | 18.548375 | 7.916622007 | 0.120857664 | 7.887528375 | 0.77 | -0.8 | -0.4 | 3.38 | 0.81 | 0 | 0 |
| KCNQ2 | 285 | P | 16.24348214 | 12.54428906 | 0.212771176 | 11.82860668 | NA | NA | NA | NA | NA | NA | NA |
| KCNQ2 | 286 | Q | 20.21840278 | 11.97115638 | 0.201387764 | 8.008490322 | NA | NA | NA | NA | NA | NA | NA |
| KCNQ2 | 287 | T | 18.66491071 | 17.52347571 | 0.311666477 | 13.55874125 | NA | NA | NA | NA | NA | NA | NA |
| KCNQ2 | 288 | W | 17.13033929 | 17.62066608 | 0.313596847 | 16.86040841 | NA | NA | NA | NA | NA | NA | NA |
| KCNQ2 | 289 | N | 15.444375 | 21.01416977 | 0.380997729 | 20.45170445 | NA | NA | NA | NA | NA | NA | NA |
| KCNQ2 | 290 | G | 13.869375 | 17.90295431 | 0.319203582 | 17.38922079 | NA | NA | NA | NA | NA | NA | NA |
| KCNQ2 | 291 | R | 15.28353409 | 13.49846939 | 0.231722856 | 13.56250851 | NA | NA | NA | NA | NA | NA | NA |
| KCNQ2 | 292 | L | 10.118375 | 18.91246221 | 0.339254164 | 17.92944875 | NA | NA | NA | NA | NA | NA | NA |
| KCNQ2 | 293 | L | 9.575875 | 17.93155409 | 0.319771623 | 17.19415605 | NA | NA | NA | NA | NA | NA | NA |
| KCNQ2 | 294 | A | 8.688225 | 14.61305286 | 0.253860421 | 13.76840066 | NA | NA | NA | NA | NA | NA | NA |

| | | | | | | | | | | | | | |
|---|---|---|---|---|---|---|---|---|---|---|---|---|---|
| KCNQ2 | 295 | A | 7.308025 | 15.6516848 | 0.274489456 | 14.61035478 | NA | NA | NA | NA | NA | NA | NA |
| KCNQ2 | 296 | T | 4.462625 | 15.6430732 | 0.274318415 | 14.56188307 | NA | NA | NA | NA | NA | NA | NA |
| KCNQ2 | 297 | F | 4.235443182 | 14.15948195 | 0.244851714 | 13.07884654 | NA | NA | NA | NA | NA | NA | NA |
| KCNQ2 | 298 | T | 4.350910714 | 10.42537203 | 0.1706858 | 9.345140034 | NA | NA | NA | NA | NA | NA | NA |
| KCNQ2 | 299 | L | 1.540875 | 12.10281393 | 0.204002712 | 10.67808663 | NA | NA | NA | NA | NA | NA | NA |
| KCNQ2 | 300 | I | 0 | 11.80104173 | 0.198008992 | 8.487443399 | NA | NA | NA | NA | NA | NA | NA |
| KCNQ2 | 301 | G | 0.199875 | 10.48660603 | 0.171902014 | 7.138611543 | NA | NA | NA | NA | NA | NA | NA |
| KCNQ2 | 302 | V | 1.100767857 | 7.309411194 | 0.108797402 | 6.066557814 | NA | NA | NA | NA | NA | NA | NA |
| KCNQ2 | 303 | S | -1.675625 | 7.993327718 | 0.122381173 | 4.433804891 | 3.63 | 0.02 | 1.8 | 0 | 0.59 | 0 | 0 |
| KCNQ2 | 304 | F | -3.211170455 | 9.866766289 | 0.159590919 | 6.337353151 | NA | NA | NA | NA | NA | NA | NA |
| KCNQ2 | 305 | F | -0.864170455 | 7.857595752 | 0.1196853 | 4.226140868 | 3.73 | -0.14 | 1.3 | 0.33 | 0.63 | 0 | 0 |
| KCNQ2 | 306 | A | -2.750625 | 4.957194368 | 0.062078287 | 1.326609737 | 3.63 | 0.02 | 1.8 | 0 | 0.59 | 0 | 0 |
| KCNQ2 | 307 | L | -6.20175 | 6.411487269 | 0.090963072 | 3.350598089 | 3.38 | -0.8 | -0.4 | 3.38 | 0.81 | 0 | 0 |
| KCNQ2 | 308 | P | -6.048482143 | 8.018877103 | 0.122888628 | 5.718487781 | NA | NA | NA | NA | NA | NA | NA |
| KCNQ2 | 309 | A | -6.122425 | 5.590868716 | 0.074664161 | 2.208402951 | 3.51 | -0.8 | -0.4 | 3.38 | 0.81 | 0 | 0 |
| KCNQ2 | 310 | G | -8.776875 | 3.793664851 | 0.038968568 | 1.724440227 | 2.15 | -0.8 | -0.4 | 3.38 | 0.81 | 0 | 0 |
| KCNQ2 | 311 | I | -10.960625 | 5.659712537 | 0.076031519 | 3.266782938 | 2.59 | -0.83 | -0.48 | 3.04 | 0.87 | 0 | 0 |
| KCNQ2 | 312 | L | -10.53575 | 7.572953057 | 0.114031801 | 5.010683303 | NA | NA | NA | NA | NA | NA | NA |
| KCNQ2 | 313 | G | -11.562125 | 5.67457038 | 0.076326622 | 3.202306249 | 2.59 | -0.83 | -0.48 | 3.04 | 0.87 | 0 | 0 |
| KCNQ2 | 314 | S | -14.30029167 | 3.384702055 | 0.030845856 | 1.322027205 | 2.07 | -0.97 | -0.8 | 1.67 | 1.11 | 0 | 0 |
| KCNQ2 | 315 | G | -15.740625 | 6.359301377 | 0.089926569 | 4.374395307 | 2.01 | -0.97 | -0.8 | 1.67 | 1.11 | 0 | 0 |
| KCNQ2 | 316 | F | -13.70598864 | 8.483681335 | 0.132120448 | 6.478378631 | NA | NA | NA | NA | NA | NA | NA |
| KCNQ2 | 317 | A | -17.141825 | 5.787489438 | 0.07856939 | 3.349457627 | 2.45 | -0.55 | 0.12 | 1.36 | 0.53 | 0 | 0 |
| KCNQ2 | 318 | L | -19.6785 | 3.174154691 | 0.02666402 | 1.274297253 | 1.92 | 1.14 | 3.8 | 0.13 | -1.79 | 0 | 0 |
| KCNQ2 | 319 | K | -20.01395833 | 8.293169539 | 0.128336553 | 6.376880713 | NA | NA | NA | NA | NA | NA | NA |
| KCNQ2 | 320 | V | -20.21019643 | 9.892414771 | 0.160100343 | 8.019707103 | NA | NA | NA | NA | NA | NA | NA |
| KCNQ2 | 321 | Q | -23.36773611 | 4.432710909 | 0.051661134 | 1.674552395 | 2.76 | -0.65 | -2.04 | 2.85 | -0.25 | 0 | 0 |
| KCNQ2 | 322 | E | -24.847625 | 8.312272012 | 0.128715961 | 5.345221862 | NA | NA | NA | NA | NA | NA | NA |
| KCNQ2 | 323 | Q | -24.07940278 | 10.96540542 | 0.181411802 | 7.922245364 | NA | NA | NA | NA | NA | NA | NA |
| KCNQ2 | 324 | H | -26.384425 | 11.68739595 | 0.195751789 | 8.691239976 | NA | NA | NA | NA | NA | NA | NA |
| KCNQ2 | 325 | R | -28.69407955 | 4.240256006 | 0.047838645 | 1.490448507 | 3.09 | -0.42 | -4.5 | 52 | 1.63 | 3 | 3 |
| KCNQ2 | 326 | Q | -29.35551389 | 12.18508194 | 0.205636698 | 8.916502007 | NA | NA | NA | NA | NA | NA | NA |
| KCNQ2 | 327 | K | -27.20518056 | 14.79996922 | 0.257572905 | 11.71623009 | NA | NA | NA | NA | NA | NA | NA |

| | | | | | | | | | | | | | |
|---|---|---|---|---|---|---|---|---|---|---|---|---|---|
| KCNQ2 | 328 | H | -31.695925 | 13.21171109 | 0.226027337 | 8.950968713 | NA | NA | NA | NA | NA | NA | NA |
| KCNQ2 | 329 | F | -33.98853409 | 11.05398155 | 0.183171078 | 7.627309686 | NA | NA | NA | NA | NA | NA | NA |
| KCNQ2 | 330 | E | -31.03129167 | 17.95670028 | 0.32027107 | 13.84768268 | NA | NA | NA | NA | NA | NA | NA |
| KCNQ2 | 331 | K | -32.45873611 | 19.5017019 | 0.350957488 | 15.57372605 | NA | NA | NA | NA | NA | NA | NA |
| KCNQ2 | 332 | R | -37.91844318 | 14.94622748 | 0.260477848 | 11.94137994 | NA | NA | NA | NA | NA | NA | NA |
| KCNQ2 | 333 | R | -35.06171591 | 15.87663595 | 0.278957377 | 12.98745092 | NA | NA | NA | NA | NA | NA | NA |
| KCNQ2 | 334 | N | -35.08125 | 23.32586474 | 0.426912009 | 20.07669737 | NA | NA | NA | NA | NA | NA | NA |
| KCNQ2 | 335 | P | -39.165625 | 23.15214275 | 0.423461589 | 20.1439876 | NA | NA | NA | NA | NA | NA | NA |
| KCNQ2 | 336 | A | -40.680625 | 22.68676365 | 0.414218351 | 19.45689823 | NA | NA | NA | NA | NA | NA | NA |
| KCNQ2 | 337 | A | -38.422625 | 25.35361603 | 0.467186675 | 22.27713233 | NA | NA | NA | NA | NA | NA | NA |
| KCNQ2 | 338 | G | -40.729125 | 27.96313525 | 0.519016262 | 24.95426705 | NA | NA | NA | NA | NA | NA | NA |
| KCNQ2 | 339 | L | -44.810125 | 25.78489876 | 0.475752699 | 22.22610912 | NA | NA | NA | NA | NA | NA | NA |
| KCNQ2 | 340 | I | -43.64625 | 27.27232293 | 0.505295529 | 23.4851868 | NA | NA | NA | NA | NA | NA | NA |
| KCNQ2 | 341 | Q | -41.39851389 | 31.28776376 | 0.585049163 | 27.49839205 | NA | NA | NA | NA | NA | NA | NA |
| KCNQ2 | 342 | S | -45.977125 | 31.44771993 | 0.588226171 | 29.98777612 | NA | NA | NA | NA | NA | NA | NA |
| KCNQ2 | 343 | A | -47.972625 | 30.59317638 | 0.571253451 | 29.40482767 | NA | NA | NA | NA | NA | NA | NA |
| KCNQ2 | 344 | W | -46.46576786 | 33.48004624 | 0.628591704 | 32.29337821 | NA | NA | NA | NA | NA | NA | NA |
| KCNQ2 | 345 | R | -45.18198864 | 35.85709308 | 0.675803986 | 34.67493255 | NA | NA | NA | NA | NA | NA | NA |
| KCNQ2 | 346 | F | -52.61753409 | 34.11075558 | 0.641118688 | 32.57989487 | NA | NA | NA | NA | NA | NA | NA |
| KCNQ2 | 347 | Y | -52.32104167 | 35.90778475 | 0.676810811 | 34.30638694 | NA | NA | NA | NA | NA | NA | NA |
| KCNQ2 | 348 | A | -50.309225 | 38.60219935 | 0.730326568 | 37.02574636 | NA | NA | NA | NA | NA | NA | NA |
| KCNQ2 | 349 | T | -52.74448214 | 38.81909955 | 0.734634583 | 37.20471037 | NA | NA | NA | NA | NA | NA | NA |
| KCNQ2 | 350 | N | -56.222375 | 41.32133069 | 0.784333243 | 39.80359583 | NA | NA | NA | NA | NA | NA | NA |
| KCNQ2 | 351 | L | -57.40425 | 43.98063975 | 0.837151744 | 42.46198414 | NA | NA | NA | NA | NA | NA | NA |
| KCNQ2 | 352 | S | -57.60529167 | 43.41743422 | 0.825965503 | 41.93934506 | NA | NA | NA | NA | NA | NA | NA |
| KCNQ2 | 353 | R | -54.52371591 | 37.8525774 | 0.715437773 | 36.10800696 | NA | NA | NA | NA | NA | NA | NA |
| KCNQ2 | 354 | T | -52.66591071 | 46.71212428 | 0.891403774 | 45.12161531 | NA | NA | NA | NA | NA | NA | NA |
| KCNQ2 | 355 | D | -50.255 | 46.04158767 | 0.878085752 | 44.48396784 | NA | NA | NA | NA | NA | NA | NA |
| KCNQ2 | 356 | L | -49.914875 | 40.88905714 | 0.775747539 | 39.78331706 | NA | NA | NA | NA | NA | NA | NA |
| KCNQ2 | 357 | H | -46.428825 | 46.10276409 | 0.879300822 | 44.91588665 | NA | NA | NA | NA | NA | NA | NA |
| KCNQ2 | 358 | S | -44.58329167 | 43.17800818 | 0.821210085 | 41.58340913 | NA | NA | NA | NA | NA | NA | NA |
| KCNQ2 | 359 | T | -46.12848214 | 40.9243274 | 0.776448068 | 39.45602895 | NA | NA | NA | NA | NA | NA | NA |
| KCNQ2 | 360 | W | -50.005125 | 45.29383758 | 0.863234135 | 44.24578052 | NA | NA | NA | NA | NA | NA | NA |

| | | | | | | | | | | | | | |
|---|---|---|---|---|---|---|---|---|---|---|---|---|---|
| KCNQ2 | 361 | Q | -43.60829167 | 47.74103085 | 0.911839647 | 44.01558281 | NA | NA | NA | NA | NA | NA | NA |
| KCNQ2 | 362 | Y | -41.32379167 | 41.73308861 | 0.792511471 | 38.36882344 | NA | NA | NA | NA | NA | NA | NA |
| KCNQ2 | 363 | Y | -45.24479167 | 40.74436537 | 0.772873709 | 37.05464868 | NA | NA | NA | NA | NA | NA | NA |
| KCNQ2 | 364 | E | -45.80518056 | 49.14253337 | 0.939675923 | 47.54262032 | NA | NA | NA | NA | NA | NA | NA |
| KCNQ2 | 365 | R | -39.19217045 | 45.71511788 | 0.871601494 | 42.64284718 | NA | NA | NA | NA | NA | NA | NA |
| KCNQ2 | 366 | T | -41.20505357 | 46.49752304 | 0.887141421 | 43.30174257 | NA | NA | NA | NA | NA | NA | NA |
| KCNQ2 | 367 | V | -43.78976786 | 50.31625807 | 0.962988136 | 46.52720132 | NA | NA | NA | NA | NA | NA | NA |
| KCNQ2 | 535 | G | -56.830125 | 47.70810512 | 0.911185685 | 46.24805734 | NA | NA | NA | NA | NA | NA | NA |
| KCNQ2 | 536 | L | -53.16 | 47.7545799 | 0.912108755 | 45.89962112 | NA | NA | NA | NA | NA | NA | NA |
| KCNQ2 | 537 | K | -51.96773611 | 45.70163702 | 0.871333741 | 43.84196276 | NA | NA | NA | NA | NA | NA | NA |
| KCNQ2 | 538 | V | -56.30119643 | 43.48383991 | 0.827284435 | 41.97533471 | NA | NA | NA | NA | NA | NA | NA |
| KCNQ2 | 539 | S | -53.777625 | 43.31159693 | 0.82386339 | 41.45094349 | NA | NA | NA | NA | NA | NA | NA |
| KCNQ2 | 540 | I | -50.24875 | 42.1979085 | 0.801743602 | 40.62454682 | NA | NA | NA | NA | NA | NA | NA |
| KCNQ2 | 541 | R | -54.68917045 | 39.77788573 | 0.753677743 | 38.15863544 | NA | NA | NA | NA | NA | NA | NA |
| KCNQ2 | 542 | A | -53.240025 | 38.71269159 | 0.732521136 | 36.81985397 | NA | NA | NA | NA | NA | NA | NA |
| KCNQ2 | 543 | V | -49.54991071 | 38.48081873 | 0.727915738 | 37.07825217 | NA | NA | NA | NA | NA | NA | NA |
| KCNQ2 | 544 | C | -48.81279167 | 36.35513668 | 0.685695998 | 35.17340116 | NA | NA | NA | NA | NA | NA | NA |
| KCNQ2 | 545 | V | -51.84791071 | 34.22010072 | 0.643290472 | 32.64185444 | NA | NA | NA | NA | NA | NA | NA |
| KCNQ2 | 546 | M | -51.487625 | 33.7726399 | 0.634403123 | 32.21448635 | NA | NA | NA | NA | NA | NA | NA |
| KCNQ2 | 547 | R | -46.05617045 | 32.81028218 | 0.615289026 | 31.63434479 | NA | NA | NA | NA | NA | NA | NA |
| KCNQ2 | 548 | F | -49.09480682 | 29.78681245 | 0.555237661 | 28.60889525 | NA | NA | NA | NA | NA | NA | NA |
| KCNQ2 | 549 | L | -51.7 | 28.78888178 | 0.535417024 | 27.23430025 | NA | NA | NA | NA | NA | NA | NA |
| KCNQ2 | 550 | V | -48.209625 | 28.66689005 | 0.532994056 | 27.48999899 | NA | NA | NA | NA | NA | NA | NA |
| KCNQ2 | 551 | S | -45.695625 | 27.02652874 | 0.500413629 | 25.42525139 | NA | NA | NA | NA | NA | NA | NA |
| KCNQ2 | 552 | K | -49.80484722 | 24.48738888 | 0.449981898 | 23.29959925 | NA | NA | NA | NA | NA | NA | NA |
| KCNQ2 | 553 | R | -50.15980682 | 23.9537308 | 0.439382521 | 22.81908548 | NA | NA | NA | NA | NA | NA | NA |
| KCNQ2 | 554 | K | -43.88295833 | 23.81352928 | 0.436597875 | 21.88920259 | NA | NA | NA | NA | NA | NA | NA |
| KCNQ2 | 555 | F | -44.35017045 | 20.85632396 | 0.377862637 | 17.09782689 | NA | NA | NA | NA | NA | NA | NA |
| KCNQ2 | 556 | K | -47.37195833 | 19.30075286 | 0.346966291 | 17.86672736 | NA | NA | NA | NA | NA | NA | NA |
| KCNQ2 | 557 | E | -43.63584722 | 20.95890162 | 0.379900007 | 17.18731362 | NA | NA | NA | NA | NA | NA | NA |
| KCNQ2 | 558 | S | -40.59145833 | 19.78904083 | 0.356664539 | 16.59889203 | NA | NA | NA | NA | NA | NA | NA |
| KCNQ2 | 559 | L | -40.6325 | 14.69838811 | 0.255555327 | 11.86333875 | NA | NA | NA | NA | NA | NA | NA |
| KCNQ2 | 560 | R | -38.50417045 | 15.08584983 | 0.26325099 | 12.14342163 | NA | NA | NA | NA | NA | NA | NA |

| | | | | | | | | | | | | | |
|---|---|---|---|---|---|---|---|---|---|---|---|---|---|
| KCNQ2 | 561 | P | -37.01248214 | 12.86037359 | 0.219049164 | 9.901145094 | NA | NA | NA | NA | NA | NA | NA |
| KCNQ2 | 562 | Y | -34.53595833 | 7.635160575 | 0.115267351 | 5.150256755 | NA | NA | NA | NA | NA | NA | NA |
| KCNQ2 | 563 | D | -35.303 | 6.412541618 | 0.090984013 | 3.102938654 | 3.31 | 1.13 | 4.2 | 0.13 | -1.3 | 0 | 0 |
| KCNQ2 | 564 | V | -37.45519643 | 4.598842246 | 0.054960791 | 1.630118197 | 3 | 1.13 | 4.2 | 0.13 | -1.3 | 0 | 0 |
| KCNQ2 | 565 | M | -39.046125 | 7.564654255 | 0.113866973 | 4.578344929 | 3.01 | 1.13 | 4.2 | 0.13 | -1.3 | 0 | 0 |
| KCNQ2 | 566 | D | -39.733 | 8.674045942 | 0.13590142 | 5.561842717 | NA | NA | NA | NA | NA | NA | NA |
| KCNQ2 | 567 | V | -41.216625 | 4.602669769 | 0.055036812 | 1.543960618 | 3.23 | 1.13 | 4.2 | 0.13 | -1.3 | 0 | 0 |
| KCNQ2 | 568 | I | -42.775 | 6.625709698 | 0.095217902 | 2.872136628 | 3.81 | 1.13 | 3.1 | 0.43 | -1.33 | 0 | 0 |
| KCNQ2 | 569 | E | -43.81651389 | 8.983610299 | 0.142049906 | 7.060029789 | NA | NA | NA | NA | NA | NA | NA |
| KCNQ2 | 570 | Q | -44.183625 | 7.888825388 | 0.120305575 | 6.324663369 | NA | NA | NA | NA | NA | NA | NA |
| KCNQ2 | 571 | Y | -47.705625 | 2.527999011 | 0.013830245 | 1.492543257 | 1.47 | 1.11 | -1.3 | 1.61 | -1.44 | 0 | 0 |
| KCNQ2 | 572 | S | -48.67429167 | 7.745849276 | 0.117465821 | 6.723258458 | NA | NA | NA | NA | NA | NA | NA |
| KCNQ2 | 573 | A | -50.220225 | 9.431320056 | 0.1509422 | 8.089194831 | NA | NA | NA | NA | NA | NA | NA |
| KCNQ2 | 574 | G | -52.036375 | 6.860594362 | 0.099883119 | 5.479644294 | 1.62 | 0.94 | -1.68 | 11.61 | -1.19 | 0 | 0 |
| KCNQ2 | 575 | H | -52.732825 | 3.107242668 | 0.025335031 | 1.488370551 | 1.62 | 0.26 | -3.2 | 51.6 | -0.2 | 0 | 0 |
| KCNQ2 | 576 | L | -53.443375 | 7.725924217 | 0.117070075 | 5.856932816 | NA | NA | NA | NA | NA | NA | NA |
| KCNQ2 | 577 | D | -56.1355 | 8.72764533 | 0.136965997 | 7.248067427 | NA | NA | NA | NA | NA | NA | NA |
| KCNQ2 | 578 | M | -57.414875 | 2.884743663 | 0.020915814 | 1.614057 | 1.49 | 1 | 1.9 | 1.43 | -0.72 | 0 | 0 |
| KCNQ2 | 579 | L | -58.376875 | 6.787597587 | 0.098433277 | 5.310873449 | 1.51 | 1 | 1.9 | 1.43 | -0.72 | 0 | 0 |
| KCNQ2 | 580 | S | -60.004625 | 9.186690862 | 0.146083439 | 7.022116884 | NA | NA | NA | NA | NA | NA | NA |
| KCNQ2 | 581 | R | -60.11744318 | 7.154656316 | 0.105723701 | 4.914579295 | 2.35 | 1.81 | 4.5 | 0.13 | -1.85 | 0 | 0 |
| KCNQ2 | 582 | I | -62.556 | 4.040139354 | 0.04386398 | 1.521702801 | 2.61 | 1.81 | 4.5 | 0.13 | -1.85 | 0 | 0 |
| KCNQ2 | 583 | K | -63.33506944 | 7.713060806 | 0.116814585 | 4.887208524 | 2.9 | 1.81 | 4.5 | 0.13 | -1.85 | 0 | 0 |
| KCNQ2 | 584 | S | -66.30579167 | 8.352359008 | 0.129512158 | 5.254068582 | NA | NA | NA | NA | NA | NA | NA |
| KCNQ2 | 585 | L | -67.51825 | 4.354372974 | 0.050105206 | 1.423804342 | 3.31 | 1.14 | 3.8 | 0.13 | -1.79 | 0 | 0 |
| KCNQ2 | 586 | Q | -66.71129167 | 6.622634748 | 0.095156828 | 3.325131676 | 3.31 | 1.14 | 3.8 | 0.13 | -1.79 | 0 | 0 |
| KCNQ2 | 587 | S | -70.201125 | 9.014342017 | 0.142660292 | 5.432338208 | NA | NA | NA | NA | NA | NA | NA |
| KCNQ2 | 588 | R | -70.38044318 | 6.742393121 | 0.097535437 | 3.29691645 | 3.52 | 1.13 | 4.2 | 0.13 | -1.3 | 0 | 0 |
| KCNQ2 | 589 | V | -72.69433929 | 4.859582492 | 0.060139546 | 1.35828686 | 3.55 | 1.27 | 4.26 | 0.13 | -1.41 | 0 | 0 |
| KCNQ2 | 590 | D | -72.89225 | 8.000493172 | 0.122523491 | 4.480245772 | 3.54 | 1.4 | 4.32 | 0.13 | -1.52 | 0 | 0 |
| KCNQ2 | 591 | Q | -75.56951389 | 8.719569542 | 0.136805598 | 5.614418961 | NA | NA | NA | NA | NA | NA | NA |
| KCNQ2 | 592 | I | -77.735375 | 3.932764295 | 0.041731325 | 1.68941913 | 2.29 | 1.81 | 4.5 | 0.13 | -1.85 | 0 | 0 |
| KCNQ2 | 593 | V | -77.57419643 | 7.302278206 | 0.108655728 | 5.106475674 | NA | NA | NA | NA | NA | NA | NA |

| | | | | | | | | | | | | | |
|---|---|---|---|---|---|---|---|---|---|---|---|---|---|
| KCNQ2 | 594 | G | -80.306125 | 9.678873591 | 0.155859044 | 7.388999777 | NA | NA | NA | NA | NA | NA | NA |
| KCNQ2 | 595 | R | -81.13544318 | 7.083373914 | 0.104307909 | 3.520911203 | 3.63 | -0.25 | 0.34 | 1.35 | 0.25 | 0 | 0 |
| KCNQ2 | 596 | G | -83.451875 | 6.85025474 | 0.099677757 | 3.224699647 | 3.63 | -0.25 | 0.34 | 1.35 | 0.25 | 0 | 0 |
| KCNQ2 | 597 | P | -84.59991071 | 8.971457017 | 0.141808521 | 5.307106743 | NA | NA | NA | NA | NA | NA | NA |
| KCNQ2 | 598 | A | -86.252025 | 9.192591474 | 0.146200636 | 5.479294964 | NA | NA | NA | NA | NA | NA | NA |
| KCNQ2 | 599 | I | -87.329625 | 5.315307517 | 0.069191037 | 1.81978935 | 3.71 | -0.79 | -0.52 | 2.69 | 0.79 | 0 | 0 |
| KCNQ2 | 600 | T | -88.33819643 | 5.324976432 | 0.069383078 | 1.613633368 | 3.71 | -0.79 | -0.52 | 2.69 | 0.79 | 0 | 0 |
| MCOLN1 | 38 | E | -37.43227778 | 37.81449083 | 0.773513082 | 31.18765417 | NA | NA | NA | NA | NA | NA | NA |
| MCOLN1 | 39 | E | -41.54905556 | 41.52484545 | 0.854309676 | 34.90026882 | NA | NA | NA | NA | NA | NA | NA |
| MCOLN1 | 40 | D | -38.08004167 | 43.84744912 | 0.904886638 | 37.211314 | NA | NA | NA | NA | NA | NA | NA |
| MCOLN1 | 41 | L | -33.65891667 | 40.80385666 | 0.838609441 | 34.13128584 | NA | NA | NA | NA | NA | NA | NA |
| MCOLN1 | 42 | R | -36.74507576 | 37.53156049 | 0.767351997 | 30.89857615 | NA | NA | NA | NA | NA | NA | NA |
| MCOLN1 | 43 | R | -35.08725758 | 39.71895031 | 0.814984546 | 33.03443831 | NA | NA | NA | NA | NA | NA | NA |
| MCOLN1 | 44 | R | -29.37225758 | 39.64710094 | 0.813419956 | 32.94870927 | NA | NA | NA | NA | NA | NA | NA |
| MCOLN1 | 45 | L | -30.74154167 | 36.42612163 | 0.743279987 | 29.72602716 | NA | NA | NA | NA | NA | NA | NA |
| MCOLN1 | 46 | K | -31.85916667 | 33.27138448 | 0.674582505 | 26.61981716 | NA | NA | NA | NA | NA | NA | NA |
| MCOLN1 | 47 | Y | -26.84133333 | 37.57733365 | 0.768348752 | 30.87959281 | NA | NA | NA | NA | NA | NA | NA |
| MCOLN1 | 48 | F | -23.79107576 | 35.96350934 | 0.733206152 | 31.36466847 | NA | NA | NA | NA | NA | NA | NA |
| MCOLN1 | 49 | F | -25.35289394 | 32.09576139 | 0.648982165 | 27.45328919 | NA | NA | NA | NA | NA | NA | NA |
| MCOLN1 | 50 | M | -26.60491667 | 33.10485697 | 0.670956206 | 28.2647166 | NA | NA | NA | NA | NA | NA | NA |
| MCOLN1 | 51 | S | -23.54666667 | 32.07800779 | 0.648595563 | 28.05800021 | NA | NA | NA | NA | NA | NA | NA |
| MCOLN1 | 52 | P | -20.6577381 | 33.78916874 | 0.685857763 | 30.01321928 | NA | NA | NA | NA | NA | NA | NA |
| MCOLN1 | 53 | C | -22.72116667 | 34.69555484 | 0.705595205 | 30.77515507 | NA | NA | NA | NA | NA | NA | NA |
| MCOLN1 | 54 | D | -26.64629167 | 33.69086568 | 0.683717118 | 28.7961844 | NA | NA | NA | NA | NA | NA | NA |
| MCOLN1 | 55 | K | -23.62561111 | 37.51017799 | 0.766886372 | 32.86103883 | NA | NA | NA | NA | NA | NA | NA |
| MCOLN1 | 56 | F | -22.34934848 | 39.86331347 | 0.818128195 | 35.84634931 | NA | NA | NA | NA | NA | NA | NA |
| MCOLN1 | 57 | R | -26.54771212 | 36.11307168 | 0.736463018 | 31.17003761 | NA | NA | NA | NA | NA | NA | NA |
| MCOLN1 | 58 | A | -28.59096667 | 41.02024722 | 0.843321557 | 34.32182928 | NA | NA | NA | NA | NA | NA | NA |
| MCOLN1 | 59 | K | -28.34461111 | 43.39179652 | 0.894964358 | 38.38234969 | NA | NA | NA | NA | NA | NA | NA |
| MCOLN1 | 60 | G | -23.90416667 | 45.82447091 | 0.947938218 | 41.72631141 | NA | NA | NA | NA | NA | NA | NA |
| MCOLN1 | 61 | R | -19.73762121 | 44.80124305 | 0.925656432 | 41.24104686 | NA | NA | NA | NA | NA | NA | NA |
| MCOLN1 | 62 | K | -16.6595 | 44.67547673 | 0.922917748 | 41.35807163 | NA | NA | NA | NA | NA | NA | NA |
| MCOLN1 | 63 | P | -16.1787381 | 41.30225691 | 0.849462593 | 37.98363609 | NA | NA | NA | NA | NA | NA | NA |

| | | | | | | | | | | | | | |
|---|---|---|---|---|---|---|---|---|---|---|---|---|---|
| MCOLN1 | 64 | C | -12.47366667 | 42.50487968 | 0.875650878 | 39.88921541 | NA | NA | NA | NA | NA | NA | NA |
| MCOLN1 | 65 | K | -13.17394444 | 37.13831329 | 0.758788655 | 34.52958702 | NA | NA | NA | NA | NA | NA | NA |
| MCOLN1 | 66 | L | -13.35616667 | 36.04825023 | 0.735051468 | 33.50349276 | NA | NA | NA | NA | NA | NA | NA |
| MCOLN1 | 67 | M | -9.206166667 | 39.07041983 | 0.800862162 | 35.45773657 | NA | NA | NA | NA | NA | NA | NA |
| MCOLN1 | 68 | L | -7.269791667 | 38.73394161 | 0.79353502 | 34.89192054 | NA | NA | NA | NA | NA | NA | NA |
| MCOLN1 | 69 | Q | -8.359388889 | 35.15031525 | 0.715498057 | 31.3009508 | NA | NA | NA | NA | NA | NA | NA |
| MCOLN1 | 70 | V | -6.944166667 | 35.38116585 | 0.720525054 | 31.43404035 | NA | NA | NA | NA | NA | NA | NA |
| MCOLN1 | 71 | V | -3.838452381 | 37.46784794 | 0.765964594 | 33.56852808 | NA | NA | NA | NA | NA | NA | NA |
| MCOLN1 | 72 | K | -3.382388889 | 35.33739071 | 0.719571808 | 33.14624564 | NA | NA | NA | NA | NA | NA | NA |
| MCOLN1 | 73 | I | -3.120041667 | 31.26870824 | 0.630972275 | 29.06681304 | NA | NA | NA | NA | NA | NA | NA |
| MCOLN1 | 74 | L | -0.332041667 | 34.45517584 | 0.700360718 | 32.69209458 | NA | NA | NA | NA | NA | NA | NA |
| MCOLN1 | 75 | V | 0.323452381 | 36.32993591 | 0.741185449 | 34.7748726 | NA | NA | NA | NA | NA | NA | NA |
| MCOLN1 | 76 | V | 0.68002381 | 33.18897306 | 0.672787916 | 32.18712061 | NA | NA | NA | NA | NA | NA | NA |
| MCOLN1 | 77 | T | 1.95902381 | 29.93735252 | 0.601980704 | 28.90703804 | NA | NA | NA | NA | NA | NA | NA |
| MCOLN1 | 78 | V | 4.845166667 | 33.43774517 | 0.678205172 | 32.32020579 | NA | NA | NA | NA | NA | NA | NA |
| MCOLN1 | 79 | Q | 6.079944444 | 34.081426 | 0.69222195 | 33.09349718 | NA | NA | NA | NA | NA | NA | NA |
| MCOLN1 | 80 | L | 5.704666667 | 28.38742021 | 0.568229415 | 27.5390613 | NA | NA | NA | NA | NA | NA | NA |
| MCOLN1 | 81 | I | 7.921791667 | 29.54118368 | 0.593353741 | 27.51922842 | NA | NA | NA | NA | NA | NA | NA |
| MCOLN1 | 82 | L | 10.13316667 | 32.95025531 | 0.667589604 | 30.14090138 | NA | NA | NA | NA | NA | NA | NA |
| MCOLN1 | 83 | F | 11.03171212 | 31.61523294 | 0.638518189 | 28.78570052 | NA | NA | NA | NA | NA | NA | NA |
| MCOLN1 | 84 | G | 12.32541667 | 28.57733661 | 0.57236503 | 25.27507486 | NA | NA | NA | NA | NA | NA | NA |
| MCOLN1 | 85 | L | 14.39679167 | 29.99340814 | 0.60320137 | 25.21769331 | NA | NA | NA | NA | NA | NA | NA |
| MCOLN1 | 86 | S | 16.55083333 | 30.93683263 | 0.62374536 | 25.74994971 | NA | NA | NA | NA | NA | NA | NA |
| MCOLN1 | 87 | N | 14.50129167 | 28.12075422 | 0.562422503 | 23.00827176 | NA | NA | NA | NA | NA | NA | NA |
| MCOLN1 | 88 | Q | 16.45727778 | 24.38166364 | 0.481000155 | 17.99056245 | NA | NA | NA | NA | NA | NA | NA |
| MCOLN1 | 89 | L | 20.36354167 | 27.97663064 | 0.559284071 | 19.32849657 | NA | NA | NA | NA | NA | NA | NA |
| MCOLN1 | 90 | A | 20.34076667 | 27.52609333 | 0.549473181 | 18.90309824 | NA | NA | NA | NA | NA | NA | NA |
| MCOLN1 | 91 | V | 19.29988095 | 23.51810794 | 0.462195387 | 14.88194791 | NA | NA | NA | NA | NA | NA | NA |
| MCOLN1 | 92 | T | 21.89645238 | 23.37436838 | 0.459065317 | 15.96105052 | NA | NA | NA | NA | NA | NA | NA |
| MCOLN1 | 93 | F | 26.10925758 | 25.22235037 | 0.499306929 | 18.91341238 | NA | NA | NA | NA | NA | NA | NA |
| MCOLN1 | 94 | R | 22.27862121 | 23.96685534 | 0.471967299 | 17.64244425 | NA | NA | NA | NA | NA | NA | NA |
| MCOLN1 | 95 | E | 23.47861111 | 17.79434219 | 0.337554799 | 11.55838627 | NA | NA | NA | NA | NA | NA | NA |
| MCOLN1 | 96 | E | 26.57972222 | 18.93774086 | 0.362453422 | 12.66950477 | NA | NA | NA | NA | NA | NA | NA |

| | | | | | | | | | | | | | |
|---|---|---|---|---|---|---|---|---|---|---|---|---|---|
| MCOLN1 | 97 | N | 29.25591667 | 22.12835943 | 0.431932256 | 16.3375372 | NA | NA | NA | NA | NA | NA | NA |
| MCOLN1 | 98 | T | 27.80845238 | 19.64570134 | 0.377869952 | 13.58377097 | NA | NA | NA | NA | NA | NA | NA |
| MCOLN1 | 99 | I | 28.49329167 | 16.1441539 | 0.301620338 | 10.43186304 | NA | NA | NA | NA | NA | NA | NA |
| MCOLN1 | 100 | A | 32.44936667 | 19.17518318 | 0.36762396 | 13.51105142 | NA | NA | NA | NA | NA | NA | NA |
| MCOLN1 | 101 | F | 32.53898485 | 19.42422696 | 0.373047131 | 13.76233139 | NA | NA | NA | NA | NA | NA | NA |
| MCOLN1 | 102 | R | 30.02680303 | 15.91744873 | 0.296683612 | 10.25899732 | NA | NA | NA | NA | NA | NA | NA |
| MCOLN1 | 103 | H | 33.94676667 | 12.37119291 | 0.219460429 | 6.692269687 | NA | NA | NA | NA | NA | NA | NA |
| MCOLN1 | 104 | L | 37.59554167 | 17.7907723 | 0.337477061 | 12.15531254 | NA | NA | NA | NA | NA | NA | NA |
| MCOLN1 | 105 | F | 37.01289394 | 18.83888503 | 0.360300739 | 13.20544159 | NA | NA | NA | NA | NA | NA | NA |
| MCOLN1 | 106 | L | 35.16741667 | 14.9013942 | 0.274558032 | 9.285038189 | NA | NA | NA | NA | NA | NA | NA |
| MCOLN1 | 107 | L | 39.77116667 | 13.46059189 | 0.243183155 | 7.867445362 | NA | NA | NA | NA | NA | NA | NA |
| MCOLN1 | 108 | G | 36.46666667 | 9.372340476 | 0.15415749 | 3.749341323 | 5.68 | -0.97 | -0.8 | 1.67 | 1.11 | 0 | 0 |
| MCOLN1 | 109 | Y | 31.62325 | 10.54116569 | 0.1796098 | 4.875773924 | 5.68 | -0.97 | -0.8 | 1.67 | 1.11 | 0 | 0 |
| MCOLN1 | 110 | S | 31.37683333 | 6.914682205 | 0.100639581 | 1.397990544 | 5.68 | -0.97 | -0.8 | 1.67 | 1.11 | 0 | 0 |
| MCOLN1 | 111 | D | 28.64679167 | 8.02986706 | 0.12492382 | 1.979528947 | 6.09 | -0.94 | -0.72 | 2.01 | 1.05 | 0 | 0 |
| MCOLN1 | 112 | G | 27.40716667 | 7.939106814 | 0.122947427 | 1.623877723 | 6.32 | -0.8 | -0.4 | 3.38 | 0.81 | 0 | 0 |
| MCOLN1 | 113 | A | 27.12456667 | 8.561936405 | 0.13651015 | 2.264981747 | 6.32 | -0.8 | -0.4 | 3.38 | 0.81 | 0 | 0 |
| MCOLN1 | 114 | D | 25.54879167 | 11.5807179 | 0.202247064 | 5.279954414 | NA | NA | NA | NA | NA | NA | NA |
| MCOLN1 | 115 | D | 24.11579167 | 11.65440488 | 0.20385167 | 5.346126862 | NA | NA | NA | NA | NA | NA | NA |
| MCOLN1 | 116 | T | 27.07702381 | 10.35975854 | 0.175659482 | 4.132679541 | 6.24 | -0.83 | -0.48 | 3.04 | 0.87 | 0 | 0 |
| MCOLN1 | 117 | F | 31.72289394 | 12.49383228 | 0.222131021 | 7.112003539 | NA | NA | NA | NA | NA | NA | NA |
| MCOLN1 | 118 | A | 32.66736667 | 14.79729012 | 0.272291065 | 9.139535284 | NA | NA | NA | NA | NA | NA | NA |
| MCOLN1 | 119 | A | 35.35536667 | 17.16237717 | 0.323793144 | 11.50455113 | NA | NA | NA | NA | NA | NA | NA |
| MCOLN1 | 120 | Y | 34.55541667 | 14.69620213 | 0.270089775 | 9.022946812 | NA | NA | NA | NA | NA | NA | NA |
| MCOLN1 | 121 | T | 40.54902381 | 18.80512063 | 0.359565487 | 13.21676852 | NA | NA | NA | NA | NA | NA | NA |
| MCOLN1 | 122 | R | 42.91880303 | 22.13369508 | 0.432048445 | 16.54297651 | NA | NA | NA | NA | NA | NA | NA |
| MCOLN1 | 123 | E | 45.53672222 | 20.98428386 | 0.407018893 | 15.42702242 | NA | NA | NA | NA | NA | NA | NA |
| MCOLN1 | 124 | Q | 40.37694444 | 16.04558815 | 0.299473973 | 10.44026533 | NA | NA | NA | NA | NA | NA | NA |
| MCOLN1 | 125 | L | 39.59054167 | 19.87511716 | 0.382865706 | 14.28395905 | NA | NA | NA | NA | NA | NA | NA |
| MCOLN1 | 126 | Y | 44.39066667 | 22.22958337 | 0.434136506 | 16.64461923 | NA | NA | NA | NA | NA | NA | NA |
| MCOLN1 | 127 | Q | 44.23527778 | 17.89020808 | 0.339642373 | 12.35634521 | NA | NA | NA | NA | NA | NA | NA |
| MCOLN1 | 128 | A | 40.05376667 | 18.73005809 | 0.357930927 | 13.12284822 | NA | NA | NA | NA | NA | NA | NA |
| MCOLN1 | 129 | I | 40.81891667 | 21.55514166 | 0.419449879 | 15.9562901 | NA | NA | NA | NA | NA | NA | NA |

| | | | | | | | | | | | | | |
|---|---|---|---|---|---|---|---|---|---|---|---|---|---|
| MCOLN1 | 130 | F | 45.36298485 | 23.01148472 | 0.451163171 | 17.43851155 | NA | NA | NA | NA | NA | NA | NA |
| MCOLN1 | 131 | H | 41.89366667 | 17.5782682 | 0.332849577 | 11.9996662 | NA | NA | NA | NA | NA | NA | NA |
| MCOLN1 | 132 | A | 40.37416667 | 21.63564307 | 0.421202876 | 16.03936969 | NA | NA | NA | NA | NA | NA | NA |
| MCOLN1 | 133 | V | 43.34745238 | 24.49938526 | 0.483563658 | 18.90281728 | NA | NA | NA | NA | NA | NA | NA |
| MCOLN1 | 134 | D | 45.73029167 | 23.54339982 | 0.462746142 | 17.99470375 | NA | NA | NA | NA | NA | NA | NA |
| MCOLN1 | 135 | Q | 42.27872222 | 20.30757556 | 0.39228291 | 14.72198536 | NA | NA | NA | NA | NA | NA | NA |
| MCOLN1 | 136 | Y | 41.33716667 | 25.23428402 | 0.499566796 | 19.63526688 | NA | NA | NA | NA | NA | NA | NA |
| MCOLN1 | 137 | L | 46.10104167 | 27.78369826 | 0.55508278 | 22.19455765 | NA | NA | NA | NA | NA | NA | NA |
| MCOLN1 | 138 | A | 45.29436667 | 25.73014876 | 0.510364735 | 20.15013463 | NA | NA | NA | NA | NA | NA | NA |
| MCOLN1 | 139 | L | 41.14291667 | 26.24800149 | 0.521641484 | 20.64067513 | NA | NA | NA | NA | NA | NA | NA |
| MCOLN1 | 140 | P | 43.47716667 | 28.39879059 | 0.568477016 | 22.79802339 | NA | NA | NA | NA | NA | NA | NA |
| MCOLN1 | 141 | D | 46.28141667 | 26.4404569 | 0.525832388 | 20.85978509 | NA | NA | NA | NA | NA | NA | NA |
| MCOLN1 | 142 | V | 42.94616667 | 23.0392106 | 0.451766929 | 17.45965852 | NA | NA | NA | NA | NA | NA | NA |
| MCOLN1 | 143 | S | 39.58116667 | 24.10631677 | 0.475004208 | 18.49084738 | NA | NA | NA | NA | NA | NA | NA |
| MCOLN1 | 144 | L | 37.49641667 | 21.39476349 | 0.415957488 | 15.75268747 | NA | NA | NA | NA | NA | NA | NA |
| MCOLN1 | 145 | G | 36.59266667 | 25.75445911 | 0.510894117 | 20.10607233 | NA | NA | NA | NA | NA | NA | NA |
| MCOLN1 | 146 | R | 37.85680303 | 28.67651691 | 0.574524778 | 23.02326595 | NA | NA | NA | NA | NA | NA | NA |
| MCOLN1 | 147 | Y | 35.76258333 | 24.94326059 | 0.493229477 | 19.26580824 | NA | NA | NA | NA | NA | NA | NA |
| MCOLN1 | 148 | A | 37.53696667 | 32.36755025 | 0.654900632 | 26.71861189 | NA | NA | NA | NA | NA | NA | NA |
| MCOLN1 | 149 | Y | 40.48625 | 29.94427374 | 0.602131421 | 24.3384764 | NA | NA | NA | NA | NA | NA | NA |
| MCOLN1 | 150 | V | 37.21888095 | 35.05971513 | 0.713525151 | 29.40039399 | NA | NA | NA | NA | NA | NA | NA |
| MCOLN1 | 151 | R | 40.72443939 | 37.30337498 | 0.762383035 | 31.67801781 | NA | NA | NA | NA | NA | NA | NA |
| MCOLN1 | 152 | G | 37.88691667 | 39.48023571 | 0.809786303 | 33.83406116 | NA | NA | NA | NA | NA | NA | NA |
| MCOLN1 | 153 | G | 34.78116667 | 40.69197083 | 0.836173018 | 35.02927461 | NA | NA | NA | NA | NA | NA | NA |
| MCOLN1 | 154 | G | 33.60091667 | 40.29133572 | 0.827448797 | 34.61144139 | NA | NA | NA | NA | NA | NA | NA |
| MCOLN1 | 155 | D | 32.24154167 | 39.36516803 | 0.807280591 | 33.69152139 | NA | NA | NA | NA | NA | NA | NA |
| MCOLN1 | 156 | P | 34.29702381 | 35.71804425 | 0.72786091 | 30.04558813 | NA | NA | NA | NA | NA | NA | NA |
| MCOLN1 | 157 | W | 35.95738095 | 33.50119461 | 0.679586845 | 27.83216991 | NA | NA | NA | NA | NA | NA | NA |
| MCOLN1 | 158 | T | 40.73045238 | 39.90776828 | 0.819096242 | 34.26485214 | NA | NA | NA | NA | NA | NA | NA |
| MCOLN1 | 159 | N | 41.42479167 | 42.91907998 | 0.884670494 | 37.28108162 | NA | NA | NA | NA | NA | NA | NA |
| MCOLN1 | 160 | G | 42.62241667 | 39.24591706 | 0.804683785 | 33.62428829 | NA | NA | NA | NA | NA | NA | NA |
| MCOLN1 | 161 | S | 41.129 | 36.86745909 | 0.752890541 | 31.23625032 | NA | NA | NA | NA | NA | NA | NA |
| MCOLN1 | 162 | G | 39.65266667 | 33.03605143 | 0.669457898 | 27.39287576 | NA | NA | NA | NA | NA | NA | NA |

| | | | | | | | | | | | | | |
|---|---|---|---|---|---|---|---|---|---|---|---|---|---|
| MCOLN1 | 163 | L | 40.41541667 | 28.41915625 | 0.568920498 | 22.78904293 | NA | NA | NA | NA | NA | NA | NA |
| MCOLN1 | 164 | A | 38.61076667 | 31.57400656 | 0.637620444 | 25.92704211 | NA | NA | NA | NA | NA | NA | NA |
| MCOLN1 | 165 | L | 40.17104167 | 29.06230662 | 0.582925726 | 23.45467602 | NA | NA | NA | NA | NA | NA | NA |
| MCOLN1 | 166 | C | 36.858 | 30.91240586 | 0.623213443 | 25.25544567 | NA | NA | NA | NA | NA | NA | NA |
| MCOLN1 | 167 | Q | 39.01516667 | 31.38120747 | 0.633422055 | 25.75142272 | NA | NA | NA | NA | NA | NA | NA |
| MCOLN1 | 168 | R | 31.74589394 | 30.7872368 | 0.620487764 | 25.43907571 | NA | NA | NA | NA | NA | NA | NA |
| MCOLN1 | 169 | Y | 37.75316667 | 34.19453064 | 0.694684914 | 28.53129068 | NA | NA | NA | NA | NA | NA | NA |
| MCOLN1 | 170 | Y | 33.24916667 | 34.28390719 | 0.696631176 | 28.72158763 | NA | NA | NA | NA | NA | NA | NA |
| MCOLN1 | 171 | H | 36.78336667 | 39.55829425 | 0.811486103 | 33.89340006 | NA | NA | NA | NA | NA | NA | NA |
| MCOLN1 | 172 | R | 36.42771212 | 40.00196171 | 0.821147396 | 34.33527663 | NA | NA | NA | NA | NA | NA | NA |
| MCOLN1 | 173 | G | 37.78116667 | 39.64410812 | 0.813354784 | 33.98766626 | NA | NA | NA | NA | NA | NA | NA |
| MCOLN1 | 174 | H | 39.97846667 | 39.41417144 | 0.808347688 | 33.76469336 | NA | NA | NA | NA | NA | NA | NA |
| MCOLN1 | 175 | V | 36.83659524 | 35.13033461 | 0.715062959 | 29.47564363 | NA | NA | NA | NA | NA | NA | NA |
| MCOLN1 | 176 | D | 38.79191667 | 37.757078 | 0.772262861 | 32.10903425 | NA | NA | NA | NA | NA | NA | NA |
| MCOLN1 | 177 | P | 35.68659524 | 34.84971944 | 0.70895229 | 29.18830215 | NA | NA | NA | NA | NA | NA | NA |
| MCOLN1 | 178 | A | 37.45016667 | 38.52165716 | 0.788912319 | 32.86666247 | NA | NA | NA | NA | NA | NA | NA |
| MCOLN1 | 179 | N | 39.39404167 | 41.30540898 | 0.849531232 | 35.65582504 | NA | NA | NA | NA | NA | NA | NA |
| MCOLN1 | 180 | D | 37.38216667 | 44.19893286 | 0.91254054 | 38.5435916 | NA | NA | NA | NA | NA | NA | NA |
| MCOLN1 | 181 | T | 36.83016667 | 43.50715503 | 0.897476402 | 37.85310552 | NA | NA | NA | NA | NA | NA | NA |
| MCOLN1 | 182 | F | 33.87698485 | 40.69455521 | 0.836229295 | 35.02775716 | NA | NA | NA | NA | NA | NA | NA |
| MCOLN1 | 183 | D | 34.75216667 | 42.80360324 | 0.882155875 | 37.13567214 | NA | NA | NA | NA | NA | NA | NA |
| MCOLN1 | 184 | I | 32.12204167 | 38.27805937 | 0.783607739 | 32.59766991 | NA | NA | NA | NA | NA | NA | NA |
| MCOLN1 | 185 | D | 31.31391667 | 40.78812854 | 0.838266946 | 35.30028236 | NA | NA | NA | NA | NA | NA | NA |
| MCOLN1 | 186 | P | 28.40788095 | 39.39600971 | 0.807952199 | 33.59615255 | NA | NA | NA | NA | NA | NA | NA |
| MCOLN1 | 187 | M | 31.93054167 | 39.77626395 | 0.816232607 | 34.34578171 | NA | NA | NA | NA | NA | NA | NA |
| MCOLN1 | 188 | V | 31.93445238 | 36.43608055 | 0.743496852 | 30.81399587 | NA | NA | NA | NA | NA | NA | NA |
| MCOLN1 | 189 | V | 36.46916667 | 36.54304099 | 0.74582602 | 30.8842025 | NA | NA | NA | NA | NA | NA | NA |
| MCOLN1 | 190 | T | 36.15245238 | 36.17374415 | 0.737784221 | 30.50865685 | NA | NA | NA | NA | NA | NA | NA |
| MCOLN1 | 191 | D | 40.83641667 | 34.21711667 | 0.695176747 | 28.58298978 | NA | NA | NA | NA | NA | NA | NA |
| MCOLN1 | 192 | C | 39.89416667 | 35.43351473 | 0.721665002 | 29.79855186 | NA | NA | NA | NA | NA | NA | NA |
| MCOLN1 | 193 | I | 43.35904167 | 33.44423433 | 0.67834648 | 27.82796472 | NA | NA | NA | NA | NA | NA | NA |
| MCOLN1 | 194 | Q | 42.60961111 | 35.44945892 | 0.722012203 | 29.84130155 | NA | NA | NA | NA | NA | NA | NA |
| MCOLN1 | 195 | V | 44.52288095 | 32.15615845 | 0.65029737 | 26.55562964 | NA | NA | NA | NA | NA | NA | NA |

| | | | | | | | | | | | | | |
|---|---|---|---|---|---|---|---|---|---|---|---|---|---|
| MCOLN1 | 196 | D | 45.36216667 | 34.9628134 | 0.711415021 | 29.36413261 | NA | NA | NA | NA | NA | NA | NA |
| MCOLN1 | 197 | P | 45.32430952 | 32.97572263 | 0.66814418 | 27.37888997 | NA | NA | NA | NA | NA | NA | NA |
| MCOLN1 | 198 | P | 48.79488095 | 34.31910579 | 0.69739766 | 28.74156765 | NA | NA | NA | NA | NA | NA | NA |
| MCOLN1 | 199 | E | 47.03683333 | 36.79242209 | 0.751256537 | 31.1822487 | NA | NA | NA | NA | NA | NA | NA |
| MCOLN1 | 200 | R | 49.05907576 | 31.78502761 | 0.642215633 | 26.20646125 | NA | NA | NA | NA | NA | NA | NA |
| MCOLN1 | 201 | P | 51.38530952 | 40.16261329 | 0.82464574 | 34.58358384 | NA | NA | NA | NA | NA | NA | NA |
| MCOLN1 | 202 | P | 54.6657381 | 40.97724492 | 0.84238514 | 35.41469001 | NA | NA | NA | NA | NA | NA | NA |
| MCOLN1 | 203 | P | 54.63216667 | 43.0609301 | 0.887759419 | 37.48579675 | NA | NA | NA | NA | NA | NA | NA |
| MCOLN1 | 204 | P | 58.78616667 | 46.66152233 | 0.96616583 | 41.09980025 | NA | NA | NA | NA | NA | NA | NA |
| MCOLN1 | 205 | P | 57.39145238 | 46.6980975 | 0.96696229 | 41.12964941 | NA | NA | NA | NA | NA | NA | NA |
| MCOLN1 | 216 | S | 51.363 | 33.54172329 | 0.680469397 | 27.98955213 | NA | NA | NA | NA | NA | NA | NA |
| MCOLN1 | 217 | S | 50.1135 | 34.27059401 | 0.696341268 | 28.69868111 | NA | NA | NA | NA | NA | NA | NA |
| MCOLN1 | 218 | Y | 47.45808333 | 28.60190057 | 0.572899934 | 23.03411217 | NA | NA | NA | NA | NA | NA | NA |
| MCOLN1 | 219 | K | 50.2375 | 27.18898801 | 0.542132384 | 21.64251872 | NA | NA | NA | NA | NA | NA | NA |
| MCOLN1 | 220 | N | 51.15541667 | 32.82283076 | 0.66481481 | 27.26254213 | NA | NA | NA | NA | NA | NA | NA |
| MCOLN1 | 221 | L | 45.68716667 | 28.95741599 | 0.58064163 | 23.3588947 | NA | NA | NA | NA | NA | NA | NA |
| MCOLN1 | 222 | T | 45.8337381 | 31.69864738 | 0.64033462 | 26.09924384 | NA | NA | NA | NA | NA | NA | NA |
| MCOLN1 | 223 | L | 41.49104167 | 26.77493113 | 0.533115891 | 21.15732576 | NA | NA | NA | NA | NA | NA | NA |
| MCOLN1 | 224 | K | 41.6695 | 31.70675565 | 0.640511185 | 26.08325463 | NA | NA | NA | NA | NA | NA | NA |
| MCOLN1 | 225 | F | 40.20434848 | 26.29787432 | 0.522727513 | 20.68617377 | NA | NA | NA | NA | NA | NA | NA |
| MCOLN1 | 226 | H | 38.90836667 | 31.18784072 | 0.629211306 | 25.55051801 | NA | NA | NA | NA | NA | NA | NA |
| MCOLN1 | 227 | K | 37.90827778 | 33.05004658 | 0.669762656 | 27.39297537 | NA | NA | NA | NA | NA | NA | NA |
| MCOLN1 | 228 | L | 35.79379167 | 27.53967992 | 0.549769043 | 21.89431719 | NA | NA | NA | NA | NA | NA | NA |
| MCOLN1 | 229 | V | 31.87288095 | 28.7054499 | 0.575154822 | 23.05219177 | NA | NA | NA | NA | NA | NA | NA |
| MCOLN1 | 230 | N | 32.66604167 | 28.04806056 | 0.560839527 | 22.3772246 | NA | NA | NA | NA | NA | NA | NA |
| MCOLN1 | 231 | V | 35.81988095 | 26.16025661 | 0.519730753 | 20.49164973 | NA | NA | NA | NA | NA | NA | NA |
| MCOLN1 | 232 | T | 33.95416667 | 28.02387254 | 0.56031281 | 22.35453418 | NA | NA | NA | NA | NA | NA | NA |
| MCOLN1 | 233 | I | 36.20741667 | 24.85857842 | 0.49138544 | 19.22806174 | NA | NA | NA | NA | NA | NA | NA |
| MCOLN1 | 234 | H | 33.75056667 | 29.10984534 | 0.583960928 | 23.44819059 | NA | NA | NA | NA | NA | NA | NA |
| MCOLN1 | 235 | F | 35.57098485 | 25.47201918 | 0.504743711 | 19.8187019 | NA | NA | NA | NA | NA | NA | NA |
| MCOLN1 | 236 | R | 33.62962121 | 31.24598326 | 0.630477416 | 25.57844783 | NA | NA | NA | NA | NA | NA | NA |
| MCOLN1 | 237 | L | 34.45516667 | 27.22001435 | 0.542808013 | 21.56172863 | NA | NA | NA | NA | NA | NA | NA |
| MCOLN1 | 238 | K | 33.71394444 | 32.39889788 | 0.655583258 | 26.73076179 | NA | NA | NA | NA | NA | NA | NA |

| | | | | | | | | | | | | | |
|---|---|---|---|---|---|---|---|---|---|---|---|---|---|
| MCOLN1 | 239 | T | 31.62845238 | 30.85492429 | 0.621961726 | 25.21505323 | NA | NA | NA | NA | NA | NA | NA |
| MCOLN1 | 240 | I | 31.66966667 | 33.04090236 | 0.669563532 | 27.55230725 | NA | NA | NA | NA | NA | NA | NA |
| MCOLN1 | 241 | N | 26.88466667 | 30.95317307 | 0.624101189 | 24.86215574 | NA | NA | NA | NA | NA | NA | NA |
| MCOLN1 | 242 | L | 29.64604167 | 32.97083831 | 0.668037819 | 26.88306505 | NA | NA | NA | NA | NA | NA | NA |
| MCOLN1 | 243 | Q | 24.61338889 | 27.11646275 | 0.540553076 | 20.83819712 | NA | NA | NA | NA | NA | NA | NA |
| MCOLN1 | 244 | S | 23.55516667 | 33.32860126 | 0.675828457 | 25.92577983 | NA | NA | NA | NA | NA | NA | NA |
| MCOLN1 | 245 | L | 25.07204167 | 35.39465727 | 0.720818843 | 29.08481209 | NA | NA | NA | NA | NA | NA | NA |
| MCOLN1 | 246 | I | 20.70566667 | 32.96017626 | 0.667805642 | 24.32968071 | NA | NA | NA | NA | NA | NA | NA |
| MCOLN1 | 247 | N | 18.76454167 | 35.13572885 | 0.715180424 | 28.62089734 | NA | NA | NA | NA | NA | NA | NA |
| MCOLN1 | 248 | N | 21.46116667 | 38.51665102 | 0.788803305 | 31.06286107 | NA | NA | NA | NA | NA | NA | NA |
| MCOLN1 | 249 | E | 21.55561111 | 37.90197478 | 0.77541813 | 30.43795853 | NA | NA | NA | NA | NA | NA | NA |
| MCOLN1 | 250 | I | 26.47704167 | 37.97518561 | 0.777012367 | 31.6727269 | NA | NA | NA | NA | NA | NA | NA |
| MCOLN1 | 251 | P | 25.14030952 | 38.90331927 | 0.797223384 | 32.57313249 | NA | NA | NA | NA | NA | NA | NA |
| MCOLN1 | 252 | D | 24.72704167 | 35.1107838 | 0.714637221 | 28.84808711 | NA | NA | NA | NA | NA | NA | NA |
| MCOLN1 | 253 | C | 28.20216667 | 35.95182613 | 0.732951739 | 29.81207679 | NA | NA | NA | NA | NA | NA | NA |
| MCOLN1 | 254 | Y | 26.12275 | 30.98926888 | 0.62488721 | 25.04831882 | NA | NA | NA | NA | NA | NA | NA |
| MCOLN1 | 255 | T | 29.46145238 | 32.26715403 | 0.652714407 | 26.43718843 | NA | NA | NA | NA | NA | NA | NA |
| MCOLN1 | 256 | F | 29.05780303 | 28.08156794 | 0.561569183 | 22.28069519 | NA | NA | NA | NA | NA | NA | NA |
| MCOLN1 | 257 | S | 29.35816667 | 28.35191685 | 0.567456295 | 22.52197298 | NA | NA | NA | NA | NA | NA | NA |
| MCOLN1 | 258 | V | 31.96230952 | 24.71251414 | 0.488204748 | 19.27874485 | NA | NA | NA | NA | NA | NA | NA |
| MCOLN1 | 259 | L | 30.00979167 | 24.3916088 | 0.481216721 | 18.92195121 | NA | NA | NA | NA | NA | NA | NA |
| MCOLN1 | 260 | I | 33.62104167 | 22.18593237 | 0.433185963 | 16.52439015 | NA | NA | NA | NA | NA | NA | NA |
| MCOLN1 | 261 | T | 30.7467381 | 22.16799339 | 0.432795324 | 16.67842547 | NA | NA | NA | NA | NA | NA | NA |
| MCOLN1 | 262 | F | 34.74880303 | 20.81985596 | 0.403438315 | 15.18032741 | NA | NA | NA | NA | NA | NA | NA |
| MCOLN1 | 263 | D | 28.95641667 | 23.43703396 | 0.460429922 | 17.6278137 | NA | NA | NA | NA | NA | NA | NA |
| MCOLN1 | 264 | N | 31.49104167 | 25.62323483 | 0.50803658 | 20.13655716 | NA | NA | NA | NA | NA | NA | NA |
| MCOLN1 | 265 | K | 26.87272222 | 27.09754839 | 0.540141197 | 20.90414711 | NA | NA | NA | NA | NA | NA | NA |
| MCOLN1 | 266 | A | 27.67096667 | 25.3330207 | 0.501716884 | 19.09296486 | NA | NA | NA | NA | NA | NA | NA |
| MCOLN1 | 267 | H | 30.75046667 | 26.93602148 | 0.536623791 | 21.44775819 | NA | NA | NA | NA | NA | NA | NA |
| MCOLN1 | 268 | S | 30.028 | 23.21111408 | 0.455510296 | 17.45718652 | NA | NA | NA | NA | NA | NA | NA |
| MCOLN1 | 269 | G | 33.25291667 | 20.91227568 | 0.405450845 | 15.24504163 | NA | NA | NA | NA | NA | NA | NA |
| MCOLN1 | 270 | R | 29.80280303 | 16.17388695 | 0.302267804 | 11.36343027 | NA | NA | NA | NA | NA | NA | NA |
| MCOLN1 | 271 | I | 33.04516667 | 19.4776822 | 0.374211172 | 13.7974472 | NA | NA | NA | NA | NA | NA | NA |

| | | | | | | | | | | | | | |
|---|---|---|---|---|---|---|---|---|---|---|---|---|---|
| MCOLN1 | 272 | P | 29.50430952 | 19.0822233 | 0.365599668 | 13.26680835 | NA | NA | NA | NA | NA | NA | NA |
| MCOLN1 | 273 | I | 32.36391667 | 16.90389597 | 0.318164464 | 11.70601486 | NA | NA | NA | NA | NA | NA | NA |
| MCOLN1 | 274 | S | 29.29033333 | 19.41444225 | 0.37283406 | 13.5861991 | NA | NA | NA | NA | NA | NA | NA |
| MCOLN1 | 275 | L | 29.93291667 | 20.24193096 | 0.390853434 | 14.4119307 | NA | NA | NA | NA | NA | NA | NA |
| MCOLN1 | 276 | E | 26.50083333 | 21.43852339 | 0.416910403 | 15.23702397 | NA | NA | NA | NA | NA | NA | NA |
| MCOLN1 | 277 | T | 26.39888095 | 24.60599555 | 0.485885201 | 18.39014715 | NA | NA | NA | NA | NA | NA | NA |
| MCOLN1 | 278 | Q | 25.21138889 | 27.90873894 | 0.557805663 | 21.58239797 | NA | NA | NA | NA | NA | NA | NA |
| MCOLN1 | 279 | A | 24.39076667 | 30.00692603 | 0.603495736 | 23.68172875 | NA | NA | NA | NA | NA | NA | NA |
| MCOLN1 | 280 | H | 25.74866667 | 33.15477899 | 0.672043307 | 26.82451692 | NA | NA | NA | NA | NA | NA | NA |
| MCOLN1 | 281 | I | 23.36541667 | 33.43426778 | 0.678129448 | 27.27465192 | NA | NA | NA | NA | NA | NA | NA |
| MCOLN1 | 282 | Q | 25.90716667 | 38.463145 | 0.787638159 | 32.13345348 | NA | NA | NA | NA | NA | NA | NA |
| MCOLN1 | 283 | E | 25.17916667 | 41.35109198 | 0.850526024 | 35.06091819 | NA | NA | NA | NA | NA | NA | NA |
| MCOLN1 | 284 | C | 30.25533333 | 40.13148127 | 0.82396781 | 34.33132031 | NA | NA | NA | NA | NA | NA | NA |
| MCOLN1 | 285 | K | 31.09638889 | 44.69515736 | 0.923346313 | 39.22287101 | NA | NA | NA | NA | NA | NA | NA |
| MCOLN1 | 286 | H | 29.60286667 | 43.91913473 | 0.906447662 | 38.08968239 | NA | NA | NA | NA | NA | NA | NA |
| MCOLN1 | 287 | P | 27.71559524 | 47.42062541 | 0.98269604 | 41.18950779 | NA | NA | NA | NA | NA | NA | NA |
| MCOLN1 | 288 | S | 26.75816667 | 48.20873222 | 0.999857836 | 41.89498215 | NA | NA | NA | NA | NA | NA | NA |
| MCOLN1 | 289 | V | 27.88745238 | 44.53371434 | 0.919830733 | 38.35395138 | NA | NA | NA | NA | NA | NA | NA |
| MCOLN1 | 290 | F | 24.15025758 | 44.15842726 | 0.911658491 | 37.84758961 | NA | NA | NA | NA | NA | NA | NA |
| MCOLN1 | 291 | Q | 21.84583333 | 43.89712812 | 0.905968447 | 36.41915882 | NA | NA | NA | NA | NA | NA | NA |
| MCOLN1 | 292 | H | 20.67916667 | 47.7625291 | 0.990141327 | 39.08854471 | NA | NA | NA | NA | NA | NA | NA |
| MCOLN1 | 293 | G | 16.25291667 | 46.35909297 | 0.959580135 | 41.1022457 | NA | NA | NA | NA | NA | NA | NA |
| MCOLN1 | 294 | D | 16.06216667 | 40.89780171 | 0.840655186 | 35.63954269 | NA | NA | NA | NA | NA | NA | NA |
| MCOLN1 | 295 | N | 16.17104167 | 45.23686613 | 0.93514255 | 39.98930217 | NA | NA | NA | NA | NA | NA | NA |
| MCOLN1 | 296 | S | 12.833 | 46.9967411 | 0.973465546 | 42.47833596 | NA | NA | NA | NA | NA | NA | NA |
| MCOLN1 | 297 | F | 11.69443939 | 44.11639841 | 0.910743272 | 41.29605266 | NA | NA | NA | NA | NA | NA | NA |
| MCOLN1 | 298 | R | 12.5395303 | 36.4485938 | 0.74376934 | 33.75889447 | NA | NA | NA | NA | NA | NA | NA |
| MCOLN1 | 299 | L | 10.13504167 | 44.31663673 | 0.915103656 | 41.49027144 | NA | NA | NA | NA | NA | NA | NA |
| MCOLN1 | 300 | L | 7.409166667 | 45.29007337 | 0.93630119 | 44.30102133 | NA | NA | NA | NA | NA | NA | NA |
| MCOLN1 | 301 | F | 5.579075758 | 38.74056594 | 0.793679271 | 37.80676299 | NA | NA | NA | NA | NA | NA | NA |
| MCOLN1 | 302 | D | 6.439041667 | 41.46689775 | 0.853047808 | 40.38394026 | NA | NA | NA | NA | NA | NA | NA |
| MCOLN1 | 303 | V | 4.065309524 | 44.25747807 | 0.913815419 | 42.67610363 | NA | NA | NA | NA | NA | NA | NA |
| MCOLN1 | 304 | V | 1.255309524 | 43.18938975 | 0.890556753 | 42.2114583 | NA | NA | NA | NA | NA | NA | NA |

| | | | | | | | | | | | | | |
|---|---|---|---|---|---|---|---|---|---|---|---|---|---|
| MCOLN1 | 305 | V | 1.22202381 | 38.84196257 | 0.795887282 | 37.81303173 | NA | NA | NA | NA | NA | NA | NA |
| MCOLN1 | 306 | I | 1.304291667 | 42.41783109 | 0.87375531 | 40.86144868 | NA | NA | NA | NA | NA | NA | NA |
| MCOLN1 | 307 | L | -0.675041667 | 44.43247638 | 0.917626178 | 42.68177825 | NA | NA | NA | NA | NA | NA | NA |
| MCOLN1 | 308 | T | -2.379309524 | 40.81000828 | 0.838743399 | 38.62532934 | NA | NA | NA | NA | NA | NA | NA |
| MCOLN1 | 309 | C | -2.521333333 | 38.65829294 | 0.791887696 | 36.48046859 | NA | NA | NA | NA | NA | NA | NA |
| MCOLN1 | 310 | S | -3.7775 | 42.88901554 | 0.884015811 | 38.92203141 | NA | NA | NA | NA | NA | NA | NA |
| MCOLN1 | 311 | L | -6.949291667 | 43.62843405 | 0.900117371 | 39.59585587 | NA | NA | NA | NA | NA | NA | NA |
| MCOLN1 | 312 | S | -7.543166667 | 39.70367585 | 0.81465193 | 35.86572413 | NA | NA | NA | NA | NA | NA | NA |
| MCOLN1 | 313 | F | -6.962984848 | 41.34350281 | 0.850360763 | 37.47766183 | NA | NA | NA | NA | NA | NA | NA |
| MCOLN1 | 314 | L | -9.808166667 | 43.92225929 | 0.906515703 | 40.05927641 | NA | NA | NA | NA | NA | NA | NA |
| MCOLN1 | 315 | L | -12.19216667 | 42.02024386 | 0.86509746 | 39.42090848 | NA | NA | NA | NA | NA | NA | NA |
| MCOLN1 | 316 | C | -12.201 | 38.43543854 | 0.787034824 | 35.80991854 | NA | NA | NA | NA | NA | NA | NA |
| MCOLN1 | 317 | A | -12.86276667 | 42.51949299 | 0.875969097 | 39.89053733 | NA | NA | NA | NA | NA | NA | NA |
| MCOLN1 | 318 | R | -15.94480303 | 43.42683032 | 0.895727253 | 40.4359724 | NA | NA | NA | NA | NA | NA | NA |
| MCOLN1 | 319 | S | -16.62716667 | 39.213788 | 0.803984143 | 35.68282383 | NA | NA | NA | NA | NA | NA | NA |
| MCOLN1 | 320 | L | -15.68991667 | 38.96729669 | 0.798616554 | 35.6652188 | NA | NA | NA | NA | NA | NA | NA |
| MCOLN1 | 321 | L | -18.32191667 | 43.51925233 | 0.897739833 | 40.20847247 | NA | NA | NA | NA | NA | NA | NA |
| MCOLN1 | 322 | R | -21.64271212 | 42.22084657 | 0.86946578 | 38.43166039 | NA | NA | NA | NA | NA | NA | NA |
| MCOLN1 | 323 | G | -21.30016667 | 40.73072389 | 0.837016904 | 36.94343635 | NA | NA | NA | NA | NA | NA | NA |
| MCOLN1 | 324 | F | -20.01225758 | 42.77202031 | 0.881468126 | 38.80558857 | NA | NA | NA | NA | NA | NA | NA |
| MCOLN1 | 325 | L | -24.61779167 | 43.92252522 | 0.906521494 | 39.83478189 | NA | NA | NA | NA | NA | NA | NA |
| MCOLN1 | 326 | L | -26.82691667 | 38.46318492 | 0.787639029 | 33.50343121 | NA | NA | NA | NA | NA | NA | NA |
| MCOLN1 | 327 | Q | -24.07961111 | 39.98690242 | 0.820819465 | 35.55354077 | NA | NA | NA | NA | NA | NA | NA |
| MCOLN1 | 328 | N | -27.33491667 | 44.28745593 | 0.914468216 | 39.28905491 | NA | NA | NA | NA | NA | NA | NA |
| MCOLN1 | 329 | E | -30.40027778 | 41.87827908 | 0.862006038 | 35.19305871 | NA | NA | NA | NA | NA | NA | NA |
| MCOLN1 | 330 | F | -29.92507576 | 40.98086982 | 0.842464075 | 34.29998839 | NA | NA | NA | NA | NA | NA | NA |
| MCOLN1 | 331 | V | -32.81102381 | 36.62512172 | 0.747613408 | 29.95789971 | NA | NA | NA | NA | NA | NA | NA |
| MCOLN1 | 332 | G | -35.27016667 | 40.57578953 | 0.833643057 | 33.90911141 | NA | NA | NA | NA | NA | NA | NA |
| MCOLN1 | 333 | F | -36.87171212 | 42.02757582 | 0.86525712 | 35.38195116 | NA | NA | NA | NA | NA | NA | NA |
| MCOLN1 | 334 | M | -36.93116667 | 39.1895966 | 0.803457352 | 32.53638227 | NA | NA | NA | NA | NA | NA | NA |
| MCOLN1 | 335 | W | -38.52938095 | 34.59660092 | 0.703440387 | 27.96422385 | NA | NA | NA | NA | NA | NA | NA |
| MCOLN1 | 336 | R | -42.79462121 | 42.31237267 | 0.87145885 | 35.68701011 | NA | NA | NA | NA | NA | NA | NA |
| MCOLN1 | 337 | Q | -43.40838889 | 38.69507019 | 0.792688557 | 32.103263 | NA | NA | NA | NA | NA | NA | NA |

| | | | | | | | | | | | | | |
|---|---|---|---|---|---|---|---|---|---|---|---|---|---|
| MCOLN1 | 338 | R | -41.68934848 | 39.58292669 | 0.812022499 | 32.96930608 | NA | NA | NA | NA | NA | NA | NA |
| MCOLN1 | 339 | G | -40.67491667 | 36.42128412 | 0.743174645 | 29.81565137 | NA | NA | NA | NA | NA | NA | NA |
| MCOLN1 | 340 | R | -37.71616667 | 29.95692611 | 0.602406938 | 23.3467095 | NA | NA | NA | NA | NA | NA | NA |
| MCOLN1 | 341 | V | -36.72502381 | 35.14779383 | 0.715443151 | 28.4956537 | NA | NA | NA | NA | NA | NA | NA |
| MCOLN1 | 342 | I | -33.24379167 | 35.03375602 | 0.712959866 | 28.37619445 | NA | NA | NA | NA | NA | NA | NA |
| MCOLN1 | 343 | S | -29.64133333 | 33.64011941 | 0.682612068 | 26.94335801 | NA | NA | NA | NA | NA | NA | NA |
| MCOLN1 | 344 | L | -27.92116667 | 36.31283175 | 0.740812989 | 31.32471038 | NA | NA | NA | NA | NA | NA | NA |
| MCOLN1 | 345 | W | -23.69616667 | 29.28506097 | 0.587776419 | 24.82171256 | NA | NA | NA | NA | NA | NA | NA |
| MCOLN1 | 346 | E | -27.45505556 | 30.47746395 | 0.613742158 | 26.10512714 | NA | NA | NA | NA | NA | NA | NA |
| MCOLN1 | 347 | R | -24.74916667 | 34.42968186 | 0.699805562 | 29.78066348 | NA | NA | NA | NA | NA | NA | NA |
| MCOLN1 | 348 | L | -21.19116667 | 34.05244466 | 0.691590853 | 30.08394152 | NA | NA | NA | NA | NA | NA | NA |
| MCOLN1 | 349 | E | -22.61361111 | 28.30591465 | 0.566454552 | 24.39635171 | NA | NA | NA | NA | NA | NA | NA |
| MCOLN1 | 350 | F | -23.34125758 | 32.78116369 | 0.663907469 | 28.79084495 | NA | NA | NA | NA | NA | NA | NA |
| MCOLN1 | 351 | V | -19.25702381 | 32.78830432 | 0.664062963 | 29.28976259 | NA | NA | NA | NA | NA | NA | NA |
| MCOLN1 | 352 | N | -17.66066667 | 29.88413474 | 0.600821835 | 26.56808486 | NA | NA | NA | NA | NA | NA | NA |
| MCOLN1 | 353 | G | -14.21741667 | 30.68797232 | 0.618326183 | 27.68339219 | NA | NA | NA | NA | NA | NA | NA |
| MCOLN1 | 354 | W | -14.20380952 | 25.14985594 | 0.497728292 | 22.72339199 | NA | NA | NA | NA | NA | NA | NA |
| MCOLN1 | 355 | Y | -14.23158333 | 32.71482485 | 0.662462876 | 30.09924098 | NA | NA | NA | NA | NA | NA | NA |
| MCOLN1 | 356 | I | -11.19041667 | 32.63771991 | 0.660783841 | 29.07917997 | NA | NA | NA | NA | NA | NA | NA |
| MCOLN1 | 357 | L | -8.294291667 | 27.64799121 | 0.552127627 | 23.80795692 | NA | NA | NA | NA | NA | NA | NA |
| MCOLN1 | 358 | L | -8.234416667 | 32.66067717 | 0.661283757 | 28.82157747 | NA | NA | NA | NA | NA | NA | NA |
| MCOLN1 | 359 | V | -7.468738095 | 35.38412126 | 0.720589411 | 31.50456515 | NA | NA | NA | NA | NA | NA | NA |
| MCOLN1 | 360 | T | -4.892166667 | 33.03444751 | 0.669422971 | 29.02945815 | NA | NA | NA | NA | NA | NA | NA |
| MCOLN1 | 361 | S | -3.386333333 | 32.20530857 | 0.651367662 | 30.02600664 | NA | NA | NA | NA | NA | NA | NA |
| MCOLN1 | 362 | D | -2.972541667 | 35.43496439 | 0.72169657 | 33.2422289 | NA | NA | NA | NA | NA | NA | NA |
| MCOLN1 | 363 | V | -1.190738095 | 37.23559546 | 0.76090707 | 35.11565001 | NA | NA | NA | NA | NA | NA | NA |
| MCOLN1 | 364 | L | 0.149291667 | 31.59009696 | 0.637970828 | 30.02672173 | NA | NA | NA | NA | NA | NA | NA |
| MCOLN1 | 365 | T | 0.567595238 | 35.28179821 | 0.718361227 | 33.74158123 | NA | NA | NA | NA | NA | NA | NA |
| MCOLN1 | 366 | I | 0.734541667 | 38.43552086 | 0.787036617 | 37.4247525 | NA | NA | NA | NA | NA | NA | NA |
| MCOLN1 | 367 | S | 3.5875 | 38.13471885 | 0.780486359 | 36.75638831 | NA | NA | NA | NA | NA | NA | NA |
| MCOLN1 | 368 | G | 5.488666667 | 36.65033134 | 0.748162372 | 35.51418362 | NA | NA | NA | NA | NA | NA | NA |
| MCOLN1 | 369 | T | 5.384738095 | 38.29407228 | 0.783956436 | 37.3412022 | NA | NA | NA | NA | NA | NA | NA |
| MCOLN1 | 370 | I | 6.641791667 | 40.87405439 | 0.840138065 | 39.8885392 | NA | NA | NA | NA | NA | NA | NA |

| | | | | | | | | | | | | | |
|---|---|---|---|---|---|---|---|---|---|---|---|---|---|
| MCOLN1 | 371 | M | 9.480416667 | 36.61317354 | 0.747353225 | 34.36407066 | NA | NA | NA | NA | NA | NA | NA |
| MCOLN1 | 372 | K | 10.50905556 | 37.41526861 | 0.764819628 | 34.58508617 | NA | NA | NA | NA | NA | NA | NA |
| MCOLN1 | 373 | I | 11.04441667 | 40.72527547 | 0.836898259 | 37.8952543 | NA | NA | NA | NA | NA | NA | NA |
| MCOLN1 | 374 | G | 13.69416667 | 41.21527515 | 0.84756848 | 36.69990043 | NA | NA | NA | NA | NA | NA | NA |
| MCOLN1 | 375 | I | 15.12991667 | 35.12104966 | 0.71486077 | 29.86145924 | NA | NA | NA | NA | NA | NA | NA |
| MCOLN1 | 376 | E | 15.84827778 | 39.36995984 | 0.807384938 | 34.11062031 | NA | NA | NA | NA | NA | NA | NA |
| MCOLN1 | 377 | A | 16.32516667 | 42.00061444 | 0.86467001 | 36.74441196 | NA | NA | NA | NA | NA | NA | NA |
| MCOLN1 | 378 | K | 16.94683333 | 42.00384635 | 0.864740388 | 35.48400789 | NA | NA | NA | NA | NA | NA | NA |
| MCOLN1 | 379 | N | 18.54691667 | 41.44630394 | 0.852599358 | 32.52222374 | NA | NA | NA | NA | NA | NA | NA |
| MCOLN1 | 380 | L | 15.46279167 | 38.86641036 | 0.796419656 | 33.67477288 | NA | NA | NA | NA | NA | NA | NA |
| MCOLN1 | 381 | A | 17.47296667 | 35.54668494 | 0.724129394 | 29.03365041 | NA | NA | NA | NA | NA | NA | NA |
| MCOLN1 | 382 | S | 14.12983333 | 33.46981186 | 0.678903455 | 28.2341087 | NA | NA | NA | NA | NA | NA | NA |
| MCOLN1 | 383 | Y | 15.44641667 | 30.801138 | 0.620790476 | 25.99314223 | NA | NA | NA | NA | NA | NA | NA |
| MCOLN1 | 384 | D | 12.34816667 | 28.94925873 | 0.580463997 | 25.61349542 | NA | NA | NA | NA | NA | NA | NA |
| MCOLN1 | 385 | V | 9.267166667 | 31.29229504 | 0.631485901 | 29.06901335 | NA | NA | NA | NA | NA | NA | NA |
| MCOLN1 | 386 | C | 8.746 | 31.97524627 | 0.64635783 | 29.72722352 | NA | NA | NA | NA | NA | NA | NA |
| MCOLN1 | 387 | S | 8.464166667 | 28.07525022 | 0.561431609 | 25.82626915 | NA | NA | NA | NA | NA | NA | NA |
| MCOLN1 | 388 | I | 6.004916667 | 27.617868 | 0.551471665 | 26.63105274 | NA | NA | NA | NA | NA | NA | NA |
| MCOLN1 | 389 | L | 4.358041667 | 30.82735493 | 0.621361376 | 29.26162808 | NA | NA | NA | NA | NA | NA | NA |
| MCOLN1 | 390 | L | 4.517791667 | 30.43987401 | 0.612923601 | 28.87417961 | NA | NA | NA | NA | NA | NA | NA |
| MCOLN1 | 391 | G | 2.765666667 | 27.17531012 | 0.541834535 | 25.74836966 | NA | NA | NA | NA | NA | NA | NA |
| MCOLN1 | 392 | T | 0.579595238 | 27.62755384 | 0.551682583 | 26.11115638 | NA | NA | NA | NA | NA | NA | NA |
| MCOLN1 | 393 | S | 0 | 30.02643126 | 0.603920481 | 28.10290377 | NA | NA | NA | NA | NA | NA | NA |
| MCOLN1 | 394 | T | 0.550738095 | 27.92237986 | 0.558102708 | 26.02121912 | NA | NA | NA | NA | NA | NA | NA |
| MCOLN1 | 395 | L | -0.959291667 | 23.04470373 | 0.451886548 | 21.29252917 | NA | NA | NA | NA | NA | NA | NA |
| MCOLN1 | 396 | L | -3.632916667 | 27.25098936 | 0.543482524 | 25.10679765 | NA | NA | NA | NA | NA | NA | NA |
| MCOLN1 | 397 | V | -3.755595238 | 29.37847508 | 0.589810603 | 25.38171378 | NA | NA | NA | NA | NA | NA | NA |
| MCOLN1 | 398 | W | -3.22902381 | 26.69800024 | 0.531440646 | 22.75829217 | NA | NA | NA | NA | NA | NA | NA |
| MCOLN1 | 399 | V | -6.943880952 | 23.75890742 | 0.46743903 | 19.73200319 | NA | NA | NA | NA | NA | NA | NA |
| MCOLN1 | 400 | G | -9.051916667 | 28.02735979 | 0.560388748 | 24.19773679 | NA | NA | NA | NA | NA | NA | NA |
| MCOLN1 | 401 | V | -9.59302381 | 27.60370923 | 0.551163343 | 23.74844397 | NA | NA | NA | NA | NA | NA | NA |
| MCOLN1 | 402 | I | -11.92541667 | 22.93935947 | 0.449592574 | 20.43513326 | NA | NA | NA | NA | NA | NA | NA |
| MCOLN1 | 403 | R | -15.36316667 | 27.52938775 | 0.549544921 | 24.62011203 | NA | NA | NA | NA | NA | NA | NA |

| | | | | | | | | | | | | | |
|---|---|---|---|---|---|---|---|---|---|---|---|---|---|
| MCOLN1 | 404 | Y | -13.78633333 | 29.43720681 | 0.591089543 | 26.42947228 | NA | NA | NA | NA | NA | NA | NA |
| MCOLN1 | 405 | L | -14.66241667 | 25.06081651 | 0.495789372 | 22.21461863 | NA | NA | NA | NA | NA | NA | NA |
| MCOLN1 | 406 | T | -19.45916667 | 27.11730425 | 0.5405714 | 23.71735384 | NA | NA | NA | NA | NA | NA | NA |
| MCOLN1 | 407 | F | -20.64716667 | 25.55667784 | 0.506587236 | 22.00226603 | NA | NA | NA | NA | NA | NA | NA |
| MCOLN1 | 408 | F | -23.02371212 | 24.07168349 | 0.474250035 | 20.09473104 | NA | NA | NA | NA | NA | NA | NA |
| MCOLN1 | 409 | H | -25.34736667 | 21.66117148 | 0.421758782 | 17.55079581 | NA | NA | NA | NA | NA | NA | NA |
| MCOLN1 | 410 | N | -21.42041667 | 19.45598324 | 0.373738656 | 15.50671896 | NA | NA | NA | NA | NA | NA | NA |
| MCOLN1 | 411 | Y | -17.49 | 19.97835223 | 0.38511375 | 16.65798548 | NA | NA | NA | NA | NA | NA | NA |
| MCOLN1 | 412 | N | -19.94716667 | 19.98591739 | 0.385278489 | 16.62667463 | NA | NA | NA | NA | NA | NA | NA |
| MCOLN1 | 413 | I | -22.09491667 | 16.15069082 | 0.301762686 | 12.19644488 | NA | NA | NA | NA | NA | NA | NA |
| MCOLN1 | 414 | L | -17.63591667 | 14.95722705 | 0.275773847 | 11.63833765 | NA | NA | NA | NA | NA | NA | NA |
| MCOLN1 | 415 | I | -16.06616667 | 18.35698714 | 0.349806942 | 14.85406245 | NA | NA | NA | NA | NA | NA | NA |
| MCOLN1 | 416 | A | -19.16896667 | 18.30857425 | 0.348752704 | 14.90828365 | NA | NA | NA | NA | NA | NA | NA |
| MCOLN1 | 417 | T | -18.49845238 | 13.46286025 | 0.243232551 | 10.06371193 | NA | NA | NA | NA | NA | NA | NA |
| MCOLN1 | 418 | L | -14.91729167 | 16.71615964 | 0.314076322 | 13.2079392 | NA | NA | NA | NA | NA | NA | NA |
| MCOLN1 | 419 | R | -17.63562121 | 19.14948075 | 0.367064264 | 15.83789116 | NA | NA | NA | NA | NA | NA | NA |
| MCOLN1 | 420 | V | -21.04930952 | 16.08753984 | 0.300387512 | 12.66780441 | NA | NA | NA | NA | NA | NA | NA |
| MCOLN1 | 421 | A | -17.73436667 | 14.85443759 | 0.273535506 | 11.55258748 | NA | NA | NA | NA | NA | NA | NA |
| MCOLN1 | 422 | L | -15.41491667 | 17.88998963 | 0.339637616 | 14.93732139 | NA | NA | NA | NA | NA | NA | NA |
| MCOLN1 | 423 | P | -18.44102381 | 19.70235877 | 0.379103723 | 16.38232674 | NA | NA | NA | NA | NA | NA | NA |
| MCOLN1 | 424 | S | -17.98016667 | 16.35909233 | 0.306300832 | 13.0688646 | NA | NA | NA | NA | NA | NA | NA |
| MCOLN1 | 425 | V | -13.84830952 | 14.97289301 | 0.276114989 | 11.98460186 | NA | NA | NA | NA | NA | NA | NA |
| MCOLN1 | 426 | M | -13.50679167 | 19.11878461 | 0.366395826 | 16.55785729 | NA | NA | NA | NA | NA | NA | NA |
| MCOLN1 | 427 | R | -17.34443939 | 20.02009955 | 0.386022839 | 17.02260395 | NA | NA | NA | NA | NA | NA | NA |
| MCOLN1 | 428 | F | -12.98616667 | 13.70949153 | 0.248603188 | 11.08459136 | NA | NA | NA | NA | NA | NA | NA |
| MCOLN1 | 429 | C | -9.681833333 | 18.12876548 | 0.344837193 | 14.44866769 | NA | NA | NA | NA | NA | NA | NA |
| MCOLN1 | 430 | C | -10.14716667 | 20.65562279 | 0.399861978 | 16.89067372 | NA | NA | NA | NA | NA | NA | NA |
| MCOLN1 | 431 | C | -10.70133333 | 20.29582198 | 0.392026964 | 16.73955002 | NA | NA | NA | NA | NA | NA | NA |
| MCOLN1 | 432 | V | -7.819595238 | 15.91223671 | 0.296570115 | 12.12501376 | NA | NA | NA | NA | NA | NA | NA |
| MCOLN1 | 433 | A | -5.372966667 | 19.46246575 | 0.373879818 | 15.09335206 | NA | NA | NA | NA | NA | NA | NA |
| MCOLN1 | 434 | V | -6.301880952 | 22.02507487 | 0.429683134 | 17.57598053 | NA | NA | NA | NA | NA | NA | NA |
| MCOLN1 | 435 | I | -5.373291667 | 19.66445608 | 0.378278355 | 15.654709 | NA | NA | NA | NA | NA | NA | NA |
| MCOLN1 | 436 | Y | -0.999166667 | 15.91173583 | 0.296559208 | 13.8767979 | NA | NA | NA | NA | NA | NA | NA |

| | | | | | | | | | | | | | |
|---|---|---|---|---|---|---|---|---|---|---|---|---|---|
| MCOLN1 | 437 | L | -0.780291667 | 21.44170837 | 0.416979759 | 19.70440306 | NA | NA | NA | NA | NA | NA | NA |
| MCOLN1 | 438 | G | -0.304666667 | 22.9432441 | 0.449677165 | 21.16413041 | NA | NA | NA | NA | NA | NA | NA |
| MCOLN1 | 439 | Y | 0.851666667 | 19.37967329 | 0.372076932 | 17.81453744 | NA | NA | NA | NA | NA | NA | NA |
| MCOLN1 | 440 | C | 2.4435 | 20.05897198 | 0.386869324 | 19.04591437 | NA | NA | NA | NA | NA | NA | NA |
| MCOLN1 | 441 | F | 2.682166667 | 23.21114924 | 0.455511061 | 21.84438402 | NA | NA | NA | NA | NA | NA | NA |
| MCOLN1 | 442 | C | 3.786333333 | 23.34585355 | 0.458444379 | 22.03399989 | NA | NA | NA | NA | NA | NA | NA |
| MCOLN1 | 443 | G | 6.309916667 | 20.73434178 | 0.401576161 | 19.7829583 | NA | NA | NA | NA | NA | NA | NA |
| MCOLN1 | 444 | W | 8.478238095 | 21.85382275 | 0.425953952 | 19.8413494 | NA | NA | NA | NA | NA | NA | NA |
| MCOLN1 | 445 | I | 8.775541667 | 24.26652254 | 0.478492845 | 22.02133782 | NA | NA | NA | NA | NA | NA | NA |
| MCOLN1 | 446 | V | 8.737166667 | 23.2685136 | 0.456760226 | 21.17045603 | NA | NA | NA | NA | NA | NA | NA |
| MCOLN1 | 447 | L | 9.473666667 | 18.92613764 | 0.36220075 | 16.39770389 | NA | NA | NA | NA | NA | NA | NA |
| MCOLN1 | 448 | G | 12.90691667 | 20.30452383 | 0.392216455 | 17.08558759 | NA | NA | NA | NA | NA | NA | NA |
| MCOLN1 | 449 | P | 14.77516667 | 21.0852148 | 0.409216763 | 15.88805672 | NA | NA | NA | NA | NA | NA | NA |
| MCOLN1 | 450 | Y | 15.1165 | 18.10264967 | 0.344268496 | 12.85780392 | NA | NA | NA | NA | NA | NA | NA |
| MCOLN1 | 451 | H | 13.59776667 | 16.26802059 | 0.304317656 | 11.39051474 | NA | NA | NA | NA | NA | NA | NA |
| MCOLN1 | 452 | V | 15.0827381 | 12.45219832 | 0.221224401 | 7.206580754 | NA | NA | NA | NA | NA | NA | NA |
| MCOLN1 | 453 | K | 10.75583333 | 8.261850217 | 0.12997548 | 5.108257501 | NA | NA | NA | NA | NA | NA | NA |
| MCOLN1 | 454 | F | 9.003348485 | 15.48417095 | 0.287248565 | 12.83130412 | NA | NA | NA | NA | NA | NA | NA |
| MCOLN1 | 455 | R | 15.55262121 | 15.30222 | 0.283286406 | 11.5911168 | NA | NA | NA | NA | NA | NA | NA |
| MCOLN1 | 456 | S | 10.58133333 | 19.51542964 | 0.375033159 | 16.73072691 | NA | NA | NA | NA | NA | NA | NA |
| MCOLN1 | 457 | L | 6.010791667 | 19.96796983 | 0.384887663 | 19.00803205 | NA | NA | NA | NA | NA | NA | NA |
| MCOLN1 | 458 | S | 7.565 | 17.56519701 | 0.332564939 | 16.57790544 | NA | NA | NA | NA | NA | NA | NA |
| MCOLN1 | 459 | M | 10.06091667 | 15.12498863 | 0.279427019 | 13.08587905 | NA | NA | NA | NA | NA | NA | NA |
| MCOLN1 | 460 | V | 5.833880952 | 15.88514167 | 0.295980094 | 15.08860529 | NA | NA | NA | NA | NA | NA | NA |
| MCOLN1 | 461 | S | 3.1135 | 15.34361603 | 0.284187845 | 13.98747331 | NA | NA | NA | NA | NA | NA | NA |
| MCOLN1 | 462 | E | 6.158833333 | 12.04420936 | 0.212340043 | 10.91625506 | NA | NA | NA | NA | NA | NA | NA |
| MCOLN1 | 463 | C | 5.218833333 | 11.12505883 | 0.192324643 | 10.02031501 | NA | NA | NA | NA | NA | NA | NA |
| MCOLN1 | 464 | L | 1.689416667 | 12.06174332 | 0.212721862 | 11.15268163 | NA | NA | NA | NA | NA | NA | NA |
| MCOLN1 | 465 | F | 0.395712121 | 9.763239524 | 0.162669698 | 8.197465851 | NA | NA | NA | NA | NA | NA | NA |
| MCOLN1 | 466 | S | 2.652333333 | 6.863656897 | 0.099528455 | 5.833992962 | 1.03 | -0.8 | -0.4 | 3.38 | 0.81 | 0 | 0 |
| MCOLN1 | 467 | L | 1.610666667 | 7.692476909 | 0.11757682 | 7.011465467 | NA | NA | NA | NA | NA | NA | NA |
| MCOLN1 | 468 | I | -1.764916667 | 7.115025931 | 0.105002262 | 4.916517717 | 2.21 | -0.8 | -0.4 | 3.38 | 0.81 | 0 | 0 |
| MCOLN1 | 469 | N | -0.840791667 | 3.357968731 | 0.023188673 | 1.604274702 | 1.79 | -0.8 | -0.4 | 3.38 | 0.81 | 0 | 0 |

| | | | | | | | | | | | | | |
|---|---|---|---|---|---|---|---|---|---|---|---|---|---|
| MCOLN1 | 470 | G | 0.982916667 | 2.448102326 | 0.003375444 | 1.532034422 | 1.57 | -0.8 | -0.4 | 3.38 | 0.81 | 0 | 0 |
| MCOLN1 | 471 | D | 4.874666667 | 2.294393602 | 2.83E-05 | 1.580723443 | 1.38 | -0.99 | -2.88 | 40.44 | 2.27 | 4 | -4 |
| MCOLN1 | 472 | D | 7.779791667 | 4.140834336 | 0.040236336 | 3.588160086 | 0.99 | -1.04 | -3.5 | 49.7 | 2.63 | 4 | -4 |
| MCOLN1 | 473 | M | 3.647666667 | 6.429175375 | 0.090067196 | 5.374366815 | 1.14 | -0.99 | -2.88 | 40.44 | 2.27 | 4 | -4 |
| MCOLN1 | 474 | F | 8.470166667 | 7.750766027 | 0.118846123 | 6.7736075 | NA | NA | NA | NA | NA | NA | NA |
| MCOLN1 | 475 | V | 10.60659524 | 5.702773097 | 0.074249077 | 2.885914681 | 2.83 | -1.04 | -3.5 | 49.7 | 2.63 | 5 | -5 |
| MCOLN1 | 476 | T | 8.29802381 | 9.903406434 | 0.165721969 | 7.785398208 | NA | NA | NA | NA | NA | NA | NA |
| MCOLN1 | 477 | F | 6.703348485 | 12.23985658 | 0.216600452 | 10.12966396 | NA | NA | NA | NA | NA | NA | NA |
| MCOLN1 | 478 | A | 12.41116667 | 10.62311677 | 0.181394365 | 7.283631282 | NA | NA | NA | NA | NA | NA | NA |
| MCOLN1 | 479 | A | 13.50296667 | 12.49936874 | 0.222251583 | 7.856990363 | NA | NA | NA | NA | NA | NA | NA |
| MCOLN1 | 480 | M | 11.52016667 | 15.30159162 | 0.283272722 | 11.96231702 | NA | NA | NA | NA | NA | NA | NA |
| MCOLN1 | 481 | Q | 13.74527778 | 15.47758098 | 0.287105062 | 10.6121367 | NA | NA | NA | NA | NA | NA | NA |
| MCOLN1 | 482 | A | 17.45416667 | 16.79702572 | 0.315837259 | 10.27751413 | NA | NA | NA | NA | NA | NA | NA |
| MCOLN1 | 483 | Q | 15.70272222 | 19.19669735 | 0.368092452 | 13.95044836 | NA | NA | NA | NA | NA | NA | NA |
| MCOLN1 | 484 | Q | 14.01183333 | 20.57180337 | 0.398036728 | 15.68565763 | NA | NA | NA | NA | NA | NA | NA |
| MCOLN1 | 485 | G | 17.92366667 | 23.02511216 | 0.451459922 | 16.54632551 | NA | NA | NA | NA | NA | NA | NA |
| MCOLN1 | 486 | R | 19.59062121 | 21.17507164 | 0.411173484 | 15.09869095 | NA | NA | NA | NA | NA | NA | NA |
| MCOLN1 | 487 | S | 14.81733333 | 25.08796494 | 0.496380555 | 19.94077137 | NA | NA | NA | NA | NA | NA | NA |
| MCOLN1 | 488 | S | 12.2915 | 25.68945605 | 0.509478612 | 22.45293682 | NA | NA | NA | NA | NA | NA | NA |
| MCOLN1 | 489 | L | 9.655416667 | 25.93092411 | 0.514736814 | 23.33483326 | NA | NA | NA | NA | NA | NA | NA |
| MCOLN1 | 490 | V | 10.51630952 | 22.54440995 | 0.440992162 | 19.76546958 | NA | NA | NA | NA | NA | NA | NA |
| MCOLN1 | 491 | W | 9.660880952 | 20.26323819 | 0.39131742 | 17.4355318 | NA | NA | NA | NA | NA | NA | NA |
| MCOLN1 | 492 | L | 6.215791667 | 23.52805 | 0.462411885 | 22.61638329 | NA | NA | NA | NA | NA | NA | NA |
| MCOLN1 | 493 | F | 4.776075758 | 21.98130108 | 0.428729917 | 20.84557708 | NA | NA | NA | NA | NA | NA | NA |
| MCOLN1 | 494 | S | 6.3355 | 18.34903131 | 0.349633696 | 17.47282678 | NA | NA | NA | NA | NA | NA | NA |
| MCOLN1 | 495 | Q | 5.090833333 | 17.81195758 | 0.337938392 | 16.83566628 | NA | NA | NA | NA | NA | NA | NA |
| MCOLN1 | 496 | L | 1.132666667 | 20.11039468 | 0.387989103 | 19.1841412 | NA | NA | NA | NA | NA | NA | NA |
| MCOLN1 | 497 | Y | 2.490333333 | 17.73993112 | 0.336369945 | 16.7715454 | NA | NA | NA | NA | NA | NA | NA |
| MCOLN1 | 498 | L | 2.965166667 | 14.46728879 | 0.265104963 | 13.50817132 | NA | NA | NA | NA | NA | NA | NA |
| MCOLN1 | 499 | Y | 1.14425 | 15.25379717 | 0.282231951 | 13.75306372 | NA | NA | NA | NA | NA | NA | NA |
| MCOLN1 | 500 | S | -1.594166667 | 16.28407796 | 0.304667321 | 14.09148047 | NA | NA | NA | NA | NA | NA | NA |
| MCOLN1 | 501 | F | -0.532166667 | 13.58579173 | 0.245909504 | 11.82818214 | NA | NA | NA | NA | NA | NA | NA |
| MCOLN1 | 502 | I | -1.062166667 | 10.3317032 | 0.17504855 | 8.305724793 | NA | NA | NA | NA | NA | NA | NA |

| | | | | | | | | | | | | | |
|---|---|---|---|---|---|---|---|---|---|---|---|---|---|
| MCOLN1 | 503 | S | -4.384666667 | 12.83433481 | 0.229545796 | 8.902244136 | NA | NA | NA | NA | NA | NA | NA |
| MCOLN1 | 504 | L | -6.431916667 | 13.17696858 | 0.237006981 | 8.799136568 | NA | NA | NA | NA | NA | NA | NA |
| MCOLN1 | 505 | F | -5.117984848 | 10.04519522 | 0.168809559 | 5.70306682 | NA | NA | NA | NA | NA | NA | NA |
| MCOLN1 | 506 | I | -4.574166667 | 6.706620684 | 0.096108839 | 2.720791198 | 4.01 | -0.41 | 0.44 | 2.73 | 0.29 | 0 | 0 |
| MCOLN1 | 507 | Y | -6.814416667 | 8.60766577 | 0.137505951 | 5.100148672 | NA | NA | NA | NA | NA | NA | NA |
| MCOLN1 | 508 | M | -10.07804167 | 11.43428682 | 0.199058384 | 7.901433355 | NA | NA | NA | NA | NA | NA | NA |
| MCOLN1 | 509 | V | -9.712166667 | 11.01965449 | 0.19002936 | 7.251254591 | NA | NA | NA | NA | NA | NA | NA |
| MCOLN1 | 510 | L | -9.042916667 | 5.42828472 | 0.068271825 | 1.603633624 | 3.83 | 1.27 | 3.94 | 0.13 | -1.8 | 0 | 0 |
| MCOLN1 | 511 | S | -12.76983333 | 7.265555588 | 0.108280192 | 4.641575362 | 2.63 | 1.81 | 4.5 | 0.13 | -1.85 | 0 | 0 |
| MCOLN1 | 512 | L | -14.67016667 | 9.306112722 | 0.152715316 | 6.433973316 | NA | NA | NA | NA | NA | NA | NA |
| MCOLN1 | 513 | F | -11.60125758 | 9.071444703 | 0.14760519 | 6.484803193 | NA | NA | NA | NA | NA | NA | NA |
| MCOLN1 | 514 | I | -13.24366667 | 4.204730788 | 0.041627744 | 1.777785527 | 2.63 | 1.81 | 4.5 | 0.13 | -1.85 | 0 | 0 |
| MCOLN1 | 515 | A | -17.35896667 | 6.964620521 | 0.101727037 | 3.629770175 | 3.43 | -0.77 | -0.7 | 1.66 | 0.77 | 0 | 0 |
| MCOLN1 | 516 | L | -17.14304167 | 9.284600638 | 0.152246869 | 5.855588242 | NA | NA | NA | NA | NA | NA | NA |
| MCOLN1 | 517 | I | -15.52991667 | 9.332929444 | 0.153299276 | 5.844304245 | NA | NA | NA | NA | NA | NA | NA |
| MCOLN1 | 518 | T | -18.3277381 | 4.907797266 | 0.056937703 | 1.669853035 | 3.32 | -0.77 | -0.7 | 1.66 | 0.77 | 0 | 0 |
| MCOLN1 | 519 | G | -21.28166667 | 7.881232201 | 0.121687151 | 4.238551305 | 3.79 | -0.82 | -1.26 | 11.27 | 1.14 | 1 | -1 |
| MCOLN1 | 520 | A | -20.96196667 | 10.50668211 | 0.178858886 | 6.738353505 | NA | NA | NA | NA | NA | NA | NA |
| MCOLN1 | 521 | Y | -20.70975 | 9.19355236 | 0.150264204 | 5.433247386 | NA | NA | NA | NA | NA | NA | NA |
| MCOLN1 | 522 | D | -23.81916667 | 5.409524009 | 0.067863292 | 1.551379624 | 4.11 | -1.04 | -3.5 | 49.7 | 2.63 | 4 | -4 |
| MCOLN1 | 523 | T | -25.55159524 | 10.28516704 | 0.174035179 | 5.386538691 | NA | NA | NA | NA | NA | NA | NA |
| MCOLN1 | 524 | I | -24.06904167 | 12.88151823 | 0.230573261 | 8.777549816 | NA | NA | NA | NA | NA | NA | NA |
| MCOLN1 | 525 | K | -25.79272222 | 7.488220149 | 0.11312893 | 2.667428321 | 4.97 | -1.04 | -3.5 | 49.7 | 2.63 | 4 | -4 |
| MCOLN1 | 526 | H | -29.47096667 | 8.257234464 | 0.129874968 | 1.566177221 | 6.7 | -0.78 | -3.44 | 50.08 | 2.06 | 4 | -4 |
| PKD2 | 215 | K | -23.43983333 | 39.40353106 | 0.822239479 | 35.32099862 | NA | NA | NA | NA | NA | NA | NA |
| PKD2 | 216 | Y | -22.85175 | 38.328787 | 0.79852453 | 34.24646342 | NA | NA | NA | NA | NA | NA | NA |
| PKD2 | 217 | L | -19.2075 | 38.10342464 | 0.793551759 | 35.30351858 | NA | NA | NA | NA | NA | NA | NA |
| PKD2 | 218 | K | -16.51905556 | 42.76678048 | 0.896451834 | 41.78401161 | NA | NA | NA | NA | NA | NA | NA |
| PKD2 | 219 | S | -17.10633333 | 40.76126502 | 0.85219879 | 39.83811046 | NA | NA | NA | NA | NA | NA | NA |
| PKD2 | 220 | V | -16.13407143 | 37.07595324 | 0.770879913 | 33.98185834 | NA | NA | NA | NA | NA | NA | NA |
| PKD2 | 221 | L | -13.230125 | 39.96574257 | 0.834645053 | 37.07311214 | NA | NA | NA | NA | NA | NA | NA |
| PKD2 | 222 | R | -10.8145 | 41.97149663 | 0.878903362 | 39.51555448 | NA | NA | NA | NA | NA | NA | NA |
| PKD2 | 223 | E | -12.10761111 | 39.35378465 | 0.821141791 | 37.28596354 | NA | NA | NA | NA | NA | NA | NA |

| | | | | | | | | | | | | | |
|---|---|---|---|---|---|---|---|---|---|---|---|---|---|
| PKD2 | 224 | L | -10.47875 | 35.25588542 | 0.730718895 | 33.12582454 | NA | NA | NA | NA | NA | NA | NA |
| PKD2 | 225 | V | -7.809785714 | 39.98883295 | 0.835154558 | 38.63098159 | NA | NA | NA | NA | NA | NA | NA |
| PKD2 | 226 | T | -6.272357143 | 39.86280841 | 0.832373742 | 37.00660816 | NA | NA | NA | NA | NA | NA | NA |
| PKD2 | 227 | Y | -6.10025 | 36.53881349 | 0.759027564 | 33.55325056 | NA | NA | NA | NA | NA | NA | NA |
| PKD2 | 228 | L | -6.046875 | 35.17734276 | 0.728985799 | 32.24754848 | NA | NA | NA | NA | NA | NA | NA |
| PKD2 | 229 | L | -1.864 | 38.91543208 | 0.811469248 | 35.67590447 | NA | NA | NA | NA | NA | NA | NA |
| PKD2 | 230 | F | -0.694681818 | 36.45084616 | 0.757086506 | 33.74193871 | NA | NA | NA | NA | NA | NA | NA |
| PKD2 | 231 | L | -1.107375 | 33.75513197 | 0.697603763 | 31.04559632 | NA | NA | NA | NA | NA | NA | NA |
| PKD2 | 232 | I | 0.615 | 36.06001808 | 0.748462622 | 33.05488201 | NA | NA | NA | NA | NA | NA | NA |
| PKD2 | 233 | V | 2.224928571 | 37.30142854 | 0.775855177 | 34.63462346 | NA | NA | NA | NA | NA | NA | NA |
| PKD2 | 234 | L | 3.290125 | 34.18463071 | 0.707080941 | 32.74898029 | NA | NA | NA | NA | NA | NA | NA |
| PKD2 | 235 | C | 4.219333333 | 32.05129036 | 0.660007355 | 30.64580327 | NA | NA | NA | NA | NA | NA | NA |
| PKD2 | 236 | I | 6.718875 | 35.39560168 | 0.733801828 | 32.97594963 | NA | NA | NA | NA | NA | NA | NA |
| PKD2 | 237 | L | 7.9055 | 34.31196592 | 0.709890678 | 31.9567861 | NA | NA | NA | NA | NA | NA | NA |
| PKD2 | 238 | T | 8.547928571 | 31.07720312 | 0.638513467 | 28.68653695 | NA | NA | NA | NA | NA | NA | NA |
| PKD2 | 239 | Y | 11.97258333 | 32.26632921 | 0.664752332 | 29.00065226 | NA | NA | NA | NA | NA | NA | NA |
| PKD2 | 240 | G | 12.5545 | 34.67411493 | 0.717881739 | 31.3942078 | NA | NA | NA | NA | NA | NA | NA |
| PKD2 | 241 | M | 12.520125 | 34.25950551 | 0.708733104 | 31.10990344 | NA | NA | NA | NA | NA | NA | NA |
| PKD2 | 242 | M | 14.542 | 30.88751639 | 0.634327902 | 27.70284116 | NA | NA | NA | NA | NA | NA | NA |
| PKD2 | 243 | S | 16.65883333 | 29.12385656 | 0.595411564 | 25.94054154 | NA | NA | NA | NA | NA | NA | NA |
| PKD2 | 244 | S | 16.15066667 | 26.20664979 | 0.531041439 | 23.0684596 | NA | NA | NA | NA | NA | NA | NA |
| PKD2 | 245 | N | 20.355125 | 26.1771694 | 0.530390935 | 20.12288944 | NA | NA | NA | NA | NA | NA | NA |
| PKD2 | 246 | V | 19.85092857 | 25.76447314 | 0.521284515 | 17.19588914 | NA | NA | NA | NA | NA | NA | NA |
| PKD2 | 247 | Y | 20.55258333 | 17.55726804 | 0.340187028 | 8.989441203 | NA | NA | NA | NA | NA | NA | NA |
| PKD2 | 248 | Y | 22.23933333 | 21.02529719 | 0.416711418 | 11.38800737 | NA | NA | NA | NA | NA | NA | NA |
| PKD2 | 249 | Y | 25.857 | 24.2176273 | 0.487152324 | 15.33718338 | NA | NA | NA | NA | NA | NA | NA |
| PKD2 | 250 | T | 23.97921429 | 21.62195296 | 0.429877028 | 12.74205115 | NA | NA | NA | NA | NA | NA | NA |
| PKD2 | 251 | R | 25.27377273 | 16.122967 | 0.308538213 | 7.206328422 | NA | NA | NA | NA | NA | NA | NA |
| PKD2 | 252 | M | 29.336625 | 20.43308787 | 0.403643922 | 14.65539806 | NA | NA | NA | NA | NA | NA | NA |
| PKD2 | 253 | M | 29.354375 | 21.82957677 | 0.434458387 | 16.05825916 | NA | NA | NA | NA | NA | NA | NA |
| PKD2 | 254 | S | 27.81716667 | 18.24056822 | 0.355264505 | 11.73575874 | NA | NA | NA | NA | NA | NA | NA |
| PKD2 | 255 | Q | 29.04061111 | 14.3356424 | 0.269099697 | 8.785858288 | NA | NA | NA | NA | NA | NA | NA |
| PKD2 | 256 | L | 33.780875 | 18.11293786 | 0.352448255 | 14.8883085 | NA | NA | NA | NA | NA | NA | NA |

| | | | | | | | | | | | | | |
|---|---|---|---|---|---|---|---|---|---|---|---|---|---|
| PKD2 | 257 | F | 33.38340909 | 18.64799952 | 0.36425475 | 15.47740789 | NA | NA | NA | NA | NA | NA | NA |
| PKD2 | 258 | L | 30.194375 | 15.20104039 | 0.288195284 | 9.768869423 | NA | NA | NA | NA | NA | NA | NA |
| PKD2 | 259 | D | 30.795625 | 11.56062481 | 0.207867072 | 8.345285425 | NA | NA | NA | NA | NA | NA | NA |
| PKD2 | 260 | T | 35.19478571 | 13.33634736 | 0.247049581 | 10.15744065 | NA | NA | NA | NA | NA | NA | NA |
| PKD2 | 261 | P | 37.5125 | 10.78609318 | 0.190776511 | 6.259867152 | NA | NA | NA | NA | NA | NA | NA |
| PKD2 | 262 | V | 41.47392857 | 10.87465788 | 0.192730751 | 7.641943411 | NA | NA | NA | NA | NA | NA | NA |
| PKD2 | 263 | S | 42.86183333 | 12.56039625 | 0.229927699 | 9.310714487 | NA | NA | NA | NA | NA | NA | NA |
| PKD2 | 264 | K | 44.27883333 | 8.971136662 | 0.150728278 | 5.740590103 | NA | NA | NA | NA | NA | NA | NA |
| PKD2 | 265 | T | 42.7695 | 4.719159459 | 0.056905548 | 1.549899688 | 3.27 | -0.77 | -0.7 | 1.66 | 0.77 | 0 | 0 |
| PKD2 | 266 | E | 40.85283333 | 8.188724137 | 0.133463821 | 4.942791608 | 3.27 | -0.77 | -0.7 | 1.66 | 0.77 | 0 | 0 |
| PKD2 | 267 | K | 35.35905556 | 4.802835933 | 0.058751926 | 1.734303753 | 4.33 | -0.48 | -3.26 | 39.93 | 1.32 | 4 | 4 |
| PKD2 | 268 | T | 35.57778571 | 9.699676489 | 0.166803998 | 6.731229011 | NA | NA | NA | NA | NA | NA | NA |
| PKD2 | 269 | N | 32.521125 | 10.82839956 | 0.19171003 | 8.039244577 | NA | NA | NA | NA | NA | NA | NA |
| PKD2 | 270 | F | 31.0915 | 14.85891847 | 0.280646134 | 11.67655822 | NA | NA | NA | NA | NA | NA | NA |
| PKD2 | 271 | K | 27.66394444 | 12.56672193 | 0.230067279 | 7.36315253 | NA | NA | NA | NA | NA | NA | NA |
| PKD2 | 272 | T | 31.06335714 | 10.83871086 | 0.191937556 | 5.073324019 | NA | NA | NA | NA | NA | NA | NA |
| PKD2 | 273 | L | 32.6695 | 15.266597 | 0.289641834 | 12.23655616 | NA | NA | NA | NA | NA | NA | NA |
| PKD2 | 274 | S | 33.82533333 | 17.61650893 | 0.341494218 | 14.39335312 | NA | NA | NA | NA | NA | NA | NA |
| PKD2 | 275 | S | 36.81216667 | 19.42058622 | 0.381302394 | 15.10268068 | NA | NA | NA | NA | NA | NA | NA |
| PKD2 | 276 | M | 37.49775 | 22.54821618 | 0.450315647 | 18.02769361 | NA | NA | NA | NA | NA | NA | NA |
| PKD2 | 277 | E | 40.8815 | 20.36443378 | 0.402129023 | 17.0961064 | NA | NA | NA | NA | NA | NA | NA |
| PKD2 | 278 | D | 38.82775 | 16.43937438 | 0.315519954 | 13.08070828 | NA | NA | NA | NA | NA | NA | NA |
| PKD2 | 279 | F | 36.53931818 | 20.51353819 | 0.405419112 | 15.98421917 | NA | NA | NA | NA | NA | NA | NA |
| PKD2 | 280 | W | 41.33264286 | 21.53192862 | 0.42789058 | 18.26770171 | NA | NA | NA | NA | NA | NA | NA |
| PKD2 | 281 | K | 42.20872222 | 15.2068936 | 0.288324439 | 11.96945419 | NA | NA | NA | NA | NA | NA | NA |
| PKD2 | 282 | F | 36.89540909 | 18.91239712 | 0.37008886 | 14.53577128 | NA | NA | NA | NA | NA | NA | NA |
| PKD2 | 283 | T | 39.73492857 | 21.7079655 | 0.431774952 | 18.2655772 | NA | NA | NA | NA | NA | NA | NA |
| PKD2 | 284 | E | 43.94705556 | 21.40807572 | 0.425157683 | 18.16483056 | NA | NA | NA | NA | NA | NA | NA |
| PKD2 | 285 | G | 42.12025 | 17.96278581 | 0.349135049 | 14.70768203 | NA | NA | NA | NA | NA | NA | NA |
| PKD2 | 286 | S | 39.12083333 | 18.52900499 | 0.361629056 | 15.19031712 | NA | NA | NA | NA | NA | NA | NA |
| PKD2 | 287 | L | 37.361125 | 21.47466177 | 0.426626949 | 16.93573605 | NA | NA | NA | NA | NA | NA | NA |
| PKD2 | 288 | L | 42.895125 | 23.4936027 | 0.471176235 | 20.2252821 | NA | NA | NA | NA | NA | NA | NA |
| PKD2 | 289 | D | 42.730625 | 19.58695206 | 0.384973368 | 16.36006628 | NA | NA | NA | NA | NA | NA | NA |

| | | | | | | | | | | | | | |
|---|---|---|---|---|---|---|---|---|---|---|---|---|---|
| PKD2 | 290 | G | 38.8845 | 22.41735143 | 0.447428029 | 17.91660116 | NA | NA | NA | NA | NA | NA | NA |
| PKD2 | 291 | L | 38.189625 | 25.1429192 | 0.50756951 | 20.60768901 | NA | NA | NA | NA | NA | NA | NA |
| PKD2 | 292 | Y | 42.07841667 | 25.68076387 | 0.519437414 | 22.43790388 | NA | NA | NA | NA | NA | NA | NA |
| PKD2 | 293 | W | 41.7095 | 23.51862502 | 0.471728369 | 20.24949519 | NA | NA | NA | NA | NA | NA | NA |
| PKD2 | 294 | K | 46.44672222 | 27.16810207 | 0.552256529 | 23.93136164 | NA | NA | NA | NA | NA | NA | NA |
| PKD2 | 295 | M | 43.6375 | 29.74458547 | 0.609108364 | 26.50269303 | NA | NA | NA | NA | NA | NA | NA |
| PKD2 | 296 | Q | 41.68194444 | 30.93946861 | 0.635474262 | 27.69940453 | NA | NA | NA | NA | NA | NA | NA |
| PKD2 | 297 | P | 46.08335714 | 32.84205392 | 0.677456084 | 29.59979822 | NA | NA | NA | NA | NA | NA | NA |
| PKD2 | 298 | S | 48.85833333 | 31.12729307 | 0.639618735 | 27.91741713 | NA | NA | NA | NA | NA | NA | NA |
| PKD2 | 299 | N | 50.169125 | 34.45523975 | 0.713052111 | 31.23961691 | NA | NA | NA | NA | NA | NA | NA |
| PKD2 | 300 | Q | 48.7125 | 37.65282073 | 0.783608881 | 34.42700484 | NA | NA | NA | NA | NA | NA | NA |
| PKD2 | 301 | T | 50.30392857 | 38.55281803 | 0.803467925 | 35.33593065 | NA | NA | NA | NA | NA | NA | NA |
| PKD2 | 302 | E | 46.9309 | 37.34539365 | 0.776825296 | 34.11353426 | NA | NA | NA | NA | NA | NA | NA |
| PKD2 | 303 | A | 47.7323 | 39.5801413 | 0.826136502 | 36.34973019 | NA | NA | NA | NA | NA | NA | NA |
| PKD2 | 304 | D | 44.262125 | 39.93807149 | 0.834034472 | 36.6816584 | NA | NA | NA | NA | NA | NA | NA |
| PKD2 | 305 | N | 41.57075 | 36.16409496 | 0.750759148 | 32.89410519 | NA | NA | NA | NA | NA | NA | NA |
| PKD2 | 306 | R | 45.63204545 | 34.17488841 | 0.706865971 | 30.93103871 | NA | NA | NA | NA | NA | NA | NA |
| PKD2 | 307 | S | 41.87333333 | 31.1015953 | 0.639051696 | 27.844913 | NA | NA | NA | NA | NA | NA | NA |
| PKD2 | 308 | F | 38.60395455 | 29.46464459 | 0.602931281 | 24.96968871 | NA | NA | NA | NA | NA | NA | NA |
| PKD2 | 309 | I | 37.077875 | 27.00548394 | 0.548668251 | 22.52339293 | NA | NA | NA | NA | NA | NA | NA |
| PKD2 | 310 | F | 36.40704545 | 24.36052857 | 0.490305536 | 19.9998141 | NA | NA | NA | NA | NA | NA | NA |
| PKD2 | 311 | Y | 37.92333333 | 26.21262995 | 0.531173396 | 21.71100305 | NA | NA | NA | NA | NA | NA | NA |
| PKD2 | 312 | E | 32.36194444 | 30.29055313 | 0.621155507 | 27.2372955 | NA | NA | NA | NA | NA | NA | NA |
| PKD2 | 313 | N | 32.359875 | 28.33950963 | 0.578104423 | 25.32677545 | NA | NA | NA | NA | NA | NA | NA |
| PKD2 | 314 | L | 34.088625 | 32.8236404 | 0.677049777 | 29.59924113 | NA | NA | NA | NA | NA | NA | NA |
| PKD2 | 315 | L | 35.984375 | 31.84015986 | 0.655348619 | 28.78556213 | NA | NA | NA | NA | NA | NA | NA |
| PKD2 | 316 | L | 33.2635 | 34.81663495 | 0.721026539 | 31.5915625 | NA | NA | NA | NA | NA | NA | NA |
| PKD2 | 317 | G | 36.059 | 35.9398 | 0.745809929 | 32.89172877 | NA | NA | NA | NA | NA | NA | NA |
| PKD2 | 318 | V | 37.1065 | 34.25396716 | 0.708610897 | 29.75699259 | NA | NA | NA | NA | NA | NA | NA |
| PKD2 | 319 | P | 36.39878571 | 30.76593366 | 0.631645097 | 27.66642781 | NA | NA | NA | NA | NA | NA | NA |
| PKD2 | 320 | R | 33.19740909 | 32.38356972 | 0.667339322 | 29.15995938 | NA | NA | NA | NA | NA | NA | NA |
| PKD2 | 321 | I | 35.715125 | 30.26994306 | 0.620700732 | 27.35163704 | NA | NA | NA | NA | NA | NA | NA |
| PKD2 | 322 | R | 29.81040909 | 32.10991725 | 0.661300997 | 28.93507763 | NA | NA | NA | NA | NA | NA | NA |

| | | | | | | | | | | | | | |
|---|---|---|---|---|---|---|---|---|---|---|---|---|---|
| PKD2 | 323 | Q | 33.17394444 | 33.27806061 | 0.687076864 | 30.09665909 | NA | NA | NA | NA | NA | NA | NA |
| PKD2 | 324 | L | 29.4425 | 34.51354082 | 0.714338564 | 28.77168979 | NA | NA | NA | NA | NA | NA | NA |
| PKD2 | 325 | R | 33.22340909 | 36.56671182 | 0.759643159 | 30.82778575 | NA | NA | NA | NA | NA | NA | NA |
| PKD2 | 326 | V | 29.78778571 | 37.25452032 | 0.774820115 | 31.4952879 | NA | NA | NA | NA | NA | NA | NA |
| PKD2 | 327 | R | 29.25386364 | 40.84816776 | 0.854116357 | 35.09093821 | NA | NA | NA | NA | NA | NA | NA |
| PKD2 | 328 | N | 34.14225 | 42.03461765 | 0.88029617 | 38.82161861 | NA | NA | NA | NA | NA | NA | NA |
| PKD2 | 329 | G | 31.0295 | 42.66216404 | 0.894143402 | 39.45543011 | NA | NA | NA | NA | NA | NA | NA |
| PKD2 | 330 | S | 27.9455 | 40.63821154 | 0.849483532 | 34.09234563 | NA | NA | NA | NA | NA | NA | NA |
| PKD2 | 331 | C | 27.61116667 | 39.89485462 | 0.833080863 | 33.23519162 | NA | NA | NA | NA | NA | NA | NA |
| PKD2 | 332 | S | 26.81716667 | 44.05010994 | 0.92476936 | 37.50847999 | NA | NA | NA | NA | NA | NA | NA |
| PKD2 | 333 | I | 29.19075 | 42.12287533 | 0.882243635 | 35.72422519 | NA | NA | NA | NA | NA | NA | NA |
| PKD2 | 334 | P | 26.88378571 | 42.8826822 | 0.899009283 | 35.97843125 | NA | NA | NA | NA | NA | NA | NA |
| PKD2 | 335 | Q | 27.1095 | 47.4595073 | 1 | 40.91123019 | NA | NA | NA | NA | NA | NA | NA |
| PKD2 | 336 | D | 28.809375 | 46.82605494 | 0.986022449 | 41.07381099 | NA | NA | NA | NA | NA | NA | NA |
| PKD2 | 337 | L | 31.35475 | 41.40556923 | 0.866415795 | 38.37551741 | NA | NA | NA | NA | NA | NA | NA |
| PKD2 | 338 | R | 31.22686364 | 46.08912021 | 0.969761489 | 42.87942466 | NA | NA | NA | NA | NA | NA | NA |
| PKD2 | 339 | D | 36.591375 | 44.89804839 | 0.943479691 | 40.56946088 | NA | NA | NA | NA | NA | NA | NA |
| PKD2 | 340 | E | 35.82505556 | 41.67904152 | 0.872450144 | 37.35103471 | NA | NA | NA | NA | NA | NA | NA |
| PKD2 | 341 | I | 35.7815 | 38.73475469 | 0.80748248 | 34.40760387 | NA | NA | NA | NA | NA | NA | NA |
| PKD2 | 342 | K | 38.3255 | 41.90388634 | 0.877411496 | 37.58615293 | NA | NA | NA | NA | NA | NA | NA |
| PKD2 | 343 | E | 36.1785 | 39.44514565 | 0.823157733 | 36.26668525 | NA | NA | NA | NA | NA | NA | NA |
| PKD2 | 344 | C | 31.115 | 39.33032127 | 0.820624056 | 36.18722756 | NA | NA | NA | NA | NA | NA | NA |
| PKD2 | 345 | Y | 32.21975 | 36.11877699 | 0.749759177 | 30.34186988 | NA | NA | NA | NA | NA | NA | NA |
| PKD2 | 346 | D | 26.0505 | 36.25377913 | 0.752738089 | 29.35457398 | NA | NA | NA | NA | NA | NA | NA |
| PKD2 | 347 | V | 23.49364286 | 33.83598633 | 0.699387869 | 24.8771329 | NA | NA | NA | NA | NA | NA | NA |
| PKD2 | 348 | Y | 24.074 | 34.87862857 | 0.72239447 | 25.89178697 | NA | NA | NA | NA | NA | NA | NA |
| PKD2 | 349 | S | 21.58866667 | 37.759152 | 0.785955152 | 27.97342735 | NA | NA | NA | NA | NA | NA | NA |
| PKD2 | 350 | V | 21.09807143 | 40.27787021 | 0.841532359 | 30.57519856 | NA | NA | NA | NA | NA | NA | NA |
| PKD2 | 351 | S | 21.8265 | 43.01276631 | 0.901879677 | 33.30500918 | NA | NA | NA | NA | NA | NA | NA |
| PKD2 | 352 | S | 24.6215 | 39.60403957 | 0.826663834 | 30.61651139 | NA | NA | NA | NA | NA | NA | NA |
| PKD2 | 353 | E | 25.18427778 | 39.56763446 | 0.825860531 | 32.77560265 | NA | NA | NA | NA | NA | NA | NA |
| PKD2 | 354 | D | 30.85 | 41.74108821 | 0.873819246 | 35.966511 | NA | NA | NA | NA | NA | NA | NA |
| PKD2 | 355 | R | 28.52995455 | 44.16231672 | 0.927245278 | 38.39428939 | NA | NA | NA | NA | NA | NA | NA |

| | | | | | | | | | | | | | |
|---|---|---|---|---|---|---|---|---|---|---|---|---|---|
| PKD2 | 356 | A | 34.2685 | 43.76232003 | 0.918419082 | 40.54737084 | NA | NA | NA | NA | NA | NA | NA |
| PKD2 | 357 | P | 37.23578571 | 44.24191186 | 0.929001598 | 39.71330425 | NA | NA | NA | NA | NA | NA | NA |
| PKD2 | 358 | F | 39.15368182 | 37.40879669 | 0.778224327 | 33.96387941 | NA | NA | NA | NA | NA | NA | NA |
| PKD2 | 359 | G | 42.702 | 40.65995604 | 0.849963339 | 37.4009352 | NA | NA | NA | NA | NA | NA | NA |
| PKD2 | 360 | P | 45.2625 | 39.62665882 | 0.827162943 | 36.38806523 | NA | NA | NA | NA | NA | NA | NA |
| PKD2 | 361 | R | 42.76722727 | 43.26867052 | 0.907526375 | 40.01990236 | NA | NA | NA | NA | NA | NA | NA |
| PKD2 | 362 | N | 45.597125 | 41.33946712 | 0.864957207 | 38.08433289 | NA | NA | NA | NA | NA | NA | NA |
| PKD2 | 363 | G | 42.42525 | 42.28027678 | 0.885716804 | 39.01524322 | NA | NA | NA | NA | NA | NA | NA |
| PKD2 | 364 | T | 39.17278571 | 39.73409767 | 0.829533653 | 36.28882992 | NA | NA | NA | NA | NA | NA | NA |
| PKD2 | 365 | A | 38.9139 | 37.21696346 | 0.773991398 | 32.68410105 | NA | NA | NA | NA | NA | NA | NA |
| PKD2 | 366 | W | 41.80485714 | 36.63091637 | 0.761059876 | 33.37676888 | NA | NA | NA | NA | NA | NA | NA |
| PKD2 | 367 | I | 37.851875 | 42.45093455 | 0.889482482 | 37.91469286 | NA | NA | NA | NA | NA | NA | NA |
| PKD2 | 368 | Y | 32.01775 | 42.76780271 | 0.89647439 | 39.70754761 | NA | NA | NA | NA | NA | NA | NA |
| PKD2 | 369 | T | 31.41478571 | 42.40432603 | 0.888454034 | 39.23630894 | NA | NA | NA | NA | NA | NA | NA |
| PKD2 | 370 | S | 27.87966667 | 43.99894242 | 0.923640314 | 37.45264936 | NA | NA | NA | NA | NA | NA | NA |
| PKD2 | 371 | E | 24.11261111 | 41.62672884 | 0.87129583 | 34.8799279 | NA | NA | NA | NA | NA | NA | NA |
| PKD2 | 372 | K | 23.30216667 | 45.26317463 | 0.951536446 | 36.28006013 | NA | NA | NA | NA | NA | NA | NA |
| PKD2 | 373 | D | 28.868875 | 45.89403922 | 0.965456897 | 40.11680635 | NA | NA | NA | NA | NA | NA | NA |
| PKD2 | 374 | L | 29.09925 | 40.54903327 | 0.847515754 | 34.79071833 | NA | NA | NA | NA | NA | NA | NA |
| PKD2 | 375 | N | 26.4985 | 42.15256411 | 0.882898738 | 35.27995365 | NA | NA | NA | NA | NA | NA | NA |
| PKD2 | 376 | G | 24.6345 | 40.06620003 | 0.836861715 | 31.08754372 | NA | NA | NA | NA | NA | NA | NA |
| PKD2 | 377 | S | 21.70116667 | 37.37940857 | 0.777575858 | 27.6710369 | NA | NA | NA | NA | NA | NA | NA |
| PKD2 | 378 | S | 20.351 | 35.29132746 | 0.731500948 | 25.60583322 | NA | NA | NA | NA | NA | NA | NA |
| PKD2 | 379 | H | 21.0085 | 28.88694584 | 0.59018397 | 22.80719973 | NA | NA | NA | NA | NA | NA | NA |
| PKD2 | 380 | W | 15.9165 | 27.27737618 | 0.554667736 | 21.56033976 | NA | NA | NA | NA | NA | NA | NA |
| PKD2 | 381 | G | 19.28 | 27.14048003 | 0.55164703 | 21.01966122 | NA | NA | NA | NA | NA | NA | NA |
| PKD2 | 382 | I | 19.1745 | 22.12155991 | 0.44090119 | 16.01367858 | NA | NA | NA | NA | NA | NA | NA |
| PKD2 | 383 | I | 20.31625 | 26.14044738 | 0.529580639 | 20.01627965 | NA | NA | NA | NA | NA | NA | NA |
| PKD2 | 384 | A | 20.6611 | 29.5838763 | 0.605562208 | 23.45607617 | NA | NA | NA | NA | NA | NA | NA |
| PKD2 | 385 | T | 19.34907143 | 32.17815932 | 0.662806804 | 26.05383478 | NA | NA | NA | NA | NA | NA | NA |
| PKD2 | 386 | Y | 23.2745 | 29.91358531 | 0.612837459 | 20.21314345 | NA | NA | NA | NA | NA | NA | NA |
| PKD2 | 387 | S | 23.3715 | 36.63309953 | 0.761108049 | 26.92453908 | NA | NA | NA | NA | NA | NA | NA |
| PKD2 | 388 | G | 26.151 | 37.49894147 | 0.780213432 | 30.6117536 | NA | NA | NA | NA | NA | NA | NA |

| | | | | | | | | | | | | | |
|---|---|---|---|---|---|---|---|---|---|---|---|---|---|
| PKD2 | 389 | A | 29.4845 | 38.82357724 | 0.809442409 | 33.0431402 | NA | NA | NA | NA | NA | NA | NA |
| PKD2 | 390 | G | 32.01175 | 38.72480825 | 0.807263005 | 32.96760236 | NA | NA | NA | NA | NA | NA | NA |
| PKD2 | 391 | Y | 35.09241667 | 37.12786946 | 0.772025479 | 33.90201724 | NA | NA | NA | NA | NA | NA | NA |
| PKD2 | 392 | Y | 35.52766667 | 37.21964332 | 0.774050531 | 34.0163241 | NA | NA | NA | NA | NA | NA | NA |
| PKD2 | 393 | L | 39.074875 | 34.25349468 | 0.708600471 | 29.83541886 | NA | NA | NA | NA | NA | NA | NA |
| PKD2 | 394 | D | 37.76275 | 35.58308876 | 0.737938857 | 31.05389206 | NA | NA | NA | NA | NA | NA | NA |
| PKD2 | 395 | L | 39.57625 | 30.03930753 | 0.615611604 | 26.79918916 | NA | NA | NA | NA | NA | NA | NA |
| PKD2 | 396 | S | 43.03283333 | 33.07065565 | 0.682500334 | 29.80802143 | NA | NA | NA | NA | NA | NA | NA |
| PKD2 | 397 | R | 41.36477273 | 33.0001602 | 0.680944805 | 29.74694616 | NA | NA | NA | NA | NA | NA | NA |
| PKD2 | 398 | T | 46.11278571 | 31.47959626 | 0.647392541 | 28.24272308 | NA | NA | NA | NA | NA | NA | NA |
| PKD2 | 399 | R | 46.67440909 | 23.03313921 | 0.461015799 | 19.82648157 | NA | NA | NA | NA | NA | NA | NA |
| PKD2 | 400 | E | 49.75883333 | 30.24294065 | 0.620104905 | 27.02970518 | NA | NA | NA | NA | NA | NA | NA |
| PKD2 | 401 | E | 48.21161111 | 32.03634678 | 0.659677615 | 28.8117281 | NA | NA | NA | NA | NA | NA | NA |
| PKD2 | 402 | T | 44.57778571 | 29.16380284 | 0.596293006 | 25.91014704 | NA | NA | NA | NA | NA | NA | NA |
| PKD2 | 403 | A | 47.4987 | 27.31046931 | 0.555397958 | 24.10045811 | NA | NA | NA | NA | NA | NA | NA |
| PKD2 | 404 | A | 49.2221 | 30.02900708 | 0.615384318 | 26.81861504 | NA | NA | NA | NA | NA | NA | NA |
| PKD2 | 405 | Q | 45.70527778 | 30.92092096 | 0.635064996 | 27.69313657 | NA | NA | NA | NA | NA | NA | NA |
| PKD2 | 406 | V | 44.93621429 | 26.42195144 | 0.535792215 | 23.18082549 | NA | NA | NA | NA | NA | NA | NA |
| PKD2 | 407 | A | 48.3803 | 28.47478804 | 0.581089432 | 25.25748645 | NA | NA | NA | NA | NA | NA | NA |
| PKD2 | 408 | S | 48.50816667 | 31.01111161 | 0.637055113 | 27.80071345 | NA | NA | NA | NA | NA | NA | NA |
| PKD2 | 409 | L | 44.489125 | 31.03718492 | 0.637630438 | 27.79854565 | NA | NA | NA | NA | NA | NA | NA |
| PKD2 | 410 | K | 45.36983333 | 24.84894859 | 0.501082852 | 21.60895422 | NA | NA | NA | NA | NA | NA | NA |
| PKD2 | 411 | K | 49.46138889 | 30.63047118 | 0.628656027 | 27.4094968 | NA | NA | NA | NA | NA | NA | NA |
| PKD2 | 412 | N | 46.2055 | 33.09800275 | 0.683103766 | 29.87026269 | NA | NA | NA | NA | NA | NA | NA |
| PKD2 | 413 | V | 43.31978571 | 30.9974707 | 0.636754117 | 27.75962663 | NA | NA | NA | NA | NA | NA | NA |
| PKD2 | 414 | W | 39.71907143 | 31.73307579 | 0.652985737 | 28.46373791 | NA | NA | NA | NA | NA | NA | NA |
| PKD2 | 415 | L | 37.94325 | 27.90875805 | 0.5685996 | 23.38002717 | NA | NA | NA | NA | NA | NA | NA |
| PKD2 | 416 | D | 38.59375 | 31.47777587 | 0.647352373 | 28.06257798 | NA | NA | NA | NA | NA | NA | NA |
| PKD2 | 417 | R | 36.02104545 | 32.85460176 | 0.67773296 | 29.75544596 | NA | NA | NA | NA | NA | NA | NA |
| PKD2 | 418 | G | 32.17175 | 33.64415683 | 0.695155023 | 30.41731957 | NA | NA | NA | NA | NA | NA | NA |
| PKD2 | 419 | T | 32.77292857 | 31.06213306 | 0.638180936 | 27.83670508 | NA | NA | NA | NA | NA | NA | NA |
| PKD2 | 420 | R | 28.97813636 | 29.46863964 | 0.603019434 | 23.68819261 | NA | NA | NA | NA | NA | NA | NA |
| PKD2 | 421 | A | 28.9533 | 30.79171585 | 0.632213998 | 25.01683984 | NA | NA | NA | NA | NA | NA | NA |

| | | | | | | | | | | | | | |
|---|---|---|---|---|---|---|---|---|---|---|---|---|---|
| PKD2 | 422 | T | 31.34678571 | 28.0883502 | 0.572562422 | 22.31959128 | NA | NA | NA | NA | NA | NA | NA |
| PKD2 | 423 | F | 28.82731818 | 29.67959024 | 0.607674201 | 24.00040762 | NA | NA | NA | NA | NA | NA | NA |
| PKD2 | 424 | I | 32.735 | 26.15698951 | 0.529945652 | 22.93779601 | NA | NA | NA | NA | NA | NA | NA |
| PKD2 | 425 | D | 30.510875 | 30.44705649 | 0.624608858 | 24.73920341 | NA | NA | NA | NA | NA | NA | NA |
| PKD2 | 426 | F | 32.39568182 | 26.66421883 | 0.541138007 | 23.46576392 | NA | NA | NA | NA | NA | NA | NA |
| PKD2 | 427 | S | 30.45616667 | 32.02275519 | 0.659377707 | 26.26383207 | NA | NA | NA | NA | NA | NA | NA |
| PKD2 | 428 | V | 30.54835714 | 30.04818758 | 0.615807548 | 24.282903 | NA | NA | NA | NA | NA | NA | NA |
| PKD2 | 429 | Y | 29.12766667 | 33.23098392 | 0.686038085 | 27.49141824 | NA | NA | NA | NA | NA | NA | NA |
| PKD2 | 430 | N | 28.215 | 32.48635746 | 0.669607403 | 26.70606 | NA | NA | NA | NA | NA | NA | NA |
| PKD2 | 431 | A | 29.7605 | 35.2462484 | 0.730506248 | 29.47335449 | NA | NA | NA | NA | NA | NA | NA |
| PKD2 | 432 | N | 28.526125 | 33.36053648 | 0.688896749 | 26.88597451 | NA | NA | NA | NA | NA | NA | NA |
| PKD2 | 433 | I | 23.943875 | 32.20769015 | 0.663458421 | 23.23806205 | NA | NA | NA | NA | NA | NA | NA |
| PKD2 | 434 | N | 23.89575 | 36.27508288 | 0.753208171 | 27.2912756 | NA | NA | NA | NA | NA | NA | NA |
| PKD2 | 435 | L | 23.7195 | 31.30594162 | 0.643560735 | 22.33269775 | NA | NA | NA | NA | NA | NA | NA |
| PKD2 | 436 | F | 25.70777273 | 33.08458297 | 0.68280765 | 26.23468983 | NA | NA | NA | NA | NA | NA | NA |
| PKD2 | 437 | C | 26.059 | 28.35423866 | 0.578429429 | 21.4552423 | NA | NA | NA | NA | NA | NA | NA |
| PKD2 | 438 | V | 25.61478571 | 28.5274199 | 0.582250789 | 21.61987411 | NA | NA | NA | NA | NA | NA | NA |
| PKD2 | 439 | V | 28.47778571 | 26.49392749 | 0.537380415 | 20.01152431 | NA | NA | NA | NA | NA | NA | NA |
| PKD2 | 440 | R | 25.52168182 | 25.87554531 | 0.523735397 | 19.35332759 | NA | NA | NA | NA | NA | NA | NA |
| PKD2 | 441 | L | 29.554125 | 23.34284698 | 0.467849709 | 17.56240385 | NA | NA | NA | NA | NA | NA | NA |
| PKD2 | 442 | L | 26.350375 | 24.03386881 | 0.483097569 | 17.48679409 | NA | NA | NA | NA | NA | NA | NA |
| PKD2 | 443 | V | 30.0185 | 23.62768823 | 0.474134922 | 17.92809983 | NA | NA | NA | NA | NA | NA | NA |
| PKD2 | 444 | E | 26.02138889 | 24.98289373 | 0.504038441 | 18.46851363 | NA | NA | NA | NA | NA | NA | NA |
| PKD2 | 445 | F | 30.82586364 | 25.3774307 | 0.512744164 | 19.82828146 | NA | NA | NA | NA | NA | NA | NA |
| PKD2 | 446 | P | 27.3265 | 25.41426656 | 0.513556972 | 18.75548643 | NA | NA | NA | NA | NA | NA | NA |
| PKD2 | 447 | A | 27.3167 | 29.92147767 | 0.613011609 | 23.3856643 | NA | NA | NA | NA | NA | NA | NA |
| PKD2 | 448 | T | 27.55892857 | 27.30777322 | 0.555338467 | 20.78040591 | NA | NA | NA | NA | NA | NA | NA |
| PKD2 | 449 | G | 31.04325 | 25.42379329 | 0.513767185 | 19.82601702 | NA | NA | NA | NA | NA | NA | NA |
| PKD2 | 450 | G | 30.2795 | 23.72740245 | 0.476335183 | 17.97362973 | NA | NA | NA | NA | NA | NA | NA |
| PKD2 | 451 | V | 29.90492857 | 20.54720835 | 0.406162067 | 14.85439093 | NA | NA | NA | NA | NA | NA | NA |
| PKD2 | 452 | I | 26.23075 | 21.31807001 | 0.423171647 | 14.75688022 | NA | NA | NA | NA | NA | NA | NA |
| PKD2 | 453 | P | 27.45535714 | 18.1764504 | 0.353849702 | 11.63528324 | NA | NA | NA | NA | NA | NA | NA |
| PKD2 | 454 | S | 25.38916667 | 21.09146135 | 0.418171375 | 14.19395695 | NA | NA | NA | NA | NA | NA | NA |

| | | | | | | | | | | | | | |
|---|---|---|---|---|---|---|---|---|---|---|---|---|---|
| PKD2 | 455 | W | 24.8085 | 16.50024051 | 0.316863006 | 9.64376999 | NA | NA | NA | NA | NA | NA | NA |
| PKD2 | 456 | Q | 22.4585 | 23.95981855 | 0.481463601 | 14.2720839 | NA | NA | NA | NA | NA | NA | NA |
| PKD2 | 457 | F | 25.06486364 | 21.71173719 | 0.431858177 | 12.93189899 | NA | NA | NA | NA | NA | NA | NA |
| PKD2 | 458 | Q | 20.78061111 | 27.22331479 | 0.553474835 | 17.51467529 | NA | NA | NA | NA | NA | NA | NA |
| PKD2 | 459 | P | 20.93692857 | 27.48289232 | 0.559202587 | 17.83918835 | NA | NA | NA | NA | NA | NA | NA |
| PKD2 | 460 | L | 20.6875 | 31.91436672 | 0.656986043 | 23.34626984 | NA | NA | NA | NA | NA | NA | NA |
| PKD2 | 461 | K | 18.2675 | 34.93408953 | 0.723618253 | 26.36863076 | NA | NA | NA | NA | NA | NA | NA |
| PKD2 | 462 | L | 21.8475 | 38.11447863 | 0.793795672 | 29.37510053 | NA | NA | NA | NA | NA | NA | NA |
| PKD2 | 463 | I | 22.186625 | 41.05444695 | 0.858668046 | 31.34558922 | NA | NA | NA | NA | NA | NA | NA |
| PKD2 | 464 | R | 16.07413636 | 41.46459287 | 0.867718191 | 35.65316678 | NA | NA | NA | NA | NA | NA | NA |
| PKD2 | 465 | Y | 18.05816667 | 43.92981153 | 0.922114895 | 37.81369731 | NA | NA | NA | NA | NA | NA | NA |
| PKD2 | 466 | V | 19.4435 | 41.97537017 | 0.878988835 | 33.41276094 | NA | NA | NA | NA | NA | NA | NA |
| PKD2 | 467 | T | 20.48664286 | 45.28132434 | 0.951936932 | 36.714213 | NA | NA | NA | NA | NA | NA | NA |
| PKD2 | 468 | T | 18.54707143 | 43.74066576 | 0.917941266 | 37.63585187 | NA | NA | NA | NA | NA | NA | NA |
| PKD2 | 469 | F | 13.83122727 | 39.55567218 | 0.825596575 | 36.45714979 | NA | NA | NA | NA | NA | NA | NA |
| PKD2 | 470 | D | 14.77825 | 42.65050449 | 0.893886126 | 39.45457167 | NA | NA | NA | NA | NA | NA | NA |
| PKD2 | 471 | F | 15.02377273 | 46.00122435 | 0.967822008 | 42.81016628 | NA | NA | NA | NA | NA | NA | NA |
| PKD2 | 472 | F | 11.04740909 | 44.59624296 | 0.936820151 | 41.32736454 | NA | NA | NA | NA | NA | NA | NA |
| PKD2 | 473 | L | 10.696875 | 39.65382451 | 0.827762372 | 36.39147148 | NA | NA | NA | NA | NA | NA | NA |
| PKD2 | 474 | A | 10.8057 | 44.22339252 | 0.928592956 | 40.94337426 | NA | NA | NA | NA | NA | NA | NA |
| PKD2 | 475 | A | 8.2365 | 46.65057516 | 0.98215037 | 44.27422379 | NA | NA | NA | NA | NA | NA | NA |
| PKD2 | 476 | C | 5.992333333 | 42.64260575 | 0.893711835 | 40.25199355 | NA | NA | NA | NA | NA | NA | NA |
| PKD2 | 477 | E | 7.825833333 | 43.21004144 | 0.906232685 | 40.79796378 | NA | NA | NA | NA | NA | NA | NA |
| PKD2 | 478 | I | 5.802625 | 45.8667355 | 0.964854421 | 43.43697688 | NA | NA | NA | NA | NA | NA | NA |
| PKD2 | 479 | I | 2.662 | 45.16459757 | 0.949361277 | 42.46784934 | NA | NA | NA | NA | NA | NA | NA |
| PKD2 | 480 | F | 2.543954545 | 38.56894354 | 0.803823746 | 35.87703861 | NA | NA | NA | NA | NA | NA | NA |
| PKD2 | 481 | C | 2.633833333 | 43.96211717 | 0.92282774 | 41.27848883 | NA | NA | NA | NA | NA | NA | NA |
| PKD2 | 482 | F | 0.2995 | 46.26000468 | 0.97353217 | 43.23979073 | NA | NA | NA | NA | NA | NA | NA |
| PKD2 | 483 | F | -1.605590909 | 40.95165966 | 0.856399975 | 38.28067154 | NA | NA | NA | NA | NA | NA | NA |
| PKD2 | 484 | I | -0.7095 | 40.63233898 | 0.84935395 | 37.93986839 | NA | NA | NA | NA | NA | NA | NA |
| PKD2 | 485 | F | -1.600863636 | 45.08512595 | 0.947607682 | 42.37616191 | NA | NA | NA | NA | NA | NA | NA |
| PKD2 | 486 | Y | -6.39775 | 44.93962159 | 0.944397031 | 41.6548383 | NA | NA | NA | NA | NA | NA | NA |
| PKD2 | 487 | Y | -6.847333333 | 41.0571646 | 0.858728013 | 38.26240197 | NA | NA | NA | NA | NA | NA | NA |

| | | | | | | | | | | | | | |
|---|---|---|---|---|---|---|---|---|---|---|---|---|---|
| PKD2 | 488 | V | -4.284642857 | 44.07279222 | 0.925269859 | 40.73578787 | NA | NA | NA | NA | NA | NA | NA |
| PKD2 | 489 | V | -7.401071429 | 46.27864299 | 0.973943437 | 43.48154851 | NA | NA | NA | NA | NA | NA | NA |
| PKD2 | 490 | E | -10.6565 | 43.2100519 | 0.906232915 | 41.9318482 | NA | NA | NA | NA | NA | NA | NA |
| PKD2 | 491 | E | -9.089722222 | 39.35713289 | 0.821215672 | 37.40638603 | NA | NA | NA | NA | NA | NA | NA |
| PKD2 | 492 | I | -9.733 | 44.14085229 | 0.926771651 | 42.02798963 | NA | NA | NA | NA | NA | NA | NA |
| PKD2 | 493 | L | -13.372 | 45.62330308 | 0.959482922 | 42.73916584 | NA | NA | NA | NA | NA | NA | NA |
| PKD2 | 494 | E | -15.7175 | 40.33804134 | 0.842860075 | 37.3804151 | NA | NA | NA | NA | NA | NA | NA |
| PKD2 | 495 | I | -13.156625 | 40.16490429 | 0.83903969 | 37.09316072 | NA | NA | NA | NA | NA | NA | NA |
| PKD2 | 496 | R | -14.92340909 | 45.18970195 | 0.949915222 | 42.09431562 | NA | NA | NA | NA | NA | NA | NA |
| PKD2 | 497 | I | -18.412125 | 45.422694 | 0.955056348 | 44.45628417 | NA | NA | NA | NA | NA | NA | NA |
| PKD2 | 498 | H | -19.8614 | 41.92137431 | 0.87779738 | 39.13639626 | NA | NA | NA | NA | NA | NA | NA |
| PKD2 | 499 | K | -18.01361111 | 40.05891424 | 0.836700949 | 39.04901603 | NA | NA | NA | NA | NA | NA | NA |
| PKD2 | 500 | L | -21.245 | 36.91847513 | 0.767405053 | 34.11689291 | NA | NA | NA | NA | NA | NA | NA |
| PKD2 | 501 | H | -20.9179 | 35.77164156 | 0.742099401 | 31.72768427 | NA | NA | NA | NA | NA | NA | NA |
| PKD2 | 502 | Y | -15.71116667 | 34.6652316 | 0.717685722 | 33.67558097 | NA | NA | NA | NA | NA | NA | NA |
| PKD2 | 503 | F | -17.33268182 | 33.77318728 | 0.698002166 | 32.80745856 | NA | NA | NA | NA | NA | NA | NA |
| PKD2 | 504 | R | -18.40986364 | 26.36700844 | 0.534579861 | 25.45493733 | NA | NA | NA | NA | NA | NA | NA |
| PKD2 | 505 | S | -15.87666667 | 29.82107632 | 0.610796186 | 26.72865608 | NA | NA | NA | NA | NA | NA | NA |
| PKD2 | 506 | F | -13.40022727 | 25.73966832 | 0.52073718 | 23.18822331 | NA | NA | NA | NA | NA | NA | NA |
| PKD2 | 507 | W | -13.87378571 | 23.79841526 | 0.477902129 | 21.34081203 | NA | NA | NA | NA | NA | NA | NA |
| PKD2 | 508 | N | -12.93725 | 30.84270578 | 0.633339126 | 28.24632792 | NA | NA | NA | NA | NA | NA | NA |
| PKD2 | 509 | C | -10.41516667 | 31.38973859 | 0.645409772 | 29.26230106 | NA | NA | NA | NA | NA | NA | NA |
| PKD2 | 510 | L | -7.79775 | 26.86514623 | 0.545571605 | 25.51122387 | NA | NA | NA | NA | NA | NA | NA |
| PKD2 | 511 | D | -9.087125 | 32.13328947 | 0.66181672 | 30.79139709 | NA | NA | NA | NA | NA | NA | NA |
| PKD2 | 512 | V | -6.684357143 | 34.84280198 | 0.721603932 | 32.04818442 | NA | NA | NA | NA | NA | NA | NA |
| PKD2 | 513 | V | -3.253928571 | 32.00610848 | 0.659010387 | 28.42709505 | NA | NA | NA | NA | NA | NA | NA |
| PKD2 | 514 | I | -2.34075 | 33.82373705 | 0.699117581 | 30.5134776 | NA | NA | NA | NA | NA | NA | NA |
| PKD2 | 515 | V | -2.373357143 | 36.51224741 | 0.758441366 | 33.20575394 | NA | NA | NA | NA | NA | NA | NA |
| PKD2 | 516 | V | -0.9955 | 36.85653062 | 0.766038205 | 34.14996375 | NA | NA | NA | NA | NA | NA | NA |
| PKD2 | 517 | L | 0.59325 | 32.13945505 | 0.661952768 | 29.12030317 | NA | NA | NA | NA | NA | NA | NA |
| PKD2 | 518 | S | 1.081833333 | 36.88606418 | 0.766689883 | 34.19562904 | NA | NA | NA | NA | NA | NA | NA |
| PKD2 | 519 | V | 1.546785714 | 39.71412198 | 0.829092876 | 37.0163819 | NA | NA | NA | NA | NA | NA | NA |
| PKD2 | 520 | V | 4.341071429 | 38.22132375 | 0.796153282 | 36.76765949 | NA | NA | NA | NA | NA | NA | NA |

| | | | | | | | | | | | | | |
|---|---|---|---|---|---|---|---|---|---|---|---|---|---|
| PKD2 | 521 | A | 6.2359 | 37.42238616 | 0.778524188 | 34.99899728 | NA | NA | NA | NA | NA | NA | NA |
| PKD2 | 522 | I | 4.89175 | 40.58727203 | 0.848359518 | 38.16671241 | NA | NA | NA | NA | NA | NA | NA |
| PKD2 | 523 | G | 8.049 | 43.19328381 | 0.905862916 | 40.7511038 | NA | NA | NA | NA | NA | NA | NA |
| PKD2 | 524 | I | 9.954625 | 38.22445232 | 0.796222316 | 34.98760615 | NA | NA | NA | NA | NA | NA | NA |
| PKD2 | 525 | N | 10.879375 | 40.8170586 | 0.853429913 | 37.53737353 | NA | NA | NA | NA | NA | NA | NA |
| PKD2 | 526 | I | 11.50975 | 43.73031964 | 0.917712972 | 40.45185717 | NA | NA | NA | NA | NA | NA | NA |
| PKD2 | 527 | Y | 10.9785 | 45.71811409 | 0.96157499 | 42.43890344 | NA | NA | NA | NA | NA | NA | NA |
| PKD2 | 528 | R | 12.83959091 | 43.04336192 | 0.902554789 | 39.88259753 | NA | NA | NA | NA | NA | NA | NA |
| PKD2 | 529 | T | 15.94707143 | 40.61571176 | 0.84898706 | 37.41769791 | NA | NA | NA | NA | NA | NA | NA |
| PKD2 | 530 | S | 16.35083333 | 44.79031005 | 0.941102372 | 41.59488575 | NA | NA | NA | NA | NA | NA | NA |
| PKD2 | 531 | N | 18.7495 | 47.31869504 | 0.996892883 | 41.27830825 | NA | NA | NA | NA | NA | NA | NA |
| PKD2 | 532 | V | 20.5135 | 42.99678076 | 0.901526945 | 34.42803681 | NA | NA | NA | NA | NA | NA | NA |
| PKD2 | 533 | E | 20.42683333 | 41.88126843 | 0.876912417 | 32.42538607 | NA | NA | NA | NA | NA | NA | NA |
| PKD2 | 534 | V | 22.42464286 | 46.99529896 | 0.989756931 | 37.28637968 | NA | NA | NA | NA | NA | NA | NA |
| PKD2 | 535 | L | 24.252375 | 46.70277805 | 0.983302262 | 37.71385283 | NA | NA | NA | NA | NA | NA | NA |
| PKD2 | 536 | L | 26.34825 | 41.94103884 | 0.878231291 | 35.28563217 | NA | NA | NA | NA | NA | NA | NA |
| PKD2 | 537 | Q | 29.13794444 | 45.84427732 | 0.964358867 | 40.09203389 | NA | NA | NA | NA | NA | NA | NA |
| PKD2 | 538 | F | 34.23077273 | 46.44279966 | 0.977565664 | 43.21876971 | NA | NA | NA | NA | NA | NA | NA |
| PKD2 | 539 | L | 30.33425 | 44.37122765 | 0.931855037 | 38.66856491 | NA | NA | NA | NA | NA | NA | NA |
| PKD2 | 540 | E | 29.82994444 | 41.40005877 | 0.866294203 | 35.74577274 | NA | NA | NA | NA | NA | NA | NA |
| PKD2 | 541 | D | 34.273625 | 42.09375291 | 0.881601029 | 38.90661206 | NA | NA | NA | NA | NA | NA | NA |
| PKD2 | 542 | Q | 37.14916667 | 41.67925226 | 0.872454794 | 37.15986821 | NA | NA | NA | NA | NA | NA | NA |
| PKD2 | 543 | N | 39.935375 | 39.17743285 | 0.817250471 | 35.74828248 | NA | NA | NA | NA | NA | NA | NA |
| PKD2 | 544 | T | 36.40135714 | 40.91571672 | 0.85560687 | 36.45949886 | NA | NA | NA | NA | NA | NA | NA |
| PKD2 | 545 | F | 32.90677273 | 36.16449372 | 0.750767947 | 33.03050591 | NA | NA | NA | NA | NA | NA | NA |
| PKD2 | 546 | P | 30.74307143 | 38.95104551 | 0.812255082 | 33.18147408 | NA | NA | NA | NA | NA | NA | NA |
| PKD2 | 547 | N | 26.78525 | 38.91039721 | 0.81135815 | 32.40038509 | NA | NA | NA | NA | NA | NA | NA |
| PKD2 | 548 | F | 24.79513636 | 36.20183229 | 0.751591848 | 29.36087851 | NA | NA | NA | NA | NA | NA | NA |
| PKD2 | 549 | E | 23.52494444 | 33.7697309 | 0.697925899 | 24.8103054 | NA | NA | NA | NA | NA | NA | NA |
| PKD2 | 550 | H | 20.3077 | 37.30757426 | 0.775990786 | 27.5991817 | NA | NA | NA | NA | NA | NA | NA |
| PKD2 | 551 | L | 21.38575 | 38.32292392 | 0.798395158 | 29.75515205 | NA | NA | NA | NA | NA | NA | NA |
| PKD2 | 552 | A | 19.7151 | 34.41118291 | 0.712079968 | 28.33197884 | NA | NA | NA | NA | NA | NA | NA |
| PKD2 | 553 | Y | 16.70766667 | 34.19898058 | 0.707397581 | 28.30898104 | NA | NA | NA | NA | NA | NA | NA |

| | | | | | | | | | | | | | |
|---|---|---|---|---|---|---|---|---|---|---|---|---|---|
| PKD2 | 554 | W | 13.93092857 | 35.02974515 | 0.725728958 | 31.8698355 | NA | NA | NA | NA | NA | NA | NA |
| PKD2 | 555 | Q | 15.47294444 | 32.83708786 | 0.677346504 | 29.67370747 | NA | NA | NA | NA | NA | NA | NA |
| PKD2 | 556 | I | 15.0465 | 29.18679669 | 0.59680038 | 26.07881915 | NA | NA | NA | NA | NA | NA | NA |
| PKD2 | 557 | Q | 12.11505556 | 32.45754254 | 0.668971582 | 29.2554262 | NA | NA | NA | NA | NA | NA | NA |
| PKD2 | 558 | F | 10.5495 | 32.82288295 | 0.677033063 | 29.58334869 | NA | NA | NA | NA | NA | NA | NA |
| PKD2 | 559 | N | 11.597125 | 29.14399164 | 0.595855858 | 25.9896943 | NA | NA | NA | NA | NA | NA | NA |
| PKD2 | 560 | N | 8.8345 | 28.32521345 | 0.577788968 | 25.93332468 | NA | NA | NA | NA | NA | NA | NA |
| PKD2 | 561 | I | 6.0315 | 30.82813269 | 0.63301756 | 28.40614068 | NA | NA | NA | NA | NA | NA | NA |
| PKD2 | 562 | A | 6.3813 | 31.56829994 | 0.649349847 | 29.14611524 | NA | NA | NA | NA | NA | NA | NA |
| PKD2 | 563 | A | 5.3117 | 28.04033017 | 0.571502827 | 25.61069139 | NA | NA | NA | NA | NA | NA | NA |
| PKD2 | 564 | V | 2.808928571 | 27.91990876 | 0.568845648 | 26.45461995 | NA | NA | NA | NA | NA | NA | NA |
| PKD2 | 565 | T | 1.192642857 | 30.18661152 | 0.618861965 | 27.5455588 | NA | NA | NA | NA | NA | NA | NA |
| PKD2 | 566 | V | 1.672928571 | 28.87628125 | 0.589948649 | 26.21118778 | NA | NA | NA | NA | NA | NA | NA |
| PKD2 | 567 | F | 0.134863636 | 22.58001198 | 0.451017243 | 19.26932347 | NA | NA | NA | NA | NA | NA | NA |
| PKD2 | 568 | F | -1.528954545 | 27.02055845 | 0.54900088 | 24.33807797 | NA | NA | NA | NA | NA | NA | NA |
| PKD2 | 569 | V | -1.7045 | 29.5907787 | 0.605714514 | 26.8834469 | NA | NA | NA | NA | NA | NA | NA |
| PKD2 | 570 | W | -1.503214286 | 26.75519839 | 0.543145532 | 23.45360627 | NA | NA | NA | NA | NA | NA | NA |
| PKD2 | 571 | I | -5.453875 | 24.16191158 | 0.485922919 | 21.30074938 | NA | NA | NA | NA | NA | NA | NA |
| PKD2 | 572 | K | -7.751388889 | 28.57564631 | 0.583314937 | 25.77790041 | NA | NA | NA | NA | NA | NA | NA |
| PKD2 | 573 | L | -7.384 | 27.86248456 | 0.567578545 | 26.56106593 | NA | NA | NA | NA | NA | NA | NA |
| PKD2 | 574 | F | -10.01822727 | 21.39952161 | 0.424968931 | 20.12910835 | NA | NA | NA | NA | NA | NA | NA |
| PKD2 | 575 | K | -12.97794444 | 27.23497997 | 0.553732235 | 24.60711153 | NA | NA | NA | NA | NA | NA | NA |
| PKD2 | 576 | F | -12.18495455 | 29.25053205 | 0.598206744 | 26.59422472 | NA | NA | NA | NA | NA | NA | NA |
| PKD2 | 577 | I | -11.746375 | 29.32848598 | 0.59992685 | 26.87237091 | NA | NA | NA | NA | NA | NA | NA |
| PKD2 | 578 | N | -13.79775 | 25.05476436 | 0.505624315 | 22.03535208 | NA | NA | NA | NA | NA | NA | NA |
| PKD2 | 579 | F | -19.26222727 | 27.81164974 | 0.56645684 | 26.89925877 | NA | NA | NA | NA | NA | NA | NA |
| PKD2 | 580 | N | -17.302125 | 25.35864912 | 0.512329736 | 24.35159094 | NA | NA | NA | NA | NA | NA | NA |
| PKD2 | 581 | R | -18.02486364 | 23.25333376 | 0.46587454 | 20.47690032 | NA | NA | NA | NA | NA | NA | NA |
| PKD2 | 582 | T | -20.9015 | 19.41065081 | 0.381083162 | 16.59337057 | NA | NA | NA | NA | NA | NA | NA |
| PKD2 | 583 | M | -16.52925 | 20.6285444 | 0.407956801 | 19.62337657 | NA | NA | NA | NA | NA | NA | NA |
| PKD2 | 584 | S | -18.4405 | 22.52957181 | 0.449904247 | 21.58487848 | NA | NA | NA | NA | NA | NA | NA |
| PKD2 | 585 | Q | -21.97305556 | 19.35152301 | 0.379778468 | 16.55552113 | NA | NA | NA | NA | NA | NA | NA |
| PKD2 | 586 | L | -18.817875 | 16.06793761 | 0.307323953 | 15.17751152 | NA | NA | NA | NA | NA | NA | NA |

| | | | | | | | | | | | | | |
|---|---|---|---|---|---|---|---|---|---|---|---|---|---|
| PKD2 | 587 | S | -16.2725 | 19.27983268 | 0.378196572 | 18.27638714 | NA | NA | NA | NA | NA | NA | NA |
| PKD2 | 588 | T | -19.54007143 | 18.03470948 | 0.350722094 | 17.11895529 | NA | NA | NA | NA | NA | NA | NA |
| PKD2 | 589 | T | -18.3945 | 14.94906639 | 0.282635309 | 14.02315596 | NA | NA | NA | NA | NA | NA | NA |
| PKD2 | 590 | M | -13.910625 | 14.58027657 | 0.274497714 | 11.70842981 | NA | NA | NA | NA | NA | NA | NA |
| PKD2 | 591 | S | -16.5775 | 17.92578149 | 0.348318524 | 16.94696911 | NA | NA | NA | NA | NA | NA | NA |
| PKD2 | 592 | R | -20.24977273 | 16.29392138 | 0.312310436 | 15.69502994 | NA | NA | NA | NA | NA | NA | NA |
| PKD2 | 593 | C | -15.72916667 | 14.11189938 | 0.264162657 | 13.24533465 | NA | NA | NA | NA | NA | NA | NA |
| PKD2 | 594 | A | -14.0601 | 16.79897797 | 0.323454849 | 13.70809198 | NA | NA | NA | NA | NA | NA | NA |
| PKD2 | 595 | K | -18.24094444 | 18.87518122 | 0.369267666 | 17.91325103 | NA | NA | NA | NA | NA | NA | NA |
| PKD2 | 596 | D | -16.405375 | 14.462052 | 0.271889009 | 13.45023774 | NA | NA | NA | NA | NA | NA | NA |
| PKD2 | 597 | L | -11.852375 | 13.72634536 | 0.255655149 | 11.04057086 | NA | NA | NA | NA | NA | NA | NA |
| PKD2 | 598 | F | -10.66722727 | 18.25702366 | 0.355627605 | 16.15431823 | NA | NA | NA | NA | NA | NA | NA |
| PKD2 | 599 | G | -12.0395 | 20.56731429 | 0.406605718 | 17.96925571 | NA | NA | NA | NA | NA | NA | NA |
| PKD2 | 600 | F | -11.65495455 | 15.2849579 | 0.29004698 | 12.746346 | NA | NA | NA | NA | NA | NA | NA |
| PKD2 | 601 | A | -7.8927 | 17.5939254 | 0.340995897 | 16.28021637 | NA | NA | NA | NA | NA | NA | NA |
| PKD2 | 602 | I | -7.724125 | 19.90862301 | 0.392071253 | 17.30621056 | NA | NA | NA | NA | NA | NA | NA |
| PKD2 | 603 | M | -9.196 | 16.90137447 | 0.325714296 | 15.6700698 | NA | NA | NA | NA | NA | NA | NA |
| PKD2 | 604 | F | -4.473045455 | 13.63483315 | 0.25363587 | 11.23462106 | NA | NA | NA | NA | NA | NA | NA |
| PKD2 | 605 | F | -2.180318182 | 18.18379677 | 0.354011805 | 14.58564377 | NA | NA | NA | NA | NA | NA | NA |
| PKD2 | 606 | I | -3.7055 | 20.63960886 | 0.408200946 | 17.03120348 | NA | NA | NA | NA | NA | NA | NA |
| PKD2 | 607 | I | -3.08875 | 17.75419441 | 0.34453234 | 14.48979313 | NA | NA | NA | NA | NA | NA | NA |
| PKD2 | 608 | F | -0.2105 | 14.70921208 | 0.277342762 | 12.2142365 | NA | NA | NA | NA | NA | NA | NA |
| PKD2 | 609 | L | 0 | 20.13441944 | 0.397053603 | 17.20949048 | NA | NA | NA | NA | NA | NA | NA |
| PKD2 | 610 | A | 0.2161 | 21.53392735 | 0.427934684 | 18.59790089 | NA | NA | NA | NA | NA | NA | NA |
| PKD2 | 611 | Y | 1.03575 | 18.19358865 | 0.354227869 | 15.61075629 | NA | NA | NA | NA | NA | NA | NA |
| PKD2 | 612 | A | 4.1117 | 19.89570059 | 0.391786111 | 18.44818894 | NA | NA | NA | NA | NA | NA | NA |
| PKD2 | 613 | Q | 4.219277778 | 22.52890554 | 0.449889545 | 21.05944927 | NA | NA | NA | NA | NA | NA | NA |
| PKD2 | 614 | L | 4.452375 | 22.44806326 | 0.448105706 | 20.99752066 | NA | NA | NA | NA | NA | NA | NA |
| PKD2 | 615 | A | 7.7919 | 19.70066062 | 0.387482423 | 17.25101945 | NA | NA | NA | NA | NA | NA | NA |
| PKD2 | 616 | Y | 10.77583333 | 22.43633702 | 0.447846959 | 19.48319049 | NA | NA | NA | NA | NA | NA | NA |
| PKD2 | 617 | L | 8.77675 | 25.09152921 | 0.506435556 | 22.09667662 | NA | NA | NA | NA | NA | NA | NA |
| PKD2 | 618 | V | 10.38207143 | 23.43334219 | 0.469846546 | 20.30792825 | NA | NA | NA | NA | NA | NA | NA |
| PKD2 | 619 | F | 11.75986364 | 19.51456597 | 0.38337612 | 16.77033261 | NA | NA | NA | NA | NA | NA | NA |

| | | | | | | | | | | | | | |
|---|---|---|---|---|---|---|---|---|---|---|---|---|---|
| PKD2 | 620 | G | 14.689 | 22.52059338 | 0.449706132 | 19.38500454 | NA | NA | NA | NA | NA | NA | NA |
| PKD2 | 621 | T | 17.28492857 | 23.10816535 | 0.462671301 | 20.11953092 | NA | NA | NA | NA | NA | NA | NA |
| PKD2 | 622 | Q | 18.37905556 | 19.06077126 | 0.373362835 | 12.9407551 | NA | NA | NA | NA | NA | NA | NA |
| PKD2 | 624 | D | 17.947125 | 13.96478475 | 0.260916474 | 7.84333246 | NA | NA | NA | NA | NA | NA | NA |
| PKD2 | 625 | D | 13.92875 | 12.53352121 | 0.229334683 | 9.585731428 | NA | NA | NA | NA | NA | NA | NA |
| PKD2 | 626 | F | 11.78559091 | 15.77291739 | 0.300814134 | 12.50391953 | NA | NA | NA | NA | NA | NA | NA |
| PKD2 | 627 | S | 14.86166667 | 19.88946611 | 0.391648543 | 16.70066195 | NA | NA | NA | NA | NA | NA | NA |
| PKD2 | 628 | T | 12.87592857 | 21.92119087 | 0.436479913 | 19.0308613 | NA | NA | NA | NA | NA | NA | NA |
| PKD2 | 629 | F | 7.228318182 | 21.93036099 | 0.436682258 | 19.5291459 | NA | NA | NA | NA | NA | NA | NA |
| PKD2 | 630 | Q | 10.51127778 | 20.68879673 | 0.409286309 | 18.28469764 | NA | NA | NA | NA | NA | NA | NA |
| PKD2 | 631 | E | 12.46827778 | 16.76023034 | 0.322599856 | 13.97995829 | NA | NA | NA | NA | NA | NA | NA |
| PKD2 | 632 | C | 8.750333333 | 17.60282091 | 0.341192182 | 15.25749365 | NA | NA | NA | NA | NA | NA | NA |
| PKD2 | 633 | I | 5.226625 | 16.52366957 | 0.317379984 | 14.5795386 | NA | NA | NA | NA | NA | NA | NA |
| PKD2 | 634 | F | 7.356318182 | 13.39389865 | 0.248319489 | 10.97687736 | NA | NA | NA | NA | NA | NA | NA |
| PKD2 | 635 | T | 8.348214286 | 11.85018291 | 0.214256365 | 9.601590302 | NA | NA | NA | NA | NA | NA | NA |
| PKD2 | 636 | Q | 3.472722222 | 12.1338695 | 0.2205161 | 10.69796796 | NA | NA | NA | NA | NA | NA | NA |
| PKD2 | 637 | F | 2.701136364 | 9.220510723 | 0.156230884 | 7.75438494 | NA | NA | NA | NA | NA | NA | NA |
| PKD2 | 638 | R | 9.552227273 | 7.190217452 | 0.111431101 | 4.762168995 | 2.43 | -0.8 | -0.4 | 3.38 | 0.81 | 0 | 0 |
| PKD2 | 639 | I | 5.88175 | 6.856200114 | 0.104060784 | 5.11502383 | NA | NA | NA | NA | NA | NA | NA |
| PKD2 | 640 | I | 1.424125 | 5.22629362 | 0.068095804 | 2.548931281 | 2.7 | -0.8 | -0.4 | 3.38 | 0.81 | 0 | 0 |
| PKD2 | 641 | L | 3.240375 | 2.95261562 | 0.017925573 | 1.499835173 | 1.47 | -0.8 | -0.4 | 3.38 | 0.81 | 0 | 0 |
| PKD2 | 642 | G | 6.5135 | 3.86349065 | 0.038024642 | 2.049535724 | 2.03 | -0.8 | -0.4 | 3.38 | 0.81 | 0 | 0 |
| PKD2 | 643 | D | 10.5475 | 4.371920516 | 0.049243488 | 1.924571148 | 3.01 | -0.8 | -0.4 | 3.38 | 0.81 | 0 | 0 |
| PKD2 | 644 | I | 11.819125 | 6.097677345 | 0.08732347 | 2.84281481 | 3.28 | -0.99 | -2.88 | 40.44 | 2.27 | 4 | -4 |
| PKD2 | 645 | N | 14.468625 | 9.023782965 | 0.151889954 | 6.012307206 | NA | NA | NA | NA | NA | NA | NA |
| PKD2 | 646 | F | 11.47977273 | 10.43067773 | 0.182934031 | 7.356791463 | NA | NA | NA | NA | NA | NA | NA |
| PKD2 | 647 | A | 16.1615 | 10.28777221 | 0.179780725 | 7.290918707 | NA | NA | NA | NA | NA | NA | NA |
| PKD2 | 648 | E | 17.25605556 | 11.38717357 | 0.204039754 | 8.722661544 | NA | NA | NA | NA | NA | NA | NA |
| PKD2 | 649 | I | 12.96175 | 13.9412621 | 0.260397431 | 11.35683294 | NA | NA | NA | NA | NA | NA | NA |
| PKD2 | 650 | E | 14.12172222 | 15.2340622 | 0.288923932 | 12.05649223 | NA | NA | NA | NA | NA | NA | NA |
| PKD2 | 651 | E | 19.0855 | 17.47567261 | 0.33838657 | 11.36980827 | NA | NA | NA | NA | NA | NA | NA |
| PKD2 | 652 | A | 16.1987 | 19.17519025 | 0.375887567 | 16.05432937 | NA | NA | NA | NA | NA | NA | NA |
| PKD2 | 653 | N | 13.006625 | 21.380213 | 0.424542873 | 18.18484398 | NA | NA | NA | NA | NA | NA | NA |

| | | | | | | | | | | | | | |
|---|---|---|---|---|---|---|---|---|---|---|---|---|---|
| PKD2 | 654 | R | 15.09095455 | 22.56389098 | 0.450661522 | 19.6777459 | NA | NA | NA | NA | NA | NA | NA |
| PKD2 | 655 | V | 9.259785714 | 23.57387471 | 0.472947491 | 20.68059091 | NA | NA | NA | NA | NA | NA | NA |
| PKD2 | 656 | L | 8.4375 | 20.74309731 | 0.410484488 | 18.3738148 | NA | NA | NA | NA | NA | NA | NA |
| PKD2 | 657 | G | 8.98625 | 17.56955002 | 0.340458038 | 14.66878292 | NA | NA | NA | NA | NA | NA | NA |
| PKD2 | 658 | P | 7.904785714 | 18.08199895 | 0.351765567 | 15.80436755 | NA | NA | NA | NA | NA | NA | NA |
| PKD2 | 659 | I | 4.10525 | 19.57851246 | 0.384787142 | 18.12346301 | NA | NA | NA | NA | NA | NA | NA |
| PKD2 | 660 | Y | 4.994666667 | 16.16995303 | 0.309574991 | 14.76383192 | NA | NA | NA | NA | NA | NA | NA |
| PKD2 | 661 | F | 5.374136364 | 11.38892247 | 0.204078344 | 9.363533244 | NA | NA | NA | NA | NA | NA | NA |
| PKD2 | 662 | T | 2.546928571 | 15.23434075 | 0.288930079 | 14.07011132 | NA | NA | NA | NA | NA | NA | NA |
| PKD2 | 663 | T | 0.1225 | 15.40564293 | 0.292709976 | 12.13326146 | NA | NA | NA | NA | NA | NA | NA |
| PKD2 | 664 | F | 1.0715 | 12.21151796 | 0.222229466 | 9.31469235 | NA | NA | NA | NA | NA | NA | NA |
| PKD2 | 665 | V | 0.6595 | 9.910629042 | 0.171458808 | 6.887601035 | NA | NA | NA | NA | NA | NA | NA |
| PKD2 | 666 | F | -2.449772727 | 12.02016888 | 0.21800722 | 9.429820055 | NA | NA | NA | NA | NA | NA | NA |
| PKD2 | 667 | F | -3.484772727 | 12.09996112 | 0.219767889 | 8.837532378 | NA | NA | NA | NA | NA | NA | NA |
| PKD2 | 668 | M | -1.9523 | 10.3277698 | 0.180663299 | 7.802807457 | NA | NA | NA | NA | NA | NA | NA |
| PKD2 | 669 | F | -1.393863636 | 4.29488568 | 0.047543663 | 1.589642717 | 2.71 | 1.35 | 2.8 | 0.35 | -1.81 | 0 | 0 |
| PKD2 | 670 | F | -4.347681818 | 9.071285135 | 0.152938121 | 6.34827067 | NA | NA | NA | NA | NA | NA | NA |
| PKD2 | 671 | I | -6.877875 | 10.29767265 | 0.179999185 | 7.73687764 | NA | NA | NA | NA | NA | NA | NA |
| PKD2 | 672 | L | -5.0995 | 6.226812828 | 0.090172931 | 3.516959303 | 2.89 | 1.14 | 3.8 | 0.13 | -1.79 | 0 | 0 |
| PKD2 | 673 | L | -7.656625 | 4.496807201 | 0.051999197 | 3.768403956 | 1.36 | 1.14 | 3.8 | 0.13 | -1.79 | 0 | 0 |
| PKD2 | 674 | N | -11.50625 | 8.215580138 | 0.134056416 | 6.186009278 | NA | NA | NA | NA | NA | NA | NA |
| PKD2 | 675 | M | -11.9165 | 9.641684811 | 0.165524373 | 7.31504968 | NA | NA | NA | NA | NA | NA | NA |
| PKD2 | 676 | F | -8.891136364 | 7.770940548 | 0.124245146 | 5.648361724 | NA | NA | NA | NA | NA | NA | NA |
| PKD2 | 677 | L | -11.421375 | 2.910278681 | 0.01699138 | 1.769126232 | 2.38 | 1.14 | 3.8 | 0.13 | -1.79 | 0 | 0 |
| PKD2 | 678 | A | -15.5915 | 5.625263549 | 0.076899343 | 2.526830316 | 3.1 | -0.39 | -2.04 | 2.73 | 0.07 | 0 | 0 |
| PKD2 | 679 | I | -15.746875 | 8.093847416 | 0.131370302 | 7.22124038 | NA | NA | NA | NA | NA | NA | NA |
| PKD2 | 680 | I | -14.001375 | 7.753376039 | 0.123857573 | 4.706061622 | 3.1 | -0.39 | -2.04 | 2.73 | 0.07 | 0 | 0 |
| PKD2 | 681 | N | -17.5495 | 2.259175292 | 0.002624347 | 1.472173528 | 1.01 | -0.77 | -3.5 | 3.38 | 0.54 | 0 | 0 |
| PKD2 | 682 | D | -19.947625 | 6.223114092 | 0.090091316 | 3.832496777 | 2.82 | -0.77 | -3.5 | 3.38 | 0.54 | 0 | 0 |
| PKD2 | 683 | T | -20.1335 | 8.867610501 | 0.148443904 | 6.106767997 | NA | NA | NA | NA | NA | NA | NA |
| PKD2 | 684 | Y | -19.60558333 | 9.303671856 | 0.15806589 | 6.483868717 | NA | NA | NA | NA | NA | NA | NA |
| PKD2 | 685 | S | -22.75383333 | 5.691235103 | 0.07835505 | 1.602305649 | 4.12 | -0.89 | -1.88 | 2.35 | 0.88 | 0 | 0 |
| PKD2 | 686 | E | -25.47027778 | 8.340763214 | 0.136818665 | 4.020579607 | 4.32 | -0.97 | -0.8 | 1.67 | 1.11 | 0 | 0 |

| | | | | | | | | | | | | | |
|---|---|---|---|---|---|---|---|---|---|---|---|---|---|
| PKD2 | 687 | V | -23.62521429 | 11.19164036 | 0.199725182 | 6.875538876 | NA | NA | NA | NA | NA | NA | NA |
| PKD2 | 688 | K | -23.68883333 | 11.58076703 | 0.208311523 | 7.28667156 | NA | NA | NA | NA | NA | NA | NA |
| PKD2 | 689 | S | -27.44366667 | 8.842784347 | 0.147896098 | 4.308573579 | 4.58 | -0.97 | -0.8 | 1.67 | 1.11 | 0 | 0 |
| PKD2 | 690 | D | -28.326875 | 12.01586851 | 0.217912329 | 4.584054337 | 7.54 | -0.97 | -0.8 | 1.67 | 1.11 | 0 | 0 |
| PKD2 | 691 | L | -26.508375 | 14.71363538 | 0.277440365 | 10.17009511 | NA | NA | NA | NA | NA | NA | NA |
| PKD2 | 692 | A | -29.8981 | 12.78081242 | 0.23479133 | 5.240845684 | NA | NA | NA | NA | NA | NA | NA |
| PKD2 | 693 | Q | -31.94405556 | 11.32451363 | 0.20265712 | 3.894218358 | 7.54 | -0.97 | -0.8 | 1.67 | 1.11 | 0 | 0 |
| PKD2 | 694 | Q | -31.16683333 | 16.03994018 | 0.306706171 | 8.589014043 | NA | NA | NA | NA | NA | NA | NA |
| PKD2 | 695 | K | -31.33327778 | 16.5422509 | 0.317789994 | 9.180146657 | NA | NA | NA | NA | NA | NA | NA |
| SCN2A | 117 | P | -16.98821429 | 57.42698453 | 0.895905829 | 54.22799851 | NA | NA | NA | NA | NA | NA | NA |
| SCN2A | 118 | I | -15.82925 | 55.62462798 | 0.866636927 | 52.62266673 | NA | NA | NA | NA | NA | NA | NA |
| SCN2A | 119 | R | -18.62522727 | 52.86413789 | 0.821808663 | 49.67473327 | NA | NA | NA | NA | NA | NA | NA |
| SCN2A | 120 | K | -18.85305556 | 53.38553619 | 0.830275776 | 50.18851993 | NA | NA | NA | NA | NA | NA | NA |
| SCN2A | 121 | L | -14.027875 | 53.53762324 | 0.832745554 | 50.54139999 | NA | NA | NA | NA | NA | NA | NA |
| SCN2A | 122 | A | -13.0509 | 50.83907952 | 0.788923252 | 49.10629623 | NA | NA | NA | NA | NA | NA | NA |
| SCN2A | 123 | I | -15.999875 | 47.52585812 | 0.735119052 | 44.32742718 | NA | NA | NA | NA | NA | NA | NA |
| SCN2A | 124 | K | -16.30361111 | 47.50364488 | 0.734758325 | 44.40023868 | NA | NA | NA | NA | NA | NA | NA |
| SCN2A | 125 | I | -11.283875 | 46.08257346 | 0.711681204 | 44.35731488 | NA | NA | NA | NA | NA | NA | NA |
| SCN2A | 126 | L | -12.9625 | 43.0511239 | 0.662452764 | 41.29783519 | NA | NA | NA | NA | NA | NA | NA |
| SCN2A | 127 | V | -15.96864286 | 42.35147342 | 0.651090972 | 39.16948507 | NA | NA | NA | NA | NA | NA | NA |
| SCN2A | 128 | H | -14.1766 | 42.72772527 | 0.657201016 | 40.00890198 | NA | NA | NA | NA | NA | NA | NA |
| SCN2A | 129 | S | -12.347 | 39.86676031 | 0.610741117 | 38.02307679 | NA | NA | NA | NA | NA | NA | NA |
| SCN2A | 130 | L | -9.012875 | 43.16733011 | 0.664339865 | 39.0270109 | NA | NA | NA | NA | NA | NA | NA |
| SCN2A | 131 | F | -8.154136364 | 41.69540053 | 0.640436846 | 37.38912212 | NA | NA | NA | NA | NA | NA | NA |
| SCN2A | 132 | N | -8.6655 | 37.30530655 | 0.569145052 | 33.19428508 | NA | NA | NA | NA | NA | NA | NA |
| SCN2A | 133 | M | -6.82775 | 38.58283927 | 0.589891213 | 33.68382318 | NA | NA | NA | NA | NA | NA | NA |
| SCN2A | 134 | L | -3.298 | 39.84698684 | 0.61042001 | 34.94906649 | NA | NA | NA | NA | NA | NA | NA |
| SCN2A | 135 | I | -3.37075 | 36.7633304 | 0.56034377 | 31.84996423 | NA | NA | NA | NA | NA | NA | NA |
| SCN2A | 136 | M | -3.4345 | 32.42585748 | 0.489906502 | 27.53298141 | NA | NA | NA | NA | NA | NA | NA |
| SCN2A | 137 | C | -0.637666667 | 35.86753252 | 0.545796693 | 31.54249976 | NA | NA | NA | NA | NA | NA | NA |
| SCN2A | 138 | T | 0.572642857 | 35.35272151 | 0.537436553 | 34.06103645 | NA | NA | NA | NA | NA | NA | NA |
| SCN2A | 139 | I | 0.6885 | 31.98078376 | 0.482678843 | 30.65169178 | NA | NA | NA | NA | NA | NA | NA |
| SCN2A | 140 | L | 1.117125 | 30.13251388 | 0.452664343 | 28.71297516 | NA | NA | NA | NA | NA | NA | NA |

| | | | | | | | | | | | | | |
|---|---|---|---|---|---|---|---|---|---|---|---|---|---|
| SCN2A | 141 | T | 4.387071429 | 32.95007176 | 0.498419344 | 31.93184717 | NA | NA | NA | NA | NA | NA | NA |
| SCN2A | 142 | N | 5.949 | 30.98444027 | 0.466498981 | 29.88633163 | NA | NA | NA | NA | NA | NA | NA |
| SCN2A | 143 | C | 5.1825 | 27.6687919 | 0.412655368 | 26.63032692 | NA | NA | NA | NA | NA | NA | NA |
| SCN2A | 144 | V | 7.9865 | 29.15541866 | 0.436797058 | 28.8589147 | NA | NA | NA | NA | NA | NA | NA |
| SCN2A | 145 | F | 10.08177273 | 30.26526978 | 0.454820198 | 29.96107042 | NA | NA | NA | NA | NA | NA | NA |
| SCN2A | 146 | M | 8.93725 | 26.84094857 | 0.399211821 | 26.48154816 | NA | NA | NA | NA | NA | NA | NA |
| SCN2A | 147 | T | 11.82135714 | 25.35880632 | 0.375142956 | 24.95081023 | NA | NA | NA | NA | NA | NA | NA |
| SCN2A | 148 | M | 14.7355 | 27.52450347 | 0.410312233 | 25.9594271 | NA | NA | NA | NA | NA | NA | NA |
| SCN2A | 149 | S | 17.84533333 | 23.98981344 | 0.35291155 | 22.43565734 | NA | NA | NA | NA | NA | NA | NA |
| SCN2A | 150 | N | 20.432375 | 27.99031867 | 0.417876719 | 26.44371646 | NA | NA | NA | NA | NA | NA | NA |
| SCN2A | 151 | P | 17.95664286 | 31.25697009 | 0.470924658 | 29.70501232 | NA | NA | NA | NA | NA | NA | NA |
| SCN2A | 152 | P | 18.88621429 | 33.38107263 | 0.505418471 | 31.82861318 | NA | NA | NA | NA | NA | NA | NA |
| SCN2A | 153 | D | 20.668875 | 37.24176112 | 0.568113122 | 35.69925032 | NA | NA | NA | NA | NA | NA | NA |
| SCN2A | 154 | W | 16.8585 | 36.36682722 | 0.55390486 | 34.80598492 | NA | NA | NA | NA | NA | NA | NA |
| SCN2A | 155 | T | 14.84992857 | 35.98643971 | 0.547727655 | 34.42330118 | NA | NA | NA | NA | NA | NA | NA |
| SCN2A | 156 | K | 15.37683333 | 39.43319428 | 0.603700333 | 37.87200494 | NA | NA | NA | NA | NA | NA | NA |
| SCN2A | 157 | N | 12.672875 | 41.96719687 | 0.644850612 | 40.40192363 | NA | NA | NA | NA | NA | NA | NA |
| SCN2A | 158 | V | 9.989214286 | 39.16107237 | 0.59928128 | 38.9001775 | NA | NA | NA | NA | NA | NA | NA |
| SCN2A | 159 | E | 10.07727778 | 35.03284061 | 0.53224193 | 34.73498498 | NA | NA | NA | NA | NA | NA | NA |
| SCN2A | 160 | Y | 10.61166667 | 41.40455495 | 0.635713734 | 41.14928447 | NA | NA | NA | NA | NA | NA | NA |
| SCN2A | 161 | T | 5.8105 | 42.03608024 | 0.645969226 | 41.93101209 | NA | NA | NA | NA | NA | NA | NA |
| SCN2A | 162 | F | 5.128409091 | 36.49187862 | 0.555935599 | 35.41517373 | NA | NA | NA | NA | NA | NA | NA |
| SCN2A | 163 | T | 4.740642857 | 40.54764835 | 0.621798222 | 39.24575691 | NA | NA | NA | NA | NA | NA | NA |
| SCN2A | 164 | G | 2.0835 | 43.45976595 | 0.669088801 | 42.11624894 | NA | NA | NA | NA | NA | NA | NA |
| SCN2A | 165 | I | 0.52075 | 42.23484729 | 0.649197052 | 40.846551 | NA | NA | NA | NA | NA | NA | NA |
| SCN2A | 166 | Y | -0.461583333 | 35.79885984 | 0.544681501 | 34.33343101 | NA | NA | NA | NA | NA | NA | NA |
| SCN2A | 167 | T | -0.493642857 | 43.73030438 | 0.67348214 | 42.1482492 | NA | NA | NA | NA | NA | NA | NA |
| SCN2A | 168 | F | -3.219045455 | 46.00650255 | 0.71044587 | 41.22184414 | NA | NA | NA | NA | NA | NA | NA |
| SCN2A | 169 | E | -4.8325 | 40.62795336 | 0.623102314 | 35.69416924 | NA | NA | NA | NA | NA | NA | NA |
| SCN2A | 170 | S | -4.438333333 | 42.51446575 | 0.653737843 | 37.56945005 | NA | NA | NA | NA | NA | NA | NA |
| SCN2A | 171 | L | -5.199 | 46.84611965 | 0.724080615 | 41.90714121 | NA | NA | NA | NA | NA | NA | NA |
| SCN2A | 172 | I | -8.43425 | 47.74154705 | 0.738621676 | 42.81054971 | NA | NA | NA | NA | NA | NA | NA |
| SCN2A | 173 | K | -9.329722222 | 40.37112223 | 0.618931571 | 35.43903055 | NA | NA | NA | NA | NA | NA | NA |

| | | | | | | | | | | | | | |
|---|---|---|---|---|---|---|---|---|---|---|---|---|---|
| SCN2A | 174 | I | -8.163375 | 47.5546608 | 0.735586785 | 42.60544692 | NA | NA | NA | NA | NA | NA | NA |
| SCN2A | 175 | L | -10.52925 | 50.09409428 | 0.776825258 | 45.15588211 | NA | NA | NA | NA | NA | NA | NA |
| SCN2A | 176 | A | -14.0369 | 47.90365017 | 0.741254108 | 46.17881485 | NA | NA | NA | NA | NA | NA | NA |
| SCN2A | 177 | R | -14.77759091 | 41.6841254 | 0.640253746 | 38.89242875 | NA | NA | NA | NA | NA | NA | NA |
| SCN2A | 178 | G | -14.75125 | 49.88659921 | 0.773455695 | 47.00579259 | NA | NA | NA | NA | NA | NA | NA |
| SCN2A | 179 | F | -16.07568182 | 53.34020373 | 0.829539611 | 51.60646555 | NA | NA | NA | NA | NA | NA | NA |
| SCN2A | 180 | C | -15.1685 | 56.7190373 | 0.884409304 | 55.02180114 | NA | NA | NA | NA | NA | NA | NA |
| SCN2A | 181 | L | -13.782625 | 55.90325863 | 0.871161677 | 54.22715451 | NA | NA | NA | NA | NA | NA | NA |
| SCN2A | 182 | E | -16.55894444 | 51.47137648 | 0.799191275 | 49.79353758 | NA | NA | NA | NA | NA | NA | NA |
| SCN2A | 183 | D | -14.201625 | 50.12512558 | 0.777329182 | 48.4242242 | NA | NA | NA | NA | NA | NA | NA |
| SCN2A | 184 | F | -12.20122727 | 45.42122829 | 0.700941461 | 43.64645997 | NA | NA | NA | NA | NA | NA | NA |
| SCN2A | 185 | T | -12.18821429 | 44.30126974 | 0.682754184 | 42.62993079 | NA | NA | NA | NA | NA | NA | NA |
| SCN2A | 186 | F | -9.344590909 | 41.61531786 | 0.639136364 | 39.81375856 | NA | NA | NA | NA | NA | NA | NA |
| SCN2A | 187 | L | -9.953375 | 41.5124031 | 0.637465106 | 39.70928253 | NA | NA | NA | NA | NA | NA | NA |
| SCN2A | 188 | R | -14.77086364 | 35.23341025 | 0.535499028 | 33.5473629 | NA | NA | NA | NA | NA | NA | NA |
| SCN2A | 189 | D | -11.759 | 36.59722663 | 0.557646371 | 34.79158102 | NA | NA | NA | NA | NA | NA | NA |
| SCN2A | 190 | P | -9.079357143 | 33.40646268 | 0.505830787 | 29.45730796 | NA | NA | NA | NA | NA | NA | NA |
| SCN2A | 191 | W | -7.179071429 | 28.96018256 | 0.433626572 | 24.01338149 | NA | NA | NA | NA | NA | NA | NA |
| SCN2A | 192 | N | -7.246375 | 37.09250121 | 0.565689254 | 32.14091035 | NA | NA | NA | NA | NA | NA | NA |
| SCN2A | 193 | W | -6.124928571 | 36.38211232 | 0.554153078 | 31.40381526 | NA | NA | NA | NA | NA | NA | NA |
| SCN2A | 194 | L | -1.8265 | 31.56321766 | 0.475897886 | 26.60323874 | NA | NA | NA | NA | NA | NA | NA |
| SCN2A | 195 | D | -2.87125 | 36.84024127 | 0.561592745 | 31.89473585 | NA | NA | NA | NA | NA | NA | NA |
| SCN2A | 196 | F | -1.404954545 | 39.49145606 | 0.60464646 | 34.5295314 | NA | NA | NA | NA | NA | NA | NA |
| SCN2A | 197 | T | 0.646214286 | 36.14144957 | 0.550244898 | 34.32273113 | NA | NA | NA | NA | NA | NA | NA |
| SCN2A | 198 | V | 1.625928571 | 35.76642809 | 0.544154833 | 34.0995736 | NA | NA | NA | NA | NA | NA | NA |
| SCN2A | 199 | I | 1.109 | 39.79515264 | 0.609578262 | 38.04817863 | NA | NA | NA | NA | NA | NA | NA |
| SCN2A | 200 | T | 2.638928571 | 41.1108943 | 0.630944908 | 39.28827516 | NA | NA | NA | NA | NA | NA | NA |
| SCN2A | 201 | F | 5.429772727 | 35.71594347 | 0.543335002 | 33.90483239 | NA | NA | NA | NA | NA | NA | NA |
| SCN2A | 202 | A | 6.5603 | 39.62818757 | 0.606866876 | 38.07516147 | NA | NA | NA | NA | NA | NA | NA |
| SCN2A | 203 | Y | 5.499583333 | 42.16004361 | 0.647982297 | 40.65985077 | NA | NA | NA | NA | NA | NA | NA |
| SCN2A | 204 | V | 9.064642857 | 40.87216899 | 0.62706819 | 39.30222415 | NA | NA | NA | NA | NA | NA | NA |
| SCN2A | 205 | T | 12.07378571 | 40.36870897 | 0.618892382 | 38.92904718 | NA | NA | NA | NA | NA | NA | NA |
| SCN2A | 206 | E | 10.9865 | 44.05056916 | 0.678682997 | 42.90903527 | NA | NA | NA | NA | NA | NA | NA |

| | | | | | | | | | | | | | |
|---|---|---|---|---|---|---|---|---|---|---|---|---|---|
| SCN2A | 207 | F | 10.5325 | 46.38646426 | 0.71661616 | 45.13231583 | NA | NA | NA | NA | NA | NA | NA |
| SCN2A | 208 | V | 15.5175 | 46.923663 | 0.72533986 | 45.36860561 | NA | NA | NA | NA | NA | NA | NA |
| SCN2A | 209 | D | 17.238875 | 46.78243424 | 0.723046412 | 45.22276578 | NA | NA | NA | NA | NA | NA | NA |
| SCN2A | 210 | L | 13.874625 | 42.73996666 | 0.657399807 | 41.6217417 | NA | NA | NA | NA | NA | NA | NA |
| SCN2A | 211 | G | 17.2345 | 40.82289555 | 0.626268027 | 39.26904458 | NA | NA | NA | NA | NA | NA | NA |
| SCN2A | 212 | N | 20.423125 | 37.62291395 | 0.574302755 | 36.07685117 | NA | NA | NA | NA | NA | NA | NA |
| SCN2A | 213 | V | 19.52121429 | 34.82587456 | 0.528880958 | 33.27403617 | NA | NA | NA | NA | NA | NA | NA |
| SCN2A | 214 | S | 16.03583333 | 33.50159275 | 0.507375627 | 31.93812979 | NA | NA | NA | NA | NA | NA | NA |
| SCN2A | 215 | A | 15.3805 | 30.83139786 | 0.464013688 | 29.25690372 | NA | NA | NA | NA | NA | NA | NA |
| SCN2A | 216 | L | 11.5955 | 30.8585307 | 0.464454305 | 28.95559515 | NA | NA | NA | NA | NA | NA | NA |
| SCN2A | 217 | R | 14.28268182 | 30.51436942 | 0.458865387 | 30.23184565 | NA | NA | NA | NA | NA | NA | NA |
| SCN2A | 218 | T | 9.855785714 | 26.38055665 | 0.391735405 | 24.88818012 | NA | NA | NA | NA | NA | NA | NA |
| SCN2A | 219 | F | 6.802409091 | 30.59565267 | 0.460185365 | 29.03591265 | NA | NA | NA | NA | NA | NA | NA |
| SCN2A | 220 | R | 9.5365 | 30.62859869 | 0.460720384 | 29.16025325 | NA | NA | NA | NA | NA | NA | NA |
| SCN2A | 221 | V | 5.3265 | 26.74251641 | 0.397613357 | 25.29709148 | NA | NA | NA | NA | NA | NA | NA |
| SCN2A | 222 | L | 2.223125 | 30.0987461 | 0.45211598 | 28.26848664 | NA | NA | NA | NA | NA | NA | NA |
| SCN2A | 223 | R | 3.243772727 | 31.64935657 | 0.477296717 | 30.00378639 | NA | NA | NA | NA | NA | NA | NA |
| SCN2A | 224 | A | 1.2047 | 28.15328333 | 0.420523141 | 26.58832381 | NA | NA | NA | NA | NA | NA | NA |
| SCN2A | 225 | L | -0.408375 | 26.38231965 | 0.391764035 | 21.42670093 | NA | NA | NA | NA | NA | NA | NA |
| SCN2A | 226 | K | -1.822833333 | 30.93292482 | 0.465662409 | 25.99398205 | NA | NA | NA | NA | NA | NA | NA |
| SCN2A | 227 | T | -2.621214286 | 27.85348405 | 0.415654628 | 22.9302871 | NA | NA | NA | NA | NA | NA | NA |
| SCN2A | 228 | I | -5.453 | 26.57375704 | 0.394872833 | 21.63828838 | NA | NA | NA | NA | NA | NA | NA |
| SCN2A | 229 | S | -7.255 | 31.22812388 | 0.470456218 | 26.89576303 | NA | NA | NA | NA | NA | NA | NA |
| SCN2A | 230 | V | -6.644642857 | 33.02158212 | 0.499580618 | 28.68423633 | NA | NA | NA | NA | NA | NA | NA |
| SCN2A | 231 | I | -7.219125 | 29.46497909 | 0.441824085 | 25.16467036 | NA | NA | NA | NA | NA | NA | NA |
| SCN2A | 232 | P | -11.40621429 | 28.91808946 | 0.432943012 | 27.1839754 | NA | NA | NA | NA | NA | NA | NA |
| SCN2A | 233 | G | -11.8125 | 25.14964723 | 0.371746371 | 23.49101033 | NA | NA | NA | NA | NA | NA | NA |
| SCN2A | 234 | L | -8.3525 | 25.86563065 | 0.383373398 | 23.05905408 | NA | NA | NA | NA | NA | NA | NA |
| SCN2A | 235 | K | -11.08805556 | 25.74813182 | 0.381465306 | 23.99568937 | NA | NA | NA | NA | NA | NA | NA |
| SCN2A | 236 | T | -13.7755 | 22.41526275 | 0.327342042 | 20.75331273 | NA | NA | NA | NA | NA | NA | NA |
| SCN2A | 237 | I | -10.458875 | 20.66907204 | 0.29898523 | 18.82824938 | NA | NA | NA | NA | NA | NA | NA |
| SCN2A | 238 | V | -8.1175 | 22.87787193 | 0.334854464 | 20.17284574 | NA | NA | NA | NA | NA | NA | NA |
| SCN2A | 239 | G | -11.6945 | 22.23721179 | 0.32445063 | 20.47483941 | NA | NA | NA | NA | NA | NA | NA |

| | | | | | | | | | | | | | |
|---|---|---|---|---|---|---|---|---|---|---|---|---|---|
| SCN2A | 240 | A | -12.2795 | 18.02736145 | 0.256085856 | 16.24127663 | NA | NA | NA | NA | NA | NA | NA |
| SCN2A | 241 | L | -8.541625 | 17.86406936 | 0.253434116 | 16.15107512 | NA | NA | NA | NA | NA | NA | NA |
| SCN2A | 242 | I | -8.808375 | 20.80402449 | 0.301176756 | 18.91774544 | NA | NA | NA | NA | NA | NA | NA |
| SCN2A | 243 | Q | -13.89683333 | 18.69909674 | 0.266994327 | 16.87533794 | NA | NA | NA | NA | NA | NA | NA |
| SCN2A | 244 | S | -10.94883333 | 14.46061026 | 0.198164523 | 12.58275775 | NA | NA | NA | NA | NA | NA | NA |
| SCN2A | 245 | V | -8.028785714 | 18.00957992 | 0.255797097 | 15.58548717 | NA | NA | NA | NA | NA | NA | NA |
| SCN2A | 246 | K | -12.65305556 | 19.19680127 | 0.275076671 | 17.27393965 | NA | NA | NA | NA | NA | NA | NA |
| SCN2A | 247 | K | -13.48661111 | 15.73354127 | 0.218835957 | 13.84723438 | NA | NA | NA | NA | NA | NA | NA |
| SCN2A | 248 | L | -8.14925 | 14.03021579 | 0.191175244 | 12.04758129 | NA | NA | NA | NA | NA | NA | NA |
| SCN2A | 249 | S | -9.043833333 | 18.82910285 | 0.269105527 | 16.80895322 | NA | NA | NA | NA | NA | NA | NA |
| SCN2A | 250 | D | -10.170375 | 18.72781132 | 0.26746063 | 16.65837291 | NA | NA | NA | NA | NA | NA | NA |
| SCN2A | 251 | V | -7.050357143 | 14.64623556 | 0.201178937 | 11.72165434 | NA | NA | NA | NA | NA | NA | NA |
| SCN2A | 252 | M | -4.046625 | 15.11738648 | 0.208830071 | 10.32334183 | NA | NA | NA | NA | NA | NA | NA |
| SCN2A | 253 | I | -5.441875 | 20.60624614 | 0.297964985 | 15.93823105 | NA | NA | NA | NA | NA | NA | NA |
| SCN2A | 254 | L | -4.892875 | 17.8942291 | 0.253923887 | 15.06300575 | NA | NA | NA | NA | NA | NA | NA |
| SCN2A | 255 | T | -1.512357143 | 15.41501739 | 0.213663371 | 10.5729447 | NA | NA | NA | NA | NA | NA | NA |
| SCN2A | 256 | V | -0.192928571 | 18.98363184 | 0.271614961 | 14.1354845 | NA | NA | NA | NA | NA | NA | NA |
| SCN2A | 257 | F | -1.602045455 | 21.15828325 | 0.306929649 | 16.31172159 | NA | NA | NA | NA | NA | NA | NA |
| SCN2A | 258 | C | -0.735 | 18.6865005 | 0.266789773 | 13.88893028 | NA | NA | NA | NA | NA | NA | NA |
| SCN2A | 259 | L | 2.659 | 15.52466918 | 0.215444033 | 10.7418308 | NA | NA | NA | NA | NA | NA | NA |
| SCN2A | 260 | S | 2.93 | 21.0087027 | 0.304500574 | 16.22083993 | NA | NA | NA | NA | NA | NA | NA |
| SCN2A | 261 | V | 1.866071429 | 22.8334851 | 0.334133656 | 18.04375999 | NA | NA | NA | NA | NA | NA | NA |
| SCN2A | 262 | F | 3.339863636 | 18.89813171 | 0.270226504 | 14.27434532 | NA | NA | NA | NA | NA | NA | NA |
| SCN2A | 263 | A | 7.2595 | 21.61815533 | 0.314397623 | 19.61786585 | NA | NA | NA | NA | NA | NA | NA |
| SCN2A | 264 | L | 6.747 | 24.29199895 | 0.357818814 | 22.28699745 | NA | NA | NA | NA | NA | NA | NA |
| SCN2A | 265 | I | 7.038125 | 24.20777305 | 0.356451049 | 22.64870335 | NA | NA | NA | NA | NA | NA | NA |
| SCN2A | 266 | G | 11.0615 | 23.0384282 | 0.337461776 | 21.00851266 | NA | NA | NA | NA | NA | NA | NA |
| SCN2A | 267 | L | 12.199875 | 24.47536715 | 0.360796574 | 22.4380501 | NA | NA | NA | NA | NA | NA | NA |
| SCN2A | 268 | Q | 11.0165 | 27.0855855 | 0.403184539 | 25.05234071 | NA | NA | NA | NA | NA | NA | NA |
| SCN2A | 269 | L | 12.52525 | 25.35162395 | 0.375026319 | 23.33904022 | NA | NA | NA | NA | NA | NA | NA |
| SCN2A | 270 | F | 14.52059091 | 21.58043153 | 0.313785017 | 19.58498098 | NA | NA | NA | NA | NA | NA | NA |
| SCN2A | 271 | M | 17.763875 | 26.49962017 | 0.393668906 | 24.50617395 | NA | NA | NA | NA | NA | NA | NA |
| SCN2A | 272 | G | 20.794 | 24.85294178 | 0.366928099 | 24.12754374 | NA | NA | NA | NA | NA | NA | NA |

| | | | | | | | | | | | | | |
|---|---|---|---|---|---|---|---|---|---|---|---|---|---|
| SCN2A | 273 | N | 20.429875 | 23.25205311 | 0.340930882 | 22.52651546 | NA | NA | NA | NA | NA | NA | NA |
| SCN2A | 274 | L | 20.973625 | 19.84229747 | 0.285559039 | 19.42265335 | NA | NA | NA | NA | NA | NA | NA |
| SCN2A | 275 | R | 25.50913636 | 22.72632566 | 0.332393468 | 22.004934 | NA | NA | NA | NA | NA | NA | NA |
| SCN2A | 276 | N | 25.867 | 21.19223386 | 0.307480981 | 20.69613734 | NA | NA | NA | NA | NA | NA | NA |
| SCN2A | 277 | K | 29.61983333 | 19.49557434 | 0.279928518 | 18.81813976 | NA | NA | NA | NA | NA | NA | NA |
| SCN2A | 278 | C | 32.2185 | 20.78734658 | 0.300905919 | 20.14091691 | NA | NA | NA | NA | NA | NA | NA |
| SCN2A | 279 | L | 36.0525 | 23.43661112 | 0.343927965 | 22.96708435 | NA | NA | NA | NA | NA | NA | NA |
| SCN2A | 280 | Q | 40.16016667 | 23.31227876 | 0.341908901 | 22.76604099 | NA | NA | NA | NA | NA | NA | NA |
| SCN2A | 281 | W | 41.26314286 | 27.64317798 | 0.412239417 | 27.02330101 | NA | NA | NA | NA | NA | NA | NA |
| SCN2A | 282 | P | 44.28935714 | 29.52574153 | 0.442810821 | 28.94260216 | NA | NA | NA | NA | NA | NA | NA |
| SCN2A | 283 | P | 46.61107143 | 27.22178431 | 0.405396304 | 26.73814746 | NA | NA | NA | NA | NA | NA | NA |
| SCN2A | 284 | D | 49.5825 | 28.55340864 | 0.427020872 | 28.18567907 | NA | NA | NA | NA | NA | NA | NA |
| SCN2A | 314 | F | 44.53522727 | 33.14880137 | 0.501646562 | 32.58865436 | NA | NA | NA | NA | NA | NA | NA |
| SCN2A | 315 | N | 42.861375 | 34.21462958 | 0.518954803 | 33.5768021 | NA | NA | NA | NA | NA | NA | NA |
| SCN2A | 316 | W | 36.99564286 | 30.42182892 | 0.4573626 | 29.80716919 | NA | NA | NA | NA | NA | NA | NA |
| SCN2A | 317 | D | 36.90675 | 29.22393126 | 0.437909651 | 28.58699142 | NA | NA | NA | NA | NA | NA | NA |
| SCN2A | 318 | E | 41.26361111 | 28.37197157 | 0.424074472 | 27.8080779 | NA | NA | NA | NA | NA | NA | NA |
| SCN2A | 319 | Y | 40.65733333 | 26.61686358 | 0.395572851 | 25.98257936 | NA | NA | NA | NA | NA | NA | NA |
| SCN2A | 320 | I | 35.585375 | 25.20553767 | 0.372653989 | 24.61441319 | NA | NA | NA | NA | NA | NA | NA |
| SCN2A | 321 | E | 35.45694444 | 24.52472864 | 0.361598167 | 23.89692457 | NA | NA | NA | NA | NA | NA | NA |
| SCN2A | 322 | D | 39.447625 | 21.02353807 | 0.30474149 | 20.50669806 | NA | NA | NA | NA | NA | NA | NA |
| SCN2A | 323 | K | 34.74472222 | 17.58741055 | 0.248941387 | 16.96962166 | NA | NA | NA | NA | NA | NA | NA |
| SCN2A | 324 | S | 39.45116667 | 19.08651183 | 0.273285654 | 18.68724159 | NA | NA | NA | NA | NA | NA | NA |
| SCN2A | 325 | H | 39.9276 | 21.14890959 | 0.306777428 | 20.55699843 | NA | NA | NA | NA | NA | NA | NA |
| SCN2A | 326 | F | 34.82904545 | 18.56944329 | 0.264888853 | 17.93547396 | NA | NA | NA | NA | NA | NA | NA |
| SCN2A | 327 | Y | 34.14158333 | 16.02758712 | 0.223611038 | 15.56423956 | NA | NA | NA | NA | NA | NA | NA |
| SCN2A | 328 | F | 33.60613636 | 12.91124727 | 0.173004044 | 12.30214673 | NA | NA | NA | NA | NA | NA | NA |
| SCN2A | 329 | L | 30.82275 | 9.939679924 | 0.124748046 | 9.286028683 | NA | NA | NA | NA | NA | NA | NA |
| SCN2A | 330 | E | 32.58627778 | 7.38997977 | 0.083342851 | 6.867848504 | 0.65 | -0.28 | 0.57 | 2.75 | 0.59 | 0 | 0 |
| SCN2A | 331 | G | 31.95925 | 6.14754455 | 0.063166646 | 5.59332278 | 0.65 | -0.28 | 0.57 | 2.75 | 0.59 | 0 | 0 |
| SCN2A | 332 | Q | 27.84516667 | 7.840745437 | 0.090662943 | 7.247331158 | 0.8 | -0.41 | 0.33 | 2.91 | 0.65 | 0 | 0 |
| SCN2A | 333 | N | 29.4805 | 8.436020211 | 0.100329754 | 7.7042699 | NA | NA | NA | NA | NA | NA | NA |
| SCN2A | 334 | D | 27.03125 | 10.84899244 | 0.139514591 | 10.42570329 | NA | NA | NA | NA | NA | NA | NA |

| | | | | | | | | | | | | | |
|---|---|---|---|---|---|---|---|---|---|---|---|---|---|
| SCN2A | 335 | A | 29.3161 | 13.81312796 | 0.187649902 | 13.14476889 | NA | NA | NA | NA | NA | NA | NA |
| SCN2A | 336 | L | 28.806125 | 15.64519009 | 0.217401201 | 15.0201225 | NA | NA | NA | NA | NA | NA | NA |
| SCN2A | 337 | L | 25.150125 | 17.28581392 | 0.244043687 | 16.57303504 | NA | NA | NA | NA | NA | NA | NA |
| SCN2A | 338 | C | 26.84816667 | 21.52404739 | 0.312869381 | 20.8610203 | NA | NA | NA | NA | NA | NA | NA |
| SCN2A | 339 | G | 25.68025 | 24.62001485 | 0.363145543 | 23.91753488 | NA | NA | NA | NA | NA | NA | NA |
| SCN2A | 340 | N | 26.6895 | 28.19269301 | 0.421163124 | 27.48832337 | NA | NA | NA | NA | NA | NA | NA |
| SCN2A | 341 | S | 22.66933333 | 28.50381911 | 0.426215576 | 27.78981871 | NA | NA | NA | NA | NA | NA | NA |
| SCN2A | 342 | S | 22.42166667 | 29.68805597 | 0.445446684 | 28.9588756 | NA | NA | NA | NA | NA | NA | NA |
| SCN2A | 343 | D | 19.49575 | 25.57334098 | 0.378626836 | 25.03020857 | NA | NA | NA | NA | NA | NA | NA |
| SCN2A | 344 | A | 22.5461 | 24.41616831 | 0.35983523 | 23.69729361 | NA | NA | NA | NA | NA | NA | NA |
| SCN2A | 345 | G | 25.155 | 22.61836796 | 0.330640317 | 21.9164335 | NA | NA | NA | NA | NA | NA | NA |
| SCN2A | 346 | Q | 28.06961111 | 21.74275682 | 0.316421056 | 21.11124119 | NA | NA | NA | NA | NA | NA | NA |
| SCN2A | 347 | C | 31.90216667 | 22.99248758 | 0.336715735 | 22.34726637 | NA | NA | NA | NA | NA | NA | NA |
| SCN2A | 348 | P | 34.54764286 | 20.87118645 | 0.302267415 | 20.21688563 | NA | NA | NA | NA | NA | NA | NA |
| SCN2A | 349 | E | 38.70794444 | 24.86895257 | 0.367188102 | 24.37639694 | NA | NA | NA | NA | NA | NA | NA |
| SCN2A | 350 | G | 38.963 | 26.6475607 | 0.396071348 | 26.07880324 | NA | NA | NA | NA | NA | NA | NA |
| SCN2A | 351 | Y | 36.90416667 | 20.80124369 | 0.301131598 | 20.168232 | NA | NA | NA | NA | NA | NA | NA |
| SCN2A | 352 | I | 34.506125 | 25.30064768 | 0.374198503 | 24.64013195 | NA | NA | NA | NA | NA | NA | NA |
| SCN2A | 353 | C | 30.24883333 | 25.63001032 | 0.379547103 | 24.98657248 | NA | NA | NA | NA | NA | NA | NA |
| SCN2A | 354 | V | 31.08535714 | 25.17017926 | 0.372079795 | 24.50785017 | NA | NA | NA | NA | NA | NA | NA |
| SCN2A | 355 | K | 26.97083333 | 26.71636835 | 0.397188733 | 26.05678358 | NA | NA | NA | NA | NA | NA | NA |
| SCN2A | 356 | A | 30.2081 | 23.31588731 | 0.341967501 | 22.63301207 | NA | NA | NA | NA | NA | NA | NA |
| SCN2A | 357 | G | 28.7625 | 22.57533402 | 0.329941478 | 21.88342638 | NA | NA | NA | NA | NA | NA | NA |
| SCN2A | 358 | R | 27.04331818 | 15.86230973 | 0.220927059 | 15.15724065 | NA | NA | NA | NA | NA | NA | NA |
| SCN2A | 359 | N | 23.325 | 17.57539783 | 0.248746309 | 16.859068 | NA | NA | NA | NA | NA | NA | NA |
| SCN2A | 360 | P | 22.59878571 | 15.48716065 | 0.214834923 | 14.76279817 | NA | NA | NA | NA | NA | NA | NA |
| SCN2A | 361 | N | 21.727625 | 11.90710679 | 0.156697565 | 11.22194845 | NA | NA | NA | NA | NA | NA | NA |
| SCN2A | 362 | Y | 23.4395 | 12.14035358 | 0.160485316 | 11.58538412 | NA | NA | NA | NA | NA | NA | NA |
| SCN2A | 363 | G | 22.08025 | 17.70896931 | 0.250915409 | 16.98324288 | NA | NA | NA | NA | NA | NA | NA |
| SCN2A | 364 | Y | 19.4675 | 16.65047702 | 0.233726297 | 15.05470102 | NA | NA | NA | NA | NA | NA | NA |
| SCN2A | 365 | T | 18.39335714 | 15.66994365 | 0.21780318 | 14.46315508 | NA | NA | NA | NA | NA | NA | NA |
| SCN2A | 366 | S | 17.214 | 19.61490589 | 0.281866372 | 17.60764334 | NA | NA | NA | NA | NA | NA | NA |
| SCN2A | 367 | F | 14.56913636 | 18.75126422 | 0.267841487 | 16.73526342 | NA | NA | NA | NA | NA | NA | NA |

| | | | | | | | | | | | | | |
|---|---|---|---|---|---|---|---|---|---|---|---|---|---|
| SCN2A | 368 | D | 18.0875 | 24.46998508 | 0.360709173 | 22.43933214 | NA | NA | NA | NA | NA | NA | NA |
| SCN2A | 369 | T | 15.36092857 | 24.38920044 | 0.359397292 | 22.32691737 | NA | NA | NA | NA | NA | NA | NA |
| SCN2A | 370 | F | 9.170227273 | 22.90281751 | 0.335259561 | 20.83568443 | NA | NA | NA | NA | NA | NA | NA |
| SCN2A | 371 | S | 11.616 | 20.82522533 | 0.301521041 | 18.7531282 | NA | NA | NA | NA | NA | NA | NA |
| SCN2A | 372 | W | 15.5625 | 19.06945754 | 0.273008705 | 17.0117612 | NA | NA | NA | NA | NA | NA | NA |
| SCN2A | 373 | A | 11.1205 | 18.94849166 | 0.271044311 | 16.88897488 | NA | NA | NA | NA | NA | NA | NA |
| SCN2A | 374 | F | 6.894227273 | 17.39092485 | 0.245750608 | 15.32178442 | NA | NA | NA | NA | NA | NA | NA |
| SCN2A | 375 | L | 11.169375 | 14.47940137 | 0.198469677 | 12.41010497 | NA | NA | NA | NA | NA | NA | NA |
| SCN2A | 376 | S | 11.2955 | 14.1408457 | 0.19297179 | 12.09244895 | NA | NA | NA | NA | NA | NA | NA |
| SCN2A | 377 | L | 7.53275 | 14.37921538 | 0.196842732 | 12.47310678 | NA | NA | NA | NA | NA | NA | NA |
| SCN2A | 378 | F | 5.583772727 | 11.68887723 | 0.153153683 | 9.623027284 | NA | NA | NA | NA | NA | NA | NA |
| SCN2A | 379 | R | 11.17713636 | 9.345789747 | 0.11510372 | 7.288611912 | NA | NA | NA | NA | NA | NA | NA |
| SCN2A | 380 | L | 7.687 | 10.50505045 | 0.133929234 | 8.982690594 | NA | NA | NA | NA | NA | NA | NA |
| SCN2A | 381 | M | 2.639875 | 8.531399651 | 0.101878643 | 3.927637823 | 4.64 | -0.35 | -1.08 | 12.03 | 0.28 | 1 | 1 |
| SCN2A | 382 | T | 5.0555 | 5.949995462 | 0.059958599 | 2.057369391 | 4.05 | -0.17 | -0.48 | 14.15 | 0.32 | 1 | 1 |
| SCN2A | 383 | Q | 5.575944444 | 4.653986033 | 0.038912389 | 2.597952306 | 2.09 | -0.25 | -1.25 | 25.78 | 0.67 | 2 | 0 |
| SCN2A | 384 | D | 10.19 | 5.052720158 | 0.045387529 | 2.979654084 | 2.09 | -0.25 | -1.25 | 25.78 | 0.67 | 2 | 0 |
| SCN2A | 385 | F | 14.8895 | 5.592969605 | 0.05416077 | 4.022863111 | 1.83 | -0.07 | -1.53 | 33.25 | 0.63 | 2 | 0 |
| SCN2A | 386 | W | 10.10857143 | 7.819925639 | 0.090324846 | 5.885990977 | 2.09 | -0.25 | -1.25 | 25.78 | 0.67 | 2 | 0 |
| SCN2A | 387 | E | 14.27394444 | 6.811571772 | 0.073949944 | 6.825611957 | 2.09 | -0.25 | -1.25 | 25.78 | 0.67 | 2 | 0 |
| SCN2A | 388 | N | 17.72925 | 10.09929473 | 0.127340069 | 7.917754023 | NA | NA | NA | NA | NA | NA | NA |
| SCN2A | 389 | L | 14.654 | 12.61427628 | 0.168181461 | 10.73646017 | NA | NA | NA | NA | NA | NA | NA |
| SCN2A | 390 | Y | 13.24075 | 13.8419624 | 0.188118152 | 11.94290323 | NA | NA | NA | NA | NA | NA | NA |
| SCN2A | 391 | Q | 17.76372222 | 12.74969604 | 0.170380575 | 10.59535331 | NA | NA | NA | NA | NA | NA | NA |
| SCN2A | 392 | L | 19.821125 | 14.86779937 | 0.204776965 | 13.40388695 | NA | NA | NA | NA | NA | NA | NA |
| SCN2A | 393 | T | 17.06335714 | 18.28146969 | 0.26021238 | 15.9241459 | NA | NA | NA | NA | NA | NA | NA |
| SCN2A | 394 | L | 16.824125 | 19.37290585 | 0.277936475 | 17.50267748 | NA | NA | NA | NA | NA | NA | NA |
| SCN2A | 395 | R | 22.89777273 | 16.4379677 | 0.230275307 | 15.73031086 | NA | NA | NA | NA | NA | NA | NA |
| SCN2A | 396 | A | 21.6109 | 21.80450965 | 0.317423875 | 21.07513631 | NA | NA | NA | NA | NA | NA | NA |
| SCN2A | 397 | A | 18.7503 | 24.15856074 | 0.355651878 | 22.0188293 | NA | NA | NA | NA | NA | NA | NA |
| SCN2A | 398 | G | 17.82625 | 23.74706506 | 0.348969501 | 21.72952592 | NA | NA | NA | NA | NA | NA | NA |
| SCN2A | 399 | K | 16.8515 | 22.93504569 | 0.335782923 | 20.77433389 | NA | NA | NA | NA | NA | NA | NA |
| SCN2A | 400 | T | 13.4115 | 25.01357188 | 0.36953661 | 23.07430745 | NA | NA | NA | NA | NA | NA | NA |

| | | | | | | | | | | | | | |
|---|---|---|---|---|---|---|---|---|---|---|---|---|---|
| SCN2A | 401 | Y | 13.55333333 | 21.32376153 | 0.309616891 | 19.35849266 | NA | NA | NA | NA | NA | NA | NA |
| SCN2A | 402 | M | 10.329875 | 20.04764053 | 0.288893654 | 18.06976391 | NA | NA | NA | NA | NA | NA | NA |
| SCN2A | 403 | I | 7.477125 | 21.13359858 | 0.306528789 | 16.5657452 | NA | NA | NA | NA | NA | NA | NA |
| SCN2A | 404 | F | 8.1985 | 18.87631092 | 0.269872151 | 14.32652258 | NA | NA | NA | NA | NA | NA | NA |
| SCN2A | 405 | F | 9.848045455 | 15.83978463 | 0.220561268 | 14.03943102 | NA | NA | NA | NA | NA | NA | NA |
| SCN2A | 406 | V | 5.0435 | 16.40697879 | 0.229772071 | 11.80658438 | NA | NA | NA | NA | NA | NA | NA |
| SCN2A | 407 | L | 2.626875 | 17.08122086 | 0.24072125 | 12.4695481 | NA | NA | NA | NA | NA | NA | NA |
| SCN2A | 408 | V | 4.342785714 | 13.97568932 | 0.190289775 | 9.369693838 | NA | NA | NA | NA | NA | NA | NA |
| SCN2A | 409 | I | 3.864 | 11.50440972 | 0.15015807 | 6.930129438 | NA | NA | NA | NA | NA | NA | NA |
| SCN2A | 410 | F | 1.024954545 | 13.96691985 | 0.190147366 | 9.349343335 | NA | NA | NA | NA | NA | NA | NA |
| SCN2A | 411 | L | -0.399125 | 14.82904667 | 0.204147651 | 10.02719006 | NA | NA | NA | NA | NA | NA | NA |
| SCN2A | 412 | G | 0.21675 | 11.5578983 | 0.151026684 | 6.758329552 | NA | NA | NA | NA | NA | NA | NA |
| SCN2A | 413 | S | 1.2425 | 8.344410824 | 0.098842087 | 3.54356874 | 4.81 | 0.09 | 0.6 | 2.15 | -0.38 | 0 | 0 |
| SCN2A | 414 | F | -2.296045455 | 8.945420001 | 0.10860202 | 5.945417197 | NA | NA | NA | NA | NA | NA | NA |
| SCN2A | 415 | Y | -4.33275 | 11.61612556 | 0.151972251 | 8.734425884 | NA | NA | NA | NA | NA | NA | NA |
| SCN2A | 416 | L | -2.200375 | 11.07289127 | 0.143150538 | 7.556986489 | NA | NA | NA | NA | NA | NA | NA |
| SCN2A | 417 | I | -1.993875 | 6.152380677 | 0.063245181 | 2.674224318 | 3.54 | 1.13 | 2.1 | 0.62 | -1.67 | 0 | 0 |
| SCN2A | 418 | N | -6.034 | 6.681827445 | 0.071842995 | 3.836775236 | 2.92 | 1.58 | 2.57 | 0.62 | -1.71 | 0 | 0 |
| SCN2A | 419 | L | -7.604 | 9.92528191 | 0.124514233 | 7.035660004 | NA | NA | NA | NA | NA | NA | NA |
| SCN2A | 420 | I | -5.280625 | 9.527130575 | 0.118048558 | 6.615170315 | NA | NA | NA | NA | NA | NA | NA |
| SCN2A | 421 | L | -6.394625 | 4.696630068 | 0.039604896 | 1.879753715 | 3.01 | 1.47 | 2.88 | 0.5 | -1.73 | 0 | 0 |
| SCN2A | 422 | A | -10.8165 | 7.021596969 | 0.077360594 | 4.95590442 | 2.24 | 1.53 | 3.9 | 0.19 | -1.83 | 0 | 0 |
| SCN2A | 423 | V | -11.38407143 | 9.124360087 | 0.111507871 | 7.08526757 | NA | NA | NA | NA | NA | NA | NA |
| SCN2A | 424 | V | -9.927071429 | 9.235608643 | 0.113314463 | 7.329798246 | NA | NA | NA | NA | NA | NA | NA |
| SCN2A | 425 | A | -12.8187 | 5.592318035 | 0.054150189 | 3.722548952 | 2.24 | 1.53 | 3.9 | 0.19 | -1.83 | 0 | 0 |
| SCN2A | 426 | M | -15.744125 | 7.890581347 | 0.091472241 | 6.308129673 | 2.09 | 1.43 | 3.7 | 0.2 | -1.82 | 0 | 0 |
| SCN2A | 427 | A | -14.9379 | 10.5356846 | 0.13442671 | 8.83145446 | NA | NA | NA | NA | NA | NA | NA |
| SCN2A | 428 | Y | -14.51766667 | 10.21370584 | 0.129198019 | 8.412670817 | NA | NA | NA | NA | NA | NA | NA |
| SCN2A | 429 | E | -18.50561111 | 4.729140091 | 0.040132834 | 1.650426575 | 3.22 | 0.57 | 1.03 | 1.29 | -1.02 | 0 | 0 |
| SCN2A | 430 | E | -19.65116667 | 10.93828967 | 0.140964711 | 7.908963534 | NA | NA | NA | NA | NA | NA | NA |
| SCN2A | 431 | Q | -17.98427778 | 13.30203281 | 0.179350105 | 10.08771665 | NA | NA | NA | NA | NA | NA | NA |
| SCN2A | 432 | N | -19.5635 | 10.74218269 | 0.137780082 | 7.53946484 | NA | NA | NA | NA | NA | NA | NA |
| SCN2A | 433 | Q | -23.73405556 | 9.939994014 | 0.124753147 | 7.033958121 | NA | NA | NA | NA | NA | NA | NA |

| | | | | | | | | | | | | | |
|---|---|---|---|---|---|---|---|---|---|---|---|---|---|
| SCN2A | 434 | A | -23.7697 | 14.75053036 | 0.202872606 | 11.63469848 | NA | NA | NA | NA | NA | NA | NA |
| SCN2A | 435 | T | -22.51278571 | 16.39593251 | 0.229592688 | 13.19466768 | NA | NA | NA | NA | NA | NA | NA |
| SCN2A | 436 | L | -25.697375 | 13.04954271 | 0.175249857 | 10.22721183 | NA | NA | NA | NA | NA | NA | NA |
| SCN2A | 437 | E | -28.43694444 | 16.04558868 | 0.22390337 | 11.67325075 | NA | NA | NA | NA | NA | NA | NA |
| SCN2A | 438 | E | -26.97394444 | 18.80744908 | 0.268753887 | 14.39699526 | NA | NA | NA | NA | NA | NA | NA |
| SCN2A | 439 | A | -27.9683 | 20.51934093 | 0.296553711 | 16.13550135 | NA | NA | NA | NA | NA | NA | NA |
| SCN2A | 440 | E | -30.37527778 | 16.62697089 | 0.233344576 | 12.43907719 | NA | NA | NA | NA | NA | NA | NA |
| SCN2A | 441 | Q | -32.30172222 | 19.98227139 | 0.287832109 | 15.78278428 | NA | NA | NA | NA | NA | NA | NA |
| SCN2A | 740 | D | -15.516625 | 63.83702369 | 1 | 60.91221518 | NA | NA | NA | NA | NA | NA | NA |
| SCN2A | 741 | C | -14.5415 | 60.9721987 | 0.953477417 | 58.04674448 | NA | NA | NA | NA | NA | NA | NA |
| SCN2A | 742 | C | -17.38066667 | 59.13231701 | 0.923599135 | 56.19589277 | NA | NA | NA | NA | NA | NA | NA |
| SCN2A | 743 | K | -18.95027778 | 60.35177686 | 0.943402237 | 57.41635352 | NA | NA | NA | NA | NA | NA | NA |
| SCN2A | 744 | P | -15.64121429 | 59.9241994 | 0.936458704 | 57.00833036 | NA | NA | NA | NA | NA | NA | NA |
| SCN2A | 745 | W | -15.34342857 | 53.56036913 | 0.83311493 | 50.62788446 | NA | NA | NA | NA | NA | NA | NA |
| SCN2A | 746 | L | -18.72725 | 54.99834007 | 0.856466487 | 52.0516631 | NA | NA | NA | NA | NA | NA | NA |
| SCN2A | 747 | K | -17.30061111 | 58.23234526 | 0.908984276 | 55.31304976 | NA | NA | NA | NA | NA | NA | NA |
| SCN2A | 748 | V | -13.05964286 | 55.59463496 | 0.866149863 | 52.69170043 | NA | NA | NA | NA | NA | NA | NA |
| SCN2A | 749 | K | -15.83572222 | 50.38809623 | 0.781599626 | 47.46538472 | NA | NA | NA | NA | NA | NA | NA |
| SCN2A | 750 | H | -18.7614 | 52.49889988 | 0.815877475 | 49.57517365 | NA | NA | NA | NA | NA | NA | NA |
| SCN2A | 751 | L | -14.6465 | 52.12244398 | 0.809764117 | 49.2159085 | NA | NA | NA | NA | NA | NA | NA |
| SCN2A | 752 | V | -13.3585 | 49.15752293 | 0.76161605 | 46.255031 | NA | NA | NA | NA | NA | NA | NA |
| SCN2A | 753 | N | -17.9695 | 47.76411403 | 0.738988147 | 44.84328079 | NA | NA | NA | NA | NA | NA | NA |
| SCN2A | 754 | L | -17.999375 | 49.59919481 | 0.768788466 | 46.68455208 | NA | NA | NA | NA | NA | NA | NA |
| SCN2A | 755 | V | -14.52892857 | 45.83487705 | 0.707658802 | 42.92788424 | NA | NA | NA | NA | NA | NA | NA |
| SCN2A | 756 | V | -13.86678571 | 42.59989692 | 0.655125181 | 40.2676694 | NA | NA | NA | NA | NA | NA | NA |
| SCN2A | 757 | M | -19.315 | 42.22262951 | 0.648998644 | 39.29239824 | NA | NA | NA | NA | NA | NA | NA |
| SCN2A | 758 | D | -18.52675 | 42.98248713 | 0.661338155 | 40.06829935 | NA | NA | NA | NA | NA | NA | NA |
| SCN2A | 759 | P | -15.07021429 | 42.48562264 | 0.653269453 | 40.10779901 | NA | NA | NA | NA | NA | NA | NA |
| SCN2A | 760 | F | -11.80686364 | 41.7421131 | 0.641195422 | 39.3119766 | NA | NA | NA | NA | NA | NA | NA |
| SCN2A | 761 | V | -10.51378571 | 41.86148331 | 0.643133904 | 39.59758143 | NA | NA | NA | NA | NA | NA | NA |
| SCN2A | 762 | D | -11.684125 | 37.89817781 | 0.578772831 | 35.44231543 | NA | NA | NA | NA | NA | NA | NA |
| SCN2A | 763 | L | -10.168375 | 36.58653989 | 0.557472827 | 34.00105311 | NA | NA | NA | NA | NA | NA | NA |
| SCN2A | 764 | A | -6.8211 | 39.84034224 | 0.610312107 | 36.46270954 | NA | NA | NA | NA | NA | NA | NA |

| | | | | | | | | | | | | | |
|---|---|---|---|---|---|---|---|---|---|---|---|---|---|
| SCN2A | 765 | I | -5.727 | 36.64131735 | 0.558362371 | 33.26125102 | NA | NA | NA | NA | NA | NA | NA |
| SCN2A | 766 | T | -6.522928571 | 34.13279356 | 0.517625849 | 30.75529869 | NA | NA | NA | NA | NA | NA | NA |
| SCN2A | 767 | I | -3.896875 | 36.35229491 | 0.553668866 | 32.97510127 | NA | NA | NA | NA | NA | NA | NA |
| SCN2A | 768 | C | -1.396 | 36.60161603 | 0.557717652 | 33.21932802 | NA | NA | NA | NA | NA | NA | NA |
| SCN2A | 769 | I | -1.70725 | 32.22112779 | 0.486581848 | 28.83951798 | NA | NA | NA | NA | NA | NA | NA |
| SCN2A | 770 | V | -1.263785714 | 31.33755206 | 0.472233248 | 27.95931539 | NA | NA | NA | NA | NA | NA | NA |
| SCN2A | 771 | L | 0.83375 | 34.75805146 | 0.527779562 | 31.37655612 | NA | NA | NA | NA | NA | NA | NA |
| SCN2A | 772 | N | 2.66975 | 32.68825939 | 0.49416771 | 31.84769423 | NA | NA | NA | NA | NA | NA | NA |
| SCN2A | 773 | T | 2.371214286 | 29.06937559 | 0.435399784 | 28.59987253 | NA | NA | NA | NA | NA | NA | NA |
| SCN2A | 774 | L | 3.7755 | 31.05656892 | 0.467670295 | 30.20860506 | NA | NA | NA | NA | NA | NA | NA |
| SCN2A | 775 | F | 6.544954545 | 31.87624772 | 0.480981257 | 29.89096618 | NA | NA | NA | NA | NA | NA | NA |
| SCN2A | 776 | M | 7.479625 | 28.25331784 | 0.422147626 | 26.27357904 | NA | NA | NA | NA | NA | NA | NA |
| SCN2A | 777 | A | 8.2493 | 27.2165078 | 0.405310618 | 25.29570168 | NA | NA | NA | NA | NA | NA | NA |
| SCN2A | 778 | M | 10.2565 | 29.36317818 | 0.440170915 | 27.21318836 | NA | NA | NA | NA | NA | NA | NA |
| SCN2A | 779 | E | 13.49527778 | 26.39727558 | 0.392006908 | 24.32385678 | NA | NA | NA | NA | NA | NA | NA |
| SCN2A | 780 | H | 16.6881 | 26.42913567 | 0.392524292 | 24.38549276 | NA | NA | NA | NA | NA | NA | NA |
| SCN2A | 781 | Y | 21.94775 | 23.5996794 | 0.34657607 | 21.69960941 | NA | NA | NA | NA | NA | NA | NA |
| SCN2A | 782 | P | 22.31692857 | 26.24551676 | 0.38954246 | 24.29705197 | NA | NA | NA | NA | NA | NA | NA |
| SCN2A | 783 | M | 19.2895 | 29.52004866 | 0.442718373 | 27.52328823 | NA | NA | NA | NA | NA | NA | NA |
| SCN2A | 784 | T | 21.69092857 | 33.01412343 | 0.499459495 | 31.02012452 | NA | NA | NA | NA | NA | NA | NA |
| SCN2A | 785 | E | 22.08461111 | 35.89970298 | 0.546319117 | 33.91029558 | NA | NA | NA | NA | NA | NA | NA |
| SCN2A | 786 | Q | 18.2775 | 38.69323531 | 0.591683961 | 36.67730156 | NA | NA | NA | NA | NA | NA | NA |
| SCN2A | 787 | F | 15.67640909 | 32.67118169 | 0.493890381 | 30.91795886 | NA | NA | NA | NA | NA | NA | NA |
| SCN2A | 788 | S | 16.9685 | 34.36122645 | 0.521335425 | 32.33869187 | NA | NA | NA | NA | NA | NA | NA |
| SCN2A | 789 | S | 15.7685 | 37.65203651 | 0.574775683 | 35.62619893 | NA | NA | NA | NA | NA | NA | NA |
| SCN2A | 790 | V | 12.50964286 | 37.82998541 | 0.577665438 | 35.75596069 | NA | NA | NA | NA | NA | NA | NA |
| SCN2A | 791 | L | 12.317875 | 32.75702729 | 0.495284449 | 30.77175095 | NA | NA | NA | NA | NA | NA | NA |
| SCN2A | 792 | S | 13.1195 | 37.12601801 | 0.566233542 | 35.14549901 | NA | NA | NA | NA | NA | NA | NA |
| SCN2A | 793 | V | 11.09992857 | 39.75217566 | 0.608880349 | 37.77454441 | NA | NA | NA | NA | NA | NA | NA |
| SCN2A | 794 | G | 8.046 | 38.69208016 | 0.591665203 | 36.72519725 | NA | NA | NA | NA | NA | NA | NA |
| SCN2A | 795 | N | 8.162125 | 34.25178973 | 0.519558256 | 32.28701383 | NA | NA | NA | NA | NA | NA | NA |
| SCN2A | 796 | L | 8.124875 | 39.58208127 | 0.606118145 | 37.62588215 | NA | NA | NA | NA | NA | NA | NA |
| SCN2A | 797 | V | 4.470357143 | 41.49208182 | 0.637135104 | 39.53955401 | NA | NA | NA | NA | NA | NA | NA |

| | | | | | | | | | | | | | |
|---|---|---|---|---|---|---|---|---|---|---|---|---|---|
| SCN2A | 798 | F | 2.413045455 | 38.22815708 | 0.584131444 | 37.34554934 | NA | NA | NA | NA | NA | NA | NA |
| SCN2A | 799 | T | 3.395785714 | 38.1228436 | 0.582421233 | 37.30831992 | NA | NA | NA | NA | NA | NA | NA |
| SCN2A | 800 | G | 1.42575 | 42.28029572 | 0.6499351 | 41.5525472 | NA | NA | NA | NA | NA | NA | NA |
| SCN2A | 801 | I | -0.375625 | 41.72252223 | 0.640877282 | 38.34062927 | NA | NA | NA | NA | NA | NA | NA |
| SCN2A | 802 | F | -0.661863636 | 37.2520498 | 0.568280202 | 33.86933567 | NA | NA | NA | NA | NA | NA | NA |
| SCN2A | 803 | T | -0.603071429 | 42.15597584 | 0.647916239 | 38.32752643 | NA | NA | NA | NA | NA | NA | NA |
| SCN2A | 804 | A | -2.5151 | 45.24856883 | 0.698137602 | 41.8704357 | NA | NA | NA | NA | NA | NA | NA |
| SCN2A | 805 | E | -5.900055556 | 40.58303776 | 0.622372919 | 37.15697605 | NA | NA | NA | NA | NA | NA | NA |
| SCN2A | 806 | M | -4.593625 | 42.08820469 | 0.646815687 | 38.66223777 | NA | NA | NA | NA | NA | NA | NA |
| SCN2A | 807 | F | -3.344045455 | 45.43579104 | 0.701177949 | 42.01034259 | NA | NA | NA | NA | NA | NA | NA |
| SCN2A | 808 | L | -7.509375 | 46.96732168 | 0.726048844 | 43.53006694 | NA | NA | NA | NA | NA | NA | NA |
| SCN2A | 809 | K | -10.14927778 | 39.55387233 | 0.605660053 | 36.19518474 | NA | NA | NA | NA | NA | NA | NA |
| SCN2A | 810 | I | -8.881 | 44.87878137 | 0.692132534 | 41.4381085 | NA | NA | NA | NA | NA | NA | NA |
| SCN2A | 811 | I | -10.8815 | 48.28484508 | 0.747444424 | 45.91337384 | NA | NA | NA | NA | NA | NA | NA |
| SCN2A | 812 | A | -14.7575 | 46.32898636 | 0.715682763 | 43.40970811 | NA | NA | NA | NA | NA | NA | NA |
| SCN2A | 813 | M | -15.1955 | 39.92725892 | 0.611723568 | 37.009961 | NA | NA | NA | NA | NA | NA | NA |
| SCN2A | 814 | D | -16.44275 | 45.32601928 | 0.699395339 | 42.39829858 | NA | NA | NA | NA | NA | NA | NA |
| SCN2A | 815 | P | -12.73692857 | 46.22744173 | 0.714033755 | 43.31490608 | NA | NA | NA | NA | NA | NA | NA |
| SCN2A | 816 | Y | -10.9525 | 43.28568647 | 0.666261882 | 40.35655878 | NA | NA | NA | NA | NA | NA | NA |
| SCN2A | 817 | Y | -17.55458333 | 40.93758877 | 0.628130558 | 38.01004655 | NA | NA | NA | NA | NA | NA | NA |
| SCN2A | 818 | Y | -10.78283333 | 39.58376295 | 0.606145454 | 36.6848233 | NA | NA | NA | NA | NA | NA | NA |
| SCN2A | 819 | F | -9.8575 | 38.74497184 | 0.592524123 | 35.82983944 | NA | NA | NA | NA | NA | NA | NA |
| SCN2A | 820 | Q | -16.13116667 | 35.15438785 | 0.534215765 | 32.22495393 | NA | NA | NA | NA | NA | NA | NA |
| SCN2A | 821 | E | -14.86761111 | 32.51830577 | 0.491407792 | 29.60047635 | NA | NA | NA | NA | NA | NA | NA |
| SCN2A | 822 | G | -10.7325 | 33.54574346 | 0.508092601 | 31.23428402 | NA | NA | NA | NA | NA | NA | NA |
| SCN2A | 823 | W | -10.09564286 | 28.22371375 | 0.421666878 | 25.12142855 | NA | NA | NA | NA | NA | NA | NA |
| SCN2A | 824 | N | -9.759125 | 35.90088418 | 0.546338298 | 32.78755985 | NA | NA | NA | NA | NA | NA | NA |
| SCN2A | 825 | I | -7.676125 | 35.76820056 | 0.544183617 | 32.32986057 | NA | NA | NA | NA | NA | NA | NA |
| SCN2A | 826 | F | -4.627136364 | 29.74469638 | 0.446366481 | 26.32291609 | NA | NA | NA | NA | NA | NA | NA |
| SCN2A | 827 | D | -4.67925 | 35.13997265 | 0.533981673 | 31.70797132 | NA | NA | NA | NA | NA | NA | NA |
| SCN2A | 828 | G | -2.87525 | 37.67812063 | 0.575199269 | 34.25018542 | NA | NA | NA | NA | NA | NA | NA |
| SCN2A | 829 | F | -0.540318182 | 32.40506426 | 0.489568836 | 28.40686703 | NA | NA | NA | NA | NA | NA | NA |
| SCN2A | 830 | I | 0.148125 | 31.97829154 | 0.482638371 | 27.67604971 | NA | NA | NA | NA | NA | NA | NA |

| | | | | | | | | | | | | | |
|---|---|---|---|---|---|---|---|---|---|---|---|---|---|
| SCN2A | 831 | V | 0.378928571 | 37.17560416 | 0.567038783 | 33.07886867 | NA | NA | NA | NA | NA | NA | NA |
| SCN2A | 832 | S | 1.439166667 | 37.45048942 | 0.571502711 | 37.18262102 | NA | NA | NA | NA | NA | NA | NA |
| SCN2A | 833 | L | 3.8885 | 32.44103699 | 0.490153006 | 32.08413138 | NA | NA | NA | NA | NA | NA | NA |
| SCN2A | 834 | S | 4.567666667 | 36.14686267 | 0.550332802 | 35.69334636 | NA | NA | NA | NA | NA | NA | NA |
| SCN2A | 835 | L | 5.17975 | 38.55688662 | 0.589469762 | 37.92111702 | NA | NA | NA | NA | NA | NA | NA |
| SCN2A | 836 | M | 6.97875 | 37.45008589 | 0.571496158 | 36.68353025 | NA | NA | NA | NA | NA | NA | NA |
| SCN2A | 837 | E | 10.06338889 | 33.863938 | 0.513259839 | 32.99700333 | NA | NA | NA | NA | NA | NA | NA |
| SCN2A | 838 | L | 10.107125 | 37.95861728 | 0.579754322 | 36.04284039 | NA | NA | NA | NA | NA | NA | NA |
| SCN2A | 839 | G | 11.27375 | 40.11362099 | 0.614749947 | 38.19777222 | NA | NA | NA | NA | NA | NA | NA |
| SCN2A | 840 | L | 13.863125 | 37.80719156 | 0.577295283 | 35.73522371 | NA | NA | NA | NA | NA | NA | NA |
| SCN2A | 841 | A | 15.9631 | 35.93433268 | 0.546881477 | 33.89177107 | NA | NA | NA | NA | NA | NA | NA |
| SCN2A | 842 | N | 18.613375 | 33.68423146 | 0.510341541 | 31.63920892 | NA | NA | NA | NA | NA | NA | NA |
| SCN2A | 843 | V | 15.8685 | 31.03085192 | 0.467252671 | 28.97837227 | NA | NA | NA | NA | NA | NA | NA |
| SCN2A | 844 | E | 20.70405556 | 28.90868025 | 0.432790214 | 26.8608021 | NA | NA | NA | NA | NA | NA | NA |
| SCN2A | 845 | G | 17.36625 | 26.36809671 | 0.391533065 | 24.29417351 | NA | NA | NA | NA | NA | NA | NA |
| SCN2A | 846 | L | 14.071375 | 28.47866939 | 0.425807164 | 26.40808223 | NA | NA | NA | NA | NA | NA | NA |
| SCN2A | 847 | S | 13.8745 | 28.29055364 | 0.422752307 | 26.22708875 | NA | NA | NA | NA | NA | NA | NA |
| SCN2A | 848 | V | 10.93021429 | 24.49826908 | 0.361168484 | 22.6112045 | NA | NA | NA | NA | NA | NA | NA |
| SCN2A | 849 | L | 8.482 | 28.78137087 | 0.430722806 | 27.87573105 | NA | NA | NA | NA | NA | NA | NA |
| SCN2A | 850 | R | 10.53986364 | 29.10425386 | 0.435966181 | 28.33615678 | NA | NA | NA | NA | NA | NA | NA |
| SCN2A | 851 | S | 7.316166667 | 25.08835144 | 0.370750974 | 24.28743469 | NA | NA | NA | NA | NA | NA | NA |
| SCN2A | 852 | F | 3.485590909 | 28.20911769 | 0.421429849 | 27.88929542 | NA | NA | NA | NA | NA | NA | NA |
| SCN2A | 853 | R | 4.3965 | 30.46037206 | 0.457988511 | 29.98850766 | NA | NA | NA | NA | NA | NA | NA |
| SCN2A | 854 | L | 3.5465 | 25.86653719 | 0.383388119 | 25.2831432 | NA | NA | NA | NA | NA | NA | NA |
| SCN2A | 855 | L | 0.3895 | 26.03858769 | 0.386182089 | 21.74581589 | NA | NA | NA | NA | NA | NA | NA |
| SCN2A | 856 | R | -0.975136364 | 30.17469465 | 0.453349327 | 26.79486568 | NA | NA | NA | NA | NA | NA | NA |
| SCN2A | 857 | V | -1.884357143 | 26.89593161 | 0.400104703 | 23.51356544 | NA | NA | NA | NA | NA | NA | NA |
| SCN2A | 858 | F | -3.844227273 | 22.73075815 | 0.332465448 | 19.31576952 | NA | NA | NA | NA | NA | NA | NA |
| SCN2A | 859 | K | -6.300055556 | 29.02742002 | 0.434718457 | 25.60138935 | NA | NA | NA | NA | NA | NA | NA |
| SCN2A | 860 | L | -6.8915 | 27.41313692 | 0.408503725 | 24.14984991 | NA | NA | NA | NA | NA | NA | NA |
| SCN2A | 861 | A | -10.1991 | 25.30557367 | 0.374278498 | 22.77434366 | NA | NA | NA | NA | NA | NA | NA |
| SCN2A | 862 | K | -12.76505556 | 27.03063236 | 0.402292142 | 24.63651412 | NA | NA | NA | NA | NA | NA | NA |
| SCN2A | 863 | S | -12.68683333 | 26.28564287 | 0.390194078 | 23.83819863 | NA | NA | NA | NA | NA | NA | NA |

| | | | | | | | | | | | | | |
|---|---|---|---|---|---|---|---|---|---|---|---|---|---|
| SCN2A | 864 | W | -9.407214286 | 22.55999699 | 0.329692416 | 19.97875544 | NA | NA | NA | NA | NA | NA | NA |
| SCN2A | 865 | P | -13.40878571 | 20.19305024 | 0.291254998 | 17.71974145 | NA | NA | NA | NA | NA | NA | NA |
| SCN2A | 866 | T | -10.38964286 | 17.92678362 | 0.254452548 | 15.34193602 | NA | NA | NA | NA | NA | NA | NA |
| SCN2A | 867 | L | -7.441125 | 21.11009922 | 0.306147177 | 18.52515519 | NA | NA | NA | NA | NA | NA | NA |
| SCN2A | 868 | N | -11.879 | 20.45990748 | 0.295588557 | 18.10044811 | NA | NA | NA | NA | NA | NA | NA |
| SCN2A | 869 | M | -12.038375 | 16.09242987 | 0.224664035 | 14.30904047 | NA | NA | NA | NA | NA | NA | NA |
| SCN2A | 870 | L | -6.57625 | 16.50596735 | 0.23137957 | 13.48126574 | NA | NA | NA | NA | NA | NA | NA |
| SCN2A | 871 | I | -8.2755 | 19.93819365 | 0.28711632 | 16.83959904 | NA | NA | NA | NA | NA | NA | NA |
| SCN2A | 872 | K | -12.78738889 | 17.8606868 | 0.253379186 | 16.61102813 | NA | NA | NA | NA | NA | NA | NA |
| SCN2A | 873 | I | -8.93425 | 13.77594418 | 0.187046066 | 10.85076212 | NA | NA | NA | NA | NA | NA | NA |
| SCN2A | 874 | I | -5.65925 | 17.80816975 | 0.252526349 | 14.69442071 | NA | NA | NA | NA | NA | NA | NA |
| SCN2A | 875 | G | -9.08675 | 20.29202964 | 0.292862348 | 17.19785294 | NA | NA | NA | NA | NA | NA | NA |
| SCN2A | 876 | N | -11.319875 | 16.22013724 | 0.226737906 | 14.35443464 | NA | NA | NA | NA | NA | NA | NA |
| SCN2A | 877 | S | -7.116333333 | 17.88489525 | 0.253772312 | 14.769483 | NA | NA | NA | NA | NA | NA | NA |
| SCN2A | 878 | V | -7.248357143 | 20.96266956 | 0.303753031 | 17.88417976 | NA | NA | NA | NA | NA | NA | NA |
| SCN2A | 879 | G | -11.0685 | 22.1950339 | 0.323765693 | 20.39244889 | NA | NA | NA | NA | NA | NA | NA |
| SCN2A | 880 | A | -13.5165 | 19.7097981 | 0.28340735 | 16.74794032 | NA | NA | NA | NA | NA | NA | NA |
| SCN2A | 881 | L | -11.01075 | 16.13798609 | 0.225403834 | 14.33936575 | NA | NA | NA | NA | NA | NA | NA |
| SCN2A | 882 | G | -8.486 | 21.3255675 | 0.309646218 | 18.56644602 | NA | NA | NA | NA | NA | NA | NA |
| SCN2A | 883 | N | -9.337125 | 21.93524983 | 0.319546997 | 19.07910352 | NA | NA | NA | NA | NA | NA | NA |
| SCN2A | 884 | L | -7.693875 | 16.53316902 | 0.231821305 | 13.68495959 | NA | NA | NA | NA | NA | NA | NA |
| SCN2A | 885 | T | -4.263785714 | 18.16571821 | 0.258332664 | 13.81997091 | NA | NA | NA | NA | NA | NA | NA |
| SCN2A | 886 | L | -4.07075 | 21.050019 | 0.30517152 | 16.69047756 | NA | NA | NA | NA | NA | NA | NA |
| SCN2A | 887 | V | -3.905071429 | 19.98378478 | 0.287856685 | 15.68875583 | NA | NA | NA | NA | NA | NA | NA |
| SCN2A | 888 | L | -0.840625 | 14.88769139 | 0.205099997 | 9.974042834 | NA | NA | NA | NA | NA | NA | NA |
| SCN2A | 889 | A | 0.2553 | 19.44084718 | 0.279039791 | 14.53782265 | NA | NA | NA | NA | NA | NA | NA |
| SCN2A | 890 | I | 0.4975 | 21.54532221 | 0.313214868 | 16.63176762 | NA | NA | NA | NA | NA | NA | NA |
| SCN2A | 891 | I | 0.881875 | 18.76463051 | 0.268058545 | 13.81160589 | NA | NA | NA | NA | NA | NA | NA |
| SCN2A | 892 | V | 3.604928571 | 16.2694085 | 0.227538034 | 14.51585414 | NA | NA | NA | NA | NA | NA | NA |
| SCN2A | 893 | F | 4.985318182 | 19.86167495 | 0.285873715 | 18.34982685 | NA | NA | NA | NA | NA | NA | NA |
| SCN2A | 894 | I | 3.926375 | 21.43222422 | 0.311378243 | 19.86377 | NA | NA | NA | NA | NA | NA | NA |
| SCN2A | 895 | F | 5.116045455 | 17.67572146 | 0.250375489 | 15.82735323 | NA | NA | NA | NA | NA | NA | NA |
| SCN2A | 896 | A | 9.1893 | 18.52016547 | 0.264088619 | 18.33855662 | NA | NA | NA | NA | NA | NA | NA |

| | | | | | | | | | | | | | |
|---|---|---|---|---|---|---|---|---|---|---|---|---|---|
| SCN2A | 897 | V | 9.328928571 | 21.29821387 | 0.309202016 | 21.07247072 | NA | NA | NA | NA | NA | NA | NA |
| SCN2A | 898 | V | 9.336214286 | 22.28322923 | 0.325197918 | 20.95731203 | NA | NA | NA | NA | NA | NA | NA |
| SCN2A | 899 | G | 12.7275 | 20.4132966 | 0.294831631 | 20.02371974 | NA | NA | NA | NA | NA | NA | NA |
| SCN2A | 900 | M | 14.4675 | 21.23083027 | 0.308107758 | 20.070852 | NA | NA | NA | NA | NA | NA | NA |
| SCN2A | 901 | Q | 14.01305556 | 23.95458977 | 0.352339545 | 22.45845518 | NA | NA | NA | NA | NA | NA | NA |
| SCN2A | 902 | L | 14.08075 | 23.69626217 | 0.3481445 | 22.12493949 | NA | NA | NA | NA | NA | NA | NA |
| SCN2A | 903 | F | 15.80631818 | 20.18841641 | 0.291179748 | 18.61605679 | NA | NA | NA | NA | NA | NA | NA |
| SCN2A | 904 | G | 19.27275 | 21.30950656 | 0.309385401 | 19.75675593 | NA | NA | NA | NA | NA | NA | NA |
| SCN2A | 905 | K | 21.86827778 | 22.95819958 | 0.336158924 | 21.41454748 | NA | NA | NA | NA | NA | NA | NA |
| SCN2A | 906 | S | 22.15616667 | 20.64951237 | 0.298667596 | 19.12382919 | NA | NA | NA | NA | NA | NA | NA |
| SCN2A | 907 | Y | 21.10741667 | 14.98705748 | 0.206713627 | 13.43702429 | NA | NA | NA | NA | NA | NA | NA |
| SCN2A | 908 | K | 26.42427778 | 19.84228361 | 0.285558814 | 18.5725177 | NA | NA | NA | NA | NA | NA | NA |
| SCN2A | 909 | E | 26.33361111 | 22.25235091 | 0.324696477 | 21.91318225 | NA | NA | NA | NA | NA | NA | NA |
| SCN2A | 910 | C | 26.584 | 18.44559679 | 0.26287768 | 17.96825419 | NA | NA | NA | NA | NA | NA | NA |
| SCN2A | 911 | V | 28.49692857 | 15.22279748 | 0.210541865 | 14.73662921 | NA | NA | NA | NA | NA | NA | NA |
| SCN2A | 912 | C | 31.10266667 | 15.0732877 | 0.20811394 | 14.63554335 | NA | NA | NA | NA | NA | NA | NA |
| SCN2A | 913 | K | 28.01683333 | 14.67004461 | 0.201565578 | 14.02121459 | NA | NA | NA | NA | NA | NA | NA |
| SCN2A | 914 | I | 25.394625 | 12.58184112 | 0.167654739 | 11.79887816 | NA | NA | NA | NA | NA | NA | NA |
| SCN2A | 915 | S | 28.99816667 | 10.63584157 | 0.136053183 | 10.22542838 | NA | NA | NA | NA | NA | NA | NA |
| SCN2A | 916 | N | 32.499625 | 7.449323929 | 0.084306555 | 7.428871046 | 0.65 | -0.28 | 0.57 | 2.75 | 0.59 | 0 | 0 |
| SCN2A | 917 | D | 34.247375 | 10.63463982 | 0.136033667 | 10.54775504 | NA | NA | NA | NA | NA | NA | NA |
| SCN2A | 918 | C | 32.55816667 | 13.37983939 | 0.180613625 | 13.24477216 | NA | NA | NA | NA | NA | NA | NA |
| SCN2A | 919 | E | 30.17516667 | 13.78492492 | 0.187191906 | 13.38248526 | NA | NA | NA | NA | NA | NA | NA |
| SCN2A | 920 | L | 25.280625 | 12.70897278 | 0.16971926 | 12.24390829 | NA | NA | NA | NA | NA | NA | NA |
| SCN2A | 921 | P | 23.41964286 | 10.67403991 | 0.136673495 | 9.172185303 | NA | NA | NA | NA | NA | NA | NA |
| SCN2A | 922 | R | 22.18759091 | 7.77115088 | 0.08953278 | 6.255923156 | 1.63 | -0.02 | -0.8 | 25.57 | 0.96 | 1 | -1 |
| SCN2A | 923 | W | 17.281 | 6.443170726 | 0.067967391 | 4.851789871 | 1.63 | -0.02 | -0.8 | 25.57 | 0.96 | 1 | -1 |
| SCN2A | 924 | H | 19.9331 | 12.30058925 | 0.163087422 | 10.74925096 | NA | NA | NA | NA | NA | NA | NA |
| SCN2A | 925 | M | 16.46025 | 15.79481595 | 0.219831011 | 14.23885481 | NA | NA | NA | NA | NA | NA | NA |
| SCN2A | 926 | H | 21.1234 | 18.10759076 | 0.257388719 | 16.55909631 | NA | NA | NA | NA | NA | NA | NA |
| SCN2A | 927 | D | 17.5545 | 18.24291441 | 0.259586272 | 16.68607142 | NA | NA | NA | NA | NA | NA | NA |
| SCN2A | 928 | F | 11.71286364 | 19.10323169 | 0.273557172 | 18.01778066 | NA | NA | NA | NA | NA | NA | NA |
| SCN2A | 929 | F | 14.06404545 | 16.66742299 | 0.234001487 | 15.17654715 | NA | NA | NA | NA | NA | NA | NA |

| | | | | | | | | | | | | | |
|---|---|---|---|---|---|---|---|---|---|---|---|---|---|
| SCN2A | 930 | H | 17.1828 | 13.76923288 | 0.186937079 | 12.2113182 | NA | NA | NA | NA | NA | NA | NA |
| SCN2A | 931 | S | 12.95183333 | 15.86353444 | 0.220946947 | 15.52792259 | NA | NA | NA | NA | NA | NA | NA |
| SCN2A | 932 | F | 9.057954545 | 13.63959757 | 0.184831901 | 13.44807888 | NA | NA | NA | NA | NA | NA | NA |
| SCN2A | 933 | L | 12.507 | 10.52717764 | 0.134288563 | 10.10539161 | NA | NA | NA | NA | NA | NA | NA |
| SCN2A | 934 | I | 12.68775 | 10.5823333 | 0.135184249 | 10.35991075 | NA | NA | NA | NA | NA | NA | NA |
| SCN2A | 935 | V | 8.131214286 | 11.47118638 | 0.149618549 | 9.958559574 | NA | NA | NA | NA | NA | NA | NA |
| SCN2A | 936 | F | 7.046954545 | 8.719020014 | 0.104925456 | 7.433688302 | NA | NA | NA | NA | NA | NA | NA |
| SCN2A | 937 | R | 11.88495455 | 5.262483349 | 0.048793924 | 5.872386391 | 0.88 | 0.11 | -0.35 | 25.13 | 0.21 | 1 | -1 |
| SCN2A | 938 | V | 7.542357143 | 7.784441598 | 0.089748612 | 5.951897405 | 1.88 | -0.25 | -1.25 | 25.78 | 0.67 | 2 | 0 |
| SCN2A | 939 | L | 3.927625 | 7.391328636 | 0.083364755 | 6.207754645 | 1.51 | 0.05 | -0.5 | 17.74 | 0.15 | 1 | 1 |
| SCN2A | 940 | C | 6.091666667 | 3.325705639 | 0.017342124 | 1.829224766 | 1.51 | 0.05 | -0.5 | 17.74 | 0.15 | 1 | 1 |
| SCN2A | 941 | G | 7.181 | 4.080211024 | 0.029594719 | 2.783495201 | 1.51 | 0.05 | -0.5 | 17.74 | 0.15 | 1 | 1 |
| SCN2A | 942 | E | 11.18661111 | 2.282564347 | 4.02E-04 | 1.684576779 | 2.09 | -0.36 | -1.08 | 21.3 | 0.7 | 2 | 0 |
| SCN2A | 943 | W | 9.287071429 | 6.958387816 | 0.076334125 | 4.917288454 | 2.09 | -0.36 | -1.08 | 21.3 | 0.7 | 2 | 0 |
| SCN2A | 944 | I | 13.333875 | 8.715170968 | 0.10486295 | 6.637190945 | NA | NA | NA | NA | NA | NA | NA |
| SCN2A | 945 | E | 16.95261111 | 4.965373198 | 0.043969081 | 3.79934956 | 1.23 | -0.15 | -1.83 | 33.54 | 1.12 | 2 | 0 |
| SCN2A | 946 | T | 15.50964286 | 9.966086644 | 0.125176871 | 8.348766293 | NA | NA | NA | NA | NA | NA | NA |
| SCN2A | 947 | M | 14.480125 | 12.69397026 | 0.169475631 | 11.8505915 | NA | NA | NA | NA | NA | NA | NA |
| SCN2A | 948 | W | 18.19814286 | 10.16663617 | 0.128433643 | 8.537784855 | NA | NA | NA | NA | NA | NA | NA |
| SCN2A | 949 | D | 20.525375 | 10.01939005 | 0.126042478 | 8.46256747 | NA | NA | NA | NA | NA | NA | NA |
| SCN2A | 950 | C | 17.7635 | 15.00059365 | 0.206933444 | 13.4268449 | NA | NA | NA | NA | NA | NA | NA |
| SCN2A | 951 | M | 17.9345 | 17.28603931 | 0.244047347 | 15.70431925 | NA | NA | NA | NA | NA | NA | NA |
| SCN2A | 952 | E | 23.41683333 | 16.74632476 | 0.235282792 | 15.1973448 | NA | NA | NA | NA | NA | NA | NA |
| SCN2A | 953 | V | 22.10521429 | 18.24445061 | 0.259611219 | 16.70411973 | NA | NA | NA | NA | NA | NA | NA |
| SCN2A | 954 | A | 19.2383 | 20.71016033 | 0.299652473 | 19.15344743 | NA | NA | NA | NA | NA | NA | NA |
| SCN2A | 955 | G | 18.16975 | 21.94088307 | 0.319638476 | 20.35217 | NA | NA | NA | NA | NA | NA | NA |
| SCN2A | 956 | Q | 16.83827778 | 21.53207417 | 0.31299973 | 20.73055443 | NA | NA | NA | NA | NA | NA | NA |
| SCN2A | 957 | T | 13.32321429 | 23.16378497 | 0.339497475 | 21.12551633 | NA | NA | NA | NA | NA | NA | NA |
| SCN2A | 958 | M | 13.84725 | 21.57788817 | 0.313743715 | 19.53732786 | NA | NA | NA | NA | NA | NA | NA |
| SCN2A | 959 | C | 13.575 | 17.75624662 | 0.251683156 | 15.72352376 | NA | NA | NA | NA | NA | NA | NA |
| SCN2A | 960 | L | 10.929125 | 18.09486076 | 0.257181993 | 16.03286056 | NA | NA | NA | NA | NA | NA | NA |
| SCN2A | 961 | T | 8.1465 | 19.74013523 | 0.283900002 | 17.71246635 | NA | NA | NA | NA | NA | NA | NA |
| SCN2A | 962 | V | 8.943928571 | 17.33534387 | 0.244848015 | 15.30138536 | NA | NA | NA | NA | NA | NA | NA |

| | | | | | | | | | | | | | |
|---|---|---|---|---|---|---|---|---|---|---|---|---|---|
| SCN2A | 963 | F | 10.55013636 | 13.93703749 | 0.189662099 | 11.99834726 | NA | NA | NA | NA | NA | NA | NA |
| SCN2A | 964 | M | 6.342 | 14.68051729 | 0.201735647 | 12.6870413 | NA | NA | NA | NA | NA | NA | NA |
| SCN2A | 965 | M | 3.5045 | 16.96454727 | 0.23882656 | 14.93555822 | NA | NA | NA | NA | NA | NA | NA |
| SCN2A | 966 | V | 4.9185 | 14.01203914 | 0.190880069 | 11.98407603 | NA | NA | NA | NA | NA | NA | NA |
| SCN2A | 967 | M | 3.971875 | 10.61252496 | 0.135674539 | 8.353727141 | NA | NA | NA | NA | NA | NA | NA |
| SCN2A | 968 | V | 0.509928571 | 14.03148292 | 0.191195821 | 9.202186199 | NA | NA | NA | NA | NA | NA | NA |
| SCN2A | 969 | I | -0.034875 | 14.87591967 | 0.204908833 | 10.19523048 | NA | NA | NA | NA | NA | NA | NA |
| SCN2A | 970 | G | 0 | 12.14882175 | 0.160622833 | 7.196501836 | NA | NA | NA | NA | NA | NA | NA |
| SCN2A | 971 | N | -0.895875 | 8.184684844 | 0.096248258 | 3.277715801 | 5.03 | 1.31 | 2.72 | 0.47 | -1.74 | 0 | 0 |
| SCN2A | 972 | L | -3.933125 | 12.1094262 | 0.159983079 | 7.769800334 | NA | NA | NA | NA | NA | NA | NA |
| SCN2A | 973 | V | -3.291071429 | 13.15107338 | 0.176898638 | 8.74846714 | NA | NA | NA | NA | NA | NA | NA |
| SCN2A | 974 | V | -1.878214286 | 9.162554829 | 0.112128124 | 4.858102352 | 4.44 | 1.3 | 2.7 | 0.5 | -1.72 | 0 | 0 |
| SCN2A | 975 | L | -5.107875 | 7.103364696 | 0.07868844 | 3.98432466 | 3.18 | 1.35 | 2.33 | 0.62 | -1.69 | 0 | 0 |
| SCN2A | 976 | N | -7.826 | 11.32286386 | 0.147209903 | 8.485953373 | NA | NA | NA | NA | NA | NA | NA |
| SCN2A | 977 | L | -6.30025 | 11.80452828 | 0.155031768 | 8.954103152 | NA | NA | NA | NA | NA | NA | NA |
| SCN2A | 978 | F | -6.091409091 | 5.325385526 | 0.049815407 | 2.282759475 | 3.18 | 1.35 | 2.33 | 0.62 | -1.69 | 0 | 0 |
| SCN2A | 979 | L | -10.02325 | 6.440563329 | 0.067925049 | 4.391156062 | 2.24 | 1.53 | 3.9 | 0.19 | -1.83 | 0 | 0 |
| SCN2A | 980 | A | -11.5445 | 10.89349448 | 0.140237271 | 9.43502259 | NA | NA | NA | NA | NA | NA | NA |
| SCN2A | 981 | L | -9.47875 | 11.3907174 | 0.148311794 | 9.586697379 | NA | NA | NA | NA | NA | NA | NA |
| SCN2A | 982 | L | -11.537875 | 5.975399066 | 0.060371134 | 4.763712597 | 2.09 | 1.43 | 3.7 | 0.2 | -1.82 | 0 | 0 |
| SCN2A | 983 | L | -15.26025 | 6.191410421 | 0.063878994 | 4.445719989 | 2.56 | 0.88 | 1.9 | 1 | -1.23 | 0 | 0 |
| SCN2A | 984 | S | -15.52716667 | 10.76467236 | 0.138145297 | 7.845284808 | NA | NA | NA | NA | NA | NA | NA |
| SCN2A | 985 | S | -14.67216667 | 11.66083955 | 0.152698372 | 9.30321579 | NA | NA | NA | NA | NA | NA | NA |
| SCN2A | 986 | F | -16.09813636 | 5.583396636 | 0.054005312 | 2.803099177 | 3.1 | 0.57 | 1.03 | 1.29 | -1.02 | 0 | 0 |
| SCN2A | 987 | S | -20.0565 | 9.428370962 | 0.116444777 | 6.428118581 | NA | NA | NA | NA | NA | NA | NA |
| SCN2A | 1191 | G | -21.81025 | 57.32317037 | 0.894219966 | 52.73251959 | NA | NA | NA | NA | NA | NA | NA |
| SCN2A | 1192 | K | -22.18727778 | 59.17455212 | 0.924285001 | 54.55659801 | NA | NA | NA | NA | NA | NA | NA |
| SCN2A | 1193 | L | -17.129125 | 56.90794234 | 0.887476978 | 54.68769345 | NA | NA | NA | NA | NA | NA | NA |
| SCN2A | 1194 | W | -15.83892857 | 53.60003715 | 0.833759109 | 51.39017519 | NA | NA | NA | NA | NA | NA | NA |
| SCN2A | 1195 | W | -21.54885714 | 49.12758925 | 0.76112995 | 45.13062018 | NA | NA | NA | NA | NA | NA | NA |
| SCN2A | 1196 | N | -19.766125 | 54.54933163 | 0.84917493 | 50.66662903 | NA | NA | NA | NA | NA | NA | NA |
| SCN2A | 1197 | L | -14.728375 | 51.22023937 | 0.795112999 | 49.00595152 | NA | NA | NA | NA | NA | NA | NA |
| SCN2A | 1198 | R | -17.29495455 | 47.20088156 | 0.729841679 | 44.97044291 | NA | NA | NA | NA | NA | NA | NA |

| | | | | | | | | | | | | | |
|---|---|---|---|---|---|---|---|---|---|---|---|---|---|
| SCN2A | 1199 | K | -19.99238889 | 47.89190201 | 0.741063327 | 43.81372815 | NA | NA | NA | NA | NA | NA | NA |
| SCN2A | 1200 | T | -15.6845 | 48.74615158 | 0.754935691 | 46.53364613 | NA | NA | NA | NA | NA | NA | NA |
| SCN2A | 1201 | C | -13.63933333 | 45.56968423 | 0.703352272 | 43.36111886 | NA | NA | NA | NA | NA | NA | NA |
| SCN2A | 1202 | Y | -18.32433333 | 41.01079707 | 0.629319405 | 39.46130352 | NA | NA | NA | NA | NA | NA | NA |
| SCN2A | 1203 | K | -17.34338889 | 44.90163499 | 0.69250366 | 42.68142255 | NA | NA | NA | NA | NA | NA | NA |
| SCN2A | 1204 | I | -12.26225 | 43.18381319 | 0.664607538 | 40.98050012 | NA | NA | NA | NA | NA | NA | NA |
| SCN2A | 1205 | V | -12.88692857 | 39.82903415 | 0.610128472 | 37.62065023 | NA | NA | NA | NA | NA | NA | NA |
| SCN2A | 1206 | E | -16.69361111 | 39.01224405 | 0.596864421 | 36.79684001 | NA | NA | NA | NA | NA | NA | NA |
| SCN2A | 1207 | H | -14.1443 | 39.80997708 | 0.609819 | 37.59592586 | NA | NA | NA | NA | NA | NA | NA |
| SCN2A | 1208 | N | -13.502 | 38.01229986 | 0.580626086 | 35.79474036 | NA | NA | NA | NA | NA | NA | NA |
| SCN2A | 1209 | W | -11.53821429 | 41.77213436 | 0.641682945 | 39.56702766 | NA | NA | NA | NA | NA | NA | NA |
| SCN2A | 1210 | F | -8.623227273 | 39.87138007 | 0.610816138 | 37.6940162 | NA | NA | NA | NA | NA | NA | NA |
| SCN2A | 1211 | E | -10.24283333 | 33.33061825 | 0.504599131 | 31.12826032 | NA | NA | NA | NA | NA | NA | NA |
| SCN2A | 1212 | T | -7.470642857 | 37.7566121 | 0.576473911 | 35.57431698 | NA | NA | NA | NA | NA | NA | NA |
| SCN2A | 1213 | F | -3.548409091 | 39.44086279 | 0.603824864 | 37.30169801 | NA | NA | NA | NA | NA | NA | NA |
| SCN2A | 1214 | I | -4.392125 | 36.60946196 | 0.557845064 | 34.58464496 | NA | NA | NA | NA | NA | NA | NA |
| SCN2A | 1215 | V | -4.323785714 | 32.90935025 | 0.497758058 | 30.75137673 | NA | NA | NA | NA | NA | NA | NA |
| SCN2A | 1216 | F | -0.635409091 | 36.56323786 | 0.557094419 | 34.22676134 | NA | NA | NA | NA | NA | NA | NA |
| SCN2A | 1217 | M | -0.203875 | 37.23283113 | 0.567968106 | 33.93728757 | NA | NA | NA | NA | NA | NA | NA |
| SCN2A | 1218 | I | -0.9815 | 32.59604511 | 0.49267022 | 29.4038934 | NA | NA | NA | NA | NA | NA | NA |
| SCN2A | 1219 | L | 0.759125 | 30.99819621 | 0.466722367 | 27.68862936 | NA | NA | NA | NA | NA | NA | NA |
| SCN2A | 1220 | L | 3.34625 | 35.80000485 | 0.544700095 | 32.39997161 | NA | NA | NA | NA | NA | NA | NA |
| SCN2A | 1221 | S | 4.1485 | 35.31099802 | 0.536758995 | 31.89806342 | NA | NA | NA | NA | NA | NA | NA |
| SCN2A | 1222 | S | 4.292166667 | 31.75360104 | 0.478989568 | 28.34258708 | NA | NA | NA | NA | NA | NA | NA |
| SCN2A | 1223 | G | 6.887 | 33.41597037 | 0.505985184 | 28.78682068 | NA | NA | NA | NA | NA | NA | NA |
| SCN2A | 1224 | A | 9.1027 | 35.21946646 | 0.535272592 | 30.58546321 | NA | NA | NA | NA | NA | NA | NA |
| SCN2A | 1225 | L | 9.15275 | 31.44779132 | 0.47402345 | 29.34204021 | NA | NA | NA | NA | NA | NA | NA |
| SCN2A | 1226 | A | 11.4703 | 30.96517634 | 0.466186149 | 28.85118558 | NA | NA | NA | NA | NA | NA | NA |
| SCN2A | 1227 | F | 13.16877273 | 32.94454152 | 0.498329537 | 30.83283484 | NA | NA | NA | NA | NA | NA | NA |
| SCN2A | 1228 | E | 14.89294444 | 30.61830173 | 0.460553169 | 28.50851846 | NA | NA | NA | NA | NA | NA | NA |
| SCN2A | 1229 | D | 18.133375 | 29.48433613 | 0.442138428 | 27.3672908 | NA | NA | NA | NA | NA | NA | NA |
| SCN2A | 1230 | I | 22.695625 | 26.81753671 | 0.39883163 | 26.16017228 | NA | NA | NA | NA | NA | NA | NA |
| SCN2A | 1231 | Y | 23.03366667 | 28.67111587 | 0.428932348 | 28.00600971 | NA | NA | NA | NA | NA | NA | NA |

| | | | | | | | | | | | | | |
|---|---|---|---|---|---|---|---|---|---|---|---|---|---|
| SCN2A | 1232 | I | 20.10075 | 34.54755462 | 0.524361253 | 32.52031648 | NA | NA | NA | NA | NA | NA | NA |
| SCN2A | 1233 | E | 23.97627778 | 36.07697845 | 0.549197936 | 35.43668623 | NA | NA | NA | NA | NA | NA | NA |
| SCN2A | 1234 | Q | 26.03661111 | 33.76371903 | 0.511632359 | 33.12840723 | NA | NA | NA | NA | NA | NA | NA |
| SCN2A | 1235 | R | 21.22222727 | 33.9379477 | 0.5144617 | 33.84711874 | NA | NA | NA | NA | NA | NA | NA |
| SCN2A | 1236 | K | 23.93138889 | 41.38332518 | 0.635368979 | 40.71882145 | NA | NA | NA | NA | NA | NA | NA |
| SCN2A | 1237 | T | 19.65278571 | 43.23294931 | 0.665405471 | 41.11213652 | NA | NA | NA | NA | NA | NA | NA |
| SCN2A | 1238 | I | 16.970625 | 39.66908885 | 0.607531082 | 37.54570666 | NA | NA | NA | NA | NA | NA | NA |
| SCN2A | 1239 | K | 19.37983333 | 40.24892443 | 0.616947172 | 38.13825735 | NA | NA | NA | NA | NA | NA | NA |
| SCN2A | 1240 | T | 17.5575 | 43.1358554 | 0.66382874 | 41.02356904 | NA | NA | NA | NA | NA | NA | NA |
| SCN2A | 1241 | M | 13.86725 | 43.01000174 | 0.661784972 | 40.9009757 | NA | NA | NA | NA | NA | NA | NA |
| SCN2A | 1242 | L | 13.33 | 37.92151602 | 0.579151826 | 35.81508494 | NA | NA | NA | NA | NA | NA | NA |
| SCN2A | 1243 | E | 14.57238889 | 42.53112485 | 0.654008374 | 40.42540523 | NA | NA | NA | NA | NA | NA | NA |
| SCN2A | 1244 | Y | 13.18058333 | 45.05063996 | 0.694923387 | 42.94599533 | NA | NA | NA | NA | NA | NA | NA |
| SCN2A | 1245 | A | 9.0129 | 42.72531283 | 0.65716184 | 38.13137117 | NA | NA | NA | NA | NA | NA | NA |
| SCN2A | 1246 | D | 9.58425 | 40.22127011 | 0.616498087 | 38.12305274 | NA | NA | NA | NA | NA | NA | NA |
| SCN2A | 1247 | K | 8.767388889 | 44.79960416 | 0.690846757 | 41.52429832 | NA | NA | NA | NA | NA | NA | NA |
| SCN2A | 1248 | V | 5.282071429 | 44.43538943 | 0.684932186 | 41.03970766 | NA | NA | NA | NA | NA | NA | NA |
| SCN2A | 1249 | F | 4.488045455 | 39.92638458 | 0.61170937 | 36.51227866 | NA | NA | NA | NA | NA | NA | NA |
| SCN2A | 1250 | T | 3.916214286 | 42.18714149 | 0.648422346 | 38.78513576 | NA | NA | NA | NA | NA | NA | NA |
| SCN2A | 1251 | Y | 4.002666667 | 44.80523127 | 0.690938137 | 41.42179989 | NA | NA | NA | NA | NA | NA | NA |
| SCN2A | 1252 | I | 0.294625 | 42.5131488 | 0.653716457 | 39.43358041 | NA | NA | NA | NA | NA | NA | NA |
| SCN2A | 1253 | F | -0.357954545 | 37.60288847 | 0.573977556 | 35.24280507 | NA | NA | NA | NA | NA | NA | NA |
| SCN2A | 1254 | I | -0.799375 | 43.13661046 | 0.663841001 | 40.76044396 | NA | NA | NA | NA | NA | NA | NA |
| SCN2A | 1255 | L | -3.025875 | 45.12405632 | 0.696115613 | 42.71131605 | NA | NA | NA | NA | NA | NA | NA |
| SCN2A | 1256 | E | -5.051833333 | 38.94102359 | 0.595707855 | 36.5297909 | NA | NA | NA | NA | NA | NA | NA |
| SCN2A | 1257 | M | -4.747875 | 41.75943991 | 0.641476797 | 39.3535327 | NA | NA | NA | NA | NA | NA | NA |
| SCN2A | 1258 | L | -5.704375 | 44.49726035 | 0.685936923 | 42.06490425 | NA | NA | NA | NA | NA | NA | NA |
| SCN2A | 1259 | L | -8.875875 | 44.33264876 | 0.683263755 | 42.14466512 | NA | NA | NA | NA | NA | NA | NA |
| SCN2A | 1260 | K | -9.959277778 | 37.36763309 | 0.570157187 | 35.17818237 | NA | NA | NA | NA | NA | NA | NA |
| SCN2A | 1261 | W | -8.294857143 | 42.68061562 | 0.656435991 | 40.52350546 | NA | NA | NA | NA | NA | NA | NA |
| SCN2A | 1262 | V | -12.59521429 | 46.05117295 | 0.711171284 | 43.8535763 | NA | NA | NA | NA | NA | NA | NA |
| SCN2A | 1263 | A | -15.0071 | 43.99360885 | 0.677758005 | 41.76617027 | NA | NA | NA | NA | NA | NA | NA |
| SCN2A | 1264 | Y | -15.03416667 | 38.41332183 | 0.587138379 | 36.197223 | NA | NA | NA | NA | NA | NA | NA |

| | | | | | | | | | | | | | |
|---|---|---|---|---|---|---|---|---|---|---|---|---|---|
| SCN2A | 1265 | G | -14.55125 | 42.68056289 | 0.656435135 | 40.4653953 | NA | NA | NA | NA | NA | NA | NA |
| SCN2A | 1266 | F | -11.94977273 | 41.6846986 | 0.640263054 | 39.46777036 | NA | NA | NA | NA | NA | NA | NA |
| SCN2A | 1267 | Q | -17.19127778 | 39.68973895 | 0.607866424 | 37.45452475 | NA | NA | NA | NA | NA | NA | NA |
| SCN2A | 1268 | V | -16.73921429 | 37.58852946 | 0.573744377 | 35.36129549 | NA | NA | NA | NA | NA | NA | NA |
| SCN2A | 1269 | Y | -10.54291667 | 37.01573098 | 0.564442564 | 34.81228033 | NA | NA | NA | NA | NA | NA | NA |
| SCN2A | 1270 | F | -10.61531818 | 36.73740451 | 0.559922754 | 34.52223153 | NA | NA | NA | NA | NA | NA | NA |
| SCN2A | 1271 | T | -15.23792857 | 33.27388375 | 0.503677805 | 31.04226148 | NA | NA | NA | NA | NA | NA | NA |
| SCN2A | 1272 | N | -13.5295 | 32.59069429 | 0.492583327 | 30.38339469 | NA | NA | NA | NA | NA | NA | NA |
| SCN2A | 1273 | A | -10.3641 | 29.38575922 | 0.440537614 | 27.18311522 | NA | NA | NA | NA | NA | NA | NA |
| SCN2A | 1274 | W | -9.331357143 | 25.01742685 | 0.369599212 | 22.84713818 | NA | NA | NA | NA | NA | NA | NA |
| SCN2A | 1275 | C | -8.835166667 | 32.94221219 | 0.498291711 | 30.75801584 | NA | NA | NA | NA | NA | NA | NA |
| SCN2A | 1276 | W | -7.79 | 33.2197183 | 0.5027982 | 30.78542356 | NA | NA | NA | NA | NA | NA | NA |
| SCN2A | 1277 | L | -4.02125 | 28.36329729 | 0.423933608 | 25.96010393 | NA | NA | NA | NA | NA | NA | NA |
| SCN2A | 1278 | D | -4.224875 | 32.93143673 | 0.498116726 | 30.52799288 | NA | NA | NA | NA | NA | NA | NA |
| SCN2A | 1279 | F | -3.424954545 | 35.81765743 | 0.544986759 | 33.40525112 | NA | NA | NA | NA | NA | NA | NA |
| SCN2A | 1280 | L | -1.128 | 33.35216086 | 0.504948966 | 30.12161701 | NA | NA | NA | NA | NA | NA | NA |
| SCN2A | 1281 | I | 0.34925 | 32.17095555 | 0.485767088 | 28.81635055 | NA | NA | NA | NA | NA | NA | NA |
| SCN2A | 1282 | V | 0.447785714 | 37.65448684 | 0.574815475 | 34.29674723 | NA | NA | NA | NA | NA | NA | NA |
| SCN2A | 1283 | D | 1.1815 | 39.29139038 | 0.601397546 | 35.92917688 | NA | NA | NA | NA | NA | NA | NA |
| SCN2A | 1284 | V | 4.949071429 | 36.14700653 | 0.550335139 | 32.75310242 | NA | NA | NA | NA | NA | NA | NA |
| SCN2A | 1285 | S | 4.989333333 | 39.02915005 | 0.597138962 | 35.6350637 | NA | NA | NA | NA | NA | NA | NA |
| SCN2A | 1286 | L | 3.745625 | 42.23802572 | 0.649248667 | 38.84925076 | NA | NA | NA | NA | NA | NA | NA |
| SCN2A | 1287 | V | 7.3655 | 42.60544573 | 0.655215289 | 39.40271767 | NA | NA | NA | NA | NA | NA | NA |
| SCN2A | 1288 | S | 10.01166667 | 40.74163891 | 0.624948481 | 38.64852975 | NA | NA | NA | NA | NA | NA | NA |
| SCN2A | 1289 | L | 8.6845 | 44.04572068 | 0.678604261 | 41.95262068 | NA | NA | NA | NA | NA | NA | NA |
| SCN2A | 1290 | T | 9.092785714 | 46.6002578 | 0.720088005 | 44.50746161 | NA | NA | NA | NA | NA | NA | NA |
| SCN2A | 1291 | A | 12.8155 | 44.83131484 | 0.691361714 | 42.73739936 | NA | NA | NA | NA | NA | NA | NA |
| SCN2A | 1292 | N | 14.587125 | 44.02143698 | 0.678209912 | 41.92340518 | NA | NA | NA | NA | NA | NA | NA |
| SCN2A | 1293 | A | 13.0767 | 48.41812017 | 0.74960871 | 46.32278912 | NA | NA | NA | NA | NA | NA | NA |
| SCN2A | 1294 | L | 13.83875 | 50.23758497 | 0.779155437 | 48.14274773 | NA | NA | NA | NA | NA | NA | NA |
| SCN2A | 1295 | G | 17.31275 | 47.99660694 | 0.742763655 | 45.89789716 | NA | NA | NA | NA | NA | NA | NA |
| SCN2A | 1296 | Y | 15.92283333 | 44.37400239 | 0.683935307 | 42.27745612 | NA | NA | NA | NA | NA | NA | NA |
| SCN2A | 1297 | S | 18.6565 | 42.00480462 | 0.645461333 | 39.90614485 | NA | NA | NA | NA | NA | NA | NA |

| | | | | | | | | | | | | | |
|---|---|---|---|---|---|---|---|---|---|---|---|---|---|
| SCN2A | 1298 | E | 21.28227778 | 41.35387144 | 0.634890673 | 39.39520583 | NA | NA | NA | NA | NA | NA | NA |
| SCN2A | 1299 | L | 17.84875 | 38.28865738 | 0.585113923 | 36.28586201 | NA | NA | NA | NA | NA | NA | NA |
| SCN2A | 1300 | G | 17.21325 | 35.70120459 | 0.543095653 | 33.70972993 | NA | NA | NA | NA | NA | NA | NA |
| SCN2A | 1301 | A | 14.0561 | 36.93503405 | 0.563132107 | 34.84095687 | NA | NA | NA | NA | NA | NA | NA |
| SCN2A | 1302 | I | 14.021 | 38.04776957 | 0.581202088 | 35.95276888 | NA | NA | NA | NA | NA | NA | NA |
| SCN2A | 1303 | K | 17.66538889 | 34.65367504 | 0.526084569 | 32.55682022 | NA | NA | NA | NA | NA | NA | NA |
| SCN2A | 1304 | S | 13.7545 | 33.07922337 | 0.500516669 | 30.9860744 | NA | NA | NA | NA | NA | NA | NA |
| SCN2A | 1305 | L | 10.41125 | 35.25460629 | 0.535843236 | 33.1621431 | NA | NA | NA | NA | NA | NA | NA |
| SCN2A | 1306 | R | 13.89359091 | 33.78436045 | 0.511967559 | 31.68944797 | NA | NA | NA | NA | NA | NA | NA |
| SCN2A | 1307 | T | 11.90821429 | 30.50245152 | 0.45867185 | 28.40934072 | NA | NA | NA | NA | NA | NA | NA |
| SCN2A | 1308 | L | 7.691375 | 31.81304013 | 0.479954814 | 29.72234045 | NA | NA | NA | NA | NA | NA | NA |
| SCN2A | 1309 | R | 9.528772727 | 32.65127581 | 0.493567125 | 30.68681222 | NA | NA | NA | NA | NA | NA | NA |
| SCN2A | 1310 | A | 6.4187 | 28.6429056 | 0.428474235 | 24.19420279 | NA | NA | NA | NA | NA | NA | NA |
| SCN2A | 1311 | L | 3.3555 | 28.3680367 | 0.424010573 | 24.9673982 | NA | NA | NA | NA | NA | NA | NA |
| SCN2A | 1312 | R | 4.215318182 | 31.26258323 | 0.471015811 | 27.87013471 | NA | NA | NA | NA | NA | NA | NA |
| SCN2A | 1313 | P | 1.670928571 | 28.16538139 | 0.420719604 | 24.79318071 | NA | NA | NA | NA | NA | NA | NA |
| SCN2A | 1314 | L | -0.4535 | 25.61217566 | 0.379257481 | 22.24795142 | NA | NA | NA | NA | NA | NA | NA |
| SCN2A | 1315 | R | -1.253409091 | 30.05206542 | 0.451357921 | 27.69414108 | NA | NA | NA | NA | NA | NA | NA |
| SCN2A | 1316 | A | -3.5053 | 27.17172153 | 0.404583323 | 24.76955489 | NA | NA | NA | NA | NA | NA | NA |
| SCN2A | 1317 | L | -5.21825 | 23.25244213 | 0.3409372 | 20.86597173 | NA | NA | NA | NA | NA | NA | NA |
| SCN2A | 1318 | S | -8.303166667 | 27.61999589 | 0.411862958 | 25.45917339 | NA | NA | NA | NA | NA | NA | NA |
| SCN2A | 1319 | R | -7.589318182 | 28.61591133 | 0.428035869 | 26.43852853 | NA | NA | NA | NA | NA | NA | NA |
| SCN2A | 1320 | F | -7.180227273 | 24.7896337 | 0.365900024 | 22.60555515 | NA | NA | NA | NA | NA | NA | NA |
| SCN2A | 1321 | E | -13.32405556 | 21.95006501 | 0.319787584 | 19.73859938 | NA | NA | NA | NA | NA | NA | NA |
| SCN2A | 1322 | G | -9.71475 | 19.68083581 | 0.282937024 | 17.50217934 | NA | NA | NA | NA | NA | NA | NA |
| SCN2A | 1323 | M | -6.50675 | 20.90738496 | 0.302855251 | 18.74236188 | NA | NA | NA | NA | NA | NA | NA |
| SCN2A | 1324 | R | -11.68904545 | 21.06818336 | 0.305466495 | 18.86493056 | NA | NA | NA | NA | NA | NA | NA |
| SCN2A | 1325 | V | -13.05835714 | 17.6595588 | 0.25011302 | 15.44135179 | NA | NA | NA | NA | NA | NA | NA |
| SCN2A | 1326 | V | -9.451928571 | 15.72703004 | 0.218730219 | 13.57464296 | NA | NA | NA | NA | NA | NA | NA |
| SCN2A | 1327 | V | -8.700928571 | 18.53589626 | 0.264344075 | 16.36347718 | NA | NA | NA | NA | NA | NA | NA |
| SCN2A | 1328 | N | -12.937625 | 18.19248573 | 0.258767348 | 15.96712069 | NA | NA | NA | NA | NA | NA | NA |
| SCN2A | 1329 | A | -12.3321 | 14.42128808 | 0.197525961 | 12.19774855 | NA | NA | NA | NA | NA | NA | NA |
| SCN2A | 1330 | L | -8.543625 | 13.96087132 | 0.190049142 | 11.51058521 | NA | NA | NA | NA | NA | NA | NA |

| | | | | | | | | | | | | | |
|---|---|---|---|---|---|---|---|---|---|---|---|---|---|
| SCN2A | 1331 | L | -9.60675 | 17.69579896 | 0.250701532 | 15.2050954 | NA | NA | NA | NA | NA | NA | NA |
| SCN2A | 1332 | G | -13.323 | 17.0672218 | 0.240493916 | 14.81356621 | NA | NA | NA | NA | NA | NA | NA |
| SCN2A | 1333 | A | -11.5555 | 13.79404861 | 0.187340068 | 11.55369034 | NA | NA | NA | NA | NA | NA | NA |
| SCN2A | 1334 | I | -8.807125 | 16.92355947 | 0.238160949 | 14.38999272 | NA | NA | NA | NA | NA | NA | NA |
| SCN2A | 1335 | P | -11.6095 | 18.16858511 | 0.258379221 | 15.6065965 | NA | NA | NA | NA | NA | NA | NA |
| SCN2A | 1336 | S | -11.32033333 | 14.98244126 | 0.206638663 | 12.40410105 | NA | NA | NA | NA | NA | NA | NA |
| SCN2A | 1337 | I | -7.11925 | 13.48781721 | 0.182367103 | 11.01314075 | NA | NA | NA | NA | NA | NA | NA |
| SCN2A | 1338 | M | -6.201875 | 17.46091372 | 0.246887174 | 14.94251115 | NA | NA | NA | NA | NA | NA | NA |
| SCN2A | 1339 | N | -9.279375 | 18.22823815 | 0.259347941 | 15.66724701 | NA | NA | NA | NA | NA | NA | NA |
| SCN2A | 1340 | V | -6.5175 | 13.86895313 | 0.188556461 | 11.46414038 | NA | NA | NA | NA | NA | NA | NA |
| SCN2A | 1341 | L | -2.48 | 16.56392955 | 0.232320832 | 13.20069739 | NA | NA | NA | NA | NA | NA | NA |
| SCN2A | 1342 | L | -3.91775 | 19.0975002 | 0.273464097 | 15.73107242 | NA | NA | NA | NA | NA | NA | NA |
| SCN2A | 1343 | V | -4.235785714 | 18.38967509 | 0.261969554 | 15.01249367 | NA | NA | NA | NA | NA | NA | NA |
| SCN2A | 1344 | C | -1.142166667 | 14.3191285 | 0.195866967 | 10.94096754 | NA | NA | NA | NA | NA | NA | NA |
| SCN2A | 1345 | L | 0.286625 | 18.86127697 | 0.269628011 | 15.46639713 | NA | NA | NA | NA | NA | NA | NA |
| SCN2A | 1346 | I | -0.567875 | 21.23365762 | 0.308153672 | 17.8516392 | NA | NA | NA | NA | NA | NA | NA |
| SCN2A | 1347 | F | -0.446681818 | 17.2868758 | 0.244060931 | 13.87044577 | NA | NA | NA | NA | NA | NA | NA |
| SCN2A | 1348 | W | 3.005 | 13.24387817 | 0.178405718 | 10.95689405 | NA | NA | NA | NA | NA | NA | NA |
| SCN2A | 1349 | L | 3.653625 | 21.46189227 | 0.311860029 | 19.24369229 | NA | NA | NA | NA | NA | NA | NA |
| SCN2A | 1350 | I | 2.895125 | 21.87841578 | 0.318624055 | 21.39889599 | NA | NA | NA | NA | NA | NA | NA |
| SCN2A | 1351 | F | 5.014045455 | 17.98233238 | 0.255354618 | 17.63049694 | NA | NA | NA | NA | NA | NA | NA |
| SCN2A | 1352 | S | 8.471833333 | 20.07202195 | 0.28928959 | 19.17225486 | NA | NA | NA | NA | NA | NA | NA |
| SCN2A | 1353 | I | 8.05575 | 22.42752619 | 0.327541191 | 20.47794862 | NA | NA | NA | NA | NA | NA | NA |
| SCN2A | 1354 | M | 8.744875 | 21.23516303 | 0.308178118 | 20.61164356 | NA | NA | NA | NA | NA | NA | NA |
| SCN2A | 1355 | G | 12.39525 | 19.55481439 | 0.280890532 | 17.59699913 | NA | NA | NA | NA | NA | NA | NA |
| SCN2A | 1356 | V | 13.74821429 | 21.23298465 | 0.308142743 | 19.25933076 | NA | NA | NA | NA | NA | NA | NA |
| SCN2A | 1357 | N | 13.130875 | 23.36332008 | 0.342737774 | 21.39402378 | NA | NA | NA | NA | NA | NA | NA |
| SCN2A | 1358 | L | 14.73375 | 20.12459841 | 0.290143392 | 18.63971162 | NA | NA | NA | NA | NA | NA | NA |
| SCN2A | 1359 | F | 16.64422727 | 16.91919171 | 0.23809002 | 14.91284879 | NA | NA | NA | NA | NA | NA | NA |
| SCN2A | 1360 | A | 18.9811 | 20.09534578 | 0.289668351 | 18.06613597 | NA | NA | NA | NA | NA | NA | NA |
| SCN2A | 1361 | G | 22.82425 | 20.55976802 | 0.297210216 | 18.59072689 | NA | NA | NA | NA | NA | NA | NA |
| SCN2A | 1362 | K | 21.99927778 | 17.0271216 | 0.23984272 | 15.07792632 | NA | NA | NA | NA | NA | NA | NA |
| SCN2A | 1363 | F | 21.5925 | 13.99656837 | 0.190628835 | 13.90330023 | NA | NA | NA | NA | NA | NA | NA |

| | | | | | | | | | | | | | |
|---|---|---|---|---|---|---|---|---|---|---|---|---|---|
| SCN2A | 1364 | Y | 26.35625 | 16.03019931 | 0.223653458 | 15.44509338 | NA | NA | NA | NA | NA | NA | NA |
| SCN2A | 1365 | H | 29.0801 | 11.06815405 | 0.143073609 | 10.14641882 | NA | NA | NA | NA | NA | NA | NA |
| SCN2A | 1366 | C | 33.6855 | 12.59630906 | 0.167889687 | 12.48963756 | NA | NA | NA | NA | NA | NA | NA |
| SCN2A | 1367 | I | 34.04025 | 10.38314042 | 0.131949508 | 10.11749194 | NA | NA | NA | NA | NA | NA | NA |
| SCN2A | 1368 | N | 38.42525 | 10.46714999 | 0.13331376 | 10.4420514 | NA | NA | NA | NA | NA | NA | NA |
| SCN2A | 1369 | Y | 38.15958333 | 12.29800004 | 0.163045375 | 12.05840725 | NA | NA | NA | NA | NA | NA | NA |
| SCN2A | 1370 | T | 39.82121429 | 11.36025119 | 0.147817045 | 11.28863862 | NA | NA | NA | NA | NA | NA | NA |
| SCN2A | 1371 | T | 36.85792857 | 6.865087763 | 0.074819003 | 6.752353062 | 0.65 | -0.28 | 0.57 | 2.75 | 0.59 | 0 | 0 |
| SCN2A | 1372 | G | 33.6395 | 6.42136037 | 0.067613207 | 6.173591233 | 0.65 | -0.28 | 0.57 | 2.75 | 0.59 | 0 | 0 |
| SCN2A | 1373 | E | 35.12394444 | 6.394111275 | 0.067170702 | 6.616293747 | 0.65 | -0.28 | 0.57 | 2.75 | 0.59 | 0 | 0 |
| SCN2A | 1374 | M | 31.674125 | 8.002581146 | 0.093291032 | 8.012707192 | 0.65 | -0.28 | 0.57 | 2.75 | 0.59 | 0 | 0 |
| SCN2A | 1375 | F | 36.66959091 | 11.37553753 | 0.148065284 | 11.43822595 | NA | NA | NA | NA | NA | NA | NA |
| SCN2A | 1376 | D | 36.138875 | 12.57010839 | 0.167464208 | 12.77565051 | NA | NA | NA | NA | NA | NA | NA |
| SCN2A | 1377 | V | 34.34378571 | 15.97132997 | 0.222697465 | 16.05290597 | NA | NA | NA | NA | NA | NA | NA |
| SCN2A | 1378 | S | 38.3865 | 18.1351732 | 0.257836636 | 18.27695797 | NA | NA | NA | NA | NA | NA | NA |
| SCN2A | 1379 | V | 39.75978571 | 17.01526177 | 0.239650125 | 17.29030706 | NA | NA | NA | NA | NA | NA | NA |
| SCN2A | 1380 | V | 36.80078571 | 17.48834623 | 0.247332658 | 17.58287807 | NA | NA | NA | NA | NA | NA | NA |
| SCN2A | 1381 | N | 34.61725 | 19.99441357 | 0.288029289 | 20.02215124 | NA | NA | NA | NA | NA | NA | NA |
| SCN2A | 1382 | N | 34.508 | 20.77473487 | 0.300701115 | 20.68002329 | NA | NA | NA | NA | NA | NA | NA |
| SCN2A | 1383 | Y | 33.94266667 | 21.96820193 | 0.320082114 | 21.35912742 | NA | NA | NA | NA | NA | NA | NA |
| SCN2A | 1384 | S | 37.47083333 | 23.99302009 | 0.352963624 | 23.86959319 | NA | NA | NA | NA | NA | NA | NA |
| SCN2A | 1385 | E | 39.35705556 | 21.92629504 | 0.319401578 | 21.98067471 | NA | NA | NA | NA | NA | NA | NA |
| SCN2A | 1386 | C | 37.83866667 | 18.44880988 | 0.262929858 | 18.35983983 | NA | NA | NA | NA | NA | NA | NA |
| SCN2A | 1387 | K | 38.76538889 | 22.43627469 | 0.32768326 | 22.36111796 | NA | NA | NA | NA | NA | NA | NA |
| SCN2A | 1388 | A | 42.4543 | 24.07895822 | 0.354359194 | 24.13706113 | NA | NA | NA | NA | NA | NA | NA |
| SCN2A | 1389 | L | 42.400875 | 18.70996927 | 0.267170888 | 18.83246426 | NA | NA | NA | NA | NA | NA | NA |
| SCN2A | 1390 | I | 40.977875 | 20.49014966 | 0.296079667 | 20.33036017 | NA | NA | NA | NA | NA | NA | NA |
| SCN2A | 1391 | E | 43.88905556 | 24.83350261 | 0.366612422 | 24.80716446 | NA | NA | NA | NA | NA | NA | NA |
| SCN2A | 1392 | S | 46.911 | 24.80619104 | 0.366168903 | 24.92347528 | NA | NA | NA | NA | NA | NA | NA |
| SCN2A | 1393 | N | 46.814 | 22.64894395 | 0.331136847 | 22.71875896 | NA | NA | NA | NA | NA | NA | NA |
| SCN2A | 1394 | Q | 44.66527778 | 19.47075212 | 0.279525424 | 19.50824778 | NA | NA | NA | NA | NA | NA | NA |
| SCN2A | 1395 | T | 41.25492857 | 15.36297136 | 0.212818183 | 15.38379411 | NA | NA | NA | NA | NA | NA | NA |
| SCN2A | 1396 | A | 38.2235 | 16.55603663 | 0.232192657 | 16.48663317 | NA | NA | NA | NA | NA | NA | NA |

| | | | | | | | | | | | | | |
|---|---|---|---|---|---|---|---|---|---|---|---|---|---|
| SCN2A | 1397 | R | 33.46695455 | 14.86469653 | 0.204726578 | 14.39711385 | NA | NA | NA | NA | NA | NA | NA |
| SCN2A | 1398 | W | 31.21421429 | 17.28778068 | 0.244075625 | 16.32484777 | NA | NA | NA | NA | NA | NA | NA |
| SCN2A | 1399 | K | 28.73283333 | 11.57070309 | 0.151234624 | 10.68081935 | NA | NA | NA | NA | NA | NA | NA |
| SCN2A | 1400 | N | 24.8025 | 13.3669051 | 0.180403582 | 12.78120993 | NA | NA | NA | NA | NA | NA | NA |
| SCN2A | 1401 | V | 22.49992857 | 11.11455393 | 0.143827108 | 11.36110144 | NA | NA | NA | NA | NA | NA | NA |
| SCN2A | 1402 | K | 21.8425 | 14.44805336 | 0.197960609 | 12.47302637 | NA | NA | NA | NA | NA | NA | NA |
| SCN2A | 1403 | V | 17.45092857 | 12.07885181 | 0.159486574 | 10.01410427 | NA | NA | NA | NA | NA | NA | NA |
| SCN2A | 1404 | N | 18.07975 | 14.8527384 | 0.204532387 | 12.96858591 | NA | NA | NA | NA | NA | NA | NA |
| SCN2A | 1405 | F | 14.58786364 | 14.79011443 | 0.203515421 | 12.80066648 | NA | NA | NA | NA | NA | NA | NA |
| SCN2A | 1406 | D | 18.324625 | 19.80334189 | 0.28492643 | 17.76522412 | NA | NA | NA | NA | NA | NA | NA |
| SCN2A | 1407 | N | 14.718875 | 20.76213578 | 0.300496515 | 18.70553952 | NA | NA | NA | NA | NA | NA | NA |
| SCN2A | 1408 | V | 9.9975 | 20.51964127 | 0.296558588 | 18.38275221 | NA | NA | NA | NA | NA | NA | NA |
| SCN2A | 1409 | G | 11.043 | 19.03119915 | 0.272387418 | 16.90715848 | NA | NA | NA | NA | NA | NA | NA |
| SCN2A | 1410 | L | 14.514125 | 16.26029766 | 0.227390081 | 14.22333664 | NA | NA | NA | NA | NA | NA | NA |
| SCN2A | 1411 | G | 11.10225 | 16.19921365 | 0.226398123 | 14.05470616 | NA | NA | NA | NA | NA | NA | NA |
| SCN2A | 1412 | Y | 6.80225 | 15.36534692 | 0.21285676 | 13.23949106 | NA | NA | NA | NA | NA | NA | NA |
| SCN2A | 1413 | L | 11.66725 | 12.31378955 | 0.163301784 | 10.19166722 | NA | NA | NA | NA | NA | NA | NA |
| SCN2A | 1414 | S | 12.42066667 | 11.18897484 | 0.145035647 | 9.039498868 | NA | NA | NA | NA | NA | NA | NA |
| SCN2A | 1415 | L | 8.393375 | 11.65817014 | 0.152655023 | 11.07869002 | NA | NA | NA | NA | NA | NA | NA |
| SCN2A | 1416 | L | 7.51175 | 9.865375512 | 0.123541399 | 9.089632553 | NA | NA | NA | NA | NA | NA | NA |
| SCN2A | 1417 | Q | 12.19716667 | 6.727081239 | 0.072577882 | 4.783130315 | 2.21 | -0.17 | -0.53 | 3.06 | -0.01 | 0 | 0 |
| SCN2A | 1418 | V | 9.572071429 | 7.103580013 | 0.078691936 | 6.739436366 | 0.86 | -0.97 | -1.95 | 26.64 | 1.52 | 1 | -1 |
| SCN2A | 1419 | A | 5.8445 | 6.884308753 | 0.075131137 | 5.72163468 | 1.51 | 0.05 | -0.5 | 17.74 | 0.15 | 1 | 1 |
| SCN2A | 1420 | T | 6.800928571 | 3.986776517 | 0.028077413 | 2.609788046 | 1.51 | 0.05 | -0.5 | 17.74 | 0.15 | 1 | 1 |
| SCN2A | 1421 | F | 7.830863636 | 2.257790956 | 0 | 1.707715229 | 0.88 | 0.11 | -0.35 | 25.13 | 0.21 | 1 | -1 |
| SCN2A | 1422 | K | 10.63494444 | 3.294892107 | 0.016841736 | 2.040122535 | 2.21 | -0.17 | -0.53 | 3.06 | -0.01 | 0 | 0 |
| SCN2A | 1423 | G | 14.8605 | 3.73562632 | 0.023998925 | 1.781095809 | 2.11 | 0.14 | -0.74 | 11.96 | -0.02 | 1 | 1 |
| SCN2A | 1424 | W | 12.32471429 | 2.932586913 | 0.010958174 | 1.659147462 | 2.03 | 0.19 | -0.73 | 14.2 | -0.04 | 1 | -1 |
| SCN2A | 1425 | M | 16.872625 | 2.785359043 | 0.008567305 | 1.506867271 | 1.27 | 0.3 | -1 | 25.47 | 0.37 | 1 | 1 |
| SCN2A | 1426 | D | 19.788625 | 3.852343702 | 0.025894326 | 1.865224281 | 2.09 | -0.38 | -2.7 | 40.47 | 1.53 | 4 | 0 |
| SCN2A | 1427 | I | 16.524625 | 7.952671186 | 0.092480532 | 5.883025095 | 2.09 | -0.38 | -2.7 | 40.47 | 1.53 | 4 | 0 |
| SCN2A | 1428 | M | 15.98525 | 8.546674733 | 0.102126699 | 6.923582948 | NA | NA | NA | NA | NA | NA | NA |
| SCN2A | 1429 | Y | 20.48483333 | 4.394670067 | 0.034701295 | 2.350937684 | 2.09 | -0.38 | -2.7 | 40.47 | 1.53 | 4 | 0 |

| | | | | | | | | | | | | | |
|---|---|---|---|---|---|---|---|---|---|---|---|---|---|
| SCN2A | 1430 | A | 21.1843 | 9.912430832 | 0.124305542 | 7.905719079 | NA | NA | NA | NA | NA | NA | NA |
| SCN2A | 1431 | A | 18.9173 | 12.34484382 | 0.163806082 | 10.3078595 | NA | NA | NA | NA | NA | NA | NA |
| SCN2A | 1432 | V | 20.07307143 | 12.90436178 | 0.172892229 | 10.84001233 | NA | NA | NA | NA | NA | NA | NA |
| SCN2A | 1433 | D | 24.050375 | 11.08117083 | 0.143284992 | 9.575756395 | NA | NA | NA | NA | NA | NA | NA |
| SCN2A | 1434 | S | 22.031 | 15.65718113 | 0.217595926 | 13.66895434 | NA | NA | NA | NA | NA | NA | NA |
| SCN2A | 1435 | R | 27.17586364 | 18.83103749 | 0.269136944 | 18.23808454 | NA | NA | NA | NA | NA | NA | NA |
| SCN2A | 1436 | N | 24.62825 | 21.31839445 | 0.309529734 | 20.68794872 | NA | NA | NA | NA | NA | NA | NA |
| SCN2A | 1437 | V | 24.9765 | 23.26827353 | 0.34119429 | 22.64528856 | NA | NA | NA | NA | NA | NA | NA |
| SCN2A | 1438 | E | 29.20461111 | 22.55388013 | 0.329593083 | 21.96285202 | NA | NA | NA | NA | NA | NA | NA |
| SCN2A | 1439 | L | 29.2785 | 19.05343762 | 0.272748554 | 18.44962894 | NA | NA | NA | NA | NA | NA | NA |
| SCN2A | 1440 | Q | 31.28372222 | 14.19424891 | 0.193839017 | 13.6267394 | NA | NA | NA | NA | NA | NA | NA |
| SCN2A | 1441 | P | 28.26178571 | 14.50769761 | 0.198929186 | 13.87903831 | NA | NA | NA | NA | NA | NA | NA |
| SCN2A | 1442 | K | 28.52683333 | 15.17747074 | 0.209805793 | 14.51943124 | NA | NA | NA | NA | NA | NA | NA |
| SCN2A | 1443 | Y | 26.18041667 | 10.95138946 | 0.141177441 | 10.36521365 | NA | NA | NA | NA | NA | NA | NA |
| SCN2A | 1444 | E | 22.67394444 | 12.85492769 | 0.172089457 | 11.39117086 | NA | NA | NA | NA | NA | NA | NA |
| SCN2A | 1445 | D | 24.297125 | 17.01083775 | 0.239578282 | 15.03262166 | NA | NA | NA | NA | NA | NA | NA |
| SCN2A | 1446 | N | 19.79925 | 16.85089395 | 0.236980916 | 14.78064558 | NA | NA | NA | NA | NA | NA | NA |
| SCN2A | 1447 | L | 18.595875 | 16.13042957 | 0.225281122 | 14.53903006 | NA | NA | NA | NA | NA | NA | NA |
| SCN2A | 1448 | Y | 15.68566667 | 18.20309155 | 0.258939579 | 16.61265133 | NA | NA | NA | NA | NA | NA | NA |
| SCN2A | 1449 | M | 15.037125 | 15.4719194 | 0.214587416 | 14.19688156 | NA | NA | NA | NA | NA | NA | NA |
| SCN2A | 1450 | Y | 14.99133333 | 10.26806472 | 0.130080766 | 10.91671589 | NA | NA | NA | NA | NA | NA | NA |
| SCN2A | 1451 | L | 10.108375 | 16.23871504 | 0.227039595 | 15.61389045 | NA | NA | NA | NA | NA | NA | NA |
| SCN2A | 1452 | Y | 10.01666667 | 15.18440809 | 0.209918451 | 14.6128588 | NA | NA | NA | NA | NA | NA | NA |
| SCN2A | 1453 | F | 11.44177273 | 11.67927729 | 0.152997787 | 11.21364223 | NA | NA | NA | NA | NA | NA | NA |
| SCN2A | 1454 | V | 7.246785714 | 12.94206494 | 0.1735045 | 12.60369426 | NA | NA | NA | NA | NA | NA | NA |
| SCN2A | 1455 | I | 4.950125 | 14.85247498 | 0.204528109 | 14.69594418 | NA | NA | NA | NA | NA | NA | NA |
| SCN2A | 1456 | F | 5.2645 | 12.25594456 | 0.162362426 | 11.89638323 | NA | NA | NA | NA | NA | NA | NA |
| SCN2A | 1457 | I | 4.8135 | 8.325405576 | 0.098533456 | 7.976711533 | 1.51 | 0.05 | -0.5 | 17.74 | 0.15 | 1 | 1 |
| SCN2A | 1458 | I | 1.285625 | 13.22105416 | 0.178035073 | 8.369943773 | NA | NA | NA | NA | NA | NA | NA |
| SCN2A | 1459 | F | 0.252409091 | 14.26581172 | 0.195001143 | 9.919980587 | NA | NA | NA | NA | NA | NA | NA |
| SCN2A | 1460 | G | 1.2155 | 12.40006863 | 0.164702891 | 8.121619001 | NA | NA | NA | NA | NA | NA | NA |
| SCN2A | 1461 | S | 1.173166667 | 8.639009492 | 0.103626146 | 3.790467071 | 5.03 | 1.31 | 2.72 | 0.47 | -1.74 | 0 | 0 |
| SCN2A | 1462 | F | -1.638045455 | 9.458675806 | 0.116936904 | 5.075834853 | NA | NA | NA | NA | NA | NA | NA |

| | | | | | | | | | | | | | |
|---|---|---|---|---|---|---|---|---|---|---|---|---|---|
| SCN2A | 1463 | F | -3.2445 | 11.95026799 | 0.15739847 | 8.741355043 | NA | NA | NA | NA | NA | NA | NA |
| SCN2A | 1464 | T | -1.810785714 | 10.87735942 | 0.13997525 | 7.495493604 | NA | NA | NA | NA | NA | NA | NA |
| SCN2A | 1465 | L | -2.90075 | 5.118018464 | 0.046447924 | 1.740233766 | 3.41 | 1.3 | 2.8 | 0.5 | -1.6 | 0 | 0 |
| SCN2A | 1466 | N | -6.65525 | 9.023297679 | 0.109866694 | 5.957138178 | NA | NA | NA | NA | NA | NA | NA |
| SCN2A | 1467 | L | -7.7495 | 10.76401858 | 0.13813468 | 7.660422765 | NA | NA | NA | NA | NA | NA | NA |
| SCN2A | 1468 | F | -5.700590909 | 8.606471867 | 0.103097759 | 5.76173479 | NA | NA | NA | NA | NA | NA | NA |
| SCN2A | 1469 | I | -7.85825 | 4.262634866 | 0.032557143 | 1.674960105 | 2.61 | 1.67 | 3.34 | 0.43 | -1.77 | 0 | 0 |
| SCN2A | 1470 | G | -11.5645 | 7.933787683 | 0.092173879 | 5.26561943 | 2.69 | 1.29 | 3.52 | 0.78 | -1.32 | 0 | 0 |
| SCN2A | 1471 | V | -11.6705 | 9.839382145 | 0.123119286 | 7.164091134 | NA | NA | NA | NA | NA | NA | NA |
| SCN2A | 1472 | I | -10.565625 | 8.06497421 | 0.094304248 | 5.506848292 | 2.61 | 1.67 | 3.34 | 0.43 | -1.77 | 0 | 0 |
| SCN2A | 1473 | I | -13.62025 | 3.730002815 | 0.023907603 | 2.039275454 | 2.09 | 1.43 | 3.7 | 0.2 | -1.82 | 0 | 0 |
| SCN2A | 1474 | D | -16.82775 | 7.475943954 | 0.084738844 | 5.057786189 | 2.56 | 0.88 | 1.9 | 1 | -1.23 | 0 | 0 |
| SCN2A | 1475 | N | -15.483375 | 9.621393298 | 0.119579313 | 7.157991484 | NA | NA | NA | NA | NA | NA | NA |
| SCN2A | 1476 | F | -15.91113636 | 6.512309421 | 0.069090151 | 4.373043721 | 2.56 | 0.88 | 1.9 | 1 | -1.23 | 0 | 0 |
| SCN2A | 1477 | N | -19.385375 | 4.615917352 | 0.038294183 | 1.644807776 | 3.1 | 0.57 | 1.03 | 1.29 | -1.02 | 0 | 0 |
| SCN2A | 1478 | Q | -21.30283333 | 9.799167771 | 0.122466235 | 6.761451135 | NA | NA | NA | NA | NA | NA | NA |
| SCN2A | 1479 | Q | -17.6535 | 11.93203377 | 0.15710236 | 9.403385667 | NA | NA | NA | NA | NA | NA | NA |
| SCN2A | 1480 | K | -20.76005556 | 5.76445765 | 0.056945606 | 1.704299604 | 4.09 | 0.24 | -0.2 | 24.97 | 0.02 | 2 | 0 |
| SCN2A | 1481 | K | -24.90738889 | 7.808032915 | 0.090131717 | 3.686114218 | 4.63 | -0.77 | -2.9 | 26.11 | 1.25 | 2 | 0 |
| SCN2A | 1482 | K | -23.77838889 | 13.25504074 | 0.178586989 | 8.802715951 | NA | NA | NA | NA | NA | NA | NA |
| SCN2A | 1483 | F | -21.83459091 | 14.84439349 | 0.204396872 | 10.32652107 | NA | NA | NA | NA | NA | NA | NA |
| SCN2A | 1484 | G | -26.94575 | 12.35022984 | 0.163893547 | 8.091809032 | NA | NA | NA | NA | NA | NA | NA |
| SCN2A | 1485 | G | -24.99725 | 10.39028017 | 0.132065452 | 5.817506599 | NA | NA | NA | NA | NA | NA | NA |
| SCN2A | 1486 | Q | -23.13816667 | 11.3638251 | 0.147875083 | 7.209797647 | NA | NA | NA | NA | NA | NA | NA |
| SCN2A | 1487 | D | -19.349 | 11.18443566 | 0.144961935 | 7.104013724 | NA | NA | NA | NA | NA | NA | NA |
| SCN2A | 1488 | I | -16.15075 | 12.31107428 | 0.163257691 | 10.03565231 | NA | NA | NA | NA | NA | NA | NA |
| SCN2A | 1489 | F | -12.30268182 | 10.57521749 | 0.135068694 | 8.358434895 | NA | NA | NA | NA | NA | NA | NA |
| SCN2A | 1490 | M | -15.796 | 15.25390619 | 0.211047047 | 12.95263133 | NA | NA | NA | NA | NA | NA | NA |
| SCN2A | 1491 | T | -16.14107143 | 19.74447411 | 0.283970462 | 17.48047826 | NA | NA | NA | NA | NA | NA | NA |
| SCN2A | 1492 | E | -19.66916667 | 22.07492464 | 0.32181521 | 18.0046189 | NA | NA | NA | NA | NA | NA | NA |
| SCN2A | 1493 | E | -17.43527778 | 23.20353863 | 0.340143044 | 19.1627906 | NA | NA | NA | NA | NA | NA | NA |
| SCN2A | 1494 | Q | -15.12672222 | 19.59128393 | 0.28148277 | 15.71841433 | NA | NA | NA | NA | NA | NA | NA |
| SCN2A | 1495 | K | -20.3835 | 17.08626887 | 0.240803226 | 13.02438293 | NA | NA | NA | NA | NA | NA | NA |

| | | | | | | | | | | | | | |
|---|---|---|---|---|---|---|---|---|---|---|---|---|---|
| SCN2A | 1496 | K | -22.16172222 | 20.8417505 | 0.301789398 | 16.82894349 | NA | NA | NA | NA | NA | NA | NA |
| SCN2A | 1497 | Y | -17.50291667 | 21.08300882 | 0.30570725 | 17.08301855 | NA | NA | NA | NA | NA | NA | NA |
| SCN2A | 1498 | Y | -18.93233333 | 12.41750526 | 0.164986049 | 8.413153358 | NA | NA | NA | NA | NA | NA | NA |
| SCN2A | 1499 | N | -23.2815 | 15.85979284 | 0.220886187 | 11.8834921 | NA | NA | NA | NA | NA | NA | NA |
| SCN2A | 1500 | A | -22.3475 | 20.19985552 | 0.29136551 | 16.2347664 | NA | NA | NA | NA | NA | NA | NA |
| SCN2A | 1501 | M | -18.240125 | 19.48465265 | 0.279751159 | 15.46724641 | NA | NA | NA | NA | NA | NA | NA |
| SCN2A | 1502 | K | -22.44538889 | 12.26242187 | 0.162467612 | 8.325861077 | NA | NA | NA | NA | NA | NA | NA |
| SCN2A | 1503 | K | -25.54427778 | 19.04665354 | 0.272638385 | 15.09580415 | NA | NA | NA | NA | NA | NA | NA |
| SCN2A | 1504 | L | -20.8405 | 21.35152247 | 0.310067707 | 17.39594375 | NA | NA | NA | NA | NA | NA | NA |
| SCN2A | 1505 | G | -20.26625 | 20.24887046 | 0.292161476 | 16.28237807 | NA | NA | NA | NA | NA | NA | NA |
| SCN2A | 1506 | S | -23.28066667 | 17.28657739 | 0.244056085 | 13.33151624 | NA | NA | NA | NA | NA | NA | NA |
| SCN2A | 1507 | K | -24.2925 | 21.93230683 | 0.319499205 | 17.98674459 | NA | NA | NA | NA | NA | NA | NA |
| SCN2A | 1508 | K | -25.04938889 | 23.50320374 | 0.345009378 | 19.56128242 | NA | NA | NA | NA | NA | NA | NA |
| SCN2A | 1509 | P | -24.68735714 | 28.98141818 | 0.433971423 | 25.036293 | NA | NA | NA | NA | NA | NA | NA |
| SCN2A | 1510 | Q | -28.2885 | 28.89871014 | 0.432628307 | 24.96974806 | NA | NA | NA | NA | NA | NA | NA |
| SCN2A | 1511 | K | -28.49227778 | 35.02646682 | 0.532138424 | 31.08721127 | NA | NA | NA | NA | NA | NA | NA |
| SCN2A | 1512 | P | -24.12778571 | 35.33692263 | 0.537179991 | 31.38832094 | NA | NA | NA | NA | NA | NA | NA |
| SCN2A | 1513 | I | -24.96575 | 39.88632233 | 0.611058789 | 35.93733805 | NA | NA | NA | NA | NA | NA | NA |
| SCN2A | 1514 | P | -24.88064286 | 42.53881468 | 0.654133251 | 38.58621191 | NA | NA | NA | NA | NA | NA | NA |
| SCN2A | 1515 | R | -24.34668182 | 45.98392068 | 0.710079158 | 42.03098232 | NA | NA | NA | NA | NA | NA | NA |
| SCN2A | 1516 | P | -20.82221429 | 47.68133577 | 0.737643891 | 43.71719147 | NA | NA | NA | NA | NA | NA | NA |
| SCN2A | 1517 | A | -24.3677 | 51.21179709 | 0.794975903 | 47.25459241 | NA | NA | NA | NA | NA | NA | NA |
| SCN2A | 1518 | N | -25.235125 | 53.67920007 | 0.835044654 | 49.72444075 | NA | NA | NA | NA | NA | NA | NA |
| SCN2A | 1519 | K | -20.16327778 | 55.53216483 | 0.865135396 | 51.57210989 | NA | NA | NA | NA | NA | NA | NA |
| SCN2A | 1520 | F | -17.32859091 | 54.54141519 | 0.849046374 | 50.56954826 | NA | NA | NA | NA | NA | NA | NA |
| SCN2A | 1521 | Q | -17.1225 | 50.99779731 | 0.791500709 | 47.02791245 | NA | NA | NA | NA | NA | NA | NA |
| SCN2A | 1522 | G | -20.10325 | 50.29477214 | 0.780084114 | 46.32892716 | NA | NA | NA | NA | NA | NA | NA |
| SCN2A | 1523 | M | -19.740375 | 51.28745308 | 0.796204499 | 47.31335417 | NA | NA | NA | NA | NA | NA | NA |
| SCN2A | 1524 | V | -15.43664286 | 48.85916418 | 0.75677093 | 44.88111153 | NA | NA | NA | NA | NA | NA | NA |
| SCN2A | 1525 | F | -19.15504545 | 44.10292989 | 0.679533295 | 40.13842262 | NA | NA | NA | NA | NA | NA | NA |
| SCN2A | 1526 | D | -19.8815 | 46.32029053 | 0.715541549 | 42.34656687 | NA | NA | NA | NA | NA | NA | NA |
| SCN2A | 1527 | F | -16.18613636 | 46.092592 | 0.711843898 | 42.10811676 | NA | NA | NA | NA | NA | NA | NA |
| SCN2A | 1528 | V | -14.09635714 | 43.36610203 | 0.66756777 | 39.38056965 | NA | NA | NA | NA | NA | NA | NA |

| | | | | | | | | | | | | | |
|---|---|---|---|---|---|---|---|---|---|---|---|---|---|
| SCN2A | 1529 | T | -17.76278571 | 41.09263921 | 0.630648459 | 37.11487226 | NA | NA | NA | NA | NA | NA | NA |
| SCN2A | 1530 | K | -17.79938889 | 41.78801968 | 0.641940911 | 37.80050325 | NA | NA | NA | NA | NA | NA | NA |
| SCN2A | 1531 | Q | -15.45516667 | 36.87419767 | 0.562144171 | 32.8784565 | NA | NA | NA | NA | NA | NA | NA |
| SCN2A | 1532 | V | -12.77907143 | 42.54940508 | 0.654305232 | 38.55582678 | NA | NA | NA | NA | NA | NA | NA |
| SCN2A | 1533 | F | -10.72604545 | 41.65583864 | 0.639794391 | 39.43752987 | NA | NA | NA | NA | NA | NA | NA |
| SCN2A | 1534 | D | -12.390125 | 36.59246477 | 0.557569042 | 34.98848466 | NA | NA | NA | NA | NA | NA | NA |
| SCN2A | 1535 | I | -10.078625 | 36.48239796 | 0.555781641 | 34.39444673 | NA | NA | NA | NA | NA | NA | NA |
| SCN2A | 1536 | S | -7.263 | 40.66475266 | 0.623699907 | 38.47803262 | NA | NA | NA | NA | NA | NA | NA |
| SCN2A | 1537 | I | -6.247 | 38.05585165 | 0.581333334 | 36.03609451 | NA | NA | NA | NA | NA | NA | NA |
| SCN2A | 1538 | M | -7.311125 | 33.71370247 | 0.510820127 | 31.50988352 | NA | NA | NA | NA | NA | NA | NA |
| SCN2A | 1539 | I | -4.175625 | 37.20624427 | 0.567536356 | 32.36069087 | NA | NA | NA | NA | NA | NA | NA |
| SCN2A | 1540 | L | -1.296625 | 37.78797556 | 0.57698323 | 32.9464038 | NA | NA | NA | NA | NA | NA | NA |
| SCN2A | 1541 | I | -1.726625 | 33.6330094 | 0.509509733 | 28.79278205 | NA | NA | NA | NA | NA | NA | NA |
| SCN2A | 1542 | C | -0.9335 | 32.94939955 | 0.498408428 | 28.10263154 | NA | NA | NA | NA | NA | NA | NA |
| SCN2A | 1543 | L | 1.051125 | 35.87069134 | 0.54584799 | 31.07904746 | NA | NA | NA | NA | NA | NA | NA |
| SCN2A | 1544 | N | 2.578875 | 33.86435704 | 0.513266643 | 29.079619 | NA | NA | NA | NA | NA | NA | NA |
| SCN2A | 1545 | M | 2.24875 | 26.77369504 | 0.398119674 | 21.98288855 | NA | NA | NA | NA | NA | NA | NA |
| SCN2A | 1546 | V | 5.040214286 | 32.78424934 | 0.495726514 | 27.99938945 | NA | NA | NA | NA | NA | NA | NA |
| SCN2A | 1547 | T | 7.308642857 | 34.36053835 | 0.521324251 | 32.29588607 | NA | NA | NA | NA | NA | NA | NA |
| SCN2A | 1548 | M | 7.005 | 30.62149072 | 0.460604956 | 28.56153708 | NA | NA | NA | NA | NA | NA | NA |
| SCN2A | 1549 | M | 7.7885 | 27.66891434 | 0.412657356 | 26.03258705 | NA | NA | NA | NA | NA | NA | NA |
| SCN2A | 1550 | V | 11.72921429 | 31.75734402 | 0.479050351 | 29.69202809 | NA | NA | NA | NA | NA | NA | NA |
| SCN2A | 1551 | E | 13.00083333 | 29.76896876 | 0.446760646 | 27.71378126 | NA | NA | NA | NA | NA | NA | NA |
| SCN2A | 1552 | T | 17.38621429 | 30.60078014 | 0.460268632 | 28.5554309 | NA | NA | NA | NA | NA | NA | NA |
| SCN2A | 1553 | D | 20.754375 | 29.05142456 | 0.435108273 | 27.41464462 | NA | NA | NA | NA | NA | NA | NA |
| SCN2A | 1554 | D | 23.61125 | 30.62462836 | 0.460655909 | 29.90776425 | NA | NA | NA | NA | NA | NA | NA |
| SCN2A | 1555 | Q | 19.58994444 | 34.93069999 | 0.530583244 | 32.90551295 | NA | NA | NA | NA | NA | NA | NA |
| SCN2A | 1556 | S | 22.1375 | 38.16153222 | 0.583049507 | 37.60293812 | NA | NA | NA | NA | NA | NA | NA |
| SCN2A | 1557 | Q | 22.29072222 | 41.36047017 | 0.634997831 | 39.35644446 | NA | NA | NA | NA | NA | NA | NA |
| SCN2A | 1558 | E | 20.1205 | 42.37021087 | 0.651395253 | 40.31439226 | NA | NA | NA | NA | NA | NA | NA |
| SCN2A | 1559 | M | 17.06225 | 37.56444652 | 0.573353288 | 35.50211069 | NA | NA | NA | NA | NA | NA | NA |
| SCN2A | 1560 | T | 17.56907143 | 39.00633505 | 0.596768463 | 36.97318696 | NA | NA | NA | NA | NA | NA | NA |
| SCN2A | 1561 | N | 17.253 | 42.37331677 | 0.651445691 | 40.32981679 | NA | NA | NA | NA | NA | NA | NA |

| | | | | | | | | | | | | | |
|---|---|---|---|---|---|---|---|---|---|---|---|---|---|
| SCN2A | 1562 | I | 13.985625 | 42.10866549 | 0.647147955 | 40.05076701 | NA | NA | NA | NA | NA | NA | NA |
| SCN2A | 1563 | L | 12.4425 | 36.65145985 | 0.558527078 | 34.59429377 | NA | NA | NA | NA | NA | NA | NA |
| SCN2A | 1564 | Y | 12.34841667 | 41.36045503 | 0.634997585 | 39.30793168 | NA | NA | NA | NA | NA | NA | NA |
| SCN2A | 1565 | W | 12.20221429 | 43.88346191 | 0.675969302 | 41.8287609 | NA | NA | NA | NA | NA | NA | NA |
| SCN2A | 1566 | I | 7.75775 | 40.6619382 | 0.623654202 | 38.60005763 | NA | NA | NA | NA | NA | NA | NA |
| SCN2A | 1567 | N | 7.633375 | 37.91893356 | 0.579109888 | 33.32806535 | NA | NA | NA | NA | NA | NA | NA |
| SCN2A | 1568 | L | 7.35375 | 43.21619424 | 0.665133381 | 38.62706264 | NA | NA | NA | NA | NA | NA | NA |
| SCN2A | 1569 | V | 4.4005 | 44.27746715 | 0.682367648 | 39.48960057 | NA | NA | NA | NA | NA | NA | NA |
| SCN2A | 1570 | F | 2.656863636 | 39.72569704 | 0.608450356 | 34.93689873 | NA | NA | NA | NA | NA | NA | NA |
| SCN2A | 1571 | I | 3.10575 | 41.29337035 | 0.633908181 | 36.51244097 | NA | NA | NA | NA | NA | NA | NA |
| SCN2A | 1572 | V | 1.559928571 | 44.9662593 | 0.693553109 | 40.17466394 | NA | NA | NA | NA | NA | NA | NA |
| SCN2A | 1573 | L | -0.809375 | 43.93786787 | 0.676852813 | 39.14216098 | NA | NA | NA | NA | NA | NA | NA |
| SCN2A | 1574 | F | -1.251227273 | 38.66857759 | 0.591283539 | 33.87266635 | NA | NA | NA | NA | NA | NA | NA |
| SCN2A | 1575 | T | -1.245214286 | 44.00603752 | 0.677959837 | 39.20831541 | NA | NA | NA | NA | NA | NA | NA |
| SCN2A | 1576 | G | -3.84775 | 46.5300546 | 0.718947958 | 41.73201321 | NA | NA | NA | NA | NA | NA | NA |
| SCN2A | 1577 | E | -6.503611111 | 41.21149658 | 0.632578613 | 38.88865667 | NA | NA | NA | NA | NA | NA | NA |
| SCN2A | 1578 | C | -6.273166667 | 42.94133357 | 0.660669853 | 40.63128988 | NA | NA | NA | NA | NA | NA | NA |
| SCN2A | 1579 | V | -6.255928571 | 46.20020298 | 0.713591418 | 43.8845451 | NA | NA | NA | NA | NA | NA | NA |
| SCN2A | 1580 | L | -9.47275 | 47.62268415 | 0.736691433 | 45.27198181 | NA | NA | NA | NA | NA | NA | NA |
| SCN2A | 1581 | K | -11.53772222 | 42.0296408 | 0.645864654 | 39.95424075 | NA | NA | NA | NA | NA | NA | NA |
| SCN2A | 1582 | L | -10.54975 | 45.40483149 | 0.700675189 | 43.07249307 | NA | NA | NA | NA | NA | NA | NA |
| SCN2A | 1583 | I | -11.681875 | 49.37040094 | 0.765073027 | 45.38432301 | NA | NA | NA | NA | NA | NA | NA |
| SCN2A | 1584 | S | -15.21 | 47.21117818 | 0.730008888 | 43.23807861 | NA | NA | NA | NA | NA | NA | NA |
| SCN2A | 1585 | L | -16.31225 | 42.46271255 | 0.652897411 | 38.49012701 | NA | NA | NA | NA | NA | NA | NA |
| SCN2A | 1586 | R | -17.15313636 | 47.14766536 | 0.728977488 | 43.17213666 | NA | NA | NA | NA | NA | NA | NA |
| SCN2A | 1587 | Y | -20.06658333 | 45.22978771 | 0.697832611 | 41.25963218 | NA | NA | NA | NA | NA | NA | NA |
| SCN2A | 1588 | Y | -18.61983333 | 39.93914163 | 0.611916534 | 35.97180232 | NA | NA | NA | NA | NA | NA | NA |
| SCN2A | 1589 | Y | -12.5345 | 39.52500196 | 0.60519122 | 35.54031782 | NA | NA | NA | NA | NA | NA | NA |
| SCN2A | 1590 | F | -13.17840909 | 37.31872466 | 0.569362952 | 33.33776737 | NA | NA | NA | NA | NA | NA | NA |
| SCN2A | 1591 | T | -18.38421429 | 34.11476329 | 0.517333051 | 30.13911032 | NA | NA | NA | NA | NA | NA | NA |
| SCN2A | 1592 | I | -16.242625 | 33.25528062 | 0.503375705 | 29.28311652 | NA | NA | NA | NA | NA | NA | NA |
| SCN2A | 1593 | G | -12.6985 | 32.55902572 | 0.492069054 | 28.5705652 | NA | NA | NA | NA | NA | NA | NA |
| SCN2A | 1594 | W | -11.12642857 | 28.06149285 | 0.419032533 | 25.70315121 | NA | NA | NA | NA | NA | NA | NA |

| | | | | | | | | | | | | | |
|---|---|---|---|---|---|---|---|---|---|---|---|---|---|
| SCN2A | 1595 | N | -11.71525 | 35.54280087 | 0.540523297 | 33.19536065 | NA | NA | NA | NA | NA | NA | NA |
| SCN2A | 1596 | I | -10.262625 | 35.72963232 | 0.543557298 | 33.42185254 | NA | NA | NA | NA | NA | NA | NA |
| SCN2A | 1597 | F | -6.032954545 | 31.01089405 | 0.46692857 | 28.74254051 | NA | NA | NA | NA | NA | NA | NA |
| SCN2A | 1598 | D | -6.230375 | 35.83341448 | 0.545242642 | 33.52642793 | NA | NA | NA | NA | NA | NA | NA |
| SCN2A | 1599 | F | -5.5175 | 38.39075001 | 0.58677183 | 36.09180314 | NA | NA | NA | NA | NA | NA | NA |
| SCN2A | 1600 | V | -3.113357143 | 36.80153819 | 0.560964236 | 34.55331216 | NA | NA | NA | NA | NA | NA | NA |
| SCN2A | 1601 | V | -1.199928571 | 34.53101011 | 0.524092583 | 29.72976349 | NA | NA | NA | NA | NA | NA | NA |
| SCN2A | 1602 | V | -1.0185 | 38.90567681 | 0.59513385 | 34.10507814 | NA | NA | NA | NA | NA | NA | NA |
| SCN2A | 1603 | I | -0.5585 | 40.59761051 | 0.622609569 | 35.8110375 | NA | NA | NA | NA | NA | NA | NA |
| SCN2A | 1604 | L | 1.653875 | 35.53361075 | 0.540374057 | 30.91599157 | NA | NA | NA | NA | NA | NA | NA |
| SCN2A | 1605 | S | 3.228166667 | 38.69946409 | 0.591785112 | 34.08247052 | NA | NA | NA | NA | NA | NA | NA |
| SCN2A | 1606 | I | 3.263375 | 41.32376171 | 0.634401713 | 36.70794206 | NA | NA | NA | NA | NA | NA | NA |
| SCN2A | 1607 | V | 5.466071429 | 41.32793134 | 0.634469425 | 36.72525171 | NA | NA | NA | NA | NA | NA | NA |
| SCN2A | 1608 | G | 8.1125 | 39.91848852 | 0.611581143 | 35.33337277 | NA | NA | NA | NA | NA | NA | NA |
| SCN2A | 1609 | M | 8.629125 | 41.92328443 | 0.644137507 | 37.33746306 | NA | NA | NA | NA | NA | NA | NA |
| SCN2A | 1610 | F | 8.1115 | 44.60656981 | 0.687712025 | 40.01971648 | NA | NA | NA | NA | NA | NA | NA |
| SCN2A | 1611 | L | 10.41475 | 42.52024692 | 0.653831725 | 38.02456234 | NA | NA | NA | NA | NA | NA | NA |
| SCN2A | 1612 | A | 13.0941 | 42.59263615 | 0.655007271 | 40.57261109 | NA | NA | NA | NA | NA | NA | NA |
| SCN2A | 1613 | E | 14.12927778 | 45.71660042 | 0.70573808 | 43.69981194 | NA | NA | NA | NA | NA | NA | NA |
| SCN2A | 1614 | L | 14.300125 | 46.52043811 | 0.718791794 | 44.52534553 | NA | NA | NA | NA | NA | NA | NA |
| SCN2A | 1615 | I | 16.1055 | 41.02551545 | 0.629558421 | 39.02519622 | NA | NA | NA | NA | NA | NA | NA |
| SCN2A | 1616 | E | 18.6135 | 45.63773768 | 0.704457409 | 43.64611042 | NA | NA | NA | NA | NA | NA | NA |
| SCN2A | 1617 | K | 18.13861111 | 48.14410389 | 0.745158894 | 46.15690995 | NA | NA | NA | NA | NA | NA | NA |
| SCN2A | 1618 | Y | 17.64925 | 46.44547552 | 0.717574458 | 44.4768806 | NA | NA | NA | NA | NA | NA | NA |
| SCN2A | 1619 | F | 18.13922727 | 43.2211235 | 0.665213429 | 41.26827524 | NA | NA | NA | NA | NA | NA | NA |
| SCN2A | 1620 | V | 19.70464286 | 41.39916712 | 0.63562624 | 39.44195869 | NA | NA | NA | NA | NA | NA | NA |
| SCN2A | 1621 | S | 17.42566667 | 37.96269507 | 0.579820542 | 35.98931363 | NA | NA | NA | NA | NA | NA | NA |
| SCN2A | 1622 | P | 18.6165 | 34.81210463 | 0.528657345 | 32.83245925 | NA | NA | NA | NA | NA | NA | NA |
| SCN2A | 1623 | T | 15.99421429 | 31.89256479 | 0.481246234 | 29.93310561 | NA | NA | NA | NA | NA | NA | NA |
| SCN2A | 1624 | L | 12.6145 | 35.01274845 | 0.531915648 | 33.03565572 | NA | NA | NA | NA | NA | NA | NA |
| SCN2A | 1625 | F | 13.98140909 | 34.91564426 | 0.53033875 | 32.90985549 | NA | NA | NA | NA | NA | NA | NA |
| SCN2A | 1626 | R | 14.72086364 | 29.38964762 | 0.440600759 | 27.3957196 | NA | NA | NA | NA | NA | NA | NA |
| SCN2A | 1627 | V | 9.630642857 | 30.96571567 | 0.466194908 | 28.96741105 | NA | NA | NA | NA | NA | NA | NA |

| | | | | | | | | | | | | | |
|---|---|---|---|---|---|---|---|---|---|---|---|---|---|
| SCN2A | 1628 | I | 7.959625 | 34.17894489 | 0.518375311 | 29.61612652 | NA | NA | NA | NA | NA | NA | NA |
| SCN2A | 1629 | R | 10.50440909 | 31.76614607 | 0.47919329 | 28.22500749 | NA | NA | NA | NA | NA | NA | NA |
| SCN2A | 1630 | L | 6.640875 | 28.62680094 | 0.428212708 | 24.0637165 | NA | NA | NA | NA | NA | NA | NA |
| SCN2A | 1631 | A | 3.1829 | 32.76412173 | 0.495399657 | 28.15986567 | NA | NA | NA | NA | NA | NA | NA |
| SCN2A | 1632 | R | 5.350590909 | 32.45614366 | 0.490398327 | 27.83593718 | NA | NA | NA | NA | NA | NA | NA |
| SCN2A | 1633 | I | 2.63275 | 27.57462143 | 0.411126111 | 22.95569602 | NA | NA | NA | NA | NA | NA | NA |
| SCN2A | 1634 | G | -0.774 | 30.78199542 | 0.46321143 | 25.98503102 | NA | NA | NA | NA | NA | NA | NA |
| SCN2A | 1635 | R | -1.0815 | 32.24684115 | 0.486999413 | 27.4489475 | NA | NA | NA | NA | NA | NA | NA |
| SCN2A | 1636 | I | -0.80125 | 28.25316975 | 0.422145221 | 23.45767418 | NA | NA | NA | NA | NA | NA | NA |
| SCN2A | 1637 | L | -3.34475 | 26.38870292 | 0.391867695 | 21.53248416 | NA | NA | NA | NA | NA | NA | NA |
| SCN2A | 1638 | R | -6.854136364 | 30.81758701 | 0.463789411 | 28.53228472 | NA | NA | NA | NA | NA | NA | NA |
| SCN2A | 1639 | L | -5.505875 | 28.74999303 | 0.430213254 | 26.48932437 | NA | NA | NA | NA | NA | NA | NA |
| SCN2A | 1640 | I | -8.1775 | 24.82171833 | 0.366421054 | 22.54640081 | NA | NA | NA | NA | NA | NA | NA |
| SCN2A | 1641 | K | -11.91627778 | 28.75748157 | 0.430334862 | 26.48357654 | NA | NA | NA | NA | NA | NA | NA |
| SCN2A | 1642 | G | -12.79675 | 27.8632171 | 0.415812686 | 25.77126417 | NA | NA | NA | NA | NA | NA | NA |
| SCN2A | 1643 | A | -11.8471 | 24.06701271 | 0.354165208 | 21.96433355 | NA | NA | NA | NA | NA | NA | NA |
| SCN2A | 1644 | K | -15.64194444 | 22.79587636 | 0.333522918 | 18.98387149 | NA | NA | NA | NA | NA | NA | NA |
| SCN2A | 1645 | G | -13.5835 | 18.59259041 | 0.265264745 | 16.50358564 | NA | NA | NA | NA | NA | NA | NA |
| SCN2A | 1646 | I | -10.065375 | 19.34858605 | 0.27754154 | 17.10022921 | NA | NA | NA | NA | NA | NA | NA |
| SCN2A | 1647 | R | -13.77759091 | 19.85935724 | 0.285836077 | 17.63195531 | NA | NA | NA | NA | NA | NA | NA |
| SCN2A | 1648 | T | -15.19592857 | 16.13779381 | 0.225400711 | 12.3408205 | NA | NA | NA | NA | NA | NA | NA |
| SCN2A | 1649 | L | -11.15325 | 14.43299092 | 0.197716006 | 12.31214865 | NA | NA | NA | NA | NA | NA | NA |
| SCN2A | 1650 | L | -9.8505 | 17.68671555 | 0.250554025 | 15.35516857 | NA | NA | NA | NA | NA | NA | NA |
| SCN2A | 1651 | F | -15.60822727 | 17.00778151 | 0.239528651 | 14.71094645 | NA | NA | NA | NA | NA | NA | NA |
| SCN2A | 1652 | A | -13.6137 | 13.11921412 | 0.176381268 | 10.87735222 | NA | NA | NA | NA | NA | NA | NA |
| SCN2A | 1653 | L | -9.382375 | 14.1578694 | 0.193248242 | 11.89933172 | NA | NA | NA | NA | NA | NA | NA |
| SCN2A | 1654 | M | -12.04125 | 16.41371226 | 0.229881417 | 14.09757689 | NA | NA | NA | NA | NA | NA | NA |
| SCN2A | 1655 | M | -15.231375 | 15.18124096 | 0.209867019 | 12.86611097 | NA | NA | NA | NA | NA | NA | NA |
| SCN2A | 1656 | S | -11.139 | 12.18157744 | 0.16115476 | 9.903617424 | NA | NA | NA | NA | NA | NA | NA |
| SCN2A | 1657 | L | -8.972375 | 15.45021029 | 0.214234877 | 13.19310626 | NA | NA | NA | NA | NA | NA | NA |
| SCN2A | 1658 | P | -12.03192857 | 16.63954137 | 0.233548711 | 14.40336237 | NA | NA | NA | NA | NA | NA | NA |
| SCN2A | 1659 | A | -10.9713 | 13.35315116 | 0.180180228 | 11.1046116 | NA | NA | NA | NA | NA | NA | NA |
| SCN2A | 1660 | L | -6.816875 | 12.60046705 | 0.16795721 | 10.43988065 | NA | NA | NA | NA | NA | NA | NA |

| | | | | | | | | | | | | | |
|---|---|---|---|---|---|---|---|---|---|---|---|---|---|
| SCN2A | 1661 | F | -6.762681818 | 16.51874587 | 0.231587083 | 14.33848961 | NA | NA | NA | NA | NA | NA | NA |
| SCN2A | 1662 | N | -9.0105 | 17.323608 | 0.244657434 | 15.16813549 | NA | NA | NA | NA | NA | NA | NA |
| SCN2A | 1663 | I | -6.20925 | 12.11306171 | 0.160042117 | 9.909928193 | NA | NA | NA | NA | NA | NA | NA |
| SCN2A | 1664 | G | -2.80375 | 16.42495239 | 0.230063949 | 14.04326423 | NA | NA | NA | NA | NA | NA | NA |
| SCN2A | 1665 | L | -3.549625 | 19.05320524 | 0.27274478 | 16.66891173 | NA | NA | NA | NA | NA | NA | NA |
| SCN2A | 1666 | L | -3.1695 | 17.42587573 | 0.246318184 | 15.06318723 | NA | NA | NA | NA | NA | NA | NA |
| SCN2A | 1667 | L | -0.303 | 13.94441759 | 0.189781946 | 10.52895513 | NA | NA | NA | NA | NA | NA | NA |
| SCN2A | 1668 | F | 1.460136364 | 19.43561579 | 0.278954837 | 16.02617908 | NA | NA | NA | NA | NA | NA | NA |
| SCN2A | 1669 | L | 0.442125 | 21.67623053 | 0.31534072 | 18.27211523 | NA | NA | NA | NA | NA | NA | NA |
| SCN2A | 1670 | V | 1.705357143 | 19.06203824 | 0.272888221 | 15.6424665 | NA | NA | NA | NA | NA | NA | NA |
| SCN2A | 1671 | M | 4.5495 | 16.42562885 | 0.230074934 | 11.80503272 | NA | NA | NA | NA | NA | NA | NA |
| SCN2A | 1672 | F | 5.302318182 | 22.79783564 | 0.333554736 | 18.15980839 | NA | NA | NA | NA | NA | NA | NA |
| SCN2A | 1673 | I | 4.650125 | 24.55454121 | 0.362082301 | 19.90452943 | NA | NA | NA | NA | NA | NA | NA |
| SCN2A | 1674 | Y | 6.536416667 | 21.05266173 | 0.305214436 | 17.02239667 | NA | NA | NA | NA | NA | NA | NA |
| SCN2A | 1675 | A | 9.9989 | 23.44136148 | 0.344005107 | 21.33993063 | NA | NA | NA | NA | NA | NA | NA |
| SCN2A | 1676 | I | 9.883125 | 25.54611176 | 0.378184654 | 23.44582456 | NA | NA | NA | NA | NA | NA | NA |
| SCN2A | 1677 | F | 9.711954545 | 24.42439835 | 0.359968879 | 22.32794188 | NA | NA | NA | NA | NA | NA | NA |
| SCN2A | 1678 | G | 14.055 | 22.4626031 | 0.328110813 | 20.36375379 | NA | NA | NA | NA | NA | NA | NA |
| SCN2A | 1679 | M | 15.179375 | 24.11443423 | 0.354935297 | 22.01197527 | NA | NA | NA | NA | NA | NA | NA |
| SCN2A | 1680 | S | 15.92666667 | 26.34589746 | 0.391172566 | 24.24358803 | NA | NA | NA | NA | NA | NA | NA |
| SCN2A | 1681 | N | 16.301375 | 24.19604224 | 0.356260549 | 22.09443665 | NA | NA | NA | NA | NA | NA | NA |
| SCN2A | 1682 | F | 17.20768182 | 19.86983973 | 0.286006304 | 17.904283 | NA | NA | NA | NA | NA | NA | NA |
| SCN2A | 1683 | A | 21.0845 | 23.72764019 | 0.348654056 | 21.6985922 | NA | NA | NA | NA | NA | NA | NA |
| SCN2A | 1684 | Y | 24.03166667 | 22.54066186 | 0.329378428 | 21.93432323 | NA | NA | NA | NA | NA | NA | NA |
| SCN2A | 1685 | V | 22.9705 | 18.66797753 | 0.266488974 | 18.01611619 | NA | NA | NA | NA | NA | NA | NA |
| SCN2A | 1686 | K | 27.47883333 | 15.4946505 | 0.214956552 | 15.8426952 | NA | NA | NA | NA | NA | NA | NA |
| SCN2A | 1687 | R | 26.80795455 | 13.95849917 | 0.19001062 | 13.58855883 | NA | NA | NA | NA | NA | NA | NA |
| SCN2A | 1688 | E | 25.66294444 | 11.73568047 | 0.153913732 | 11.11844028 | NA | NA | NA | NA | NA | NA | NA |
| SCN2A | 1689 | V | 22.75335714 | 7.904198947 | 0.09169338 | 7.184108881 | 0.75 | -0.67 | -1.77 | 19.59 | 1.08 | 1 | 1 |
| SCN2A | 1690 | G | 20.3965 | 11.19080632 | 0.145065389 | 10.47310999 | NA | NA | NA | NA | NA | NA | NA |
| SCN2A | 1691 | I | 21.74075 | 13.72805886 | 0.186268445 | 13.0208187 | NA | NA | NA | NA | NA | NA | NA |
| SCN2A | 1692 | D | 23.163625 | 15.0990803 | 0.208532792 | 14.3851602 | NA | NA | NA | NA | NA | NA | NA |
| SCN2A | 1693 | D | 25.409875 | 18.18981322 | 0.258723949 | 17.53141571 | NA | NA | NA | NA | NA | NA | NA |

| | | | | | | | | | | | | | |
|---|---|---|---|---|---|---|---|---|---|---|---|---|---|
| SCN2A | 1694 | M | 21.477375 | 19.34090807 | 0.277416856 | 18.62275888 | NA | NA | NA | NA | NA | NA | NA |
| SCN2A | 1695 | F | 18.77877273 | 16.66830909 | 0.234015877 | 14.61390615 | NA | NA | NA | NA | NA | NA | NA |
| SCN2A | 1696 | N | 19.551125 | 18.73347904 | 0.267552669 | 16.69488178 | NA | NA | NA | NA | NA | NA | NA |
| SCN2A | 1697 | F | 16.3605 | 17.54536286 | 0.248258564 | 15.43928652 | NA | NA | NA | NA | NA | NA | NA |
| SCN2A | 1698 | E | 21.07772222 | 23.06970266 | 0.33796965 | 21.12324747 | NA | NA | NA | NA | NA | NA | NA |
| SCN2A | 1699 | T | 17.27864286 | 24.56359652 | 0.362229352 | 22.44563222 | NA | NA | NA | NA | NA | NA | NA |
| SCN2A | 1700 | F | 11.41986364 | 24.91883547 | 0.367998163 | 22.79917875 | NA | NA | NA | NA | NA | NA | NA |
| SCN2A | 1701 | G | 13.875 | 22.96782284 | 0.336315199 | 20.83766721 | NA | NA | NA | NA | NA | NA | NA |
| SCN2A | 1702 | N | 16.68825 | 20.1152037 | 0.289990829 | 17.99124329 | NA | NA | NA | NA | NA | NA | NA |
| SCN2A | 1703 | S | 13.162 | 20.20027309 | 0.291372291 | 18.08769455 | NA | NA | NA | NA | NA | NA | NA |
| SCN2A | 1704 | M | 9.3365 | 19.5198302 | 0.280322415 | 17.61755278 | NA | NA | NA | NA | NA | NA | NA |
| SCN2A | 1705 | I | 12.749375 | 16.64995892 | 0.233717884 | 14.79947968 | NA | NA | NA | NA | NA | NA | NA |
| SCN2A | 1706 | C | 12.72766667 | 15.32698069 | 0.212233722 | 13.21447804 | NA | NA | NA | NA | NA | NA | NA |
| SCN2A | 1707 | L | 8.895125 | 15.63239825 | 0.217193471 | 13.37148113 | NA | NA | NA | NA | NA | NA | NA |
| SCN2A | 1708 | F | 6.739136364 | 13.549031 | 0.183361168 | 9.514779328 | NA | NA | NA | NA | NA | NA | NA |
| SCN2A | 1709 | Q | 11.18005556 | 10.8128794 | 0.138928143 | 9.00279763 | NA | NA | NA | NA | NA | NA | NA |
| SCN2A | 1710 | I | 8.8585 | 11.30740947 | 0.146958936 | 9.010416979 | NA | NA | NA | NA | NA | NA | NA |
| SCN2A | 1711 | T | 4.5015 | 9.954433937 | 0.12498764 | 5.313928722 | NA | NA | NA | NA | NA | NA | NA |
| SCN2A | 1712 | T | 5.611214286 | 6.825127691 | 0.074170082 | 2.860889685 | 4.05 | -0.17 | -0.48 | 14.15 | 0.32 | 1 | 1 |
| SCN2A | 1713 | S | 7.336166667 | 6.349321224 | 0.066443346 | 4.036855386 | 2.32 | -0.17 | -0.48 | 14.15 | 0.32 | 1 | 1 |
| SCN2A | 1714 | A | 10.3443 | 7.376939338 | 0.083131084 | 5.805849968 | 1.83 | -0.07 | -1.53 | 33.25 | 0.63 | 2 | 0 |
| SCN2A | 1715 | G | 13.31875 | 7.012010339 | 0.077204914 | 4.898571681 | 2.21 | -0.17 | -0.53 | 3.06 | -0.01 | 0 | 0 |
| SCN2A | 1716 | W | 11.214 | 8.341103284 | 0.098788375 | 6.242545356 | 2.21 | -0.17 | -0.53 | 3.06 | -0.01 | 0 | 0 |
| SCN2A | 1717 | D | 15.384375 | 4.441698436 | 0.035465 | 2.347062466 | 2.21 | -0.17 | -0.53 | 3.06 | -0.01 | 0 | 0 |
| SCN2A | 1718 | G | 17.887 | 8.25496705 | 0.097389588 | 6.229001755 | 2.11 | 0.14 | -0.74 | 11.96 | -0.02 | 1 | 1 |
| SCN2A | 1719 | L | 16.56025 | 10.84286798 | 0.139415135 | 8.818770754 | NA | NA | NA | NA | NA | NA | NA |
| SCN2A | 1720 | L | 16.149 | 12.17390685 | 0.161030196 | 10.16208854 | NA | NA | NA | NA | NA | NA | NA |
| SCN2A | 1721 | A | 20.3575 | 11.27326479 | 0.146404452 | 9.353368076 | NA | NA | NA | NA | NA | NA | NA |
| SCN2A | 1722 | P | 21.89735714 | 13.50517434 | 0.182648969 | 12.81664913 | NA | NA | NA | NA | NA | NA | NA |
| SCN2A | 1723 | I | 19.1945 | 16.49234168 | 0.231158299 | 14.42417825 | NA | NA | NA | NA | NA | NA | NA |
| SCN2A | 1724 | L | 19.773375 | 14.53195641 | 0.199323131 | 12.46638345 | NA | NA | NA | NA | NA | NA | NA |
| SCN2A | 1725 | N | 24.25675 | 15.63426452 | 0.217223778 | 15.05150596 | NA | NA | NA | NA | NA | NA | NA |
| SCN2A | 1726 | S | 24.87616667 | 20.47574248 | 0.295845705 | 19.90417389 | NA | NA | NA | NA | NA | NA | NA |

| | | | | | | | | | | | | | |
|---|---|---|---|---|---|---|---|---|---|---|---|---|---|
| SCN2A | 1727 | G | 28.33 | 20.4527834 | 0.295472867 | 19.48993798 | NA | NA | NA | NA | NA | NA | NA |
| SCN2A | 1728 | P | 31.59292857 | 21.18837743 | 0.307418356 | 20.20110265 | NA | NA | NA | NA | NA | NA | NA |
| SCN2A | 1729 | P | 32.70621429 | 18.05232309 | 0.256491214 | 17.40725554 | NA | NA | NA | NA | NA | NA | NA |
| SCN2A | 1730 | D | 29.592 | 15.79172812 | 0.219780867 | 14.82197683 | NA | NA | NA | NA | NA | NA | NA |
| SCN2A | 1731 | C | 29.80033333 | 18.91505646 | 0.270501349 | 17.92476646 | NA | NA | NA | NA | NA | NA | NA |
| SCN2A | 1732 | D | 32.528375 | 21.73610975 | 0.316313113 | 21.01130705 | NA | NA | NA | NA | NA | NA | NA |
| SCN2A | 1733 | P | 30.77935714 | 25.03435905 | 0.369874178 | 24.03446323 | NA | NA | NA | NA | NA | NA | NA |
| SCN2A | 1734 | D | 33.43675 | 27.058862 | 0.402750569 | 26.33224648 | NA | NA | NA | NA | NA | NA | NA |
| SCN2A | 1735 | K | 32.46227778 | 22.10709239 | 0.322337589 | 21.41202747 | NA | NA | NA | NA | NA | NA | NA |
| SCN2A | 1736 | D | 34.304375 | 28.4192727 | 0.424842607 | 27.7166863 | NA | NA | NA | NA | NA | NA | NA |
| SCN2A | 1737 | H | 29.632 | 27.46811193 | 0.409396478 | 26.79064055 | NA | NA | NA | NA | NA | NA | NA |
| SCN2A | 1738 | P | 32.5455 | 30.56933912 | 0.459758053 | 29.88579315 | NA | NA | NA | NA | NA | NA | NA |
| SCN2A | 1739 | G | 29.90575 | 33.1595547 | 0.501821188 | 32.49609192 | NA | NA | NA | NA | NA | NA | NA |
| SCN2A | 1740 | S | 28.26383333 | 32.61503915 | 0.492978669 | 32.06170242 | NA | NA | NA | NA | NA | NA | NA |
| SCN2A | 1741 | S | 27.50066667 | 34.0341091 | 0.516023288 | 33.37621696 | NA | NA | NA | NA | NA | NA | NA |
| SCN2A | 1742 | V | 26.41607143 | 29.96735557 | 0.449982297 | 29.35382225 | NA | NA | NA | NA | NA | NA | NA |
| SCN2A | 1743 | K | 30.04483333 | 28.66638636 | 0.428855545 | 27.97387632 | NA | NA | NA | NA | NA | NA | NA |
| SCN2A | 1744 | G | 28.33925 | 25.28543955 | 0.373951535 | 24.30464209 | NA | NA | NA | NA | NA | NA | NA |
| SCN2A | 1745 | D | 28.322625 | 20.51685846 | 0.296513397 | 19.6454195 | NA | NA | NA | NA | NA | NA | NA |
| SCN2A | 1746 | C | 25.53066667 | 22.22331145 | 0.324224899 | 21.63761136 | NA | NA | NA | NA | NA | NA | NA |
| SCN2A | 1747 | G | 22.1265 | 21.61208916 | 0.314299113 | 20.99427226 | NA | NA | NA | NA | NA | NA | NA |
| SCN2A | 1748 | N | 18.486125 | 21.21594047 | 0.307865959 | 19.19611727 | NA | NA | NA | NA | NA | NA | NA |
| SCN2A | 1749 | P | 18.28392857 | 20.07622589 | 0.289357859 | 18.06720534 | NA | NA | NA | NA | NA | NA | NA |
| SCN2A | 1750 | S | 14.869 | 22.21054493 | 0.32401758 | 20.11121427 | NA | NA | NA | NA | NA | NA | NA |
| SCN2A | 1751 | V | 13.9735 | 21.2002125 | 0.307610548 | 19.10732954 | NA | NA | NA | NA | NA | NA | NA |
| SCN2A | 1752 | G | 13.7955 | 17.47683226 | 0.247145679 | 15.38389175 | NA | NA | NA | NA | NA | NA | NA |
| SCN2A | 1753 | I | 12.486875 | 17.56653264 | 0.248602345 | 15.46473542 | NA | NA | NA | NA | NA | NA | NA |
| SCN2A | 1754 | F | 8.082136364 | 19.87759415 | 0.28613223 | 17.78395307 | NA | NA | NA | NA | NA | NA | NA |
| SCN2A | 1755 | F | 10.52186364 | 18.29732079 | 0.26046979 | 16.20687671 | NA | NA | NA | NA | NA | NA | NA |
| SCN2A | 1756 | F | 11.56831818 | 14.0139488 | 0.19091108 | 11.92186949 | NA | NA | NA | NA | NA | NA | NA |
| SCN2A | 1757 | V | 7.385785714 | 15.79868529 | 0.219893846 | 13.70180386 | NA | NA | NA | NA | NA | NA | NA |
| SCN2A | 1758 | S | 4.732166667 | 17.7209006 | 0.251109164 | 13.75586175 | NA | NA | NA | NA | NA | NA | NA |
| SCN2A | 1759 | Y | 5.262666667 | 14.88352747 | 0.205032378 | 10.84962167 | NA | NA | NA | NA | NA | NA | NA |

| | | | | | | | | | | | | | |
|---|---|---|---|---|---|---|---|---|---|---|---|---|---|
| SCN2A | 1760 | I | 5.043125 | 11.66426697 | 0.152754031 | 7.887111336 | NA | NA | NA | NA | NA | NA | NA |
| SCN2A | 1761 | I | 1.432375 | 13.17300364 | 0.177254769 | 9.761833284 | NA | NA | NA | NA | NA | NA | NA |
| SCN2A | 1762 | I | 0.338375 | 13.56606343 | 0.183637762 | 10.17144012 | NA | NA | NA | NA | NA | NA | NA |
| SCN2A | 1763 | S | 0.968 | 10.70554053 | 0.137185041 | 7.284347353 | NA | NA | NA | NA | NA | NA | NA |
| SCN2A | 1764 | F | -0.387227273 | 6.651505995 | 0.071350597 | 3.298807748 | 3.43 | 1.13 | 2.23 | 0.62 | -1.51 | 0 | 0 |
| SCN2A | 1765 | L | -3.093625 | 10.4433577 | 0.132927391 | 7.315324832 | NA | NA | NA | NA | NA | NA | NA |
| SCN2A | 1766 | V | -2.861928571 | 10.86098748 | 0.139709382 | 7.535167053 | NA | NA | NA | NA | NA | NA | NA |
| SCN2A | 1767 | V | -1.855357143 | 7.482924228 | 0.084852198 | 4.14947198 | 3.43 | 1.13 | 2.23 | 0.62 | -1.51 | 0 | 0 |
| SCN2A | 1768 | V | -5.039357143 | 5.031861286 | 0.045048797 | 1.702054677 | 3.41 | 1.3 | 2.8 | 0.5 | -1.6 | 0 | 0 |
| SCN2A | 1769 | N | -7.60525 | 7.752138415 | 0.089224032 | 5.409792149 | 2.61 | 1.67 | 3.34 | 0.43 | -1.77 | 0 | 0 |
| SCN2A | 1770 | M | -5.245625 | 7.508639557 | 0.085269796 | 5.125375315 | 2.58 | 1.64 | 3.05 | 0.5 | -1.75 | 0 | 0 |
| SCN2A | 1771 | Y | -4.648 | 4.335937038 | 0.033747515 | 1.873381416 | 2.58 | 1.64 | 3.05 | 0.5 | -1.75 | 0 | 0 |
| SCN2A | 1772 | I | -10.471625 | 3.884883653 | 0.02642275 | 1.740456588 | 2.36 | 1.64 | 4.33 | 0.13 | -1.84 | 0 | 0 |
| SCN2A | 1773 | A | -11.3385 | 6.432347239 | 0.067791625 | 4.226641517 | 2.36 | 1.64 | 4.33 | 0.13 | -1.84 | 0 | 0 |
| SCN2A | 1774 | V | -10.06992857 | 7.121272779 | 0.078979253 | 4.8081531 | 2.36 | 1.64 | 4.33 | 0.13 | -1.84 | 0 | 0 |
| SCN2A | 1775 | I | -11.05125 | 3.837526417 | 0.025653705 | 1.667220206 | 2.36 | 1.64 | 4.33 | 0.13 | -1.84 | 0 | 0 |
| SCN2A | 1776 | L | -15.219125 | 3.676047062 | 0.023031403 | 1.782424314 | 2.09 | 1.43 | 3.7 | 0.2 | -1.82 | 0 | 0 |
| SCN2A | 1777 | E | -15.43705556 | 8.276984837 | 0.09774714 | 5.979037128 | 2.3 | 1.59 | 4.27 | 0.13 | -1.83 | 0 | 0 |
| SCN2A | 1778 | N | -14.002125 | 8.86915182 | 0.107363482 | 6.736399737 | NA | NA | NA | NA | NA | NA | NA |
| SCN2A | 1779 | F | -16.97677273 | 3.822514487 | 0.025409922 | 1.575594103 | 3.89 | 0.15 | 0.6 | 13.01 | -0.15 | 1 | -1 |
| SCN2A | 1780 | S | -20.20116667 | 7.325160681 | 0.082290238 | 3.316112017 | 4.1 | 0.03 | -0.3 | 20.31 | 0.17 | 2 | 0 |
| SCN2A | 1781 | V | -19.30178571 | 9.639222479 | 0.119868845 | 5.646619641 | NA | NA | NA | NA | NA | NA | NA |
| SCN2A | 1782 | A | -20.5939 | 9.559514318 | 0.118574445 | 5.577146181 | NA | NA | NA | NA | NA | NA | NA |
| SCN2A | 1783 | T | -23.58621429 | 5.690488028 | 0.055744395 | 1.725282281 | 4.1 | 0.03 | -0.3 | 20.31 | 0.17 | 2 | 0 |
| SCN2A | 1784 | E | -25.1125 | 9.707083496 | 0.120970857 | 5.70789686 | NA | NA | NA | NA | NA | NA | NA |
| SCN2A | 1785 | E | -22.82327778 | 12.25259311 | 0.162308001 | 8.305796988 | NA | NA | NA | NA | NA | NA | NA |
| SCN4A | 119 | V | -19.74389286 | 55.15482975 | 0.910140769 | 52.06315055 | NA | NA | NA | NA | NA | NA | NA |
| SCN4A | 120 | V | -15.73975 | 52.28919733 | 0.860913147 | 49.20296547 | NA | NA | NA | NA | NA | NA | NA |
| SCN4A | 121 | R | -19.15347727 | 48.40991654 | 0.794272439 | 45.58832378 | NA | NA | NA | NA | NA | NA | NA |
| SCN4A | 122 | R | -22.00247727 | 50.53224679 | 0.830731153 | 47.43850882 | NA | NA | NA | NA | NA | NA | NA |
| SCN4A | 123 | G | -17.531 | 49.51453724 | 0.813248303 | 46.42982254 | NA | NA | NA | NA | NA | NA | NA |
| SCN4A | 124 | A | -15.57795 | 46.3797181 | 0.759396423 | 43.29698578 | NA | NA | NA | NA | NA | NA | NA |
| SCN4A | 125 | I | -18.989 | 44.76706631 | 0.731693284 | 41.67766909 | NA | NA | NA | NA | NA | NA | NA |

| | | | | | | | | | | | | | |
|---|---|---|---|---|---|---|---|---|---|---|---|---|---|
| SCN4A | 126 | K | -19.15252778 | 45.85664436 | 0.750410736 | 42.77261844 | NA | NA | NA | NA | NA | NA | NA |
| SCN4A | 127 | V | -14.32103571 | 43.73693481 | 0.713997042 | 40.75497461 | NA | NA | NA | NA | NA | NA | NA |
| SCN4A | 128 | L | -15.09825 | 40.85849623 | 0.664549427 | 37.778842 | NA | NA | NA | NA | NA | NA | NA |
| SCN4A | 129 | I | -18.561625 | 40.05126098 | 0.650682236 | 36.95785421 | NA | NA | NA | NA | NA | NA | NA |
| SCN4A | 130 | H | -15.89675 | 41.25141604 | 0.671299248 | 38.1735928 | NA | NA | NA | NA | NA | NA | NA |
| SCN4A | 131 | A | -15.15695 | 38.74004165 | 0.628157291 | 35.65804582 | NA | NA | NA | NA | NA | NA | NA |
| SCN4A | 132 | L | -11.549 | 40.46679127 | 0.657820474 | 37.95353208 | NA | NA | NA | NA | NA | NA | NA |
| SCN4A | 133 | F | -10.25320455 | 39.13048888 | 0.634864638 | 36.61794091 | NA | NA | NA | NA | NA | NA | NA |
| SCN4A | 134 | S | -10.99908333 | 35.16887856 | 0.566809622 | 32.62230267 | NA | NA | NA | NA | NA | NA | NA |
| SCN4A | 135 | M | -8.9725 | 35.30814396 | 0.56920201 | 31.8892518 | NA | NA | NA | NA | NA | NA | NA |
| SCN4A | 136 | F | -5.140659091 | 37.60549822 | 0.608667396 | 34.50369924 | NA | NA | NA | NA | NA | NA | NA |
| SCN4A | 137 | I | -5.80025 | 34.1535993 | 0.549368521 | 30.33058724 | NA | NA | NA | NA | NA | NA | NA |
| SCN4A | 138 | M | -5.936125 | 30.59269032 | 0.488197004 | 27.13191418 | NA | NA | NA | NA | NA | NA | NA |
| SCN4A | 139 | I | -2.752125 | 34.8058614 | 0.560573487 | 30.96856494 | NA | NA | NA | NA | NA | NA | NA |
| SCN4A | 140 | T | -1.156607143 | 34.20797394 | 0.550302603 | 30.32460892 | NA | NA | NA | NA | NA | NA | NA |
| SCN4A | 141 | I | -1.348375 | 29.83791147 | 0.475230941 | 25.98545831 | NA | NA | NA | NA | NA | NA | NA |
| SCN4A | 142 | L | -1.219 | 28.40715695 | 0.450652547 | 24.54009644 | NA | NA | NA | NA | NA | NA | NA |
| SCN4A | 143 | T | 2.764892857 | 32.69590143 | 0.524327277 | 28.98750264 | NA | NA | NA | NA | NA | NA | NA |
| SCN4A | 144 | N | 3.91275 | 31.10258145 | 0.496956232 | 27.29567738 | NA | NA | NA | NA | NA | NA | NA |
| SCN4A | 145 | C | 3.524416667 | 27.72809119 | 0.438987131 | 24.02661122 | NA | NA | NA | NA | NA | NA | NA |
| SCN4A | 146 | V | 6.55775 | 30.21925196 | 0.481781846 | 26.39688932 | NA | NA | NA | NA | NA | NA | NA |
| SCN4A | 147 | F | 8.664295455 | 31.71104377 | 0.507408777 | 27.80726593 | NA | NA | NA | NA | NA | NA | NA |
| SCN4A | 148 | M | 7.408125 | 28.03723817 | 0.444297851 | 24.26599665 | NA | NA | NA | NA | NA | NA | NA |
| SCN4A | 149 | T | 9.243035714 | 26.31808572 | 0.414765177 | 24.19027303 | NA | NA | NA | NA | NA | NA | NA |
| SCN4A | 150 | M | 13.20725 | 28.8455282 | 0.458183162 | 27.06609041 | NA | NA | NA | NA | NA | NA | NA |
| SCN4A | 151 | S | 16.57475 | 27.43493496 | 0.43395111 | 25.61693671 | NA | NA | NA | NA | NA | NA | NA |
| SCN4A | 152 | D | 19.65 | 31.81327443 | 0.509164959 | 30.05223243 | NA | NA | NA | NA | NA | NA | NA |
| SCN4A | 153 | P | 16.60460714 | 34.08771553 | 0.548236729 | 32.27519993 | NA | NA | NA | NA | NA | NA | NA |
| SCN4A | 154 | P | 17.45732143 | 36.99915246 | 0.59825121 | 35.1898054 | NA | NA | NA | NA | NA | NA | NA |
| SCN4A | 155 | P | 18.88389286 | 40.80353037 | 0.663605189 | 38.98996242 | NA | NA | NA | NA | NA | NA | NA |
| SCN4A | 156 | W | 15.94439286 | 41.75509578 | 0.679951774 | 39.97378993 | NA | NA | NA | NA | NA | NA | NA |
| SCN4A | 157 | S | 13.22658333 | 38.74834682 | 0.628299963 | 36.96742212 | NA | NA | NA | NA | NA | NA | NA |
| SCN4A | 158 | K | 14.91952778 | 41.73416947 | 0.679592288 | 38.02561232 | NA | NA | NA | NA | NA | NA | NA |

| | | | | | | | | | | | | | |
|---|---|---|---|---|---|---|---|---|---|---|---|---|---|
| SCN4A | 159 | N | 10.731 | 42.88921222 | 0.699434334 | 39.06901901 | NA | NA | NA | NA | NA | NA | NA |
| SCN4A | 160 | V | 8.634321429 | 39.23177715 | 0.636604631 | 35.39147878 | NA | NA | NA | NA | NA | NA | NA |
| SCN4A | 161 | E | 9.230305556 | 37.74287338 | 0.611027312 | 33.87442197 | NA | NA | NA | NA | NA | NA | NA |
| SCN4A | 162 | Y | 8.862333333 | 41.18835243 | 0.670215902 | 37.38334285 | NA | NA | NA | NA | NA | NA | NA |
| SCN4A | 163 | T | 4.629321429 | 41.15599604 | 0.669660064 | 37.43620837 | NA | NA | NA | NA | NA | NA | NA |
| SCN4A | 164 | F | 3.452204545 | 36.07169102 | 0.582318698 | 32.13706017 | NA | NA | NA | NA | NA | NA | NA |
| SCN4A | 165 | T | 2.933321429 | 39.28827576 | 0.637575199 | 35.37752298 | NA | NA | NA | NA | NA | NA | NA |
| SCN4A | 166 | G | 0.928 | 41.89098615 | 0.682286183 | 38.00680149 | NA | NA | NA | NA | NA | NA | NA |
| SCN4A | 167 | I | -1.039125 | 39.79624407 | 0.646301396 | 35.95219903 | NA | NA | NA | NA | NA | NA | NA |
| SCN4A | 168 | Y | -1.105416667 | 33.6242097 | 0.540274336 | 29.75314759 | NA | NA | NA | NA | NA | NA | NA |
| SCN4A | 169 | T | -1.620321429 | 40.59321286 | 0.659992223 | 36.75372125 | NA | NA | NA | NA | NA | NA | NA |
| SCN4A | 170 | F | -5.588113636 | 42.79329458 | 0.697786601 | 38.96704519 | NA | NA | NA | NA | NA | NA | NA |
| SCN4A | 171 | E | -6.897083333 | 36.65383108 | 0.592319064 | 32.87903151 | NA | NA | NA | NA | NA | NA | NA |
| SCN4A | 172 | S | -6.597416667 | 40.76854974 | 0.66300427 | 36.96941277 | NA | NA | NA | NA | NA | NA | NA |
| SCN4A | 173 | L | -6.864 | 43.55827934 | 0.710927986 | 39.7660315 | NA | NA | NA | NA | NA | NA | NA |
| SCN4A | 174 | I | -10.5105 | 43.16720313 | 0.704209835 | 39.39700137 | NA | NA | NA | NA | NA | NA | NA |
| SCN4A | 175 | K | -11.66730556 | 38.40665579 | 0.622430181 | 34.66461432 | NA | NA | NA | NA | NA | NA | NA |
| SCN4A | 176 | I | -11.078875 | 43.53727467 | 0.710567155 | 39.77184817 | NA | NA | NA | NA | NA | NA | NA |
| SCN4A | 177 | L | -12.3215 | 46.77861697 | 0.766248957 | 43.02427339 | NA | NA | NA | NA | NA | NA | NA |
| SCN4A | 178 | A | -16.05375 | 45.75098492 | 0.748595652 | 42.67169689 | NA | NA | NA | NA | NA | NA | NA |
| SCN4A | 179 | R | -17.21793182 | 38.62518012 | 0.626184128 | 35.53304096 | NA | NA | NA | NA | NA | NA | NA |
| SCN4A | 180 | G | -16.5805 | 47.4436972 | 0.77767412 | 44.35017211 | NA | NA | NA | NA | NA | NA | NA |
| SCN4A | 181 | F | -15.98138636 | 49.20399441 | 0.807913605 | 46.86128652 | NA | NA | NA | NA | NA | NA | NA |
| SCN4A | 182 | C | -13.09058333 | 48.91211352 | 0.802899493 | 45.15983833 | NA | NA | NA | NA | NA | NA | NA |
| SCN4A | 183 | V | -16.20803571 | 47.25400259 | 0.774415428 | 43.52112467 | NA | NA | NA | NA | NA | NA | NA |
| SCN4A | 184 | D | -15.521625 | 44.48008405 | 0.726763323 | 41.95563883 | NA | NA | NA | NA | NA | NA | NA |
| SCN4A | 185 | D | -10.947875 | 42.84257856 | 0.698633232 | 39.09102461 | NA | NA | NA | NA | NA | NA | NA |
| SCN4A | 186 | F | -13.98211364 | 41.70149037 | 0.679030906 | 37.97183469 | NA | NA | NA | NA | NA | NA | NA |
| SCN4A | 187 | T | -16.39232143 | 40.57241149 | 0.659634884 | 37.58850659 | NA | NA | NA | NA | NA | NA | NA |
| SCN4A | 188 | F | -11.61238636 | 38.64886815 | 0.626591056 | 34.91711775 | NA | NA | NA | NA | NA | NA | NA |
| SCN4A | 189 | L | -10.635125 | 37.44966917 | 0.605990467 | 33.71445328 | NA | NA | NA | NA | NA | NA | NA |
| SCN4A | 190 | R | -16.48756818 | 35.82864921 | 0.578143574 | 33.42689845 | NA | NA | NA | NA | NA | NA | NA |
| SCN4A | 191 | D | -14.260375 | 33.29245305 | 0.534575213 | 30.88721059 | NA | NA | NA | NA | NA | NA | NA |

| | | | | | | | | | | | | | |
|---|---|---|---|---|---|---|---|---|---|---|---|---|---|
| SCN4A | 192 | P | -11.76232143 | 30.88002362 | 0.493132994 | 27.14200957 | NA | NA | NA | NA | NA | NA | NA |
| SCN4A | 193 | W | -10.30603571 | 25.59739381 | 0.402384682 | 21.8796656 | NA | NA | NA | NA | NA | NA | NA |
| SCN4A | 194 | N | -9.811625 | 33.46553938 | 0.537548598 | 29.72473413 | NA | NA | NA | NA | NA | NA | NA |
| SCN4A | 195 | W | -8.352964286 | 32.39610824 | 0.519177242 | 28.63877799 | NA | NA | NA | NA | NA | NA | NA |
| SCN4A | 196 | L | -3.98375 | 27.65065968 | 0.437656965 | 23.85299889 | NA | NA | NA | NA | NA | NA | NA |
| SCN4A | 197 | D | -5.494125 | 32.71233524 | 0.524609587 | 28.92999707 | NA | NA | NA | NA | NA | NA | NA |
| SCN4A | 198 | F | -3.852204545 | 35.47823064 | 0.572123865 | 31.67883181 | NA | NA | NA | NA | NA | NA | NA |
| SCN4A | 199 | S | -0.89075 | 32.84099304 | 0.526819751 | 28.99209734 | NA | NA | NA | NA | NA | NA | NA |
| SCN4A | 200 | V | 0.537892857 | 32.46839342 | 0.520419002 | 28.62147565 | NA | NA | NA | NA | NA | NA | NA |
| SCN4A | 201 | I | 0.927375 | 36.7452702 | 0.593889862 | 32.89042685 | NA | NA | NA | NA | NA | NA | NA |
| SCN4A | 202 | M | 1.2525 | 38.27677284 | 0.62019897 | 34.41367332 | NA | NA | NA | NA | NA | NA | NA |
| SCN4A | 203 | M | 4.14525 | 34.93412043 | 0.5627768 | 31.03110548 | NA | NA | NA | NA | NA | NA | NA |
| SCN4A | 204 | A | 5.72855 | 37.35114736 | 0.604297998 | 33.62333842 | NA | NA | NA | NA | NA | NA | NA |
| SCN4A | 205 | Y | 4.287 | 40.06817077 | 0.650972723 | 36.33270314 | NA | NA | NA | NA | NA | NA | NA |
| SCN4A | 206 | L | 7.58325 | 39.30142107 | 0.637801017 | 35.54820891 | NA | NA | NA | NA | NA | NA | NA |
| SCN4A | 207 | T | 10.58460714 | 37.67838431 | 0.609919479 | 33.82888363 | NA | NA | NA | NA | NA | NA | NA |
| SCN4A | 208 | E | 10.10597222 | 41.88456225 | 0.682175829 | 38.05983046 | NA | NA | NA | NA | NA | NA | NA |
| SCN4A | 209 | F | 10.20838636 | 44.34380401 | 0.72442222 | 40.52314886 | NA | NA | NA | NA | NA | NA | NA |
| SCN4A | 210 | V | 14.13460714 | 42.62543813 | 0.694903058 | 38.75598758 | NA | NA | NA | NA | NA | NA | NA |
| SCN4A | 211 | D | 14.445 | 38.60351857 | 0.625812013 | 34.69019556 | NA | NA | NA | NA | NA | NA | NA |
| SCN4A | 212 | L | 14.933 | 42.2777341 | 0.688929981 | 38.37428073 | NA | NA | NA | NA | NA | NA | NA |
| SCN4A | 213 | G | 15.9185 | 38.47109796 | 0.623537209 | 36.64046614 | NA | NA | NA | NA | NA | NA | NA |
| SCN4A | 214 | N | 17.77 | 37.51263827 | 0.60707219 | 35.68175101 | NA | NA | NA | NA | NA | NA | NA |
| SCN4A | 215 | I | 17.42 | 33.26868972 | 0.534166992 | 31.45914351 | NA | NA | NA | NA | NA | NA | NA |
| SCN4A | 216 | S | 15.82058333 | 31.6840502 | 0.506945065 | 29.86180237 | NA | NA | NA | NA | NA | NA | NA |
| SCN4A | 217 | A | 12.45675 | 29.99653603 | 0.477955893 | 28.16990031 | NA | NA | NA | NA | NA | NA | NA |
| SCN4A | 218 | L | 10.73475 | 32.221292 | 0.51617414 | 28.33636312 | NA | NA | NA | NA | NA | NA | NA |
| SCN4A | 219 | R | 12.25029545 | 30.13171767 | 0.480278128 | 26.26254768 | NA | NA | NA | NA | NA | NA | NA |
| SCN4A | 220 | T | 8.230178571 | 26.99628228 | 0.426415661 | 23.10167422 | NA | NA | NA | NA | NA | NA | NA |
| SCN4A | 221 | F | 5.672386364 | 29.22578656 | 0.464715477 | 25.40789837 | NA | NA | NA | NA | NA | NA | NA |
| SCN4A | 222 | R | 6.579840909 | 29.65610092 | 0.472107686 | 25.89455865 | NA | NA | NA | NA | NA | NA | NA |
| SCN4A | 223 | V | 4.377321429 | 24.91735026 | 0.390702469 | 21.12586589 | NA | NA | NA | NA | NA | NA | NA |
| SCN4A | 224 | L | 1.039 | 25.95836308 | 0.408585637 | 22.09266261 | NA | NA | NA | NA | NA | NA | NA |

| | | | | | | | | | | | | | |
|---|---|---|---|---|---|---|---|---|---|---|---|---|---|
| SCN4A | 225 | R | 1.969022727 | 28.81641036 | 0.457682957 | 24.96204737 | NA | NA | NA | NA | NA | NA | NA |
| SCN4A | 226 | A | -0.72415 | 25.56193128 | 0.401775484 | 21.73263576 | NA | NA | NA | NA | NA | NA | NA |
| SCN4A | 227 | L | -2.796125 | 24.0749161 | 0.376230609 | 20.27402012 | NA | NA | NA | NA | NA | NA | NA |
| SCN4A | 228 | K | -4.471305556 | 28.21075752 | 0.447278675 | 24.4067362 | NA | NA | NA | NA | NA | NA | NA |
| SCN4A | 229 | T | -5.397892857 | 25.72866709 | 0.404639776 | 21.92764919 | NA | NA | NA | NA | NA | NA | NA |
| SCN4A | 230 | I | -7.986125 | 23.33493113 | 0.363518685 | 21.59847058 | NA | NA | NA | NA | NA | NA | NA |
| SCN4A | 231 | T | -9.41475 | 28.7580866 | 0.456681036 | 26.41107397 | NA | NA | NA | NA | NA | NA | NA |
| SCN4A | 232 | V | -9.33575 | 30.6217892 | 0.488696882 | 28.05889928 | NA | NA | NA | NA | NA | NA | NA |
| SCN4A | 233 | I | -9.916875 | 26.88085328 | 0.42443275 | 24.34409505 | NA | NA | NA | NA | NA | NA | NA |
| SCN4A | 234 | P | -13.98003571 | 26.32629045 | 0.414906123 | 23.2525545 | NA | NA | NA | NA | NA | NA | NA |
| SCN4A | 235 | G | -14.40125 | 23.01972635 | 0.3581039 | 19.94511452 | NA | NA | NA | NA | NA | NA | NA |
| SCN4A | 236 | L | -10.662875 | 22.62986277 | 0.351406581 | 20.08256657 | NA | NA | NA | NA | NA | NA | NA |
| SCN4A | 237 | K | -13.52097222 | 23.80535782 | 0.371599968 | 21.39391503 | NA | NA | NA | NA | NA | NA | NA |
| SCN4A | 238 | T | -16.09532143 | 20.34164342 | 0.31209812 | 17.25420501 | NA | NA | NA | NA | NA | NA | NA |
| SCN4A | 239 | I | -12.582875 | 18.02698036 | 0.272335393 | 15.63585597 | NA | NA | NA | NA | NA | NA | NA |
| SCN4A | 240 | V | -10.38489286 | 20.49109158 | 0.314665434 | 18.13543811 | NA | NA | NA | NA | NA | NA | NA |
| SCN4A | 241 | G | -13.9785 | 20.65127028 | 0.317417083 | 18.25988897 | NA | NA | NA | NA | NA | NA | NA |
| SCN4A | 242 | A | -14.08695 | 16.5427835 | 0.246838933 | 13.65477357 | NA | NA | NA | NA | NA | NA | NA |
| SCN4A | 243 | L | -10.167125 | 14.54747277 | 0.212562239 | 12.21791365 | NA | NA | NA | NA | NA | NA | NA |
| SCN4A | 244 | I | -10.743 | 18.20036013 | 0.275313819 | 15.80073554 | NA | NA | NA | NA | NA | NA | NA |
| SCN4A | 245 | Q | -15.50963889 | 17.27253432 | 0.259375048 | 14.1964567 | NA | NA | NA | NA | NA | NA | NA |
| SCN4A | 246 | S | -12.56825 | 13.51140692 | 0.194764053 | 11.12207198 | NA | NA | NA | NA | NA | NA | NA |
| SCN4A | 247 | V | -10.10375 | 15.79525251 | 0.233997379 | 13.41110034 | NA | NA | NA | NA | NA | NA | NA |
| SCN4A | 248 | K | -12.93663889 | 17.79410827 | 0.268334971 | 15.42450706 | NA | NA | NA | NA | NA | NA | NA |
| SCN4A | 249 | K | -15.44930556 | 15.31719198 | 0.225784956 | 12.16832664 | NA | NA | NA | NA | NA | NA | NA |
| SCN4A | 250 | L | -9.755375 | 12.38121161 | 0.175348853 | 9.994145508 | NA | NA | NA | NA | NA | NA | NA |
| SCN4A | 251 | S | -10.63575 | 17.69590964 | 0.266648053 | 15.32102138 | NA | NA | NA | NA | NA | NA | NA |
| SCN4A | 252 | D | -11.694375 | 17.98718508 | 0.271651765 | 15.60861972 | NA | NA | NA | NA | NA | NA | NA |
| SCN4A | 253 | V | -8.456464286 | 13.68824441 | 0.197801878 | 11.26125726 | NA | NA | NA | NA | NA | NA | NA |
| SCN4A | 254 | M | -5.407125 | 12.56471731 | 0.178501228 | 8.750302914 | NA | NA | NA | NA | NA | NA | NA |
| SCN4A | 255 | I | -6.668 | 18.64109063 | 0.282884963 | 14.77271564 | NA | NA | NA | NA | NA | NA | NA |
| SCN4A | 256 | L | -6.1355 | 17.55442446 | 0.264217533 | 13.59933128 | NA | NA | NA | NA | NA | NA | NA |
| SCN4A | 257 | T | -2.711464286 | 13.15273652 | 0.188602589 | 9.165167805 | NA | NA | NA | NA | NA | NA | NA |

| | | | | | | | | | | | | | |
|---|---|---|---|---|---|---|---|---|---|---|---|---|---|
| SCN4A | 258 | V | -0.911607143 | 16.94773902 | 0.253795511 | 12.9651508 | NA | NA | NA | NA | NA | NA | NA |
| SCN4A | 259 | F | -2.581840909 | 19.41216675 | 0.296130989 | 15.40511374 | NA | NA | NA | NA | NA | NA | NA |
| SCN4A | 260 | C | -1.265416667 | 17.69126233 | 0.266568219 | 13.61841472 | NA | NA | NA | NA | NA | NA | NA |
| SCN4A | 261 | L | 2.244625 | 14.17597944 | 0.206180495 | 10.28849585 | NA | NA | NA | NA | NA | NA | NA |
| SCN4A | 262 | S | 2.494083333 | 19.47862749 | 0.297272694 | 15.56143831 | NA | NA | NA | NA | NA | NA | NA |
| SCN4A | 263 | V | 1.855464286 | 21.61577928 | 0.333986021 | 17.69851058 | NA | NA | NA | NA | NA | NA | NA |
| SCN4A | 264 | F | 2.931568182 | 17.52161813 | 0.263653965 | 15.01293146 | NA | NA | NA | NA | NA | NA | NA |
| SCN4A | 265 | A | 7.15955 | 20.24182556 | 0.310383386 | 18.0777799 | NA | NA | NA | NA | NA | NA | NA |
| SCN4A | 266 | L | 6.711625 | 22.8095112 | 0.354492693 | 20.66751148 | NA | NA | NA | NA | NA | NA | NA |
| SCN4A | 267 | V | 7.406178571 | 22.94151935 | 0.356760412 | 21.12381842 | NA | NA | NA | NA | NA | NA | NA |
| SCN4A | 268 | G | 10.5845 | 20.91342442 | 0.321920531 | 18.97625329 | NA | NA | NA | NA | NA | NA | NA |
| SCN4A | 269 | L | 12.48825 | 22.23195147 | 0.344571012 | 20.29559531 | NA | NA | NA | NA | NA | NA | NA |
| SCN4A | 270 | Q | 11.17475 | 24.86180478 | 0.389748274 | 22.90235705 | NA | NA | NA | NA | NA | NA | NA |
| SCN4A | 271 | L | 12.492875 | 23.2502586 | 0.362064127 | 21.30654095 | NA | NA | NA | NA | NA | NA | NA |
| SCN4A | 272 | F | 14.69765909 | 19.78473811 | 0.302531253 | 17.95678504 | NA | NA | NA | NA | NA | NA | NA |
| SCN4A | 273 | M | 17.201625 | 23.81882084 | 0.371831245 | 22.07469248 | NA | NA | NA | NA | NA | NA | NA |
| SCN4A | 274 | G | 21.033 | 22.13057326 | 0.342829474 | 18.53420476 | NA | NA | NA | NA | NA | NA | NA |
| SCN4A | 275 | N | 20.497625 | 18.73842648 | 0.284557058 | 15.18657132 | NA | NA | NA | NA | NA | NA | NA |
| SCN4A | 276 | L | 20.95125 | 15.96090082 | 0.236842989 | 12.54513731 | NA | NA | NA | NA | NA | NA | NA |
| SCN4A | 277 | R | 24.70147727 | 18.63866784 | 0.282843342 | 15.05517629 | NA | NA | NA | NA | NA | NA | NA |
| SCN4A | 278 | Q | 25.49008333 | 16.17775497 | 0.240568245 | 14.17141415 | NA | NA | NA | NA | NA | NA | NA |
| SCN4A | 279 | K | 29.54152778 | 14.92040261 | 0.218968661 | 12.80511979 | NA | NA | NA | NA | NA | NA | NA |
| SCN4A | 280 | C | 32.37808333 | 12.11204252 | 0.170724898 | 9.804674188 | NA | NA | NA | NA | NA | NA | NA |
| SCN4A | 281 | V | 35.65403571 | 14.23559328 | 0.207204579 | 11.53787055 | NA | NA | NA | NA | NA | NA | NA |
| SCN4A | 282 | R | 41.20129545 | 12.76090863 | 0.181871525 | 10.50981854 | NA | NA | NA | NA | NA | NA | NA |
| SCN4A | 283 | W | 40.10667857 | 15.39489022 | 0.227119705 | 14.89486653 | NA | NA | NA | NA | NA | NA | NA |
| SCN4A | 284 | P | 44.34403571 | 18.05641664 | 0.272841068 | 17.71970445 | NA | NA | NA | NA | NA | NA | NA |
| SCN4A | 285 | P | 48.10646429 | 17.67916924 | 0.266360476 | 16.77100858 | NA | NA | NA | NA | NA | NA | NA |
| SCN4A | 286 | P | 48.54846429 | 20.6318116 | 0.31708281 | 19.72483904 | NA | NA | NA | NA | NA | NA | NA |
| SCN4A | 336 | F | 44.62420455 | 26.81193482 | 0.423248825 | 24.20540905 | NA | NA | NA | NA | NA | NA | NA |
| SCN4A | 337 | D | 43.51125 | 24.76330691 | 0.388056216 | 22.15298398 | NA | NA | NA | NA | NA | NA | NA |
| SCN4A | 338 | W | 37.97103571 | 22.38025078 | 0.34711859 | 19.72070907 | NA | NA | NA | NA | NA | NA | NA |
| SCN4A | 339 | D | 37.4015 | 24.27899701 | 0.379736438 | 21.59112882 | NA | NA | NA | NA | NA | NA | NA |

| | | | | | | | | | | | | | |
|---|---|---|---|---|---|---|---|---|---|---|---|---|---|
| SCN4A | 340 | A | 40.61075 | 21.88720377 | 0.338648721 | 19.23608604 | NA | NA | NA | NA | NA | NA | NA |
| SCN4A | 341 | Y | 40.66591667 | 16.87108515 | 0.252478703 | 14.32324978 | NA | NA | NA | NA | NA | NA | NA |
| SCN4A | 342 | I | 35.863625 | 19.15198898 | 0.291661493 | 16.44802689 | NA | NA | NA | NA | NA | NA | NA |
| SCN4A | 343 | S | 36.35408333 | 19.67463192 | 0.30063978 | 16.96158971 | NA | NA | NA | NA | NA | NA | NA |
| SCN4A | 344 | D | 39.432 | 16.47241515 | 0.245630101 | 13.77672118 | NA | NA | NA | NA | NA | NA | NA |
| SCN4A | 345 | E | 35.91063889 | 13.20103742 | 0.189432332 | 10.49120073 | NA | NA | NA | NA | NA | NA | NA |
| SCN4A | 346 | G | 39.7835 | 12.24207674 | 0.172958707 | 9.641604188 | NA | NA | NA | NA | NA | NA | NA |
| SCN4A | 347 | N | 39.246 | 12.97284283 | 0.185512263 | 10.29904718 | NA | NA | NA | NA | NA | NA | NA |
| SCN4A | 348 | F | 34.64729545 | 10.27003062 | 0.139081667 | 7.567724464 | NA | NA | NA | NA | NA | NA | NA |
| SCN4A | 349 | Y | 34.0165 | 3.891557657 | 0.029508277 | 1.557044166 | 2.58 | -0.41 | -1.4 | 14.57 | 0.6 | 1 | -1 |
| SCN4A | 350 | F | 33.68620455 | 5.324872017 | 0.054130646 | 2.987632525 | 2.39 | -0.28 | -1.73 | 18.3 | 0.53 | 1 | -1 |
| SCN4A | 351 | L | 30.786 | 3.869032308 | 0.029121322 | 2.027316093 | 2.22 | -1.14 | -3.5 | 49.9 | 2.23 | 2 | -2 |
| SCN4A | 352 | E | 33.99230556 | 3.847323615 | 0.028748397 | 1.85407143 | 2.23 | -0.38 | -1.07 | 33.31 | 0.89 | 2 | -2 |
| SCN4A | 353 | G | 30.66325 | 5.418435752 | 0.055737942 | 1.936814915 | 3.68 | -0.32 | 0.65 | 2.57 | 0.16 | 0 | 0 |
| SCN4A | 354 | S | 29.55658333 | 5.884087525 | 0.063737199 | 2.234672188 | 3.68 | -0.32 | 0.65 | 2.57 | 0.16 | 0 | 0 |
| SCN4A | 355 | N | 28.453375 | 8.214653127 | 0.10377311 | 4.6457481 | 3.8 | -0.86 | -0.53 | 2.81 | 0.91 | 0 | 0 |
| SCN4A | 356 | D | 27.25675 | 9.601535346 | 0.127597838 | 5.959750435 | NA | NA | NA | NA | NA | NA | NA |
| SCN4A | 357 | A | 29.24595 | 11.08280411 | 0.153043998 | 9.274235468 | NA | NA | NA | NA | NA | NA | NA |
| SCN4A | 358 | L | 28.91625 | 7.343902573 | 0.088814813 | 5.356952818 | 2.22 | -1.14 | -3.5 | 49.9 | 2.23 | 2 | -2 |
| SCN4A | 359 | L | 25.10525 | 11.73019667 | 0.164165311 | 8.416862976 | NA | NA | NA | NA | NA | NA | NA |
| SCN4A | 360 | C | 26.84125 | 12.78169234 | 0.18222856 | 10.71432997 | NA | NA | NA | NA | NA | NA | NA |
| SCN4A | 361 | G | 25.8195 | 16.55709727 | 0.247084823 | 14.49949243 | NA | NA | NA | NA | NA | NA | NA |
| SCN4A | 362 | N | 26.1885 | 19.89285631 | 0.304388575 | 17.80370683 | NA | NA | NA | NA | NA | NA | NA |
| SCN4A | 363 | S | 23.42191667 | 19.87016593 | 0.303998785 | 17.7525005 | NA | NA | NA | NA | NA | NA | NA |
| SCN4A | 364 | S | 23.42741667 | 20.44124823 | 0.313809194 | 18.34598317 | NA | NA | NA | NA | NA | NA | NA |
| SCN4A | 365 | D | 20.175 | 17.89217857 | 0.270019684 | 14.35866844 | NA | NA | NA | NA | NA | NA | NA |
| SCN4A | 366 | A | 22.74895 | 16.56850639 | 0.247280816 | 13.2023152 | NA | NA | NA | NA | NA | NA | NA |
| SCN4A | 367 | G | 25.46125 | 14.22413301 | 0.207007707 | 12.17218079 | NA | NA | NA | NA | NA | NA | NA |
| SCN4A | 368 | H | 28.23295 | 11.20791524 | 0.155193235 | 9.126448755 | NA | NA | NA | NA | NA | NA | NA |
| SCN4A | 369 | C | 32.06108333 | 10.97031932 | 0.151111664 | 8.665730307 | NA | NA | NA | NA | NA | NA | NA |
| SCN4A | 370 | P | 34.95760714 | 7.822715321 | 0.097040157 | 5.153433504 | 2.72 | -0.48 | -1.82 | 12.33 | 0.59 | 1 | -1 |
| SCN4A | 371 | E | 37.56563889 | 8.277274491 | 0.104848858 | 7.852647985 | NA | NA | NA | NA | NA | NA | NA |
| SCN4A | 372 | G | 39.2005 | 11.16442462 | 0.154446126 | 10.84585987 | NA | NA | NA | NA | NA | NA | NA |

| | | | | | | | | | | | | | |
|---|---|---|---|---|---|---|---|---|---|---|---|---|---|
| SCN4A | 373 | Y | 37.1035 | 8.397249728 | 0.106909868 | 5.812987487 | NA | NA | NA | NA | NA | NA | NA |
| SCN4A | 374 | E | 34.31008333 | 14.67484749 | 0.214750362 | 12.01001032 | NA | NA | NA | NA | NA | NA | NA |
| SCN4A | 375 | C | 30.46458333 | 16.10440787 | 0.239308242 | 13.87904299 | NA | NA | NA | NA | NA | NA | NA |
| SCN4A | 376 | I | 31.869625 | 18.17653141 | 0.274904474 | 15.84585601 | NA | NA | NA | NA | NA | NA | NA |
| SCN4A | 377 | K | 26.91297222 | 20.85475807 | 0.320912724 | 18.56284592 | NA | NA | NA | NA | NA | NA | NA |
| SCN4A | 378 | T | 30.71775 | 19.3288025 | 0.294698906 | 16.99538003 | NA | NA | NA | NA | NA | NA | NA |
| SCN4A | 379 | G | 28.80025 | 20.01868749 | 0.306550181 | 17.7281094 | NA | NA | NA | NA | NA | NA | NA |
| SCN4A | 380 | R | 27.44611364 | 14.16811819 | 0.206045449 | 11.1186459 | NA | NA | NA | NA | NA | NA | NA |
| SCN4A | 381 | N | 24.015 | 15.09153786 | 0.221908529 | 10.84030154 | NA | NA | NA | NA | NA | NA | NA |
| SCN4A | 382 | P | 22.40417857 | 13.24802906 | 0.190239584 | 8.935590103 | NA | NA | NA | NA | NA | NA | NA |
| SCN4A | 383 | N | 21.736625 | 10.42090668 | 0.141673511 | 6.130285565 | NA | NA | NA | NA | NA | NA | NA |
| SCN4A | 384 | Y | 23.33466667 | 12.74769944 | 0.181644609 | 8.472356269 | NA | NA | NA | NA | NA | NA | NA |
| SCN4A | 385 | G | 21.30925 | 16.39058437 | 0.244224361 | 12.08723785 | NA | NA | NA | NA | NA | NA | NA |
| SCN4A | 386 | Y | 18.89283333 | 15.20730144 | 0.223897188 | 13.2871287 | NA | NA | NA | NA | NA | NA | NA |
| SCN4A | 387 | T | 17.57403571 | 13.50234254 | 0.19460834 | 11.97243891 | NA | NA | NA | NA | NA | NA | NA |
| SCN4A | 388 | S | 17.83225 | 17.51839873 | 0.26359866 | 15.7995449 | NA | NA | NA | NA | NA | NA | NA |
| SCN4A | 389 | Y | 14.303 | 15.5209924 | 0.229285967 | 13.64407521 | NA | NA | NA | NA | NA | NA | NA |
| SCN4A | 390 | D | 17.554875 | 22.16562668 | 0.343431643 | 20.27988875 | NA | NA | NA | NA | NA | NA | NA |
| SCN4A | 391 | T | 15.21132143 | 22.47169598 | 0.348689493 | 20.57197954 | NA | NA | NA | NA | NA | NA | NA |
| SCN4A | 392 | F | 9.086477273 | 21.39030764 | 0.330112729 | 19.49836644 | NA | NA | NA | NA | NA | NA | NA |
| SCN4A | 393 | S | 11.61558333 | 19.52107643 | 0.298001908 | 17.62337362 | NA | NA | NA | NA | NA | NA | NA |
| SCN4A | 394 | W | 15.22296429 | 17.30493805 | 0.259931699 | 15.4726134 | NA | NA | NA | NA | NA | NA | NA |
| SCN4A | 395 | A | 10.68455 | 17.88965883 | 0.269976398 | 16.00716994 | NA | NA | NA | NA | NA | NA | NA |
| SCN4A | 396 | F | 6.404022727 | 15.75662048 | 0.233333733 | 13.60259287 | NA | NA | NA | NA | NA | NA | NA |
| SCN4A | 397 | L | 10.584875 | 12.98488325 | 0.185719101 | 11.08749937 | NA | NA | NA | NA | NA | NA | NA |
| SCN4A | 398 | A | 10.90715 | 12.72349217 | 0.181228762 | 10.90058754 | NA | NA | NA | NA | NA | NA | NA |
| SCN4A | 399 | L | 7.4725 | 13.07356856 | 0.187242592 | 11.67218802 | NA | NA | NA | NA | NA | NA | NA |
| SCN4A | 400 | F | 5.281113636 | 9.751527214 | 0.130174492 | 7.598805899 | NA | NA | NA | NA | NA | NA | NA |
| SCN4A | 401 | R | 10.69547727 | 7.894089181 | 0.098266262 | 5.99746487 | 1.91 | -0.79 | -2.05 | 26.54 | 1.33 | 2 | 0 |
| SCN4A | 402 | L | 6.821 | 9.038658584 | 0.11792839 | 6.894370594 | NA | NA | NA | NA | NA | NA | NA |
| SCN4A | 403 | M | 1.955875 | 7.429341895 | 0.090282543 | 4.935692557 | 2.52 | -0.8 | -0.4 | 3.38 | 0.81 | 0 | 0 |
| SCN4A | 404 | T | 4.024607143 | 3.950359224 | 0.030518407 | 1.691654397 | 2.52 | -0.8 | -0.4 | 3.38 | 0.81 | 0 | 0 |
| SCN4A | 405 | Q | 5.028305556 | 3.566566136 | 0.023925369 | 2.015091332 | 2.17 | -0.8 | -0.4 | 3.38 | 0.81 | 0 | 0 |

| | | | | | | | | | | | | | |
|---|---|---|---|---|---|---|---|---|---|---|---|---|---|
| SCN4A | 406 | D | 9.4115 | 4.009768821 | 0.031538982 | 2.113323637 | 1.91 | -0.79 | -2.05 | 26.54 | 1.33 | 2 | 0 |
| SCN4A | 407 | Y | 14.35458333 | 3.564460829 | 0.023889203 | 1.54154579 | 2 | -0.78 | -2.6 | 34.26 | 1.5 | 2 | 0 |
| SCN4A | 408 | W | 9.39675 | 5.409556266 | 0.055585404 | 3.54239072 | 2 | -0.78 | -2.6 | 34.26 | 1.5 | 2 | 0 |
| SCN4A | 409 | E | 13.62052778 | 4.066806364 | 0.032518808 | 2.985872864 | 2.07 | -0.15 | -2.9 | 33.67 | 0.75 | 2 | 0 |
| SCN4A | 410 | N | 16.9555 | 7.385447989 | 0.089528506 | 4.951194999 | 2.54 | -0.36 | -2.77 | 33.74 | 1.14 | 2 | -2 |
| SCN4A | 411 | L | 13.954125 | 9.486483753 | 0.12562141 | 7.816282854 | NA | NA | NA | NA | NA | NA | NA |
| SCN4A | 412 | F | 13.06529545 | 10.74224427 | 0.147193648 | 9.082260893 | NA | NA | NA | NA | NA | NA | NA |
| SCN4A | 413 | Q | 17.44930556 | 7.33148607 | 0.088601515 | 3.675121087 | 3.77 | -0.47 | -2.18 | 26.15 | 1.06 | 2 | -2 |
| SCN4A | 414 | L | 19.2895 | 11.32504857 | 0.157205424 | 7.884884583 | NA | NA | NA | NA | NA | NA | NA |
| SCN4A | 415 | T | 16.32932143 | 14.99919658 | 0.220322233 | 11.24996101 | NA | NA | NA | NA | NA | NA | NA |
| SCN4A | 416 | L | 16.558 | 15.76236559 | 0.233432426 | 12.0906207 | NA | NA | NA | NA | NA | NA | NA |
| SCN4A | 417 | R | 22.60756818 | 9.242764359 | 0.121434646 | 6.030681304 | NA | NA | NA | NA | NA | NA | NA |
| SCN4A | 418 | A | 21.03255 | 16.13725243 | 0.239872467 | 12.84867758 | NA | NA | NA | NA | NA | NA | NA |
| SCN4A | 419 | A | 18.37555 | 19.02427894 | 0.28946761 | 15.7357672 | NA | NA | NA | NA | NA | NA | NA |
| SCN4A | 420 | G | 17.565 | 19.45027851 | 0.296785697 | 16.1584644 | NA | NA | NA | NA | NA | NA | NA |
| SCN4A | 421 | K | 16.77963889 | 16.72193165 | 0.249916451 | 14.39834499 | NA | NA | NA | NA | NA | NA | NA |
| SCN4A | 422 | T | 13.72489286 | 21.82891051 | 0.337647323 | 18.17556796 | NA | NA | NA | NA | NA | NA | NA |
| SCN4A | 423 | Y | 13.42433333 | 19.01419746 | 0.289294425 | 17.36601026 | NA | NA | NA | NA | NA | NA | NA |
| SCN4A | 424 | M | 9.69325 | 17.861519 | 0.269492994 | 16.18361767 | NA | NA | NA | NA | NA | NA | NA |
| SCN4A | 425 | I | 7.2665 | 20.40682011 | 0.313217766 | 18.55519889 | NA | NA | NA | NA | NA | NA | NA |
| SCN4A | 426 | F | 8.017840909 | 17.68094005 | 0.266390896 | 15.71295316 | NA | NA | NA | NA | NA | NA | NA |
| SCN4A | 427 | F | 8.986840909 | 13.86932868 | 0.200912657 | 12.02125323 | NA | NA | NA | NA | NA | NA | NA |
| SCN4A | 428 | V | 4.271607143 | 17.08233901 | 0.256107754 | 15.21797394 | NA | NA | NA | NA | NA | NA | NA |
| SCN4A | 429 | V | 2.281321429 | 17.00625682 | 0.254800767 | 14.87823792 | NA | NA | NA | NA | NA | NA | NA |
| SCN4A | 430 | I | 3.7515 | 13.67638084 | 0.197598078 | 11.54817662 | NA | NA | NA | NA | NA | NA | NA |
| SCN4A | 431 | I | 3.346625 | 10.85513712 | 0.149132992 | 8.864460871 | NA | NA | NA | NA | NA | NA | NA |
| SCN4A | 432 | F | -1.900295455 | 14.06530284 | 0.204279223 | 11.92860873 | NA | NA | NA | NA | NA | NA | NA |
| SCN4A | 433 | L | -1.790625 | 15.04990455 | 0.221193326 | 10.95931469 | NA | NA | NA | NA | NA | NA | NA |
| SCN4A | 434 | G | 0 | 11.51773459 | 0.160515505 | 7.430920823 | NA | NA | NA | NA | NA | NA | NA |
| SCN4A | 435 | S | 0.756916667 | 8.631862429 | 0.110940191 | 6.148702796 | NA | NA | NA | NA | NA | NA | NA |
| SCN4A | 436 | F | -3.296477273 | 10.59211504 | 0.144614635 | 6.241737852 | NA | NA | NA | NA | NA | NA | NA |
| SCN4A | 437 | Y | -6.683083333 | 11.81973426 | 0.165703444 | 9.325297005 | NA | NA | NA | NA | NA | NA | NA |
| SCN4A | 438 | L | -3.747125 | 10.71067267 | 0.146651292 | 6.643753466 | NA | NA | NA | NA | NA | NA | NA |

| | | | | | | | | | | | | | |
|---|---|---|---|---|---|---|---|---|---|---|---|---|---|
| SCN4A | 439 | I | -3.395 | 6.084899177 | 0.067186867 | 1.740292383 | 4.37 | -0.27 | -0.04 | 2.73 | 0.22 | 0 | 0 |
| SCN4A | 440 | N | -7.179875 | 7.239087857 | 0.08701424 | 4.67766447 | 2.61 | 1.21 | 3.47 | 0.2 | -1.8 | 0 | 0 |
| SCN4A | 441 | L | -9.1375 | 9.613800341 | 0.127808534 | 7.175850238 | NA | NA | NA | NA | NA | NA | NA |
| SCN4A | 442 | I | -6.76275 | 7.901713738 | 0.098397241 | 5.485660059 | 2.61 | 1.21 | 3.47 | 0.2 | -1.8 | 0 | 0 |
| SCN4A | 443 | L | -8.1805 | 4.389983599 | 0.038070549 | 1.916516891 | 2.61 | 1.21 | 3.47 | 0.2 | -1.8 | 0 | 0 |
| SCN4A | 444 | A | -12.54935 | 6.752530044 | 0.078655846 | 4.08491737 | 3.1 | 1.64 | 4.33 | 0.13 | -1.84 | 0 | 0 |
| SCN4A | 445 | V | -12.89275 | 8.952862056 | 0.116454523 | 5.929290781 | NA | NA | NA | NA | NA | NA | NA |
| SCN4A | 446 | V | -11.74332143 | 8.444204699 | 0.10771649 | 6.077929534 | NA | NA | NA | NA | NA | NA | NA |
| SCN4A | 447 | A | -14.75815 | 4.956778793 | 0.04780731 | 1.733407473 | 3.25 | 1.66 | 3.93 | 0.2 | -1.84 | 0 | 0 |
| SCN4A | 448 | M | -17.357875 | 7.615641339 | 0.093482911 | 4.601700511 | 3.14 | 1.36 | 3.73 | 0.19 | -1.81 | 0 | 0 |
| SCN4A | 449 | A | -16.67755 | 10.08247023 | 0.135859638 | 6.96393458 | NA | NA | NA | NA | NA | NA | NA |
| SCN4A | 450 | Y | -16.29775 | 8.240577346 | 0.104218452 | 5.144472605 | NA | NA | NA | NA | NA | NA | NA |
| SCN4A | 451 | A | -19.85795 | 6.528963853 | 0.074815287 | 3.664802041 | 2.93 | 1.5 | 3.37 | 0.28 | -1.82 | 0 | 0 |
| SCN4A | 452 | E | -21.80108333 | 9.272683161 | 0.12194861 | 3.909965129 | 5.39 | 0.49 | 0.98 | 1.02 | -0.62 | 0 | 0 |
| SCN4A | 453 | Q | -20.11497222 | 11.77696633 | 0.16496875 | 6.388908648 | NA | NA | NA | NA | NA | NA | NA |
| SCN4A | 454 | N | -21.196 | 9.67808044 | 0.128912778 | 4.66316212 | 5.39 | 0.49 | 0.98 | 1.02 | -0.62 | 0 | 0 |
| SCN4A | 455 | E | -25.75375 | 7.965627784 | 0.099495197 | 2.083936362 | 5.91 | -0.04 | 0.1 | 1.35 | 0.02 | 0 | 0 |
| SCN4A | 456 | A | -25.12335 | 11.83451744 | 0.165957399 | 5.977603298 | NA | NA | NA | NA | NA | NA | NA |
| SCN4A | 457 | T | -24.45960714 | 14.02329747 | 0.203557629 | 8.109433262 | NA | NA | NA | NA | NA | NA | NA |
| SCN4A | 458 | L | -26.853625 | 10.84564078 | 0.148969858 | 4.153582138 | 6.92 | -0.35 | -1.76 | 20.96 | 0.64 | 2 | 0 |
| SCN4A | 459 | A | -29.90995 | 13.13928118 | 0.188371445 | 6.386278177 | NA | NA | NA | NA | NA | NA | NA |
| SCN4A | 460 | E | -29.35286111 | 15.82076992 | 0.234435733 | 9.05627059 | NA | NA | NA | NA | NA | NA | NA |
| SCN4A | 461 | D | -29.065625 | 17.03958506 | 0.2553733 | 10.17441659 | NA | NA | NA | NA | NA | NA | NA |
| SCN4A | 462 | K | -33.30363889 | 12.23414247 | 0.172822407 | 5.924992576 | 6.92 | -0.35 | -1.76 | 20.96 | 0.64 | 2 | 0 |
| SCN4A | 463 | E | -34.35952778 | 16.93536625 | 0.253582964 | 10.0596248 | NA | NA | NA | NA | NA | NA | NA |
| SCN4A | 559 | N | -14.424875 | 60.38570441 | 1 | 57.29696291 | NA | NA | NA | NA | NA | NA | NA |
| SCN4A | 560 | C | -17.31308333 | 60.18924994 | 0.996625182 | 57.09536839 | NA | NA | NA | NA | NA | NA | NA |
| SCN4A | 561 | C | -19.63158333 | 58.00511769 | 0.959104795 | 54.90709716 | NA | NA | NA | NA | NA | NA | NA |
| SCN4A | 562 | A | -22.25755 | 59.19002587 | 0.979459887 | 56.08817951 | NA | NA | NA | NA | NA | NA | NA |
| SCN4A | 563 | P | -19.78260714 | 58.24689092 | 0.963258127 | 55.15050195 | NA | NA | NA | NA | NA | NA | NA |
| SCN4A | 564 | W | -17.24103571 | 52.7957309 | 0.869614696 | 49.70656326 | NA | NA | NA | NA | NA | NA | NA |
| SCN4A | 565 | L | -21.964875 | 53.39760701 | 0.8799541 | 50.29411979 | NA | NA | NA | NA | NA | NA | NA |
| SCN4A | 566 | K | -20.12108333 | 55.43458586 | 0.914946594 | 52.33466878 | NA | NA | NA | NA | NA | NA | NA |

| | | | | | | | | | | | | | |
|---|---|---|---|---|---|---|---|---|---|---|---|---|---|
| SCN4A | 567 | F | -16.22720455 | 52.58095131 | 0.865925079 | 49.48483627 | NA | NA | NA | NA | NA | NA | NA |
| SCN4A | 568 | K | -19.46852778 | 49.92878257 | 0.820364468 | 46.82862546 | NA | NA | NA | NA | NA | NA | NA |
| SCN4A | 569 | N | -22.71975 | 50.49601403 | 0.830108724 | 47.39004391 | NA | NA | NA | NA | NA | NA | NA |
| SCN4A | 570 | I | -19.249375 | 50.11931661 | 0.823637581 | 47.01859027 | NA | NA | NA | NA | NA | NA | NA |
| SCN4A | 571 | I | -16.6325 | 46.8384087 | 0.767276097 | 43.74025823 | NA | NA | NA | NA | NA | NA | NA |
| SCN4A | 572 | H | -21.44165 | 43.97214434 | 0.718037618 | 40.86458252 | NA | NA | NA | NA | NA | NA | NA |
| SCN4A | 573 | L | -21.633 | 46.19646664 | 0.756248415 | 43.08978793 | NA | NA | NA | NA | NA | NA | NA |
| SCN4A | 574 | I | -16.990375 | 44.03571998 | 0.71912976 | 40.93542129 | NA | NA | NA | NA | NA | NA | NA |
| SCN4A | 575 | V | -17.70017857 | 41.12429153 | 0.669115424 | 38.02198498 | NA | NA | NA | NA | NA | NA | NA |
| SCN4A | 576 | M | -22.3015 | 39.82324994 | 0.64676532 | 36.99057326 | NA | NA | NA | NA | NA | NA | NA |
| SCN4A | 577 | D | -21.374125 | 42.02561433 | 0.68459891 | 38.92046932 | NA | NA | NA | NA | NA | NA | NA |
| SCN4A | 578 | P | -19.50975 | 40.35376448 | 0.65587883 | 37.24594662 | NA | NA | NA | NA | NA | NA | NA |
| SCN4A | 579 | F | -16.58584091 | 41.22429967 | 0.670833426 | 38.12098859 | NA | NA | NA | NA | NA | NA | NA |
| SCN4A | 580 | V | -13.63889286 | 39.94543104 | 0.648864223 | 36.85549854 | NA | NA | NA | NA | NA | NA | NA |
| SCN4A | 581 | D | -14.7655 | 35.36961781 | 0.570258046 | 32.26987168 | NA | NA | NA | NA | NA | NA | NA |
| SCN4A | 582 | L | -13.39825 | 36.22559126 | 0.584962493 | 33.12712592 | NA | NA | NA | NA | NA | NA | NA |
| SCN4A | 583 | G | -10.12175 | 39.52229621 | 0.641595349 | 36.43136005 | NA | NA | NA | NA | NA | NA | NA |
| SCN4A | 584 | I | -9.0855 | 36.14039736 | 0.583498979 | 33.88004046 | NA | NA | NA | NA | NA | NA | NA |
| SCN4A | 585 | T | -9.837464286 | 33.52872404 | 0.538634024 | 30.44212664 | NA | NA | NA | NA | NA | NA | NA |
| SCN4A | 586 | I | -7.526125 | 36.15144062 | 0.583688687 | 33.85562017 | NA | NA | NA | NA | NA | NA | NA |
| SCN4A | 587 | C | -4.633583333 | 36.30151524 | 0.586266762 | 33.00049655 | NA | NA | NA | NA | NA | NA | NA |
| SCN4A | 588 | I | -4.8165 | 31.59692677 | 0.505448404 | 28.31353629 | NA | NA | NA | NA | NA | NA | NA |
| SCN4A | 589 | V | -3.696321429 | 31.86246533 | 0.510009991 | 28.64727124 | NA | NA | NA | NA | NA | NA | NA |
| SCN4A | 590 | L | -1.4055 | 34.95898791 | 0.56320399 | 31.59821136 | NA | NA | NA | NA | NA | NA | NA |
| SCN4A | 591 | N | 0.738875 | 33.75464795 | 0.542515086 | 30.29388357 | NA | NA | NA | NA | NA | NA | NA |
| SCN4A | 592 | T | 0.75775 | 29.03507179 | 0.46143926 | 25.57389075 | NA | NA | NA | NA | NA | NA | NA |
| SCN4A | 593 | L | 1.488125 | 31.01874691 | 0.49551607 | 27.58338219 | NA | NA | NA | NA | NA | NA | NA |
| SCN4A | 594 | F | 5.184386364 | 33.65341879 | 0.540776108 | 30.91199056 | NA | NA | NA | NA | NA | NA | NA |
| SCN4A | 595 | M | 5.425 | 30.00551913 | 0.47811021 | 27.27308618 | NA | NA | NA | NA | NA | NA | NA |
| SCN4A | 596 | A | 6.51635 | 28.42465662 | 0.450953167 | 25.70238004 | NA | NA | NA | NA | NA | NA | NA |
| SCN4A | 597 | M | 8.397375 | 30.61099711 | 0.488511489 | 27.88653918 | NA | NA | NA | NA | NA | NA | NA |
| SCN4A | 598 | E | 11.65619444 | 27.67222788 | 0.438027477 | 24.97528306 | NA | NA | NA | NA | NA | NA | NA |
| SCN4A | 599 | H | 14.95865 | 27.92413891 | 0.442354961 | 25.25272283 | NA | NA | NA | NA | NA | NA | NA |

| | | | | | | | | | | | | | |
|---|---|---|---|---|---|---|---|---|---|---|---|---|---|
| SCN4A | 600 | Y | 20.3055 | 25.93886969 | 0.408250767 | 22.10393508 | NA | NA | NA | NA | NA | NA | NA |
| SCN4A | 601 | P | 19.94646429 | 27.26464196 | 0.431025711 | 23.4753223 | NA | NA | NA | NA | NA | NA | NA |
| SCN4A | 602 | M | 17.75425 | 31.40414406 | 0.502136662 | 27.59917474 | NA | NA | NA | NA | NA | NA | NA |
| SCN4A | 603 | T | 19.62346429 | 34.79113756 | 0.560320552 | 30.99412004 | NA | NA | NA | NA | NA | NA | NA |
| SCN4A | 604 | E | 20.85497222 | 37.57806702 | 0.608196166 | 33.76042084 | NA | NA | NA | NA | NA | NA | NA |
| SCN4A | 605 | H | 16.71025 | 39.75393968 | 0.645574665 | 35.95392517 | NA | NA | NA | NA | NA | NA | NA |
| SCN4A | 606 | F | 13.80475 | 34.33966016 | 0.552564791 | 31.65553708 | NA | NA | NA | NA | NA | NA | NA |
| SCN4A | 607 | D | 15.637875 | 35.43852843 | 0.571441836 | 32.7551195 | NA | NA | NA | NA | NA | NA | NA |
| SCN4A | 608 | N | 15.045125 | 39.21994417 | 0.636401356 | 36.53878728 | NA | NA | NA | NA | NA | NA | NA |
| SCN4A | 609 | V | 10.96360714 | 39.50387208 | 0.641278847 | 36.7982975 | NA | NA | NA | NA | NA | NA | NA |
| SCN4A | 610 | L | 10.063375 | 33.90129682 | 0.545034312 | 31.18769206 | NA | NA | NA | NA | NA | NA | NA |
| SCN4A | 611 | T | 11.30832143 | 38.45564138 | 0.623271686 | 35.75534949 | NA | NA | NA | NA | NA | NA | NA |
| SCN4A | 612 | V | 8.752892857 | 41.09795287 | 0.668662962 | 38.38254091 | NA | NA | NA | NA | NA | NA | NA |
| SCN4A | 613 | G | 5.77 | 40.16598698 | 0.65265307 | 37.42674125 | NA | NA | NA | NA | NA | NA | NA |
| SCN4A | 614 | N | 5.659125 | 35.60041823 | 0.57422288 | 32.86405621 | NA | NA | NA | NA | NA | NA | NA |
| SCN4A | 615 | L | 5.955375 | 40.92463085 | 0.665685528 | 38.18270613 | NA | NA | NA | NA | NA | NA | NA |
| SCN4A | 616 | V | 2.289892857 | 42.54418581 | 0.693507255 | 39.08450717 | NA | NA | NA | NA | NA | NA | NA |
| SCN4A | 617 | F | 0.627022727 | 38.88362337 | 0.630623828 | 35.46467052 | NA | NA | NA | NA | NA | NA | NA |
| SCN4A | 618 | T | 1.128464286 | 39.32265155 | 0.638165728 | 35.84220907 | NA | NA | NA | NA | NA | NA | NA |
| SCN4A | 619 | G | -0.91075 | 42.52009982 | 0.693093491 | 39.07210531 | NA | NA | NA | NA | NA | NA | NA |
| SCN4A | 620 | I | -3.20275 | 41.5458697 | 0.676357557 | 38.23478719 | NA | NA | NA | NA | NA | NA | NA |
| SCN4A | 621 | F | -4.284659091 | 36.54957632 | 0.59052811 | 33.24404134 | NA | NA | NA | NA | NA | NA | NA |
| SCN4A | 622 | T | -3.161892857 | 41.45331674 | 0.674767625 | 38.08710183 | NA | NA | NA | NA | NA | NA | NA |
| SCN4A | 623 | A | -5.80355 | 44.08516373 | 0.719979135 | 40.74637953 | NA | NA | NA | NA | NA | NA | NA |
| SCN4A | 624 | E | -8.911527778 | 39.4318131 | 0.640040973 | 36.27278563 | NA | NA | NA | NA | NA | NA | NA |
| SCN4A | 625 | M | -8.19725 | 40.34211063 | 0.655678632 | 37.1022269 | NA | NA | NA | NA | NA | NA | NA |
| SCN4A | 626 | V | -8.117892857 | 44.01072327 | 0.718700351 | 40.78098629 | NA | NA | NA | NA | NA | NA | NA |
| SCN4A | 627 | L | -11.234875 | 45.54292189 | 0.745021415 | 42.45809288 | NA | NA | NA | NA | NA | NA | NA |
| SCN4A | 628 | K | -13.58941667 | 40.30679801 | 0.65507201 | 37.2148853 | NA | NA | NA | NA | NA | NA | NA |
| SCN4A | 629 | L | -12.12725 | 43.65241827 | 0.712545164 | 40.58074269 | NA | NA | NA | NA | NA | NA | NA |
| SCN4A | 630 | I | -13.850125 | 46.97092835 | 0.769552602 | 43.88537322 | NA | NA | NA | NA | NA | NA | NA |
| SCN4A | 631 | A | -17.29355 | 45.33357432 | 0.741425112 | 42.23950667 | NA | NA | NA | NA | NA | NA | NA |
| SCN4A | 632 | M | -19.2895 | 40.68232724 | 0.661523086 | 37.57949902 | NA | NA | NA | NA | NA | NA | NA |

| | | | | | | | | | | | | | |
|---|---|---|---|---|---|---|---|---|---|---|---|---|---|
| SCN4A | 633 | D | -18.07575 | 43.75624967 | 0.714328844 | 40.6655441 | NA | NA | NA | NA | NA | NA | NA |
| SCN4A | 634 | P | -15.12489286 | 43.41953048 | 0.708544472 | 40.33398074 | NA | NA | NA | NA | NA | NA | NA |
| SCN4A | 635 | Y | -19.78691667 | 41.02786373 | 0.667458927 | 37.93442045 | NA | NA | NA | NA | NA | NA | NA |
| SCN4A | 636 | E | -20.89297222 | 38.61228465 | 0.625962602 | 35.50594703 | NA | NA | NA | NA | NA | NA | NA |
| SCN4A | 637 | Y | -14.29741667 | 37.61302596 | 0.608796712 | 34.52532504 | NA | NA | NA | NA | NA | NA | NA |
| SCN4A | 638 | F | -13.37920455 | 36.37540132 | 0.587536023 | 33.28978941 | NA | NA | NA | NA | NA | NA | NA |
| SCN4A | 639 | Q | -19.51997222 | 33.7122197 | 0.541786227 | 30.61865636 | NA | NA | NA | NA | NA | NA | NA |
| SCN4A | 640 | Q | -17.77808333 | 32.90058512 | 0.527843461 | 29.81008967 | NA | NA | NA | NA | NA | NA | NA |
| SCN4A | 641 | G | -14.647 | 30.85285389 | 0.492666256 | 27.76646035 | NA | NA | NA | NA | NA | NA | NA |
| SCN4A | 642 | W | -13.13110714 | 26.61421013 | 0.419852187 | 23.5351275 | NA | NA | NA | NA | NA | NA | NA |
| SCN4A | 643 | N | -12.961125 | 33.82334209 | 0.543695157 | 30.74161133 | NA | NA | NA | NA | NA | NA | NA |
| SCN4A | 644 | I | -10.9275 | 33.49673448 | 0.538084487 | 31.1877349 | NA | NA | NA | NA | NA | NA | NA |
| SCN4A | 645 | F | -7.573022727 | 28.05303957 | 0.444569297 | 25.86278338 | NA | NA | NA | NA | NA | NA | NA |
| SCN4A | 646 | D | -8.1705 | 33.24762431 | 0.533805117 | 30.1238888 | NA | NA | NA | NA | NA | NA | NA |
| SCN4A | 647 | S | -6.554416667 | 35.53119365 | 0.573033697 | 32.26755397 | NA | NA | NA | NA | NA | NA | NA |
| SCN4A | 648 | I | -3.925625 | 31.91809946 | 0.510965709 | 28.56994355 | NA | NA | NA | NA | NA | NA | NA |
| SCN4A | 649 | I | -2.361875 | 31.04125484 | 0.495902725 | 27.67579927 | NA | NA | NA | NA | NA | NA | NA |
| SCN4A | 650 | V | -2.286178571 | 36.1744327 | 0.584083659 | 32.75633871 | NA | NA | NA | NA | NA | NA | NA |
| SCN4A | 651 | T | -1.156607143 | 37.67131111 | 0.609797971 | 34.19197518 | NA | NA | NA | NA | NA | NA | NA |
| SCN4A | 652 | L | 1.34225 | 32.12206043 | 0.514469478 | 28.63600641 | NA | NA | NA | NA | NA | NA | NA |
| SCN4A | 653 | S | 2.28075 | 36.47463833 | 0.589240779 | 32.99134112 | NA | NA | NA | NA | NA | NA | NA |
| SCN4A | 654 | L | 2.7025 | 39.14680689 | 0.635144959 | 35.66367768 | NA | NA | NA | NA | NA | NA | NA |
| SCN4A | 655 | V | 4.939464286 | 38.56517317 | 0.625153291 | 34.38184184 | NA | NA | NA | NA | NA | NA | NA |
| SCN4A | 656 | E | 7.840083333 | 35.50642966 | 0.572608286 | 32.75264318 | NA | NA | NA | NA | NA | NA | NA |
| SCN4A | 657 | L | 7.340625 | 39.51271225 | 0.641430709 | 36.75812749 | NA | NA | NA | NA | NA | NA | NA |
| SCN4A | 658 | G | 8.6025 | 41.81958633 | 0.681059632 | 39.06633777 | NA | NA | NA | NA | NA | NA | NA |
| SCN4A | 659 | L | 10.99775 | 37.46090546 | 0.606183491 | 34.71041819 | NA | NA | NA | NA | NA | NA | NA |
| SCN4A | 660 | A | 13.77835 | 40.79700382 | 0.663493072 | 38.06233973 | NA | NA | NA | NA | NA | NA | NA |
| SCN4A | 661 | N | 16.52575 | 38.87318959 | 0.63044459 | 36.15975802 | NA | NA | NA | NA | NA | NA | NA |
| SCN4A | 662 | V | 15.49546429 | 35.61836203 | 0.57453113 | 32.88595211 | NA | NA | NA | NA | NA | NA | NA |
| SCN4A | 663 | Q | 18.70830556 | 31.86066707 | 0.5099791 | 29.14984591 | NA | NA | NA | NA | NA | NA | NA |
| SCN4A | 664 | G | 15.94625 | 30.64148709 | 0.489035265 | 27.90624095 | NA | NA | NA | NA | NA | NA | NA |
| SCN4A | 665 | L | 11.9745 | 31.91609403 | 0.510931259 | 29.17554458 | NA | NA | NA | NA | NA | NA | NA |

| | | | | | | | | | | | | | |
|---|---|---|---|---|---|---|---|---|---|---|---|---|---|
| SCN4A | 666 | S | 12.01208333 | 28.96712752 | 0.460272071 | 26.22395945 | NA | NA | NA | NA | NA | NA | NA |
| SCN4A | 667 | V | 8.723607143 | 26.82718688 | 0.423510835 | 24.06822574 | NA | NA | NA | NA | NA | NA | NA |
| SCN4A | 668 | L | 6.456625 | 30.86715181 | 0.492911874 | 28.10448393 | NA | NA | NA | NA | NA | NA | NA |
| SCN4A | 669 | R | 9.011022727 | 31.31964561 | 0.500685095 | 28.57681268 | NA | NA | NA | NA | NA | NA | NA |
| SCN4A | 670 | S | 5.401083333 | 27.68744082 | 0.438288814 | 24.92911533 | NA | NA | NA | NA | NA | NA | NA |
| SCN4A | 671 | F | 2.511022727 | 27.56460341 | 0.436178636 | 24.12746009 | NA | NA | NA | NA | NA | NA | NA |
| SCN4A | 672 | R | 2.240477273 | 30.8999404 | 0.493475137 | 27.42647877 | NA | NA | NA | NA | NA | NA | NA |
| SCN4A | 673 | L | 1.69425 | 27.33476676 | 0.432230359 | 23.87801387 | NA | NA | NA | NA | NA | NA | NA |
| SCN4A | 674 | L | -2.201625 | 25.00556718 | 0.392217914 | 21.56543935 | NA | NA | NA | NA | NA | NA | NA |
| SCN4A | 675 | R | -2.847931818 | 29.27274734 | 0.465522199 | 25.89279748 | NA | NA | NA | NA | NA | NA | NA |
| SCN4A | 676 | V | -4.492607143 | 26.01783306 | 0.409607249 | 22.69973809 | NA | NA | NA | NA | NA | NA | NA |
| SCN4A | 677 | F | -6.569204545 | 21.77959788 | 0.3368002 | 19.58016281 | NA | NA | NA | NA | NA | NA | NA |
| SCN4A | 678 | K | -8.682416667 | 28.35220508 | 0.449708549 | 26.06418662 | NA | NA | NA | NA | NA | NA | NA |
| SCN4A | 679 | L | -9.72475 | 26.51486168 | 0.418145517 | 23.42935059 | NA | NA | NA | NA | NA | NA | NA |
| SCN4A | 680 | A | -12.87275 | 24.34726393 | 0.38090917 | 21.25426003 | NA | NA | NA | NA | NA | NA | NA |
| SCN4A | 681 | K | -15.47097222 | 26.49317342 | 0.417772943 | 23.38830934 | NA | NA | NA | NA | NA | NA | NA |
| SCN4A | 682 | S | -15.30091667 | 26.12645452 | 0.411473217 | 23.02132491 | NA | NA | NA | NA | NA | NA | NA |
| SCN4A | 683 | W | -12.72553571 | 22.54800656 | 0.350000404 | 19.46364774 | NA | NA | NA | NA | NA | NA | NA |
| SCN4A | 684 | P | -16.37360714 | 19.06571339 | 0.290179397 | 15.96207821 | NA | NA | NA | NA | NA | NA | NA |
| SCN4A | 685 | T | -13.16403571 | 17.80696869 | 0.268555895 | 14.71995298 | NA | NA | NA | NA | NA | NA | NA |
| SCN4A | 686 | L | -10.44625 | 20.3430407 | 0.312122123 | 17.2592855 | NA | NA | NA | NA | NA | NA | NA |
| SCN4A | 687 | N | -14.987625 | 18.99553953 | 0.288973907 | 15.90917069 | NA | NA | NA | NA | NA | NA | NA |
| SCN4A | 688 | M | -14.67525 | 15.03309053 | 0.220904484 | 11.93024112 | NA | NA | NA | NA | NA | NA | NA |
| SCN4A | 689 | L | -9.264375 | 15.04344857 | 0.221082421 | 12.64392905 | NA | NA | NA | NA | NA | NA | NA |
| SCN4A | 690 | I | -11.104125 | 18.96342443 | 0.288422214 | 15.89490812 | NA | NA | NA | NA | NA | NA | NA |
| SCN4A | 691 | K | -15.64130556 | 16.23375003 | 0.241530163 | 13.13285151 | NA | NA | NA | NA | NA | NA | NA |
| SCN4A | 692 | I | -11.395125 | 12.72719015 | 0.181292288 | 9.664843388 | NA | NA | NA | NA | NA | NA | NA |
| SCN4A | 693 | I | -8.505 | 16.42213814 | 0.244766411 | 14.03256361 | NA | NA | NA | NA | NA | NA | NA |
| SCN4A | 694 | G | -12.081 | 18.68271353 | 0.283599987 | 15.74127186 | NA | NA | NA | NA | NA | NA | NA |
| SCN4A | 695 | N | -14.0545 | 14.42952605 | 0.210536077 | 11.34661262 | NA | NA | NA | NA | NA | NA | NA |
| SCN4A | 696 | S | -9.661416667 | 15.67311115 | 0.231899158 | 13.24517296 | NA | NA | NA | NA | NA | NA | NA |
| SCN4A | 697 | V | -9.93375 | 19.03274087 | 0.289612975 | 16.59960387 | NA | NA | NA | NA | NA | NA | NA |
| SCN4A | 698 | G | -13.69575 | 20.15346583 | 0.308865488 | 17.07861164 | NA | NA | NA | NA | NA | NA | NA |

| | | | | | | | | | | | | | |
|---|---|---|---|---|---|---|---|---|---|---|---|---|---|
| SCN4A | 699 | A | -16.08175 | 17.90697442 | 0.270273856 | 14.81694775 | NA | NA | NA | NA | NA | NA | NA |
| SCN4A | 700 | L | -13.497875 | 14.16355016 | 0.205966977 | 11.08547564 | NA | NA | NA | NA | NA | NA | NA |
| SCN4A | 701 | G | -11.007 | 19.13941135 | 0.291445427 | 16.79494494 | NA | NA | NA | NA | NA | NA | NA |
| SCN4A | 702 | N | -11.694875 | 19.33706488 | 0.294840843 | 16.82812816 | NA | NA | NA | NA | NA | NA | NA |
| SCN4A | 703 | L | -10.0525 | 13.71974821 | 0.19834307 | 11.22942152 | NA | NA | NA | NA | NA | NA | NA |
| SCN4A | 704 | T | -6.647464286 | 16.49175712 | 0.24596237 | 13.06726688 | NA | NA | NA | NA | NA | NA | NA |
| SCN4A | 705 | L | -6.359375 | 19.00022718 | 0.289054434 | 15.59224987 | NA | NA | NA | NA | NA | NA | NA |
| SCN4A | 706 | V | -6.073321429 | 17.85571654 | 0.269393316 | 14.52988006 | NA | NA | NA | NA | NA | NA | NA |
| SCN4A | 707 | L | -3.114625 | 12.82951012 | 0.183050004 | 8.702199571 | NA | NA | NA | NA | NA | NA | NA |
| SCN4A | 708 | A | -1.35415 | 17.44153471 | 0.262278242 | 13.14655559 | NA | NA | NA | NA | NA | NA | NA |
| SCN4A | 709 | I | -1.94625 | 19.40159254 | 0.295949339 | 15.55866982 | NA | NA | NA | NA | NA | NA | NA |
| SCN4A | 710 | I | -1.278625 | 15.37464074 | 0.226771847 | 11.53375488 | NA | NA | NA | NA | NA | NA | NA |
| SCN4A | 711 | V | 2.223321429 | 14.33560093 | 0.208922572 | 10.60454369 | NA | NA | NA | NA | NA | NA | NA |
| SCN4A | 712 | F | 3.446931818 | 18.82666216 | 0.286072826 | 15.05704679 | NA | NA | NA | NA | NA | NA | NA |
| SCN4A | 713 | I | 2.629 | 20.2243257 | 0.310082763 | 16.40939564 | NA | NA | NA | NA | NA | NA | NA |
| SCN4A | 714 | F | 4.113022727 | 15.61197861 | 0.230848985 | 11.90477171 | NA | NA | NA | NA | NA | NA | NA |
| SCN4A | 715 | A | 8.03495 | 19.2158922 | 0.292759263 | 15.29695954 | NA | NA | NA | NA | NA | NA | NA |
| SCN4A | 716 | V | 8.128892857 | 21.88473233 | 0.338606265 | 17.96579698 | NA | NA | NA | NA | NA | NA | NA |
| SCN4A | 717 | V | 8.059321429 | 22.04634473 | 0.341382543 | 18.15044363 | NA | NA | NA | NA | NA | NA | NA |
| SCN4A | 718 | G | 11.34925 | 20.62578895 | 0.316979349 | 18.82509022 | NA | NA | NA | NA | NA | NA | NA |
| SCN4A | 719 | M | 13.2185 | 22.14248304 | 0.343034067 | 20.33726993 | NA | NA | NA | NA | NA | NA | NA |
| SCN4A | 720 | Q | 12.28808333 | 24.82445935 | 0.389106731 | 23.01388554 | NA | NA | NA | NA | NA | NA | NA |
| SCN4A | 721 | L | 13.019875 | 23.55232296 | 0.367253178 | 21.74125182 | NA | NA | NA | NA | NA | NA | NA |
| SCN4A | 722 | F | 14.86193182 | 19.8842554 | 0.304240823 | 18.14714168 | NA | NA | NA | NA | NA | NA | NA |
| SCN4A | 723 | G | 18.2755 | 23.94588737 | 0.374014073 | 22.09883945 | NA | NA | NA | NA | NA | NA | NA |
| SCN4A | 724 | K | 19.75697222 | 26.33538094 | 0.415062285 | 24.45606245 | NA | NA | NA | NA | NA | NA | NA |
| SCN4A | 725 | S | 21.43941667 | 24.12320891 | 0.377060213 | 19.82333823 | NA | NA | NA | NA | NA | NA | NA |
| SCN4A | 726 | Y | 20.34958333 | 17.42321305 | 0.261963501 | 14.78691997 | NA | NA | NA | NA | NA | NA | NA |
| SCN4A | 727 | K | 25.70163889 | 23.95978894 | 0.374252883 | 19.65967053 | NA | NA | NA | NA | NA | NA | NA |
| SCN4A | 728 | E | 25.94475 | 24.94339392 | 0.391149863 | 20.68323678 | NA | NA | NA | NA | NA | NA | NA |
| SCN4A | 729 | C | 26.03591667 | 20.69556525 | 0.31817801 | 16.42963686 | NA | NA | NA | NA | NA | NA | NA |
| SCN4A | 730 | V | 27.52960714 | 18.37087317 | 0.278242998 | 14.13335243 | NA | NA | NA | NA | NA | NA | NA |
| SCN4A | 731 | C | 30.48208333 | 18.43431661 | 0.279332869 | 14.26942403 | NA | NA | NA | NA | NA | NA | NA |

| | | | | | | | | | | | | | |
|---|---|---|---|---|---|---|---|---|---|---|---|---|---|
| SCN4A | 732 | K | 27.17830556 | 16.47204329 | 0.245623713 | 12.23692477 | NA | NA | NA | NA | NA | NA | NA |
| SCN4A | 733 | I | 25.32575 | 13.58317345 | 0.195996903 | 9.36449195 | NA | NA | NA | NA | NA | NA | NA |
| SCN4A | 734 | A | 28.54275 | 14.52460416 | 0.212169388 | 10.32748651 | NA | NA | NA | NA | NA | NA | NA |
| SCN4A | 735 | L | 31.76525 | 12.07319225 | 0.170057504 | 8.062503412 | NA | NA | NA | NA | NA | NA | NA |
| SCN4A | 736 | D | 32.23575 | 15.70863247 | 0.232509365 | 11.61433785 | NA | NA | NA | NA | NA | NA | NA |
| SCN4A | 737 | C | 31.58141667 | 17.98635007 | 0.27163742 | 13.84740726 | NA | NA | NA | NA | NA | NA | NA |
| SCN4A | 738 | N | 29.28575 | 17.61481754 | 0.265255003 | 13.3701463 | NA | NA | NA | NA | NA | NA | NA |
| SCN4A | 739 | L | 24.584125 | 16.7426371 | 0.250272142 | 12.45714057 | NA | NA | NA | NA | NA | NA | NA |
| SCN4A | 740 | P | 22.75946429 | 14.67478855 | 0.214749349 | 10.36648049 | NA | NA | NA | NA | NA | NA | NA |
| SCN4A | 741 | R | 21.10965909 | 13.02853998 | 0.186469063 | 8.738682015 | NA | NA | NA | NA | NA | NA | NA |
| SCN4A | 742 | W | 16.14239286 | 10.8478986 | 0.149008644 | 8.086275611 | NA | NA | NA | NA | NA | NA | NA |
| SCN4A | 743 | H | 18.69335 | 16.82060421 | 0.25161151 | 14.04013992 | NA | NA | NA | NA | NA | NA | NA |
| SCN4A | 744 | M | 15.270625 | 17.07957851 | 0.256060332 | 15.34805712 | NA | NA | NA | NA | NA | NA | NA |
| SCN4A | 745 | H | 20.00955 | 22.2729008 | 0.345274465 | 20.3892798 | NA | NA | NA | NA | NA | NA | NA |
| SCN4A | 746 | D | 16.31375 | 22.44585543 | 0.348245588 | 20.71033604 | NA | NA | NA | NA | NA | NA | NA |
| SCN4A | 747 | F | 10.68356818 | 21.00017079 | 0.323410714 | 19.20333593 | NA | NA | NA | NA | NA | NA | NA |
| SCN4A | 748 | F | 12.21056818 | 18.79187548 | 0.285475239 | 16.97888736 | NA | NA | NA | NA | NA | NA | NA |
| SCN4A | 749 | H | 15.58735 | 16.85088024 | 0.252131611 | 15.10227141 | NA | NA | NA | NA | NA | NA | NA |
| SCN4A | 750 | S | 11.59541667 | 16.88984707 | 0.252801007 | 15.10685034 | NA | NA | NA | NA | NA | NA | NA |
| SCN4A | 751 | F | 7.736931818 | 15.23963244 | 0.22445259 | 13.49745905 | NA | NA | NA | NA | NA | NA | NA |
| SCN4A | 752 | L | 11.184875 | 12.6578721 | 0.180101499 | 10.84650629 | NA | NA | NA | NA | NA | NA | NA |
| SCN4A | 753 | I | 11.441625 | 11.57793704 | 0.1615497 | 9.791274253 | NA | NA | NA | NA | NA | NA | NA |
| SCN4A | 754 | V | 6.867464286 | 11.83012113 | 0.165881876 | 9.745880574 | NA | NA | NA | NA | NA | NA | NA |
| SCN4A | 755 | F | 5.747113636 | 9.923776449 | 0.133133497 | 7.704144931 | NA | NA | NA | NA | NA | NA | NA |
| SCN4A | 756 | R | 10.71129545 | 6.897145497 | 0.081140141 | 5.176011658 | 1.96 | -0.79 | -2.05 | 26.54 | 1.33 | 2 | 0 |
| SCN4A | 757 | I | 6.588875 | 6.800180953 | 0.079474424 | 5.063601166 | 2.3 | -0.7 | -1.28 | 14.91 | 0.97 | 1 | 1 |
| SCN4A | 758 | L | 2.5375 | 5.627109649 | 0.059322672 | 1.720683177 | 3.92 | -0.79 | -1.02 | 3.38 | 0.76 | 0 | 0 |
| SCN4A | 759 | C | 4.74475 | 3.907792471 | 0.029787168 | 1.691281118 | 2.52 | -0.8 | -0.4 | 3.38 | 0.81 | 0 | 0 |
| SCN4A | 760 | G | 5.9405 | 3.701208721 | 0.026238344 | 1.807375852 | 2.17 | -0.8 | -0.4 | 3.38 | 0.81 | 0 | 0 |
| SCN4A | 761 | E | 9.864416667 | 3.50513238 | 0.022870022 | 1.823997865 | 1.96 | -0.79 | -2.05 | 26.54 | 1.33 | 2 | 0 |
| SCN4A | 762 | W | 8.614178571 | 6.47141561 | 0.073826687 | 4.703260242 | 1.9 | -0.78 | -2.6 | 34.26 | 1.5 | 2 | 0 |
| SCN4A | 763 | I | 12.645125 | 7.417006135 | 0.090070632 | 5.633231169 | 1.91 | -0.79 | -2.05 | 26.54 | 1.33 | 2 | 0 |
| SCN4A | 764 | E | 16.06919444 | 6.100920422 | 0.06746209 | 5.006427826 | 2 | -0.78 | -2.6 | 34.26 | 1.5 | 2 | 0 |

| | | | | | | | | | | | | | |
|---|---|---|---|---|---|---|---|---|---|---|---|---|---|
| SCN4A | 765 | T | 14.78446429 | 10.54746268 | 0.143847569 | 8.543304626 | NA | NA | NA | NA | NA | NA | NA |
| SCN4A | 766 | M | 13.741875 | 12.31690209 | 0.174244103 | 10.41857535 | NA | NA | NA | NA | NA | NA | NA |
| SCN4A | 767 | W | 17.47889286 | 10.11545619 | 0.136426291 | 8.59527814 | NA | NA | NA | NA | NA | NA | NA |
| SCN4A | 768 | D | 19.86475 | 13.45819724 | 0.193849984 | 9.155911452 | NA | NA | NA | NA | NA | NA | NA |
| SCN4A | 769 | C | 17.12908333 | 16.91963847 | 0.253312783 | 15.06599566 | NA | NA | NA | NA | NA | NA | NA |
| SCN4A | 770 | M | 17.483125 | 16.23130127 | 0.241488096 | 14.38900907 | NA | NA | NA | NA | NA | NA | NA |
| SCN4A | 771 | E | 22.79963889 | 18.00006972 | 0.271873105 | 13.73575855 | NA | NA | NA | NA | NA | NA | NA |
| SCN4A | 772 | V | 21.57717857 | 19.82812896 | 0.303276648 | 15.52384856 | NA | NA | NA | NA | NA | NA | NA |
| SCN4A | 773 | A | 19.25015 | 22.73150855 | 0.353152715 | 20.85093564 | NA | NA | NA | NA | NA | NA | NA |
| SCN4A | 774 | G | 17.287 | 21.44882759 | 0.331118021 | 19.54704562 | NA | NA | NA | NA | NA | NA | NA |
| SCN4A | 775 | Q | 16.37352778 | 21.00726839 | 0.323532641 | 19.13899335 | NA | NA | NA | NA | NA | NA | NA |
| SCN4A | 776 | A | 12.94555 | 22.1394974 | 0.342982778 | 20.35201118 | NA | NA | NA | NA | NA | NA | NA |
| SCN4A | 777 | M | 12.793 | 20.13747603 | 0.308590805 | 18.29218587 | NA | NA | NA | NA | NA | NA | NA |
| SCN4A | 778 | C | 12.86141667 | 16.66374175 | 0.248916829 | 14.87047567 | NA | NA | NA | NA | NA | NA | NA |
| SCN4A | 779 | L | 10.419375 | 16.40364609 | 0.244448743 | 14.54274322 | NA | NA | NA | NA | NA | NA | NA |
| SCN4A | 780 | T | 7.736035714 | 18.03718321 | 0.272510664 | 14.13182347 | NA | NA | NA | NA | NA | NA | NA |
| SCN4A | 781 | V | 7.978178571 | 15.72750409 | 0.232833554 | 11.81066654 | NA | NA | NA | NA | NA | NA | NA |
| SCN4A | 782 | F | 9.906568182 | 12.4683055 | 0.176845006 | 9.958847598 | NA | NA | NA | NA | NA | NA | NA |
| SCN4A | 783 | L | 5.445875 | 13.21921306 | 0.189744565 | 9.390353081 | NA | NA | NA | NA | NA | NA | NA |
| SCN4A | 784 | M | 2.859625 | 13.9196269 | 0.201776711 | 10.20447718 | NA | NA | NA | NA | NA | NA | NA |
| SCN4A | 785 | V | 3.773607143 | 11.08582667 | 0.153095921 | 7.346098441 | NA | NA | NA | NA | NA | NA | NA |
| SCN4A | 786 | M | 2.898875 | 8.130241386 | 0.102323032 | 4.211855949 | 3.92 | -0.79 | -1.02 | 3.38 | 0.76 | 0 | 0 |
| SCN4A | 787 | V | -0.675892857 | 10.41236193 | 0.141526724 | 6.502865228 | NA | NA | NA | NA | NA | NA | NA |
| SCN4A | 788 | I | -1.467 | 11.65048192 | 0.162795922 | 7.858670993 | NA | NA | NA | NA | NA | NA | NA |
| SCN4A | 789 | G | -0.8615 | 9.336955125 | 0.123052714 | 5.538542149 | NA | NA | NA | NA | NA | NA | NA |
| SCN4A | 790 | N | -1.363875 | 5.568291928 | 0.058312265 | 1.644032369 | 4.1 | -0.79 | -1.43 | 3.38 | 0.72 | 0 | 0 |
| SCN4A | 791 | L | -5.734375 | 9.040364926 | 0.117957702 | 5.332890156 | NA | NA | NA | NA | NA | NA | NA |
| SCN4A | 792 | V | -5.279892857 | 10.87495738 | 0.149473477 | 7.347465809 | NA | NA | NA | NA | NA | NA | NA |
| SCN4A | 793 | V | -3.930892857 | 7.364911473 | 0.089175717 | 3.894145855 | 3.54 | 1.21 | 3.47 | 0.2 | -1.8 | 0 | 0 |
| SCN4A | 794 | L | -7.28425 | 4.97672774 | 0.048150006 | 2.589748648 | 2.61 | 1.25 | 3.3 | 0.24 | -1.8 | 0 | 0 |
| SCN4A | 795 | N | -9.965875 | 8.691345465 | 0.111962028 | 6.447801392 | NA | NA | NA | NA | NA | NA | NA |
| SCN4A | 796 | L | -8.558375 | 9.302808071 | 0.122466115 | 6.859830554 | NA | NA | NA | NA | NA | NA | NA |
| SCN4A | 797 | F | -8.344386364 | 4.060165144 | 0.032404721 | 1.771047375 | 2.61 | 1.25 | 3.3 | 0.24 | -1.8 | 0 | 0 |

| | | | | | | | | | | | | | |
|---|---|---|---|---|---|---|---|---|---|---|---|---|---|
| SCN4A | 798 | L | -12.193125 | 4.421901514 | 0.038618854 | 2.02138753 | 2.61 | 1.21 | 3.47 | 0.2 | -1.8 | 0 | 0 |
| SCN4A | 799 | A | -13.71755 | 8.82552633 | 0.114267071 | 5.775910647 | NA | NA | NA | NA | NA | NA | NA |
| SCN4A | 800 | L | -12.078625 | 8.936661289 | 0.116176217 | 5.873949356 | NA | NA | NA | NA | NA | NA | NA |
| SCN4A | 801 | L | -14.111 | 4.6444455 | 0.042441854 | 1.817359457 | 3.11 | 1.36 | 3.73 | 0.19 | -1.81 | 0 | 0 |
| SCN4A | 802 | L | -17.887 | 4.697591936 | 0.043354837 | 1.594644275 | 3.11 | 1.36 | 3.73 | 0.19 | -1.81 | 0 | 0 |
| SCN4A | 803 | S | -17.76425 | 9.157974503 | 0.119978073 | 6.054102156 | NA | NA | NA | NA | NA | NA | NA |
| SCN4A | 804 | S | -17.23791667 | 9.5977364 | 0.127532578 | 6.504373805 | NA | NA | NA | NA | NA | NA | NA |
| SCN4A | 805 | F | -19.68993182 | 4.313275553 | 0.03675281 | 1.37760692 | 2.93 | 1.5 | 3.37 | 0.28 | -1.82 | 0 | 0 |
| SCN4A | 806 | S | -23.66208333 | 10.16689977 | 0.137310021 | 6.933246512 | NA | NA | NA | NA | NA | NA | NA |
| SCN4A | 1014 | G | -22.747 | 58.72015671 | 0.971388182 | 55.77863869 | NA | NA | NA | NA | NA | NA | NA |
| SCN4A | 1015 | K | -26.23508333 | 58.21840763 | 0.962768823 | 55.29140536 | NA | NA | NA | NA | NA | NA | NA |
| SCN4A | 1016 | K | -20.96597222 | 57.81439636 | 0.955828465 | 54.85968657 | NA | NA | NA | NA | NA | NA | NA |
| SCN4A | 1017 | W | -18.42682143 | 53.6696818 | 0.884627971 | 50.69279414 | NA | NA | NA | NA | NA | NA | NA |
| SCN4A | 1018 | W | -23.75296429 | 49.50980076 | 0.813166937 | 46.58525266 | NA | NA | NA | NA | NA | NA | NA |
| SCN4A | 1019 | T | -22.03675 | 53.66145803 | 0.884486698 | 50.71609377 | NA | NA | NA | NA | NA | NA | NA |
| SCN4A | 1020 | L | -17.5195 | 52.48518056 | 0.864279869 | 49.51818265 | NA | NA | NA | NA | NA | NA | NA |
| SCN4A | 1021 | R | -18.87129545 | 47.927002 | 0.785976632 | 44.95662897 | NA | NA | NA | NA | NA | NA | NA |
| SCN4A | 1022 | R | -23.69411364 | 49.06269024 | 0.805486193 | 46.11807754 | NA | NA | NA | NA | NA | NA | NA |
| SCN4A | 1023 | A | -18.83575 | 49.89886144 | 0.819850464 | 46.93505132 | NA | NA | NA | NA | NA | NA | NA |
| SCN4A | 1024 | C | -16.03158333 | 47.40089647 | 0.776938863 | 44.42424879 | NA | NA | NA | NA | NA | NA | NA |
| SCN4A | 1025 | F | -19.66765909 | 42.10710097 | 0.685998738 | 39.16464825 | NA | NA | NA | NA | NA | NA | NA |
| SCN4A | 1026 | K | -20.04141667 | 45.90043983 | 0.751163082 | 42.93617588 | NA | NA | NA | NA | NA | NA | NA |
| SCN4A | 1027 | I | -14.462625 | 44.81835756 | 0.732574397 | 41.98871736 | NA | NA | NA | NA | NA | NA | NA |
| SCN4A | 1028 | V | -14.39560714 | 41.58594639 | 0.67704602 | 38.57701393 | NA | NA | NA | NA | NA | NA | NA |
| SCN4A | 1029 | E | -18.27741667 | 38.23988087 | 0.619565217 | 35.29228632 | NA | NA | NA | NA | NA | NA | NA |
| SCN4A | 1030 | H | -15.99245 | 41.82259322 | 0.681111287 | 39.01075655 | NA | NA | NA | NA | NA | NA | NA |
| SCN4A | 1031 | N | -14.659875 | 38.93365534 | 0.631483308 | 35.91816992 | NA | NA | NA | NA | NA | NA | NA |
| SCN4A | 1032 | W | -11.53303571 | 42.96928261 | 0.700809833 | 39.96888294 | NA | NA | NA | NA | NA | NA | NA |
| SCN4A | 1033 | F | -10.13893182 | 41.67894398 | 0.67864359 | 38.71515908 | NA | NA | NA | NA | NA | NA | NA |
| SCN4A | 1034 | E | -10.60619444 | 35.94343516 | 0.580115439 | 32.97424265 | NA | NA | NA | NA | NA | NA | NA |
| SCN4A | 1035 | T | -8.710321429 | 39.37296153 | 0.639029984 | 36.39159504 | NA | NA | NA | NA | NA | NA | NA |
| SCN4A | 1036 | F | -4.902931818 | 41.32959144 | 0.672642194 | 38.37761143 | NA | NA | NA | NA | NA | NA | NA |
| SCN4A | 1037 | I | -5.535375 | 38.8771029 | 0.630511815 | 36.00119173 | NA | NA | NA | NA | NA | NA | NA |

| | | | | | | | | | | | | | |
|---|---|---|---|---|---|---|---|---|---|---|---|---|---|
| SCN4A | 1038 | V | -5.405607143 | 34.98234535 | 0.563605238 | 32.12480445 | NA | NA | NA | NA | NA | NA | NA |
| SCN4A | 1039 | F | -1.333113636 | 38.50062364 | 0.624044419 | 35.62315646 | NA | NA | NA | NA | NA | NA | NA |
| SCN4A | 1040 | M | -0.589125 | 38.67315498 | 0.62700827 | 36.90829863 | NA | NA | NA | NA | NA | NA | NA |
| SCN4A | 1041 | I | -1.575875 | 34.88353156 | 0.561907753 | 31.76470032 | NA | NA | NA | NA | NA | NA | NA |
| SCN4A | 1042 | L | 1.01125 | 33.4472148 | 0.537233807 | 31.68988583 | NA | NA | NA | NA | NA | NA | NA |
| SCN4A | 1043 | L | 3.43475 | 35.42429377 | 0.571197304 | 31.91903804 | NA | NA | NA | NA | NA | NA | NA |
| SCN4A | 1044 | S | 3.707583333 | 33.72337077 | 0.541977787 | 30.22975608 | NA | NA | NA | NA | NA | NA | NA |
| SCN4A | 1045 | S | 4.023416667 | 30.66517031 | 0.48944211 | 27.16698852 | NA | NA | NA | NA | NA | NA | NA |
| SCN4A | 1046 | G | 6.85475 | 31.52482382 | 0.504209775 | 28.00375395 | NA | NA | NA | NA | NA | NA | NA |
| SCN4A | 1047 | A | 8.85235 | 31.58527616 | 0.505248263 | 28.01334165 | NA | NA | NA | NA | NA | NA | NA |
| SCN4A | 1048 | L | 8.921 | 27.98067233 | 0.443326128 | 24.46359954 | NA | NA | NA | NA | NA | NA | NA |
| SCN4A | 1049 | A | 11.38755 | 26.15088058 | 0.411892823 | 22.44999385 | NA | NA | NA | NA | NA | NA | NA |
| SCN4A | 1050 | F | 13.18920455 | 27.40418443 | 0.433422859 | 23.85816376 | NA | NA | NA | NA | NA | NA | NA |
| SCN4A | 1051 | E | 15.06486111 | 24.91925723 | 0.390735228 | 21.36391087 | NA | NA | NA | NA | NA | NA | NA |
| SCN4A | 1052 | D | 18.120875 | 22.54031987 | 0.349868357 | 18.99124946 | NA | NA | NA | NA | NA | NA | NA |
| SCN4A | 1053 | I | 22.2425 | 18.03130336 | 0.272409656 | 14.5191056 | NA | NA | NA | NA | NA | NA | NA |
| SCN4A | 1054 | Y | 22.672 | 18.15139504 | 0.274472666 | 14.65341421 | NA | NA | NA | NA | NA | NA | NA |
| SCN4A | 1055 | I | 20.417625 | 25.31677442 | 0.397564026 | 21.80327908 | NA | NA | NA | NA | NA | NA | NA |
| SCN4A | 1056 | E | 25.48086111 | 26.31567886 | 0.414723831 | 22.82570115 | NA | NA | NA | NA | NA | NA | NA |
| SCN4A | 1057 | Q | 25.39441667 | 23.63739861 | 0.36871466 | 23.49336299 | NA | NA | NA | NA | NA | NA | NA |
| SCN4A | 1058 | R | 20.68375 | 25.8551767 | 0.406813037 | 22.33878099 | NA | NA | NA | NA | NA | NA | NA |
| SCN4A | 1059 | R | 24.39975 | 29.1959037 | 0.464202131 | 28.07756001 | NA | NA | NA | NA | NA | NA | NA |
| SCN4A | 1060 | V | 19.26103571 | 34.17815645 | 0.549790379 | 30.6765591 | NA | NA | NA | NA | NA | NA | NA |
| SCN4A | 1061 | I | 16.56075 | 32.03569144 | 0.512985778 | 28.51239413 | NA | NA | NA | NA | NA | NA | NA |
| SCN4A | 1062 | R | 19.81147727 | 31.06633788 | 0.496333617 | 27.56759345 | NA | NA | NA | NA | NA | NA | NA |
| SCN4A | 1063 | T | 17.43560714 | 35.4161347 | 0.571057142 | 31.89691163 | NA | NA | NA | NA | NA | NA | NA |
| SCN4A | 1064 | I | 13.7665 | 37.15510066 | 0.600930185 | 33.62837548 | NA | NA | NA | NA | NA | NA | NA |
| SCN4A | 1065 | L | 12.419375 | 32.77549788 | 0.525694635 | 29.22546021 | NA | NA | NA | NA | NA | NA | NA |
| SCN4A | 1066 | E | 15.02308333 | 34.78224826 | 0.560167845 | 31.25707784 | NA | NA | NA | NA | NA | NA | NA |
| SCN4A | 1067 | Y | 13.23133333 | 39.39404781 | 0.639392217 | 35.85292893 | NA | NA | NA | NA | NA | NA | NA |
| SCN4A | 1068 | A | 8.53915 | 39.45711232 | 0.640475579 | 35.90028112 | NA | NA | NA | NA | NA | NA | NA |
| SCN4A | 1069 | D | 8.4875 | 36.64041124 | 0.592088529 | 33.07363325 | NA | NA | NA | NA | NA | NA | NA |
| SCN4A | 1070 | K | 8.87475 | 40.54924168 | 0.659236859 | 37.00373106 | NA | NA | NA | NA | NA | NA | NA |

| | | | | | | | | | | | | | |
|---|---|---|---|---|---|---|---|---|---|---|---|---|---|
| SCN4A | 1071 | V | 4.723464286 | 43.36949451 | 0.707684923 | 39.80519295 | NA | NA | NA | NA | NA | NA | NA |
| SCN4A | 1072 | F | 3.656022727 | 38.96067877 | 0.631947533 | 35.45409127 | NA | NA | NA | NA | NA | NA | NA |
| SCN4A | 1073 | T | 3.245035714 | 40.94441492 | 0.666025392 | 37.4530896 | NA | NA | NA | NA | NA | NA | NA |
| SCN4A | 1074 | Y | 0.707 | 45.02898392 | 0.736192668 | 41.53899371 | NA | NA | NA | NA | NA | NA | NA |
| SCN4A | 1075 | I | -0.928 | 44.81613214 | 0.732536167 | 41.69726402 | NA | NA | NA | NA | NA | NA | NA |
| SCN4A | 1076 | F | -1.326204545 | 40.21564827 | 0.653506183 | 37.39303371 | NA | NA | NA | NA | NA | NA | NA |
| SCN4A | 1077 | I | -1.901875 | 45.806173 | 0.749543707 | 42.92348772 | NA | NA | NA | NA | NA | NA | NA |
| SCN4A | 1078 | M | -4.2295 | 47.39084227 | 0.776766145 | 44.55029221 | NA | NA | NA | NA | NA | NA | NA |
| SCN4A | 1079 | E | -6.615083333 | 40.86119032 | 0.664595707 | 37.92432144 | NA | NA | NA | NA | NA | NA | NA |
| SCN4A | 1080 | M | -6.280125 | 44.13020455 | 0.720752875 | 41.2003661 | NA | NA | NA | NA | NA | NA | NA |
| SCN4A | 1081 | L | -7.161375 | 46.83710097 | 0.767253631 | 43.88824768 | NA | NA | NA | NA | NA | NA | NA |
| SCN4A | 1082 | L | -10.5845 | 46.71472952 | 0.765151458 | 43.73492151 | NA | NA | NA | NA | NA | NA | NA |
| SCN4A | 1083 | K | -11.62619444 | 40.45739983 | 0.657659142 | 37.47271077 | NA | NA | NA | NA | NA | NA | NA |
| SCN4A | 1084 | W | -10.06667857 | 44.6305717 | 0.729348494 | 41.66412966 | NA | NA | NA | NA | NA | NA | NA |
| SCN4A | 1085 | V | -14.19132143 | 48.0853874 | 0.788697475 | 45.09053466 | NA | NA | NA | NA | NA | NA | NA |
| SCN4A | 1086 | A | -16.39995 | 45.76946766 | 0.74891316 | 42.79366729 | NA | NA | NA | NA | NA | NA | NA |
| SCN4A | 1087 | Y | -16.52925 | 39.37170118 | 0.639008333 | 36.39099527 | NA | NA | NA | NA | NA | NA | NA |
| SCN4A | 1088 | G | -16.18925 | 44.697758 | 0.730502662 | 41.71659652 | NA | NA | NA | NA | NA | NA | NA |
| SCN4A | 1089 | F | -13.70529545 | 44.5579805 | 0.728101477 | 41.60534619 | NA | NA | NA | NA | NA | NA | NA |
| SCN4A | 1090 | K | -17.15575 | 41.00827519 | 0.667122423 | 38.02635617 | NA | NA | NA | NA | NA | NA | NA |
| SCN4A | 1091 | V | -18.13175 | 39.20893513 | 0.636212236 | 36.22636868 | NA | NA | NA | NA | NA | NA | NA |
| SCN4A | 1092 | Y | -11.83625 | 38.8886643 | 0.630710424 | 35.88891603 | NA | NA | NA | NA | NA | NA | NA |
| SCN4A | 1093 | F | -11.81593182 | 38.61838268 | 0.626067358 | 35.61460225 | NA | NA | NA | NA | NA | NA | NA |
| SCN4A | 1094 | T | -16.56275 | 35.63628585 | 0.574839036 | 32.65433231 | NA | NA | NA | NA | NA | NA | NA |
| SCN4A | 1095 | N | -14.417625 | 32.90428887 | 0.527907087 | 29.89349051 | NA | NA | NA | NA | NA | NA | NA |
| SCN4A | 1096 | A | -11.41835 | 32.01580616 | 0.512644176 | 29.05188471 | NA | NA | NA | NA | NA | NA | NA |
| SCN4A | 1097 | W | -10.87746429 | 27.31636901 | 0.431914311 | 24.35436579 | NA | NA | NA | NA | NA | NA | NA |
| SCN4A | 1098 | C | -10.19791667 | 35.25010845 | 0.56820504 | 32.27904858 | NA | NA | NA | NA | NA | NA | NA |
| SCN4A | 1099 | W | -8.82875 | 35.82074082 | 0.578007719 | 32.86478287 | NA | NA | NA | NA | NA | NA | NA |
| SCN4A | 1100 | L | -5.243125 | 30.96539402 | 0.49459954 | 28.08783557 | NA | NA | NA | NA | NA | NA | NA |
| SCN4A | 1101 | D | -5.31625 | 35.23866608 | 0.568008476 | 32.3764759 | NA | NA | NA | NA | NA | NA | NA |
| SCN4A | 1102 | F | -4.727659091 | 38.58476343 | 0.625489825 | 35.71637774 | NA | NA | NA | NA | NA | NA | NA |
| SCN4A | 1103 | L | -2.029375 | 36.60899798 | 0.591548893 | 33.48127604 | NA | NA | NA | NA | NA | NA | NA |

| | | | | | | | | | | | | | |
|---|---|---|---|---|---|---|---|---|---|---|---|---|---|
| SCN4A | 1104 | I | 0.539125 | 35.13369741 | 0.566205258 | 32.55714887 | NA | NA | NA | NA | NA | NA | NA |
| SCN4A | 1105 | V | 0.656321429 | 39.97231743 | 0.649326094 | 37.25028177 | NA | NA | NA | NA | NA | NA | NA |
| SCN4A | 1106 | D | 1.149 | 40.84175816 | 0.66426189 | 37.37455336 | NA | NA | NA | NA | NA | NA | NA |
| SCN4A | 1107 | V | 3.948321429 | 36.14406282 | 0.583561946 | 32.6711672 | NA | NA | NA | NA | NA | NA | NA |
| SCN4A | 1108 | S | 4.340083333 | 39.22048271 | 0.636410608 | 35.74445468 | NA | NA | NA | NA | NA | NA | NA |
| SCN4A | 1109 | I | 4.074 | 41.608024 | 0.677425283 | 38.12708573 | NA | NA | NA | NA | NA | NA | NA |
| SCN4A | 1110 | I | 6.408375 | 40.60604204 | 0.660212611 | 37.11982565 | NA | NA | NA | NA | NA | NA | NA |
| SCN4A | 1111 | S | 9.114416667 | 37.96387011 | 0.614823732 | 34.40635015 | NA | NA | NA | NA | NA | NA | NA |
| SCN4A | 1112 | L | 8.299875 | 41.0742981 | 0.668256606 | 37.51842819 | NA | NA | NA | NA | NA | NA | NA |
| SCN4A | 1113 | V | 8.549607143 | 43.57475051 | 0.711210938 | 40.02259734 | NA | NA | NA | NA | NA | NA | NA |
| SCN4A | 1114 | A | 11.94235 | 41.34030365 | 0.672826215 | 37.80695742 | NA | NA | NA | NA | NA | NA | NA |
| SCN4A | 1115 | N | 14.023125 | 38.75558605 | 0.628424323 | 35.21471034 | NA | NA | NA | NA | NA | NA | NA |
| SCN4A | 1116 | W | 11.47625 | 43.14950497 | 0.703905805 | 39.61461798 | NA | NA | NA | NA | NA | NA | NA |
| SCN4A | 1117 | L | 13.304875 | 44.44958085 | 0.72623932 | 40.92056927 | NA | NA | NA | NA | NA | NA | NA |
| SCN4A | 1118 | G | 16.04975 | 42.73631376 | 0.696807749 | 39.22036035 | NA | NA | NA | NA | NA | NA | NA |
| SCN4A | 1119 | Y | 12.97033333 | 40.95950182 | 0.666284564 | 37.43364214 | NA | NA | NA | NA | NA | NA | NA |
| SCN4A | 1120 | S | 16.64825 | 37.64699287 | 0.609380217 | 34.11781066 | NA | NA | NA | NA | NA | NA | NA |
| SCN4A | 1121 | E | 18.64397222 | 35.0447132 | 0.564676632 | 31.53439897 | NA | NA | NA | NA | NA | NA | NA |
| SCN4A | 1122 | L | 19.01975 | 34.6731925 | 0.558294418 | 31.1698747 | NA | NA | NA | NA | NA | NA | NA |
| SCN4A | 1123 | G | 16.02075 | 33.02757689 | 0.530025005 | 29.48830759 | NA | NA | NA | NA | NA | NA | NA |
| SCN4A | 1124 | P | 12.82189286 | 35.1729643 | 0.56687981 | 31.5002695 | NA | NA | NA | NA | NA | NA | NA |
| SCN4A | 1125 | I | 12.945125 | 34.34612972 | 0.552675929 | 30.79834042 | NA | NA | NA | NA | NA | NA | NA |
| SCN4A | 1126 | K | 16.43119444 | 26.83501761 | 0.423645356 | 23.30232093 | NA | NA | NA | NA | NA | NA | NA |
| SCN4A | 1127 | S | 12.05108333 | 31.12356978 | 0.497316782 | 27.8993168 | NA | NA | NA | NA | NA | NA | NA |
| SCN4A | 1128 | L | 8.9925 | 33.30180395 | 0.534735849 | 29.80662617 | NA | NA | NA | NA | NA | NA | NA |
| SCN4A | 1129 | R | 12.51638636 | 30.91104891 | 0.493665966 | 27.49656643 | NA | NA | NA | NA | NA | NA | NA |
| SCN4A | 1130 | T | 10.88775 | 27.6328834 | 0.437351592 | 24.12966245 | NA | NA | NA | NA | NA | NA | NA |
| SCN4A | 1131 | L | 6.518875 | 30.3818807 | 0.484575584 | 26.9020576 | NA | NA | NA | NA | NA | NA | NA |
| SCN4A | 1132 | R | 8.977931818 | 30.33429496 | 0.483758127 | 27.31914245 | NA | NA | NA | NA | NA | NA | NA |
| SCN4A | 1133 | A | 6.05375 | 27.27863536 | 0.431266098 | 23.80501251 | NA | NA | NA | NA | NA | NA | NA |
| SCN4A | 1134 | L | 2.525 | 29.20758117 | 0.464402734 | 26.46683959 | NA | NA | NA | NA | NA | NA | NA |
| SCN4A | 1135 | R | 3.612568182 | 32.34921283 | 0.518371644 | 29.67566259 | NA | NA | NA | NA | NA | NA | NA |
| SCN4A | 1136 | P | 1.47975 | 29.75272147 | 0.473767494 | 27.97102248 | NA | NA | NA | NA | NA | NA | NA |

| | | | | | | | | | | | | | |
|---|---|---|---|---|---|---|---|---|---|---|---|---|---|
| SCN4A | 1137 | L | -1.6275 | 28.62459359 | 0.454387809 | 25.50626479 | NA | NA | NA | NA | NA | NA | NA |
| SCN4A | 1138 | R | -2.206022727 | 32.8198392 | 0.526456357 | 29.75297576 | NA | NA | NA | NA | NA | NA | NA |
| SCN4A | 1139 | A | -4.51635 | 29.9107536 | 0.476482269 | 27.04516439 | NA | NA | NA | NA | NA | NA | NA |
| SCN4A | 1140 | L | -6.497875 | 26.2523437 | 0.41363582 | 23.39250689 | NA | NA | NA | NA | NA | NA | NA |
| SCN4A | 1141 | S | -9.127916667 | 29.95796744 | 0.477293338 | 27.02237311 | NA | NA | NA | NA | NA | NA | NA |
| SCN4A | 1142 | R | -8.82275 | 30.9081365 | 0.493615935 | 27.94551738 | NA | NA | NA | NA | NA | NA | NA |
| SCN4A | 1143 | F | -7.942295455 | 27.46035726 | 0.434387831 | 24.48782011 | NA | NA | NA | NA | NA | NA | NA |
| SCN4A | 1144 | E | -14.76319444 | 25.01285148 | 0.392343048 | 22.00266728 | NA | NA | NA | NA | NA | NA | NA |
| SCN4A | 1145 | G | -11.44425 | 22.23870448 | 0.344687019 | 19.26581979 | NA | NA | NA | NA | NA | NA | NA |
| SCN4A | 1146 | M | -8.143875 | 23.35501432 | 0.363863687 | 20.40169099 | NA | NA | NA | NA | NA | NA | NA |
| SCN4A | 1147 | R | -13.35620455 | 23.25309141 | 0.362112791 | 20.24340089 | NA | NA | NA | NA | NA | NA | NA |
| SCN4A | 1148 | V | -14.87717857 | 19.94438129 | 0.305273703 | 16.91497843 | NA | NA | NA | NA | NA | NA | NA |
| SCN4A | 1149 | V | -11.30960714 | 18.16141674 | 0.274644825 | 15.22177862 | NA | NA | NA | NA | NA | NA | NA |
| SCN4A | 1150 | V | -10.67289286 | 20.84898117 | 0.320813485 | 17.88110485 | NA | NA | NA | NA | NA | NA | NA |
| SCN4A | 1151 | N | -14.96675 | 20.46013903 | 0.314133711 | 17.46517229 | NA | NA | NA | NA | NA | NA | NA |
| SCN4A | 1152 | A | -14.26315 | 16.79134599 | 0.251108894 | 13.76537533 | NA | NA | NA | NA | NA | NA | NA |
| SCN4A | 1153 | L | -10.381 | 16.23119299 | 0.241486236 | 13.29419482 | NA | NA | NA | NA | NA | NA | NA |
| SCN4A | 1154 | L | -12.13175 | 19.57888138 | 0.298994917 | 16.59085017 | NA | NA | NA | NA | NA | NA | NA |
| SCN4A | 1155 | G | -15.419 | 17.78774823 | 0.268225714 | 14.93621406 | NA | NA | NA | NA | NA | NA | NA |
| SCN4A | 1156 | A | -13.28355 | 15.26696601 | 0.224922143 | 12.25242913 | NA | NA | NA | NA | NA | NA | NA |
| SCN4A | 1157 | I | -10.932 | 18.33149511 | 0.277566537 | 15.32734103 | NA | NA | NA | NA | NA | NA | NA |
| SCN4A | 1158 | P | -14.36603571 | 19.54492371 | 0.298411571 | 16.49596381 | NA | NA | NA | NA | NA | NA | NA |
| SCN4A | 1159 | S | -13.72958333 | 15.44917972 | 0.228052324 | 12.49083022 | NA | NA | NA | NA | NA | NA | NA |
| SCN4A | 1160 | I | -9.696125 | 15.06653739 | 0.221479055 | 12.16119526 | NA | NA | NA | NA | NA | NA | NA |
| SCN4A | 1161 | M | -8.76725 | 18.80873297 | 0.285764827 | 15.88738003 | NA | NA | NA | NA | NA | NA | NA |
| SCN4A | 1162 | N | -11.91625 | 18.81235397 | 0.285827031 | 15.8437771 | NA | NA | NA | NA | NA | NA | NA |
| SCN4A | 1163 | V | -8.68275 | 14.51123337 | 0.211939696 | 11.3881233 | NA | NA | NA | NA | NA | NA | NA |
| SCN4A | 1164 | L | -4.8175 | 17.72654484 | 0.267174324 | 15.35689271 | NA | NA | NA | NA | NA | NA | NA |
| SCN4A | 1165 | L | -6.528625 | 20.14697977 | 0.308754066 | 17.82249462 | NA | NA | NA | NA | NA | NA | NA |
| SCN4A | 1166 | V | -6.391178571 | 18.34500747 | 0.277798661 | 16.05302556 | NA | NA | NA | NA | NA | NA | NA |
| SCN4A | 1167 | C | -3.034083333 | 15.16517066 | 0.223173439 | 11.92017231 | NA | NA | NA | NA | NA | NA | NA |
| SCN4A | 1168 | L | -1.119 | 19.48113962 | 0.297315849 | 16.08446097 | NA | NA | NA | NA | NA | NA | NA |
| SCN4A | 1169 | I | -2.9455 | 21.58142278 | 0.333395824 | 18.20121023 | NA | NA | NA | NA | NA | NA | NA |

| | | | | | | | | | | | | | |
|---|---|---|---|---|---|---|---|---|---|---|---|---|---|
| SCN4A | 1170 | F | -2.167659091 | 16.82472814 | 0.251682354 | 13.39097307 | NA | NA | NA | NA | NA | NA | NA |
| SCN4A | 1171 | W | 1.251321429 | 14.30147478 | 0.208336332 | 10.85988782 | NA | NA | NA | NA | NA | NA | NA |
| SCN4A | 1172 | L | 1.795375 | 22.06058904 | 0.341627241 | 18.58150435 | NA | NA | NA | NA | NA | NA | NA |
| SCN4A | 1173 | I | 1.64425 | 23.44332084 | 0.365380671 | 19.96474043 | NA | NA | NA | NA | NA | NA | NA |
| SCN4A | 1174 | F | 3.197840909 | 18.60180704 | 0.282210124 | 14.48731809 | NA | NA | NA | NA | NA | NA | NA |
| SCN4A | 1175 | S | 6.43225 | 20.83941108 | 0.320649084 | 18.07078305 | NA | NA | NA | NA | NA | NA | NA |
| SCN4A | 1176 | I | 6.327375 | 23.99620524 | 0.374878465 | 21.23830999 | NA | NA | NA | NA | NA | NA | NA |
| SCN4A | 1177 | M | 6.893625 | 23.45233204 | 0.365535471 | 20.70179262 | NA | NA | NA | NA | NA | NA | NA |
| SCN4A | 1178 | G | 10.60675 | 21.49516087 | 0.331913963 | 18.75591666 | NA | NA | NA | NA | NA | NA | NA |
| SCN4A | 1179 | V | 11.78203571 | 23.01077758 | 0.357950173 | 20.28495565 | NA | NA | NA | NA | NA | NA | NA |
| SCN4A | 1180 | N | 11.002875 | 25.41728872 | 0.399290724 | 22.69407333 | NA | NA | NA | NA | NA | NA | NA |
| SCN4A | 1181 | L | 12.849625 | 22.99472535 | 0.357674418 | 20.24628875 | NA | NA | NA | NA | NA | NA | NA |
| SCN4A | 1182 | F | 14.75138636 | 19.16429352 | 0.291872868 | 16.43370386 | NA | NA | NA | NA | NA | NA | NA |
| SCN4A | 1183 | A | 16.98355 | 23.35276348 | 0.36382502 | 20.67419644 | NA | NA | NA | NA | NA | NA | NA |
| SCN4A | 1184 | G | 20.735 | 23.81811827 | 0.371819175 | 19.97277326 | NA | NA | NA | NA | NA | NA | NA |
| SCN4A | 1185 | K | 19.95619444 | 20.86458706 | 0.321081572 | 17.08181461 | NA | NA | NA | NA | NA | NA | NA |
| SCN4A | 1186 | F | 20.08193182 | 16.79887618 | 0.251238253 | 13.17008217 | NA | NA | NA | NA | NA | NA | NA |
| SCN4A | 1187 | Y | 24.965 | 20.02076727 | 0.306585909 | 16.25537651 | NA | NA | NA | NA | NA | NA | NA |
| SCN4A | 1188 | Y | 27.59591667 | 13.01481721 | 0.186233325 | 9.284292316 | NA | NA | NA | NA | NA | NA | NA |
| SCN4A | 1189 | C | 32.48925 | 18.63301481 | 0.282746231 | 14.90127319 | NA | NA | NA | NA | NA | NA | NA |
| SCN4A | 1190 | I | 33.18925 | 13.85046887 | 0.200588671 | 10.19426463 | NA | NA | NA | NA | NA | NA | NA |
| SCN4A | 1191 | N | 37.7585 | 15.25618409 | 0.224736925 | 12.34094955 | NA | NA | NA | NA | NA | NA | NA |
| SCN4A | 1192 | T | 37.34917857 | 12.8959135 | 0.184190723 | 10.98123925 | NA | NA | NA | NA | NA | NA | NA |
| SCN4A | 1193 | T | 39.64003571 | 12.19814314 | 0.172203988 | 10.02003083 | NA | NA | NA | NA | NA | NA | NA |
| SCN4A | 1194 | T | 37.79232143 | 12.58347702 | 0.178823494 | 10.65312951 | NA | NA | NA | NA | NA | NA | NA |
| SCN4A | 1195 | S | 33.70441667 | 10.57655374 | 0.144347313 | 7.471579514 | NA | NA | NA | NA | NA | NA | NA |
| SCN4A | 1196 | E | 34.24352778 | 13.921636 | 0.201811224 | 10.19107922 | NA | NA | NA | NA | NA | NA | NA |
| SCN4A | 1197 | R | 29.74584091 | 15.31143383 | 0.225686039 | 11.60292587 | NA | NA | NA | NA | NA | NA | NA |
| SCN4A | 1198 | F | 35.53747727 | 18.84433445 | 0.286376412 | 15.13905925 | NA | NA | NA | NA | NA | NA | NA |
| SCN4A | 1199 | D | 33.53025 | 21.3354322 | 0.329170044 | 17.27324741 | NA | NA | NA | NA | NA | NA | NA |
| SCN4A | 1200 | I | 31.89 | 23.5826582 | 0.367774295 | 19.43460422 | NA | NA | NA | NA | NA | NA | NA |
| SCN4A | 1201 | S | 36.34641667 | 26.07881819 | 0.41065489 | 21.95466626 | NA | NA | NA | NA | NA | NA | NA |
| SCN4A | 1202 | E | 37.70252778 | 24.02643765 | 0.375397816 | 20.34463578 | NA | NA | NA | NA | NA | NA | NA |

| | | | | | | | | | | | | | |
|---|---|---|---|---|---|---|---|---|---|---|---|---|---|
| SCN4A | 1203 | V | 35.19317857 | 24.65702857 | 0.3862305 | 20.93550829 | NA | NA | NA | NA | NA | NA | NA |
| SCN4A | 1204 | N | 32.305125 | 26.39114353 | 0.41602021 | 22.19477155 | NA | NA | NA | NA | NA | NA | NA |
| SCN4A | 1205 | N | 31.658375 | 26.36239583 | 0.415526364 | 22.29801418 | NA | NA | NA | NA | NA | NA | NA |
| SCN4A | 1206 | K | 30.85997222 | 27.00648805 | 0.426590982 | 23.27656166 | NA | NA | NA | NA | NA | NA | NA |
| SCN4A | 1207 | S | 34.50808333 | 29.22807782 | 0.464754838 | 25.49899945 | NA | NA | NA | NA | NA | NA | NA |
| SCN4A | 1208 | E | 36.78597222 | 28.10435891 | 0.445450893 | 24.37459353 | NA | NA | NA | NA | NA | NA | NA |
| SCN4A | 1209 | C | 35.58808333 | 23.3400057 | 0.363605859 | 19.67129395 | NA | NA | NA | NA | NA | NA | NA |
| SCN4A | 1210 | E | 35.85719444 | 27.25339146 | 0.430832443 | 24.07004633 | NA | NA | NA | NA | NA | NA | NA |
| SCN4A | 1211 | S | 40.17725 | 28.80899047 | 0.457555494 | 25.79132436 | NA | NA | NA | NA | NA | NA | NA |
| SCN4A | 1212 | L | 40.566625 | 24.55475044 | 0.384473503 | 21.62272166 | NA | NA | NA | NA | NA | NA | NA |
| SCN4A | 1213 | M | 39.1565 | 23.18214123 | 0.360893965 | 20.05406604 | NA | NA | NA | NA | NA | NA | NA |
| SCN4A | 1214 | H | 44.64795 | 25.559799 | 0.401738854 | 24.24118357 | NA | NA | NA | NA | NA | NA | NA |
| SCN4A | 1215 | T | 44.21032143 | 21.76117692 | 0.336483753 | 20.36018938 | NA | NA | NA | NA | NA | NA | NA |
| SCN4A | 1216 | G | 41.81925 | 19.32934603 | 0.294708243 | 17.80257768 | NA | NA | NA | NA | NA | NA | NA |
| SCN4A | 1217 | Q | 41.78686111 | 18.19499022 | 0.275221571 | 16.59971562 | NA | NA | NA | NA | NA | NA | NA |
| SCN4A | 1218 | V | 37.88646429 | 19.86431547 | 0.303898282 | 16.96353694 | NA | NA | NA | NA | NA | NA | NA |
| SCN4A | 1219 | R | 32.42884091 | 14.18685381 | 0.206367301 | 10.90060004 | NA | NA | NA | NA | NA | NA | NA |
| SCN4A | 1220 | W | 30.08996429 | 20.22092802 | 0.310024395 | 16.60782721 | NA | NA | NA | NA | NA | NA | NA |
| SCN4A | 1221 | L | 28.231375 | 14.58436416 | 0.213195982 | 10.93586615 | NA | NA | NA | NA | NA | NA | NA |
| SCN4A | 1222 | N | 23.516 | 15.35701433 | 0.226469049 | 11.41454813 | NA | NA | NA | NA | NA | NA | NA |
| SCN4A | 1223 | V | 21.30646429 | 13.26018684 | 0.190448438 | 9.415082372 | NA | NA | NA | NA | NA | NA | NA |
| SCN4A | 1224 | K | 20.81419444 | 12.24715796 | 0.173045995 | 8.601565013 | NA | NA | NA | NA | NA | NA | NA |
| SCN4A | 1225 | V | 16.04860714 | 12.49326451 | 0.177273767 | 8.872829676 | NA | NA | NA | NA | NA | NA | NA |
| SCN4A | 1226 | N | 16.652 | 16.07068057 | 0.238728854 | 13.62168736 | NA | NA | NA | NA | NA | NA | NA |
| SCN4A | 1227 | Y | 12.88016667 | 16.20863307 | 0.241098688 | 13.46440295 | NA | NA | NA | NA | NA | NA | NA |
| SCN4A | 1228 | D | 16.562125 | 21.41744303 | 0.330578877 | 18.75721311 | NA | NA | NA | NA | NA | NA | NA |
| SCN4A | 1229 | N | 12.761375 | 22.23269127 | 0.344583721 | 19.52884915 | NA | NA | NA | NA | NA | NA | NA |
| SCN4A | 1230 | V | 8.072035714 | 22.03833027 | 0.341244866 | 19.30381266 | NA | NA | NA | NA | NA | NA | NA |
| SCN4A | 1231 | G | 9.4155 | 19.94016535 | 0.305201279 | 17.22596719 | NA | NA | NA | NA | NA | NA | NA |
| SCN4A | 1232 | L | 12.948125 | 17.50786635 | 0.263417728 | 14.81025791 | NA | NA | NA | NA | NA | NA | NA |
| SCN4A | 1233 | G | 9.529 | 17.29949736 | 0.259838236 | 14.55439808 | NA | NA | NA | NA | NA | NA | NA |
| SCN4A | 1234 | Y | 5.137916667 | 16.59685539 | 0.247767813 | 13.83183277 | NA | NA | NA | NA | NA | NA | NA |
| SCN4A | 1235 | L | 10.356875 | 13.26813393 | 0.190584958 | 10.54037229 | NA | NA | NA | NA | NA | NA | NA |

| | | | | | | | | | | | | | |
|---|---|---|---|---|---|---|---|---|---|---|---|---|---|
| SCN4A | 1236 | S | 10.95191667 | 12.924098 | 0.184674894 | 10.17247009 | NA | NA | NA | NA | NA | NA | NA |
| SCN4A | 1237 | L | 6.861 | 13.78052916 | 0.199387203 | 11.00929128 | NA | NA | NA | NA | NA | NA | NA |
| SCN4A | 1238 | L | 6.292 | 11.07524293 | 0.152914107 | 8.387242515 | NA | NA | NA | NA | NA | NA | NA |
| SCN4A | 1239 | Q | 10.76886111 | 8.37216137 | 0.106478885 | 5.5868701 | NA | NA | NA | NA | NA | NA | NA |
| SCN4A | 1240 | V | 8.07375 | 9.715408483 | 0.129554022 | 7.655731324 | NA | NA | NA | NA | NA | NA | NA |
| SCN4A | 1241 | A | 4.54575 | 8.791679532 | 0.113685629 | 6.567531631 | NA | NA | NA | NA | NA | NA | NA |
| SCN4A | 1242 | T | 5.696178571 | 6.058940666 | 0.066740935 | 3.884778258 | 2.3 | -0.7 | -1.28 | 14.91 | 0.97 | 1 | 1 |
| SCN4A | 1243 | F | 6.479659091 | 4.547013965 | 0.040768114 | 3.241414354 | 1.96 | -0.79 | -2.05 | 26.54 | 1.33 | 2 | 0 |
| SCN4A | 1244 | K | 9.606527778 | 3.583302667 | 0.02421288 | 1.520968251 | 2.76 | -0.4 | -3.05 | 37.73 | 1.12 | 3 | -1 |
| SCN4A | 1245 | G | 13.58475 | 4.60660059 | 0.041791731 | 1.881051835 | 2.79 | -0.29 | -2.28 | 26.05 | 0.87 | 2 | 0 |
| SCN4A | 1246 | W | 11.13125 | 5.804863995 | 0.062376247 | 3.022761106 | 2.79 | -0.29 | -2.28 | 26.05 | 0.87 | 2 | 0 |
| SCN4A | 1247 | M | 15.53125 | 5.819213864 | 0.062622758 | 3.086962826 | 2.79 | -0.29 | -2.28 | 26.05 | 0.87 | 2 | 0 |
| SCN4A | 1248 | D | 18.584875 | 6.891969095 | 0.081051218 | 4.990600959 | 2.54 | -0.36 | -2.77 | 33.74 | 1.14 | 2 | -2 |
| SCN4A | 1249 | I | 15.12675 | 10.6216342 | 0.145121733 | 7.963749763 | NA | NA | NA | NA | NA | NA | NA |
| SCN4A | 1250 | M | 14.48725 | 11.20136585 | 0.155080725 | 8.432989259 | NA | NA | NA | NA | NA | NA | NA |
| SCN4A | 1251 | Y | 19.14283333 | 9.894657447 | 0.132633273 | 7.177399459 | NA | NA | NA | NA | NA | NA | NA |
| SCN4A | 1252 | A | 19.76855 | 14.64788043 | 0.214287105 | 10.97191645 | NA | NA | NA | NA | NA | NA | NA |
| SCN4A | 1253 | A | 17.19655 | 16.31737605 | 0.242966743 | 13.64160907 | NA | NA | NA | NA | NA | NA | NA |
| SCN4A | 1254 | V | 18.41146429 | 17.24037851 | 0.258822655 | 14.51295749 | NA | NA | NA | NA | NA | NA | NA |
| SCN4A | 1255 | D | 22.45025 | 16.10715894 | 0.239355502 | 11.79261125 | NA | NA | NA | NA | NA | NA | NA |
| SCN4A | 1256 | S | 20.68408333 | 20.92208711 | 0.322069344 | 16.9581479 | NA | NA | NA | NA | NA | NA | NA |
| SCN4A | 1257 | R | 24.94256818 | 23.95098132 | 0.37410158 | 19.98438206 | NA | NA | NA | NA | NA | NA | NA |
| SCN4A | 1258 | E | 22.30052778 | 25.86291803 | 0.406946022 | 21.89043787 | NA | NA | NA | NA | NA | NA | NA |
| SCN4A | 1259 | K | 21.49230556 | 26.74413225 | 0.42208407 | 22.94162687 | NA | NA | NA | NA | NA | NA | NA |
| SCN4A | 1260 | E | 26.84586111 | 26.79958255 | 0.42303663 | 22.55768913 | NA | NA | NA | NA | NA | NA | NA |
| SCN4A | 1261 | E | 26.75452778 | 23.73808899 | 0.370444382 | 19.44477175 | NA | NA | NA | NA | NA | NA | NA |
| SCN4A | 1262 | Q | 29.43308333 | 19.87267393 | 0.304041869 | 15.739549 | NA | NA | NA | NA | NA | NA | NA |
| SCN4A | 1263 | P | 26.47732143 | 19.77259467 | 0.302322645 | 15.46320166 | NA | NA | NA | NA | NA | NA | NA |
| SCN4A | 1264 | Q | 27.04286111 | 20.16913717 | 0.3091347 | 15.85594134 | NA | NA | NA | NA | NA | NA | NA |
| SCN4A | 1265 | Y | 25.01991667 | 15.88880062 | 0.235604406 | 11.59285494 | NA | NA | NA | NA | NA | NA | NA |
| SCN4A | 1266 | E | 21.87441667 | 18.30792659 | 0.277161663 | 14.01642453 | NA | NA | NA | NA | NA | NA | NA |
| SCN4A | 1267 | V | 22.36417857 | 22.18645407 | 0.343789429 | 18.27655719 | NA | NA | NA | NA | NA | NA | NA |
| SCN4A | 1268 | N | 18.104 | 21.1625924 | 0.326200894 | 18.43494765 | NA | NA | NA | NA | NA | NA | NA |

| | | | | | | | | | | | | | |
|---|---|---|---|---|---|---|---|---|---|---|---|---|---|
| SCN4A | 1269 | L | 17.008875 | 19.92766198 | 0.304986488 | 17.1785007 | NA | NA | NA | NA | NA | NA | NA |
| SCN4A | 1270 | Y | 13.77041667 | 21.48960895 | 0.331818589 | 18.72500365 | NA | NA | NA | NA | NA | NA | NA |
| SCN4A | 1271 | M | 13.23675 | 18.99160172 | 0.288906261 | 16.22280156 | NA | NA | NA | NA | NA | NA | NA |
| SCN4A | 1272 | Y | 13.46516667 | 14.3331259 | 0.208880055 | 11.57281647 | NA | NA | NA | NA | NA | NA | NA |
| SCN4A | 1273 | L | 8.1695 | 18.91128893 | 0.287526598 | 17.11377515 | NA | NA | NA | NA | NA | NA | NA |
| SCN4A | 1274 | Y | 8.175166667 | 17.83596092 | 0.269053942 | 15.45107697 | NA | NA | NA | NA | NA | NA | NA |
| SCN4A | 1275 | F | 9.76975 | 14.18950686 | 0.206412877 | 12.45654258 | NA | NA | NA | NA | NA | NA | NA |
| SCN4A | 1276 | V | 5.601464286 | 15.27002295 | 0.224974657 | 13.17363697 | NA | NA | NA | NA | NA | NA | NA |
| SCN4A | 1277 | I | 3.174375 | 16.26300799 | 0.242032774 | 11.83841346 | NA | NA | NA | NA | NA | NA | NA |
| SCN4A | 1278 | F | 3.430568182 | 14.13743064 | 0.205518279 | 9.711408869 | NA | NA | NA | NA | NA | NA | NA |
| SCN4A | 1279 | I | 3.092 | 10.6980279 | 0.146434072 | 6.048512881 | NA | NA | NA | NA | NA | NA | NA |
| SCN4A | 1280 | I | -0.75025 | 12.36014781 | 0.174987005 | 8.369578541 | NA | NA | NA | NA | NA | NA | NA |
| SCN4A | 1281 | F | -1.696568182 | 13.67336718 | 0.197546308 | 10.22086499 | NA | NA | NA | NA | NA | NA | NA |
| SCN4A | 1282 | G | -0.3105 | 12.66919248 | 0.180295968 | 9.190544921 | NA | NA | NA | NA | NA | NA | NA |
| SCN4A | 1283 | S | -1.092416667 | 8.77133479 | 0.113336135 | 5.312765972 | NA | NA | NA | NA | NA | NA | NA |
| SCN4A | 1284 | F | -4.044931818 | 7.481573698 | 0.091179813 | 4.070890002 | 3.54 | 1.21 | 3.47 | 0.2 | -1.8 | 0 | 0 |
| SCN4A | 1285 | F | -5.788295455 | 10.55091138 | 0.143906812 | 7.208633091 | NA | NA | NA | NA | NA | NA | NA |
| SCN4A | 1286 | T | -4.791321429 | 10.56386378 | 0.144129317 | 7.459213951 | NA | NA | NA | NA | NA | NA | NA |
| SCN4A | 1287 | L | -5.256875 | 4.870678905 | 0.046328233 | 3.179896999 | 2.61 | 1.25 | 3.3 | 0.24 | -1.8 | 0 | 0 |
| SCN4A | 1288 | N | -9.368875 | 8.284848399 | 0.104978968 | 5.950173441 | NA | NA | NA | NA | NA | NA | NA |
| SCN4A | 1289 | L | -10.0835 | 10.5348345 | 0.143630634 | 7.555019759 | NA | NA | NA | NA | NA | NA | NA |
| SCN4A | 1290 | F | -8.350022727 | 9.415541726 | 0.124402724 | 6.327967116 | NA | NA | NA | NA | NA | NA | NA |
| SCN4A | 1291 | I | -10.45425 | 4.613158788 | 0.041904391 | 1.630363256 | 3.01 | 1.45 | 3.88 | 0.17 | -1.82 | 0 | 0 |
| SCN4A | 1292 | G | -14.243 | 7.6539659 | 0.094141274 | 4.569493122 | 3.11 | 1.36 | 3.73 | 0.19 | -1.81 | 0 | 0 |
| SCN4A | 1293 | V | -14.27775 | 9.628823604 | 0.128066613 | 6.535694776 | NA | NA | NA | NA | NA | NA | NA |
| SCN4A | 1294 | I | -13.344875 | 8.587455094 | 0.110177335 | 5.565218123 | NA | NA | NA | NA | NA | NA | NA |
| SCN4A | 1295 | I | -16.383125 | 4.243750228 | 0.03555846 | 1.335213612 | 2.93 | 1.5 | 3.37 | 0.28 | -1.82 | 0 | 0 |
| SCN4A | 1296 | D | -19.94275 | 7.702491155 | 0.094974871 | 4.953354957 | 3.06 | 0.94 | 1.65 | 1.05 | -1.23 | 0 | 0 |
| SCN4A | 1297 | N | -17.711375 | 10.09439711 | 0.136064525 | 7.207770848 | NA | NA | NA | NA | NA | NA | NA |
| SCN4A | 1298 | F | -16.55820455 | 9.440753254 | 0.124835823 | 6.460207219 | NA | NA | NA | NA | NA | NA | NA |
| SCN4A | 1299 | N | -21.731 | 4.804064217 | 0.045183884 | 1.856954548 | 3.06 | 0.94 | 1.65 | 1.05 | -1.23 | 0 | 0 |
| SCN4A | 1300 | Q | -22.71741667 | 10.30210221 | 0.139632613 | 7.390901095 | NA | NA | NA | NA | NA | NA | NA |
| SCN4A | 1301 | Q | -20.04697222 | 12.5680117 | 0.178557821 | 9.629608374 | NA | NA | NA | NA | NA | NA | NA |

| | | | | | | | | | | | | | |
|---|---|---|---|---|---|---|---|---|---|---|---|---|---|
| SCN4A | 1302 | K | -22.55930556 | 9.912170146 | 0.132934117 | 6.955796424 | NA | NA | NA | NA | NA | NA | NA |
| SCN4A | 1303 | K | -27.65363889 | 8.716917976 | 0.112401329 | 1.762931197 | 7.05 | -0.77 | -2.9 | 26.11 | 1.25 | 2 | 0 |
| SCN4A | 1304 | K | -25.61486111 | 14.32603525 | 0.208758247 | 7.344461697 | NA | NA | NA | NA | NA | NA | NA |
| SCN4A | 1305 | L | -24.05025 | 16.50441059 | 0.246179739 | 13.7001428 | NA | NA | NA | NA | NA | NA | NA |
| SCN4A | 1306 | G | -26.52725 | 19.04153245 | 0.289764002 | 12.06845519 | NA | NA | NA | NA | NA | NA | NA |
| SCN4A | 1307 | G | -24.398 | 16.56702541 | 0.247255375 | 13.40193558 | NA | NA | NA | NA | NA | NA | NA |
| SCN4A | 1308 | K | -26.01808333 | 13.65338306 | 0.197203008 | 10.41515136 | NA | NA | NA | NA | NA | NA | NA |
| SCN4A | 1309 | D | -21.60825 | 13.55228357 | 0.195466258 | 10.75073253 | NA | NA | NA | NA | NA | NA | NA |
| SCN4A | 1310 | I | -18.02960714 | 14.1994334 | 0.206583402 | 11.22571868 | NA | NA | NA | NA | NA | NA | NA |
| SCN4A | 1311 | F | -14.22029545 | 12.93687126 | 0.184894321 | 9.917158997 | NA | NA | NA | NA | NA | NA | NA |
| SCN4A | 1312 | M | -17.352875 | 17.64903864 | 0.265842874 | 14.64494188 | NA | NA | NA | NA | NA | NA | NA |
| SCN4A | 1313 | T | -17.53589286 | 21.78085995 | 0.33682188 | 18.79466167 | NA | NA | NA | NA | NA | NA | NA |
| SCN4A | 1314 | E | -22.09886111 | 23.01002601 | 0.357937262 | 20.16088706 | NA | NA | NA | NA | NA | NA | NA |
| SCN4A | 1315 | E | -17.83141667 | 25.87272127 | 0.407114428 | 22.93605893 | NA | NA | NA | NA | NA | NA | NA |
| SCN4A | 1316 | Q | -16.79063889 | 22.25320804 | 0.34493617 | 19.50748654 | NA | NA | NA | NA | NA | NA | NA |
| SCN4A | 1317 | K | -21.89275 | 19.03607412 | 0.289670235 | 15.90271621 | NA | NA | NA | NA | NA | NA | NA |
| SCN4A | 1318 | K | -23.42108333 | 24.34559496 | 0.380880499 | 21.07368248 | NA | NA | NA | NA | NA | NA | NA |
| SCN4A | 1319 | Y | -17.1575 | 24.12340967 | 0.377063662 | 21.19092052 | NA | NA | NA | NA | NA | NA | NA |
| SCN4A | 1320 | Y | -20.40725 | 15.41620774 | 0.227485911 | 12.11102349 | NA | NA | NA | NA | NA | NA | NA |
| SCN4A | 1321 | N | -24.146875 | 19.73218166 | 0.301628405 | 14.37345618 | NA | NA | NA | NA | NA | NA | NA |
| SCN4A | 1322 | A | -22.21695 | 24.29167629 | 0.379954251 | 18.90330448 | NA | NA | NA | NA | NA | NA | NA |
| SCN4A | 1323 | M | -18.741375 | 22.97932464 | 0.357409855 | 19.56333531 | NA | NA | NA | NA | NA | NA | NA |
| SCN4A | 1324 | K | -23.34319444 | 16.86208943 | 0.252324169 | 11.63825723 | NA | NA | NA | NA | NA | NA | NA |
| SCN4A | 1325 | K | -25.67630556 | 22.92301451 | 0.356442524 | 17.23059236 | NA | NA | NA | NA | NA | NA | NA |
| SCN4A | 1326 | L | -20.3295 | 24.98449867 | 0.391855986 | 19.63220995 | NA | NA | NA | NA | NA | NA | NA |
| SCN4A | 1327 | G | -21.3125 | 22.28466473 | 0.345476553 | 16.96319693 | NA | NA | NA | NA | NA | NA | NA |
| SCN4A | 1328 | S | -24.99841667 | 21.74709206 | 0.336241794 | 16.12321101 | NA | NA | NA | NA | NA | NA | NA |
| SCN4A | 1329 | K | -23.67552778 | 25.16989889 | 0.395040907 | 19.51719824 | NA | NA | NA | NA | NA | NA | NA |
| SCN4A | 1330 | K | -26.13119444 | 23.70551168 | 0.369884749 | 18.11727772 | NA | NA | NA | NA | NA | NA | NA |
| SCN4A | 1331 | P | -25.61189286 | 30.88904966 | 0.493288049 | 25.26317236 | NA | NA | NA | NA | NA | NA | NA |
| SCN4A | 1332 | Q | -28.35597222 | 29.99008219 | 0.477845025 | 16.51505155 | NA | NA | NA | NA | NA | NA | NA |
| SCN4A | 1333 | K | -29.95963889 | 35.05857169 | 0.564914702 | 21.57707935 | NA | NA | NA | NA | NA | NA | NA |
| SCN4A | 1334 | P | -25.99489286 | 36.92596069 | 0.596993876 | 23.53904493 | NA | NA | NA | NA | NA | NA | NA |

| | | | | | | | | | | | | | |
|---|---|---|---|---|---|---|---|---|---|---|---|---|---|
| SCN4A | 1335 | I | -24.33 | 40.46867013 | 0.657852751 | 27.12857769 | NA | NA | NA | NA | NA | NA | NA |
| SCN4A | 1336 | P | -26.25103571 | 42.50888488 | 0.692900833 | 25.93970188 | NA | NA | NA | NA | NA | NA | NA |
| SCN4A | 1337 | R | -22.07902273 | 42.30568981 | 0.689410221 | 36.64470492 | NA | NA | NA | NA | NA | NA | NA |
| SCN4A | 1338 | P | -22.12146429 | 49.45103054 | 0.812157345 | 43.79329802 | NA | NA | NA | NA | NA | NA | NA |
| SCN4A | 1339 | Q | -24.58797222 | 52.08747666 | 0.857447863 | 35.48455291 | NA | NA | NA | NA | NA | NA | NA |
| SCN4A | 1340 | N | -20.330625 | 55.34621398 | 0.913428487 | 49.78229564 | NA | NA | NA | NA | NA | NA | NA |
| SCN4A | 1341 | K | -19.76986111 | 57.99385842 | 0.958911376 | 52.33489792 | NA | NA | NA | NA | NA | NA | NA |
| SCN4A | 1342 | I | -15.782875 | 57.1672582 | 0.944711522 | 51.4973026 | NA | NA | NA | NA | NA | NA | NA |
| SCN4A | 1343 | Q | -16.45863889 | 52.54812177 | 0.865361112 | 46.8747107 | NA | NA | NA | NA | NA | NA | NA |
| SCN4A | 1344 | G | -18.58075 | 52.64572133 | 0.867037738 | 46.98877296 | NA | NA | NA | NA | NA | NA | NA |
| SCN4A | 1345 | M | -16.022875 | 52.87706592 | 0.87101192 | 47.21126544 | NA | NA | NA | NA | NA | NA | NA |
| SCN4A | 1346 | V | -14.26717857 | 50.019206 | 0.821917818 | 44.6779159 | NA | NA | NA | NA | NA | NA | NA |
| SCN4A | 1347 | Y | -19.38633333 | 47.27222772 | 0.77472851 | 41.6099805 | NA | NA | NA | NA | NA | NA | NA |
| SCN4A | 1348 | D | -18.881875 | 48.06392428 | 0.788328768 | 42.40782558 | NA | NA | NA | NA | NA | NA | NA |
| SCN4A | 1349 | L | -15.043875 | 46.19920053 | 0.75629538 | 40.53241885 | NA | NA | NA | NA | NA | NA | NA |
| SCN4A | 1350 | V | -14.64089286 | 43.34896368 | 0.707332231 | 40.2877101 | NA | NA | NA | NA | NA | NA | NA |
| SCN4A | 1351 | T | -18.79103571 | 42.0528487 | 0.685066759 | 36.3953416 | NA | NA | NA | NA | NA | NA | NA |
| SCN4A | 1352 | K | -18.30397222 | 42.55295207 | 0.693657847 | 36.88733304 | NA | NA | NA | NA | NA | NA | NA |
| SCN4A | 1353 | Q | -16.75430556 | 37.37652586 | 0.604733966 | 31.72049653 | NA | NA | NA | NA | NA | NA | NA |
| SCN4A | 1354 | A | -13.05975 | 42.79751148 | 0.697859041 | 39.65045687 | NA | NA | NA | NA | NA | NA | NA |
| SCN4A | 1355 | F | -11.00302273 | 41.16664001 | 0.669842913 | 37.20417104 | NA | NA | NA | NA | NA | NA | NA |
| SCN4A | 1356 | D | -11.343625 | 35.9327589 | 0.579932036 | 32.41583269 | NA | NA | NA | NA | NA | NA | NA |
| SCN4A | 1357 | I | -10.460125 | 38.11629996 | 0.617442267 | 34.16439112 | NA | NA | NA | NA | NA | NA | NA |
| SCN4A | 1358 | T | -7.290607143 | 39.07881555 | 0.633976961 | 35.08885665 | NA | NA | NA | NA | NA | NA | NA |
| SCN4A | 1359 | I | -6.313625 | 36.38407374 | 0.587685004 | 32.37631075 | NA | NA | NA | NA | NA | NA | NA |
| SCN4A | 1360 | M | -7.088 | 31.90128225 | 0.510676813 | 27.94906643 | NA | NA | NA | NA | NA | NA | NA |
| SCN4A | 1361 | I | -3.803125 | 34.59067436 | 0.55687687 | 30.62285863 | NA | NA | NA | NA | NA | NA | NA |
| SCN4A | 1362 | L | -1.80625 | 36.02283311 | 0.581479387 | 31.98456452 | NA | NA | NA | NA | NA | NA | NA |
| SCN4A | 1363 | I | -2.05325 | 31.87665149 | 0.51025369 | 27.8214875 | NA | NA | NA | NA | NA | NA | NA |
| SCN4A | 1364 | C | -1.181083333 | 30.46051511 | 0.485926415 | 26.46212257 | NA | NA | NA | NA | NA | NA | NA |
| SCN4A | 1365 | L | 1.66875 | 33.57678951 | 0.539459723 | 29.52886535 | NA | NA | NA | NA | NA | NA | NA |
| SCN4A | 1366 | N | 2.7115 | 32.44802029 | 0.52006902 | 28.36767549 | NA | NA | NA | NA | NA | NA | NA |
| SCN4A | 1367 | M | 2.588 | 25.20429122 | 0.39563172 | 21.32739419 | NA | NA | NA | NA | NA | NA | NA |

| | | | | | | | | | | | | | |
|---|---|---|---|---|---|---|---|---|---|---|---|---|---|
| SCN4A | 1368 | V | 5.473464286 | 30.5819744 | 0.488012919 | 26.68929186 | NA | NA | NA | NA | NA | NA | NA |
| SCN4A | 1369 | T | 7.626321429 | 32.81797669 | 0.526424362 | 30.32017761 | NA | NA | NA | NA | NA | NA | NA |
| SCN4A | 1370 | M | 6.99975 | 29.15074244 | 0.463426323 | 27.14652752 | NA | NA | NA | NA | NA | NA | NA |
| SCN4A | 1371 | M | 7.544375 | 25.69435745 | 0.404050383 | 23.98288527 | NA | NA | NA | NA | NA | NA | NA |
| SCN4A | 1372 | V | 11.58832143 | 30.03570119 | 0.478628697 | 28.14857589 | NA | NA | NA | NA | NA | NA | NA |
| SCN4A | 1373 | E | 13.53852778 | 28.18040479 | 0.446757257 | 26.36119189 | NA | NA | NA | NA | NA | NA | NA |
| SCN4A | 1374 | T | 17.15717857 | 28.57577101 | 0.453549105 | 26.60001961 | NA | NA | NA | NA | NA | NA | NA |
| SCN4A | 1375 | D | 20.75475 | 25.89183783 | 0.407442825 | 25.41355013 | NA | NA | NA | NA | NA | NA | NA |
| SCN4A | 1376 | N | 23.9985 | 29.23176502 | 0.464818179 | 25.58424963 | NA | NA | NA | NA | NA | NA | NA |
| SCN4A | 1377 | Q | 19.92441667 | 32.51087114 | 0.521148711 | 28.20527462 | NA | NA | NA | NA | NA | NA | NA |
| SCN4A | 1378 | S | 22.32525 | 35.47782994 | 0.572116982 | 31.17603569 | NA | NA | NA | NA | NA | NA | NA |
| SCN4A | 1379 | Q | 23.51030556 | 38.26826709 | 0.620052853 | 34.62158888 | NA | NA | NA | NA | NA | NA | NA |
| SCN4A | 1380 | L | 20.695125 | 40.59229881 | 0.659976521 | 38.62067007 | NA | NA | NA | NA | NA | NA | NA |
| SCN4A | 1381 | K | 17.43086111 | 32.73938384 | 0.525074245 | 30.82485519 | NA | NA | NA | NA | NA | NA | NA |
| SCN4A | 1382 | V | 18.10917857 | 36.60799112 | 0.591531596 | 34.71213904 | NA | NA | NA | NA | NA | NA | NA |
| SCN4A | 1383 | D | 18.32825 | 40.4066203 | 0.65678682 | 38.43688108 | NA | NA | NA | NA | NA | NA | NA |
| SCN4A | 1384 | I | 14.77225 | 40.41971513 | 0.657011771 | 38.58142227 | NA | NA | NA | NA | NA | NA | NA |
| SCN4A | 1385 | L | 12.702375 | 34.57090998 | 0.556537345 | 32.72872511 | NA | NA | NA | NA | NA | NA | NA |
| SCN4A | 1386 | Y | 13.25258333 | 39.33657846 | 0.638404973 | 37.42631462 | NA | NA | NA | NA | NA | NA | NA |
| SCN4A | 1387 | N | 12.1255 | 41.8414195 | 0.681434696 | 40.00518552 | NA | NA | NA | NA | NA | NA | NA |
| SCN4A | 1388 | I | 8.622625 | 39.59707023 | 0.642879863 | 37.51038854 | NA | NA | NA | NA | NA | NA | NA |
| SCN4A | 1389 | N | 7.624375 | 36.56329504 | 0.590763779 | 34.47515459 | NA | NA | NA | NA | NA | NA | NA |
| SCN4A | 1390 | M | 8.246625 | 41.42273311 | 0.67424224 | 39.27321312 | NA | NA | NA | NA | NA | NA | NA |
| SCN4A | 1391 | I | 4.939625 | 42.68933039 | 0.696000639 | 38.77157807 | NA | NA | NA | NA | NA | NA | NA |
| SCN4A | 1392 | F | 2.793477273 | 38.36966152 | 0.62179467 | 34.28880729 | NA | NA | NA | NA | NA | NA | NA |
| SCN4A | 1393 | I | 3.26375 | 40.76368856 | 0.662920761 | 36.75785784 | NA | NA | NA | NA | NA | NA | NA |
| SCN4A | 1394 | I | 2.099 | 43.61663335 | 0.711930428 | 39.52558357 | NA | NA | NA | NA | NA | NA | NA |
| SCN4A | 1395 | I | -1.095375 | 42.37974533 | 0.690682393 | 38.32509765 | NA | NA | NA | NA | NA | NA | NA |
| SCN4A | 1396 | F | -1.653022727 | 36.90234016 | 0.596588108 | 32.82375942 | NA | NA | NA | NA | NA | NA | NA |
| SCN4A | 1397 | T | -1.862607143 | 42.86701774 | 0.699053063 | 38.78649587 | NA | NA | NA | NA | NA | NA | NA |
| SCN4A | 1398 | G | -4.36625 | 45.19977357 | 0.739126599 | 41.12294606 | NA | NA | NA | NA | NA | NA | NA |
| SCN4A | 1399 | E | -6.664083333 | 40.19098633 | 0.653082525 | 36.14363145 | NA | NA | NA | NA | NA | NA | NA |
| SCN4A | 1400 | C | -7.001416667 | 43.50838598 | 0.710070887 | 39.44808136 | NA | NA | NA | NA | NA | NA | NA |

| | | | | | | | | | | | | | |
|---|---|---|---|---|---|---|---|---|---|---|---|---|---|
| SCN4A | 1401 | V | -7.197035714 | 46.44218696 | 0.760469552 | 42.38263188 | NA | NA | NA | NA | NA | NA | NA |
| SCN4A | 1402 | L | -9.355375 | 47.75403663 | 0.783005325 | 43.87234723 | NA | NA | NA | NA | NA | NA | NA |
| SCN4A | 1403 | K | -12.02463889 | 40.4609136 | 0.657719504 | 37.51717037 | NA | NA | NA | NA | NA | NA | NA |
| SCN4A | 1404 | M | -10.920375 | 46.41240778 | 0.759957987 | 43.37283189 | NA | NA | NA | NA | NA | NA | NA |
| SCN4A | 1405 | L | -12.527875 | 49.09689202 | 0.806073733 | 46.02238232 | NA | NA | NA | NA | NA | NA | NA |
| SCN4A | 1406 | A | -15.45195 | 48.69055811 | 0.799093475 | 43.3440304 | NA | NA | NA | NA | NA | NA | NA |
| SCN4A | 1407 | L | -16.857625 | 44.33937529 | 0.72434614 | 39.0367711 | NA | NA | NA | NA | NA | NA | NA |
| SCN4A | 1408 | R | -19.21547727 | 47.98777898 | 0.787020697 | 42.6237722 | NA | NA | NA | NA | NA | NA | NA |
| SCN4A | 1409 | Q | -18.25786111 | 44.09099296 | 0.720079274 | 38.72085918 | NA | NA | NA | NA | NA | NA | NA |
| SCN4A | 1410 | Y | -19.25475 | 37.38617223 | 0.604899677 | 32.10263976 | NA | NA | NA | NA | NA | NA | NA |
| SCN4A | 1411 | Y | -12.20733333 | 40.9656905 | 0.666390877 | 37.88315097 | NA | NA | NA | NA | NA | NA | NA |
| SCN4A | 1412 | F | -12.91629545 | 40.05077351 | 0.650673862 | 36.97269574 | NA | NA | NA | NA | NA | NA | NA |
| SCN4A | 1413 | T | -17.77446429 | 37.44595192 | 0.60592661 | 32.05828158 | NA | NA | NA | NA | NA | NA | NA |
| SCN4A | 1414 | V | -16.27503571 | 35.1395034 | 0.566304997 | 29.75164704 | NA | NA | NA | NA | NA | NA | NA |
| SCN4A | 1415 | G | -13.574 | 33.57577241 | 0.53944225 | 30.51157954 | NA | NA | NA | NA | NA | NA | NA |
| SCN4A | 1416 | W | -12.25117857 | 28.47684061 | 0.451849616 | 25.45987143 | NA | NA | NA | NA | NA | NA | NA |
| SCN4A | 1417 | N | -11.752125 | 36.0441992 | 0.581846427 | 32.97012833 | NA | NA | NA | NA | NA | NA | NA |
| SCN4A | 1418 | I | -10.7425 | 36.79198267 | 0.594692318 | 33.76483512 | NA | NA | NA | NA | NA | NA | NA |
| SCN4A | 1419 | F | -6.407204545 | 31.09807007 | 0.496878732 | 29.05116429 | NA | NA | NA | NA | NA | NA | NA |
| SCN4A | 1420 | D | -6.91275 | 36.32936451 | 0.586745174 | 32.32868158 | NA | NA | NA | NA | NA | NA | NA |
| SCN4A | 1421 | F | -6.471022727 | 38.89829909 | 0.630875936 | 34.80965464 | NA | NA | NA | NA | NA | NA | NA |
| SCN4A | 1422 | V | -3.572607143 | 37.15161286 | 0.60087027 | 33.06316848 | NA | NA | NA | NA | NA | NA | NA |
| SCN4A | 1423 | V | -1.697607143 | 34.48068179 | 0.554987348 | 30.39802385 | NA | NA | NA | NA | NA | NA | NA |
| SCN4A | 1424 | V | -1.970464286 | 38.87938432 | 0.630551007 | 34.82562476 | NA | NA | NA | NA | NA | NA | NA |
| SCN4A | 1425 | I | -1.302625 | 40.97434854 | 0.66653961 | 36.92452301 | NA | NA | NA | NA | NA | NA | NA |
| SCN4A | 1426 | L | 1.97625 | 36.18803152 | 0.584317268 | 33.67321848 | NA | NA | NA | NA | NA | NA | NA |
| SCN4A | 1427 | S | 2.97325 | 37.38259859 | 0.604838287 | 34.91938948 | NA | NA | NA | NA | NA | NA | NA |
| SCN4A | 1428 | I | 3.748875 | 41.17166548 | 0.669929243 | 39.0235049 | NA | NA | NA | NA | NA | NA | NA |
| SCN4A | 1429 | V | 5.64475 | 40.50078108 | 0.658404372 | 38.48823586 | NA | NA | NA | NA | NA | NA | NA |
| SCN4A | 1430 | G | 8.05725 | 38.53488605 | 0.624633 | 36.59004146 | NA | NA | NA | NA | NA | NA | NA |
| SCN4A | 1431 | L | 9.408625 | 40.43824597 | 0.657330105 | 38.5098933 | NA | NA | NA | NA | NA | NA | NA |
| SCN4A | 1432 | A | 10.40035 | 42.90163089 | 0.699647669 | 40.95027156 | NA | NA | NA | NA | NA | NA | NA |
| SCN4A | 1433 | L | 11.341 | 40.22458583 | 0.653659718 | 38.29439481 | NA | NA | NA | NA | NA | NA | NA |

| | | | | | | | | | | | | | |
|---|---|---|---|---|---|---|---|---|---|---|---|---|---|
| SCN4A | 1434 | S | 14.44108333 | 39.43651104 | 0.640121677 | 37.49013235 | NA | NA | NA | NA | NA | NA | NA |
| SCN4A | 1435 | D | 16.801625 | 42.74623268 | 0.696978142 | 40.66540818 | NA | NA | NA | NA | NA | NA | NA |
| SCN4A | 1436 | L | 15.241125 | 40.90631124 | 0.665370823 | 38.81254524 | NA | NA | NA | NA | NA | NA | NA |
| SCN4A | 1437 | I | 16.957375 | 36.72196837 | 0.593489569 | 34.77676009 | NA | NA | NA | NA | NA | NA | NA |
| SCN4A | 1438 | Q | 21.38419444 | 38.78627733 | 0.628951557 | 36.66630887 | NA | NA | NA | NA | NA | NA | NA |
| SCN4A | 1439 | K | 19.31230556 | 40.22116608 | 0.653600971 | 38.09345634 | NA | NA | NA | NA | NA | NA | NA |
| SCN4A | 1440 | Y | 17.90566667 | 37.10157421 | 0.600010675 | 35.05910486 | NA | NA | NA | NA | NA | NA | NA |
| SCN4A | 1441 | F | 20.48265909 | 32.33478862 | 0.518123856 | 30.28764712 | NA | NA | NA | NA | NA | NA | NA |
| SCN4A | 1442 | V | 22.88389286 | 29.63529985 | 0.471750352 | 27.5142018 | NA | NA | NA | NA | NA | NA | NA |
| SCN4A | 1443 | S | 20.12075 | 30.89034399 | 0.493310284 | 28.87432462 | NA | NA | NA | NA | NA | NA | NA |
| SCN4A | 1444 | P | 19.97946429 | 27.33838582 | 0.432292529 | 24.10102729 | NA | NA | NA | NA | NA | NA | NA |
| SCN4A | 1445 | T | 17.20503571 | 27.49359718 | 0.434958847 | 24.20188821 | NA | NA | NA | NA | NA | NA | NA |
| SCN4A | 1446 | L | 14.3865 | 31.29107139 | 0.50019423 | 27.99791583 | NA | NA | NA | NA | NA | NA | NA |
| SCN4A | 1447 | F | 15.07993182 | 32.79280496 | 0.525991946 | 30.9849879 | NA | NA | NA | NA | NA | NA | NA |
| SCN4A | 1448 | R | 14.93702273 | 27.62973652 | 0.437297533 | 25.94520963 | NA | NA | NA | NA | NA | NA | NA |
| SCN4A | 1449 | V | 10.19175 | 28.76336936 | 0.456771786 | 26.96252881 | NA | NA | NA | NA | NA | NA | NA |
| SCN4A | 1450 | I | 8.966125 | 32.33479632 | 0.518123988 | 30.38561625 | NA | NA | NA | NA | NA | NA | NA |
| SCN4A | 1451 | R | 10.42011364 | 31.44213285 | 0.502789258 | 29.54411143 | NA | NA | NA | NA | NA | NA | NA |
| SCN4A | 1452 | L | 6.836875 | 26.67108579 | 0.420829233 | 24.86616243 | NA | NA | NA | NA | NA | NA | NA |
| SCN4A | 1453 | A | 3.56735 | 31.368772 | 0.501529019 | 29.24286697 | NA | NA | NA | NA | NA | NA | NA |
| SCN4A | 1454 | R | 5.871386364 | 31.80170458 | 0.508966205 | 29.28245993 | NA | NA | NA | NA | NA | NA | NA |
| SCN4A | 1455 | I | 3.08725 | 26.89399779 | 0.424658554 | 24.37904136 | NA | NA | NA | NA | NA | NA | NA |
| SCN4A | 1456 | G | -0.863 | 30.05325478 | 0.478930243 | 25.9878298 | NA | NA | NA | NA | NA | NA | NA |
| SCN4A | 1457 | R | -1.605022727 | 31.44224588 | 0.502791199 | 27.36357104 | NA | NA | NA | NA | NA | NA | NA |
| SCN4A | 1458 | V | -1.280892857 | 27.24549625 | 0.430696814 | 23.16830461 | NA | NA | NA | NA | NA | NA | NA |
| SCN4A | 1459 | L | -3.839625 | 25.88286943 | 0.40728876 | 21.82710676 | NA | NA | NA | NA | NA | NA | NA |
| SCN4A | 1460 | R | -6.445113636 | 30.3359986 | 0.483787393 | 26.29287147 | NA | NA | NA | NA | NA | NA | NA |
| SCN4A | 1461 | L | -5.672 | 28.16953237 | 0.446570483 | 24.15953478 | NA | NA | NA | NA | NA | NA | NA |
| SCN4A | 1462 | I | -8.158 | 25.09916489 | 0.393825794 | 21.11766695 | NA | NA | NA | NA | NA | NA | NA |
| SCN4A | 1463 | R | -12.74002273 | 28.87749421 | 0.458732294 | 26.1820965 | NA | NA | NA | NA | NA | NA | NA |
| SCN4A | 1464 | G | -12.965 | 28.59614804 | 0.453899154 | 25.47767452 | NA | NA | NA | NA | NA | NA | NA |
| SCN4A | 1465 | A | -13.03295 | 24.80220625 | 0.388724453 | 21.66355932 | NA | NA | NA | NA | NA | NA | NA |
| SCN4A | 1466 | K | -18.02641667 | 22.42614978 | 0.347907072 | 19.27571887 | NA | NA | NA | NA | NA | NA | NA |

| | | | | | | | | | | | | | |
|---|---|---|---|---|---|---|---|---|---|---|---|---|---|
| SCN4A | 1467 | G | -14.73775 | 19.68017208 | 0.300734952 | 16.53685034 | NA | NA | NA | NA | NA | NA | NA |
| SCN4A | 1468 | I | -11.21375 | 20.7911693 | 0.319820356 | 17.75752187 | NA | NA | NA | NA | NA | NA | NA |
| SCN4A | 1469 | R | -15.40829545 | 21.65697167 | 0.33469365 | 18.51888658 | NA | NA | NA | NA | NA | NA | NA |
| SCN4A | 1470 | T | -16.25146429 | 17.67663981 | 0.266317024 | 14.53833284 | NA | NA | NA | NA | NA | NA | NA |
| SCN4A | 1471 | L | -12.368125 | 15.43142181 | 0.227747268 | 12.44808974 | NA | NA | NA | NA | NA | NA | NA |
| SCN4A | 1472 | L | -10.6815 | 18.85519401 | 0.286562964 | 15.78231976 | NA | NA | NA | NA | NA | NA | NA |
| SCN4A | 1473 | F | -16.53238636 | 18.3701595 | 0.278230738 | 15.24071665 | NA | NA | NA | NA | NA | NA | NA |
| SCN4A | 1474 | A | -14.23375 | 14.75520532 | 0.216130799 | 11.62484996 | NA | NA | NA | NA | NA | NA | NA |
| SCN4A | 1475 | L | -10.253 | 16.36546247 | 0.243792801 | 13.30256848 | NA | NA | NA | NA | NA | NA | NA |
| SCN4A | 1476 | M | -13.2575 | 18.63866505 | 0.282843294 | 15.58817034 | NA | NA | NA | NA | NA | NA | NA |
| SCN4A | 1477 | M | -16.328875 | 17.62432328 | 0.265418298 | 14.47965587 | NA | NA | NA | NA | NA | NA | NA |
| SCN4A | 1478 | S | -12.71341667 | 14.00371561 | 0.203221239 | 10.98684765 | NA | NA | NA | NA | NA | NA | NA |
| SCN4A | 1479 | L | -10.337625 | 17.83496027 | 0.269036752 | 14.80920811 | NA | NA | NA | NA | NA | NA | NA |
| SCN4A | 1480 | P | -13.18975 | 19.38613149 | 0.295683739 | 16.37025812 | NA | NA | NA | NA | NA | NA | NA |
| SCN4A | 1481 | A | -12.38855 | 16.09707905 | 0.239182343 | 13.05844379 | NA | NA | NA | NA | NA | NA | NA |
| SCN4A | 1482 | L | -8.409125 | 14.37284641 | 0.209562399 | 11.42470711 | NA | NA | NA | NA | NA | NA | NA |
| SCN4A | 1483 | F | -7.684022727 | 18.91316994 | 0.287558911 | 16.03558235 | NA | NA | NA | NA | NA | NA | NA |
| SCN4A | 1484 | N | -10.20175 | 19.80445036 | 0.302869882 | 16.87752644 | NA | NA | NA | NA | NA | NA | NA |
| SCN4A | 1485 | I | -7.403375 | 15.35477105 | 0.226430513 | 12.50957631 | NA | NA | NA | NA | NA | NA | NA |
| SCN4A | 1486 | G | -3.9855 | 19.02255469 | 0.28943799 | 15.90583073 | NA | NA | NA | NA | NA | NA | NA |
| SCN4A | 1487 | L | -4.6795 | 21.7258021 | 0.335876062 | 18.61034884 | NA | NA | NA | NA | NA | NA | NA |
| SCN4A | 1488 | L | -4.448125 | 20.37493355 | 0.312669999 | 17.39905944 | NA | NA | NA | NA | NA | NA | NA |
| SCN4A | 1489 | L | -0.800375 | 16.9992784 | 0.254680887 | 13.76659005 | NA | NA | NA | NA | NA | NA | NA |
| SCN4A | 1490 | F | 1.261204545 | 21.92893787 | 0.339365655 | 19.82806254 | NA | NA | NA | NA | NA | NA | NA |
| SCN4A | 1491 | L | -0.869375 | 24.24345029 | 0.379125794 | 21.98620905 | NA | NA | NA | NA | NA | NA | NA |
| SCN4A | 1492 | V | 1.13875 | 21.64160274 | 0.334429633 | 19.55511145 | NA | NA | NA | NA | NA | NA | NA |
| SCN4A | 1493 | M | 3.93775 | 17.43519317 | 0.262169303 | 15.68077092 | NA | NA | NA | NA | NA | NA | NA |
| SCN4A | 1494 | F | 4.772295455 | 22.80273688 | 0.35437632 | 21.02488437 | NA | NA | NA | NA | NA | NA | NA |
| SCN4A | 1495 | I | 3.987 | 23.87786862 | 0.372845604 | 21.14037593 | NA | NA | NA | NA | NA | NA | NA |
| SCN4A | 1496 | Y | 5.752666667 | 20.18587848 | 0.309422293 | 17.45931522 | NA | NA | NA | NA | NA | NA | NA |
| SCN4A | 1497 | S | 9.240583333 | 20.59612085 | 0.316469692 | 17.11384613 | NA | NA | NA | NA | NA | NA | NA |
| SCN4A | 1498 | I | 9.428875 | 22.91969009 | 0.356385415 | 19.43410385 | NA | NA | NA | NA | NA | NA | NA |
| SCN4A | 1499 | F | 10.04220455 | 21.82014688 | 0.337496776 | 18.33453417 | NA | NA | NA | NA | NA | NA | NA |

| | | | | | | | | | | | | | |
|---|---|---|---|---|---|---|---|---|---|---|---|---|---|
| SCN4A | 1500 | G | 13.3295 | 19.36998495 | 0.295406364 | 15.8538745 | NA | NA | NA | NA | NA | NA | NA |
| SCN4A | 1501 | M | 14.640375 | 20.69141554 | 0.318106724 | 17.26781438 | NA | NA | NA | NA | NA | NA | NA |
| SCN4A | 1502 | S | 15.14441667 | 22.33847869 | 0.346401003 | 18.649996 | NA | NA | NA | NA | NA | NA | NA |
| SCN4A | 1503 | N | 15.378375 | 19.42114338 | 0.296285196 | 15.72235033 | NA | NA | NA | NA | NA | NA | NA |
| SCN4A | 1504 | F | 16.21193182 | 15.73407624 | 0.232946454 | 12.01895387 | NA | NA | NA | NA | NA | NA | NA |
| SCN4A | 1505 | A | 20.40395 | 16.43669133 | 0.245016415 | 12.89005336 | NA | NA | NA | NA | NA | NA | NA |
| SCN4A | 1506 | Y | 23.61608333 | 14.52898696 | 0.212244678 | 11.01993881 | NA | NA | NA | NA | NA | NA | NA |
| SCN4A | 1507 | V | 22.64175 | 11.48016786 | 0.15987016 | 7.948588611 | NA | NA | NA | NA | NA | NA | NA |
| SCN4A | 1508 | K | 27.04475 | 8.750801163 | 0.112983395 | 5.305076698 | NA | NA | NA | NA | NA | NA | NA |
| SCN4A | 1509 | K | 25.91397222 | 5.279677452 | 0.053354265 | 1.773241242 | 3.58 | -0.4 | -0.13 | 14.2 | 0.52 | 1 | -1 |
| SCN4A | 1510 | E | 25.10830556 | 5.201307624 | 0.052007979 | 1.593421477 | 3.67 | -0.32 | 0.65 | 2.57 | 0.16 | 0 | 0 |
| SCN4A | 1511 | S | 22.08391667 | 5.087601891 | 0.050054671 | 1.637086573 | 3.47 | -0.15 | 1 | 2.3 | -0.06 | 0 | 0 |
| SCN4A | 1512 | G | 19.718 | 6.045107609 | 0.066503302 | 2.419710227 | 3.75 | -0.95 | -1.95 | 26.59 | 1.62 | 2 | -2 |
| SCN4A | 1513 | I | 21.020125 | 6.912653977 | 0.081406555 | 3.534122982 | 3.5 | -0.32 | 0.65 | 2.57 | 0.16 | 0 | 0 |
| SCN4A | 1514 | D | 22.903625 | 5.575200176 | 0.058430939 | 2.320568822 | 3.67 | -0.32 | 0.65 | 2.57 | 0.16 | 0 | 0 |
| SCN4A | 1515 | D | 24.5905 | 8.548972043 | 0.109516249 | 5.05533118 | NA | NA | NA | NA | NA | NA | NA |
| SCN4A | 1516 | M | 20.772875 | 11.28271661 | 0.156478219 | 7.72294049 | NA | NA | NA | NA | NA | NA | NA |
| SCN4A | 1517 | F | 18.29802273 | 11.13427541 | 0.153928204 | 7.646374249 | NA | NA | NA | NA | NA | NA | NA |
| SCN4A | 1518 | N | 19.30725 | 11.67813042 | 0.163270885 | 8.110655852 | NA | NA | NA | NA | NA | NA | NA |
| SCN4A | 1519 | F | 15.77447727 | 14.088482 | 0.204677409 | 11.40854554 | NA | NA | NA | NA | NA | NA | NA |
| SCN4A | 1520 | E | 20.94308333 | 13.70265055 | 0.198049356 | 10.18204317 | NA | NA | NA | NA | NA | NA | NA |
| SCN4A | 1521 | T | 17.10246429 | 18.67834481 | 0.283524938 | 15.1117533 | NA | NA | NA | NA | NA | NA | NA |
| SCN4A | 1522 | F | 11.36847727 | 21.24873229 | 0.327680659 | 17.71600214 | NA | NA | NA | NA | NA | NA | NA |
| SCN4A | 1523 | G | 13.90775 | 18.82463854 | 0.286038063 | 15.27246643 | NA | NA | NA | NA | NA | NA | NA |
| SCN4A | 1524 | N | 16.58575 | 15.818041 | 0.234388853 | 12.2769814 | NA | NA | NA | NA | NA | NA | NA |
| SCN4A | 1525 | S | 12.73808333 | 17.40701321 | 0.26168521 | 13.88519114 | NA | NA | NA | NA | NA | NA | NA |
| SCN4A | 1526 | I | 9.33825 | 17.60222375 | 0.265038659 | 14.45527354 | NA | NA | NA | NA | NA | NA | NA |
| SCN4A | 1527 | I | 12.29575 | 14.05760783 | 0.204147033 | 10.66447084 | NA | NA | NA | NA | NA | NA | NA |
| SCN4A | 1528 | C | 12.36558333 | 12.64173857 | 0.179824347 | 9.117578591 | NA | NA | NA | NA | NA | NA | NA |
| SCN4A | 1529 | L | 8.5265 | 14.89576628 | 0.218545442 | 12.5575194 | NA | NA | NA | NA | NA | NA | NA |
| SCN4A | 1530 | F | 6.700659091 | 12.84389481 | 0.183297113 | 11.10890569 | NA | NA | NA | NA | NA | NA | NA |
| SCN4A | 1531 | E | 10.39252778 | 9.290391757 | 0.122252819 | 7.533794511 | NA | NA | NA | NA | NA | NA | NA |
| SCN4A | 1532 | I | 8.0025 | 10.79546595 | 0.148107923 | 9.000072662 | NA | NA | NA | NA | NA | NA | NA |

| | | | | | | | | | | | | | |
|---|---|---|---|---|---|---|---|---|---|---|---|---|---|
| SCN4A | 1533 | T | 3.659892857 | 11.30814304 | 0.156915011 | 9.916949501 | NA | NA | NA | NA | NA | NA | NA |
| SCN4A | 1534 | T | 5.064607143 | 7.311766202 | 0.088262754 | 5.086120698 | 2.36 | -0.8 | -0.4 | 3.38 | 0.81 | 0 | 0 |
| SCN4A | 1535 | S | 6.102083333 | 6.739721433 | 0.078435812 | 5.165283222 | 2 | -0.78 | -2.6 | 34.26 | 1.5 | 2 | 0 |
| SCN4A | 1536 | A | 9.43335 | 5.607833628 | 0.058991537 | 3.854572473 | 2.07 | -0.15 | -2.9 | 33.67 | 0.75 | 2 | 0 |
| SCN4A | 1537 | G | 12.705 | 5.344892889 | 0.054474577 | 2.623627264 | 2.79 | -0.37 | -3.05 | 37.68 | 1.22 | 3 | -1 |
| SCN4A | 1538 | W | 10.49317857 | 7.12844927 | 0.085113622 | 4.384845693 | 2.76 | -0.4 | -3.05 | 37.73 | 1.12 | 3 | -1 |
| SCN4A | 1539 | D | 14.67025 | 4.022645025 | 0.031760177 | 1.561160551 | 2.54 | -0.36 | -2.77 | 33.74 | 1.14 | 2 | -2 |
| SCN4A | 1540 | G | 17.2765 | 5.372905173 | 0.054955789 | 1.885079306 | 3.75 | -0.95 | -1.95 | 26.59 | 1.62 | 2 | -2 |
| SCN4A | 1541 | L | 15.704375 | 7.36552768 | 0.089186303 | 3.887923878 | 3.75 | -0.95 | -1.95 | 26.59 | 1.62 | 2 | -2 |
| SCN4A | 1542 | L | 15.201125 | 9.495993524 | 0.125784775 | 6.01684261 | NA | NA | NA | NA | NA | NA | NA |
| SCN4A | 1543 | N | 19.836125 | 7.692299721 | 0.094799797 | 4.086352757 | 3.75 | -0.95 | -1.95 | 26.59 | 1.62 | 2 | -2 |
| SCN4A | 1544 | P | 21.12732143 | 9.46126741 | 0.125188228 | 6.304192695 | NA | NA | NA | NA | NA | NA | NA |
| SCN4A | 1545 | I | 18.282875 | 12.91529407 | 0.184523654 | 9.353088902 | NA | NA | NA | NA | NA | NA | NA |
| SCN4A | 1546 | L | 18.680125 | 13.14770524 | 0.188516159 | 9.615095868 | NA | NA | NA | NA | NA | NA | NA |
| SCN4A | 1547 | N | 23.474125 | 11.88416821 | 0.166810331 | 8.228817059 | NA | NA | NA | NA | NA | NA | NA |
| SCN4A | 1548 | S | 24.09025 | 17.08391071 | 0.256134754 | 13.38913357 | NA | NA | NA | NA | NA | NA | NA |
| SCN4A | 1549 | G | 27.703 | 17.55191537 | 0.26417443 | 14.0576747 | NA | NA | NA | NA | NA | NA | NA |
| SCN4A | 1550 | P | 31.00832143 | 17.95063481 | 0.271023881 | 14.67616924 | NA | NA | NA | NA | NA | NA | NA |
| SCN4A | 1551 | P | 31.42203571 | 14.92424115 | 0.219034602 | 11.65638258 | NA | NA | NA | NA | NA | NA | NA |
| SCN4A | 1552 | D | 28.577375 | 11.39580405 | 0.158420906 | 7.878664018 | NA | NA | NA | NA | NA | NA | NA |
| SCN4A | 1553 | C | 28.54458333 | 13.62692607 | 0.196748513 | 10.14900128 | NA | NA | NA | NA | NA | NA | NA |
| SCN4A | 1554 | D | 31.4155 | 15.11504767 | 0.222312395 | 11.86452172 | NA | NA | NA | NA | NA | NA | NA |
| SCN4A | 1555 | P | 30.54232143 | 18.59071984 | 0.282019662 | 15.32573095 | NA | NA | NA | NA | NA | NA | NA |
| SCN4A | 1556 | N | 32.308875 | 20.03787838 | 0.306879854 | 16.77431601 | NA | NA | NA | NA | NA | NA | NA |
| SCN4A | 1557 | L | 31.848375 | 14.62139453 | 0.213832113 | 11.36358591 | NA | NA | NA | NA | NA | NA | NA |
| SCN4A | 1558 | E | 34.36597222 | 16.95817871 | 0.253974851 | 16.29685756 | NA | NA | NA | NA | NA | NA | NA |
| SCN4A | 1559 | N | 29.568125 | 16.32331109 | 0.243068699 | 14.33712397 | NA | NA | NA | NA | NA | NA | NA |
| SCN4A | 1560 | P | 31.95360714 | 15.98969077 | 0.23733756 | 15.01973636 | NA | NA | NA | NA | NA | NA | NA |
| SCN4A | 1561 | G | 29.4655 | 20.39427422 | 0.313002245 | 19.41741684 | NA | NA | NA | NA | NA | NA | NA |
| SCN4A | 1562 | T | 27.67889286 | 21.10329408 | 0.325182231 | 17.86711618 | NA | NA | NA | NA | NA | NA | NA |
| SCN4A | 1563 | S | 27.09641667 | 24.41619501 | 0.382093311 | 21.19434669 | NA | NA | NA | NA | NA | NA | NA |
| SCN4A | 1564 | V | 25.43975 | 20.76398531 | 0.319353373 | 17.28950271 | NA | NA | NA | NA | NA | NA | NA |
| SCN4A | 1565 | K | 29.61175 | 20.87949707 | 0.321337706 | 17.66406366 | NA | NA | NA | NA | NA | NA | NA |

| | | | | | | | | | | | | | |
|---|---|---|---|---|---|---|---|---|---|---|---|---|---|
| SCN4A | 1566 | G | 27.826 | 17.170947 | 0.257629917 | 13.69745467 | NA | NA | NA | NA | NA | NA | NA |
| SCN4A | 1567 | D | 27.25475 | 12.43408839 | 0.176257203 | 8.978231473 | NA | NA | NA | NA | NA | NA | NA |
| SCN4A | 1568 | C | 24.50691667 | 16.18654287 | 0.240719209 | 12.67410612 | NA | NA | NA | NA | NA | NA | NA |
| SCN4A | 1569 | G | 21.18975 | 16.11324437 | 0.239460041 | 12.43207565 | NA | NA | NA | NA | NA | NA | NA |
| SCN4A | 1570 | N | 17.620375 | 18.71607496 | 0.28417309 | 15.04794069 | NA | NA | NA | NA | NA | NA | NA |
| SCN4A | 1571 | P | 17.15475 | 18.64759665 | 0.282996727 | 15.12226268 | NA | NA | NA | NA | NA | NA | NA |
| SCN4A | 1572 | S | 13.77291667 | 20.61239693 | 0.316749293 | 17.10243224 | NA | NA | NA | NA | NA | NA | NA |
| SCN4A | 1573 | I | 12.9315 | 19.55435445 | 0.298573579 | 16.08356551 | NA | NA | NA | NA | NA | NA | NA |
| SCN4A | 1574 | G | 12.748 | 16.45782744 | 0.245379504 | 12.99547703 | NA | NA | NA | NA | NA | NA | NA |
| SCN4A | 1575 | I | 11.255 | 17.14268465 | 0.257144409 | 14.9408722 | NA | NA | NA | NA | NA | NA | NA |
| SCN4A | 1576 | C | 8.366583333 | 19.62333562 | 0.29975858 | 16.97829972 | NA | NA | NA | NA | NA | NA | NA |
| SCN4A | 1577 | F | 9.480477273 | 16.90143062 | 0.252999997 | 14.59664795 | NA | NA | NA | NA | NA | NA | NA |
| SCN4A | 1578 | F | 10.29929545 | 12.63306131 | 0.179675284 | 10.33754925 | NA | NA | NA | NA | NA | NA | NA |
| SCN4A | 1579 | C | 6.165916667 | 15.94212194 | 0.236520393 | 13.20058286 | NA | NA | NA | NA | NA | NA | NA |
| SCN4A | 1580 | S | 3.824083333 | 18.22344394 | 0.275710367 | 15.50148898 | NA | NA | NA | NA | NA | NA | NA |
| SCN4A | 1581 | Y | 4.21975 | 15.38454968 | 0.226942069 | 12.88126838 | NA | NA | NA | NA | NA | NA | NA |
| SCN4A | 1582 | I | 4.06925 | 12.23937686 | 0.172912327 | 10.42426143 | NA | NA | NA | NA | NA | NA | NA |
| SCN4A | 1583 | I | 0.843625 | 15.69357671 | 0.232250728 | 11.67779871 | NA | NA | NA | NA | NA | NA | NA |
| SCN4A | 1584 | I | -0.44325 | 16.47265449 | 0.245634213 | 12.97352012 | NA | NA | NA | NA | NA | NA | NA |
| SCN4A | 1585 | S | -0.629083333 | 13.6370598 | 0.196922597 | 9.483029822 | NA | NA | NA | NA | NA | NA | NA |
| SCN4A | 1586 | F | -2.051840909 | 8.815193985 | 0.114089575 | 5.327406496 | NA | NA | NA | NA | NA | NA | NA |
| SCN4A | 1587 | L | -5.103875 | 13.27580009 | 0.190716652 | 10.15889865 | NA | NA | NA | NA | NA | NA | NA |
| SCN4A | 1588 | I | -4.116125 | 13.25070236 | 0.190285507 | 10.13432851 | NA | NA | NA | NA | NA | NA | NA |
| SCN4A | 1589 | V | -3.230464286 | 10.37984961 | 0.140968207 | 7.391029149 | NA | NA | NA | NA | NA | NA | NA |
| SCN4A | 1590 | V | -6.47575 | 8.485927468 | 0.10843323 | 5.370066764 | NA | NA | NA | NA | NA | NA | NA |
| SCN4A | 1591 | N | -8.9565 | 10.6452984 | 0.145528252 | 7.783024788 | NA | NA | NA | NA | NA | NA | NA |
| SCN4A | 1592 | M | -6.64375 | 10.07422627 | 0.135718018 | 6.95551749 | NA | NA | NA | NA | NA | NA | NA |
| SCN4A | 1593 | Y | -6.173583333 | 6.600905998 | 0.076051154 | 3.498970933 | 3.12 | 1.45 | 3.88 | 0.17 | -1.82 | 0 | 0 |
| SCN4A | 1594 | I | -12.115125 | 5.421009685 | 0.055782158 | 2.479502083 | 3.1 | 1.64 | 4.33 | 0.13 | -1.84 | 0 | 0 |
| SCN4A | 1595 | A | -12.43595 | 9.180854699 | 0.120371123 | 6.195614987 | NA | NA | NA | NA | NA | NA | NA |
| SCN4A | 1596 | I | -10.752375 | 9.103160001 | 0.119036435 | 6.059511807 | NA | NA | NA | NA | NA | NA | NA |
| SCN4A | 1597 | I | -12.702375 | 4.640234585 | 0.042369516 | 1.626459608 | 3.1 | 1.64 | 4.33 | 0.13 | -1.84 | 0 | 0 |
| SCN4A | 1598 | L | -16.768 | 5.912097174 | 0.064218366 | 2.817964916 | 3.19 | 1.43 | 3.7 | 0.2 | -1.82 | 0 | 0 |

| | | | | | | | | | | | | | |
|---|---|---|---|---|---|---|---|---|---|---|---|---|---|
| SCN4A | 1599 | E | -16.94475 | 9.858547002 | 0.132012945 | 6.712879856 | NA | NA | NA | NA | NA | NA | NA |
| SCN4A | 1600 | N | -15.22625 | 10.20774338 | 0.138011658 | 7.069330882 | NA | NA | NA | NA | NA | NA | NA |
| SCN4A | 1601 | F | -18.13075 | 4.320202542 | 0.036871806 | 1.773931599 | 3.06 | 0.94 | 1.65 | 1.05 | -1.23 | 0 | 0 |
| SCN4A | 1602 | N | -21.588375 | 9.385315392 | 0.123883477 | 5.997222618 | NA | NA | NA | NA | NA | NA | NA |
| SCN4A | 1603 | V | -20.20717857 | 11.8769118 | 0.166685676 | 6.554704212 | NA | NA | NA | NA | NA | NA | NA |
| SCN4A | 1604 | A | -21.94735 | 10.85337722 | 0.149102759 | 5.574744951 | NA | NA | NA | NA | NA | NA | NA |
| SCN4A | 1605 | T | -24.71717857 | 7.37106919 | 0.089281498 | 1.694938734 | 5.73 | 0.29 | 0.35 | 1.44 | -0.58 | 0 | 0 |
| SCN4A | 1606 | E | -26.10841667 | 11.33114879 | 0.157310218 | 5.688709867 | 5.73 | 0.29 | 0.35 | 1.44 | -0.58 | 0 | 0 |
| SCN4A | 1607 | E | -25.00408333 | 13.36384424 | 0.192229129 | 7.755311885 | 5.73 | 0.29 | 0.35 | 1.44 | -0.58 | 0 | 0 |
| SCN9A | 114 | P | -18.77778571 | 57.15867813 | 0.888634281 | 53.87890618 | NA | NA | NA | NA | NA | NA | NA |
| SCN9A | 115 | L | -14.8425 | 54.42020461 | 0.843424556 | 51.18364512 | NA | NA | NA | NA | NA | NA | NA |
| SCN9A | 116 | R | -18.62504545 | 51.29216367 | 0.791783424 | 48.00244645 | NA | NA | NA | NA | NA | NA | NA |
| SCN9A | 117 | R | -21.22468182 | 53.05209822 | 0.820838355 | 49.72728476 | NA | NA | NA | NA | NA | NA | NA |
| SCN9A | 118 | I | -16.2135 | 51.76334247 | 0.79956216 | 48.48351679 | NA | NA | NA | NA | NA | NA | NA |
| SCN9A | 119 | S | -14.83566667 | 48.91714101 | 0.752573944 | 45.64696139 | NA | NA | NA | NA | NA | NA | NA |
| SCN9A | 120 | I | -18.438125 | 48.45221904 | 0.744898502 | 45.11258605 | NA | NA | NA | NA | NA | NA | NA |
| SCN9A | 121 | K | -18.23683333 | 48.71728224 | 0.749274456 | 45.3732392 | NA | NA | NA | NA | NA | NA | NA |
| SCN9A | 122 | I | -13.43625 | 46.13160814 | 0.706587311 | 42.84274971 | NA | NA | NA | NA | NA | NA | NA |
| SCN9A | 123 | L | -15.037375 | 43.42965702 | 0.661980537 | 40.12314165 | NA | NA | NA | NA | NA | NA | NA |
| SCN9A | 124 | V | -17.88635714 | 44.02606012 | 0.671826614 | 40.64275387 | NA | NA | NA | NA | NA | NA | NA |
| SCN9A | 125 | H | -15.235 | 42.55398779 | 0.647524028 | 39.18976455 | NA | NA | NA | NA | NA | NA | NA |
| SCN9A | 126 | S | -14.99483333 | 41.13576501 | 0.624110451 | 37.77136985 | NA | NA | NA | NA | NA | NA | NA |
| SCN9A | 127 | L | -11.339125 | 41.96537647 | 0.637806587 | 38.68159317 | NA | NA | NA | NA | NA | NA | NA |
| SCN9A | 128 | F | -10.04668182 | 40.5887506 | 0.615079736 | 37.35365885 | NA | NA | NA | NA | NA | NA | NA |
| SCN9A | 129 | S | -10.74283333 | 37.11711095 | 0.5577661 | 33.79267588 | NA | NA | NA | NA | NA | NA | NA |
| SCN9A | 130 | M | -9.11425 | 38.16715904 | 0.575101446 | 34.94657819 | NA | NA | NA | NA | NA | NA | NA |
| SCN9A | 131 | L | -5.764375 | 39.49303982 | 0.596990541 | 36.3718213 | NA | NA | NA | NA | NA | NA | NA |
| SCN9A | 132 | I | -5.804625 | 36.84052216 | 0.553199869 | 33.75879819 | NA | NA | NA | NA | NA | NA | NA |
| SCN9A | 133 | M | -5.218875 | 32.90833331 | 0.48828298 | 29.64884938 | NA | NA | NA | NA | NA | NA | NA |
| SCN9A | 134 | C | -2.536833333 | 36.29474467 | 0.544189575 | 32.83357494 | NA | NA | NA | NA | NA | NA | NA |
| SCN9A | 135 | T | -0.497214286 | 36.30091606 | 0.544291459 | 31.56926528 | NA | NA | NA | NA | NA | NA | NA |
| SCN9A | 136 | I | -0.891625 | 31.47986051 | 0.464700184 | 26.85646964 | NA | NA | NA | NA | NA | NA | NA |
| SCN9A | 137 | L | -0.967875 | 30.50518479 | 0.448609168 | 25.90388334 | NA | NA | NA | NA | NA | NA | NA |

| | | | | | | | | | | | | | |
|---|---|---|---|---|---|---|---|---|---|---|---|---|---|
| SCN9A | 138 | T | 2.495357143 | 34.41150649 | 0.513099013 | 29.622615 | NA | NA | NA | NA | NA | NA | NA |
| SCN9A | 139 | N | 3.727625 | 33.12831322 | 0.49191465 | 30.63824949 | NA | NA | NA | NA | NA | NA | NA |
| SCN9A | 140 | C | 3.7795 | 29.74725174 | 0.436096377 | 27.50878889 | NA | NA | NA | NA | NA | NA | NA |
| SCN9A | 141 | I | 6.30775 | 31.21006562 | 0.460246114 | 28.70203892 | NA | NA | NA | NA | NA | NA | NA |
| SCN9A | 142 | F | 8.644409091 | 32.72274053 | 0.485219011 | 30.8462975 | NA | NA | NA | NA | NA | NA | NA |
| SCN9A | 143 | M | 7.581625 | 29.63720945 | 0.434279678 | 27.82593358 | NA | NA | NA | NA | NA | NA | NA |
| SCN9A | 144 | T | 10.44535714 | 28.03054512 | 0.407755101 | 26.21904379 | NA | NA | NA | NA | NA | NA | NA |
| SCN9A | 145 | M | 13.52675 | 30.51316773 | 0.448740959 | 28.95438798 | NA | NA | NA | NA | NA | NA | NA |
| SCN9A | 146 | N | 16.70225 | 26.80315836 | 0.387492053 | 26.3046011 | NA | NA | NA | NA | NA | NA | NA |
| SCN9A | 147 | N | 18.614625 | 31.09823926 | 0.458399962 | 27.82126256 | NA | NA | NA | NA | NA | NA | NA |
| SCN9A | 148 | P | 16.18292857 | 34.45908742 | 0.513884531 | 31.25073912 | NA | NA | NA | NA | NA | NA | NA |
| SCN9A | 149 | P | 17.38107143 | 37.39196642 | 0.562303716 | 34.16366961 | NA | NA | NA | NA | NA | NA | NA |
| SCN9A | 150 | D | 19.0565 | 40.43389682 | 0.61252324 | 37.17467087 | NA | NA | NA | NA | NA | NA | NA |
| SCN9A | 151 | W | 15.15 | 40.59510652 | 0.615184667 | 37.4448403 | NA | NA | NA | NA | NA | NA | NA |
| SCN9A | 152 | T | 13.1195 | 38.88713525 | 0.586987603 | 37.07211162 | NA | NA | NA | NA | NA | NA | NA |
| SCN9A | 153 | K | 13.49783333 | 41.93275838 | 0.637268092 | 40.11934429 | NA | NA | NA | NA | NA | NA | NA |
| SCN9A | 154 | N | 11.39925 | 44.12124851 | 0.673398088 | 42.3088834 | NA | NA | NA | NA | NA | NA | NA |
| SCN9A | 155 | V | 8.445071429 | 40.73611357 | 0.617512566 | 38.92495438 | NA | NA | NA | NA | NA | NA | NA |
| SCN9A | 156 | E | 8.973277778 | 37.35790662 | 0.561741419 | 35.49862198 | NA | NA | NA | NA | NA | NA | NA |
| SCN9A | 157 | Y | 8.819666667 | 42.87018927 | 0.652744229 | 40.38157826 | NA | NA | NA | NA | NA | NA | NA |
| SCN9A | 158 | T | 3.980642857 | 43.09958388 | 0.656531327 | 40.63437014 | NA | NA | NA | NA | NA | NA | NA |
| SCN9A | 159 | F | 3.4055 | 38.05898824 | 0.573315644 | 35.57037252 | NA | NA | NA | NA | NA | NA | NA |
| SCN9A | 160 | T | 2.770928571 | 41.60390871 | 0.63183908 | 39.14810353 | NA | NA | NA | NA | NA | NA | NA |
| SCN9A | 161 | G | 0.68225 | 44.02725554 | 0.671846349 | 39.29651567 | NA | NA | NA | NA | NA | NA | NA |
| SCN9A | 162 | I | -1.0445 | 41.74038228 | 0.634092136 | 37.17522321 | NA | NA | NA | NA | NA | NA | NA |
| SCN9A | 163 | Y | -1.017666667 | 35.96088232 | 0.538677809 | 32.5727336 | NA | NA | NA | NA | NA | NA | NA |
| SCN9A | 164 | T | -1.8965 | 43.07943339 | 0.656198661 | 39.71735028 | NA | NA | NA | NA | NA | NA | NA |
| SCN9A | 165 | F | -5.375590909 | 45.1371792 | 0.690170187 | 41.75865272 | NA | NA | NA | NA | NA | NA | NA |
| SCN9A | 166 | E | -6.031277778 | 38.52933928 | 0.581080715 | 35.48925104 | NA | NA | NA | NA | NA | NA | NA |
| SCN9A | 167 | S | -6.256666667 | 40.73992166 | 0.617575434 | 37.75285626 | NA | NA | NA | NA | NA | NA | NA |
| SCN9A | 168 | L | -7.128125 | 45.05455839 | 0.688806192 | 42.03026176 | NA | NA | NA | NA | NA | NA | NA |
| SCN9A | 169 | V | -10.31535714 | 45.50468099 | 0.696237309 | 42.34119473 | NA | NA | NA | NA | NA | NA | NA |
| SCN9A | 170 | K | -11.28216667 | 39.03886767 | 0.589492568 | 35.90142385 | NA | NA | NA | NA | NA | NA | NA |

| | | | | | | | | | | | | | |
|---|---|---|---|---|---|---|---|---|---|---|---|---|---|
| SCN9A | 171 | I | -10.40625 | 44.49802327 | 0.679618299 | 41.44085573 | NA | NA | NA | NA | NA | NA | NA |
| SCN9A | 172 | L | -12.3595 | 48.18762096 | 0.740530226 | 45.06204984 | NA | NA | NA | NA | NA | NA | NA |
| SCN9A | 173 | A | -15.8129 | 48.81405324 | 0.750872058 | 45.565002 | NA | NA | NA | NA | NA | NA | NA |
| SCN9A | 174 | R | -17.04022727 | 42.25158475 | 0.642531627 | 38.92212016 | NA | NA | NA | NA | NA | NA | NA |
| SCN9A | 175 | G | -16.08075 | 48.36139748 | 0.74339912 | 45.15887902 | NA | NA | NA | NA | NA | NA | NA |
| SCN9A | 176 | F | -18.91622727 | 51.27984784 | 0.791580101 | 48.07741519 | NA | NA | NA | NA | NA | NA | NA |
| SCN9A | 177 | C | -14.81566667 | 52.90853393 | 0.818468238 | 49.74830989 | NA | NA | NA | NA | NA | NA | NA |
| SCN9A | 178 | V | -17.29878571 | 50.82835926 | 0.784126432 | 47.65792035 | NA | NA | NA | NA | NA | NA | NA |
| SCN9A | 179 | G | -13.78325 | 47.60849484 | 0.730969377 | 44.50853432 | NA | NA | NA | NA | NA | NA | NA |
| SCN9A | 180 | E | -12.08705556 | 45.25747553 | 0.69215617 | 42.10438888 | NA | NA | NA | NA | NA | NA | NA |
| SCN9A | 181 | F | -17.52495455 | 44.27164212 | 0.675880951 | 41.03958031 | NA | NA | NA | NA | NA | NA | NA |
| SCN9A | 182 | T | -17.39378571 | 42.53986776 | 0.647290919 | 39.28578634 | NA | NA | NA | NA | NA | NA | NA |
| SCN9A | 183 | F | -11.51422727 | 40.79450925 | 0.618476626 | 37.63119749 | NA | NA | NA | NA | NA | NA | NA |
| SCN9A | 184 | L | -11.590125 | 39.83132901 | 0.60257539 | 36.68010677 | NA | NA | NA | NA | NA | NA | NA |
| SCN9A | 185 | R | -16.96022727 | 38.33188695 | 0.577820955 | 35.09140448 | NA | NA | NA | NA | NA | NA | NA |
| SCN9A | 186 | D | -14.80775 | 36.42070275 | 0.546269029 | 33.21506225 | NA | NA | NA | NA | NA | NA | NA |
| SCN9A | 187 | P | -12.02964286 | 32.96488723 | 0.489216635 | 29.72800197 | NA | NA | NA | NA | NA | NA | NA |
| SCN9A | 188 | W | -9.711071429 | 27.65995005 | 0.401636911 | 24.51118598 | NA | NA | NA | NA | NA | NA | NA |
| SCN9A | 189 | N | -9.671125 | 35.60217062 | 0.532755803 | 32.47876554 | NA | NA | NA | NA | NA | NA | NA |
| SCN9A | 190 | W | -8.852714286 | 35.26262628 | 0.527150232 | 32.18548298 | NA | NA | NA | NA | NA | NA | NA |
| SCN9A | 191 | L | -4.145 | 30.76129419 | 0.452837303 | 27.42735243 | NA | NA | NA | NA | NA | NA | NA |
| SCN9A | 192 | D | -4.65925 | 35.68387312 | 0.534104637 | 32.35237321 | NA | NA | NA | NA | NA | NA | NA |
| SCN9A | 193 | F | -3.879954545 | 38.24363909 | 0.576364062 | 34.91933695 | NA | NA | NA | NA | NA | NA | NA |
| SCN9A | 194 | V | -0.920071429 | 35.43125441 | 0.52993413 | 30.85256679 | NA | NA | NA | NA | NA | NA | NA |
| SCN9A | 195 | V | 0.178071429 | 34.68549657 | 0.517622341 | 32.54708871 | NA | NA | NA | NA | NA | NA | NA |
| SCN9A | 196 | I | -0.76525 | 39.26273665 | 0.593188443 | 34.6714081 | NA | NA | NA | NA | NA | NA | NA |
| SCN9A | 197 | V | 1.181642857 | 41.05657252 | 0.622803054 | 38.63282366 | NA | NA | NA | NA | NA | NA | NA |
| SCN9A | 198 | F | 3.564681818 | 35.04847863 | 0.523614848 | 32.62483415 | NA | NA | NA | NA | NA | NA | NA |
| SCN9A | 199 | A | 4.9235 | 39.7097757 | 0.600568654 | 37.26866002 | NA | NA | NA | NA | NA | NA | NA |
| SCN9A | 200 | Y | 3.122083333 | 42.36919754 | 0.644473308 | 39.92577144 | NA | NA | NA | NA | NA | NA | NA |
| SCN9A | 201 | L | 6.153875 | 41.00503228 | 0.621952171 | 38.56950698 | NA | NA | NA | NA | NA | NA | NA |
| SCN9A | 202 | T | 9.596785714 | 40.27193001 | 0.609849315 | 38.41841307 | NA | NA | NA | NA | NA | NA | NA |
| SCN9A | 203 | E | 9.786722222 | 43.89946712 | 0.669736677 | 42.06024919 | NA | NA | NA | NA | NA | NA | NA |

| | | | | | | | | | | | | | |
|---|---|---|---|---|---|---|---|---|---|---|---|---|---|
| SCN9A | 204 | F | 8.415045455 | 44.84462191 | 0.68534033 | 42.98198953 | NA | NA | NA | NA | NA | NA | NA |
| SCN9A | 205 | V | 11.38378571 | 44.49089998 | 0.6795007 | 42.69777894 | NA | NA | NA | NA | NA | NA | NA |
| SCN9A | 206 | N | 12.75275 | 41.63419496 | 0.632339079 | 39.81859089 | NA | NA | NA | NA | NA | NA | NA |
| SCN9A | 207 | L | 12.91175 | 36.67400803 | 0.550450871 | 34.86712769 | NA | NA | NA | NA | NA | NA | NA |
| SCN9A | 208 | G | 16.159 | 37.53446648 | 0.564656263 | 35.72320593 | NA | NA | NA | NA | NA | NA | NA |
| SCN9A | 209 | N | 14.51625 | 33.30555253 | 0.494840711 | 31.49502972 | NA | NA | NA | NA | NA | NA | NA |
| SCN9A | 210 | V | 14.77107143 | 33.91947952 | 0.504976092 | 32.11214978 | NA | NA | NA | NA | NA | NA | NA |
| SCN9A | 211 | S | 14.12116667 | 30.59903665 | 0.450158577 | 28.78654446 | NA | NA | NA | NA | NA | NA | NA |
| SCN9A | 212 | A | 11.4111 | 31.0373583 | 0.457394872 | 29.22572015 | NA | NA | NA | NA | NA | NA | NA |
| SCN9A | 213 | L | 9.7615 | 33.10548604 | 0.491537794 | 31.28950048 | NA | NA | NA | NA | NA | NA | NA |
| SCN9A | 214 | R | 10.86913636 | 32.12059087 | 0.475278064 | 30.34013448 | NA | NA | NA | NA | NA | NA | NA |
| SCN9A | 215 | T | 8.172071429 | 27.40532859 | 0.39743334 | 25.5391247 | NA | NA | NA | NA | NA | NA | NA |
| SCN9A | 216 | F | 5.110136364 | 31.0446042 | 0.457514495 | 28.58901419 | NA | NA | NA | NA | NA | NA | NA |
| SCN9A | 217 | R | 6.566681818 | 31.59439857 | 0.466591104 | 29.14235032 | NA | NA | NA | NA | NA | NA | NA |
| SCN9A | 218 | V | 3.967928571 | 26.7110924 | 0.385972127 | 24.26239605 | NA | NA | NA | NA | NA | NA | NA |
| SCN9A | 219 | L | 0.692875 | 29.99625845 | 0.440207253 | 27.61856678 | NA | NA | NA | NA | NA | NA | NA |
| SCN9A | 220 | R | 1.873772727 | 31.129245 | 0.458911838 | 26.4948977 | NA | NA | NA | NA | NA | NA | NA |
| SCN9A | 221 | A | 0.5345 | 27.98698758 | 0.407036005 | 23.26037217 | NA | NA | NA | NA | NA | NA | NA |
| SCN9A | 222 | L | -2.141875 | 25.76743777 | 0.370393241 | 22.4336161 | NA | NA | NA | NA | NA | NA | NA |
| SCN9A | 223 | K | -4.038277778 | 30.19611482 | 0.443506702 | 26.83107807 | NA | NA | NA | NA | NA | NA | NA |
| SCN9A | 224 | T | -4.219214286 | 27.64804691 | 0.401440401 | 24.59720635 | NA | NA | NA | NA | NA | NA | NA |
| SCN9A | 225 | I | -6.8995 | 24.56041734 | 0.350466424 | 21.43562965 | NA | NA | NA | NA | NA | NA | NA |
| SCN9A | 226 | S | -9.3035 | 29.96067521 | 0.439619806 | 26.77679485 | NA | NA | NA | NA | NA | NA | NA |
| SCN9A | 227 | V | -8.621357143 | 32.07889202 | 0.474589653 | 28.8559732 | NA | NA | NA | NA | NA | NA | NA |
| SCN9A | 228 | I | -9.306125 | 28.03692597 | 0.407860443 | 24.78784223 | NA | NA | NA | NA | NA | NA | NA |
| SCN9A | 229 | P | -13.26107143 | 29.04867558 | 0.424563516 | 25.65599604 | NA | NA | NA | NA | NA | NA | NA |
| SCN9A | 230 | G | -14.117 | 26.35551269 | 0.380101827 | 22.98202408 | NA | NA | NA | NA | NA | NA | NA |
| SCN9A | 231 | L | -10.331 | 24.86213645 | 0.355447534 | 21.50083143 | NA | NA | NA | NA | NA | NA | NA |
| SCN9A | 232 | K | -13.07172222 | 26.41602745 | 0.381100871 | 23.04050525 | NA | NA | NA | NA | NA | NA | NA |
| SCN9A | 233 | T | -15.62735714 | 23.6988282 | 0.336242365 | 21.02963479 | NA | NA | NA | NA | NA | NA | NA |
| SCN9A | 234 | I | -12.31475 | 20.74203618 | 0.287428398 | 17.34633652 | NA | NA | NA | NA | NA | NA | NA |
| SCN9A | 235 | V | -10.32864286 | 22.2530746 | 0.312374278 | 18.92712045 | NA | NA | NA | NA | NA | NA | NA |
| SCN9A | 236 | G | -13.4245 | 22.42200885 | 0.31516323 | 19.76711486 | NA | NA | NA | NA | NA | NA | NA |

| | | | | | | | | | | | | | |
|---|---|---|---|---|---|---|---|---|---|---|---|---|---|
| SCN9A | 237 | A | -14.0207 | 18.98444466 | 0.258412148 | 16.32358238 | NA | NA | NA | NA | NA | NA | NA |
| SCN9A | 238 | L | -9.896125 | 17.04105598 | 0.226328556 | 13.68469024 | NA | NA | NA | NA | NA | NA | NA |
| SCN9A | 239 | I | -10.953625 | 20.66837045 | 0.286212243 | 17.32746557 | NA | NA | NA | NA | NA | NA | NA |
| SCN9A | 240 | Q | -15.37161111 | 19.0753022 | 0.259912124 | 16.40873561 | NA | NA | NA | NA | NA | NA | NA |
| SCN9A | 241 | S | -12.57066667 | 15.84197431 | 0.2065328 | 13.18278505 | NA | NA | NA | NA | NA | NA | NA |
| SCN9A | 242 | V | -9.733642857 | 17.74951681 | 0.238024605 | 15.22722328 | NA | NA | NA | NA | NA | NA | NA |
| SCN9A | 243 | K | -13.92238889 | 19.37711751 | 0.264894823 | 16.72323257 | NA | NA | NA | NA | NA | NA | NA |
| SCN9A | 244 | K | -15.29494444 | 15.19634986 | 0.195874123 | 13.71061849 | NA | NA | NA | NA | NA | NA | NA |
| SCN9A | 245 | L | -9.56075 | 13.70898497 | 0.171319071 | 11.16560862 | NA | NA | NA | NA | NA | NA | NA |
| SCN9A | 246 | S | -10.67083333 | 18.70107711 | 0.253734006 | 16.1612132 | NA | NA | NA | NA | NA | NA | NA |
| SCN9A | 247 | D | -11.403 | 18.34059546 | 0.24778278 | 15.70776463 | NA | NA | NA | NA | NA | NA | NA |
| SCN9A | 248 | V | -8.135357143 | 14.19397217 | 0.179325772 | 11.64227006 | NA | NA | NA | NA | NA | NA | NA |
| SCN9A | 249 | M | -5.389 | 14.88559082 | 0.190743772 | 11.96450982 | NA | NA | NA | NA | NA | NA | NA |
| SCN9A | 250 | I | -6.831 | 19.94460356 | 0.274263505 | 16.84863691 | NA | NA | NA | NA | NA | NA | NA |
| SCN9A | 251 | L | -6.030125 | 17.02533788 | 0.226069064 | 14.44526254 | NA | NA | NA | NA | NA | NA | NA |
| SCN9A | 252 | T | -2.616928571 | 15.01513623 | 0.19288245 | 11.6161774 | NA | NA | NA | NA | NA | NA | NA |
| SCN9A | 253 | V | -0.9635 | 18.72572914 | 0.254140989 | 16.31087257 | NA | NA | NA | NA | NA | NA | NA |
| SCN9A | 254 | F | -2.831681818 | 21.02265126 | 0.292061099 | 18.43150397 | NA | NA | NA | NA | NA | NA | NA |
| SCN9A | 255 | C | -1.167166667 | 18.35983077 | 0.248100337 | 15.78403867 | NA | NA | NA | NA | NA | NA | NA |
| SCN9A | 256 | L | 1.457375 | 14.79945036 | 0.189321671 | 12.38426939 | NA | NA | NA | NA | NA | NA | NA |
| SCN9A | 257 | S | 1.760666667 | 19.94122318 | 0.274207698 | 17.51736784 | NA | NA | NA | NA | NA | NA | NA |
| SCN9A | 258 | V | 1.2295 | 21.88368161 | 0.306275933 | 19.44544133 | NA | NA | NA | NA | NA | NA | NA |
| SCN9A | 259 | F | 2.461227273 | 17.84356497 | 0.239577255 | 15.38736546 | NA | NA | NA | NA | NA | NA | NA |
| SCN9A | 260 | A | 6.2809 | 20.19740048 | 0.278436954 | 18.40579776 | NA | NA | NA | NA | NA | NA | NA |
| SCN9A | 261 | L | 5.59675 | 22.84739771 | 0.322186016 | 20.98568149 | NA | NA | NA | NA | NA | NA | NA |
| SCN9A | 262 | I | 5.931875 | 22.91415277 | 0.323288082 | 21.17700027 | NA | NA | NA | NA | NA | NA | NA |
| SCN9A | 263 | G | 9.688 | 20.9752789 | 0.291279025 | 19.09356602 | NA | NA | NA | NA | NA | NA | NA |
| SCN9A | 264 | L | 11.275125 | 22.38353629 | 0.314528083 | 20.49667817 | NA | NA | NA | NA | NA | NA | NA |
| SCN9A | 265 | Q | 11.06416667 | 25.10987338 | 0.359537447 | 23.20982294 | NA | NA | NA | NA | NA | NA | NA |
| SCN9A | 266 | L | 11.50975 | 23.44224597 | 0.332006424 | 21.21213017 | NA | NA | NA | NA | NA | NA | NA |
| SCN9A | 267 | F | 13.5925 | 19.54811633 | 0.267717859 | 17.33973394 | NA | NA | NA | NA | NA | NA | NA |
| SCN9A | 268 | M | 16.089875 | 23.71951975 | 0.336583964 | 21.56066706 | NA | NA | NA | NA | NA | NA | NA |
| SCN9A | 269 | G | 20.028 | 23.95120135 | 0.340408818 | 21.87284977 | NA | NA | NA | NA | NA | NA | NA |

| | | | | | | | | | | | | | |
|---|---|---|---|---|---|---|---|---|---|---|---|---|---|
| SCN9A | 270 | N | 19.413125 | 21.58516382 | 0.301347674 | 19.52876684 | NA | NA | NA | NA | NA | NA | NA |
| SCN9A | 271 | L | 20.167375 | 17.03077382 | 0.226158807 | 14.99063699 | NA | NA | NA | NA | NA | NA | NA |
| SCN9A | 272 | K | 23.8015 | 21.294308 | 0.296545907 | 15.51997205 | NA | NA | NA | NA | NA | NA | NA |
| SCN9A | 273 | H | 24.6647 | 19.31976296 | 0.263947951 | 13.5798622 | NA | NA | NA | NA | NA | NA | NA |
| SCN9A | 274 | K | 28.84127778 | 18.55889789 | 0.251386756 | 16.16993668 | NA | NA | NA | NA | NA | NA | NA |
| SCN9A | 275 | C | 32.02116667 | 18.72754199 | 0.254170917 | 16.29179605 | NA | NA | NA | NA | NA | NA | NA |
| SCN9A | 276 | F | 36.00259091 | 20.93049968 | 0.29053976 | 18.49036842 | NA | NA | NA | NA | NA | NA | NA |
| SCN9A | 277 | R | 40.47813636 | 20.44772129 | 0.282569525 | 18.15172185 | NA | NA | NA | NA | NA | NA | NA |
| SCN9A | 278 | N | 39.314375 | 23.93067498 | 0.340069946 | 21.57077782 | NA | NA | NA | NA | NA | NA | NA |
| SCN9A | 279 | S | 43.57916667 | 24.87630065 | 0.355681372 | 22.56372128 | NA | NA | NA | NA | NA | NA | NA |
| SCN9A | 280 | L | 43.439 | 24.8614714 | 0.355436554 | 22.51562849 | NA | NA | NA | NA | NA | NA | NA |
| SCN9A | 281 | E | 48.5905 | 25.86065281 | 0.371932138 | 23.61209741 | NA | NA | NA | NA | NA | NA | NA |
| SCN9A | 282 | N | 49.478125 | 28.61456482 | 0.417396739 | 26.33343931 | NA | NA | NA | NA | NA | NA | NA |
| SCN9A | 283 | N | 46.864 | 31.61122158 | 0.466868837 | 29.27670321 | NA | NA | NA | NA | NA | NA | NA |
| SCN9A | 284 | E | 44.37627778 | 25.93643285 | 0.373183198 | 23.6182008 | NA | NA | NA | NA | NA | NA | NA |
| SCN9A | 285 | T | 40.53178571 | 27.79431391 | 0.403855137 | 25.41035112 | NA | NA | NA | NA | NA | NA | NA |
| SCN9A | 286 | L | 36.014 | 25.28859951 | 0.362488054 | 22.86349681 | NA | NA | NA | NA | NA | NA | NA |
| SCN9A | 287 | E | 35.53405556 | 28.69030695 | 0.418647173 | 26.28567917 | NA | NA | NA | NA | NA | NA | NA |
| SCN9A | 288 | S | 40.2005 | 27.76928811 | 0.403441983 | 25.39211992 | NA | NA | NA | NA | NA | NA | NA |
| SCN9A | 289 | I | 39.442625 | 23.31065628 | 0.329833997 | 20.93798303 | NA | NA | NA | NA | NA | NA | NA |
| SCN9A | 290 | M | 34.852625 | 22.53311117 | 0.316997429 | 20.09796903 | NA | NA | NA | NA | NA | NA | NA |
| SCN9A | 291 | N | 37.478 | 26.26132828 | 0.378546928 | 23.86892342 | NA | NA | NA | NA | NA | NA | NA |
| SCN9A | 292 | T | 40.14392857 | 24.68844479 | 0.352580041 | 22.32667862 | NA | NA | NA | NA | NA | NA | NA |
| SCN9A | 293 | L | 38.112375 | 19.28595409 | 0.263389797 | 16.93146791 | NA | NA | NA | NA | NA | NA | NA |
| SCN9A | 294 | E | 34.79083333 | 18.27446421 | 0.246691012 | 15.86672214 | NA | NA | NA | NA | NA | NA | NA |
| SCN9A | 295 | S | 36.71916667 | 15.67656423 | 0.203802029 | 13.30182338 | NA | NA | NA | NA | NA | NA | NA |
| SCN9A | 296 | E | 34.15427778 | 9.448135954 | 0.100976301 | 7.029613032 | NA | NA | NA | NA | NA | NA | NA |
| SCN9A | 297 | E | 38.75938889 | 10.59898415 | 0.119975765 | 8.295962786 | NA | NA | NA | NA | NA | NA | NA |
| SCN9A | 298 | D | 40.01325 | 15.62760087 | 0.202993688 | 13.29865867 | NA | NA | NA | NA | NA | NA | NA |
| SCN9A | 299 | F | 34.83077273 | 15.16785084 | 0.19540363 | 12.79518397 | NA | NA | NA | NA | NA | NA | NA |
| SCN9A | 300 | R | 35.85104545 | 7.065781839 | 0.061645785 | 4.661316789 | 2.47 | -0.88 | -1.43 | 18.82 | 1.42 | 1 | -1 |
| SCN9A | 301 | K | 41.57672222 | 14.8993369 | 0.190970707 | 12.57880889 | NA | NA | NA | NA | NA | NA | NA |
| SCN9A | 302 | Y | 39.90133333 | 17.26956774 | 0.230101079 | 14.92818748 | NA | NA | NA | NA | NA | NA | NA |

| | | | | | | | | | | | | | |
|---|---|---|---|---|---|---|---|---|---|---|---|---|---|
| SCN9A | 303 | F | 34.25740909 | 14.79934208 | 0.189319883 | 12.41198347 | NA | NA | NA | NA | NA | NA | NA |
| SCN9A | 304 | Y | 33.57433333 | 12.29229124 | 0.147930737 | 9.904817683 | NA | NA | NA | NA | NA | NA | NA |
| SCN9A | 305 | Y | 32.42766667 | 10.11257578 | 0.111945602 | 7.714961386 | NA | NA | NA | NA | NA | NA | NA |
| SCN9A | 306 | L | 29.809125 | 7.066545266 | 0.061658388 | 4.463581675 | 2.59 | -0.8 | -0.4 | 3.38 | 0.81 | 0 | 0 |
| SCN9A | 307 | E | 33.22505556 | 4.145954534 | 0.013442072 | 1.762239382 | 2.47 | -0.88 | -1.43 | 18.82 | 1.42 | 1 | -1 |
| SCN9A | 308 | G | 30.93025 | 4.073012153 | 0.012237859 | 1.619495774 | 2.47 | -0.88 | -1.43 | 18.82 | 1.42 | 1 | -1 |
| SCN9A | 309 | S | 28.82366667 | 5.689453225 | 0.038923842 | 3.256845477 | 2.59 | -0.8 | -0.4 | 3.38 | 0.81 | 0 | 0 |
| SCN9A | 310 | K | 28.03283333 | 7.643566772 | 0.071184492 | 5.336496566 | 2.59 | -0.8 | -0.4 | 3.38 | 0.81 | 0 | 0 |
| SCN9A | 311 | D | 26.69075 | 10.51445229 | 0.118580221 | 8.00392796 | NA | NA | NA | NA | NA | NA | NA |
| SCN9A | 312 | A | 28.6455 | 13.00366706 | 0.159674909 | 10.60357509 | NA | NA | NA | NA | NA | NA | NA |
| SCN9A | 313 | L | 27.95975 | 13.32339221 | 0.164953283 | 11.04928414 | NA | NA | NA | NA | NA | NA | NA |
| SCN9A | 314 | L | 24.2875 | 15.48255544 | 0.200599119 | 10.33346743 | NA | NA | NA | NA | NA | NA | NA |
| SCN9A | 315 | C | 26.053 | 18.72738583 | 0.254168339 | 13.55045917 | NA | NA | NA | NA | NA | NA | NA |
| SCN9A | 316 | G | 24.83925 | 21.57890463 | 0.30124434 | 16.82078912 | NA | NA | NA | NA | NA | NA | NA |
| SCN9A | 317 | F | 25.01159091 | 25.14269319 | 0.360079272 | 20.38862297 | NA | NA | NA | NA | NA | NA | NA |
| SCN9A | 318 | S | 22.07416667 | 25.53472344 | 0.366551338 | 20.80199904 | NA | NA | NA | NA | NA | NA | NA |
| SCN9A | 319 | T | 22.53121429 | 25.08401646 | 0.359110573 | 20.36749148 | NA | NA | NA | NA | NA | NA | NA |
| SCN9A | 320 | D | 19.387375 | 22.79820383 | 0.32137387 | 18.09234697 | NA | NA | NA | NA | NA | NA | NA |
| SCN9A | 321 | S | 21.8095 | 21.37165768 | 0.29782288 | 16.64975264 | NA | NA | NA | NA | NA | NA | NA |
| SCN9A | 322 | G | 24.49725 | 19.12673244 | 0.260761191 | 14.39151038 | NA | NA | NA | NA | NA | NA | NA |
| SCN9A | 323 | Q | 27.49194444 | 18.49507202 | 0.250333048 | 13.31338707 | NA | NA | NA | NA | NA | NA | NA |
| SCN9A | 324 | C | 31.0915 | 19.90576331 | 0.273622288 | 17.29725834 | NA | NA | NA | NA | NA | NA | NA |
| SCN9A | 325 | P | 34.19535714 | 17.56178359 | 0.234925299 | 15.21795137 | NA | NA | NA | NA | NA | NA | NA |
| SCN9A | 326 | E | 36.91527778 | 21.29893915 | 0.296622363 | 19.00291388 | NA | NA | NA | NA | NA | NA | NA |
| SCN9A | 327 | G | 38.53 | 22.30800545 | 0.313281137 | 19.98088541 | NA | NA | NA | NA | NA | NA | NA |
| SCN9A | 328 | Y | 36.50275 | 16.63261967 | 0.219585641 | 14.30695432 | NA | NA | NA | NA | NA | NA | NA |
| SCN9A | 329 | T | 33.94407143 | 22.6186084 | 0.318408911 | 20.17055415 | NA | NA | NA | NA | NA | NA | NA |
| SCN9A | 330 | C | 29.92083333 | 23.7554687 | 0.337177448 | 21.3003453 | NA | NA | NA | NA | NA | NA | NA |
| SCN9A | 331 | V | 30.42007143 | 23.20076397 | 0.328019774 | 20.75770287 | NA | NA | NA | NA | NA | NA | NA |
| SCN9A | 332 | K | 26.10516667 | 24.72129582 | 0.353122382 | 22.17439917 | NA | NA | NA | NA | NA | NA | NA |
| SCN9A | 333 | I | 29.43625 | 21.7586023 | 0.304210987 | 19.32438062 | NA | NA | NA | NA | NA | NA | NA |
| SCN9A | 334 | G | 27.34825 | 21.88543612 | 0.306304899 | 19.47270668 | NA | NA | NA | NA | NA | NA | NA |
| SCN9A | 335 | R | 26.09031818 | 14.75434611 | 0.18857704 | 9.154366075 | NA | NA | NA | NA | NA | NA | NA |

| | | | | | | | | | | | | | |
|---|---|---|---|---|---|---|---|---|---|---|---|---|---|
| SCN9A | 336 | N | 21.902125 | 16.96661961 | 0.225099679 | 11.18905144 | NA | NA | NA | NA | NA | NA | NA |
| SCN9A | 337 | P | 21.77392857 | 14.12584847 | 0.178201111 | 8.345677602 | NA | NA | NA | NA | NA | NA | NA |
| SCN9A | 338 | D | 20.393375 | 11.11176003 | 0.128441232 | 5.908798665 | NA | NA | NA | NA | NA | NA | NA |
| SCN9A | 339 | Y | 22.79716667 | 12.3496757 | 0.148878103 | 6.57971328 | NA | NA | NA | NA | NA | NA | NA |
| SCN9A | 340 | G | 20.66925 | 17.40705894 | 0.232370934 | 12.03362012 | NA | NA | NA | NA | NA | NA | NA |
| SCN9A | 341 | Y | 18.40808333 | 16.07598883 | 0.210396168 | 14.03803251 | NA | NA | NA | NA | NA | NA | NA |
| SCN9A | 342 | T | 17.49564286 | 12.9945903 | 0.15952506 | 10.97061114 | NA | NA | NA | NA | NA | NA | NA |
| SCN9A | 343 | S | 16.88416667 | 17.03500857 | 0.226228719 | 14.94163699 | NA | NA | NA | NA | NA | NA | NA |
| SCN9A | 344 | F | 13.47868182 | 16.83918148 | 0.22299579 | 14.65720129 | NA | NA | NA | NA | NA | NA | NA |
| SCN9A | 345 | D | 17.169125 | 21.91517577 | 0.306795873 | 19.7768992 | NA | NA | NA | NA | NA | NA | NA |
| SCN9A | 346 | T | 13.72564286 | 22.16077208 | 0.310850447 | 20.36819397 | NA | NA | NA | NA | NA | NA | NA |
| SCN9A | 347 | F | 7.877590909 | 21.45505978 | 0.299199774 | 19.62149989 | NA | NA | NA | NA | NA | NA | NA |
| SCN9A | 348 | S | 10.642 | 19.46160132 | 0.266289574 | 17.63139801 | NA | NA | NA | NA | NA | NA | NA |
| SCN9A | 349 | W | 14.39428571 | 17.4879751 | 0.233706787 | 15.6946767 | NA | NA | NA | NA | NA | NA | NA |
| SCN9A | 350 | A | 10.1647 | 17.1575479 | 0.228251732 | 15.34238576 | NA | NA | NA | NA | NA | NA | NA |
| SCN9A | 351 | F | 5.808136364 | 15.93028691 | 0.207990761 | 14.12401041 | NA | NA | NA | NA | NA | NA | NA |
| SCN9A | 352 | L | 9.999875 | 13.21616332 | 0.163183031 | 11.37696501 | NA | NA | NA | NA | NA | NA | NA |
| SCN9A | 353 | A | 10.0411 | 12.33466355 | 0.148630265 | 10.53421119 | NA | NA | NA | NA | NA | NA | NA |
| SCN9A | 354 | L | 6.388375 | 12.43701319 | 0.150319966 | 10.87071776 | NA | NA | NA | NA | NA | NA | NA |
| SCN9A | 355 | F | 4.692045455 | 10.15535342 | 0.112651822 | 7.699542364 | NA | NA | NA | NA | NA | NA | NA |
| SCN9A | 356 | R | 10.22040909 | 7.894460463 | 0.075326521 | 6.121648736 | 1.82 | -0.79 | -2.05 | 26.54 | 1.33 | 2 | 0 |
| SCN9A | 357 | L | 6.7265 | 8.787046717 | 0.090062314 | 7.036266686 | 1.89 | -0.67 | -1.57 | 18.75 | 1.03 | 1 | 1 |
| SCN9A | 358 | M | 1.82825 | 7.976196775 | 0.076675914 | 5.578796082 | 2.58 | -0.8 | -0.4 | 3.38 | 0.81 | 0 | 0 |
| SCN9A | 359 | T | 3.702642857 | 4.208643249 | 0.014477006 | 1.773790342 | 2.58 | -0.8 | -0.4 | 3.38 | 0.81 | 0 | 0 |
| SCN9A | 360 | Q | 4.592277778 | 3.368329259 | 6.04E-04 | 1.560203636 | 1.89 | -0.67 | -1.57 | 18.75 | 1.03 | 1 | 1 |
| SCN9A | 361 | D | 9.158125 | 4.118424092 | 0.012987569 | 2.312501781 | 1.85 | -0.79 | -1.72 | 21.91 | 1.22 | 2 | 0 |
| SCN9A | 362 | Y | 13.95291667 | 3.541069189 | 0.003455961 | 1.607634914 | 2.24 | -0.87 | -2.83 | 38.17 | 1.68 | 3 | -1 |
| SCN9A | 363 | W | 8.929571429 | 5.371246503 | 0.033670536 | 3.393266908 | 1.98 | -0.78 | -2.6 | 34.26 | 1.5 | 2 | 0 |
| SCN9A | 364 | E | 13.19083333 | 3.9330642 | 0.009927445 | 1.89756268 | 2.24 | -0.87 | -2.83 | 38.17 | 1.68 | 3 | -1 |
| SCN9A | 365 | N | 16.42575 | 7.293572581 | 0.065406404 | 5.272965753 | 2.24 | -0.39 | -2.77 | 33.8 | 1.01 | 2 | -2 |
| SCN9A | 366 | L | 13.552875 | 9.455265464 | 0.101094002 | 7.351797753 | NA | NA | NA | NA | NA | NA | NA |
| SCN9A | 367 | Y | 12.36425 | 10.88980161 | 0.124776899 | 8.731734013 | NA | NA | NA | NA | NA | NA | NA |
| SCN9A | 368 | Q | 16.89216667 | 10.14314852 | 0.112450331 | 8.34211038 | NA | NA | NA | NA | NA | NA | NA |

| | | | | | | | | | | | | | |
|---|---|---|---|---|---|---|---|---|---|---|---|---|---|
| SCN9A | 369 | Q | 18.79716667 | 12.23906308 | 0.147051988 | 10.61300934 | NA | NA | NA | NA | NA | NA | NA |
| SCN9A | 370 | T | 15.90892857 | 15.15285221 | 0.195156016 | 13.0482331 | NA | NA | NA | NA | NA | NA | NA |
| SCN9A | 371 | L | 15.8085 | 16.33146821 | 0.214613902 | 14.18265306 | NA | NA | NA | NA | NA | NA | NA |
| SCN9A | 372 | R | 21.98977273 | 14.16793076 | 0.178895852 | 12.78267903 | NA | NA | NA | NA | NA | NA | NA |
| SCN9A | 373 | A | 20.6193 | 19.32486833 | 0.264032236 | 13.74169623 | NA | NA | NA | NA | NA | NA | NA |
| SCN9A | 374 | A | 17.8659 | 20.73816393 | 0.28736447 | 18.64109306 | NA | NA | NA | NA | NA | NA | NA |
| SCN9A | 375 | G | 17.0535 | 20.72473674 | 0.2871428 | 18.56739693 | NA | NA | NA | NA | NA | NA | NA |
| SCN9A | 376 | K | 16.22494444 | 20.56307319 | 0.28447388 | 18.4634313 | NA | NA | NA | NA | NA | NA | NA |
| SCN9A | 377 | T | 12.8465 | 22.44631794 | 0.315564551 | 20.24597214 | NA | NA | NA | NA | NA | NA | NA |
| SCN9A | 378 | Y | 12.762 | 19.14284477 | 0.261027191 | 16.92383096 | NA | NA | NA | NA | NA | NA | NA |
| SCN9A | 379 | M | 9.664125 | 17.52843595 | 0.234374759 | 15.33495993 | NA | NA | NA | NA | NA | NA | NA |
| SCN9A | 380 | I | 6.768375 | 19.74990162 | 0.271049152 | 17.85413218 | NA | NA | NA | NA | NA | NA | NA |
| SCN9A | 381 | F | 7.3765 | 17.5098173 | 0.234067382 | 15.6313165 | NA | NA | NA | NA | NA | NA | NA |
| SCN9A | 382 | F | 8.784045455 | 13.80664423 | 0.172931338 | 11.83142767 | NA | NA | NA | NA | NA | NA | NA |
| SCN9A | 383 | V | 4.1215 | 16.30629388 | 0.214198297 | 14.71871673 | NA | NA | NA | NA | NA | NA | NA |
| SCN9A | 384 | V | 1.976071429 | 16.89328512 | 0.223888992 | 14.07654797 | NA | NA | NA | NA | NA | NA | NA |
| SCN9A | 385 | V | 3.198928571 | 13.92698564 | 0.174918066 | 11.43711092 | NA | NA | NA | NA | NA | NA | NA |
| SCN9A | 386 | I | 3.15175 | 10.4553196 | 0.117603993 | 7.851528318 | NA | NA | NA | NA | NA | NA | NA |
| SCN9A | 387 | F | 0.087772727 | 13.87754265 | 0.174101807 | 9.131832214 | NA | NA | NA | NA | NA | NA | NA |
| SCN9A | 388 | L | -1.955625 | 14.35611288 | 0.182002569 | 9.500682478 | NA | NA | NA | NA | NA | NA | NA |
| SCN9A | 389 | G | 0 | 12.46050577 | 0.150707807 | 8.477311578 | NA | NA | NA | NA | NA | NA | NA |
| SCN9A | 390 | S | 0.543666667 | 8.354956972 | 0.082928902 | 4.293562222 | 4.11 | -0.35 | -1.08 | 12.03 | 0.28 | 1 | 1 |
| SCN9A | 391 | F | -1.009409091 | 7.984553651 | 0.076813878 | 3.236258764 | 4.84 | 0.04 | -0.22 | 2.41 | -0.31 | 0 | 0 |
| SCN9A | 392 | Y | -6.19375 | 10.77606106 | 0.122899145 | 8.81644581 | NA | NA | NA | NA | NA | NA | NA |
| SCN9A | 393 | L | -3.509625 | 10.69939947 | 0.121633532 | 7.445054821 | NA | NA | NA | NA | NA | NA | NA |
| SCN9A | 394 | I | -2.834875 | 5.78865943 | 0.040561647 | 2.656656386 | 3.4 | 1.13 | 2.1 | 0.62 | -1.67 | 0 | 0 |
| SCN9A | 395 | N | -6.7185 | 6.047449462 | 0.044834037 | 3.779189984 | 2.65 | 1.47 | 2.88 | 0.5 | -1.73 | 0 | 0 |
| SCN9A | 396 | L | -8.867875 | 8.972038174 | 0.093116356 | 6.395889495 | 2.65 | 1.47 | 2.88 | 0.5 | -1.73 | 0 | 0 |
| SCN9A | 397 | I | -6.7315 | 8.906619729 | 0.092036357 | 6.35358551 | 2.65 | 1.47 | 2.88 | 0.5 | -1.73 | 0 | 0 |
| SCN9A | 398 | L | -8.18925 | 3.670732216 | 0.00559658 | 1.741863055 | 2.65 | 1.47 | 2.88 | 0.5 | -1.73 | 0 | 0 |
| SCN9A | 399 | A | -12.2791 | 6.832414873 | 0.057793107 | 4.170107807 | 2.74 | 1.54 | 3.2 | 0.43 | -1.76 | 0 | 0 |
| SCN9A | 400 | V | -12.56164286 | 9.439280958 | 0.100830113 | 6.752322777 | NA | NA | NA | NA | NA | NA | NA |
| SCN9A | 401 | V | -11.28164286 | 10.00902328 | 0.110236044 | 7.331501134 | NA | NA | NA | NA | NA | NA | NA |

| | | | | | | | | | | | | | |
|---|---|---|---|---|---|---|---|---|---|---|---|---|---|
| SCN9A | 402 | A | -14.0841 | 6.998204484 | 0.060530144 | 4.277387719 | 2.74 | 1.54 | 3.2 | 0.43 | -1.76 | 0 | 0 |
| SCN9A | 403 | M | -17.6135 | 8.810161009 | 0.09044391 | 6.162140112 | 2.77 | 0.4 | 0.85 | 1.81 | -0.58 | 0 | 0 |
| SCN9A | 404 | A | -16.5305 | 11.84601617 | 0.140563138 | 9.181703785 | NA | NA | NA | NA | NA | NA | NA |
| SCN9A | 405 | Y | -16.16683333 | 12.20491352 | 0.14648821 | 9.492627011 | NA | NA | NA | NA | NA | NA | NA |
| SCN9A | 407 | E | -21.25038889 | 12.86816514 | 0.157437895 | 7.029933972 | NA | NA | NA | NA | NA | NA | NA |
| SCN9A | 408 | Q | -19.44072222 | 15.67838021 | 0.203832009 | 13.05278591 | NA | NA | NA | NA | NA | NA | NA |
| SCN9A | 409 | N | -20.598375 | 14.97324494 | 0.192190862 | 12.32833733 | NA | NA | NA | NA | NA | NA | NA |
| SCN9A | 410 | Q | -25.15938889 | 10.92530091 | 0.125362961 | 4.967123252 | 6.19 | -0.15 | -1.33 | 13.72 | 0.24 | 1 | 1 |
| SCN9A | 411 | A | -25.4007 | 15.26488601 | 0.197005593 | 8.519182681 | NA | NA | NA | NA | NA | NA | NA |
| SCN9A | 412 | N | -23.833625 | 18.02359626 | 0.242549409 | 11.26104161 | NA | NA | NA | NA | NA | NA | NA |
| SCN9A | 413 | I | -25.869 | 15.64753399 | 0.203322766 | 8.825102157 | NA | NA | NA | NA | NA | NA | NA |
| SCN9A | 414 | E | -29.82538889 | 15.17337741 | 0.195494869 | 8.37404293 | NA | NA | NA | NA | NA | NA | NA |
| SCN9A | 415 | E | -29.26527778 | 19.10797943 | 0.260451596 | 12.29706808 | NA | NA | NA | NA | NA | NA | NA |
| SCN9A | 416 | A | -30.2265 | 21.22637767 | 0.295424439 | 14.41358999 | NA | NA | NA | NA | NA | NA | NA |
| SCN9A | 417 | K | -31.31161111 | 19.04032171 | 0.259334628 | 12.2632783 | NA | NA | NA | NA | NA | NA | NA |
| SCN9A | 726 | C | -15.05683333 | 61.9259329 | 0.967337354 | 59.69738422 | NA | NA | NA | NA | NA | NA | NA |
| SCN9A | 727 | S | -17.77016667 | 61.12426874 | 0.954102603 | 58.92704954 | NA | NA | NA | NA | NA | NA | NA |
| SCN9A | 728 | P | -19.22492857 | 61.97318243 | 0.968117401 | 60.09647981 | NA | NA | NA | NA | NA | NA | NA |
| SCN9A | 729 | Y | -15.68658333 | 59.90522123 | 0.933977229 | 57.72839637 | NA | NA | NA | NA | NA | NA | NA |
| SCN9A | 730 | W | -15.55578571 | 53.68945953 | 0.831360614 | 51.45421144 | NA | NA | NA | NA | NA | NA | NA |
| SCN9A | 731 | I | -20.420625 | 57.17381858 | 0.888884237 | 55.32776908 | NA | NA | NA | NA | NA | NA | NA |
| SCN9A | 732 | K | -19.57116667 | 57.55375022 | 0.895156565 | 55.70178007 | NA | NA | NA | NA | NA | NA | NA |
| SCN9A | 733 | F | -14.37986364 | 54.76474666 | 0.849112634 | 52.92233357 | NA | NA | NA | NA | NA | NA | NA |
| SCN9A | 734 | K | -18.24216667 | 52.49596764 | 0.811657141 | 50.63999437 | NA | NA | NA | NA | NA | NA | NA |
| SCN9A | 735 | K | -21.5335 | 52.44854823 | 0.810874289 | 50.94623084 | NA | NA | NA | NA | NA | NA | NA |
| SCN9A | 736 | C | -17.7105 | 51.9531196 | 0.802695209 | 50.10858896 | NA | NA | NA | NA | NA | NA | NA |
| SCN9A | 737 | I | -15.23 | 48.80093603 | 0.750655504 | 46.94063371 | NA | NA | NA | NA | NA | NA | NA |
| SCN9A | 738 | Y | -20.22266667 | 44.86854739 | 0.685735318 | 43.60453533 | NA | NA | NA | NA | NA | NA | NA |
| SCN9A | 739 | F | -20.31440909 | 48.44583665 | 0.744793134 | 46.7440014 | NA | NA | NA | NA | NA | NA | NA |
| SCN9A | 740 | I | -15.198625 | 46.3041746 | 0.709436227 | 44.44447268 | NA | NA | NA | NA | NA | NA | NA |
| SCN9A | 741 | V | -16.83478571 | 43.4744705 | 0.662720367 | 41.64147539 | NA | NA | NA | NA | NA | NA | NA |
| SCN9A | 742 | M | -17.026625 | 42.84759367 | 0.652371196 | 41.1032654 | NA | NA | NA | NA | NA | NA | NA |
| SCN9A | 743 | D | -18.051 | 38.22042914 | 0.575980887 | 36.5581033 | NA | NA | NA | NA | NA | NA | NA |

| | | | | | | | | | | | | | |
|---|---|---|---|---|---|---|---|---|---|---|---|---|---|
| SCN9A | 744 | P | -18.41864286 | 40.49786864 | 0.613579357 | 38.83465787 | NA | NA | NA | NA | NA | NA | NA |
| SCN9A | 745 | F | -15.61786364 | 41.81247976 | 0.6352824 | 40.30758244 | NA | NA | NA | NA | NA | NA | NA |
| SCN9A | 746 | V | -12.6655 | 41.40377125 | 0.628534992 | 39.60243192 | NA | NA | NA | NA | NA | NA | NA |
| SCN9A | 747 | D | -13.098875 | 37.37234554 | 0.561979792 | 35.51148896 | NA | NA | NA | NA | NA | NA | NA |
| SCN9A | 748 | L | -12.173125 | 36.95934166 | 0.555161471 | 35.12894218 | NA | NA | NA | NA | NA | NA | NA |
| SCN9A | 749 | A | -9.2317 | 39.21819531 | 0.592453106 | 37.59080166 | NA | NA | NA | NA | NA | NA | NA |
| SCN9A | 750 | I | -7.272125 | 36.95278838 | 0.555053283 | 34.7947896 | NA | NA | NA | NA | NA | NA | NA |
| SCN9A | 751 | T | -7.924071429 | 34.20935998 | 0.509761756 | 32.10854414 | NA | NA | NA | NA | NA | NA | NA |
| SCN9A | 752 | I | -6.01975 | 35.74658959 | 0.535140029 | 33.69278335 | NA | NA | NA | NA | NA | NA | NA |
| SCN9A | 753 | C | -3.026333333 | 36.43111622 | 0.546440946 | 34.20878641 | NA | NA | NA | NA | NA | NA | NA |
| SCN9A | 754 | I | -3.2595 | 32.37928057 | 0.479548797 | 30.15689954 | NA | NA | NA | NA | NA | NA | NA |
| SCN9A | 755 | V | -2.3605 | 30.85415142 | 0.454370292 | 28.70316134 | NA | NA | NA | NA | NA | NA | NA |
| SCN9A | 756 | L | -0.38375 | 34.46368763 | 0.513960476 | 32.25810897 | NA | NA | NA | NA | NA | NA | NA |
| SCN9A | 757 | N | 1.433125 | 34.17174665 | 0.509140794 | 31.87288372 | NA | NA | NA | NA | NA | NA | NA |
| SCN9A | 758 | T | 1.502071429 | 29.37072205 | 0.429880212 | 27.0658334 | NA | NA | NA | NA | NA | NA | NA |
| SCN9A | 759 | L | 2.942375 | 31.11598345 | 0.458692902 | 28.8536892 | NA | NA | NA | NA | NA | NA | NA |
| SCN9A | 760 | F | 6.259318182 | 33.82683082 | 0.503446545 | 30.82341875 | NA | NA | NA | NA | NA | NA | NA |
| SCN9A | 761 | M | 6.10925 | 30.38966517 | 0.446702043 | 27.38643104 | NA | NA | NA | NA | NA | NA | NA |
| SCN9A | 762 | A | 7.5011 | 28.69820404 | 0.418777547 | 25.69712129 | NA | NA | NA | NA | NA | NA | NA |
| SCN9A | 763 | M | 9.529625 | 30.58354666 | 0.449902851 | 26.99204988 | NA | NA | NA | NA | NA | NA | NA |
| SCN9A | 764 | E | 12.66972222 | 27.27882014 | 0.3953448 | 23.69199881 | NA | NA | NA | NA | NA | NA | NA |
| SCN9A | 765 | H | 16.2193 | 26.85333424 | 0.388320412 | 23.29398302 | NA | NA | NA | NA | NA | NA | NA |
| SCN9A | 766 | H | 21.6146 | 23.79602189 | 0.337846945 | 20.7726253 | NA | NA | NA | NA | NA | NA | NA |
| SCN9A | 767 | P | 20.92792857 | 27.04049404 | 0.391410251 | 22.30744243 | NA | NA | NA | NA | NA | NA | NA |
| SCN9A | 768 | M | 18.920375 | 30.76286726 | 0.452863273 | 25.2549144 | NA | NA | NA | NA | NA | NA | NA |
| SCN9A | 769 | T | 20.68092857 | 34.06336263 | 0.507351472 | 29.32205488 | NA | NA | NA | NA | NA | NA | NA |
| SCN9A | 770 | E | 23.05905556 | 36.73646378 | 0.551481959 | 31.23392143 | NA | NA | NA | NA | NA | NA | NA |
| SCN9A | 771 | E | 18.24094444 | 39.32220525 | 0.594170216 | 33.80538927 | NA | NA | NA | NA | NA | NA | NA |
| SCN9A | 772 | F | 15.2725 | 34.78674365 | 0.519293839 | 31.19822379 | NA | NA | NA | NA | NA | NA | NA |
| SCN9A | 773 | K | 16.80283333 | 34.6843882 | 0.517604043 | 31.16179138 | NA | NA | NA | NA | NA | NA | NA |
| SCN9A | 774 | N | 16.43375 | 38.53235163 | 0.581130446 | 34.98897303 | NA | NA | NA | NA | NA | NA | NA |
| SCN9A | 775 | V | 12.74592857 | 39.58830931 | 0.598563354 | 36.01747422 | NA | NA | NA | NA | NA | NA | NA |
| SCN9A | 776 | L | 11.328375 | 34.7068272 | 0.517974491 | 31.12691548 | NA | NA | NA | NA | NA | NA | NA |

| | | | | | | | | | | | | | |
|---|---|---|---|---|---|---|---|---|---|---|---|---|---|
| SCN9A | 777 | A | 11.6807 | 38.50567954 | 0.580690114 | 34.94922739 | NA | NA | NA | NA | NA | NA | NA |
| SCN9A | 778 | I | 10.45225 | 41.27746274 | 0.626449752 | 37.71905993 | NA | NA | NA | NA | NA | NA | NA |
| SCN9A | 779 | G | 7.0745 | 40.85962541 | 0.619551635 | 37.29829936 | NA | NA | NA | NA | NA | NA | NA |
| SCN9A | 780 | N | 6.765125 | 36.6049186 | 0.549310267 | 33.60022884 | NA | NA | NA | NA | NA | NA | NA |
| SCN9A | 781 | L | 6.61475 | 41.7114847 | 0.633615063 | 38.72348355 | NA | NA | NA | NA | NA | NA | NA |
| SCN9A | 782 | V | 3.404785714 | 43.21426636 | 0.658424631 | 40.90188518 | NA | NA | NA | NA | NA | NA | NA |
| SCN9A | 783 | F | 1.557681818 | 38.8537667 | 0.586436719 | 36.56110217 | NA | NA | NA | NA | NA | NA | NA |
| SCN9A | 784 | T | 2.511642857 | 40.30917308 | 0.610464164 | 38.06037929 | NA | NA | NA | NA | NA | NA | NA |
| SCN9A | 785 | G | 0.65675 | 43.33444423 | 0.66040866 | 41.01068928 | NA | NA | NA | NA | NA | NA | NA |
| SCN9A | 786 | I | -1.389375 | 42.57891584 | 0.647935568 | 40.31274024 | NA | NA | NA | NA | NA | NA | NA |
| SCN9A | 787 | F | -2.032227273 | 36.91486369 | 0.55442718 | 34.65933576 | NA | NA | NA | NA | NA | NA | NA |
| SCN9A | 788 | A | -2.1463 | 42.23167378 | 0.642202915 | 39.93637772 | NA | NA | NA | NA | NA | NA | NA |
| SCN9A | 789 | A | -3.9963 | 44.8178644 | 0.684898587 | 42.53644628 | NA | NA | NA | NA | NA | NA | NA |
| SCN9A | 790 | E | -6.882388889 | 39.13852543 | 0.591137828 | 36.92039815 | NA | NA | NA | NA | NA | NA | NA |
| SCN9A | 791 | M | -6.262875 | 41.21966544 | 0.625495571 | 38.94373812 | NA | NA | NA | NA | NA | NA | NA |
| SCN9A | 792 | V | -6.055928571 | 45.28188673 | 0.692559177 | 43.01433044 | NA | NA | NA | NA | NA | NA | NA |
| SCN9A | 793 | L | -9.390125 | 47.10524741 | 0.722661216 | 44.87946546 | NA | NA | NA | NA | NA | NA | NA |
| SCN9A | 794 | K | -11.40372222 | 41.54513396 | 0.630868762 | 39.40635819 | NA | NA | NA | NA | NA | NA | NA |
| SCN9A | 795 | L | -10.397875 | 44.58529343 | 0.681059051 | 42.33080666 | NA | NA | NA | NA | NA | NA | NA |
| SCN9A | 796 | I | -11.94325 | 48.8632789 | 0.751684729 | 46.64261234 | NA | NA | NA | NA | NA | NA | NA |
| SCN9A | 797 | A | -15.7271 | 48.63793299 | 0.747964471 | 46.78117436 | NA | NA | NA | NA | NA | NA | NA |
| SCN9A | 798 | M | -16.443625 | 42.64815475 | 0.64907864 | 40.77080423 | NA | NA | NA | NA | NA | NA | NA |
| SCN9A | 799 | D | -16.23825 | 46.6799222 | 0.715639481 | 44.46279076 | NA | NA | NA | NA | NA | NA | NA |
| SCN9A | 800 | P | -13.15735714 | 44.97477392 | 0.687489022 | 42.72239176 | NA | NA | NA | NA | NA | NA | NA |
| SCN9A | 801 | Y | -16.72825 | 44.0812175 | 0.672737212 | 41.85566889 | NA | NA | NA | NA | NA | NA | NA |
| SCN9A | 802 | E | -18.46805556 | 42.72440634 | 0.650337485 | 40.84056055 | NA | NA | NA | NA | NA | NA | NA |
| SCN9A | 803 | Y | -11.9165 | 40.77992396 | 0.618235836 | 38.56038446 | NA | NA | NA | NA | NA | NA | NA |
| SCN9A | 804 | F | -11.21804545 | 39.33716517 | 0.594417191 | 37.08592308 | NA | NA | NA | NA | NA | NA | NA |
| SCN9A | 805 | Q | -17.59594444 | 37.82962683 | 0.569529094 | 35.93675434 | NA | NA | NA | NA | NA | NA | NA |
| SCN9A | 806 | V | -14.87407143 | 36.16105438 | 0.541982469 | 34.30152369 | NA | NA | NA | NA | NA | NA | NA |
| SCN9A | 807 | G | -12.28425 | 33.39673777 | 0.496346097 | 31.1480844 | NA | NA | NA | NA | NA | NA | NA |
| SCN9A | 808 | W | -10.92442857 | 28.04761091 | 0.408036842 | 25.93396325 | NA | NA | NA | NA | NA | NA | NA |
| SCN9A | 809 | N | -10.465 | 35.47419685 | 0.530643071 | 33.25715345 | NA | NA | NA | NA | NA | NA | NA |

| | | | | | | | | | | | | | |
|---|---|---|---|---|---|---|---|---|---|---|---|---|---|
| SCN9A | 810 | I | -8.76175 | 33.62234562 | 0.500070679 | 31.31279581 | NA | NA | NA | NA | NA | NA | NA |
| SCN9A | 811 | F | -6.135318182 | 29.0705146 | 0.424924058 | 26.76794748 | NA | NA | NA | NA | NA | NA | NA |
| SCN9A | 812 | D | -5.772125 | 34.30338307 | 0.511313993 | 32.02223867 | NA | NA | NA | NA | NA | NA | NA |
| SCN9A | 813 | S | -4.769833333 | 36.74433645 | 0.551611929 | 34.44415061 | NA | NA | NA | NA | NA | NA | NA |
| SCN9A | 814 | L | -2.616875 | 34.4819737 | 0.514262363 | 32.14354247 | NA | NA | NA | NA | NA | NA | NA |
| SCN9A | 815 | I | -0.761125 | 32.51988299 | 0.481870016 | 30.17738335 | NA | NA | NA | NA | NA | NA | NA |
| SCN9A | 816 | V | -0.988214286 | 37.39025094 | 0.562275394 | 35.05442909 | NA | NA | NA | NA | NA | NA | NA |
| SCN9A | 817 | T | 0.602642857 | 39.03033217 | 0.589351655 | 35.68483447 | NA | NA | NA | NA | NA | NA | NA |
| SCN9A | 818 | L | 3.314 | 35.90055569 | 0.537681871 | 30.99716735 | NA | NA | NA | NA | NA | NA | NA |
| SCN9A | 819 | S | 3.799666667 | 38.21835619 | 0.575946665 | 33.31523064 | NA | NA | NA | NA | NA | NA | NA |
| SCN9A | 820 | L | 3.472625 | 41.71216269 | 0.633626256 | 36.8084033 | NA | NA | NA | NA | NA | NA | NA |
| SCN9A | 821 | V | 6.256785714 | 41.84694828 | 0.635851444 | 36.94390869 | NA | NA | NA | NA | NA | NA | NA |
| SCN9A | 822 | E | 9.4795 | 36.39658432 | 0.545870856 | 32.90030757 | NA | NA | NA | NA | NA | NA | NA |
| SCN9A | 823 | L | 10.1835 | 42.05822217 | 0.639339385 | 38.57274241 | NA | NA | NA | NA | NA | NA | NA |
| SCN9A | 824 | F | 12.29731818 | 43.37480934 | 0.661075051 | 39.97522973 | NA | NA | NA | NA | NA | NA | NA |
| SCN9A | 825 | L | 10.981375 | 41.16515618 | 0.624595672 | 37.77552882 | NA | NA | NA | NA | NA | NA | NA |
| SCN9A | 831 | L | 12.358375 | 30.72151137 | 0.452180524 | 27.35437643 | NA | NA | NA | NA | NA | NA | NA |
| SCN9A | 832 | S | 12.3615 | 31.76441767 | 0.469397966 | 28.3004672 | NA | NA | NA | NA | NA | NA | NA |
| SCN9A | 833 | V | 10.13678571 | 27.70730469 | 0.402418693 | 24.2484683 | NA | NA | NA | NA | NA | NA | NA |
| SCN9A | 834 | L | 7.575375 | 31.27651996 | 0.461343215 | 28.31671765 | NA | NA | NA | NA | NA | NA | NA |
| SCN9A | 835 | R | 9.018136364 | 32.95361504 | 0.489030541 | 29.95343521 | NA | NA | NA | NA | NA | NA | NA |
| SCN9A | 836 | S | 7.193666667 | 28.54724722 | 0.416285386 | 25.54197335 | NA | NA | NA | NA | NA | NA | NA |
| SCN9A | 837 | F | 3.246409091 | 27.12333735 | 0.392777919 | 22.22005401 | NA | NA | NA | NA | NA | NA | NA |
| SCN9A | 838 | R | 2.474590909 | 31.6547982 | 0.467588247 | 26.84645622 | NA | NA | NA | NA | NA | NA | NA |
| SCN9A | 839 | L | 2.02125 | 27.91217564 | 0.405800928 | 23.06716106 | NA | NA | NA | NA | NA | NA | NA |
| SCN9A | 840 | L | -0.864375 | 26.26766017 | 0.378651462 | 23.92621031 | NA | NA | NA | NA | NA | NA | NA |
| SCN9A | 841 | R | -2.213954545 | 29.75620994 | 0.436244269 | 27.45179445 | NA | NA | NA | NA | NA | NA | NA |
| SCN9A | 842 | V | -2.757214286 | 27.50699905 | 0.399111828 | 25.27584066 | NA | NA | NA | NA | NA | NA | NA |
| SCN9A | 843 | F | -5.1265 | 22.44514732 | 0.315545225 | 20.19804527 | NA | NA | NA | NA | NA | NA | NA |
| SCN9A | 844 | K | -8.135388889 | 28.6604964 | 0.418155028 | 26.48202086 | NA | NA | NA | NA | NA | NA | NA |
| SCN9A | 845 | L | -8.22525 | 27.34545249 | 0.39644484 | 25.49105248 | NA | NA | NA | NA | NA | NA | NA |
| SCN9A | 846 | A | -11.0961 | 25.24319651 | 0.361738491 | 23.38322597 | NA | NA | NA | NA | NA | NA | NA |
| SCN9A | 847 | K | -13.92516667 | 27.45029342 | 0.398175669 | 25.68892399 | NA | NA | NA | NA | NA | NA | NA |

| | | | | | | | | | | | | | |
|---|---|---|---|---|---|---|---|---|---|---|---|---|---|
| SCN9A | 848 | S | -14.38833333 | 26.97102679 | 0.39026341 | 25.27884096 | NA | NA | NA | NA | NA | NA | NA |
| SCN9A | 849 | W | -10.8525 | 22.35948027 | 0.31413094 | 20.57365173 | NA | NA | NA | NA | NA | NA | NA |
| SCN9A | 850 | P | -14.97892857 | 20.56219893 | 0.284459447 | 18.79937835 | NA | NA | NA | NA | NA | NA | NA |
| SCN9A | 851 | T | -12.05107143 | 17.6909651 | 0.237057969 | 15.7778064 | NA | NA | NA | NA | NA | NA | NA |
| SCN9A | 852 | L | -9.542375 | 20.72125626 | 0.28708534 | 18.91665308 | NA | NA | NA | NA | NA | NA | NA |
| SCN9A | 853 | N | -13.985125 | 20.71012038 | 0.286901497 | 18.56201692 | NA | NA | NA | NA | NA | NA | NA |
| SCN9A | 854 | M | -13.3605 | 15.30543511 | 0.197675022 | 13.23054931 | NA | NA | NA | NA | NA | NA | NA |
| SCN9A | 855 | L | -8.292875 | 15.87784749 | 0.207125034 | 14.3722108 | NA | NA | NA | NA | NA | NA | NA |
| SCN9A | 856 | I | -10.15175 | 19.65903347 | 0.269549001 | 17.75953222 | NA | NA | NA | NA | NA | NA | NA |
| SCN9A | 857 | K | -14.35061111 | 17.87703124 | 0.240129753 | 16.71907068 | NA | NA | NA | NA | NA | NA | NA |
| SCN9A | 858 | I | -10.357375 | 13.4160357 | 0.166482743 | 11.56583221 | NA | NA | NA | NA | NA | NA | NA |
| SCN9A | 859 | I | -7.51025 | 17.50323193 | 0.233958663 | 15.27865444 | NA | NA | NA | NA | NA | NA | NA |
| SCN9A | 860 | G | -10.889 | 19.83716615 | 0.272489811 | 17.58540617 | NA | NA | NA | NA | NA | NA | NA |
| SCN9A | 861 | N | -13.024875 | 17.13675451 | 0.227908452 | 15.44480274 | NA | NA | NA | NA | NA | NA | NA |
| SCN9A | 862 | S | -8.728833333 | 17.14001251 | 0.227962239 | 14.77678819 | NA | NA | NA | NA | NA | NA | NA |
| SCN9A | 863 | V | -8.9395 | 20.2419793 | 0.27917291 | 17.8800848 | NA | NA | NA | NA | NA | NA | NA |
| SCN9A | 864 | G | -12.792 | 22.105607 | 0.309939721 | 20.23499024 | NA | NA | NA | NA | NA | NA | NA |
| SCN9A | 865 | A | -15.0565 | 21.3958428 | 0.298222155 | 18.68347796 | NA | NA | NA | NA | NA | NA | NA |
| SCN9A | 866 | L | -12.38125 | 15.81777699 | 0.206133324 | 13.86162817 | NA | NA | NA | NA | NA | NA | NA |
| SCN9A | 867 | G | -10.132 | 21.2258176 | 0.295415192 | 18.00400424 | NA | NA | NA | NA | NA | NA | NA |
| SCN9A | 868 | N | -11.216 | 21.41981307 | 0.298617882 | 18.02544596 | NA | NA | NA | NA | NA | NA | NA |
| SCN9A | 869 | L | -9.385375 | 16.43916747 | 0.21639192 | 13.1190228 | NA | NA | NA | NA | NA | NA | NA |
| SCN9A | 870 | T | -6.418071429 | 18.01699825 | 0.242440482 | 14.62774536 | NA | NA | NA | NA | NA | NA | NA |
| SCN9A | 871 | L | -5.585375 | 20.14566199 | 0.277582798 | 16.78717601 | NA | NA | NA | NA | NA | NA | NA |
| SCN9A | 872 | V | -5.4915 | 19.11378657 | 0.260547467 | 15.95644174 | NA | NA | NA | NA | NA | NA | NA |
| SCN9A | 873 | L | -2.267875 | 13.68087665 | 0.170855028 | 10.52809178 | NA | NA | NA | NA | NA | NA | NA |
| SCN9A | 874 | A | -0.8337 | 18.7114466 | 0.253905197 | 15.2500388 | NA | NA | NA | NA | NA | NA | NA |
| SCN9A | 875 | I | -1.629875 | 21.02144736 | 0.292041224 | 17.58341336 | NA | NA | NA | NA | NA | NA | NA |
| SCN9A | 876 | I | -0.978375 | 17.84312747 | 0.239570032 | 13.22698973 | NA | NA | NA | NA | NA | NA | NA |
| SCN9A | 877 | V | 2.351071429 | 16.53291045 | 0.217939532 | 11.63499231 | NA | NA | NA | NA | NA | NA | NA |
| SCN9A | 878 | F | 3.549318182 | 20.83809919 | 0.289014311 | 18.38221386 | NA | NA | NA | NA | NA | NA | NA |
| SCN9A | 879 | I | 2.419875 | 22.3175783 | 0.313439176 | 19.86021919 | NA | NA | NA | NA | NA | NA | NA |
| SCN9A | 880 | F | 3.899954545 | 18.01858826 | 0.242466731 | 15.59927816 | NA | NA | NA | NA | NA | NA | NA |

| | | | | | | | | | | | | | |
|---|---|---|---|---|---|---|---|---|---|---|---|---|---|
| SCN9A | 881 | A | 7.9237 | 20.37096115 | 0.281302284 | 18.58184105 | NA | NA | NA | NA | NA | NA | NA |
| SCN9A | 882 | V | 7.992928571 | 22.95641599 | 0.323985809 | 21.07108151 | NA | NA | NA | NA | NA | NA | NA |
| SCN9A | 883 | V | 7.8045 | 23.41123561 | 0.331494471 | 21.56690715 | NA | NA | NA | NA | NA | NA | NA |
| SCN9A | 884 | G | 11.11 | 21.51780928 | 0.300235711 | 19.70372484 | NA | NA | NA | NA | NA | NA | NA |
| SCN9A | 885 | M | 12.840875 | 22.83935168 | 0.322053183 | 21.03394946 | NA | NA | NA | NA | NA | NA | NA |
| SCN9A | 886 | Q | 12.11783333 | 25.59101889 | 0.367480725 | 23.7899141 | NA | NA | NA | NA | NA | NA | NA |
| SCN9A | 887 | L | 12.668125 | 24.63030672 | 0.351620234 | 22.81390029 | NA | NA | NA | NA | NA | NA | NA |
| SCN9A | 888 | F | 14.69131818 | 20.53361347 | 0.283987527 | 18.72583919 | NA | NA | NA | NA | NA | NA | NA |
| SCN9A | 889 | G | 18.1195 | 24.22527694 | 0.344933558 | 20.94880499 | NA | NA | NA | NA | NA | NA | NA |
| SCN9A | 890 | K | 20.73216667 | 25.45883047 | 0.365298414 | 20.11829881 | NA | NA | NA | NA | NA | NA | NA |
| SCN9A | 891 | S | 21.07116667 | 22.71226561 | 0.319955107 | 16.96222857 | NA | NA | NA | NA | NA | NA | NA |
| SCN9A | 892 | Y | 20.00866667 | 16.78946259 | 0.222174976 | 11.97885635 | NA | NA | NA | NA | NA | NA | NA |
| SCN9A | 893 | K | 25.22494444 | 22.42960343 | 0.31528861 | 16.69078984 | NA | NA | NA | NA | NA | NA | NA |
| SCN9A | 894 | E | 25.62683333 | 22.84389308 | 0.322128158 | 17.4948367 | NA | NA | NA | NA | NA | NA | NA |
| SCN9A | 895 | C | 25.83383333 | 19.83573936 | 0.272466256 | 14.39004997 | NA | NA | NA | NA | NA | NA | NA |
| SCN9A | 896 | V | 26.84392857 | 16.62555831 | 0.219469064 | 10.87106913 | NA | NA | NA | NA | NA | NA | NA |
| SCN9A | 897 | C | 29.47316667 | 15.65945267 | 0.203519532 | 13.23895505 | NA | NA | NA | NA | NA | NA | NA |
| SCN9A | 898 | K | 26.40116667 | 15.6802595 | 0.203863034 | 13.45185653 | NA | NA | NA | NA | NA | NA | NA |
| SCN9A | 899 | I | 24.510125 | 12.57910382 | 0.152665754 | 6.830211153 | NA | NA | NA | NA | NA | NA | NA |
| SCN9A | 900 | N | 27.9755 | 12.18371479 | 0.146138238 | 9.76281443 | NA | NA | NA | NA | NA | NA | NA |
| SCN9A | 901 | D | 30.896 | 8.551667264 | 0.086176412 | 6.132512847 | 2.47 | -0.88 | -1.43 | 18.82 | 1.42 | 1 | -1 |
| SCN9A | 902 | D | 31.998625 | 11.72668308 | 0.138593057 | 9.310057342 | NA | NA | NA | NA | NA | NA | NA |
| SCN9A | 903 | C | 31.452 | 14.53496182 | 0.184955203 | 12.11672048 | NA | NA | NA | NA | NA | NA | NA |
| SCN9A | 904 | T | 28.22092857 | 14.70953609 | 0.187837267 | 12.2848688 | NA | NA | NA | NA | NA | NA | NA |
| SCN9A | 905 | L | 24.288625 | 14.76408175 | 0.188737767 | 9.086788745 | NA | NA | NA | NA | NA | NA | NA |
| SCN9A | 906 | P | 22.35578571 | 12.83906632 | 0.1569575 | 7.123199806 | NA | NA | NA | NA | NA | NA | NA |
| SCN9A | 907 | R | 21.14459091 | 10.53953367 | 0.118994292 | 7.295566132 | NA | NA | NA | NA | NA | NA | NA |
| SCN9A | 908 | W | 15.99685714 | 9.395290043 | 0.100103862 | 6.501208926 | NA | NA | NA | NA | NA | NA | NA |
| SCN9A | 909 | H | 18.4109 | 15.29805625 | 0.197553204 | 12.1335848 | NA | NA | NA | NA | NA | NA | NA |
| SCN9A | 910 | M | 14.68625 | 17.69582487 | 0.237138199 | 16.03478668 | NA | NA | NA | NA | NA | NA | NA |
| SCN9A | 911 | N | 18.820125 | 20.92208866 | 0.290400902 | 17.72381464 | NA | NA | NA | NA | NA | NA | NA |
| SCN9A | 912 | D | 16.563625 | 22.33447302 | 0.313718092 | 19.33129027 | NA | NA | NA | NA | NA | NA | NA |
| SCN9A | 913 | F | 10.22131818 | 21.8014589 | 0.30491851 | 19.98862586 | NA | NA | NA | NA | NA | NA | NA |

| | | | | | | | | | | | | | |
|---|---|---|---|---|---|---|---|---|---|---|---|---|---|
| SCN9A | 914 | F | 12.38195455 | 19.72270248 | 0.270600119 | 17.92666599 | NA | NA | NA | NA | NA | NA | NA |
| SCN9A | 915 | H | 15.7266 | 17.16407778 | 0.228359535 | 14.87748826 | NA | NA | NA | NA | NA | NA | NA |
| SCN9A | 916 | S | 11.56083333 | 17.8063722 | 0.238963236 | 15.99281114 | NA | NA | NA | NA | NA | NA | NA |
| SCN9A | 917 | F | 7.943227273 | 16.46460403 | 0.216811855 | 14.68982207 | NA | NA | NA | NA | NA | NA | NA |
| SCN9A | 918 | L | 11.1265 | 13.66444049 | 0.170583682 | 11.8513466 | NA | NA | NA | NA | NA | NA | NA |
| SCN9A | 919 | I | 11.550625 | 12.68036612 | 0.154337503 | 10.86596879 | NA | NA | NA | NA | NA | NA | NA |
| SCN9A | 920 | V | 6.928214286 | 12.78094054 | 0.155997896 | 11.00588052 | NA | NA | NA | NA | NA | NA | NA |
| SCN9A | 921 | F | 5.641590909 | 10.76993682 | 0.12279804 | 8.967017225 | NA | NA | NA | NA | NA | NA | NA |
| SCN9A | 922 | R | 10.60186364 | 6.425339368 | 0.051072658 | 6.386152818 | 1.82 | -0.79 | -2.05 | 26.54 | 1.33 | 2 | 0 |
| SCN9A | 923 | V | 6.334357143 | 8.308809421 | 0.082167048 | 6.802112519 | 1.89 | -0.67 | -1.57 | 18.75 | 1.03 | 1 | 1 |
| SCN9A | 924 | L | 2.5785 | 7.277700805 | 0.065144376 | 4.8927658 | 2.58 | -0.8 | -0.4 | 3.38 | 0.81 | 0 | 0 |
| SCN9A | 925 | C | 4.8 | 5.497239489 | 0.035750567 | 2.988537685 | 2.58 | -0.8 | -0.4 | 3.38 | 0.81 | 0 | 0 |
| SCN9A | 926 | G | 5.98925 | 3.333580807 | 3.05E-05 | 1.637715722 | 2.58 | -0.8 | -0.4 | 3.38 | 0.81 | 0 | 0 |
| SCN9A | 927 | E | 9.762722222 | 3.370339152 | 6.37E-04 | 1.822255763 | 1.82 | -0.79 | -2.05 | 26.54 | 1.33 | 2 | 0 |
| SCN9A | 928 | W | 8.565642857 | 7.15566971 | 0.063129752 | 5.343035569 | 1.82 | -0.79 | -2.05 | 26.54 | 1.33 | 2 | 0 |
| SCN9A | 929 | I | 12.432125 | 7.504735838 | 0.068892519 | 5.759628234 | 1.85 | -0.79 | -1.72 | 21.91 | 1.22 | 2 | 0 |
| SCN9A | 930 | E | 15.60438889 | 4.678693942 | 0.022237119 | 3.431384165 | 2.24 | -0.39 | -2.77 | 33.8 | 1.01 | 2 | -2 |
| SCN9A | 931 | T | 14.57735714 | 10.4253727 | 0.117109597 | 8.753712739 | NA | NA | NA | NA | NA | NA | NA |
| SCN9A | 932 | M | 13.578125 | 12.39577303 | 0.149639128 | 10.61826108 | NA | NA | NA | NA | NA | NA | NA |
| SCN9A | 933 | W | 17.25057143 | 9.958219821 | 0.109397325 | 7.882243957 | NA | NA | NA | NA | NA | NA | NA |
| SCN9A | 934 | D | 19.479625 | 12.45266763 | 0.150578406 | 9.15833746 | NA | NA | NA | NA | NA | NA | NA |
| SCN9A | 935 | C | 16.78483333 | 16.57526024 | 0.218638689 | 15.24063103 | NA | NA | NA | NA | NA | NA | NA |
| SCN9A | 936 | M | 16.966625 | 17.29482541 | 0.23051806 | 15.51496289 | NA | NA | NA | NA | NA | NA | NA |
| SCN9A | 937 | E | 22.15983333 | 17.0376212 | 0.226271851 | 11.64231058 | NA | NA | NA | NA | NA | NA | NA |
| SCN9A | 938 | V | 21.26721429 | 19.50028864 | 0.266928267 | 14.11909925 | NA | NA | NA | NA | NA | NA | NA |
| SCN9A | 939 | A | 18.0545 | 22.0450445 | 0.308939889 | 20.79589286 | NA | NA | NA | NA | NA | NA | NA |
| SCN9A | 940 | G | 17.3115 | 21.80743593 | 0.305017186 | 20.011324 | NA | NA | NA | NA | NA | NA | NA |
| SCN9A | 941 | Q | 15.91372222 | 20.97393881 | 0.291256901 | 19.15812924 | NA | NA | NA | NA | NA | NA | NA |
| SCN9A | 942 | A | 12.4333 | 22.55977584 | 0.317437639 | 20.74168893 | NA | NA | NA | NA | NA | NA | NA |
| SCN9A | 943 | M | 12.8275 | 21.19021069 | 0.294827354 | 19.37983981 | NA | NA | NA | NA | NA | NA | NA |
| SCN9A | 944 | C | 12.60083333 | 17.78964165 | 0.238687029 | 15.98650798 | NA | NA | NA | NA | NA | NA | NA |
| SCN9A | 945 | L | 10.141 | 17.37327212 | 0.231813144 | 15.57155697 | NA | NA | NA | NA | NA | NA | NA |
| SCN9A | 946 | I | 7.267 | 19.15820119 | 0.261280712 | 17.45146463 | NA | NA | NA | NA | NA | NA | NA |

| | | | | | | | | | | | | | |
|---|---|---|---|---|---|---|---|---|---|---|---|---|---|
| SCN9A | 947 | V | 7.7925 | 17.22105697 | 0.22930021 | 15.46811005 | NA | NA | NA | NA | NA | NA | NA |
| SCN9A | 948 | Y | 9.605916667 | 13.37195517 | 0.165755014 | 11.79742146 | NA | NA | NA | NA | NA | NA | NA |
| SCN9A | 949 | M | 5.34375 | 14.64779205 | 0.186817929 | 12.20864141 | NA | NA | NA | NA | NA | NA | NA |
| SCN9A | 950 | M | 2.59025 | 16.15953372 | 0.211775419 | 13.78714714 | NA | NA | NA | NA | NA | NA | NA |
| SCN9A | 951 | V | 3.642642857 | 13.52801423 | 0.168331408 | 11.14707858 | NA | NA | NA | NA | NA | NA | NA |
| SCN9A | 952 | M | 2.634375 | 7.678492105 | 0.071761078 | 5.297498903 | 2.58 | -0.8 | -0.4 | 3.38 | 0.81 | 0 | 0 |
| SCN9A | 953 | V | -0.530357143 | 13.30793335 | 0.164698071 | 8.947450323 | NA | NA | NA | NA | NA | NA | NA |
| SCN9A | 954 | I | -1.304 | 14.40222754 | 0.18276388 | 11.09508351 | NA | NA | NA | NA | NA | NA | NA |
| SCN9A | 955 | G | -0.3685 | 11.96278387 | 0.142490868 | 7.516659967 | NA | NA | NA | NA | NA | NA | NA |
| SCN9A | 956 | N | -1.052375 | 7.939701128 | 0.076073403 | 4.688755313 | 3.73 | 1.3 | 2.7 | 0.5 | -1.72 | 0 | 0 |
| SCN9A | 957 | L | -5.398375 | 11.34307657 | 0.132260059 | 8.420729804 | NA | NA | NA | NA | NA | NA | NA |
| SCN9A | 958 | V | -4.761214286 | 12.19067225 | 0.146253099 | 8.958859793 | NA | NA | NA | NA | NA | NA | NA |
| SCN9A | 959 | V | -3.237642857 | 8.326129293 | 0.082452984 | 5.23387756 | 3.4 | 1.13 | 2.1 | 0.62 | -1.67 | 0 | 0 |
| SCN9A | 960 | L | -6.421375 | 7.645997515 | 0.071224622 | 4.252075763 | 3.4 | 1.13 | 2.1 | 0.62 | -1.67 | 0 | 0 |
| SCN9A | 961 | N | -9.344 | 10.88206033 | 0.124649098 | 7.487008187 | NA | NA | NA | NA | NA | NA | NA |
| SCN9A | 962 | L | -7.727875 | 11.35946909 | 0.132530685 | 8.152289439 | NA | NA | NA | NA | NA | NA | NA |
| SCN9A | 963 | F | -7.400863636 | 5.411146459 | 0.034329248 | 3.310666624 | 2.48 | 1.3 | 2.7 | 0.5 | -1.72 | 0 | 0 |
| SCN9A | 964 | L | -10.902125 | 5.898725371 | 0.042378736 | 3.211075122 | 2.74 | 1.54 | 3.2 | 0.43 | -1.76 | 0 | 0 |
| SCN9A | 965 | A | -12.8357 | 10.83760255 | 0.12391514 | 8.149061873 | NA | NA | NA | NA | NA | NA | NA |
| SCN9A | 966 | L | -11.126625 | 11.94176281 | 0.142143829 | 9.214966028 | NA | NA | NA | NA | NA | NA | NA |
| SCN9A | 967 | L | -12.6995 | 6.223880542 | 0.047746754 | 3.489443316 | 2.74 | 1.54 | 3.2 | 0.43 | -1.76 | 0 | 0 |
| SCN9A | 968 | L | -16.17125 | 7.761261431 | 0.073127525 | 6.057374057 | 1.99 | 1.81 | 4.5 | 0.13 | -1.85 | 0 | 0 |
| SCN9A | 969 | S | -16.9875 | 12.78926253 | 0.156135284 | 10.14428847 | NA | NA | NA | NA | NA | NA | NA |
| SCN9A | 970 | S | -15.79416667 | 13.75530785 | 0.17208382 | 11.20319797 | NA | NA | NA | NA | NA | NA | NA |
| SCN9A | 971 | F | -18.48868182 | 8.660635889 | 0.087975385 | 7.646070685 | 1.99 | 1.81 | 4.5 | 0.13 | -1.85 | 0 | 0 |
| SCN9A | 972 | S | -21.33766667 | 12.54768648 | 0.152147082 | 10.0011107 | NA | NA | NA | NA | NA | NA | NA |
| SCN9A | 1175 | G | -24.2045 | 57.36242197 | 0.891997908 | 54.75114637 | NA | NA | NA | NA | NA | NA | NA |
| SCN9A | 1176 | K | -24.06772222 | 52.36837023 | 0.809550623 | 49.75660975 | NA | NA | NA | NA | NA | NA | NA |
| SCN9A | 1177 | I | -20.23025 | 53.52332103 | 0.828617818 | 50.90994875 | NA | NA | NA | NA | NA | NA | NA |
| SCN9A | 1178 | W | -17.3915 | 48.92027631 | 0.752625705 | 46.30070763 | NA | NA | NA | NA | NA | NA | NA |
| SCN9A | 1179 | W | -22.90657143 | 46.1287926 | 0.706540829 | 43.51626908 | NA | NA | NA | NA | NA | NA | NA |
| SCN9A | 1180 | N | -21.488625 | 49.33956285 | 0.759547747 | 46.72857491 | NA | NA | NA | NA | NA | NA | NA |
| SCN9A | 1181 | I | -16.7025 | 47.84341797 | 0.734847746 | 45.22966321 | NA | NA | NA | NA | NA | NA | NA |

| | | | | | | | | | | | | | |
|---|---|---|---|---|---|---|---|---|---|---|---|---|---|
| SCN9A | 1182 | R | -18.21586364 | 43.28005146 | 0.659510684 | 40.66279737 | NA | NA | NA | NA | NA | NA | NA |
| SCN9A | 1183 | K | -22.16238889 | 44.56312864 | 0.68069313 | 41.95320683 | NA | NA | NA | NA | NA | NA | NA |
| SCN9A | 1184 | T | -17.58192857 | 44.97054096 | 0.68741914 | 42.35958709 | NA | NA | NA | NA | NA | NA | NA |
| SCN9A | 1185 | C | -14.83083333 | 42.4711157 | 0.646155885 | 39.85782746 | NA | NA | NA | NA | NA | NA | NA |
| SCN9A | 1186 | Y | -18.928 | 36.84509859 | 0.553275421 | 34.23625095 | NA | NA | NA | NA | NA | NA | NA |
| SCN9A | 1187 | K | -18.86716667 | 41.32449431 | 0.627226201 | 38.71569776 | NA | NA | NA | NA | NA | NA | NA |
| SCN9A | 1188 | I | -13.581 | 40.51266416 | 0.613823618 | 37.90102071 | NA | NA | NA | NA | NA | NA | NA |
| SCN9A | 1189 | V | -13.65978571 | 37.17258066 | 0.558681855 | 34.560406 | NA | NA | NA | NA | NA | NA | NA |
| SCN9A | 1190 | E | -17.0745 | 35.92715124 | 0.538120939 | 33.31668977 | NA | NA | NA | NA | NA | NA | NA |
| SCN9A | 1191 | H | -15.0152 | 37.79934383 | 0.569029149 | 35.18984283 | NA | NA | NA | NA | NA | NA | NA |
| SCN9A | 1192 | S | -13.53416667 | 36.20449078 | 0.542699565 | 33.59865765 | NA | NA | NA | NA | NA | NA | NA |
| SCN9A | 1193 | W | -11.74878571 | 38.8751524 | 0.586789777 | 36.26684177 | NA | NA | NA | NA | NA | NA | NA |
| SCN9A | 1194 | F | -9.499045455 | 37.7441612 | 0.568118133 | 35.12915381 | NA | NA | NA | NA | NA | NA | NA |
| SCN9A | 1195 | E | -10.33227778 | 32.60603283 | 0.483292272 | 29.99056721 | NA | NA | NA | NA | NA | NA | NA |
| SCN9A | 1196 | S | -8.144666667 | 35.6510617 | 0.53356295 | 33.03847287 | NA | NA | NA | NA | NA | NA | NA |
| SCN9A | 1197 | F | -4.534227273 | 37.64459438 | 0.566474375 | 35.03027251 | NA | NA | NA | NA | NA | NA | NA |
| SCN9A | 1198 | I | -5.311875 | 35.78896463 | 0.535839603 | 33.17010092 | NA | NA | NA | NA | NA | NA | NA |
| SCN9A | 1199 | V | -4.974928571 | 31.84205943 | 0.470679761 | 29.22525921 | NA | NA | NA | NA | NA | NA | NA |
| SCN9A | 1200 | L | -1.566875 | 35.52969973 | 0.531559374 | 32.91257279 | NA | NA | NA | NA | NA | NA | NA |
| SCN9A | 1201 | M | -0.340125 | 36.99630561 | 0.555771713 | 34.38137432 | NA | NA | NA | NA | NA | NA | NA |
| SCN9A | 1202 | I | -1.243875 | 32.50992284 | 0.481705583 | 30.03385041 | NA | NA | NA | NA | NA | NA | NA |
| SCN9A | 1203 | L | 0.875625 | 31.1743539 | 0.459656546 | 29.24392076 | NA | NA | NA | NA | NA | NA | NA |
| SCN9A | 1204 | L | 3.088375 | 36.26075627 | 0.543628457 | 34.36065136 | NA | NA | NA | NA | NA | NA | NA |
| SCN9A | 1205 | S | 3.594 | 35.06555237 | 0.52389672 | 32.98373073 | NA | NA | NA | NA | NA | NA | NA |
| SCN9A | 1206 | S | 3.830333333 | 31.64962221 | 0.467502796 | 29.56238368 | NA | NA | NA | NA | NA | NA | NA |
| SCN9A | 1207 | G | 6.74725 | 32.44433072 | 0.480622717 | 30.36340221 | NA | NA | NA | NA | NA | NA | NA |
| SCN9A | 1208 | A | 8.9481 | 33.59042323 | 0.499543669 | 31.53266279 | NA | NA | NA | NA | NA | NA | NA |
| SCN9A | 1209 | L | 8.8335 | 30.32969911 | 0.445712058 | 28.29442879 | NA | NA | NA | NA | NA | NA | NA |
| SCN9A | 1210 | A | 11.1917 | 29.12364582 | 0.425801207 | 27.0843635 | NA | NA | NA | NA | NA | NA | NA |
| SCN9A | 1211 | F | 13.10786364 | 31.20176759 | 0.460109121 | 29.18895456 | NA | NA | NA | NA | NA | NA | NA |
| SCN9A | 1212 | E | 15.00827778 | 29.05905769 | 0.424734915 | 25.8816654 | NA | NA | NA | NA | NA | NA | NA |
| SCN9A | 1213 | D | 18.24175 | 27.63386652 | 0.401206295 | 22.97682064 | NA | NA | NA | NA | NA | NA | NA |
| SCN9A | 1214 | I | 22.21675 | 23.97668555 | 0.340829539 | 19.29946799 | NA | NA | NA | NA | NA | NA | NA |

| | | | | | | | | | | | | | |
|---|---|---|---|---|---|---|---|---|---|---|---|---|---|
| SCN9A | 1215 | Y | 22.55183333 | 24.15197982 | 0.343723489 | 19.46227552 | NA | NA | NA | NA | NA | NA | NA |
| SCN9A | 1216 | I | 20.193125 | 30.74144917 | 0.452509679 | 26.07164661 | NA | NA | NA | NA | NA | NA | NA |
| SCN9A | 1217 | E | 24.54827778 | 32.15573947 | 0.475858336 | 27.47676318 | NA | NA | NA | NA | NA | NA | NA |
| SCN9A | 1218 | R | 24.69768182 | 30.27736118 | 0.444848006 | 25.58741066 | NA | NA | NA | NA | NA | NA | NA |
| SCN9A | 1219 | K | 20.64494444 | 32.39740261 | 0.479847976 | 27.72762317 | NA | NA | NA | NA | NA | NA | NA |
| SCN9A | 1220 | K | 24.18872222 | 37.75260122 | 0.568257471 | 33.07350522 | NA | NA | NA | NA | NA | NA | NA |
| SCN9A | 1221 | T | 19.73392857 | 40.16724962 | 0.608121136 | 35.49141383 | NA | NA | NA | NA | NA | NA | NA |
| SCN9A | 1222 | I | 16.721125 | 37.66172748 | 0.566757227 | 33.11029209 | NA | NA | NA | NA | NA | NA | NA |
| SCN9A | 1223 | K | 18.89938889 | 36.64604586 | 0.54998924 | 31.98101846 | NA | NA | NA | NA | NA | NA | NA |
| SCN9A | 1224 | I | 17.537 | 40.78340786 | 0.618293352 | 36.11920365 | NA | NA | NA | NA | NA | NA | NA |
| SCN9A | 1225 | I | 13.79075 | 41.78924858 | 0.634898874 | 39.7711663 | NA | NA | NA | NA | NA | NA | NA |
| SCN9A | 1226 | L | 12.5725 | 36.35964545 | 0.545261028 | 34.32173901 | NA | NA | NA | NA | NA | NA | NA |
| SCN9A | 1227 | E | 14.64894444 | 40.86187558 | 0.619588783 | 37.85633508 | NA | NA | NA | NA | NA | NA | NA |
| SCN9A | 1228 | Y | 13.21541667 | 43.42287465 | 0.661868566 | 40.44668419 | NA | NA | NA | NA | NA | NA | NA |
| SCN9A | 1229 | A | 8.6917 | 41.86384428 | 0.636130382 | 39.80597311 | NA | NA | NA | NA | NA | NA | NA |
| SCN9A | 1230 | D | 7.90975 | 39.1755767 | 0.591749511 | 37.1343679 | NA | NA | NA | NA | NA | NA | NA |
| SCN9A | 1231 | K | 8.658944444 | 43.47598861 | 0.66274543 | 41.43373197 | NA | NA | NA | NA | NA | NA | NA |
| SCN9A | 1232 | I | 4.48475 | 44.78481392 | 0.684352954 | 42.70967322 | NA | NA | NA | NA | NA | NA | NA |
| SCN9A | 1233 | F | 3.662136364 | 39.99298335 | 0.605244157 | 39.57767844 | NA | NA | NA | NA | NA | NA | NA |
| SCN9A | 1234 | T | 3.420071429 | 42.39815364 | 0.644951347 | 41.61445578 | NA | NA | NA | NA | NA | NA | NA |
| SCN9A | 1235 | Y | 0.803083333 | 45.27721446 | 0.692482042 | 42.80687987 | NA | NA | NA | NA | NA | NA | NA |
| SCN9A | 1236 | I | -0.52325 | 42.24534203 | 0.642428565 | 39.62527799 | NA | NA | NA | NA | NA | NA | NA |
| SCN9A | 1237 | F | -0.855590909 | 37.68371432 | 0.56712021 | 35.09324985 | NA | NA | NA | NA | NA | NA | NA |
| SCN9A | 1238 | I | -1.125625 | 42.65777553 | 0.64923747 | 40.18565871 | NA | NA | NA | NA | NA | NA | NA |
| SCN9A | 1239 | L | -3.694375 | 43.6580425 | 0.665750975 | 41.03656292 | NA | NA | NA | NA | NA | NA | NA |
| SCN9A | 1240 | E | -6.028277778 | 36.66849495 | 0.550359855 | 34.04828095 | NA | NA | NA | NA | NA | NA | NA |
| SCN9A | 1241 | M | -5.49375 | 40.61682713 | 0.615543255 | 38.00241189 | NA | NA | NA | NA | NA | NA | NA |
| SCN9A | 1242 | L | -6.625375 | 42.87850653 | 0.65288154 | 40.26149465 | NA | NA | NA | NA | NA | NA | NA |
| SCN9A | 1243 | L | -9.92275 | 42.00690987 | 0.638492266 | 39.38690014 | NA | NA | NA | NA | NA | NA | NA |
| SCN9A | 1244 | K | -10.72838889 | 36.48484172 | 0.547327906 | 33.86537575 | NA | NA | NA | NA | NA | NA | NA |
| SCN9A | 1245 | W | -9.182642857 | 40.93022268 | 0.620717132 | 38.32532074 | NA | NA | NA | NA | NA | NA | NA |
| SCN9A | 1246 | I | -13.279125 | 43.61416498 | 0.665026597 | 40.99474072 | NA | NA | NA | NA | NA | NA | NA |
| SCN9A | 1247 | A | -15.8607 | 40.90762681 | 0.620344095 | 38.2919051 | NA | NA | NA | NA | NA | NA | NA |

| | | | | | | | | | | | | | |
|---|---|---|---|---|---|---|---|---|---|---|---|---|---|
| SCN9A | 1248 | Y | -15.66416667 | 35.24192234 | 0.526808429 | 32.62647669 | NA | NA | NA | NA | NA | NA | NA |
| SCN9A | 1249 | G | -15.168 | 39.96266192 | 0.604743578 | 37.33961686 | NA | NA | NA | NA | NA | NA | NA |
| SCN9A | 1250 | Y | -12.22625 | 39.09486745 | 0.590417074 | 36.50100573 | NA | NA | NA | NA | NA | NA | NA |
| SCN9A | 1251 | K | -18.15794444 | 36.89420591 | 0.554086139 | 34.29728265 | NA | NA | NA | NA | NA | NA | NA |
| SCN9A | 1252 | T | -17.12521429 | 34.8192243 | 0.519830066 | 32.19690587 | NA | NA | NA | NA | NA | NA | NA |
| SCN9A | 1253 | Y | -11.02316667 | 34.92660698 | 0.521602857 | 32.3197878 | NA | NA | NA | NA | NA | NA | NA |
| SCN9A | 1254 | F | -10.68440909 | 34.20609922 | 0.509707924 | 31.61856283 | NA | NA | NA | NA | NA | NA | NA |
| SCN9A | 1255 | T | -15.57335714 | 30.94905667 | 0.455937092 | 28.3554042 | NA | NA | NA | NA | NA | NA | NA |
| SCN9A | 1256 | N | -13.23125 | 29.30982088 | 0.428874789 | 26.7143226 | NA | NA | NA | NA | NA | NA | NA |
| SCN9A | 1257 | A | -10.5983 | 27.24756617 | 0.394828825 | 24.66225158 | NA | NA | NA | NA | NA | NA | NA |
| SCN9A | 1258 | W | -9.732285714 | 23.721637 | 0.336618918 | 21.1321948 | NA | NA | NA | NA | NA | NA | NA |
| SCN9A | 1259 | C | -9.0165 | 31.32302894 | 0.462111036 | 28.7339621 | NA | NA | NA | NA | NA | NA | NA |
| SCN9A | 1260 | W | -7.199071429 | 32.64307628 | 0.483903826 | 30.22995729 | NA | NA | NA | NA | NA | NA | NA |
| SCN9A | 1261 | L | -4.156875 | 28.16526409 | 0.409979189 | 25.76268173 | NA | NA | NA | NA | NA | NA | NA |
| SCN9A | 1262 | D | -4.594 | 32.63232638 | 0.483726355 | 30.21456153 | NA | NA | NA | NA | NA | NA | NA |
| SCN9A | 1263 | F | -3.866318182 | 35.38468926 | 0.529165382 | 32.9670864 | NA | NA | NA | NA | NA | NA | NA |
| SCN9A | 1264 | L | -1.277375 | 35.04797371 | 0.523606512 | 33.03796013 | NA | NA | NA | NA | NA | NA | NA |
| SCN9A | 1265 | I | 0.395125 | 32.5171486 | 0.481824874 | 30.48402225 | NA | NA | NA | NA | NA | NA | NA |
| SCN9A | 1266 | V | 0.442642857 | 37.94153041 | 0.571376521 | 35.90999036 | NA | NA | NA | NA | NA | NA | NA |
| SCN9A | 1267 | D | 0.914 | 40.02181098 | 0.605720075 | 37.97480478 | NA | NA | NA | NA | NA | NA | NA |
| SCN9A | 1268 | V | 4.403785714 | 37.0985859 | 0.557460268 | 34.05774271 | NA | NA | NA | NA | NA | NA | NA |
| SCN9A | 1269 | S | 4.785666667 | 39.65023827 | 0.599585745 | 36.71122425 | NA | NA | NA | NA | NA | NA | NA |
| SCN9A | 1270 | L | 4.0055 | 42.3246622 | 0.64373807 | 39.38069086 | NA | NA | NA | NA | NA | NA | NA |
| SCN9A | 1271 | V | 6.768357143 | 42.69423927 | 0.649839454 | 39.54823719 | NA | NA | NA | NA | NA | NA | NA |
| SCN9A | 1272 | T | 9.393785714 | 39.23718152 | 0.592766551 | 36.09054905 | NA | NA | NA | NA | NA | NA | NA |
| SCN9A | 1273 | L | 8.522875 | 43.32644037 | 0.660276523 | 40.1769569 | NA | NA | NA | NA | NA | NA | NA |
| SCN9A | 1274 | V | 8.822785714 | 46.05606446 | 0.705340153 | 42.90795868 | NA | NA | NA | NA | NA | NA | NA |
| SCN9A | 1275 | A | 12.1395 | 44.32267331 | 0.67672343 | 41.1544346 | NA | NA | NA | NA | NA | NA | NA |
| SCN9A | 1276 | N | 13.727125 | 41.61674568 | 0.632051007 | 38.47226808 | NA | NA | NA | NA | NA | NA | NA |
| SCN9A | 1277 | T | 12.97078571 | 46.92867866 | 0.719746225 | 42.27866423 | NA | NA | NA | NA | NA | NA | NA |
| SCN9A | 1278 | L | 14.569625 | 48.46996118 | 0.745191408 | 43.81481479 | NA | NA | NA | NA | NA | NA | NA |
| SCN9A | 1279 | G | 16.76025 | 45.85438295 | 0.702010573 | 41.19945595 | NA | NA | NA | NA | NA | NA | NA |
| SCN9A | 1280 | Y | 17.17525 | 43.81040841 | 0.668266399 | 39.15493251 | NA | NA | NA | NA | NA | NA | NA |

| | | | | | | | | | | | | | |
|---|---|---|---|---|---|---|---|---|---|---|---|---|---|
| SCN9A | 1281 | S | 18.047 | 40.60598462 | 0.615364255 | 35.94803621 | NA | NA | NA | NA | NA | NA | NA |
| SCN9A | 1282 | D | 19.53 | 36.91966786 | 0.554506493 | 32.2591101 | NA | NA | NA | NA | NA | NA | NA |
| SCN9A | 1283 | L | 17.942625 | 38.07732607 | 0.573618385 | 33.41979728 | NA | NA | NA | NA | NA | NA | NA |
| SCN9A | 1284 | G | 16.68125 | 36.01060356 | 0.539498662 | 31.355277 | NA | NA | NA | NA | NA | NA | NA |
| SCN9A | 1285 | P | 13.77478571 | 37.0475173 | 0.556617171 | 33.94019608 | NA | NA | NA | NA | NA | NA | NA |
| SCN9A | 1286 | I | 13.07625 | 37.34773158 | 0.561573438 | 34.18522721 | NA | NA | NA | NA | NA | NA | NA |
| SCN9A | 1287 | K | 15.89238889 | 32.8215026 | 0.486849484 | 28.17205486 | NA | NA | NA | NA | NA | NA | NA |
| SCN9A | 1288 | S | 12.91466667 | 32.50966358 | 0.481701303 | 29.34703279 | NA | NA | NA | NA | NA | NA | NA |
| SCN9A | 1289 | L | 9.592875 | 34.78737594 | 0.519304278 | 31.64227906 | NA | NA | NA | NA | NA | NA | NA |
| SCN9A | 1290 | R | 12.96877273 | 33.46099044 | 0.497406851 | 30.31247053 | NA | NA | NA | NA | NA | NA | NA |
| SCN9A | 1291 | T | 11.32592857 | 29.01279962 | 0.423971236 | 25.88535913 | NA | NA | NA | NA | NA | NA | NA |
| SCN9A | 1292 | L | 6.958 | 31.29625211 | 0.461668975 | 28.14762057 | NA | NA | NA | NA | NA | NA | NA |
| SCN9A | 1293 | R | 8.684409091 | 31.92316891 | 0.472018805 | 29.05949866 | NA | NA | NA | NA | NA | NA | NA |
| SCN9A | 1294 | A | 6.2809 | 28.27353726 | 0.411766682 | 26.24017437 | NA | NA | NA | NA | NA | NA | NA |
| SCN9A | 1295 | L | 2.77575 | 28.91733852 | 0.422395259 | 24.36568334 | NA | NA | NA | NA | NA | NA | NA |
| SCN9A | 1296 | R | 3.982954545 | 32.1159653 | 0.4752017 | 27.52286538 | NA | NA | NA | NA | NA | NA | NA |
| SCN9A | 1297 | P | 1.588071429 | 28.94736658 | 0.422890995 | 27.10183401 | NA | NA | NA | NA | NA | NA | NA |
| SCN9A | 1298 | L | -0.867375 | 26.37869341 | 0.38048452 | 24.22113106 | NA | NA | NA | NA | NA | NA | NA |
| SCN9A | 1299 | R | -1.690136364 | 29.65977217 | 0.434652169 | 27.30268467 | NA | NA | NA | NA | NA | NA | NA |
| SCN9A | 1300 | A | -3.7499 | 26.9805267 | 0.390420245 | 24.52423866 | NA | NA | NA | NA | NA | NA | NA |
| SCN9A | 1301 | L | -5.544875 | 22.86112434 | 0.32241263 | 20.42229572 | NA | NA | NA | NA | NA | NA | NA |
| SCN9A | 1302 | S | -8.9475 | 26.35915031 | 0.380161881 | 23.74013043 | NA | NA | NA | NA | NA | NA | NA |
| SCN9A | 1303 | R | -8.134409091 | 26.98273919 | 0.390456771 | 24.36545301 | NA | NA | NA | NA | NA | NA | NA |
| SCN9A | 1304 | F | -7.334318182 | 23.22666868 | 0.328447437 | 20.60928208 | NA | NA | NA | NA | NA | NA | NA |
| SCN9A | 1305 | E | -12.54605556 | 20.51075316 | 0.283610124 | 17.89731185 | NA | NA | NA | NA | NA | NA | NA |
| SCN9A | 1306 | G | -10.835 | 18.38133961 | 0.248455429 | 15.76427618 | NA | NA | NA | NA | NA | NA | NA |
| SCN9A | 1307 | M | -7.378125 | 19.40028085 | 0.265277229 | 16.78444039 | NA | NA | NA | NA | NA | NA | NA |
| SCN9A | 1308 | R | -12.43822727 | 19.00446437 | 0.258742656 | 16.38296157 | NA | NA | NA | NA | NA | NA | NA |
| SCN9A | 1309 | V | -13.5075 | 15.64387145 | 0.203262301 | 13.02303915 | NA | NA | NA | NA | NA | NA | NA |
| SCN9A | 1310 | V | -10.08264286 | 15.19165985 | 0.195796695 | 12.62001794 | NA | NA | NA | NA | NA | NA | NA |
| SCN9A | 1311 | V | -9.430071429 | 17.27535126 | 0.230196559 | 14.90551725 | NA | NA | NA | NA | NA | NA | NA |
| SCN9A | 1312 | N | -13.679625 | 16.14170527 | 0.211481087 | 13.55329404 | NA | NA | NA | NA | NA | NA | NA |
| SCN9A | 1313 | A | -12.8763 | 12.66286385 | 0.154048556 | 10.07579177 | NA | NA | NA | NA | NA | NA | NA |

| | | | | | | | | | | | | | |
|---|---|---|---|---|---|---|---|---|---|---|---|---|---|
| SCN9A | 1314 | L | -9.18875 | 13.4523453 | 0.167082182 | 11.12534714 | NA | NA | NA | NA | NA | NA | NA |
| SCN9A | 1315 | I | -11.2575 | 16.79143654 | 0.222207564 | 14.40880791 | NA | NA | NA | NA | NA | NA | NA |
| SCN9A | 1316 | G | -14.0185 | 15.20529756 | 0.196021842 | 12.83189532 | NA | NA | NA | NA | NA | NA | NA |
| SCN9A | 1317 | A | -11.8485 | 12.62415898 | 0.153409574 | 10.37570641 | NA | NA | NA | NA | NA | NA | NA |
| SCN9A | 1318 | I | -9.565875 | 16.11219879 | 0.210993962 | 14.339601 | NA | NA | NA | NA | NA | NA | NA |
| SCN9A | 1319 | P | -12.1325 | 17.7065515 | 0.237315286 | 15.59625217 | NA | NA | NA | NA | NA | NA | NA |
| SCN9A | 1320 | S | -12.05616667 | 13.30115848 | 0.164586224 | 11.61316093 | NA | NA | NA | NA | NA | NA | NA |
| SCN9A | 1321 | I | -8.219 | 12.67912872 | 0.154317074 | 10.5401693 | NA | NA | NA | NA | NA | NA | NA |
| SCN9A | 1322 | M | -7.251375 | 17.57959459 | 0.235219342 | 15.45230496 | NA | NA | NA | NA | NA | NA | NA |
| SCN9A | 1323 | N | -10.361875 | 18.02129435 | 0.242511406 | 16.14427753 | NA | NA | NA | NA | NA | NA | NA |
| SCN9A | 1324 | V | -7.506214286 | 13.63850483 | 0.170155508 | 11.75356353 | NA | NA | NA | NA | NA | NA | NA |
| SCN9A | 1325 | L | -3.577125 | 16.30070419 | 0.214106016 | 14.78469674 | NA | NA | NA | NA | NA | NA | NA |
| SCN9A | 1326 | L | -5.115375 | 19.08488287 | 0.260070293 | 17.39020274 | NA | NA | NA | NA | NA | NA | NA |
| SCN9A | 1327 | V | -5.739928571 | 18.23643608 | 0.246063202 | 16.19404284 | NA | NA | NA | NA | NA | NA | NA |
| SCN9A | 1328 | C | -2.175333333 | 14.02770441 | 0.176580842 | 11.86251427 | NA | NA | NA | NA | NA | NA | NA |
| SCN9A | 1329 | L | -0.32875 | 18.71925429 | 0.254034095 | 16.52509376 | NA | NA | NA | NA | NA | NA | NA |
| SCN9A | 1330 | I | -1.878125 | 21.16827912 | 0.294465284 | 18.96371035 | NA | NA | NA | NA | NA | NA | NA |
| SCN9A | 1331 | F | -1.374772727 | 17.39235631 | 0.232128207 | 15.08408137 | NA | NA | NA | NA | NA | NA | NA |
| SCN9A | 1332 | W | 1.750357143 | 13.77751026 | 0.172450362 | 8.927820575 | NA | NA | NA | NA | NA | NA | NA |
| SCN9A | 1333 | L | 2.67175 | 21.81004961 | 0.305060335 | 16.93508163 | NA | NA | NA | NA | NA | NA | NA |
| SCN9A | 1334 | I | 1.986 | 23.54664717 | 0.333729993 | 18.64243612 | NA | NA | NA | NA | NA | NA | NA |
| SCN9A | 1335 | F | 3.677590909 | 18.83012974 | 0.255864548 | 13.92648981 | NA | NA | NA | NA | NA | NA | NA |
| SCN9A | 1336 | S | 7.2145 | 21.81634729 | 0.305164304 | 18.80132471 | NA | NA | NA | NA | NA | NA | NA |
| SCN9A | 1337 | I | 6.707 | 24.14976824 | 0.343686978 | 21.13656537 | NA | NA | NA | NA | NA | NA | NA |
| SCN9A | 1338 | M | 7.136375 | 23.91761253 | 0.339854297 | 20.90972433 | NA | NA | NA | NA | NA | NA | NA |
| SCN9A | 1339 | G | 10.94175 | 21.73219779 | 0.303775072 | 18.73275788 | NA | NA | NA | NA | NA | NA | NA |
| SCN9A | 1340 | V | 12.29435714 | 22.87484693 | 0.322639178 | 19.30294079 | NA | NA | NA | NA | NA | NA | NA |
| SCN9A | 1341 | N | 12.20725 | 25.09767045 | 0.359335988 | 21.5472688 | NA | NA | NA | NA | NA | NA | NA |
| SCN9A | 1342 | L | 13.605625 | 22.95003534 | 0.323880471 | 19.48464869 | NA | NA | NA | NA | NA | NA | NA |
| SCN9A | 1343 | F | 15.3625 | 19.0195649 | 0.258991952 | 15.51360442 | NA | NA | NA | NA | NA | NA | NA |
| SCN9A | 1344 | A | 17.5075 | 22.38208875 | 0.314504185 | 18.87881879 | NA | NA | NA | NA | NA | NA | NA |
| SCN9A | 1345 | G | 21.15 | 21.89286827 | 0.306427597 | 18.55804734 | NA | NA | NA | NA | NA | NA | NA |
| SCN9A | 1346 | K | 20.80338889 | 18.77716872 | 0.25499021 | 13.15757001 | NA | NA | NA | NA | NA | NA | NA |

| | | | | | | | | | | | | | |
|---|---|---|---|---|---|---|---|---|---|---|---|---|---|
| SCN9A | 1347 | F | 20.3175 | 15.35176075 | 0.198439816 | 9.865993677 | NA | NA | NA | NA | NA | NA | NA |
| SCN9A | 1348 | Y | 25.5045 | 16.99753544 | 0.225610071 | 11.46103974 | NA | NA | NA | NA | NA | NA | NA |
| SCN9A | 1349 | E | 27.84116667 | 12.0465752 | 0.143874187 | 6.513453515 | NA | NA | NA | NA | NA | NA | NA |
| SCN9A | 1350 | C | 32.56666667 | 13.92858959 | 0.174944545 | 11.51046124 | NA | NA | NA | NA | NA | NA | NA |
| SCN9A | 1351 | I | 31.977875 | 10.35768961 | 0.11599221 | 8.471695583 | NA | NA | NA | NA | NA | NA | NA |
| SCN9A | 1352 | N | 35.930125 | 8.484770592 | 0.085072008 | 6.218361417 | 2.47 | -0.88 | -1.43 | 18.82 | 1.42 | 1 | -1 |
| SCN9A | 1353 | T | 34.98607143 | 8.211467652 | 0.080560023 | 6.020329293 | 2.47 | -0.88 | -1.43 | 18.82 | 1.42 | 1 | -1 |
| SCN9A | 1354 | T | 36.6655 | 5.608981904 | 0.037595333 | 3.496075048 | 2.47 | -0.88 | -1.43 | 18.82 | 1.42 | 1 | -1 |
| SCN9A | 1355 | D | 34.10275 | 4.145574025 | 0.01343579 | 1.765592837 | 2.47 | -0.88 | -1.43 | 18.82 | 1.42 | 1 | -1 |
| SCN9A | 1356 | G | 31.16 | 6.03117816 | 0.044565412 | 3.919873087 | 2.59 | -0.8 | -0.4 | 3.38 | 0.81 | 0 | 0 |
| SCN9A | 1357 | S | 32.66 | 7.293342444 | 0.065402605 | 4.887383566 | 2.47 | -0.88 | -1.43 | 18.82 | 1.42 | 1 | -1 |
| SCN9A | 1358 | R | 29.30777273 | 10.21630564 | 0.113658089 | 7.755235858 | NA | NA | NA | NA | NA | NA | NA |
| SCN9A | 1359 | F | 35.32659091 | 13.24315586 | 0.163628653 | 10.81077775 | NA | NA | NA | NA | NA | NA | NA |
| SCN9A | 1360 | P | 34.6235 | 15.39304473 | 0.199121378 | 12.97699285 | NA | NA | NA | NA | NA | NA | NA |
| SCN9A | 1361 | A | 33.6885 | 18.16861365 | 0.244943515 | 15.73545246 | NA | NA | NA | NA | NA | NA | NA |
| SCN9A | 1362 | S | 36.6755 | 20.32870616 | 0.280604692 | 17.91083688 | NA | NA | NA | NA | NA | NA | NA |
| SCN9A | 1363 | Q | 38.46583333 | 17.12595159 | 0.227730106 | 14.79777709 | NA | NA | NA | NA | NA | NA | NA |
| SCN9A | 1364 | V | 35.94292857 | 18.20461582 | 0.245537878 | 15.82588726 | NA | NA | NA | NA | NA | NA | NA |
| SCN9A | 1365 | P | 34.12578571 | 22.14090895 | 0.310522524 | 19.73159112 | NA | NA | NA | NA | NA | NA | NA |
| SCN9A | 1366 | N | 33.527625 | 22.75288766 | 0.320625741 | 20.36746315 | NA | NA | NA | NA | NA | NA | NA |
| SCN9A | 1367 | R | 31.87440909 | 23.29814469 | 0.329627442 | 20.7104341 | NA | NA | NA | NA | NA | NA | NA |
| SCN9A | 1368 | S | 35.88066667 | 25.17322784 | 0.360583372 | 22.80428384 | NA | NA | NA | NA | NA | NA | NA |
| SCN9A | 1369 | E | 38.31572222 | 22.72609174 | 0.320183364 | 20.41860688 | NA | NA | NA | NA | NA | NA | NA |
| SCN9A | 1370 | C | 36.267 | 18.99358063 | 0.258562975 | 16.66654308 | NA | NA | NA | NA | NA | NA | NA |
| SCN9A | 1371 | F | 36.44068182 | 22.38265299 | 0.3145135 | 20.08160032 | NA | NA | NA | NA | NA | NA | NA |
| SCN9A | 1372 | A | 41.4041 | 23.27811165 | 0.329296714 | 21.0335009 | NA | NA | NA | NA | NA | NA | NA |
| SCN9A | 1373 | L | 41.066875 | 17.85549154 | 0.239774152 | 15.58539706 | NA | NA | NA | NA | NA | NA | NA |
| SCN9A | 1374 | M | 39.443375 | 19.31633754 | 0.2638914 | 17.07912278 | NA | NA | NA | NA | NA | NA | NA |
| SCN9A | 1375 | N | 44.456 | 22.67961536 | 0.319416081 | 20.47596863 | NA | NA | NA | NA | NA | NA | NA |
| SCN9A | 1376 | V | 45.90221429 | 21.5172254 | 0.300226072 | 19.3590492 | NA | NA | NA | NA | NA | NA | NA |
| SCN9A | 1377 | S | 43.305 | 18.40570306 | 0.248857647 | 16.22753627 | NA | NA | NA | NA | NA | NA | NA |
| SCN9A | 1378 | Q | 42.77616667 | 16.72265353 | 0.221072019 | 14.5793202 | NA | NA | NA | NA | NA | NA | NA |
| SCN9A | 1379 | N | 39.290125 | 12.77030043 | 0.155822237 | 10.66378385 | NA | NA | NA | NA | NA | NA | NA |

| | | | | | | | | | | | | | |
|---|---|---|---|---|---|---|---|---|---|---|---|---|---|
| SCN9A | 1380 | V | 37.47507143 | 14.88794956 | 0.190782712 | 12.65979009 | NA | NA | NA | NA | NA | NA | NA |
| SCN9A | 1381 | R | 32.06604545 | 13.25194827 | 0.163773808 | 12.23426222 | NA | NA | NA | NA | NA | NA | NA |
| SCN9A | 1382 | W | 30.56778571 | 17.04331696 | 0.226365883 | 12.26778975 | NA | NA | NA | NA | NA | NA | NA |
| SCN9A | 1383 | K | 27.76272222 | 10.41668479 | 0.116966168 | 5.652996151 | NA | NA | NA | NA | NA | NA | NA |
| SCN9A | 1384 | N | 23.5355 | 13.66714414 | 0.170628317 | 8.928399268 | NA | NA | NA | NA | NA | NA | NA |
| SCN9A | 1385 | L | 21.1695 | 11.71624765 | 0.138420777 | 6.644252303 | NA | NA | NA | NA | NA | NA | NA |
| SCN9A | 1386 | K | 20.59194444 | 11.43093776 | 0.133710568 | 9.322270929 | NA | NA | NA | NA | NA | NA | NA |
| SCN9A | 1387 | V | 16.38421429 | 12.6130857 | 0.153226764 | 9.015711684 | NA | NA | NA | NA | NA | NA | NA |
| SCN9A | 1388 | N | 16.85475 | 15.41406371 | 0.199468382 | 11.82795866 | NA | NA | NA | NA | NA | NA | NA |
| SCN9A | 1389 | F | 13.3755 | 16.82219228 | 0.222715314 | 13.25072357 | NA | NA | NA | NA | NA | NA | NA |
| SCN9A | 1390 | D | 17.4915 | 20.47510532 | 0.28302161 | 16.90313483 | NA | NA | NA | NA | NA | NA | NA |
| SCN9A | 1391 | N | 13.463875 | 22.15931716 | 0.310826427 | 18.5584802 | NA | NA | NA | NA | NA | NA | NA |
| SCN9A | 1392 | V | 9.230214286 | 22.23170641 | 0.312021509 | 19.22577423 | NA | NA | NA | NA | NA | NA | NA |
| SCN9A | 1393 | G | 10.006 | 20.3531585 | 0.281008378 | 17.34833002 | NA | NA | NA | NA | NA | NA | NA |
| SCN9A | 1394 | L | 13.419625 | 17.70962693 | 0.237366059 | 14.12715267 | NA | NA | NA | NA | NA | NA | NA |
| SCN9A | 1395 | G | 10.198 | 17.59251869 | 0.235432707 | 14.58365469 | NA | NA | NA | NA | NA | NA | NA |
| SCN9A | 1396 | Y | 5.671333333 | 16.7079409 | 0.220829126 | 13.68762261 | NA | NA | NA | NA | NA | NA | NA |
| SCN9A | 1397 | L | 10.594875 | 13.64544653 | 0.170270109 | 10.63450426 | NA | NA | NA | NA | NA | NA | NA |
| SCN9A | 1398 | S | 11.3135 | 12.85144517 | 0.157161863 | 9.840634988 | NA | NA | NA | NA | NA | NA | NA |
| SCN9A | 1399 | L | 7.326625 | 13.62043065 | 0.169857119 | 10.58640808 | NA | NA | NA | NA | NA | NA | NA |
| SCN9A | 1400 | L | 6.53 | 11.6187795 | 0.136811666 | 8.57584343 | NA | NA | NA | NA | NA | NA | NA |
| SCN9A | 1401 | Q | 10.85405556 | 8.773243072 | 0.089834428 | 5.734489584 | 3.05 | -0.47 | -1.2 | 12.29 | 0.72 | 1 | -1 |
| SCN9A | 1402 | V | 8.271214286 | 10.04817287 | 0.110882368 | 7.546124772 | NA | NA | NA | NA | NA | NA | NA |
| SCN9A | 1403 | A | 4.7577 | 8.231619768 | 0.080892717 | 4.144106224 | 4.11 | -0.35 | -1.08 | 12.03 | 0.28 | 1 | 1 |
| SCN9A | 1404 | T | 5.932214286 | 5.851320022 | 0.041596117 | 2.800714898 | 3.6 | -0.78 | -2.05 | 14.95 | 0.8 | 1 | 1 |
| SCN9A | 1405 | F | 6.663954545 | 4.805134337 | 0.024324535 | 3.741236307 | 1.82 | -0.79 | -2.05 | 26.54 | 1.33 | 2 | 0 |
| SCN9A | 1406 | K | 9.595055556 | 3.331732432 | 0 | 1.823745757 | 3.05 | -0.47 | -1.2 | 12.29 | 0.72 | 1 | -1 |
| SCN9A | 1407 | G | 13.825 | 4.605628187 | 0.021030869 | 1.785238729 | 3.1 | -0.47 | -2.18 | 26.15 | 1.06 | 2 | -2 |
| SCN9A | 1408 | W | 11.27 | 6.144172279 | 0.046430843 | 3.138655672 | 3.05 | -0.47 | -1.2 | 12.29 | 0.72 | 1 | -1 |
| SCN9A | 1409 | T | 15.58835714 | 5.454556536 | 0.03504591 | 2.150023687 | 3.67 | 0.19 | -0.95 | 25.34 | 0.39 | 2 | -2 |
| SCN9A | 1410 | I | 18.4405 | 5.245967118 | 0.031602287 | 1.715941292 | 3.67 | 0.19 | -0.95 | 25.34 | 0.39 | 2 | -2 |
| SCN9A | 1411 | I | 15.647875 | 9.17960887 | 0.096543161 | 5.716589677 | 3.67 | 0.19 | -0.95 | 25.34 | 0.39 | 2 | -2 |
| SCN9A | 1412 | M | 14.850125 | 11.0668496 | 0.127699802 | 7.638774887 | NA | NA | NA | NA | NA | NA | NA |

| | | | | | | | | | | | | | |
|---|---|---|---|---|---|---|---|---|---|---|---|---|---|
| SCN9A | 1413 | Y | 19.29666667 | 7.670654275 | 0.071631683 | 4.333065246 | 3.63 | 0.25 | -0.25 | 13.33 | -0.07 | 1 | -1 |
| SCN9A | 1414 | A | 19.8641 | 12.4322792 | 0.150241812 | 9.004210164 | NA | NA | NA | NA | NA | NA | NA |
| SCN9A | 1415 | A | 17.5949 | 14.81368914 | 0.18955674 | 11.52878169 | NA | NA | NA | NA | NA | NA | NA |
| SCN9A | 1416 | V | 18.6905 | 15.60596581 | 0.202636513 | 12.27702746 | NA | NA | NA | NA | NA | NA | NA |
| SCN9A | 1417 | D | 22.8625 | 13.86168915 | 0.17384008 | 8.257608578 | NA | NA | NA | NA | NA | NA | NA |
| SCN9A | 1418 | S | 21.27933333 | 18.07230486 | 0.243353544 | 12.47834897 | NA | NA | NA | NA | NA | NA | NA |
| SCN9A | 1419 | V | 23.89078571 | 21.17024832 | 0.294497794 | 15.56258555 | NA | NA | NA | NA | NA | NA | NA |
| SCN9A | 1420 | N | 23.8795 | 22.98184312 | 0.324405588 | 17.37206396 | NA | NA | NA | NA | NA | NA | NA |
| SCN9A | 1421 | V | 23.78378571 | 23.51078769 | 0.333137986 | 18.00950805 | NA | NA | NA | NA | NA | NA | NA |
| SCN9A | 1422 | D | 28.25275 | 24.33740079 | 0.346784622 | 18.81044023 | NA | NA | NA | NA | NA | NA | NA |
| SCN9A | 1423 | K | 28.60672222 | 21.28495671 | 0.296391526 | 15.81626813 | NA | NA | NA | NA | NA | NA | NA |
| SCN9A | 1424 | Q | 30.09505556 | 16.39072381 | 0.215592159 | 11.22673232 | NA | NA | NA | NA | NA | NA | NA |
| SCN9A | 1425 | P | 26.88792857 | 17.00429316 | 0.225721635 | 11.40363937 | NA | NA | NA | NA | NA | NA | NA |
| SCN9A | 1426 | K | 27.57761111 | 17.78146394 | 0.238552023 | 12.17650012 | NA | NA | NA | NA | NA | NA | NA |
| SCN9A | 1427 | Y | 25.182 | 14.00894964 | 0.176271217 | 8.429952965 | NA | NA | NA | NA | NA | NA | NA |
| SCN9A | 1428 | E | 21.58672222 | 16.07208471 | 0.210331715 | 10.7707816 | NA | NA | NA | NA | NA | NA | NA |
| SCN9A | 1429 | Y | 23.14716667 | 19.96170406 | 0.274545819 | 14.38804003 | NA | NA | NA | NA | NA | NA | NA |
| SCN9A | 1430 | S | 18.42433333 | 20.99642617 | 0.291628147 | 17.6892305 | NA | NA | NA | NA | NA | NA | NA |
| SCN9A | 1431 | L | 17.086875 | 19.39975654 | 0.265268573 | 16.2301464 | NA | NA | NA | NA | NA | NA | NA |
| SCN9A | 1432 | Y | 14.07458333 | 21.3247328 | 0.297048193 | 17.97682484 | NA | NA | NA | NA | NA | NA | NA |
| SCN9A | 1433 | M | 13.873 | 18.71340987 | 0.253937609 | 15.3222928 | NA | NA | NA | NA | NA | NA | NA |
| SCN9A | 1434 | Y | 13.62508333 | 13.21303375 | 0.163131365 | 11.34000323 | NA | NA | NA | NA | NA | NA | NA |
| SCN9A | 1435 | I | 8.808625 | 19.39030418 | 0.265112523 | 16.46165141 | NA | NA | NA | NA | NA | NA | NA |
| SCN9A | 1436 | Y | 8.622083333 | 18.60574892 | 0.252160224 | 15.59316037 | NA | NA | NA | NA | NA | NA | NA |
| SCN9A | 1437 | F | 10.20786364 | 14.60234372 | 0.186067618 | 11.61576435 | NA | NA | NA | NA | NA | NA | NA |
| SCN9A | 1438 | V | 6.008785714 | 15.54871943 | 0.201691427 | 14.29556844 | NA | NA | NA | NA | NA | NA | NA |
| SCN9A | 1439 | V | 3.542357143 | 16.66471254 | 0.220115465 | 11.76240262 | NA | NA | NA | NA | NA | NA | NA |
| SCN9A | 1440 | F | 4.116681818 | 14.34798076 | 0.181868315 | 9.445592567 | NA | NA | NA | NA | NA | NA | NA |
| SCN9A | 1441 | I | 3.729 | 10.97987386 | 0.126263911 | 6.096337555 | NA | NA | NA | NA | NA | NA | NA |
| SCN9A | 1442 | I | 0.610125 | 13.15487085 | 0.162171147 | 8.401135599 | NA | NA | NA | NA | NA | NA | NA |
| SCN9A | 1443 | F | -0.702681818 | 14.1478874 | 0.178564954 | 10.87657885 | NA | NA | NA | NA | NA | NA | NA |
| SCN9A | 1444 | G | 0.03125 | 12.52681388 | 0.151802494 | 7.732291321 | NA | NA | NA | NA | NA | NA | NA |
| SCN9A | 1445 | S | 0.649666667 | 8.248894108 | 0.0811779 | 3.416053668 | 4.86 | 1.35 | 2.45 | 0.56 | -1.72 | 0 | 0 |

| | | | | | | | | | | | | | |
|---|---|---|---|---|---|---|---|---|---|---|---|---|---|
| SCN9A | 1446 | F | -2.785136364 | 8.770252505 | 0.089785057 | 5.373694991 | 3.4 | 1.13 | 2.1 | 0.62 | -1.67 | 0 | 0 |
| SCN9A | 1447 | F | -4.567227273 | 11.10221532 | 0.128283658 | 8.839382733 | NA | NA | NA | NA | NA | NA | NA |
| SCN9A | 1448 | T | -3.3345 | 10.09750766 | 0.111696841 | 7.945467221 | NA | NA | NA | NA | NA | NA | NA |
| SCN9A | 1449 | L | -3.79425 | 4.500551966 | 0.019296155 | 2.203670318 | 2.48 | 1.3 | 2.7 | 0.5 | -1.72 | 0 | 0 |
| SCN9A | 1450 | N | -7.775125 | 8.107437141 | 0.078842574 | 6.556419323 | 2.09 | 1.35 | 2.33 | 0.62 | -1.69 | 0 | 0 |
| SCN9A | 1451 | L | -8.632125 | 10.0459124 | 0.11084505 | 8.15163702 | NA | NA | NA | NA | NA | NA | NA |
| SCN9A | 1452 | F | -5.4725 | 8.374550615 | 0.083252376 | 6.535993943 | 2.09 | 1.35 | 2.33 | 0.62 | -1.69 | 0 | 0 |
| SCN9A | 1453 | I | -8.254625 | 3.677029644 | 0.005700545 | 1.791195055 | 2.09 | 1.35 | 2.33 | 0.62 | -1.69 | 0 | 0 |
| SCN9A | 1454 | G | -11.9655 | 7.465946022 | 0.068252134 | 5.4760401 | 2.17 | 1.64 | 3.05 | 0.5 | -1.75 | 0 | 0 |
| SCN9A | 1455 | V | -12.32878571 | 9.275216116 | 0.09812155 | 7.501575316 | 1.99 | 1.81 | 4.5 | 0.13 | -1.85 | 0 | 0 |
| SCN9A | 1456 | I | -10.518125 | 7.352657275 | 0.066381839 | 5.455039376 | 1.99 | 1.81 | 4.5 | 0.13 | -1.85 | 0 | 0 |
| SCN9A | 1457 | I | -13.662625 | 3.540083756 | 0.003439692 | 1.645467516 | 1.99 | 1.81 | 4.5 | 0.13 | -1.85 | 0 | 0 |
| SCN9A | 1458 | D | -17.102625 | 6.339942034 | 0.049662825 | 4.312381434 | 2.42 | 1.81 | 4.5 | 0.13 | -1.85 | 0 | 0 |
| SCN9A | 1459 | N | -15.72025 | 8.575818853 | 0.086575133 | 6.369787748 | 2.42 | 1.81 | 4.5 | 0.13 | -1.85 | 0 | 0 |
| SCN9A | 1460 | F | -14.22695455 | 6.565131986 | 0.053380508 | 4.147645424 | 2.42 | 1.81 | 4.5 | 0.13 | -1.85 | 0 | 0 |
| SCN9A | 1461 | N | -19.529375 | 4.239092945 | 0.014979703 | 1.62563369 | 2.65 | 0.51 | 1.28 | 1.76 | -0.59 | 0 | 0 |
| SCN9A | 1462 | Q | -21.65627778 | 9.090356209 | 0.09506968 | 6.445890804 | 2.65 | 0.51 | 1.28 | 1.76 | -0.59 | 0 | 0 |
| SCN9A | 1463 | Q | -18.37527778 | 10.5706207 | 0.11950751 | 7.904899002 | NA | NA | NA | NA | NA | NA | NA |
| SCN9A | 1464 | K | -21.56727778 | 5.287183182 | 0.032282727 | 1.97833645 | 3.35 | 0.33 | -0.2 | 13.34 | -0.4 | 1 | 1 |
| SCN9A | 1465 | K | -25.55038889 | 8.142561268 | 0.079422441 | 5.211668551 | 3.8 | 0.06 | -1.53 | 17.74 | 0.06 | 1 | 1 |
| SCN9A | 1466 | K | -25.01572222 | 11.83982268 | 0.140460889 | 8.215150172 | NA | NA | NA | NA | NA | NA | NA |
| SCN9A | 1467 | L | -23.223375 | 12.44027596 | 0.150373831 | 9.124273646 | NA | NA | NA | NA | NA | NA | NA |
| SCN9A | 1468 | G | -26.972 | 11.09180747 | 0.128111834 | 4.721671237 | 6.53 | -0.53 | -3.77 | 34.13 | 1.15 | 2 | 2 |
| SCN9A | 1469 | G | -25.41 | 8.876191526 | 0.091534014 | 5.096685219 | 3.8 | 0.06 | -1.53 | 17.74 | 0.06 | 1 | 1 |
| SCN9A | 1470 | Q | -23.90083333 | 10.0002498 | 0.110091202 | 6.844609915 | NA | NA | NA | NA | NA | NA | NA |
| SCN9A | 1471 | D | -18.82275 | 8.585926916 | 0.086742008 | 5.583644029 | 3.35 | 0.33 | -0.2 | 13.34 | -0.4 | 1 | 1 |
| SCN9A | 1472 | I | -17.1225 | 9.264204499 | 0.097939758 | 6.644637787 | 2.63 | -0.14 | 0.05 | 2.57 | 0.08 | 0 | 0 |
| SCN9A | 1473 | F | -13.41995455 | 8.280850077 | 0.081705465 | 5.662096554 | 2.63 | -0.14 | 0.05 | 2.57 | 0.08 | 0 | 0 |
| SCN9A | 1474 | M | -16.301 | 13.41795711 | 0.166514464 | 10.81901197 | NA | NA | NA | NA | NA | NA | NA |
| SCN9A | 1475 | T | -16.22321429 | 17.27282753 | 0.230154895 | 14.66999916 | NA | NA | NA | NA | NA | NA | NA |
| SCN9A | 1476 | E | -21.10905556 | 18.76971481 | 0.254867153 | 15.51247788 | NA | NA | NA | NA | NA | NA | NA |
| SCN9A | 1477 | E | -18.98505556 | 20.92289249 | 0.290414172 | 17.63741586 | NA | NA | NA | NA | NA | NA | NA |
| SCN9A | 1478 | Q | -15.75216667 | 17.88038372 | 0.240185099 | 15.27439846 | NA | NA | NA | NA | NA | NA | NA |

| | | | | | | | | | | | | | |
|---|---|---|---|---|---|---|---|---|---|---|---|---|---|
| SCN9A | 1479 | K | -20.89994444 | 14.99769919 | 0.19259458 | 11.69067221 | NA | NA | NA | NA | NA | NA | NA |
| SCN9A | 1480 | K | -22.83494444 | 18.93561047 | 0.25760594 | 15.6396453 | NA | NA | NA | NA | NA | NA | NA |
| SCN9A | 1481 | Y | -18.33158333 | 18.86917372 | 0.256509129 | 15.57048713 | NA | NA | NA | NA | NA | NA | NA |
| SCN9A | 1482 | Y | -19.55408333 | 10.40765709 | 0.116817128 | 7.120318543 | NA | NA | NA | NA | NA | NA | NA |
| SCN9A | 1483 | N | -23.548875 | 15.10281725 | 0.194329984 | 11.88871404 | NA | NA | NA | NA | NA | NA | NA |
| SCN9A | 1484 | A | -21.7979 | 18.98801006 | 0.25847101 | 15.71172325 | NA | NA | NA | NA | NA | NA | NA |
| SCN9A | 1485 | M | -18.367625 | 17.85377744 | 0.239745853 | 14.5476574 | NA | NA | NA | NA | NA | NA | NA |
| SCN9A | 1486 | K | -23.20261111 | 12.34701008 | 0.148834096 | 9.179181678 | NA | NA | NA | NA | NA | NA | NA |
| SCN9A | 1487 | K | -25.23694444 | 18.44248145 | 0.249464825 | 14.78803548 | NA | NA | NA | NA | NA | NA | NA |
| SCN9A | 1488 | L | -20.608 | 20.20770789 | 0.27860712 | 16.92648441 | NA | NA | NA | NA | NA | NA | NA |
| SCN9A | 1489 | G | -22.789 | 18.66436725 | 0.253127959 | 15.38358418 | NA | NA | NA | NA | NA | NA | NA |
| SCN9A | 1490 | S | -25.62716667 | 20.01640752 | 0.275448924 | 16.1608536 | NA | NA | NA | NA | NA | NA | NA |
| SCN9A | 1491 | K | -27.24183333 | 22.27746633 | 0.312776963 | 16.15077963 | NA | NA | NA | NA | NA | NA | NA |
| SCN9A | 1492 | K | -29.4385 | 24.5442907 | 0.350200188 | 18.40442926 | NA | NA | NA | NA | NA | NA | NA |
| SCN9A | 1493 | P | -29.17492857 | 27.85681836 | 0.404887029 | 21.72165579 | NA | NA | NA | NA | NA | NA | NA |
| SCN9A | 1494 | Q | -25.3565 | 30.35536055 | 0.446135705 | 24.28553506 | NA | NA | NA | NA | NA | NA | NA |
| SCN9A | 1495 | K | -28.8475 | 33.88034677 | 0.504330045 | 27.74077981 | NA | NA | NA | NA | NA | NA | NA |
| SCN9A | 1496 | P | -27.44092857 | 36.51923048 | 0.547895633 | 30.38062577 | NA | NA | NA | NA | NA | NA | NA |
| SCN9A | 1497 | I | -25.216875 | 38.17559091 | 0.575240648 | 34.39099961 | NA | NA | NA | NA | NA | NA | NA |
| SCN9A | 1498 | P | -26.72792857 | 42.70466168 | 0.650011518 | 36.57561159 | NA | NA | NA | NA | NA | NA | NA |
| SCN9A | 1499 | R | -25.88213636 | 45.35821202 | 0.693819239 | 41.60745033 | NA | NA | NA | NA | NA | NA | NA |
| SCN9A | 1500 | P | -23.70292857 | 47.69766775 | 0.732441541 | 43.94088173 | NA | NA | NA | NA | NA | NA | NA |
| SCN9A | 1501 | G | -25.8365 | 51.17872841 | 0.78991071 | 47.382904 | NA | NA | NA | NA | NA | NA | NA |
| SCN9A | 1502 | N | -22.887875 | 52.46627646 | 0.811166966 | 48.68579728 | NA | NA | NA | NA | NA | NA | NA |
| SCN9A | 1503 | K | -22.38072222 | 52.9989066 | 0.819960209 | 49.26254123 | NA | NA | NA | NA | NA | NA | NA |
| SCN9A | 1504 | I | -18.888375 | 52.35217533 | 0.809283259 | 49.03250069 | NA | NA | NA | NA | NA | NA | NA |
| SCN9A | 1505 | Q | -19.40138889 | 49.27873259 | 0.758543494 | 45.96032956 | NA | NA | NA | NA | NA | NA | NA |
| SCN9A | 1506 | G | -21.22075 | 48.43242264 | 0.744571681 | 44.67668861 | NA | NA | NA | NA | NA | NA | NA |
| SCN9A | 1507 | C | -19.441 | 48.49837418 | 0.745660481 | 45.17504108 | NA | NA | NA | NA | NA | NA | NA |
| SCN9A | 1508 | I | -16.3245 | 46.21137118 | 0.707904126 | 42.8960764 | NA | NA | NA | NA | NA | NA | NA |
| SCN9A | 1509 | F | -21.18668182 | 42.83418255 | 0.652149791 | 39.54619741 | NA | NA | NA | NA | NA | NA | NA |
| SCN9A | 1510 | D | -21.24375 | 44.74012228 | 0.683615135 | 41.41666595 | NA | NA | NA | NA | NA | NA | NA |
| SCN9A | 1511 | L | -16.070625 | 43.77493288 | 0.66768073 | 40.46676643 | NA | NA | NA | NA | NA | NA | NA |

| | | | | | | | | | | | | | |
|---|---|---|---|---|---|---|---|---|---|---|---|---|---|
| SCN9A | 1512 | V | -16.4515 | 40.69135076 | 0.616773573 | 37.38277339 | NA | NA | NA | NA | NA | NA | NA |
| SCN9A | 1513 | T | -20.31778571 | 39.28695203 | 0.593588217 | 35.96242276 | NA | NA | NA | NA | NA | NA | NA |
| SCN9A | 1514 | N | -18.4555 | 40.04789359 | 0.606150675 | 36.73850261 | NA | NA | NA | NA | NA | NA | NA |
| SCN9A | 1515 | Q | -18.05938889 | 36.73283194 | 0.551422 | 33.40721863 | NA | NA | NA | NA | NA | NA | NA |
| SCN9A | 1516 | A | -15.3083 | 39.79741933 | 0.602015571 | 37.08628288 | NA | NA | NA | NA | NA | NA | NA |
| SCN9A | 1517 | F | -12.94631818 | 40.27759289 | 0.609942804 | 37.56296567 | NA | NA | NA | NA | NA | NA | NA |
| SCN9A | 1518 | D | -13.2615 | 34.45502171 | 0.51381741 | 31.7412009 | NA | NA | NA | NA | NA | NA | NA |
| SCN9A | 1519 | I | -12.176125 | 35.95157977 | 0.538524232 | 33.22875668 | NA | NA | NA | NA | NA | NA | NA |
| SCN9A | 1520 | S | -8.956833333 | 39.42184412 | 0.595815164 | 36.73039266 | NA | NA | NA | NA | NA | NA | NA |
| SCN9A | 1521 | I | -8.16725 | 37.40580226 | 0.562532133 | 35.06471892 | NA | NA | NA | NA | NA | NA | NA |
| SCN9A | 1522 | M | -8.247125 | 32.42471898 | 0.480298944 | 29.85013198 | NA | NA | NA | NA | NA | NA | NA |
| SCN9A | 1523 | V | -5.472071429 | 36.49943317 | 0.547568797 | 33.30281359 | NA | NA | NA | NA | NA | NA | NA |
| SCN9A | 1524 | L | -3.331125 | 36.71023292 | 0.551048911 | 34.29960282 | NA | NA | NA | NA | NA | NA | NA |
| SCN9A | 1525 | I | -3.52725 | 33.18985645 | 0.492930673 | 30.78999985 | NA | NA | NA | NA | NA | NA | NA |
| SCN9A | 1526 | C | -1.9975 | 32.00267865 | 0.47333144 | 29.6883197 | NA | NA | NA | NA | NA | NA | NA |
| SCN9A | 1527 | L | -0.337375 | 34.6973945 | 0.517818766 | 32.29015311 | NA | NA | NA | NA | NA | NA | NA |
| SCN9A | 1528 | N | 1.050625 | 33.15849606 | 0.492412942 | 30.70960395 | NA | NA | NA | NA | NA | NA | NA |
| SCN9A | 1529 | M | 1.072625 | 26.01422413 | 0.374467461 | 23.59484883 | NA | NA | NA | NA | NA | NA | NA |
| SCN9A | 1530 | V | 3.661642857 | 31.38856762 | 0.463193021 | 28.9487447 | NA | NA | NA | NA | NA | NA | NA |
| SCN9A | 1531 | T | 5.923071429 | 32.5716713 | 0.482724994 | 30.72379223 | NA | NA | NA | NA | NA | NA | NA |
| SCN9A | 1532 | M | 5.786125 | 29.27111949 | 0.428235864 | 27.46043989 | NA | NA | NA | NA | NA | NA | NA |
| SCN9A | 1533 | M | 6.379125 | 26.51923558 | 0.382804745 | 25.31282695 | NA | NA | NA | NA | NA | NA | NA |
| SCN9A | 1534 | V | 10.15392857 | 30.22441968 | 0.443973989 | 28.39510692 | NA | NA | NA | NA | NA | NA | NA |
| SCN9A | 1535 | E | 11.42772222 | 28.18717073 | 0.410340848 | 26.39547792 | NA | NA | NA | NA | NA | NA | NA |
| SCN9A | 1536 | K | 16.4735 | 28.6862458 | 0.418580127 | 26.79626213 | NA | NA | NA | NA | NA | NA | NA |
| SCN9A | 1537 | E | 19.67272222 | 26.75610983 | 0.386715325 | 24.65029539 | NA | NA | NA | NA | NA | NA | NA |
| SCN9A | 1538 | G | 21.043 | 30.40911332 | 0.447023115 | 28.30221988 | NA | NA | NA | NA | NA | NA | NA |
| SCN9A | 1539 | Q | 17.80483333 | 32.73062482 | 0.485349174 | 30.609972 | NA | NA | NA | NA | NA | NA | NA |
| SCN9A | 1540 | S | 20.649 | 35.68828511 | 0.534177475 | 33.56737828 | NA | NA | NA | NA | NA | NA | NA |
| SCN9A | 1541 | Q | 21.41016667 | 39.08969761 | 0.590331725 | 36.97739431 | NA | NA | NA | NA | NA | NA | NA |
| SCN9A | 1542 | H | 18.1413 | 40.91126664 | 0.620404185 | 38.75672669 | NA | NA | NA | NA | NA | NA | NA |
| SCN9A | 1543 | M | 15.831125 | 35.7303507 | 0.53487194 | 33.886645 | NA | NA | NA | NA | NA | NA | NA |
| SCN9A | 1544 | T | 16.20321429 | 36.69455792 | 0.550790131 | 34.9315015 | NA | NA | NA | NA | NA | NA | NA |

| | | | | | | | | | | | | | |
|---|---|---|---|---|---|---|---|---|---|---|---|---|---|
| SCN9A | 1545 | E | 15.39161111 | 40.25668458 | 0.609597626 | 38.35675226 | NA | NA | NA | NA | NA | NA | NA |
| SCN9A | 1546 | V | 12.27664286 | 40.30614313 | 0.610414142 | 38.5073942 | NA | NA | NA | NA | NA | NA | NA |
| SCN9A | 1547 | L | 10.743125 | 34.42090501 | 0.513254174 | 32.62354957 | NA | NA | NA | NA | NA | NA | NA |
| SCN9A | 1548 | Y | 10.781 | 39.14187664 | 0.591193154 | 37.35669483 | NA | NA | NA | NA | NA | NA | NA |
| SCN9A | 1549 | W | 10.27357143 | 41.66341293 | 0.632821441 | 39.87417391 | NA | NA | NA | NA | NA | NA | NA |
| SCN9A | 1550 | I | 6.533 | 39.71426829 | 0.600642823 | 38.00237157 | NA | NA | NA | NA | NA | NA | NA |
| SCN9A | 1551 | N | 5.8375 | 36.12209047 | 0.54133921 | 34.38631566 | NA | NA | NA | NA | NA | NA | NA |
| SCN9A | 1552 | V | 5.049642857 | 41.53704905 | 0.630735288 | 39.71290644 | NA | NA | NA | NA | NA | NA | NA |
| SCN9A | 1553 | V | 2.349642857 | 42.91430575 | 0.653472553 | 40.45965959 | NA | NA | NA | NA | NA | NA | NA |
| SCN9A | 1554 | F | 0.958590909 | 37.84748375 | 0.569823895 | 35.40145417 | NA | NA | NA | NA | NA | NA | NA |
| SCN9A | 1555 | I | 1.2255 | 40.483692 | 0.613345314 | 38.00832806 | NA | NA | NA | NA | NA | NA | NA |
| SCN9A | 1556 | I | -0.77375 | 43.74120344 | 0.667123887 | 41.28415918 | NA | NA | NA | NA | NA | NA | NA |
| SCN9A | 1557 | L | -2.38725 | 43.66071812 | 0.665795147 | 41.23357252 | NA | NA | NA | NA | NA | NA | NA |
| SCN9A | 1558 | F | -3.413409091 | 37.5295304 | 0.564574773 | 35.10337731 | NA | NA | NA | NA | NA | NA | NA |
| SCN9A | 1559 | T | -3.774071429 | 43.38403114 | 0.661227295 | 41.05416783 | NA | NA | NA | NA | NA | NA | NA |
| SCN9A | 1560 | G | -6.34825 | 45.48414549 | 0.695898287 | 42.79943181 | NA | NA | NA | NA | NA | NA | NA |
| SCN9A | 1561 | E | -8.660388889 | 38.81894061 | 0.585861771 | 36.15707802 | NA | NA | NA | NA | NA | NA | NA |
| SCN9A | 1562 | C | -9.04 | 42.01331005 | 0.638597927 | 39.34399414 | NA | NA | NA | NA | NA | NA | NA |
| SCN9A | 1563 | V | -9.411928571 | 44.68046945 | 0.682630321 | 42.00251515 | NA | NA | NA | NA | NA | NA | NA |
| SCN9A | 1564 | L | -12.077875 | 44.89811896 | 0.686223518 | 42.22423099 | NA | NA | NA | NA | NA | NA | NA |
| SCN9A | 1565 | K | -13.80472222 | 37.78712463 | 0.568827421 | 34.63855073 | NA | NA | NA | NA | NA | NA | NA |
| SCN9A | 1566 | L | -13.822875 | 42.35874093 | 0.644300679 | 39.10804395 | NA | NA | NA | NA | NA | NA | NA |
| SCN9A | 1567 | I | -14.927375 | 46.24222811 | 0.708413547 | 42.96376777 | NA | NA | NA | NA | NA | NA | NA |
| SCN9A | 1568 | S | -18.35816667 | 44.97946155 | 0.68756641 | 41.66263965 | NA | NA | NA | NA | NA | NA | NA |
| SCN9A | 1569 | L | -19.041875 | 39.48518834 | 0.59686092 | 36.16938042 | NA | NA | NA | NA | NA | NA | NA |
| SCN9A | 1570 | R | -20.84959091 | 42.28493927 | 0.64308228 | 38.97311552 | NA | NA | NA | NA | NA | NA | NA |
| SCN9A | 1571 | H | -21.4718 | 39.861048 | 0.603066024 | 36.55104168 | NA | NA | NA | NA | NA | NA | NA |
| SCN9A | 1572 | Y | -23.09833333 | 37.60450373 | 0.565812515 | 34.29300079 | NA | NA | NA | NA | NA | NA | NA |
| SCN9A | 1573 | Y | -15.16516667 | 36.87144729 | 0.553710414 | 33.57262906 | NA | NA | NA | NA | NA | NA | NA |
| SCN9A | 1574 | F | -15.48759091 | 35.46469536 | 0.53048621 | 32.16283887 | NA | NA | NA | NA | NA | NA | NA |
| SCN9A | 1575 | T | -20.19935714 | 32.73268277 | 0.485383148 | 29.42111446 | NA | NA | NA | NA | NA | NA | NA |
| SCN9A | 1576 | V | -17.85921429 | 31.68920215 | 0.468156226 | 28.3869356 | NA | NA | NA | NA | NA | NA | NA |
| SCN9A | 1577 | G | -15.21425 | 30.36960741 | 0.446370908 | 27.11250956 | NA | NA | NA | NA | NA | NA | NA |

| | | | | | | | | | | | | | |
|---|---|---|---|---|---|---|---|---|---|---|---|---|---|
| SCN9A | 1578 | W | -12.857 | 26.74284134 | 0.386496274 | 24.07425115 | NA | NA | NA | NA | NA | NA | NA |
| SCN9A | 1579 | N | -13.96775 | 33.15367102 | 0.492333285 | 30.48901064 | NA | NA | NA | NA | NA | NA | NA |
| SCN9A | 1580 | I | -12.0155 | 34.00622833 | 0.506408236 | 31.35420797 | NA | NA | NA | NA | NA | NA | NA |
| SCN9A | 1581 | F | -7.951136364 | 30.72205197 | 0.452189449 | 28.11092528 | NA | NA | NA | NA | NA | NA | NA |
| SCN9A | 1582 | D | -8.394125 | 34.21865779 | 0.509915255 | 31.67592058 | NA | NA | NA | NA | NA | NA | NA |
| SCN9A | 1583 | F | -8.173136364 | 36.89945437 | 0.554172786 | 34.37397694 | NA | NA | NA | NA | NA | NA | NA |
| SCN9A | 1584 | V | -5.728642857 | 35.605517 | 0.532811048 | 32.99766724 | NA | NA | NA | NA | NA | NA | NA |
| SCN9A | 1585 | V | -3.769357143 | 34.40021123 | 0.512912538 | 31.79863338 | NA | NA | NA | NA | NA | NA | NA |
| SCN9A | 1586 | V | -3.069214286 | 38.79371435 | 0.585445308 | 36.25083217 | NA | NA | NA | NA | NA | NA | NA |
| SCN9A | 1587 | I | -2.228 | 40.38172881 | 0.611661994 | 37.6620215 | NA | NA | NA | NA | NA | NA | NA |
| SCN9A | 1588 | I | -0.630375 | 35.3599471 | 0.528756911 | 32.87767001 | NA | NA | NA | NA | NA | NA | NA |
| SCN9A | 1589 | S | 1.163833333 | 38.00423863 | 0.572411777 | 35.5275375 | NA | NA | NA | NA | NA | NA | NA |
| SCN9A | 1590 | I | 1.509 | 40.60753867 | 0.615389911 | 38.11862447 | NA | NA | NA | NA | NA | NA | NA |
| SCN9A | 1591 | V | 3.2615 | 40.4223142 | 0.612332022 | 38.02452908 | NA | NA | NA | NA | NA | NA | NA |
| SCN9A | 1592 | G | 5.856 | 38.22272378 | 0.57601877 | 36.37635564 | NA | NA | NA | NA | NA | NA | NA |
| SCN9A | 1593 | M | 6.76625 | 39.46673892 | 0.596556336 | 37.61555679 | NA | NA | NA | NA | NA | NA | NA |
| SCN9A | 1594 | F | 6.749045455 | 41.98038039 | 0.638054288 | 40.11447452 | NA | NA | NA | NA | NA | NA | NA |
| SCN9A | 1595 | L | 8.391875 | 40.37743459 | 0.6115911 | 38.43867467 | NA | NA | NA | NA | NA | NA | NA |
| SCN9A | 1596 | A | 11.5789 | 38.37366816 | 0.578510725 | 36.16042547 | NA | NA | NA | NA | NA | NA | NA |
| SCN9A | 1597 | D | 12.175625 | 41.35810089 | 0.627781015 | 39.1499973 | NA | NA | NA | NA | NA | NA | NA |
| SCN9A | 1598 | L | 12.66775 | 43.26989477 | 0.659343006 | 41.07452587 | NA | NA | NA | NA | NA | NA | NA |
| SCN9A | 1599 | I | 14.4415 | 38.00800481 | 0.572473953 | 35.80517183 | NA | NA | NA | NA | NA | NA | NA |
| SCN9A | 1600 | E | 16.7005 | 39.59554023 | 0.59868273 | 37.41205715 | NA | NA | NA | NA | NA | NA | NA |
| SCN9A | 1601 | T | 17.37835714 | 44.02893199 | 0.671874026 | 41.85325738 | NA | NA | NA | NA | NA | NA | NA |
| SCN9A | 1602 | Y | 17.29275 | 43.73707336 | 0.667055703 | 41.57429243 | NA | NA | NA | NA | NA | NA | NA |
| SCN9A | 1603 | F | 20.65586364 | 40.87420915 | 0.619792399 | 38.71221822 | NA | NA | NA | NA | NA | NA | NA |
| SCN9A | 1604 | V | 20.54264286 | 38.38945534 | 0.578771357 | 36.23823681 | NA | NA | NA | NA | NA | NA | NA |
| SCN9A | 1605 | S | 18.404 | 34.85642748 | 0.520444257 | 32.69444495 | NA | NA | NA | NA | NA | NA | NA |
| SCN9A | 1606 | P | 18.3765 | 32.66854991 | 0.484324373 | 30.49573904 | NA | NA | NA | NA | NA | NA | NA |
| SCN9A | 1607 | T | 15.63864286 | 29.25682474 | 0.42799987 | 27.07968863 | NA | NA | NA | NA | NA | NA | NA |
| SCN9A | 1608 | L | 12.564 | 32.65200796 | 0.48405128 | 30.46632623 | NA | NA | NA | NA | NA | NA | NA |
| SCN9A | 1609 | F | 13.27504545 | 32.98041936 | 0.489473056 | 30.77824635 | NA | NA | NA | NA | NA | NA | NA |
| SCN9A | 1610 | R | 13.50313636 | 26.08723926 | 0.375672875 | 23.91918314 | NA | NA | NA | NA | NA | NA | NA |

| | | | | | | | | | | | | | |
|---|---|---|---|---|---|---|---|---|---|---|---|---|---|
| SCN9A | 1611 | V | 9.128357143 | 29.24191121 | 0.427753661 | 27.02488731 | NA | NA | NA | NA | NA | NA | NA |
| SCN9A | 1612 | I | 8.172125 | 32.56222801 | 0.482569094 | 30.59677732 | NA | NA | NA | NA | NA | NA | NA |
| SCN9A | 1613 | R | 9.445772727 | 32.01841915 | 0.473591301 | 30.16366021 | NA | NA | NA | NA | NA | NA | NA |
| SCN9A | 1614 | L | 5.79125 | 27.02216411 | 0.391107641 | 25.16528267 | NA | NA | NA | NA | NA | NA | NA |
| SCN9A | 1615 | A | 2.7317 | 32.11199868 | 0.475136215 | 29.60506748 | NA | NA | NA | NA | NA | NA | NA |
| SCN9A | 1616 | R | 4.825136364 | 31.6677872 | 0.467802684 | 29.44514513 | NA | NA | NA | NA | NA | NA | NA |
| SCN9A | 1617 | I | 1.815875 | 26.5454679 | 0.383237817 | 24.05144759 | NA | NA | NA | NA | NA | NA | NA |
| SCN9A | 1618 | G | -1.16075 | 30.88023973 | 0.454800986 | 28.4138033 | NA | NA | NA | NA | NA | NA | NA |
| SCN9A | 1619 | R | -1.504590909 | 32.2073454 | 0.476710303 | 29.78268549 | NA | NA | NA | NA | NA | NA | NA |
| SCN9A | 1620 | I | -1.348625 | 27.83961113 | 0.404602953 | 25.41816959 | NA | NA | NA | NA | NA | NA | NA |
| SCN9A | 1621 | L | -3.803125 | 26.93249459 | 0.389627278 | 24.90420931 | NA | NA | NA | NA | NA | NA | NA |
| SCN9A | 1622 | R | -6.7135 | 30.44719513 | 0.447651811 | 27.85663183 | NA | NA | NA | NA | NA | NA | NA |
| SCN9A | 1623 | L | -6.364375 | 28.42891894 | 0.414331893 | 25.85112413 | NA | NA | NA | NA | NA | NA | NA |
| SCN9A | 1624 | V | -9.2325 | 24.60635954 | 0.351224888 | 22.25723119 | NA | NA | NA | NA | NA | NA | NA |
| SCN9A | 1625 | K | -12.58227778 | 27.8135566 | 0.404172816 | 25.08670148 | NA | NA | NA | NA | NA | NA | NA |
| SCN9A | 1626 | G | -12.5105 | 28.14569072 | 0.409656051 | 25.45447894 | NA | NA | NA | NA | NA | NA | NA |
| SCN9A | 1627 | A | -12.3481 | 24.13054838 | 0.343369675 | 21.43014847 | NA | NA | NA | NA | NA | NA | NA |
| SCN9A | 1628 | K | -16.93583333 | 21.32567274 | 0.297063711 | 18.61538826 | NA | NA | NA | NA | NA | NA | NA |
| SCN9A | 1629 | G | -13.93425 | 19.33577025 | 0.264212217 | 16.61821511 | NA | NA | NA | NA | NA | NA | NA |
| SCN9A | 1630 | I | -10.36175 | 19.80537442 | 0.271964958 | 17.0475716 | NA | NA | NA | NA | NA | NA | NA |
| SCN9A | 1631 | R | -14.53568182 | 19.01406022 | 0.258901075 | 16.33441779 | NA | NA | NA | NA | NA | NA | NA |
| SCN9A | 1632 | T | -15.79507143 | 15.72358703 | 0.204578333 | 13.01648471 | NA | NA | NA | NA | NA | NA | NA |
| SCN9A | 1633 | L | -12.034 | 14.77212679 | 0.188870583 | 12.02674357 | NA | NA | NA | NA | NA | NA | NA |
| SCN9A | 1634 | L | -10.341 | 17.56938272 | 0.235050753 | 14.97163741 | NA | NA | NA | NA | NA | NA | NA |
| SCN9A | 1635 | F | -16.16177273 | 15.67028261 | 0.203698325 | 13.01518993 | NA | NA | NA | NA | NA | NA | NA |
| SCN9A | 1636 | A | -13.6983 | 12.76359268 | 0.155711498 | 10.01528613 | NA | NA | NA | NA | NA | NA | NA |
| SCN9A | 1637 | L | -9.686125 | 14.50103265 | 0.184395064 | 11.8912664 | NA | NA | NA | NA | NA | NA | NA |
| SCN9A | 1638 | M | -12.931125 | 16.08240794 | 0.210502142 | 13.47999711 | NA | NA | NA | NA | NA | NA | NA |
| SCN9A | 1639 | M | -15.6365 | 13.38991464 | 0.166051508 | 10.99406337 | NA | NA | NA | NA | NA | NA | NA |
| SCN9A | 1640 | S | -11.89066667 | 11.36232327 | 0.132577805 | 8.753351266 | NA | NA | NA | NA | NA | NA | NA |
| SCN9A | 1641 | L | -9.639875 | 14.68501338 | 0.187432419 | 12.07741825 | NA | NA | NA | NA | NA | NA | NA |
| SCN9A | 1642 | P | -12.53478571 | 15.22233392 | 0.196303096 | 12.61479123 | NA | NA | NA | NA | NA | NA | NA |
| SCN9A | 1643 | A | -11.4317 | 11.59574931 | 0.136431458 | 8.986712926 | NA | NA | NA | NA | NA | NA | NA |

| | | | | | | | | | | | | | |
|---|---|---|---|---|---|---|---|---|---|---|---|---|---|
| SCN9A | 1644 | L | -8.024 | 11.92453077 | 0.141859343 | 9.322968519 | NA | NA | NA | NA | NA | NA | NA |
| SCN9A | 1645 | F | -7.150772727 | 16.26447998 | 0.213507987 | 13.64961638 | NA | NA | NA | NA | NA | NA | NA |
| SCN9A | 1646 | N | -9.34525 | 15.94908728 | 0.208301138 | 13.33836679 | NA | NA | NA | NA | NA | NA | NA |
| SCN9A | 1647 | I | -6.57225 | 11.83734366 | 0.140419963 | 9.684821143 | NA | NA | NA | NA | NA | NA | NA |
| SCN9A | 1648 | G | -3.403 | 16.82367169 | 0.222739737 | 14.65646585 | NA | NA | NA | NA | NA | NA | NA |
| SCN9A | 1649 | L | -3.913875 | 19.54216984 | 0.267619688 | 17.5777551 | NA | NA | NA | NA | NA | NA | NA |
| SCN9A | 1650 | L | -3.850875 | 18.3188058 | 0.247423052 | 16.15761209 | NA | NA | NA | NA | NA | NA | NA |
| SCN9A | 1651 | L | -0.48825 | 15.68146211 | 0.203882888 | 13.87941182 | NA | NA | NA | NA | NA | NA | NA |
| SCN9A | 1652 | F | 1.112954545 | 20.21471073 | 0.278722731 | 15.71983805 | NA | NA | NA | NA | NA | NA | NA |
| SCN9A | 1653 | L | 0.58275 | 22.6228919 | 0.318479628 | 20.67331915 | NA | NA | NA | NA | NA | NA | NA |
| SCN9A | 1654 | V | 1.043928571 | 19.92334801 | 0.273912595 | 15.15747264 | NA | NA | NA | NA | NA | NA | NA |
| SCN9A | 1655 | M | 4.05075 | 17.74392364 | 0.237932266 | 13.65815604 | NA | NA | NA | NA | NA | NA | NA |
| SCN9A | 1656 | F | 4.870409091 | 22.78194597 | 0.321105467 | 20.6568381 | NA | NA | NA | NA | NA | NA | NA |
| SCN9A | 1657 | I | 3.946625 | 24.15879273 | 0.343835964 | 22.08994222 | NA | NA | NA | NA | NA | NA | NA |
| SCN9A | 1658 | Y | 5.828 | 20.30501679 | 0.280213602 | 18.25455435 | NA | NA | NA | NA | NA | NA | NA |
| SCN9A | 1659 | A | 9.4767 | 21.84343283 | 0.305611462 | 19.78844444 | NA | NA | NA | NA | NA | NA | NA |
| SCN9A | 1660 | I | 9.551875 | 24.18191004 | 0.34421761 | 22.14241453 | NA | NA | NA | NA | NA | NA | NA |
| SCN9A | 1661 | F | 8.976590909 | 23.85800926 | 0.338870301 | 20.75712928 | NA | NA | NA | NA | NA | NA | NA |
| SCN9A | 1662 | G | 13.22275 | 21.85438439 | 0.305792263 | 18.75347268 | NA | NA | NA | NA | NA | NA | NA |
| SCN9A | 1663 | M | 14.611625 | 23.35169863 | 0.330511569 | 20.15505267 | NA | NA | NA | NA | NA | NA | NA |
| SCN9A | 1664 | S | 15.33933333 | 25.47369565 | 0.365543824 | 22.30151398 | NA | NA | NA | NA | NA | NA | NA |
| SCN9A | 1665 | N | 15.17075 | 22.70578488 | 0.319848116 | 19.76558044 | NA | NA | NA | NA | NA | NA | NA |
| SCN9A | 1666 | F | 16.13359091 | 19.32257286 | 0.26399434 | 16.14066642 | NA | NA | NA | NA | NA | NA | NA |
| SCN9A | 1667 | A | 20.1019 | 21.08883783 | 0.29315378 | 16.43382317 | NA | NA | NA | NA | NA | NA | NA |
| SCN9A | 1668 | Y | 23.4305 | 19.13098944 | 0.260831471 | 14.46231658 | NA | NA | NA | NA | NA | NA | NA |
| SCN9A | 1669 | V | 22.32578571 | 16.19702797 | 0.212394415 | 11.53271276 | NA | NA | NA | NA | NA | NA | NA |
| SCN9A | 1670 | K | 26.51872222 | 11.63545831 | 0.137087018 | 6.929927916 | NA | NA | NA | NA | NA | NA | NA |
| SCN9A | 1671 | K | 25.31816667 | 10.39219775 | 0.116561909 | 5.706621354 | NA | NA | NA | NA | NA | NA | NA |
| SCN9A | 1672 | E | 24.89372222 | 7.06052668 | 0.061559027 | 2.397789846 | 4.79 | -0.88 | -1.43 | 18.82 | 1.42 | 1 | -1 |
| SCN9A | 1673 | D | 21.46925 | 6.355697365 | 0.049922931 | 1.722456184 | 4.79 | -0.88 | -1.43 | 18.82 | 1.42 | 1 | -1 |
| SCN9A | 1674 | G | 19.25 | 8.747872256 | 0.089415579 | 5.352441364 | 3.67 | 0.19 | -0.95 | 25.34 | 0.39 | 2 | -2 |
| SCN9A | 1675 | I | 20.778375 | 10.87565626 | 0.124543372 | 6.213680224 | NA | NA | NA | NA | NA | NA | NA |
| SCN9A | 1676 | N | 22.512 | 10.63709989 | 0.120605022 | 5.954755393 | NA | NA | NA | NA | NA | NA | NA |

| | | | | | | | | | | | | | |
|---|---|---|---|---|---|---|---|---|---|---|---|---|---|
| SCN9A | 1677 | D | 24.281625 | 14.35082959 | 0.181915347 | 9.639242538 | NA | NA | NA | NA | NA | NA | NA |
| SCN9A | 1678 | M | 20.470375 | 15.46123633 | 0.20024716 | 10.7888403 | NA | NA | NA | NA | NA | NA | NA |
| SCN9A | 1679 | F | 17.59486364 | 14.08655739 | 0.177552451 | 10.92289863 | NA | NA | NA | NA | NA | NA | NA |
| SCN9A | 1680 | N | 19.1285 | 16.5988046 | 0.219027385 | 13.45857779 | NA | NA | NA | NA | NA | NA | NA |
| SCN9A | 1681 | F | 15.77104545 | 16.5573404 | 0.218342848 | 13.36406221 | NA | NA | NA | NA | NA | NA | NA |
| SCN9A | 1682 | E | 20.98694444 | 19.20591771 | 0.262068469 | 14.52940498 | NA | NA | NA | NA | NA | NA | NA |
| SCN9A | 1683 | T | 17.07607143 | 22.86914493 | 0.322545043 | 19.72714279 | NA | NA | NA | NA | NA | NA | NA |
| SCN9A | 1684 | F | 11.18822727 | 22.67162868 | 0.319284228 | 20.60770456 | NA | NA | NA | NA | NA | NA | NA |
| SCN9A | 1685 | G | 13.39525 | 20.73280138 | 0.28727594 | 18.65484337 | NA | NA | NA | NA | NA | NA | NA |
| SCN9A | 1686 | N | 16.141125 | 18.20938275 | 0.245616576 | 16.13760559 | NA | NA | NA | NA | NA | NA | NA |
| SCN9A | 1687 | S | 12.6705 | 17.92565963 | 0.240932563 | 15.94695478 | NA | NA | NA | NA | NA | NA | NA |
| SCN9A | 1688 | M | 9.046125 | 17.25095563 | 0.229793809 | 15.136031 | NA | NA | NA | NA | NA | NA | NA |
| SCN9A | 1689 | I | 12.261625 | 14.38241085 | 0.182436725 | 12.27836572 | NA | NA | NA | NA | NA | NA | NA |
| SCN9A | 1690 | C | 12.1615 | 12.93143101 | 0.158482357 | 10.86580531 | NA | NA | NA | NA | NA | NA | NA |
| SCN9A | 1691 | L | 8.341875 | 14.04570999 | 0.176878097 | 11.91793299 | NA | NA | NA | NA | NA | NA | NA |
| SCN9A | 1692 | F | 6.507590909 | 12.39228635 | 0.149581566 | 10.21492614 | NA | NA | NA | NA | NA | NA | NA |
| SCN9A | 1693 | Q | 10.62527778 | 8.675744118 | 0.088224809 | 6.66775223 | 2.24 | -0.87 | -2.83 | 38.17 | 1.68 | 3 | -1 |
| SCN9A | 1694 | I | 8.493875 | 10.44220326 | 0.117387454 | 8.342184199 | NA | NA | NA | NA | NA | NA | NA |
| SCN9A | 1695 | T | 3.9405 | 10.20344324 | 0.113445742 | 6.751255611 | NA | NA | NA | NA | NA | NA | NA |
| SCN9A | 1696 | T | 4.706928571 | 6.775277559 | 0.056849821 | 3.194246086 | 3.6 | -0.78 | -2.05 | 14.95 | 0.8 | 1 | 1 |
| SCN9A | 1697 | S | 6.425666667 | 6.375701687 | 0.050253184 | 4.416999306 | 1.98 | -0.78 | -2.6 | 34.26 | 1.5 | 2 | 0 |
| SCN9A | 1698 | A | 9.5005 | 5.209712564 | 0.031003757 | 3.031255492 | 2.24 | -0.87 | -2.83 | 38.17 | 1.68 | 3 | -1 |
| SCN9A | 1699 | G | 12.5535 | 5.252465612 | 0.031709571 | 2.503015133 | 3.11 | -0.47 | -2.18 | 26.15 | 1.06 | 2 | -2 |
| SCN9A | 1700 | W | 10.1835 | 7.456551884 | 0.068097045 | 4.432214104 | 3.05 | -0.47 | -1.2 | 12.29 | 0.72 | 1 | -1 |
| SCN9A | 1701 | D | 14.315 | 4.623649749 | 0.021328389 | 1.624425535 | 3.05 | -0.47 | -1.2 | 12.29 | 0.72 | 1 | -1 |
| SCN9A | 1702 | G | 16.71 | 7.081269025 | 0.061901464 | 3.871159963 | 3.24 | -0.36 | -2.77 | 33.74 | 1.14 | 2 | -2 |
| SCN9A | 1703 | L | 15.481625 | 9.377356024 | 0.099807788 | 6.212696695 | 3.24 | -0.36 | -2.77 | 33.74 | 1.14 | 2 | -2 |
| SCN9A | 1704 | L | 15.119625 | 11.40133369 | 0.133221832 | 8.241696278 | NA | NA | NA | NA | NA | NA | NA |
| SCN9A | 1705 | A | 19.0771 | 10.07469558 | 0.111320234 | 6.499276012 | NA | NA | NA | NA | NA | NA | NA |
| SCN9A | 1706 | P | 20.57164286 | 11.70176367 | 0.13818166 | 7.064836253 | NA | NA | NA | NA | NA | NA | NA |
| SCN9A | 1707 | I | 18.0515 | 14.77529837 | 0.188922943 | 10.13156457 | NA | NA | NA | NA | NA | NA | NA |
| SCN9A | 1708 | L | 18.57025 | 13.99255195 | 0.176000506 | 10.66353612 | NA | NA | NA | NA | NA | NA | NA |
| SCN9A | 1709 | N | 23.427875 | 12.81054351 | 0.156486614 | 8.134023361 | NA | NA | NA | NA | NA | NA | NA |

| | | | | | | | | | | | | | |
|---|---|---|---|---|---|---|---|---|---|---|---|---|---|
| SCN9A | 1710 | S | 23.79116667 | 17.77082004 | 0.238376302 | 13.08957225 | NA | NA | NA | NA | NA | NA | NA |
| SCN9A | 1711 | K | 26.69305556 | 18.65628867 | 0.252994589 | 13.94608155 | NA | NA | NA | NA | NA | NA | NA |
| SCN9A | 1712 | P | 30.86035714 | 19.42013741 | 0.265605042 | 14.68925747 | NA | NA | NA | NA | NA | NA | NA |
| SCN9A | 1713 | P | 31.54907143 | 15.65854265 | 0.203504509 | 10.92216924 | NA | NA | NA | NA | NA | NA | NA |
| SCN9A | 1714 | D | 27.613875 | 13.60298004 | 0.169569026 | 8.885906032 | NA | NA | NA | NA | NA | NA | NA |
| SCN9A | 1715 | C | 28.553 | 16.47595254 | 0.216999208 | 11.76759285 | NA | NA | NA | NA | NA | NA | NA |
| SCN9A | 1716 | D | 31.306625 | 18.76855727 | 0.254848043 | 14.06112567 | NA | NA | NA | NA | NA | NA | NA |
| SCN9A | 1717 | P | 29.3345 | 22.04411318 | 0.308924514 | 17.34197614 | NA | NA | NA | NA | NA | NA | NA |
| SCN9A | 1718 | K | 32.36972222 | 24.24200283 | 0.345209688 | 19.54139391 | NA | NA | NA | NA | NA | NA | NA |
| SCN9A | 1719 | K | 31.40738889 | 20.00182364 | 0.275208157 | 15.29543311 | NA | NA | NA | NA | NA | NA | NA |
| SCN9A | 1720 | V | 33.21935714 | 24.85015262 | 0.355249692 | 20.14115651 | NA | NA | NA | NA | NA | NA | NA |
| SCN9A | 1721 | H | 29.0726 | 21.29247334 | 0.296515618 | 16.60750945 | NA | NA | NA | NA | NA | NA | NA |
| SCN9A | 1722 | P | 32.07878571 | 25.0857681 | 0.359139491 | 20.37289715 | NA | NA | NA | NA | NA | NA | NA |
| SCN9A | 1723 | G | 29.51925 | 28.94006346 | 0.422770428 | 24.23851704 | NA | NA | NA | NA | NA | NA | NA |
| SCN9A | 1724 | S | 27.64966667 | 28.14738746 | 0.409684062 | 23.46610489 | NA | NA | NA | NA | NA | NA | NA |
| SCN9A | 1725 | S | 26.7975 | 30.0050722 | 0.44035276 | 25.32539677 | NA | NA | NA | NA | NA | NA | NA |
| SCN9A | 1726 | V | 25.71235714 | 26.04466611 | 0.374970031 | 21.37260008 | NA | NA | NA | NA | NA | NA | NA |
| SCN9A | 1727 | E | 29.03416667 | 25.27635128 | 0.362285847 | 20.59390138 | NA | NA | NA | NA | NA | NA | NA |
| SCN9A | 1728 | G | 27.07575 | 22.03247787 | 0.308732425 | 17.35267681 | NA | NA | NA | NA | NA | NA | NA |
| SCN9A | 1729 | D | 26.74075 | 17.82106041 | 0.239205725 | 13.12814277 | NA | NA | NA | NA | NA | NA | NA |
| SCN9A | 1730 | C | 23.66083333 | 19.63684267 | 0.269182651 | 14.96819564 | NA | NA | NA | NA | NA | NA | NA |
| SCN9A | 1731 | G | 20.461 | 18.90082019 | 0.257031584 | 14.24199397 | NA | NA | NA | NA | NA | NA | NA |
| SCN9A | 1732 | N | 16.92575 | 20.58947556 | 0.28490976 | 16.37633539 | NA | NA | NA | NA | NA | NA | NA |
| SCN9A | 1733 | P | 17.01735714 | 20.00322337 | 0.275231265 | 16.43077168 | NA | NA | NA | NA | NA | NA | NA |
| SCN9A | 1734 | S | 13.1365 | 22.10185696 | 0.309877811 | 18.5281626 | NA | NA | NA | NA | NA | NA | NA |
| SCN9A | 1735 | V | 12.83564286 | 21.25936492 | 0.295969028 | 18.19545408 | NA | NA | NA | NA | NA | NA | NA |
| SCN9A | 1736 | G | 12.83325 | 17.81315073 | 0.239075143 | 14.78765549 | NA | NA | NA | NA | NA | NA | NA |
| SCN9A | 1737 | I | 11.436875 | 18.0117161 | 0.242353278 | 14.98733031 | NA | NA | NA | NA | NA | NA | NA |
| SCN9A | 1738 | F | 6.771772727 | 19.97693733 | 0.274797306 | 16.9398474 | NA | NA | NA | NA | NA | NA | NA |
| SCN9A | 1739 | Y | 9.440333333 | 18.10583928 | 0.243907167 | 15.06474058 | NA | NA | NA | NA | NA | NA | NA |
| SCN9A | 1740 | F | 10.6945 | 13.93646458 | 0.175074554 | 10.90143189 | NA | NA | NA | NA | NA | NA | NA |
| SCN9A | 1741 | V | 6.2725 | 16.28978447 | 0.213925741 | 13.25149655 | NA | NA | NA | NA | NA | NA | NA |
| SCN9A | 1742 | S | 3.808833333 | 17.29263907 | 0.230481965 | 12.47987397 | NA | NA | NA | NA | NA | NA | NA |

| | | | | | | | | | | | | | |
|---|---|---|---|---|---|---|---|---|---|---|---|---|---|
| SCN9A | 1743 | Y | 4.580833333 | 14.79762988 | 0.189291616 | 10.76766562 | NA | NA | NA | NA | NA | NA | NA |
| SCN9A | 1744 | I | 4.341875 | 11.7276493 | 0.138609008 | 7.659811124 | NA | NA | NA | NA | NA | NA | NA |
| SCN9A | 1745 | I | 0.79525 | 13.17978452 | 0.16258245 | 9.173782555 | NA | NA | NA | NA | NA | NA | NA |
| SCN9A | 1746 | I | -0.141 | 13.97325037 | 0.175681855 | 11.91368426 | NA | NA | NA | NA | NA | NA | NA |
| SCN9A | 1747 | S | 0.6395 | 11.44406388 | 0.133927269 | 7.171389588 | NA | NA | NA | NA | NA | NA | NA |
| SCN9A | 1748 | F | -1.441772727 | 6.708089818 | 0.055740612 | 4.523319191 | 2.48 | 1.3 | 2.7 | 0.5 | -1.72 | 0 | 0 |
| SCN9A | 1749 | L | -4.222125 | 10.46367512 | 0.117741935 | 8.487424797 | NA | NA | NA | NA | NA | NA | NA |
| SCN9A | 1750 | V | -4.017785714 | 10.44166663 | 0.117378595 | 8.598398584 | NA | NA | NA | NA | NA | NA | NA |
| SCN9A | 1751 | V | -3.093071429 | 7.343565006 | 0.066231734 | 5.248733907 | 2.09 | 1.35 | 2.33 | 0.62 | -1.69 | 0 | 0 |
| SCN9A | 1752 | V | -6.497642857 | 5.888568756 | 0.04221106 | 4.827810181 | 2.09 | 1.35 | 2.33 | 0.62 | -1.69 | 0 | 0 |
| SCN9A | 1753 | N | -8.462125 | 7.914528603 | 0.075657828 | 5.92871857 | 2.17 | 1.64 | 3.05 | 0.5 | -1.75 | 0 | 0 |
| SCN9A | 1754 | M | -6.19125 | 7.444046816 | 0.067890598 | 5.277413368 | 2.17 | 1.64 | 3.05 | 0.5 | -1.75 | 0 | 0 |
| SCN9A | 1755 | Y | -6.375916667 | 3.678490723 | 0.005724666 | 2.076941754 | 2.17 | 1.64 | 3.05 | 0.5 | -1.75 | 0 | 0 |
| SCN9A | 1756 | I | -11.93425 | 3.509163576 | 0.002929228 | 1.43527137 | 1.99 | 1.81 | 4.5 | 0.13 | -1.85 | 0 | 0 |
| SCN9A | 1757 | A | -12.2269 | 6.298101381 | 0.048972073 | 3.753635581 | 2.63 | -0.14 | 0.05 | 2.57 | 0.08 | 0 | 0 |
| SCN9A | 1758 | V | -10.91307143 | 6.964796479 | 0.059978608 | 5.066737678 | 1.99 | 1.81 | 4.5 | 0.13 | -1.85 | 0 | 0 |
| SCN9A | 1759 | I | -13.398625 | 3.581851197 | 0.004129235 | 1.625464998 | 2.49 | 0.94 | 2.87 | 1.21 | -0.96 | 0 | 0 |
| SCN9A | 1760 | L | -17.330875 | 4.177398832 | 0.013961189 | 1.653085391 | 2.63 | 0.73 | 1.6 | 1.21 | -1.03 | 0 | 0 |
| SCN9A | 1761 | E | -16.17105556 | 6.654483752 | 0.054855625 | 4.104229523 | 2.77 | 0.57 | 1.03 | 1.29 | -1.02 | 0 | 0 |
| SCN9A | 1762 | N | -15.343625 | 7.930478989 | 0.075921154 | 5.181573618 | 2.77 | 0.4 | 0.85 | 1.81 | -0.58 | 0 | 0 |
| SCN9A | 1763 | F | -18.22631818 | 4.372220717 | 0.017177522 | 1.687227102 | 2.77 | 0.57 | 1.03 | 1.29 | -1.02 | 0 | 0 |
| SCN9A | 1764 | S | -20.86783333 | 6.961869505 | 0.059930286 | 3.679382086 | 3.35 | 0.33 | -0.2 | 13.34 | -0.4 | 1 | 1 |
| SCN9A | 1765 | V | -19.04835714 | 9.839955589 | 0.107444889 | 6.544178522 | NA | NA | NA | NA | NA | NA | NA |
| SCN9A | 1766 | A | -19.9691 | 11.13055731 | 0.128751558 | 7.853253065 | NA | NA | NA | NA | NA | NA | NA |
| SCN9A | 1767 | T | -23.45878571 | 7.894055105 | 0.075319829 | 4.363310731 | 3.8 | 0.06 | -1.53 | 17.74 | 0.06 | 1 | 1 |
| SCN9A | 1768 | E | -24.98883333 | 11.25083237 | 0.130737191 | 5.233561915 | NA | NA | NA | NA | NA | NA | NA |
| TRPA1 | 446 | K | -95.7302 | 22.27680581 | 0.397982259 | 19.90012454 | NA | NA | NA | NA | NA | NA | NA |
| TRPA1 | 447 | K | -92.1904 | 22.57851798 | 0.403935858 | 20.18762052 | NA | NA | NA | NA | NA | NA | NA |
| TRPA1 | 448 | S | -90.85666667 | 19.32834949 | 0.33980122 | 16.22900336 | NA | NA | NA | NA | NA | NA | NA |
| TRPA1 | 449 | P | -92.98214286 | 17.85191359 | 0.310667136 | 15.46013315 | NA | NA | NA | NA | NA | NA | NA |
| TRPA1 | 450 | L | -91.214375 | 19.95462801 | 0.352159394 | 17.56668319 | NA | NA | NA | NA | NA | NA | NA |
| TRPA1 | 451 | H | -87.6696 | 18.83309069 | 0.330028421 | 16.21568632 | NA | NA | NA | NA | NA | NA | NA |
| TRPA1 | 452 | F | -87.904 | 12.09434087 | 0.197054611 | 9.445233418 | NA | NA | NA | NA | NA | NA | NA |

| | | | | | | | | | | | | | |
|---|---|---|---|---|---|---|---|---|---|---|---|---|---|
| TRPA1 | 453 | A | -89.0656 | 16.37696327 | 0.281562366 | 13.74207402 | NA | NA | NA | NA | NA | NA | NA |
| TRPA1 | 454 | A | -86.3568 | 18.22190797 | 0.317968128 | 15.54816374 | NA | NA | NA | NA | NA | NA | NA |
| TRPA1 | 455 | S | -83.473 | 15.64664351 | 0.267151176 | 13.76877039 | NA | NA | NA | NA | NA | NA | NA |
| TRPA1 | 456 | Y | -83.63 | 12.21130746 | 0.19936268 | 10.18708123 | NA | NA | NA | NA | NA | NA | NA |
| TRPA1 | 457 | G | -86.143 | 15.33407144 | 0.260983282 | 12.84985762 | NA | NA | NA | NA | NA | NA | NA |
| TRPA1 | 458 | R | -89.65681818 | 8.575367222 | 0.127615718 | 5.987299912 | NA | NA | NA | NA | NA | NA | NA |
| TRPA1 | 459 | I | -90.710125 | 16.02982395 | 0.274712366 | 15.0401478 | NA | NA | NA | NA | NA | NA | NA |
| TRPA1 | 460 | N | -94.412125 | 13.57781223 | 0.226327524 | 12.68035779 | NA | NA | NA | NA | NA | NA | NA |
| TRPA1 | 461 | T | -94.25514286 | 13.2999715 | 0.220844973 | 12.32416312 | NA | NA | NA | NA | NA | NA | NA |
| TRPA1 | 462 | C | -92.80183333 | 17.56377297 | 0.30498134 | 16.57401311 | NA | NA | NA | NA | NA | NA | NA |
| TRPA1 | 463 | Q | -95.55877778 | 19.20125165 | 0.337293235 | 18.24324213 | NA | NA | NA | NA | NA | NA | NA |
| TRPA1 | 464 | R | -98.69454545 | 13.445072 | 0.2237082 | 12.57965079 | NA | NA | NA | NA | NA | NA | NA |
| TRPA1 | 465 | L | -96.90425 | 18.3113876 | 0.319733804 | 17.34720196 | NA | NA | NA | NA | NA | NA | NA |
| TRPA1 | 466 | L | -95.97825 | 22.3625387 | 0.399674002 | 21.3915012 | NA | NA | NA | NA | NA | NA | NA |
| TRPA1 | 467 | Q | -101.8025556 | 22.00341983 | 0.392587612 | 21.10449738 | NA | NA | NA | NA | NA | NA | NA |
| TRPA1 | 468 | D | -101.0836 | 25.81539335 | 0.467808188 | 24.87429727 | NA | NA | NA | NA | NA | NA | NA |
| TRPA1 | 469 | I | -99.7816 | 28.88566186 | 0.528392912 | 27.93917658 | NA | NA | NA | NA | NA | NA | NA |
| TRPA1 | 470 | S | -100.8386 | 31.15977429 | 0.573267318 | 30.20066075 | NA | NA | NA | NA | NA | NA | NA |
| TRPA1 | 471 | D | -98.4452 | 30.5070238 | 0.56038678 | 29.52808647 | NA | NA | NA | NA | NA | NA | NA |
| TRPA1 | 472 | T | -94.73685714 | 32.11525402 | 0.592121524 | 30.83762991 | NA | NA | NA | NA | NA | NA | NA |
| TRPA1 | 473 | R | -93.848 | 31.34959387 | 0.577012973 | 30.13806223 | NA | NA | NA | NA | NA | NA | NA |
| TRPA1 | 474 | L | -93.961 | 25.63880881 | 0.464323696 | 23.30075519 | NA | NA | NA | NA | NA | NA | NA |
| TRPA1 | 475 | L | -91.294125 | 27.72321428 | 0.505454669 | 26.43908818 | NA | NA | NA | NA | NA | NA | NA |
| TRPA1 | 476 | N | -88.94 | 30.41817355 | 0.558633524 | 27.86262659 | NA | NA | NA | NA | NA | NA | NA |
| TRPA1 | 477 | E | -88.611 | 29.18542001 | 0.534307953 | 26.60787579 | NA | NA | NA | NA | NA | NA | NA |
| TRPA1 | 478 | G | -86.627 | 28.31985514 | 0.517228011 | 25.83243494 | NA | NA | NA | NA | NA | NA | NA |
| TRPA1 | 479 | D | -84.00175 | 24.41807965 | 0.440235399 | 22.3109629 | NA | NA | NA | NA | NA | NA | NA |
| TRPA1 | 480 | L | -82.290375 | 27.21309631 | 0.495388658 | 25.3347539 | NA | NA | NA | NA | NA | NA | NA |
| TRPA1 | 481 | H | -78.7455 | 25.34698069 | 0.458565136 | 23.05826321 | NA | NA | NA | NA | NA | NA | NA |
| TRPA1 | 482 | G | -81.21925 | 29.04239169 | 0.531485617 | 27.04938478 | NA | NA | NA | NA | NA | NA | NA |
| TRPA1 | 483 | M | -80.69475 | 24.33647892 | 0.438625195 | 22.48476403 | NA | NA | NA | NA | NA | NA | NA |
| TRPA1 | 484 | T | -83.898 | 25.20856025 | 0.455833725 | 23.12426835 | NA | NA | NA | NA | NA | NA | NA |
| TRPA1 | 485 | P | -86.095 | 23.48764043 | 0.42187531 | 21.06724052 | NA | NA | NA | NA | NA | NA | NA |

| | | | | | | | | | | | | | |
|---|---|---|---|---|---|---|---|---|---|---|---|---|---|
| TRPA1 | 486 | L | -83.148 | 24.81748053 | 0.448116662 | 22.95078656 | NA | NA | NA | NA | NA | NA | NA |
| TRPA1 | 487 | H | -80.1857 | 23.04249056 | 0.413091296 | 20.85867176 | NA | NA | NA | NA | NA | NA | NA |
| TRPA1 | 488 | L | -81.924125 | 19.89861435 | 0.351054092 | 17.8998879 | NA | NA | NA | NA | NA | NA | NA |
| TRPA1 | 489 | A | -81.97 | 20.69773169 | 0.366822844 | 18.93655784 | NA | NA | NA | NA | NA | NA | NA |
| TRPA1 | 490 | A | -78.578 | 21.84566527 | 0.389474687 | 19.67653321 | NA | NA | NA | NA | NA | NA | NA |
| TRPA1 | 491 | K | -76.46166667 | 18.92826141 | 0.331906397 | 18.00073518 | NA | NA | NA | NA | NA | NA | NA |
| TRPA1 | 492 | N | -78.246625 | 16.02594662 | 0.274635855 | 13.80936007 | NA | NA | NA | NA | NA | NA | NA |
| TRPA1 | 493 | G | -78.40025 | 18.30043448 | 0.319517669 | 16.07128607 | NA | NA | NA | NA | NA | NA | NA |
| TRPA1 | 494 | H | -81.7358 | 14.90638685 | 0.252543905 | 12.80221253 | NA | NA | NA | NA | NA | NA | NA |
| TRPA1 | 495 | D | -81.88175 | 21.0261487 | 0.373303403 | 19.01658087 | NA | NA | NA | NA | NA | NA | NA |
| TRPA1 | 496 | K | -85.79844444 | 21.12857224 | 0.375324497 | 19.08723432 | NA | NA | NA | NA | NA | NA | NA |
| TRPA1 | 497 | V | -86.37457143 | 19.29074612 | 0.339059203 | 16.79067052 | NA | NA | NA | NA | NA | NA | NA |
| TRPA1 | 498 | V | -83.83942857 | 23.41453715 | 0.420432784 | 21.5162472 | NA | NA | NA | NA | NA | NA | NA |
| TRPA1 | 499 | Q | -86.12377778 | 25.69256262 | 0.465384404 | 23.23368911 | NA | NA | NA | NA | NA | NA | NA |
| TRPA1 | 500 | L | -90.291125 | 23.47667796 | 0.421658991 | 22.51657938 | NA | NA | NA | NA | NA | NA | NA |
| TRPA1 | 501 | L | -88.299625 | 24.11038718 | 0.434163792 | 23.28361734 | NA | NA | NA | NA | NA | NA | NA |
| TRPA1 | 502 | L | -86.185125 | 28.78761987 | 0.526458278 | 26.13859383 | NA | NA | NA | NA | NA | NA | NA |
| TRPA1 | 503 | K | -91.36933333 | 28.89082323 | 0.52849476 | 27.98660474 | NA | NA | NA | NA | NA | NA | NA |
| TRPA1 | 504 | K | -93.58177778 | 26.37686822 | 0.47888761 | 25.43947392 | NA | NA | NA | NA | NA | NA | NA |
| TRPA1 | 505 | G | -90.4065 | 31.25075265 | 0.575062567 | 30.26917005 | NA | NA | NA | NA | NA | NA | NA |
| TRPA1 | 506 | A | -87.3334 | 30.64863457 | 0.563181145 | 27.9997689 | NA | NA | NA | NA | NA | NA | NA |
| TRPA1 | 507 | L | -85.856125 | 32.61450648 | 0.601973129 | 30.52891139 | NA | NA | NA | NA | NA | NA | NA |
| TRPA1 | 508 | F | -81.10745455 | 32.12119092 | 0.592238675 | 29.98872047 | NA | NA | NA | NA | NA | NA | NA |
| TRPA1 | 509 | L | -82.150125 | 32.74852586 | 0.604617695 | 30.63576537 | NA | NA | NA | NA | NA | NA | NA |
| TRPA1 | 510 | S | -78.58216667 | 32.70506588 | 0.603760111 | 30.48054493 | NA | NA | NA | NA | NA | NA | NA |
| TRPA1 | 511 | D | -76.7195 | 28.17590421 | 0.514387469 | 26.17698052 | NA | NA | NA | NA | NA | NA | NA |
| TRPA1 | 512 | H | -77.4881 | 31.8191184 | 0.586277965 | 31.08889873 | NA | NA | NA | NA | NA | NA | NA |
| TRPA1 | 513 | N | -72.4155 | 29.5617134 | 0.541733242 | 28.63468688 | NA | NA | NA | NA | NA | NA | NA |
| TRPA1 | 514 | G | -73.428 | 32.65596347 | 0.602791188 | 31.71443956 | NA | NA | NA | NA | NA | NA | NA |
| TRPA1 | 515 | W | -72.24485714 | 25.83097369 | 0.46811563 | 25.40745823 | NA | NA | NA | NA | NA | NA | NA |
| TRPA1 | 516 | T | -75.72614286 | 28.11592981 | 0.513204011 | 27.15774943 | NA | NA | NA | NA | NA | NA | NA |
| TRPA1 | 517 | A | -77.1408 | 26.51348221 | 0.481583374 | 24.478069 | NA | NA | NA | NA | NA | NA | NA |
| TRPA1 | 518 | L | -73.788875 | 27.43660261 | 0.499799043 | 26.55931864 | NA | NA | NA | NA | NA | NA | NA |

| | | | | | | | | | | | | | |
|---|---|---|---|---|---|---|---|---|---|---|---|---|---|
| TRPA1 | 519 | H | -71.6219 | 25.66327047 | 0.464806391 | 24.92077323 | NA | NA | NA | NA | NA | NA | NA |
| TRPA1 | 520 | H | -74.5765 | 22.66467291 | 0.405635929 | 21.75281423 | NA | NA | NA | NA | NA | NA | NA |
| TRPA1 | 521 | A | -73.3542 | 23.1317455 | 0.414852538 | 22.37229302 | NA | NA | NA | NA | NA | NA | NA |
| TRPA1 | 522 | S | -69.84666667 | 23.17649956 | 0.415735657 | 22.06117454 | NA | NA | NA | NA | NA | NA | NA |
| TRPA1 | 523 | M | -68.836125 | 20.26523639 | 0.358288539 | 18.79767613 | NA | NA | NA | NA | NA | NA | NA |
| TRPA1 | 524 | G | -70.3595 | 18.53674011 | 0.32418062 | 17.53227672 | NA | NA | NA | NA | NA | NA | NA |
| TRPA1 | 525 | G | -70.17275 | 20.47859678 | 0.362498718 | 19.19046501 | NA | NA | NA | NA | NA | NA | NA |
| TRPA1 | 526 | Y | -73.66141667 | 15.58839613 | 0.266001797 | 14.92031813 | NA | NA | NA | NA | NA | NA | NA |
| TRPA1 | 527 | T | -72.68885714 | 22.81025109 | 0.408508581 | 22.05009571 | NA | NA | NA | NA | NA | NA | NA |
| TRPA1 | 528 | Q | -76.67955556 | 23.08366888 | 0.413903856 | 22.14597151 | NA | NA | NA | NA | NA | NA | NA |
| TRPA1 | 529 | T | -77.62257143 | 22.45423811 | 0.40148348 | 20.22518593 | NA | NA | NA | NA | NA | NA | NA |
| TRPA1 | 530 | M | -73.73025 | 25.57797287 | 0.463123238 | 24.96315444 | NA | NA | NA | NA | NA | NA | NA |
| TRPA1 | 531 | K | -76.26877778 | 28.14700986 | 0.513817305 | 27.18845623 | NA | NA | NA | NA | NA | NA | NA |
| TRPA1 | 532 | V | -80.03857143 | 27.47015621 | 0.500461147 | 25.28356327 | NA | NA | NA | NA | NA | NA | NA |
| TRPA1 | 533 | I | -78.384625 | 27.92080309 | 0.509353632 | 25.69184101 | NA | NA | NA | NA | NA | NA | NA |
| TRPA1 | 534 | L | -75.348625 | 31.83555399 | 0.586602284 | 30.88868098 | NA | NA | NA | NA | NA | NA | NA |
| TRPA1 | 535 | D | -78.866375 | 32.57363544 | 0.601166633 | 30.34485544 | NA | NA | NA | NA | NA | NA | NA |
| TRPA1 | 536 | T | -81.56 | 32.7315854 | 0.604283414 | 30.66202391 | NA | NA | NA | NA | NA | NA | NA |
| TRPA1 | 537 | N | -80.183125 | 35.90657094 | 0.666934489 | 33.68158931 | NA | NA | NA | NA | NA | NA | NA |
| TRPA1 | 538 | L | -75.9308 | 38.37873247 | 0.71571694 | 37.52726741 | NA | NA | NA | NA | NA | NA | NA |
| TRPA1 | 539 | K | -76.359 | 38.6814023 | 0.721689437 | 37.72279782 | NA | NA | NA | NA | NA | NA | NA |
| TRPA1 | 540 | C | -75.3355 | 33.84571196 | 0.626268152 | 32.88830902 | NA | NA | NA | NA | NA | NA | NA |
| TRPA1 | 541 | T | -71.49671429 | 35.2758554 | 0.654488761 | 34.24959787 | NA | NA | NA | NA | NA | NA | NA |
| TRPA1 | 542 | D | -70.4448 | 37.39575116 | 0.696320053 | 35.93981738 | NA | NA | NA | NA | NA | NA | NA |
| TRPA1 | 543 | R | -73.15109091 | 35.18256719 | 0.652647931 | 34.24911437 | NA | NA | NA | NA | NA | NA | NA |
| TRPA1 | 544 | L | -68.76375 | 35.86914027 | 0.66619588 | 34.39159445 | NA | NA | NA | NA | NA | NA | NA |
| TRPA1 | 545 | D | -68.69875 | 31.59336915 | 0.58182332 | 30.16119911 | NA | NA | NA | NA | NA | NA | NA |
| TRPA1 | 546 | E | -68.89588889 | 32.84146472 | 0.606451631 | 31.3638612 | NA | NA | NA | NA | NA | NA | NA |
| TRPA1 | 547 | D | -64.331625 | 32.27058695 | 0.595186664 | 30.52256094 | NA | NA | NA | NA | NA | NA | NA |
| TRPA1 | 548 | G | -64.9105 | 35.16218055 | 0.652245648 | 33.33231525 | NA | NA | NA | NA | NA | NA | NA |
| TRPA1 | 549 | N | -64.884875 | 31.18580738 | 0.573781021 | 29.38113265 | NA | NA | NA | NA | NA | NA | NA |
| TRPA1 | 550 | T | -66.317 | 30.35539232 | 0.55739468 | 28.51725904 | NA | NA | NA | NA | NA | NA | NA |
| TRPA1 | 551 | A | -67.9358 | 28.19382959 | 0.514741185 | 26.47666515 | NA | NA | NA | NA | NA | NA | NA |

| | | | | | | | | | | | | | |
|---|---|---|---|---|---|---|---|---|---|---|---|---|---|
| TRPA1 | 552 | L | -64.189625 | 28.60699839 | 0.522894126 | 26.84670082 | NA | NA | NA | NA | NA | NA | NA |
| TRPA1 | 553 | H | -62.6543 | 26.9877751 | 0.490942459 | 25.74672801 | NA | NA | NA | NA | NA | NA | NA |
| TRPA1 | 554 | F | -66.15245455 | 24.07296045 | 0.433425261 | 22.25478785 | NA | NA | NA | NA | NA | NA | NA |
| TRPA1 | 555 | A | -64.6726 | 23.99169752 | 0.431821723 | 22.25800101 | NA | NA | NA | NA | NA | NA | NA |
| TRPA1 | 556 | A | -61.0732 | 23.73826889 | 0.426820889 | 22.34080521 | NA | NA | NA | NA | NA | NA | NA |
| TRPA1 | 557 | R | -61.95736364 | 21.36333764 | 0.379957055 | 19.95604822 | NA | NA | NA | NA | NA | NA | NA |
| TRPA1 | 558 | E | -63.46766667 | 18.73628368 | 0.328118156 | 17.60601093 | NA | NA | NA | NA | NA | NA | NA |
| TRPA1 | 559 | G | -61.75325 | 19.4901951 | 0.342994873 | 18.1420745 | NA | NA | NA | NA | NA | NA | NA |
| TRPA1 | 560 | H | -65.1129 | 18.17586188 | 0.317059514 | 15.88135129 | NA | NA | NA | NA | NA | NA | NA |
| TRPA1 | 561 | A | -62.9242 | 22.51440661 | 0.402670767 | 21.27436421 | NA | NA | NA | NA | NA | NA | NA |
| TRPA1 | 562 | K | -66.37411111 | 22.68247081 | 0.40598713 | 20.8454841 | NA | NA | NA | NA | NA | NA | NA |
| TRPA1 | 563 | A | -67.0184 | 24.76878348 | 0.447155737 | 22.9331605 | NA | NA | NA | NA | NA | NA | NA |
| TRPA1 | 564 | V | -63.64428571 | 26.52526196 | 0.481815821 | 24.16636156 | NA | NA | NA | NA | NA | NA | NA |
| TRPA1 | 565 | A | -64.853 | 28.57027245 | 0.522169424 | 26.74389923 | NA | NA | NA | NA | NA | NA | NA |
| TRPA1 | 566 | L | -69.233625 | 28.70964941 | 0.524919709 | 27.27269158 | NA | NA | NA | NA | NA | NA | NA |
| TRPA1 | 567 | L | -67.693875 | 29.55541949 | 0.541609046 | 27.74350584 | NA | NA | NA | NA | NA | NA | NA |
| TRPA1 | 568 | L | -63.408125 | 32.59220836 | 0.601533127 | 30.75762258 | NA | NA | NA | NA | NA | NA | NA |
| TRPA1 | 569 | S | -66.81416667 | 34.16569276 | 0.632582241 | 32.33039257 | NA | NA | NA | NA | NA | NA | NA |
| TRPA1 | 570 | H | -69.8718 | 34.53817521 | 0.63993233 | 32.99587244 | NA | NA | NA | NA | NA | NA | NA |
| TRPA1 | 571 | N | -67.458875 | 37.80942588 | 0.704482977 | 35.97533046 | NA | NA | NA | NA | NA | NA | NA |
| TRPA1 | 572 | A | -65.7048 | 35.35469052 | 0.656044391 | 33.52214479 | NA | NA | NA | NA | NA | NA | NA |
| TRPA1 | 573 | D | -64.6864 | 38.12562503 | 0.710722444 | 36.29728851 | NA | NA | NA | NA | NA | NA | NA |
| TRPA1 | 574 | I | -60.684 | 36.52951696 | 0.679226903 | 35.14002618 | NA | NA | NA | NA | NA | NA | NA |
| TRPA1 | 575 | V | -61.44742857 | 36.1677544 | 0.672088347 | 34.76218129 | NA | NA | NA | NA | NA | NA | NA |
| TRPA1 | 576 | L | -58.1825 | 36.80200963 | 0.684603923 | 35.58029752 | NA | NA | NA | NA | NA | NA | NA |
| TRPA1 | 577 | N | -59.896625 | 32.24036957 | 0.594590393 | 30.83292541 | NA | NA | NA | NA | NA | NA | NA |
| TRPA1 | 578 | K | -60.70466667 | 36.34420016 | 0.6755701 | 34.93622581 | NA | NA | NA | NA | NA | NA | NA |
| TRPA1 | 579 | Q | -56.49288889 | 31.98411965 | 0.589533887 | 30.29411788 | NA | NA | NA | NA | NA | NA | NA |
| TRPA1 | 580 | Q | -55.28055556 | 35.60447838 | 0.660973383 | 33.91910274 | NA | NA | NA | NA | NA | NA | NA |
| TRPA1 | 581 | A | -56.4868 | 32.06554185 | 0.591140568 | 30.37601748 | NA | NA | NA | NA | NA | NA | NA |
| TRPA1 | 582 | S | -56.52316667 | 30.83829541 | 0.566923668 | 29.13872591 | NA | NA | NA | NA | NA | NA | NA |
| TRPA1 | 583 | F | -58.95327273 | 28.30882873 | 0.51701043 | 27.06859339 | NA | NA | NA | NA | NA | NA | NA |
| TRPA1 | 584 | L | -54.5145 | 27.81519567 | 0.507269711 | 26.14036971 | NA | NA | NA | NA | NA | NA | NA |

| | | | | | | | | | | | | | |
|---|---|---|---|---|---|---|---|---|---|---|---|---|---|
| TRPA1 | 585 | H | -52.7814 | 27.02143477 | 0.491606656 | 25.17589106 | NA | NA | NA | NA | NA | NA | NA |
| TRPA1 | 586 | L | -57.52325 | 24.74260192 | 0.446639104 | 23.04433643 | NA | NA | NA | NA | NA | NA | NA |
| TRPA1 | 587 | A | -55.749 | 22.96589839 | 0.411579925 | 21.2889868 | NA | NA | NA | NA | NA | NA | NA |
| TRPA1 | 588 | L | -51.732875 | 22.91830177 | 0.410640714 | 20.98228624 | NA | NA | NA | NA | NA | NA | NA |
| TRPA1 | 589 | H | -52.0197 | 21.82261976 | 0.389019937 | 19.92133112 | NA | NA | NA | NA | NA | NA | NA |
| TRPA1 | 590 | N | -55.61375 | 18.60885246 | 0.325603592 | 16.75960856 | NA | NA | NA | NA | NA | NA | NA |
| TRPA1 | 591 | K | -51.44911111 | 16.74691389 | 0.288862495 | 15.10979868 | NA | NA | NA | NA | NA | NA | NA |
| TRPA1 | 592 | R | -57.20072727 | 16.38699744 | 0.281760367 | 14.86821969 | NA | NA | NA | NA | NA | NA | NA |
| TRPA1 | 593 | K | -53.17666667 | 17.91622921 | 0.311936257 | 16.4969708 | NA | NA | NA | NA | NA | NA | NA |
| TRPA1 | 594 | E | -57.41788889 | 20.88649978 | 0.370547751 | 19.2009247 | NA | NA | NA | NA | NA | NA | NA |
| TRPA1 | 595 | V | -58.14328571 | 22.99413799 | 0.412137169 | 21.46437281 | NA | NA | NA | NA | NA | NA | NA |
| TRPA1 | 596 | V | -54.20957143 | 24.40151686 | 0.43990857 | 22.78575501 | NA | NA | NA | NA | NA | NA | NA |
| TRPA1 | 597 | L | -54.137875 | 24.05412881 | 0.433053662 | 22.51685106 | NA | NA | NA | NA | NA | NA | NA |
| TRPA1 | 598 | T | -58.229 | 26.75499935 | 0.486349162 | 25.24030043 | NA | NA | NA | NA | NA | NA | NA |
| TRPA1 | 599 | I | -56.61675 | 28.82847705 | 0.527264501 | 27.12963232 | NA | NA | NA | NA | NA | NA | NA |
| TRPA1 | 600 | I | -52.65925 | 28.53311159 | 0.521436139 | 27.32232976 | NA | NA | NA | NA | NA | NA | NA |
| TRPA1 | 601 | R | -54.79263636 | 27.75157005 | 0.506014205 | 26.18623046 | NA | NA | NA | NA | NA | NA | NA |
| TRPA1 | 602 | S | -57.54366667 | 32.92294297 | 0.608059417 | 31.4041879 | NA | NA | NA | NA | NA | NA | NA |
| TRPA1 | 603 | K | -58.43744444 | 36.30433368 | 0.674783426 | 34.60680392 | NA | NA | NA | NA | NA | NA | NA |
| TRPA1 | 604 | R | -59.65618182 | 37.17552191 | 0.691974333 | 35.47947125 | NA | NA | NA | NA | NA | NA | NA |
| TRPA1 | 605 | W | -52.29478571 | 31.33143339 | 0.576654617 | 29.88699972 | NA | NA | NA | NA | NA | NA | NA |
| TRPA1 | 606 | D | -51.918625 | 37.97482635 | 0.707746777 | 36.07134213 | NA | NA | NA | NA | NA | NA | NA |
| TRPA1 | 607 | E | -56.55344444 | 38.62576854 | 0.720591632 | 36.93983251 | NA | NA | NA | NA | NA | NA | NA |
| TRPA1 | 608 | C | -54.79966667 | 34.90904215 | 0.64725054 | 33.2100248 | NA | NA | NA | NA | NA | NA | NA |
| TRPA1 | 609 | L | -49.94475 | 35.15249212 | 0.652054468 | 32.78581679 | NA | NA | NA | NA | NA | NA | NA |
| TRPA1 | 610 | K | -51.95288889 | 38.87278486 | 0.725465934 | 36.93637229 | NA | NA | NA | NA | NA | NA | NA |
| TRPA1 | 611 | I | -54.010375 | 37.08493611 | 0.690186829 | 35.23995698 | NA | NA | NA | NA | NA | NA | NA |
| TRPA1 | 612 | F | -49.46827273 | 36.8659877 | 0.685866383 | 34.97010329 | NA | NA | NA | NA | NA | NA | NA |
| TRPA1 | 613 | S | -50.024 | 37.78658182 | 0.704032202 | 34.88298914 | NA | NA | NA | NA | NA | NA | NA |
| TRPA1 | 614 | H | -46.4975 | 40.36689764 | 0.754948831 | 38.53311366 | NA | NA | NA | NA | NA | NA | NA |
| TRPA1 | 615 | N | -47.36275 | 39.35258671 | 0.734933725 | 37.1314742 | NA | NA | NA | NA | NA | NA | NA |
| TRPA1 | 616 | S | -48.4705 | 36.05308565 | 0.669825621 | 33.17750994 | NA | NA | NA | NA | NA | NA | NA |
| TRPA1 | 617 | P | -46.56085714 | 32.62697684 | 0.602219203 | 31.26627251 | NA | NA | NA | NA | NA | NA | NA |

| | | | | | | | | | | | | | |
|---|---|---|---|---|---|---|---|---|---|---|---|---|---|
| TRPA1 | 618 | G | -48.28275 | 30.57792599 | 0.561785873 | 27.67085321 | NA | NA | NA | NA | NA | NA | NA |
| TRPA1 | 619 | N | -50.178625 | 32.59356384 | 0.601559874 | 29.68852858 | NA | NA | NA | NA | NA | NA | NA |
| TRPA1 | 620 | K | -45.61566667 | 29.99842936 | 0.550350833 | 28.52939564 | NA | NA | NA | NA | NA | NA | NA |
| TRPA1 | 621 | C | -45.753 | 31.87924812 | 0.587464488 | 30.5173728 | NA | NA | NA | NA | NA | NA | NA |
| TRPA1 | 622 | P | -48.09471429 | 29.79629524 | 0.546362178 | 26.89858802 | NA | NA | NA | NA | NA | NA | NA |
| TRPA1 | 623 | I | -44.738875 | 28.89261984 | 0.528530212 | 27.55491842 | NA | NA | NA | NA | NA | NA | NA |
| TRPA1 | 624 | T | -43.14114286 | 27.32789983 | 0.497654043 | 25.09831664 | NA | NA | NA | NA | NA | NA | NA |
| TRPA1 | 625 | E | -46.73066667 | 25.44494282 | 0.460498194 | 24.09052752 | NA | NA | NA | NA | NA | NA | NA |
| TRPA1 | 626 | M | -46.71025 | 24.21073169 | 0.436143862 | 22.86231832 | NA | NA | NA | NA | NA | NA | NA |
| TRPA1 | 627 | I | -41.89075 | 22.98403711 | 0.411937851 | 20.75709579 | NA | NA | NA | NA | NA | NA | NA |
| TRPA1 | 628 | E | -43.57455556 | 21.37820044 | 0.380250338 | 19.16123396 | NA | NA | NA | NA | NA | NA | NA |
| TRPA1 | 629 | Y | -47.109 | 19.78160097 | 0.348745101 | 18.43847192 | NA | NA | NA | NA | NA | NA | NA |
| TRPA1 | 630 | L | -48.22875 | 19.5834318 | 0.344834686 | 17.93318954 | NA | NA | NA | NA | NA | NA | NA |
| TRPA1 | 631 | P | -44.01642857 | 19.70510068 | 0.347235543 | 18.3796593 | NA | NA | NA | NA | NA | NA | NA |
| TRPA1 | 632 | E | -46.77555556 | 17.65993516 | 0.306878881 | 16.5737871 | NA | NA | NA | NA | NA | NA | NA |
| TRPA1 | 633 | C | -48.08766667 | 23.06221204 | 0.413480454 | 20.141724 | NA | NA | NA | NA | NA | NA | NA |
| TRPA1 | 634 | M | -43.990125 | 25.4030548 | 0.45967163 | 24.11189399 | NA | NA | NA | NA | NA | NA | NA |
| TRPA1 | 635 | K | -43.08544444 | 22.43225225 | 0.401049639 | 20.98334889 | NA | NA | NA | NA | NA | NA | NA |
| TRPA1 | 636 | V | -47.147 | 25.56878505 | 0.462941937 | 24.3668161 | NA | NA | NA | NA | NA | NA | NA |
| TRPA1 | 637 | L | -46.176 | 28.92890191 | 0.529246156 | 27.56847587 | NA | NA | NA | NA | NA | NA | NA |
| TRPA1 | 638 | L | -41.87 | 28.75345758 | 0.525784163 | 27.05620996 | NA | NA | NA | NA | NA | NA | NA |
| TRPA1 | 639 | D | -43.896375 | 27.57537296 | 0.502537359 | 26.09786129 | NA | NA | NA | NA | NA | NA | NA |
| TRPA1 | 640 | F | -47.87309091 | 31.32379029 | 0.576503798 | 30.35766702 | NA | NA | NA | NA | NA | NA | NA |
| TRPA1 | 641 | C | -42.553 | 34.55511554 | 0.640266609 | 33.07513731 | NA | NA | NA | NA | NA | NA | NA |
| TRPA1 | 642 | M | -40.813375 | 32.80328619 | 0.605698265 | 31.18903596 | NA | NA | NA | NA | NA | NA | NA |
| TRPA1 | 643 | L | -42.452125 | 37.94883538 | 0.707233905 | 36.24229408 | NA | NA | NA | NA | NA | NA | NA |
| TRPA1 | 644 | H | -41.6207 | 35.74517762 | 0.663749761 | 34.04121051 | NA | NA | NA | NA | NA | NA | NA |
| TRPA1 | 645 | S | -41.658 | 42.41483723 | 0.795360233 | 40.71005804 | NA | NA | NA | NA | NA | NA | NA |
| TRPA1 | 646 | T | -42.2988 | 44.0927265 | 0.828469538 | 42.6224185 | NA | NA | NA | NA | NA | NA | NA |
| TRPA1 | 647 | E | -42.6734 | 41.59728739 | 0.779227757 | 40.11748253 | NA | NA | NA | NA | NA | NA | NA |
| TRPA1 | 648 | D | -39.4636 | 43.02393996 | 0.807379482 | 41.32177775 | NA | NA | NA | NA | NA | NA | NA |
| TRPA1 | 649 | K | -36.212 | 40.54036343 | 0.758371781 | 38.30881435 | NA | NA | NA | NA | NA | NA | NA |
| TRPA1 | 650 | S | -34.9108 | 43.87510495 | 0.824175275 | 38.04831763 | NA | NA | NA | NA | NA | NA | NA |

| | | | | | | | | | | | | | |
|---|---|---|---|---|---|---|---|---|---|---|---|---|---|
| TRPA1 | 651 | C | -35.5315 | 45.96905282 | 0.865494545 | 43.83578151 | NA | NA | NA | NA | NA | NA | NA |
| TRPA1 | 652 | R | -32.276 | 47.7512016 | 0.900661174 | 40.60597721 | NA | NA | NA | NA | NA | NA | NA |
| TRPA1 | 653 | D | -34.9404 | 47.87799817 | 0.903163214 | 40.7767946 | NA | NA | NA | NA | NA | NA | NA |
| TRPA1 | 654 | Y | -37.00016667 | 40.15795042 | 0.750825735 | 38.07548976 | NA | NA | NA | NA | NA | NA | NA |
| TRPA1 | 655 | Y | -37.47925 | 42.60715523 | 0.799155189 | 40.40071223 | NA | NA | NA | NA | NA | NA | NA |
| TRPA1 | 656 | I | -36.538625 | 37.54636753 | 0.699292123 | 35.32294303 | NA | NA | NA | NA | NA | NA | NA |
| TRPA1 | 657 | E | -39.07544444 | 39.10713102 | 0.730090219 | 37.04276592 | NA | NA | NA | NA | NA | NA | NA |
| TRPA1 | 658 | Y | -36.10416667 | 35.09260396 | 0.650872713 | 32.89473743 | NA | NA | NA | NA | NA | NA | NA |
| TRPA1 | 659 | N | -40.3995 | 37.16196155 | 0.69170675 | 34.94846135 | NA | NA | NA | NA | NA | NA | NA |
| TRPA1 | 660 | F | -36.575 | 36.70667902 | 0.682722791 | 35.05356927 | NA | NA | NA | NA | NA | NA | NA |
| TRPA1 | 661 | K | -43.19377778 | 38.25622924 | 0.713299619 | 36.03577488 | NA | NA | NA | NA | NA | NA | NA |
| TRPA1 | 662 | Y | -44.22175 | 34.55307596 | 0.640226362 | 32.84529998 | NA | NA | NA | NA | NA | NA | NA |
| TRPA1 | 663 | L | -38.045375 | 33.5976378 | 0.621372976 | 31.66614089 | NA | NA | NA | NA | NA | NA | NA |
| TRPA1 | 664 | Q | -37.17722222 | 36.75516526 | 0.683679556 | 34.53607342 | NA | NA | NA | NA | NA | NA | NA |
| TRPA1 | 665 | C | -38.86016667 | 38.73605931 | 0.722767968 | 36.51473339 | NA | NA | NA | NA | NA | NA | NA |
| TRPA1 | 666 | P | -39.89257143 | 42.13821721 | 0.78990177 | 39.91504018 | NA | NA | NA | NA | NA | NA | NA |
| TRPA1 | 667 | L | -35.171125 | 43.30808095 | 0.812986354 | 41.24470687 | NA | NA | NA | NA | NA | NA | NA |
| TRPA1 | 668 | E | -38.04133333 | 46.49548083 | 0.875882398 | 44.3145391 | NA | NA | NA | NA | NA | NA | NA |
| TRPA1 | 669 | F | -40.13690909 | 43.45732281 | 0.815931301 | 41.26119369 | NA | NA | NA | NA | NA | NA | NA |
| TRPA1 | 670 | T | -34.20214286 | 42.09844459 | 0.789116948 | 34.87896392 | NA | NA | NA | NA | NA | NA | NA |
| TRPA1 | 671 | K | -31.70355556 | 45.14808284 | 0.849294581 | 37.94221316 | NA | NA | NA | NA | NA | NA | NA |
| TRPA1 | 672 | K | -33.74177778 | 46.68781063 | 0.879677586 | 39.4708426 | NA | NA | NA | NA | NA | NA | NA |
| TRPA1 | 673 | T | -36.21214286 | 49.2073167 | 0.929394274 | 42.03008334 | NA | NA | NA | NA | NA | NA | NA |
| TRPA1 | 674 | P | -36.62871429 | 52.45149808 | 0.993410771 | 50.36843367 | NA | NA | NA | NA | NA | NA | NA |
| TRPA1 | 675 | T | -40.41528571 | 51.00146418 | 0.964797669 | 48.77146494 | NA | NA | NA | NA | NA | NA | NA |
| TRPA1 | 676 | Q | -40.17344444 | 47.38541988 | 0.893443309 | 45.15908874 | NA | NA | NA | NA | NA | NA | NA |
| TRPA1 | 677 | D | -40.2854 | 45.67502203 | 0.85969252 | 43.44630326 | NA | NA | NA | NA | NA | NA | NA |
| TRPA1 | 678 | V | -40.86885714 | 43.0502294 | 0.807898243 | 40.82025655 | NA | NA | NA | NA | NA | NA | NA |
| TRPA1 | 679 | I | -36.703875 | 40.26179124 | 0.752874797 | 38.03610302 | NA | NA | NA | NA | NA | NA | NA |
| TRPA1 | 680 | Y | -39.12683333 | 37.91852365 | 0.706635772 | 36.14099586 | NA | NA | NA | NA | NA | NA | NA |
| TRPA1 | 681 | E | -33.65155556 | 32.93025244 | 0.608203653 | 25.74390918 | NA | NA | NA | NA | NA | NA | NA |
| TRPA1 | 682 | P | -32.01142857 | 35.56053083 | 0.660106179 | 28.3389217 | NA | NA | NA | NA | NA | NA | NA |
| TRPA1 | 683 | L | -32.38575 | 33.21658783 | 0.613853827 | 26.03705857 | NA | NA | NA | NA | NA | NA | NA |

| | | | | | | | | | | | | | |
|---|---|---|---|---|---|---|---|---|---|---|---|---|---|
| TRPA1 | 684 | T | -37.05785714 | 31.91087547 | 0.588088581 | 29.70865534 | NA | NA | NA | NA | NA | NA | NA |
| TRPA1 | 685 | A | -37.5622 | 29.32636462 | 0.537089173 | 27.86481551 | NA | NA | NA | NA | NA | NA | NA |
| TRPA1 | 686 | L | -33.856375 | 28.18434754 | 0.514554079 | 21.42271172 | NA | NA | NA | NA | NA | NA | NA |
| TRPA1 | 687 | N | -33.294875 | 27.90666996 | 0.509074747 | 20.76491186 | NA | NA | NA | NA | NA | NA | NA |
| TRPA1 | 688 | A | -37.22 | 25.93111937 | 0.470091776 | 23.76495881 | NA | NA | NA | NA | NA | NA | NA |
| TRPA1 | 689 | M | -36.42375 | 23.67598792 | 0.425591917 | 21.46491471 | NA | NA | NA | NA | NA | NA | NA |
| TRPA1 | 690 | V | -32.45542857 | 23.06770272 | 0.4135888 | 15.87122334 | NA | NA | NA | NA | NA | NA | NA |
| TRPA1 | 691 | Q | -34.61588889 | 23.25094918 | 0.41720475 | 16.19905782 | NA | NA | NA | NA | NA | NA | NA |
| TRPA1 | 692 | N | -37.828625 | 20.05247957 | 0.35409027 | 17.89680459 | NA | NA | NA | NA | NA | NA | NA |
| TRPA1 | 693 | N | -34.42175 | 18.48721042 | 0.323203265 | 16.29863823 | NA | NA | NA | NA | NA | NA | NA |
| TRPA1 | 694 | R | -38.50572727 | 18.74414346 | 0.328273251 | 16.93837218 | NA | NA | NA | NA | NA | NA | NA |
| TRPA1 | 695 | I | -33.23225 | 18.00764851 | 0.313740208 | 11.21726429 | NA | NA | NA | NA | NA | NA | NA |
| TRPA1 | 696 | E | -37.31488889 | 16.73726148 | 0.288672027 | 14.61100542 | NA | NA | NA | NA | NA | NA | NA |
| TRPA1 | 697 | L | -38.083875 | 22.64043275 | 0.405157605 | 20.54822062 | NA | NA | NA | NA | NA | NA | NA |
| TRPA1 | 698 | L | -33.0615 | 24.00304591 | 0.432045658 | 17.25815281 | NA | NA | NA | NA | NA | NA | NA |
| TRPA1 | 699 | N | -32.5525 | 21.25563899 | 0.377831869 | 14.44177061 | NA | NA | NA | NA | NA | NA | NA |
| TRPA1 | 700 | H | -37.1205 | 26.20216787 | 0.475440298 | 23.99332555 | NA | NA | NA | NA | NA | NA | NA |
| TRPA1 | 701 | P | -34.83657143 | 28.99009422 | 0.530453646 | 22.33651368 | NA | NA | NA | NA | NA | NA | NA |
| TRPA1 | 702 | V | -36.29314286 | 31.65152295 | 0.582970852 | 29.42066518 | NA | NA | NA | NA | NA | NA | NA |
| TRPA1 | 703 | C | -33.779 | 29.38658172 | 0.538277419 | 22.68501619 | NA | NA | NA | NA | NA | NA | NA |
| TRPA1 | 704 | K | -30.46355556 | 26.54260087 | 0.482157964 | 19.28214912 | NA | NA | NA | NA | NA | NA | NA |
| TRPA1 | 705 | E | -31.09944444 | 33.2517934 | 0.614548528 | 26.02388935 | NA | NA | NA | NA | NA | NA | NA |
| TRPA1 | 706 | Y | -32.58566667 | 34.57856391 | 0.640729308 | 27.36515481 | NA | NA | NA | NA | NA | NA | NA |
| TRPA1 | 707 | L | -28.31125 | 30.04762563 | 0.551321608 | 23.00457028 | NA | NA | NA | NA | NA | NA | NA |
| TRPA1 | 708 | L | -26.396 | 34.53251497 | 0.639820638 | 27.54291209 | NA | NA | NA | NA | NA | NA | NA |
| TRPA1 | 709 | M | -28.80025 | 37.3406298 | 0.695232359 | 30.35060913 | NA | NA | NA | NA | NA | NA | NA |
| TRPA1 | 710 | K | -27.87744444 | 36.96373722 | 0.687795246 | 29.974111 | NA | NA | NA | NA | NA | NA | NA |
| TRPA1 | 711 | W | -21.45864286 | 35.0355637 | 0.649747154 | 30.52988934 | NA | NA | NA | NA | NA | NA | NA |
| TRPA1 | 712 | L | -23.244375 | 38.84535109 | 0.724924591 | 34.59858304 | NA | NA | NA | NA | NA | NA | NA |
| TRPA1 | 713 | A | -24.2694 | 41.19543611 | 0.771298142 | 36.53713292 | NA | NA | NA | NA | NA | NA | NA |
| TRPA1 | 714 | Y | -23.91491667 | 39.42043202 | 0.736272497 | 35.38775703 | NA | NA | NA | NA | NA | NA | NA |
| TRPA1 | 715 | G | -20.2275 | 37.96729278 | 0.707598119 | 33.62571486 | NA | NA | NA | NA | NA | NA | NA |
| TRPA1 | 716 | F | -16.81463636 | 37.6859909 | 0.70204727 | 36.44788457 | NA | NA | NA | NA | NA | NA | NA |

| | | | | | | | | | | | | | |
|---|---|---|---|---|---|---|---|---|---|---|---|---|---|
| TRPA1 | 717 | R | -16.65554545 | 40.71994996 | 0.761915511 | 39.65275823 | NA | NA | NA | NA | NA | NA | NA |
| TRPA1 | 718 | A | -16.4446 | 39.36473206 | 0.735173386 | 38.33726803 | NA | NA | NA | NA | NA | NA | NA |
| TRPA1 | 719 | H | -14.8293 | 32.75409985 | 0.604727685 | 31.63863828 | NA | NA | NA | NA | NA | NA | NA |
| TRPA1 | 720 | M | -12.06 | 37.12330797 | 0.69094401 | 35.29441559 | NA | NA | NA | NA | NA | NA | NA |
| TRPA1 | 721 | M | -11.977125 | 40.36418494 | 0.754895302 | 38.53497241 | NA | NA | NA | NA | NA | NA | NA |
| TRPA1 | 722 | N | -11.611 | 36.7406004 | 0.683392152 | 34.92575163 | NA | NA | NA | NA | NA | NA | NA |
| TRPA1 | 723 | L | -8.851 | 33.52152201 | 0.619871005 | 31.91198222 | NA | NA | NA | NA | NA | NA | NA |
| TRPA1 | 724 | G | -6.59 | 38.44564529 | 0.717037312 | 36.77779534 | NA | NA | NA | NA | NA | NA | NA |
| TRPA1 | 725 | S | -6.506833333 | 39.5734196 | 0.739291357 | 36.20596317 | NA | NA | NA | NA | NA | NA | NA |
| TRPA1 | 726 | Y | -4.341416667 | 31.77191293 | 0.585346473 | 28.44049865 | NA | NA | NA | NA | NA | NA | NA |
| TRPA1 | 727 | C | -2.797666667 | 35.69174934 | 0.662695476 | 31.54617074 | NA | NA | NA | NA | NA | NA | NA |
| TRPA1 | 728 | L | -1.099875 | 38.64248711 | 0.720921534 | 33.95626912 | NA | NA | NA | NA | NA | NA | NA |
| TRPA1 | 729 | G | -0.0015 | 36.56147921 | 0.679857605 | 31.7763257 | NA | NA | NA | NA | NA | NA | NA |
| TRPA1 | 730 | L | -0.314125 | 31.63397659 | 0.582624615 | 26.84925883 | NA | NA | NA | NA | NA | NA | NA |
| TRPA1 | 731 | I | 0.98475 | 35.2832459 | 0.654634595 | 31.63635697 | NA | NA | NA | NA | NA | NA | NA |
| TRPA1 | 732 | P | 3.669857143 | 36.72246057 | 0.683034204 | 32.84567942 | NA | NA | NA | NA | NA | NA | NA |
| TRPA1 | 733 | M | 4.994875 | 33.56411861 | 0.620711552 | 29.71480765 | NA | NA | NA | NA | NA | NA | NA |
| TRPA1 | 734 | T | 4.916857143 | 31.40148035 | 0.578036834 | 27.52442531 | NA | NA | NA | NA | NA | NA | NA |
| TRPA1 | 735 | I | 7.137125 | 34.16559268 | 0.632580266 | 31.2288001 | NA | NA | NA | NA | NA | NA | NA |
| TRPA1 | 736 | L | 9.850625 | 34.1587479 | 0.6324452 | 30.7015742 | NA | NA | NA | NA | NA | NA | NA |
| TRPA1 | 737 | V | 9.936285714 | 29.45298107 | 0.539587659 | 26.00183435 | NA | NA | NA | NA | NA | NA | NA |
| TRPA1 | 738 | V | 10.419 | 29.62924996 | 0.543065922 | 26.22307608 | NA | NA | NA | NA | NA | NA | NA |
| TRPA1 | 739 | N | 12.80175 | 32.78795837 | 0.605395805 | 28.60884095 | NA | NA | NA | NA | NA | NA | NA |
| TRPA1 | 740 | I | 15.009 | 30.94304119 | 0.568990586 | 26.05306478 | NA | NA | NA | NA | NA | NA | NA |
| TRPA1 | 741 | K | 19.08188889 | 27.33288543 | 0.497752422 | 22.53839933 | NA | NA | NA | NA | NA | NA | NA |
| TRPA1 | 742 | P | 16.39057143 | 27.63726025 | 0.503758563 | 22.74736747 | NA | NA | NA | NA | NA | NA | NA |
| TRPA1 | 743 | G | 18.8365 | 29.83890005 | 0.547202887 | 24.97527253 | NA | NA | NA | NA | NA | NA | NA |
| TRPA1 | 744 | M | 20.3425 | 31.89633868 | 0.587801731 | 27.10326201 | NA | NA | NA | NA | NA | NA | NA |
| TRPA1 | 745 | A | 19.5188 | 35.6084817 | 0.66105238 | 30.7659956 | NA | NA | NA | NA | NA | NA | NA |
| TRPA1 | 746 | F | 15.37318182 | 36.24285611 | 0.673570307 | 31.41128878 | NA | NA | NA | NA | NA | NA | NA |
| TRPA1 | 747 | N | 17.340375 | 41.16600318 | 0.77071735 | 36.29454789 | NA | NA | NA | NA | NA | NA | NA |
| TRPA1 | 748 | S | 15.29116667 | 44.33948951 | 0.833338842 | 39.45493416 | NA | NA | NA | NA | NA | NA | NA |
| TRPA1 | 749 | T | 19.20471429 | 44.51192464 | 0.836741455 | 39.65401113 | NA | NA | NA | NA | NA | NA | NA |

| | | | | | | | | | | | | | |
|---|---|---|---|---|---|---|---|---|---|---|---|---|---|
| TRPA1 | 750 | G | 19.1205 | 41.05565185 | 0.768539819 | 36.20366985 | NA | NA | NA | NA | NA | NA | NA |
| TRPA1 | 751 | I | 20.524125 | 36.51782324 | 0.678996154 | 31.68247656 | NA | NA | NA | NA | NA | NA | NA |
| TRPA1 | 752 | I | 22.949 | 37.19305244 | 0.692320258 | 32.38597798 | NA | NA | NA | NA | NA | NA | NA |
| TRPA1 | 753 | N | 26.36325 | 34.46286142 | 0.638446185 | 29.7060781 | NA | NA | NA | NA | NA | NA | NA |
| TRPA1 | 761 | I | 28.281 | 39.20674158 | 0.732055805 | 34.44299718 | NA | NA | NA | NA | NA | NA | NA |
| TRPA1 | 762 | L | 24.854125 | 42.10384981 | 0.789223608 | 37.30200386 | NA | NA | NA | NA | NA | NA | NA |
| TRPA1 | 763 | D | 22.633 | 37.79956864 | 0.704288467 | 32.98084667 | NA | NA | NA | NA | NA | NA | NA |
| TRPA1 | 764 | T | 21.75557143 | 41.44102588 | 0.776144294 | 36.60083921 | NA | NA | NA | NA | NA | NA | NA |
| TRPA1 | 765 | T | 19.61228571 | 41.26766487 | 0.772723411 | 36.42372568 | NA | NA | NA | NA | NA | NA | NA |
| TRPA1 | 766 | N | 16.178625 | 38.41984645 | 0.71652823 | 33.5463089 | NA | NA | NA | NA | NA | NA | NA |
| TRPA1 | 767 | S | 15.746 | 43.38824253 | 0.814568159 | 38.50642359 | NA | NA | NA | NA | NA | NA | NA |
| TRPA1 | 768 | Y | 11.47091667 | 45.20690604 | 0.850455323 | 41.02715273 | NA | NA | NA | NA | NA | NA | NA |
| TRPA1 | 769 | L | 10.728625 | 39.63269249 | 0.740460972 | 35.83683622 | NA | NA | NA | NA | NA | NA | NA |
| TRPA1 | 770 | I | 12.7005 | 39.54274368 | 0.738686038 | 35.41242323 | NA | NA | NA | NA | NA | NA | NA |
| TRPA1 | 771 | K | 11.297 | 43.85012089 | 0.823682272 | 40.05401452 | NA | NA | NA | NA | NA | NA | NA |
| TRPA1 | 772 | T | 7.602285714 | 44.15359349 | 0.829670609 | 41.14205257 | NA | NA | NA | NA | NA | NA | NA |
| TRPA1 | 773 | C | 6.943666667 | 40.27003241 | 0.753037417 | 36.64529387 | NA | NA | NA | NA | NA | NA | NA |
| TRPA1 | 774 | M | 9.011625 | 41.37053548 | 0.774753327 | 38.43782295 | NA | NA | NA | NA | NA | NA | NA |
| TRPA1 | 775 | I | 5.878625 | 44.53903888 | 0.837276492 | 40.66650814 | NA | NA | NA | NA | NA | NA | NA |
| TRPA1 | 776 | L | 2.4555 | 42.75998937 | 0.802171021 | 39.06077453 | NA | NA | NA | NA | NA | NA | NA |
| TRPA1 | 777 | V | 3.270571429 | 39.02712477 | 0.728511479 | 35.33663382 | NA | NA | NA | NA | NA | NA | NA |
| TRPA1 | 778 | F | 3.865363636 | 42.85459169 | 0.804037781 | 38.97553056 | NA | NA | NA | NA | NA | NA | NA |
| TRPA1 | 779 | L | 0.7265 | 44.74567216 | 0.841353927 | 39.9599307 | NA | NA | NA | NA | NA | NA | NA |
| TRPA1 | 780 | S | -0.3715 | 40.39765027 | 0.755555663 | 35.71572383 | NA | NA | NA | NA | NA | NA | NA |
| TRPA1 | 781 | S | -0.098666667 | 40.33186616 | 0.754257565 | 35.54631877 | NA | NA | NA | NA | NA | NA | NA |
| TRPA1 | 782 | I | -0.330625 | 44.18423506 | 0.830275251 | 39.3981614 | NA | NA | NA | NA | NA | NA | NA |
| TRPA1 | 783 | F | -3.341181818 | 44.33157323 | 0.833182632 | 40.18369702 | NA | NA | NA | NA | NA | NA | NA |
| TRPA1 | 784 | G | -4.394 | 42.1319618 | 0.789778333 | 37.98457863 | NA | NA | NA | NA | NA | NA | NA |
| TRPA1 | 785 | Y | -1.784416667 | 42.07053475 | 0.788566211 | 38.04783404 | NA | NA | NA | NA | NA | NA | NA |
| TRPA1 | 786 | C | -6.038 | 45.75561407 | 0.86128282 | 42.38817212 | NA | NA | NA | NA | NA | NA | NA |
| TRPA1 | 787 | K | -9.074777778 | 39.74564175 | 0.742689767 | 38.07535566 | NA | NA | NA | NA | NA | NA | NA |
| TRPA1 | 788 | E | -8.067444444 | 39.30216979 | 0.733938863 | 37.6320725 | NA | NA | NA | NA | NA | NA | NA |
| TRPA1 | 789 | A | -8.7804 | 44.78000453 | 0.842031398 | 43.11049103 | NA | NA | NA | NA | NA | NA | NA |

| | | | | | | | | | | | | | |
|---|---|---|---|---|---|---|---|---|---|---|---|---|---|
| TRPA1 | 790 | G | -11.93675 | 45.60201949 | 0.858251982 | 43.77748813 | NA | NA | NA | NA | NA | NA | NA |
| TRPA1 | 791 | Q | -12.68755556 | 39.49027402 | 0.737650669 | 37.66676992 | NA | NA | NA | NA | NA | NA | NA |
| TRPA1 | 792 | I | -10.985125 | 43.80270536 | 0.822746635 | 42.02765166 | NA | NA | NA | NA | NA | NA | NA |
| TRPA1 | 793 | F | -12.39163636 | 46.60744943 | 0.878091842 | 45.52989897 | NA | NA | NA | NA | NA | NA | NA |
| TRPA1 | 794 | Q | -16.8916 | 44.97358116 | 0.84585119 | 43.91154957 | NA | NA | NA | NA | NA | NA | NA |
| TRPA1 | 795 | Q | -17.2404 | 41.48520065 | 0.777015982 | 40.22401721 | NA | NA | NA | NA | NA | NA | NA |
| TRPA1 | 796 | K | -17.199 | 43.40981098 | 0.814993763 | 42.14653814 | NA | NA | NA | NA | NA | NA | NA |
| TRPA1 | 797 | R | -15.6916 | 40.32989984 | 0.754218764 | 39.20571047 | NA | NA | NA | NA | NA | NA | NA |
| TRPA1 | 798 | N | -17.2896 | 36.88423162 | 0.686226385 | 35.61341054 | NA | NA | NA | NA | NA | NA | NA |
| TRPA1 | 799 | Y | -12.84975 | 36.46684919 | 0.677990298 | 35.43195789 | NA | NA | NA | NA | NA | NA | NA |
| TRPA1 | 800 | F | -11.965 | 35.22914396 | 0.653567017 | 33.54497698 | NA | NA | NA | NA | NA | NA | NA |
| TRPA1 | 801 | M | -13.8012 | 32.1439842 | 0.592688448 | 30.51518472 | NA | NA | NA | NA | NA | NA | NA |
| TRPA1 | 802 | D | -12.0605 | 31.5394285 | 0.580758925 | 29.85042374 | NA | NA | NA | NA | NA | NA | NA |
| TRPA1 | 803 | I | -8.596625 | 27.22206473 | 0.495565629 | 25.6428314 | NA | NA | NA | NA | NA | NA | NA |
| TRPA1 | 804 | S | -9.118 | 31.13853569 | 0.572848222 | 29.52987006 | NA | NA | NA | NA | NA | NA | NA |
| TRPA1 | 805 | N | -9.101 | 33.82111101 | 0.625782709 | 32.1487614 | NA | NA | NA | NA | NA | NA | NA |
| TRPA1 | 806 | V | -5.279142857 | 32.97846379 | 0.609154994 | 29.6103055 | NA | NA | NA | NA | NA | NA | NA |
| TRPA1 | 807 | L | -4.234 | 28.519829 | 0.521174038 | 24.45384574 | NA | NA | NA | NA | NA | NA | NA |
| TRPA1 | 808 | E | -5.560777778 | 33.93453219 | 0.628020816 | 30.56527883 | NA | NA | NA | NA | NA | NA | NA |
| TRPA1 | 809 | W | -3.636 | 36.18388411 | 0.67240663 | 31.50481825 | NA | NA | NA | NA | NA | NA | NA |
| TRPA1 | 810 | I | -0.136375 | 31.77404521 | 0.585388549 | 26.98558359 | NA | NA | NA | NA | NA | NA | NA |
| TRPA1 | 811 | I | -0.274125 | 32.05745898 | 0.590981071 | 27.27770506 | NA | NA | NA | NA | NA | NA | NA |
| TRPA1 | 812 | Y | -1.815916667 | 36.80483353 | 0.684659646 | 32.01743182 | NA | NA | NA | NA | NA | NA | NA |
| TRPA1 | 813 | T | 1.679142857 | 37.55021442 | 0.699368032 | 33.95648732 | NA | NA | NA | NA | NA | NA | NA |
| TRPA1 | 814 | T | 4.017285714 | 34.01565653 | 0.629621619 | 30.1394434 | NA | NA | NA | NA | NA | NA | NA |
| TRPA1 | 815 | G | 4.31 | 36.7595936 | 0.683766939 | 32.88048544 | NA | NA | NA | NA | NA | NA | NA |
| TRPA1 | 816 | I | 4.5905 | 39.47228447 | 0.737295687 | 35.59357618 | NA | NA | NA | NA | NA | NA | NA |
| TRPA1 | 817 | I | 7.87275 | 38.89692658 | 0.725942315 | 35.90223825 | NA | NA | NA | NA | NA | NA | NA |
| TRPA1 | 818 | F | 9.324545455 | 34.37177623 | 0.636648827 | 30.90266254 | NA | NA | NA | NA | NA | NA | NA |
| TRPA1 | 819 | V | 8.743571429 | 39.68871964 | 0.74156654 | 36.24447518 | NA | NA | NA | NA | NA | NA | NA |
| TRPA1 | 820 | L | 10.088375 | 42.16816006 | 0.790492623 | 38.70029768 | NA | NA | NA | NA | NA | NA | NA |
| TRPA1 | 821 | P | 13.99214286 | 41.13811807 | 0.770167101 | 37.04632681 | NA | NA | NA | NA | NA | NA | NA |
| TRPA1 | 822 | L | 11.686125 | 44.79629862 | 0.842352925 | 40.62056112 | NA | NA | NA | NA | NA | NA | NA |

| | | | | | | | | | | | | | |
|---|---|---|---|---|---|---|---|---|---|---|---|---|---|
| TRPA1 | 823 | F | 10.89518182 | 47.13527042 | 0.888507181 | 42.95761224 | NA | NA | NA | NA | NA | NA | NA |
| TRPA1 | 824 | V | 14.13471429 | 44.63420556 | 0.839154389 | 40.54270786 | NA | NA | NA | NA | NA | NA | NA |
| TRPA1 | 825 | E | 19.04344444 | 43.67342323 | 0.820195548 | 38.79927705 | NA | NA | NA | NA | NA | NA | NA |
| TRPA1 | 826 | I | 15.6125 | 39.72041887 | 0.742192051 | 34.85304271 | NA | NA | NA | NA | NA | NA | NA |
| TRPA1 | 827 | P | 18.08728571 | 37.09285309 | 0.690343053 | 32.21705574 | NA | NA | NA | NA | NA | NA | NA |
| TRPA1 | 828 | A | 17.8188 | 33.62714158 | 0.621955166 | 28.74920876 | NA | NA | NA | NA | NA | NA | NA |
| TRPA1 | 829 | H | 16.7907 | 30.12351503 | 0.552819112 | 25.25779289 | NA | NA | NA | NA | NA | NA | NA |
| TRPA1 | 830 | L | 13.159875 | 34.55970036 | 0.64035708 | 30.49380179 | NA | NA | NA | NA | NA | NA | NA |
| TRPA1 | 831 | Q | 13.56888889 | 33.93820327 | 0.628093256 | 29.85466188 | NA | NA | NA | NA | NA | NA | NA |
| TRPA1 | 832 | W | 13.97757143 | 30.2594036 | 0.555500562 | 26.08954061 | NA | NA | NA | NA | NA | NA | NA |
| TRPA1 | 833 | Q | 11.16322222 | 30.7282053 | 0.564751291 | 27.01133684 | NA | NA | NA | NA | NA | NA | NA |
| TRPA1 | 834 | C | 8.397666667 | 32.42094221 | 0.598153581 | 29.41807194 | NA | NA | NA | NA | NA | NA | NA |
| TRPA1 | 835 | G | 8.0935 | 30.48542002 | 0.559960479 | 27.47436927 | NA | NA | NA | NA | NA | NA | NA |
| TRPA1 | 836 | A | 7.5014 | 27.63820692 | 0.503777243 | 24.63734199 | NA | NA | NA | NA | NA | NA | NA |
| TRPA1 | 837 | I | 4.821375 | 29.16209451 | 0.533847678 | 25.28650364 | NA | NA | NA | NA | NA | NA | NA |
| TRPA1 | 838 | A | 3.4212 | 30.62256847 | 0.56266679 | 26.74696259 | NA | NA | NA | NA | NA | NA | NA |
| TRPA1 | 839 | V | 4.139571429 | 27.62495332 | 0.503515714 | 23.7674978 | NA | NA | NA | NA | NA | NA | NA |
| TRPA1 | 840 | Y | 1.21325 | 21.84456328 | 0.389452942 | 18.57426908 | NA | NA | NA | NA | NA | NA | NA |
| TRPA1 | 841 | F | 0 | 27.33900287 | 0.497873136 | 22.55064619 | NA | NA | NA | NA | NA | NA | NA |
| TRPA1 | 842 | Y | -0.155333333 | 29.07169046 | 0.532063761 | 24.28748572 | NA | NA | NA | NA | NA | NA | NA |
| TRPA1 | 843 | W | -0.370214286 | 25.57125357 | 0.462990648 | 20.8857232 | NA | NA | NA | NA | NA | NA | NA |
| TRPA1 | 844 | M | -2.329375 | 21.88872815 | 0.390324434 | 17.21479308 | NA | NA | NA | NA | NA | NA | NA |
| TRPA1 | 845 | N | -5.192 | 27.37997087 | 0.498681546 | 24.03129081 | NA | NA | NA | NA | NA | NA | NA |
| TRPA1 | 846 | F | -5.257 | 27.56214173 | 0.502276271 | 24.2424327 | NA | NA | NA | NA | NA | NA | NA |
| TRPA1 | 847 | L | -7.077875 | 22.02255775 | 0.392965256 | 20.48796295 | NA | NA | NA | NA | NA | NA | NA |
| TRPA1 | 848 | L | -9.9625 | 26.19430467 | 0.475285136 | 24.57889177 | NA | NA | NA | NA | NA | NA | NA |
| TRPA1 | 849 | Y | -9.775333333 | 28.71656423 | 0.525056157 | 26.89312436 | NA | NA | NA | NA | NA | NA | NA |
| TRPA1 | 850 | L | -10.89025 | 25.25522087 | 0.456754466 | 23.63602044 | NA | NA | NA | NA | NA | NA | NA |
| TRPA1 | 851 | Q | -15.34777778 | 24.76031199 | 0.446988571 | 23.65654499 | NA | NA | NA | NA | NA | NA | NA |
| TRPA1 | 852 | R | -15.70909091 | 28.03727467 | 0.511651932 | 26.98320974 | NA | NA | NA | NA | NA | NA | NA |
| TRPA1 | 853 | F | -14.74036364 | 26.06714511 | 0.472775933 | 25.01391821 | NA | NA | NA | NA | NA | NA | NA |
| TRPA1 | 854 | E | -20.05677778 | 23.58983641 | 0.423891914 | 21.80423756 | NA | NA | NA | NA | NA | NA | NA |
| TRPA1 | 855 | N | -16.167375 | 22.34170211 | 0.399262839 | 21.33112004 | NA | NA | NA | NA | NA | NA | NA |

| | | | | | | | | | | | | | |
|---|---|---|---|---|---|---|---|---|---|---|---|---|---|
| TRPA1 | 856 | C | -13.69783333 | 21.15704809 | 0.375886403 | 20.06152908 | NA | NA | NA | NA | NA | NA | NA |
| TRPA1 | 857 | G | -15.78175 | 20.87423623 | 0.370305758 | 19.75292444 | NA | NA | NA | NA | NA | NA | NA |
| TRPA1 | 858 | I | -17.3285 | 17.27729996 | 0.29932845 | 16.05688337 | NA | NA | NA | NA | NA | NA | NA |
| TRPA1 | 859 | F | -13.144 | 16.20362243 | 0.278141881 | 14.51841272 | NA | NA | NA | NA | NA | NA | NA |
| TRPA1 | 860 | I | -12.573 | 19.36773433 | 0.34057839 | 17.54209586 | NA | NA | NA | NA | NA | NA | NA |
| TRPA1 | 861 | V | -16.37585714 | 17.46126908 | 0.30295866 | 16.39483191 | NA | NA | NA | NA | NA | NA | NA |
| TRPA1 | 862 | M | -14.967625 | 13.27302294 | 0.220313205 | 12.17797149 | NA | NA | NA | NA | NA | NA | NA |
| TRPA1 | 863 | L | -10.274375 | 16.07375426 | 0.27557923 | 14.24402626 | NA | NA | NA | NA | NA | NA | NA |
| TRPA1 | 864 | E | -14.01188889 | 17.92880936 | 0.312184498 | 16.10291776 | NA | NA | NA | NA | NA | NA | NA |
| TRPA1 | 865 | V | -15.28128571 | 14.72483266 | 0.248961348 | 13.68945059 | NA | NA | NA | NA | NA | NA | NA |
| TRPA1 | 866 | I | -11.621375 | 11.4214014 | 0.183775691 | 9.737505988 | NA | NA | NA | NA | NA | NA | NA |
| TRPA1 | 867 | L | -8.95725 | 15.98690671 | 0.273865492 | 14.61185143 | NA | NA | NA | NA | NA | NA | NA |
| TRPA1 | 868 | K | -13.26055556 | 18.02649697 | 0.314112139 | 16.34259449 | NA | NA | NA | NA | NA | NA | NA |
| TRPA1 | 869 | T | -12.704 | 14.42153296 | 0.242976423 | 12.75466702 | NA | NA | NA | NA | NA | NA | NA |
| TRPA1 | 870 | L | -7.648625 | 14.6842455 | 0.248160454 | 13.26148268 | NA | NA | NA | NA | NA | NA | NA |
| TRPA1 | 871 | L | -8.939125 | 18.44964455 | 0.322461988 | 16.77247443 | NA | NA | NA | NA | NA | NA | NA |
| TRPA1 | 872 | R | -13.51118182 | 19.04077168 | 0.33412653 | 17.60074614 | NA | NA | NA | NA | NA | NA | NA |
| TRPA1 | 873 | S | -8.497166667 | 16.17478872 | 0.277572913 | 14.58061353 | NA | NA | NA | NA | NA | NA | NA |
| TRPA1 | 874 | T | -5.591428571 | 19.32118203 | 0.339659786 | 18.01418366 | NA | NA | NA | NA | NA | NA | NA |
| TRPA1 | 875 | V | -6.472142857 | 21.52817006 | 0.383209646 | 20.16208523 | NA | NA | NA | NA | NA | NA | NA |
| TRPA1 | 876 | V | -6.286 | 18.91248437 | 0.331595073 | 17.56272326 | NA | NA | NA | NA | NA | NA | NA |
| TRPA1 | 877 | F | -3.856545455 | 14.37278178 | 0.24201443 | 11.09896968 | NA | NA | NA | NA | NA | NA | NA |
| TRPA1 | 878 | I | -1.050625 | 20.57798447 | 0.364459907 | 15.78782839 | NA | NA | NA | NA | NA | NA | NA |
| TRPA1 | 879 | F | -2.770272727 | 22.3287621 | 0.399007498 | 17.65994804 | NA | NA | NA | NA | NA | NA | NA |
| TRPA1 | 880 | L | -1.34375 | 17.3183108 | 0.300137705 | 12.67498983 | NA | NA | NA | NA | NA | NA | NA |
| TRPA1 | 881 | L | 0.968625 | 16.94596332 | 0.29279028 | 13.03671664 | NA | NA | NA | NA | NA | NA | NA |
| TRPA1 | 882 | L | 1.92225 | 21.52812646 | 0.383208785 | 17.8419009 | NA | NA | NA | NA | NA | NA | NA |
| TRPA1 | 883 | A | 1.8432 | 22.36213641 | 0.399666063 | 18.65882521 | NA | NA | NA | NA | NA | NA | NA |
| TRPA1 | 884 | F | 2.222181818 | 18.50284113 | 0.323511701 | 14.89986849 | NA | NA | NA | NA | NA | NA | NA |
| TRPA1 | 885 | G | 6.349 | 20.39351284 | 0.360819781 | 17.24665706 | NA | NA | NA | NA | NA | NA | NA |
| TRPA1 | 886 | L | 6.730625 | 22.86989777 | 0.409685572 | 19.6173942 | NA | NA | NA | NA | NA | NA | NA |
| TRPA1 | 887 | S | 7.260333333 | 22.50574516 | 0.402499853 | 19.33948026 | NA | NA | NA | NA | NA | NA | NA |
| TRPA1 | 888 | F | 8.656454545 | 16.92018706 | 0.292281644 | 13.98890271 | NA | NA | NA | NA | NA | NA | NA |

| | | | | | | | | | | | | | |
|---|---|---|---|---|---|---|---|---|---|---|---|---|---|
| TRPA1 | 889 | Y | 12.63208333 | 21.75729581 | 0.387730918 | 18.30510527 | NA | NA | NA | NA | NA | NA | NA |
| TRPA1 | 890 | I | 11.69 | 24.12168338 | 0.434386697 | 20.32709985 | NA | NA | NA | NA | NA | NA | NA |
| TRPA1 | 891 | L | 12.0045 | 22.06937068 | 0.393889001 | 18.21653557 | NA | NA | NA | NA | NA | NA | NA |
| TRPA1 | 892 | L | 14.335375 | 17.18147575 | 0.297437579 | 12.53209832 | NA | NA | NA | NA | NA | NA | NA |
| TRPA1 | 893 | N | 17.551375 | 21.32496108 | 0.379199781 | 16.50750906 | NA | NA | NA | NA | NA | NA | NA |
| TRPA1 | 894 | L | 20.9155 | 19.34478728 | 0.340125582 | 14.59518598 | NA | NA | NA | NA | NA | NA | NA |
| TRPA1 | 895 | Q | 18.08144444 | 14.79455508 | 0.250337161 | 9.938163322 | NA | NA | NA | NA | NA | NA | NA |
| TRPA1 | 896 | D | 20.0715 | 15.8982828 | 0.272116702 | 11.14162082 | NA | NA | NA | NA | NA | NA | NA |
| TRPA1 | 897 | P | 15.55285714 | 13.33969677 | 0.221628861 | 8.470962133 | NA | NA | NA | NA | NA | NA | NA |
| TRPA1 | 898 | F | 13.55372727 | 15.39881466 | 0.262260841 | 10.63788338 | NA | NA | NA | NA | NA | NA | NA |
| TRPA1 | 899 | S | 17.35233333 | 19.93605573 | 0.351792912 | 15.07727771 | NA | NA | NA | NA | NA | NA | NA |
| TRPA1 | 900 | S | 14.66033333 | 22.00906343 | 0.392698976 | 17.27339763 | NA | NA | NA | NA | NA | NA | NA |
| TRPA1 | 901 | P | 11.27371429 | 22.54727258 | 0.403319302 | 19.13598459 | NA | NA | NA | NA | NA | NA | NA |
| TRPA1 | 902 | L | 9.64925 | 20.88070911 | 0.370433485 | 17.48515346 | NA | NA | NA | NA | NA | NA | NA |
| TRPA1 | 903 | L | 12.763125 | 17.89714117 | 0.311559598 | 14.12585525 | NA | NA | NA | NA | NA | NA | NA |
| TRPA1 | 904 | S | 9.799166667 | 17.98610694 | 0.313315134 | 14.75102734 | NA | NA | NA | NA | NA | NA | NA |
| TRPA1 | 905 | I | 6.34025 | 17.70217526 | 0.307712392 | 14.47329268 | NA | NA | NA | NA | NA | NA | NA |
| TRPA1 | 906 | I | 7.9525 | 15.11673936 | 0.25669473 | 12.14538721 | NA | NA | NA | NA | NA | NA | NA |
| TRPA1 | 907 | Q | 9.580222222 | 11.68496187 | 0.188976454 | 8.751620655 | NA | NA | NA | NA | NA | NA | NA |
| TRPA1 | 908 | T | 5.478142857 | 13.41000391 | 0.223016211 | 10.4100975 | NA | NA | NA | NA | NA | NA | NA |
| TRPA1 | 909 | F | 2.675909091 | 12.30023715 | 0.201117503 | 8.434365235 | NA | NA | NA | NA | NA | NA | NA |
| TRPA1 | 910 | S | 6.0255 | 9.265910155 | 0.141242003 | 5.656018651 | NA | NA | NA | NA | NA | NA | NA |
| TRPA1 | 911 | M | 5.897625 | 9.417037007 | 0.144224145 | 5.826160008 | NA | NA | NA | NA | NA | NA | NA |
| TRPA1 | 912 | M | 0.682125 | 8.690561835 | 0.12988882 | 5.162384366 | NA | NA | NA | NA | NA | NA | NA |
| TRPA1 | 913 | L | 1.130625 | 4.798559055 | 0.05308905 | 0.892311277 | 3.91 | -0.8 | -0.4 | 3.38 | 0.81 | 0 | 0 |
| TRPA1 | 914 | G | 3.71275 | 5.239117292 | 0.061782459 | 1.941685563 | 3.61 | -0.8 | -0.4 | 3.38 | 0.81 | 0 | 0 |
| TRPA1 | 915 | D | 7.55725 | 4.424495112 | 0.045707754 | 2.058244662 | 3.26 | -1.04 | -3.5 | 49.7 | 2.63 | 4 | -4 |
| TRPA1 | 916 | I | 7.346125 | 7.159226983 | 0.099671431 | 4.267518115 | 3.02 | -1.04 | -3.5 | 49.7 | 2.63 | 4 | -4 |
| TRPA1 | 917 | N | 11.72375 | 5.595934953 | 0.068823439 | 2.183237483 | 3.47 | -0.99 | -3.5 | 40.44 | 2.21 | 4 | -4 |
| TRPA1 | 918 | Y | 11.41725 | 9.033419397 | 0.13665433 | 5.564440362 | NA | NA | NA | NA | NA | NA | NA |
| TRPA1 | 919 | R | 14.70754545 | 8.543317681 | 0.126983294 | 3.692739502 | 4.86 | -0.92 | -3.5 | 21.99 | 1.22 | 2 | -2 |
| TRPA1 | 920 | E | 16.993 | 6.410521352 | 0.084897438 | 1.601166021 | 4.89 | -0.99 | -3.5 | 31.29 | 1.55 | 3 | -3 |
| TRPA1 | 921 | S | 15.054 | 10.93548499 | 0.174187242 | 6.067702118 | NA | NA | NA | NA | NA | NA | NA |

| | | | | | | | | | | | | | |
|---|---|---|---|---|---|---|---|---|---|---|---|---|---|
| TRPA1 | 922 | F | 12.659 | 13.33775243 | 0.221590494 | 8.682176194 | NA | NA | NA | NA | NA | NA | NA |
| TRPA1 | 923 | L | 16.288625 | 13.85552099 | 0.231807471 | 9.010499722 | NA | NA | NA | NA | NA | NA | NA |
| TRPA1 | 924 | E | 19.85866667 | 10.05820859 | 0.1568762 | 5.370331302 | NA | NA | NA | NA | NA | NA | NA |
| TRPA1 | 925 | P | 19.53828571 | 14.16754107 | 0.237964474 | 9.384553972 | NA | NA | NA | NA | NA | NA | NA |
| TRPA1 | 926 | Y | 17.7845 | 17.47317959 | 0.303193687 | 12.68098827 | NA | NA | NA | NA | NA | NA | NA |
| TRPA1 | 927 | L | 21.0235 | 17.06668717 | 0.295172489 | 12.30503295 | NA | NA | NA | NA | NA | NA | NA |
| TRPA1 | 928 | R | 24.14254545 | 14.61275973 | 0.246749845 | 9.944660011 | NA | NA | NA | NA | NA | NA | NA |
| TRPA1 | 929 | N | 23.510875 | 21.35987811 | 0.379888789 | 16.63451455 | NA | NA | NA | NA | NA | NA | NA |
| TRPA1 | 930 | E | 22.91344444 | 17.6465215 | 0.306614193 | 12.96057537 | NA | NA | NA | NA | NA | NA | NA |
| TRPA1 | 931 | L | 17.97475 | 18.90484396 | 0.331444307 | 14.05570611 | NA | NA | NA | NA | NA | NA | NA |
| TRPA1 | 932 | A | 17.5178 | 23.45460198 | 0.421223372 | 18.63242623 | NA | NA | NA | NA | NA | NA | NA |
| TRPA1 | 933 | H | 14.6645 | 23.88648463 | 0.429745588 | 19.02565035 | NA | NA | NA | NA | NA | NA | NA |
| TRPA1 | 934 | P | 14.42928571 | 22.52257687 | 0.402831989 | 17.97858333 | NA | NA | NA | NA | NA | NA | NA |
| TRPA1 | 935 | V | 10.75528571 | 23.94159496 | 0.430833064 | 20.61707497 | NA | NA | NA | NA | NA | NA | NA |
| TRPA1 | 936 | L | 9.229625 | 22.91609899 | 0.410597247 | 19.90082126 | NA | NA | NA | NA | NA | NA | NA |
| TRPA1 | 937 | S | 10.63216667 | 19.52975742 | 0.343775545 | 16.08287999 | NA | NA | NA | NA | NA | NA | NA |
| TRPA1 | 938 | F | 10.40118182 | 17.94736053 | 0.312550562 | 15.19062173 | NA | NA | NA | NA | NA | NA | NA |
| TRPA1 | 939 | A | 5.6124 | 20.67318928 | 0.366338556 | 17.50278071 | NA | NA | NA | NA | NA | NA | NA |
| TRPA1 | 940 | Q | 5.412555556 | 18.2499057 | 0.318520599 | 15.05343648 | NA | NA | NA | NA | NA | NA | NA |
| TRPA1 | 941 | L | 6.424875 | 14.10799954 | 0.236789558 | 10.88588166 | NA | NA | NA | NA | NA | NA | NA |
| TRPA1 | 942 | V | 2.888142857 | 16.72988302 | 0.28852643 | 13.07913358 | NA | NA | NA | NA | NA | NA | NA |
| TRPA1 | 943 | S | 0.846 | 17.04718208 | 0.2947876 | 13.18164881 | NA | NA | NA | NA | NA | NA | NA |
| TRPA1 | 944 | F | 2.452 | 12.63175229 | 0.207659196 | 9.025087206 | NA | NA | NA | NA | NA | NA | NA |
| TRPA1 | 945 | T | 1.371142857 | 11.86194984 | 0.192468906 | 7.981904997 | NA | NA | NA | NA | NA | NA | NA |
| TRPA1 | 946 | I | -0.88975 | 13.640677 | 0.227568017 | 9.3858104 | NA | NA | NA | NA | NA | NA | NA |
| TRPA1 | 947 | F | -2.223909091 | 13.34203736 | 0.221675047 | 8.589642407 | NA | NA | NA | NA | NA | NA | NA |
| TRPA1 | 948 | V | -0.442714286 | 10.78521697 | 0.171222046 | 6.079909892 | NA | NA | NA | NA | NA | NA | NA |
| TRPA1 | 949 | P | -0.467857143 | 7.632010155 | 0.109000725 | 3.016114611 | 4.67 | 0.24 | 1.56 | 2.08 | -0.25 | 0 | 0 |
| TRPA1 | 950 | I | -3.180375 | 9.048229551 | 0.136946574 | 5.898829258 | NA | NA | NA | NA | NA | NA | NA |
| TRPA1 | 951 | V | -6.178571429 | 10.6646872 | 0.168843667 | 9.34105001 | NA | NA | NA | NA | NA | NA | NA |
| TRPA1 | 952 | L | -5.226 | 10.27900384 | 0.161233089 | 8.900205753 | NA | NA | NA | NA | NA | NA | NA |
| TRPA1 | 953 | M | -4.43175 | 6.720869735 | 0.091021453 | 5.458313931 | 1.37 | 1.81 | 4.5 | 0.13 | -1.85 | 0 | 0 |
| TRPA1 | 954 | N | -7.95425 | 5.648681351 | 0.069864269 | 4.160234346 | 1.62 | 1.81 | 4.5 | 0.13 | -1.85 | 0 | 0 |

| | | | | | | | | | | | | | |
|---|---|---|---|---|---|---|---|---|---|---|---|---|---|
| TRPA1 | 955 | L | -10.32275 | 8.85901648 | 0.133212887 | 7.498124825 | NA | NA | NA | NA | NA | NA | NA |
| TRPA1 | 956 | L | -8.4265 | 7.865824114 | 0.113614508 | 6.41674314 | NA | NA | NA | NA | NA | NA | NA |
| TRPA1 | 957 | I | -9.774125 | 2.946654035 | 0.016545942 | 1.527256196 | 1.42 | 1.81 | 4.5 | 0.13 | -1.85 | 0 | 0 |
| TRPA1 | 958 | G | -13.7935 | 5.848535714 | 0.073807937 | 4.410470132 | 1.6 | 1.27 | 4.26 | 0.13 | -1.41 | 0 | 0 |
| TRPA1 | 959 | L | -13.938375 | 8.014643972 | 0.116551128 | 6.569504219 | NA | NA | NA | NA | NA | NA | NA |
| TRPA1 | 960 | A | -13.4842 | 7.278102088 | 0.102017159 | 5.756062998 | NA | NA | NA | NA | NA | NA | NA |
| TRPA1 | 961 | V | -15.73028571 | 2.922356925 | 0.016066494 | 1.98068917 | 1.07 | 1.13 | 4.2 | 0.13 | -1.3 | 0 | 0 |
| TRPA1 | 962 | G | -18.51425 | 6.259932108 | 0.081925904 | 4.604991213 | 1.81 | 1.13 | 4.2 | 0.13 | -1.3 | 0 | 0 |
| TRPA1 | 963 | D | -17.42 | 8.918152555 | 0.134379802 | 7.150372756 | NA | NA | NA | NA | NA | NA | NA |
| TRPA1 | 964 | I | -16.819 | 8.98737075 | 0.135745665 | 7.727326209 | NA | NA | NA | NA | NA | NA | NA |
| TRPA1 | 965 | A | -21.0296 | 6.070045964 | 0.078178935 | 1.672016962 | 4.41 | 0.69 | 3.24 | 0.08 | -0.54 | 0 | 0 |
| TRPA1 | 966 | E | -22.80477778 | 8.995944586 | 0.13591485 | 4.767390648 | 4.26 | 0.91 | 3.72 | 0.1 | -0.92 | 0 | 0 |
| TRPA1 | 967 | V | -20.86071429 | 11.64073099 | 0.188103658 | 7.233290666 | NA | NA | NA | NA | NA | NA | NA |
| TRPA1 | 968 | Q | -20.98355556 | 10.58515229 | 0.167274228 | 6.471509863 | NA | NA | NA | NA | NA | NA | NA |
| TRPA1 | 969 | K | -26.98366667 | 8.596537559 | 0.128033466 | 3.757524955 | 5.96 | 0.02 | 1.8 | 0 | 0.59 | 0 | 0 |
| TRPA1 | 970 | H | -27.1963 | 12.67470923 | 0.208506853 | 5.70351194 | NA | NA | NA | NA | NA | NA | NA |
| TRPA1 | 971 | A | -24.0696 | 15.5640603 | 0.265521585 | 10.96559613 | NA | NA | NA | NA | NA | NA | NA |
| TRPA1 | 972 | S | -26.71883333 | 17.68913149 | 0.307455003 | 10.67085434 | NA | NA | NA | NA | NA | NA | NA |
| TRPA1 | 973 | L | -28.728125 | 19.03137646 | 0.333941137 | 11.98836907 | NA | NA | NA | NA | NA | NA | NA |
| TRPA1 | 974 | K | -23.79455556 | 19.69226777 | 0.346982315 | 13.76507675 | NA | NA | NA | NA | NA | NA | NA |
| TRPA1 | 975 | R | -22.88972727 | 20.11536791 | 0.355331228 | 15.58850001 | NA | NA | NA | NA | NA | NA | NA |
| TRPA1 | 976 | I | -27.051625 | 23.96921594 | 0.431378101 | 16.94238174 | NA | NA | NA | NA | NA | NA | NA |
| TRPA1 | 977 | A | -26.4384 | 24.10021792 | 0.433963125 | 17.05421242 | NA | NA | NA | NA | NA | NA | NA |
| TRPA1 | 978 | M | -22.28875 | 23.02288596 | 0.412704444 | 18.53667335 | NA | NA | NA | NA | NA | NA | NA |
| TRPA1 | 979 | Q | -23.28944444 | 27.8528852 | 0.508013428 | 23.1924276 | NA | NA | NA | NA | NA | NA | NA |
| TRPA1 | 980 | V | -27.74471429 | 28.55011275 | 0.521771618 | 21.57919579 | NA | NA | NA | NA | NA | NA | NA |
| TRPA1 | 981 | E | -25.37033333 | 29.66013987 | 0.543675464 | 22.63320227 | NA | NA | NA | NA | NA | NA | NA |
| TRPA1 | 982 | L | -21.909375 | 31.59747878 | 0.581904414 | 27.36136462 | NA | NA | NA | NA | NA | NA | NA |
| TRPA1 | 983 | H | -24.9186 | 33.64580042 | 0.622323355 | 26.62086553 | NA | NA | NA | NA | NA | NA | NA |
| TRPA1 | 984 | T | -27.652 | 33.54280683 | 0.620291012 | 26.55463216 | NA | NA | NA | NA | NA | NA | NA |
| TRPA1 | 985 | S | -23.85666667 | 35.38423334 | 0.656627351 | 29.46690296 | NA | NA | NA | NA | NA | NA | NA |
| TRPA1 | 986 | L | -22.324875 | 37.46742298 | 0.697734333 | 33.2196976 | NA | NA | NA | NA | NA | NA | NA |
| TRPA1 | 987 | E | -27.65911111 | 38.97988348 | 0.72757928 | 31.98472077 | NA | NA | NA | NA | NA | NA | NA |

| | | | | | | | | | | | | | |
|---|---|---|---|---|---|---|---|---|---|---|---|---|---|
| TRPA1 | 988 | K | -27.18955556 | 37.38931952 | 0.696193139 | 30.37690474 | NA | NA | NA | NA | NA | NA | NA |
| TRPA1 | 989 | K | -22.10744444 | 40.63615015 | 0.760261913 | 36.17641108 | NA | NA | NA | NA | NA | NA | NA |
| TRPA1 | 990 | L | -24.083 | 43.71739316 | 0.821063194 | 39.05753771 | NA | NA | NA | NA | NA | NA | NA |
| TRPA1 | 991 | P | -25.41214286 | 48.14202284 | 0.908373137 | 41.10925872 | NA | NA | NA | NA | NA | NA | NA |
| TRPA1 | 992 | L | -30.756875 | 49.31041306 | 0.931428645 | 42.05383377 | NA | NA | NA | NA | NA | NA | NA |
| TRPA1 | 993 | W | -26.44485714 | 51.16297437 | 0.967984703 | 44.00309504 | NA | NA | NA | NA | NA | NA | NA |
| TRPA1 | 994 | F | -24.93063636 | 46.45440518 | 0.875071864 | 39.51596203 | NA | NA | NA | NA | NA | NA | NA |
| TRPA1 | 995 | L | -29.09925 | 43.90189465 | 0.824703909 | 36.74191929 | NA | NA | NA | NA | NA | NA | NA |
| TRPA1 | 996 | R | -33.32672727 | 48.62710161 | 0.917945057 | 41.42231809 | NA | NA | NA | NA | NA | NA | NA |
| TRPA1 | 997 | K | -27.41155556 | 49.53285303 | 0.935817988 | 42.37679196 | NA | NA | NA | NA | NA | NA | NA |
| TRPA1 | 998 | V | -27.39957143 | 45.57637556 | 0.857745958 | 38.59275938 | NA | NA | NA | NA | NA | NA | NA |
| TRPA1 | 999 | D | -31.64675 | 44.58915609 | 0.838265441 | 37.34926381 | NA | NA | NA | NA | NA | NA | NA |
| TRPA1 | 1000 | Q | -32.11077778 | 43.22144619 | 0.811276815 | 36.04398536 | NA | NA | NA | NA | NA | NA | NA |
| TRPA1 | 1001 | K | -37.51233333 | 41.92293851 | 0.785653737 | 39.72730379 | NA | NA | NA | NA | NA | NA | NA |
| TRPA1 | 1002 | S | -36.36933333 | 42.73586941 | 0.801695068 | 40.50491861 | NA | NA | NA | NA | NA | NA | NA |
| TRPA1 | 1003 | T | -33.86614286 | 39.81613795 | 0.744080848 | 32.99992379 | NA | NA | NA | NA | NA | NA | NA |
| TRPA1 | 1004 | I | -34.005125 | 42.95249299 | 0.805969639 | 36.1386113 | NA | NA | NA | NA | NA | NA | NA |
| TRPA1 | 1005 | V | -32.25442857 | 39.61517424 | 0.740115289 | 32.47394691 | NA | NA | NA | NA | NA | NA | NA |
| TRPA1 | 1006 | Y | -31.793 | 42.40086123 | 0.795084448 | 35.26840748 | NA | NA | NA | NA | NA | NA | NA |
| TRPA1 | 1007 | P | -31.22228571 | 39.80843988 | 0.743928945 | 32.66454891 | NA | NA | NA | NA | NA | NA | NA |
| TRPA1 | 1008 | N | -29.4 | 42.67870379 | 0.800567036 | 35.81316912 | NA | NA | NA | NA | NA | NA | NA |
| TRPA1 | 1009 | K | -28.02888889 | 43.28239713 | 0.812479542 | 36.30472522 | NA | NA | NA | NA | NA | NA | NA |
| TRPA1 | 1010 | P | -23.85557143 | 43.23847272 | 0.811612795 | 38.57996912 | NA | NA | NA | NA | NA | NA | NA |
| TRPA1 | 1011 | R | -22.0666 | 41.36201881 | 0.77458527 | 37.10295039 | NA | NA | NA | NA | NA | NA | NA |
| TRPA1 | 1012 | S | -19.161 | 43.41978957 | 0.815190668 | 39.10137872 | NA | NA | NA | NA | NA | NA | NA |
| TRPA1 | 1015 | M | -12.3418 | 39.9590857 | 0.746901595 | 38.12992461 | NA | NA | NA | NA | NA | NA | NA |
| TRPA1 | 1016 | L | -12.177 | 38.5996807 | 0.720076848 | 36.77074091 | NA | NA | NA | NA | NA | NA | NA |
| TRPA1 | 1017 | F | -15.4592 | 37.28870502 | 0.694207742 | 36.16716872 | NA | NA | NA | NA | NA | NA | NA |
| TRPA1 | 1018 | H | -14.3284 | 35.64076371 | 0.661689391 | 34.53831065 | NA | NA | NA | NA | NA | NA | NA |
| TRPA1 | 1019 | I | -11.7374 | 33.95272815 | 0.628379872 | 32.12351775 | NA | NA | NA | NA | NA | NA | NA |
| TRPA1 | 1020 | F | -14.0588 | 32.90937139 | 0.607791613 | 31.27090106 | NA | NA | NA | NA | NA | NA | NA |
| TRPA1 | 1021 | C | -16.1782 | 31.47322344 | 0.57945252 | 30.41093097 | NA | NA | NA | NA | NA | NA | NA |
| TRPA1 | 1022 | F | -13.4118 | 29.64217834 | 0.543321034 | 27.95706899 | NA | NA | NA | NA | NA | NA | NA |

| | | | | | | | | | | | | | |
|---|---|---|---|---|---|---|---|---|---|---|---|---|---|
| TRPA1 | 1023 | L | -12.5334 | 28.4432755 | 0.519663429 | 26.61591793 | NA | NA | NA | NA | NA | NA | NA |
| TRPA1 | 1024 | F | -16.1476 | 28.0646352 | 0.51219183 | 26.97915233 | NA | NA | NA | NA | NA | NA | NA |
| TRPA1 | 1025 | C | -16.1674 | 25.34334086 | 0.458493312 | 24.28969319 | NA | NA | NA | NA | NA | NA | NA |
| TRPA1 | 1039 | S | -40.79083333 | 8.232383009 | 0.120847709 | 6.840569207 | NA | NA | NA | NA | NA | NA | NA |
| TRPA1 | 1040 | L | -42.143375 | 3.432939848 | 0.02614168 | 1.965353276 | 1.72 | 1.27 | 3.94 | 0.13 | -1.8 | 0 | 0 |
| TRPA1 | 1041 | E | -42.65666667 | 6.888321203 | 0.094325725 | 5.422912184 | 1.48 | 1.54 | 4.22 | 0.13 | -1.83 | 0 | 0 |
| TRPA1 | 1042 | M | -45.355 | 9.261101122 | 0.141147107 | 7.982454196 | NA | NA | NA | NA | NA | NA | NA |
| TRPA1 | 1043 | E | -47.76 | 6.973645532 | 0.096009405 | 5.232411311 | 1.9 | 1.81 | 4.5 | 0.13 | -1.85 | 0 | 0 |
| TRPA1 | 1044 | I | -47.54575 | 2.890133907 | 0.015430646 | 1.810328692 | 2.14 | 1.81 | 4.5 | 0.13 | -1.85 | 0 | 0 |
| TRPA1 | 1045 | L | -48.96825 | 7.842193124 | 0.113148205 | 4.932835125 | 2.92 | 0.65 | 1.3 | 1.49 | -1.06 | 0 | 0 |
| TRPA1 | 1046 | K | -51.47911111 | 8.465140341 | 0.125440643 | 6.589223794 | NA | NA | NA | NA | NA | NA | NA |
| TRPA1 | 1047 | Q | -52.82555556 | 2.928196202 | 0.016181719 | 1.733629718 | 1.73 | -1.1 | -3.5 | 3.53 | 0.13 | 0 | 0 |
| TRPA1 | 1048 | K | -53.04166667 | 6.778905 | 0.092166646 | 5.152250699 | 1.69 | -1.1 | -3.5 | 3.53 | 0.13 | 0 | 0 |
| TRPA1 | 1049 | Y | -53.92258333 | 9.103038284 | 0.138028099 | 7.272446533 | NA | NA | NA | NA | NA | NA | NA |
| TRPA1 | 1050 | R | -57.56818182 | 7.0048995 | 0.096626131 | 5.627017528 | 1.53 | 1.14 | 3.8 | 0.13 | -1.79 | 0 | 0 |
| TRPA1 | 1051 | L | -58.233625 | 2.881307516 | 0.015256478 | 1.742332977 | 1.27 | 1.14 | 3.8 | 0.13 | -1.79 | 0 | 0 |
| TRPA1 | 1052 | K | -58.66377778 | 7.762229126 | 0.111570298 | 6.469410625 | NA | NA | NA | NA | NA | NA | NA |
| TRPA1 | 1053 | D | -62.259625 | 8.590321356 | 0.127910804 | 7.69829466 | NA | NA | NA | NA | NA | NA | NA |
| TRPA1 | 1054 | L | -63.111375 | 2.64746728 | 0.010642176 | 1.527176641 | 1.57 | 1.14 | 3.8 | 0.13 | -1.79 | 0 | 0 |
| TRPA1 | 1055 | T | -63.711 | 6.553665997 | 0.08772207 | 4.287144113 | 2.42 | 1.14 | 3.8 | 0.13 | -1.79 | 0 | 0 |
| TRPA1 | 1056 | F | -66.31127273 | 8.841947636 | 0.132876072 | 6.384001802 | NA | NA | NA | NA | NA | NA | NA |
| TRPA1 | 1057 | L | -68.2765 | 7.013389623 | 0.096793664 | 5.468675462 | 1.73 | 1.14 | 3.8 | 0.13 | -1.79 | 0 | 0 |
| TRPA1 | 1058 | L | -68.59375 | 2.684738162 | 0.011377631 | 1.302465845 | 1.48 | 1.14 | 3.8 | 0.13 | -1.79 | 0 | 0 |
| TRPA1 | 1059 | E | -69.769 | 7.634716039 | 0.109054119 | 6.319075699 | NA | NA | NA | NA | NA | NA | NA |
| TRPA1 | 1060 | K | -71.89644444 | 8.637426584 | 0.128840317 | 7.964848894 | NA | NA | NA | NA | NA | NA | NA |
| TRPA1 | 1061 | Q | -73.27266667 | 2.108150848 | 0 | 1.453744163 | 0.72 | -1.1 | -3.5 | 3.53 | 0.13 | 0 | 0 |
| TRPA1 | 1062 | H | -73.9781 | 6.429966485 | 0.085281143 | 5.550216754 | 0.96 | 0.06 | -0.3 | 2.17 | -0.66 | 0 | 0 |
| TRPA1 | 1063 | E | -76.41144444 | 8.659172997 | 0.129269433 | 7.763466207 | NA | NA | NA | NA | NA | NA | NA |
| TRPA1 | 1064 | L | -78.236 | 6.769419768 | 0.091979477 | 4.489587158 | 2.29 | 1.81 | 4.5 | 0.13 | -1.85 | 0 | 0 |
| TRPA1 | 1065 | I | -78.5675 | 2.640614512 | 0.010506952 | 1.639243584 | 2.29 | 1.81 | 4.5 | 0.13 | -1.85 | 0 | 0 |
| TRPA1 | 1066 | K | -78.945 | 7.600832915 | 0.108385513 | 5.323025312 | NA | NA | NA | NA | NA | NA | NA |
| TRPA1 | 1067 | L | -82.912375 | 8.244021834 | 0.121077375 | 6.367636086 | NA | NA | NA | NA | NA | NA | NA |
| TRPA1 | 1068 | I | -83.623625 | 3.392960065 | 0.02535277 | 1.310031754 | 2.11 | 1.81 | 4.5 | 0.13 | -1.85 | 0 | 0 |

| | | | | | | | | | | | | | |
|---|---|---|---|---|---|---|---|---|---|---|---|---|---|
| TRPA1 | 1069 | I | -83.97925 | 6.677128874 | 0.090158327 | 4.574043774 | 2.11 | 1.81 | 4.5 | 0.13 | -1.85 | 0 | 0 |
| TRPA1 | 1070 | Q | -85.65222222 | 9.17063591 | 0.139361984 | 6.687423758 | NA | NA | NA | NA | NA | NA | NA |
| TRPA1 | 1071 | K | -88.49177778 | 7.234332105 | 0.101153458 | 4.803462426 | 2.64 | 1 | 1.9 | 1.43 | -0.72 | 0 | 0 |
| TRPA1 | 1072 | M | -88.984 | 4.25831105 | 0.042428492 | 1.539687959 | 2.79 | 1 | 1.9 | 1.43 | -0.72 | 0 | 0 |
| TRPA1 | 1073 | E | -94.31655556 | 4.48832452 | 0.046967282 | 1.721594886 | 2.78 | -0.91 | -1.43 | 18.89 | 1.28 | 1 | -1 |
| TRPA1 | 1074 | I | -92.938375 | 6.545228644 | 0.087555578 | 3.79316101 | 2.78 | -0.91 | -1.43 | 18.89 | 1.28 | 1 | -1 |
| TRPA1 | 1075 | I | -97.2645 | 7.46721213 | 0.105748813 | 4.776138211 | 2.77 | -0.91 | -1.43 | 18.89 | 1.28 | 1 | -1 |
| TRPA1 | 1076 | S | -96.7585 | 10.2462113 | 0.160586003 | 7.561025351 | NA | NA | NA | NA | NA | NA | NA |
| TRPA1 | 1077 | E | -92.81566667 | 10.21702941 | 0.160010165 | 7.460134451 | NA | NA | NA | NA | NA | NA | NA |
| TRPA1 | 1078 | T | -94.90071429 | 12.21970986 | 0.199528481 | 9.489218305 | NA | NA | NA | NA | NA | NA | NA |
| TRPA1 | 1079 | E | -93.9156 | 15.84979549 | 0.271159916 | 13.10027392 | NA | NA | NA | NA | NA | NA | NA |
| TRPC6 | 85 | A | -59.0282 | 40.30822797 | 0.637457816 | 38.46272452 | NA | NA | NA | NA | NA | NA | NA |
| TRPC6 | 86 | Y | -57.70908333 | 36.93882225 | 0.580488071 | 35.08954763 | NA | NA | NA | NA | NA | NA | NA |
| TRPC6 | 87 | M | -61.024375 | 33.83066591 | 0.527935517 | 32.13648959 | NA | NA | NA | NA | NA | NA | NA |
| TRPC6 | 88 | F | -59.90245455 | 30.38572207 | 0.469688577 | 28.58729689 | NA | NA | NA | NA | NA | NA | NA |
| TRPC6 | 89 | S | -63.48183333 | 34.19700678 | 0.53412959 | 33.08544186 | NA | NA | NA | NA | NA | NA | NA |
| TRPC6 | 90 | D | -64.659 | 31.920623 | 0.495640606 | 30.79529206 | NA | NA | NA | NA | NA | NA | NA |
| TRPC6 | 91 | R | -69.85145455 | 32.76859762 | 0.509978119 | 31.09411567 | NA | NA | NA | NA | NA | NA | NA |
| TRPC6 | 92 | S | -67.5656 | 36.32698249 | 0.570143114 | 34.88737074 | NA | NA | NA | NA | NA | NA | NA |
| TRPC6 | 93 | T | -70.843 | 35.74699021 | 0.560336633 | 33.76755253 | NA | NA | NA | NA | NA | NA | NA |
| TRPC6 | 94 | S | -73.89716667 | 34.89253711 | 0.545889583 | 33.14339776 | NA | NA | NA | NA | NA | NA | NA |
| TRPC6 | 95 | L | -75.4515 | 29.44904852 | 0.453851347 | 27.02336907 | NA | NA | NA | NA | NA | NA | NA |
| TRPC6 | 96 | S | -79.7415 | 30.61402951 | 0.473548788 | 28.67533281 | NA | NA | NA | NA | NA | NA | NA |
| TRPC6 | 97 | I | -82.14125 | 28.29176877 | 0.434284118 | 26.4108834 | NA | NA | NA | NA | NA | NA | NA |
| TRPC6 | 98 | E | -82.8124 | 28.23276726 | 0.433286523 | 26.45579803 | NA | NA | NA | NA | NA | NA | NA |
| TRPC6 | 99 | E | -79.22611111 | 26.97375421 | 0.411999193 | 25.17587832 | NA | NA | NA | NA | NA | NA | NA |
| TRPC6 | 100 | E | -77.901 | 24.46090264 | 0.369512022 | 22.30926607 | NA | NA | NA | NA | NA | NA | NA |
| TRPC6 | 101 | R | -82.69227273 | 22.33217119 | 0.333519536 | 20.51673233 | NA | NA | NA | NA | NA | NA | NA |
| TRPC6 | 102 | F | -79.96654545 | 22.75483951 | 0.340665991 | 20.94489417 | NA | NA | NA | NA | NA | NA | NA |
| TRPC6 | 103 | L | -75.62425 | 21.65767783 | 0.322115236 | 19.25100595 | NA | NA | NA | NA | NA | NA | NA |
| TRPC6 | 104 | D | -76.454625 | 18.73841832 | 0.27275654 | 16.24078782 | NA | NA | NA | NA | NA | NA | NA |
| TRPC6 | 105 | A | -78.93 | 17.85708425 | 0.257854987 | 15.92315957 | NA | NA | NA | NA | NA | NA | NA |
| TRPC6 | 106 | A | -76.4302 | 18.52610456 | 0.26916675 | 16.46054443 | NA | NA | NA | NA | NA | NA | NA |

| | | | | | | | | | | | | | |
|---|---|---|---|---|---|---|---|---|---|---|---|---|---|
| TRPC6 | 107 | E | -72.55966667 | 16.22487242 | 0.23025763 | 14.45560518 | NA | NA | NA | NA | NA | NA | NA |
| TRPC6 | 108 | Y | -73.99783333 | 13.49755626 | 0.184144304 | 11.45587444 | NA | NA | NA | NA | NA | NA | NA |
| TRPC6 | 109 | G | -78.1855 | 13.41263889 | 0.182708525 | 11.48568745 | NA | NA | NA | NA | NA | NA | NA |
| TRPC6 | 110 | N | -81.712625 | 15.03792293 | 0.210188747 | 13.29322353 | NA | NA | NA | NA | NA | NA | NA |
| TRPC6 | 111 | I | -82.6865 | 13.29364453 | 0.180696574 | 11.41113409 | NA | NA | NA | NA | NA | NA | NA |
| TRPC6 | 112 | P | -86.19157143 | 13.51921666 | 0.184510536 | 11.55156478 | NA | NA | NA | NA | NA | NA | NA |
| TRPC6 | 113 | V | -84.98285714 | 17.35793288 | 0.249415361 | 15.40071659 | NA | NA | NA | NA | NA | NA | NA |
| TRPC6 | 114 | V | -83.25542857 | 19.41003993 | 0.284112285 | 17.57945218 | NA | NA | NA | NA | NA | NA | NA |
| TRPC6 | 115 | R | -87.37481818 | 14.15387452 | 0.1952413 | 12.01781411 | NA | NA | NA | NA | NA | NA | NA |
| TRPC6 | 116 | K | -88.99377778 | 20.6340963 | 0.30480857 | 17.0586348 | NA | NA | NA | NA | NA | NA | NA |
| TRPC6 | 117 | M | -85.69625 | 22.96536588 | 0.344225561 | 20.83047429 | NA | NA | NA | NA | NA | NA | NA |
| TRPC6 | 118 | L | -85.247 | 24.51483753 | 0.370423951 | 22.58210209 | NA | NA | NA | NA | NA | NA | NA |
| TRPC6 | 119 | E | -90.0302 | 24.8625549 | 0.376303139 | 21.25325819 | NA | NA | NA | NA | NA | NA | NA |
| TRPC6 | 120 | E | -91.2222 | 25.27862126 | 0.383337969 | 21.63560238 | NA | NA | NA | NA | NA | NA | NA |
| TRPC6 | 121 | C | -88.46 | 27.6593236 | 0.423590766 | 24.08763059 | NA | NA | NA | NA | NA | NA | NA |
| TRPC6 | 122 | H | -90.4756 | 30.99639295 | 0.48001377 | 27.69513297 | NA | NA | NA | NA | NA | NA | NA |
| TRPC6 | 123 | S | -86.3594 | 33.32062288 | 0.519311734 | 31.19875962 | NA | NA | NA | NA | NA | NA | NA |
| TRPC6 | 124 | L | -83.504625 | 28.13548409 | 0.431641664 | 26.31922074 | NA | NA | NA | NA | NA | NA | NA |
| TRPC6 | 125 | N | -79.508625 | 30.760058 | 0.47601783 | 28.76949334 | NA | NA | NA | NA | NA | NA | NA |
| TRPC6 | 126 | V | -79.15685714 | 29.0639037 | 0.447339337 | 27.21177382 | NA | NA | NA | NA | NA | NA | NA |
| TRPC6 | 127 | N | -75.778125 | 31.36289835 | 0.486210625 | 28.89970662 | NA | NA | NA | NA | NA | NA | NA |
| TRPC6 | 128 | C | -75.36633333 | 29.29102567 | 0.451179504 | 27.41643036 | NA | NA | NA | NA | NA | NA | NA |
| TRPC6 | 129 | V | -71.45414286 | 29.71444884 | 0.458338722 | 27.8484951 | NA | NA | NA | NA | NA | NA | NA |
| TRPC6 | 130 | D | -69.548625 | 25.24750946 | 0.382811932 | 23.34858686 | NA | NA | NA | NA | NA | NA | NA |
| TRPC6 | 131 | Y | -66.06008333 | 23.89626356 | 0.359965133 | 22.02725017 | NA | NA | NA | NA | NA | NA | NA |
| TRPC6 | 132 | M | -64.07025 | 26.40086249 | 0.402312768 | 25.30737349 | NA | NA | NA | NA | NA | NA | NA |
| TRPC6 | 133 | G | -67.16425 | 29.047517 | 0.447062271 | 27.61148244 | NA | NA | NA | NA | NA | NA | NA |
| TRPC6 | 134 | Q | -67.63444444 | 24.51049271 | 0.370350489 | 22.61988969 | NA | NA | NA | NA | NA | NA | NA |
| TRPC6 | 135 | N | -72.09475 | 25.06161633 | 0.37966886 | 23.20582532 | NA | NA | NA | NA | NA | NA | NA |
| TRPC6 | 136 | A | -74.6236 | 22.81542658 | 0.341690394 | 20.73835433 | NA | NA | NA | NA | NA | NA | NA |
| TRPC6 | 137 | L | -73.062875 | 22.95835162 | 0.344106964 | 21.25920732 | NA | NA | NA | NA | NA | NA | NA |
| TRPC6 | 138 | Q | -68.876 | 22.20853847 | 0.33142916 | 20.33626764 | NA | NA | NA | NA | NA | NA | NA |
| TRPC6 | 139 | L | -71.09075 | 19.02532862 | 0.277607605 | 17.16103142 | NA | NA | NA | NA | NA | NA | NA |

| | | | | | | | | | | | | | |
|---|---|---|---|---|---|---|---|---|---|---|---|---|---|
| TRPC6 | 140 | A | -72.791 | 18.14356225 | 0.262698743 | 16.41911126 | NA | NA | NA | NA | NA | NA | NA |
| TRPC6 | 141 | V | -69.80228571 | 18.28154044 | 0.265031672 | 16.41791943 | NA | NA | NA | NA | NA | NA | NA |
| TRPC6 | 142 | A | -67.5892 | 16.29084194 | 0.23137304 | 14.22042089 | NA | NA | NA | NA | NA | NA | NA |
| TRPC6 | 143 | N | -69.91325 | 13.44654491 | 0.183281806 | 11.57309326 | NA | NA | NA | NA | NA | NA | NA |
| TRPC6 | 144 | E | -71.30222222 | 12.15129791 | 0.161381834 | 10.3089043 | NA | NA | NA | NA | NA | NA | NA |
| TRPC6 | 145 | H | -74.5392 | 13.22643164 | 0.179560142 | 11.14339329 | NA | NA | NA | NA | NA | NA | NA |
| TRPC6 | 146 | L | -76.125125 | 16.12874725 | 0.228632351 | 13.64931473 | NA | NA | NA | NA | NA | NA | NA |
| TRPC6 | 147 | E | -79.32977778 | 13.39783673 | 0.182458251 | 11.4586624 | NA | NA | NA | NA | NA | NA | NA |
| TRPC6 | 148 | I | -78.51675 | 16.9631508 | 0.242740405 | 15.03185579 | NA | NA | NA | NA | NA | NA | NA |
| TRPC6 | 149 | T | -76.21657143 | 20.05846445 | 0.295075815 | 17.57222905 | NA | NA | NA | NA | NA | NA | NA |
| TRPC6 | 150 | E | -79.479 | 19.99032599 | 0.293923733 | 18.07484309 | NA | NA | NA | NA | NA | NA | NA |
| TRPC6 | 151 | L | -82.51125 | 19.91162088 | 0.292592991 | 18.11665014 | NA | NA | NA | NA | NA | NA | NA |
| TRPC6 | 152 | L | -79.113 | 23.34376784 | 0.350623562 | 21.62491673 | NA | NA | NA | NA | NA | NA | NA |
| TRPC6 | 153 | L | -77.80225 | 25.50206084 | 0.387115874 | 23.74237586 | NA | NA | NA | NA | NA | NA | NA |
| TRPC6 | 154 | K | -83.35355556 | 23.74386255 | 0.357388344 | 22.15425449 | NA | NA | NA | NA | NA | NA | NA |
| TRPC6 | 155 | K | -82.727 | 28.70723292 | 0.441308765 | 26.91156511 | NA | NA | NA | NA | NA | NA | NA |
| TRPC6 | 156 | E | -80.8564 | 32.08876794 | 0.498483592 | 30.00124848 | NA | NA | NA | NA | NA | NA | NA |
| TRPC6 | 157 | N | -79.515125 | 33.51364179 | 0.522575289 | 31.58973559 | NA | NA | NA | NA | NA | NA | NA |
| TRPC6 | 158 | L | -76.37625 | 28.30734027 | 0.4345474 | 25.80756428 | NA | NA | NA | NA | NA | NA | NA |
| TRPC6 | 159 | S | -72.8255 | 30.62401512 | 0.473717624 | 28.86998724 | NA | NA | NA | NA | NA | NA | NA |
| TRPC6 | 160 | R | -69.23790909 | 28.86378738 | 0.44395578 | 26.96905412 | NA | NA | NA | NA | NA | NA | NA |
| TRPC6 | 161 | V | -70.94814286 | 28.89935918 | 0.444557226 | 26.95811672 | NA | NA | NA | NA | NA | NA | NA |
| TRPC6 | 162 | G | -66.93825 | 29.16765479 | 0.449093555 | 27.768084 | NA | NA | NA | NA | NA | NA | NA |
| TRPC6 | 163 | D | -65.336625 | 26.31643154 | 0.400885213 | 25.19630793 | NA | NA | NA | NA | NA | NA | NA |
| TRPC6 | 164 | A | -68.0016 | 24.33443065 | 0.36737364 | 22.1624058 | NA | NA | NA | NA | NA | NA | NA |
| TRPC6 | 165 | L | -66.675125 | 25.18456029 | 0.38174759 | 23.81842576 | NA | NA | NA | NA | NA | NA | NA |
| TRPC6 | 166 | L | -62.69225 | 24.07264927 | 0.362947454 | 23.05903251 | NA | NA | NA | NA | NA | NA | NA |
| TRPC6 | 167 | L | -64.332 | 20.89251974 | 0.309177981 | 19.78253007 | NA | NA | NA | NA | NA | NA | NA |
| TRPC6 | 168 | A | -65.591 | 20.50846991 | 0.302684485 | 19.37803282 | NA | NA | NA | NA | NA | NA | NA |
| TRPC6 | 169 | I | -62.7455 | 21.24846173 | 0.31519623 | 19.85219655 | NA | NA | NA | NA | NA | NA | NA |
| TRPC6 | 170 | S | -60.33833333 | 18.80096551 | 0.273814085 | 17.09409816 | NA | NA | NA | NA | NA | NA | NA |
| TRPC6 | 171 | K | -62.93177778 | 16.02114506 | 0.226813018 | 14.61725067 | NA | NA | NA | NA | NA | NA | NA |
| TRPC6 | 172 | G | -63.86925 | 17.21525658 | 0.247002997 | 16.01904963 | NA | NA | NA | NA | NA | NA | NA |

| | | | | | | | | | | | | | |
|---|---|---|---|---|---|---|---|---|---|---|---|---|---|
| TRPC6 | 173 | Y | -66.88241667 | 14.15096127 | 0.195192043 | 12.76039102 | NA | NA | NA | NA | NA | NA | NA |
| TRPC6 | 174 | V | -68.169 | 18.87221463 | 0.275018762 | 16.69442061 | NA | NA | NA | NA | NA | NA | NA |
| TRPC6 | 175 | R | -73.49163636 | 15.04362642 | 0.210285181 | 13.74801051 | NA | NA | NA | NA | NA | NA | NA |
| TRPC6 | 176 | I | -71.257625 | 18.07102534 | 0.261472293 | 16.20099677 | NA | NA | NA | NA | NA | NA | NA |
| TRPC6 | 177 | V | -68.925 | 22.28608294 | 0.332740278 | 20.3859993 | NA | NA | NA | NA | NA | NA | NA |
| TRPC6 | 178 | E | -72.3892 | 23.97924749 | 0.361368221 | 22.11189368 | NA | NA | NA | NA | NA | NA | NA |
| TRPC6 | 179 | A | -74.5818 | 23.10555732 | 0.346595911 | 21.00558292 | NA | NA | NA | NA | NA | NA | NA |
| TRPC6 | 180 | I | -72.09775 | 24.5423824 | 0.370889678 | 22.91132669 | NA | NA | NA | NA | NA | NA | NA |
| TRPC6 | 181 | L | -70.9275 | 27.85768831 | 0.426944707 | 25.99889775 | NA | NA | NA | NA | NA | NA | NA |
| TRPC6 | 182 | S | -76.12633333 | 28.18178699 | 0.432424551 | 25.9298757 | NA | NA | NA | NA | NA | NA | NA |
| TRPC6 | 183 | H | -75.6885 | 30.30185283 | 0.46827052 | 27.8142857 | NA | NA | NA | NA | NA | NA | NA |
| TRPC6 | 184 | P | -76.99028571 | 33.09057886 | 0.515422161 | 30.62565222 | NA | NA | NA | NA | NA | NA | NA |
| TRPC6 | 185 | A | -72.6064 | 35.00143878 | 0.547730887 | 33.24075515 | NA | NA | NA | NA | NA | NA | NA |
| TRPC6 | 186 | F | -70.48936364 | 31.33658502 | 0.48576572 | 29.4753755 | NA | NA | NA | NA | NA | NA | NA |
| TRPC6 | 187 | A | -73.412 | 36.34291466 | 0.570412495 | 34.58251017 | NA | NA | NA | NA | NA | NA | NA |
| TRPC6 | 188 | E | -70.7954 | 38.3095043 | 0.603663495 | 36.44936354 | NA | NA | NA | NA | NA | NA | NA |
| TRPC6 | 189 | G | -67.94825 | 36.8493305 | 0.578974949 | 34.66997065 | NA | NA | NA | NA | NA | NA | NA |
| TRPC6 | 190 | K | -67.3214 | 37.75793481 | 0.594337585 | 35.68283837 | NA | NA | NA | NA | NA | NA | NA |
| TRPC6 | 191 | R | -66.71445455 | 34.8718099 | 0.545539128 | 33.74808662 | NA | NA | NA | NA | NA | NA | NA |
| TRPC6 | 192 | L | -64.353125 | 32.46958831 | 0.504922484 | 31.36824441 | NA | NA | NA | NA | NA | NA | NA |
| TRPC6 | 193 | A | -60.7052 | 35.97854818 | 0.564251804 | 34.21207643 | NA | NA | NA | NA | NA | NA | NA |
| TRPC6 | 194 | T | -58.36185714 | 37.54870541 | 0.590799945 | 35.6996004 | NA | NA | NA | NA | NA | NA | NA |
| TRPC6 | 203 | Q | -60.945 | 41.47710139 | 0.65722107 | 39.7259582 | NA | NA | NA | NA | NA | NA | NA |
| TRPC6 | 204 | D | -64.322 | 40.51063172 | 0.640880049 | 39.38038911 | NA | NA | NA | NA | NA | NA | NA |
| TRPC6 | 205 | D | -64.886 | 37.46241815 | 0.589341004 | 36.37225205 | NA | NA | NA | NA | NA | NA | NA |
| TRPC6 | 206 | F | -67.27845455 | 34.07934345 | 0.532140144 | 32.64250586 | NA | NA | NA | NA | NA | NA | NA |
| TRPC6 | 207 | Y | -63.6075 | 32.02500039 | 0.497405413 | 30.90322572 | NA | NA | NA | NA | NA | NA | NA |
| TRPC6 | 208 | A | -64.3138 | 32.4405198 | 0.504430995 | 31.3242952 | NA | NA | NA | NA | NA | NA | NA |
| TRPC6 | 209 | Y | -62.77625 | 28.47163905 | 0.437325355 | 27.48044486 | NA | NA | NA | NA | NA | NA | NA |
| TRPC6 | 210 | D | -63.597875 | 32.42950175 | 0.504244703 | 31.3096092 | NA | NA | NA | NA | NA | NA | NA |
| TRPC6 | 211 | E | -64.4004 | 35.63496544 | 0.558442524 | 34.50567944 | NA | NA | NA | NA | NA | NA | NA |
| TRPC6 | 212 | D | -59.994875 | 35.20150017 | 0.551113515 | 33.45777884 | NA | NA | NA | NA | NA | NA | NA |
| TRPC6 | 213 | G | -59.92375 | 34.00448748 | 0.530874483 | 32.24701666 | NA | NA | NA | NA | NA | NA | NA |

| | | | | | | | | | | | | | |
|---|---|---|---|---|---|---|---|---|---|---|---|---|---|
| TRPC6 | 214 | T | -59.02071429 | 30.84414526 | 0.477439573 | 28.99877743 | NA | NA | NA | NA | NA | NA | NA |
| TRPC6 | 215 | R | -60.69009091 | 24.23432821 | 0.365681113 | 22.66517075 | NA | NA | NA | NA | NA | NA | NA |
| TRPC6 | 216 | F | -57.50918182 | 24.16667611 | 0.364537255 | 22.30274419 | NA | NA | NA | NA | NA | NA | NA |
| TRPC6 | 217 | S | -54.258 | 30.16751785 | 0.46599919 | 27.08295556 | NA | NA | NA | NA | NA | NA | NA |
| TRPC6 | 218 | H | -54.589 | 33.72219241 | 0.526101452 | 31.63796205 | NA | NA | NA | NA | NA | NA | NA |
| TRPC6 | 219 | D | -53.652125 | 32.53822196 | 0.506082938 | 29.47254275 | NA | NA | NA | NA | NA | NA | NA |
| TRPC6 | 220 | V | -56.34714286 | 29.03382918 | 0.446830838 | 26.94301446 | NA | NA | NA | NA | NA | NA | NA |
| TRPC6 | 221 | T | -58.39857143 | 29.97344034 | 0.462717738 | 28.12440742 | NA | NA | NA | NA | NA | NA | NA |
| TRPC6 | 222 | P | -61.32728571 | 28.19090387 | 0.432578699 | 26.44552931 | NA | NA | NA | NA | NA | NA | NA |
| TRPC6 | 223 | I | -58.56525 | 28.77161523 | 0.442397338 | 26.96037426 | NA | NA | NA | NA | NA | NA | NA |
| TRPC6 | 224 | I | -55.124875 | 27.12584697 | 0.41457077 | 24.90202862 | NA | NA | NA | NA | NA | NA | NA |
| TRPC6 | 225 | L | -57.742 | 24.24580269 | 0.365875123 | 22.3203573 | NA | NA | NA | NA | NA | NA | NA |
| TRPC6 | 226 | A | -58.382 | 24.23835791 | 0.365749247 | 22.41566226 | NA | NA | NA | NA | NA | NA | NA |
| TRPC6 | 227 | A | -54.8676 | 24.38629625 | 0.368250581 | 22.31538812 | NA | NA | NA | NA | NA | NA | NA |
| TRPC6 | 228 | H | -52.7951 | 21.50395333 | 0.31951607 | 18.45841876 | NA | NA | NA | NA | NA | NA | NA |
| TRPC6 | 229 | C | -56.113 | 19.25534734 | 0.281496751 | 17.04375757 | NA | NA | NA | NA | NA | NA | NA |
| TRPC6 | 230 | Q | -54.32888889 | 20.04219936 | 0.294800806 | 17.83943499 | NA | NA | NA | NA | NA | NA | NA |
| TRPC6 | 231 | E | -59.69266667 | 18.5885691 | 0.270222897 | 16.7688628 | NA | NA | NA | NA | NA | NA | NA |
| TRPC6 | 232 | Y | -57.84508333 | 21.7344677 | 0.323413595 | 20.05951873 | NA | NA | NA | NA | NA | NA | NA |
| TRPC6 | 233 | E | -65.057 | 21.28499314 | 0.315813902 | 20.21163438 | NA | NA | NA | NA | NA | NA | NA |
| TRPC6 | 234 | I | -63.414875 | 21.9569624 | 0.327175525 | 20.8447063 | NA | NA | NA | NA | NA | NA | NA |
| TRPC6 | 235 | V | -60.22742857 | 26.14672308 | 0.398015791 | 24.37710374 | NA | NA | NA | NA | NA | NA | NA |
| TRPC6 | 236 | H | -63.6718 | 27.93116888 | 0.428187113 | 26.52131392 | NA | NA | NA | NA | NA | NA | NA |
| TRPC6 | 237 | T | -66.27042857 | 26.88922496 | 0.410569976 | 25.85727498 | NA | NA | NA | NA | NA | NA | NA |
| TRPC6 | 238 | L | -63.7515 | 28.67232929 | 0.440718616 | 27.61052132 | NA | NA | NA | NA | NA | NA | NA |
| TRPC6 | 239 | L | -61.406375 | 31.91111897 | 0.495479912 | 30.50894714 | NA | NA | NA | NA | NA | NA | NA |
| TRPC6 | 240 | R | -67.61890909 | 29.85633261 | 0.460737686 | 29.03232367 | NA | NA | NA | NA | NA | NA | NA |
| TRPC6 | 241 | K | -67.72688889 | 31.29754765 | 0.485105678 | 29.94586958 | NA | NA | NA | NA | NA | NA | NA |
| TRPC6 | 242 | G | -63.59425 | 36.54153366 | 0.573770735 | 35.41138292 | NA | NA | NA | NA | NA | NA | NA |
| TRPC6 | 243 | A | -61.1204 | 34.52040008 | 0.539597508 | 32.94235297 | NA | NA | NA | NA | NA | NA | NA |
| TRPC6 | 244 | R | -58.24645455 | 35.7596399 | 0.560550514 | 33.96050453 | NA | NA | NA | NA | NA | NA | NA |
| TRPC6 | 245 | I | -53.97775 | 31.28353848 | 0.484868812 | 28.19627866 | NA | NA | NA | NA | NA | NA | NA |
| TRPC6 | 246 | E | -52.3556 | 37.08005961 | 0.582876105 | 33.99807276 | NA | NA | NA | NA | NA | NA | NA |

| | | | | | | | | | | | | | |
|---|---|---|---|---|---|---|---|---|---|---|---|---|---|
| TRPC6 | 247 | R | -47.67209091 | 36.44597786 | 0.572155082 | 32.89756514 | NA | NA | NA | NA | NA | NA | NA |
| TRPC6 | 248 | P | -46.625 | 36.49408238 | 0.572968431 | 30.67905516 | NA | NA | NA | NA | NA | NA | NA |
| TRPC6 | 249 | H | -45.7685 | 40.19839656 | 0.635600792 | 34.46376876 | NA | NA | NA | NA | NA | NA | NA |
| TRPC6 | 250 | D | -40.937 | 41.60969516 | 0.659462959 | 30.26018542 | NA | NA | NA | NA | NA | NA | NA |
| TRPC6 | 251 | Y | -37.4595 | 38.07635143 | 0.599721358 | 28.11502221 | NA | NA | NA | NA | NA | NA | NA |
| TRPC6 | 252 | F | -35.58036364 | 41.04021907 | 0.649834285 | 32.22558219 | NA | NA | NA | NA | NA | NA | NA |
| TRPC6 | 253 | C | -41.01933333 | 44.94549052 | 0.715864422 | 33.60984445 | NA | NA | NA | NA | NA | NA | NA |
| TRPC6 | 254 | K | -41.5968 | 48.06960005 | 0.768686712 | 36.51989106 | NA | NA | NA | NA | NA | NA | NA |
| TRPC6 | 255 | C | -45.368 | 47.4322601 | 0.757910599 | 41.61157931 | NA | NA | NA | NA | NA | NA | NA |
| TRPC6 | 256 | N | -47.8736 | 50.21839668 | 0.805018459 | 44.46025958 | NA | NA | NA | NA | NA | NA | NA |
| TRPC6 | 257 | D | -48.602 | 45.896612 | 0.731945937 | 42.22500604 | NA | NA | NA | NA | NA | NA | NA |
| TRPC6 | 258 | C | -45.33783333 | 45.05615502 | 0.717735532 | 39.25055896 | NA | NA | NA | NA | NA | NA | NA |
| TRPC6 | 259 | N | -44.3776 | 47.21719253 | 0.754274247 | 41.39396654 | NA | NA | NA | NA | NA | NA | NA |
| TRPC6 | 260 | Q | -47.3656 | 46.43377607 | 0.741028281 | 40.63775653 | NA | NA | NA | NA | NA | NA | NA |
| TRPC6 | 261 | K | -48.29422222 | 42.2802686 | 0.670800982 | 36.72563473 | NA | NA | NA | NA | NA | NA | NA |
| TRPC6 | 262 | Q | -42.03111111 | 43.20061895 | 0.68636222 | 34.87849732 | NA | NA | NA | NA | NA | NA | NA |
| TRPC6 | 263 | K | -44.4284 | 45.88925326 | 0.731821516 | 40.08556888 | NA | NA | NA | NA | NA | NA | NA |
| TRPC6 | 264 | H | -46.9944 | 44.11324331 | 0.701792827 | 38.29144333 | NA | NA | NA | NA | NA | NA | NA |
| TRPC6 | 265 | D | -46.1015 | 39.66589096 | 0.626597214 | 33.87705893 | NA | NA | NA | NA | NA | NA | NA |
| TRPC6 | 266 | S | -42.19483333 | 38.06599283 | 0.599546215 | 29.64692886 | NA | NA | NA | NA | NA | NA | NA |
| TRPC6 | 267 | F | -43.71281818 | 32.67929754 | 0.508468237 | 24.28855226 | NA | NA | NA | NA | NA | NA | NA |
| TRPC6 | 268 | S | -47.42033333 | 36.05938244 | 0.565618546 | 30.28154437 | NA | NA | NA | NA | NA | NA | NA |
| TRPC6 | 269 | H | -45.2864 | 36.6378012 | 0.575398422 | 30.8051147 | NA | NA | NA | NA | NA | NA | NA |
| TRPC6 | 270 | S | -43.244 | 33.36704227 | 0.520096591 | 25.0245511 | NA | NA | NA | NA | NA | NA | NA |
| TRPC6 | 271 | R | -46.92563636 | 28.81218451 | 0.443083281 | 23.10366781 | NA | NA | NA | NA | NA | NA | NA |
| TRPC6 | 272 | S | -48.32466667 | 33.83671241 | 0.528037751 | 30.1621738 | NA | NA | NA | NA | NA | NA | NA |
| TRPC6 | 273 | R | -44.06445455 | 33.08541259 | 0.51533481 | 27.25565636 | NA | NA | NA | NA | NA | NA | NA |
| TRPC6 | 274 | I | -44.821875 | 28.72799948 | 0.441659885 | 23.10014048 | NA | NA | NA | NA | NA | NA | NA |
| TRPC6 | 275 | N | -49.327625 | 28.63158789 | 0.440029763 | 24.9438051 | NA | NA | NA | NA | NA | NA | NA |
| TRPC6 | 276 | A | -49.2566 | 31.82126002 | 0.493960581 | 28.10902451 | NA | NA | NA | NA | NA | NA | NA |
| TRPC6 | 277 | Y | -45.08458333 | 29.45178035 | 0.453897536 | 23.64017017 | NA | NA | NA | NA | NA | NA | NA |
| TRPC6 | 278 | K | -47.96533333 | 24.87762724 | 0.376557981 | 19.152864 | NA | NA | NA | NA | NA | NA | NA |
| TRPC6 | 279 | G | -50.92425 | 28.55716313 | 0.438771392 | 24.81438189 | NA | NA | NA | NA | NA | NA | NA |

| | | | | | | | | | | | | | |
|---|---|---|---|---|---|---|---|---|---|---|---|---|---|
| TRPC6 | 280 | L | -49.37 | 30.11409972 | 0.465095999 | 26.42567709 | NA | NA | NA | NA | NA | NA | NA |
| TRPC6 | 281 | A | -47.3094 | 26.07981045 | 0.396884436 | 20.28567034 | NA | NA | NA | NA | NA | NA | NA |
| TRPC6 | 282 | S | -50.57716667 | 25.4177403 | 0.385690186 | 21.65394824 | NA | NA | NA | NA | NA | NA | NA |
| TRPC6 | 283 | P | -51.01185714 | 23.52553774 | 0.353696919 | 19.8117411 | NA | NA | NA | NA | NA | NA | NA |
| TRPC6 | 284 | A | -54.85 | 24.69861053 | 0.373531176 | 22.34936985 | NA | NA | NA | NA | NA | NA | NA |
| TRPC6 | 285 | Y | -54.13725 | 27.65795772 | 0.423567672 | 24.57009514 | NA | NA | NA | NA | NA | NA | NA |
| TRPC6 | 286 | L | -50.397875 | 28.97664565 | 0.445863982 | 25.41013082 | NA | NA | NA | NA | NA | NA | NA |
| TRPC6 | 287 | S | -54.009 | 27.52090774 | 0.421250438 | 24.43203975 | NA | NA | NA | NA | NA | NA | NA |
| TRPC6 | 288 | L | -57.562 | 27.84373197 | 0.426708734 | 26.00765995 | NA | NA | NA | NA | NA | NA | NA |
| TRPC6 | 289 | S | -55.21216667 | 32.63924719 | 0.507791068 | 30.41040553 | NA | NA | NA | NA | NA | NA | NA |
| TRPC6 | 290 | S | -53.341 | 34.79692593 | 0.544272993 | 31.81975383 | NA | NA | NA | NA | NA | NA | NA |
| TRPC6 | 291 | E | -53.84088889 | 37.51444278 | 0.590220634 | 34.46718259 | NA | NA | NA | NA | NA | NA | NA |
| TRPC6 | 292 | D | -48.941375 | 36.23142793 | 0.568527482 | 32.51333502 | NA | NA | NA | NA | NA | NA | NA |
| TRPC6 | 293 | P | -49.30885714 | 32.69784228 | 0.508781791 | 28.95052965 | NA | NA | NA | NA | NA | NA | NA |
| TRPC6 | 294 | V | -45.32371429 | 33.38347397 | 0.520374418 | 27.55634129 | NA | NA | NA | NA | NA | NA | NA |
| TRPC6 | 295 | M | -43.72575 | 35.76427302 | 0.56062885 | 29.94115407 | NA | NA | NA | NA | NA | NA | NA |
| TRPC6 | 296 | T | -46.49228571 | 34.20040213 | 0.534186999 | 28.38539847 | NA | NA | NA | NA | NA | NA | NA |
| TRPC6 | 297 | A | -44.102 | 30.93932388 | 0.479048849 | 25.1186172 | NA | NA | NA | NA | NA | NA | NA |
| TRPC6 | 298 | L | -40.62675 | 32.30488059 | 0.502137614 | 21.29779122 | NA | NA | NA | NA | NA | NA | NA |
| TRPC6 | 299 | E | -40.87888889 | 34.97171143 | 0.547228258 | 23.92189956 | NA | NA | NA | NA | NA | NA | NA |
| TRPC6 | 300 | L | -43.310375 | 31.66179508 | 0.491264356 | 23.4809864 | NA | NA | NA | NA | NA | NA | NA |
| TRPC6 | 301 | S | -39.40216667 | 29.97483338 | 0.462741291 | 19.51585772 | NA | NA | NA | NA | NA | NA | NA |
| TRPC6 | 302 | N | -36.67025 | 32.57598381 | 0.506721414 | 23.55851787 | NA | NA | NA | NA | NA | NA | NA |
| TRPC6 | 303 | E | -40.22411111 | 33.71041299 | 0.525902286 | 23.72417663 | NA | NA | NA | NA | NA | NA | NA |
| TRPC6 | 304 | L | -39.835875 | 29.66071978 | 0.457430274 | 19.83475755 | NA | NA | NA | NA | NA | NA | NA |
| TRPC6 | 305 | A | -35.083 | 30.31721207 | 0.468530213 | 23.09030014 | NA | NA | NA | NA | NA | NA | NA |
| TRPC6 | 306 | V | -34.45614286 | 32.90642797 | 0.512308547 | 27.28048393 | NA | NA | NA | NA | NA | NA | NA |
| TRPC6 | 307 | L | -37.30875 | 32.60091635 | 0.507142972 | 25.36058168 | NA | NA | NA | NA | NA | NA | NA |
| TRPC6 | 308 | A | -34.2676 | 29.14918429 | 0.448781257 | 21.91930599 | NA | NA | NA | NA | NA | NA | NA |
| TRPC6 | 309 | N | -30.99725 | 30.08475544 | 0.464599848 | 27.11611772 | NA | NA | NA | NA | NA | NA | NA |
| TRPC6 | 310 | I | -33.0955 | 34.35743973 | 0.536842183 | 28.7541131 | NA | NA | NA | NA | NA | NA | NA |
| TRPC6 | 311 | E | -36.26433333 | 31.92494011 | 0.495713599 | 25.51576768 | NA | NA | NA | NA | NA | NA | NA |
| TRPC6 | 312 | K | -32.5538 | 29.94008532 | 0.462153772 | 27.00607712 | NA | NA | NA | NA | NA | NA | NA |

| | | | | | | | | | | | | | |
|---|---|---|---|---|---|---|---|---|---|---|---|---|---|
| TRPC6 | 313 | E | -35.42466667 | 28.85321031 | 0.443776943 | 21.63262252 | NA | NA | NA | NA | NA | NA | NA |
| TRPC6 | 314 | F | -39.10445455 | 28.26722494 | 0.433869132 | 19.23794545 | NA | NA | NA | NA | NA | NA | NA |
| TRPC6 | 315 | K | -33.68655556 | 24.79577879 | 0.375174092 | 17.79843688 | NA | NA | NA | NA | NA | NA | NA |
| TRPC6 | 316 | N | -35.72275 | 21.96947805 | 0.327387139 | 14.80563757 | NA | NA | NA | NA | NA | NA | NA |
| TRPC6 | 317 | D | -40.243125 | 23.62812657 | 0.355431486 | 13.18619896 | NA | NA | NA | NA | NA | NA | NA |
| TRPC6 | 318 | Y | -39.52416667 | 26.5573297 | 0.404958308 | 16.57654914 | NA | NA | NA | NA | NA | NA | NA |
| TRPC6 | 319 | K | -35.45266667 | 23.79724602 | 0.358290949 | 14.77472498 | NA | NA | NA | NA | NA | NA | NA |
| TRPC6 | 320 | K | -40.33622222 | 19.75004684 | 0.289861105 | 9.308181197 | NA | NA | NA | NA | NA | NA | NA |
| TRPC6 | 321 | L | -42.782875 | 24.48198572 | 0.369868494 | 16.07217733 | NA | NA | NA | NA | NA | NA | NA |
| TRPC6 | 322 | S | -39.36066667 | 25.67934941 | 0.39011346 | 15.22010204 | NA | NA | NA | NA | NA | NA | NA |
| TRPC6 | 323 | M | -39.0905 | 21.05828877 | 0.311980795 | 11.08127967 | NA | NA | NA | NA | NA | NA | NA |
| TRPC6 | 324 | Q | -44.49933333 | 22.01628954 | 0.328178625 | 13.92286481 | NA | NA | NA | NA | NA | NA | NA |
| TRPC6 | 325 | C | -43.17783333 | 25.55678125 | 0.388041084 | 17.16077935 | NA | NA | NA | NA | NA | NA | NA |
| TRPC6 | 326 | K | -39.088 | 25.31077723 | 0.38388166 | 13.97103491 | NA | NA | NA | NA | NA | NA | NA |
| TRPC6 | 327 | D | -42.566375 | 19.90378858 | 0.292460563 | 11.48463687 | NA | NA | NA | NA | NA | NA | NA |
| TRPC6 | 328 | F | -47.231 | 24.72686292 | 0.374008866 | 18.9234659 | NA | NA | NA | NA | NA | NA | NA |
| TRPC6 | 329 | V | -43.498 | 27.1377416 | 0.414771883 | 21.34863741 | NA | NA | NA | NA | NA | NA | NA |
| TRPC6 | 330 | V | -42.004 | 24.09181739 | 0.363271547 | 15.70828817 | NA | NA | NA | NA | NA | NA | NA |
| TRPC6 | 331 | G | -46.03475 | 24.25163654 | 0.365973762 | 18.42084126 | NA | NA | NA | NA | NA | NA | NA |
| TRPC6 | 332 | L | -47.746375 | 26.51573525 | 0.404255031 | 20.69224692 | NA | NA | NA | NA | NA | NA | NA |
| TRPC6 | 333 | L | -43.488125 | 28.45673463 | 0.437073352 | 22.63350983 | NA | NA | NA | NA | NA | NA | NA |
| TRPC6 | 334 | D | -45.341375 | 23.38851985 | 0.351380227 | 17.6339483 | NA | NA | NA | NA | NA | NA | NA |
| TRPC6 | 335 | L | -49.636125 | 25.24012381 | 0.382687055 | 21.48412853 | NA | NA | NA | NA | NA | NA | NA |
| TRPC6 | 336 | C | -47.45416667 | 29.97786699 | 0.462792583 | 24.21280324 | NA | NA | NA | NA | NA | NA | NA |
| TRPC6 | 337 | R | -51.39336364 | 29.08837087 | 0.447753027 | 25.47584376 | NA | NA | NA | NA | NA | NA | NA |
| TRPC6 | 338 | N | -49.33025 | 35.22457151 | 0.551503604 | 29.519402 | NA | NA | NA | NA | NA | NA | NA |
| TRPC6 | 339 | T | -45.23528571 | 38.04647932 | 0.599216282 | 32.22334825 | NA | NA | NA | NA | NA | NA | NA |
| TRPC6 | 340 | E | -49.20744444 | 38.75866284 | 0.611257845 | 32.96553123 | NA | NA | NA | NA | NA | NA | NA |
| TRPC6 | 341 | E | -49.20033333 | 33.6255514 | 0.524467451 | 28.13973813 | NA | NA | NA | NA | NA | NA | NA |
| TRPC6 | 342 | V | -43.88628571 | 34.45989768 | 0.538574536 | 28.67397608 | NA | NA | NA | NA | NA | NA | NA |
| TRPC6 | 343 | E | -43.27911111 | 37.85780317 | 0.596026155 | 29.48707406 | NA | NA | NA | NA | NA | NA | NA |
| TRPC6 | 344 | A | -46.219 | 37.30427855 | 0.586667188 | 31.47249154 | NA | NA | NA | NA | NA | NA | NA |
| TRPC6 | 345 | I | -44.90825 | 32.67395816 | 0.508377959 | 26.88152084 | NA | NA | NA | NA | NA | NA | NA |

| | | | | | | | | | | | | | |
|---|---|---|---|---|---|---|---|---|---|---|---|---|---|
| TRPC6 | 346 | L | -40.46775 | 33.59415635 | 0.523936625 | 22.26969021 | NA | NA | NA | NA | NA | NA | NA |
| TRPC6 | 347 | N | -41.172125 | 38.295694 | 0.603429991 | 26.74566136 | NA | NA | NA | NA | NA | NA | NA |
| TRPC6 | 348 | G | -44.1315 | 40.85363164 | 0.646679474 | 32.43739891 | NA | NA | NA | NA | NA | NA | NA |
| TRPC6 | 349 | D | -47.035 | 42.93663602 | 0.68189881 | 37.10854398 | NA | NA | NA | NA | NA | NA | NA |
| TRPC6 | 350 | V | -44.88257143 | 44.03688347 | 0.700501739 | 38.22834686 | NA | NA | NA | NA | NA | NA | NA |
| TRPC6 | 351 | E | -47.94388889 | 46.68008229 | 0.745192814 | 40.95539276 | NA | NA | NA | NA | NA | NA | NA |
| TRPC6 | 352 | T | -46.097 | 47.59420471 | 0.760648751 | 41.77138754 | NA | NA | NA | NA | NA | NA | NA |
| TRPC6 | 353 | L | -42.005625 | 44.52881948 | 0.708819369 | 36.11287485 | NA | NA | NA | NA | NA | NA | NA |
| TRPC6 | 354 | Q | -38.9456 | 48.40654874 | 0.774383824 | 38.41863723 | NA | NA | NA | NA | NA | NA | NA |
| TRPC6 | 355 | S | -37.629 | 48.64047827 | 0.778339092 | 39.61916113 | NA | NA | NA | NA | NA | NA | NA |
| TRPC6 | 356 | G | -34.40425 | 48.63365708 | 0.77822376 | 42.2241176 | NA | NA | NA | NA | NA | NA | NA |
| TRPC6 | 357 | D | -31.2724 | 49.09756625 | 0.786067514 | 46.1241448 | NA | NA | NA | NA | NA | NA | NA |
| TRPC6 | 358 | H | -30.1672 | 49.46943401 | 0.792355035 | 46.48452113 | NA | NA | NA | NA | NA | NA | NA |
| TRPC6 | 359 | G | -34.377 | 47.71863017 | 0.76275253 | 42.08276243 | NA | NA | NA | NA | NA | NA | NA |
| TRPC6 | 360 | R | -32.215 | 44.88183799 | 0.714788188 | 38.93730725 | NA | NA | NA | NA | NA | NA | NA |
| TRPC6 | 361 | P | -36.69457143 | 42.37976751 | 0.672483304 | 33.35464958 | NA | NA | NA | NA | NA | NA | NA |
| TRPC6 | 362 | N | -37.414875 | 42.06374843 | 0.667140069 | 33.03333169 | NA | NA | NA | NA | NA | NA | NA |
| TRPC6 | 363 | L | -34.146625 | 39.95666239 | 0.631513563 | 32.73396459 | NA | NA | NA | NA | NA | NA | NA |
| TRPC6 | 364 | S | -36.17783333 | 40.06550244 | 0.633353825 | 32.82563443 | NA | NA | NA | NA | NA | NA | NA |
| TRPC6 | 365 | R | -40.43854545 | 36.46749635 | 0.572518916 | 27.44383788 | NA | NA | NA | NA | NA | NA | NA |
| TRPC6 | 366 | L | -35.198625 | 34.37588931 | 0.537154127 | 27.13583558 | NA | NA | NA | NA | NA | NA | NA |
| TRPC6 | 367 | K | -31.88177778 | 36.45835853 | 0.572364414 | 33.47889653 | NA | NA | NA | NA | NA | NA | NA |
| TRPC6 | 368 | L | -35.5165 | 34.70316081 | 0.542687617 | 28.29334983 | NA | NA | NA | NA | NA | NA | NA |
| TRPC6 | 369 | A | -34.8342 | 31.35202666 | 0.486026807 | 24.14366356 | NA | NA | NA | NA | NA | NA | NA |
| TRPC6 | 370 | I | -30.753875 | 32.25685679 | 0.50132563 | 29.28408773 | NA | NA | NA | NA | NA | NA | NA |
| TRPC6 | 371 | K | -31.36411111 | 33.3495009 | 0.519800003 | 30.36091505 | NA | NA | NA | NA | NA | NA | NA |
| TRPC6 | 372 | Y | -35.07741667 | 30.38740191 | 0.469716979 | 24.74794117 | NA | NA | NA | NA | NA | NA | NA |
| TRPC6 | 373 | E | -30.991 | 26.95213687 | 0.411633688 | 24.14821034 | NA | NA | NA | NA | NA | NA | NA |
| TRPC6 | 374 | V | -35.60157143 | 27.57429374 | 0.422153085 | 20.34593563 | NA | NA | NA | NA | NA | NA | NA |
| TRPC6 | 375 | K | -31.90255556 | 23.57256579 | 0.354492067 | 18.6480786 | NA | NA | NA | NA | NA | NA | NA |
| TRPC6 | 376 | K | -36.551 | 19.65777803 | 0.288301029 | 12.43375545 | NA | NA | NA | NA | NA | NA | NA |
| TRPC6 | 377 | F | -38.59363636 | 26.80607204 | 0.409164031 | 17.77869151 | NA | NA | NA | NA | NA | NA | NA |
| TRPC6 | 378 | V | -33.81042857 | 28.68187722 | 0.440880052 | 23.04257283 | NA | NA | NA | NA | NA | NA | NA |

| | | | | | | | | | | | | | |
|---|---|---|---|---|---|---|---|---|---|---|---|---|---|
| TRPC6 | 379 | A | -34.003 | 25.28192336 | 0.3833938 | 19.67091049 | NA | NA | NA | NA | NA | NA | NA |
| TRPC6 | 380 | H | -38.6947 | 25.51847623 | 0.387393424 | 16.73183376 | NA | NA | NA | NA | NA | NA | NA |
| TRPC6 | 381 | P | -37.537 | 26.4245543 | 0.402713348 | 17.39412366 | NA | NA | NA | NA | NA | NA | NA |
| TRPC6 | 382 | N | -40.471375 | 29.81432592 | 0.460027439 | 20.04268555 | NA | NA | NA | NA | NA | NA | NA |
| TRPC6 | 383 | C | -36.25966667 | 31.6180102 | 0.490524043 | 22.58907328 | NA | NA | NA | NA | NA | NA | NA |
| TRPC6 | 384 | Q | -33.11822222 | 29.01394516 | 0.44649464 | 23.55771211 | NA | NA | NA | NA | NA | NA | NA |
| TRPC6 | 385 | Q | -37.252 | 33.40131801 | 0.520676124 | 26.16601434 | NA | NA | NA | NA | NA | NA | NA |
| TRPC6 | 386 | Q | -36.46422222 | 35.78909253 | 0.561048497 | 28.54881749 | NA | NA | NA | NA | NA | NA | NA |
| TRPC6 | 387 | L | -32.066875 | 34.0682511 | 0.531952595 | 31.08198638 | NA | NA | NA | NA | NA | NA | NA |
| TRPC6 | 388 | L | -30.57025 | 33.70672856 | 0.52583999 | 30.71837982 | NA | NA | NA | NA | NA | NA | NA |
| TRPC6 | 389 | S | -32.6635 | 38.81263841 | 0.612170462 | 35.82909137 | NA | NA | NA | NA | NA | NA | NA |
| TRPC6 | 390 | I | -30.6925 | 40.60382478 | 0.642455752 | 37.61400233 | NA | NA | NA | NA | NA | NA | NA |
| TRPC6 | 391 | W | -25.14771429 | 36.52826376 | 0.573546368 | 34.66383191 | NA | NA | NA | NA | NA | NA | NA |
| TRPC6 | 392 | Y | -26.65641667 | 36.1496377 | 0.567144577 | 34.16336617 | NA | NA | NA | NA | NA | NA | NA |
| TRPC6 | 393 | E | -28.32766667 | 43.34698963 | 0.688837048 | 40.8725323 | NA | NA | NA | NA | NA | NA | NA |
| TRPC6 | 394 | N | -26.72525 | 46.21410959 | 0.737314171 | 44.16478988 | NA | NA | NA | NA | NA | NA | NA |
| TRPC6 | 395 | L | -25.077 | 43.42491439 | 0.690154596 | 41.68822583 | NA | NA | NA | NA | NA | NA | NA |
| TRPC6 | 396 | S | -28.11716667 | 43.94708023 | 0.69898335 | 41.55841392 | NA | NA | NA | NA | NA | NA | NA |
| TRPC6 | 397 | G | -25.27475 | 42.76007026 | 0.678913444 | 40.9947517 | NA | NA | NA | NA | NA | NA | NA |
| TRPC6 | 398 | L | -22.4315 | 39.60179413 | 0.625513468 | 37.58928956 | NA | NA | NA | NA | NA | NA | NA |
| TRPC6 | 399 | R | -28.96718182 | 37.28623372 | 0.586362087 | 35.67811567 | NA | NA | NA | NA | NA | NA | NA |
| TRPC6 | 400 | Q | -26.8536 | 38.92200558 | 0.614019636 | 36.88006814 | NA | NA | NA | NA | NA | NA | NA |
| TRPC6 | 401 | Q | -22.88522222 | 36.87062431 | 0.579334983 | 35.07081898 | NA | NA | NA | NA | NA | NA | NA |
| TRPC6 | 402 | T | -21.35471429 | 34.83673748 | 0.544946125 | 33.05285879 | NA | NA | NA | NA | NA | NA | NA |
| TRPC6 | 403 | M | -18.787125 | 31.86108579 | 0.494633954 | 30.76309696 | NA | NA | NA | NA | NA | NA | NA |
| TRPC6 | 404 | A | -15.5658 | 33.94610366 | 0.529887333 | 30.75897072 | NA | NA | NA | NA | NA | NA | NA |
| TRPC6 | 405 | V | -17.07 | 36.95745906 | 0.580803181 | 33.77460974 | NA | NA | NA | NA | NA | NA | NA |
| TRPC6 | 406 | K | -18.62988889 | 35.76689327 | 0.560673153 | 35.04077449 | NA | NA | NA | NA | NA | NA | NA |
| TRPC6 | 407 | F | -11.60690909 | 31.98183863 | 0.496675636 | 28.79027758 | NA | NA | NA | NA | NA | NA | NA |
| TRPC6 | 408 | L | -12.1925 | 36.3428809 | 0.570411924 | 33.25193822 | NA | NA | NA | NA | NA | NA | NA |
| TRPC6 | 409 | V | -14.41457143 | 39.44293147 | 0.622827426 | 36.25609337 | NA | NA | NA | NA | NA | NA | NA |
| TRPC6 | 410 | V | -12.10071429 | 36.06908744 | 0.565782638 | 32.8923892 | NA | NA | NA | NA | NA | NA | NA |
| TRPC6 | 411 | L | -8.527125 | 35.0372919 | 0.54833709 | 31.84030195 | NA | NA | NA | NA | NA | NA | NA |

| | | | | | | | | | | | | | |
|---|---|---|---|---|---|---|---|---|---|---|---|---|---|
| TRPC6 | 412 | A | -9.9724 | 40.26742993 | 0.636768005 | 37.07356072 | NA | NA | NA | NA | NA | NA | NA |
| TRPC6 | 413 | V | -10.87557143 | 41.84652751 | 0.663467309 | 38.6610272 | NA | NA | NA | NA | NA | NA | NA |
| TRPC6 | 414 | A | -6.9572 | 38.58730736 | 0.608360575 | 35.39627663 | NA | NA | NA | NA | NA | NA | NA |
| TRPC6 | 415 | I | -5.28375 | 40.59837594 | 0.642363624 | 39.11039322 | NA | NA | NA | NA | NA | NA | NA |
| TRPC6 | 416 | G | -7.5785 | 44.26995131 | 0.704442439 | 41.0727509 | NA | NA | NA | NA | NA | NA | NA |
| TRPC6 | 417 | L | -7.837 | 42.81311717 | 0.679810359 | 39.63738909 | NA | NA | NA | NA | NA | NA | NA |
| TRPC6 | 418 | P | -5.562428571 | 46.34130006 | 0.739464701 | 43.1500814 | NA | NA | NA | NA | NA | NA | NA |
| TRPC6 | 419 | F | -6.479727273 | 48.51119506 | 0.776153178 | 45.57652689 | NA | NA | NA | NA | NA | NA | NA |
| TRPC6 | 420 | L | -11.635 | 46.83620986 | 0.747832611 | 43.65662052 | NA | NA | NA | NA | NA | NA | NA |
| TRPC6 | 421 | A | -9.7826 | 49.42458126 | 0.791596667 | 46.25180869 | NA | NA | NA | NA | NA | NA | NA |
| TRPC6 | 422 | L | -8.33125 | 52.1721884 | 0.838053073 | 48.99511713 | NA | NA | NA | NA | NA | NA | NA |
| TRPC6 | 423 | I | -13.036 | 53.35903628 | 0.858120238 | 50.17733184 | NA | NA | NA | NA | NA | NA | NA |
| TRPC6 | 424 | Y | -15.48633333 | 52.28208326 | 0.83991117 | 48.813865 | NA | NA | NA | NA | NA | NA | NA |
| TRPC6 | 425 | W | -9.614642857 | 51.69504124 | 0.829985492 | 48.06742539 | NA | NA | NA | NA | NA | NA | NA |
| TRPC6 | 426 | F | -10.38872727 | 57.42201812 | 0.926816934 | 53.78677572 | NA | NA | NA | NA | NA | NA | NA |
| TRPC6 | 427 | A | -15.6154 | 57.71319447 | 0.93174013 | 54.06979809 | NA | NA | NA | NA | NA | NA | NA |
| TRPC6 | 428 | P | -18.12785714 | 57.26858949 | 0.924222771 | 56.19857604 | NA | NA | NA | NA | NA | NA | NA |
| TRPC6 | 429 | C | -21.828 | 57.37722589 | 0.92605959 | 55.71337088 | NA | NA | NA | NA | NA | NA | NA |
| TRPC6 | 430 | S | -20.987 | 55.81476182 | 0.899641524 | 54.15466855 | NA | NA | NA | NA | NA | NA | NA |
| TRPC6 | 431 | K | -21.3658 | 53.43426685 | 0.859392233 | 51.76494582 | NA | NA | NA | NA | NA | NA | NA |
| TRPC6 | 432 | M | -18.13325 | 50.62975908 | 0.811973754 | 49.93365068 | NA | NA | NA | NA | NA | NA | NA |
| TRPC6 | 433 | G | -20.02575 | 52.77360854 | 0.848221855 | 51.69618597 | NA | NA | NA | NA | NA | NA | NA |
| TRPC6 | 434 | K | -23.068 | 50.17539483 | 0.804291386 | 48.03163151 | NA | NA | NA | NA | NA | NA | NA |
| TRPC6 | 435 | I | -21.654125 | 47.43992976 | 0.758040278 | 45.7699112 | NA | NA | NA | NA | NA | NA | NA |
| TRPC6 | 436 | M | -17.842375 | 47.63058163 | 0.76126381 | 46.53204016 | NA | NA | NA | NA | NA | NA | NA |
| TRPC6 | 437 | R | -21.69827273 | 46.96947939 | 0.750085926 | 45.32770782 | NA | NA | NA | NA | NA | NA | NA |
| TRPC6 | 438 | G | -22.64875 | 44.29208879 | 0.704816738 | 42.51708513 | NA | NA | NA | NA | NA | NA | NA |
| TRPC6 | 439 | P | -23.48114286 | 40.74069035 | 0.644769869 | 38.76542292 | NA | NA | NA | NA | NA | NA | NA |
| TRPC6 | 440 | F | -21.15481818 | 37.64939636 | 0.592502423 | 35.98959353 | NA | NA | NA | NA | NA | NA | NA |
| TRPC6 | 441 | M | -18.684375 | 42.47341339 | 0.674066664 | 41.38738891 | NA | NA | NA | NA | NA | NA | NA |
| TRPC6 | 442 | K | -18.77444444 | 41.351926 | 0.655104611 | 40.34076175 | NA | NA | NA | NA | NA | NA | NA |
| TRPC6 | 443 | F | -17.61627273 | 34.15695544 | 0.533452404 | 32.9973768 | NA | NA | NA | NA | NA | NA | NA |
| TRPC6 | 444 | V | -15.46528571 | 39.50137438 | 0.623815575 | 37.67401343 | NA | NA | NA | NA | NA | NA | NA |

| | | | | | | | | | | | | | |
|---|---|---|---|---|---|---|---|---|---|---|---|---|---|
| TRPC6 | 445 | A | -13.1242 | 40.5382944 | 0.641347768 | 36.88103439 | NA | NA | NA | NA | NA | NA | NA |
| TRPC6 | 446 | H | -13.3796 | 37.62012913 | 0.592007574 | 33.93538976 | NA | NA | NA | NA | NA | NA | NA |
| TRPC6 | 447 | A | -11.56 | 35.57303638 | 0.557395431 | 31.93223118 | NA | NA | NA | NA | NA | NA | NA |
| TRPC6 | 448 | A | -9.2106 | 37.61829313 | 0.591976531 | 34.44095321 | NA | NA | NA | NA | NA | NA | NA |
| TRPC6 | 449 | S | -7.524666667 | 37.59607025 | 0.591600787 | 33.95917883 | NA | NA | NA | NA | NA | NA | NA |
| TRPC6 | 450 | F | -7.077545455 | 30.58709777 | 0.473093427 | 26.91418392 | NA | NA | NA | NA | NA | NA | NA |
| TRPC6 | 451 | T | -5.987428571 | 33.8360275 | 0.52802617 | 30.65785131 | NA | NA | NA | NA | NA | NA | NA |
| TRPC6 | 452 | I | -3.454875 | 35.92309398 | 0.563314187 | 32.7386099 | NA | NA | NA | NA | NA | NA | NA |
| TRPC6 | 453 | F | -1.738636364 | 33.86506994 | 0.528517218 | 30.68529179 | NA | NA | NA | NA | NA | NA | NA |
| TRPC6 | 454 | L | -2.362875 | 29.30528406 | 0.451420584 | 26.13147449 | NA | NA | NA | NA | NA | NA | NA |
| TRPC6 | 455 | G | -0.445 | 32.78082693 | 0.510184891 | 31.09883686 | NA | NA | NA | NA | NA | NA | NA |
| TRPC6 | 456 | L | 0.909875 | 33.82285881 | 0.527803515 | 32.13518468 | NA | NA | NA | NA | NA | NA | NA |
| TRPC6 | 457 | L | 1.245 | 30.44729244 | 0.470729605 | 28.76118374 | NA | NA | NA | NA | NA | NA | NA |
| TRPC6 | 458 | V | 2.428857143 | 28.45502421 | 0.437044433 | 26.7671134 | NA | NA | NA | NA | NA | NA | NA |
| TRPC6 | 459 | M | 4.4235 | 31.56214825 | 0.489579532 | 29.95654608 | NA | NA | NA | NA | NA | NA | NA |
| TRPC6 | 460 | N | 7.07975 | 33.18268832 | 0.516979544 | 31.48705248 | NA | NA | NA | NA | NA | NA | NA |
| TRPC6 | 461 | A | 8.716 | 29.84491402 | 0.460544621 | 28.13685006 | NA | NA | NA | NA | NA | NA | NA |
| TRPC6 | 462 | A | 9.5382 | 31.67043096 | 0.491410371 | 29.95839367 | NA | NA | NA | NA | NA | NA | NA |
| TRPC6 | 463 | D | 13.2544 | 32.0608899 | 0.498012231 | 31.0531255 | NA | NA | NA | NA | NA | NA | NA |
| TRPC6 | 464 | R | 12.80127273 | 27.63215252 | 0.423131359 | 26.21677783 | NA | NA | NA | NA | NA | NA | NA |
| TRPC6 | 465 | F | 9.139818182 | 25.198977 | 0.381991347 | 24.15980107 | NA | NA | NA | NA | NA | NA | NA |
| TRPC6 | 466 | E | 14.3496 | 26.46032747 | 0.403318199 | 25.42160249 | NA | NA | NA | NA | NA | NA | NA |
| TRPC6 | 467 | G | 16.28175 | 27.35625789 | 0.418466546 | 25.96296068 | NA | NA | NA | NA | NA | NA | NA |
| TRPC6 | 468 | T | 17.80414286 | 29.64018195 | 0.457083021 | 28.26335606 | NA | NA | NA | NA | NA | NA | NA |
| TRPC6 | 469 | K | 21.1288 | 31.9177374 | 0.495591816 | 30.55294678 | NA | NA | NA | NA | NA | NA | NA |
| TRPC6 | 470 | L | 21.535125 | 33.39432676 | 0.520557916 | 32.06582167 | NA | NA | NA | NA | NA | NA | NA |
| TRPC6 | 471 | L | 24.125875 | 31.8057018 | 0.493697524 | 30.50460835 | NA | NA | NA | NA | NA | NA | NA |
| TRPC6 | 472 | P | 23.22085714 | 31.30422029 | 0.485218499 | 29.65905505 | NA | NA | NA | NA | NA | NA | NA |
| TRPC6 | 473 | N | 27.3986 | 34.9540404 | 0.546929477 | 33.52086534 | NA | NA | NA | NA | NA | NA | NA |
| TRPC6 | 474 | E | 27.0822 | 36.95090198 | 0.580692314 | 35.67442332 | NA | NA | NA | NA | NA | NA | NA |
| TRPC6 | 475 | T | 28.33314286 | 39.96835155 | 0.631711202 | 38.67296074 | NA | NA | NA | NA | NA | NA | NA |
| TRPC6 | 476 | S | 25.8434 | 40.89501545 | 0.64737919 | 39.5540871 | NA | NA | NA | NA | NA | NA | NA |
| TRPC6 | 477 | T | 26.5566 | 44.2416985 | 0.703964741 | 42.90244665 | NA | NA | NA | NA | NA | NA | NA |

| | | | | | | | | | | | | | |
|---|---|---|---|---|---|---|---|---|---|---|---|---|---|
| TRPC6 | 478 | D | 25.8564 | 45.14063267 | 0.719163876 | 43.76559364 | NA | NA | NA | NA | NA | NA | NA |
| TRPC6 | 479 | N | 27.8822 | 48.36521147 | 0.773684895 | 47.00126923 | NA | NA | NA | NA | NA | NA | NA |
| TRPC6 | 480 | A | 31.3472 | 50.27110763 | 0.805909693 | 48.92162972 | NA | NA | NA | NA | NA | NA | NA |
| TRPC6 | 481 | K | 30.2306 | 52.13043208 | 0.837347059 | 50.55440567 | NA | NA | NA | NA | NA | NA | NA |
| TRPC6 | 482 | Q | 26.34488889 | 47.23064617 | 0.754501721 | 45.51699777 | NA | NA | NA | NA | NA | NA | NA |
| TRPC6 | 483 | L | 26.9945 | 46.67960202 | 0.745184694 | 44.96893802 | NA | NA | NA | NA | NA | NA | NA |
| TRPC6 | 484 | F | 24.62809091 | 39.55178503 | 0.624667916 | 37.83906053 | NA | NA | NA | NA | NA | NA | NA |
| TRPC6 | 485 | R | 21.98545455 | 41.70305339 | 0.661041455 | 40.01036363 | NA | NA | NA | NA | NA | NA | NA |
| TRPC6 | 486 | M | 21.602625 | 44.92722908 | 0.715555659 | 43.22550307 | NA | NA | NA | NA | NA | NA | NA |
| TRPC6 | 487 | K | 20.4986 | 41.29036486 | 0.654063738 | 39.58168634 | NA | NA | NA | NA | NA | NA | NA |
| TRPC6 | 488 | T | 18.54557143 | 39.61465528 | 0.625730923 | 37.91713287 | NA | NA | NA | NA | NA | NA | NA |
| TRPC6 | 489 | S | 16.1345 | 41.70430789 | 0.661062666 | 40.00724508 | NA | NA | NA | NA | NA | NA | NA |
| TRPC6 | 490 | C | 15.46 | 41.95095028 | 0.665232884 | 40.25487039 | NA | NA | NA | NA | NA | NA | NA |
| TRPC6 | 491 | F | 11.04736364 | 42.271709 | 0.670656256 | 40.59073053 | NA | NA | NA | NA | NA | NA | NA |
| TRPC6 | 492 | S | 10.506 | 40.69308031 | 0.64396488 | 38.99440595 | NA | NA | NA | NA | NA | NA | NA |
| TRPC6 | 493 | W | 7.506214286 | 40.97973424 | 0.648811611 | 39.27617655 | NA | NA | NA | NA | NA | NA | NA |
| TRPC6 | 494 | M | 5.84175 | 36.65636842 | 0.575712355 | 34.96011711 | NA | NA | NA | NA | NA | NA | NA |
| TRPC6 | 495 | E | 6.7376 | 39.42239014 | 0.622480114 | 37.7353307 | NA | NA | NA | NA | NA | NA | NA |
| TRPC6 | 496 | M | 5.68 | 41.83894591 | 0.663339119 | 40.15466773 | NA | NA | NA | NA | NA | NA | NA |
| TRPC6 | 497 | L | 1.742125 | 41.06082203 | 0.650182639 | 39.3834398 | NA | NA | NA | NA | NA | NA | NA |
| TRPC6 | 498 | I | 1.932125 | 36.05332946 | 0.565516202 | 34.37263435 | NA | NA | NA | NA | NA | NA | NA |
| TRPC6 | 499 | I | 2.744875 | 40.89327205 | 0.647349713 | 39.22001852 | NA | NA | NA | NA | NA | NA | NA |
| TRPC6 | 500 | S | 0.1355 | 43.04048789 | 0.683654732 | 39.40245063 | NA | NA | NA | NA | NA | NA | NA |
| TRPC6 | 501 | W | -2.211 | 39.06823346 | 0.61649205 | 35.43179955 | NA | NA | NA | NA | NA | NA | NA |
| TRPC6 | 502 | V | -0.723714286 | 38.02658068 | 0.598879836 | 34.3676386 | NA | NA | NA | NA | NA | NA | NA |
| TRPC6 | 503 | I | -1.29425 | 42.41730142 | 0.673117926 | 38.74286782 | NA | NA | NA | NA | NA | NA | NA |
| TRPC6 | 504 | G | -5.06575 | 43.00122469 | 0.682990872 | 39.30022924 | NA | NA | NA | NA | NA | NA | NA |
| TRPC6 | 505 | M | -5.569375 | 36.74565868 | 0.577222071 | 33.04595608 | NA | NA | NA | NA | NA | NA | NA |
| TRPC6 | 506 | I | -4.342375 | 41.19087318 | 0.652381538 | 37.48268216 | NA | NA | NA | NA | NA | NA | NA |
| TRPC6 | 507 | W | -7.948428571 | 44.2655571 | 0.704368141 | 40.23554236 | NA | NA | NA | NA | NA | NA | NA |
| TRPC6 | 508 | A | -10.1198 | 42.52545225 | 0.674946535 | 39.5339132 | NA | NA | NA | NA | NA | NA | NA |
| TRPC6 | 509 | E | -9.877 | 37.65159868 | 0.592539659 | 34.60335179 | NA | NA | NA | NA | NA | NA | NA |
| TRPC6 | 510 | C | -9.644833333 | 43.82024712 | 0.696838862 | 40.75262479 | NA | NA | NA | NA | NA | NA | NA |

| | | | | | | | | | | | | | |
|---|---|---|---|---|---|---|---|---|---|---|---|---|---|
| TRPC6 | 511 | K | -13.33533333 | 45.41020535 | 0.723721798 | 43.7952911 | NA | NA | NA | NA | NA | NA | NA |
| TRPC6 | 512 | E | -15.31444444 | 39.17317441 | 0.618266387 | 37.05036261 | NA | NA | NA | NA | NA | NA | NA |
| TRPC6 | 513 | I | -12.876625 | 42.38652844 | 0.672597618 | 40.80229057 | NA | NA | NA | NA | NA | NA | NA |
| TRPC6 | 514 | W | -12.96521429 | 46.10001537 | 0.735385071 | 43.89540998 | NA | NA | NA | NA | NA | NA | NA |
| TRPC6 | 515 | T | -18.44971429 | 45.40491101 | 0.723632281 | 44.34073939 | NA | NA | NA | NA | NA | NA | NA |
| TRPC6 | 516 | Q | -19.3234 | 41.80906027 | 0.662833815 | 40.24236192 | NA | NA | NA | NA | NA | NA | NA |
| TRPC6 | 517 | G | -17.67025 | 42.36298957 | 0.672199624 | 41.3102462 | NA | NA | NA | NA | NA | NA | NA |
| TRPC6 | 518 | P | -15.41442857 | 41.83693202 | 0.663305069 | 39.87710483 | NA | NA | NA | NA | NA | NA | NA |
| TRPC6 | 519 | K | -18.0238 | 39.06524345 | 0.616441495 | 37.92156375 | NA | NA | NA | NA | NA | NA | NA |
| TRPC6 | 520 | E | -19.4882 | 37.6131628 | 0.591889787 | 35.90813797 | NA | NA | NA | NA | NA | NA | NA |
| TRPC6 | 521 | Y | -14.06275 | 36.51226676 | 0.573275892 | 35.40030913 | NA | NA | NA | NA | NA | NA | NA |
| TRPC6 | 522 | L | -14.287125 | 34.57091588 | 0.540451627 | 32.36518476 | NA | NA | NA | NA | NA | NA | NA |
| TRPC6 | 523 | F | -18.12 | 32.14405597 | 0.499418399 | 31.07470554 | NA | NA | NA | NA | NA | NA | NA |
| TRPC6 | 524 | E | -16.2838 | 31.82974236 | 0.494104 | 30.71910548 | NA | NA | NA | NA | NA | NA | NA |
| TRPC6 | 525 | L | -13.720625 | 27.7996573 | 0.425963521 | 26.21126224 | NA | NA | NA | NA | NA | NA | NA |
| TRPC6 | 526 | W | -12.2025 | 25.05488226 | 0.379555 | 23.48471496 | NA | NA | NA | NA | NA | NA | NA |
| TRPC6 | 527 | N | -12.9085 | 32.46845092 | 0.504903253 | 30.84993901 | NA | NA | NA | NA | NA | NA | NA |
| TRPC6 | 528 | M | -9.2304 | 32.41931306 | 0.504072433 | 29.35143407 | NA | NA | NA | NA | NA | NA | NA |
| TRPC6 | 529 | L | -7.9055 | 26.68123207 | 0.407053243 | 23.61576495 | NA | NA | NA | NA | NA | NA | NA |
| TRPC6 | 530 | D | -8.000375 | 32.17375143 | 0.499920489 | 26.09458616 | NA | NA | NA | NA | NA | NA | NA |
| TRPC6 | 531 | F | -5.923090909 | 34.66686294 | 0.542073895 | 28.62911266 | NA | NA | NA | NA | NA | NA | NA |
| TRPC6 | 532 | G | -3.20475 | 33.408944 | 0.520805064 | 26.78175779 | NA | NA | NA | NA | NA | NA | NA |
| TRPC6 | 533 | M | -2.7 | 32.00840093 | 0.497124751 | 28.30049076 | NA | NA | NA | NA | NA | NA | NA |
| TRPC6 | 534 | L | -2.84925 | 35.15240768 | 0.550283461 | 31.4535584 | NA | NA | NA | NA | NA | NA | NA |
| TRPC6 | 535 | A | -0.5646 | 36.79020841 | 0.577975315 | 32.791415 | NA | NA | NA | NA | NA | NA | NA |
| TRPC6 | 536 | I | 0.729125 | 31.5353608 | 0.489126612 | 27.83632528 | NA | NA | NA | NA | NA | NA | NA |
| TRPC6 | 537 | F | 0.862272727 | 33.44225375 | 0.521368263 | 29.79788794 | NA | NA | NA | NA | NA | NA | NA |
| TRPC6 | 538 | A | 2.1776 | 38.30541206 | 0.603594304 | 34.61319599 | NA | NA | NA | NA | NA | NA | NA |
| TRPC6 | 539 | A | 4.679 | 38.1516507 | 0.600994514 | 34.45112444 | NA | NA | NA | NA | NA | NA | NA |
| TRPC6 | 540 | S | 6.4025 | 35.79592844 | 0.561164079 | 34.12623562 | NA | NA | NA | NA | NA | NA | NA |
| TRPC6 | 541 | F | 5.403818182 | 38.04399071 | 0.599174204 | 36.37277427 | NA | NA | NA | NA | NA | NA | NA |
| TRPC6 | 542 | I | 7.093375 | 41.73956686 | 0.661658823 | 40.07287474 | NA | NA | NA | NA | NA | NA | NA |
| TRPC6 | 543 | A | 10.5718 | 39.81267921 | 0.629079102 | 38.14254172 | NA | NA | NA | NA | NA | NA | NA |

| | | | | | | | | | | | | | |
|---|---|---|---|---|---|---|---|---|---|---|---|---|---|
| TRPC6 | 544 | R | 11.80827273 | 37.88371461 | 0.596464264 | 36.19483023 | NA | NA | NA | NA | NA | NA | NA |
| TRPC6 | 545 | F | 11.25281818 | 43.14043833 | 0.685344689 | 41.46388075 | NA | NA | NA | NA | NA | NA | NA |
| TRPC6 | 546 | M | 13.3005 | 44.42886947 | 0.707129419 | 42.75527046 | NA | NA | NA | NA | NA | NA | NA |
| TRPC6 | 547 | A | 16.161 | 42.44827642 | 0.67364165 | 40.76421061 | NA | NA | NA | NA | NA | NA | NA |
| TRPC6 | 548 | F | 16.19618182 | 44.86834083 | 0.714559979 | 43.18314887 | NA | NA | NA | NA | NA | NA | NA |
| TRPC6 | 549 | W | 16.51485714 | 47.58316095 | 0.760462023 | 45.89922141 | NA | NA | NA | NA | NA | NA | NA |
| TRPC6 | 550 | H | 18.9296 | 47.08660023 | 0.752066199 | 45.40527278 | NA | NA | NA | NA | NA | NA | NA |
| TRPC6 | 551 | A | 21.262 | 46.8737129 | 0.748466711 | 45.17977231 | NA | NA | NA | NA | NA | NA | NA |
| TRPC6 | 552 | S | 20.82683333 | 49.81932249 | 0.798270932 | 48.12733091 | NA | NA | NA | NA | NA | NA | NA |
| TRPC6 | 553 | K | 21.87577778 | 52.21566586 | 0.838788188 | 50.52560278 | NA | NA | NA | NA | NA | NA | NA |
| TRPC6 | 554 | A | 25.4054 | 51.34546441 | 0.824074864 | 49.6503378 | NA | NA | NA | NA | NA | NA | NA |
| TRPC6 | 555 | Q | 25.66433333 | 51.86382725 | 0.832839318 | 50.16349446 | NA | NA | NA | NA | NA | NA | NA |
| TRPC6 | 556 | S | 26.089 | 54.88189783 | 0.883868706 | 53.18432844 | NA | NA | NA | NA | NA | NA | NA |
| TRPC6 | 557 | I | 29.43175 | 56.82222252 | 0.91667562 | 55.12374724 | NA | NA | NA | NA | NA | NA | NA |
| TRPC6 | 558 | I | 30.3806 | 55.194884 | 0.889160661 | 53.48799131 | NA | NA | NA | NA | NA | NA | NA |
| TRPC6 | 559 | D | 30.574 | 57.43460542 | 0.92702976 | 55.72784994 | NA | NA | NA | NA | NA | NA | NA |
| TRPC6 | 569 | K | 39.3582 | 53.05758052 | 0.853023239 | 51.78403708 | NA | NA | NA | NA | NA | NA | NA |
| TRPC6 | 570 | V | 39.4066 | 54.10237413 | 0.870688558 | 52.66166518 | NA | NA | NA | NA | NA | NA | NA |
| TRPC6 | 571 | T | 38.9516 | 55.21042233 | 0.889423382 | 53.59450471 | NA | NA | NA | NA | NA | NA | NA |
| TRPC6 | 572 | L | 35.133125 | 51.95790268 | 0.834429941 | 50.24491598 | NA | NA | NA | NA | NA | NA | NA |
| TRPC6 | 573 | G | 34.58775 | 53.88001834 | 0.866928977 | 52.17489519 | NA | NA | NA | NA | NA | NA | NA |
| TRPC6 | 574 | D | 32.8442 | 54.23904647 | 0.872999407 | 52.53948257 | NA | NA | NA | NA | NA | NA | NA |
| TRPC6 | 575 | N | 29.7338 | 52.37807046 | 0.841534117 | 50.68122958 | NA | NA | NA | NA | NA | NA | NA |
| TRPC6 | 576 | V | 29.92928571 | 49.92603434 | 0.800075211 | 48.22110558 | NA | NA | NA | NA | NA | NA | NA |
| TRPC6 | 577 | K | 30.8788 | 47.81821022 | 0.764436225 | 46.1116241 | NA | NA | NA | NA | NA | NA | NA |
| TRPC6 | 578 | Y | 25.54216667 | 41.93713597 | 0.664999312 | 40.24454223 | NA | NA | NA | NA | NA | NA | NA |
| TRPC6 | 579 | Y | 25.26191667 | 46.28536534 | 0.738518959 | 44.58848501 | NA | NA | NA | NA | NA | NA | NA |
| TRPC6 | 580 | N | 29.133125 | 45.09659077 | 0.718419218 | 43.38536021 | NA | NA | NA | NA | NA | NA | NA |
| TRPC6 | 581 | L | 27.894375 | 41.23326658 | 0.653098323 | 39.52436418 | NA | NA | NA | NA | NA | NA | NA |
| TRPC6 | 582 | A | 25.9384 | 38.54546077 | 0.607653035 | 36.83488186 | NA | NA | NA | NA | NA | NA | NA |
| TRPC6 | 583 | R | 20.61890909 | 33.45632834 | 0.521606236 | 31.76003396 | NA | NA | NA | NA | NA | NA | NA |
| TRPC6 | 584 | I | 26.04675 | 35.70740909 | 0.559667398 | 34.07550539 | NA | NA | NA | NA | NA | NA | NA |
| TRPC6 | 585 | K | 27.87266667 | 38.47404617 | 0.606445561 | 36.76571187 | NA | NA | NA | NA | NA | NA | NA |

| | | | | | | | | | | | | | |
|---|---|---|---|---|---|---|---|---|---|---|---|---|---|
| TRPC6 | 586 | W | 21.89142857 | 36.90875541 | 0.579979702 | 35.2115799 | NA | NA | NA | NA | NA | NA | NA |
| TRPC6 | 587 | D | 21.251 | 37.98674349 | 0.598206271 | 36.29781595 | NA | NA | NA | NA | NA | NA | NA |
| TRPC6 | 588 | P | 19.136 | 36.21621667 | 0.568270291 | 34.53587594 | NA | NA | NA | NA | NA | NA | NA |
| TRPC6 | 589 | S | 16.81216667 | 38.41990221 | 0.605530097 | 36.73846234 | NA | NA | NA | NA | NA | NA | NA |
| TRPC6 | 590 | D | 17.83975 | 38.11884533 | 0.600439843 | 36.43867299 | NA | NA | NA | NA | NA | NA | NA |
| TRPC6 | 591 | P | 14.91571429 | 38.34445052 | 0.604254364 | 36.66642838 | NA | NA | NA | NA | NA | NA | NA |
| TRPC6 | 592 | Q | 15.49633333 | 35.80880682 | 0.561381826 | 34.12378329 | NA | NA | NA | NA | NA | NA | NA |
| TRPC6 | 593 | I | 14.788125 | 33.52335915 | 0.522739589 | 31.84585108 | NA | NA | NA | NA | NA | NA | NA |
| TRPC6 | 594 | I | 11.17275 | 34.54553185 | 0.540022435 | 32.87597828 | NA | NA | NA | NA | NA | NA | NA |
| TRPC6 | 595 | S | 10.07583333 | 33.94802576 | 0.529919831 | 32.27482225 | NA | NA | NA | NA | NA | NA | NA |
| TRPC6 | 596 | E | 11.89055556 | 30.44096837 | 0.470622678 | 28.76390377 | NA | NA | NA | NA | NA | NA | NA |
| TRPC6 | 597 | G | 8.122 | 29.91271853 | 0.461691056 | 28.24715225 | NA | NA | NA | NA | NA | NA | NA |
| TRPC6 | 598 | L | 5.578125 | 31.57511623 | 0.489798794 | 30.05328083 | NA | NA | NA | NA | NA | NA | NA |
| TRPC6 | 599 | Y | 5.295333333 | 30.34384714 | 0.468980557 | 28.676242 | NA | NA | NA | NA | NA | NA | NA |
| TRPC6 | 600 | A | 4.35 | 27.11937848 | 0.414461401 | 25.51224303 | NA | NA | NA | NA | NA | NA | NA |
| TRPC6 | 601 | I | 1.93975 | 27.91677254 | 0.4279437 | 24.22012881 | NA | NA | NA | NA | NA | NA | NA |
| TRPC6 | 602 | A | 0.4954 | 29.76475374 | 0.459189275 | 26.09162953 | NA | NA | NA | NA | NA | NA | NA |
| TRPC6 | 603 | V | 1.034857143 | 27.30109613 | 0.417533874 | 23.66358131 | NA | NA | NA | NA | NA | NA | NA |
| TRPC6 | 604 | V | -0.400714286 | 23.55865894 | 0.354256931 | 19.87323658 | NA | NA | NA | NA | NA | NA | NA |
| TRPC6 | 605 | L | -2.51575 | 26.85254357 | 0.409949769 | 23.15777576 | NA | NA | NA | NA | NA | NA | NA |
| TRPC6 | 606 | S | -2.999 | 28.73310392 | 0.441746191 | 25.04030119 | NA | NA | NA | NA | NA | NA | NA |
| TRPC6 | 607 | F | -3.286 | 25.6379547 | 0.38941356 | 21.97452691 | NA | NA | NA | NA | NA | NA | NA |
| TRPC6 | 608 | S | -7.601333333 | 26.51914861 | 0.404312744 | 22.47627841 | NA | NA | NA | NA | NA | NA | NA |
| TRPC6 | 609 | R | -8.787 | 28.97394709 | 0.445818355 | 24.94032233 | NA | NA | NA | NA | NA | NA | NA |
| TRPC6 | 610 | I | -10.37975 | 26.41979097 | 0.40263281 | 22.74762613 | NA | NA | NA | NA | NA | NA | NA |
| TRPC6 | 611 | A | -13.7616 | 26.88860184 | 0.410559441 | 25.26811851 | NA | NA | NA | NA | NA | NA | NA |
| TRPC6 | 612 | Y | -14.15075 | 29.82846729 | 0.46026654 | 28.16929976 | NA | NA | NA | NA | NA | NA | NA |
| TRPC6 | 613 | I | -14.504875 | 30.22601411 | 0.466988242 | 28.27520282 | NA | NA | NA | NA | NA | NA | NA |
| TRPC6 | 614 | L | -15.752125 | 24.35696428 | 0.367754638 | 22.54310458 | NA | NA | NA | NA | NA | NA | NA |
| TRPC6 | 615 | P | -19.67714286 | 27.30213578 | 0.417551452 | 25.59462457 | NA | NA | NA | NA | NA | NA | NA |
| TRPC6 | 616 | A | -20.8022 | 29.32717508 | 0.451790716 | 27.68504741 | NA | NA | NA | NA | NA | NA | NA |
| TRPC6 | 617 | N | -20.033125 | 26.38763623 | 0.402089139 | 24.83930417 | NA | NA | NA | NA | NA | NA | NA |
| TRPC6 | 618 | E | -22.9832 | 23.91534106 | 0.360287694 | 21.89752432 | NA | NA | NA | NA | NA | NA | NA |

| | | | | | | | | | | | | | |
|---|---|---|---|---|---|---|---|---|---|---|---|---|---|
| TRPC6 | 619 | S | -19.89566667 | 21.77350654 | 0.324073662 | 20.21017871 | NA | NA | NA | NA | NA | NA | NA |
| TRPC6 | 620 | F | -16.59981818 | 22.77785165 | 0.34105508 | 21.86440695 | NA | NA | NA | NA | NA | NA | NA |
| TRPC6 | 621 | G | -20.3165 | 22.10426009 | 0.329666027 | 20.51508141 | NA | NA | NA | NA | NA | NA | NA |
| TRPC6 | 622 | P | -21.84142857 | 18.46430857 | 0.268121906 | 16.73221571 | NA | NA | NA | NA | NA | NA | NA |
| TRPC6 | 623 | L | -17.93325 | 17.64588978 | 0.254284122 | 16.57664487 | NA | NA | NA | NA | NA | NA | NA |
| TRPC6 | 624 | Q | -17.13511111 | 20.28667713 | 0.298934424 | 19.13104443 | NA | NA | NA | NA | NA | NA | NA |
| TRPC6 | 625 | I | -21.08675 | 18.47584147 | 0.268316904 | 16.88721228 | NA | NA | NA | NA | NA | NA | NA |
| TRPC6 | 626 | S | -19.298 | 15.05191629 | 0.210425346 | 13.46242591 | NA | NA | NA | NA | NA | NA | NA |
| TRPC6 | 627 | L | -14.93275 | 16.78653326 | 0.239754164 | 14.93138333 | NA | NA | NA | NA | NA | NA | NA |
| TRPC6 | 628 | G | -17.13175 | 18.54555068 | 0.269495544 | 17.38907558 | NA | NA | NA | NA | NA | NA | NA |
| TRPC6 | 629 | R | -21.31690909 | 15.01004004 | 0.209717305 | 14.21518909 | NA | NA | NA | NA | NA | NA | NA |
| TRPC6 | 630 | T | -16.145 | 13.01453741 | 0.175977445 | 11.86028913 | NA | NA | NA | NA | NA | NA | NA |
| TRPC6 | 631 | V | -13.81342857 | 16.62702192 | 0.237057154 | 14.45629833 | NA | NA | NA | NA | NA | NA | NA |
| TRPC6 | 632 | K | -17.76377778 | 18.06143596 | 0.261310156 | 16.90202428 | NA | NA | NA | NA | NA | NA | NA |
| TRPC6 | 633 | D | -17.73825 | 13.75359986 | 0.188473476 | 12.84073805 | NA | NA | NA | NA | NA | NA | NA |
| TRPC6 | 634 | I | -12.539 | 13.66452539 | 0.186967409 | 12.04463673 | NA | NA | NA | NA | NA | NA | NA |
| TRPC6 | 635 | F | -12.34345455 | 18.03699848 | 0.260896968 | 16.41889304 | NA | NA | NA | NA | NA | NA | NA |
| TRPC6 | 636 | K | -16.23177778 | 17.81687936 | 0.257175205 | 16.41655893 | NA | NA | NA | NA | NA | NA | NA |
| TRPC6 | 637 | F | -13.75390909 | 14.16895218 | 0.195496233 | 12.96359554 | NA | NA | NA | NA | NA | NA | NA |
| TRPC6 | 638 | M | -8.776 | 17.19249444 | 0.246618136 | 14.14643409 | NA | NA | NA | NA | NA | NA | NA |
| TRPC6 | 639 | V | -10.12057143 | 20.29442598 | 0.299065441 | 17.25416298 | NA | NA | NA | NA | NA | NA | NA |
| TRPC6 | 640 | I | -10.2135 | 19.3178095 | 0.282552858 | 16.25418221 | NA | NA | NA | NA | NA | NA | NA |
| TRPC6 | 641 | F | -5.750090909 | 13.33224321 | 0.181349199 | 7.27384112 | NA | NA | NA | NA | NA | NA | NA |
| TRPC6 | 642 | I | -4.944875 | 19.73260295 | 0.289566165 | 13.1047153 | NA | NA | NA | NA | NA | NA | NA |
| TRPC6 | 643 | M | -6.687125 | 21.87757226 | 0.3258332 | 15.17955011 | NA | NA | NA | NA | NA | NA | NA |
| TRPC6 | 644 | V | -3.976142857 | 18.51094849 | 0.268910492 | 11.88276483 | NA | NA | NA | NA | NA | NA | NA |
| TRPC6 | 645 | F | -0.462090909 | 17.37650742 | 0.249729418 | 10.80183414 | NA | NA | NA | NA | NA | NA | NA |
| TRPC6 | 646 | V | -0.472571429 | 22.18946013 | 0.331106585 | 15.58504897 | NA | NA | NA | NA | NA | NA | NA |
| TRPC6 | 647 | A | -0.4946 | 23.74964393 | 0.357486095 | 17.14465342 | NA | NA | NA | NA | NA | NA | NA |
| TRPC6 | 648 | F | 0.549272727 | 20.14960077 | 0.296616743 | 13.5444846 | NA | NA | NA | NA | NA | NA | NA |
| TRPC6 | 649 | M | 2.93975 | 22.77750544 | 0.341049226 | 16.20615316 | NA | NA | NA | NA | NA | NA | NA |
| TRPC6 | 650 | I | 3.00525 | 25.61270804 | 0.388986691 | 19.04268336 | NA | NA | NA | NA | NA | NA | NA |
| TRPC6 | 651 | G | 4.26975 | 26.48780266 | 0.403782748 | 19.91807703 | NA | NA | NA | NA | NA | NA | NA |

| | | | | | | | | | | | | | |
|---|---|---|---|---|---|---|---|---|---|---|---|---|---|
| TRPC6 | 652 | M | 6.026875 | 20.63857936 | 0.304884369 | 14.09734882 | NA | NA | NA | NA | NA | NA | NA |
| TRPC6 | 653 | F | 8.887363636 | 26.72526095 | 0.407797681 | 20.18386289 | NA | NA | NA | NA | NA | NA | NA |
| TRPC6 | 654 | N | 7.785625 | 29.39329592 | 0.452908684 | 22.84808644 | NA | NA | NA | NA | NA | NA | NA |
| TRPC6 | 655 | L | 9.856375 | 27.39764769 | 0.419166363 | 20.86485149 | NA | NA | NA | NA | NA | NA | NA |
| TRPC6 | 656 | Y | 12.00116667 | 25.72431313 | 0.390873705 | 19.21133737 | NA | NA | NA | NA | NA | NA | NA |
| TRPC6 | 657 | S | 14.05183333 | 31.23035115 | 0.483969523 | 24.71112305 | NA | NA | NA | NA | NA | NA | NA |
| TRPC6 | 658 | Y | 16.16908333 | 33.8243371 | 0.52782851 | 27.31001499 | NA | NA | NA | NA | NA | NA | NA |
| TRPC6 | 659 | Y | 17.52058333 | 31.92952607 | 0.495791138 | 25.43830027 | NA | NA | NA | NA | NA | NA | NA |
| TRPC6 | 660 | I | 19.02675 | 31.88771152 | 0.49508414 | 30.1916441 | NA | NA | NA | NA | NA | NA | NA |
| TRPC6 | 661 | G | 22.0795 | 33.13241171 | 0.516129469 | 31.42509057 | NA | NA | NA | NA | NA | NA | NA |
| TRPC6 | 662 | A | 21.212 | 31.87551656 | 0.494877948 | 30.16943844 | NA | NA | NA | NA | NA | NA | NA |
| TRPC6 | 663 | K | 17.23388889 | 25.92362409 | 0.394243644 | 24.65965784 | NA | NA | NA | NA | NA | NA | NA |
| TRPC6 | 664 | Q | 18.83155556 | 24.62976957 | 0.372367216 | 22.91548649 | NA | NA | NA | NA | NA | NA | NA |
| TRPC6 | 665 | N | 16.0705 | 21.76206056 | 0.323880134 | 20.06268985 | NA | NA | NA | NA | NA | NA | NA |
| TRPC6 | 666 | E | 16.77211111 | 25.63337639 | 0.389336151 | 23.93724134 | NA | NA | NA | NA | NA | NA | NA |
| TRPC6 | 667 | A | 12.5924 | 23.33422075 | 0.350462141 | 16.84384573 | NA | NA | NA | NA | NA | NA | NA |
| TRPC6 | 668 | F | 9.366 | 20.98014797 | 0.310659595 | 14.46499084 | NA | NA | NA | NA | NA | NA | NA |
| TRPC6 | 669 | T | 13.778 | 27.69133251 | 0.424131971 | 21.18263099 | NA | NA | NA | NA | NA | NA | NA |
| TRPC6 | 670 | T | 12.13785714 | 27.40605207 | 0.419308463 | 20.88272006 | NA | NA | NA | NA | NA | NA | NA |
| TRPC6 | 671 | V | 7.965142857 | 25.81095293 | 0.392338606 | 19.27192929 | NA | NA | NA | NA | NA | NA | NA |
| TRPC6 | 672 | E | 9.792111111 | 24.18581109 | 0.364860788 | 17.66189749 | NA | NA | NA | NA | NA | NA | NA |
| TRPC6 | 673 | E | 11.239 | 22.82805653 | 0.341903941 | 16.31603519 | NA | NA | NA | NA | NA | NA | NA |
| TRPC6 | 674 | S | 6.901833333 | 21.69696168 | 0.322779445 | 15.1618621 | NA | NA | NA | NA | NA | NA | NA |
| TRPC6 | 675 | F | 3.800818182 | 19.89100023 | 0.292244338 | 13.35118345 | NA | NA | NA | NA | NA | NA | NA |
| TRPC6 | 676 | K | 7.696333333 | 18.28670998 | 0.265119078 | 11.77329516 | NA | NA | NA | NA | NA | NA | NA |
| TRPC6 | 677 | T | 6.957428571 | 17.10710502 | 0.245174376 | 10.5969805 | NA | NA | NA | NA | NA | NA | NA |
| TRPC6 | 678 | L | 2.831125 | 15.59414621 | 0.219593343 | 9.0474918 | NA | NA | NA | NA | NA | NA | NA |
| TRPC6 | 679 | F | 1.096909091 | 14.03921023 | 0.193302562 | 7.500334793 | NA | NA | NA | NA | NA | NA | NA |
| TRPC6 | 680 | W | 4.2465 | 12.92510402 | 0.174465309 | 6.429846273 | NA | NA | NA | NA | NA | NA | NA |
| TRPC6 | 681 | A | 1.9156 | 10.97006244 | 0.141409563 | 4.452358197 | 6.64 | -0.41 | 0.52 | 2.73 | 0.39 | 0 | 0 |
| TRPC6 | 682 | I | -0.4915 | 8.202390444 | 0.094613902 | 1.664382417 | 6.64 | -0.41 | 0.52 | 2.73 | 0.39 | 0 | 0 |
| TRPC6 | 683 | F | 0.303090909 | 8.76986773 | 0.104208779 | 2.446728965 | 6.64 | -0.41 | 0.52 | 2.73 | 0.39 | 0 | 0 |
| TRPC6 | 684 | G | 3.019 | 9.617428658 | 0.118539298 | 3.13363359 | 6.64 | -0.41 | 0.52 | 2.73 | 0.39 | 0 | 0 |

| | | | | | | | | | | | | | |
|---|---|---|---|---|---|---|---|---|---|---|---|---|---|
| TRPC6 | 685 | L | 5.356375 | 10.00229304 | 0.125046566 | 8.332944624 | NA | NA | NA | NA | NA | NA | NA |
| TRPC6 | 686 | S | 6.738833333 | 14.18262867 | 0.195727474 | 12.5088019 | NA | NA | NA | NA | NA | NA | NA |
| TRPC6 | 687 | E | 10.07722222 | 14.45213593 | 0.200284289 | 12.74457001 | NA | NA | NA | NA | NA | NA | NA |
| TRPC6 | 688 | V | 9.329857143 | 16.76884176 | 0.239455037 | 15.26252596 | NA | NA | NA | NA | NA | NA | NA |
| TRPC6 | 689 | K | 14.35877778 | 19.71698458 | 0.28930209 | 18.00900336 | NA | NA | NA | NA | NA | NA | NA |
| TRPC6 | 690 | S | 11.35866667 | 20.09446476 | 0.295684506 | 18.96254865 | NA | NA | NA | NA | NA | NA | NA |
| TRPC6 | 691 | V | 11.02042857 | 21.62103009 | 0.321495598 | 15.11820808 | NA | NA | NA | NA | NA | NA | NA |
| TRPC6 | 692 | V | 15.23314286 | 24.49676121 | 0.370118317 | 22.78568334 | NA | NA | NA | NA | NA | NA | NA |
| TRPC6 | 693 | I | 17.342625 | 28.57303089 | 0.439039684 | 26.86664166 | NA | NA | NA | NA | NA | NA | NA |
| TRPC6 | 694 | N | 21.70875 | 29.69531993 | 0.458015291 | 27.98062208 | NA | NA | NA | NA | NA | NA | NA |
| TRPC6 | 695 | Y | 22.34941667 | 33.54400395 | 0.523088651 | 31.83178139 | NA | NA | NA | NA | NA | NA | NA |
| TRPC6 | 696 | N | 20.811625 | 35.69381857 | 0.55943761 | 34.07767482 | NA | NA | NA | NA | NA | NA | NA |
| TRPC6 | 697 | H | 18.5062 | 32.36160183 | 0.503096654 | 25.85444053 | NA | NA | NA | NA | NA | NA | NA |
| TRPC6 | 698 | K | 15.74011111 | 30.52673533 | 0.472072822 | 24.00826314 | NA | NA | NA | NA | NA | NA | NA |
| TRPC6 | 699 | F | 11.24509091 | 30.23085154 | 0.467070033 | 23.70055406 | NA | NA | NA | NA | NA | NA | NA |
| TRPC6 | 700 | I | 12.528125 | 28.2189509 | 0.433052917 | 21.69717151 | NA | NA | NA | NA | NA | NA | NA |
| TRPC6 | 701 | E | 13.01711111 | 25.91081822 | 0.394027123 | 19.39505572 | NA | NA | NA | NA | NA | NA | NA |
| TRPC6 | 702 | N | 9.62125 | 25.57684865 | 0.388380382 | 19.0397241 | NA | NA | NA | NA | NA | NA | NA |
| TRPC6 | 703 | I | 7.25375 | 24.85769501 | 0.376220968 | 18.31401259 | NA | NA | NA | NA | NA | NA | NA |
| TRPC6 | 704 | G | 7.40475 | 22.16424386 | 0.330680229 | 15.63605422 | NA | NA | NA | NA | NA | NA | NA |
| TRPC6 | 705 | Y | 8.458 | 20.54459428 | 0.303295274 | 14.00804462 | NA | NA | NA | NA | NA | NA | NA |
| TRPC6 | 706 | V | 3.339571429 | 20.86857496 | 0.308773124 | 14.30463258 | NA | NA | NA | NA | NA | NA | NA |
| TRPC6 | 707 | L | 2.98025 | 19.70101901 | 0.289032145 | 13.13908312 | NA | NA | NA | NA | NA | NA | NA |
| TRPC6 | 708 | Y | 4.23375 | 15.89678939 | 0.224710419 | 9.398812138 | NA | NA | NA | NA | NA | NA | NA |
| TRPC6 | 709 | G | 0.75575 | 16.02631352 | 0.226900406 | 9.450127299 | NA | NA | NA | NA | NA | NA | NA |
| TRPC6 | 710 | V | -0.498285714 | 16.96668494 | 0.24280016 | 10.36327521 | NA | NA | NA | NA | NA | NA | NA |
| TRPC6 | 711 | Y | 0 | 14.71777595 | 0.204775718 | 8.123586317 | NA | NA | NA | NA | NA | NA | NA |
| TRPC6 | 712 | N | -0.77075 | 11.87134883 | 0.156648469 | 5.270430279 | NA | NA | NA | NA | NA | NA | NA |
| TRPC6 | 713 | V | -3.557857143 | 13.35233792 | 0.181688959 | 6.717908758 | NA | NA | NA | NA | NA | NA | NA |
| TRPC6 | 714 | T | -4.974857143 | 14.20792698 | 0.196155217 | 7.648386431 | NA | NA | NA | NA | NA | NA | NA |
| TRPC6 | 715 | M | -4.238125 | 11.0444975 | 0.142668107 | 4.478757907 | 6.64 | -0.41 | 0.52 | 2.73 | 0.39 | 0 | 0 |
| TRPC6 | 716 | V | -6.106857143 | 8.862056251 | 0.105767498 | 2.95831422 | 6.06 | 0.75 | 2.96 | 0.78 | -1.27 | 0 | 0 |
| TRPC6 | 717 | I | -9.149625 | 11.69993949 | 0.153750288 | 8.691129502 | NA | NA | NA | NA | NA | NA | NA |

| | | | | | | | | | | | | | |
|---|---|---|---|---|---|---|---|---|---|---|---|---|---|
| TRPC6 | 718 | V | -9.472571429 | 11.77633054 | 0.155041904 | 8.801514379 | NA | NA | NA | NA | NA | NA | NA |
| TRPC6 | 719 | L | -8.611 | 8.30895908 | 0.096415759 | 5.576628802 | 3.07 | 1.14 | 3.8 | 0.13 | -1.79 | 0 | 0 |
| TRPC6 | 720 | L | -12.1145 | 7.609703082 | 0.084592773 | 6.135497985 | 1.62 | 1.14 | 3.8 | 0.13 | -1.79 | 0 | 0 |
| TRPC6 | 721 | N | -14.748 | 10.0014491 | 0.125032296 | 7.948616009 | NA | NA | NA | NA | NA | NA | NA |
| TRPC6 | 722 | M | -13.27225 | 9.36553581 | 0.114280306 | 7.185578426 | NA | NA | NA | NA | NA | NA | NA |
| TRPC6 | 723 | L | -14.4935 | 3.436170252 | 0.014026886 | 1.566581809 | 2.18 | 1.27 | 3.94 | 0.13 | -1.8 | 0 | 0 |
| TRPC6 | 724 | I | -17.6255 | 6.651470063 | 0.068391016 | 5.505500806 | 1.16 | 1.81 | 4.5 | 0.13 | -1.85 | 0 | 0 |
| TRPC6 | 725 | A | -19.1712 | 8.860045429 | 0.1057335 | 7.800174493 | NA | NA | NA | NA | NA | NA | NA |
| TRPC6 | 726 | M | -17.768125 | 7.980285083 | 0.090858555 | 6.891106369 | 1.1 | 1.81 | 4.5 | 0.13 | -1.85 | 0 | 0 |
| TRPC6 | 727 | I | -19.95525 | 2.663991179 | 9.71E-04 | 1.684200178 | 1.59 | 1.81 | 4.5 | 0.13 | -1.85 | 0 | 0 |
| TRPC6 | 728 | N | -23.891875 | 6.564998248 | 0.066928955 | 4.62838648 | 2.02 | 1.35 | 2.8 | 0.35 | -1.81 | 0 | 0 |
| TRPC6 | 729 | S | -23.025 | 8.861361126 | 0.105755745 | 7.071771638 | NA | NA | NA | NA | NA | NA | NA |
| TRPC6 | 730 | S | -24.17266667 | 8.149824231 | 0.093725115 | 6.598670728 | 1.8 | 1.35 | 2.8 | 0.35 | -1.81 | 0 | 0 |
| TRPC6 | 731 | F | -26.671 | 3.292278239 | 0.011593967 | 1.695070959 | 2.11 | 1.35 | 2.8 | 0.35 | -1.81 | 0 | 0 |
| TRPC6 | 732 | Q | -28.96244444 | 8.076483084 | 0.092485066 | 5.555258562 | 2.54 | 1.35 | 2.8 | 0.35 | -1.81 | 0 | 0 |
| TRPC6 | 733 | E | -29.168 | 10.08462662 | 0.126438658 | 7.589005925 | NA | NA | NA | NA | NA | NA | NA |
| TRPC6 | 734 | I | -31.5624 | 11.69870668 | 0.153729444 | 8.727303799 | NA | NA | NA | NA | NA | NA | NA |
| TRPC6 | 735 | E | -31.0378 | 11.81188084 | 0.155642987 | 8.825297615 | NA | NA | NA | NA | NA | NA | NA |
| TRPC6 | 736 | D | -32.9324 | 14.34400819 | 0.198456071 | 8.808975744 | NA | NA | NA | NA | NA | NA | NA |
| TRPC6 | 737 | D | -30.757 | 16.29452966 | 0.231435392 | 13.31546807 | NA | NA | NA | NA | NA | NA | NA |
| TRPC6 | 738 | A | -27.8494 | 16.08659945 | 0.227919718 | 13.75104189 | NA | NA | NA | NA | NA | NA | NA |
| TRPC6 | 739 | D | -29.503875 | 18.80398067 | 0.273865065 | 16.26461085 | NA | NA | NA | NA | NA | NA | NA |
| TRPC6 | 740 | V | -31.27914286 | 20.81420796 | 0.307853889 | 17.84079656 | NA | NA | NA | NA | NA | NA | NA |
| TRPC6 | 741 | E | -27.4014 | 21.08168034 | 0.312376299 | 18.86435116 | NA | NA | NA | NA | NA | NA | NA |
| TRPC6 | 742 | W | -25.71842857 | 22.52594062 | 0.33679578 | 20.79029211 | NA | NA | NA | NA | NA | NA | NA |
| TRPC6 | 743 | K | -29.46077778 | 23.52452924 | 0.353679867 | 21.20634267 | NA | NA | NA | NA | NA | NA | NA |
| TRPC6 | 744 | F | -28.87818182 | 26.27693759 | 0.400217452 | 23.7380991 | NA | NA | NA | NA | NA | NA | NA |
| TRPC6 | 745 | A | -24.6356 | 26.41528796 | 0.402556673 | 24.69150051 | NA | NA | NA | NA | NA | NA | NA |
| TRPC6 | 746 | R | -25.97427273 | 28.3629132 | 0.435487024 | 26.59460384 | NA | NA | NA | NA | NA | NA | NA |
| TRPC6 | 747 | A | -27.5206 | 30.5108893 | 0.471804898 | 28.40709257 | NA | NA | NA | NA | NA | NA | NA |
| TRPC6 | 748 | K | -25.17488889 | 29.21241272 | 0.44985032 | 27.46033286 | NA | NA | NA | NA | NA | NA | NA |
| TRPC6 | 749 | L | -21.999125 | 31.88362548 | 0.495015054 | 29.78572009 | NA | NA | NA | NA | NA | NA | NA |
| TRPC6 | 750 | W | -26.30228571 | 32.7224151 | 0.509197267 | 31.08980737 | NA | NA | NA | NA | NA | NA | NA |

| | | | | | | | | | | | | | |
|---|---|---|---|---|---|---|---|---|---|---|---|---|---|
| TRPC6 | 751 | F | -28.22381818 | 35.88391318 | 0.56265172 | 33.97158554 | NA | NA | NA | NA | NA | NA | NA |
| TRPC6 | 752 | S | -22.294 | 36.89807026 | 0.579799038 | 34.98721765 | NA | NA | NA | NA | NA | NA | NA |
| TRPC6 | 753 | Y | -20.70566667 | 36.98004415 | 0.581185048 | 35.30197058 | NA | NA | NA | NA | NA | NA | NA |
| TRPC6 | 754 | F | -27.13018182 | 38.6454857 | 0.609344252 | 37.0022308 | NA | NA | NA | NA | NA | NA | NA |
| TRPC6 | 755 | E | -24.12566667 | 41.53207923 | 0.658150633 | 39.80381959 | NA | NA | NA | NA | NA | NA | NA |
| TRPC6 | 756 | E | -23.5498 | 46.74638265 | 0.746313817 | 44.95270411 | NA | NA | NA | NA | NA | NA | NA |
| TRPC6 | 757 | G | -19.71575 | 47.96930661 | 0.766990955 | 46.21213899 | NA | NA | NA | NA | NA | NA | NA |
| TRPC6 | 758 | R | -17.09409091 | 41.40455596 | 0.655994476 | 40.25130109 | NA | NA | NA | NA | NA | NA | NA |
| TRPC6 | 759 | T | -16.62228571 | 46.41026818 | 0.740630811 | 44.55645507 | NA | NA | NA | NA | NA | NA | NA |
| TRPC6 | 760 | L | -12.4035 | 46.09135414 | 0.735238628 | 42.39087285 | NA | NA | NA | NA | NA | NA | NA |
| TRPC6 | 761 | P | -10.56914286 | 42.9414746 | 0.68198062 | 39.27105268 | NA | NA | NA | NA | NA | NA | NA |
| TRPC6 | 762 | V | -7.560714286 | 40.26805172 | 0.636778518 | 36.59298041 | NA | NA | NA | NA | NA | NA | NA |
| TRPC6 | 763 | P | -5.369 | 42.89067079 | 0.681121632 | 39.24334902 | NA | NA | NA | NA | NA | NA | NA |
| TRPC6 | 764 | F | -7.166727273 | 45.83274295 | 0.730866042 | 42.21398457 | NA | NA | NA | NA | NA | NA | NA |
| TRPC6 | 765 | N | -8.77875 | 45.55096918 | 0.726101825 | 41.8414688 | NA | NA | NA | NA | NA | NA | NA |
| TRPC6 | 766 | L | -5.3295 | 47.47360549 | 0.758609665 | 43.80347431 | NA | NA | NA | NA | NA | NA | NA |
| TRPC6 | 767 | V | -6.864714286 | 51.73657331 | 0.830687714 | 48.06105019 | NA | NA | NA | NA | NA | NA | NA |
| TRPC6 | 853 | R | -53.94027273 | 44.03148393 | 0.700410444 | 41.90582029 | NA | NA | NA | NA | NA | NA | NA |
| TRPC6 | 854 | Q | -53.75766667 | 41.69997282 | 0.660989369 | 38.628581 | NA | NA | NA | NA | NA | NA | NA |
| TRPC6 | 855 | Y | -50.16425 | 38.99657822 | 0.615280507 | 35.3978985 | NA | NA | NA | NA | NA | NA | NA |
| TRPC6 | 856 | Q | -54.75088889 | 38.91043395 | 0.613823984 | 35.84192664 | NA | NA | NA | NA | NA | NA | NA |
| TRPC6 | 857 | K | -57.18566667 | 37.5420451 | 0.590687333 | 35.39477946 | NA | NA | NA | NA | NA | NA | NA |
| TRPC6 | 858 | I | -52.588125 | 35.58699483 | 0.557631439 | 32.51937866 | NA | NA | NA | NA | NA | NA | NA |
| TRPC6 | 859 | M | -52.16275 | 34.26428435 | 0.535267116 | 31.17668317 | NA | NA | NA | NA | NA | NA | NA |
| TRPC6 | 860 | K | -58.52166667 | 33.12830154 | 0.516059975 | 31.19557789 | NA | NA | NA | NA | NA | NA | NA |
| TRPC6 | 861 | R | -55.73209091 | 31.41927994 | 0.487163922 | 29.48772113 | NA | NA | NA | NA | NA | NA | NA |
| TRPC6 | 862 | L | -52.3535 | 29.63296479 | 0.456960994 | 26.56405192 | NA | NA | NA | NA | NA | NA | NA |
| TRPC6 | 863 | I | -54.913625 | 28.54359072 | 0.438541911 | 26.2075207 | NA | NA | NA | NA | NA | NA | NA |
| TRPC6 | 864 | K | -59.48211111 | 27.06913161 | 0.413611829 | 25.23777985 | NA | NA | NA | NA | NA | NA | NA |
| TRPC6 | 865 | R | -54.47654545 | 25.29285682 | 0.383578663 | 23.07134236 | NA | NA | NA | NA | NA | NA | NA |
| TRPC6 | 866 | Y | -52.1135 | 23.57055021 | 0.354457987 | 20.75527428 | NA | NA | NA | NA | NA | NA | NA |
| TRPC6 | 867 | V | -57.32914286 | 22.57308366 | 0.337592872 | 20.65805549 | NA | NA | NA | NA | NA | NA | NA |
| TRPC6 | 868 | L | -58.94675 | 21.00794138 | 0.311129524 | 19.18572143 | NA | NA | NA | NA | NA | NA | NA |

| | | | | | | | | | | | | | |
|---|---|---|---|---|---|---|---|---|---|---|---|---|---|
| TRPC6 | 869 | Q | -54.05088889 | 19.48070784 | 0.285307135 | 17.24301427 | NA | NA | NA | NA | NA | NA | NA |
| TRPC6 | 870 | A | -54.8328 | 18.18042876 | 0.26332208 | 15.10838733 | NA | NA | NA | NA | NA | NA | NA |
| TRPC6 | 871 | Q | -59.13866667 | 16.45201586 | 0.234098161 | 14.61280675 | NA | NA | NA | NA | NA | NA | NA |
| TRPC6 | 872 | I | -56.99875 | 15.07840064 | 0.210873142 | 13.14096406 | NA | NA | NA | NA | NA | NA | NA |
| TRPC6 | 873 | D | -53.0875 | 13.71815301 | 0.187874142 | 10.64693635 | NA | NA | NA | NA | NA | NA | NA |
| TRPC6 | 874 | K | -56.49344444 | 12.42835749 | 0.166066344 | 10.24426222 | NA | NA | NA | NA | NA | NA | NA |
| TRPC6 | 875 | E | -59.57022222 | 11.42057704 | 0.149026842 | 9.596974491 | NA | NA | NA | NA | NA | NA | NA |
| TRPC6 | 876 | S | -54.73933333 | 9.064148333 | 0.109184461 | 7.054314265 | NA | NA | NA | NA | NA | NA | NA |
| TRPC6 | 877 | D | -54.032375 | 7.429229099 | 0.081541328 | 4.392507551 | 3.09 | -1.07 | -2.88 | 40.6 | 1.95 | 4 | -4 |
| TRPC6 | 878 | E | -54.7388 | 4.056424287 | 0.024514111 | 1.821462188 | 2.36 | -1.07 | -2.88 | 40.6 | 1.95 | 4 | -4 |
| TRPC6 | 879 | V | -58.24285714 | 2.632556742 | 4.39E-04 | 0.802177407 | 1.85 | 1.13 | 4.2 | 0.13 | -1.3 | 0 | 0 |
| TRPC6 | 880 | N | -57.897125 | 5.707795546 | 0.052435414 | 3.943870159 | 1.94 | 1.13 | 4.2 | 0.13 | -1.3 | 0 | 0 |
| TRPC6 | 881 | E | -59.47655556 | 7.437903334 | 0.081687991 | 5.589066474 | 1.85 | 1.13 | 4.2 | 0.13 | -1.3 | 0 | 0 |
| TRPC6 | 882 | G | -61.77825 | 9.251630397 | 0.112354399 | 7.528680135 | NA | NA | NA | NA | NA | NA | NA |
| TRPC6 | 883 | E | -61.91533333 | 6.807064419 | 0.071021798 | 5.841871643 | 1.41 | 1.14 | 3.88 | 0.13 | -1.69 | 0 | 0 |
| TRPC6 | 884 | L | -64.773 | 2.713211013 | 0.001803127 | 1.581020841 | 1.13 | 1.14 | 3.8 | 0.13 | -1.79 | 0 | 0 |
| TRPC6 | 885 | K | -66.35488889 | 7.935896295 | 0.090108032 | 6.70735497 | 1.44 | 1.27 | 3.94 | 0.13 | -1.8 | 0 | 0 |
| TRPC6 | 886 | E | -68.03966667 | 8.187294181 | 0.094358655 | 6.077840536 | 2.18 | 1.68 | 4.36 | 0.13 | -1.84 | 0 | 0 |
| TRPC6 | 887 | I | -69.305875 | 3.431905593 | 0.013954779 | 1.533701466 | 1.9 | 1.81 | 4.5 | 0.13 | -1.85 | 0 | 0 |
| TRPC6 | 888 | K | -70.428 | 6.445480122 | 0.064908149 | 4.533559346 | 1.98 | 1.81 | 4.5 | 0.13 | -1.85 | 0 | 0 |
| TRPC6 | 889 | Q | -72.07155556 | 8.811328277 | 0.104909792 | 6.941763951 | NA | NA | NA | NA | NA | NA | NA |
| TRPC6 | 890 | D | -73.6655 | 6.806798513 | 0.071017302 | 5.204181676 | 1.77 | 1.81 | 4.5 | 0.13 | -1.85 | 0 | 0 |
| TRPC6 | 891 | I | -75.061 | 2.919920204 | 0.005298156 | 1.7908925 | 1.98 | 1.81 | 4.5 | 0.13 | -1.85 | 0 | 0 |
| TRPC6 | 892 | S | -76.69983333 | 7.663836768 | 0.085508063 | 5.17857423 | 2.52 | 1.27 | 3.94 | 0.13 | -1.8 | 0 | 0 |
| TRPC6 | 893 | S | -78.70866667 | 8.853557759 | 0.105623806 | 6.932080365 | NA | NA | NA | NA | NA | NA | NA |
| TRPC6 | 894 | L | -80.178625 | 3.253270508 | 0.010934426 | 1.614376634 | 1.89 | 1.14 | 3.8 | 0.13 | -1.79 | 0 | 0 |
| TRPC6 | 895 | R | -81.21654545 | 6.567024745 | 0.066963219 | 4.725575709 | 1.87 | 1.14 | 3.8 | 0.13 | -1.79 | 0 | 0 |
| TRPC6 | 896 | Y | -81.45825 | 8.975484388 | 0.107685336 | 7.087241224 | NA | NA | NA | NA | NA | NA | NA |
| TRPC6 | 897 | E | -84.24188889 | 7.964459241 | 0.090590972 | 6.226025012 | 1.83 | 1.14 | 3.8 | 0.13 | -1.79 | 0 | 0 |
| TRPC6 | 898 | L | -85.466375 | 2.632235932 | 4.34E-04 | 1.740593753 | 1.97 | 1.14 | 3.8 | 0.13 | -1.79 | 0 | 0 |
| TRPC6 | 899 | L | -86.628125 | 6.951717845 | 0.073467591 | 4.563693135 | 2.46 | 1.14 | 3.8 | 0.13 | -1.79 | 0 | 0 |
| TRPC6 | 900 | E | -89.41055556 | 9.137841047 | 0.110430454 | 5.625114439 | NA | NA | NA | NA | NA | NA | NA |
| TRPC6 | 901 | E | -90.6882 | 6.510683067 | 0.066010597 | 3.183129903 | 3.62 | -0.41 | -3.9 | 49.5 | 1.46 | 4 | 4 |

| | | | | | | | | | | | | | |
|---|---|---|---|---|---|---|---|---|---|---|---|---|---|
| TRPC6 | 902 | K | -91.9342 | 5.15417782 | 0.043074873 | 1.520497165 | 3.66 | -0.41 | -3.9 | 49.5 | 1.46 | 4 | 4 |
| TRPC6 | 903 | S | -93.19416667 | 7.769595356 | 0.087296224 | 3.898623826 | 3.92 | -0.78 | -2.88 | 3.38 | 0.59 | 0 | 0 |
| TRPC6 | 904 | Q | -95.15077778 | 8.4157254 | 0.098220958 | 4.945779711 | 3.7 | -0.78 | -2.88 | 3.38 | 0.59 | 0 | 0 |
| TRPC6 | 905 | N | -96.5874 | 4.900633633 | 0.03878796 | 1.370806188 | 3.57 | -0.77 | -3.5 | 3.38 | 0.54 | 0 | 0 |
| TRPC6 | 906 | T | -97.9242 | 6.116820171 | 0.059351182 | 2.895871088 | 3.29 | -0.77 | -3.5 | 3.38 | 0.54 | 0 | 0 |
| TRPC6 | 907 | E | -99.4338 | 8.505988361 | 0.09974712 | 5.711591525 | 2.83 | 0.75 | 2.96 | 0.78 | -1.27 | 0 | 0 |
| TRPC6 | 908 | D | -101.1126 | 7.529519108 | 0.083237026 | 5.07539015 | 2.63 | 1.14 | 3.8 | 0.13 | -1.79 | 0 | 0 |
| TRPC6 | 909 | L | -102.5226 | 4.127703599 | 0.025719298 | 1.667001937 | 2.47 | 1.14 | 3.8 | 0.13 | -1.79 | 0 | 0 |
| TRPC6 | 910 | A | -103.8156 | 6.905415411 | 0.072684712 | 4.226994025 | 2.76 | 1.81 | 4.5 | 0.13 | -1.85 | 0 | 0 |
| TRPC6 | 911 | E | -105.463 | 9.063059362 | 0.109166049 | 6.223595396 | NA | NA | NA | NA | NA | NA | NA |
| TRPC6 | 912 | L | -106.61475 | 6.123934193 | 0.059471466 | 4.906547383 | 1.57 | 1.81 | 4.5 | 0.13 | -1.85 | 0 | 0 |
| TRPC6 | 913 | I | -108.0245 | 2.631560184 | 4.23E-04 | 1.282100629 | 1.57 | 1.81 | 4.5 | 0.13 | -1.85 | 0 | 0 |
| TRPC6 | 914 | R | -109.5992727 | 7.929881273 | 0.09000633 | 6.578552833 | 1.45 | 1.68 | 4.36 | 0.13 | -1.84 | 0 | 0 |
| TRPC6 | 915 | E | -111.4776 | 8.590646425 | 0.101178514 | 6.83423197 | NA | NA | NA | NA | NA | NA | NA |
| TRPC6 | 916 | L | -113.187625 | 2.743926748 | 0.002322467 | 1.630979663 | 1.11 | 1.14 | 3.8 | 0.13 | -1.79 | 0 | 0 |
| TRPC6 | 917 | G | -114.049 | 6.703204756 | 0.069265744 | 5.59081926 | 1.11 | 1.14 | 3.8 | 0.13 | -1.79 | 0 | 0 |
| TRPC6 | 918 | E | -115.271 | 9.026540256 | 0.108548586 | 7.457458982 | NA | NA | NA | NA | NA | NA | NA |
| TRPC6 | 919 | K | -117.6742222 | 6.02918701 | 0.057869485 | 3.940013781 | 2.22 | 1.14 | 3.8 | 0.13 | -1.79 | 0 | 0 |
| TRPC6 | 920 | L | -118.74 | 3.721271423 | 0.018847363 | 1.575997441 | 2.26 | 1.14 | 3.8 | 0.13 | -1.79 | 0 | 0 |
| TRPM4 | 7 | E | -53.5572 | 53.33753306 | 0.739366873 | 47.88671865 | NA | NA | NA | NA | NA | NA | NA |
| TRPM4 | 8 | Q | -56.85355556 | 52.8021307 | 0.731613666 | 48.45499787 | NA | NA | NA | NA | NA | NA | NA |
| TRPM4 | 9 | S | -56.781 | 58.06303224 | 0.807797224 | 54.56146405 | NA | NA | NA | NA | NA | NA | NA |
| TRPM4 | 10 | W | -59.80785714 | 54.84550309 | 0.761203914 | 51.47658054 | NA | NA | NA | NA | NA | NA | NA |
| TRPM4 | 11 | I | -62.356625 | 54.82561626 | 0.760915931 | 52.70787269 | NA | NA | NA | NA | NA | NA | NA |
| TRPM4 | 12 | P | -62.02357143 | 60.33786122 | 0.840739214 | 57.99403821 | NA | NA | NA | NA | NA | NA | NA |
| TRPM4 | 13 | K | -62.347 | 62.66472428 | 0.874434714 | 60.54526267 | NA | NA | NA | NA | NA | NA | NA |
| TRPM4 | 14 | I | -64.83775 | 60.34631556 | 0.840861642 | 58.2926704 | NA | NA | NA | NA | NA | NA | NA |
| TRPM4 | 15 | F | -67.16581818 | 56.41044113 | 0.783865913 | 53.77112876 | NA | NA | NA | NA | NA | NA | NA |
| TRPM4 | 16 | K | -69.94755556 | 61.70398712 | 0.860522198 | 58.86304318 | NA | NA | NA | NA | NA | NA | NA |
| TRPM4 | 17 | K | -71.13811111 | 62.93317712 | 0.878322202 | 60.52319178 | NA | NA | NA | NA | NA | NA | NA |
| TRPM4 | 18 | K | -76.40455556 | 64.1233803 | 0.895557634 | 62.39844537 | NA | NA | NA | NA | NA | NA | NA |
| TRPM4 | 19 | T | -75.49942857 | 67.31784662 | 0.94181697 | 65.55791603 | NA | NA | NA | NA | NA | NA | NA |
| TRPM4 | 66 | H | -81.8324 | 71.33571085 | 1 | 68.9140954 | NA | NA | NA | NA | NA | NA | NA |

| | | | | | | | | | | | | | |
|---|---|---|---|---|---|---|---|---|---|---|---|---|---|
| TRPM4 | 67 | T | -78.40814286 | 68.6632708 | 0.961300171 | 65.48410006 | NA | NA | NA | NA | NA | NA | NA |
| TRPM4 | 68 | T | -74.8886 | 68.62891503 | 0.960802662 | 66.87452097 | NA | NA | NA | NA | NA | NA | NA |
| TRPM4 | 69 | E | -71.666 | 68.76696234 | 0.962801737 | 66.8906816 | NA | NA | NA | NA | NA | NA | NA |
| TRPM4 | 70 | K | -69.05222222 | 65.7448706 | 0.919038572 | 63.01554004 | NA | NA | NA | NA | NA | NA | NA |
| TRPM4 | 71 | P | -65.95185714 | 64.51329775 | 0.901204062 | 61.94449614 | NA | NA | NA | NA | NA | NA | NA |
| TRPM4 | 72 | T | -66.58071429 | 60.01695101 | 0.836092086 | 57.45750228 | NA | NA | NA | NA | NA | NA | NA |
| TRPM4 | 73 | D | -65.256125 | 59.06012817 | 0.822236254 | 57.05885226 | NA | NA | NA | NA | NA | NA | NA |
| TRPM4 | 74 | A | -66.1314 | 56.08513273 | 0.779155095 | 53.51553198 | NA | NA | NA | NA | NA | NA | NA |
| TRPM4 | 75 | Y | -62.54566667 | 54.30502141 | 0.753377153 | 52.25258968 | NA | NA | NA | NA | NA | NA | NA |
| TRPM4 | 76 | G | -62.72925 | 50.36450517 | 0.696314206 | 48.26427654 | NA | NA | NA | NA | NA | NA | NA |
| TRPM4 | 77 | E | -61.7984 | 47.68869997 | 0.657565646 | 45.587682 | NA | NA | NA | NA | NA | NA | NA |
| TRPM4 | 78 | L | -64.9935 | 45.30789751 | 0.623089045 | 43.24922157 | NA | NA | NA | NA | NA | NA | NA |
| TRPM4 | 79 | D | -62.26975 | 43.53699926 | 0.597444519 | 41.18007512 | NA | NA | NA | NA | NA | NA | NA |
| TRPM4 | 80 | F | -66.63809091 | 42.4131881 | 0.581170515 | 40.35538367 | NA | NA | NA | NA | NA | NA | NA |
| TRPM4 | 81 | T | -62.95342857 | 42.28035938 | 0.579247011 | 39.93899244 | NA | NA | NA | NA | NA | NA | NA |
| TRPM4 | 82 | G | -63.4835 | 38.79442283 | 0.52876687 | 36.73711236 | NA | NA | NA | NA | NA | NA | NA |
| TRPM4 | 83 | A | -64.0036 | 35.19128712 | 0.476589559 | 33.13128704 | NA | NA | NA | NA | NA | NA | NA |
| TRPM4 | 84 | G | -61.918 | 36.40247538 | 0.494128878 | 34.0517408 | NA | NA | NA | NA | NA | NA | NA |
| TRPM4 | 85 | R | -60.3256 | 34.72237994 | 0.469799275 | 31.54239406 | NA | NA | NA | NA | NA | NA | NA |
| TRPM4 | 86 | K | -61.5046 | 38.18243253 | 0.519904587 | 35.52300047 | NA | NA | NA | NA | NA | NA | NA |
| TRPM4 | 87 | H | -60.3462 | 41.78852545 | 0.572124722 | 39.67674677 | NA | NA | NA | NA | NA | NA | NA |
| TRPM4 | 88 | S | -65.22816667 | 43.77443805 | 0.60088289 | 41.74449913 | NA | NA | NA | NA | NA | NA | NA |
| TRPM4 | 89 | N | -64.737 | 45.79886099 | 0.630198729 | 43.75523594 | NA | NA | NA | NA | NA | NA | NA |
| TRPM4 | 90 | F | -66.57981818 | 49.19202919 | 0.679335483 | 46.57540281 | NA | NA | NA | NA | NA | NA | NA |
| TRPM4 | 91 | L | -68.344875 | 51.97849583 | 0.719686541 | 49.32859173 | NA | NA | NA | NA | NA | NA | NA |
| TRPM4 | 92 | R | -71.26127273 | 55.29248827 | 0.767676744 | 52.16866513 | NA | NA | NA | NA | NA | NA | NA |
| TRPM4 | 93 | L | -72.456125 | 54.80773953 | 0.760657057 | 52.39782453 | NA | NA | NA | NA | NA | NA | NA |
| TRPM4 | 94 | S | -74.566 | 59.19749077 | 0.824225414 | 57.44248865 | NA | NA | NA | NA | NA | NA | NA |
| TRPM4 | 95 | D | -78.52875 | 58.5464587 | 0.814797764 | 56.29890447 | NA | NA | NA | NA | NA | NA | NA |
| TRPM4 | 96 | R | -77.3738 | 60.26043911 | 0.839618058 | 56.81043596 | NA | NA | NA | NA | NA | NA | NA |
| TRPM4 | 97 | T | -74.17 | 58.30820068 | 0.811347529 | 56.54835187 | NA | NA | NA | NA | NA | NA | NA |
| TRPM4 | 98 | D | -72.9078 | 57.34014807 | 0.797329078 | 55.4687924 | NA | NA | NA | NA | NA | NA | NA |
| TRPM4 | 99 | P | -73.65471429 | 53.48221265 | 0.74146199 | 51.61523285 | NA | NA | NA | NA | NA | NA | NA |

| | | | | | | | | | | | | | |
|---|---|---|---|---|---|---|---|---|---|---|---|---|---|
| TRPM4 | 100 | A | -70.161 | 54.1550944 | 0.751206048 | 51.50102331 | NA | NA | NA | NA | NA | NA | NA |
| TRPM4 | 101 | A | -68.1098 | 54.14742773 | 0.751095026 | 51.509576 | NA | NA | NA | NA | NA | NA | NA |
| TRPM4 | 102 | V | -69.64871429 | 50.94151123 | 0.704669879 | 48.09190135 | NA | NA | NA | NA | NA | NA | NA |
| TRPM4 | 103 | Y | -69.10066667 | 46.53848099 | 0.640909228 | 43.67928301 | NA | NA | NA | NA | NA | NA | NA |
| TRPM4 | 104 | S | -65.38933333 | 50.81408697 | 0.702824638 | 48.3430354 | NA | NA | NA | NA | NA | NA | NA |
| TRPM4 | 105 | L | -64.6245 | 49.38956908 | 0.682196075 | 47.33093632 | NA | NA | NA | NA | NA | NA | NA |
| TRPM4 | 106 | V | -66.31328571 | 46.00407186 | 0.633170404 | 43.4384625 | NA | NA | NA | NA | NA | NA | NA |
| TRPM4 | 107 | T | -64.29885714 | 45.973896 | 0.632733425 | 43.95034856 | NA | NA | NA | NA | NA | NA | NA |
| TRPM4 | 108 | R | -60.8122 | 47.46416088 | 0.654314076 | 44.84248907 | NA | NA | NA | NA | NA | NA | NA |
| TRPM4 | 109 | T | -59.82885714 | 47.17407676 | 0.650113344 | 43.80866529 | NA | NA | NA | NA | NA | NA | NA |
| TRPM4 | 110 | W | -63.03292857 | 45.10022242 | 0.620081685 | 42.44047728 | NA | NA | NA | NA | NA | NA | NA |
| TRPM4 | 111 | G | -61.02075 | 40.82604965 | 0.558187029 | 37.65864872 | NA | NA | NA | NA | NA | NA | NA |
| TRPM4 | 112 | F | -64.819 | 41.24381734 | 0.564236758 | 39.19639339 | NA | NA | NA | NA | NA | NA | NA |
| TRPM4 | 113 | R | -64.9854 | 39.13725793 | 0.533731494 | 37.07778985 | NA | NA | NA | NA | NA | NA | NA |
| TRPM4 | 114 | A | -67.0036 | 40.11603177 | 0.5479052 | 37.5531296 | NA | NA | NA | NA | NA | NA | NA |
| TRPM4 | 115 | P | -70.44885714 | 40.05928948 | 0.547083511 | 37.02539703 | NA | NA | NA | NA | NA | NA | NA |
| TRPM4 | 116 | N | -71.57975 | 36.55989877 | 0.496408539 | 34.21291077 | NA | NA | NA | NA | NA | NA | NA |
| TRPM4 | 117 | L | -76.246375 | 35.06744534 | 0.474796196 | 33.37822732 | NA | NA | NA | NA | NA | NA | NA |
| TRPM4 | 118 | V | -77.52214286 | 39.4587404 | 0.538386909 | 37.18905412 | NA | NA | NA | NA | NA | NA | NA |
| TRPM4 | 119 | V | -79.91414286 | 36.21459106 | 0.491408109 | 33.08659939 | NA | NA | NA | NA | NA | NA | NA |
| TRPM4 | 120 | S | -82.54483333 | 39.61897972 | 0.540707348 | 37.1856105 | NA | NA | NA | NA | NA | NA | NA |
| TRPM4 | 121 | V | -83.60428571 | 36.925367 | 0.501700915 | 34.82920459 | NA | NA | NA | NA | NA | NA | NA |
| TRPM4 | 122 | L | -87.009125 | 40.38982358 | 0.551870003 | 38.28403427 | NA | NA | NA | NA | NA | NA | NA |
| TRPM4 | 123 | G | -88.49675 | 41.05346325 | 0.561480225 | 38.8738986 | NA | NA | NA | NA | NA | NA | NA |
| TRPM4 | 124 | G | -91.00225 | 41.0202864 | 0.560999788 | 39.06832186 | NA | NA | NA | NA | NA | NA | NA |
| TRPM4 | 125 | S | -91.615 | 38.44283025 | 0.523675428 | 36.56535729 | NA | NA | NA | NA | NA | NA | NA |
| TRPM4 | 126 | G | -91.26675 | 41.75861158 | 0.571691537 | 39.85998573 | NA | NA | NA | NA | NA | NA | NA |
| TRPM4 | 127 | G | -89.5845 | 40.59083893 | 0.554780923 | 38.41618363 | NA | NA | NA | NA | NA | NA | NA |
| TRPM4 | 128 | P | -91.18442857 | 37.45342032 | 0.5093477 | 35.49209317 | NA | NA | NA | NA | NA | NA | NA |
| TRPM4 | 129 | V | -89.43385714 | 33.70093608 | 0.45500766 | 31.53339432 | NA | NA | NA | NA | NA | NA | NA |
| TRPM4 | 130 | L | -89.617125 | 31.22714236 | 0.419184445 | 29.2581561 | NA | NA | NA | NA | NA | NA | NA |
| TRPM4 | 131 | Q | -92.75133333 | 28.41944225 | 0.378525903 | 26.66490756 | NA | NA | NA | NA | NA | NA | NA |
| TRPM4 | 132 | T | -89.396 | 24.73462856 | 0.325165806 | 22.59089006 | NA | NA | NA | NA | NA | NA | NA |

| | | | | | | | | | | | | | |
|---|---|---|---|---|---|---|---|---|---|---|---|---|---|
| TRPM4 | 133 | W | -91.68821429 | 24.57149922 | 0.322803517 | 22.17566107 | NA | NA | NA | NA | NA | NA | NA |
| TRPM4 | 134 | L | -89.223875 | 27.66282484 | 0.367569263 | 25.42903602 | NA | NA | NA | NA | NA | NA | NA |
| TRPM4 | 135 | Q | -85.19888889 | 27.31019496 | 0.3624628 | 25.20339461 | NA | NA | NA | NA | NA | NA | NA |
| TRPM4 | 136 | D | -85.017625 | 21.5650207 | 0.279266448 | 19.46656317 | NA | NA | NA | NA | NA | NA | NA |
| TRPM4 | 137 | L | -85.799125 | 26.32838761 | 0.348245165 | 24.22179816 | NA | NA | NA | NA | NA | NA | NA |
| TRPM4 | 138 | L | -83.206 | 28.75302913 | 0.383356603 | 26.50618256 | NA | NA | NA | NA | NA | NA | NA |
| TRPM4 | 139 | R | -79.49 | 22.98326306 | 0.299804135 | 19.98196219 | NA | NA | NA | NA | NA | NA | NA |
| TRPM4 | 140 | R | -80.64663636 | 19.4567304 | 0.248736119 | 16.32131655 | NA | NA | NA | NA | NA | NA | NA |
| TRPM4 | 141 | G | -81.006 | 26.48200759 | 0.350469749 | 23.46653802 | NA | NA | NA | NA | NA | NA | NA |
| TRPM4 | 142 | L | -80.600875 | 29.25041868 | 0.390559343 | 26.12683859 | NA | NA | NA | NA | NA | NA | NA |
| TRPM4 | 143 | V | -76.38271429 | 28.4183285 | 0.378509775 | 26.19841479 | NA | NA | NA | NA | NA | NA | NA |
| TRPM4 | 144 | R | -76.64736364 | 22.72593822 | 0.296077793 | 20.98231001 | NA | NA | NA | NA | NA | NA | NA |
| TRPM4 | 145 | A | -76.407 | 27.66933995 | 0.367663609 | 25.92993398 | NA | NA | NA | NA | NA | NA | NA |
| TRPM4 | 146 | A | -74.4414 | 30.18933522 | 0.404155872 | 28.42848672 | NA | NA | NA | NA | NA | NA | NA |
| TRPM4 | 147 | Q | -70.98977778 | 25.54360323 | 0.336880637 | 23.14516404 | NA | NA | NA | NA | NA | NA | NA |
| TRPM4 | 148 | S | -72.066 | 27.38311454 | 0.363518755 | 25.0144287 | NA | NA | NA | NA | NA | NA | NA |
| TRPM4 | 149 | T | -72.53257143 | 30.220948 | 0.40461366 | 27.82021038 | NA | NA | NA | NA | NA | NA | NA |
| TRPM4 | 150 | G | -70.59725 | 33.22678481 | 0.448141436 | 30.18833426 | NA | NA | NA | NA | NA | NA | NA |
| TRPM4 | 151 | A | -72.6234 | 35.14603143 | 0.475934207 | 33.28631905 | NA | NA | NA | NA | NA | NA | NA |
| TRPM4 | 152 | W | -72.27757143 | 37.61982016 | 0.51175735 | 35.74942041 | NA | NA | NA | NA | NA | NA | NA |
| TRPM4 | 153 | I | -76.643375 | 35.03418939 | 0.474314613 | 33.29193306 | NA | NA | NA | NA | NA | NA | NA |
| TRPM4 | 154 | V | -78.44257143 | 40.77502169 | 0.557448089 | 37.32570613 | NA | NA | NA | NA | NA | NA | NA |
| TRPM4 | 155 | T | -79.44328571 | 39.96096226 | 0.545659626 | 36.83660475 | NA | NA | NA | NA | NA | NA | NA |
| TRPM4 | 156 | G | -80.4245 | 43.92516065 | 0.603065516 | 40.79677397 | NA | NA | NA | NA | NA | NA | NA |
| TRPM4 | 157 | G | -77.67525 | 44.88148706 | 0.616914159 | 42.60175015 | NA | NA | NA | NA | NA | NA | NA |
| TRPM4 | 158 | L | -79.454 | 43.87119835 | 0.602284084 | 40.68229725 | NA | NA | NA | NA | NA | NA | NA |
| TRPM4 | 159 | H | -79.2657 | 41.78754878 | 0.572110579 | 38.60320143 | NA | NA | NA | NA | NA | NA | NA |
| TRPM4 | 160 | T | -83.18242857 | 43.57519627 | 0.597997653 | 41.29679618 | NA | NA | NA | NA | NA | NA | NA |
| TRPM4 | 161 | G | -84.4395 | 40.74881601 | 0.557068602 | 38.64147889 | NA | NA | NA | NA | NA | NA | NA |
| TRPM4 | 162 | I | -85.112875 | 38.30201322 | 0.521636245 | 36.23139716 | NA | NA | NA | NA | NA | NA | NA |
| TRPM4 | 163 | G | -81.0755 | 38.61411768 | 0.526155856 | 35.50027776 | NA | NA | NA | NA | NA | NA | NA |
| TRPM4 | 164 | R | -81.09936364 | 36.79696535 | 0.49984152 | 33.66878437 | NA | NA | NA | NA | NA | NA | NA |
| TRPM4 | 165 | H | -83.6341 | 33.78081854 | 0.456164445 | 31.38240831 | NA | NA | NA | NA | NA | NA | NA |

| | | | | | | | | | | | | | |
|---|---|---|---|---|---|---|---|---|---|---|---|---|---|
| TRPM4 | 166 | V | -79.67914286 | 34.20363438 | 0.462287276 | 31.14894477 | NA | NA | NA | NA | NA | NA | NA |
| TRPM4 | 167 | G | -76.94925 | 34.29903467 | 0.463668776 | 32.07567792 | NA | NA | NA | NA | NA | NA | NA |
| TRPM4 | 168 | V | -78.373 | 31.36506313 | 0.421181687 | 27.93003849 | NA | NA | NA | NA | NA | NA | NA |
| TRPM4 | 169 | A | -77.7766 | 29.6025747 | 0.395658944 | 26.37896208 | NA | NA | NA | NA | NA | NA | NA |
| TRPM4 | 170 | V | -74.35828571 | 30.51788304 | 0.408913601 | 28.87073328 | NA | NA | NA | NA | NA | NA | NA |
| TRPM4 | 171 | R | -74.16527273 | 29.75811535 | 0.397911341 | 27.8834834 | NA | NA | NA | NA | NA | NA | NA |
| TRPM4 | 172 | D | -74.6104 | 26.48235131 | 0.350474727 | 24.80453338 | NA | NA | NA | NA | NA | NA | NA |
| TRPM4 | 173 | H | -73.3491 | 24.53051457 | 0.322210014 | 22.67183971 | NA | NA | NA | NA | NA | NA | NA |
| TRPM4 | 174 | Q | -69.456 | 26.47807546 | 0.350412808 | 23.46877066 | NA | NA | NA | NA | NA | NA | NA |
| TRPM4 | 175 | M | -70.258 | 24.90449578 | 0.327625668 | 21.86557468 | NA | NA | NA | NA | NA | NA | NA |
| TRPM4 | 176 | A | -71.1676 | 21.59407118 | 0.279687131 | 19.23443228 | NA | NA | NA | NA | NA | NA | NA |
| TRPM4 | 177 | S | -68.745 | 20.50404511 | 0.263902371 | 17.77553776 | NA | NA | NA | NA | NA | NA | NA |
| TRPM4 | 178 | T | -65.152 | 21.89834491 | 0.284093344 | 19.42359648 | NA | NA | NA | NA | NA | NA | NA |
| TRPM4 | 179 | G | -65.68025 | 23.30110163 | 0.304406782 | 20.74702291 | NA | NA | NA | NA | NA | NA | NA |
| TRPM4 | 180 | G | -65.1055 | 25.40866642 | 0.334926605 | 23.3637868 | NA | NA | NA | NA | NA | NA | NA |
| TRPM4 | 181 | T | -65.65628571 | 28.81021043 | 0.38418465 | 26.75553777 | NA | NA | NA | NA | NA | NA | NA |
| TRPM4 | 182 | K | -66.9542 | 30.9578659 | 0.41528503 | 28.40084681 | NA | NA | NA | NA | NA | NA | NA |
| TRPM4 | 183 | V | -70.79614286 | 31.17614519 | 0.418445951 | 28.19304882 | NA | NA | NA | NA | NA | NA | NA |
| TRPM4 | 184 | V | -70.06985714 | 35.43131094 | 0.480065364 | 32.39177462 | NA | NA | NA | NA | NA | NA | NA |
| TRPM4 | 185 | A | -72.394 | 38.59800246 | 0.52592249 | 36.74105179 | NA | NA | NA | NA | NA | NA | NA |
| TRPM4 | 186 | M | -73.3245 | 41.61834901 | 0.569660383 | 39.74111212 | NA | NA | NA | NA | NA | NA | NA |
| TRPM4 | 187 | G | -74.902 | 44.58775842 | 0.61266065 | 42.85075599 | NA | NA | NA | NA | NA | NA | NA |
| TRPM4 | 188 | V | -77.682 | 46.93968276 | 0.646719065 | 44.67254695 | NA | NA | NA | NA | NA | NA | NA |
| TRPM4 | 189 | A | -77.7608 | 49.72820996 | 0.687099962 | 46.43320969 | NA | NA | NA | NA | NA | NA | NA |
| TRPM4 | 190 | P | -79.77 | 51.84731497 | 0.7177869 | 48.6827691 | NA | NA | NA | NA | NA | NA | NA |
| TRPM4 | 191 | W | -74.82857143 | 55.38426939 | 0.769005834 | 53.10459778 | NA | NA | NA | NA | NA | NA | NA |
| TRPM4 | 192 | G | -79.415 | 56.69626335 | 0.788004929 | 53.5137874 | NA | NA | NA | NA | NA | NA | NA |
| TRPM4 | 193 | V | -80.30028571 | 52.76146158 | 0.731024733 | 49.63377904 | NA | NA | NA | NA | NA | NA | NA |
| TRPM4 | 194 | V | -76.09057143 | 52.5965886 | 0.728637194 | 50.83616523 | NA | NA | NA | NA | NA | NA | NA |
| TRPM4 | 195 | R | -76.93781818 | 51.56930288 | 0.713760983 | 49.95729755 | NA | NA | NA | NA | NA | NA | NA |
| TRPM4 | 196 | N | -72.458375 | 54.08889761 | 0.750247446 | 52.21291555 | NA | NA | NA | NA | NA | NA | NA |
| TRPM4 | 197 | R | -73.28372727 | 57.27534786 | 0.796390701 | 54.86923944 | NA | NA | NA | NA | NA | NA | NA |
| TRPM4 | 198 | D | -67.7726 | 59.03088657 | 0.821812804 | 56.39203569 | NA | NA | NA | NA | NA | NA | NA |

| | | | | | | | | | | | | | |
|---|---|---|---|---|---|---|---|---|---|---|---|---|---|
| TRPM4 | 199 | T | -66.03742857 | 54.94870168 | 0.762698341 | 52.40825045 | NA | NA | NA | NA | NA | NA | NA |
| TRPM4 | 200 | L | -66.725125 | 52.9766608 | 0.734141051 | 50.40691749 | NA | NA | NA | NA | NA | NA | NA |
| TRPM4 | 201 | I | -66.226375 | 58.39567016 | 0.812614182 | 55.83287715 | NA | NA | NA | NA | NA | NA | NA |
| TRPM4 | 202 | N | -62.2394 | 60.04326918 | 0.836473202 | 57.92865327 | NA | NA | NA | NA | NA | NA | NA |
| TRPM4 | 203 | P | -60.77828571 | 62.2230395 | 0.86803864 | 59.57292588 | NA | NA | NA | NA | NA | NA | NA |
| TRPM4 | 204 | K | -58.301 | 58.76118517 | 0.817907236 | 55.23228824 | NA | NA | NA | NA | NA | NA | NA |
| TRPM4 | 205 | G | -59.03975 | 56.66918839 | 0.787612854 | 53.30193031 | NA | NA | NA | NA | NA | NA | NA |
| TRPM4 | 206 | S | -59.722 | 52.9224086 | 0.733355421 | 49.55364131 | NA | NA | NA | NA | NA | NA | NA |
| TRPM4 | 207 | F | -56.96381818 | 49.51922159 | 0.684073583 | 46.00091013 | NA | NA | NA | NA | NA | NA | NA |
| TRPM4 | 208 | P | -58.13785714 | 49.30096959 | 0.680913058 | 45.96296248 | NA | NA | NA | NA | NA | NA | NA |
| TRPM4 | 209 | A | -62.2876 | 49.61324231 | 0.685435105 | 47.50833556 | NA | NA | NA | NA | NA | NA | NA |
| TRPM4 | 210 | R | -63.1774 | 50.46311193 | 0.697742139 | 48.10603595 | NA | NA | NA | NA | NA | NA | NA |
| TRPM4 | 211 | Y | -68.7035 | 49.20082204 | 0.679462813 | 46.71138469 | NA | NA | NA | NA | NA | NA | NA |
| TRPM4 | 212 | R | -67.06545455 | 50.98345152 | 0.70527722 | 48.38936765 | NA | NA | NA | NA | NA | NA | NA |
| TRPM4 | 213 | W | -72.50328571 | 43.81180793 | 0.601424046 | 41.93431192 | NA | NA | NA | NA | NA | NA | NA |
| TRPM4 | 214 | R | -72.26018182 | 47.80410118 | 0.65923678 | 45.95611907 | NA | NA | NA | NA | NA | NA | NA |
| TRPM4 | 215 | G | -72.818 | 55.01838734 | 0.763707465 | 52.62197291 | NA | NA | NA | NA | NA | NA | NA |
| TRPM4 | 221 | V | -85.15271429 | 53.80862806 | 0.746188839 | 51.70193134 | NA | NA | NA | NA | NA | NA | NA |
| TRPM4 | 222 | Q | -80.57822222 | 51.9305223 | 0.718991832 | 48.91914111 | NA | NA | NA | NA | NA | NA | NA |
| TRPM4 | 223 | F | -81.63163636 | 48.87744971 | 0.674780031 | 45.84901615 | NA | NA | NA | NA | NA | NA | NA |
| TRPM4 | 224 | P | -77.17657143 | 47.60459912 | 0.656347774 | 45.87144565 | NA | NA | NA | NA | NA | NA | NA |
| TRPM4 | 225 | L | -74.018875 | 45.51306729 | 0.626060126 | 43.78851938 | NA | NA | NA | NA | NA | NA | NA |
| TRPM4 | 226 | D | -74.697375 | 40.12809242 | 0.548079852 | 38.37494313 | NA | NA | NA | NA | NA | NA | NA |
| TRPM4 | 227 | Y | -71.87733333 | 38.98600735 | 0.531541222 | 36.57735129 | NA | NA | NA | NA | NA | NA | NA |
| TRPM4 | 228 | N | -71.966125 | 35.35103247 | 0.478902845 | 32.99653745 | NA | NA | NA | NA | NA | NA | NA |
| TRPM4 | 229 | Y | -73.2905 | 39.53627233 | 0.539509655 | 37.3031977 | NA | NA | NA | NA | NA | NA | NA |
| TRPM4 | 230 | S | -67.8955 | 40.4732748 | 0.553078467 | 37.82492683 | NA | NA | NA | NA | NA | NA | NA |
| TRPM4 | 231 | A | -70.0198 | 43.61483017 | 0.598571594 | 40.75484376 | NA | NA | NA | NA | NA | NA | NA |
| TRPM4 | 232 | F | -71.24236364 | 44.96041288 | 0.618057091 | 42.55120705 | NA | NA | NA | NA | NA | NA | NA |
| TRPM4 | 233 | F | -72.33718182 | 49.23497978 | 0.679957454 | 47.36164915 | NA | NA | NA | NA | NA | NA | NA |
| TRPM4 | 234 | L | -73.611125 | 51.73569295 | 0.716170492 | 49.87376037 | NA | NA | NA | NA | NA | NA | NA |
| TRPM4 | 235 | V | -77.15242857 | 53.92122734 | 0.747819399 | 52.186746 | NA | NA | NA | NA | NA | NA | NA |
| TRPM4 | 236 | D | -76.99325 | 57.77924209 | 0.803687635 | 55.5079068 | NA | NA | NA | NA | NA | NA | NA |

| | | | | | | | | | | | | | |
|---|---|---|---|---|---|---|---|---|---|---|---|---|---|
| TRPM4 | 237 | D | -80.387 | 60.23601867 | 0.839264424 | 57.11417256 | NA | NA | NA | NA | NA | NA | NA |
| TRPM4 | 238 | G | -80.88675 | 62.34402886 | 0.869790697 | 59.21565783 | NA | NA | NA | NA | NA | NA | NA |
| TRPM4 | 239 | T | -82.152 | 60.28906801 | 0.840032636 | 57.85081578 | NA | NA | NA | NA | NA | NA | NA |
| TRPM4 | 240 | H | -81.5541 | 60.40243873 | 0.841674367 | 57.96851833 | NA | NA | NA | NA | NA | NA | NA |
| TRPM4 | 241 | G | -84.37225 | 57.52023872 | 0.799936986 | 55.49290264 | NA | NA | NA | NA | NA | NA | NA |
| TRPM4 | 242 | C | -85.4582 | 55.90441282 | 0.776538075 | 53.79520675 | NA | NA | NA | NA | NA | NA | NA |
| TRPM4 | 243 | L | -86.8296 | 52.72535381 | 0.730501854 | 50.63297207 | NA | NA | NA | NA | NA | NA | NA |
| TRPM4 | 244 | G | -87.2035 | 52.70393595 | 0.7301917 | 50.3481422 | NA | NA | NA | NA | NA | NA | NA |
| TRPM4 | 245 | G | -85.2055 | 54.49772334 | 0.756167686 | 52.391457 | NA | NA | NA | NA | NA | NA | NA |
| TRPM4 | 246 | E | -82.35533333 | 51.76337785 | 0.716571399 | 49.73048676 | NA | NA | NA | NA | NA | NA | NA |
| TRPM4 | 247 | N | -85.3525 | 50.88709519 | 0.703881876 | 48.78169134 | NA | NA | NA | NA | NA | NA | NA |
| TRPM4 | 248 | R | -84.61654545 | 54.43751407 | 0.755295791 | 52.35264031 | NA | NA | NA | NA | NA | NA | NA |
| TRPM4 | 249 | F | -77.983 | 52.475229 | 0.726879775 | 49.35531394 | NA | NA | NA | NA | NA | NA | NA |
| TRPM4 | 250 | R | -81.64790909 | 44.34254898 | 0.609109752 | 41.27378039 | NA | NA | NA | NA | NA | NA | NA |
| TRPM4 | 251 | L | -83.692625 | 49.81295629 | 0.68832718 | 47.59345754 | NA | NA | NA | NA | NA | NA | NA |
| TRPM4 | 252 | R | -79.55927273 | 52.33925113 | 0.724910668 | 49.22974991 | NA | NA | NA | NA | NA | NA | NA |
| TRPM4 | 253 | L | -76.7475 | 48.70873837 | 0.672336908 | 46.48298276 | NA | NA | NA | NA | NA | NA | NA |
| TRPM4 | 254 | E | -79.93 | 44.29960377 | 0.608487859 | 41.14127267 | NA | NA | NA | NA | NA | NA | NA |
| TRPM4 | 255 | S | -80.28433333 | 49.56568626 | 0.684746442 | 46.43702079 | NA | NA | NA | NA | NA | NA | NA |
| TRPM4 | 256 | Y | -74.521 | 50.29727497 | 0.69534064 | 48.04432761 | NA | NA | NA | NA | NA | NA | NA |
| TRPM4 | 257 | I | -75.380625 | 45.53225478 | 0.626337982 | 43.77639123 | NA | NA | NA | NA | NA | NA | NA |
| TRPM4 | 258 | S | -78.2545 | 46.89715153 | 0.646103166 | 43.48254434 | NA | NA | NA | NA | NA | NA | NA |
| TRPM4 | 259 | Q | -77.05277778 | 49.4050457 | 0.682420193 | 47.65324876 | NA | NA | NA | NA | NA | NA | NA |
| TRPM4 | 260 | Q | -71.548 | 46.97669991 | 0.647255113 | 45.09719422 | NA | NA | NA | NA | NA | NA | NA |
| TRPM4 | 261 | K | -71.04155556 | 46.07285553 | 0.634166467 | 43.37272039 | NA | NA | NA | NA | NA | NA | NA |
| TRPM4 | 262 | T | -69.33842857 | 41.42685605 | 0.566887357 | 38.56703613 | NA | NA | NA | NA | NA | NA | NA |
| TRPM4 | 263 | G | -65.838 | 42.56081077 | 0.583308251 | 39.99811719 | NA | NA | NA | NA | NA | NA | NA |
| TRPM4 | 264 | V | -62.25585714 | 42.42416191 | 0.581329427 | 40.07903877 | NA | NA | NA | NA | NA | NA | NA |
| TRPM4 | 265 | G | -64.47775 | 46.02483883 | 0.633471133 | 43.9705492 | NA | NA | NA | NA | NA | NA | NA |
| TRPM4 | 266 | G | -66.98275 | 48.61911145 | 0.671039013 | 46.05351562 | NA | NA | NA | NA | NA | NA | NA |
| TRPM4 | 267 | T | -68.6776 | 46.3172457 | 0.637705501 | 43.64860346 | NA | NA | NA | NA | NA | NA | NA |
| TRPM4 | 268 | G | -71.32425 | 44.40227273 | 0.609974616 | 41.6920525 | NA | NA | NA | NA | NA | NA | NA |
| TRPM4 | 269 | I | -74.11575 | 40.5938181 | 0.554824064 | 38.77677346 | NA | NA | NA | NA | NA | NA | NA |

| | | | | | | | | | | | | | |
|---|---|---|---|---|---|---|---|---|---|---|---|---|---|
| TRPM4 | 270 | D | -76.4825 | 43.26478293 | 0.593502531 | 41.50788083 | NA | NA | NA | NA | NA | NA | NA |
| TRPM4 | 271 | I | -74.979625 | 42.16518234 | 0.577579122 | 40.41358645 | NA | NA | NA | NA | NA | NA | NA |
| TRPM4 | 272 | P | -78.16842857 | 39.51905719 | 0.539260361 | 36.34141432 | NA | NA | NA | NA | NA | NA | NA |
| TRPM4 | 273 | V | -80.90857143 | 41.42615524 | 0.566877209 | 38.3253808 | NA | NA | NA | NA | NA | NA | NA |
| TRPM4 | 274 | L | -83.2865 | 36.77916047 | 0.499583686 | 34.67990785 | NA | NA | NA | NA | NA | NA | NA |
| TRPM4 | 275 | L | -86.05625 | 39.19201283 | 0.534524404 | 37.14431372 | NA | NA | NA | NA | NA | NA | NA |
| TRPM4 | 276 | L | -86.59375 | 34.12944203 | 0.461212891 | 31.87890353 | NA | NA | NA | NA | NA | NA | NA |
| TRPM4 | 277 | L | -90.780625 | 36.30729547 | 0.49275057 | 33.91738081 | NA | NA | NA | NA | NA | NA | NA |
| TRPM4 | 278 | I | -90.273375 | 34.3320151 | 0.464146368 | 32.37830619 | NA | NA | NA | NA | NA | NA | NA |
| TRPM4 | 279 | D | -94.156125 | 36.82969374 | 0.500315463 | 34.91328341 | NA | NA | NA | NA | NA | NA | NA |
| TRPM4 | 280 | G | -94.2705 | 40.14798244 | 0.548367881 | 38.23516799 | NA | NA | NA | NA | NA | NA | NA |
| TRPM4 | 281 | D | -96.0584 | 43.37663955 | 0.595122336 | 42.23176184 | NA | NA | NA | NA | NA | NA | NA |
| TRPM4 | 282 | E | -98.516 | 45.9361099 | 0.632186242 | 44.21880198 | NA | NA | NA | NA | NA | NA | NA |
| TRPM4 | 283 | K | -95.4432 | 47.77209647 | 0.658773318 | 46.62975259 | NA | NA | NA | NA | NA | NA | NA |
| TRPM4 | 284 | M | -93.236375 | 43.86674352 | 0.602219573 | 42.28750598 | NA | NA | NA | NA | NA | NA | NA |
| TRPM4 | 285 | L | -97.17675 | 43.94538433 | 0.603358377 | 42.85822803 | NA | NA | NA | NA | NA | NA | NA |
| TRPM4 | 286 | T | -94.64871429 | 48.69544965 | 0.672144473 | 47.54627478 | NA | NA | NA | NA | NA | NA | NA |
| TRPM4 | 287 | R | -89.71472727 | 47.41643504 | 0.653622954 | 45.452271 | NA | NA | NA | NA | NA | NA | NA |
| TRPM4 | 288 | I | -92.697875 | 43.24594754 | 0.593229774 | 41.33973393 | NA | NA | NA | NA | NA | NA | NA |
| TRPM4 | 289 | E | -93.8378 | 47.95640755 | 0.661442342 | 46.36988475 | NA | NA | NA | NA | NA | NA | NA |
| TRPM4 | 290 | N | -91.07575 | 50.32175221 | 0.695695097 | 48.37176523 | NA | NA | NA | NA | NA | NA | NA |
| TRPM4 | 291 | A | -88.8446 | 47.75497064 | 0.658525317 | 45.52240691 | NA | NA | NA | NA | NA | NA | NA |
| TRPM4 | 292 | T | -91.3182 | 46.48252593 | 0.640098938 | 44.52428502 | NA | NA | NA | NA | NA | NA | NA |
| TRPM4 | 293 | Q | -92.72211111 | 49.93388074 | 0.690078297 | 48.02872004 | NA | NA | NA | NA | NA | NA | NA |
| TRPM4 | 294 | A | -88.3106 | 51.92914112 | 0.718971831 | 49.69885163 | NA | NA | NA | NA | NA | NA | NA |
| TRPM4 | 295 | Q | -85.6586 | 49.33476902 | 0.68140251 | 47.22590241 | NA | NA | NA | NA | NA | NA | NA |
| TRPM4 | 296 | L | -87.551875 | 46.40030481 | 0.638908287 | 44.10736999 | NA | NA | NA | NA | NA | NA | NA |
| TRPM4 | 297 | P | -86.38742857 | 43.77430587 | 0.600880975 | 41.69224804 | NA | NA | NA | NA | NA | NA | NA |
| TRPM4 | 298 | C | -86.8688 | 40.45532186 | 0.552818489 | 38.12141188 | NA | NA | NA | NA | NA | NA | NA |
| TRPM4 | 299 | L | -89.862375 | 37.93382756 | 0.516304518 | 35.75582966 | NA | NA | NA | NA | NA | NA | NA |
| TRPM4 | 300 | L | -89.301625 | 33.37708689 | 0.450317973 | 31.00093792 | NA | NA | NA | NA | NA | NA | NA |
| TRPM4 | 301 | V | -93.20671429 | 33.29967787 | 0.449197006 | 31.39551251 | NA | NA | NA | NA | NA | NA | NA |
| TRPM4 | 302 | A | -92.5858 | 32.33619257 | 0.435244694 | 30.44501064 | NA | NA | NA | NA | NA | NA | NA |

| | | | | | | | | | | | | | |
|---|---|---|---|---|---|---|---|---|---|---|---|---|---|
| TRPM4 | 303 | G | -94.854 | 30.42118665 | 0.407513332 | 29.29621417 | NA | NA | NA | NA | NA | NA | NA |
| TRPM4 | 304 | S | -96.7395 | 29.04646297 | 0.387605843 | 27.89652457 | NA | NA | NA | NA | NA | NA | NA |
| TRPM4 | 305 | G | -97.06775 | 32.77538131 | 0.441604623 | 31.6329434 | NA | NA | NA | NA | NA | NA | NA |
| TRPM4 | 306 | G | -98.69025 | 35.85527855 | 0.486204875 | 34.14900474 | NA | NA | NA | NA | NA | NA | NA |
| TRPM4 | 307 | A | -98.0514 | 38.23994059 | 0.520737366 | 36.75378867 | NA | NA | NA | NA | NA | NA | NA |
| TRPM4 | 308 | A | -97.5132 | 36.98544922 | 0.502570971 | 35.82008592 | NA | NA | NA | NA | NA | NA | NA |
| TRPM4 | 309 | D | -99.77925 | 33.17727249 | 0.447424443 | 31.50849028 | NA | NA | NA | NA | NA | NA | NA |
| TRPM4 | 310 | C | -102.5101667 | 37.1167564 | 0.504472441 | 35.83668356 | NA | NA | NA | NA | NA | NA | NA |
| TRPM4 | 311 | L | -99.613875 | 37.36661752 | 0.508090701 | 35.70476768 | NA | NA | NA | NA | NA | NA | NA |
| TRPM4 | 312 | A | -100.6616 | 33.79349355 | 0.456347992 | 32.12458702 | NA | NA | NA | NA | NA | NA | NA |
| TRPM4 | 313 | E | -104.8563333 | 32.84308943 | 0.44258511 | 29.92906774 | NA | NA | NA | NA | NA | NA | NA |
| TRPM4 | 314 | T | -104.1262857 | 36.97033841 | 0.50235215 | 34.07479655 | NA | NA | NA | NA | NA | NA | NA |
| TRPM4 | 315 | L | -100.8285 | 35.98719979 | 0.488115237 | 34.69693251 | NA | NA | NA | NA | NA | NA | NA |
| TRPM4 | 316 | E | -104.0406 | 32.56671241 | 0.438582872 | 31.29576309 | NA | NA | NA | NA | NA | NA | NA |
| TRPM4 | 317 | D | -106.7196 | 35.16644039 | 0.476229751 | 31.96972268 | NA | NA | NA | NA | NA | NA | NA |
| TRPM4 | 327 | R | -115.4494 | 41.63557162 | 0.569909785 | 38.03568937 | NA | NA | NA | NA | NA | NA | NA |
| TRPM4 | 328 | Q | -112.582 | 37.38364483 | 0.508337275 | 33.87100498 | NA | NA | NA | NA | NA | NA | NA |
| TRPM4 | 329 | G | -110.601 | 41.54429313 | 0.568587973 | 38.18314765 | NA | NA | NA | NA | NA | NA | NA |
| TRPM4 | 330 | E | -113.2052222 | 41.69209177 | 0.570728258 | 38.29256317 | NA | NA | NA | NA | NA | NA | NA |
| TRPM4 | 331 | A | -112.0956 | 38.17355033 | 0.519775963 | 34.79344014 | NA | NA | NA | NA | NA | NA | NA |
| TRPM4 | 332 | R | -107.9352727 | 37.00434058 | 0.502844539 | 33.51915372 | NA | NA | NA | NA | NA | NA | NA |
| TRPM4 | 333 | D | -107.3345 | 40.890368 | 0.559118428 | 37.70892429 | NA | NA | NA | NA | NA | NA | NA |
| TRPM4 | 334 | R | -109.3024 | 39.6498709 | 0.541154686 | 36.1614212 | NA | NA | NA | NA | NA | NA | NA |
| TRPM4 | 335 | I | -108.111 | 36.35858965 | 0.493493365 | 33.00091048 | NA | NA | NA | NA | NA | NA | NA |
| TRPM4 | 336 | R | -104.4522 | 36.07219641 | 0.48934608 | 32.8642577 | NA | NA | NA | NA | NA | NA | NA |
| TRPM4 | 337 | R | -104.137 | 38.82883658 | 0.529265219 | 37.59614455 | NA | NA | NA | NA | NA | NA | NA |
| TRPM4 | 338 | F | -104.4478 | 41.83191965 | 0.572753117 | 38.64899349 | NA | NA | NA | NA | NA | NA | NA |
| TRPM4 | 339 | F | -108.1118 | 42.58511802 | 0.583660246 | 39.21951364 | NA | NA | NA | NA | NA | NA | NA |
| TRPM4 | 340 | P | -107.1528571 | 45.32651701 | 0.623358676 | 42.12713213 | NA | NA | NA | NA | NA | NA | NA |
| TRPM4 | 341 | K | -106.2522 | 47.48347664 | 0.65459379 | 44.19480052 | NA | NA | NA | NA | NA | NA | NA |
| TRPM4 | 342 | G | -108.219 | 49.49892689 | 0.683779694 | 46.13086946 | NA | NA | NA | NA | NA | NA | NA |
| TRPM4 | 343 | D | -110.6895 | 47.10657045 | 0.649135779 | 43.75541747 | NA | NA | NA | NA | NA | NA | NA |
| TRPM4 | 344 | L | -108.7996 | 51.3395387 | 0.710433749 | 47.83154432 | NA | NA | NA | NA | NA | NA | NA |

| | | | | | | | | | | | | | |
|---|---|---|---|---|---|---|---|---|---|---|---|---|---|
| TRPM4 | 345 | E | -105.6918 | 50.73680536 | 0.701705516 | 47.45689995 | NA | NA | NA | NA | NA | NA | NA |
| TRPM4 | 346 | V | -104.8798571 | 46.7990125 | 0.644682007 | 43.59006082 | NA | NA | NA | NA | NA | NA | NA |
| TRPM4 | 347 | L | -108.621375 | 46.02407907 | 0.63346013 | 42.99880205 | NA | NA | NA | NA | NA | NA | NA |
| TRPM4 | 348 | Q | -106.2212 | 49.29648609 | 0.680848132 | 46.09659132 | NA | NA | NA | NA | NA | NA | NA |
| TRPM4 | 349 | A | -102.8258 | 49.66366286 | 0.68616525 | 48.37774909 | NA | NA | NA | NA | NA | NA | NA |
| TRPM4 | 350 | Q | -101.297 | 46.20200219 | 0.63603665 | 44.93732996 | NA | NA | NA | NA | NA | NA | NA |
| TRPM4 | 351 | V | -103.0544286 | 43.99091119 | 0.604017655 | 42.74036903 | NA | NA | NA | NA | NA | NA | NA |
| TRPM4 | 352 | E | -102.9572 | 47.48858493 | 0.654667763 | 46.20776029 | NA | NA | NA | NA | NA | NA | NA |
| TRPM4 | 353 | R | -100.7479091 | 48.4189873 | 0.668140998 | 46.75067134 | NA | NA | NA | NA | NA | NA | NA |
| TRPM4 | 354 | I | -98.351625 | 43.75811698 | 0.600646543 | 42.27296158 | NA | NA | NA | NA | NA | NA | NA |
| TRPM4 | 355 | M | -101.509 | 40.96513481 | 0.560201133 | 39.67442463 | NA | NA | NA | NA | NA | NA | NA |
| TRPM4 | 356 | T | -100.0654286 | 46.82919032 | 0.645119014 | 45.17191093 | NA | NA | NA | NA | NA | NA | NA |
| TRPM4 | 357 | R | -96.354 | 43.97892832 | 0.60384413 | 42.81212893 | NA | NA | NA | NA | NA | NA | NA |
| TRPM4 | 358 | K | -97.07 | 40.40078044 | 0.55202867 | 39.30398693 | NA | NA | NA | NA | NA | NA | NA |
| TRPM4 | 359 | E | -93.7504 | 41.18675736 | 0.563410468 | 39.61929114 | NA | NA | NA | NA | NA | NA | NA |
| TRPM4 | 360 | L | -91.671 | 40.148246 | 0.548371697 | 38.25481512 | NA | NA | NA | NA | NA | NA | NA |
| TRPM4 | 361 | L | -94.430875 | 38.17589195 | 0.519809872 | 36.59084903 | NA | NA | NA | NA | NA | NA | NA |
| TRPM4 | 362 | T | -92.62957143 | 34.16494341 | 0.461726989 | 32.2738112 | NA | NA | NA | NA | NA | NA | NA |
| TRPM4 | 363 | V | -95.33314286 | 31.16399584 | 0.418270015 | 30.08022505 | NA | NA | NA | NA | NA | NA | NA |
| TRPM4 | 364 | Y | -90.9595 | 26.06190219 | 0.344386168 | 24.49790996 | NA | NA | NA | NA | NA | NA | NA |
| TRPM4 | 365 | S | -93.3845 | 24.63808858 | 0.323767803 | 23.05806493 | NA | NA | NA | NA | NA | NA | NA |
| TRPM4 | 366 | S | -95.20616667 | 23.21069465 | 0.303097591 | 22.09635342 | NA | NA | NA | NA | NA | NA | NA |
| TRPM4 | 367 | E | -95.082 | 20.09025289 | 0.257910211 | 19.03579091 | NA | NA | NA | NA | NA | NA | NA |
| TRPM4 | 368 | D | -91.7676 | 21.78687509 | 0.28247914 | 19.88099994 | NA | NA | NA | NA | NA | NA | NA |
| TRPM4 | 369 | G | -88.93 | 21.54017263 | 0.278906621 | 19.1256775 | NA | NA | NA | NA | NA | NA | NA |
| TRPM4 | 370 | S | -85.6714 | 21.10274612 | 0.272572211 | 18.99912263 | NA | NA | NA | NA | NA | NA | NA |
| TRPM4 | 371 | E | -83.1762 | 19.81105613 | 0.25386714 | 17.54201861 | NA | NA | NA | NA | NA | NA | NA |
| TRPM4 | 372 | E | -84.7 | 22.98929055 | 0.29989142 | 20.90770243 | NA | NA | NA | NA | NA | NA | NA |
| TRPM4 | 373 | F | -83.63227273 | 25.64080625 | 0.338288242 | 23.54626243 | NA | NA | NA | NA | NA | NA | NA |
| TRPM4 | 374 | E | -80.54022222 | 25.24616575 | 0.332573419 | 22.37043059 | NA | NA | NA | NA | NA | NA | NA |
| TRPM4 | 375 | T | -83.93571429 | 26.1527836 | 0.345702229 | 24.05493517 | NA | NA | NA | NA | NA | NA | NA |
| TRPM4 | 376 | I | -87.073875 | 27.50589962 | 0.365296816 | 25.58986864 | NA | NA | NA | NA | NA | NA | NA |
| TRPM4 | 377 | V | -83.77685714 | 31.34593476 | 0.420904688 | 29.23672097 | NA | NA | NA | NA | NA | NA | NA |

| | | | | | | | | | | | | | |
|---|---|---|---|---|---|---|---|---|---|---|---|---|---|
| TRPM4 | 378 | L | -82.586625 | 32.03982177 | 0.430952924 | 29.90967866 | NA | NA | NA | NA | NA | NA | NA |
| TRPM4 | 379 | K | -88.12211111 | 29.69716837 | 0.397028763 | 28.14639788 | NA | NA | NA | NA | NA | NA | NA |
| TRPM4 | 380 | A | -87.6902 | 34.47482625 | 0.466214429 | 32.12021437 | NA | NA | NA | NA | NA | NA | NA |
| TRPM4 | 381 | L | -83.790125 | 36.74565968 | 0.499098559 | 34.64083453 | NA | NA | NA | NA | NA | NA | NA |
| TRPM4 | 382 | V | -85.28271429 | 37.03663359 | 0.503312176 | 34.97016484 | NA | NA | NA | NA | NA | NA | NA |
| TRPM4 | 383 | K | -90.83411111 | 36.2457861 | 0.491859847 | 34.48605699 | NA | NA | NA | NA | NA | NA | NA |
| TRPM4 | 384 | A | -88.5708 | 40.26370035 | 0.550043602 | 38.08542223 | NA | NA | NA | NA | NA | NA | NA |
| TRPM4 | 385 | C | -86.494 | 42.46488646 | 0.581919163 | 40.40064757 | NA | NA | NA | NA | NA | NA | NA |
| TRPM4 | 386 | G | -89.37725 | 44.19381459 | 0.606955917 | 41.96289836 | NA | NA | NA | NA | NA | NA | NA |
| TRPM4 | 393 | Y | -79.737 | 44.56535621 | 0.612336242 | 41.45160307 | NA | NA | NA | NA | NA | NA | NA |
| TRPM4 | 394 | L | -80.5305 | 40.95998243 | 0.560126521 | 37.84450891 | NA | NA | NA | NA | NA | NA | NA |
| TRPM4 | 395 | D | -79.870375 | 38.64902176 | 0.526661305 | 35.52700059 | NA | NA | NA | NA | NA | NA | NA |
| TRPM4 | 396 | E | -79.93755556 | 31.89875551 | 0.428910131 | 28.77961387 | NA | NA | NA | NA | NA | NA | NA |
| TRPM4 | 397 | L | -75.8365 | 35.57378171 | 0.482128495 | 33.8155573 | NA | NA | NA | NA | NA | NA | NA |
| TRPM4 | 398 | R | -74.26509091 | 37.97599745 | 0.516915183 | 36.21615478 | NA | NA | NA | NA | NA | NA | NA |
| TRPM4 | 399 | L | -76.41525 | 35.27137368 | 0.477749299 | 33.5227982 | NA | NA | NA | NA | NA | NA | NA |
| TRPM4 | 400 | A | -73.9634 | 32.89227683 | 0.443297397 | 31.13841368 | NA | NA | NA | NA | NA | NA | NA |
| TRPM4 | 401 | V | -70.60085714 | 34.80786661 | 0.471037214 | 32.13557848 | NA | NA | NA | NA | NA | NA | NA |
| TRPM4 | 402 | A | -71.2278 | 35.77035232 | 0.484975051 | 33.36261429 | NA | NA | NA | NA | NA | NA | NA |
| TRPM4 | 403 | W | -74.31735714 | 32.17099406 | 0.432852441 | 30.11044969 | NA | NA | NA | NA | NA | NA | NA |
| TRPM4 | 404 | N | -68.47375 | 28.41065043 | 0.378398588 | 25.55087269 | NA | NA | NA | NA | NA | NA | NA |
| TRPM4 | 405 | R | -73.60081818 | 24.43195934 | 0.320782828 | 22.41508829 | NA | NA | NA | NA | NA | NA | NA |
| TRPM4 | 406 | V | -71.37142857 | 26.03347439 | 0.343974502 | 23.64474402 | NA | NA | NA | NA | NA | NA | NA |
| TRPM4 | 407 | D | -75.434 | 23.38419391 | 0.305610049 | 21.62851676 | NA | NA | NA | NA | NA | NA | NA |
| TRPM4 | 408 | I | -77.15975 | 26.87097767 | 0.356102458 | 24.64413124 | NA | NA | NA | NA | NA | NA | NA |
| TRPM4 | 409 | A | -75.1928 | 28.84791322 | 0.384730628 | 27.08755476 | NA | NA | NA | NA | NA | NA | NA |
| TRPM4 | 410 | Q | -74.664 | 23.91348887 | 0.313274813 | 22.25749562 | NA | NA | NA | NA | NA | NA | NA |
| TRPM4 | 411 | S | -79.53583333 | 25.86189717 | 0.341489878 | 22.69410259 | NA | NA | NA | NA | NA | NA | NA |
| TRPM4 | 412 | E | -79.5622 | 28.81611124 | 0.3842701 | 25.69820165 | NA | NA | NA | NA | NA | NA | NA |
| TRPM4 | 413 | L | -77.7442 | 30.20266674 | 0.404348927 | 27.92776866 | NA | NA | NA | NA | NA | NA | NA |
| TRPM4 | 414 | F | -79.117 | 26.79837592 | 0.355051106 | 23.50097773 | NA | NA | NA | NA | NA | NA | NA |
| TRPM4 | 415 | R | -82.2624 | 28.16348208 | 0.374819323 | 25.73248903 | NA | NA | NA | NA | NA | NA | NA |
| TRPM4 | 416 | G | -84.899 | 29.33305393 | 0.391755991 | 27.26125215 | NA | NA | NA | NA | NA | NA | NA |

| | | | | | | | | | | | | | |
|---|---|---|---|---|---|---|---|---|---|---|---|---|---|
| TRPM4 | 417 | D | -87.1382 | 30.73322362 | 0.412031966 | 28.37309426 | NA | NA | NA | NA | NA | NA | NA |
| TRPM4 | 418 | I | -85.0616 | 33.37874306 | 0.450341956 | 31.27678337 | NA | NA | NA | NA | NA | NA | NA |
| TRPM4 | 419 | Q | -83.5534 | 36.30344762 | 0.492694849 | 34.23271667 | NA | NA | NA | NA | NA | NA | NA |
| TRPM4 | 420 | W | -80.06214286 | 33.73577574 | 0.455512176 | 30.6083648 | NA | NA | NA | NA | NA | NA | NA |
| TRPM4 | 421 | R | -80.48781818 | 40.22579706 | 0.549494721 | 37.04733148 | NA | NA | NA | NA | NA | NA | NA |
| TRPM4 | 422 | S | -75.46883333 | 42.90754509 | 0.58832934 | 41.18557484 | NA | NA | NA | NA | NA | NA | NA |
| TRPM4 | 423 | F | -76.54745455 | 43.86797864 | 0.602237459 | 42.10718895 | NA | NA | NA | NA | NA | NA | NA |
| TRPM4 | 424 | H | -77.6579 | 40.45423461 | 0.552802744 | 38.9050116 | NA | NA | NA | NA | NA | NA | NA |
| TRPM4 | 425 | L | -73.170125 | 37.33173409 | 0.507585551 | 35.46436045 | NA | NA | NA | NA | NA | NA | NA |
| TRPM4 | 426 | E | -71.0908 | 42.38579707 | 0.580773863 | 39.98183074 | NA | NA | NA | NA | NA | NA | NA |
| TRPM4 | 427 | A | -71.5108 | 42.8160361 | 0.58700419 | 40.4081525 | NA | NA | NA | NA | NA | NA | NA |
| TRPM4 | 428 | S | -70.947 | 38.90810045 | 0.530413045 | 36.50299266 | NA | NA | NA | NA | NA | NA | NA |
| TRPM4 | 429 | L | -66.878125 | 38.34827527 | 0.52230617 | 35.76767824 | NA | NA | NA | NA | NA | NA | NA |
| TRPM4 | 430 | M | -65.746375 | 41.87443957 | 0.573368852 | 39.30533834 | NA | NA | NA | NA | NA | NA | NA |
| TRPM4 | 431 | D | -67.822125 | 39.3363817 | 0.536615022 | 36.77167795 | NA | NA | NA | NA | NA | NA | NA |
| TRPM4 | 432 | A | -65.5546 | 36.91391525 | 0.501535082 | 34.34502656 | NA | NA | NA | NA | NA | NA | NA |
| TRPM4 | 433 | L | -62.193375 | 38.45514283 | 0.523853728 | 36.10369975 | NA | NA | NA | NA | NA | NA | NA |
| TRPM4 | 434 | L | -62.271625 | 39.85714477 | 0.544156236 | 37.73850163 | NA | NA | NA | NA | NA | NA | NA |
| TRPM4 | 435 | N | -63.62125 | 36.13626835 | 0.490273911 | 33.77718385 | NA | NA | NA | NA | NA | NA | NA |
| TRPM4 | 436 | D | -60.8794 | 33.32805954 | 0.449608003 | 30.37469164 | NA | NA | NA | NA | NA | NA | NA |
| TRPM4 | 437 | R | -64.32136364 | 32.36601262 | 0.435676521 | 30.3182612 | NA | NA | NA | NA | NA | NA | NA |
| TRPM4 | 438 | P | -61.45971429 | 29.99655384 | 0.401364189 | 27.35158701 | NA | NA | NA | NA | NA | NA | NA |
| TRPM4 | 439 | E | -64.7944 | 27.57590463 | 0.366310564 | 25.52800914 | NA | NA | NA | NA | NA | NA | NA |
| TRPM4 | 440 | F | -68.18972727 | 29.82402902 | 0.398865843 | 27.39598207 | NA | NA | NA | NA | NA | NA | NA |
| TRPM4 | 441 | V | -64.30285714 | 32.61245465 | 0.439245269 | 30.57886441 | NA | NA | NA | NA | NA | NA | NA |
| TRPM4 | 442 | R | -63.30881818 | 27.34132131 | 0.362913543 | 25.42854827 | NA | NA | NA | NA | NA | NA | NA |
| TRPM4 | 443 | L | -69.448875 | 28.09793336 | 0.373870106 | 25.23966524 | NA | NA | NA | NA | NA | NA | NA |
| TRPM4 | 444 | L | -69.40475 | 33.46562657 | 0.451600123 | 30.60836787 | NA | NA | NA | NA | NA | NA | NA |
| TRPM4 | 445 | I | -65.99 | 35.08858485 | 0.475102319 | 32.58315321 | NA | NA | NA | NA | NA | NA | NA |
| TRPM4 | 446 | S | -68.26366667 | 31.33849275 | 0.420796919 | 28.68901348 | NA | NA | NA | NA | NA | NA | NA |
| TRPM4 | 447 | H | -72.7005 | 31.60663876 | 0.424679964 | 29.25403185 | NA | NA | NA | NA | NA | NA | NA |
| TRPM4 | 448 | G | -71.0215 | 36.28330451 | 0.492403154 | 33.87589937 | NA | NA | NA | NA | NA | NA | NA |
| TRPM4 | 449 | L | -69.193875 | 39.5935397 | 0.540338949 | 36.55591613 | NA | NA | NA | NA | NA | NA | NA |

| | | | | | | | | | | | | | |
|---|---|---|---|---|---|---|---|---|---|---|---|---|---|
| TRPM4 | 450 | S | -67.1732 | 42.13925563 | 0.577203675 | 39.59947242 | NA | NA | NA | NA | NA | NA | NA |
| TRPM4 | 451 | L | -63.127375 | 41.75758195 | 0.571676627 | 39.72627748 | NA | NA | NA | NA | NA | NA | NA |
| TRPM4 | 452 | G | -64.31275 | 45.276259 | 0.622630885 | 43.21585593 | NA | NA | NA | NA | NA | NA | NA |
| TRPM4 | 453 | H | -68.2594 | 46.95560301 | 0.646949607 | 44.39215968 | NA | NA | NA | NA | NA | NA | NA |
| TRPM4 | 454 | F | -65.40336364 | 43.74080645 | 0.600395868 | 41.20274552 | NA | NA | NA | NA | NA | NA | NA |
| TRPM4 | 455 | L | -61.246375 | 46.0643246 | 0.634042929 | 43.83681609 | NA | NA | NA | NA | NA | NA | NA |
| TRPM4 | 456 | T | -62.93071429 | 49.41672521 | 0.682589325 | 47.05777999 | NA | NA | NA | NA | NA | NA | NA |
| TRPM4 | 457 | P | -59.97428571 | 52.3817177 | 0.72552563 | 49.02093545 | NA | NA | NA | NA | NA | NA | NA |
| TRPM4 | 458 | M | -62.042 | 52.09252133 | 0.721337754 | 49.97425912 | NA | NA | NA | NA | NA | NA | NA |
| TRPM4 | 459 | R | -64.32036364 | 46.57937134 | 0.641501365 | 44.43759233 | NA | NA | NA | NA | NA | NA | NA |
| TRPM4 | 460 | L | -57.94125 | 46.84324166 | 0.645322493 | 43.54475691 | NA | NA | NA | NA | NA | NA | NA |
| TRPM4 | 461 | A | -58.3524 | 50.89667353 | 0.704020581 | 47.36728435 | NA | NA | NA | NA | NA | NA | NA |
| TRPM4 | 462 | Q | -60.6446 | 48.80203014 | 0.673687874 | 45.84551062 | NA | NA | NA | NA | NA | NA | NA |
| TRPM4 | 463 | L | -58.923 | 43.87170822 | 0.602291467 | 40.50547494 | NA | NA | NA | NA | NA | NA | NA |
| TRPM4 | 464 | Y | -54.379 | 46.22438885 | 0.636360833 | 41.92552297 | NA | NA | NA | NA | NA | NA | NA |
| TRPM4 | 465 | S | -57.685 | 49.60203649 | 0.685272833 | 46.08766147 | NA | NA | NA | NA | NA | NA | NA |
| TRPM4 | 466 | A | -57.6914 | 46.40325189 | 0.638950964 | 42.88709431 | NA | NA | NA | NA | NA | NA | NA |
| TRPM4 | 467 | A | -55.2288 | 45.95503265 | 0.632460263 | 41.59530612 | NA | NA | NA | NA | NA | NA | NA |
| TRPM4 | 468 | P | -55.54857143 | 42.51562382 | 0.582653895 | 38.15991609 | NA | NA | NA | NA | NA | NA | NA |
| TRPM4 | 469 | S | -52.3808 | 43.32042518 | 0.594308291 | 36.47693141 | NA | NA | NA | NA | NA | NA | NA |
| TRPM4 | 470 | N | -49.8804 | 45.95472187 | 0.632455763 | 37.34590868 | NA | NA | NA | NA | NA | NA | NA |
| TRPM4 | 471 | S | -46.4618 | 45.48215971 | 0.625612551 | 35.52633622 | NA | NA | NA | NA | NA | NA | NA |
| TRPM4 | 472 | L | -46.036875 | 40.59811658 | 0.554886311 | 30.63370044 | NA | NA | NA | NA | NA | NA | NA |
| TRPM4 | 473 | I | -48.580125 | 45.1917023 | 0.621406412 | 35.21259656 | NA | NA | NA | NA | NA | NA | NA |
| TRPM4 | 474 | R | -50.05563636 | 47.67565656 | 0.657376763 | 37.69816882 | NA | NA | NA | NA | NA | NA | NA |
| TRPM4 | 475 | N | -43.72325 | 48.48326773 | 0.669071848 | 37.63203042 | NA | NA | NA | NA | NA | NA | NA |
| TRPM4 | 476 | L | -43.39725 | 46.58969612 | 0.641650879 | 35.73192934 | NA | NA | NA | NA | NA | NA | NA |
| TRPM4 | 477 | L | -47.352375 | 50.278189 | 0.695064254 | 40.33029725 | NA | NA | NA | NA | NA | NA | NA |
| TRPM4 | 478 | D | -44.7604 | 52.85368471 | 0.732360224 | 41.96010863 | NA | NA | NA | NA | NA | NA | NA |
| TRPM4 | 479 | Q | -42.5146 | 53.88036197 | 0.747227624 | 37.46022506 | NA | NA | NA | NA | NA | NA | NA |
| TRPM4 | 480 | A | -45.4598 | 54.6506173 | 0.758381756 | 43.75251854 | NA | NA | NA | NA | NA | NA | NA |
| TRPM4 | 558 | P | -56.675 | 43.94724003 | 0.60338525 | 39.59008533 | NA | NA | NA | NA | NA | NA | NA |
| TRPM4 | 559 | W | -57.28607143 | 37.84257119 | 0.514983026 | 33.66376462 | NA | NA | NA | NA | NA | NA | NA |

| | | | | | | | | | | | | | |
|---|---|---|---|---|---|---|---|---|---|---|---|---|---|
| TRPM4 | 560 | S | -51.68166667 | 42.03401857 | 0.575679729 | 35.43589291 | NA | NA | NA | NA | NA | NA | NA |
| TRPM4 | 561 | D | -53.3802 | 43.53796395 | 0.597458488 | 38.08221442 | NA | NA | NA | NA | NA | NA | NA |
| TRPM4 | 562 | L | -55.883125 | 40.03074311 | 0.546670128 | 35.70419613 | NA | NA | NA | NA | NA | NA | NA |
| TRPM4 | 563 | L | -51.10675 | 36.519905 | 0.495829386 | 29.93527109 | NA | NA | NA | NA | NA | NA | NA |
| TRPM4 | 564 | L | -49.05425 | 40.85948327 | 0.558671184 | 31.66488266 | NA | NA | NA | NA | NA | NA | NA |
| TRPM4 | 565 | W | -54.66607143 | 39.87203272 | 0.54437183 | 33.62963999 | NA | NA | NA | NA | NA | NA | NA |
| TRPM4 | 566 | A | -52.1814 | 37.07485372 | 0.503865645 | 31.65710282 | NA | NA | NA | NA | NA | NA | NA |
| TRPM4 | 567 | L | -48.283 | 35.35293302 | 0.478930367 | 26.18644487 | NA | NA | NA | NA | NA | NA | NA |
| TRPM4 | 568 | L | -48.126375 | 39.62324447 | 0.540769106 | 29.65445197 | NA | NA | NA | NA | NA | NA | NA |
| TRPM4 | 569 | L | -51.568 | 37.7237672 | 0.513262616 | 29.12376432 | NA | NA | NA | NA | NA | NA | NA |
| TRPM4 | 570 | N | -49.779125 | 33.40336146 | 0.450698457 | 24.20518792 | NA | NA | NA | NA | NA | NA | NA |
| TRPM4 | 571 | R | -54.93263636 | 32.87943488 | 0.443111432 | 27.42011441 | NA | NA | NA | NA | NA | NA | NA |
| TRPM4 | 572 | A | -52.7056 | 29.35350925 | 0.392052206 | 22.50261102 | NA | NA | NA | NA | NA | NA | NA |
| TRPM4 | 573 | Q | -56.15466667 | 26.07674853 | 0.344601159 | 21.7606532 | NA | NA | NA | NA | NA | NA | NA |
| TRPM4 | 574 | M | -56.96475 | 31.87452509 | 0.428559249 | 27.51998026 | NA | NA | NA | NA | NA | NA | NA |
| TRPM4 | 575 | A | -52.8652 | 32.36267195 | 0.435628145 | 26.90284269 | NA | NA | NA | NA | NA | NA | NA |
| TRPM4 | 576 | M | -52.583125 | 28.28533921 | 0.376583946 | 22.83874758 | NA | NA | NA | NA | NA | NA | NA |
| TRPM4 | 577 | Y | -58.79316667 | 30.91738999 | 0.414698895 | 26.66322157 | NA | NA | NA | NA | NA | NA | NA |
| TRPM4 | 578 | F | -56.19981818 | 33.73828598 | 0.455548527 | 29.37874967 | NA | NA | NA | NA | NA | NA | NA |
| TRPM4 | 579 | W | -50.42664286 | 30.05553323 | 0.402218275 | 23.51335323 | NA | NA | NA | NA | NA | NA | NA |
| TRPM4 | 580 | E | -54.8722 | 30.79244154 | 0.412889506 | 26.43302494 | NA | NA | NA | NA | NA | NA | NA |
| TRPM4 | 581 | M | -57.67075 | 32.82376994 | 0.442305343 | 28.85404543 | NA | NA | NA | NA | NA | NA | NA |
| TRPM4 | 582 | G | -53.78475 | 36.25553263 | 0.492000987 | 30.82182278 | NA | NA | NA | NA | NA | NA | NA |
| TRPM4 | 583 | S | -51.5156 | 38.95531169 | 0.531096715 | 32.35956842 | NA | NA | NA | NA | NA | NA | NA |
| TRPM4 | 584 | N | -47.91325 | 37.91344114 | 0.5160093 | 27.93293982 | NA | NA | NA | NA | NA | NA | NA |
| TRPM4 | 585 | A | -47.1238 | 34.93049135 | 0.472812953 | 24.98812796 | NA | NA | NA | NA | NA | NA | NA |
| TRPM4 | 586 | V | -43.43485714 | 33.8660341 | 0.457398458 | 23.0337447 | NA | NA | NA | NA | NA | NA | NA |
| TRPM4 | 587 | S | -42.46983333 | 36.66552442 | 0.497938113 | 20.28714074 | NA | NA | NA | NA | NA | NA | NA |
| TRPM4 | 588 | S | -45.4285 | 35.77315186 | 0.485015591 | 24.85651251 | NA | NA | NA | NA | NA | NA | NA |
| TRPM4 | 589 | A | -43.7008 | 32.25629708 | 0.434087721 | 21.32837385 | NA | NA | NA | NA | NA | NA | NA |
| TRPM4 | 590 | L | -39.871625 | 31.67523176 | 0.425673265 | 15.31836671 | NA | NA | NA | NA | NA | NA | NA |
| TRPM4 | 591 | G | -41.46225 | 36.14789998 | 0.49044235 | 19.72785294 | NA | NA | NA | NA | NA | NA | NA |
| TRPM4 | 592 | A | -43.3676 | 34.69836039 | 0.469451445 | 23.82612925 | NA | NA | NA | NA | NA | NA | NA |

| | | | | | | | | | | | | | |
|---|---|---|---|---|---|---|---|---|---|---|---|---|---|
| TRPM4 | 593 | C | -40.2415 | 31.02573321 | 0.416267822 | 14.67552234 | NA | NA | NA | NA | NA | NA | NA |
| TRPM4 | 594 | L | -37.983625 | 34.99860199 | 0.473799269 | 19.11854571 | NA | NA | NA | NA | NA | NA | NA |
| TRPM4 | 595 | L | -41.7255 | 37.18449006 | 0.505453298 | 20.77326354 | NA | NA | NA | NA | NA | NA | NA |
| TRPM4 | 596 | L | -42.310625 | 32.73222951 | 0.440979739 | 16.42265345 | NA | NA | NA | NA | NA | NA | NA |
| TRPM4 | 597 | R | -37.14872727 | 28.17989274 | 0.375056967 | 12.38573552 | NA | NA | NA | NA | NA | NA | NA |
| TRPM4 | 598 | V | -37.356 | 37.05578984 | 0.50358958 | 21.20736138 | NA | NA | NA | NA | NA | NA | NA |
| TRPM4 | 599 | M | -40.4296 | 38.34104731 | 0.522201501 | 22.47140593 | NA | NA | NA | NA | NA | NA | NA |
| TRPM4 | 600 | A | -38.939 | 34.98875276 | 0.473656642 | 19.10974048 | NA | NA | NA | NA | NA | NA | NA |
| TRPM4 | 601 | R | -34.33327273 | 35.20701718 | 0.476817347 | 20.27531149 | NA | NA | NA | NA | NA | NA | NA |
| TRPM4 | 602 | L | -38.1365 | 40.5255424 | 0.553835358 | 24.66947332 | NA | NA | NA | NA | NA | NA | NA |
| TRPM4 | 603 | E | -40.2396 | 39.55029459 | 0.539712713 | 23.18737095 | NA | NA | NA | NA | NA | NA | NA |
| TRPM4 | 604 | P | -39.95871429 | 38.95902798 | 0.531150531 | 23.08191829 | NA | NA | NA | NA | NA | NA | NA |
| TRPM4 | 605 | D | -41.2832 | 36.45833344 | 0.494937763 | 20.04491553 | NA | NA | NA | NA | NA | NA | NA |
| TRPM4 | 606 | A | -41.1074 | 32.50187935 | 0.437644018 | 16.09205436 | NA | NA | NA | NA | NA | NA | NA |
| TRPM4 | 607 | E | -44.6656 | 32.55739838 | 0.438447994 | 21.67674498 | NA | NA | NA | NA | NA | NA | NA |
| TRPM4 | 608 | E | -43.331 | 33.18596443 | 0.447550312 | 22.25417403 | NA | NA | NA | NA | NA | NA | NA |
| TRPM4 | 609 | A | -41.391 | 30.66553458 | 0.411051755 | 14.24749945 | NA | NA | NA | NA | NA | NA | NA |
| TRPM4 | 610 | A | -43.1428 | 27.94235382 | 0.371617146 | 11.95221343 | NA | NA | NA | NA | NA | NA | NA |
| TRPM4 | 611 | R | -45.3918 | 29.65282347 | 0.396386601 | 18.76261466 | NA | NA | NA | NA | NA | NA | NA |
| TRPM4 | 612 | R | -43.95345455 | 30.5944031 | 0.410021694 | 19.71949133 | NA | NA | NA | NA | NA | NA | NA |
| TRPM4 | 613 | K | -39.58866667 | 27.39580809 | 0.363702571 | 10.97700268 | NA | NA | NA | NA | NA | NA | NA |
| TRPM4 | 614 | D | -44.500125 | 23.65068274 | 0.309469096 | 12.73416508 | NA | NA | NA | NA | NA | NA | NA |
| TRPM4 | 615 | L | -46.235875 | 28.05757459 | 0.373285667 | 17.15187471 | NA | NA | NA | NA | NA | NA | NA |
| TRPM4 | 616 | A | -42.1876 | 28.15637054 | 0.37471634 | 12.1014619 | NA | NA | NA | NA | NA | NA | NA |
| TRPM4 | 617 | F | -40.26654545 | 24.33530117 | 0.319383113 | 8.050295593 | NA | NA | NA | NA | NA | NA | NA |
| TRPM4 | 618 | K | -46.55077778 | 23.05558913 | 0.300851495 | 12.48636608 | NA | NA | NA | NA | NA | NA | NA |
| TRPM4 | 619 | F | -45.99554545 | 27.64932298 | 0.367373741 | 16.74778026 | NA | NA | NA | NA | NA | NA | NA |
| TRPM4 | 620 | E | -39.86866667 | 26.51997244 | 0.351019521 | 10.1075805 | NA | NA | NA | NA | NA | NA | NA |
| TRPM4 | 621 | G | -43.17025 | 24.37609011 | 0.319973781 | 8.342416827 | NA | NA | NA | NA | NA | NA | NA |
| TRPM4 | 622 | M | -46.805125 | 24.83185416 | 0.326573738 | 14.89556753 | NA | NA | NA | NA | NA | NA | NA |
| TRPM4 | 623 | G | -43.34075 | 28.00165256 | 0.372475856 | 17.07647438 | NA | NA | NA | NA | NA | NA | NA |
| TRPM4 | 624 | V | -40.98028571 | 24.4494774 | 0.321036508 | 8.059453454 | NA | NA | NA | NA | NA | NA | NA |
| TRPM4 | 625 | D | -44.340125 | 22.03488482 | 0.28607059 | 11.14001693 | NA | NA | NA | NA | NA | NA | NA |

| | | | | | | | | | | | | | |
|---|---|---|---|---|---|---|---|---|---|---|---|---|---|
| TRPM4 | 626 | L | -45.9775 | 27.24111306 | 0.361462419 | 16.34529037 | NA | NA | NA | NA | NA | NA | NA |
| TRPM4 | 627 | F | -40.53618182 | 29.10682836 | 0.388479999 | 12.70022213 | NA | NA | NA | NA | NA | NA | NA |
| TRPM4 | 628 | G | -42.442 | 27.03085818 | 0.358417701 | 10.62137101 | NA | NA | NA | NA | NA | NA | NA |
| TRPM4 | 629 | E | -45.5728 | 27.35235498 | 0.363073323 | 16.58313627 | NA | NA | NA | NA | NA | NA | NA |
| TRPM4 | 630 | C | -44.27383333 | 30.1132124 | 0.403053531 | 19.19425424 | NA | NA | NA | NA | NA | NA | NA |
| TRPM4 | 631 | Y | -38.58575 | 29.8083076 | 0.398638179 | 13.55065649 | NA | NA | NA | NA | NA | NA | NA |
| TRPM4 | 632 | R | -43.08927273 | 24.87189279 | 0.327153541 | 8.512315029 | NA | NA | NA | NA | NA | NA | NA |
| TRPM4 | 633 | S | -43.92416667 | 32.54228869 | 0.43822919 | 21.63149573 | NA | NA | NA | NA | NA | NA | NA |
| TRPM4 | 634 | S | -41.701 | 35.81589738 | 0.485634592 | 19.40966841 | NA | NA | NA | NA | NA | NA | NA |
| TRPM4 | 635 | E | -37.41811111 | 32.42498336 | 0.436530481 | 16.65393486 | NA | NA | NA | NA | NA | NA | NA |
| TRPM4 | 636 | V | -38.07071429 | 38.50942459 | 0.524639787 | 22.64402088 | NA | NA | NA | NA | NA | NA | NA |
| TRPM4 | 637 | R | -42.69854545 | 38.63515583 | 0.526460511 | 22.21796287 | NA | NA | NA | NA | NA | NA | NA |
| TRPM4 | 638 | A | -39.319 | 35.12076984 | 0.475568392 | 19.24206964 | NA | NA | NA | NA | NA | NA | NA |
| TRPM4 | 639 | A | -36.4606 | 37.57011633 | 0.511037585 | 21.72448859 | NA | NA | NA | NA | NA | NA | NA |
| TRPM4 | 640 | R | -39.30045455 | 40.17398011 | 0.548744355 | 24.32182575 | NA | NA | NA | NA | NA | NA | NA |
| TRPM4 | 641 | L | -41.171 | 36.75625729 | 0.499252023 | 20.37602363 | NA | NA | NA | NA | NA | NA | NA |
| TRPM4 | 642 | L | -36.926625 | 34.8363427 | 0.471449579 | 19.18171829 | NA | NA | NA | NA | NA | NA | NA |
| TRPM4 | 643 | L | -34.4715 | 39.07722772 | 0.532862192 | 29.3826965 | NA | NA | NA | NA | NA | NA | NA |
| TRPM4 | 644 | R | -40.33354545 | 42.15158022 | 0.577382149 | 26.32366773 | NA | NA | NA | NA | NA | NA | NA |
| TRPM4 | 645 | R | -37.04372727 | 44.7017271 | 0.61431104 | 28.83196713 | NA | NA | NA | NA | NA | NA | NA |
| TRPM4 | 646 | C | -41.76266667 | 42.89004815 | 0.588075965 | 26.51884327 | NA | NA | NA | NA | NA | NA | NA |
| TRPM4 | 647 | P | -43.63985714 | 46.38524271 | 0.638690172 | 29.98949723 | NA | NA | NA | NA | NA | NA | NA |
| TRPM4 | 648 | L | -44.6956 | 44.82481431 | 0.616093477 | 33.94886376 | NA | NA | NA | NA | NA | NA | NA |
| TRPM4 | 649 | W | -42.41264286 | 39.7689726 | 0.542879407 | 23.39538819 | NA | NA | NA | NA | NA | NA | NA |
| TRPM4 | 650 | G | -39.46 | 46.62747665 | 0.642197982 | 30.74805623 | NA | NA | NA | NA | NA | NA | NA |
| TRPM4 | 651 | D | -36.2132 | 46.82747619 | 0.645094192 | 31.06353036 | NA | NA | NA | NA | NA | NA | NA |
| TRPM4 | 652 | A | -36.5248 | 43.55068662 | 0.597642726 | 34.00714546 | NA | NA | NA | NA | NA | NA | NA |
| TRPM4 | 653 | T | -35.32628571 | 40.22911241 | 0.549542731 | 30.68444595 | NA | NA | NA | NA | NA | NA | NA |
| TRPM4 | 654 | C | -37.04683333 | 35.95381711 | 0.48763182 | 20.08050008 | NA | NA | NA | NA | NA | NA | NA |
| TRPM4 | 655 | L | -32.819875 | 34.60774868 | 0.46813929 | 24.90954086 | NA | NA | NA | NA | NA | NA | NA |
| TRPM4 | 656 | Q | -31.84677778 | 37.82953616 | 0.514794265 | 28.13107021 | NA | NA | NA | NA | NA | NA | NA |
| TRPM4 | 657 | L | -36.210625 | 35.81353857 | 0.485600434 | 26.20282762 | NA | NA | NA | NA | NA | NA | NA |
| TRPM4 | 658 | A | -34.174 | 32.85112054 | 0.44270141 | 23.15397105 | NA | NA | NA | NA | NA | NA | NA |

| | | | | | | | | | | | | | |
|---|---|---|---|---|---|---|---|---|---|---|---|---|---|
| TRPM4 | 659 | M | -29.717125 | 34.26101022 | 0.463118141 | 24.63885212 | NA | NA | NA | NA | NA | NA | NA |
| TRPM4 | 660 | Q | -31.58033333 | 35.87847993 | 0.486540856 | 26.17958703 | NA | NA | NA | NA | NA | NA | NA |
| TRPM4 | 661 | A | -34.0232 | 32.06113049 | 0.431261497 | 22.39071074 | NA | NA | NA | NA | NA | NA | NA |
| TRPM4 | 662 | D | -32.2268 | 28.99910957 | 0.386920115 | 19.30161567 | NA | NA | NA | NA | NA | NA | NA |
| TRPM4 | 663 | A | -33.9898 | 27.86787966 | 0.370538679 | 18.18062737 | NA | NA | NA | NA | NA | NA | NA |
| TRPM4 | 664 | R | -30.98536364 | 24.62518262 | 0.32358091 | 14.93396185 | NA | NA | NA | NA | NA | NA | NA |
| TRPM4 | 665 | A | -35.5776 | 23.51561634 | 0.307513188 | 14.12250365 | NA | NA | NA | NA | NA | NA | NA |
| TRPM4 | 666 | F | -36.20072727 | 26.38444134 | 0.349056884 | 16.82673804 | NA | NA | NA | NA | NA | NA | NA |
| TRPM4 | 667 | F | -31.26063636 | 26.14040149 | 0.345522922 | 16.48299091 | NA | NA | NA | NA | NA | NA | NA |
| TRPM4 | 668 | A | -32.5476 | 22.72998841 | 0.296136444 | 13.0329082 | NA | NA | NA | NA | NA | NA | NA |
| TRPM4 | 669 | Q | -35.50255556 | 24.38918929 | 0.320163471 | 14.7406853 | NA | NA | NA | NA | NA | NA | NA |
| TRPM4 | 670 | D | -33.737375 | 25.4262629 | 0.335181421 | 15.75295987 | NA | NA | NA | NA | NA | NA | NA |
| TRPM4 | 671 | G | -33.7505 | 28.08982346 | 0.373752666 | 18.42857187 | NA | NA | NA | NA | NA | NA | NA |
| TRPM4 | 672 | V | -32.33242857 | 29.89217749 | 0.399852706 | 20.1972886 | NA | NA | NA | NA | NA | NA | NA |
| TRPM4 | 673 | Q | -28.69922222 | 26.84665642 | 0.355750259 | 18.9356742 | NA | NA | NA | NA | NA | NA | NA |
| TRPM4 | 674 | S | -30.12383333 | 30.33378885 | 0.406247718 | 20.74854838 | NA | NA | NA | NA | NA | NA | NA |
| TRPM4 | 675 | L | -31.252125 | 32.98133601 | 0.444587071 | 23.35172493 | NA | NA | NA | NA | NA | NA | NA |
| TRPM4 | 676 | L | -27.176375 | 32.01112346 | 0.430537341 | 24.03152603 | NA | NA | NA | NA | NA | NA | NA |
| TRPM4 | 677 | T | -25.21785714 | 31.26562608 | 0.419741731 | 24.25600982 | NA | NA | NA | NA | NA | NA | NA |
| TRPM4 | 678 | Q | -27.65555556 | 35.60512415 | 0.482582368 | 27.59486936 | NA | NA | NA | NA | NA | NA | NA |
| TRPM4 | 679 | K | -25.72677778 | 37.47310995 | 0.509632827 | 30.4718184 | NA | NA | NA | NA | NA | NA | NA |
| TRPM4 | 680 | W | -19.97714286 | 32.53369973 | 0.438104812 | 27.7043594 | NA | NA | NA | NA | NA | NA | NA |
| TRPM4 | 681 | W | -20.46485714 | 35.05816558 | 0.474661815 | 30.04273209 | NA | NA | NA | NA | NA | NA | NA |
| TRPM4 | 682 | G | -22.6835 | 40.3467509 | 0.551246264 | 35.31788613 | NA | NA | NA | NA | NA | NA | NA |
| TRPM4 | 683 | D | -22.5252 | 43.91376499 | 0.602900495 | 38.88294771 | NA | NA | NA | NA | NA | NA | NA |
| TRPM4 | 684 | M | -23.3574 | 44.15136698 | 0.606341229 | 38.55349087 | NA | NA | NA | NA | NA | NA | NA |
| TRPM4 | 685 | A | -22.795 | 40.55713998 | 0.554292925 | 35.5682858 | NA | NA | NA | NA | NA | NA | NA |
| TRPM4 | 686 | S | -23.7146 | 38.51374916 | 0.524702411 | 32.91223875 | NA | NA | NA | NA | NA | NA | NA |
| TRPM4 | 687 | T | -22.44428571 | 35.0945267 | 0.475188363 | 30.11037613 | NA | NA | NA | NA | NA | NA | NA |
| TRPM4 | 688 | T | -18.60057143 | 36.4367279 | 0.494624891 | 33.92852024 | NA | NA | NA | NA | NA | NA | NA |
| TRPM4 | 689 | P | -16.10628571 | 35.64472585 | 0.483155843 | 33.5254174 | NA | NA | NA | NA | NA | NA | NA |
| TRPM4 | 690 | I | -12.956125 | 32.51710415 | 0.43786449 | 30.28764577 | NA | NA | NA | NA | NA | NA | NA |
| TRPM4 | 691 | W | -11.7475 | 34.2679933 | 0.463219263 | 33.19872668 | NA | NA | NA | NA | NA | NA | NA |

| | | | | | | | | | | | | | |
|---|---|---|---|---|---|---|---|---|---|---|---|---|---|
| TRPM4 | 692 | A | -12.0694 | 38.5981848 | 0.525925131 | 37.34837023 | NA | NA | NA | NA | NA | NA | NA |
| TRPM4 | 693 | L | -12.119375 | 37.75235496 | 0.513676598 | 36.5345017 | NA | NA | NA | NA | NA | NA | NA |
| TRPM4 | 694 | V | -7.866714286 | 36.0369676 | 0.488835929 | 34.95845278 | NA | NA | NA | NA | NA | NA | NA |
| TRPM4 | 695 | L | -7.5264 | 40.13589055 | 0.548192777 | 39.04775091 | NA | NA | NA | NA | NA | NA | NA |
| TRPM4 | 696 | A | -9.0978 | 42.60381073 | 0.583930937 | 41.1806352 | NA | NA | NA | NA | NA | NA | NA |
| TRPM4 | 697 | F | -6.942636364 | 40.81185467 | 0.55798147 | 39.75151516 | NA | NA | NA | NA | NA | NA | NA |
| TRPM4 | 698 | F | -4.062 | 41.94997774 | 0.574462726 | 38.60671255 | NA | NA | NA | NA | NA | NA | NA |
| TRPM4 | 699 | C | -4.998666667 | 44.93407096 | 0.617675631 | 42.41473659 | NA | NA | NA | NA | NA | NA | NA |
| TRPM4 | 700 | P | -6.325857143 | 48.02915979 | 0.662495873 | 46.98440788 | NA | NA | NA | NA | NA | NA | NA |
| TRPM4 | 701 | P | -6.559857143 | 50.45443741 | 0.697616522 | 49.37850758 | NA | NA | NA | NA | NA | NA | NA |
| TRPM4 | 702 | L | -8.882 | 46.27480245 | 0.637090877 | 44.92414649 | NA | NA | NA | NA | NA | NA | NA |
| TRPM4 | 703 | I | -12.491125 | 47.28789956 | 0.651761621 | 46.03775445 | NA | NA | NA | NA | NA | NA | NA |
| TRPM4 | 704 | Y | -12.73641667 | 51.27382344 | 0.70948212 | 49.06366139 | NA | NA | NA | NA | NA | NA | NA |
| TRPM4 | 705 | T | -13.8122 | 50.15132036 | 0.693227059 | 47.94151135 | NA | NA | NA | NA | NA | NA | NA |
| TRPM4 | 706 | R | -15.6558 | 47.6627151 | 0.657189357 | 45.56162803 | NA | NA | NA | NA | NA | NA | NA |
| TRPM4 | 707 | L | -14.7792 | 44.48879258 | 0.611227518 | 41.52722438 | NA | NA | NA | NA | NA | NA | NA |
| TRPM4 | 708 | I | -16.7356 | 45.60519384 | 0.627394218 | 43.49338772 | NA | NA | NA | NA | NA | NA | NA |
| TRPM4 | 709 | T | -20.2318 | 43.99288535 | 0.604046243 | 41.54504202 | NA | NA | NA | NA | NA | NA | NA |
| TRPM4 | 710 | F | -21.59854545 | 46.12481267 | 0.634918862 | 41.09791358 | NA | NA | NA | NA | NA | NA | NA |
| TRPM4 | 766 | R | -19.689 | 59.85136969 | 0.833694289 | 57.40972643 | NA | NA | NA | NA | NA | NA | NA |
| TRPM4 | 767 | R | -19.1006 | 56.41793543 | 0.783974439 | 53.91007459 | NA | NA | NA | NA | NA | NA | NA |
| TRPM4 | 768 | C | -15.7926 | 55.55461503 | 0.771472624 | 53.43544664 | NA | NA | NA | NA | NA | NA | NA |
| TRPM4 | 769 | L | -15.116125 | 56.30238579 | 0.782301155 | 54.19016023 | NA | NA | NA | NA | NA | NA | NA |
| TRPM4 | 770 | R | -18.8522 | 53.209938 | 0.737519158 | 50.70381561 | NA | NA | NA | NA | NA | NA | NA |
| TRPM4 | 771 | R | -17.698 | 49.75713482 | 0.687518825 | 47.24978604 | NA | NA | NA | NA | NA | NA | NA |
| TRPM4 | 772 | W | -12.26885714 | 50.95825465 | 0.704912342 | 47.95944862 | NA | NA | NA | NA | NA | NA | NA |
| TRPM4 | 773 | F | -16.8858 | 50.10610897 | 0.692572349 | 48.0272898 | NA | NA | NA | NA | NA | NA | NA |
| TRPM4 | 774 | H | -20.5279 | 47.20299753 | 0.650532148 | 44.69504139 | NA | NA | NA | NA | NA | NA | NA |
| TRPM4 | 775 | F | -14.79363636 | 44.25818626 | 0.607888088 | 42.20148761 | NA | NA | NA | NA | NA | NA | NA |
| TRPM4 | 776 | W | -11.86671429 | 45.61589246 | 0.627549146 | 42.90846696 | NA | NA | NA | NA | NA | NA | NA |
| TRPM4 | 777 | G | -17.56075 | 44.77733401 | 0.61540591 | 42.26822546 | NA | NA | NA | NA | NA | NA | NA |
| TRPM4 | 778 | A | -17.718 | 42.11677236 | 0.576878093 | 39.61010921 | NA | NA | NA | NA | NA | NA | NA |
| TRPM4 | 779 | P | -17.96557143 | 38.93325699 | 0.530777339 | 36.51672796 | NA | NA | NA | NA | NA | NA | NA |

| | | | | | | | | | | | | | |
|---|---|---|---|---|---|---|---|---|---|---|---|---|---|
| TRPM4 | 780 | V | -15.75028571 | 36.58276376 | 0.496739649 | 34.46471265 | NA | NA | NA | NA | NA | NA | NA |
| TRPM4 | 781 | T | -12.98685714 | 40.0585232 | 0.547072414 | 37.83152292 | NA | NA | NA | NA | NA | NA | NA |
| TRPM4 | 782 | I | -12.737 | 39.87239002 | 0.544377004 | 38.63800415 | NA | NA | NA | NA | NA | NA | NA |
| TRPM4 | 783 | F | -12.12527273 | 32.45131054 | 0.436911727 | 31.20137883 | NA | NA | NA | NA | NA | NA | NA |
| TRPM4 | 784 | M | -9.05375 | 35.34767956 | 0.478854291 | 33.91983896 | NA | NA | NA | NA | NA | NA | NA |
| TRPM4 | 785 | G | -7.27425 | 39.44774776 | 0.538227723 | 38.36981153 | NA | NA | NA | NA | NA | NA | NA |
| TRPM4 | 786 | N | -7.72225 | 37.3256519 | 0.507497475 | 36.26732467 | NA | NA | NA | NA | NA | NA | NA |
| TRPM4 | 787 | V | -6.155714286 | 33.24937693 | 0.448468594 | 30.74633947 | NA | NA | NA | NA | NA | NA | NA |
| TRPM4 | 788 | V | -3.200142857 | 36.67501472 | 0.498075543 | 32.57890343 | NA | NA | NA | NA | NA | NA | NA |
| TRPM4 | 789 | S | -2.535666667 | 38.12156076 | 0.519023098 | 33.88061528 | NA | NA | NA | NA | NA | NA | NA |
| TRPM4 | 790 | Y | -2.491583333 | 32.85413525 | 0.442745066 | 28.60468961 | NA | NA | NA | NA | NA | NA | NA |
| TRPM4 | 791 | L | -0.743625 | 31.19145 | 0.418667581 | 25.73867435 | NA | NA | NA | NA | NA | NA | NA |
| TRPM4 | 792 | L | 0.754375 | 36.11823354 | 0.490012748 | 31.30340189 | NA | NA | NA | NA | NA | NA | NA |
| TRPM4 | 793 | F | 0.722636364 | 35.94218548 | 0.487463381 | 31.30306597 | NA | NA | NA | NA | NA | NA | NA |
| TRPM4 | 794 | L | 1.396875 | 30.85672042 | 0.413820334 | 26.77103971 | NA | NA | NA | NA | NA | NA | NA |
| TRPM4 | 795 | L | 5.09075 | 33.38373805 | 0.450414289 | 31.00474042 | NA | NA | NA | NA | NA | NA | NA |
| TRPM4 | 796 | L | 6.432875 | 35.79924907 | 0.485393507 | 33.57419189 | NA | NA | NA | NA | NA | NA | NA |
| TRPM4 | 797 | F | 5.781090909 | 33.52351299 | 0.452438381 | 31.29534615 | NA | NA | NA | NA | NA | NA | NA |
| TRPM4 | 798 | S | 8.277166667 | 30.01937265 | 0.40169463 | 28.22002041 | NA | NA | NA | NA | NA | NA | NA |
| TRPM4 | 799 | R | 12.37909091 | 33.5254947 | 0.452467079 | 31.52054557 | NA | NA | NA | NA | NA | NA | NA |
| TRPM4 | 800 | V | 11.00557143 | 34.9315091 | 0.472827691 | 32.32187589 | NA | NA | NA | NA | NA | NA | NA |
| TRPM4 | 801 | L | 11.23825 | 30.58482202 | 0.40988295 | 28.05931184 | NA | NA | NA | NA | NA | NA | NA |
| TRPM4 | 802 | L | 13.293375 | 27.83632312 | 0.370081706 | 24.95334758 | NA | NA | NA | NA | NA | NA | NA |
| TRPM4 | 803 | V | 16.66357143 | 31.98979315 | 0.430228455 | 27.57558633 | NA | NA | NA | NA | NA | NA | NA |
| TRPM4 | 804 | D | 15.955875 | 34.81303137 | 0.471112005 | 30.40154485 | NA | NA | NA | NA | NA | NA | NA |
| TRPM4 | 805 | F | 14.67754545 | 35.92229061 | 0.487175282 | 31.37273407 | NA | NA | NA | NA | NA | NA | NA |
| TRPM4 | 806 | Q | 19.143 | 37.73225348 | 0.513385506 | 30.12146678 | NA | NA | NA | NA | NA | NA | NA |
| TRPM4 | 807 | P | 21.37471429 | 37.5219879 | 0.510340633 | 29.94997903 | NA | NA | NA | NA | NA | NA | NA |
| TRPM4 | 808 | A | 19.9456 | 41.80450355 | 0.572356102 | 34.22553027 | NA | NA | NA | NA | NA | NA | NA |
| TRPM4 | 809 | P | 19.33628571 | 44.8068737 | 0.615833677 | 38.00769422 | NA | NA | NA | NA | NA | NA | NA |
| TRPM4 | 810 | P | 15.27371429 | 43.85095238 | 0.6019909 | 39.29161713 | NA | NA | NA | NA | NA | NA | NA |
| TRPM4 | 811 | G | 14.2055 | 45.88271134 | 0.631412973 | 41.39400289 | NA | NA | NA | NA | NA | NA | NA |
| TRPM4 | 812 | S | 12.42216667 | 45.35104642 | 0.623713888 | 42.48109917 | NA | NA | NA | NA | NA | NA | NA |

| | | | | | | | | | | | | | |
|---|---|---|---|---|---|---|---|---|---|---|---|---|---|
| TRPM4 | 813 | L | 11.74725 | 41.43227114 | 0.566965774 | 38.84555342 | NA | NA | NA | NA | NA | NA | NA |
| TRPM4 | 814 | E | 11.46344444 | 40.23502487 | 0.549628349 | 38.32773199 | NA | NA | NA | NA | NA | NA | NA |
| TRPM4 | 815 | L | 9.396 | 45.21111822 | 0.621687576 | 43.27354747 | NA | NA | NA | NA | NA | NA | NA |
| TRPM4 | 816 | L | 6.058875 | 43.65328581 | 0.599128474 | 41.71521889 | NA | NA | NA | NA | NA | NA | NA |
| TRPM4 | 817 | L | 5.57225 | 39.19182715 | 0.534521715 | 37.06855422 | NA | NA | NA | NA | NA | NA | NA |
| TRPM4 | 818 | Y | 6.740916667 | 43.19459294 | 0.592486104 | 41.14559959 | NA | NA | NA | NA | NA | NA | NA |
| TRPM4 | 819 | F | 1.904727273 | 46.07624301 | 0.634215521 | 42.94580382 | NA | NA | NA | NA | NA | NA | NA |
| TRPM4 | 820 | W | 0.651571429 | 41.19301631 | 0.563501104 | 36.58214753 | NA | NA | NA | NA | NA | NA | NA |
| TRPM4 | 821 | A | 0.3304 | 42.40214485 | 0.581010596 | 37.63376552 | NA | NA | NA | NA | NA | NA | NA |
| TRPM4 | 822 | F | 0 | 45.1981355 | 0.621499572 | 39.84294892 | NA | NA | NA | NA | NA | NA | NA |
| TRPM4 | 823 | T | -1.711285714 | 44.94273602 | 0.617801111 | 40.14154871 | NA | NA | NA | NA | NA | NA | NA |
| TRPM4 | 824 | L | -2.856625 | 39.64416739 | 0.541072093 | 35.42017417 | NA | NA | NA | NA | NA | NA | NA |
| TRPM4 | 825 | L | -3.474375 | 43.79705494 | 0.601210407 | 39.68354159 | NA | NA | NA | NA | NA | NA | NA |
| TRPM4 | 826 | C | -5.891 | 46.15143886 | 0.635304438 | 43.63431557 | NA | NA | NA | NA | NA | NA | NA |
| TRPM4 | 827 | E | -8.0432 | 43.91123804 | 0.602863902 | 42.86280244 | NA | NA | NA | NA | NA | NA | NA |
| TRPM4 | 828 | E | -8.6272 | 41.99678176 | 0.575140499 | 40.91032632 | NA | NA | NA | NA | NA | NA | NA |
| TRPM4 | 829 | L | -9.313 | 44.97403941 | 0.618254418 | 43.55661417 | NA | NA | NA | NA | NA | NA | NA |
| TRPM4 | 830 | R | -12.61590909 | 45.89787688 | 0.631632586 | 44.65555126 | NA | NA | NA | NA | NA | NA | NA |
| TRPM4 | 831 | Q | -13.529 | 38.65947691 | 0.526812707 | 36.42980946 | NA | NA | NA | NA | NA | NA | NA |
| TRPM4 | 832 | G | -14.31175 | 43.84013331 | 0.601834228 | 41.61803637 | NA | NA | NA | NA | NA | NA | NA |
| TRPM4 | 833 | L | -14.95175 | 46.2922858 | 0.637344055 | 44.28664344 | NA | NA | NA | NA | NA | NA | NA |
| TRPM4 | 834 | S | -18.02533333 | 46.30657541 | 0.637550984 | 43.79918536 | NA | NA | NA | NA | NA | NA | NA |
| TRPM4 | 835 | G | -19.626 | 42.44849116 | 0.581681741 | 39.99450084 | NA | NA | NA | NA | NA | NA | NA |
| TRPM4 | 838 | G | -25.52725 | 38.07434183 | 0.518339317 | 31.31495242 | NA | NA | NA | NA | NA | NA | NA |
| TRPM4 | 839 | S | -27.85583333 | 35.31998661 | 0.478453267 | 27.35105506 | NA | NA | NA | NA | NA | NA | NA |
| TRPM4 | 840 | L | -31.581375 | 37.92524747 | 0.516180269 | 28.23886295 | NA | NA | NA | NA | NA | NA | NA |
| TRPM4 | 841 | A | -31.8102 | 34.23653693 | 0.462763741 | 24.54602116 | NA | NA | NA | NA | NA | NA | NA |
| TRPM4 | 842 | S | -29.5672 | 36.1303415 | 0.490188084 | 26.51736443 | NA | NA | NA | NA | NA | NA | NA |
| TRPM4 | 843 | G | -29.914 | 38.71117123 | 0.527561297 | 29.08899178 | NA | NA | NA | NA | NA | NA | NA |
| TRPM4 | 844 | G | -28.67475 | 40.73103772 | 0.556811153 | 32.80046931 | NA | NA | NA | NA | NA | NA | NA |
| TRPM4 | 845 | P | -27.32385714 | 43.62733025 | 0.598752609 | 35.61878053 | NA | NA | NA | NA | NA | NA | NA |
| TRPM4 | 846 | G | -28.3035 | 44.19732044 | 0.607006685 | 36.23427913 | NA | NA | NA | NA | NA | NA | NA |
| TRPM4 | 847 | P | -29.08385714 | 44.25391551 | 0.607826243 | 36.3851235 | NA | NA | NA | NA | NA | NA | NA |

| | | | | | | | | | | | | | |
|---|---|---|---|---|---|---|---|---|---|---|---|---|---|
| TRPM4 | 848 | G | -27.4325 | 48.46449449 | 0.668799991 | 40.49014674 | NA | NA | NA | NA | NA | NA | NA |
| TRPM4 | 849 | H | -26.8532 | 50.42427214 | 0.697179697 | 42.42301749 | NA | NA | NA | NA | NA | NA | NA |
| TRPM4 | 850 | A | -24.2174 | 48.62185241 | 0.671078705 | 42.72240012 | NA | NA | NA | NA | NA | NA | NA |
| TRPM4 | 851 | S | -21.8212 | 49.50623497 | 0.683885523 | 44.50516844 | NA | NA | NA | NA | NA | NA | NA |
| TRPM4 | 852 | L | -17.71925 | 48.99881514 | 0.676537534 | 46.49345519 | NA | NA | NA | NA | NA | NA | NA |
| TRPM4 | 853 | S | -18.99733333 | 47.67039227 | 0.65730053 | 45.17768572 | NA | NA | NA | NA | NA | NA | NA |
| TRPM4 | 854 | Q | -22.10055556 | 44.91524384 | 0.617402994 | 39.93194729 | NA | NA | NA | NA | NA | NA | NA |
| TRPM4 | 855 | R | -19.44618182 | 44.29094619 | 0.608362487 | 41.78892873 | NA | NA | NA | NA | NA | NA | NA |
| TRPM4 | 856 | L | -15.0665 | 43.63277312 | 0.598831428 | 41.51321407 | NA | NA | NA | NA | NA | NA | NA |
| TRPM4 | 857 | R | -18.14145455 | 40.30844921 | 0.550691614 | 37.79913753 | NA | NA | NA | NA | NA | NA | NA |
| TRPM4 | 858 | L | -19.631125 | 39.43972141 | 0.538111493 | 36.93029209 | NA | NA | NA | NA | NA | NA | NA |
| TRPM4 | 859 | Y | -13.82108333 | 38.98790333 | 0.531568678 | 36.88071266 | NA | NA | NA | NA | NA | NA | NA |
| TRPM4 | 860 | L | -13.182 | 37.96848663 | 0.516806419 | 35.73876531 | NA | NA | NA | NA | NA | NA | NA |
| TRPM4 | 861 | A | -16.4164 | 34.37445496 | 0.464760944 | 32.2644989 | NA | NA | NA | NA | NA | NA | NA |
| TRPM4 | 862 | D | -15.43275 | 32.42976677 | 0.43659975 | 30.31296964 | NA | NA | NA | NA | NA | NA | NA |
| TRPM4 | 863 | S | -12.00516667 | 30.51597629 | 0.408885989 | 29.27224248 | NA | NA | NA | NA | NA | NA | NA |
| TRPM4 | 864 | W | -10.97335714 | 26.92953089 | 0.356950372 | 25.73169431 | NA | NA | NA | NA | NA | NA | NA |
| TRPM4 | 865 | N | -11.019125 | 34.80379538 | 0.470978258 | 33.61431478 | NA | NA | NA | NA | NA | NA | NA |
| TRPM4 | 866 | Q | -9.363111111 | 34.94243965 | 0.472985978 | 33.55129901 | NA | NA | NA | NA | NA | NA | NA |
| TRPM4 | 867 | C | -6.033166667 | 31.21265367 | 0.418974633 | 30.22127073 | NA | NA | NA | NA | NA | NA | NA |
| TRPM4 | 868 | D | -5.3554 | 34.94453557 | 0.473016329 | 32.44477926 | NA | NA | NA | NA | NA | NA | NA |
| TRPM4 | 869 | L | -5.182375 | 37.58972441 | 0.511321531 | 35.0753944 | NA | NA | NA | NA | NA | NA | NA |
| TRPM4 | 870 | V | -2.679142857 | 35.11781706 | 0.475525633 | 30.87267475 | NA | NA | NA | NA | NA | NA | NA |
| TRPM4 | 871 | A | -0.7128 | 34.95673121 | 0.473192935 | 29.5135743 | NA | NA | NA | NA | NA | NA | NA |
| TRPM4 | 872 | L | -0.782 | 38.04963931 | 0.517981597 | 32.59662037 | NA | NA | NA | NA | NA | NA | NA |
| TRPM4 | 873 | T | -0.451714286 | 39.54290665 | 0.539605727 | 34.1861871 | NA | NA | NA | NA | NA | NA | NA |
| TRPM4 | 874 | C | 1.98 | 35.29980272 | 0.478160982 | 31.13424659 | NA | NA | NA | NA | NA | NA | NA |
| TRPM4 | 875 | F | 3.370090909 | 38.23039579 | 0.520599147 | 35.09206482 | NA | NA | NA | NA | NA | NA | NA |
| TRPM4 | 876 | L | 2.696375 | 40.93490864 | 0.559763425 | 37.78558447 | NA | NA | NA | NA | NA | NA | NA |
| TRPM4 | 877 | L | 4.8205 | 40.16094684 | 0.548555619 | 38.07822758 | NA | NA | NA | NA | NA | NA | NA |
| TRPM4 | 878 | G | 7.99025 | 39.12739384 | 0.533588651 | 37.33255664 | NA | NA | NA | NA | NA | NA | NA |
| TRPM4 | 879 | V | 7.830428571 | 40.78285116 | 0.557561468 | 38.98842621 | NA | NA | NA | NA | NA | NA | NA |
| TRPM4 | 880 | G | 9.42875 | 43.31542942 | 0.594235947 | 41.38137594 | NA | NA | NA | NA | NA | NA | NA |

| | | | | | | | | | | | | | |
|---|---|---|---|---|---|---|---|---|---|---|---|---|---|
| TRPM4 | 881 | C | 11.5405 | 40.16026301 | 0.548545717 | 38.15432401 | NA | NA | NA | NA | NA | NA | NA |
| TRPM4 | 882 | R | 14.26981818 | 40.42394124 | 0.552364063 | 37.52816284 | NA | NA | NA | NA | NA | NA | NA |
| TRPM4 | 883 | L | 14.0394 | 44.15226124 | 0.606354179 | 41.25288778 | NA | NA | NA | NA | NA | NA | NA |
| TRPM4 | 884 | T | 15.04028571 | 43.97154968 | 0.60373728 | 39.42730627 | NA | NA | NA | NA | NA | NA | NA |
| TRPM4 | 885 | P | 18.94528571 | 44.31309694 | 0.608683254 | 36.69446309 | NA | NA | NA | NA | NA | NA | NA |
| TRPM4 | 886 | G | 19.4645 | 41.1197773 | 0.562440524 | 33.56572874 | NA | NA | NA | NA | NA | NA | NA |
| TRPM4 | 887 | L | 16.047125 | 39.49156757 | 0.538862282 | 35.10152227 | NA | NA | NA | NA | NA | NA | NA |
| TRPM4 | 888 | Y | 17.98875 | 35.99477036 | 0.488224867 | 31.64826559 | NA | NA | NA | NA | NA | NA | NA |
| TRPM4 | 889 | H | 17.983 | 33.07832459 | 0.445991571 | 28.75154022 | NA | NA | NA | NA | NA | NA | NA |
| TRPM4 | 890 | L | 12.445875 | 35.45662914 | 0.480431999 | 32.62625292 | NA | NA | NA | NA | NA | NA | NA |
| TRPM4 | 891 | G | 11.488 | 35.5446427 | 0.481706531 | 32.95918772 | NA | NA | NA | NA | NA | NA | NA |
| TRPM4 | 892 | R | 14.00981818 | 31.53127016 | 0.423588545 | 29.11800622 | NA | NA | NA | NA | NA | NA | NA |
| TRPM4 | 893 | T | 10.98285714 | 30.89542688 | 0.414380845 | 28.29120252 | NA | NA | NA | NA | NA | NA | NA |
| TRPM4 | 894 | V | 7.424571429 | 32.9497273 | 0.444129342 | 31.2243329 | NA | NA | NA | NA | NA | NA | NA |
| TRPM4 | 895 | L | 7.55075 | 32.34438345 | 0.435363307 | 30.38833087 | NA | NA | NA | NA | NA | NA | NA |
| TRPM4 | 896 | C | 7.9055 | 28.82782853 | 0.38443978 | 27.12801139 | NA | NA | NA | NA | NA | NA | NA |
| TRPM4 | 897 | I | 4.828 | 27.32796291 | 0.362720099 | 24.98814046 | NA | NA | NA | NA | NA | NA | NA |
| TRPM4 | 898 | D | 2.169375 | 31.32153515 | 0.420551355 | 28.18455338 | NA | NA | NA | NA | NA | NA | NA |
| TRPM4 | 899 | F | 3.375181818 | 27.9522305 | 0.371760171 | 24.81855499 | NA | NA | NA | NA | NA | NA | NA |
| TRPM4 | 900 | M | 0.956625 | 23.6925908 | 0.31007597 | 19.09910817 | NA | NA | NA | NA | NA | NA | NA |
| TRPM4 | 901 | V | -0.475714286 | 29.23691148 | 0.390363744 | 23.80961338 | NA | NA | NA | NA | NA | NA | NA |
| TRPM4 | 902 | F | -0.044727273 | 30.68574532 | 0.411344429 | 25.22756931 | NA | NA | NA | NA | NA | NA | NA |
| TRPM4 | 903 | T | -0.390285714 | 26.93229344 | 0.356990376 | 21.49061215 | NA | NA | NA | NA | NA | NA | NA |
| TRPM4 | 904 | V | -3.491857143 | 24.8396911 | 0.326687226 | 20.60700295 | NA | NA | NA | NA | NA | NA | NA |
| TRPM4 | 905 | R | -6.230272727 | 28.7984515 | 0.384014368 | 26.28831873 | NA | NA | NA | NA | NA | NA | NA |
| TRPM4 | 906 | L | -5.092375 | 27.38741539 | 0.363581036 | 24.93466022 | NA | NA | NA | NA | NA | NA | NA |
| TRPM4 | 907 | L | -8.371875 | 23.00481447 | 0.300116223 | 21.92078046 | NA | NA | NA | NA | NA | NA | NA |
| TRPM4 | 908 | H | -11.9322 | 27.9984866 | 0.372430009 | 26.80373378 | NA | NA | NA | NA | NA | NA | NA |
| TRPM4 | 909 | I | -10.43075 | 29.94628012 | 0.400636171 | 28.74838544 | NA | NA | NA | NA | NA | NA | NA |
| TRPM4 | 910 | F | -10.33727273 | 26.49433938 | 0.350648327 | 25.28495298 | NA | NA | NA | NA | NA | NA | NA |
| TRPM4 | 911 | T | -15.28757143 | 26.57440063 | 0.351807701 | 23.67286786 | NA | NA | NA | NA | NA | NA | NA |
| TRPM4 | 912 | V | -16.98757143 | 28.82488737 | 0.384397188 | 26.39035748 | NA | NA | NA | NA | NA | NA | NA |
| TRPM4 | 913 | N | -15.76025 | 26.12263758 | 0.345265682 | 24.07149546 | NA | NA | NA | NA | NA | NA | NA |

| | | | | | | | | | | | | | |
|---|---|---|---|---|---|---|---|---|---|---|---|---|---|
| TRPM4 | 914 | K | -20.08955556 | 23.62609415 | 0.309113026 | 21.12200202 | NA | NA | NA | NA | NA | NA | NA |
| TRPM4 | 915 | Q | -16.67388889 | 21.30224749 | 0.275461207 | 19.18336531 | NA | NA | NA | NA | NA | NA | NA |
| TRPM4 | 916 | L | -13.718625 | 21.79565769 | 0.282606322 | 18.84789907 | NA | NA | NA | NA | NA | NA | NA |
| TRPM4 | 917 | G | -17.414 | 22.05824082 | 0.28640881 | 19.55418336 | NA | NA | NA | NA | NA | NA | NA |
| TRPM4 | 918 | P | -19.18842857 | 18.05823917 | 0.22848445 | 15.54849333 | NA | NA | NA | NA | NA | NA | NA |
| TRPM4 | 919 | K | -15.89388889 | 14.57762268 | 0.17808135 | 12.57261659 | NA | NA | NA | NA | NA | NA | NA |
| TRPM4 | 920 | I | -13.328375 | 20.03781418 | 0.257150842 | 17.81835159 | NA | NA | NA | NA | NA | NA | NA |
| TRPM4 | 921 | V | -17.08042857 | 19.67420329 | 0.251885362 | 17.58102713 | NA | NA | NA | NA | NA | NA | NA |
| TRPM4 | 922 | I | -16.975 | 14.76900118 | 0.180852718 | 12.74294661 | NA | NA | NA | NA | NA | NA | NA |
| TRPM4 | 923 | V | -12.41614286 | 17.0054468 | 0.213238875 | 15.75504767 | NA | NA | NA | NA | NA | NA | NA |
| TRPM4 | 924 | S | -14.20033333 | 19.23350311 | 0.245503546 | 17.00950391 | NA | NA | NA | NA | NA | NA | NA |
| TRPM4 | 925 | K | -18.08988889 | 16.97252916 | 0.212762192 | 14.87959583 | NA | NA | NA | NA | NA | NA | NA |
| TRPM4 | 926 | M | -14.050375 | 11.57846868 | 0.134650348 | 9.375602562 | NA | NA | NA | NA | NA | NA | NA |
| TRPM4 | 927 | M | -11.305 | 17.57892775 | 0.221543501 | 16.3605126 | NA | NA | NA | NA | NA | NA | NA |
| TRPM4 | 928 | K | -15.35066667 | 17.29496022 | 0.217431343 | 15.07758252 | NA | NA | NA | NA | NA | NA | NA |
| TRPM4 | 929 | D | -12.4605 | 12.97884899 | 0.154929373 | 11.95272538 | NA | NA | NA | NA | NA | NA | NA |
| TRPM4 | 930 | V | -7.933142857 | 15.09954668 | 0.185639375 | 14.02329433 | NA | NA | NA | NA | NA | NA | NA |
| TRPM4 | 931 | F | -8.451181818 | 18.21059939 | 0.230690791 | 17.14748985 | NA | NA | NA | NA | NA | NA | NA |
| TRPM4 | 932 | F | -10.02745455 | 15.50061796 | 0.191447322 | 14.10659227 | NA | NA | NA | NA | NA | NA | NA |
| TRPM4 | 933 | F | -5.123545455 | 12.06451035 | 0.141688758 | 11.47282597 | NA | NA | NA | NA | NA | NA | NA |
| TRPM4 | 934 | L | -3.725375 | 17.34489576 | 0.218154464 | 14.03935785 | NA | NA | NA | NA | NA | NA | NA |
| TRPM4 | 935 | F | -6.345272727 | 19.81365968 | 0.253904842 | 17.29482262 | NA | NA | NA | NA | NA | NA | NA |
| TRPM4 | 936 | F | -5.691 | 17.53555229 | 0.220915377 | 15.01936439 | NA | NA | NA | NA | NA | NA | NA |
| TRPM4 | 937 | L | -0.959375 | 15.04612325 | 0.184865746 | 10.28003878 | NA | NA | NA | NA | NA | NA | NA |
| TRPM4 | 938 | G | -0.21275 | 19.49858072 | 0.249342157 | 14.17612537 | NA | NA | NA | NA | NA | NA | NA |
| TRPM4 | 939 | V | -1.751571429 | 21.85594246 | 0.283479311 | 17.04667009 | NA | NA | NA | NA | NA | NA | NA |
| TRPM4 | 940 | W | -0.364357143 | 17.97869122 | 0.227332509 | 12.64760455 | NA | NA | NA | NA | NA | NA | NA |
| TRPM4 | 941 | L | 2.04225 | 17.06364853 | 0.214081699 | 12.90298806 | NA | NA | NA | NA | NA | NA | NA |
| TRPM4 | 942 | V | 2.623857143 | 21.79841063 | 0.282646187 | 18.6748111 | NA | NA | NA | NA | NA | NA | NA |
| TRPM4 | 943 | A | 2.9432 | 23.40795722 | 0.305954167 | 20.26935514 | NA | NA | NA | NA | NA | NA | NA |
| TRPM4 | 944 | Y | 5.74925 | 16.70861152 | 0.208940378 | 14.66838193 | NA | NA | NA | NA | NA | NA | NA |
| TRPM4 | 945 | G | 7.5225 | 21.31894932 | 0.275703068 | 19.36844335 | NA | NA | NA | NA | NA | NA | NA |
| TRPM4 | 946 | V | 7.637285714 | 23.84179964 | 0.312236676 | 22.06432029 | NA | NA | NA | NA | NA | NA | NA |

| | | | | | | | | | | | | | |
|---|---|---|---|---|---|---|---|---|---|---|---|---|---|
| TRPM4 | 947 | A | 8.1822 | 24.37198763 | 0.319914372 | 22.58937633 | NA | NA | NA | NA | NA | NA | NA |
| TRPM4 | 948 | T | 10.62314286 | 20.07322383 | 0.257663612 | 18.06336959 | NA | NA | NA | NA | NA | NA | NA |
| TRPM4 | 949 | E | 13.24066667 | 20.49922694 | 0.263832599 | 17.60673175 | NA | NA | NA | NA | NA | NA | NA |
| TRPM4 | 950 | G | 12.87875 | 25.85415205 | 0.34137772 | 23.24862179 | NA | NA | NA | NA | NA | NA | NA |
| TRPM4 | 951 | L | 12.80475 | 25.35096907 | 0.334091085 | 22.46707678 | NA | NA | NA | NA | NA | NA | NA |
| TRPM4 | 952 | L | 16.304 | 20.03615919 | 0.257126876 | 15.66363121 | NA | NA | NA | NA | NA | NA | NA |
| TRPM4 | 953 | R | 18.71909091 | 25.39583826 | 0.33474084 | 18.64658791 | NA | NA | NA | NA | NA | NA | NA |
| TRPM4 | 954 | P | 19.40142857 | 23.08262539 | 0.301243009 | 15.62743427 | NA | NA | NA | NA | NA | NA | NA |
| TRPM4 | 955 | R | 22.77672727 | 25.17803594 | 0.331586826 | 17.45365564 | NA | NA | NA | NA | NA | NA | NA |
| TRPM4 | 956 | D | 22.8738 | 23.79869757 | 0.311612511 | 15.99863167 | NA | NA | NA | NA | NA | NA | NA |
| TRPM4 | 957 | S | 20.9446 | 26.34436331 | 0.348476511 | 18.79924397 | NA | NA | NA | NA | NA | NA | NA |
| TRPM4 | 958 | D | 20.04925 | 26.57117002 | 0.351760918 | 19.10721242 | NA | NA | NA | NA | NA | NA | NA |
| TRPM4 | 959 | F | 14.378 | 26.82416882 | 0.355424615 | 22.49875417 | NA | NA | NA | NA | NA | NA | NA |
| TRPM4 | 960 | P | 16.30085714 | 23.52488081 | 0.307647347 | 18.97959844 | NA | NA | NA | NA | NA | NA | NA |
| TRPM4 | 961 | S | 17.0805 | 22.16994511 | 0.288026409 | 17.80157847 | NA | NA | NA | NA | NA | NA | NA |
| TRPM4 | 962 | I | 14.533125 | 23.58163105 | 0.308469152 | 19.1271515 | NA | NA | NA | NA | NA | NA | NA |
| TRPM4 | 963 | L | 11.08425 | 22.3935806 | 0.291264894 | 19.81226403 | NA | NA | NA | NA | NA | NA | NA |
| TRPM4 | 964 | R | 13.85318182 | 16.55376181 | 0.206697987 | 13.75808125 | NA | NA | NA | NA | NA | NA | NA |
| TRPM4 | 965 | R | 14.91045455 | 19.32388338 | 0.24681235 | 16.4852876 | NA | NA | NA | NA | NA | NA | NA |
| TRPM4 | 966 | V | 9.377714286 | 20.04956668 | 0.257321031 | 18.01941327 | NA | NA | NA | NA | NA | NA | NA |
| TRPM4 | 967 | F | 6.394909091 | 17.85200048 | 0.22549789 | 16.10679324 | NA | NA | NA | NA | NA | NA | NA |
| TRPM4 | 968 | Y | 9.936583333 | 14.25155925 | 0.173359598 | 12.41776115 | NA | NA | NA | NA | NA | NA | NA |
| TRPM4 | 969 | R | 11.54918182 | 13.33555653 | 0.160094885 | 11.25797873 | NA | NA | NA | NA | NA | NA | NA |
| TRPM4 | 970 | P | 6.690571429 | 13.416831 | 0.161271828 | 11.54804073 | NA | NA | NA | NA | NA | NA | NA |
| TRPM4 | 971 | Y | 3.5 | 11.94797326 | 0.140001175 | 9.572641128 | NA | NA | NA | NA | NA | NA | NA |
| TRPM4 | 972 | L | 7.448875 | 8.860141139 | 0.095286018 | 6.925015074 | 1.96 | -0.8 | -0.4 | 3.38 | 0.81 | 0 | 0 |
| TRPM4 | 973 | Q | 6.683444444 | 8.746061113 | 0.093634016 | 6.607478782 | 2.23 | -0.8 | -0.4 | 3.38 | 0.81 | 0 | 0 |
| TRPM4 | 974 | I | 1.672875 | 7.069787267 | 0.069359753 | 4.133125237 | 3.15 | -0.8 | -0.4 | 3.38 | 0.81 | 0 | 0 |
| TRPM4 | 975 | F | 2.634636364 | 4.773569733 | 0.036108034 | 2.088119614 | 3.15 | -0.8 | -0.4 | 3.38 | 0.81 | 0 | 0 |
| TRPM4 | 976 | G | 6.32775 | 3.225259059 | 0.013686817 | 1.603754973 | 2.23 | -0.8 | -0.4 | 3.38 | 0.81 | 0 | 0 |
| TRPM4 | 977 | Q | 9.512555556 | 3.559842272 | 0.018531944 | 1.671165889 | 1.95 | -0.86 | -1.02 | 3.41 | 0.67 | 0 | 0 |
| TRPM4 | 978 | I | 9.50925 | 6.735683633 | 0.06452157 | 4.626422043 | 2.11 | -0.86 | -1.02 | 3.41 | 0.67 | 0 | 0 |
| TRPM4 | 979 | P | 13.38485714 | 8.56777795 | 0.091052282 | 5.722030175 | 2.9 | -1.1 | -3.5 | 3.53 | 0.13 | 0 | 0 |

| | | | | | | | | | | | | | |
|---|---|---|---|---|---|---|---|---|---|---|---|---|---|
| TRPM4 | 980 | Q | 14.72555556 | 9.917665098 | 0.110600112 | 5.358831539 | NA | NA | NA | NA | NA | NA | NA |
| TRPM4 | 981 | E | 19.51011111 | 8.997600013 | 0.097276572 | 2.461551053 | 6.8 | -1.11 | -3.5 | 12.8 | 0.55 | 1 | -1 |
| TRPM4 | 982 | D | 16.9565 | 13.40087273 | 0.161040735 | 6.616957732 | NA | NA | NA | NA | NA | NA | NA |
| TRPM4 | 983 | M | 14.463375 | 14.91162399 | 0.182918051 | 10.36802144 | NA | NA | NA | NA | NA | NA | NA |
| TRPM4 | 984 | D | 16.704125 | 15.49871298 | 0.191419736 | 11.15156622 | NA | NA | NA | NA | NA | NA | NA |
| TRPM4 | 985 | V | 20.53557143 | 18.37802133 | 0.233115242 | 10.87445126 | NA | NA | NA | NA | NA | NA | NA |
| TRPM4 | 986 | A | 20.7432 | 16.76288716 | 0.209726348 | 9.207928125 | NA | NA | NA | NA | NA | NA | NA |
| TRPM4 | 987 | L | 20.909 | 12.77781284 | 0.152018152 | 5.616114678 | NA | NA | NA | NA | NA | NA | NA |
| TRPM4 | 988 | M | 23.60525 | 15.23521821 | 0.187604046 | 7.50360273 | NA | NA | NA | NA | NA | NA | NA |
| TRPM4 | 989 | E | 27.7772 | 16.05235441 | 0.199437064 | 6.737562366 | NA | NA | NA | NA | NA | NA | NA |
| TRPM4 | 990 | H | 27.4476 | 19.17512467 | 0.244658163 | 9.857806997 | NA | NA | NA | NA | NA | NA | NA |
| TRPM4 | 991 | S | 30.327 | 20.08950109 | 0.257899325 | 10.77727419 | NA | NA | NA | NA | NA | NA | NA |
| TRPM4 | 992 | N | 31.6878 | 23.47192374 | 0.306880472 | 14.15919558 | NA | NA | NA | NA | NA | NA | NA |
| TRPM4 | 993 | C | 31.5424 | 23.41953702 | 0.306121855 | 14.10799957 | NA | NA | NA | NA | NA | NA | NA |
| TRPM4 | 994 | S | 29.36533333 | 26.24589791 | 0.347050625 | 16.93108632 | NA | NA | NA | NA | NA | NA | NA |
| TRPM4 | 995 | S | 30.0902 | 29.62537792 | 0.395989159 | 20.30932083 | NA | NA | NA | NA | NA | NA | NA |
| TRPM4 | 996 | E | 27.0336 | 31.21698751 | 0.419037392 | 21.90030309 | NA | NA | NA | NA | NA | NA | NA |
| TRPM4 | 997 | P | 26.43714286 | 32.68467049 | 0.440291032 | 23.3128367 | NA | NA | NA | NA | NA | NA | NA |
| TRPM4 | 998 | G | 25.90675 | 30.1560325 | 0.403673613 | 22.39042985 | NA | NA | NA | NA | NA | NA | NA |
| TRPM4 | 999 | F | 25.53527273 | 26.65494585 | 0.352974083 | 17.4471917 | NA | NA | NA | NA | NA | NA | NA |
| TRPM4 | 1000 | W | 25.27314286 | 23.11133144 | 0.301658704 | 13.88108177 | NA | NA | NA | NA | NA | NA | NA |
| TRPM4 | 1001 | A | 25.0384 | 19.95488815 | 0.255949983 | 10.73188823 | NA | NA | NA | NA | NA | NA | NA |
| TRPM4 | 1002 | H | 27.5197 | 15.40113408 | 0.190006687 | 6.261146997 | NA | NA | NA | NA | NA | NA | NA |
| TRPM4 | 1003 | P | 24.43771429 | 14.19278782 | 0.172508524 | 6.404043467 | NA | NA | NA | NA | NA | NA | NA |
| TRPM4 | 1004 | P | 24.48314286 | 10.72384954 | 0.122274536 | 2.98342887 | 7.8 | -1.14 | -3.5 | 49.9 | 2.23 | 4 | -4 |
| TRPM4 | 1005 | G | 23.50725 | 11.50369845 | 0.133567594 | 3.889738222 | 7.65 | -1.14 | -3.5 | 49.9 | 2.23 | 4 | -4 |
| TRPM4 | 1006 | A | 21.975 | 12.78332738 | 0.152098008 | 5.146565975 | NA | NA | NA | NA | NA | NA | NA |
| TRPM4 | 1007 | Q | 19.39088889 | 16.15016566 | 0.200853477 | 8.60130806 | NA | NA | NA | NA | NA | NA | NA |
| TRPM4 | 1008 | A | 20.5664 | 16.46762357 | 0.205450611 | 8.914508442 | NA | NA | NA | NA | NA | NA | NA |
| TRPM4 | 1009 | G | 23.208 | 18.33381823 | 0.232475133 | 10.83709196 | NA | NA | NA | NA | NA | NA | NA |
| TRPM4 | 1010 | T | 26.517 | 19.66406212 | 0.251738507 | 10.45263098 | NA | NA | NA | NA | NA | NA | NA |
| TRPM4 | 1011 | C | 24.84533333 | 21.33575361 | 0.275946412 | 13.54949654 | NA | NA | NA | NA | NA | NA | NA |
| TRPM4 | 1012 | V | 21.553 | 19.94739765 | 0.255841513 | 12.32101021 | NA | NA | NA | NA | NA | NA | NA |

| | | | | | | | | | | | | | |
|---|---|---|---|---|---|---|---|---|---|---|---|---|---|
| TRPM4 | 1013 | S | 20.33333333 | 23.55674725 | 0.308108808 | 15.98983845 | NA | NA | NA | NA | NA | NA | NA |
| TRPM4 | 1014 | Q | 21.42977778 | 22.77750752 | 0.296824572 | 15.44223989 | NA | NA | NA | NA | NA | NA | NA |
| TRPM4 | 1015 | Y | 17.15958333 | 27.24455764 | 0.361512301 | 20.45482558 | NA | NA | NA | NA | NA | NA | NA |
| TRPM4 | 1016 | A | 14.7216 | 26.11076355 | 0.345093733 | 21.55851671 | NA | NA | NA | NA | NA | NA | NA |
| TRPM4 | 1017 | N | 15.347875 | 20.98823282 | 0.270913934 | 16.54034535 | NA | NA | NA | NA | NA | NA | NA |
| TRPM4 | 1018 | W | 10.85271429 | 22.98706308 | 0.299859164 | 20.43225864 | NA | NA | NA | NA | NA | NA | NA |
| TRPM4 | 1019 | L | 9.158875 | 22.8246364 | 0.297507049 | 20.74813638 | NA | NA | NA | NA | NA | NA | NA |
| TRPM4 | 1020 | V | 11.00471429 | 19.48676079 | 0.249170992 | 17.50783243 | NA | NA | NA | NA | NA | NA | NA |
| TRPM4 | 1021 | V | 9.754 | 18.57979884 | 0.236037199 | 16.47294358 | NA | NA | NA | NA | NA | NA | NA |
| TRPM4 | 1022 | L | 5.63575 | 20.13268919 | 0.258524735 | 18.27496706 | NA | NA | NA | NA | NA | NA | NA |
| TRPM4 | 1023 | L | 5.5515 | 18.47503294 | 0.234520075 | 16.35430729 | NA | NA | NA | NA | NA | NA | NA |
| TRPM4 | 1024 | L | 6.8315 | 14.15467432 | 0.171956599 | 12.23042237 | NA | NA | NA | NA | NA | NA | NA |
| TRPM4 | 1025 | V | 3.740428571 | 15.91986774 | 0.197518513 | 13.77867401 | NA | NA | NA | NA | NA | NA | NA |
| TRPM4 | 1026 | I | 0.83725 | 16.78721317 | 0.210078615 | 12.71427968 | NA | NA | NA | NA | NA | NA | NA |
| TRPM4 | 1027 | F | 2.031272727 | 14.08159061 | 0.170898267 | 10.0132125 | NA | NA | NA | NA | NA | NA | NA |
| TRPM4 | 1028 | L | 2.048875 | 10.20077674 | 0.114699875 | 6.136370583 | NA | NA | NA | NA | NA | NA | NA |
| TRPM4 | 1029 | L | -0.6975 | 13.13113224 | 0.1571346 | 7.9516608 | NA | NA | NA | NA | NA | NA | NA |
| TRPM4 | 1030 | V | -1.867571429 | 13.05389923 | 0.156016182 | 8.326546968 | NA | NA | NA | NA | NA | NA | NA |
| TRPM4 | 1031 | A | -0.9382 | 10.38816841 | 0.11741351 | 5.244420548 | NA | NA | NA | NA | NA | NA | NA |
| TRPM4 | 1032 | N | -0.58625 | 6.877742798 | 0.066578741 | 1.651812622 | 5.36 | -0.39 | -1.96 | 2.73 | 0.17 | 0 | 0 |
| TRPM4 | 1033 | I | -3.6305 | 8.903369138 | 0.095912007 | 4.85098415 | 4.13 | 1.27 | 4.26 | 0.13 | -1.41 | 0 | 0 |
| TRPM4 | 1034 | L | -6.946375 | 10.52486166 | 0.119392976 | 8.328623574 | NA | NA | NA | NA | NA | NA | NA |
| TRPM4 | 1035 | L | -5.77725 | 9.908047689 | 0.110460841 | 7.497554236 | NA | NA | NA | NA | NA | NA | NA |
| TRPM4 | 1036 | V | -5.489571429 | 5.652784624 | 0.048840019 | 3.157105378 | 2.52 | 1.29 | 2.9 | 0.78 | -1.37 | 0 | 0 |
| TRPM4 | 1037 | N | -8.14575 | 5.071401088 | 0.040420955 | 4.066951667 | 1.09 | 1.81 | 4.5 | 0.13 | -1.85 | 0 | 0 |
| TRPM4 | 1038 | L | -11.17375 | 8.478182175 | 0.089754838 | 7.292374307 | 1.2 | 1.81 | 4.5 | 0.13 | -1.85 | 0 | 0 |
| TRPM4 | 1039 | L | -9.64025 | 7.876296147 | 0.081038876 | 6.699807817 | 1.2 | 1.81 | 4.5 | 0.13 | -1.85 | 0 | 0 |
| TRPM4 | 1040 | I | -10.83125 | 2.82530724 | 0.007895081 | 1.812165044 | 1.25 | 1.81 | 4.5 | 0.13 | -1.85 | 0 | 0 |
| TRPM4 | 1041 | A | -14.6304 | 5.947066252 | 0.053101536 | 3.246794148 | 2.84 | 1.25 | 3.44 | 0.44 | -1.26 | 0 | 0 |
| TRPM4 | 1042 | M | -14.62225 | 8.149140016 | 0.084989951 | 5.404093497 | 2.77 | -0.97 | -0.8 | 1.67 | 1.11 | 0 | 0 |
| TRPM4 | 1043 | F | -12.91709091 | 7.787353145 | 0.079750885 | 5.540857166 | 2.32 | 1.25 | 3.44 | 0.44 | -1.26 | 0 | 0 |
| TRPM4 | 1044 | S | -17.34216667 | 3.51122799 | 0.017827957 | 2.033420269 | 2.12 | -0.97 | -0.8 | 1.67 | 1.11 | 0 | 0 |
| TRPM4 | 1045 | Y | -21.31316667 | 6.693858454 | 0.063915896 | 4.179288994 | 2.51 | -0.97 | -0.8 | 1.67 | 1.11 | 0 | 0 |

| | | | | | | | | | | | | | |
|---|---|---|---|---|---|---|---|---|---|---|---|---|---|
| TRPM4 | 1046 | T | -18.99757143 | 9.349428271 | 0.102371426 | 6.863872852 | NA | NA | NA | NA | NA | NA | NA |
| TRPM4 | 1047 | F | -18.72018182 | 9.340654046 | 0.102244366 | 6.837058107 | NA | NA | NA | NA | NA | NA | NA |
| TRPM4 | 1048 | G | -22.67575 | 7.887674816 | 0.081203652 | 2.863074575 | 5.03 | -0.97 | -0.8 | 1.67 | 1.11 | 0 | 0 |
| TRPM4 | 1049 | K | -24.616 | 9.41385224 | 0.103304355 | 3.836850215 | 5.61 | -0.97 | -0.8 | 1.67 | 1.11 | 0 | 0 |
| TRPM4 | 1050 | V | -22.39814286 | 12.29487592 | 0.145024702 | 7.279348124 | NA | NA | NA | NA | NA | NA | NA |
| TRPM4 | 1051 | Q | -23.27777778 | 12.17197622 | 0.143244981 | 7.264573244 | NA | NA | NA | NA | NA | NA | NA |
| TRPM4 | 1052 | G | -26.62025 | 13.61012307 | 0.164070906 | 5.641734929 | NA | NA | NA | NA | NA | NA | NA |
| TRPM4 | 1053 | N | -26.466375 | 15.87911691 | 0.196928397 | 7.868373453 | NA | NA | NA | NA | NA | NA | NA |
| TRPM4 | 1054 | S | -22.77616667 | 17.70854308 | 0.223420471 | 12.77372279 | NA | NA | NA | NA | NA | NA | NA |
| TRPM4 | 1055 | D | -24.83275 | 17.93386088 | 0.226683317 | 11.8961856 | NA | NA | NA | NA | NA | NA | NA |
| TRPM4 | 1056 | L | -28.353875 | 19.85520169 | 0.254506415 | 11.84475022 | NA | NA | NA | NA | NA | NA | NA |
| TRPM4 | 1057 | Y | -23.89341667 | 21.95172296 | 0.284866316 | 14.94698097 | NA | NA | NA | NA | NA | NA | NA |
| TRPM4 | 1058 | W | -21.93092857 | 22.9699493 | 0.299611338 | 17.94278514 | NA | NA | NA | NA | NA | NA | NA |
| TRPM4 | 1059 | K | -26.71677778 | 21.87100537 | 0.283697438 | 14.96436766 | NA | NA | NA | NA | NA | NA | NA |
| TRPM4 | 1060 | A | -26.7368 | 26.09388185 | 0.344849268 | 18.08368878 | NA | NA | NA | NA | NA | NA | NA |
| TRPM4 | 1061 | Q | -23.67188889 | 28.30755311 | 0.376905627 | 22.71075903 | NA | NA | NA | NA | NA | NA | NA |
| TRPM4 | 1062 | R | -23.30381818 | 28.74659801 | 0.383263473 | 23.13761479 | NA | NA | NA | NA | NA | NA | NA |
| TRPM4 | 1063 | Y | -26.60266667 | 32.03134073 | 0.430830109 | 25.02736326 | NA | NA | NA | NA | NA | NA | NA |
| TRPM4 | 1064 | R | -22.24445455 | 32.75753745 | 0.441346225 | 27.33085021 | NA | NA | NA | NA | NA | NA | NA |
| TRPM4 | 1065 | L | -19.4525 | 30.37854218 | 0.406895794 | 27.87279056 | NA | NA | NA | NA | NA | NA | NA |
| TRPM4 | 1066 | I | -21.39125 | 34.20090546 | 0.462247759 | 29.20965567 | NA | NA | NA | NA | NA | NA | NA |
| TRPM4 | 1067 | R | -23.803 | 37.35551266 | 0.507929891 | 32.32772833 | NA | NA | NA | NA | NA | NA | NA |
| TRPM4 | 1068 | E | -17.92822222 | 38.02243718 | 0.517587681 | 35.51130564 | NA | NA | NA | NA | NA | NA | NA |
| TRPM4 | 1069 | F | -16.01736364 | 35.41266426 | 0.47979534 | 33.10304906 | NA | NA | NA | NA | NA | NA | NA |
| TRPM4 | 1070 | H | -20.2102 | 41.4105723 | 0.566651551 | 38.90309208 | NA | NA | NA | NA | NA | NA | NA |
| TRPM4 | 1071 | S | -18.06416667 | 43.66970206 | 0.599366199 | 41.16310496 | NA | NA | NA | NA | NA | NA | NA |
| TRPM4 | 1072 | R | -14.8052 | 42.73643198 | 0.585851436 | 39.7420858 | NA | NA | NA | NA | NA | NA | NA |
| TRPM4 | 1073 | P | -12.06885714 | 45.17678122 | 0.621190339 | 43.92740726 | NA | NA | NA | NA | NA | NA | NA |
| TRPM4 | 1074 | A | -11.7908 | 46.78933516 | 0.644541868 | 45.53934223 | NA | NA | NA | NA | NA | NA | NA |
| TRPM4 | 1075 | L | -8.2832 | 45.55206449 | 0.626624848 | 44.46453626 | NA | NA | NA | NA | NA | NA | NA |
| TRPM4 | 1076 | A | -5.1372 | 44.76723071 | 0.615259604 | 42.25078605 | NA | NA | NA | NA | NA | NA | NA |
| TRPM4 | 1077 | P | -3.245571429 | 42.54020246 | 0.58300982 | 38.42375207 | NA | NA | NA | NA | NA | NA | NA |
| TRPM4 | 1078 | P | -0.863428571 | 44.43670718 | 0.610473265 | 38.98990006 | NA | NA | NA | NA | NA | NA | NA |

| | | | | | | | | | | | | | |
|---|---|---|---|---|---|---|---|---|---|---|---|---|---|
| TRPM4 | 1079 | F | -3.8688 | 48.09511371 | 0.663450957 | 43.98473112 | NA | NA | NA | NA | NA | NA | NA |
| TRPM4 | 1080 | I | -4.801 | 47.60164973 | 0.656305064 | 45.08233185 | NA | NA | NA | NA | NA | NA | NA |
| TRPM4 | 1081 | V | -3.4104 | 50.67080772 | 0.700749799 | 46.42756996 | NA | NA | NA | NA | NA | NA | NA |
| TRPM4 | 1082 | I | -5.553 | 52.98287246 | 0.734231003 | 50.4680891 | NA | NA | NA | NA | NA | NA | NA |
| TRPM4 | 1083 | S | -8.4066 | 50.57057822 | 0.699298367 | 49.4889996 | NA | NA | NA | NA | NA | NA | NA |
| TRPM4 | 1084 | H | -7.7646 | 52.40491311 | 0.725861525 | 51.31716406 | NA | NA | NA | NA | NA | NA | NA |
| TRPM4 | 1085 | L | -7.6368 | 55.09188915 | 0.764771851 | 54.00721298 | NA | NA | NA | NA | NA | NA | NA |
| TRPM4 | 1086 | R | -10.941 | 55.75564113 | 0.7743837 | 54.56073558 | NA | NA | NA | NA | NA | NA | NA |
| TRPM4 | 1087 | L | -12.0802 | 53.72410515 | 0.744964856 | 52.47389491 | NA | NA | NA | NA | NA | NA | NA |
| TRPM4 | 1088 | L | -10.2944 | 56.6549095 | 0.78740608 | 55.46013211 | NA | NA | NA | NA | NA | NA | NA |
| TRPM4 | 1089 | L | -11.6714 | 59.22356702 | 0.824603026 | 57.98027793 | NA | NA | NA | NA | NA | NA | NA |
| TRPM4 | 1090 | R | -15.2332 | 60.26252499 | 0.839648264 | 57.28747878 | NA | NA | NA | NA | NA | NA | NA |
| TRPM4 | 1105 | A | -25.0604 | 45.15907575 | 0.620933945 | 39.11966474 | NA | NA | NA | NA | NA | NA | NA |
| TRPM4 | 1106 | L | -26.714875 | 47.30148443 | 0.651958345 | 39.31538403 | NA | NA | NA | NA | NA | NA | NA |
| TRPM4 | 1107 | E | -25.6508 | 43.98753111 | 0.603968708 | 36.98028953 | NA | NA | NA | NA | NA | NA | NA |
| TRPM4 | 1108 | H | -28.0656 | 41.64430973 | 0.570036322 | 33.7044749 | NA | NA | NA | NA | NA | NA | NA |
| TRPM4 | 1109 | F | -30.037 | 43.6624867 | 0.599261713 | 34.06131859 | NA | NA | NA | NA | NA | NA | NA |
| TRPM4 | 1110 | R | -32.099 | 45.4700854 | 0.625437701 | 35.79332647 | NA | NA | NA | NA | NA | NA | NA |
| TRPM4 | 1111 | V | -34.87814286 | 45.01818595 | 0.618893708 | 35.32686503 | NA | NA | NA | NA | NA | NA | NA |
| TRPM4 | 1112 | Y | -35.35533333 | 49.02599835 | 0.676931176 | 33.2774213 | NA | NA | NA | NA | NA | NA | NA |
| TRPM4 | 1113 | L | -40.069125 | 46.19506693 | 0.63593622 | 30.33361252 | NA | NA | NA | NA | NA | NA | NA |
| TRPM4 | 1114 | S | -43.23466667 | 50.40865951 | 0.696953609 | 34.03896818 | NA | NA | NA | NA | NA | NA | NA |
| TRPM4 | 1115 | K | -45.98188889 | 50.15007008 | 0.693208953 | 40.16968146 | NA | NA | NA | NA | NA | NA | NA |
| TRPM4 | 1116 | E | -47.8808 | 48.88356511 | 0.674868588 | 38.90704134 | NA | NA | NA | NA | NA | NA | NA |
| TRPM4 | 1117 | A | -45.6422 | 46.15268904 | 0.635322542 | 36.20667419 | NA | NA | NA | NA | NA | NA | NA |
| TRPM4 | 1118 | E | -45.93077778 | 45.16577876 | 0.621031011 | 35.18522754 | NA | NA | NA | NA | NA | NA | NA |
| TRPM4 | 1119 | R | -51.54454545 | 45.1773388 | 0.621198413 | 36.00607686 | NA | NA | NA | NA | NA | NA | NA |
| TRPM4 | 1120 | K | -49.14666667 | 42.95600044 | 0.589031026 | 33.74953336 | NA | NA | NA | NA | NA | NA | NA |
| TRPM4 | 1121 | L | -46.081875 | 40.50605715 | 0.553553191 | 30.52574087 | NA | NA | NA | NA | NA | NA | NA |
| TRPM4 | 1122 | L | -48.9485 | 39.67632457 | 0.541537763 | 30.49094391 | NA | NA | NA | NA | NA | NA | NA |
| TRPM4 | 1123 | T | -52.40757143 | 38.74332164 | 0.52802687 | 32.14529583 | NA | NA | NA | NA | NA | NA | NA |
| TRPM4 | 1124 | W | -47.68492857 | 36.20349567 | 0.491247436 | 28.51955178 | NA | NA | NA | NA | NA | NA | NA |
| TRPM4 | 1125 | E | -48.88688889 | 34.76031192 | 0.470348571 | 25.56092756 | NA | NA | NA | NA | NA | NA | NA |

| | | | | | | | | | | | | | |
|---|---|---|---|---|---|---|---|---|---|---|---|---|---|
| TRPM4 | 1126 | S | -52.8265 | 34.50239473 | 0.46661365 | 27.94613607 | NA | NA | NA | NA | NA | NA | NA |
| TRPM4 | 1127 | V | -53.652 | 32.67721953 | 0.440183134 | 27.22482052 | NA | NA | NA | NA | NA | NA | NA |
| TRPM4 | 1128 | H | -50.0341 | 30.18961949 | 0.404159989 | 22.93912516 | NA | NA | NA | NA | NA | NA | NA |
| TRPM4 | 1129 | K | -52.13677778 | 29.62914898 | 0.396043768 | 23.06277811 | NA | NA | NA | NA | NA | NA | NA |
| TRPM4 | 1130 | E | -55.6606 | 29.25133682 | 0.390572639 | 24.91107438 | NA | NA | NA | NA | NA | NA | NA |
| TRPM4 | 1131 | N | -54.62825 | 26.64349979 | 0.352808331 | 22.29298133 | NA | NA | NA | NA | NA | NA | NA |
| TRPM4 | 1132 | F | -51.01427273 | 24.12987486 | 0.316408317 | 17.10179231 | NA | NA | NA | NA | NA | NA | NA |
| TRPM4 | 1133 | L | -54.723875 | 23.98774031 | 0.314350055 | 19.63536133 | NA | NA | NA | NA | NA | NA | NA |
| TRPM4 | 1134 | L | -58.4185 | 22.8244513 | 0.297504369 | 19.29663621 | NA | NA | NA | NA | NA | NA | NA |
| TRPM4 | 1135 | A | -55.3408 | 20.5263356 | 0.264225162 | 16.16582774 | NA | NA | NA | NA | NA | NA | NA |
| TRPM4 | 1136 | R | -53.88918182 | 18.97926308 | 0.241821875 | 13.68945609 | NA | NA | NA | NA | NA | NA | NA |
| TRPM4 | 1137 | A | -58.1556 | 18.60010271 | 0.236331221 | 15.08628335 | NA | NA | NA | NA | NA | NA | NA |
| TRPM4 | 1138 | R | -60.22636364 | 16.96148375 | 0.212602242 | 13.60494238 | NA | NA | NA | NA | NA | NA | NA |
| TRPM4 | 1139 | D | -55.454125 | 14.54727064 | 0.177641819 | 10.22913821 | NA | NA | NA | NA | NA | NA | NA |
| TRPM4 | 1140 | K | -58.09733333 | 13.81526685 | 0.167041611 | 10.29841922 | NA | NA | NA | NA | NA | NA | NA |
| TRPM4 | 1141 | R | -63.54827273 | 13.38305724 | 0.160782747 | 10.4946284 | NA | NA | NA | NA | NA | NA | NA |
| TRPM4 | 1142 | E | -60.1536 | 12.15533373 | 0.143003979 | 9.180897645 | NA | NA | NA | NA | NA | NA | NA |
| TRPM4 | 1143 | S | -58.843 | 9.36702402 | 0.102626232 | 5.863414602 | NA | NA | NA | NA | NA | NA | NA |
| TRPM4 | 1144 | D | -58.543625 | 4.995254648 | 0.039318272 | 1.545315141 | 3.53 | -0.41 | 0.44 | 2.73 | 0.29 | 0 | 0 |
| TRPM4 | 1145 | S | -59.77483333 | 7.000653255 | 0.068358618 | 3.630735098 | 3.37 | -0.41 | 0.44 | 2.73 | 0.29 | 0 | 0 |
| TRPM4 | 1146 | E | -62.31844444 | 8.748068015 | 0.093663078 | 6.436804731 | 2.36 | 1.14 | 3.8 | 0.13 | -1.79 | 0 | 0 |
| TRPM4 | 1147 | R | -64.21963636 | 6.636558747 | 0.063086135 | 4.787663999 | 2.06 | 1.14 | 3.8 | 0.13 | -1.79 | 0 | 0 |
| TRPM4 | 1148 | L | -64.836875 | 2.93854692 | 0.009534914 | 1.079214751 | 2.06 | 1.14 | 3.8 | 0.13 | -1.79 | 0 | 0 |
| TRPM4 | 1149 | K | -66.09866667 | 7.532663938 | 0.076062709 | 5.099846524 | 2.57 | 0.76 | 2.9 | 0.44 | -1.28 | 0 | 0 |
| TRPM4 | 1150 | R | -67.75163636 | 8.194770162 | 0.085650725 | 5.332758754 | 2.93 | -0.77 | -0.7 | 1.66 | 0.77 | 0 | 0 |
| TRPM4 | 1151 | T | -69.83614286 | 4.346815041 | 0.029928163 | 1.600401533 | 3.08 | -0.77 | -0.7 | 1.66 | 0.77 | 0 | 0 |
| TRPM4 | 1152 | S | -70.54383333 | 6.241110318 | 0.057359613 | 3.043708781 | 3.23 | -0.39 | 0.28 | 1.35 | 0.36 | 0 | 0 |
| TRPM4 | 1153 | Q | -71.70477778 | 8.647505883 | 0.092206829 | 6.895392598 | 1.88 | 1.13 | 4.2 | 0.13 | -1.3 | 0 | 0 |
| TRPM4 | 1154 | K | -74.19211111 | 6.677160325 | 0.063674089 | 4.964229718 | 1.76 | 1.13 | 4.2 | 0.13 | -1.3 | 0 | 0 |
| TRPM4 | 1155 | V | -75.24585714 | 3.061503879 | 0.011315464 | 1.697303431 | 1.76 | 1.13 | 4.2 | 0.13 | -1.3 | 0 | 0 |
| TRPM4 | 1156 | D | -76.6282 | 7.349216693 | 0.073406194 | 5.588719361 | 1.76 | 1.13 | 4.2 | 0.13 | -1.3 | 0 | 0 |
| TRPM4 | 1157 | L | -79.133875 | 8.321560731 | 0.08748679 | 5.123803194 | 3.3 | 0.24 | 2.28 | 0.03 | 0.21 | 0 | 0 |
| TRPM4 | 1158 | A | -80.4088 | 4.787979741 | 0.036316706 | 1.692627426 | 3.13 | 0.24 | 2.2 | 0.03 | 0.11 | 0 | 0 |

| | | | | | | | | | | | | | |
|---|---|---|---|---|---|---|---|---|---|---|---|---|---|
| TRPM4 | 1159 | L | -80.606375 | 6.204783074 | 0.056833555 | 3.112435095 | 3.14 | 0.92 | 3.4 | 0.1 | -1.31 | 0 | 0 |
| TRPM4 | 1160 | K | -82.00044444 | 8.687451237 | 0.092785281 | 6.287729635 | 2.44 | 1.14 | 3.8 | 0.13 | -1.79 | 0 | 0 |
| TRPM4 | 1161 | Q | -85.07255556 | 7.23422933 | 0.071741053 | 5.138947907 | 2.11 | 1.14 | 3.8 | 0.13 | -1.79 | 0 | 0 |
| TRPM4 | 1162 | L | -85.422375 | 2.775726211 | 0.007177094 | 0.743611039 | 2.11 | 1.14 | 3.8 | 0.13 | -1.79 | 0 | 0 |
| TRPM4 | 1163 | G | -87.14175 | 7.77939265 | 0.079635608 | 5.68671025 | 2.11 | 1.14 | 3.8 | 0.13 | -1.79 | 0 | 0 |
| TRPM4 | 1164 | H | -89.2271 | 8.117066958 | 0.084525499 | 5.763511352 | 2.43 | 1.81 | 4.5 | 0.13 | -1.85 | 0 | 0 |
| TRPM4 | 1165 | I | -91.029125 | 3.261202386 | 0.014207315 | 1.375101035 | 1.98 | 1.81 | 4.5 | 0.13 | -1.85 | 0 | 0 |
| TRPM4 | 1166 | R | -90.831 | 7.077267481 | 0.069468075 | 5.252014278 | 1.91 | 1.22 | 2.9 | 10.08 | -1.03 | 1 | -1 |
| TRPM4 | 1167 | E | -94.14011111 | 9.036296642 | 0.097836941 | 7.451975176 | 1.59 | -0.55 | -1.9 | 39.95 | 1.41 | 4 | -4 |
| TRPM4 | 1168 | Y | -96.10725 | 6.418233636 | 0.05992455 | 5.307640268 | 1.15 | -1.14 | -3.5 | 49.9 | 2.23 | 4 | -4 |
| TRPM4 | 1169 | E | -97.197 | 2.344240602 | 9.29E-04 | 1.959307517 | 1.17 | -0.68 | -2.04 | 39.95 | 1.43 | 4 | -4 |
| TRPM4 | 1170 | Q | -99.0842 | 8.43107502 | 0.089072676 | 6.963271682 | 1.49 | -1.14 | -3.5 | 49.9 | 2.23 | 4 | -4 |
| TRPM4 | 1171 | R | -101.2474 | 8.650707832 | 0.092253197 | 7.368493128 | 1.29 | 1.14 | 3.8 | 0.13 | -1.79 | 0 | 0 |
| TRPM4 | 1172 | L | -102.3415 | 2.819525847 | 0.00781136 | 1.68734987 | 1.29 | 1.14 | 3.8 | 0.13 | -1.79 | 0 | 0 |
| TRPM4 | 1173 | K | -103.3972 | 6.874215737 | 0.066527665 | 5.584550113 | 1.29 | 1.14 | 3.8 | 0.13 | -1.79 | 0 | 0 |
| TRPM4 | 1174 | V | -105.293 | 9.208690841 | 0.100333396 | 6.400424338 | NA | NA | NA | NA | NA | NA | NA |
| TRPM4 | 1175 | L | -107.12 | 6.981736245 | 0.068084679 | 3.893103128 | 3.2 | -1.14 | -3.5 | 49.9 | 2.23 | 4 | -4 |
| TRPM4 | 1176 | E | -107.9856 | 4.889246772 | 0.037783163 | 1.778868294 | 3.37 | -1.14 | -3.5 | 49.9 | 2.23 | 4 | -4 |
| TRPM4 | 1177 | R | -109.322 | 7.834146795 | 0.080428508 | 4.387630949 | 3.49 | -0.69 | -1.96 | 39.95 | 1.52 | 4 | -4 |
| TRPM4 | 1178 | E | -111.447 | 8.446365905 | 0.089294104 | 5.046337131 | 3.4 | 1.13 | 4.2 | 0.13 | -1.3 | 0 | 0 |
| TRPM4 | 1179 | V | -112.7908 | 4.876894914 | 0.037604295 | 1.480311304 | 3.4 | 1.13 | 4.2 | 0.13 | -1.3 | 0 | 0 |
| TRPM4 | 1180 | Q | -113.5846 | 6.389447785 | 0.0595077 | 3.019340967 | 3.54 | 1.13 | 4.2 | 0.13 | -1.3 | 0 | 0 |
| TRPM4 | 1181 | Q | -115.3884 | 8.812758535 | 0.094599867 | 5.260090039 | 3.6 | -0.94 | -0.72 | 2.01 | 1.05 | 0 | 0 |
| TRPM4 | 1182 | C | -117.3674 | 7.174894773 | 0.070881824 | 3.489484261 | 3.69 | -0.97 | -0.8 | 1.67 | 1.11 | 0 | 0 |
| TRPM4 | 1183 | S | -118.328 | 4.681016343 | 0.03476776 | 1.018305985 | 3.69 | -0.97 | -0.8 | 1.67 | 1.11 | 0 | 0 |
| TRPM4 | 1184 | R | -119.4344 | 7.664492155 | 0.077971725 | 3.767068705 | 4.18 | 0.3 | 1.96 | 0.75 | -0.63 | 0 | 0 |
| TRPM4 | 1185 | V | -121.4826 | 9.036157646 | 0.097834928 | 4.820416097 | 4.26 | 0.72 | 2.88 | 0.44 | -1.21 | 0 | 0 |
| TRPM4 | 1186 | L | -123.096 | 5.796213074 | 0.050917018 | 1.509745012 | 4.3 | 0.75 | 2.96 | 0.78 | -1.27 | 0 | 0 |
| TRPM4 | 1187 | G | -124.18725 | 6.519187756 | 0.061386475 | 2.244662435 | 4.3 | 0.75 | 2.96 | 0.78 | -1.27 | 0 | 0 |
| TRPM4 | 1188 | W | -125.346 | 9.161303292 | 0.099647173 | 4.89468254 | 4.3 | 0.75 | 2.96 | 0.78 | -1.27 | 0 | 0 |
| TRPM4 | 1189 | V | -127.5646 | 10.28930474 | 0.115981857 | 6.04235801 | NA | NA | NA | NA | NA | NA | NA |
| TRPM4 | 1190 | A | -128.618 | 7.54385717 | 0.076224799 | 3.338547513 | 4.3 | 0.75 | 2.96 | 0.78 | -1.27 | 0 | 0 |
| TRPM4 | 1191 | E | -128.6332 | 9.80754271 | 0.10900542 | 5.590646142 | NA | NA | NA | NA | NA | NA | NA |

| | | | | | | | | | | | | | |
|---|---|---|---|---|---|---|---|---|---|---|---|---|---|
| TRPM4 | 1192 | A | -128.8924 | 12.16960595 | 0.143210657 | 7.941023071 | NA | NA | NA | NA | NA | NA | NA |
| TRPV3 | 114 | R | -63.52065 | 57.2060909 | 0.725999931 | 46.63169709 | NA | NA | NA | NA | NA | NA | NA |
| TRPV3 | 115 | K | -66.56405 | 56.3090293 | 0.714113645 | 45.62834919 | NA | NA | NA | NA | NA | NA | NA |
| TRPV3 | 116 | K | -65.23825 | 54.48958962 | 0.690005623 | 43.81362403 | NA | NA | NA | NA | NA | NA | NA |
| TRPV3 | 117 | R | -62.46985 | 52.38029799 | 0.662056991 | 41.80690367 | NA | NA | NA | NA | NA | NA | NA |
| TRPV3 | 118 | R | -64.46165 | 51.61067264 | 0.651859266 | 40.93020082 | NA | NA | NA | NA | NA | NA | NA |
| TRPV3 | 119 | L | -67.610875 | 50.66275584 | 0.639299136 | 39.98204281 | NA | NA | NA | NA | NA | NA | NA |
| TRPV3 | 120 | K | -62.89380556 | 48.32544944 | 0.608329252 | 37.64516999 | NA | NA | NA | NA | NA | NA | NA |
| TRPV3 | 121 | K | -62.86865 | 46.76243626 | 0.587618944 | 36.08359039 | NA | NA | NA | NA | NA | NA | NA |
| TRPV3 | 122 | R | -66.06665 | 46.46626077 | 0.583694546 | 35.78701323 | NA | NA | NA | NA | NA | NA | NA |
| TRPV3 | 123 | I | -66.877625 | 44.79347878 | 0.561529774 | 34.11448881 | NA | NA | NA | NA | NA | NA | NA |
| TRPV3 | 124 | F | -62.75443182 | 42.28278319 | 0.528262442 | 31.60198851 | NA | NA | NA | NA | NA | NA | NA |
| TRPV3 | 125 | A | -64.12485 | 41.73904831 | 0.521057821 | 31.05905943 | NA | NA | NA | NA | NA | NA | NA |
| TRPV3 | 126 | A | -67.34525 | 41.34728535 | 0.515866866 | 30.67693848 | NA | NA | NA | NA | NA | NA | NA |
| TRPV3 | 127 | V | -65.87553571 | 38.76090435 | 0.481596683 | 28.08593417 | NA | NA | NA | NA | NA | NA | NA |
| TRPV3 | 128 | S | -63.43005 | 36.90210212 | 0.456967097 | 26.22264806 | NA | NA | NA | NA | NA | NA | NA |
| TRPV3 | 129 | E | -66.01485 | 37.2848319 | 0.46203836 | 26.61894008 | NA | NA | NA | NA | NA | NA | NA |
| TRPV3 | 130 | G | -69.108 | 38.22897705 | 0.474548515 | 27.57202457 | NA | NA | NA | NA | NA | NA | NA |
| TRPV3 | 131 | C | -71.29105 | 39.27842139 | 0.48845391 | 28.63821379 | NA | NA | NA | NA | NA | NA | NA |
| TRPV3 | 132 | V | -73.78465 | 41.59441911 | 0.519141449 | 30.58089359 | NA | NA | NA | NA | NA | NA | NA |
| TRPV3 | 133 | E | -75.49485 | 42.19287112 | 0.527071085 | 31.16016991 | NA | NA | NA | NA | NA | NA | NA |
| TRPV3 | 134 | E | -77.48885 | 44.81136663 | 0.561766793 | 33.77748853 | NA | NA | NA | NA | NA | NA | NA |
| TRPV3 | 135 | L | -74.359875 | 44.55277142 | 0.558340343 | 33.55388464 | NA | NA | NA | NA | NA | NA | NA |
| TRPV3 | 136 | V | -73.25396429 | 43.44304599 | 0.543636208 | 32.79121256 | NA | NA | NA | NA | NA | NA | NA |
| TRPV3 | 137 | E | -77.04125 | 47.24032922 | 0.593951143 | 36.21922403 | NA | NA | NA | NA | NA | NA | NA |
| TRPV3 | 138 | L | -77.11425 | 49.89229071 | 0.629090284 | 38.87143455 | NA | NA | NA | NA | NA | NA | NA |
| TRPV3 | 139 | L | -72.78925 | 48.06629395 | 0.604895379 | 37.39578542 | NA | NA | NA | NA | NA | NA | NA |
| TRPV3 | 140 | V | -74.28625 | 48.71697145 | 0.613517015 | 38.06295604 | NA | NA | NA | NA | NA | NA | NA |
| TRPV3 | 141 | E | -77.42565 | 52.26958761 | 0.660590051 | 41.26275844 | NA | NA | NA | NA | NA | NA | NA |
| TRPV3 | 142 | L | -75.40605 | 54.30223831 | 0.687523172 | 43.65397344 | NA | NA | NA | NA | NA | NA | NA |
| TRPV3 | 143 | Q | -73.38605 | 53.55989039 | 0.67768688 | 42.89153152 | NA | NA | NA | NA | NA | NA | NA |
| TRPV3 | 144 | E | -76.15145 | 55.04101353 | 0.697312125 | 44.38612174 | NA | NA | NA | NA | NA | NA | NA |
| TRPV3 | 145 | L | -77.12225 | 57.56801975 | 0.730795577 | 46.91602692 | NA | NA | NA | NA | NA | NA | NA |

| | | | | | | | | | | | | | |
|---|---|---|---|---|---|---|---|---|---|---|---|---|---|
| TRPV3 | 146 | C | -73.85845 | 58.65495147 | 0.745197689 | 47.99024414 | NA | NA | NA | NA | NA | NA | NA |
| TRPV3 | 147 | R | -74.15605 | 58.47010681 | 0.742748452 | 47.80308244 | NA | NA | NA | NA | NA | NA | NA |
| TRPV3 | 148 | R | -76.54665 | 61.24049021 | 0.779456712 | 50.57739948 | NA | NA | NA | NA | NA | NA | NA |
| TRPV3 | 149 | R | -74.03945 | 62.90845147 | 0.801557608 | 52.23714711 | NA | NA | NA | NA | NA | NA | NA |
| TRPV3 | 150 | H | -70.90345 | 60.95271181 | 0.775643578 | 50.27281184 | NA | NA | NA | NA | NA | NA | NA |
| TRPV3 | 151 | D | -73.42125 | 65.1484938 | 0.831238719 | 54.47330954 | NA | NA | NA | NA | NA | NA | NA |
| TRPV3 | 152 | E | -70.31765 | 63.70905947 | 0.812165861 | 53.02723573 | NA | NA | NA | NA | NA | NA | NA |
| TRPV3 | 153 | D | -67.93825 | 60.43314832 | 0.768759234 | 49.75055726 | NA | NA | NA | NA | NA | NA | NA |
| TRPV3 | 154 | V | -70.62810714 | 57.93534635 | 0.735662745 | 47.25497718 | NA | NA | NA | NA | NA | NA | NA |
| TRPV3 | 155 | P | -69.04482143 | 54.95827731 | 0.69621585 | 44.27671814 | NA | NA | NA | NA | NA | NA | NA |
| TRPV3 | 156 | D | -64.986125 | 55.25636746 | 0.700165617 | 44.57725951 | NA | NA | NA | NA | NA | NA | NA |
| TRPV3 | 157 | F | -66.21961364 | 55.49945053 | 0.703386528 | 44.81786233 | NA | NA | NA | NA | NA | NA | NA |
| TRPV3 | 158 | L | -68.782 | 52.68382151 | 0.666078752 | 42.00273132 | NA | NA | NA | NA | NA | NA | NA |
| TRPV3 | 159 | M | -64.31925 | 49.79035394 | 0.627739597 | 39.10729355 | NA | NA | NA | NA | NA | NA | NA |
| TRPV3 | 160 | H | -61.41195 | 52.06748616 | 0.657912157 | 41.409409 | NA | NA | NA | NA | NA | NA | NA |
| TRPV3 | 161 | K | -64.85805 | 51.12985596 | 0.645488327 | 40.44892652 | NA | NA | NA | NA | NA | NA | NA |
| TRPV3 | 162 | L | -65.738875 | 47.44174847 | 0.596619998 | 36.76546666 | NA | NA | NA | NA | NA | NA | NA |
| TRPV3 | 163 | T | -61.35010714 | 47.02810607 | 0.591139134 | 36.38174558 | NA | NA | NA | NA | NA | NA | NA |
| TRPV3 | 164 | A | -58.05185 | 45.76385355 | 0.574387478 | 35.19861007 | NA | NA | NA | NA | NA | NA | NA |
| TRPV3 | 165 | S | -55.87685 | 49.48132753 | 0.623644921 | 38.9176618 | NA | NA | NA | NA | NA | NA | NA |
| TRPV3 | 166 | D | -52.431375 | 47.25366565 | 0.594127854 | 36.70440368 | NA | NA | NA | NA | NA | NA | NA |
| TRPV3 | 167 | T | -53.05325 | 43.26767686 | 0.541312524 | 32.7107362 | NA | NA | NA | NA | NA | NA | NA |
| TRPV3 | 168 | G | -55.70275 | 44.37690709 | 0.556010097 | 33.81176526 | NA | NA | NA | NA | NA | NA | NA |
| TRPV3 | 169 | K | -57.17947222 | 38.85383464 | 0.482828032 | 28.29339927 | NA | NA | NA | NA | NA | NA | NA |
| TRPV3 | 170 | T | -61.32010714 | 41.94431196 | 0.523777615 | 31.27295975 | NA | NA | NA | NA | NA | NA | NA |
| TRPV3 | 171 | C | -64.33941667 | 40.87331704 | 0.509586669 | 30.19274966 | NA | NA | NA | NA | NA | NA | NA |
| TRPV3 | 172 | L | -64.966125 | 38.26158048 | 0.474980519 | 27.58135868 | NA | NA | NA | NA | NA | NA | NA |
| TRPV3 | 173 | M | -60.263625 | 36.94721579 | 0.457564864 | 26.27222065 | NA | NA | NA | NA | NA | NA | NA |
| TRPV3 | 174 | K | -61.24385 | 37.32188246 | 0.462529289 | 26.64119403 | NA | NA | NA | NA | NA | NA | NA |
| TRPV3 | 175 | A | -63.76605 | 35.07723406 | 0.432787147 | 24.39813559 | NA | NA | NA | NA | NA | NA | NA |
| TRPV3 | 176 | L | -62.847 | 32.41350644 | 0.397492102 | 21.73230285 | NA | NA | NA | NA | NA | NA | NA |
| TRPV3 | 177 | L | -58.671125 | 32.07205793 | 0.392967826 | 21.40208437 | NA | NA | NA | NA | NA | NA | NA |
| TRPV3 | 178 | N | -61.348875 | 33.24167918 | 0.408465595 | 22.56114598 | NA | NA | NA | NA | NA | NA | NA |

| | | | | | | | | | | | | | |
|---|---|---|---|---|---|---|---|---|---|---|---|---|---|
| TRPV3 | 179 | I | -63.130375 | 28.56312408 | 0.346473591 | 17.89727282 | NA | NA | NA | NA | NA | NA | NA |
| TRPV3 | 180 | N | -66.360375 | 31.30875438 | 0.382853867 | 20.65571814 | NA | NA | NA | NA | NA | NA | NA |
| TRPV3 | 181 | P | -69.54582143 | 29.62348158 | 0.360523588 | 18.5934732 | NA | NA | NA | NA | NA | NA | NA |
| TRPV3 | 182 | N | -71.44265 | 33.88614385 | 0.41700491 | 22.85500526 | NA | NA | NA | NA | NA | NA | NA |
| TRPV3 | 183 | T | -69.00453571 | 33.31529081 | 0.409440967 | 22.33347981 | NA | NA | NA | NA | NA | NA | NA |
| TRPV3 | 184 | K | -72.17005 | 34.6428573 | 0.427031548 | 23.60976065 | NA | NA | NA | NA | NA | NA | NA |
| TRPV3 | 185 | E | -74.30145 | 37.19429577 | 0.460838734 | 26.1564321 | NA | NA | NA | NA | NA | NA | NA |
| TRPV3 | 186 | I | -71.51425 | 38.71577066 | 0.48099865 | 27.68920588 | NA | NA | NA | NA | NA | NA | NA |
| TRPV3 | 187 | V | -70.55853571 | 36.75081304 | 0.454962479 | 25.75384047 | NA | NA | NA | NA | NA | NA | NA |
| TRPV3 | 188 | R | -74.97225 | 38.52964075 | 0.478532383 | 27.50347815 | NA | NA | NA | NA | NA | NA | NA |
| TRPV3 | 189 | I | -75.15125 | 42.4565908 | 0.530565435 | 31.42441565 | NA | NA | NA | NA | NA | NA | NA |
| TRPV3 | 190 | L | -71.18475 | 41.54492852 | 0.518485686 | 30.88338841 | NA | NA | NA | NA | NA | NA | NA |
| TRPV3 | 191 | L | -72.1375 | 40.39371008 | 0.503231759 | 29.38232164 | NA | NA | NA | NA | NA | NA | NA |
| TRPV3 | 192 | A | -76.28165 | 45.37534314 | 0.56923962 | 34.34495667 | NA | NA | NA | NA | NA | NA | NA |
| TRPV3 | 193 | F | -74.47088636 | 47.56090691 | 0.598198876 | 36.53857522 | NA | NA | NA | NA | NA | NA | NA |
| TRPV3 | 194 | A | -72.78585 | 47.34735931 | 0.595369318 | 36.6939662 | NA | NA | NA | NA | NA | NA | NA |
| TRPV3 | 195 | E | -75.57805 | 48.83381014 | 0.615065156 | 37.81536208 | NA | NA | NA | NA | NA | NA | NA |
| TRPV3 | 196 | E | -77.56625 | 51.03140923 | 0.644183884 | 40.00516147 | NA | NA | NA | NA | NA | NA | NA |
| TRPV3 | 197 | N | -74.862875 | 52.64923333 | 0.66562045 | 42.00987976 | NA | NA | NA | NA | NA | NA | NA |
| TRPV3 | 198 | D | -73.22845 | 52.87220133 | 0.668574831 | 42.228443 | NA | NA | NA | NA | NA | NA | NA |
| TRPV3 | 199 | I | -69.9615 | 49.19652782 | 0.619871255 | 38.52232096 | NA | NA | NA | NA | NA | NA | NA |
| TRPV3 | 200 | L | -70.292375 | 44.82711194 | 0.561975422 | 34.18240593 | NA | NA | NA | NA | NA | NA | NA |
| TRPV3 | 201 | G | -66.819 | 47.29491004 | 0.594674352 | 36.61930619 | NA | NA | NA | NA | NA | NA | NA |
| TRPV3 | 202 | R | -65.20445 | 48.23309492 | 0.607105532 | 37.5495611 | NA | NA | NA | NA | NA | NA | NA |
| TRPV3 | 203 | F | -66.09552273 | 43.44588875 | 0.543673876 | 32.77361606 | NA | NA | NA | NA | NA | NA | NA |
| TRPV3 | 204 | I | -65.209625 | 41.31552086 | 0.515445978 | 30.63686002 | NA | NA | NA | NA | NA | NA | NA |
| TRPV3 | 205 | N | -62.099875 | 43.26479265 | 0.541274308 | 32.58051512 | NA | NA | NA | NA | NA | NA | NA |
| TRPV3 | 206 | A | -59.92845 | 42.65627481 | 0.533211297 | 32.0025038 | NA | NA | NA | NA | NA | NA | NA |
| TRPV3 | 207 | E | -56.35285 | 42.33483424 | 0.528952131 | 31.76611543 | NA | NA | NA | NA | NA | NA | NA |
| TRPV3 | 208 | Y | -53.89466667 | 37.87286593 | 0.469829956 | 27.31015909 | NA | NA | NA | NA | NA | NA | NA |
| TRPV3 | 209 | T | -51.22225 | 45.39029718 | 0.569437765 | 34.83516942 | NA | NA | NA | NA | NA | NA | NA |
| TRPV3 | 210 | E | -48.50025 | 43.18731622 | 0.540247726 | 32.64204894 | NA | NA | NA | NA | NA | NA | NA |
| TRPV3 | 211 | E | -45.15185 | 42.48058058 | 0.530883305 | 31.94122303 | NA | NA | NA | NA | NA | NA | NA |

| | | | | | | | | | | | | | |
|---|---|---|---|---|---|---|---|---|---|---|---|---|---|
| TRPV3 | 212 | A | -46.14585 | 38.64494924 | 0.480060249 | 28.10862051 | NA | NA | NA | NA | NA | NA | NA |
| TRPV3 | 213 | Y | -49.63875 | 35.73950247 | 0.441562366 | 25.1922925 | NA | NA | NA | NA | NA | NA | NA |
| TRPV3 | 214 | E | -49.00380556 | 40.49500148 | 0.504573895 | 29.94174923 | NA | NA | NA | NA | NA | NA | NA |
| TRPV3 | 215 | G | -53.18725 | 39.75927396 | 0.494825325 | 29.19335902 | NA | NA | NA | NA | NA | NA | NA |
| TRPV3 | 216 | Q | -54.31936111 | 34.35126114 | 0.423167828 | 23.78712292 | NA | NA | NA | NA | NA | NA | NA |
| TRPV3 | 217 | T | -58.10882143 | 36.38663307 | 0.450137005 | 25.81065578 | NA | NA | NA | NA | NA | NA | NA |
| TRPV3 | 218 | A | -61.12165 | 34.95232254 | 0.431132039 | 24.26913656 | NA | NA | NA | NA | NA | NA | NA |
| TRPV3 | 219 | L | -61.086125 | 32.96582359 | 0.40481044 | 22.28134786 | NA | NA | NA | NA | NA | NA | NA |
| TRPV3 | 220 | N | -56.776875 | 31.59366093 | 0.386628948 | 21.01945394 | NA | NA | NA | NA | NA | NA | NA |
| TRPV3 | 221 | I | -58.43 | 30.61633829 | 0.373679183 | 19.94193438 | NA | NA | NA | NA | NA | NA | NA |
| TRPV3 | 222 | A | -61.02285 | 28.97464944 | 0.351926403 | 18.29377825 | NA | NA | NA | NA | NA | NA | NA |
| TRPV3 | 223 | I | -58.90675 | 27.07918368 | 0.326811017 | 16.40133194 | NA | NA | NA | NA | NA | NA | NA |
| TRPV3 | 224 | E | -56.36205 | 26.14220174 | 0.314395776 | 15.56973912 | NA | NA | NA | NA | NA | NA | NA |
| TRPV3 | 225 | R | -58.48225 | 26.12218969 | 0.314130611 | 15.44017159 | NA | NA | NA | NA | NA | NA | NA |
| TRPV3 | 226 | R | -59.91152273 | 19.65338284 | 0.228417333 | 8.991495132 | NA | NA | NA | NA | NA | NA | NA |
| TRPV3 | 227 | Q | -64.37436111 | 26.14340942 | 0.314411778 | 15.51310057 | NA | NA | NA | NA | NA | NA | NA |
| TRPV3 | 228 | G | -65.97075 | 27.76885143 | 0.335949284 | 16.74069273 | NA | NA | NA | NA | NA | NA | NA |
| TRPV3 | 229 | D | -69.51175 | 29.44712337 | 0.358186799 | 18.41226353 | NA | NA | NA | NA | NA | NA | NA |
| TRPV3 | 230 | I | -67.5185 | 31.50658106 | 0.385475119 | 20.50690107 | NA | NA | NA | NA | NA | NA | NA |
| TRPV3 | 231 | A | -66.10765 | 31.98081495 | 0.391758834 | 21.33011292 | NA | NA | NA | NA | NA | NA | NA |
| TRPV3 | 232 | A | -69.15365 | 33.26682964 | 0.408798844 | 22.22443321 | NA | NA | NA | NA | NA | NA | NA |
| TRPV3 | 233 | L | -71.404125 | 35.46794114 | 0.437964112 | 24.43025267 | NA | NA | NA | NA | NA | NA | NA |
| TRPV3 | 234 | L | -67.51125 | 35.96076798 | 0.444494188 | 24.98832734 | NA | NA | NA | NA | NA | NA | NA |
| TRPV3 | 235 | I | -67.451 | 35.10122613 | 0.433105048 | 24.10295041 | NA | NA | NA | NA | NA | NA | NA |
| TRPV3 | 236 | A | -71.74665 | 39.09140362 | 0.485975879 | 28.05437466 | NA | NA | NA | NA | NA | NA | NA |
| TRPV3 | 237 | A | -70.85485 | 41.22709932 | 0.514274371 | 30.19739061 | NA | NA | NA | NA | NA | NA | NA |
| TRPV3 | 238 | G | -68.11875 | 42.62124462 | 0.532747139 | 31.98026304 | NA | NA | NA | NA | NA | NA | NA |
| TRPV3 | 239 | A | -65.34105 | 39.5547854 | 0.492115801 | 28.88453948 | NA | NA | NA | NA | NA | NA | NA |
| TRPV3 | 240 | D | -61.1395 | 39.24650382 | 0.488030995 | 28.56097299 | NA | NA | NA | NA | NA | NA | NA |
| TRPV3 | 241 | V | -60.87882143 | 37.51734132 | 0.465119167 | 26.83502926 | NA | NA | NA | NA | NA | NA | NA |
| TRPV3 | 242 | N | -57.589125 | 39.06280903 | 0.485596994 | 28.47886507 | NA | NA | NA | NA | NA | NA | NA |
| TRPV3 | 243 | A | -55.94685 | 37.92009723 | 0.470455782 | 27.34830584 | NA | NA | NA | NA | NA | NA | NA |
| TRPV3 | 244 | H | -52.38985 | 38.75374074 | 0.481501763 | 28.1877097 | NA | NA | NA | NA | NA | NA | NA |

| | | | | | | | | | | | | | |
|---|---|---|---|---|---|---|---|---|---|---|---|---|---|
| TRPV3 | 245 | A | -50.53605 | 36.3695781 | 0.449911023 | 25.81114354 | NA | NA | NA | NA | NA | NA | NA |
| TRPV3 | 246 | K | -48.03569444 | 39.48601956 | 0.491204637 | 28.93727061 | NA | NA | NA | NA | NA | NA | NA |
| TRPV3 | 247 | G | -45.11525 | 38.2144196 | 0.474355625 | 27.67621299 | NA | NA | NA | NA | NA | NA | NA |
| TRPV3 | 248 | A | -42.65245 | 36.64356271 | 0.453541386 | 26.17490071 | NA | NA | NA | NA | NA | NA | NA |
| TRPV3 | 249 | F | -46.17588636 | 31.17189261 | 0.381040414 | 20.72164347 | NA | NA | NA | NA | NA | NA | NA |
| TRPV3 | 250 | F | -47.86379545 | 32.53745007 | 0.399134386 | 22.08859993 | NA | NA | NA | NA | NA | NA | NA |
| TRPV3 | 251 | N | -42.19245 | 35.84756988 | 0.442994286 | 25.37321853 | NA | NA | NA | NA | NA | NA | NA |
| TRPV3 | 252 | P | -40.52625 | 32.58511656 | 0.399765979 | 22.68622535 | NA | NA | NA | NA | NA | NA | NA |
| TRPV3 | 253 | K | -36.61385 | 34.00577661 | 0.418590073 | 24.2368363 | NA | NA | NA | NA | NA | NA | NA |
| TRPV3 | 254 | Y | -35.23958333 | 30.43323284 | 0.371252991 | 20.76949879 | NA | NA | NA | NA | NA | NA | NA |
| TRPV3 | 255 | Q | -38.76185 | 28.49542256 | 0.345576529 | 18.6863042 | NA | NA | NA | NA | NA | NA | NA |
| TRPV3 | 256 | H | -38.83705 | 29.80013064 | 0.362864232 | 19.98480478 | NA | NA | NA | NA | NA | NA | NA |
| TRPV3 | 257 | E | -41.37665 | 30.76576206 | 0.375659085 | 20.42237926 | NA | NA | NA | NA | NA | NA | NA |
| TRPV3 | 258 | G | -43.32 | 29.70592342 | 0.361615963 | 19.23603811 | NA | NA | NA | NA | NA | NA | NA |
| TRPV3 | 259 | F | -46.88770455 | 25.05004772 | 0.299924467 | 14.5828454 | NA | NA | NA | NA | NA | NA | NA |
| TRPV3 | 260 | Y | -44.54425 | 29.01237755 | 0.35242631 | 18.53899997 | NA | NA | NA | NA | NA | NA | NA |
| TRPV3 | 261 | F | -49.68215909 | 28.65549631 | 0.347697546 | 18.10224149 | NA | NA | NA | NA | NA | NA | NA |
| TRPV3 | 262 | G | -48.38025 | 32.29288338 | 0.395893817 | 21.73920576 | NA | NA | NA | NA | NA | NA | NA |
| TRPV3 | 263 | E | -48.77236111 | 33.22456123 | 0.408238777 | 22.66759651 | NA | NA | NA | NA | NA | NA | NA |
| TRPV3 | 264 | T | -53.33196429 | 32.42465149 | 0.397639777 | 21.85186635 | NA | NA | NA | NA | NA | NA | NA |
| TRPV3 | 265 | P | -56.81953571 | 30.8686022 | 0.377021742 | 20.28413348 | NA | NA | NA | NA | NA | NA | NA |
| TRPV3 | 266 | L | -56.851875 | 29.55379431 | 0.359600215 | 18.88942018 | NA | NA | NA | NA | NA | NA | NA |
| TRPV3 | 267 | A | -53.19585 | 28.02511866 | 0.339344887 | 17.45222956 | NA | NA | NA | NA | NA | NA | NA |
| TRPV3 | 268 | L | -54.24025 | 26.48382408 | 0.318922356 | 15.90872323 | NA | NA | NA | NA | NA | NA | NA |
| TRPV3 | 269 | A | -56.77305 | 24.95536844 | 0.298669943 | 14.27038019 | NA | NA | NA | NA | NA | NA | NA |
| TRPV3 | 270 | A | -54.66945 | 23.69984643 | 0.282033969 | 13.11706393 | NA | NA | NA | NA | NA | NA | NA |
| TRPV3 | 271 | C | -51.81391667 | 22.20472594 | 0.262223255 | 11.63756684 | NA | NA | NA | NA | NA | NA | NA |
| TRPV3 | 272 | T | -54.74225 | 21.3396391 | 0.250760642 | 10.75535401 | NA | NA | NA | NA | NA | NA | NA |
| TRPV3 | 273 | N | -56.855 | 17.14925392 | 0.19523701 | 6.466999108 | NA | NA | NA | NA | NA | NA | NA |
| TRPV3 | 274 | Q | -59.51558333 | 22.9264007 | 0.271785623 | 12.24361342 | NA | NA | NA | NA | NA | NA | NA |
| TRPV3 | 275 | P | -61.26653571 | 21.70233061 | 0.255566394 | 10.65692687 | NA | NA | NA | NA | NA | NA | NA |
| TRPV3 | 276 | E | -64.48285 | 24.90610909 | 0.298017245 | 13.86146875 | NA | NA | NA | NA | NA | NA | NA |
| TRPV3 | 277 | I | -62.826 | 27.05447057 | 0.326483562 | 16.02259641 | NA | NA | NA | NA | NA | NA | NA |

| | | | | | | | | | | | | | |
|---|---|---|---|---|---|---|---|---|---|---|---|---|---|
| TRPV3 | 278 | V | -60.63910714 | 26.85563622 | 0.323848958 | 16.16811016 | NA | NA | NA | NA | NA | NA | NA |
| TRPV3 | 279 | Q | -64.58391667 | 26.8048646 | 0.323176221 | 15.85940637 | NA | NA | NA | NA | NA | NA | NA |
| TRPV3 | 280 | L | -66.284125 | 30.93988636 | 0.377966275 | 19.89541384 | NA | NA | NA | NA | NA | NA | NA |
| TRPV3 | 281 | L | -62.289625 | 31.62326455 | 0.387021203 | 20.93965631 | NA | NA | NA | NA | NA | NA | NA |
| TRPV3 | 282 | M | -62.166125 | 29.98754123 | 0.365347468 | 18.95019378 | NA | NA | NA | NA | NA | NA | NA |
| TRPV3 | 283 | E | -66.53725 | 35.01834856 | 0.4320069 | 23.973751 | NA | NA | NA | NA | NA | NA | NA |
| TRPV3 | 284 | H | -66.72835 | 36.75396765 | 0.455004279 | 25.72195532 | NA | NA | NA | NA | NA | NA | NA |
| TRPV3 | 285 | E | -65.42825 | 41.40858486 | 0.516679099 | 30.37061853 | NA | NA | NA | NA | NA | NA | NA |
| TRPV3 | 286 | Q | -64.38036111 | 41.76551748 | 0.521408544 | 31.08279364 | NA | NA | NA | NA | NA | NA | NA |
| TRPV3 | 287 | T | -61.19753571 | 36.47899257 | 0.451360791 | 25.79290054 | NA | NA | NA | NA | NA | NA | NA |
| TRPV3 | 288 | D | -56.76375 | 38.59244606 | 0.479364569 | 27.92946994 | NA | NA | NA | NA | NA | NA | NA |
| TRPV3 | 289 | I | -56.334 | 34.9562128 | 0.431183586 | 24.26351105 | NA | NA | NA | NA | NA | NA | NA |
| TRPV3 | 290 | T | -53.16825 | 38.84880164 | 0.482761343 | 28.26398936 | NA | NA | NA | NA | NA | NA | NA |
| TRPV3 | 291 | S | -52.42125 | 38.37443412 | 0.476475857 | 27.8009269 | NA | NA | NA | NA | NA | NA | NA |
| TRPV3 | 292 | R | -47.06243182 | 39.23363538 | 0.487860485 | 28.66628138 | NA | NA | NA | NA | NA | NA | NA |
| TRPV3 | 293 | D | -46.640125 | 36.53874559 | 0.452152534 | 25.98552477 | NA | NA | NA | NA | NA | NA | NA |
| TRPV3 | 294 | S | -44.55391667 | 40.05010318 | 0.498678884 | 29.57918309 | NA | NA | NA | NA | NA | NA | NA |
| TRPV3 | 295 | R | -41.58915909 | 33.63949216 | 0.413736714 | 23.34577445 | NA | NA | NA | NA | NA | NA | NA |
| TRPV3 | 296 | G | -44.277 | 39.23450759 | 0.487872042 | 28.7470945 | NA | NA | NA | NA | NA | NA | NA |
| TRPV3 | 297 | N | -44.59825 | 34.82439903 | 0.429437021 | 24.33642189 | NA | NA | NA | NA | NA | NA | NA |
| TRPV3 | 298 | N | -48.426375 | 33.78895102 | 0.415717081 | 23.22348213 | NA | NA | NA | NA | NA | NA | NA |
| TRPV3 | 299 | I | -51.335375 | 30.48455133 | 0.371932974 | 19.91400772 | NA | NA | NA | NA | NA | NA | NA |
| TRPV3 | 300 | L | -48.87075 | 30.20760244 | 0.368263333 | 19.6415237 | NA | NA | NA | NA | NA | NA | NA |
| TRPV3 | 301 | H | -44.68295 | 30.6263082 | 0.373811287 | 20.17878003 | NA | NA | NA | NA | NA | NA | NA |
| TRPV3 | 302 | A | -46.70485 | 28.22975505 | 0.34205637 | 17.74214639 | NA | NA | NA | NA | NA | NA | NA |
| TRPV3 | 303 | L | -48.68525 | 25.45057227 | 0.305231516 | 14.97093825 | NA | NA | NA | NA | NA | NA | NA |
| TRPV3 | 304 | V | -43.86967857 | 26.93304251 | 0.32487461 | 17.01040328 | NA | NA | NA | NA | NA | NA | NA |
| TRPV3 | 305 | T | -43.19767857 | 26.64048012 | 0.320998087 | 16.77220417 | NA | NA | NA | NA | NA | NA | NA |
| TRPV3 | 306 | V | -45.45082143 | 22.30963711 | 0.263613354 | 12.40483338 | NA | NA | NA | NA | NA | NA | NA |
| TRPV3 | 307 | A | -44.13225 | 21.47021928 | 0.252490862 | 11.54482642 | NA | NA | NA | NA | NA | NA | NA |
| TRPV3 | 308 | E | -42.13845 | 19.57532482 | 0.227383045 | 9.758676394 | NA | NA | NA | NA | NA | NA | NA |
| TRPV3 | 309 | D | -43.16305 | 16.0176104 | 0.180242456 | 6.311898156 | NA | NA | NA | NA | NA | NA | NA |
| TRPV3 | 310 | F | -46.77625 | 10.87546431 | 0.112107758 | 1.010017184 | 10.05 | 0.76 | 0.57 | 16.73 | -0.72 | 1 | 1 |

| | | | | | | | | | | | | | |
|---|---|---|---|---|---|---|---|---|---|---|---|---|---|
| TRPV3 | 311 | K | -44.53065 | 11.14656328 | 0.115699886 | 1.347880764 | 10.05 | 0.76 | 0.57 | 16.73 | -0.72 | 1 | 1 |
| TRPV3 | 312 | T | -45.14839286 | 13.57917416 | 0.147932577 | 3.787617976 | 10.06 | 0.63 | 1.73 | 1.36 | -0.94 | 0 | 0 |
| TRPV3 | 313 | Q | -47.84336111 | 14.53316559 | 0.160573198 | 4.043909581 | 10.55 | 0.28 | 1.13 | 1.44 | -0.51 | 0 | 0 |
| TRPV3 | 314 | N | -50.50005 | 12.38486476 | 0.132107684 | 1.897309545 | 10.55 | 0.28 | 1.13 | 1.44 | -0.51 | 0 | 0 |
| TRPV3 | 315 | D | -50.67225 | 15.56859172 | 0.174292848 | 5.064771055 | NA | NA | NA | NA | NA | NA | NA |
| TRPV3 | 316 | F | -52.32243182 | 16.60172946 | 0.187982177 | 6.026982304 | NA | NA | NA | NA | NA | NA | NA |
| TRPV3 | 317 | V | -50.46639286 | 19.71285451 | 0.229205347 | 9.135312291 | NA | NA | NA | NA | NA | NA | NA |
| TRPV3 | 318 | K | -50.82658333 | 19.77635505 | 0.230046745 | 9.176614937 | NA | NA | NA | NA | NA | NA | NA |
| TRPV3 | 319 | R | -55.14525 | 15.86709176 | 0.178248047 | 5.16790429 | NA | NA | NA | NA | NA | NA | NA |
| TRPV3 | 320 | M | -55.4375 | 22.38847625 | 0.264657992 | 11.69879488 | NA | NA | NA | NA | NA | NA | NA |
| TRPV3 | 321 | Y | -50.79633333 | 24.99109623 | 0.299143346 | 14.39669195 | NA | NA | NA | NA | NA | NA | NA |
| TRPV3 | 322 | D | -55.10045 | 25.67949055 | 0.308264739 | 14.99624723 | NA | NA | NA | NA | NA | NA | NA |
| TRPV3 | 323 | M | -58.59375 | 24.42877076 | 0.291692395 | 13.37879842 | NA | NA | NA | NA | NA | NA | NA |
| TRPV3 | 324 | I | -56.10775 | 28.44280176 | 0.344879291 | 17.74711987 | NA | NA | NA | NA | NA | NA | NA |
| TRPV3 | 325 | L | -54.435625 | 30.33415689 | 0.36994021 | 19.63671758 | NA | NA | NA | NA | NA | NA | NA |
| TRPV3 | 326 | L | -58.9495 | 28.4324209 | 0.344741742 | 17.37923682 | NA | NA | NA | NA | NA | NA | NA |
| TRPV3 | 327 | R | -60.26705 | 32.15093854 | 0.394013013 | 21.10847615 | NA | NA | NA | NA | NA | NA | NA |
| TRPV3 | 328 | S | -57.43975 | 34.45632797 | 0.424559989 | 23.76863476 | NA | NA | NA | NA | NA | NA | NA |
| TRPV3 | 329 | G | -57.57125 | 37.10338223 | 0.459634108 | 26.41186075 | NA | NA | NA | NA | NA | NA | NA |
| TRPV3 | 330 | N | -55.345375 | 37.9378462 | 0.47069096 | 27.23809523 | NA | NA | NA | NA | NA | NA | NA |
| TRPV3 | 331 | W | -50.26767857 | 37.0219023 | 0.458554479 | 26.42815169 | NA | NA | NA | NA | NA | NA | NA |
| TRPV3 | 332 | E | -50.79145 | 40.06462304 | 0.498871275 | 29.47613833 | NA | NA | NA | NA | NA | NA | NA |
| TRPV3 | 333 | L | -50.525625 | 34.70460062 | 0.427849663 | 24.11425107 | NA | NA | NA | NA | NA | NA | NA |
| TRPV3 | 334 | E | -47.23691667 | 33.26053871 | 0.408715488 | 22.74409029 | NA | NA | NA | NA | NA | NA | NA |
| TRPV3 | 335 | T | -45.68845 | 39.48312407 | 0.491166271 | 28.97328252 | NA | NA | NA | NA | NA | NA | NA |
| TRPV3 | 336 | T | -45.57625 | 38.35789425 | 0.4762567 | 27.86144514 | NA | NA | NA | NA | NA | NA | NA |
| TRPV3 | 337 | R | -40.60206818 | 40.16554607 | 0.50020853 | 30.25008969 | NA | NA | NA | NA | NA | NA | NA |
| TRPV3 | 338 | N | -40.180125 | 37.02199244 | 0.458555673 | 27.17371722 | NA | NA | NA | NA | NA | NA | NA |
| TRPV3 | 339 | N | -39.28125 | 41.01842089 | 0.511509331 | 31.1211669 | NA | NA | NA | NA | NA | NA | NA |
| TRPV3 | 340 | D | -36.23145 | 39.90826952 | 0.496799553 | 30.08784528 | NA | NA | NA | NA | NA | NA | NA |
| TRPV3 | 341 | G | -37.2675 | 41.14926834 | 0.513243092 | 31.29220498 | NA | NA | NA | NA | NA | NA | NA |
| TRPV3 | 342 | L | -38.029 | 36.51915111 | 0.451892902 | 26.61087527 | NA | NA | NA | NA | NA | NA | NA |
| TRPV3 | 343 | T | -41.25167857 | 35.43118619 | 0.4374771 | 25.59381622 | NA | NA | NA | NA | NA | NA | NA |

| | | | | | | | | | | | | | |
|---|---|---|---|---|---|---|---|---|---|---|---|---|---|
| TRPV3 | 344 | P | -43.60096429 | 31.98152288 | 0.391768214 | 22.12500963 | NA | NA | NA | NA | NA | NA | NA |
| TRPV3 | 345 | L | -41.233375 | 31.40591651 | 0.384141289 | 21.43531791 | NA | NA | NA | NA | NA | NA | NA |
| TRPV3 | 346 | Q | -37.78647222 | 33.83291006 | 0.416299549 | 23.88907789 | NA | NA | NA | NA | NA | NA | NA |
| TRPV3 | 347 | L | -39.744 | 31.19226906 | 0.381310407 | 21.29246968 | NA | NA | NA | NA | NA | NA | NA |
| TRPV3 | 348 | A | -39.48125 | 28.00939701 | 0.339136572 | 18.08726124 | NA | NA | NA | NA | NA | NA | NA |
| TRPV3 | 349 | A | -36.28605 | 30.07183082 | 0.366464326 | 20.10536862 | NA | NA | NA | NA | NA | NA | NA |
| TRPV3 | 350 | K | -34.38510714 | 31.38443594 | 0.383856666 | 21.43645219 | NA | NA | NA | NA | NA | NA | NA |
| TRPV3 | 351 | M | -37.06975 | 27.34657187 | 0.330353976 | 17.52374542 | NA | NA | NA | NA | NA | NA | NA |
| TRPV3 | 352 | G | -36.82975 | 23.64718554 | 0.281336199 | 13.69533043 | NA | NA | NA | NA | NA | NA | NA |
| TRPV3 | 353 | K | -40.19136111 | 23.67055342 | 0.281645829 | 13.75220188 | NA | NA | NA | NA | NA | NA | NA |
| TRPV3 | 354 | A | -40.69505 | 21.32567879 | 0.250575665 | 11.3653123 | NA | NA | NA | NA | NA | NA | NA |
| TRPV3 | 355 | E | -44.13585 | 20.0801307 | 0.234071847 | 10.13019585 | NA | NA | NA | NA | NA | NA | NA |
| TRPV3 | 356 | I | -45.124625 | 22.90897346 | 0.271554708 | 13.08658728 | NA | NA | NA | NA | NA | NA | NA |
| TRPV3 | 357 | L | -41.539375 | 25.11604724 | 0.300798977 | 15.14655061 | NA | NA | NA | NA | NA | NA | NA |
| TRPV3 | 358 | K | -43.85958333 | 19.95568556 | 0.232422918 | 10.14665292 | NA | NA | NA | NA | NA | NA | NA |
| TRPV3 | 359 | Y | -49.28008333 | 24.04604606 | 0.286621199 | 14.17073102 | NA | NA | NA | NA | NA | NA | NA |
| TRPV3 | 360 | I | -45.778125 | 27.30885842 | 0.329854263 | 17.36003798 | NA | NA | NA | NA | NA | NA | NA |
| TRPV3 | 361 | L | -42.7995 | 29.02877774 | 0.352643616 | 19.01770236 | NA | NA | NA | NA | NA | NA | NA |
| TRPV3 | 362 | S | -46.63408333 | 26.16498594 | 0.314697672 | 16.16762566 | NA | NA | NA | NA | NA | NA | NA |
| TRPV3 | 363 | R | -49.86188636 | 28.8269468 | 0.349969307 | 18.83165817 | NA | NA | NA | NA | NA | NA | NA |
| TRPV3 | 364 | E | -52.96525 | 29.38116097 | 0.357312781 | 18.33316533 | NA | NA | NA | NA | NA | NA | NA |
| TRPV3 | 365 | I | -55.914 | 30.62973725 | 0.373856723 | 18.90538981 | NA | NA | NA | NA | NA | NA | NA |
| TRPV3 | 366 | K | -59.58785 | 30.68966663 | 0.374650802 | 17.24846395 | NA | NA | NA | NA | NA | NA | NA |
| TRPV3 | 367 | E | -60.52965 | 32.39335997 | 0.397225157 | 18.94347036 | NA | NA | NA | NA | NA | NA | NA |
| TRPV3 | 368 | K | -60.88725 | 36.260488 | 0.448465552 | 22.81455953 | NA | NA | NA | NA | NA | NA | NA |
| TRPV3 | 369 | R | -61.58753571 | 36.8564196 | 0.456361792 | 23.40902965 | NA | NA | NA | NA | NA | NA | NA |
| TRPV3 | 370 | L | -58.509375 | 33.86004812 | 0.416659135 | 22.14048076 | NA | NA | NA | NA | NA | NA | NA |
| TRPV3 | 371 | R | -55.71115909 | 36.11684034 | 0.446562185 | 24.39905982 | NA | NA | NA | NA | NA | NA | NA |
| TRPV3 | 372 | S | -53.74745 | 38.26298582 | 0.47499914 | 27.55832538 | NA | NA | NA | NA | NA | NA | NA |
| TRPV3 | 373 | L | -53.393 | 35.16150853 | 0.433903805 | 24.55080634 | NA | NA | NA | NA | NA | NA | NA |
| TRPV3 | 374 | S | -51.18275 | 33.61567686 | 0.413421156 | 22.9996008 | NA | NA | NA | NA | NA | NA | NA |
| TRPV3 | 375 | R | -45.81979545 | 33.19553008 | 0.407854108 | 23.21310742 | NA | NA | NA | NA | NA | NA | NA |
| TRPV3 | 376 | K | -47.99191667 | 28.85245315 | 0.350307272 | 19.10391866 | NA | NA | NA | NA | NA | NA | NA |

| | | | | | | | | | | | | | |
|---|---|---|---|---|---|---|---|---|---|---|---|---|---|
| TRPV3 | 377 | F | -49.91243182 | 34.76446739 | 0.428642912 | 24.76223558 | NA | NA | NA | NA | NA | NA | NA |
| TRPV3 | 378 | T | -51.04253571 | 32.93505071 | 0.404402692 | 22.2523568 | NA | NA | NA | NA | NA | NA | NA |
| TRPV3 | 379 | D | -49.37475 | 37.70817751 | 0.467647794 | 27.05441479 | NA | NA | NA | NA | NA | NA | NA |
| TRPV3 | 380 | W | -48.81667857 | 36.9451999 | 0.457538153 | 22.40706233 | NA | NA | NA | NA | NA | NA | NA |
| TRPV3 | 381 | A | -52.54805 | 34.78673788 | 0.428938001 | 20.25264191 | NA | NA | NA | NA | NA | NA | NA |
| TRPV3 | 382 | Y | -53.25466667 | 37.1144564 | 0.459780843 | 22.60570166 | NA | NA | NA | NA | NA | NA | NA |
| TRPV3 | 383 | G | -52.45075 | 33.15121806 | 0.407266962 | 17.72185866 | NA | NA | NA | NA | NA | NA | NA |
| TRPV3 | 384 | P | -50.08839286 | 30.16101515 | 0.36764604 | 14.7309625 | NA | NA | NA | NA | NA | NA | NA |
| TRPV3 | 385 | V | -48.18296429 | 32.53506264 | 0.399102752 | 17.74485798 | NA | NA | NA | NA | NA | NA | NA |
| TRPV3 | 386 | S | -47.88908333 | 30.99459277 | 0.378691148 | 16.45978429 | NA | NA | NA | NA | NA | NA | NA |
| TRPV3 | 387 | S | -46.53725 | 34.98334068 | 0.431543037 | 20.44010345 | NA | NA | NA | NA | NA | NA | NA |
| TRPV3 | 388 | S | -45.41708333 | 31.39822055 | 0.384039315 | 16.88485412 | NA | NA | NA | NA | NA | NA | NA |
| TRPV3 | 389 | L | -45.63 | 34.90109703 | 0.430453288 | 24.22582132 | NA | NA | NA | NA | NA | NA | NA |
| TRPV3 | 390 | Y | -43.57941667 | 29.29598013 | 0.356184114 | 18.64426944 | NA | NA | NA | NA | NA | NA | NA |
| TRPV3 | 391 | D | -43.803375 | 35.756192 | 0.441783506 | 25.807198 | NA | NA | NA | NA | NA | NA | NA |
| TRPV3 | 392 | L | -39.0565 | 36.98902134 | 0.458118798 | 26.34328144 | NA | NA | NA | NA | NA | NA | NA |
| TRPV3 | 393 | T | -41.06125 | 39.38791318 | 0.489904703 | 29.38480833 | NA | NA | NA | NA | NA | NA | NA |
| TRPV3 | 394 | N | -39.467 | 38.09729695 | 0.472803722 | 28.10351353 | NA | NA | NA | NA | NA | NA | NA |
| TRPV3 | 395 | V | -36.96382143 | 35.0673782 | 0.432656555 | 25.19287839 | NA | NA | NA | NA | NA | NA | NA |
| TRPV3 | 396 | D | -34.48725 | 39.45759293 | 0.490827977 | 28.87207757 | NA | NA | NA | NA | NA | NA | NA |
| TRPV3 | 397 | T | -30.73867857 | 39.44151707 | 0.490614968 | 32.51166116 | NA | NA | NA | NA | NA | NA | NA |
| TRPV3 | 398 | T | -33.29439286 | 43.46187328 | 0.543885675 | 36.44397576 | NA | NA | NA | NA | NA | NA | NA |
| TRPV3 | 399 | T | -33.75745 | 42.7199671 | 0.534055236 | 32.74071302 | NA | NA | NA | NA | NA | NA | NA |
| TRPV3 | 400 | D | -32.18945 | 43.76136586 | 0.547854025 | 33.80168 | NA | NA | NA | NA | NA | NA | NA |
| TRPV3 | 401 | N | -33.3755 | 39.57962922 | 0.492444988 | 29.60731171 | NA | NA | NA | NA | NA | NA | NA |
| TRPV3 | 402 | S | -33.43041667 | 37.50259674 | 0.464923798 | 27.56755701 | NA | NA | NA | NA | NA | NA | NA |
| TRPV3 | 403 | V | -35.59610714 | 32.4697943 | 0.39823793 | 22.50578821 | NA | NA | NA | NA | NA | NA | NA |
| TRPV3 | 404 | L | -31.6285 | 33.08654302 | 0.406410002 | 26.08622581 | NA | NA | NA | NA | NA | NA | NA |
| TRPV3 | 405 | E | -29.39213889 | 35.27526072 | 0.435411049 | 31.1623011 | NA | NA | NA | NA | NA | NA | NA |
| TRPV3 | 406 | I | -32.252375 | 33.63603753 | 0.413690939 | 23.68693912 | NA | NA | NA | NA | NA | NA | NA |
| TRPV3 | 407 | T | -31.60525 | 30.5749592 | 0.3731309 | 23.5743059 | NA | NA | NA | NA | NA | NA | NA |
| TRPV3 | 408 | V | -27.56967857 | 30.79108785 | 0.375994658 | 26.70172996 | NA | NA | NA | NA | NA | NA | NA |
| TRPV3 | 409 | Y | -25.41041667 | 32.99146805 | 0.405150236 | 30.51358427 | NA | NA | NA | NA | NA | NA | NA |

| | | | | | | | | | | | | | |
|---|---|---|---|---|---|---|---|---|---|---|---|---|---|
| TRPV3 | 410 | N | -28.546125 | 30.3255077 | 0.369825607 | 27.84506474 | NA | NA | NA | NA | NA | NA | NA |
| TRPV3 | 411 | T | -25.93905 | 28.61550828 | 0.347167695 | 26.11308327 | NA | NA | NA | NA | NA | NA | NA |
| TRPV3 | 412 | N | -28.204625 | 29.78037394 | 0.362602451 | 27.37332214 | NA | NA | NA | NA | NA | NA | NA |
| TRPV3 | 413 | I | -31.376875 | 26.17582291 | 0.314841265 | 22.07923799 | NA | NA | NA | NA | NA | NA | NA |
| TRPV3 | 414 | D | -29.74945 | 22.88629688 | 0.271254238 | 18.78474201 | NA | NA | NA | NA | NA | NA | NA |
| TRPV3 | 415 | N | -33.47325 | 21.50149341 | 0.252905252 | 15.62755778 | NA | NA | NA | NA | NA | NA | NA |
| TRPV3 | 416 | R | -29.05425 | 24.82046736 | 0.296882471 | 20.77605321 | NA | NA | NA | NA | NA | NA | NA |
| TRPV3 | 417 | H | -31.40525 | 18.63400059 | 0.214910268 | 11.62468929 | NA | NA | NA | NA | NA | NA | NA |
| TRPV3 | 418 | E | -35.34265 | 21.45813592 | 0.252330754 | 12.33604916 | NA | NA | NA | NA | NA | NA | NA |
| TRPV3 | 419 | M | -35.44675 | 24.64175126 | 0.294514439 | 14.07959359 | NA | NA | NA | NA | NA | NA | NA |
| TRPV3 | 420 | L | -32.335125 | 25.83604511 | 0.310339125 | 18.81887207 | NA | NA | NA | NA | NA | NA | NA |
| TRPV3 | 421 | T | -35.13710714 | 21.2721722 | 0.249866689 | 11.84113027 | NA | NA | NA | NA | NA | NA | NA |
| TRPV3 | 422 | L | -38.1705 | 23.91920494 | 0.284940523 | 13.31468774 | NA | NA | NA | NA | NA | NA | NA |
| TRPV3 | 423 | E | -38.57145 | 27.04388068 | 0.326343243 | 12.67430605 | NA | NA | NA | NA | NA | NA | NA |
| TRPV3 | 424 | P | -38.28239286 | 29.1204483 | 0.353858274 | 18.45650573 | NA | NA | NA | NA | NA | NA | NA |
| TRPV3 | 425 | L | -34.999375 | 29.71833262 | 0.361780388 | 20.57750894 | NA | NA | NA | NA | NA | NA | NA |
| TRPV3 | 426 | H | -32.92715 | 25.52979769 | 0.306281272 | 16.33670887 | NA | NA | NA | NA | NA | NA | NA |
| TRPV3 | 427 | T | -34.74939286 | 30.1771042 | 0.367859224 | 20.79062365 | NA | NA | NA | NA | NA | NA | NA |
| TRPV3 | 428 | L | -33.305 | 33.12002431 | 0.406853638 | 26.16270483 | NA | NA | NA | NA | NA | NA | NA |
| TRPV3 | 429 | L | -29.46975 | 29.65484466 | 0.360939157 | 25.58034283 | NA | NA | NA | NA | NA | NA | NA |
| TRPV3 | 430 | H | -29.39515 | 30.18170686 | 0.36792021 | 26.06464374 | NA | NA | NA | NA | NA | NA | NA |
| TRPV3 | 431 | M | -30.679 | 34.05910394 | 0.419296673 | 30.0236573 | NA | NA | NA | NA | NA | NA | NA |
| TRPV3 | 432 | K | -28.16447222 | 35.73232849 | 0.441467309 | 31.62207116 | NA | NA | NA | NA | NA | NA | NA |
| TRPV3 | 433 | W | -23.40210714 | 29.74276014 | 0.362104059 | 28.00017345 | NA | NA | NA | NA | NA | NA | NA |
| TRPV3 | 434 | K | -25.64985 | 35.06504644 | 0.432625658 | 32.59799556 | NA | NA | NA | NA | NA | NA | NA |
| TRPV3 | 435 | K | -26.70758333 | 37.7224056 | 0.467836319 | 35.24009058 | NA | NA | NA | NA | NA | NA | NA |
| TRPV3 | 436 | F | -23.22752273 | 38.47757854 | 0.477842546 | 35.88513098 | NA | NA | NA | NA | NA | NA | NA |
| TRPV3 | 437 | A | -20.95545 | 35.04773428 | 0.432396268 | 32.46858276 | NA | NA | NA | NA | NA | NA | NA |
| TRPV3 | 438 | K | -19.36545 | 33.8801881 | 0.416925994 | 31.26885298 | NA | NA | NA | NA | NA | NA | NA |
| TRPV3 | 439 | H | -18.32455 | 36.91297341 | 0.457111144 | 34.29335338 | NA | NA | NA | NA | NA | NA | NA |
| TRPV3 | 440 | M | -17.291125 | 37.18016103 | 0.460651446 | 34.54322444 | NA | NA | NA | NA | NA | NA | NA |
| TRPV3 | 441 | F | -15.15161364 | 28.96195106 | 0.351758146 | 26.90025996 | NA | NA | NA | NA | NA | NA | NA |
| TRPV3 | 442 | F | -12.66715909 | 33.53614338 | 0.412367317 | 31.46988131 | NA | NA | NA | NA | NA | NA | NA |

| | | | | | | | | | | | | | |
|---|---|---|---|---|---|---|---|---|---|---|---|---|---|
| TRPV3 | 443 | L | -11.87075 | 36.97204666 | 0.457893879 | 33.73262345 | NA | NA | NA | NA | NA | NA | NA |
| TRPV3 | 444 | S | -10.72975 | 35.08613371 | 0.43290507 | 31.7610423 | NA | NA | NA | NA | NA | NA | NA |
| TRPV3 | 445 | F | -8.996068182 | 28.35737467 | 0.343747361 | 24.58152399 | NA | NA | NA | NA | NA | NA | NA |
| TRPV3 | 446 | C | -7.441083333 | 34.83298128 | 0.429550738 | 31.06327979 | NA | NA | NA | NA | NA | NA | NA |
| TRPV3 | 447 | F | -7.020068182 | 36.23711978 | 0.448155918 | 32.17336926 | NA | NA | NA | NA | NA | NA | NA |
| TRPV3 | 448 | Y | -4.661166667 | 28.90740972 | 0.35103546 | 24.85014724 | NA | NA | NA | NA | NA | NA | NA |
| TRPV3 | 449 | F | -3.149886364 | 29.77547909 | 0.362537593 | 25.71700081 | NA | NA | NA | NA | NA | NA | NA |
| TRPV3 | 450 | F | -1.890068182 | 35.05127874 | 0.432443233 | 30.99294044 | NA | NA | NA | NA | NA | NA | NA |
| TRPV3 | 451 | Y | -0.70025 | 34.38591604 | 0.423627014 | 30.33726745 | NA | NA | NA | NA | NA | NA | NA |
| TRPV3 | 452 | N | -0.251 | 28.48444253 | 0.345431041 | 23.93159246 | NA | NA | NA | NA | NA | NA | NA |
| TRPV3 | 453 | I | 1.171875 | 31.60833797 | 0.386823423 | 28.82632244 | NA | NA | NA | NA | NA | NA | NA |
| TRPV3 | 454 | T | 2.92825 | 35.03846884 | 0.432273499 | 32.78977113 | NA | NA | NA | NA | NA | NA | NA |
| TRPV3 | 455 | L | 3.631 | 31.72664606 | 0.388391034 | 29.51644522 | NA | NA | NA | NA | NA | NA | NA |
| TRPV3 | 456 | T | 5.162678571 | 28.71466904 | 0.348481599 | 26.60291678 | NA | NA | NA | NA | NA | NA | NA |
| TRPV3 | 457 | L | 7.73105 | 32.80333456 | 0.402657421 | 30.77313711 | NA | NA | NA | NA | NA | NA | NA |
| TRPV3 | 458 | V | 8.68525 | 33.98335231 | 0.418292945 | 32.00831805 | NA | NA | NA | NA | NA | NA | NA |
| TRPV3 | 459 | S | 9.429416667 | 29.70590231 | 0.361615683 | 27.67232619 | NA | NA | NA | NA | NA | NA | NA |
| TRPV3 | 460 | Y | 12.18783333 | 26.53398672 | 0.319587023 | 23.74681393 | NA | NA | NA | NA | NA | NA | NA |
| TRPV3 | 461 | Y | 13.26958333 | 32.35983636 | 0.396780961 | 29.56524053 | NA | NA | NA | NA | NA | NA | NA |
| TRPV3 | 462 | R | 13.80425 | 32.57568636 | 0.399641026 | 29.78959816 | NA | NA | NA | NA | NA | NA | NA |
| TRPV3 | 484 | L | 8.73425 | 43.55938808 | 0.54517777 | 41.5356418 | NA | NA | NA | NA | NA | NA | NA |
| TRPV3 | 485 | L | 5.56165 | 44.00202596 | 0.55104283 | 41.85576501 | NA | NA | NA | NA | NA | NA | NA |
| TRPV3 | 486 | G | 4.4965 | 40.81614805 | 0.508829166 | 38.52709904 | NA | NA | NA | NA | NA | NA | NA |
| TRPV3 | 487 | R | 5.38085 | 40.95976019 | 0.510732062 | 38.81214751 | NA | NA | NA | NA | NA | NA | NA |
| TRPV3 | 488 | M | 3.24105 | 44.05099773 | 0.551691718 | 41.75809497 | NA | NA | NA | NA | NA | NA | NA |
| TRPV3 | 489 | F | 0.710886364 | 39.84798843 | 0.496000813 | 37.40080719 | NA | NA | NA | NA | NA | NA | NA |
| TRPV3 | 490 | V | 1.075107143 | 38.14658349 | 0.47345678 | 35.34710867 | NA | NA | NA | NA | NA | NA | NA |
| TRPV3 | 491 | L | 0.86925 | 42.5457158 | 0.531746363 | 39.73823718 | NA | NA | NA | NA | NA | NA | NA |
| TRPV3 | 492 | I | -0.70425 | 44.28873968 | 0.554841857 | 40.23264419 | NA | NA | NA | NA | NA | NA | NA |
| TRPV3 | 493 | W | -4.714321429 | 37.59945176 | 0.466207151 | 33.53560286 | NA | NA | NA | NA | NA | NA | NA |
| TRPV3 | 494 | A | -2.17425 | 39.83237691 | 0.495793957 | 35.77596317 | NA | NA | NA | NA | NA | NA | NA |
| TRPV3 | 495 | M | -4.07625 | 42.81533977 | 0.535318947 | 38.75711607 | NA | NA | NA | NA | NA | NA | NA |
| TRPV3 | 496 | C | -7.13725 | 41.7004909 | 0.520546926 | 37.63670296 | NA | NA | NA | NA | NA | NA | NA |

| | | | | | | | | | | | | | |
|---|---|---|---|---|---|---|---|---|---|---|---|---|---|
| TRPV3 | 497 | I | -6.528125 | 36.20258291 | 0.447698296 | 32.13849523 | NA | NA | NA | NA | NA | NA | NA |
| TRPV3 | 498 | S | -5.815083333 | 39.72484595 | 0.494369145 | 35.6605004 | NA | NA | NA | NA | NA | NA | NA |
| TRPV3 | 499 | V | -8.756821429 | 42.43032285 | 0.530217378 | 38.36611472 | NA | NA | NA | NA | NA | NA | NA |
| TRPV3 | 500 | K | -11.24641667 | 39.99677231 | 0.497972236 | 36.22110551 | NA | NA | NA | NA | NA | NA | NA |
| TRPV3 | 501 | E | -10.03945 | 37.67155967 | 0.467162598 | 33.89762019 | NA | NA | NA | NA | NA | NA | NA |
| TRPV3 | 502 | G | -10.7015 | 40.45968692 | 0.504105969 | 36.68386593 | NA | NA | NA | NA | NA | NA | NA |
| TRPV3 | 503 | I | -13.72245 | 41.57330468 | 0.518861677 | 39.53306632 | NA | NA | NA | NA | NA | NA | NA |
| TRPV3 | 504 | A | -14.62605 | 37.28637728 | 0.462058837 | 35.24165845 | NA | NA | NA | NA | NA | NA | NA |
| TRPV3 | 505 | I | -12.299 | 36.77619693 | 0.455298822 | 34.81048196 | NA | NA | NA | NA | NA | NA | NA |
| TRPV3 | 506 | F | -15.16825 | 41.1906757 | 0.513791749 | 39.15607335 | NA | NA | NA | NA | NA | NA | NA |
| TRPV3 | 507 | L | -18.12465 | 39.70092697 | 0.494052213 | 36.86186378 | NA | NA | NA | NA | NA | NA | NA |
| TRPV3 | 508 | L | -17.44905 | 36.82002534 | 0.455879559 | 33.96974328 | NA | NA | NA | NA | NA | NA | NA |
| TRPV3 | 518 | S | -16.76385 | 25.32534908 | 0.303572278 | 22.48786814 | NA | NA | NA | NA | NA | NA | NA |
| TRPV3 | 519 | D | -17.41205 | 26.83586006 | 0.323586919 | 24.01022331 | NA | NA | NA | NA | NA | NA | NA |
| TRPV3 | 520 | A | -14.90165 | 29.72937924 | 0.361926758 | 27.72837057 | NA | NA | NA | NA | NA | NA | NA |
| TRPV3 | 521 | W | -10.19610714 | 23.13880667 | 0.274600054 | 21.86757778 | NA | NA | NA | NA | NA | NA | NA |
| TRPV3 | 522 | F | -11.79915909 | 23.76128511 | 0.282848046 | 20.8477906 | NA | NA | NA | NA | NA | NA | NA |
| TRPV3 | 523 | H | -12.65465 | 31.24914786 | 0.382064065 | 28.11172372 | NA | NA | NA | NA | NA | NA | NA |
| TRPV3 | 524 | F | -8.56005 | 31.97572976 | 0.391691454 | 28.20728047 | NA | NA | NA | NA | NA | NA | NA |
| TRPV3 | 525 | V | -6.489678571 | 27.05175519 | 0.326447582 | 22.98940865 | NA | NA | NA | NA | NA | NA | NA |
| TRPV3 | 526 | F | -7.402977273 | 29.84835414 | 0.363503205 | 25.80185344 | NA | NA | NA | NA | NA | NA | NA |
| TRPV3 | 527 | F | -5.68025 | 33.85787899 | 0.416630393 | 29.79341662 | NA | NA | NA | NA | NA | NA | NA |
| TRPV3 | 528 | I | -2.63105 | 32.47830939 | 0.398350757 | 28.42848663 | NA | NA | NA | NA | NA | NA | NA |
| TRPV3 | 529 | Q | -1.586027778 | 30.88915554 | 0.377294079 | 26.82511144 | NA | NA | NA | NA | NA | NA | NA |
| TRPV3 | 530 | A | -1.29725 | 34.95842602 | 0.431212911 | 30.89215805 | NA | NA | NA | NA | NA | NA | NA |
| TRPV3 | 531 | V | -0.707535714 | 36.65747906 | 0.453725781 | 31.74191645 | NA | NA | NA | NA | NA | NA | NA |
| TRPV3 | 532 | L | 1.462625 | 31.48533552 | 0.38519361 | 28.67897385 | NA | NA | NA | NA | NA | NA | NA |
| TRPV3 | 533 | V | 2.061964286 | 34.42780307 | 0.424182027 | 31.63072452 | NA | NA | NA | NA | NA | NA | NA |
| TRPV3 | 534 | I | 2.254964286 | 38.5188394 | 0.478389263 | 35.71219799 | NA | NA | NA | NA | NA | NA | NA |
| TRPV3 | 535 | L | 4.329875 | 37.32439474 | 0.462562578 | 35.06001303 | NA | NA | NA | NA | NA | NA | NA |
| TRPV3 | 536 | S | 6.80625 | 35.20731045 | 0.434510691 | 33.17352298 | NA | NA | NA | NA | NA | NA | NA |
| TRPV3 | 537 | V | 7.037678571 | 38.81003615 | 0.482247691 | 36.78660444 | NA | NA | NA | NA | NA | NA | NA |
| TRPV3 | 538 | F | 8.159083333 | 41.16115486 | 0.513400591 | 39.12557682 | NA | NA | NA | NA | NA | NA | NA |

| | | | | | | | | | | | | | |
|---|---|---|---|---|---|---|---|---|---|---|---|---|---|
| TRPV3 | 539 | L | 10.498625 | 37.03833476 | 0.458772213 | 34.22120966 | NA | NA | NA | NA | NA | NA | NA |
| TRPV3 | 540 | Y | 12.55566667 | 37.86922757 | 0.469781746 | 35.06576 | NA | NA | NA | NA | NA | NA | NA |
| TRPV3 | 541 | L | 12.624125 | 42.33615385 | 0.528969616 | 39.52638987 | NA | NA | NA | NA | NA | NA | NA |
| TRPV3 | 542 | F | 13.21370455 | 41.29288149 | 0.515146001 | 37.18361141 | NA | NA | NA | NA | NA | NA | NA |
| TRPV3 | 543 | A | 17.88405 | 40.5547987 | 0.505366223 | 36.43380958 | NA | NA | NA | NA | NA | NA | NA |
| TRPV3 | 544 | Y | 15.26758333 | 37.25232807 | 0.461607677 | 33.13864671 | NA | NA | NA | NA | NA | NA | NA |
| TRPV3 | 545 | K | 16.02605 | 33.04061396 | 0.405801431 | 28.93053308 | NA | NA | NA | NA | NA | NA | NA |
| TRPV3 | 546 | E | 12.55385 | 32.87845405 | 0.403652772 | 30.06997984 | NA | NA | NA | NA | NA | NA | NA |
| TRPV3 | 547 | Y | 12.75425 | 33.66888149 | 0.41412613 | 30.85109213 | NA | NA | NA | NA | NA | NA | NA |
| TRPV3 | 548 | L | 11.507 | 28.31264551 | 0.343154689 | 25.50594733 | NA | NA | NA | NA | NA | NA | NA |
| TRPV3 | 549 | A | 9.16085 | 29.16243565 | 0.354414617 | 26.34423502 | NA | NA | NA | NA | NA | NA | NA |
| TRPV3 | 550 | C | 6.33925 | 31.15143558 | 0.380769354 | 29.13381555 | NA | NA | NA | NA | NA | NA | NA |
| TRPV3 | 551 | L | 6.684 | 30.42867616 | 0.371192614 | 28.28513243 | NA | NA | NA | NA | NA | NA | NA |
| TRPV3 | 552 | V | 6.95625 | 26.58125366 | 0.320213322 | 24.54679399 | NA | NA | NA | NA | NA | NA | NA |
| TRPV3 | 553 | L | 3.337 | 26.22982244 | 0.315556772 | 23.98139277 | NA | NA | NA | NA | NA | NA | NA |
| TRPV3 | 554 | A | 1.59225 | 28.92256055 | 0.351236213 | 26.14815671 | NA | NA | NA | NA | NA | NA | NA |
| TRPV3 | 555 | M | 2.64125 | 26.62364971 | 0.32077508 | 23.85462816 | NA | NA | NA | NA | NA | NA | NA |
| TRPV3 | 556 | A | 0.97225 | 23.57912846 | 0.280434426 | 20.77419329 | NA | NA | NA | NA | NA | NA | NA |
| TRPV3 | 557 | L | -1.284 | 25.95992854 | 0.311980611 | 20.4622547 | NA | NA | NA | NA | NA | NA | NA |
| TRPV3 | 558 | G | -1.52325 | 27.35509622 | 0.330466925 | 23.29114235 | NA | NA | NA | NA | NA | NA | NA |
| TRPV3 | 559 | W | -0.688892857 | 23.58472105 | 0.280508529 | 19.5205115 | NA | NA | NA | NA | NA | NA | NA |
| TRPV3 | 560 | A | -3.99125 | 22.66677469 | 0.268345515 | 18.61184887 | NA | NA | NA | NA | NA | NA | NA |
| TRPV3 | 561 | N | -5.626125 | 25.56451281 | 0.306741256 | 21.50292505 | NA | NA | NA | NA | NA | NA | NA |
| TRPV3 | 562 | M | -6.74525 | 25.32737023 | 0.303599059 | 21.27028563 | NA | NA | NA | NA | NA | NA | NA |
| TRPV3 | 563 | L | -9.124625 | 19.72311023 | 0.229341238 | 15.97568307 | NA | NA | NA | NA | NA | NA | NA |
| TRPV3 | 564 | Y | -12.23991667 | 25.23496806 | 0.302374707 | 21.88920836 | NA | NA | NA | NA | NA | NA | NA |
| TRPV3 | 565 | Y | -10.59666667 | 26.83502512 | 0.323575856 | 23.63868244 | NA | NA | NA | NA | NA | NA | NA |
| TRPV3 | 566 | T | -12.03925 | 22.00954395 | 0.259637046 | 19.96391841 | NA | NA | NA | NA | NA | NA | NA |
| TRPV3 | 567 | R | -16.37288636 | 23.5688023 | 0.280297602 | 21.50055467 | NA | NA | NA | NA | NA | NA | NA |
| TRPV3 | 568 | G | -17.31825 | 26.03409566 | 0.312963344 | 23.41699339 | NA | NA | NA | NA | NA | NA | NA |
| TRPV3 | 569 | F | -17.21385 | 22.97109327 | 0.27237781 | 20.33699236 | NA | NA | NA | NA | NA | NA | NA |
| TRPV3 | 570 | Q | -21.43980556 | 20.01625015 | 0.233225414 | 17.40476215 | NA | NA | NA | NA | NA | NA | NA |
| TRPV3 | 571 | S | -17.54941667 | 18.53869807 | 0.213647486 | 15.8772178 | NA | NA | NA | NA | NA | NA | NA |

| | | | | | | | | | | | | | |
|---|---|---|---|---|---|---|---|---|---|---|---|---|---|
| TRPV3 | 572 | M | -15.05485 | 19.77709235 | 0.230056514 | 17.70531897 | NA | NA | NA | NA | NA | NA | NA |
| TRPV3 | 573 | G | -17.28975 | 19.9395435 | 0.232209032 | 17.30199404 | NA | NA | NA | NA | NA | NA | NA |
| TRPV3 | 574 | M | -18.07845 | 16.46677582 | 0.186194008 | 13.81791914 | NA | NA | NA | NA | NA | NA | NA |
| TRPV3 | 575 | Y | -12.45566667 | 14.41756238 | 0.159041427 | 12.36789732 | NA | NA | NA | NA | NA | NA | NA |
| TRPV3 | 576 | S | -14.14275 | 18.67398653 | 0.215440092 | 16.61111968 | NA | NA | NA | NA | NA | NA | NA |
| TRPV3 | 577 | V | -17.27539286 | 17.66506238 | 0.202071599 | 15.00917288 | NA | NA | NA | NA | NA | NA | NA |
| TRPV3 | 578 | M | -15.32425 | 14.39738608 | 0.158774086 | 12.33731761 | NA | NA | NA | NA | NA | NA | NA |
| TRPV3 | 579 | I | -11.620125 | 15.2950879 | 0.170668855 | 13.26490043 | NA | NA | NA | NA | NA | NA | NA |
| TRPV3 | 580 | Q | -13.77525 | 18.03865693 | 0.207021818 | 15.99606374 | NA | NA | NA | NA | NA | NA | NA |
| TRPV3 | 581 | K | -15.76805 | 16.54573498 | 0.187240236 | 13.73558217 | NA | NA | NA | NA | NA | NA | NA |
| TRPV3 | 582 | V | -13.08239286 | 11.64577331 | 0.122314541 | 9.629754359 | NA | NA | NA | NA | NA | NA | NA |
| TRPV3 | 583 | I | -10.125 | 16.12866275 | 0.181713927 | 14.13852988 | NA | NA | NA | NA | NA | NA | NA |
| TRPV3 | 584 | L | -13.39965 | 18.10259233 | 0.207868978 | 16.11327233 | NA | NA | NA | NA | NA | NA | NA |
| TRPV3 | 585 | H | -16.17365 | 16.0936357 | 0.18124981 | 14.03612911 | NA | NA | NA | NA | NA | NA | NA |
| TRPV3 | 586 | D | -12.00225 | 13.22599565 | 0.143252875 | 11.22272944 | NA | NA | NA | NA | NA | NA | NA |
| TRPV3 | 587 | V | -8.46525 | 13.22840588 | 0.143284811 | 9.88066798 | NA | NA | NA | NA | NA | NA | NA |
| TRPV3 | 588 | L | -8.726625 | 17.65113713 | 0.201887086 | 13.88864926 | NA | NA | NA | NA | NA | NA | NA |
| TRPV3 | 589 | K | -10.95369444 | 16.90869247 | 0.192049512 | 15.02699415 | NA | NA | NA | NA | NA | NA | NA |
| TRPV3 | 590 | F | -5.25125 | 11.79293789 | 0.124264508 | 7.77069619 | NA | NA | NA | NA | NA | NA | NA |
| TRPV3 | 591 | L | -3.866 | 15.45939698 | 0.172845991 | 11.39137868 | NA | NA | NA | NA | NA | NA | NA |
| TRPV3 | 592 | F | -4.951795455 | 19.40331296 | 0.225103846 | 15.32018202 | NA | NA | NA | NA | NA | NA | NA |
| TRPV3 | 593 | V | -4.644678571 | 17.80694081 | 0.203951523 | 13.74130717 | NA | NA | NA | NA | NA | NA | NA |
| TRPV3 | 594 | Y | -0.7515 | 13.54320926 | 0.147456033 | 8.419095104 | NA | NA | NA | NA | NA | NA | NA |
| TRPV3 | 595 | I | 0 | 19.47169045 | 0.226009864 | 14.00998955 | NA | NA | NA | NA | NA | NA | NA |
| TRPV3 | 596 | V | -1.015678571 | 21.66837737 | 0.255116505 | 16.17626827 | NA | NA | NA | NA | NA | NA | NA |
| TRPV3 | 597 | F | 1.49225 | 18.76088409 | 0.216591506 | 13.62301079 | NA | NA | NA | NA | NA | NA | NA |
| TRPV3 | 598 | L | 2.77125 | 15.9543265 | 0.179403929 | 13.14763898 | NA | NA | NA | NA | NA | NA | NA |
| TRPV3 | 599 | L | 3.349875 | 22.05571316 | 0.260248799 | 19.2455111 | NA | NA | NA | NA | NA | NA | NA |
| TRPV3 | 600 | G | 3.3145 | 23.41742484 | 0.278291813 | 20.61949353 | NA | NA | NA | NA | NA | NA | NA |
| TRPV3 | 601 | F | 3.443340909 | 20.02553914 | 0.233348495 | 17.68355163 | NA | NA | NA | NA | NA | NA | NA |
| TRPV3 | 602 | G | 7.98025 | 20.83074067 | 0.244017613 | 18.7002026 | NA | NA | NA | NA | NA | NA | NA |
| TRPV3 | 603 | V | 8.012964286 | 23.90952666 | 0.284812283 | 21.50559126 | NA | NA | NA | NA | NA | NA | NA |
| TRPV3 | 604 | A | 9.22705 | 23.85852512 | 0.2841365 | 21.25336258 | NA | NA | NA | NA | NA | NA | NA |

| | | | | | | | | | | | | | |
|---|---|---|---|---|---|---|---|---|---|---|---|---|---|
| TRPV3 | 605 | L | 10.527375 | 17.80110078 | 0.203874141 | 15.20210757 | NA | NA | NA | NA | NA | NA | NA |
| TRPV3 | 606 | A | 13.43285 | 22.29438014 | 0.263411195 | 19.40850286 | NA | NA | NA | NA | NA | NA | NA |
| TRPV3 | 607 | S | 13.36391667 | 24.89452659 | 0.297863774 | 21.84565072 | NA | NA | NA | NA | NA | NA | NA |
| TRPV3 | 608 | L | 14.067625 | 23.07168752 | 0.273710709 | 19.7849883 | NA | NA | NA | NA | NA | NA | NA |
| TRPV3 | 609 | I | 17.20765 | 21.18116741 | 0.248660854 | 17.32949484 | NA | NA | NA | NA | NA | NA | NA |
| TRPV3 | 610 | E | 18.88265 | 24.60329224 | 0.294004847 | 20.74862488 | NA | NA | NA | NA | NA | NA | NA |
| TRPV3 | 611 | K | 21.26305 | 23.74072581 | 0.282575631 | 19.69439032 | NA | NA | NA | NA | NA | NA | NA |
| TRPV3 | 612 | C | 24.52325 | 24.58902166 | 0.293815759 | 20.54609393 | NA | NA | NA | NA | NA | NA | NA |
| TRPV3 | 613 | P | 25.53482143 | 20.82902864 | 0.243994928 | 16.68508052 | NA | NA | NA | NA | NA | NA | NA |
| TRPV3 | 614 | K | 29.22225 | 24.69488368 | 0.295218456 | 20.54849187 | NA | NA | NA | NA | NA | NA | NA |
| TRPV3 | 615 | D | 28.83565 | 22.66808792 | 0.268362915 | 18.51308348 | NA | NA | NA | NA | NA | NA | NA |
| TRPV3 | 616 | N | 26.38705 | 19.40449785 | 0.225119546 | 15.25373761 | NA | NA | NA | NA | NA | NA | NA |
| TRPV3 | 617 | K | 23.37765 | 16.54454853 | 0.187224515 | 12.39154168 | NA | NA | NA | NA | NA | NA | NA |
| TRPV3 | 618 | D | 23.43165 | 14.57987222 | 0.161192072 | 10.42704787 | NA | NA | NA | NA | NA | NA | NA |
| TRPV3 | 619 | C | 23.93475 | 16.70402284 | 0.189337589 | 12.56340249 | NA | NA | NA | NA | NA | NA | NA |
| TRPV3 | 620 | S | 20.84058333 | 17.85517869 | 0.204590687 | 13.80698381 | NA | NA | NA | NA | NA | NA | NA |
| TRPV3 | 621 | S | 17.86958333 | 14.0650711 | 0.154370831 | 10.01634853 | NA | NA | NA | NA | NA | NA | NA |
| TRPV3 | 622 | Y | 13.67891667 | 12.92847191 | 0.139310613 | 9.747450698 | NA | NA | NA | NA | NA | NA | NA |
| TRPV3 | 623 | G | 16.88475 | 18.53582461 | 0.213609412 | 14.48355697 | NA | NA | NA | NA | NA | NA | NA |
| TRPV3 | 624 | S | 15.37825 | 20.16492604 | 0.235195406 | 16.31725499 | NA | NA | NA | NA | NA | NA | NA |
| TRPV3 | 625 | F | 10.15534091 | 20.60839467 | 0.241071474 | 18.00579517 | NA | NA | NA | NA | NA | NA | NA |
| TRPV3 | 626 | S | 11.68241667 | 18.45161188 | 0.212493573 | 15.90394456 | NA | NA | NA | NA | NA | NA | NA |
| TRPV3 | 627 | D | 13.92208333 | 16.10049797 | 0.181340737 | 13.04248361 | NA | NA | NA | NA | NA | NA | NA |
| TRPV3 | 628 | A | 11.13545 | 16.30431762 | 0.184041397 | 13.71742652 | NA | NA | NA | NA | NA | NA | NA |
| TRPV3 | 629 | V | 8.081964286 | 15.49378801 | 0.173301681 | 13.36562653 | NA | NA | NA | NA | NA | NA | NA |
| TRPV3 | 630 | L | 10.64175 | 12.82585985 | 0.137950978 | 10.28783614 | NA | NA | NA | NA | NA | NA | NA |
| TRPV3 | 631 | E | 11.91269444 | 11.02654352 | 0.114109595 | 8.637011861 | NA | NA | NA | NA | NA | NA | NA |
| TRPV3 | 632 | L | 6.870625 | 11.67879403 | 0.122752074 | 9.551012338 | NA | NA | NA | NA | NA | NA | NA |
| TRPV3 | 633 | F | 5.444613636 | 9.749673738 | 0.097190757 | 7.613271317 | 2.15 | -0.8 | -0.4 | 3.38 | 0.81 | 0 | 0 |
| TRPV3 | 634 | K | 9.871694444 | 7.132400788 | 0.062511248 | 5.988674237 | 1.38 | -0.8 | -0.4 | 3.38 | 0.81 | 0 | 0 |
| TRPV3 | 635 | L | 6.691875 | 7.261990567 | 0.064228344 | 5.184244777 | 2.15 | -0.8 | -0.4 | 3.38 | 0.81 | 0 | 0 |
| TRPV3 | 636 | T | 3.138392857 | 6.770661711 | 0.057718116 | 3.96640251 | 2.81 | -0.8 | -0.4 | 3.38 | 0.81 | 0 | 0 |
| TRPV3 | 637 | I | 4.522375 | 3.860045207 | 0.019151733 | 1.559574318 | 2.3 | -0.8 | -0.4 | 3.38 | 0.81 | 0 | 0 |

| | | | | | | | | | | | | | |
|---|---|---|---|---|---|---|---|---|---|---|---|---|---|
| TRPV3 | 638 | G | 7.312 | 3.245109551 | 0.011003685 | 1.48868667 | 2.15 | -0.8 | -0.4 | 3.38 | 0.81 | 0 | 0 |
| TRPV3 | 639 | L | 9.873125 | 3.842239061 | 0.018915797 | 1.461567266 | 2.55 | -0.8 | -0.4 | 3.38 | 0.81 | 0 | 0 |
| TRPV3 | 640 | G | 11.79 | 4.8878483 | 0.032770376 | 2.299713819 | 2.62 | -0.8 | -0.4 | 3.38 | 0.81 | 0 | 0 |
| TRPV3 | 641 | D | 14.84065 | 5.588487183 | 0.042054014 | 2.359543934 | 3.32 | -0.8 | -0.4 | 3.38 | 0.81 | 0 | 0 |
| TRPV3 | 642 | L | 12.9795 | 7.679553698 | 0.069761158 | 4.632269835 | 3.09 | -0.8 | -0.4 | 3.38 | 0.81 | 0 | 0 |
| TRPV3 | 643 | N | 17.158625 | 10.42339283 | 0.1061177 | 6.733486156 | NA | NA | NA | NA | NA | NA | NA |
| TRPV3 | 644 | I | 16.9635 | 14.25706583 | 0.156914808 | 10.53831807 | NA | NA | NA | NA | NA | NA | NA |
| TRPV3 | 645 | Q | 18.75447222 | 13.69068256 | 0.149410091 | 9.851468419 | NA | NA | NA | NA | NA | NA | NA |
| TRPV3 | 646 | Q | 22.13165 | 17.83613562 | 0.204338361 | 13.79821588 | NA | NA | NA | NA | NA | NA | NA |
| TRPV3 | 647 | N | 21.38825 | 19.31045895 | 0.223873507 | 15.26894425 | NA | NA | NA | NA | NA | NA | NA |
| TRPV3 | 648 | S | 19.11091667 | 20.62664987 | 0.24131336 | 16.7795367 | NA | NA | NA | NA | NA | NA | NA |
| TRPV3 | 649 | K | 18.14125 | 24.76385856 | 0.296132391 | 20.91194006 | NA | NA | NA | NA | NA | NA | NA |
| TRPV3 | 650 | Y | 15.19391667 | 22.88092859 | 0.271183106 | 19.57215416 | NA | NA | NA | NA | NA | NA | NA |
| TRPV3 | 651 | P | 15.13325 | 20.67257763 | 0.241921914 | 17.36281278 | NA | NA | NA | NA | NA | NA | NA |
| TRPV3 | 652 | I | 11.955125 | 22.46425877 | 0.265662129 | 19.71367018 | NA | NA | NA | NA | NA | NA | NA |
| TRPV3 | 653 | L | 9.573875 | 22.50280276 | 0.266172846 | 19.78119402 | NA | NA | NA | NA | NA | NA | NA |
| TRPV3 | 654 | F | 11.79425 | 15.92198508 | 0.178975397 | 13.16821617 | NA | NA | NA | NA | NA | NA | NA |
| TRPV3 | 655 | L | 9.512 | 16.14754449 | 0.181964114 | 13.56231769 | NA | NA | NA | NA | NA | NA | NA |
| TRPV3 | 656 | F | 4.803886364 | 20.14963769 | 0.234992832 | 17.80829102 | NA | NA | NA | NA | NA | NA | NA |
| TRPV3 | 657 | L | 5.77325 | 18.96104011 | 0.219243622 | 16.61783748 | NA | NA | NA | NA | NA | NA | NA |
| TRPV3 | 658 | L | 6.93925 | 13.33106605 | 0.144645084 | 11.4099948 | NA | NA | NA | NA | NA | NA | NA |
| TRPV3 | 659 | I | 4.21 | 15.57008343 | 0.174312614 | 13.20932731 | NA | NA | NA | NA | NA | NA | NA |
| TRPV3 | 660 | T | 1.508535714 | 17.27824001 | 0.196946108 | 12.73969957 | NA | NA | NA | NA | NA | NA | NA |
| TRPV3 | 661 | Y | 2.32425 | 13.69513202 | 0.149469047 | 10.90879798 | NA | NA | NA | NA | NA | NA | NA |
| TRPV3 | 662 | V | 1.65425 | 10.77654685 | 0.110797077 | 6.271002186 | NA | NA | NA | NA | NA | NA | NA |
| TRPV3 | 663 | I | -0.597375 | 13.59104981 | 0.148089932 | 8.074492884 | NA | NA | NA | NA | NA | NA | NA |
| TRPV3 | 664 | L | -1.828125 | 14.51604275 | 0.160346316 | 9.257219111 | NA | NA | NA | NA | NA | NA | NA |
| TRPV3 | 665 | T | -0.907107143 | 10.72120278 | 0.110063755 | 5.227607304 | NA | NA | NA | NA | NA | NA | NA |
| TRPV3 | 666 | F | -3.326522727 | 6.609630096 | 0.055584408 | 1.458215413 | 5.5 | -0.02 | 1.28 | 2.08 | -0.23 | 0 | 0 |
| TRPV3 | 667 | V | -6.020107143 | 11.85077095 | 0.12503081 | 7.851967854 | NA | NA | NA | NA | NA | NA | NA |
| TRPV3 | 668 | L | -5.96975 | 11.88401439 | 0.125471294 | 7.816629597 | NA | NA | NA | NA | NA | NA | NA |
| TRPV3 | 669 | L | -5.528125 | 5.985011362 | 0.047308056 | 1.993625667 | 4.09 | 1.14 | 3.8 | 0.13 | -1.79 | 0 | 0 |
| TRPV3 | 670 | L | -8.8075 | 6.8570245 | 0.058862444 | 4.891366628 | 2.17 | 1.14 | 3.8 | 0.13 | -1.79 | 0 | 0 |

| | | | | | | | | | | | | | |
|---|---|---|---|---|---|---|---|---|---|---|---|---|---|
| TRPV3 | 671 | N | -11.440625 | 10.48159668 | 0.106888915 | 8.468453965 | NA | NA | NA | NA | NA | NA | NA |
| TRPV3 | 672 | M | -10.64625 | 9.200383688 | 0.089912529 | 7.199592008 | 2.03 | 1.14 | 3.8 | 0.13 | -1.79 | 0 | 0 |
| TRPV3 | 673 | L | -11.47525 | 2.724352767 | 0.00410353 | 0.746329983 | 2.03 | 1.14 | 3.8 | 0.13 | -1.79 | 0 | 0 |
| TRPV3 | 674 | I | -14.4175 | 7.386951536 | 0.065884108 | 5.406416254 | 2.08 | 1.07 | 2.85 | 0.78 | -1.26 | 0 | 0 |
| TRPV3 | 675 | A | -15.73025 | 9.9160357 | 0.099395094 | 7.514821977 | 2.42 | 1.1 | 3.63 | 0.46 | -1.16 | 0 | 0 |
| TRPV3 | 676 | L | -15.15375 | 6.57889246 | 0.055177127 | 4.540352506 | 2.09 | 1.14 | 3.8 | 0.13 | -1.79 | 0 | 0 |
| TRPV3 | 677 | M | -16.753125 | 3.318815451 | 0.011980306 | 1.177771537 | 2.68 | 1.09 | 3.43 | 0.56 | -1.11 | 0 | 0 |
| TRPV3 | 678 | G | -20.4745 | 5.946209633 | 0.046793924 | 3.693791761 | 2.27 | 0.36 | 2.36 | 1.43 | -0.46 | 0 | 0 |
| TRPV3 | 679 | E | -21.27065 | 7.787837569 | 0.071195946 | 5.87132585 | 1.94 | 0.65 | 3.05 | 0.94 | -0.77 | 0 | 0 |
| TRPV3 | 680 | T | -20.27325 | 5.804137404 | 0.044911432 | 3.014958172 | 2.85 | 1.07 | 3.05 | 0.78 | -1.01 | 0 | 0 |
| TRPV3 | 681 | V | -23.46696429 | 2.414658154 | 0 | 0.99180389 | 1.8 | 1.13 | 4.2 | 0.13 | -1.3 | 0 | 0 |
| TRPV3 | 682 | E | -25.94865 | 6.559706548 | 0.054922909 | 4.242338423 | 2.51 | 1.13 | 4.2 | 0.13 | -1.3 | 0 | 0 |
| TRPV3 | 683 | N | -24.72165 | 9.228962834 | 0.090291209 | 6.833003718 | 2.51 | 1.13 | 4.2 | 0.13 | -1.3 | 0 | 0 |
| TRPV3 | 684 | V | -21.60253571 | 9.762021768 | 0.097354371 | 6.988574813 | 2.85 | 1.07 | 3.05 | 0.78 | -1.01 | 0 | 0 |
| TRPV3 | 685 | S | -24.35541667 | 10.55263019 | 0.107830127 | 9.022896387 | NA | NA | NA | NA | NA | NA | NA |
| TRPV3 | 686 | K | -26.55265 | 12.44683341 | 0.132928784 | 9.945058259 | NA | NA | NA | NA | NA | NA | NA |
| TRPV3 | 687 | E | -24.33865 | 14.39071235 | 0.158685657 | 11.91558514 | NA | NA | NA | NA | NA | NA | NA |
| TRPV3 | 688 | S | -22.20541667 | 14.8202582 | 0.164377245 | 12.08088104 | NA | NA | NA | NA | NA | NA | NA |
| TRPV3 | 689 | E | -24.56902778 | 16.47511011 | 0.186304439 | 14.01690258 | NA | NA | NA | NA | NA | NA | NA |
| TRPV3 | 690 | R | -28.18879545 | 16.18130189 | 0.182411408 | 14.5188626 | NA | NA | NA | NA | NA | NA | NA |
| TRPV3 | 691 | I | -22.4965 | 19.26829149 | 0.223314778 | 17.8450666 | NA | NA | NA | NA | NA | NA | NA |
| TRPV3 | 692 | W | -22.36975 | 20.96789293 | 0.245834914 | 18.22311655 | NA | NA | NA | NA | NA | NA | NA |
| TRPV3 | 693 | R | -25.48625 | 22.85182439 | 0.270797469 | 20.36575871 | NA | NA | NA | NA | NA | NA | NA |
| TRPV3 | 694 | L | -25.25825 | 24.46688572 | 0.292197428 | 21.97119199 | NA | NA | NA | NA | NA | NA | NA |
| TRPV3 | 695 | Q | -20.58958333 | 24.33489355 | 0.290448499 | 21.52022742 | NA | NA | NA | NA | NA | NA | NA |
| TRPV3 | 696 | R | -21.80743182 | 26.15491606 | 0.314564244 | 23.58005877 | NA | NA | NA | NA | NA | NA | NA |
| TRPV3 | 697 | A | -25.16145 | 28.89710202 | 0.350898881 | 26.40628041 | NA | NA | NA | NA | NA | NA | NA |
| TRPV3 | 698 | R | -21.78806818 | 30.4264273 | 0.371162816 | 27.96813665 | NA | NA | NA | NA | NA | NA | NA |
| TRPV3 | 699 | T | -20.41425 | 30.19327425 | 0.368073481 | 27.58255031 | NA | NA | NA | NA | NA | NA | NA |
| TRPV3 | 700 | I | -23.63710714 | 32.9018122 | 0.403962274 | 30.33232937 | NA | NA | NA | NA | NA | NA | NA |
| TRPV3 | 701 | L | -25.183875 | 34.37594236 | 0.42349486 | 31.88143292 | NA | NA | NA | NA | NA | NA | NA |
| TRPV3 | 702 | E | -20.73725 | 35.98623215 | 0.444831595 | 33.22137291 | NA | NA | NA | NA | NA | NA | NA |
| TRPV3 | 703 | F | -18.85370455 | 34.01704659 | 0.418739403 | 31.39782053 | NA | NA | NA | NA | NA | NA | NA |

| | | | | | | | | | | | | | |
|---|---|---|---|---|---|---|---|---|---|---|---|---|---|
| TRPV3 | 704 | E | -25.10547222 | 38.91914795 | 0.483693449 | 36.34411703 | NA | NA | NA | NA | NA | NA | NA |
| TRPV3 | 705 | K | -21.82708333 | 40.62350653 | 0.506276619 | 37.86331793 | NA | NA | NA | NA | NA | NA | NA |
| TRPV3 | 706 | M | -18.10875 | 40.69230628 | 0.507188232 | 38.06917779 | NA | NA | NA | NA | NA | NA | NA |
| TRPV3 | 707 | L | -21.169125 | 42.7468801 | 0.534411839 | 40.16213741 | NA | NA | NA | NA | NA | NA | NA |
| TRPV3 | 708 | P | -21.66710714 | 47.41994928 | 0.596331153 | 44.82384874 | NA | NA | NA | NA | NA | NA | NA |
| TRPV3 | 709 | E | -25.40205 | 49.92897251 | 0.629576327 | 47.42843162 | NA | NA | NA | NA | NA | NA | NA |
| TRPV3 | 710 | W | -23.06567857 | 50.76293142 | 0.640626487 | 48.27872354 | NA | NA | NA | NA | NA | NA | NA |
| TRPV3 | 711 | L | -23.311625 | 47.29509444 | 0.594676796 | 44.71186087 | NA | NA | NA | NA | NA | NA | NA |
| TRPV3 | 712 | R | -26.87388636 | 44.29595688 | 0.554937487 | 41.88923384 | NA | NA | NA | NA | NA | NA | NA |
| TRPV3 | 713 | S | -29.25825 | 48.66309449 | 0.612803132 | 44.54734159 | NA | NA | NA | NA | NA | NA | NA |
| TRPV3 | 714 | R | -27.17343182 | 47.35720145 | 0.595499729 | 43.25790823 | NA | NA | NA | NA | NA | NA | NA |
| TRPV3 | 715 | F | -26.51597727 | 43.27278594 | 0.541380221 | 39.17289057 | NA | NA | NA | NA | NA | NA | NA |
| TRPV3 | 716 | R | -33.04143182 | 43.9397179 | 0.550217233 | 36.92364977 | NA | NA | NA | NA | NA | NA | NA |
| TRPV3 | 717 | M | -33.09039286 | 39.15833206 | 0.486862697 | 29.80321018 | NA | NA | NA | NA | NA | NA | NA |
| TRPV3 | 718 | G | -36.88225 | 41.369923 | 0.51616682 | 30.94830321 | NA | NA | NA | NA | NA | NA | NA |
| TRPV3 | 719 | E | -40.65247222 | 42.61014364 | 0.532600048 | 28.07593036 | NA | NA | NA | NA | NA | NA | NA |
| TRPV3 | 720 | L | -42.8355 | 45.88342434 | 0.57597182 | 35.22500037 | NA | NA | NA | NA | NA | NA | NA |
| TRPV3 | 721 | C | -45.79758333 | 42.47013172 | 0.530744856 | 31.80579449 | NA | NA | NA | NA | NA | NA | NA |
| TRPV3 | 722 | K | -51.07569444 | 44.55568001 | 0.558378882 | 34.50745934 | NA | NA | NA | NA | NA | NA | NA |
| TRPV3 | 723 | V | -51.14910714 | 40.31726799 | 0.502218883 | 29.77375838 | NA | NA | NA | NA | NA | NA | NA |
| TRPV3 | 724 | A | -54.19725 | 42.04463058 | 0.525106861 | 30.99227097 | NA | NA | NA | NA | NA | NA | NA |
| TRPV3 | 725 | E | -55.66945 | 45.87809424 | 0.575901195 | 34.82088705 | NA | NA | NA | NA | NA | NA | NA |
| TRPV3 | 726 | D | -53.12745 | 48.41099621 | 0.609462768 | 37.72736645 | NA | NA | NA | NA | NA | NA | NA |
| TRPV3 | 727 | D | -50.429375 | 43.46135773 | 0.543878844 | 32.84700126 | NA | NA | NA | NA | NA | NA | NA |
| TRPV3 | 728 | F | -47.98825 | 46.53907417 | 0.584659342 | 36.50376728 | NA | NA | NA | NA | NA | NA | NA |
| TRPV3 | 729 | R | -46.54652273 | 39.0116601 | 0.484919258 | 29.07306825 | NA | NA | NA | NA | NA | NA | NA |
| TRPV3 | 730 | L | -41.024875 | 41.12590142 | 0.512933474 | 30.5606473 | NA | NA | NA | NA | NA | NA | NA |
| TRPV3 | 731 | C | -42.23208333 | 39.55479161 | 0.492115884 | 28.8845838 | NA | NA | NA | NA | NA | NA | NA |
| TRPV3 | 732 | L | -39.62175 | 34.76216393 | 0.428612391 | 20.23196214 | NA | NA | NA | NA | NA | NA | NA |
| TRPV3 | 733 | R | -42.10561364 | 35.90175971 | 0.443712314 | 21.370185 | NA | NA | NA | NA | NA | NA | NA |
| TRPV3 | 734 | I | -41.269 | 30.6879069 | 0.374627485 | 15.8919042 | NA | NA | NA | NA | NA | NA | NA |
| TRPV3 | 735 | N | -42.167625 | 31.63848059 | 0.38722282 | 16.84968917 | NA | NA | NA | NA | NA | NA | NA |
| TRPV3 | 736 | E | -44.78236111 | 26.32533901 | 0.31682239 | 11.56112409 | NA | NA | NA | NA | NA | NA | NA |

| | | | | | | | | | | | | | |
|---|---|---|---|---|---|---|---|---|---|---|---|---|---|
| TRPV3 | 737 | V | -45.26482143 | 26.6120083 | 0.320620828 | 11.82216998 | NA | NA | NA | NA | NA | NA | NA |
| TRPV3 | 738 | K | -48.65280556 | 25.19883223 | 0.301895898 | 10.49192264 | NA | NA | NA | NA | NA | NA | NA |
| TRPV3 | 739 | W | -48.60246429 | 25.07973909 | 0.300317885 | 9.649080252 | NA | NA | NA | NA | NA | NA | NA |
| TRPV3 | 740 | T | -51.37510714 | 20.86761723 | 0.244506237 | 5.439134673 | NA | NA | NA | NA | NA | NA | NA |
| TRPV3 | 741 | E | -53.59525 | 24.78730716 | 0.29644309 | 9.36042528 | NA | NA | NA | NA | NA | NA | NA |
| TRPV3 | 742 | W | -55.37396429 | 26.42668398 | 0.318165235 | 12.98132661 | NA | NA | NA | NA | NA | NA | NA |
| TRPV3 | 743 | K | -56.99191667 | 27.887763 | 0.337524891 | 14.43866506 | NA | NA | NA | NA | NA | NA | NA |
| TRPV3 | 744 | T | -60.73725 | 31.31460226 | 0.382931352 | 17.87665558 | NA | NA | NA | NA | NA | NA | NA |
| TRPV3 | 745 | H | -56.33245 | 31.93710646 | 0.391179686 | 18.67740105 | NA | NA | NA | NA | NA | NA | NA |
| TRPV3 | 746 | V | -60.30025 | 33.40660593 | 0.410650914 | 19.9662626 | NA | NA | NA | NA | NA | NA | NA |
| TRPV3 | 747 | S | -56.80475 | 33.65300608 | 0.413915777 | 20.21123062 | NA | NA | NA | NA | NA | NA | NA |
| TRPV3 | 748 | F | -56.32288636 | 38.14794801 | 0.473474861 | 25.11856844 | NA | NA | NA | NA | NA | NA | NA |
| TRPV3 | 749 | L | -52.45025 | 39.49231102 | 0.491288 | 26.60455888 | NA | NA | NA | NA | NA | NA | NA |
| TRPV3 | 750 | N | -52.049875 | 42.30540885 | 0.528562237 | 31.62294872 | NA | NA | NA | NA | NA | NA | NA |
| TRPV3 | 751 | E | -56.79180556 | 43.90413876 | 0.549745801 | 30.45438067 | NA | NA | NA | NA | NA | NA | NA |
| TRPV3 | 752 | D | -53.054375 | 44.48288176 | 0.557414288 | 29.93852257 | NA | NA | NA | NA | NA | NA | NA |
| TRPV3 | 753 | P | -50.44510714 | 40.85929441 | 0.509400866 | 26.3176649 | NA | NA | NA | NA | NA | NA | NA |
| TRPV3 | 754 | G | -47.50175 | 44.04020865 | 0.55154876 | 29.25133742 | NA | NA | NA | NA | NA | NA | NA |
| TRPV3 | 755 | P | -44.10510714 | 42.33332707 | 0.52893216 | 27.54419827 | NA | NA | NA | NA | NA | NA | NA |
| TRPV3 | 756 | V | -44.17625 | 45.78239134 | 0.574633108 | 30.99024174 | NA | NA | NA | NA | NA | NA | NA |
| SCN1A | 116 | P | -16.02146429 | 56.45688565 | 0.827115036 | 54.06549154 | NA | NA | NA | NA | NA | NA | NA |
| SCN1A | 117 | L | -14.006375 | 54.7186621 | 0.800986926 | 52.40197244 | NA | NA | NA | NA | NA | NA | NA |
| SCN1A | 118 | R | -17.65038636 | 51.9091376 | 0.758755555 | 49.49957267 | NA | NA | NA | NA | NA | NA | NA |
| SCN1A | 119 | K | -18.19419444 | 51.69629483 | 0.755556208 | 49.25682937 | NA | NA | NA | NA | NA | NA | NA |
| SCN1A | 120 | I | -13.490875 | 50.62356335 | 0.739431444 | 48.2292668 | NA | NA | NA | NA | NA | NA | NA |
| SCN1A | 121 | A | -12.98755 | 48.07065356 | 0.701057376 | 45.76045247 | NA | NA | NA | NA | NA | NA | NA |
| SCN1A | 122 | I | -16.51975 | 46.43075986 | 0.676407312 | 43.99277317 | NA | NA | NA | NA | NA | NA | NA |
| SCN1A | 123 | K | -17.22941667 | 46.27824838 | 0.674114835 | 43.83456394 | NA | NA | NA | NA | NA | NA | NA |
| SCN1A | 124 | I | -11.706625 | 44.65107696 | 0.649656006 | 42.26373484 | NA | NA | NA | NA | NA | NA | NA |
| SCN1A | 125 | L | -13.061125 | 41.86289051 | 0.607745377 | 39.55501147 | NA | NA | NA | NA | NA | NA | NA |
| SCN1A | 126 | V | -16.30589286 | 41.70193975 | 0.605326046 | 39.22439136 | NA | NA | NA | NA | NA | NA | NA |
| SCN1A | 127 | H | -14.54735 | 41.48308235 | 0.60203629 | 39.07553565 | NA | NA | NA | NA | NA | NA | NA |
| SCN1A | 128 | S | -12.94208333 | 39.37651183 | 0.570371373 | 36.91536945 | NA | NA | NA | NA | NA | NA | NA |

| | | | | | | | | | | | | | |
|---|---|---|---|---|---|---|---|---|---|---|---|---|---|
| SCN1A | 129 | L | -9.863375 | 41.83621992 | 0.607344478 | 39.52337462 | NA | NA | NA | NA | NA | NA | NA |
| SCN1A | 130 | F | -8.137931818 | 41.05924836 | 0.595665429 | 38.2848765 | NA | NA | NA | NA | NA | NA | NA |
| SCN1A | 131 | S | -9.028916667 | 37.79619321 | 0.546616811 | 35.01781747 | NA | NA | NA | NA | NA | NA | NA |
| SCN1A | 132 | M | -7.9075 | 35.9211449 | 0.51843202 | 33.40297292 | NA | NA | NA | NA | NA | NA | NA |
| SCN1A | 133 | L | -4.05275 | 40.24169068 | 0.58337631 | 37.45771564 | NA | NA | NA | NA | NA | NA | NA |
| SCN1A | 134 | I | -3.929875 | 37.71999403 | 0.545471423 | 34.94290388 | NA | NA | NA | NA | NA | NA | NA |
| SCN1A | 135 | M | -4.436875 | 33.34147917 | 0.47965577 | 30.55099334 | NA | NA | NA | NA | NA | NA | NA |
| SCN1A | 136 | C | -1.467416667 | 37.40748334 | 0.540773918 | 34.61779306 | NA | NA | NA | NA | NA | NA | NA |
| SCN1A | 137 | T | 0.26075 | 38.37563727 | 0.555326725 | 35.60021114 | NA | NA | NA | NA | NA | NA | NA |
| SCN1A | 138 | I | 0.713125 | 33.89992735 | 0.488050085 | 31.12881894 | NA | NA | NA | NA | NA | NA | NA |
| SCN1A | 139 | L | 0.9075 | 31.86425943 | 0.45745094 | 27.90429886 | NA | NA | NA | NA | NA | NA | NA |
| SCN1A | 140 | T | 3.704464286 | 36.46004417 | 0.526532485 | 32.11564036 | NA | NA | NA | NA | NA | NA | NA |
| SCN1A | 141 | N | 5.075875 | 35.35645532 | 0.509943887 | 30.98873935 | NA | NA | NA | NA | NA | NA | NA |
| SCN1A | 142 | C | 4.350083333 | 32.15920601 | 0.46188443 | 27.80507115 | NA | NA | NA | NA | NA | NA | NA |
| SCN1A | 143 | V | 6.753464286 | 33.48454638 | 0.481806285 | 29.46770376 | NA | NA | NA | NA | NA | NA | NA |
| SCN1A | 144 | F | 9.212113636 | 35.44426942 | 0.511263865 | 31.40201529 | NA | NA | NA | NA | NA | NA | NA |
| SCN1A | 145 | M | 8.4835 | 32.5612595 | 0.467927897 | 28.53921157 | NA | NA | NA | NA | NA | NA | NA |
| SCN1A | 146 | T | 10.66432143 | 30.61375812 | 0.438654028 | 26.55271888 | NA | NA | NA | NA | NA | NA | NA |
| SCN1A | 147 | M | 13.7485 | 33.70062934 | 0.485054337 | 30.75411114 | NA | NA | NA | NA | NA | NA | NA |
| SCN1A | 148 | S | 17.54691667 | 32.04739136 | 0.460203688 | 29.29179758 | NA | NA | NA | NA | NA | NA | NA |
| SCN1A | 149 | N | 19.8565 | 34.34125255 | 0.494683866 | 31.56386179 | NA | NA | NA | NA | NA | NA | NA |
| SCN1A | 150 | P | 16.77389286 | 38.00560609 | 0.549764601 | 35.26301055 | NA | NA | NA | NA | NA | NA | NA |
| SCN1A | 151 | P | 18.28889286 | 39.92728303 | 0.578650291 | 37.16058637 | NA | NA | NA | NA | NA | NA | NA |
| SCN1A | 152 | D | 19.17625 | 44.02086438 | 0.640182965 | 41.24706256 | NA | NA | NA | NA | NA | NA | NA |
| SCN1A | 153 | W | 15.70132143 | 42.26494337 | 0.613788835 | 39.49723297 | NA | NA | NA | NA | NA | NA | NA |
| SCN1A | 154 | T | 13.94517857 | 41.07478841 | 0.595899019 | 37.00186044 | NA | NA | NA | NA | NA | NA | NA |
| SCN1A | 155 | K | 14.46597222 | 44.74205453 | 0.651023535 | 40.66636667 | NA | NA | NA | NA | NA | NA | NA |
| SCN1A | 156 | N | 12.361875 | 46.99154027 | 0.684836683 | 42.94410811 | NA | NA | NA | NA | NA | NA | NA |
| SCN1A | 157 | V | 9.736607143 | 42.97062975 | 0.624396361 | 38.9315541 | NA | NA | NA | NA | NA | NA | NA |
| SCN1A | 158 | E | 9.366305556 | 42.32009284 | 0.614617815 | 37.93491607 | NA | NA | NA | NA | NA | NA | NA |
| SCN1A | 159 | Y | 10.67491667 | 45.8241179 | 0.667288572 | 41.44557795 | NA | NA | NA | NA | NA | NA | NA |
| SCN1A | 160 | T | 6.069464286 | 45.82535833 | 0.667307218 | 41.46281773 | NA | NA | NA | NA | NA | NA | NA |
| SCN1A | 161 | F | 4.993204545 | 40.53059568 | 0.587718986 | 36.16614425 | NA | NA | NA | NA | NA | NA | NA |

| | | | | | | | | | | | | | |
|---|---|---|---|---|---|---|---|---|---|---|---|---|---|
| SCN1A | 162 | T | 4.81775 | 44.55981322 | 0.648284175 | 40.18142379 | NA | NA | NA | NA | NA | NA | NA |
| SCN1A | 163 | G | 2.20425 | 47.04441256 | 0.685631433 | 44.28553896 | NA | NA | NA | NA | NA | NA | NA |
| SCN1A | 164 | I | 0.5955 | 43.79007747 | 0.636713891 | 41.01788006 | NA | NA | NA | NA | NA | NA | NA |
| SCN1A | 165 | Y | -0.511833333 | 38.31330739 | 0.554389813 | 35.5504038 | NA | NA | NA | NA | NA | NA | NA |
| SCN1A | 166 | T | -0.618035714 | 45.3981898 | 0.660886234 | 42.63864631 | NA | NA | NA | NA | NA | NA | NA |
| SCN1A | 167 | F | -3.269931818 | 46.61142744 | 0.679123017 | 43.85192835 | NA | NA | NA | NA | NA | NA | NA |
| SCN1A | 168 | E | -4.54175 | 40.46573428 | 0.586744021 | 37.71073573 | NA | NA | NA | NA | NA | NA | NA |
| SCN1A | 169 | S | -4.357083333 | 44.30745622 | 0.64449087 | 41.55602436 | NA | NA | NA | NA | NA | NA | NA |
| SCN1A | 170 | L | -4.70725 | 46.82524171 | 0.682336966 | 44.0720672 | NA | NA | NA | NA | NA | NA | NA |
| SCN1A | 171 | I | -8.150375 | 46.74407135 | 0.681116854 | 44.07643942 | NA | NA | NA | NA | NA | NA | NA |
| SCN1A | 172 | K | -9.323083333 | 40.69051065 | 0.590122748 | 38.3166025 | NA | NA | NA | NA | NA | NA | NA |
| SCN1A | 173 | I | -7.944625 | 46.61264872 | 0.679141375 | 43.87213518 | NA | NA | NA | NA | NA | NA | NA |
| SCN1A | 174 | I | -10.030125 | 49.13494257 | 0.717055238 | 46.7669951 | NA | NA | NA | NA | NA | NA | NA |
| SCN1A | 175 | A | -13.42575 | 46.43675404 | 0.676497413 | 44.11445886 | NA | NA | NA | NA | NA | NA | NA |
| SCN1A | 176 | R | -14.64929545 | 41.65479926 | 0.604617453 | 39.36856493 | NA | NA | NA | NA | NA | NA | NA |
| SCN1A | 177 | G | -12.35525 | 48.69244255 | 0.710403798 | 46.3496406 | NA | NA | NA | NA | NA | NA | NA |
| SCN1A | 178 | F | -15.49820455 | 50.55498565 | 0.738400618 | 48.26432885 | NA | NA | NA | NA | NA | NA | NA |
| SCN1A | 179 | C | -14.40675 | 54.97015171 | 0.804767193 | 52.63660102 | NA | NA | NA | NA | NA | NA | NA |
| SCN1A | 180 | L | -12.632 | 54.95119694 | 0.804482274 | 52.60450822 | NA | NA | NA | NA | NA | NA | NA |
| SCN1A | 181 | E | -16.24052778 | 50.90844363 | 0.743713622 | 48.56166092 | NA | NA | NA | NA | NA | NA | NA |
| SCN1A | 182 | D | -14.2195 | 50.9943815 | 0.745005397 | 48.60916716 | NA | NA | NA | NA | NA | NA | NA |
| SCN1A | 183 | F | -13.09584091 | 45.88024426 | 0.668132236 | 43.52854667 | NA | NA | NA | NA | NA | NA | NA |
| SCN1A | 184 | T | -9.898321429 | 44.98977303 | 0.654747116 | 42.60957119 | NA | NA | NA | NA | NA | NA | NA |
| SCN1A | 185 | F | -8.95375 | 41.24425462 | 0.598446351 | 39.09368459 | NA | NA | NA | NA | NA | NA | NA |
| SCN1A | 186 | L | -9.857125 | 40.49219829 | 0.587141815 | 38.10628553 | NA | NA | NA | NA | NA | NA | NA |
| SCN1A | 187 | R | -14.90329545 | 35.10553411 | 0.506172165 | 32.78546268 | NA | NA | NA | NA | NA | NA | NA |
| SCN1A | 188 | D | -10.897875 | 33.72692473 | 0.485449596 | 31.36548866 | NA | NA | NA | NA | NA | NA | NA |
| SCN1A | 189 | P | -8.969321429 | 32.07452213 | 0.460611504 | 29.3475235 | NA | NA | NA | NA | NA | NA | NA |
| SCN1A | 190 | W | -7.251178571 | 27.8098329 | 0.396506822 | 25.06235117 | NA | NA | NA | NA | NA | NA | NA |
| SCN1A | 191 | N | -7.619 | 35.8746421 | 0.517733013 | 33.13482692 | NA | NA | NA | NA | NA | NA | NA |
| SCN1A | 192 | W | -6.350392857 | 35.54510266 | 0.51277954 | 32.80722897 | NA | NA | NA | NA | NA | NA | NA |
| SCN1A | 193 | L | -1.943375 | 32.75406921 | 0.470826117 | 30.00344905 | NA | NA | NA | NA | NA | NA | NA |
| SCN1A | 194 | D | -2.496875 | 36.94731735 | 0.533856932 | 34.19914369 | NA | NA | NA | NA | NA | NA | NA |

| | | | | | | | | | | | | | |
|---|---|---|---|---|---|---|---|---|---|---|---|---|---|
| SCN1A | 195 | F | -1.705204545 | 39.65808465 | 0.574603835 | 36.91071654 | NA | NA | NA | NA | NA | NA | NA |
| SCN1A | 196 | T | 0.632464286 | 38.18800834 | 0.552506381 | 35.44168422 | NA | NA | NA | NA | NA | NA | NA |
| SCN1A | 197 | V | 1.591607143 | 36.99772613 | 0.534614652 | 34.24957216 | NA | NA | NA | NA | NA | NA | NA |
| SCN1A | 198 | I | 1.082125 | 41.43762298 | 0.601352968 | 38.68748492 | NA | NA | NA | NA | NA | NA | NA |
| SCN1A | 199 | T | 2.60875 | 43.5120129 | 0.632534163 | 40.76325489 | NA | NA | NA | NA | NA | NA | NA |
| SCN1A | 200 | F | 7.150477273 | 40.12282865 | 0.581589635 | 35.59673998 | NA | NA | NA | NA | NA | NA | NA |
| SCN1A | 201 | A | 6.54675 | 43.45586232 | 0.631690135 | 39.06663747 | NA | NA | NA | NA | NA | NA | NA |
| SCN1A | 202 | Y | 5.697833333 | 46.23022 | 0.673392897 | 41.84244899 | NA | NA | NA | NA | NA | NA | NA |
| SCN1A | 203 | V | 8.21475 | 46.37380339 | 0.675551171 | 41.84067916 | NA | NA | NA | NA | NA | NA | NA |
| SCN1A | 204 | T | 11.19875 | 43.82883568 | 0.637296485 | 39.29920614 | NA | NA | NA | NA | NA | NA | NA |
| SCN1A | 205 | E | 10.69797222 | 47.71601637 | 0.695726647 | 43.18778671 | NA | NA | NA | NA | NA | NA | NA |
| SCN1A | 206 | F | 10.88084091 | 49.75387713 | 0.726358754 | 45.22772963 | NA | NA | NA | NA | NA | NA | NA |
| SCN1A | 207 | V | 16.78960714 | 50.13041777 | 0.732018725 | 45.60904038 | NA | NA | NA | NA | NA | NA | NA |
| SCN1A | 208 | D | 17.58575 | 49.1167482 | 0.71678175 | 44.5934652 | NA | NA | NA | NA | NA | NA | NA |
| SCN1A | 209 | L | 14.194375 | 45.04866021 | 0.655632279 | 40.52629037 | NA | NA | NA | NA | NA | NA | NA |
| SCN1A | 210 | G | 18.93225 | 46.18449097 | 0.672705521 | 41.66604952 | NA | NA | NA | NA | NA | NA | NA |
| SCN1A | 211 | N | 21.292125 | 44.48088965 | 0.647097835 | 41.7427313 | NA | NA | NA | NA | NA | NA | NA |
| SCN1A | 212 | V | 21.78932143 | 41.22872568 | 0.598212928 | 38.49308274 | NA | NA | NA | NA | NA | NA | NA |
| SCN1A | 213 | S | 18.17041667 | 39.6073331 | 0.573840963 | 35.11411149 | NA | NA | NA | NA | NA | NA | NA |
| SCN1A | 214 | A | 15.32415 | 36.81760355 | 0.531907139 | 32.29721154 | NA | NA | NA | NA | NA | NA | NA |
| SCN1A | 215 | L | 12.473125 | 37.90305321 | 0.548223077 | 33.376713 | NA | NA | NA | NA | NA | NA | NA |
| SCN1A | 216 | R | 13.21075 | 35.78600981 | 0.516400736 | 31.26167707 | NA | NA | NA | NA | NA | NA | NA |
| SCN1A | 217 | T | 10.33346429 | 32.94363363 | 0.473675555 | 28.41968125 | NA | NA | NA | NA | NA | NA | NA |
| SCN1A | 218 | F | 7.785386364 | 35.09909926 | 0.506075439 | 30.57003769 | NA | NA | NA | NA | NA | NA | NA |
| SCN1A | 219 | R | 8.81475 | 35.23633649 | 0.508138321 | 30.74251676 | NA | NA | NA | NA | NA | NA | NA |
| SCN1A | 220 | V | 5.827321429 | 31.06387674 | 0.445419986 | 26.66982184 | NA | NA | NA | NA | NA | NA | NA |
| SCN1A | 221 | L | 2.490375 | 33.02966633 | 0.474968755 | 28.63997273 | NA | NA | NA | NA | NA | NA | NA |
| SCN1A | 222 | R | 3.39375 | 34.43897181 | 0.496152733 | 30.27563838 | NA | NA | NA | NA | NA | NA | NA |
| SCN1A | 223 | A | 1.39875 | 30.44161789 | 0.436066502 | 26.45699494 | NA | NA | NA | NA | NA | NA | NA |
| SCN1A | 224 | L | -0.766625 | 28.58684813 | 0.408186527 | 25.84012654 | NA | NA | NA | NA | NA | NA | NA |
| SCN1A | 225 | K | -3.089638889 | 31.92741783 | 0.458400305 | 29.16591915 | NA | NA | NA | NA | NA | NA | NA |
| SCN1A | 226 | T | -2.971607143 | 28.60000465 | 0.408384289 | 25.83400202 | NA | NA | NA | NA | NA | NA | NA |
| SCN1A | 227 | I | -5.684625 | 25.76382524 | 0.365752255 | 23.00215888 | NA | NA | NA | NA | NA | NA | NA |

| | | | | | | | | | | | | | |
|---|---|---|---|---|---|---|---|---|---|---|---|---|---|
| SCN1A | 228 | S | -7.597416667 | 30.27765676 | 0.43360192 | 27.51966612 | NA | NA | NA | NA | NA | NA | NA |
| SCN1A | 229 | V | -7.902178571 | 31.72173339 | 0.455308559 | 28.95759232 | NA | NA | NA | NA | NA | NA | NA |
| SCN1A | 230 | I | -7.76375 | 27.93003833 | 0.39831369 | 25.14743742 | NA | NA | NA | NA | NA | NA | NA |
| SCN1A | 231 | P | -11.81275 | 27.44872638 | 0.391078849 | 25.13483814 | NA | NA | NA | NA | NA | NA | NA |
| SCN1A | 232 | G | -12.2695 | 23.60261649 | 0.333266042 | 21.2672771 | NA | NA | NA | NA | NA | NA | NA |
| SCN1A | 233 | L | -9.092 | 23.6609754 | 0.334143264 | 21.30331805 | NA | NA | NA | NA | NA | NA | NA |
| SCN1A | 234 | K | -11.75841667 | 23.93799597 | 0.338307299 | 21.57229196 | NA | NA | NA | NA | NA | NA | NA |
| SCN1A | 235 | T | -14.18617857 | 20.75446036 | 0.290453979 | 18.42359713 | NA | NA | NA | NA | NA | NA | NA |
| SCN1A | 236 | I | -10.841 | 19.34083509 | 0.269205069 | 16.96083736 | NA | NA | NA | NA | NA | NA | NA |
| SCN1A | 237 | V | -8.800892857 | 20.78677354 | 0.290939695 | 18.05745361 | NA | NA | NA | NA | NA | NA | NA |
| SCN1A | 238 | G | -12.177 | 20.52685804 | 0.287032775 | 18.14378424 | NA | NA | NA | NA | NA | NA | NA |
| SCN1A | 239 | A | -12.83195 | 16.56506668 | 0.227481102 | 14.16461448 | NA | NA | NA | NA | NA | NA | NA |
| SCN1A | 240 | L | -8.866625 | 16.68224784 | 0.229242511 | 13.88617929 | NA | NA | NA | NA | NA | NA | NA |
| SCN1A | 241 | I | -9.57325 | 18.92672864 | 0.262980427 | 15.86976864 | NA | NA | NA | NA | NA | NA | NA |
| SCN1A | 242 | Q | -14.50919444 | 16.73537839 | 0.230041143 | 14.29282887 | NA | NA | NA | NA | NA | NA | NA |
| SCN1A | 243 | S | -11.07558333 | 13.9241836 | 0.187784664 | 10.87662737 | NA | NA | NA | NA | NA | NA | NA |
| SCN1A | 244 | V | -8.238178571 | 16.04355986 | 0.219642072 | 13.04308476 | NA | NA | NA | NA | NA | NA | NA |
| SCN1A | 245 | K | -11.89886111 | 17.76612715 | 0.245534844 | 14.69854622 | NA | NA | NA | NA | NA | NA | NA |
| SCN1A | 246 | K | -13.38486111 | 13.88317907 | 0.187168305 | 11.26492973 | NA | NA | NA | NA | NA | NA | NA |
| SCN1A | 247 | L | -7.848625 | 13.2044552 | 0.176966066 | 10.35643281 | NA | NA | NA | NA | NA | NA | NA |
| SCN1A | 248 | S | -8.85675 | 18.02070692 | 0.249361561 | 14.91875067 | NA | NA | NA | NA | NA | NA | NA |
| SCN1A | 249 | D | -9.51425 | 18.42557467 | 0.255447331 | 15.32738753 | NA | NA | NA | NA | NA | NA | NA |
| SCN1A | 250 | V | -6.734178571 | 14.23729237 | 0.192491159 | 11.37371914 | NA | NA | NA | NA | NA | NA | NA |
| SCN1A | 251 | M | -3.635125 | 16.80106369 | 0.231028491 | 13.99356041 | NA | NA | NA | NA | NA | NA | NA |
| SCN1A | 252 | I | -5.132 | 20.35502319 | 0.284449839 | 17.50437996 | NA | NA | NA | NA | NA | NA | NA |
| SCN1A | 253 | L | -4.549625 | 17.84044938 | 0.246652019 | 15.04155571 | NA | NA | NA | NA | NA | NA | NA |
| SCN1A | 254 | T | -0.981892857 | 16.98382495 | 0.233775668 | 14.1846899 | NA | NA | NA | NA | NA | NA | NA |
| SCN1A | 255 | V | 0.098464286 | 20.87067301 | 0.29220083 | 17.32027284 | NA | NA | NA | NA | NA | NA | NA |
| SCN1A | 256 | F | -1.242113636 | 23.0764463 | 0.325356914 | 19.97177636 | NA | NA | NA | NA | NA | NA | NA |
| SCN1A | 257 | C | 0.772083333 | 20.11961814 | 0.280911347 | 16.81804512 | NA | NA | NA | NA | NA | NA | NA |
| SCN1A | 258 | L | 3.255625 | 18.47978934 | 0.256262258 | 13.94324372 | NA | NA | NA | NA | NA | NA | NA |
| SCN1A | 259 | S | 3.48175 | 23.38342873 | 0.329971321 | 18.84448825 | NA | NA | NA | NA | NA | NA | NA |
| SCN1A | 260 | V | 2.797464286 | 25.02842071 | 0.35469802 | 20.48942857 | NA | NA | NA | NA | NA | NA | NA |

| | | | | | | | | | | | | | |
|---|---|---|---|---|---|---|---|---|---|---|---|---|---|
| SCN1A | 261 | F | 3.986113636 | 21.27209597 | 0.29823482 | 16.75626509 | NA | NA | NA | NA | NA | NA | NA |
| SCN1A | 262 | A | 8.02675 | 23.69143683 | 0.334601145 | 19.15950056 | NA | NA | NA | NA | NA | NA | NA |
| SCN1A | 263 | L | 7.605125 | 26.4239217 | 0.375674495 | 21.89332999 | NA | NA | NA | NA | NA | NA | NA |
| SCN1A | 264 | I | 7.787 | 25.92212406 | 0.368131724 | 21.38485192 | NA | NA | NA | NA | NA | NA | NA |
| SCN1A | 265 | G | 11.61275 | 25.00875729 | 0.35440245 | 24.60016226 | NA | NA | NA | NA | NA | NA | NA |
| SCN1A | 266 | L | 12.903125 | 26.81656438 | 0.381576504 | 26.41493679 | NA | NA | NA | NA | NA | NA | NA |
| SCN1A | 267 | Q | 12.99330556 | 29.45827466 | 0.421285376 | 29.05379176 | NA | NA | NA | NA | NA | NA | NA |
| SCN1A | 268 | L | 13.356 | 26.81084775 | 0.381490575 | 26.3952109 | NA | NA | NA | NA | NA | NA | NA |
| SCN1A | 269 | F | 15.20047727 | 23.54551078 | 0.332407658 | 24.37918007 | NA | NA | NA | NA | NA | NA | NA |
| SCN1A | 270 | M | 17.818875 | 28.75848866 | 0.410766542 | 29.4947049 | NA | NA | NA | NA | NA | NA | NA |
| SCN1A | 271 | G | 21.26075 | 27.72386954 | 0.395214663 | 28.03364176 | NA | NA | NA | NA | NA | NA | NA |
| SCN1A | 272 | N | 20.831125 | 24.78902759 | 0.351099582 | 25.09125811 | NA | NA | NA | NA | NA | NA | NA |
| SCN1A | 273 | L | 20.7745 | 20.94487322 | 0.29331617 | 21.38636727 | NA | NA | NA | NA | NA | NA | NA |
| SCN1A | 274 | R | 25.41502273 | 25.3833485 | 0.360033118 | 23.74016608 | NA | NA | NA | NA | NA | NA | NA |
| SCN1A | 275 | N | 25.825125 | 23.77007817 | 0.335783243 | 21.9331101 | NA | NA | NA | NA | NA | NA | NA |
| SCN1A | 276 | K | 29.66330556 | 24.49729299 | 0.346714373 | 20.81307107 | NA | NA | NA | NA | NA | NA | NA |
| SCN1A | 277 | C | 32.66808333 | 23.04222468 | 0.324842512 | 19.42717082 | NA | NA | NA | NA | NA | NA | NA |
| SCN1A | 278 | I | 36.073125 | 25.82336837 | 0.366647277 | 22.22068424 | NA | NA | NA | NA | NA | NA | NA |
| SCN1A | 279 | Q | 40.07386111 | 24.22187941 | 0.342574494 | 20.73245224 | NA | NA | NA | NA | NA | NA | NA |
| SCN1A | 280 | W | 41.32710714 | 28.66110685 | 0.409302747 | 24.89030053 | NA | NA | NA | NA | NA | NA | NA |
| SCN1A | 281 | P | 45.23875 | 31.5610502 | 0.45289325 | 27.77560195 | NA | NA | NA | NA | NA | NA | NA |
| SCN1A | 282 | P | 48.87975 | 31.87966363 | 0.457682488 | 30.70117335 | NA | NA | NA | NA | NA | NA | NA |
| SCN1A | 312 | F | 46.44965909 | 42.20368484 | 0.612868028 | 38.62046668 | NA | NA | NA | NA | NA | NA | NA |
| SCN1A | 313 | D | 43.7155 | 38.96355966 | 0.564164081 | 35.37357649 | NA | NA | NA | NA | NA | NA | NA |
| SCN1A | 314 | W | 37.64625 | 34.67550076 | 0.499708118 | 31.07523506 | NA | NA | NA | NA | NA | NA | NA |
| SCN1A | 315 | K | 35.15230556 | 35.67044735 | 0.514663659 | 32.02107252 | NA | NA | NA | NA | NA | NA | NA |
| SCN1A | 316 | S | 39.25125 | 33.03586095 | 0.47506187 | 29.4288967 | NA | NA | NA | NA | NA | NA | NA |
| SCN1A | 317 | Y | 40.75558333 | 29.86812508 | 0.427446044 | 26.38934718 | NA | NA | NA | NA | NA | NA | NA |
| SCN1A | 318 | I | 35.218375 | 30.84958655 | 0.442198883 | 27.20530698 | NA | NA | NA | NA | NA | NA | NA |
| SCN1A | 319 | Q | 34.82797222 | 28.45171666 | 0.406155298 | 24.82796527 | NA | NA | NA | NA | NA | NA | NA |
| SCN1A | 320 | D | 38.308125 | 26.01324272 | 0.369501374 | 22.39009001 | NA | NA | NA | NA | NA | NA | NA |
| SCN1A | 321 | S | 35.40325 | 22.99204117 | 0.324088178 | 19.3757398 | NA | NA | NA | NA | NA | NA | NA |
| SCN1A | 322 | R | 41.07138636 | 22.03247716 | 0.309664491 | 19.37334184 | NA | NA | NA | NA | NA | NA | NA |

| | | | | | | | | | | | | | |
|---|---|---|---|---|---|---|---|---|---|---|---|---|---|
| SCN1A | 323 | Y | 40.3535 | 24.51244594 | 0.346942145 | 20.92052878 | NA | NA | NA | NA | NA | NA | NA |
| SCN1A | 324 | H | 34.98095 | 20.93169979 | 0.293118154 | 17.37216267 | NA | NA | NA | NA | NA | NA | NA |
| SCN1A | 325 | Y | 34.27175 | 15.53726031 | 0.212031629 | 12.06316651 | NA | NA | NA | NA | NA | NA | NA |
| SCN1A | 326 | F | 33.71820455 | 14.78465062 | 0.200718776 | 11.25529652 | NA | NA | NA | NA | NA | NA | NA |
| SCN1A | 327 | L | 30.56275 | 11.1357975 | 0.145871035 | 7.548579267 | NA | NA | NA | NA | NA | NA | NA |
| SCN1A | 328 | E | 34.09908333 | 9.151790863 | 0.116048436 | 5.561806626 | NA | NA | NA | NA | NA | NA | NA |
| SCN1A | 329 | G | 31.43625 | 10.17359022 | 0.131407615 | 6.489462969 | NA | NA | NA | NA | NA | NA | NA |
| SCN1A | 330 | F | 28.03438636 | 9.502136654 | 0.121314659 | 5.805036127 | NA | NA | NA | NA | NA | NA | NA |
| SCN1A | 331 | L | 28.93575 | 14.52998369 | 0.196890749 | 10.8395083 | NA | NA | NA | NA | NA | NA | NA |
| SCN1A | 332 | D | 26.70625 | 16.94910369 | 0.233253755 | 13.33914387 | NA | NA | NA | NA | NA | NA | NA |
| SCN1A | 333 | A | 29.22475 | 18.58159444 | 0.257792542 | 14.89232117 | NA | NA | NA | NA | NA | NA | NA |
| SCN1A | 334 | L | 28.912 | 17.44035008 | 0.240637926 | 13.80003369 | NA | NA | NA | NA | NA | NA | NA |
| SCN1A | 335 | L | 25.757375 | 19.34625527 | 0.269286542 | 17.50702513 | NA | NA | NA | NA | NA | NA | NA |
| SCN1A | 336 | C | 27.30508333 | 22.63204111 | 0.318676837 | 20.79468944 | NA | NA | NA | NA | NA | NA | NA |
| SCN1A | 337 | G | 26.6975 | 25.10166036 | 0.355798922 | 23.31210869 | NA | NA | NA | NA | NA | NA | NA |
| SCN1A | 338 | N | 27.4995 | 28.14697323 | 0.401574547 | 28.35099099 | NA | NA | NA | NA | NA | NA | NA |
| SCN1A | 339 | S | 23.74575 | 26.98632806 | 0.384128307 | 27.32385607 | NA | NA | NA | NA | NA | NA | NA |
| SCN1A | 340 | S | 23.09058333 | 26.44419963 | 0.375979303 | 26.96893739 | NA | NA | NA | NA | NA | NA | NA |
| SCN1A | 341 | D | 20.27975 | 22.41235733 | 0.31537466 | 23.03783426 | NA | NA | NA | NA | NA | NA | NA |
| SCN1A | 342 | A | 23.25075 | 22.56353698 | 0.317647117 | 22.85971523 | NA | NA | NA | NA | NA | NA | NA |
| SCN1A | 343 | G | 25.7 | 21.52662598 | 0.302060788 | 19.80782893 | NA | NA | NA | NA | NA | NA | NA |
| SCN1A | 344 | Q | 28.62375 | 22.00377154 | 0.309233002 | 20.46553478 | NA | NA | NA | NA | NA | NA | NA |
| SCN1A | 345 | C | 32.05708333 | 22.77125495 | 0.32076943 | 19.23682537 | NA | NA | NA | NA | NA | NA | NA |
| SCN1A | 346 | P | 35.11532143 | 20.62869026 | 0.288563466 | 16.90663605 | NA | NA | NA | NA | NA | NA | NA |
| SCN1A | 347 | E | 39.61897222 | 22.56260229 | 0.317633067 | 18.78880461 | NA | NA | NA | NA | NA | NA | NA |
| SCN1A | 348 | G | 39.14 | 26.24449293 | 0.372977412 | 22.47856928 | NA | NA | NA | NA | NA | NA | NA |
| SCN1A | 349 | Y | 37.1015 | 21.79766451 | 0.306134904 | 18.01821185 | NA | NA | NA | NA | NA | NA | NA |
| SCN1A | 350 | M | 34.99075 | 27.34436251 | 0.389510103 | 23.72799193 | NA | NA | NA | NA | NA | NA | NA |
| SCN1A | 351 | C | 30.73758333 | 27.60902805 | 0.393488423 | 24.44736829 | NA | NA | NA | NA | NA | NA | NA |
| SCN1A | 352 | V | 31.75375 | 28.87659719 | 0.412541891 | 25.20684608 | NA | NA | NA | NA | NA | NA | NA |
| SCN1A | 353 | K | 27.24797222 | 29.87965694 | 0.427619385 | 28.04559277 | NA | NA | NA | NA | NA | NA | NA |
| SCN1A | 354 | A | 30.52175 | 28.073663 | 0.400472584 | 24.39390819 | NA | NA | NA | NA | NA | NA | NA |
| SCN1A | 355 | G | 28.8035 | 28.11036967 | 0.401024341 | 24.61935341 | NA | NA | NA | NA | NA | NA | NA |

| | | | | | | | | | | | | | |
|---|---|---|---|---|---|---|---|---|---|---|---|---|---|
| SCN1A | 356 | R | 27.16629545 | 21.7428706 | 0.305311269 | 20.07726947 | NA | NA | NA | NA | NA | NA | NA |
| SCN1A | 357 | N | 23.473875 | 22.34153193 | 0.314310048 | 20.56765116 | NA | NA | NA | NA | NA | NA | NA |
| SCN1A | 358 | P | 22.36475 | 19.67869513 | 0.274283613 | 17.83953069 | NA | NA | NA | NA | NA | NA | NA |
| SCN1A | 359 | N | 21.75975 | 16.95703011 | 0.233372901 | 14.93832962 | NA | NA | NA | NA | NA | NA | NA |
| SCN1A | 360 | Y | 23.85466667 | 19.05810919 | 0.264955274 | 17.23447896 | NA | NA | NA | NA | NA | NA | NA |
| SCN1A | 361 | G | 22.10275 | 23.30705975 | 0.328823381 | 21.2888474 | NA | NA | NA | NA | NA | NA | NA |
| SCN1A | 362 | Y | 19.37241667 | 22.91621046 | 0.322948329 | 23.32370178 | NA | NA | NA | NA | NA | NA | NA |
| SCN1A | 363 | T | 18.59189286 | 19.60714411 | 0.273208093 | 19.91881793 | NA | NA | NA | NA | NA | NA | NA |
| SCN1A | 364 | S | 17.75691667 | 23.32885359 | 0.329150975 | 23.7970467 | NA | NA | NA | NA | NA | NA | NA |
| SCN1A | 365 | F | 14.87893182 | 20.94472294 | 0.293313911 | 20.62063611 | NA | NA | NA | NA | NA | NA | NA |
| SCN1A | 366 | D | 17.7515 | 27.67003424 | 0.394405438 | 27.33705339 | NA | NA | NA | NA | NA | NA | NA |
| SCN1A | 367 | T | 15.87689286 | 28.7232509 | 0.410236866 | 28.341689 | NA | NA | NA | NA | NA | NA | NA |
| SCN1A | 368 | F | 9.809659091 | 26.52263452 | 0.377158297 | 22.00508061 | NA | NA | NA | NA | NA | NA | NA |
| SCN1A | 369 | S | 12.06358333 | 24.94209434 | 0.353400405 | 20.42360766 | NA | NA | NA | NA | NA | NA | NA |
| SCN1A | 370 | W | 15.63110714 | 23.02277753 | 0.324550192 | 22.64103065 | NA | NA | NA | NA | NA | NA | NA |
| SCN1A | 371 | A | 11.35595 | 22.04016227 | 0.309780009 | 19.23029659 | NA | NA | NA | NA | NA | NA | NA |
| SCN1A | 372 | F | 6.993931818 | 20.45635874 | 0.285973064 | 15.9328629 | NA | NA | NA | NA | NA | NA | NA |
| SCN1A | 373 | L | 11.362 | 18.73063686 | 0.260032873 | 14.33922179 | NA | NA | NA | NA | NA | NA | NA |
| SCN1A | 374 | S | 11.71458333 | 17.25467177 | 0.237846902 | 16.87280937 | NA | NA | NA | NA | NA | NA | NA |
| SCN1A | 375 | L | 7.974625 | 16.50991121 | 0.226652033 | 11.98404442 | NA | NA | NA | NA | NA | NA | NA |
| SCN1A | 376 | F | 5.817931818 | 14.49458716 | 0.196358686 | 9.971205871 | NA | NA | NA | NA | NA | NA | NA |
| SCN1A | 377 | R | 11.45920455 | 12.64267788 | 0.168521709 | 9.926522928 | NA | NA | NA | NA | NA | NA | NA |
| SCN1A | 378 | L | 8.175125 | 12.43019686 | 0.1653278 | 9.576741684 | NA | NA | NA | NA | NA | NA | NA |
| SCN1A | 379 | M | 2.95075 | 10.73405431 | 0.139832231 | 6.197125282 | NA | NA | NA | NA | NA | NA | NA |
| SCN1A | 380 | T | 5.114321429 | 8.408523414 | 0.10487601 | 3.896258747 | 4.54 | 0.44 | 0.12 | 10.65 | -0.49 | 1 | 1 |
| SCN1A | 381 | Q | 5.696194444 | 7.100386116 | 0.085212743 | 4.28412318 | 2.96 | -0.21 | -0.58 | 13.73 | 0.39 | 1 | 1 |
| SCN1A | 382 | D | 9.98 | 9.678222616 | 0.123961496 | 9.302317245 | NA | NA | NA | NA | NA | NA | NA |
| SCN1A | 383 | F | 14.81720455 | 9.352166273 | 0.119060379 | 9.506666628 | NA | NA | NA | NA | NA | NA | NA |
| SCN1A | 384 | W | 10.17953571 | 8.297582961 | 0.103208409 | 7.998296229 | NA | NA | NA | NA | NA | NA | NA |
| SCN1A | 385 | E | 14.25763889 | 6.970065279 | 0.083253825 | 7.515811173 | -0.01 | -1.04 | -3.5 | 49.7 | 2.63 | 1 | -1 |
| SCN1A | 386 | N | 17.58775 | 11.34617513 | 0.149033326 | 12.18268301 | NA | NA | NA | NA | NA | NA | NA |
| SCN1A | 387 | L | 14.851875 | 14.18922186 | 0.191768587 | 14.78103243 | NA | NA | NA | NA | NA | NA | NA |
| SCN1A | 388 | Y | 14.027 | 12.81321353 | 0.171085116 | 13.48354483 | NA | NA | NA | NA | NA | NA | NA |

| | | | | | | | | | | | | | |
|---|---|---|---|---|---|---|---|---|---|---|---|---|---|
| SCN1A | 389 | Q | 18.26875 | 9.469600942 | 0.120825599 | 10.07406245 | NA | NA | NA | NA | NA | NA | NA |
| SCN1A | 390 | L | 20.290875 | 15.47830818 | 0.21114549 | 15.98479504 | NA | NA | NA | NA | NA | NA | NA |
| SCN1A | 391 | T | 17.29646429 | 17.91034511 | 0.247702657 | 18.54736756 | NA | NA | NA | NA | NA | NA | NA |
| SCN1A | 392 | L | 17.165 | 17.31434296 | 0.23874385 | 18.40433621 | NA | NA | NA | NA | NA | NA | NA |
| SCN1A | 393 | R | 23.52020455 | 15.80447111 | 0.216048209 | 16.17971962 | NA | NA | NA | NA | NA | NA | NA |
| SCN1A | 394 | A | 21.91455 | 20.97508579 | 0.293770311 | 21.38224638 | NA | NA | NA | NA | NA | NA | NA |
| SCN1A | 395 | A | 19.13295 | 22.97495388 | 0.323831331 | 23.55695131 | NA | NA | NA | NA | NA | NA | NA |
| SCN1A | 396 | G | 18.34425 | 22.07754074 | 0.310341864 | 22.97868003 | NA | NA | NA | NA | NA | NA | NA |
| SCN1A | 397 | K | 17.73163889 | 19.67429391 | 0.274217456 | 20.43114132 | NA | NA | NA | NA | NA | NA | NA |
| SCN1A | 398 | T | 14.31275 | 23.02257433 | 0.324547138 | 23.83989998 | NA | NA | NA | NA | NA | NA | NA |
| SCN1A | 399 | Y | 14.11483333 | 21.49002608 | 0.301510636 | 22.2494215 | NA | NA | NA | NA | NA | NA | NA |
| SCN1A | 400 | M | 10.860625 | 18.17706362 | 0.251711836 | 17.8648701 | NA | NA | NA | NA | NA | NA | NA |
| SCN1A | 401 | I | 8.244625 | 20.81175259 | 0.291315167 | 20.32809715 | NA | NA | NA | NA | NA | NA | NA |
| SCN1A | 402 | F | 8.717113636 | 19.57930167 | 0.27278958 | 19.11983711 | NA | NA | NA | NA | NA | NA | NA |
| SCN1A | 403 | F | 10.32484091 | 16.18910269 | 0.221829799 | 15.71810668 | NA | NA | NA | NA | NA | NA | NA |
| SCN1A | 404 | V | 5.52275 | 16.85121628 | 0.23178236 | 16.41852906 | NA | NA | NA | NA | NA | NA | NA |
| SCN1A | 405 | L | 3.485625 | 17.75422032 | 0.245355867 | 13.37117026 | NA | NA | NA | NA | NA | NA | NA |
| SCN1A | 406 | V | 5.029178571 | 15.58550923 | 0.212756883 | 11.11846072 | NA | NA | NA | NA | NA | NA | NA |
| SCN1A | 407 | I | 4.288 | 11.75064449 | 0.155113108 | 7.816705898 | NA | NA | NA | NA | NA | NA | NA |
| SCN1A | 408 | F | 0.844386364 | 14.33279599 | 0.193926722 | 10.95251805 | NA | NA | NA | NA | NA | NA | NA |
| SCN1A | 409 | L | -0.043875 | 14.69360929 | 0.199350288 | 11.11461052 | NA | NA | NA | NA | NA | NA | NA |
| SCN1A | 410 | G | 0.312 | 12.68097197 | 0.169097327 | 9.195277288 | NA | NA | NA | NA | NA | NA | NA |
| SCN1A | 411 | S | 1.714416667 | 9.176176164 | 0.116414984 | 4.721763433 | 4.59 | 0.44 | 0.12 | 10.65 | -0.49 | 1 | 1 |
| SCN1A | 412 | F | -2.09575 | 8.592760965 | 0.107645377 | 5.383550186 | NA | NA | NA | NA | NA | NA | NA |
| SCN1A | 413 | Y | -3.9775 | 11.61710399 | 0.153105794 | 8.530883438 | NA | NA | NA | NA | NA | NA | NA |
| SCN1A | 414 | L | -1.952625 | 11.58435639 | 0.152613548 | 8.482908083 | NA | NA | NA | NA | NA | NA | NA |
| SCN1A | 415 | I | -2.092 | 7.171790502 | 0.086286058 | 4.274743155 | 3.15 | 1.35 | 2.33 | 0.62 | -1.69 | 0 | 0 |
| SCN1A | 416 | N | -5.741125 | 6.97109353 | 0.083269281 | 3.863065745 | 3.14 | 1.35 | 2.33 | 0.62 | -1.69 | 0 | 0 |
| SCN1A | 417 | L | -7.719625 | 10.03909782 | 0.129385992 | 7.118934636 | NA | NA | NA | NA | NA | NA | NA |
| SCN1A | 418 | I | -5.390125 | 8.928738657 | 0.112695626 | 6.093351212 | NA | NA | NA | NA | NA | NA | NA |
| SCN1A | 419 | L | -6.80325 | 4.618720819 | 0.047909587 | 2.096244961 | 3.01 | 1.41 | 4.08 | 0.13 | -1.81 | 0 | 0 |
| SCN1A | 420 | A | -10.98535 | 6.665429469 | 0.078674691 | 3.555007842 | 3.13 | 1.09 | 3.34 | 0.15 | -1.33 | 0 | 0 |
| SCN1A | 421 | V | -11.30789286 | 8.721054638 | 0.109573824 | 5.612019846 | NA | NA | NA | NA | NA | NA | NA |

| | | | | | | | | | | | | | |
|---|---|---|---|---|---|---|---|---|---|---|---|---|---|
| SCN1A | 422 | V | -10.00075 | 8.248483255 | 0.102470366 | 5.189742182 | NA | NA | NA | NA | NA | NA | NA |
| SCN1A | 423 | A | -12.97255 | 4.729764265 | 0.049578737 | 1.692115816 | 3.13 | 1.09 | 3.34 | 0.15 | -1.33 | 0 | 0 |
| SCN1A | 424 | M | -16.39675 | 7.345948067 | 0.088903907 | 4.851764581 | 2.64 | 1.36 | 3.73 | 0.19 | -1.81 | 0 | 0 |
| SCN1A | 425 | A | -15.22755 | 9.769365742 | 0.125331514 | 7.340931768 | NA | NA | NA | NA | NA | NA | NA |
| SCN1A | 426 | Y | -13.75366667 | 8.555949801 | 0.10709205 | 6.203516339 | NA | NA | NA | NA | NA | NA | NA |
| SCN1A | 427 | E | -18.31808333 | 4.346178206 | 0.043812863 | 1.695025409 | 2.73 | 0.79 | 1.9 | 12.63 | -0.81 | 1 | -1 |
| SCN1A | 428 | E | -20.09397222 | 9.983449805 | 0.128549519 | 7.329084657 | NA | NA | NA | NA | NA | NA | NA |
| SCN1A | 429 | Q | -17.58352778 | 12.17919131 | 0.16155481 | 9.541406506 | NA | NA | NA | NA | NA | NA | NA |
| SCN1A | 430 | N | -19.296625 | 11.37628437 | 0.149485913 | 8.728158281 | NA | NA | NA | NA | NA | NA | NA |
| SCN1A | 431 | Q | -23.50086111 | 8.648541958 | 0.108483849 | 1.710925859 | 6.97 | -0.95 | -2.8 | 14.62 | 0.92 | 1 | -1 |
| SCN1A | 432 | A | -23.94235 | 14.05636514 | 0.189771551 | 7.120785949 | NA | NA | NA | NA | NA | NA | NA |
| SCN1A | 433 | T | -22.70946429 | 15.67261717 | 0.214066246 | 13.02270786 | NA | NA | NA | NA | NA | NA | NA |
| SCN1A | 434 | L | -25.962125 | 12.59720405 | 0.167838169 | 5.644658272 | NA | NA | NA | NA | NA | NA | NA |
| SCN1A | 435 | E | -28.61452778 | 14.59718254 | 0.197900849 | 7.637410311 | NA | NA | NA | NA | NA | NA | NA |
| SCN1A | 436 | E | -27.66608333 | 17.53032458 | 0.241990378 | 10.57304161 | NA | NA | NA | NA | NA | NA | NA |
| SCN1A | 437 | A | -28.22075 | 19.2185309 | 0.267366653 | 12.26183115 | NA | NA | NA | NA | NA | NA | NA |
| SCN1A | 438 | E | -31.25797222 | 13.75380471 | 0.185223614 | 6.848000937 | 6.97 | -0.95 | -2.8 | 14.62 | 0.92 | 1 | -1 |
| SCN1A | 439 | Q | -32.56208333 | 18.86414339 | 0.262039677 | 6.068596279 | NA | NA | NA | NA | NA | NA | NA |
| SCN1A | 749 | D | -16.596875 | 61.74851309 | 0.906656142 | 59.07102825 | NA | NA | NA | NA | NA | NA | NA |
| SCN1A | 750 | C | -14.62258333 | 58.60283985 | 0.85937195 | 55.92149546 | NA | NA | NA | NA | NA | NA | NA |
| SCN1A | 751 | S | -17.84191667 | 56.76153418 | 0.831694361 | 54.08253273 | NA | NA | NA | NA | NA | NA | NA |
| SCN1A | 752 | P | -19.56032143 | 58.73667585 | 0.861383706 | 56.10231676 | NA | NA | NA | NA | NA | NA | NA |
| SCN1A | 753 | Y | -17.185 | 58.26646647 | 0.854315753 | 55.61061406 | NA | NA | NA | NA | NA | NA | NA |
| SCN1A | 754 | W | -15.79203571 | 51.35913661 | 0.750488214 | 48.67722266 | NA | NA | NA | NA | NA | NA | NA |
| SCN1A | 755 | L | -19.894375 | 54.29519453 | 0.794621573 | 51.65146551 | NA | NA | NA | NA | NA | NA | NA |
| SCN1A | 756 | K | -19.73975 | 56.89903975 | 0.833761276 | 54.29169492 | NA | NA | NA | NA | NA | NA | NA |
| SCN1A | 757 | V | -15.28403571 | 54.43713706 | 0.796755182 | 51.80277837 | NA | NA | NA | NA | NA | NA | NA |
| SCN1A | 758 | K | -17.60708333 | 51.13020846 | 0.74704708 | 48.49571535 | NA | NA | NA | NA | NA | NA | NA |
| SCN1A | 759 | H | -21.23315 | 52.13878988 | 0.762207573 | 49.57921291 | NA | NA | NA | NA | NA | NA | NA |
| SCN1A | 760 | V | -17.75632143 | 51.76750597 | 0.756626619 | 49.19527769 | NA | NA | NA | NA | NA | NA | NA |
| SCN1A | 761 | V | -15.27560714 | 48.62691494 | 0.70941882 | 46.02412843 | NA | NA | NA | NA | NA | NA | NA |
| SCN1A | 762 | N | -19.10775 | 46.66113444 | 0.679870188 | 44.10757558 | NA | NA | NA | NA | NA | NA | NA |
| SCN1A | 763 | L | -20.9475 | 49.44904729 | 0.721776704 | 46.90137105 | NA | NA | NA | NA | NA | NA | NA |

| | | | | | | | | | | | | | |
|---|---|---|---|---|---|---|---|---|---|---|---|---|---|
| SCN1A | 764 | V | -16.49117857 | 46.7227254 | 0.680795992 | 44.18270875 | NA | NA | NA | NA | NA | NA | NA |
| SCN1A | 765 | V | -15.57560714 | 43.23503144 | 0.628370716 | 40.62666883 | NA | NA | NA | NA | NA | NA | NA |
| SCN1A | 766 | M | -19.6165 | 39.25940804 | 0.568611127 | 36.77946182 | NA | NA | NA | NA | NA | NA | NA |
| SCN1A | 767 | D | -19.91875 | 42.77292038 | 0.621424493 | 39.33532148 | NA | NA | NA | NA | NA | NA | NA |
| SCN1A | 768 | P | -16.70189286 | 41.72882138 | 0.605730117 | 38.55403614 | NA | NA | NA | NA | NA | NA | NA |
| SCN1A | 769 | F | -13.91793182 | 41.38609625 | 0.600578444 | 38.87116898 | NA | NA | NA | NA | NA | NA | NA |
| SCN1A | 770 | V | -12.28332143 | 40.95434664 | 0.594088599 | 38.32295902 | NA | NA | NA | NA | NA | NA | NA |
| SCN1A | 771 | D | -12.67225 | 36.69966428 | 0.530134335 | 34.08061484 | NA | NA | NA | NA | NA | NA | NA |
| SCN1A | 772 | L | -11.534875 | 36.29475506 | 0.524047941 | 33.73754142 | NA | NA | NA | NA | NA | NA | NA |
| SCN1A | 773 | A | -8.28795 | 38.6678402 | 0.559718974 | 35.39571035 | NA | NA | NA | NA | NA | NA | NA |
| SCN1A | 774 | I | -7.6505 | 36.20404073 | 0.522684369 | 32.74129071 | NA | NA | NA | NA | NA | NA | NA |
| SCN1A | 775 | T | -7.999464286 | 33.15495622 | 0.476852051 | 29.64022848 | NA | NA | NA | NA | NA | NA | NA |
| SCN1A | 776 | I | -5.767375 | 34.62186691 | 0.498901921 | 33.45007916 | NA | NA | NA | NA | NA | NA | NA |
| SCN1A | 777 | C | -2.965083333 | 34.35712948 | 0.49492252 | 33.19450419 | NA | NA | NA | NA | NA | NA | NA |
| SCN1A | 778 | I | -3.3675 | 31.04900876 | 0.445196498 | 29.88439303 | NA | NA | NA | NA | NA | NA | NA |
| SCN1A | 779 | V | -2.416321429 | 30.0492942 | 0.430169287 | 28.88238202 | NA | NA | NA | NA | NA | NA | NA |
| SCN1A | 780 | L | -0.511125 | 31.73966745 | 0.455578135 | 30.5834408 | NA | NA | NA | NA | NA | NA | NA |
| SCN1A | 781 | N | 1.5 | 30.03992179 | 0.430028406 | 28.89006641 | NA | NA | NA | NA | NA | NA | NA |
| SCN1A | 782 | T | 1.136464286 | 26.25819678 | 0.373183401 | 25.09601315 | NA | NA | NA | NA | NA | NA | NA |
| SCN1A | 783 | L | 2.548875 | 27.59799922 | 0.393322643 | 26.4276053 | NA | NA | NA | NA | NA | NA | NA |
| SCN1A | 784 | F | 5.985659091 | 29.38103247 | 0.42012431 | 28.81737828 | NA | NA | NA | NA | NA | NA | NA |
| SCN1A | 785 | M | 5.53675 | 25.94275643 | 0.368441859 | 25.3807317 | NA | NA | NA | NA | NA | NA | NA |
| SCN1A | 786 | A | 6.82915 | 24.14372318 | 0.341399688 | 23.58004946 | NA | NA | NA | NA | NA | NA | NA |
| SCN1A | 787 | M | 8.6845 | 26.32684284 | 0.374215254 | 25.76233944 | NA | NA | NA | NA | NA | NA | NA |
| SCN1A | 788 | E | 12.04141667 | 23.48539621 | 0.331504045 | 20.68403954 | NA | NA | NA | NA | NA | NA | NA |
| SCN1A | 789 | H | 15.23495 | 22.03034859 | 0.309632495 | 19.30139877 | NA | NA | NA | NA | NA | NA | NA |
| SCN1A | 790 | Y | 20.80883333 | 18.56117779 | 0.257485649 | 15.97984014 | NA | NA | NA | NA | NA | NA | NA |
| SCN1A | 791 | P | 20.45117857 | 20.45695258 | 0.285981991 | 17.46188275 | NA | NA | NA | NA | NA | NA | NA |
| SCN1A | 792 | M | 17.883 | 24.59128872 | 0.34812727 | 21.95849454 | NA | NA | NA | NA | NA | NA | NA |
| SCN1A | 793 | T | 19.92146429 | 28.08001141 | 0.40056801 | 25.05041079 | NA | NA | NA | NA | NA | NA | NA |
| SCN1A | 794 | D | 19.92625 | 30.89750809 | 0.442919216 | 27.8305736 | NA | NA | NA | NA | NA | NA | NA |
| SCN1A | 795 | H | 18.55825 | 33.00032365 | 0.474527691 | 30.37787667 | NA | NA | NA | NA | NA | NA | NA |
| SCN1A | 796 | F | 14.03193182 | 29.27117377 | 0.418472969 | 26.52965066 | NA | NA | NA | NA | NA | NA | NA |

| | | | | | | | | | | | | | |
|---|---|---|---|---|---|---|---|---|---|---|---|---|---|
| SCN1A | 797 | N | 15.543 | 29.39255814 | 0.420297558 | 26.7109691 | NA | NA | NA | NA | NA | NA | NA |
| SCN1A | 798 | N | 14.827875 | 33.84726135 | 0.487258436 | 31.16693248 | NA | NA | NA | NA | NA | NA | NA |
| SCN1A | 799 | V | 10.90017857 | 34.95449309 | 0.503901791 | 32.20855893 | NA | NA | NA | NA | NA | NA | NA |
| SCN1A | 800 | L | 10.644 | 30.48973982 | 0.436789846 | 27.70717618 | NA | NA | NA | NA | NA | NA | NA |
| SCN1A | 801 | T | 11.06017857 | 34.31762591 | 0.494328722 | 31.57209429 | NA | NA | NA | NA | NA | NA | NA |
| SCN1A | 802 | V | 8.78975 | 37.06479958 | 0.535622867 | 34.31674981 | NA | NA | NA | NA | NA | NA | NA |
| SCN1A | 803 | G | 5.83225 | 36.66254871 | 0.529576432 | 36.10131257 | NA | NA | NA | NA | NA | NA | NA |
| SCN1A | 804 | N | 6.34875 | 32.64177903 | 0.469138227 | 32.07520732 | NA | NA | NA | NA | NA | NA | NA |
| SCN1A | 805 | L | 6.38775 | 37.93733128 | 0.548738328 | 37.3729233 | NA | NA | NA | NA | NA | NA | NA |
| SCN1A | 806 | V | 2.893892857 | 39.90395093 | 0.578299574 | 38.8455265 | NA | NA | NA | NA | NA | NA | NA |
| SCN1A | 807 | F | 0.831931818 | 35.35776024 | 0.509963502 | 34.1934502 | NA | NA | NA | NA | NA | NA | NA |
| SCN1A | 808 | T | 1.71875 | 37.29589421 | 0.539096566 | 36.13824744 | NA | NA | NA | NA | NA | NA | NA |
| SCN1A | 809 | G | 0.583 | 41.02090352 | 0.595089049 | 39.86256768 | NA | NA | NA | NA | NA | NA | NA |
| SCN1A | 810 | I | -2.211625 | 40.42934208 | 0.586196992 | 39.27408785 | NA | NA | NA | NA | NA | NA | NA |
| SCN1A | 811 | F | -2.74975 | 36.12400579 | 0.521481323 | 34.96142049 | NA | NA | NA | NA | NA | NA | NA |
| SCN1A | 812 | T | -2.00575 | 41.61681539 | 0.604046499 | 40.46282655 | NA | NA | NA | NA | NA | NA | NA |
| SCN1A | 813 | A | -4.35015 | 44.31000314 | 0.644529154 | 43.15531274 | NA | NA | NA | NA | NA | NA | NA |
| SCN1A | 814 | E | -7.83575 | 40.89362564 | 0.593175871 | 37.40655545 | NA | NA | NA | NA | NA | NA | NA |
| SCN1A | 815 | M | -7.0235 | 40.50905602 | 0.587395212 | 37.03458144 | NA | NA | NA | NA | NA | NA | NA |
| SCN1A | 816 | F | -5.335931818 | 45.56124403 | 0.663337184 | 41.54412704 | NA | NA | NA | NA | NA | NA | NA |
| SCN1A | 817 | L | -9.358 | 46.98251202 | 0.684700975 | 44.29566989 | NA | NA | NA | NA | NA | NA | NA |
| SCN1A | 818 | K | -12.00875 | 39.11972196 | 0.566511435 | 36.43067233 | NA | NA | NA | NA | NA | NA | NA |
| SCN1A | 819 | I | -11.162875 | 44.83661651 | 0.652444944 | 42.12903535 | NA | NA | NA | NA | NA | NA | NA |
| SCN1A | 820 | I | -13.14675 | 47.11402884 | 0.686677871 | 44.42754618 | NA | NA | NA | NA | NA | NA | NA |
| SCN1A | 821 | A | -17.08675 | 45.62852097 | 0.664348457 | 43.01798872 | NA | NA | NA | NA | NA | NA | NA |
| SCN1A | 822 | M | -17.247125 | 41.290566 | 0.599142481 | 38.64444421 | NA | NA | NA | NA | NA | NA | NA |
| SCN1A | 823 | D | -17.603 | 44.6531514 | 0.649687188 | 41.97257655 | NA | NA | NA | NA | NA | NA | NA |
| SCN1A | 824 | P | -13.92146429 | 45.2226677 | 0.658247873 | 42.5289118 | NA | NA | NA | NA | NA | NA | NA |
| SCN1A | 825 | Y | -16.46883333 | 42.10815466 | 0.611432066 | 39.42235012 | NA | NA | NA | NA | NA | NA | NA |
| SCN1A | 826 | Y | -20.04666667 | 41.94216635 | 0.608937012 | 39.28293641 | NA | NA | NA | NA | NA | NA | NA |
| SCN1A | 827 | Y | -13.38683333 | 38.95643939 | 0.564057053 | 36.26630372 | NA | NA | NA | NA | NA | NA | NA |
| SCN1A | 828 | F | -12.00402273 | 37.78431852 | 0.546438317 | 35.0910399 | NA | NA | NA | NA | NA | NA | NA |
| SCN1A | 829 | Q | -17.32141667 | 34.38943021 | 0.495408048 | 31.71471908 | NA | NA | NA | NA | NA | NA | NA |

| | | | | | | | | | | | | | |
|---|---|---|---|---|---|---|---|---|---|---|---|---|---|
| SCN1A | 830 | E | -16.56452778 | 31.70219429 | 0.455014858 | 29.02369917 | NA | NA | NA | NA | NA | NA | NA |
| SCN1A | 831 | G | -12.6395 | 32.18866523 | 0.462327246 | 29.48923973 | NA | NA | NA | NA | NA | NA | NA |
| SCN1A | 832 | W | -11.30132143 | 27.64247704 | 0.393991212 | 24.9316766 | NA | NA | NA | NA | NA | NA | NA |
| SCN1A | 833 | N | -11.8025 | 34.71184814 | 0.500254474 | 32.00680021 | NA | NA | NA | NA | NA | NA | NA |
| SCN1A | 834 | I | -9.147 | 33.46688992 | 0.481540882 | 30.82185266 | NA | NA | NA | NA | NA | NA | NA |
| SCN1A | 835 | F | -5.944113636 | 30.23265794 | 0.43292552 | 27.58912027 | NA | NA | NA | NA | NA | NA | NA |
| SCN1A | 836 | D | -6.495 | 35.05338938 | 0.505388351 | 31.97336255 | NA | NA | NA | NA | NA | NA | NA |
| SCN1A | 837 | G | -4.864 | 37.70051948 | 0.545178691 | 33.7789657 | NA | NA | NA | NA | NA | NA | NA |
| SCN1A | 838 | F | -1.898295455 | 32.72349031 | 0.47036647 | 28.71191424 | NA | NA | NA | NA | NA | NA | NA |
| SCN1A | 839 | I | -1.337 | 31.83489661 | 0.457009573 | 27.81893899 | NA | NA | NA | NA | NA | NA | NA |
| SCN1A | 840 | V | -1.534607143 | 37.23603874 | 0.538196848 | 33.21720296 | NA | NA | NA | NA | NA | NA | NA |
| SCN1A | 841 | T | -0.89675 | 38.7263414 | 0.560598335 | 34.71111971 | NA | NA | NA | NA | NA | NA | NA |
| SCN1A | 842 | L | 2.1765 | 33.97470437 | 0.489174096 | 32.82811344 | NA | NA | NA | NA | NA | NA | NA |
| SCN1A | 843 | S | 2.623916667 | 35.52262461 | 0.512441661 | 34.36735183 | NA | NA | NA | NA | NA | NA | NA |
| SCN1A | 844 | L | 2.41725 | 39.33521314 | 0.569750591 | 38.17978116 | NA | NA | NA | NA | NA | NA | NA |
| SCN1A | 845 | V | 5.025178571 | 39.68159326 | 0.574957205 | 38.52468741 | NA | NA | NA | NA | NA | NA | NA |
| SCN1A | 846 | E | 7.612416667 | 34.87932749 | 0.50277194 | 34.30758443 | NA | NA | NA | NA | NA | NA | NA |
| SCN1A | 847 | L | 7.886375 | 38.75659659 | 0.561053116 | 35.99093138 | NA | NA | NA | NA | NA | NA | NA |
| SCN1A | 848 | G | 10.1725 | 40.76582295 | 0.591254805 | 37.98749882 | NA | NA | NA | NA | NA | NA | NA |
| SCN1A | 849 | L | 12.213625 | 36.64322849 | 0.52928602 | 33.88740012 | NA | NA | NA | NA | NA | NA | NA |
| SCN1A | 850 | A | 15.29015 | 36.13607392 | 0.521662725 | 33.45867561 | NA | NA | NA | NA | NA | NA | NA |
| SCN1A | 851 | N | 16.848625 | 34.03801866 | 0.490125805 | 30.92631137 | NA | NA | NA | NA | NA | NA | NA |
| SCN1A | 852 | V | 14.98017857 | 31.25499618 | 0.448292798 | 28.56318869 | NA | NA | NA | NA | NA | NA | NA |
| SCN1A | 853 | E | 18.68452778 | 28.36476367 | 0.404848264 | 25.21689226 | NA | NA | NA | NA | NA | NA | NA |
| SCN1A | 854 | G | 16.5525 | 26.99009835 | 0.384184981 | 24.32164135 | NA | NA | NA | NA | NA | NA | NA |
| SCN1A | 855 | L | 12.97925 | 29.16195907 | 0.416831308 | 26.38115612 | NA | NA | NA | NA | NA | NA | NA |
| SCN1A | 856 | S | 12.83591667 | 28.51598836 | 0.407121398 | 25.76778995 | NA | NA | NA | NA | NA | NA | NA |
| SCN1A | 857 | V | 9.525892857 | 26.90110325 | 0.382847251 | 24.10026583 | NA | NA | NA | NA | NA | NA | NA |
| SCN1A | 858 | L | 7.683625 | 30.36381928 | 0.434897072 | 29.78598036 | NA | NA | NA | NA | NA | NA | NA |
| SCN1A | 859 | R | 9.923295455 | 29.53441303 | 0.42242985 | 28.96690612 | NA | NA | NA | NA | NA | NA | NA |
| SCN1A | 860 | S | 6.231916667 | 25.91359601 | 0.368003534 | 24.86862148 | NA | NA | NA | NA | NA | NA | NA |
| SCN1A | 861 | F | 3.510204545 | 29.27938461 | 0.41859639 | 28.1237167 | NA | NA | NA | NA | NA | NA | NA |
| SCN1A | 862 | R | 3.614477273 | 30.09400937 | 0.430841424 | 28.94579471 | NA | NA | NA | NA | NA | NA | NA |

| | | | | | | | | | | | | | |
|---|---|---|---|---|---|---|---|---|---|---|---|---|---|
| SCN1A | 863 | L | 1.920125 | 24.07932096 | 0.340431626 | 22.93393581 | NA | NA | NA | NA | NA | NA | NA |
| SCN1A | 864 | L | -0.73275 | 27.39915008 | 0.390333642 | 23.38445249 | NA | NA | NA | NA | NA | NA | NA |
| SCN1A | 865 | R | -1.47375 | 29.85818518 | 0.427296632 | 25.83469155 | NA | NA | NA | NA | NA | NA | NA |
| SCN1A | 866 | V | -2.666035714 | 26.57702079 | 0.377975805 | 22.58083019 | NA | NA | NA | NA | NA | NA | NA |
| SCN1A | 867 | F | -4.691295455 | 23.04017747 | 0.324811739 | 19.62982177 | NA | NA | NA | NA | NA | NA | NA |
| SCN1A | 868 | K | -7.099194444 | 29.35469709 | 0.41972845 | 25.84701032 | NA | NA | NA | NA | NA | NA | NA |
| SCN1A | 869 | L | -7.94625 | 27.61439337 | 0.393569072 | 24.47556356 | NA | NA | NA | NA | NA | NA | NA |
| SCN1A | 870 | A | -11.21775 | 25.73568971 | 0.365329335 | 23.07946682 | NA | NA | NA | NA | NA | NA | NA |
| SCN1A | 871 | K | -13.58386111 | 27.59420057 | 0.393265544 | 25.03866469 | NA | NA | NA | NA | NA | NA | NA |
| SCN1A | 872 | S | -13.96125 | 25.9940669 | 0.369213132 | 23.46621521 | NA | NA | NA | NA | NA | NA | NA |
| SCN1A | 873 | W | -9.538535714 | 22.76911505 | 0.320737264 | 20.87825895 | NA | NA | NA | NA | NA | NA | NA |
| SCN1A | 874 | P | -14.15875 | 20.0275121 | 0.279526855 | 17.42418896 | NA | NA | NA | NA | NA | NA | NA |
| SCN1A | 875 | T | -11.29303571 | 17.91202817 | 0.247727956 | 15.45430563 | NA | NA | NA | NA | NA | NA | NA |
| SCN1A | 876 | L | -8.371875 | 20.83482964 | 0.29166205 | 17.92776008 | NA | NA | NA | NA | NA | NA | NA |
| SCN1A | 877 | N | -12.994 | 20.03027182 | 0.279568338 | 17.32637051 | NA | NA | NA | NA | NA | NA | NA |
| SCN1A | 878 | M | -13.094375 | 16.26700252 | 0.22300075 | 13.70843274 | NA | NA | NA | NA | NA | NA | NA |
| SCN1A | 879 | L | -7.617 | 16.75453962 | 0.230329165 | 14.10216992 | NA | NA | NA | NA | NA | NA | NA |
| SCN1A | 880 | I | -9.153125 | 19.79282852 | 0.275999209 | 16.71016646 | NA | NA | NA | NA | NA | NA | NA |
| SCN1A | 881 | K | -13.56797222 | 16.60794355 | 0.228125606 | 13.93791855 | NA | NA | NA | NA | NA | NA | NA |
| SCN1A | 882 | I | -10.018125 | 13.25630552 | 0.177745454 | 10.61258242 | NA | NA | NA | NA | NA | NA | NA |
| SCN1A | 883 | I | -6.327375 | 17.62283167 | 0.243380898 | 14.62681369 | NA | NA | NA | NA | NA | NA | NA |
| SCN1A | 884 | G | -9.53525 | 19.80621385 | 0.27620041 | 16.86072049 | NA | NA | NA | NA | NA | NA | NA |
| SCN1A | 885 | N | -11.9085 | 14.99235769 | 0.203840925 | 12.35729234 | NA | NA | NA | NA | NA | NA | NA |
| SCN1A | 886 | S | -7.56875 | 17.57641431 | 0.242683175 | 14.68739451 | NA | NA | NA | NA | NA | NA | NA |
| SCN1A | 887 | V | -7.701892857 | 20.0752124 | 0.280243863 | 17.19525263 | NA | NA | NA | NA | NA | NA | NA |
| SCN1A | 888 | G | -11.64675 | 20.71650796 | 0.289883498 | 18.18279759 | NA | NA | NA | NA | NA | NA | NA |
| SCN1A | 889 | A | -13.89055 | 18.2999826 | 0.253559493 | 15.66104877 | NA | NA | NA | NA | NA | NA | NA |
| SCN1A | 890 | L | -11.01075 | 15.07457243 | 0.205076736 | 12.75198597 | NA | NA | NA | NA | NA | NA | NA |
| SCN1A | 891 | G | -8.63 | 20.09877524 | 0.280598047 | 17.35008149 | NA | NA | NA | NA | NA | NA | NA |
| SCN1A | 892 | N | -9.657125 | 20.78988179 | 0.290986417 | 17.9991473 | NA | NA | NA | NA | NA | NA | NA |
| SCN1A | 893 | L | -8.035875 | 15.25295778 | 0.207758135 | 12.46923784 | NA | NA | NA | NA | NA | NA | NA |
| SCN1A | 894 | T | -4.686321429 | 18.41422146 | 0.255276675 | 15.60967843 | NA | NA | NA | NA | NA | NA | NA |
| SCN1A | 895 | L | -4.305125 | 20.98258521 | 0.293883038 | 18.17966377 | NA | NA | NA | NA | NA | NA | NA |

| | | | | | | | | | | | | | |
|---|---|---|---|---|---|---|---|---|---|---|---|---|---|
| SCN1A | 896 | V | -4.525464286 | 20.48959575 | 0.286472666 | 17.72568705 | NA | NA | NA | NA | NA | NA | NA |
| SCN1A | 897 | L | -1.339375 | 16.51830806 | 0.22677825 | 13.79217253 | NA | NA | NA | NA | NA | NA | NA |
| SCN1A | 898 | A | -0.24395 | 21.11466204 | 0.295868351 | 16.97301442 | NA | NA | NA | NA | NA | NA | NA |
| SCN1A | 899 | I | -0.538 | 23.57770381 | 0.332891567 | 19.34930456 | NA | NA | NA | NA | NA | NA | NA |
| SCN1A | 900 | I | 0.746625 | 20.39964764 | 0.285120611 | 16.24222458 | NA | NA | NA | NA | NA | NA | NA |
| SCN1A | 901 | V | 3.32775 | 18.70669808 | 0.259673038 | 14.34817446 | NA | NA | NA | NA | NA | NA | NA |
| SCN1A | 902 | F | 4.776204545 | 23.18656531 | 0.327012168 | 18.82069486 | NA | NA | NA | NA | NA | NA | NA |
| SCN1A | 903 | I | 3.364375 | 25.1828772 | 0.357019733 | 20.8094311 | NA | NA | NA | NA | NA | NA | NA |
| SCN1A | 904 | F | 4.812568182 | 20.84440724 | 0.291806016 | 16.35073635 | NA | NA | NA | NA | NA | NA | NA |
| SCN1A | 905 | A | 8.81155 | 23.60279265 | 0.33326869 | 19.54010284 | NA | NA | NA | NA | NA | NA | NA |
| SCN1A | 906 | V | 8.899892857 | 26.277094 | 0.373467455 | 21.76246126 | NA | NA | NA | NA | NA | NA | NA |
| SCN1A | 907 | V | 8.83175 | 27.01106272 | 0.384500106 | 22.48474873 | NA | NA | NA | NA | NA | NA | NA |
| SCN1A | 908 | G | 12.17525 | 26.04044965 | 0.369910335 | 21.52515199 | NA | NA | NA | NA | NA | NA | NA |
| SCN1A | 909 | M | 13.948375 | 27.35095686 | 0.389609226 | 24.39844694 | NA | NA | NA | NA | NA | NA | NA |
| SCN1A | 910 | Q | 13.56652778 | 30.10576334 | 0.431018103 | 27.15142524 | NA | NA | NA | NA | NA | NA | NA |
| SCN1A | 911 | L | 13.3995 | 28.76366522 | 0.410844354 | 24.24231563 | NA | NA | NA | NA | NA | NA | NA |
| SCN1A | 912 | F | 15.37847727 | 25.88256374 | 0.367537073 | 22.94331648 | NA | NA | NA | NA | NA | NA | NA |
| SCN1A | 913 | G | 18.8025 | 28.8440397 | 0.412052503 | 26.09997283 | NA | NA | NA | NA | NA | NA | NA |
| SCN1A | 914 | K | 21.06675 | 30.54208477 | 0.43757667 | 27.76821237 | NA | NA | NA | NA | NA | NA | NA |
| SCN1A | 915 | S | 21.80025 | 28.13230024 | 0.40135399 | 25.358956 | NA | NA | NA | NA | NA | NA | NA |
| SCN1A | 916 | Y | 20.43175 | 22.85555237 | 0.322036547 | 20.15274707 | NA | NA | NA | NA | NA | NA | NA |
| SCN1A | 917 | K | 26.02319444 | 26.28994028 | 0.373660554 | 23.70179192 | NA | NA | NA | NA | NA | NA | NA |
| SCN1A | 918 | D | 26.80875 | 29.63969286 | 0.424012364 | 26.84172268 | NA | NA | NA | NA | NA | NA | NA |
| SCN1A | 919 | C | 26.73458333 | 26.05054554 | 0.370062091 | 23.24267602 | NA | NA | NA | NA | NA | NA | NA |
| SCN1A | 920 | V | 27.87946429 | 23.22268434 | 0.327555092 | 19.6761516 | NA | NA | NA | NA | NA | NA | NA |
| SCN1A | 921 | C | 30.95891667 | 23.42689843 | 0.330624736 | 19.90946675 | NA | NA | NA | NA | NA | NA | NA |
| SCN1A | 922 | K | 27.59363889 | 22.86331013 | 0.322153158 | 19.31807543 | NA | NA | NA | NA | NA | NA | NA |
| SCN1A | 923 | I | 25.33375 | 20.30232543 | 0.283657712 | 17.48090469 | NA | NA | NA | NA | NA | NA | NA |
| SCN1A | 924 | A | 29.18255 | 18.53263095 | 0.257056547 | 15.00584272 | NA | NA | NA | NA | NA | NA | NA |
| SCN1A | 925 | S | 32.44908333 | 15.85932313 | 0.216872717 | 12.17347222 | NA | NA | NA | NA | NA | NA | NA |
| SCN1A | 926 | D | 32.571 | 17.01642295 | 0.234265665 | 13.33233089 | NA | NA | NA | NA | NA | NA | NA |
| SCN1A | 927 | C | 32.04308333 | 20.84370984 | 0.291795533 | 17.15439083 | NA | NA | NA | NA | NA | NA | NA |
| SCN1A | 928 | Q | 29.68397222 | 19.6101629 | 0.27325347 | 16.06190266 | NA | NA | NA | NA | NA | NA | NA |

| | | | | | | | | | | | | | |
|---|---|---|---|---|---|---|---|---|---|---|---|---|---|
| SCN1A | 929 | L | 24.8495 | 19.17494514 | 0.266711494 | 16.37407955 | NA | NA | NA | NA | NA | NA | NA |
| SCN1A | 930 | P | 23.09303571 | 17.37909957 | 0.239717239 | 14.58621139 | NA | NA | NA | NA | NA | NA | NA |
| SCN1A | 931 | R | 21.59665909 | 12.90168427 | 0.172414964 | 10.11444711 | NA | NA | NA | NA | NA | NA | NA |
| SCN1A | 932 | W | 16.44025 | 13.66036559 | 0.183819083 | 10.99449782 | NA | NA | NA | NA | NA | NA | NA |
| SCN1A | 933 | H | 19.16075 | 19.95501686 | 0.278437143 | 17.24742394 | NA | NA | NA | NA | NA | NA | NA |
| SCN1A | 934 | M | 15.531375 | 22.67774543 | 0.319363842 | 20.85050342 | NA | NA | NA | NA | NA | NA | NA |
| SCN1A | 935 | N | 19.860875 | 25.53574869 | 0.362323922 | 22.80788187 | NA | NA | NA | NA | NA | NA | NA |
| SCN1A | 936 | D | 17.02075 | 25.00909361 | 0.354407505 | 22.26243384 | NA | NA | NA | NA | NA | NA | NA |
| SCN1A | 937 | F | 11.04347727 | 25.01120799 | 0.354439287 | 22.08848976 | NA | NA | NA | NA | NA | NA | NA |
| SCN1A | 938 | F | 13.19593182 | 22.2794151 | 0.313376339 | 22.01968002 | NA | NA | NA | NA | NA | NA | NA |
| SCN1A | 939 | H | 16.42325 | 20.11580078 | 0.280853967 | 17.36860889 | NA | NA | NA | NA | NA | NA | NA |
| SCN1A | 940 | S | 12.28025 | 21.60239295 | 0.303199679 | 18.66180261 | NA | NA | NA | NA | NA | NA | NA |
| SCN1A | 941 | F | 8.668022727 | 19.0762208 | 0.265227519 | 16.12238371 | NA | NA | NA | NA | NA | NA | NA |
| SCN1A | 942 | L | 11.796125 | 16.76645302 | 0.230508241 | 16.20701109 | NA | NA | NA | NA | NA | NA | NA |
| SCN1A | 943 | I | 12.222875 | 16.58197787 | 0.227735303 | 16.18313574 | NA | NA | NA | NA | NA | NA | NA |
| SCN1A | 944 | V | 7.59575 | 15.61530288 | 0.213204726 | 12.75292158 | NA | NA | NA | NA | NA | NA | NA |
| SCN1A | 945 | F | 6.465022727 | 13.01367089 | 0.174098291 | 10.06834155 | NA | NA | NA | NA | NA | NA | NA |
| SCN1A | 946 | R | 11.61747727 | 11.65498799 | 0.153675247 | 8.753930405 | NA | NA | NA | NA | NA | NA | NA |
| SCN1A | 947 | V | 7.421464286 | 12.02820386 | 0.159285242 | 9.274250058 | NA | NA | NA | NA | NA | NA | NA |
| SCN1A | 948 | L | 3.6085 | 9.887660896 | 0.127109668 | 5.395327186 | NA | NA | NA | NA | NA | NA | NA |
| SCN1A | 949 | C | 5.586916667 | 7.271573282 | 0.087785943 | 4.314323956 | 2.96 | -0.21 | -0.58 | 13.73 | 0.39 | 1 | 1 |
| SCN1A | 950 | G | 6.96775 | 9.060631932 | 0.11467818 | 6.108514228 | NA | NA | NA | NA | NA | NA | NA |
| SCN1A | 951 | E | 11.01063889 | 8.211794018 | 0.101918872 | 8.070162302 | NA | NA | NA | NA | NA | NA | NA |
| SCN1A | 952 | W | 9.229607143 | 12.62007076 | 0.16818189 | 9.735118265 | NA | NA | NA | NA | NA | NA | NA |
| SCN1A | 953 | I | 13.46125 | 13.84006376 | 0.186520217 | 13.5013798 | NA | NA | NA | NA | NA | NA | NA |
| SCN1A | 954 | E | 15.66530556 | 11.21887659 | 0.147119838 | 11.83960044 | NA | NA | NA | NA | NA | NA | NA |
| SCN1A | 955 | T | 15.58346429 | 17.00273143 | 0.23405986 | 15.33725533 | NA | NA | NA | NA | NA | NA | NA |
| SCN1A | 956 | M | 14.085 | 18.75105354 | 0.260339767 | 18.34082875 | NA | NA | NA | NA | NA | NA | NA |
| SCN1A | 957 | W | 17.95775 | 15.95897042 | 0.218370565 | 13.96758548 | NA | NA | NA | NA | NA | NA | NA |
| SCN1A | 958 | D | 20.4375 | 17.73405935 | 0.245052817 | 15.01470813 | NA | NA | NA | NA | NA | NA | NA |
| SCN1A | 959 | C | 17.39208333 | 22.4431654 | 0.315837751 | 20.62627336 | NA | NA | NA | NA | NA | NA | NA |
| SCN1A | 960 | M | 17.711625 | 23.01564133 | 0.324442924 | 22.57749103 | NA | NA | NA | NA | NA | NA | NA |
| SCN1A | 961 | E | 23.17686111 | 23.89431426 | 0.337650698 | 21.18112738 | NA | NA | NA | NA | NA | NA | NA |

| | | | | | | | | | | | | | |
|---|---|---|---|---|---|---|---|---|---|---|---|---|---|
| SCN1A | 962 | V | 22.06717857 | 25.65596098 | 0.364130893 | 22.94244247 | NA | NA | NA | NA | NA | NA | NA |
| SCN1A | 963 | A | 18.92495 | 28.14214222 | 0.40150193 | 25.45939188 | NA | NA | NA | NA | NA | NA | NA |
| SCN1A | 964 | G | 17.98025 | 28.30940639 | 0.404016161 | 27.8920754 | NA | NA | NA | NA | NA | NA | NA |
| SCN1A | 965 | Q | 16.91130556 | 26.92421821 | 0.383194703 | 24.04773003 | NA | NA | NA | NA | NA | NA | NA |
| SCN1A | 966 | A | 13.09835 | 27.51631823 | 0.392094855 | 22.99663066 | NA | NA | NA | NA | NA | NA | NA |
| SCN1A | 967 | M | 13.6035 | 26.64724087 | 0.379031318 | 22.12936969 | NA | NA | NA | NA | NA | NA | NA |
| SCN1A | 968 | C | 13.39841667 | 23.27673712 | 0.328367586 | 18.76178698 | NA | NA | NA | NA | NA | NA | NA |
| SCN1A | 969 | L | 11.089125 | 22.60893222 | 0.318329476 | 18.08585468 | NA | NA | NA | NA | NA | NA | NA |
| SCN1A | 970 | T | 7.994178571 | 23.32926808 | 0.329157205 | 18.79911206 | NA | NA | NA | NA | NA | NA | NA |
| SCN1A | 971 | V | 8.700178571 | 21.56934413 | 0.302702906 | 17.04259352 | NA | NA | NA | NA | NA | NA | NA |
| SCN1A | 972 | F | 10.20575 | 18.42346903 | 0.25541568 | 13.89811314 | NA | NA | NA | NA | NA | NA | NA |
| SCN1A | 973 | M | 6.170625 | 18.78738486 | 0.260885881 | 14.2613341 | NA | NA | NA | NA | NA | NA | NA |
| SCN1A | 974 | M | 3.476125 | 19.65784815 | 0.273970251 | 15.11954015 | NA | NA | NA | NA | NA | NA | NA |
| SCN1A | 975 | V | 4.763464286 | 16.98686448 | 0.233821356 | 12.4496346 | NA | NA | NA | NA | NA | NA | NA |
| SCN1A | 976 | M | 4.157 | 13.93943729 | 0.18801395 | 9.410383302 | NA | NA | NA | NA | NA | NA | NA |
| SCN1A | 977 | V | 0.573607143 | 15.98624599 | 0.218780558 | 11.78464793 | NA | NA | NA | NA | NA | NA | NA |
| SCN1A | 978 | I | -0.222625 | 16.54553574 | 0.227187523 | 13.74748325 | NA | NA | NA | NA | NA | NA | NA |
| SCN1A | 979 | G | 0 | 14.46131688 | 0.195858584 | 10.28901707 | NA | NA | NA | NA | NA | NA | NA |
| SCN1A | 980 | N | -0.892625 | 10.49323477 | 0.136212352 | 7.177990964 | NA | NA | NA | NA | NA | NA | NA |
| SCN1A | 981 | L | -4.04425 | 12.2165947 | 0.162117039 | 9.471549199 | NA | NA | NA | NA | NA | NA | NA |
| SCN1A | 982 | V | -3.63375 | 13.00665687 | 0.17399286 | 10.24706879 | NA | NA | NA | NA | NA | NA | NA |
| SCN1A | 983 | V | -2.288178571 | 10.3671933 | 0.13431776 | 7.527988987 | NA | NA | NA | NA | NA | NA | NA |
| SCN1A | 984 | L | -5.643625 | 6.251097024 | 0.072446652 | 3.512468383 | 3.01 | 1.41 | 4.08 | 0.13 | -1.81 | 0 | 0 |
| SCN1A | 985 | N | -8.216625 | 10.20919835 | 0.131942858 | 7.451417287 | NA | NA | NA | NA | NA | NA | NA |
| SCN1A | 986 | L | -6.6185 | 10.66611387 | 0.138810984 | 7.856314119 | NA | NA | NA | NA | NA | NA | NA |
| SCN1A | 987 | F | -6.67675 | 5.291471062 | 0.058022033 | 2.485556061 | 3.01 | 1.41 | 4.08 | 0.13 | -1.81 | 0 | 0 |
| SCN1A | 988 | L | -10.503125 | 4.619210863 | 0.047916953 | 1.593864811 | 3.14 | 1.48 | 4.15 | 0.13 | -1.82 | 0 | 0 |
| SCN1A | 989 | A | -11.84815 | 9.561776665 | 0.122211138 | 7.116603301 | NA | NA | NA | NA | NA | NA | NA |
| SCN1A | 990 | L | -10.22375 | 10.01802406 | 0.129069222 | 7.597502611 | NA | NA | NA | NA | NA | NA | NA |
| SCN1A | 991 | L | -12.2565 | 4.33811353 | 0.043691638 | 1.751780549 | 2.64 | 1.36 | 3.73 | 0.19 | -1.81 | 0 | 0 |
| SCN1A | 992 | L | -15.65175 | 4.251455868 | 0.042389044 | 1.77417607 | 2.73 | 0.79 | 1.9 | 12.63 | -0.81 | 1 | -1 |
| SCN1A | 993 | S | -16.36575 | 9.261039143 | 0.117690602 | 6.59090939 | NA | NA | NA | NA | NA | NA | NA |
| SCN1A | 994 | S | -15.83308333 | 10.44075677 | 0.135423529 | 7.745425189 | NA | NA | NA | NA | NA | NA | NA |

| | | | | | | | | | | | | | |
|---|---|---|---|---|---|---|---|---|---|---|---|---|---|
| SCN1A | 995 | F | -17.18284091 | 4.908825623 | 0.052270298 | 2.375505147 | 2.73 | 0.79 | 1.9 | 12.63 | -0.81 | 1 | -1 |
| SCN1A | 996 | S | -20.90375 | 9.707519611 | 0.124401874 | 7.2521638 | NA | NA | NA | NA | NA | NA | NA |
| SCN1A | 1201 | G | -21.09075 | 57.50641488 | 0.842891036 | 54.15376456 | NA | NA | NA | NA | NA | NA | NA |
| SCN1A | 1202 | K | -21.42619444 | 59.02181998 | 0.86566985 | 55.65914287 | NA | NA | NA | NA | NA | NA | NA |
| SCN1A | 1203 | Q | -16.11663889 | 56.70903386 | 0.830905202 | 53.34251578 | NA | NA | NA | NA | NA | NA | NA |
| SCN1A | 1204 | W | -15.59232143 | 53.07765195 | 0.77632008 | 49.71578185 | NA | NA | NA | NA | NA | NA | NA |
| SCN1A | 1205 | W | -21.19410714 | 49.12331346 | 0.716880435 | 45.78596299 | NA | NA | NA | NA | NA | NA | NA |
| SCN1A | 1206 | N | -18.628 | 53.97615431 | 0.789825919 | 50.60915912 | NA | NA | NA | NA | NA | NA | NA |
| SCN1A | 1207 | L | -14.336625 | 51.81047273 | 0.757272474 | 48.44627847 | NA | NA | NA | NA | NA | NA | NA |
| SCN1A | 1208 | R | -15.86993182 | 47.77249612 | 0.696575623 | 44.41063989 | NA | NA | NA | NA | NA | NA | NA |
| SCN1A | 1209 | R | -20.37147727 | 49.07022292 | 0.716082405 | 45.70672462 | NA | NA | NA | NA | NA | NA | NA |
| SCN1A | 1210 | T | -14.82389286 | 49.45355514 | 0.721844464 | 46.10937079 | NA | NA | NA | NA | NA | NA | NA |
| SCN1A | 1211 | C | -12.90658333 | 46.70311348 | 0.680501196 | 43.34888151 | NA | NA | NA | NA | NA | NA | NA |
| SCN1A | 1212 | F | -16.91484091 | 42.21925183 | 0.613102023 | 38.85124782 | NA | NA | NA | NA | NA | NA | NA |
| SCN1A | 1213 | R | -17.45647727 | 45.74593418 | 0.666113354 | 42.43762441 | NA | NA | NA | NA | NA | NA | NA |
| SCN1A | 1214 | I | -11.499 | 44.79389079 | 0.651802712 | 41.56146631 | NA | NA | NA | NA | NA | NA | NA |
| SCN1A | 1215 | V | -11.92803571 | 41.45481547 | 0.601611397 | 38.46766202 | NA | NA | NA | NA | NA | NA | NA |
| SCN1A | 1216 | E | -15.92941667 | 40.04427386 | 0.580408838 | 36.67940046 | NA | NA | NA | NA | NA | NA | NA |
| SCN1A | 1217 | H | -12.70165 | 41.86609834 | 0.607793596 | 38.60274263 | NA | NA | NA | NA | NA | NA | NA |
| SCN1A | 1218 | N | -12.368375 | 39.35758241 | 0.570086835 | 36.15206112 | NA | NA | NA | NA | NA | NA | NA |
| SCN1A | 1219 | W | -9.04375 | 42.05920633 | 0.610696299 | 38.9766282 | NA | NA | NA | NA | NA | NA | NA |
| SCN1A | 1220 | F | -7.602386364 | 40.70178523 | 0.590292222 | 37.61513761 | NA | NA | NA | NA | NA | NA | NA |
| SCN1A | 1221 | E | -9.408861111 | 36.88350344 | 0.532897713 | 33.81749073 | NA | NA | NA | NA | NA | NA | NA |
| SCN1A | 1222 | T | -6.513464286 | 38.12235348 | 0.55151949 | 35.04473579 | NA | NA | NA | NA | NA | NA | NA |
| SCN1A | 1223 | F | -2.350386364 | 39.23229122 | 0.56820352 | 36.83858216 | NA | NA | NA | NA | NA | NA | NA |
| SCN1A | 1224 | I | -3.305125 | 36.3649934 | 0.525103729 | 35.20635116 | NA | NA | NA | NA | NA | NA | NA |
| SCN1A | 1225 | V | -3.017464286 | 33.18287265 | 0.477271676 | 30.09136517 | NA | NA | NA | NA | NA | NA | NA |
| SCN1A | 1226 | F | -0.473204545 | 35.55472613 | 0.512924195 | 34.51367454 | NA | NA | NA | NA | NA | NA | NA |
| SCN1A | 1227 | M | 0.992 | 34.78474995 | 0.501350297 | 33.78612398 | NA | NA | NA | NA | NA | NA | NA |
| SCN1A | 1228 | I | 0.74325 | 31.38032253 | 0.450176642 | 30.23756019 | NA | NA | NA | NA | NA | NA | NA |
| SCN1A | 1229 | L | 1.221625 | 30.04455873 | 0.430098106 | 28.93722463 | NA | NA | NA | NA | NA | NA | NA |
| SCN1A | 1230 | L | 5.068625 | 32.72510428 | 0.47039073 | 32.75814485 | NA | NA | NA | NA | NA | NA | NA |
| SCN1A | 1231 | S | 5.150916667 | 30.78950992 | 0.441295841 | 30.49210882 | NA | NA | NA | NA | NA | NA | NA |

| | | | | | | | | | | | | | |
|---|---|---|---|---|---|---|---|---|---|---|---|---|---|
| SCN1A | 1232 | S | 5.267416667 | 27.8458245 | 0.397047829 | 27.87107993 | NA | NA | NA | NA | NA | NA | NA |
| SCN1A | 1233 | G | 8.10475 | 28.93247556 | 0.413381826 | 28.88707715 | NA | NA | NA | NA | NA | NA | NA |
| SCN1A | 1234 | A | 10.23955 | 29.72167191 | 0.425244632 | 30.18583082 | NA | NA | NA | NA | NA | NA | NA |
| SCN1A | 1235 | L | 10.2335 | 25.9001466 | 0.367801369 | 26.4354453 | NA | NA | NA | NA | NA | NA | NA |
| SCN1A | 1236 | A | 12.49535 | 25.24514989 | 0.357955785 | 25.91679334 | NA | NA | NA | NA | NA | NA | NA |
| SCN1A | 1237 | F | 14.10938636 | 27.25509486 | 0.388168276 | 27.88781348 | NA | NA | NA | NA | NA | NA | NA |
| SCN1A | 1238 | E | 16.46063889 | 24.17429683 | 0.341859256 | 24.81756548 | NA | NA | NA | NA | NA | NA | NA |
| SCN1A | 1239 | D | 18.97175 | 24.40546926 | 0.345334125 | 25.00968808 | NA | NA | NA | NA | NA | NA | NA |
| SCN1A | 1240 | I | 23.503125 | 21.39181743 | 0.300034413 | 21.67139171 | NA | NA | NA | NA | NA | NA | NA |
| SCN1A | 1241 | Y | 23.64675 | 23.85979809 | 0.337131868 | 24.11637036 | NA | NA | NA | NA | NA | NA | NA |
| SCN1A | 1242 | I | 21.11075 | 28.57331787 | 0.407983147 | 28.95522451 | NA | NA | NA | NA | NA | NA | NA |
| SCN1A | 1243 | D | 25.53 | 28.5160671 | 0.407122582 | 28.8309732 | NA | NA | NA | NA | NA | NA | NA |
| SCN1A | 1244 | Q | 27.34608333 | 30.3717334 | 0.435016033 | 26.64033378 | NA | NA | NA | NA | NA | NA | NA |
| SCN1A | 1245 | R | 22.69711364 | 29.49756466 | 0.421875964 | 30.05541003 | NA | NA | NA | NA | NA | NA | NA |
| SCN1A | 1246 | K | 24.85330556 | 34.94297825 | 0.503728706 | 35.80615234 | NA | NA | NA | NA | NA | NA | NA |
| SCN1A | 1247 | T | 20.88989286 | 37.85295457 | 0.547470019 | 38.35576038 | NA | NA | NA | NA | NA | NA | NA |
| SCN1A | 1248 | I | 17.801625 | 33.7025204 | 0.485082762 | 34.38021274 | NA | NA | NA | NA | NA | NA | NA |
| SCN1A | 1249 | K | 20.34397222 | 34.10085838 | 0.49107038 | 34.47737161 | NA | NA | NA | NA | NA | NA | NA |
| SCN1A | 1250 | T | 18.79675 | 37.11367931 | 0.536357602 | 37.6995899 | NA | NA | NA | NA | NA | NA | NA |
| SCN1A | 1251 | M | 15.553875 | 37.55632593 | 0.543011246 | 38.18491585 | NA | NA | NA | NA | NA | NA | NA |
| SCN1A | 1252 | L | 14.252125 | 31.75732852 | 0.455843608 | 32.41012439 | NA | NA | NA | NA | NA | NA | NA |
| SCN1A | 1253 | E | 15.39497222 | 36.09766675 | 0.521085408 | 36.71262251 | NA | NA | NA | NA | NA | NA | NA |
| SCN1A | 1254 | Y | 14.31308333 | 38.65445073 | 0.55951771 | 39.22144571 | NA | NA | NA | NA | NA | NA | NA |
| SCN1A | 1255 | A | 9.98835 | 37.15471577 | 0.536974442 | 37.61087292 | NA | NA | NA | NA | NA | NA | NA |
| SCN1A | 1256 | D | 10.30675 | 33.15870668 | 0.476908426 | 33.53796956 | NA | NA | NA | NA | NA | NA | NA |
| SCN1A | 1257 | K | 10.39275 | 38.13884037 | 0.551767312 | 38.51493705 | NA | NA | NA | NA | NA | NA | NA |
| SCN1A | 1258 | V | 6.659321429 | 40.26921716 | 0.583790074 | 39.94971777 | NA | NA | NA | NA | NA | NA | NA |
| SCN1A | 1259 | F | 5.468022727 | 35.74924963 | 0.515848176 | 35.44094494 | NA | NA | NA | NA | NA | NA | NA |
| SCN1A | 1260 | T | 5.186464286 | 36.9275057 | 0.533559133 | 36.57032315 | NA | NA | NA | NA | NA | NA | NA |
| SCN1A | 1261 | Y | 5.274333333 | 40.68841207 | 0.590091203 | 40.32002207 | NA | NA | NA | NA | NA | NA | NA |
| SCN1A | 1262 | I | 0.73275 | 39.99820387 | 0.579716337 | 39.9811146 | NA | NA | NA | NA | NA | NA | NA |
| SCN1A | 1263 | F | 0.429113636 | 35.34752583 | 0.509809664 | 34.27648254 | NA | NA | NA | NA | NA | NA | NA |
| SCN1A | 1264 | I | 0.904625 | 40.36101898 | 0.585169993 | 39.2882415 | NA | NA | NA | NA | NA | NA | NA |

| | | | | | | | | | | | | | |
|---|---|---|---|---|---|---|---|---|---|---|---|---|---|
| SCN1A | 1265 | L | -1.4265 | 42.98442156 | 0.624603673 | 41.8830555 | NA | NA | NA | NA | NA | NA | NA |
| SCN1A | 1266 | E | -3.74575 | 38.02997457 | 0.550130896 | 36.82451954 | NA | NA | NA | NA | NA | NA | NA |
| SCN1A | 1267 | M | -4.018625 | 40.02838585 | 0.580170018 | 38.80173883 | NA | NA | NA | NA | NA | NA | NA |
| SCN1A | 1268 | L | -4.553625 | 43.56913694 | 0.633392823 | 42.34796758 | NA | NA | NA | NA | NA | NA | NA |
| SCN1A | 1269 | L | -7.70675 | 45.15507824 | 0.657231902 | 42.06258622 | NA | NA | NA | NA | NA | NA | NA |
| SCN1A | 1270 | K | -8.922527778 | 38.0607783 | 0.550593922 | 34.97101678 | NA | NA | NA | NA | NA | NA | NA |
| SCN1A | 1271 | W | -7.299035714 | 42.32301429 | 0.614661729 | 39.79674908 | NA | NA | NA | NA | NA | NA | NA |
| SCN1A | 1272 | V | -10.99289286 | 46.79459257 | 0.681876264 | 43.70102728 | NA | NA | NA | NA | NA | NA | NA |
| SCN1A | 1273 | A | -13.70815 | 44.55812906 | 0.648258859 | 41.19225604 | NA | NA | NA | NA | NA | NA | NA |
| SCN1A | 1274 | Y | -14.05391667 | 39.41241462 | 0.570911045 | 36.05236389 | NA | NA | NA | NA | NA | NA | NA |
| SCN1A | 1275 | G | -13.8435 | 42.77749933 | 0.621493321 | 39.41651476 | NA | NA | NA | NA | NA | NA | NA |
| SCN1A | 1276 | Y | -10.88875 | 41.76549316 | 0.606281349 | 38.40762701 | NA | NA | NA | NA | NA | NA | NA |
| SCN1A | 1277 | Q | -16.64286111 | 39.25253201 | 0.56850777 | 35.9204558 | NA | NA | NA | NA | NA | NA | NA |
| SCN1A | 1278 | T | -16.10017857 | 37.93189357 | 0.548656591 | 34.57083998 | NA | NA | NA | NA | NA | NA | NA |
| SCN1A | 1279 | Y | -9.819666667 | 37.59446249 | 0.543584495 | 34.28309863 | NA | NA | NA | NA | NA | NA | NA |
| SCN1A | 1280 | F | -9.793386364 | 37.0072907 | 0.534758422 | 33.90828338 | NA | NA | NA | NA | NA | NA | NA |
| SCN1A | 1281 | T | -14.65917857 | 33.76775335 | 0.486063311 | 30.41117931 | NA | NA | NA | NA | NA | NA | NA |
| SCN1A | 1282 | N | -12.1575 | 32.22227115 | 0.462832394 | 28.85865902 | NA | NA | NA | NA | NA | NA | NA |
| SCN1A | 1283 | A | -9.49835 | 29.63609996 | 0.423958358 | 26.53948739 | NA | NA | NA | NA | NA | NA | NA |
| SCN1A | 1284 | W | -8.52975 | 25.6901629 | 0.364644999 | 22.59475934 | NA | NA | NA | NA | NA | NA | NA |
| SCN1A | 1285 | C | -8.15325 | 33.6127423 | 0.483733263 | 30.51586416 | NA | NA | NA | NA | NA | NA | NA |
| SCN1A | 1286 | W | -6.550964286 | 33.11424631 | 0.476240119 | 31.92064523 | NA | NA | NA | NA | NA | NA | NA |
| SCN1A | 1287 | L | -3.361 | 27.86120134 | 0.397278966 | 26.71283483 | NA | NA | NA | NA | NA | NA | NA |
| SCN1A | 1288 | D | -2.9015 | 32.49835307 | 0.466982319 | 31.30686991 | NA | NA | NA | NA | NA | NA | NA |
| SCN1A | 1289 | F | -2.632022727 | 34.79538604 | 0.501510174 | 33.57416143 | NA | NA | NA | NA | NA | NA | NA |
| SCN1A | 1290 | L | -0.973125 | 31.13095373 | 0.446428254 | 29.8809546 | NA | NA | NA | NA | NA | NA | NA |
| SCN1A | 1291 | I | 1.207 | 28.99670645 | 0.414347313 | 27.81754473 | NA | NA | NA | NA | NA | NA | NA |
| SCN1A | 1292 | V | 0.914178571 | 34.26302447 | 0.49350798 | 34.14326974 | NA | NA | NA | NA | NA | NA | NA |
| SCN1A | 1293 | D | 1.819375 | 35.35952094 | 0.509989968 | 34.92144719 | NA | NA | NA | NA | NA | NA | NA |
| SCN1A | 1294 | V | 5.027178571 | 31.00454247 | 0.444528103 | 30.50734329 | NA | NA | NA | NA | NA | NA | NA |
| SCN1A | 1295 | S | 5.744416667 | 34.11381235 | 0.491265098 | 33.6442423 | NA | NA | NA | NA | NA | NA | NA |
| SCN1A | 1296 | L | 4.710375 | 36.88456748 | 0.532913707 | 36.40223155 | NA | NA | NA | NA | NA | NA | NA |
| SCN1A | 1297 | V | 7.705607143 | 35.85557883 | 0.517446463 | 34.92462579 | NA | NA | NA | NA | NA | NA | NA |

| | | | | | | | | | | | | | |
|---|---|---|---|---|---|---|---|---|---|---|---|---|---|
| SCN1A | 1298 | S | 10.74775 | 34.21761663 | 0.492825432 | 34.46975544 | NA | NA | NA | NA | NA | NA | NA |
| SCN1A | 1299 | L | 9.458125 | 37.44134863 | 0.541282964 | 37.65639356 | NA | NA | NA | NA | NA | NA | NA |
| SCN1A | 1300 | T | 9.88175 | 40.02562966 | 0.580128588 | 40.2293466 | NA | NA | NA | NA | NA | NA | NA |
| SCN1A | 1301 | A | 13.47115 | 37.39576131 | 0.540597718 | 37.6664988 | NA | NA | NA | NA | NA | NA | NA |
| SCN1A | 1302 | N | 14.887125 | 35.92632748 | 0.518509922 | 36.32087193 | NA | NA | NA | NA | NA | NA | NA |
| SCN1A | 1303 | A | 13.92135 | 41.28803656 | 0.599104459 | 41.65128646 | NA | NA | NA | NA | NA | NA | NA |
| SCN1A | 1304 | L | 13.84325 | 43.31580686 | 0.629584891 | 43.65459155 | NA | NA | NA | NA | NA | NA | NA |
| SCN1A | 1305 | G | 18.10875 | 41.32863585 | 0.599714728 | 41.57054711 | NA | NA | NA | NA | NA | NA | NA |
| SCN1A | 1306 | Y | 16.61116667 | 37.8186937 | 0.546955027 | 38.11119087 | NA | NA | NA | NA | NA | NA | NA |
| SCN1A | 1307 | S | 19.42691667 | 35.51031463 | 0.512256624 | 35.75627192 | NA | NA | NA | NA | NA | NA | NA |
| SCN1A | 1308 | E | 20.94719444 | 33.05268735 | 0.475314796 | 29.59924129 | NA | NA | NA | NA | NA | NA | NA |
| SCN1A | 1309 | L | 18.847875 | 31.36915399 | 0.450008762 | 28.7843047 | NA | NA | NA | NA | NA | NA | NA |
| SCN1A | 1310 | G | 17.95475 | 29.24848943 | 0.418131989 | 26.86837876 | NA | NA | NA | NA | NA | NA | NA |
| SCN1A | 1311 | A | 14.90815 | 30.13096623 | 0.431396941 | 30.38270008 | NA | NA | NA | NA | NA | NA | NA |
| SCN1A | 1312 | I | 14.81175 | 31.05186426 | 0.445239421 | 31.40787345 | NA | NA | NA | NA | NA | NA | NA |
| SCN1A | 1313 | K | 18.39863889 | 25.09454156 | 0.355691916 | 25.37625866 | NA | NA | NA | NA | NA | NA | NA |
| SCN1A | 1314 | S | 14.68608333 | 26.59940939 | 0.378312339 | 26.81916937 | NA | NA | NA | NA | NA | NA | NA |
| SCN1A | 1315 | L | 11.412375 | 28.71782417 | 0.410155294 | 28.97342099 | NA | NA | NA | NA | NA | NA | NA |
| SCN1A | 1316 | R | 15.09565909 | 27.46063005 | 0.391257779 | 27.78356113 | NA | NA | NA | NA | NA | NA | NA |
| SCN1A | 1317 | T | 12.32132143 | 24.04873015 | 0.3399718 | 24.35083428 | NA | NA | NA | NA | NA | NA | NA |
| SCN1A | 1318 | L | 8.371375 | 26.40411 | 0.375376696 | 25.57270918 | NA | NA | NA | NA | NA | NA | NA |
| SCN1A | 1319 | R | 10.30365909 | 26.9878152 | 0.384150661 | 26.52881881 | NA | NA | NA | NA | NA | NA | NA |
| SCN1A | 1320 | A | 7.52675 | 23.30257209 | 0.328755924 | 22.84413223 | NA | NA | NA | NA | NA | NA | NA |
| SCN1A | 1321 | L | 4.22075 | 23.39706838 | 0.330176346 | 23.266461 | NA | NA | NA | NA | NA | NA | NA |
| SCN1A | 1322 | R | 4.994386364 | 27.01212983 | 0.384516147 | 26.94237211 | NA | NA | NA | NA | NA | NA | NA |
| SCN1A | 1323 | P | 2.497321429 | 25.10656577 | 0.355872658 | 24.05098539 | NA | NA | NA | NA | NA | NA | NA |
| SCN1A | 1324 | L | -0.383125 | 23.61141938 | 0.333398363 | 22.42111546 | NA | NA | NA | NA | NA | NA | NA |
| SCN1A | 1325 | R | -1.14875 | 28.63641226 | 0.40893155 | 27.45740913 | NA | NA | NA | NA | NA | NA | NA |
| SCN1A | 1326 | A | -2.80695 | 27.95607789 | 0.398705103 | 27.41368194 | NA | NA | NA | NA | NA | NA | NA |
| SCN1A | 1327 | L | -4.4425 | 23.74437159 | 0.335396834 | 20.6417509 | NA | NA | NA | NA | NA | NA | NA |
| SCN1A | 1328 | S | -7.546416667 | 28.41864988 | 0.405658255 | 25.32604095 | NA | NA | NA | NA | NA | NA | NA |
| SCN1A | 1329 | R | -6.804477273 | 29.6770615 | 0.424574071 | 26.59564857 | NA | NA | NA | NA | NA | NA | NA |
| SCN1A | 1330 | F | -6.141295455 | 25.94681478 | 0.368502862 | 22.86738688 | NA | NA | NA | NA | NA | NA | NA |

| | | | | | | | | | | | | | |
|---|---|---|---|---|---|---|---|---|---|---|---|---|---|
| SCN1A | 1331 | E | -12.26152778 | 23.49115585 | 0.331590621 | 20.39838825 | NA | NA | NA | NA | NA | NA | NA |
| SCN1A | 1332 | G | -8.7065 | 20.75766102 | 0.29050209 | 17.66264995 | NA | NA | NA | NA | NA | NA | NA |
| SCN1A | 1333 | M | -5.66525 | 21.40738545 | 0.300268424 | 18.32088474 | NA | NA | NA | NA | NA | NA | NA |
| SCN1A | 1334 | R | -10.83320455 | 22.34663362 | 0.314386734 | 19.26835628 | NA | NA | NA | NA | NA | NA | NA |
| SCN1A | 1335 | V | -12.01589286 | 19.49164688 | 0.271471997 | 16.39617612 | NA | NA | NA | NA | NA | NA | NA |
| SCN1A | 1336 | V | -8.510892857 | 16.57653335 | 0.227653463 | 13.47385636 | NA | NA | NA | NA | NA | NA | NA |
| SCN1A | 1337 | V | -8.088464286 | 19.63597803 | 0.27364151 | 16.54938635 | NA | NA | NA | NA | NA | NA | NA |
| SCN1A | 1338 | N | -12.310875 | 19.47776183 | 0.271263283 | 16.65546229 | NA | NA | NA | NA | NA | NA | NA |
| SCN1A | 1339 | A | -11.82515 | 16.2522879 | 0.222779567 | 13.24327317 | NA | NA | NA | NA | NA | NA | NA |
| SCN1A | 1340 | L | -8.21525 | 14.88029855 | 0.202156508 | 11.78721557 | NA | NA | NA | NA | NA | NA | NA |
| SCN1A | 1341 | L | -9.77525 | 18.69245674 | 0.259458969 | 15.59869889 | NA | NA | NA | NA | NA | NA | NA |
| SCN1A | 1342 | G | -13.46875 | 17.20243079 | 0.237061642 | 13.85152736 | NA | NA | NA | NA | NA | NA | NA |
| SCN1A | 1343 | A | -11.73715 | 15.12314008 | 0.20580678 | 11.77822005 | NA | NA | NA | NA | NA | NA | NA |
| SCN1A | 1344 | I | -8.496875 | 16.8804787 | 0.232222218 | 13.79849061 | NA | NA | NA | NA | NA | NA | NA |
| SCN1A | 1345 | P | -11.35532143 | 18.75542111 | 0.260405418 | 15.67387168 | NA | NA | NA | NA | NA | NA | NA |
| SCN1A | 1346 | S | -11.69291667 | 15.95485484 | 0.218308702 | 13.1553402 | NA | NA | NA | NA | NA | NA | NA |
| SCN1A | 1347 | I | -7.624375 | 13.07689374 | 0.175048626 | 9.993298333 | NA | NA | NA | NA | NA | NA | NA |
| SCN1A | 1348 | M | -6.689375 | 17.33126773 | 0.238998254 | 13.84568493 | NA | NA | NA | NA | NA | NA | NA |
| SCN1A | 1349 | N | -10.092 | 18.49631826 | 0.256510713 | 15.46387928 | NA | NA | NA | NA | NA | NA | NA |
| SCN1A | 1350 | V | -7.625178571 | 14.16030402 | 0.191333909 | 10.68189765 | NA | NA | NA | NA | NA | NA | NA |
| SCN1A | 1351 | L | -3.281625 | 15.32023463 | 0.208769407 | 12.50326806 | NA | NA | NA | NA | NA | NA | NA |
| SCN1A | 1352 | L | -4.72375 | 18.80903179 | 0.261211267 | 15.31533406 | NA | NA | NA | NA | NA | NA | NA |
| SCN1A | 1353 | V | -5.337035714 | 18.24935939 | 0.25279855 | 14.87015617 | NA | NA | NA | NA | NA | NA | NA |
| SCN1A | 1354 | C | -2.187916667 | 14.15356651 | 0.191232634 | 10.93127407 | NA | NA | NA | NA | NA | NA | NA |
| SCN1A | 1355 | L | -0.249125 | 16.12466787 | 0.220861247 | 14.92949514 | NA | NA | NA | NA | NA | NA | NA |
| SCN1A | 1356 | I | -1.109 | 19.69301978 | 0.274498934 | 18.55286691 | NA | NA | NA | NA | NA | NA | NA |
| SCN1A | 1357 | F | -0.998568182 | 17.08023568 | 0.235224866 | 13.06555033 | NA | NA | NA | NA | NA | NA | NA |
| SCN1A | 1358 | W | 2.394107143 | 10.02558851 | 0.129182927 | 8.849457156 | NA | NA | NA | NA | NA | NA | NA |
| SCN1A | 1359 | L | 2.958625 | 18.29553978 | 0.253492711 | 17.13343621 | NA | NA | NA | NA | NA | NA | NA |
| SCN1A | 1360 | I | 2.0975 | 20.31848087 | 0.283900553 | 19.17055449 | NA | NA | NA | NA | NA | NA | NA |
| SCN1A | 1361 | F | 4.066659091 | 17.14167043 | 0.236148322 | 15.98668968 | NA | NA | NA | NA | NA | NA | NA |
| SCN1A | 1362 | S | 7.630083333 | 17.25877948 | 0.237908647 | 16.34312247 | NA | NA | NA | NA | NA | NA | NA |
| SCN1A | 1363 | I | 7.094125 | 20.44465219 | 0.285797097 | 19.89142467 | NA | NA | NA | NA | NA | NA | NA |

| | | | | | | | | | | | | | |
|---|---|---|---|---|---|---|---|---|---|---|---|---|---|
| SCN1A | 1364 | M | 7.578875 | 20.9200446 | 0.292942959 | 20.34892066 | NA | NA | NA | NA | NA | NA | NA |
| SCN1A | 1365 | G | 11.503 | 18.85376504 | 0.261883675 | 18.27524274 | NA | NA | NA | NA | NA | NA | NA |
| SCN1A | 1366 | V | 12.48532143 | 19.80778506 | 0.276224028 | 17.28436813 | NA | NA | NA | NA | NA | NA | NA |
| SCN1A | 1367 | N | 12.1495 | 22.2879543 | 0.313504696 | 19.46247491 | NA | NA | NA | NA | NA | NA | NA |
| SCN1A | 1368 | L | 13.424375 | 20.34325797 | 0.28427299 | 17.58760149 | NA | NA | NA | NA | NA | NA | NA |
| SCN1A | 1369 | F | 15.39193182 | 17.08560994 | 0.235305649 | 14.49222366 | NA | NA | NA | NA | NA | NA | NA |
| SCN1A | 1370 | A | 17.64015 | 17.20916721 | 0.2371629 | 14.43606114 | NA | NA | NA | NA | NA | NA | NA |
| SCN1A | 1371 | G | 21.52 | 17.06617573 | 0.235013523 | 14.04933029 | NA | NA | NA | NA | NA | NA | NA |
| SCN1A | 1372 | K | 21.02208333 | 14.54986856 | 0.197189649 | 11.40011207 | NA | NA | NA | NA | NA | NA | NA |
| SCN1A | 1373 | F | 20.49647727 | 10.73331291 | 0.139821087 | 7.578938854 | NA | NA | NA | NA | NA | NA | NA |
| SCN1A | 1374 | Y | 25.2105 | 12.58708187 | 0.167686018 | 9.398983352 | NA | NA | NA | NA | NA | NA | NA |
| SCN1A | 1375 | H | 27.93935 | 7.566109568 | 0.092213265 | 7.179674529 | 1.56 | -0.8 | -0.4 | 3.38 | 0.81 | 0 | 0 |
| SCN1A | 1376 | C | 32.72525 | 8.289853376 | 0.103092221 | 6.77343605 | NA | NA | NA | NA | NA | NA | NA |
| SCN1A | 1377 | I | 32.899875 | 3.766519215 | 0.035099718 | 2.257337235 | 1.56 | -0.8 | -0.4 | 3.38 | 0.81 | 0 | 0 |
| SCN1A | 1378 | N | 36.924625 | 3.714688278 | 0.034320621 | 2.549305385 | 1.56 | -0.8 | -0.4 | 3.38 | 0.81 | 0 | 0 |
| SCN1A | 1379 | T | 36.53675 | 3.174239751 | 0.026196868 | 1.868353482 | 1.56 | -0.8 | -0.4 | 3.38 | 0.81 | 0 | 0 |
| SCN1A | 1380 | T | 39.01017857 | 4.533967137 | 0.046635612 | 3.407389002 | 1.56 | -0.8 | -0.4 | 3.38 | 0.81 | 0 | 0 |
| SCN1A | 1381 | T | 35.93132143 | 4.311401164 | 0.043290111 | 3.19890188 | 1.56 | -0.8 | -0.4 | 3.38 | 0.81 | 0 | 0 |
| SCN1A | 1382 | G | 32.7375 | 3.174840153 | 0.026205893 | 1.886619436 | 1.56 | -0.8 | -0.4 | 3.38 | 0.81 | 0 | 0 |
| SCN1A | 1383 | D | 33.90875 | 5.115800719 | 0.055381444 | 3.853405289 | 1.56 | -0.8 | -0.4 | 3.38 | 0.81 | 0 | 0 |
| SCN1A | 1384 | R | 29.72847727 | 8.289977684 | 0.10309409 | 6.91595129 | NA | NA | NA | NA | NA | NA | NA |
| SCN1A | 1385 | F | 35.82665909 | 9.89947837 | 0.127287302 | 8.603618702 | NA | NA | NA | NA | NA | NA | NA |
| SCN1A | 1386 | D | 34.90525 | 13.26891514 | 0.177934996 | 11.88595892 | NA | NA | NA | NA | NA | NA | NA |
| SCN1A | 1387 | I | 33.060625 | 16.65156851 | 0.228781355 | 15.22269314 | NA | NA | NA | NA | NA | NA | NA |
| SCN1A | 1388 | E | 37.62419444 | 18.57818533 | 0.257741298 | 17.19076097 | NA | NA | NA | NA | NA | NA | NA |
| SCN1A | 1389 | D | 38.732125 | 15.07841669 | 0.205134521 | 13.79399541 | NA | NA | NA | NA | NA | NA | NA |
| SCN1A | 1390 | V | 35.64746429 | 13.8134436 | 0.186120076 | 12.33981017 | NA | NA | NA | NA | NA | NA | NA |
| SCN1A | 1391 | N | 33.3005 | 17.74398132 | 0.245201959 | 16.2583641 | NA | NA | NA | NA | NA | NA | NA |
| SCN1A | 1392 | N | 32.9725 | 16.5958907 | 0.227944433 | 15.07385132 | NA | NA | NA | NA | NA | NA | NA |
| SCN1A | 1393 | H | 32.24195 | 16.94301145 | 0.233162179 | 15.44218816 | NA | NA | NA | NA | NA | NA | NA |
| SCN1A | 1394 | T | 36.11817857 | 19.09578168 | 0.265521548 | 17.56918498 | NA | NA | NA | NA | NA | NA | NA |
| SCN1A | 1395 | D | 38.031625 | 16.67308973 | 0.229104851 | 15.18128918 | NA | NA | NA | NA | NA | NA | NA |
| SCN1A | 1396 | C | 36.54691667 | 12.06495342 | 0.159837643 | 10.54235855 | NA | NA | NA | NA | NA | NA | NA |

| | | | | | | | | | | | | | |
|---|---|---|---|---|---|---|---|---|---|---|---|---|---|
| SCN1A | 1397 | L | 37.20925 | 15.81415221 | 0.21619373 | 14.28753095 | NA | NA | NA | NA | NA | NA | NA |
| SCN1A | 1398 | K | 41.61741667 | 17.65466596 | 0.243859415 | 16.16717814 | NA | NA | NA | NA | NA | NA | NA |
| SCN1A | 1399 | L | 41.298625 | 13.6539606 | 0.183722807 | 12.25837275 | NA | NA | NA | NA | NA | NA | NA |
| SCN1A | 1400 | I | 39.898625 | 12.39678188 | 0.164825523 | 10.86298257 | NA | NA | NA | NA | NA | NA | NA |
| SCN1A | 1401 | E | 43.92208333 | 16.86532466 | 0.23199443 | 15.35106543 | NA | NA | NA | NA | NA | NA | NA |
| SCN1A | 1402 | R | 46.14647727 | 17.32080977 | 0.238841055 | 15.90924191 | NA | NA | NA | NA | NA | NA | NA |
| SCN1A | 1403 | N | 46.106125 | 14.8341993 | 0.201463567 | 13.4403298 | NA | NA | NA | NA | NA | NA | NA |
| SCN1A | 1404 | E | 43.41808333 | 11.52779233 | 0.151763305 | 10.09392043 | NA | NA | NA | NA | NA | NA | NA |
| SCN1A | 1405 | T | 40.23317857 | 7.123944062 | 0.085566854 | 5.793139202 | 1.56 | -0.8 | -0.4 | 3.38 | 0.81 | 0 | 0 |
| SCN1A | 1406 | A | 37.24415 | 8.309376691 | 0.103385686 | 6.787105452 | NA | NA | NA | NA | NA | NA | NA |
| SCN1A | 1407 | R | 32.47429545 | 6.003855928 | 0.068730248 | 4.492460508 | 1.56 | -0.8 | -0.4 | 3.38 | 0.81 | 0 | 0 |
| SCN1A | 1408 | W | 30.66975 | 9.26206532 | 0.117706027 | 7.753447105 | NA | NA | NA | NA | NA | NA | NA |
| SCN1A | 1409 | K | 27.75241667 | 4.898095446 | 0.052109007 | 4.357346419 | 1.93 | 0.07 | 1.23 | 2.3 | -0.08 | 0 | 0 |
| SCN1A | 1410 | N | 23.832375 | 7.134334797 | 0.085723042 | 3.964419415 | 3.33 | -0.37 | -1.68 | 37.31 | 1.26 | 3 | -1 |
| SCN1A | 1411 | V | 21.60503571 | 5.88446786 | 0.066935666 | 3.125525917 | 2.92 | -0.26 | -0.25 | 24.93 | 1.04 | 2 | -2 |
| SCN1A | 1412 | K | 21.77697222 | 6.87518807 | 0.081827678 | 4.254068218 | 2.92 | -0.26 | -0.25 | 24.93 | 1.04 | 2 | -2 |
| SCN1A | 1413 | V | 16.95803571 | 7.367537513 | 0.089228429 | 4.480317656 | 2.89 | -0.23 | -0.25 | 24.88 | 1.14 | 2 | -2 |
| SCN1A | 1414 | N | 17.07825 | 10.96817346 | 0.143351394 | 8.312136681 | NA | NA | NA | NA | NA | NA | NA |
| SCN1A | 1415 | F | 13.35365909 | 14.05382475 | 0.189733365 | 11.40422949 | NA | NA | NA | NA | NA | NA | NA |
| SCN1A | 1416 | D | 17.341625 | 14.66856319 | 0.198973807 | 11.98999117 | NA | NA | NA | NA | NA | NA | NA |
| SCN1A | 1417 | N | 13.815125 | 17.91852977 | 0.247825684 | 15.1484199 | NA | NA | NA | NA | NA | NA | NA |
| SCN1A | 1418 | V | 9.029178571 | 17.09777322 | 0.235488481 | 16.53912999 | NA | NA | NA | NA | NA | NA | NA |
| SCN1A | 1419 | G | 10.31225 | 14.40887247 | 0.195070266 | 13.84015714 | NA | NA | NA | NA | NA | NA | NA |
| SCN1A | 1420 | F | 14.41356818 | 12.61811396 | 0.168152477 | 12.03729195 | NA | NA | NA | NA | NA | NA | NA |
| SCN1A | 1421 | G | 10.0575 | 13.00555155 | 0.173976245 | 12.45192789 | NA | NA | NA | NA | NA | NA | NA |
| SCN1A | 1422 | Y | 5.6995 | 11.31106012 | 0.148505495 | 10.53723293 | NA | NA | NA | NA | NA | NA | NA |
| SCN1A | 1423 | L | 11.02125 | 8.292463205 | 0.103131451 | 7.423708566 | NA | NA | NA | NA | NA | NA | NA |
| SCN1A | 1424 | S | 11.50041667 | 8.877357602 | 0.111923292 | 8.319090893 | NA | NA | NA | NA | NA | NA | NA |
| SCN1A | 1425 | L | 7.4205 | 8.773946546 | 0.110368869 | 7.61752907 | NA | NA | NA | NA | NA | NA | NA |
| SCN1A | 1426 | L | 6.75525 | 5.372467683 | 0.059239534 | 4.213554448 | 1.25 | -0.07 | -1.48 | 14.16 | 0.14 | 1 | 1 |
| SCN1A | 1427 | Q | 11.41919444 | 4.868763395 | 0.051668103 | 4.382661905 | 0.81 | 0.17 | -1.73 | 18.33 | -0.07 | 1 | 1 |
| SCN1A | 1428 | V | 8.924464286 | 7.255015644 | 0.087537057 | 6.09050518 | 1.25 | -0.07 | -1.48 | 14.16 | 0.14 | 1 | 1 |
| SCN1A | 1429 | A | 5.21495 | 5.777095983 | 0.065321705 | 5.087006043 | 1.24 | 0.18 | -1.83 | 17.75 | -0.08 | 1 | 1 |

| | | | | | | | | | | | | |
|---|---|---|---|---|---|---|---|---|---|---|---|---|
| SCN1A | 1430 | T | 6.474321429 | 2.5533807 | 0.016864424 | 1.947789581 | 0.69 | -0.73 | -1.7 | 18.18 | 1.13 | 1 | 1 |
| SCN1A | 1431 | F | 7.085931818 | 4.591732788 | 0.047503917 | 3.875550554 | 0.81 | -0.72 | -1.8 | 17.61 | 1.11 | 1 | 1 |
| SCN1A | 1432 | K | 10.12652778 | 1.973879682 | 0.008153654 | 1.655360812 | 0.5 | 0.65 | -2.4 | 25.8 | -0.51 | 1 | 1 |
| SCN1A | 1433 | G | 14.2945 | 3.876587804 | 0.036754214 | 3.225936961 | 0.67 | -0.92 | -1.95 | 26.54 | 1.72 | 1 | -1 |
| SCN1A | 1434 | W | 12.15275 | 6.251202604 | 0.072448239 | 5.615909678 | 0.67 | -0.92 | -1.95 | 26.54 | 1.72 | 1 | -1 |
| SCN1A | 1435 | M | 16.42025 | 7.616344596 | 0.092968373 | 4.938921892 | 2.89 | -0.23 | -0.25 | 24.88 | 1.14 | 2 | -2 |
| SCN1A | 1436 | D | 19.779375 | 4.17346331 | 0.041216698 | 1.622801809 | 2.89 | -0.23 | -0.25 | 24.88 | 1.14 | 2 | -2 |
| SCN1A | 1437 | I | 15.885625 | 7.586412195 | 0.092518444 | 4.821100245 | 2.89 | -0.23 | -0.25 | 24.88 | 1.14 | 2 | -2 |
| SCN1A | 1438 | M | 15.223 | 10.27001212 | 0.132856981 | 7.518771841 | NA | NA | NA | NA | NA | NA | NA |
| SCN1A | 1439 | Y | 19.88733333 | 9.199904836 | 0.116771661 | 6.412256215 | NA | NA | NA | NA | NA | NA | NA |
| SCN1A | 1440 | A | 20.45855 | 10.27778678 | 0.132973845 | 7.684121141 | NA | NA | NA | NA | NA | NA | NA |
| SCN1A | 1441 | A | 17.91995 | 12.87336463 | 0.171989278 | 10.09095314 | NA | NA | NA | NA | NA | NA | NA |
| SCN1A | 1442 | V | 18.87746429 | 15.00195271 | 0.203985152 | 12.2129353 | NA | NA | NA | NA | NA | NA | NA |
| SCN1A | 1443 | D | 22.88125 | 13.46257397 | 0.180845979 | 10.26604386 | NA | NA | NA | NA | NA | NA | NA |
| SCN1A | 1444 | S | 20.73525 | 16.5290468 | 0.226939669 | 13.38477033 | NA | NA | NA | NA | NA | NA | NA |
| SCN1A | 1445 | R | 25.92175 | 18.43608887 | 0.255605375 | 15.26592294 | NA | NA | NA | NA | NA | NA | NA |
| SCN1A | 1446 | N | 23.476375 | 19.3150933 | 0.268818131 | 16.14080014 | NA | NA | NA | NA | NA | NA | NA |
| SCN1A | 1447 | V | 23.19975 | 20.38387333 | 0.2848835 | 17.22985281 | NA | NA | NA | NA | NA | NA | NA |
| SCN1A | 1448 | E | 27.74875 | 19.11335664 | 0.265785726 | 17.59073756 | NA | NA | NA | NA | NA | NA | NA |
| SCN1A | 1449 | L | 27.7085 | 17.34648325 | 0.239226966 | 15.82187485 | NA | NA | NA | NA | NA | NA | NA |
| SCN1A | 1450 | Q | 30.06763889 | 12.3058187 | 0.16345821 | 10.78485681 | NA | NA | NA | NA | NA | NA | NA |
| SCN1A | 1451 | P | 27.06632143 | 13.91602562 | 0.187662038 | 10.75086312 | NA | NA | NA | NA | NA | NA | NA |
| SCN1A | 1452 | K | 28.01619444 | 16.91484118 | 0.232738738 | 13.62678568 | NA | NA | NA | NA | NA | NA | NA |
| SCN1A | 1453 | Y | 25.93891667 | 15.52014108 | 0.211774301 | 12.16704166 | NA | NA | NA | NA | NA | NA | NA |
| SCN1A | 1454 | E | 22.37908333 | 17.32897729 | 0.238963826 | 14.44957157 | NA | NA | NA | NA | NA | NA | NA |
| SCN1A | 1455 | E | 24.09419444 | 19.82878799 | 0.276539734 | 16.6355837 | NA | NA | NA | NA | NA | NA | NA |
| SCN1A | 1456 | S | 19.16858333 | 19.72132333 | 0.274924379 | 16.93124424 | NA | NA | NA | NA | NA | NA | NA |
| SCN1A | 1457 | L | 18.095625 | 20.04329963 | 0.279764166 | 17.27043954 | NA | NA | NA | NA | NA | NA | NA |
| SCN1A | 1458 | Y | 14.583 | 21.64512927 | 0.30384207 | 18.88058268 | NA | NA | NA | NA | NA | NA | NA |
| SCN1A | 1459 | M | 13.890875 | 18.94964931 | 0.263324959 | 16.21001522 | NA | NA | NA | NA | NA | NA | NA |
| SCN1A | 1460 | Y | 14.21791667 | 15.0230225 | 0.204301863 | 12.27257589 | NA | NA | NA | NA | NA | NA | NA |
| SCN1A | 1461 | L | 9.267375 | 19.53408705 | 0.272109936 | 18.36839331 | NA | NA | NA | NA | NA | NA | NA |
| SCN1A | 1462 | Y | 9.097416667 | 16.99146027 | 0.233890438 | 15.84590729 | NA | NA | NA | NA | NA | NA | NA |

| | | | | | | | | | | | | | |
|---|---|---|---|---|---|---|---|---|---|---|---|---|---|
| SCN1A | 1463 | F | 10.61893182 | 13.55871804 | 0.182291168 | 12.82936235 | NA | NA | NA | NA | NA | NA | NA |
| SCN1A | 1464 | V | 6.560178571 | 15.43413911 | 0.210481563 | 12.5130164 | NA | NA | NA | NA | NA | NA | NA |
| SCN1A | 1465 | I | 4.163875 | 16.30250054 | 0.223534339 | 13.45552986 | NA | NA | NA | NA | NA | NA | NA |
| SCN1A | 1466 | F | 4.627295455 | 11.46809561 | 0.150865974 | 9.566664663 | NA | NA | NA | NA | NA | NA | NA |
| SCN1A | 1467 | I | 4.508875 | 10.85048681 | 0.141582387 | 7.905431506 | NA | NA | NA | NA | NA | NA | NA |
| SCN1A | 1468 | I | 0.874 | 12.81058055 | 0.171045538 | 8.835314914 | NA | NA | NA | NA | NA | NA | NA |
| SCN1A | 1469 | F | -0.16875 | 13.77125644 | 0.18548594 | 9.795122326 | NA | NA | NA | NA | NA | NA | NA |
| SCN1A | 1470 | G | 0.209 | 11.43379368 | 0.150350365 | 7.40811184 | NA | NA | NA | NA | NA | NA | NA |
| SCN1A | 1471 | S | 0.681583333 | 7.835234968 | 0.096258624 | 3.814096433 | 4.1 | 0.72 | 2.15 | 0.63 | -1.08 | 0 | 0 |
| SCN1A | 1472 | F | -2.401022727 | 9.899217191 | 0.127283376 | 6.657139325 | NA | NA | NA | NA | NA | NA | NA |
| SCN1A | 1473 | F | -4.120204545 | 11.89838149 | 0.157333817 | 8.47350543 | NA | NA | NA | NA | NA | NA | NA |
| SCN1A | 1474 | T | -3.168464286 | 10.80794023 | 0.140942848 | 7.333434334 | NA | NA | NA | NA | NA | NA | NA |
| SCN1A | 1475 | L | -3.711 | 5.13946398 | 0.055737139 | 1.600283346 | 3.58 | 1.13 | 2.1 | 0.62 | -1.67 | 0 | 0 |
| SCN1A | 1476 | N | -7.739375 | 8.157492813 | 0.101102643 | 5.206155301 | NA | NA | NA | NA | NA | NA | NA |
| SCN1A | 1477 | L | -8.685875 | 10.59704605 | 0.137772791 | 7.580532043 | NA | NA | NA | NA | NA | NA | NA |
| SCN1A | 1478 | F | -6.877659091 | 7.879373389 | 0.096922091 | 5.318207313 | 3.02 | 1.59 | 4.27 | 0.13 | -1.83 | 0 | 0 |
| SCN1A | 1479 | I | -8.907125 | 4.630793237 | 0.048091054 | 1.801371574 | 3.02 | 1.59 | 4.27 | 0.13 | -1.83 | 0 | 0 |
| SCN1A | 1480 | G | -12.50775 | 8.226096644 | 0.102133862 | 5.590860039 | NA | NA | NA | NA | NA | NA | NA |
| SCN1A | 1481 | V | -12.37075 | 10.09272059 | 0.130192023 | 7.440288136 | NA | NA | NA | NA | NA | NA | NA |
| SCN1A | 1482 | I | -11.4825 | 8.204575796 | 0.101810371 | 6.095278535 | NA | NA | NA | NA | NA | NA | NA |
| SCN1A | 1483 | I | -14.77725 | 4.350934382 | 0.043884355 | 1.634668145 | 2.76 | 0.67 | 1.27 | 16.79 | -0.48 | 1 | -1 |
| SCN1A | 1484 | D | -17.16775 | 7.944711197 | 0.097904216 | 4.611580335 | 3.37 | 0.17 | -0.72 | 20.65 | 0.11 | 2 | 0 |
| SCN1A | 1485 | N | -15.342875 | 9.818872593 | 0.126075676 | 6.494343412 | NA | NA | NA | NA | NA | NA | NA |
| SCN1A | 1486 | F | -16.30693182 | 6.604006208 | 0.077751407 | 3.254976499 | 3.37 | 0.17 | -0.72 | 20.65 | 0.11 | 2 | 0 |
| SCN1A | 1487 | N | -20.70825 | 5.31592071 | 0.058389548 | 2.002120928 | 3.51 | -0.13 | -1.6 | 25.73 | 0.6 | 2 | 0 |
| SCN1A | 1488 | Q | -21.49986111 | 8.847275456 | 0.111471112 | 5.388490701 | NA | NA | NA | NA | NA | NA | NA |
| SCN1A | 1489 | Q | -18.17641667 | 10.78483222 | 0.1405955 | 7.320034107 | NA | NA | NA | NA | NA | NA | NA |
| SCN1A | 1490 | K | -21.02775 | 5.243654642 | 0.057303281 | 1.859000097 | 3.51 | -0.13 | -1.6 | 25.73 | 0.6 | 2 | 0 |
| SCN1A | 1491 | K | -24.92708333 | 8.506598556 | 0.106350227 | 4.621556686 | 3.96 | -0.77 | -3.63 | 34.26 | 1.41 | 2 | 0 |
| SCN1A | 1492 | K | -24.42441667 | 12.49688257 | 0.166330187 | 8.629053574 | NA | NA | NA | NA | NA | NA | NA |
| SCN1A | 1493 | F | -21.66656818 | 13.88003383 | 0.187121027 | 10.26984549 | NA | NA | NA | NA | NA | NA | NA |
| SCN1A | 1494 | G | -26.1675 | 12.62514182 | 0.168258116 | 6.446792436 | NA | NA | NA | NA | NA | NA | NA |
| SCN1A | 1495 | G | -24.48075 | 11.94830185 | 0.158084195 | 8.067835569 | NA | NA | NA | NA | NA | NA | NA |

| | | | | | | | | | | | | | |
|---|---|---|---|---|---|---|---|---|---|---|---|---|---|
| SCN1A | 1496 | Q | -21.95030556 | 10.15163898 | 0.131077654 | 6.356177742 | NA | NA | NA | NA | NA | NA | NA |
| SCN1A | 1497 | D | -18.508625 | 12.52108917 | 0.166694048 | 9.346412375 | NA | NA | NA | NA | NA | NA | NA |
| SCN1A | 1498 | I | -15.77775 | 12.60976693 | 0.168027008 | 9.269530384 | NA | NA | NA | NA | NA | NA | NA |
| SCN1A | 1499 | F | -11.58956818 | 12.86689232 | 0.171891989 | 10.41409296 | NA | NA | NA | NA | NA | NA | NA |
| SCN1A | 1500 | M | -14.80175 | 17.33487892 | 0.239052536 | 14.28842894 | NA | NA | NA | NA | NA | NA | NA |
| SCN1A | 1501 | T | -14.63232143 | 21.10797702 | 0.295767865 | 17.95466802 | NA | NA | NA | NA | NA | NA | NA |
| SCN1A | 1502 | E | -18.72641667 | 22.88828174 | 0.322528518 | 19.64561352 | NA | NA | NA | NA | NA | NA | NA |
| SCN1A | 1503 | E | -15.85597222 | 24.60531278 | 0.348338073 | 21.52161296 | NA | NA | NA | NA | NA | NA | NA |
| SCN1A | 1504 | Q | -13.97163889 | 21.57784 | 0.302830611 | 18.52878955 | NA | NA | NA | NA | NA | NA | NA |
| SCN1A | 1505 | K | -19.09130556 | 18.73717295 | 0.260131121 | 15.53733198 | NA | NA | NA | NA | NA | NA | NA |
| SCN1A | 1506 | K | -20.70908333 | 22.67698404 | 0.319352397 | 19.02159567 | NA | NA | NA | NA | NA | NA | NA |
| SCN1A | 1507 | Y | -16.714 | 22.88597433 | 0.322493834 | 19.91309203 | NA | NA | NA | NA | NA | NA | NA |
| SCN1A | 1508 | Y | -18.47133333 | 14.29203565 | 0.193314033 | 11.34874237 | NA | NA | NA | NA | NA | NA | NA |
| SCN1A | 1509 | N | -22.27475 | 18.65374027 | 0.258877002 | 12.49482494 | NA | NA | NA | NA | NA | NA | NA |
| SCN1A | 1510 | A | -20.90375 | 22.76720866 | 0.320708608 | 16.61404723 | NA | NA | NA | NA | NA | NA | NA |
| SCN1A | 1511 | M | -17.41775 | 21.6856281 | 0.304450828 | 17.9586418 | NA | NA | NA | NA | NA | NA | NA |
| SCN1A | 1512 | K | -21.88708333 | 13.66938817 | 0.183954706 | 7.556031555 | NA | NA | NA | NA | NA | NA | NA |
| SCN1A | 1513 | K | -23.85619444 | 21.03324666 | 0.294644556 | 14.92910683 | NA | NA | NA | NA | NA | NA | NA |
| SCN1A | 1514 | L | -19.267125 | 23.36240292 | 0.329655271 | 17.25148325 | NA | NA | NA | NA | NA | NA | NA |
| SCN1A | 1515 | G | -20.23175 | 21.93133776 | 0.308144214 | 15.82281714 | NA | NA | NA | NA | NA | NA | NA |
| SCN1A | 1516 | S | -23.70058333 | 19.88674626 | 0.277410934 | 13.24385297 | NA | NA | NA | NA | NA | NA | NA |
| SCN1A | 1517 | K | -23.02430556 | 23.66234993 | 0.334163925 | 17.0088499 | NA | NA | NA | NA | NA | NA | NA |
| SCN1A | 1518 | K | -24.02263889 | 23.96673816 | 0.338739338 | 17.31861846 | NA | NA | NA | NA | NA | NA | NA |
| SCN1A | 1519 | P | -22.83089286 | 30.50323822 | 0.436992748 | 23.83513932 | NA | NA | NA | NA | NA | NA | NA |
| SCN1A | 1520 | Q | -26.76363889 | 29.24658823 | 0.418103411 | 20.76428912 | NA | NA | NA | NA | NA | NA | NA |
| SCN1A | 1521 | K | -25.94397222 | 35.76334505 | 0.516060051 | 27.27251403 | NA | NA | NA | NA | NA | NA | NA |
| SCN1A | 1522 | P | -22.59532143 | 36.5322685 | 0.527618125 | 28.07048245 | NA | NA | NA | NA | NA | NA | NA |
| SCN1A | 1523 | I | -22.381875 | 40.79923972 | 0.591757109 | 34.25644231 | NA | NA | NA | NA | NA | NA | NA |
| SCN1A | 1524 | P | -24.44717857 | 43.69206677 | 0.635240643 | 35.19934701 | NA | NA | NA | NA | NA | NA | NA |
| SCN1A | 1525 | R | -23.87675 | 46.58652483 | 0.678748693 | 38.09463537 | NA | NA | NA | NA | NA | NA | NA |
| SCN1A | 1526 | P | -20.33803571 | 48.89032336 | 0.713378244 | 42.21761522 | NA | NA | NA | NA | NA | NA | NA |
| SCN1A | 1527 | G | -23.09675 | 52.21848922 | 0.763405574 | 43.72444051 | NA | NA | NA | NA | NA | NA | NA |
| SCN1A | 1528 | N | -22.82225 | 54.90229221 | 0.803747163 | 46.41097651 | NA | NA | NA | NA | NA | NA | NA |

| | | | | | | | | | | | | | |
|---|---|---|---|---|---|---|---|---|---|---|---|---|---|
| SCN1A | 1529 | K | -18.43386111 | 56.54294403 | 0.828408623 | 49.86119057 | NA | NA | NA | NA | NA | NA | NA |
| SCN1A | 1530 | F | -15.36620455 | 55.42302058 | 0.811574492 | 48.75270466 | NA | NA | NA | NA | NA | NA | NA |
| SCN1A | 1531 | Q | -15.95630556 | 52.13464401 | 0.762145255 | 45.4651565 | NA | NA | NA | NA | NA | NA | NA |
| SCN1A | 1532 | G | -18.75 | 51.5326565 | 0.753096479 | 44.84609286 | NA | NA | NA | NA | NA | NA | NA |
| SCN1A | 1533 | M | -16.796375 | 52.17704969 | 0.762782676 | 45.49507218 | NA | NA | NA | NA | NA | NA | NA |
| SCN1A | 1534 | V | -14.07075 | 49.94093684 | 0.729170542 | 46.8133551 | NA | NA | NA | NA | NA | NA | NA |
| SCN1A | 1535 | F | -18.65129545 | 45.43311574 | 0.661411223 | 38.75242789 | NA | NA | NA | NA | NA | NA | NA |
| SCN1A | 1536 | D | -18.650625 | 47.98122427 | 0.69971312 | 41.28916473 | NA | NA | NA | NA | NA | NA | NA |
| SCN1A | 1537 | F | -14.39684091 | 47.30869049 | 0.689603927 | 44.18430315 | NA | NA | NA | NA | NA | NA | NA |
| SCN1A | 1538 | V | -13.15875 | 44.14886651 | 0.642107029 | 41.02330126 | NA | NA | NA | NA | NA | NA | NA |
| SCN1A | 1539 | T | -16.70375 | 42.08305721 | 0.611054813 | 36.00134707 | NA | NA | NA | NA | NA | NA | NA |
| SCN1A | 1540 | R | -16.78756818 | 42.07521788 | 0.610936976 | 35.8878738 | NA | NA | NA | NA | NA | NA | NA |
| SCN1A | 1541 | Q | -15.68619444 | 36.31560217 | 0.524361305 | 33.20696682 | NA | NA | NA | NA | NA | NA | NA |
| SCN1A | 1542 | V | -12.62960714 | 41.84180854 | 0.607428484 | 38.72514319 | NA | NA | NA | NA | NA | NA | NA |
| SCN1A | 1543 | F | -9.792386364 | 41.1903635 | 0.597636286 | 38.07570709 | NA | NA | NA | NA | NA | NA | NA |
| SCN1A | 1544 | D | -11.3745 | 36.52034181 | 0.527438849 | 33.39531957 | NA | NA | NA | NA | NA | NA | NA |
| SCN1A | 1545 | I | -9.611125 | 35.63844702 | 0.514182646 | 32.52388398 | NA | NA | NA | NA | NA | NA | NA |
| SCN1A | 1546 | S | -6.756416667 | 39.99707139 | 0.579699315 | 36.88595485 | NA | NA | NA | NA | NA | NA | NA |
| SCN1A | 1547 | I | -5.755 | 38.14843526 | 0.551911538 | 35.05294367 | NA | NA | NA | NA | NA | NA | NA |
| SCN1A | 1548 | M | -6.779875 | 34.53166034 | 0.49754598 | 31.41396287 | NA | NA | NA | NA | NA | NA | NA |
| SCN1A | 1549 | I | -3.687125 | 36.87015788 | 0.53269711 | 33.7699279 | NA | NA | NA | NA | NA | NA | NA |
| SCN1A | 1550 | L | -1.5035 | 38.80958702 | 0.561849642 | 35.90964813 | NA | NA | NA | NA | NA | NA | NA |
| SCN1A | 1551 | I | -1.395125 | 34.66752893 | 0.499588289 | 31.55544697 | NA | NA | NA | NA | NA | NA | NA |
| SCN1A | 1552 | C | -0.62775 | 33.55675944 | 0.482891756 | 30.68538331 | NA | NA | NA | NA | NA | NA | NA |
| SCN1A | 1553 | L | 1.62275 | 37.37558204 | 0.540294394 | 33.96915083 | NA | NA | NA | NA | NA | NA | NA |
| SCN1A | 1554 | N | 3.258 | 36.46189323 | 0.526560279 | 33.02520493 | NA | NA | NA | NA | NA | NA | NA |
| SCN1A | 1555 | M | 2.81 | 29.17170208 | 0.41697776 | 24.73448349 | NA | NA | NA | NA | NA | NA | NA |
| SCN1A | 1556 | V | 5.229035714 | 35.14034609 | 0.506695441 | 30.60135347 | NA | NA | NA | NA | NA | NA | NA |
| SCN1A | 1557 | T | 7.770178571 | 36.98294507 | 0.53439247 | 32.44813886 | NA | NA | NA | NA | NA | NA | NA |
| SCN1A | 1558 | M | 7.346625 | 33.48310946 | 0.481784686 | 28.9526252 | NA | NA | NA | NA | NA | NA | NA |
| SCN1A | 1559 | M | 7.90425 | 30.74233807 | 0.440586777 | 26.20306474 | NA | NA | NA | NA | NA | NA | NA |
| SCN1A | 1560 | V | 12.16946429 | 34.92158562 | 0.503407143 | 30.39283094 | NA | NA | NA | NA | NA | NA | NA |
| SCN1A | 1561 | E | 13.05975 | 33.55340352 | 0.482841312 | 29.21111299 | NA | NA | NA | NA | NA | NA | NA |

| | | | | | | | | | | | | | |
|---|---|---|---|---|---|---|---|---|---|---|---|---|---|
| SCN1A | 1562 | T | 17.51103571 | 34.38735944 | 0.495376921 | 34.01547493 | NA | NA | NA | NA | NA | NA | NA |
| SCN1A | 1563 | D | 21.092 | 32.10378459 | 0.461051363 | 32.41189336 | NA | NA | NA | NA | NA | NA | NA |
| SCN1A | 1564 | D | 24.25125 | 35.39766588 | 0.510563344 | 35.78914393 | NA | NA | NA | NA | NA | NA | NA |
| SCN1A | 1565 | Q | 20.24919444 | 38.12775266 | 0.551600647 | 38.52721311 | NA | NA | NA | NA | NA | NA | NA |
| SCN1A | 1566 | S | 22.96741667 | 41.69430238 | 0.605211245 | 42.08340005 | NA | NA | NA | NA | NA | NA | NA |
| SCN1A | 1567 | E | 22.53286111 | 44.53786276 | 0.647954226 | 44.950373 | NA | NA | NA | NA | NA | NA | NA |
| SCN1A | 1568 | Y | 20.41225 | 46.2566691 | 0.673790466 | 45.87418485 | NA | NA | NA | NA | NA | NA | NA |
| SCN1A | 1569 | V | 18.26560714 | 42.12761267 | 0.611724549 | 37.77478879 | NA | NA | NA | NA | NA | NA | NA |
| SCN1A | 1570 | T | 18.11989286 | 41.7431529 | 0.605945541 | 41.38239793 | NA | NA | NA | NA | NA | NA | NA |
| SCN1A | 1571 | T | 17.56703571 | 45.34590813 | 0.660100362 | 41.12723093 | NA | NA | NA | NA | NA | NA | NA |
| SCN1A | 1572 | I | 14.578375 | 45.25614953 | 0.658751155 | 40.72822504 | NA | NA | NA | NA | NA | NA | NA |
| SCN1A | 1573 | L | 12.97175 | 39.8555424 | 0.577571921 | 35.33086279 | NA | NA | NA | NA | NA | NA | NA |
| SCN1A | 1574 | S | 13.10308333 | 43.66924847 | 0.63489765 | 39.14155214 | NA | NA | NA | NA | NA | NA | NA |
| SCN1A | 1575 | R | 13.20738636 | 45.97019505 | 0.669484331 | 41.43995957 | NA | NA | NA | NA | NA | NA | NA |
| SCN1A | 1576 | I | 8.532875 | 43.25018359 | 0.628598475 | 38.71609608 | NA | NA | NA | NA | NA | NA | NA |
| SCN1A | 1577 | N | 8.163625 | 39.58524345 | 0.573508922 | 35.04886919 | NA | NA | NA | NA | NA | NA | NA |
| SCN1A | 1578 | L | 8.01225 | 44.72385119 | 0.650749912 | 40.18705214 | NA | NA | NA | NA | NA | NA | NA |
| SCN1A | 1579 | V | 5.337464286 | 46.0837223 | 0.671190816 | 41.54502494 | NA | NA | NA | NA | NA | NA | NA |
| SCN1A | 1580 | F | 3.214113636 | 41.79708478 | 0.606756218 | 38.28158005 | NA | NA | NA | NA | NA | NA | NA |
| SCN1A | 1581 | I | 3.634375 | 42.78116559 | 0.62154843 | 39.97436456 | NA | NA | NA | NA | NA | NA | NA |
| SCN1A | 1582 | V | 2.193321429 | 46.25838806 | 0.673816305 | 43.16434881 | NA | NA | NA | NA | NA | NA | NA |
| SCN1A | 1583 | L | 0.455375 | 45.30641388 | 0.659506704 | 42.19764849 | NA | NA | NA | NA | NA | NA | NA |
| SCN1A | 1584 | F | -0.787386364 | 39.49633616 | 0.572172512 | 36.38482808 | NA | NA | NA | NA | NA | NA | NA |
| SCN1A | 1585 | T | -0.892607143 | 44.89833712 | 0.653372697 | 41.79196142 | NA | NA | NA | NA | NA | NA | NA |
| SCN1A | 1586 | G | -3.402 | 47.28006944 | 0.68917371 | 44.17476093 | NA | NA | NA | NA | NA | NA | NA |
| SCN1A | 1587 | E | -5.565305556 | 40.42780332 | 0.586173862 | 37.32144139 | NA | NA | NA | NA | NA | NA | NA |
| SCN1A | 1588 | C | -5.804416667 | 43.89438598 | 0.638281804 | 40.91385838 | NA | NA | NA | NA | NA | NA | NA |
| SCN1A | 1589 | V | -5.820178571 | 46.99511516 | 0.684890419 | 43.8952647 | NA | NA | NA | NA | NA | NA | NA |
| SCN1A | 1590 | L | -8.678875 | 48.31446941 | 0.704722295 | 45.19632006 | NA | NA | NA | NA | NA | NA | NA |
| SCN1A | 1591 | K | -11.16075 | 41.18010453 | 0.597482079 | 38.05350617 | NA | NA | NA | NA | NA | NA | NA |
| SCN1A | 1592 | L | -9.82525 | 45.82674035 | 0.667327992 | 42.70987209 | NA | NA | NA | NA | NA | NA | NA |
| SCN1A | 1593 | I | -11.973125 | 49.59819292 | 0.724018587 | 46.47397925 | NA | NA | NA | NA | NA | NA | NA |
| SCN1A | 1594 | S | -15.30541667 | 47.39882726 | 0.690958818 | 41.31749348 | NA | NA | NA | NA | NA | NA | NA |

| | | | | | | | | | | | | | |
|---|---|---|---|---|---|---|---|---|---|---|---|---|---|
| SCN1A | 1595 | L | -15.717375 | 43.25934915 | 0.628736247 | 37.15347887 | NA | NA | NA | NA | NA | NA | NA |
| SCN1A | 1596 | R | -15.30820455 | 48.42443563 | 0.706375252 | 42.33735776 | NA | NA | NA | NA | NA | NA | NA |
| SCN1A | 1597 | H | -19.34335 | 46.69602418 | 0.680394633 | 40.6155718 | NA | NA | NA | NA | NA | NA | NA |
| SCN1A | 1598 | Y | -17.93666667 | 41.99025464 | 0.609659851 | 35.91928719 | NA | NA | NA | NA | NA | NA | NA |
| SCN1A | 1599 | Y | -11.75516667 | 41.4581242 | 0.601661132 | 38.39118428 | NA | NA | NA | NA | NA | NA | NA |
| SCN1A | 1600 | F | -12.56038636 | 39.65188062 | 0.574510579 | 37.27452175 | NA | NA | NA | NA | NA | NA | NA |
| SCN1A | 1601 | T | -17.33075 | 36.03759377 | 0.520182421 | 29.97237913 | NA | NA | NA | NA | NA | NA | NA |
| SCN1A | 1602 | I | -15.375 | 34.81395699 | 0.501789323 | 28.70675523 | NA | NA | NA | NA | NA | NA | NA |
| SCN1A | 1603 | G | -11.92375 | 34.02379344 | 0.489911978 | 30.94966946 | NA | NA | NA | NA | NA | NA | NA |
| SCN1A | 1604 | W | -10.31339286 | 28.71147178 | 0.410059808 | 25.59435352 | NA | NA | NA | NA | NA | NA | NA |
| SCN1A | 1605 | N | -10.885 | 36.33316393 | 0.524625284 | 33.21161762 | NA | NA | NA | NA | NA | NA | NA |
| SCN1A | 1606 | I | -9.414125 | 36.94344375 | 0.533798706 | 34.08507072 | NA | NA | NA | NA | NA | NA | NA |
| SCN1A | 1607 | F | -5.162295455 | 31.53748034 | 0.452538959 | 28.43633391 | NA | NA | NA | NA | NA | NA | NA |
| SCN1A | 1608 | D | -5.396375 | 36.44829995 | 0.526355951 | 33.3511094 | NA | NA | NA | NA | NA | NA | NA |
| SCN1A | 1609 | F | -5.170659091 | 39.13306855 | 0.566712055 | 36.03579776 | NA | NA | NA | NA | NA | NA | NA |
| SCN1A | 1610 | V | -2.80375 | 36.57757018 | 0.528299077 | 33.47904287 | NA | NA | NA | NA | NA | NA | NA |
| SCN1A | 1611 | V | -0.972035714 | 34.72573397 | 0.500463199 | 31.64167022 | NA | NA | NA | NA | NA | NA | NA |
| SCN1A | 1612 | V | -0.770178571 | 39.40180561 | 0.570751576 | 36.31824629 | NA | NA | NA | NA | NA | NA | NA |
| SCN1A | 1613 | I | 0.589625 | 41.42389495 | 0.601146615 | 38.33997231 | NA | NA | NA | NA | NA | NA | NA |
| SCN1A | 1614 | L | 1.8535 | 36.15443151 | 0.521938668 | 33.06555191 | NA | NA | NA | NA | NA | NA | NA |
| SCN1A | 1615 | S | 3.53525 | 40.36374232 | 0.585210929 | 36.87377725 | NA | NA | NA | NA | NA | NA | NA |
| SCN1A | 1616 | I | 3.448 | 43.12775554 | 0.626758198 | 39.59429241 | NA | NA | NA | NA | NA | NA | NA |
| SCN1A | 1617 | V | 5.834607143 | 43.0181923 | 0.625111298 | 38.47847729 | NA | NA | NA | NA | NA | NA | NA |
| SCN1A | 1618 | G | 8.475 | 41.5176643 | 0.602556109 | 36.98255061 | NA | NA | NA | NA | NA | NA | NA |
| SCN1A | 1619 | M | 8.5095 | 43.29732392 | 0.629307065 | 38.76208643 | NA | NA | NA | NA | NA | NA | NA |
| SCN1A | 1620 | F | 9.165477273 | 45.83111989 | 0.667393823 | 41.29636768 | NA | NA | NA | NA | NA | NA | NA |
| SCN1A | 1621 | L | 11.972125 | 42.63046697 | 0.619283204 | 42.19540497 | NA | NA | NA | NA | NA | NA | NA |
| SCN1A | 1622 | A | 14.41555 | 44.48514706 | 0.64716183 | 44.06330856 | NA | NA | NA | NA | NA | NA | NA |
| SCN1A | 1623 | E | 15.16375 | 47.06070311 | 0.685876305 | 46.65205587 | NA | NA | NA | NA | NA | NA | NA |
| SCN1A | 1624 | L | 15.7505 | 46.42316855 | 0.676293203 | 46.36830609 | NA | NA | NA | NA | NA | NA | NA |
| SCN1A | 1625 | I | 18.052875 | 42.13133393 | 0.611780485 | 42.84771477 | NA | NA | NA | NA | NA | NA | NA |
| SCN1A | 1626 | E | 20.70275 | 46.3033295 | 0.674491842 | 47.07697783 | NA | NA | NA | NA | NA | NA | NA |
| SCN1A | 1627 | K | 20.54608333 | 48.45324101 | 0.70680824 | 49.21032215 | NA | NA | NA | NA | NA | NA | NA |

| | | | | | | | | | | | | | |
|---|---|---|---|---|---|---|---|---|---|---|---|---|---|
| SCN1A | 1628 | Y | 18.85025 | 45.70520238 | 0.665501094 | 46.48296212 | NA | NA | NA | NA | NA | NA | NA |
| SCN1A | 1629 | F | 18.96084091 | 40.39197136 | 0.585635254 | 41.23791039 | NA | NA | NA | NA | NA | NA | NA |
| SCN1A | 1630 | V | 21.48060714 | 40.22326761 | 0.583099383 | 40.93840388 | NA | NA | NA | NA | NA | NA | NA |
| SCN1A | 1631 | S | 19.49308333 | 36.17708362 | 0.522279163 | 36.96843281 | NA | NA | NA | NA | NA | NA | NA |
| SCN1A | 1632 | P | 20.41517857 | 35.34050677 | 0.509704157 | 36.11048765 | NA | NA | NA | NA | NA | NA | NA |
| SCN1A | 1633 | T | 18.23089286 | 31.41727125 | 0.450732036 | 32.13877323 | NA | NA | NA | NA | NA | NA | NA |
| SCN1A | 1634 | L | 14.98525 | 34.78160116 | 0.501302966 | 35.57356437 | NA | NA | NA | NA | NA | NA | NA |
| SCN1A | 1635 | F | 15.73893182 | 36.35183612 | 0.524905955 | 35.98730806 | NA | NA | NA | NA | NA | NA | NA |
| SCN1A | 1636 | R | 16.34220455 | 30.6038782 | 0.438505518 | 31.21102425 | NA | NA | NA | NA | NA | NA | NA |
| SCN1A | 1637 | V | 11.51560714 | 31.67094665 | 0.454545159 | 31.30682488 | NA | NA | NA | NA | NA | NA | NA |
| SCN1A | 1638 | I | 10.8395 | 35.24767757 | 0.508308794 | 34.81835377 | NA | NA | NA | NA | NA | NA | NA |
| SCN1A | 1639 | R | 11.56856818 | 35.46798912 | 0.511620408 | 31.02955122 | NA | NA | NA | NA | NA | NA | NA |
| SCN1A | 1640 | L | 8.10675 | 30.15877081 | 0.431814885 | 25.90279595 | NA | NA | NA | NA | NA | NA | NA |
| SCN1A | 1641 | A | 4.86115 | 34.67649955 | 0.499723131 | 30.14021026 | NA | NA | NA | NA | NA | NA | NA |
| SCN1A | 1642 | R | 6.950295455 | 34.7572868 | 0.500937485 | 30.21822751 | NA | NA | NA | NA | NA | NA | NA |
| SCN1A | 1643 | I | 3.748375 | 29.83025407 | 0.426876785 | 25.24151651 | NA | NA | NA | NA | NA | NA | NA |
| SCN1A | 1644 | G | 0.546 | 32.72536663 | 0.470394674 | 29.63304816 | NA | NA | NA | NA | NA | NA | NA |
| SCN1A | 1645 | R | -1.666022727 | 33.69554377 | 0.484977893 | 30.6044363 | NA | NA | NA | NA | NA | NA | NA |
| SCN1A | 1646 | I | -0.7705 | 30.25721106 | 0.43329459 | 27.15894367 | NA | NA | NA | NA | NA | NA | NA |
| SCN1A | 1647 | L | -2.088875 | 26.94238184 | 0.38346773 | 23.83663348 | NA | NA | NA | NA | NA | NA | NA |
| SCN1A | 1648 | R | -4.897295455 | 31.45342085 | 0.451275419 | 28.34802807 | NA | NA | NA | NA | NA | NA | NA |
| SCN1A | 1649 | L | -4.409 | 28.78175681 | 0.411116297 | 25.68038202 | NA | NA | NA | NA | NA | NA | NA |
| SCN1A | 1650 | I | -7.47875 | 25.43916312 | 0.360872095 | 22.33272592 | NA | NA | NA | NA | NA | NA | NA |
| SCN1A | 1651 | K | -10.96452778 | 29.40919633 | 0.420547655 | 26.29713578 | NA | NA | NA | NA | NA | NA | NA |
| SCN1A | 1652 | G | -11.56525 | 28.75231158 | 0.410673691 | 25.62991821 | NA | NA | NA | NA | NA | NA | NA |
| SCN1A | 1653 | A | -10.75575 | 24.64587763 | 0.348947823 | 21.52586887 | NA | NA | NA | NA | NA | NA | NA |
| SCN1A | 1654 | K | -14.48363889 | 24.01837815 | 0.339515564 | 20.90473635 | NA | NA | NA | NA | NA | NA | NA |
| SCN1A | 1655 | G | -12.8925 | 19.61130801 | 0.273270683 | 16.48692725 | NA | NA | NA | NA | NA | NA | NA |
| SCN1A | 1656 | I | -9.015875 | 20.63093253 | 0.28859717 | 17.51791883 | NA | NA | NA | NA | NA | NA | NA |
| SCN1A | 1657 | R | -11.79220455 | 21.30931984 | 0.29879435 | 18.18511517 | NA | NA | NA | NA | NA | NA | NA |
| SCN1A | 1658 | T | -14.00903571 | 18.07339506 | 0.250153542 | 14.95359533 | NA | NA | NA | NA | NA | NA | NA |
| SCN1A | 1659 | L | -10.684375 | 15.55175116 | 0.212249448 | 12.42679475 | NA | NA | NA | NA | NA | NA | NA |
| SCN1A | 1660 | L | -8.541375 | 18.95892534 | 0.263464392 | 15.84489392 | NA | NA | NA | NA | NA | NA | NA |

| | | | | | | | | | | | | | |
|---|---|---|---|---|---|---|---|---|---|---|---|---|---|
| SCN1A | 1661 | F | -14.44693182 | 18.90170807 | 0.262604331 | 15.82740107 | NA | NA | NA | NA | NA | NA | NA |
| SCN1A | 1662 | A | -12.25495 | 15.28405381 | 0.208225555 | 12.24864709 | NA | NA | NA | NA | NA | NA | NA |
| SCN1A | 1663 | L | -8.396625 | 15.65146233 | 0.213748257 | 12.55889118 | NA | NA | NA | NA | NA | NA | NA |
| SCN1A | 1664 | M | -11.298375 | 19.27350682 | 0.268193024 | 16.18875249 | NA | NA | NA | NA | NA | NA | NA |
| SCN1A | 1665 | M | -14.299875 | 16.90185274 | 0.232543502 | 13.91432666 | NA | NA | NA | NA | NA | NA | NA |
| SCN1A | 1666 | S | -10.31975 | 13.78202924 | 0.185647871 | 10.76391055 | NA | NA | NA | NA | NA | NA | NA |
| SCN1A | 1667 | L | -8.0235 | 17.53858472 | 0.24211454 | 14.45573069 | NA | NA | NA | NA | NA | NA | NA |
| SCN1A | 1668 | P | -10.80717857 | 19.04811012 | 0.264804973 | 15.96549495 | NA | NA | NA | NA | NA | NA | NA |
| SCN1A | 1669 | A | -10.20575 | 15.99320484 | 0.21888516 | 12.90713321 | NA | NA | NA | NA | NA | NA | NA |
| SCN1A | 1670 | L | -6.561875 | 12.46717426 | 0.165883626 | 9.377120269 | NA | NA | NA | NA | NA | NA | NA |
| SCN1A | 1671 | F | -5.85075 | 17.8884458 | 0.247373477 | 14.81779076 | NA | NA | NA | NA | NA | NA | NA |
| SCN1A | 1672 | N | -7.6145 | 18.52567737 | 0.256952024 | 15.44455878 | NA | NA | NA | NA | NA | NA | NA |
| SCN1A | 1673 | I | -5.255375 | 13.45543853 | 0.180738722 | 10.38414983 | NA | NA | NA | NA | NA | NA | NA |
| SCN1A | 1674 | G | -1.79525 | 17.01894336 | 0.23430355 | 13.92740761 | NA | NA | NA | NA | NA | NA | NA |
| SCN1A | 1675 | L | -2.139125 | 19.71936525 | 0.274894946 | 16.6241318 | NA | NA | NA | NA | NA | NA | NA |
| SCN1A | 1676 | L | -2.374375 | 17.45532873 | 0.240863077 | 14.40142336 | NA | NA | NA | NA | NA | NA | NA |
| SCN1A | 1677 | L | 0.59 | 14.00654294 | 0.189022649 | 13.49363824 | NA | NA | NA | NA | NA | NA | NA |
| SCN1A | 1678 | F | 2.399386364 | 18.61376974 | 0.258276185 | 18.00121287 | NA | NA | NA | NA | NA | NA | NA |
| SCN1A | 1679 | L | 1.01125 | 19.98909138 | 0.278949334 | 18.84072059 | NA | NA | NA | NA | NA | NA | NA |
| SCN1A | 1680 | V | 2.31275 | 15.90849239 | 0.217611805 | 14.74743953 | NA | NA | NA | NA | NA | NA | NA |
| SCN1A | 1681 | M | 5.616375 | 14.82944753 | 0.20139214 | 13.75241745 | NA | NA | NA | NA | NA | NA | NA |
| SCN1A | 1682 | F | 6.354931818 | 18.91592453 | 0.262818025 | 18.98852134 | NA | NA | NA | NA | NA | NA | NA |
| SCN1A | 1683 | I | 5.338 | 19.09288595 | 0.265478021 | 19.05783248 | NA | NA | NA | NA | NA | NA | NA |
| SCN1A | 1684 | Y | 6.976 | 14.4150974 | 0.195163836 | 14.13278883 | NA | NA | NA | NA | NA | NA | NA |
| SCN1A | 1685 | A | 10.76555 | 17.42251101 | 0.240369778 | 17.04949291 | NA | NA | NA | NA | NA | NA | NA |
| SCN1A | 1686 | I | 10.670375 | 19.67901603 | 0.274288436 | 19.99732191 | NA | NA | NA | NA | NA | NA | NA |
| SCN1A | 1687 | F | 10.25056818 | 18.01352242 | 0.249253567 | 18.37594657 | NA | NA | NA | NA | NA | NA | NA |
| SCN1A | 1688 | G | 14.652 | 16.16002082 | 0.221392655 | 16.65447694 | NA | NA | NA | NA | NA | NA | NA |
| SCN1A | 1689 | M | 15.768625 | 17.93656729 | 0.248096815 | 18.5365886 | NA | NA | NA | NA | NA | NA | NA |
| SCN1A | 1690 | S | 16.17825 | 20.04015758 | 0.279716936 | 20.70319597 | NA | NA | NA | NA | NA | NA | NA |
| SCN1A | 1691 | N | 16.380125 | 17.54749105 | 0.242248415 | 17.82374264 | NA | NA | NA | NA | NA | NA | NA |
| SCN1A | 1692 | F | 16.90702273 | 13.2196335 | 0.177194219 | 13.95377478 | NA | NA | NA | NA | NA | NA | NA |
| SCN1A | 1693 | A | 20.88475 | 17.12681532 | 0.235925028 | 17.36576969 | NA | NA | NA | NA | NA | NA | NA |

| | | | | | | | | | | | | | |
|---|---|---|---|---|---|---|---|---|---|---|---|---|---|
| SCN1A | 1694 | Y | 24.11041667 | 16.09435171 | 0.220405549 | 15.0343608 | NA | NA | NA | NA | NA | NA | NA |
| SCN1A | 1695 | V | 23.23917857 | 12.64314933 | 0.168528796 | 10.64524178 | NA | NA | NA | NA | NA | NA | NA |
| SCN1A | 1696 | K | 27.62708333 | 6.399512325 | 0.074677557 | 2.634602831 | 3.8 | -0.23 | 0.05 | 14.2 | 0.5 | 1 | -1 |
| SCN1A | 1697 | R | 27.11965909 | 9.285853057 | 0.118063592 | 5.647964086 | NA | NA | NA | NA | NA | NA | NA |
| SCN1A | 1698 | E | 25.82119444 | 4.691992008 | 0.049010963 | 1.654557883 | 3.78 | -0.14 | -0.83 | 25.73 | 0.66 | 2 | 0 |
| SCN1A | 1699 | V | 23.48889286 | 3.724650588 | 0.034470369 | 2.059225758 | 1.84 | -0.61 | -0.7 | 17.69 | 1.34 | 1 | -1 |
| SCN1A | 1700 | G | 20.84775 | 6.08327831 | 0.069924085 | 4.277601981 | 1.84 | -0.61 | -0.7 | 17.69 | 1.34 | 1 | -1 |
| SCN1A | 1701 | I | 21.889875 | 8.452963149 | 0.105544006 | 6.784610594 | NA | NA | NA | NA | NA | NA | NA |
| SCN1A | 1702 | D | 23.632625 | 11.7180137 | 0.154622618 | 10.00631127 | NA | NA | NA | NA | NA | NA | NA |
| SCN1A | 1703 | D | 25.691 | 15.18704715 | 0.206767399 | 13.41492284 | NA | NA | NA | NA | NA | NA | NA |
| SCN1A | 1704 | M | 21.82 | 15.445907 | 0.210658452 | 15.8258159 | NA | NA | NA | NA | NA | NA | NA |
| SCN1A | 1705 | F | 19.16193182 | 11.92335343 | 0.157709183 | 12.22073498 | NA | NA | NA | NA | NA | NA | NA |
| SCN1A | 1706 | N | 20.221 | 13.55110453 | 0.182176726 | 13.94874667 | NA | NA | NA | NA | NA | NA | NA |
| SCN1A | 1707 | F | 16.80093182 | 11.37417443 | 0.149454198 | 12.16999751 | NA | NA | NA | NA | NA | NA | NA |
| SCN1A | 1708 | E | 21.72363889 | 17.42685003 | 0.240435 | 17.87531267 | NA | NA | NA | NA | NA | NA | NA |
| SCN1A | 1709 | T | 18.14960714 | 19.00188701 | 0.26411017 | 19.8620391 | NA | NA | NA | NA | NA | NA | NA |
| SCN1A | 1710 | F | 12.30147727 | 19.43339654 | 0.270596406 | 20.07952324 | NA | NA | NA | NA | NA | NA | NA |
| SCN1A | 1711 | G | 14.75175 | 18.0882715 | 0.250377158 | 18.82298006 | NA | NA | NA | NA | NA | NA | NA |
| SCN1A | 1712 | N | 17.486125 | 14.92635609 | 0.202848821 | 15.97061329 | NA | NA | NA | NA | NA | NA | NA |
| SCN1A | 1713 | S | 14.00225 | 14.77099255 | 0.200513474 | 15.2911185 | NA | NA | NA | NA | NA | NA | NA |
| SCN1A | 1714 | M | 10.105 | 15.47907278 | 0.211156983 | 15.31539736 | NA | NA | NA | NA | NA | NA | NA |
| SCN1A | 1715 | I | 13.59175 | 13.14096652 | 0.176011736 | 13.7034756 | NA | NA | NA | NA | NA | NA | NA |
| SCN1A | 1716 | C | 13.52525 | 10.62861407 | 0.138247306 | 10.99795984 | NA | NA | NA | NA | NA | NA | NA |
| SCN1A | 1717 | L | 9.63475 | 11.20157016 | 0.146859696 | 11.32322922 | NA | NA | NA | NA | NA | NA | NA |
| SCN1A | 1718 | F | 7.60275 | 10.9735533 | 0.143432261 | 10.41652535 | NA | NA | NA | NA | NA | NA | NA |
| SCN1A | 1719 | Q | 11.79308333 | 7.162513037 | 0.086146604 | 8.037929041 | 0.3 | 0.17 | -1.73 | 18.33 | -0.07 | 1 | 1 |
| SCN1A | 1720 | I | 9.44625 | 6.742431164 | 0.079832142 | 6.960681116 | 0.3 | 0.17 | -1.73 | 18.33 | -0.07 | 1 | 1 |
| SCN1A | 1721 | T | 5.162607143 | 8.209786234 | 0.101888692 | 7.581356609 | NA | NA | NA | NA | NA | NA | NA |
| SCN1A | 1722 | T | 5.685178571 | 4.299572769 | 0.043112313 | 3.752749229 | 0.69 | -0.73 | -1.7 | 18.18 | 1.13 | 1 | 1 |
| SCN1A | 1723 | S | 7.08475 | 2.214839723 | 0.011775645 | 2.062253389 | 0.81 | -0.72 | -1.8 | 17.61 | 1.11 | 1 | 1 |
| SCN1A | 1724 | A | 10.20555 | 4.439381714 | 0.045213851 | 4.680701315 | 0.3 | 0.17 | -1.73 | 18.33 | -0.07 | 1 | 1 |
| SCN1A | 1725 | G | 13.433 | 3.439386864 | 0.030182427 | 3.783515035 | -0.01 | -1.04 | -3.5 | 49.7 | 2.63 | 1 | -1 |
| SCN1A | 1726 | W | 11.40196429 | 1.911777445 | 0.007220164 | 2.824833581 | -0.01 | -1.04 | -3.5 | 49.7 | 2.63 | 1 | -1 |

| | | | | | | | | | | | | | |
|---|---|---|---|---|---|---|---|---|---|---|---|---|---|
| SCN1A | 1727 | D | 15.348875 | 1.43144193 | 0 | 1.516748112 | -0.01 | -1.04 | -3.5 | 49.7 | 2.63 | 1 | -1 |
| SCN1A | 1728 | G | 18.011 | 1.471687807 | 6.05E-04 | 1.859097162 | -0.03 | -0.92 | -1.95 | 26.54 | 1.72 | 1 | -1 |
| SCN1A | 1729 | L | 16.665375 | 4.137853308 | 0.040681427 | 4.48973487 | -0.03 | -0.92 | -1.95 | 26.54 | 1.72 | 1 | -1 |
| SCN1A | 1730 | L | 16.1875 | 5.293456432 | 0.058051877 | 5.605761405 | -0.03 | -0.92 | -1.95 | 26.54 | 1.72 | 1 | -1 |
| SCN1A | 1731 | A | 20.25575 | 3.74644338 | 0.034797948 | 1.923811385 | 1.84 | -0.61 | -0.7 | 17.69 | 1.34 | 1 | -1 |
| SCN1A | 1732 | P | 21.68632143 | 6.4893794 | 0.076028394 | 4.328552031 | 2.92 | -0.26 | -0.25 | 24.93 | 1.04 | 2 | -2 |
| SCN1A | 1733 | I | 19.076 | 9.19977228 | 0.116769669 | 7.538873211 | NA | NA | NA | NA | NA | NA | NA |
| SCN1A | 1734 | L | 19.369625 | 8.350440168 | 0.104002932 | 5.893983071 | NA | NA | NA | NA | NA | NA | NA |
| SCN1A | 1735 | N | 24.126 | 7.50257036 | 0.091258176 | 4.951499806 | 2.92 | -0.26 | -0.25 | 24.93 | 1.04 | 2 | -2 |
| SCN1A | 1736 | S | 24.61375 | 12.06020904 | 0.159766328 | 9.475826483 | NA | NA | NA | NA | NA | NA | NA |
| SCN1A | 1737 | K | 27.69252778 | 11.07411527 | 0.144943858 | 7.313782771 | NA | NA | NA | NA | NA | NA | NA |
| SCN1A | 1738 | P | 31.30660714 | 11.11343628 | 0.145534912 | 7.468982096 | NA | NA | NA | NA | NA | NA | NA |
| SCN1A | 1739 | P | 32.69817857 | 7.229431305 | 0.087152486 | 5.038024057 | 2.46 | 0.08 | 1.13 | 1.72 | -0.09 | 0 | 0 |
| SCN1A | 1740 | D | 29.36225 | 6.655696733 | 0.078528394 | 2.886533713 | 3.8 | -0.23 | 0.05 | 14.2 | 0.5 | 1 | -1 |
| SCN1A | 1741 | C | 29.70725 | 9.515230213 | 0.121511475 | 5.749421942 | NA | NA | NA | NA | NA | NA | NA |
| SCN1A | 1742 | D | 32.660125 | 12.22977068 | 0.162315094 | 8.474267595 | NA | NA | NA | NA | NA | NA | NA |
| SCN1A | 1743 | P | 30.36889286 | 15.26281645 | 0.207906326 | 11.52568093 | NA | NA | NA | NA | NA | NA | NA |
| SCN1A | 1744 | N | 32.8935 | 17.6246769 | 0.243408635 | 13.87691417 | NA | NA | NA | NA | NA | NA | NA |
| SCN1A | 1745 | K | 32.19641667 | 13.51596856 | 0.181648579 | 9.754692819 | NA | NA | NA | NA | NA | NA | NA |
| SCN1A | 1746 | V | 34.00832143 | 20.3029305 | 0.283666808 | 16.54375293 | NA | NA | NA | NA | NA | NA | NA |
| SCN1A | 1747 | N | 30.225375 | 19.10286183 | 0.265627974 | 15.35334793 | NA | NA | NA | NA | NA | NA | NA |
| SCN1A | 1748 | P | 31.44203571 | 21.42143795 | 0.300479654 | 17.65604563 | NA | NA | NA | NA | NA | NA | NA |
| SCN1A | 1749 | G | 28.40725 | 25.55647474 | 0.362635465 | 21.80322446 | NA | NA | NA | NA | NA | NA | NA |
| SCN1A | 1750 | S | 27.44175 | 25.59941704 | 0.363280953 | 21.85727089 | NA | NA | NA | NA | NA | NA | NA |
| SCN1A | 1751 | S | 27.43741667 | 25.54112531 | 0.36240474 | 21.77370029 | NA | NA | NA | NA | NA | NA | NA |
| SCN1A | 1752 | V | 25.75032143 | 22.70956574 | 0.319842148 | 18.95070335 | NA | NA | NA | NA | NA | NA | NA |
| SCN1A | 1753 | K | 29.13452778 | 20.27032743 | 0.283176734 | 16.50326614 | NA | NA | NA | NA | NA | NA | NA |
| SCN1A | 1754 | G | 27.4535 | 17.06673926 | 0.235021994 | 13.30967959 | NA | NA | NA | NA | NA | NA | NA |
| SCN1A | 1755 | D | 28.10075 | 12.09965929 | 0.160359325 | 8.334268985 | NA | NA | NA | NA | NA | NA | NA |
| SCN1A | 1756 | C | 24.58025 | 12.93142475 | 0.172862008 | 9.216619932 | NA | NA | NA | NA | NA | NA | NA |
| SCN1A | 1757 | G | 21.19825 | 14.32080113 | 0.193746421 | 11.73041439 | NA | NA | NA | NA | NA | NA | NA |
| SCN1A | 1758 | N | 18.761875 | 14.8839787 | 0.202211826 | 12.846287 | NA | NA | NA | NA | NA | NA | NA |
| SCN1A | 1759 | P | 17.82903571 | 13.78370172 | 0.185673011 | 13.99895203 | NA | NA | NA | NA | NA | NA | NA |

| | | | | | | | | | | | | | |
|---|---|---|---|---|---|---|---|---|---|---|---|---|---|
| SCN1A | 1760 | S | 14.35208333 | 15.57863043 | 0.212653484 | 15.60498338 | NA | NA | NA | NA | NA | NA | NA |
| SCN1A | 1761 | V | 13.70760714 | 14.72747395 | 0.199859324 | 14.78789524 | NA | NA | NA | NA | NA | NA | NA |
| SCN1A | 1762 | G | 13.6695 | 11.33419838 | 0.148853298 | 11.36768827 | NA | NA | NA | NA | NA | NA | NA |
| SCN1A | 1763 | I | 12.22 | 11.5059001 | 0.151434232 | 10.53968477 | NA | NA | NA | NA | NA | NA | NA |
| SCN1A | 1764 | F | 7.835113636 | 13.36726999 | 0.179413417 | 12.41975697 | NA | NA | NA | NA | NA | NA | NA |
| SCN1A | 1765 | F | 10.24902273 | 11.87456648 | 0.156975842 | 10.9696428 | NA | NA | NA | NA | NA | NA | NA |
| SCN1A | 1766 | F | 11.45720455 | 8.464812874 | 0.105722125 | 7.696672641 | NA | NA | NA | NA | NA | NA | NA |
| SCN1A | 1767 | V | 6.968035714 | 9.824513067 | 0.126160461 | 9.527638402 | NA | NA | NA | NA | NA | NA | NA |
| SCN1A | 1768 | S | 4.682916667 | 11.75343312 | 0.155155025 | 10.53752781 | NA | NA | NA | NA | NA | NA | NA |
| SCN1A | 1769 | Y | 5.447666667 | 9.81941317 | 0.126083802 | 8.576782734 | NA | NA | NA | NA | NA | NA | NA |
| SCN1A | 1770 | I | 5.07425 | 5.428112563 | 0.06007596 | 4.277516674 | 1.29 | -0.07 | -1.48 | 14.16 | 0.14 | 1 | 1 |
| SCN1A | 1771 | I | 1.303 | 9.251974168 | 0.117554342 | 8.079626476 | NA | NA | NA | NA | NA | NA | NA |
| SCN1A | 1772 | I | 0.464125 | 11.52255067 | 0.151684516 | 10.43325249 | NA | NA | NA | NA | NA | NA | NA |
| SCN1A | 1773 | S | 1.284916667 | 8.50244653 | 0.106287816 | 8.350694125 | NA | NA | NA | NA | NA | NA | NA |
| SCN1A | 1774 | F | 0.636022727 | 5.24307591 | 0.057294582 | 2.054446954 | 3.53 | 1.2 | 1.77 | 0.7 | -1.68 | 0 | 0 |
| SCN1A | 1775 | L | -2.93925 | 9.941278942 | 0.127915627 | 6.786982567 | NA | NA | NA | NA | NA | NA | NA |
| SCN1A | 1776 | V | -2.377321429 | 10.74795743 | 0.140041216 | 7.274502633 | NA | NA | NA | NA | NA | NA | NA |
| SCN1A | 1777 | V | -1.56475 | 7.036978542 | 0.084259632 | 4.020011342 | 3.55 | 1.13 | 1.25 | 0.87 | -1.62 | 0 | 0 |
| SCN1A | 1778 | V | -4.99675 | 5.742622746 | 0.064803521 | 2.719738779 | 3.15 | 1.35 | 2.33 | 0.62 | -1.69 | 0 | 0 |
| SCN1A | 1779 | N | -6.7965 | 8.373818305 | 0.10435434 | 5.279732148 | NA | NA | NA | NA | NA | NA | NA |
| SCN1A | 1780 | M | -4.904375 | 8.384000537 | 0.104507395 | 5.313633849 | NA | NA | NA | NA | NA | NA | NA |
| SCN1A | 1781 | Y | -4.45675 | 4.734384226 | 0.049648182 | 1.640674995 | 3.14 | 1.35 | 2.33 | 0.62 | -1.69 | 0 | 0 |
| SCN1A | 1782 | I | -10.582 | 5.826001545 | 0.066056829 | 3.408914771 | 3.14 | 1.35 | 2.33 | 0.62 | -1.69 | 0 | 0 |
| SCN1A | 1783 | A | -10.57215 | 9.189261287 | 0.116611673 | 6.111251474 | NA | NA | NA | NA | NA | NA | NA |
| SCN1A | 1784 | V | -9.38275 | 9.376915644 | 0.1194324 | 6.526531803 | NA | NA | NA | NA | NA | NA | NA |
| SCN1A | 1785 | I | -11.3065 | 4.674631536 | 0.048750009 | 1.628790421 | 3.05 | 1.43 | 3.7 | 0.2 | -1.82 | 0 | 0 |
| SCN1A | 1786 | L | -15.024 | 7.457660558 | 0.090583114 | 4.36458325 | 3.34 | 0.4 | -0.03 | 24.97 | 0.01 | 2 | 0 |
| SCN1A | 1787 | E | -14.54786111 | 9.62390534 | 0.123145025 | 6.570685916 | NA | NA | NA | NA | NA | NA | NA |
| SCN1A | 1788 | N | -13.232875 | 9.424093909 | 0.12014156 | 6.388107103 | NA | NA | NA | NA | NA | NA | NA |
| SCN1A | 1789 | F | -16.28693182 | 4.330688052 | 0.043580022 | 1.931187736 | 3.34 | 0.4 | -0.03 | 24.97 | 0.01 | 2 | 0 |
| SCN1A | 1790 | S | -19.74558333 | 8.693751147 | 0.109163411 | 5.627770163 | NA | NA | NA | NA | NA | NA | NA |
| SCN1A | 1791 | V | -18.13946429 | 11.34397219 | 0.149000213 | 8.395130105 | NA | NA | NA | NA | NA | NA | NA |
| SCN1A | 1792 | A | -19.19175 | 11.83403874 | 0.156366649 | 8.887135176 | NA | NA | NA | NA | NA | NA | NA |

| | | | | | | | | | | | | | |
|---|---|---|---|---|---|---|---|---|---|---|---|---|---|
| SCN1A | 1793 | T | -22.09889286 | 7.956172698 | 0.0980765 | 1.943299849 | 6.19 | -0.77 | -2.9 | 26.11 | 1.25 | 2 | 0 |
| SCN1A | 1794 | E | -23.67475 | 10.4212078 | 0.135129679 | 4.327910489 | 6.19 | -0.77 | -2.9 | 26.11 | 1.25 | 2 | 0 |
| SCN1A | 1795 | E | -21.82997222 | 13.05139701 | 0.174665371 | 6.378198129 | NA | NA | NA | NA | NA | NA | NA |
| SCN5A | 119 | P | -19.77424675 | 55.22623838 | 0.91690573 | 52.10869109 | NA | NA | NA | NA | NA | NA | NA |
| SCN5A | 120 | I | -16.44719318 | 52.65913912 | 0.871888692 | 49.61608157 | NA | NA | NA | NA | NA | NA | NA |
| SCN5A | 121 | R | -19.03863636 | 47.39461852 | 0.779569268 | 44.29932783 | NA | NA | NA | NA | NA | NA | NA |
| SCN5A | 122 | R | -21.64727273 | 49.19700151 | 0.811176127 | 46.06207145 | NA | NA | NA | NA | NA | NA | NA |
| SCN5A | 123 | A | -16.79021818 | 48.88032113 | 0.805622772 | 45.90583038 | NA | NA | NA | NA | NA | NA | NA |
| SCN5A | 124 | A | -15.23761818 | 46.15645924 | 0.757856721 | 43.69098155 | NA | NA | NA | NA | NA | NA | NA |
| SCN5A | 125 | V | -18.25196104 | 44.41996047 | 0.727405218 | 41.26779463 | NA | NA | NA | NA | NA | NA | NA |
| SCN5A | 126 | K | -17.49726263 | 44.94975925 | 0.736695849 | 41.78702622 | NA | NA | NA | NA | NA | NA | NA |
| SCN5A | 127 | I | -13.63294318 | 43.37305525 | 0.70904653 | 40.93865171 | NA | NA | NA | NA | NA | NA | NA |
| SCN5A | 128 | L | -14.71231818 | 40.41861107 | 0.657236954 | 37.96108804 | NA | NA | NA | NA | NA | NA | NA |
| SCN5A | 129 | V | -17.58481818 | 39.60968344 | 0.643051477 | 36.47154158 | NA | NA | NA | NA | NA | NA | NA |
| SCN5A | 130 | H | -15.09231818 | 39.4729074 | 0.640652952 | 36.75007609 | NA | NA | NA | NA | NA | NA | NA |
| SCN5A | 131 | S | -13.02065152 | 38.06243468 | 0.615918691 | 35.8654111 | NA | NA | NA | NA | NA | NA | NA |
| SCN5A | 132 | L | -10.68544318 | 40.97789801 | 0.667044693 | 39.02064218 | NA | NA | NA | NA | NA | NA | NA |
| SCN5A | 133 | F | -9.740090909 | 39.47135075 | 0.640625654 | 37.30267552 | NA | NA | NA | NA | NA | NA | NA |
| SCN5A | 134 | N | -10.04581818 | 34.97023349 | 0.561693386 | 33.20704244 | NA | NA | NA | NA | NA | NA | NA |
| SCN5A | 135 | M | -6.596943182 | 37.21162672 | 0.600998794 | 35.24549047 | NA | NA | NA | NA | NA | NA | NA |
| SCN5A | 136 | L | -4.636693182 | 38.22525643 | 0.618773957 | 33.36424259 | NA | NA | NA | NA | NA | NA | NA |
| SCN5A | 137 | I | -5.210443182 | 35.40101312 | 0.569247602 | 30.53867836 | NA | NA | NA | NA | NA | NA | NA |
| SCN5A | 138 | M | -4.090943182 | 30.84268732 | 0.489312116 | 25.98636322 | NA | NA | NA | NA | NA | NA | NA |
| SCN5A | 139 | C | -1.257318182 | 35.55280369 | 0.571909424 | 30.69310019 | NA | NA | NA | NA | NA | NA | NA |
| SCN5A | 140 | T | -0.351103896 | 35.58577439 | 0.572487604 | 30.71645866 | NA | NA | NA | NA | NA | NA | NA |
| SCN5A | 141 | I | -0.726443182 | 31.46732644 | 0.500265882 | 26.59997969 | NA | NA | NA | NA | NA | NA | NA |
| SCN5A | 142 | L | 0.687068182 | 29.64878176 | 0.468375609 | 24.7844803 | NA | NA | NA | NA | NA | NA | NA |
| SCN5A | 143 | T | 3.539818182 | 34.13651831 | 0.547073231 | 29.2677693 | NA | NA | NA | NA | NA | NA | NA |
| SCN5A | 144 | N | 4.364443182 | 32.27117364 | 0.514362267 | 29.81335901 | NA | NA | NA | NA | NA | NA | NA |
| SCN5A | 145 | C | 4.270984848 | 29.22457374 | 0.460936633 | 26.77450308 | NA | NA | NA | NA | NA | NA | NA |
| SCN5A | 146 | V | 6.910818182 | 30.22606438 | 0.478498924 | 29.44564205 | NA | NA | NA | NA | NA | NA | NA |
| SCN5A | 147 | F | 8.815818182 | 30.58296608 | 0.484757606 | 26.68826985 | NA | NA | NA | NA | NA | NA | NA |
| SCN5A | 148 | M | 7.741943182 | 27.58612553 | 0.43220456 | 23.68650916 | NA | NA | NA | NA | NA | NA | NA |

| | | | | | | | | | | | | | |
|---|---|---|---|---|---|---|---|---|---|---|---|---|---|
| SCN5A | 149 | A | 10.60341818 | 26.04864701 | 0.405243105 | 22.13411014 | NA | NA | NA | NA | NA | NA | NA |
| SCN5A | 150 | Q | 13.54526263 | 26.21435715 | 0.408149023 | 22.37185913 | NA | NA | NA | NA | NA | NA | NA |
| SCN5A | 151 | H | 17.94131818 | 22.90174862 | 0.350058623 | 16.79102955 | NA | NA | NA | NA | NA | NA | NA |
| SCN5A | 152 | D | 19.44681818 | 26.06036091 | 0.405448522 | 19.94979679 | NA | NA | NA | NA | NA | NA | NA |
| SCN5A | 153 | P | 16.56053247 | 29.24775114 | 0.461343076 | 23.13715827 | NA | NA | NA | NA | NA | NA | NA |
| SCN5A | 154 | P | 17.55438961 | 32.00449834 | 0.509685809 | 25.89627007 | NA | NA | NA | NA | NA | NA | NA |
| SCN5A | 155 | P | 18.75624675 | 34.59094237 | 0.55504208 | 28.4961881 | NA | NA | NA | NA | NA | NA | NA |
| SCN5A | 156 | W | 16.66638961 | 35.72117956 | 0.574862089 | 29.62252724 | NA | NA | NA | NA | NA | NA | NA |
| SCN5A | 157 | T | 12.84881818 | 35.31302203 | 0.567704577 | 29.20273128 | NA | NA | NA | NA | NA | NA | NA |
| SCN5A | 158 | K | 13.05615152 | 39.63949488 | 0.643574255 | 33.52967971 | NA | NA | NA | NA | NA | NA | NA |
| SCN5A | 159 | Y | 9.163234848 | 41.93464664 | 0.683822381 | 35.97192009 | NA | NA | NA | NA | NA | NA | NA |
| SCN5A | 160 | V | 8.585961039 | 37.66725424 | 0.608988747 | 33.77718747 | NA | NA | NA | NA | NA | NA | NA |
| SCN5A | 161 | E | 8.974707071 | 37.33955862 | 0.603242227 | 33.5268344 | NA | NA | NA | NA | NA | NA | NA |
| SCN5A | 162 | Y | 8.203484848 | 41.54690886 | 0.677022953 | 37.68232479 | NA | NA | NA | NA | NA | NA | NA |
| SCN5A | 163 | T | 4.269961039 | 42.35980232 | 0.691277975 | 38.46088868 | NA | NA | NA | NA | NA | NA | NA |
| SCN5A | 164 | F | 3.465272727 | 37.61191568 | 0.608018322 | 35.14729098 | NA | NA | NA | NA | NA | NA | NA |
| SCN5A | 165 | T | 2.552961039 | 41.50716738 | 0.676326041 | 39.04129187 | NA | NA | NA | NA | NA | NA | NA |
| SCN5A | 166 | A | 0.713418182 | 44.06295453 | 0.721144708 | 39.35384954 | NA | NA | NA | NA | NA | NA | NA |
| SCN5A | 167 | I | -0.502693182 | 41.50895297 | 0.676357353 | 36.64210664 | NA | NA | NA | NA | NA | NA | NA |
| SCN5A | 168 | Y | -0.498068182 | 35.16269832 | 0.565068478 | 30.29138058 | NA | NA | NA | NA | NA | NA | NA |
| SCN5A | 169 | T | -1.56738961 | 43.00549109 | 0.70260087 | 38.13369835 | NA | NA | NA | NA | NA | NA | NA |
| SCN5A | 170 | F | -5.078636364 | 44.66797892 | 0.731754506 | 39.80216382 | NA | NA | NA | NA | NA | NA | NA |
| SCN5A | 171 | E | -5.984707071 | 37.74610462 | 0.610371479 | 32.88265842 | NA | NA | NA | NA | NA | NA | NA |
| SCN5A | 172 | S | -6.465151515 | 42.56987388 | 0.694961821 | 37.70045066 | NA | NA | NA | NA | NA | NA | NA |
| SCN5A | 173 | L | -7.511318182 | 45.20673865 | 0.741202278 | 40.33973754 | NA | NA | NA | NA | NA | NA | NA |
| SCN5A | 174 | V | -10.41481818 | 44.62482302 | 0.730997718 | 42.50569823 | NA | NA | NA | NA | NA | NA | NA |
| SCN5A | 175 | K | -10.68415152 | 37.69619287 | 0.609496219 | 35.58981916 | NA | NA | NA | NA | NA | NA | NA |
| SCN5A | 176 | I | -11.22681818 | 44.43416472 | 0.727654305 | 42.03114125 | NA | NA | NA | NA | NA | NA | NA |
| SCN5A | 177 | L | -12.62581818 | 46.96912288 | 0.772107713 | 44.5459564 | NA | NA | NA | NA | NA | NA | NA |
| SCN5A | 178 | A | -16.21501818 | 44.60805799 | 0.730703724 | 42.1205227 | NA | NA | NA | NA | NA | NA | NA |
| SCN5A | 179 | R | -18.56472727 | 42.7591771 | 0.698281471 | 39.62183218 | NA | NA | NA | NA | NA | NA | NA |
| SCN5A | 180 | G | -15.71481818 | 46.55744574 | 0.76488848 | 44.06235916 | NA | NA | NA | NA | NA | NA | NA |
| SCN5A | 181 | F | -15.96209091 | 49.42667715 | 0.815203753 | 46.95218707 | NA | NA | NA | NA | NA | NA | NA |

| | | | | | | | | | | | | | |
|---|---|---|---|---|---|---|---|---|---|---|---|---|---|
| SCN5A | 182 | C | -16.11248485 | 48.10206471 | 0.791975151 | 45.58695997 | NA | NA | NA | NA | NA | NA | NA |
| SCN5A | 183 | L | -17.65894318 | 46.17203553 | 0.758129869 | 43.62557039 | NA | NA | NA | NA | NA | NA | NA |
| SCN5A | 184 | H | -13.57011818 | 43.21007611 | 0.706188504 | 40.70786225 | NA | NA | NA | NA | NA | NA | NA |
| SCN5A | 185 | A | -17.31781818 | 40.33516564 | 0.655773642 | 37.83333826 | NA | NA | NA | NA | NA | NA | NA |
| SCN5A | 186 | F | -21.06790909 | 40.70184642 | 0.662203811 | 37.6170441 | NA | NA | NA | NA | NA | NA | NA |
| SCN5A | 187 | T | -17.6061039 | 39.7851334 | 0.646128194 | 36.78973073 | NA | NA | NA | NA | NA | NA | NA |
| SCN5A | 188 | F | -12.48009091 | 40.00739958 | 0.650025887 | 37.61352566 | NA | NA | NA | NA | NA | NA | NA |
| SCN5A | 189 | L | -11.48806818 | 37.88665891 | 0.61283626 | 35.42350969 | NA | NA | NA | NA | NA | NA | NA |
| SCN5A | 190 | R | -16.44172727 | 34.43195705 | 0.552254089 | 31.8745938 | NA | NA | NA | NA | NA | NA | NA |
| SCN5A | 191 | D | -13.99081818 | 33.28914313 | 0.532213533 | 30.75715024 | NA | NA | NA | NA | NA | NA | NA |
| SCN5A | 192 | P | -10.75838961 | 30.47423371 | 0.482850858 | 28.02824583 | NA | NA | NA | NA | NA | NA | NA |
| SCN5A | 193 | W | -8.54288961 | 26.74943012 | 0.417532143 | 24.82120403 | NA | NA | NA | NA | NA | NA | NA |
| SCN5A | 194 | N | -9.486818182 | 34.66097139 | 0.556270119 | 32.85077179 | NA | NA | NA | NA | NA | NA | NA |
| SCN5A | 195 | W | -7.867032468 | 33.6320096 | 0.538226091 | 29.86416808 | NA | NA | NA | NA | NA | NA | NA |
| SCN5A | 196 | L | -3.229943182 | 30.40547309 | 0.481645062 | 25.53154321 | NA | NA | NA | NA | NA | NA | NA |
| SCN5A | 197 | D | -3.971943182 | 35.63684607 | 0.573383204 | 30.76406133 | NA | NA | NA | NA | NA | NA | NA |
| SCN5A | 198 | F | -3.897454545 | 38.23001347 | 0.618857377 | 33.35569761 | NA | NA | NA | NA | NA | NA | NA |
| SCN5A | 199 | S | -0.406984848 | 35.71060261 | 0.57467661 | 33.15008575 | NA | NA | NA | NA | NA | NA | NA |
| SCN5A | 200 | V | 0.069961039 | 34.93973449 | 0.561158551 | 32.45144354 | NA | NA | NA | NA | NA | NA | NA |
| SCN5A | 201 | I | -0.856693182 | 39.40347884 | 0.639435442 | 36.837112 | NA | NA | NA | NA | NA | NA | NA |
| SCN5A | 202 | I | 0.849443182 | 40.86658531 | 0.665092697 | 38.31573635 | NA | NA | NA | NA | NA | NA | NA |
| SCN5A | 203 | M | 4.442193182 | 35.09132327 | 0.563816835 | 32.62529152 | NA | NA | NA | NA | NA | NA | NA |
| SCN5A | 204 | A | 4.386418182 | 39.66386137 | 0.644001549 | 37.19681112 | NA | NA | NA | NA | NA | NA | NA |
| SCN5A | 205 | Y | 3.138484848 | 42.34837848 | 0.691077645 | 39.88266918 | NA | NA | NA | NA | NA | NA | NA |
| SCN5A | 206 | T | 6.16738961 | 41.57777736 | 0.677564268 | 37.84701225 | NA | NA | NA | NA | NA | NA | NA |
| SCN5A | 207 | T | 9.506818182 | 38.05050435 | 0.615709478 | 34.25568165 | NA | NA | NA | NA | NA | NA | NA |
| SCN5A | 208 | E | 9.637373737 | 40.08971882 | 0.65146945 | 34.29154809 | NA | NA | NA | NA | NA | NA | NA |
| SCN5A | 209 | F | 10.88763636 | 43.82889099 | 0.717040134 | 37.72268435 | NA | NA | NA | NA | NA | NA | NA |
| SCN5A | 210 | V | 11.40938961 | 40.72212771 | 0.662559467 | 34.61656716 | NA | NA | NA | NA | NA | NA | NA |
| SCN5A | 211 | D | 15.37619318 | 36.76230277 | 0.593119382 | 30.65849418 | NA | NA | NA | NA | NA | NA | NA |
| SCN5A | 212 | L | 15.35706818 | 38.17408013 | 0.617876522 | 32.072296 | NA | NA | NA | NA | NA | NA | NA |
| SCN5A | 213 | G | 14.51581818 | 35.15316882 | 0.564901367 | 29.04844797 | NA | NA | NA | NA | NA | NA | NA |
| SCN5A | 214 | N | 18.18956818 | 32.99824556 | 0.527112309 | 26.89789074 | NA | NA | NA | NA | NA | NA | NA |

| | | | | | | | | | | | | | |
|---|---|---|---|---|---|---|---|---|---|---|---|---|---|
| SCN5A | 215 | V | 16.3231039 | 29.20580167 | 0.460607444 | 23.10073634 | NA | NA | NA | NA | NA | NA | NA |
| SCN5A | 216 | S | 16.11948485 | 29.28275993 | 0.461956995 | 23.20030475 | NA | NA | NA | NA | NA | NA | NA |
| SCN5A | 217 | A | 12.83741818 | 28.48698764 | 0.448002213 | 24.76192853 | NA | NA | NA | NA | NA | NA | NA |
| SCN5A | 218 | L | 9.693193182 | 31.36795569 | 0.498523301 | 27.60064438 | NA | NA | NA | NA | NA | NA | NA |
| SCN5A | 219 | R | 12.65572727 | 27.28184917 | 0.426868724 | 23.80182289 | NA | NA | NA | NA | NA | NA | NA |
| SCN5A | 220 | T | 9.17538961 | 26.63916692 | 0.415598551 | 22.91292068 | NA | NA | NA | NA | NA | NA | NA |
| SCN5A | 221 | F | 5.607363636 | 30.52155065 | 0.483680615 | 28.31116519 | NA | NA | NA | NA | NA | NA | NA |
| SCN5A | 222 | R | 7.157545455 | 31.02220555 | 0.492460175 | 28.55661635 | NA | NA | NA | NA | NA | NA | NA |
| SCN5A | 223 | V | 4.488246753 | 26.12283832 | 0.406544135 | 23.65574594 | NA | NA | NA | NA | NA | NA | NA |
| SCN5A | 224 | L | 0.981443182 | 28.59799115 | 0.449948787 | 26.0350164 | NA | NA | NA | NA | NA | NA | NA |
| SCN5A | 225 | R | 1.660636364 | 31.29421196 | 0.49723012 | 26.41941054 | NA | NA | NA | NA | NA | NA | NA |
| SCN5A | 226 | A | 0.426818182 | 27.64724634 | 0.433276383 | 22.77645455 | NA | NA | NA | NA | NA | NA | NA |
| SCN5A | 227 | L | -1.838818182 | 25.68196914 | 0.398812987 | 20.81103511 | NA | NA | NA | NA | NA | NA | NA |
| SCN5A | 228 | K | -3.735151515 | 29.86247259 | 0.472122924 | 24.99327573 | NA | NA | NA | NA | NA | NA | NA |
| SCN5A | 229 | T | -4.037103896 | 28.24105938 | 0.443689578 | 23.38121245 | NA | NA | NA | NA | NA | NA | NA |
| SCN5A | 230 | I | -6.600068182 | 24.3526716 | 0.375502226 | 22.21034193 | NA | NA | NA | NA | NA | NA | NA |
| SCN5A | 231 | S | -9.451151515 | 29.10263474 | 0.458798293 | 26.9930628 | NA | NA | NA | NA | NA | NA | NA |
| SCN5A | 232 | V | -8.61338961 | 30.87251755 | 0.489835224 | 28.83054418 | NA | NA | NA | NA | NA | NA | NA |
| SCN5A | 233 | I | -9.448818182 | 26.98626032 | 0.421685233 | 24.89990594 | NA | NA | NA | NA | NA | NA | NA |
| SCN5A | 234 | S | -13.58831818 | 25.77511084 | 0.400446334 | 23.27943277 | NA | NA | NA | NA | NA | NA | NA |
| SCN5A | 235 | G | -12.29331818 | 22.09229606 | 0.335863941 | 19.95838006 | NA | NA | NA | NA | NA | NA | NA |
| SCN5A | 236 | L | -8.745193182 | 22.35667623 | 0.340500151 | 20.4993659 | NA | NA | NA | NA | NA | NA | NA |
| SCN5A | 237 | K | -12.46926263 | 22.77917826 | 0.34790921 | 20.70113993 | NA | NA | NA | NA | NA | NA | NA |
| SCN5A | 238 | T | -14.29496104 | 19.44814564 | 0.289495722 | 16.95817082 | NA | NA | NA | NA | NA | NA | NA |
| SCN5A | 239 | I | -10.89544318 | 17.85430853 | 0.261545955 | 15.76143481 | NA | NA | NA | NA | NA | NA | NA |
| SCN5A | 240 | V | -9.193532468 | 19.85445182 | 0.296620768 | 17.78726113 | NA | NA | NA | NA | NA | NA | NA |
| SCN5A | 241 | G | -12.58256818 | 18.99802379 | 0.281602318 | 16.49558814 | NA | NA | NA | NA | NA | NA | NA |
| SCN5A | 242 | A | -12.64261818 | 15.60463678 | 0.222095373 | 13.14043337 | NA | NA | NA | NA | NA | NA | NA |
| SCN5A | 243 | L | -8.635568182 | 15.11370127 | 0.213486254 | 13.00844469 | NA | NA | NA | NA | NA | NA | NA |
| SCN5A | 244 | I | -9.889068182 | 18.19316825 | 0.26748825 | 15.81960959 | NA | NA | NA | NA | NA | NA | NA |
| SCN5A | 245 | Q | -14.35559596 | 16.33847759 | 0.234964116 | 13.76753209 | NA | NA | NA | NA | NA | NA | NA |
| SCN5A | 246 | S | -10.74015152 | 13.30703908 | 0.181804355 | 10.87954236 | NA | NA | NA | NA | NA | NA | NA |
| SCN5A | 247 | V | -8.036818182 | 15.77606988 | 0.22510165 | 13.73094047 | NA | NA | NA | NA | NA | NA | NA |

| | | | | | | | | | | | | | |
|---|---|---|---|---|---|---|---|---|---|---|---|---|---|
| SCN5A | 248 | K | -12.36126263 | 17.75820061 | 0.259860592 | 15.29032544 | NA | NA | NA | NA | NA | NA | NA |
| SCN5A | 249 | K | -13.23915152 | 14.54055188 | 0.203435421 | 11.93114502 | NA | NA | NA | NA | NA | NA | NA |
| SCN5A | 250 | L | -7.567818182 | 13.16270892 | 0.17927336 | 10.78970251 | NA | NA | NA | NA | NA | NA | NA |
| SCN5A | 251 | A | -8.444018182 | 17.97379965 | 0.263641369 | 15.47071885 | NA | NA | NA | NA | NA | NA | NA |
| SCN5A | 252 | D | -9.606693182 | 18.39529627 | 0.271032797 | 15.85703517 | NA | NA | NA | NA | NA | NA | NA |
| SCN5A | 253 | V | -6.107961039 | 14.33671601 | 0.199860924 | 12.30609391 | NA | NA | NA | NA | NA | NA | NA |
| SCN5A | 254 | M | -3.585068182 | 17.99920801 | 0.264086934 | 13.4977918 | NA | NA | NA | NA | NA | NA | NA |
| SCN5A | 255 | V | -4.235103896 | 20.62474429 | 0.310128733 | 16.0075771 | NA | NA | NA | NA | NA | NA | NA |
| SCN5A | 256 | L | -4.124318182 | 17.89094388 | 0.262188398 | 14.41789513 | NA | NA | NA | NA | NA | NA | NA |
| SCN5A | 257 | T | -0.775532468 | 15.02340341 | 0.211902778 | 11.95213724 | NA | NA | NA | NA | NA | NA | NA |
| SCN5A | 258 | V | 0.043961039 | 18.7143029 | 0.276626945 | 15.63942791 | NA | NA | NA | NA | NA | NA | NA |
| SCN5A | 259 | F | -1.152454545 | 20.92137285 | 0.315330456 | 17.84819103 | NA | NA | NA | NA | NA | NA | NA |
| SCN5A | 260 | C | 0.638151515 | 18.7743349 | 0.277679676 | 15.7222202 | NA | NA | NA | NA | NA | NA | NA |
| SCN5A | 261 | L | 3.511068182 | 14.64318411 | 0.205235195 | 11.99139499 | NA | NA | NA | NA | NA | NA | NA |
| SCN5A | 262 | S | 3.535318182 | 19.98258537 | 0.298867737 | 17.33033274 | NA | NA | NA | NA | NA | NA | NA |
| SCN5A | 263 | V | 2.779103896 | 21.94284223 | 0.333243096 | 19.30141497 | NA | NA | NA | NA | NA | NA | NA |
| SCN5A | 264 | F | 3.866545455 | 17.66848406 | 0.258287309 | 15.15363504 | NA | NA | NA | NA | NA | NA | NA |
| SCN5A | 265 | A | 8.011618182 | 20.08167229 | 0.30060534 | 17.56064652 | NA | NA | NA | NA | NA | NA | NA |
| SCN5A | 266 | L | 7.495818182 | 22.70693498 | 0.346642341 | 20.19545502 | NA | NA | NA | NA | NA | NA | NA |
| SCN5A | 267 | I | 7.775193182 | 22.99983204 | 0.351778628 | 20.78197511 | NA | NA | NA | NA | NA | NA | NA |
| SCN5A | 268 | G | 11.59031818 | 21.10217854 | 0.318501091 | 19.41485127 | NA | NA | NA | NA | NA | NA | NA |
| SCN5A | 269 | L | 12.72694318 | 22.40575647 | 0.34136083 | 20.7220634 | NA | NA | NA | NA | NA | NA | NA |
| SCN5A | 270 | Q | 11.81070707 | 25.11360422 | 0.388846055 | 23.43292748 | NA | NA | NA | NA | NA | NA | NA |
| SCN5A | 271 | L | 13.29269318 | 23.70808936 | 0.364198735 | 22.05432837 | NA | NA | NA | NA | NA | NA | NA |
| SCN5A | 272 | F | 15.44781818 | 19.89206601 | 0.297280376 | 18.203741 | NA | NA | NA | NA | NA | NA | NA |
| SCN5A | 273 | M | 17.75219318 | 23.74479482 | 0.364842407 | 22.11356793 | NA | NA | NA | NA | NA | NA | NA |
| SCN5A | 274 | G | 21.42856818 | 23.36583208 | 0.35819686 | 21.85399505 | NA | NA | NA | NA | NA | NA | NA |
| SCN5A | 275 | N | 20.60994318 | 20.78595001 | 0.31295566 | 19.21881359 | NA | NA | NA | NA | NA | NA | NA |
| SCN5A | 276 | L | 20.94381818 | 16.37245409 | 0.235559933 | 11.6581915 | NA | NA | NA | NA | NA | NA | NA |
| SCN5A | 277 | R | 25.34963636 | 20.07667893 | 0.300517776 | 14.17645656 | NA | NA | NA | NA | NA | NA | NA |
| SCN5A | 278 | H | 25.86241818 | 19.09487706 | 0.283300751 | 13.17856503 | NA | NA | NA | NA | NA | NA | NA |
| SCN5A | 279 | K | 29.74248485 | 17.30310435 | 0.251879956 | 11.38633701 | NA | NA | NA | NA | NA | NA | NA |
| SCN5A | 280 | C | 32.80281818 | 18.46090404 | 0.272183305 | 11.51087203 | NA | NA | NA | NA | NA | NA | NA |

| | | | | | | | | | | | | | |
|---|---|---|---|---|---|---|---|---|---|---|---|---|---|
| SCN5A | 281 | V | 35.51438961 | 19.82214782 | 0.29605428 | 12.35743078 | NA | NA | NA | NA | NA | NA | NA |
| SCN5A | 282 | R | 41.02781818 | 15.49281959 | 0.22013453 | 8.139342419 | NA | NA | NA | NA | NA | NA | NA |
| SCN5A | 283 | N | 40.97881818 | 21.64459997 | 0.328013075 | 14.23344421 | NA | NA | NA | NA | NA | NA | NA |
| SCN5A | 284 | F | 41.124 | 23.79850155 | 0.365784217 | 16.4011175 | NA | NA | NA | NA | NA | NA | NA |
| SCN5A | 285 | T | 45.83553247 | 26.30626355 | 0.409760708 | 18.94715986 | NA | NA | NA | NA | NA | NA | NA |
| SCN5A | 286 | A | 45.61661818 | 29.18472808 | 0.460237894 | 21.83227152 | NA | NA | NA | NA | NA | NA | NA |
| SCN5A | 287 | L | 49.76894318 | 27.49502188 | 0.430606952 | 20.19091637 | NA | NA | NA | NA | NA | NA | NA |
| SCN5A | 288 | N | 49.61831818 | 32.09015541 | 0.511187904 | 24.7661938 | NA | NA | NA | NA | NA | NA | NA |
| SCN5A | 289 | G | 51.11256818 | 32.06738243 | 0.510788554 | 24.77185771 | NA | NA | NA | NA | NA | NA | NA |
| SCN5A | 290 | T | 48.80538961 | 30.34886387 | 0.480652354 | 23.04710324 | NA | NA | NA | NA | NA | NA | NA |
| SCN5A | 291 | N | 47.00806818 | 33.23495562 | 0.531263293 | 25.91293886 | NA | NA | NA | NA | NA | NA | NA |
| SCN5A | 292 | G | 44.69506818 | 31.01060654 | 0.492256773 | 23.67681579 | NA | NA | NA | NA | NA | NA | NA |
| SCN5A | 293 | S | 42.28281818 | 27.92200047 | 0.438094513 | 20.57383516 | NA | NA | NA | NA | NA | NA | NA |
| SCN5A | 294 | V | 45.6351039 | 26.78937618 | 0.418232643 | 19.45205728 | NA | NA | NA | NA | NA | NA | NA |
| SCN5A | 295 | E | 43.7900404 | 23.74369043 | 0.364823041 | 16.39466522 | NA | NA | NA | NA | NA | NA | NA |
| SCN5A | 296 | A | 46.97941818 | 23.14902013 | 0.354394813 | 15.81632763 | NA | NA | NA | NA | NA | NA | NA |
| SCN5A | 297 | D | 45.58244318 | 18.96255223 | 0.280980283 | 11.59373913 | NA | NA | NA | NA | NA | NA | NA |
| SCN5A | 298 | G | 49.99931818 | 20.27992372 | 0.304081907 | 12.98423546 | NA | NA | NA | NA | NA | NA | NA |
| SCN5A | 299 | L | 49.62806818 | 17.50441556 | 0.255410179 | 10.22583358 | NA | NA | NA | NA | NA | NA | NA |
| SCN5A | 300 | V | 49.14124675 | 21.33812206 | 0.322638632 | 14.04266824 | NA | NA | NA | NA | NA | NA | NA |
| SCN5A | 301 | W | 44.79753247 | 16.85100558 | 0.243951884 | 9.524710518 | NA | NA | NA | NA | NA | NA | NA |
| SCN5A | 302 | E | 45.3430404 | 23.81050302 | 0.365994676 | 16.48383138 | NA | NA | NA | NA | NA | NA | NA |
| SCN5A | 303 | S | 40.67898485 | 25.28926422 | 0.391926455 | 17.91628569 | NA | NA | NA | NA | NA | NA | NA |
| SCN5A | 304 | L | 38.57669318 | 23.95720831 | 0.368567322 | 16.56447088 | NA | NA | NA | NA | NA | NA | NA |
| SCN5A | 305 | D | 35.81669318 | 22.71365288 | 0.346760147 | 15.30437282 | NA | NA | NA | NA | NA | NA | NA |
| SCN5A | 306 | L | 39.92481818 | 20.03708075 | 0.299823377 | 12.65805551 | NA | NA | NA | NA | NA | NA | NA |
| SCN5A | 307 | Y | 40.37665152 | 19.16408219 | 0.284514343 | 11.766604 | NA | NA | NA | NA | NA | NA | NA |
| SCN5A | 308 | L | 35.76656818 | 19.35285408 | 0.287824675 | 11.9109245 | NA | NA | NA | NA | NA | NA | NA |
| SCN5A | 309 | S | 34.95148485 | 17.63778027 | 0.257748883 | 10.19602506 | NA | NA | NA | NA | NA | NA | NA |
| SCN5A | 310 | D | 38.47881818 | 14.39389381 | 0.200863603 | 7.013971088 | NA | NA | NA | NA | NA | NA | NA |
| SCN5A | 311 | P | 36.35724675 | 12.10981528 | 0.160809658 | 4.65042905 | 7.51 | -0.55 | -1.68 | 37.41 | 1.33 | 3 | -3 |
| SCN5A | 312 | E | 40.16237374 | 9.124560757 | 0.108459787 | 1.778954944 | 7.51 | -0.55 | -1.68 | 37.41 | 1.33 | 3 | -3 |
| SCN5A | 313 | N | 38.81981818 | 14.61033928 | 0.204659223 | 7.193838843 | NA | NA | NA | NA | NA | NA | NA |

| | | | | | | | | | | | | | |
|---|---|---|---|---|---|---|---|---|---|---|---|---|---|
| SCN5A | 314 | Y | 34.38698485 | 14.00334453 | 0.194014872 | 6.515959895 | NA | NA | NA | NA | NA | NA | NA |
| SCN5A | 315 | L | 34.33569318 | 13.96788878 | 0.193393115 | 7.272078429 | NA | NA | NA | NA | NA | NA | NA |
| SCN5A | 316 | L | 33.58281818 | 8.3170511 | 0.094299176 | 1.962347809 | 6.5 | -0.5 | -1 | 25.35 | 1.05 | 2 | -2 |
| SCN5A | 317 | K | 28.43870707 | 12.42470273 | 0.166331572 | 6.523403139 | NA | NA | NA | NA | NA | NA | NA |
| SCN5A | 318 | N | 31.27456818 | 12.36284458 | 0.165246818 | 6.459272082 | NA | NA | NA | NA | NA | NA | NA |
| SCN5A | 319 | G | 30.47006818 | 9.569855224 | 0.116268538 | 3.671704939 | 5.93 | -0.33 | -2.53 | 26.25 | 0.98 | 2 | 0 |
| SCN5A | 320 | T | 28.5531039 | 8.516398946 | 0.09779497 | 2.611740666 | 5.93 | -0.33 | -2.53 | 26.25 | 0.98 | 2 | 0 |
| SCN5A | 321 | S | 30.00648485 | 7.37386832 | 0.077759381 | 1.631651237 | 5.93 | -0.33 | -2.53 | 26.25 | 0.98 | 2 | 0 |
| SCN5A | 322 | D | 27.46244318 | 7.62138288 | 0.082099834 | 1.708154172 | 5.93 | -0.33 | -2.53 | 26.25 | 0.98 | 2 | 0 |
| SCN5A | 323 | V | 29.62296104 | 12.13661172 | 0.161279565 | 6.226912477 | NA | NA | NA | NA | NA | NA | NA |
| SCN5A | 324 | L | 29.71306818 | 15.08279195 | 0.212944224 | 9.174428794 | NA | NA | NA | NA | NA | NA | NA |
| SCN5A | 325 | L | 25.87094318 | 15.42528097 | 0.218950163 | 9.508120214 | NA | NA | NA | NA | NA | NA | NA |
| SCN5A | 326 | C | 27.98898485 | 20.20121001 | 0.302701572 | 14.28740454 | NA | NA | NA | NA | NA | NA | NA |
| SCN5A | 327 | G | 25.40181818 | 23.29865599 | 0.35701885 | 17.38337665 | NA | NA | NA | NA | NA | NA | NA |
| SCN5A | 328 | N | 27.49756818 | 26.17225397 | 0.407410696 | 20.25733585 | NA | NA | NA | NA | NA | NA | NA |
| SCN5A | 329 | S | 23.95465152 | 27.57807091 | 0.432063313 | 25.95956145 | NA | NA | NA | NA | NA | NA | NA |
| SCN5A | 330 | S | 23.50348485 | 29.07539621 | 0.458320634 | 27.44601609 | NA | NA | NA | NA | NA | NA | NA |
| SCN5A | 331 | D | 21.11081818 | 25.17021452 | 0.389838781 | 23.54203201 | NA | NA | NA | NA | NA | NA | NA |
| SCN5A | 332 | A | 22.94041818 | 23.35503299 | 0.358007485 | 21.73390937 | NA | NA | NA | NA | NA | NA | NA |
| SCN5A | 333 | G | 25.78756818 | 22.84284895 | 0.349025749 | 16.92899781 | NA | NA | NA | NA | NA | NA | NA |
| SCN5A | 334 | T | 29.42767532 | 23.21605074 | 0.355570272 | 17.30810929 | NA | NA | NA | NA | NA | NA | NA |
| SCN5A | 335 | C | 32.21681818 | 23.87619949 | 0.367146739 | 17.02839785 | NA | NA | NA | NA | NA | NA | NA |
| SCN5A | 336 | P | 34.83524675 | 21.01797543 | 0.317024493 | 14.12163676 | NA | NA | NA | NA | NA | NA | NA |
| SCN5A | 337 | E | 39.43626263 | 23.66520429 | 0.363446696 | 16.16804826 | NA | NA | NA | NA | NA | NA | NA |
| SCN5A | 338 | G | 39.76206818 | 20.56600078 | 0.309098598 | 13.12331427 | NA | NA | NA | NA | NA | NA | NA |
| SCN5A | 339 | Y | 37.63565152 | 15.57680574 | 0.221607324 | 8.118337811 | NA | NA | NA | NA | NA | NA | NA |
| SCN5A | 340 | R | 35.89872727 | 22.71227391 | 0.346735966 | 15.23153775 | NA | NA | NA | NA | NA | NA | NA |
| SCN5A | 341 | C | 31.30698485 | 24.5521004 | 0.378999439 | 17.6272975 | NA | NA | NA | NA | NA | NA | NA |
| SCN5A | 342 | L | 31.85881818 | 23.3977753 | 0.358757021 | 17.19399225 | NA | NA | NA | NA | NA | NA | NA |
| SCN5A | 343 | K | 27.81470707 | 24.31709946 | 0.374878427 | 18.39975645 | NA | NA | NA | NA | NA | NA | NA |
| SCN5A | 344 | A | 30.82581818 | 20.91243444 | 0.31517371 | 15.22311071 | NA | NA | NA | NA | NA | NA | NA |
| SCN5A | 345 | G | 29.22806818 | 19.86247298 | 0.296761428 | 13.94188383 | NA | NA | NA | NA | NA | NA | NA |
| SCN5A | 346 | E | 27.00092929 | 17.79542784 | 0.260513414 | 11.88187458 | NA | NA | NA | NA | NA | NA | NA |

| | | | | | | | | | | | | | |
|---|---|---|---|---|---|---|---|---|---|---|---|---|---|
| SCN5A | 347 | N | 24.28394318 | 14.16561365 | 0.196860448 | 8.261793375 | NA | NA | NA | NA | NA | NA | NA |
| SCN5A | 348 | P | 23.30081818 | 12.19928215 | 0.162378563 | 6.326697407 | NA | NA | NA | NA | NA | NA | NA |
| SCN5A | 349 | D | 21.70469318 | 9.080339091 | 0.107684309 | 3.197923557 | 5.96 | -0.49 | -2.72 | 30.98 | 1.23 | 3 | -1 |
| SCN5A | 350 | H | 24.21421818 | 8.353349807 | 0.094935716 | 2.49021906 | 5.96 | -0.49 | -2.72 | 30.98 | 1.23 | 3 | -1 |
| SCN5A | 351 | G | 21.96931818 | 14.03228224 | 0.194522328 | 8.212499468 | NA | NA | NA | NA | NA | NA | NA |
| SCN5A | 352 | Y | 19.58931818 | 13.20289453 | 0.17997806 | 8.827584678 | NA | NA | NA | NA | NA | NA | NA |
| SCN5A | 353 | T | 18.50938961 | 13.09106516 | 0.178017004 | 11.52469819 | NA | NA | NA | NA | NA | NA | NA |
| SCN5A | 354 | S | 18.76481818 | 16.84459154 | 0.243839406 | 15.25052289 | NA | NA | NA | NA | NA | NA | NA |
| SCN5A | 355 | F | 14.90181818 | 16.34205694 | 0.235026884 | 14.68832157 | NA | NA | NA | NA | NA | NA | NA |
| SCN5A | 356 | D | 18.67344318 | 21.25940086 | 0.321258166 | 19.63886936 | NA | NA | NA | NA | NA | NA | NA |
| SCN5A | 357 | S | 15.36631818 | 21.61028609 | 0.327411342 | 19.94720557 | NA | NA | NA | NA | NA | NA | NA |
| SCN5A | 358 | F | 9.592545455 | 21.0869029 | 0.318233215 | 19.39010515 | NA | NA | NA | NA | NA | NA | NA |
| SCN5A | 359 | A | 12.05661818 | 18.64108243 | 0.27534294 | 16.96749086 | NA | NA | NA | NA | NA | NA | NA |
| SCN5A | 360 | W | 15.88788961 | 16.49391388 | 0.23768987 | 14.81919861 | NA | NA | NA | NA | NA | NA | NA |
| SCN5A | 361 | A | 11.59221818 | 16.99546089 | 0.246485074 | 15.30911573 | NA | NA | NA | NA | NA | NA | NA |
| SCN5A | 362 | F | 7.058454545 | 15.81621971 | 0.225805723 | 14.38418217 | NA | NA | NA | NA | NA | NA | NA |
| SCN5A | 363 | L | 11.31994318 | 12.93138573 | 0.175216841 | 11.24783493 | NA | NA | NA | NA | NA | NA | NA |
| SCN5A | 364 | A | 11.46461818 | 12.29340189 | 0.164029061 | 10.61196754 | NA | NA | NA | NA | NA | NA | NA |
| SCN5A | 365 | L | 7.647943182 | 12.61219572 | 0.169619477 | 10.11106094 | NA | NA | NA | NA | NA | NA | NA |
| SCN5A | 366 | F | 6.096 | 9.978374016 | 0.123432384 | 7.467028608 | NA | NA | NA | NA | NA | NA | NA |
| SCN5A | 367 | R | 11.63945455 | 7.662379917 | 0.082818764 | 5.751218375 | 2.27 | -0.79 | -1.72 | 21.91 | 1.22 | 2 | 0 |
| SCN5A | 368 | L | 8.387193182 | 8.88019493 | 0.104174551 | 6.465958599 | NA | NA | NA | NA | NA | NA | NA |
| SCN5A | 369 | M | 3.303943182 | 7.951393337 | 0.087886947 | 5.511567138 | 2.68 | -0.72 | -1.1 | 12.6 | 0.94 | 1 | 1 |
| SCN5A | 370 | T | 5.43838961 | 4.287523411 | 0.023636773 | 1.635299971 | 2.68 | -0.72 | -1.1 | 12.6 | 0.94 | 1 | 1 |
| SCN5A | 371 | Q | 6.069040404 | 3.885404226 | 0.016585151 | 1.77197087 | 2.68 | -0.72 | -1.1 | 12.6 | 0.94 | 1 | 1 |
| SCN5A | 372 | D | 10.38431818 | 3.44359231 | 0.008837471 | 1.883318774 | 2.27 | -0.79 | -1.72 | 21.91 | 1.22 | 2 | 0 |
| SCN5A | 373 | C | 13.59931818 | 3.332687204 | 0.006892622 | 1.749202849 | 1.69 | -0.47 | -1.5 | 33.69 | 1.67 | 2 | -2 |
| SCN5A | 374 | W | 10.62017532 | 5.766692466 | 0.049575704 | 4.038256641 | 1.74 | -0.99 | -2.47 | 34.33 | 1.89 | 2 | -2 |
| SCN5A | 375 | E | 15.21170707 | 6.407980727 | 0.060821431 | 4.821029585 | 1.69 | -0.47 | -1.5 | 33.69 | 1.67 | 2 | -2 |
| SCN5A | 376 | R | 19.27354545 | 7.464101419 | 0.079341723 | 6.320492132 | 1.69 | -0.47 | -1.5 | 33.69 | 1.67 | 2 | -2 |
| SCN5A | 377 | L | 15.34669318 | 9.614062669 | 0.117043766 | 8.002620788 | NA | NA | NA | NA | NA | NA | NA |
| SCN5A | 378 | Y | 14.39273485 | 11.48584912 | 0.149867694 | 9.818549548 | NA | NA | NA | NA | NA | NA | NA |
| SCN5A | 379 | Q | 18.81748485 | 11.75263051 | 0.154546013 | 10.11494131 | NA | NA | NA | NA | NA | NA | NA |

| | | | | | | | | | | | | | |
|---|---|---|---|---|---|---|---|---|---|---|---|---|---|
| SCN5A | 380 | Q | 20.42648485 | 11.61840092 | 0.152192143 | 10.62507759 | NA | NA | NA | NA | NA | NA | NA |
| SCN5A | 381 | T | 17.49538961 | 15.45016032 | 0.219386451 | 13.81209471 | NA | NA | NA | NA | NA | NA | NA |
| SCN5A | 382 | L | 17.29169318 | 17.25084131 | 0.250963463 | 15.60730522 | NA | NA | NA | NA | NA | NA | NA |
| SCN5A | 383 | R | 23.69309091 | 17.35442056 | 0.252779845 | 16.74465222 | NA | NA | NA | NA | NA | NA | NA |
| SCN5A | 384 | S | 22.11698485 | 20.15833309 | 0.301949676 | 18.57296181 | NA | NA | NA | NA | NA | NA | NA |
| SCN5A | 385 | A | 19.28901818 | 21.57707506 | 0.326828948 | 19.96314927 | NA | NA | NA | NA | NA | NA | NA |
| SCN5A | 386 | G | 17.88181818 | 21.65047886 | 0.328116168 | 19.96746009 | NA | NA | NA | NA | NA | NA | NA |
| SCN5A | 387 | K | 17.79826263 | 21.78756517 | 0.330520134 | 20.19643595 | NA | NA | NA | NA | NA | NA | NA |
| SCN5A | 388 | I | 13.71669318 | 23.37155068 | 0.358297142 | 21.2567584 | NA | NA | NA | NA | NA | NA | NA |
| SCN5A | 389 | Y | 14.16765152 | 19.86456524 | 0.296798119 | 17.7627733 | NA | NA | NA | NA | NA | NA | NA |
| SCN5A | 390 | M | 11.01344318 | 18.54257625 | 0.273615521 | 16.4859779 | NA | NA | NA | NA | NA | NA | NA |
| SCN5A | 391 | I | 8.253318182 | 19.86407909 | 0.296789593 | 17.89138949 | NA | NA | NA | NA | NA | NA | NA |
| SCN5A | 392 | F | 8.719 | 17.72094617 | 0.259207293 | 15.63752707 | NA | NA | NA | NA | NA | NA | NA |
| SCN5A | 393 | F | 10.49572727 | 13.97808889 | 0.193571985 | 11.86585619 | NA | NA | NA | NA | NA | NA | NA |
| SCN5A | 394 | M | 5.789943182 | 16.30447856 | 0.234367904 | 13.27184148 | NA | NA | NA | NA | NA | NA | NA |
| SCN5A | 395 | L | 3.696318182 | 16.28570898 | 0.234038758 | 13.24740838 | NA | NA | NA | NA | NA | NA | NA |
| SCN5A | 396 | V | 4.948103896 | 13.29479075 | 0.181589566 | 10.23168319 | NA | NA | NA | NA | NA | NA | NA |
| SCN5A | 397 | I | 4.683443182 | 10.10723251 | 0.125692066 | 7.071198055 | NA | NA | NA | NA | NA | NA | NA |
| SCN5A | 398 | F | 0.715545455 | 13.27349106 | 0.181216052 | 9.428491079 | NA | NA | NA | NA | NA | NA | NA |
| SCN5A | 399 | L | -0.139943182 | 14.51095269 | 0.202916365 | 10.67953217 | NA | NA | NA | NA | NA | NA | NA |
| SCN5A | 400 | G | 0.297568182 | 12.08011813 | 0.160288885 | 8.275200941 | NA | NA | NA | NA | NA | NA | NA |
| SCN5A | 401 | S | 1.466984848 | 7.926199846 | 0.08744515 | 4.170902964 | 3.86 | -0.35 | -1.08 | 12.03 | 0.28 | 1 | 1 |
| SCN5A | 402 | F | -0.701727273 | 7.389630573 | 0.078035791 | 2.890770914 | 4.66 | 0.34 | 0.36 | 1.8 | -0.72 | 0 | 0 |
| SCN5A | 403 | Y | -4.281651515 | 11.54856957 | 0.150967569 | 9.962061571 | NA | NA | NA | NA | NA | NA | NA |
| SCN5A | 404 | L | -1.811568182 | 11.13944936 | 0.143793176 | 6.498873713 | NA | NA | NA | NA | NA | NA | NA |
| SCN5A | 405 | V | -1.458818182 | 6.297094012 | 0.058876905 | 1.658833595 | 4.66 | 0.7 | 1.33 | 1.37 | -0.94 | 0 | 0 |
| SCN5A | 406 | N | -4.413693182 | 6.766551411 | 0.067109381 | 4.868701928 | 1.86 | 1.43 | 3.7 | 0.2 | -1.82 | 0 | 0 |
| SCN5A | 407 | L | -7.138193182 | 9.557988125 | 0.116060434 | 7.479420144 | NA | NA | NA | NA | NA | NA | NA |
| SCN5A | 408 | I | -4.898818182 | 8.730567794 | 0.101550667 | 6.599943859 | NA | NA | NA | NA | NA | NA | NA |
| SCN5A | 409 | L | -6.203068182 | 3.48128324 | 0.009498425 | 1.800475343 | 2.19 | 1.45 | 3.88 | 0.17 | -1.82 | 0 | 0 |
| SCN5A | 410 | A | -10.28881818 | 6.34126336 | 0.059651466 | 3.785982324 | 2.76 | 1.2 | 3.65 | 0.1 | -1.23 | 0 | 0 |
| SCN5A | 411 | V | -10.79453247 | 8.326041376 | 0.094456831 | 5.767489792 | 2.76 | 1.2 | 3.65 | 0.1 | -1.23 | 0 | 0 |
| SCN5A | 412 | V | -9.691246753 | 7.779651535 | 0.084875257 | 5.319791279 | 2.76 | 1.2 | 3.65 | 0.1 | -1.23 | 0 | 0 |

| | | | | | | | | | | | | | |
|---|---|---|---|---|---|---|---|---|---|---|---|---|---|
| SCN5A | 413 | A | -12.53461818 | 4.16384858 | 0.021467993 | 1.629229416 | 2.76 | 1.2 | 3.65 | 0.1 | -1.23 | 0 | 0 |
| SCN5A | 414 | M | -15.51931818 | 6.788849755 | 0.067500408 | 4.032125013 | 2.84 | 1.08 | 3.23 | 0.15 | -1.22 | 0 | 0 |
| SCN5A | 415 | A | -14.97441818 | 9.449376117 | 0.114155798 | 6.721704695 | NA | NA | NA | NA | NA | NA | NA |
| SCN5A | 416 | Y | -14.09765152 | 8.903408842 | 0.104581634 | 6.251906797 | NA | NA | NA | NA | NA | NA | NA |
| SCN5A | 417 | E | -18.6750404 | 4.996711018 | 0.036073194 | 1.937830919 | 3.29 | 0.39 | 0.48 | 10.82 | -0.53 | 1 | -1 |
| SCN5A | 418 | E | -19.99937374 | 8.692019961 | 0.100874686 | 5.602026331 | NA | NA | NA | NA | NA | NA | NA |
| SCN5A | 419 | Q | -18.04026263 | 11.38189097 | 0.148044668 | 8.307214475 | NA | NA | NA | NA | NA | NA | NA |
| SCN5A | 420 | N | -19.54694318 | 10.01533449 | 0.124080529 | 6.912848931 | NA | NA | NA | NA | NA | NA | NA |
| SCN5A | 421 | Q | -23.96526263 | 8.470656999 | 0.096992832 | 5.0619382 | 3.51 | -0.19 | -1.4 | 17.88 | 0.32 | 1 | -1 |
| SCN5A | 422 | A | -23.85381818 | 12.41141672 | 0.166098587 | 9.222654208 | NA | NA | NA | NA | NA | NA | NA |
| SCN5A | 423 | T | -22.72838961 | 14.2668439 | 0.198635637 | 11.25375193 | NA | NA | NA | NA | NA | NA | NA |
| SCN5A | 424 | I | -24.85506818 | 12.37809703 | 0.165514288 | 9.480725022 | NA | NA | NA | NA | NA | NA | NA |
| SCN5A | 425 | A | -28.08981818 | 14.42592905 | 0.201425378 | 6.557038556 | NA | NA | NA | NA | NA | NA | NA |
| SCN5A | 426 | E | -26.76748485 | 17.32149102 | 0.252202387 | 9.598051676 | NA | NA | NA | NA | NA | NA | NA |
| SCN5A | 427 | T | -27.21896104 | 17.39038591 | 0.253410538 | 14.5162897 | NA | NA | NA | NA | NA | NA | NA |
| SCN5A | 428 | E | -31.80248485 | 16.59409684 | 0.239446694 | 8.791675448 | NA | NA | NA | NA | NA | NA | NA |
| SCN5A | 429 | E | -33.75470707 | 19.87065145 | 0.296904847 | 11.36082691 | NA | NA | NA | NA | NA | NA | NA |
| SCN5A | 699 | C | -17.13365152 | 59.96469402 | 1 | 56.94220332 | NA | NA | NA | NA | NA | NA | NA |
| SCN5A | 700 | C | -20.45548485 | 58.98144682 | 0.982757629 | 55.93498254 | NA | NA | NA | NA | NA | NA | NA |
| SCN5A | 701 | P | -20.14867532 | 59.35366127 | 0.989284838 | 56.30773696 | NA | NA | NA | NA | NA | NA | NA |
| SCN5A | 702 | L | -16.01006818 | 57.60291541 | 0.958583496 | 54.58065414 | NA | NA | NA | NA | NA | NA | NA |
| SCN5A | 703 | W | -17.39496104 | 53.26546972 | 0.882521396 | 50.23898429 | NA | NA | NA | NA | NA | NA | NA |
| SCN5A | 704 | M | -21.24719318 | 54.54976111 | 0.905042923 | 51.49517863 | NA | NA | NA | NA | NA | NA | NA |
| SCN5A | 705 | S | -19.07715152 | 54.85085846 | 0.910323012 | 51.80853027 | NA | NA | NA | NA | NA | NA | NA |
| SCN5A | 706 | I | -15.38669318 | 52.38562821 | 0.867092364 | 49.36409734 | NA | NA | NA | NA | NA | NA | NA |
| SCN5A | 707 | K | -18.65470707 | 50.09247766 | 0.826879331 | 47.05116634 | NA | NA | NA | NA | NA | NA | NA |
| SCN5A | 708 | Q | -21.25970707 | 50.38688104 | 0.832042033 | 47.33085795 | NA | NA | NA | NA | NA | NA | NA |
| SCN5A | 709 | G | -17.59081818 | 50.14061212 | 0.827723424 | 47.10780953 | NA | NA | NA | NA | NA | NA | NA |
| SCN5A | 710 | V | -15.14896104 | 47.22131805 | 0.776530245 | 44.19810373 | NA | NA | NA | NA | NA | NA | NA |
| SCN5A | 711 | K | -19.2660404 | 43.8143993 | 0.716786006 | 40.73517005 | NA | NA | NA | NA | NA | NA | NA |
| SCN5A | 712 | L | -19.54069318 | 46.25846661 | 0.759645537 | 43.20792732 | NA | NA | NA | NA | NA | NA | NA |
| SCN5A | 713 | V | -15.07067532 | 45.28095792 | 0.742503798 | 42.2636071 | NA | NA | NA | NA | NA | NA | NA |
| SCN5A | 714 | V | -14.24167532 | 42.26361047 | 0.68959114 | 39.25545156 | NA | NA | NA | NA | NA | NA | NA |

| | | | | | | | | | | | | | |
|---|---|---|---|---|---|---|---|---|---|---|---|---|---|
| SCN5A | 715 | M | -17.32394318 | 39.85335431 | 0.647324526 | 36.80819878 | NA | NA | NA | NA | NA | NA | NA |
| SCN5A | 716 | D | -15.92631818 | 41.0138131 | 0.667674506 | 37.99834878 | NA | NA | NA | NA | NA | NA | NA |
| SCN5A | 717 | P | -15.91524675 | 38.43142205 | 0.622389308 | 35.38679638 | NA | NA | NA | NA | NA | NA | NA |
| SCN5A | 718 | F | -13.99363636 | 40.58915879 | 0.660227704 | 37.57170391 | NA | NA | NA | NA | NA | NA | NA |
| SCN5A | 719 | T | -11.0111039 | 39.72191493 | 0.645019585 | 37.85625076 | NA | NA | NA | NA | NA | NA | NA |
| SCN5A | 720 | D | -11.47181818 | 34.70695819 | 0.557076551 | 33.24083651 | NA | NA | NA | NA | NA | NA | NA |
| SCN5A | 721 | L | -10.18531818 | 35.85090062 | 0.577136897 | 33.9832184 | NA | NA | NA | NA | NA | NA | NA |
| SCN5A | 722 | T | -6.672961039 | 38.38111725 | 0.621507156 | 36.72245318 | NA | NA | NA | NA | NA | NA | NA |
| SCN5A | 723 | I | -6.259068182 | 35.69326158 | 0.574372515 | 34.02031859 | NA | NA | NA | NA | NA | NA | NA |
| SCN5A | 724 | T | -6.314103896 | 32.26265713 | 0.51421292 | 30.60467334 | NA | NA | NA | NA | NA | NA | NA |
| SCN5A | 725 | M | -3.734068182 | 35.24028279 | 0.566429011 | 33.59044979 | NA | NA | NA | NA | NA | NA | NA |
| SCN5A | 726 | C | -1.381984848 | 35.65051075 | 0.57362283 | 32.55139474 | NA | NA | NA | NA | NA | NA | NA |
| SCN5A | 727 | I | -2.055318182 | 31.54062339 | 0.501551228 | 28.52141694 | NA | NA | NA | NA | NA | NA | NA |
| SCN5A | 728 | V | -0.958818182 | 30.71001501 | 0.486985555 | 27.78208674 | NA | NA | NA | NA | NA | NA | NA |
| SCN5A | 729 | L | 1.390068182 | 33.87579888 | 0.542501216 | 30.78127096 | NA | NA | NA | NA | NA | NA | NA |
| SCN5A | 730 | N | 2.495068182 | 33.08337277 | 0.528605113 | 29.95563318 | NA | NA | NA | NA | NA | NA | NA |
| SCN5A | 731 | T | 2.13938961 | 29.52320228 | 0.466173429 | 26.38605074 | NA | NA | NA | NA | NA | NA | NA |
| SCN5A | 732 | L | 3.714068182 | 29.56840254 | 0.466966067 | 26.44362737 | NA | NA | NA | NA | NA | NA | NA |
| SCN5A | 733 | F | 6.654454545 | 33.37712576 | 0.53375641 | 30.28057626 | NA | NA | NA | NA | NA | NA | NA |
| SCN5A | 734 | M | 6.891068182 | 30.79449965 | 0.48846709 | 26.2723946 | NA | NA | NA | NA | NA | NA | NA |
| SCN5A | 735 | A | 8.624418182 | 29.29647926 | 0.46219758 | 25.40829497 | NA | NA | NA | NA | NA | NA | NA |
| SCN5A | 736 | L | 10.64794318 | 31.88975201 | 0.5076736 | 29.5728283 | NA | NA | NA | NA | NA | NA | NA |
| SCN5A | 737 | E | 13.44437374 | 28.59559767 | 0.449906815 | 26.30441332 | NA | NA | NA | NA | NA | NA | NA |
| SCN5A | 738 | H | 16.92151818 | 28.26933804 | 0.444185477 | 26.00803291 | NA | NA | NA | NA | NA | NA | NA |
| SCN5A | 739 | Y | 22.02115152 | 26.20027139 | 0.407902013 | 23.9932609 | NA | NA | NA | NA | NA | NA | NA |
| SCN5A | 740 | N | 22.82356818 | 29.35328956 | 0.463193814 | 27.10885716 | NA | NA | NA | NA | NA | NA | NA |
| SCN5A | 741 | M | 19.08006818 | 32.29349563 | 0.514753709 | 30.04679666 | NA | NA | NA | NA | NA | NA | NA |
| SCN5A | 742 | T | 21.24467532 | 35.5649631 | 0.572122654 | 33.31270655 | NA | NA | NA | NA | NA | NA | NA |
| SCN5A | 743 | S | 22.23331818 | 38.6410834 | 0.626065961 | 36.39966989 | NA | NA | NA | NA | NA | NA | NA |
| SCN5A | 744 | E | 19.37570707 | 40.58234815 | 0.660108271 | 38.33808827 | NA | NA | NA | NA | NA | NA | NA |
| SCN5A | 745 | F | 15.82254545 | 35.62274292 | 0.573135889 | 33.336611 | NA | NA | NA | NA | NA | NA | NA |
| SCN5A | 746 | E | 17.71215152 | 36.73008746 | 0.59255445 | 34.48432774 | NA | NA | NA | NA | NA | NA | NA |
| SCN5A | 747 | E | 16.09281818 | 39.74888489 | 0.645492535 | 37.49631149 | NA | NA | NA | NA | NA | NA | NA |

| | | | | | | | | | | | | | |
|---|---|---|---|---|---|---|---|---|---|---|---|---|---|
| SCN5A | 748 | M | 12.46056818 | 39.93716863 | 0.648794307 | 37.6411547 | NA | NA | NA | NA | NA | NA | NA |
| SCN5A | 749 | L | 11.61156818 | 35.01970337 | 0.562560897 | 32.74099824 | NA | NA | NA | NA | NA | NA | NA |
| SCN5A | 750 | Q | 12.72926263 | 38.41285682 | 0.622063746 | 36.17514245 | NA | NA | NA | NA | NA | NA | NA |
| SCN5A | 751 | V | 9.970675325 | 42.10784043 | 0.686859533 | 39.83821314 | NA | NA | NA | NA | NA | NA | NA |
| SCN5A | 752 | G | 6.980318182 | 40.62977598 | 0.660939973 | 36.73852132 | NA | NA | NA | NA | NA | NA | NA |
| SCN5A | 753 | N | 6.175318182 | 36.32981038 | 0.58553513 | 32.42075713 | NA | NA | NA | NA | NA | NA | NA |
| SCN5A | 754 | L | 6.538318182 | 41.51656277 | 0.676490799 | 37.61459052 | NA | NA | NA | NA | NA | NA | NA |
| SCN5A | 755 | V | 3.694246753 | 42.49682589 | 0.693680841 | 39.36569419 | NA | NA | NA | NA | NA | NA | NA |
| SCN5A | 756 | F | 2.049090909 | 37.83128539 | 0.611865222 | 34.69475945 | NA | NA | NA | NA | NA | NA | NA |
| SCN5A | 757 | T | 1.997675325 | 40.11021769 | 0.651828921 | 37.07120527 | NA | NA | NA | NA | NA | NA | NA |
| SCN5A | 758 | G | 0.512318182 | 43.26081377 | 0.707078247 | 40.13078516 | NA | NA | NA | NA | NA | NA | NA |
| SCN5A | 759 | I | -1.031818182 | 41.49411985 | 0.676097237 | 38.46972301 | NA | NA | NA | NA | NA | NA | NA |
| SCN5A | 760 | F | -1.653363636 | 36.0595491 | 0.580795788 | 32.97222616 | NA | NA | NA | NA | NA | NA | NA |
| SCN5A | 761 | T | -1.810246753 | 42.4796931 | 0.693380398 | 39.39393363 | NA | NA | NA | NA | NA | NA | NA |
| SCN5A | 762 | A | -4.381218182 | 44.73160517 | 0.732870266 | 43.04583969 | NA | NA | NA | NA | NA | NA | NA |
| SCN5A | 763 | E | -7.42159596 | 40.08003525 | 0.651299637 | 38.34051714 | NA | NA | NA | NA | NA | NA | NA |
| SCN5A | 764 | M | -6.430943182 | 41.51438403 | 0.676452593 | 39.76997928 | NA | NA | NA | NA | NA | NA | NA |
| SCN5A | 765 | T | -7.08038961 | 45.21874447 | 0.741412814 | 43.48472206 | NA | NA | NA | NA | NA | NA | NA |
| SCN5A | 766 | F | -8.642909091 | 46.01668169 | 0.755405561 | 44.04056432 | NA | NA | NA | NA | NA | NA | NA |
| SCN5A | 767 | K | -12.21137374 | 37.9507978 | 0.613961009 | 36.36963778 | NA | NA | NA | NA | NA | NA | NA |
| SCN5A | 768 | I | -11.09806818 | 44.95679114 | 0.736819161 | 42.99871997 | NA | NA | NA | NA | NA | NA | NA |
| SCN5A | 769 | I | -12.90894318 | 47.39064982 | 0.779499672 | 44.41006945 | NA | NA | NA | NA | NA | NA | NA |
| SCN5A | 770 | A | -16.45901818 | 45.08765403 | 0.739113992 | 42.06624196 | NA | NA | NA | NA | NA | NA | NA |
| SCN5A | 771 | L | -18.30269318 | 43.10276754 | 0.704306725 | 40.04363743 | NA | NA | NA | NA | NA | NA | NA |
| SCN5A | 772 | D | -17.96106818 | 44.1957119 | 0.723472761 | 41.17053931 | NA | NA | NA | NA | NA | NA | NA |
| SCN5A | 773 | P | -15.14824675 | 42.43804018 | 0.692649966 | 39.42884543 | NA | NA | NA | NA | NA | NA | NA |
| SCN5A | 774 | Y | -19.17415152 | 41.2668499 | 0.672111797 | 38.22231622 | NA | NA | NA | NA | NA | NA | NA |
| SCN5A | 775 | Y | -21.26031818 | 39.25513268 | 0.636834022 | 36.22165319 | NA | NA | NA | NA | NA | NA | NA |
| SCN5A | 776 | Y | -13.52948485 | 39.410895 | 0.639565493 | 36.41978849 | NA | NA | NA | NA | NA | NA | NA |
| SCN5A | 777 | F | -12.505 | 37.71605124 | 0.609844458 | 34.72365158 | NA | NA | NA | NA | NA | NA | NA |
| SCN5A | 778 | Q | -17.6430404 | 35.10201366 | 0.564004303 | 32.0868406 | NA | NA | NA | NA | NA | NA | NA |
| SCN5A | 779 | Q | -17.39759596 | 30.38253317 | 0.481242784 | 27.31725904 | NA | NA | NA | NA | NA | NA | NA |
| SCN5A | 780 | G | -13.08506818 | 30.7514022 | 0.487711327 | 27.80958276 | NA | NA | NA | NA | NA | NA | NA |

| | | | | | | | | | | | | | |
|---|---|---|---|---|---|---|---|---|---|---|---|---|---|
| SCN5A | 781 | W | -11.05846104 | 27.29616843 | 0.427119828 | 25.03336737 | NA | NA | NA | NA | NA | NA | NA |
| SCN5A | 782 | N | -11.08531818 | 34.53246213 | 0.554016562 | 32.67911266 | NA | NA | NA | NA | NA | NA | NA |
| SCN5A | 783 | I | -9.693693182 | 33.66537969 | 0.538811274 | 31.73222557 | NA | NA | NA | NA | NA | NA | NA |
| SCN5A | 784 | F | -5.854909091 | 28.90864724 | 0.455396499 | 27.2686794 | NA | NA | NA | NA | NA | NA | NA |
| SCN5A | 785 | D | -6.765443182 | 33.21741817 | 0.530955753 | 31.45960334 | NA | NA | NA | NA | NA | NA | NA |
| SCN5A | 786 | S | -5.609818182 | 36.79479004 | 0.593689084 | 35.04227809 | NA | NA | NA | NA | NA | NA | NA |
| SCN5A | 787 | I | -3.334818182 | 34.30893814 | 0.550096811 | 31.36212767 | NA | NA | NA | NA | NA | NA | NA |
| SCN5A | 788 | I | -1.377068182 | 31.70441652 | 0.504423529 | 28.6246254 | NA | NA | NA | NA | NA | NA | NA |
| SCN5A | 789 | V | -1.353818182 | 37.21973812 | 0.601141037 | 34.09330005 | NA | NA | NA | NA | NA | NA | NA |
| SCN5A | 790 | I | -1.023943182 | 39.06455583 | 0.633492037 | 34.21728754 | NA | NA | NA | NA | NA | NA | NA |
| SCN5A | 791 | L | 1.339568182 | 34.00010979 | 0.544681151 | 29.14830418 | NA | NA | NA | NA | NA | NA | NA |
| SCN5A | 792 | S | 2.558984848 | 37.55784634 | 0.607070154 | 32.70277519 | NA | NA | NA | NA | NA | NA | NA |
| SCN5A | 793 | L | 2.168068182 | 40.28437399 | 0.654882952 | 35.42872263 | NA | NA | NA | NA | NA | NA | NA |
| SCN5A | 794 | M | 4.768568182 | 41.0883128 | 0.668980944 | 36.23103393 | NA | NA | NA | NA | NA | NA | NA |
| SCN5A | 795 | E | 7.156373737 | 37.6652007 | 0.608952736 | 32.80390931 | NA | NA | NA | NA | NA | NA | NA |
| SCN5A | 796 | L | 7.176568182 | 40.89549233 | 0.665599615 | 38.65409011 | NA | NA | NA | NA | NA | NA | NA |
| SCN5A | 797 | G | 8.924818182 | 43.41200369 | 0.709729536 | 41.16669283 | NA | NA | NA | NA | NA | NA | NA |
| SCN5A | 798 | L | 10.99519318 | 39.09823918 | 0.634082714 | 36.87514442 | NA | NA | NA | NA | NA | NA | NA |
| SCN5A | 799 | S | 13.44715152 | 40.79221918 | 0.663788602 | 38.57528033 | NA | NA | NA | NA | NA | NA | NA |
| SCN5A | 800 | R | 16.15927273 | 42.28069619 | 0.689890758 | 40.12149084 | NA | NA | NA | NA | NA | NA | NA |
| SCN5A | 801 | M | 15.80744318 | 37.85877581 | 0.612347298 | 34.52772339 | NA | NA | NA | NA | NA | NA | NA |
| SCN5A | 802 | S | 17.73331818 | 34.7710501 | 0.558200476 | 31.41047302 | NA | NA | NA | NA | NA | NA | NA |
| SCN5A | 803 | N | 13.76944318 | 32.84367971 | 0.524401819 | 30.62082755 | NA | NA | NA | NA | NA | NA | NA |
| SCN5A | 804 | L | 15.65706818 | 28.97801223 | 0.456612894 | 26.76345861 | NA | NA | NA | NA | NA | NA | NA |
| SCN5A | 805 | S | 13.41331818 | 27.58041325 | 0.432104388 | 25.36664486 | NA | NA | NA | NA | NA | NA | NA |
| SCN5A | 806 | V | 9.922675325 | 28.92201105 | 0.455630849 | 26.68143125 | NA | NA | NA | NA | NA | NA | NA |
| SCN5A | 807 | L | 10.02619318 | 31.99250383 | 0.509475471 | 29.75238519 | NA | NA | NA | NA | NA | NA | NA |
| SCN5A | 808 | R | 10.53636364 | 31.02617685 | 0.492529816 | 28.76140613 | NA | NA | NA | NA | NA | NA | NA |
| SCN5A | 809 | S | 5.754818182 | 28.31192261 | 0.444932246 | 23.44923214 | NA | NA | NA | NA | NA | NA | NA |
| SCN5A | 810 | F | 5.417090909 | 31.79304007 | 0.505977645 | 26.93115203 | NA | NA | NA | NA | NA | NA | NA |
| SCN5A | 811 | R | 6.200454545 | 31.74222316 | 0.505086512 | 26.88646688 | NA | NA | NA | NA | NA | NA | NA |
| SCN5A | 812 | L | 3.946943182 | 26.26818945 | 0.409093035 | 21.40312422 | NA | NA | NA | NA | NA | NA | NA |
| SCN5A | 813 | L | 0.297193182 | 28.53744475 | 0.448887036 | 24.63348372 | NA | NA | NA | NA | NA | NA | NA |

| | | | | | | | | | | | | | |
|---|---|---|---|---|---|---|---|---|---|---|---|---|---|
| SCN5A | 814 | R | -1.153545455 | 29.80559186 | 0.471125455 | 26.66873334 | NA | NA | NA | NA | NA | NA | NA |
| SCN5A | 815 | V | -0.781818182 | 25.72714809 | 0.399605252 | 22.59111377 | NA | NA | NA | NA | NA | NA | NA |
| SCN5A | 816 | F | -2.394363636 | 23.28494101 | 0.356778342 | 21.63970582 | NA | NA | NA | NA | NA | NA | NA |
| SCN5A | 817 | K | -4.916929293 | 28.58340179 | 0.449692946 | 26.87686668 | NA | NA | NA | NA | NA | NA | NA |
| SCN5A | 818 | L | -5.677943182 | 25.90354968 | 0.402698657 | 24.22916897 | NA | NA | NA | NA | NA | NA | NA |
| SCN5A | 819 | A | -8.591618182 | 24.74665054 | 0.3824111 | 22.81332441 | NA | NA | NA | NA | NA | NA | NA |
| SCN5A | 820 | K | -10.95637374 | 26.45082647 | 0.412295785 | 24.46156006 | NA | NA | NA | NA | NA | NA | NA |
| SCN5A | 821 | S | -12.37415152 | 26.83842672 | 0.419092801 | 23.86498437 | NA | NA | NA | NA | NA | NA | NA |
| SCN5A | 822 | W | -8.934246753 | 22.00126128 | 0.334267541 | 20.35992753 | NA | NA | NA | NA | NA | NA | NA |
| SCN5A | 823 | P | -13.5771039 | 19.01565476 | 0.281911497 | 16.02994915 | NA | NA | NA | NA | NA | NA | NA |
| SCN5A | 824 | T | -9.982961039 | 17.95681876 | 0.26334359 | 15.84984986 | NA | NA | NA | NA | NA | NA | NA |
| SCN5A | 825 | L | -7.669443182 | 20.31667783 | 0.304726433 | 18.32202555 | NA | NA | NA | NA | NA | NA | NA |
| SCN5A | 826 | N | -12.43769318 | 19.24042105 | 0.285853032 | 16.96388712 | NA | NA | NA | NA | NA | NA | NA |
| SCN5A | 827 | T | -11.67667532 | 14.71074063 | 0.206419876 | 12.41750627 | NA | NA | NA | NA | NA | NA | NA |
| SCN5A | 828 | L | -6.932693182 | 16.31899868 | 0.234622531 | 14.60497899 | NA | NA | NA | NA | NA | NA | NA |
| SCN5A | 829 | I | -8.876193182 | 18.62958287 | 0.275141282 | 16.62733395 | NA | NA | NA | NA | NA | NA | NA |
| SCN5A | 830 | K | -13.28792929 | 15.53742363 | 0.220916713 | 14.57791342 | NA | NA | NA | NA | NA | NA | NA |
| SCN5A | 831 | I | -9.133068182 | 12.1933846 | 0.162275143 | 9.952412636 | NA | NA | NA | NA | NA | NA | NA |
| SCN5A | 832 | I | -6.240568182 | 16.55737431 | 0.238802722 | 14.56313906 | NA | NA | NA | NA | NA | NA | NA |
| SCN5A | 833 | G | -9.653568182 | 18.88634006 | 0.279643815 | 16.75833698 | NA | NA | NA | NA | NA | NA | NA |
| SCN5A | 834 | N | -11.72856818 | 14.08387965 | 0.195427148 | 12.09083277 | NA | NA | NA | NA | NA | NA | NA |
| SCN5A | 835 | S | -7.355484848 | 15.88224518 | 0.226963556 | 13.77713331 | NA | NA | NA | NA | NA | NA | NA |
| SCN5A | 836 | V | -6.876675325 | 18.82335918 | 0.278539373 | 16.86619322 | NA | NA | NA | NA | NA | NA | NA |
| SCN5A | 837 | G | -10.83806818 | 20.89314534 | 0.314835454 | 18.61447928 | NA | NA | NA | NA | NA | NA | NA |
| SCN5A | 838 | A | -12.84141818 | 18.6799886 | 0.276025205 | 15.75813297 | NA | NA | NA | NA | NA | NA | NA |
| SCN5A | 839 | L | -11.10894318 | 15.04461462 | 0.212274741 | 12.86455179 | NA | NA | NA | NA | NA | NA | NA |
| SCN5A | 840 | G | -8.165318182 | 19.09240367 | 0.283257377 | 16.92401746 | NA | NA | NA | NA | NA | NA | NA |
| SCN5A | 841 | N | -9.291818182 | 19.37357339 | 0.288188012 | 17.20329344 | NA | NA | NA | NA | NA | NA | NA |
| SCN5A | 842 | L | -6.870943182 | 14.1237985 | 0.196127171 | 11.94835386 | NA | NA | NA | NA | NA | NA | NA |
| SCN5A | 843 | T | -3.822246753 | 17.07068183 | 0.247804159 | 12.22211569 | NA | NA | NA | NA | NA | NA | NA |
| SCN5A | 844 | L | -3.710568182 | 19.5027547 | 0.290453355 | 14.65254886 | NA | NA | NA | NA | NA | NA | NA |
| SCN5A | 845 | V | -3.914246753 | 18.79653545 | 0.278068988 | 13.93587302 | NA | NA | NA | NA | NA | NA | NA |
| SCN5A | 846 | L | -0.507568182 | 13.59540647 | 0.186861209 | 8.729139379 | NA | NA | NA | NA | NA | NA | NA |

| | | | | | | | | | | | | | |
|---|---|---|---|---|---|---|---|---|---|---|---|---|---|
| SCN5A | 847 | A | 0.190818182 | 18.2805257 | 0.269020163 | 13.41724157 | NA | NA | NA | NA | NA | NA | NA |
| SCN5A | 848 | I | 0.687068182 | 20.7115041 | 0.311650166 | 15.84482995 | NA | NA | NA | NA | NA | NA | NA |
| SCN5A | 849 | I | 0.814443182 | 17.74500758 | 0.259629237 | 12.86562238 | NA | NA | NA | NA | NA | NA | NA |
| SCN5A | 850 | V | 3.780818182 | 16.29832611 | 0.234260014 | 13.98125763 | NA | NA | NA | NA | NA | NA | NA |
| SCN5A | 851 | F | 4.8 | 20.56965566 | 0.30916269 | 18.10713727 | NA | NA | NA | NA | NA | NA | NA |
| SCN5A | 852 | I | 3.336818182 | 21.86467603 | 0.331872362 | 19.35511113 | NA | NA | NA | NA | NA | NA | NA |
| SCN5A | 853 | F | 4.949636364 | 18.08671949 | 0.265621549 | 15.66501713 | NA | NA | NA | NA | NA | NA | NA |
| SCN5A | 854 | A | 8.986618182 | 19.58500248 | 0.291895664 | 18.3495203 | NA | NA | NA | NA | NA | NA | NA |
| SCN5A | 855 | V | 8.955818182 | 21.94943596 | 0.333358725 | 20.8207562 | NA | NA | NA | NA | NA | NA | NA |
| SCN5A | 856 | V | 8.904103896 | 22.0811319 | 0.335668165 | 20.83035156 | NA | NA | NA | NA | NA | NA | NA |
| SCN5A | 857 | G | 12.28731818 | 19.66798398 | 0.29335084 | 15.82286341 | NA | NA | NA | NA | NA | NA | NA |
| SCN5A | 858 | M | 13.95631818 | 20.47021529 | 0.307418889 | 16.58691322 | NA | NA | NA | NA | NA | NA | NA |
| SCN5A | 859 | Q | 13.12092929 | 22.83389923 | 0.348868805 | 18.99118622 | NA | NA | NA | NA | NA | NA | NA |
| SCN5A | 860 | L | 13.73431818 | 21.52295284 | 0.325879853 | 17.78479948 | NA | NA | NA | NA | NA | NA | NA |
| SCN5A | 861 | F | 15.88318182 | 17.77145503 | 0.260093024 | 14.10106818 | NA | NA | NA | NA | NA | NA | NA |
| SCN5A | 862 | G | 19.38306818 | 17.91159557 | 0.262550549 | 11.80426628 | NA | NA | NA | NA | NA | NA | NA |
| SCN5A | 863 | K | 21.6820404 | 18.10284201 | 0.265904276 | 12.01143197 | NA | NA | NA | NA | NA | NA | NA |
| SCN5A | 864 | N | 22.02769318 | 15.5398805 | 0.220959797 | 9.447046261 | NA | NA | NA | NA | NA | NA | NA |
| SCN5A | 865 | Y | 21.40073485 | 12.28794226 | 0.16393332 | 6.458547068 | NA | NA | NA | NA | NA | NA | NA |
| SCN5A | 866 | S | 26.16231818 | 14.36452648 | 0.200348613 | 8.325563526 | NA | NA | NA | NA | NA | NA | NA |
| SCN5A | 867 | E | 28.4600404 | 13.83351669 | 0.191036745 | 8.260737778 | NA | NA | NA | NA | NA | NA | NA |
| SCN5A | 868 | L | 27.42744318 | 11.35830806 | 0.147631115 | 5.33125482 | NA | NA | NA | NA | NA | NA | NA |
| SCN5A | 869 | R | 25.63954545 | 8.23087711 | 0.092788016 | 2.183739216 | 6.08 | -0.22 | -0.63 | 2.63 | 0.06 | 0 | 0 |
| SCN5A | 870 | D | 29.58269318 | 7.427979133 | 0.078708277 | 1.644983408 | 5.89 | 0.27 | 0.43 | 1.82 | -0.59 | 0 | 0 |
| SCN5A | 871 | S | 32.80065152 | 7.19170119 | 0.074564871 | 1.80148124 | 5.8 | -0.4 | 0.45 | 1.71 | 0.31 | 0 | 0 |
| SCN5A | 872 | D | 36.46706818 | 7.316870233 | 0.076759854 | 1.722775645 | 5.69 | -0.46 | -0.33 | 13.29 | 0.77 | 1 | -1 |
| SCN5A | 873 | S | 34.76115152 | 8.268327582 | 0.093444753 | 2.772462739 | 5.69 | -0.46 | -0.33 | 13.29 | 0.77 | 1 | -1 |
| SCN5A | 874 | G | 31.71956818 | 10.60603116 | 0.134439074 | 5.081200468 | NA | NA | NA | NA | NA | NA | NA |
| SCN5A | 875 | L | 29.70556818 | 8.596084341 | 0.099192345 | 2.704761709 | 5.89 | 0.27 | 0.43 | 1.82 | -0.59 | 0 | 0 |
| SCN5A | 876 | L | 25.53744318 | 8.638266609 | 0.09993206 | 2.576386138 | 6.08 | -0.22 | -0.63 | 2.63 | 0.06 | 0 | 0 |
| SCN5A | 877 | P | 23.81738961 | 7.72068857 | 0.083841273 | 1.611484669 | 6.12 | -0.04 | -0.93 | 2.18 | -0.47 | 0 | 0 |
| SCN5A | 878 | R | 22.04436364 | 7.137449264 | 0.073613501 | 3.238651983 | 3.98 | -0.15 | -3.1 | 34.5 | 0.81 | 2 | 0 |
| SCN5A | 879 | W | 17.12981818 | 6.596940655 | 0.064135061 | 2.85947033 | 3.98 | -0.15 | -3.1 | 34.5 | 0.81 | 2 | 0 |

| | | | | | | | | | | | | | |
|---|---|---|---|---|---|---|---|---|---|---|---|---|---|
| SCN5A | 880 | H | 19.59701818 | 11.40479851 | 0.148446379 | 7.571167054 | NA | NA | NA | NA | NA | NA | NA |
| SCN5A | 881 | M | 15.93519318 | 14.73425455 | 0.20683222 | 10.9307511 | NA | NA | NA | NA | NA | NA | NA |
| SCN5A | 882 | M | 20.15394318 | 15.760839 | 0.224834559 | 9.655345515 | NA | NA | NA | NA | NA | NA | NA |
| SCN5A | 883 | D | 17.41456818 | 17.89518198 | 0.262262718 | 14.04481107 | NA | NA | NA | NA | NA | NA | NA |
| SCN5A | 884 | F | 11.10036364 | 19.82911347 | 0.296176431 | 15.92244967 | NA | NA | NA | NA | NA | NA | NA |
| SCN5A | 885 | F | 13.39118182 | 16.72840572 | 0.241801954 | 13.00965542 | NA | NA | NA | NA | NA | NA | NA |
| SCN5A | 886 | H | 16.73461818 | 13.41103695 | 0.183628077 | 9.488761237 | NA | NA | NA | NA | NA | NA | NA |
| SCN5A | 887 | A | 12.51341818 | 15.86177449 | 0.226604579 | 12.40105728 | NA | NA | NA | NA | NA | NA | NA |
| SCN5A | 888 | F | 8.910909091 | 15.24974262 | 0.215871896 | 14.01446437 | NA | NA | NA | NA | NA | NA | NA |
| SCN5A | 889 | L | 12.21106818 | 11.88119556 | 0.156800549 | 9.941948931 | NA | NA | NA | NA | NA | NA | NA |
| SCN5A | 890 | I | 12.69344318 | 11.29504697 | 0.146521759 | 10.20971173 | NA | NA | NA | NA | NA | NA | NA |
| SCN5A | 891 | I | 8.101068182 | 11.92287797 | 0.157531498 | 9.461471761 | NA | NA | NA | NA | NA | NA | NA |
| SCN5A | 892 | F | 7.066181818 | 9.900901424 | 0.122073813 | 7.43731433 | NA | NA | NA | NA | NA | NA | NA |
| SCN5A | 893 | R | 12.15381818 | 3.940622413 | 0.017553465 | 4.88336045 | 2.38 | -0.79 | -2.05 | 26.54 | 1.33 | 2 | 0 |
| SCN5A | 894 | I | 7.987943182 | 7.911387615 | 0.0871854 | 5.458047252 | 2.51 | -0.67 | -1.57 | 18.75 | 1.03 | 1 | 1 |
| SCN5A | 895 | L | 4.136443182 | 7.490975504 | 0.07981299 | 4.95553621 | 2.58 | -0.7 | -1.28 | 14.91 | 0.97 | 1 | 1 |
| SCN5A | 896 | C | 6.332484848 | 4.219855566 | 0.02245014 | 1.673166585 | 2.58 | -0.7 | -1.28 | 14.91 | 0.97 | 1 | 1 |
| SCN5A | 897 | G | 7.675318182 | 3.993993115 | 0.018489382 | 1.58290674 | 2.51 | -0.67 | -1.57 | 18.75 | 1.03 | 1 | 1 |
| SCN5A | 898 | E | 11.55181818 | 2.976573533 | 6.48E-04 | 1.719864991 | 1.41 | -1.11 | -3.5 | 49.83 | 2.36 | 3 | -3 |
| SCN5A | 899 | W | 9.796032468 | 5.969187298 | 0.053126684 | 4.681981737 | 1.41 | -1.11 | -3.5 | 49.83 | 2.36 | 3 | -3 |
| SCN5A | 900 | I | 13.98844318 | 5.94161165 | 0.052643113 | 4.594876312 | 1.41 | -1.11 | -3.5 | 49.83 | 2.36 | 3 | -3 |
| SCN5A | 901 | E | 16.52326263 | 2.939635011 | 0 | 2.073672446 | 2.63 | -0.19 | -2.03 | 34.44 | 1.05 | 2 | 0 |
| SCN5A | 902 | T | 15.78896104 | 8.120259725 | 0.090848213 | 4.186887227 | 3.98 | -0.15 | -3.1 | 34.5 | 0.81 | 2 | 0 |
| SCN5A | 903 | M | 14.62794318 | 10.2277852 | 0.127806096 | 8.973581984 | NA | NA | NA | NA | NA | NA | NA |
| SCN5A | 904 | W | 18.26296104 | 8.399672196 | 0.095748032 | 4.585249748 | 3.98 | -0.15 | -3.1 | 34.5 | 0.81 | 2 | 0 |
| SCN5A | 905 | D | 20.64294318 | 6.229768615 | 0.057696277 | 2.215506443 | 4.5 | -0.36 | -2.77 | 33.74 | 1.14 | 2 | -2 |
| SCN5A | 906 | C | 18.21631818 | 11.54837984 | 0.150964242 | 7.878058069 | NA | NA | NA | NA | NA | NA | NA |
| SCN5A | 907 | M | 17.96831818 | 13.84664418 | 0.191266951 | 9.68976835 | NA | NA | NA | NA | NA | NA | NA |
| SCN5A | 908 | E | 23.60081818 | 9.276941522 | 0.111131959 | 3.173290885 | 6.12 | -0.04 | -0.93 | 2.18 | -0.47 | 0 | 0 |
| SCN5A | 909 | V | 22.60953247 | 13.1241634 | 0.17859742 | 7.023008718 | NA | NA | NA | NA | NA | NA | NA |
| SCN5A | 910 | S | 19.23365152 | 16.98814793 | 0.246356833 | 10.88480577 | NA | NA | NA | NA | NA | NA | NA |
| SCN5A | 911 | G | 18.19306818 | 18.81549561 | 0.278401476 | 14.65428097 | NA | NA | NA | NA | NA | NA | NA |
| SCN5A | 912 | Q | 16.87337374 | 19.28503218 | 0.28663534 | 15.79810395 | NA | NA | NA | NA | NA | NA | NA |

| | | | | | | | | | | | | | |
|---|---|---|---|---|---|---|---|---|---|---|---|---|---|
| SCN5A | 913 | S | 13.31398485 | 20.77987283 | 0.31284909 | 19.46993121 | NA | NA | NA | NA | NA | NA | NA |
| SCN5A | 914 | L | 12.48919318 | 19.4791582 | 0.290039563 | 18.17533218 | NA | NA | NA | NA | NA | NA | NA |
| SCN5A | 915 | C | 13.87381818 | 15.59344224 | 0.221899064 | 14.63076951 | NA | NA | NA | NA | NA | NA | NA |
| SCN5A | 916 | L | 11.08406818 | 15.5927462 | 0.221886858 | 14.29036411 | NA | NA | NA | NA | NA | NA | NA |
| SCN5A | 917 | L | 7.653193182 | 18.26731141 | 0.268788435 | 17.24169349 | NA | NA | NA | NA | NA | NA | NA |
| SCN5A | 918 | V | 8.818532468 | 16.4049466 | 0.236129727 | 15.35857749 | NA | NA | NA | NA | NA | NA | NA |
| SCN5A | 919 | F | 10.86109091 | 12.94147364 | 0.175393744 | 11.83442961 | NA | NA | NA | NA | NA | NA | NA |
| SCN5A | 920 | L | 6.462568182 | 13.69256167 | 0.188564937 | 11.13679459 | NA | NA | NA | NA | NA | NA | NA |
| SCN5A | 921 | L | 3.467318182 | 15.60276418 | 0.222062535 | 13.06059362 | NA | NA | NA | NA | NA | NA | NA |
| SCN5A | 922 | V | 4.971961039 | 13.13187378 | 0.17873263 | 10.58233783 | NA | NA | NA | NA | NA | NA | NA |
| SCN5A | 923 | M | 4.240318182 | 7.046881651 | 0.072025294 | 4.531216362 | 2.58 | -0.7 | -1.28 | 14.91 | 0.97 | 1 | 1 |
| SCN5A | 924 | V | 0.708818182 | 12.76366417 | 0.172275651 | 10.12540381 | NA | NA | NA | NA | NA | NA | NA |
| SCN5A | 925 | I | -0.010818182 | 13.60493657 | 0.18702833 | 8.724244393 | NA | NA | NA | NA | NA | NA | NA |
| SCN5A | 926 | G | 0.101568182 | 10.9570428 | 0.140594467 | 6.078808856 | NA | NA | NA | NA | NA | NA | NA |
| SCN5A | 927 | N | 1.074693182 | 7.101657694 | 0.072985855 | 2.743424151 | 4.89 | 1.25 | 3.3 | 0.24 | -1.8 | 0 | 0 |
| SCN5A | 928 | L | -3.320568182 | 10.87066893 | 0.139079802 | 5.988375372 | NA | NA | NA | NA | NA | NA | NA |
| SCN5A | 929 | V | -3.085103896 | 11.55098355 | 0.151009901 | 6.69173419 | NA | NA | NA | NA | NA | NA | NA |
| SCN5A | 930 | V | -1.527532468 | 7.532720956 | 0.080545045 | 2.672426061 | 4.89 | 1.25 | 3.3 | 0.24 | -1.8 | 0 | 0 |
| SCN5A | 931 | L | -4.535568182 | 7.132190267 | 0.073521279 | 5.539542192 | 1.86 | 1.43 | 3.7 | 0.2 | -1.82 | 0 | 0 |
| SCN5A | 932 | N | -7.863443182 | 9.290672473 | 0.111372747 | 7.116629451 | NA | NA | NA | NA | NA | NA | NA |
| SCN5A | 933 | L | -5.870318182 | 9.312733755 | 0.111759617 | 7.160858194 | NA | NA | NA | NA | NA | NA | NA |
| SCN5A | 934 | F | -5.597727273 | 3.404565464 | 0.00815309 | 1.49896825 | 2.03 | 1.36 | 3.73 | 0.19 | -1.81 | 0 | 0 |
| SCN5A | 935 | L | -9.350443182 | 3.692093986 | 0.013195234 | 1.650614932 | 2.4 | 1.36 | 4.03 | 0.13 | -1.81 | 0 | 0 |
| SCN5A | 936 | A | -11.12321818 | 7.986673901 | 0.088505632 | 5.792919295 | 2.45 | 1.03 | 3.48 | 0.1 | -1.21 | 0 | 0 |
| SCN5A | 937 | L | -9.497693182 | 9.12211675 | 0.108416928 | 6.723838048 | NA | NA | NA | NA | NA | NA | NA |
| SCN5A | 938 | L | -10.91431818 | 3.937197099 | 0.017493399 | 1.656475936 | 2.4 | 1.36 | 4.03 | 0.13 | -1.81 | 0 | 0 |
| SCN5A | 939 | L | -15.19644318 | 4.712490743 | 0.031089064 | 1.85046259 | 3.08 | 1.36 | 3.54 | 0.22 | -1.81 | 0 | 0 |
| SCN5A | 940 | S | -15.16931818 | 8.475833587 | 0.09708361 | 5.515339094 | 3.12 | 1.41 | 3.48 | 0.24 | -1.82 | 0 | 0 |
| SCN5A | 941 | S | -14.13765152 | 10.17372675 | 0.126858119 | 7.175008472 | NA | NA | NA | NA | NA | NA | NA |
| SCN5A | 942 | F | -16.321 | 4.690081982 | 0.030696101 | 1.69125872 | 3.12 | 1.41 | 3.48 | 0.24 | -1.82 | 0 | 0 |
| SCN5A | 943 | S | -19.29831818 | 5.635439379 | 0.04727403 | 2.45287776 | 3.29 | 0.39 | 0.48 | 10.82 | -0.53 | 1 | -1 |
| SCN5A | 944 | A | -19.41201818 | 10.27173822 | 0.128576863 | 7.042778546 | NA | NA | NA | NA | NA | NA | NA |
| SCN5A | 1188 | G | -18.02681818 | 58.72076897 | 0.978186343 | 53.00695181 | NA | NA | NA | NA | NA | NA | NA |

| | | | | | | | | | | | | | |
|---|---|---|---|---|---|---|---|---|---|---|---|---|---|
| SCN5A | 1189 | K | -21.20215152 | 57.18039089 | 0.951174042 | 51.47699164 | NA | NA | NA | NA | NA | NA | NA |
| SCN5A | 1190 | V | -16.71867532 | 56.82729839 | 0.944982159 | 51.0378341 | NA | NA | NA | NA | NA | NA | NA |
| SCN5A | 1191 | W | -15.21417532 | 52.39391165 | 0.867237623 | 46.59399542 | NA | NA | NA | NA | NA | NA | NA |
| SCN5A | 1192 | W | -20.47674675 | 48.05181905 | 0.791094035 | 42.37781302 | NA | NA | NA | NA | NA | NA | NA |
| SCN5A | 1193 | R | -18.20027273 | 53.20036174 | 0.881379653 | 47.39207689 | NA | NA | NA | NA | NA | NA | NA |
| SCN5A | 1194 | L | -13.22381818 | 52.29293484 | 0.865466879 | 46.48797988 | NA | NA | NA | NA | NA | NA | NA |
| SCN5A | 1195 | R | -16.40445455 | 45.41826489 | 0.744911634 | 39.64842049 | NA | NA | NA | NA | NA | NA | NA |
| SCN5A | 1196 | K | -18.31815152 | 45.44829842 | 0.745438306 | 39.71503024 | NA | NA | NA | NA | NA | NA | NA |
| SCN5A | 1197 | T | -14.4351039 | 50.36555859 | 0.831668119 | 44.59370113 | NA | NA | NA | NA | NA | NA | NA |
| SCN5A | 1198 | C | -11.84581818 | 47.83467594 | 0.787286181 | 42.08874171 | NA | NA | NA | NA | NA | NA | NA |
| SCN5A | 1199 | Y | -15.87156818 | 40.90532427 | 0.665772029 | 35.08839027 | NA | NA | NA | NA | NA | NA | NA |
| SCN5A | 1200 | H | -15.50011818 | 45.94879439 | 0.754215079 | 40.19432991 | NA | NA | NA | NA | NA | NA | NA |
| SCN5A | 1201 | I | -10.69744318 | 44.88061926 | 0.735483399 | 39.61253942 | NA | NA | NA | NA | NA | NA | NA |
| SCN5A | 1202 | V | -11.68024675 | 41.56160654 | 0.677280694 | 36.29830464 | NA | NA | NA | NA | NA | NA | NA |
| SCN5A | 1203 | E | -16.01548485 | 41.11733518 | 0.669489884 | 35.38913628 | NA | NA | NA | NA | NA | NA | NA |
| SCN5A | 1204 | H | -12.25821818 | 42.55703587 | 0.694736692 | 37.30696025 | NA | NA | NA | NA | NA | NA | NA |
| SCN5A | 1205 | S | -11.18448485 | 39.97774912 | 0.649505932 | 34.72973289 | NA | NA | NA | NA | NA | NA | NA |
| SCN5A | 1206 | W | -8.512103896 | 42.87394887 | 0.700294126 | 37.60507846 | NA | NA | NA | NA | NA | NA | NA |
| SCN5A | 1207 | F | -6.745909091 | 41.84940704 | 0.682327607 | 39.92876951 | NA | NA | NA | NA | NA | NA | NA |
| SCN5A | 1208 | E | -7.466373737 | 36.14083758 | 0.582221275 | 34.05143413 | NA | NA | NA | NA | NA | NA | NA |
| SCN5A | 1209 | T | -5.520818182 | 39.50700432 | 0.641250881 | 37.93894699 | NA | NA | NA | NA | NA | NA | NA |
| SCN5A | 1210 | F | -1.795909091 | 40.89684377 | 0.665623314 | 39.33620553 | NA | NA | NA | NA | NA | NA | NA |
| SCN5A | 1211 | I | -2.941443182 | 38.01622604 | 0.615108369 | 36.46512432 | NA | NA | NA | NA | NA | NA | NA |
| SCN5A | 1212 | I | -2.795568182 | 34.81907987 | 0.559042733 | 33.26931256 | NA | NA | NA | NA | NA | NA | NA |
| SCN5A | 1213 | F | -0.912636364 | 38.03368882 | 0.615414599 | 35.21679166 | NA | NA | NA | NA | NA | NA | NA |
| SCN5A | 1214 | M | 1.551068182 | 38.33289926 | 0.620661598 | 34.39650538 | NA | NA | NA | NA | NA | NA | NA |
| SCN5A | 1215 | I | 0.790943182 | 34.03843553 | 0.545353237 | 31.09774168 | NA | NA | NA | NA | NA | NA | NA |
| SCN5A | 1216 | L | 1.677318182 | 33.44348209 | 0.534920044 | 29.70107454 | NA | NA | NA | NA | NA | NA | NA |
| SCN5A | 1217 | L | 5.138068182 | 36.55586195 | 0.589499205 | 32.6096059 | NA | NA | NA | NA | NA | NA | NA |
| SCN5A | 1218 | S | 5.530818182 | 35.11584538 | 0.564246858 | 31.20979872 | NA | NA | NA | NA | NA | NA | NA |
| SCN5A | 1219 | S | 5.791318182 | 32.00276922 | 0.509655487 | 28.07620642 | NA | NA | NA | NA | NA | NA | NA |
| SCN5A | 1220 | G | 8.618818182 | 33.25682503 | 0.531646798 | 29.43580867 | NA | NA | NA | NA | NA | NA | NA |
| SCN5A | 1221 | A | 10.72661818 | 34.98687654 | 0.561985241 | 31.27999371 | NA | NA | NA | NA | NA | NA | NA |

| | | | | | | | | | | | | | |
|---|---|---|---|---|---|---|---|---|---|---|---|---|---|
| SCN5A | 1222 | L | 10.67094318 | 31.71057119 | 0.504531458 | 29.79224711 | NA | NA | NA | NA | NA | NA | NA |
| SCN5A | 1223 | A | 13.11981818 | 30.62978265 | 0.485578588 | 28.69675744 | NA | NA | NA | NA | NA | NA | NA |
| SCN5A | 1224 | F | 15.04509091 | 32.72163691 | 0.522261659 | 30.77155291 | NA | NA | NA | NA | NA | NA | NA |
| SCN5A | 1225 | E | 16.00237374 | 30.96333768 | 0.491427859 | 28.99884147 | NA | NA | NA | NA | NA | NA | NA |
| SCN5A | 1226 | D | 19.85081818 | 30.53220773 | 0.4838675 | 28.52051592 | NA | NA | NA | NA | NA | NA | NA |
| SCN5A | 1227 | I | 23.96881818 | 29.79301571 | 0.470904917 | 27.40734418 | NA | NA | NA | NA | NA | NA | NA |
| SCN5A | 1228 | Y | 24.18081818 | 29.60057462 | 0.467530242 | 28.03361506 | NA | NA | NA | NA | NA | NA | NA |
| SCN5A | 1229 | L | 21.53281818 | 35.72579004 | 0.574942939 | 34.02684554 | NA | NA | NA | NA | NA | NA | NA |
| SCN5A | 1230 | E | 26.1030404 | 37.91202903 | 0.613281154 | 35.56340081 | NA | NA | NA | NA | NA | NA | NA |
| SCN5A | 1231 | E | 27.9040404 | 37.09035996 | 0.598872242 | 34.70502964 | NA | NA | NA | NA | NA | NA | NA |
| SCN5A | 1232 | R | 22.76409091 | 34.58122336 | 0.554871646 | 32.9110324 | NA | NA | NA | NA | NA | NA | NA |
| SCN5A | 1233 | K | 26.38681818 | 42.27715059 | 0.689828582 | 40.58195581 | NA | NA | NA | NA | NA | NA | NA |
| SCN5A | 1234 | T | 22.09853247 | 43.82640261 | 0.716996498 | 41.79566166 | NA | NA | NA | NA | NA | NA | NA |
| SCN5A | 1235 | I | 18.54019318 | 39.77623709 | 0.645972187 | 37.78040594 | NA | NA | NA | NA | NA | NA | NA |
| SCN5A | 1236 | K | 21.06315152 | 41.46925058 | 0.675661126 | 39.4829771 | NA | NA | NA | NA | NA | NA | NA |
| SCN5A | 1237 | V | 19.2031039 | 44.15767721 | 0.722805779 | 42.18718333 | NA | NA | NA | NA | NA | NA | NA |
| SCN5A | 1238 | L | 15.41169318 | 43.49344269 | 0.711157663 | 41.55294916 | NA | NA | NA | NA | NA | NA | NA |
| SCN5A | 1239 | L | 14.87056818 | 38.13342451 | 0.617163579 | 36.18360745 | NA | NA | NA | NA | NA | NA | NA |
| SCN5A | 1240 | E | 16.39837374 | 42.81387823 | 0.699240718 | 40.88946573 | NA | NA | NA | NA | NA | NA | NA |
| SCN5A | 1241 | Y | 14.92406818 | 45.25019984 | 0.74196442 | 43.33924372 | NA | NA | NA | NA | NA | NA | NA |
| SCN5A | 1242 | A | 10.48381818 | 42.6423554 | 0.696232868 | 38.85001744 | NA | NA | NA | NA | NA | NA | NA |
| SCN5A | 1243 | D | 10.47919318 | 40.25177702 | 0.654311327 | 36.38068442 | NA | NA | NA | NA | NA | NA | NA |
| SCN5A | 1244 | K | 11.10948485 | 44.34840833 | 0.726150468 | 40.46668461 | NA | NA | NA | NA | NA | NA | NA |
| SCN5A | 1245 | M | 6.649068182 | 44.85112018 | 0.734966099 | 40.94742873 | NA | NA | NA | NA | NA | NA | NA |
| SCN5A | 1246 | F | 5.818181818 | 40.00380153 | 0.649962791 | 36.09139129 | NA | NA | NA | NA | NA | NA | NA |
| SCN5A | 1247 | T | 5.393961039 | 42.19029734 | 0.68830551 | 38.31721671 | NA | NA | NA | NA | NA | NA | NA |
| SCN5A | 1248 | Y | 5.668901515 | 45.3814554 | 0.744266137 | 42.42332839 | NA | NA | NA | NA | NA | NA | NA |
| SCN5A | 1249 | V | 1.107675325 | 43.6998421 | 0.714777114 | 40.73740138 | NA | NA | NA | NA | NA | NA | NA |
| SCN5A | 1250 | F | 0.793818182 | 38.48821122 | 0.623385172 | 35.53437085 | NA | NA | NA | NA | NA | NA | NA |
| SCN5A | 1251 | V | 0.699532468 | 43.78579364 | 0.716284373 | 40.87397582 | NA | NA | NA | NA | NA | NA | NA |
| SCN5A | 1252 | L | -1.078068182 | 46.07362215 | 0.756404077 | 44.55772271 | NA | NA | NA | NA | NA | NA | NA |
| SCN5A | 1253 | E | -3.87259596 | 40.70833765 | 0.662317642 | 39.19367529 | NA | NA | NA | NA | NA | NA | NA |
| SCN5A | 1254 | M | -3.356318182 | 42.02038827 | 0.685325959 | 40.52141227 | NA | NA | NA | NA | NA | NA | NA |

| | | | | | | | | | | | | | |
|---|---|---|---|---|---|---|---|---|---|---|---|---|---|
| SCN5A | 1255 | L | -3.701068182 | 45.24180914 | 0.741817279 | 43.73951404 | NA | NA | NA | NA | NA | NA | NA |
| SCN5A | 1256 | L | -7.240693182 | 46.03879392 | 0.755793324 | 44.54600913 | NA | NA | NA | NA | NA | NA | NA |
| SCN5A | 1257 | K | -8.353040404 | 39.36258471 | 0.638718317 | 37.26842577 | NA | NA | NA | NA | NA | NA | NA |
| SCN5A | 1258 | W | -6.89588961 | 42.96478205 | 0.70188699 | 41.24850434 | NA | NA | NA | NA | NA | NA | NA |
| SCN5A | 1259 | V | -10.27738961 | 47.6133662 | 0.78340526 | 45.59633517 | NA | NA | NA | NA | NA | NA | NA |
| SCN5A | 1260 | A | -12.98221818 | 44.99581236 | 0.737503443 | 39.18122718 | NA | NA | NA | NA | NA | NA | NA |
| SCN5A | 1261 | Y | -13.38798485 | 40.15443284 | 0.652604284 | 34.39437369 | NA | NA | NA | NA | NA | NA | NA |
| SCN5A | 1262 | G | -12.89431818 | 45.44086946 | 0.745308031 | 39.94740644 | NA | NA | NA | NA | NA | NA | NA |
| SCN5A | 1263 | F | -10.99890909 | 42.38024225 | 0.691636413 | 39.29880272 | NA | NA | NA | NA | NA | NA | NA |
| SCN5A | 1264 | K | -16.68092929 | 38.9477962 | 0.631444523 | 36.83923752 | NA | NA | NA | NA | NA | NA | NA |
| SCN5A | 1265 | K | -15.74548485 | 38.26140329 | 0.619407834 | 32.62191638 | NA | NA | NA | NA | NA | NA | NA |
| SCN5A | 1266 | Y | -9.120318182 | 37.94559993 | 0.613869859 | 35.09031645 | NA | NA | NA | NA | NA | NA | NA |
| SCN5A | 1267 | F | -9.174181818 | 37.629399 | 0.608324912 | 35.48945225 | NA | NA | NA | NA | NA | NA | NA |
| SCN5A | 1268 | T | -13.9001039 | 34.43671443 | 0.552337516 | 31.34720331 | NA | NA | NA | NA | NA | NA | NA |
| SCN5A | 1269 | N | -12.11944318 | 32.41875582 | 0.51695029 | 29.37721513 | NA | NA | NA | NA | NA | NA | NA |
| SCN5A | 1270 | A | -9.303018182 | 30.749239 | 0.487673393 | 28.88562782 | NA | NA | NA | NA | NA | NA | NA |
| SCN5A | 1271 | W | -8.226032468 | 26.58045111 | 0.414568902 | 24.65475514 | NA | NA | NA | NA | NA | NA | NA |
| SCN5A | 1272 | C | -7.102484848 | 34.21338948 | 0.548421256 | 32.73272752 | NA | NA | NA | NA | NA | NA | NA |
| SCN5A | 1273 | W | -5.911461039 | 34.32414563 | 0.550363492 | 32.83476332 | NA | NA | NA | NA | NA | NA | NA |
| SCN5A | 1274 | L | -2.414693182 | 29.70835381 | 0.469420274 | 28.20653296 | NA | NA | NA | NA | NA | NA | NA |
| SCN5A | 1275 | D | -2.790443182 | 33.61396191 | 0.537909604 | 32.10779717 | NA | NA | NA | NA | NA | NA | NA |
| SCN5A | 1276 | F | -1.980727273 | 36.80854163 | 0.593930234 | 35.31173347 | NA | NA | NA | NA | NA | NA | NA |
| SCN5A | 1277 | L | -0.635443182 | 33.15644693 | 0.529886552 | 30.44161354 | NA | NA | NA | NA | NA | NA | NA |
| SCN5A | 1278 | I | 1.347193182 | 32.75324599 | 0.52281596 | 29.8176776 | NA | NA | NA | NA | NA | NA | NA |
| SCN5A | 1279 | V | 0.964103896 | 38.01914819 | 0.615159612 | 35.08238638 | NA | NA | NA | NA | NA | NA | NA |
| SCN5A | 1280 | D | 1.865693182 | 39.59581443 | 0.642808268 | 36.64430644 | NA | NA | NA | NA | NA | NA | NA |
| SCN5A | 1281 | V | 5.748818182 | 36.82498447 | 0.594218578 | 33.80391879 | NA | NA | NA | NA | NA | NA | NA |
| SCN5A | 1282 | S | 5.778484848 | 39.73664904 | 0.645277965 | 35.90252179 | NA | NA | NA | NA | NA | NA | NA |
| SCN5A | 1283 | L | 5.025068182 | 42.44620017 | 0.692793061 | 39.43948897 | NA | NA | NA | NA | NA | NA | NA |
| SCN5A | 1284 | V | 8.277961039 | 42.64551137 | 0.696288212 | 38.82282694 | NA | NA | NA | NA | NA | NA | NA |
| SCN5A | 1285 | S | 10.73065152 | 40.30379406 | 0.655223505 | 36.46578777 | NA | NA | NA | NA | NA | NA | NA |
| SCN5A | 1286 | L | 10.23831818 | 44.63348098 | 0.731149545 | 40.79299199 | NA | NA | NA | NA | NA | NA | NA |
| SCN5A | 1287 | V | 10.07838961 | 46.76500329 | 0.768528241 | 42.93073159 | NA | NA | NA | NA | NA | NA | NA |

| | | | | | | | | | | | | | |
|---|---|---|---|---|---|---|---|---|---|---|---|---|---|
| SCN5A | 1288 | A | 13.84061818 | 45.56804492 | 0.747538199 | 41.75398546 | NA | NA | NA | NA | NA | NA | NA |
| SCN5A | 1289 | N | 15.62369318 | 45.27119192 | 0.74233254 | 43.39643683 | NA | NA | NA | NA | NA | NA | NA |
| SCN5A | 1290 | T | 15.54024675 | 48.95670297 | 0.806962216 | 47.08324796 | NA | NA | NA | NA | NA | NA | NA |
| SCN5A | 1291 | L | 14.15144318 | 51.11606656 | 0.84482914 | 47.45290974 | NA | NA | NA | NA | NA | NA | NA |
| SCN5A | 1292 | G | 17.35856818 | 50.81952788 | 0.839628993 | 48.94093388 | NA | NA | NA | NA | NA | NA | NA |
| SCN5A | 1293 | F | 20.01172727 | 48.92687107 | 0.806439079 | 47.01834493 | NA | NA | NA | NA | NA | NA | NA |
| SCN5A | 1294 | A | 19.52661818 | 46.31470993 | 0.760631829 | 44.41398306 | NA | NA | NA | NA | NA | NA | NA |
| SCN5A | 1295 | E | 20.00259596 | 42.77573288 | 0.698571796 | 40.88418485 | NA | NA | NA | NA | NA | NA | NA |
| SCN5A | 1296 | M | 20.30781818 | 42.39549344 | 0.69190386 | 39.96895838 | NA | NA | NA | NA | NA | NA | NA |
| SCN5A | 1297 | G | 18.87481818 | 39.39225773 | 0.639238667 | 37.04756324 | NA | NA | NA | NA | NA | NA | NA |
| SCN5A | 1298 | P | 15.4021039 | 38.9645036 | 0.631737507 | 37.11427082 | NA | NA | NA | NA | NA | NA | NA |
| SCN5A | 1299 | I | 15.40394318 | 39.42732461 | 0.639853605 | 37.56557327 | NA | NA | NA | NA | NA | NA | NA |
| SCN5A | 1300 | K | 17.70548485 | 34.67685431 | 0.556548645 | 32.91285675 | NA | NA | NA | NA | NA | NA | NA |
| SCN5A | 1301 | S | 14.26998485 | 34.82263652 | 0.559105103 | 32.98502945 | NA | NA | NA | NA | NA | NA | NA |
| SCN5A | 1302 | L | 11.14969318 | 36.23739741 | 0.583914563 | 32.41038989 | NA | NA | NA | NA | NA | NA | NA |
| SCN5A | 1303 | R | 14.50318182 | 34.69724712 | 0.556906256 | 32.82947608 | NA | NA | NA | NA | NA | NA | NA |
| SCN5A | 1304 | T | 13.37124675 | 31.21556929 | 0.495851031 | 29.37205844 | NA | NA | NA | NA | NA | NA | NA |
| SCN5A | 1305 | L | 8.735568182 | 31.27399896 | 0.496875662 | 27.46063024 | NA | NA | NA | NA | NA | NA | NA |
| SCN5A | 1306 | R | 10.87590909 | 32.64338883 | 0.520889488 | 28.78550008 | NA | NA | NA | NA | NA | NA | NA |
| SCN5A | 1307 | A | 7.898618182 | 28.97060229 | 0.456482952 | 25.09887915 | NA | NA | NA | NA | NA | NA | NA |
| SCN5A | 1308 | L | 4.404568182 | 28.51243667 | 0.448448491 | 24.66755173 | NA | NA | NA | NA | NA | NA | NA |
| SCN5A | 1309 | R | 5.487545455 | 31.89116525 | 0.507698383 | 28.012845 | NA | NA | NA | NA | NA | NA | NA |
| SCN5A | 1310 | P | 2.706246753 | 28.77055024 | 0.452974809 | 24.87045023 | NA | NA | NA | NA | NA | NA | NA |
| SCN5A | 1311 | L | 0.497318182 | 26.63039363 | 0.415444702 | 23.69991009 | NA | NA | NA | NA | NA | NA | NA |
| SCN5A | 1312 | R | -0.824909091 | 31.06117443 | 0.493143539 | 29.5346488 | NA | NA | NA | NA | NA | NA | NA |
| SCN5A | 1313 | A | -1.954818182 | 28.8245153 | 0.453921149 | 27.28420197 | NA | NA | NA | NA | NA | NA | NA |
| SCN5A | 1314 | L | -3.642443182 | 24.43745596 | 0.376989017 | 22.91117421 | NA | NA | NA | NA | NA | NA | NA |
| SCN5A | 1315 | S | -6.666818182 | 29.09306392 | 0.458630458 | 27.5805889 | NA | NA | NA | NA | NA | NA | NA |
| SCN5A | 1316 | R | -5.812545455 | 30.40928195 | 0.481711855 | 28.39484289 | NA | NA | NA | NA | NA | NA | NA |
| SCN5A | 1317 | F | -5.087545455 | 26.61333797 | 0.415145611 | 24.42360172 | NA | NA | NA | NA | NA | NA | NA |
| SCN5A | 1318 | E | -10.70348485 | 24.78470766 | 0.383078475 | 22.49616262 | NA | NA | NA | NA | NA | NA | NA |
| SCN5A | 1319 | G | -8.546818182 | 22.14636921 | 0.336812176 | 19.86471843 | NA | NA | NA | NA | NA | NA | NA |
| SCN5A | 1320 | M | -5.170068182 | 22.71012316 | 0.34669825 | 20.66759564 | NA | NA | NA | NA | NA | NA | NA |

| | | | | | | | | | | | | | |
|---|---|---|---|---|---|---|---|---|---|---|---|---|---|
| SCN5A | 1321 | R | -10.33972727 | 23.29811488 | 0.357009361 | 21.01165939 | NA | NA | NA | NA | NA | NA | NA |
| SCN5A | 1322 | V | -11.5271039 | 20.14599695 | 0.301733347 | 17.27692648 | NA | NA | NA | NA | NA | NA | NA |
| SCN5A | 1323 | V | -7.95338961 | 17.79880976 | 0.26057272 | 15.81984256 | NA | NA | NA | NA | NA | NA | NA |
| SCN5A | 1324 | V | -7.262532468 | 20.58095578 | 0.309360851 | 18.44545986 | NA | NA | NA | NA | NA | NA | NA |
| SCN5A | 1325 | N | -11.65344318 | 19.87943226 | 0.297058829 | 17.63417007 | NA | NA | NA | NA | NA | NA | NA |
| SCN5A | 1326 | A | -11.00441818 | 16.48019566 | 0.237449305 | 14.2362344 | NA | NA | NA | NA | NA | NA | NA |
| SCN5A | 1327 | L | -6.946068182 | 16.1573206 | 0.23178732 | 14.90196044 | NA | NA | NA | NA | NA | NA | NA |
| SCN5A | 1328 | V | -8.883675325 | 19.15040796 | 0.284274549 | 17.04470069 | NA | NA | NA | NA | NA | NA | NA |
| SCN5A | 1329 | G | -11.88031818 | 18.16280116 | 0.266955728 | 15.13242463 | NA | NA | NA | NA | NA | NA | NA |
| SCN5A | 1330 | A | -10.06021818 | 14.74565173 | 0.207032082 | 12.53588596 | NA | NA | NA | NA | NA | NA | NA |
| SCN5A | 1331 | I | -7.855568182 | 18.00230569 | 0.264141255 | 16.25155293 | NA | NA | NA | NA | NA | NA | NA |
| SCN5A | 1332 | P | -10.47467532 | 18.57055546 | 0.274106169 | 16.3906239 | NA | NA | NA | NA | NA | NA | NA |
| SCN5A | 1333 | S | -9.947151515 | 14.92658464 | 0.210204949 | 12.75043205 | NA | NA | NA | NA | NA | NA | NA |
| SCN5A | 1334 | I | -6.106568182 | 12.57750548 | 0.169011144 | 11.08725364 | NA | NA | NA | NA | NA | NA | NA |
| SCN5A | 1335 | M | -5.394568182 | 17.53531919 | 0.25595211 | 16.01968168 | NA | NA | NA | NA | NA | NA | NA |
| SCN5A | 1336 | N | -8.268943182 | 17.76334034 | 0.259950723 | 16.19015851 | NA | NA | NA | NA | NA | NA | NA |
| SCN5A | 1337 | V | -5.599675325 | 13.62627256 | 0.187402481 | 12.06892893 | NA | NA | NA | NA | NA | NA | NA |
| SCN5A | 1338 | L | -1.868193182 | 14.86224458 | 0.209076672 | 13.33416163 | NA | NA | NA | NA | NA | NA | NA |
| SCN5A | 1339 | L | -2.991568182 | 18.65977302 | 0.275670701 | 17.0851908 | NA | NA | NA | NA | NA | NA | NA |
| SCN5A | 1340 | V | -3.745532468 | 16.67803067 | 0.24091857 | 15.0324308 | NA | NA | NA | NA | NA | NA | NA |
| SCN5A | 1341 | C | -0.933818182 | 13.77132768 | 0.18994619 | 10.81113333 | NA | NA | NA | NA | NA | NA | NA |
| SCN5A | 1342 | L | 0.666443182 | 17.91913756 | 0.262682807 | 14.82325564 | NA | NA | NA | NA | NA | NA | NA |
| SCN5A | 1343 | I | 0.412693182 | 20.38087834 | 0.305852263 | 17.26901311 | NA | NA | NA | NA | NA | NA | NA |
| SCN5A | 1344 | F | 0 | 16.56588464 | 0.23895196 | 13.51324748 | NA | NA | NA | NA | NA | NA | NA |
| SCN5A | 1345 | W | 3.432103896 | 12.82050081 | 0.173272347 | 8.92768609 | NA | NA | NA | NA | NA | NA | NA |
| SCN5A | 1346 | L | 4.003193182 | 20.76871058 | 0.312653347 | 16.8827448 | NA | NA | NA | NA | NA | NA | NA |
| SCN5A | 1347 | I | 3.000943182 | 22.57310796 | 0.344295531 | 18.67713622 | NA | NA | NA | NA | NA | NA | NA |
| SCN5A | 1348 | F | 5.035636364 | 17.73162951 | 0.259394637 | 12.8698055 | NA | NA | NA | NA | NA | NA | NA |
| SCN5A | 1349 | S | 8.724151515 | 21.77007244 | 0.330213379 | 16.89491447 | NA | NA | NA | NA | NA | NA | NA |
| SCN5A | 1350 | I | 8.113943182 | 24.54669014 | 0.378904564 | 22.40588639 | NA | NA | NA | NA | NA | NA | NA |
| SCN5A | 1351 | M | 9.410068182 | 23.49610627 | 0.360481368 | 21.21867997 | NA | NA | NA | NA | NA | NA | NA |
| SCN5A | 1352 | G | 12.37656818 | 21.44461296 | 0.324506073 | 19.1696574 | NA | NA | NA | NA | NA | NA | NA |
| SCN5A | 1353 | V | 13.64681818 | 23.07747657 | 0.353140214 | 20.80387445 | NA | NA | NA | NA | NA | NA | NA |

| | | | | | | | | | | | | | |
|---|---|---|---|---|---|---|---|---|---|---|---|---|---|
| SCN5A | 1354 | N | 13.38806818 | 25.32076004 | 0.39247877 | 23.06194549 | NA | NA | NA | NA | NA | NA | NA |
| SCN5A | 1355 | L | 14.69581818 | 22.38319356 | 0.340965163 | 20.14056171 | NA | NA | NA | NA | NA | NA | NA |
| SCN5A | 1356 | F | 16.57954545 | 18.51826571 | 0.273189208 | 16.2817693 | NA | NA | NA | NA | NA | NA | NA |
| SCN5A | 1357 | A | 18.91481818 | 23.1984473 | 0.355261575 | 20.96941708 | NA | NA | NA | NA | NA | NA | NA |
| SCN5A | 1358 | G | 22.59706818 | 23.13952415 | 0.35422829 | 19.80320995 | NA | NA | NA | NA | NA | NA | NA |
| SCN5A | 1359 | K | 21.92526263 | 19.04546111 | 0.282434186 | 15.69117805 | NA | NA | NA | NA | NA | NA | NA |
| SCN5A | 1360 | F | 21.49163636 | 15.83295468 | 0.22609919 | 12.83830308 | NA | NA | NA | NA | NA | NA | NA |
| SCN5A | 1361 | G | 27.16256818 | 19.49969013 | 0.290399614 | 13.48054382 | NA | NA | NA | NA | NA | NA | NA |
| SCN5A | 1362 | R | 27.68345455 | 12.53806337 | 0.168319481 | 6.709781117 | NA | NA | NA | NA | NA | NA | NA |
| SCN5A | 1363 | C | 33.14065152 | 15.34125057 | 0.217476593 | 9.66431816 | NA | NA | NA | NA | NA | NA | NA |
| SCN5A | 1364 | I | 33.85594318 | 14.06800359 | 0.195148743 | 8.031570196 | NA | NA | NA | NA | NA | NA | NA |
| SCN5A | 1365 | N | 38.89856818 | 12.38902656 | 0.165705949 | 6.412443445 | NA | NA | NA | NA | NA | NA | NA |
| SCN5A | 1366 | Q | 37.04681818 | 13.72163128 | 0.189074706 | 7.826481382 | NA | NA | NA | NA | NA | NA | NA |
| SCN5A | 1367 | T | 40.66881818 | 11.31116939 | 0.146804484 | 5.059708246 | NA | NA | NA | NA | NA | NA | NA |
| SCN5A | 1368 | E | 39.45370707 | 7.901932485 | 0.087019594 | 2.091995826 | 6.14 | -0.5 | -1 | 25.35 | 1.05 | 2 | -2 |
| SCN5A | 1369 | G | 34.62931818 | 10.04318804 | 0.124568973 | 4.045067163 | 6.14 | -0.5 | -1 | 25.35 | 1.05 | 2 | -2 |
| SCN5A | 1370 | D | 30.21881818 | 10.21485022 | 0.127579267 | 5.052706624 | NA | NA | NA | NA | NA | NA | NA |
| SCN5A | 1371 | L | 31.98069318 | 7.326942814 | 0.076936489 | 1.580001725 | 5.8 | -0.4 | 0.45 | 1.71 | 0.31 | 0 | 0 |
| SCN5A | 1372 | P | 31.4911039 | 11.48729272 | 0.149893009 | 6.146647017 | NA | NA | NA | NA | NA | NA | NA |
| SCN5A | 1373 | L | 35.96031818 | 13.25846933 | 0.180952629 | 7.89401199 | NA | NA | NA | NA | NA | NA | NA |
| SCN5A | 1374 | N | 35.64294318 | 13.78507087 | 0.190187192 | 8.162074874 | NA | NA | NA | NA | NA | NA | NA |
| SCN5A | 1375 | Y | 32.84790152 | 16.41374168 | 0.236283958 | 10.71751636 | NA | NA | NA | NA | NA | NA | NA |
| SCN5A | 1376 | T | 37.0551039 | 17.58905916 | 0.256894502 | 11.94057007 | NA | NA | NA | NA | NA | NA | NA |
| SCN5A | 1377 | I | 39.32544318 | 15.58416575 | 0.22173639 | 10.07979055 | NA | NA | NA | NA | NA | NA | NA |
| SCN5A | 1378 | V | 36.64367532 | 19.13056019 | 0.283926496 | 13.482848 | NA | NA | NA | NA | NA | NA | NA |
| SCN5A | 1379 | N | 33.94831818 | 21.25309058 | 0.321147507 | 15.63819403 | NA | NA | NA | NA | NA | NA | NA |
| SCN5A | 1380 | N | 33.05619318 | 23.58036285 | 0.361958904 | 17.845643 | NA | NA | NA | NA | NA | NA | NA |
| SCN5A | 1381 | K | 32.43992929 | 24.96607656 | 0.386258987 | 19.48225939 | NA | NA | NA | NA | NA | NA | NA |
| SCN5A | 1382 | S | 36.34448485 | 26.72738418 | 0.417145542 | 21.17114609 | NA | NA | NA | NA | NA | NA | NA |
| SCN5A | 1383 | Q | 38.19292929 | 23.65027224 | 0.363184845 | 17.61764549 | NA | NA | NA | NA | NA | NA | NA |
| SCN5A | 1384 | C | 36.36731818 | 20.78111751 | 0.312870917 | 14.76665972 | NA | NA | NA | NA | NA | NA | NA |
| SCN5A | 1385 | E | 36.79670707 | 24.62443427 | 0.380267897 | 18.59255808 | NA | NA | NA | NA | NA | NA | NA |
| SCN5A | 1386 | S | 41.35631818 | 25.8312832 | 0.401431381 | 19.82484929 | NA | NA | NA | NA | NA | NA | NA |

| | | | | | | | | | | | | | |
|---|---|---|---|---|---|---|---|---|---|---|---|---|---|
| SCN5A | 1387 | L | 42.37044318 | 21.96090387 | 0.333559828 | 15.93656926 | NA | NA | NA | NA | NA | NA | NA |
| SCN5A | 1388 | N | 41.72494318 | 23.48789563 | 0.360337385 | 17.50026144 | NA | NA | NA | NA | NA | NA | NA |
| SCN5A | 1389 | L | 46.88444318 | 23.58561483 | 0.362051003 | 17.2798098 | NA | NA | NA | NA | NA | NA | NA |
| SCN5A | 1390 | T | 44.88924675 | 19.53222757 | 0.290970195 | 13.23984054 | NA | NA | NA | NA | NA | NA | NA |
| SCN5A | 1391 | G | 42.07106818 | 19.92956003 | 0.297937877 | 14.04479108 | NA | NA | NA | NA | NA | NA | NA |
| SCN5A | 1392 | E | 41.21426263 | 16.09222051 | 0.230645715 | 10.25720775 | NA | NA | NA | NA | NA | NA | NA |
| SCN5A | 1393 | L | 38.60056818 | 17.64370451 | 0.257852771 | 11.63917089 | NA | NA | NA | NA | NA | NA | NA |
| SCN5A | 1394 | Y | 35.67615152 | 19.01100242 | 0.281829913 | 12.98902843 | NA | NA | NA | NA | NA | NA | NA |
| SCN5A | 1395 | W | 30.66253247 | 21.14158298 | 0.319192094 | 15.60379434 | NA | NA | NA | NA | NA | NA | NA |
| SCN5A | 1396 | T | 29.25296104 | 17.66545728 | 0.258234231 | 11.80091756 | NA | NA | NA | NA | NA | NA | NA |
| SCN5A | 1397 | K | 25.27715152 | 17.68914981 | 0.258649707 | 14.33163818 | NA | NA | NA | NA | NA | NA | NA |
| SCN5A | 1398 | V | 22.6021039 | 13.82728538 | 0.190927472 | 10.51528922 | NA | NA | NA | NA | NA | NA | NA |
| SCN5A | 1399 | K | 22.04692929 | 15.28445109 | 0.216480549 | 13.1236062 | NA | NA | NA | NA | NA | NA | NA |
| SCN5A | 1400 | V | 17.64953247 | 14.03086334 | 0.194497446 | 11.69905573 | NA | NA | NA | NA | NA | NA | NA |
| SCN5A | 1401 | N | 18.39906818 | 16.64095322 | 0.240268374 | 14.36245647 | NA | NA | NA | NA | NA | NA | NA |
| SCN5A | 1402 | F | 14.86636364 | 16.48645802 | 0.237559123 | 14.2136416 | NA | NA | NA | NA | NA | NA | NA |
| SCN5A | 1403 | D | 18.41156818 | 21.47805808 | 0.325092571 | 19.23190042 | NA | NA | NA | NA | NA | NA | NA |
| SCN5A | 1404 | N | 14.66094318 | 23.13271938 | 0.354108961 | 20.81993291 | NA | NA | NA | NA | NA | NA | NA |
| SCN5A | 1405 | V | 10.1541039 | 23.67281656 | 0.363580186 | 21.31671808 | NA | NA | NA | NA | NA | NA | NA |
| SCN5A | 1406 | G | 11.77306818 | 22.01447326 | 0.334499229 | 19.65032353 | NA | NA | NA | NA | NA | NA | NA |
| SCN5A | 1407 | A | 13.80321818 | 18.73087013 | 0.276917471 | 16.39392891 | NA | NA | NA | NA | NA | NA | NA |
| SCN5A | 1408 | G | 11.41656818 | 18.48781518 | 0.272655223 | 16.13509304 | NA | NA | NA | NA | NA | NA | NA |
| SCN5A | 1409 | Y | 7.084568182 | 17.20929278 | 0.250234862 | 13.49466012 | NA | NA | NA | NA | NA | NA | NA |
| SCN5A | 1410 | L | 12.24081818 | 14.98280271 | 0.211190798 | 12.58680899 | NA | NA | NA | NA | NA | NA | NA |
| SCN5A | 1411 | A | 12.15281818 | 13.65039966 | 0.187825578 | 11.28525462 | NA | NA | NA | NA | NA | NA | NA |
| SCN5A | 1412 | L | 8.383318182 | 13.35343143 | 0.182617898 | 11.62514398 | NA | NA | NA | NA | NA | NA | NA |
| SCN5A | 1413 | L | 7.754943182 | 11.32780371 | 0.147096186 | 7.612614134 | NA | NA | NA | NA | NA | NA | NA |
| SCN5A | 1414 | Q | 12.41115152 | 9.266735671 | 0.110952988 | 7.394170424 | NA | NA | NA | NA | NA | NA | NA |
| SCN5A | 1415 | V | 9.771532468 | 9.51249184 | 0.115262605 | 7.71172398 | NA | NA | NA | NA | NA | NA | NA |
| SCN5A | 1416 | A | 6.025818182 | 7.89002839 | 0.086810842 | 4.165851304 | 3.98 | -0.24 | -1.25 | 14.19 | 0.15 | 1 | 1 |
| SCN5A | 1417 | T | 7.39538961 | 6.138914318 | 0.056103042 | 2.855454953 | 3.98 | -0.24 | -1.25 | 14.19 | 0.15 | 1 | 1 |
| SCN5A | 1418 | F | 7.909636364 | 4.398169278 | 0.025577076 | 2.661020808 | 2.31 | -0.79 | -1.72 | 21.91 | 1.22 | 2 | 0 |
| SCN5A | 1419 | K | 11.16926263 | 3.665305171 | 0.012725461 | 1.949294266 | 2.12 | -0.78 | -2.6 | 34.26 | 1.5 | 2 | 0 |

| | | | | | | | | | | | | | |
|---|---|---|---|---|---|---|---|---|---|---|---|---|---|
| SCN5A | 1420 | G | 15.30606818 | 6.479652923 | 0.062078286 | 4.053392348 | 2.47 | -0.43 | -1.7 | 33.74 | 1.25 | 2 | -2 |
| SCN5A | 1421 | W | 12.63538961 | 5.589405872 | 0.04646678 | 3.221797255 | 2.47 | -0.43 | -1.7 | 33.74 | 1.25 | 2 | -2 |
| SCN5A | 1422 | M | 16.77606818 | 3.791153518 | 0.014932356 | 1.59733972 | 2.47 | -0.43 | -1.7 | 33.74 | 1.25 | 2 | -2 |
| SCN5A | 1423 | D | 19.84094318 | 6.581059109 | 0.06385656 | 4.093188658 | 2.63 | -0.19 | -2.03 | 34.44 | 1.05 | 2 | 0 |
| SCN5A | 1424 | I | 16.53244318 | 9.884630241 | 0.121788479 | 7.658394036 | NA | NA | NA | NA | NA | NA | NA |
| SCN5A | 1425 | M | 15.97794318 | 9.094994283 | 0.107941305 | 6.860785932 | NA | NA | NA | NA | NA | NA | NA |
| SCN5A | 1426 | Y | 20.32290152 | 5.358426915 | 0.042416298 | 1.9457353 | 3.63 | 0.14 | -1.85 | 26.24 | 0.43 | 2 | 0 |
| SCN5A | 1427 | A | 21.18441818 | 11.84122764 | 0.156099665 | 8.503952178 | NA | NA | NA | NA | NA | NA | NA |
| SCN5A | 1428 | A | 18.97421818 | 14.35976702 | 0.20026515 | 11.031013 | NA | NA | NA | NA | NA | NA | NA |
| SCN5A | 1429 | V | 19.50653247 | 13.86011955 | 0.191503257 | 10.49728471 | NA | NA | NA | NA | NA | NA | NA |
| SCN5A | 1430 | D | 23.52469318 | 12.11835467 | 0.160959407 | 6.08504077 | NA | NA | NA | NA | NA | NA | NA |
| SCN5A | 1431 | S | 22.12648485 | 17.37712683 | 0.253178025 | 14.00219327 | NA | NA | NA | NA | NA | NA | NA |
| SCN5A | 1432 | R | 27.236 | 20.12287243 | 0.301327832 | 14.26760866 | NA | NA | NA | NA | NA | NA | NA |
| SCN5A | 1433 | G | 24.24656818 | 22.93226125 | 0.350593697 | 16.95096057 | NA | NA | NA | NA | NA | NA | NA |
| SCN5A | 1434 | Y | 24.10406818 | 25.39328338 | 0.39375055 | 22.03172237 | NA | NA | NA | NA | NA | NA | NA |
| SCN5A | 1435 | E | 27.87115152 | 24.55614058 | 0.379070288 | 18.72012526 | NA | NA | NA | NA | NA | NA | NA |
| SCN5A | 1436 | E | 28.33015152 | 20.70164293 | 0.311477239 | 15.09703894 | NA | NA | NA | NA | NA | NA | NA |
| SCN5A | 1437 | Q | 30.85226263 | 16.90667898 | 0.244928181 | 11.30913607 | NA | NA | NA | NA | NA | NA | NA |
| SCN5A | 1438 | P | 27.87853247 | 15.93054943 | 0.227810626 | 9.955766101 | NA | NA | NA | NA | NA | NA | NA |
| SCN5A | 1439 | Q | 27.81881818 | 15.12835288 | 0.213743187 | 9.464587818 | NA | NA | NA | NA | NA | NA | NA |
| SCN5A | 1440 | W | 26.39474675 | 7.643639513 | 0.082490129 | 1.954624443 | 5.89 | 0.27 | 0.43 | 1.82 | -0.59 | 0 | 0 |
| SCN5A | 1441 | E | 22.70881818 | 13.29884168 | 0.181660604 | 7.300919429 | NA | NA | NA | NA | NA | NA | NA |
| SCN5A | 1442 | Y | 24.45640152 | 17.62101711 | 0.257454922 | 11.50761266 | NA | NA | NA | NA | NA | NA | NA |
| SCN5A | 1443 | N | 19.97694318 | 17.91641834 | 0.262635122 | 14.24333554 | NA | NA | NA | NA | NA | NA | NA |
| SCN5A | 1444 | L | 18.52069318 | 16.93166085 | 0.245366267 | 13.02108439 | NA | NA | NA | NA | NA | NA | NA |
| SCN5A | 1445 | Y | 15.60165152 | 19.01935764 | 0.281976431 | 15.62103004 | NA | NA | NA | NA | NA | NA | NA |
| SCN5A | 1446 | M | 15.27831818 | 17.02251891 | 0.246959567 | 14.80160412 | NA | NA | NA | NA | NA | NA | NA |
| SCN5A | 1447 | Y | 14.96098485 | 10.65018671 | 0.135213393 | 9.300158302 | NA | NA | NA | NA | NA | NA | NA |
| SCN5A | 1448 | I | 9.730693182 | 17.54021123 | 0.256037898 | 15.32089969 | NA | NA | NA | NA | NA | NA | NA |
| SCN5A | 1449 | Y | 9.943401515 | 17.53366548 | 0.25592311 | 15.26821648 | NA | NA | NA | NA | NA | NA | NA |
| SCN5A | 1450 | F | 11.45045455 | 13.86284437 | 0.19155104 | 11.59825775 | NA | NA | NA | NA | NA | NA | NA |
| SCN5A | 1451 | V | 7.28338961 | 15.12499131 | 0.213684238 | 13.32281293 | NA | NA | NA | NA | NA | NA | NA |
| SCN5A | 1452 | I | 4.831818182 | 15.78914934 | 0.225331013 | 10.92402402 | NA | NA | NA | NA | NA | NA | NA |

| | | | | | | | | | | | | | |
|---|---|---|---|---|---|---|---|---|---|---|---|---|---|
| SCN5A | 1453 | F | 5.653727273 | 13.51110073 | 0.185382811 | 8.642719932 | NA | NA | NA | NA | NA | NA | NA |
| SCN5A | 1454 | I | 5.315818182 | 10.65073993 | 0.135223094 | 5.767122727 | NA | NA | NA | NA | NA | NA | NA |
| SCN5A | 1455 | I | 1.290443182 | 12.30598131 | 0.164249656 | 7.454157168 | NA | NA | NA | NA | NA | NA | NA |
| SCN5A | 1456 | F | 0.259636364 | 12.72497155 | 0.171597131 | 7.88778501 | NA | NA | NA | NA | NA | NA | NA |
| SCN5A | 1457 | G | 0.721818182 | 11.43570291 | 0.148988323 | 6.693803237 | NA | NA | NA | NA | NA | NA | NA |
| SCN5A | 1458 | S | 1.459984848 | 7.958625886 | 0.088013778 | 3.085531941 | 4.89 | 1.25 | 3.3 | 0.24 | -1.8 | 0 | 0 |
| SCN5A | 1459 | F | -1.091454545 | 7.407214389 | 0.078344143 | 2.567414413 | 4.89 | 1.25 | 3.3 | 0.24 | -1.8 | 0 | 0 |
| SCN5A | 1460 | F | -2.944272727 | 10.35823199 | 0.130093631 | 7.380014863 | NA | NA | NA | NA | NA | NA | NA |
| SCN5A | 1461 | T | -1.458532468 | 9.371005656 | 0.112781482 | 6.419305915 | NA | NA | NA | NA | NA | NA | NA |
| SCN5A | 1462 | L | -1.586318182 | 5.602654192 | 0.046699104 | 2.501638452 | 3.15 | 1.35 | 2.45 | 0.56 | -1.72 | 0 | 0 |
| SCN5A | 1463 | N | -6.164568182 | 7.152493691 | 0.073877323 | 5.417965737 | 1.86 | 1.43 | 3.7 | 0.2 | -1.82 | 0 | 0 |
| SCN5A | 1464 | L | -6.804318182 | 8.99627534 | 0.106210155 | 7.140812627 | NA | NA | NA | NA | NA | NA | NA |
| SCN5A | 1465 | F | -4.348 | 7.738945729 | 0.084161434 | 6.22816571 | 1.86 | 1.43 | 3.7 | 0.2 | -1.82 | 0 | 0 |
| SCN5A | 1466 | I | -6.542068182 | 3.40068052 | 0.008084963 | 1.495271053 | 2.03 | 1.36 | 3.73 | 0.19 | -1.81 | 0 | 0 |
| SCN5A | 1467 | G | -10.30756818 | 6.462841945 | 0.061783486 | 4.294505513 | 2.33 | 1.48 | 4.15 | 0.13 | -1.82 | 0 | 0 |
| SCN5A | 1468 | V | -10.33996104 | 8.815999603 | 0.103048812 | 6.659454394 | NA | NA | NA | NA | NA | NA | NA |
| SCN5A | 1469 | I | -8.882818182 | 8.539012941 | 0.098191532 | 6.353071441 | 2.33 | 1.48 | 4.15 | 0.13 | -1.82 | 0 | 0 |
| SCN5A | 1470 | I | -12.46819318 | 4.592850096 | 0.028991028 | 1.642686799 | 2.97 | 1.53 | 3.9 | 0.19 | -1.83 | 0 | 0 |
| SCN5A | 1471 | D | -15.42631818 | 7.428044292 | 0.078709419 | 4.439131355 | 3.12 | 1.41 | 3.48 | 0.24 | -1.82 | 0 | 0 |
| SCN5A | 1472 | N | -13.89431818 | 10.0283364 | 0.124308532 | 6.989954082 | NA | NA | NA | NA | NA | NA | NA |
| SCN5A | 1473 | F | -12.27709091 | 10.00091766 | 0.123827713 | 6.891653873 | NA | NA | NA | NA | NA | NA | NA |
| SCN5A | 1474 | N | -17.85756818 | 5.016261058 | 0.036416026 | 1.950681875 | 3.17 | 0.77 | 1.48 | 1.05 | -1.22 | 0 | 0 |
| SCN5A | 1475 | Q | -18.80359596 | 10.01442095 | 0.124064509 | 6.783944904 | NA | NA | NA | NA | NA | NA | NA |
| SCN5A | 1476 | Q | -16.73692929 | 12.55949278 | 0.168695271 | 9.39831824 | NA | NA | NA | NA | NA | NA | NA |
| SCN5A | 1477 | K | -19.1990404 | 7.289590112 | 0.076281466 | 2.14713295 | 5.35 | -0.33 | -1.23 | 13.82 | 0.43 | 1 | -1 |
| SCN5A | 1478 | K | -23.12659596 | 8.86151759 | 0.103847022 | 3.289887587 | 5.83 | -0.77 | -2.9 | 26.11 | 1.25 | 2 | 0 |
| SCN5A | 1479 | K | -22.1810404 | 12.335258 | 0.164763056 | 6.624093232 | NA | NA | NA | NA | NA | NA | NA |
| SCN5A | 1480 | L | -21.91981818 | 13.15466967 | 0.179132382 | 7.5804487 | NA | NA | NA | NA | NA | NA | NA |
| SCN5A | 1481 | G | -24.36281818 | 12.87318593 | 0.174196241 | 5.931142761 | NA | NA | NA | NA | NA | NA | NA |
| SCN5A | 1482 | G | -22.39556818 | 14.07624911 | 0.195293338 | 8.276170427 | NA | NA | NA | NA | NA | NA | NA |
| SCN5A | 1483 | Q | -21.84048485 | 9.812122757 | 0.120516977 | 3.990288194 | 5.83 | -0.77 | -2.9 | 26.11 | 1.25 | 2 | 0 |
| SCN5A | 1484 | D | -16.78681818 | 12.66387105 | 0.170525664 | 7.383975679 | NA | NA | NA | NA | NA | NA | NA |
| SCN5A | 1485 | I | -14.63844318 | 13.87924324 | 0.191838613 | 9.170865999 | NA | NA | NA | NA | NA | NA | NA |

| | | | | | | | | | | | | | |
|---|---|---|---|---|---|---|---|---|---|---|---|---|---|
| SCN5A | 1486 | F | -11.101 | 13.10019439 | 0.178177095 | 10.21501363 | NA | NA | NA | NA | NA | NA | NA |
| SCN5A | 1487 | M | -14.03906818 | 17.1403423 | 0.249025736 | 11.826675 | NA | NA | NA | NA | NA | NA | NA |
| SCN5A | 1488 | T | -14.11681818 | 21.5743761 | 0.326781619 | 16.30432476 | NA | NA | NA | NA | NA | NA | NA |
| SCN5A | 1489 | E | -19.06115152 | 21.13567737 | 0.319088532 | 16.00509823 | NA | NA | NA | NA | NA | NA | NA |
| SCN5A | 1490 | E | -14.67070707 | 23.72574507 | 0.364508348 | 18.5010583 | NA | NA | NA | NA | NA | NA | NA |
| SCN5A | 1491 | Q | -13.20681818 | 20.62333748 | 0.310104063 | 15.36604593 | NA | NA | NA | NA | NA | NA | NA |
| SCN5A | 1492 | K | -18.14148485 | 17.69651276 | 0.258778825 | 12.44013671 | NA | NA | NA | NA | NA | NA | NA |
| SCN5A | 1493 | K | -20.11859596 | 21.14606928 | 0.319270766 | 15.96914627 | NA | NA | NA | NA | NA | NA | NA |
| SCN5A | 1494 | Y | -15.52873485 | 21.48087617 | 0.32514199 | 16.26350673 | NA | NA | NA | NA | NA | NA | NA |
| SCN5A | 1495 | Y | -17.28073485 | 13.15518183 | 0.179141363 | 7.86878871 | NA | NA | NA | NA | NA | NA | NA |
| SCN5A | 1496 | N | -21.35319318 | 16.38715088 | 0.235817658 | 11.2878277 | NA | NA | NA | NA | NA | NA | NA |
| SCN5A | 1497 | A | -20.00161818 | 20.6675993 | 0.310880245 | 15.53403182 | NA | NA | NA | NA | NA | NA | NA |
| SCN5A | 1498 | M | -16.37594318 | 19.94479383 | 0.298205019 | 14.7423895 | NA | NA | NA | NA | NA | NA | NA |
| SCN5A | 1499 | K | -21.56481818 | 16.4522083 | 0.236958515 | 11.33502997 | NA | NA | NA | NA | NA | NA | NA |
| SCN5A | 1500 | K | -22.73581818 | 19.50288722 | 0.290455679 | 14.371045 | NA | NA | NA | NA | NA | NA | NA |
| SCN5A | 1501 | L | -18.12831818 | 21.55617304 | 0.326462407 | 16.39491482 | NA | NA | NA | NA | NA | NA | NA |
| SCN5A | 1502 | G | -19.68606818 | 19.03106016 | 0.282181648 | 13.84374316 | NA | NA | NA | NA | NA | NA | NA |
| SCN5A | 1503 | S | -22.60865152 | 20.39337711 | 0.306071443 | 15.25836277 | NA | NA | NA | NA | NA | NA | NA |
| SCN5A | 1504 | K | -25.35859596 | 14.55615117 | 0.203708972 | 10.13302614 | NA | NA | NA | NA | NA | NA | NA |
| SCN5A | 1505 | K | -25.78759596 | 22.13864641 | 0.336676748 | 11.06091382 | NA | NA | NA | NA | NA | NA | NA |
| SCN5A | 1506 | P | -23.29781818 | 24.76090055 | 0.38266099 | 14.22636344 | NA | NA | NA | NA | NA | NA | NA |
| SCN5A | 1507 | Q | -25.81226263 | 27.61120772 | 0.432644405 | 16.54755177 | NA | NA | NA | NA | NA | NA | NA |
| SCN5A | 1508 | K | -25.94092929 | 30.23391832 | 0.478636652 | 19.18098737 | NA | NA | NA | NA | NA | NA | NA |
| SCN5A | 1509 | P | -23.92053247 | 33.54880748 | 0.536767046 | 22.48913991 | NA | NA | NA | NA | NA | NA | NA |
| SCN5A | 1510 | I | -26.46106818 | 35.86680333 | 0.577415769 | 24.82812495 | NA | NA | NA | NA | NA | NA | NA |
| SCN5A | 1511 | P | -23.69967532 | 38.98955949 | 0.63217689 | 27.93038424 | NA | NA | NA | NA | NA | NA | NA |
| SCN5A | 1512 | R | -25.24090909 | 41.99763371 | 0.684926932 | 30.94891182 | NA | NA | NA | NA | NA | NA | NA |
| SCN5A | 1513 | P | -24.45467532 | 44.62022193 | 0.730917033 | 33.57431922 | NA | NA | NA | NA | NA | NA | NA |
| SCN5A | 1514 | L | -21.92781818 | 46.6896132 | 0.767206189 | 35.72554837 | NA | NA | NA | NA | NA | NA | NA |
| SCN5A | 1515 | N | -22.68019318 | 50.74890987 | 0.838390625 | 39.69736815 | NA | NA | NA | NA | NA | NA | NA |
| SCN5A | 1516 | K | -21.49737374 | 54.01640687 | 0.895689943 | 42.96624977 | NA | NA | NA | NA | NA | NA | NA |
| SCN5A | 1517 | Y | -17.00990152 | 53.12457053 | 0.880050567 | 47.90903603 | NA | NA | NA | NA | NA | NA | NA |
| SCN5A | 1518 | Q | -17.42181818 | 49.42519521 | 0.815177766 | 44.21731829 | NA | NA | NA | NA | NA | NA | NA |

| | | | | | | | | | | | | | |
|---|---|---|---|---|---|---|---|---|---|---|---|---|---|
| SCN5A | 1519 | G | -19.69181818 | 49.90559089 | 0.823602057 | 44.69793509 | NA | NA | NA | NA | NA | NA | NA |
| SCN5A | 1520 | F | -18.61290909 | 51.27517406 | 0.847619273 | 46.03889607 | NA | NA | NA | NA | NA | NA | NA |
| SCN5A | 1521 | I | -14.94106818 | 48.70507418 | 0.802549615 | 43.45901678 | NA | NA | NA | NA | NA | NA | NA |
| SCN5A | 1522 | F | -19.39363636 | 44.94560892 | 0.736623068 | 39.75586979 | NA | NA | NA | NA | NA | NA | NA |
| SCN5A | 1523 | D | -19.03194318 | 46.75684628 | 0.768385198 | 41.51844937 | NA | NA | NA | NA | NA | NA | NA |
| SCN5A | 1524 | I | -14.80669318 | 44.68965998 | 0.732134709 | 39.42007388 | NA | NA | NA | NA | NA | NA | NA |
| SCN5A | 1525 | V | -14.3331039 | 41.86465899 | 0.682595067 | 36.59473089 | NA | NA | NA | NA | NA | NA | NA |
| SCN5A | 1526 | T | -18.01767532 | 40.01705589 | 0.650195222 | 34.77494323 | NA | NA | NA | NA | NA | NA | NA |
| SCN5A | 1527 | K | -17.04215152 | 39.98063868 | 0.649556604 | 34.70226869 | NA | NA | NA | NA | NA | NA | NA |
| SCN5A | 1528 | Q | -17.98237374 | 35.27978658 | 0.567121755 | 30.01685505 | NA | NA | NA | NA | NA | NA | NA |
| SCN5A | 1529 | A | -13.04641818 | 39.83699965 | 0.647037728 | 34.54519886 | NA | NA | NA | NA | NA | NA | NA |
| SCN5A | 1530 | F | -10.62618182 | 40.3994066 | 0.656900181 | 35.10325181 | NA | NA | NA | NA | NA | NA | NA |
| SCN5A | 1531 | D | -11.94906818 | 35.53286127 | 0.571559711 | 30.24181534 | NA | NA | NA | NA | NA | NA | NA |
| SCN5A | 1532 | V | -9.879961039 | 36.59756146 | 0.590230454 | 34.10179057 | NA | NA | NA | NA | NA | NA | NA |
| SCN5A | 1533 | T | -7.288532468 | 39.5681813 | 0.642323689 | 36.891891 | NA | NA | NA | NA | NA | NA | NA |
| SCN5A | 1534 | I | -6.206818182 | 37.55633648 | 0.607043676 | 35.47000004 | NA | NA | NA | NA | NA | NA | NA |
| SCN5A | 1535 | M | -6.354318182 | 33.34984409 | 0.533277994 | 30.67993336 | NA | NA | NA | NA | NA | NA | NA |
| SCN5A | 1536 | F | -4.5 | 35.58320832 | 0.572442605 | 32.51415315 | NA | NA | NA | NA | NA | NA | NA |
| SCN5A | 1537 | L | -1.737068182 | 36.67391426 | 0.591569388 | 34.03016808 | NA | NA | NA | NA | NA | NA | NA |
| SCN5A | 1538 | I | -1.398818182 | 33.19509031 | 0.530564209 | 30.56485335 | NA | NA | NA | NA | NA | NA | NA |
| SCN5A | 1539 | C | -0.529984848 | 31.16014528 | 0.494879107 | 28.52497749 | NA | NA | NA | NA | NA | NA | NA |
| SCN5A | 1540 | L | 1.698193182 | 34.44885944 | 0.552550492 | 31.80743077 | NA | NA | NA | NA | NA | NA | NA |
| SCN5A | 1541 | N | 2.690443182 | 32.82405894 | 0.524057747 | 30.18901972 | NA | NA | NA | NA | NA | NA | NA |
| SCN5A | 1542 | M | 2.632693182 | 25.79725817 | 0.400834713 | 23.1447611 | NA | NA | NA | NA | NA | NA | NA |
| SCN5A | 1543 | V | 4.927246753 | 31.25285665 | 0.496504907 | 28.74634378 | NA | NA | NA | NA | NA | NA | NA |
| SCN5A | 1544 | T | 7.258675325 | 32.81204725 | 0.523847108 | 30.28832977 | NA | NA | NA | NA | NA | NA | NA |
| SCN5A | 1545 | M | 6.755193182 | 29.37753844 | 0.463619046 | 27.06615313 | NA | NA | NA | NA | NA | NA | NA |
| SCN5A | 1546 | M | 7.728568182 | 26.48527398 | 0.412899862 | 25.35911173 | NA | NA | NA | NA | NA | NA | NA |
| SCN5A | 1547 | V | 11.68496104 | 29.82102592 | 0.471396109 | 28.13245941 | NA | NA | NA | NA | NA | NA | NA |
| SCN5A | 1548 | E | 13.38759596 | 27.60965248 | 0.432617132 | 25.94976545 | NA | NA | NA | NA | NA | NA | NA |
| SCN5A | 1549 | T | 17.57824675 | 27.57588274 | 0.432024941 | 25.94333732 | NA | NA | NA | NA | NA | NA | NA |
| SCN5A | 1550 | D | 21.25956818 | 26.29688976 | 0.409596328 | 24.98412857 | NA | NA | NA | NA | NA | NA | NA |
| SCN5A | 1551 | D | 23.60919318 | 28.9569994 | 0.45624441 | 23.06267377 | NA | NA | NA | NA | NA | NA | NA |

| | | | | | | | | | | | | | |
|---|---|---|---|---|---|---|---|---|---|---|---|---|---|
| SCN5A | 1552 | Q | 19.37426263 | 31.88990163 | 0.507676224 | 30.27797929 | NA | NA | NA | NA | NA | NA | NA |
| SCN5A | 1553 | S | 21.78398485 | 34.73431897 | 0.557556353 | 28.98395806 | NA | NA | NA | NA | NA | NA | NA |
| SCN5A | 1554 | P | 23.23996104 | 37.65754288 | 0.608818447 | 31.77325148 | NA | NA | NA | NA | NA | NA | NA |
| SCN5A | 1555 | E | 21.0600404 | 40.21815637 | 0.65372175 | 38.59709876 | NA | NA | NA | NA | NA | NA | NA |
| SCN5A | 1556 | K | 17.51215152 | 32.2888785 | 0.514672742 | 30.63608139 | NA | NA | NA | NA | NA | NA | NA |
| SCN5A | 1557 | I | 17.94181818 | 36.22885717 | 0.5837648 | 34.60461364 | NA | NA | NA | NA | NA | NA | NA |
| SCN5A | 1558 | N | 17.56331818 | 40.09319102 | 0.651530339 | 38.46638337 | NA | NA | NA | NA | NA | NA | NA |
| SCN5A | 1559 | I | 14.41244318 | 40.12974677 | 0.652171386 | 38.46440167 | NA | NA | NA | NA | NA | NA | NA |
| SCN5A | 1560 | L | 12.75131818 | 34.23394498 | 0.54878172 | 32.58167791 | NA | NA | NA | NA | NA | NA | NA |
| SCN5A | 1561 | A | 11.99641818 | 39.09392327 | 0.634007029 | 37.43205776 | NA | NA | NA | NA | NA | NA | NA |
| SCN5A | 1562 | K | 12.02981818 | 41.69357098 | 0.679594842 | 40.0259378 | NA | NA | NA | NA | NA | NA | NA |
| SCN5A | 1563 | I | 7.966818182 | 39.01164936 | 0.632564262 | 37.33543591 | NA | NA | NA | NA | NA | NA | NA |
| SCN5A | 1564 | N | 7.307943182 | 37.95036324 | 0.613953389 | 35.45461852 | NA | NA | NA | NA | NA | NA | NA |
| SCN5A | 1565 | L | 7.371568182 | 41.49458589 | 0.67610541 | 38.88167185 | NA | NA | NA | NA | NA | NA | NA |
| SCN5A | 1566 | L | 4.428193182 | 43.00030505 | 0.702509927 | 40.354734 | NA | NA | NA | NA | NA | NA | NA |
| SCN5A | 1567 | F | 2.668545455 | 38.27256745 | 0.61960361 | 35.63299969 | NA | NA | NA | NA | NA | NA | NA |
| SCN5A | 1568 | V | 2.190818182 | 41.14739249 | 0.670016974 | 38.49333394 | NA | NA | NA | NA | NA | NA | NA |
| SCN5A | 1569 | A | 0.975618182 | 43.77624682 | 0.716116958 | 41.12498118 | NA | NA | NA | NA | NA | NA | NA |
| SCN5A | 1570 | I | -0.635693182 | 43.72362443 | 0.715194164 | 41.06945003 | NA | NA | NA | NA | NA | NA | NA |
| SCN5A | 1571 | F | -1.119454545 | 37.64650632 | 0.608624908 | 34.59699267 | NA | NA | NA | NA | NA | NA | NA |
| SCN5A | 1572 | T | -1.452246753 | 43.82849148 | 0.717033128 | 40.75699348 | NA | NA | NA | NA | NA | NA | NA |
| SCN5A | 1573 | G | -4.264068182 | 46.30529837 | 0.760466786 | 43.23279263 | NA | NA | NA | NA | NA | NA | NA |
| SCN5A | 1574 | E | -6.10359596 | 41.42560061 | 0.674895673 | 39.08727458 | NA | NA | NA | NA | NA | NA | NA |
| SCN5A | 1575 | C | -6.710484848 | 44.12213556 | 0.722182515 | 39.04429437 | NA | NA | NA | NA | NA | NA | NA |
| SCN5A | 1576 | I | -7.159318182 | 46.79723927 | 0.769093536 | 41.50913736 | NA | NA | NA | NA | NA | NA | NA |
| SCN5A | 1577 | V | -9.859961039 | 46.43297859 | 0.762705806 | 41.16111926 | NA | NA | NA | NA | NA | NA | NA |
| SCN5A | 1578 | K | -11.5870404 | 39.28756344 | 0.637402732 | 33.99985741 | NA | NA | NA | NA | NA | NA | NA |
| SCN5A | 1579 | L | -10.79556818 | 44.4895913 | 0.728626274 | 39.22691252 | NA | NA | NA | NA | NA | NA | NA |
| SCN5A | 1580 | A | -13.23261818 | 47.43905323 | 0.780348482 | 42.19039679 | NA | NA | NA | NA | NA | NA | NA |
| SCN5A | 1581 | A | -15.93461818 | 44.80588061 | 0.734172771 | 39.57165533 | NA | NA | NA | NA | NA | NA | NA |
| SCN5A | 1582 | L | -17.50981818 | 42.38697641 | 0.691754504 | 37.19352296 | NA | NA | NA | NA | NA | NA | NA |
| SCN5A | 1583 | R | -18.96754545 | 44.83635475 | 0.73470717 | 39.63693526 | NA | NA | NA | NA | NA | NA | NA |
| SCN5A | 1584 | H | -18.70761818 | 41.18658794 | 0.670704311 | 36.01055918 | NA | NA | NA | NA | NA | NA | NA |

| | | | | | | | | | | | | | |
|---|---|---|---|---|---|---|---|---|---|---|---|---|---|
| SCN5A | 1585 | Y | -21.08306818 | 39.17870341 | 0.635493747 | 33.98417998 | NA | NA | NA | NA | NA | NA | NA |
| SCN5A | 1586 | Y | -13.10656818 | 38.51316114 | 0.623822697 | 33.28213195 | NA | NA | NA | NA | NA | NA | NA |
| SCN5A | 1587 | F | -14.04609091 | 36.87206706 | 0.595044225 | 31.66885061 | NA | NA | NA | NA | NA | NA | NA |
| SCN5A | 1588 | T | -18.21196104 | 33.13496953 | 0.529509922 | 27.93904589 | NA | NA | NA | NA | NA | NA | NA |
| SCN5A | 1589 | N | -15.75894318 | 31.40913232 | 0.499245381 | 26.16868893 | NA | NA | NA | NA | NA | NA | NA |
| SCN5A | 1590 | S | -13.68198485 | 30.07679087 | 0.475881241 | 24.83185014 | NA | NA | NA | NA | NA | NA | NA |
| SCN5A | 1591 | W | -10.71174675 | 28.60742575 | 0.450114234 | 23.31626336 | NA | NA | NA | NA | NA | NA | NA |
| SCN5A | 1592 | N | -11.41856818 | 35.05857912 | 0.563242628 | 29.78328811 | NA | NA | NA | NA | NA | NA | NA |
| SCN5A | 1593 | I | -10.27906818 | 36.08269315 | 0.581201646 | 30.83887534 | NA | NA | NA | NA | NA | NA | NA |
| SCN5A | 1594 | F | -5.942545455 | 32.17445549 | 0.512666203 | 30.08854896 | NA | NA | NA | NA | NA | NA | NA |
| SCN5A | 1595 | D | -6.934818182 | 35.90520806 | 0.57808924 | 33.17365564 | NA | NA | NA | NA | NA | NA | NA |
| SCN5A | 1596 | F | -6.034363636 | 39.68188676 | 0.644317645 | 36.93932679 | NA | NA | NA | NA | NA | NA | NA |
| SCN5A | 1597 | V | -3.670818182 | 37.4081702 | 0.60444541 | 33.6174081 | NA | NA | NA | NA | NA | NA | NA |
| SCN5A | 1598 | V | -1.361246753 | 34.98258775 | 0.561910032 | 31.93726934 | NA | NA | NA | NA | NA | NA | NA |
| SCN5A | 1599 | V | -1.426532468 | 39.17689484 | 0.635462031 | 36.11078532 | NA | NA | NA | NA | NA | NA | NA |
| SCN5A | 1600 | I | -1.003318182 | 40.97384887 | 0.666973687 | 37.89633575 | NA | NA | NA | NA | NA | NA | NA |
| SCN5A | 1601 | L | 0.787568182 | 35.91496437 | 0.578260329 | 32.84946858 | NA | NA | NA | NA | NA | NA | NA |
| SCN5A | 1602 | S | 2.382318182 | 38.79064245 | 0.628688651 | 36.18578768 | NA | NA | NA | NA | NA | NA | NA |
| SCN5A | 1603 | I | 2.316693182 | 41.39804233 | 0.674412407 | 38.78988145 | NA | NA | NA | NA | NA | NA | NA |
| SCN5A | 1604 | V | 4.793246753 | 41.5259953 | 0.67665621 | 38.87715757 | NA | NA | NA | NA | NA | NA | NA |
| SCN5A | 1605 | G | 7.380318182 | 39.61579393 | 0.643158632 | 36.96057862 | NA | NA | NA | NA | NA | NA | NA |
| SCN5A | 1606 | T | 7.409961039 | 40.98606774 | 0.667187959 | 38.33385917 | NA | NA | NA | NA | NA | NA | NA |
| SCN5A | 1607 | V | 8.537818182 | 43.59295404 | 0.712902709 | 41.36521028 | NA | NA | NA | NA | NA | NA | NA |
| SCN5A | 1608 | L | 11.11444318 | 40.47582743 | 0.658240308 | 38.8658075 | NA | NA | NA | NA | NA | NA | NA |
| SCN5A | 1609 | S | 12.82015152 | 40.11392528 | 0.651893938 | 38.44121627 | NA | NA | NA | NA | NA | NA | NA |
| SCN5A | 1610 | D | 15.78906818 | 42.28253855 | 0.689923066 | 40.60718888 | NA | NA | NA | NA | NA | NA | NA |
| SCN5A | 1611 | I | 14.72894318 | 43.88725787 | 0.718063665 | 42.21132729 | NA | NA | NA | NA | NA | NA | NA |
| SCN5A | 1612 | I | 17.19469318 | 39.2955306 | 0.637542446 | 37.61774393 | NA | NA | NA | NA | NA | NA | NA |
| SCN5A | 1613 | Q | 20.18392929 | 43.55311384 | 0.712204065 | 41.89748032 | NA | NA | NA | NA | NA | NA | NA |
| SCN5A | 1614 | K | 22.22881818 | 44.39452161 | 0.726959118 | 42.7419411 | NA | NA | NA | NA | NA | NA | NA |
| SCN5A | 1615 | Y | 18.55231818 | 41.37905906 | 0.674079514 | 39.72840435 | NA | NA | NA | NA | NA | NA | NA |
| SCN5A | 1616 | F | 20.37845455 | 36.3375602 | 0.585671032 | 34.66618605 | NA | NA | NA | NA | NA | NA | NA |
| SCN5A | 1617 | F | 22.08436364 | 32.70666606 | 0.521999128 | 31.06610998 | NA | NA | NA | NA | NA | NA | NA |

| | | | | | | | | | | | | | |
|---|---|---|---|---|---|---|---|---|---|---|---|---|---|
| SCN5A | 1618 | S | 18.81981818 | 33.98149991 | 0.544354805 | 32.32637015 | NA | NA | NA | NA | NA | NA | NA |
| SCN5A | 1619 | P | 18.8361039 | 30.19065246 | 0.477877935 | 28.52082095 | NA | NA | NA | NA | NA | NA | NA |
| SCN5A | 1620 | T | 16.06453247 | 30.73123157 | 0.487357612 | 29.04359611 | NA | NA | NA | NA | NA | NA | NA |
| SCN5A | 1621 | L | 14.24181818 | 33.26919411 | 0.531863704 | 31.58583437 | NA | NA | NA | NA | NA | NA | NA |
| SCN5A | 1622 | F | 14.76345455 | 33.60278633 | 0.537713627 | 31.91912845 | NA | NA | NA | NA | NA | NA | NA |
| SCN5A | 1623 | R | 15.27363636 | 27.01123674 | 0.422123223 | 25.37325235 | NA | NA | NA | NA | NA | NA | NA |
| SCN5A | 1624 | V | 10.48681818 | 29.93440813 | 0.473384396 | 27.8415723 | NA | NA | NA | NA | NA | NA | NA |
| SCN5A | 1625 | I | 10.07156818 | 33.76836681 | 0.540617271 | 31.68578803 | NA | NA | NA | NA | NA | NA | NA |
| SCN5A | 1626 | R | 11.31145455 | 31.62086433 | 0.502958345 | 29.95860176 | NA | NA | NA | NA | NA | NA | NA |
| SCN5A | 1627 | L | 7.659943182 | 27.5245413 | 0.431124609 | 24.98771993 | NA | NA | NA | NA | NA | NA | NA |
| SCN5A | 1628 | A | 4.267418182 | 32.11865081 | 0.511687604 | 29.46891004 | NA | NA | NA | NA | NA | NA | NA |
| SCN5A | 1629 | R | 6.514090909 | 31.63473042 | 0.503201503 | 29.40042329 | NA | NA | NA | NA | NA | NA | NA |
| SCN5A | 1630 | I | 2.984193182 | 27.12433347 | 0.424106505 | 24.46976611 | NA | NA | NA | NA | NA | NA | NA |
| SCN5A | 1631 | G | -0.381068182 | 31.00998368 | 0.49224585 | 27.93554859 | NA | NA | NA | NA | NA | NA | NA |
| SCN5A | 1632 | R | -0.997363636 | 31.83082976 | 0.506640331 | 28.75709962 | NA | NA | NA | NA | NA | NA | NA |
| SCN5A | 1633 | I | -0.656193182 | 27.41238992 | 0.429157906 | 24.35580616 | NA | NA | NA | NA | NA | NA | NA |
| SCN5A | 1634 | L | -1.674693182 | 26.06604387 | 0.405548179 | 23.0078982 | NA | NA | NA | NA | NA | NA | NA |
| SCN5A | 1635 | R | -4.873636364 | 30.79096265 | 0.488405065 | 27.04821724 | NA | NA | NA | NA | NA | NA | NA |
| SCN5A | 1636 | L | -4.155818182 | 28.5136299 | 0.448469416 | 25.43948958 | NA | NA | NA | NA | NA | NA | NA |
| SCN5A | 1637 | I | -7.321318182 | 26.04615945 | 0.405199483 | 23.48974347 | NA | NA | NA | NA | NA | NA | NA |
| SCN5A | 1638 | R | -11.12218182 | 29.21305662 | 0.460734668 | 26.4965545 | NA | NA | NA | NA | NA | NA | NA |
| SCN5A | 1639 | G | -12.01731818 | 27.77760805 | 0.435562426 | 22.50050023 | NA | NA | NA | NA | NA | NA | NA |
| SCN5A | 1640 | A | -10.84061818 | 23.95269187 | 0.368488121 | 21.22611455 | NA | NA | NA | NA | NA | NA | NA |
| SCN5A | 1641 | K | -15.00759596 | 21.31514743 | 0.322235746 | 16.07641191 | NA | NA | NA | NA | NA | NA | NA |
| SCN5A | 1642 | G | -11.67356818 | 19.29360798 | 0.286785726 | 16.60052933 | NA | NA | NA | NA | NA | NA | NA |
| SCN5A | 1643 | I | -8.876693182 | 20.90455503 | 0.315035536 | 18.18412287 | NA | NA | NA | NA | NA | NA | NA |
| SCN5A | 1644 | R | -12.97654545 | 21.10239463 | 0.318504881 | 18.34500855 | NA | NA | NA | NA | NA | NA | NA |
| SCN5A | 1645 | T | -13.6011039 | 17.66050965 | 0.258147469 | 12.74756222 | NA | NA | NA | NA | NA | NA | NA |
| SCN5A | 1646 | L | -9.908818182 | 16.02982873 | 0.229551603 | 13.33960194 | NA | NA | NA | NA | NA | NA | NA |
| SCN5A | 1647 | L | -8.193068182 | 19.02032071 | 0.28199332 | 16.28379165 | NA | NA | NA | NA | NA | NA | NA |
| SCN5A | 1648 | F | -13.96181818 | 18.27950645 | 0.26900229 | 15.58811744 | NA | NA | NA | NA | NA | NA | NA |
| SCN5A | 1649 | A | -11.34821818 | 14.87613535 | 0.209320263 | 12.1201737 | NA | NA | NA | NA | NA | NA | NA |
| SCN5A | 1650 | L | -7.566568182 | 16.90864575 | 0.24496267 | 14.21751833 | NA | NA | NA | NA | NA | NA | NA |

| | | | | | | | | | | | | | |
|---|---|---|---|---|---|---|---|---|---|---|---|---|---|
| SCN5A | 1651 | M | -10.83019318 | 19.48199779 | 0.290089358 | 16.72655127 | NA | NA | NA | NA | NA | NA | NA |
| SCN5A | 1652 | M | -14.38856818 | 14.53298005 | 0.20330264 | 9.283608089 | NA | NA | NA | NA | NA | NA | NA |
| SCN5A | 1653 | S | -10.25698485 | 14.76014421 | 0.207286225 | 12.01183116 | NA | NA | NA | NA | NA | NA | NA |
| SCN5A | 1654 | L | -7.364568182 | 18.69228691 | 0.27624087 | 16.44651484 | NA | NA | NA | NA | NA | NA | NA |
| SCN5A | 1655 | P | -9.867675325 | 19.75270093 | 0.294836449 | 17.49140945 | NA | NA | NA | NA | NA | NA | NA |
| SCN5A | 1656 | A | -8.722418182 | 16.44712571 | 0.236869386 | 14.16947791 | NA | NA | NA | NA | NA | NA | NA |
| SCN5A | 1657 | L | -5.519693182 | 14.6962124 | 0.206165107 | 12.67524459 | NA | NA | NA | NA | NA | NA | NA |
| SCN5A | 1658 | F | -4.747727273 | 19.43277556 | 0.28922619 | 17.814134 | NA | NA | NA | NA | NA | NA | NA |
| SCN5A | 1659 | N | -6.816193182 | 19.63559485 | 0.292782859 | 17.71624798 | NA | NA | NA | NA | NA | NA | NA |
| SCN5A | 1660 | I | -4.199443182 | 14.31631995 | 0.199503256 | 12.76906871 | NA | NA | NA | NA | NA | NA | NA |
| SCN5A | 1661 | G | -0.919568182 | 18.40046551 | 0.271123446 | 15.62851237 | NA | NA | NA | NA | NA | NA | NA |
| SCN5A | 1662 | L | -1.805568182 | 21.01414221 | 0.316957273 | 19.39803542 | NA | NA | NA | NA | NA | NA | NA |
| SCN5A | 1663 | L | -1.541443182 | 18.8641778 | 0.279255174 | 16.03793905 | NA | NA | NA | NA | NA | NA | NA |
| SCN5A | 1664 | L | 1.037068182 | 14.99989287 | 0.211490493 | 11.07014143 | NA | NA | NA | NA | NA | NA | NA |
| SCN5A | 1665 | F | 3.143454545 | 20.211514 | 0.302882264 | 16.26312209 | NA | NA | NA | NA | NA | NA | NA |
| SCN5A | 1666 | L | 1.320193182 | 22.50153435 | 0.343040405 | 18.56053654 | NA | NA | NA | NA | NA | NA | NA |
| SCN5A | 1667 | V | 3.060961039 | 19.15538405 | 0.284361811 | 15.27376704 | NA | NA | NA | NA | NA | NA | NA |
| SCN5A | 1668 | M | 6.125818182 | 16.63286761 | 0.240126584 | 12.73641047 | NA | NA | NA | NA | NA | NA | NA |
| SCN5A | 1669 | F | 6.939 | 22.69637557 | 0.34645717 | 18.78819665 | NA | NA | NA | NA | NA | NA | NA |
| SCN5A | 1670 | I | 5.947943182 | 24.25338003 | 0.373761034 | 20.38614809 | NA | NA | NA | NA | NA | NA | NA |
| SCN5A | 1671 | Y | 7.497484848 | 20.19480344 | 0.302589225 | 16.37256786 | NA | NA | NA | NA | NA | NA | NA |
| SCN5A | 1672 | S | 11.20431818 | 22.78148013 | 0.347949576 | 20.92254296 | NA | NA | NA | NA | NA | NA | NA |
| SCN5A | 1673 | I | 11.23831818 | 25.35568719 | 0.393091258 | 23.47924959 | NA | NA | NA | NA | NA | NA | NA |
| SCN5A | 1674 | F | 10.744 | 25.31394248 | 0.392359216 | 23.4303973 | NA | NA | NA | NA | NA | NA | NA |
| SCN5A | 1675 | G | 15.23281818 | 23.64438136 | 0.363081542 | 21.7308729 | NA | NA | NA | NA | NA | NA | NA |
| SCN5A | 1676 | M | 16.43481818 | 25.18586084 | 0.390113157 | 23.25072819 | NA | NA | NA | NA | NA | NA | NA |
| SCN5A | 1677 | A | 16.90481818 | 27.72812163 | 0.434694625 | 25.79794752 | NA | NA | NA | NA | NA | NA | NA |
| SCN5A | 1678 | N | 17.08206818 | 26.32265431 | 0.410048139 | 24.36705432 | NA | NA | NA | NA | NA | NA | NA |
| SCN5A | 1679 | F | 17.523 | 21.49733844 | 0.325430675 | 19.82354901 | NA | NA | NA | NA | NA | NA | NA |
| SCN5A | 1680 | A | 21.79981818 | 25.45426958 | 0.394820014 | 22.99839224 | NA | NA | NA | NA | NA | NA | NA |
| SCN5A | 1681 | Y | 25.30273485 | 25.33933906 | 0.392804575 | 22.96231716 | NA | NA | NA | NA | NA | NA | NA |
| SCN5A | 1682 | V | 24.2061039 | 21.43936988 | 0.324414129 | 19.00649332 | NA | NA | NA | NA | NA | NA | NA |
| SCN5A | 1683 | K | 28.41681818 | 19.2319616 | 0.285704686 | 13.50608934 | NA | NA | NA | NA | NA | NA | NA |

| | | | | | | | | | | | | | |
|---|---|---|---|---|---|---|---|---|---|---|---|---|---|
| SCN5A | 1684 | W | 27.36267532 | 17.38391676 | 0.253297094 | 11.47651614 | NA | NA | NA | NA | NA | NA | NA |
| SCN5A | 1685 | E | 25.82570707 | 14.47256981 | 0.202243277 | 8.66510074 | NA | NA | NA | NA | NA | NA | NA |
| SCN5A | 1686 | A | 23.39341818 | 11.05708578 | 0.142348836 | 8.747532768 | NA | NA | NA | NA | NA | NA | NA |
| SCN5A | 1687 | G | 20.82156818 | 12.6393948 | 0.170096445 | 10.3355415 | NA | NA | NA | NA | NA | NA | NA |
| SCN5A | 1688 | I | 22.21981818 | 15.05631897 | 0.21247999 | 12.69739312 | NA | NA | NA | NA | NA | NA | NA |
| SCN5A | 1689 | D | 23.46081818 | 15.78161497 | 0.225198889 | 10.33307948 | NA | NA | NA | NA | NA | NA | NA |
| SCN5A | 1690 | D | 25.99894318 | 19.50735415 | 0.290534011 | 13.59579706 | NA | NA | NA | NA | NA | NA | NA |
| SCN5A | 1691 | M | 22.09031818 | 19.80767139 | 0.29580042 | 18.17077656 | NA | NA | NA | NA | NA | NA | NA |
| SCN5A | 1692 | F | 19.15718182 | 16.4364526 | 0.23668222 | 14.75328914 | NA | NA | NA | NA | NA | NA | NA |
| SCN5A | 1693 | N | 20.83019318 | 19.69340829 | 0.293796685 | 18.02548948 | NA | NA | NA | NA | NA | NA | NA |
| SCN5A | 1694 | F | 17.49709091 | 18.65555132 | 0.275596669 | 17.21723032 | NA | NA | NA | NA | NA | NA | NA |
| SCN5A | 1695 | Q | 22.63748485 | 24.48259678 | 0.377780613 | 22.75384698 | NA | NA | NA | NA | NA | NA | NA |
| SCN5A | 1696 | T | 18.4071039 | 24.78414449 | 0.383068599 | 22.77968934 | NA | NA | NA | NA | NA | NA | NA |
| SCN5A | 1697 | F | 12.90081818 | 23.80983958 | 0.365983042 | 21.82788435 | NA | NA | NA | NA | NA | NA | NA |
| SCN5A | 1698 | A | 15.39401818 | 21.69929506 | 0.328972216 | 19.66862842 | NA | NA | NA | NA | NA | NA | NA |
| SCN5A | 1699 | N | 18.34919318 | 20.02840448 | 0.299671228 | 18.37831153 | NA | NA | NA | NA | NA | NA | NA |
| SCN5A | 1700 | S | 14.71815152 | 19.31766539 | 0.287207601 | 17.36129063 | NA | NA | NA | NA | NA | NA | NA |
| SCN5A | 1701 | M | 10.72606818 | 18.34691699 | 0.270184411 | 16.39591557 | NA | NA | NA | NA | NA | NA | NA |
| SCN5A | 1702 | L | 13.95906818 | 15.69165657 | 0.223621365 | 13.69010611 | NA | NA | NA | NA | NA | NA | NA |
| SCN5A | 1703 | C | 14.13781818 | 14.3683941 | 0.200416436 | 12.42298324 | NA | NA | NA | NA | NA | NA | NA |
| SCN5A | 1704 | L | 10.39381818 | 14.97136831 | 0.210990282 | 13.11195527 | NA | NA | NA | NA | NA | NA | NA |
| SCN5A | 1705 | F | 8.656545455 | 13.09600508 | 0.178103631 | 11.20235187 | NA | NA | NA | NA | NA | NA | NA |
| SCN5A | 1706 | Q | 12.36892929 | 10.17217823 | 0.126830964 | 8.273817212 | NA | NA | NA | NA | NA | NA | NA |
| SCN5A | 1707 | I | 10.19169318 | 11.25253718 | 0.145776301 | 9.541980573 | NA | NA | NA | NA | NA | NA | NA |
| SCN5A | 1708 | T | 5.679675325 | 10.03077589 | 0.124351312 | 6.271433303 | NA | NA | NA | NA | NA | NA | NA |
| SCN5A | 1709 | T | 6.313532468 | 6.762374583 | 0.067036135 | 2.931494388 | 3.86 | -0.35 | -1.08 | 12.03 | 0.28 | 1 | 1 |
| SCN5A | 1710 | S | 7.916984848 | 6.906535528 | 0.069564163 | 3.890655034 | 3.86 | -0.35 | -1.08 | 12.03 | 0.28 | 1 | 1 |
| SCN5A | 1711 | A | 10.89761818 | 6.611365668 | 0.06438802 | 4.728299854 | 2.12 | -0.78 | -2.6 | 34.26 | 1.5 | 2 | 0 |
| SCN5A | 1712 | G | 14.10831818 | 6.905310058 | 0.069542673 | 5.450857558 | 1.74 | -0.99 | -2.47 | 34.33 | 1.89 | 2 | -2 |
| SCN5A | 1713 | W | 11.92646104 | 9.30346629 | 0.111597101 | 7.362010022 | NA | NA | NA | NA | NA | NA | NA |
| SCN5A | 1714 | D | 15.79744318 | 7.283281266 | 0.076170833 | 4.87102439 | 2.47 | -0.43 | -1.7 | 33.74 | 1.25 | 2 | -2 |
| SCN5A | 1715 | G | 18.38581818 | 9.998194037 | 0.123779951 | 7.552931396 | NA | NA | NA | NA | NA | NA | NA |
| SCN5A | 1716 | L | 17.14844318 | 12.42281824 | 0.166298525 | 9.966134809 | NA | NA | NA | NA | NA | NA | NA |

| | | | | | | | | | | | | | |
|---|---|---|---|---|---|---|---|---|---|---|---|---|---|
| SCN5A | 1717 | L | 16.72544318 | 14.25374365 | 0.198405908 | 11.81116142 | NA | NA | NA | NA | NA | NA | NA |
| SCN5A | 1718 | S | 20.99531818 | 13.01539246 | 0.176689996 | 10.65423819 | NA | NA | NA | NA | NA | NA | NA |
| SCN5A | 1719 | P | 22.36924675 | 15.22761505 | 0.215483864 | 12.82387834 | NA | NA | NA | NA | NA | NA | NA |
| SCN5A | 1720 | I | 19.91131818 | 18.26345348 | 0.268720782 | 15.86948259 | NA | NA | NA | NA | NA | NA | NA |
| SCN5A | 1721 | L | 20.42731818 | 16.83510624 | 0.243673071 | 14.47931711 | NA | NA | NA | NA | NA | NA | NA |
| SCN5A | 1722 | N | 24.93719318 | 18.73091765 | 0.276918304 | 16.62182884 | NA | NA | NA | NA | NA | NA | NA |
| SCN5A | 1723 | T | 26.66867532 | 23.3310285 | 0.357586539 | 20.90870244 | NA | NA | NA | NA | NA | NA | NA |
| SCN5A | 1724 | G | 30.13681818 | 25.08071494 | 0.388269303 | 19.16878634 | NA | NA | NA | NA | NA | NA | NA |
| SCN5A | 1725 | P | 33.38267532 | 24.09736326 | 0.3710251 | 18.07980523 | NA | NA | NA | NA | NA | NA | NA |
| SCN5A | 1726 | P | 32.48781818 | 20.48761526 | 0.307724017 | 14.46581966 | NA | NA | NA | NA | NA | NA | NA |
| SCN5A | 1727 | Y | 29.20548485 | 16.28339286 | 0.233998142 | 10.41622959 | NA | NA | NA | NA | NA | NA | NA |
| SCN5A | 1728 | C | 30.82048485 | 22.14448579 | 0.336779148 | 16.23543054 | NA | NA | NA | NA | NA | NA | NA |
| SCN5A | 1729 | D | 33.69469318 | 25.23191001 | 0.390920683 | 19.23116839 | NA | NA | NA | NA | NA | NA | NA |
| SCN5A | 1730 | P | 32.13253247 | 28.52789643 | 0.448719596 | 22.75566495 | NA | NA | NA | NA | NA | NA | NA |
| SCN5A | 1731 | T | 33.8891039 | 31.52563271 | 0.501288349 | 25.61570436 | NA | NA | NA | NA | NA | NA | NA |
| SCN5A | 1732 | L | 32.95144318 | 28.15569124 | 0.442192549 | 22.22992213 | NA | NA | NA | NA | NA | NA | NA |
| SCN5A | 1733 | P | 33.80938961 | 31.63449861 | 0.503197438 | 25.35347598 | NA | NA | NA | NA | NA | NA | NA |
| SCN5A | 1734 | N | 29.57644318 | 30.07873039 | 0.475915253 | 24.45372928 | NA | NA | NA | NA | NA | NA | NA |
| SCN5A | 1735 | S | 28.59731818 | 34.10201622 | 0.546468197 | 31.84737652 | NA | NA | NA | NA | NA | NA | NA |
| SCN5A | 1736 | N | 25.22669318 | 34.13949251 | 0.547125387 | 31.68394405 | NA | NA | NA | NA | NA | NA | NA |
| SCN5A | 1737 | G | 25.85206818 | 32.27786601 | 0.514479625 | 29.85964927 | NA | NA | NA | NA | NA | NA | NA |
| SCN5A | 1738 | S | 28.79381818 | 33.47677838 | 0.535503933 | 31.07472094 | NA | NA | NA | NA | NA | NA | NA |
| SCN5A | 1739 | R | 31.26527273 | 31.58790702 | 0.502380401 | 26.06944068 | NA | NA | NA | NA | NA | NA | NA |
| SCN5A | 1740 | G | 28.52206818 | 28.14149804 | 0.441943655 | 26.00563293 | NA | NA | NA | NA | NA | NA | NA |
| SCN5A | 1741 | D | 29.13981818 | 23.86692647 | 0.366984126 | 18.08307266 | NA | NA | NA | NA | NA | NA | NA |
| SCN5A | 1742 | C | 25.92998485 | 23.68317116 | 0.363761766 | 21.44238331 | NA | NA | NA | NA | NA | NA | NA |
| SCN5A | 1743 | G | 22.55556818 | 23.85657817 | 0.366802657 | 21.46180003 | NA | NA | NA | NA | NA | NA | NA |
| SCN5A | 1744 | S | 19.67315152 | 23.66390014 | 0.363423826 | 21.25863332 | NA | NA | NA | NA | NA | NA | NA |
| SCN5A | 1745 | P | 18.58167532 | 22.42903828 | 0.341769103 | 20.02414902 | NA | NA | NA | NA | NA | NA | NA |
| SCN5A | 1746 | A | 15.25301818 | 24.44383534 | 0.377100887 | 22.05166237 | NA | NA | NA | NA | NA | NA | NA |
| SCN5A | 1747 | V | 14.5151039 | 22.80895828 | 0.348431437 | 20.95717637 | NA | NA | NA | NA | NA | NA | NA |
| SCN5A | 1748 | G | 14.42706818 | 19.53761406 | 0.291064654 | 17.74085412 | NA | NA | NA | NA | NA | NA | NA |
| SCN5A | 1749 | I | 13.13344318 | 19.28471905 | 0.286629849 | 17.47443554 | NA | NA | NA | NA | NA | NA | NA |

| | | | | | | | | | | | | | |
|---|---|---|---|---|---|---|---|---|---|---|---|---|---|
| SCN5A | 1750 | L | 9.606943182 | 20.15867654 | 0.301955699 | 16.38930981 | NA | NA | NA | NA | NA | NA | NA |
| SCN5A | 1751 | F | 10.908 | 18.66901703 | 0.275832806 | 17.15305998 | NA | NA | NA | NA | NA | NA | NA |
| SCN5A | 1752 | F | 12.07345455 | 15.50851124 | 0.220409701 | 13.75578156 | NA | NA | NA | NA | NA | NA | NA |
| SCN5A | 1753 | T | 7.899961039 | 15.45481006 | 0.219467989 | 11.68625395 | NA | NA | NA | NA | NA | NA | NA |
| SCN5A | 1754 | T | 5.242961039 | 16.62965129 | 0.240070182 | 12.82901373 | NA | NA | NA | NA | NA | NA | NA |
| SCN5A | 1755 | Y | 6.330234848 | 14.14068563 | 0.196423306 | 10.31256898 | NA | NA | NA | NA | NA | NA | NA |
| SCN5A | 1756 | I | 5.693193182 | 10.99779737 | 0.141309145 | 7.232219328 | NA | NA | NA | NA | NA | NA | NA |
| SCN5A | 1757 | I | 1.978193182 | 12.50589549 | 0.167755381 | 9.474909127 | NA | NA | NA | NA | NA | NA | NA |
| SCN5A | 1758 | I | 1.033318182 | 13.34604694 | 0.182488403 | 10.35238348 | NA | NA | NA | NA | NA | NA | NA |
| SCN5A | 1759 | S | 1.925984848 | 10.8009975 | 0.137858033 | 6.834826559 | NA | NA | NA | NA | NA | NA | NA |
| SCN5A | 1760 | F | 0.367818182 | 6.025638887 | 0.054116627 | 3.020899218 | 3.15 | 1.35 | 2.45 | 0.56 | -1.72 | 0 | 0 |
| SCN5A | 1761 | L | -2.045943182 | 10.5117236 | 0.132785283 | 7.667393375 | NA | NA | NA | NA | NA | NA | NA |
| SCN5A | 1762 | I | -1.496068182 | 11.13015054 | 0.14363011 | 9.570195646 | NA | NA | NA | NA | NA | NA | NA |
| SCN5A | 1763 | V | -1.119246753 | 7.794015012 | 0.085127137 | 4.757313682 | 3.15 | 1.35 | 2.45 | 0.56 | -1.72 | 0 | 0 |
| SCN5A | 1764 | V | -4.508532468 | 6.338620907 | 0.059605127 | 4.84055853 | 1.86 | 1.43 | 3.7 | 0.2 | -1.82 | 0 | 0 |
| SCN5A | 1765 | N | -6.740068182 | 9.340111616 | 0.112239719 | 7.826938467 | NA | NA | NA | NA | NA | NA | NA |
| SCN5A | 1766 | M | -4.337693182 | 9.460562933 | 0.114351971 | 7.890642312 | NA | NA | NA | NA | NA | NA | NA |
| SCN5A | 1767 | Y | -4.175318182 | 4.749690306 | 0.031741402 | 3.867048251 | 2.32 | 1.41 | 4.08 | 0.13 | -1.81 | 0 | 0 |
| SCN5A | 1768 | I | -9.874443182 | 5.954551285 | 0.052870025 | 3.6814079 | 2.32 | 1.41 | 4.08 | 0.13 | -1.81 | 0 | 0 |
| SCN5A | 1769 | A | -10.11681818 | 9.649641237 | 0.117667677 | 7.182658604 | NA | NA | NA | NA | NA | NA | NA |
| SCN5A | 1770 | I | -8.424693182 | 9.134422259 | 0.10863272 | 7.156334569 | NA | NA | NA | NA | NA | NA | NA |
| SCN5A | 1771 | I | -11.35806818 | 3.948008612 | 0.017682991 | 1.633629472 | 2.76 | 1.2 | 3.65 | 0.1 | -1.23 | 0 | 0 |
| SCN5A | 1772 | L | -15.04331818 | 7.699440824 | 0.08346867 | 4.822676669 | 2.92 | 1.23 | 3.48 | 0.15 | -1.34 | 0 | 0 |
| SCN5A | 1773 | E | -14.15548485 | 10.3484939 | 0.129922862 | 7.464461344 | NA | NA | NA | NA | NA | NA | NA |
| SCN5A | 1774 | N | -13.57781818 | 9.007228708 | 0.106402234 | 6.192312055 | NA | NA | NA | NA | NA | NA | NA |
| SCN5A | 1775 | F | -16.19390909 | 4.582943486 | 0.028817304 | 2.031226507 | 3.51 | -0.19 | -1.4 | 17.88 | 0.32 | 1 | -1 |
| SCN5A | 1776 | S | -18.60765152 | 8.710041447 | 0.101190714 | 3.461561279 | 5.35 | -0.33 | -1.23 | 13.82 | 0.43 | 1 | -1 |
| SCN5A | 1777 | V | -17.08096104 | 11.03786596 | 0.142011794 | 5.744273154 | NA | NA | NA | NA | NA | NA | NA |
| SCN5A | 1778 | A | -18.22761818 | 11.13310752 | 0.143681965 | 5.836350046 | NA | NA | NA | NA | NA | NA | NA |
| SCN5A | 1779 | T | -21.17096104 | 7.074403014 | 0.072507913 | 1.794528094 | 5.35 | -0.33 | -1.23 | 13.82 | 0.43 | 1 | -1 |
| SCN5A | 1780 | E | -22.77981818 | 10.0567278 | 0.124806408 | 4.822599397 | 5.35 | -0.33 | -1.23 | 13.82 | 0.43 | 1 | -1 |
| SCN5A | 1781 | E | -22.40937374 | 12.56812961 | 0.168846728 | 7.353231677 | NA | NA | NA | NA | NA | NA | NA |
| GRIN2B | 34 | S | 102.3868 | 56.13122786 | 0.860063251 | 55.18126945 | NA | NA | NA | NA | NA | NA | NA |

| | | | | | | | | | | | | | |
|---|---|---|---|---|---|---|---|---|---|---|---|---|---|
| GRIN2B | 35 | I | 100.673925 | 55.60620585 | 0.851621221 | 54.65524901 | NA | NA | NA | NA | NA | NA | NA |
| GRIN2B | 36 | G | 103.5293 | 52.15889803 | 0.796190636 | 51.20444327 | NA | NA | NA | NA | NA | NA | NA |
| GRIN2B | 37 | I | 104.152925 | 49.9925971 | 0.761357853 | 49.04033565 | NA | NA | NA | NA | NA | NA | NA |
| GRIN2B | 38 | A | 107.0916 | 45.96049689 | 0.696524167 | 45.00502117 | NA | NA | NA | NA | NA | NA | NA |
| GRIN2B | 39 | V | 107.2859429 | 44.4477717 | 0.672200479 | 43.4929421 | NA | NA | NA | NA | NA | NA | NA |
| GRIN2B | 40 | I | 110.47055 | 39.79950285 | 0.597459181 | 38.84840182 | NA | NA | NA | NA | NA | NA | NA |
| GRIN2B | 41 | L | 110.39505 | 39.11405011 | 0.586437524 | 38.15639562 | NA | NA | NA | NA | NA | NA | NA |
| GRIN2B | 42 | V | 112.8290857 | 34.36874134 | 0.510135884 | 33.41298017 | NA | NA | NA | NA | NA | NA | NA |
| GRIN2B | 45 | S | 110.4173 | 39.8413472 | 0.598132013 | 38.89154461 | NA | NA | NA | NA | NA | NA | NA |
| GRIN2B | 46 | D | 110.9993 | 43.28565351 | 0.653514335 | 42.33447284 | NA | NA | NA | NA | NA | NA | NA |
| GRIN2B | 47 | E | 107.8744 | 45.3874612 | 0.687310107 | 44.43746687 | NA | NA | NA | NA | NA | NA | NA |
| GRIN2B | 48 | V | 107.1732 | 48.18444764 | 0.732283924 | 47.2335037 | NA | NA | NA | NA | NA | NA | NA |
| GRIN2B | 49 | A | 103.6194 | 50.48099695 | 0.769211021 | 49.53654261 | NA | NA | NA | NA | NA | NA | NA |
| GRIN2B | 50 | I | 104.2282 | 48.5004995 | 0.737365843 | 47.55204456 | NA | NA | NA | NA | NA | NA | NA |
| GRIN2B | 51 | K | 105.6572 | 50.96045613 | 0.776920429 | 50.00656452 | NA | NA | NA | NA | NA | NA | NA |
| GRIN2B | 52 | D | 103.3178 | 53.65839314 | 0.820301593 | 52.70543967 | NA | NA | NA | NA | NA | NA | NA |
| GRIN2B | 53 | A | 100.139 | 54.13559281 | 0.82797467 | 53.20322277 | NA | NA | NA | NA | NA | NA | NA |
| GRIN2B | 54 | H | 102.8381 | 53.03442886 | 0.810268632 | 52.07674222 | NA | NA | NA | NA | NA | NA | NA |
| GRIN2B | 55 | E | 99.511 | 56.57782921 | 0.867244326 | 55.62490028 | NA | NA | NA | NA | NA | NA | NA |
| GRIN2B | 56 | K | 96.3156 | 58.45683502 | 0.897457581 | 57.50875475 | NA | NA | NA | NA | NA | NA | NA |
| GRIN2B | 57 | D | 94.9052 | 58.68045501 | 0.901053252 | 56.73413386 | NA | NA | NA | NA | NA | NA | NA |
| GRIN2B | 58 | D | 92.576 | 61.46024365 | 0.94575054 | 52.97546959 | NA | NA | NA | NA | NA | NA | NA |
| GRIN2B | 59 | F | 93.7982 | 63.44456472 | 0.9776572 | 62.4900199 | NA | NA | NA | NA | NA | NA | NA |
| GRIN2B | 60 | H | 90.4926 | 64.83409564 | 1 | 56.4071848 | NA | NA | NA | NA | NA | NA | NA |
| GRIN2B | 61 | H | 89.6394 | 61.28939505 | 0.9430034 | 52.85757985 | NA | NA | NA | NA | NA | NA | NA |
| GRIN2B | 62 | L | 92.8094 | 62.2854944 | 0.959020063 | 61.33322729 | NA | NA | NA | NA | NA | NA | NA |
| GRIN2B | 63 | S | 93.567 | 63.94546898 | 0.985711431 | 62.991676 | NA | NA | NA | NA | NA | NA | NA |
| GRIN2B | 64 | V | 96.9166 | 61.38910621 | 0.944606694 | 60.43226072 | NA | NA | NA | NA | NA | NA | NA |
| GRIN2B | 65 | V | 99.294 | 60.27314232 | 0.926662682 | 59.32183728 | NA | NA | NA | NA | NA | NA | NA |
| GRIN2B | 66 | P | 100.2808 | 57.65012446 | 0.884486172 | 56.69340454 | NA | NA | NA | NA | NA | NA | NA |
| GRIN2B | 67 | R | 104.3768 | 54.26709552 | 0.830089152 | 53.3161703 | NA | NA | NA | NA | NA | NA | NA |
| GRIN2B | 68 | V | 105.7912286 | 53.38037799 | 0.815831281 | 52.43250407 | NA | NA | NA | NA | NA | NA | NA |
| GRIN2B | 69 | E | 108.7062 | 49.41483542 | 0.752067801 | 48.4653491 | NA | NA | NA | NA | NA | NA | NA |

| | | | | | | | | | | | | | |
|---|---|---|---|---|---|---|---|---|---|---|---|---|---|
| GRIN2B | 70 | L | 111.1344 | 47.19898766 | 0.716438335 | 46.25433446 | NA | NA | NA | NA | NA | NA | NA |
| GRIN2B | 71 | V | 113.1729429 | 43.21602317 | 0.652394722 | 42.28429209 | NA | NA | NA | NA | NA | NA | NA |
| GRIN2B | 72 | A | 115.387 | 41.05252291 | 0.617606971 | 40.11489833 | NA | NA | NA | NA | NA | NA | NA |
| GRIN2B | 73 | M | 116.2744 | 37.51282279 | 0.560690776 | 36.56950677 | NA | NA | NA | NA | NA | NA | NA |
| GRIN2B | 74 | N | 117.685425 | 33.62171049 | 0.498124088 | 32.69197589 | NA | NA | NA | NA | NA | NA | NA |
| GRIN2B | 75 | E | 118.136 | 31.59671231 | 0.465563366 | 30.6523459 | NA | NA | NA | NA | NA | NA | NA |
| GRIN2B | 76 | T | 118.0116571 | 27.63793505 | 0.401908668 | 26.69150417 | NA | NA | NA | NA | NA | NA | NA |
| GRIN2B | 77 | D | 118.9822 | 29.43077418 | 0.430736416 | 28.5200542 | NA | NA | NA | NA | NA | NA | NA |
| GRIN2B | 78 | P | 118.1966571 | 29.06628898 | 0.424875719 | 28.14377204 | NA | NA | NA | NA | NA | NA | NA |
| GRIN2B | 79 | K | 120.186 | 32.24461799 | 0.47598129 | 31.33961187 | NA | NA | NA | NA | NA | NA | NA |
| GRIN2B | 80 | S | 118.3283 | 34.12620292 | 0.506236016 | 33.2181848 | NA | NA | NA | NA | NA | NA | NA |
| GRIN2B | 81 | I | 114.788425 | 33.84642553 | 0.501737368 | 32.90058923 | NA | NA | NA | NA | NA | NA | NA |
| GRIN2B | 82 | I | 117.320425 | 35.78661265 | 0.532934381 | 34.87191129 | NA | NA | NA | NA | NA | NA | NA |
| GRIN2B | 83 | T | 118.3569429 | 37.96084637 | 0.567894718 | 37.05926847 | NA | NA | NA | NA | NA | NA | NA |
| GRIN2B | 84 | R | 114.5502 | 39.9279602 | 0.599524696 | 39.01081887 | NA | NA | NA | NA | NA | NA | NA |
| GRIN2B | 85 | I | 113.1264 | 40.969195 | 0.61626711 | 40.03568635 | NA | NA | NA | NA | NA | NA | NA |
| GRIN2B | 86 | C | 115.8283 | 42.50635109 | 0.640983632 | 41.59115144 | NA | NA | NA | NA | NA | NA | NA |
| GRIN2B | 87 | D | 115.905675 | 43.54845725 | 0.657740057 | 42.64874969 | NA | NA | NA | NA | NA | NA | NA |
| GRIN2B | 88 | L | 111.2174 | 45.72354268 | 0.69271409 | 44.79946819 | NA | NA | NA | NA | NA | NA | NA |
| GRIN2B | 89 | M | 111.9176 | 47.46601683 | 0.720732 | 46.54222736 | NA | NA | NA | NA | NA | NA | NA |
| GRIN2B | 90 | S | 115.0939667 | 47.37383246 | 0.719249732 | 46.47821259 | NA | NA | NA | NA | NA | NA | NA |
| GRIN2B | 91 | D | 114.62055 | 49.62683111 | 0.755476561 | 48.72220167 | NA | NA | NA | NA | NA | NA | NA |
| GRIN2B | 92 | R | 110.772 | 51.25417436 | 0.781643237 | 50.32705606 | NA | NA | NA | NA | NA | NA | NA |
| GRIN2B | 93 | K | 107.3478 | 53.11391408 | 0.811546706 | 52.1718597 | NA | NA | NA | NA | NA | NA | NA |
| GRIN2B | 94 | I | 107.0173 | 49.28637533 | 0.750002242 | 48.34559521 | NA | NA | NA | NA | NA | NA | NA |
| GRIN2B | 95 | Q | 104.379 | 49.80263998 | 0.75830346 | 48.85077074 | NA | NA | NA | NA | NA | NA | NA |
| GRIN2B | 96 | G | 105.04055 | 47.20918128 | 0.716602242 | 46.25503438 | NA | NA | NA | NA | NA | NA | NA |
| GRIN2B | 97 | V | 106.5440857 | 43.09637141 | 0.650470795 | 42.14276738 | NA | NA | NA | NA | NA | NA | NA |
| GRIN2B | 98 | V | 105.1466571 | 42.83686914 | 0.646298159 | 41.87935983 | NA | NA | NA | NA | NA | NA | NA |
| GRIN2B | 99 | F | 108.5498909 | 35.50777133 | 0.528450784 | 34.54970117 | NA | NA | NA | NA | NA | NA | NA |
| GRIN2B | 100 | A | 108.0854 | 37.18282214 | 0.555384568 | 36.23126695 | NA | NA | NA | NA | NA | NA | NA |
| GRIN2B | 101 | D | 109.917425 | 31.95310486 | 0.471293939 | 31.01530683 | NA | NA | NA | NA | NA | NA | NA |
| GRIN2B | 102 | D | 110.0392 | 32.08863476 | 0.473473176 | 31.25889342 | NA | NA | NA | NA | NA | NA | NA |

| | | | | | | | | | | | | | |
|---|---|---|---|---|---|---|---|---|---|---|---|---|---|
| GRIN2B | 103 | T | 111.2000857 | 27.87367455 | 0.405699214 | 26.96868555 | NA | NA | NA | NA | NA | NA | NA |
| GRIN2B | 104 | D | 108.2154 | 29.20685551 | 0.427135942 | 28.33567026 | NA | NA | NA | NA | NA | NA | NA |
| GRIN2B | 105 | Q | 109.2994 | 27.78166226 | 0.404219713 | 26.88475012 | NA | NA | NA | NA | NA | NA | NA |
| GRIN2B | 106 | E | 107.0562 | 29.17788474 | 0.42667011 | 28.36015487 | NA | NA | NA | NA | NA | NA | NA |
| GRIN2B | 107 | A | 109.5354 | 28.70401312 | 0.419050546 | 27.76303026 | NA | NA | NA | NA | NA | NA | NA |
| GRIN2B | 108 | I | 108.345 | 31.60159464 | 0.465641871 | 30.66886163 | NA | NA | NA | NA | NA | NA | NA |
| GRIN2B | 109 | A | 105.4946 | 34.16721389 | 0.506895447 | 33.24457979 | NA | NA | NA | NA | NA | NA | NA |
| GRIN2B | 110 | Q | 107.4322 | 32.68305458 | 0.48303108 | 31.74187277 | NA | NA | NA | NA | NA | NA | NA |
| GRIN2B | 111 | I | 110.522925 | 31.18251095 | 0.458903264 | 30.22814494 | NA | NA | NA | NA | NA | NA | NA |
| GRIN2B | 112 | L | 106.855 | 36.26777917 | 0.540671242 | 35.31346239 | NA | NA | NA | NA | NA | NA | NA |
| GRIN2B | 113 | D | 105.4982 | 38.05622021 | 0.569428271 | 37.10545976 | NA | NA | NA | NA | NA | NA | NA |
| GRIN2B | 114 | F | 110.3019818 | 36.04895741 | 0.537152723 | 35.10311092 | NA | NA | NA | NA | NA | NA | NA |
| GRIN2B | 115 | I | 109.56755 | 37.91994223 | 0.567237005 | 36.97621065 | NA | NA | NA | NA | NA | NA | NA |
| GRIN2B | 116 | S | 105.1651333 | 41.20674135 | 0.620086709 | 40.25231541 | NA | NA | NA | NA | NA | NA | NA |
| GRIN2B | 117 | A | 106.2226 | 41.41657413 | 0.623460691 | 40.46147819 | NA | NA | NA | NA | NA | NA | NA |
| GRIN2B | 118 | Q | 109.9981333 | 39.75635347 | 0.596765366 | 38.8129054 | NA | NA | NA | NA | NA | NA | NA |
| GRIN2B | 119 | T | 108.1872286 | 43.26118503 | 0.653120897 | 42.32043332 | NA | NA | NA | NA | NA | NA | NA |
| GRIN2B | 120 | L | 104.4614 | 45.98237568 | 0.696875965 | 45.03343302 | NA | NA | NA | NA | NA | NA | NA |
| GRIN2B | 121 | T | 104.7558 | 45.47338859 | 0.688691766 | 44.52293534 | NA | NA | NA | NA | NA | NA | NA |
| GRIN2B | 122 | P | 102.0505143 | 45.8129646 | 0.694151939 | 44.85540891 | NA | NA | NA | NA | NA | NA | NA |
| GRIN2B | 123 | I | 103.45255 | 41.49346938 | 0.624697119 | 40.5422631 | NA | NA | NA | NA | NA | NA | NA |
| GRIN2B | 124 | L | 102.389 | 42.47676311 | 0.640507876 | 41.53176285 | NA | NA | NA | NA | NA | NA | NA |
| GRIN2B | 125 | G | 101.55355 | 41.83575021 | 0.630200783 | 40.92382779 | NA | NA | NA | NA | NA | NA | NA |
| GRIN2B | 126 | I | 102.634925 | 41.1584288 | 0.619309873 | 40.31596044 | NA | NA | NA | NA | NA | NA | NA |
| GRIN2B | 127 | H | 103.8922 | 37.34301998 | 0.557960451 | 36.46589701 | NA | NA | NA | NA | NA | NA | NA |
| GRIN2B | 128 | G | 104.9338 | 34.71562831 | 0.515713613 | 33.90674222 | NA | NA | NA | NA | NA | NA | NA |
| GRIN2B | 129 | G | 105.57505 | 33.68903718 | 0.49920666 | 32.83040854 | NA | NA | NA | NA | NA | NA | NA |
| GRIN2B | 130 | S | 103.5801333 | 35.98978836 | 0.536201321 | 35.11015083 | NA | NA | NA | NA | NA | NA | NA |
| GRIN2B | 131 | S | 100.8226333 | 36.24453799 | 0.540297538 | 34.28452528 | NA | NA | NA | NA | NA | NA | NA |
| GRIN2B | 132 | M | 101.5714 | 33.55043691 | 0.496978053 | 31.58542913 | NA | NA | NA | NA | NA | NA | NA |
| GRIN2B | 133 | I | 99.4528 | 32.8706097 | 0.486046851 | 30.9053282 | NA | NA | NA | NA | NA | NA | NA |
| GRIN2B | 134 | M | 100.9242 | 34.62244899 | 0.514215347 | 32.66236388 | NA | NA | NA | NA | NA | NA | NA |
| GRIN2B | 135 | A | 100.5846 | 32.88459113 | 0.486271664 | 30.88179363 | NA | NA | NA | NA | NA | NA | NA |

| | | | | | | | | | | | | | |
|---|---|---|---|---|---|---|---|---|---|---|---|---|---|
| GRIN2B | 136 | D | 99.7396 | 35.82610765 | 0.533569436 | 33.82293991 | NA | NA | NA | NA | NA | NA | NA |
| GRIN2B | 137 | K | 98.7806 | 38.1363224 | 0.570716265 | 36.18942626 | NA | NA | NA | NA | NA | NA | NA |
| GRIN2B | 138 | D | 96.926 | 40.31834148 | 0.605801787 | 39.23091081 | NA | NA | NA | NA | NA | NA | NA |
| GRIN2B | 139 | E | 98.6452 | 42.08032878 | 0.634133456 | 41.15241746 | NA | NA | NA | NA | NA | NA | NA |
| GRIN2B | 140 | S | 99.86013333 | 44.63084061 | 0.675144114 | 43.68759227 | NA | NA | NA | NA | NA | NA | NA |
| GRIN2B | 141 | S | 101.3581333 | 41.79497254 | 0.629545104 | 40.84667573 | NA | NA | NA | NA | NA | NA | NA |
| GRIN2B | 142 | M | 100.1052 | 44.25431525 | 0.669089818 | 43.30699613 | NA | NA | NA | NA | NA | NA | NA |
| GRIN2B | 143 | F | 100.5600727 | 39.53066124 | 0.593136374 | 38.62756413 | NA | NA | NA | NA | NA | NA | NA |
| GRIN2B | 144 | F | 96.67152727 | 42.82901515 | 0.646171871 | 41.15652238 | NA | NA | NA | NA | NA | NA | NA |
| GRIN2B | 145 | Q | 97.4958 | 39.74181282 | 0.596531561 | 37.78256383 | NA | NA | NA | NA | NA | NA | NA |
| GRIN2B | 146 | F | 96.411 | 38.38049434 | 0.574642399 | 36.44496299 | NA | NA | NA | NA | NA | NA | NA |
| GRIN2B | 147 | G | 95.80605 | 35.97402411 | 0.535947841 | 34.14496408 | NA | NA | NA | NA | NA | NA | NA |
| GRIN2B | 148 | P | 93.9158 | 33.08383926 | 0.489475451 | 31.93960622 | NA | NA | NA | NA | NA | NA | NA |
| GRIN2B | 149 | S | 93.17 | 31.27875197 | 0.46045076 | 30.16239766 | NA | NA | NA | NA | NA | NA | NA |
| GRIN2B | 150 | I | 93.118425 | 27.7035828 | 0.402964244 | 26.59513672 | NA | NA | NA | NA | NA | NA | NA |
| GRIN2B | 151 | E | 89.3228 | 29.44202136 | 0.430917264 | 26.94420699 | NA | NA | NA | NA | NA | NA | NA |
| GRIN2B | 152 | Q | 89.2596 | 29.88624891 | 0.438060169 | 27.39331176 | NA | NA | NA | NA | NA | NA | NA |
| GRIN2B | 153 | Q | 90.5052 | 26.70888259 | 0.386970077 | 24.21352143 | NA | NA | NA | NA | NA | NA | NA |
| GRIN2B | 154 | A | 87.9614 | 25.32525881 | 0.36472226 | 22.83582979 | NA | NA | NA | NA | NA | NA | NA |
| GRIN2B | 155 | S | 85.15346667 | 26.50887163 | 0.383754024 | 18.02178053 | NA | NA | NA | NA | NA | NA | NA |
| GRIN2B | 156 | V | 86.99051429 | 26.26805339 | 0.379881816 | 17.76384613 | NA | NA | NA | NA | NA | NA | NA |
| GRIN2B | 157 | M | 86.6218 | 22.92757504 | 0.326168983 | 20.61230154 | NA | NA | NA | NA | NA | NA | NA |
| GRIN2B | 158 | L | 83.2324 | 22.18109125 | 0.314165984 | 13.69623219 | NA | NA | NA | NA | NA | NA | NA |
| GRIN2B | 159 | N | 82.2224 | 24.05226116 | 0.344253243 | 15.55992217 | NA | NA | NA | NA | NA | NA | NA |
| GRIN2B | 160 | I | 83.8272 | 22.45526384 | 0.31857451 | 13.93922244 | NA | NA | NA | NA | NA | NA | NA |
| GRIN2B | 161 | M | 83.08055 | 18.92835928 | 0.261864059 | 10.59832186 | NA | NA | NA | NA | NA | NA | NA |
| GRIN2B | 162 | E | 78.1778 | 19.40969023 | 0.269603564 | 10.92683052 | NA | NA | NA | NA | NA | NA | NA |
| GRIN2B | 163 | E | 79.38891111 | 22.14467087 | 0.313580367 | 13.63097154 | NA | NA | NA | NA | NA | NA | NA |
| GRIN2B | 164 | Y | 80.7674 | 18.62286831 | 0.256951953 | 10.08423813 | NA | NA | NA | NA | NA | NA | NA |
| GRIN2B | 165 | D | 77.635175 | 15.63425396 | 0.208896876 | 7.107805642 | NA | NA | NA | NA | NA | NA | NA |
| GRIN2B | 166 | W | 80.9324 | 15.47532116 | 0.206341335 | 6.967087155 | NA | NA | NA | NA | NA | NA | NA |
| GRIN2B | 167 | Y | 78.95688333 | 12.62932421 | 0.160579458 | 4.150912805 | 8.72 | -0.21 | -1.78 | 37.26 | 0.96 | 3 | 1 |
| GRIN2B | 168 | I | 84.473425 | 12.53251136 | 0.159022767 | 7.430893607 | NA | NA | NA | NA | NA | NA | NA |

| | | | | | | | | | | | | | |
|---|---|---|---|---|---|---|---|---|---|---|---|---|---|
| GRIN2B | 169 | F | 83.751 | 10.56388238 | 0.127368427 | 4.992096874 | 5.75 | -0.9 | -1.43 | 3.43 | 0.58 | 0 | 0 |
| GRIN2B | 170 | S | 87.3608 | 11.13380712 | 0.136532466 | 8.624202142 | NA | NA | NA | NA | NA | NA | NA |
| GRIN2B | 171 | I | 89.261675 | 13.86039953 | 0.180374391 | 11.34188114 | NA | NA | NA | NA | NA | NA | NA |
| GRIN2B | 172 | V | 93.35822857 | 13.7252039 | 0.178200528 | 12.59952509 | NA | NA | NA | NA | NA | NA | NA |
| GRIN2B | 173 | T | 95.4598 | 17.21521461 | 0.234317749 | 16.09202658 | NA | NA | NA | NA | NA | NA | NA |
| GRIN2B | 174 | T | 99.138 | 18.19507123 | 0.250073239 | 16.22764309 | NA | NA | NA | NA | NA | NA | NA |
| GRIN2B | 175 | Y | 101.1729667 | 20.7156341 | 0.290602336 | 18.81792697 | NA | NA | NA | NA | NA | NA | NA |
| GRIN2B | 176 | F | 100.336 | 23.44024125 | 0.33441234 | 21.45855603 | NA | NA | NA | NA | NA | NA | NA |
| GRIN2B | 177 | P | 100.8842286 | 26.21730425 | 0.379065801 | 24.41942618 | NA | NA | NA | NA | NA | NA | NA |
| GRIN2B | 178 | G | 97.07255 | 25.41852974 | 0.366221999 | 24.31446397 | NA | NA | NA | NA | NA | NA | NA |
| GRIN2B | 179 | Y | 95.4024 | 24.21517437 | 0.346872787 | 23.10630613 | NA | NA | NA | NA | NA | NA | NA |
| GRIN2B | 180 | Q | 93.8308 | 26.88319821 | 0.389772965 | 25.75758002 | NA | NA | NA | NA | NA | NA | NA |
| GRIN2B | 181 | D | 92.461925 | 27.66209063 | 0.402297075 | 25.14345168 | NA | NA | NA | NA | NA | NA | NA |
| GRIN2B | 182 | F | 91.37452727 | 21.95267212 | 0.310493145 | 19.43909721 | NA | NA | NA | NA | NA | NA | NA |
| GRIN2B | 183 | V | 89.2088 | 22.10057142 | 0.312871275 | 19.57694838 | NA | NA | NA | NA | NA | NA | NA |
| GRIN2B | 184 | N | 86.9678 | 25.71124715 | 0.370928715 | 23.19052891 | NA | NA | NA | NA | NA | NA | NA |
| GRIN2B | 185 | K | 86.5014 | 25.52057621 | 0.367862843 | 23.42242107 | NA | NA | NA | NA | NA | NA | NA |
| GRIN2B | 186 | I | 85.8153 | 21.08335635 | 0.296515083 | 15.47877008 | NA | NA | NA | NA | NA | NA | NA |
| GRIN2B | 187 | R | 82.767 | 22.79644964 | 0.324060567 | 14.31316091 | NA | NA | NA | NA | NA | NA | NA |
| GRIN2B | 188 | S | 81.5923 | 24.70572796 | 0.354760586 | 16.27710158 | NA | NA | NA | NA | NA | NA | NA |
| GRIN2B | 189 | T | 81.59437143 | 22.64155679 | 0.321569986 | 14.18925025 | NA | NA | NA | NA | NA | NA | NA |
| GRIN2B | 190 | I | 79.94005 | 18.97135277 | 0.262555368 | 10.5510529 | NA | NA | NA | NA | NA | NA | NA |
| GRIN2B | 191 | E | 77.65424444 | 21.72460416 | 0.306825953 | 13.24411641 | NA | NA | NA | NA | NA | NA | NA |
| GRIN2B | 192 | N | 76.810175 | 23.82389613 | 0.340581274 | 15.32222565 | NA | NA | NA | NA | NA | NA | NA |
| GRIN2B | 193 | S | 75.68646667 | 20.61827796 | 0.28903691 | 12.12905943 | NA | NA | NA | NA | NA | NA | NA |
| GRIN2B | 194 | F | 72.039 | 19.3757622 | 0.269058022 | 10.79557383 | NA | NA | NA | NA | NA | NA | NA |
| GRIN2B | 195 | V | 73.8166 | 17.4172079 | 0.237565676 | 8.877564962 | NA | NA | NA | NA | NA | NA | NA |
| GRIN2B | 196 | G | 74.92055 | 14.56319587 | 0.191674922 | 6.059940918 | NA | NA | NA | NA | NA | NA | NA |
| GRIN2B | 197 | W | 78.88922857 | 13.77123749 | 0.17894072 | 5.328121206 | NA | NA | NA | NA | NA | NA | NA |
| GRIN2B | 198 | E | 79.8382 | 10.59510462 | 0.127870461 | 2.111450403 | 8.72 | -0.21 | -1.78 | 37.26 | 0.96 | 3 | 1 |
| GRIN2B | 199 | L | 82.1243 | 12.33436926 | 0.155836765 | 4.167857516 | 8.21 | -0.88 | -1.18 | 3.42 | 0.64 | 0 | 0 |
| GRIN2B | 200 | E | 84.6896 | 10.76644965 | 0.130625584 | 5.018743595 | NA | NA | NA | NA | NA | NA | NA |
| GRIN2B | 215 | K | 99.0506 | 10.21674552 | 0.12178668 | 9.186833723 | NA | NA | NA | NA | NA | NA | NA |

| | | | | | | | | | | | | | |
|---|---|---|---|---|---|---|---|---|---|---|---|---|---|
| GRIN2B | 216 | I | 99.453175 | 8.684243951 | 0.097145 | 7.512461822 | 1.52 | -0.26 | 0.1 | 2.35 | -0.12 | 0 | 0 |
| GRIN2B | 217 | Q | 97.6428 | 5.393901093 | 0.044238315 | 4.373188368 | 1.36 | 0.39 | 1.63 | 1.26 | -0.82 | 0 | 0 |
| GRIN2B | 218 | N | 95.829425 | 6.348237393 | 0.059583454 | 5.370957269 | 1.16 | 0.36 | 0.15 | 24.82 | 0.08 | 2 | 2 |
| GRIN2B | 219 | Q | 93.4948 | 7.959709103 | 0.085494926 | 6.846516681 | 1.2 | 0.1 | -1.2 | 33.04 | 0.54 | 2 | 2 |
| GRIN2B | 220 | L | 93.5198 | 5.886168023 | 0.052153664 | 4.759580229 | 1.16 | 0.36 | 0.15 | 24.82 | 0.08 | 2 | 2 |
| GRIN2B | 221 | K | 91.9306 | 2.642656618 | 0 | 1.591578876 | 1.2 | 0.1 | -1.2 | 33.04 | 0.54 | 2 | 2 |
| GRIN2B | 222 | K | 89.294 | 4.607724927 | 0.031597087 | 3.323094954 | 1.74 | -0.9 | -1.43 | 3.43 | 0.58 | 0 | 0 |
| GRIN2B | 223 | L | 88.751 | 6.347212301 | 0.059566972 | 4.239317474 | 2.13 | -0.92 | -1.64 | 3.44 | 0.54 | 0 | 0 |
| GRIN2B | 224 | Q | 87.2868 | 4.4947654 | 0.029780767 | 2.406262365 | 2.13 | -0.92 | -1.64 | 3.44 | 0.54 | 0 | 0 |
| GRIN2B | 225 | S | 86.3043 | 7.514320329 | 0.078333349 | 5.021453721 | 2.67 | -0.92 | -1.64 | 3.44 | 0.54 | 0 | 0 |
| GRIN2B | 226 | P | 85.4468 | 9.96834886 | 0.117792615 | 7.630893603 | NA | NA | NA | NA | NA | NA | NA |
| GRIN2B | 227 | I | 86.67955 | 12.5901633 | 0.159949775 | 10.10600391 | NA | NA | NA | NA | NA | NA | NA |
| GRIN2B | 228 | I | 90.784175 | 11.75454695 | 0.146513579 | 9.890169857 | NA | NA | NA | NA | NA | NA | NA |
| GRIN2B | 229 | L | 90.669925 | 16.18542681 | 0.217759396 | 15.06051079 | NA | NA | NA | NA | NA | NA | NA |
| GRIN2B | 230 | L | 96.113675 | 15.60545078 | 0.208433739 | 14.84822421 | NA | NA | NA | NA | NA | NA | NA |
| GRIN2B | 231 | Y | 97.64121667 | 19.92823773 | 0.277941488 | 18.20772813 | NA | NA | NA | NA | NA | NA | NA |
| GRIN2B | 232 | C | 101.0061333 | 18.323006 | 0.252130352 | 16.31363116 | NA | NA | NA | NA | NA | NA | NA |
| GRIN2B | 233 | T | 104.7290857 | 19.07450424 | 0.26421398 | 12.04172542 | NA | NA | NA | NA | NA | NA | NA |
| GRIN2B | 234 | K | 105.7442 | 17.30282023 | 0.235726393 | 10.25611199 | NA | NA | NA | NA | NA | NA | NA |
| GRIN2B | 235 | E | 107.973 | 14.91558182 | 0.197341071 | 7.774567148 | NA | NA | NA | NA | NA | NA | NA |
| GRIN2B | 236 | E | 105.2576 | 14.60758724 | 0.192388708 | 7.676220381 | NA | NA | NA | NA | NA | NA | NA |
| GRIN2B | 237 | A | 103.0474 | 14.37719211 | 0.188684097 | 11.79017935 | NA | NA | NA | NA | NA | NA | NA |
| GRIN2B | 238 | T | 105.1394 | 11.91410127 | 0.149079114 | 9.173023112 | NA | NA | NA | NA | NA | NA | NA |
| GRIN2B | 239 | Y | 104.6734 | 9.639194417 | 0.112500015 | 6.918774822 | 2.69 | -0.99 | -1.7 | 17.68 | 1.62 | 1 | -1 |
| GRIN2B | 240 | I | 100.9488 | 9.849187479 | 0.115876574 | 7.833839259 | NA | NA | NA | NA | NA | NA | NA |
| GRIN2B | 241 | F | 100.9898 | 9.596146831 | 0.111807836 | 7.634753727 | NA | NA | NA | NA | NA | NA | NA |
| GRIN2B | 242 | E | 102.8446 | 6.886597999 | 0.068239961 | 4.766418088 | 2.22 | -1.01 | -1.7 | 2.29 | 0.78 | 0 | 0 |
| GRIN2B | 243 | V | 100.8950857 | 4.994740233 | 0.037820055 | 3.030479861 | 2.04 | -0.48 | -0.23 | 1.75 | 0.26 | 0 | 0 |
| GRIN2B | 244 | A | 97.7894 | 6.715794666 | 0.065493549 | 4.851192508 | 1.97 | -0.31 | -0.03 | 1.78 | -0.02 | 0 | 0 |
| GRIN2B | 245 | N | 99.1052 | 6.339734616 | 0.059446735 | 4.397058991 | 1.97 | -0.31 | -0.03 | 1.78 | -0.02 | 0 | 0 |
| GRIN2B | 246 | S | 100.0471333 | 3.357816701 | 0.011499333 | 1.446869445 | 1.97 | -0.31 | -0.03 | 1.78 | -0.02 | 0 | 0 |
| GRIN2B | 247 | V | 96.49522857 | 2.68015466 | 6.03E-04 | 1.688442446 | 1.27 | 0.39 | 1.63 | 1.26 | -0.82 | 0 | 0 |
| GRIN2B | 248 | G | 94.35855 | 4.728995031 | 0.033547036 | 3.451591058 | 1.37 | -0.26 | 0.1 | 2.35 | -0.12 | 0 | 0 |

| | | | | | | | | | | | | | |
|---|---|---|---|---|---|---|---|---|---|---|---|---|---|
| GRIN2B | 253 | G | 87.7943 | 12.36863849 | 0.156387793 | 10.47336947 | NA | NA | NA | NA | NA | NA | NA |
| GRIN2B | 254 | Y | 89.5492 | 11.65615528 | 0.144931502 | 9.561946125 | NA | NA | NA | NA | NA | NA | NA |
| GRIN2B | 255 | T | 89.1968 | 14.75174613 | 0.194706694 | 13.77509563 | NA | NA | NA | NA | NA | NA | NA |
| GRIN2B | 256 | W | 93.5978 | 11.56652139 | 0.143490244 | 10.40898894 | NA | NA | NA | NA | NA | NA | NA |
| GRIN2B | 257 | I | 90.669175 | 18.49614138 | 0.254914262 | 17.35058162 | NA | NA | NA | NA | NA | NA | NA |
| GRIN2B | 258 | V | 95.2618 | 18.7573375 | 0.259114134 | 17.60035368 | NA | NA | NA | NA | NA | NA | NA |
| GRIN2B | 259 | P | 95.08265714 | 23.11725384 | 0.329218901 | 21.9725803 | NA | NA | NA | NA | NA | NA | NA |
| GRIN2B | 260 | S | 98.90196667 | 24.00144279 | 0.343436115 | 22.04174621 | NA | NA | NA | NA | NA | NA | NA |
| GRIN2B | 261 | L | 100.3312 | 20.32899611 | 0.284385436 | 18.36470285 | NA | NA | NA | NA | NA | NA | NA |
| GRIN2B | 262 | V | 100.7052 | 20.03431898 | 0.279647209 | 18.07222297 | NA | NA | NA | NA | NA | NA | NA |
| GRIN2B | 263 | A | 102.9688 | 22.10899882 | 0.313006782 | 20.14651927 | NA | NA | NA | NA | NA | NA | NA |
| GRIN2B | 264 | G | 105.0258 | 24.2813339 | 0.347936591 | 17.24587239 | NA | NA | NA | NA | NA | NA | NA |
| GRIN2B | 265 | D | 105.3074 | 26.46560048 | 0.383058251 | 23.76469367 | NA | NA | NA | NA | NA | NA | NA |
| GRIN2B | 266 | T | 103.1408 | 27.92235916 | 0.406482033 | 25.96293269 | NA | NA | NA | NA | NA | NA | NA |
| GRIN2B | 267 | D | 106.094 | 28.31474844 | 0.41279141 | 25.57990558 | NA | NA | NA | NA | NA | NA | NA |
| GRIN2B | 268 | T | 107.0184 | 24.73929459 | 0.355300317 | 17.69915586 | NA | NA | NA | NA | NA | NA | NA |
| GRIN2B | 269 | V | 104.6082286 | 21.62799681 | 0.305272566 | 19.08661643 | NA | NA | NA | NA | NA | NA | NA |
| GRIN2B | 270 | P | 105.7200857 | 18.47728622 | 0.254611082 | 15.77826697 | NA | NA | NA | NA | NA | NA | NA |
| GRIN2B | 271 | A | 104.9754 | 16.23692144 | 0.218587398 | 13.89942465 | NA | NA | NA | NA | NA | NA | NA |
| GRIN2B | 272 | E | 104.4402444 | 11.74823821 | 0.146412139 | 8.988654586 | NA | NA | NA | NA | NA | NA | NA |
| GRIN2B | 273 | F | 101.7224364 | 15.35601543 | 0.204422972 | 13.39820673 | NA | NA | NA | NA | NA | NA | NA |
| GRIN2B | 274 | P | 97.83565714 | 14.10276416 | 0.184271464 | 12.16838276 | NA | NA | NA | NA | NA | NA | NA |
| GRIN2B | 275 | T | 95.83022857 | 17.25132044 | 0.234898308 | 15.99351925 | NA | NA | NA | NA | NA | NA | NA |
| GRIN2B | 276 | G | 92.66005 | 19.33914592 | 0.268469255 | 18.07266456 | NA | NA | NA | NA | NA | NA | NA |
| GRIN2B | 277 | L | 93.6982 | 19.38785274 | 0.269252431 | 18.08027186 | NA | NA | NA | NA | NA | NA | NA |
| GRIN2B | 278 | I | 92.7312 | 21.79607336 | 0.307975134 | 20.63253035 | NA | NA | NA | NA | NA | NA | NA |
| GRIN2B | 279 | S | 94.53396667 | 24.10368903 | 0.345080171 | 23.3234719 | NA | NA | NA | NA | NA | NA | NA |
| GRIN2B | 280 | V | 93.70394286 | 27.47391006 | 0.399271248 | 26.31443409 | NA | NA | NA | NA | NA | NA | NA |
| GRIN2B | 281 | S | 95.3066 | 30.92400016 | 0.454746569 | 28.99241181 | NA | NA | NA | NA | NA | NA | NA |
| GRIN2B | 282 | Y | 96.2924 | 33.9797118 | 0.503880529 | 32.03757086 | NA | NA | NA | NA | NA | NA | NA |
| GRIN2B | 283 | D | 96.4486 | 36.56053514 | 0.545378577 | 34.62107795 | NA | NA | NA | NA | NA | NA | NA |
| GRIN2B | 284 | E | 98.5978 | 34.03675333 | 0.504797721 | 32.08655386 | NA | NA | NA | NA | NA | NA | NA |
| GRIN2B | 285 | W | 97.484 | 36.86283756 | 0.550239414 | 34.91512243 | NA | NA | NA | NA | NA | NA | NA |

| | | | | | | | | | | | | | |
|---|---|---|---|---|---|---|---|---|---|---|---|---|---|
| GRIN2B | 286 | D | 95.1024 | 39.20029451 | 0.587824281 | 37.25761335 | NA | NA | NA | NA | NA | NA | NA |
| GRIN2B | 287 | Y | 96.7756 | 41.41583481 | 0.623448803 | 39.47318925 | NA | NA | NA | NA | NA | NA | NA |
| GRIN2B | 288 | G | 98.9023 | 44.21266211 | 0.668420062 | 42.26348671 | NA | NA | NA | NA | NA | NA | NA |
| GRIN2B | 289 | L | 101.9004 | 44.16147615 | 0.667597023 | 43.29757098 | NA | NA | NA | NA | NA | NA | NA |
| GRIN2B | 290 | P | 101.3815143 | 47.3353051 | 0.718630236 | 46.45703101 | NA | NA | NA | NA | NA | NA | NA |
| GRIN2B | 291 | A | 97.6856 | 48.19908258 | 0.732519245 | 46.25127895 | NA | NA | NA | NA | NA | NA | NA |
| GRIN2B | 292 | R | 98.7876 | 45.3325377 | 0.686426971 | 43.37701468 | NA | NA | NA | NA | NA | NA | NA |
| GRIN2B | 293 | V | 101.4523714 | 46.21994499 | 0.700695933 | 45.28816579 | NA | NA | NA | NA | NA | NA | NA |
| GRIN2B | 294 | R | 98.12 | 50.38482246 | 0.767664595 | 49.46660857 | NA | NA | NA | NA | NA | NA | NA |
| GRIN2B | 295 | D | 95.933 | 49.16779481 | 0.748095541 | 47.22307898 | NA | NA | NA | NA | NA | NA | NA |
| GRIN2B | 296 | G | 97.9958 | 49.56977015 | 0.754559056 | 48.65280323 | NA | NA | NA | NA | NA | NA | NA |
| GRIN2B | 297 | I | 98.8554 | 50.79740687 | 0.774298698 | 49.86145287 | NA | NA | NA | NA | NA | NA | NA |
| GRIN2B | 298 | A | 95.4206 | 53.57623191 | 0.818980492 | 52.04735328 | NA | NA | NA | NA | NA | NA | NA |
| GRIN2B | 299 | I | 94.8842 | 51.11698476 | 0.779437313 | 49.97177747 | NA | NA | NA | NA | NA | NA | NA |
| GRIN2B | 300 | I | 98.436675 | 48.92044792 | 0.744118355 | 47.97075009 | NA | NA | NA | NA | NA | NA | NA |
| GRIN2B | 301 | T | 96.6238 | 54.57989781 | 0.83511882 | 53.63139434 | NA | NA | NA | NA | NA | NA | NA |
| GRIN2B | 302 | T | 92.83008571 | 54.85442766 | 0.839533091 | 53.89560698 | NA | NA | NA | NA | NA | NA | NA |
| GRIN2B | 303 | A | 94.1182 | 54.39254968 | 0.832106378 | 53.26080832 | NA | NA | NA | NA | NA | NA | NA |
| GRIN2B | 304 | A | 96.1512 | 55.99438212 | 0.857862856 | 55.03834053 | NA | NA | NA | NA | NA | NA | NA |
| GRIN2B | 305 | S | 93.38813333 | 58.75546522 | 0.90225937 | 57.8033126 | NA | NA | NA | NA | NA | NA | NA |
| GRIN2B | 306 | D | 91.3128 | 57.35696501 | 0.879772349 | 48.93771539 | NA | NA | NA | NA | NA | NA | NA |
| GRIN2B | 307 | M | 93.6228 | 57.69080699 | 0.885140322 | 56.73809427 | NA | NA | NA | NA | NA | NA | NA |
| GRIN2B | 308 | L | 94.218 | 60.3306215 | 0.927586912 | 59.3747934 | NA | NA | NA | NA | NA | NA | NA |
| GRIN2B | 309 | S | 90.6023 | 61.31409083 | 0.943400493 | 52.88209004 | NA | NA | NA | NA | NA | NA | NA |
| GRIN2B | 310 | E | 90.1624 | 60.02378423 | 0.922653158 | 51.58908477 | NA | NA | NA | NA | NA | NA | NA |
| GRIN2B | 311 | H | 93.1748 | 61.48345404 | 0.946123749 | 60.52594164 | NA | NA | NA | NA | NA | NA | NA |
| GRIN2B | 312 | S | 93.60863333 | 64.45119432 | 0.993843183 | 63.49435282 | NA | NA | NA | NA | NA | NA | NA |
| GRIN2B | 313 | F | 97.0352 | 60.56893981 | 0.931418924 | 59.61682133 | NA | NA | NA | NA | NA | NA | NA |
| GRIN2B | 314 | I | 99.6298 | 58.83485849 | 0.903535965 | 57.88623125 | NA | NA | NA | NA | NA | NA | NA |
| GRIN2B | 315 | P | 100.9652286 | 55.56076218 | 0.850890515 | 54.61344386 | NA | NA | NA | NA | NA | NA | NA |
| GRIN2B | 316 | E | 105.2916 | 54.23917623 | 0.829640227 | 53.3019011 | NA | NA | NA | NA | NA | NA | NA |
| GRIN2B | 317 | P | 108.1103714 | 50.72657627 | 0.773159785 | 49.79386836 | NA | NA | NA | NA | NA | NA | NA |
| GRIN2B | 318 | K | 110.8492 | 52.38462862 | 0.799820245 | 51.48342475 | NA | NA | NA | NA | NA | NA | NA |

| | | | | | | | | | | | | | |
|---|---|---|---|---|---|---|---|---|---|---|---|---|---|
| GRIN2B | 319 | S | 112.5004 | 49.05946909 | 0.74635373 | 48.14863608 | NA | NA | NA | NA | NA | NA | NA |
| GRIN2B | 320 | S | 115.4638 | 46.42085261 | 0.703926403 | 45.52815835 | NA | NA | NA | NA | NA | NA | NA |
| GRIN2B | 321 | C | 117.3304667 | 43.62539381 | 0.658977149 | 42.72680036 | NA | NA | NA | NA | NA | NA | NA |
| GRIN2B | 322 | Y | 117.6086 | 42.11176402 | 0.634638915 | 41.233148 | NA | NA | NA | NA | NA | NA | NA |
| GRIN2B | 323 | N | 115.9146 | 45.50660472 | 0.689225861 | 44.62137436 | NA | NA | NA | NA | NA | NA | NA |
| GRIN2B | 324 | T | 112.3220857 | 46.29188718 | 0.701852719 | 45.37668652 | NA | NA | NA | NA | NA | NA | NA |
| GRIN2B | 325 | H | 112.6416 | 44.6009315 | 0.674663194 | 43.67723553 | NA | NA | NA | NA | NA | NA | NA |
| GRIN2B | 326 | E | 109.602 | 45.27923628 | 0.685569917 | 44.34976146 | NA | NA | NA | NA | NA | NA | NA |
| GRIN2B | 327 | K | 108.0178 | 48.08019798 | 0.730607654 | 47.1526582 | NA | NA | NA | NA | NA | NA | NA |
| GRIN2B | 328 | R | 104.5126 | 50.57442497 | 0.770713286 | 49.63261154 | NA | NA | NA | NA | NA | NA | NA |
| GRIN2B | 329 | I | 102.0194 | 52.05357952 | 0.79449718 | 51.10670717 | NA | NA | NA | NA | NA | NA | NA |
| GRIN2B | 330 | Y | 99.062 | 55.31508772 | 0.846940221 | 54.36648049 | NA | NA | NA | NA | NA | NA | NA |
| GRIN2B | 331 | Q | 98.2426 | 55.05821665 | 0.842809893 | 54.10240123 | NA | NA | NA | NA | NA | NA | NA |
| GRIN2B | 332 | S | 99.0248 | 50.362474 | 0.767305245 | 49.40569788 | NA | NA | NA | NA | NA | NA | NA |
| GRIN2B | 333 | N | 95.257175 | 53.53913105 | 0.818383933 | 52.58362079 | NA | NA | NA | NA | NA | NA | NA |
| GRIN2B | 334 | M | 94.0938 | 55.69866715 | 0.853107942 | 54.74672363 | NA | NA | NA | NA | NA | NA | NA |
| GRIN2B | 335 | L | 97.02655 | 52.07556308 | 0.794850662 | 51.11763811 | NA | NA | NA | NA | NA | NA | NA |
| GRIN2B | 336 | N | 96.0703 | 49.10724698 | 0.747121969 | 48.15895328 | NA | NA | NA | NA | NA | NA | NA |
| GRIN2B | 337 | R | 91.5716 | 51.60057583 | 0.78721316 | 43.13518013 | NA | NA | NA | NA | NA | NA | NA |
| GRIN2B | 338 | Y | 92.0104 | 52.53300809 | 0.802206096 | 44.06246621 | NA | NA | NA | NA | NA | NA | NA |
| GRIN2B | 339 | L | 94.668925 | 49.0488303 | 0.746182665 | 48.72978187 | NA | NA | NA | NA | NA | NA | NA |
| GRIN2B | 340 | I | 91.12105 | 47.1101998 | 0.715010681 | 44.97971517 | NA | NA | NA | NA | NA | NA | NA |
| GRIN2B | 341 | N | 87.52805 | 48.60656303 | 0.73907128 | 40.1671325 | NA | NA | NA | NA | NA | NA | NA |
| GRIN2B | 342 | V | 89.6278 | 49.43099172 | 0.752327585 | 41.01591961 | NA | NA | NA | NA | NA | NA | NA |
| GRIN2B | 343 | T | 87.7902 | 51.15511311 | 0.780050394 | 42.73027991 | NA | NA | NA | NA | NA | NA | NA |
| GRIN2B | 344 | F | 89.2592 | 50.56014723 | 0.770483709 | 42.08606303 | NA | NA | NA | NA | NA | NA | NA |
| GRIN2B | 345 | E | 88.063 | 51.57238985 | 0.786759946 | 43.10054137 | NA | NA | NA | NA | NA | NA | NA |
| GRIN2B | 346 | G | 84.6893 | 50.47430191 | 0.769103369 | 42.01913943 | NA | NA | NA | NA | NA | NA | NA |
| GRIN2B | 347 | R | 85.4104 | 47.6172359 | 0.723163509 | 39.18228674 | NA | NA | NA | NA | NA | NA | NA |
| GRIN2B | 348 | N | 84.158925 | 45.54541858 | 0.689849964 | 37.11625632 | NA | NA | NA | NA | NA | NA | NA |
| GRIN2B | 349 | L | 88.911425 | 45.7040081 | 0.692399986 | 37.22579429 | NA | NA | NA | NA | NA | NA | NA |
| GRIN2B | 350 | S | 87.2342 | 43.14013219 | 0.651174441 | 34.72794021 | NA | NA | NA | NA | NA | NA | NA |
| GRIN2B | 351 | F | 91.3918 | 43.30303372 | 0.653793798 | 34.82882072 | NA | NA | NA | NA | NA | NA | NA |

| | | | | | | | | | | | | | |
|---|---|---|---|---|---|---|---|---|---|---|---|---|---|
| GRIN2B | 352 | S | 90.17613333 | 40.04089893 | 0.601340681 | 31.5611902 | NA | NA | NA | NA | NA | NA | NA |
| GRIN2B | 353 | E | 93.1734 | 41.79837377 | 0.629599793 | 39.28757588 | NA | NA | NA | NA | NA | NA | NA |
| GRIN2B | 354 | D | 92.905925 | 37.34989807 | 0.558071046 | 36.80688555 | NA | NA | NA | NA | NA | NA | NA |
| GRIN2B | 355 | G | 94.22055 | 40.03872718 | 0.601305761 | 38.90760625 | NA | NA | NA | NA | NA | NA | NA |
| GRIN2B | 356 | Y | 89.95796667 | 35.12997801 | 0.5223761 | 32.97316826 | NA | NA | NA | NA | NA | NA | NA |
| GRIN2B | 357 | Q | 90.88391111 | 38.97180923 | 0.584150378 | 36.70251033 | NA | NA | NA | NA | NA | NA | NA |
| GRIN2B | 358 | M | 85.556175 | 38.59905768 | 0.578156763 | 30.13058858 | NA | NA | NA | NA | NA | NA | NA |
| GRIN2B | 359 | H | 87.595 | 40.23440802 | 0.604452188 | 31.75440937 | NA | NA | NA | NA | NA | NA | NA |
| GRIN2B | 360 | P | 89.27237143 | 37.10026365 | 0.554057079 | 34.61072131 | NA | NA | NA | NA | NA | NA | NA |
| GRIN2B | 361 | K | 91.891 | 37.02201496 | 0.552798888 | 35.86329465 | NA | NA | NA | NA | NA | NA | NA |
| GRIN2B | 362 | L | 90.1283 | 33.74986987 | 0.500184812 | 33.06790418 | NA | NA | NA | NA | NA | NA | NA |
| GRIN2B | 363 | V | 92.178 | 31.32748702 | 0.461234389 | 30.16645407 | NA | NA | NA | NA | NA | NA | NA |
| GRIN2B | 364 | I | 90.02855 | 29.40090917 | 0.430256205 | 28.58532169 | NA | NA | NA | NA | NA | NA | NA |
| GRIN2B | 365 | I | 93.5756 | 26.10811619 | 0.377310124 | 24.80466368 | NA | NA | NA | NA | NA | NA | NA |
| GRIN2B | 366 | L | 91.600925 | 25.29463433 | 0.364229837 | 24.01341997 | NA | NA | NA | NA | NA | NA | NA |
| GRIN2B | 367 | L | 95.68055 | 20.87121137 | 0.293103923 | 19.88209006 | NA | NA | NA | NA | NA | NA | NA |
| GRIN2B | 368 | N | 95.34605 | 22.58166212 | 0.320606917 | 21.3455419 | NA | NA | NA | NA | NA | NA | NA |
| GRIN2B | 369 | K | 97.6404 | 22.4320208 | 0.318200776 | 20.48600309 | NA | NA | NA | NA | NA | NA | NA |
| GRIN2B | 370 | E | 100.2862 | 24.40827821 | 0.349977777 | 22.46438252 | NA | NA | NA | NA | NA | NA | NA |
| GRIN2B | 371 | R | 100.2078 | 22.67181587 | 0.322056533 | 20.71887898 | NA | NA | NA | NA | NA | NA | NA |
| GRIN2B | 372 | K | 98.5858 | 25.9557122 | 0.374859562 | 24.00830396 | NA | NA | NA | NA | NA | NA | NA |
| GRIN2B | 373 | W | 98.73172857 | 28.10391725 | 0.409401375 | 26.15797808 | NA | NA | NA | NA | NA | NA | NA |
| GRIN2B | 374 | E | 94.1302 | 29.84554597 | 0.437405691 | 28.56935241 | NA | NA | NA | NA | NA | NA | NA |
| GRIN2B | 375 | R | 93.3374 | 33.1252053 | 0.490140591 | 31.83737501 | NA | NA | NA | NA | NA | NA | NA |
| GRIN2B | 376 | V | 90.0488 | 32.95907119 | 0.487469257 | 31.69255598 | NA | NA | NA | NA | NA | NA | NA |
| GRIN2B | 377 | G | 88.3118 | 34.08231126 | 0.505530265 | 31.59706842 | NA | NA | NA | NA | NA | NA | NA |
| GRIN2B | 378 | K | 86.0926 | 34.67646781 | 0.515083936 | 26.17005343 | NA | NA | NA | NA | NA | NA | NA |
| GRIN2B | 379 | W | 85.06851429 | 30.53627721 | 0.448512223 | 22.04683626 | NA | NA | NA | NA | NA | NA | NA |
| GRIN2B | 380 | K | 83.4462 | 37.19106686 | 0.555517138 | 28.70137728 | NA | NA | NA | NA | NA | NA | NA |
| GRIN2B | 381 | D | 81.1168 | 36.75266899 | 0.548467971 | 28.28530789 | NA | NA | NA | NA | NA | NA | NA |
| GRIN2B | 382 | K | 80.85 | 33.27593017 | 0.492564154 | 24.81218977 | NA | NA | NA | NA | NA | NA | NA |
| GRIN2B | 383 | S | 80.47313333 | 33.03682734 | 0.488719528 | 24.5920198 | NA | NA | NA | NA | NA | NA | NA |
| GRIN2B | 384 | L | 81.022925 | 27.14329446 | 0.393955152 | 18.71567503 | NA | NA | NA | NA | NA | NA | NA |

| | | | | | | | | | | | | | |
|---|---|---|---|---|---|---|---|---|---|---|---|---|---|
| GRIN2B | 385 | Q | 81.93113333 | 31.75561503 | 0.468118424 | 23.25621552 | NA | NA | NA | NA | NA | NA | NA |
| GRIN2B | 386 | M | 83.174675 | 27.18325266 | 0.394597656 | 18.6614121 | NA | NA | NA | NA | NA | NA | NA |
| GRIN2B | 387 | K | 83.9958 | 31.99303829 | 0.471936044 | 23.43670926 | NA | NA | NA | NA | NA | NA | NA |
| GRIN2B | 388 | Y | 84.81713333 | 27.55381986 | 0.400556148 | 19.16534789 | NA | NA | NA | NA | NA | NA | NA |
| GRIN2B | 389 | Y | 80.788 | 24.87121768 | 0.357421559 | 16.31972981 | NA | NA | NA | NA | NA | NA | NA |
| GRIN2B | 390 | V | 80.95208571 | 23.65418479 | 0.33785242 | 15.06450282 | NA | NA | NA | NA | NA | NA | NA |
| GRIN2B | 391 | W | 85.17222857 | 20.02908198 | 0.279563002 | 15.06592476 | NA | NA | NA | NA | NA | NA | NA |
| GRIN2B | 392 | P | 86.0298 | 20.13406685 | 0.281251094 | 15.24367314 | NA | NA | NA | NA | NA | NA | NA |
| GRIN2B | 406 | L | 77.6424 | 25.63050105 | 0.369630367 | 16.62454444 | NA | NA | NA | NA | NA | NA | NA |
| GRIN2B | 407 | S | 74.10713333 | 26.70226846 | 0.386863726 | 18.85419432 | NA | NA | NA | NA | NA | NA | NA |
| GRIN2B | 408 | I | 70.6983 | 22.15114196 | 0.313684418 | 14.31000685 | NA | NA | NA | NA | NA | NA | NA |
| GRIN2B | 409 | V | 67.47465714 | 26.70449996 | 0.386899607 | 18.88796592 | NA | NA | NA | NA | NA | NA | NA |
| GRIN2B | 410 | T | 65.0188 | 23.29639994 | 0.33209946 | 20.28128927 | NA | NA | NA | NA | NA | NA | NA |
| GRIN2B | 411 | L | 61.32 | 23.26526183 | 0.331598779 | 20.24635093 | NA | NA | NA | NA | NA | NA | NA |
| GRIN2B | 412 | E | 58.7316 | 21.3642133 | 0.30103109 | 18.36516242 | NA | NA | NA | NA | NA | NA | NA |
| GRIN2B | 413 | E | 56.0356 | 20.5671279 | 0.288214448 | 17.71367467 | NA | NA | NA | NA | NA | NA | NA |
| GRIN2B | 414 | A | 54.7184 | 18.95116181 | 0.26223071 | 16.07265538 | NA | NA | NA | NA | NA | NA | NA |
| GRIN2B | 415 | P | 54.70265714 | 15.51200973 | 0.206931264 | 12.63372618 | NA | NA | NA | NA | NA | NA | NA |
| GRIN2B | 416 | F | 57.30443636 | 16.95416282 | 0.230120197 | 14.11383403 | NA | NA | NA | NA | NA | NA | NA |
| GRIN2B | 417 | V | 60.05265714 | 18.57931177 | 0.25625159 | 15.5667836 | NA | NA | NA | NA | NA | NA | NA |
| GRIN2B | 418 | I | 58.3474 | 16.99220342 | 0.230731866 | 13.98414105 | NA | NA | NA | NA | NA | NA | NA |
| GRIN2B | 419 | V | 61.3266 | 18.61284677 | 0.256790812 | 15.71367102 | NA | NA | NA | NA | NA | NA | NA |
| GRIN2B | 420 | E | 61.68 | 21.87859378 | 0.309302011 | 18.86222407 | NA | NA | NA | NA | NA | NA | NA |
| GRIN2B | 421 | S | 62.9208 | 22.59578611 | 0.320834022 | 18.41702847 | NA | NA | NA | NA | NA | NA | NA |
| GRIN2B | 422 | V | 66.58065714 | 24.86414545 | 0.357307841 | 20.69509714 | NA | NA | NA | NA | NA | NA | NA |
| GRIN2B | 423 | D | 67.1542 | 24.05504729 | 0.344298042 | 19.92484568 | NA | NA | NA | NA | NA | NA | NA |
| GRIN2B | 424 | P | 67.69108571 | 27.49810388 | 0.399660269 | 23.47591034 | NA | NA | NA | NA | NA | NA | NA |
| GRIN2B | 425 | L | 69.4724 | 27.40135008 | 0.398104528 | 23.58233135 | NA | NA | NA | NA | NA | NA | NA |
| GRIN2B | 426 | S | 72.3552 | 25.9857582 | 0.375342683 | 21.77236782 | NA | NA | NA | NA | NA | NA | NA |
| GRIN2B | 427 | G | 72.81055 | 26.15787583 | 0.378110228 | 18.44661731 | NA | NA | NA | NA | NA | NA | NA |
| GRIN2B | 428 | T | 71.7462 | 23.06688245 | 0.32840896 | 15.30783266 | NA | NA | NA | NA | NA | NA | NA |
| GRIN2B | 429 | C | 70.2398 | 19.73628146 | 0.27485495 | 11.87196011 | NA | NA | NA | NA | NA | NA | NA |
| GRIN2B | 430 | M | 68.8168 | 16.63152504 | 0.22493238 | 12.47285639 | NA | NA | NA | NA | NA | NA | NA |

| | | | | | | | | | | | | | |
|---|---|---|---|---|---|---|---|---|---|---|---|---|---|
| GRIN2B | 431 | R | 69.8424 | 12.72077777 | 0.162049975 | 4.8658198 | 7.9 | -0.39 | -1.68 | 37.36 | 1.16 | 3 | -1 |
| GRIN2B | 432 | N | 67.8336 | 12.65536606 | 0.160998195 | 8.372523258 | NA | NA | NA | NA | NA | NA | NA |
| GRIN2B | 433 | T | 68.0476 | 15.89585525 | 0.213103264 | 11.60781192 | NA | NA | NA | NA | NA | NA | NA |
| GRIN2B | 434 | V | 69.72294286 | 16.83004234 | 0.228124416 | 8.986022808 | NA | NA | NA | NA | NA | NA | NA |
| GRIN2B | 435 | P | 71.70751429 | 20.66218365 | 0.289742886 | 12.80492609 | NA | NA | NA | NA | NA | NA | NA |
| GRIN2B | 436 | C | 68.89646667 | 23.78387269 | 0.339937721 | 15.94370648 | NA | NA | NA | NA | NA | NA | NA |
| GRIN2B | 437 | Q | 70.347 | 26.80682902 | 0.388544996 | 18.96430327 | NA | NA | NA | NA | NA | NA | NA |
| GRIN2B | 438 | K | 68.0268 | 29.97402799 | 0.439471603 | 22.13660154 | NA | NA | NA | NA | NA | NA | NA |
| GRIN2B | 451 | G | 72.8488 | 32.3220056 | 0.477225635 | 24.46932906 | NA | NA | NA | NA | NA | NA | NA |
| GRIN2B | 452 | Y | 75.0654 | 29.38716994 | 0.430035287 | 21.52400541 | NA | NA | NA | NA | NA | NA | NA |
| GRIN2B | 453 | I | 72.522 | 30.23438744 | 0.443658022 | 22.37573621 | NA | NA | NA | NA | NA | NA | NA |
| GRIN2B | 454 | K | 69.5398 | 27.49778384 | 0.399655123 | 19.65070011 | NA | NA | NA | NA | NA | NA | NA |
| GRIN2B | 455 | K | 66.9786 | 24.83868414 | 0.356898439 | 20.97665612 | NA | NA | NA | NA | NA | NA | NA |
| GRIN2B | 456 | C | 66.59846667 | 21.55019596 | 0.304021577 | 17.6954164 | NA | NA | NA | NA | NA | NA | NA |
| GRIN2B | 457 | C | 66.04363333 | 19.12736561 | 0.265063958 | 15.25527992 | NA | NA | NA | NA | NA | NA | NA |
| GRIN2B | 458 | K | 63.3068 | 16.55174979 | 0.223649644 | 12.52062324 | NA | NA | NA | NA | NA | NA | NA |
| GRIN2B | 459 | G | 62.03955 | 14.75453615 | 0.194751556 | 11.76046784 | NA | NA | NA | NA | NA | NA | NA |
| GRIN2B | 460 | F | 60.08470909 | 12.1202361 | 0.152393635 | 9.096143402 | NA | NA | NA | NA | NA | NA | NA |
| GRIN2B | 461 | C | 63.05963333 | 16.36764867 | 0.220689411 | 13.35223696 | NA | NA | NA | NA | NA | NA | NA |
| GRIN2B | 462 | I | 65.9053 | 15.05037491 | 0.199508461 | 11.34288257 | NA | NA | NA | NA | NA | NA | NA |
| GRIN2B | 463 | D | 64.7528 | 11.78413658 | 0.146989362 | 7.568929028 | NA | NA | NA | NA | NA | NA | NA |
| GRIN2B | 464 | I | 65.2753 | 14.56473522 | 0.191699674 | 9.543243077 | NA | NA | NA | NA | NA | NA | NA |
| GRIN2B | 465 | L | 68.9303 | 14.91397921 | 0.197315302 | 7.076544412 | NA | NA | NA | NA | NA | NA | NA |
| GRIN2B | 466 | K | 70.3026 | 11.59279535 | 0.143912713 | 3.746453357 | 7.9 | -0.39 | -1.68 | 37.36 | 1.16 | 3 | -1 |
| GRIN2B | 467 | K | 68.01602222 | 9.633158672 | 0.112402964 | 2.351590292 | 7.9 | -0.39 | -1.68 | 37.36 | 1.16 | 3 | -1 |
| GRIN2B | 468 | I | 71.105425 | 13.45080317 | 0.173788334 | 5.617686511 | NA | NA | NA | NA | NA | NA | NA |
| GRIN2B | 469 | S | 74.66096667 | 13.2205155 | 0.17008545 | 5.403498756 | NA | NA | NA | NA | NA | NA | NA |
| GRIN2B | 470 | K | 74.5774 | 9.624586744 | 0.112265132 | 1.656016994 | 8 | -0.21 | -1.78 | 37.26 | 0.96 | 3 | 1 |
| GRIN2B | 471 | S | 74.34696667 | 11.98857715 | 0.150276641 | 3.198598428 | 9.02 | -0.61 | -2.15 | 26.44 | 1.14 | 2 | 2 |
| GRIN2B | 472 | V | 76.34094286 | 14.26434667 | 0.186869612 | 5.277896668 | NA | NA | NA | NA | NA | NA | NA |
| GRIN2B | 473 | K | 79.6424 | 14.59064899 | 0.192116352 | 5.709023768 | NA | NA | NA | NA | NA | NA | NA |
| GRIN2B | 474 | F | 77.0578 | 17.18278548 | 0.233796309 | 8.177790677 | NA | NA | NA | NA | NA | NA | NA |
| GRIN2B | 475 | T | 78.7914 | 20.08793023 | 0.280509245 | 11.08164797 | NA | NA | NA | NA | NA | NA | NA |

| | | | | | | | | | | | | | |
|---|---|---|---|---|---|---|---|---|---|---|---|---|---|
| GRIN2B | 476 | Y | 74.5848 | 16.30980484 | 0.219759318 | 8.450955427 | NA | NA | NA | NA | NA | NA | NA |
| GRIN2B | 477 | D | 74.3904 | 23.31337474 | 0.332372405 | 15.45232425 | NA | NA | NA | NA | NA | NA | NA |
| GRIN2B | 478 | L | 71.0488 | 23.19576869 | 0.330481372 | 15.36029206 | NA | NA | NA | NA | NA | NA | NA |
| GRIN2B | 479 | Y | 70.48313333 | 26.46091595 | 0.382982927 | 18.624594 | NA | NA | NA | NA | NA | NA | NA |
| GRIN2B | 480 | L | 66.7398 | 28.81441828 | 0.420825793 | 26.39481879 | NA | NA | NA | NA | NA | NA | NA |
| GRIN2B | 481 | V | 63.84408571 | 30.07842679 | 0.441150271 | 27.05989534 | NA | NA | NA | NA | NA | NA | NA |
| GRIN2B | 482 | T | 65.1138 | 33.68485596 | 0.499139429 | 30.66515837 | NA | NA | NA | NA | NA | NA | NA |
| GRIN2B | 483 | N | 62.5982 | 34.86945053 | 0.518186979 | 31.86035686 | NA | NA | NA | NA | NA | NA | NA |
| GRIN2B | 484 | G | 59.66005 | 32.45020151 | 0.479286946 | 29.57156833 | NA | NA | NA | NA | NA | NA | NA |
| GRIN2B | 485 | K | 58.1566 | 29.45431778 | 0.431114983 | 26.57383393 | NA | NA | NA | NA | NA | NA | NA |
| GRIN2B | 486 | H | 57.6747 | 26.57668378 | 0.384844402 | 23.72281737 | NA | NA | NA | NA | NA | NA | NA |
| GRIN2B | 487 | G | 58.2413 | 32.29893333 | 0.476854647 | 29.42519886 | NA | NA | NA | NA | NA | NA | NA |
| GRIN2B | 488 | K | 57.85 | 35.71674875 | 0.531811012 | 32.8560977 | NA | NA | NA | NA | NA | NA | NA |
| GRIN2B | 489 | K | 57.4 | 38.73547648 | 0.580350293 | 35.90568795 | NA | NA | NA | NA | NA | NA | NA |
| GRIN2B | 490 | I | 58.6988 | 41.29232318 | 0.621462812 | 38.43781357 | NA | NA | NA | NA | NA | NA | NA |
| GRIN2B | 491 | N | 59.554 | 44.51349662 | 0.673257295 | 41.66221844 | NA | NA | NA | NA | NA | NA | NA |
| GRIN2B | 492 | G | 61.55505 | 43.80846468 | 0.661920816 | 40.92538686 | NA | NA | NA | NA | NA | NA | NA |
| GRIN2B | 493 | T | 62.68708571 | 41.01938835 | 0.617074188 | 38.13744136 | NA | NA | NA | NA | NA | NA | NA |
| GRIN2B | 494 | W | 61.6713 | 36.57728357 | 0.545647881 | 33.72404611 | NA | NA | NA | NA | NA | NA | NA |
| GRIN2B | 495 | N | 60.955175 | 34.66415454 | 0.514885946 | 31.78307475 | NA | NA | NA | NA | NA | NA | NA |
| GRIN2B | 496 | G | 63.17105 | 32.93359517 | 0.487059618 | 30.06907842 | NA | NA | NA | NA | NA | NA | NA |
| GRIN2B | 497 | M | 64.1626 | 29.82904427 | 0.437140354 | 26.80778459 | NA | NA | NA | NA | NA | NA | NA |
| GRIN2B | 498 | I | 63.426425 | 32.0349322 | 0.472609672 | 29.02725201 | NA | NA | NA | NA | NA | NA | NA |
| GRIN2B | 499 | G | 65.96655 | 35.39848211 | 0.526693481 | 32.3805721 | NA | NA | NA | NA | NA | NA | NA |
| GRIN2B | 500 | E | 68.3584 | 33.39715537 | 0.494513381 | 25.57031445 | NA | NA | NA | NA | NA | NA | NA |
| GRIN2B | 501 | V | 68.73422857 | 31.57341477 | 0.465188756 | 23.74389174 | NA | NA | NA | NA | NA | NA | NA |
| GRIN2B | 502 | V | 68.49265714 | 35.45199437 | 0.527553925 | 27.62354328 | NA | NA | NA | NA | NA | NA | NA |
| GRIN2B | 503 | M | 71.6 | 37.02846167 | 0.552902547 | 29.18623341 | NA | NA | NA | NA | NA | NA | NA |
| GRIN2B | 504 | K | 74.1566 | 34.18746536 | 0.507221078 | 26.33435473 | NA | NA | NA | NA | NA | NA | NA |
| GRIN2B | 505 | R | 73.2644 | 31.75338801 | 0.468082615 | 23.90299082 | NA | NA | NA | NA | NA | NA | NA |
| GRIN2B | 506 | A | 71.8528 | 29.6290435 | 0.433924465 | 21.7904862 | NA | NA | NA | NA | NA | NA | NA |
| GRIN2B | 507 | Y | 73.9048 | 23.68540067 | 0.338354352 | 15.82457626 | NA | NA | NA | NA | NA | NA | NA |
| GRIN2B | 508 | M | 70.2312 | 25.0144586 | 0.359724784 | 17.17816034 | NA | NA | NA | NA | NA | NA | NA |

| | | | | | | | | | | | | | |
|---|---|---|---|---|---|---|---|---|---|---|---|---|---|
| GRIN2B | 509 | A | 66.4342 | 25.88815859 | 0.373773341 | 18.07582899 | NA | NA | NA | NA | NA | NA | NA |
| GRIN2B | 510 | V | 64.92637143 | 21.98466381 | 0.311007552 | 18.97160834 | NA | NA | NA | NA | NA | NA | NA |
| GRIN2B | 511 | G | 60.8478 | 22.81191358 | 0.324309218 | 19.80965795 | NA | NA | NA | NA | NA | NA | NA |
| GRIN2B | 512 | S | 57.09946667 | 23.28834329 | 0.331969914 | 20.43874871 | NA | NA | NA | NA | NA | NA | NA |
| GRIN2B | 513 | L | 59.8483 | 25.06521608 | 0.360540933 | 22.19198431 | NA | NA | NA | NA | NA | NA | NA |
| GRIN2B | 514 | T | 55.58522857 | 26.00276293 | 0.375616109 | 23.16350505 | NA | NA | NA | NA | NA | NA | NA |
| GRIN2B | 515 | I | 57.74155 | 26.91359989 | 0.390261805 | 24.01833894 | NA | NA | NA | NA | NA | NA | NA |
| GRIN2B | 516 | N | 56.748175 | 30.74165451 | 0.451814564 | 27.873794 | NA | NA | NA | NA | NA | NA | NA |
| GRIN2B | 517 | E | 57.5978 | 34.03371768 | 0.50474891 | 31.22553421 | NA | NA | NA | NA | NA | NA | NA |
| GRIN2B | 518 | E | 58.6224 | 36.11634727 | 0.53823631 | 33.24754255 | NA | NA | NA | NA | NA | NA | NA |
| GRIN2B | 519 | R | 60.6834 | 32.76601369 | 0.484365011 | 29.91518142 | NA | NA | NA | NA | NA | NA | NA |
| GRIN2B | 520 | S | 61.9968 | 32.60760864 | 0.481817956 | 29.77563852 | NA | NA | NA | NA | NA | NA | NA |
| GRIN2B | 521 | E | 63.2926 | 36.78362811 | 0.548965775 | 33.91586773 | NA | NA | NA | NA | NA | NA | NA |
| GRIN2B | 522 | V | 65.01051429 | 36.89868151 | 0.550815762 | 29.0767904 | NA | NA | NA | NA | NA | NA | NA |
| GRIN2B | 523 | V | 65.87094286 | 32.30735379 | 0.476990043 | 24.48452915 | NA | NA | NA | NA | NA | NA | NA |
| GRIN2B | 524 | D | 66.7568 | 30.48256592 | 0.447648579 | 22.66228718 | NA | NA | NA | NA | NA | NA | NA |
| GRIN2B | 525 | F | 64.7632 | 28.4742241 | 0.415355681 | 20.65860233 | NA | NA | NA | NA | NA | NA | NA |
| GRIN2B | 526 | S | 64.3823 | 24.65959923 | 0.354018864 | 16.84955939 | NA | NA | NA | NA | NA | NA | NA |
| GRIN2B | 527 | V | 63.18851429 | 22.80372972 | 0.324177627 | 19.77507203 | NA | NA | NA | NA | NA | NA | NA |
| GRIN2B | 528 | P | 59.63808571 | 20.99247434 | 0.295053757 | 18.1066666 | NA | NA | NA | NA | NA | NA | NA |
| GRIN2B | 529 | F | 60.05689091 | 16.62249274 | 0.224787147 | 13.85809867 | NA | NA | NA | NA | NA | NA | NA |
| GRIN2B | 530 | I | 55.55105 | 14.76471337 | 0.1949152 | 12.04144908 | NA | NA | NA | NA | NA | NA | NA |
| GRIN2B | 531 | E | 52.5204 | 17.69580575 | 0.242045358 | 13.86566643 | NA | NA | NA | NA | NA | NA | NA |
| GRIN2B | 532 | T | 51.4618 | 19.9398838 | 0.27812875 | 16.10530181 | NA | NA | NA | NA | NA | NA | NA |
| GRIN2B | 533 | G | 48.8983 | 22.06983638 | 0.312377074 | 17.31694749 | NA | NA | NA | NA | NA | NA | NA |
| GRIN2B | 534 | I | 45.8038 | 24.21471197 | 0.346865352 | 19.33623321 | NA | NA | NA | NA | NA | NA | NA |
| GRIN2B | 535 | S | 43.0743 | 21.04818306 | 0.295949519 | 16.18061517 | NA | NA | NA | NA | NA | NA | NA |
| GRIN2B | 536 | V | 39.26851429 | 22.45902569 | 0.318634998 | 17.62979147 | NA | NA | NA | NA | NA | NA | NA |
| GRIN2B | 537 | M | 39.142 | 21.23009863 | 0.298874609 | 16.41948691 | NA | NA | NA | NA | NA | NA | NA |
| GRIN2B | 538 | V | 35.03937143 | 22.91372877 | 0.325946344 | 17.77272568 | NA | NA | NA | NA | NA | NA | NA |
| GRIN2B | 539 | S | 33.6623 | 20.2392112 | 0.28294175 | 14.04339357 | NA | NA | NA | NA | NA | NA | NA |
| GRIN2B | 540 | R | 30.908 | 24.44013265 | 0.350489977 | 17.98962612 | NA | NA | NA | NA | NA | NA | NA |
| GRIN2B | 541 | S | 28.5263 | 22.44485478 | 0.318407139 | 16.26015667 | NA | NA | NA | NA | NA | NA | NA |

| | | | | | | | | | | | | | |
|---|---|---|---|---|---|---|---|---|---|---|---|---|---|
| GRIN2B | 542 | N | 26.222425 | 18.91961154 | 0.261723401 | 13.14069125 | NA | NA | NA | NA | NA | NA | NA |
| GRIN2B | 543 | G | 25.39205 | 17.86918907 | 0.244833255 | 16.5550965 | NA | NA | NA | NA | NA | NA | NA |
| GRIN2B | 544 | T | 22.8858 | 16.67763766 | 0.225673843 | 15.02672187 | NA | NA | NA | NA | NA | NA | NA |
| GRIN2B | 545 | V | 21.5656 | 14.00954949 | 0.18277263 | 12.30638208 | NA | NA | NA | NA | NA | NA | NA |
| GRIN2B | 546 | S | 19.93513333 | 16.55177163 | 0.223649995 | 14.88018197 | NA | NA | NA | NA | NA | NA | NA |
| GRIN2B | 547 | P | 16.56508571 | 18.2286988 | 0.25061395 | 16.05619583 | NA | NA | NA | NA | NA | NA | NA |
| GRIN2B | 548 | S | 16.1298 | 15.28582 | 0.203294273 | 13.19958982 | NA | NA | NA | NA | NA | NA | NA |
| GRIN2B | 549 | A | 13.2118 | 12.88153997 | 0.164634932 | 10.2644796 | NA | NA | NA | NA | NA | NA | NA |
| GRIN2B | 550 | F | 10.91907273 | 14.64687987 | 0.19302051 | 11.9838604 | NA | NA | NA | NA | NA | NA | NA |
| GRIN2B | 551 | L | 12.2418 | 16.16882126 | 0.217492389 | 13.55153873 | NA | NA | NA | NA | NA | NA | NA |
| GRIN2B | 552 | E | 14.7448 | 13.33178049 | 0.171874522 | 11.40366243 | NA | NA | NA | NA | NA | NA | NA |
| GRIN2B | 553 | P | 18.0488 | 16.9733036 | 0.230427969 | 14.94737206 | NA | NA | NA | NA | NA | NA | NA |
| GRIN2B | 554 | F | 16.7618 | 15.0965604 | 0.200251095 | 12.98558442 | NA | NA | NA | NA | NA | NA | NA |
| GRIN2B | 555 | S | 13.6808 | 15.12360989 | 0.200686034 | 12.4275221 | NA | NA | NA | NA | NA | NA | NA |
| GRIN2B | 556 | A | 12.6872 | 18.55834438 | 0.255914448 | 15.85297194 | NA | NA | NA | NA | NA | NA | NA |
| GRIN2B | 557 | D | 11.5398 | 20.37335782 | 0.285098745 | 17.74162504 | NA | NA | NA | NA | NA | NA | NA |
| GRIN2B | 558 | V | 9.737942857 | 16.028107 | 0.21522979 | 13.42610219 | NA | NA | NA | NA | NA | NA | NA |
| GRIN2B | 559 | W | 5.977728571 | 14.2894798 | 0.187273737 | 11.66507388 | NA | NA | NA | NA | NA | NA | NA |
| GRIN2B | 560 | V | 5.868657143 | 20.60445297 | 0.288814612 | 17.99610051 | NA | NA | NA | NA | NA | NA | NA |
| GRIN2B | 561 | M | 5.4114 | 20.99284026 | 0.29505964 | 18.35332313 | NA | NA | NA | NA | NA | NA | NA |
| GRIN2B | 562 | M | 3.3654 | 17.52881642 | 0.239360273 | 14.90439517 | NA | NA | NA | NA | NA | NA | NA |
| GRIN2B | 563 | F | 1.2466 | 20.21621693 | 0.282572016 | 17.58937755 | NA | NA | NA | NA | NA | NA | NA |
| GRIN2B | 564 | V | 1.0958 | 22.98321303 | 0.327063608 | 20.35679728 | NA | NA | NA | NA | NA | NA | NA |
| GRIN2B | 565 | M | 0.832 | 21.74184928 | 0.307103244 | 19.11675064 | NA | NA | NA | NA | NA | NA | NA |
| GRIN2B | 566 | L | -1.51055 | 18.56714146 | 0.256055899 | 15.975412 | NA | NA | NA | NA | NA | NA | NA |
| GRIN2B | 567 | L | -3.6494 | 23.85072169 | 0.341012612 | 21.19150997 | NA | NA | NA | NA | NA | NA | NA |
| GRIN2B | 568 | I | -3.6736 | 25.95626032 | 0.374868375 | 23.33442738 | NA | NA | NA | NA | NA | NA | NA |
| GRIN2B | 569 | V | -4.528514286 | 22.30684884 | 0.316188088 | 19.62743468 | NA | NA | NA | NA | NA | NA | NA |
| GRIN2B | 570 | S | -7.7054 | 24.68828666 | 0.35448014 | 22.00217323 | NA | NA | NA | NA | NA | NA | NA |
| GRIN2B | 571 | A | -8.6598 | 27.38021837 | 0.397764743 | 24.69444488 | NA | NA | NA | NA | NA | NA | NA |
| GRIN2B | 572 | V | -8.3318 | 28.55180495 | 0.416603133 | 25.86023256 | NA | NA | NA | NA | NA | NA | NA |
| GRIN2B | 573 | A | -10.9896 | 25.95433615 | 0.374837436 | 24.55769168 | NA | NA | NA | NA | NA | NA | NA |
| GRIN2B | 574 | V | -13.2668 | 28.65795607 | 0.418309977 | 27.27195986 | NA | NA | NA | NA | NA | NA | NA |

| | | | | | | | | | | | | | |
|---|---|---|---|---|---|---|---|---|---|---|---|---|---|
| GRIN2B | 575 | F | -13.5344 | 31.4007315 | 0.462412116 | 30.00370036 | NA | NA | NA | NA | NA | NA | NA |
| GRIN2B | 576 | V | -13.63322857 | 29.67074482 | 0.434594996 | 28.27711536 | NA | NA | NA | NA | NA | NA | NA |
| GRIN2B | 577 | F | -16.8172 | 30.21004244 | 0.443266569 | 28.94003434 | NA | NA | NA | NA | NA | NA | NA |
| GRIN2B | 578 | E | -18.5442 | 33.23257754 | 0.491867071 | 31.9372403 | NA | NA | NA | NA | NA | NA | NA |
| GRIN2B | 579 | Y | -18.3806 | 35.33396646 | 0.525656109 | 34.0339369 | NA | NA | NA | NA | NA | NA | NA |
| GRIN2B | 600 | F | -17.5618 | 27.4777161 | 0.399332446 | 26.32290983 | NA | NA | NA | NA | NA | NA | NA |
| GRIN2B | 601 | T | -16.2178 | 26.79518162 | 0.388357713 | 25.55118117 | NA | NA | NA | NA | NA | NA | NA |
| GRIN2B | 602 | I | -12.806 | 25.54591658 | 0.368270301 | 24.24437492 | NA | NA | NA | NA | NA | NA | NA |
| GRIN2B | 603 | G | -12.46555 | 23.25053733 | 0.331362018 | 22.03520698 | NA | NA | NA | NA | NA | NA | NA |
| GRIN2B | 604 | K | -12.9986 | 21.92233158 | 0.310005288 | 20.71407587 | NA | NA | NA | NA | NA | NA | NA |
| GRIN2B | 605 | A | -10.2776 | 21.45934633 | 0.302560771 | 20.11472921 | NA | NA | NA | NA | NA | NA | NA |
| GRIN2B | 606 | I | -8.134 | 19.66515095 | 0.273711215 | 18.31463208 | NA | NA | NA | NA | NA | NA | NA |
| GRIN2B | 607 | W | -9.331157143 | 16.86434716 | 0.228676017 | 15.8412965 | NA | NA | NA | NA | NA | NA | NA |
| GRIN2B | 608 | L | -8.49855 | 16.76689217 | 0.227109 | 15.43834437 | NA | NA | NA | NA | NA | NA | NA |
| GRIN2B | 609 | L | -4.8788 | 16.08523354 | 0.21614835 | 14.74167025 | NA | NA | NA | NA | NA | NA | NA |
| GRIN2B | 610 | W | -2.983442857 | 13.7191803 | 0.178103672 | 12.32055829 | NA | NA | NA | NA | NA | NA | NA |
| GRIN2B | 611 | G | -4.31605 | 11.85897491 | 0.148192717 | 10.5344391 | NA | NA | NA | NA | NA | NA | NA |
| GRIN2B | 612 | L | -2.749 | 12.21716223 | 0.153952148 | 10.93682379 | NA | NA | NA | NA | NA | NA | NA |
| GRIN2B | 613 | V | -0.4584 | 10.7082739 | 0.129690154 | 8.043809942 | NA | NA | NA | NA | NA | NA | NA |
| GRIN2B | 614 | F | 0 | 8.376862539 | 0.092202496 | 5.702204626 | 2.71 | 0.5 | 1.5 | 1.21 | -0.85 | 0 | 0 |
| GRIN2B | 615 | N | -0.921675 | 5.642966773 | 0.048243138 | 4.249465672 | 1.4 | 0.18 | 0.35 | 1.76 | -0.38 | 0 | 0 |
| GRIN2B | 616 | N | -2.57505 | 8.31516446 | 0.091210429 | 7.363944584 | 1.4 | 0.18 | 0.35 | 1.76 | -0.38 | 0 | 0 |
| GRIN2B | 617 | S | -5.8148 | 9.904391046 | 0.116764213 | 8.890507052 | NA | NA | NA | NA | NA | NA | NA |
| GRIN2B | 618 | V | -8.4942 | 11.7791175 | 0.146908659 | 10.86512666 | NA | NA | NA | NA | NA | NA | NA |
| GRIN2B | 619 | P | -9.116514286 | 13.99421348 | 0.182526036 | 12.8925746 | NA | NA | NA | NA | NA | NA | NA |
| GRIN2B | 620 | V | -12.9574 | 18.07659949 | 0.248168287 | 17.06225972 | NA | NA | NA | NA | NA | NA | NA |
| GRIN2B | 621 | Q | -14.1088 | 20.37183826 | 0.285074311 | 19.32674779 | NA | NA | NA | NA | NA | NA | NA |
| GRIN2B | 622 | N | -14.9906 | 22.65626723 | 0.321806521 | 21.52832251 | NA | NA | NA | NA | NA | NA | NA |
| GRIN2B | 623 | P | -14.58237143 | 23.86110167 | 0.341179516 | 22.64434861 | NA | NA | NA | NA | NA | NA | NA |
| GRIN2B | 624 | K | -18.1516 | 27.50118723 | 0.399709848 | 26.34955893 | NA | NA | NA | NA | NA | NA | NA |
| GRIN2B | 625 | G | -16.21955 | 26.87053786 | 0.389569395 | 25.65412605 | NA | NA | NA | NA | NA | NA | NA |
| GRIN2B | 626 | T | -13.6816 | 27.19415866 | 0.394773017 | 25.85445147 | NA | NA | NA | NA | NA | NA | NA |
| GRIN2B | 627 | T | -12.3558 | 28.31734629 | 0.412833182 | 26.92333497 | NA | NA | NA | NA | NA | NA | NA |

| | | | | | | | | | | | | | |
|---|---|---|---|---|---|---|---|---|---|---|---|---|---|
| GRIN2B | 628 | S | -11.5078 | 25.36512844 | 0.365363339 | 23.96874844 | NA | NA | NA | NA | NA | NA | NA |
| GRIN2B | 629 | K | -9.3658 | 22.57607092 | 0.320517013 | 21.18220724 | NA | NA | NA | NA | NA | NA | NA |
| GRIN2B | 630 | I | -6.4408 | 22.90429078 | 0.325794587 | 20.20723378 | NA | NA | NA | NA | NA | NA | NA |
| GRIN2B | 631 | M | -5.6008 | 21.89192399 | 0.309516353 | 19.2005121 | NA | NA | NA | NA | NA | NA | NA |
| GRIN2B | 632 | V | -6.502657143 | 18.82255535 | 0.260162797 | 16.11725492 | NA | NA | NA | NA | NA | NA | NA |
| GRIN2B | 633 | S | -3.967133333 | 17.99635013 | 0.246877927 | 15.3074179 | NA | NA | NA | NA | NA | NA | NA |
| GRIN2B | 634 | V | -1.075085714 | 18.73630551 | 0.258775953 | 16.18856475 | NA | NA | NA | NA | NA | NA | NA |
| GRIN2B | 635 | W | -0.994014286 | 16.89182376 | 0.229117823 | 14.23645527 | NA | NA | NA | NA | NA | NA | NA |
| GRIN2B | 636 | A | -1.0558 | 13.9082789 | 0.181144261 | 11.24655067 | NA | NA | NA | NA | NA | NA | NA |
| GRIN2B | 637 | F | 2.507254545 | 14.19151542 | 0.18569853 | 12.16014958 | NA | NA | NA | NA | NA | NA | NA |
| GRIN2B | 638 | F | 3.067 | 14.22677058 | 0.186265411 | 11.6106941 | NA | NA | NA | NA | NA | NA | NA |
| GRIN2B | 639 | A | 2.5544 | 11.43855013 | 0.141432545 | 8.814530217 | NA | NA | NA | NA | NA | NA | NA |
| GRIN2B | 640 | V | 3.6678 | 9.472066564 | 0.109812702 | 6.862158022 | NA | NA | NA | NA | NA | NA | NA |
| GRIN2B | 641 | I | 7.05155 | 10.11956723 | 0.120224113 | 8.261869298 | NA | NA | NA | NA | NA | NA | NA |
| GRIN2B | 642 | F | 7.3368 | 10.15641083 | 0.120816536 | 7.73406102 | NA | NA | NA | NA | NA | NA | NA |
| GRIN2B | 643 | L | 7.2008 | 6.752525231 | 0.066084154 | 4.334951381 | 2.65 | -0.3 | 0.53 | 1.28 | 0.25 | 0 | 0 |
| GRIN2B | 644 | A | 9.7732 | 5.473432378 | 0.045517129 | 4.132569305 | 2.41 | -0.77 | -0.7 | 1.66 | 0.77 | 0 | 0 |
| GRIN2B | 645 | S | 11.70646667 | 7.648128268 | 0.080484898 | 4.972510465 | 2.75 | -0.38 | 0.63 | 1.26 | 0.69 | 0 | 0 |
| GRIN2B | 646 | Y | 10.83913333 | 7.153032993 | 0.072524072 | 4.542374421 | 2.75 | -0.38 | 0.63 | 1.26 | 0.69 | 0 | 0 |
| GRIN2B | 647 | T | 11.82294286 | 3.950099999 | 0.021022884 | 1.587011347 | 2.75 | -0.38 | 0.63 | 1.26 | 0.69 | 0 | 0 |
| GRIN2B | 648 | A | 15.4086 | 4.436554294 | 0.028844769 | 2.288161241 | 2.19 | -0.77 | -0.7 | 1.66 | 0.77 | 0 | 0 |
| GRIN2B | 649 | N | 16.549425 | 6.671812198 | 0.064786338 | 4.686037885 | 2.12 | -0.31 | 0.92 | 1.35 | 0.68 | 0 | 0 |
| GRIN2B | 650 | L | 15.475925 | 6.681437196 | 0.064941102 | 4.581845958 | 2.12 | -0.31 | 0.92 | 1.35 | 0.68 | 0 | 0 |
| GRIN2B | 651 | A | 18.0662 | 3.004554043 | 0.005819088 | 1.074975746 | 2.25 | 0.02 | 1.8 | 0 | 0.59 | 0 | 0 |
| GRIN2B | 652 | A | 20.697 | 5.143919906 | 0.040218772 | 2.476792703 | 2.75 | 0.3 | 2.4 | 0.03 | 0.12 | 0 | 0 |
| GRIN2B | 653 | F | 20.69452727 | 7.521043611 | 0.078441455 | 5.773088905 | 1.76 | 1.58 | 4.4 | 0.13 | -1.67 | 0 | 0 |
| GRIN2B | 654 | M | 20.8754 | 7.330577944 | 0.075378885 | 4.574907348 | 2.81 | 0.27 | 1.94 | 1.4 | -0.19 | 0 | 0 |
| GRIN2B | 655 | I | 24.327675 | 3.268078487 | 0.010056398 | 1.878855331 | 1.49 | 1.58 | 4.4 | 0.13 | -1.67 | 0 | 0 |
| GRIN2B | 656 | Q | 25.4466 | 7.590240905 | 0.079554105 | 6.253152078 | 1.49 | 1.58 | 4.4 | 0.13 | -1.67 | 0 | 0 |
| GRIN2B | 657 | E | 28.8656 | 9.688934152 | 0.113299799 | 3.509302639 | 6.21 | 0.3 | 1.4 | 2.08 | -0.24 | 0 | 0 |
| GRIN2B | 658 | E | 29.077 | 12.61756213 | 0.160390331 | 6.448811177 | NA | NA | NA | NA | NA | NA | NA |
| GRIN2B | 659 | Y | 29.2646 | 14.926102 | 0.197510229 | 8.753907999 | NA | NA | NA | NA | NA | NA | NA |
| GRIN2B | 660 | V | 27.8062 | 17.82860048 | 0.244180616 | 11.65907321 | NA | NA | NA | NA | NA | NA | NA |

| | | | | | | | | | | | | | |
|---|---|---|---|---|---|---|---|---|---|---|---|---|---|
| GRIN2B | 661 | D | 29.416 | 20.4870637 | 0.286927065 | 14.31967413 | NA | NA | NA | NA | NA | NA | NA |
| GRIN2B | 662 | Q | 28.915 | 23.98676164 | 0.343200051 | 17.81551123 | NA | NA | NA | NA | NA | NA | NA |
| GRIN2B | 663 | V | 32.36622857 | 22.91865144 | 0.326025497 | 16.36257444 | NA | NA | NA | NA | NA | NA | NA |
| GRIN2B | 664 | S | 33.15963333 | 26.71547158 | 0.387076024 | 20.16577133 | NA | NA | NA | NA | NA | NA | NA |
| GRIN2B | 665 | G | 36.08455 | 27.45859931 | 0.39902506 | 22.76429787 | NA | NA | NA | NA | NA | NA | NA |
| GRIN2B | 666 | L | 38.98455 | 29.06619826 | 0.42487426 | 24.20367719 | NA | NA | NA | NA | NA | NA | NA |
| GRIN2B | 667 | S | 35.76146667 | 30.73560471 | 0.451717287 | 25.87338945 | NA | NA | NA | NA | NA | NA | NA |
| GRIN2B | 668 | D | 32.2493 | 31.14917458 | 0.458367235 | 24.66485182 | NA | NA | NA | NA | NA | NA | NA |
| GRIN2B | 669 | K | 31.3816 | 35.09576833 | 0.52182603 | 28.56906526 | NA | NA | NA | NA | NA | NA | NA |
| GRIN2B | 670 | K | 31.2566 | 34.08965233 | 0.505648305 | 27.53520431 | NA | NA | NA | NA | NA | NA | NA |
| GRIN2B | 671 | F | 34.61861818 | 28.91349489 | 0.422418884 | 24.49266061 | NA | NA | NA | NA | NA | NA | NA |
| GRIN2B | 672 | Q | 35.404 | 35.21010392 | 0.523664476 | 30.36276603 | NA | NA | NA | NA | NA | NA | NA |
| GRIN2B | 673 | R | 32.4496 | 37.93381295 | 0.567460038 | 33.09058542 | NA | NA | NA | NA | NA | NA | NA |
| GRIN2B | 674 | P | 32.20751429 | 37.32380341 | 0.55765146 | 32.01705553 | NA | NA | NA | NA | NA | NA | NA |
| GRIN2B | 675 | N | 29.7578 | 40.36711802 | 0.606586083 | 35.0822087 | NA | NA | NA | NA | NA | NA | NA |
| GRIN2B | 676 | D | 27.7406 | 41.38200721 | 0.622904876 | 34.93688198 | NA | NA | NA | NA | NA | NA | NA |
| GRIN2B | 677 | F | 26.848 | 37.73962578 | 0.564337628 | 31.65949711 | NA | NA | NA | NA | NA | NA | NA |
| GRIN2B | 678 | S | 27.03696667 | 36.44618617 | 0.543539916 | 30.45973983 | NA | NA | NA | NA | NA | NA | NA |
| GRIN2B | 679 | P | 27.30522857 | 39.08046265 | 0.585897458 | 33.22805922 | NA | NA | NA | NA | NA | NA | NA |
| GRIN2B | 680 | P | 27.47665714 | 35.27382324 | 0.524689043 | 29.09125389 | NA | NA | NA | NA | NA | NA | NA |
| GRIN2B | 681 | F | 31.5676 | 35.21779553 | 0.523788152 | 30.01343202 | NA | NA | NA | NA | NA | NA | NA |
| GRIN2B | 682 | R | 35.2164 | 32.433485 | 0.479018155 | 27.62606177 | NA | NA | NA | NA | NA | NA | NA |
| GRIN2B | 683 | F | 38.3168 | 30.74460741 | 0.451862045 | 25.91757927 | NA | NA | NA | NA | NA | NA | NA |
| GRIN2B | 684 | G | 41.02005 | 30.34061446 | 0.445366087 | 25.51215876 | NA | NA | NA | NA | NA | NA | NA |
| GRIN2B | 685 | T | 43.68122857 | 31.46471519 | 0.463440934 | 26.61547533 | NA | NA | NA | NA | NA | NA | NA |
| GRIN2B | 686 | V | 47.18594286 | 29.75931184 | 0.436019099 | 25.84000017 | NA | NA | NA | NA | NA | NA | NA |
| GRIN2B | 687 | P | 48.76508571 | 32.68935194 | 0.483132338 | 28.93458034 | NA | NA | NA | NA | NA | NA | NA |
| GRIN2B | 688 | N | 53.1838 | 31.80919969 | 0.468980032 | 28.93684204 | NA | NA | NA | NA | NA | NA | NA |
| GRIN2B | 689 | G | 52.8113 | 31.36301253 | 0.461805618 | 28.48899691 | NA | NA | NA | NA | NA | NA | NA |
| GRIN2B | 690 | S | 53.40013333 | 30.02091706 | 0.44022555 | 27.15981512 | NA | NA | NA | NA | NA | NA | NA |
| GRIN2B | 691 | T | 49.48165714 | 27.2869631 | 0.396265256 | 23.4122317 | NA | NA | NA | NA | NA | NA | NA |
| GRIN2B | 692 | E | 48.6962 | 31.65808044 | 0.466550128 | 27.77760304 | NA | NA | NA | NA | NA | NA | NA |
| GRIN2B | 693 | R | 50.5026 | 34.11901797 | 0.506120486 | 30.24455692 | NA | NA | NA | NA | NA | NA | NA |

| | | | | | | | | | | | | | |
|---|---|---|---|---|---|---|---|---|---|---|---|---|---|
| GRIN2B | 694 | N | 48.8353 | 31.57616813 | 0.465233028 | 27.84293219 | NA | NA | NA | NA | NA | NA | NA |
| GRIN2B | 695 | I | 45.15755 | 31.48393008 | 0.463749897 | 26.61526847 | NA | NA | NA | NA | NA | NA | NA |
| GRIN2B | 696 | R | 46.419 | 36.09522092 | 0.537896611 | 31.2565568 | NA | NA | NA | NA | NA | NA | NA |
| GRIN2B | 697 | N | 47.5238 | 37.76003434 | 0.564665785 | 32.9408833 | NA | NA | NA | NA | NA | NA | NA |
| GRIN2B | 698 | N | 44.6466 | 35.49013893 | 0.528167266 | 30.63196059 | NA | NA | NA | NA | NA | NA | NA |
| GRIN2B | 699 | Y | 41.22688333 | 35.36676492 | 0.526183488 | 30.47955484 | NA | NA | NA | NA | NA | NA | NA |
| GRIN2B | 700 | A | 41.8346 | 40.48199201 | 0.608433186 | 35.60678306 | NA | NA | NA | NA | NA | NA | NA |
| GRIN2B | 701 | E | 38.6496 | 39.94409307 | 0.599784103 | 35.07561937 | NA | NA | NA | NA | NA | NA | NA |
| GRIN2B | 702 | M | 39.918 | 36.36958441 | 0.542308207 | 31.49564257 | NA | NA | NA | NA | NA | NA | NA |
| GRIN2B | 703 | H | 42.4018 | 38.00447708 | 0.568596273 | 33.12885087 | NA | NA | NA | NA | NA | NA | NA |
| GRIN2B | 704 | A | 40.0594 | 40.21978166 | 0.604217005 | 35.35207451 | NA | NA | NA | NA | NA | NA | NA |
| GRIN2B | 705 | Y | 38.0356 | 38.36831044 | 0.57444649 | 33.5280784 | NA | NA | NA | NA | NA | NA | NA |
| GRIN2B | 706 | M | 40.8472 | 35.42992269 | 0.527199026 | 30.57885611 | NA | NA | NA | NA | NA | NA | NA |
| GRIN2B | 707 | G | 41.81155 | 38.39438344 | 0.574865727 | 33.54842524 | NA | NA | NA | NA | NA | NA | NA |
| GRIN2B | 708 | K | 39.1546 | 38.90921139 | 0.583143843 | 34.09451087 | NA | NA | NA | NA | NA | NA | NA |
| GRIN2B | 709 | F | 38.12516364 | 34.43022373 | 0.511124483 | 29.61349437 | NA | NA | NA | NA | NA | NA | NA |
| GRIN2B | 710 | N | 43.8968 | 35.08976448 | 0.521729492 | 31.22538261 | NA | NA | NA | NA | NA | NA | NA |
| GRIN2B | 711 | Q | 45.1022 | 31.68021334 | 0.466906011 | 27.84008223 | NA | NA | NA | NA | NA | NA | NA |
| GRIN2B | 712 | R | 47.5398 | 30.80667723 | 0.452860089 | 27.95021714 | NA | NA | NA | NA | NA | NA | NA |
| GRIN2B | 713 | G | 49.12355 | 28.14969375 | 0.410137433 | 25.28345164 | NA | NA | NA | NA | NA | NA | NA |
| GRIN2B | 714 | V | 48.80965714 | 23.43420585 | 0.334315294 | 20.58846681 | NA | NA | NA | NA | NA | NA | NA |
| GRIN2B | 715 | D | 47.133 | 25.37133363 | 0.365463115 | 22.50718222 | NA | NA | NA | NA | NA | NA | NA |
| GRIN2B | 716 | D | 45.0796 | 27.81384141 | 0.404737134 | 24.02699285 | NA | NA | NA | NA | NA | NA | NA |
| GRIN2B | 717 | A | 43.5104 | 27.08299915 | 0.392985641 | 23.34571584 | NA | NA | NA | NA | NA | NA | NA |
| GRIN2B | 718 | L | 42.6966 | 23.73526722 | 0.339156175 | 18.97229375 | NA | NA | NA | NA | NA | NA | NA |
| GRIN2B | 719 | L | 41.1156 | 26.32449773 | 0.380789406 | 21.57686482 | NA | NA | NA | NA | NA | NA | NA |
| GRIN2B | 720 | S | 39.04713333 | 28.70818291 | 0.419117594 | 23.95984574 | NA | NA | NA | NA | NA | NA | NA |
| GRIN2B | 721 | L | 37.896 | 26.19148635 | 0.378650665 | 20.94817247 | NA | NA | NA | NA | NA | NA | NA |
| GRIN2B | 722 | K | 36.9632 | 24.94449208 | 0.358599766 | 19.63013954 | NA | NA | NA | NA | NA | NA | NA |
| GRIN2B | 723 | T | 35.60137143 | 27.80347586 | 0.404570462 | 22.55125698 | NA | NA | NA | NA | NA | NA | NA |
| GRIN2B | 724 | G | 33.65555 | 29.10969661 | 0.425573687 | 23.68263573 | NA | NA | NA | NA | NA | NA | NA |
| GRIN2B | 725 | K | 35.1788 | 31.45978032 | 0.463361584 | 26.22110905 | NA | NA | NA | NA | NA | NA | NA |
| GRIN2B | 726 | L | 37.747925 | 28.90572585 | 0.422293963 | 24.11474648 | NA | NA | NA | NA | NA | NA | NA |

| | | | | | | | | | | | | | |
|---|---|---|---|---|---|---|---|---|---|---|---|---|---|
| GRIN2B | 727 | D | 35.033 | 28.04942798 | 0.408525221 | 22.78469407 | NA | NA | NA | NA | NA | NA | NA |
| GRIN2B | 728 | A | 37.907 | 27.65552202 | 0.402191456 | 22.84756797 | NA | NA | NA | NA | NA | NA | NA |
| GRIN2B | 729 | F | 42.97061818 | 24.50070991 | 0.351464022 | 19.68797083 | NA | NA | NA | NA | NA | NA | NA |
| GRIN2B | 730 | I | 42.657675 | 26.53547539 | 0.384181796 | 21.66961039 | NA | NA | NA | NA | NA | NA | NA |
| GRIN2B | 731 | Y | 48.43055 | 24.20127585 | 0.346649307 | 19.3480279 | NA | NA | NA | NA | NA | NA | NA |
| GRIN2B | 732 | D | 48.6426 | 21.22702831 | 0.29882524 | 17.37821832 | NA | NA | NA | NA | NA | NA | NA |
| GRIN2B | 733 | A | 47.719 | 17.92447031 | 0.245722143 | 13.08531658 | NA | NA | NA | NA | NA | NA | NA |
| GRIN2B | 734 | A | 50.0468 | 16.66580787 | 0.225483627 | 12.77636242 | NA | NA | NA | NA | NA | NA | NA |
| GRIN2B | 735 | V | 49.28965714 | 18.6063652 | 0.256686593 | 14.6838994 | NA | NA | NA | NA | NA | NA | NA |
| GRIN2B | 736 | L | 45.014425 | 17.37720921 | 0.236922522 | 12.55738069 | NA | NA | NA | NA | NA | NA | NA |
| GRIN2B | 737 | N | 46.3252 | 14.10220458 | 0.184262467 | 9.285637004 | NA | NA | NA | NA | NA | NA | NA |
| GRIN2B | 738 | Y | 49.85155 | 13.54551236 | 0.175311199 | 10.28622823 | NA | NA | NA | NA | NA | NA | NA |
| GRIN2B | 739 | M | 45.2994 | 16.91558695 | 0.229499921 | 13.01652962 | NA | NA | NA | NA | NA | NA | NA |
| GRIN2B | 740 | A | 42.9706 | 14.53349431 | 0.19119734 | 9.789809155 | NA | NA | NA | NA | NA | NA | NA |
| GRIN2B | 741 | G | 45.0853 | 12.37871019 | 0.156549739 | 7.654744806 | NA | NA | NA | NA | NA | NA | NA |
| GRIN2B | 742 | R | 44.9616 | 14.45946354 | 0.190006971 | 10.74923684 | NA | NA | NA | NA | NA | NA | NA |
| GRIN2B | 743 | D | 43.0228 | 16.46182472 | 0.222203704 | 11.76219023 | NA | NA | NA | NA | NA | NA | NA |
| GRIN2B | 744 | E | 41.7118 | 18.72396542 | 0.258577532 | 13.50235821 | NA | NA | NA | NA | NA | NA | NA |
| GRIN2B | 745 | G | 40.1313 | 16.248467 | 0.218773043 | 10.97266233 | NA | NA | NA | NA | NA | NA | NA |
| GRIN2B | 746 | C | 37.5413 | 15.20510204 | 0.201996378 | 9.753588281 | NA | NA | NA | NA | NA | NA | NA |
| GRIN2B | 747 | K | 35.4462 | 18.16599777 | 0.249605756 | 12.74396429 | NA | NA | NA | NA | NA | NA | NA |
| GRIN2B | 748 | L | 38.366675 | 18.86761039 | 0.260887254 | 13.53242196 | NA | NA | NA | NA | NA | NA | NA |
| GRIN2B | 749 | V | 37.7878 | 15.41506516 | 0.205372456 | 10.27936587 | NA | NA | NA | NA | NA | NA | NA |
| GRIN2B | 750 | T | 40.1678 | 16.38386191 | 0.22095011 | 11.55778688 | NA | NA | NA | NA | NA | NA | NA |
| GRIN2B | 751 | I | 38.178175 | 18.7192007 | 0.258500918 | 13.38821202 | NA | NA | NA | NA | NA | NA | NA |
| GRIN2B | 752 | G | 39.2043 | 16.40870677 | 0.2213496 | 11.03544379 | NA | NA | NA | NA | NA | NA | NA |
| GRIN2B | 753 | S | 40.5188 | 18.1562329 | 0.249448743 | 13.30737717 | NA | NA | NA | NA | NA | NA | NA |
| GRIN2B | 754 | G | 43.77955 | 17.50937423 | 0.239047654 | 12.60429036 | NA | NA | NA | NA | NA | NA | NA |
| GRIN2B | 755 | K | 43.2482 | 19.90090345 | 0.27750197 | 15.00003199 | NA | NA | NA | NA | NA | NA | NA |
| GRIN2B | 756 | V | 45.24108571 | 20.55032744 | 0.287944307 | 15.67091585 | NA | NA | NA | NA | NA | NA | NA |
| GRIN2B | 757 | F | 43.06443636 | 24.91838735 | 0.358180018 | 20.04279247 | NA | NA | NA | NA | NA | NA | NA |
| GRIN2B | 758 | A | 48.6692 | 25.68707274 | 0.370540005 | 20.91824149 | NA | NA | NA | NA | NA | NA | NA |
| GRIN2B | 759 | S | 50.2263 | 23.78231519 | 0.339912678 | 18.98861535 | NA | NA | NA | NA | NA | NA | NA |

| | | | | | | | | | | | | | |
|---|---|---|---|---|---|---|---|---|---|---|---|---|---|
| GRIN2B | 760 | T | 53.3588 | 22.15119667 | 0.313685298 | 18.32239683 | NA | NA | NA | NA | NA | NA | NA |
| GRIN2B | 761 | G | 56.33005 | 22.4724949 | 0.318851575 | 19.62601596 | NA | NA | NA | NA | NA | NA | NA |
| GRIN2B | 762 | Y | 56.59355 | 19.82010179 | 0.276202729 | 17.1914402 | NA | NA | NA | NA | NA | NA | NA |
| GRIN2B | 763 | G | 61.59505 | 23.83635455 | 0.340781597 | 20.95624157 | NA | NA | NA | NA | NA | NA | NA |
| GRIN2B | 764 | I | 65.0648 | 23.88170582 | 0.341510818 | 16.06874748 | NA | NA | NA | NA | NA | NA | NA |
| GRIN2B | 765 | A | 66.2962 | 27.23908446 | 0.395495397 | 19.42585652 | NA | NA | NA | NA | NA | NA | NA |
| GRIN2B | 766 | I | 69.30905 | 26.3126649 | 0.380599141 | 18.47962374 | NA | NA | NA | NA | NA | NA | NA |
| GRIN2B | 767 | Q | 71.6102 | 30.74158971 | 0.451813522 | 22.90446199 | NA | NA | NA | NA | NA | NA | NA |
| GRIN2B | 768 | K | 73.1256 | 31.15196793 | 0.458412151 | 23.30789531 | NA | NA | NA | NA | NA | NA | NA |
| GRIN2B | 769 | D | 76.3768 | 32.64187179 | 0.482368886 | 23.81417294 | NA | NA | NA | NA | NA | NA | NA |
| GRIN2B | 770 | S | 78.44513333 | 29.49710865 | 0.431803034 | 20.51958823 | NA | NA | NA | NA | NA | NA | NA |
| GRIN2B | 771 | G | 76.21705 | 27.67382057 | 0.402485685 | 18.70420444 | NA | NA | NA | NA | NA | NA | NA |
| GRIN2B | 772 | W | 76.08658571 | 21.79741349 | 0.307996682 | 12.80242469 | NA | NA | NA | NA | NA | NA | NA |
| GRIN2B | 773 | K | 71.6552 | 24.83797435 | 0.356887026 | 16.99608909 | NA | NA | NA | NA | NA | NA | NA |
| GRIN2B | 774 | R | 72.6484 | 23.39860034 | 0.33374278 | 15.54696252 | NA | NA | NA | NA | NA | NA | NA |
| GRIN2B | 775 | Q | 73.6672 | 20.34533435 | 0.284648145 | 11.75646544 | NA | NA | NA | NA | NA | NA | NA |
| GRIN2B | 776 | V | 70.52494286 | 20.11252908 | 0.280904779 | 12.28029289 | NA | NA | NA | NA | NA | NA | NA |
| GRIN2B | 777 | D | 68.3148 | 20.97842251 | 0.294827812 | 13.14906039 | NA | NA | NA | NA | NA | NA | NA |
| GRIN2B | 778 | L | 70.696425 | 18.51440545 | 0.255207937 | 10.67278004 | NA | NA | NA | NA | NA | NA | NA |
| GRIN2B | 779 | A | 69.888 | 15.96888055 | 0.214277466 | 8.140063532 | NA | NA | NA | NA | NA | NA | NA |
| GRIN2B | 780 | I | 66.104675 | 17.49459336 | 0.238809987 | 9.684746534 | NA | NA | NA | NA | NA | NA | NA |
| GRIN2B | 781 | L | 65.557675 | 17.75427526 | 0.242985512 | 9.934797117 | NA | NA | NA | NA | NA | NA | NA |
| GRIN2B | 782 | Q | 68.66613333 | 14.23141557 | 0.1863401 | 6.376216904 | NA | NA | NA | NA | NA | NA | NA |
| GRIN2B | 783 | L | 66.392175 | 11.80662954 | 0.147351035 | 3.989095283 | 7.9 | -0.39 | -1.68 | 37.36 | 1.16 | 3 | -1 |
| GRIN2B | 784 | F | 60.84343636 | 15.38747055 | 0.204928751 | 10.64768241 | NA | NA | NA | NA | NA | NA | NA |
| GRIN2B | 785 | G | 63.35305 | 15.29018365 | 0.203364438 | 10.23222018 | NA | NA | NA | NA | NA | NA | NA |
| GRIN2B | 786 | D | 63.8312 | 11.84281609 | 0.147932893 | 6.901943527 | NA | NA | NA | NA | NA | NA | NA |
| GRIN2B | 787 | G | 61.8198 | 9.275317191 | 0.106649093 | 4.602067531 | 4.94 | -1.14 | -3.5 | 49.9 | 2.23 | 2 | -2 |
| GRIN2B | 788 | E | 62.9138 | 9.008751911 | 0.102362888 | 4.181135016 | 4.94 | -1.14 | -3.5 | 49.9 | 2.23 | 2 | -2 |
| GRIN2B | 789 | M | 60.4886 | 10.15723535 | 0.120829794 | 6.61687339 | NA | NA | NA | NA | NA | NA | NA |
| GRIN2B | 790 | E | 57.7594 | 7.42477232 | 0.076893472 | 4.068529484 | 3.55 | -0.11 | 1.2 | 1.72 | 0.11 | 0 | 0 |
| GRIN2B | 791 | E | 59.85913333 | 5.280717186 | 0.042418388 | 1.757888976 | 3.55 | -0.11 | 1.2 | 1.72 | 0.11 | 0 | 0 |
| GRIN2B | 792 | L | 59.952925 | 7.371836135 | 0.076042291 | 4.117566847 | 3.27 | 0.17 | 1.7 | 1.76 | -0.49 | 0 | 0 |

| | | | | | | | | | | | | | |
|---|---|---|---|---|---|---|---|---|---|---|---|---|---|
| GRIN2B | 793 | E | 54.70957778 | 7.496404538 | 0.078045274 | 4.78800089 | 2.89 | -0.52 | -0.7 | 2.25 | 0.65 | 0 | 0 |
| GRIN2B | 794 | A | 54.3274 | 4.291826418 | 0.026517634 | 1.382015019 | 2.92 | -0.38 | -0.85 | 1.69 | 0.57 | 0 | 0 |
| GRIN2B | 795 | L | 56.67205 | 4.912882148 | 0.036503827 | 2.043982572 | 2.89 | -0.52 | -0.7 | 2.25 | 0.65 | 0 | 0 |
| GRIN2B | 796 | W | 57.41315714 | 7.329037045 | 0.075354108 | 4.452766421 | 2.89 | -0.52 | -0.7 | 2.25 | 0.65 | 0 | 0 |
| GRIN2B | 797 | L | 53.2343 | 9.02778295 | 0.102668895 | 6.346434625 | NA | NA | NA | NA | NA | NA | NA |
| GRIN2B | 798 | T | 50.46194286 | 5.83322698 | 0.051302405 | 2.018938946 | 3.92 | -0.51 | -0.8 | 1.68 | 0.63 | 0 | 0 |
| GRIN2B | 799 | G | 47.10455 | 7.786048035 | 0.082702563 | 4.205875171 | 3.92 | -0.51 | -0.8 | 1.68 | 0.63 | 0 | 0 |
| GRIN2B | 800 | I | 44.2216 | 9.683606043 | 0.113214126 | 4.98576362 | 4.91 | -0.77 | -1.63 | 2.23 | 0.69 | 0 | 0 |
| GRIN2B | 801 | C | 40.7518 | 9.585504942 | 0.111636721 | 4.195952381 | 5.48 | -0.78 | -0.6 | 2.23 | 0.78 | 0 | 0 |
| GRIN2B | 802 | H | 41.1158 | 7.518407345 | 0.078399066 | 2.436901473 | 5.48 | -0.78 | -0.6 | 2.23 | 0.78 | 0 | 0 |
| GRIN2B | 803 | N | 38.527925 | 8.150377169 | 0.088560751 | 1.910205719 | 6.47 | -0.86 | -1.7 | 12.34 | 1.03 | 1 | -1 |
| GRIN2B | 809 | M | 29.1706 | 7.668044927 | 0.080805146 | 1.50776795 | 6.21 | 0.3 | 1.4 | 2.08 | -0.24 | 0 | 0 |
| GRIN2B | 810 | S | 29.0393 | 10.28203156 | 0.12283644 | 4.116081346 | 6.21 | 0.3 | 1.4 | 2.08 | -0.24 | 0 | 0 |
| GRIN2B | 811 | S | 25.0642 | 10.37410415 | 0.12431691 | 8.679078556 | NA | NA | NA | NA | NA | NA | NA |
| GRIN2B | 812 | Q | 23.0028 | 13.61428346 | 0.176416996 | 11.89072501 | NA | NA | NA | NA | NA | NA | NA |
| GRIN2B | 813 | L | 19.7738 | 15.61830269 | 0.20864039 | 12.82426083 | NA | NA | NA | NA | NA | NA | NA |
| GRIN2B | 814 | D | 18.1808 | 17.66015207 | 0.241472069 | 14.86099472 | NA | NA | NA | NA | NA | NA | NA |
| GRIN2B | 815 | I | 14.880175 | 19.97495297 | 0.27869264 | 17.46897292 | NA | NA | NA | NA | NA | NA | NA |
| GRIN2B | 816 | D | 16.7928 | 20.2588009 | 0.28325674 | 17.45291466 | NA | NA | NA | NA | NA | NA | NA |
| GRIN2B | 817 | N | 16.5266 | 16.66418465 | 0.225457527 | 13.8607741 | NA | NA | NA | NA | NA | NA | NA |
| GRIN2B | 818 | M | 12.8748 | 16.00692469 | 0.214889192 | 13.43650426 | NA | NA | NA | NA | NA | NA | NA |
| GRIN2B | 819 | A | 12.1712 | 18.73095291 | 0.258689886 | 16.18090733 | NA | NA | NA | NA | NA | NA | NA |
| GRIN2B | 820 | G | 11.06755 | 17.95001663 | 0.246132912 | 15.33274306 | NA | NA | NA | NA | NA | NA | NA |
| GRIN2B | 821 | V | 8.642371429 | 13.93411644 | 0.181559713 | 11.36950696 | NA | NA | NA | NA | NA | NA | NA |
| GRIN2B | 822 | F | 7.665981818 | 17.12816598 | 0.232918061 | 14.69809213 | NA | NA | NA | NA | NA | NA | NA |
| GRIN2B | 823 | Y | 8.558133333 | 20.50630715 | 0.287236488 | 17.94581022 | NA | NA | NA | NA | NA | NA | NA |
| GRIN2B | 824 | M | 5.845 | 18.43398847 | 0.253914881 | 15.88412128 | NA | NA | NA | NA | NA | NA | NA |
| GRIN2B | 825 | L | 2.534675 | 17.9949938 | 0.246856118 | 15.29907508 | NA | NA | NA | NA | NA | NA | NA |
| GRIN2B | 826 | G | 2.6528 | 21.74821301 | 0.307205569 | 18.96806824 | NA | NA | NA | NA | NA | NA | NA |
| GRIN2B | 827 | A | 2.8468 | 23.58191351 | 0.336690342 | 21.02150706 | NA | NA | NA | NA | NA | NA | NA |
| GRIN2B | 828 | A | 0.8594 | 21.00167455 | 0.295201691 | 18.3760912 | NA | NA | NA | NA | NA | NA | NA |
| GRIN2B | 829 | M | -1.2144 | 22.95965202 | 0.326684761 | 21.58600138 | NA | NA | NA | NA | NA | NA | NA |
| GRIN2B | 830 | A | -0.9556 | 25.71803884 | 0.371037921 | 24.34307218 | NA | NA | NA | NA | NA | NA | NA |

| | | | | | | | | | | | | | |
|---|---|---|---|---|---|---|---|---|---|---|---|---|---|
| GRIN2B | 831 | L | -1.8972 | 26.35235728 | 0.38123737 | 25.07573569 | NA | NA | NA | NA | NA | NA | NA |
| GRIN2B | 832 | S | -4.732633333 | 24.3007624 | 0.348248989 | 23.02215043 | NA | NA | NA | NA | NA | NA | NA |
| GRIN2B | 833 | L | -5.7968 | 27.29809938 | 0.39644432 | 26.01730556 | NA | NA | NA | NA | NA | NA | NA |
| GRIN2B | 834 | I | -5.4872 | 29.92952056 | 0.438755951 | 28.61808039 | NA | NA | NA | NA | NA | NA | NA |
| GRIN2B | 835 | T | -7.241371429 | 28.95279762 | 0.423050848 | 27.58266457 | NA | NA | NA | NA | NA | NA | NA |
| GRIN2B | 836 | F | -10.1854 | 29.49946942 | 0.431840993 | 28.31850158 | NA | NA | NA | NA | NA | NA | NA |
| GRIN2B | 837 | I | -9.9142 | 32.2992472 | 0.476859694 | 31.08036206 | NA | NA | NA | NA | NA | NA | NA |
